# A Co-Fractionation Mass Spectrometry-based Prediction of Protein Complex Assemblies in the Developing Rice Aleurone-subaleurone

**DOI:** 10.1101/2021.06.16.448567

**Authors:** Youngwoo Lee, Thomas W. Okita, Daniel B. Szymanski

## Abstract

Multiprotein complexes execute and coordinate diverse cellular processes such as organelle biogenesis, vesicle trafficking, cell signaling, and metabolism. Knowledge about their composition and localization provides useful clues about the mechanisms of cellular homeostasis and systems-level control. This is of great biological importance and practical significance in heterotrophic rice endosperm and aleurone-subaleurone tissues that are a primary source of seed vitamins and stored energy. Dozens of protein complexes have been implicated in the synthesis, transport, and storage of seed proteins, lipids, vitamins, and minerals. Mutations in protein complexes that control RNA transport result in aberrant endosperm with shrunken and floury phenotypes, significantly reducing seed yield and quality. The purpose of this research is to broadly predict protein complex composition in the aleurone-subaleurone layers of developing rice seeds using co-fractionation mass spectrometry. Following orthogonal chromatographic separations of biological replicates, thousands of protein elution profiles were subjected to distance-based clustering to enable a large-scale determination of multimerization state and complex composition. Predictions included evolutionarily conserved proteins across diverse functional categories, including novel heteromeric RNA binding protein complexes that influence seed quality. This effective and open-ended proteomics pipeline provides useful clues about systems-level controls in the early stage of rice seed development.

**One-sentence summary:** A co-fractionation mass spectrometry pipeline predicts compositions of cytosolic protein complexes present in the early stages of rice seed development.

## Introduction

Rice is one of the three world’s major food crops (FAO, 2003), and its aleurone tissue and endosperm are an essential source of human nutrition. The endosperm development consists of coenocytic nuclear division, cytokinesis, and differentiation into the starchy endosperm and aleurone layer (Olsen, 2004; Wu et al., 2016). Like most cereal grains, rice has a single cell layer aleurone differentiated from the endosperm epidermis at 5 days after flowering (DAF) and is completely formed at 7 DAF (Krishnan and Dayanandan, 2003; Wu et al., 2016). During the grain filling stage, the aleurone and subaleurone tissues store proteins, lipids, vitamins, and minerals, whereas the endosperm mainly accumulates starch (Krishnan and Dayanandan, 2003; Becraft and Yi, 2010; Wu et al., 2016). The subaleurone tissue consists of 4-6 cell layers that are biochemically distinct from the rest of the starchy endosperm. These cells are much lower in starch content and accumulate the bulk of storage proteins, especially glutelins and prolamines. The aleurone also protects the endosperm during the grain filling stage, and its induced desiccation tolerance maintains stored starch in the endosperm and ensures seed survival (Fath et al., 2000; Young and Gallie, 2000; Bethke et al., 2001). Given the importance of the aleurone-subaleurone for human nutrition and seed development, we focused on the tissue in this high-throughput analysis of protein complex formation and composition using quantitative proteomics.

Data on protein multimerization and binding partner identity are some of the most valuable to analyze regulatory pathways and interactions among them. Genetic data indicate that subunits of complex and interacting protein complexes have similar phenotypes and function as part of signaling input-output modules (Yanagisawa et al., 2018). Protein multimerization is the cornerstone of cellular complexity, enabling mechanical tasks and the flow of genetic information that could never be achieved by individual proteins (Alberts, 1998; Marsh and Teichmann, 2015). For example, multiprotein complexes coordinate gene expression (Burd and Dreyfuss, 1994), organelle biogenesis (Li et al., 2019), vesicle trafficking (Kaksonen et al., 2005), metabolism (Weng et al., 2012), and signal transduction (Basu et al., 2008). Recent proteomic profiling studies indicate that more than 1/3 of all proteins exist as parts of stable protein complexes (Aryal et al., 2014; Aryal et al., 2017; McBride et al., 2017; McBride et al., 2019; Lee and Szymanski, 2021). Because of the widespread occurrence and crucial roles of protein multimerization, numerous large-scale projects to characterize protein-protein interaction networks have been conducted by yeast two-hybrid system (Van Leene et al., 2007; Arabidopsis Interactome Mapping Consortium, 2011; Jones et al., 2014). However, comparisons of global protein-protein interaction studies have shown that combinations of approaches are needed. Overlap among interactome dataset types is low, and technical bias and limitations of individual methods determine the extent to which they capture the full spectrum of physical interactions that vary greatly in terms of affinity, cell type, or subcellular localization (Wodak et al., 2009; Rattray and Foster, 2019; Salas et al., 2020).

Protein correlation profiling, also known as co-fractionation mass spectrometry (CF-MS), is gaining momentum as an effective approach to broadly analyze the multimerization behaviors of endogenous proteins (Kristensen et al., 2012; Aryal et al., 2014; McWhite et al., 2020; Salas et al., 2020). The CF-MS has the major advantage that it analyzes native protein complexes in an unbiased way with no requirement for transformation or gene cloning. It is a “guilt by association” protein chromatography method based on the expected indistinguishable elution profiles of subunits of stable protein complexes regardless of the separation method. This MS-based profiling of cell lysates separated by size exclusion chromatography (SEC) provides broad information on whether or not a protein is likely to multimerize (Aryal et al., 2014; Aryal et al., 2017; Gilbert and Schulze, 2019), form distinct complexes at different subcellular locations (McBride et al., 2017), or evolve unique properties in diverse species (Lee and Szymanski, 2021). Chance co-elution of non-interacting proteins is a confounding factor and increases the rate of false positive predictions. One approach to improve accuracy is to carry out multiple fractionations and use presumed “golden standards” of evolutionarily conserved protein complexes in combination with machine learning-based predictions (Havugimana et al., 2012; Wan et al., 2015; McWhite et al., 2020). There is uncertainty about the extent to which golden standard complexes exist as stable, fully assembled complexes (Aryal et al., 2014; McBride et al., 2017; Lee and Szymanski, 2021); however, the increasing number of validated known complexes will further empower these approaches. Our group predicts multimerization and complex composition based on experimental profile data alone (Aryal et al., 2014; Aryal et al., 2017; McBride et al., 2017; McBride et al., 2019). In this workflow, biological replicates and automated peak detection algorithms (McBride et al., 2017; McBride et al., 2019) are used to remove unreliable profiles. The filtered data is used in distance-based clustering analyses to group proteins with the most similar elution profiles. This overall approach has been validated repeatedly using known complexes or subcomplexes, co-immunoprecipitation, and profiling a mutant in which expression of a predicted novel subunit was knocked out (Aryal et al., 2014; McBride et al., 2017; McBride et al., 2019). This approach generates a valuable data resource for the community to develop and test hypotheses about protein interactions and systems-level controls.

Here we adopt this technology to a dissected tissue with great developmental and agronomic importance, the aleurone-subaleurone cell layers of developing rice seeds. This pipeline begins with 1 mg of soluble protein extracts whereupon 2,610 proteins were reproducibly profiled across two SEC and two ion exchange chromatography (IEX) column separations. This combination of orthogonal separation strategies greatly reduces chance coelution and can generate reliable protein complex composition predictions. The clustering analysis and systematic classification method predicted 771 protein complexes (proteins of your interest can be searched in Supplemental Table S3A), 170 of which correspond to self-interacting proteins. Numerous novel protein complexes involved in translation, cellular homeostasis, and tissue-specific physiology were predicted. These protein complexes could play an essential role in determining the fate of aleurone, regulating endosperm development, and the biosynthesis of seed reserves. These new findings will facilitate a deeper understanding of seed development and quality.

## Results and Discussion

### A high-quality co-fractionation mass spectrometry dataset

The CF-MS approach, coupled with biological replicates of SEC and IEX fractionations, was carried out to profile endogenous protein complexes in the aleurone-subaleurone layers of developing rice seeds (Figure 1). Expressed proteome in the developing seeds at 10 DAF was resolved under native condition through SEC and orthogonal IEX separations to reduce chance co-elution and false-positive predictions (McBride et al., 2019). The biological duplicates of 24 SEC and 71 IEX fractions were subjected to label-free shotgun proteomics to obtain abundance profiles of thousands of resolved proteins. The abundance profiles of all peptides (Supplemental Table S1, A and B) and proteins (Supplemental Table S1, C and D) are provided. These elution profiles were subjected to the Gaussian fitting to smooth the raw data and deconvolve multiple peaks, which likely reflect an individual protein being present in multiple protein complexes (McBride et al., 2017). With 1 % of false discovery rate (FDR) for both the peptide and protein levels, 3746 and 3633 endogenous rice aleurone-subaleurone proteins were reproducibly identified from SEC and IEX fractions, respectively (Figure 2, A and B). Overall, the SEC and IEX profile data were highly reproducible based on Pearson correlation coefficients (PCC) falling into diagonal across SEC and IEX fractions (Figure 2, C and D). The elution peak locations were also reproducible between replicates (Figure 2, E and F). When the peak locations for individual proteins were compared, about 88 % and 73 % of the total protein peaks fell within two SEC fractions (representing about 40 % size difference) and within four IEX fractions. The reproducible peak locations and their apparent mass (*M*_app_) values are provided (Supplemental Table S2, A and B). To eliminate noisy profile data, the set of 2,610 proteins identified with reproducible peaks between both SEC and IEX replicates was chosen for the protein complex prediction (Figure 2G).

**Figure 1.**
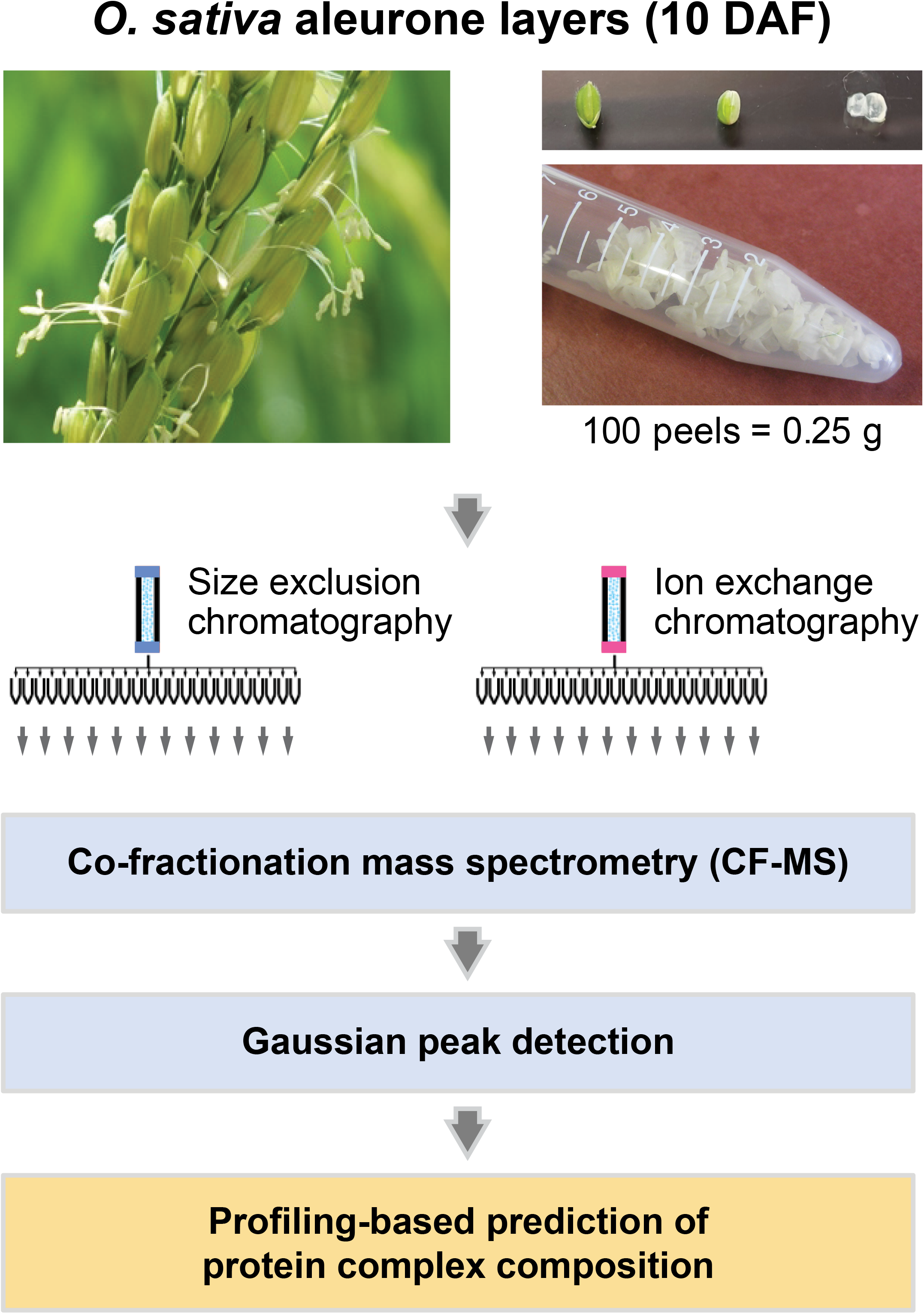
The CF-MS pipeline composed of SEC and IEX separations to predict protein complex composition in rice seed aleurone-subaleurone layers. Soluble cell fraction enriched from the isolated aleurone-subaleurone layers at 10 DAF is separated on a sizing column and a mixed-bed ion exchange column under a non-denaturing condition. Each column fraction is analyzed by LC-MS/MS for protein identification and quantification. Gaussian fitting is applied to choose reproducible peaks in SEC and IEX datasets. *M*_app_ values are calculated for the reproducible SEC peaks. Profile-based clustering analysis is conducted to predict protein complex composition using concatenated SEC and IEX datasets.

**Figure 2.**
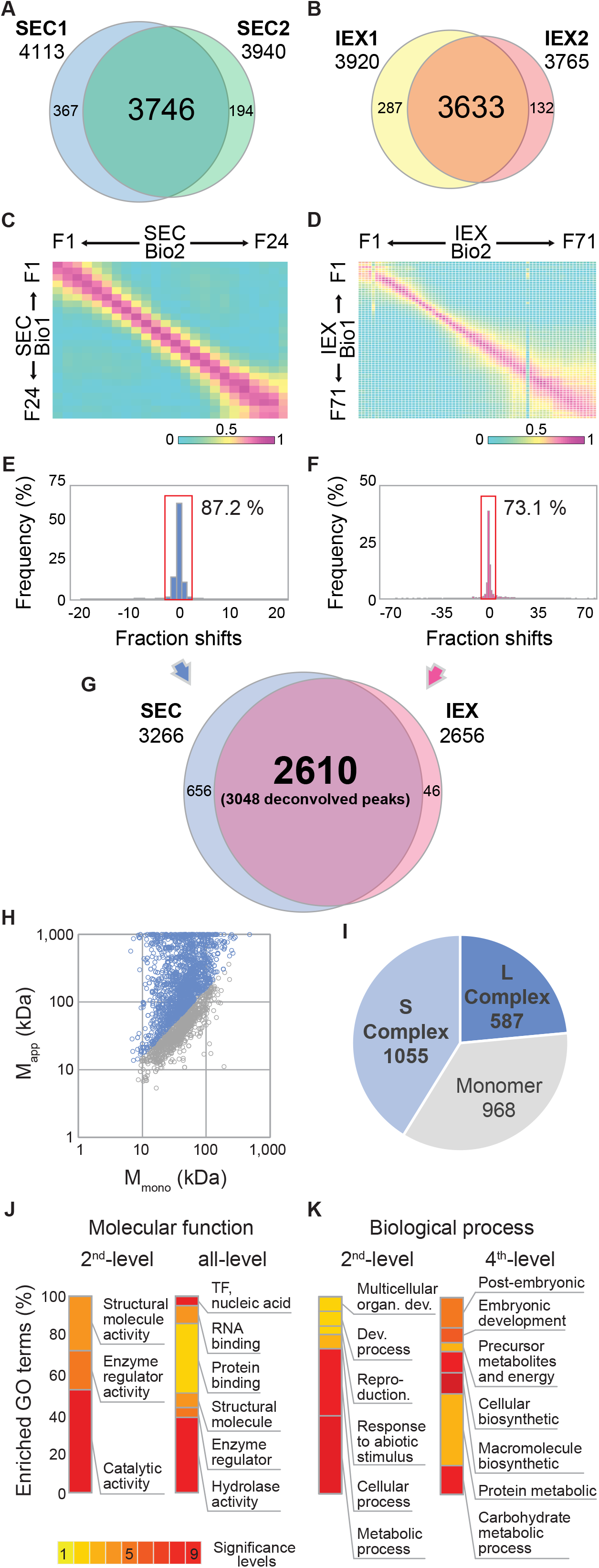
The CF-MS pipeline generated a highly reproducible dataset of protein complexes on rice aleurone-subaleurone proteome. **A to G,** Reproducibility of CF-MS datasets. Protein overlap between two biological replicates in SEC (A) and IEX (B). PCC of protein abundances across SEC (C) and IEX (D) fractions. Column fraction shifts were measured by the distances between the elution peaks of the replicates for all proteins in SEC (E) and IEX (F) fractions. Reproducible proteins present within 2 or 4-fraction shifts were boxed with the ratio of reproducibility, respectively. (G) Reproducible proteins overlapped between SEC and IEX datasets were selected for the prediction of protein complex composition. **H and I,** Protein complexes are common in the rice aleurone-subaleurone proteome. The quantification of protein multimerization was performed using *M*_app_ and *R*_app_. (H) A scatter plot of the *M*_mono_ and *M*_app_ of the reproducible proteins. Open circles rendered with light blue are proteins with *R*_app_ ≥ 1.6, while those with grey are proteins with *R*_app_ < 1.6. (I) Distribution of protein multimeric states. Proteins are classified as M (monomer: 0.62 ≤ *R*_app_ < 1.6), S (small complex: 1.6 ≤ *R*_app_ < 5), or L (large complex: *R*_app_ ≥ 5). **J and K,** Aleuronesubaleurone proteome shows functions in diverse processes. Overrepresented GO terms are highlighted according to the significant levels. Molecular function GO terms (J) and biological process GO terms (K) are visualized. Full GO analysis results are provided in Supplemental Figure S1.

### Protein multimerization in rice aleurone-subaleurone proteome

To estimate the proportion of proteins eluting at larger than expected masses, the distribution of expected monomeric masses (*M*_mono_) was compared with the distribution of *M*_app_ (Figure 2H). The scatter plot of *M*_app_ and *M*_mono_ was strongly skewed toward elevated *M*_app_ values, suggesting widespread multimerization. Using *R*_app_ (*R*_app_ = *M*_app_ / *M*_mono_) as a diagnostic for multimerization of individual proteins, more than half of the proteins fell into this class with 37.1 % detected as relatively small complexes and 22.5 % as large with *R*_app_ values of ≥ 5 (Figure 2I). Similar distributions of stable protein complexes were reported in different tissues and plant species (Aryal et al., 2014; McBride et al., 2017; McBride et al., 2019), although the types and sizes of complexes can vary within an orthologous group (Lee and Szymanski, 2021).

To gain insights into the types of proteins in the rice aleurone-subaleurone proteome, gene ontology (GO) enrichment analysis revealed 70 significantly enriched terms at a 5% FDR (Figure 2, J and K; Supplemental Figure S1). At molecular function GO hierarchy, enzyme activities including “catalytic activity” and “enzyme regulator activity” were overrepresented in the developing aleurone-subaleurone layer cells, reflecting many interesting enzymes, lipid binding, RNA binding, and signaling proteins present in the tissue (Figure 2J). For biological process, four GO terms in “cellular biosynthetic process,” “macromolecule biosynthetic process,” “carbohydrate metabolic process,” and “response to abiotic stimulus” were the most highly enriched (Figure 2K). These overrepresenting terms reflect that proteins in the aleurone-subaleurone layers promote the rapid biosynthesis of carbohydrates, proteins, and other seed storage substances. A similar pattern of GO term distribution was reported in a gene expression analysis for the developing aleurone of wheat (Gillies et al., 2012). Moreover, two other highly enriched GO terms include proteins that regulate “embryonic development” and “post-embryonic development” (Supplemental Figure S1). During wheat seed development, these terms were highly enriched in aleurone layers compared to starchy endosperm cells (Gillies et al., 2012). These analyses indicate that a broad range of protein types with aleurone-subaleurone-enriched functions is captured in our dataset.

### Profiling-based clustering and protein complex composition prediction

To begin distance-based clustering analysis, elution profiles with multiple peaks were deconvolved into separate profiles as previously described (McBride et al., 2019). Three hundred seventy-four proteins had 2 IEX peaks, and 32 had three, and these multipeak proteins were annotated with a peak number suffix on the locus ID. Among these 406 multiple peak proteins, 57 had multiple SEC peaks. These proteins were assigned the same SEC peak with the largest *M*_app_, enabling a protein to reside in multiple distinct protein complexes. In the end, the four SEC and IEX elution profile datasets were concatenated, and a total of 3,048 reproducible profile entries were used for further analysis.

As the intrinsic test of the enhanced resolution afforded by combining SEC and IEX, the mean distances between clusters were analyzed as a function of the numbers of clusters in the individual and combined datasets (Figure 3). When the elution profiles of the proteins in a cluster are similar to each other, the average distance among them is low. In the box plot of mean distance within a cluster for SEC or IEX alone, the third quartile approached zero at ~530 or ~630 clusters, respectively (Figure 3, A and B). The combination of SEC and IEX fractionations increased the resolution so that the third quartile approached zero at 1,000 clusters (Figure 3C). These data demonstrate the utility of the combined SEC and IEX datasets and indicate that the dendrogram loses resolution beyond ~1000 clusters. As a representative clustering result, dendrograms at a 1,000 cluster-cut were plotted in Supplemental Data Set 1, and a heatmap with the color code representing the relative protein abundance is shown in Supplemental Figure S2.

**Figure 3.**
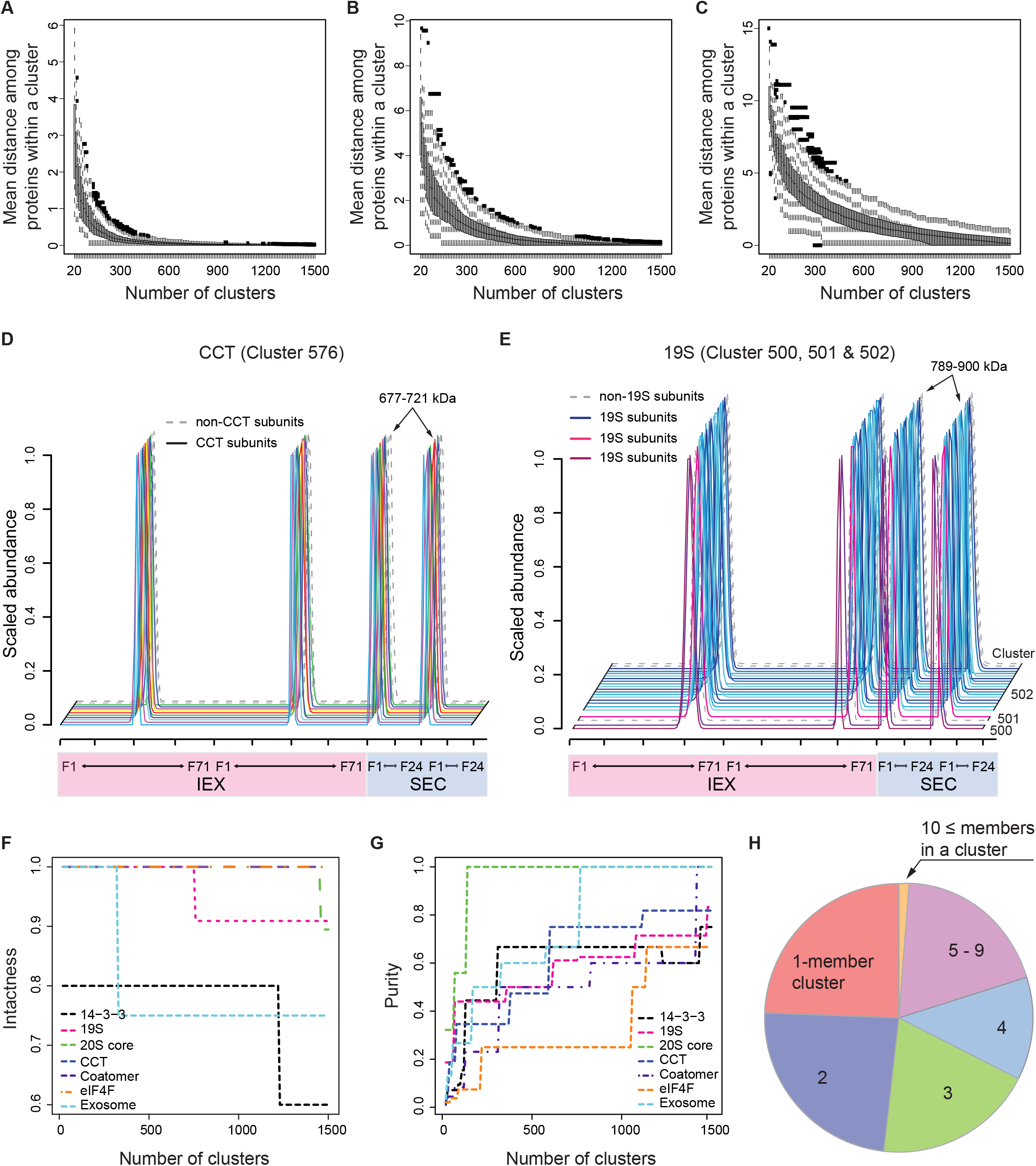
The strong resolving power of the clustering analysis evaluated by the intrinsic and extrinsic tests. **A to C,** Intrinsic tests evaluated the resolving power for independent and concatenated protein profile datasets. Box-and-whisker plots visualize the distributions of average distances of elution profiles within the clusters in the SEC only (A), IEX only (B), and combined SEC and IEX (C) datasets as a function of ascending cluster number. **D and E,** The concatenated profiles of known protein complexes are visualized over the two IEX (pink) and two SEC (light blue) separations. Solid lines indicate elution profiles of CCT (D) and 19S proteasome complex (E) subunits. Dashed lines represent elution profiles of proteins that are not subunits of the known complexes. Numbers shown above the SEC peaks show *M*_app_ values of subunits in a given complex. Numbers on the right-handed side of profiles are cluster numbers when multiple clusters are plotted. **F and G,** Extrinsic tests evaluated the resolving power for the independent and combined protein profile datasets as a function of ascending cluster number. (F) The intactness of known complexes was measured. (G) The purities of the known complexes were evaluated. Individual intactness and purity plots are provided in Supplemental Figure S3. **H,** The distribution of cluster size in the resulting clustering. The pie chart shows the distribution of predicted protein complexes by the number of members in each cluster.

### Validation of protein complex prediction using known complexes

In the concatenated SEC and IEX separations, subunits of stable known complexes should co-elute and can serve as a validation of the prediction. Known chaperonin containing TCP1 folding complex (CCT) subunits co-eluted and were assigned to cluster 576 with *M*_app_ of 677-721 kDa corresponding to the fully assembled complex (Figure 3D). Another validating known is 19S proteasome cap complex. Seventeen different 19S subunits with *M*_app_ of 789-900 kDa were also clustered into cluster 502 (14 of 19S subunits / 16 of cluster members) and neighboring clusters 500 (2 / 2) and 501 (1 / 2) (Figure 3E). The 19S subunits in clusters 500 and 501 also had similar *M*_app_ to other subunits, but a slight shift in the IEX fractions assigned these three subunits into the adjacent clusters. Further discussion on the CCT and 19S complexes is provided in Supplemental Data Set 2.

To further inform the decision of where to divide the dendrogram to generate a specific protein complex prediction using extrinsic data, the intactness and purity of seven known complexes were evaluated as a function of increasing cluster number (Figure 3, F and G). The intactness of exosome and 19S proteasome subunits dropped in intactness at around 300 and 750 clusters but remained stable until 1,500 clusters. The 14-3-3 had stable intactness until at around 1,250. Coatomer, CCT, eIFs, and 20S proteasome maintained perfect intactness until around 1,450 clusters. Because purity can be utilized to detect false positives (McBride et al., 2019), the purity of knowns was observed as cluster numbers increased to 1,400 clusters. The purity of most knowns became stable after 1,100 clusters. The intactness and purity indices indicated that splitting the dendrogram into 1,000 clusters based on the intrinsic resolution test would be appropriate. Approximately 20 % of clusters were predicted large complexes with 5 to 19 subunits, about 55 % of clusters were 2 to 4-meric, and 224 proteins were assigned into single-entry clusters (Figure 3H). The reduced number of singletons compared to the predicted number of 400 (40 % monomeric × 1000 clusters, estimation from Figure 2I) indicates common occurrences of false positives. The baseline complex composition prediction from this study is provided in Supplemental Table S3A, with the caveat that true interactors may be located in nearby clusters, and clusters will also contain false positives due to chance co-elution. A classification system to guide the user is provided below, and we encourage readers to comment on the paper online as clusters are validated or refuted over time.

### Protein complex heterogeneities and cross-validation using multiple peak entries

A small number of proteins displayed multiple peaks, and this could reflect the existence of heteromeric complexes with differing assembly states on the IEX column. It is also possible that functionally interchangeable orthologs/paralogs could assemble into homomers or ortholog-selective multimers with differing subunit stoichiometries and resolvable peaks on the IEX column. In the scenarios above, an accurate clustering result would place interacting proteins within the same cluster in two separate instances. There were 8 cases in which a pair of multiple peak proteins that co-occurred in two distinct clusters (Supplemental Table S3B). In the first three cases listed, two proteins in ribosomal protein S2 cluster, three in a DnaK family cluster, and two in another DnaK family cluster had two IEX peaks and two SEC peaks neither of which correspond to an expected monomer. This could be explained by protein assembly into two complexes with partial subunit overlap that were resolved on both columns, supporting true protein-protein interactions between these multiple peak proteins.

Other instances in which pairs of proteins had a single high mass SEC peak and multiple IEX peaks could reflect a complex that partially dissociated during high salt elution. Two proteins of the 19S proteasome cap complex [RPN12 (LOC_Os07g25420.1) and RPN9 (LOC_Os01g32800.2 and LOC_Os03g11570.1) in cluster 502] were also assigned into cluster 518. A solved structure (Lander et al., 2012), native MS analysis (Sharon et al., 2006) and a CF-MS prediction (Drew et al., 2017) did not show a direct interaction between RPN9 and RPN12. However, RPN9 and RPN12 possess significant sequence homologies with two interacting subunits of the COP9 signalosome, CSN7 and CSN8 subunits, respectively (Kapelari et al., 2000; Fu et al., 2001), and the adjacency of RPN9 and RPN12 subunits in the lid structure could enable a stable physical interaction in some species (da Fonseca et al., 2012; Lander et al., 2012). In another example, two eIFiso4F subunits, showed a similar behavior as the 19S pair did. A large subunit eIFiso4G (LOC_Os04g42140.1) and a small cap-binding protein eIFiso4E (LOC_Os10g32970.1) co-occurred in two clusters with a single SEC peak. This result is consistent with their known direct physical interaction (Mayberry et al., 2011), further supporting the accuracy of the predictions in this study.

Two homologous pyruvate kinases (OsPKpα1; LOC_Os07g08340.1 and OsPKpß2; LOC_Os10g42100.1) had two IEX peaks, a single SEC peak, and co-occurred in two clusters. Mammalian pyruvate kinases are known homotetramers (Larsen et al., 1998; Christofk et al., 2008), while plant orthologs are obligate heterooligomers of ancient paralogs (Negm et al., 1995; Andre et al., 2007; Cai et al., 2018), and the clustering pattern here could reflect partial disassembly of a multimeric form during high salt elution. Arabidopsis 14-3-3 paralogs showed IEX-resolved heteromerization patterns (McBride et al., 2019). Taken together these coclustering pairs provide clear cross validations for protein complex predictions.

### Cross-species comparisons with published co-fractionation MS datasets

We made cross-comparisons of our rice predictions with previously published CF-MS datasets from McWhite et al. (2020) and Arabidopsis leaf data from McBride et al. (2019). The clustering result here and that of McBride operated on protein groups defined by unique peptides. Both predictions were based on the profiled data from ~ 200 fractions generated from duplicates of SEC and IEX separations. The McWhite prediction was generated from ~2,000 fractions from 13 plant species and diverse tissues using four different separation methods. Orthologs and paralogs were merged into an averaged profile of a single ortholog group. Ortholog averaging obscures the commonly observed multimerization variability among paralogs and orthologs (Lee and Szymanski, 2021). The protein coverage of our rice dataset was three times greater than previous studies because of the increased LC/MS sensitivity and the use of protein groups. The key parameters for the three CF-MS studies are summarized in Supplemental Table S4A. The overlapping protein hits discussed here are summarized in Supplemental Table S4, B and C.

Differences in protein definitions and complexity make direct comparisons among studies difficult. In this comparison, we looked for two distinct protein/ortholog group members present within a single cluster in both the rice and the comparison studies. If the interaction was reproduced in both studies, they should fall within a single cluster in both instances. There were 31 rice pairs that were also present in the McBride Arabidopsis dataset, and 29 % fell into a single cluster in both cases. There were 149 rice pairs that were in the McWhite Arabidopsis dataset, and 24 % fell into a single cluster in both cases. Of the 144 rice pairs that were in the McWhite rice prediction, 25% co-occurred in the same cluster. Similar results are expected with the McWhite prediction regardless of species due to the ortholog merging. Pairs present across all three datasets were subunits in 20S, 19S, CCT, and coatomer complexes, showing highly conserved protein-protein interactions. The McBride prediction assigned subunits of exon junction complex, CCT, and HSP70 into a single cluster, while these complexes were resolved into three distinct clusters in our rice prediction and in the McWhite datasets. Our clustering had more prediction resolution and protein coverage than both published CF-MS datasets, suggesting a better overall complex prediction.

### Systematic classification of rice protein complex prediction

A systematic cluster classification was adopted based on the sum of *M*_mono_ values of all proteins within one cluster (*M*_calc_) and the measured *M*_app_ (Figure 4; Supplemental Table S3A). A reliable class contains proteins in the classes “homomer” and “possible homomer or heteromer / high subunit stoichiometry” categories. These proteins had elution profiles that placed them into small clusters of one to three proteins, yet on the SEC column these proteins had very large *M*_app_ values (Figure 4A area highlighted as light yellow). These data suggest a high subunit stoichiometry and/or homomers. Known homomers with *M*_app_ values corresponding to their expected stoichiometries and lots of novel homomers were identified from the single-entry clusters (Figure 4, A–G). An IAA-amino acid hydrolase ILR1-like 6 (ILL6) was assigned into a single-member cluster with R_app_ of 13.7 (Figure 4F). The ILL6 was identified as a member of *Arabidopsis* amidohydrolase family, which hydrolyzes IAA conjugates, but has very little activity on IAA-aa *in vitro* (LeClere et al., 2002; Zhang et al., 2016). The ILL6 was also characterized to have a catalytic activity of jasmonoyl-isoleucine conjugate, a bioactive form of JA, cleaving the JA-Ile signal upon wounding (Widemann et al., 2013). The ILL6 potentially has cross-talking ability on both IAA and JA hormone signaling pathways, but mainly on the JA signaling pathway (Zhang et al., 2016). Perhaps the ILL6 multimerization might imply a sensing mechanism that switch form IAA to JA hormone signaling during rice seed development in the aleurone-subaleurone cells. Discussion of a subset of these interesting homomers, GCN5-related N-acetyltransferases (GNAT) and autophagy protein 5 (ATG5), is provided in Supplemental Data Set 2. This systematic, reliable classification method of the prediction has the power to distinguish self-interaction proteins from single-entry clusters that resemble monomers in the hierarchical clustering.

**Figure 4.**
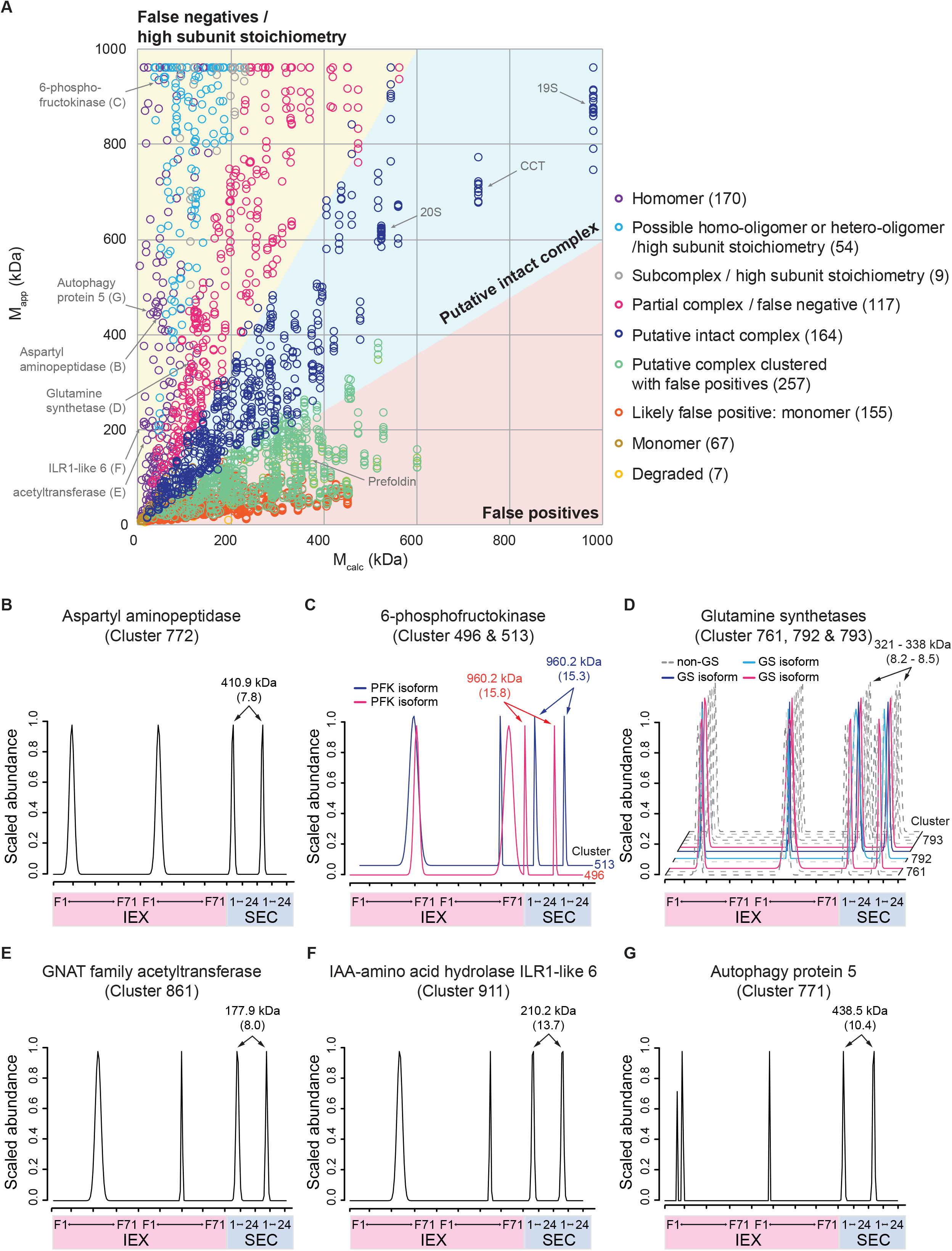
The systematic classification of the resulting clustering analysis assessed the stoichiometry of subunits in the predicted protein complexes. **A,** The clustering analysis was evaluated by comparing *M*_calc_ and *M*_app_. The most reliable predictions are present in the light blue area, where there are less than 2-fold size differences between *M*_calc_ and *M*_app_. Other reliable predictions are on the light yellow area where *M*_app_ is 2-fold greater than *M*_calc_, including “false negatives” and “proteins with high subunit stoichiometry.” Proteins present in “putative complexes with false positives” in the clustering were saved from the false positives in the light pink region where *M*_calc_ is 2-fold greater than *M*_app_. Number in parenthesis next to each category indicates number of clusters that belong to the category. **B to G,** Many of self-interacting homomers were assigned into single entry clusters and classified as homomers. B to D, The profiles of known homomers: Aspartyl aminopeptidase (B), 6-phosphofructokinase isoforms (C), and glutamine synthetase isoforms (D). Dashed lines represent elution profiles of proteins that are not subunits of the known complexes. E to G, The profiles of novel homomers: GNAT family acetyltransferase (E), IAA-amino acid hydrolase ILR1-like 6 (F), and ATG5 (G). Profiles of homomers are visualized over the combined two IEX (pink) and two SEC (light blue) separations. Numbers shown above the SEC peaks show *M*_app_ values of homomers. Numbers in parenthesis indicate *R*_app_ values of homomers. Numbers on the right-handed side of profiles are cluster numbers when multiple clusters are plotted.

Under the assumption of 1:1 subunit stoichiometries, another reliable predicted class is “putative intact complex” because the ratios of *M*_calc_ to *M*_app_ for these proteins are close to 1:1. One hundred sixty-four protein complexes fell into an interval along this diagonal (Figure 4A graph sector highlighted as light blue; Supplemental Table S3A). As an example of the “putative intact complex” class, three subunits of the VPS35-VPS29-VPS26 trimeric cargo recognition core complex were assigned into cluster 917 with *M*_app_ of 213-231 kDa (Figure 5B). *M*_calc_ of the complex was present within 40 % of *M*_app_ of the VPS cluster. This trimeric complex assembles with a membrane-associated sorting nexin dimer subcomplex into a retromer complex, functioning in retrograde transport from endosome to Golgi apparatus (Seaman et al., 1998). The ratio of *M*_calc_ to *M*_app_ and the known complex assembly support reliability of this clustering analysis. In Supplemental Data Set 2, other examples of known and novel heteromeric complexes in Figure 5 are discussed.

**Figure 5.**
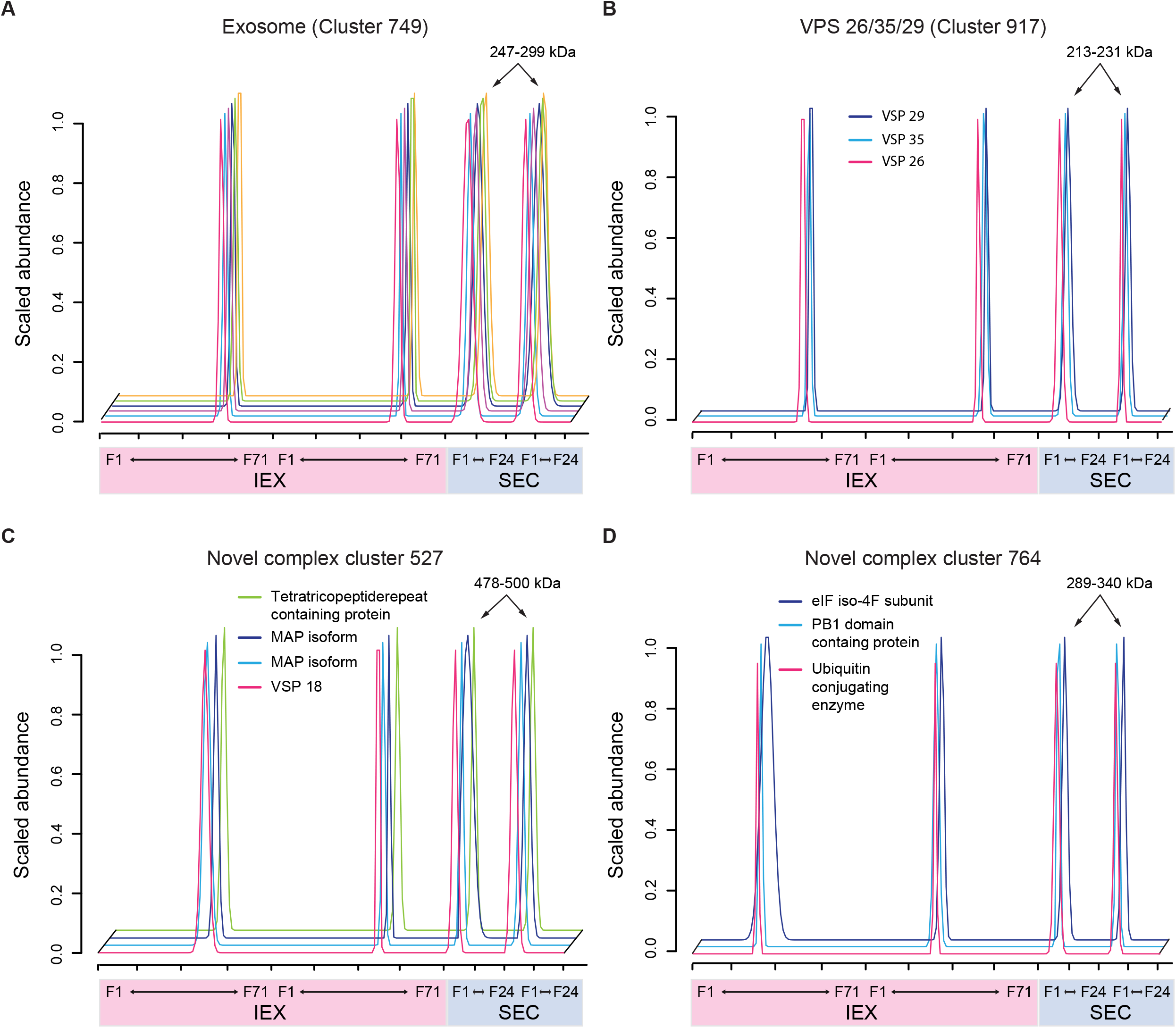
The elution profiling-based clustering result placed known or novel protein complex subunits into the “putative intact complex” clusters. The profiles of known and novel protein complexes are overlapped each other across two IEX (pink) and two SEC (light blue) separations. The clusters were classified as “putative intact complexes.”**A and B,** Known protein complexes. Elution profiles of five exosome subunits (A) and a heterotrimeric VPS complex (B) are visualized. **C and D,** Novel protein complexes. Elution profiles of a MAP containing heterotetrameric complex (C) and a heterotrimeric novel complex (D). Numbers shown above the SEC peaks show *M*_app_ values of subunits in a given complex.

About 22 % of the rice aleurone-subaleurone protein clusters were classified into either “homomer” and “possible homomer or heteromer/high subunit stoichiometry.” About 17 % fell into the “putative intact complex” category, and 12.6 % of the clusters were present in “partial complex/false negatives” and “subcomplex or high subunit stoichiometry.” Collectively about 51 % of the clusters were annotated as reliable complex prediction classes (Figure 4A; Supplemental Table S3C). The least reliable clusters contain false positives due to co-elution by chance (Figure 4A light pink area). About 26 % of the clusters were in “putative complex clustered with false positives,” and about 23 % of the proteins were filtered from clusters where they were as “likely false positive: monomer,” “monomeric,” and “degraded” in this clustering analysis. A total of 657 out of 3,048 profiles (or 229 out of 1,000 clusters) were filtered out as monomeric proteins from this clustering, and the rest of them (70 % of total profiles) were predicted in 771 different protein complexes with different reliabilities.

### Predicted subunits of RPB-associated complexes

During the grain filling stage, putative *trans-acting* factors that recognize *cis*-acting elements on mRNAs of storage proteins (two major rice storage proteins: glutelin and prolamine) maintain the restricted transportation of messenger ribonucleoprotein (mRNP) complexes to specific subdomains of the ER (Okita et al., 1994; Choi et al., 2000; Crofts et al., 2004; Washida et al., 2012; Tian et al., 2020). In rice, 257 RNA binding proteins (RBPs) were experimentally identified as putative *trans*-acting factors expressed from at least 221 distinct genes (Hamada et al., 2003; Doroshenk et al., 2009; Morris et al., 2011). Even though their dynamic regulation of mRNP complex assembly and disassembly is critical for seed productivity, there is less information on whether they are associated with multiprotein complexes or arranged in several smaller complexes. Our profiling identified 133 out of the 250 RBPs with 161 reproducible resolved peaks (27 RBPs exhibited multiple peaks) across the concatenated SEC and IEX separations (Supplemental Table S3A). The clustering analysis and classification strategy assigned 92 cytosolic RBPs (with 114 peaks) into 93 distinct protein complex clusters.

The scaffolding-nuclease protein, Tudor-SN (LOC_Os02g32350.2), a central player in RNA storage and processing (Sami-Subbu et al., 2001; Gutierrez-Beltran et al., 2016), was clustered with chorismate synthase (CS; LOC_Os03g14990.1) and aspartyl/glutamyl-tRNA amidotransferase subunit B (LOC_Os11g34210.2) (Figure 6C). In tobacco BY-2 cells, the same co-elution of Tudor-SN with CS was detected in a combination of ion exchange and gel filtration chromatography (Shan, 2018), and a two-hybrid interaction between rice Tudor-SN and isochorismate synthase (ICS) had been reported (Chou et al., 2017). Chorismate is a key metabolic precursor for salicylic acid (SA), phylloquinone (vitamin K_1_), tetrahydrofolate (vitamin B_9_), and aromatic amino acids (Tzin and Galili, 2010; Maeda and Dudareva, 2012). ICS converts chorismate to isochorismate an route to the synthesis of SA and vitamin K_1_ (Tzin and Galili, 2010). Chorismate and SA metabolism are compartmentalized among the plastid and cytosol with CS and ICS activities thought to reside solely in the plastid (Mousdale and Coggins, 1986; Strawn et al., 2007; Garcion et al., 2008). However, TUDOR-SN functions are cytosolic. Perhaps TUDOR-SN complexes are related to chorismate mediate feedback control of chorismate-dependent metabolites on mRNA processing (Lin et al., 2020). TUDOR-SN complexes may also affect the subcellular localization and activity of CS and other enzymes that dictate the flux of chorismate. In addition to the Tudor-SN complex, a subset of novel RBP complexes in Figure 6, RBP-Q associated putative complex, RBP-T associated putative EJC, and RBP-149 (eIF2α) associated complex, are discussed in Supplemental Data Set 2.

**Figure 6.**
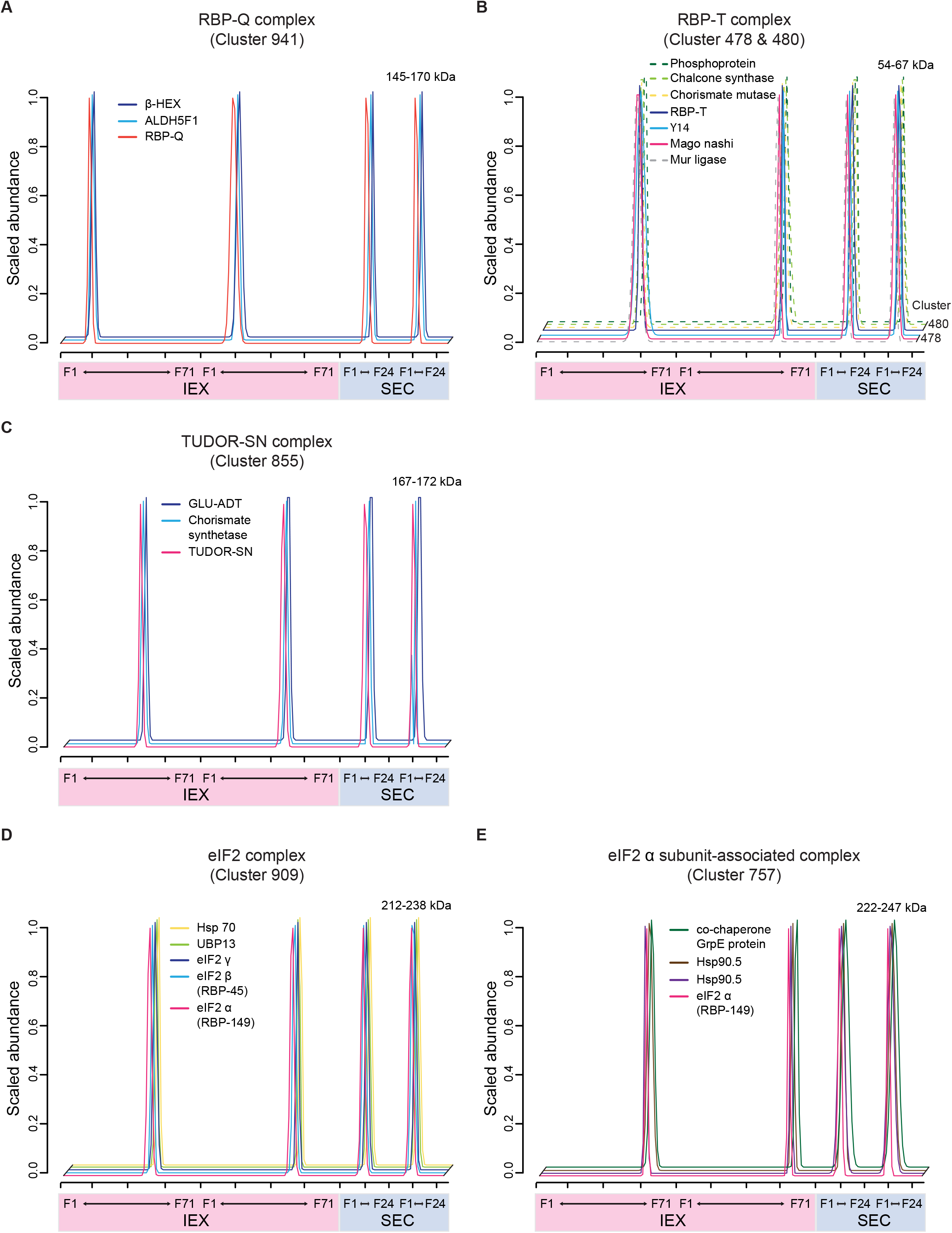
Association of RBPs into novel protein complexes. The elution profiles of subunits in RBP-associated novel protein complexes: RBP-Q associated putative complex (A), RBP-T associated putative EJC (C), TUDOR-SN associated putative complex (C), eIF2 complex (D), and eIF2α associated complex (E). Dashed lines represent elution profiles of proteins that are not subunits of the known complexes. Numbers shown above the SEC peaks show *M*_app_ values of subunits in a given complex. Numbers on the right-handed side of profiles are cluster numbers when multiple clusters are plotted.

## Conclusion

Multiprotein complexes coordinate metabolism and cell/tissue structure in the rice aleurone-subaleurone. Here we provide systems-level protein complex prediction using a robust CF-MS approach that utilizes biological replicates, reproducibility filters, and orthogonal separations that increase reliability. Using a simple classification system, more than 700 novel protein complex predictions were made. Self-interaction was common, and this type of interaction can provide clues about allosteric control (Llorca et al., 2006) and paths to neofunctionalization over evolutionary time scales (Lee and Szymanski, 2021). Predicted novel heteromeric protein complexes are associated with protein translation, metabolism, signaling, and vesicle trafficking, all of which are crucial for seed development and quality. The data provided here can be broadly leveraged by the research community to generate testable hypotheses about the functional relevance of specific protein-protein interactions. This method can also be further developed to analyze how systems of protein complex assemblies change during development or in response to any desired experimental manipulation.

## Materials and methods

### Plant growth conditions and soluble protein extraction

The rice (*Oryza sativa* L. ssp. *japonica*) cultivar Kitaake was grown in a Conviron®E15 growth chamber (Conviron) with a day/night setting of 26/22 °C and 12/12 h. Seed peels containing purified aleurone-subaleurone cell layers were collected as described previously (Yang et al., 2018). Developing seeds at 10-12 days after flowering (DAF) were pooled from panicles from 10 different plants. Seeds were dehulled, cut open, and the outer seed layers (pericarp and nucellus) were stripped away. The inner peels were washed in phosphate buffered saline buffer to remove milky starchy endosperm and embryo. This left a semi-translucent tissue containing primarily aleurone and 4-6 layers of subaleurone cells. About 250 mg of the fresh sub- and aleurone layers were prepared for cell fractionation in this project as described previously (Aryal et al., 2014; Aryal et al., 2017; McBride et al., 2017; McBride et al., 2019). The collected tissue was disrupted by a Polytron homogenizer (Brinkman Instruments) under 1 mL of ice-cold microsome isolation buffer solution [50 mM Hepes/KOH (pH 7.5), 250 mM sorbitol, 50 mM KOAc, 2 mM Mg(OAc)_2_, 1 mM EDTA, 1 mM EGTA, 1 mM dithiothreitol (DTT), 2 mM PMSF and 1 % (v/v) protein inhibitor cocktail (160 mg/mL benzamidine-HCl, 100 mg/mL leupeptin, 12 mg/mL phenanthroline, 0.1 mg/mL pepstatin A, and 0.1 mg/mL aprotinin)]. To remove debris, the homogenate was centrifuged at 1,000 *g* using a Beckman Avanti 30 (Beckman) for 10 minutes at 4°C. A soluble fraction was obtained by ultracentrifugation at 200,000 *g* for 20 minutes at 4°C on a Beckman Optima Ultracentrifuge (Beckman). Two biological replicates for SEC and IEX column separations were prepared in this manner.

### Size exclusion chromatography (SEC)

From the soluble sample, endogenous proteins were fractionated by an ÄKTA FPLC system (Amersham Biosciences) as described previously (Aryal et al., 2014; Aryal et al., 2017; McBride et al., 2019). Approximately 1 mg of total cytosolic proteins was injected onto a Superdex^®^ 200 10/300 GL column (GE Healthcare). The SEC elution was performed with the mobile phase [50 mM Hepes/KOH, (pH 7.5), 100 mM NaCl, 10 mM MgCl_2_ 5% glycerol, and 1mM DTT] at a flow rate of 0.65 mL/min. Elution chromatogram was monitored by absorption at 280 nm. The column was calibrated using a Gel Filtration Markers Kit MWGF1000 (Sigma-Aldrich), determining a mass range from 669 to 29 kDa. SEC fractions of 500 μL were collected, and proteins were precipitated by cold acetone for LC-MS/MS Analysis.

### Ion exchange chromatography (IEX)

One milligram of the cytosolic proteins was fractionated on a PolyCATWAX 204CTWX0510 column (200 x 4.6 mm id, 5 μm, 1000 A) using an UltiMate 3000 Standard HPLC System (Thermo Fisher Scientific Inc.) as described previously (McBride et al., 2019). IEX separation was over a 100-minute linear gradient elution program (from 0.0 to 1.5 M NaCl) at a flow rate of 1.0 mL/min. The absorbances of 214 nm and 280 nm were set to monitor protein elution. One hundred and seven sample fractions were collected every 22 sec (approximately 367 mL) between 3 min and 40 min. Each fraction was used for protein precipitation by a cold acetone method.

### Sample preparation for LC-MS/MS analysis

Fractionated protein samples were digested for LC-MS/MS analysis using trypsin as described previously (McBride et al., 2017; McBride et al., 2019). Protein pellets were dissolved and denatured in 8 M urea for 1 hour at room temperature, reduced in 10 mM DTT for 45 minutes at 60 °C, and then alkylated with 20 mM iodoacetamide for 45 minutes at room temperature under dark. The urea concentration in the peptide solution was brought to 1.5 M by additional ammonium bicarbonate for trypsin digestion. The digested peptides were purified using a Pierce® C18 Spin Columns (Thermo Fisher Scientific Inc.), and all samples were adjusted to an equal volume. Peptide concentrations were measured by bicinchoninic acid assay following the manufacturer’s protocol (Thermo Fisher Scientific Inc.). The most concentrated sample had a peptide concentration of 0.2 μg/μL, and 5 μL of each sample was injected onto LC-MS/MS.

### LC-MS/MS data acquisition

LC-MS/MS analysis was carried out as described previously (McBride et al., 2017). In brief, a Q-Exactive™ HF Hybrid Quadrupole-Orbitrap™ mass spectrometer in conjunction with reverse-phase HPLC-ESI-MS/MS using a Dionex UltiMate 3000 RSLC nano System (Thermo Fisher Scientific Inc.) was used. Peptides were resolved over a 125-min gradient with a flow rate of 300 nL/min. An MS survey scan was obtained from 350 to 1600 mass/charge ratio range. MS/MS spectra were acquired by selecting the 20 most abundant precursor ions for sequencing with high-energy collisional dissociation normalized collision energy of 27%. A 15-second dynamic exclusion window was applied to reduce the number of times the same ion was sequenced.

### Peptide identification and quantification

MaxQuant (version 1.6.14.0) was used for relative protein abundance quantification and protein identification (Cox et al., 2014). The search was conducted as described (McBride et al., 2019). Raw files of total cytosolic, SEC, and IEX fractions were searched on MaxQuant together against the rice proteome Osativa_323_v7.0.protein.fa (Ouyang et al., 2007). The search parameters were as follows: Cysteine carbamidomethylation was a fixed modification; oxidation on methionine and acetylation on protein N terminus were variable modifications; up to two missed trypsin cleavages were accepted; 1 % FDR at the protein and peptide level was chosen using a reverse decoy database; peptide abundance was calculated using the extracted ion current for both unique and razer, and protein level signals were aggregated from peptide intensities using MaxQuant “razor peptide” signal allocation among protein groups; the “match between runs” function was set with a maximum matching time window of 0.7 min as default; all other parameters were set as default.

### Reproducibility, Gaussian peak fitting, and *R*_app_ calculations

Reproducibility between two biological replicates was determined as described before (McBride et al., 2017; McBride et al., 2019). Pearson correlation coefficients (PCC) were estimated based on protein abundances in each fraction between duplicates and visualized by Data Analysis and Extension Tool (DAnTE). An optimized Gaussian fitting algorithm was applied to fit the chromatography resolution, and the Bayesian information criterion was utilized to prevent overfitting (McBride et al., 2017). The algorithm identified protein peaks when they had more than three non-zero fractions, with two being adjacent. Multiple reproducible peaks from a protein were split into multiple entries by labeling with a “_peak number” on their locus identifications (IDs). The reproducible peaks were selected from the two replicates if they were present within two or four fraction shifts considering the increment rate between fractions in SEC or IEX column, respectively. All non-reproducible peaks were not used for the following analysis. The fraction locations of the fitted peaks were used to determine the apparent mass (*M*_app_) values of proteins using the SEC calibration curve obtained above. The protein multimerization state (*R*_app_) was defined as the ratio of the *M*_app_ of a protein to the theoretical monomer mass (*M*_mono_) of the protein. The *R*_app_ of ≥ 1.6 thresholds was applied to determine whether a protein is present in a complex.

### Hierarchical clustering analysis

Hierarchical clustering was conducted as described previously (McBride et al., 2019). Briefly, a set of profiles that reflect compositions of a protein complex was clustered based on protein elution similarity. This cluster analysis was carried out with IEX only, SEC only, and combined IEX and SEC datasets. The Euclidean distance was used as a metric for measuring similarity in profiles of a pair of proteins. A series of dendrograms over a wide range of cluster numbers were generated to determine an optimal cluster number value for the prediction.

### Distance within clusters and purity and intactness of known protein complexes

To minimize false positives and negatives, the clustering results were evaluated by distance within clusters and intactness and purity as described previously (McBride et al., 2019). The distance within a cluster indicates how much similar or dissimilar protein elutions are in the given cluster. A cluster center was first calculated as the average elution profiles of all proteins in a cluster. Then a mean distance of each protein from the cluster center was calculated in the cluster.

The behavior of known protein complexes is that members of a known are co-migrating. This characteristic of known complexes was applied for the determination of the final cluster number to predict protein complexes. Rice orthologs were searched against the CORUM database (Ruepp et al., 2009), which provides annotated protein complexes from mammalian organisms at the purity and intactness tests. Purity was calculated by the ratio of the highest number of subunits for a known protein complex in a given group to the total number of proteins assigned to the group. Intactness is measured by the ratio of how many identified subunits of a known complex are assigned into one group to the total number of identified subunits in the known protein complex.

### Validation and complex heterogeneity using external datasets and multiple peak proteins

To validate the resulting clustering, external datasets for Arabidopsis and rice cofractionation-based protein complex predictions were downloaded from the supplemental table 2 in McBride et al. (2019) and the Table S4 in McWhite et al. (2020), respectively. To facilitate comparisons among the studies, key parameters including the number of proteins, the number of clusters, and the definition of protein group and ortholog group used in the McBride et al.’s and McWhite et al.’s studies were summarized in Supplemental Table S4A. The Arabidopsis orthologs of rice proteins were searched using the Phytozome ortholog database (Goodstein et al., 2011) to map the McBride Arabidopsis complex prediction onto our rice dataset (column F in Supplemental Table S4B). For the McWhite prediction, their ortholog groups containing rice and Arabidopsis were similarly mapped onto our rice clustering dataset (columns G–J in Supplemental Table S4C). For the rice data here, only protein groups in a cluster with two or more members but not classified as a “false positive” or “degraded” were used for comparisons. When the same protein pair was present in McBride and McWhite datasets it was scored as “within a single cluster in both studies.”

Another method to validate the clustering result using the rice data here was to test for co-occurrence of multiple peak proteins in two distinct clusters (Supplemental Table S3B). Interpretations of the multiple IEX peaks were based on whether or not the proteins had either one or two distinct SEC peaks and the *M*_app_ values of the peaks.

### Systematic classification of clustering results

The systematic classification was adopted similar to what was described previously (McBride et al., 2019) and is based on *M*_app_, the sum of *M*_mono_ of all proteins within one cluster (*M*_calc_), the average of *M*_app_ within one cluster (*M*_app-avg_), and *R*_app_ of all proteins in the given cluster. In the assumed scenario of 1:1 subunit stoichiometry, *M*_calc_ should be similar to the *M*_app-avg_ of its members. These proteins were classified as “**putative intact complex**.” In addition, clusters with likely high subunit stoichiometry contain reliable predictions. Single entry clusters are classified as “**homomer**” when the *R*_app_ ≥ 1.6. Clusters with 2 or 3 protein members were defined as “**possible homomer or heteromer / high subunit stoichiometry**” if the protein had *M*_app_ ≥ (4 * *M*_calc_). Cluster members were classified as “**subcomplex or high subunit stoichiometry**” when *M*_app_ ≥ (4 * *M*_calc_) in the clusters with more than 3 protein members and as “**partial complex/false negatives**” when *M*_app_ > 1.4 * *M*_calc_. When a protein within a cluster had *M*_calc_ > 1.4 * *M*_app_ and *R*_app_ ≥ 1.6, the protein was defined as “**putative complex clustered with false positives**.” If *R*_app_ < 1.6 and *M*_calc_ > 1.4 * *M*_app_, the protein was flagged as “**likely false positive: monomer**.” Single proteins were classified as “degraded” when the *R*_app_ < 0.5, and “monomeric” when 0.5 ≤ the *R*_app_ < 1.6.

### GO-term analysis

Singular Enrichment Analysis (SEA) tool in AgriGO v2.0 was used for gene ontology (GO) enrichment analysis (Tian et al., 2017). The enrichment was analyzed using Fisher’s exact test at the 5 % FDR level as Hochberg corrected correction against all proteins in the MSU7.0 nonTE transcript ID (TIGR) background.

### Data analysis

Gaussian fitting was applied using MATLAB_R2016a. Clustering analysis was performed using R version 3.5.1 (R Core Team, 2018) on RStudio 1.1.463 (RStudio Team, 2018).

## Supporting information

Supplemental Data Set 1

Supplemental Data Set 2

## Data and materials availability

The MS raw files have been deposited to the ProteomeXchange Consortium via the PRIDE with accession codes PXD022357. The mass spectra are available at the Protein Prospector with search key uij64faovq. The Gaussian fitting code and the clustering analysis code described in McBride et al., 2017 are available at (https://github.com/dlchenstat/Gaussian-fitting) and at (https://github.com/dlchenstat/ProteinComplexPredict), respectively.

## Supplemental Data files

Supplemental Table S1. Raw profiles of peptides and proteins (.xlsx).

Supplemental Table S2. Lists of reproducible protein peaks in SEC and IEX (.xlsx).

Supplemental Table S3. Clustering result and systematic classification (.xlsx).

Supplemental Table S4. Clustering result comparisons among plant CF-MS predictions (.xlsx).

Supplemental Data Set 1. Elution profiles of clustered proteins and dendrogram of clusters (.pdf).

Supplemental Data Set 2. Supplemental texts for discussions on the predicted novel and known protein complexes (.pdf).

## Acknowledgments

This work was supported by the National Science Foundation IOS-PGRP grant nos. 1444610 to T.W.O. and D.B.S. and 1951819 to D.B.S. Y.L. was mainly supported by the above NSF funds and partially by Bilsland Graduate Dissertation Fellowship from the Graduate School at Purdue University and CPB Graduate Research Award from the Center for Plant Biology at Purdue University. We thank the Purdue Proteomics Facility and Dr. Uma Aryal for running the samples.

## Author contributions

Y.L. and D.B.S. designed the project; Y.L. performed experiments; Y.L., T.W.O., and D.B.S. analyzed the data and wrote the paper.

