## Supplemental Figures for "A Co-Fractionation Mass Spectrometry-based Prediction of Protein Complex Assemblies in the Developing Rice Aleurone-subaleurone"

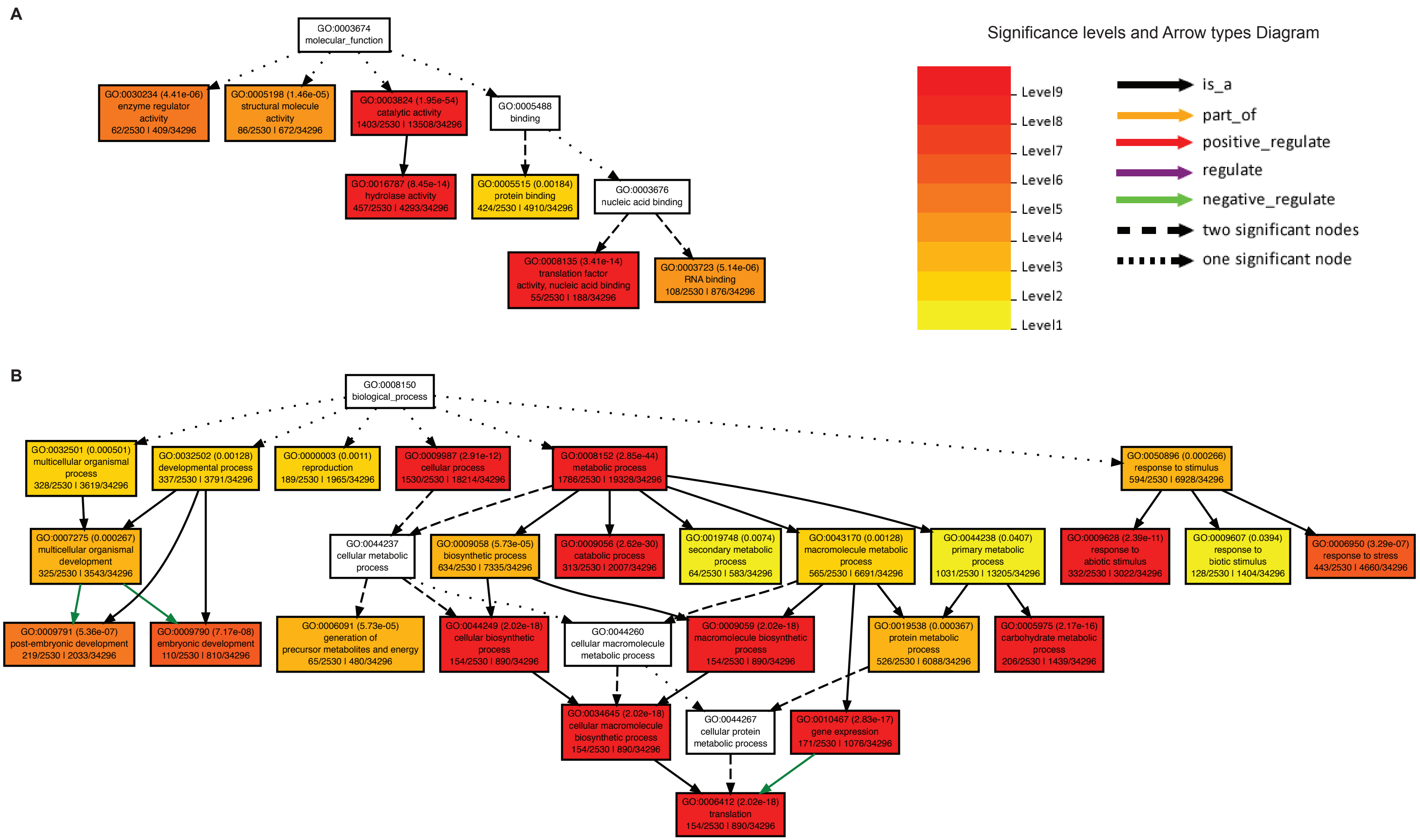


**Supplemental Figure S1. GO term enrichment analysis showed overrepresented GO terms in the sink type aleurone-subaleurone layers.**

GO terms of rice subaleurone proteins were enriched using SEA tool in AgriGO v2. Overrepresented GO terms were highlighted with their significance levels. Molecular function GO terms (A) and biological process GO terms (B) were visualized.

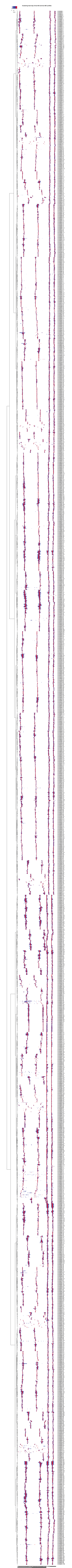


**Supplemental Figure S2. Clustering heatmap of rice aleurone and subaleurone proteome.** Hierarchical clustering analysis of elution profiles was performed, based on the elution similarity in the concatenated SEC and IEX separations.

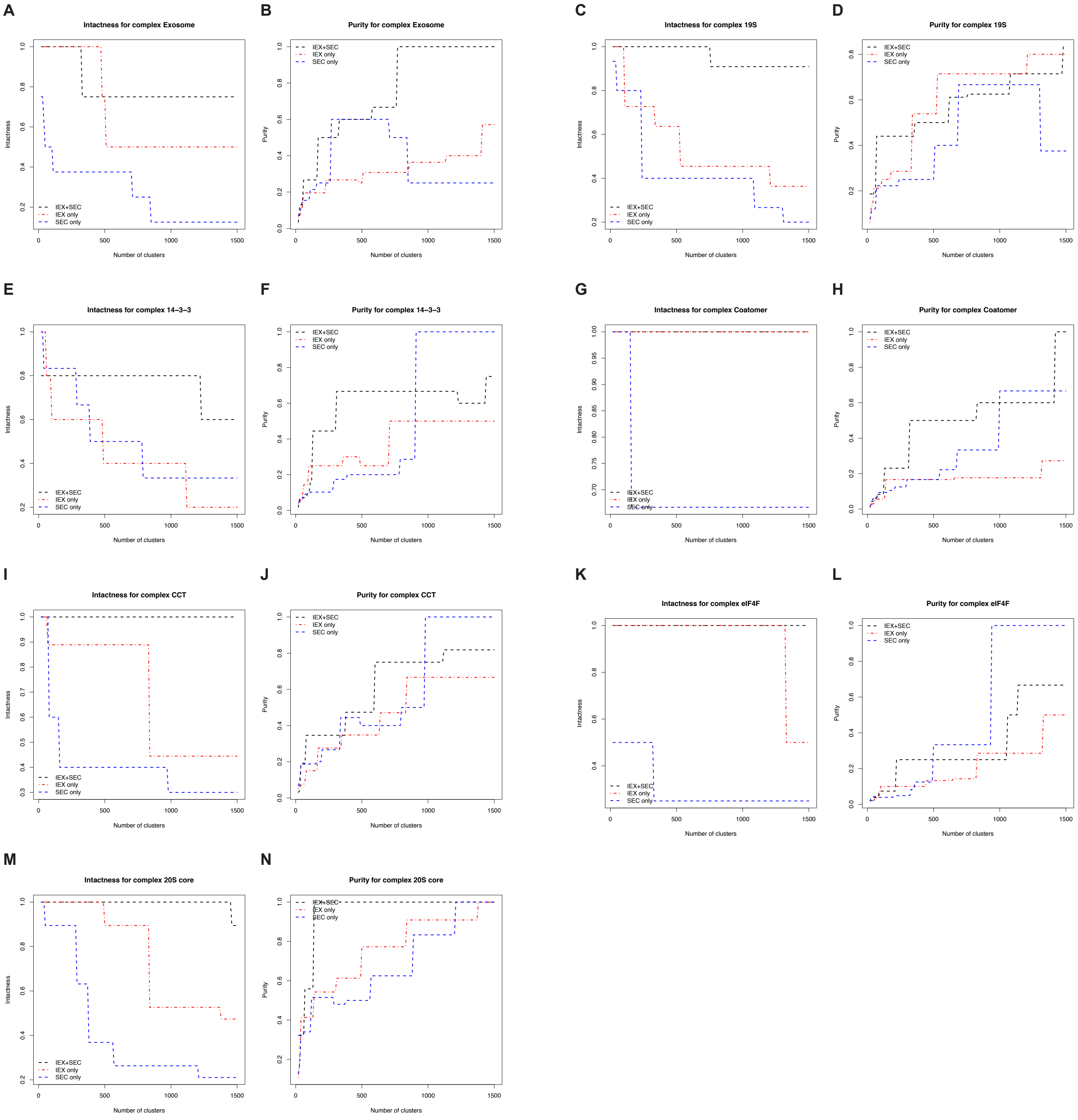


**Supplemental Figure S3. Extrinsic tests evaluated the resolving power for independent and combined protein profile datasets as a function of ascending cluster numbers.**
The intactness and purity of know complexes were plotted for IEX only (red), SEC only (blue), and concatenated IEX and SEC (black) profiles. (A) & (B) Exosome complex. (C) & (D) 19S complex. (E) & (F) 14-3-3 complex. (G) & (H) Coatomer complex. (I) & (J) CCT complex. (K) & (L) eIF4F complex. (M) & (N) 20S core complex.

**Supplemental Table S1. Raw profiles of peptides and proteins (.xlsx).**
1. Supplemental Table S1A. The abundance of peptides across the SEC fractions for both biological replicates was obtained from rice aleurone and subaleurone layers.
2. Supplemental Table S1B. The abundance of peptides across the IEX fractions for both biological replicates was obtained from rice aleurone and subaleurone layers.
3. Supplemental Table S1C. The abundance of proteins across the SEC fractions for both biological replicates was obtained from rice aleurone and subaleurone layers.
4. Supplemental Table S1D. The abundance of proteins across the IEX fractions for both biological replicates was obtained from rice aleurone and subaleurone layers.

**Supplemental Table S2. Lists of reproducible protein peaks in SEC and IEX (.xlsx).**
1. Supplemental Table S2A. The apparent masses ($M_{app}$) and multimeric states ($R_{app}$) of reproducible proteins between SEC replicates.
2. Supplemental Table S2B. The reproducible proteins between IEX replicates deconvoluted by Gaussian fitting algorism.

**Supplemental Table S3. Clustering result and systematic classification (.xlsx).**
1. Supplemental Table S3A. Clustering result predicted by the concatenated SEC and IEX datasets.
2. Supplemental Table S3B. Co-occurrence of multiple peak proteins in the same cluster.
3. Supplemental Table S3C. The systematic classification of protein complex prediction.

**Supplemental Table S4. Clustering result comparisons among plant CF-MS predictions (.xlsx).**
1. Supplemental Table S4A. Clustering result comparisons with prior multidimensional CF-MS predictions: McBride MCP 2019, 18(8): 1588-1606 and McWhite Cell 2020, 181(2): 460-474.
2. Supplemental Table S4B. List of rice clusters that were hit by ortholog pairs present in both the rice dataset and the Arabidopsis complex prediction dataset (McBride et al., 2019).
3. Supplemental Table S4C. List of rice clusters that were hit by ortholog pairs present in both the rice dataset and the interaction clustering dataset (McWhite et al., 2020).

**Supplemental Data Set 1. Elution profiles of clustered proteins and dendrogram of clusters (.pdf).**
Provided separately

**Supplemental Data Set 2. Supplemental texts for discussions on the predicted novel and known protein complexes (.pdf).**
Provided separately
1. Supplemental text 1. Discussion on two known protein complexes: CCT and 19S proteasome complexes.

2. Supplemental text 2. Discussion on predicted homomers: One known homomer and three novel homomers are discussed.
3. Supplemental text 3. Discussion on predicted heteromers: Two known and two novel heteromers are discussed.
4. Supplemental text 4. Discussion on predicted RPB-associated novel heteromeric complexes: Three novel RBP complexes are discussed.
