## Supplemental Data Set 1 for "A Co-Fractionation Mass Spectrometry-based Prediction of Protein Complex Assemblies in the Developing Rice Aleurone-subaleurone"

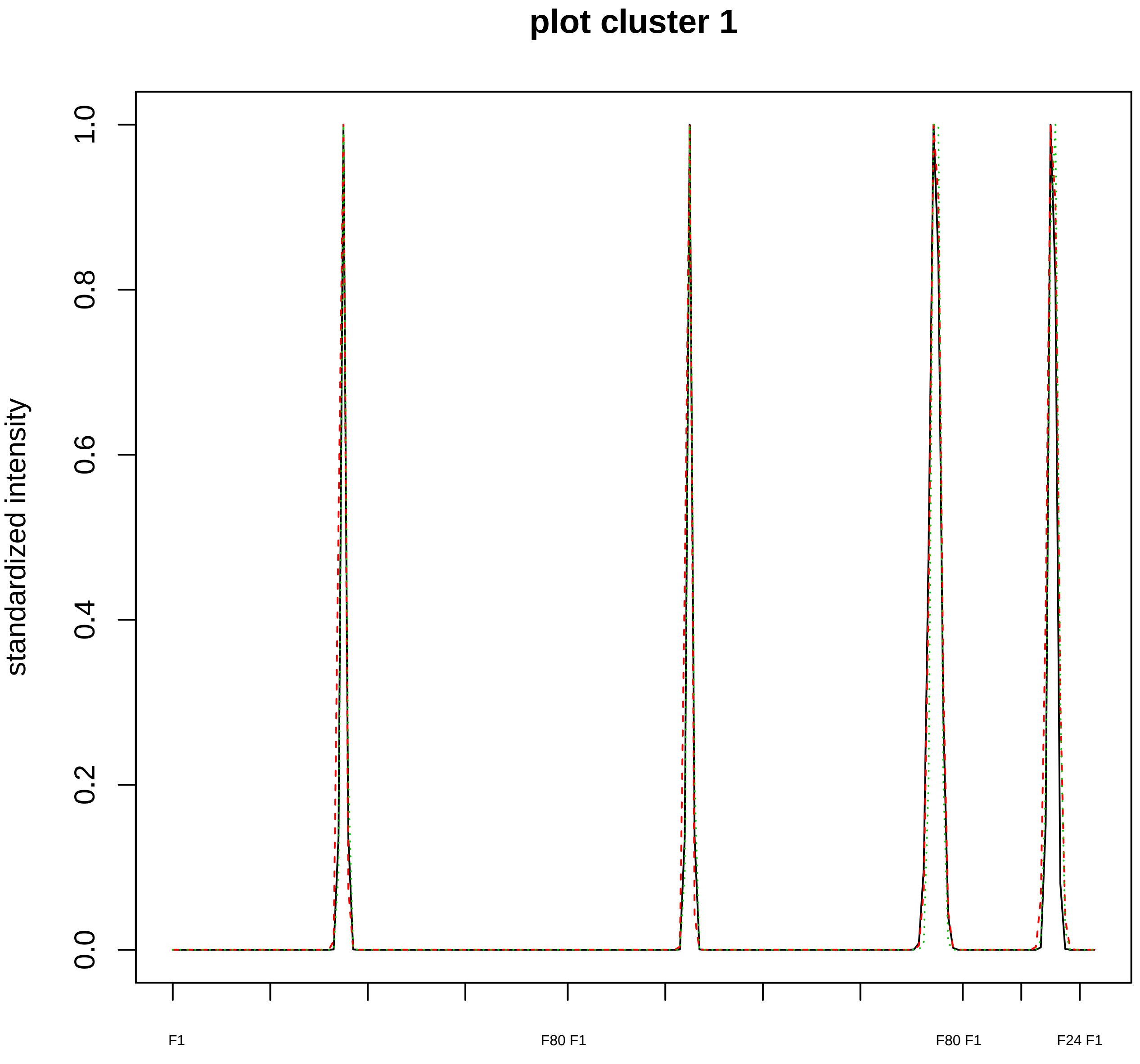

plot cluster 1
standardized intensity
0.0
0.2
0.4
0.6
0.8
1.0
F1
F80 F1
F80 F1
F24 F1

### Dendrogram of cluster 1

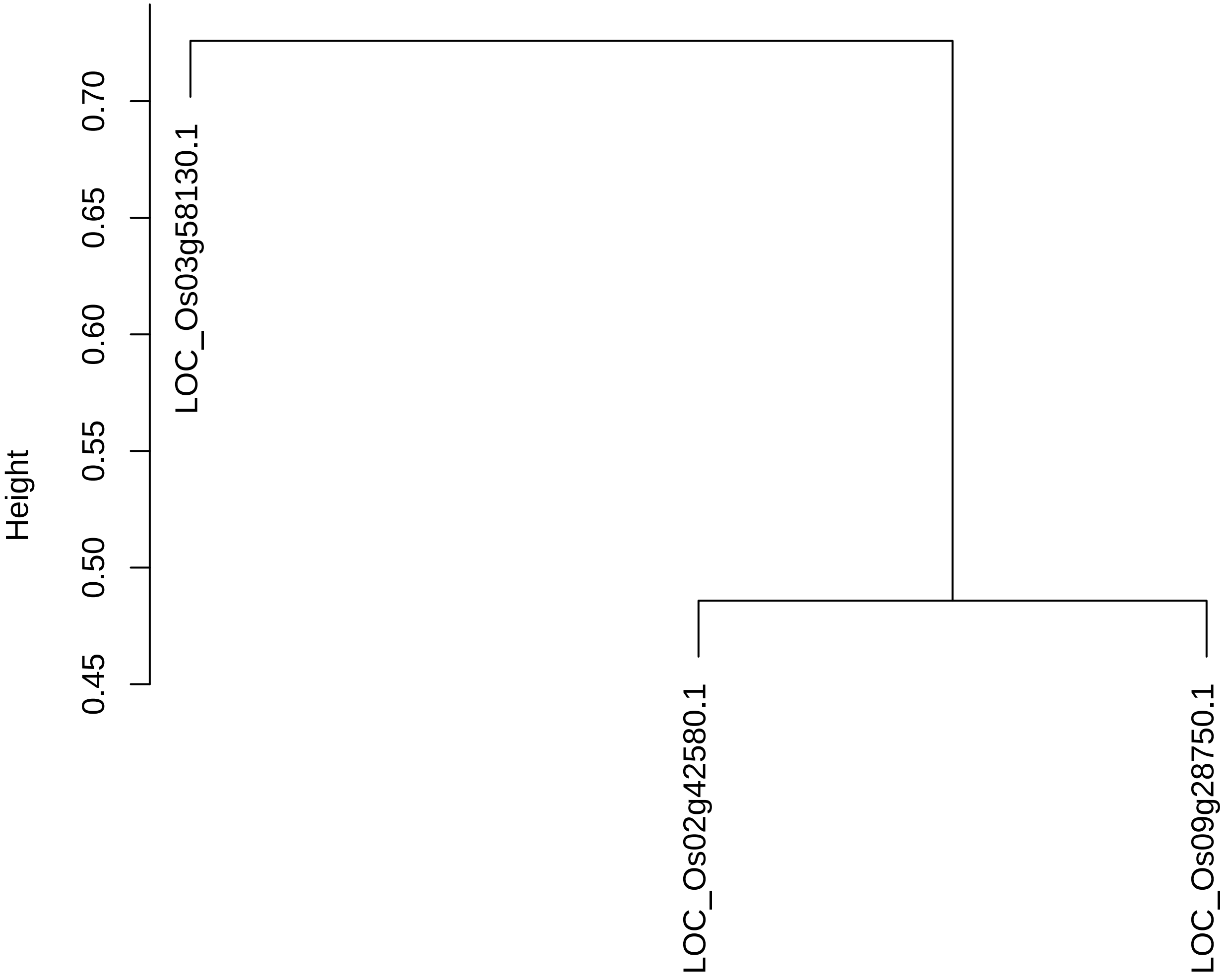


dist(spfitiex12sec12[lab[, 1] == i, ])
hclust (*, "ward.D2")

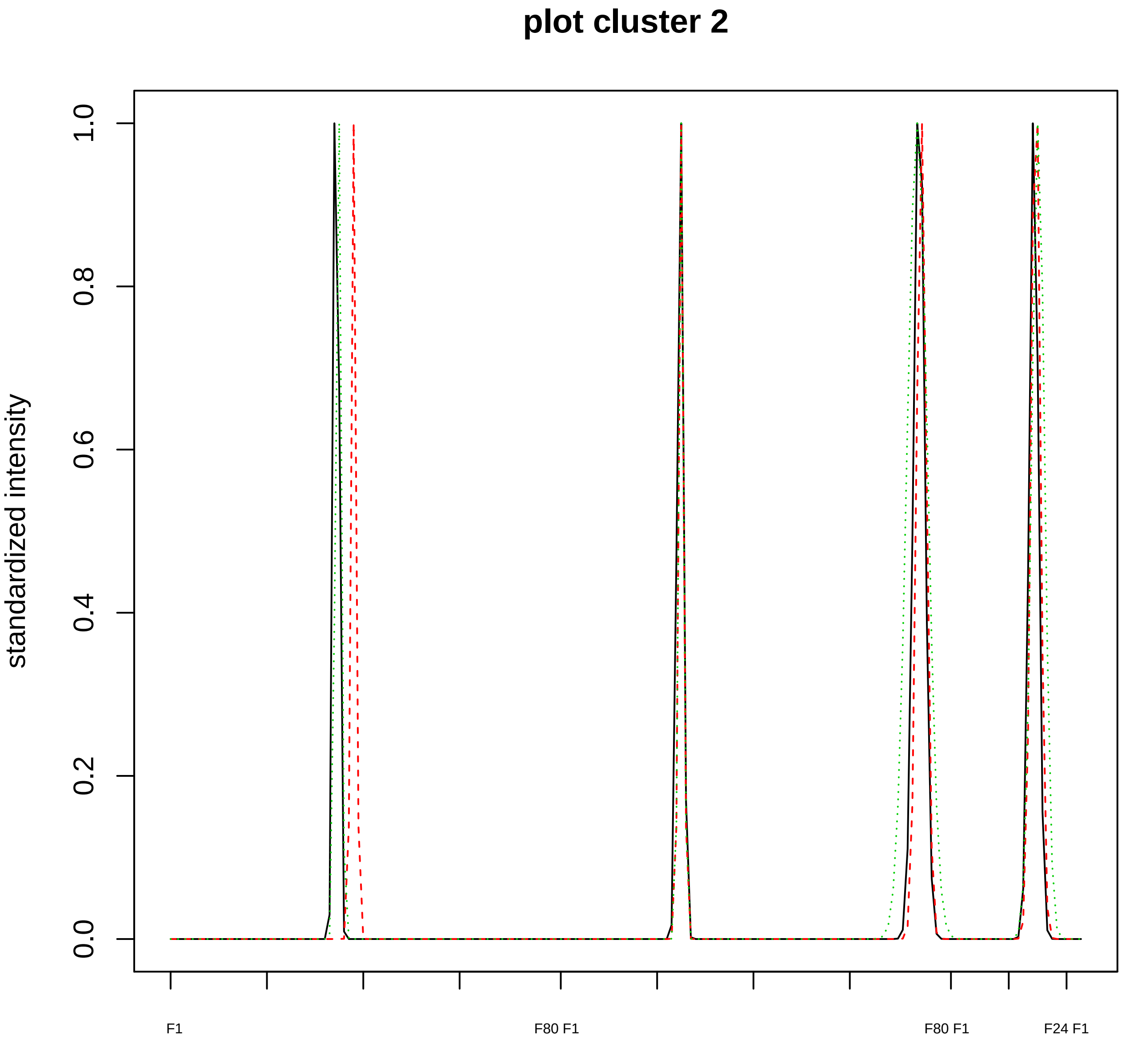

plot cluster 2
standardized intensity
0.0
0.2
0.4
0.6
0.8
1.0
F1
F80 F1
F80 F1
F24 F1

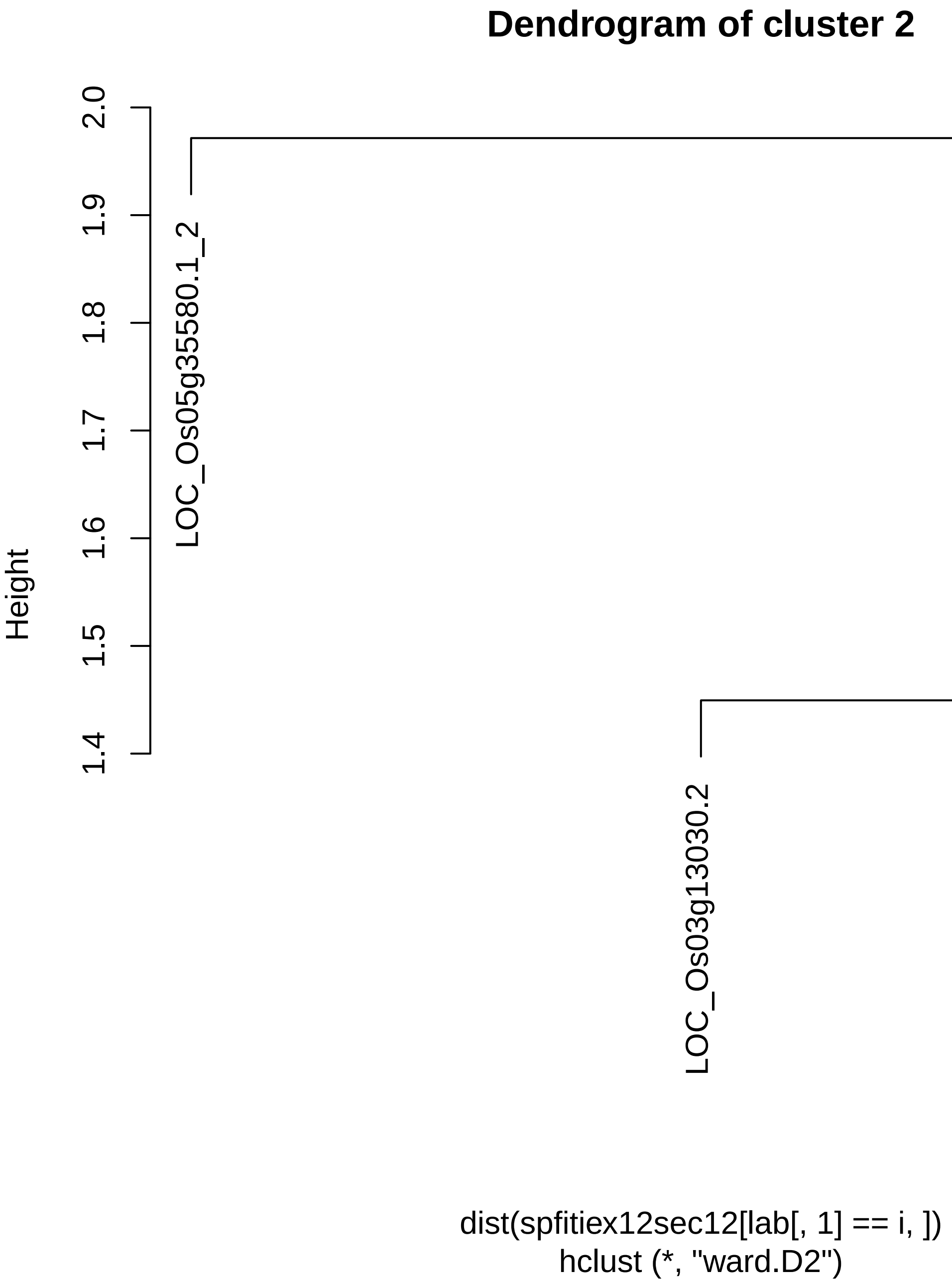
Dendrogram of cluster 2
Height
2.0
1.9
1.8
1.7
1.6
1.5
1.4
LOC_Os05g35580.1_2
LOC_Os03g13030.2
LOC_Os07g42924.1
dist(spfitiex12sec12[lab[, 1] == i, ])
hclust (*, "ward.D2")

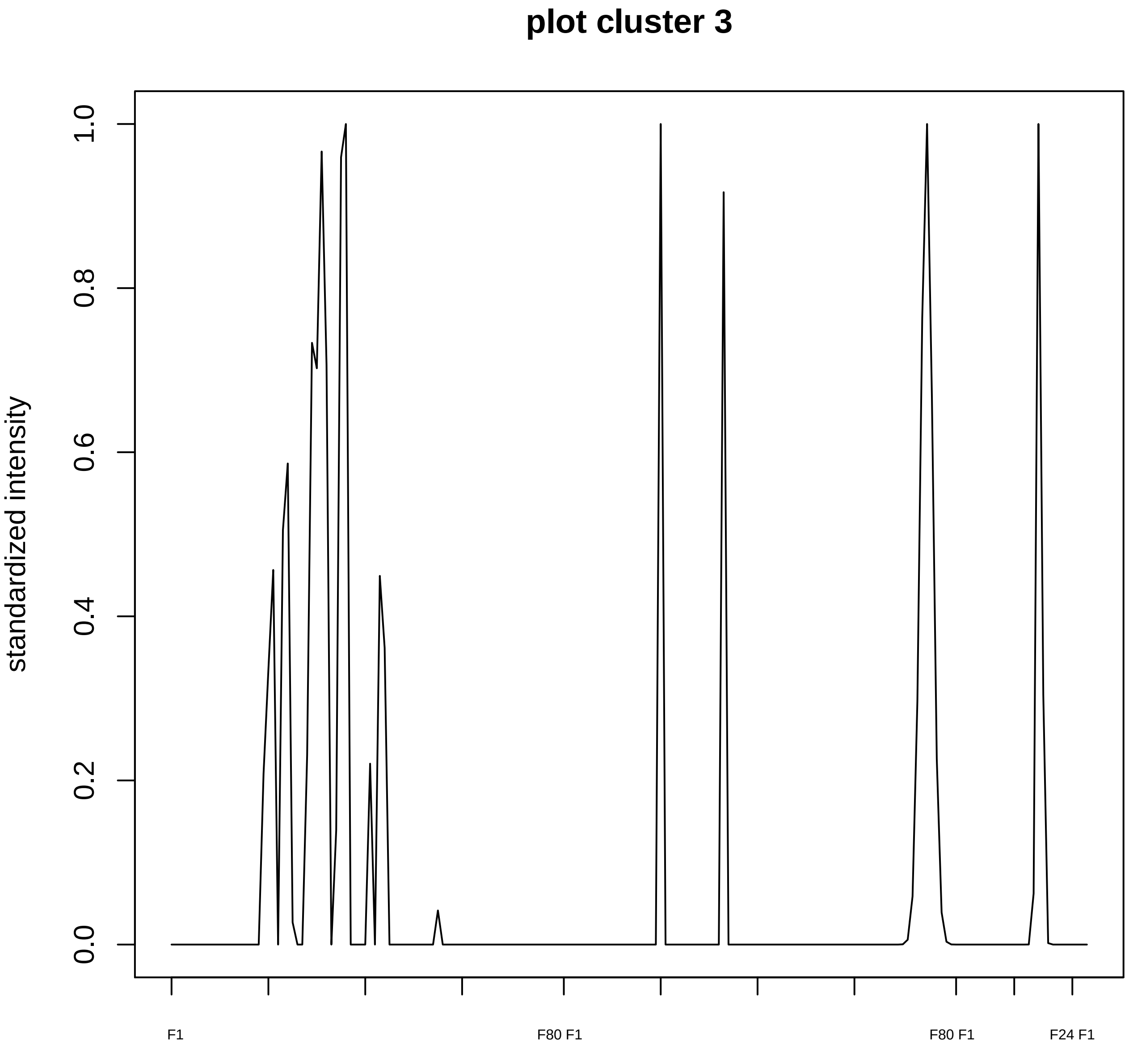

plot cluster 3
standardized intensity
0.0
0.2
0.4
0.6
0.8
1.0
F1
F80 F1
F80 F1
F24 F1

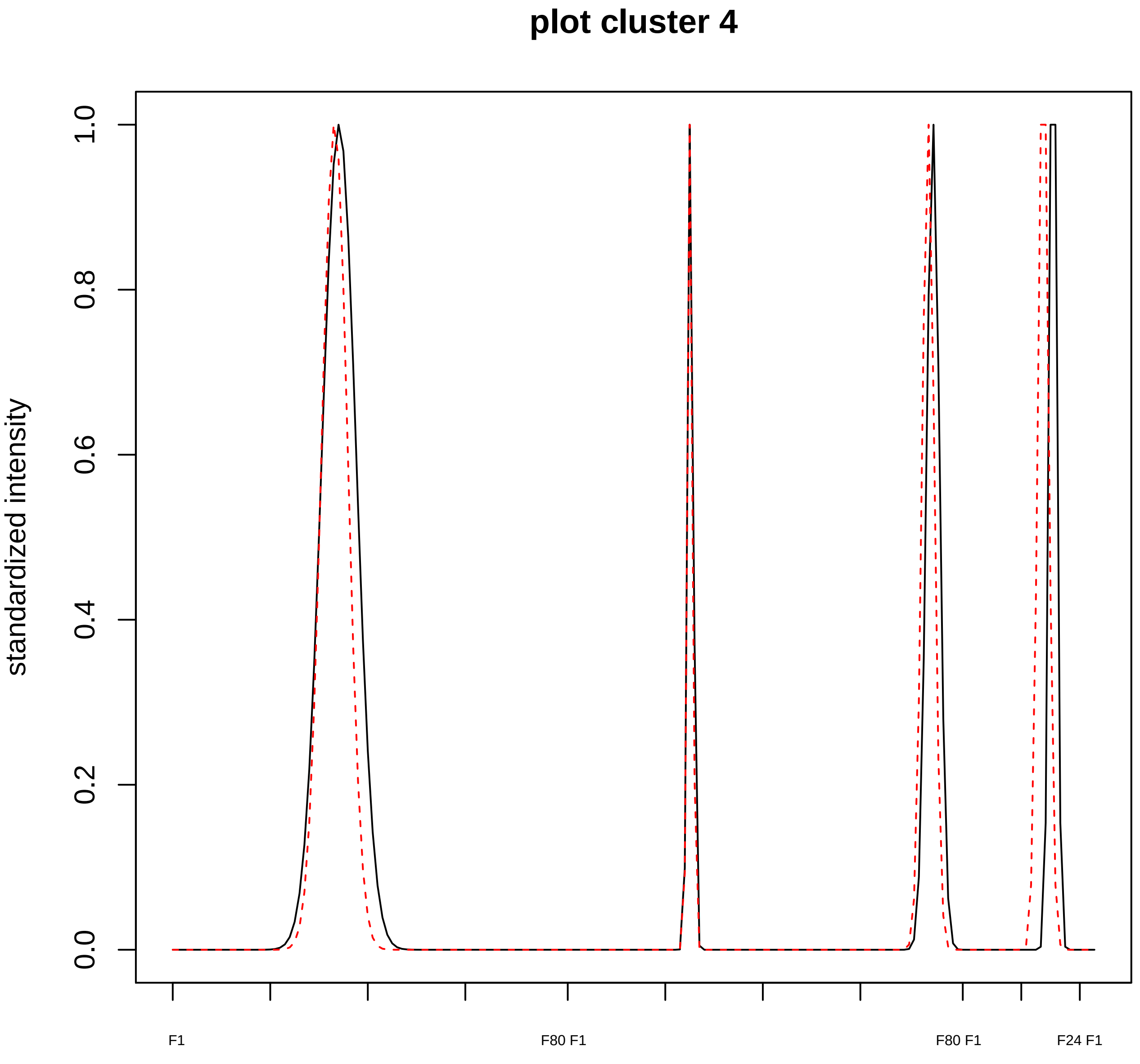
plot cluster 4
standardized intensity
0.0
0.2
0.4
0.6
0.8
1.0
F1
F80 F1
F80 F1
F24 F1

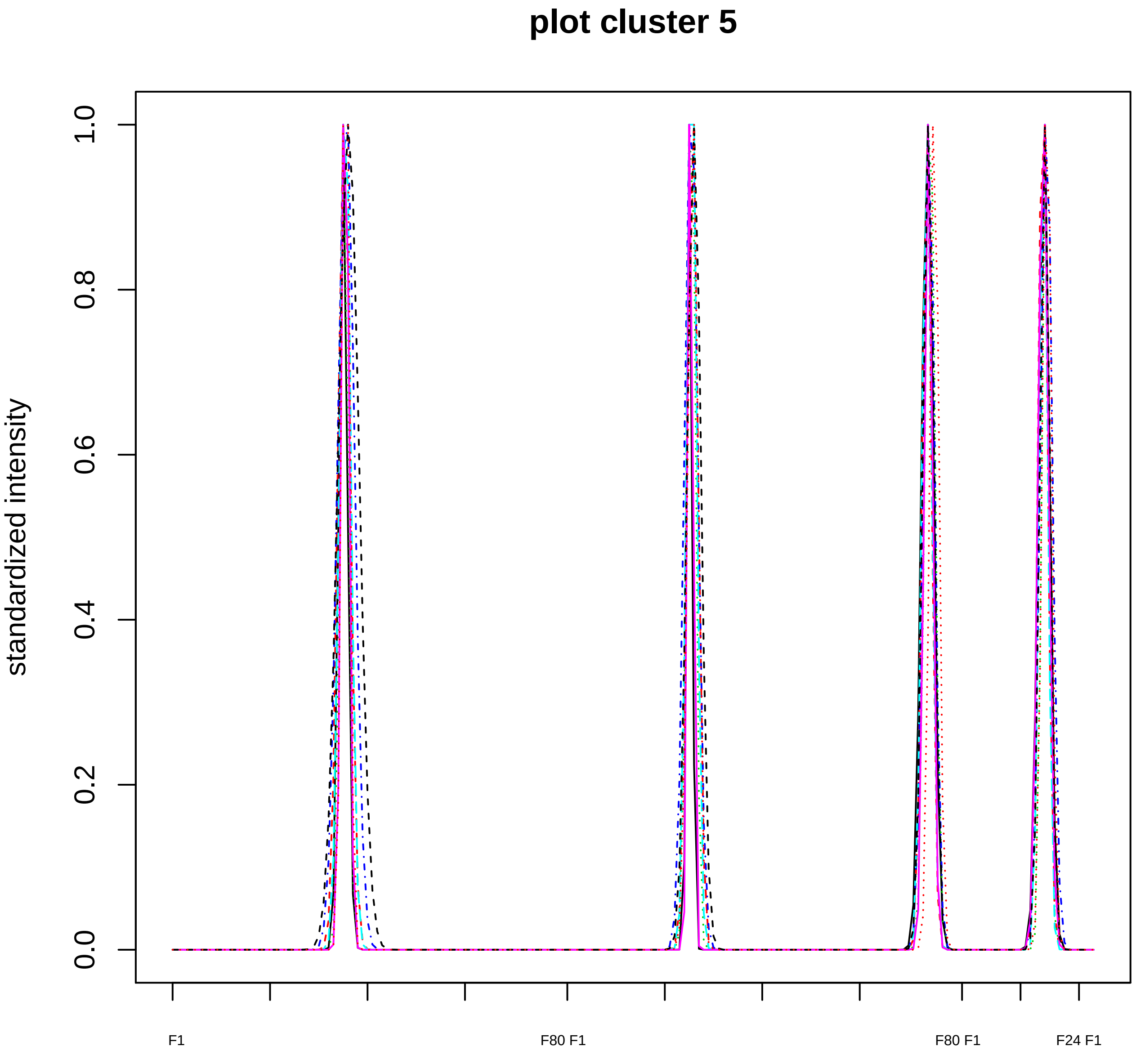
plot cluster 5
standardized intensity
0.0
0.2
0.4
0.6
0.8
1.0
F1
F80 F1
F80 F1
F24 F1

### Dendrogram of cluster 5

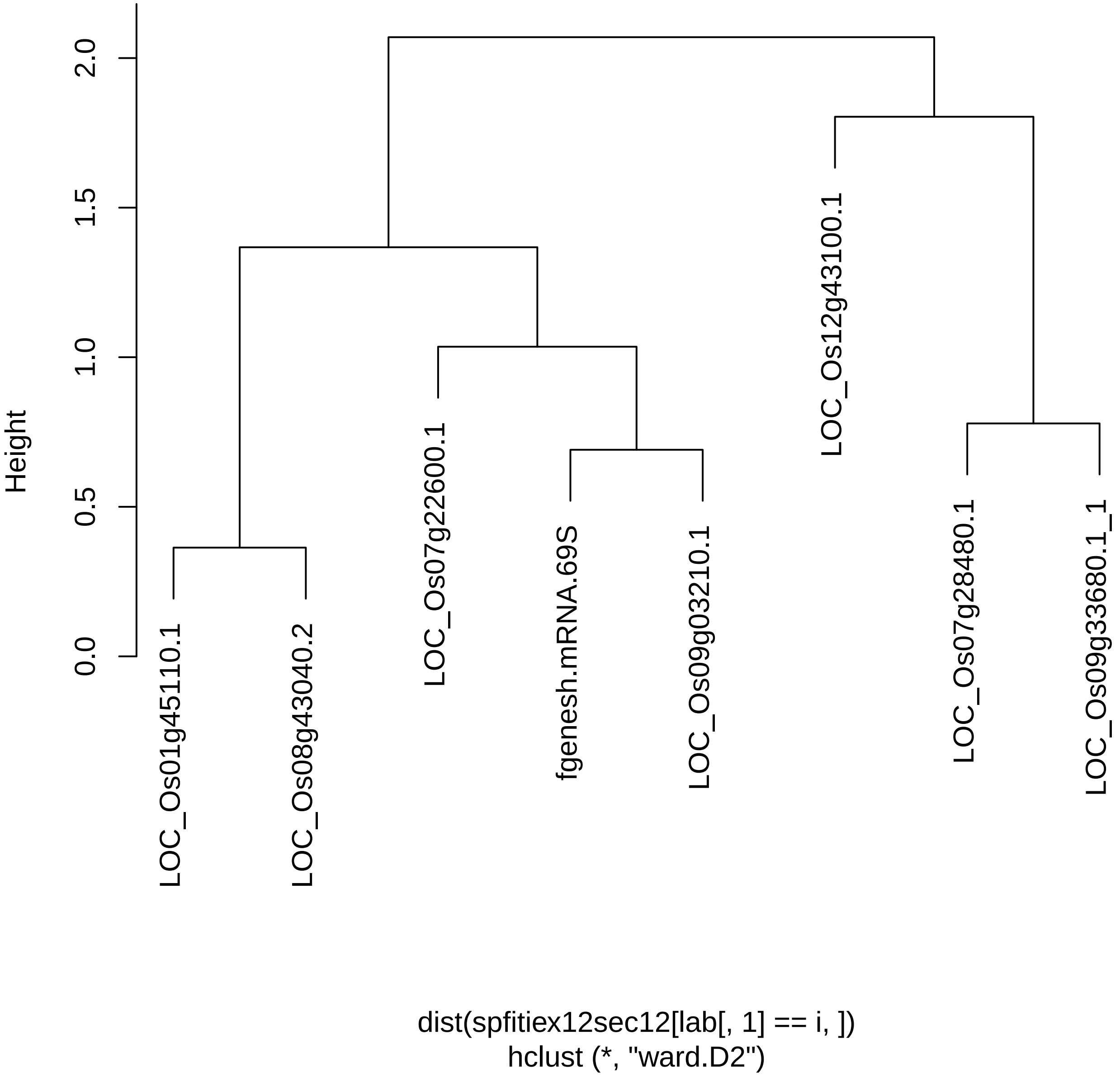

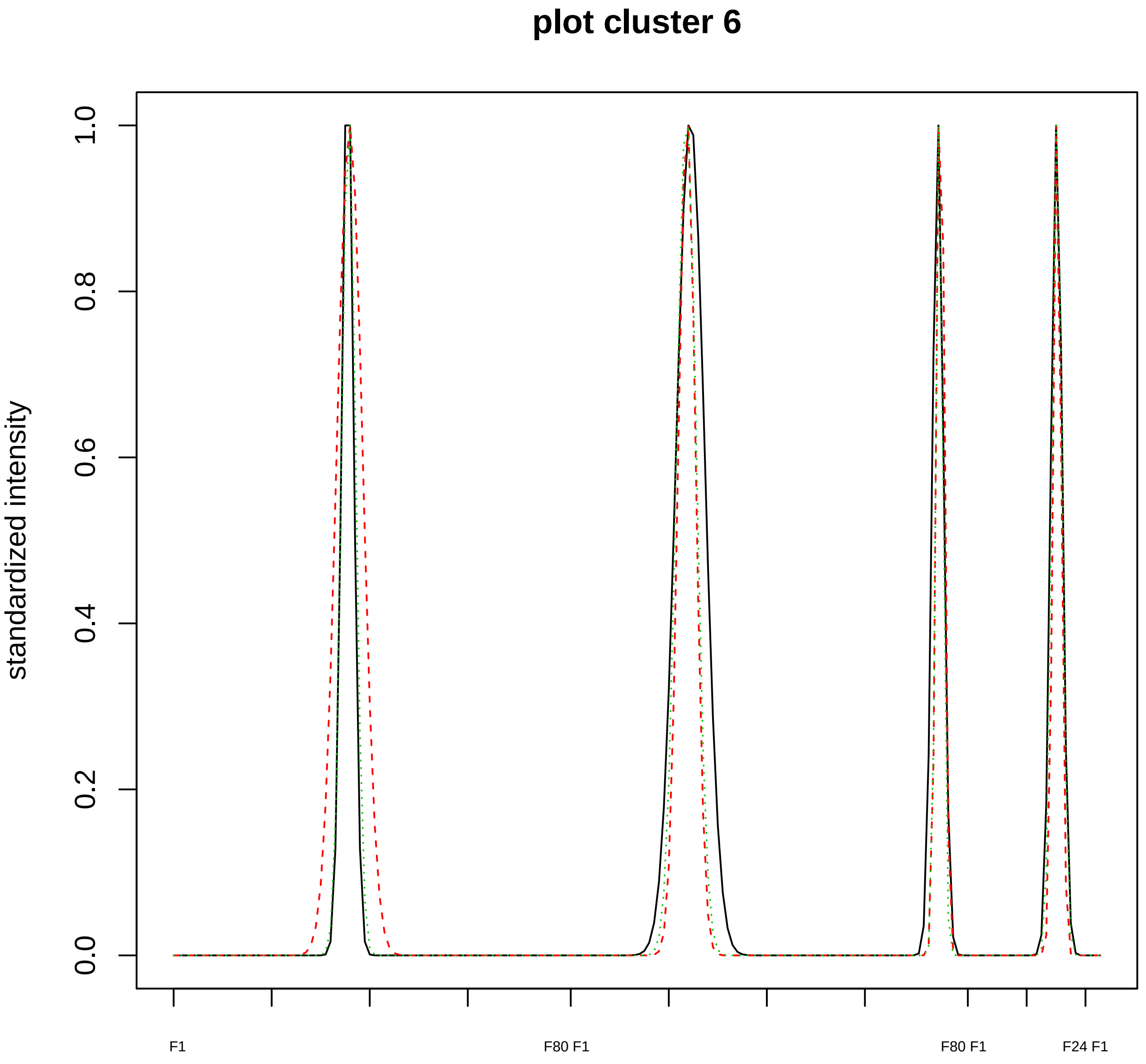
plot cluster 6
standardized intensity
0.0
0.2
0.4
0.6
0.8
1.0
F1
F80 F1
F80 F1
F24 F1

#### Dendrogram of cluster 6

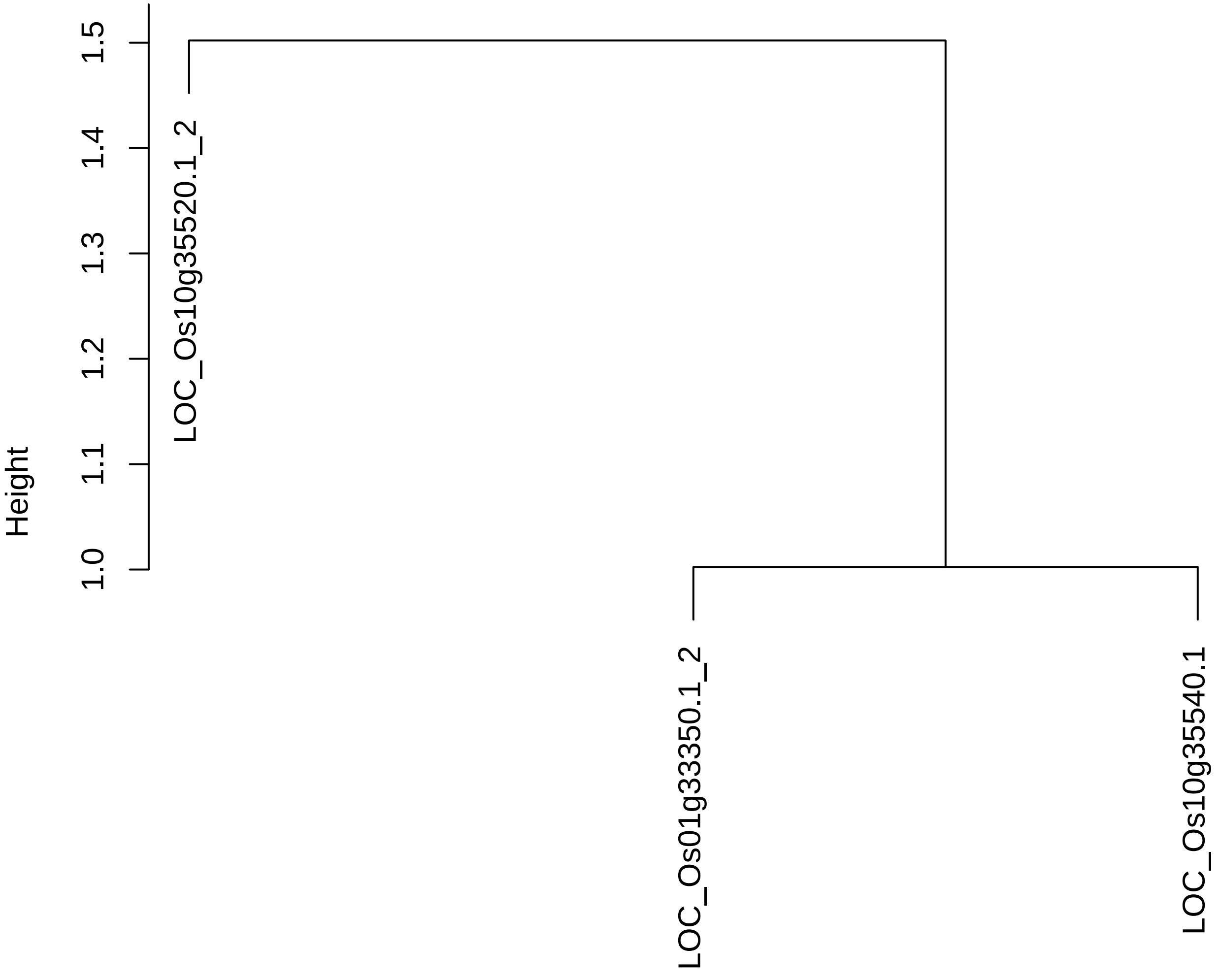


dist(spfitiex12sec12[lab[, 1] == i, ])
hclust (*, "ward.D2")

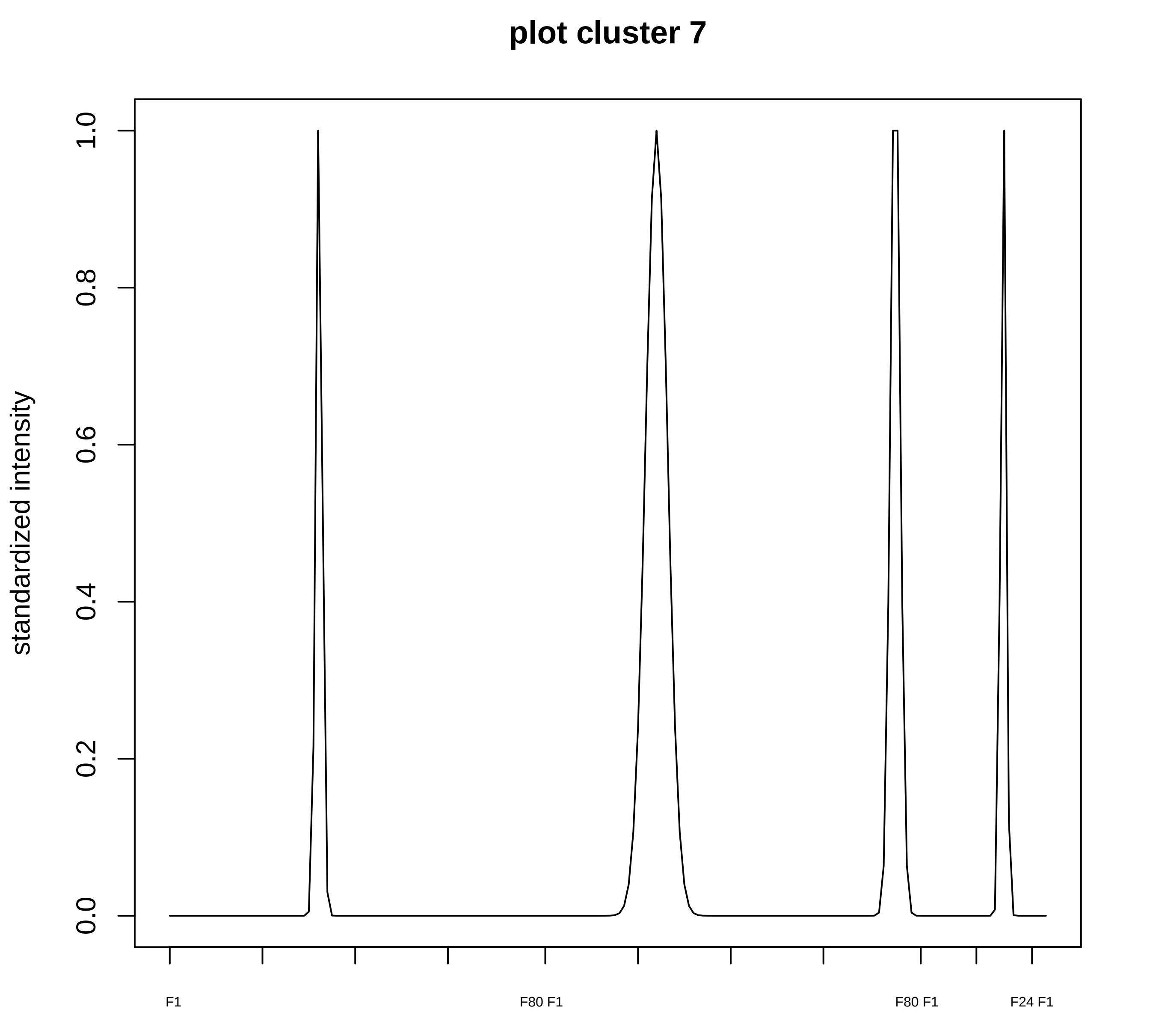

plot cluster 7
standardized intensity
0.0
0.2
0.4
0.6
0.8
1.0
F1
F80 F1
F80 F1
F24 F1

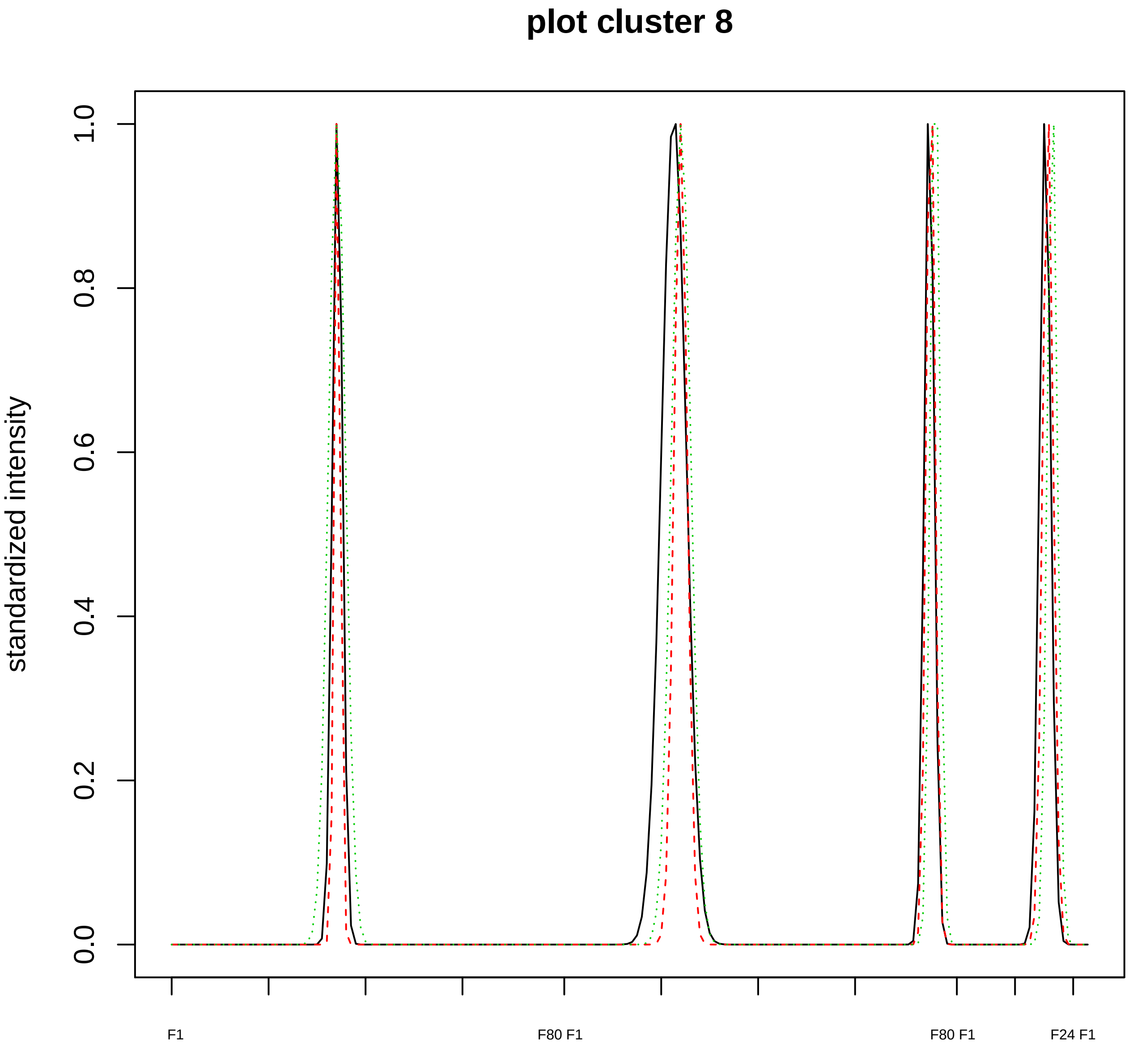
plot cluster 8
standardized intensity
0.0
0.2
0.4
0.6
0.8
1.0
F1
F80 F1
F80 F1
F24 F1

### Dendrogram of cluster 8

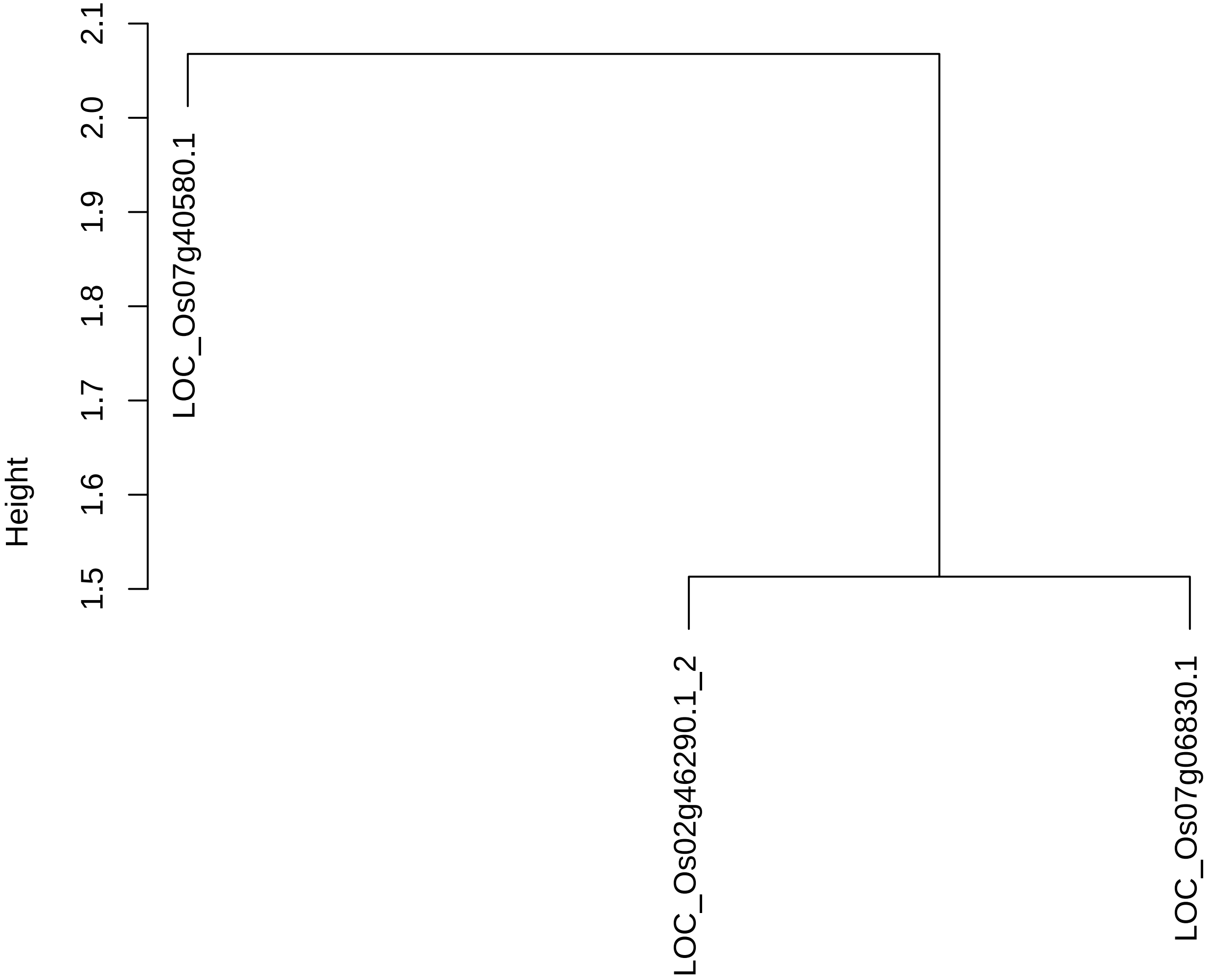


dist(spfitiex12sec12[lab[, 1] == i, ])
hclust (*, "ward.D2")

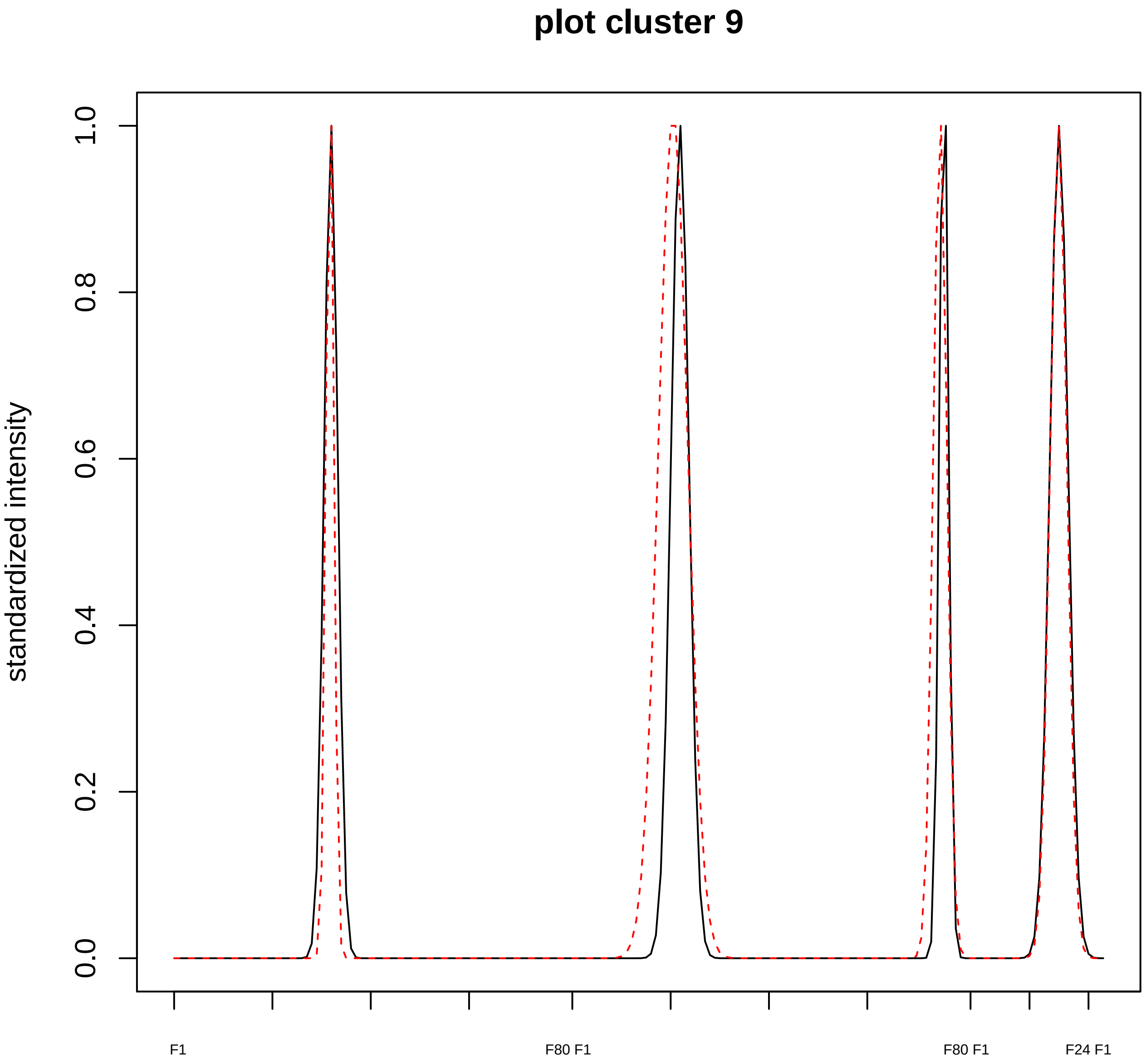

plot cluster 9
standardized intensity
0.0
0.2
0.4
0.6
0.8
1.0
F1
F80 F1
F80 F1
F24 F1

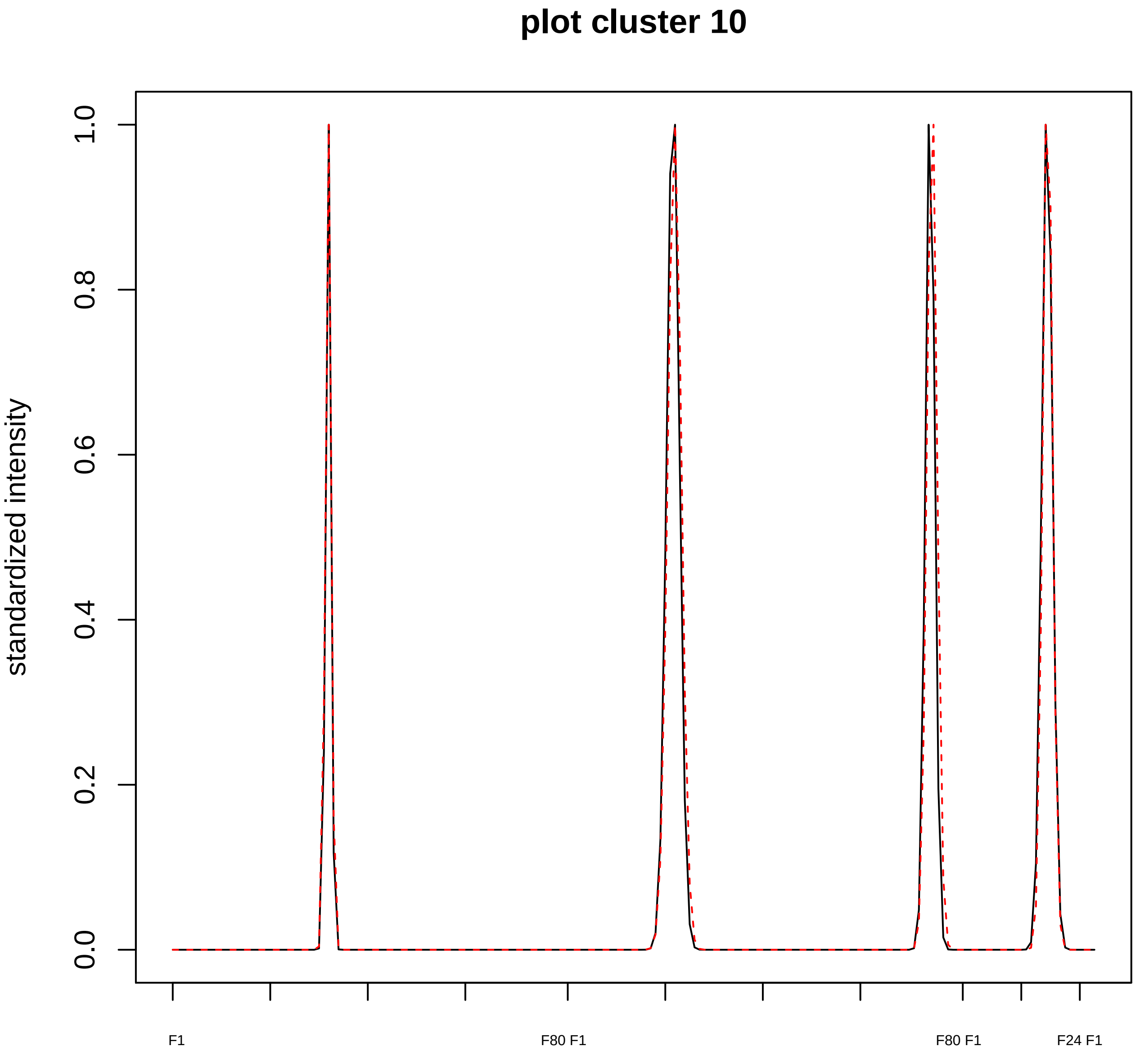
plot cluster 10
standardized intensity
0.0
0.2
0.4
0.6
0.8
1.0
F1
F80 F1
F80 F1
F24 F1

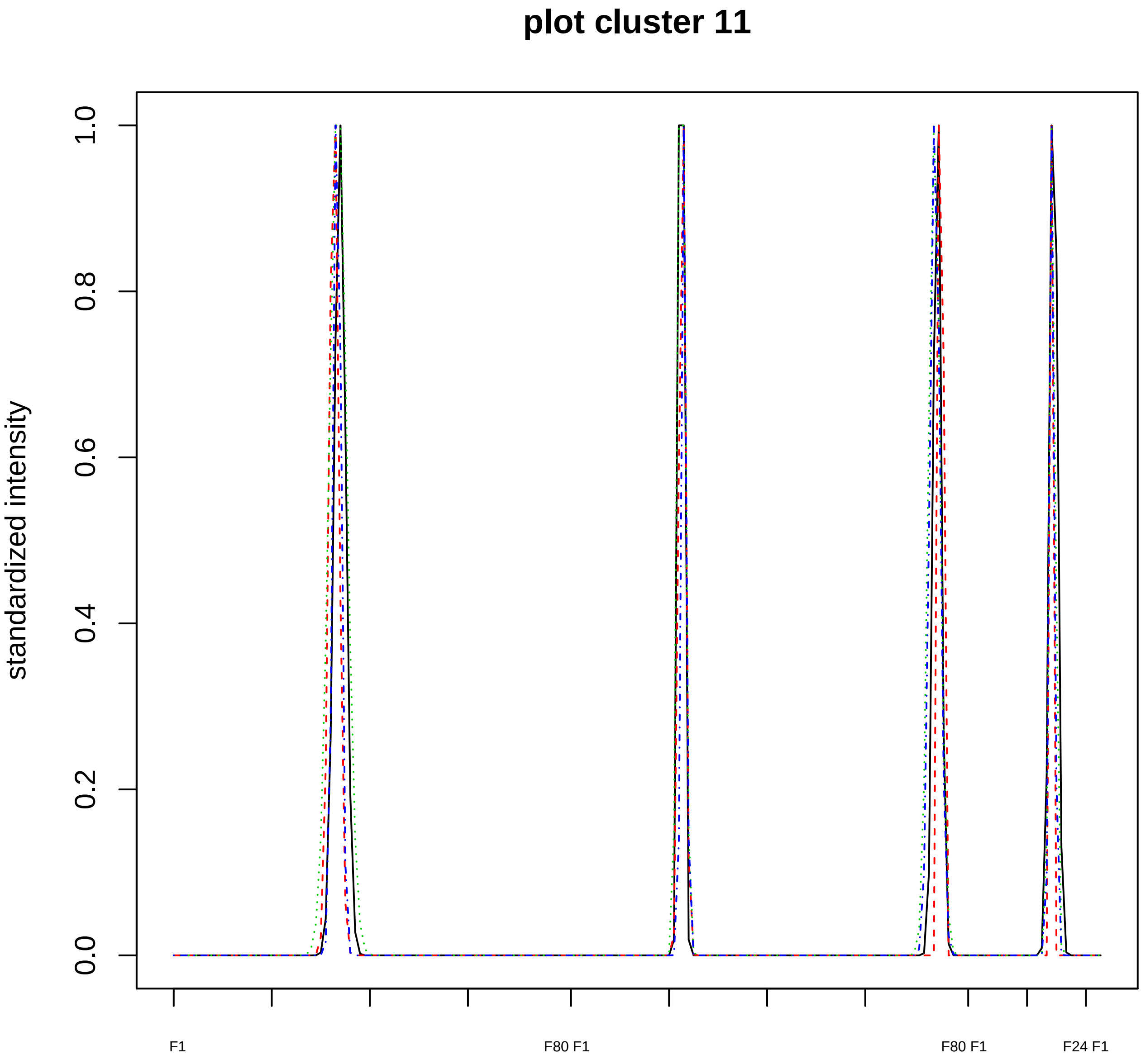

plot cluster 11
standardized intensity
0.0
0.2
0.4
0.6
0.8
1.0
F1
F80 F1
F80 F1
F24 F1

#### Dendrogram of cluster 11

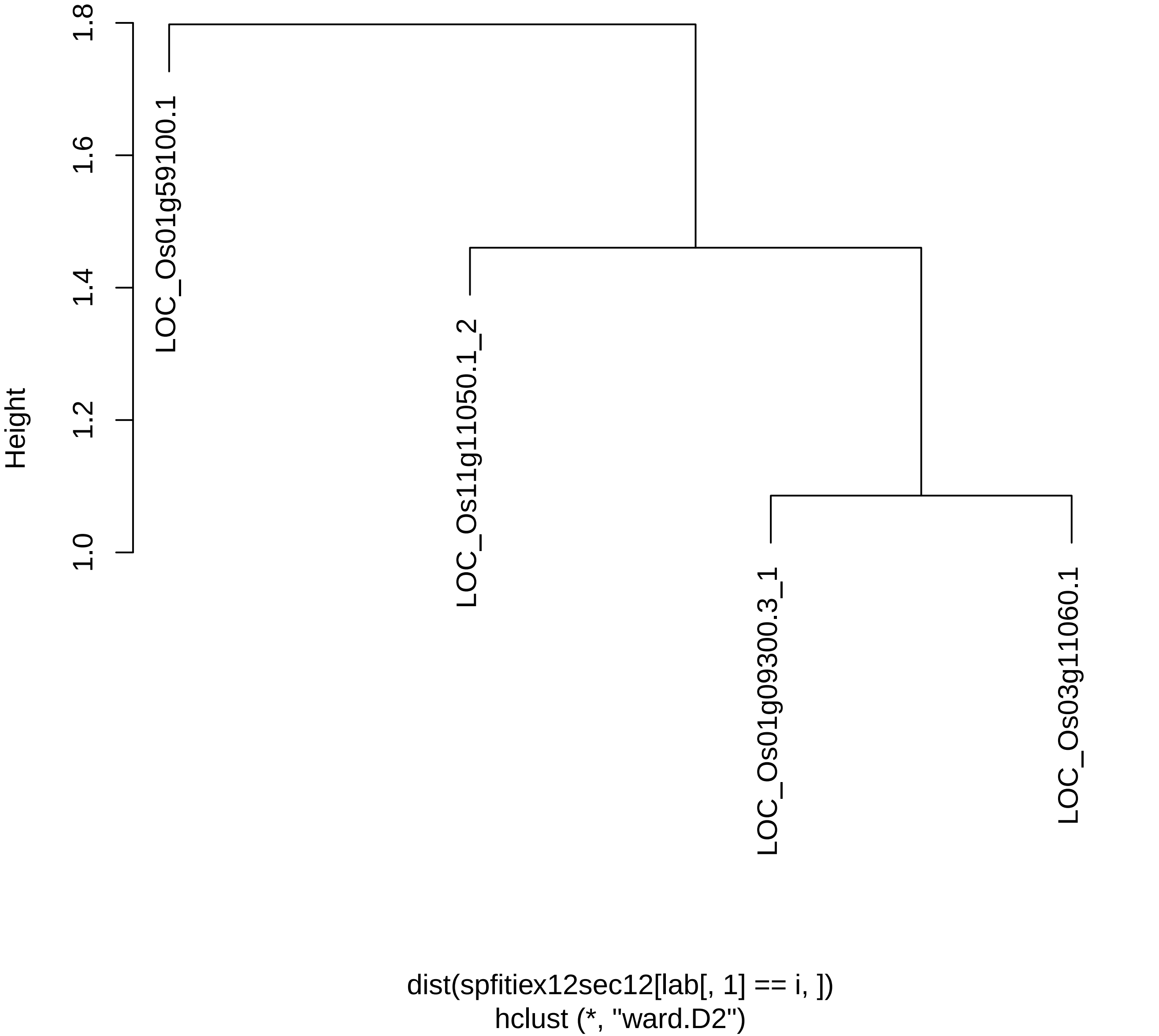

plot cluster 12

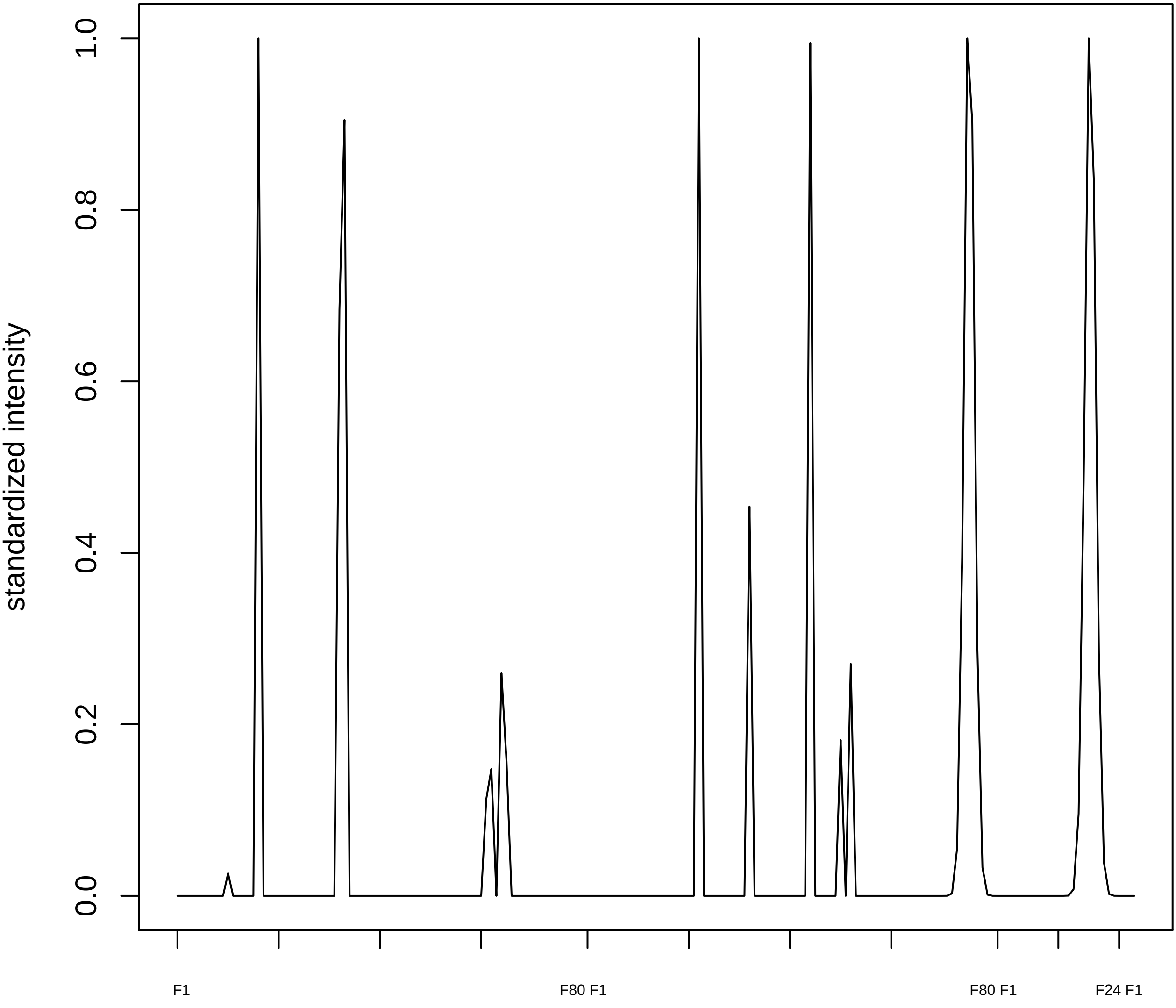

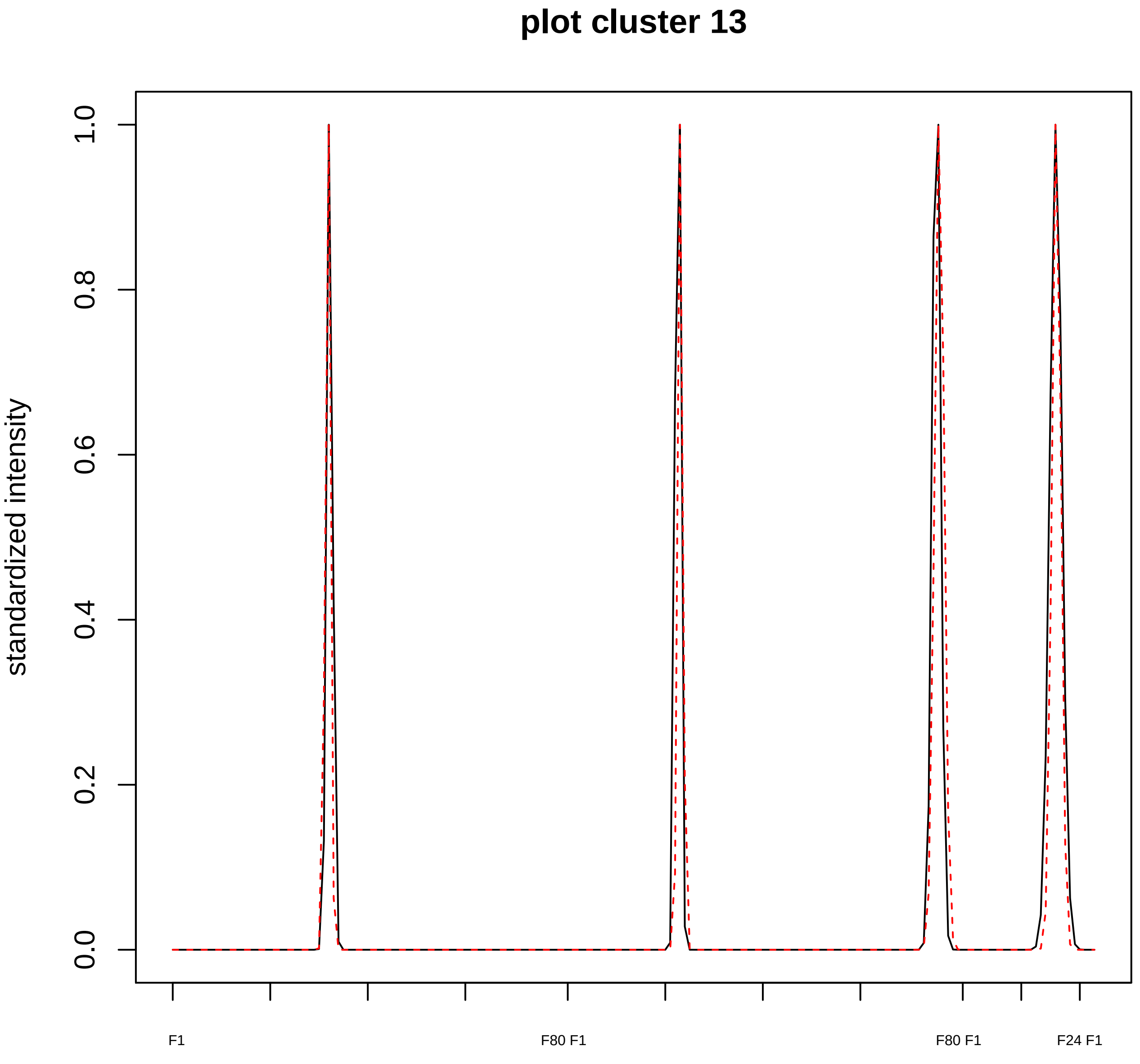
plot cluster 13
standardized intensity
0.0
0.2
0.4
0.6
0.8
1.0
F1
F80 F1
F80 F1
F24 F1

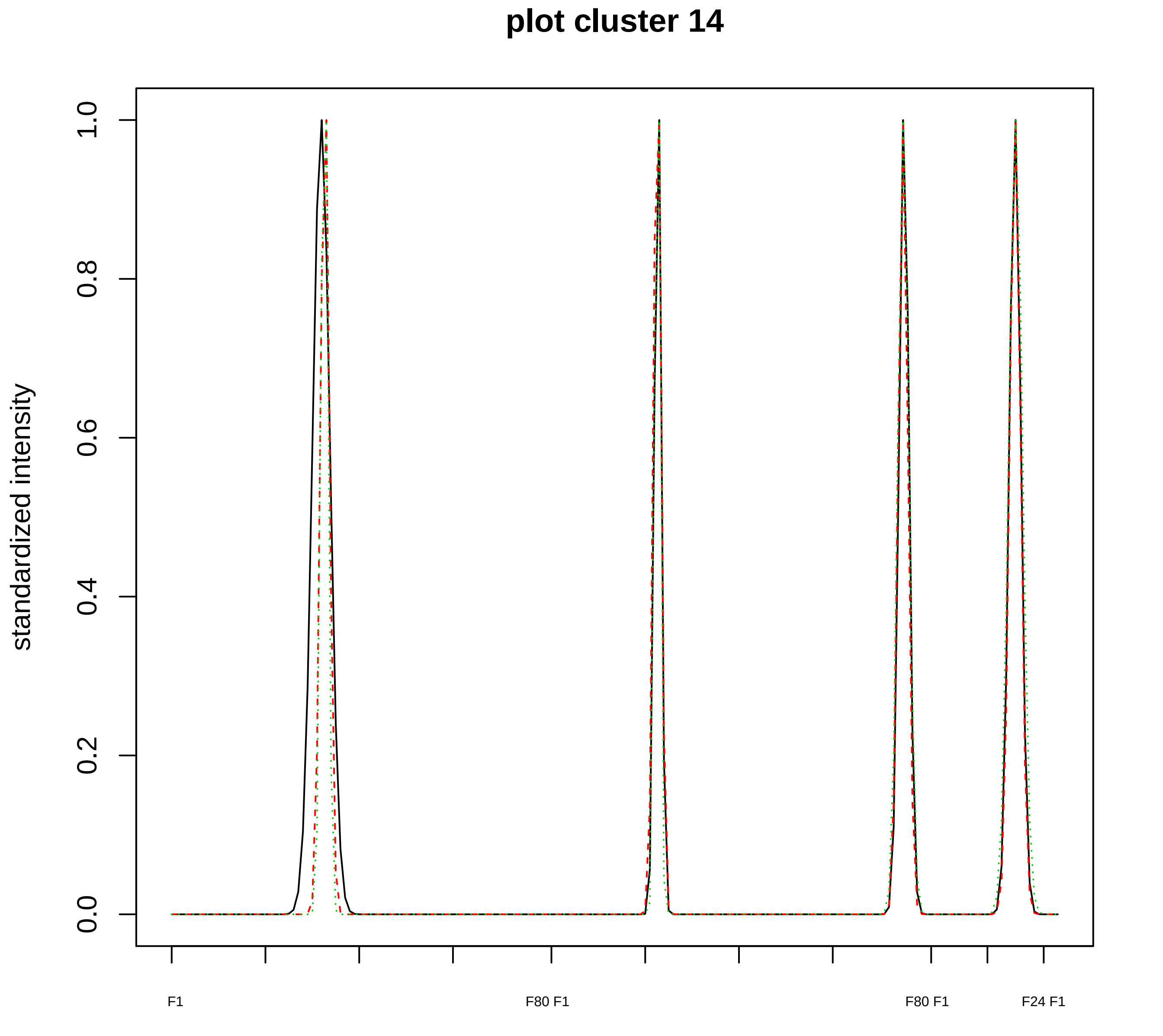

plot cluster 14
standardized intensity
0.0
0.2
0.4
0.6
0.8
1.0
F1
F80 F1
F80 F1
F24 F1

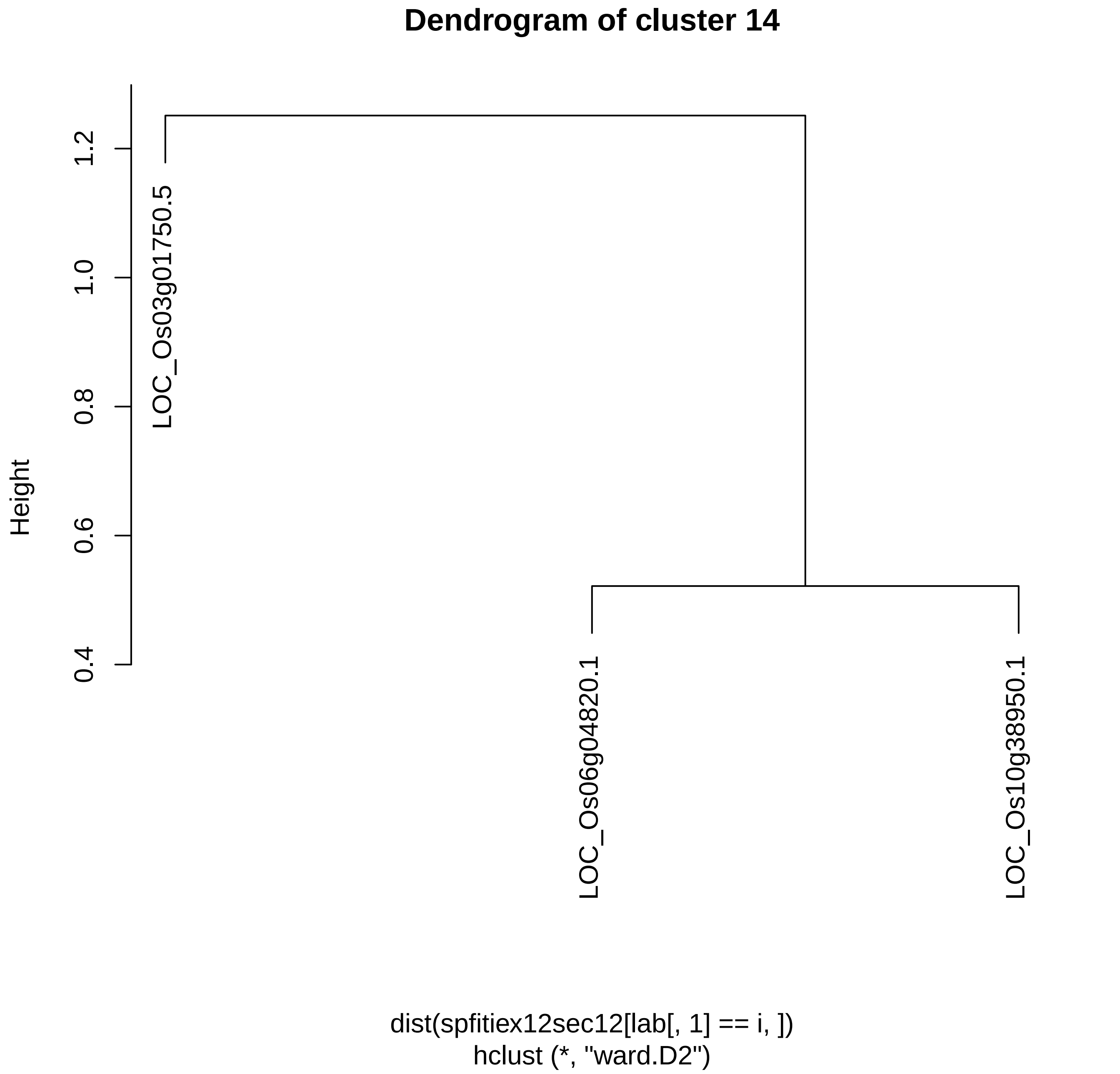
Dendrogram of cluster 14
Height
0.4
0.6
0.8
1.0
1.2
LOC_Os03g01750.5
LOC_Os06g04820.1
LOC_Os10g38950.1
dist(spfitiex12sec12[lab[, 1] == i, ])
hclust (*, "ward.D2")

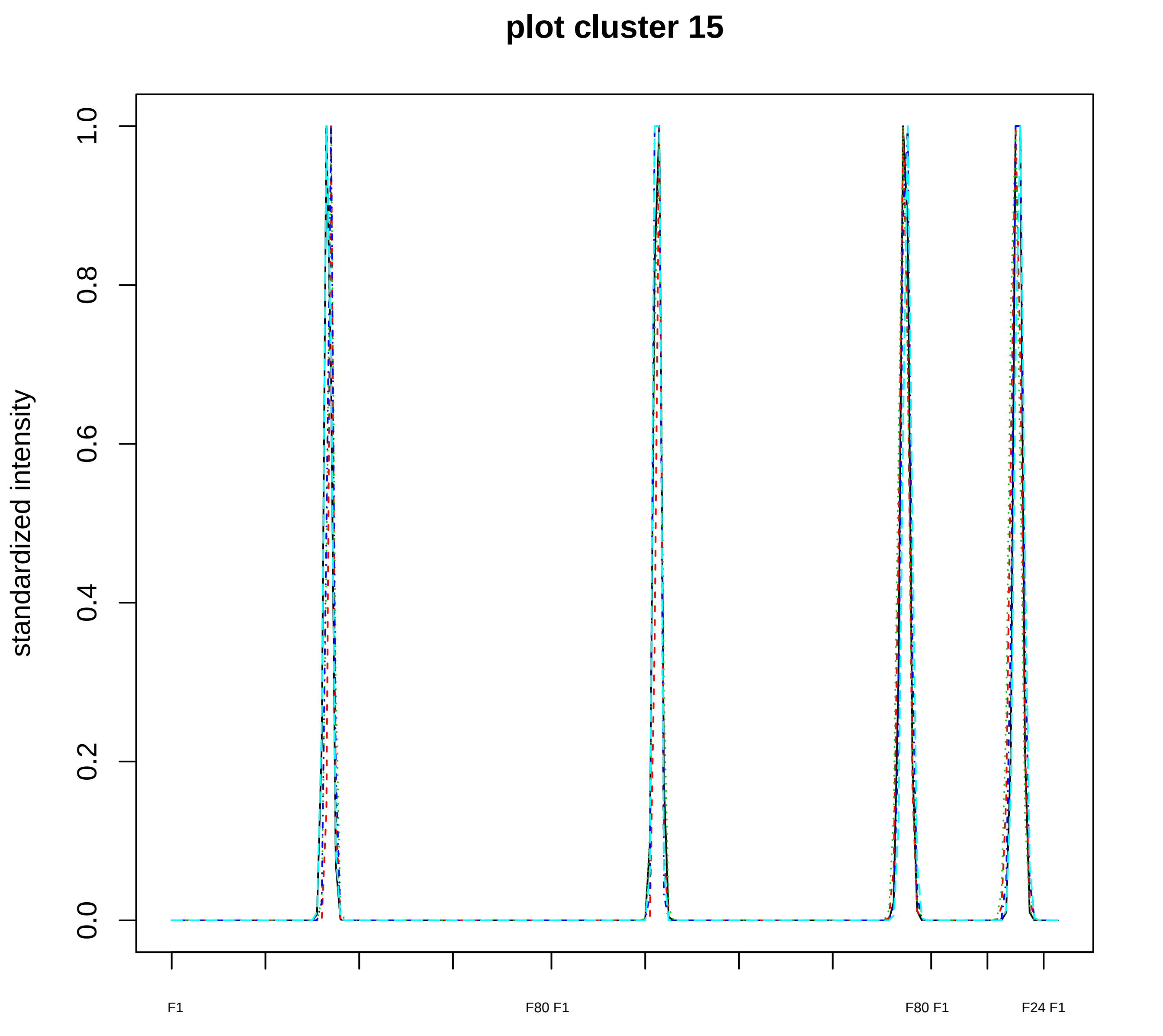

plot cluster 15
standardized intensity
0.0
0.2
0.4
0.6
0.8
1.0
F1
F80 F1
F80 F1
F24 F1

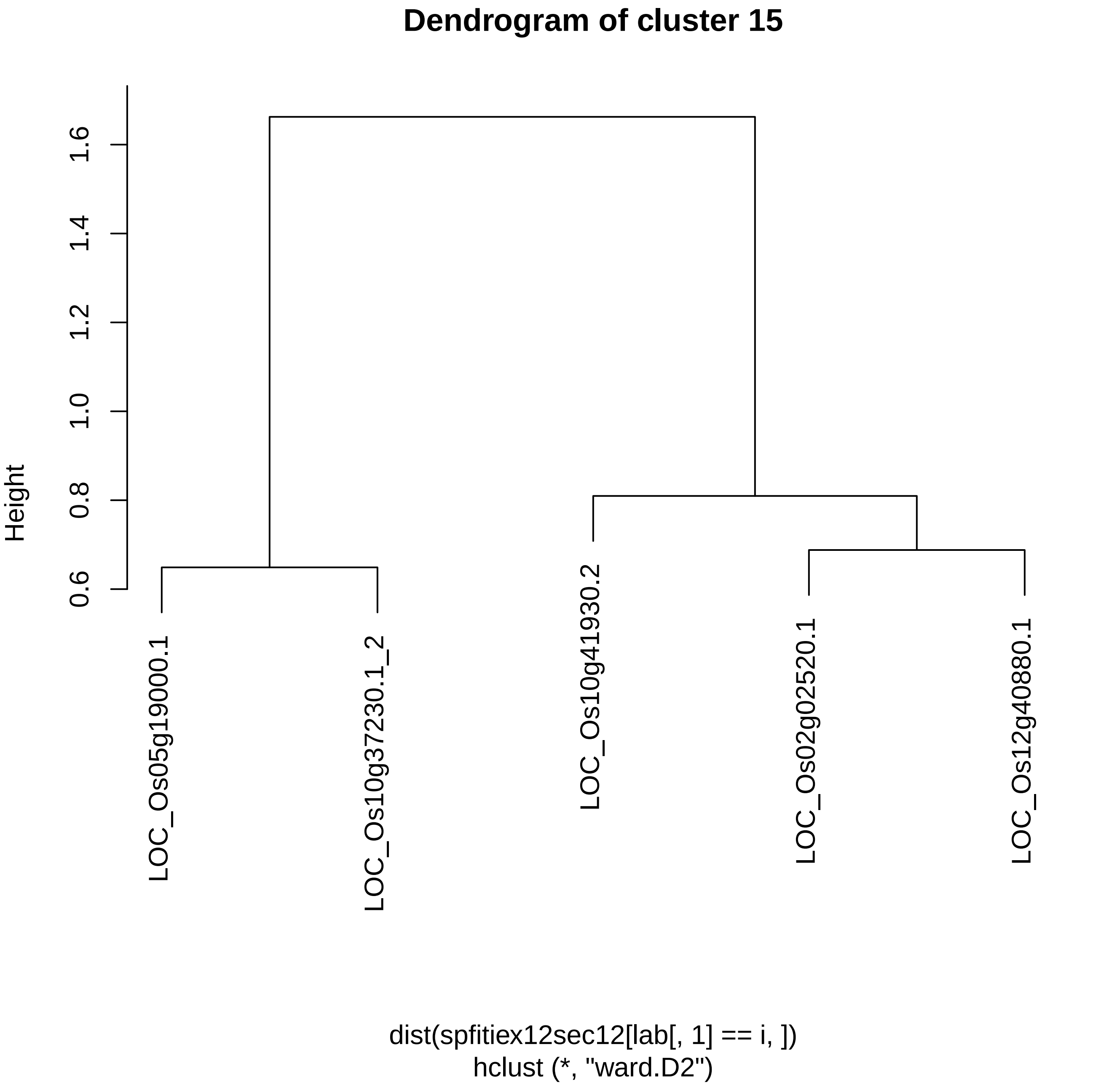
Dendrogram of cluster 15
Height
0.6
0.8
1.0
1.2
1.4
1.6
LOC_Os05g19000.1
LOC_Os10g37230.1_2
LOC_Os10g41930.2
LOC_Os02g02520.1
LOC_Os12g40880.1
dist(spfitiex12sec12[lab[, 1] == i, ])
hclust (*, "ward.D2")

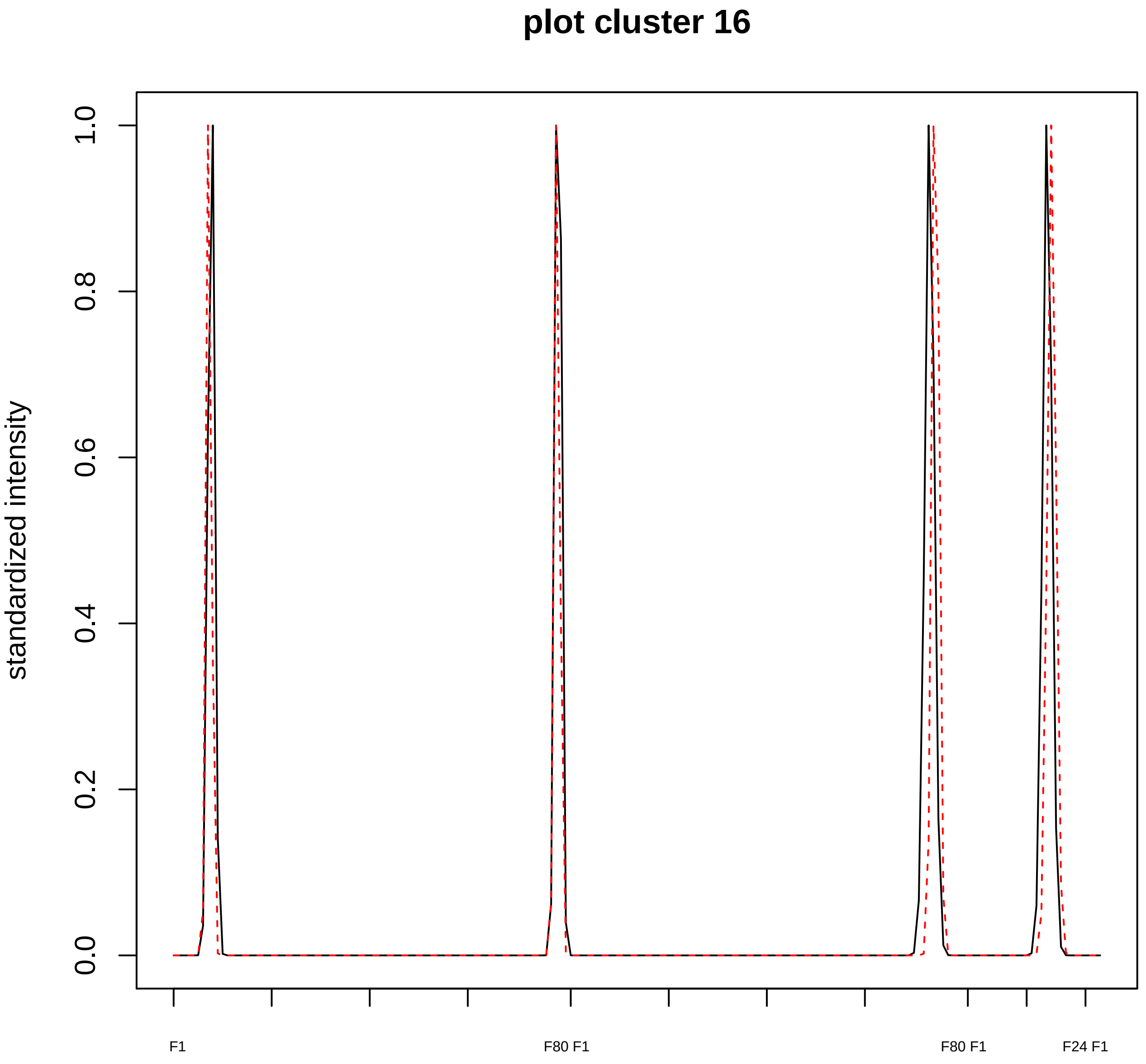

plot cluster 16
standardized intensity
0.0
0.2
0.4
0.6
0.8
1.0
F1
F80 F1
F80 F1
F24 F1

#### plot cluster 17

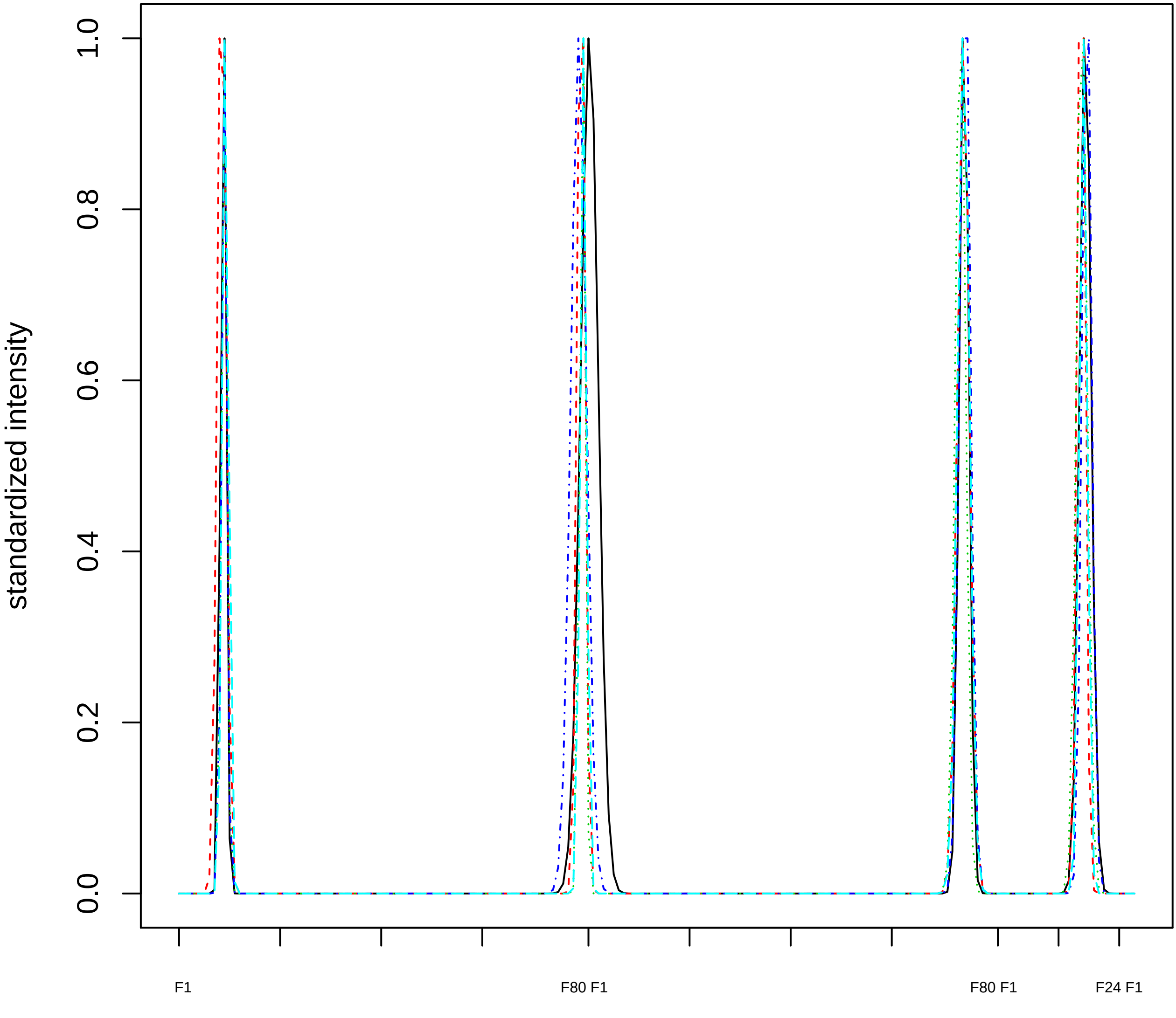

#### Dendrogram of cluster 17

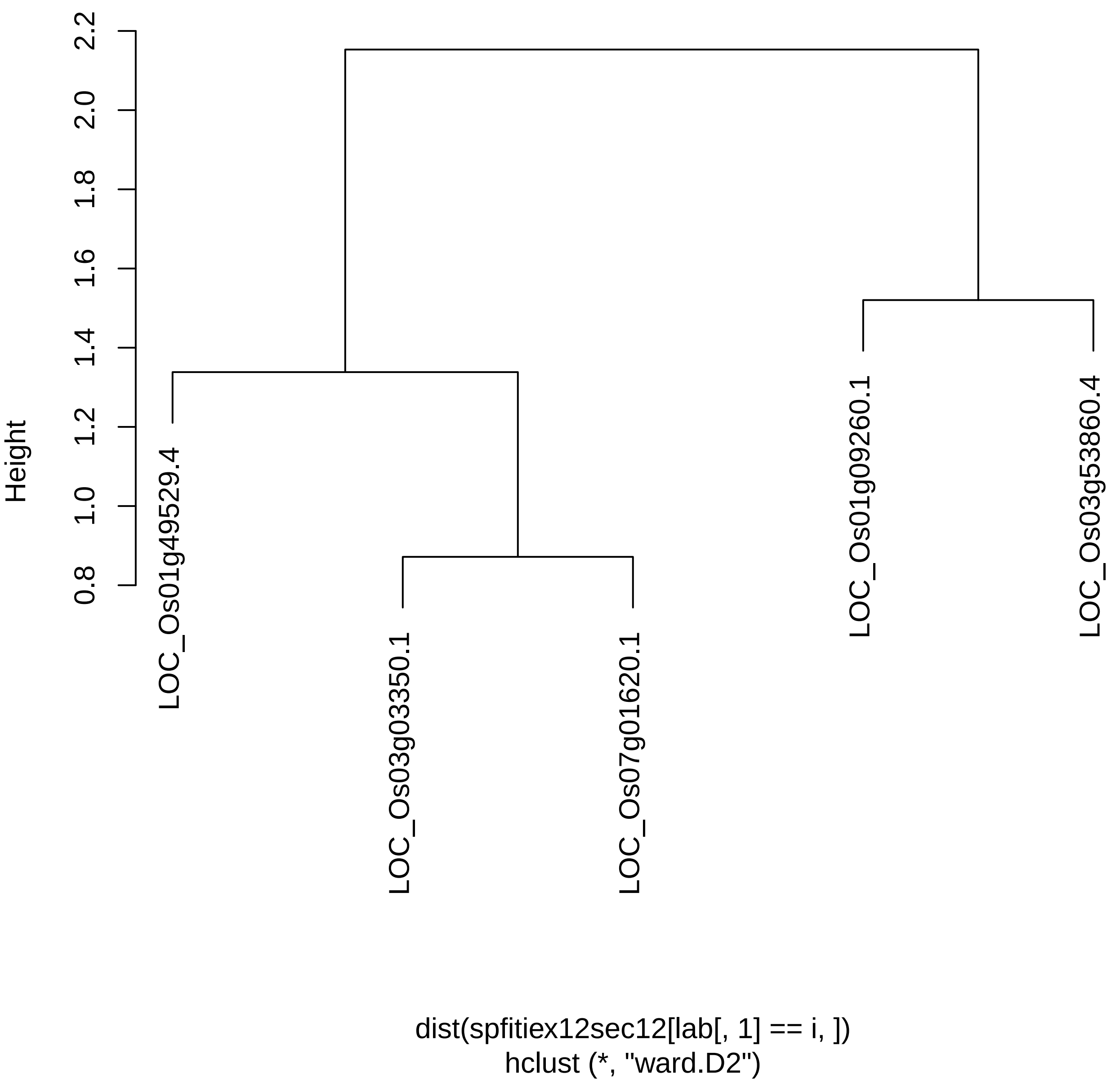

### plot cluster 18

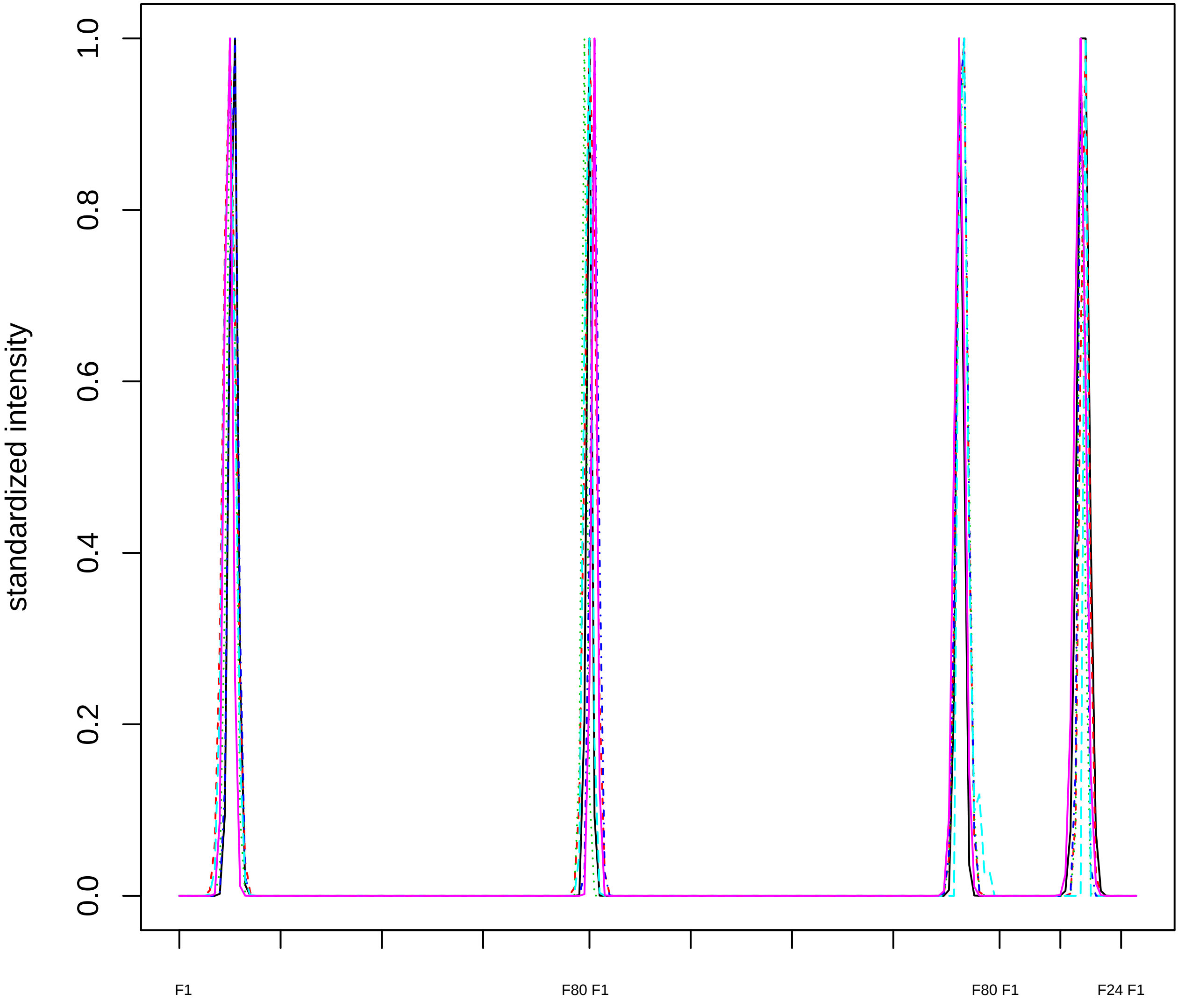

### Dendrogram of cluster 18

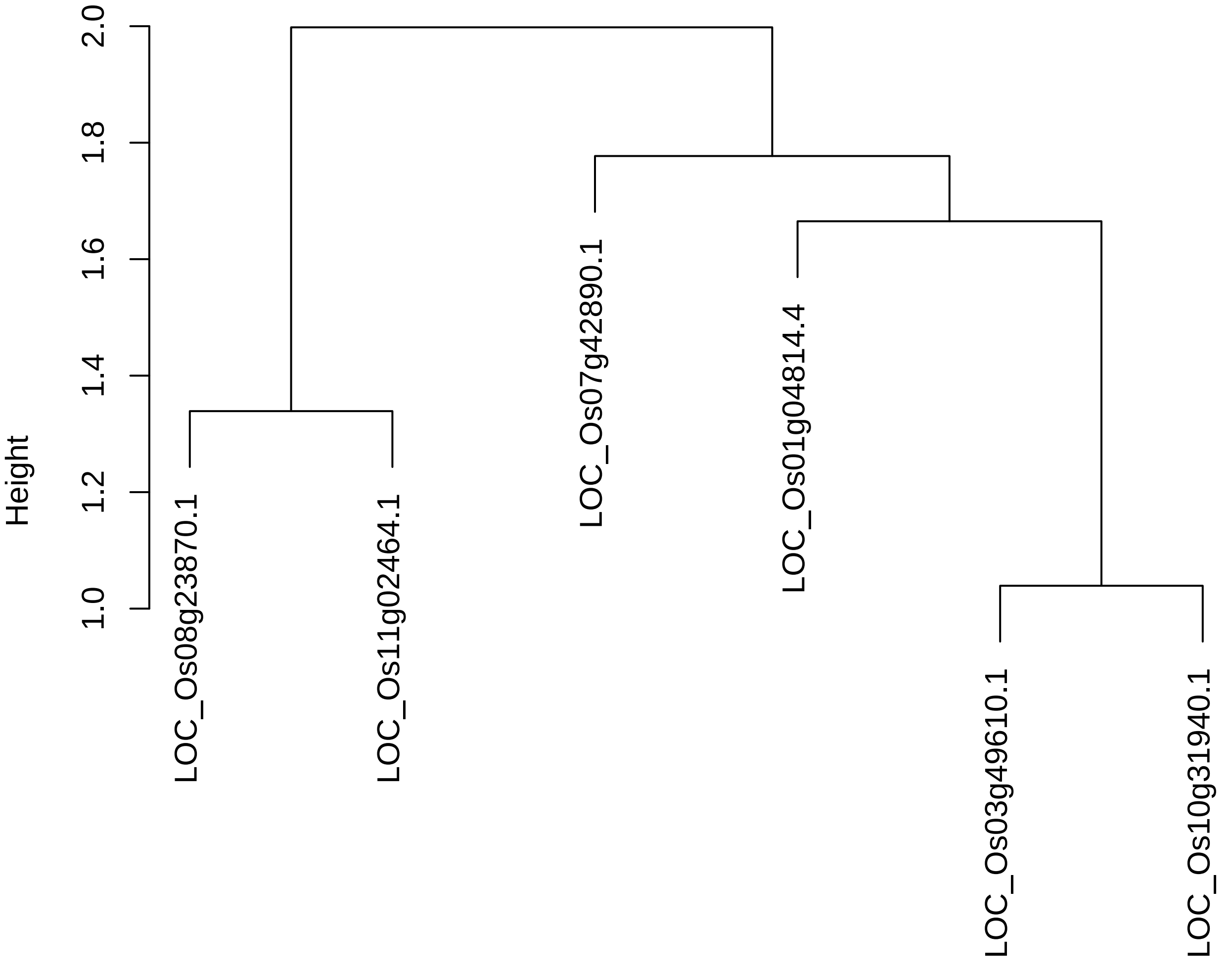


dist(spfitiex12sec12[lab[, 1] == i, ])
hclust (*, "ward.D2")

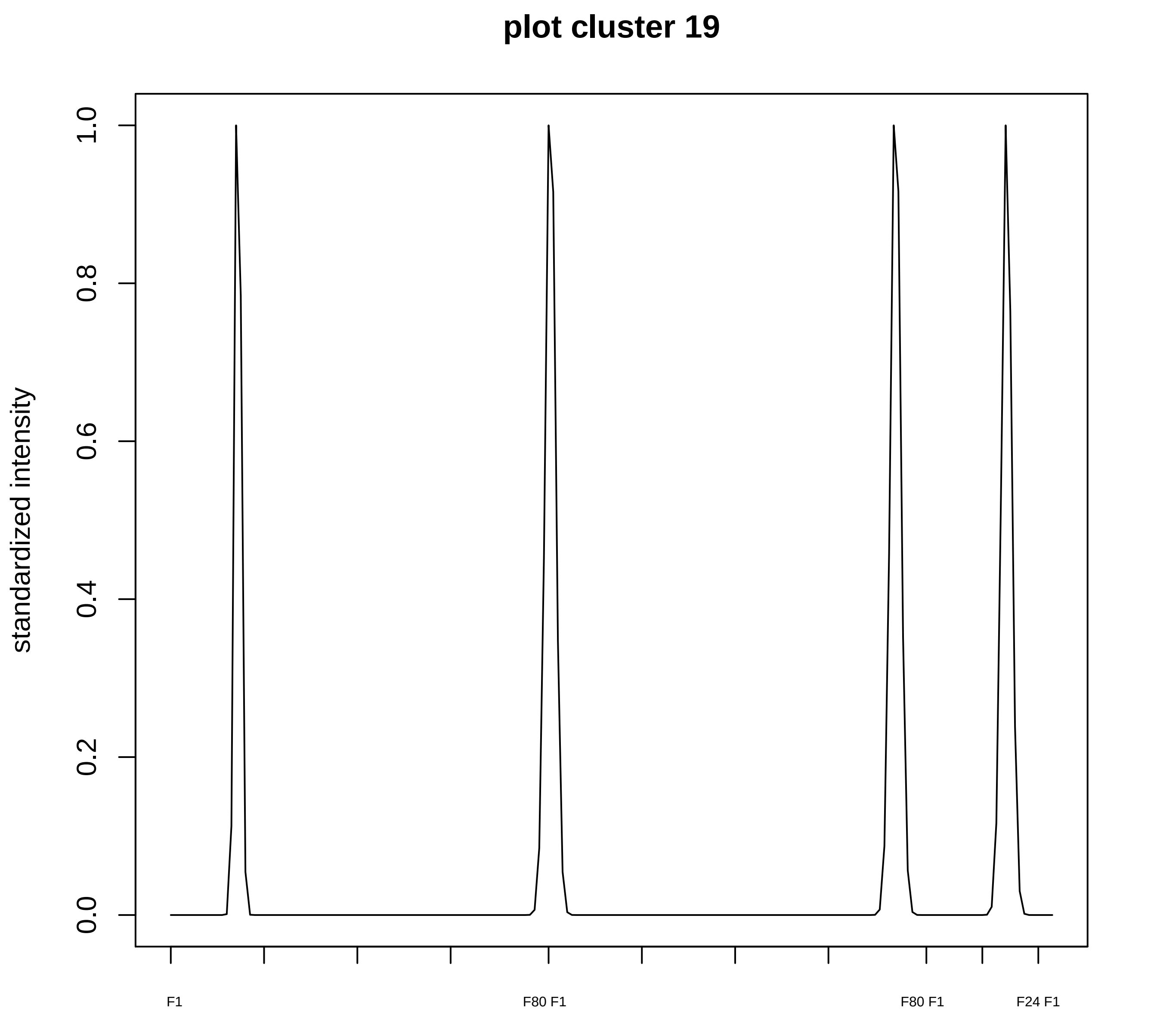
plot cluster 19
standardized intensity
0.0
0.2
0.4
0.6
0.8
1.0
F1
F80 F1
F80 F1
F24 F1

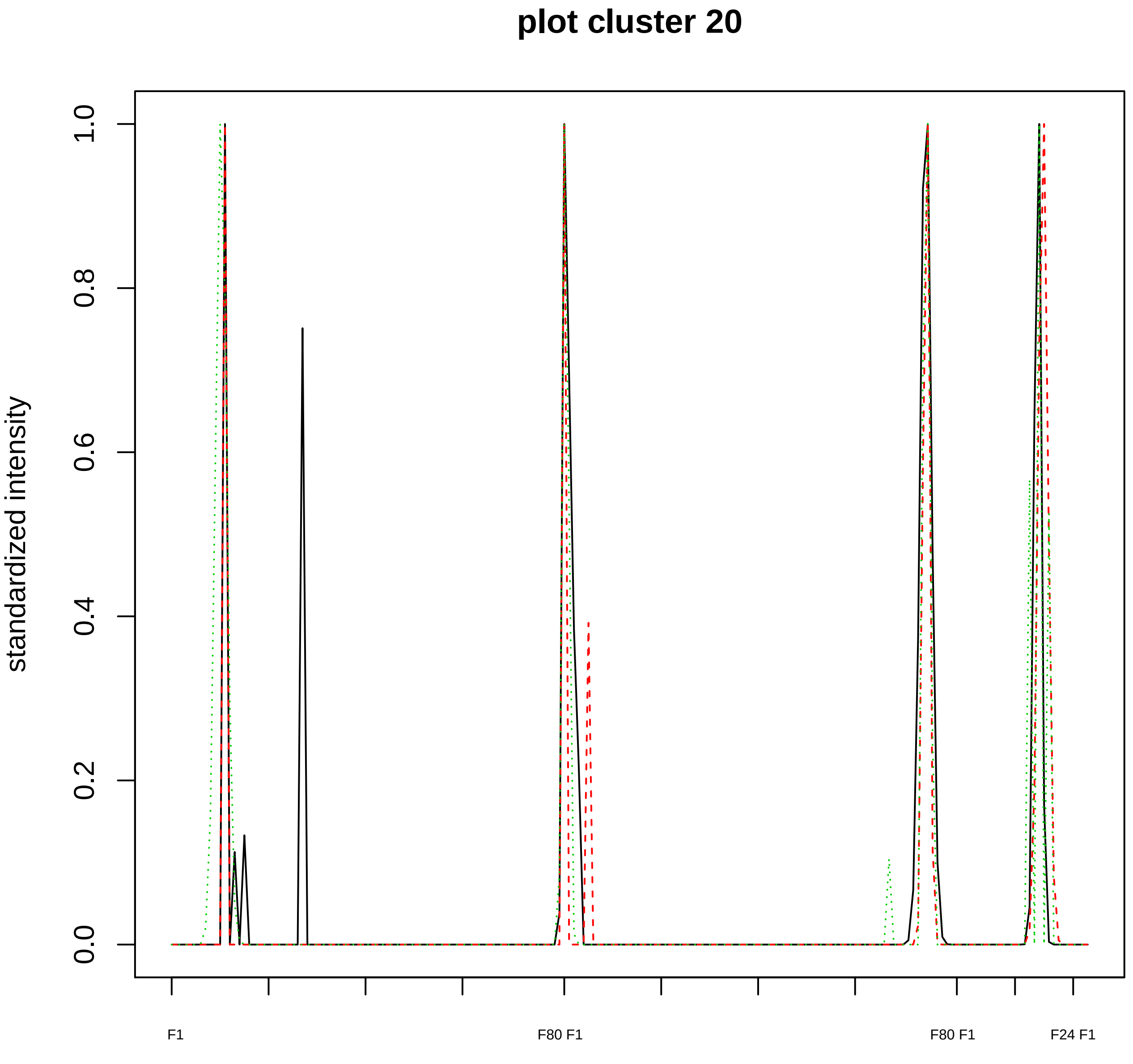
plot cluster 20
standardized intensity
0.0
0.2
0.4
0.6
0.8
1.0
F1
F80 F1
F80 F1
F24 F1

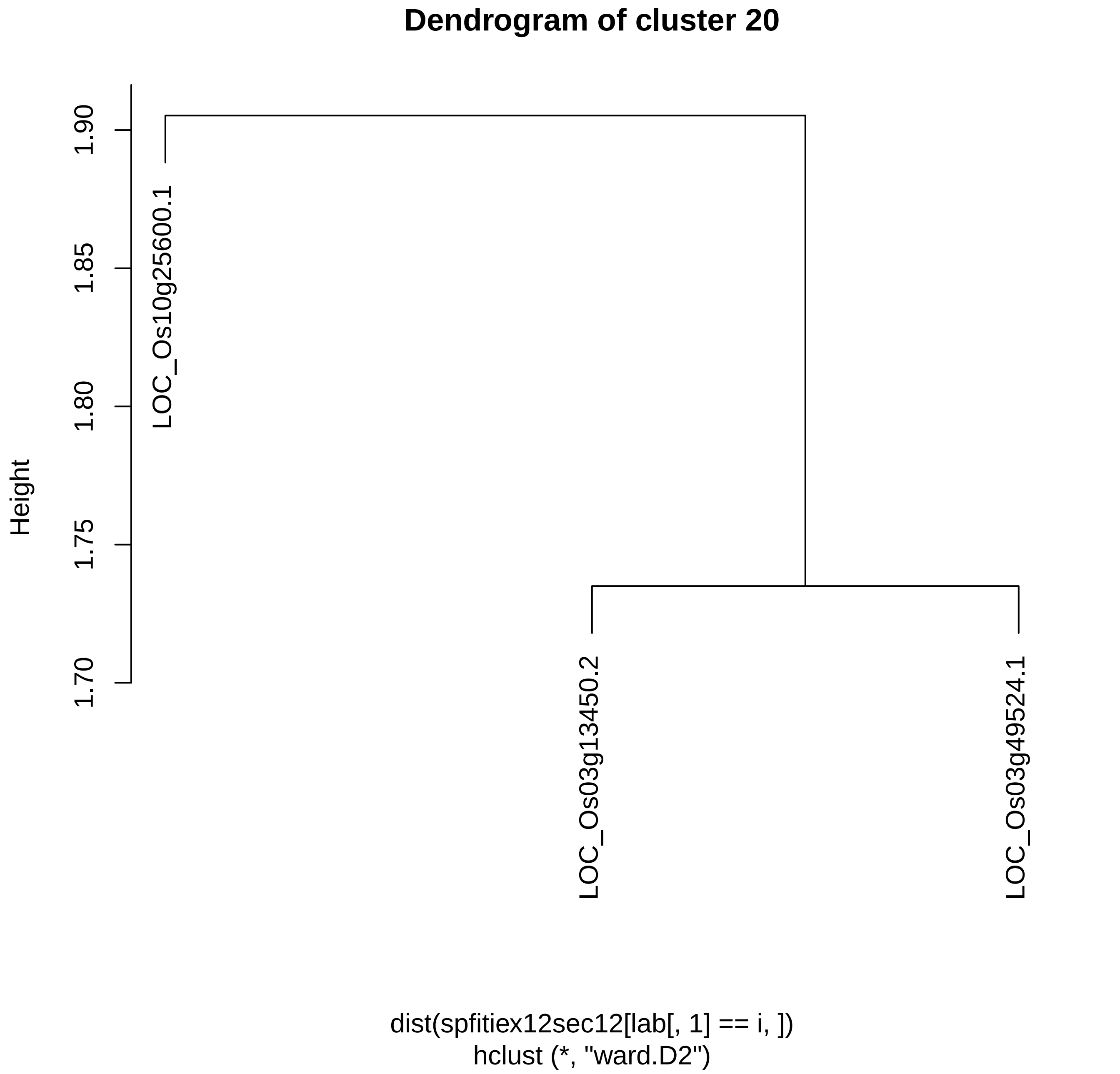
Dendrogram of cluster 20
Height
1.70
1.75
1.80
1.85
1.90
LOC_Os10g25600.1
LOC_Os03g13450.2
LOC_Os03g49524.1
dist(spfitiex12sec12[lab[, 1] == i, ])
hclust (*, "ward.D2")

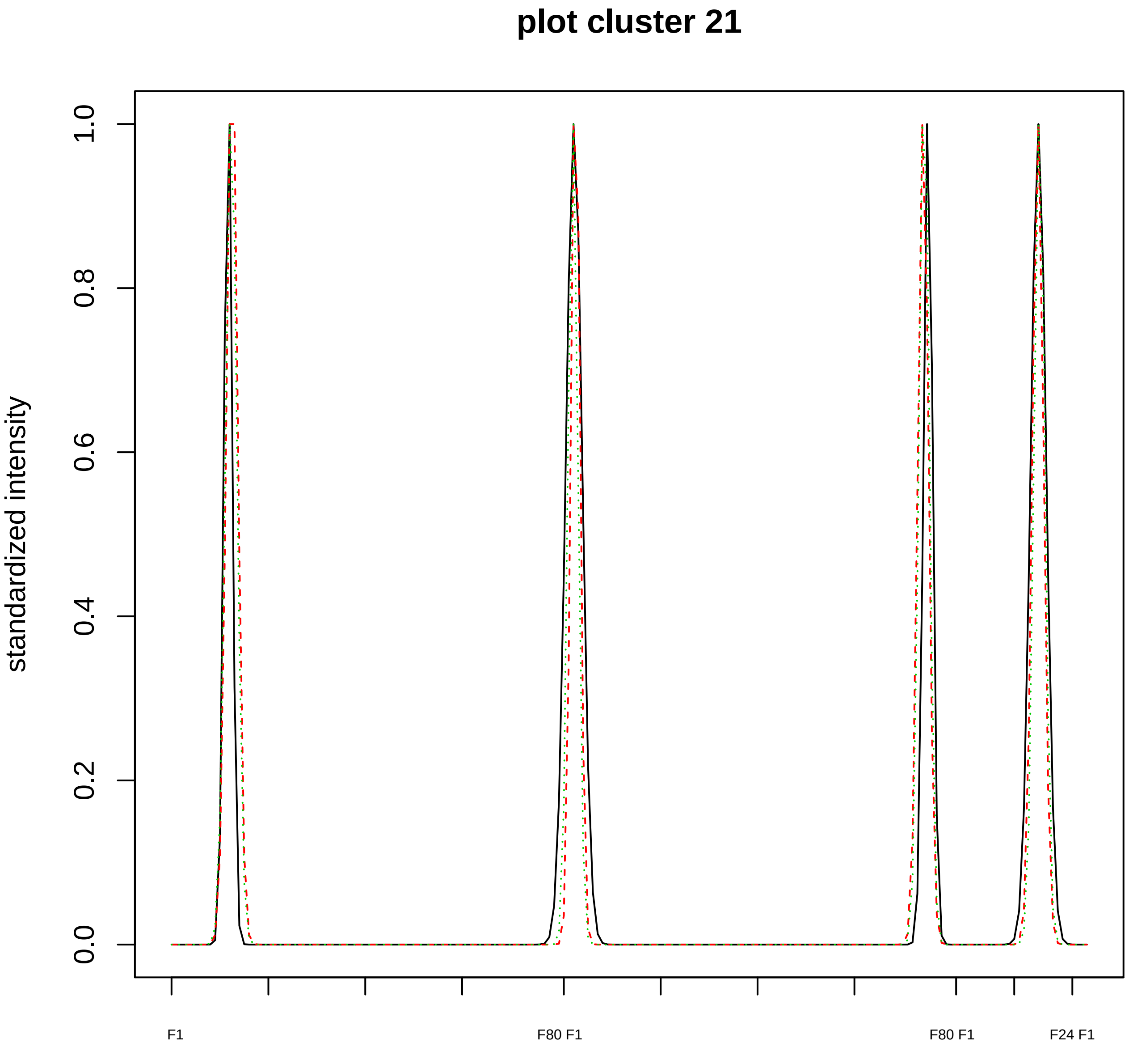
plot cluster 21
standardized intensity
0.0
0.2
0.4
0.6
0.8
1.0
F1
F80 F1
F80 F1
F24 F1

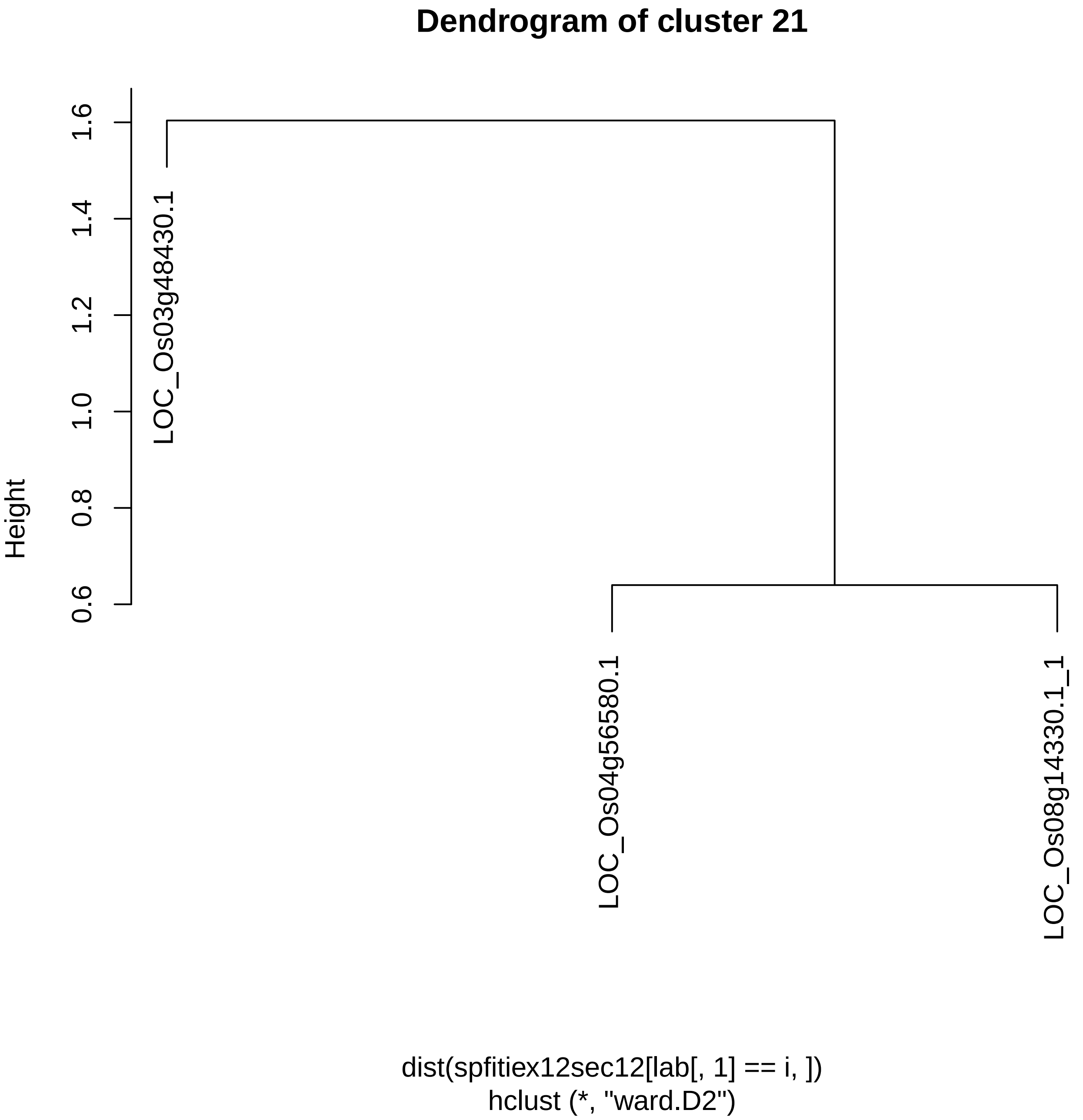
Dendrogram of cluster 21
Height
0.6
0.8
1.0
1.2
1.4
1.6
LOC_Os03g48430.1
LOC_Os04g56580.1
LOC_Os08g14330.1_1
dist(spfitiex12sec12[lab[, 1] == i, ])
hclust (*, "ward.D2")

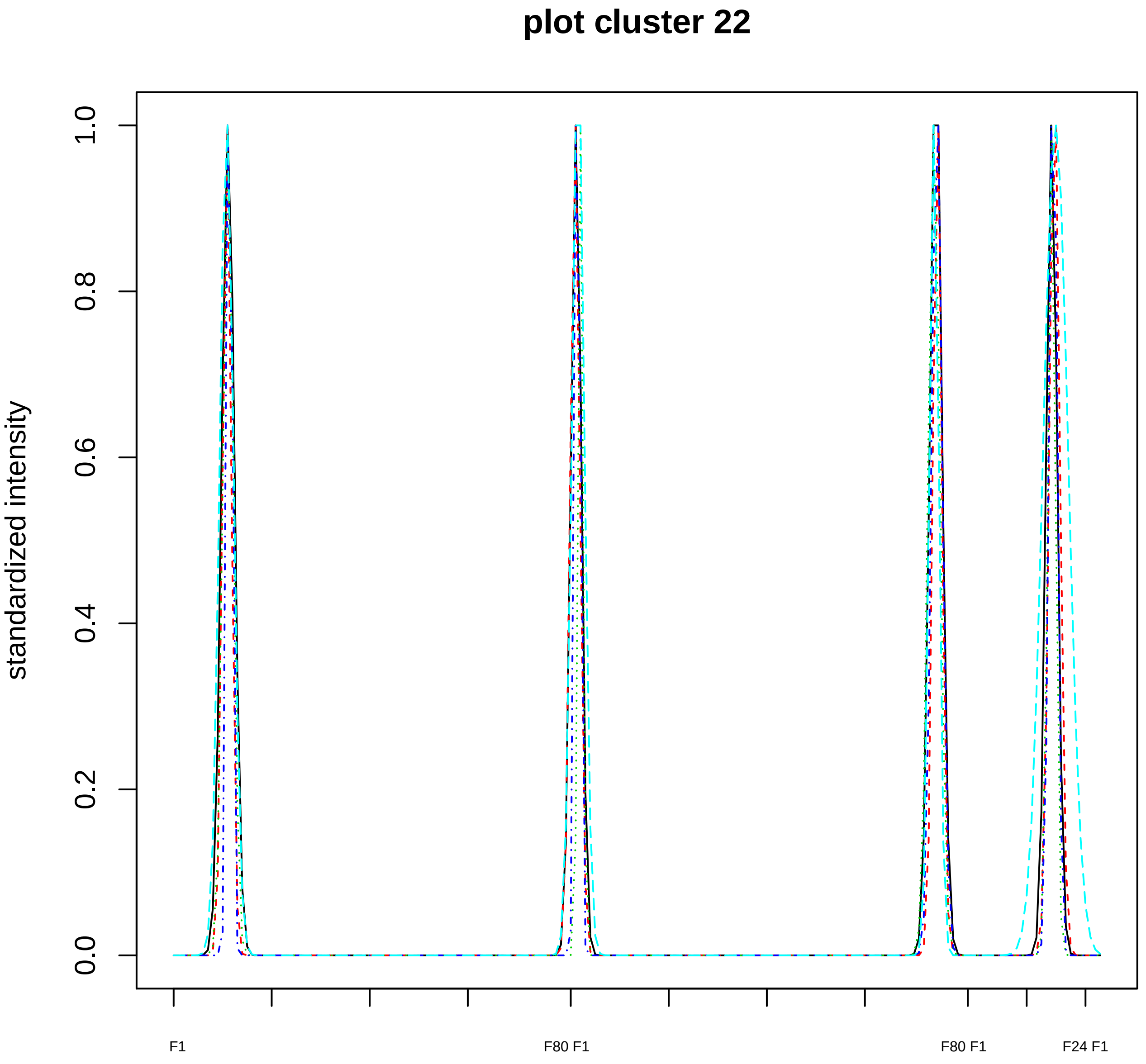
plot cluster 22
standardized intensity
0.0
0.2
0.4
0.6
0.8
1.0
F1
F80 F1
F80 F1
F24 F1

### Dendrogram of cluster 22

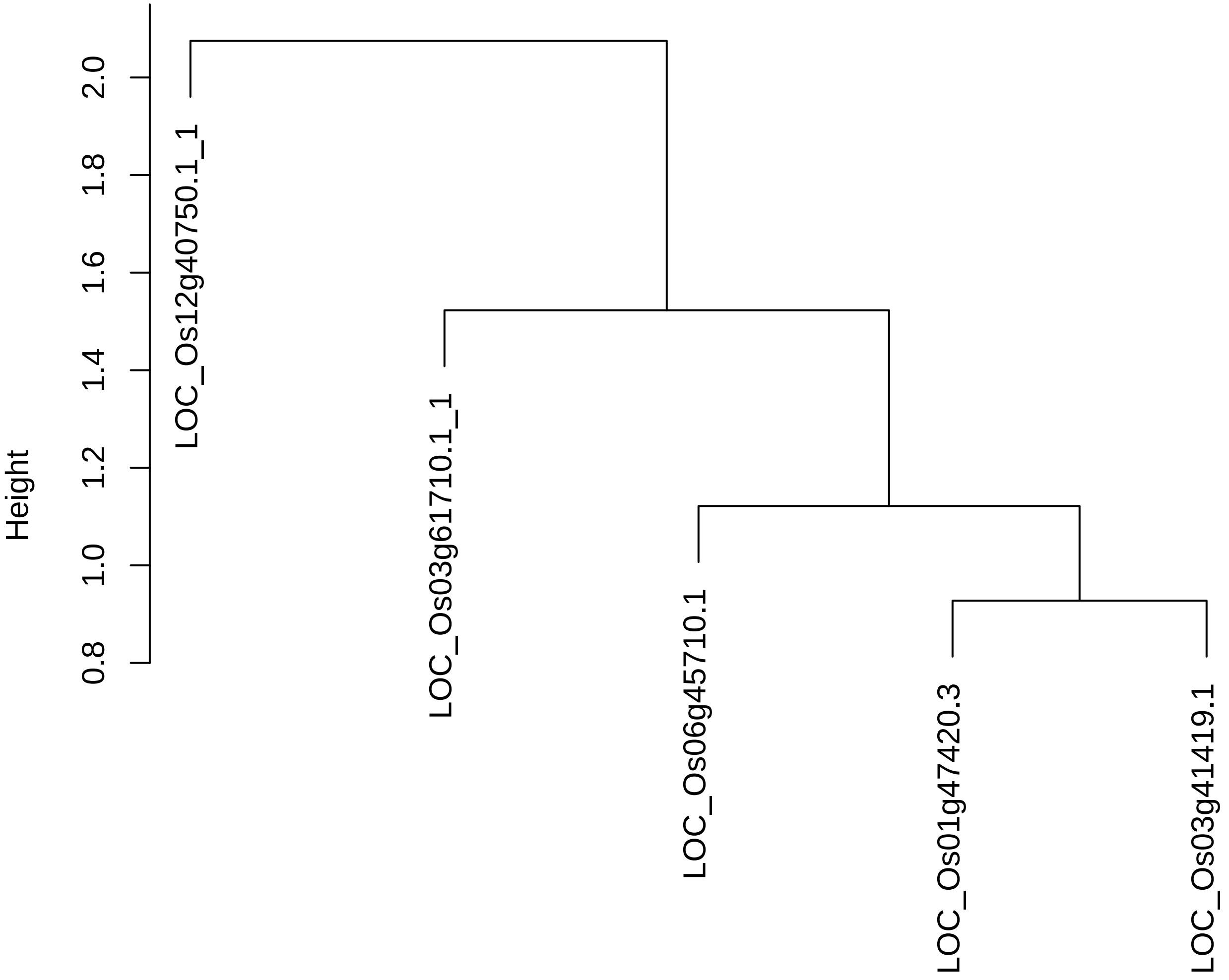


dist(spfitiex12sec12[lab[, 1] == i, ])
hclust (*, "ward.D2")

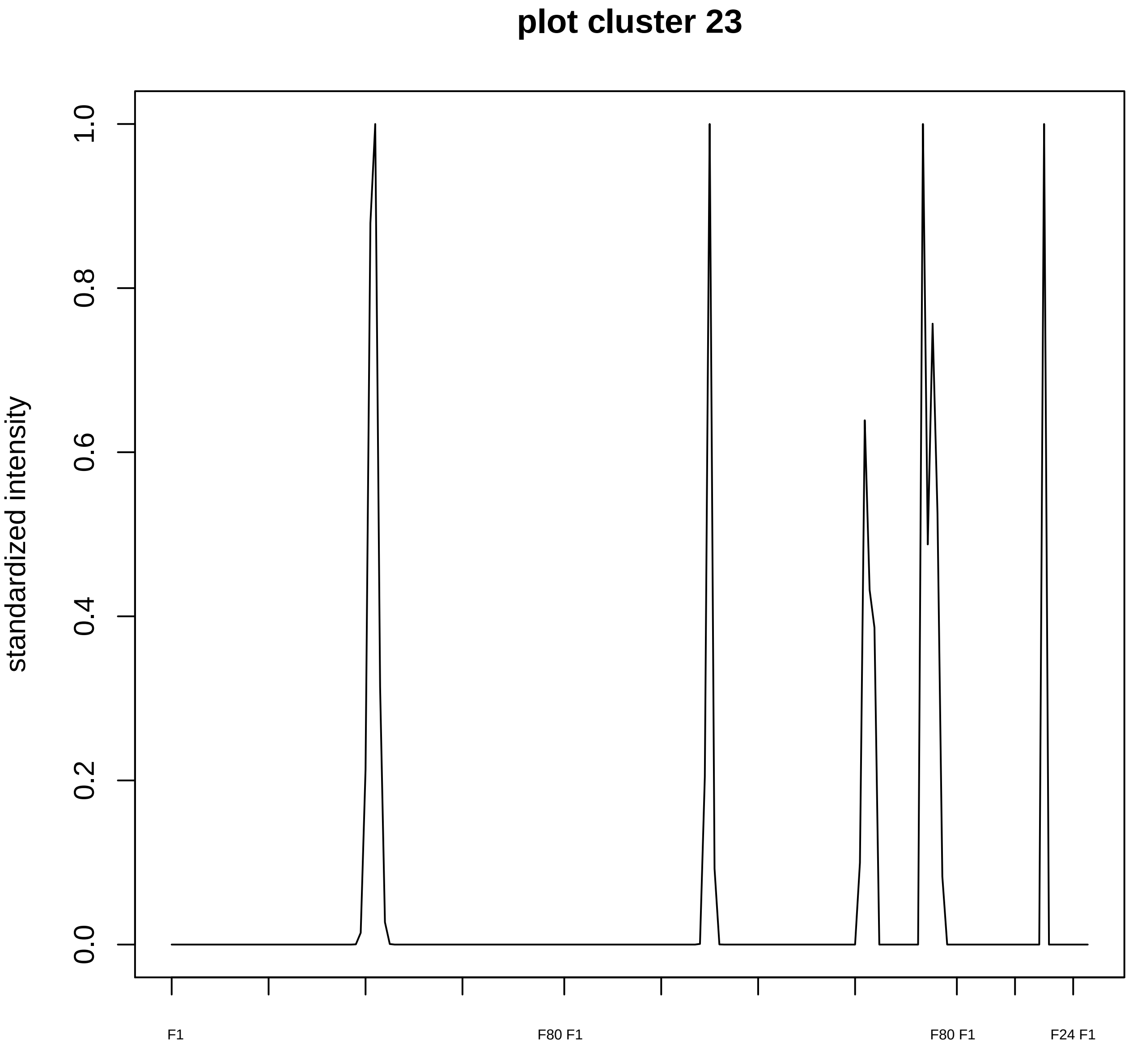

plot cluster 23
standardized intensity
0.0
0.2
0.4
0.6
0.8
1.0
F1
F80 F1
F80 F1
F24 F1

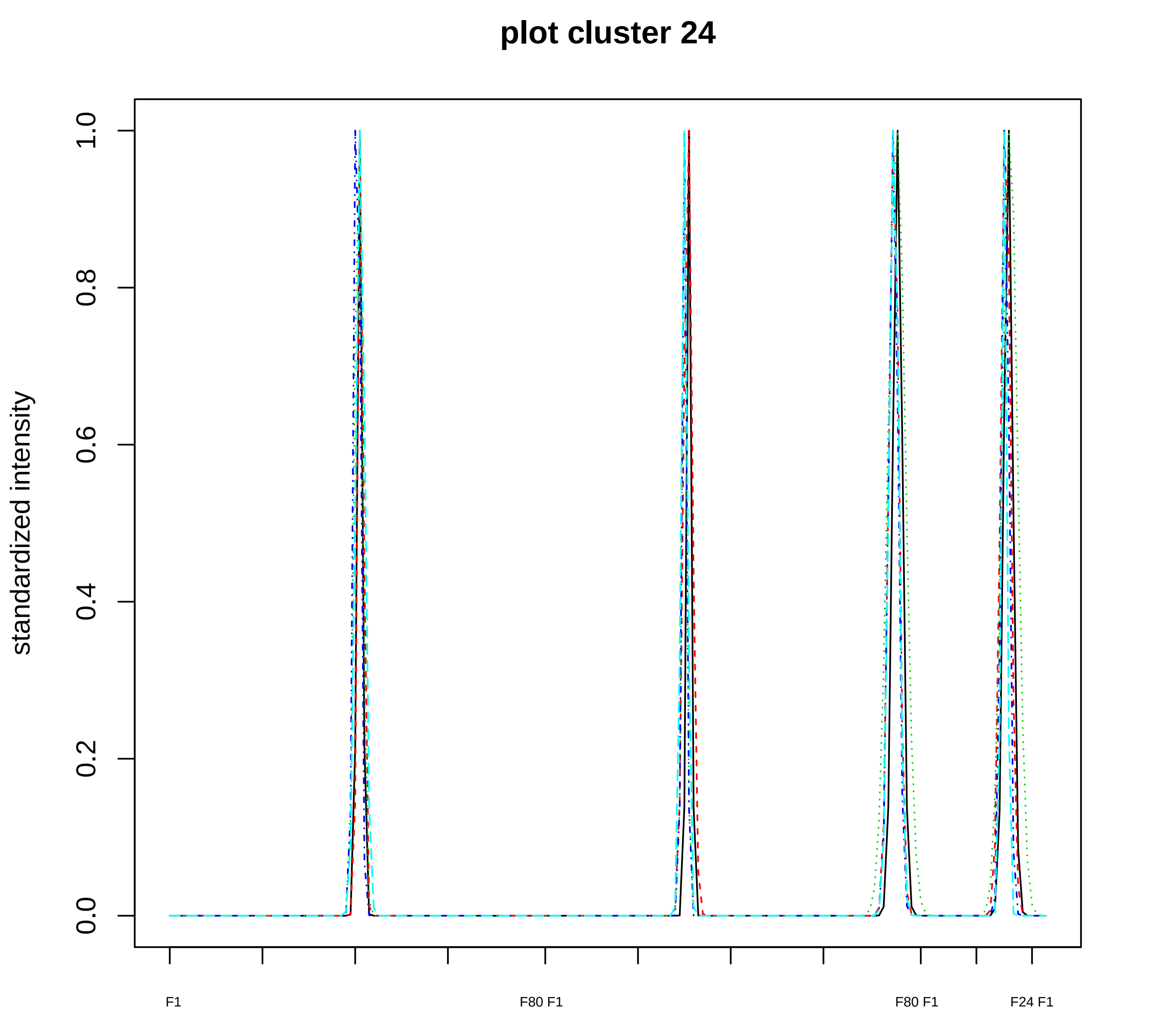

plot cluster 24
standardized intensity
0.0
0.2
0.4
0.6
0.8
1.0
F1
F80 F1
F80 F1
F24 F1

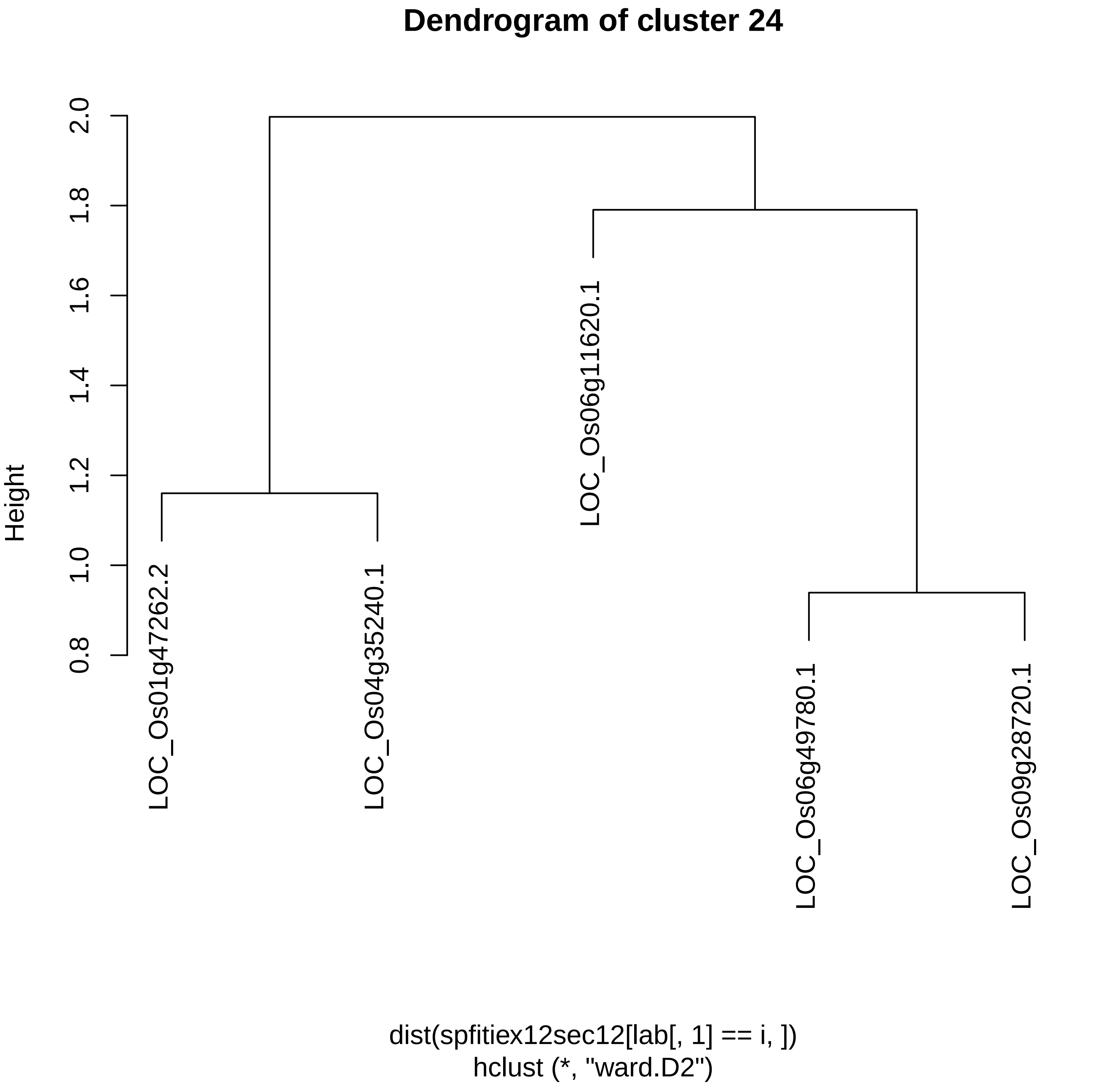
Dendrogram of cluster 24
Height
2.0
1.8
1.6
1.4
1.2
1.0
0.8
LOC_Os01g47262.2
LOC_Os04g35240.1
LOC_Os06g11620.1
LOC_Os06g49780.1
LOC_Os09g28720.1
dist(spfitiex12sec12[lab[, 1] == i, ])
hclust (*, "ward.D2")

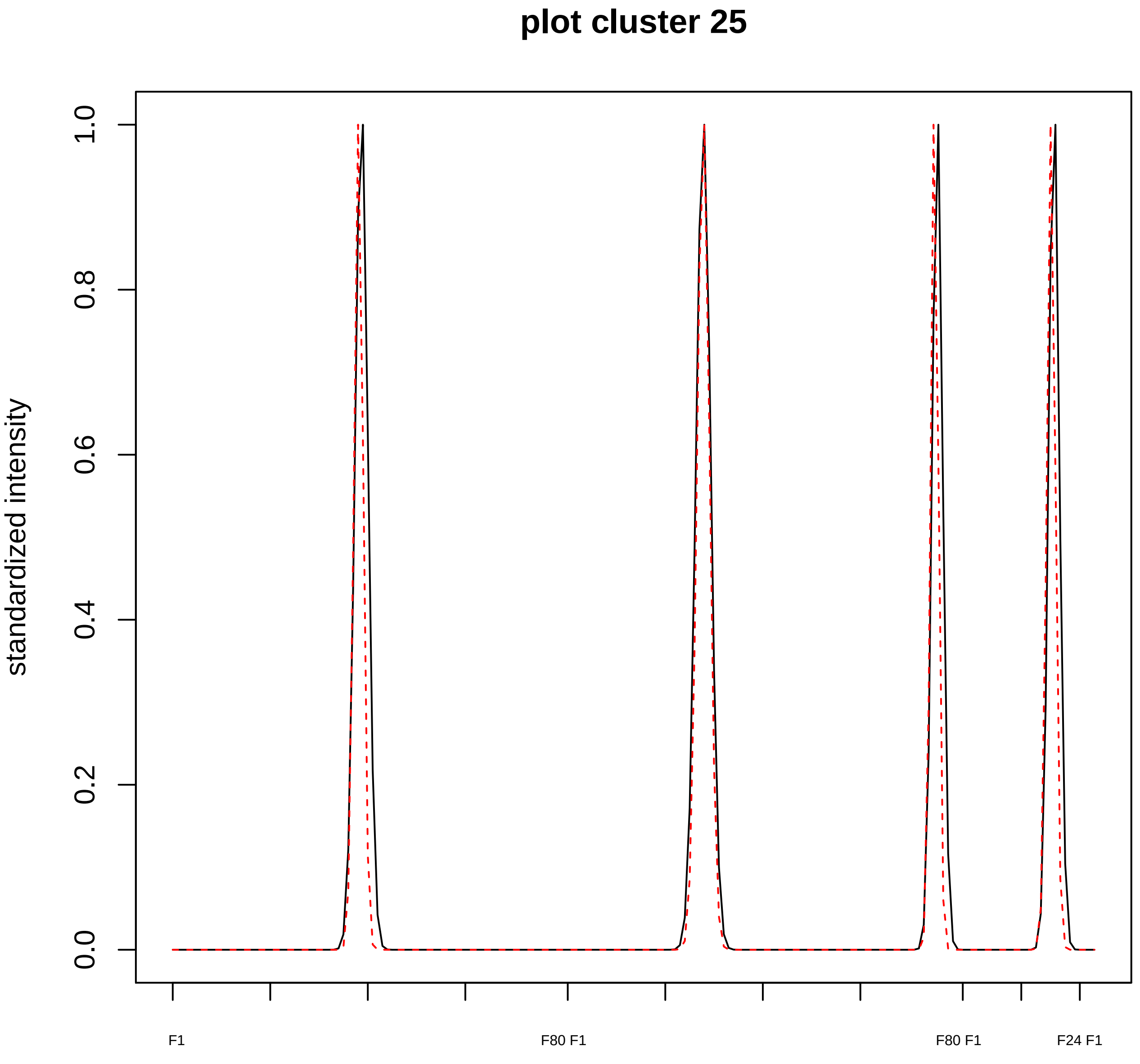
plot cluster 25
standardized intensity
0.0
0.2
0.4
0.6
0.8
1.0
F1
F80 F1
F80 F1
F24 F1

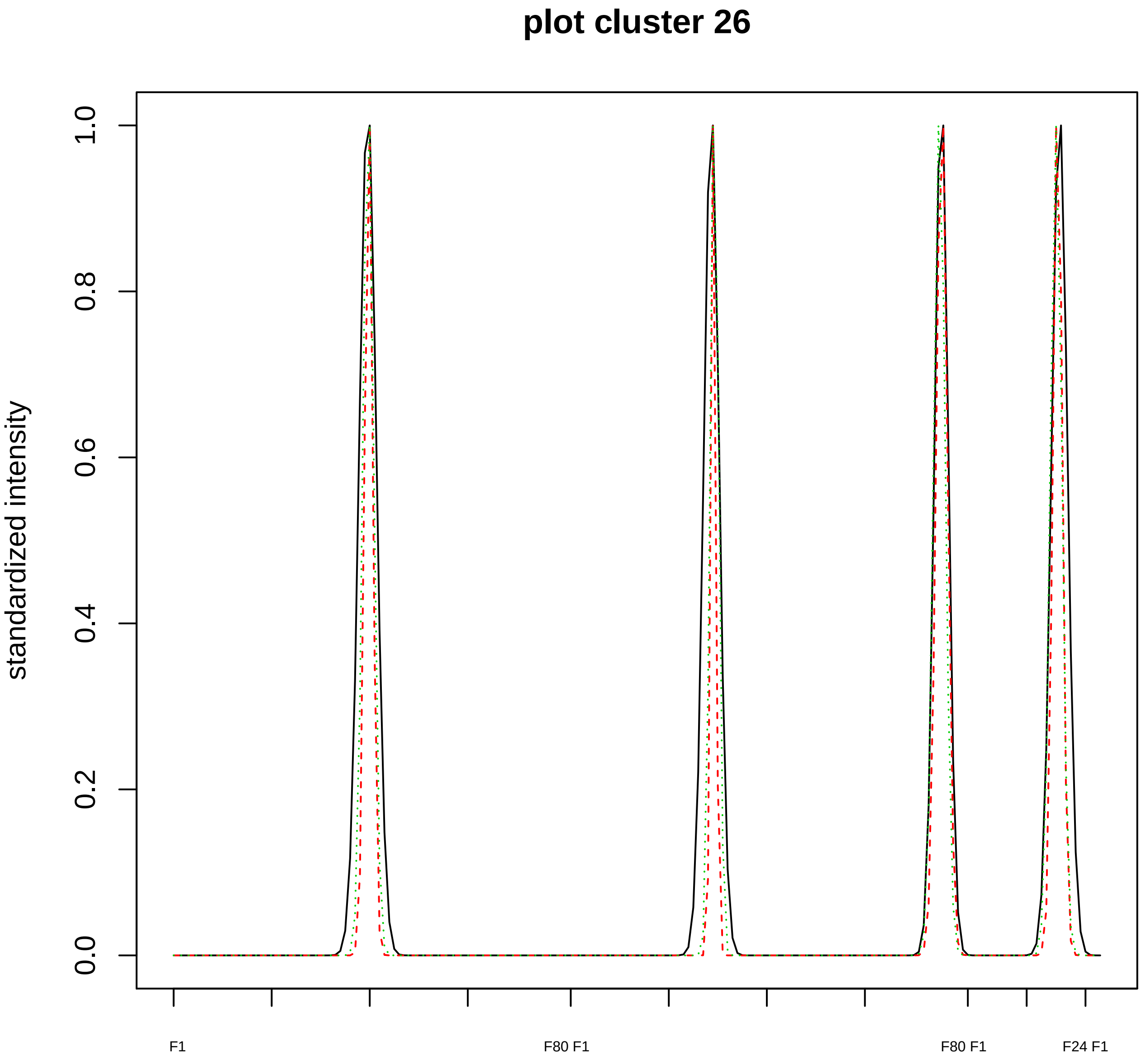
plot cluster 26
standardized intensity
0.0
0.2
0.4
0.6
0.8
1.0
F1
F80 F1
F80 F1
F24 F1

#### Dendrogram of cluster 26

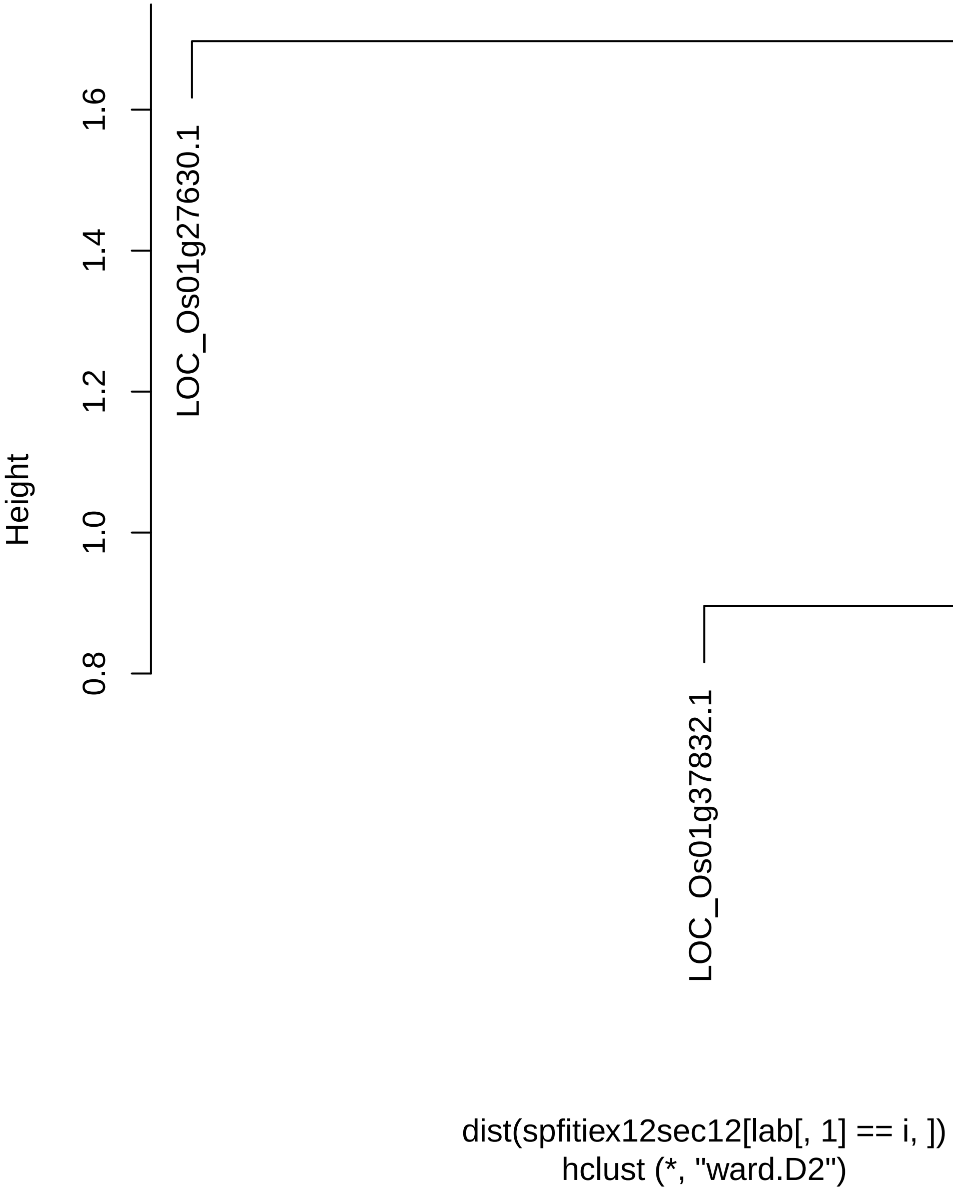

### plot cluster 27

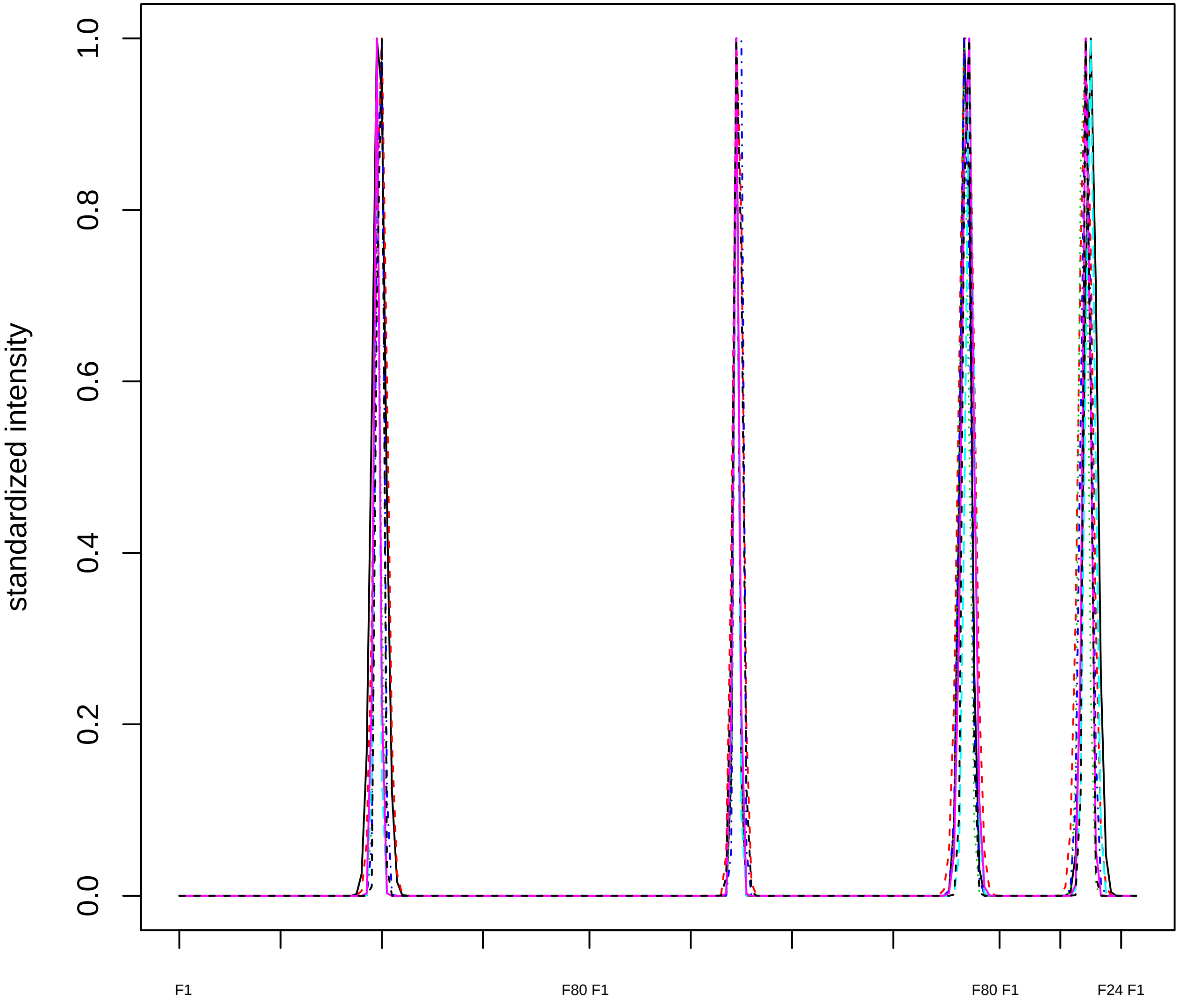

#### Dendrogram of cluster 27

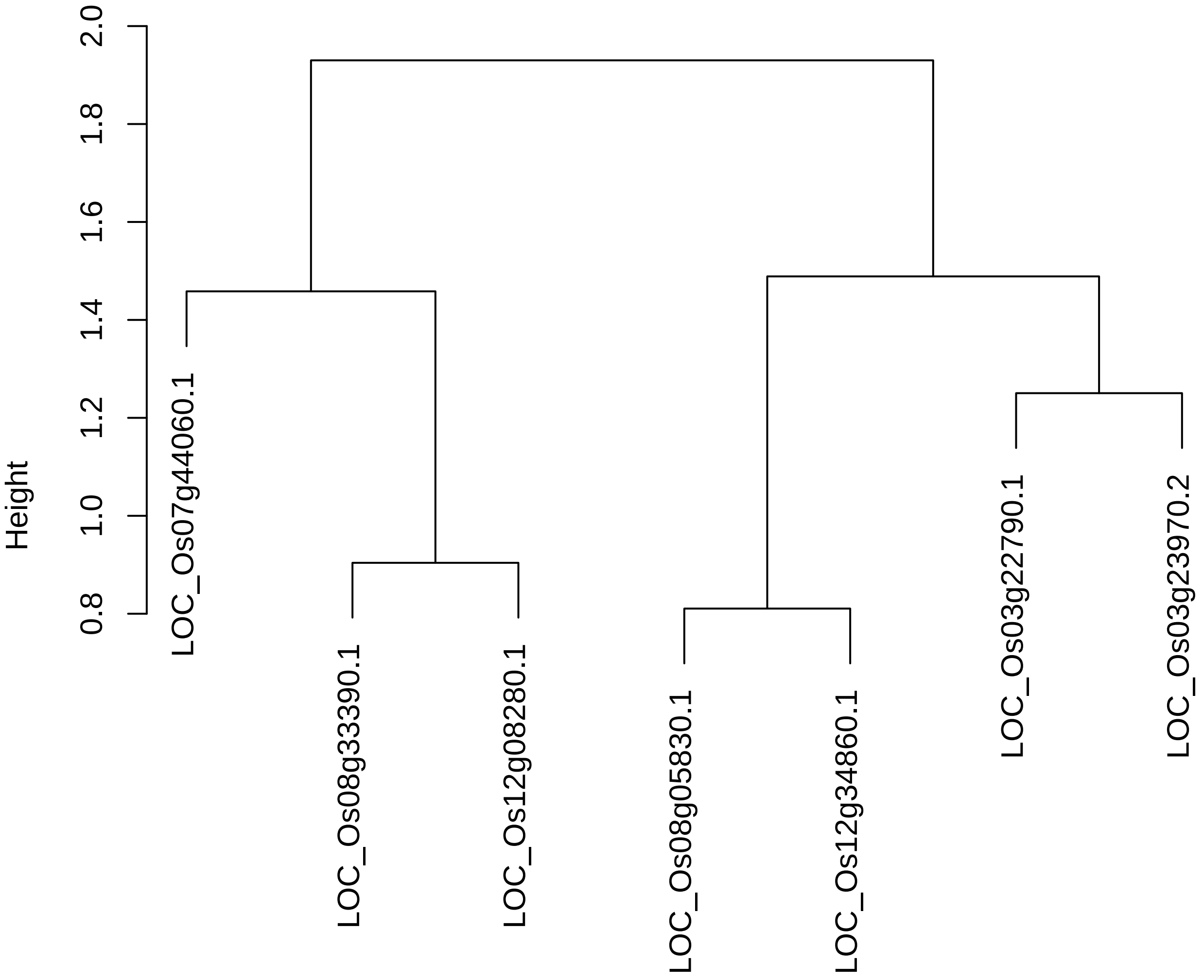


dist(spfitiex12sec12[lab[, 1] == i, ])
hclust (*, "ward.D2")

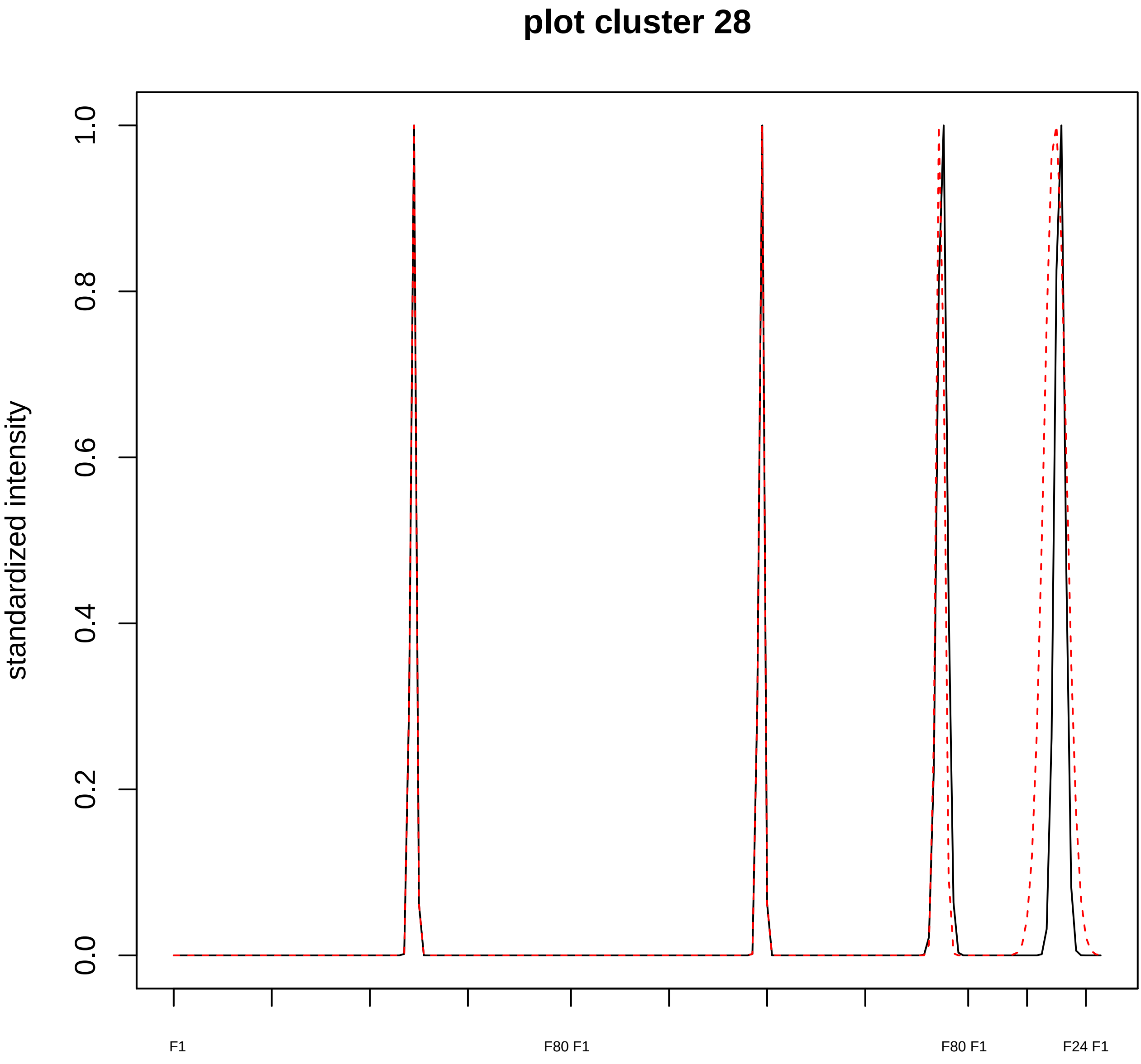

plot cluster 28
standardized intensity
0.0
0.2
0.4
0.6
0.8
1.0
F1
F80 F1
F80 F1
F24 F1

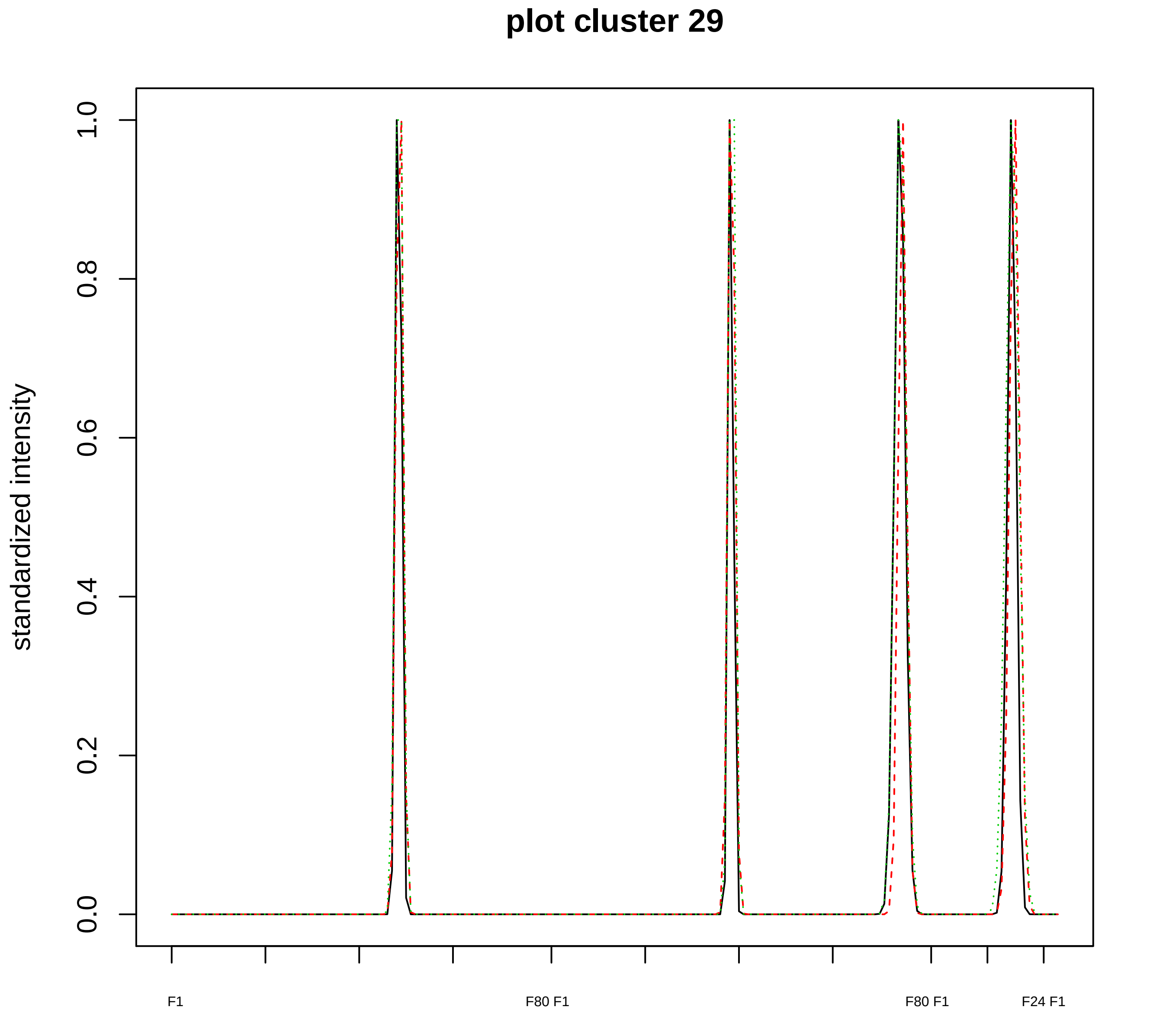
plot cluster 29
standardized intensity
0.0
0.2
0.4
0.6
0.8
1.0
F1
F80 F1
F80 F1
F24 F1

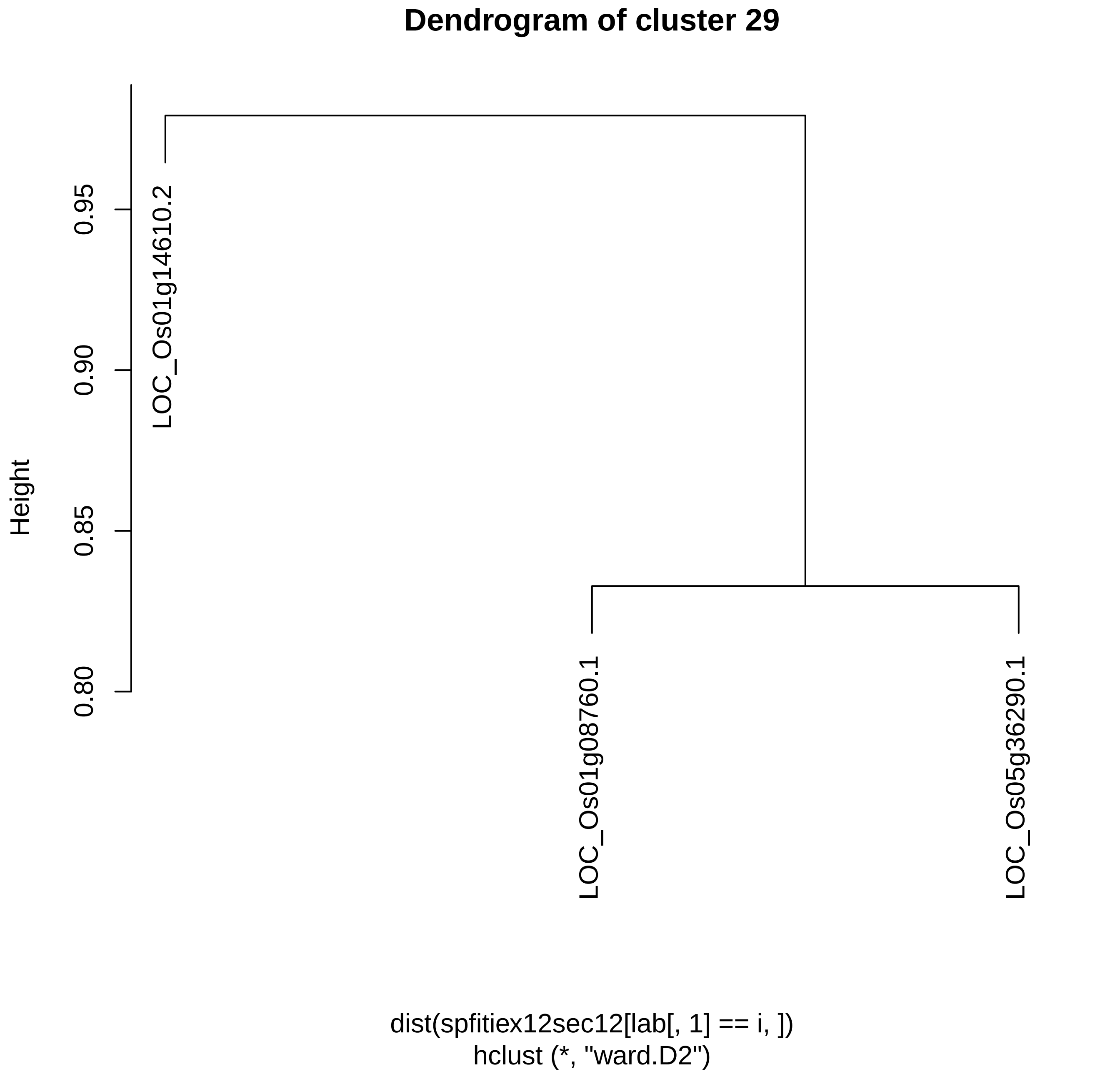
Dendrogram of cluster 29
Height
0.80
0.85
0.90
0.95
LOC_Os01g14610.2
LOC_Os01g08760.1
LOC_Os05g36290.1
dist(spfitiex12sec12[lab[, 1] == i, ])
hclust (*, "ward.D2")

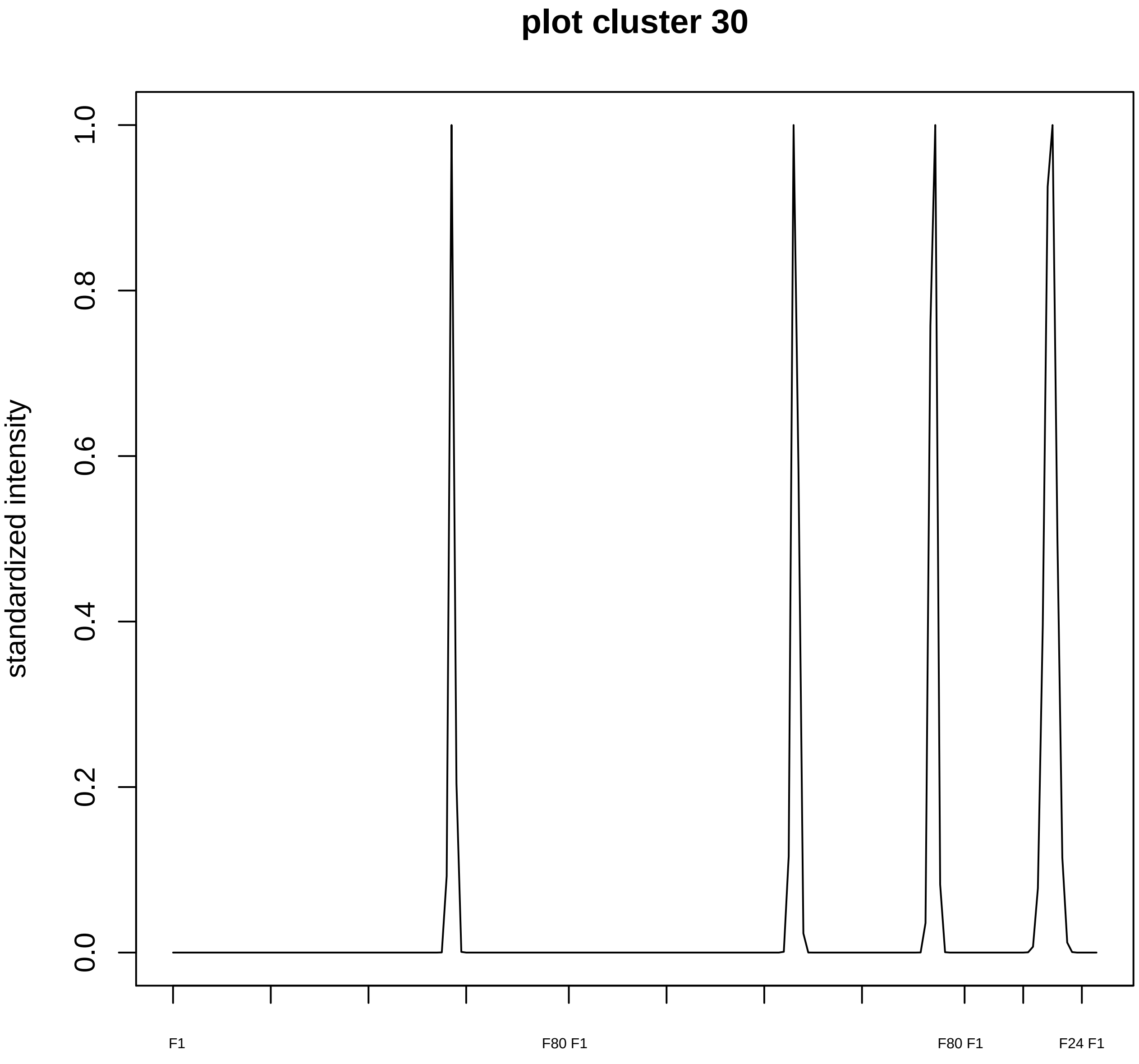

plot cluster 30
standardized intensity
0.0
0.2
0.4
0.6
0.8
1.0
F1
F80 F1
F80 F1
F24 F1

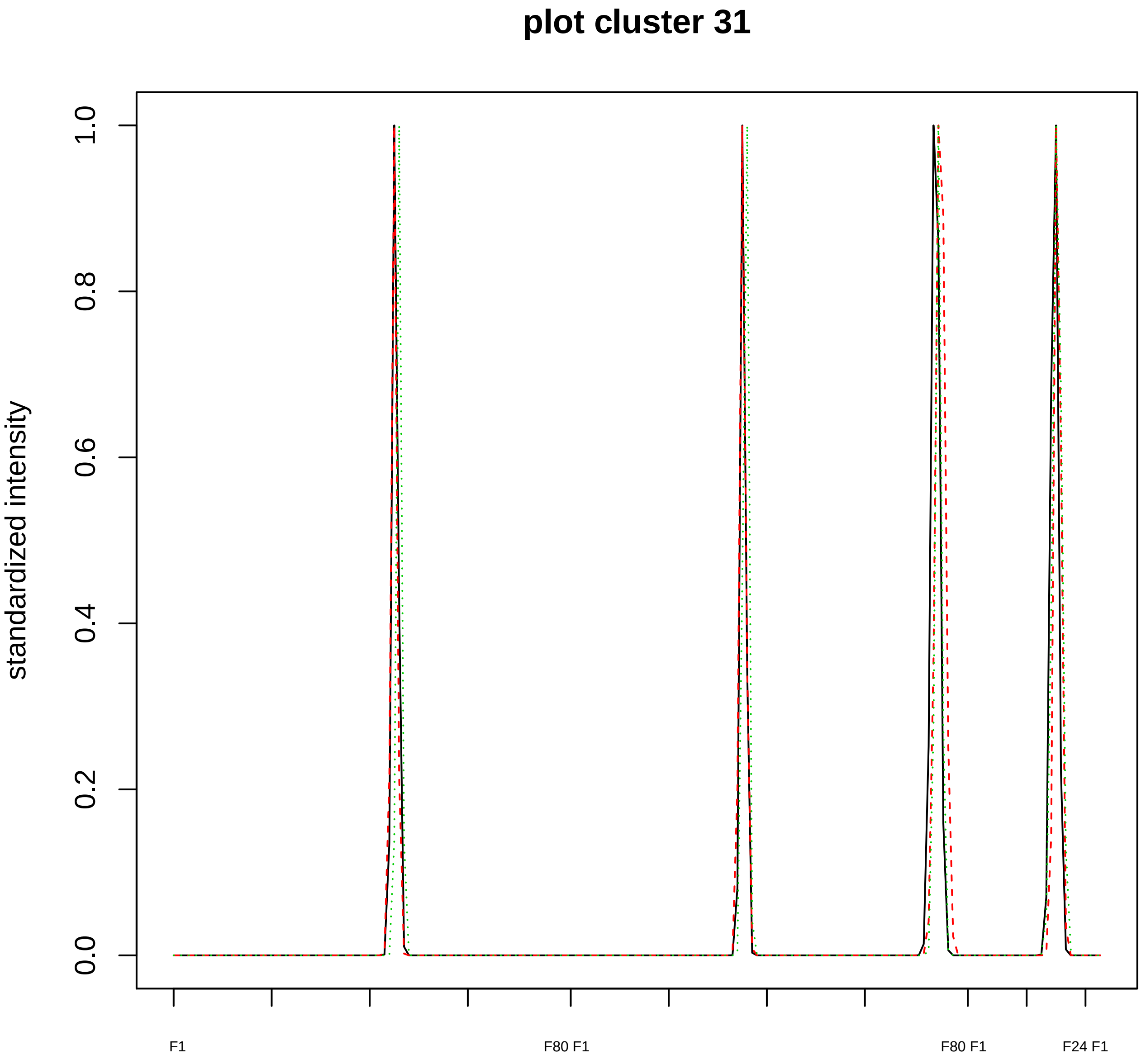
plot cluster 31
standardized intensity
1.0
0.8
0.6
0.4
0.2
0.0
F1
F80 F1
F80 F1
F24 F1

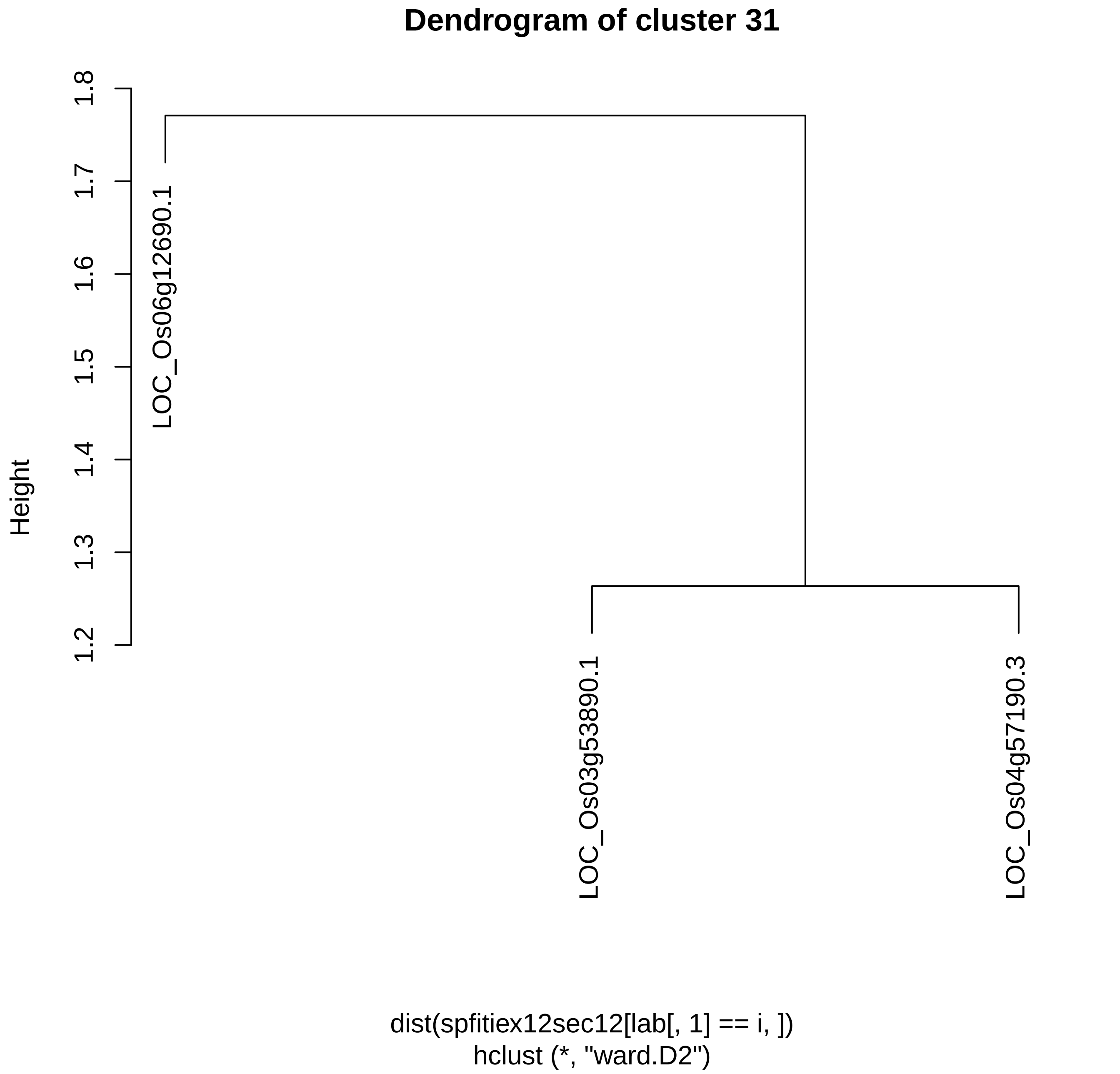
Dendrogram of cluster 31
Height
1.2
1.3
1.4
1.5
1.6
1.7
1.8
LOC_Os06g12690.1
LOC_Os03g53890.1
LOC_Os04g57190.3
dist(spfitiex12sec12[lab[, 1] == i, ])
hclust (*, "ward.D2")

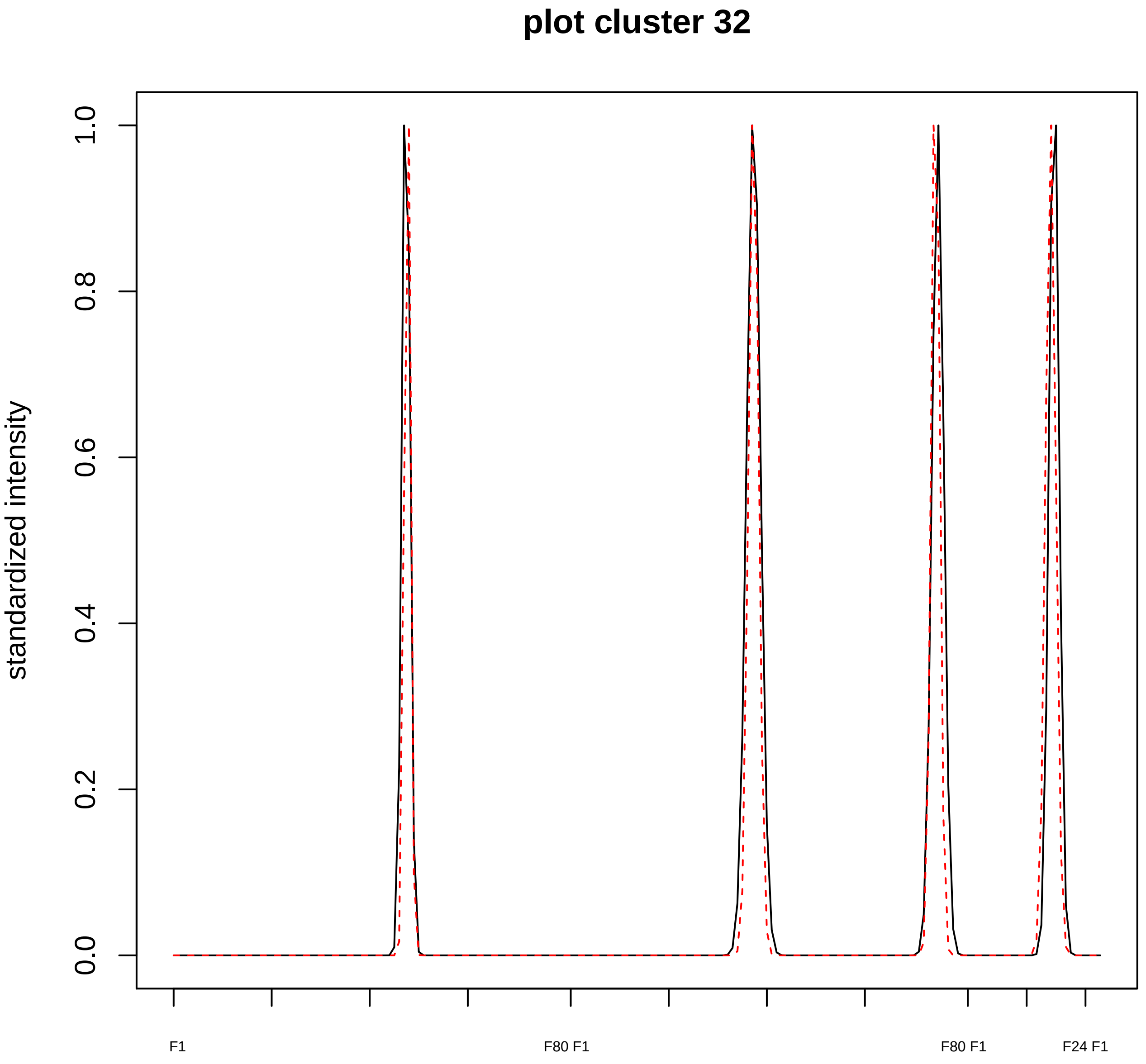

plot cluster 32
standardized intensity
0.0
0.2
0.4
0.6
0.8
1.0
F1
F80 F1
F80 F1
F24 F1

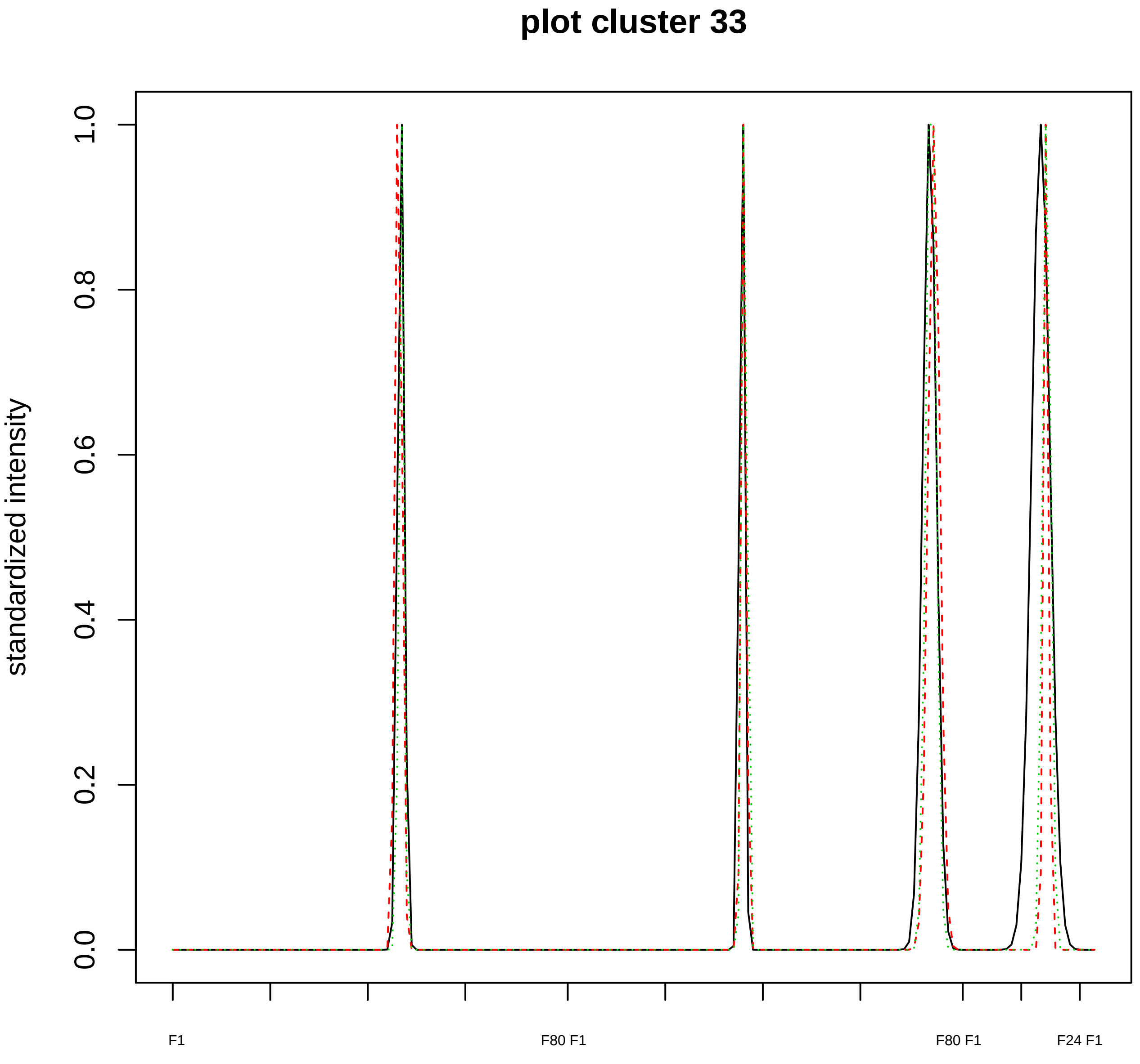
plot cluster 33
standardized intensity
0.0
0.2
0.4
0.6
0.8
1.0
F1
F80 F1
F80 F1
F24 F1

**Dendrogram of cluster 33**

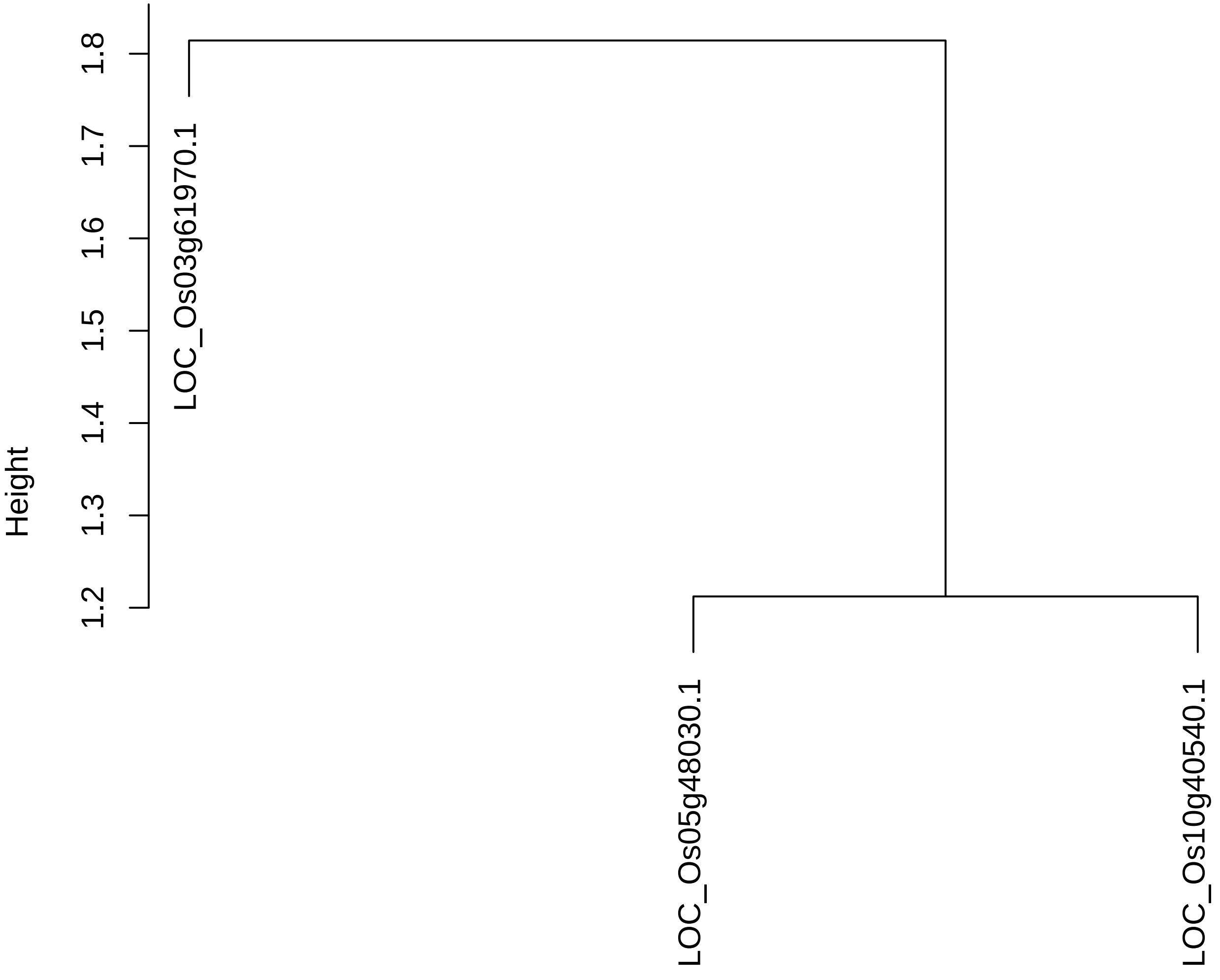


dist(spfitiex12sec12[lab[, 1] == i, ])
hclust (*, "ward.D2")

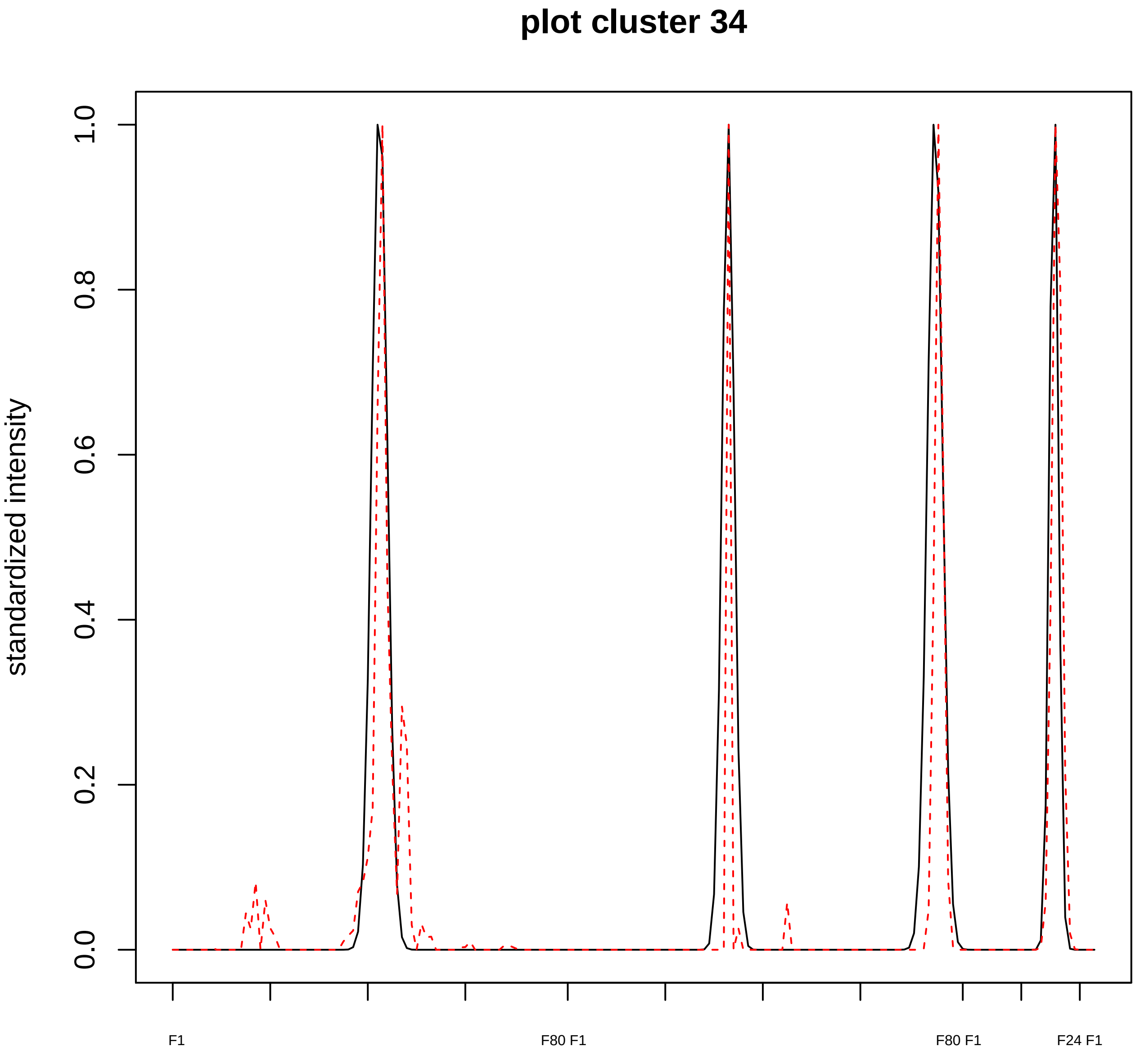
plot cluster 34
standardized intensity
0.0
0.2
0.4
0.6
0.8
1.0
F1
F80 F1
F80 F1
F24 F1

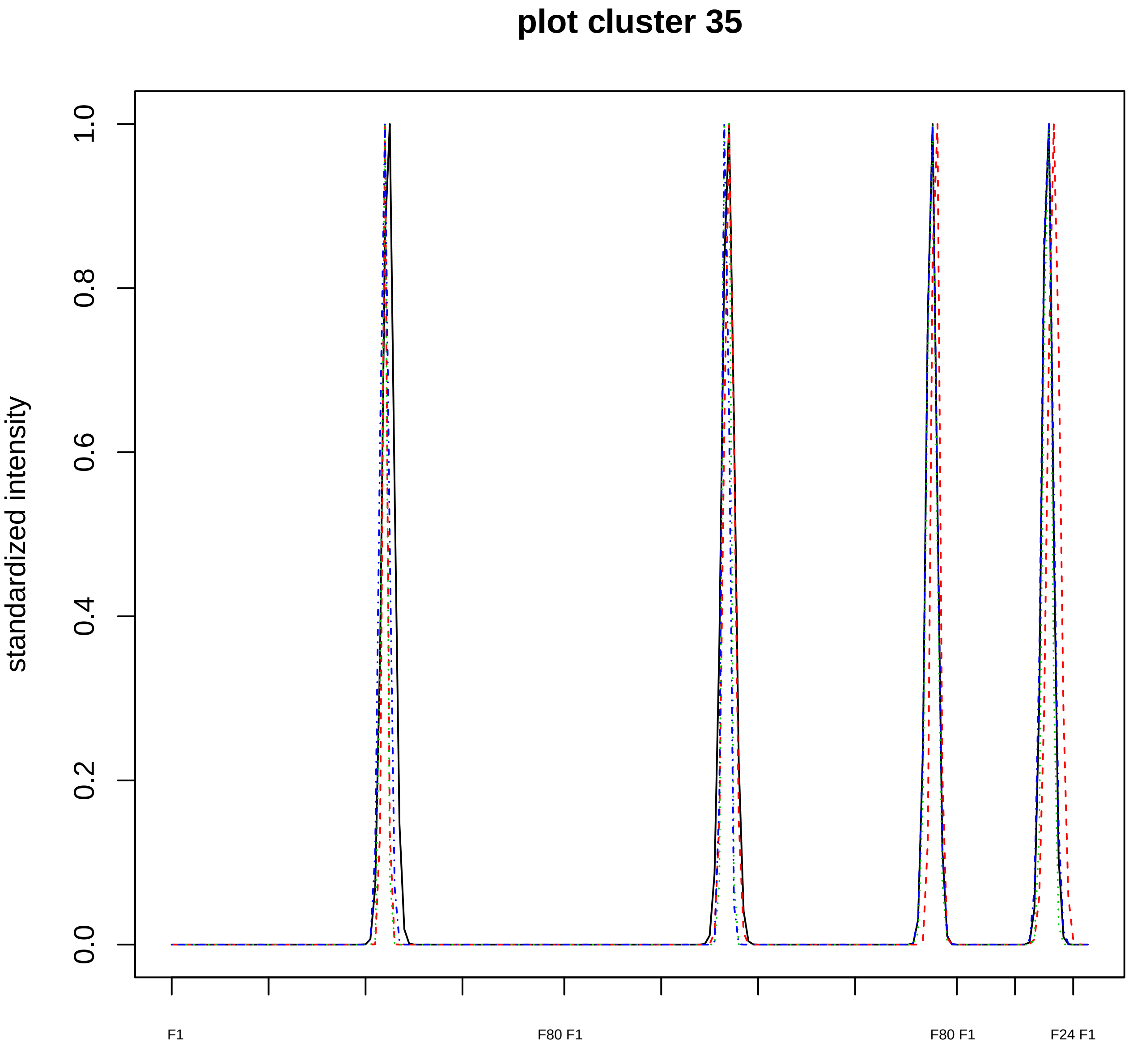
plot cluster 35
standardized intensity
0.0
0.2
0.4
0.6
0.8
1.0
F1
F80 F1
F80 F1
F24 F1

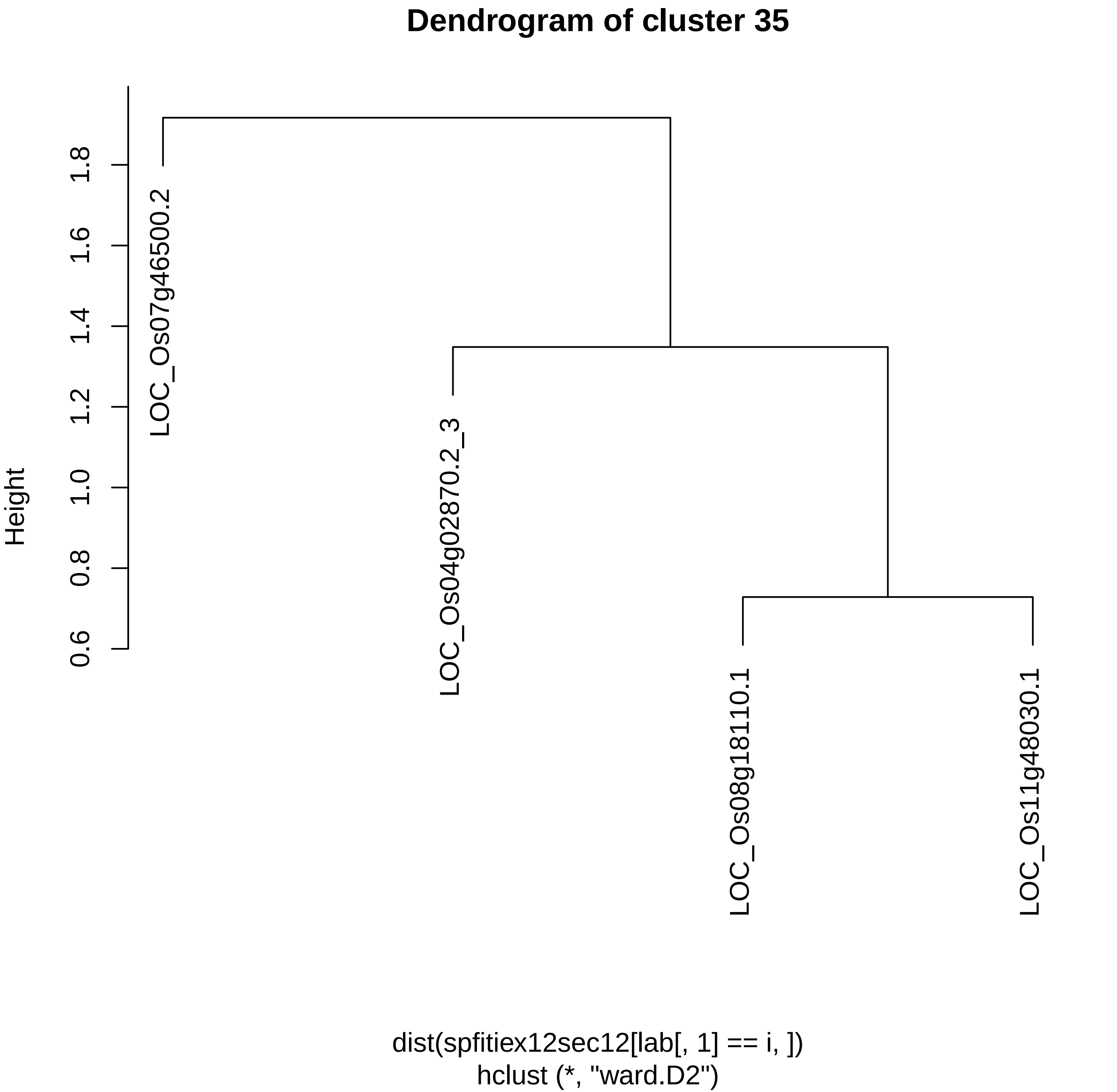
Dendrogram of cluster 35
Height
0.6
0.8
1.0
1.2
1.4
1.6
1.8
LOC_Os07g46500.2
LOC_Os04g02870.2_3
LOC_Os08g18110.1
LOC_Os11g48030.1
dist(spfitiex12sec12[lab[, 1] == i, ])
hclust (*, "ward.D2")

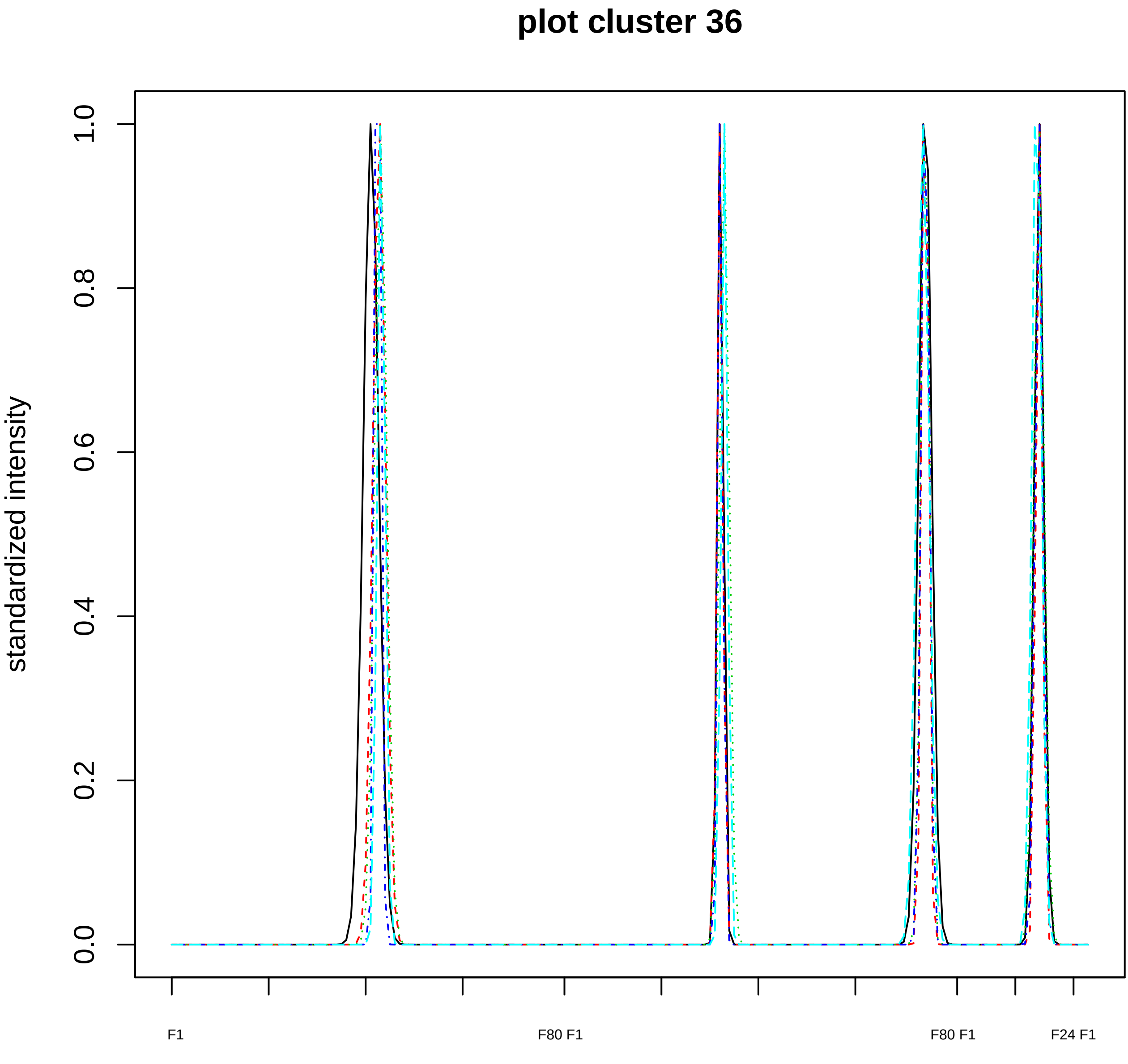

plot cluster 36
standardized intensity
0.0
0.2
0.4
0.6
0.8
1.0
F1
F80 F1
F80 F1
F24 F1

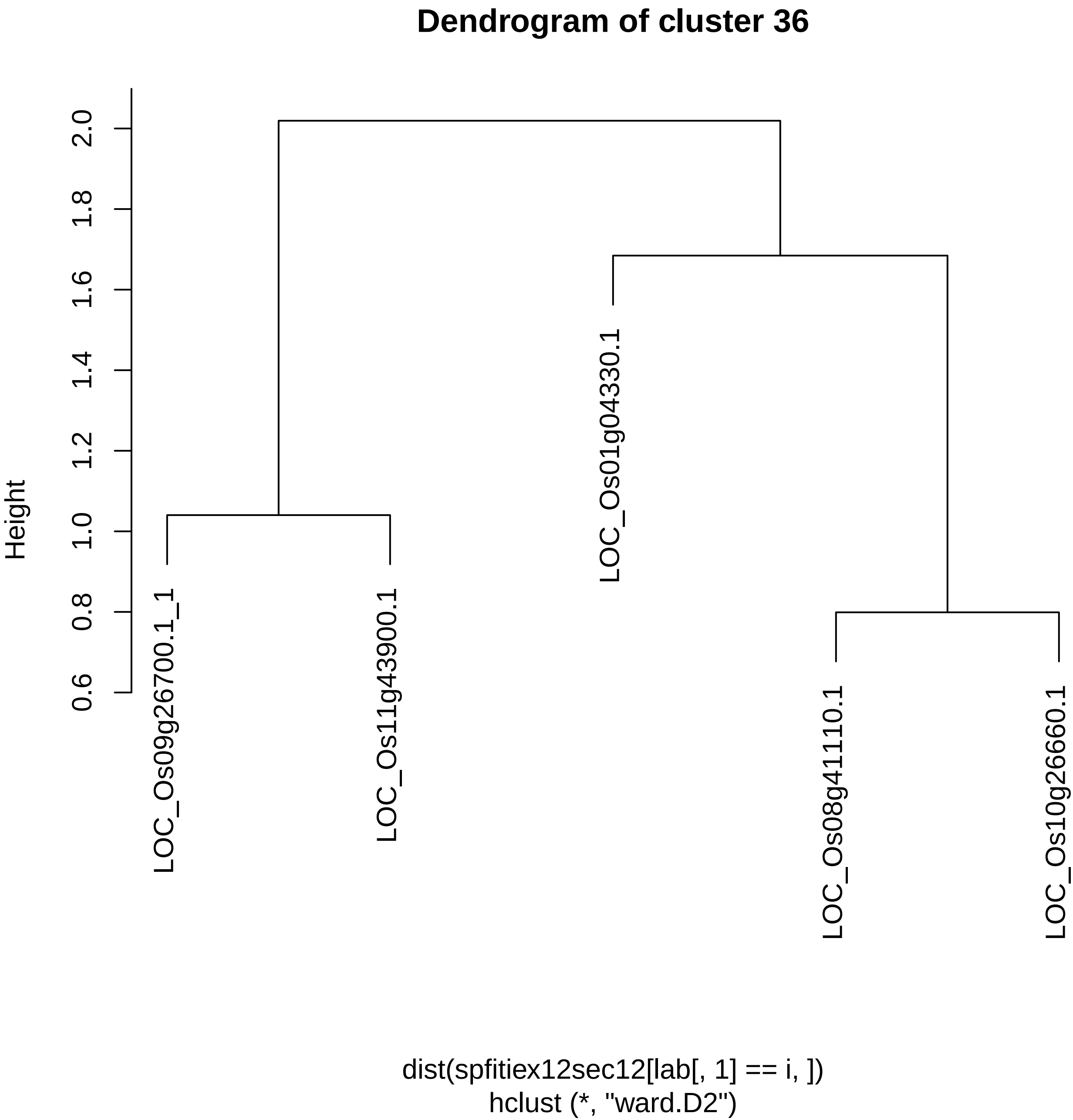
Dendrogram of cluster 36
Height
2.0
1.8
1.6
1.4
1.2
1.0
0.8
0.6
LOC_Os09g26700.1_1
LOC_Os11g43900.1
LOC_Os01g04330.1
LOC_Os08g41110.1
LOC_Os10g26660.1
dist(spfitiex12sec12[lab[, 1] == i, ])
hclust (*, "ward.D2")

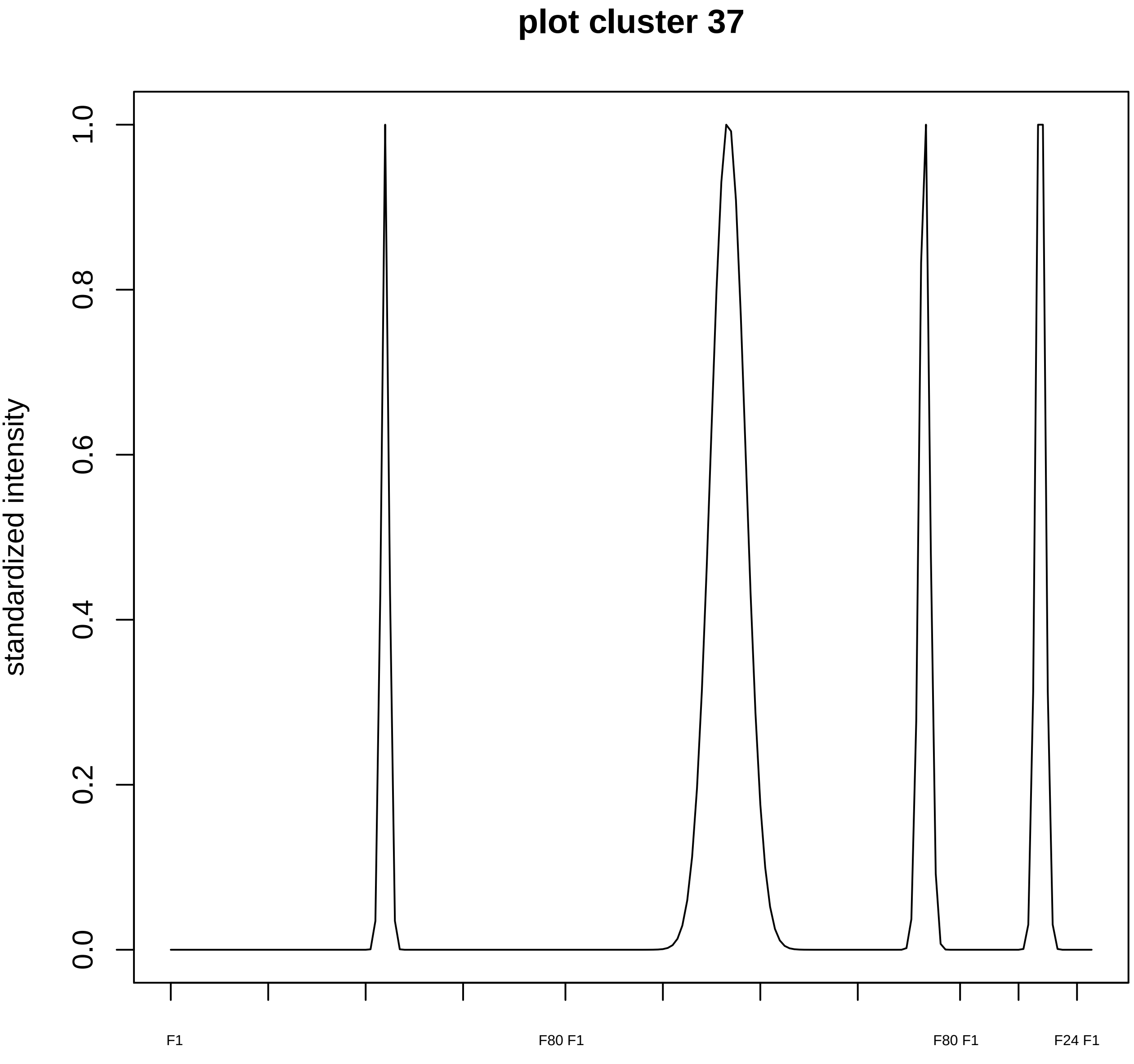
plot cluster 37
standardized intensity
1.0
0.8
0.6
0.4
0.2
0.0
F1
F80 F1
F80 F1
F24 F1

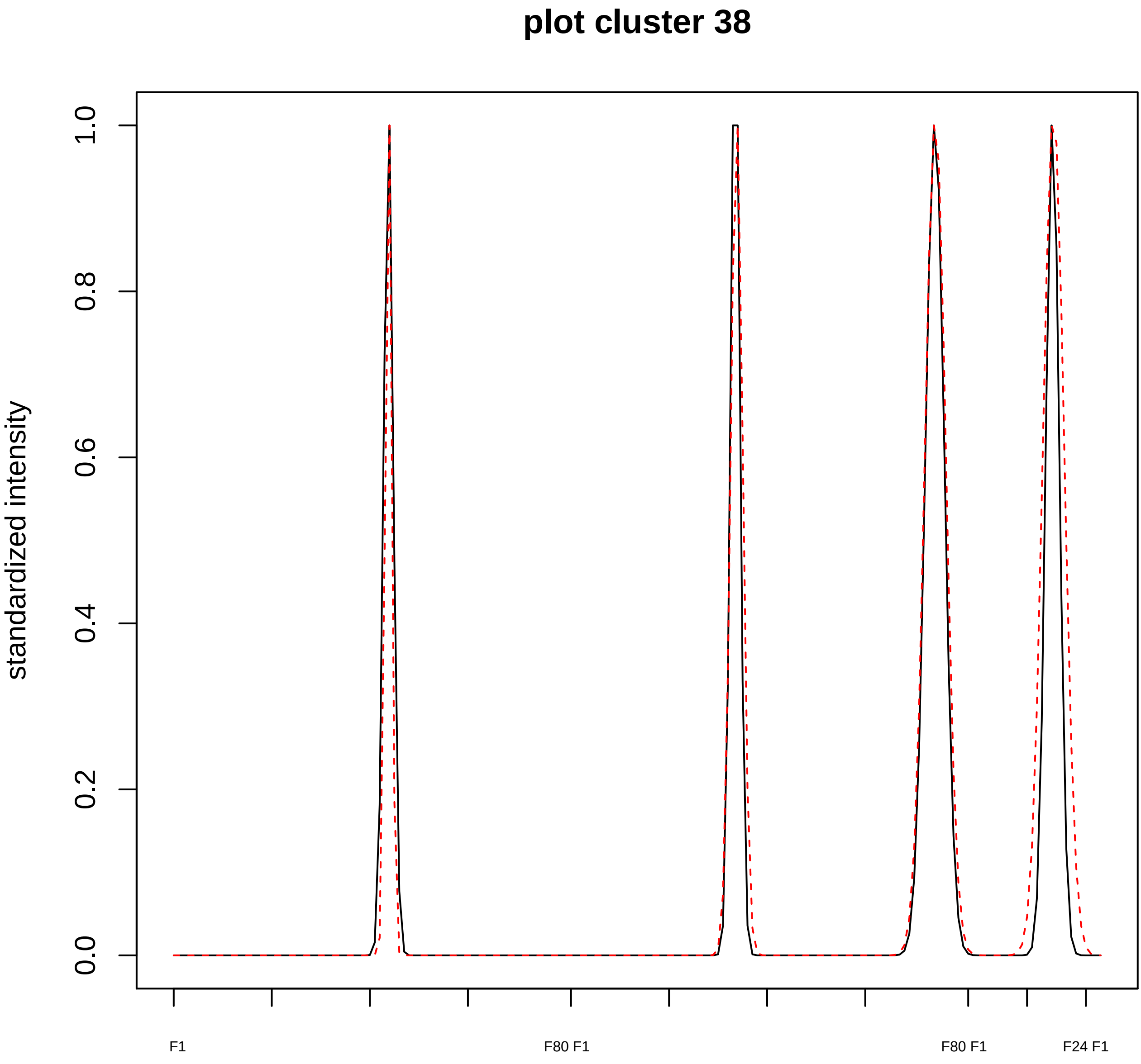
plot cluster 38
standardized intensity
0.0
0.2
0.4
0.6
0.8
1.0
F1
F80 F1
F80 F1
F24 F1

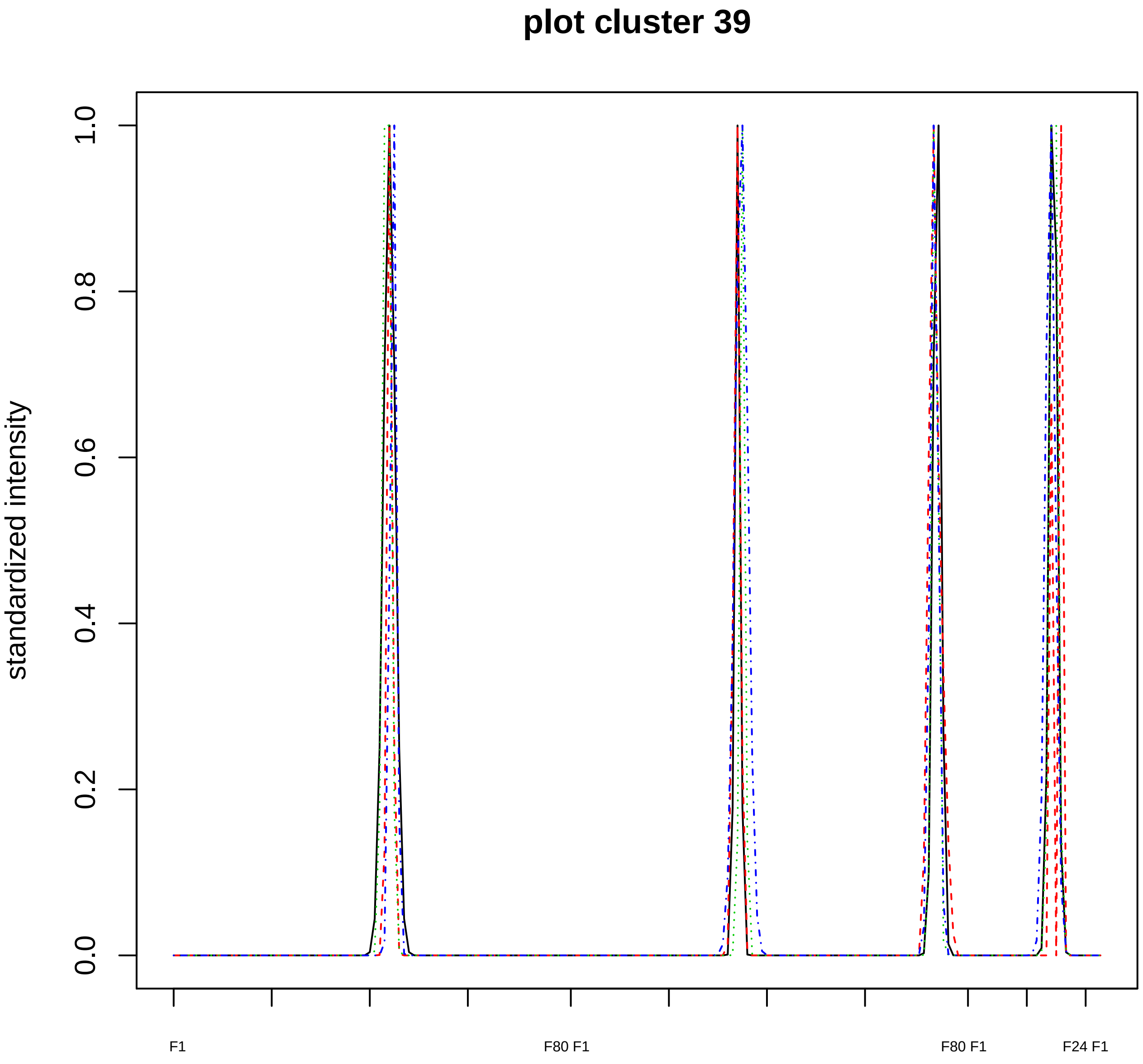
plot cluster 39
standardized intensity
0.0
0.2
0.4
0.6
0.8
1.0
F1
F80 F1
F80 F1
F24 F1

**Dendrogram of cluster 39**

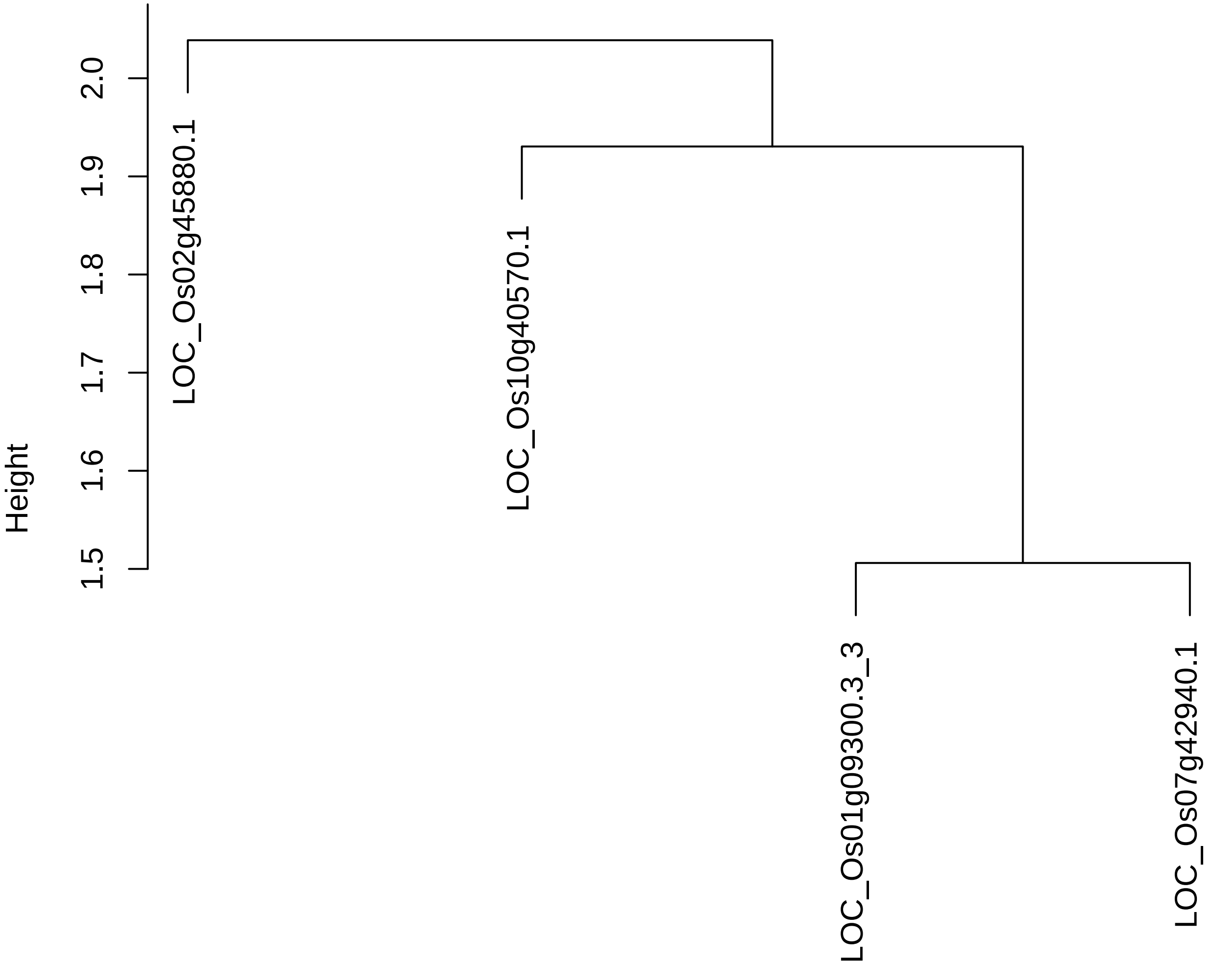


dist(spfitiex12sec12[lab[, 1] == i, ])
hclust (*, "ward.D2")

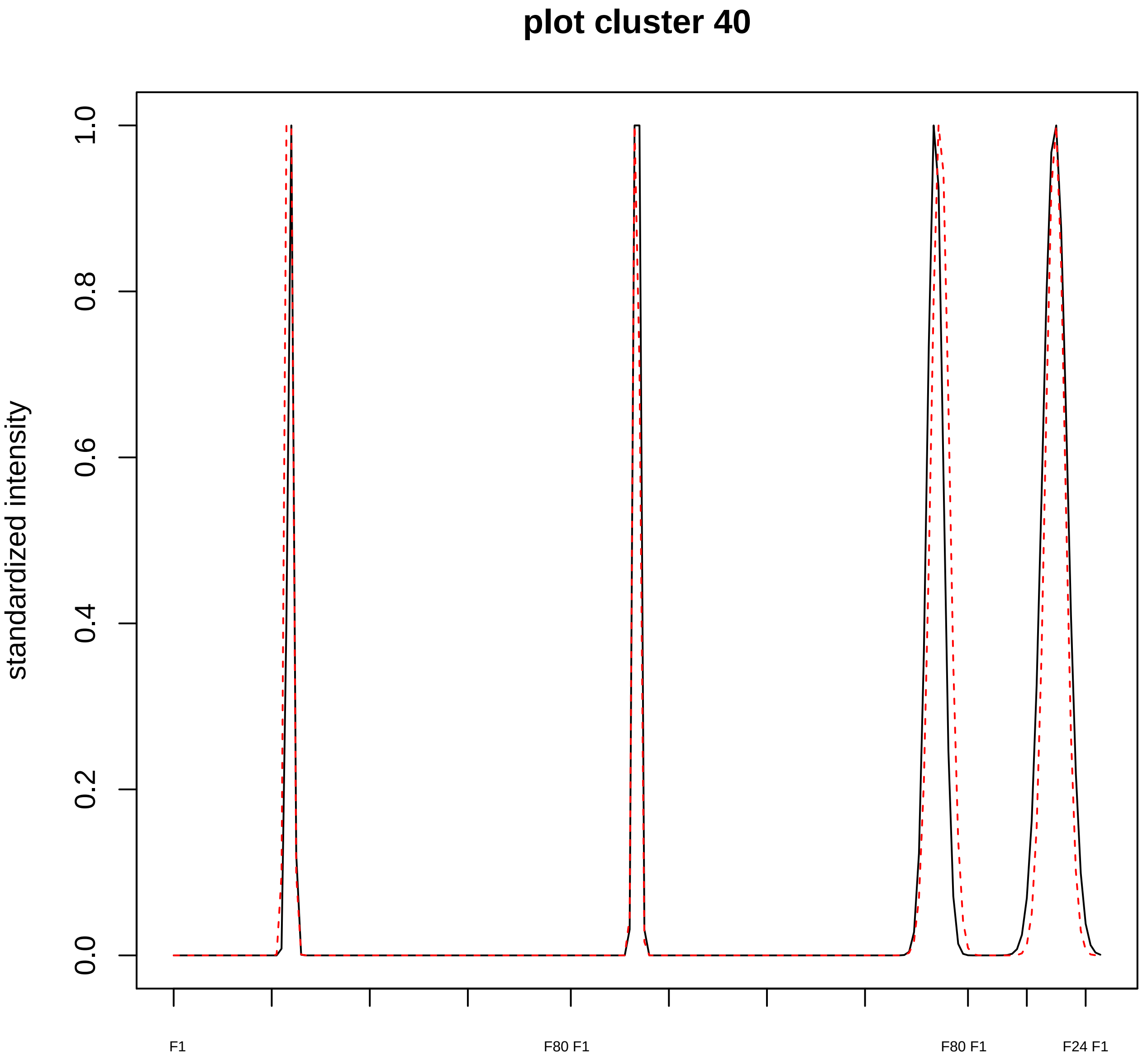
plot cluster 40
standardized intensity
0.0
0.2
0.4
0.6
0.8
1.0
F1
F80 F1
F80 F1
F24 F1

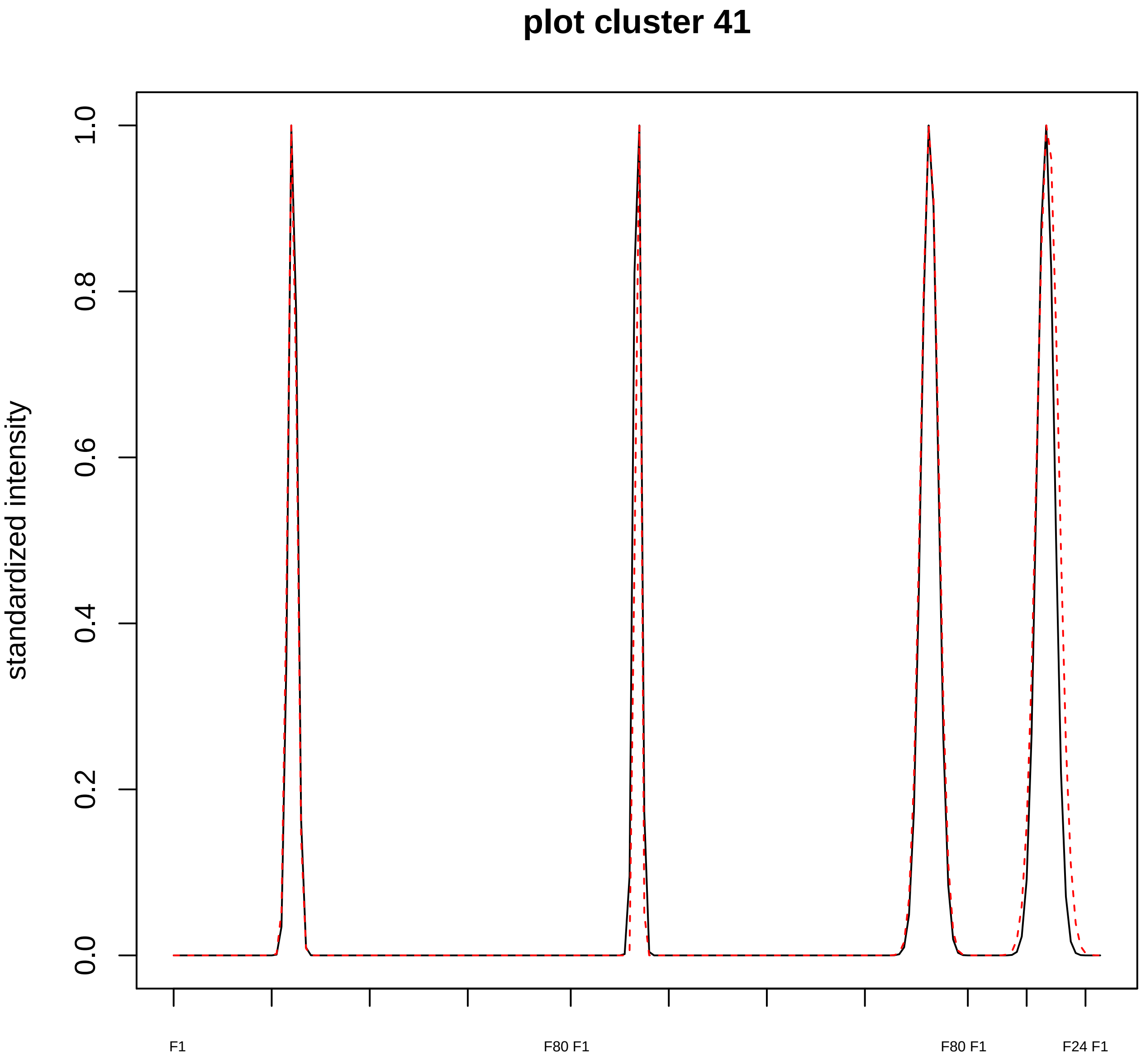
plot cluster 41
standardized intensity
0.0
0.2
0.4
0.6
0.8
1.0
F1
F80 F1
F80 F1
F24 F1

#### plot cluster 42

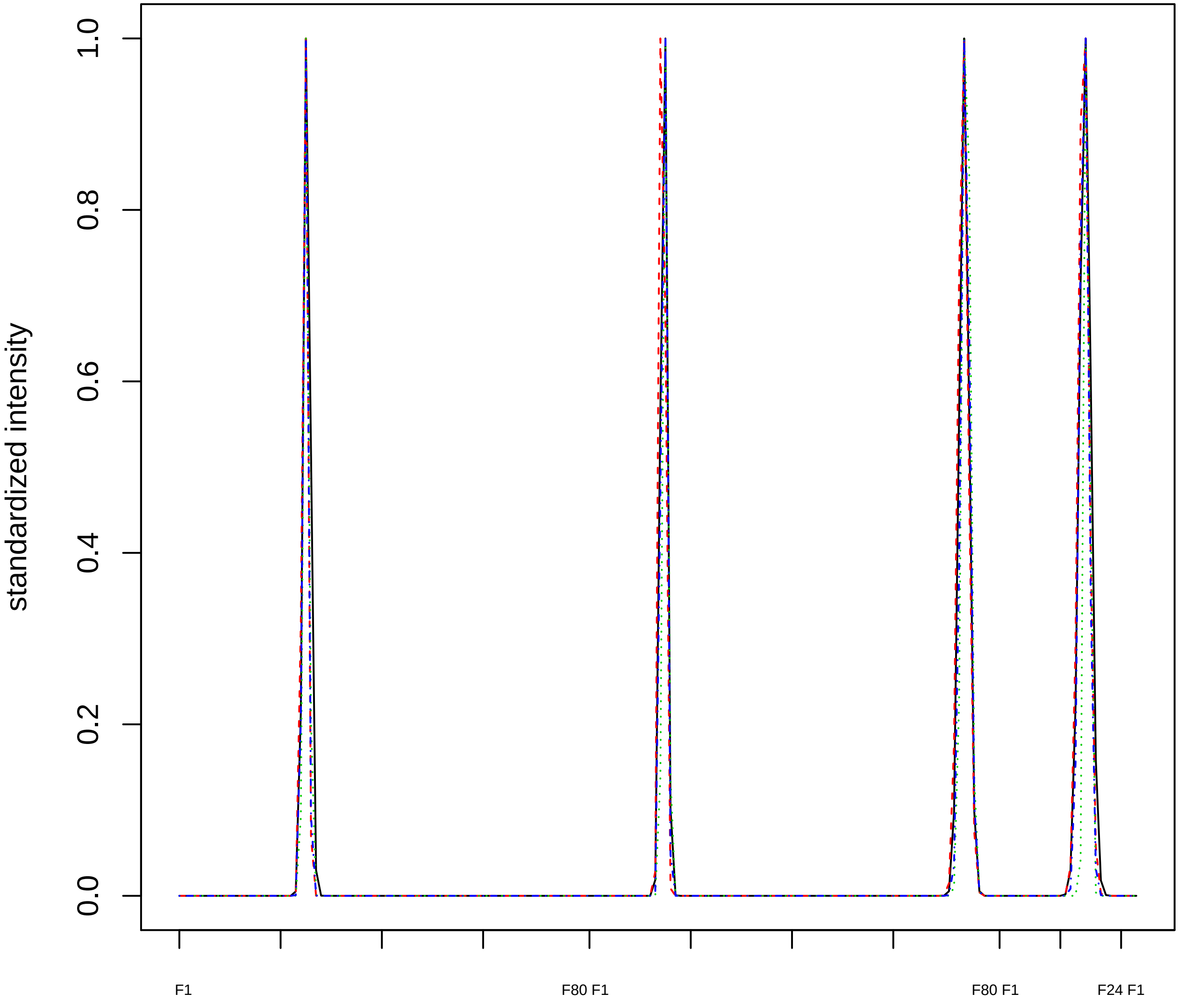

### Dendrogram of cluster 42

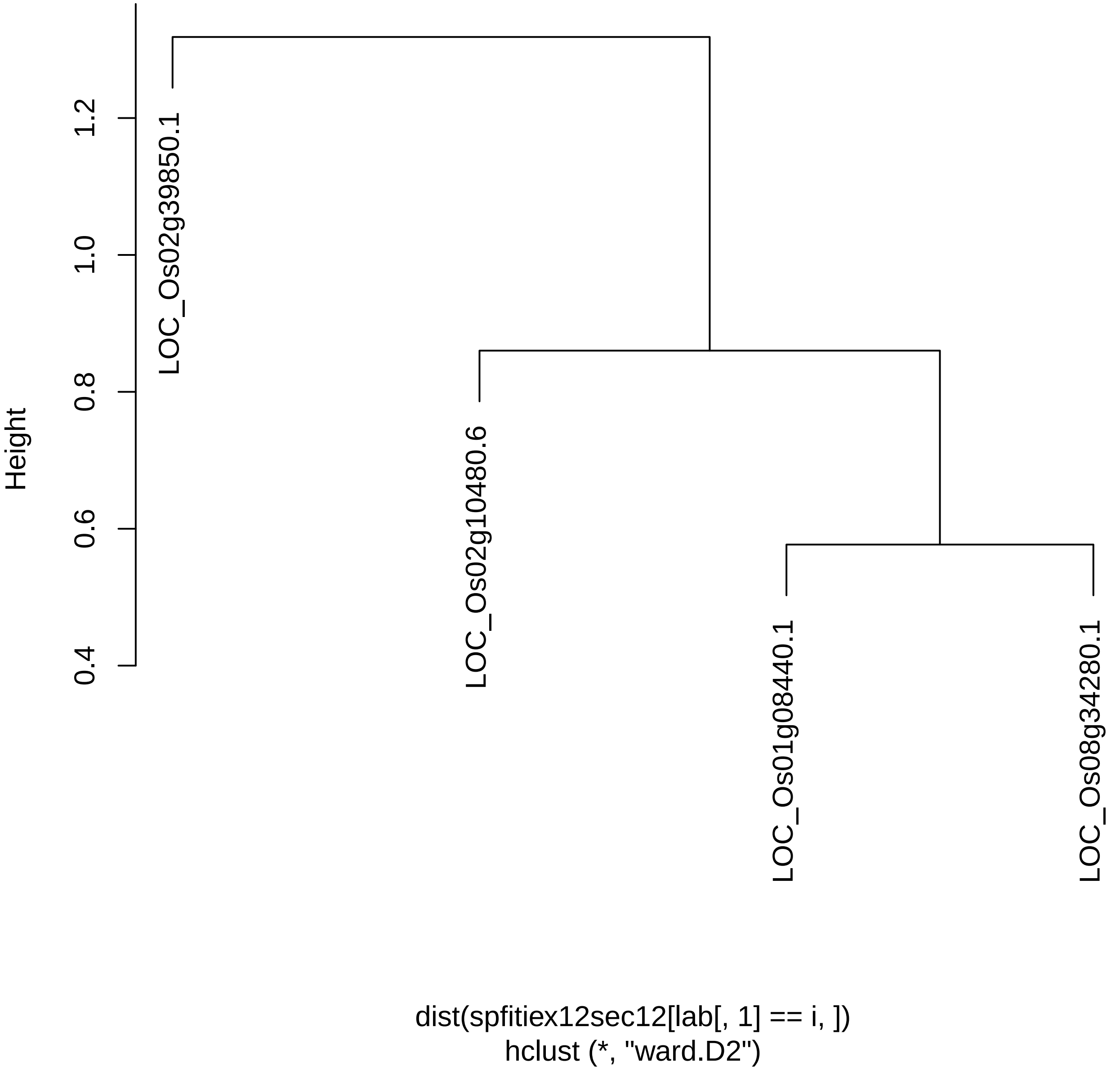

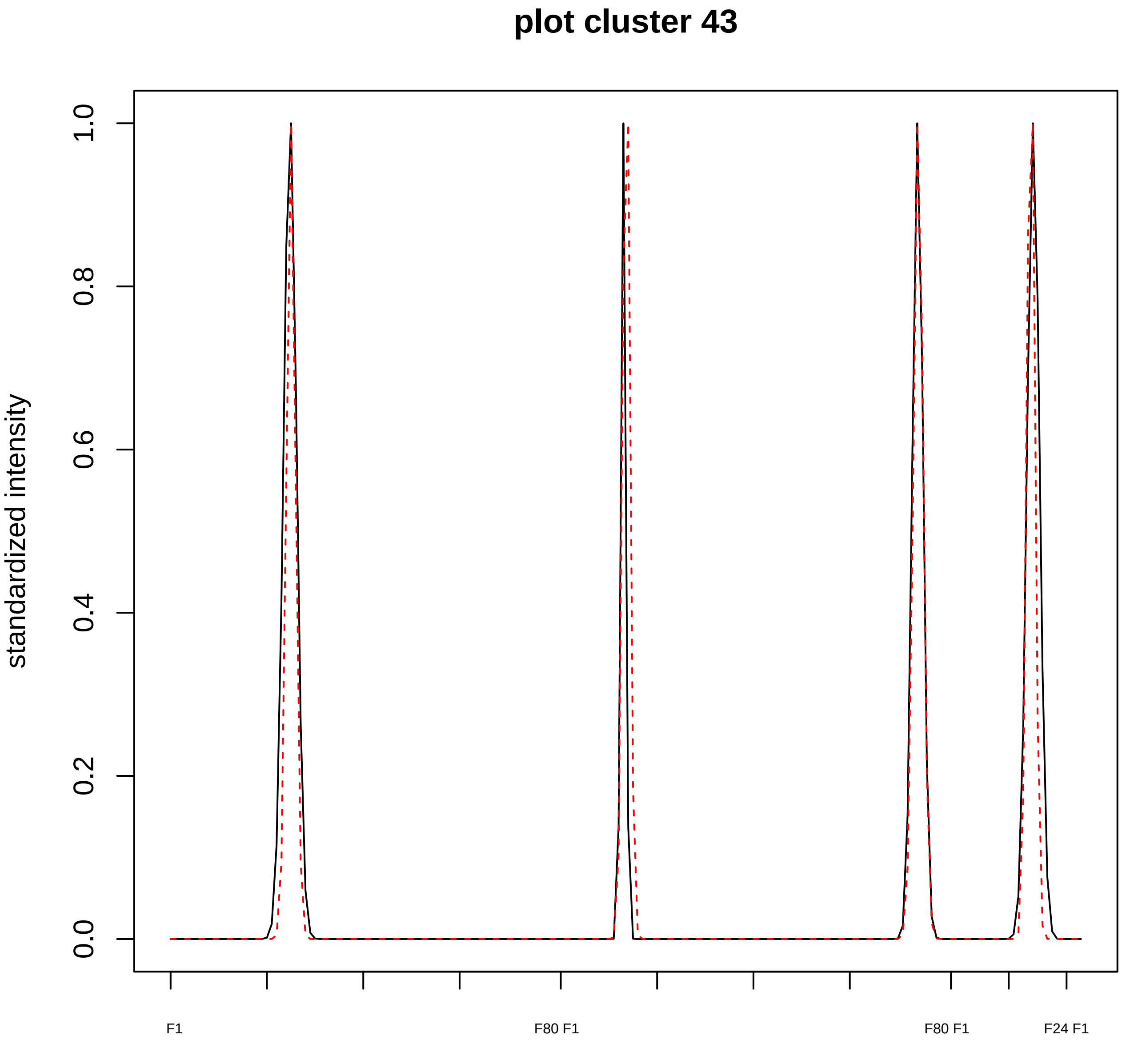
plot cluster 43
standardized intensity
0.0
0.2
0.4
0.6
0.8
1.0
F1
F80 F1
F80 F1
F24 F1

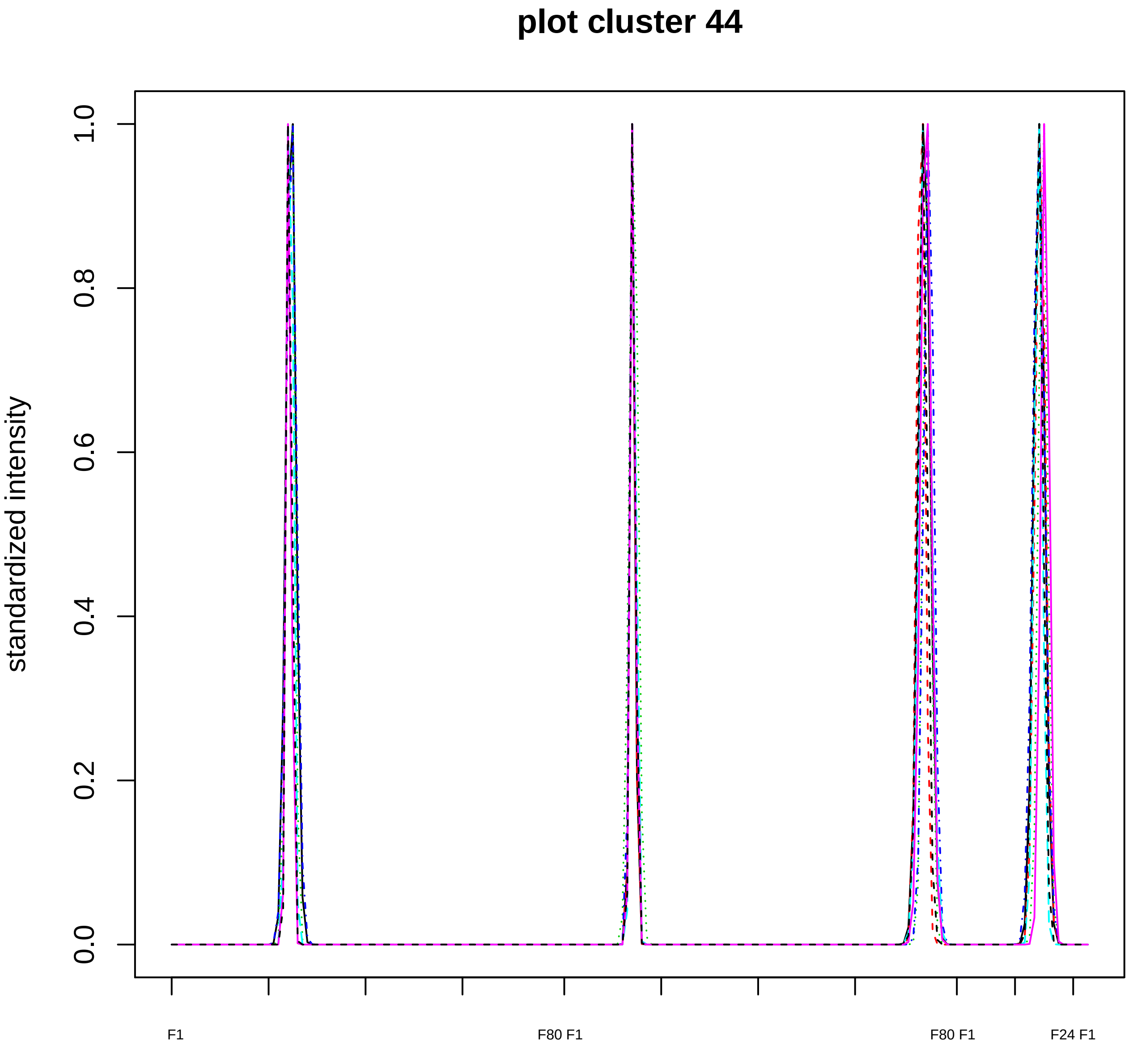

plot cluster 44
standardized intensity
0.0
0.2
0.4
0.6
0.8
1.0
F1
F80 F1
F80 F1
F24 F1

#### Dendrogram of cluster 44

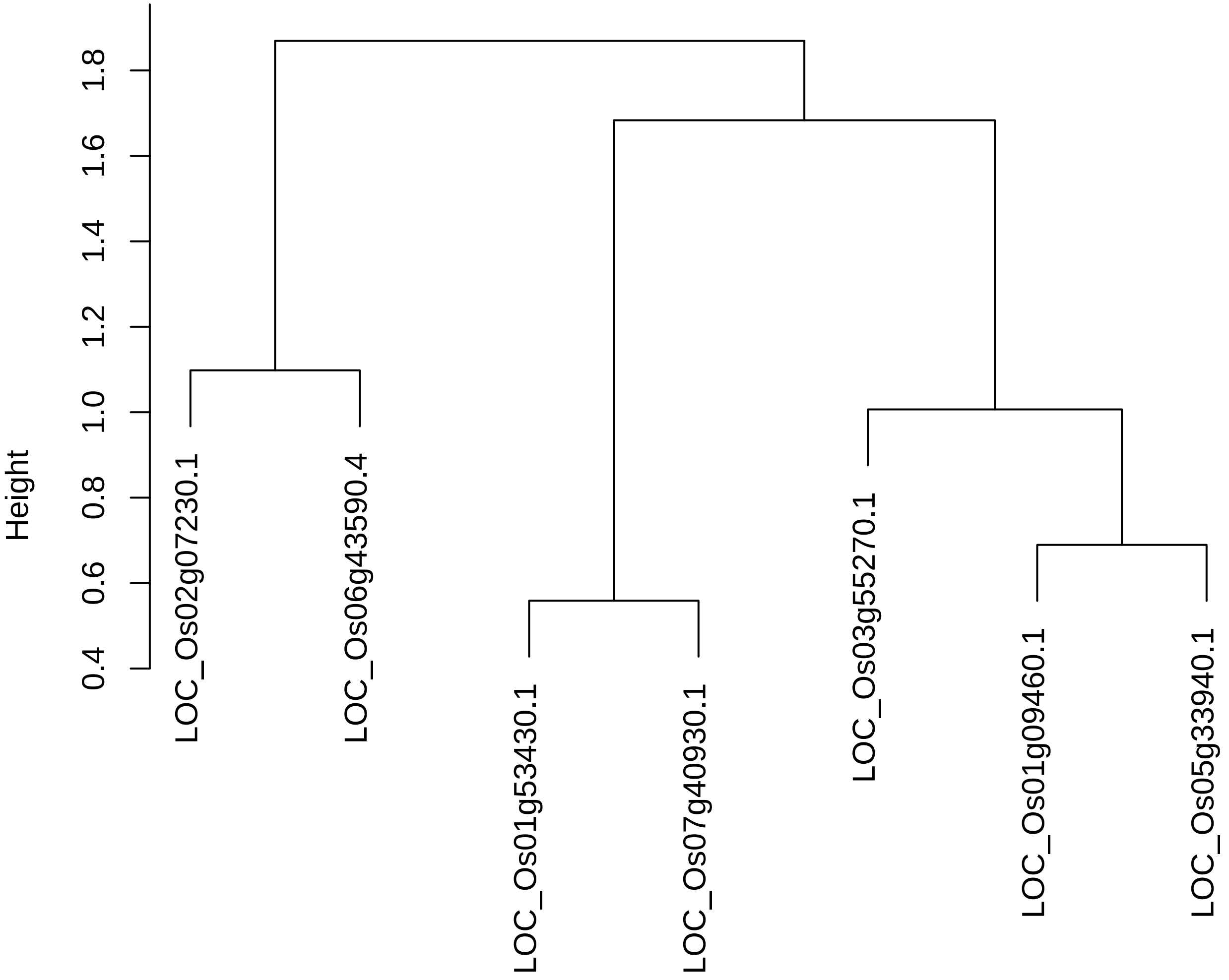


dist(spfitiex12sec12[lab[, 1] == i, ])
hclust (*, "ward.D2")

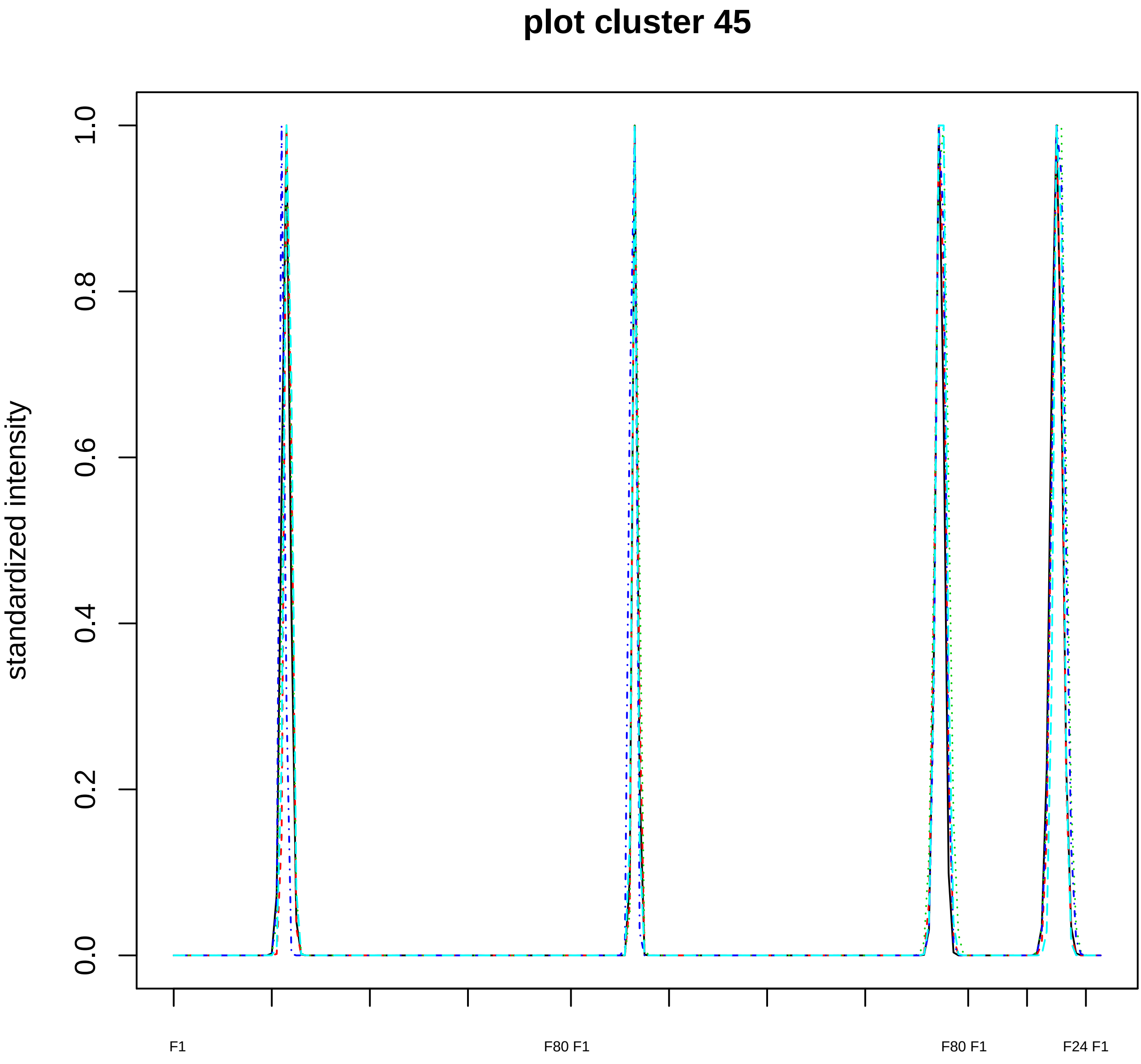
plot cluster 45
standardized intensity
0.0
0.2
0.4
0.6
0.8
1.0
F1
F80 F1
F80 F1
F24 F1

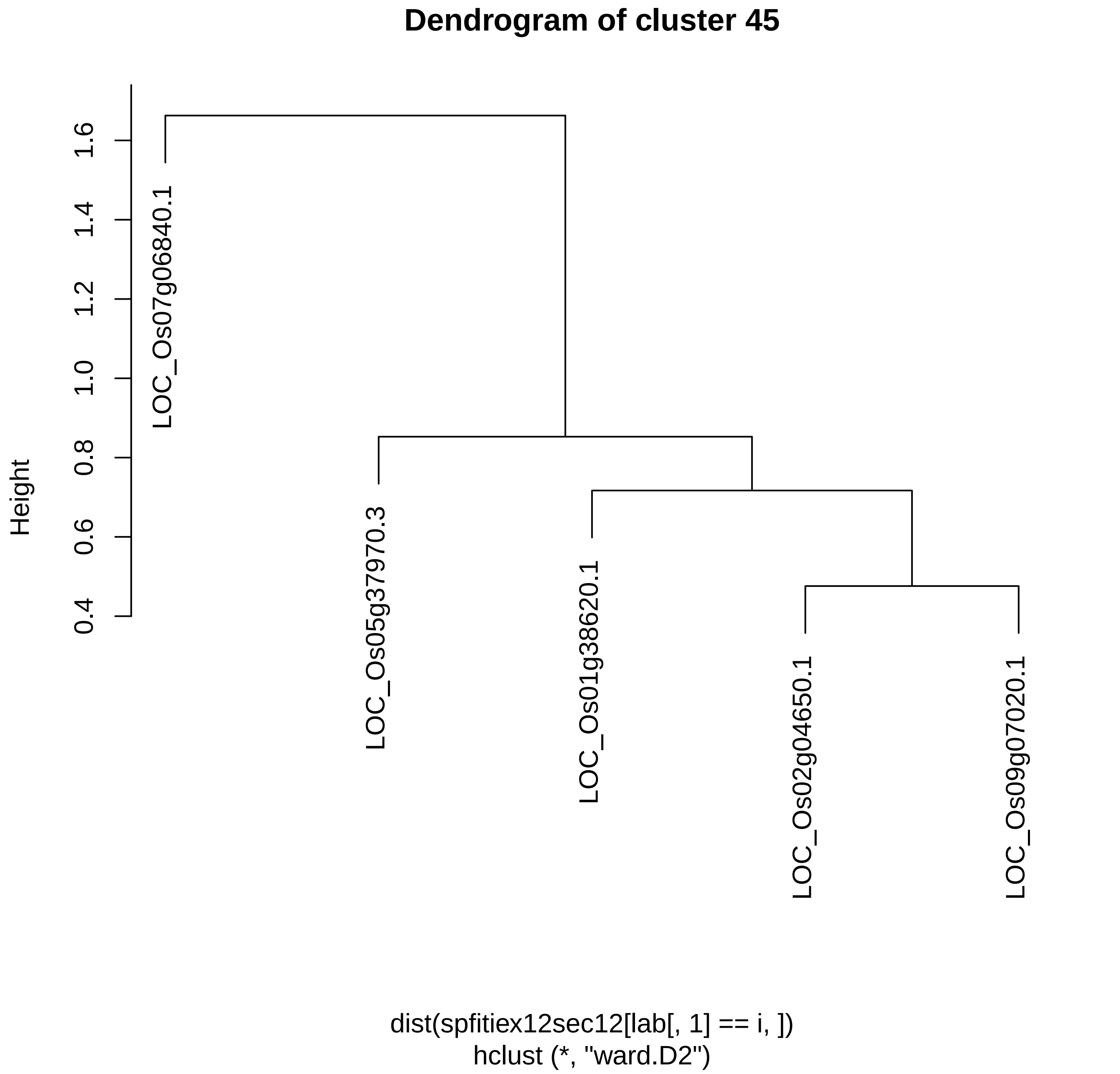
Dendrogram of cluster 45
Height
0.4
0.6
0.8
1.0
1.2
1.4
1.6
LOC_Os07g06840.1
LOC_Os05g37970.3
LOC_Os01g38620.1
LOC_Os02g04650.1
LOC_Os09g07020.1
dist(spfitiex12sec12[lab[, 1] == i, ])
hclust (*, "ward.D2")

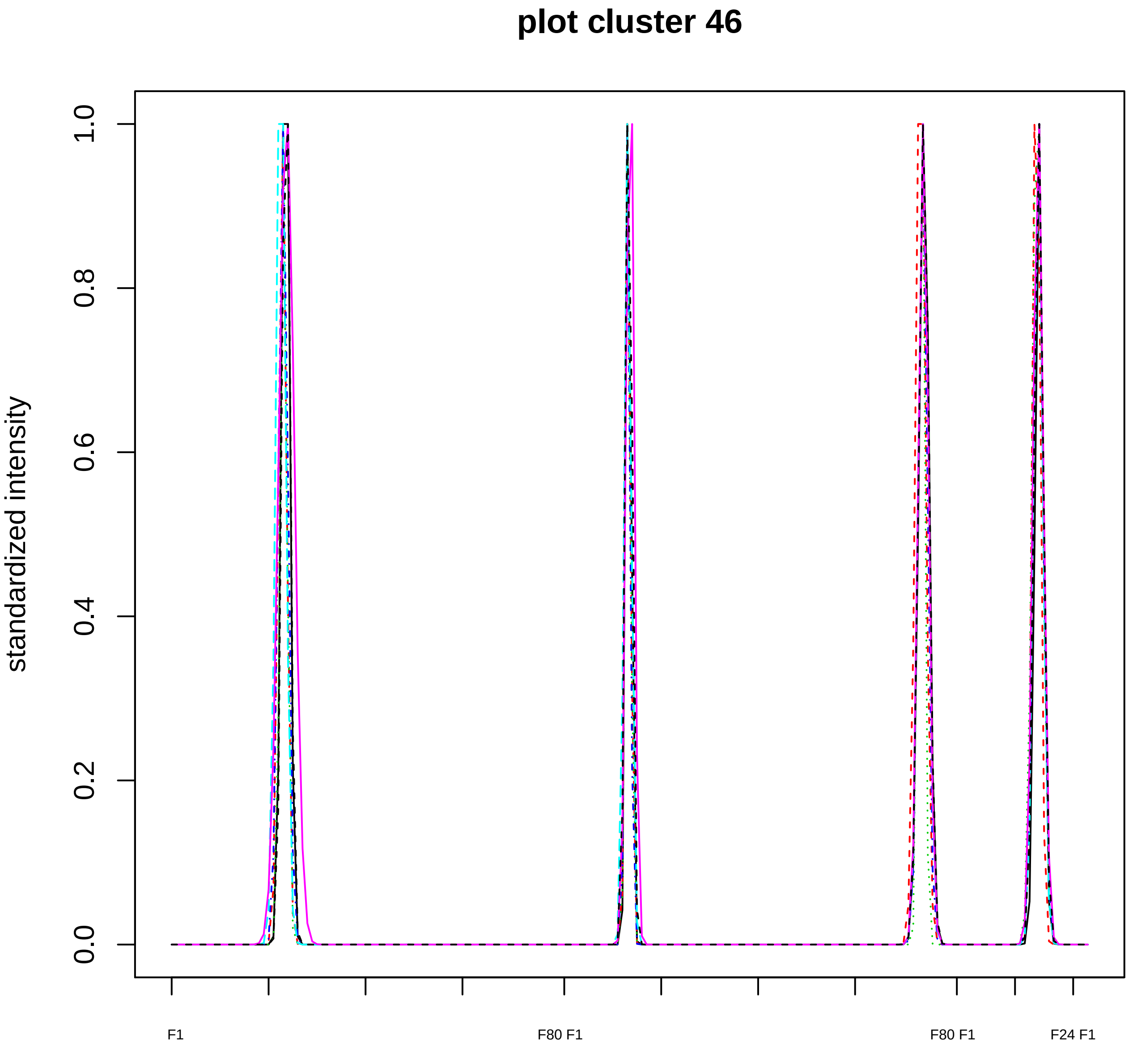

plot cluster 46
standardized intensity
0.0
0.2
0.4
0.6
0.8
1.0
F1
F80 F1
F80 F1
F24 F1

### Dendrogram of cluster 46

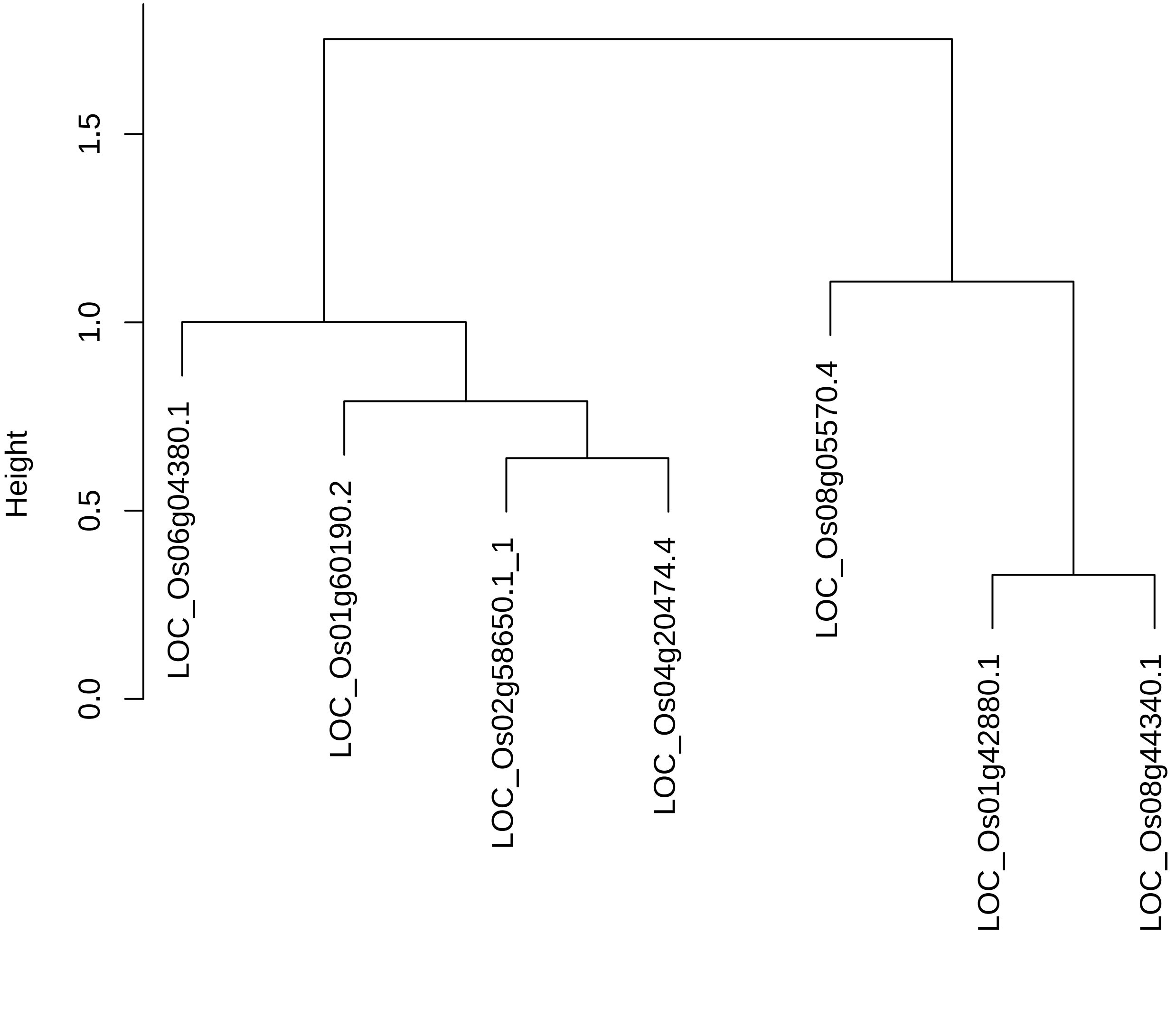


dist(spfitiex12sec12[lab[, 1] == i, ])
hclust (*, "ward.D2")

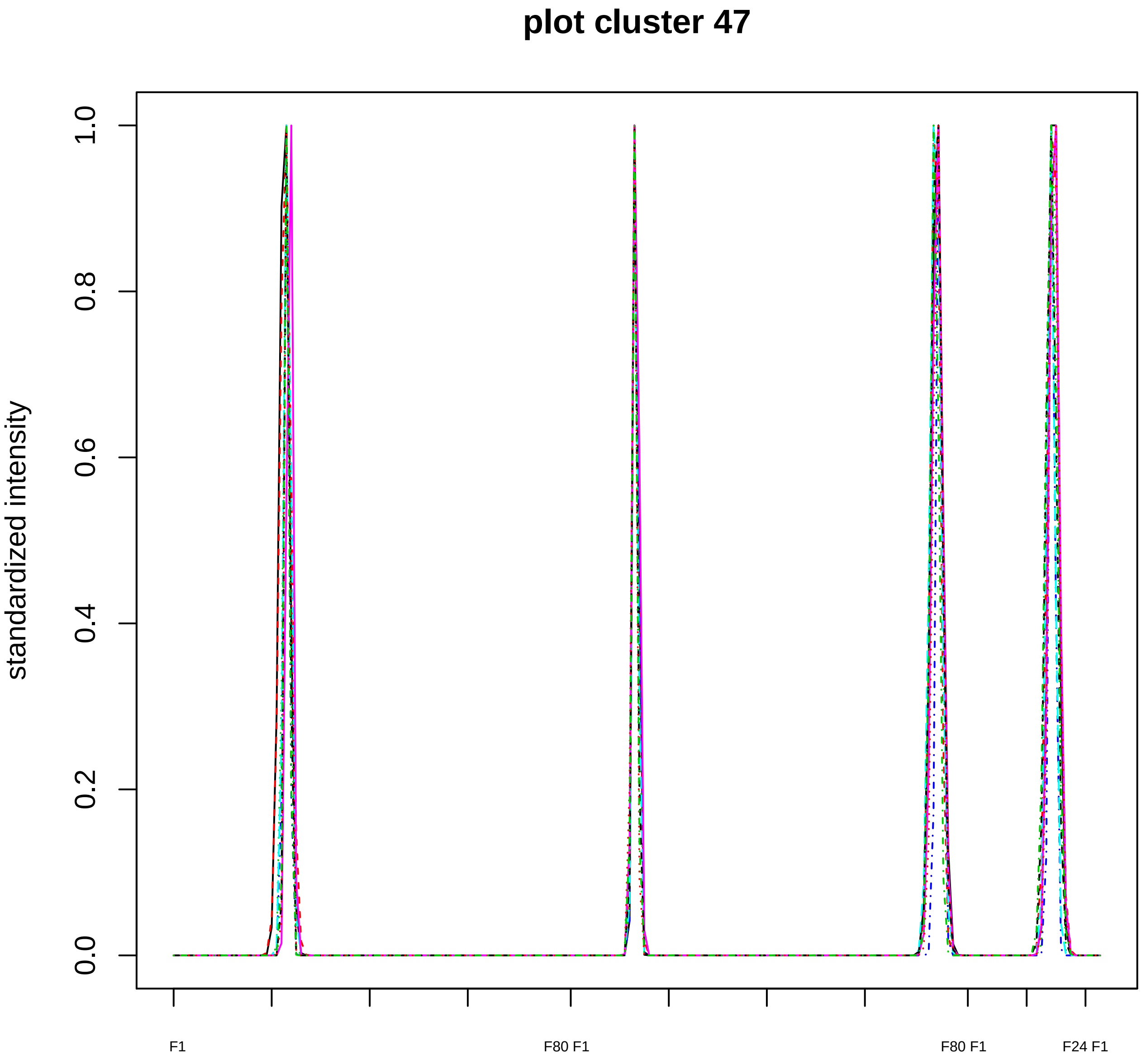
plot cluster 47
standardized intensity
0.0
0.2
0.4
0.6
0.8
1.0
F1
F80 F1
F80 F1
F24 F1

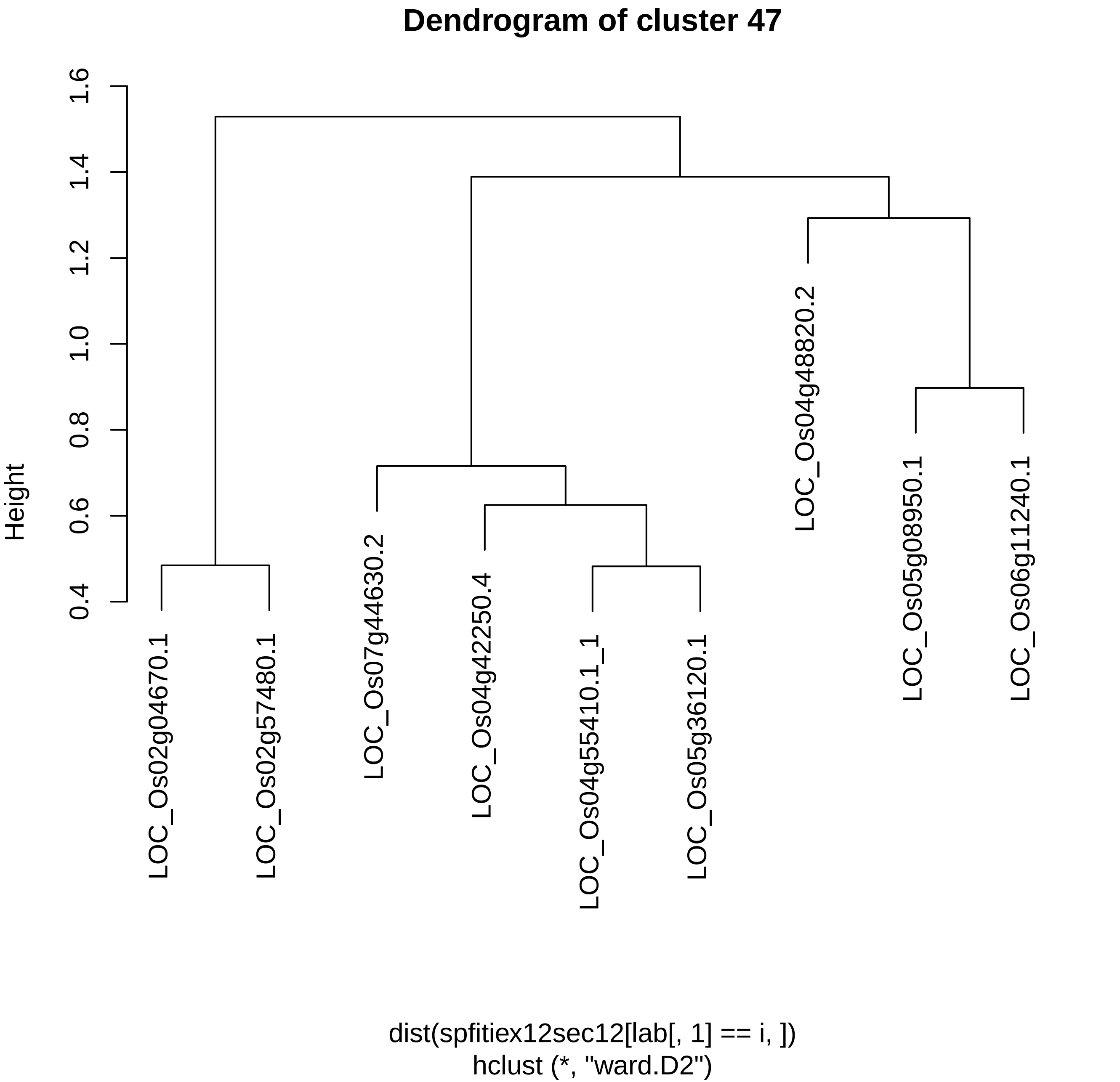
Dendrogram of cluster 47
Height
1.6
1.4
1.2
1.0
0.8
0.6
0.4
LOC_Os02g04670.1
LOC_Os02g57480.1
LOC_Os07g44630.2
LOC_Os04g42250.4
LOC_Os04g55410.1_1
LOC_Os05g36120.1
LOC_Os04g48820.2
LOC_Os05g08950.1
LOC_Os06g11240.1
dist(spfitiex12sec12[lab[, 1] == i, ])
hclust (*, "ward.D2")

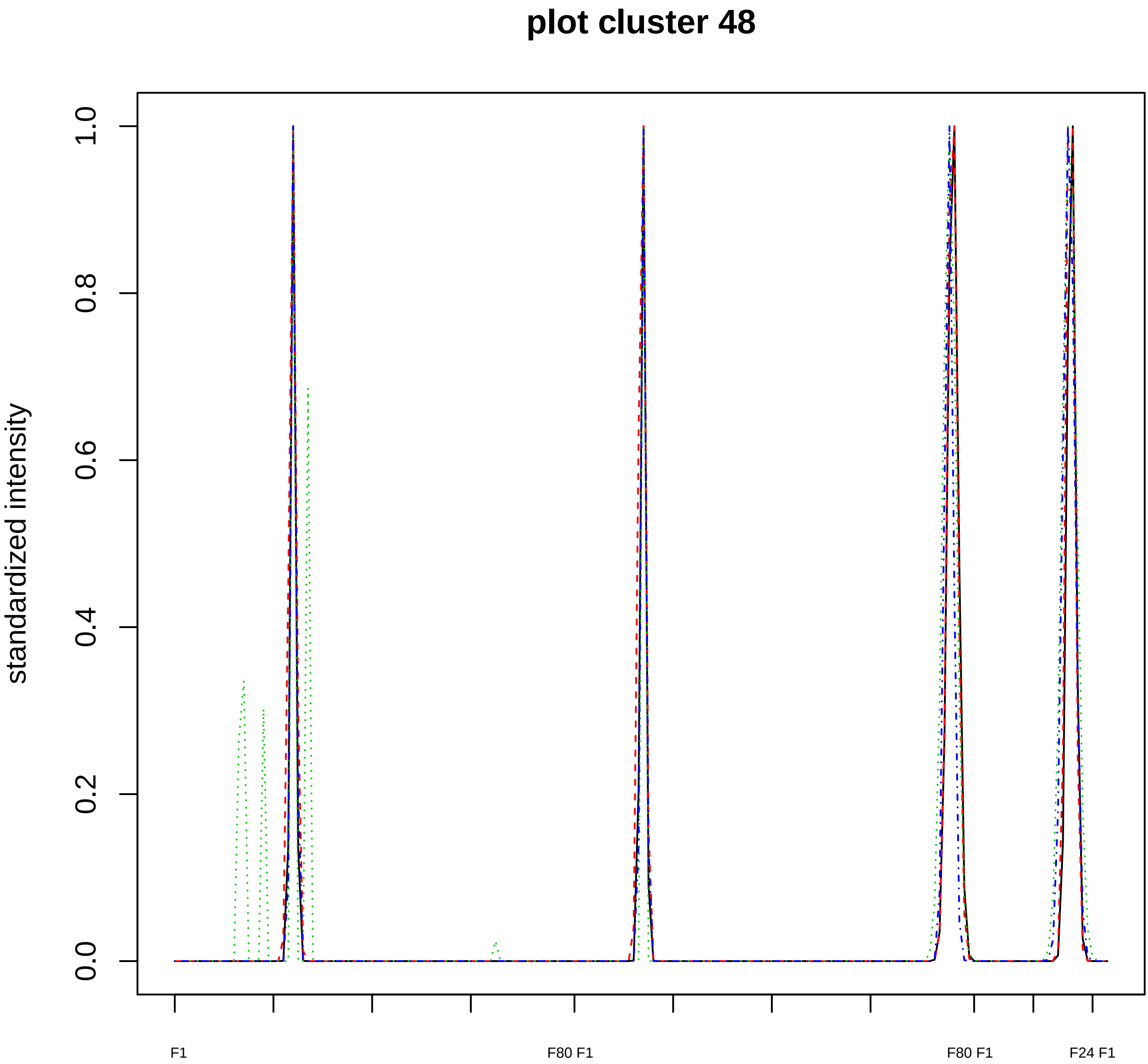

plot cluster 48
standardized intensity
0.0
0.2
0.4
0.6
0.8
1.0
F1
F80 F1
F80 F1
F24 F1

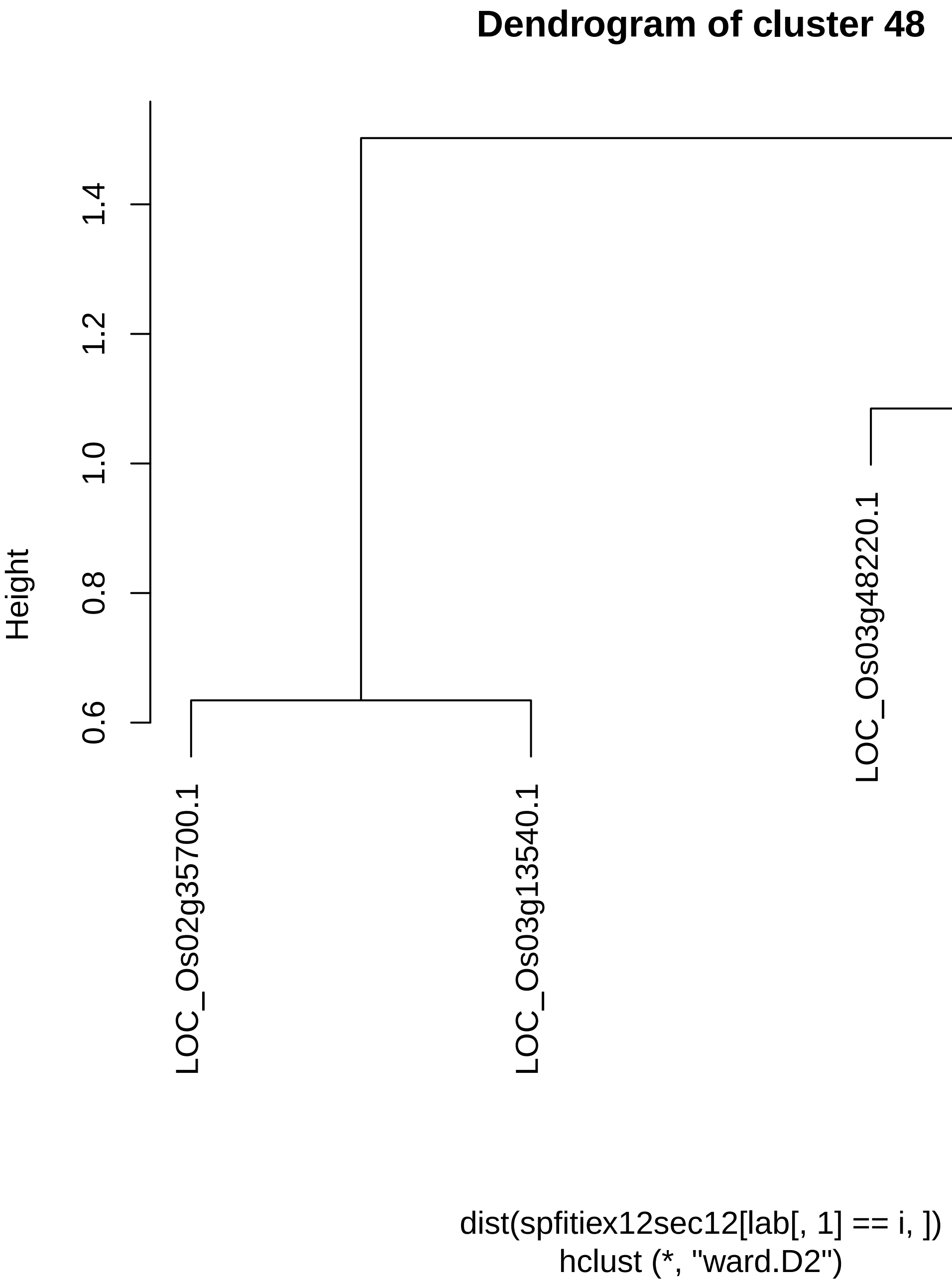
Dendrogram of cluster 48
Height
0.6
0.8
1.0
1.2
1.4
LOC_Os02g35700.1
LOC_Os03g13540.1
LOC_Os03g48220.1
LOC_Os05g33260.3
dist(spfitiex12sec12[lab[, 1] == i, ])
hclust (*, "ward.D2")

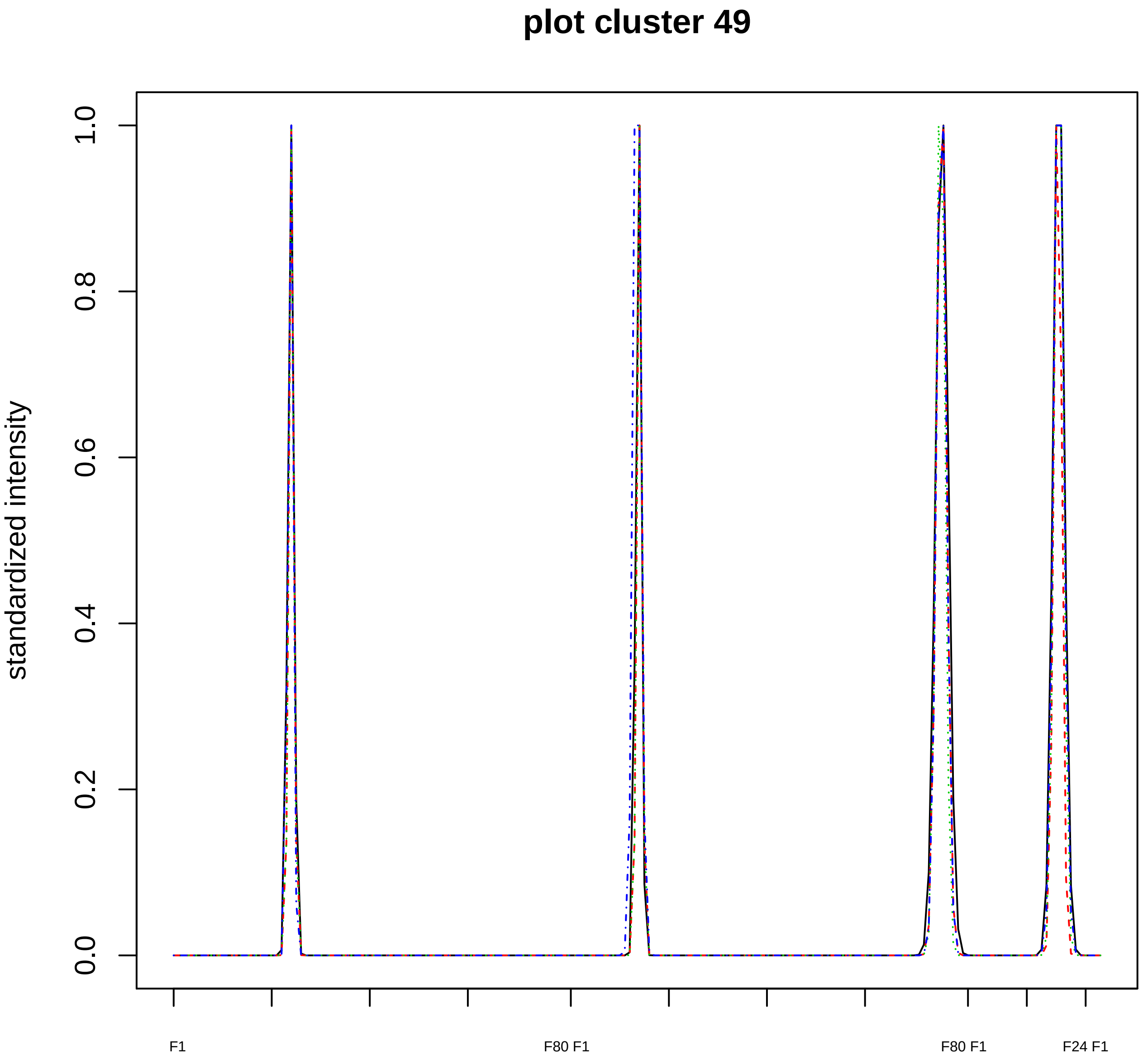

plot cluster 49
standardized intensity
0.0
0.2
0.4
0.6
0.8
1.0
F1
F80 F1
F80 F1
F24 F1

### Dendrogram of cluster 49

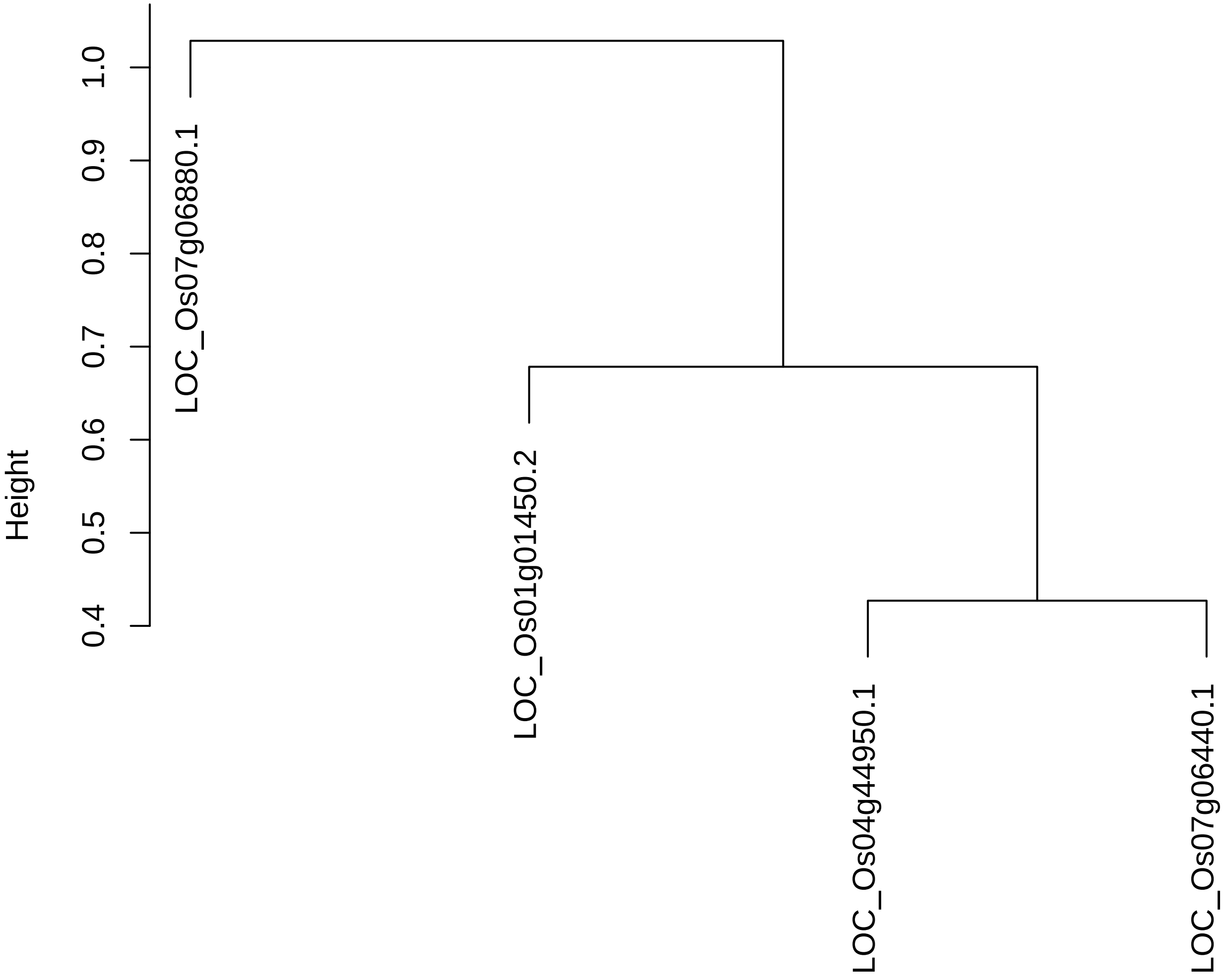


dist(spfitiex12sec12[lab[, 1] == i, ])
hclust (*, "ward.D2")

#### plot cluster 50

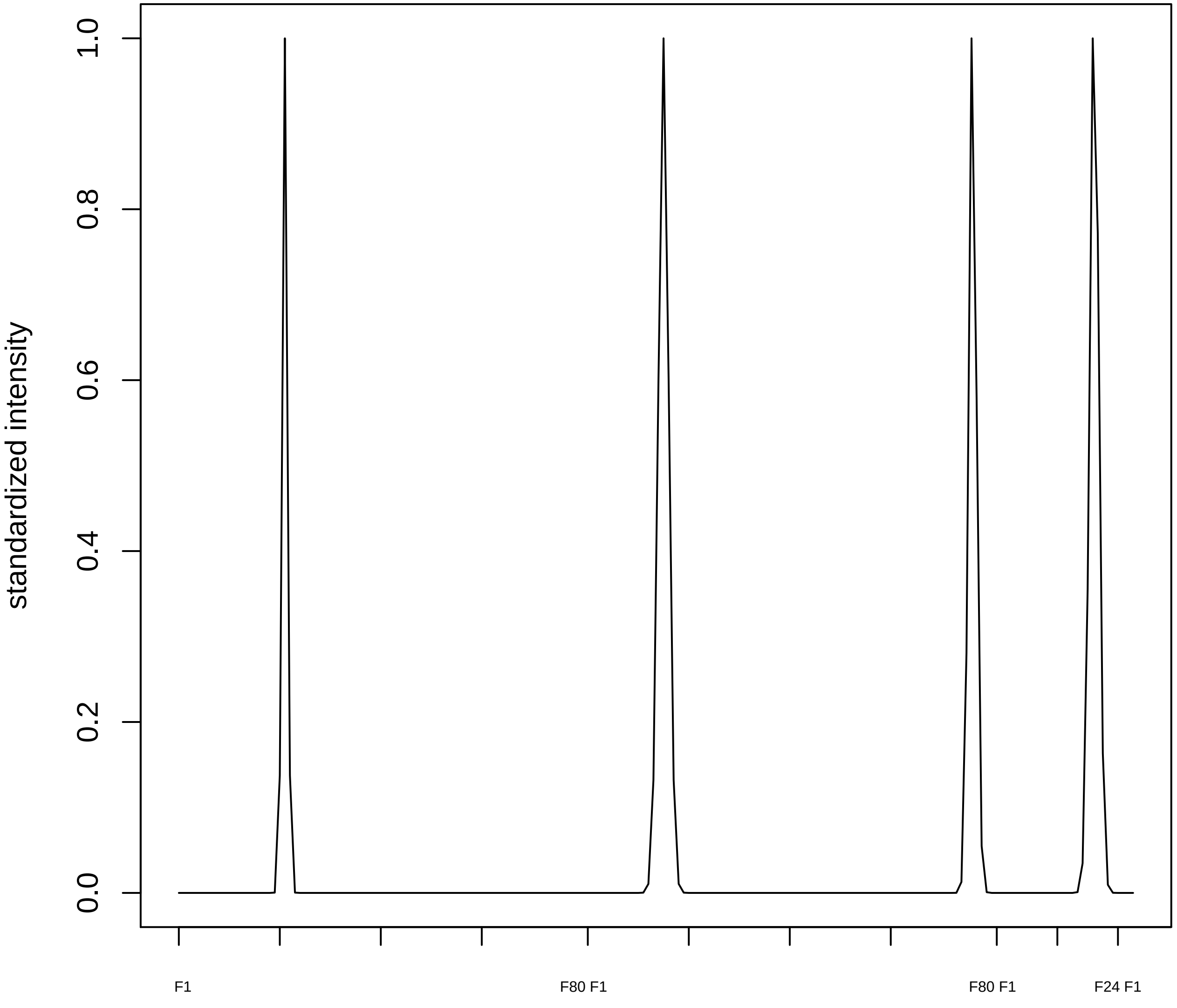

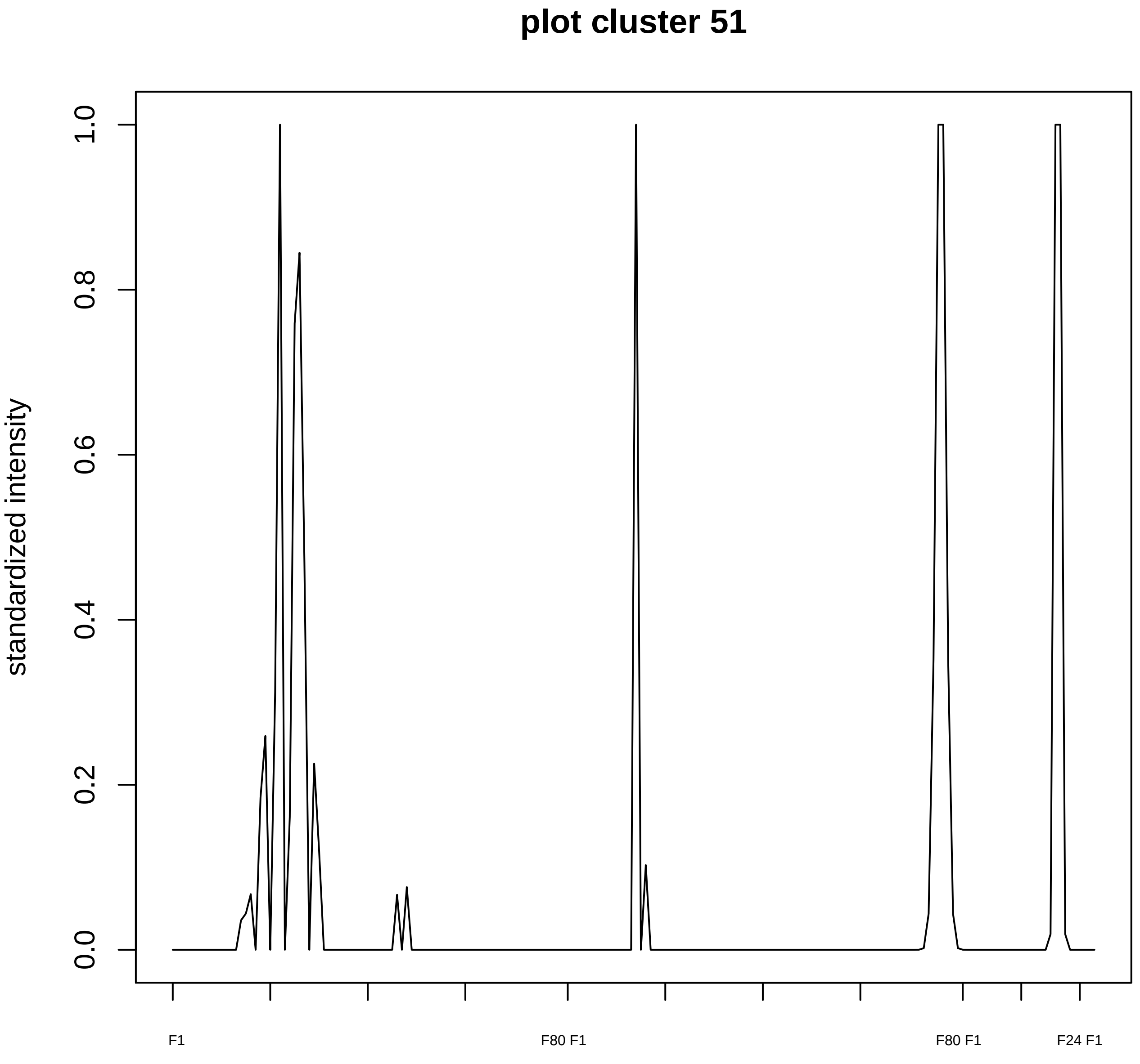

plot cluster 51
standardized intensity
0.0
0.2
0.4
0.6
0.8
1.0
F1
F80 F1
F80 F1
F24 F1

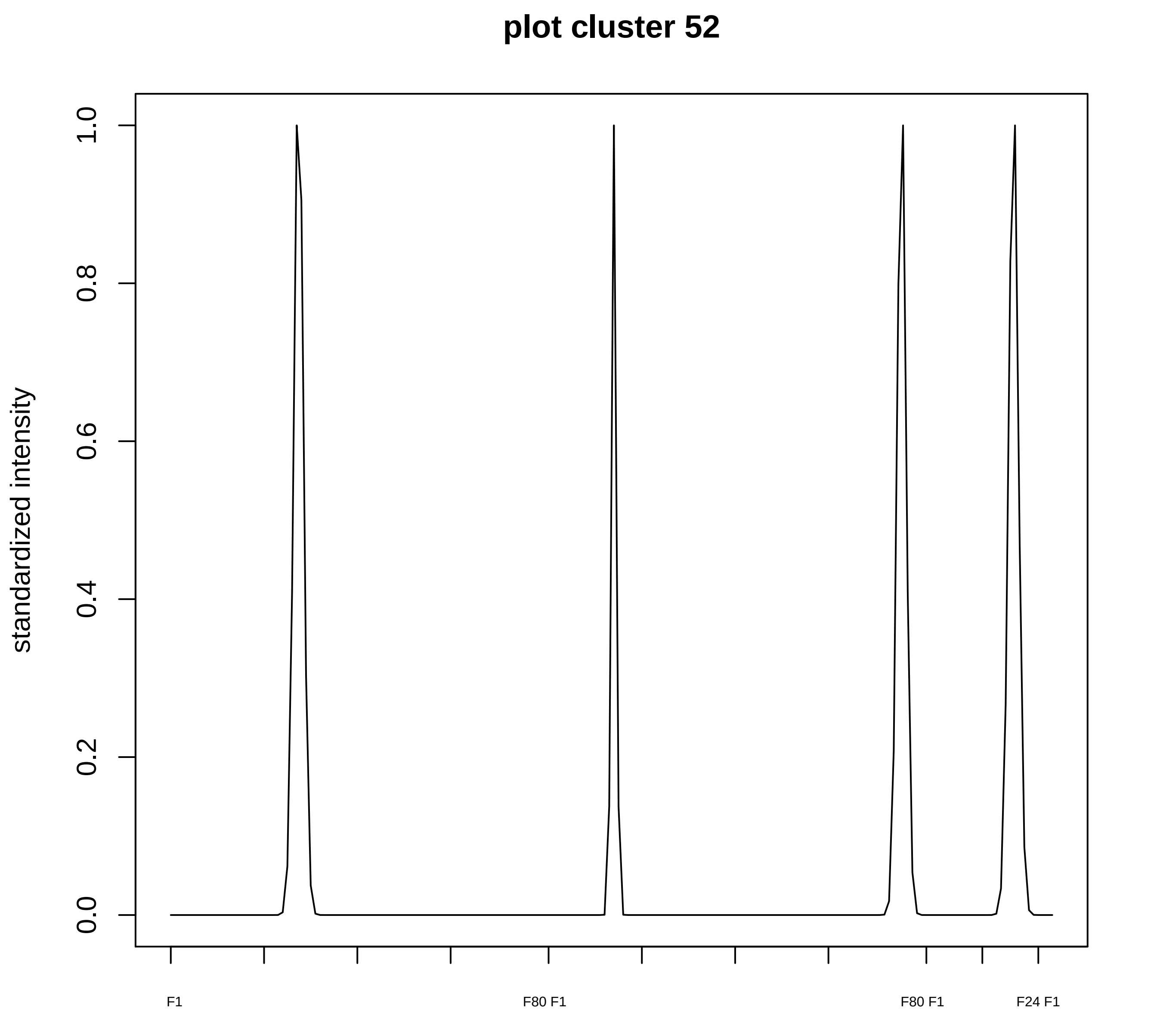

plot cluster 52
standardized intensity
0.0
0.2
0.4
0.6
0.8
1.0
F1
F80 F1
F80 F1
F24 F1

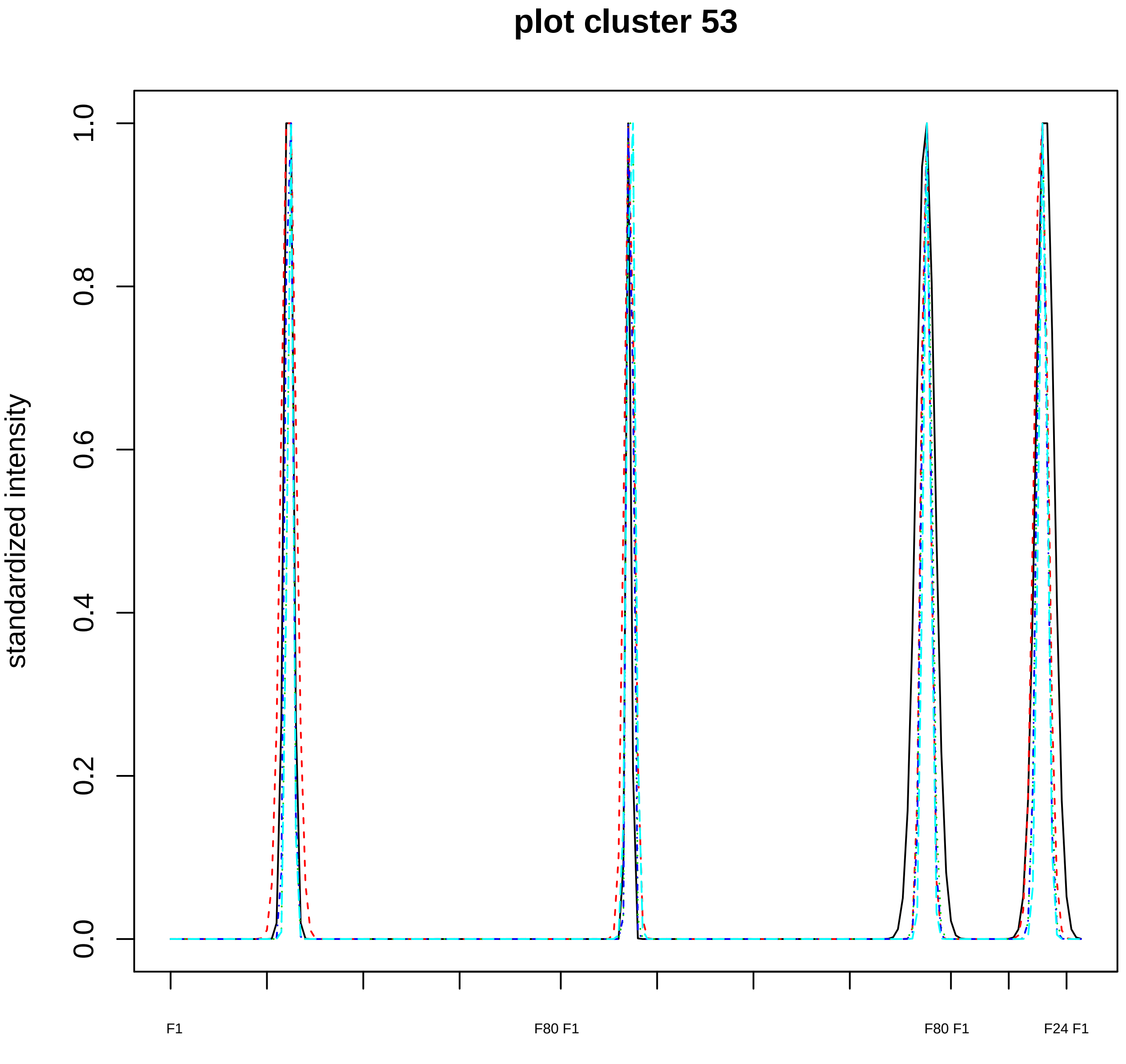
plot cluster 53
standardized intensity
0.0
0.2
0.4
0.6
0.8
1.0
F1
F80 F1
F80 F1
F24 F1

### Dendrogram of cluster 53

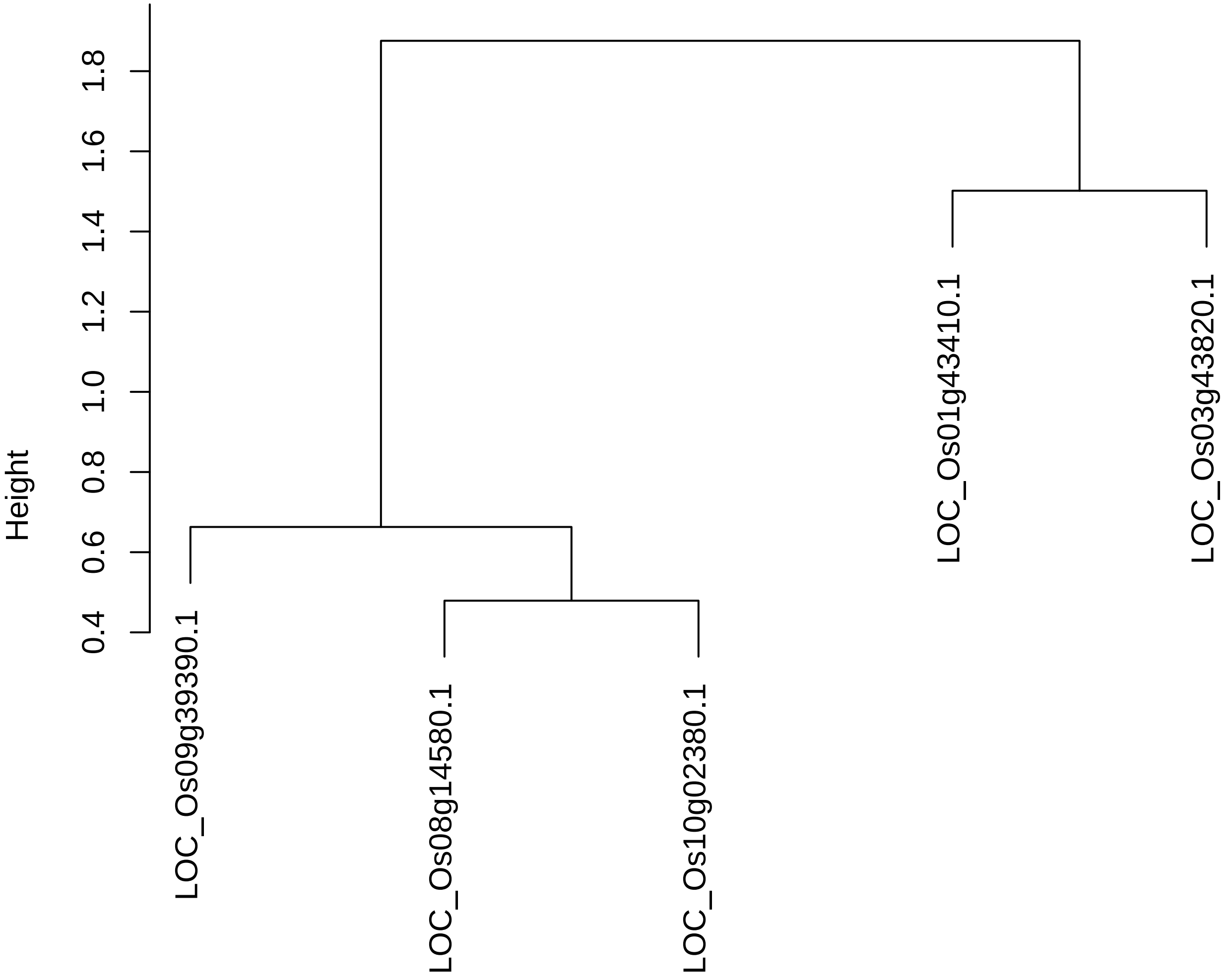


dist(spfitiex12sec12[lab[, 1] == i, ])
hclust (*, "ward.D2")

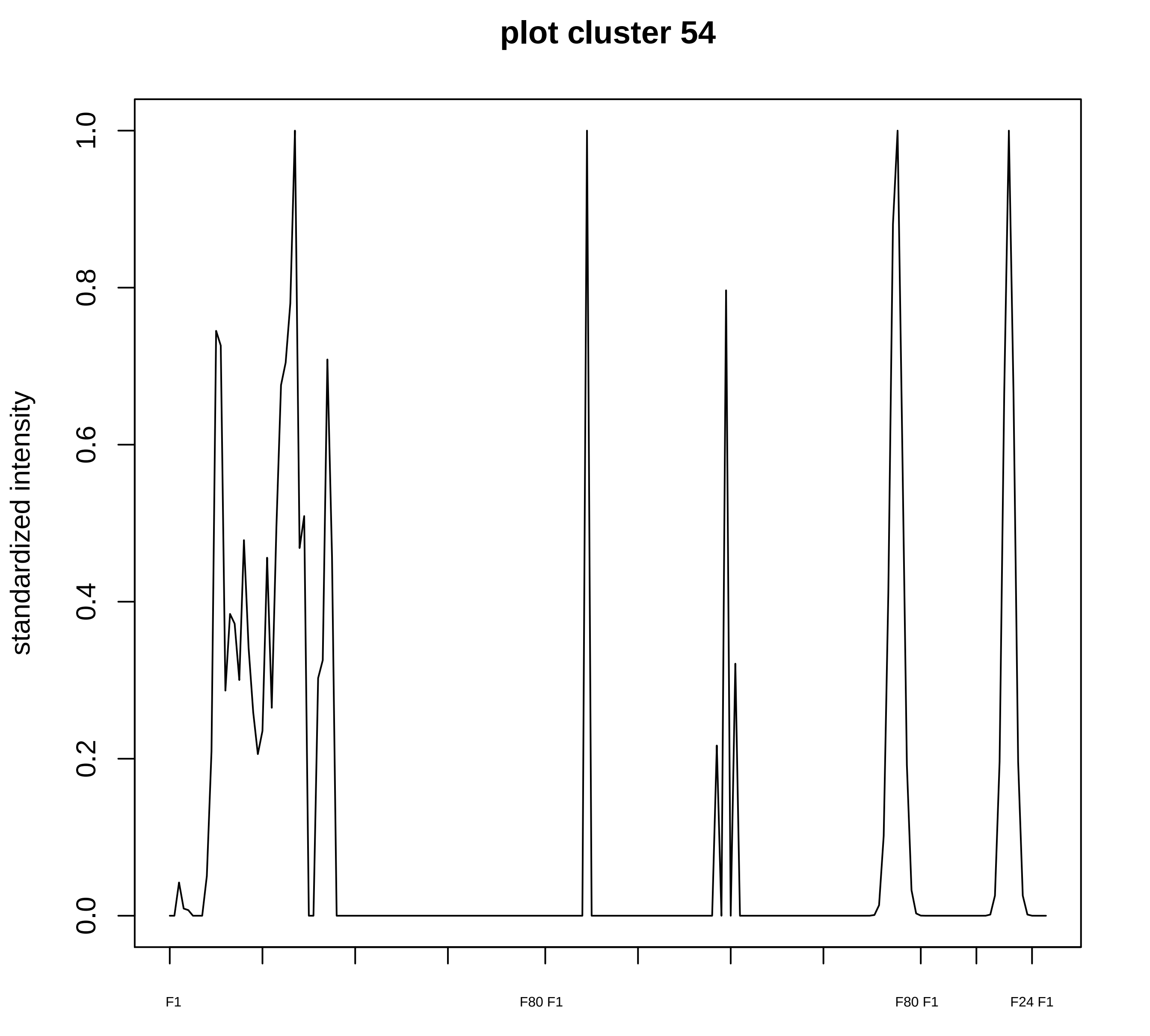
plot cluster 54
standardized intensity
0.0
0.2
0.4
0.6
0.8
1.0
F1
F80 F1
F80 F1
F24 F1

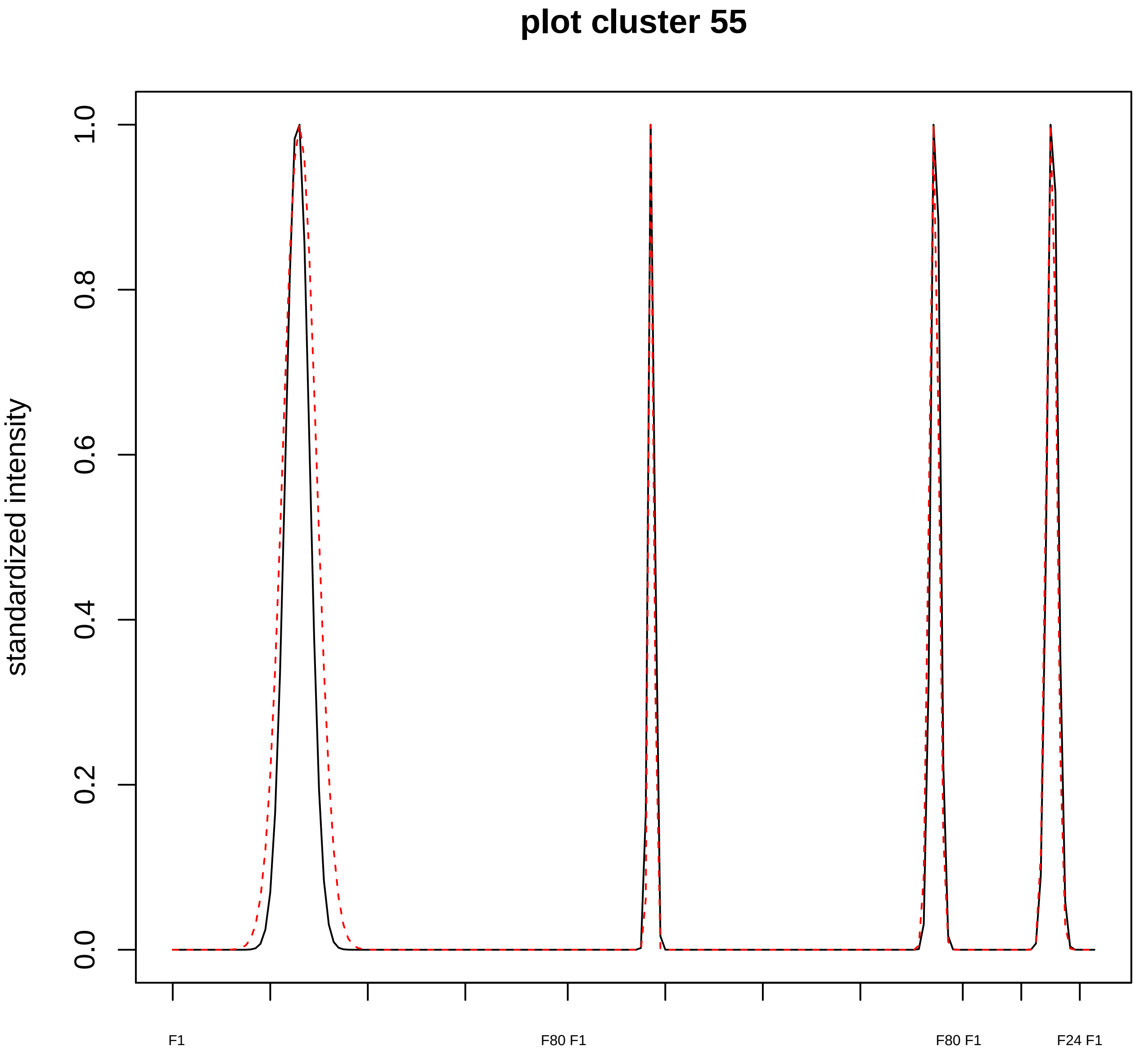
plot cluster 55
standardized intensity
0.0
0.2
0.4
0.6
0.8
1.0
F1
F80 F1
F80 F1
F24 F1

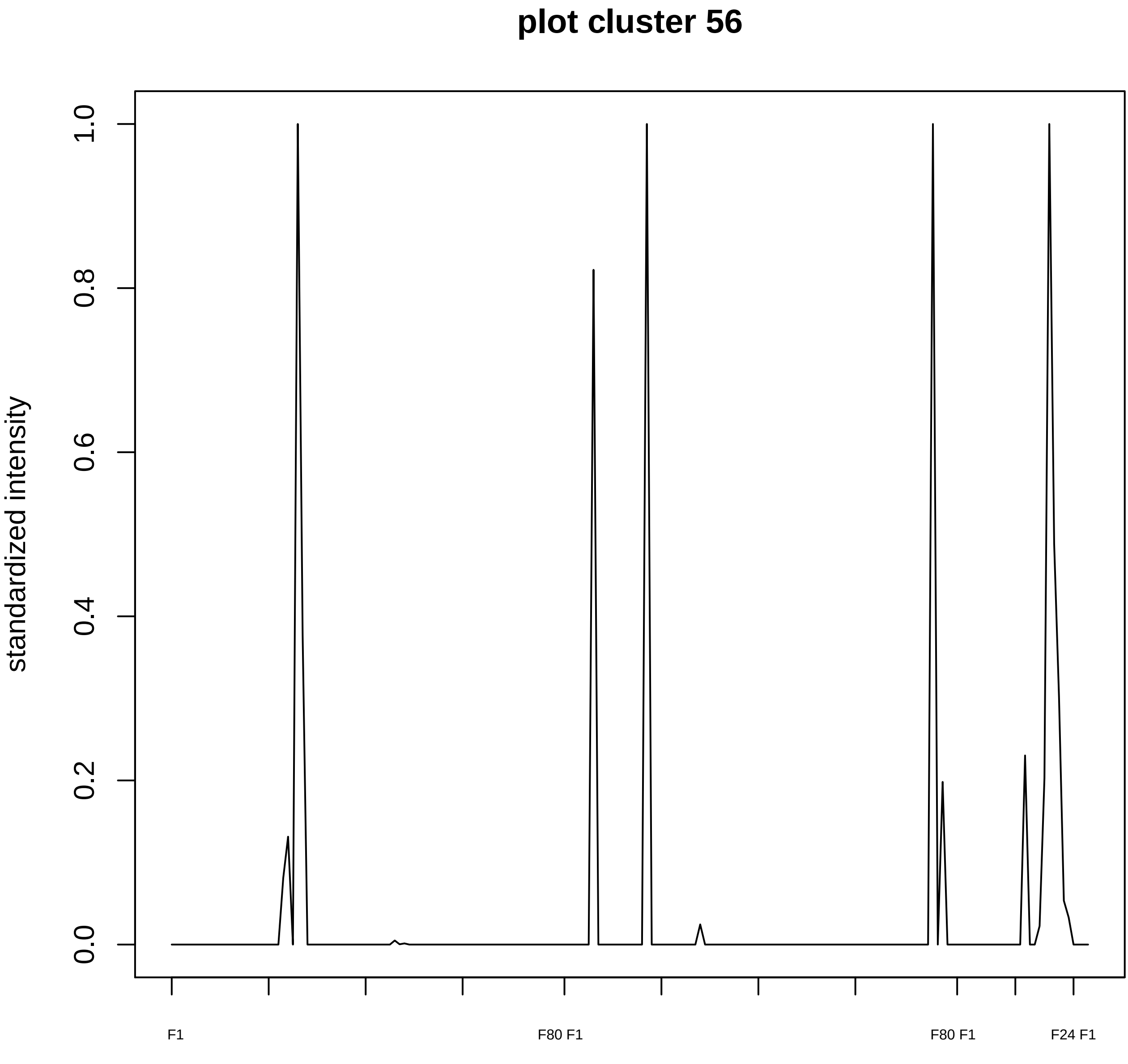
plot cluster 56
standardized intensity
0.0
0.2
0.4
0.6
0.8
1.0
F1
F80 F1
F80 F1
F24 F1

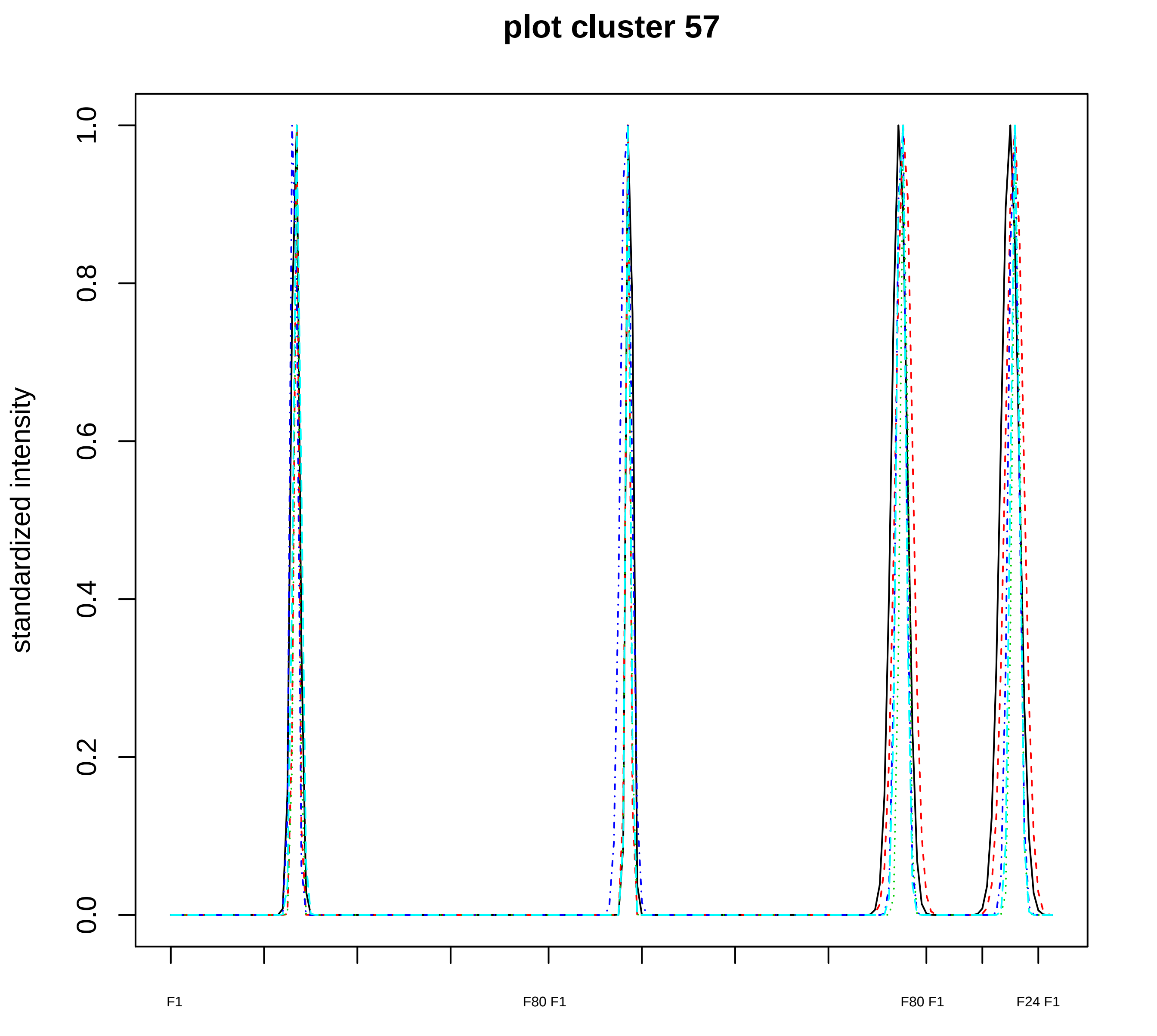

plot cluster 57
standardized intensity
0.0
0.2
0.4
0.6
0.8
1.0
F1
F80 F1
F80 F1
F24 F1

#### Dendrogram of cluster 57

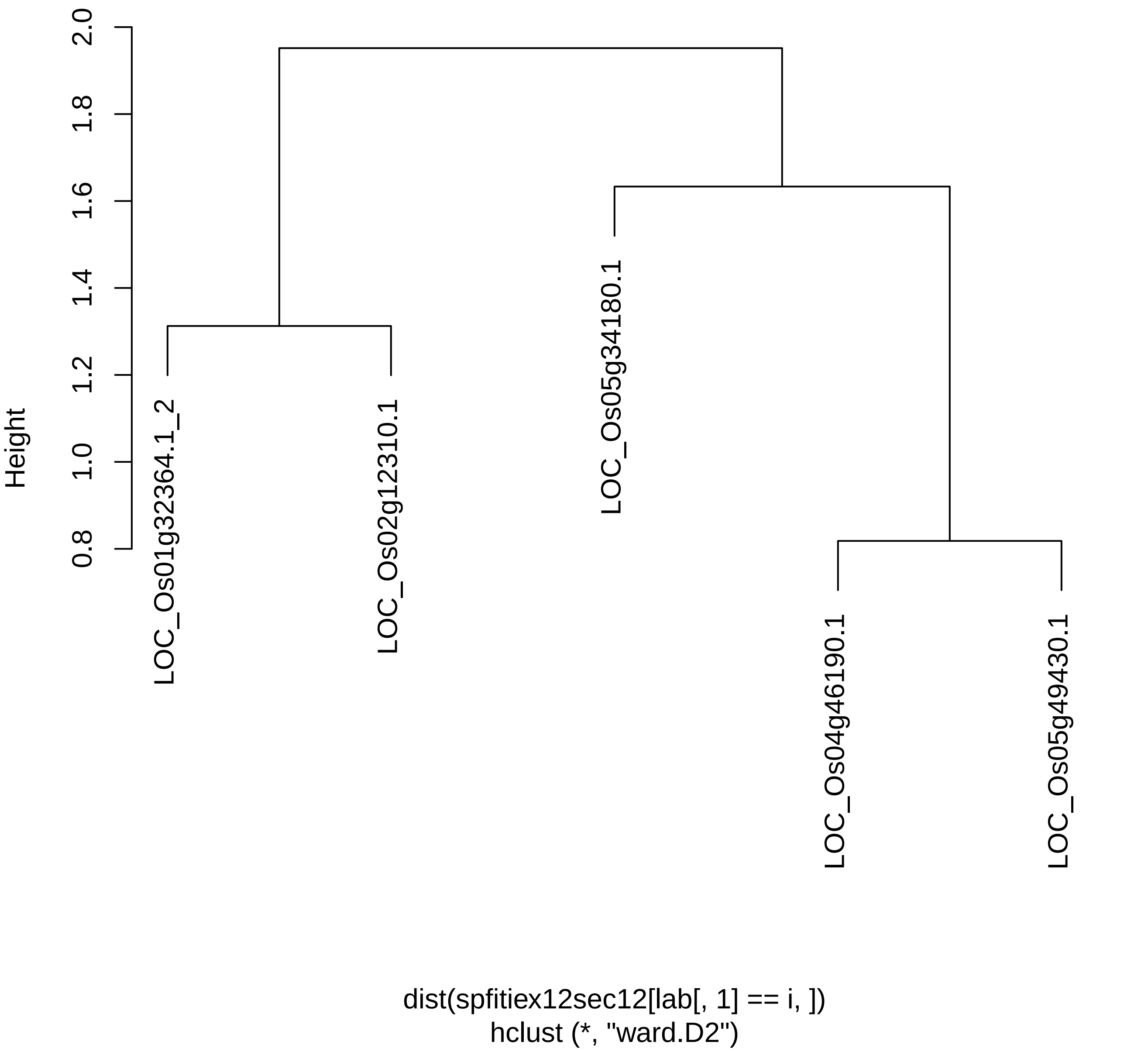

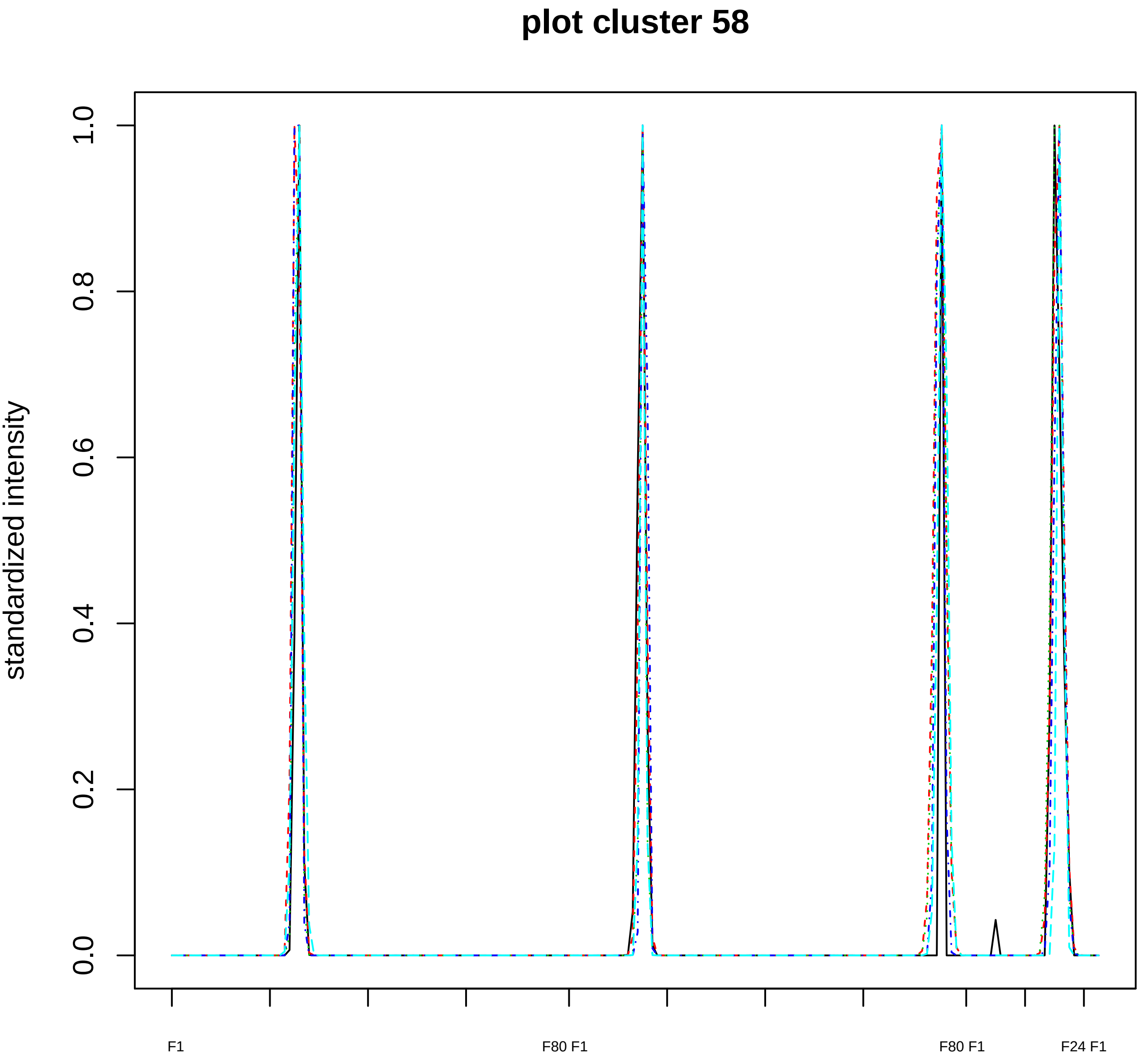
plot cluster 58
standardized intensity
0.0
0.2
0.4
0.6
0.8
1.0
F1
F80 F1
F80 F1
F24 F1

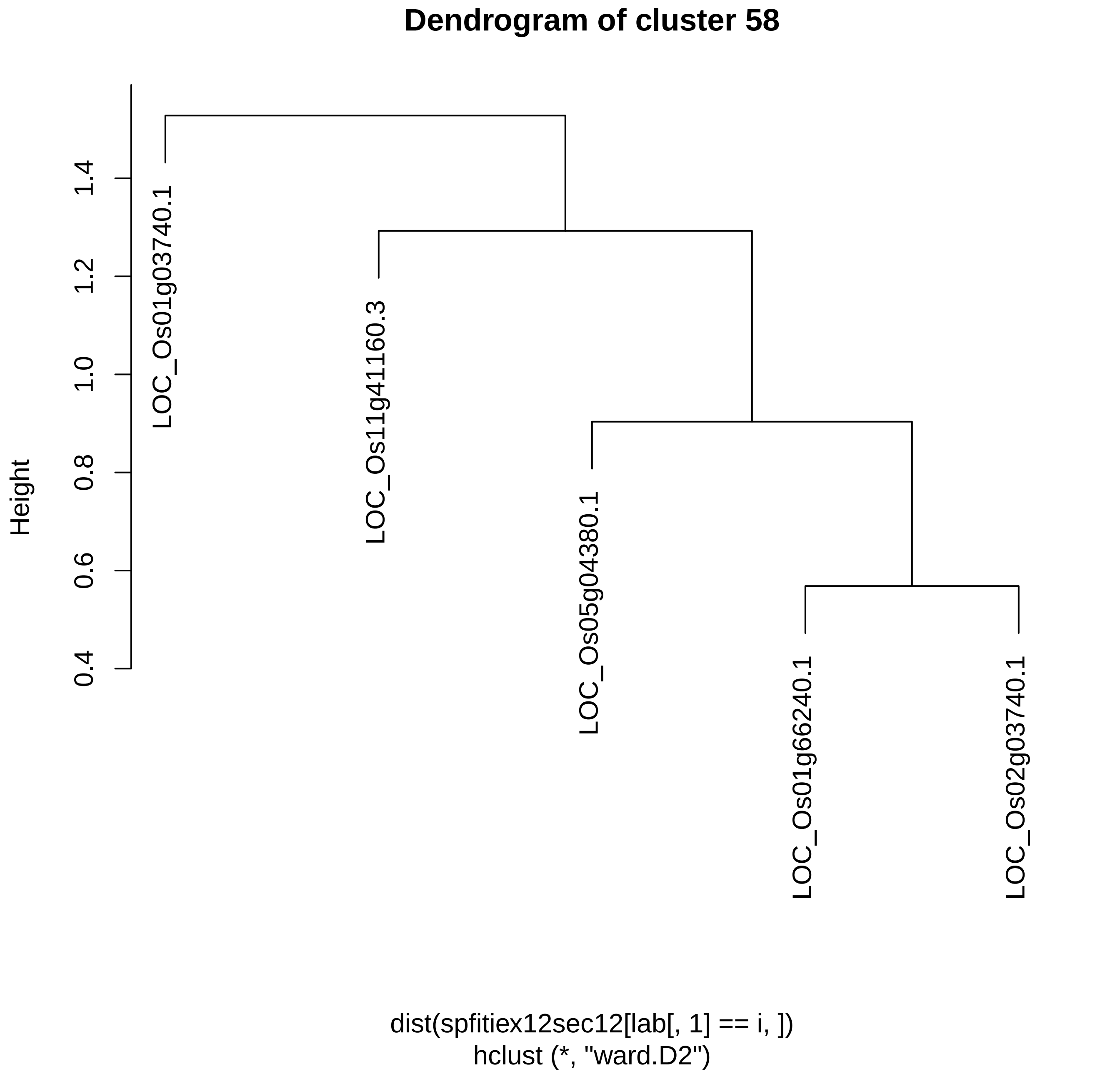
Dendrogram of cluster 58
Height
1.4
1.2
1.0
0.8
0.6
0.4
LOC_Os01g03740.1
LOC_Os11g41160.3
LOC_Os05g04380.1
LOC_Os01g66240.1
LOC_Os02g03740.1
dist(spfitiex12sec12[lab[, 1] == i, ])
hclust (*, "ward.D2")

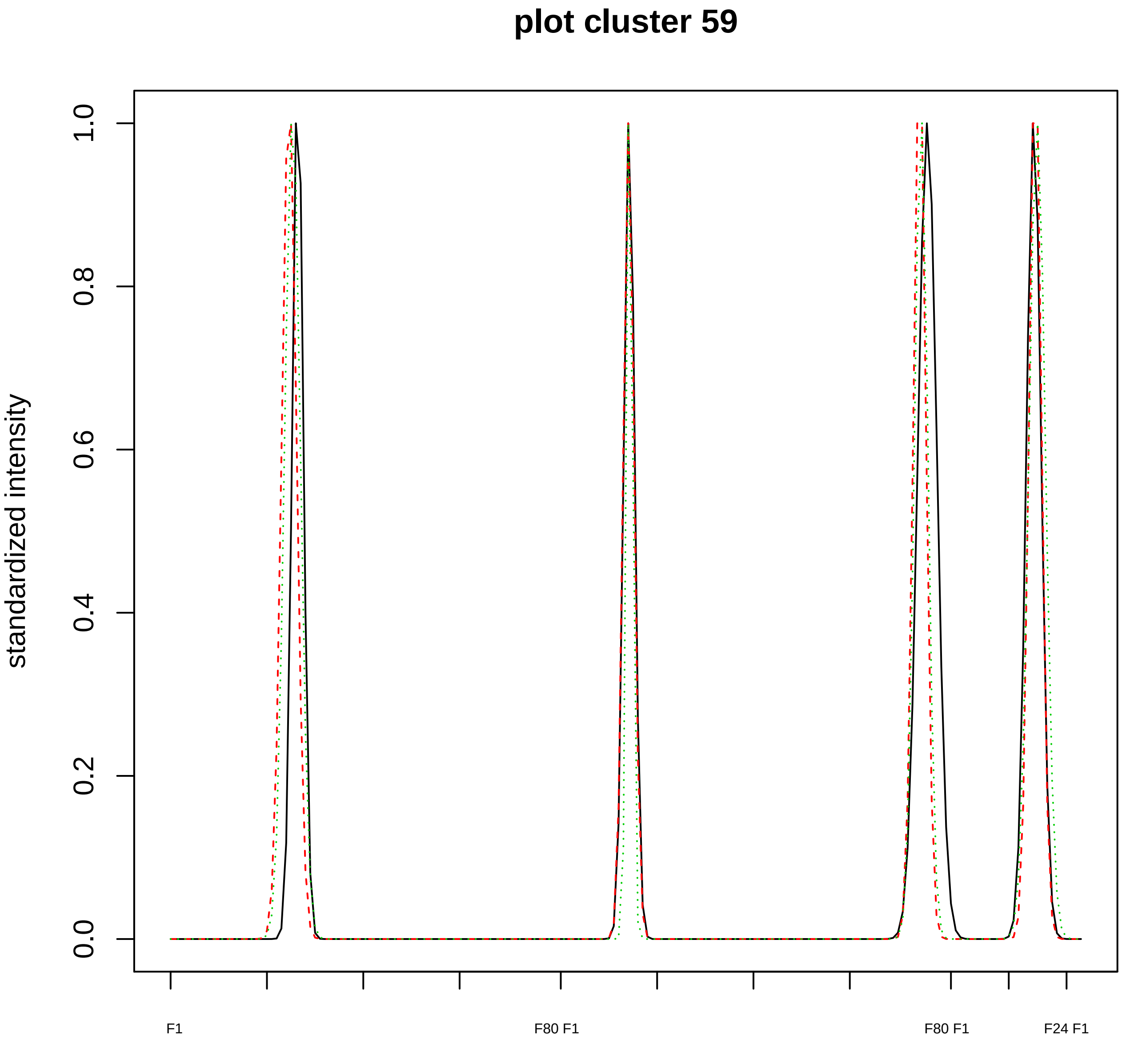
plot cluster 59
standardized intensity
0.0
0.2
0.4
0.6
0.8
1.0
F1
F80 F1
F80 F1
F24 F1

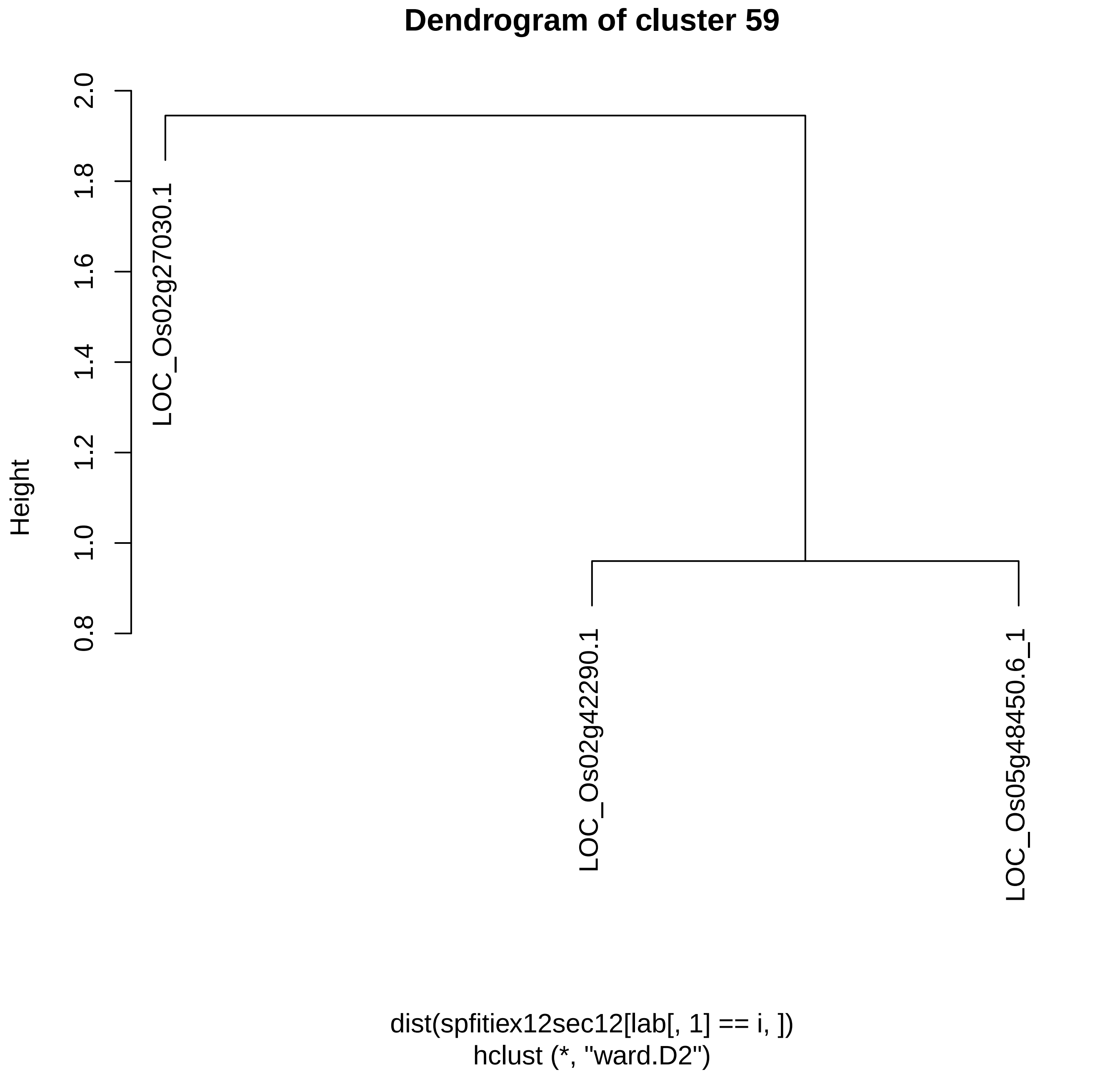
Dendrogram of cluster 59
Height
2.0
1.8
1.6
1.4
1.2
1.0
0.8
LOC_Os02g27030.1
LOC_Os02g42290.1
LOC_Os05g48450.6_1
dist(spfitiex12sec12[lab[, 1] == i, ])
hclust (*, "ward.D2")

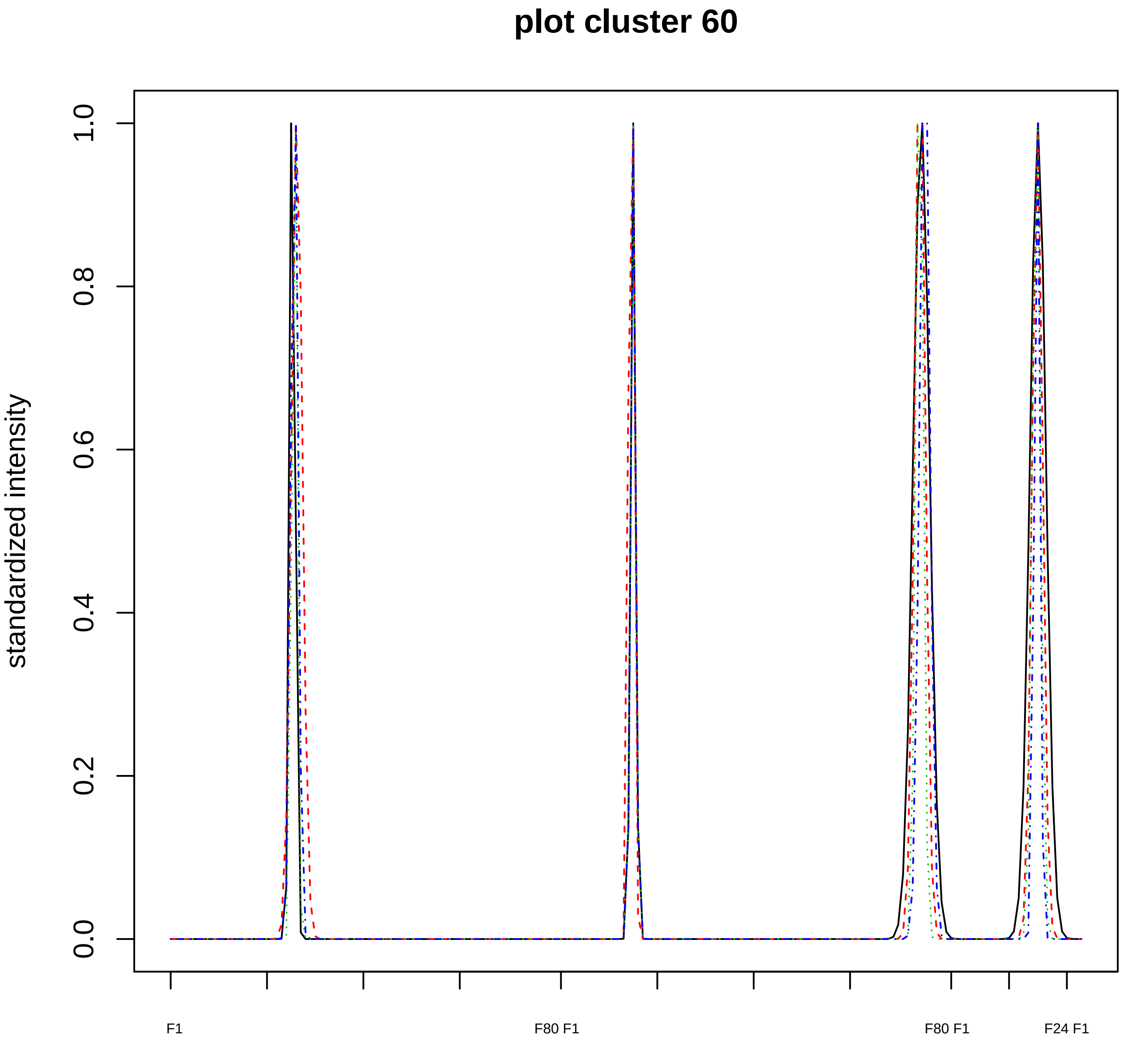

plot cluster 60
standardized intensity
0.0
0.2
0.4
0.6
0.8
1.0
F1
F80 F1
F80 F1
F24 F1

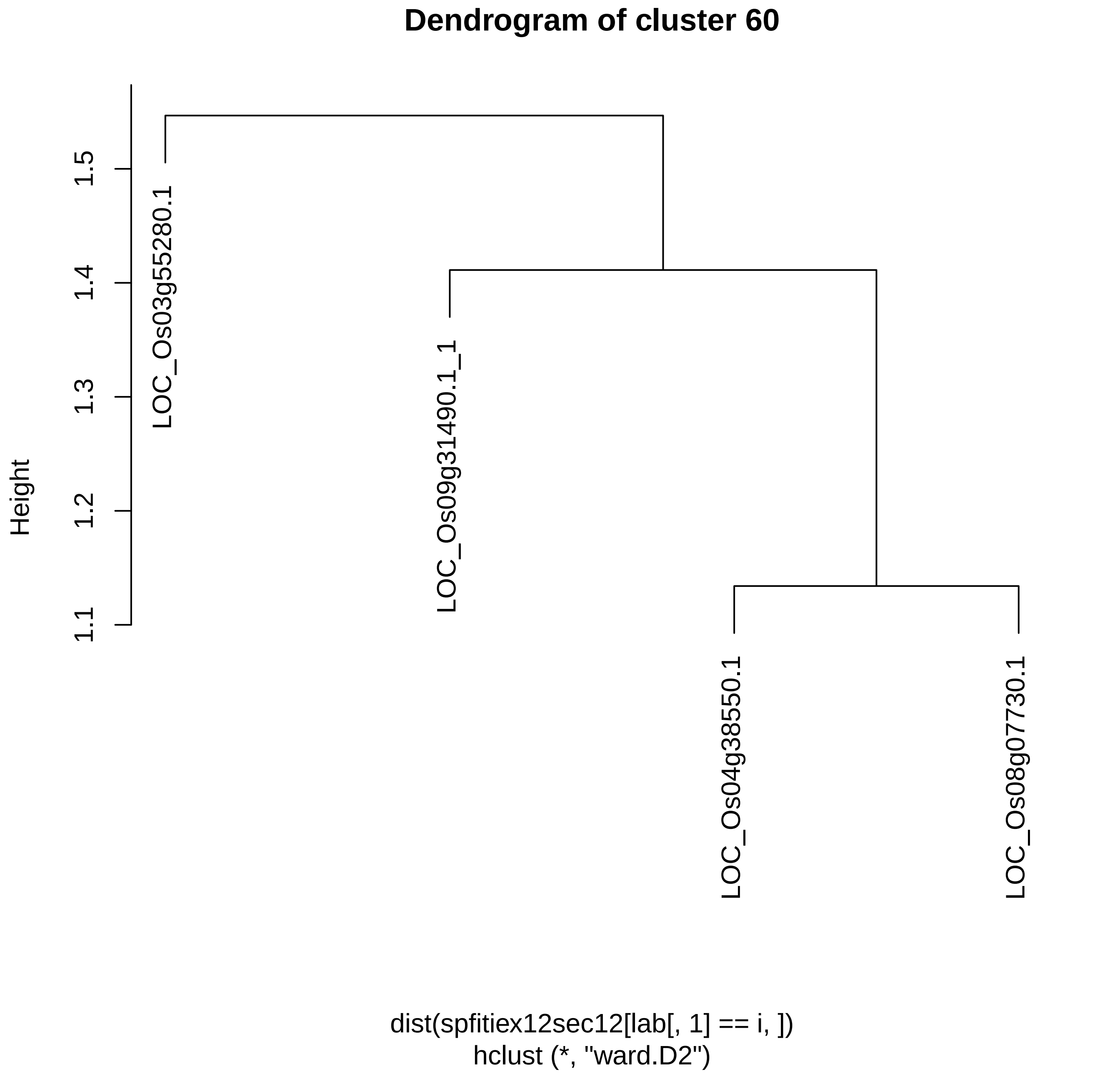
Dendrogram of cluster 60
Height
1.1
1.2
1.3
1.4
1.5
LOC_Os03g55280.1
LOC_Os09g31490.1_1
LOC_Os04g38550.1
LOC_Os08g07730.1
dist(spfitiex12sec12[lab[, 1] == i, ])
hclust (*, "ward.D2")

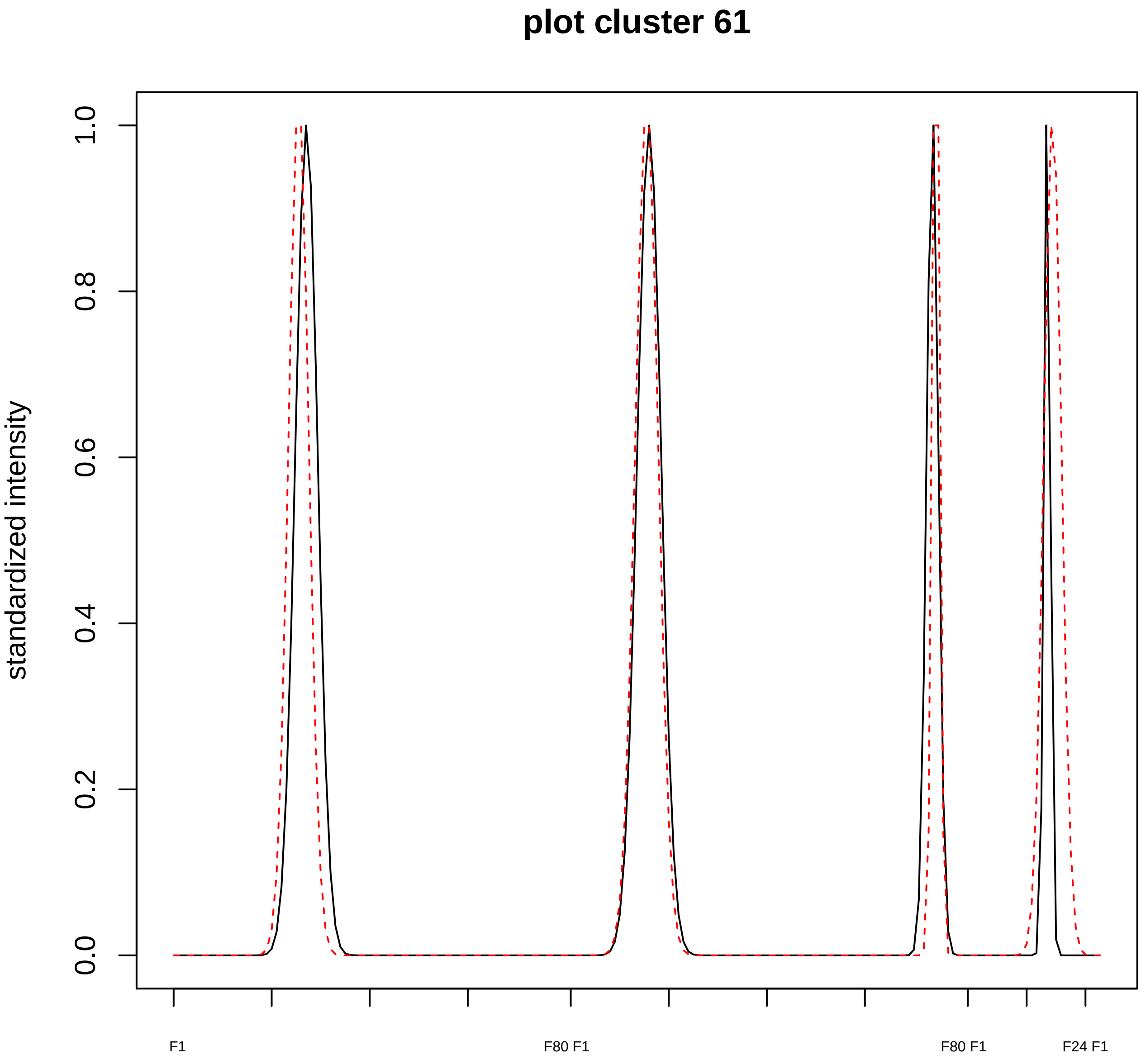

plot cluster 61
standardized intensity
0.0
0.2
0.4
0.6
0.8
1.0
F1
F80 F1
F80 F1
F24 F1

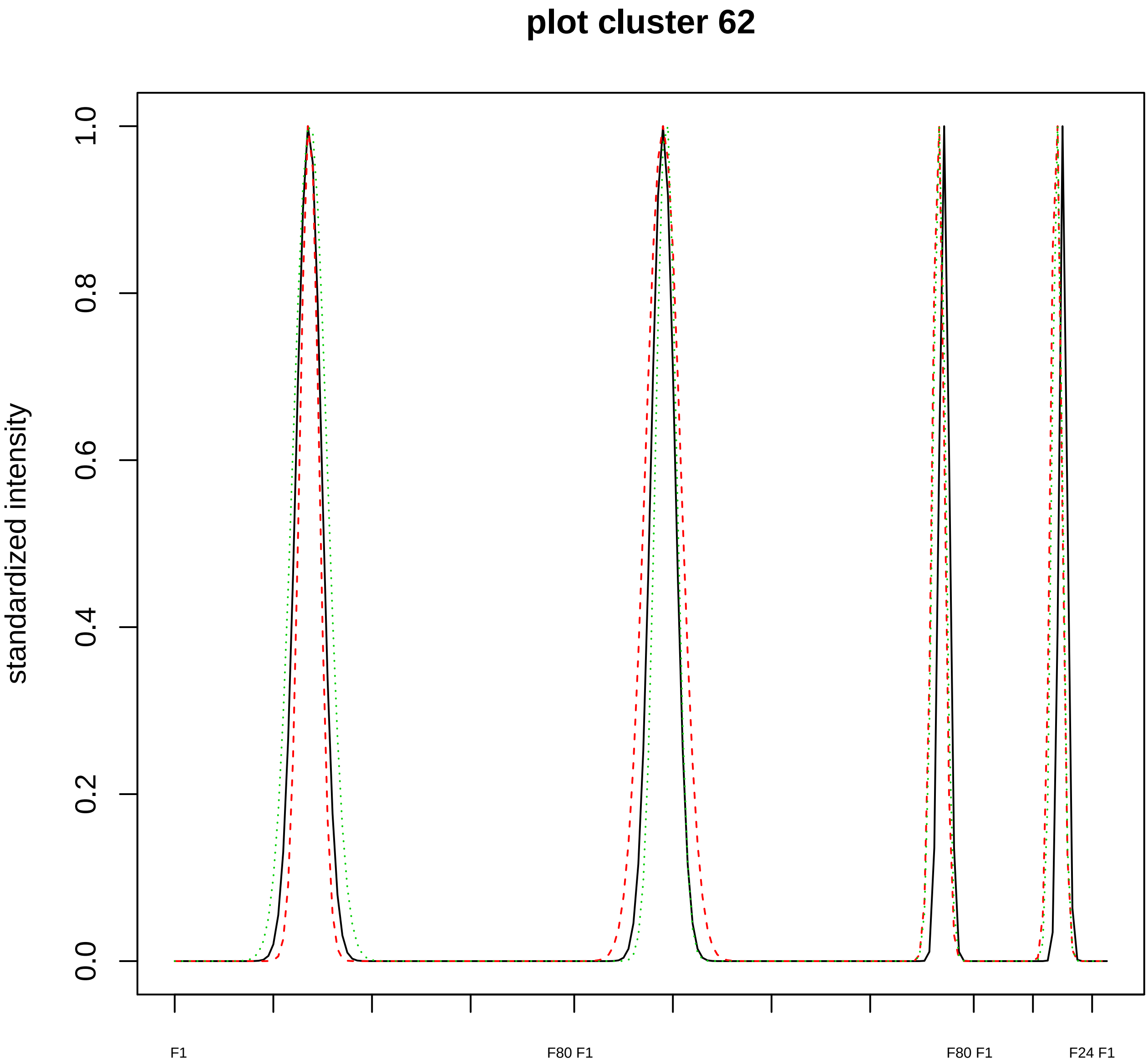
plot cluster 62
standardized intensity
1.0
0.8
0.6
0.4
0.2
0.0
F1
F80 F1
F80 F1
F24 F1

#### Dendrogram of cluster 62

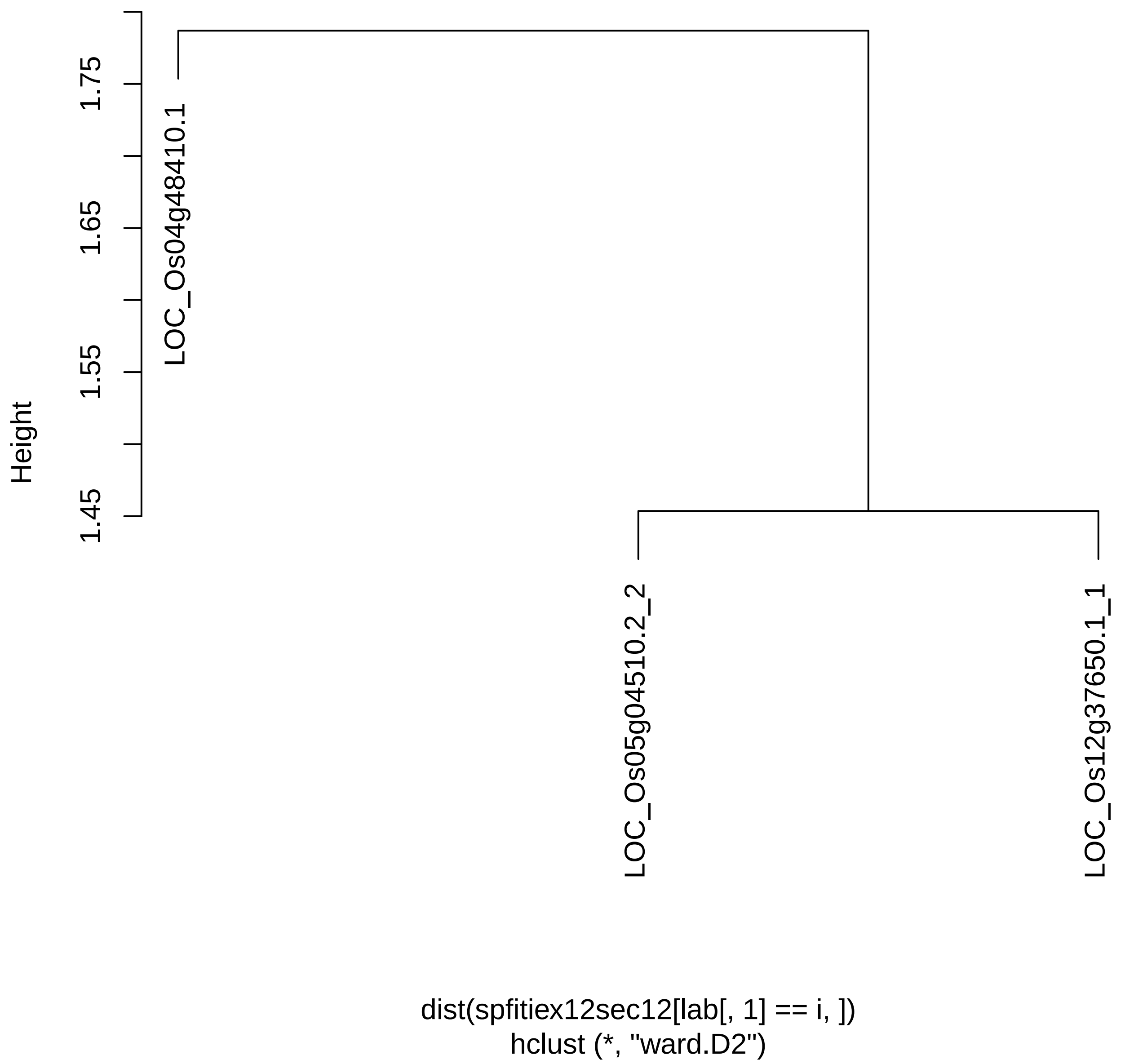

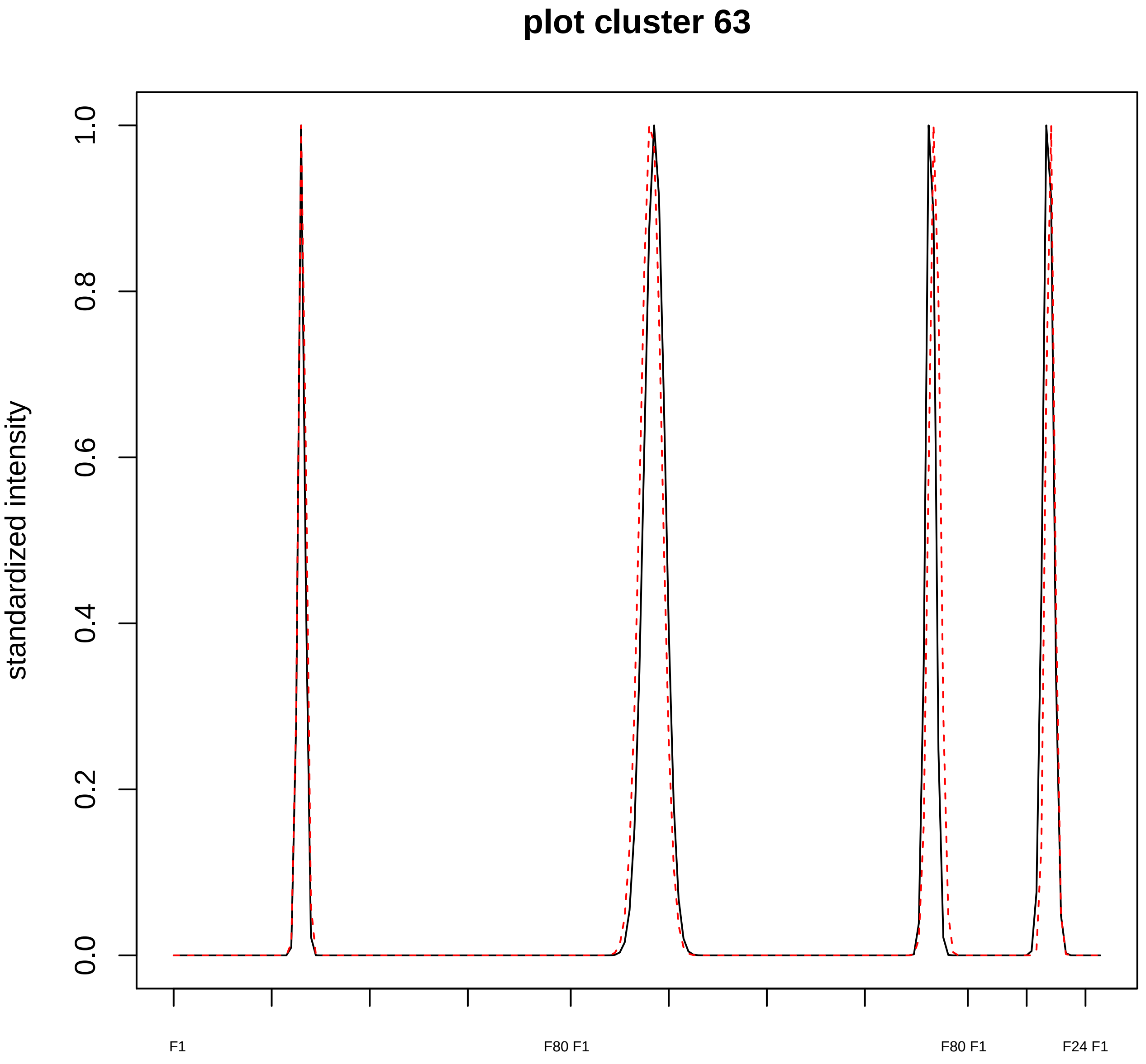
plot cluster 63
standardized intensity
0.0
0.2
0.4
0.6
0.8
1.0
F1
F80 F1
F80 F1
F24 F1

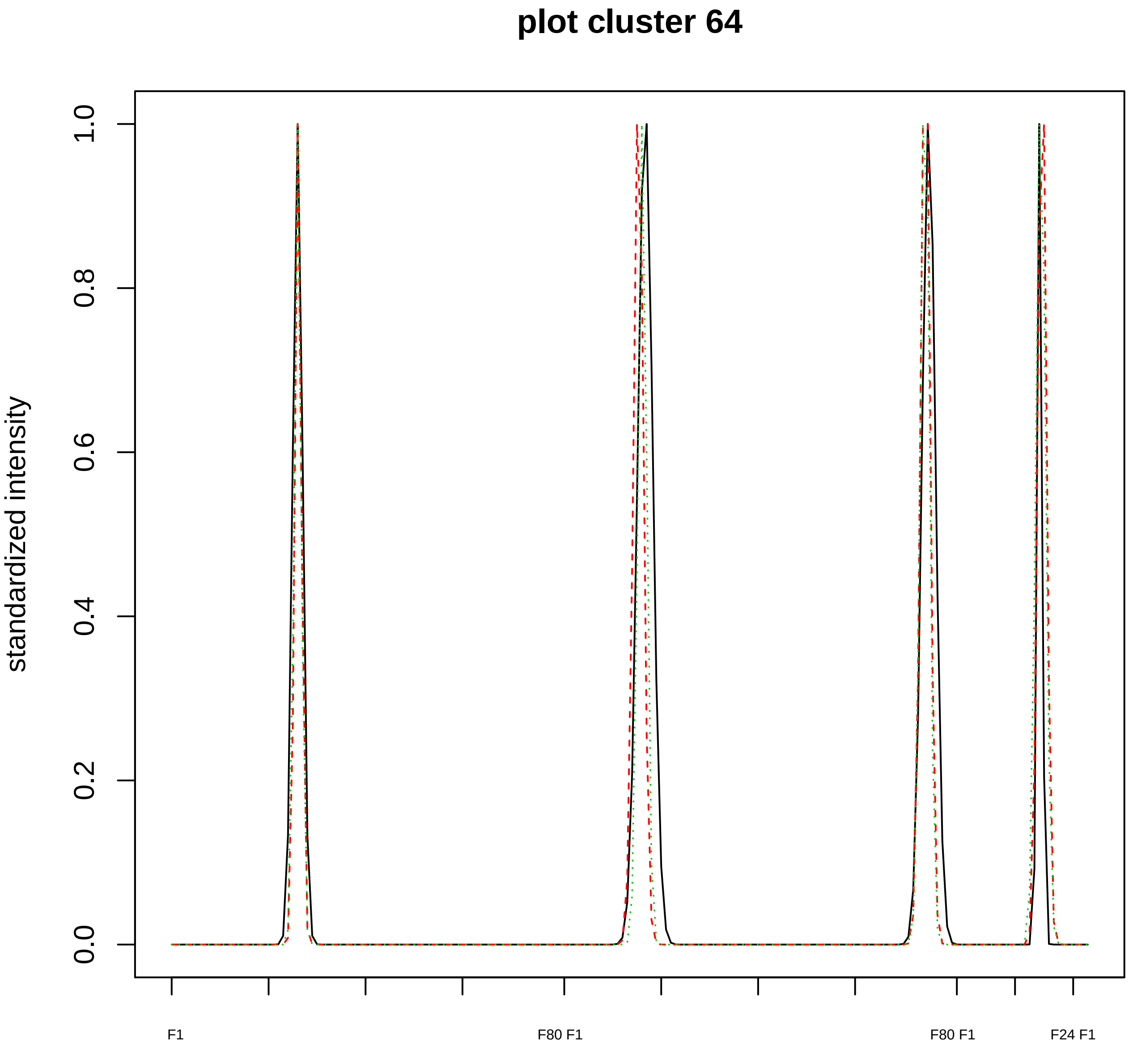
plot cluster 64
standardized intensity
0.0
0.2
0.4
0.6
0.8
1.0
F1
F80 F1
F80 F1
F24 F1

### Dendrogram of cluster 64

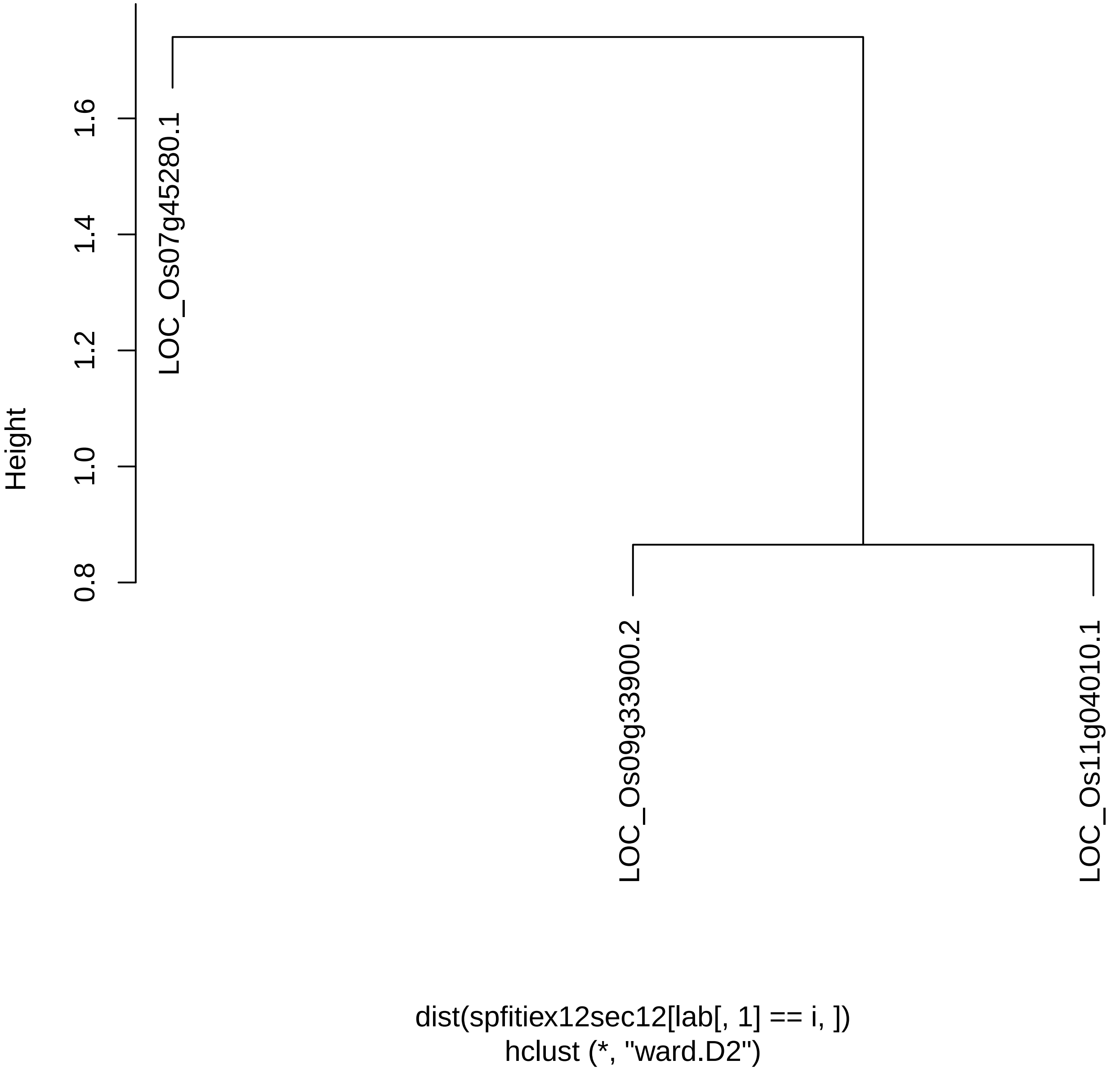

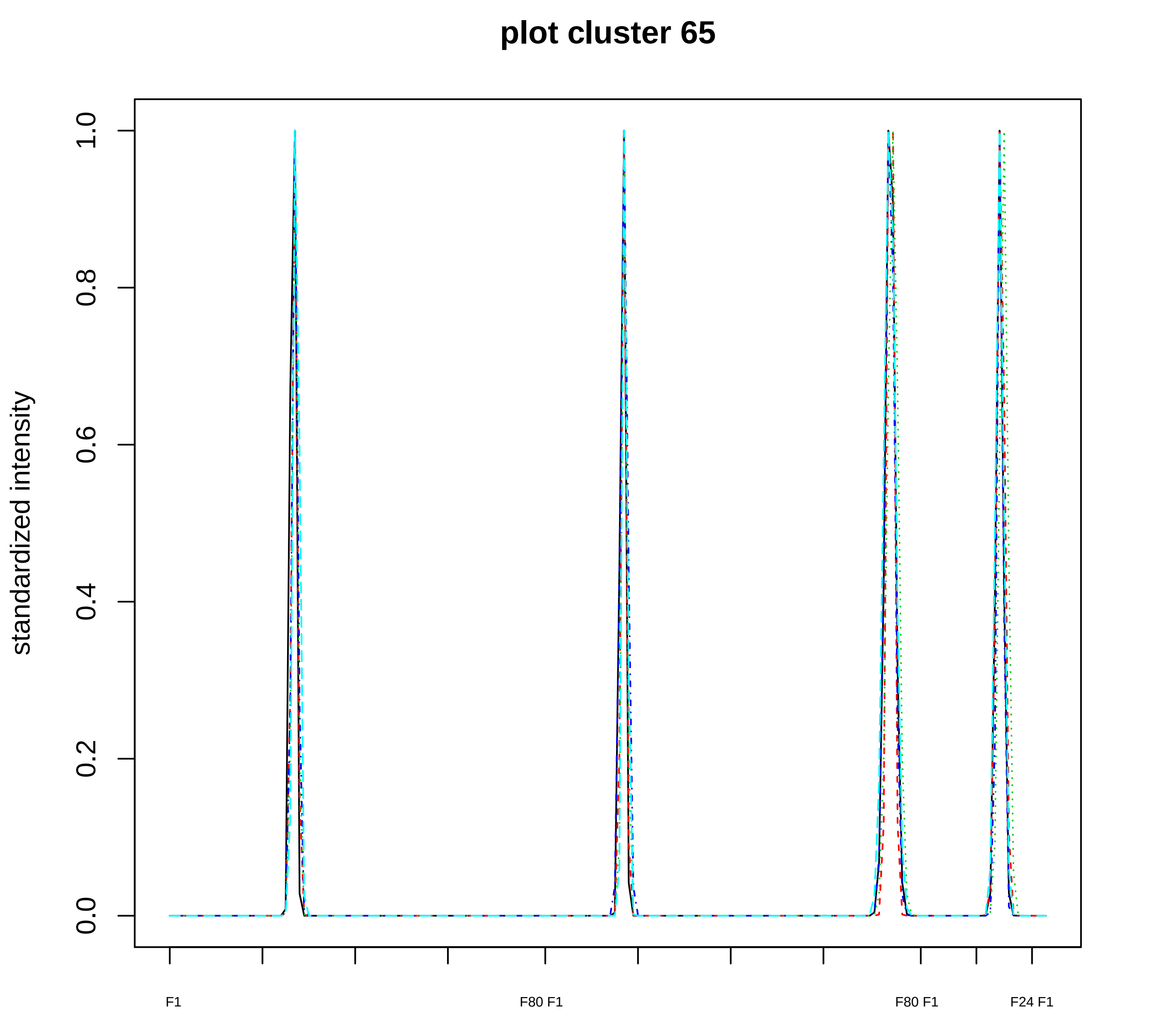
plot cluster 65
standardized intensity
0.0
0.2
0.4
0.6
0.8
1.0
F1
F80 F1
F80 F1
F24 F1

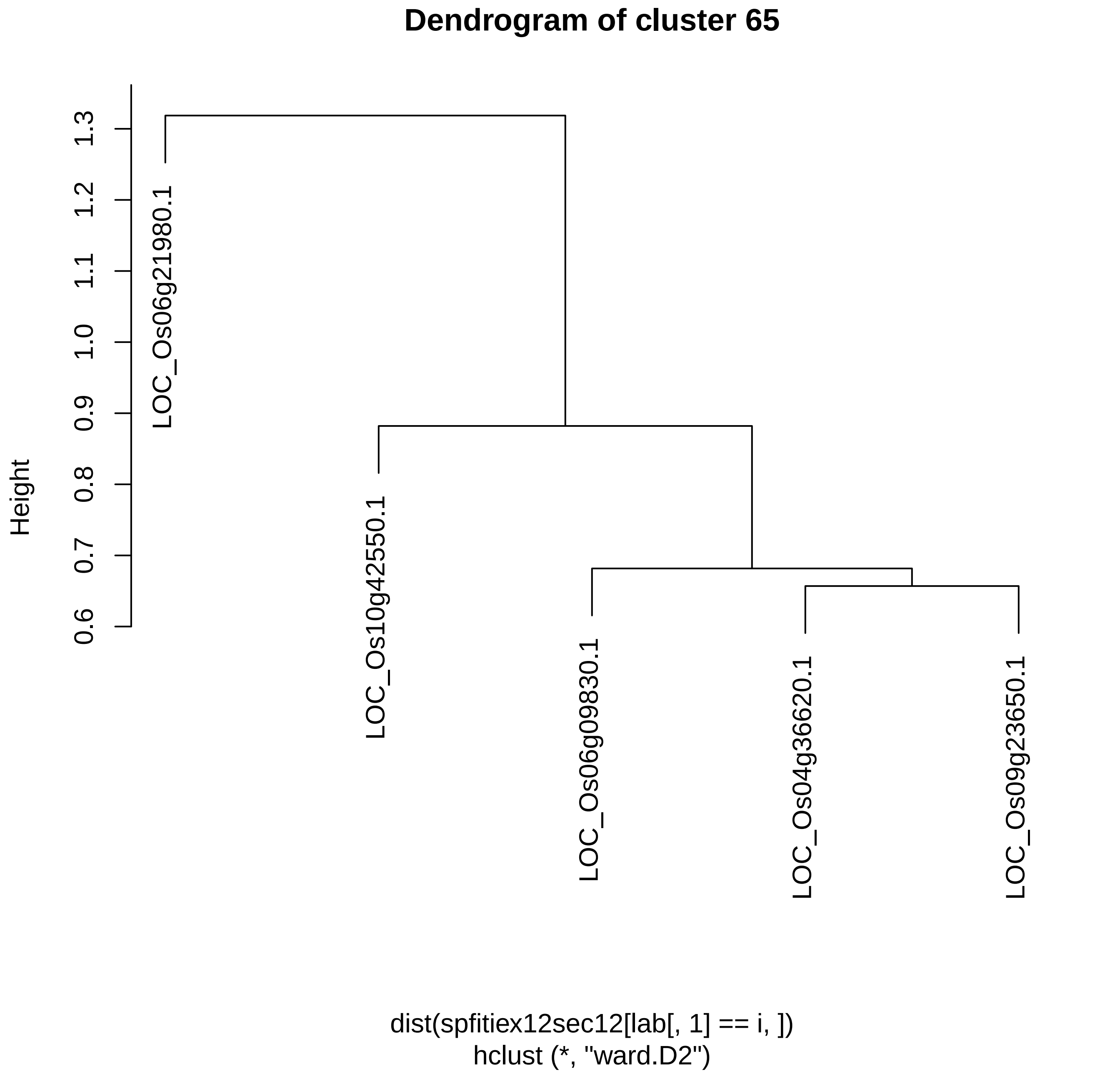
Dendrogram of cluster 65
Height
0.6
0.7
0.8
0.9
1.0
1.1
1.2
1.3
LOC_Os06g21980.1
LOC_Os10g42550.1
LOC_Os06g09830.1
LOC_Os04g36620.1
LOC_Os09g23650.1
dist(spfitiex12sec12[lab[, 1] == i, ])
hclust (*, "ward.D2")

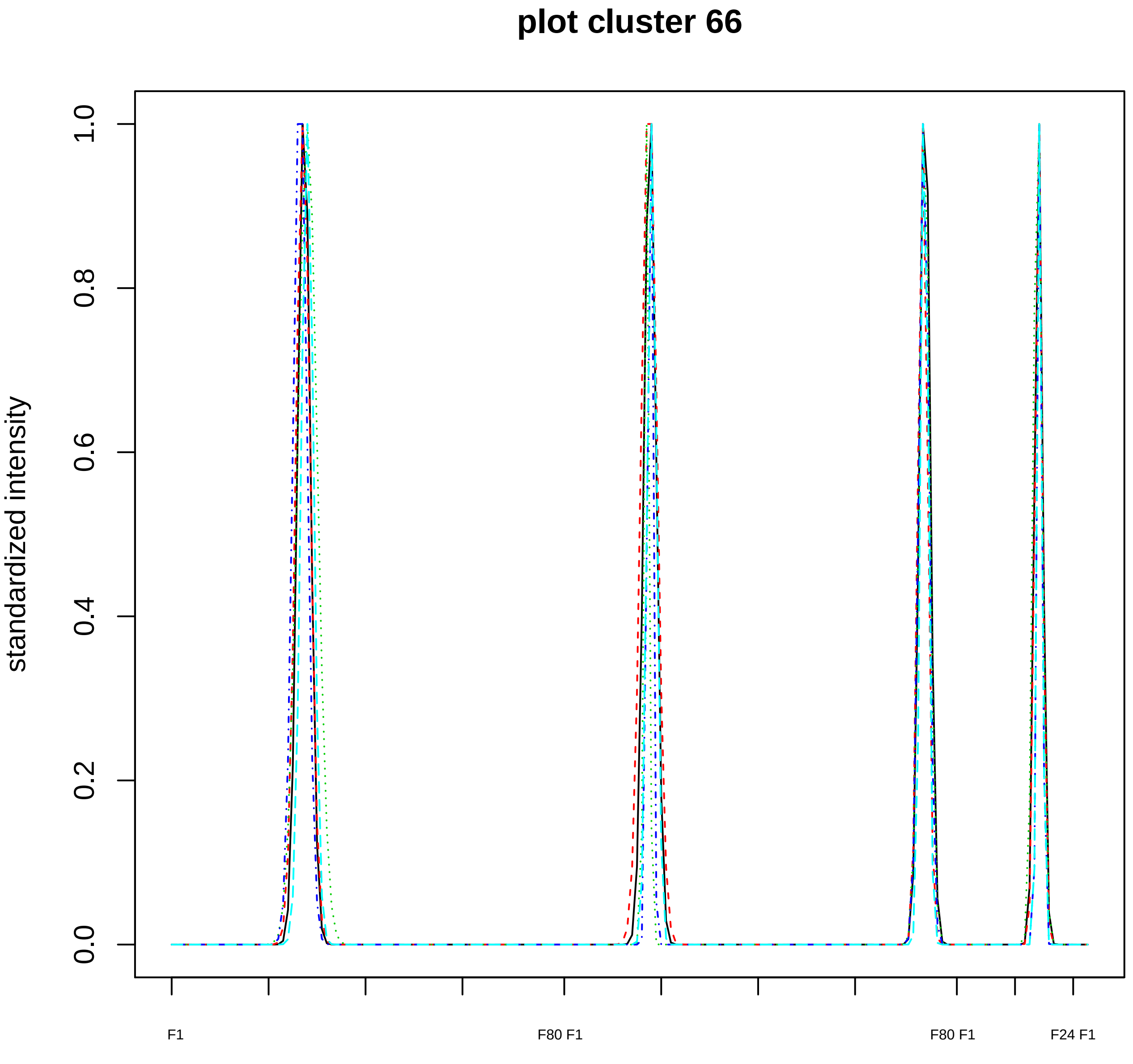

plot cluster 66
standardized intensity
0.0
0.2
0.4
0.6
0.8
1.0
F1
F80 F1
F80 F1
F24 F1

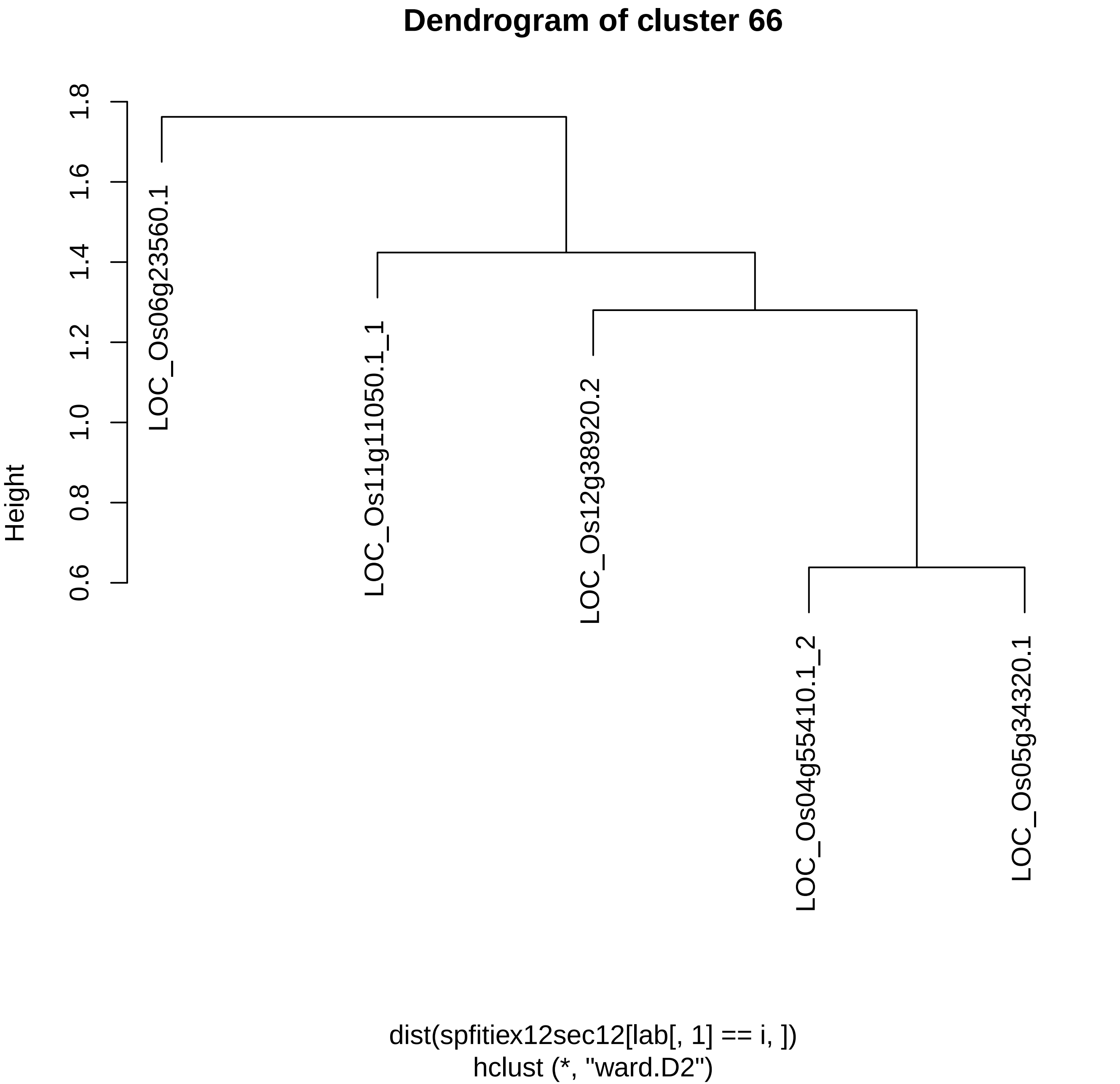
Dendrogram of cluster 66
Height
0.6
0.8
1.0
1.2
1.4
1.6
1.8
LOC_Os06g23560.1
LOC_Os11g11050.1_1
LOC_Os12g38920.2
LOC_Os04g55410.1_2
LOC_Os05g34320.1
dist(spfitiex12sec12[lab[, 1] == i, ])
hclust (*, "ward.D2")

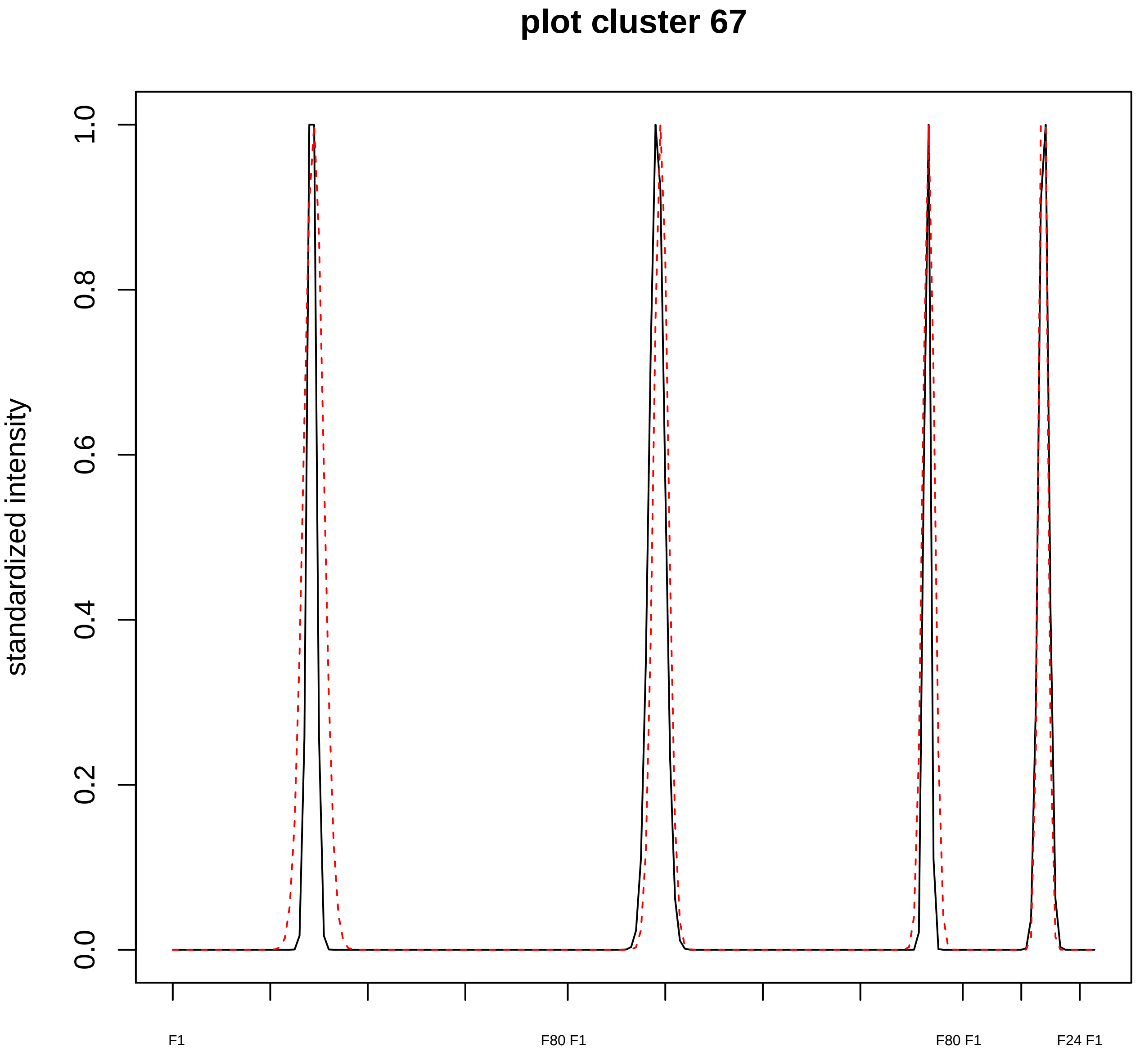
plot cluster 67
standardized intensity
0.0
0.2
0.4
0.6
0.8
1.0
F1
F80 F1
F80 F1
F24 F1

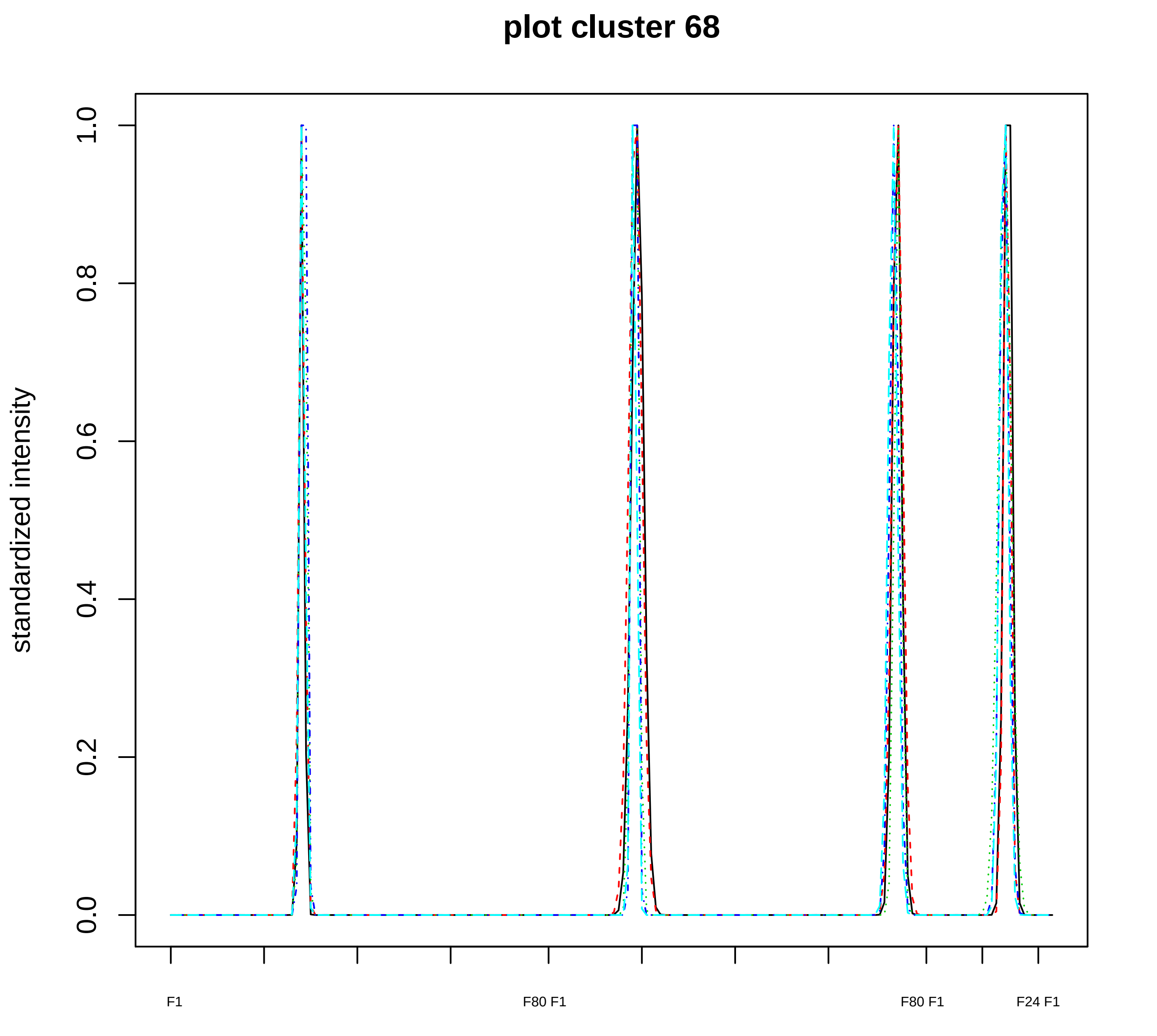
plot cluster 68
standardized intensity
0.0
0.2
0.4
0.6
0.8
1.0
F1
F80 F1
F80 F1
F24 F1

### Dendrogram of cluster 68

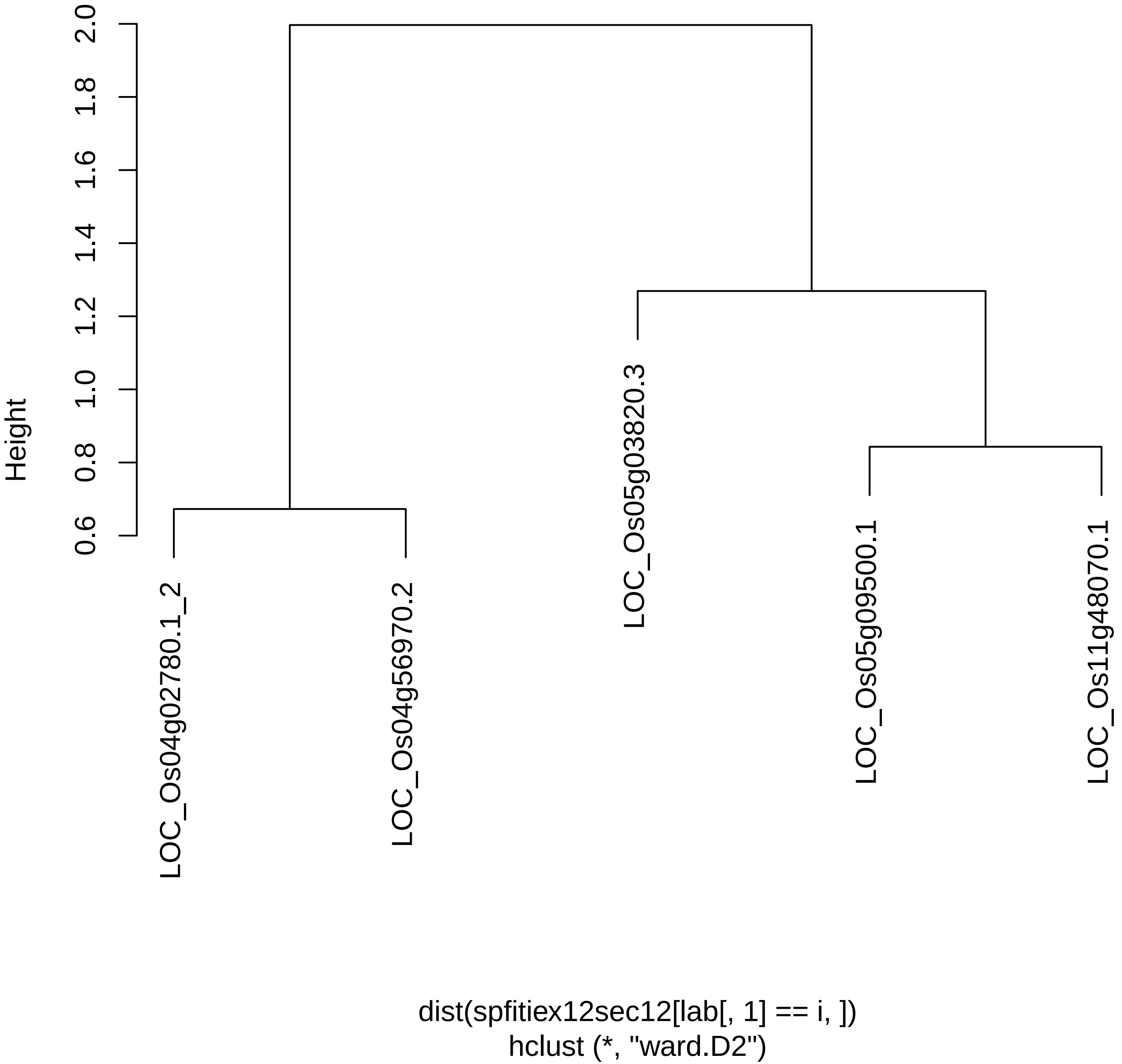

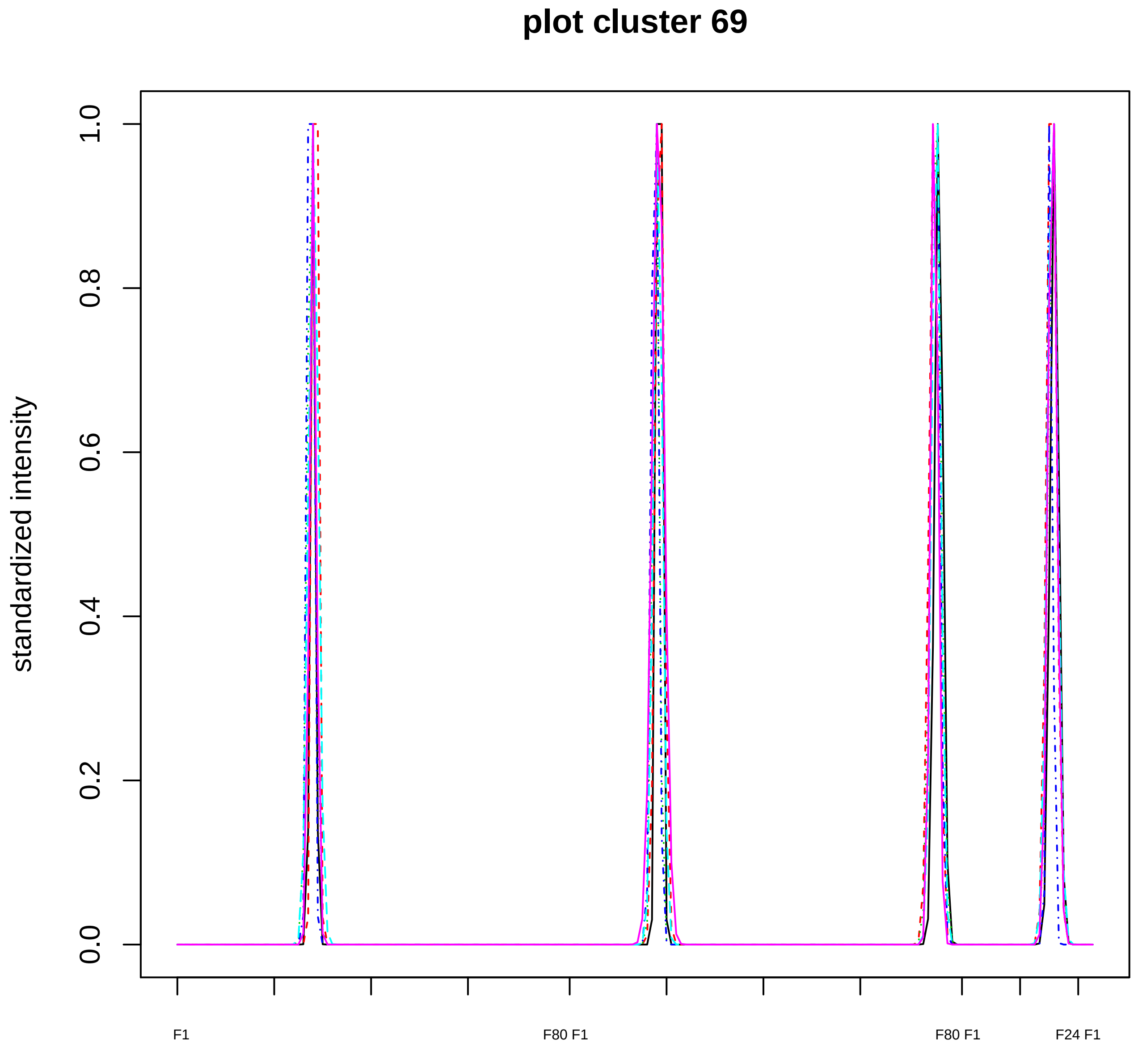
plot cluster 69
standardized intensity
0.0
0.2
0.4
0.6
0.8
1.0
F1
F80 F1
F80 F1
F24 F1

### Dendrogram of cluster 69

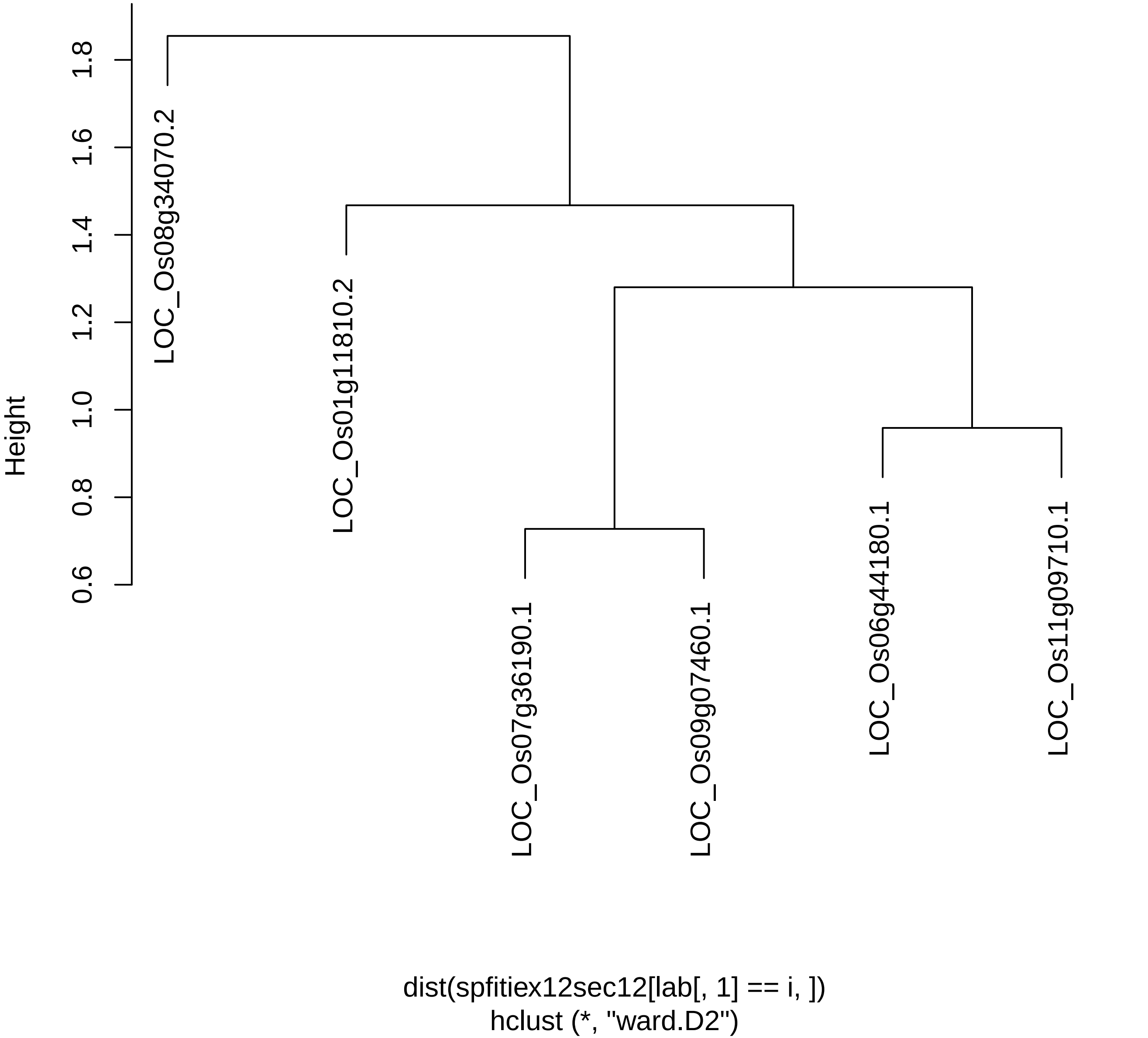

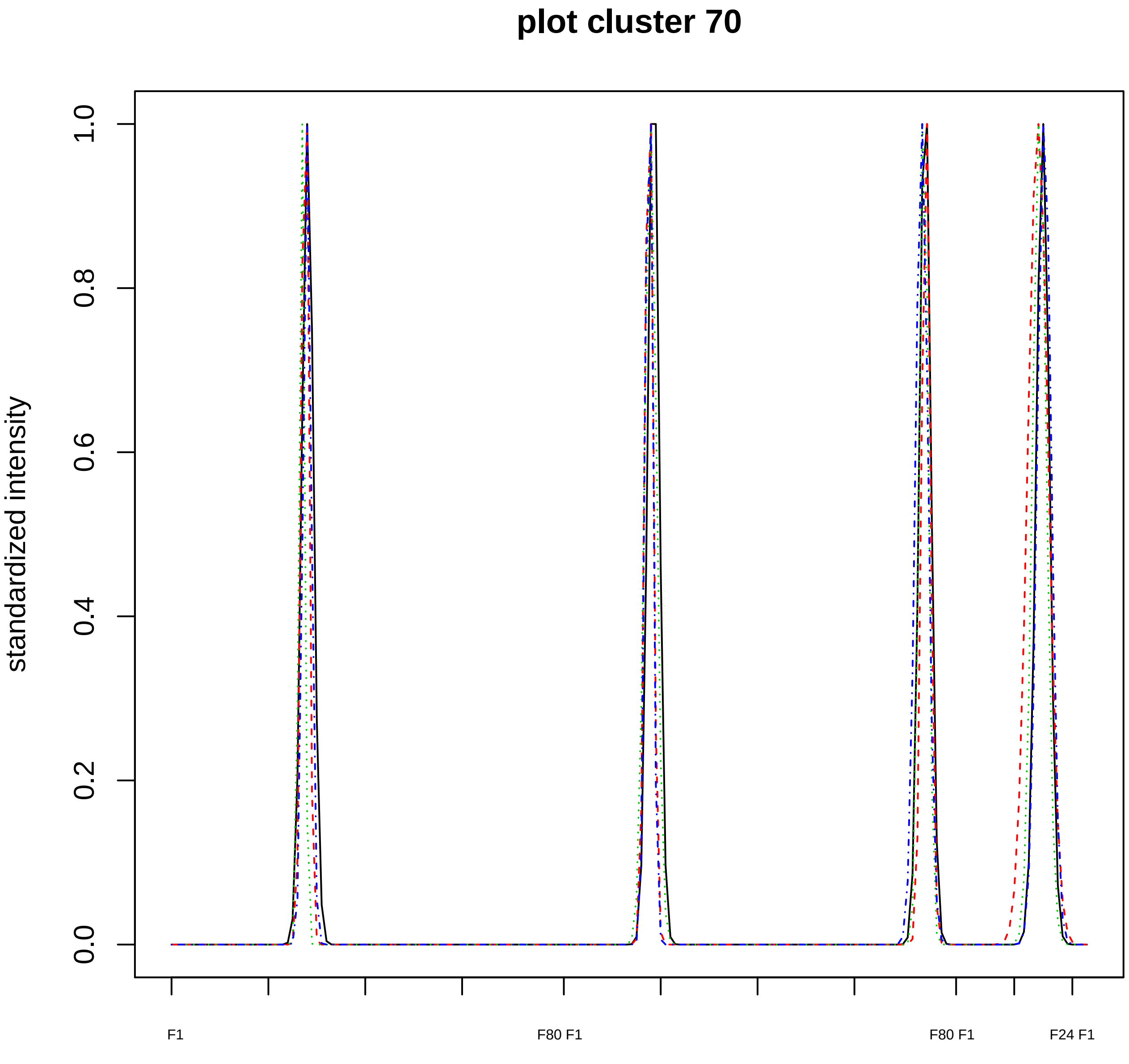
plot cluster 70
standardized intensity
0.0
0.2
0.4
0.6
0.8
1.0
F1
F80 F1
F80 F1
F24 F1

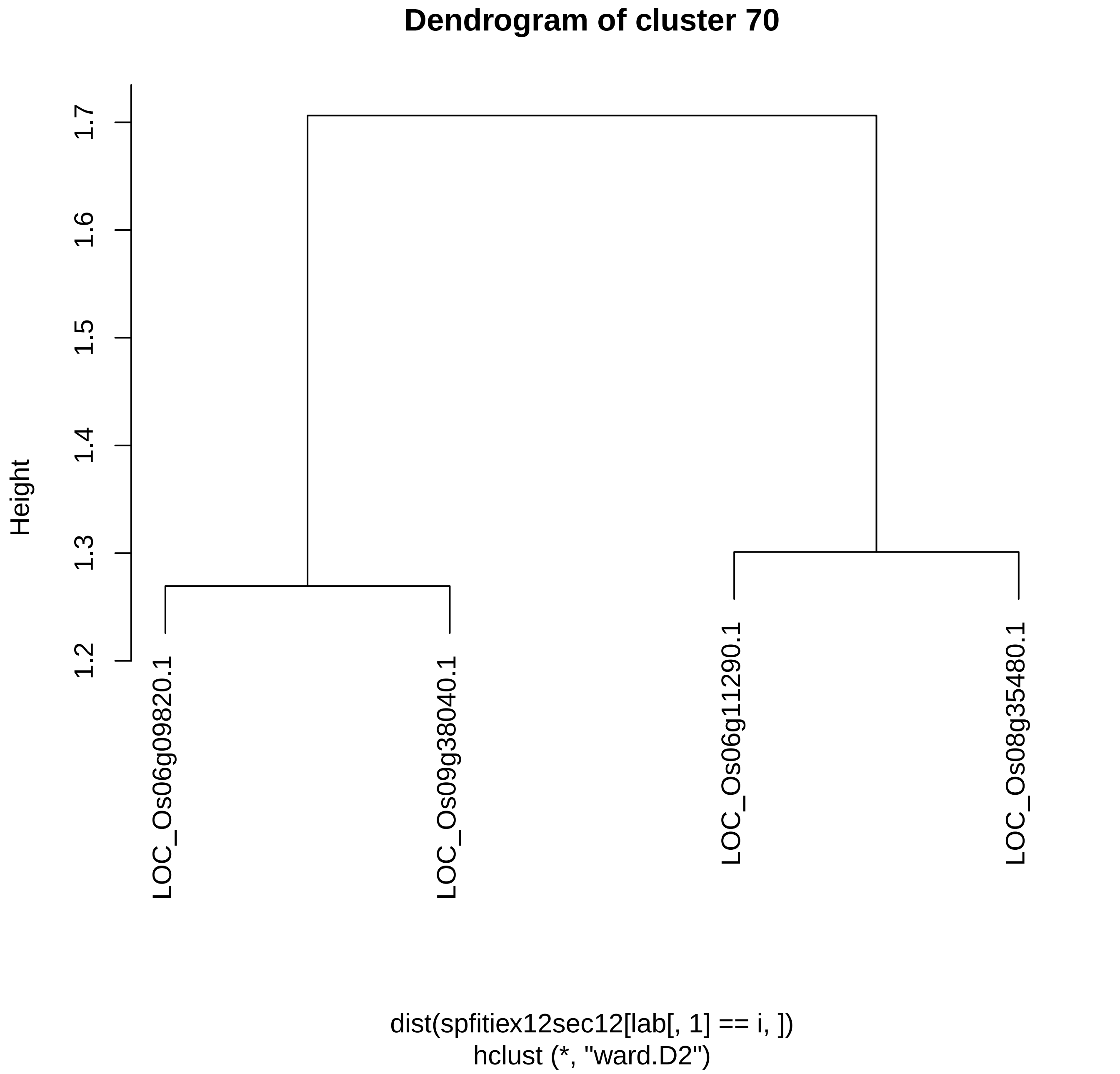
Dendrogram of cluster 70
Height
1.2
1.3
1.4
1.5
1.6
1.7
LOC_Os06g09820.1
LOC_Os09g38040.1
LOC_Os06g11290.1
LOC_Os08g35480.1
dist(spfitiex12sec12[lab[, 1] == i, ])
hclust (*, "ward.D2")

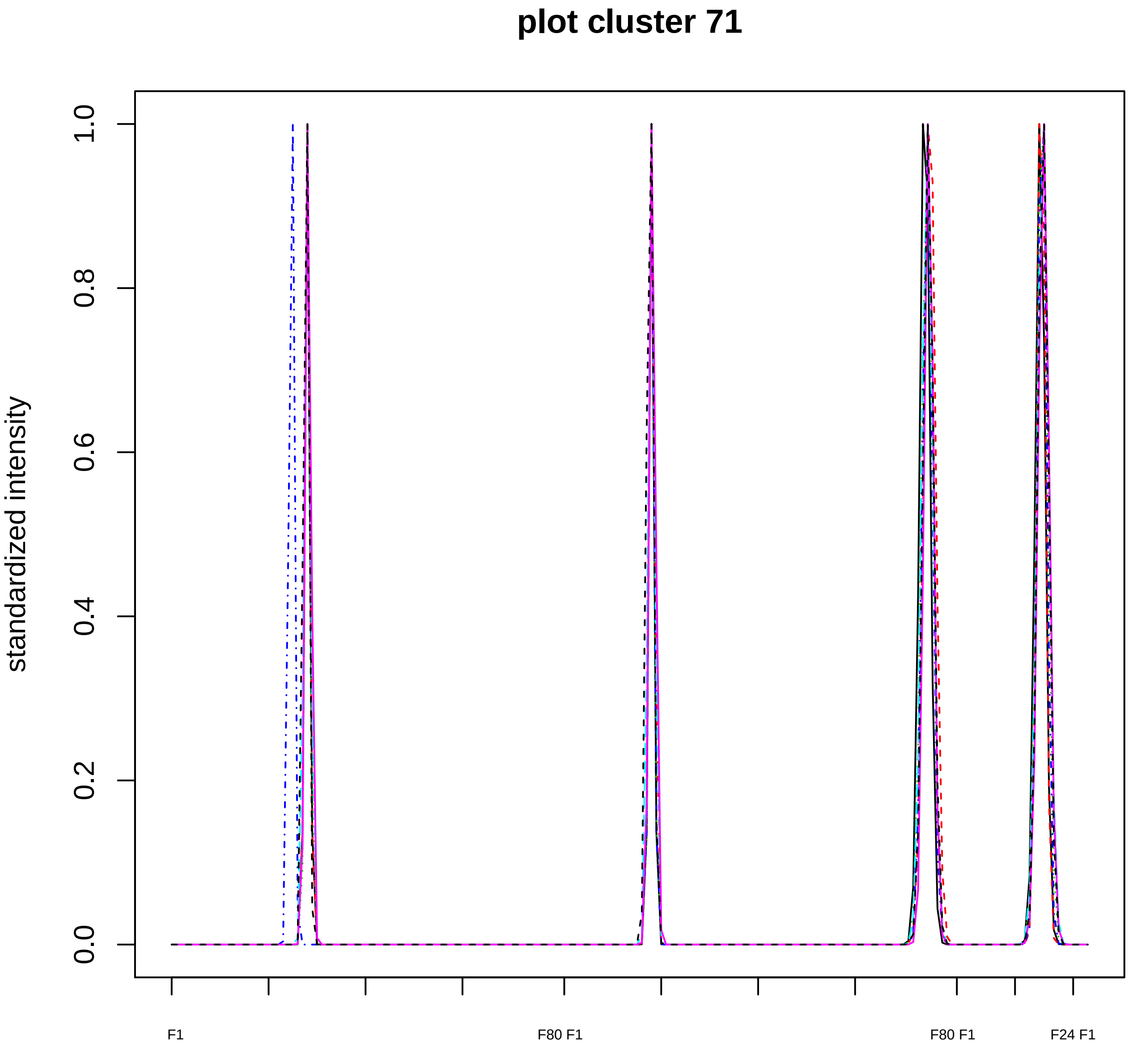

plot cluster 71
standardized intensity
0.0
0.2
0.4
0.6
0.8
1.0
F1
F80 F1
F80 F1
F24 F1

### Dendrogram of cluster 71

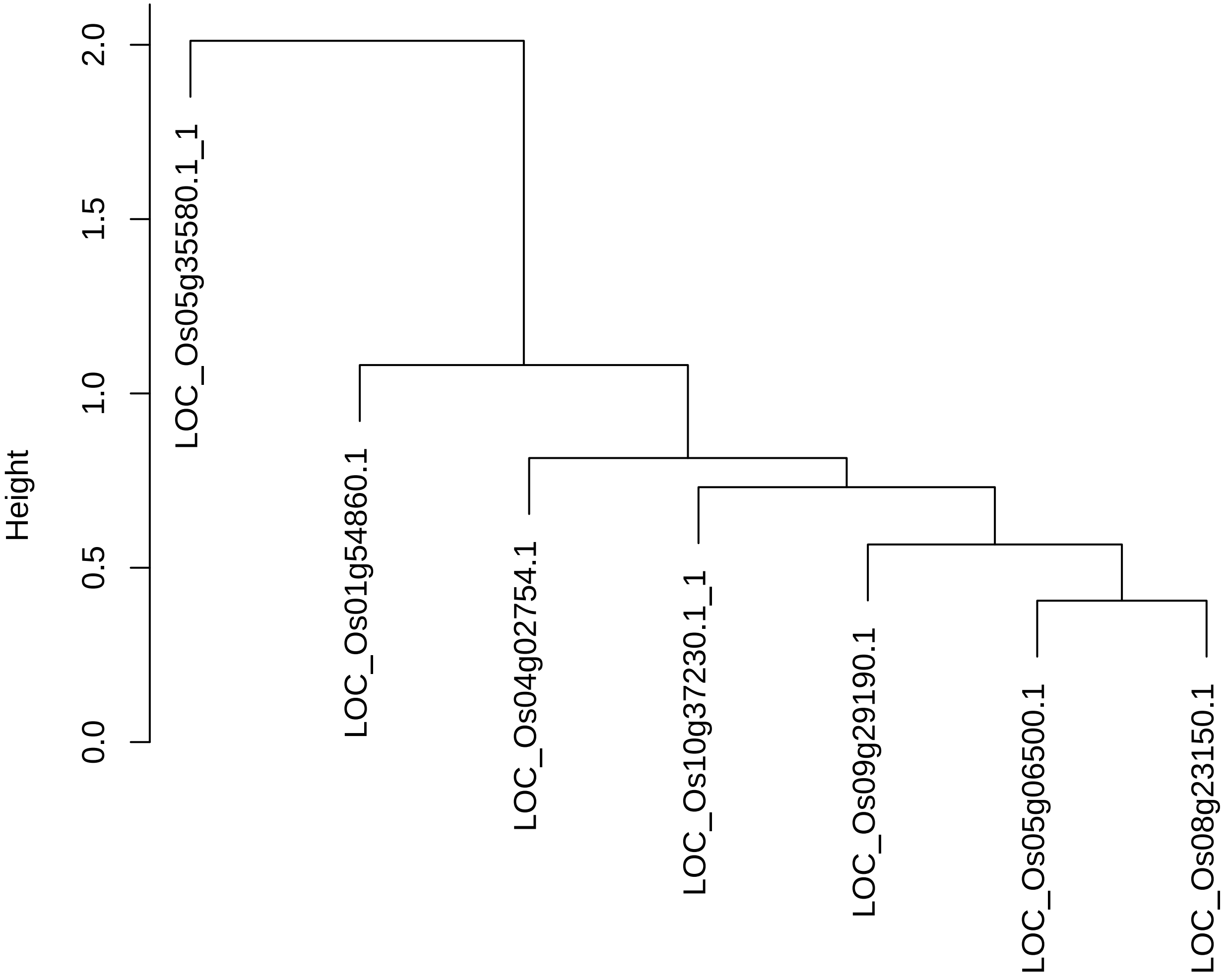


dist(spfitiex12sec12[lab[, 1] == i, ])
hclust (*, "ward.D2")

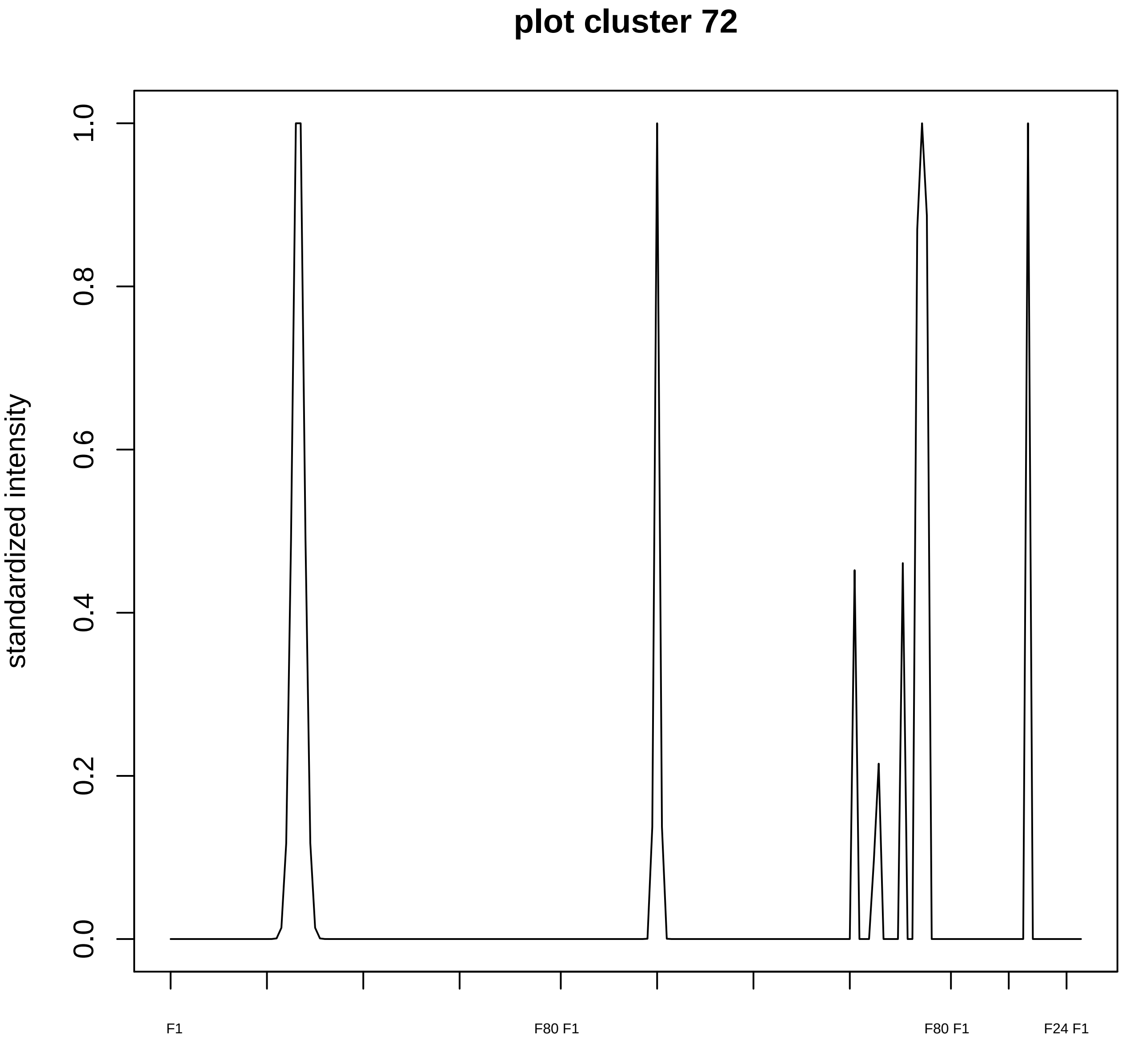
plot cluster 72
standardized intensity
0.0
0.2
0.4
0.6
0.8
1.0
F1
F80 F1
F80 F1
F24 F1

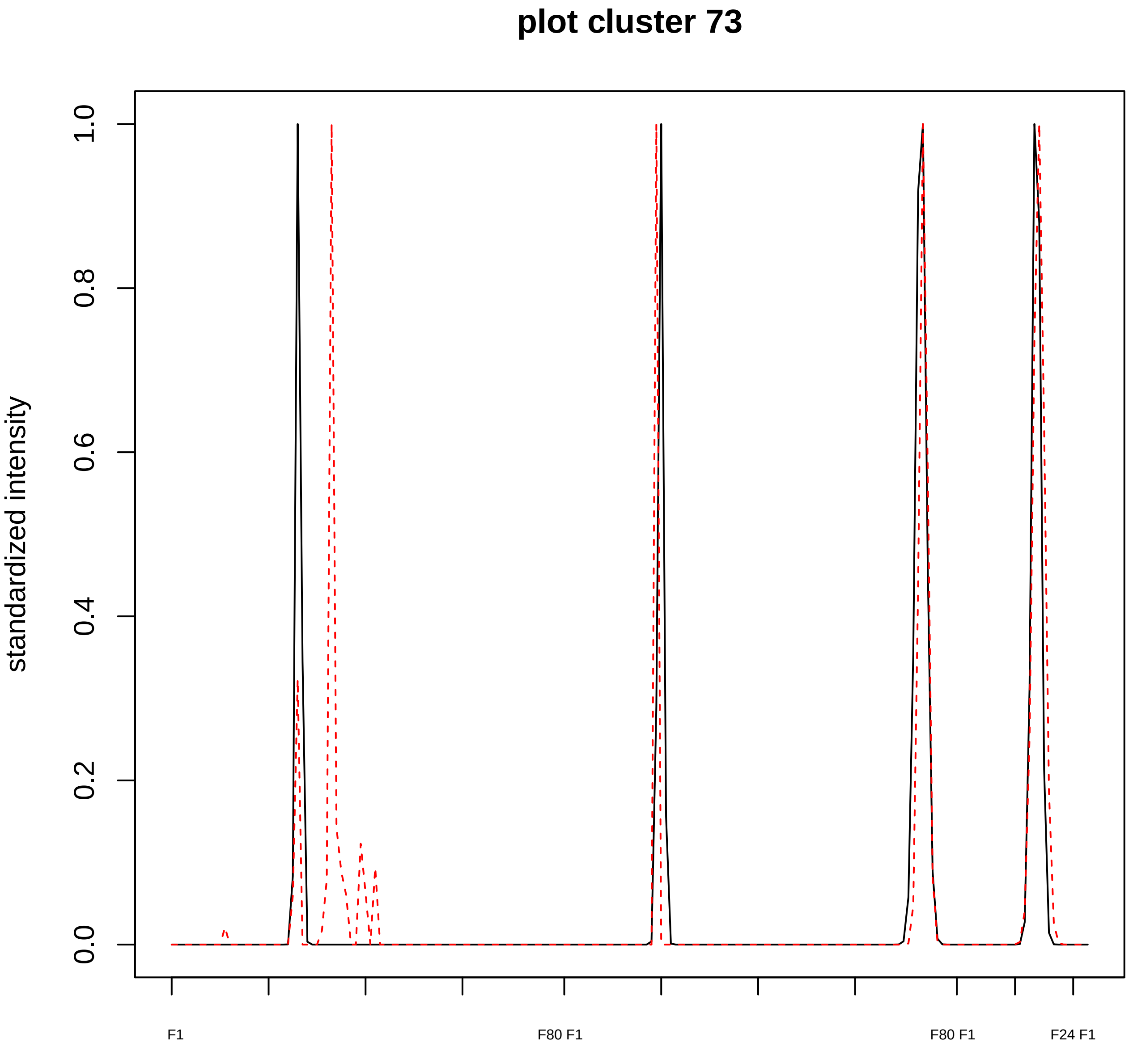
plot cluster 73
standardized intensity
0.0
0.2
0.4
0.6
0.8
1.0
F1
F80 F1
F80 F1
F24 F1

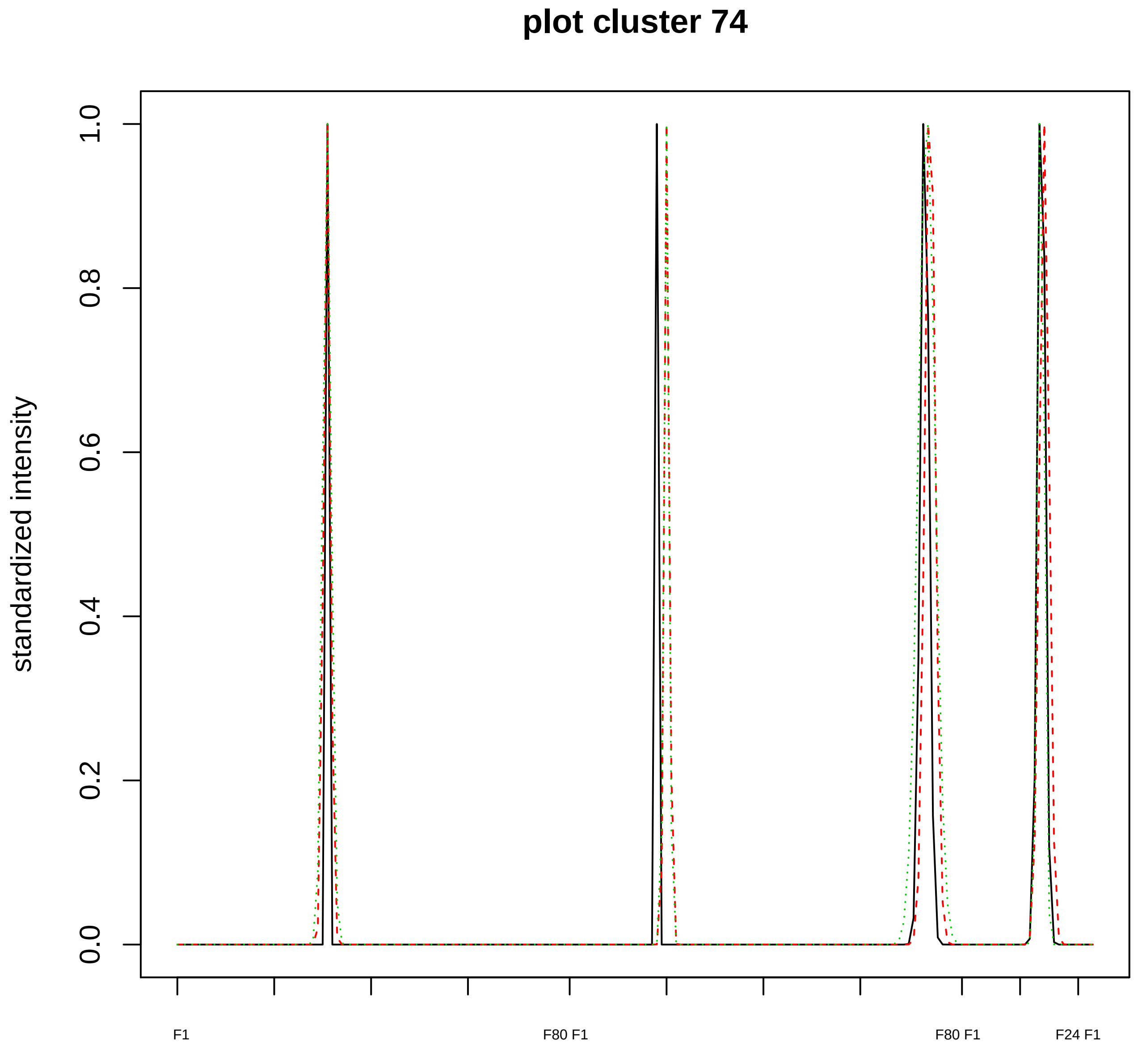

plot cluster 74
standardized intensity
0.0
0.2
0.4
0.6
0.8
1.0
F1
F80 F1
F80 F1
F24 F1

#### Dendrogram of cluster 74

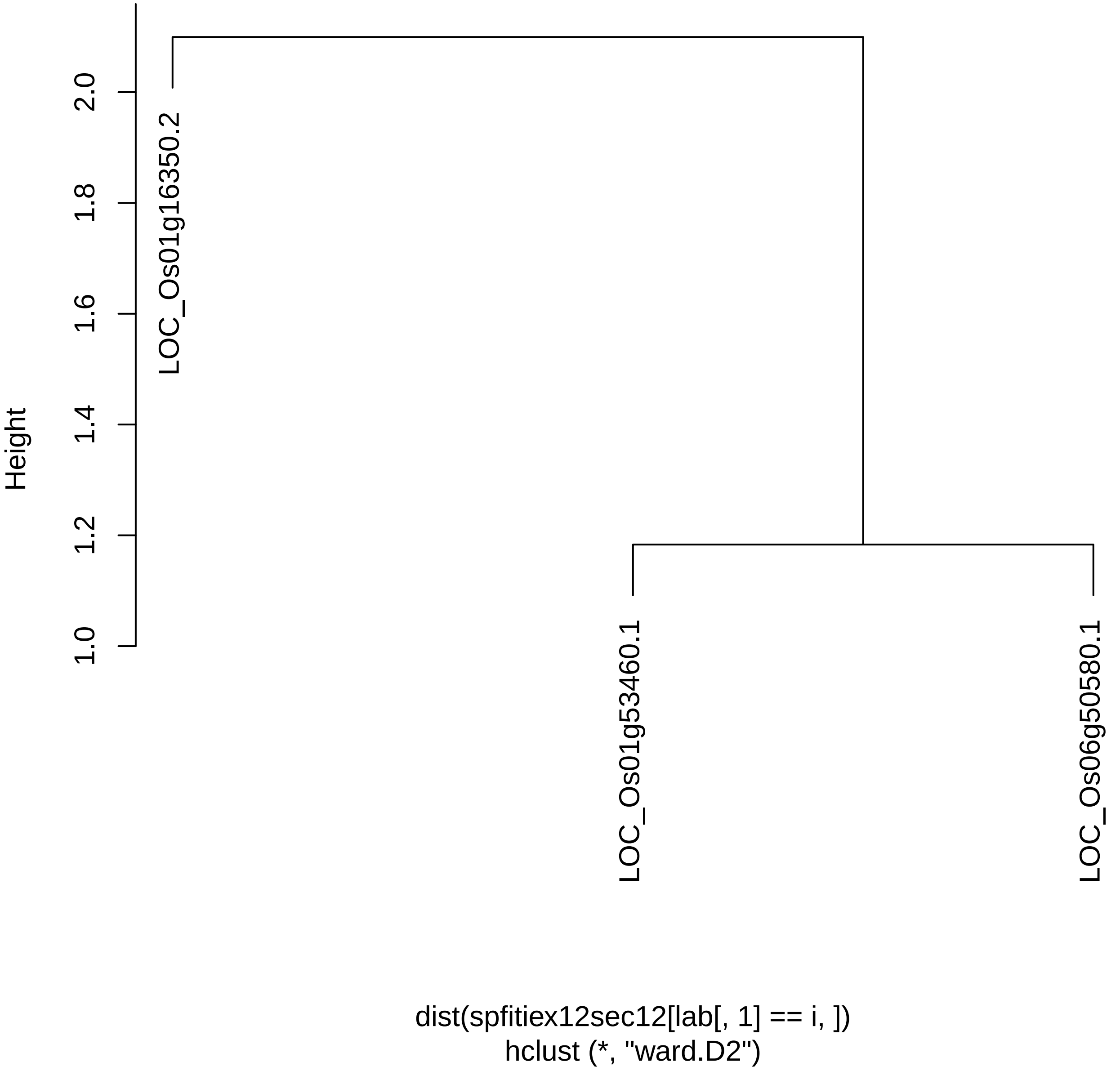

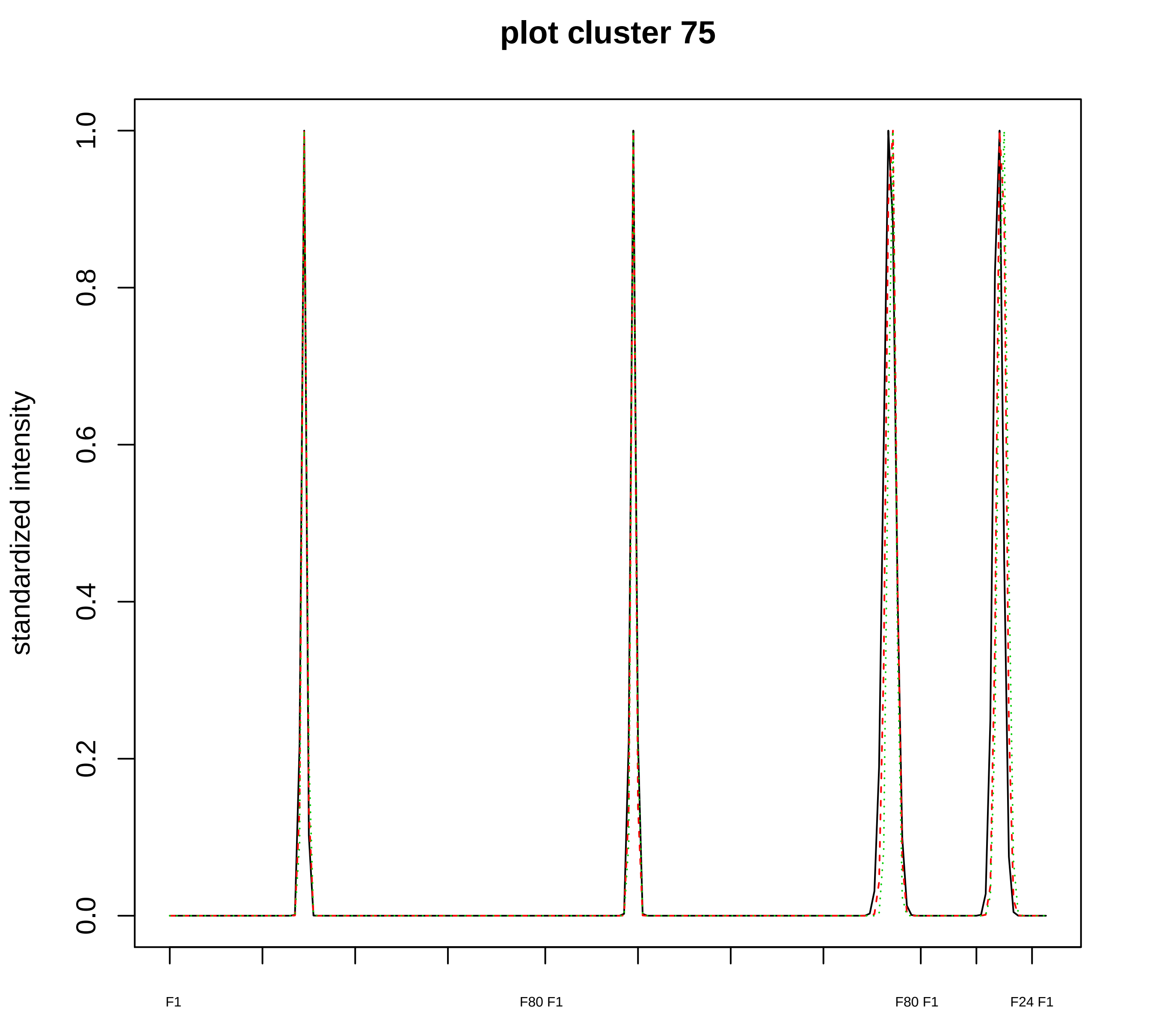
plot cluster 75
standardized intensity
0.0
0.2
0.4
0.6
0.8
1.0
F1
F80 F1
F80 F1
F24 F1

### Dendrogram of cluster 75

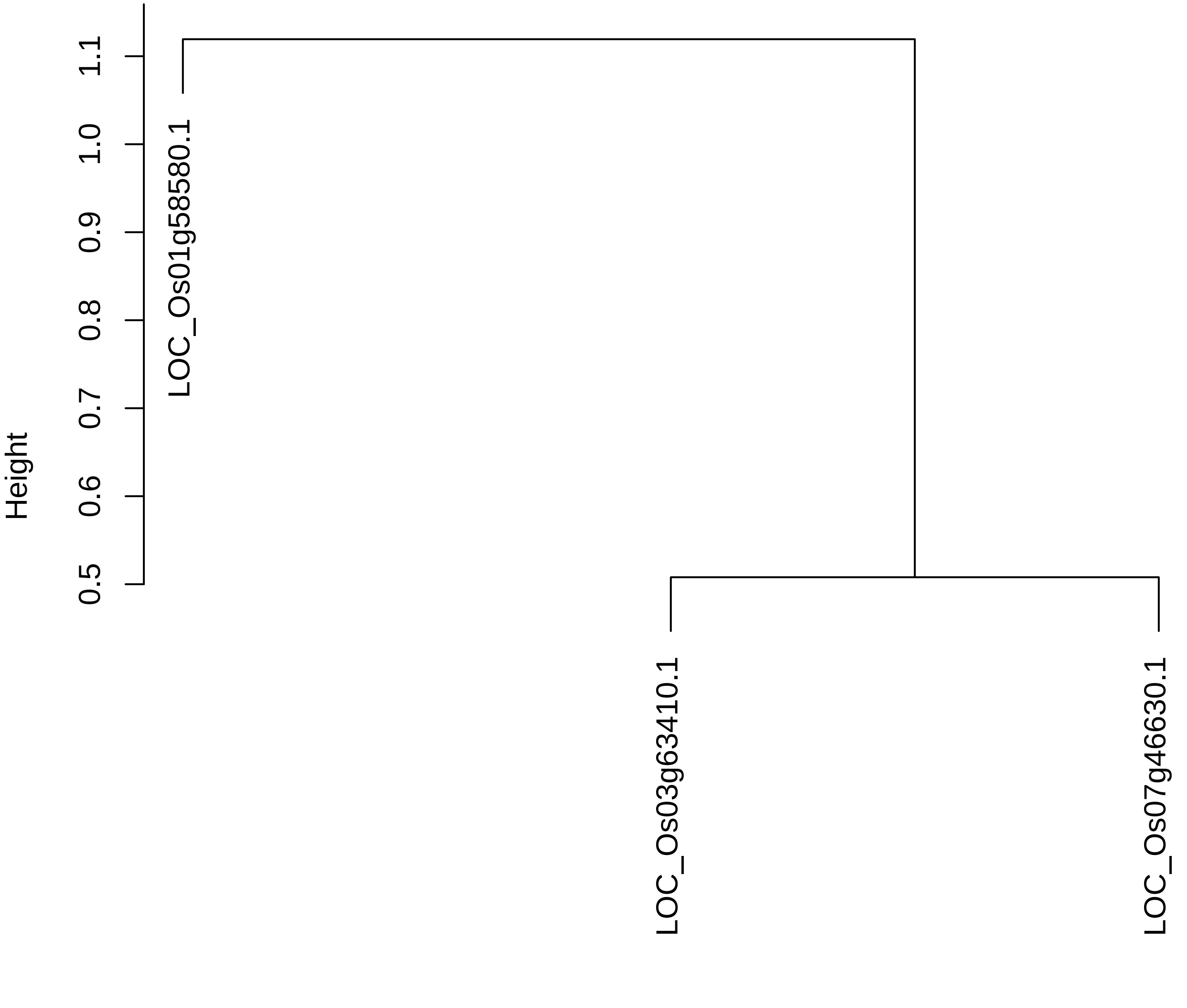


dist(spfitiex12sec12[lab[, 1] == i, ])
hclust (*, "ward.D2")

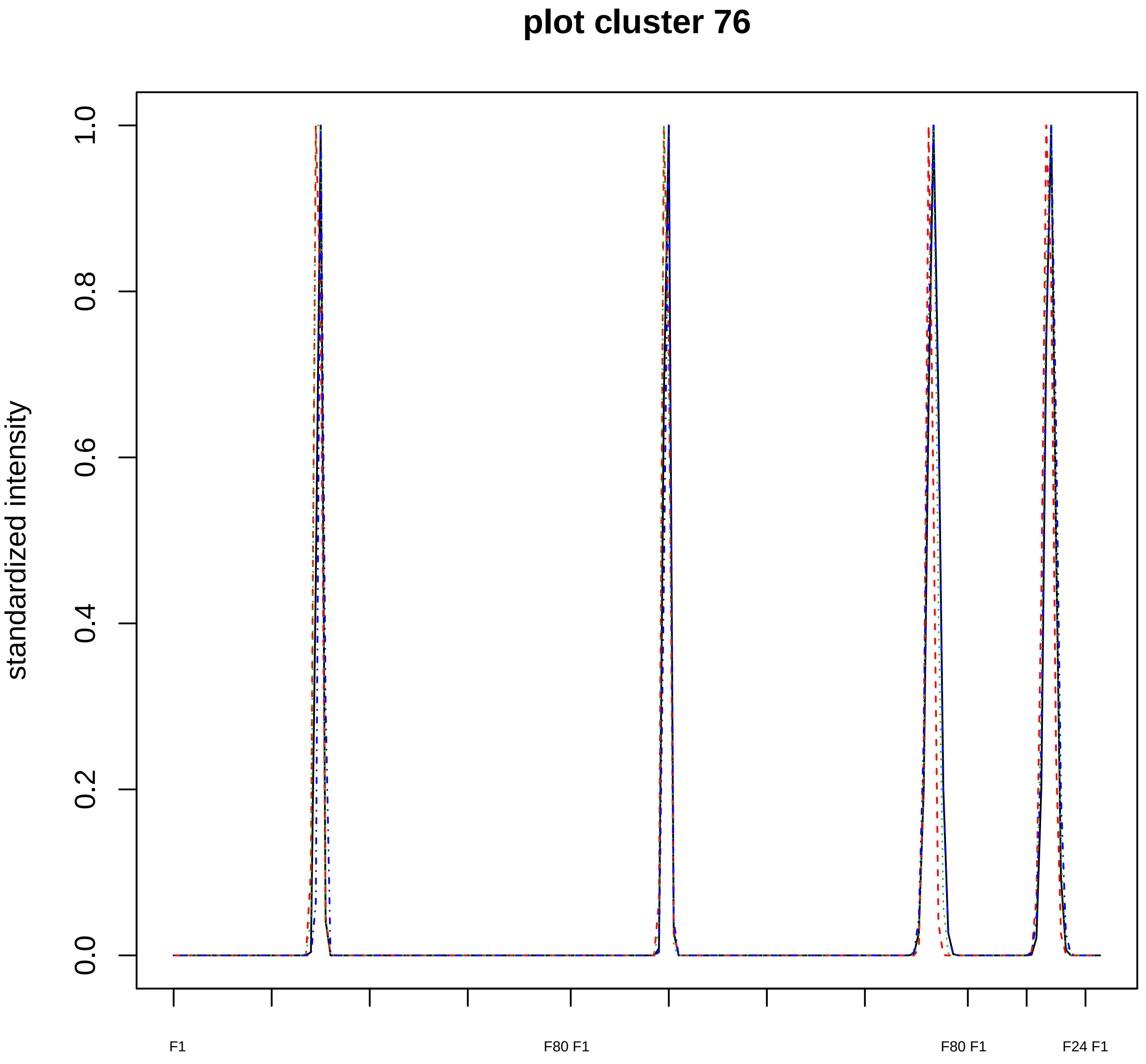
plot cluster 76
standardized intensity
0.0
0.2
0.4
0.6
0.8
1.0
F1
F80 F1
F80 F1
F24 F1

### Dendrogram of cluster 76

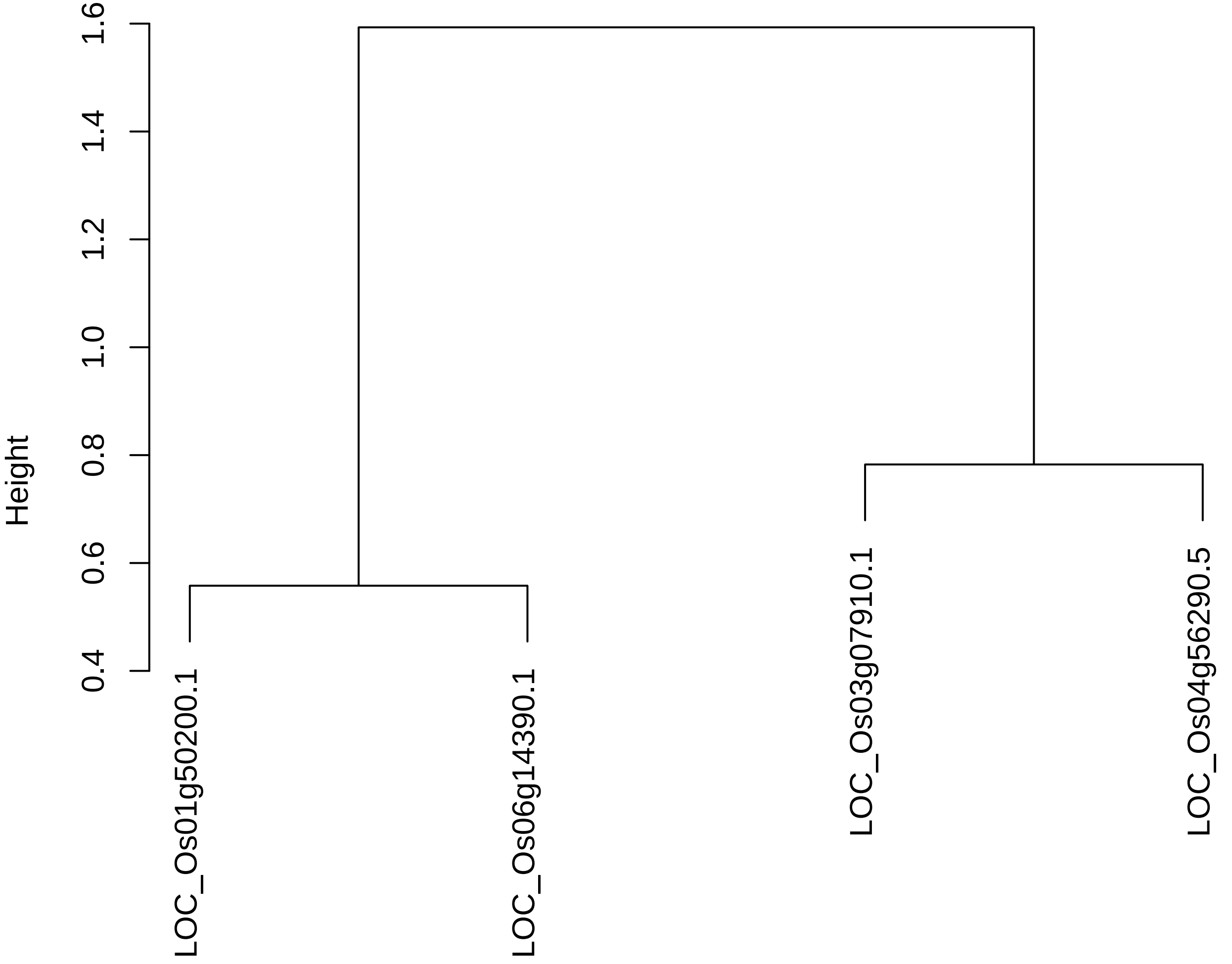


dist(spfitiex12sec12[lab[, 1] == i, ])
hclust (*, "ward.D2")

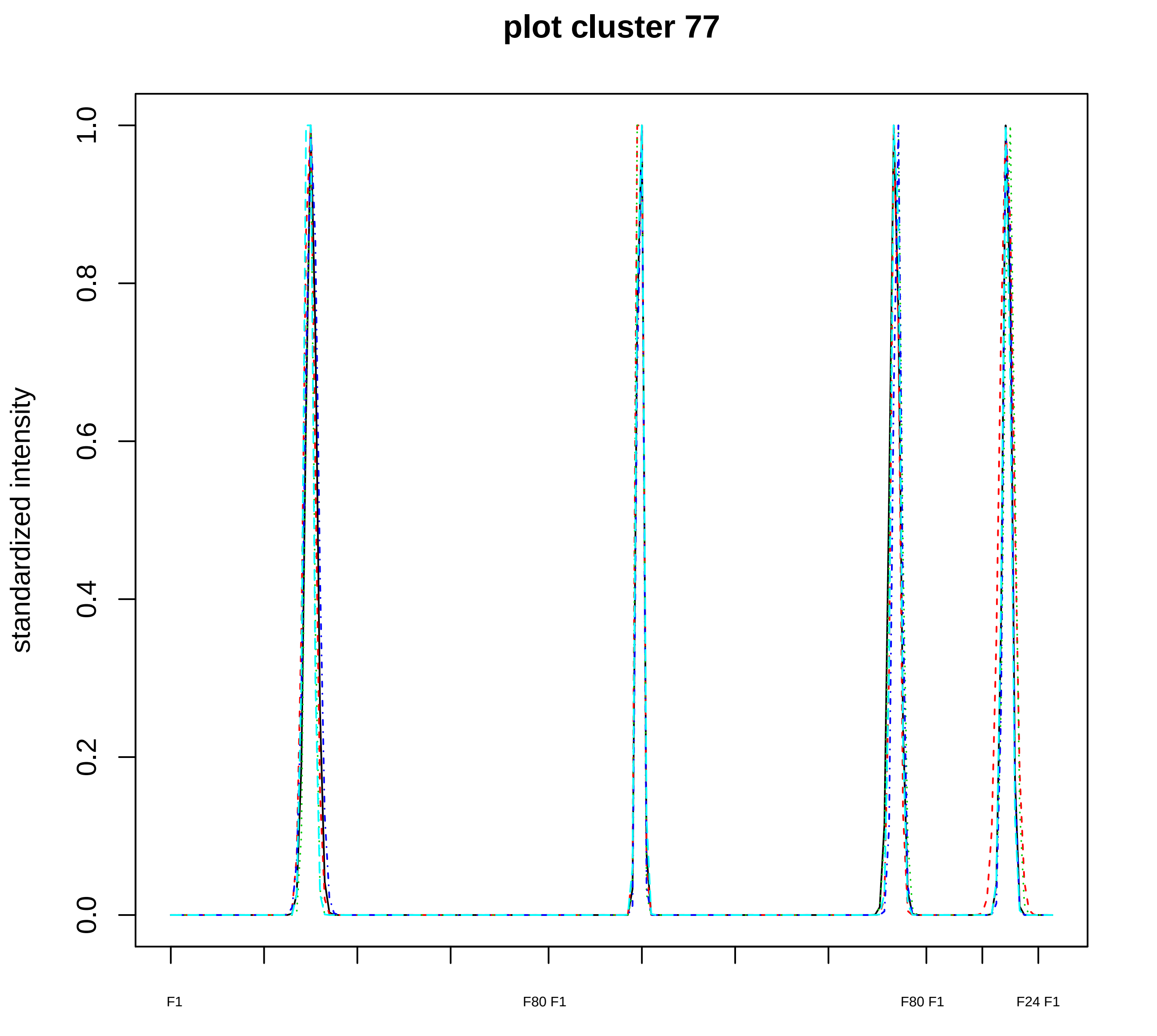
plot cluster 77
standardized intensity
0.0
0.2
0.4
0.6
0.8
1.0
F1
F80 F1
F80 F1
F24 F1

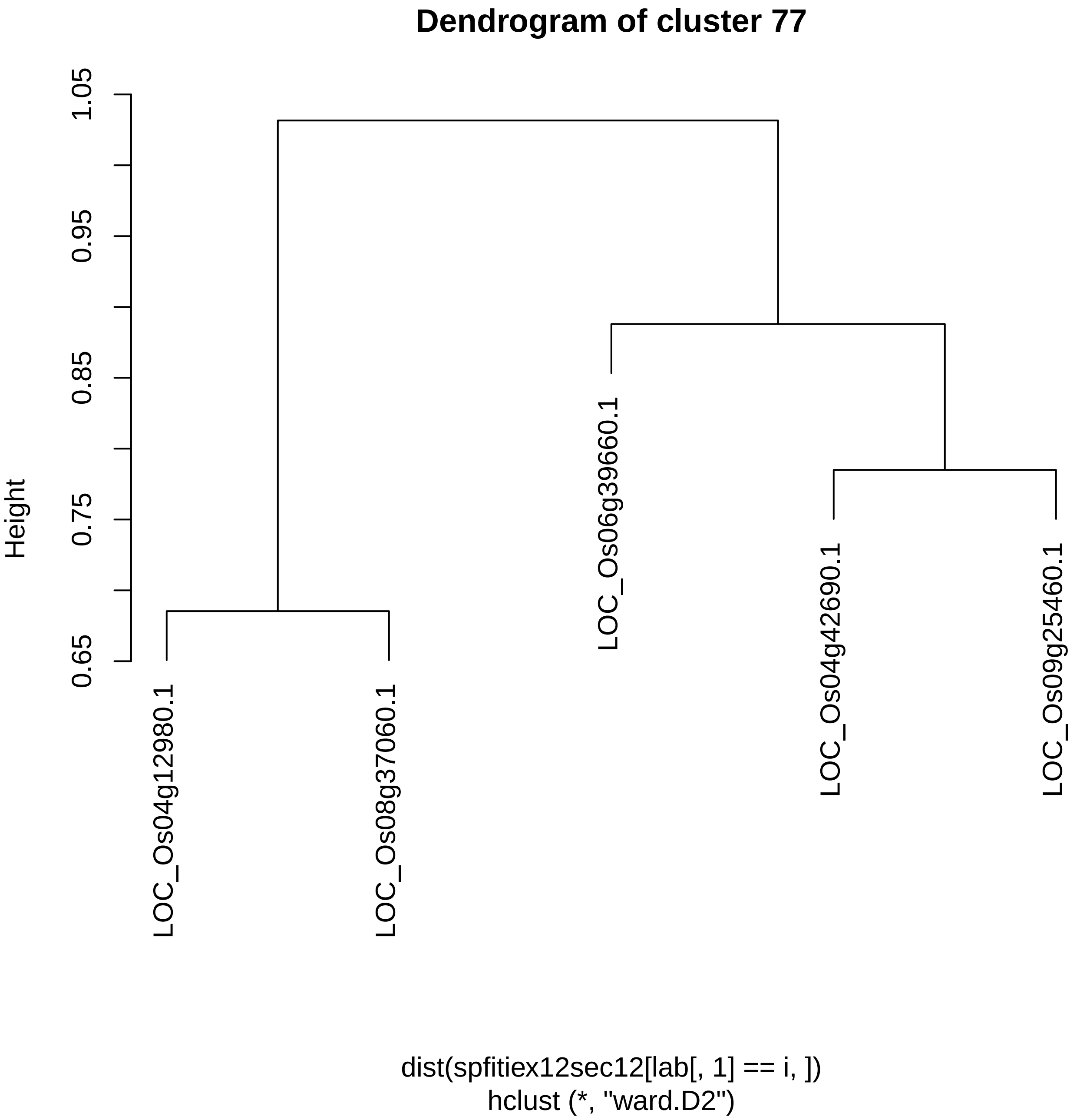
Dendrogram of cluster 77
Height
1.05
0.95
0.85
0.75
0.65
LOC_Os04g12980.1
LOC_Os08g37060.1
LOC_Os06g39660.1
LOC_Os04g42690.1
LOC_Os09g25460.1
dist(spfitiex12sec12[lab[, 1] == i, ])
hclust (*, "ward.D2")

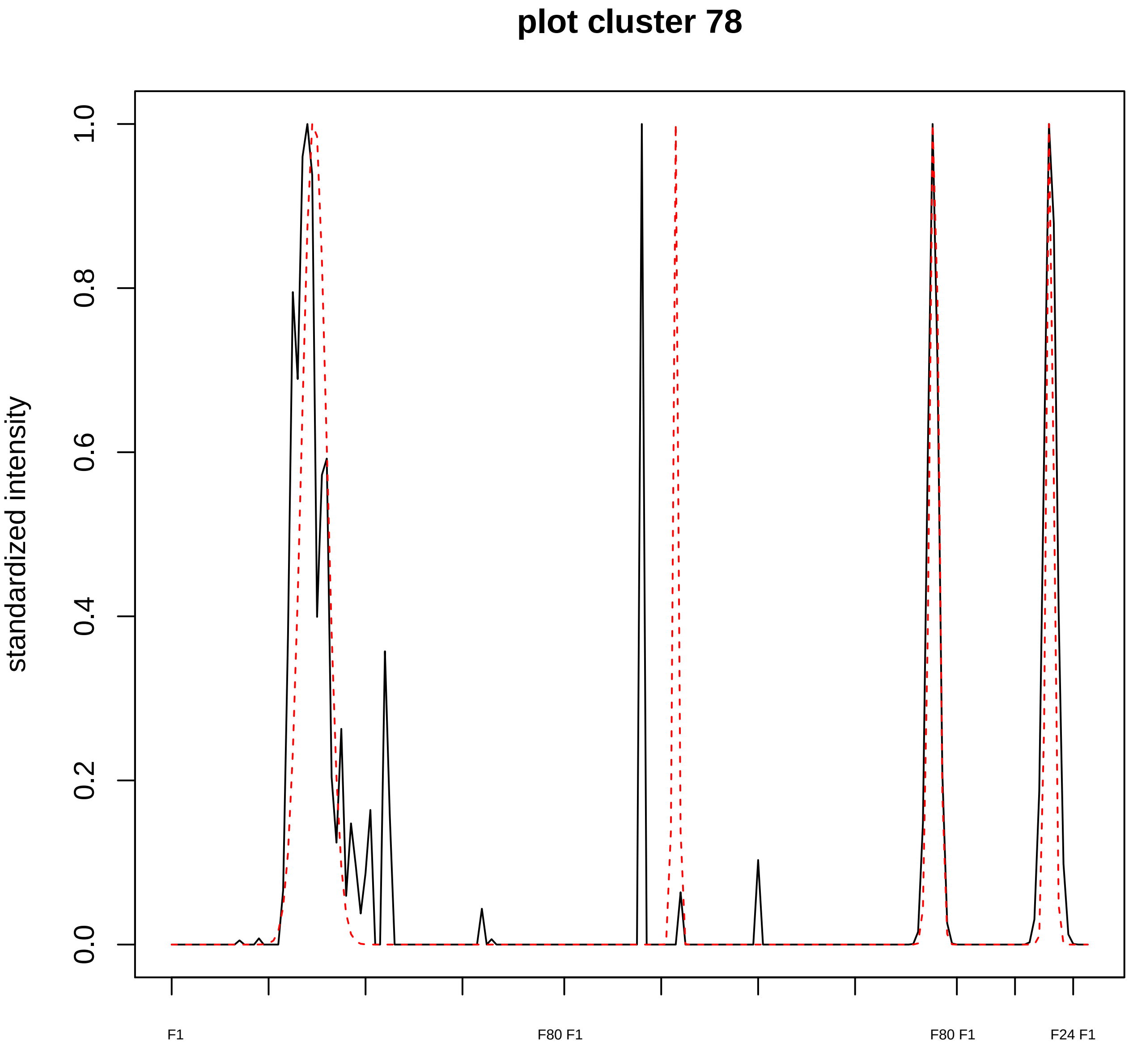
plot cluster 78
standardized intensity
1.0
0.8
0.6
0.4
0.2
0.0
F1
F80 F1
F80 F1
F24 F1

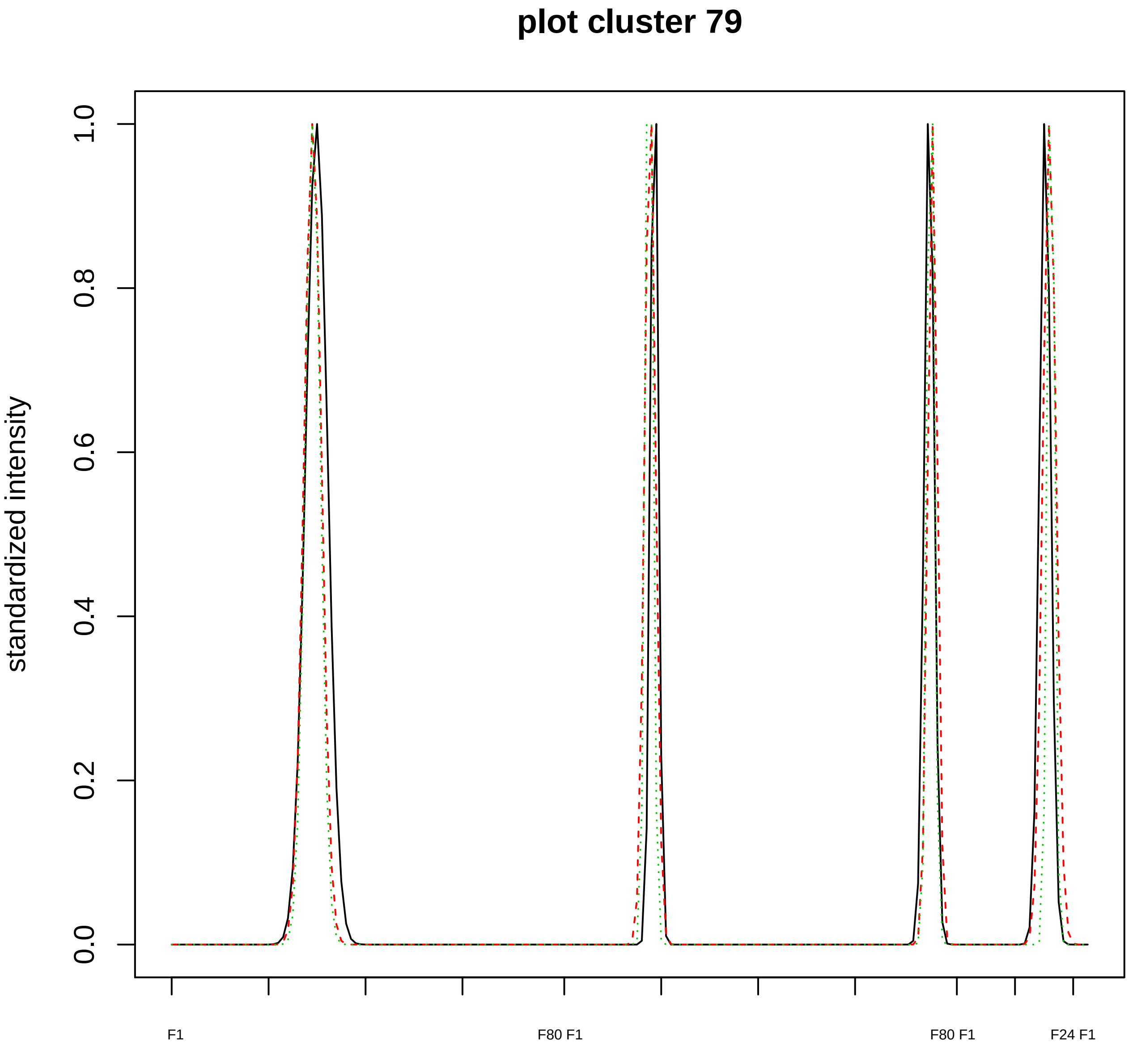
plot cluster 79
standardized intensity
0.0
0.2
0.4
0.6
0.8
1.0
F1
F80 F1
F80 F1
F24 F1

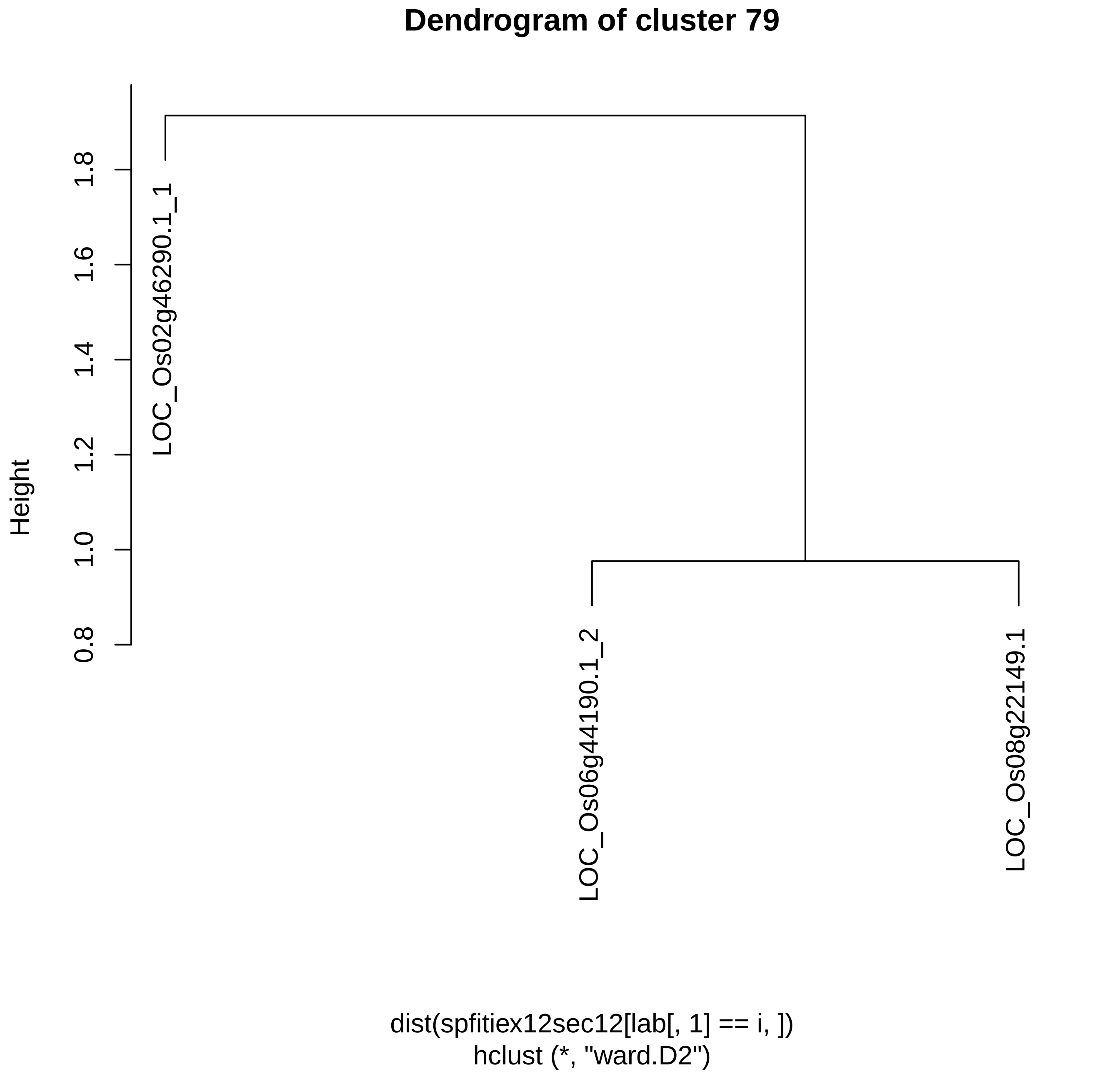
Dendrogram of cluster 79
Height
0.8
1.0
1.2
1.4
1.6
1.8
LOC_Os02g46290.1_1
LOC_Os06g44190.1_2
LOC_Os08g22149.1
dist(spfitiex12sec12[lab[, 1] == i, ])
hclust (*, "ward.D2")

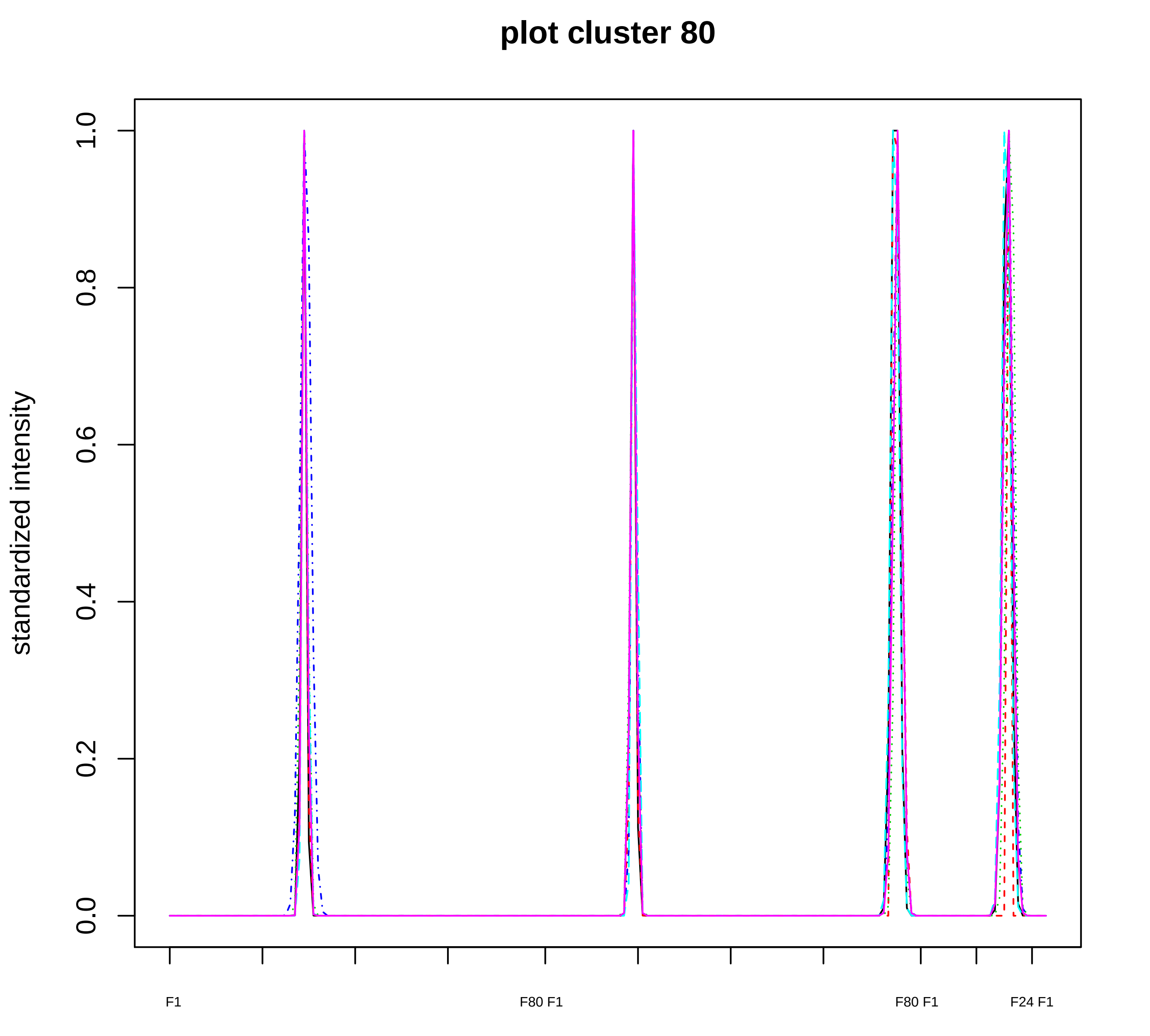

plot cluster 80
standardized intensity
0.0
0.2
0.4
0.6
0.8
1.0
F1
F80 F1
F80 F1
F24 F1

### Dendrogram of cluster 80

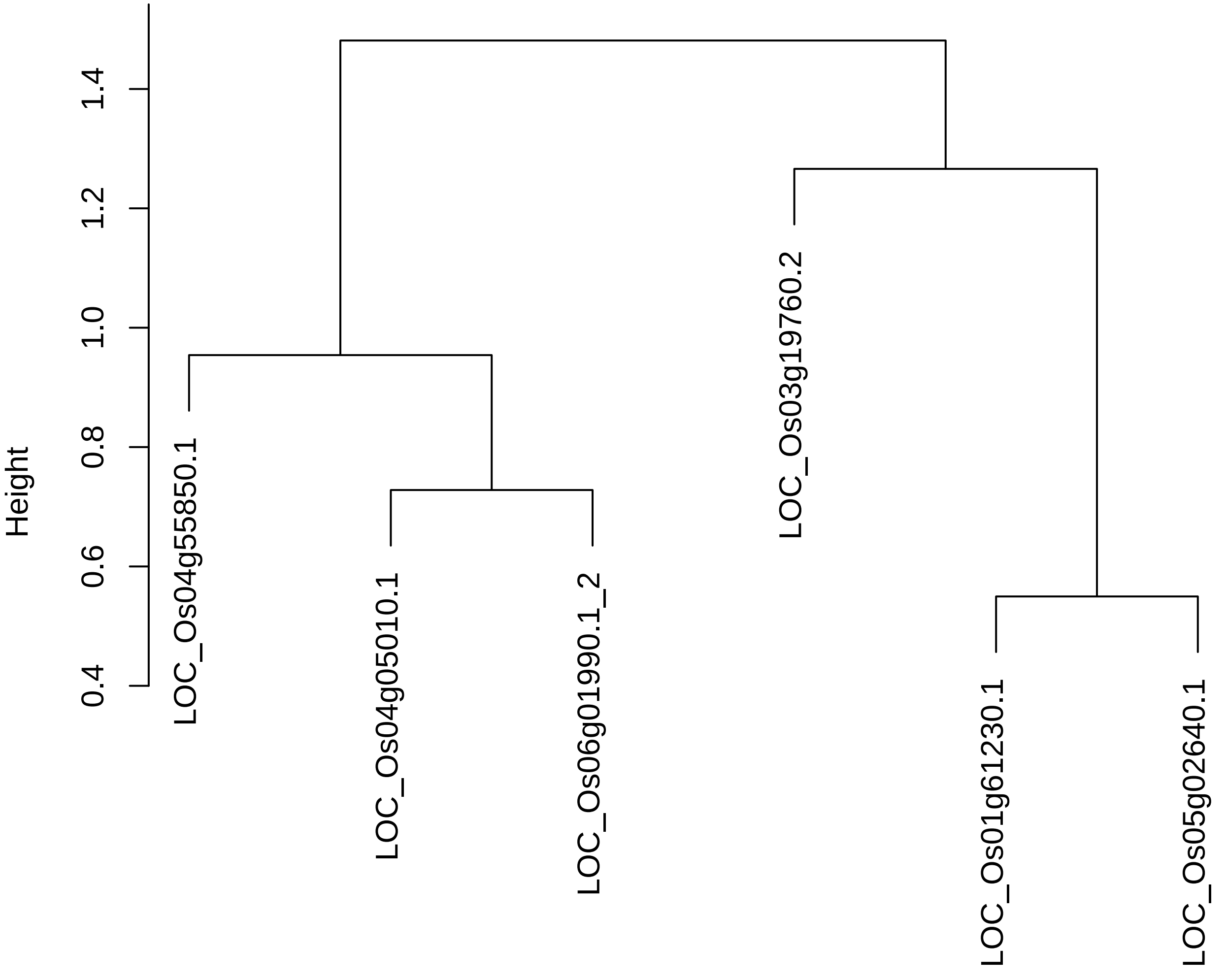


dist(spfitiex12sec12[lab[, 1] == i, ])
hclust (*, "ward.D2")

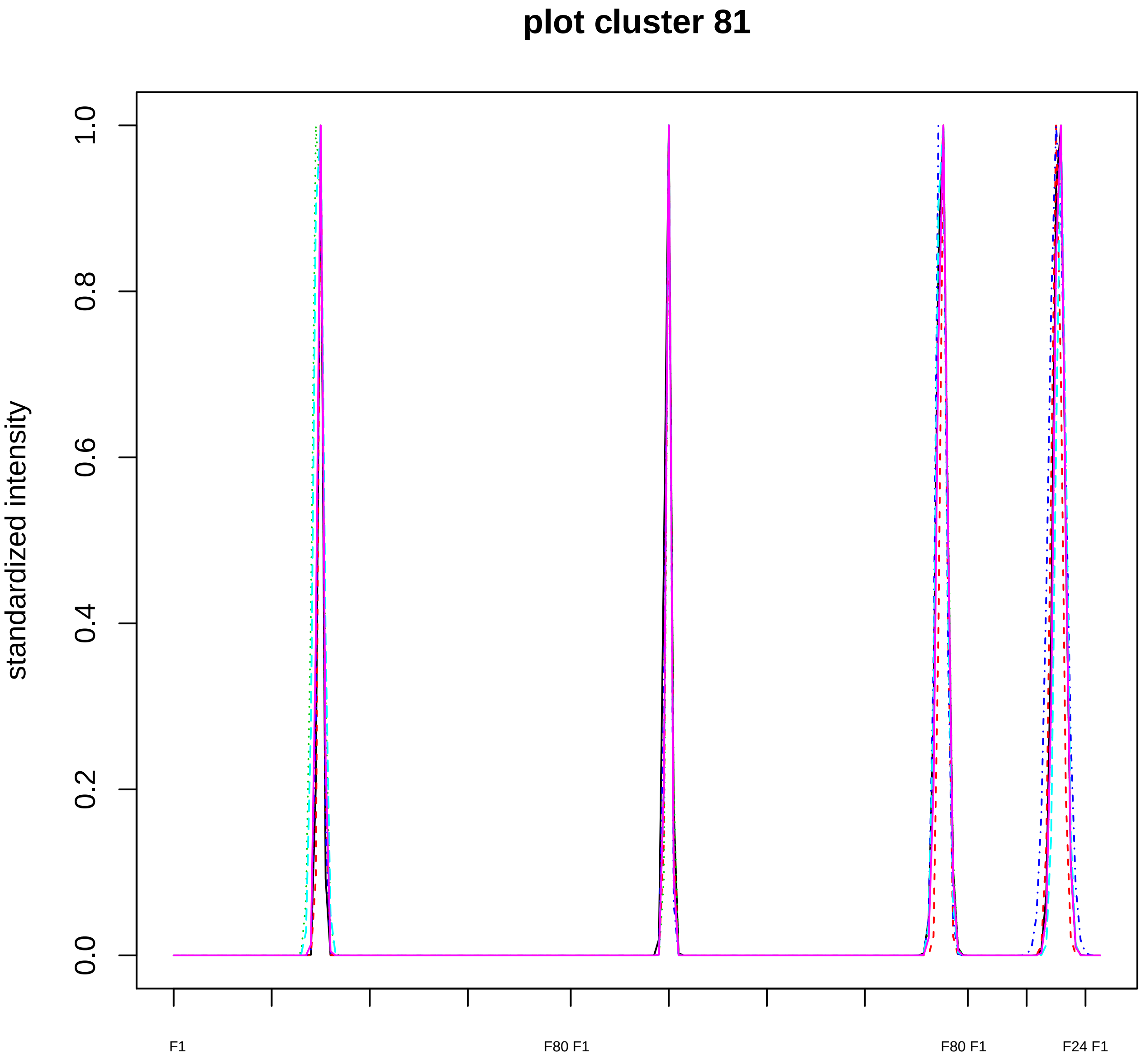
plot cluster 81
standardized intensity
0.0
0.2
0.4
0.6
0.8
1.0
F1
F80 F1
F80 F1
F24 F1

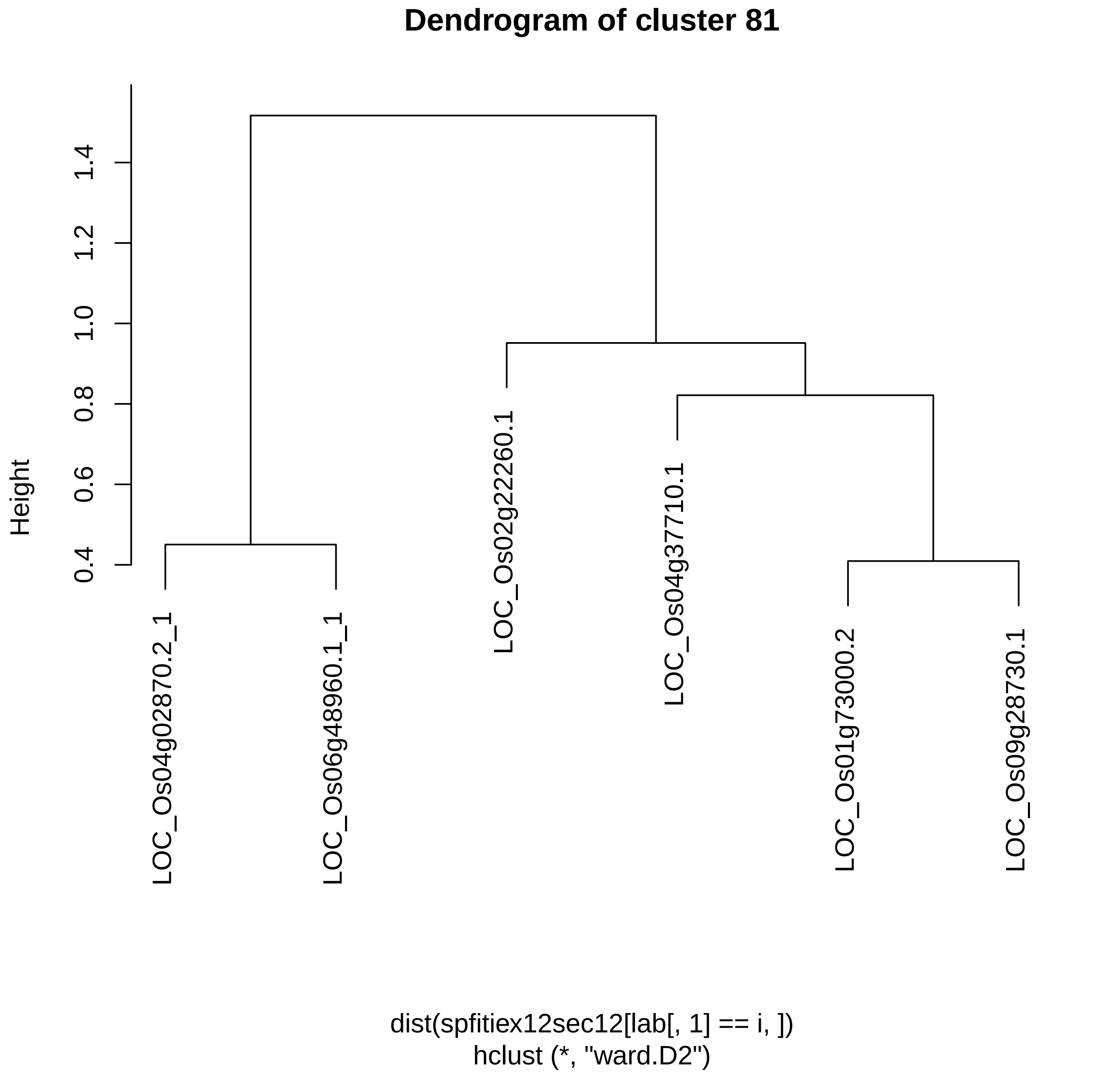
Dendrogram of cluster 81
Height
0.4
0.6
0.8
1.0
1.2
1.4
LOC_Os04g02870.2_1
LOC_Os06g48960.1_1
LOC_Os02g22260.1
LOC_Os04g37710.1
LOC_Os01g73000.2
LOC_Os09g28730.1
dist(spfitiex12sec12[lab[, 1] == i, ])
hclust (*, "ward.D2")

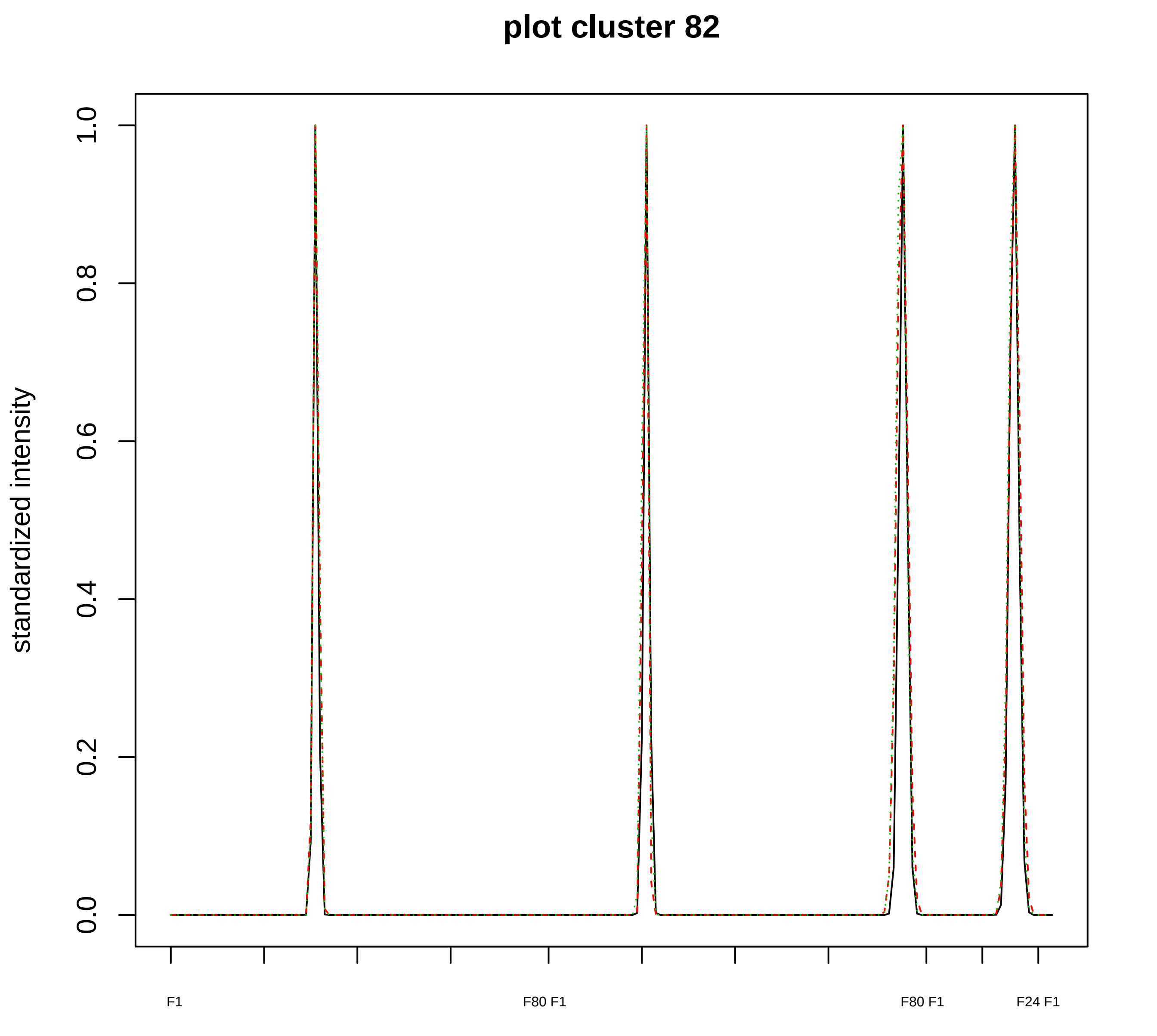

plot cluster 82
standardized intensity
0.0
0.2
0.4
0.6
0.8
1.0
F1
F80 F1
F80 F1
F24 F1

### Dendrogram of cluster 82

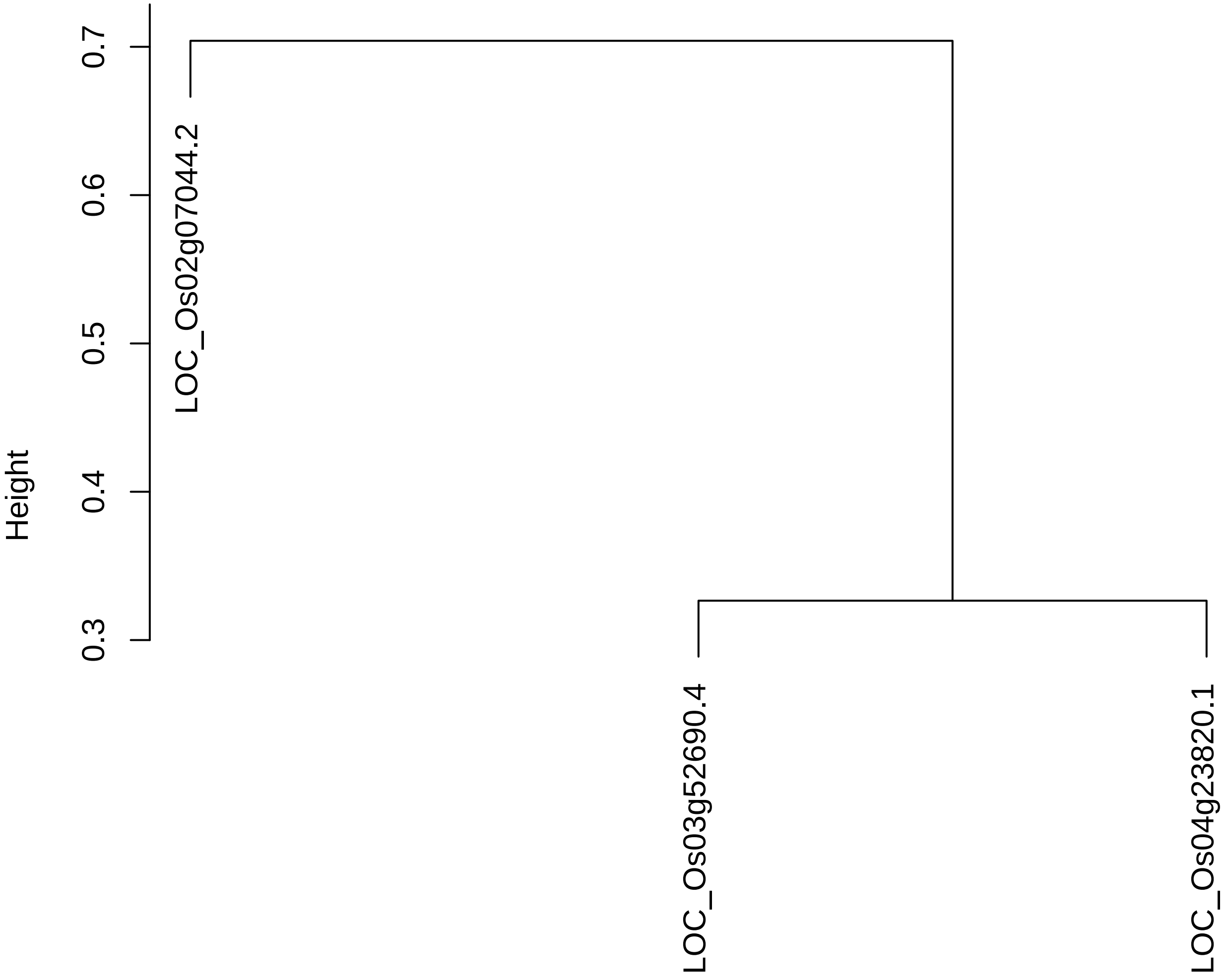


dist(spfitiex12sec12[lab[, 1] == i, ])
hclust (*, "ward.D2")

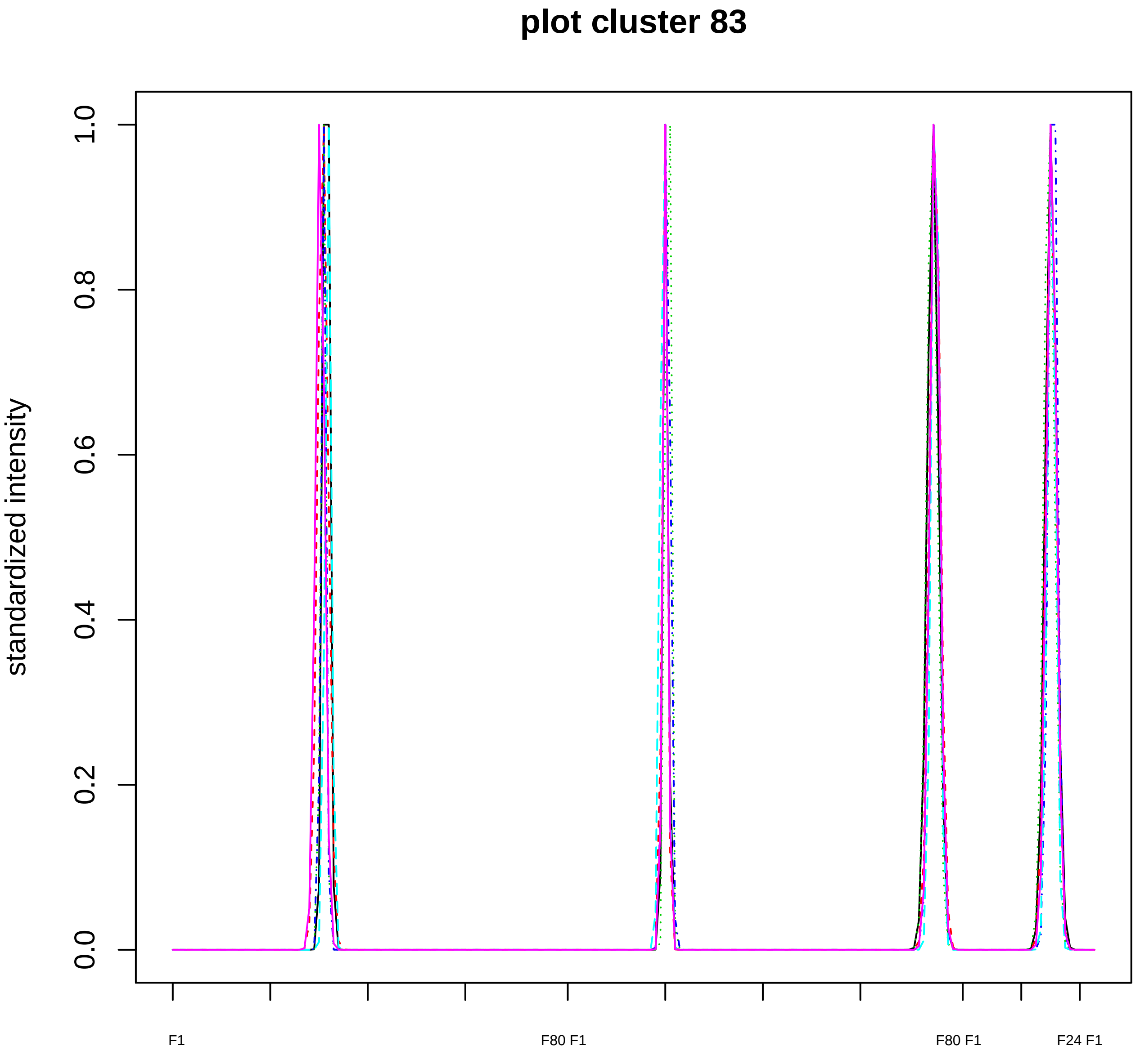

plot cluster 83
standardized intensity
0.0
0.2
0.4
0.6
0.8
1.0
F1
F80 F1
F80 F1
F24 F1

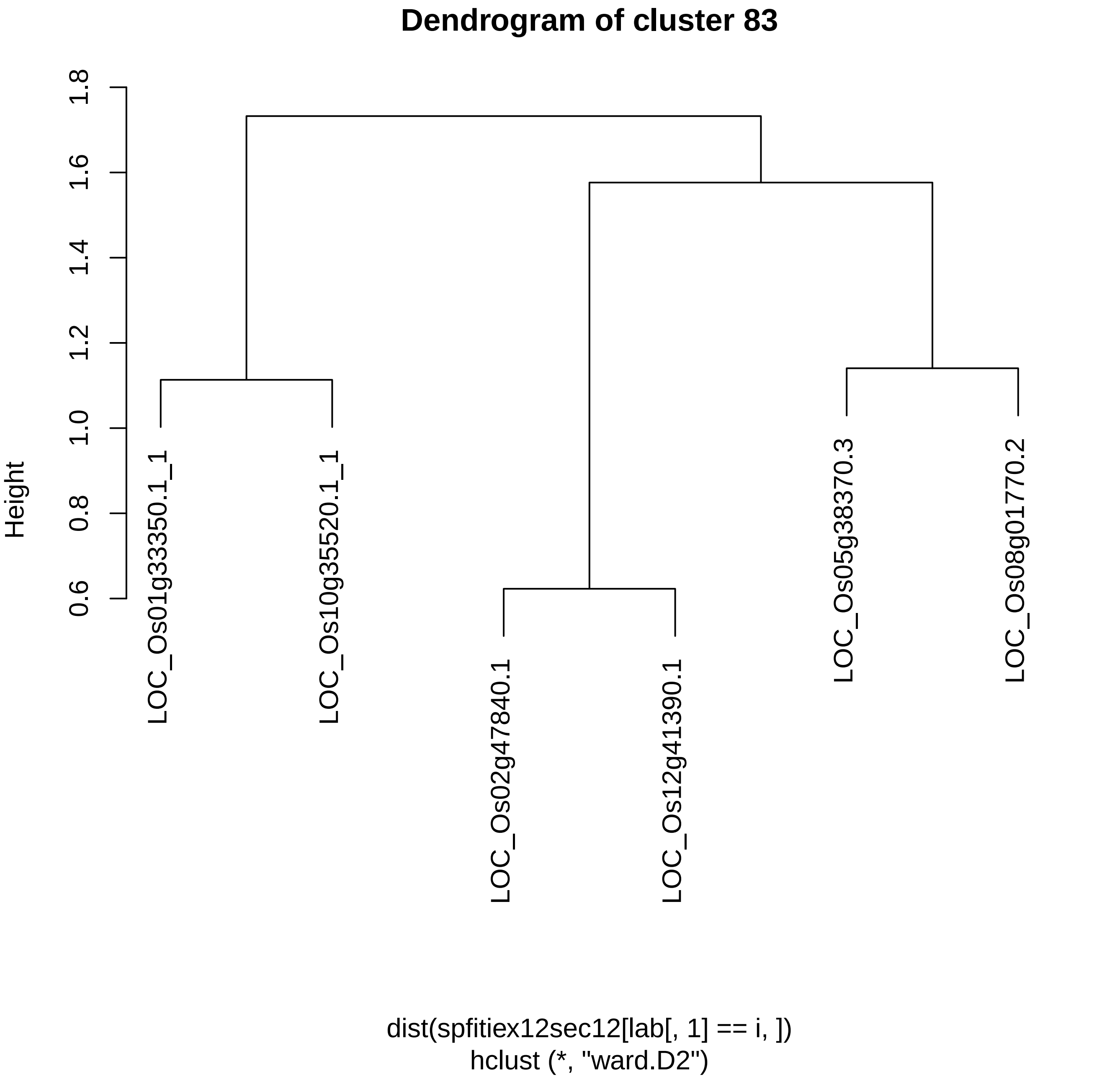
Dendrogram of cluster 83
Height
0.6
0.8
1.0
1.2
1.4
1.6
1.8
LOC_Os01g33350.1_1
LOC_Os10g35520.1_1
LOC_Os02g47840.1
LOC_Os12g41390.1
LOC_Os05g38370.3
LOC_Os08g01770.2
dist(spfitiex12sec12[lab[, 1] == i, ])
hclust (*, "ward.D2")

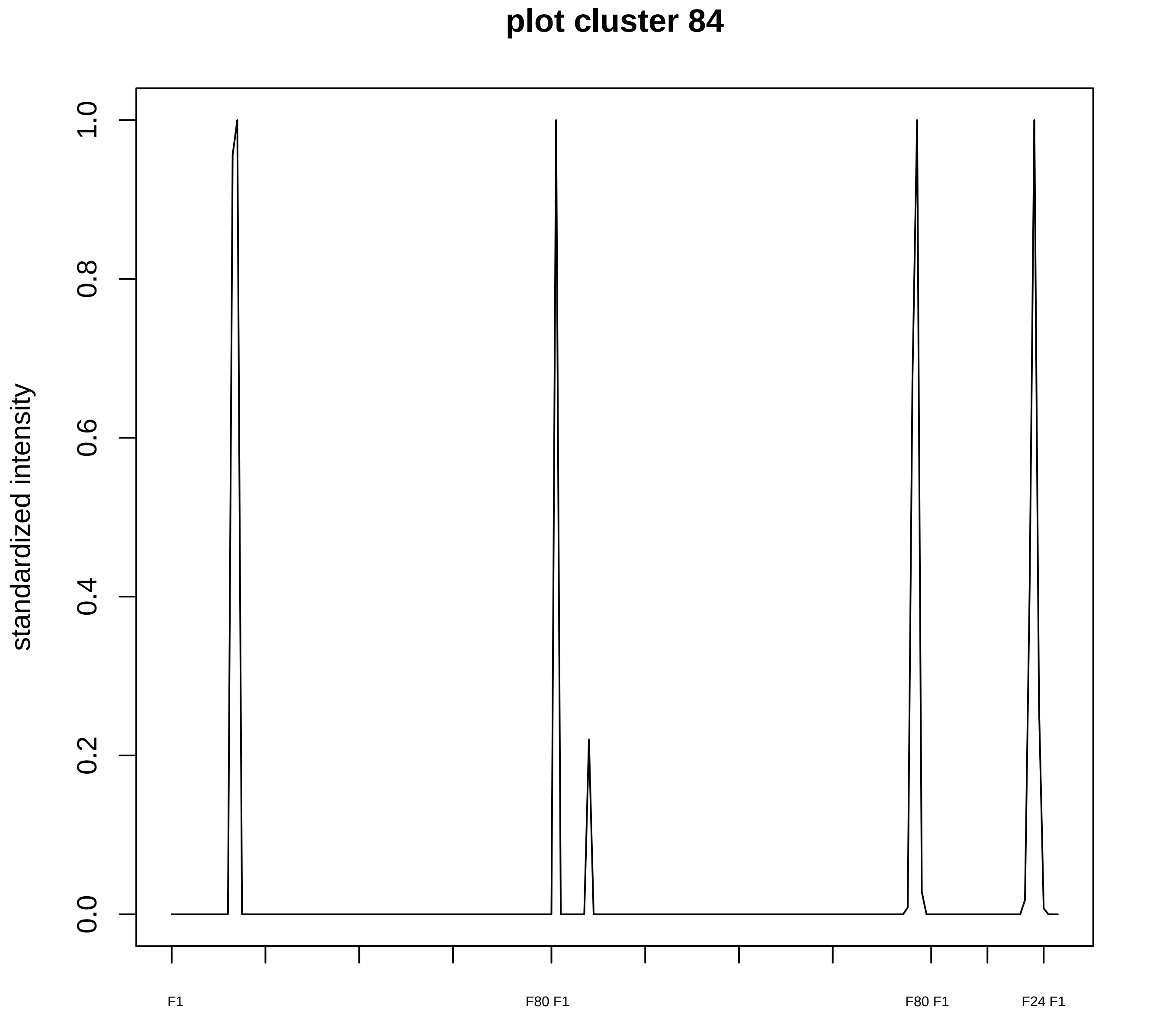
plot cluster 84
standardized intensity
0.0
0.2
0.4
0.6
0.8
1.0
F1
F80 F1
F80 F1
F24 F1

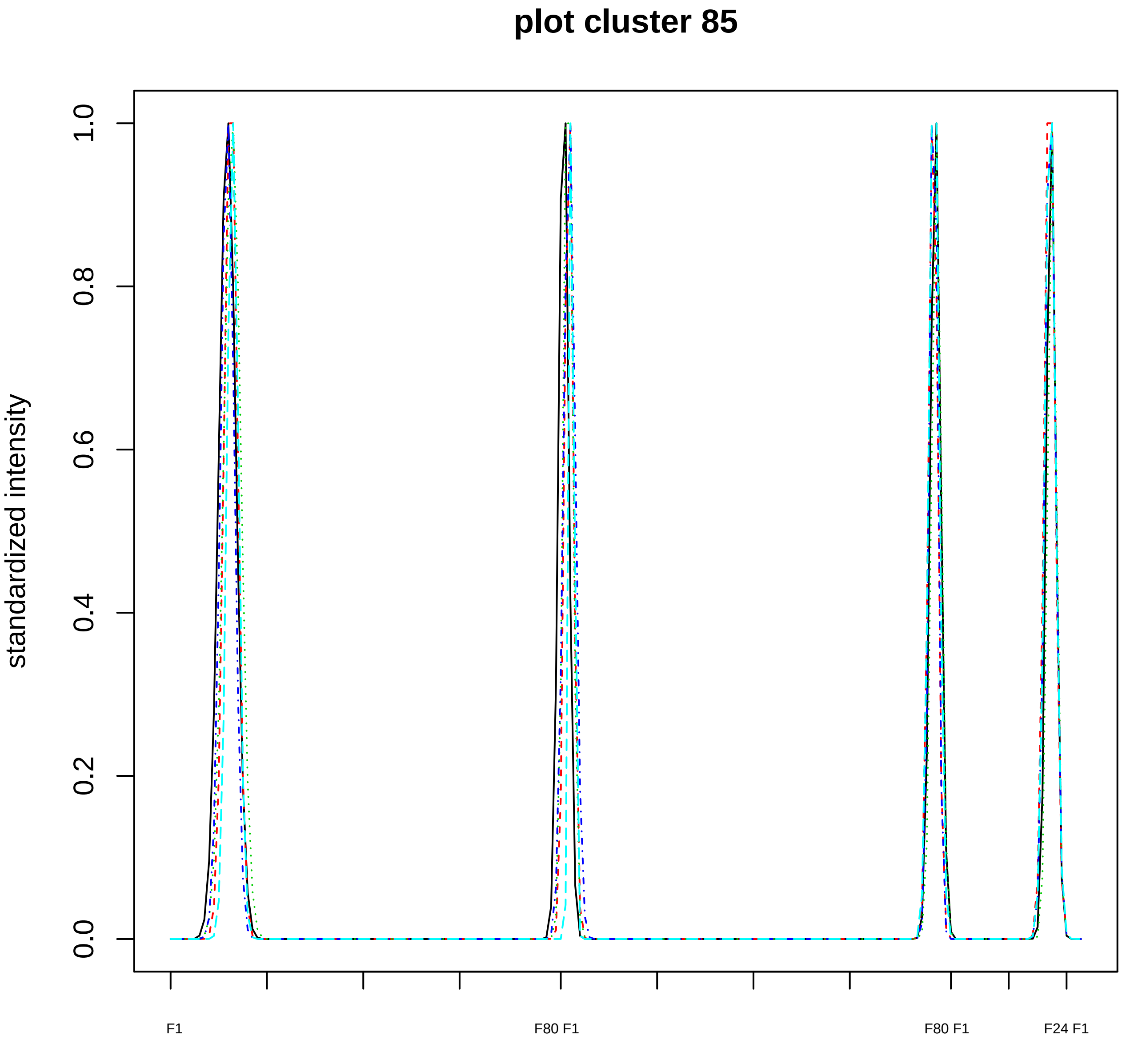
plot cluster 85
standardized intensity
0.0
0.2
0.4
0.6
0.8
1.0
F1
F80 F1
F80 F1
F24 F1

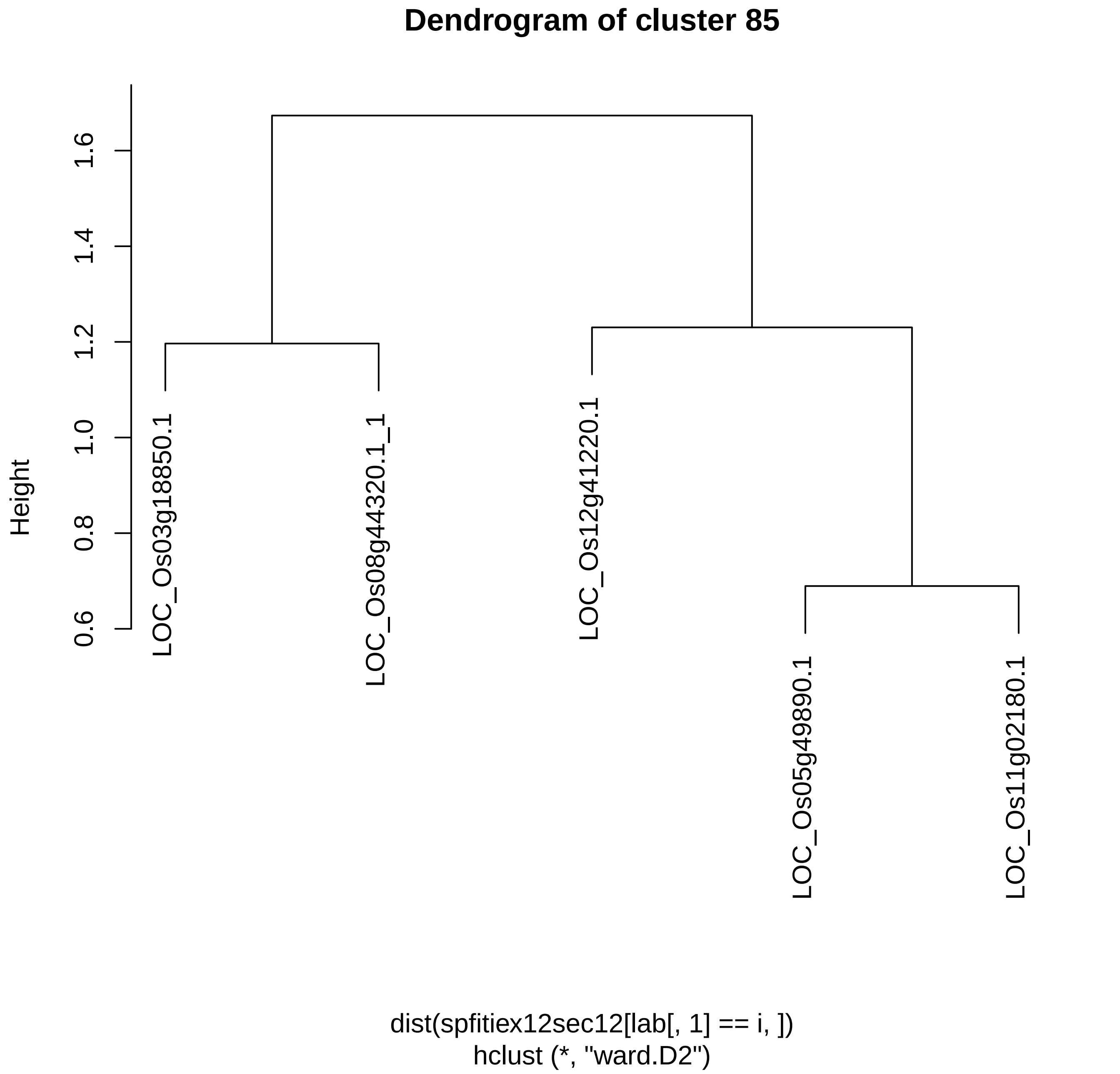
Dendrogram of cluster 85
Height
0.6
0.8
1.0
1.2
1.4
1.6
LOC_Os03g18850.1
LOC_Os08g44320.1_1
LOC_Os12g41220.1
LOC_Os05g49890.1
LOC_Os11g02180.1
dist(spfitiex12sec12[lab[, 1] == i, ])
hclust (*, "ward.D2")

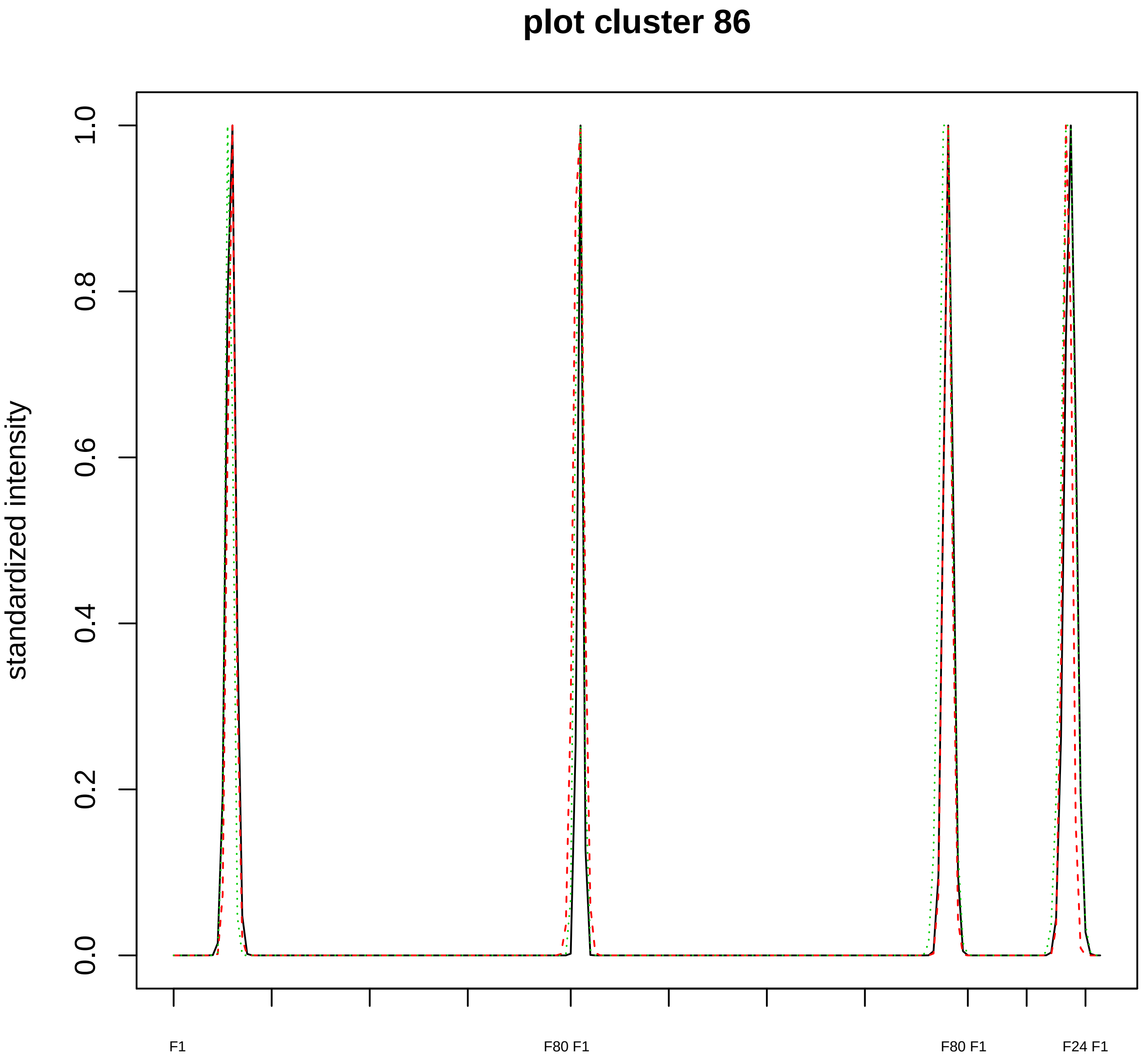

plot cluster 86
standardized intensity
0.0
0.2
0.4
0.6
0.8
1.0
F1
F80 F1
F80 F1
F24 F1

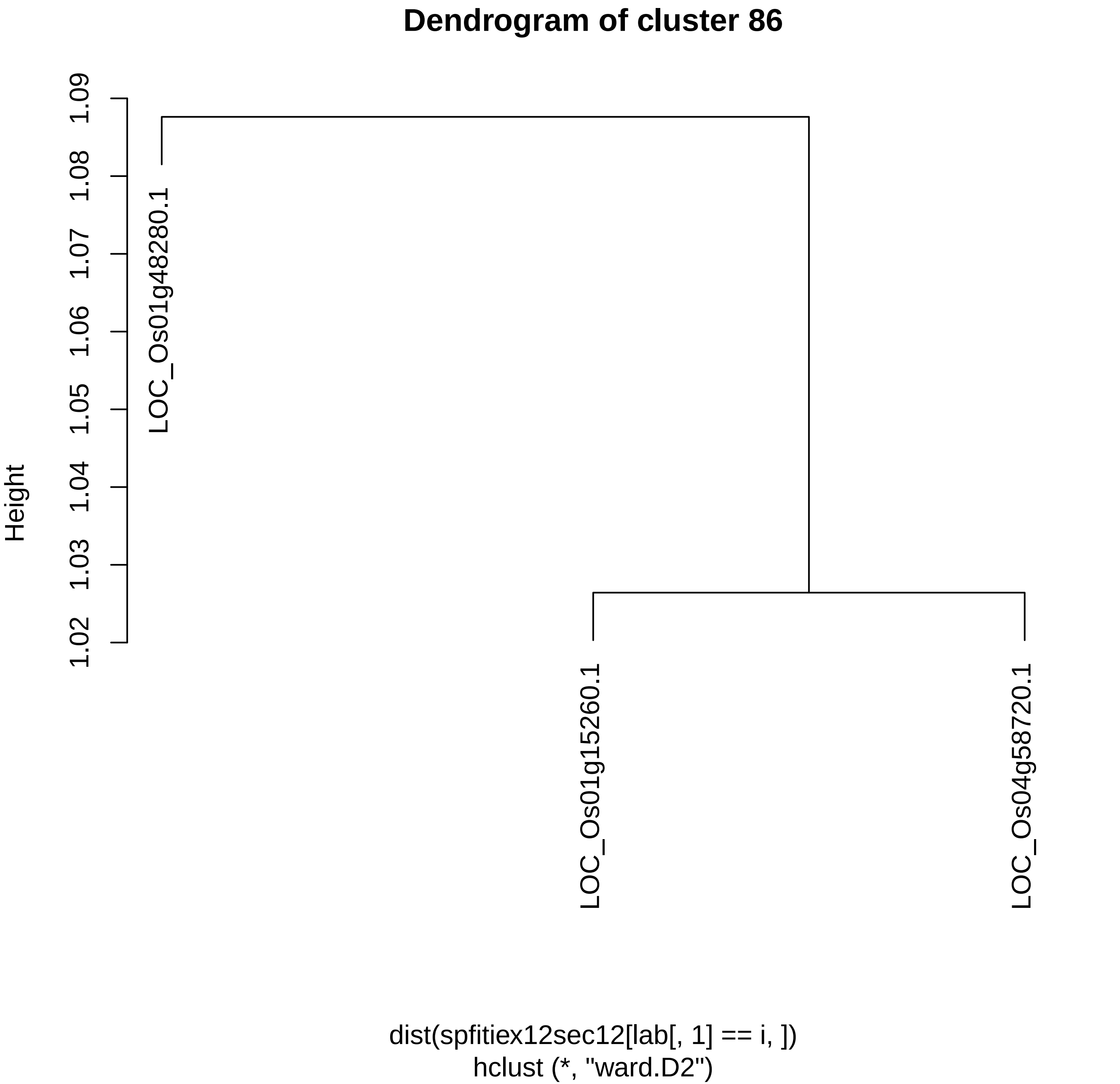
Dendrogram of cluster 86
Height
1.02
1.03
1.04
1.05
1.06
1.07
1.08
1.09
LOC_Os01g48280.1
LOC_Os01g15260.1
LOC_Os04g58720.1
dist(spfitiex12sec12[lab[, 1] == i, ])
hclust (*, "ward.D2")

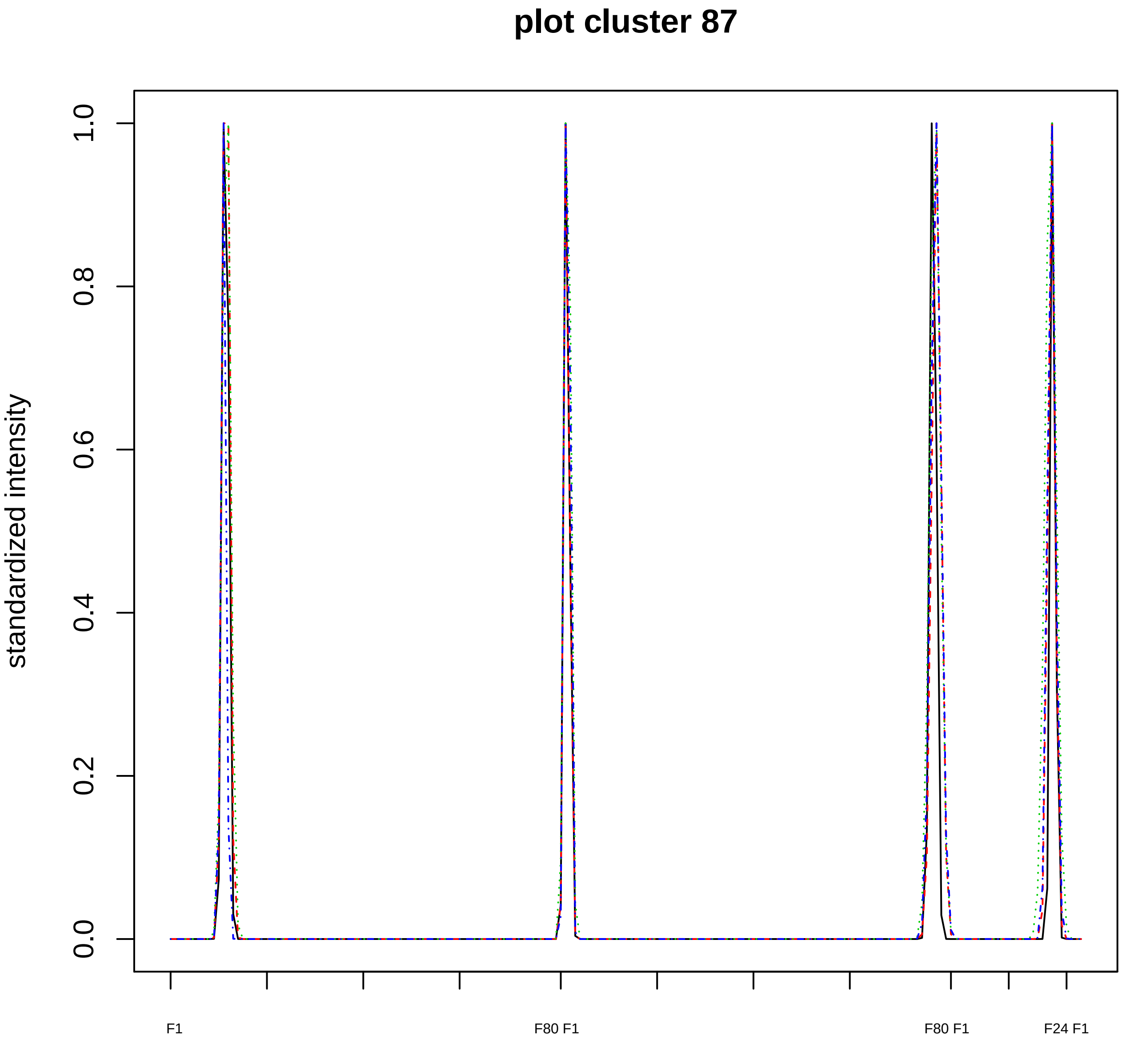
plot cluster 87
standardized intensity
0.0
0.2
0.4
0.6
0.8
1.0
F1
F80 F1
F80 F1
F24 F1

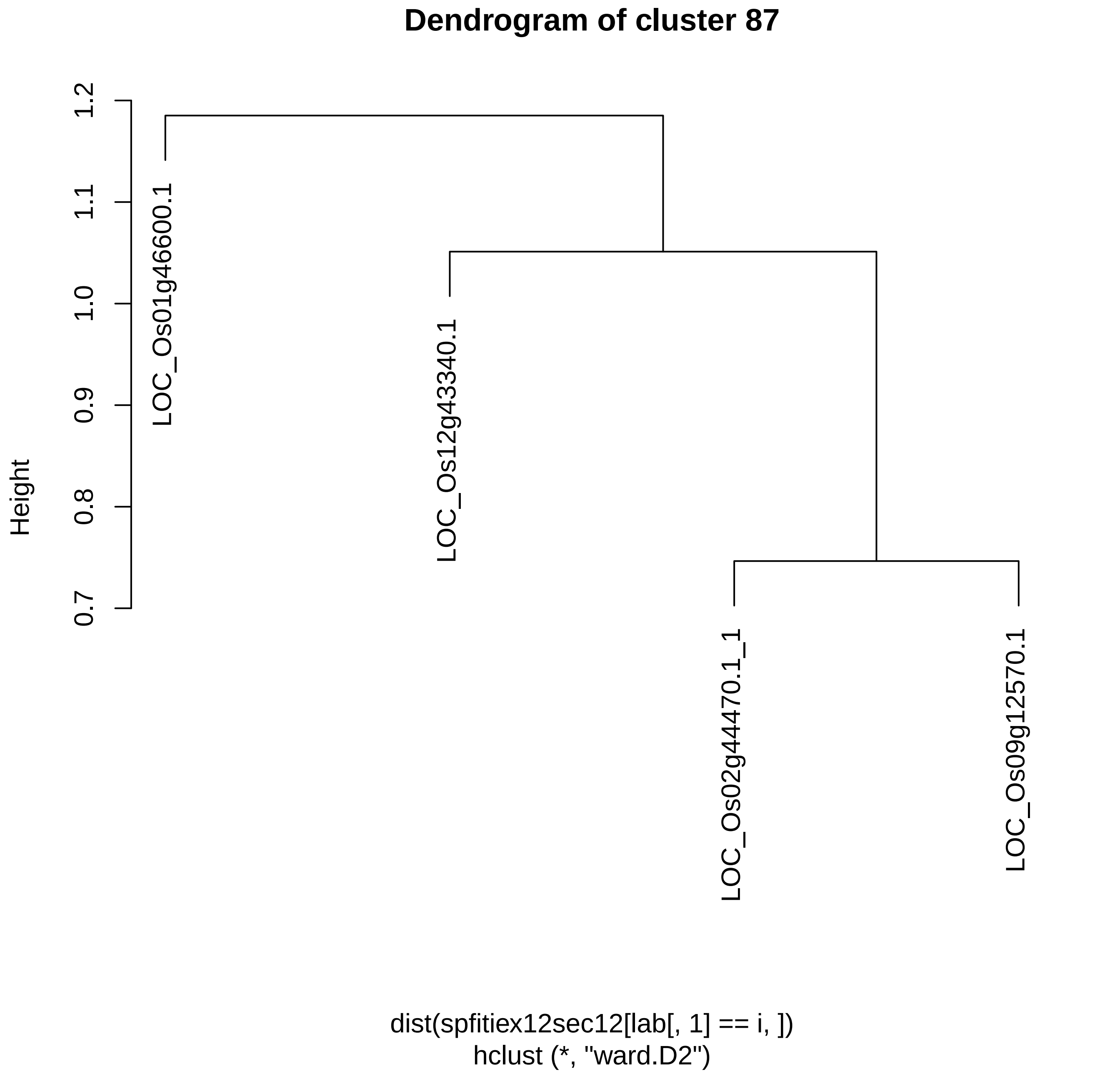
Dendrogram of cluster 87
Height
0.7
0.8
0.9
1.0
1.1
1.2
LOC_Os01g46600.1
LOC_Os12g43340.1
LOC_Os02g44470.1_1
LOC_Os09g12570.1
dist(spfitiex12sec12[lab[, 1] == i, ])
hclust (*, "ward.D2")

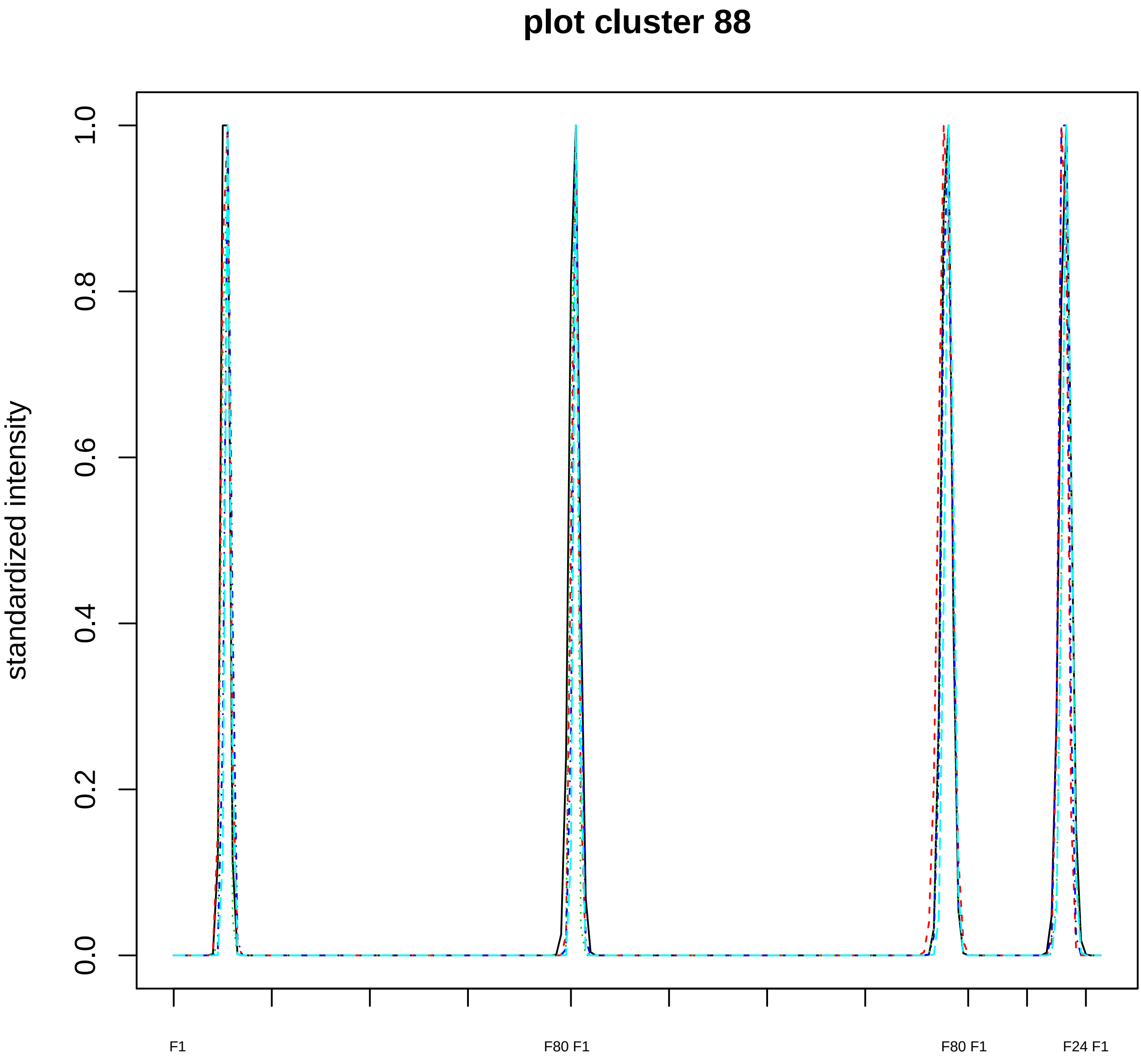
plot cluster 88
standardized intensity
0.0
0.2
0.4
0.6
0.8
1.0
F1
F80 F1
F80 F1
F24 F1

### Dendrogram of cluster 88

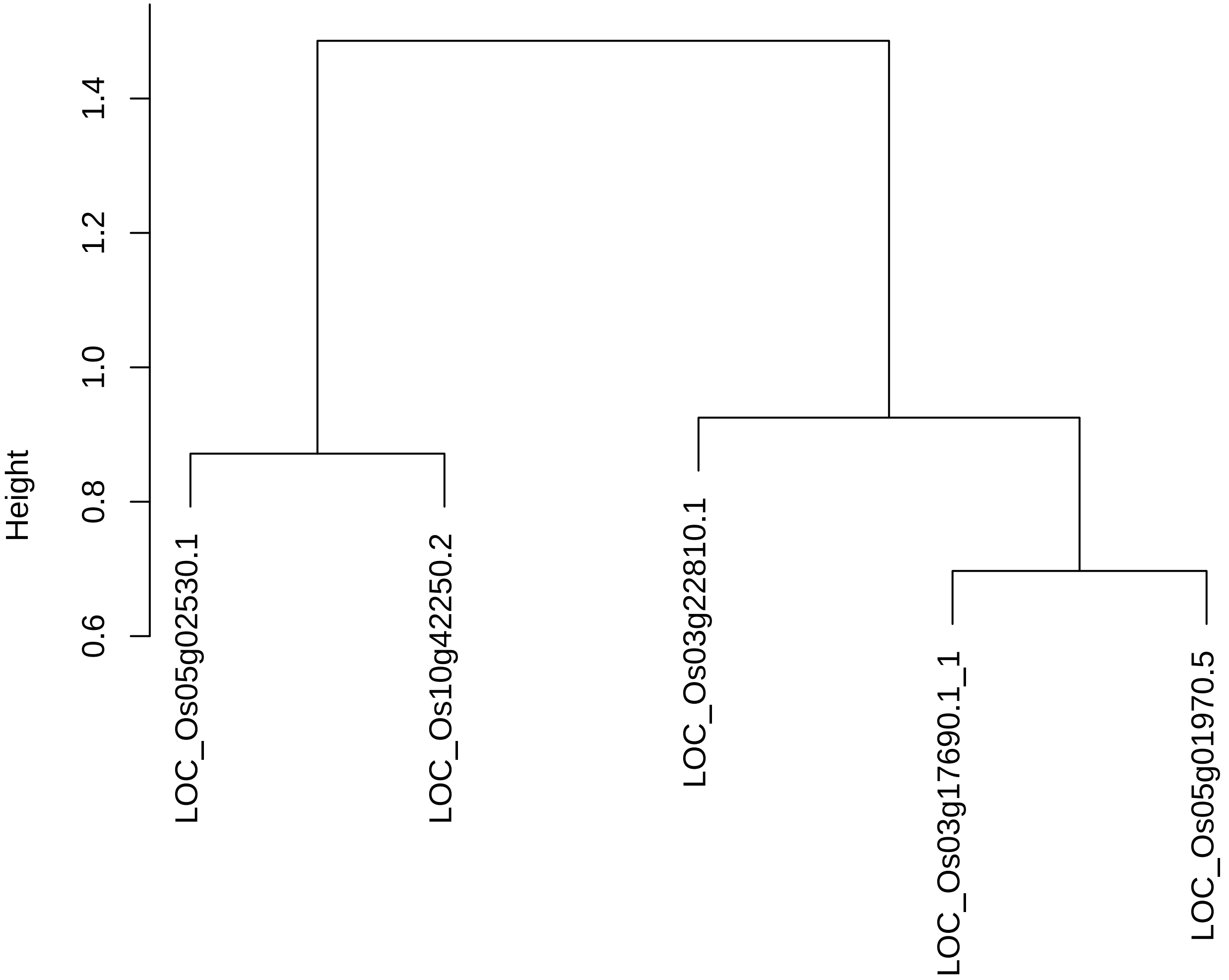


dist(spfitiex12sec12[lab[, 1] == i, ])
hclust (*, "ward.D2")

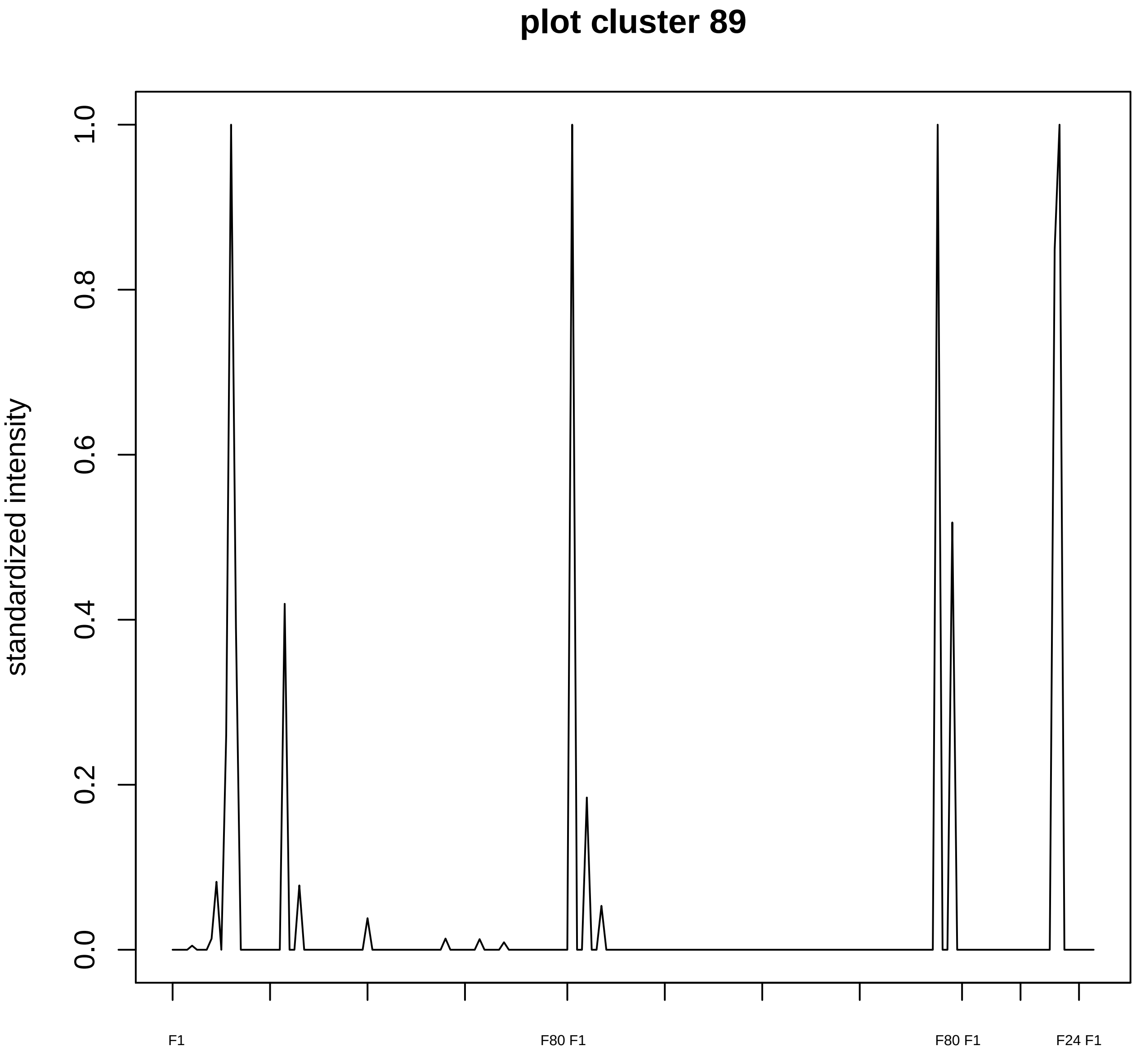

plot cluster 89
standardized intensity
0.0
0.2
0.4
0.6
0.8
1.0
F1
F80 F1
F80 F1
F24 F1

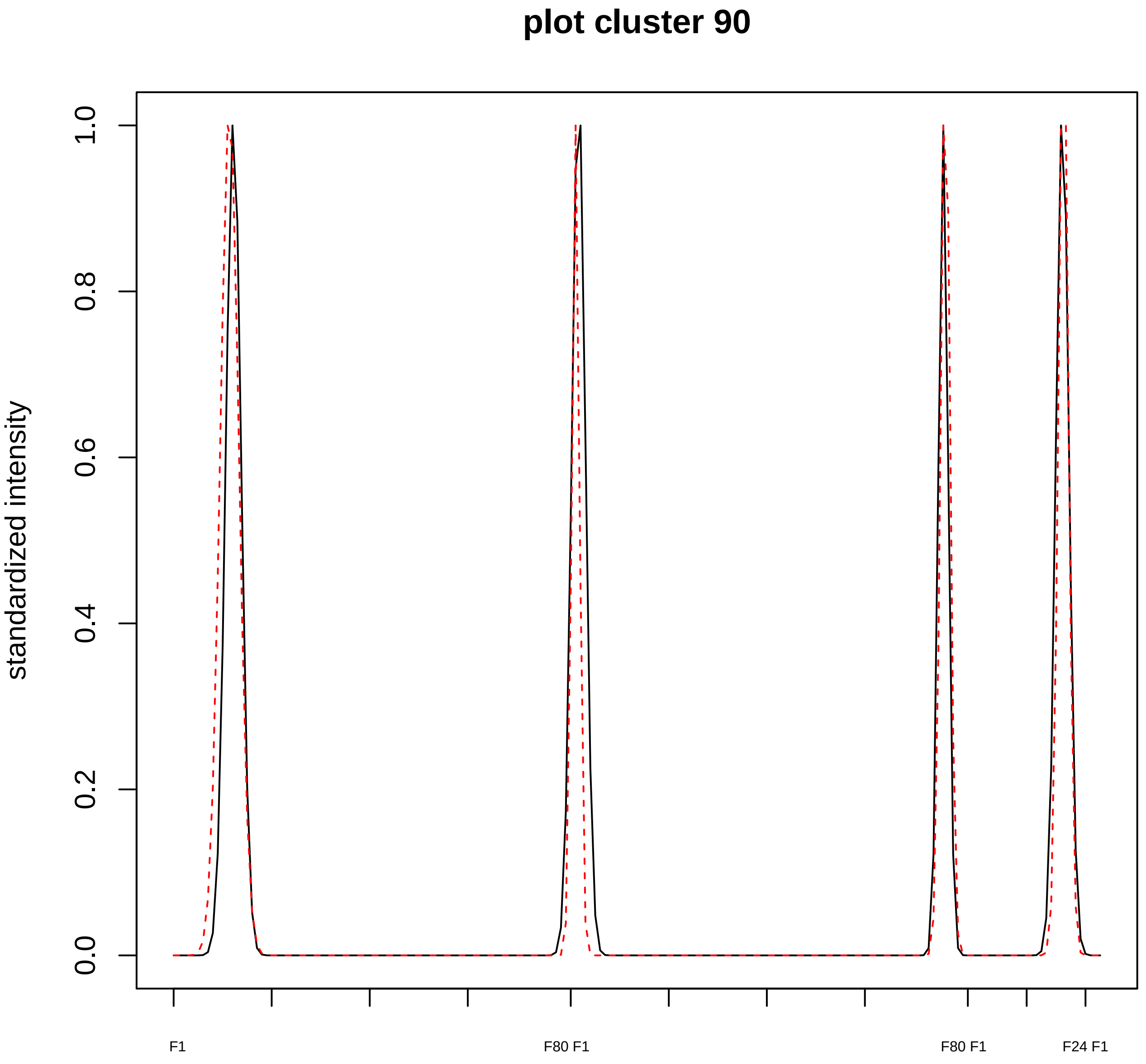

plot cluster 90
standardized intensity
0.0
0.2
0.4
0.6
0.8
1.0
F1
F80 F1
F80 F1
F24 F1

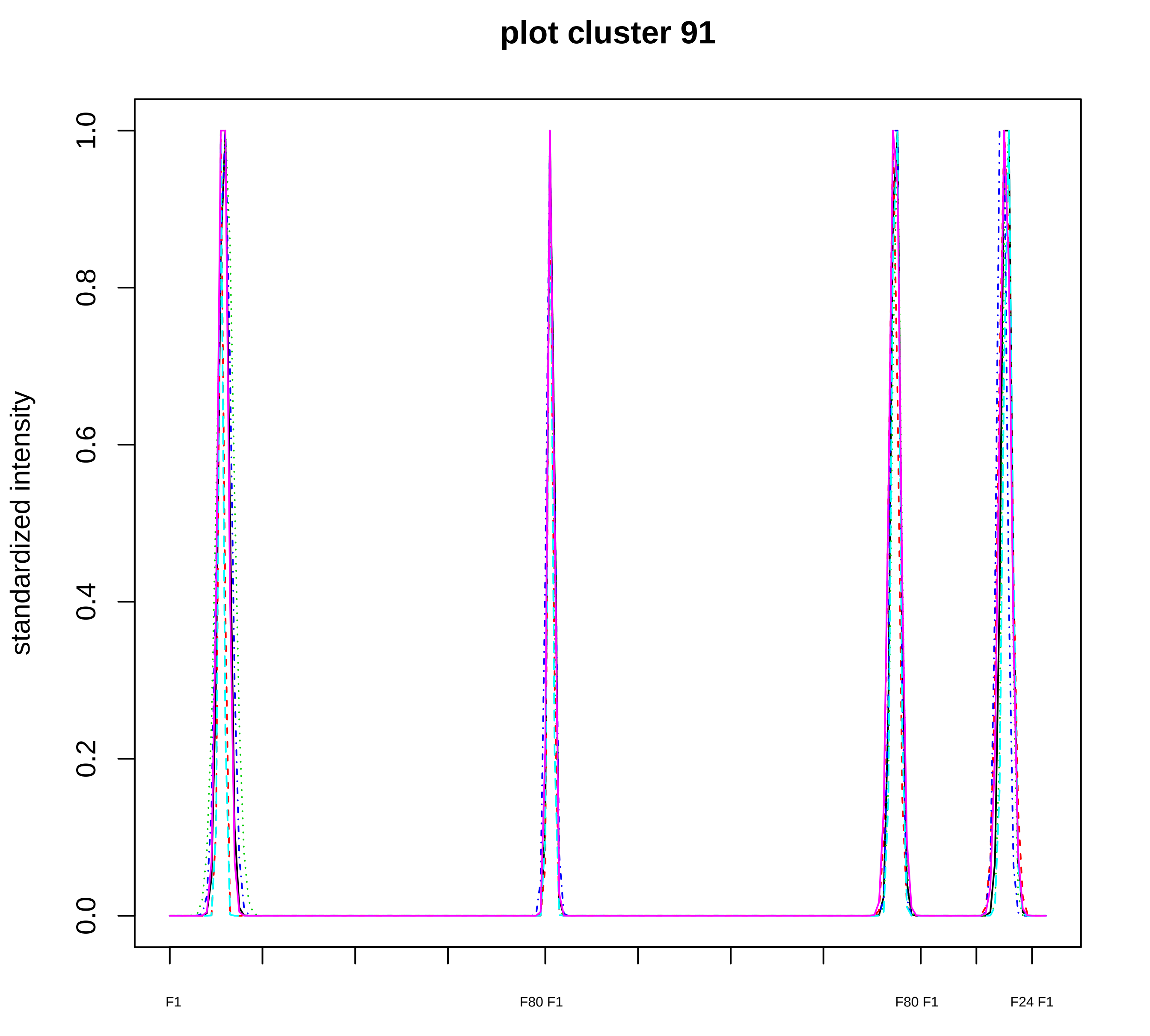
plot cluster 91
standardized intensity
0.0
0.2
0.4
0.6
0.8
1.0
F1
F80 F1
F80 F1
F24 F1

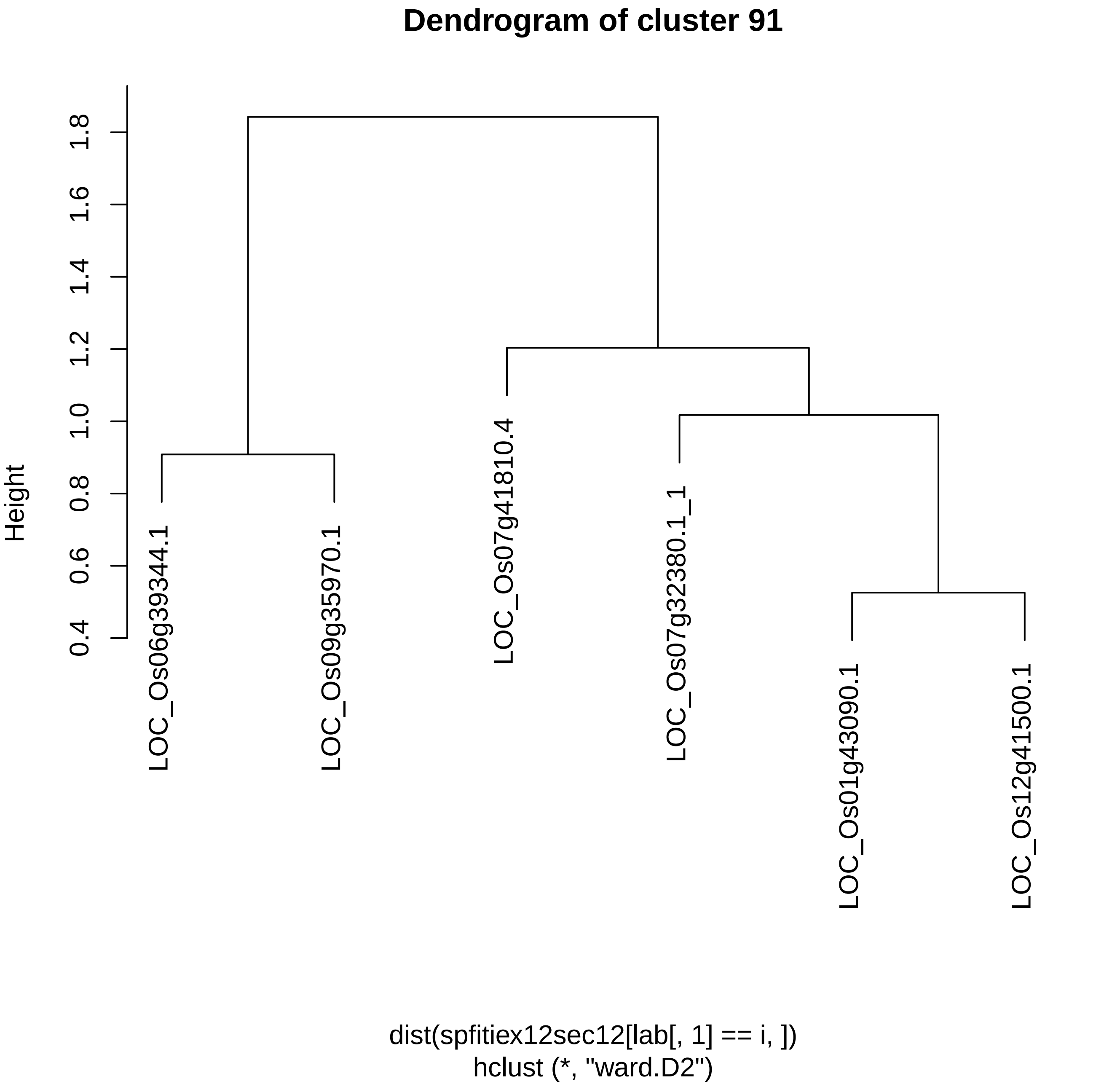
Dendrogram of cluster 91
Height
0.4
0.6
0.8
1.0
1.2
1.4
1.6
1.8
LOC_Os06g39344.1
LOC_Os09g35970.1
LOC_Os07g41810.4
LOC_Os07g32380.1_1
LOC_Os01g43090.1
LOC_Os12g41500.1
dist(spfitiex12sec12[lab[, 1] == i, ])
hclust (*, "ward.D2")

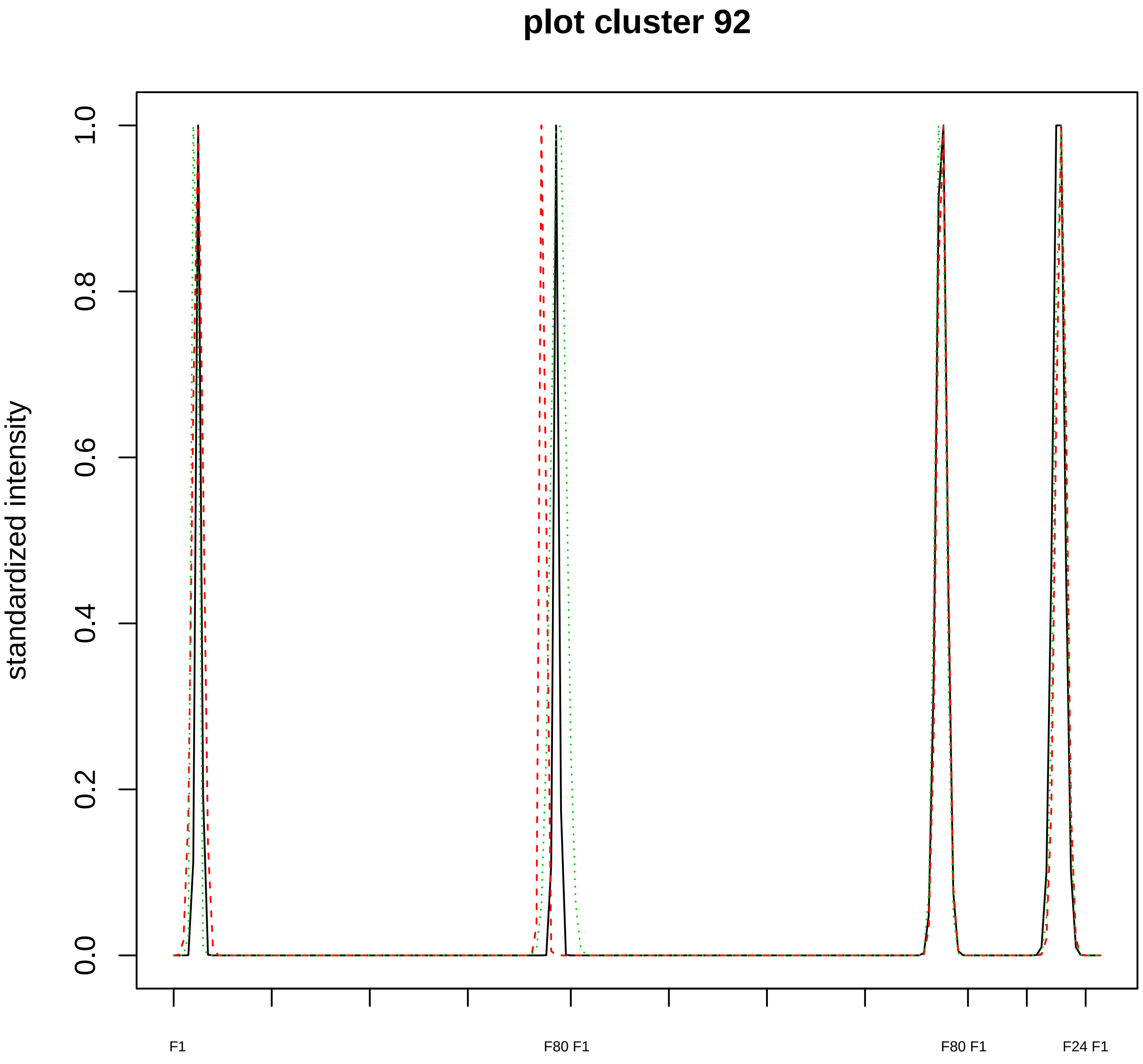
plot cluster 92
standardized intensity
0.0
0.2
0.4
0.6
0.8
1.0
F1
F80 F1
F80 F1
F24 F1

#### Dendrogram of cluster 92

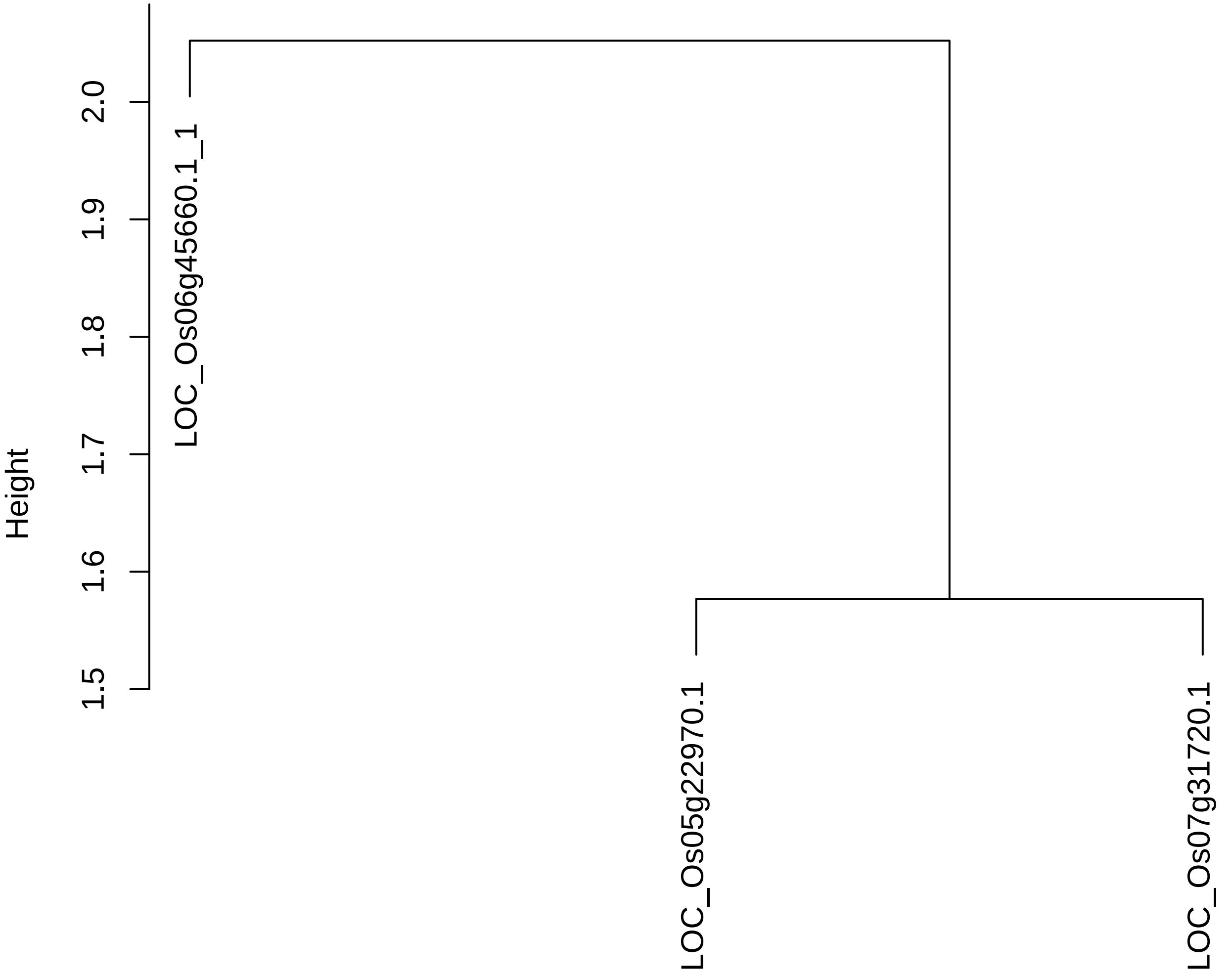


dist(spfitiex12sec12[lab[, 1] == i, ])
hclust (*, "ward.D2")

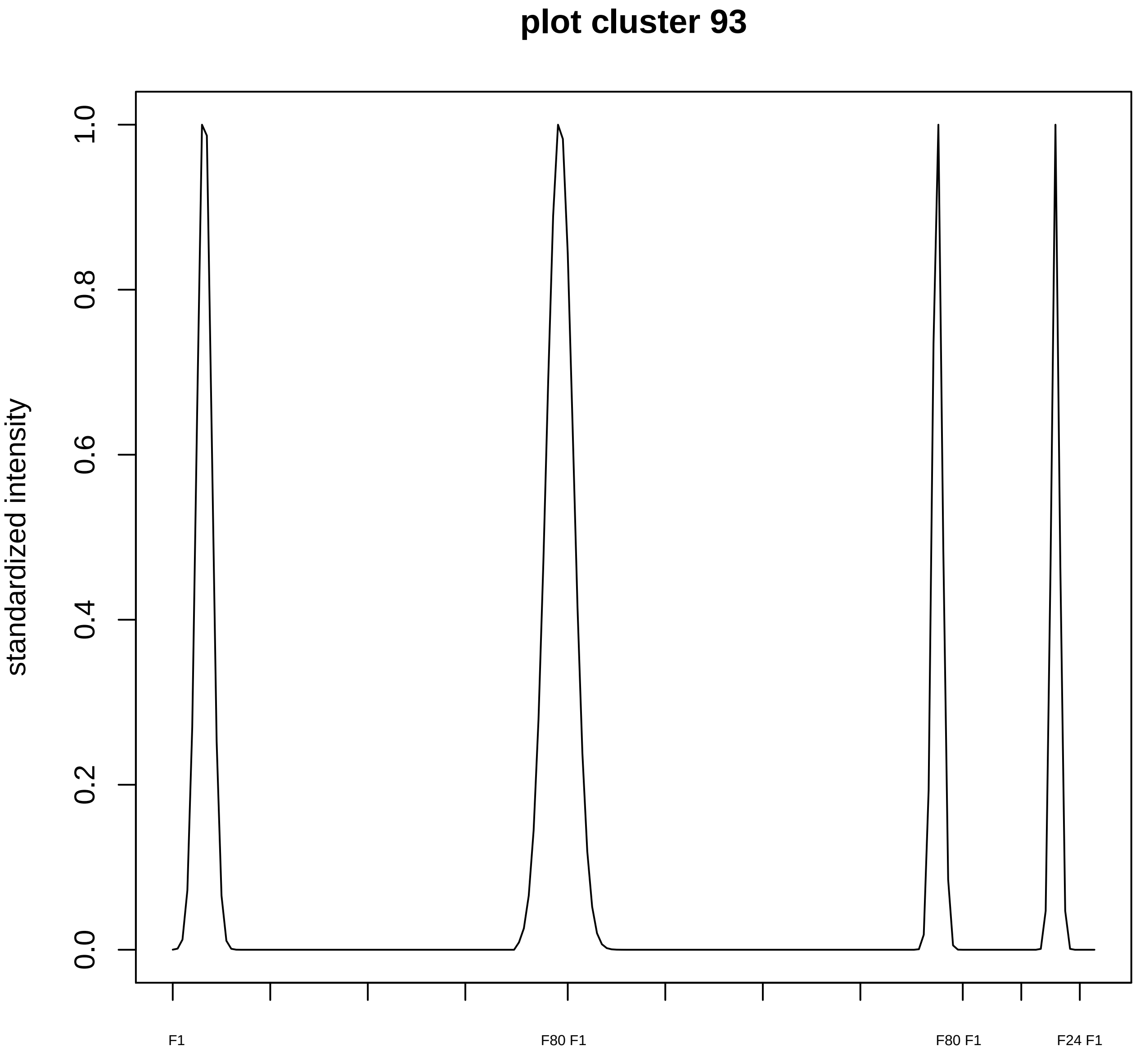
plot cluster 93
standardized intensity
0.0
0.2
0.4
0.6
0.8
1.0
F1
F80 F1
F80 F1
F24 F1

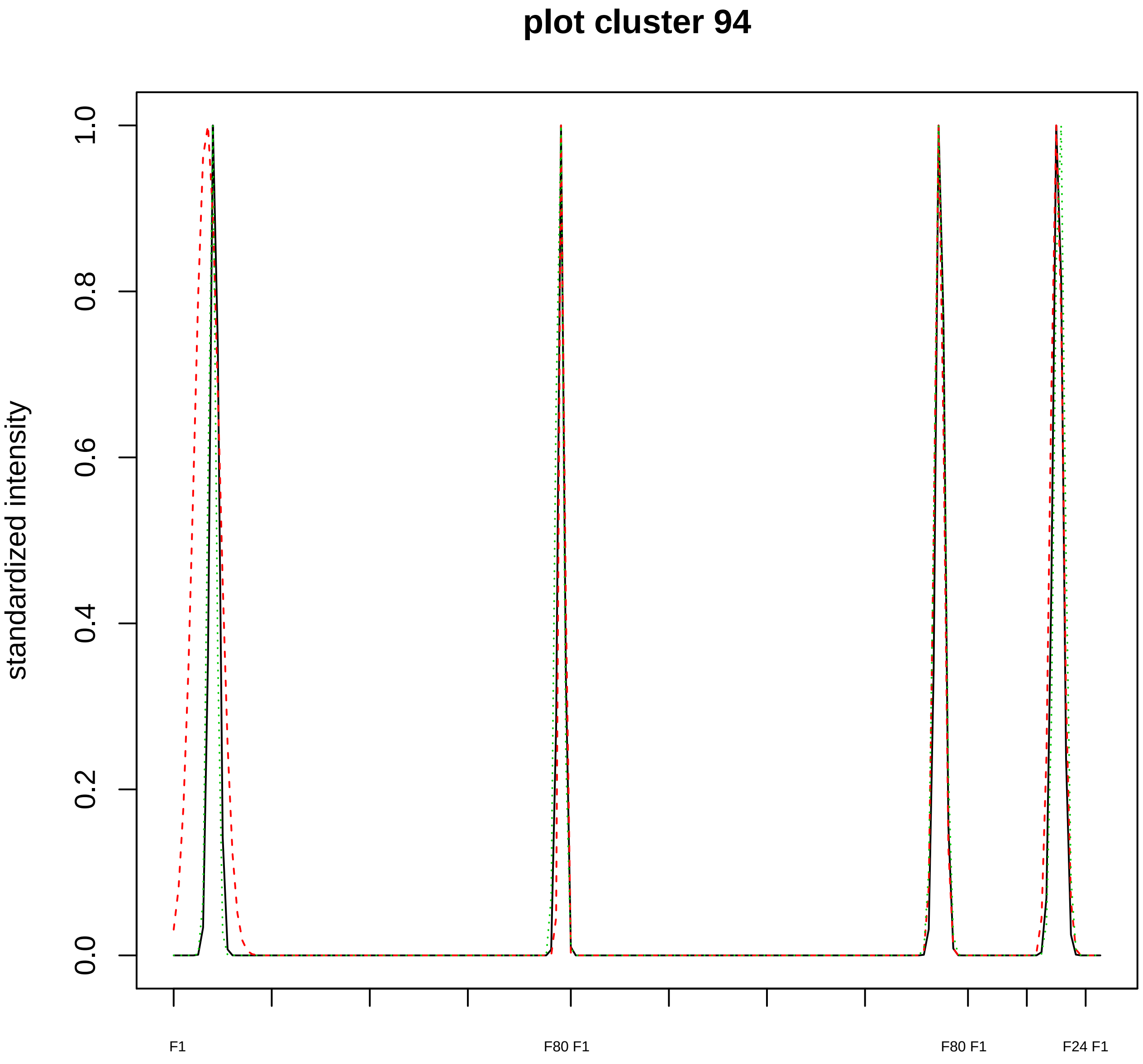
plot cluster 94
standardized intensity
0.0
0.2
0.4
0.6
0.8
1.0
F1
F80 F1
F80 F1
F24 F1

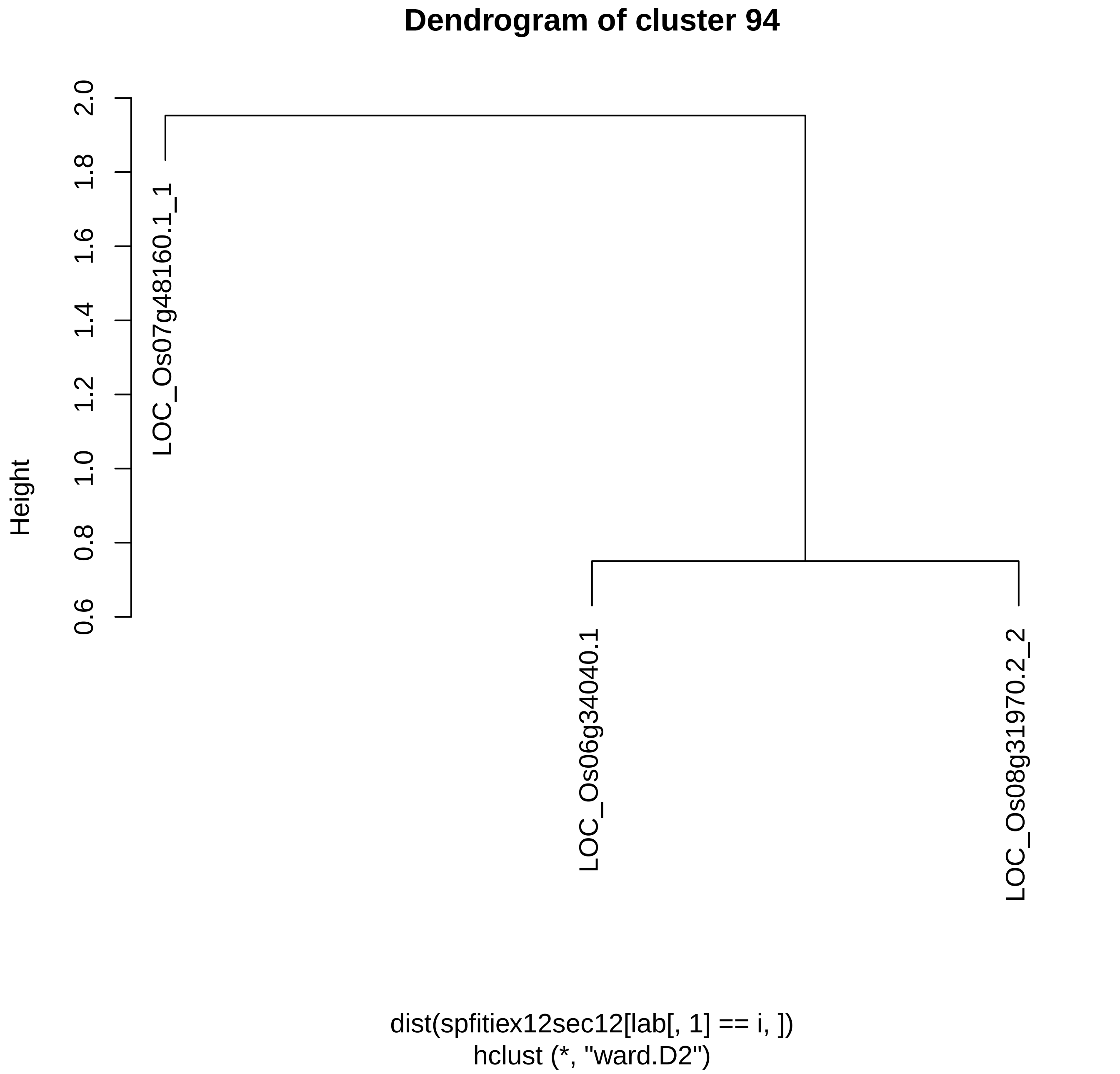
Dendrogram of cluster 94
Height
2.0
1.8
1.6
1.4
1.2
1.0
0.8
0.6
LOC_Os07g48160.1_1
LOC_Os06g34040.1
LOC_Os08g31970.2_2
dist(spfitiex12sec12[lab[, 1] == i, ])
hclust (*, "ward.D2")

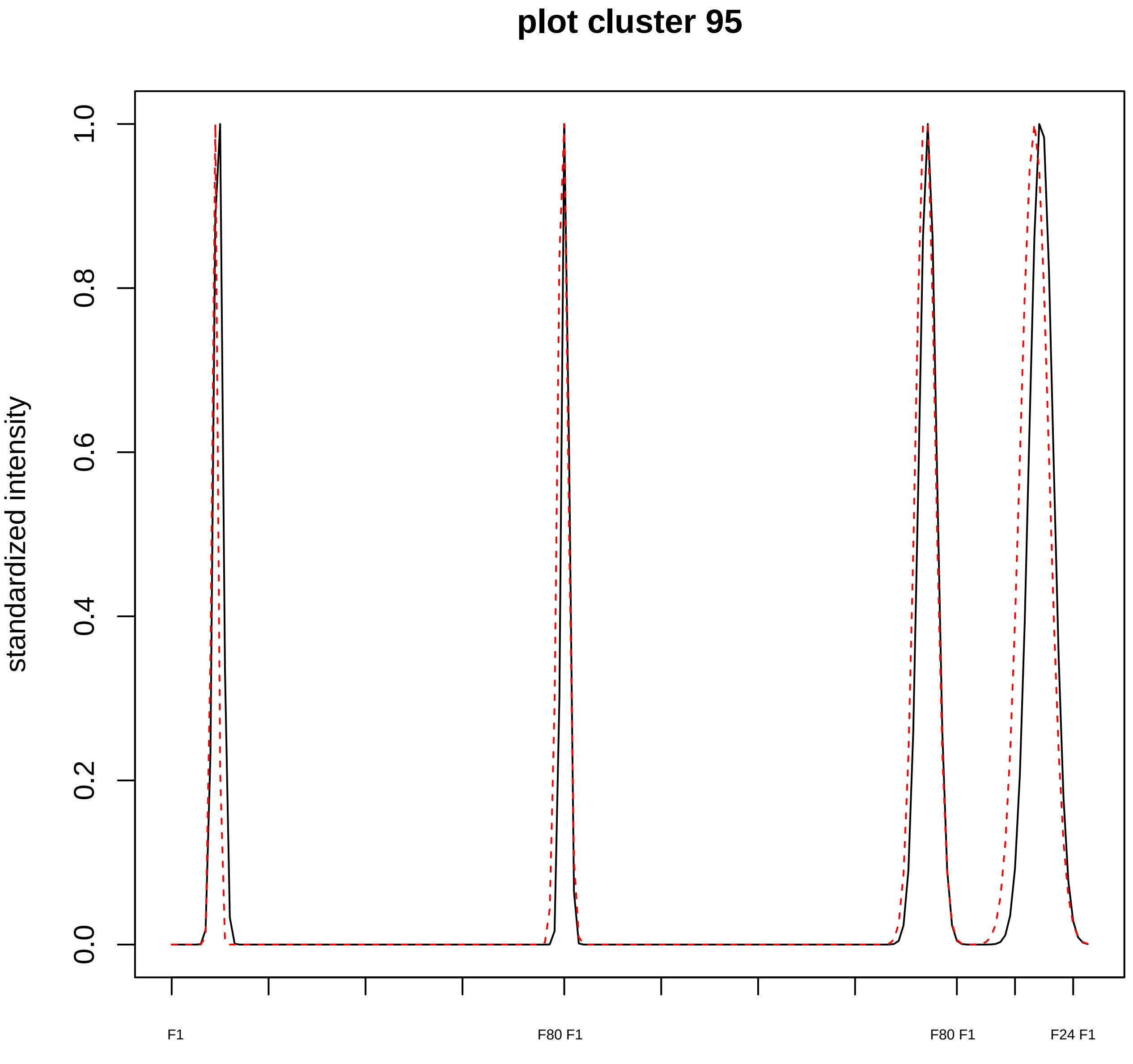
plot cluster 95
standardized intensity
0.0
0.2
0.4
0.6
0.8
1.0
F1
F80 F1
F80 F1
F24 F1

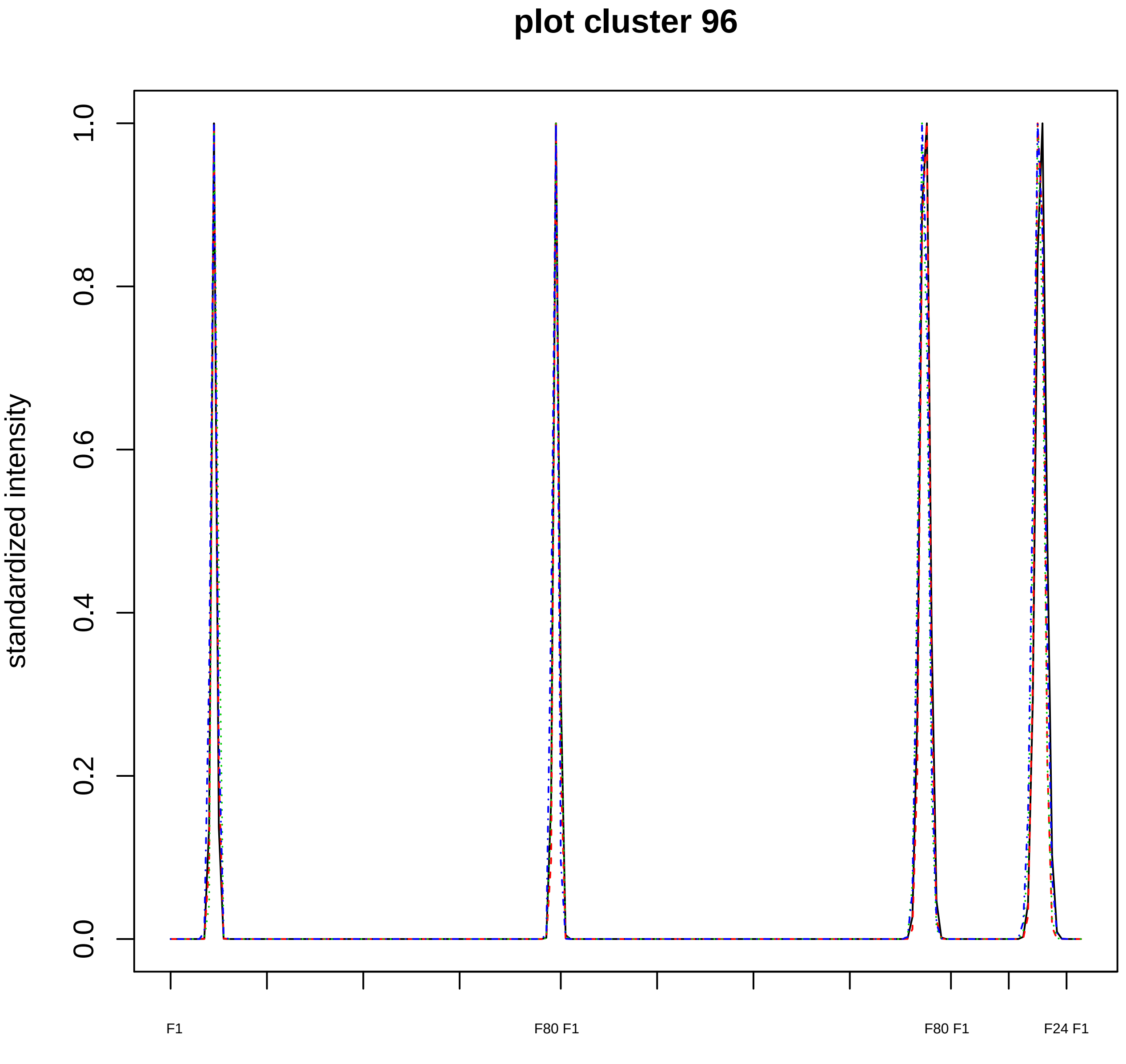
plot cluster 96
standardized intensity
0.0
0.2
0.4
0.6
0.8
1.0
F1
F80 F1
F80 F1
F24 F1

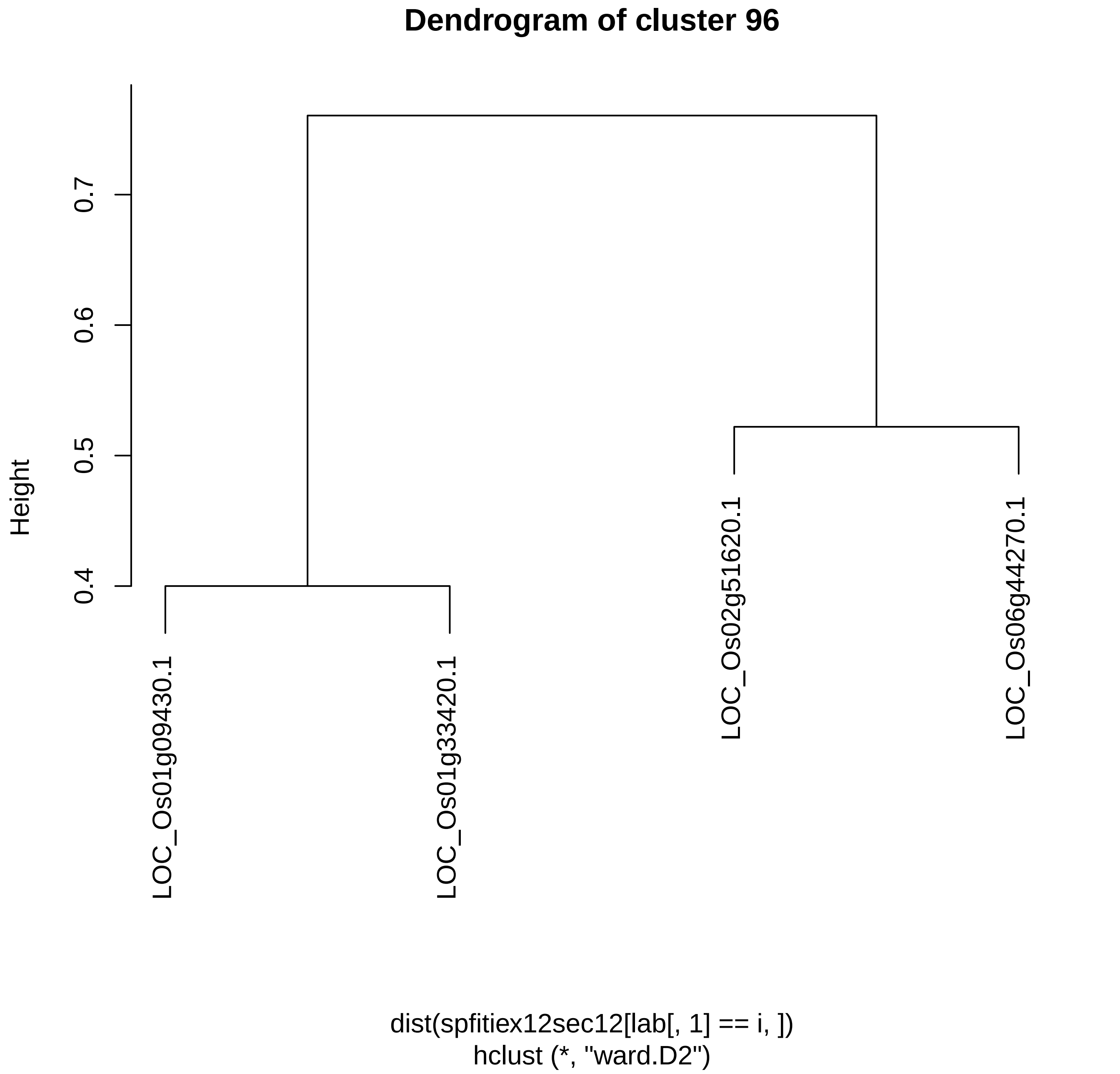
Dendrogram of cluster 96
Height
0.4
0.5
0.6
0.7
LOC_Os01g09430.1
LOC_Os01g33420.1
LOC_Os02g51620.1
LOC_Os06g44270.1
dist(spfitiex12sec12[lab[, 1] == i, ])
hclust (*, "ward.D2")

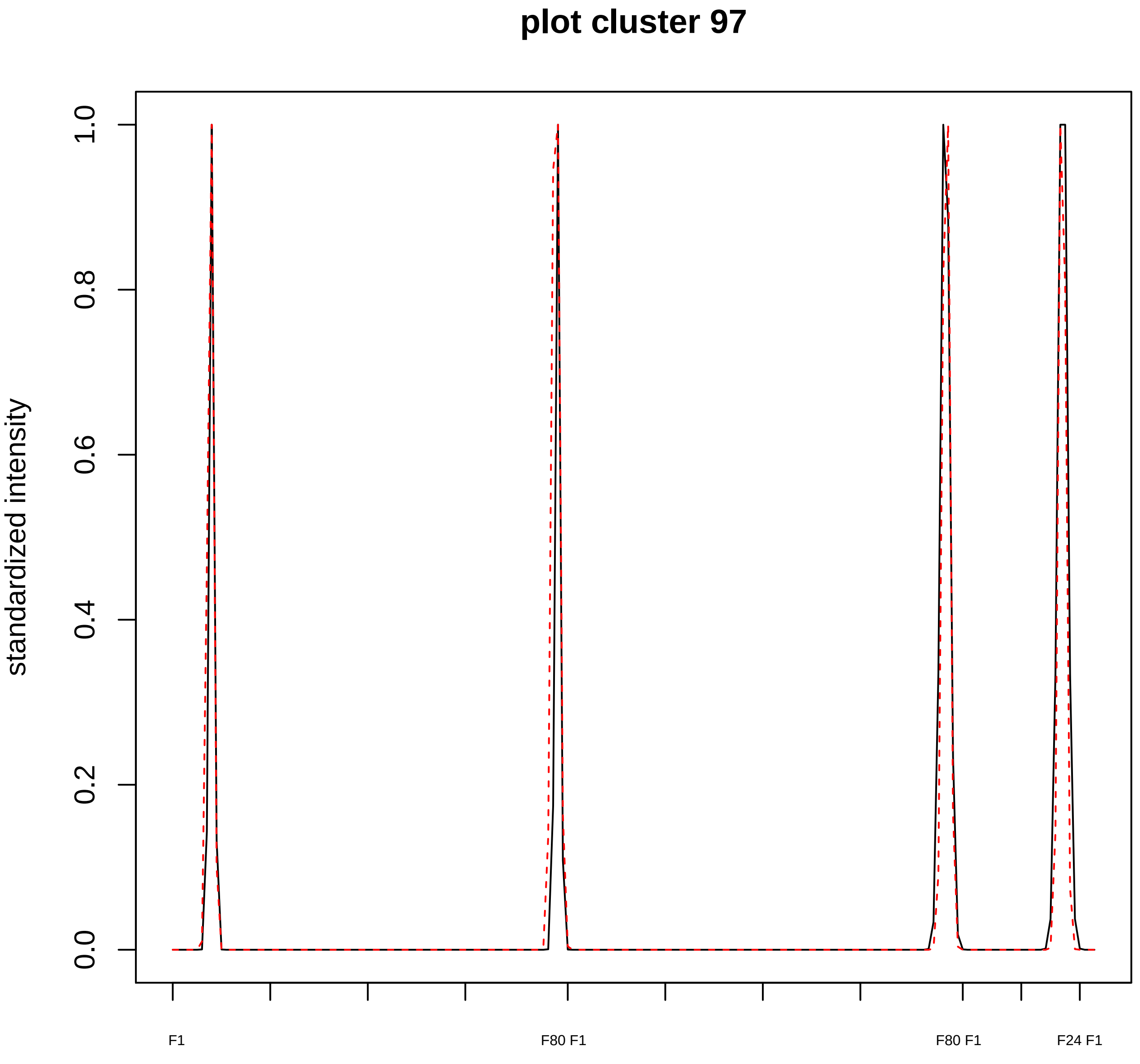
plot cluster 97
standardized intensity
0.0
0.2
0.4
0.6
0.8
1.0
F1
F80 F1
F80 F1
F24 F1

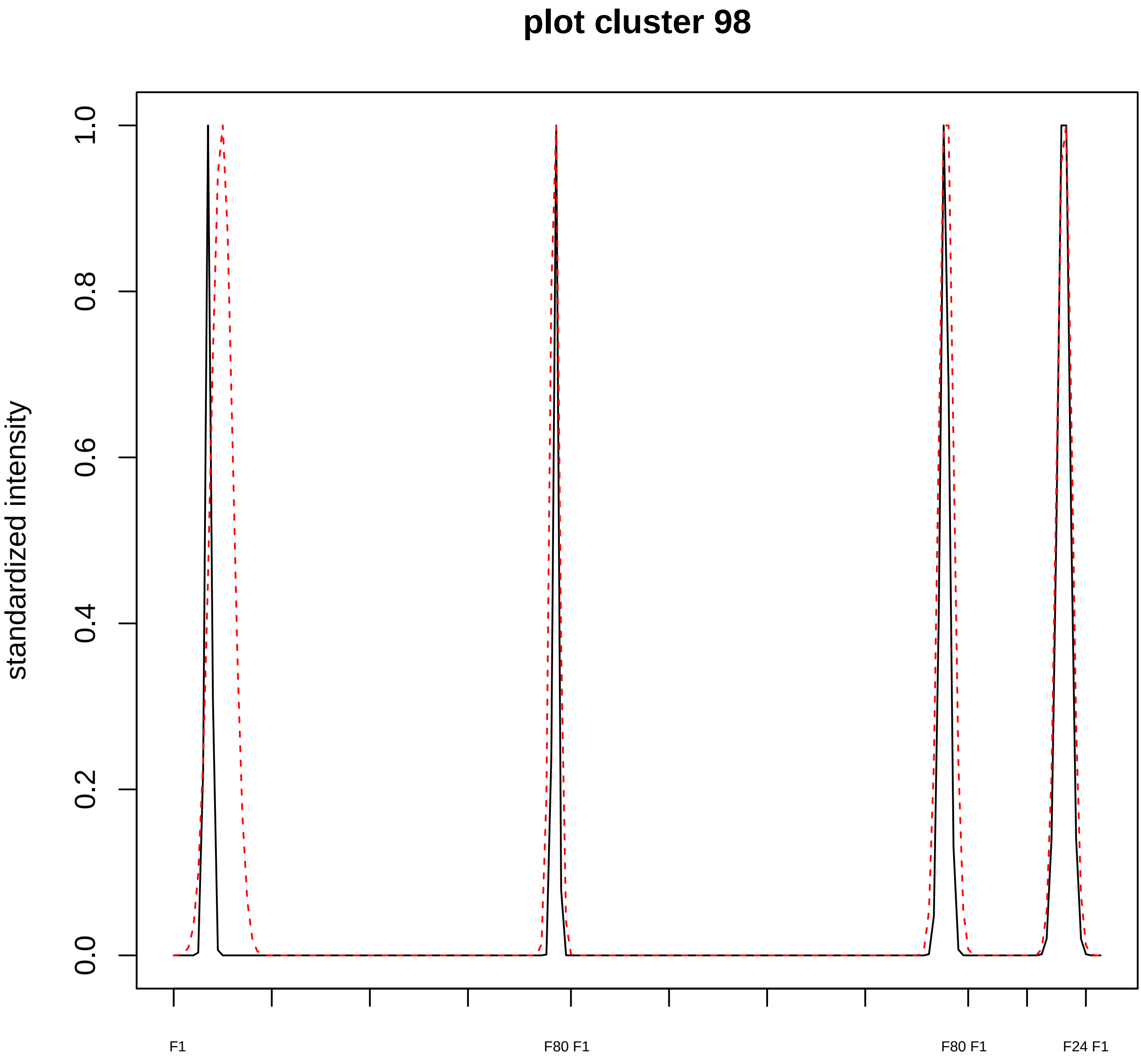
plot cluster 98
standardized intensity
0.0
0.2
0.4
0.6
0.8
1.0
F1
F80 F1
F80 F1
F24 F1

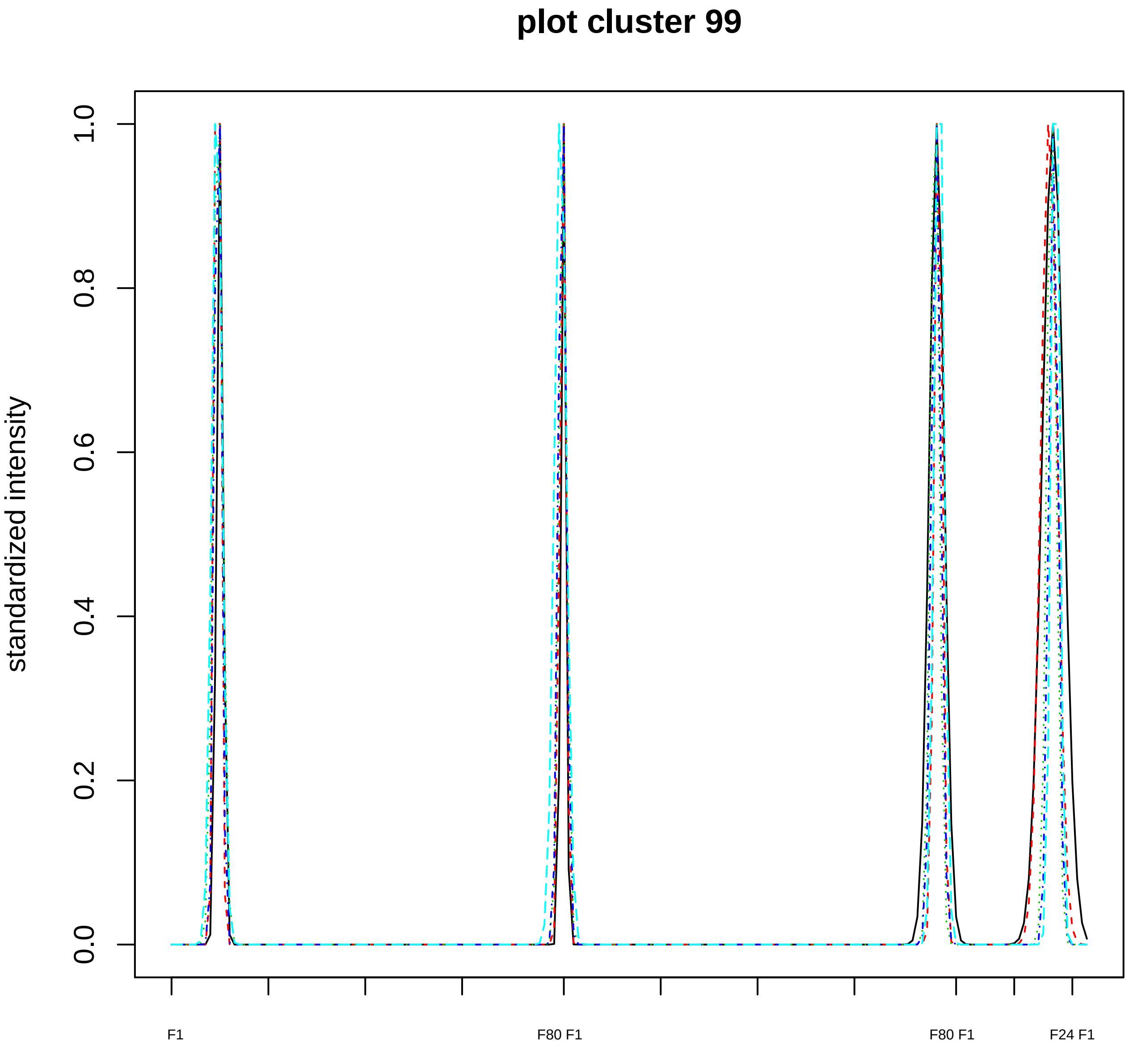

plot cluster 99
standardized intensity
0.0
0.2
0.4
0.6
0.8
1.0
F1
F80 F1
F80 F1
F24 F1

### Dendrogram of cluster 99

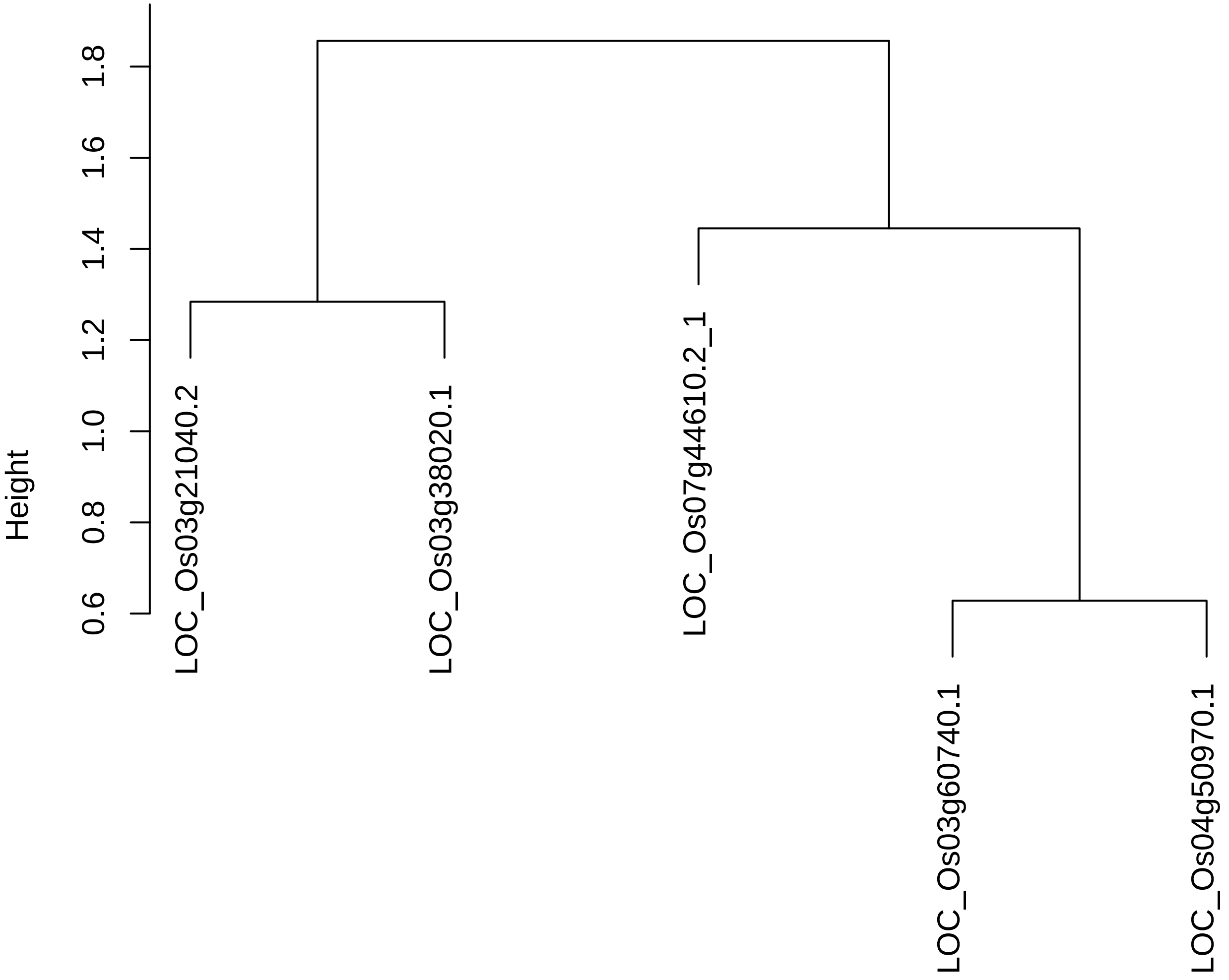


dist(spfitiex12sec12[lab[, 1] == i, ])
hclust (*, "ward.D2")

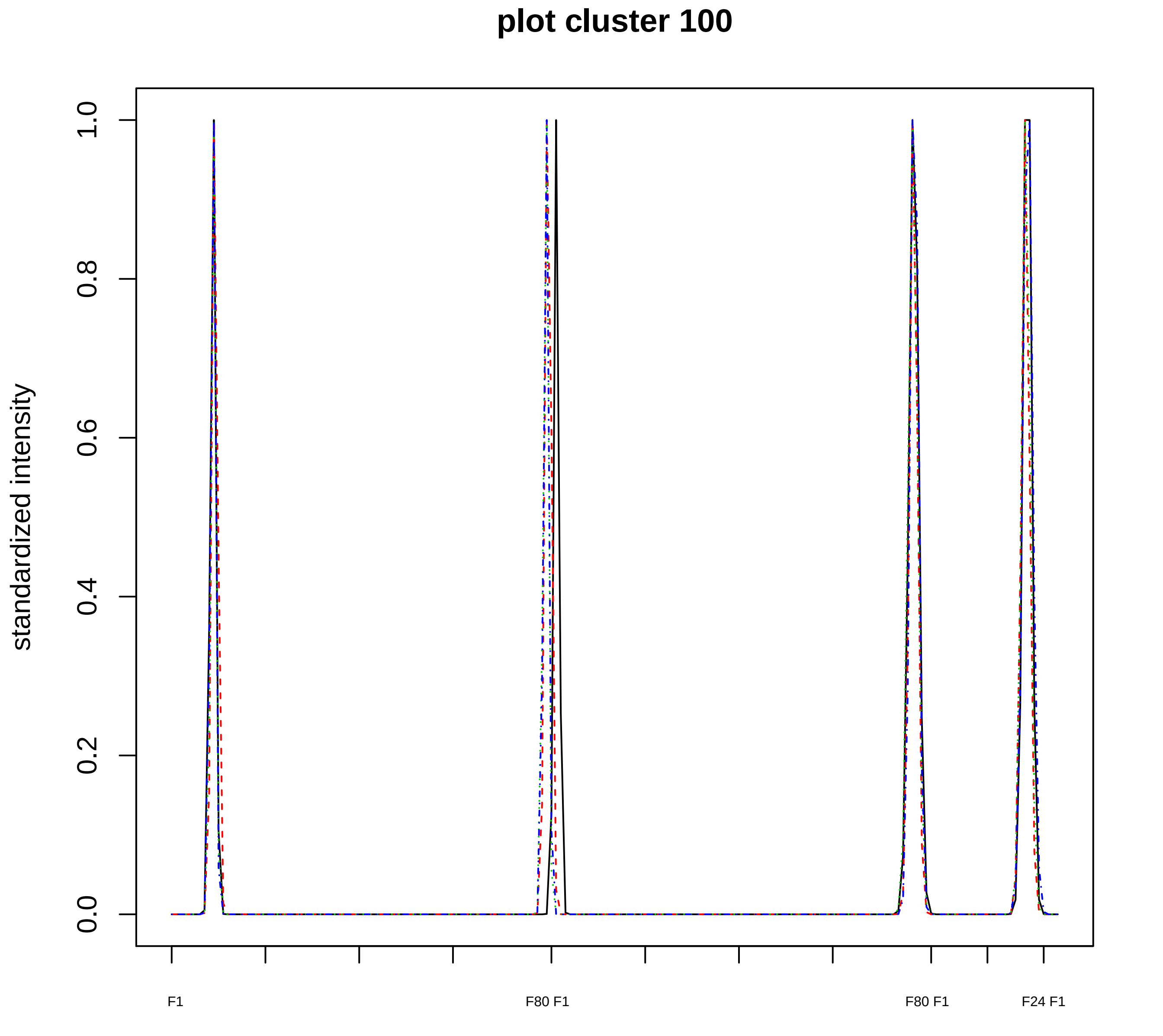
plot cluster 100
standardized intensity
0.0
0.2
0.4
0.6
0.8
1.0
F1
F80 F1
F80 F1
F24 F1

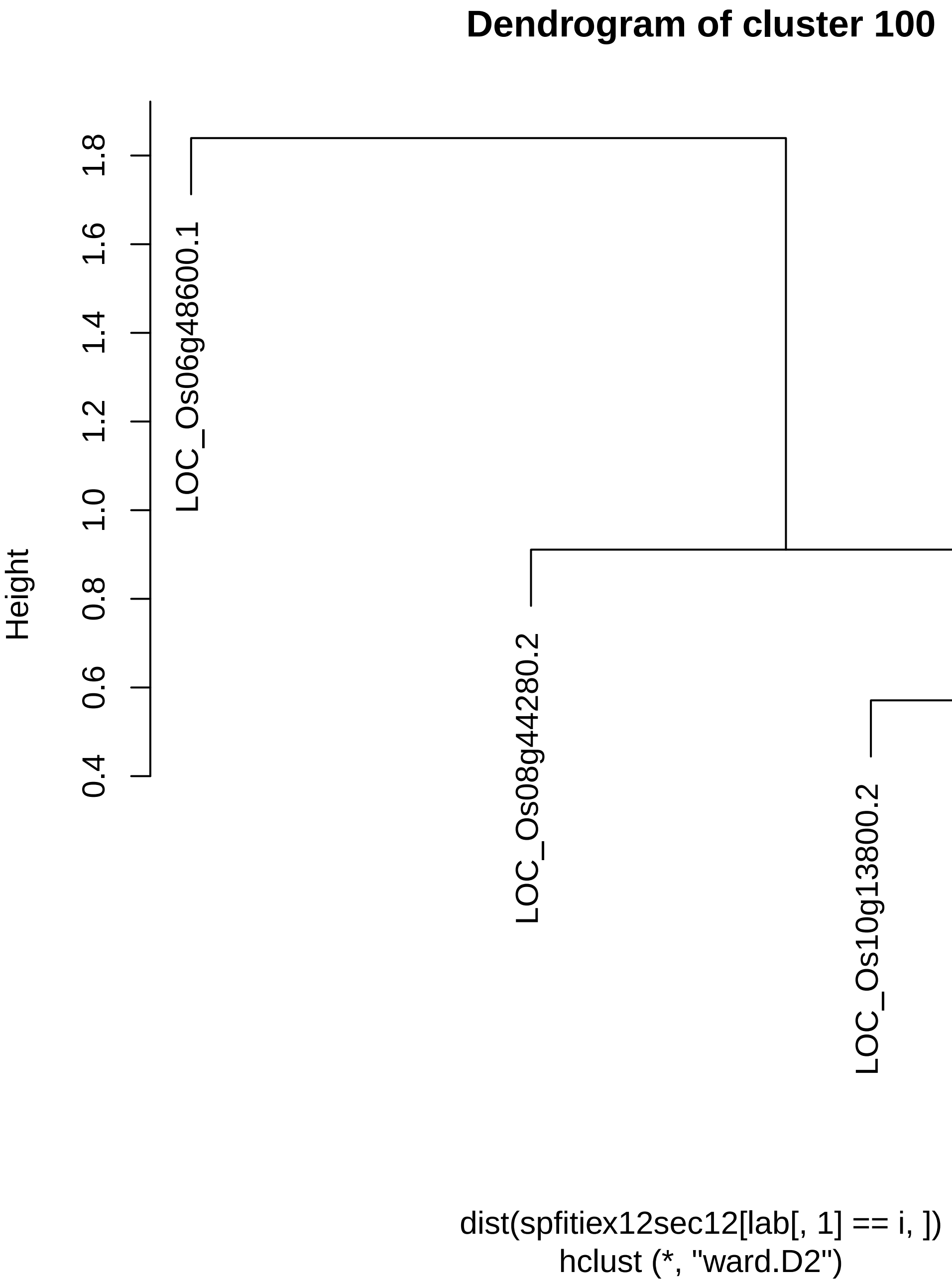
Dendrogram of cluster 100
Height
1.8
1.6
1.4
1.2
1.0
0.8
0.6
0.4
LOC_Os06g48600.1
LOC_Os08g44280.2
LOC_Os10g13800.2
LOC_Os12g01680.1
dist(spfitiex12sec12[lab[, 1] == i, ])
hclust (*, "ward.D2")

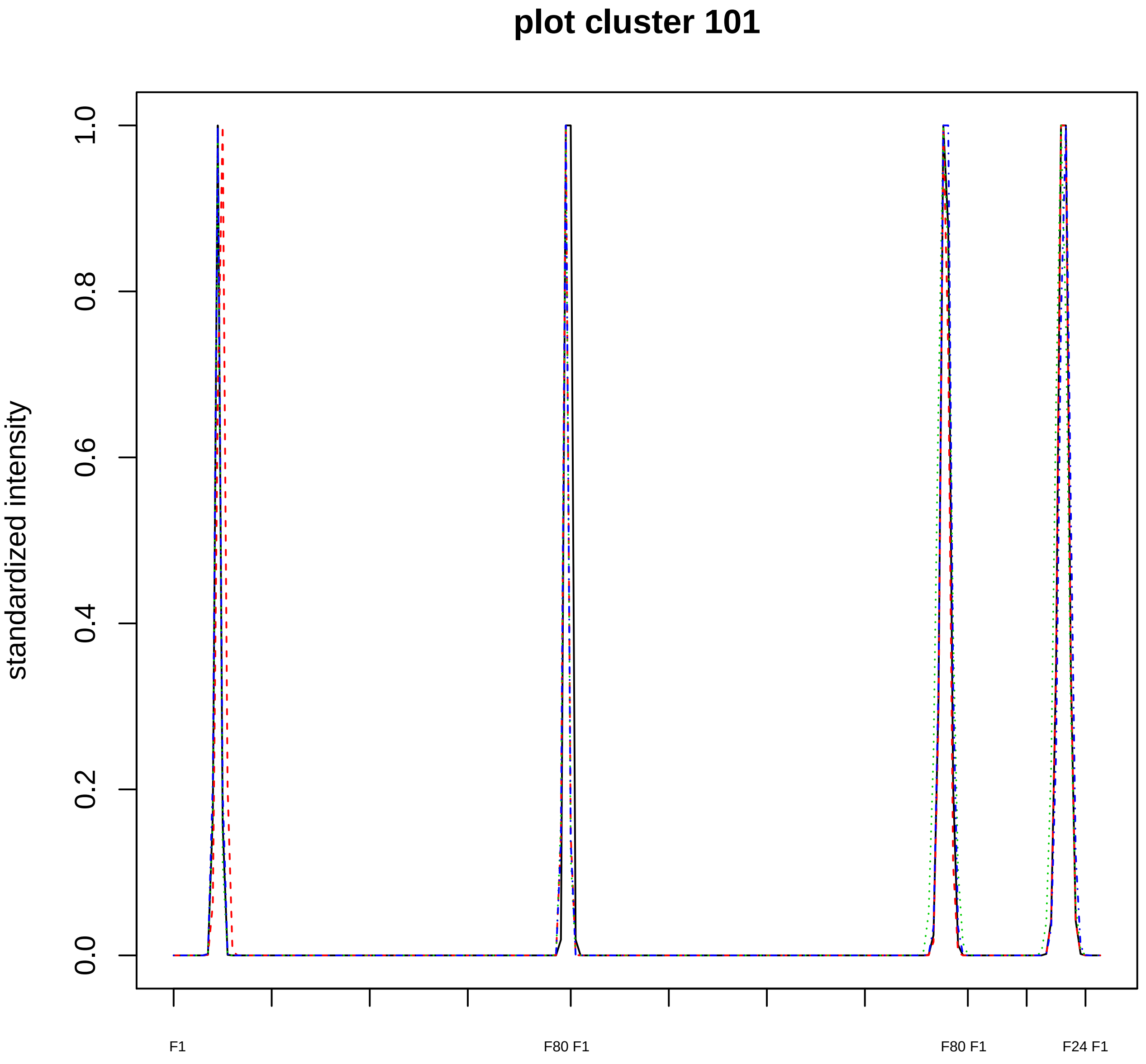
plot cluster 101
standardized intensity
1.0
0.8
0.6
0.4
0.2
0.0
F1
F80 F1
F80 F1
F24 F1

### Dendrogram of cluster 101

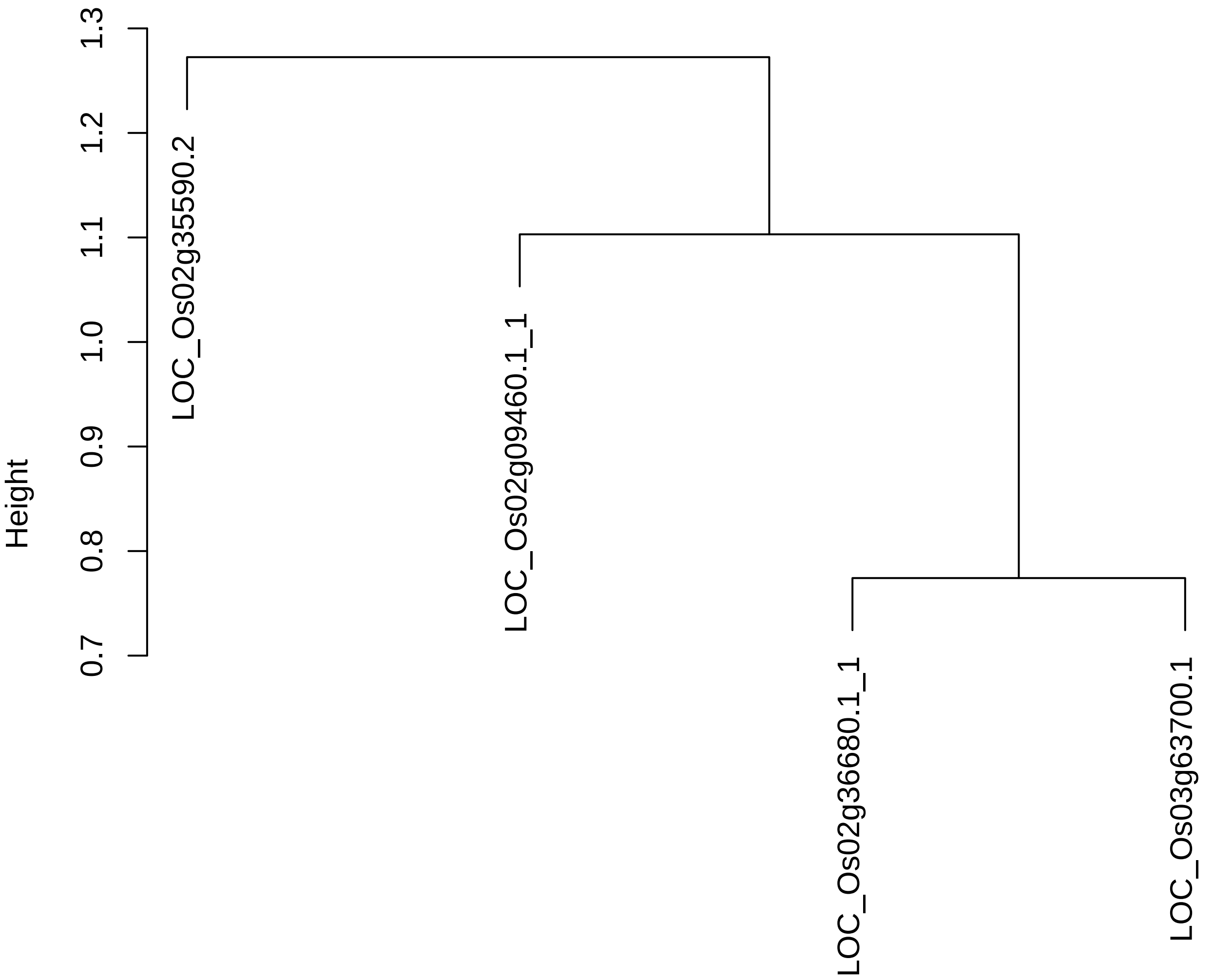


dist(spfitiex12sec12[lab[, 1] == i, ])
hclust (*, "ward.D2")

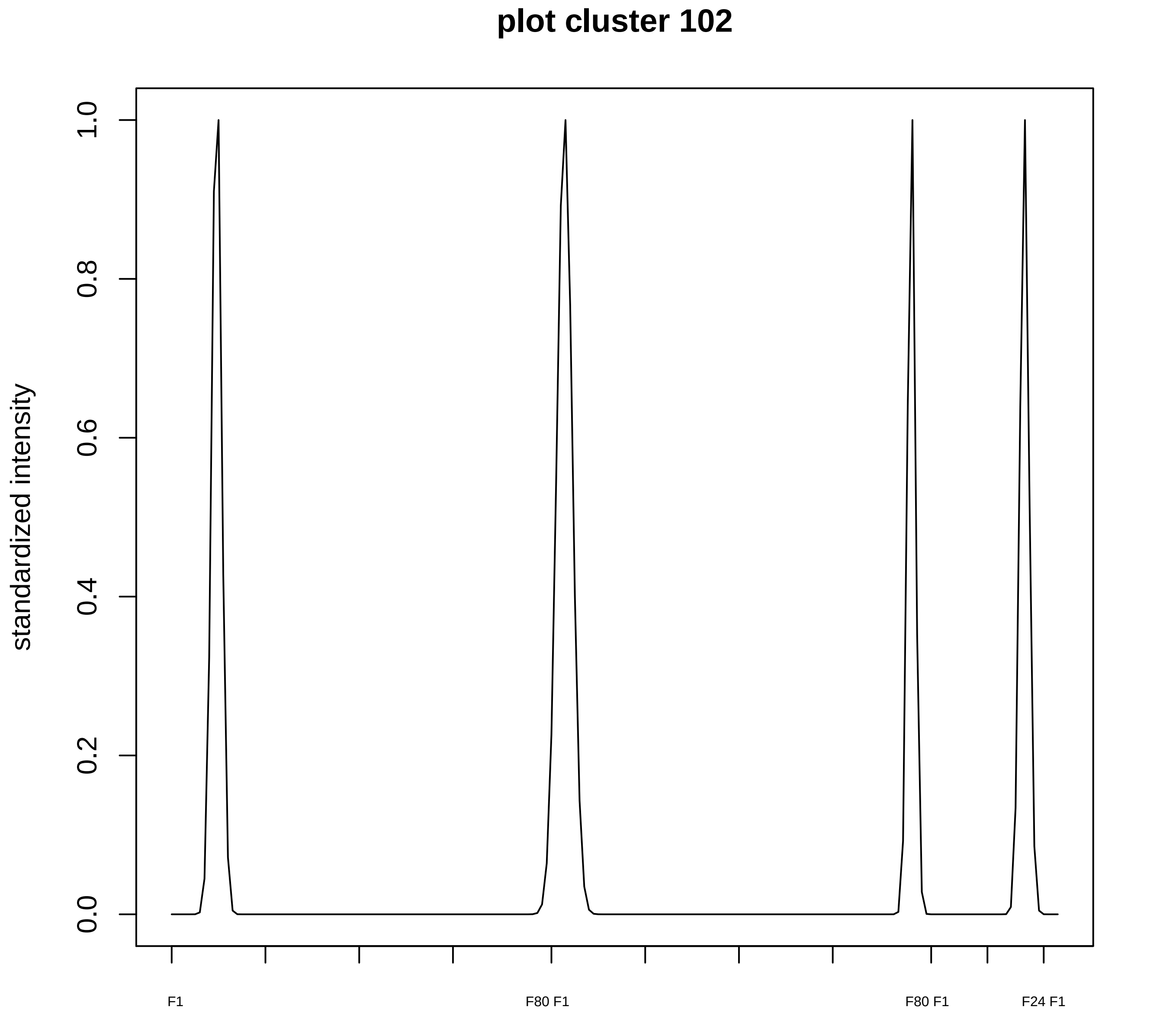
plot cluster 102
standardized intensity
0.0
0.2
0.4
0.6
0.8
1.0
F1
F80 F1
F80 F1
F24 F1

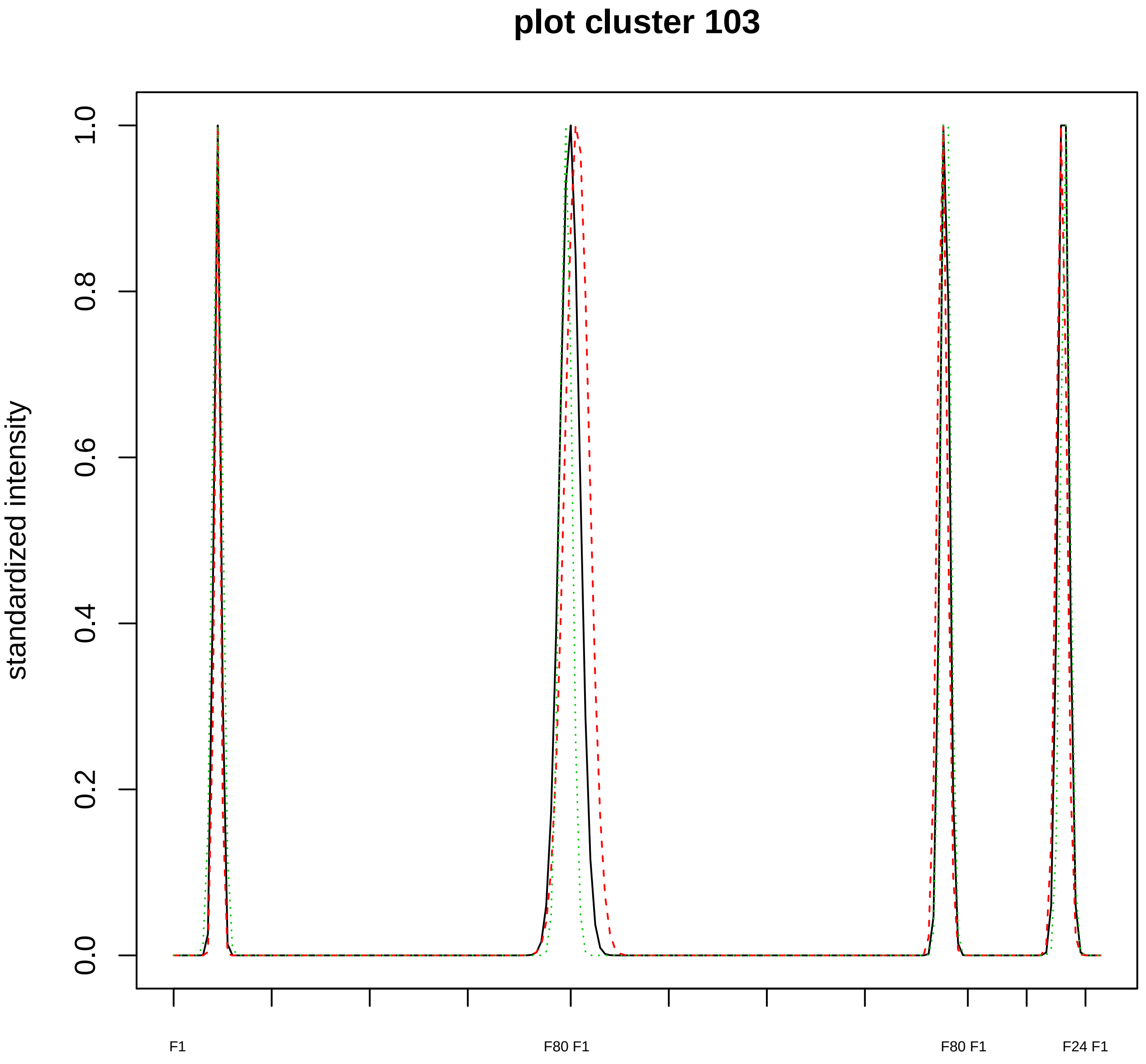
plot cluster 103
standardized intensity
0.0
0.2
0.4
0.6
0.8
1.0
F1
F80 F1
F80 F1
F24 F1

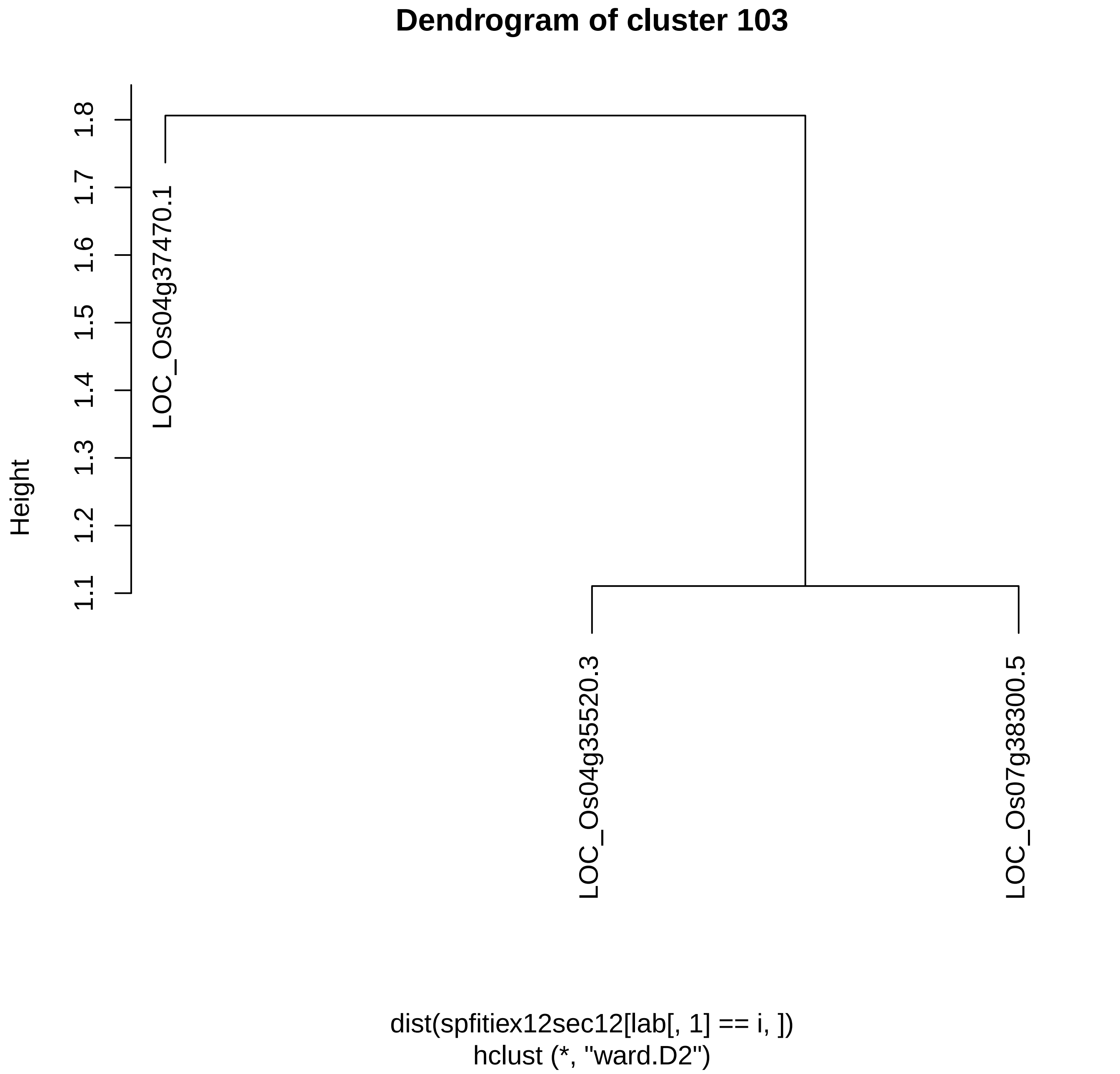
Dendrogram of cluster 103
Height
1.1
1.2
1.3
1.4
1.5
1.6
1.7
1.8
LOC_Os04g37470.1
LOC_Os04g35520.3
LOC_Os07g38300.5
dist(spfitiex12sec12[lab[, 1] == i, ])
hclust (*, "ward.D2")

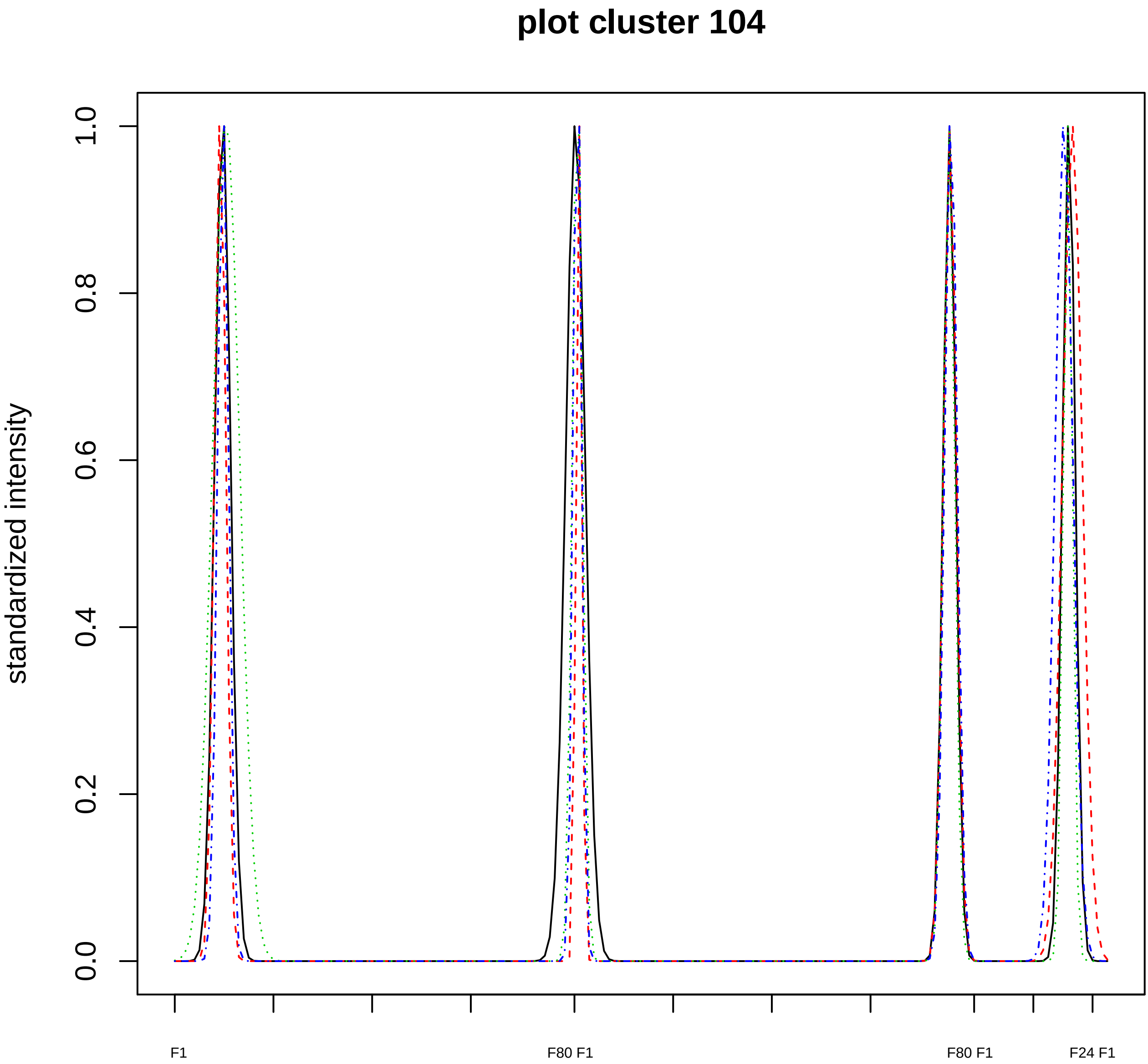
plot cluster 104
standardized intensity
0.0
0.2
0.4
0.6
0.8
1.0
F1
F80 F1
F80 F1
F24 F1

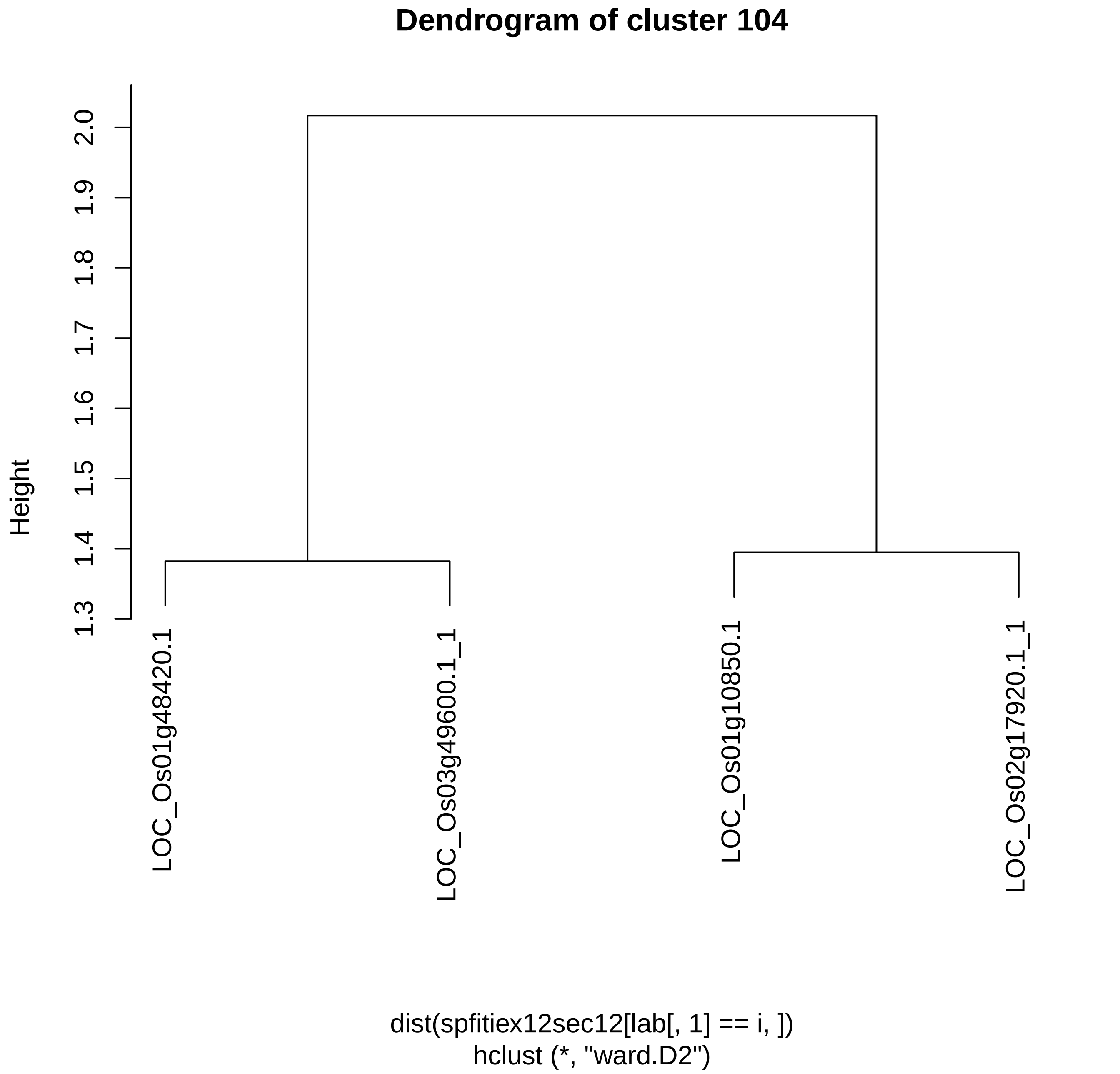

Dendrogram of cluster 104
Height
2.0
1.9
1.8
1.7
1.6
1.5
1.4
1.3
LOC_Os01g48420.1
LOC_Os03g49600.1_1
LOC_Os01g10850.1
LOC_Os02g17920.1_1
dist(spfitiex12sec12[lab[, 1] == i, ])
hclust (*, "ward.D2")

#### plot cluster 105

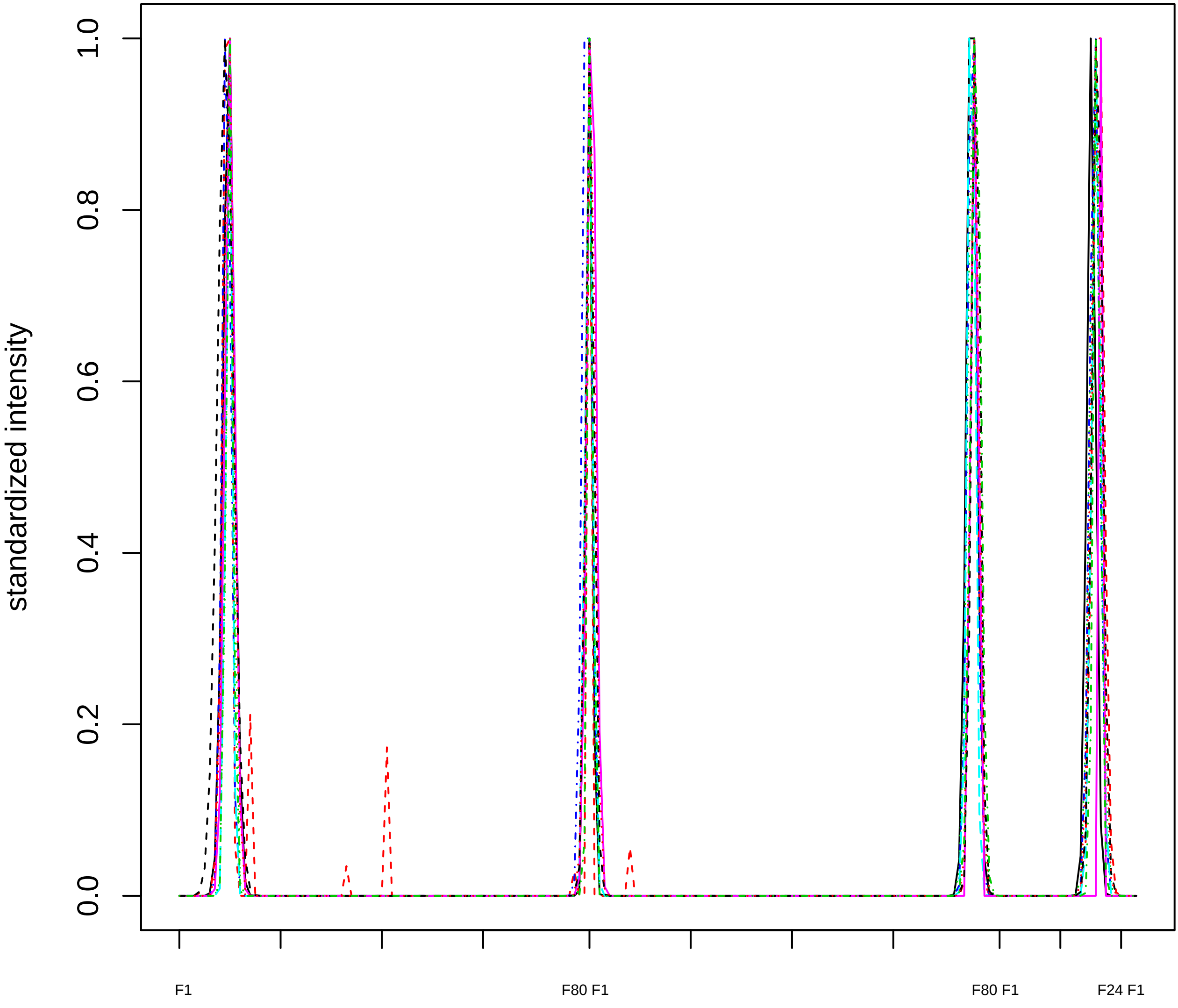

### Dendrogram of cluster 105

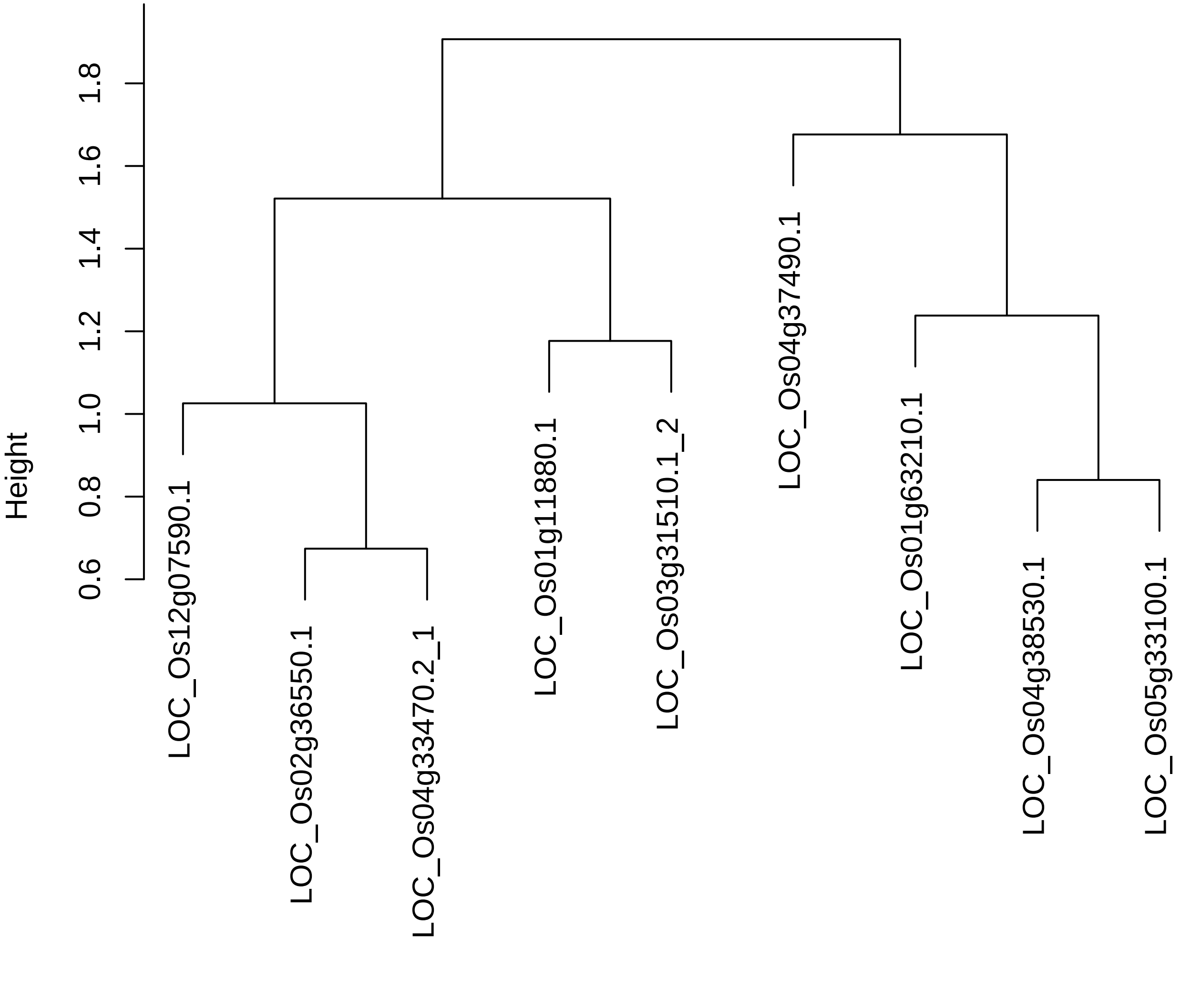


dist(spfitiex12sec12[lab[, 1] == i, ])
hclust (*, "ward.D2")

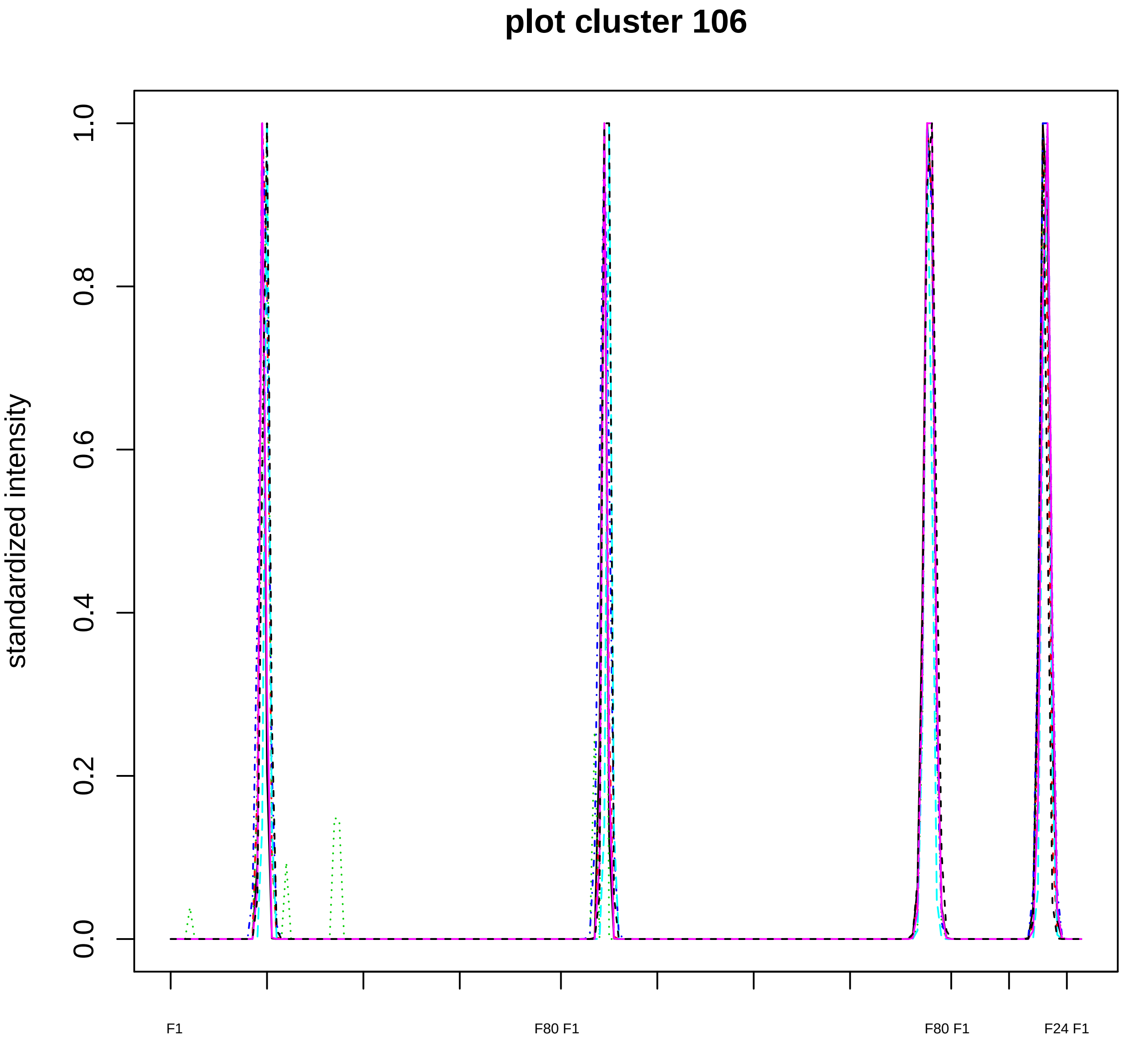
plot cluster 106
standardized intensity
0.0
0.2
0.4
0.6
0.8
1.0
F1
F80 F1
F80 F1
F24 F1

#### Dendrogram of cluster 106

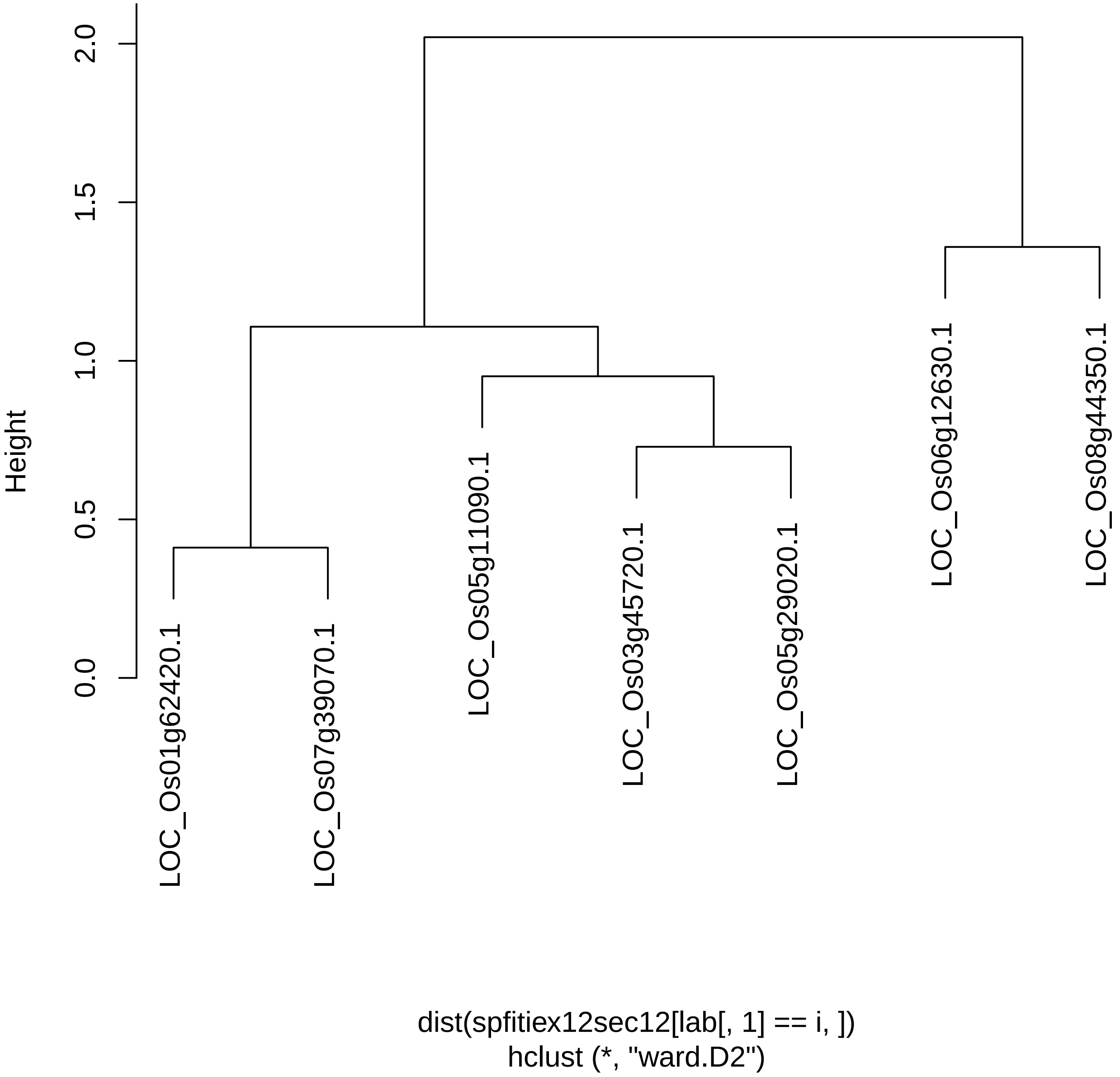

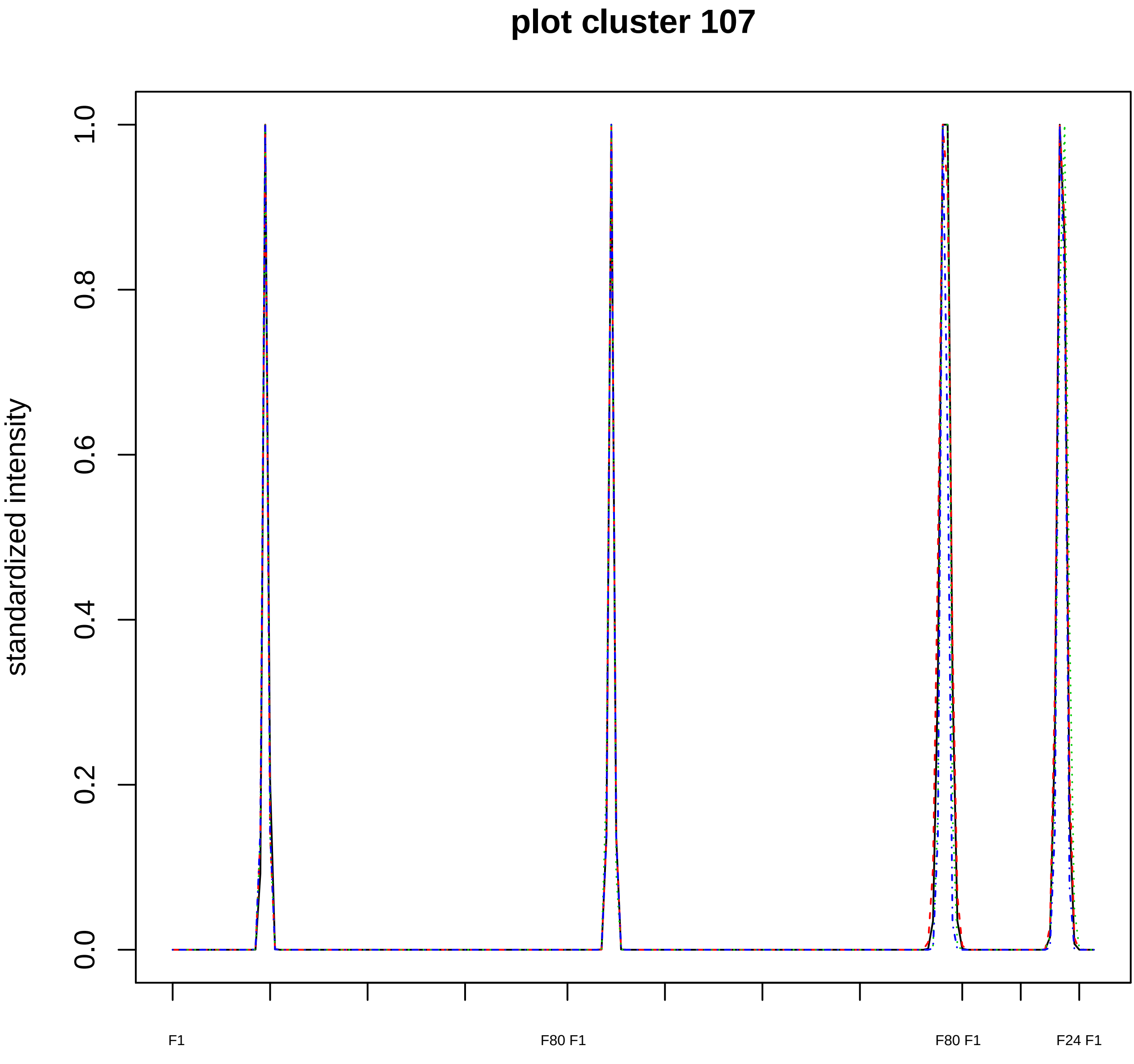

plot cluster 107
standardized intensity
0.0
0.2
0.4
0.6
0.8
1.0
F1
F80 F1
F80 F1
F24 F1

### Dendrogram of cluster 107

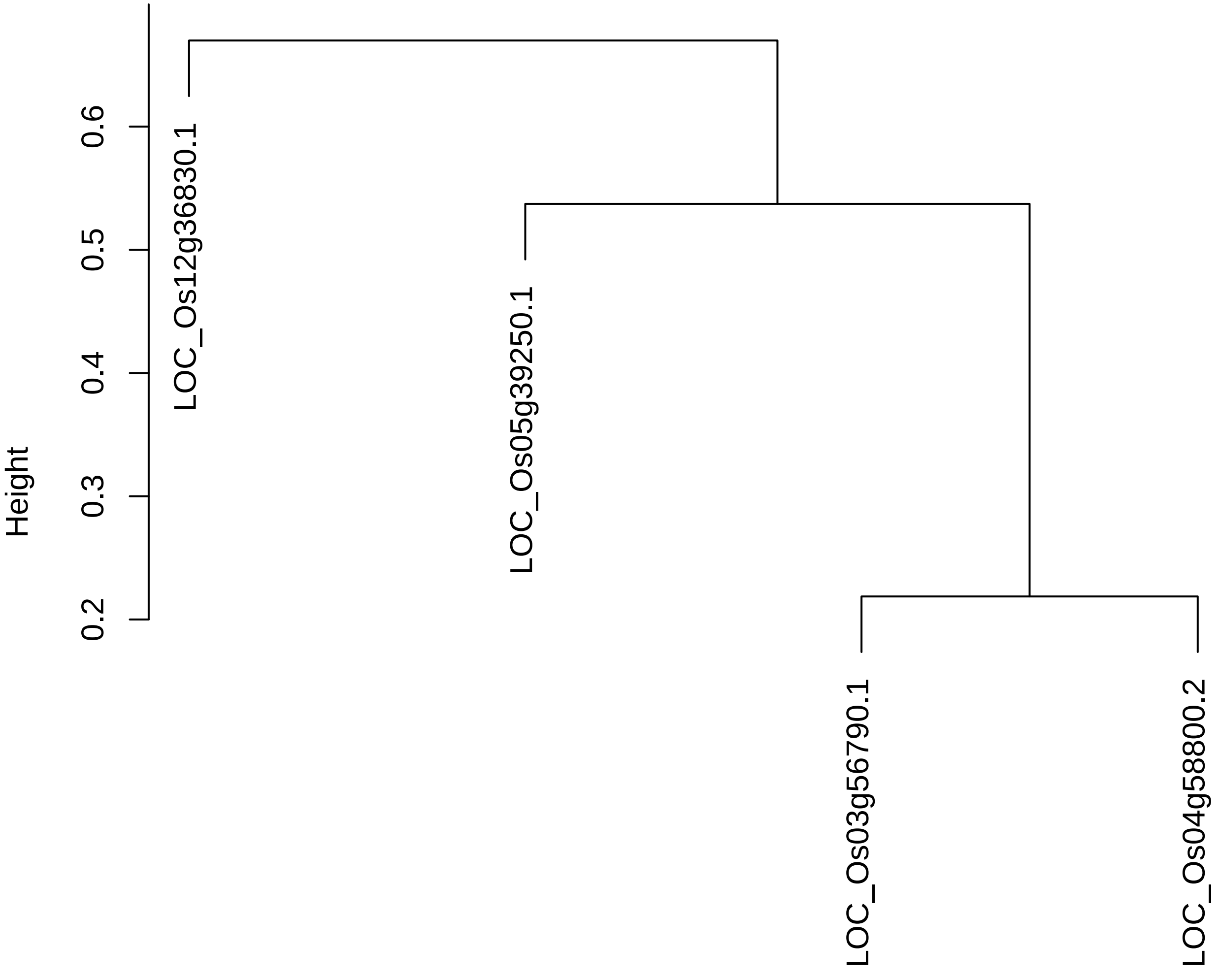


dist(spfitiex12sec12[lab[, 1] == i, ])
hclust (*, "ward.D2")

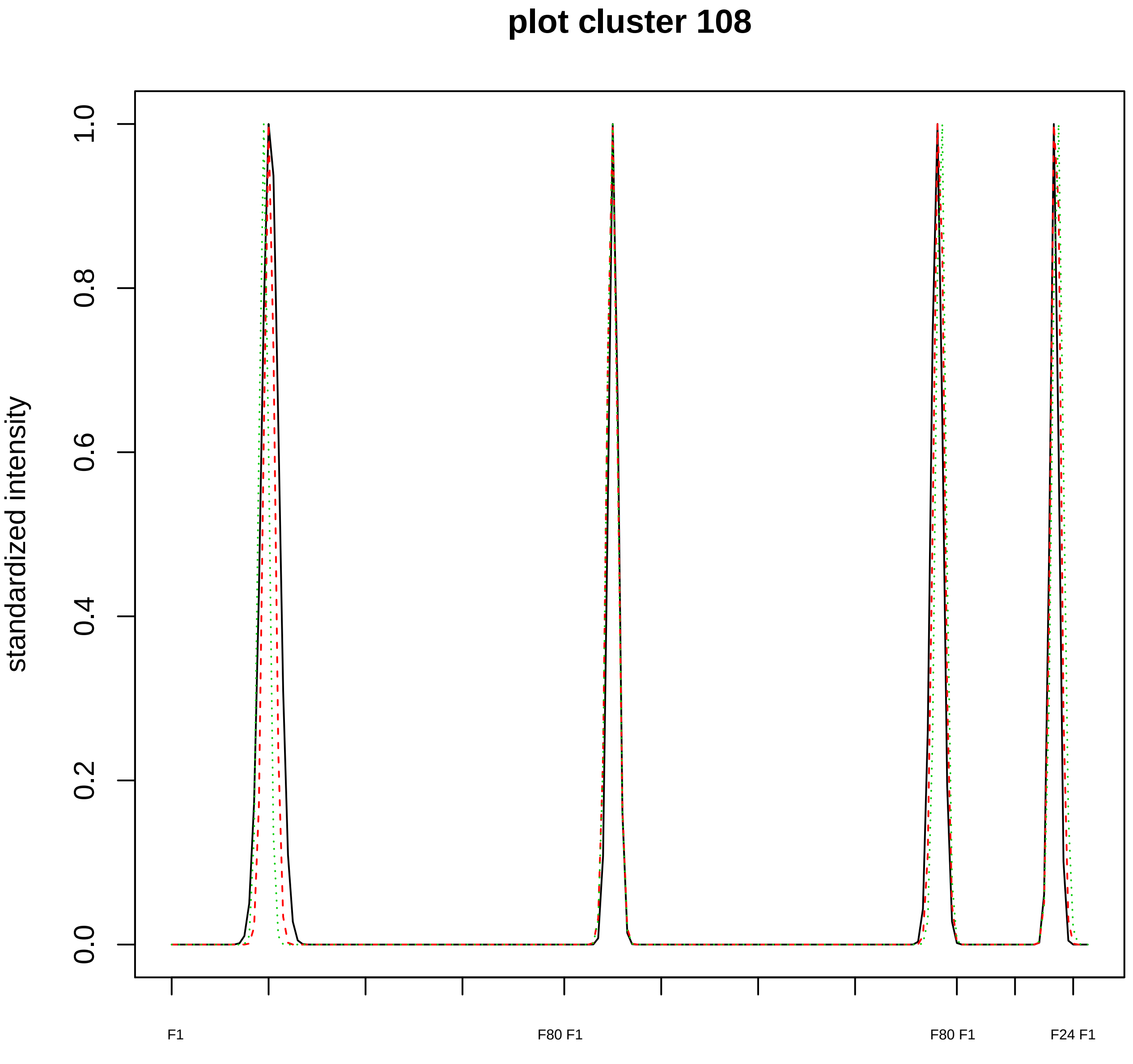

plot cluster 108
standardized intensity
0.0
0.2
0.4
0.6
0.8
1.0
F1
F80 F1
F80 F1
F24 F1

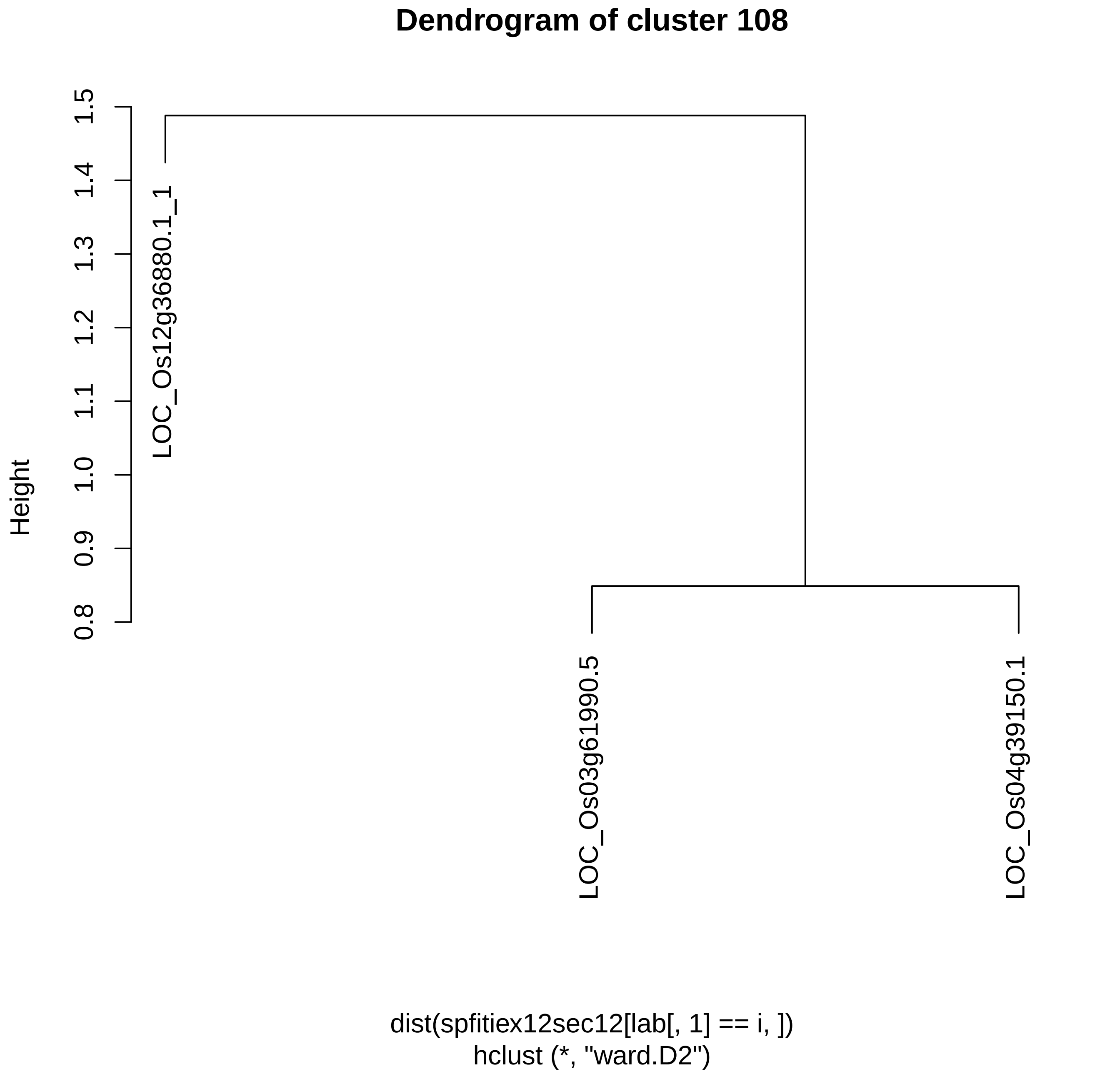
Dendrogram of cluster 108
Height
1.5
1.4
1.3
1.2
1.1
1.0
0.9
0.8
LOC_Os12g36880.1_1
LOC_Os03g61990.5
LOC_Os04g39150.1
dist(spfitiex12sec12[lab[, 1] == i, ])
hclust (*, "ward.D2")

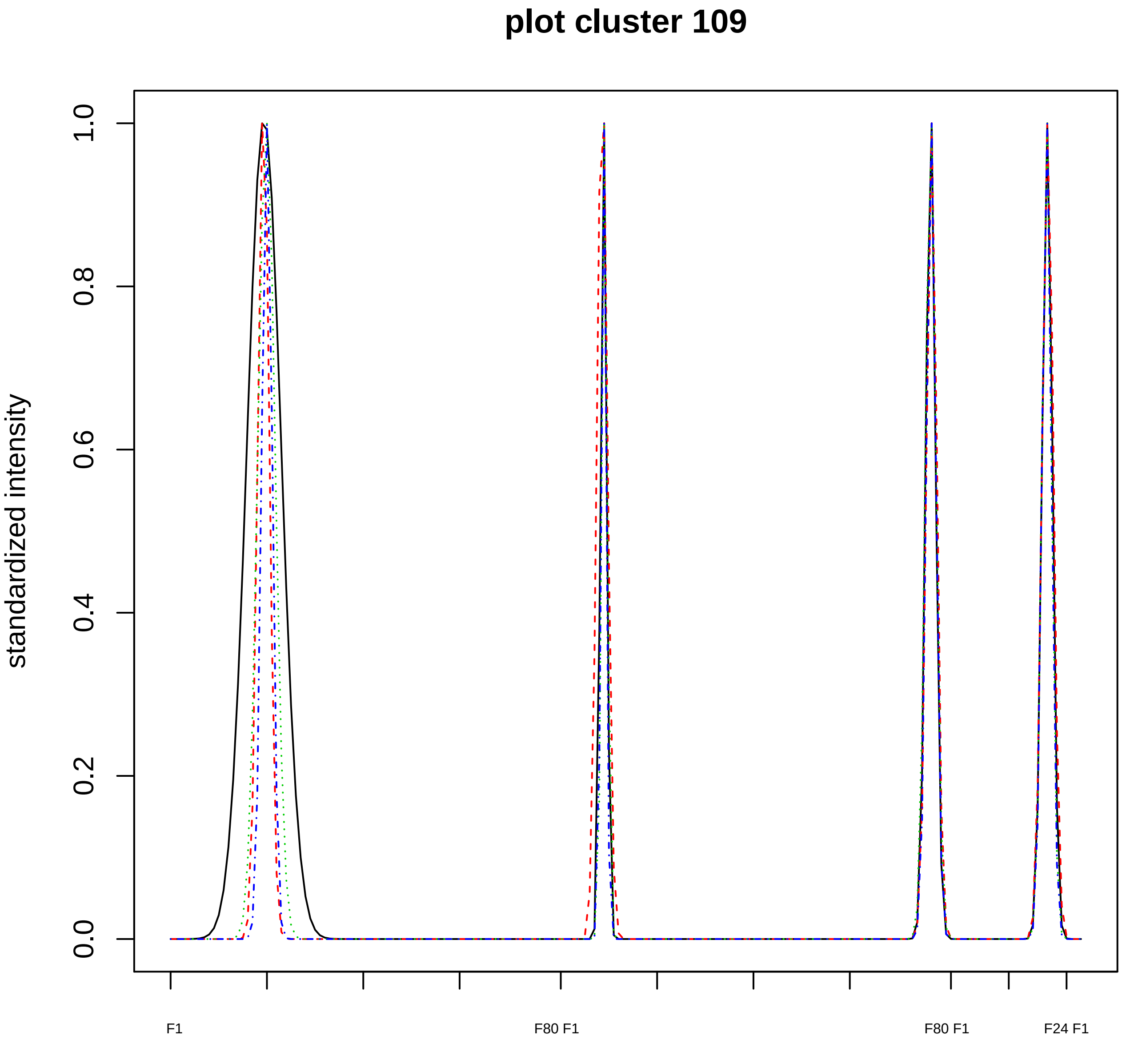
plot cluster 109
standardized intensity
0.0
0.2
0.4
0.6
0.8
1.0
F1
F80 F1
F80 F1
F24 F1

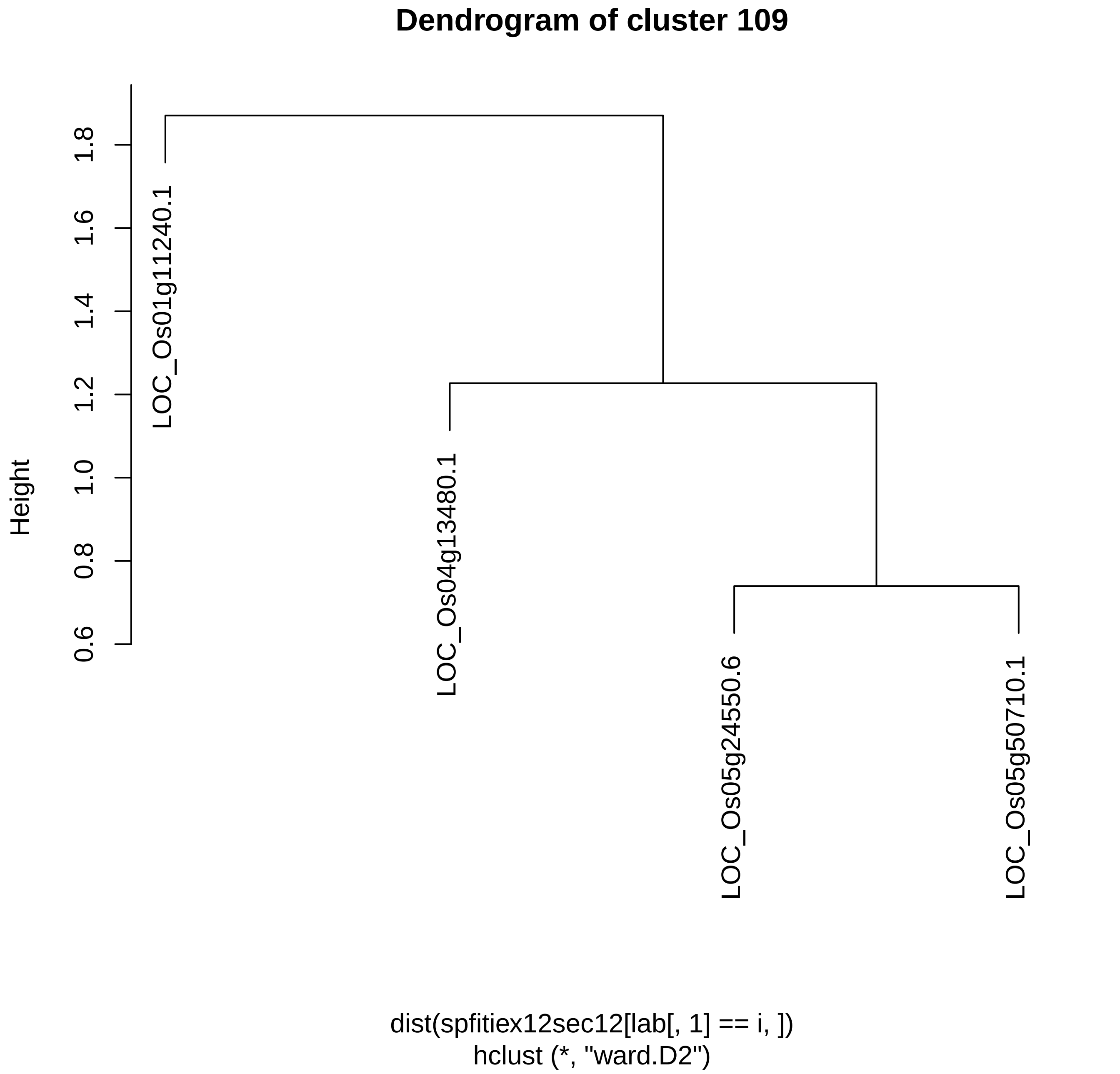

Dendrogram of cluster 109
Height
0.6
0.8
1.0
1.2
1.4
1.6
1.8
LOC_Os01g11240.1
LOC_Os04g13480.1
LOC_Os05g24550.6
LOC_Os05g50710.1
dist(spfitiex12sec12[lab[, 1] == i, ])
hclust (*, "ward.D2")

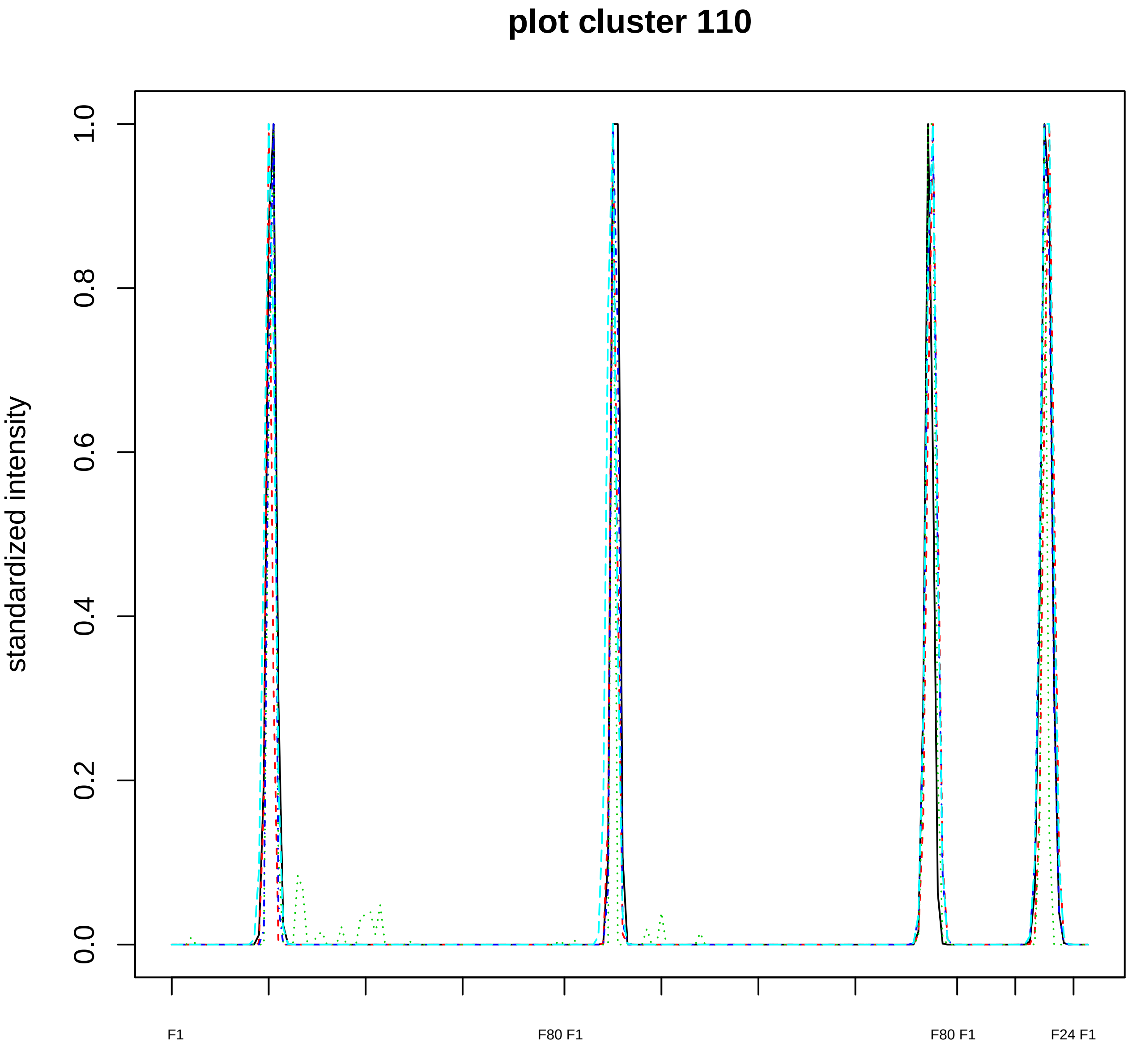
plot cluster 110
standardized intensity
0.0
0.2
0.4
0.6
0.8
1.0
F1
F80 F1
F80 F1
F24 F1

#### Dendrogram of cluster 110

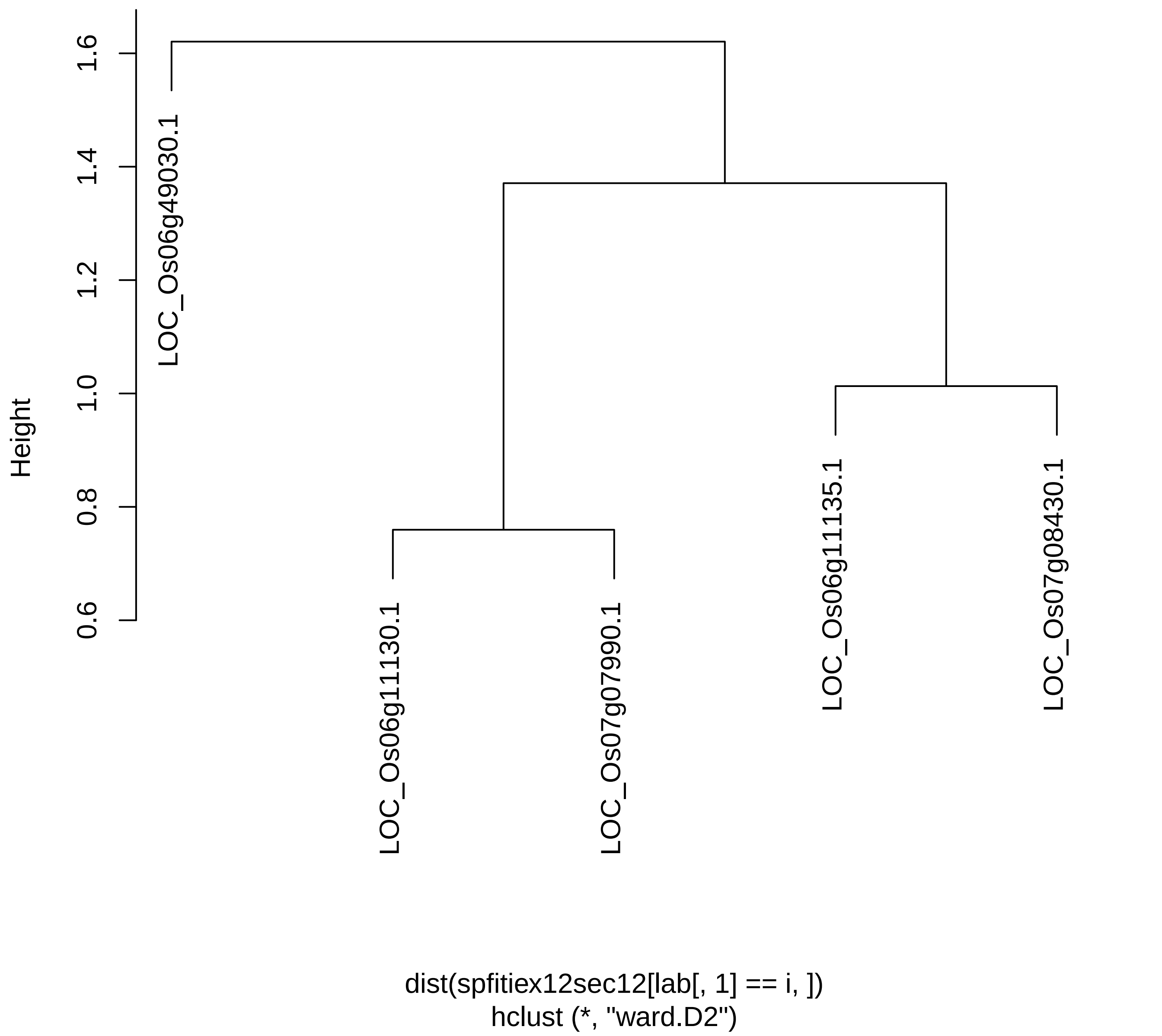

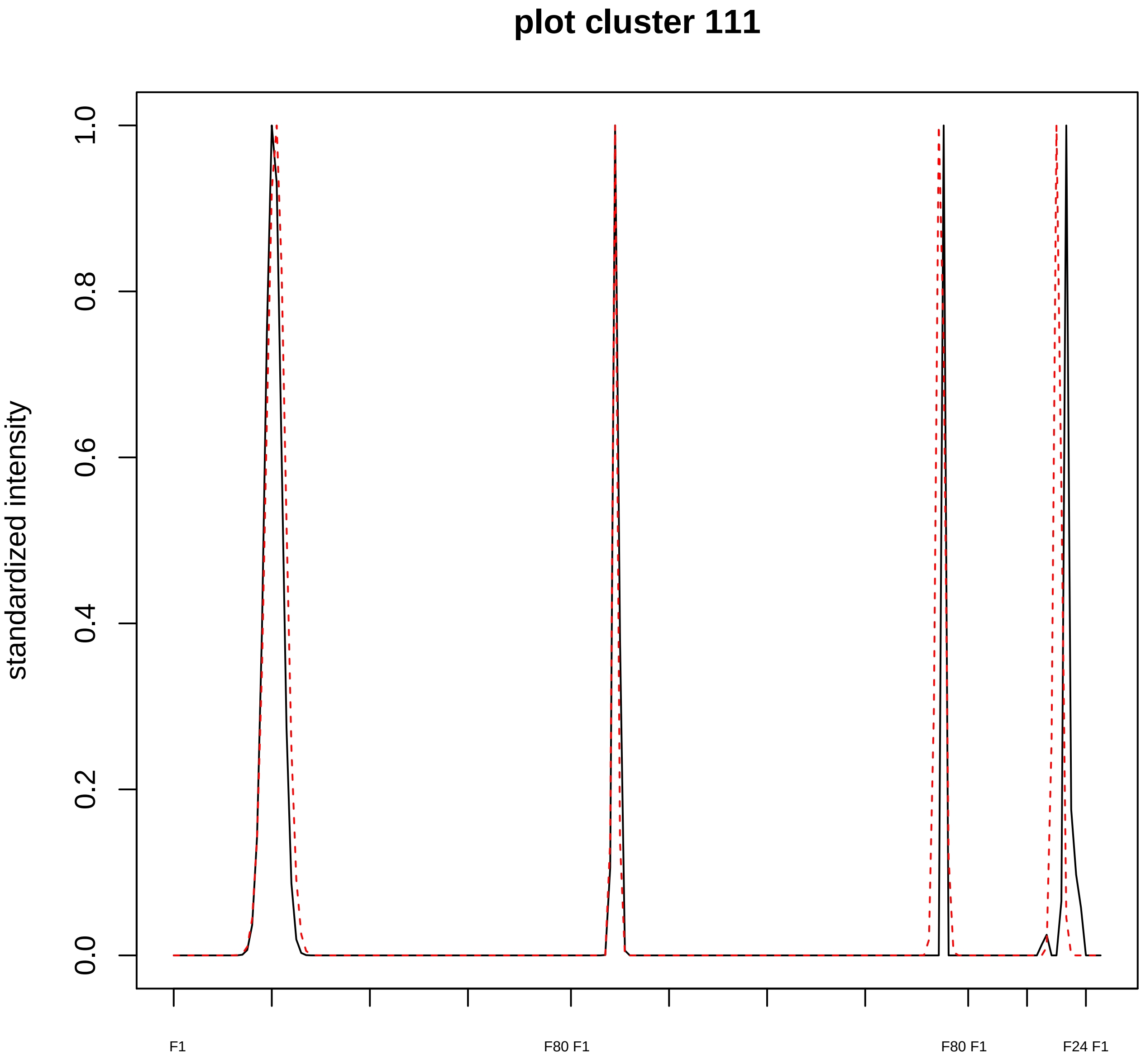
plot cluster 111
standardized intensity
0.0
0.2
0.4
0.6
0.8
1.0
F1
F80 F1
F80 F1
F24 F1

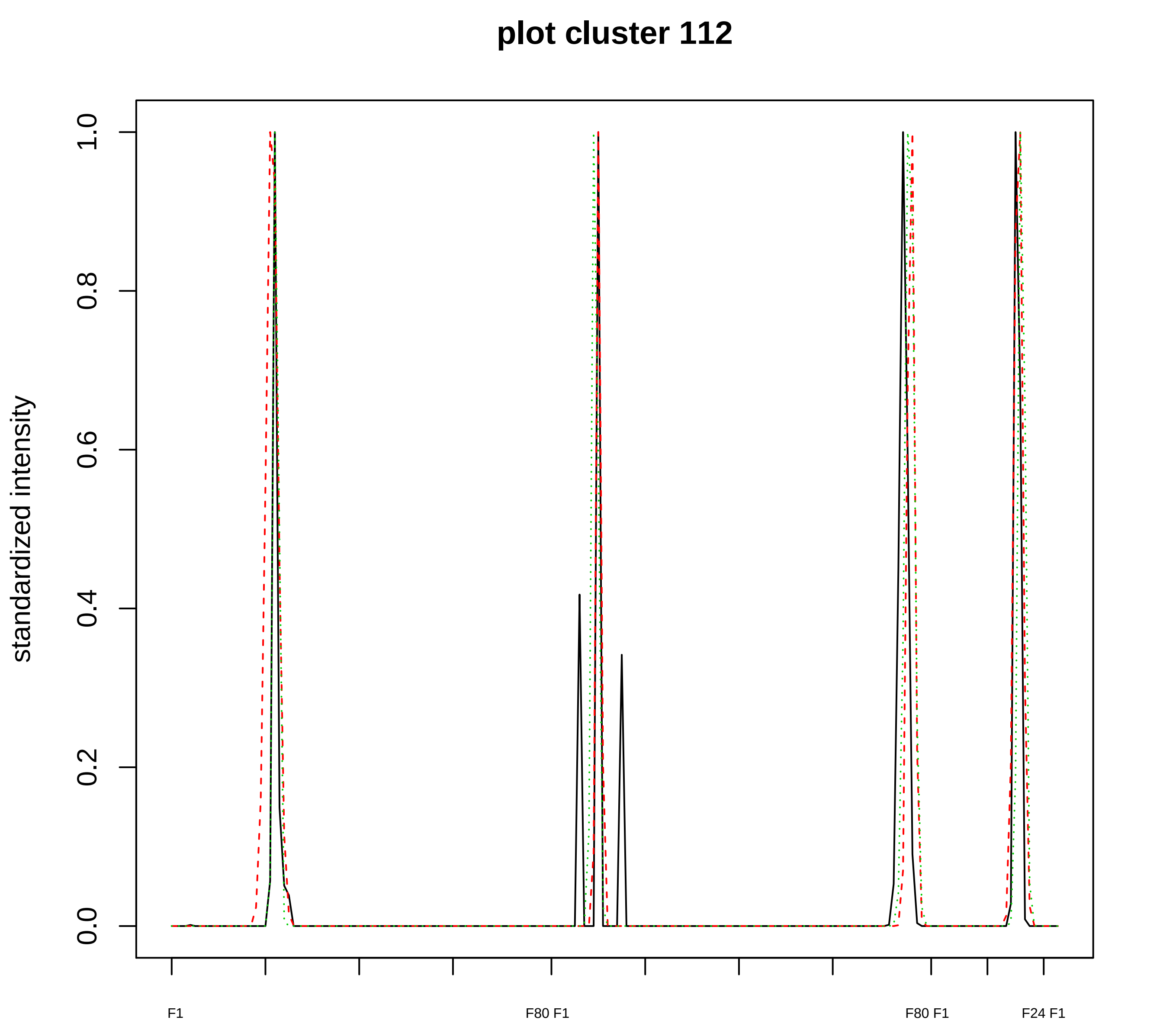
plot cluster 112
standardized intensity
0.0
0.2
0.4
0.6
0.8
1.0
F1
F80 F1
F80 F1
F24 F1

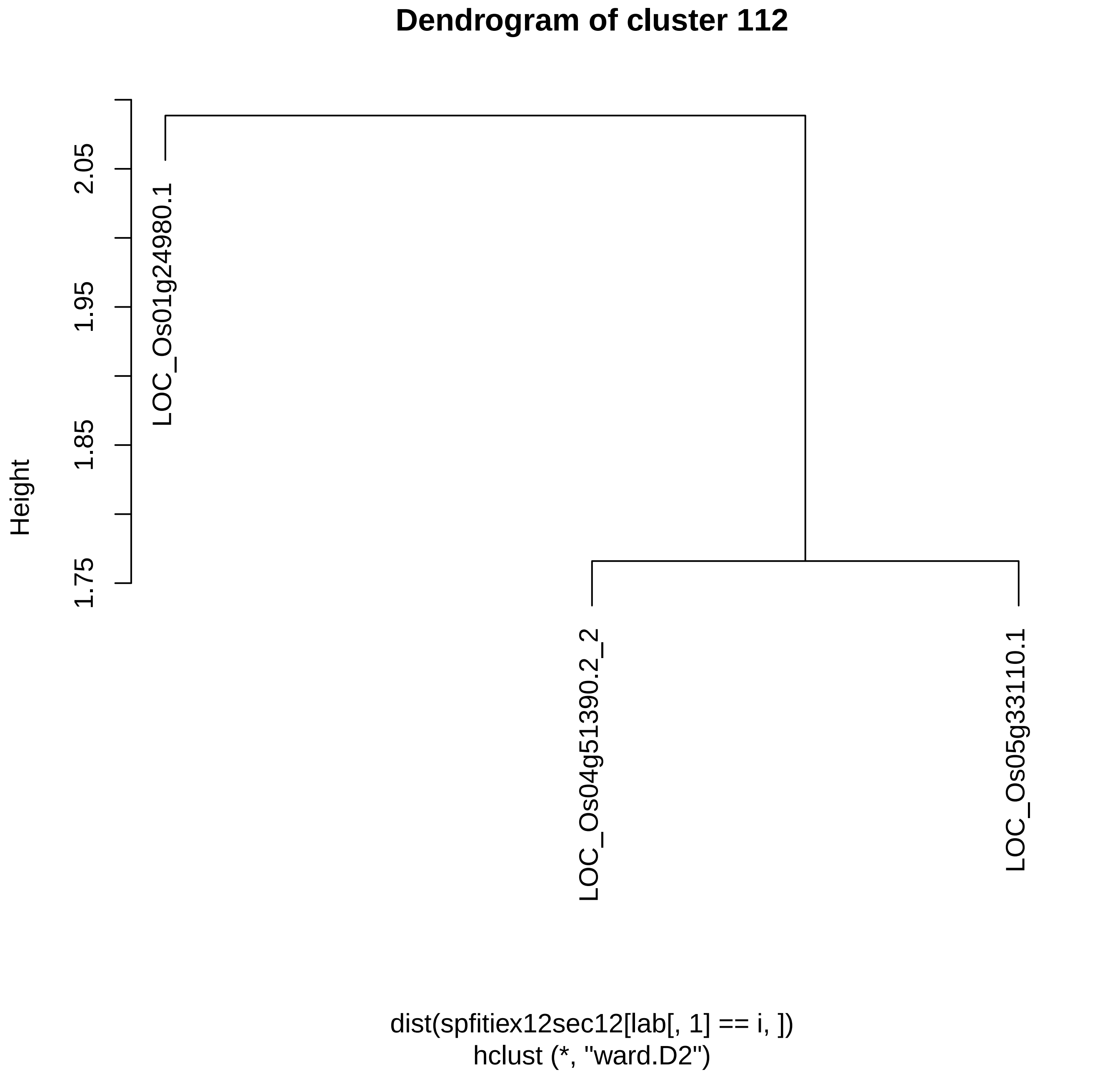
Dendrogram of cluster 112
Height
2.05
1.95
1.85
1.75
LOC_Os01g24980.1
LOC_Os04g51390.2_2
LOC_Os05g33110.1
dist(spfitiex12sec12[lab[, 1] == i, ])
hclust (*, "ward.D2")

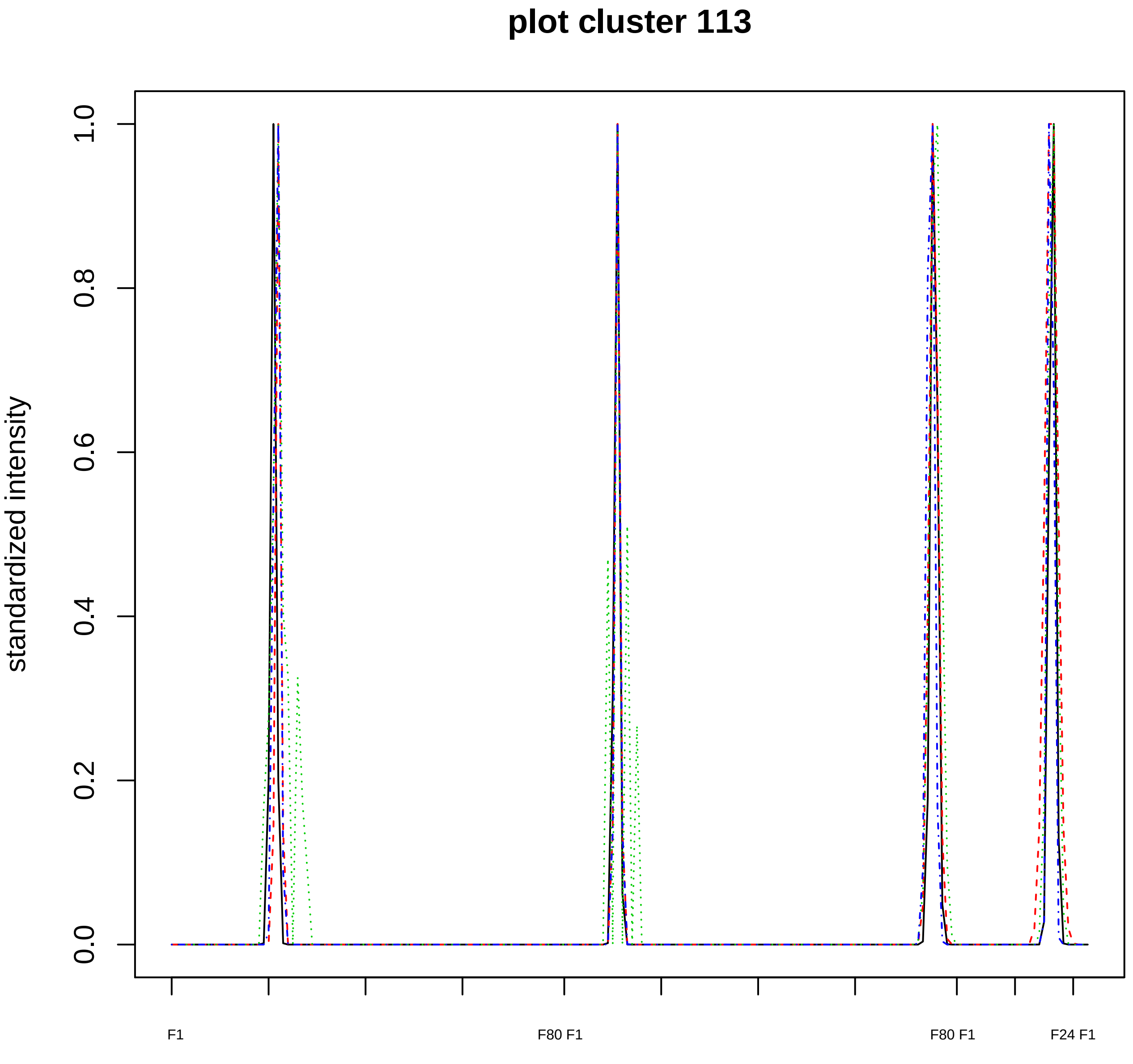
plot cluster 113
standardized intensity
0.0
0.2
0.4
0.6
0.8
1.0
F1
F80 F1
F80 F1
F24 F1

### Dendrogram of cluster 113

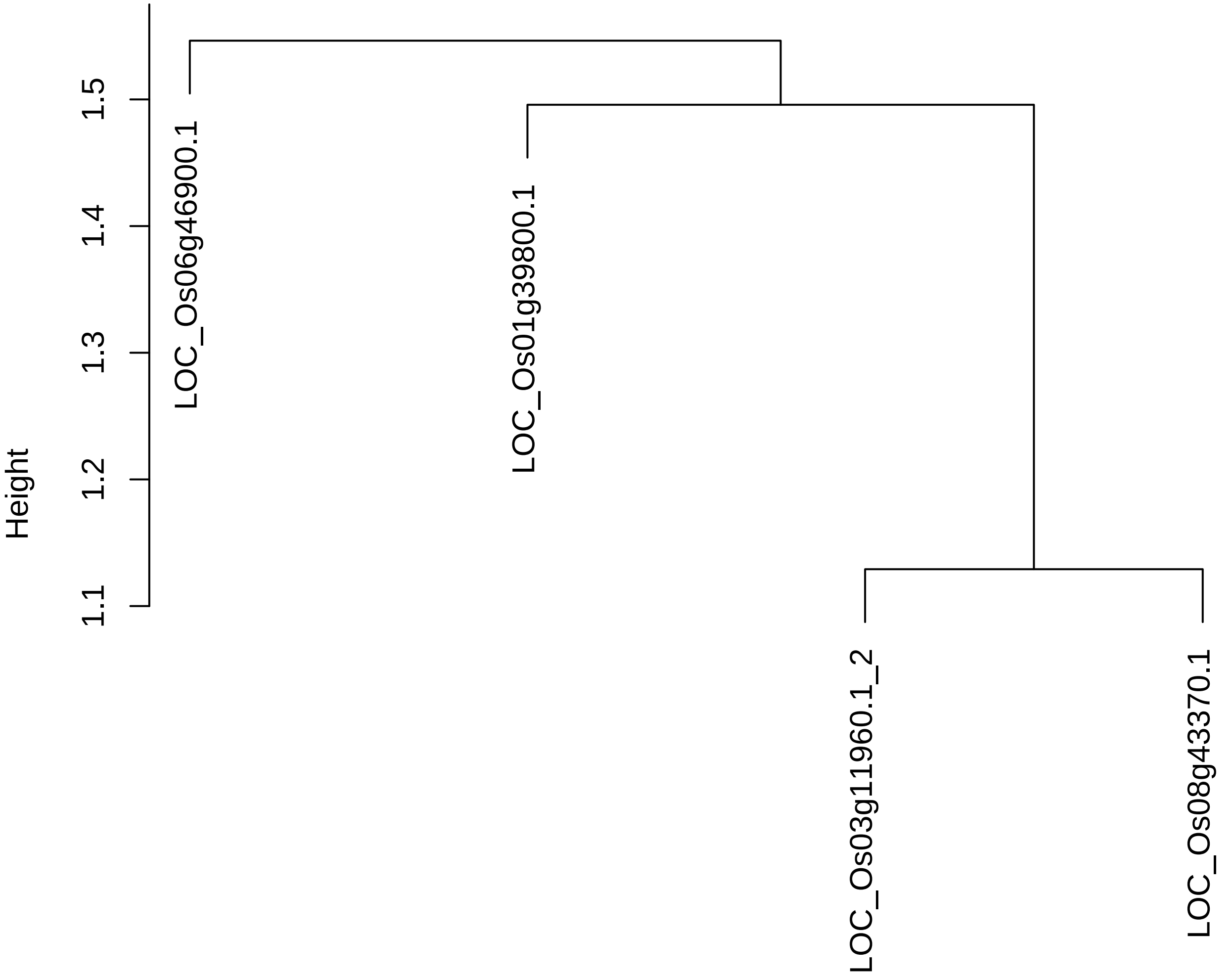


dist(spfitiex12sec12[lab[, 1] == i, ])
hclust (*, "ward.D2")

#### plot cluster 114

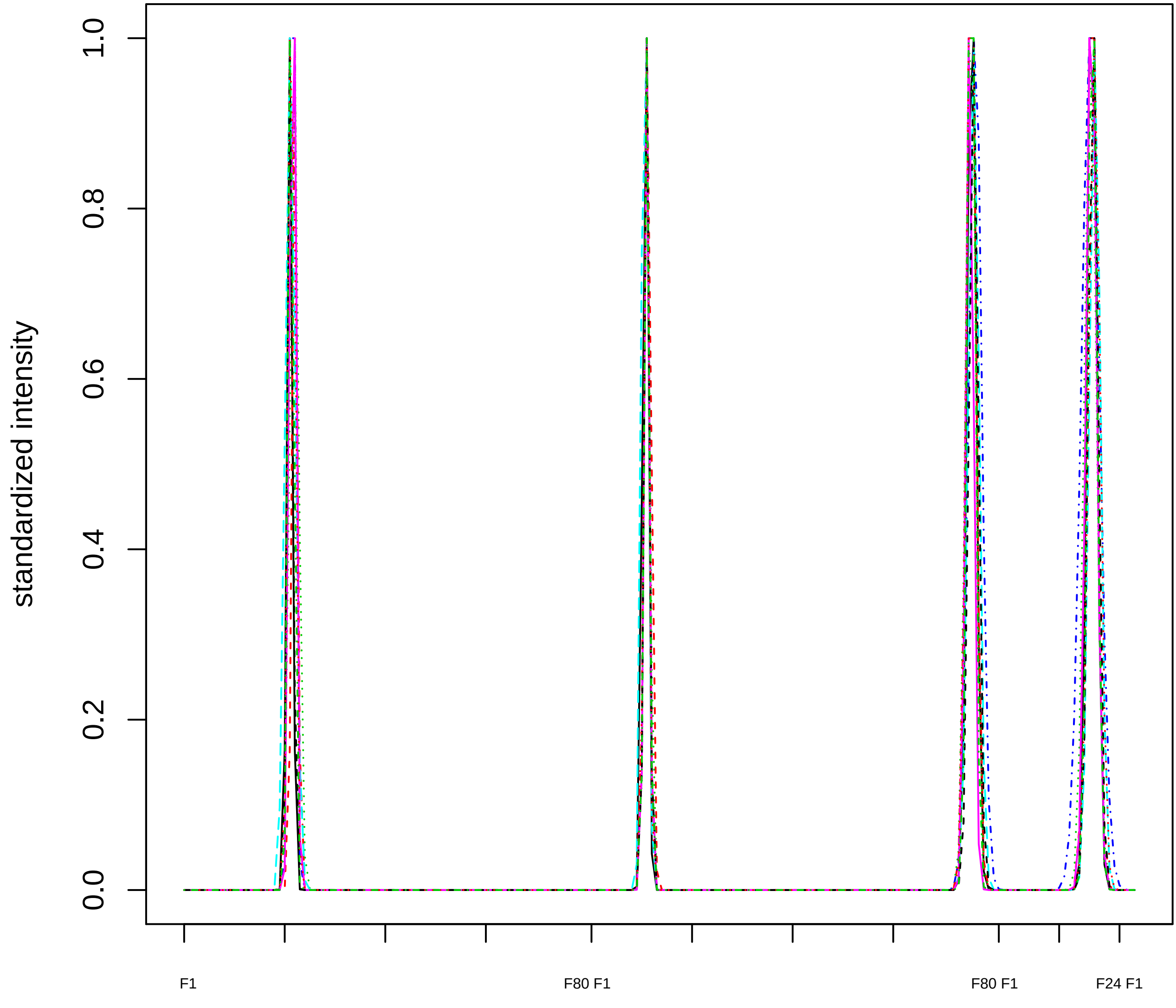

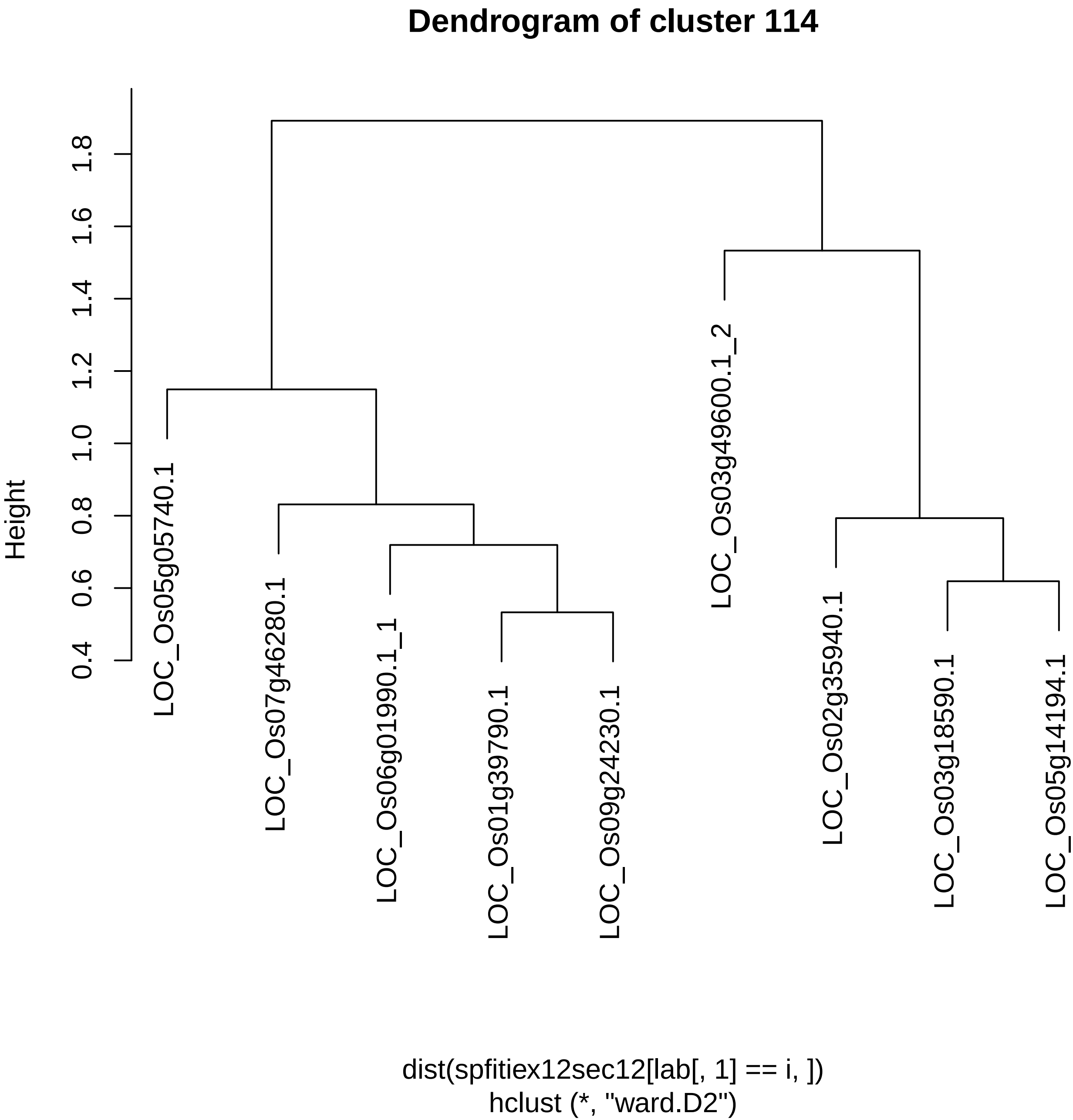
Dendrogram of cluster 114
Height
1.8
1.6
1.4
1.2
1.0
0.8
0.6
0.4
LOC_Os05g05740.1
LOC_Os07g46280.1
LOC_Os06g01990.1_1
LOC_Os01g39790.1
LOC_Os09g24230.1
LOC_Os03g49600.1_2
LOC_Os02g35940.1
LOC_Os03g18590.1
LOC_Os05g14194.1
dist(spfitiex12sec12[lab[, 1] == i, ])
hclust (*, "ward.D2")

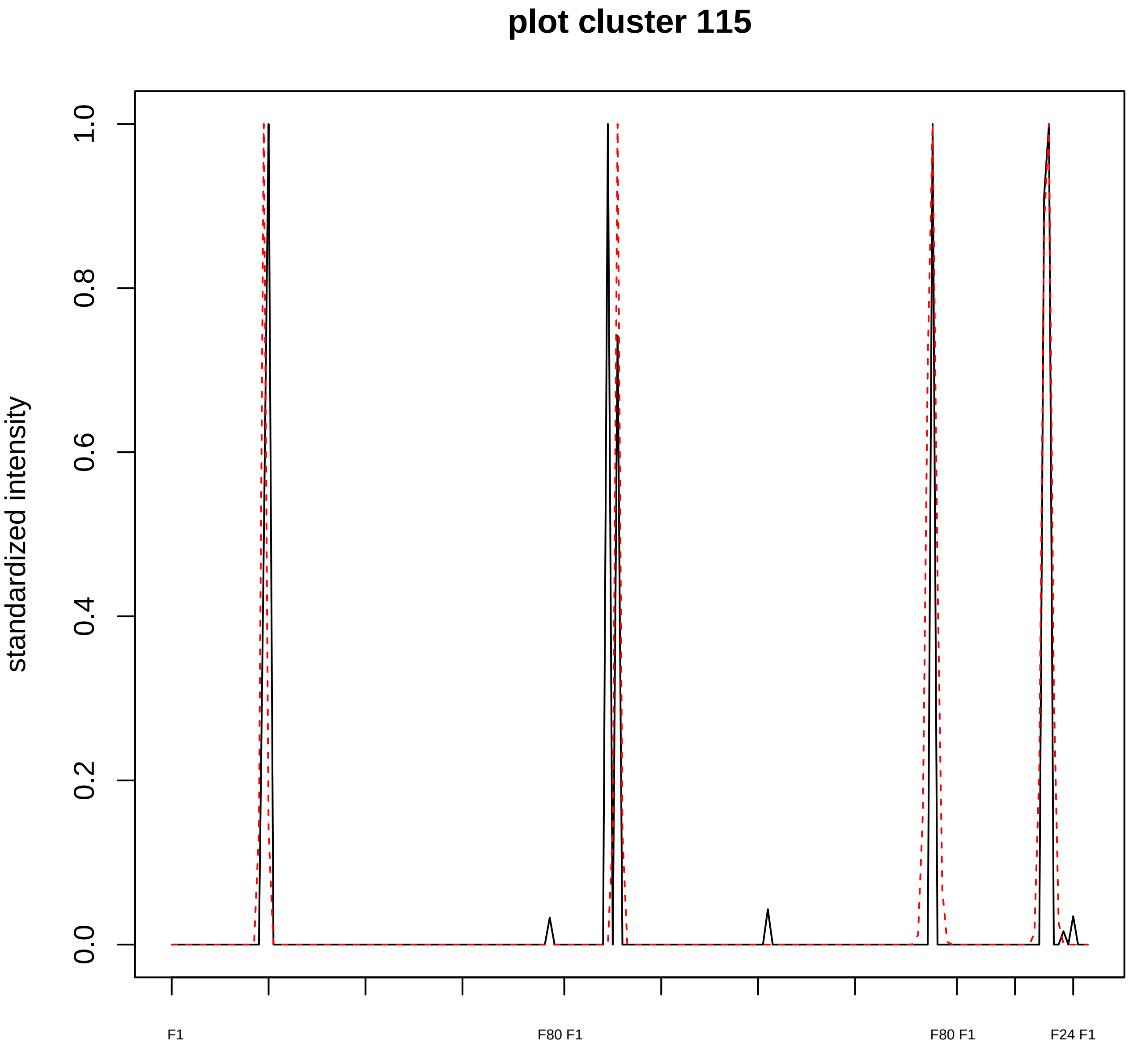
plot cluster 115
standardized intensity
0.0
0.2
0.4
0.6
0.8
1.0
F1
F80 F1
F80 F1
F24 F1

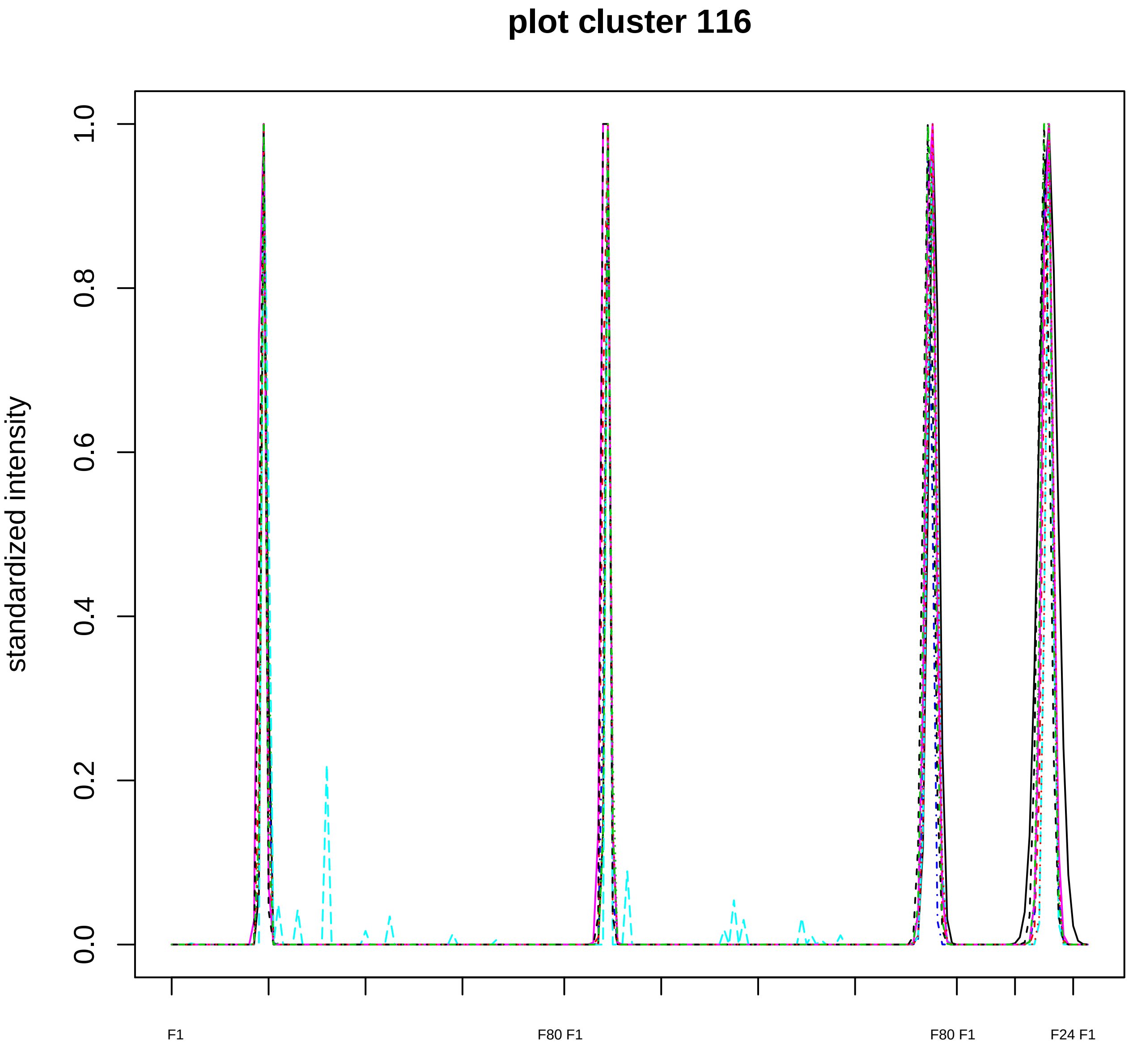
plot cluster 116
standardized intensity
0.0
0.2
0.4
0.6
0.8
1.0
F1
F80 F1
F80 F1
F24 F1

### Dendrogram of cluster 116

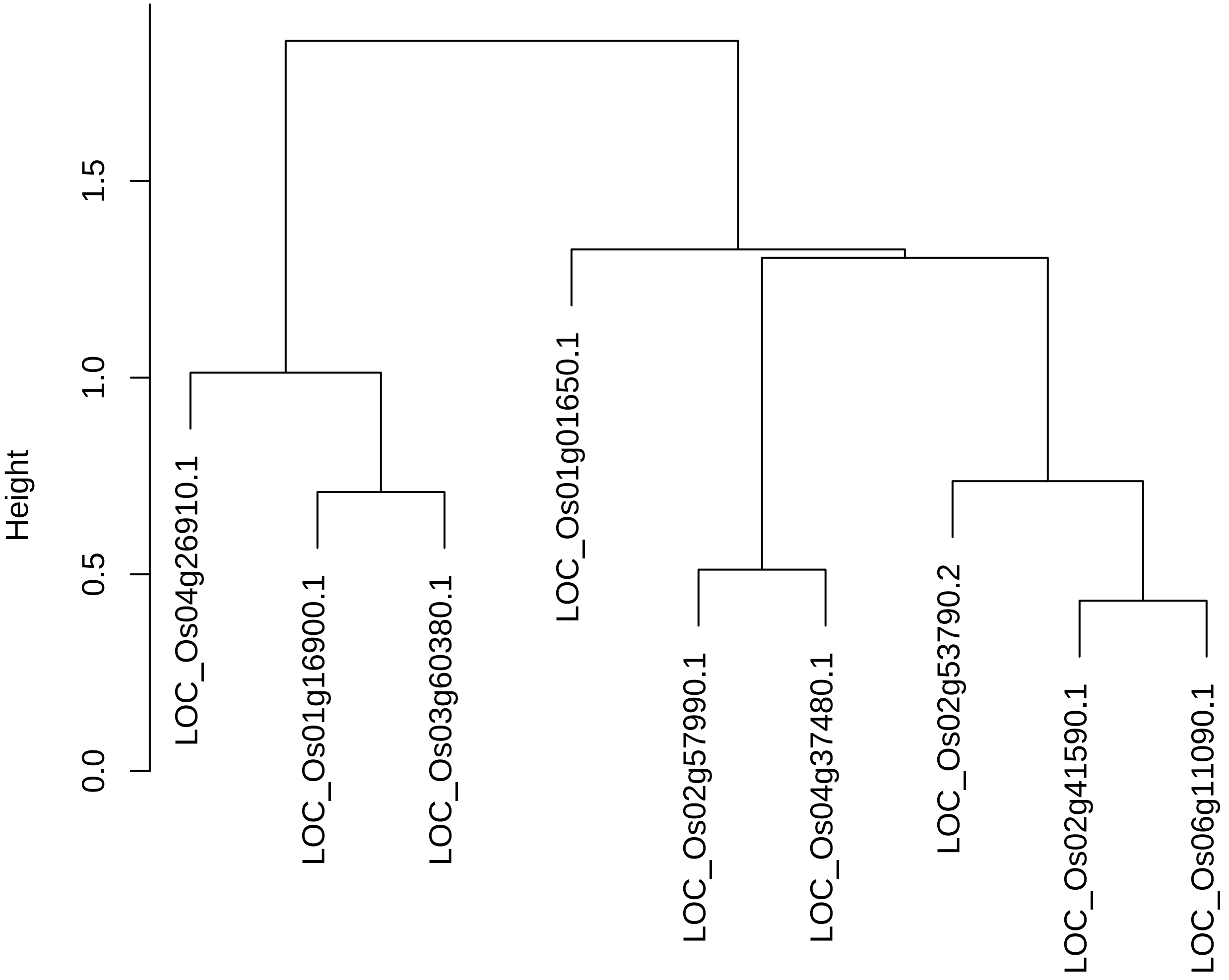


dist(spfitiex12sec12[lab[, 1] == i, ])
hclust (*, "ward.D2")

### plot cluster 117

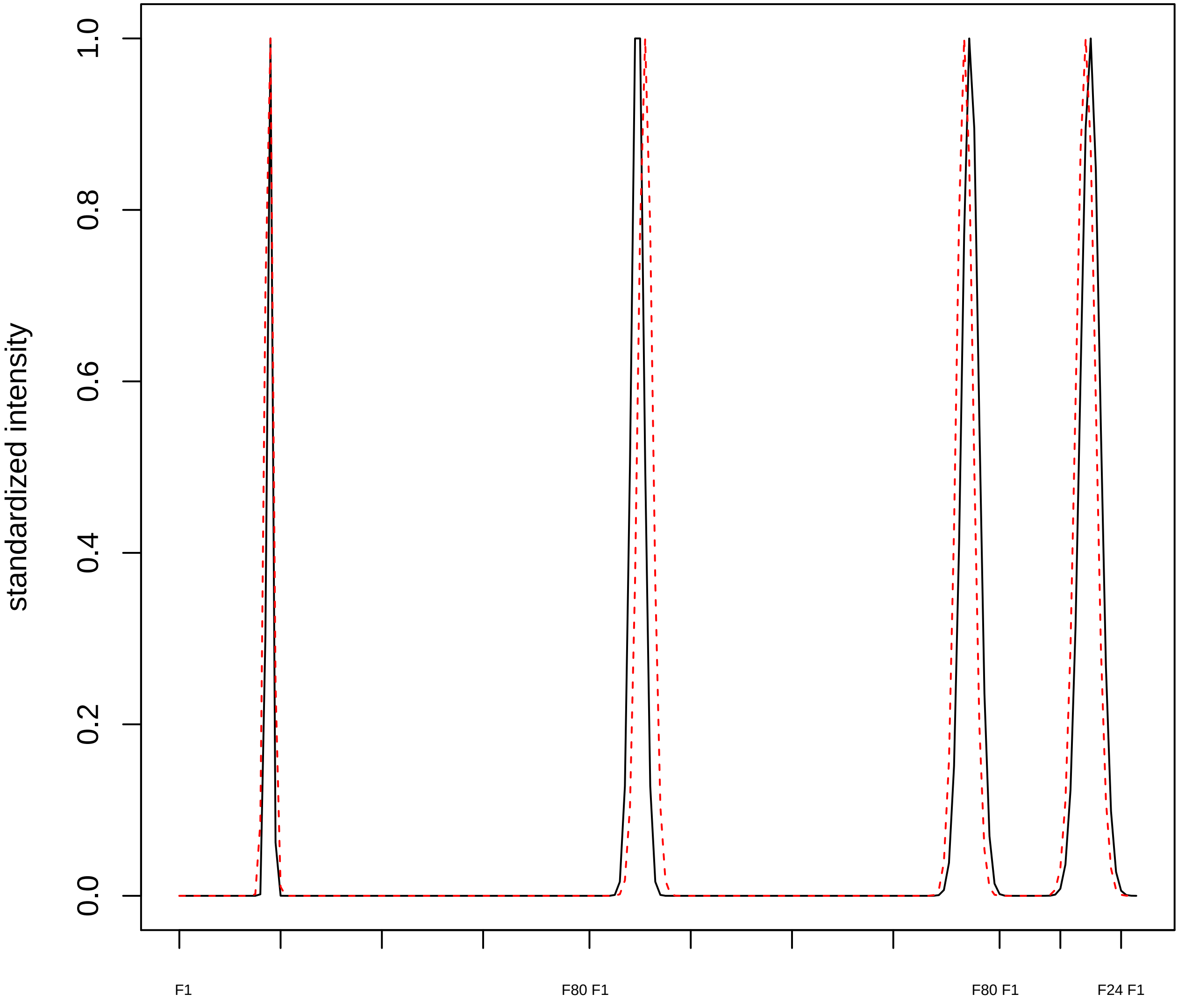

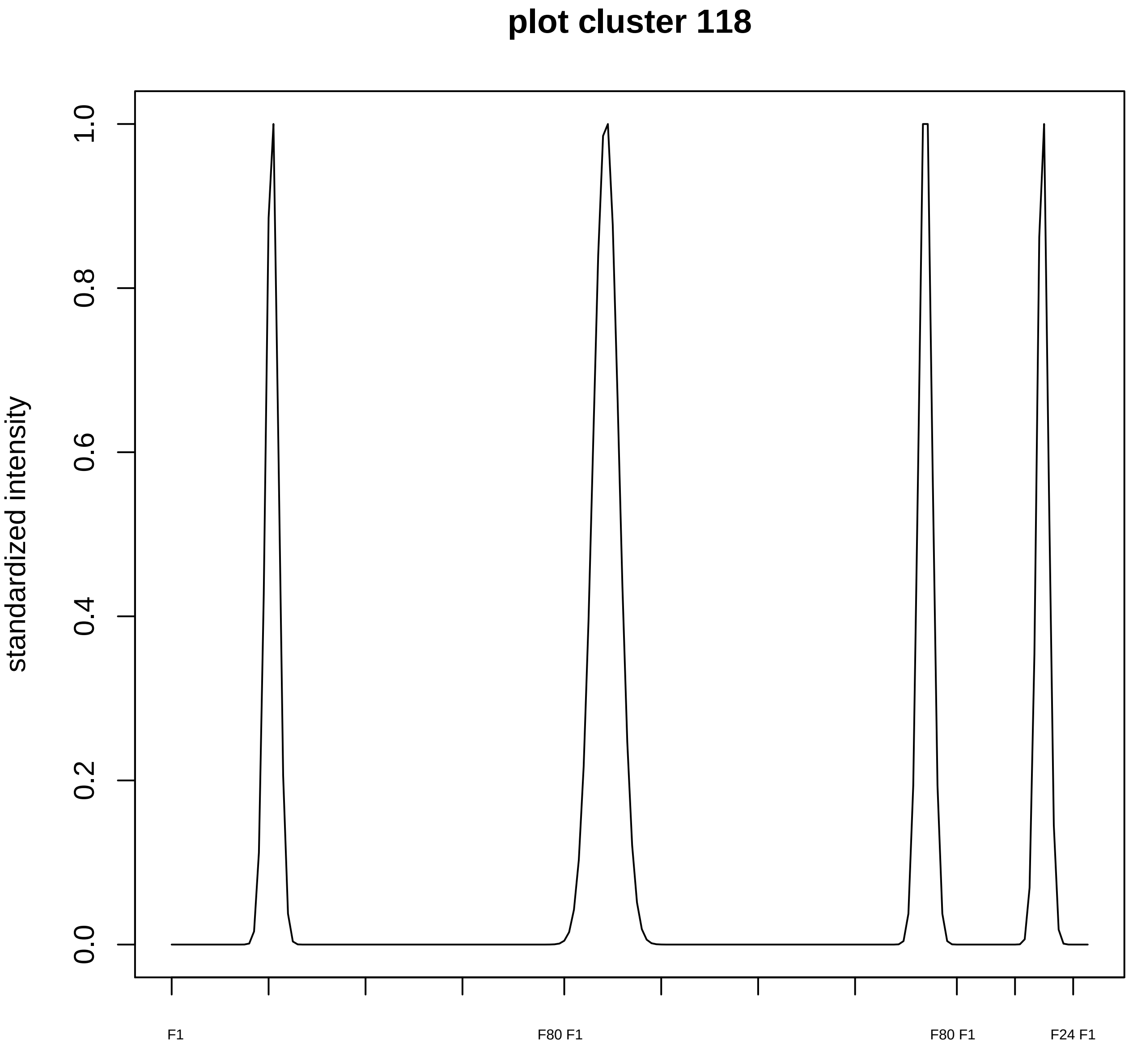
plot cluster 118
standardized intensity
0.0
0.2
0.4
0.6
0.8
1.0
F1
F80 F1
F80 F1
F24 F1

### plot cluster 119

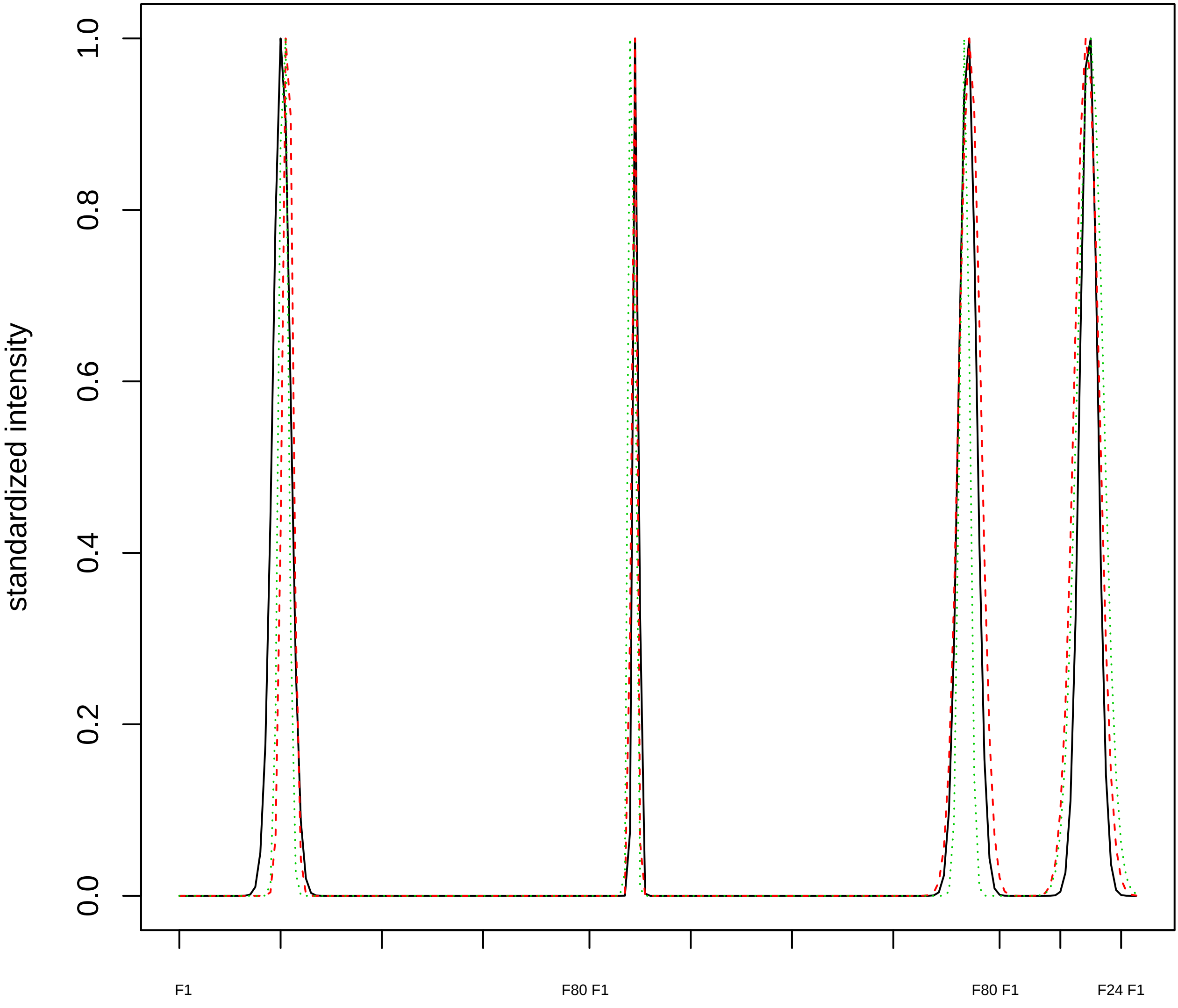

#### Dendrogram of cluster 119

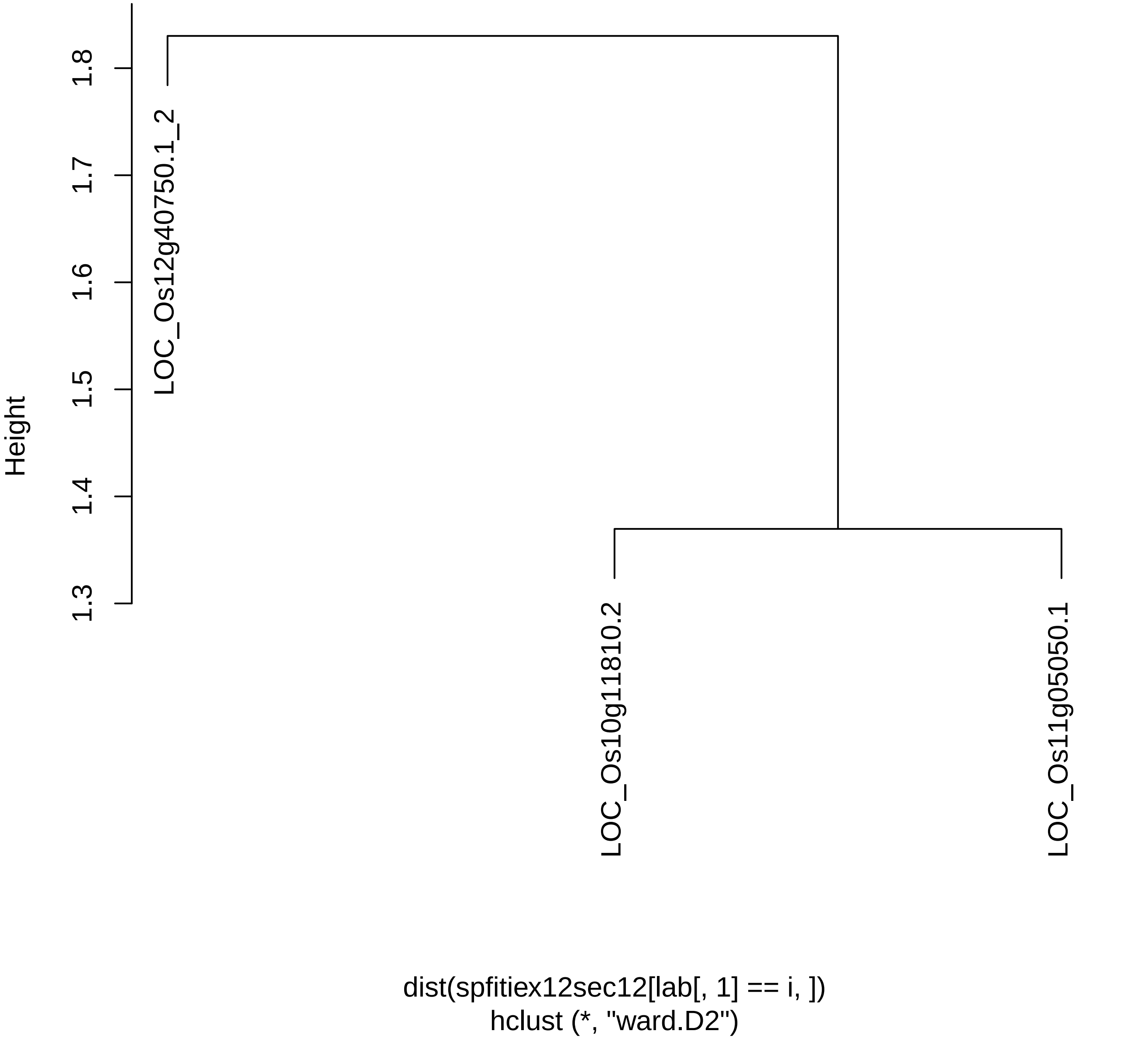

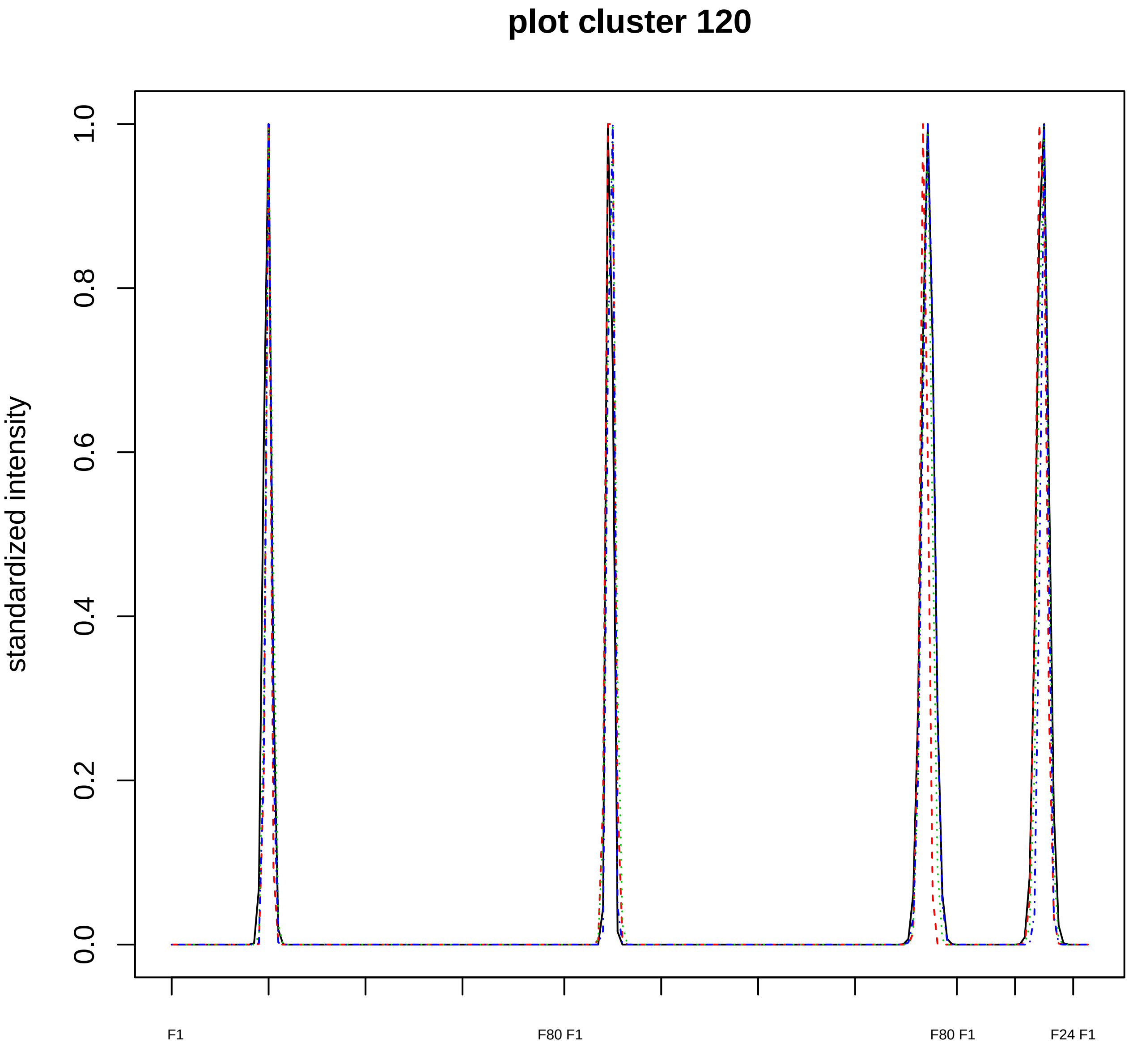

plot cluster 120
standardized intensity
0.0
0.2
0.4
0.6
0.8
1.0
F1
F80 F1
F80 F1
F24 F1

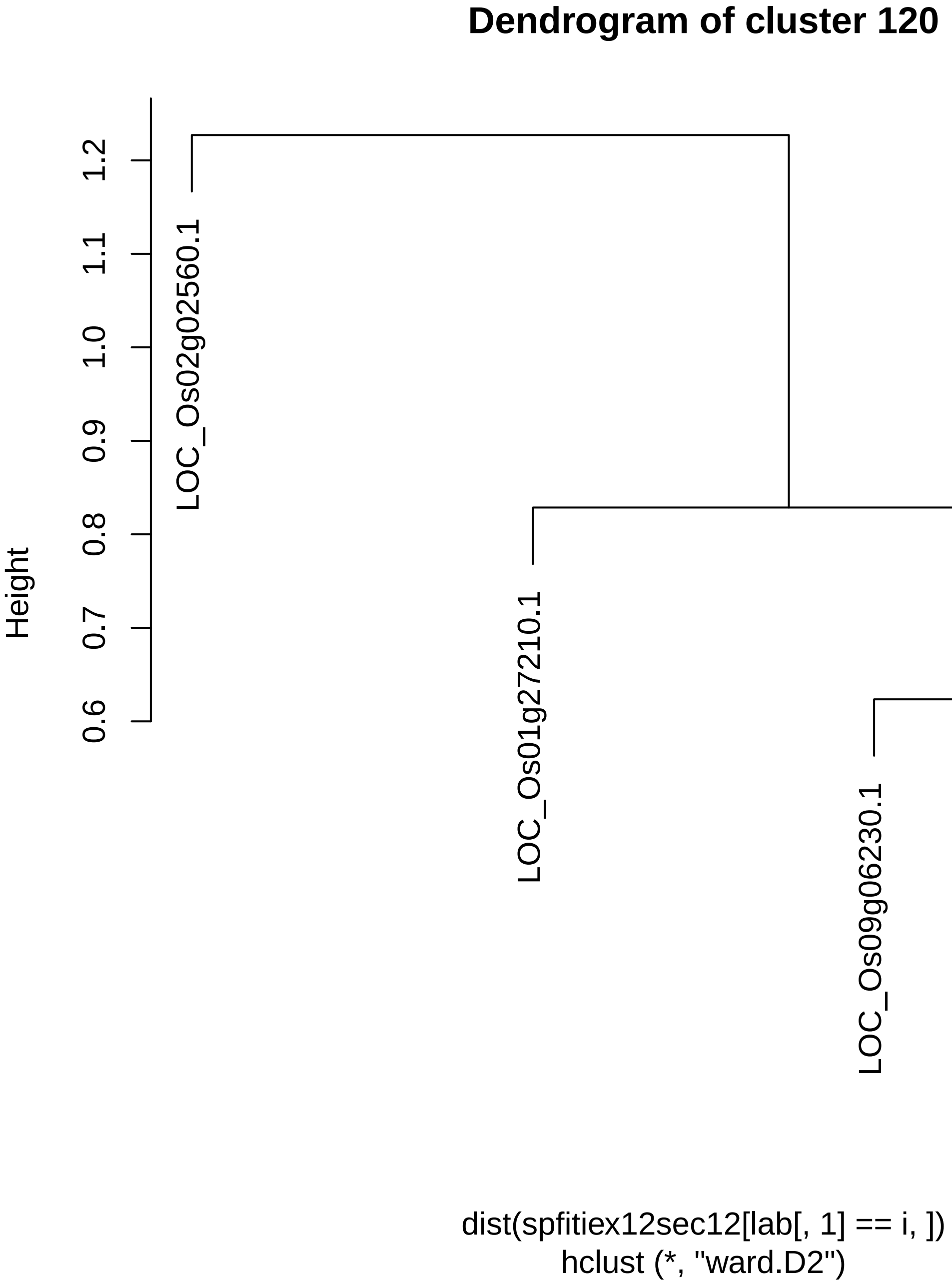

Dendrogram of cluster 120
Height
1.2
1.1
1.0
0.9
0.8
0.7
0.6
LOC_Os02g02560.1
LOC_Os01g27210.1
LOC_Os09g06230.1
LOC_Os12g43520.1
dist(spfitiex12sec12[lab[, 1] == i, ])
hclust (*, "ward.D2")

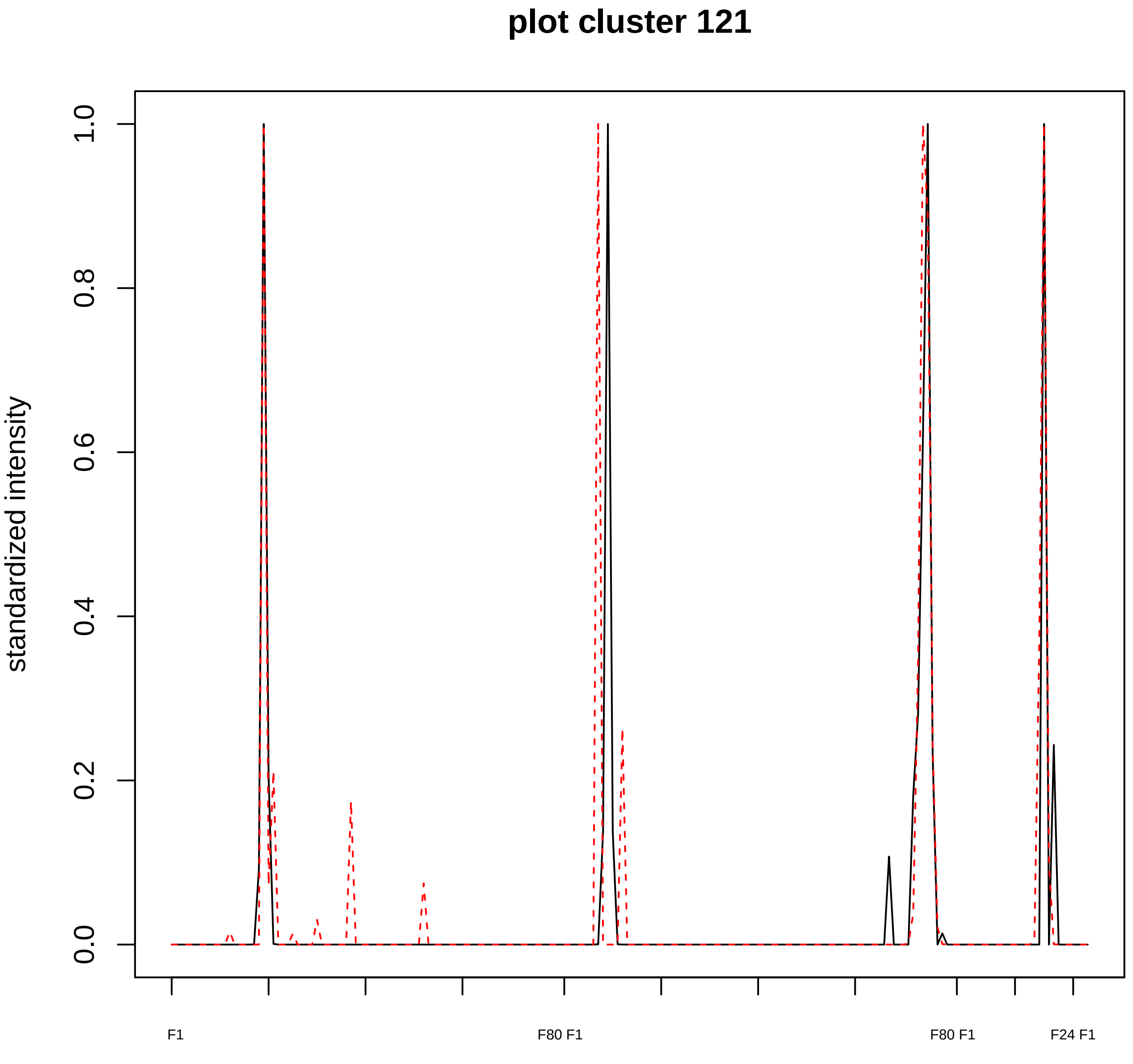
plot cluster 121
standardized intensity
0.0
0.2
0.4
0.6
0.8
1.0
F1
F80 F1
F80 F1
F24 F1

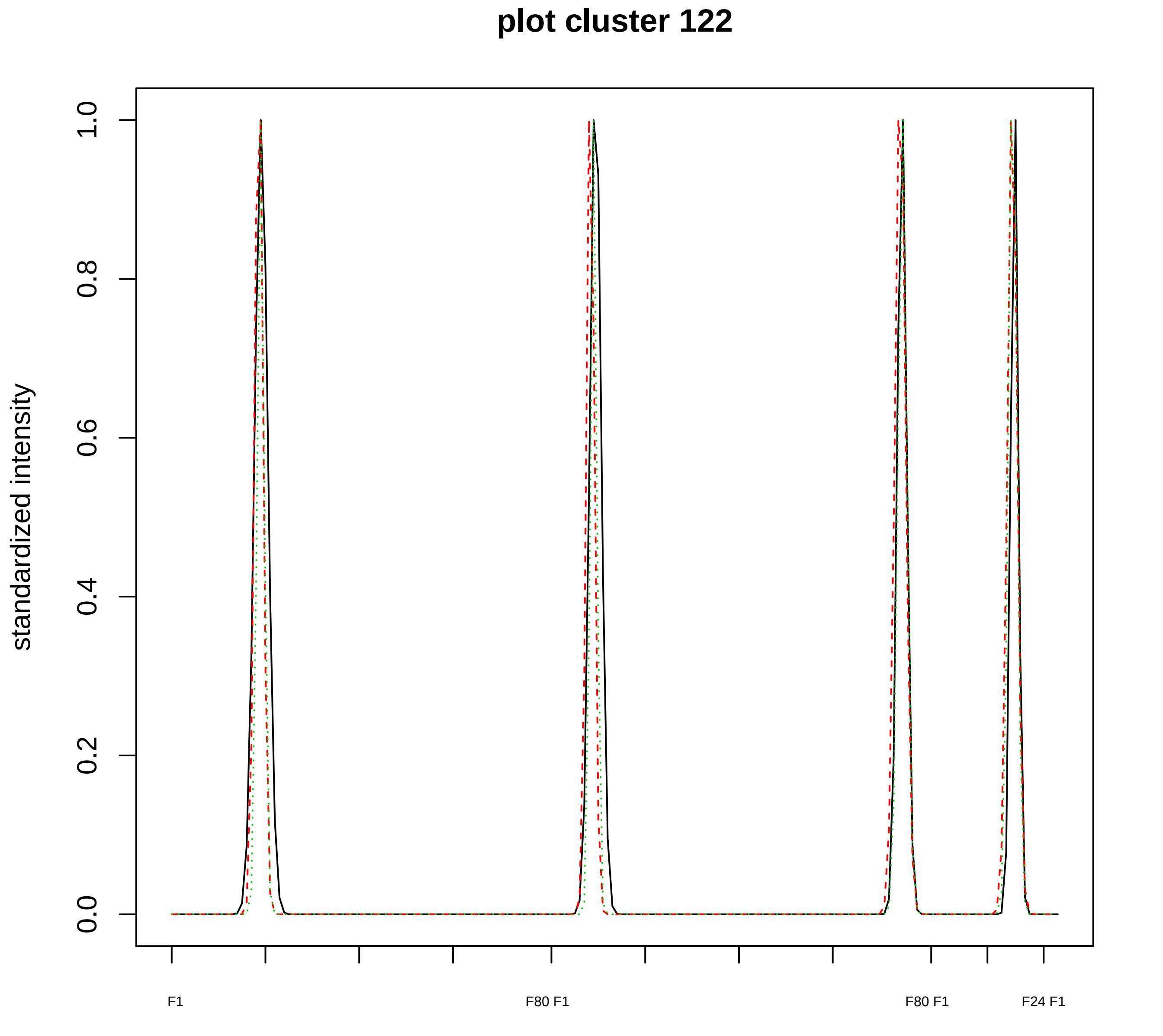
plot cluster 122
standardized intensity
0.0
0.2
0.4
0.6
0.8
1.0
F1
F80 F1
F80 F1
F24 F1

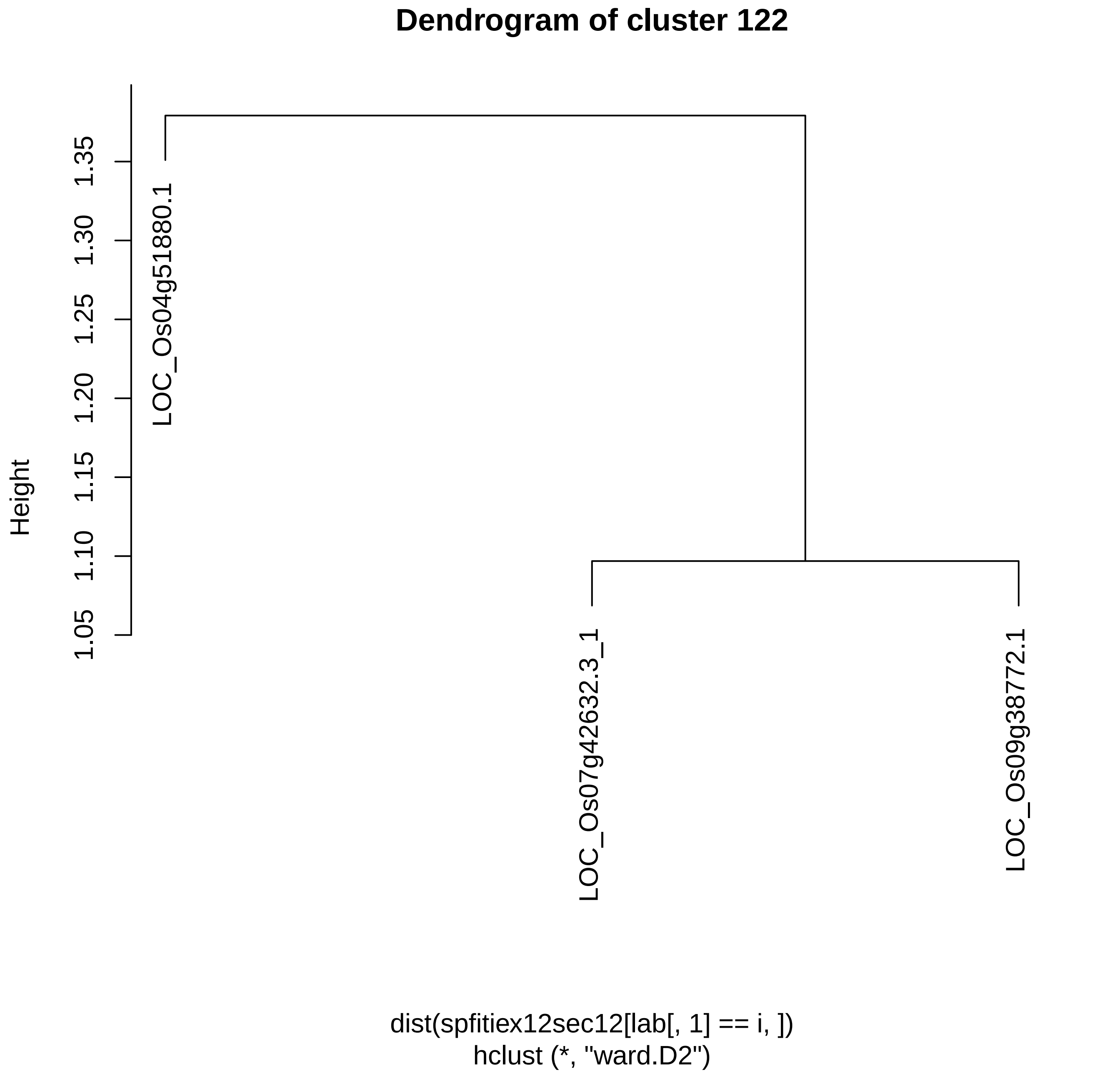
Dendrogram of cluster 122
Height
1.05
1.10
1.15
1.20
1.25
1.30
1.35
LOC_Os04g51880.1
LOC_Os07g42632.3_1
LOC_Os09g38772.1
dist(spfitiex12sec12[lab[, 1] == i, ])
hclust (*, "ward.D2")

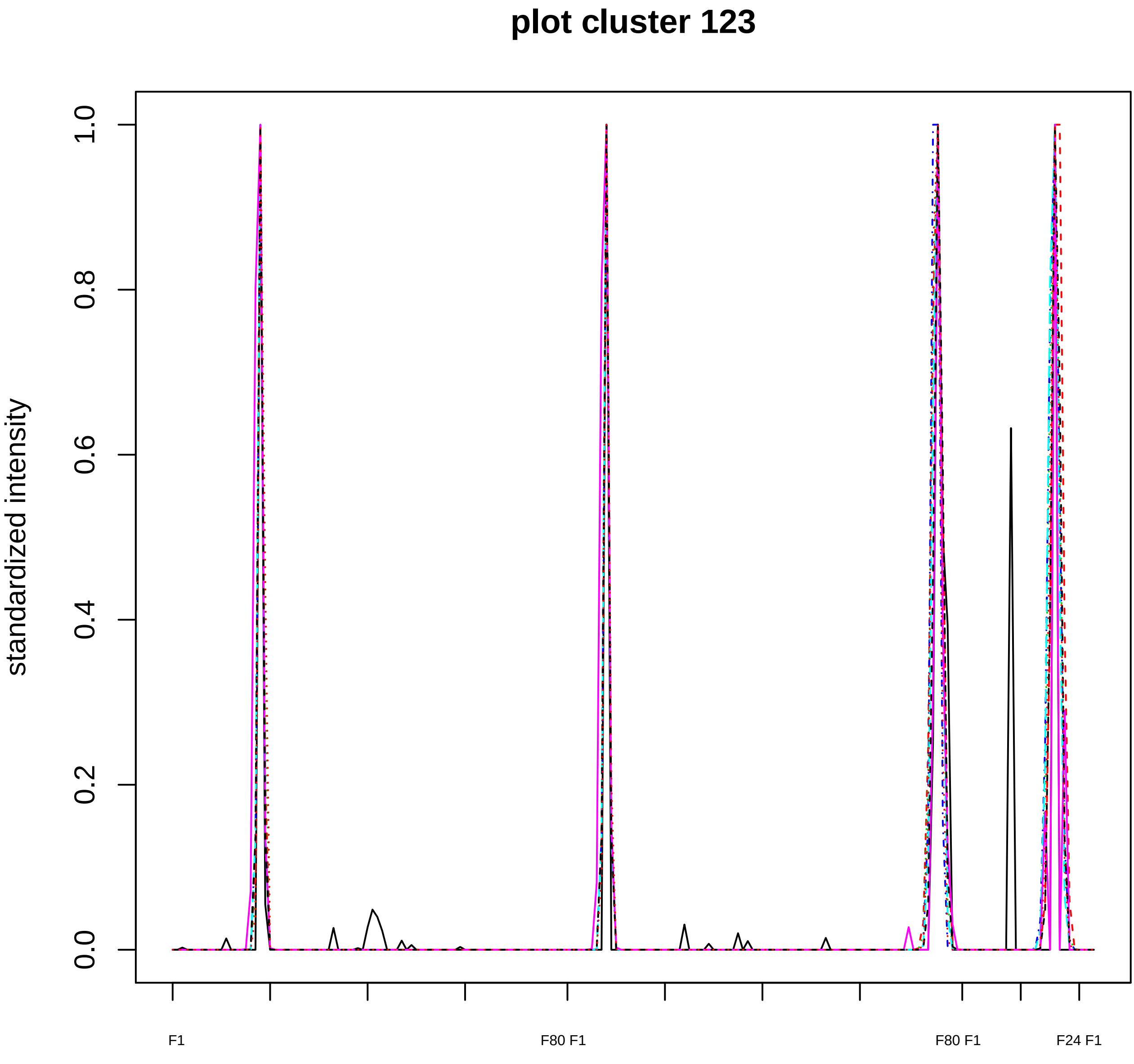
plot cluster 123
standardized intensity
0.0
0.2
0.4
0.6
0.8
1.0
F1
F80 F1
F80 F1
F24 F1

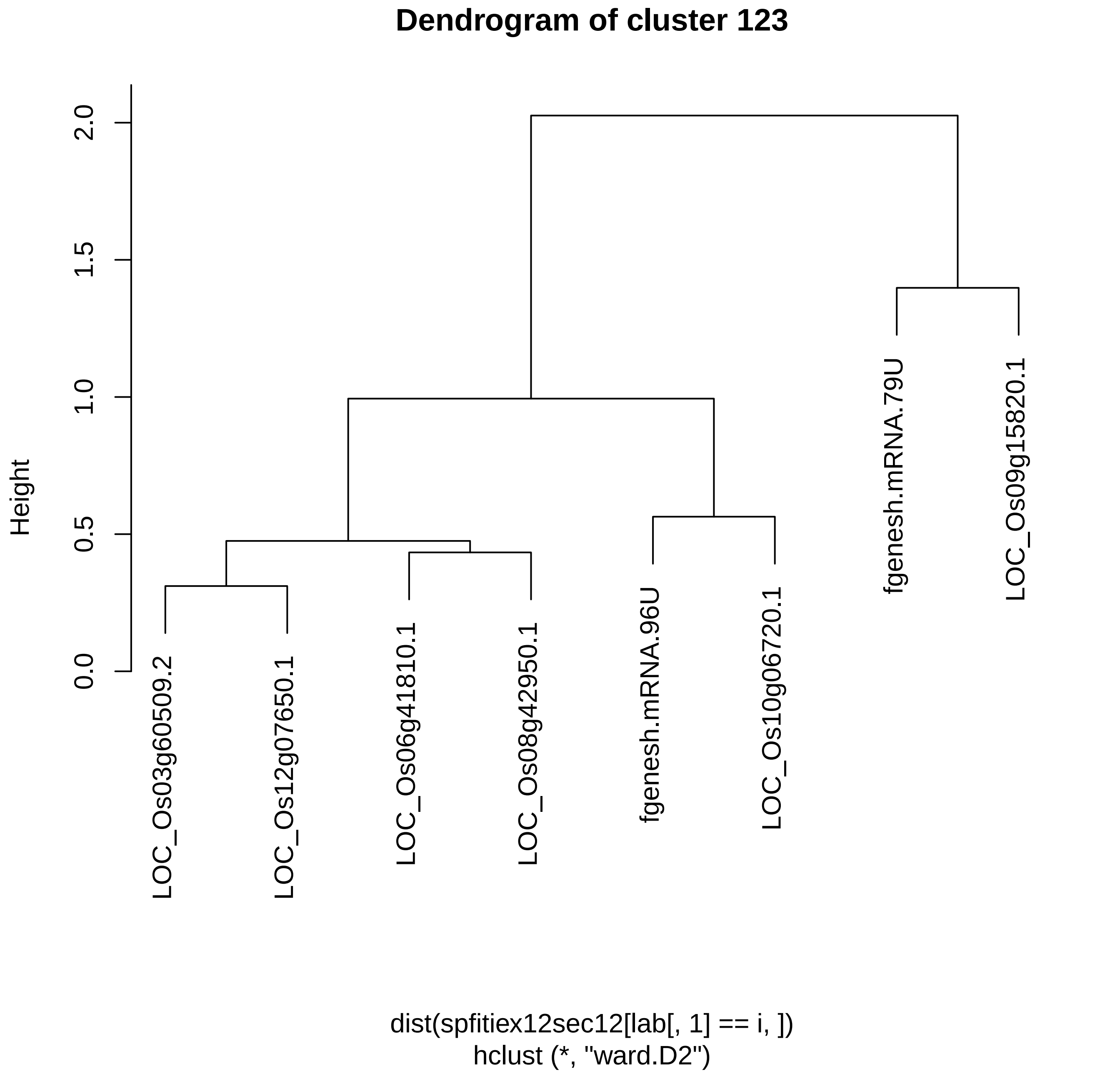
Dendrogram of cluster 123
Height
0.0
0.5
1.0
1.5
2.0
LOC_Os03g60509.2
LOC_Os12g07650.1
LOC_Os06g41810.1
LOC_Os08g42950.1
fgenesh.mRNA.96U
LOC_Os10g06720.1
fgenesh.mRNA.79U
LOC_Os09g15820.1
dist(spfitiex12sec12[lab[, 1] == i, ])
hclust (*, "ward.D2")

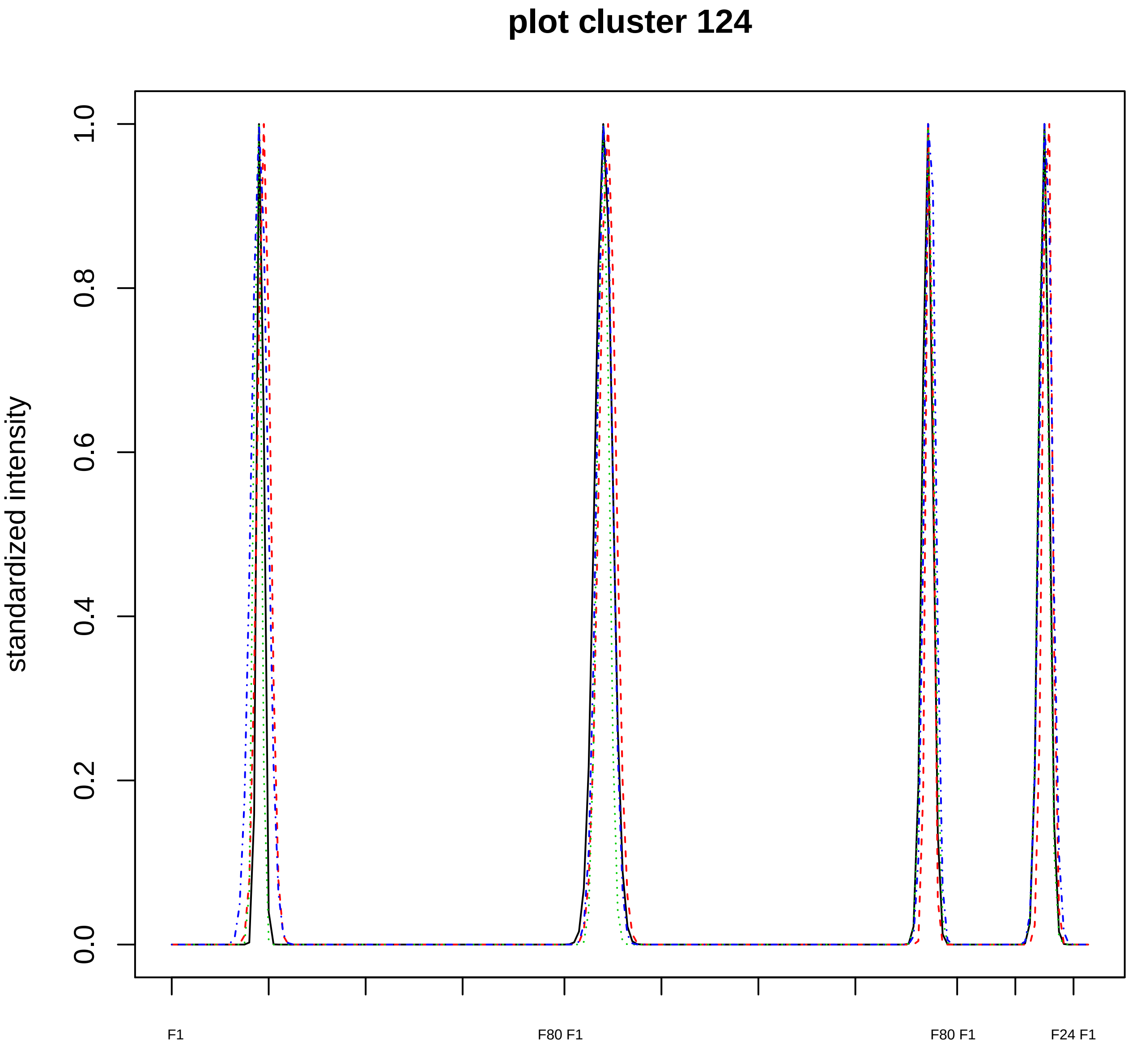
plot cluster 124
standardized intensity
0.0
0.2
0.4
0.6
0.8
1.0
F1
F80 F1
F80 F1
F24 F1

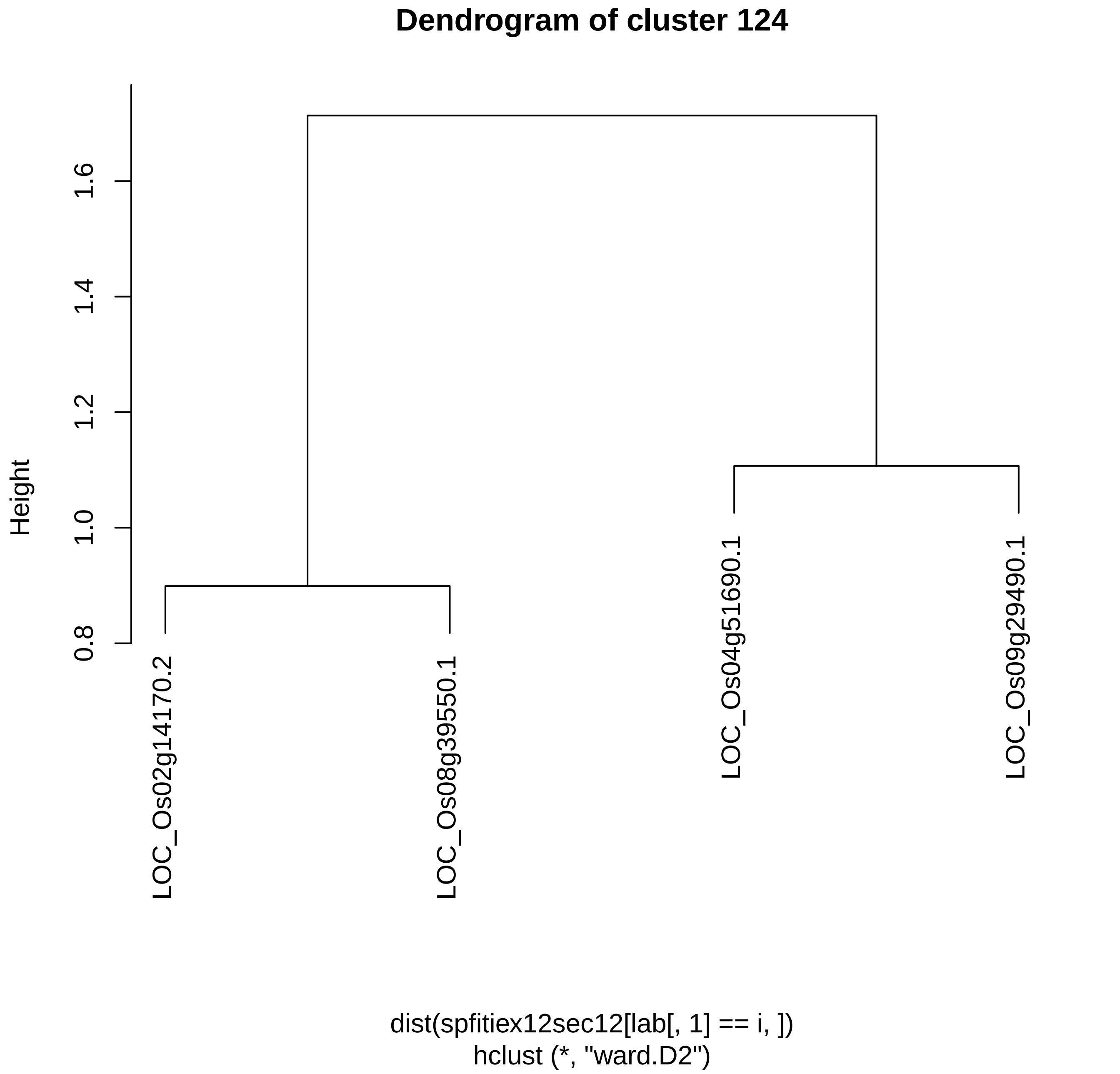

Dendrogram of cluster 124
Height
0.8
1.0
1.2
1.4
1.6
LOC_Os02g14170.2
LOC_Os08g39550.1
LOC_Os04g51690.1
LOC_Os09g29490.1
dist(spfitiex12sec12[lab[, 1] == i, ])
hclust (*, "ward.D2")

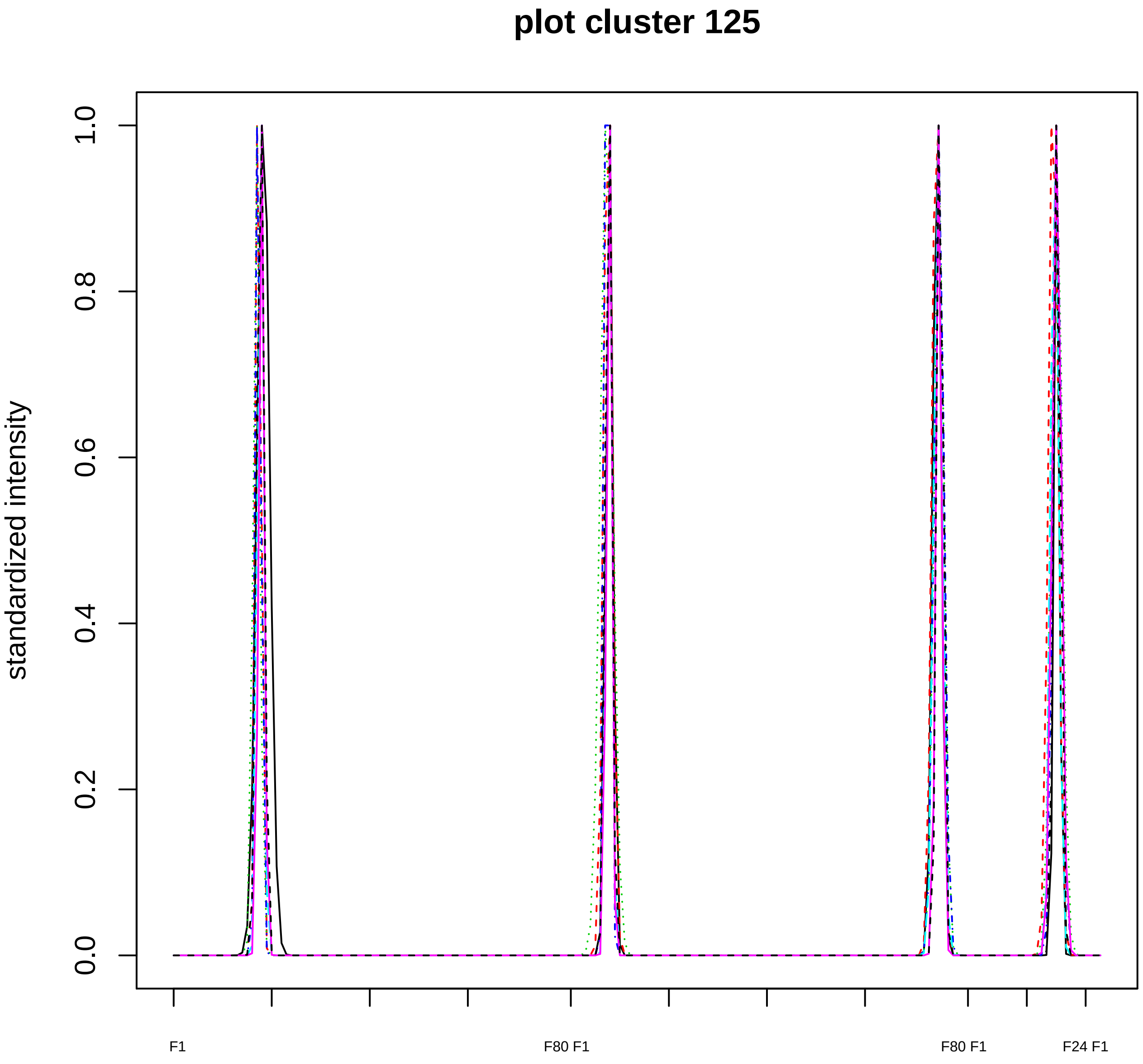
plot cluster 125
standardized intensity
0.0
0.2
0.4
0.6
0.8
1.0
F1
F80 F1
F80 F1
F24 F1

### Dendrogram of cluster 125

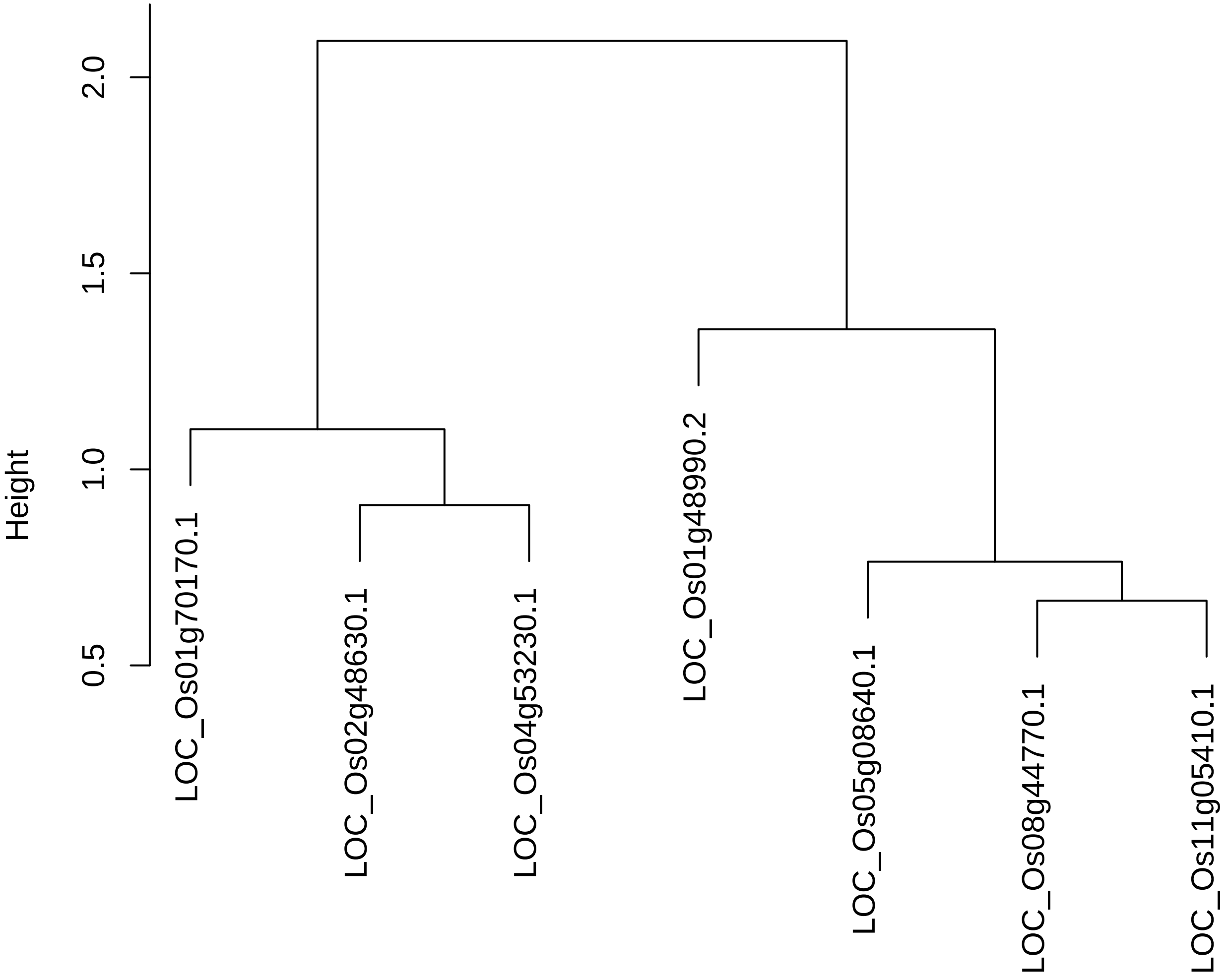


dist(spfitiex12sec12[lab[, 1] == i, ])
hclust (*, "ward.D2")

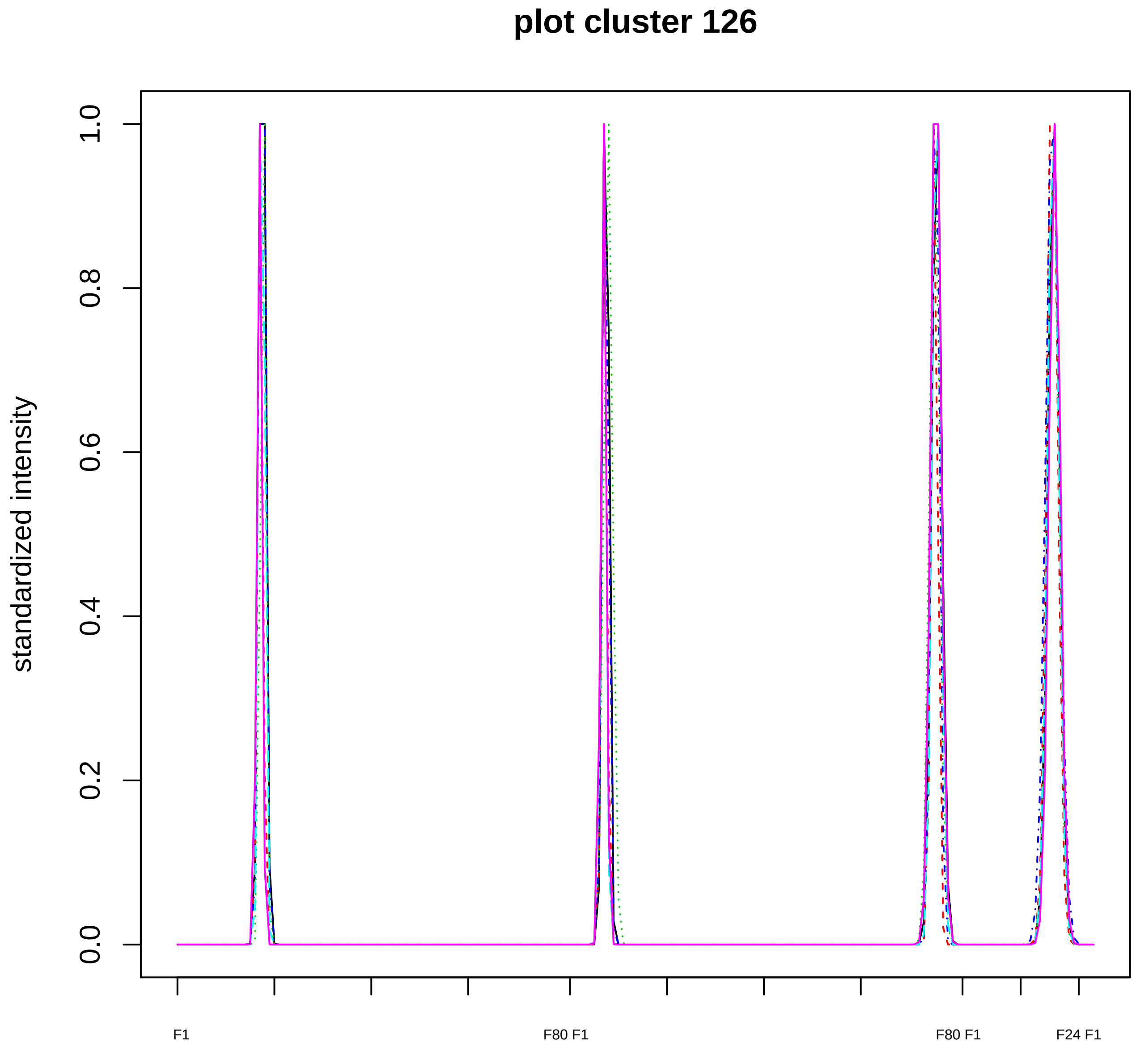
plot cluster 126
standardized intensity
0.0
0.2
0.4
0.6
0.8
1.0
F1
F80 F1
F80 F1
F24 F1

### Dendrogram of cluster 126

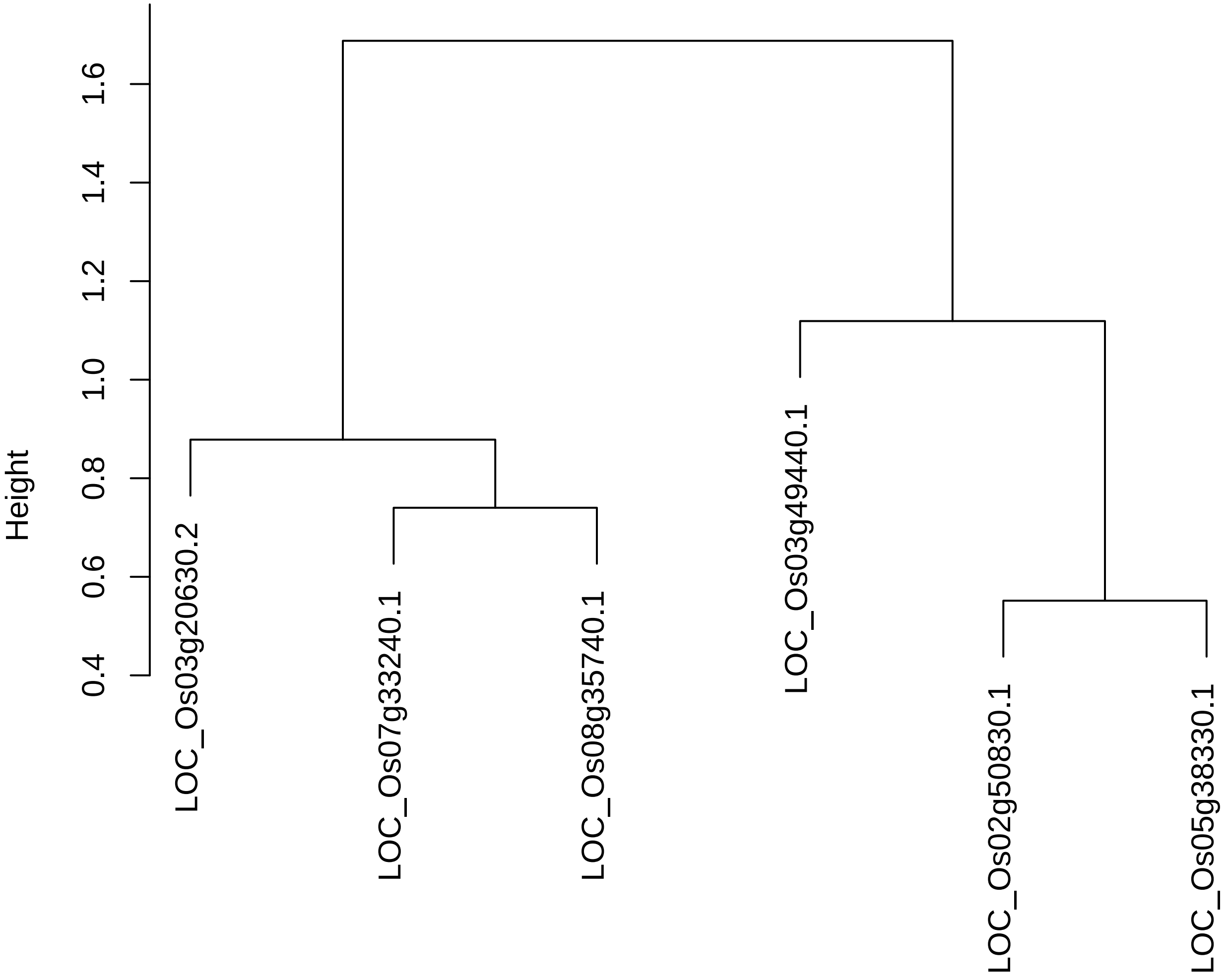


dist(spfitiex12sec12[lab[, 1] == i, ])
hclust (*, "ward.D2")

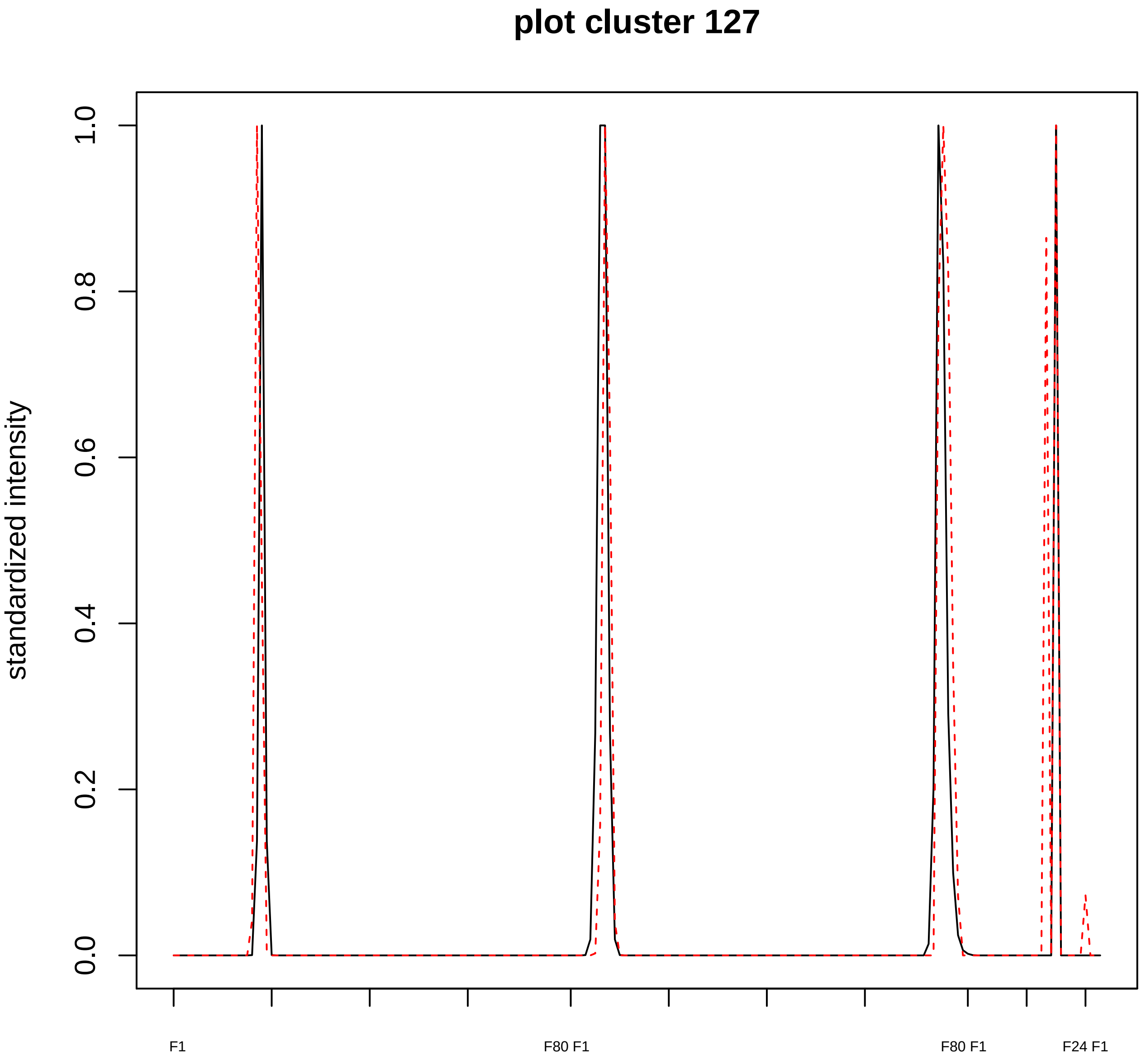
plot cluster 127
standardized intensity
0.0
0.2
0.4
0.6
0.8
1.0
F1
F80 F1
F80 F1
F24 F1

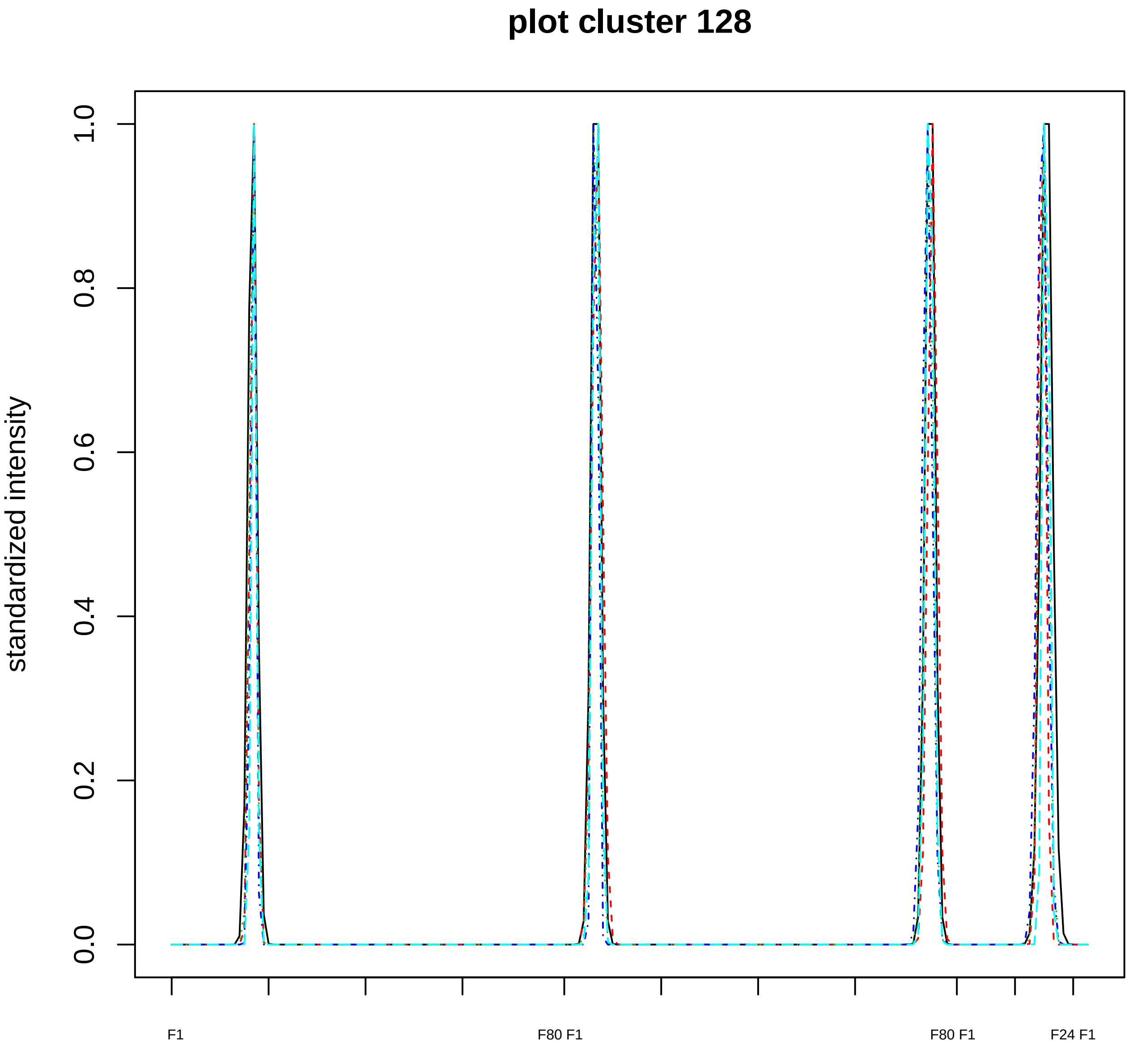
plot cluster 128
standardized intensity
0.0
0.2
0.4
0.6
0.8
1.0
F1
F80 F1
F80 F1
F24 F1

#### Dendrogram of cluster 128

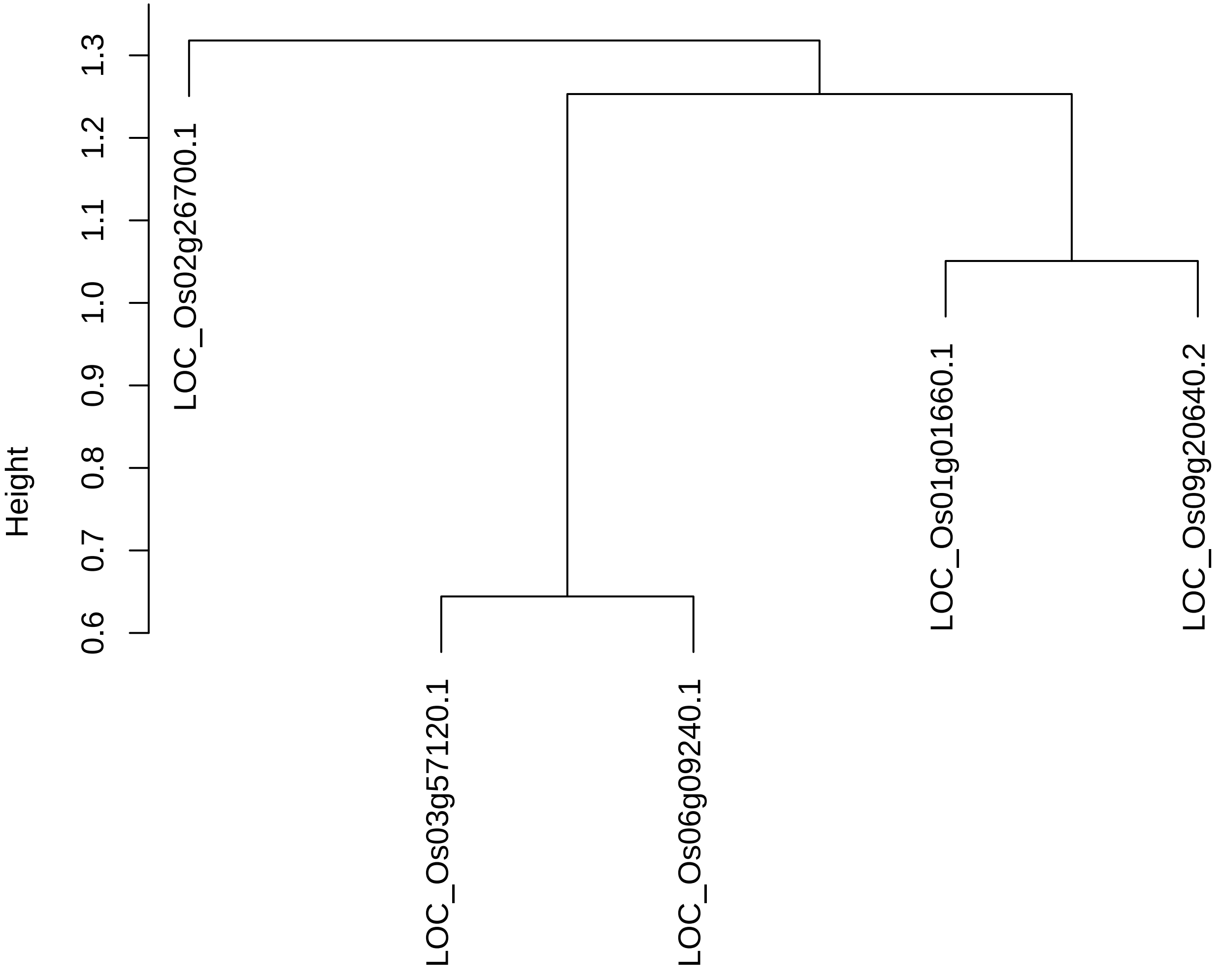


dist(spfitiex12sec12[lab[, 1] == i, ])
hclust (*, "ward.D2")

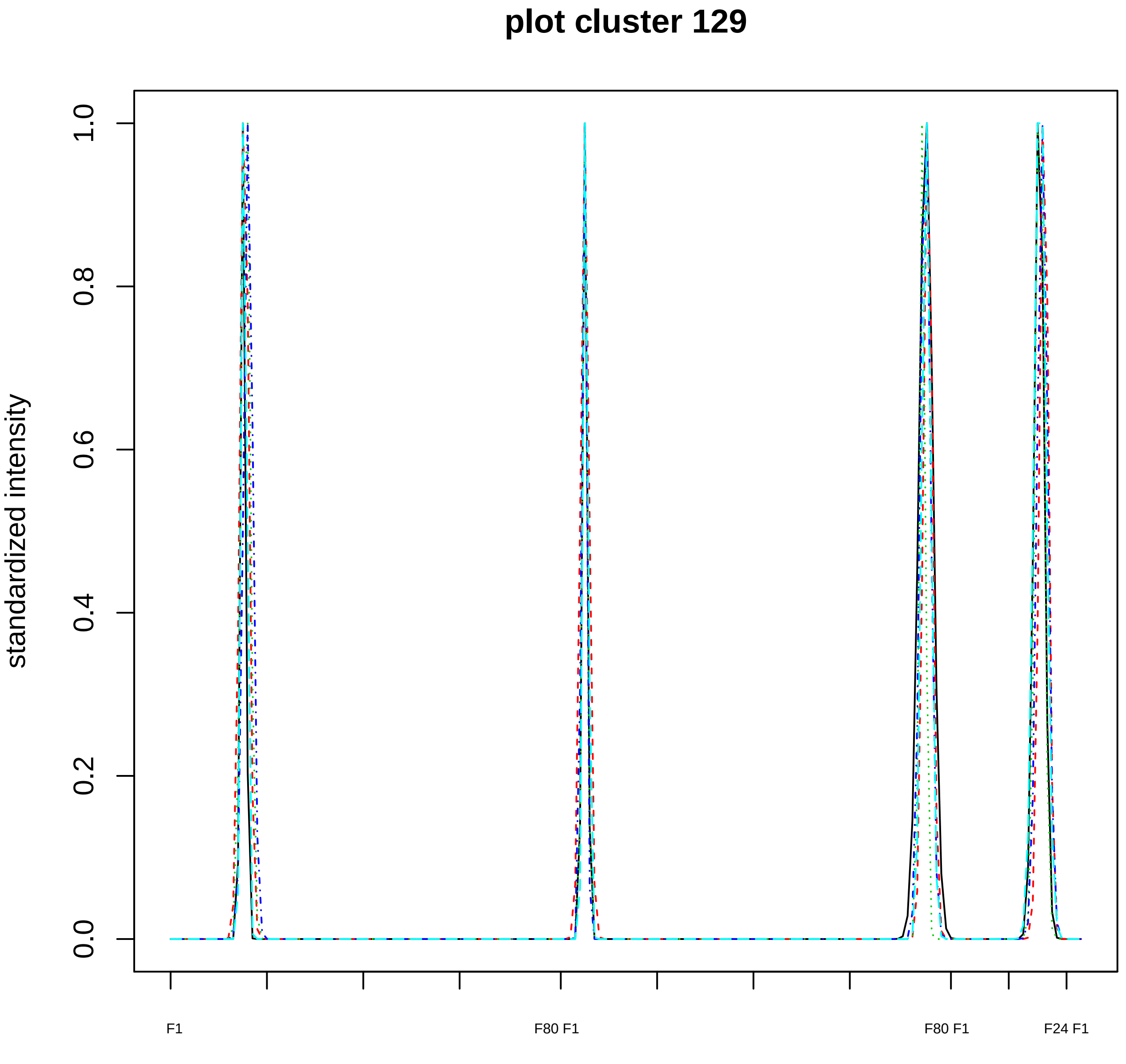
plot cluster 129
standardized intensity
0.0
0.2
0.4
0.6
0.8
1.0
F1
F80 F1
F80 F1
F24 F1

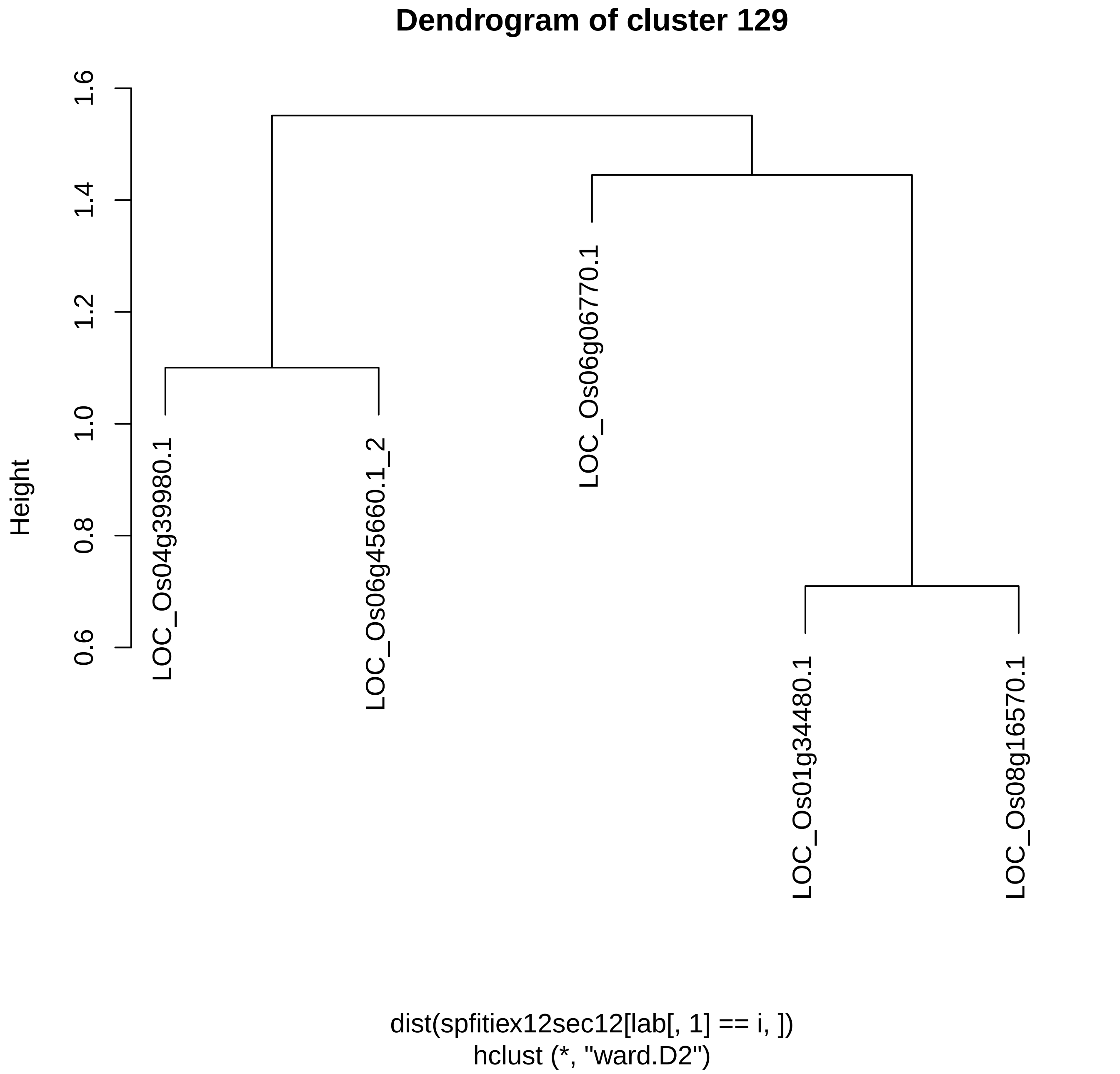
Dendrogram of cluster 129
Height
0.6
0.8
1.0
1.2
1.4
1.6
LOC_Os04g39980.1
LOC_Os06g45660.1_2
LOC_Os06g06770.1
LOC_Os01g34480.1
LOC_Os08g16570.1
dist(spfitiex12sec12[lab[, 1] == i, ])
hclust (*, "ward.D2")

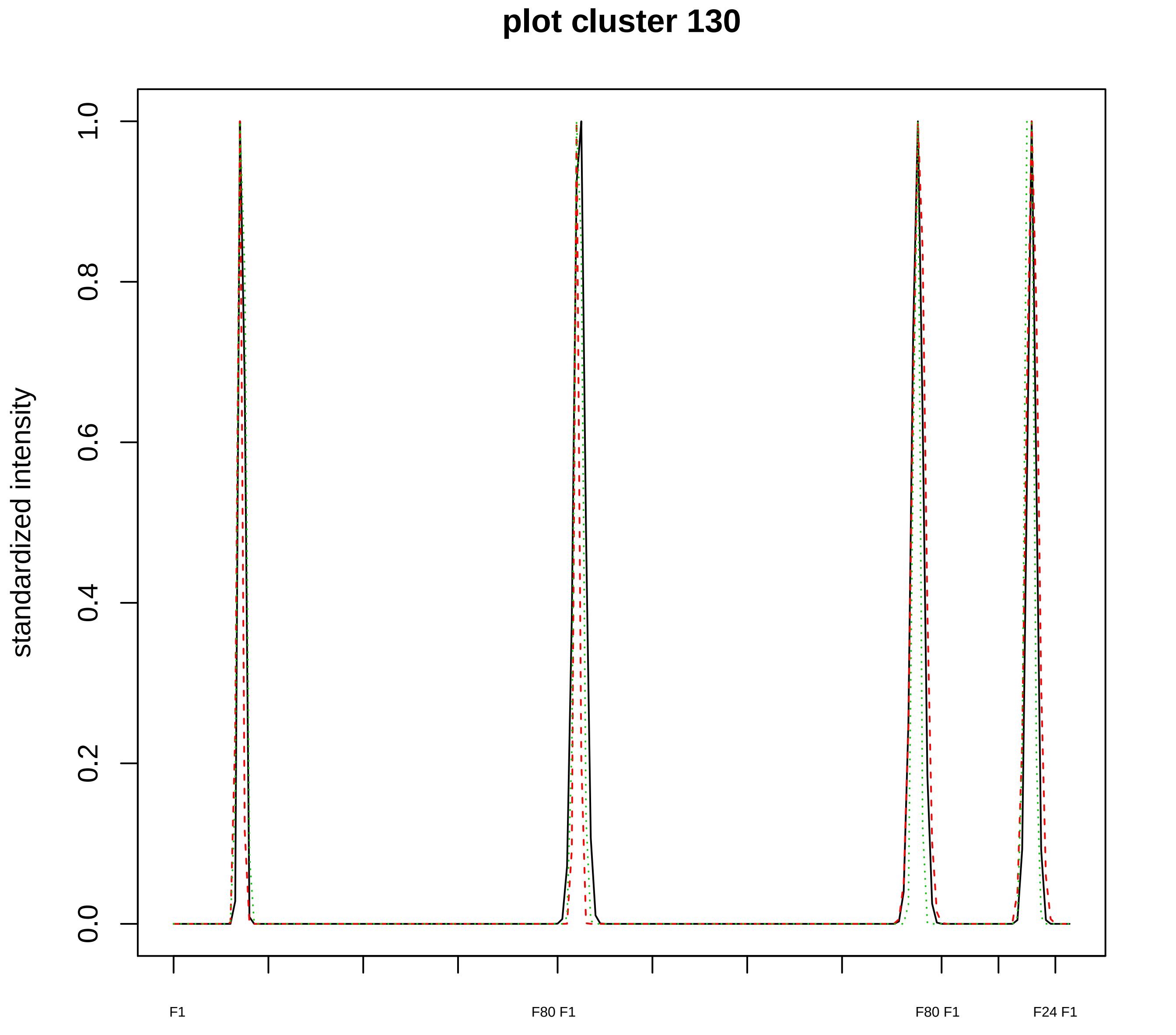
plot cluster 130
standardized intensity
0.0
0.2
0.4
0.6
0.8
1.0
F1
F80 F1
F80 F1
F24 F1

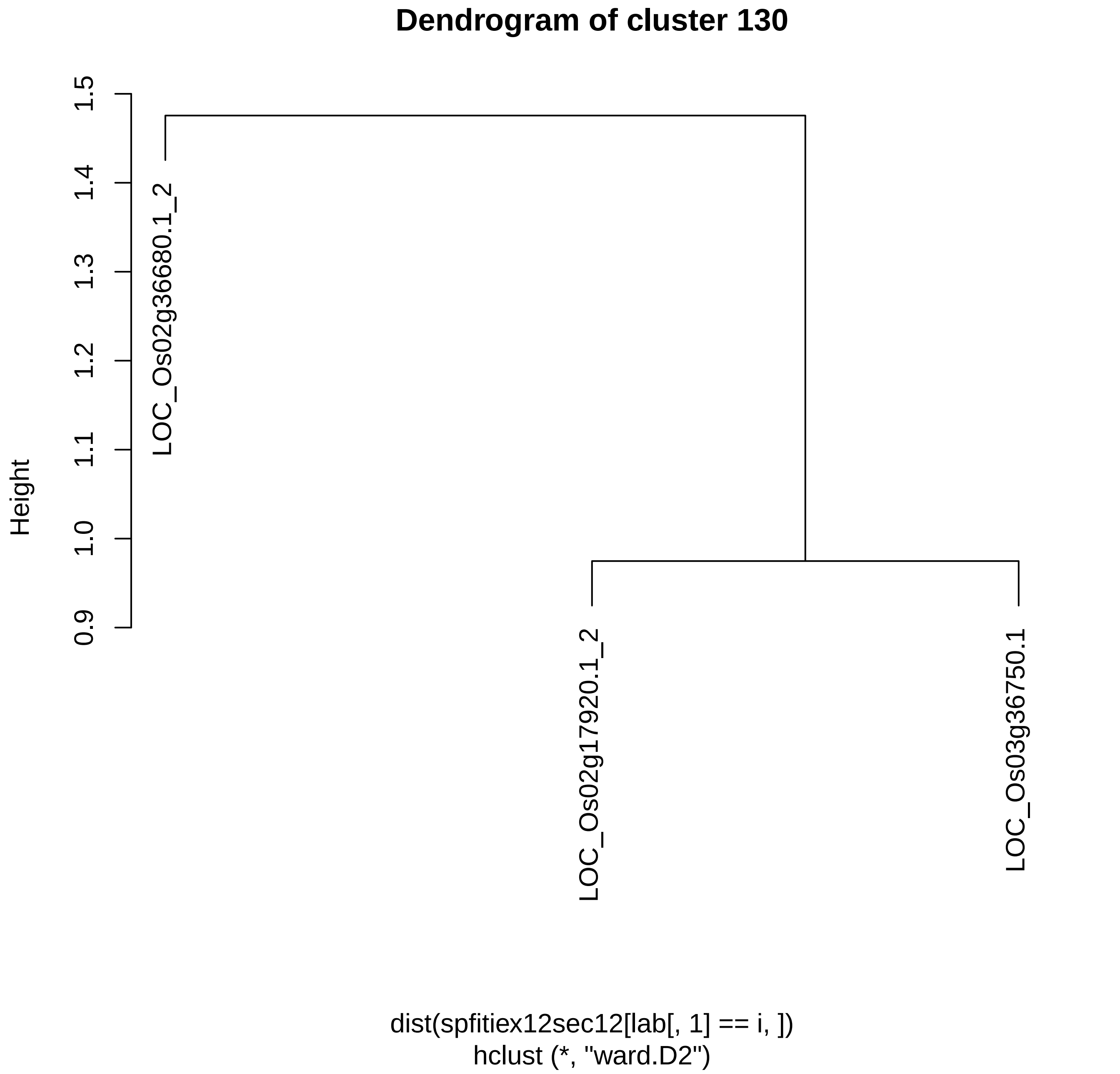
Dendrogram of cluster 130
Height
0.9
1.0
1.1
1.2
1.3
1.4
1.5
LOC_Os02g36680.1_2
LOC_Os02g17920.1_2
LOC_Os03g36750.1
dist(spfitiex12sec12[lab[, 1] == i, ])
hclust (*, "ward.D2")

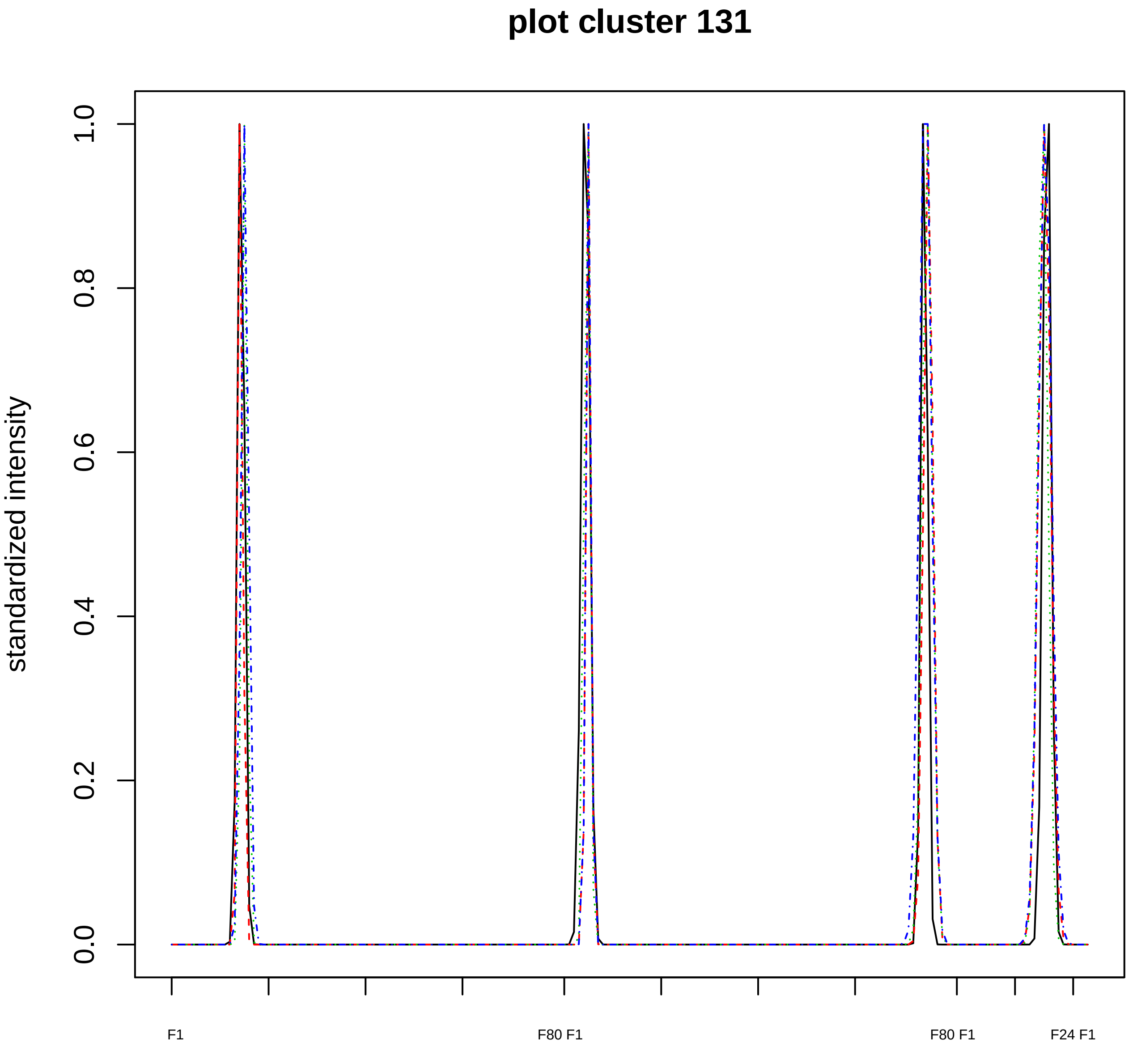
plot cluster 131
standardized intensity
0.0
0.2
0.4
0.6
0.8
1.0
F1
F80 F1
F80 F1
F24 F1

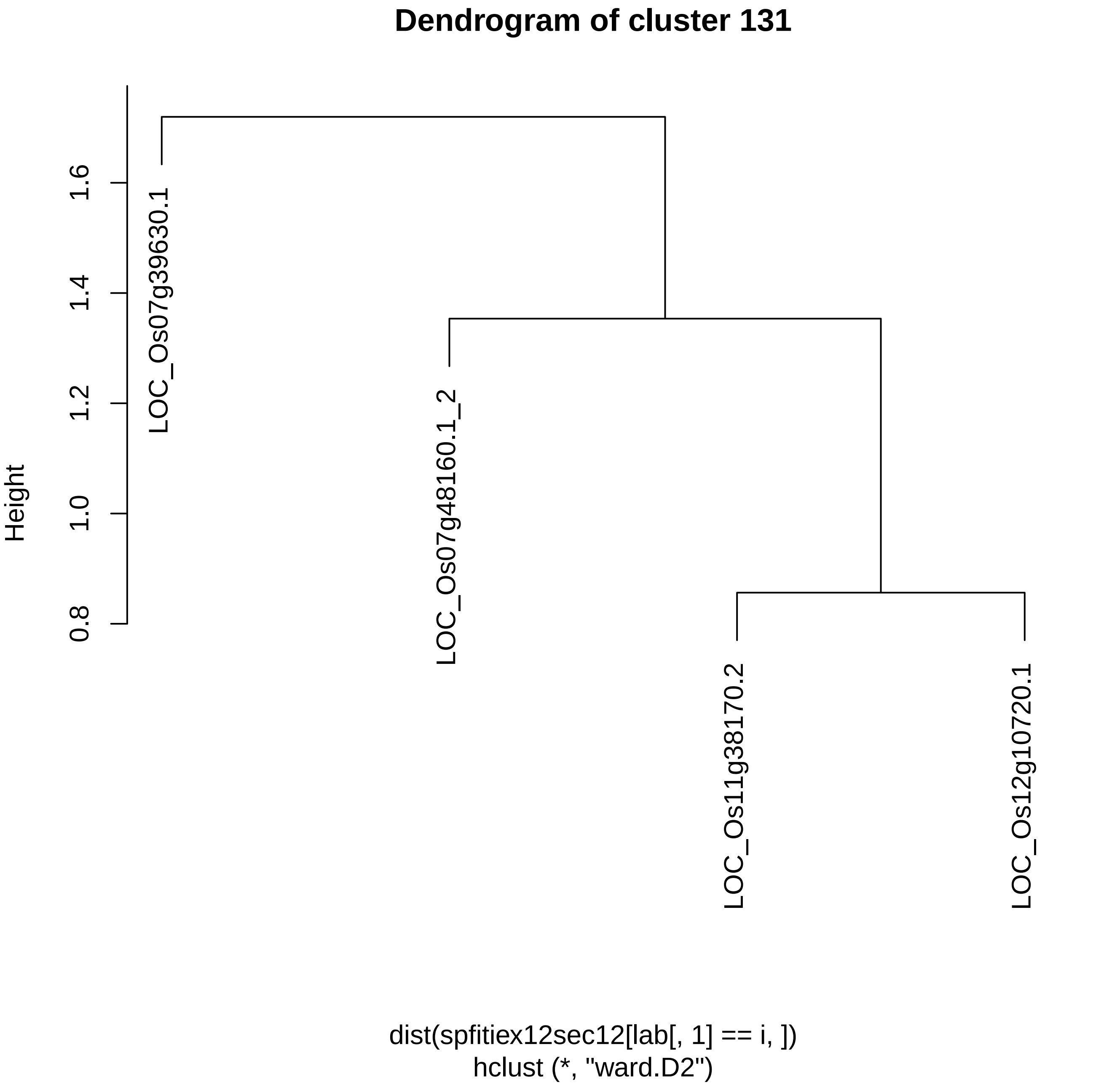
Dendrogram of cluster 131
Height
0.8
1.0
1.2
1.4
1.6
LOC_Os07g39630.1
LOC_Os07g48160.1_2
LOC_Os11g38170.2
LOC_Os12g10720.1
dist(spfitiex12sec12[lab[, 1] == i, ])
hclust (*, "ward.D2")

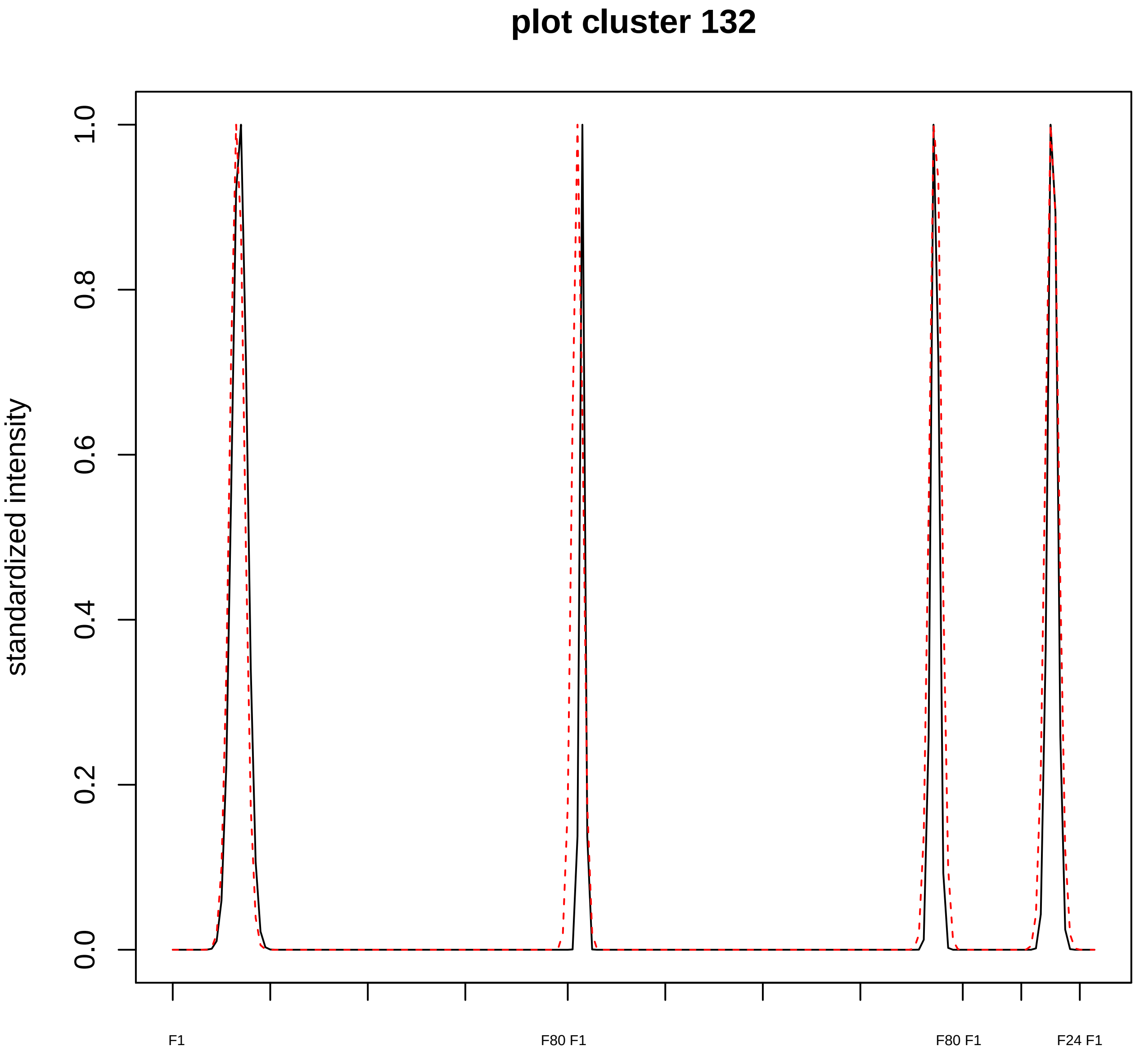
plot cluster 132
standardized intensity
0.0
0.2
0.4
0.6
0.8
1.0
F1
F80 F1
F80 F1
F24 F1

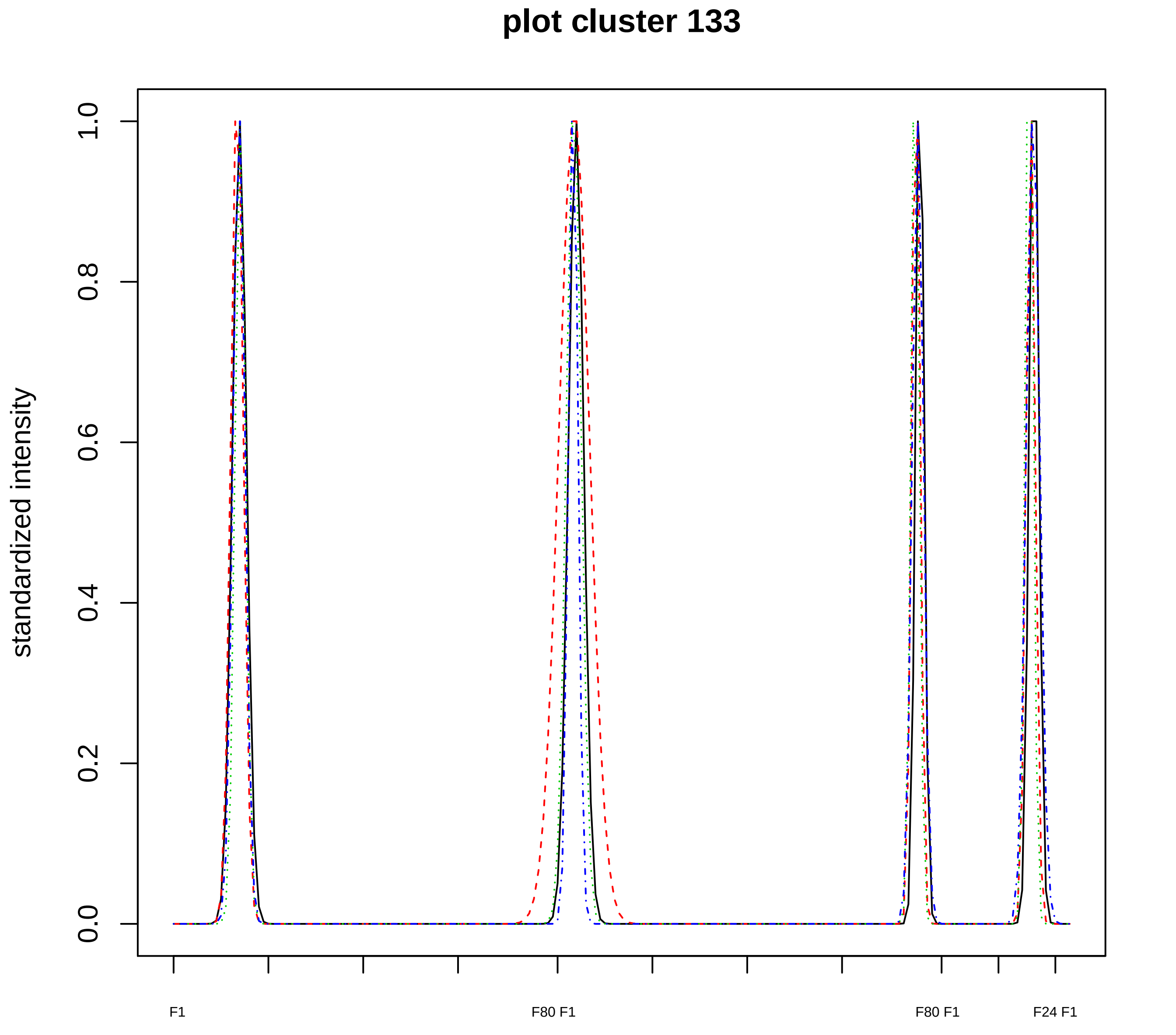

plot cluster 133
standardized intensity
0.0
0.2
0.4
0.6
0.8
1.0
F1
F80 F1
F80 F1
F24 F1

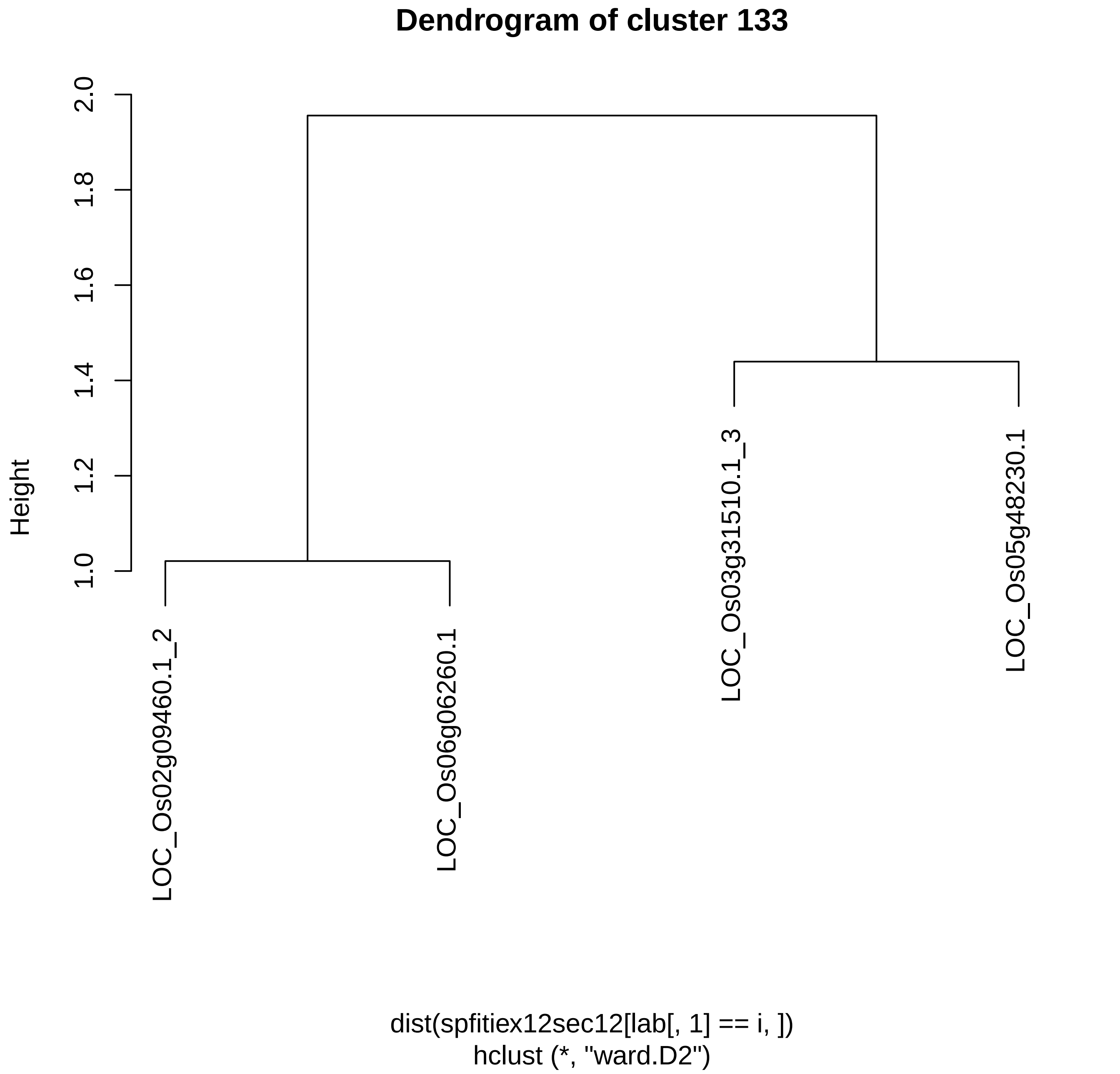
Dendrogram of cluster 133
Height
2.0
1.8
1.6
1.4
1.2
1.0
LOC_Os02g09460.1_2
LOC_Os06g06260.1
LOC_Os03g31510.1_3
LOC_Os05g48230.1
dist(spfitiex12sec12[lab[, 1] == i, ])
hclust (*, "ward.D2")

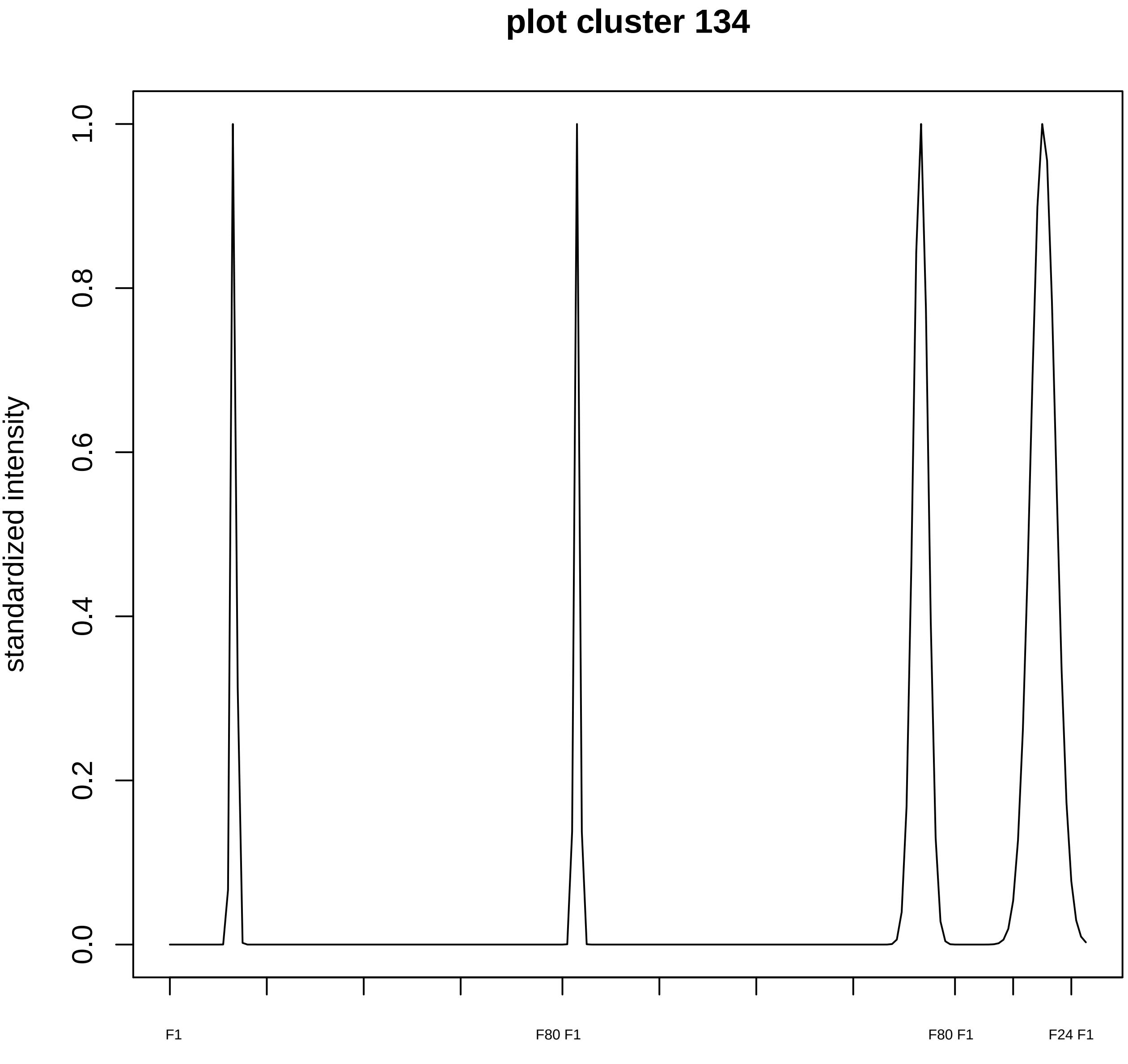

plot cluster 134
standardized intensity
0.0
0.2
0.4
0.6
0.8
1.0
F1
F80 F1
F80 F1
F24 F1

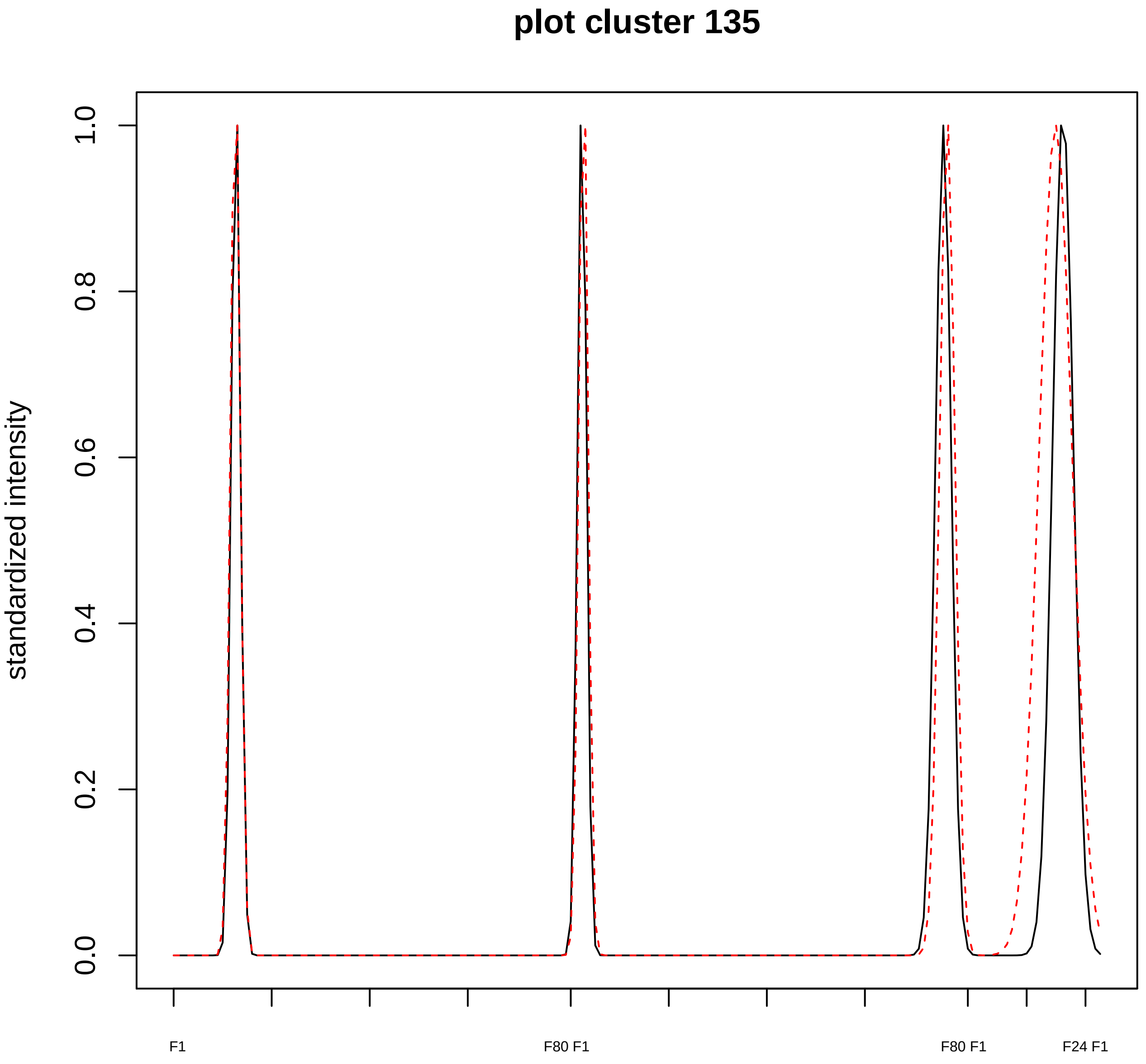
plot cluster 135
standardized intensity
0.0
0.2
0.4
0.6
0.8
1.0
F1
F80 F1
F80 F1
F24 F1

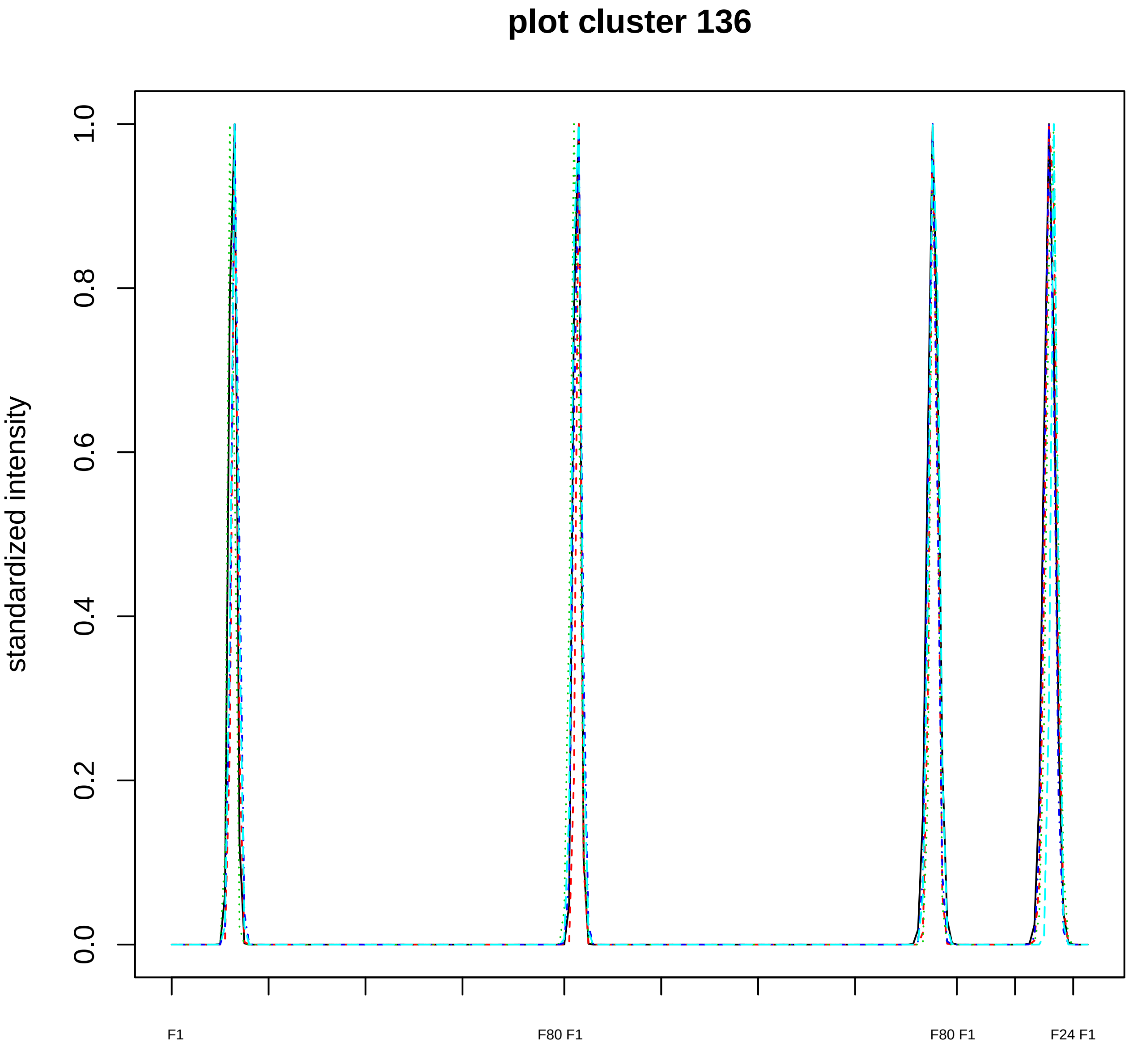
plot cluster 136
standardized intensity
0.0
0.2
0.4
0.6
0.8
1.0
F1
F80 F1
F80 F1
F24 F1

### Dendrogram of cluster 136

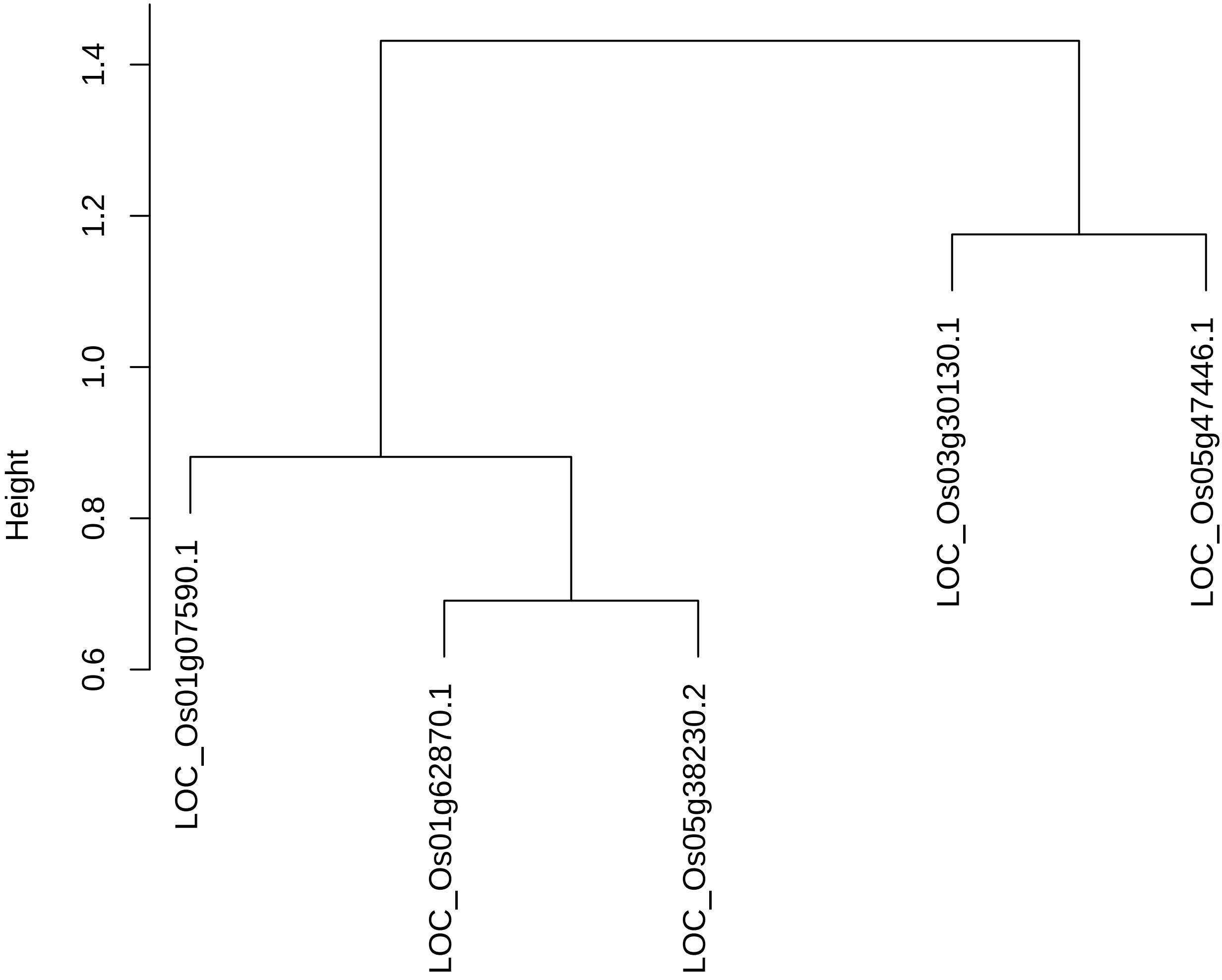


dist(spfitiex12sec12[lab[, 1] == i, ])
hclust (*, "ward.D2")

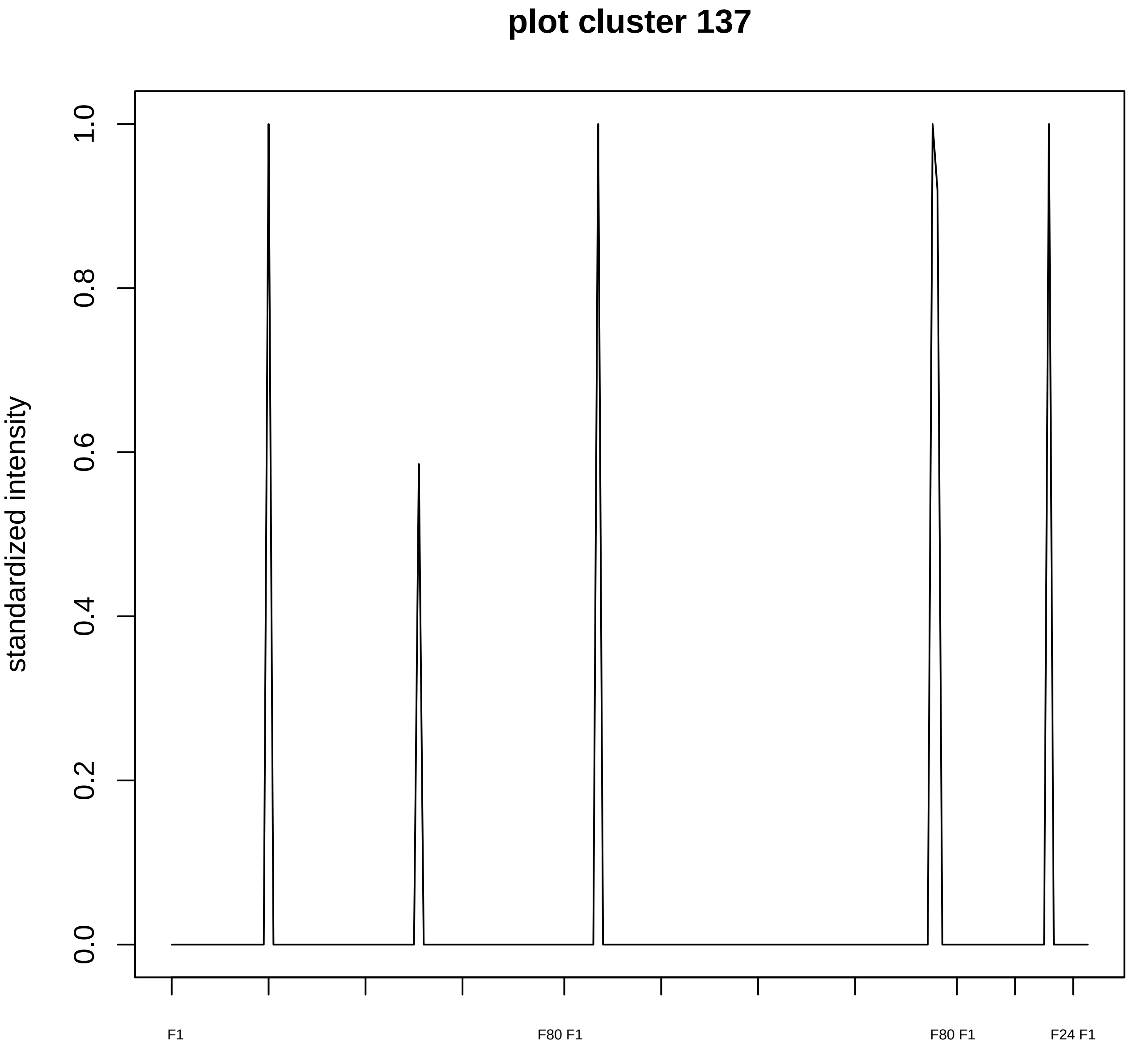

plot cluster 137
standardized intensity
0.0
0.2
0.4
0.6
0.8
1.0
F1
F80 F1
F80 F1
F24 F1

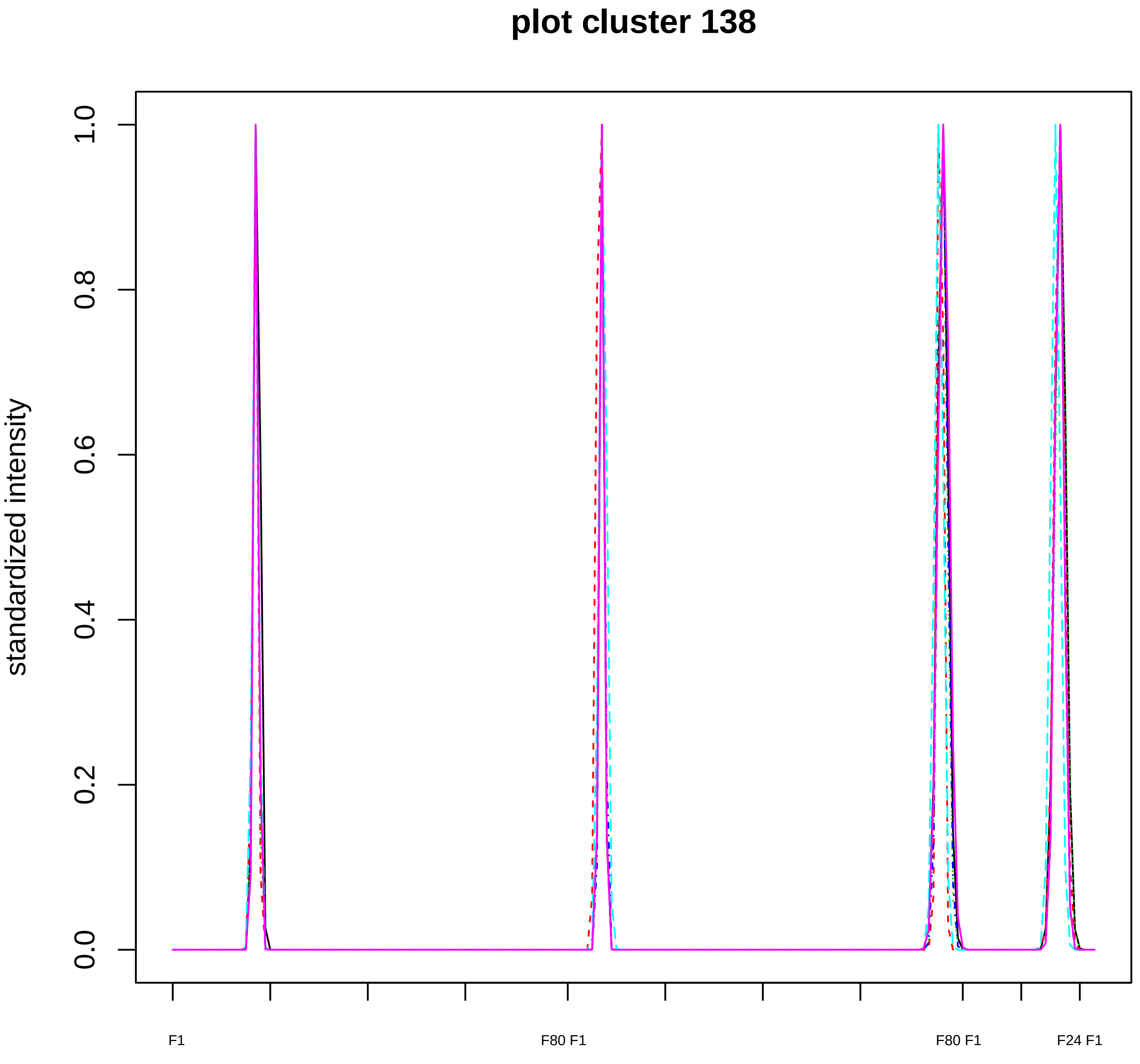

plot cluster 138
standardized intensity
0.0
0.2
0.4
0.6
0.8
1.0
F1
F80 F1
F80 F1
F24 F1

### Dendrogram of cluster 138

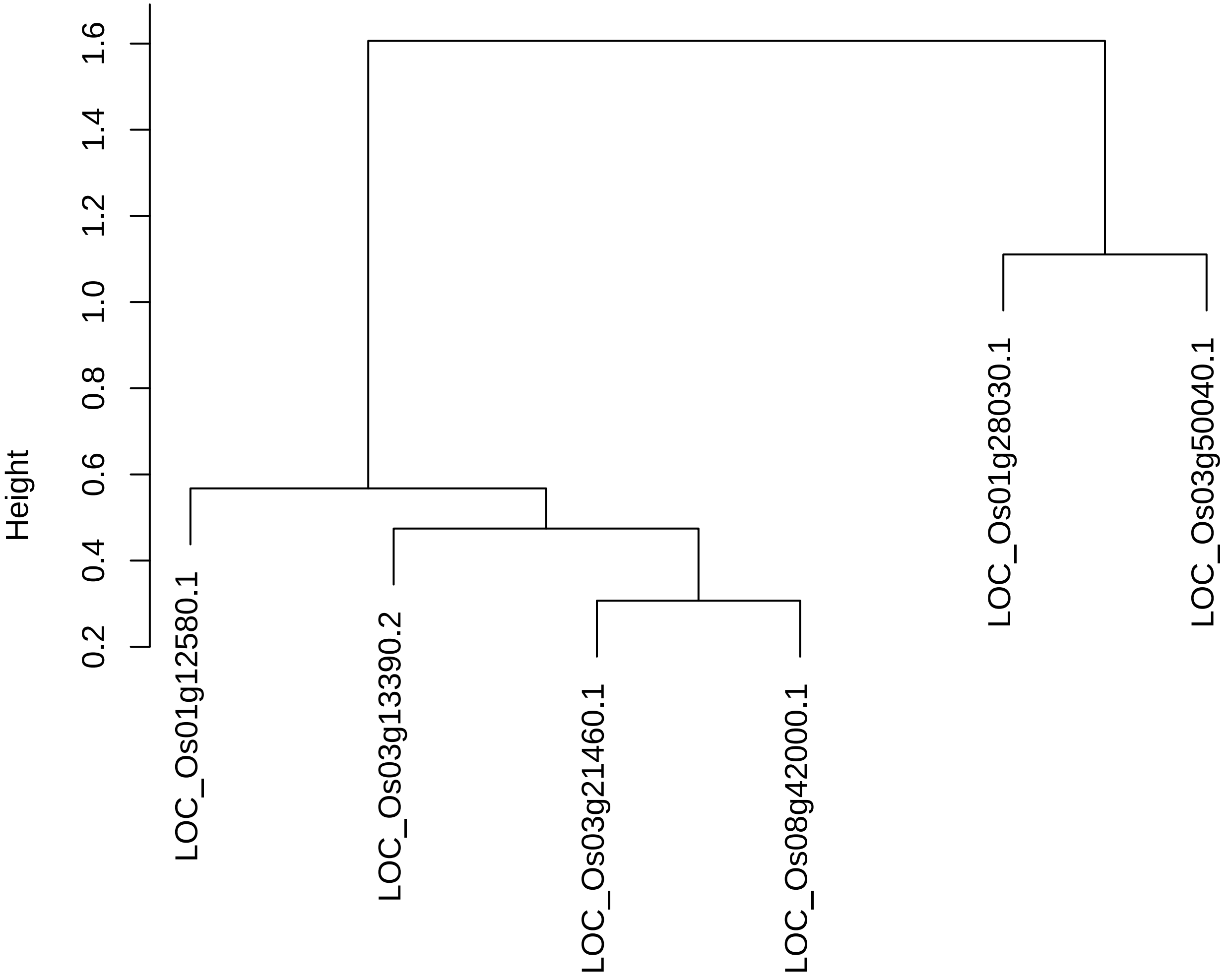


dist(spfitiex12sec12[lab[, 1] == i, ])
hclust (*, "ward.D2")

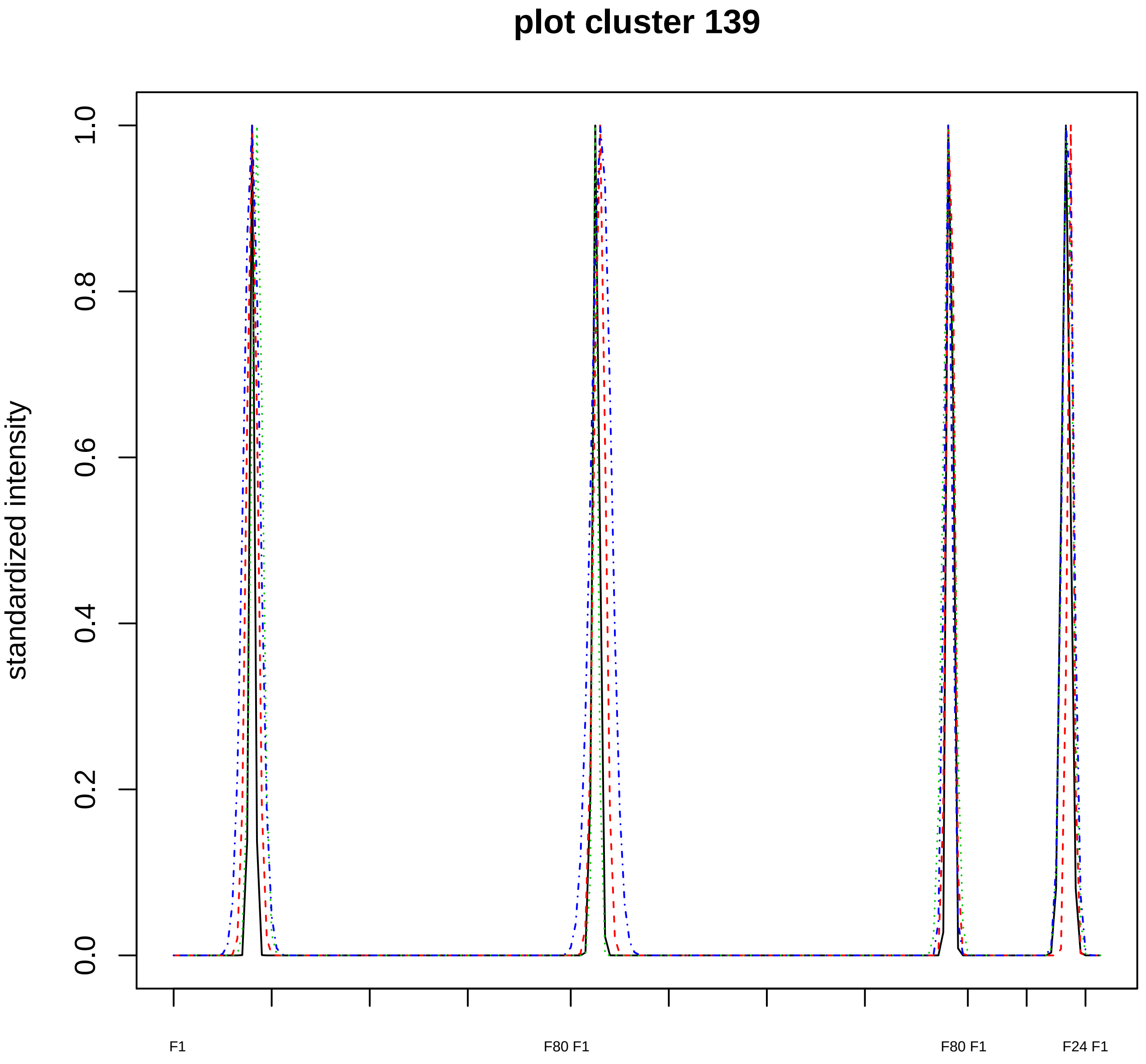
plot cluster 139
standardized intensity
1.0
0.8
0.6
0.4
0.2
0.0
F1
F80 F1
F80 F1
F24 F1

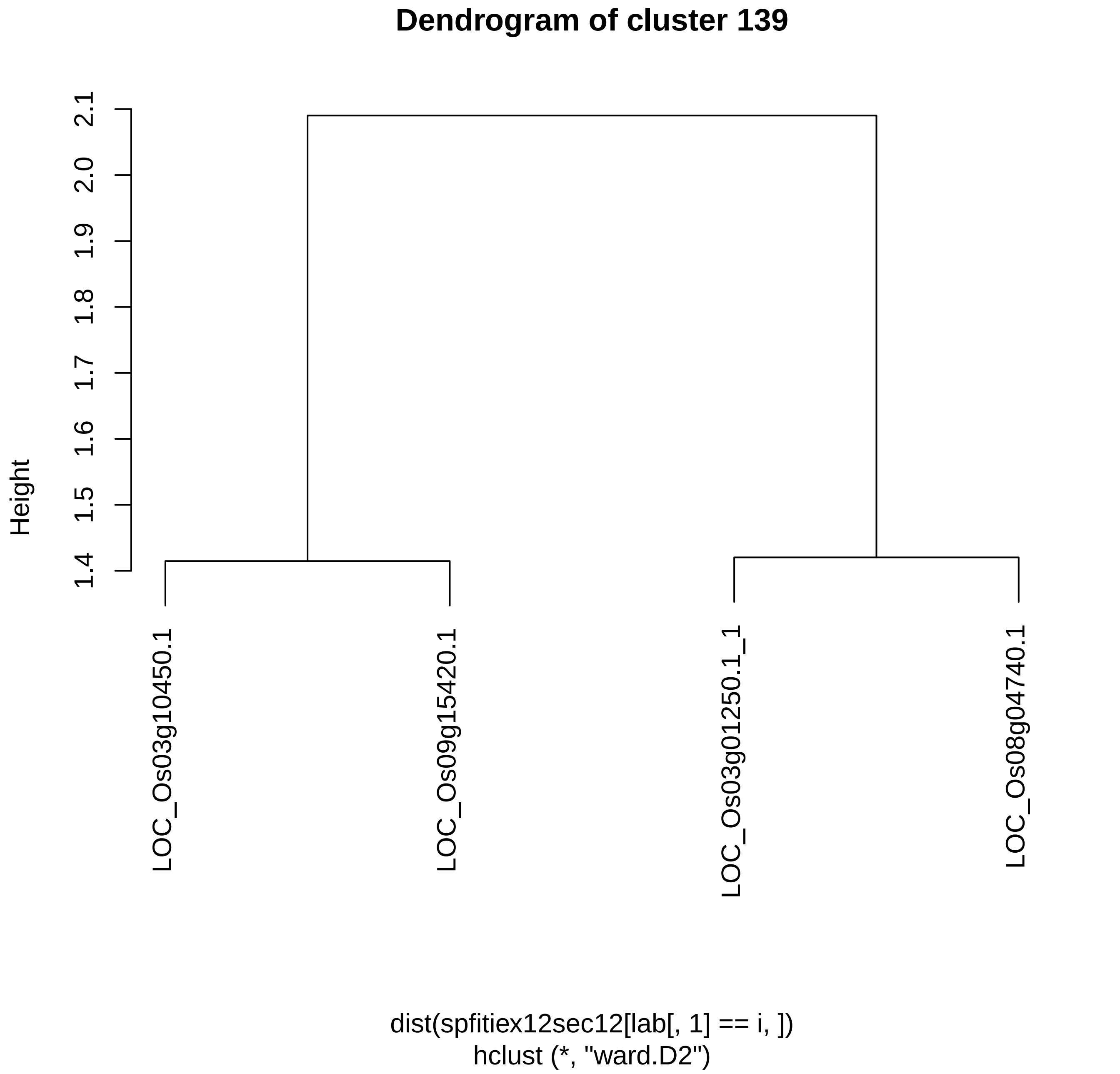

Dendrogram of cluster 139
Height
2.1
2.0
1.9
1.8
1.7
1.6
1.5
1.4
LOC_Os03g10450.1
LOC_Os09g15420.1
LOC_Os03g01250.1_1
LOC_Os08g04740.1
dist(spfitiex12sec12[lab[, 1] == i, ])
hclust (*, "ward.D2")

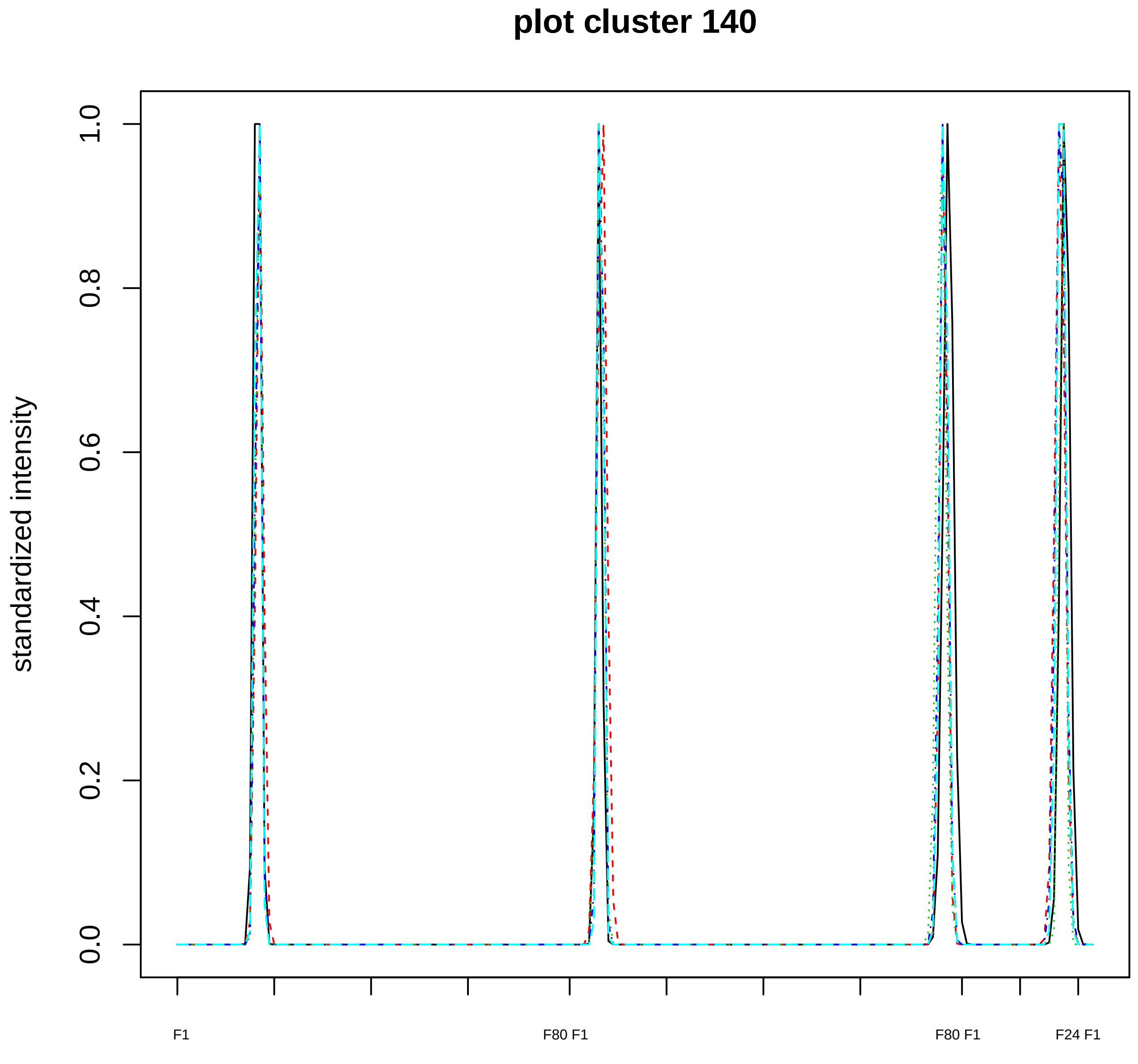
plot cluster 140
standardized intensity
0.0
0.2
0.4
0.6
0.8
1.0
F1
F80 F1
F80 F1
F24 F1

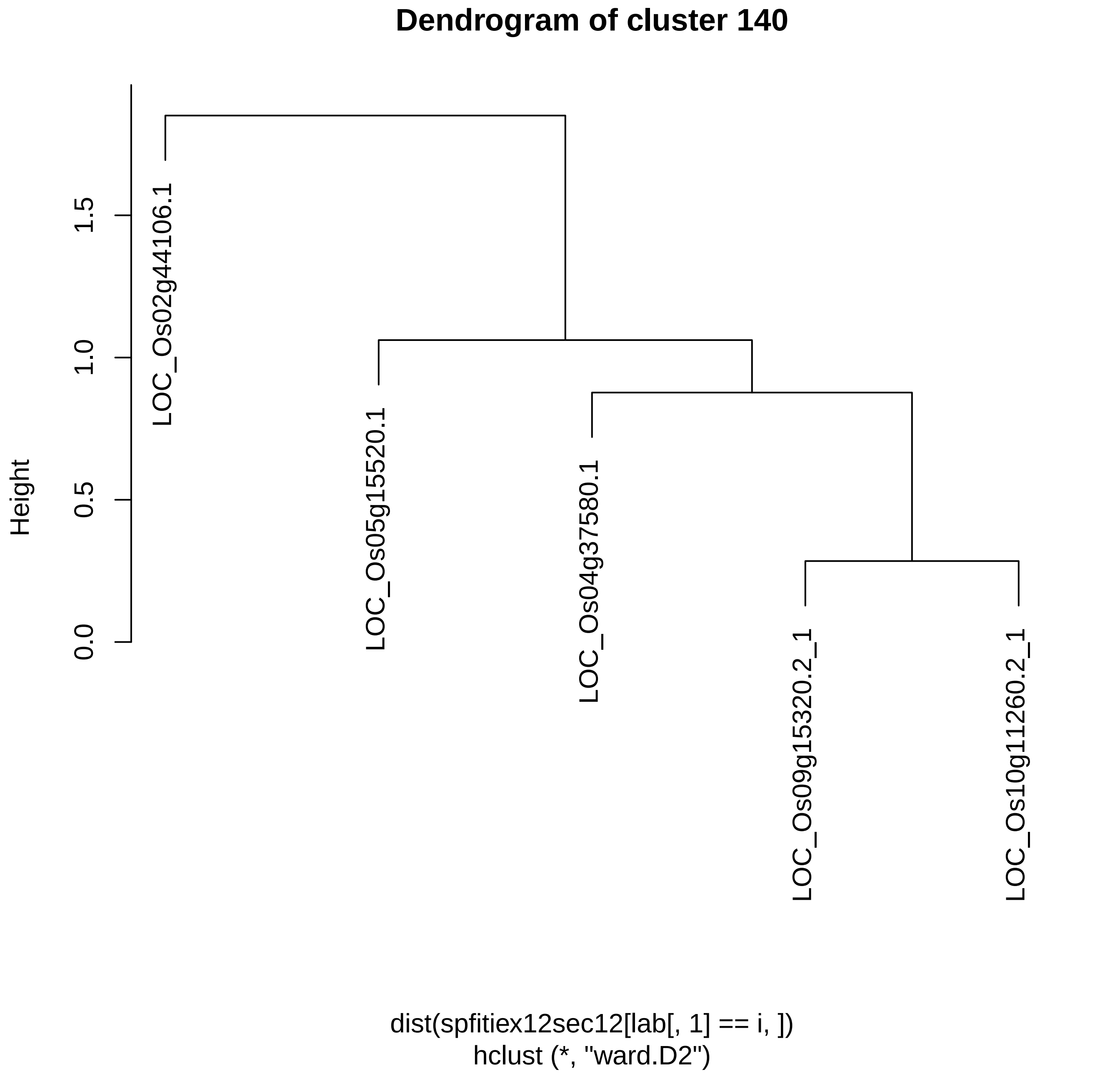

Dendrogram of cluster 140
Height
0.0
0.5
1.0
1.5
LOC_Os02g44106.1
LOC_Os05g15520.1
LOC_Os04g37580.1
LOC_Os09g15320.2_1
LOC_Os10g11260.2_1
dist(spfitiex12sec12[lab[, 1] == i, ])
hclust (*, "ward.D2")

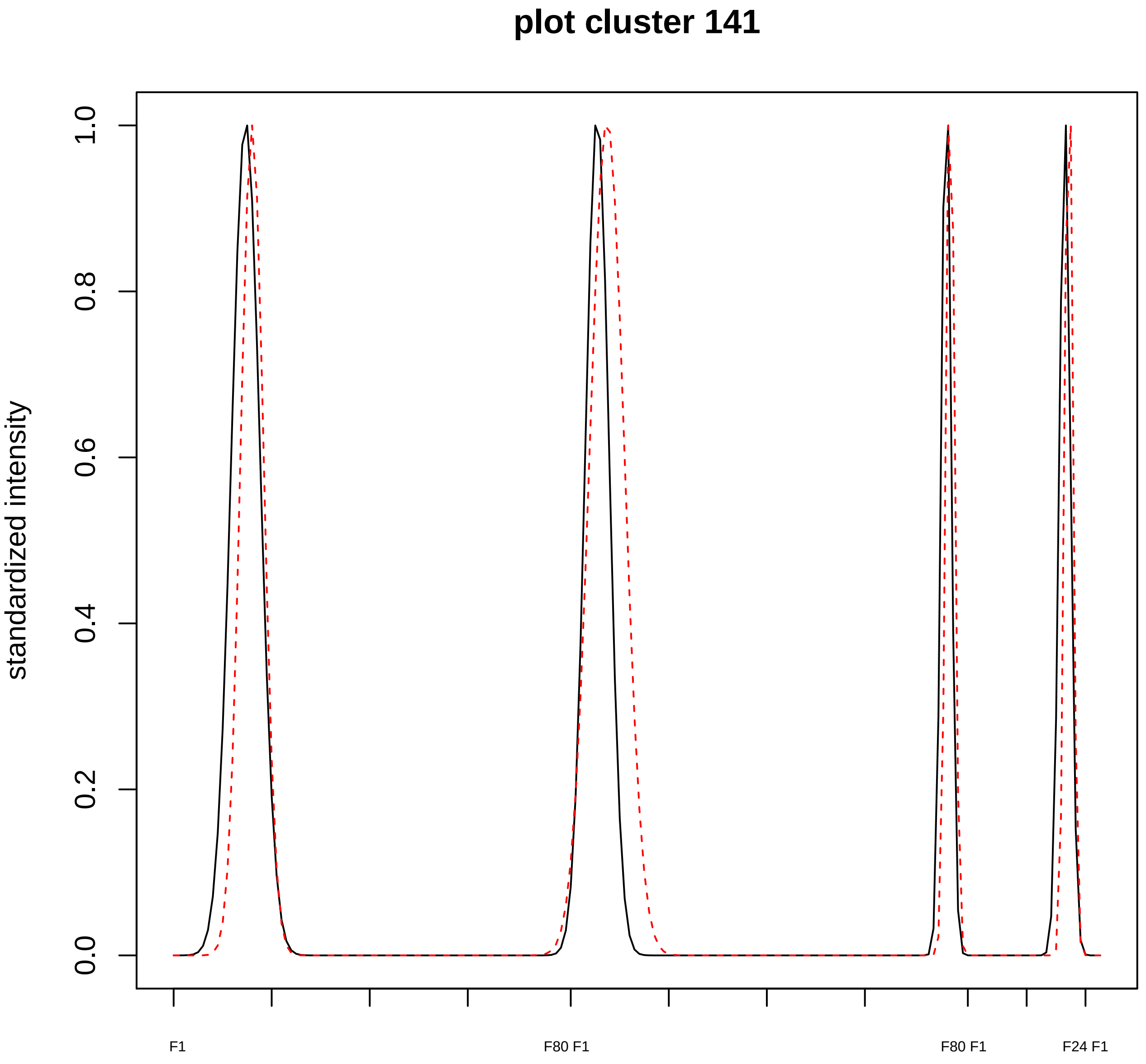
plot cluster 141
standardized intensity
0.0
0.2
0.4
0.6
0.8
1.0
F1
F80 F1
F80 F1
F24 F1

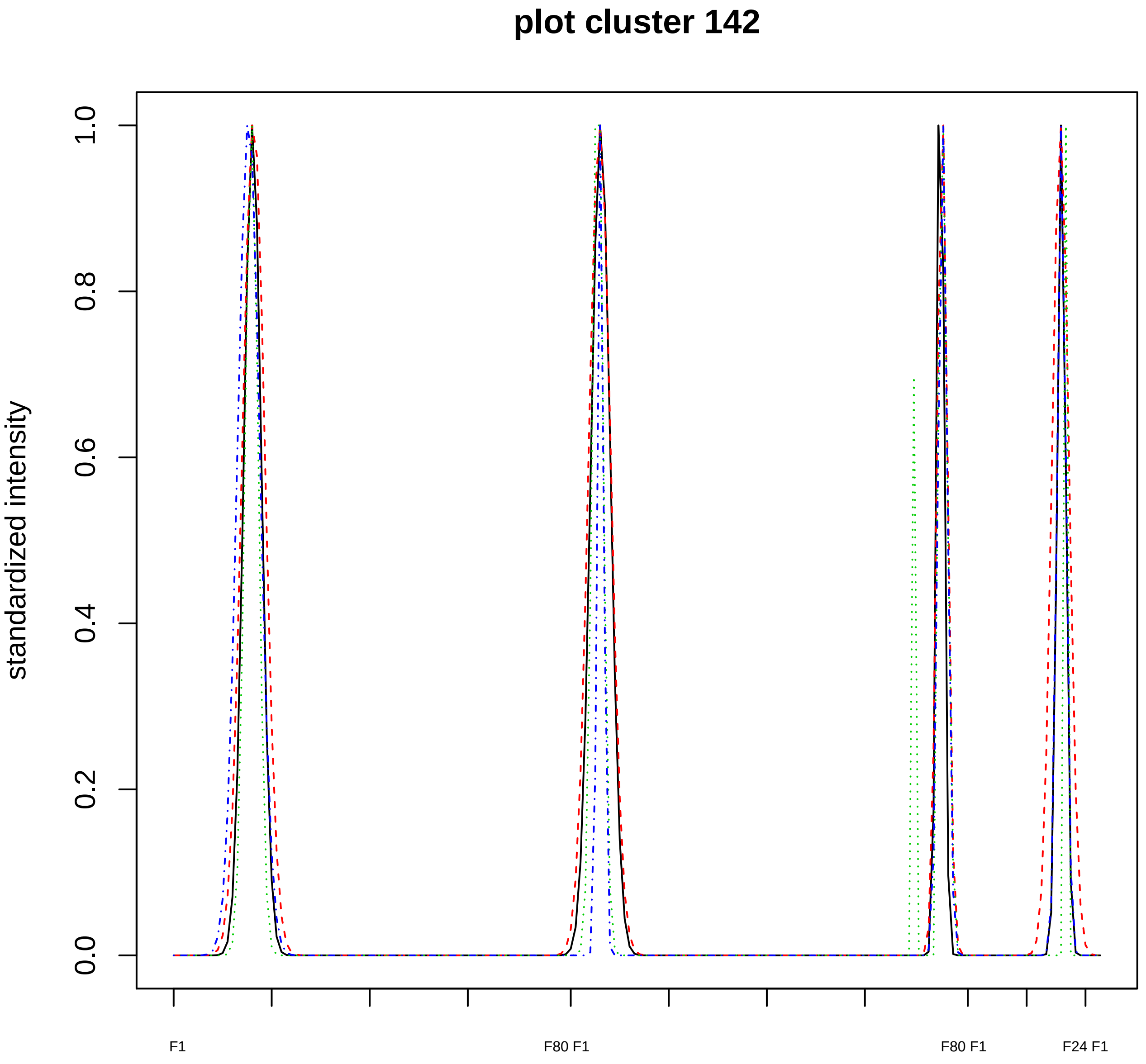
plot cluster 142
standardized intensity
1.0
0.8
0.6
0.4
0.2
0.0
F1
F80 F1
F80 F1
F24 F1

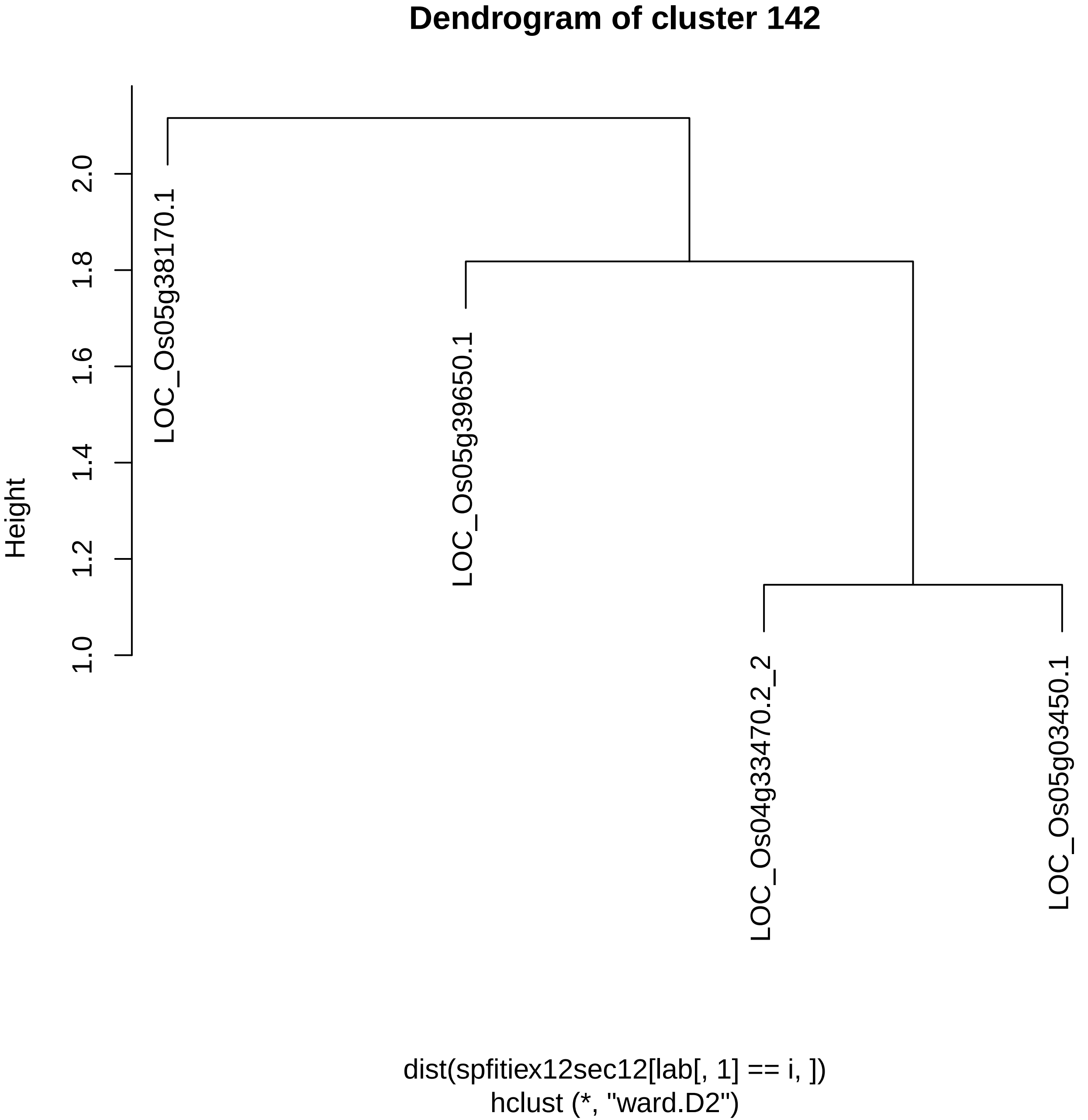

Dendrogram of cluster 142
Height
2.0
1.8
1.6
1.4
1.2
1.0
LOC_Os05g38170.1
LOC_Os05g39650.1
LOC_Os04g33470.2_2
LOC_Os05g03450.1
dist(spfitiex12sec12[lab[, 1] == i, ])
hclust (*, "ward.D2")

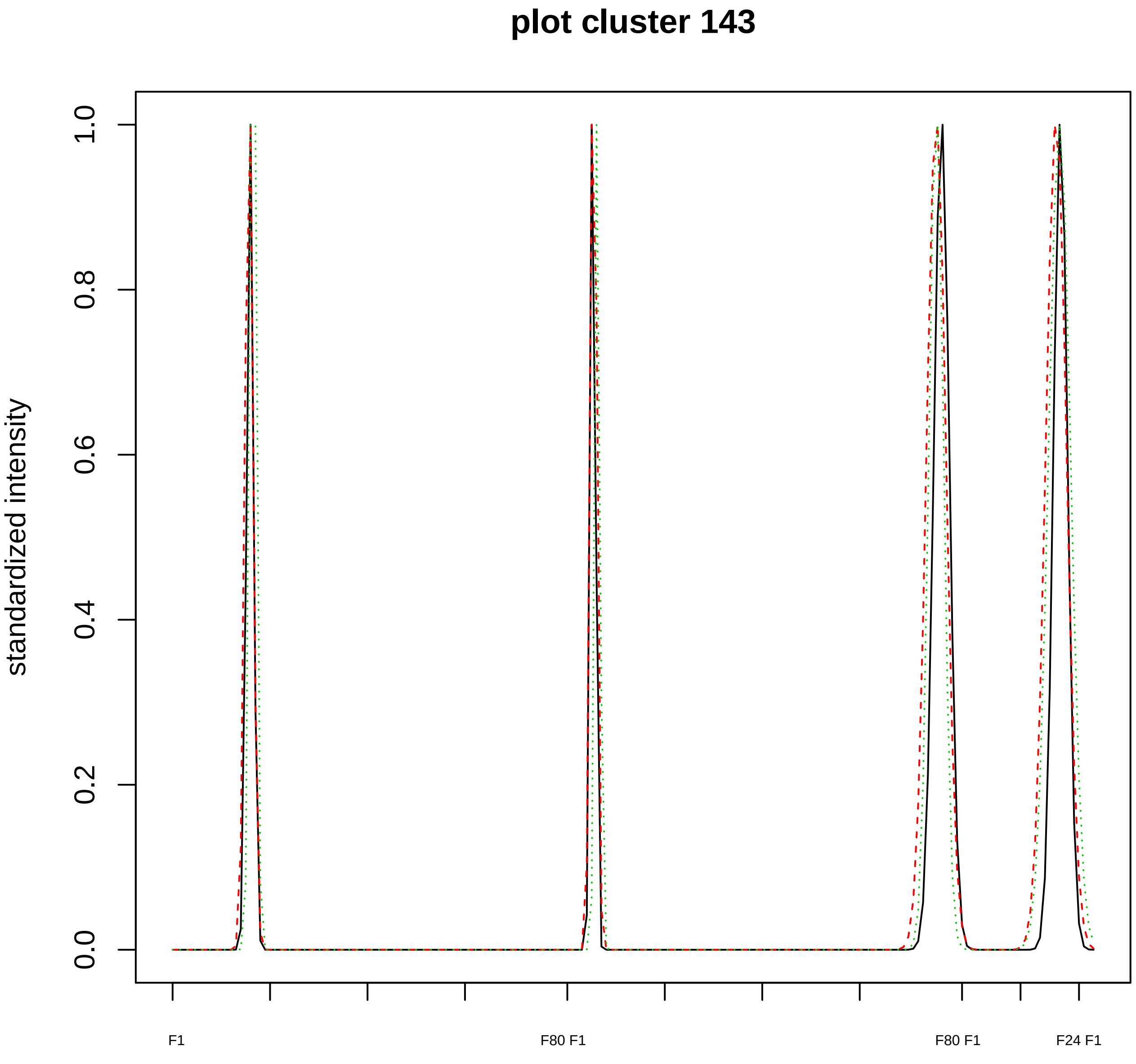
plot cluster 143
standardized intensity
0.0
0.2
0.4
0.6
0.8
1.0
F1
F80 F1
F80 F1
F24 F1

#### Dendrogram of cluster 143

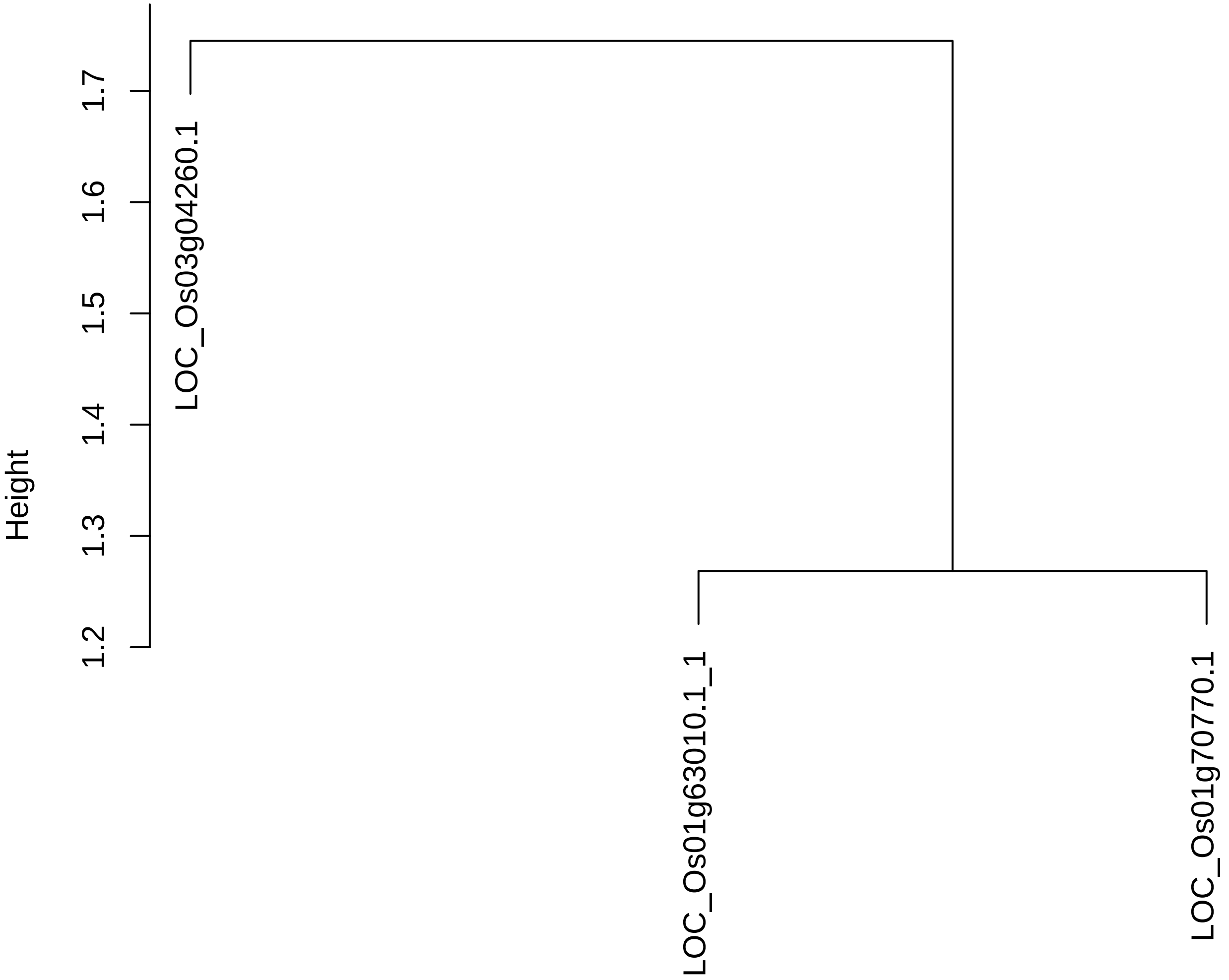


dist(spfitiex12sec12[lab[, 1] == i, ])
hclust (*, "ward.D2")

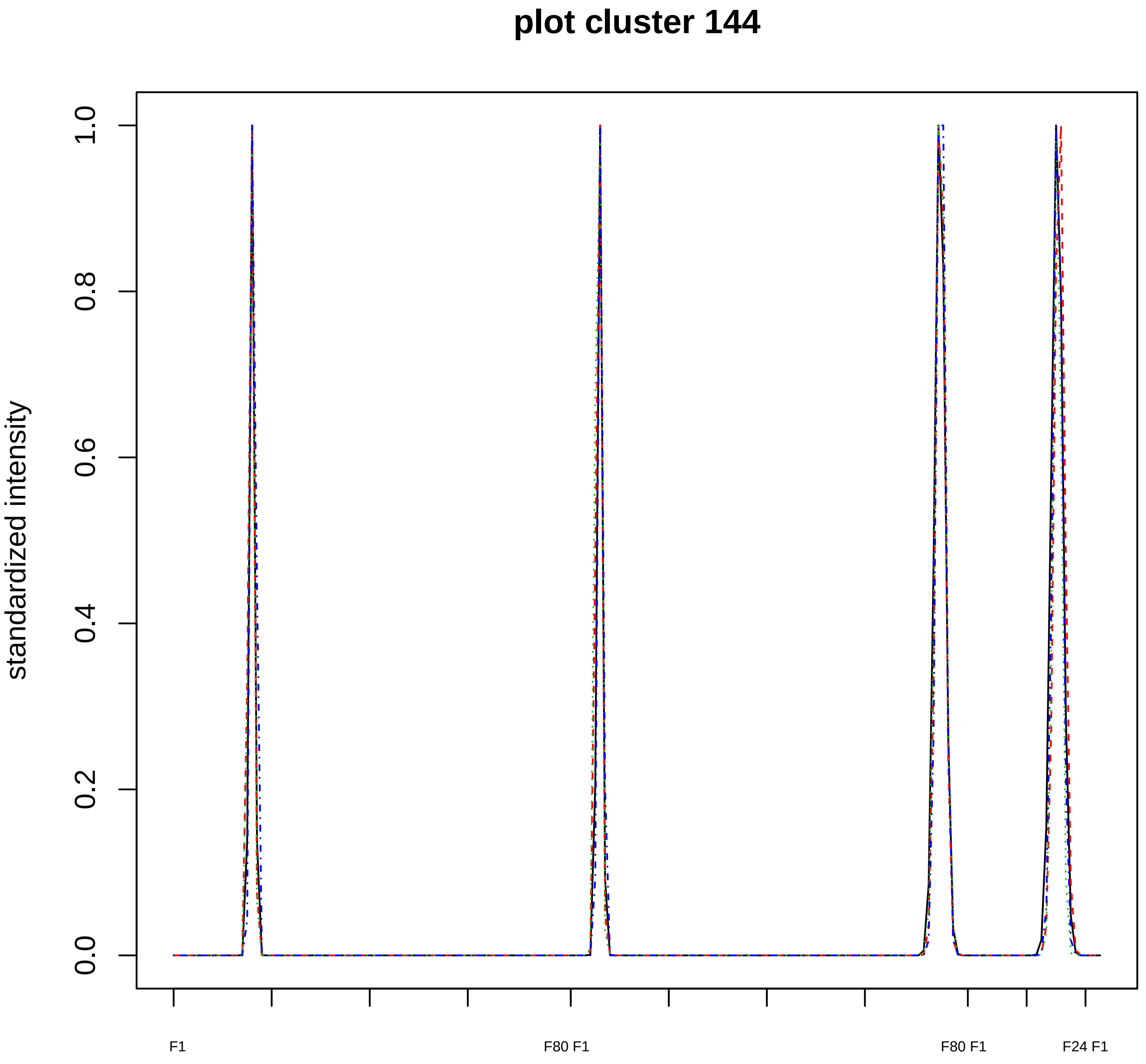
plot cluster 144
standardized intensity
0.0
0.2
0.4
0.6
0.8
1.0
F1
F80 F1
F80 F1
F24 F1

#### Dendrogram of cluster 144

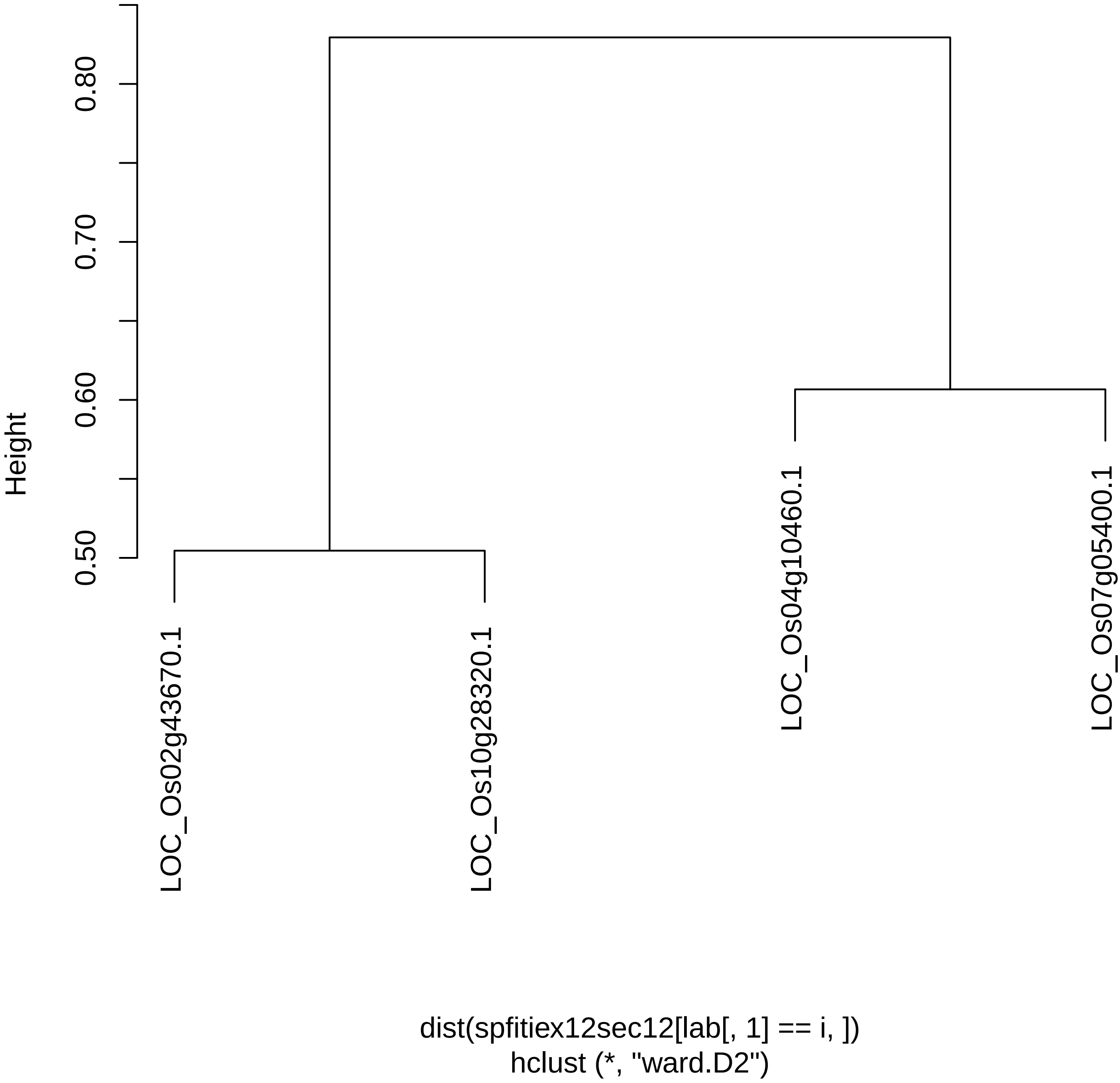

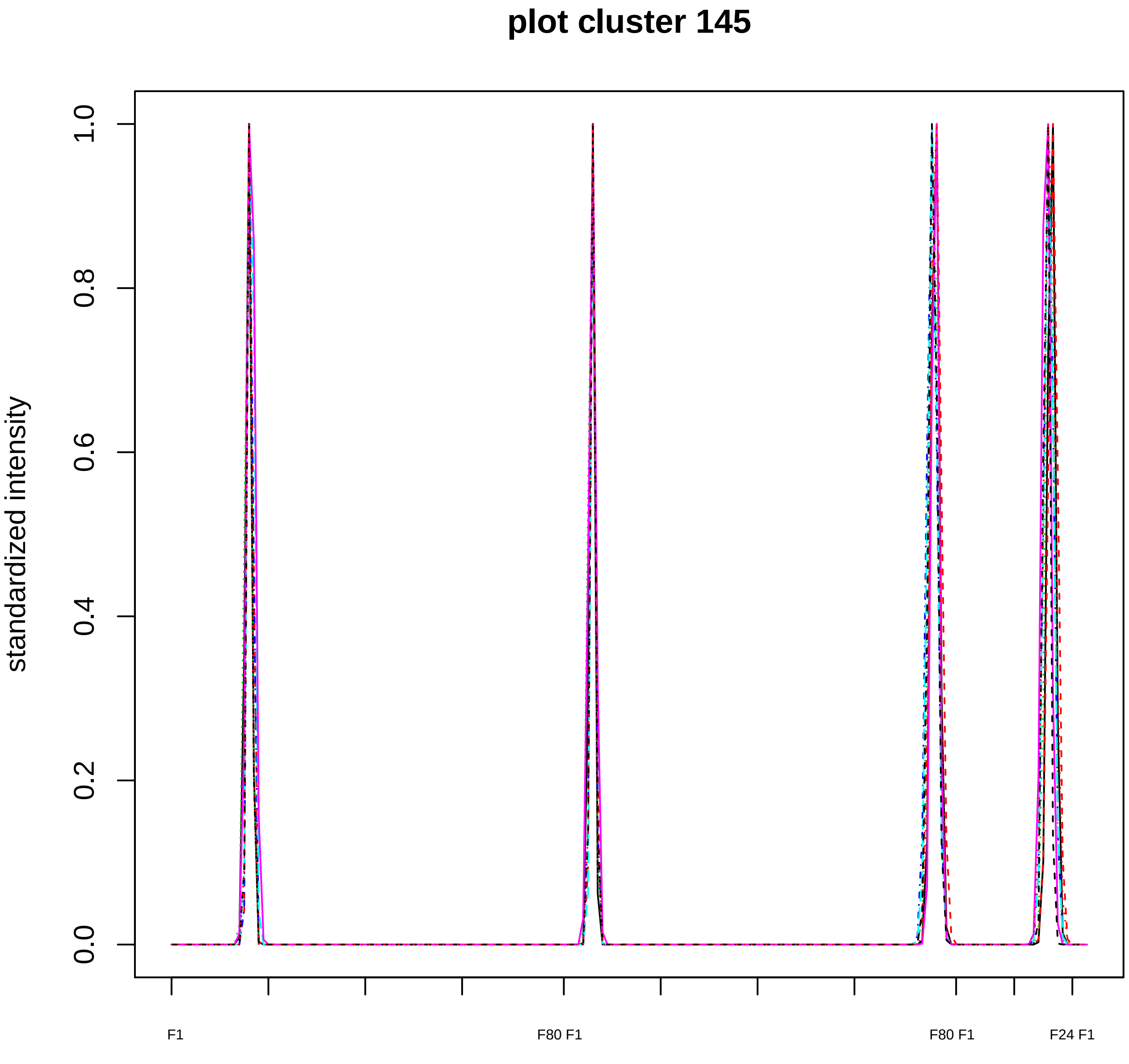

plot cluster 145
standardized intensity
0.0
0.2
0.4
0.6
0.8
1.0
F1
F80 F1
F80 F1
F24 F1

#### Dendrogram of cluster 145

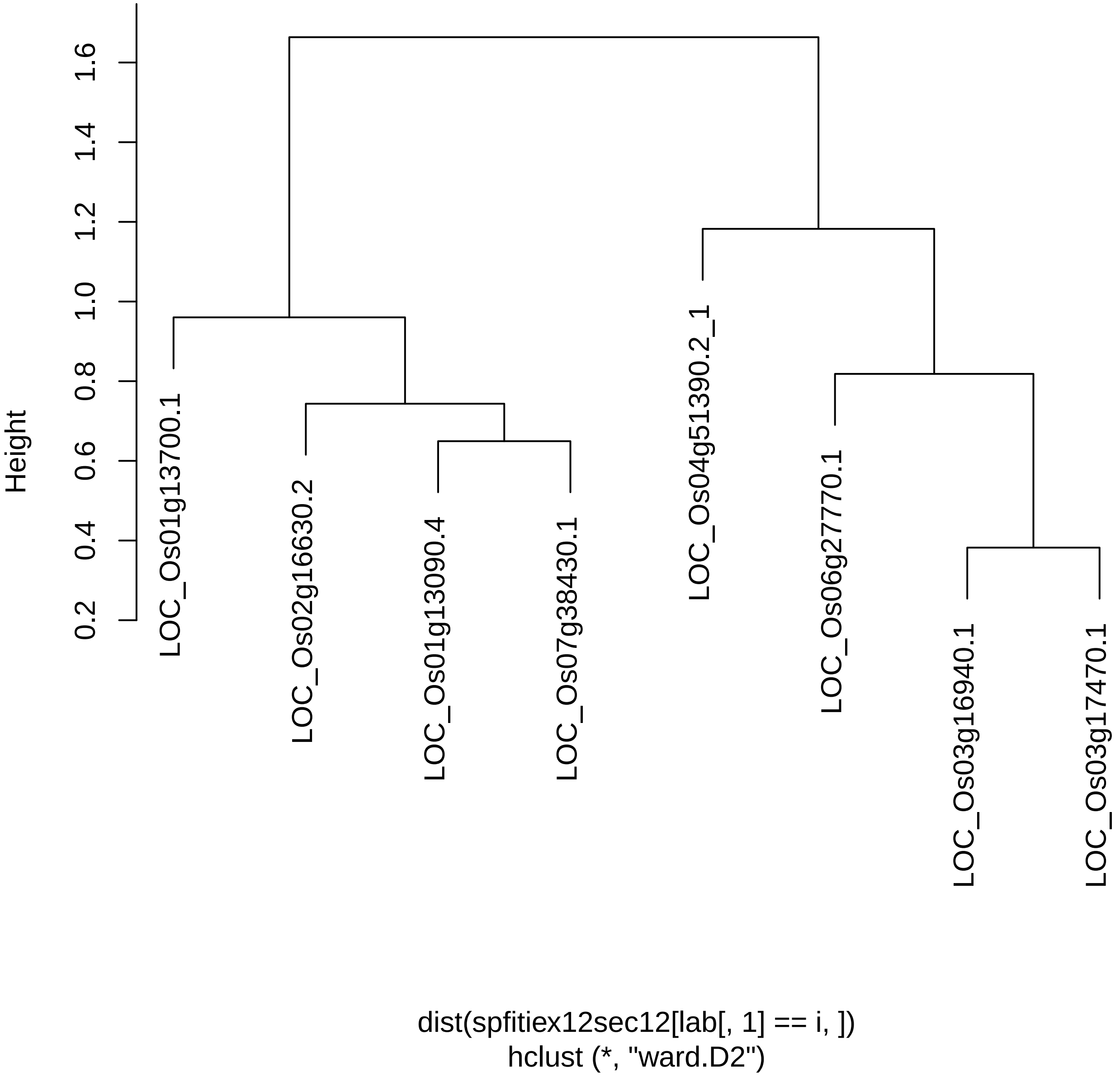

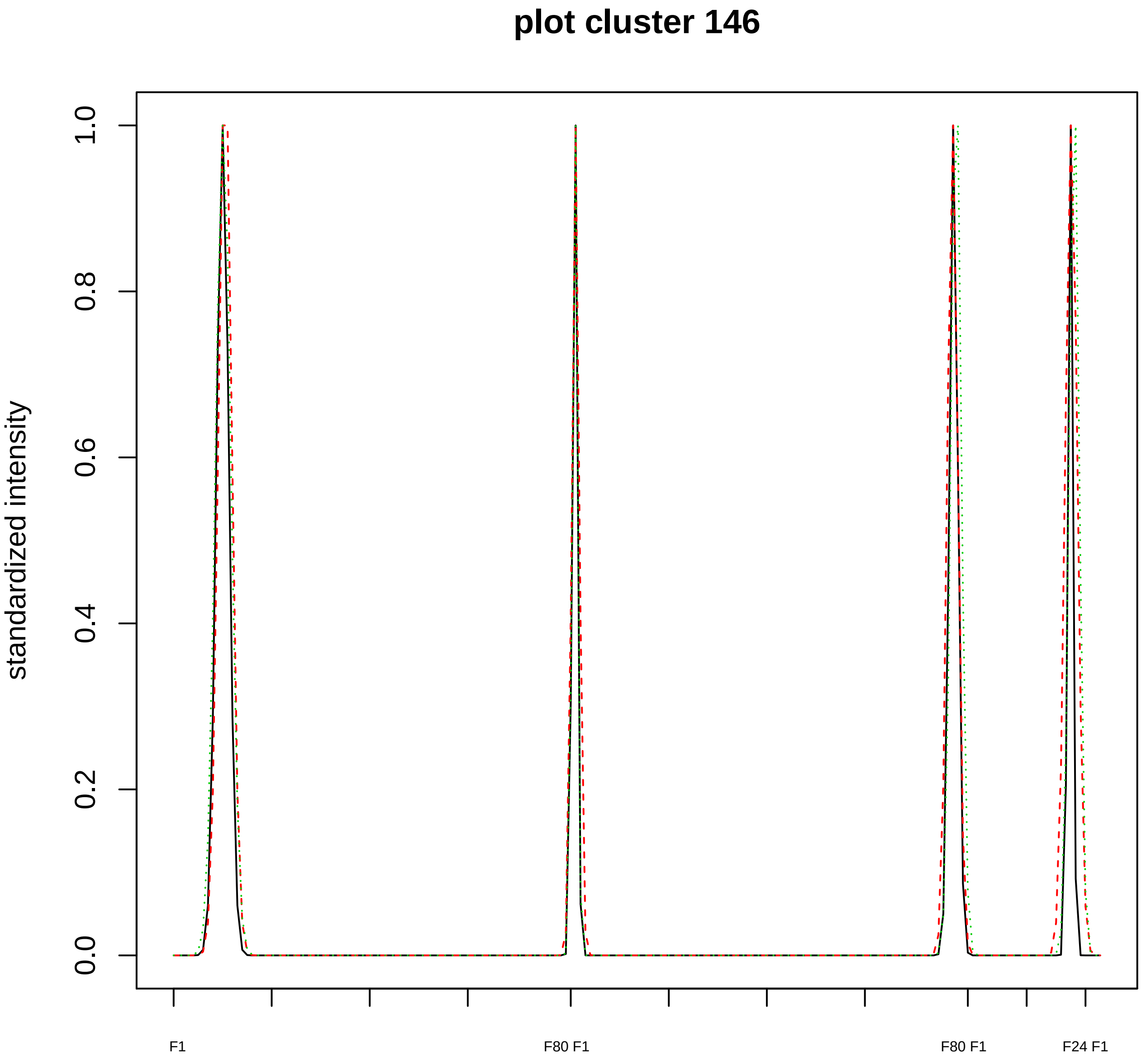

plot cluster 146
standardized intensity
0.0
0.2
0.4
0.6
0.8
1.0
F1
F80 F1
F80 F1
F24 F1

#### Dendrogram of cluster 146

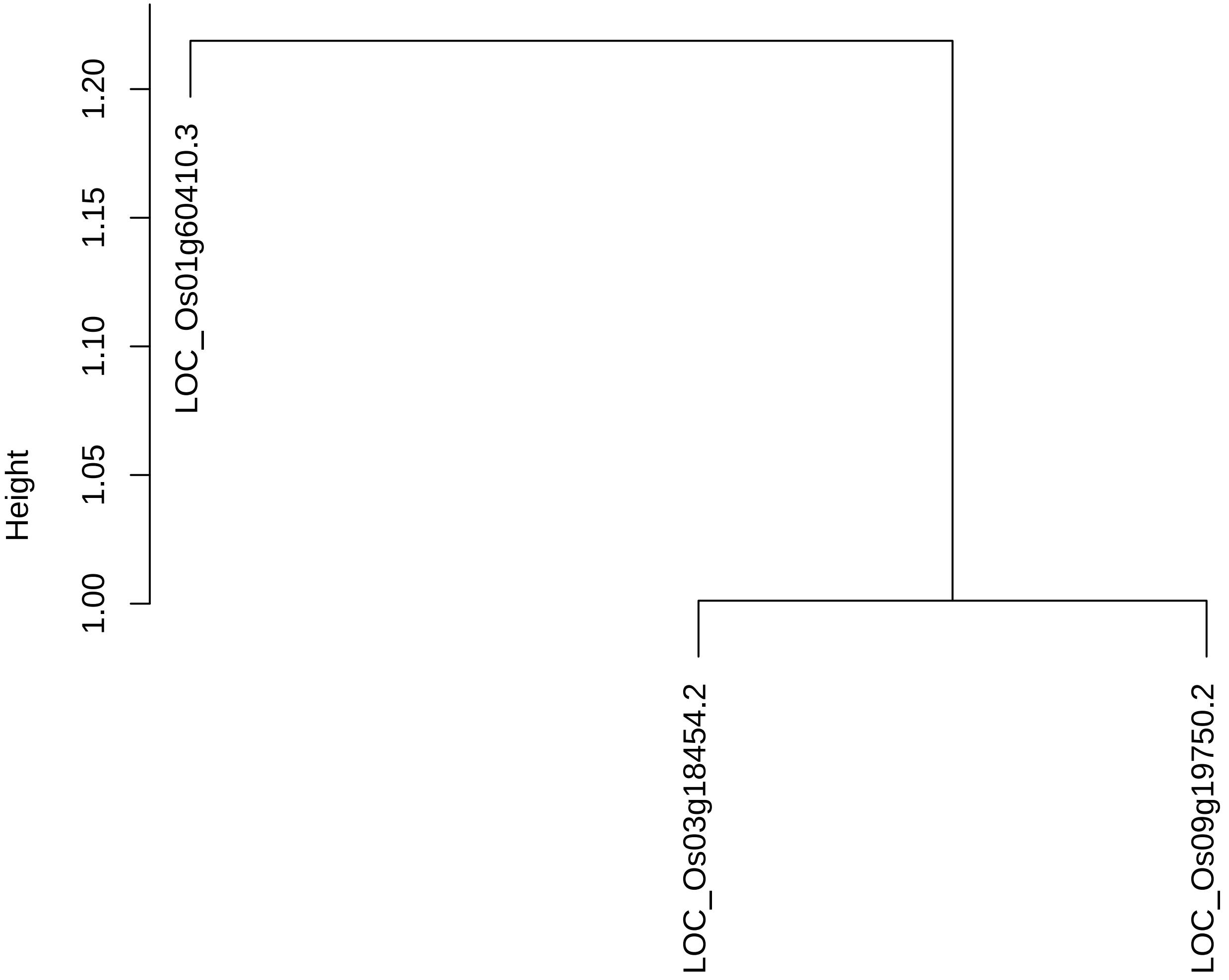


dist(spfitiex12sec12[lab[, 1] == i, ])
hclust (*, "ward.D2")

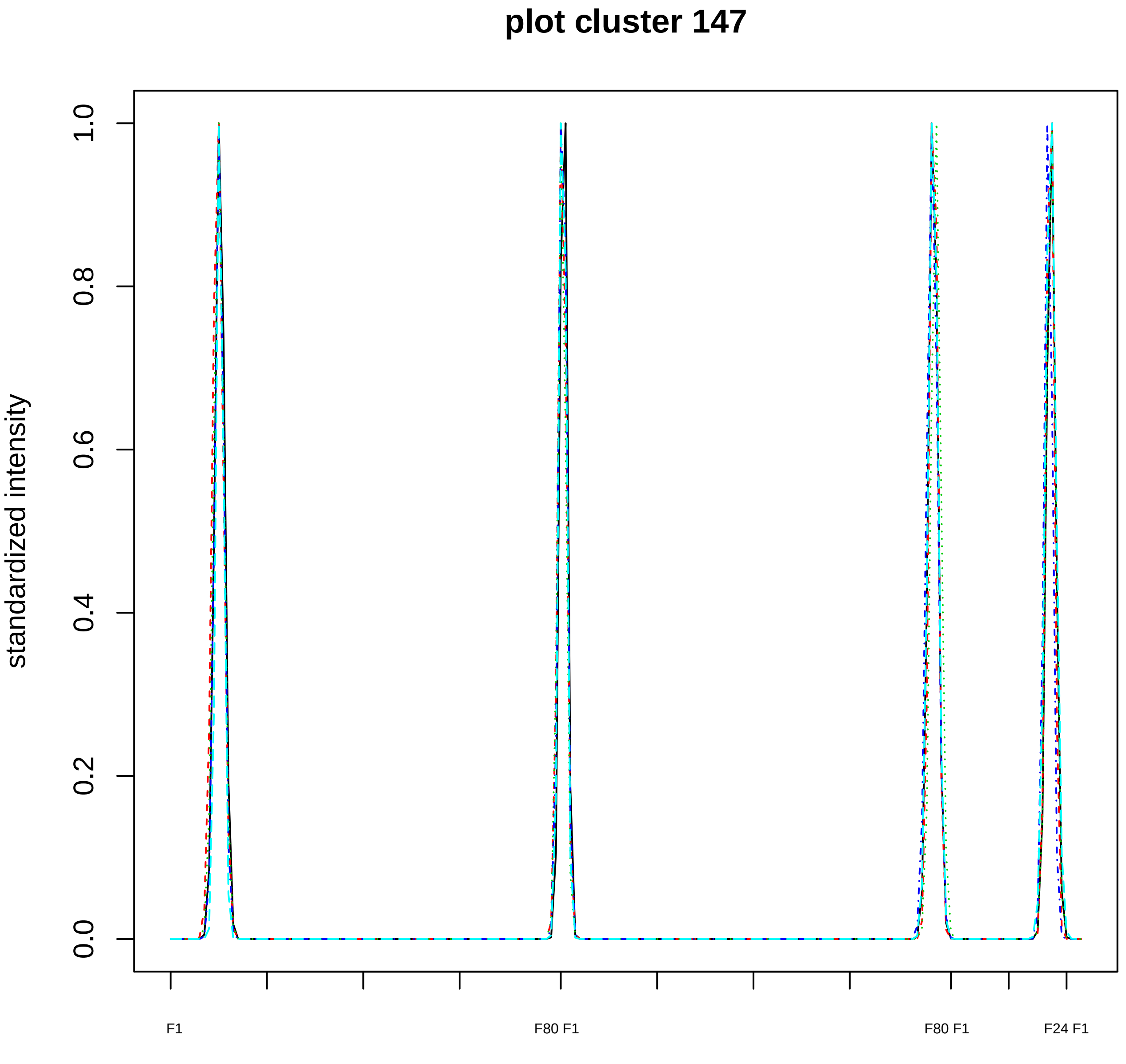

plot cluster 147
standardized intensity
0.0
0.2
0.4
0.6
0.8
1.0
F1
F80 F1
F80 F1
F24 F1

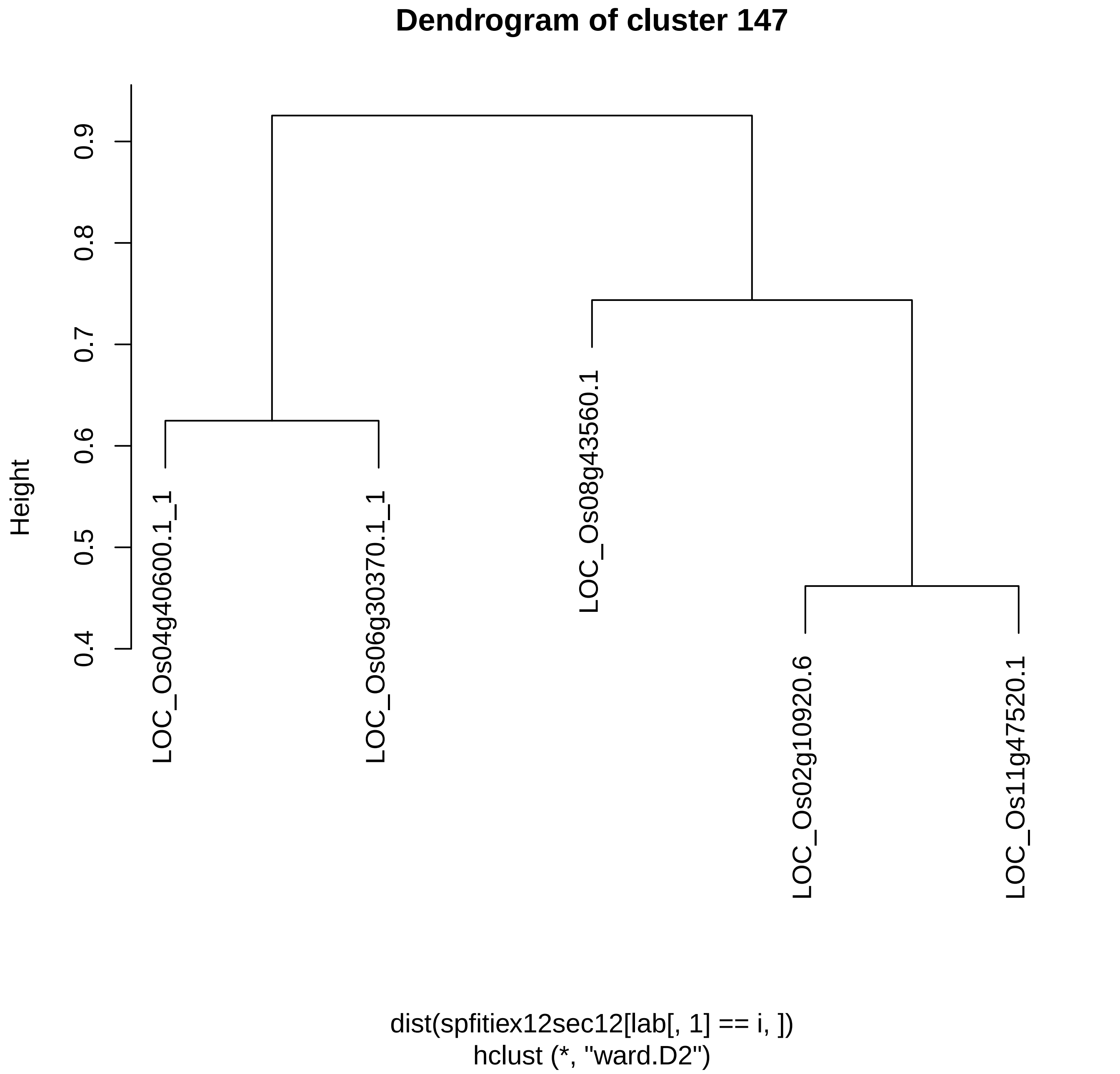
Dendrogram of cluster 147
Height
0.4
0.5
0.6
0.7
0.8
0.9
LOC_Os04g40600.1_1
LOC_Os06g30370.1_1
LOC_Os08g43560.1
LOC_Os02g10920.6
LOC_Os11g47520.1
dist(spfitiex12sec12[lab[, 1] == i, ])
hclust (*, "ward.D2")

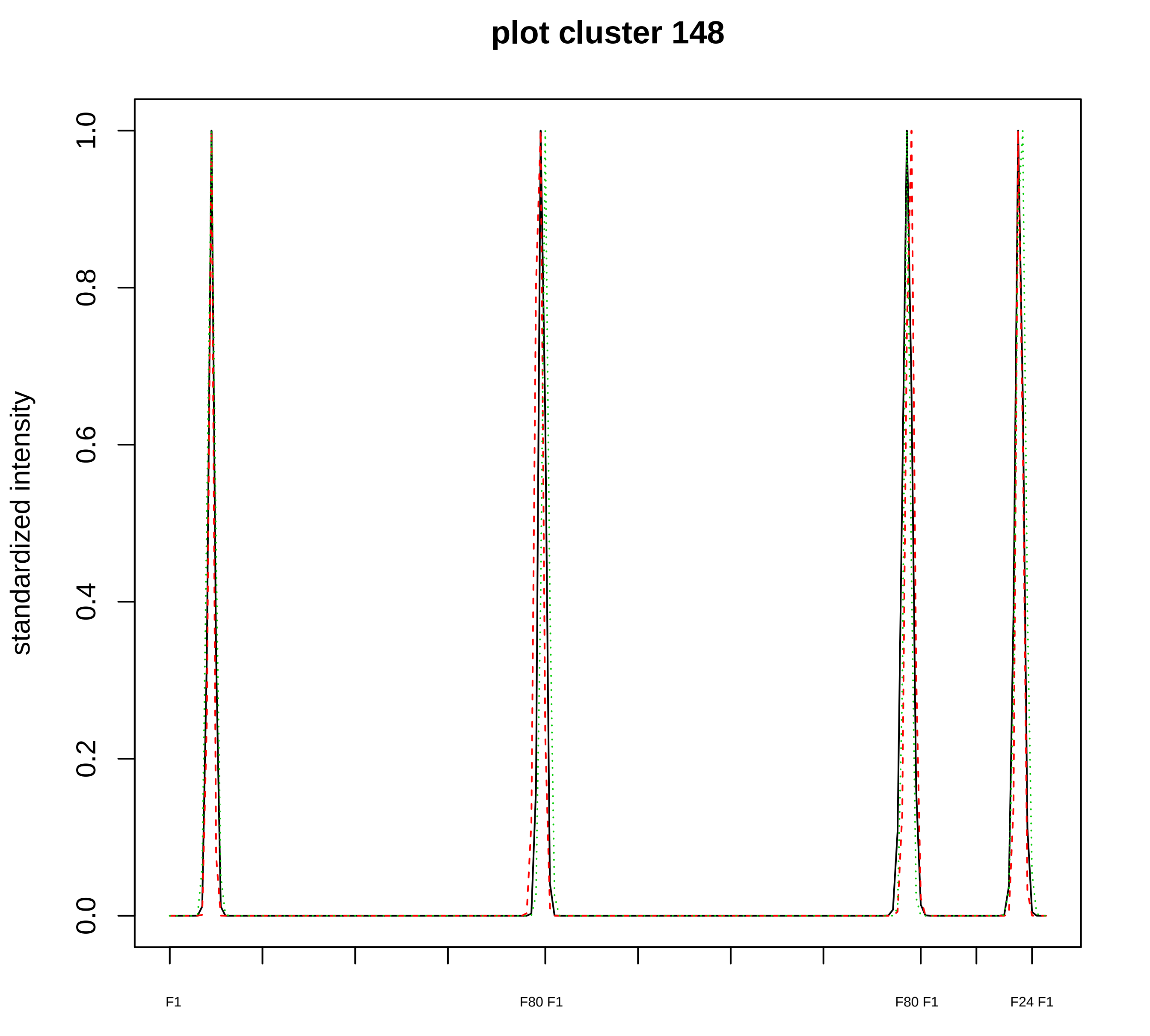
plot cluster 148
standardized intensity
0.0
0.2
0.4
0.6
0.8
1.0
F1
F80 F1
F80 F1
F24 F1

#### Dendrogram of cluster 148

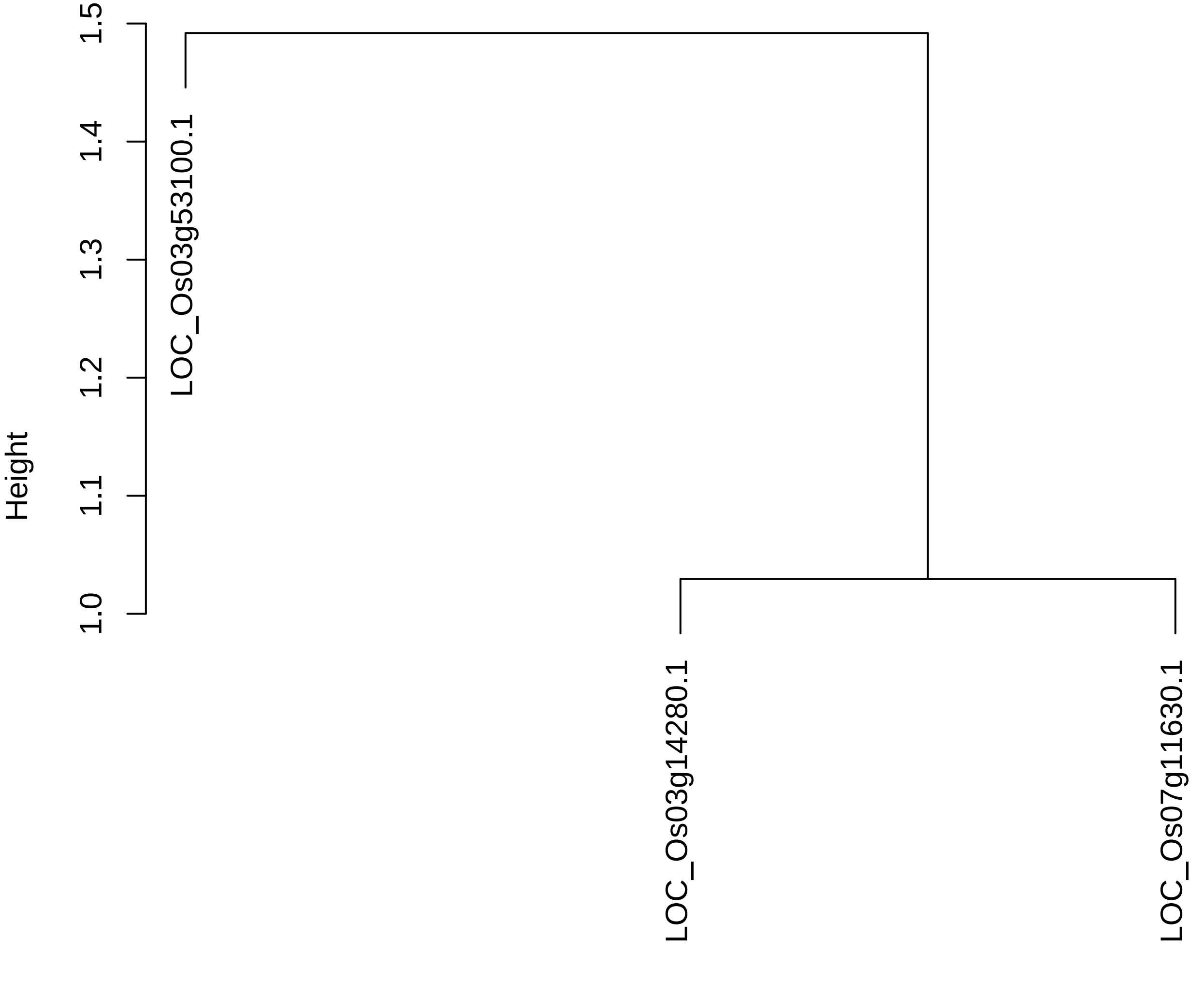


dist(spfitiex12sec12[lab[, 1] == i, ])
hclust (*, "ward.D2")

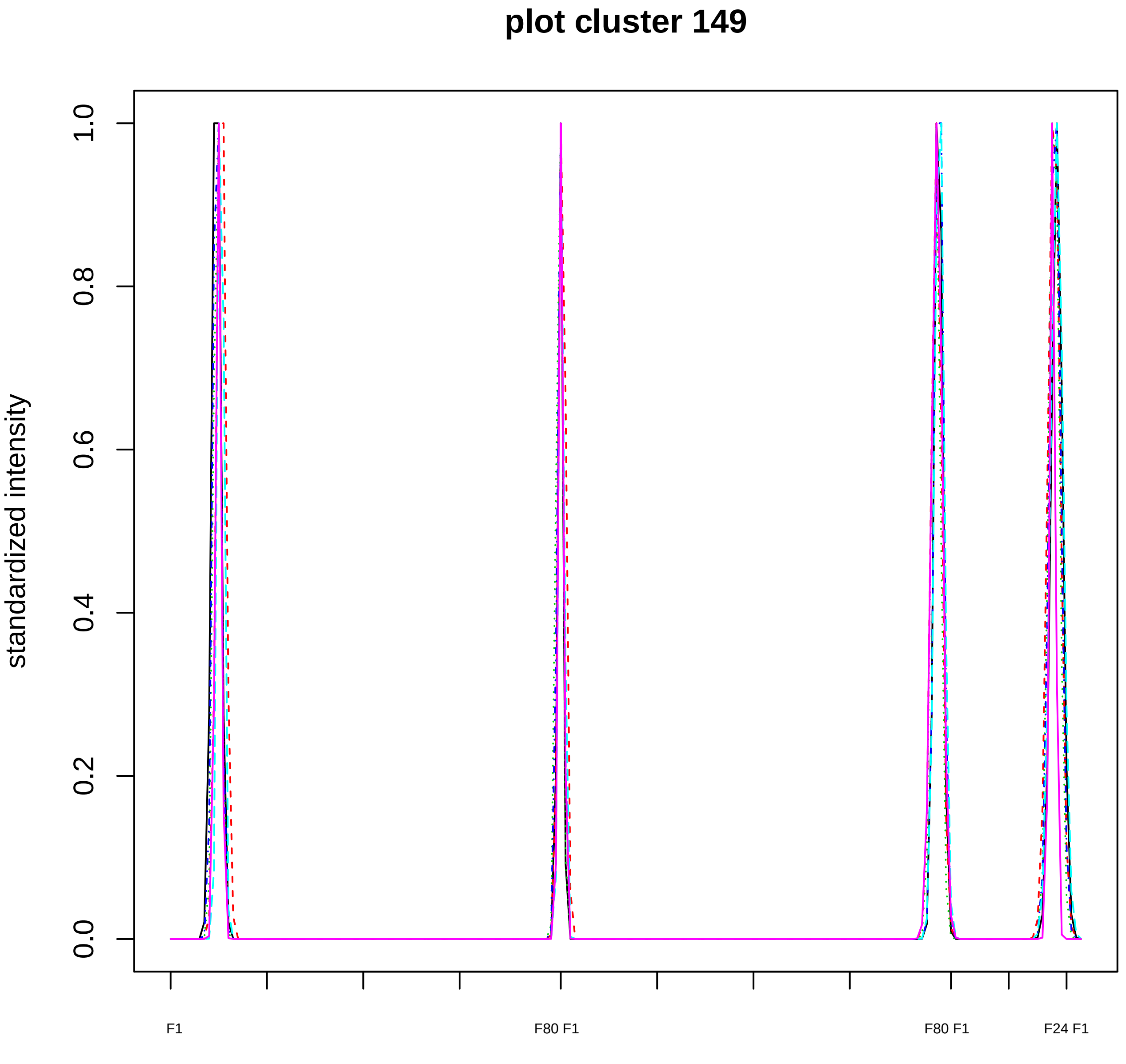

plot cluster 149
standardized intensity
0.0
0.2
0.4
0.6
0.8
1.0
F1
F80 F1
F80 F1
F24 F1

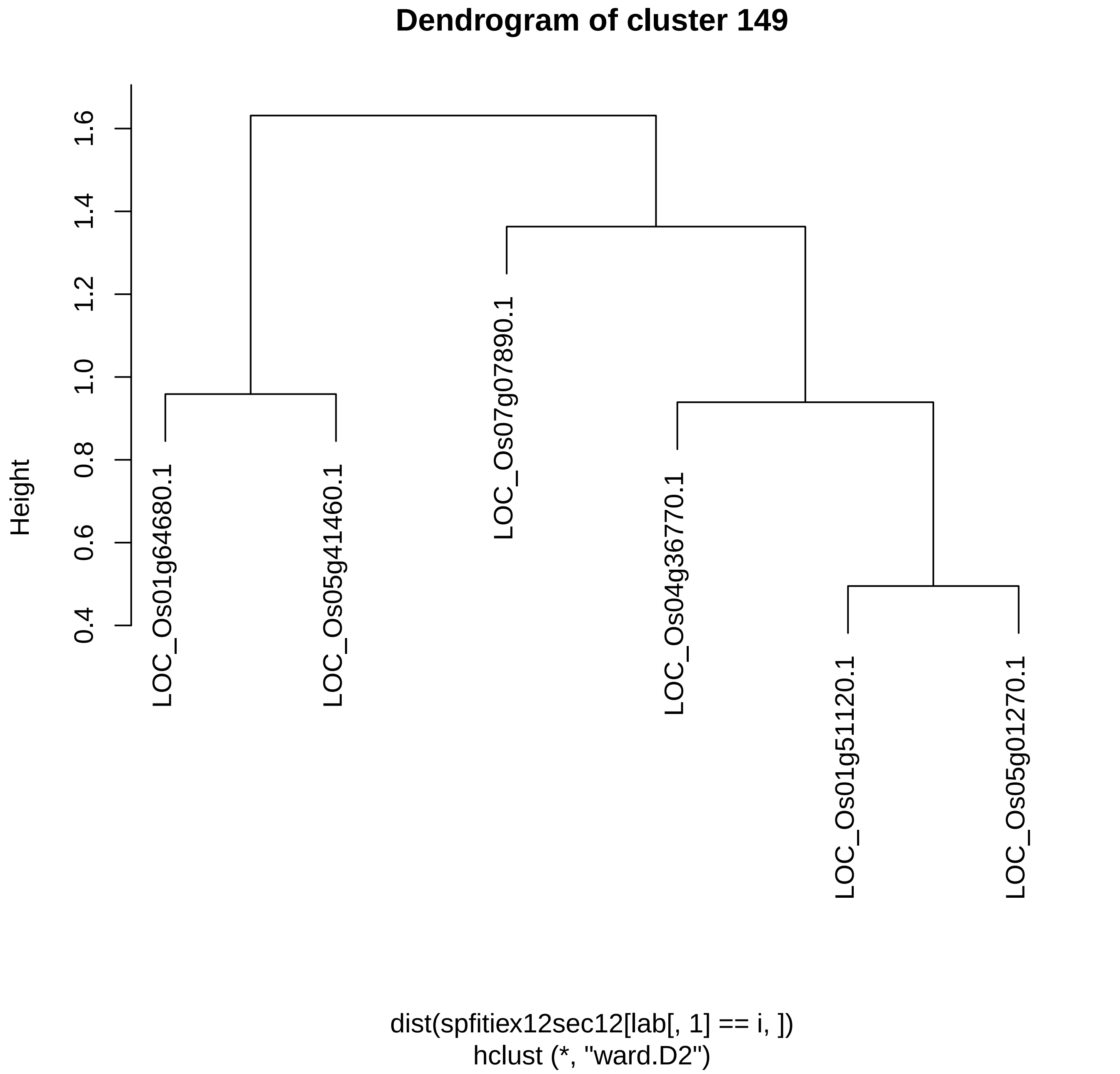
Dendrogram of cluster 149
Height
1.6
1.4
1.2
1.0
0.8
0.6
0.4
LOC_Os01g64680.1
LOC_Os05g41460.1
LOC_Os07g07890.1
LOC_Os04g36770.1
LOC_Os01g51120.1
LOC_Os05g01270.1
dist(spfitiex12sec12[lab[, 1] == i, ])
hclust (*, "ward.D2")

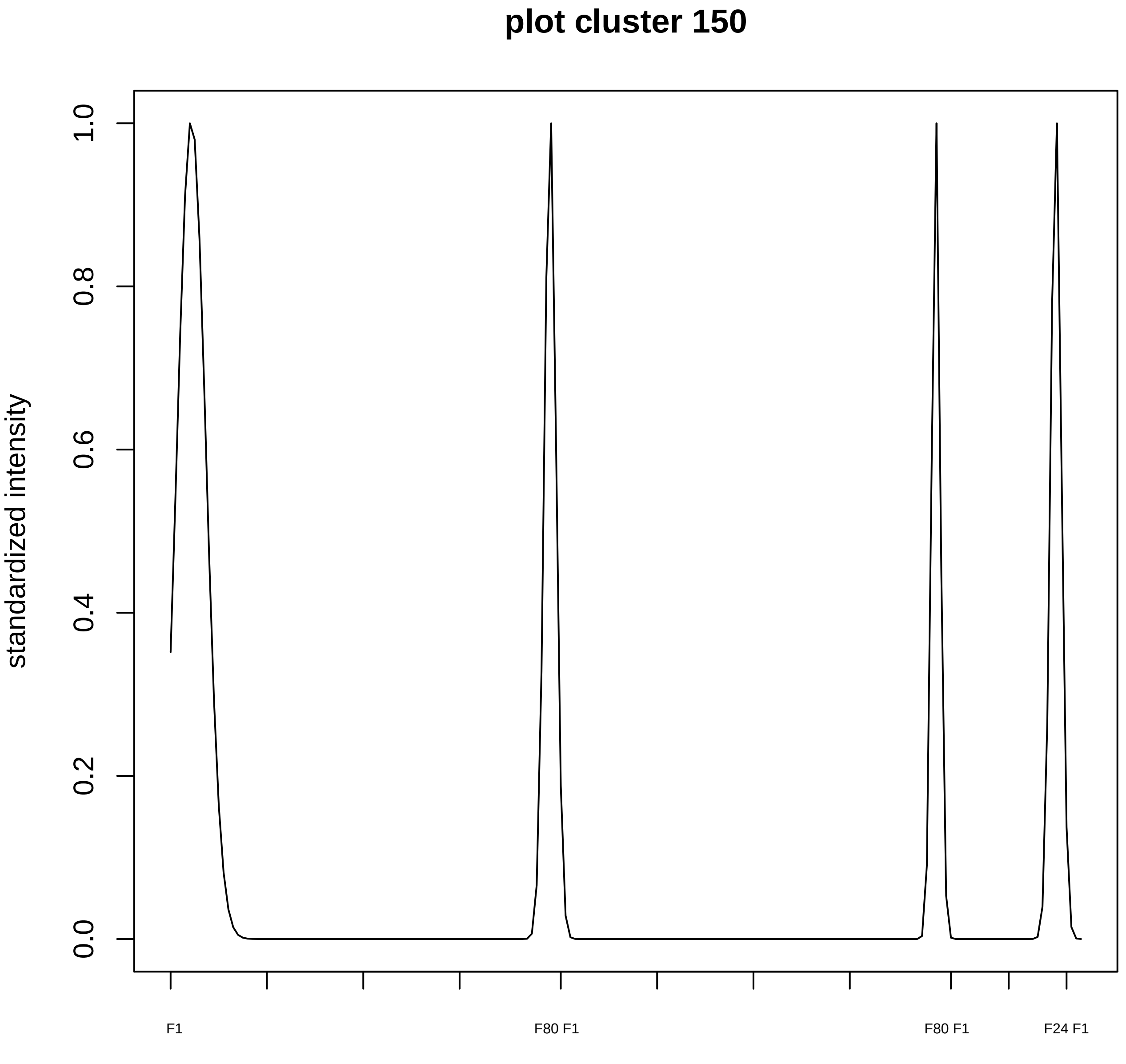

plot cluster 150
standardized intensity
0.0
0.2
0.4
0.6
0.8
1.0
F1
F80 F1
F80 F1
F24 F1

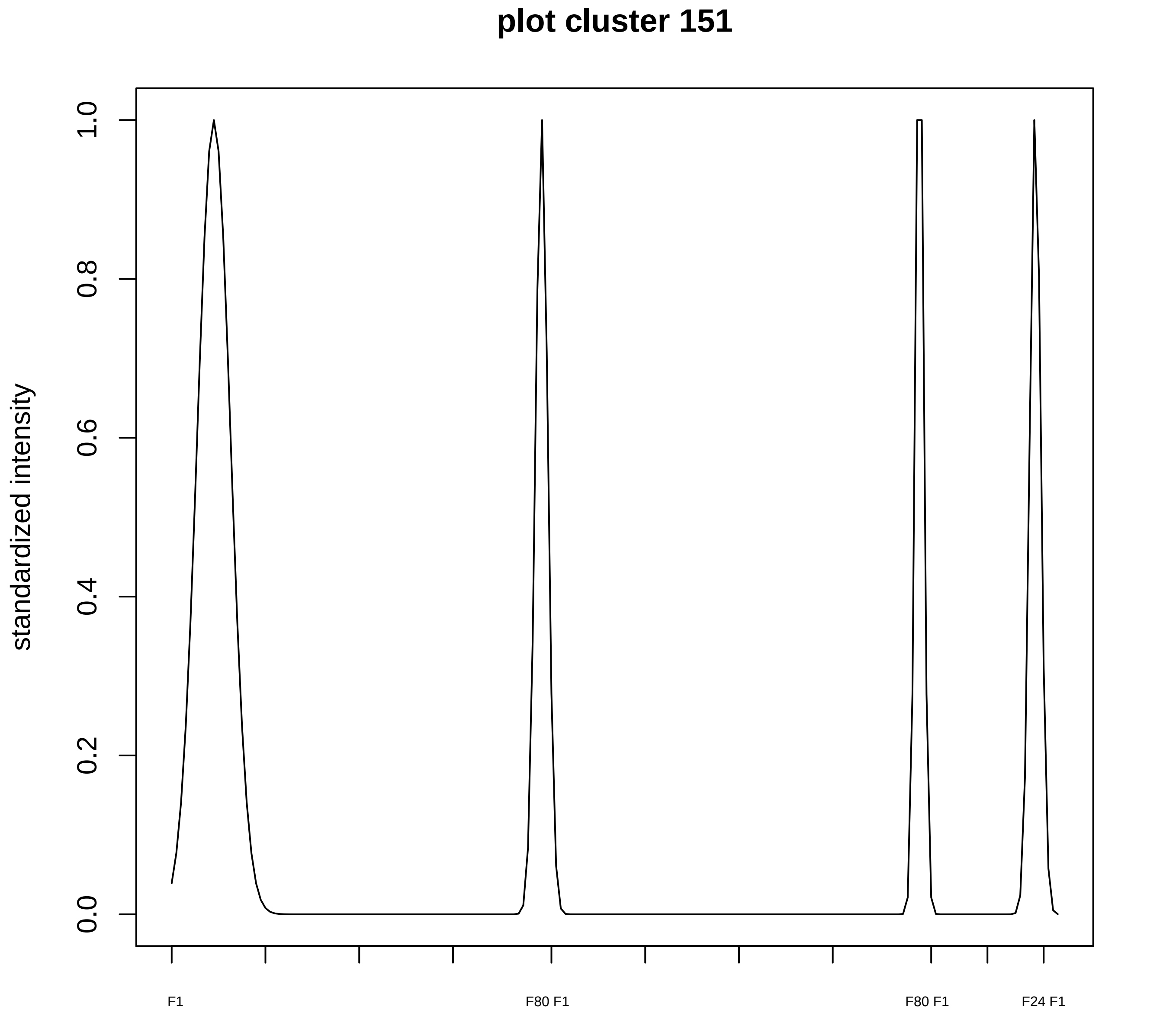

plot cluster 151
standardized intensity
0.0
0.2
0.4
0.6
0.8
1.0
F1
F80 F1
F80 F1
F24 F1

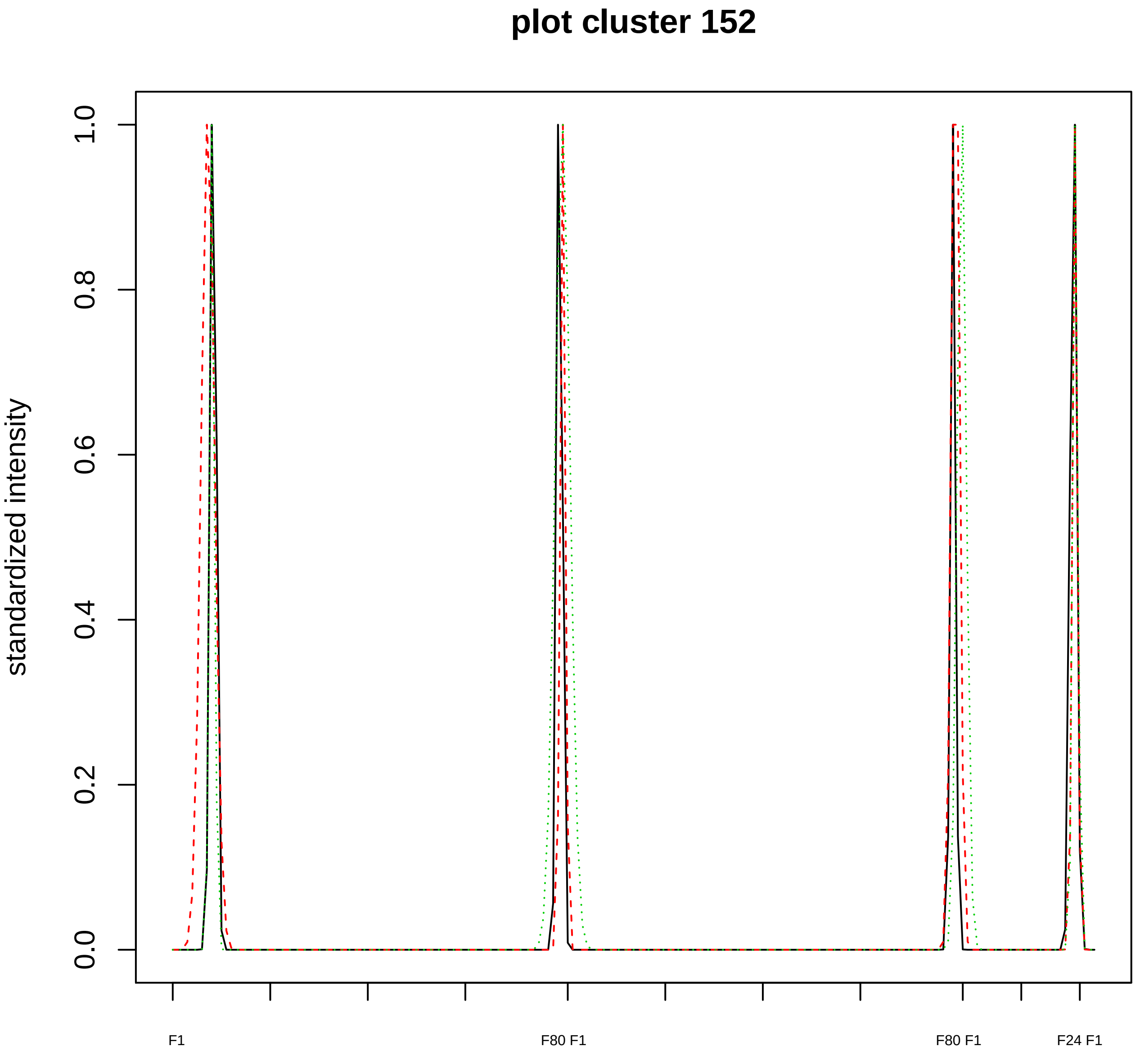
plot cluster 152
standardized intensity
0.0
0.2
0.4
0.6
0.8
1.0
F1
F80 F1
F80 F1
F24 F1

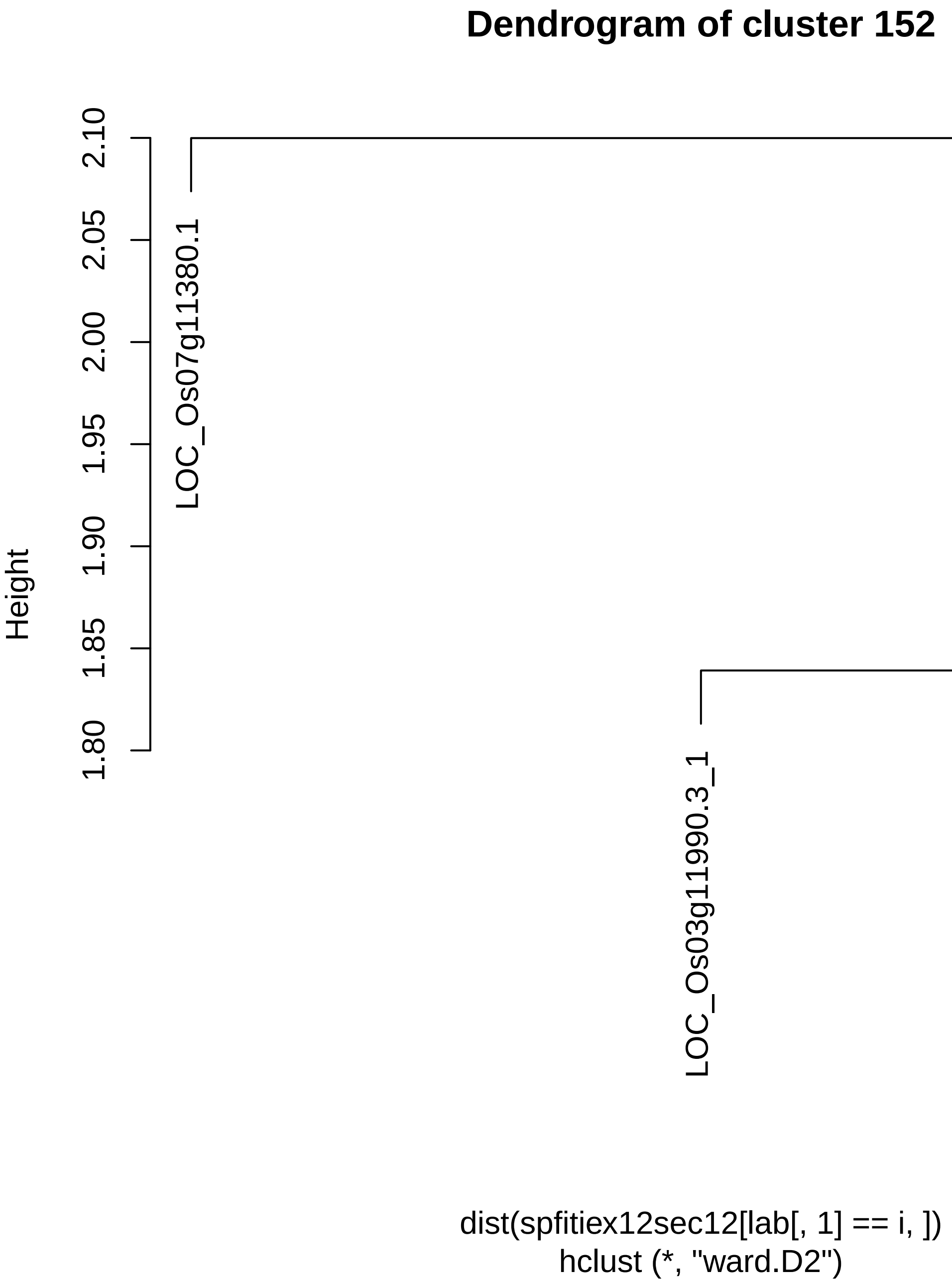
Dendrogram of cluster 152
Height
2.10
2.05
2.00
1.95
1.90
1.85
1.80
LOC_Os07g11380.1
LOC_Os03g11990.3_1
LOC_Os04g36760.1_1
dist(spfitiex12sec12[lab[, 1] == i, ])
hclust (*, "ward.D2")

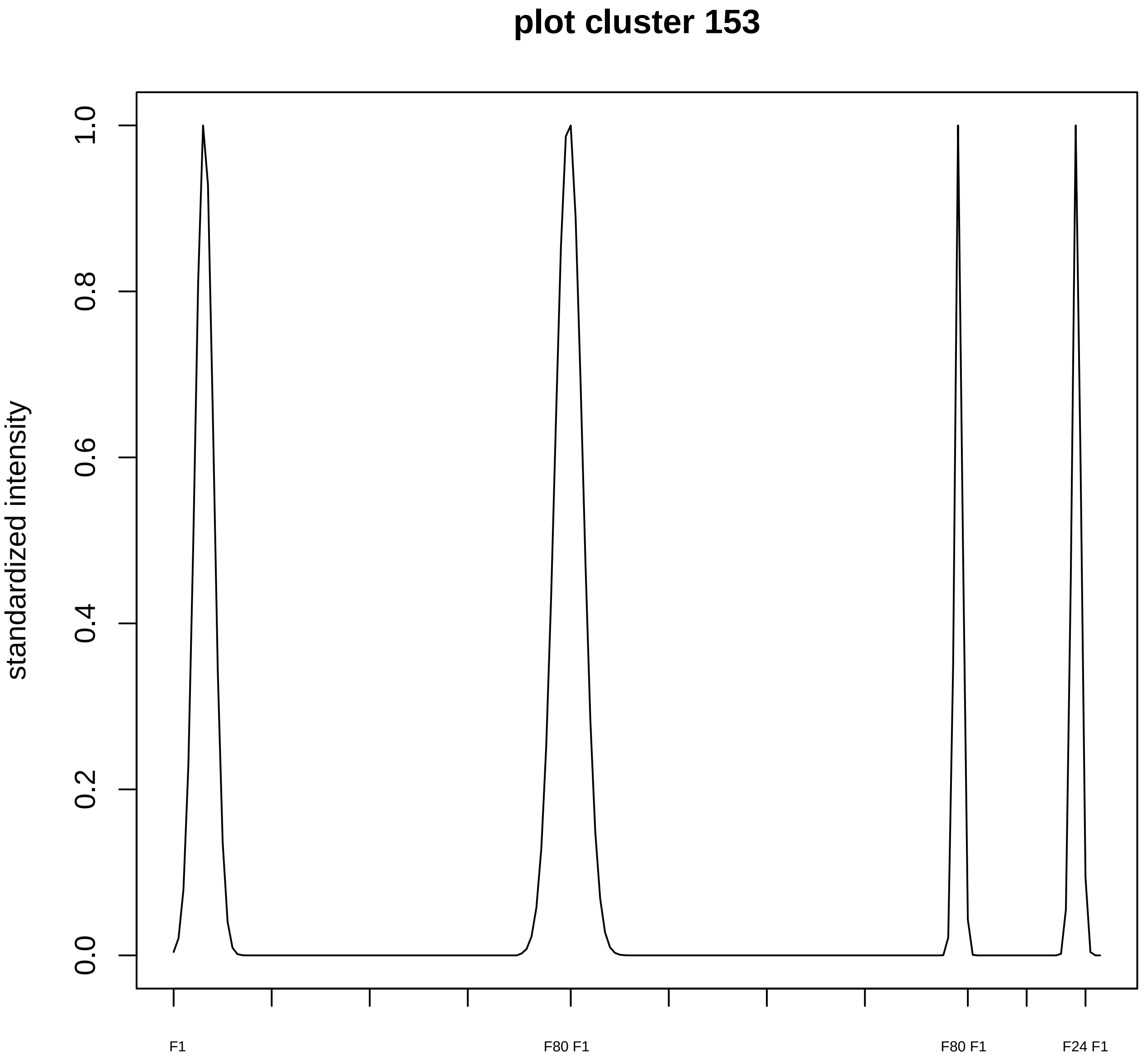
plot cluster 153
standardized intensity
0.0
0.2
0.4
0.6
0.8
1.0
F1
F80 F1
F80 F1
F24 F1

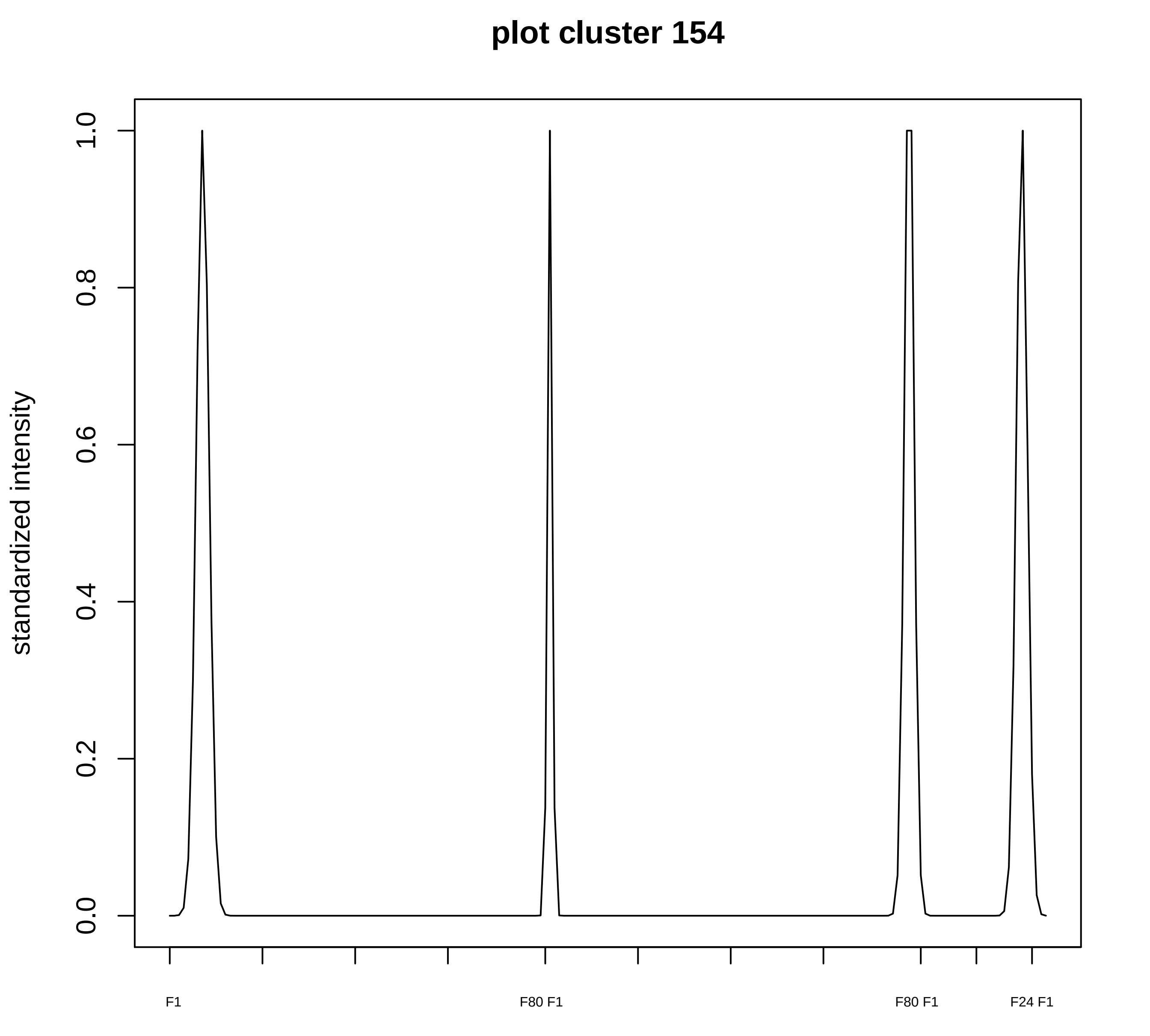
plot cluster 154
standardized intensity
0.0
0.2
0.4
0.6
0.8
1.0
F1
F80 F1
F80 F1
F24 F1

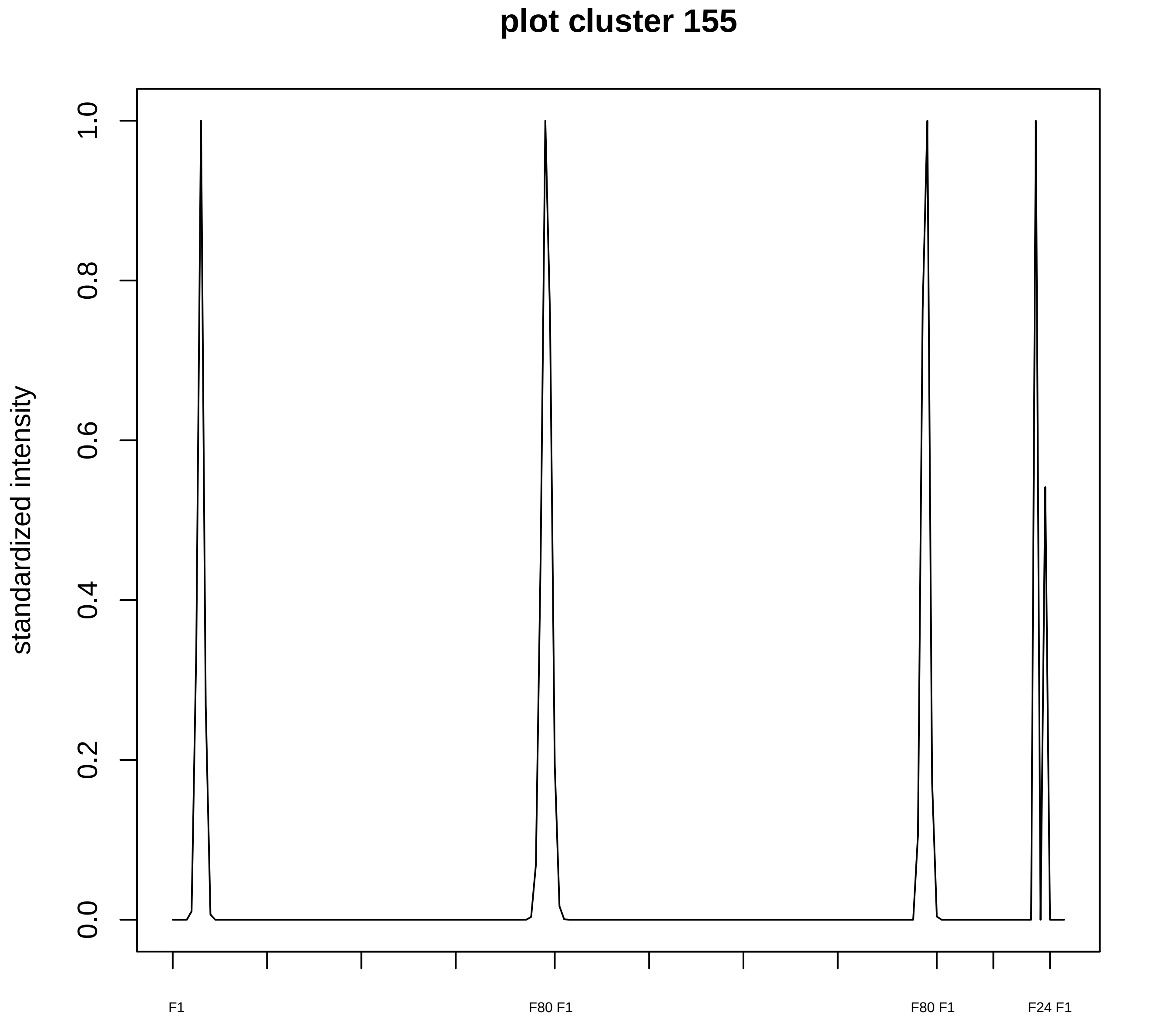
plot cluster 155
standardized intensity
0.0
0.2
0.4
0.6
0.8
1.0
F1
F80 F1
F80 F1
F24 F1

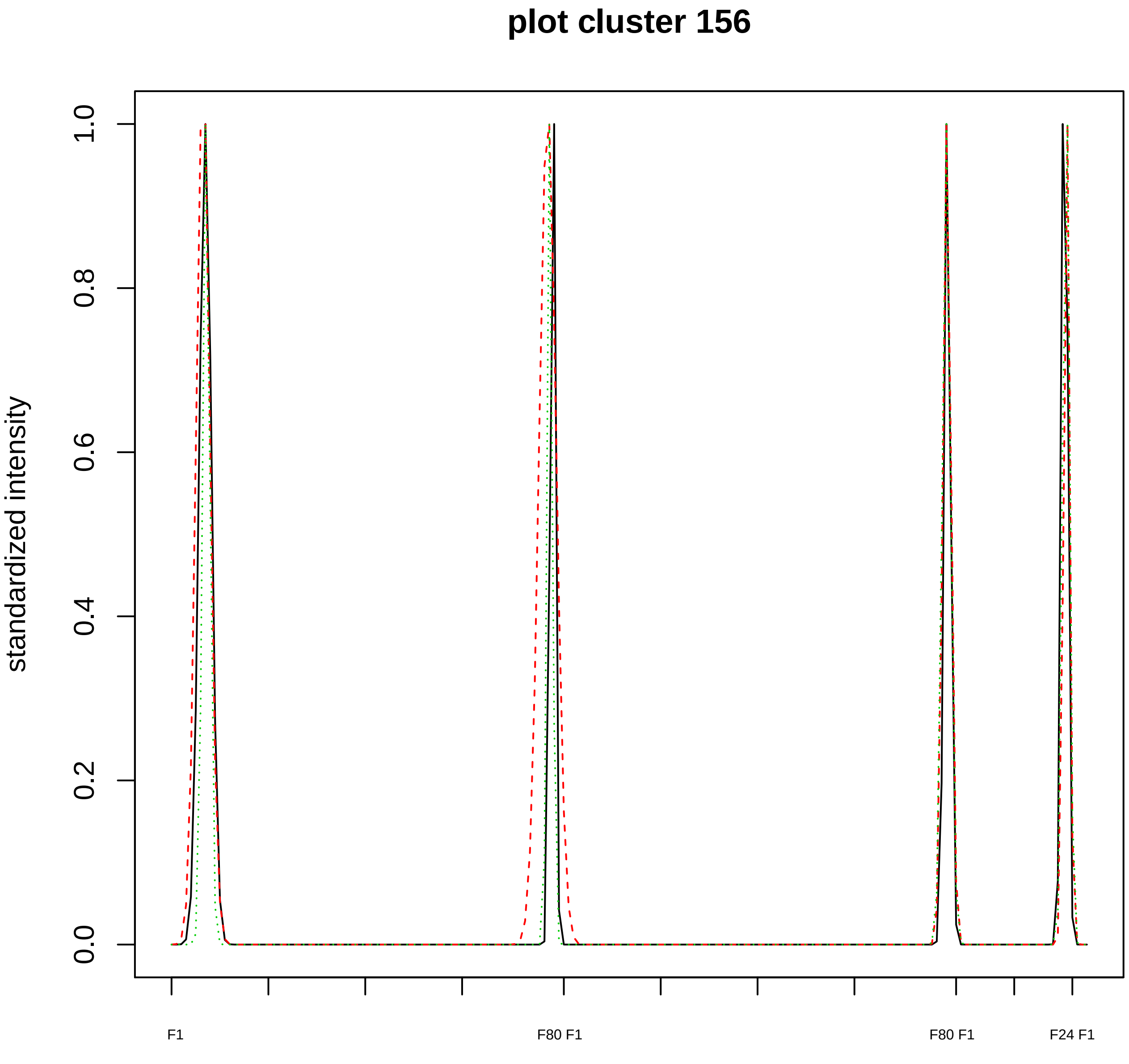
plot cluster 156
standardized intensity
0.0
0.2
0.4
0.6
0.8
1.0
F1
F80 F1
F80 F1
F24 F1

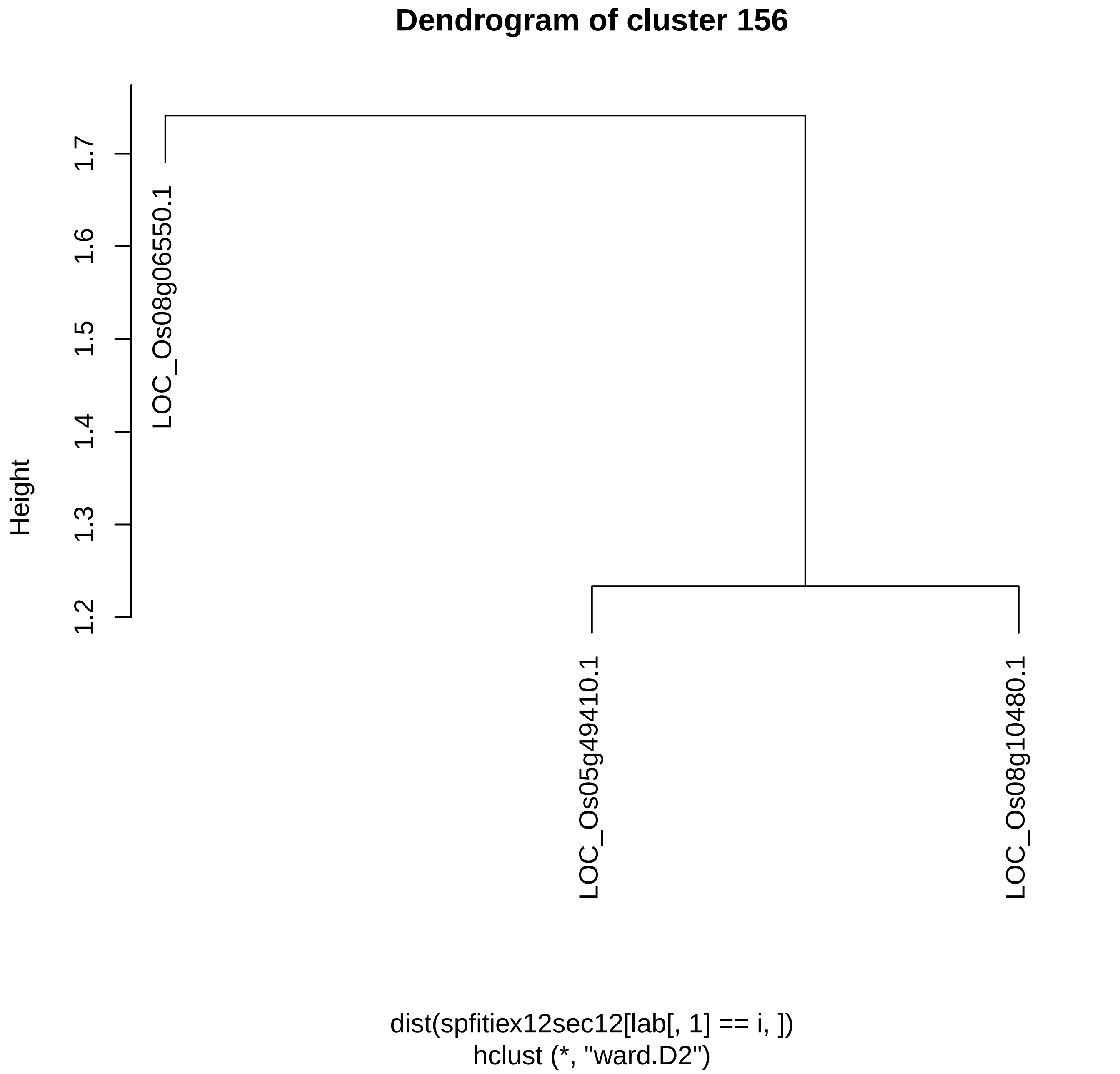
Dendrogram of cluster 156
Height
1.2
1.3
1.4
1.5
1.6
1.7
LOC_Os08g06550.1
LOC_Os05g49410.1
LOC_Os08g10480.1
dist(spfitiex12sec12[lab[, 1] == i, ])
hclust (*, "ward.D2")

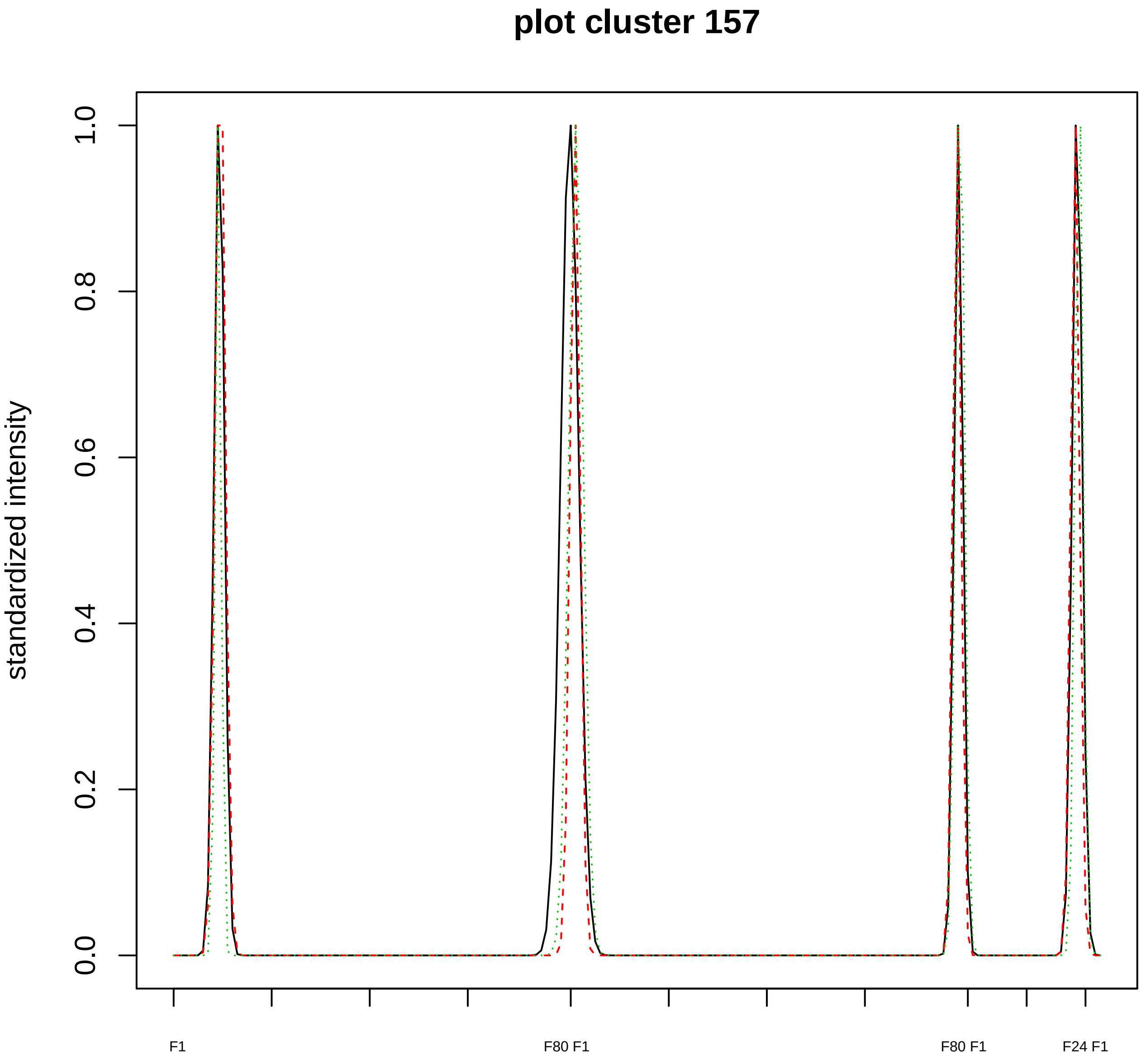
plot cluster 157
standardized intensity
0.0
0.2
0.4
0.6
0.8
1.0
F1
F80 F1
F80 F1
F24 F1

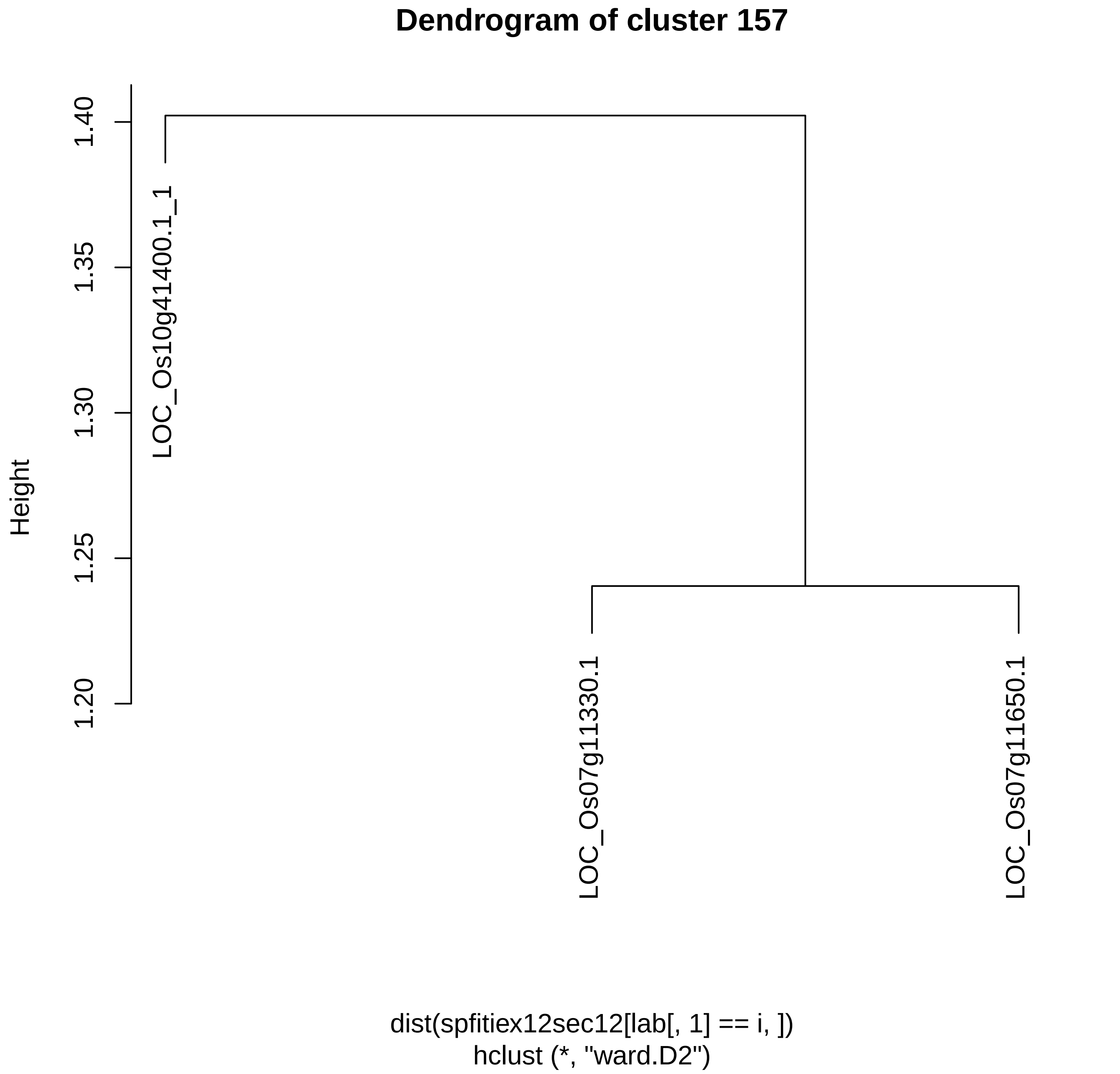

Dendrogram of cluster 157
Height
1.20
1.25
1.30
1.35
1.40
LOC_Os10g41400.1_1
LOC_Os07g11330.1
LOC_Os07g11650.1
dist(spfitiex12sec12[lab[, 1] == i, ])
hclust (*, "ward.D2")

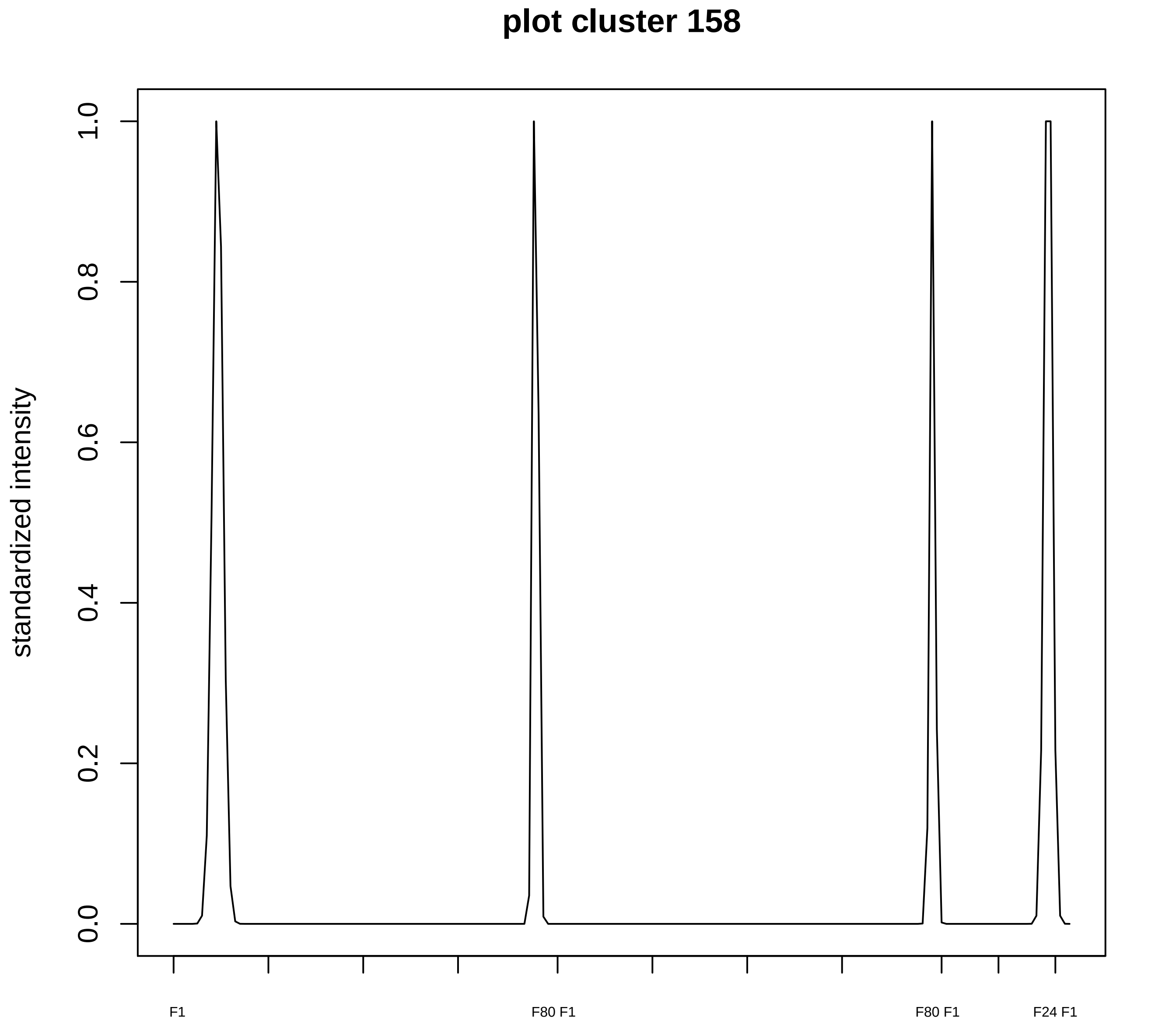

plot cluster 158
standardized intensity
0.0
0.2
0.4
0.6
0.8
1.0
F1
F80 F1
F80 F1
F24 F1

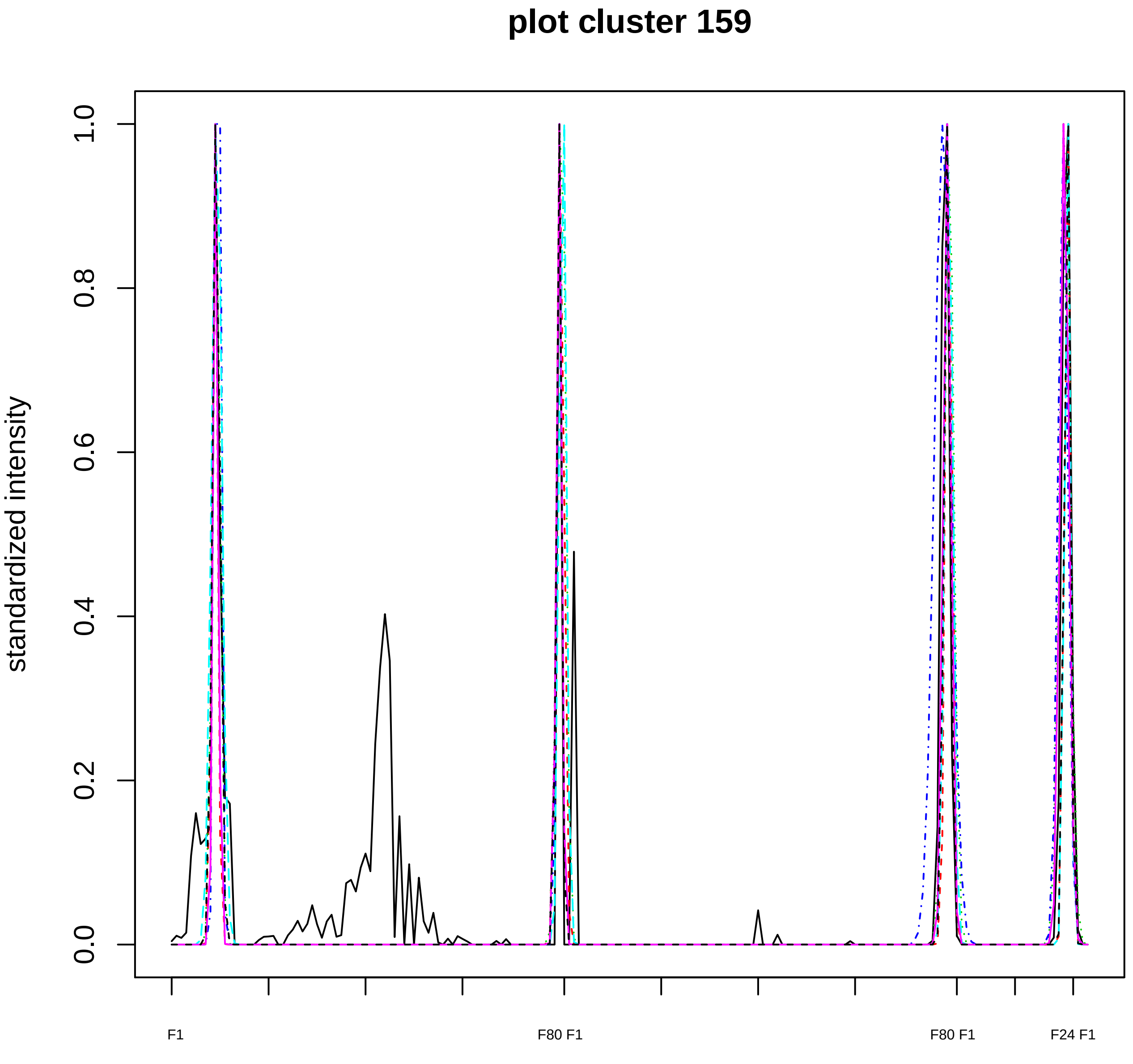

plot cluster 159
standardized intensity
1.0
0.8
0.6
0.4
0.2
0.0
F1
F80 F1
F80 F1
F24 F1

### Dendrogram of cluster 159

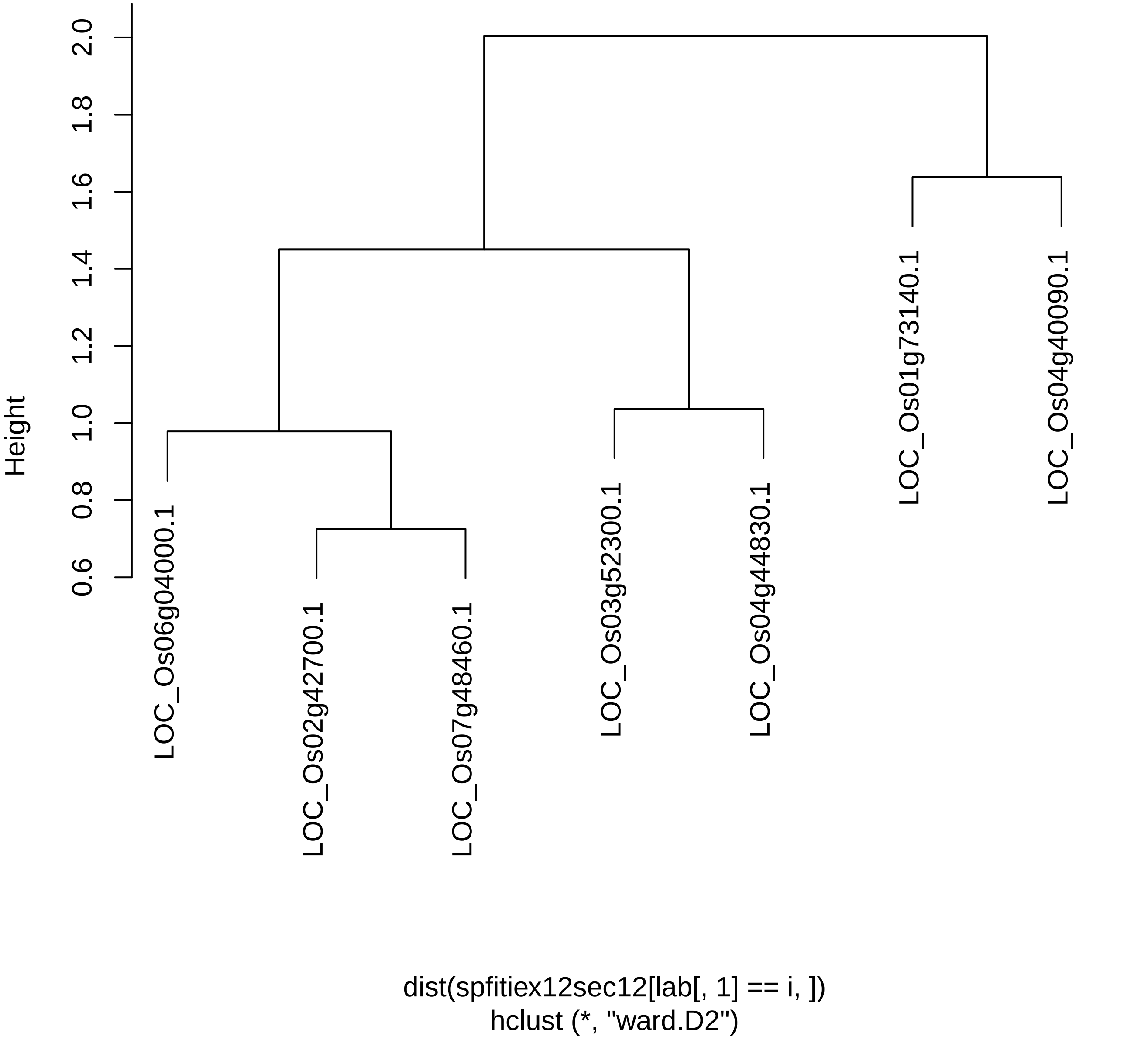

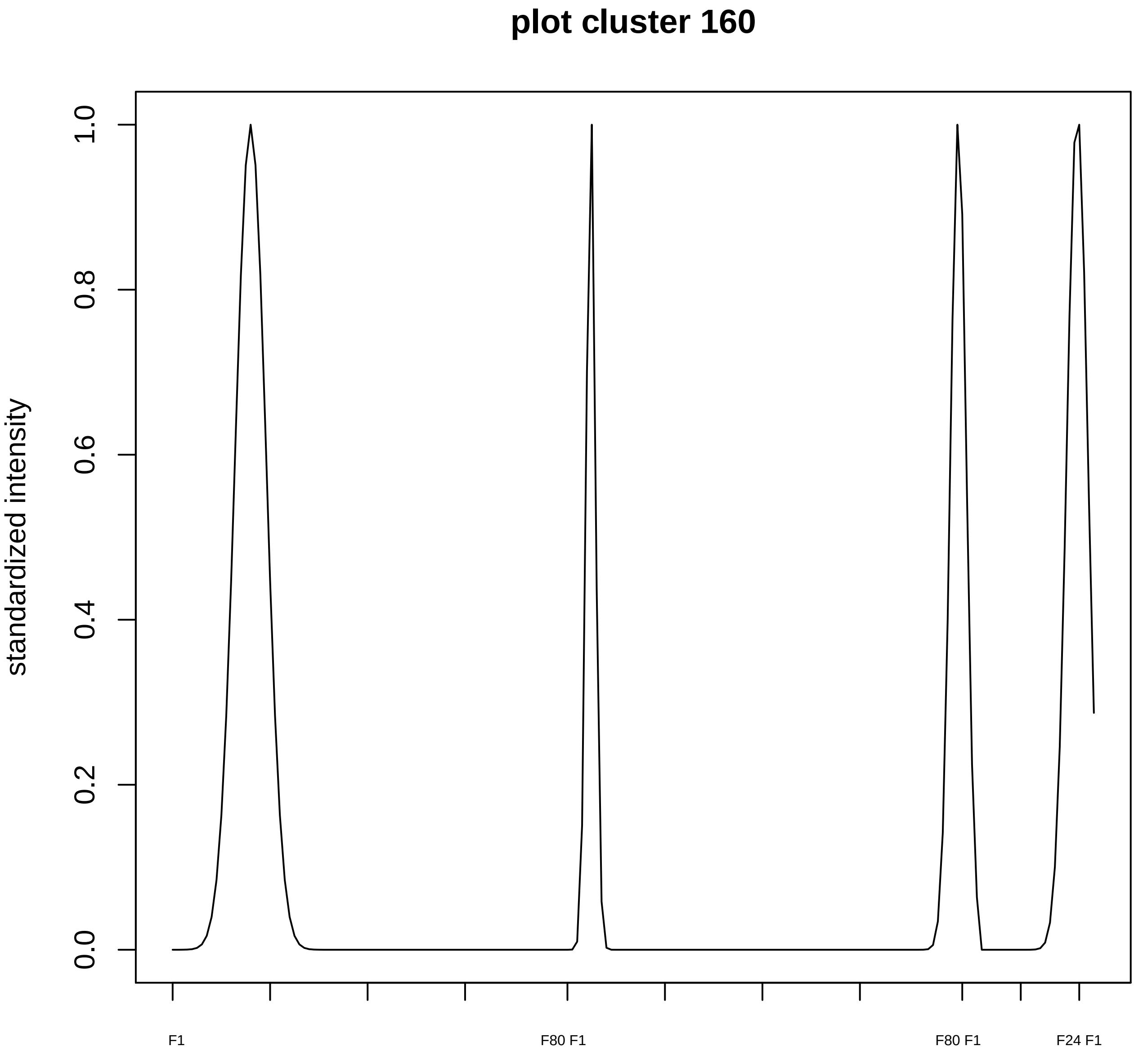
plot cluster 160
standardized intensity
0.0
0.2
0.4
0.6
0.8
1.0
F1
F80 F1
F80 F1
F24 F1

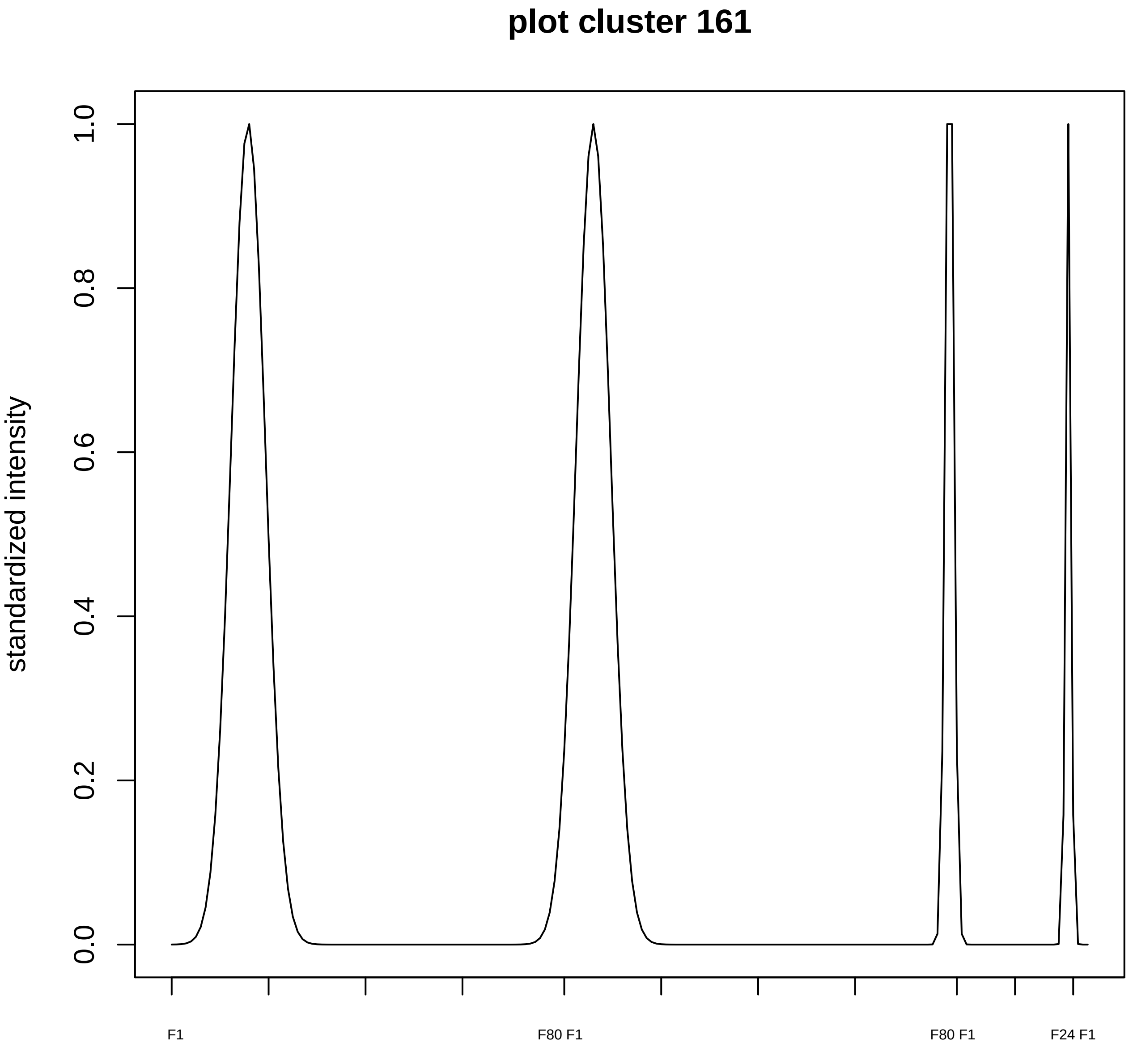

plot cluster 161
standardized intensity
0.0
0.2
0.4
0.6
0.8
1.0
F1
F80 F1
F80 F1
F24 F1

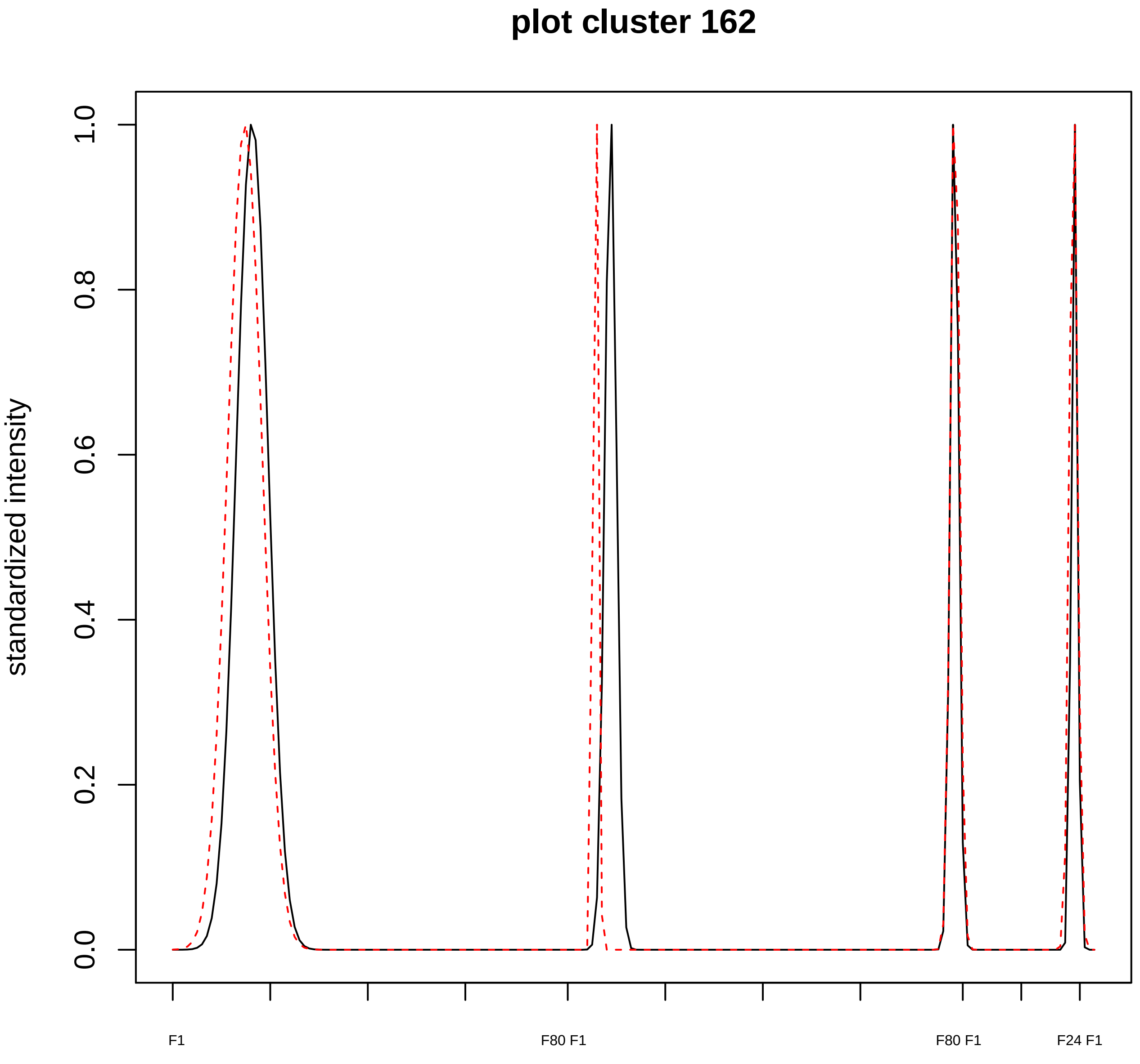
plot cluster 162
standardized intensity
0.0
0.2
0.4
0.6
0.8
1.0
F1
F80 F1
F80 F1
F24 F1

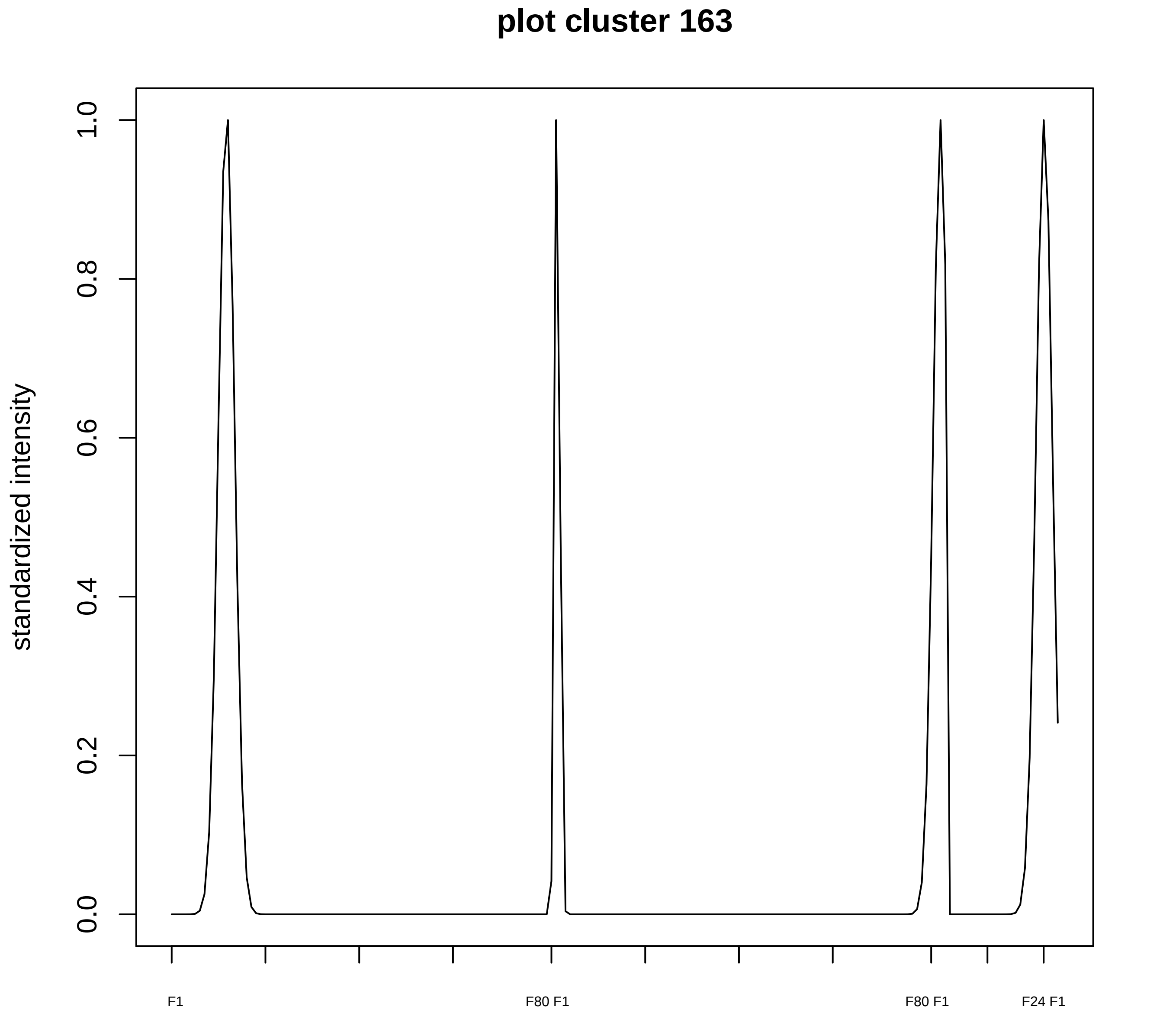
plot cluster 163
standardized intensity
0.0
0.2
0.4
0.6
0.8
1.0
F1
F80 F1
F80 F1
F24 F1

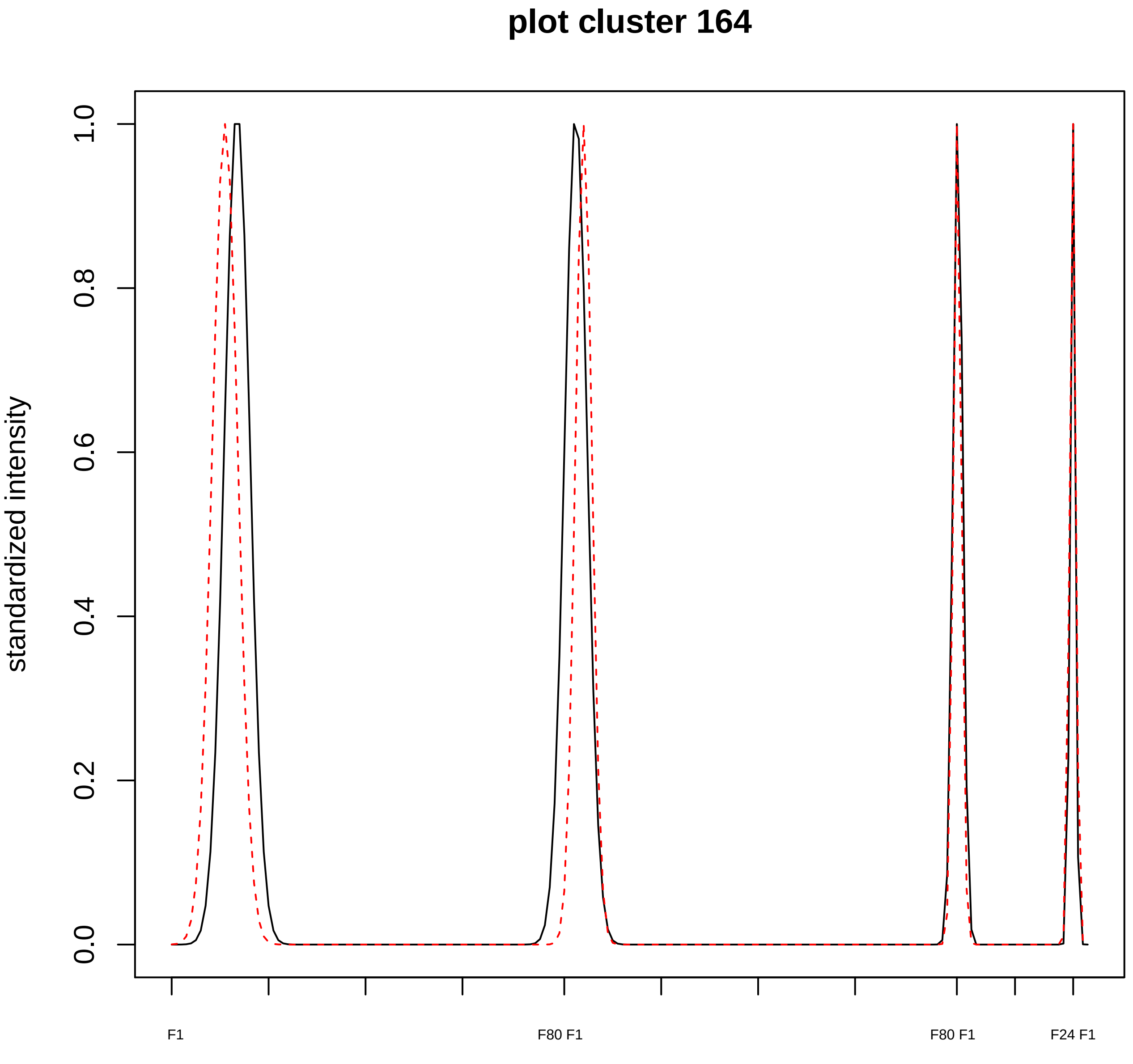
plot cluster 164
standardized intensity
0.0
0.2
0.4
0.6
0.8
1.0
F1
F80 F1
F80 F1
F24 F1

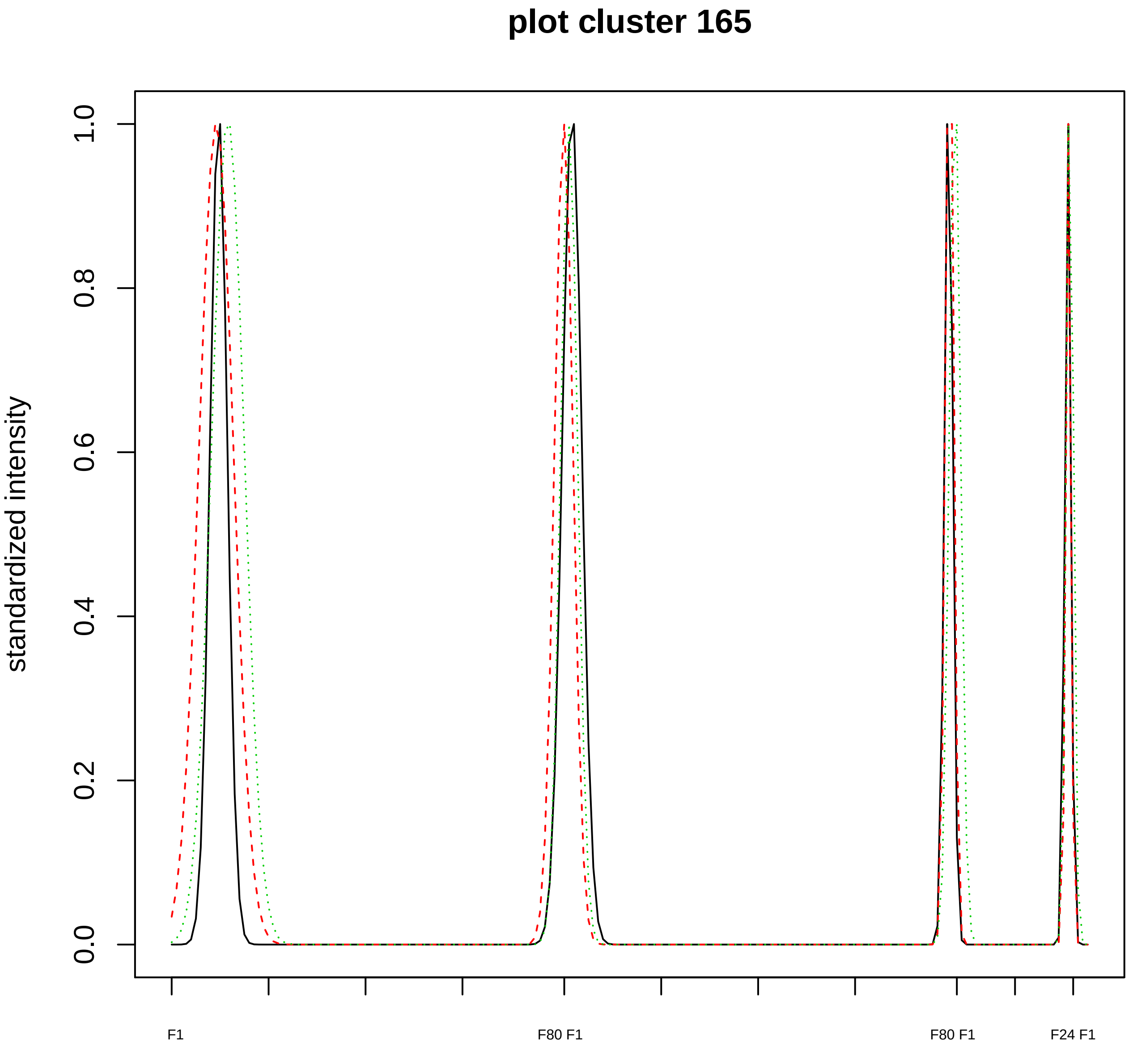
plot cluster 165
standardized intensity
0.0
0.2
0.4
0.6
0.8
1.0
F1
F80 F1
F80 F1
F24 F1

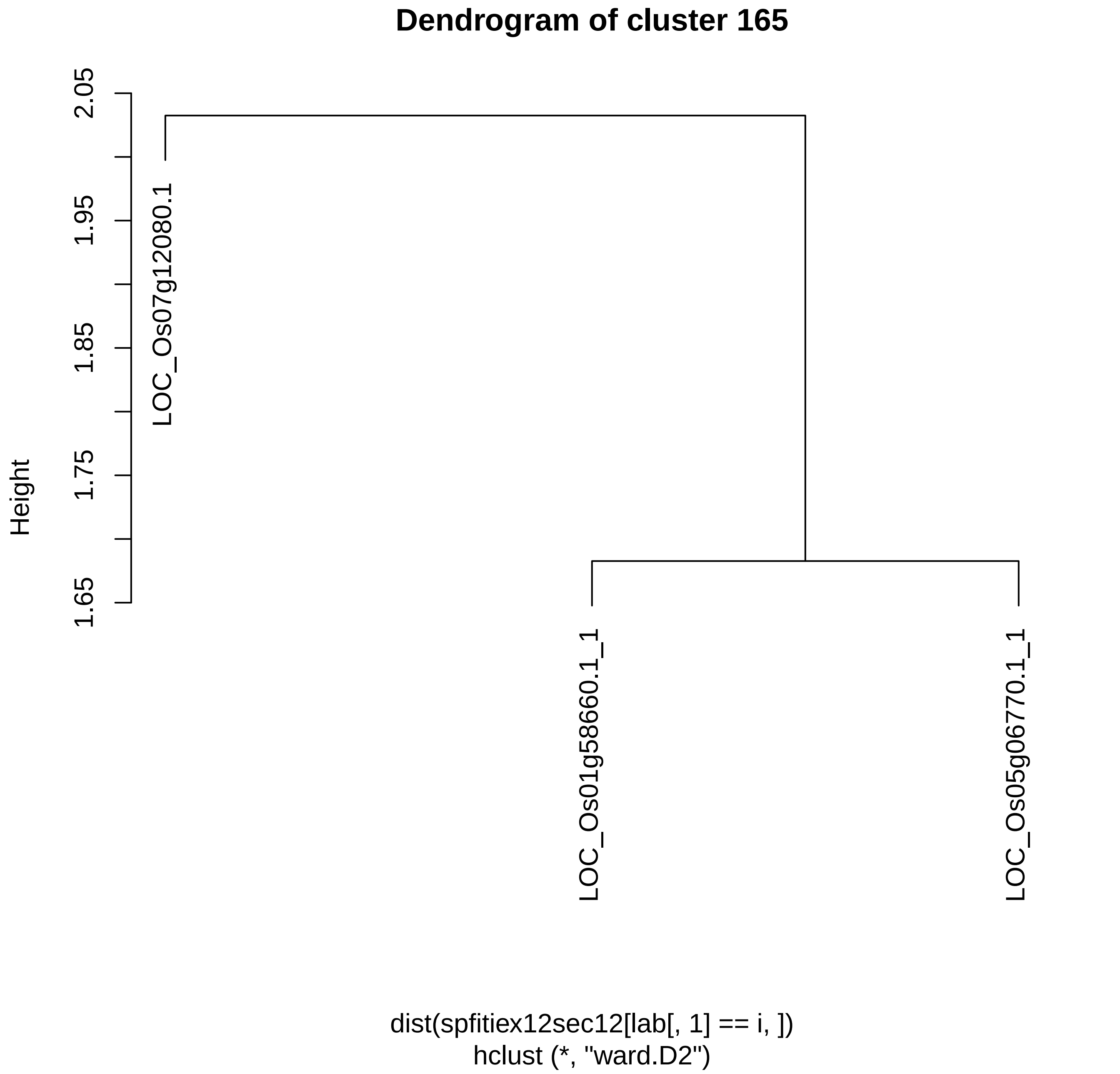
Dendrogram of cluster 165
Height
2.05
1.95
1.85
1.75
1.65
LOC_Os07g12080.1
LOC_Os01g58660.1_1
LOC_Os05g06770.1_1
dist(spfitiex12sec12[lab[, 1] == i, ])
hclust (*, "ward.D2")

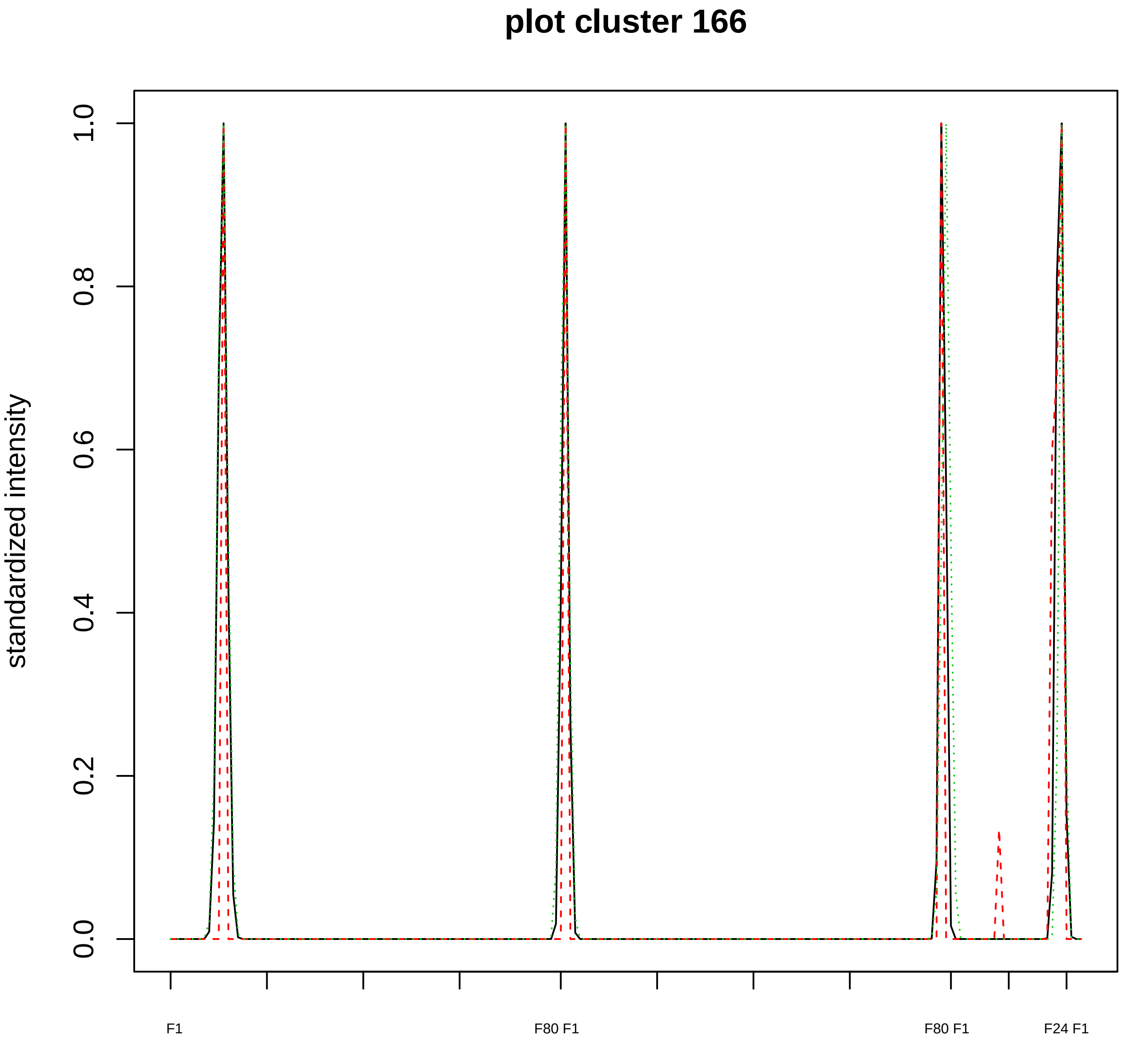
plot cluster 166
standardized intensity
0.0
0.2
0.4
0.6
0.8
1.0
F1
F80 F1
F80 F1
F24 F1

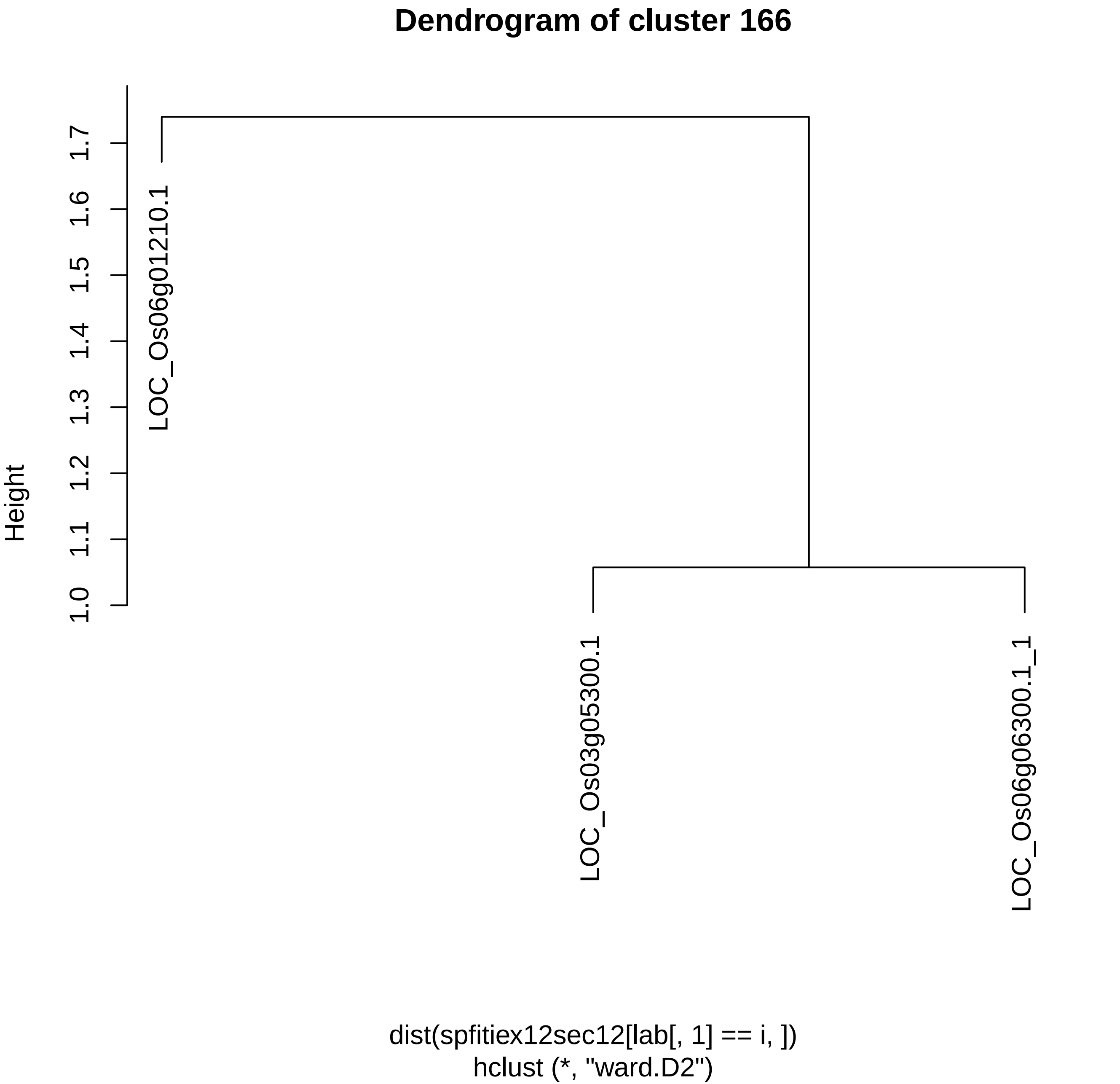
Dendrogram of cluster 166
Height
1.0
1.1
1.2
1.3
1.4
1.5
1.6
1.7
LOC_Os06g01210.1
LOC_Os03g05300.1
LOC_Os06g06300.1_1
dist(spfitiex12sec12[lab[, 1] == i, ])
hclust (*, "ward.D2")

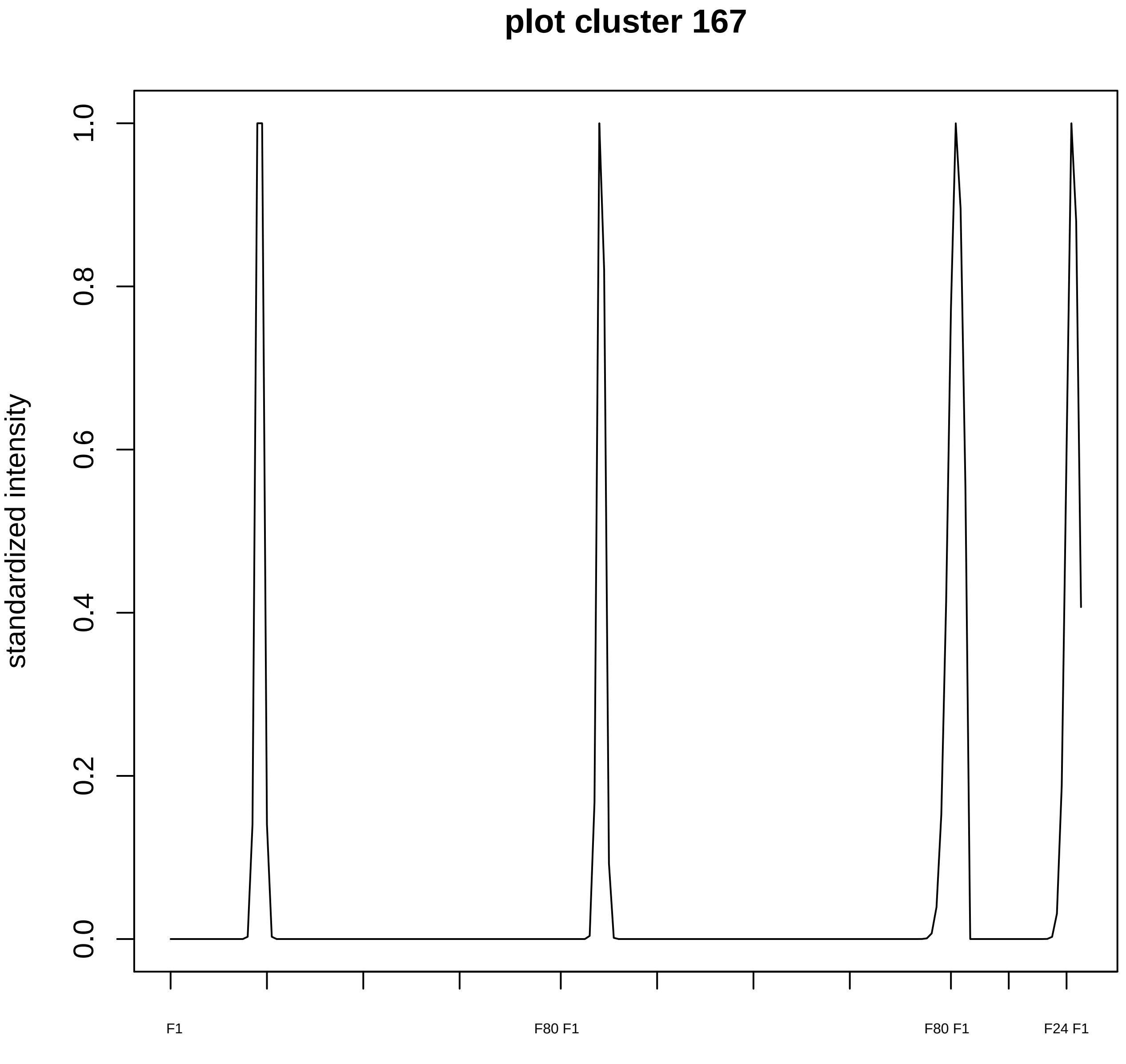

plot cluster 167
standardized intensity
0.0
0.2
0.4
0.6
0.8
1.0
F1
F80 F1
F80 F1
F24 F1

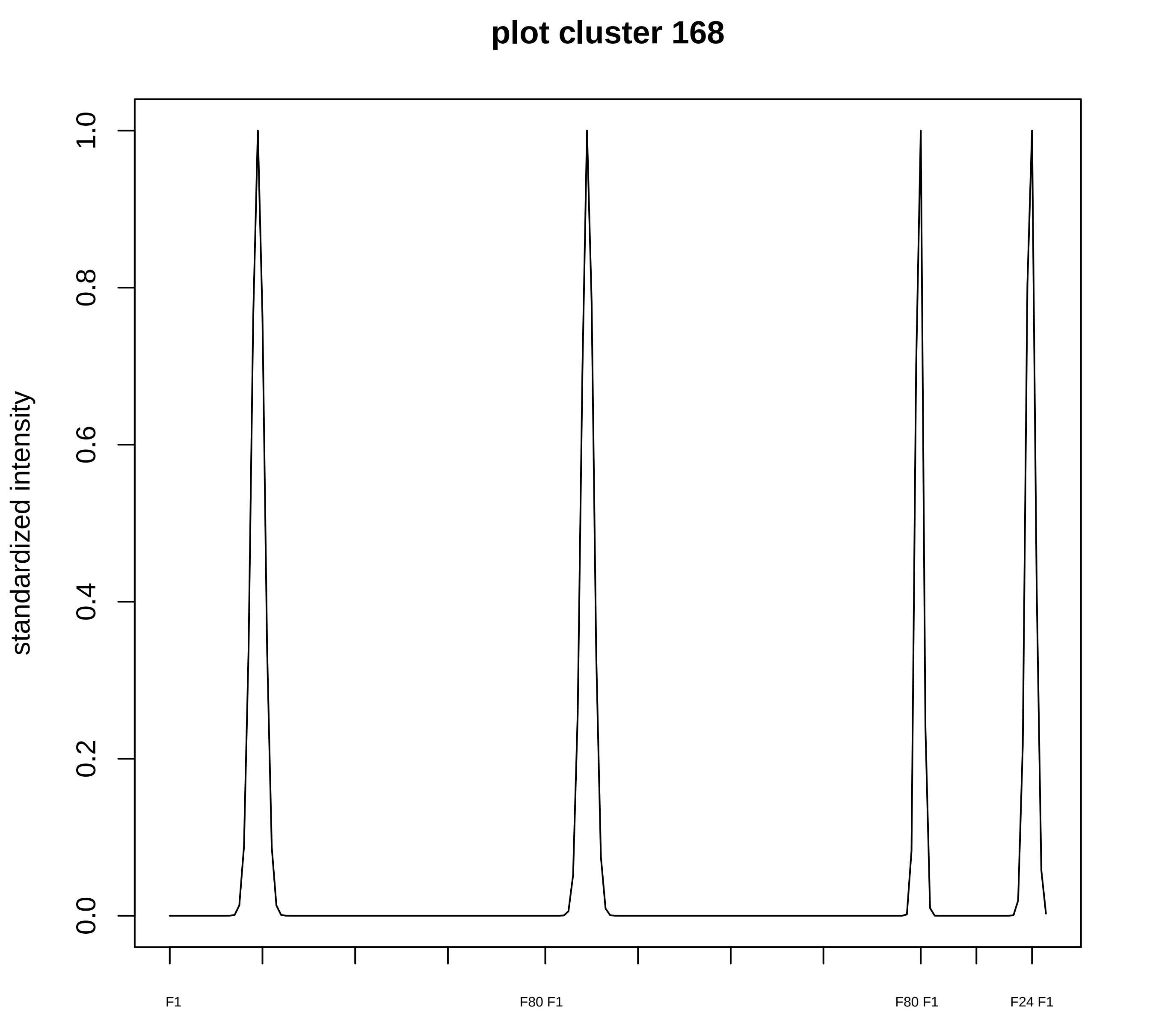

plot cluster 168
standardized intensity
0.0
0.2
0.4
0.6
0.8
1.0
F1
F80 F1
F80 F1
F24 F1

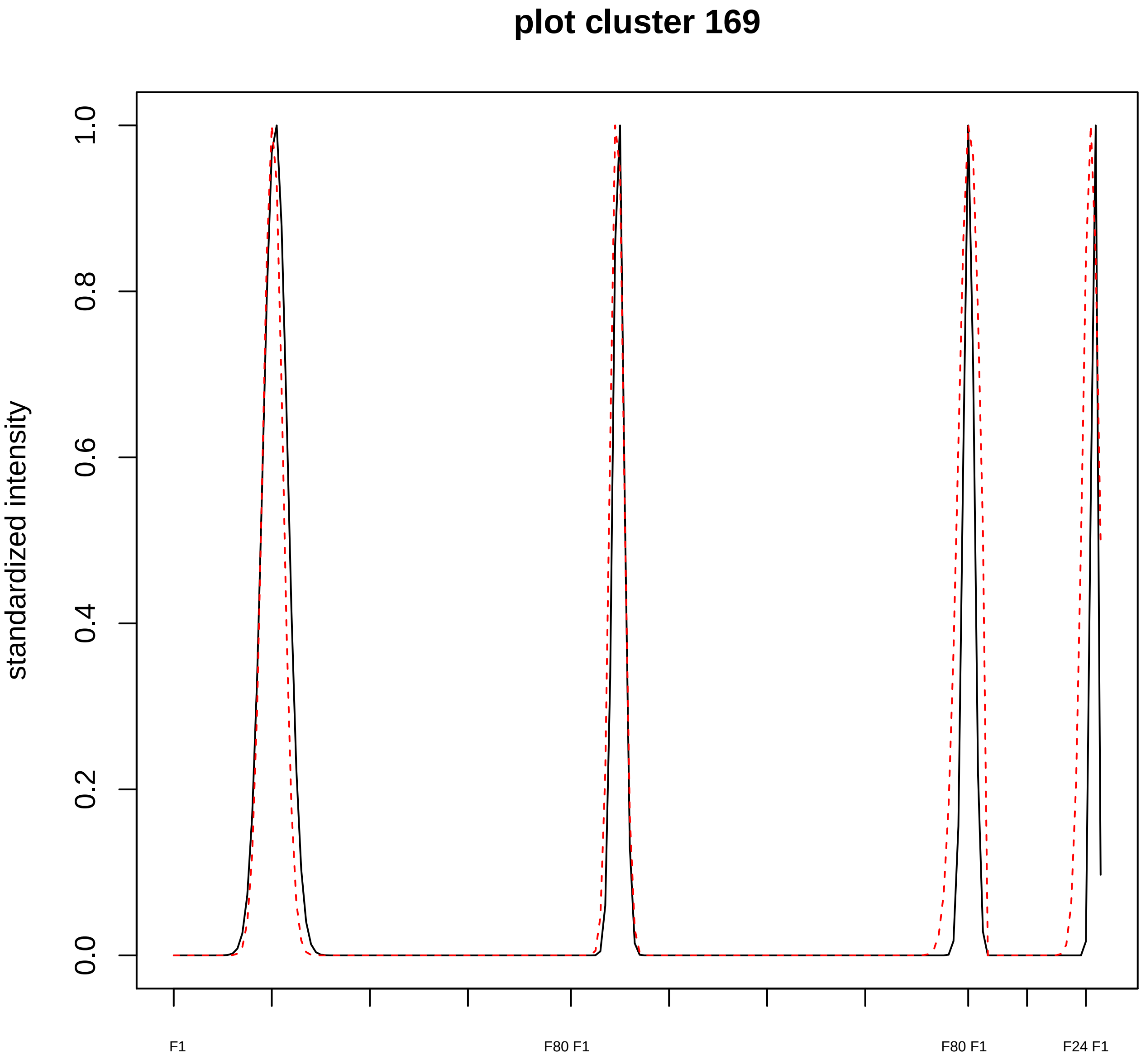

plot cluster 169
standardized intensity
0.0
0.2
0.4
0.6
0.8
1.0
F1
F80 F1
F80 F1
F24 F1

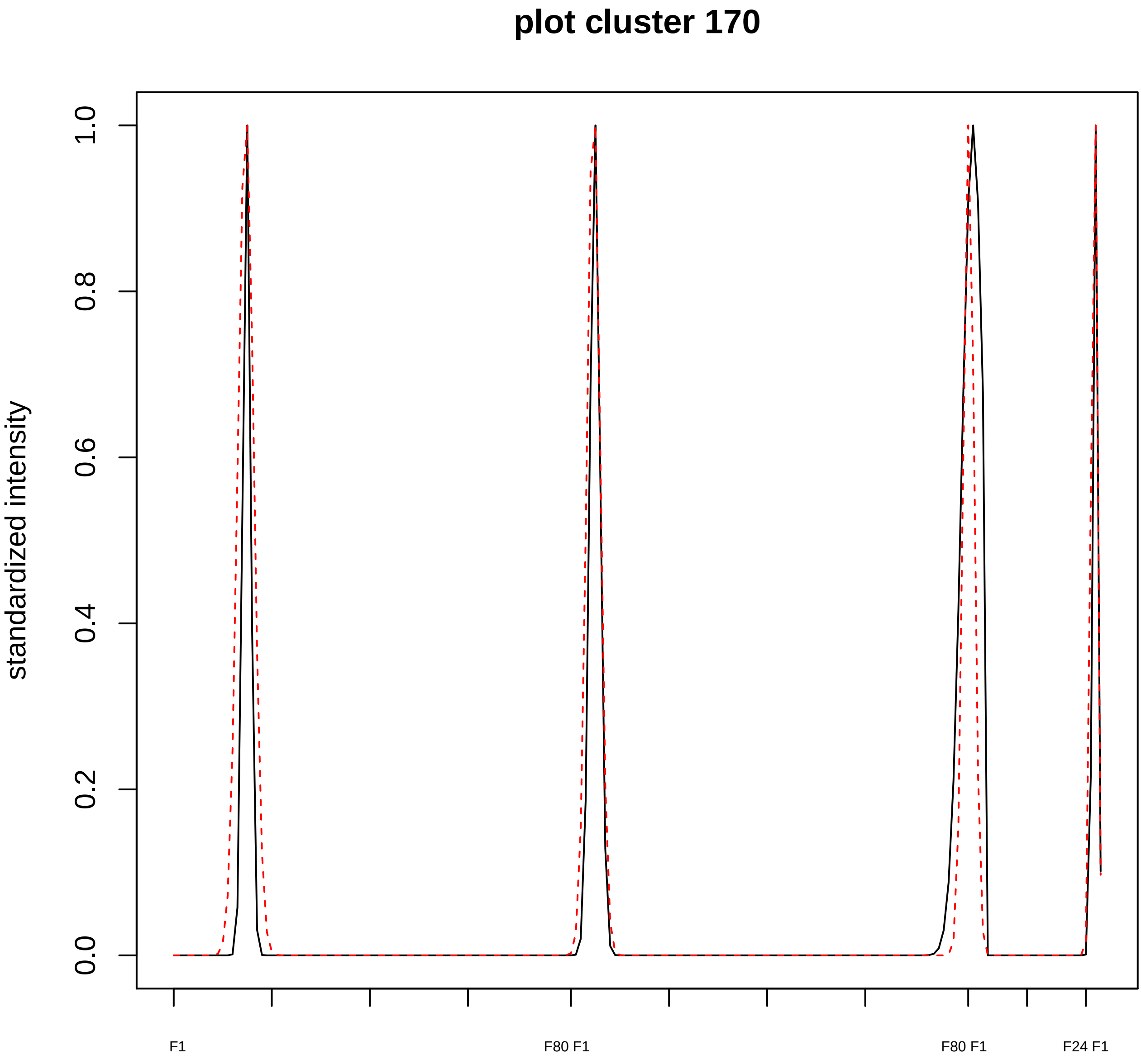
plot cluster 170
standardized intensity
0.0
0.2
0.4
0.6
0.8
1.0
F1
F80 F1
F80 F1
F24 F1

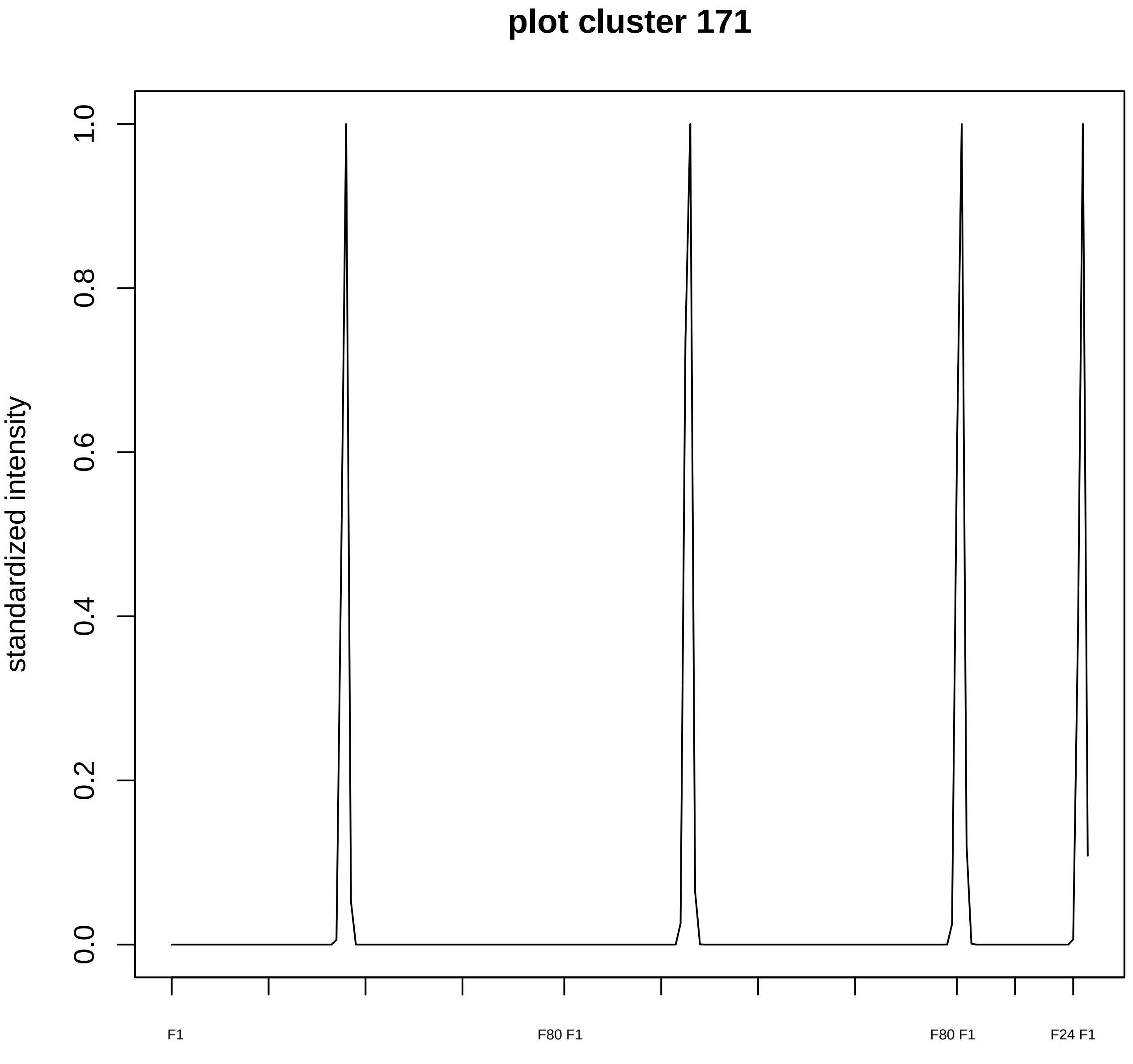

plot cluster 171
standardized intensity
0.0
0.2
0.4
0.6
0.8
1.0
F1
F80 F1
F80 F1
F24 F1

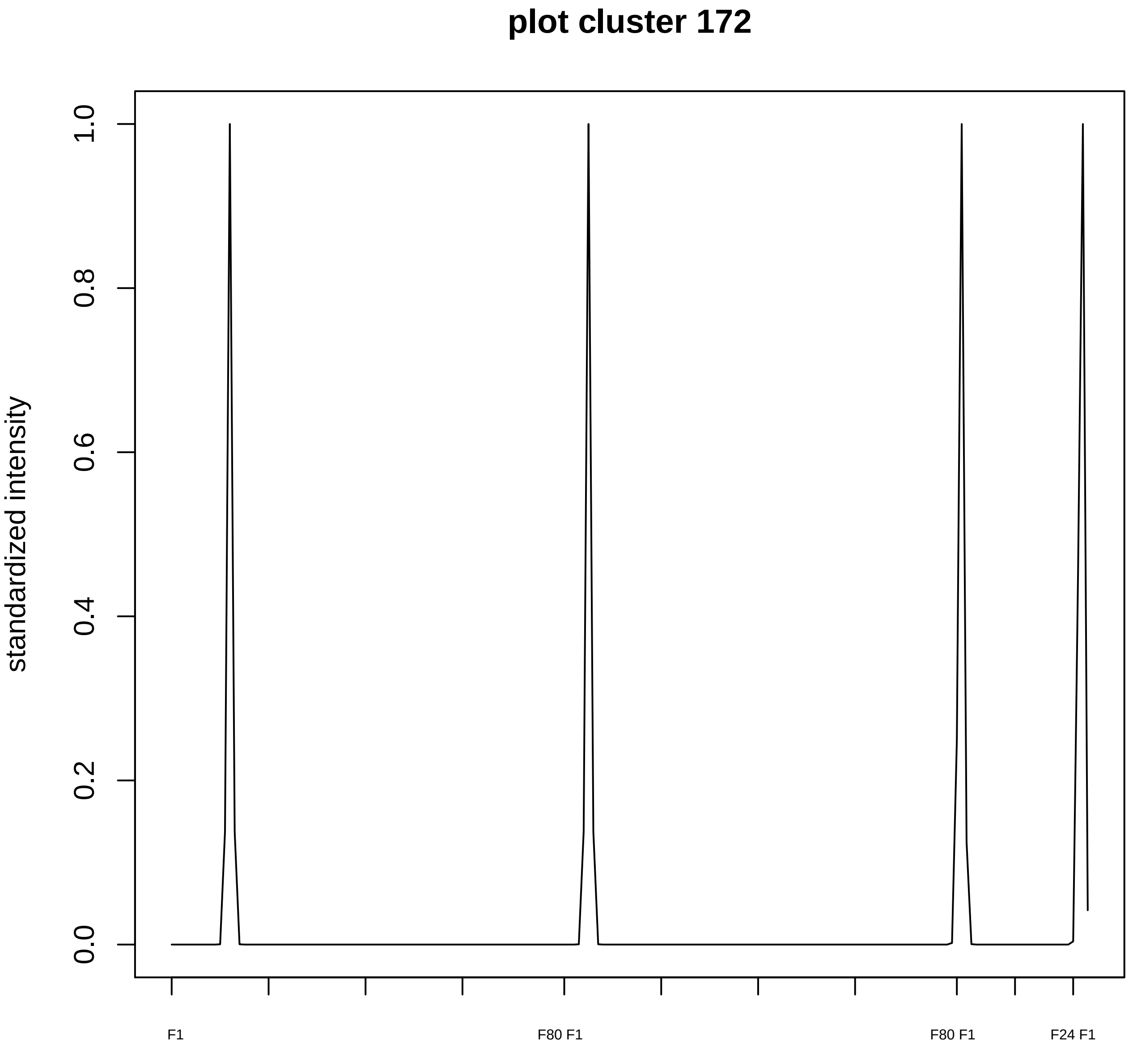

plot cluster 172
standardized intensity
1.0
0.8
0.6
0.4
0.2
0.0
F1
F80 F1
F80 F1
F24 F1

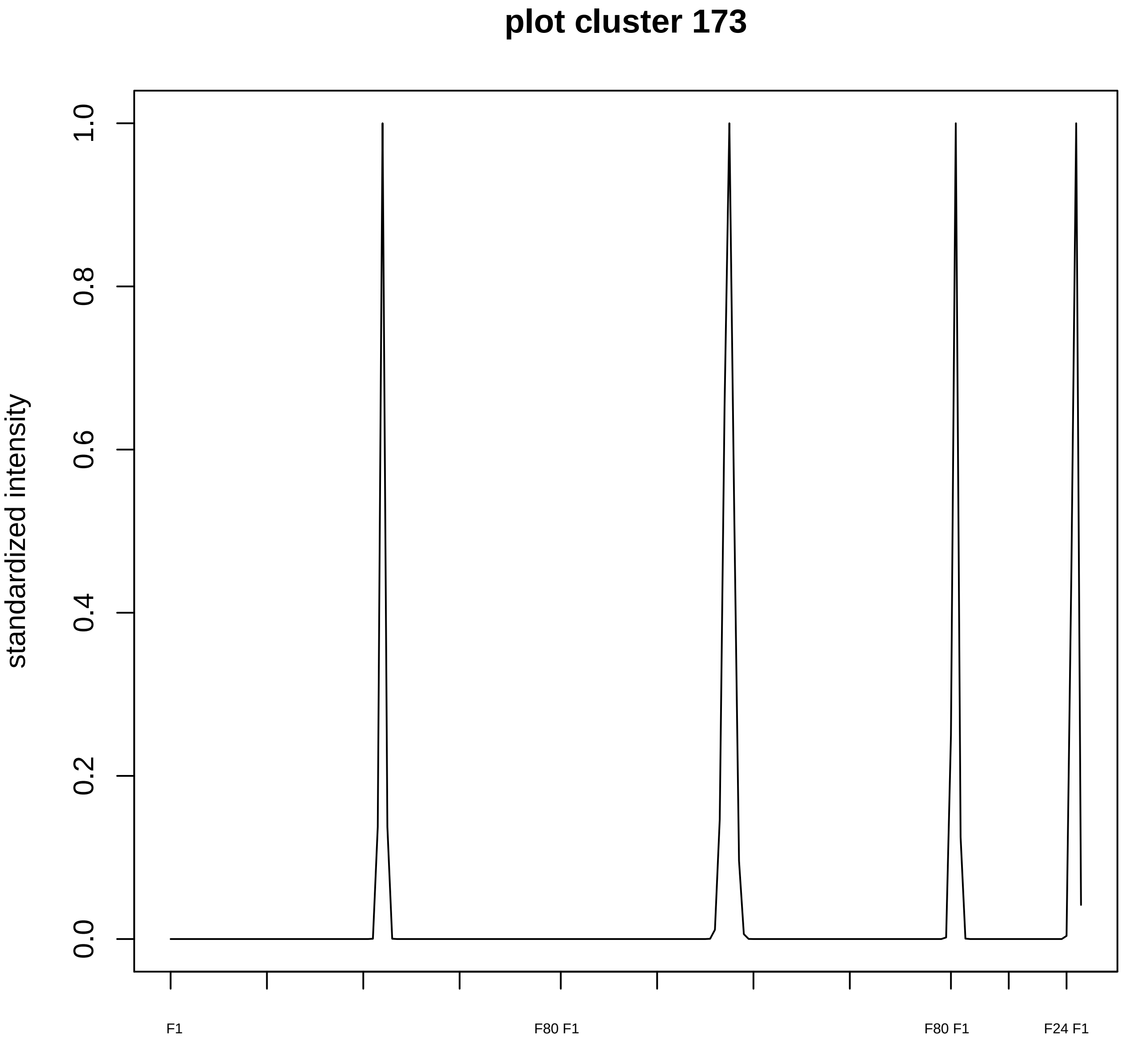
plot cluster 173
standardized intensity
0.0
0.2
0.4
0.6
0.8
1.0
F1
F80 F1
F80 F1
F24 F1

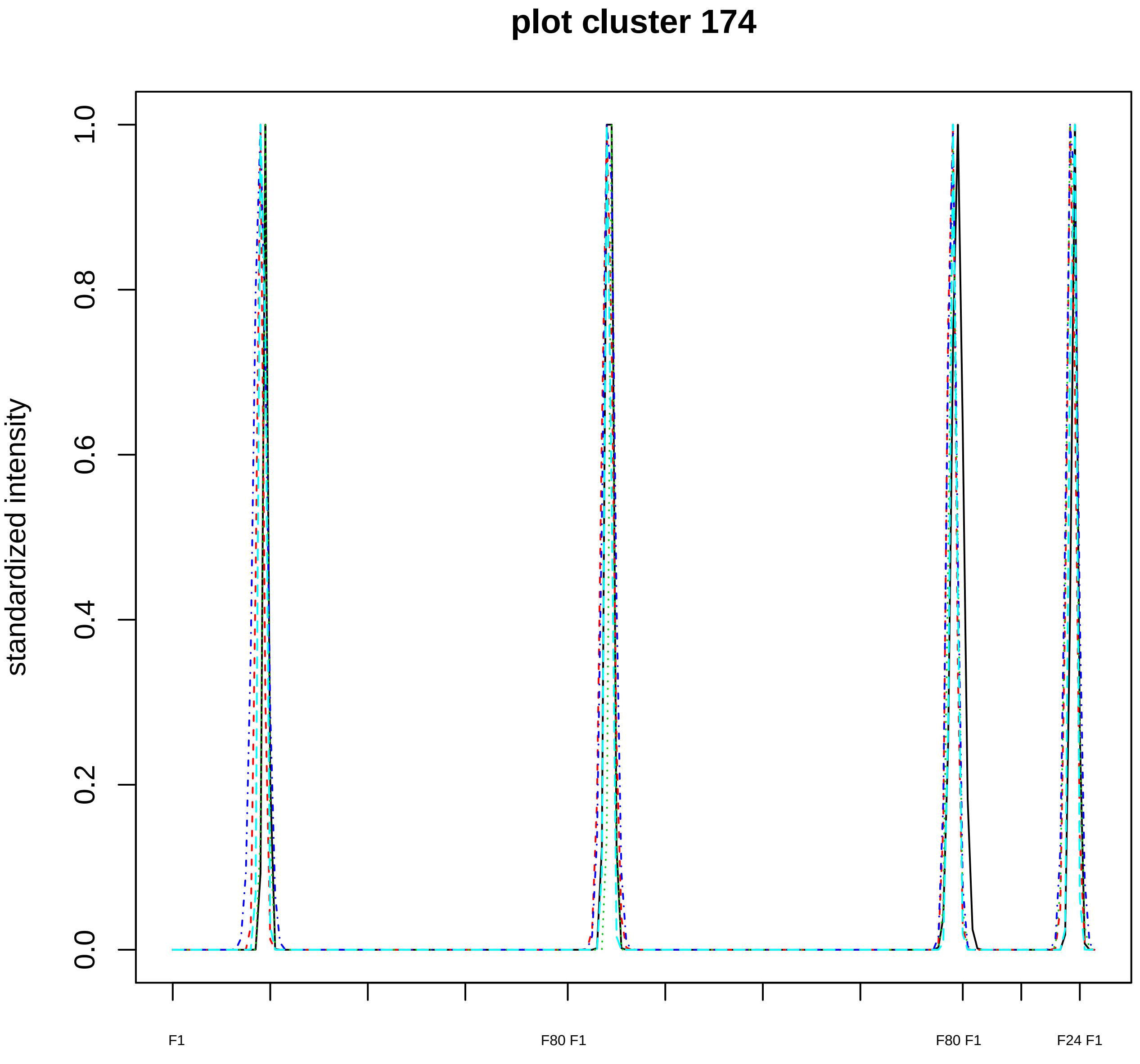
plot cluster 174
standardized intensity
0.0
0.2
0.4
0.6
0.8
1.0
F1
F80 F1
F80 F1
F24 F1

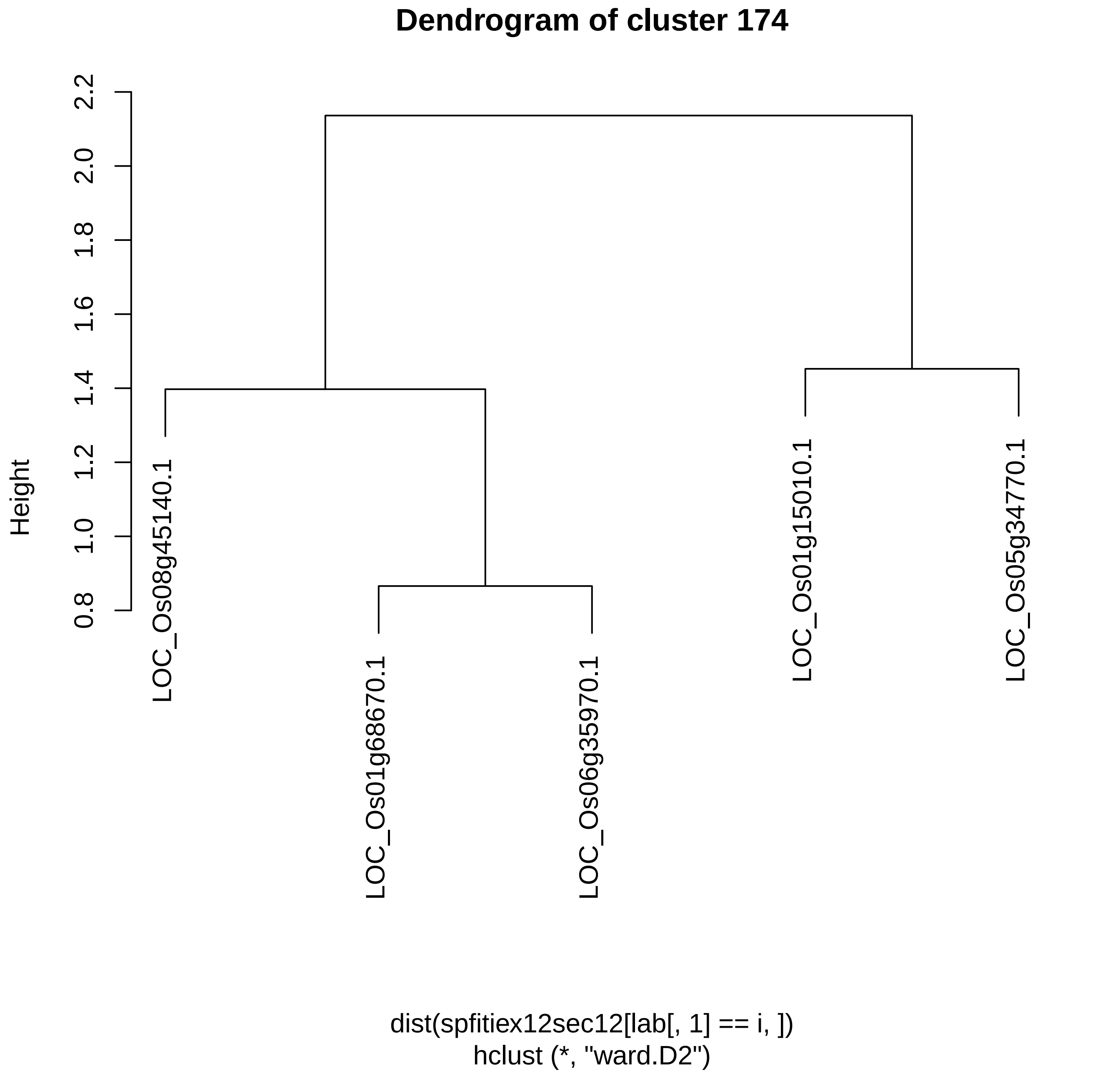
Dendrogram of cluster 174
Height
0.8
1.0
1.2
1.4
1.6
1.8
2.0
2.2
LOC_Os08g45140.1
LOC_Os01g68670.1
LOC_Os06g35970.1
LOC_Os01g15010.1
LOC_Os05g34770.1
dist(spfitiex12sec12[lab[, 1] == i, ])
hclust (*, "ward.D2")

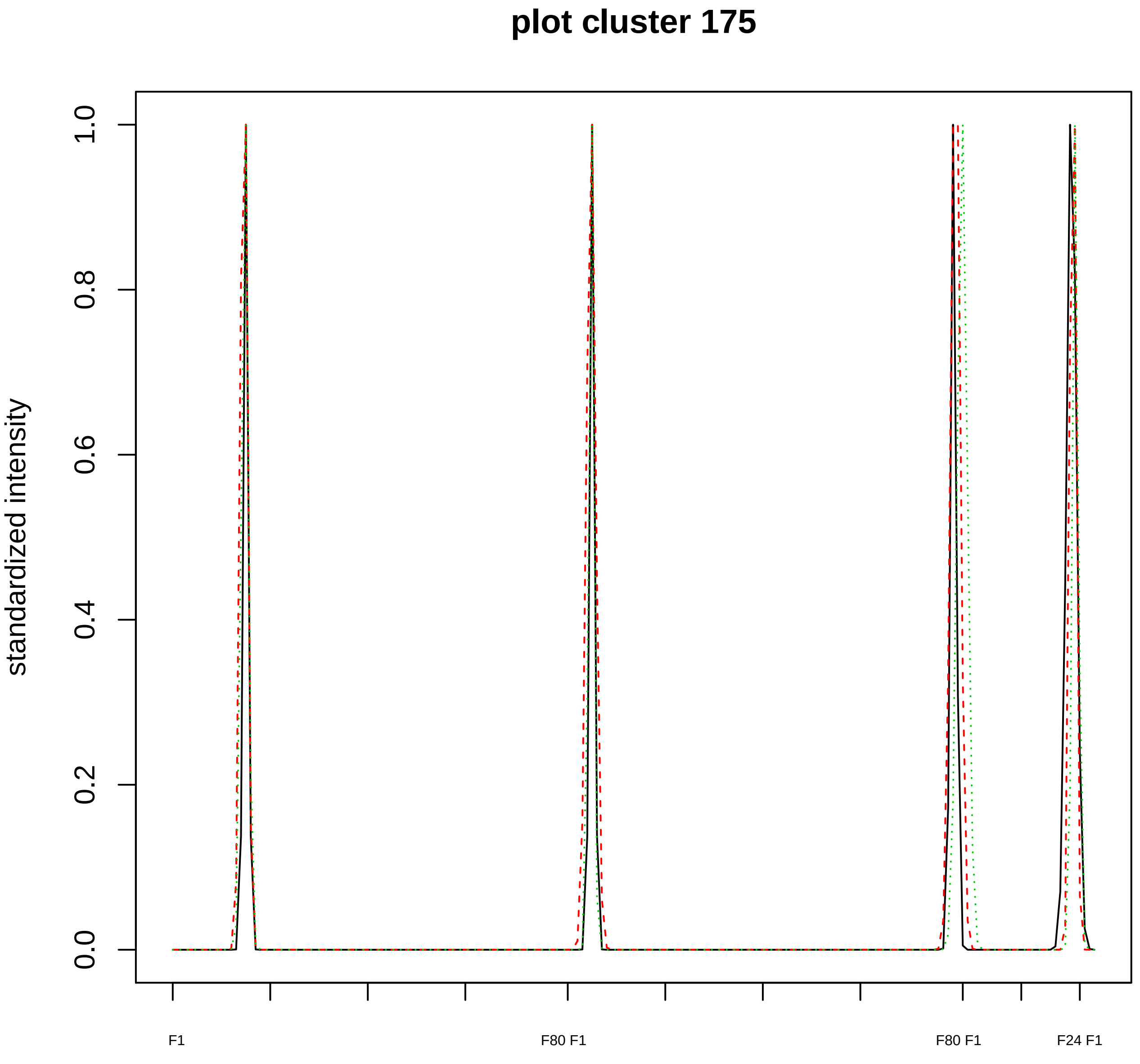

plot cluster 175
standardized intensity
0.0
0.2
0.4
0.6
0.8
1.0
F1
F80 F1
F80 F1
F24 F1

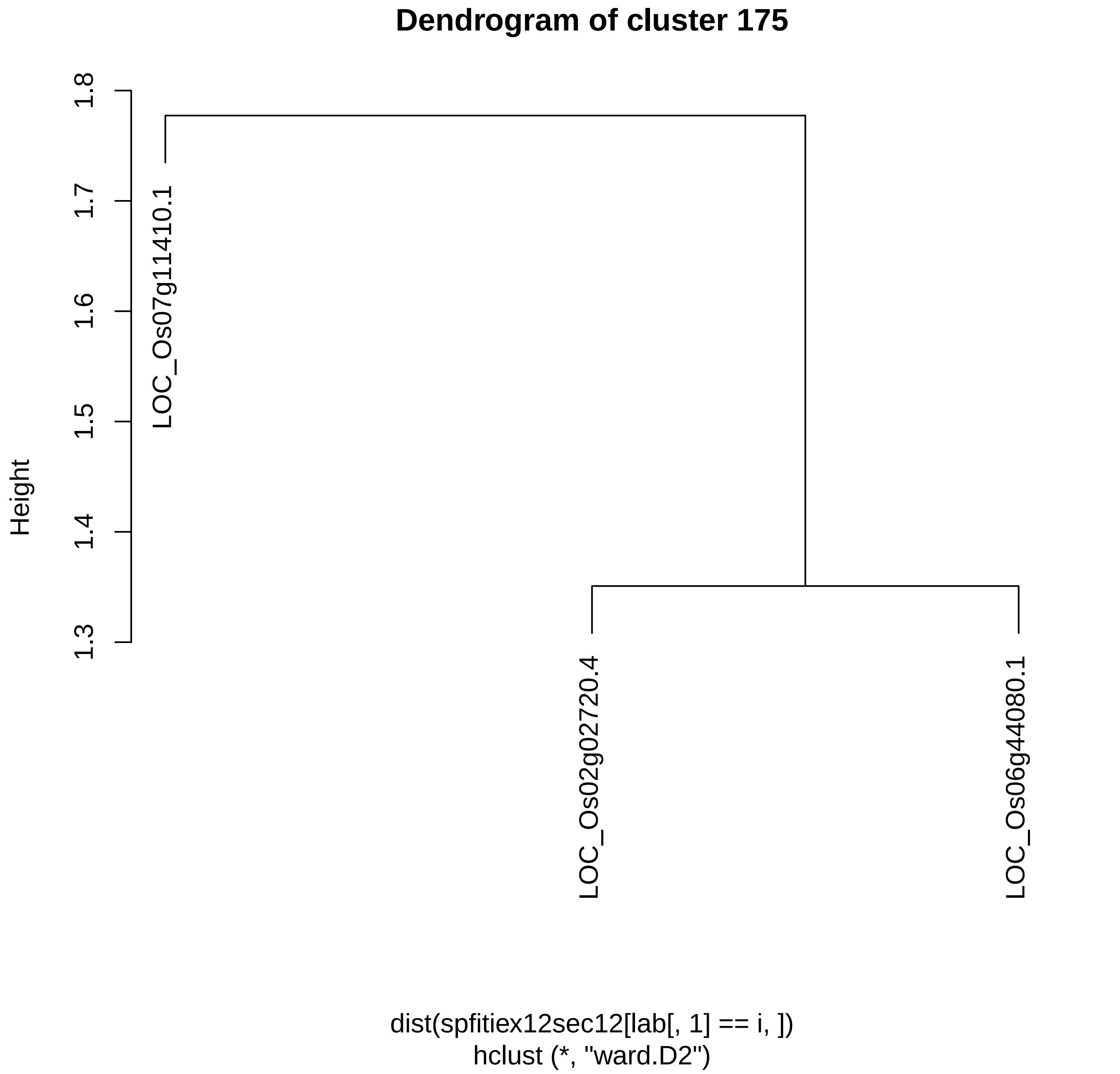
Dendrogram of cluster 175
Height
1.8
1.7
1.6
1.5
1.4
1.3
LOC_Os07g11410.1
LOC_Os02g02720.4
LOC_Os06g44080.1
dist(spfitiex12sec12[lab[, 1] == i, ])
hclust (*, "ward.D2")

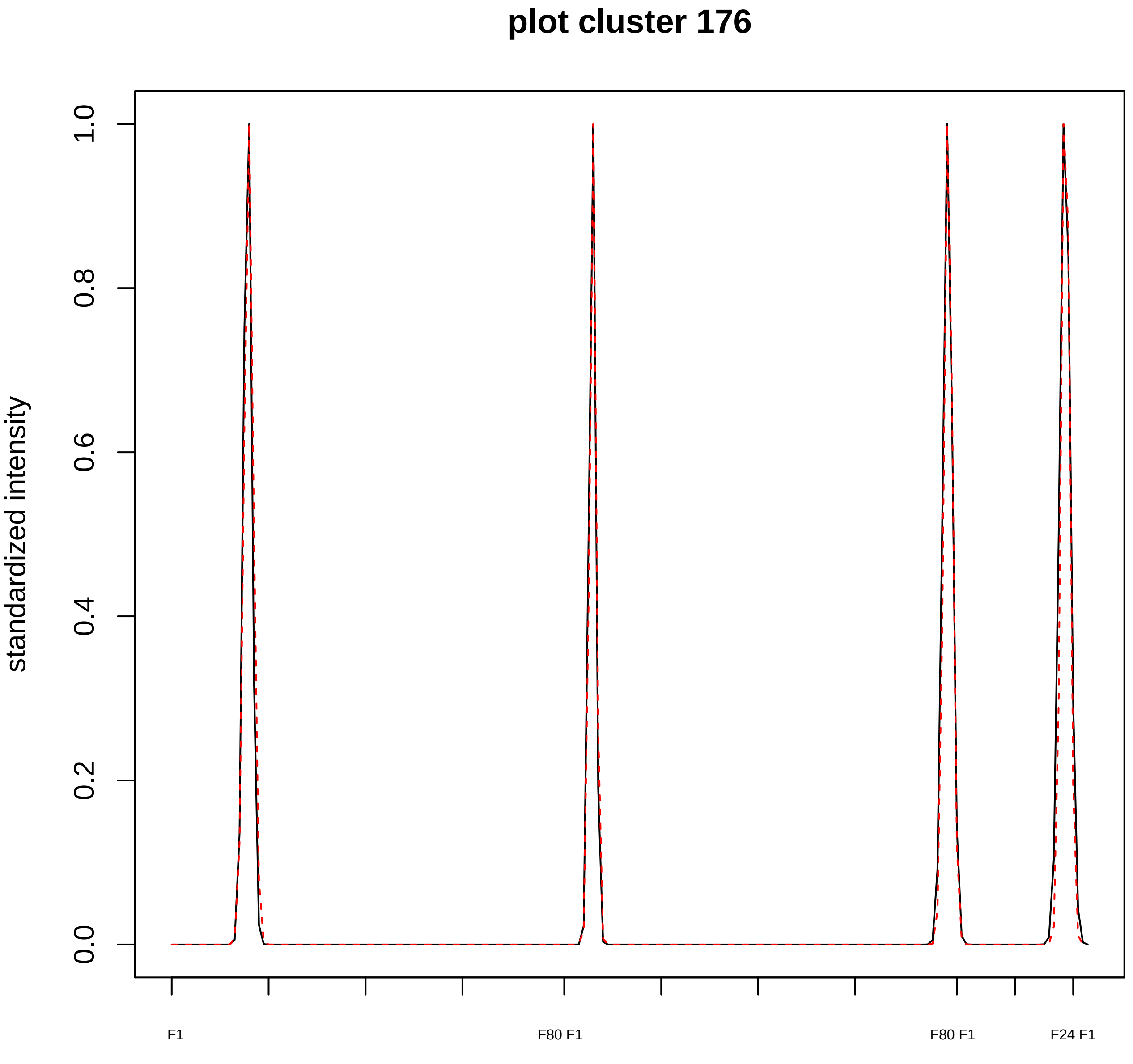
plot cluster 176
standardized intensity
0.0
0.2
0.4
0.6
0.8
1.0
F1
F80 F1
F80 F1
F24 F1

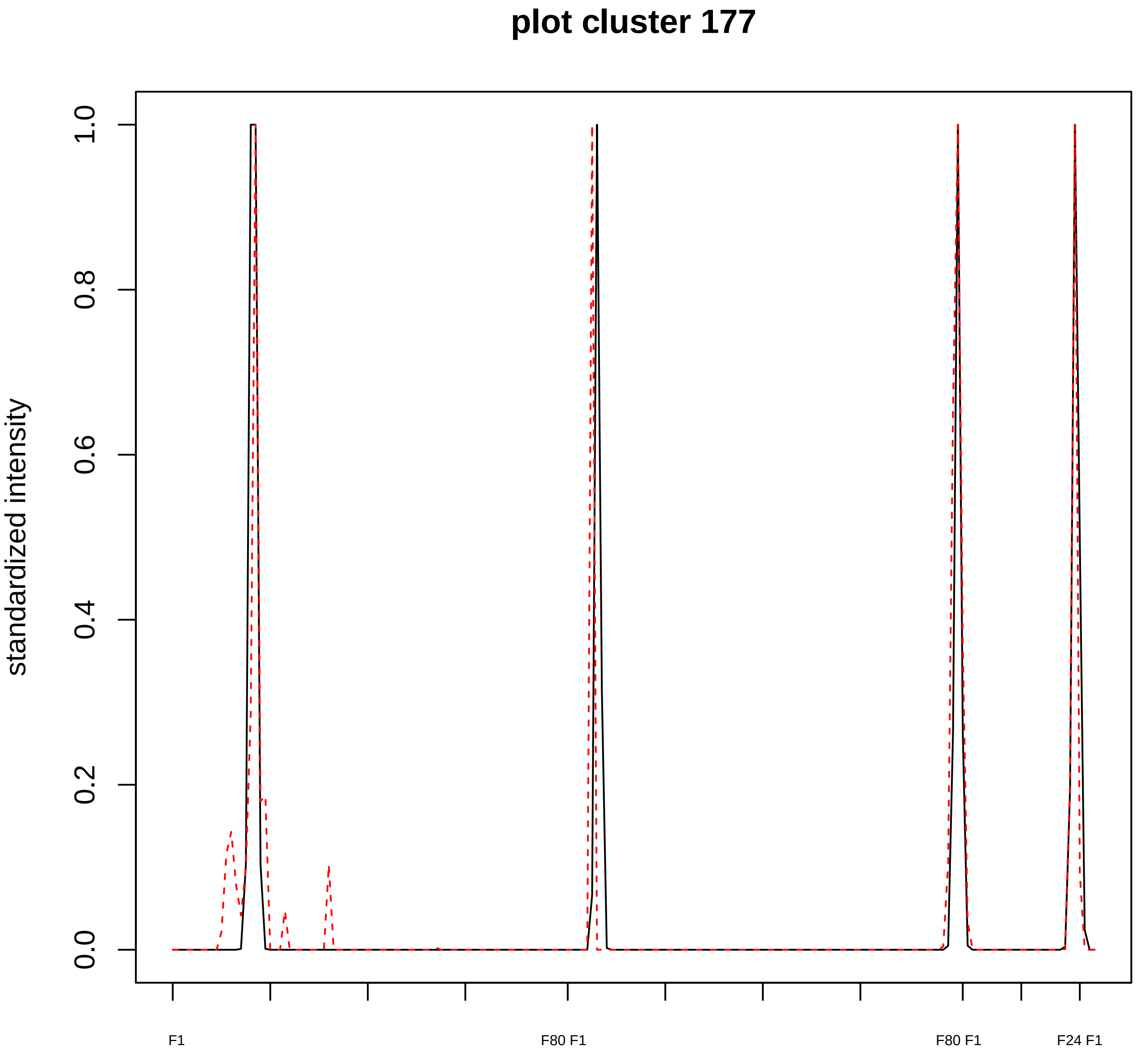
plot cluster 177
standardized intensity
0.0
0.2
0.4
0.6
0.8
1.0
F1
F80 F1
F80 F1
F24 F1

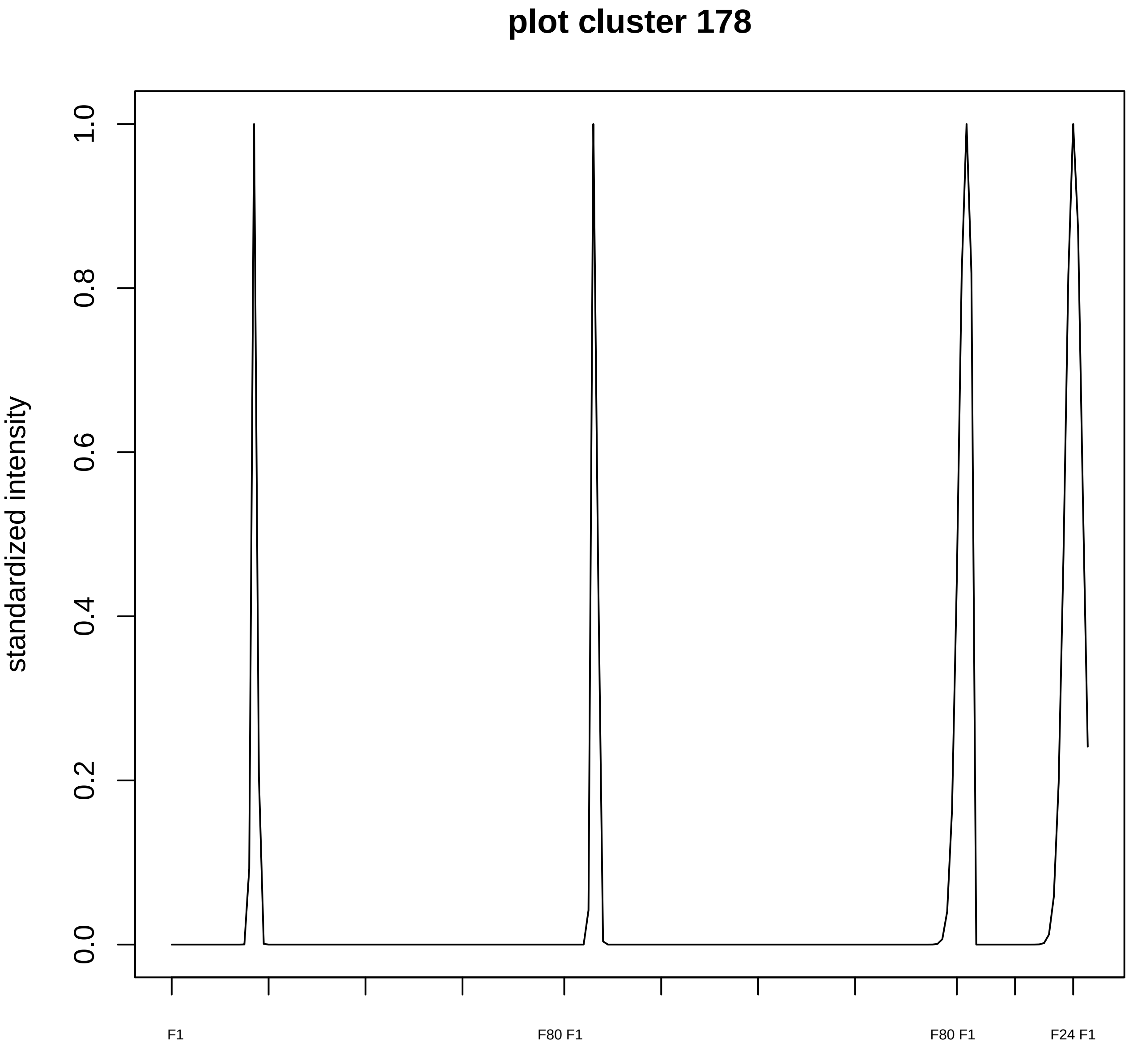

plot cluster 178
standardized intensity
0.0
0.2
0.4
0.6
0.8
1.0
F1
F80 F1
F80 F1
F24 F1

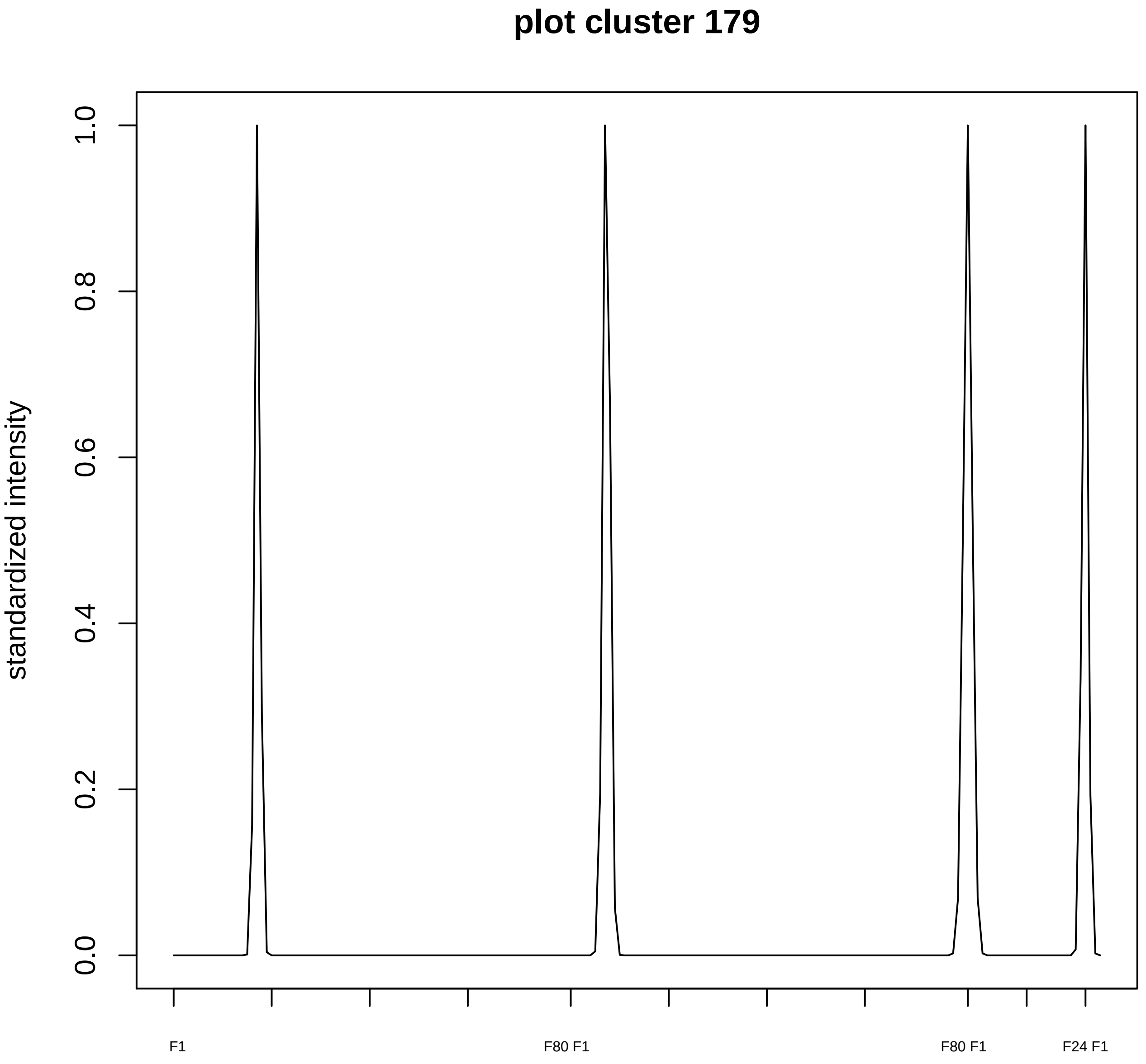

plot cluster 179
standardized intensity
0.0
0.2
0.4
0.6
0.8
1.0
F1
F80 F1
F80 F1
F24 F1

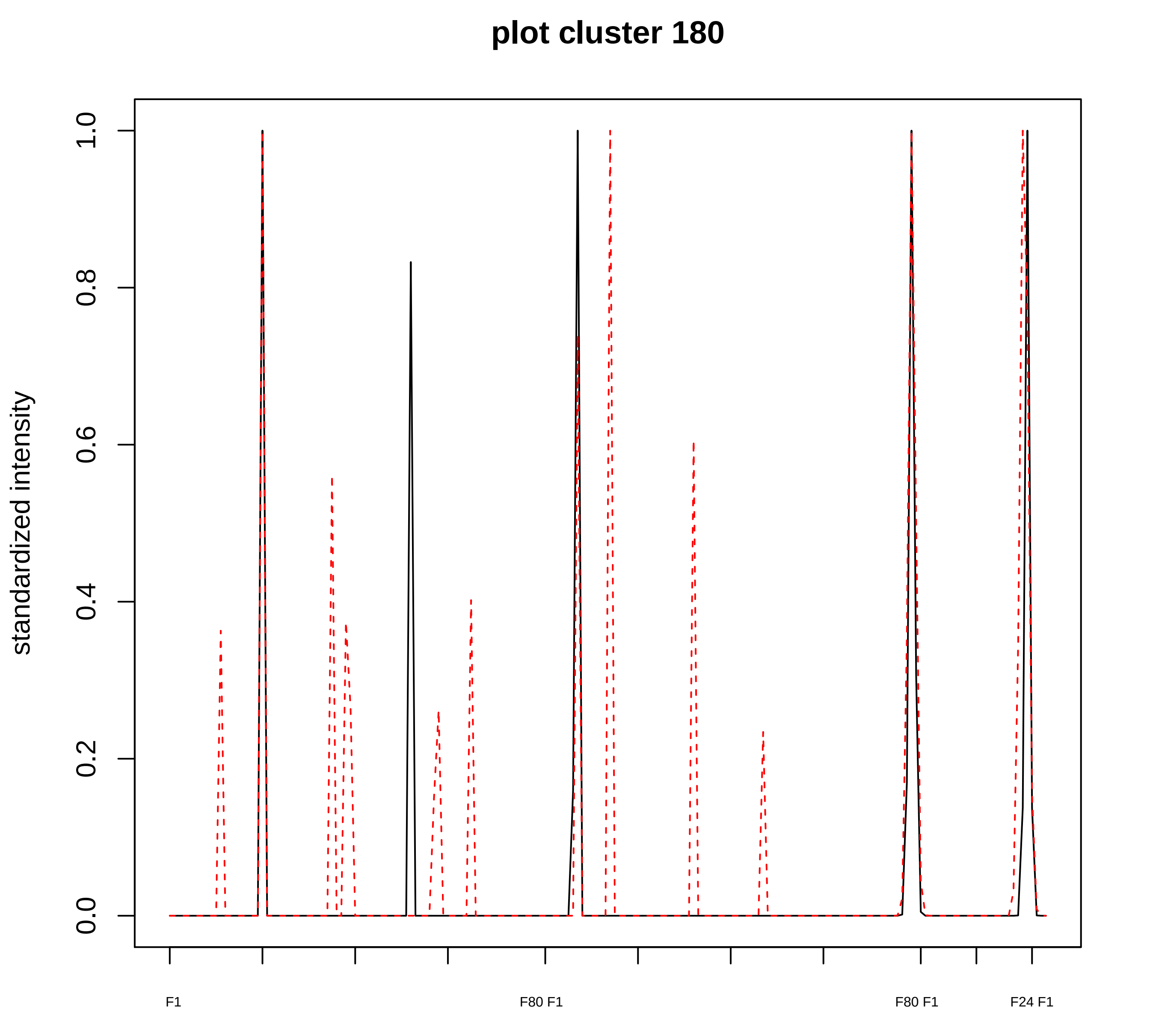

plot cluster 180
standardized intensity
0.0
0.2
0.4
0.6
0.8
1.0
F1
F80 F1
F80 F1
F24 F1

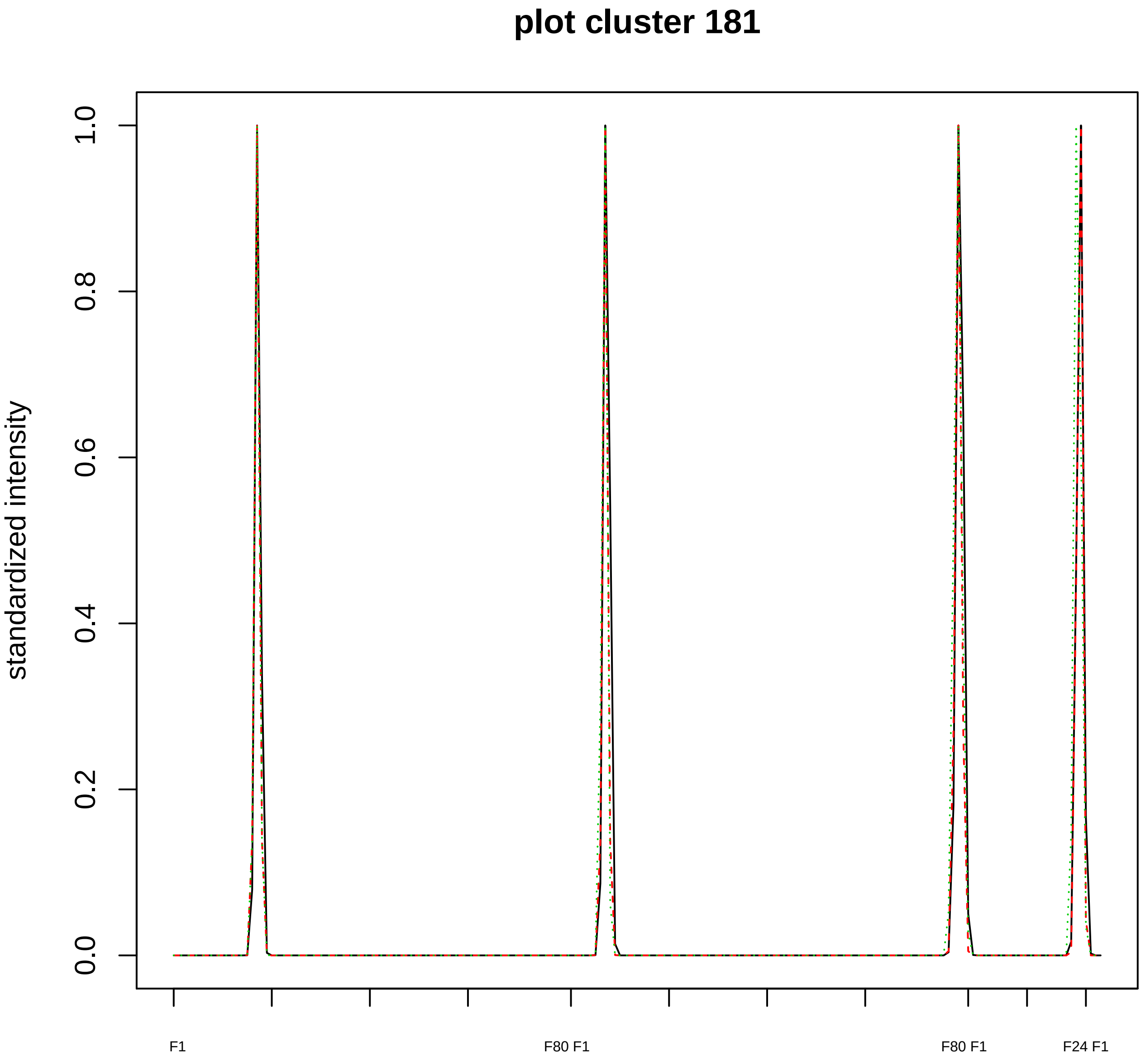

plot cluster 181
standardized intensity
0.0
0.2
0.4
0.6
0.8
1.0
F1
F80 F1
F80 F1
F24 F1

#### Dendrogram of cluster 181

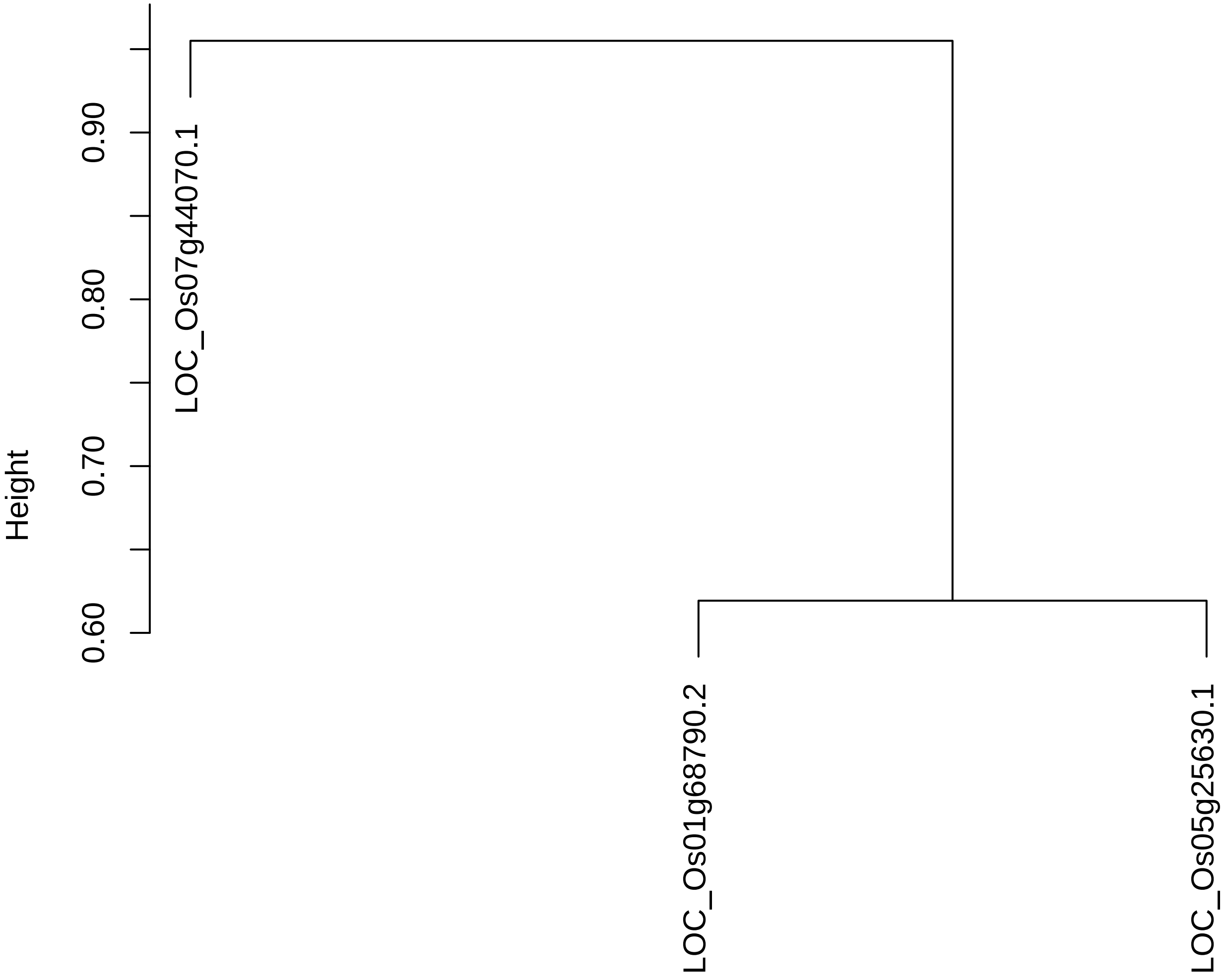


dist(spfitiex12sec12[lab[, 1] == i, ])
hclust (*, "ward.D2")

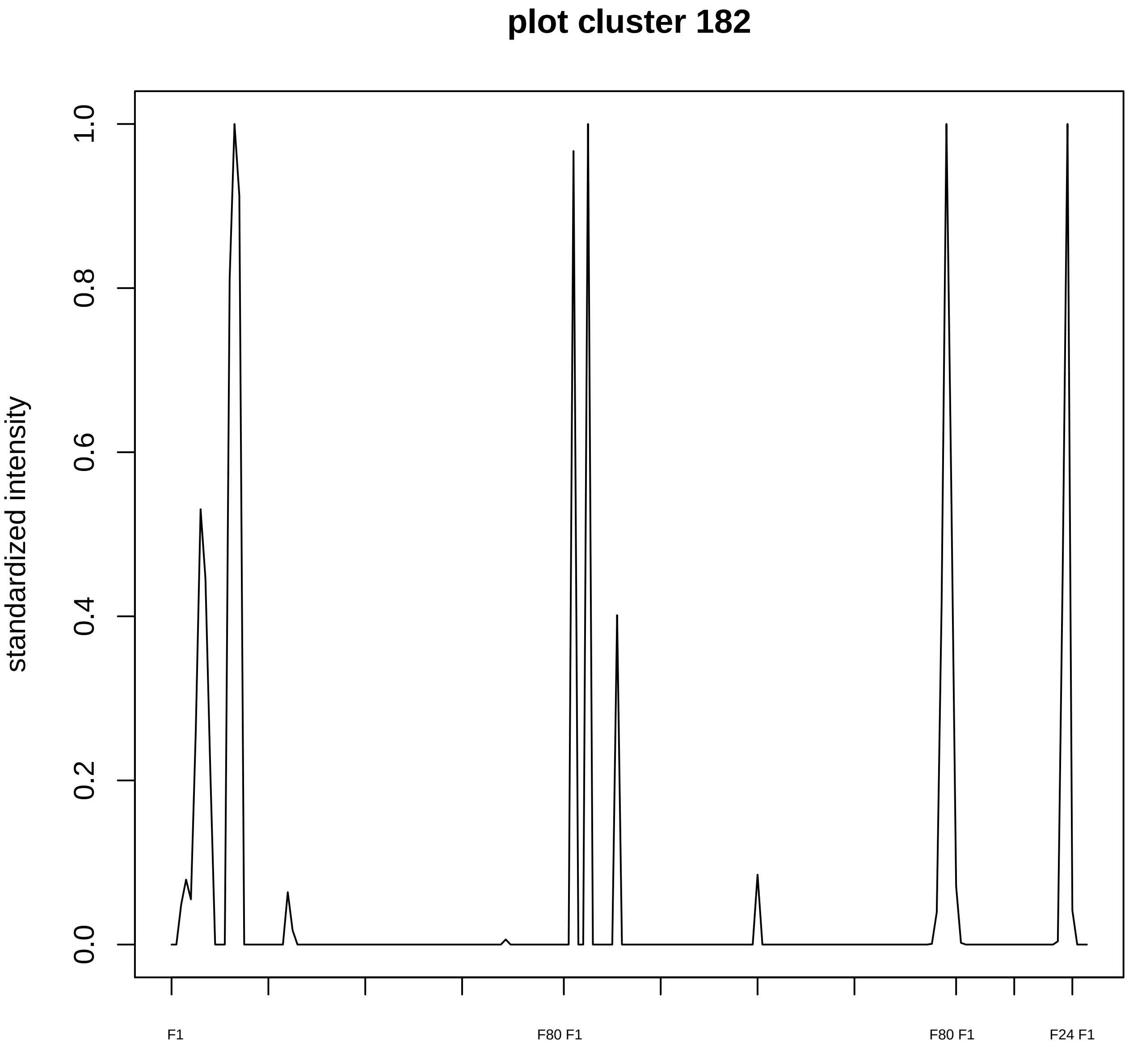

plot cluster 182
standardized intensity
0.0
0.2
0.4
0.6
0.8
1.0
F1
F80 F1
F80 F1
F24 F1

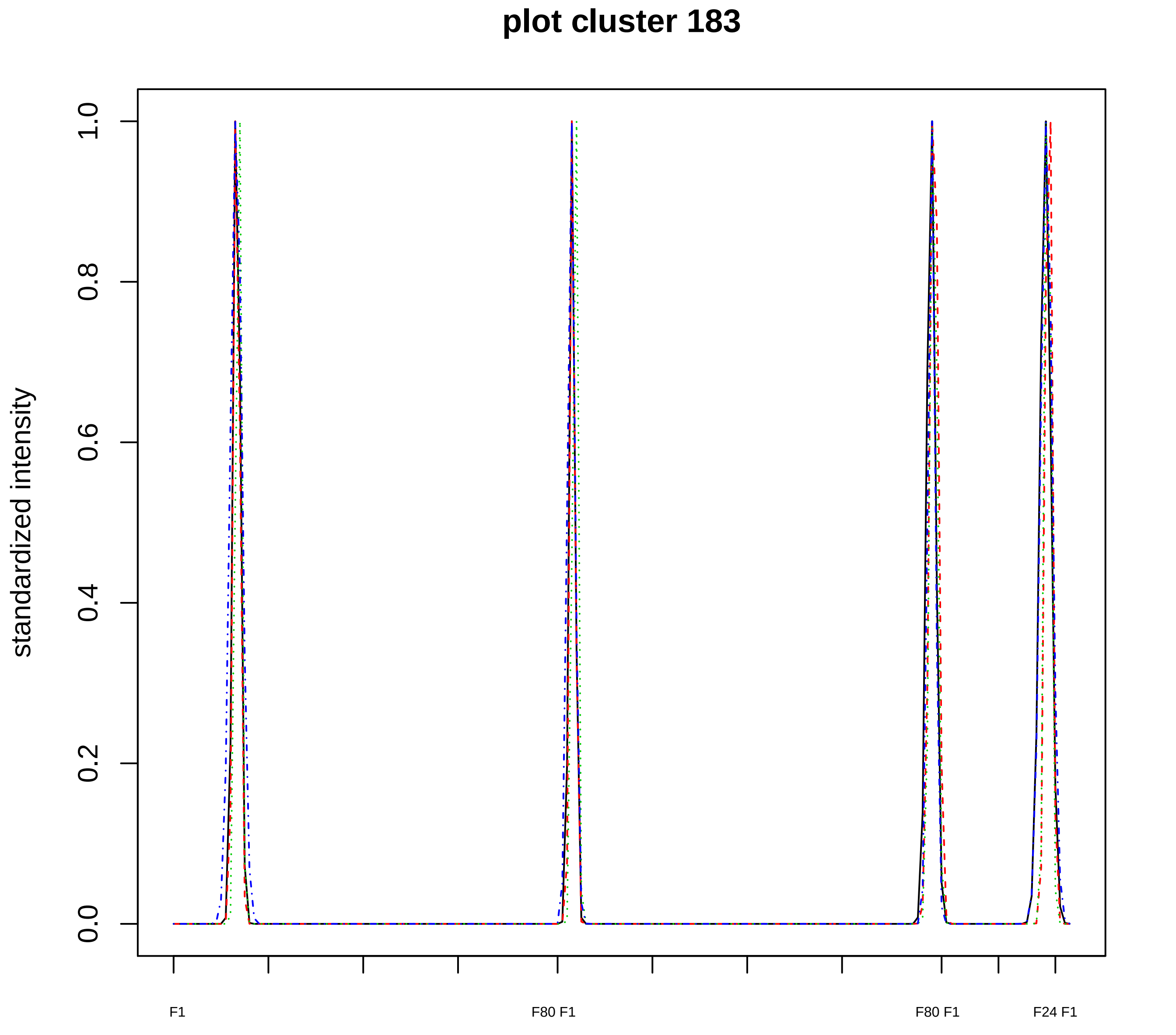

plot cluster 183
standardized intensity
1.0
0.8
0.6
0.4
0.2
0.0
F1
F80 F1
F80 F1
F24 F1

### Dendrogram of cluster 183

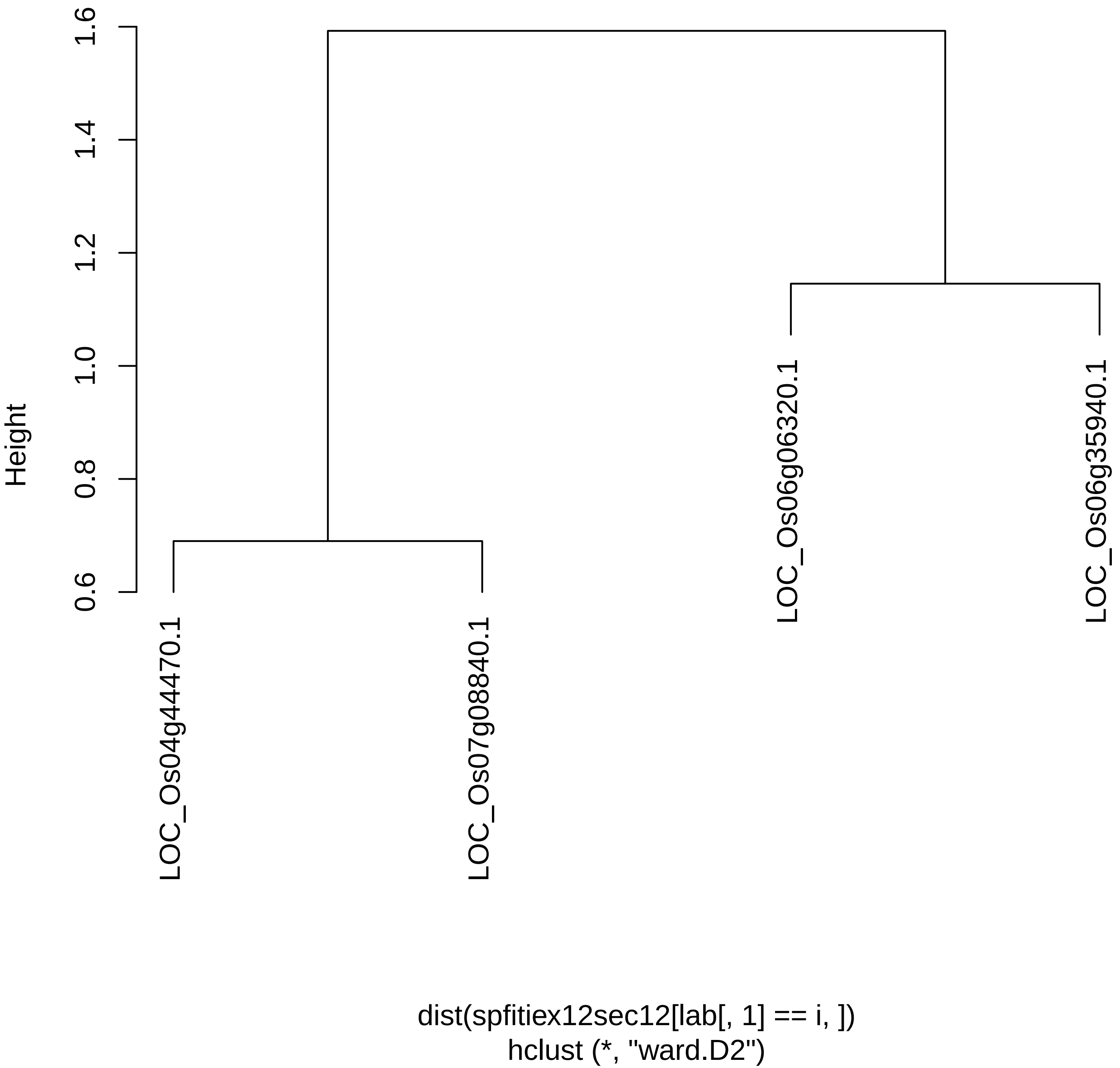

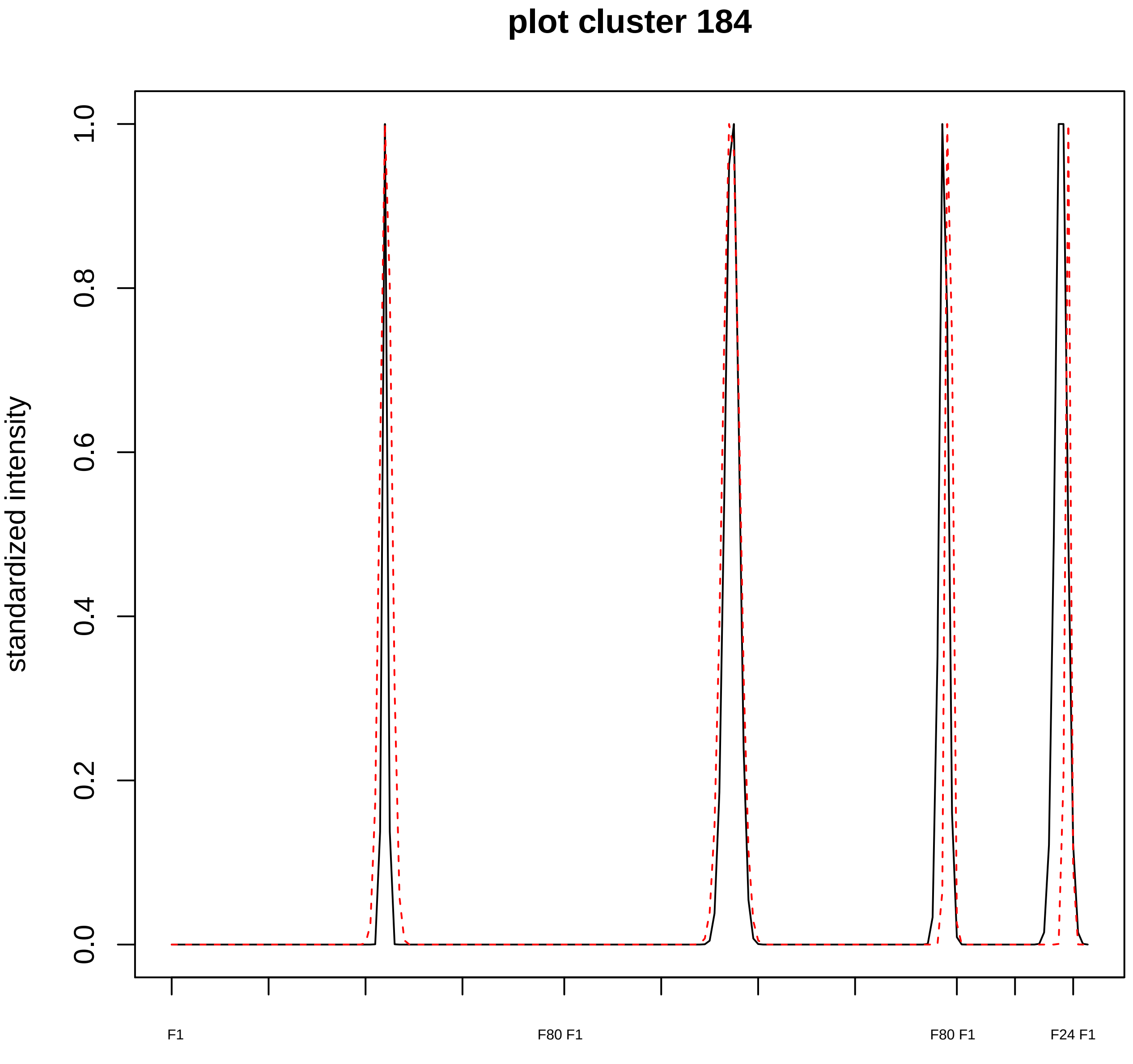
plot cluster 184
standardized intensity
0.0
0.2
0.4
0.6
0.8
1.0
F1
F80 F1
F80 F1
F24 F1

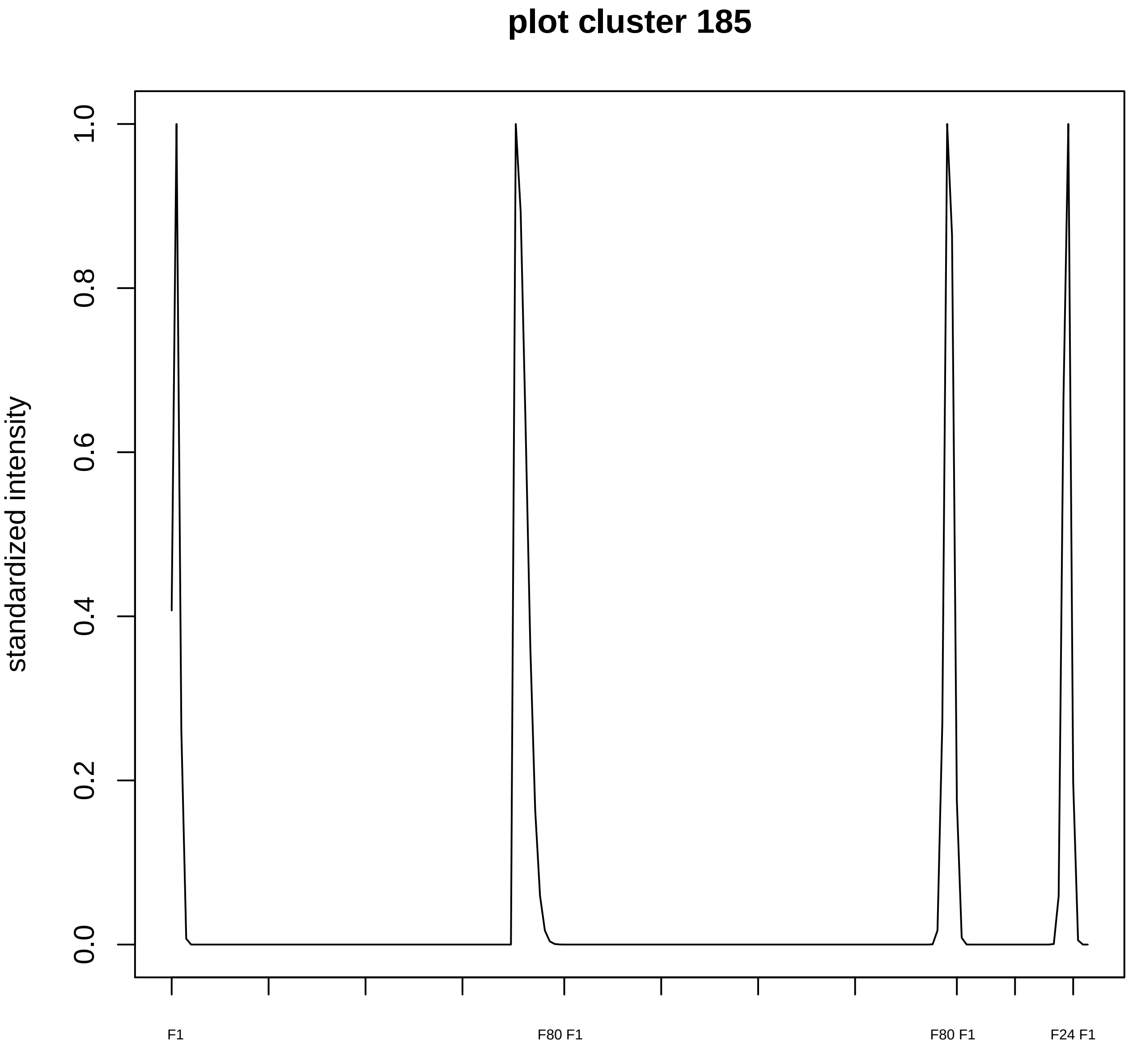

plot cluster 185
standardized intensity
0.0
0.2
0.4
0.6
0.8
1.0
F1
F80 F1
F80 F1
F24 F1

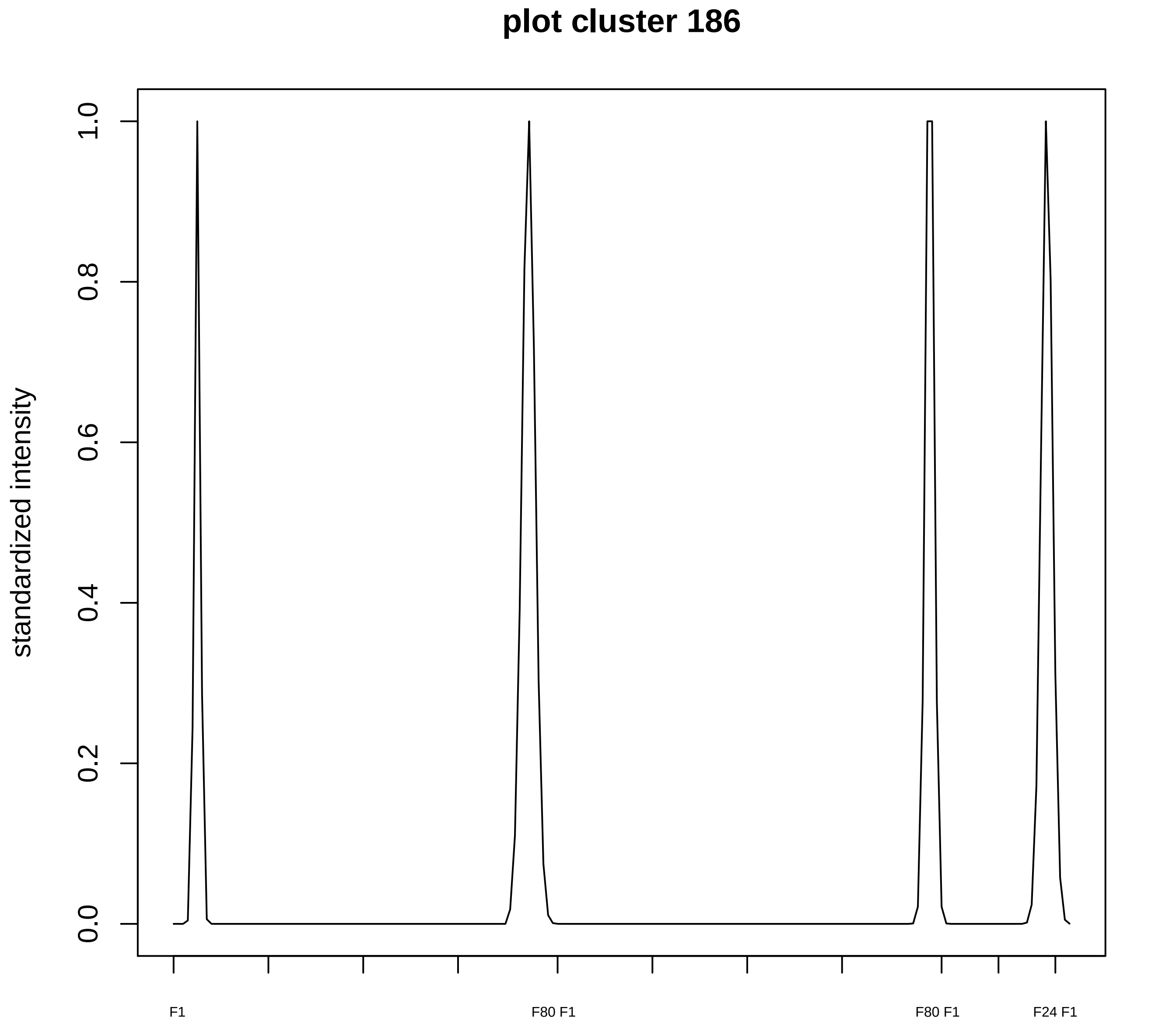
plot cluster 186
standardized intensity
0.0
0.2
0.4
0.6
0.8
1.0
F1
F80 F1
F80 F1
F24 F1

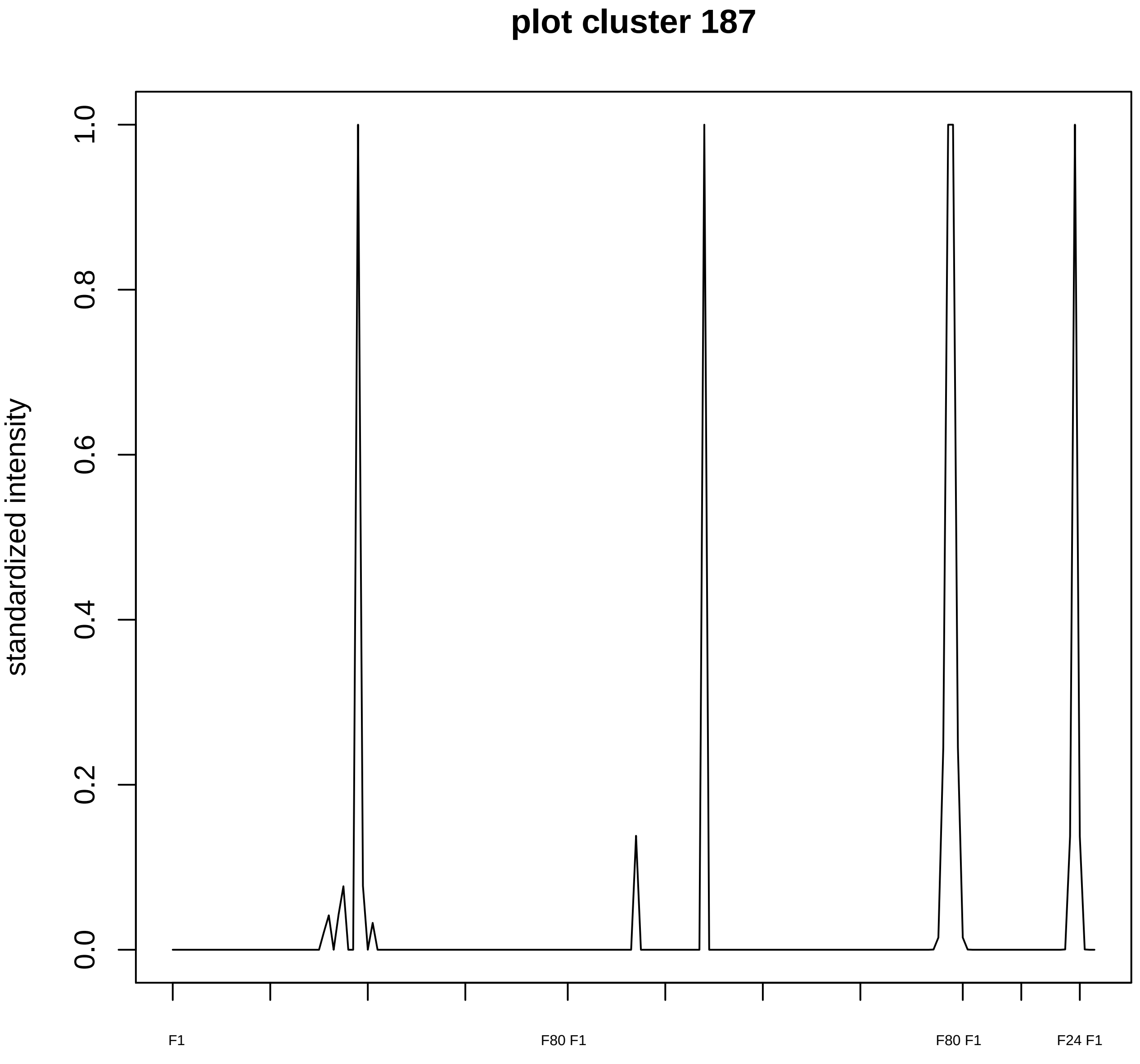

plot cluster 187
standardized intensity
0.0
0.2
0.4
0.6
0.8
1.0
F1
F80 F1
F80 F1
F24 F1

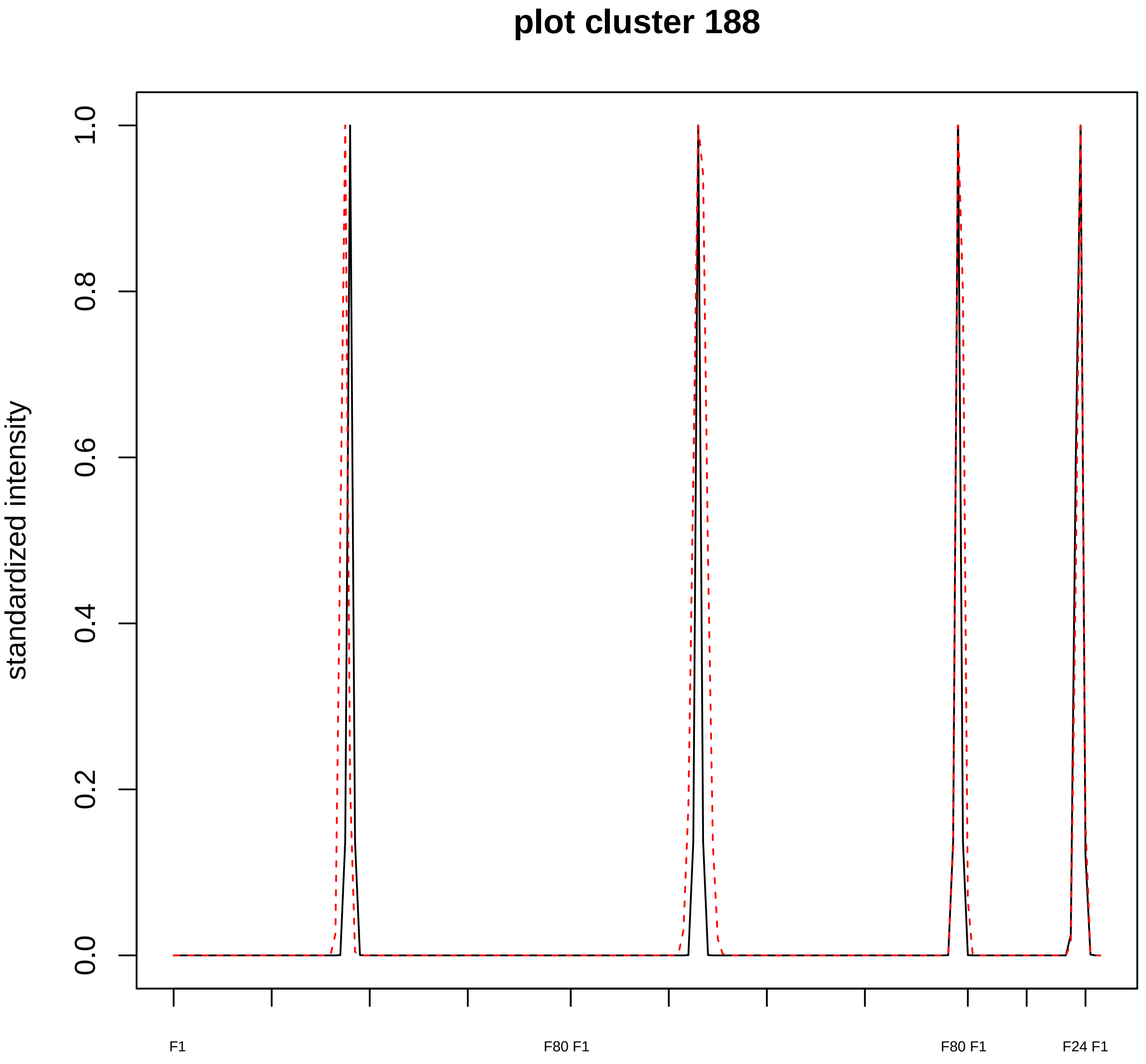
plot cluster 188
standardized intensity
0.0
0.2
0.4
0.6
0.8
1.0
F1
F80 F1
F80 F1
F24 F1

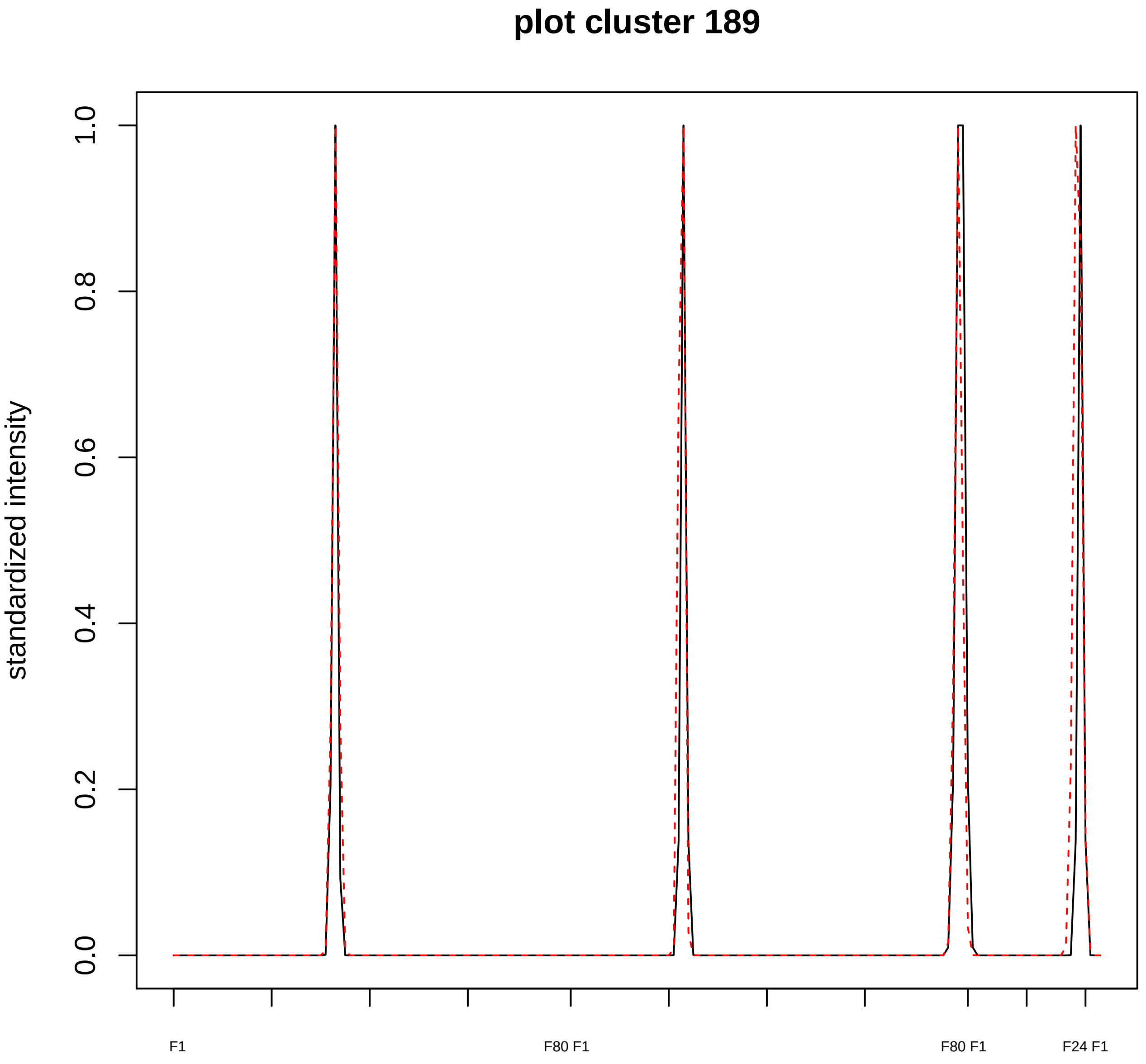
plot cluster 189
standardized intensity
0.0
0.2
0.4
0.6
0.8
1.0
F1
F80 F1
F80 F1
F24 F1

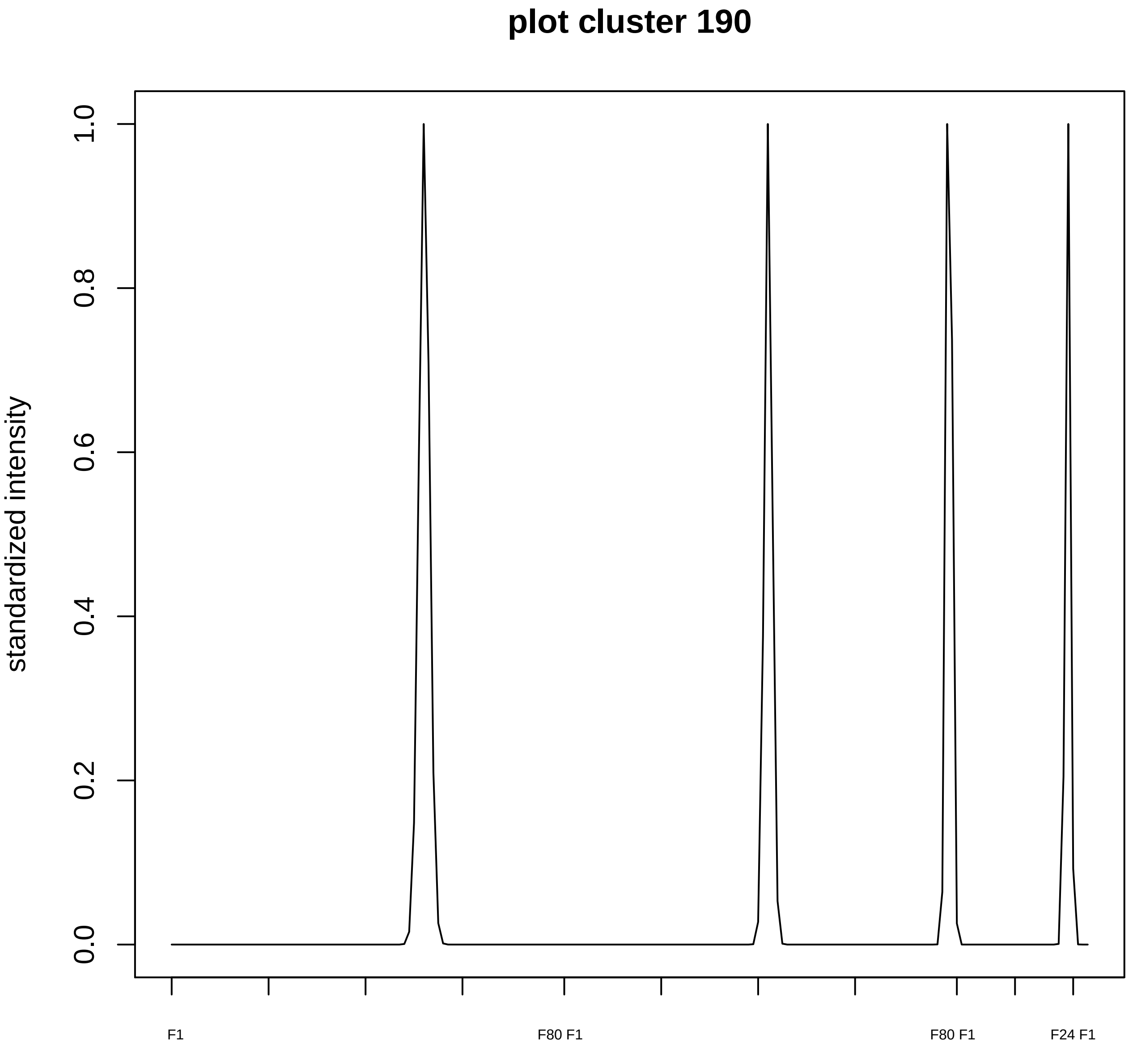
plot cluster 190
standardized intensity
0.0
0.2
0.4
0.6
0.8
1.0
F1
F80 F1
F80 F1
F24 F1

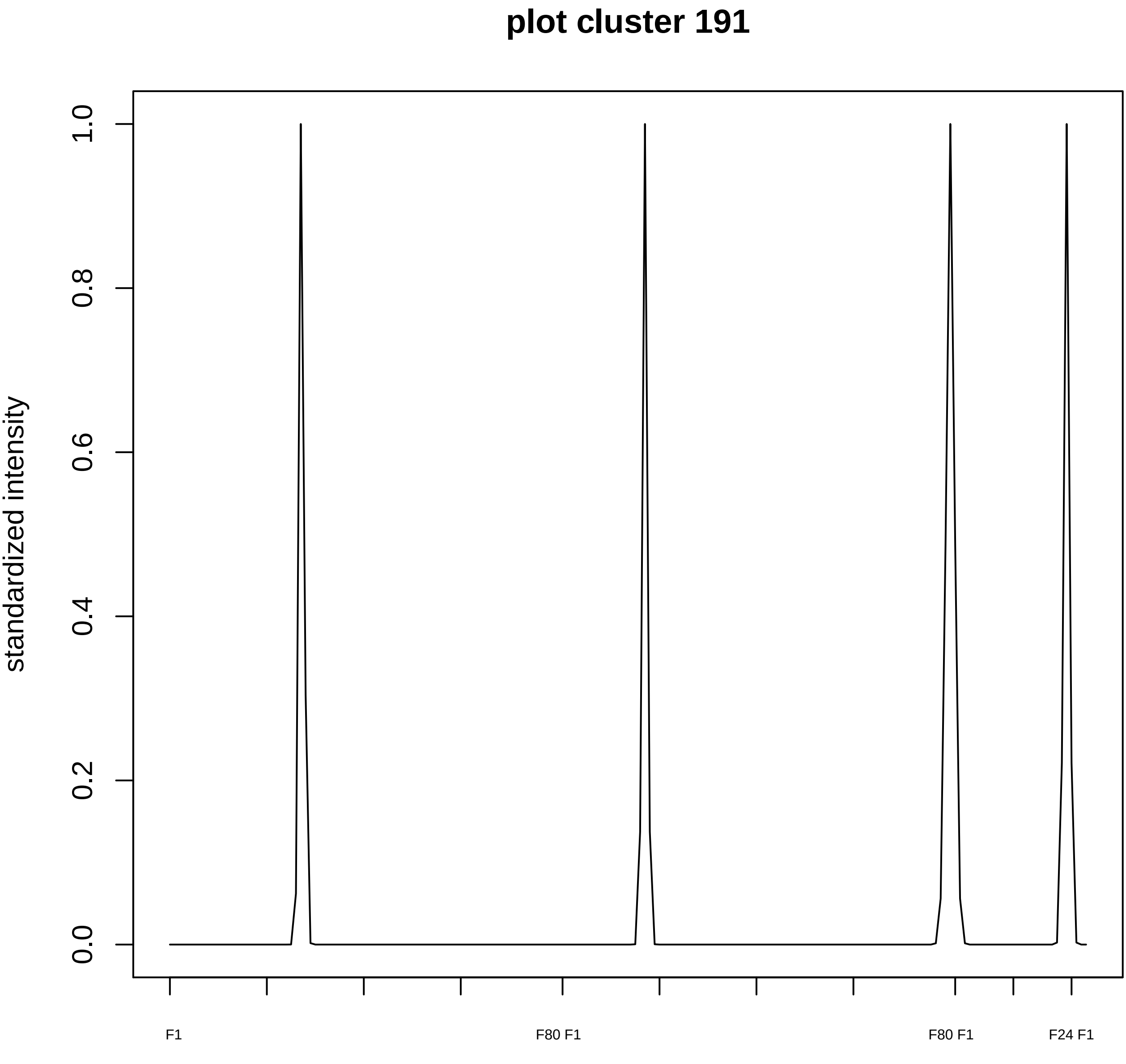

plot cluster 191
standardized intensity
0.0
0.2
0.4
0.6
0.8
1.0
F1
F80 F1
F80 F1
F24 F1

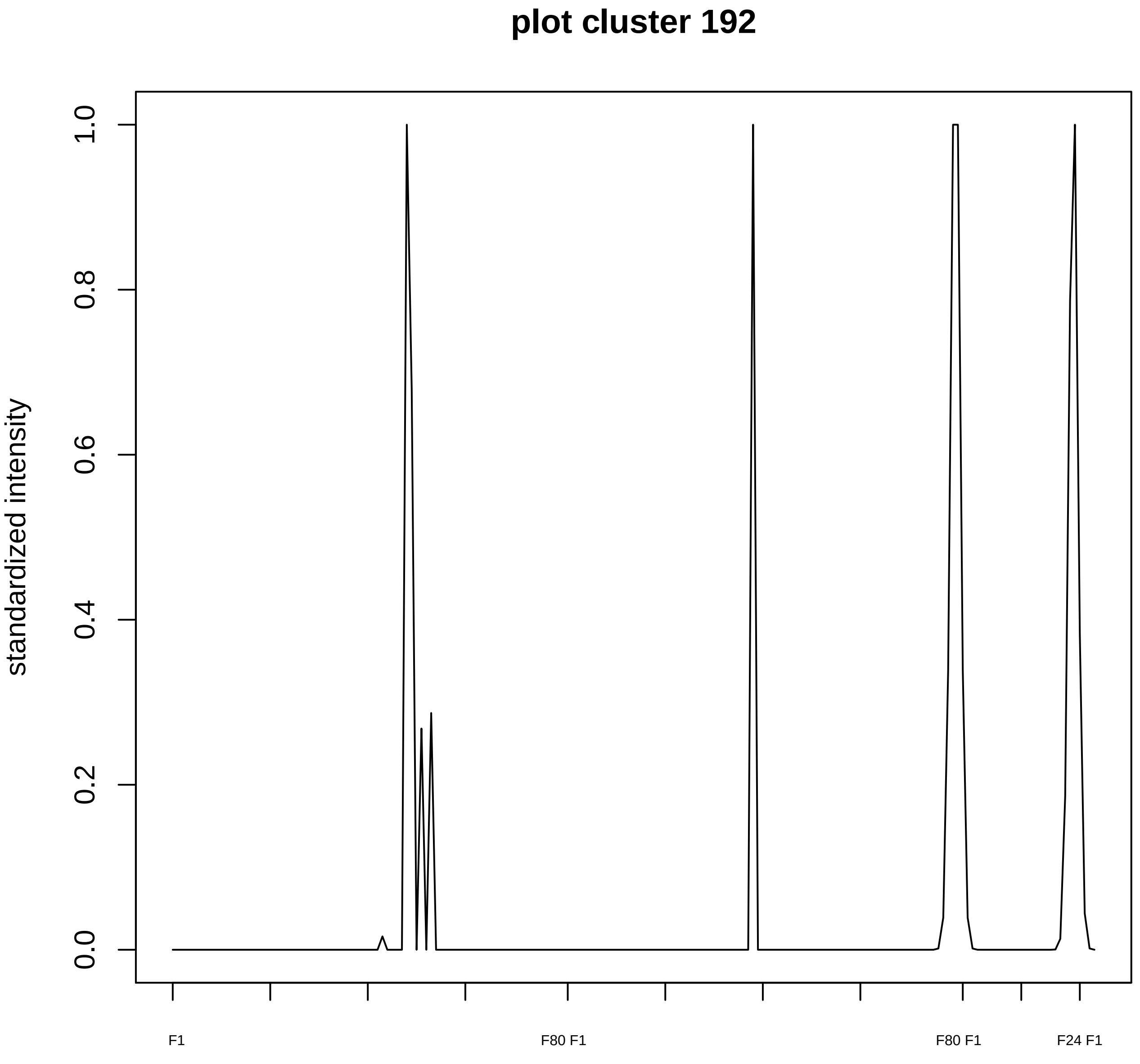

plot cluster 192
standardized intensity
1.0
0.8
0.6
0.4
0.2
0.0
F1
F80 F1
F80 F1
F24 F1

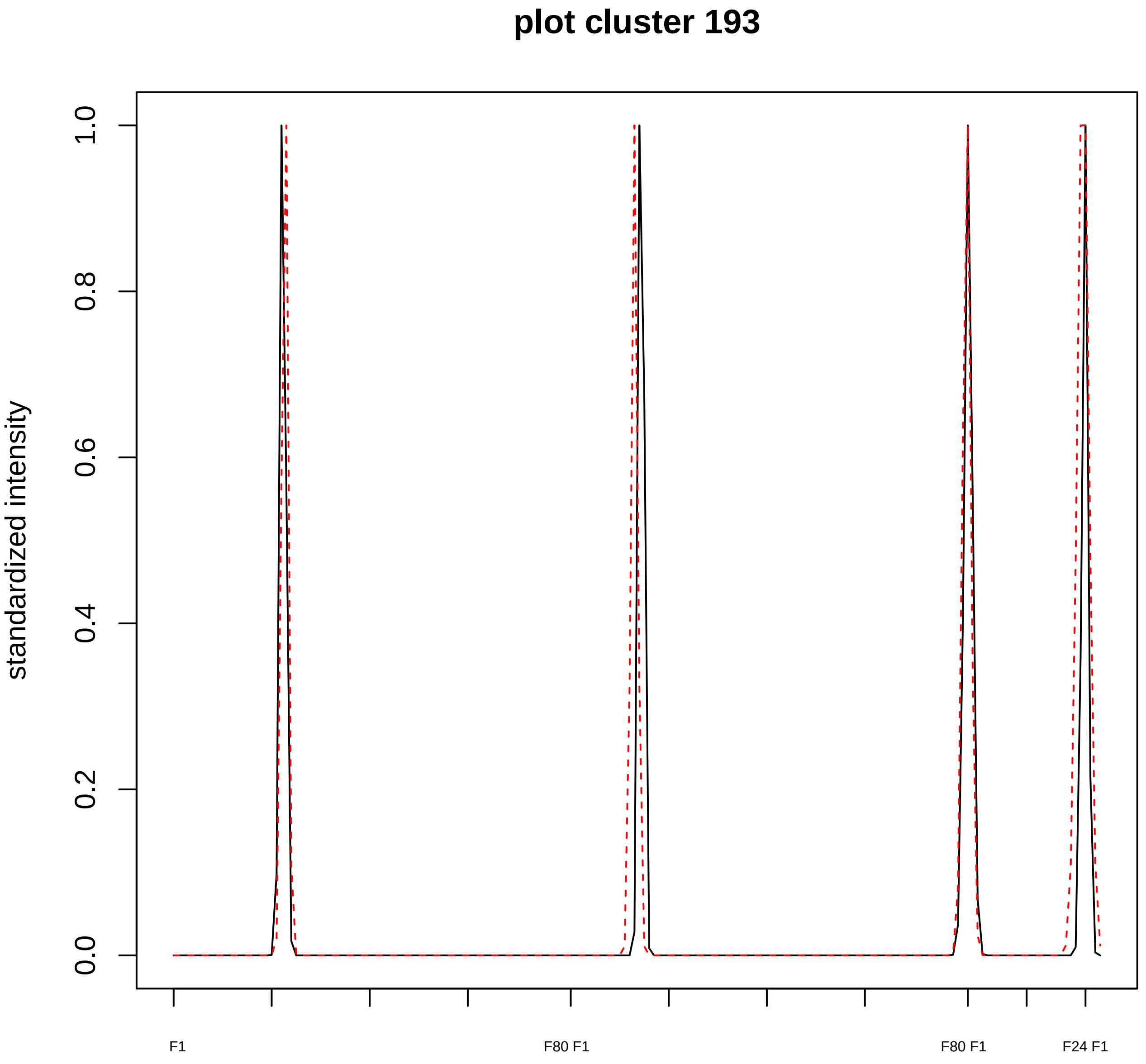
plot cluster 193
standardized intensity
0.0
0.2
0.4
0.6
0.8
1.0
F1
F80 F1
F80 F1
F24 F1

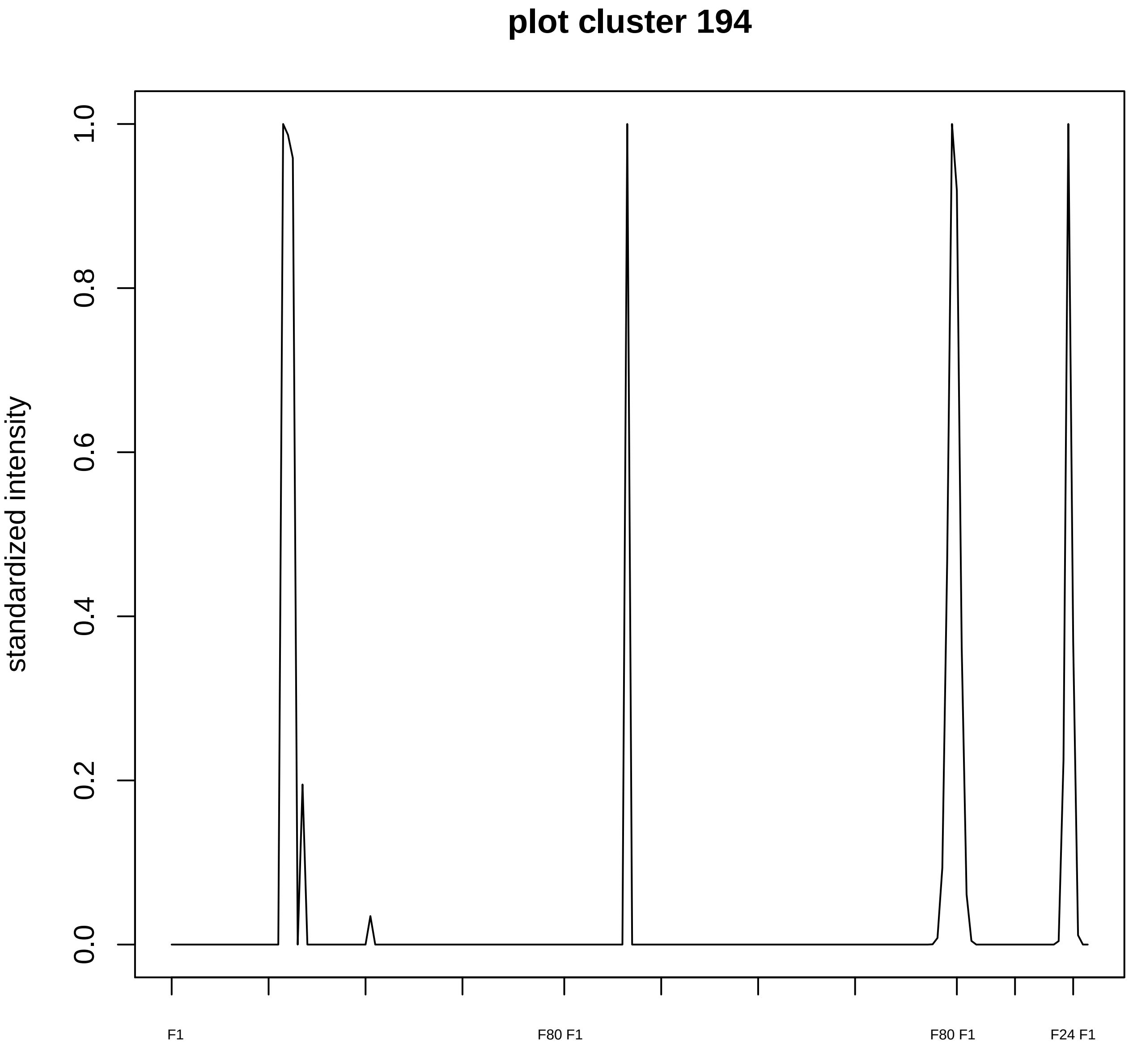

plot cluster 194
standardized intensity
0.0
0.2
0.4
0.6
0.8
1.0
F1
F80 F1
F80 F1
F24 F1

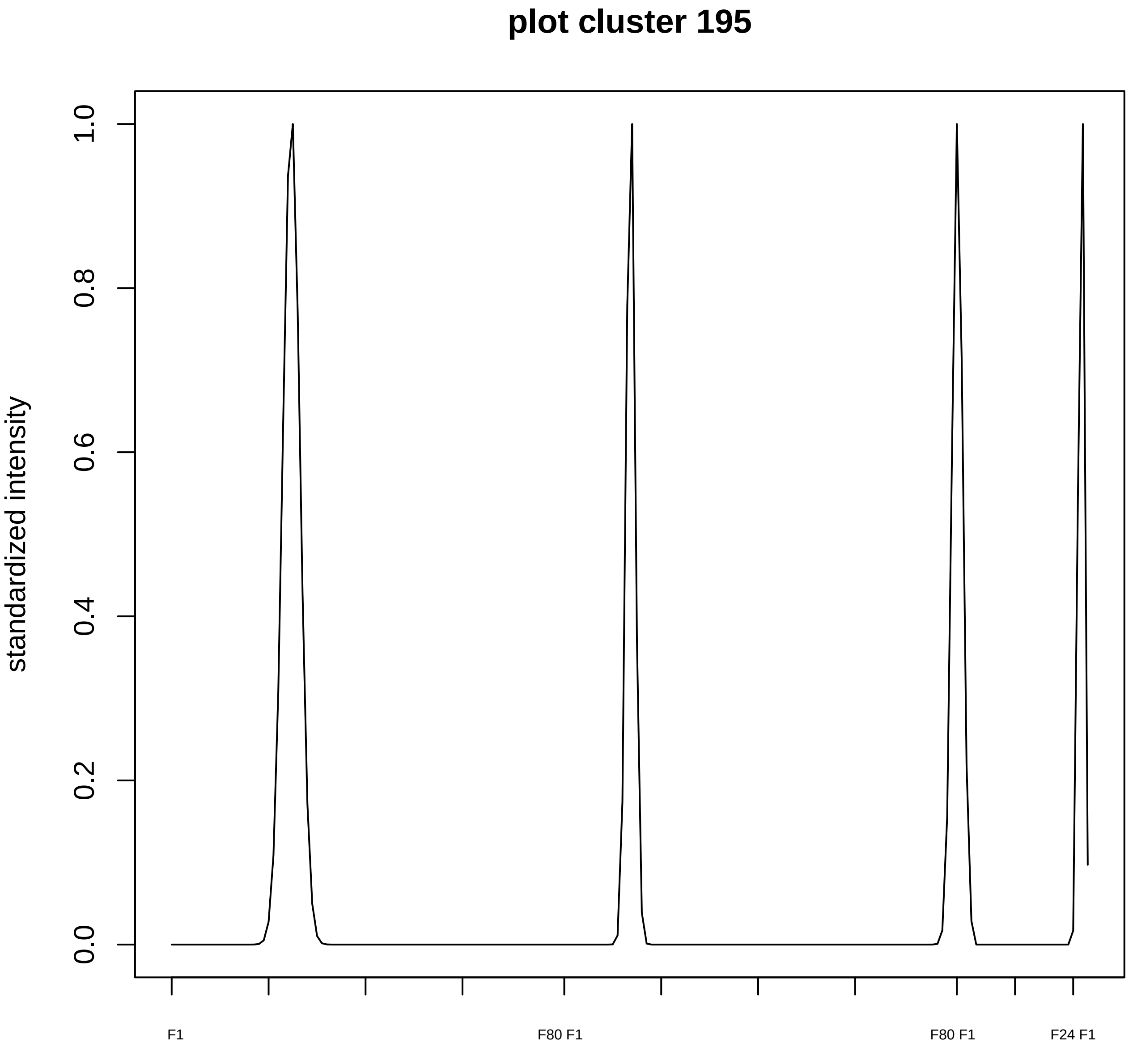

plot cluster 195
standardized intensity
0.0
0.2
0.4
0.6
0.8
1.0
F1
F80 F1
F80 F1
F24 F1

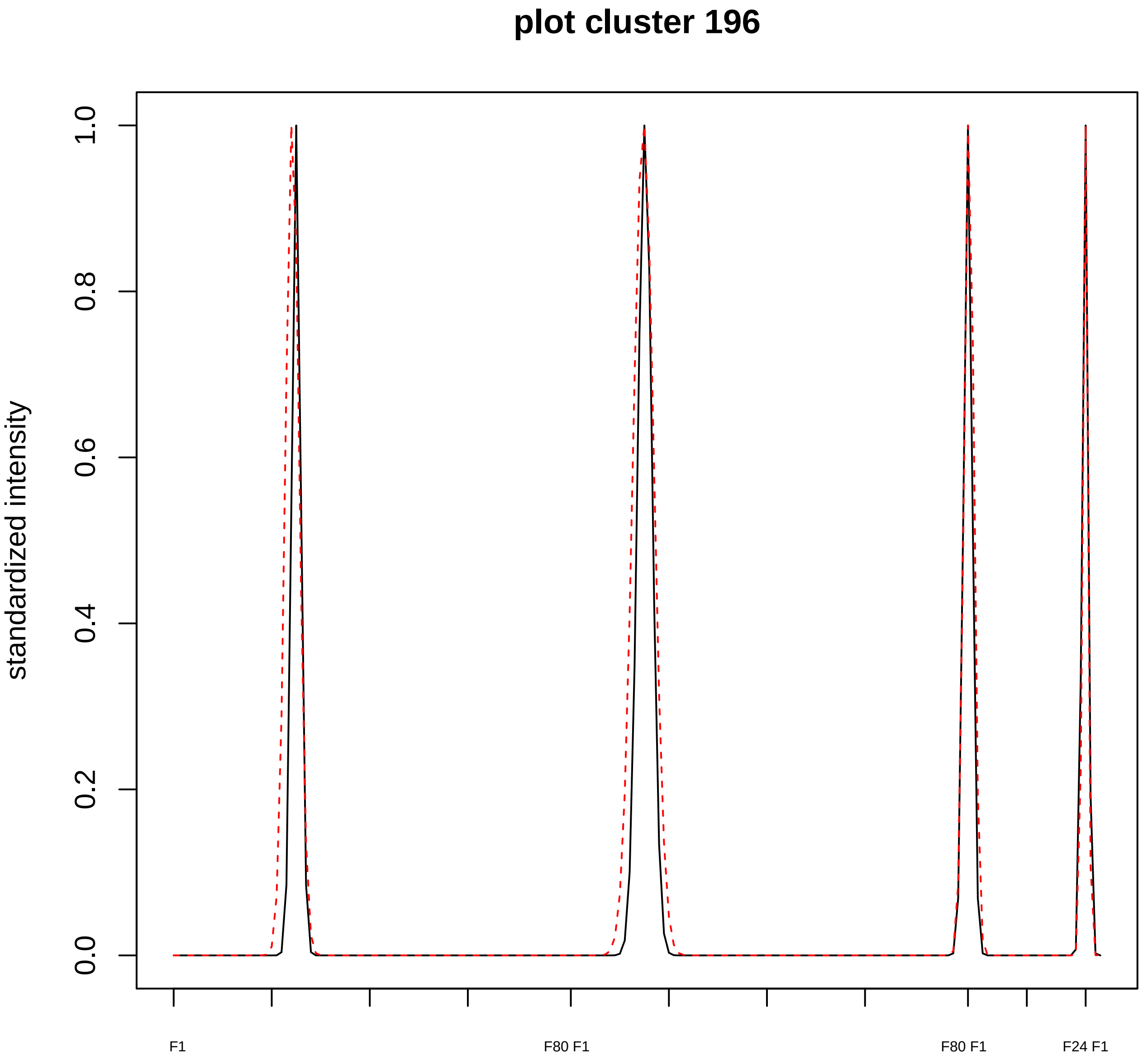

plot cluster 196
standardized intensity
0.0
0.2
0.4
0.6
0.8
1.0
F1
F80 F1
F80 F1
F24 F1

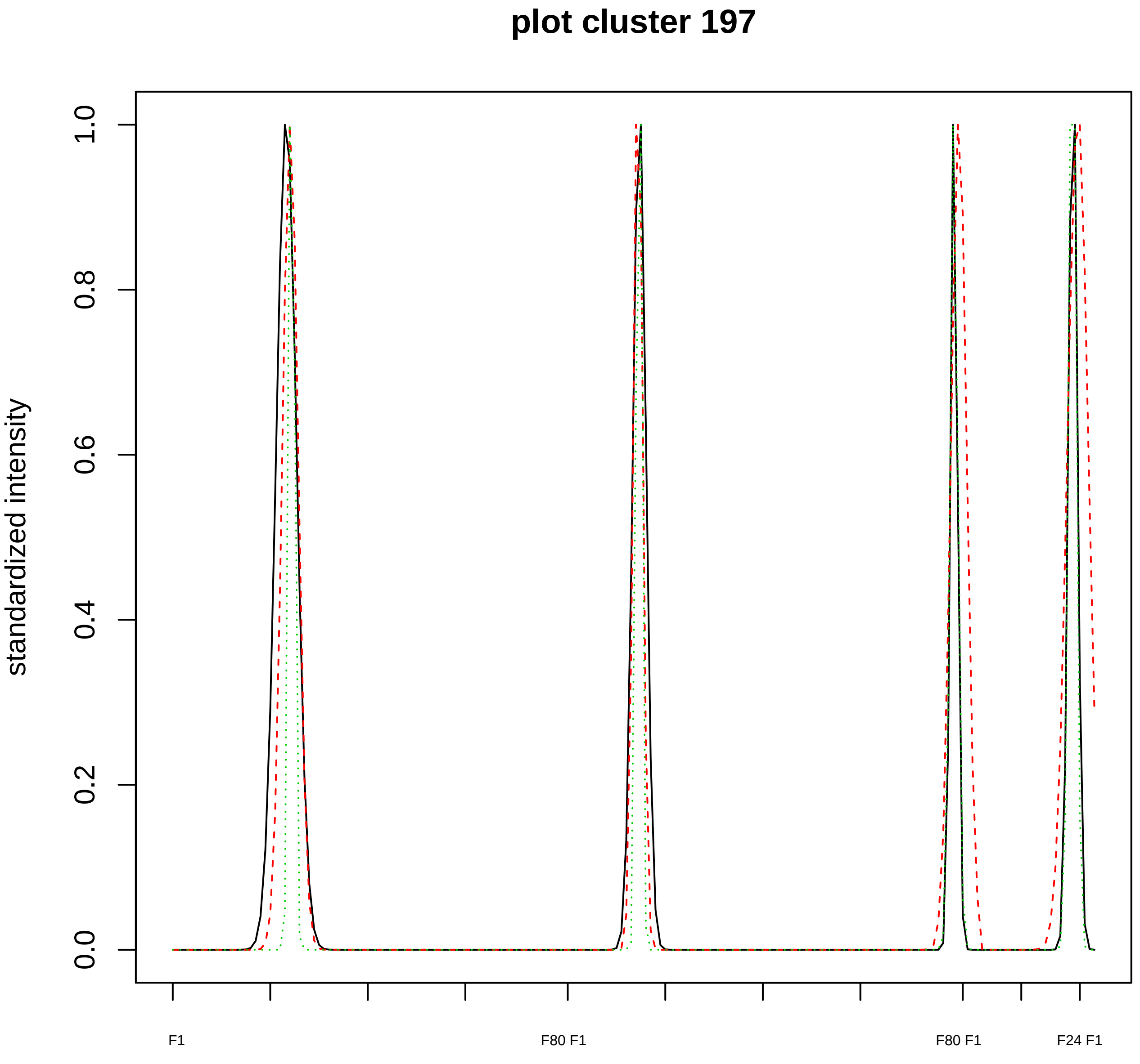

plot cluster 197
standardized intensity
0.0
0.2
0.4
0.6
0.8
1.0
F1
F80 F1
F80 F1
F24 F1

### Dendrogram of cluster 197

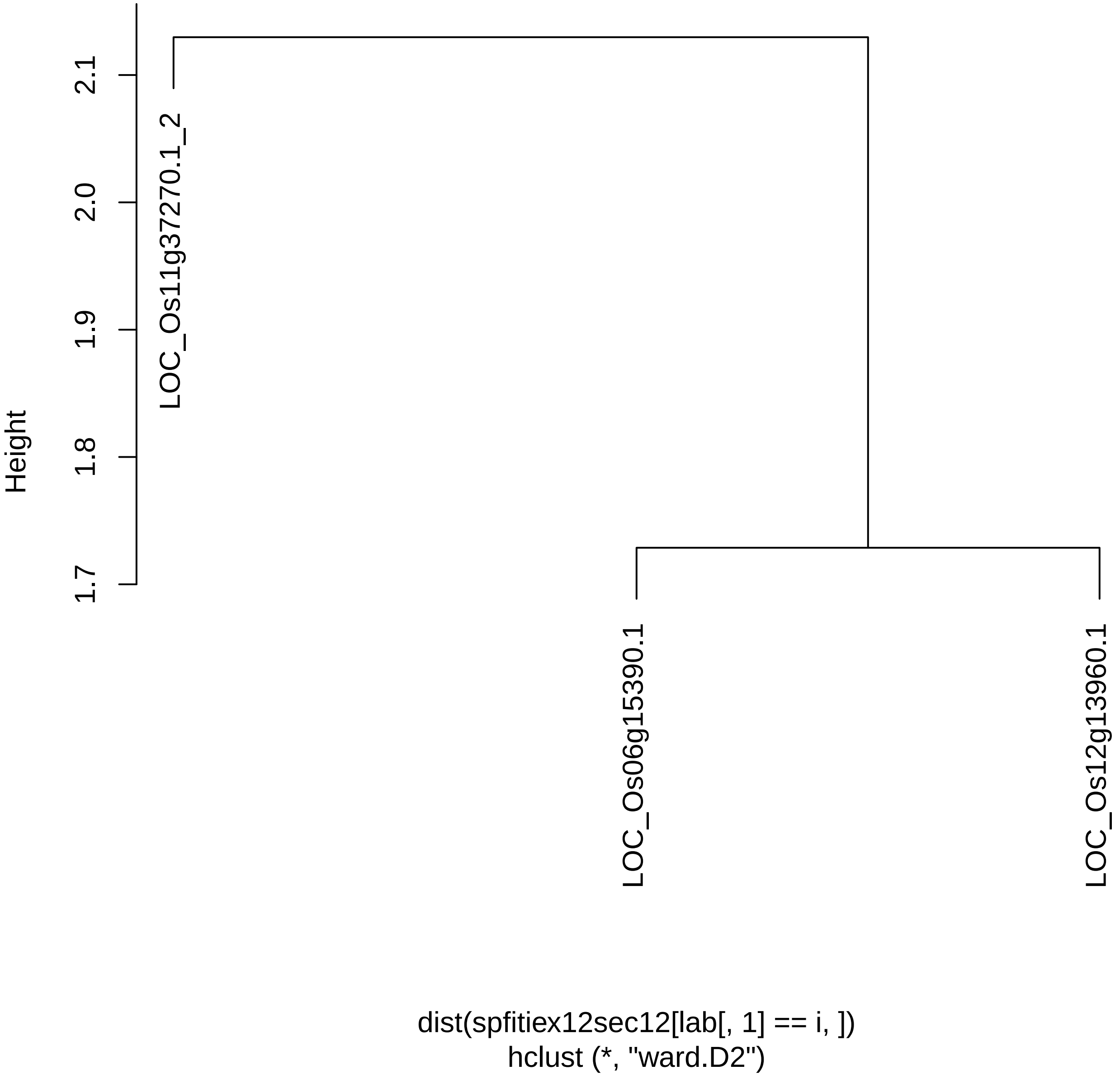

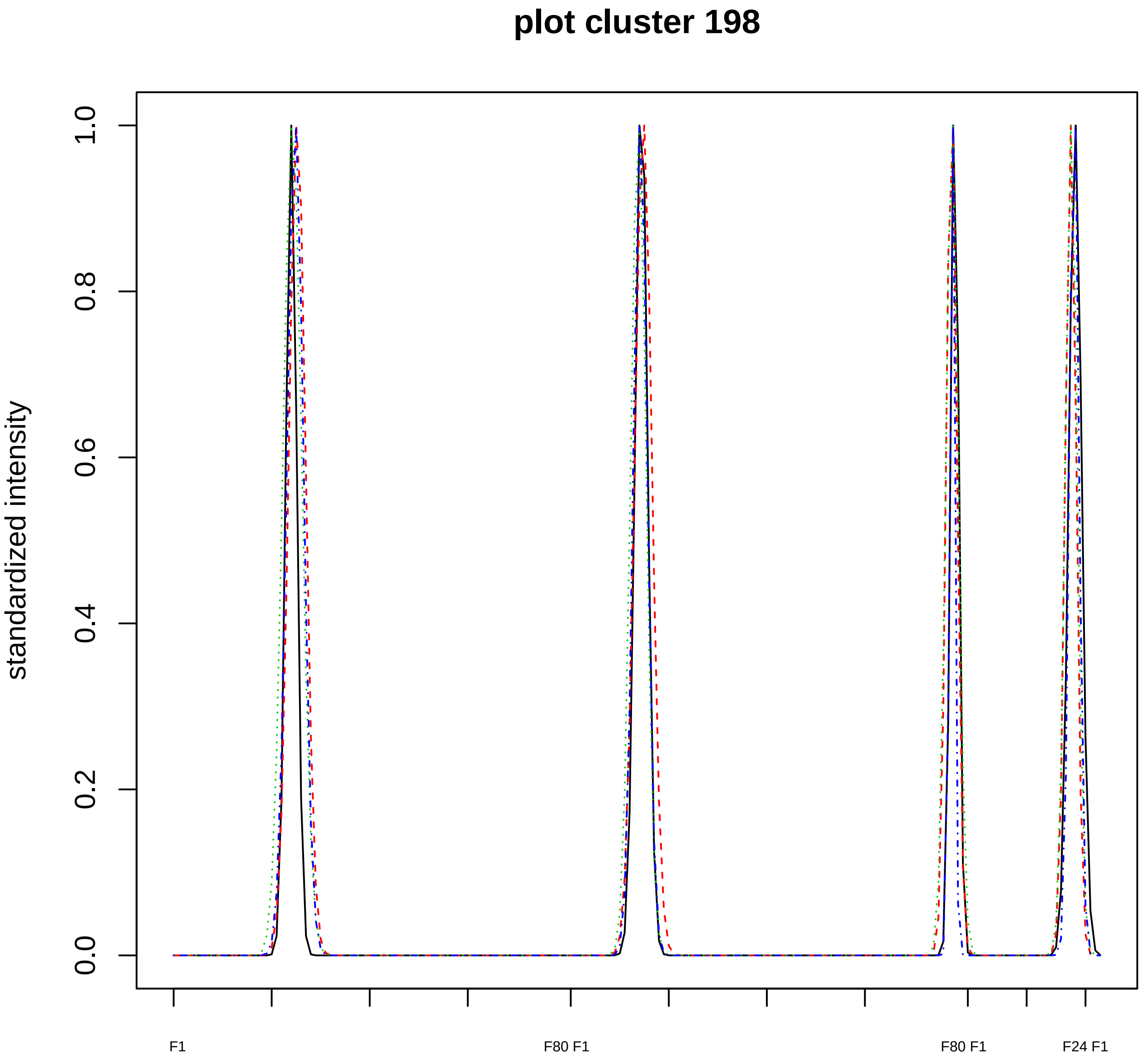

plot cluster 198
standardized intensity
0.0
0.2
0.4
0.6
0.8
1.0
F1
F80 F1
F80 F1
F24 F1

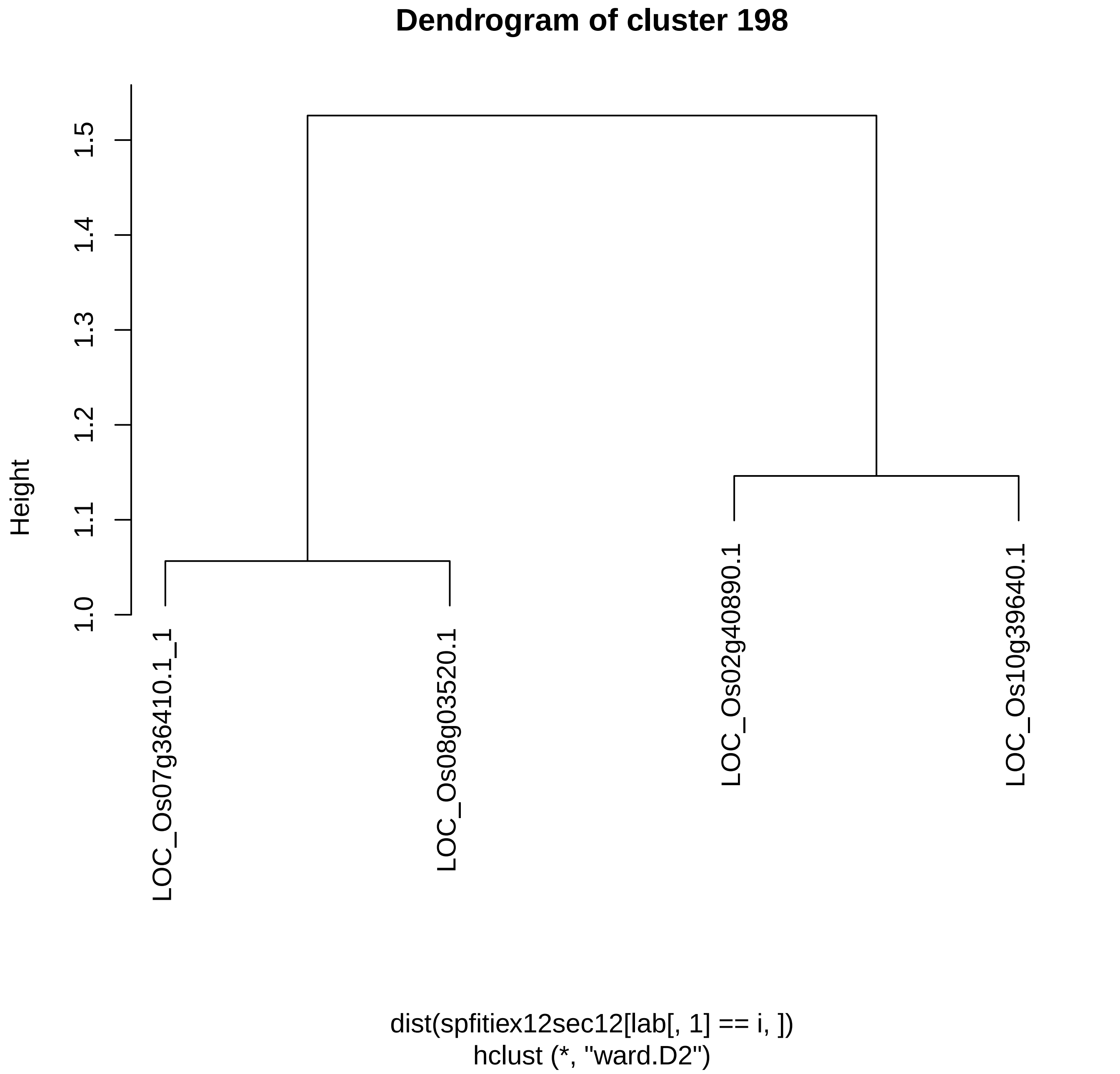
Dendrogram of cluster 198
Height
1.0
1.1
1.2
1.3
1.4
1.5
LOC_Os07g36410.1_1_
LOC_Os08g03520.1
LOC_Os02g40890.1
LOC_Os10g39640.1
dist(spfitiex12sec12[lab[, 1] == i, ])
hclust (*, "ward.D2")

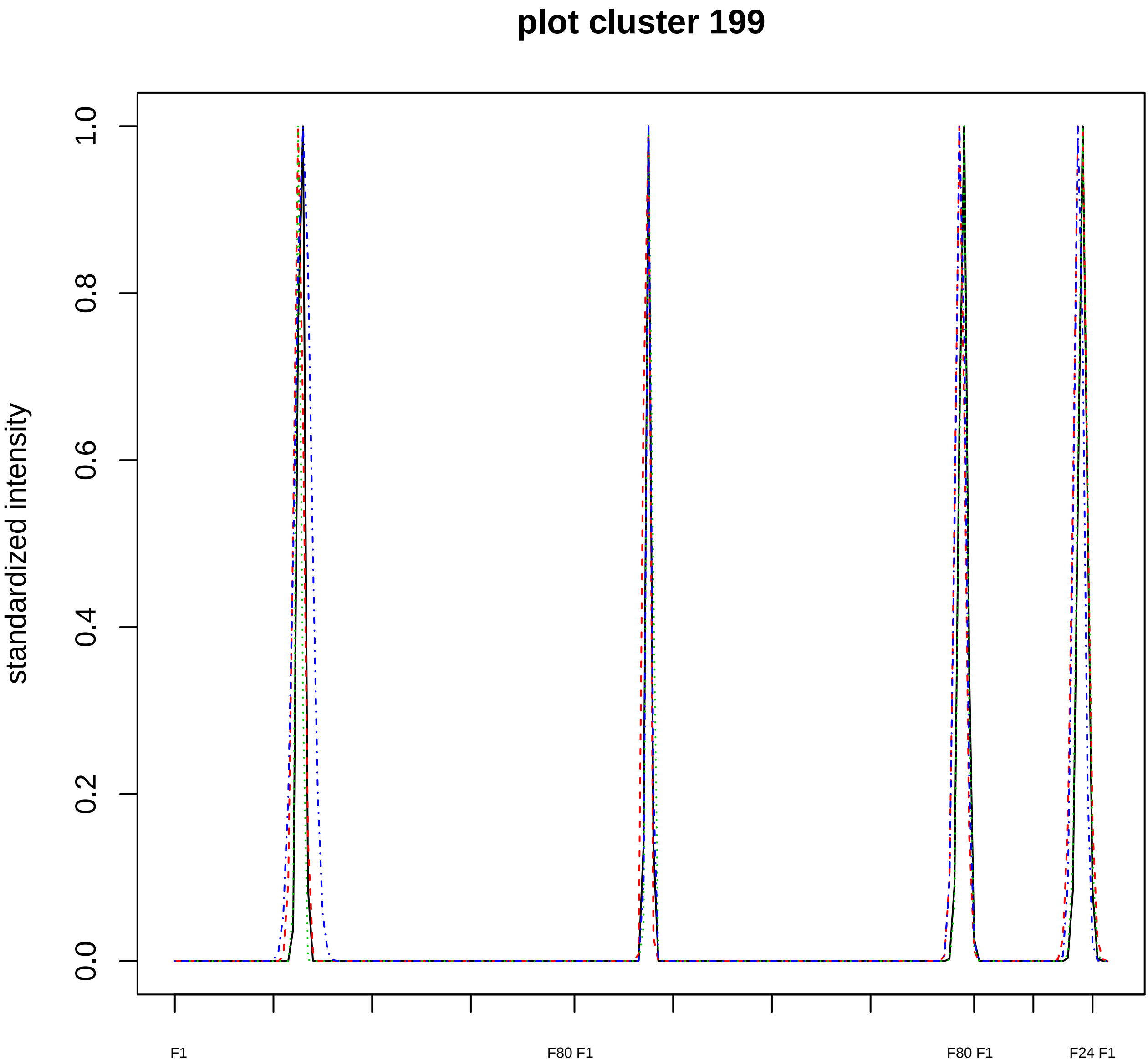

plot cluster 199
standardized intensity
0.0
0.2
0.4
0.6
0.8
1.0
F1
F80 F1
F80 F1
F24 F1

### Dendrogram of cluster 199

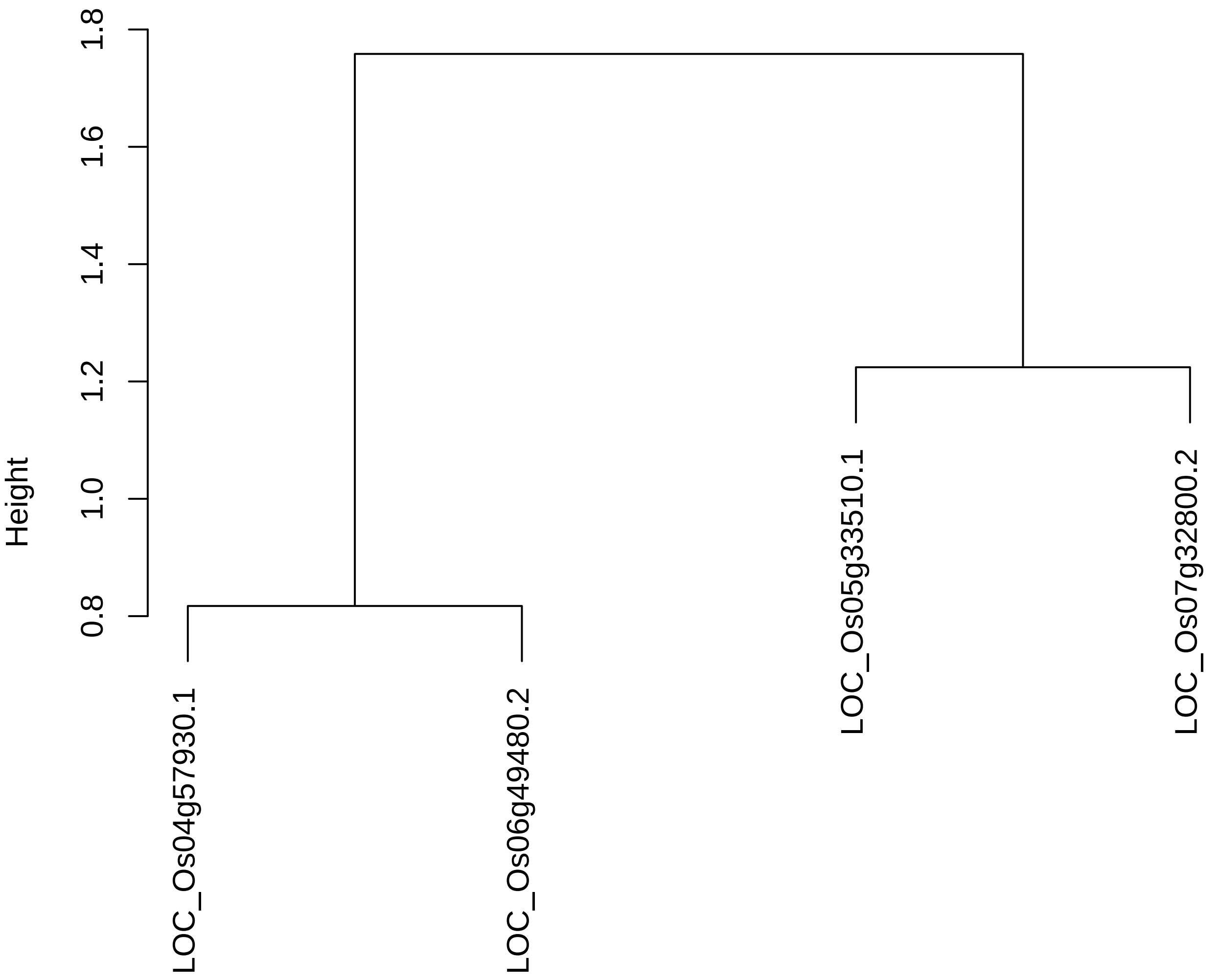


dist(spfitiex12sec12[lab[, 1] == i, ])
hclust (*, "ward.D2")

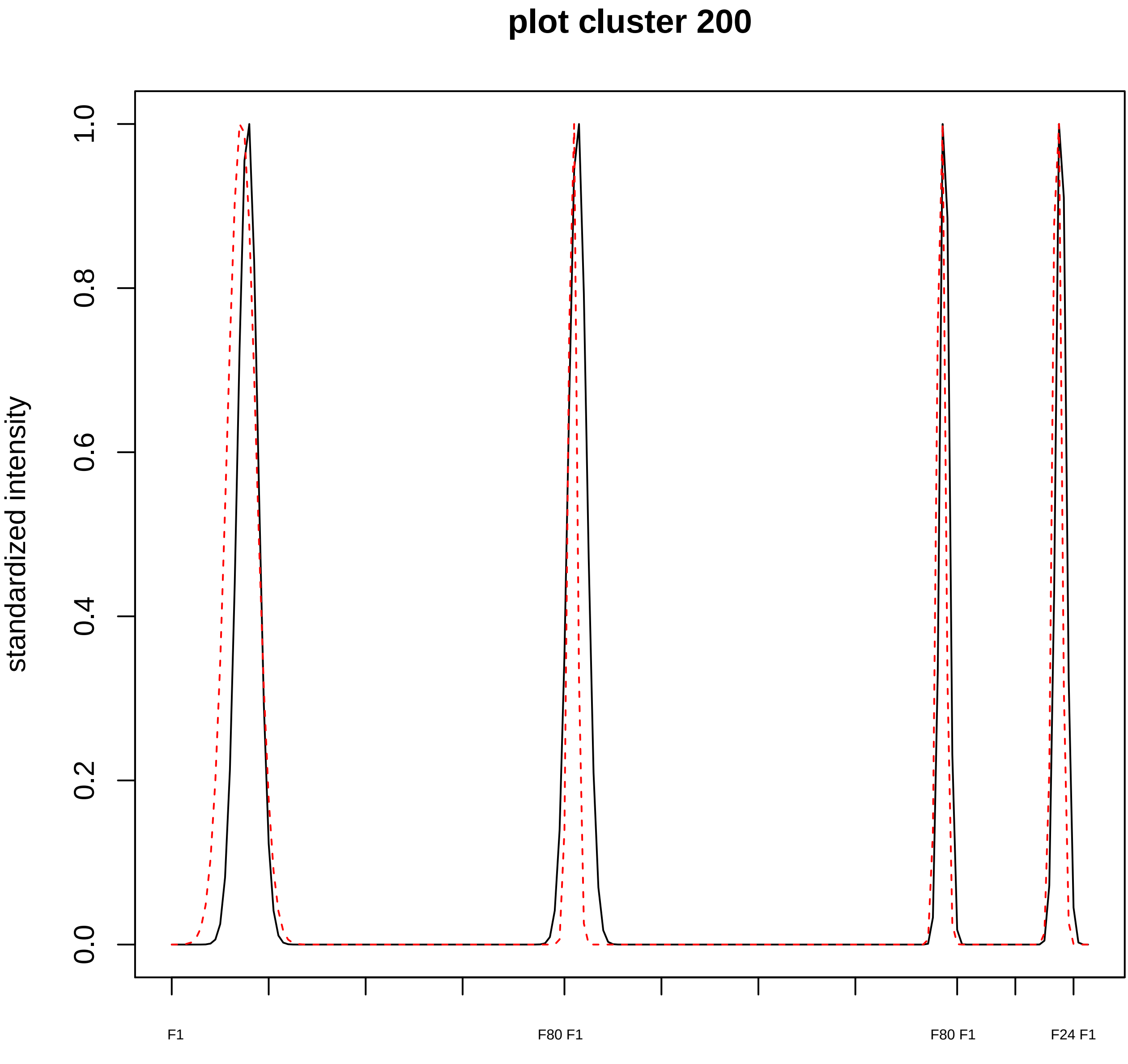
plot cluster 200
standardized intensity
0.0
0.2
0.4
0.6
0.8
1.0
F1
F80 F1
F80 F1
F24 F1

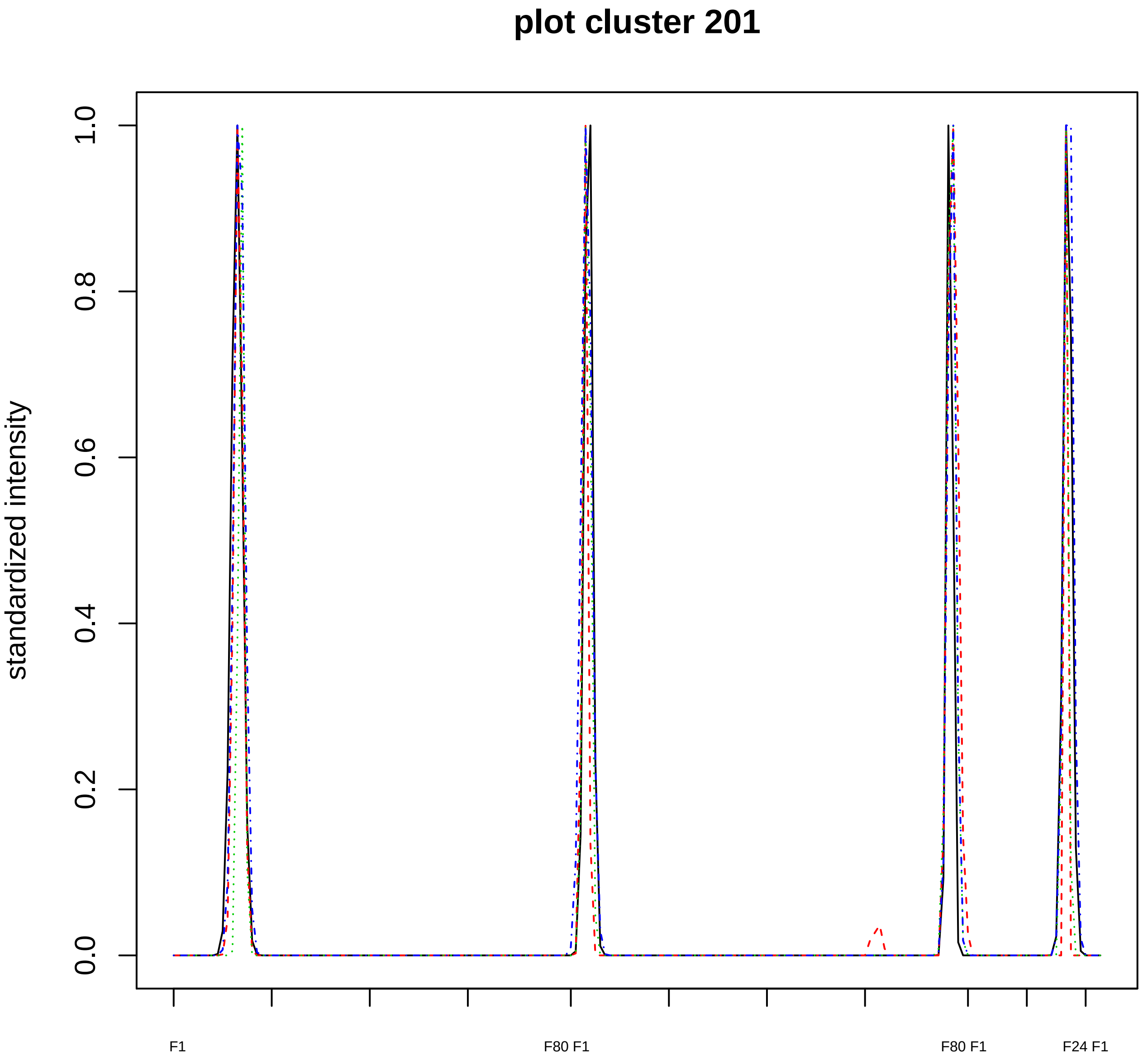
plot cluster 201
standardized intensity
0.0
0.2
0.4
0.6
0.8
1.0
F1
F80 F1
F80 F1
F24 F1

### Dendrogram of cluster 201

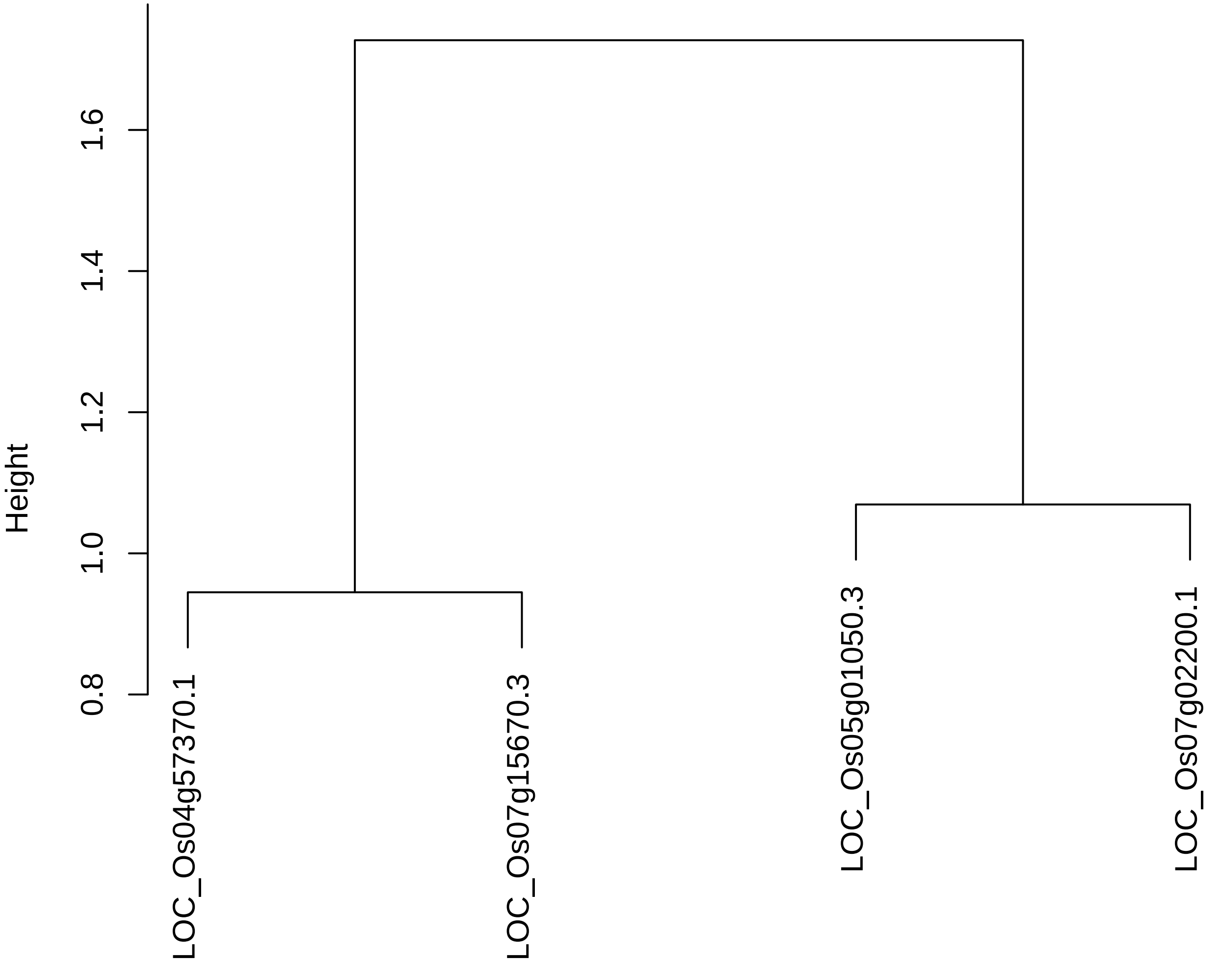


dist(spfitiex12sec12[lab[, 1] == i, ])
hclust (*, "ward.D2")

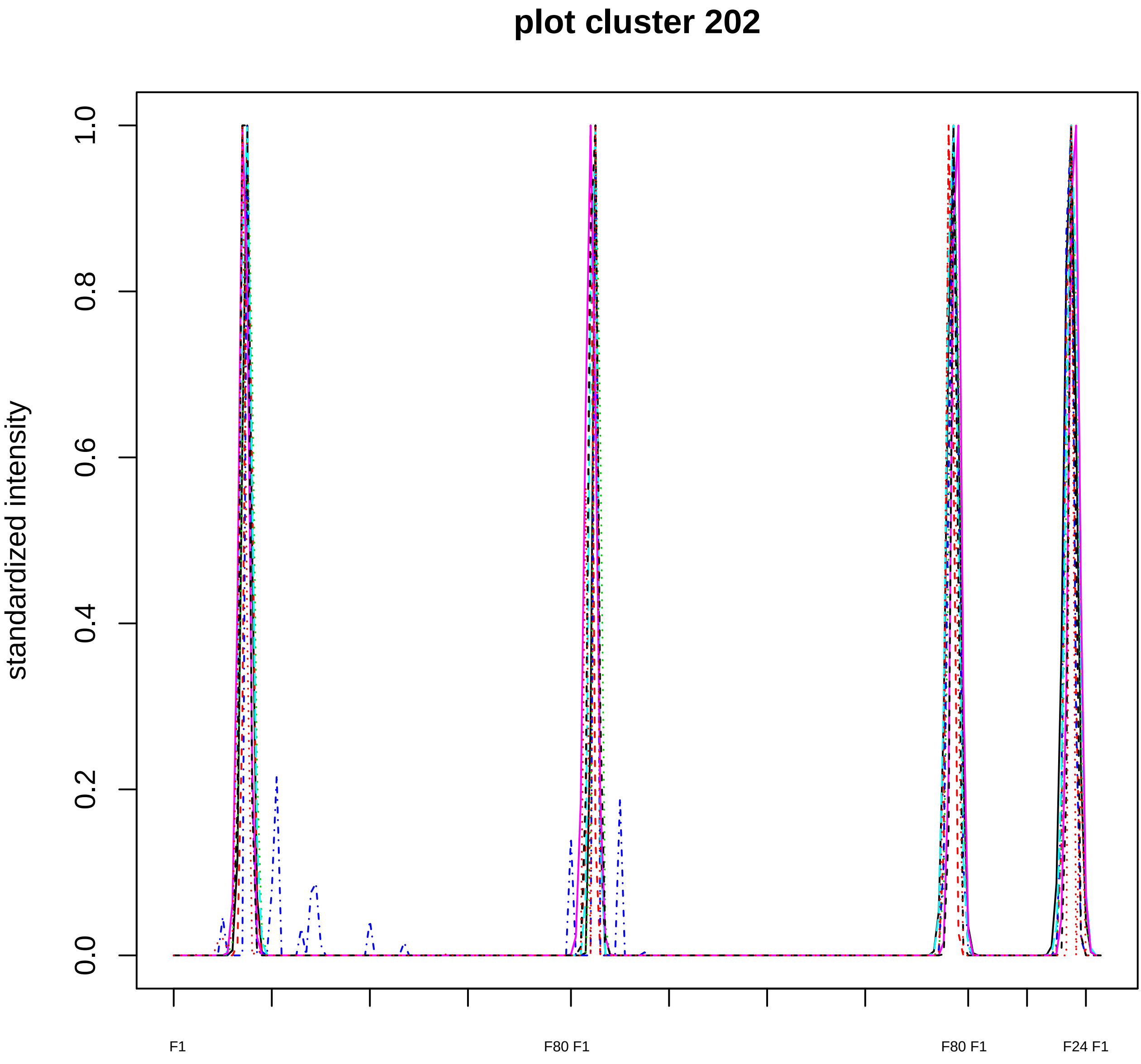
plot cluster 202
standardized intensity
0.0
0.2
0.4
0.6
0.8
1.0
F1
F80 F1
F80 F1
F24 F1

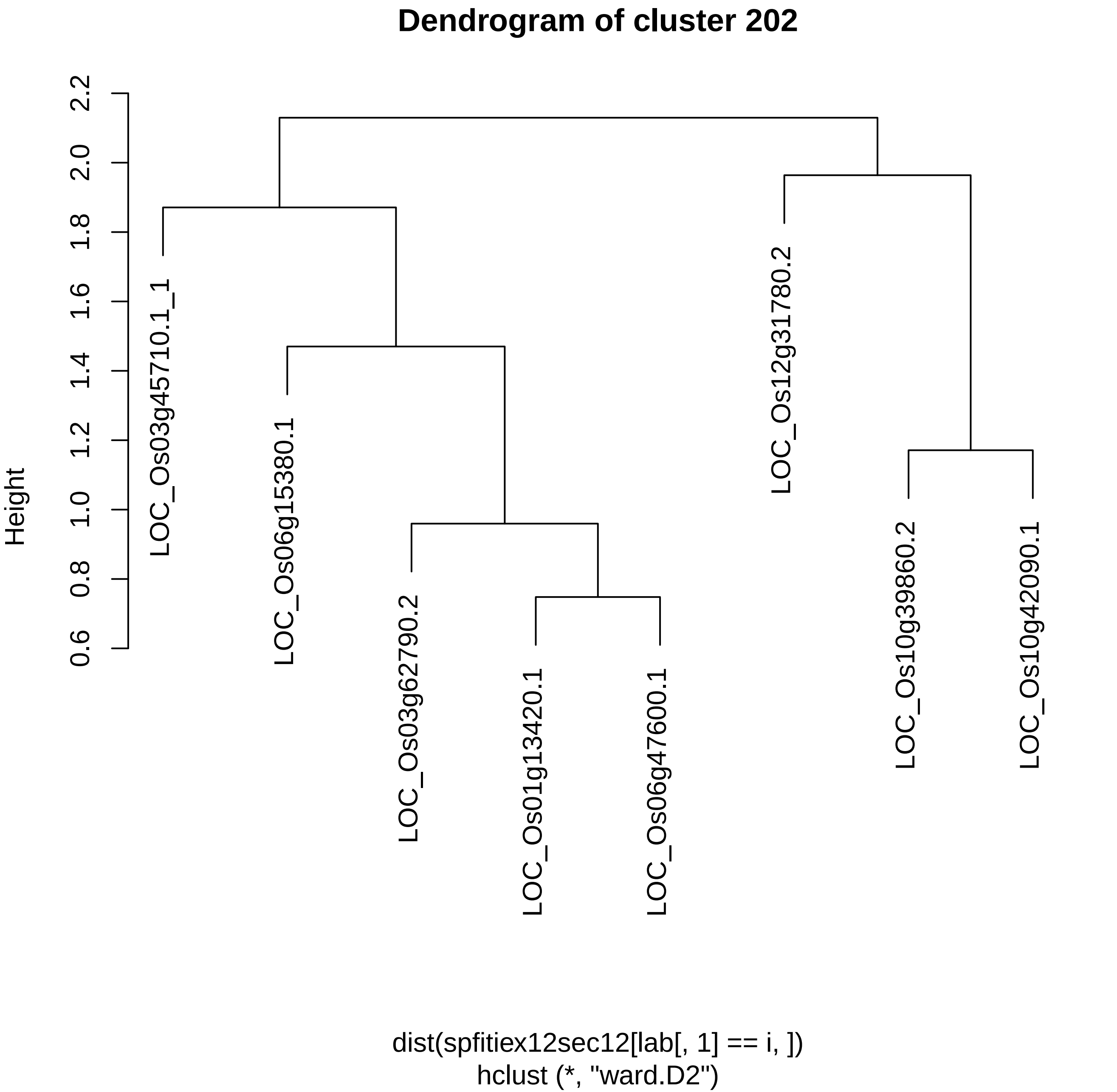
Dendrogram of cluster 202
Height
0.6
0.8
1.0
1.2
1.4
1.6
1.8
2.0
2.2
LOC_Os03g45710.1_1_
LOC_Os06g15380.1
LOC_Os03g62790.2
LOC_Os01g13420.1
LOC_Os06g47600.1
LOC_Os12g31780.2
LOC_Os10g39860.2
LOC_Os10g42090.1
dist(spfitiex12sec12[lab[, 1] == i, ])
hclust (*, "ward.D2")

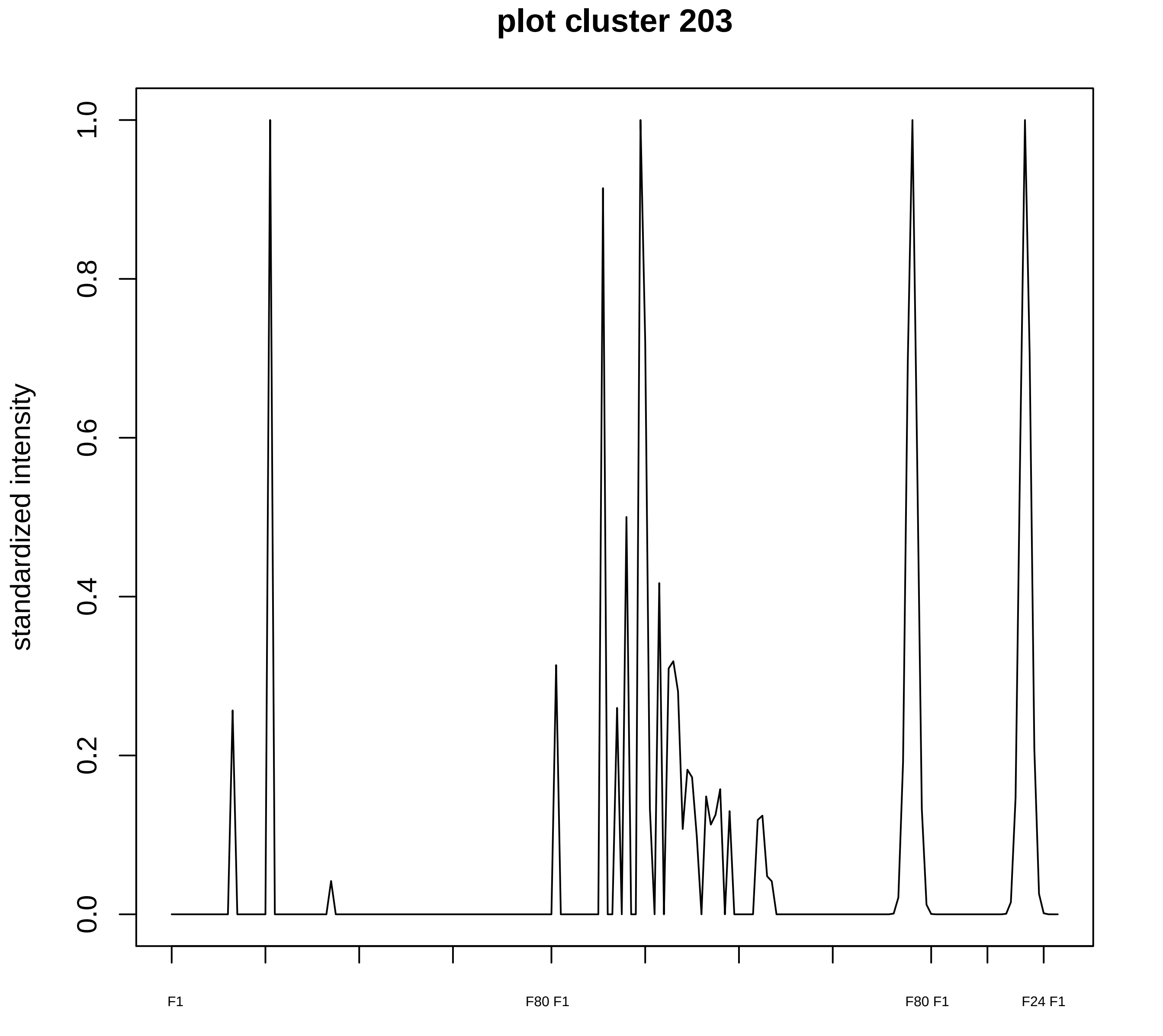
plot cluster 203
standardized intensity
0.0
0.2
0.4
0.6
0.8
1.0
F1
F80 F1
F80 F1
F24 F1

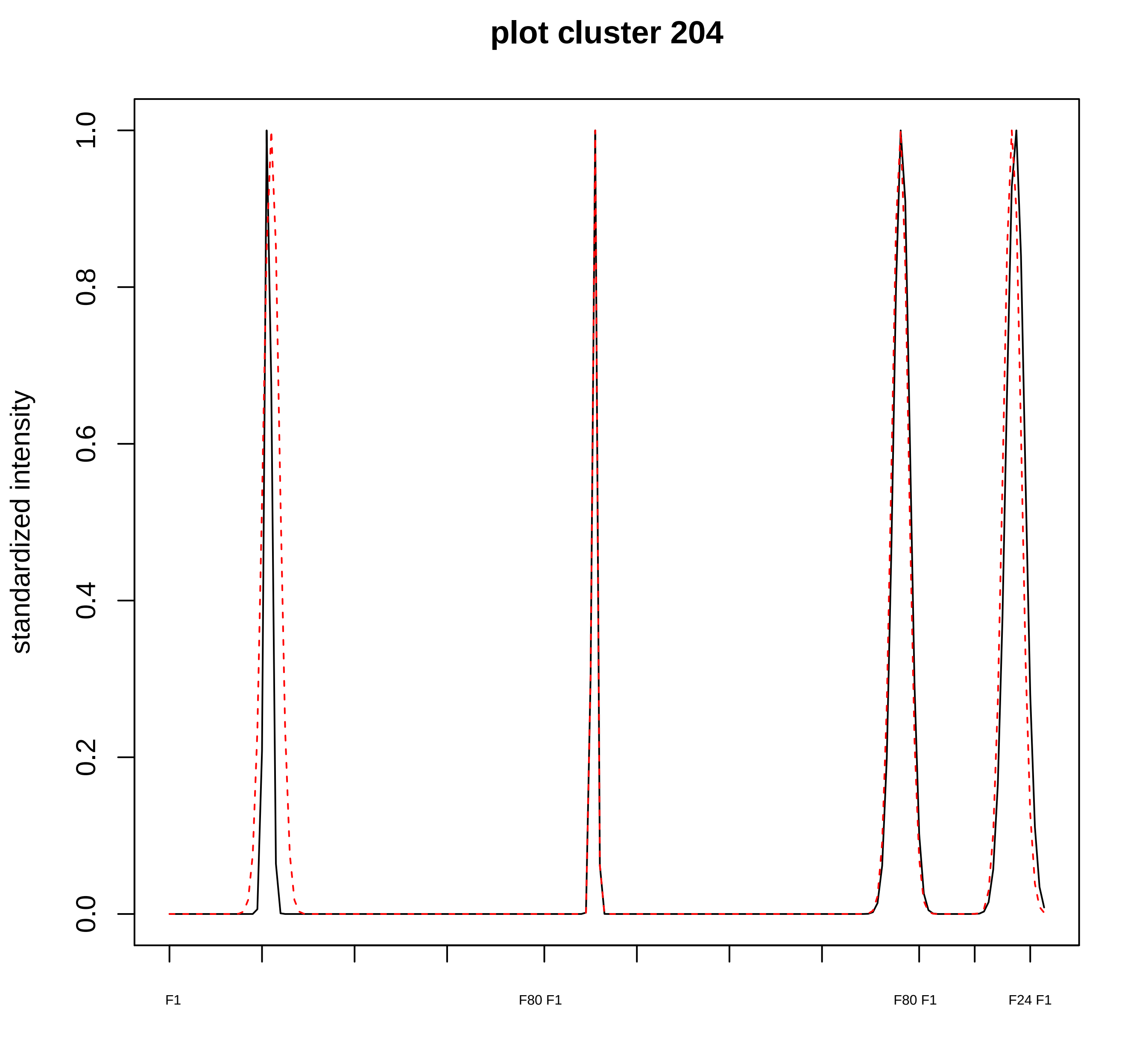
plot cluster 204
standardized intensity
0.0
0.2
0.4
0.6
0.8
1.0
F1
F80 F1
F80 F1
F24 F1

#### plot cluster 205

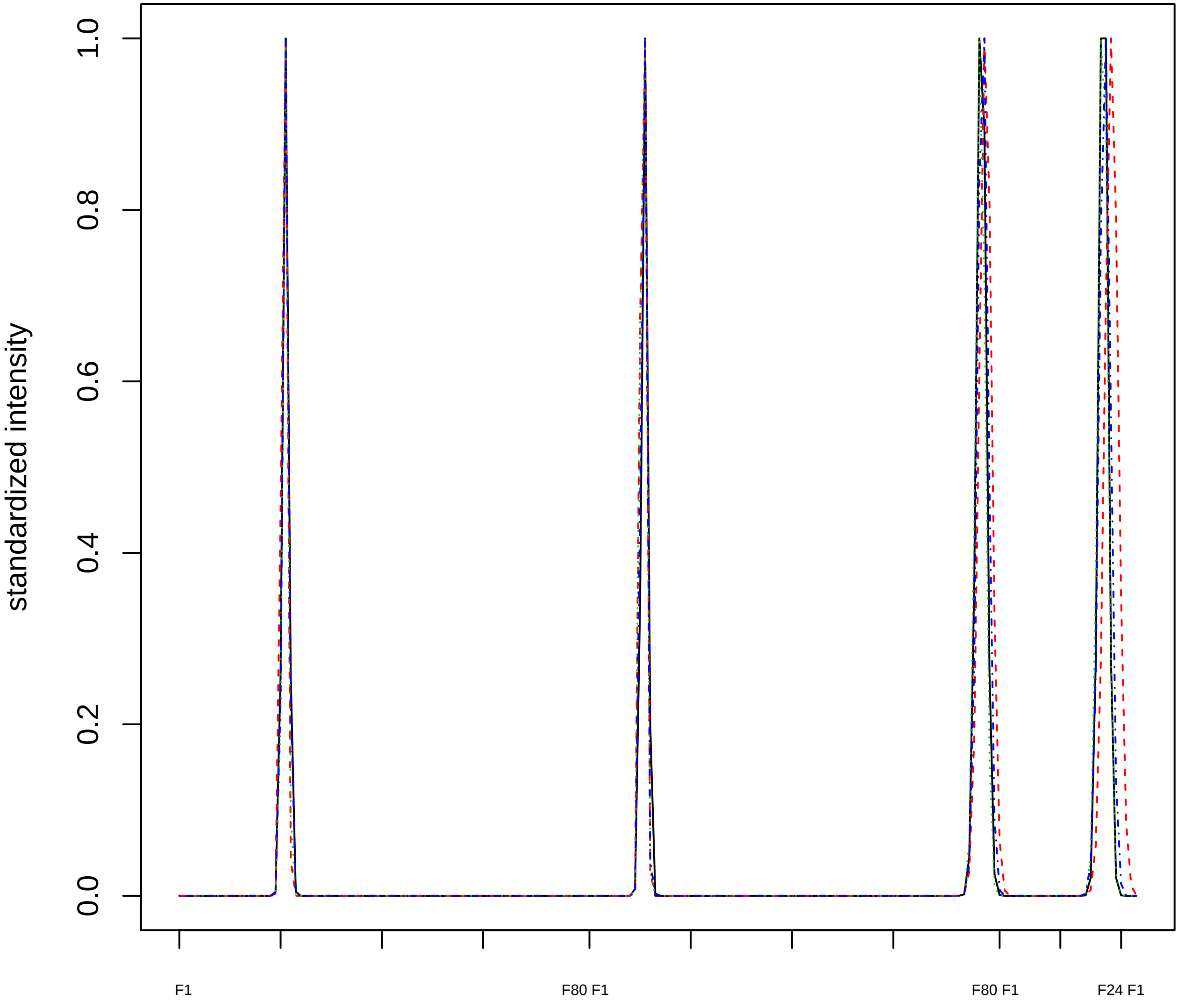

### Dendrogram of cluster 205

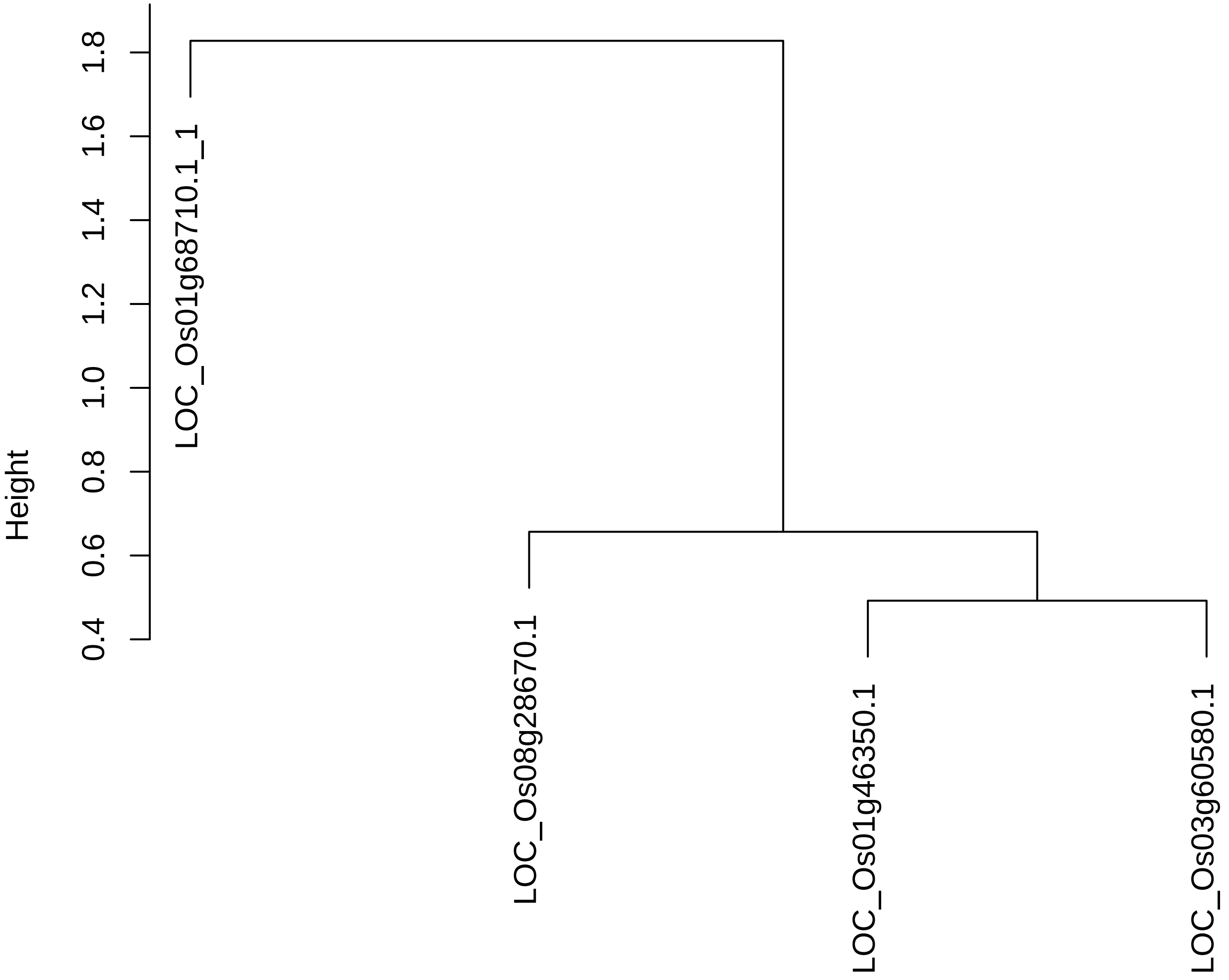


dist(spfitiex12sec12[lab[, 1] == i, ])
hclust (*, "ward.D2")

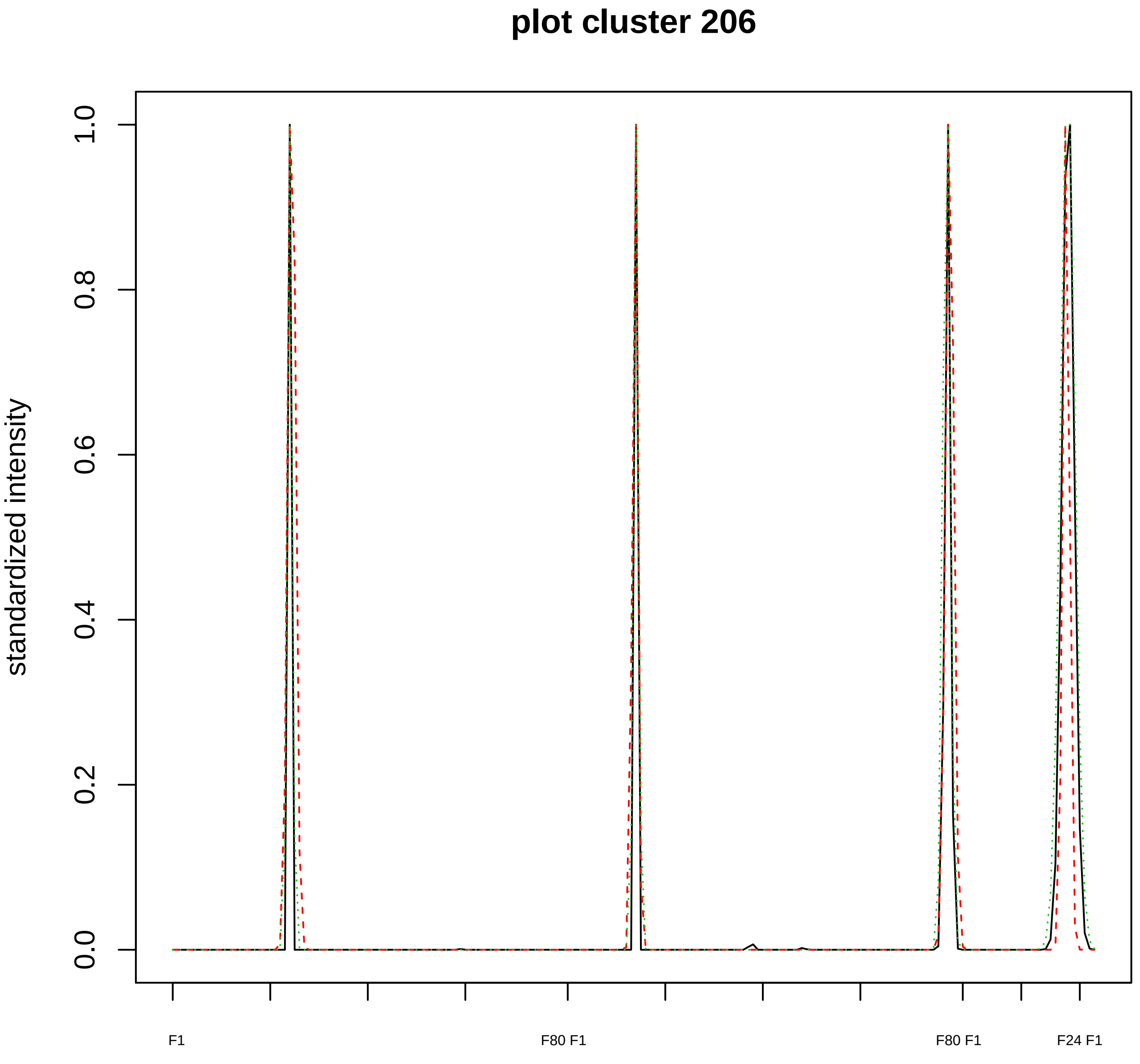
plot cluster 206
standardized intensity
0.0
0.2
0.4
0.6
0.8
1.0
F1
F80 F1
F80 F1
F24 F1

### Dendrogram of cluster 206

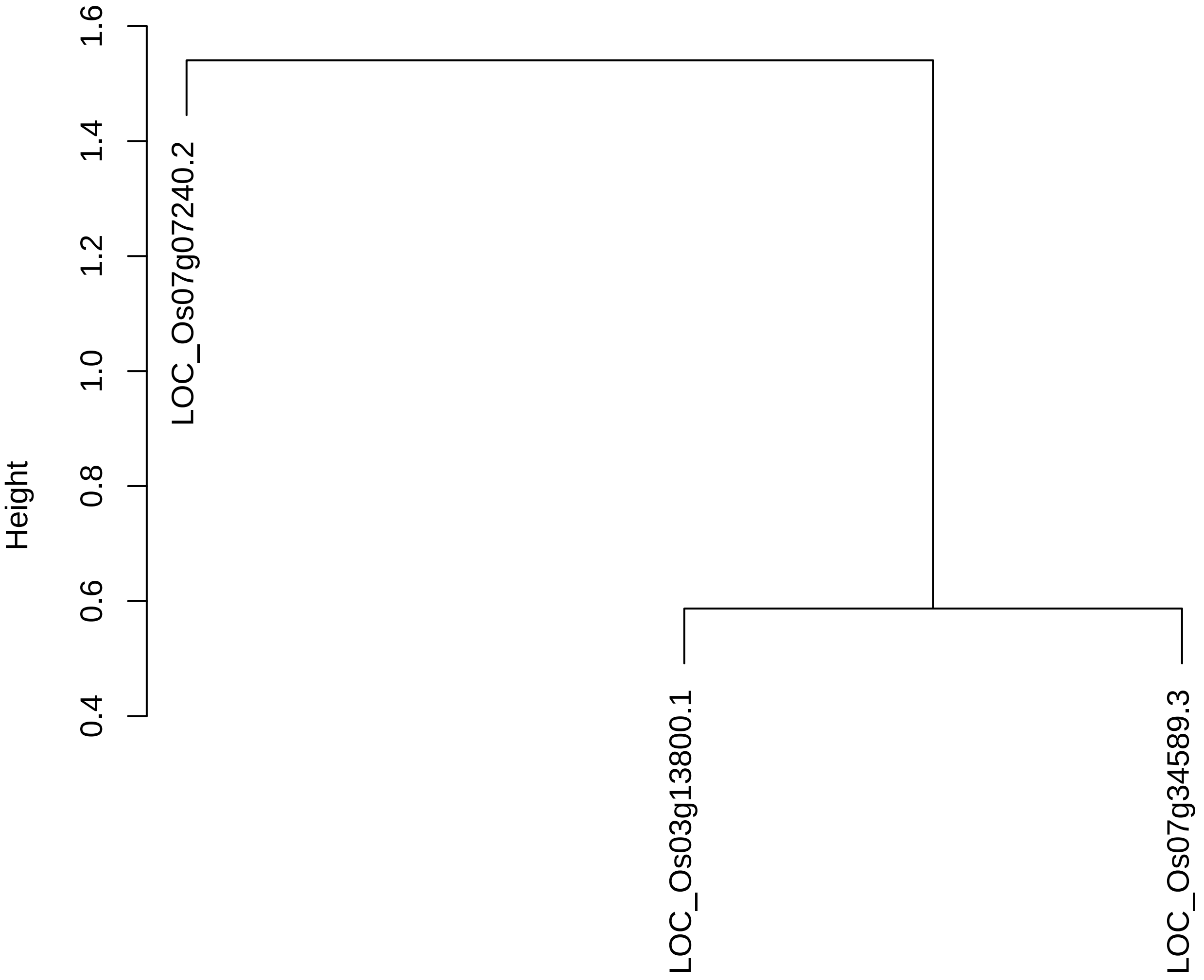


dist(spfitiex12sec12[lab[, 1] == i, ])
hclust (*, "ward.D2")

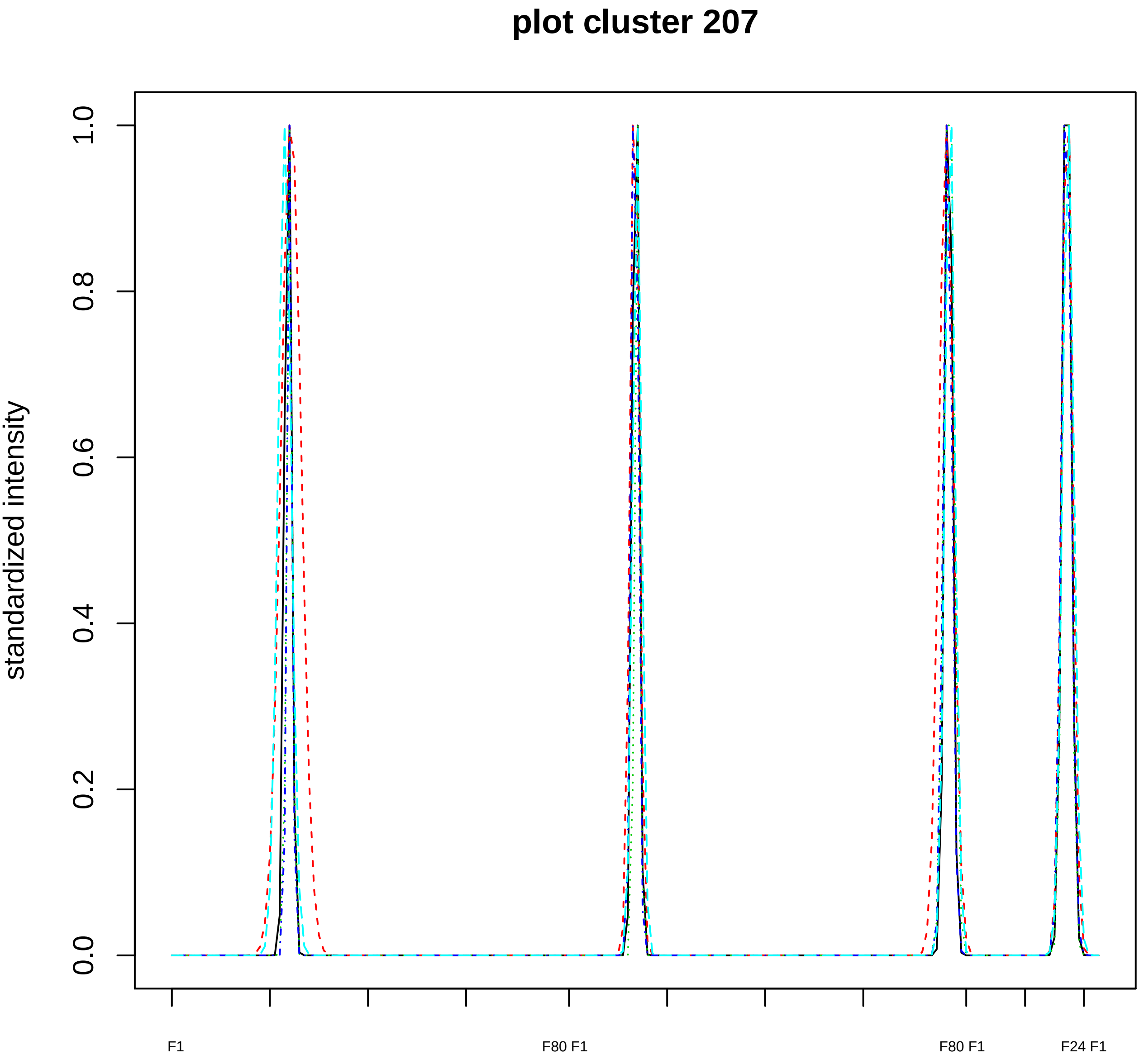

plot cluster 207
standardized intensity
0.0
0.2
0.4
0.6
0.8
1.0
F1
F80 F1
F80 F1
F24 F1

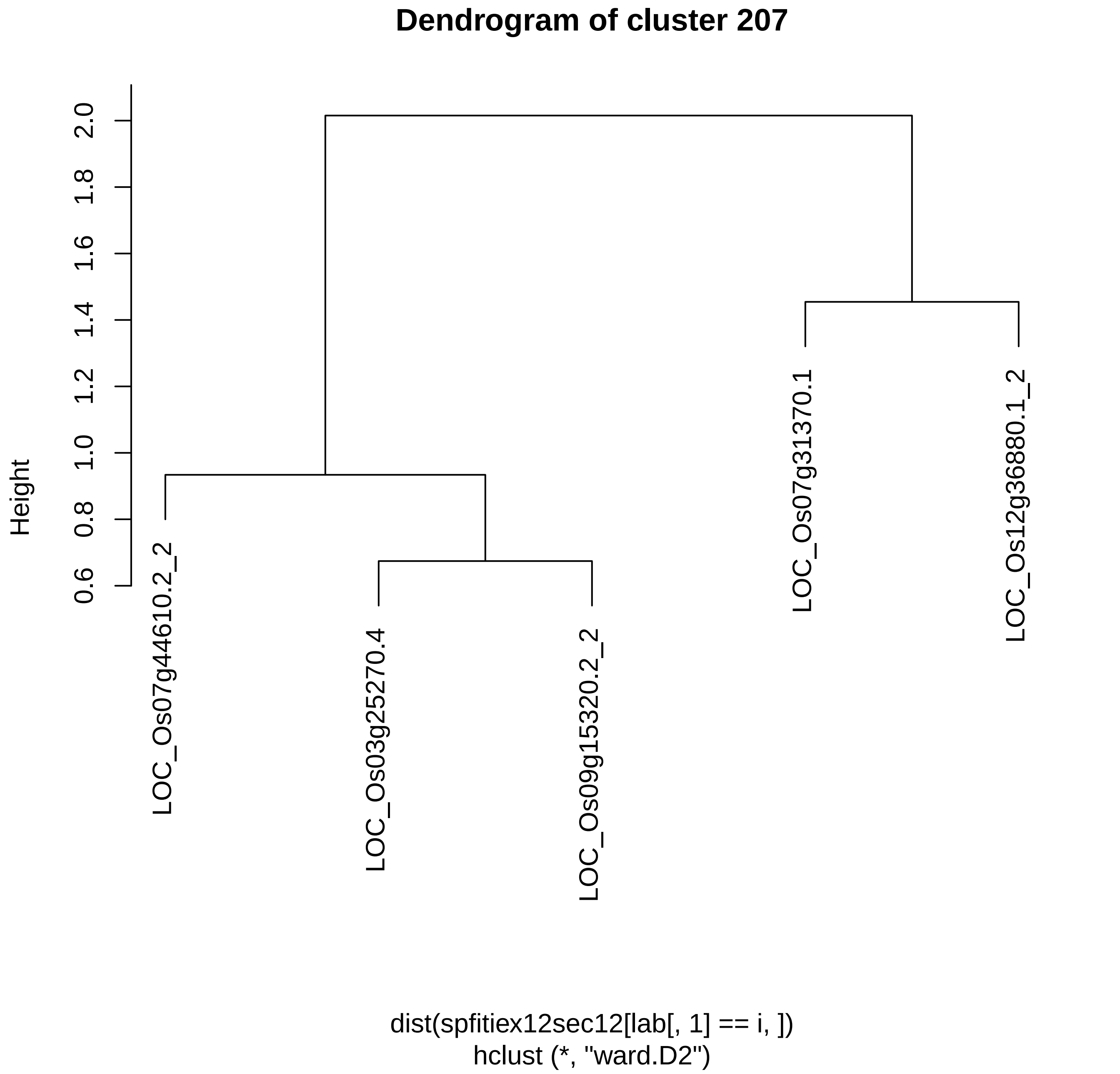
Dendrogram of cluster 207
Height
0.6
0.8
1.0
1.2
1.4
1.6
1.8
2.0
LOC_Os07g44610.2_2
LOC_Os03g25270.4
LOC_Os09g15320.2_2
LOC_Os07g31370.1
LOC_Os12g36880.1_2
dist(spfitiex12sec12[lab[, 1] == i, ])
hclust (*, "ward.D2")

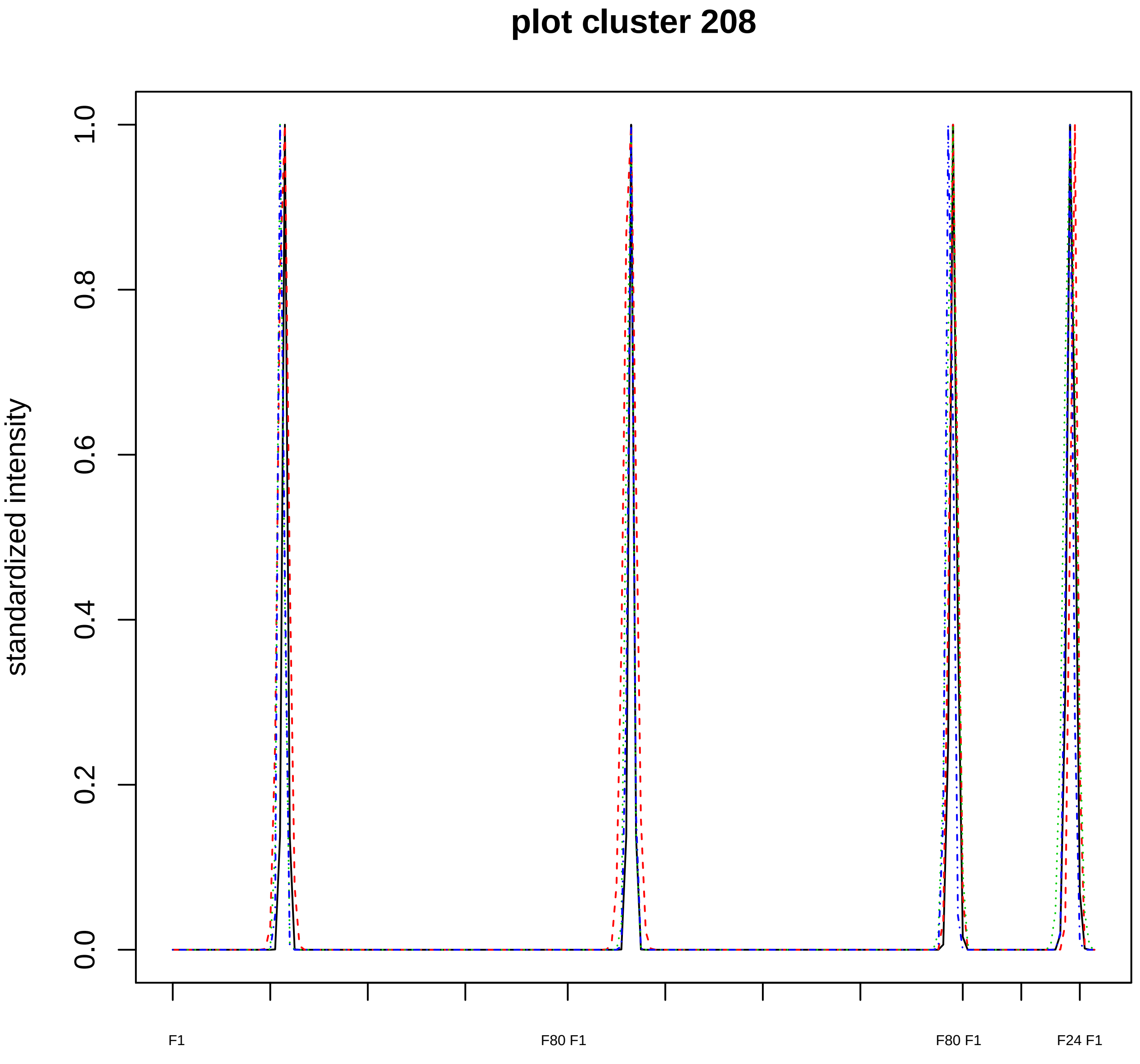
plot cluster 208
standardized intensity
0.0
0.2
0.4
0.6
0.8
1.0
F1
F80 F1
F80 F1
F24 F1

#### Dendrogram of cluster 208

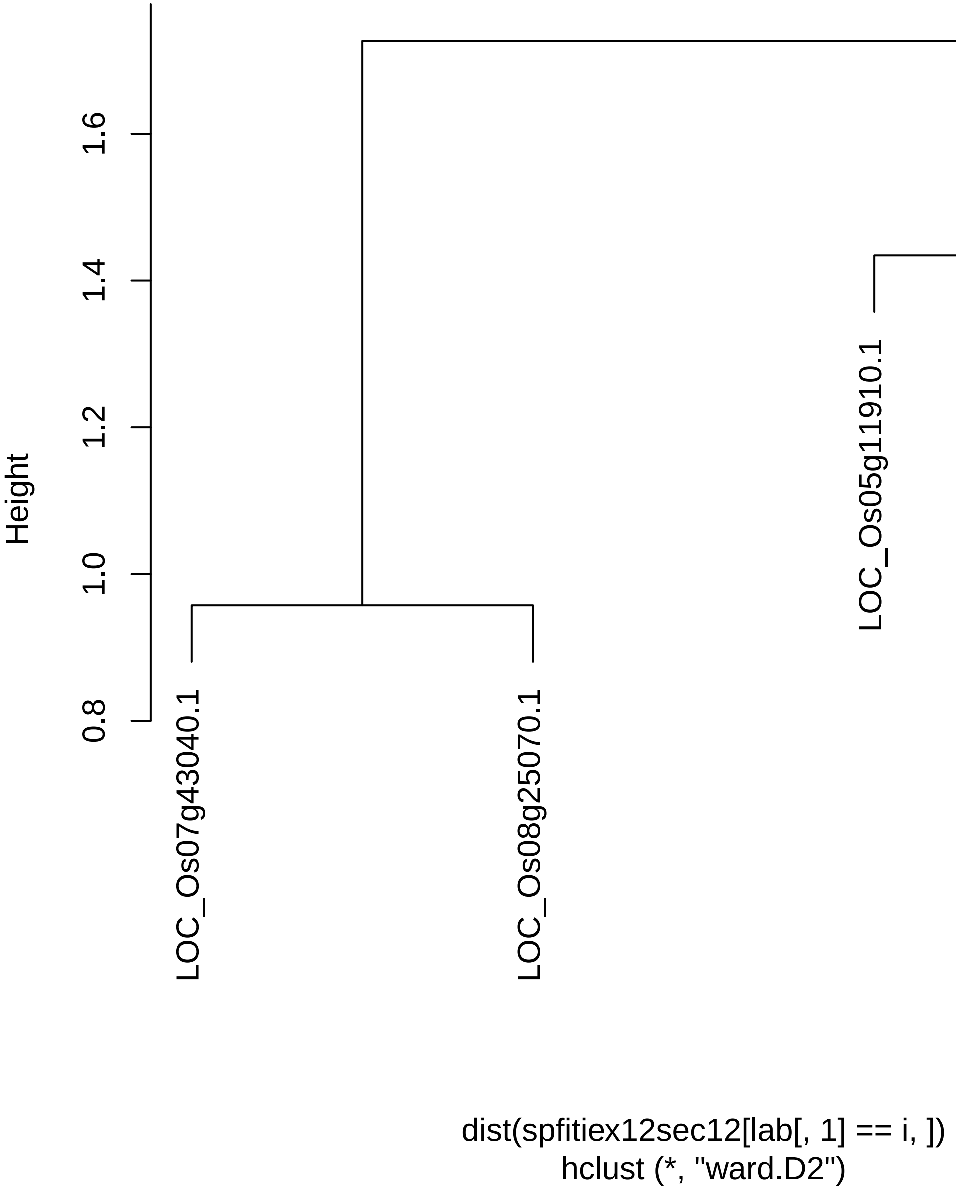

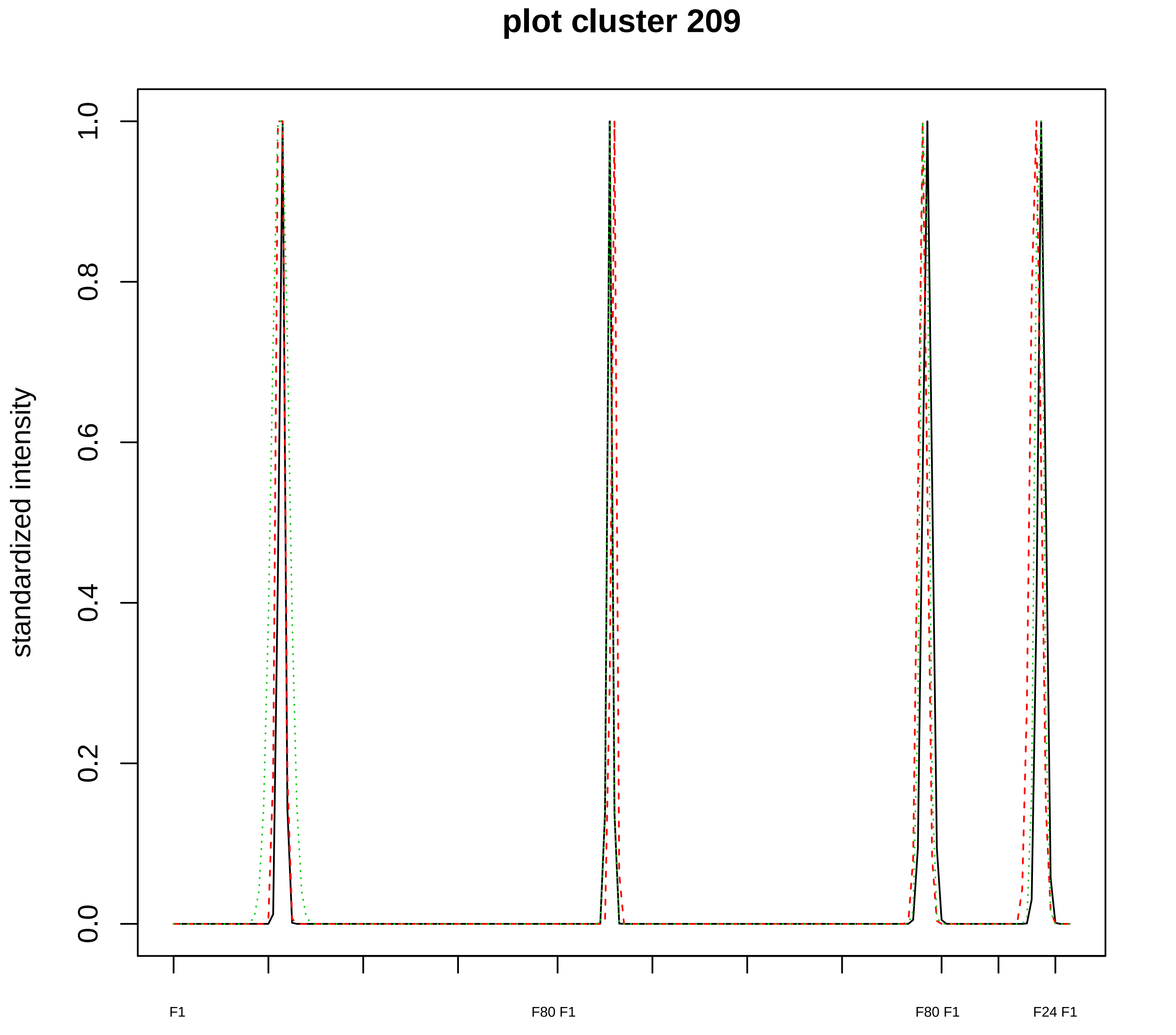
plot cluster 209
standardized intensity
0.0
0.2
0.4
0.6
0.8
1.0
F1
F80 F1
F80 F1
F24 F1

#### Dendrogram of cluster 209

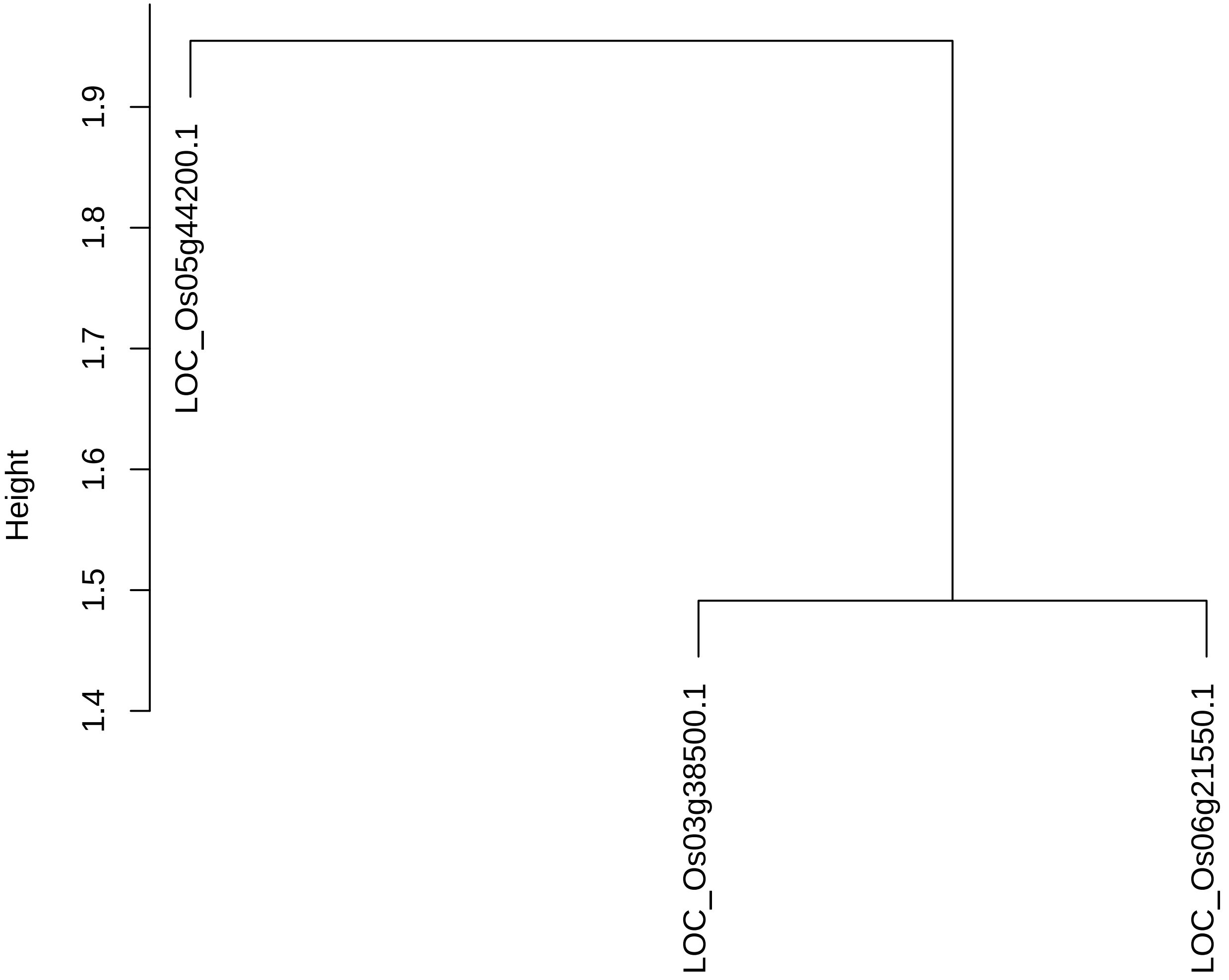


dist(spfitiex12sec12[lab[, 1] == i, ])
hclust (*, "ward.D2")

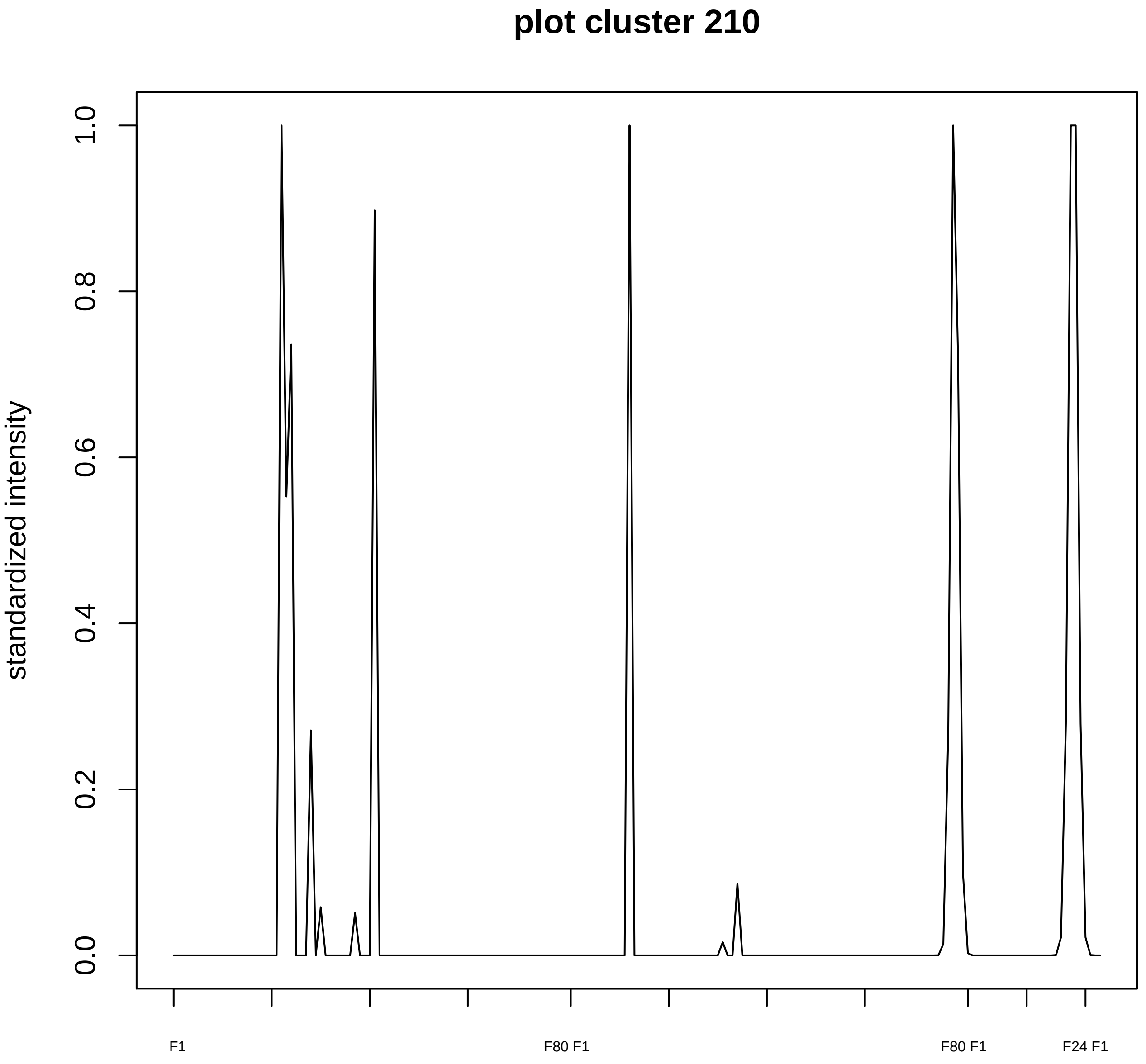
plot cluster 210
standardized intensity
0.0
0.2
0.4
0.6
0.8
1.0
F1
F80 F1
F80 F1
F24 F1

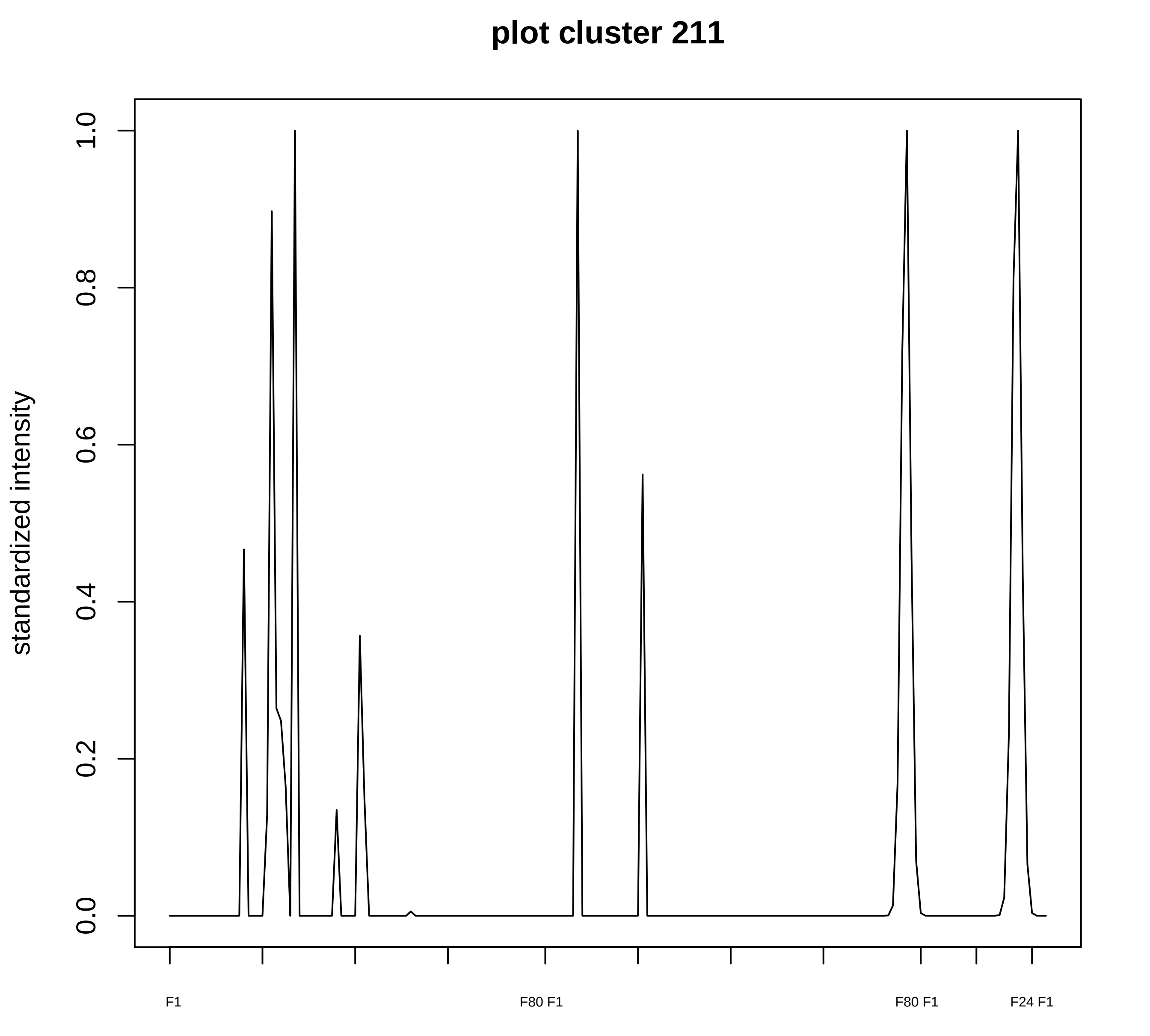

plot cluster 211
standardized intensity
0.0
0.2
0.4
0.6
0.8
1.0
F1
F80 F1
F80 F1
F24 F1

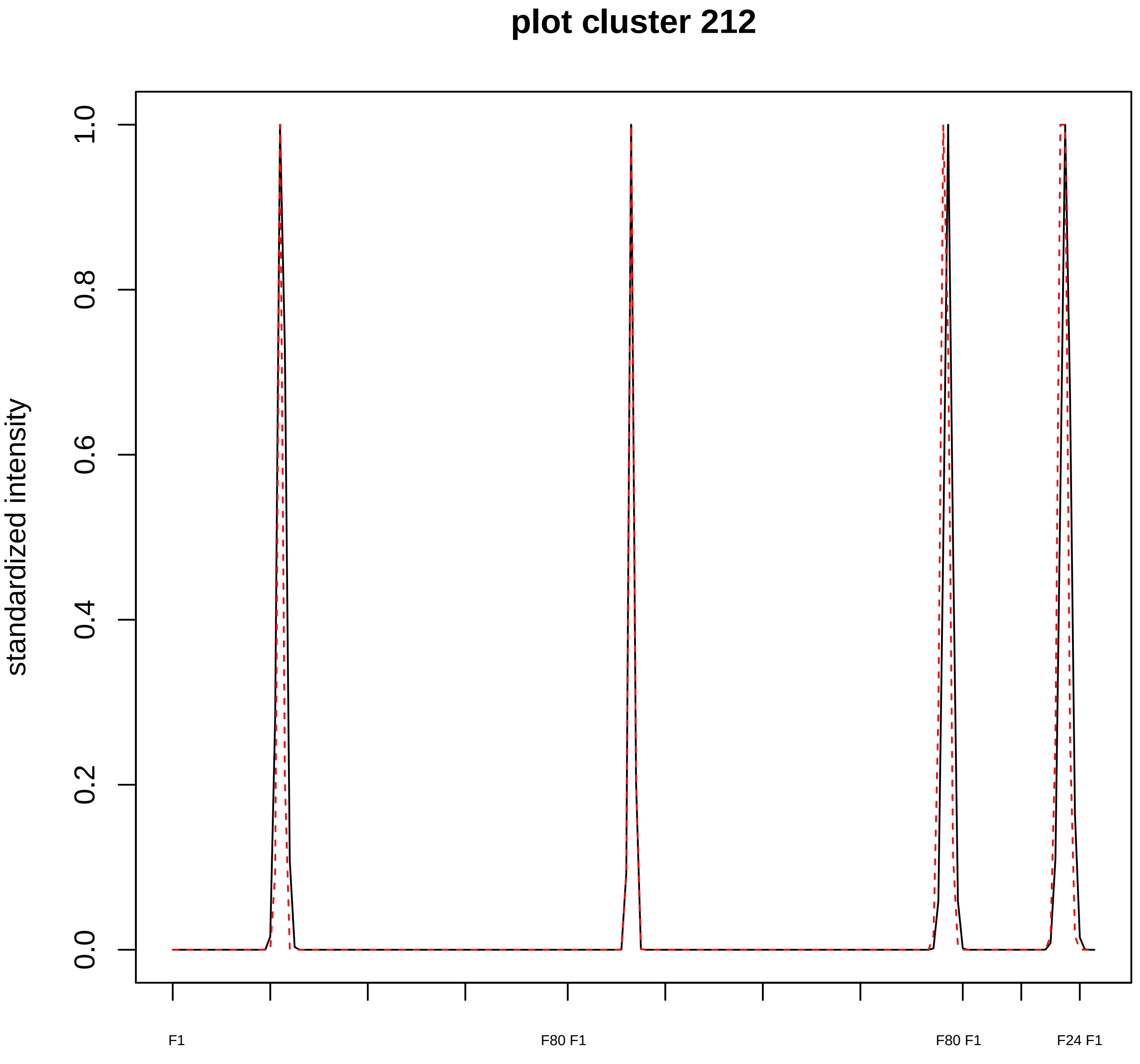
plot cluster 212
standardized intensity
0.0
0.2
0.4
0.6
0.8
1.0
F1
F80 F1
F80 F1
F24 F1

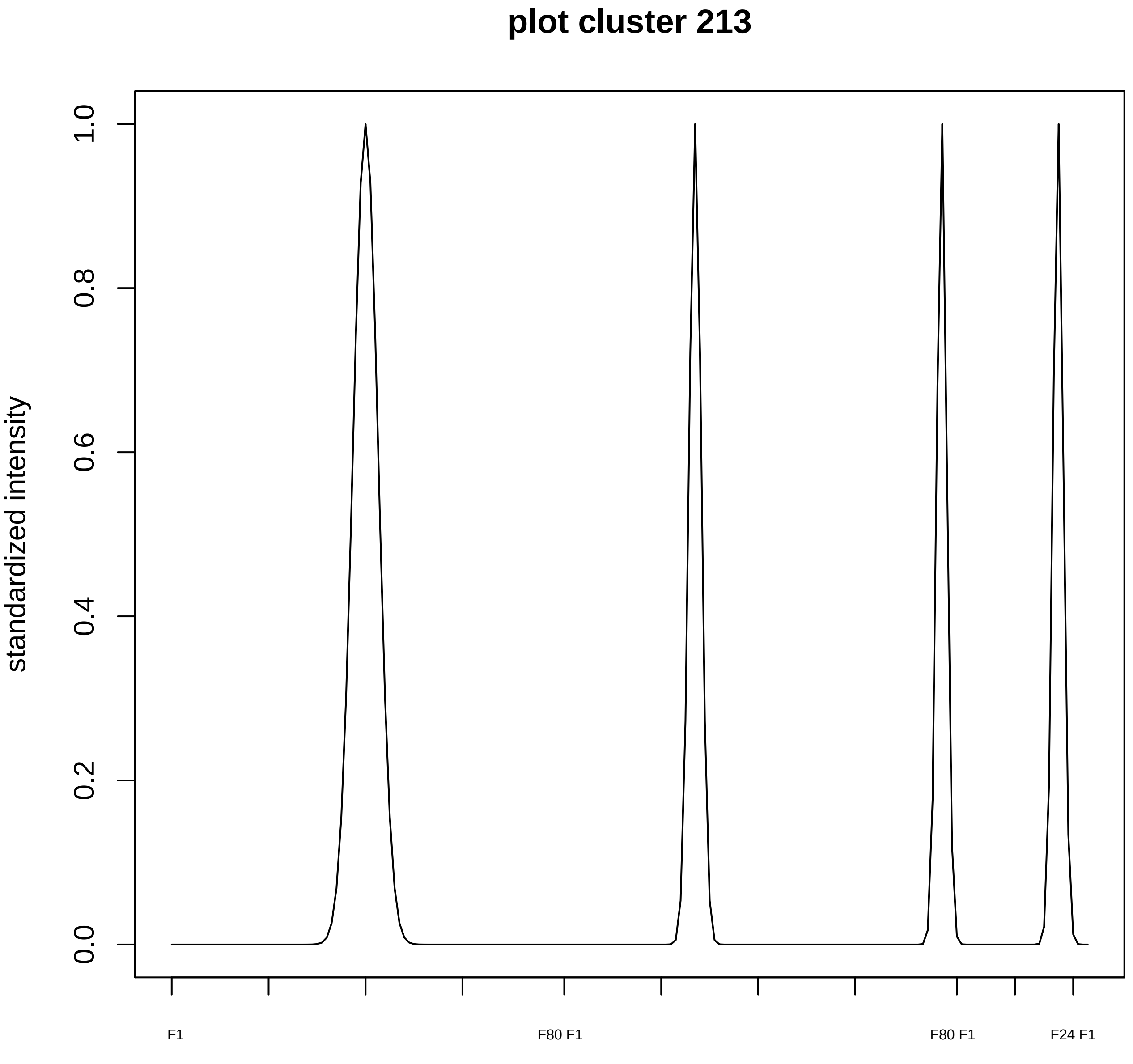

plot cluster 213
standardized intensity
0.0
0.2
0.4
0.6
0.8
1.0
F1
F80 F1
F80 F1
F24 F1

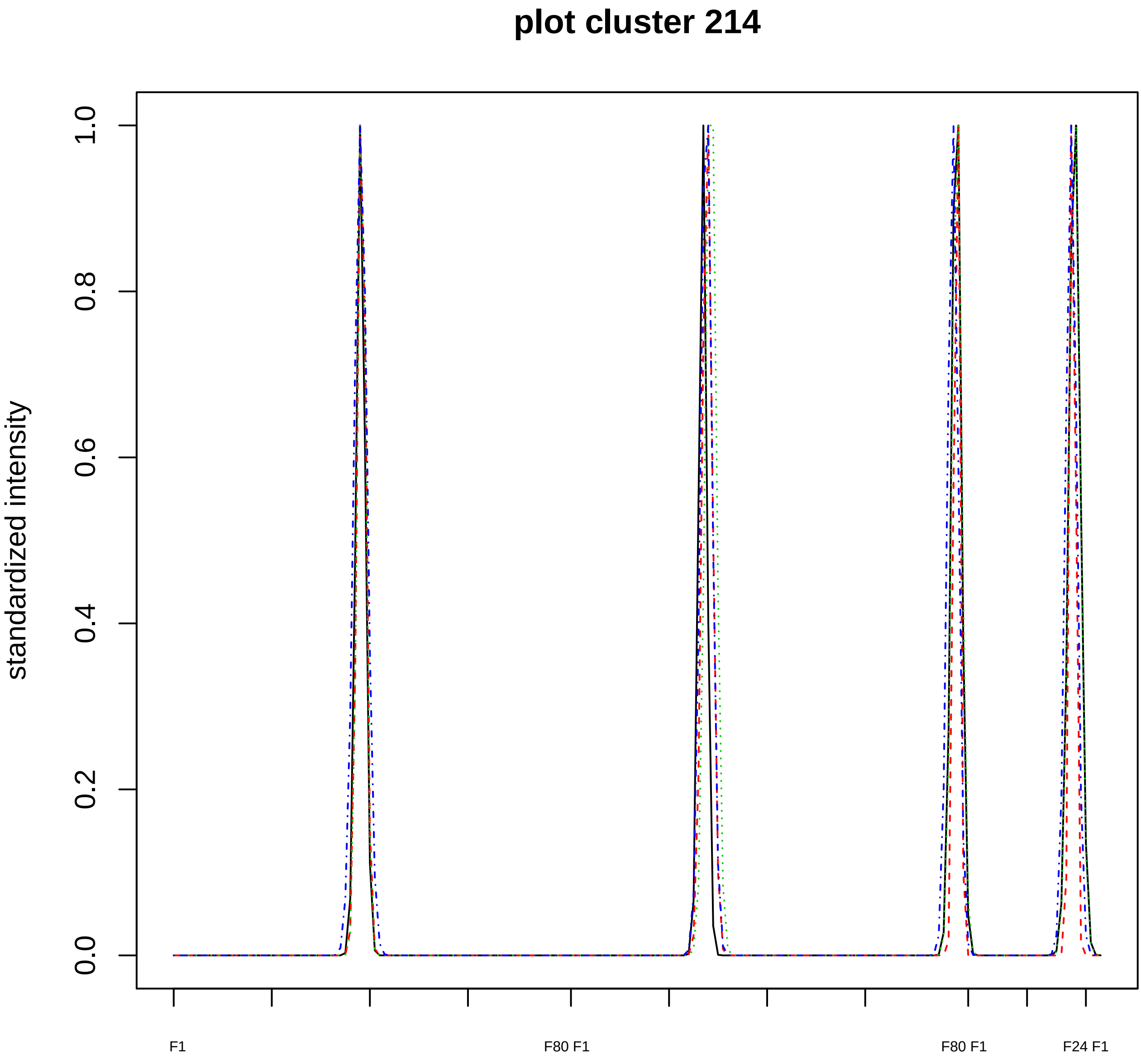
plot cluster 214
standardized intensity
0.0
0.2
0.4
0.6
0.8
1.0
F1
F80 F1
F80 F1
F24 F1

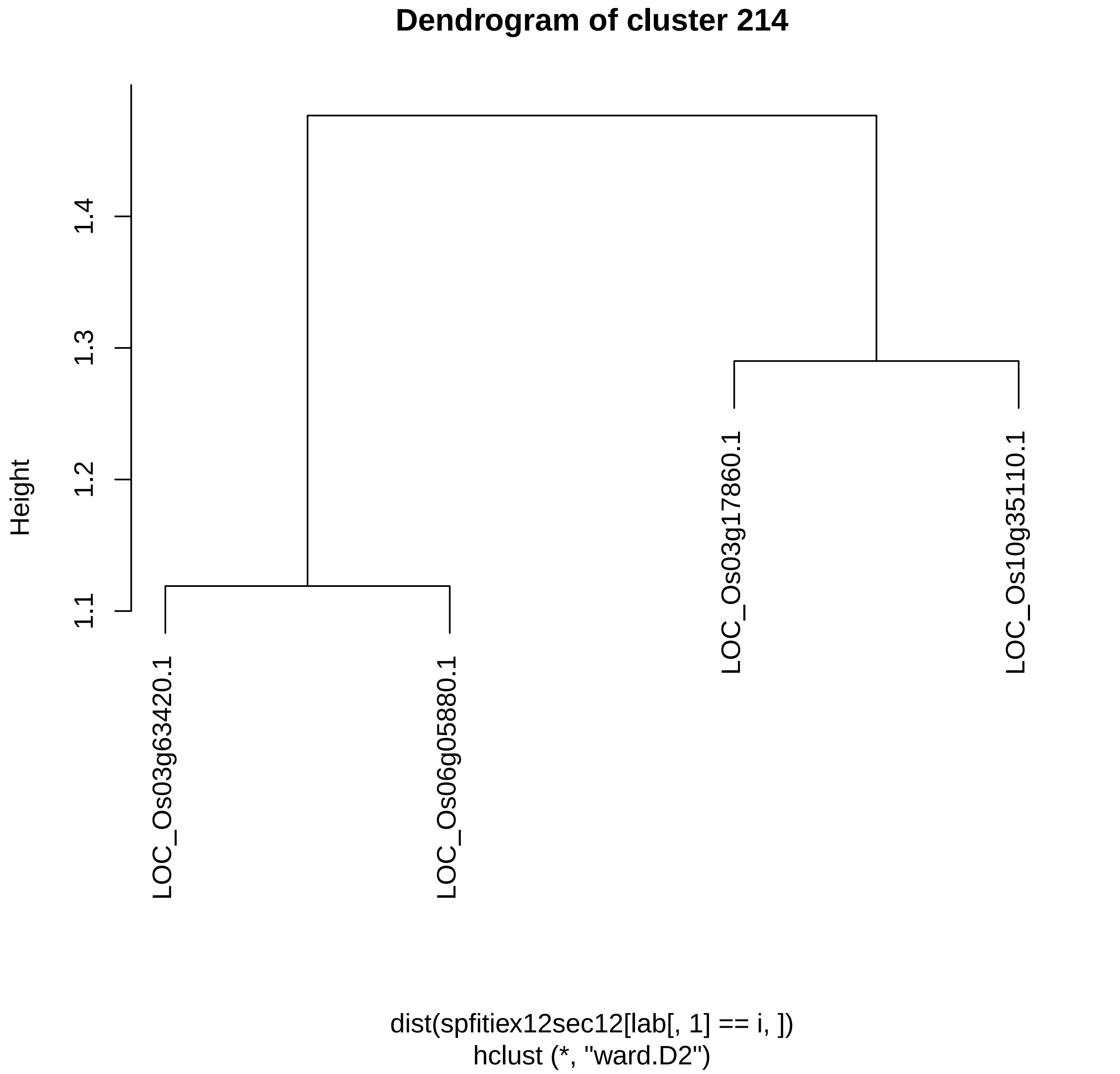
Dendrogram of cluster 214
Height
1.1
1.2
1.3
1.4
LOC_Os03g63420.1
LOC_Os06g05880.1
LOC_Os03g17860.1
LOC_Os10g35110.1
dist(spfitiex12sec12[lab[, 1] == i, ])
hclust (*, "ward.D2")

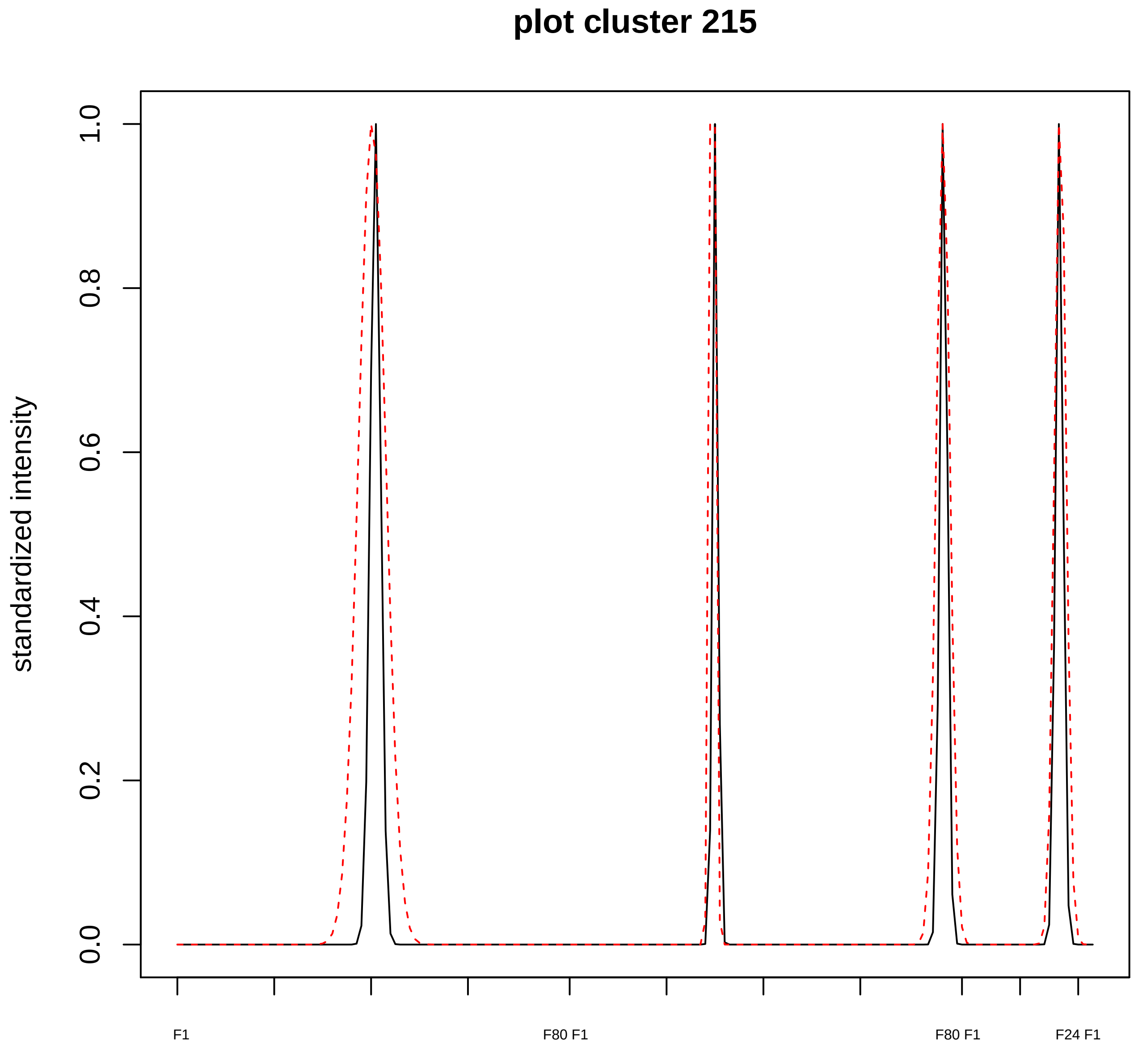
plot cluster 215
standardized intensity
0.0
0.2
0.4
0.6
0.8
1.0
F1
F80 F1
F80 F1
F24 F1

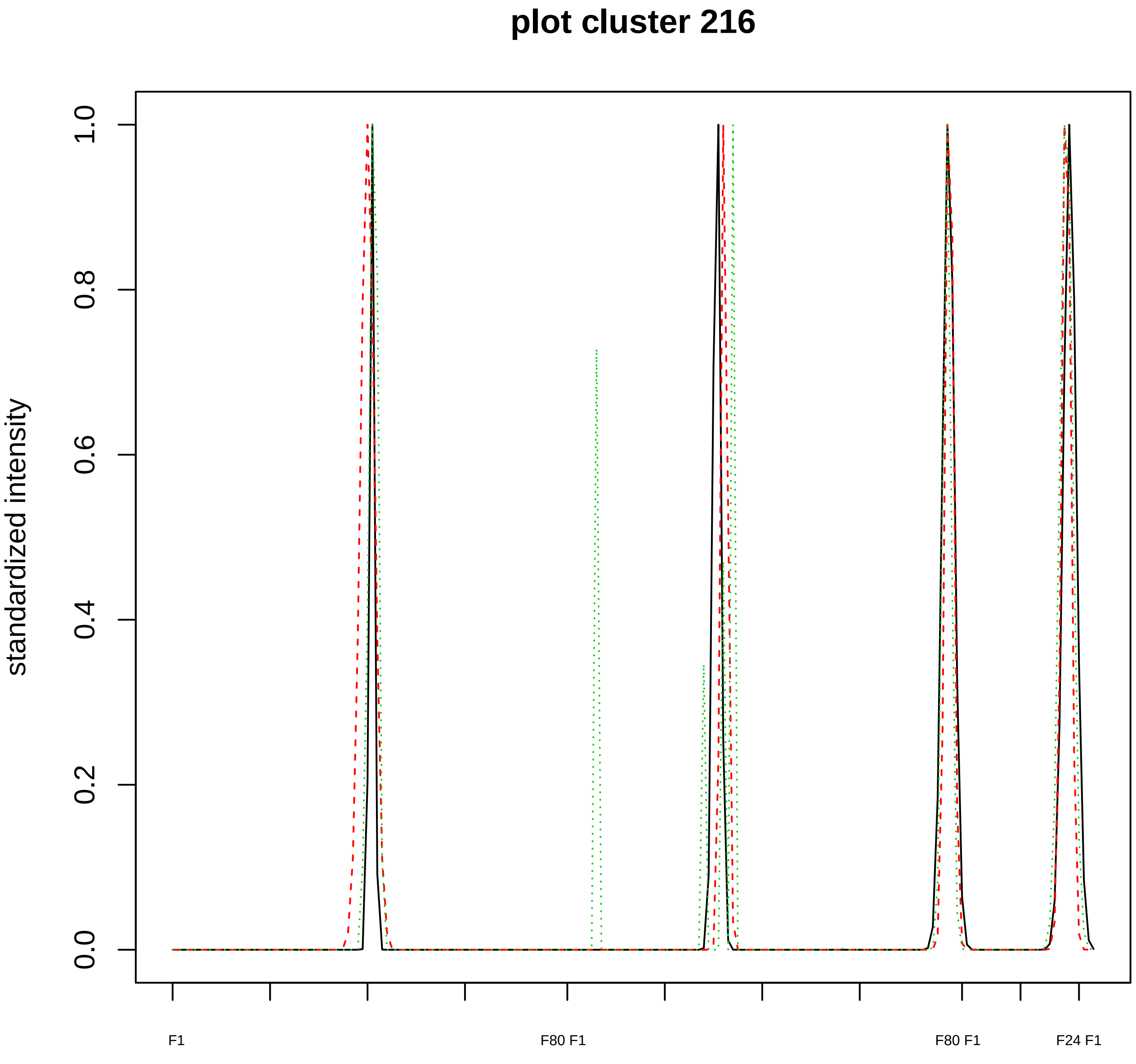
plot cluster 216
standardized intensity
0.0
0.2
0.4
0.6
0.8
1.0
F1
F80 F1
F80 F1
F24 F1

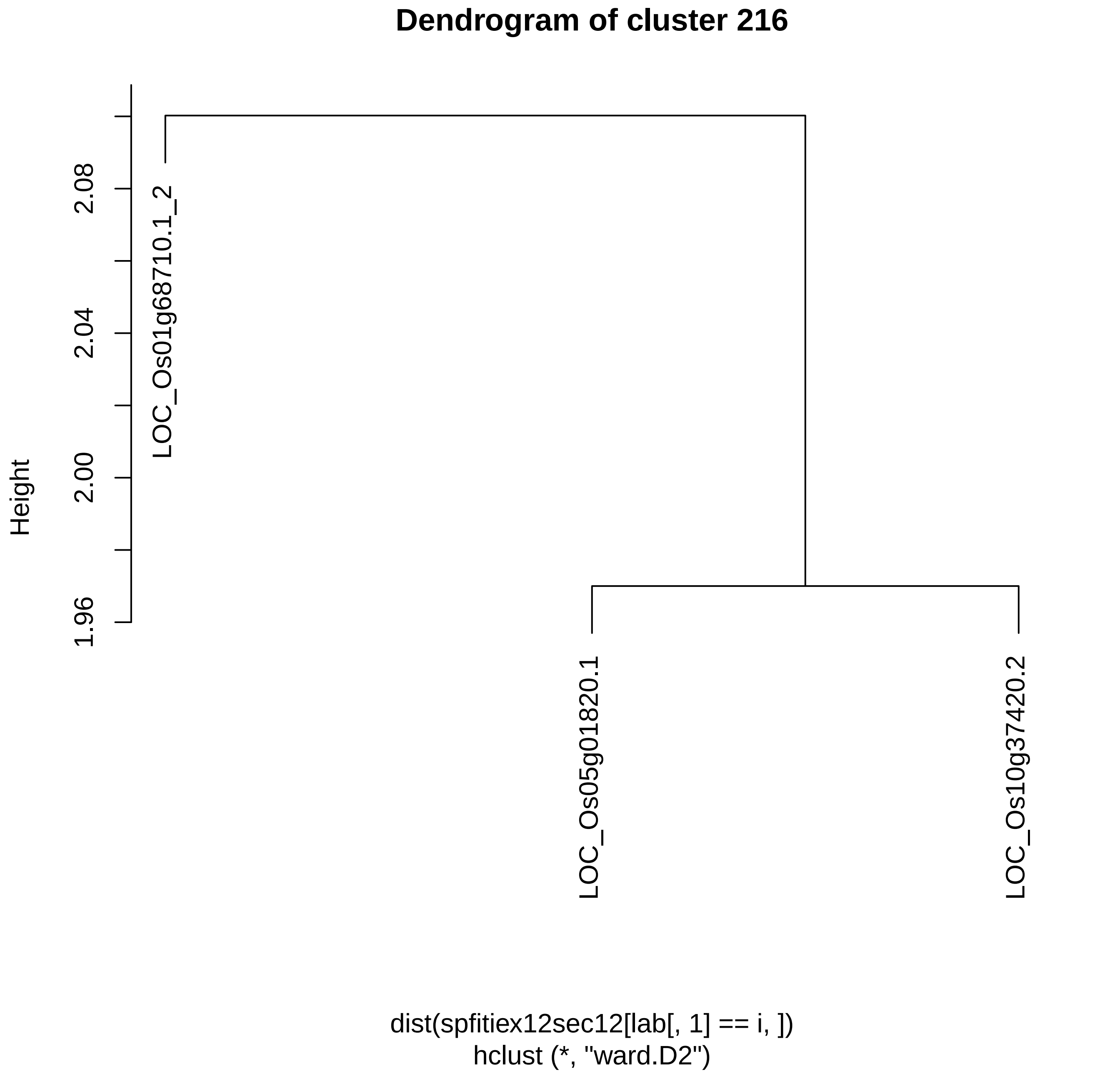
Dendrogram of cluster 216
Height
1.96
2.00
2.04
2.08
LOC_Os01g68710.1_2
LOC_Os05g01820.1
LOC_Os10g37420.2
dist(spfitiex12sec12[lab[, 1] == i, ])
hclust (*, "ward.D2")

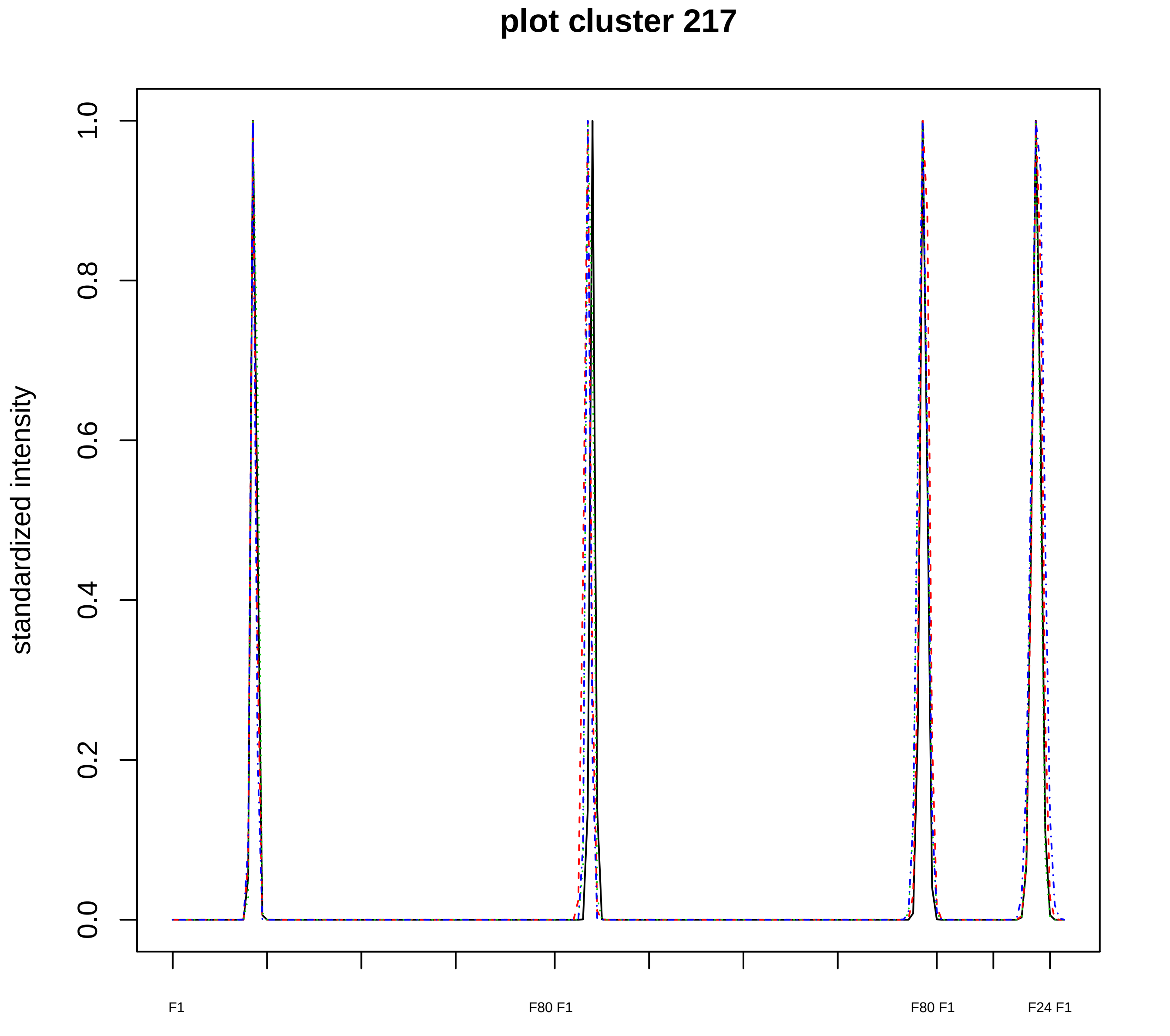
plot cluster 217
standardized intensity
1.0
0.8
0.6
0.4
0.2
0.0
F1
F80 F1
F80 F1
F24 F1

### Dendrogram of cluster 217

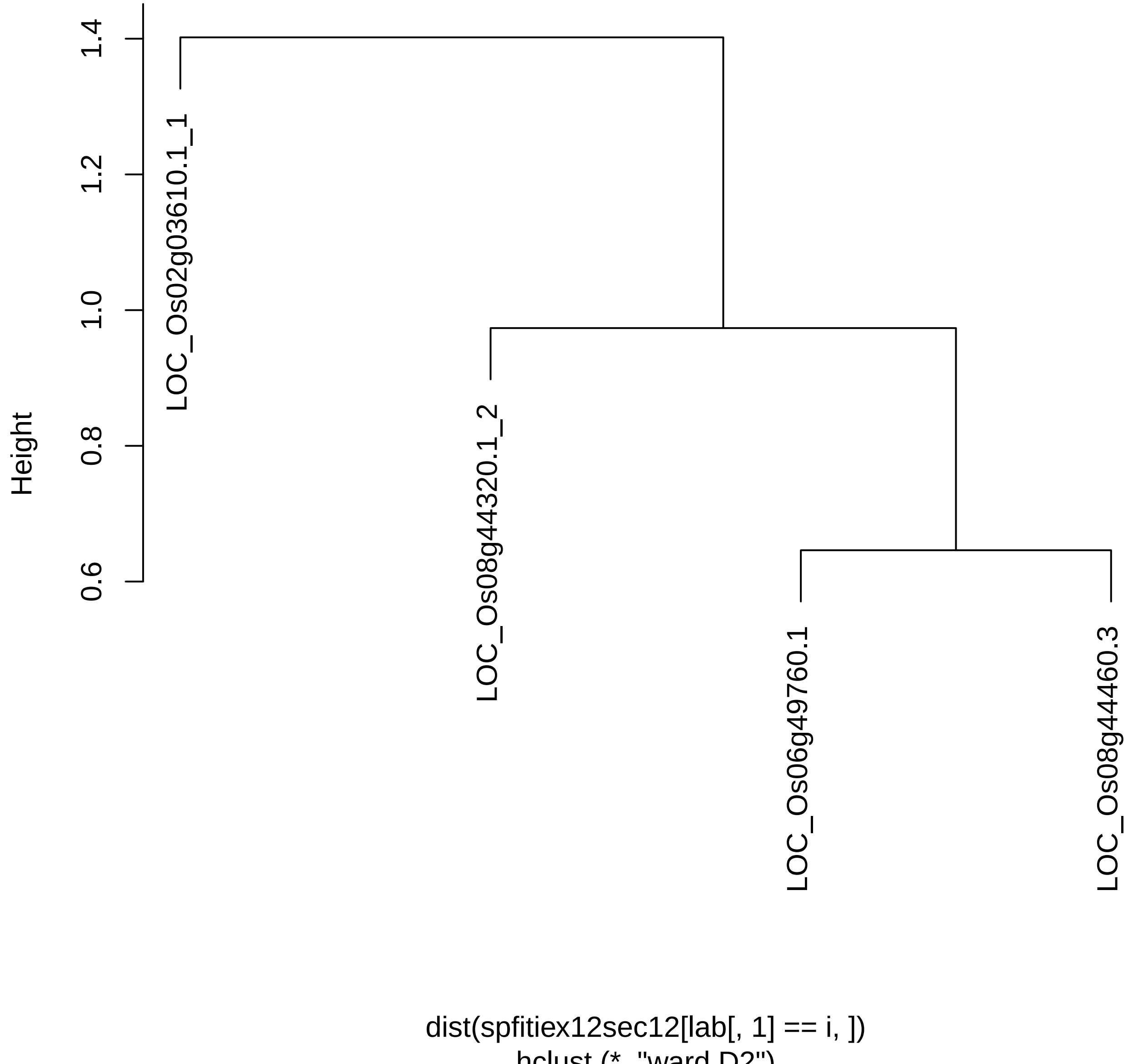

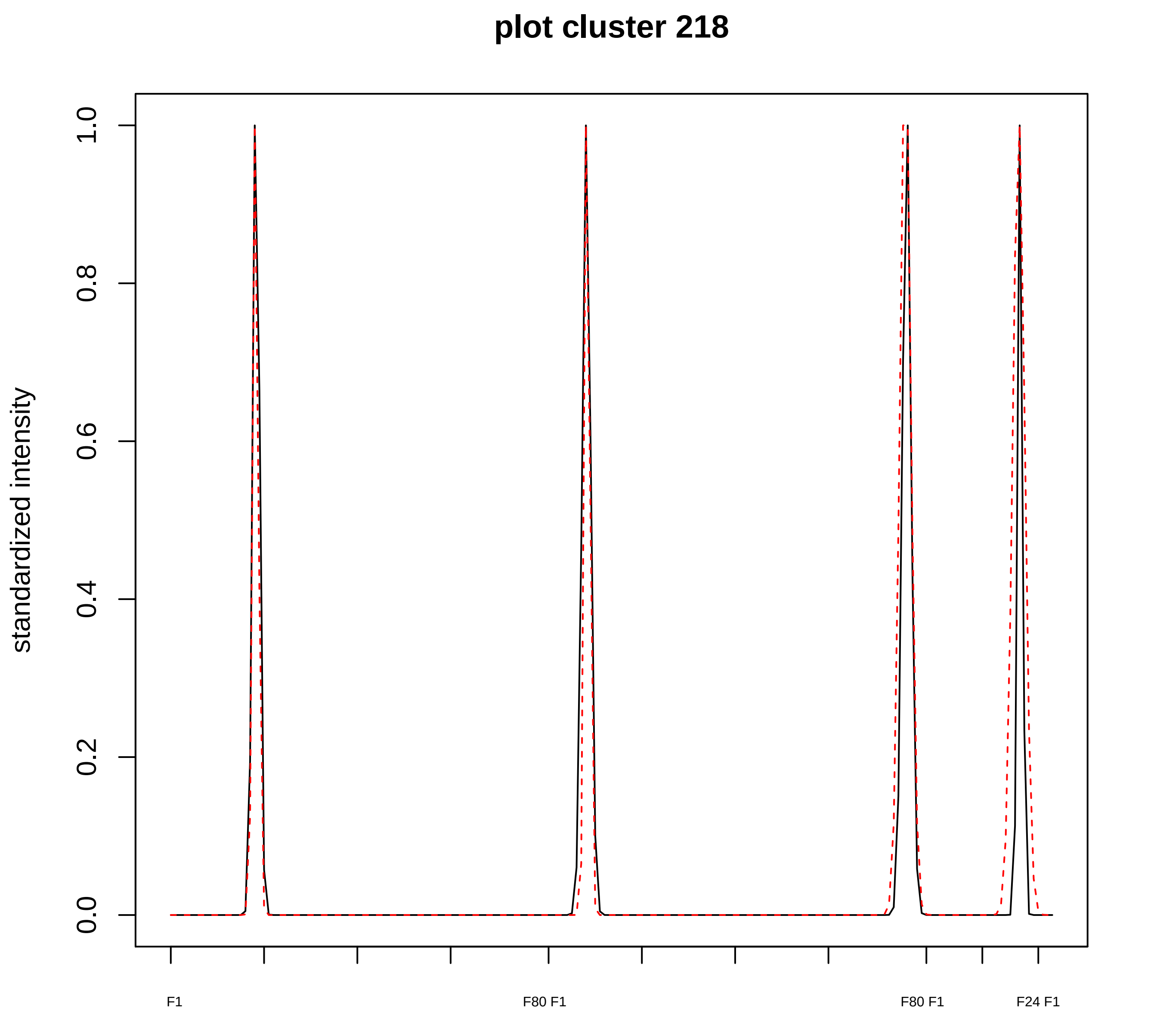

plot cluster 218
standardized intensity
0.0
0.2
0.4
0.6
0.8
1.0
F1
F80 F1
F80 F1
F24 F1

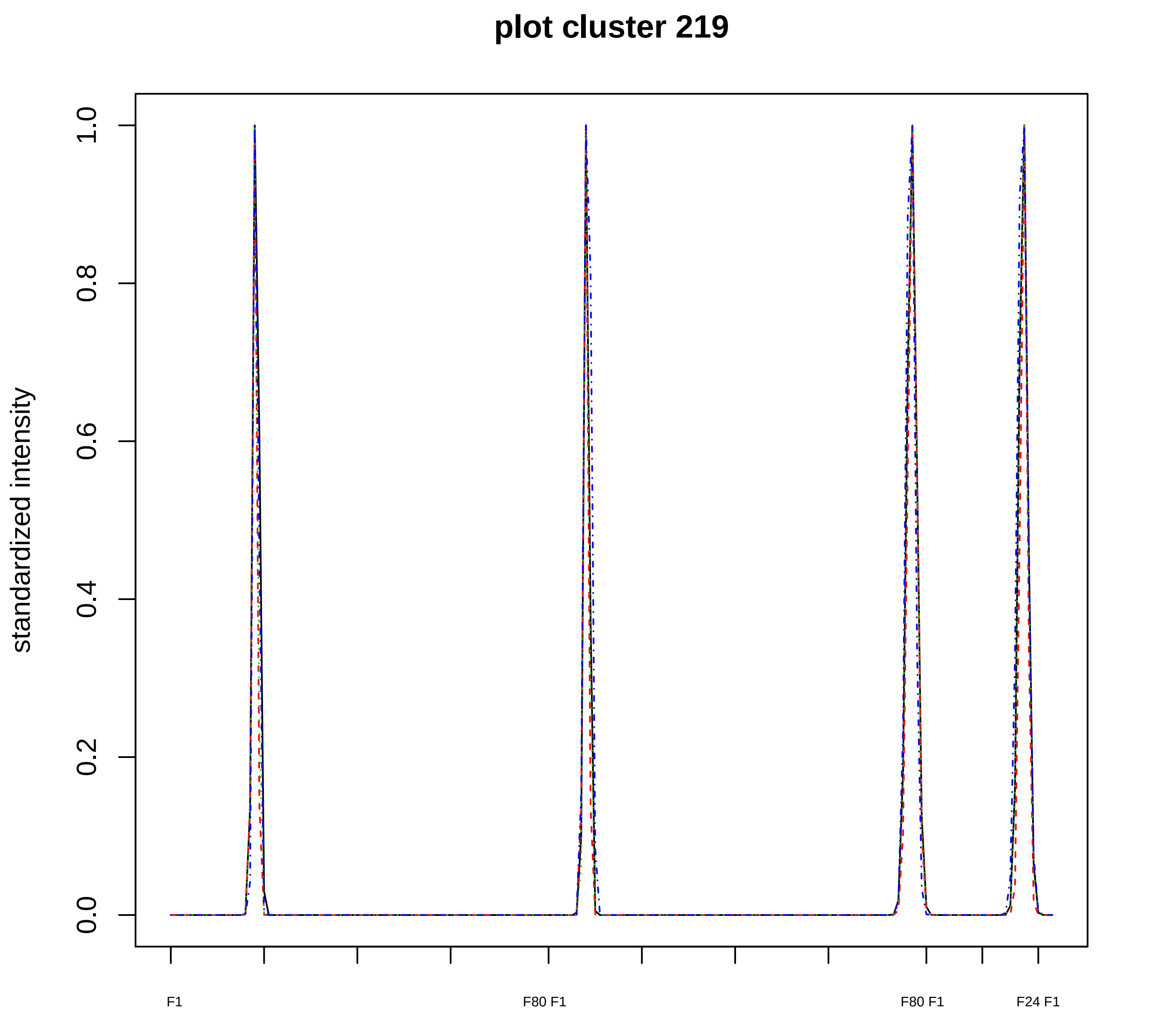
plot cluster 219
standardized intensity
0.0
0.2
0.4
0.6
0.8
1.0
F1
F80 F1
F80 F1
F24 F1

#### Dendrogram of cluster 219

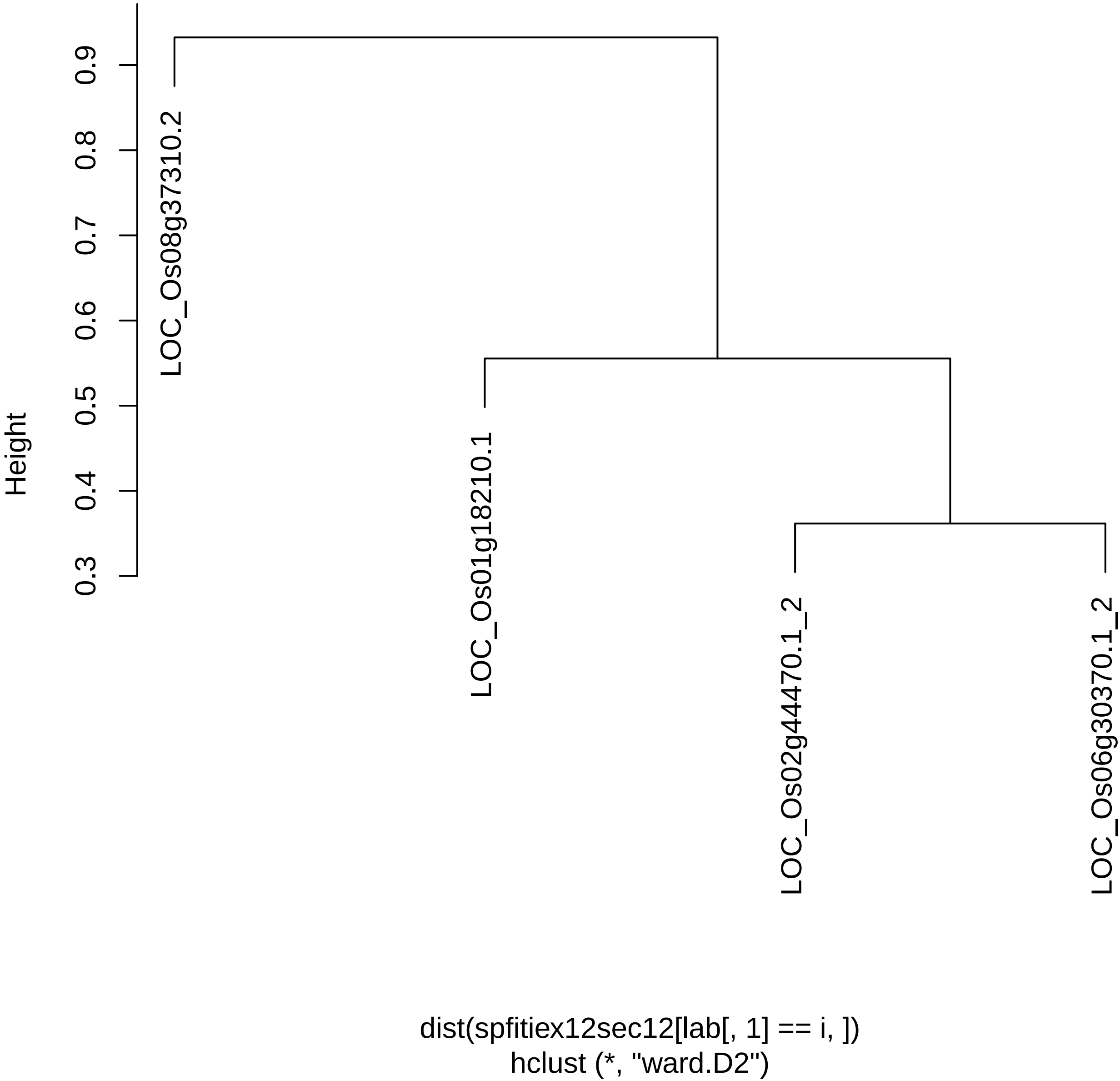

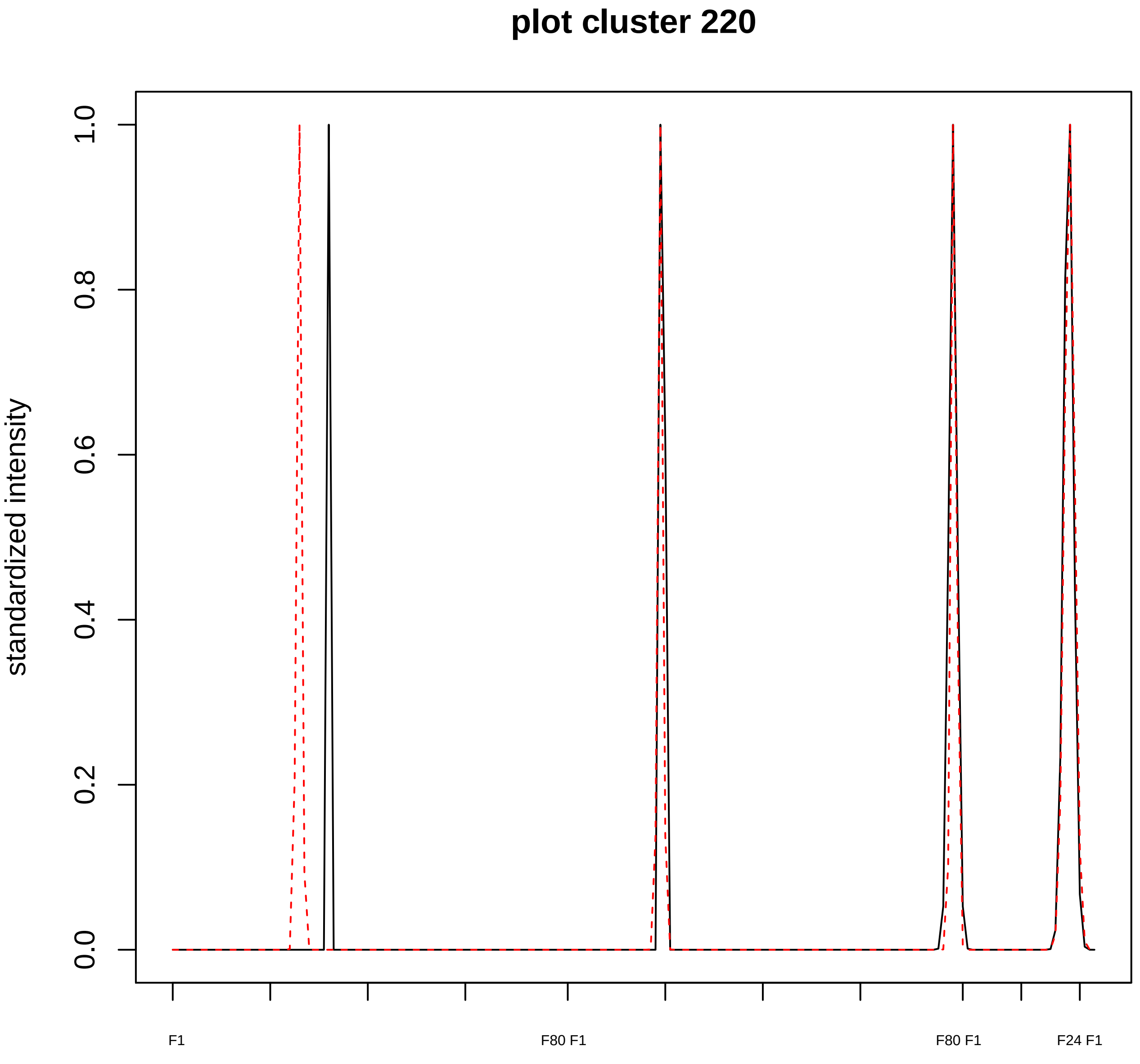
plot cluster 220
standardized intensity
0.0
0.2
0.4
0.6
0.8
1.0
F1
F80 F1
F80 F1
F24 F1

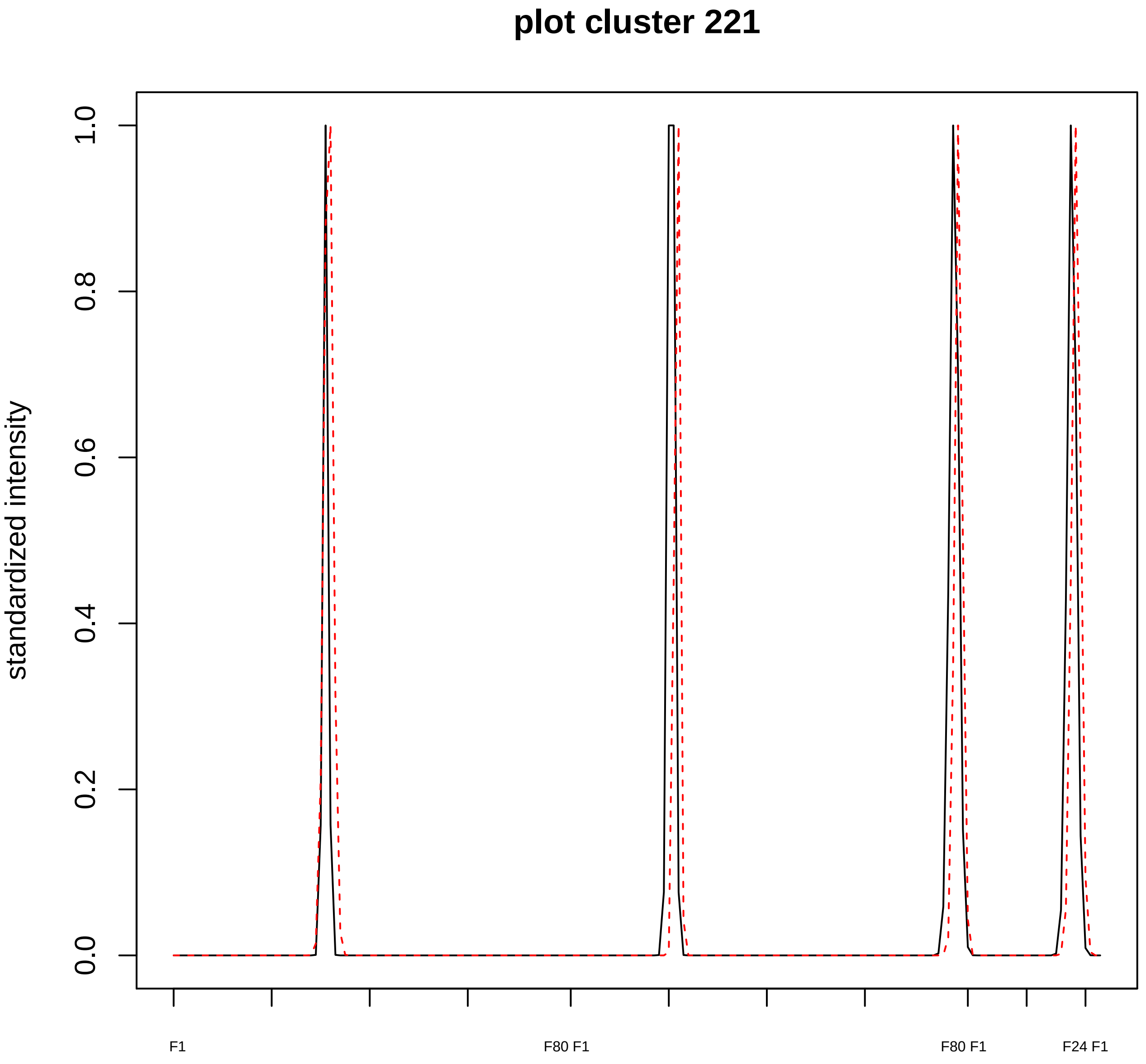
plot cluster 221
standardized intensity
0.0
0.2
0.4
0.6
0.8
1.0
F1
F80 F1
F80 F1
F24 F1

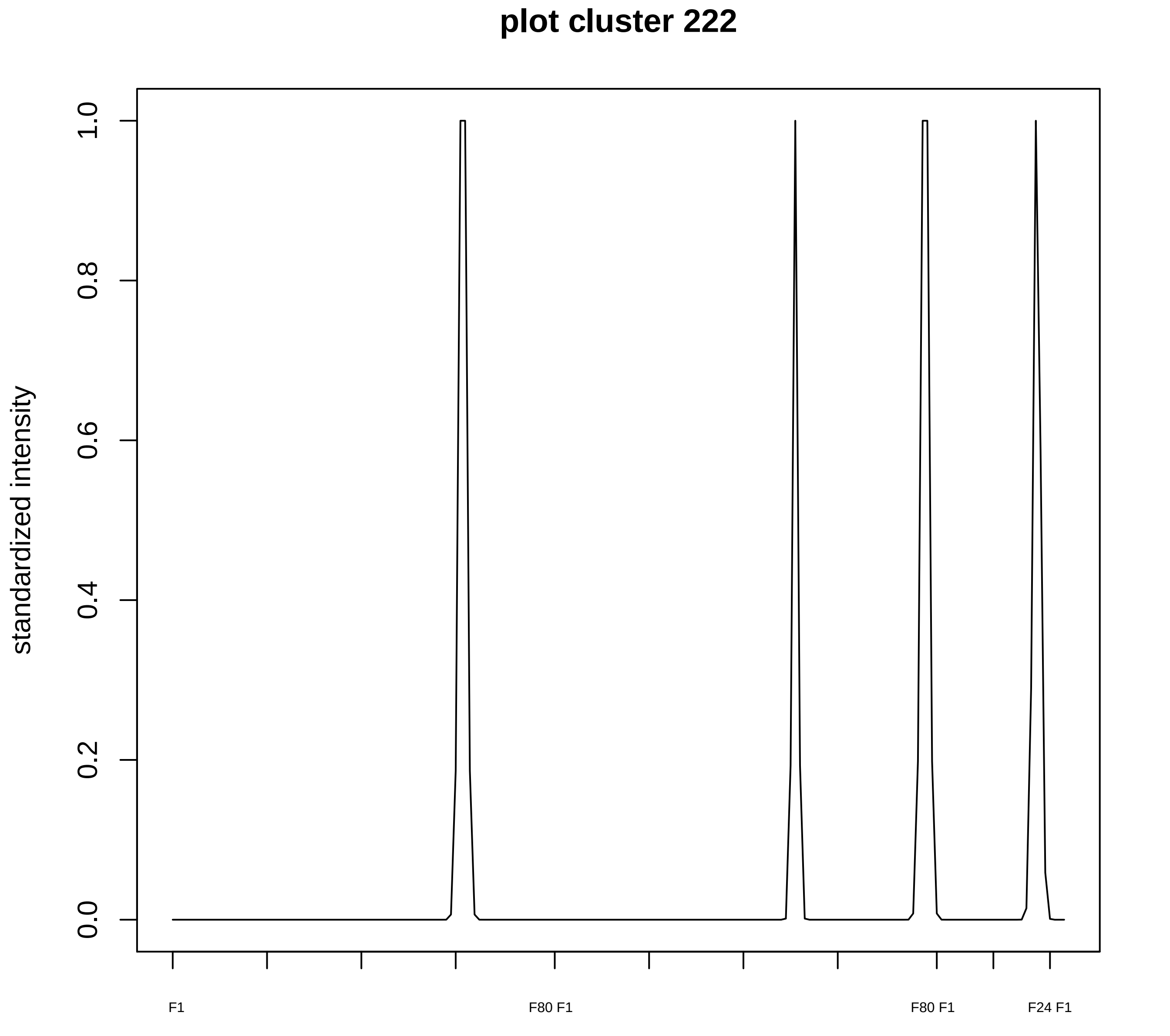

plot cluster 222
standardized intensity
0.0
0.2
0.4
0.6
0.8
1.0
F1
F80 F1
F80 F1
F24 F1

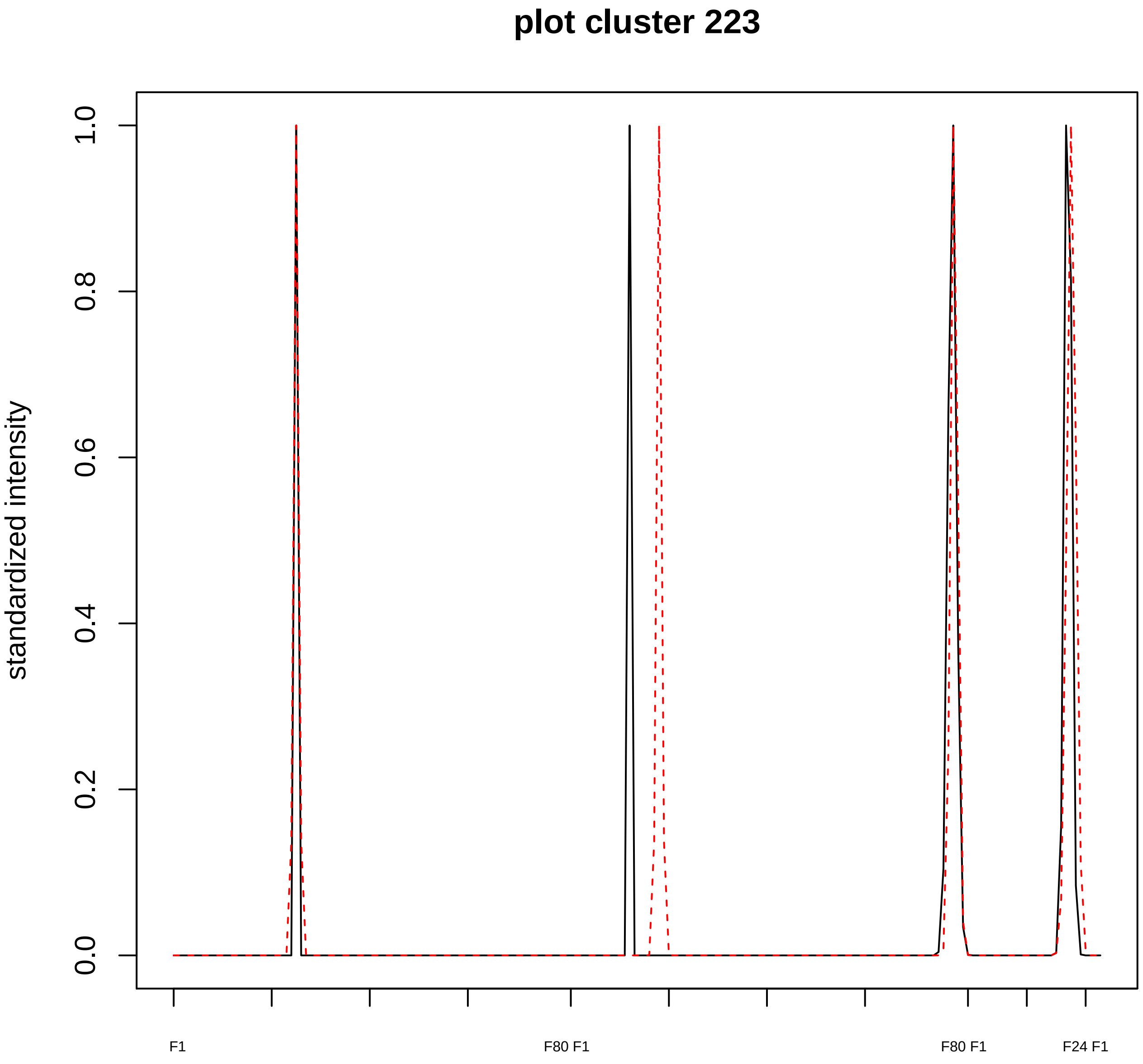

plot cluster 223
standardized intensity
0.0
0.2
0.4
0.6
0.8
1.0
F1
F80 F1
F80 F1
F24 F1

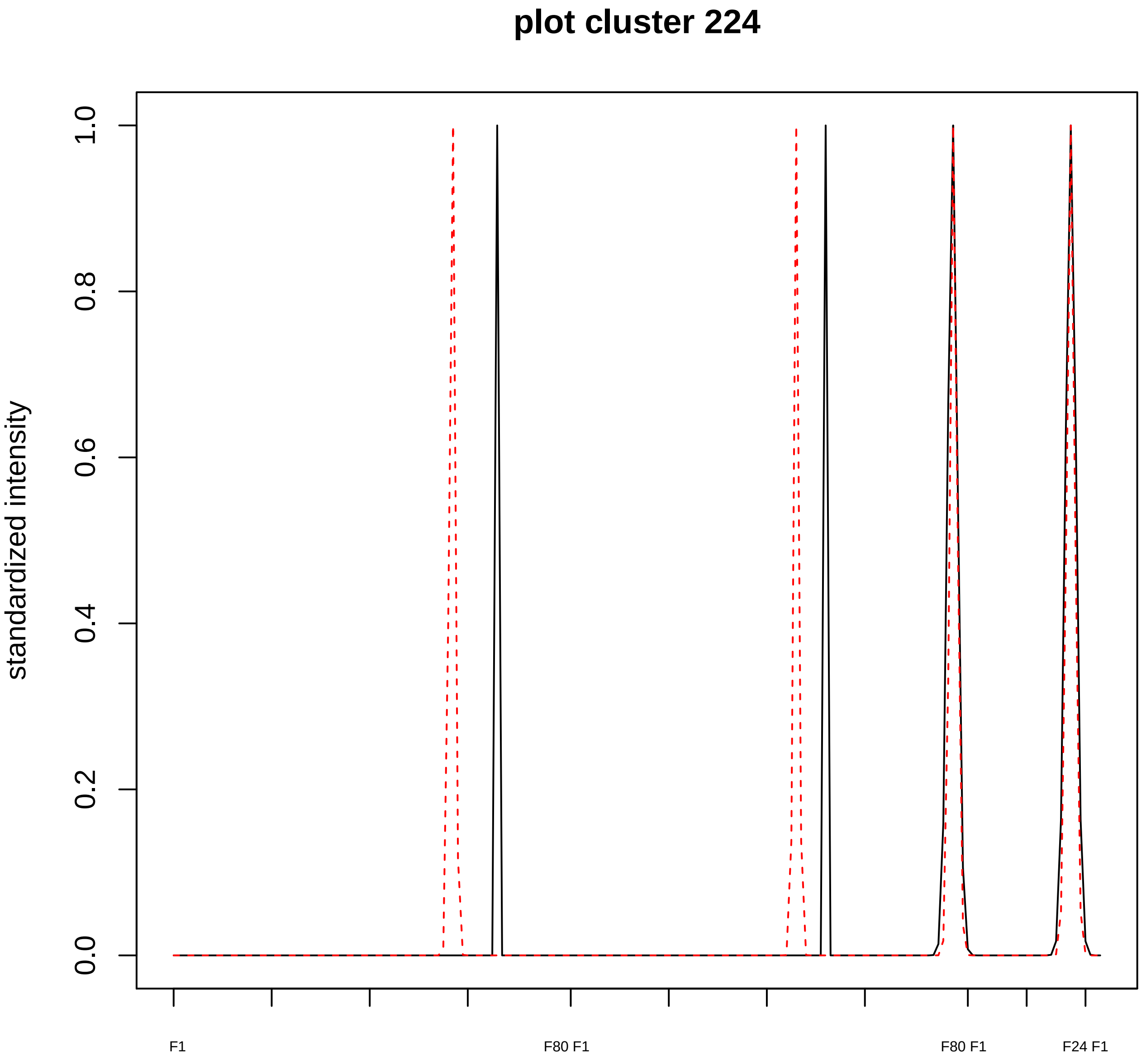
plot cluster 224
standardized intensity
0.0
0.2
0.4
0.6
0.8
1.0
F1
F80 F1
F80 F1
F24 F1

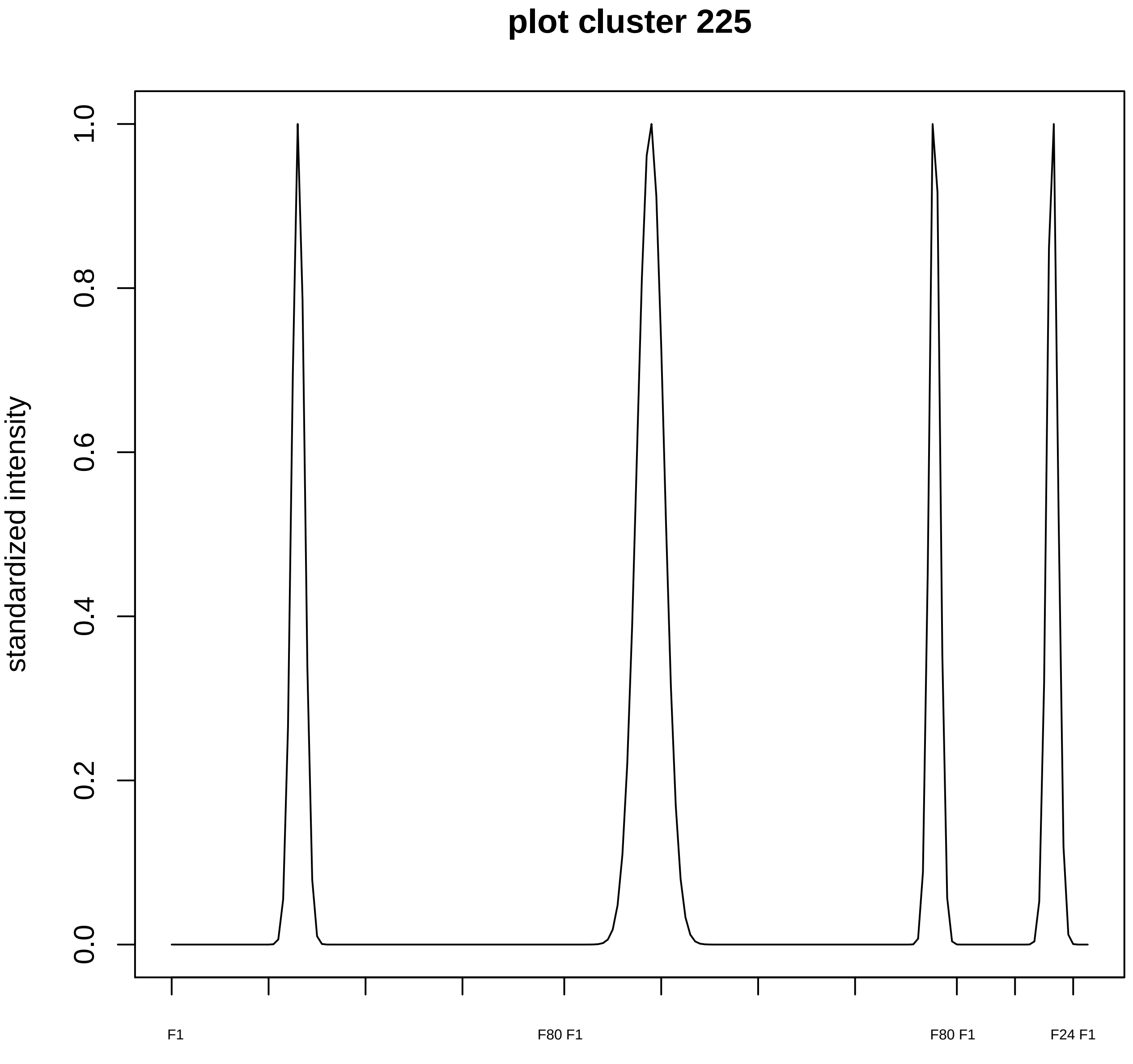
plot cluster 225
standardized intensity
0.0
0.2
0.4
0.6
0.8
1.0
F1
F80 F1
F80 F1
F24 F1

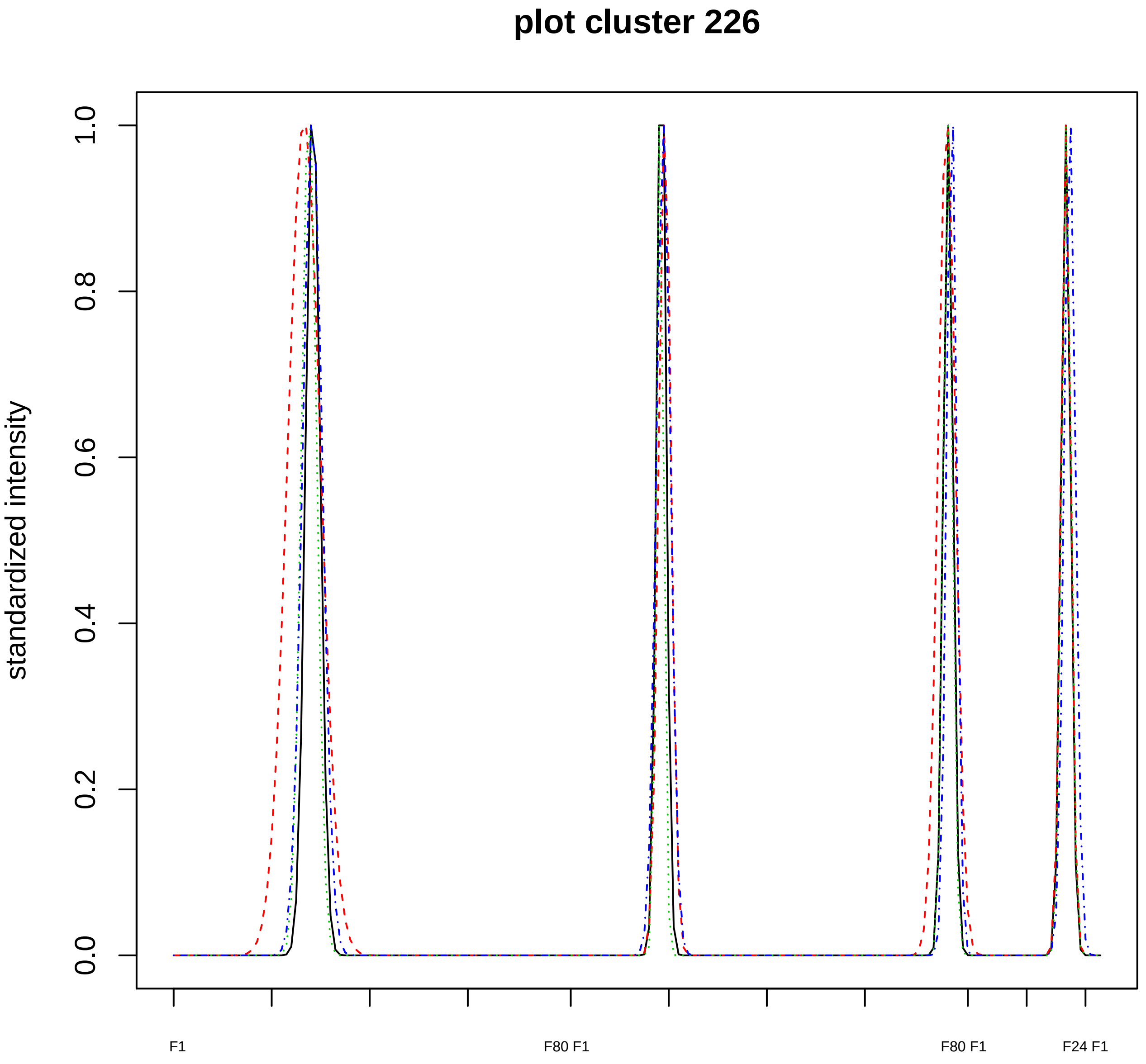

plot cluster 226
standardized intensity
0.0
0.2
0.4
0.6
0.8
1.0
F1
F80 F1
F80 F1
F24 F1

#### Dendrogram of cluster 226

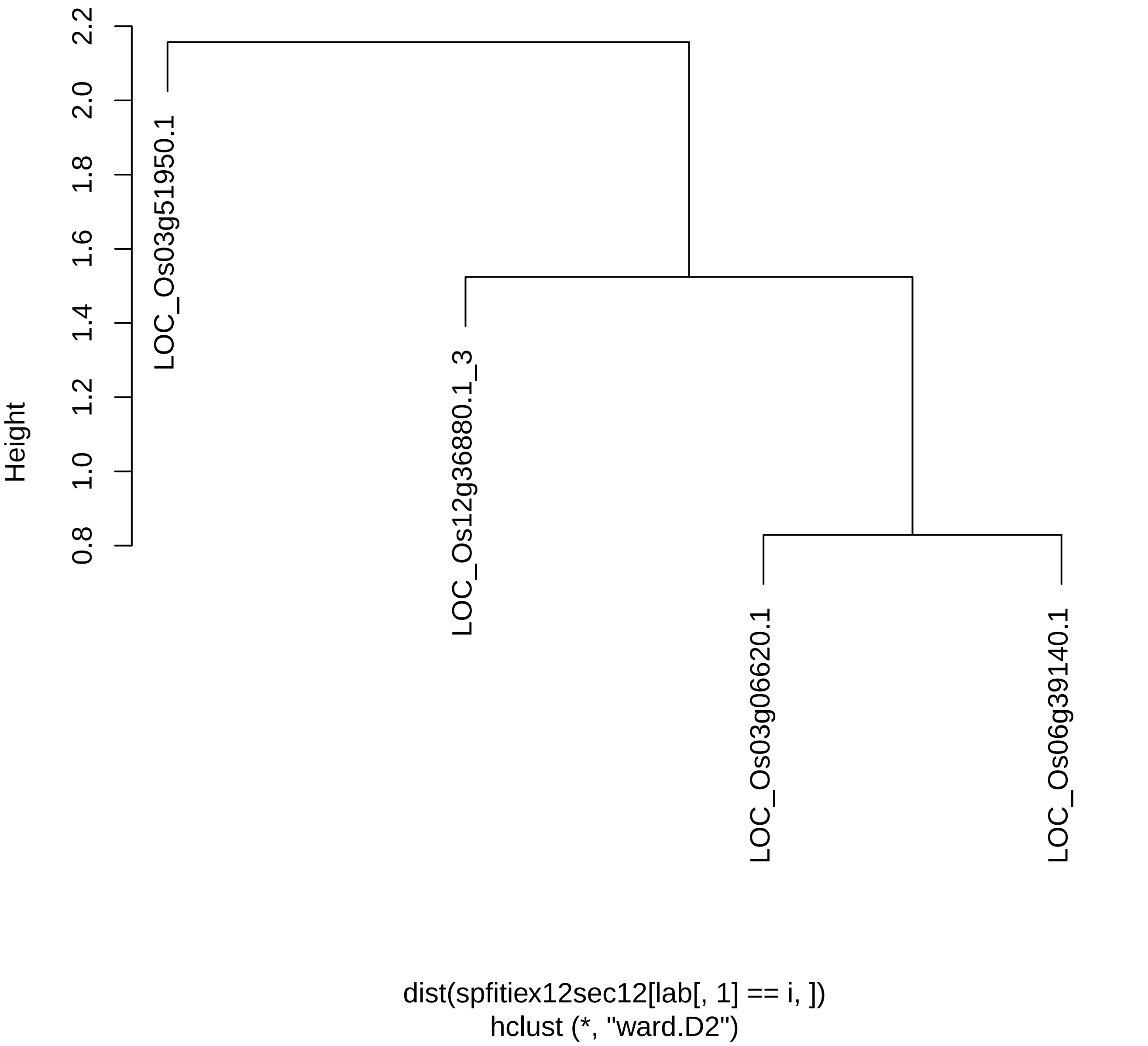

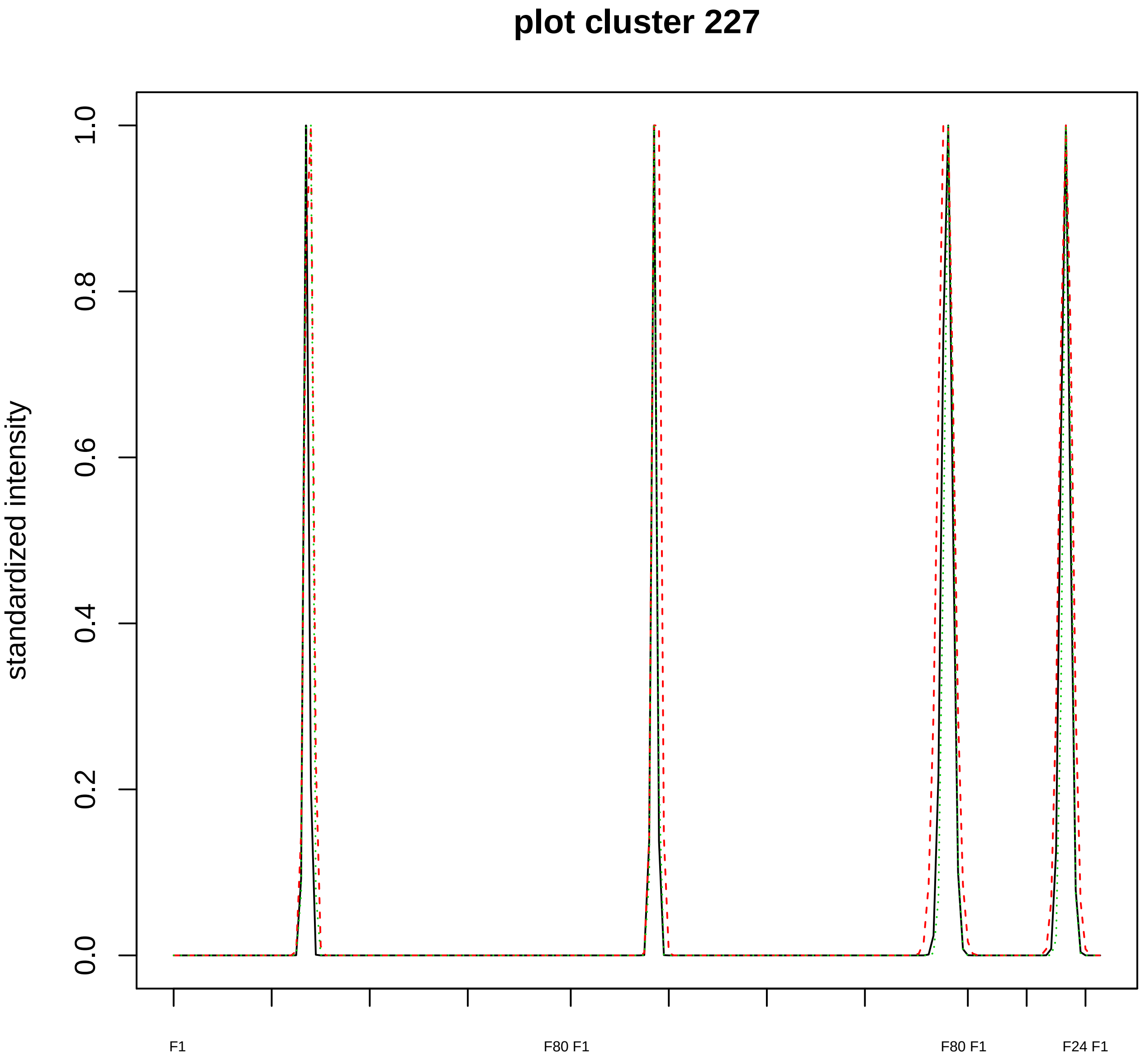

plot cluster 227
standardized intensity
0.0
0.2
0.4
0.6
0.8
1.0
F1
F80 F1
F80 F1
F24 F1

### Dendrogram of cluster 227

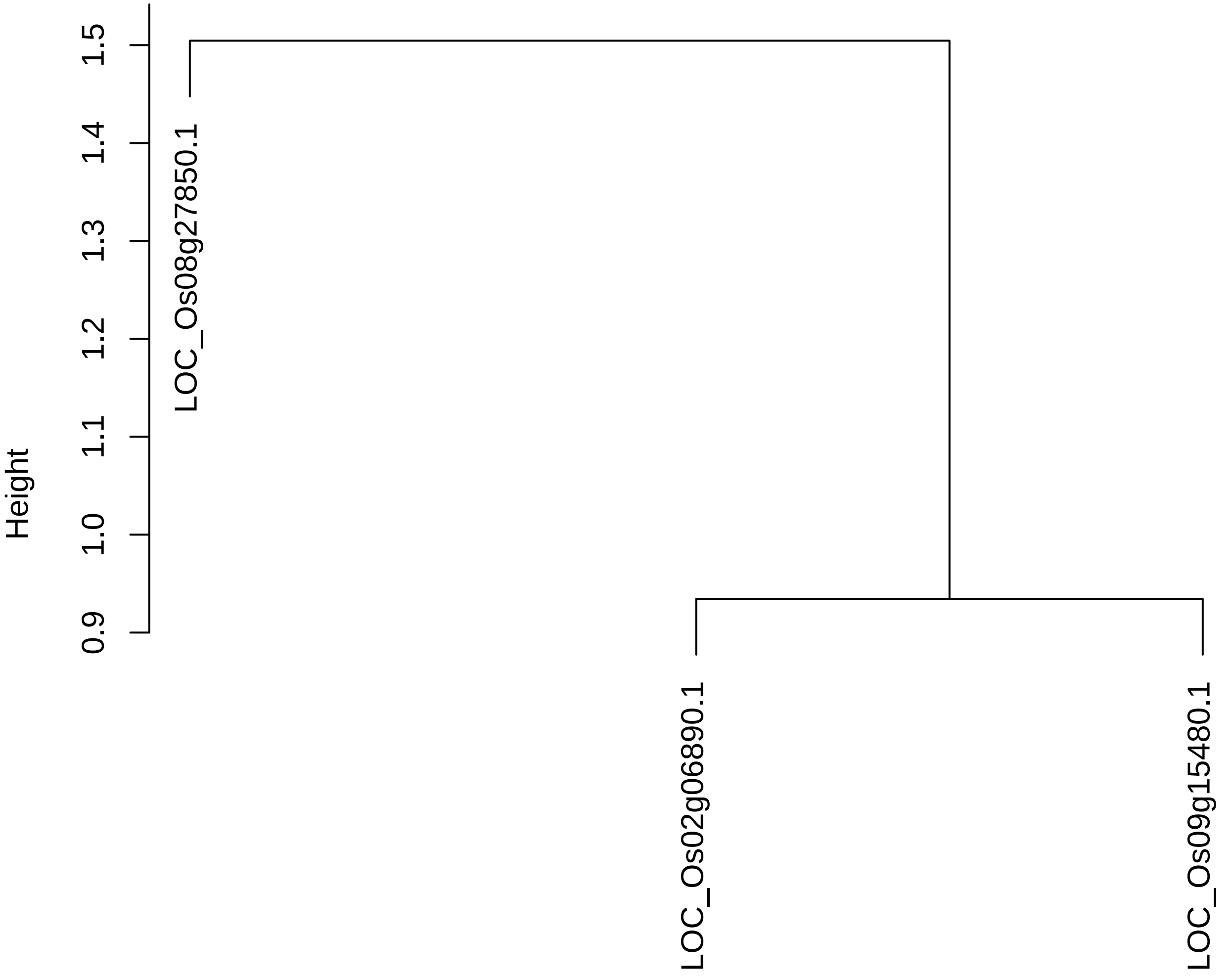


dist(spfitiex12sec12[lab[, 1] == i, ])
hclust (*, "ward.D2")

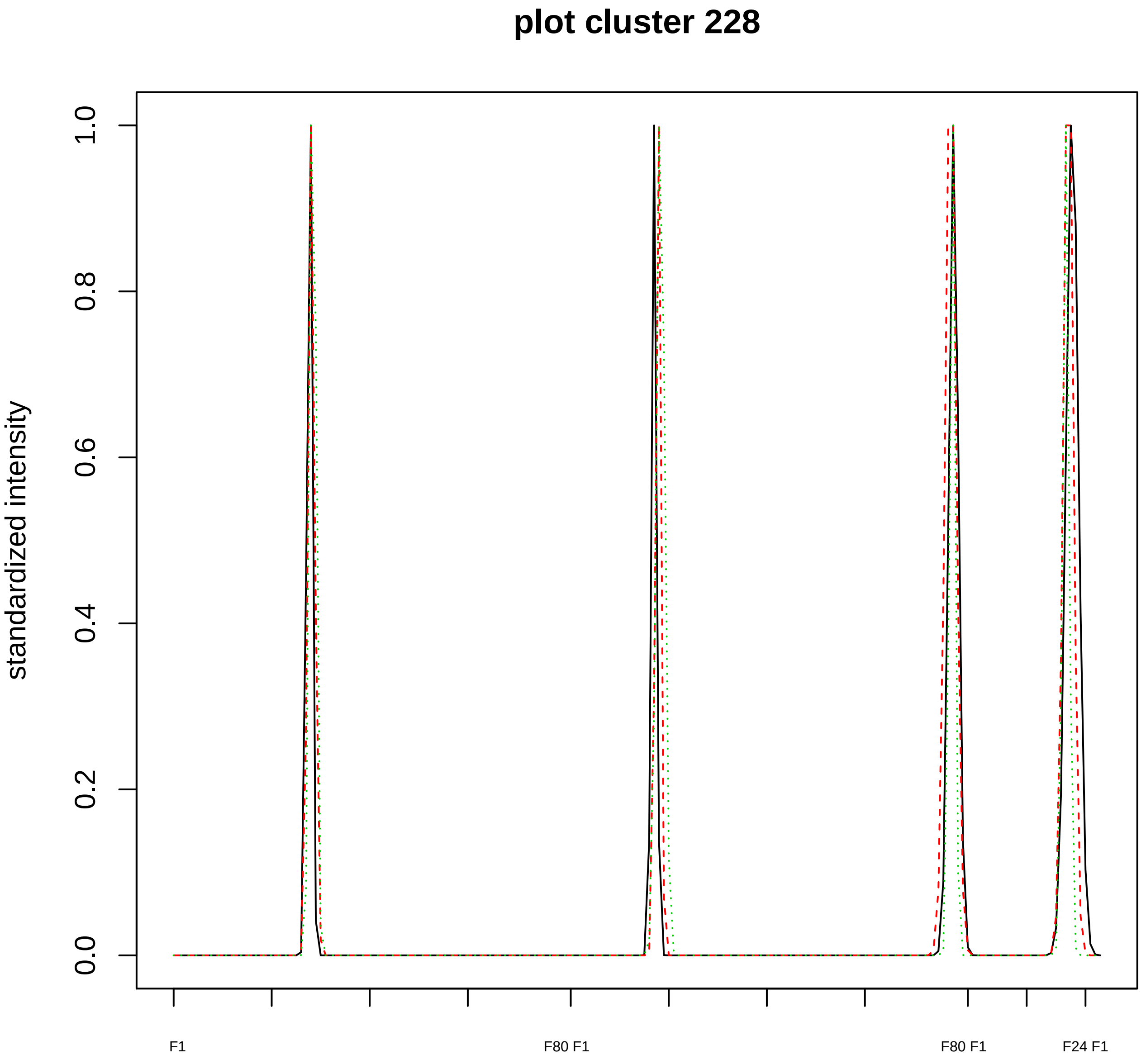
plot cluster 228
standardized intensity
0.0
0.2
0.4
0.6
0.8
1.0
F1
F80 F1
F80 F1
F24 F1

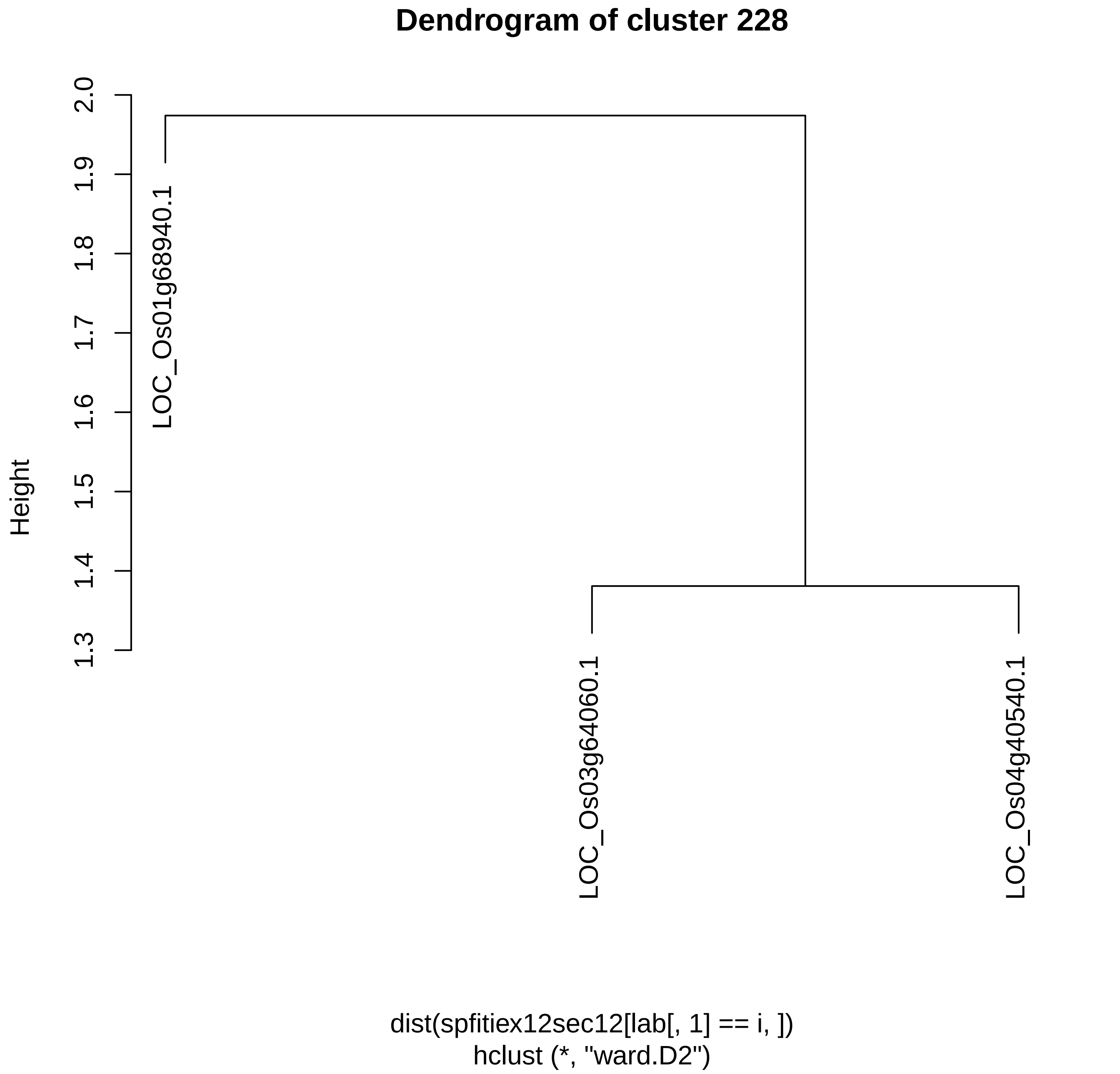
Dendrogram of cluster 228
Height
2.0
1.9
1.8
1.7
1.6
1.5
1.4
1.3
LOC_Os01g68940.1
LOC_Os03g64060.1
LOC_Os04g40540.1
dist(spfitiex12sec12[lab[, 1] == i, ])
hclust (*, "ward.D2")

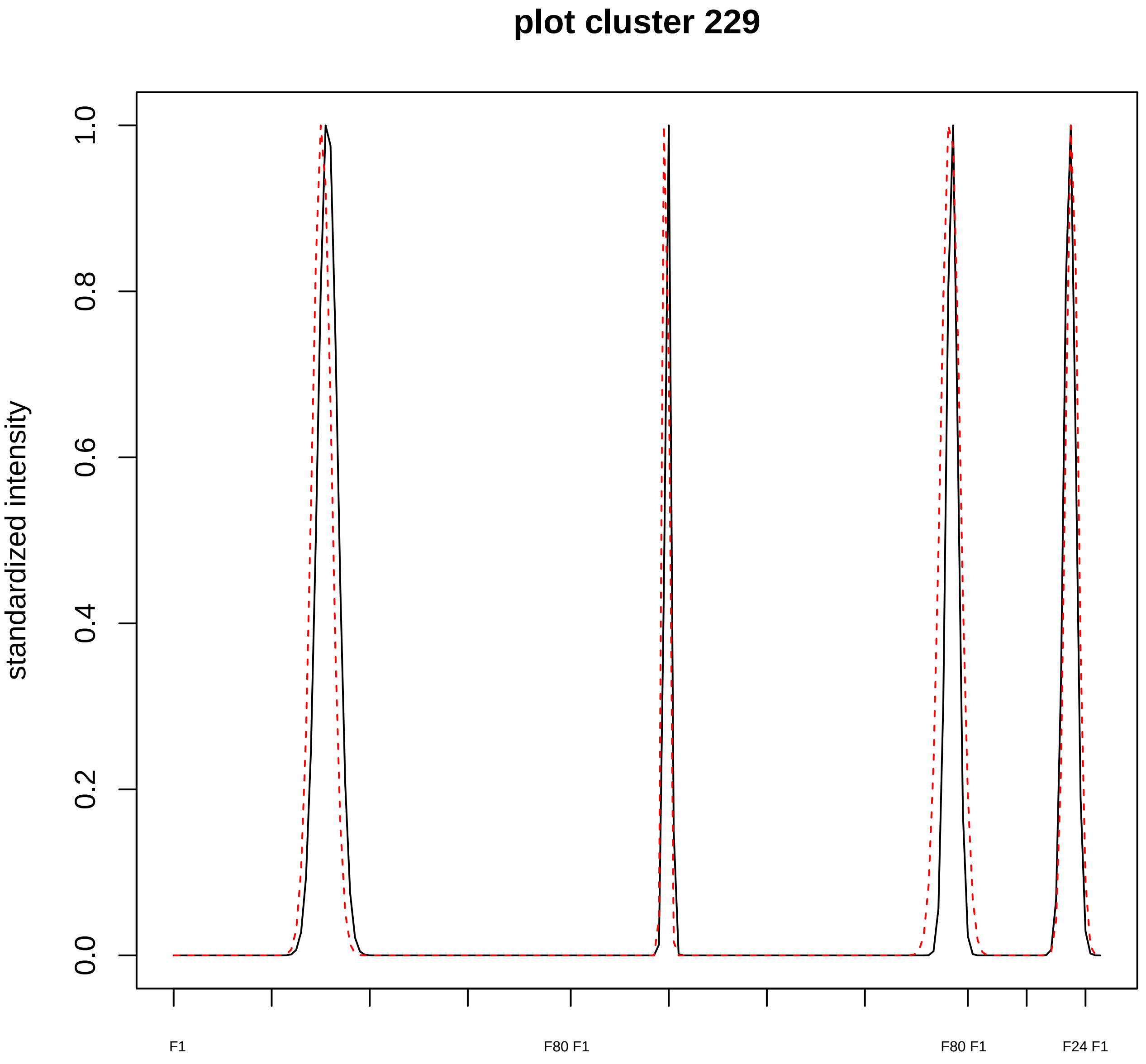

plot cluster 229
standardized intensity
0.0
0.2
0.4
0.6
0.8
1.0
F1
F80 F1
F80 F1
F24 F1

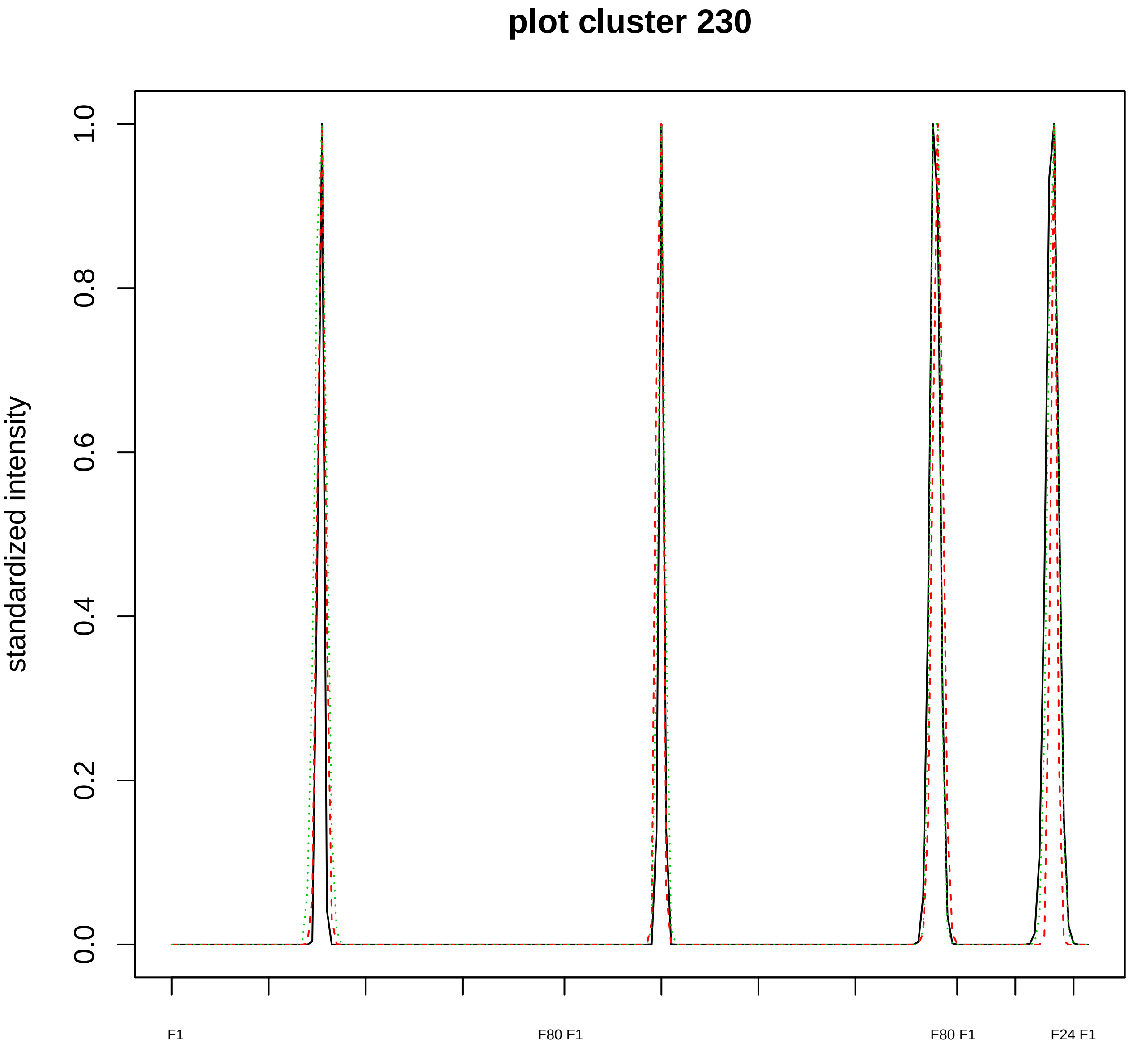
plot cluster 230
standardized intensity
0.0
0.2
0.4
0.6
0.8
1.0
F1
F80 F1
F80 F1
F24 F1

### Dendrogram of cluster 230

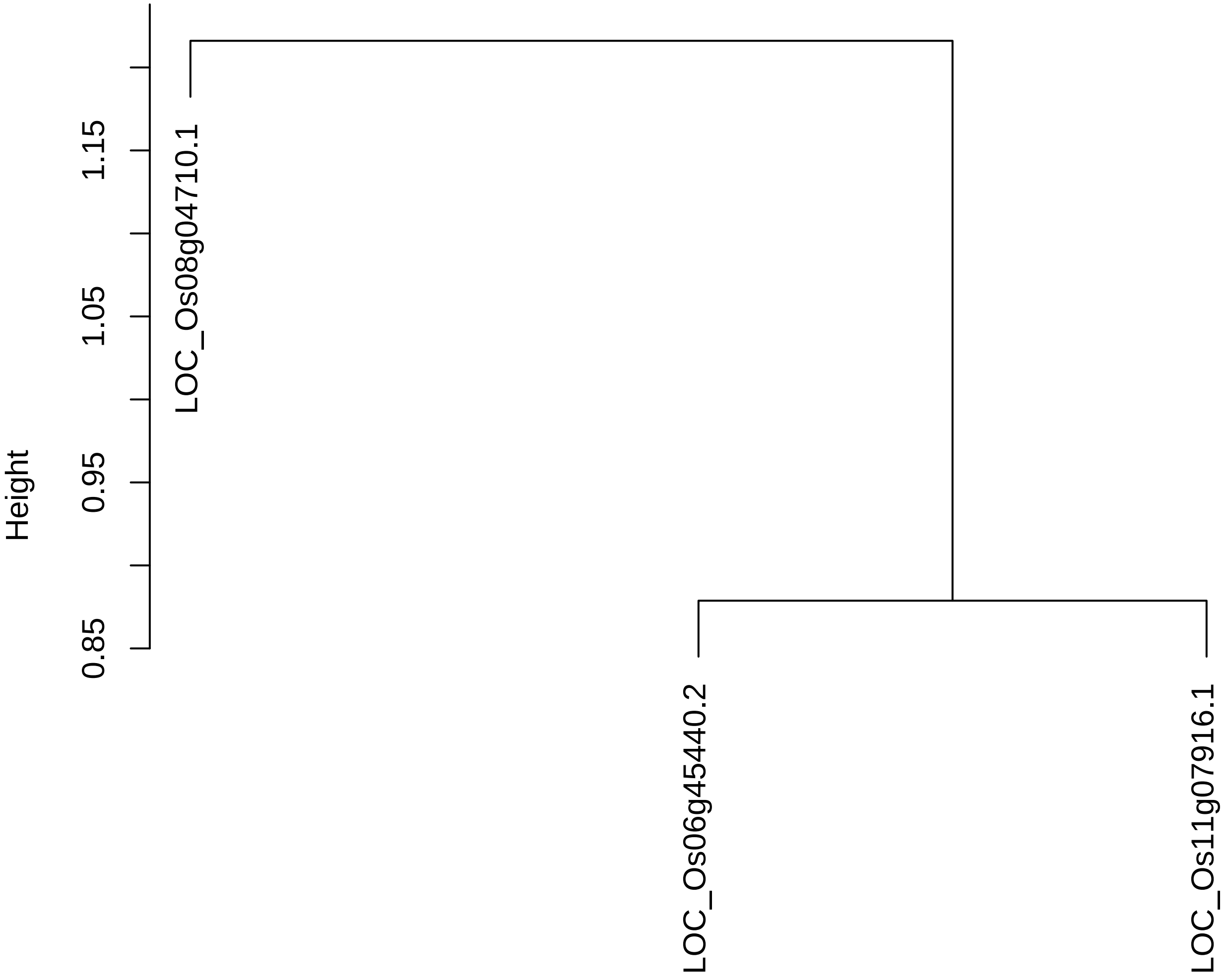


dist(spfitiex12sec12[lab[, 1] == i, ])
hclust (*, "ward.D2")

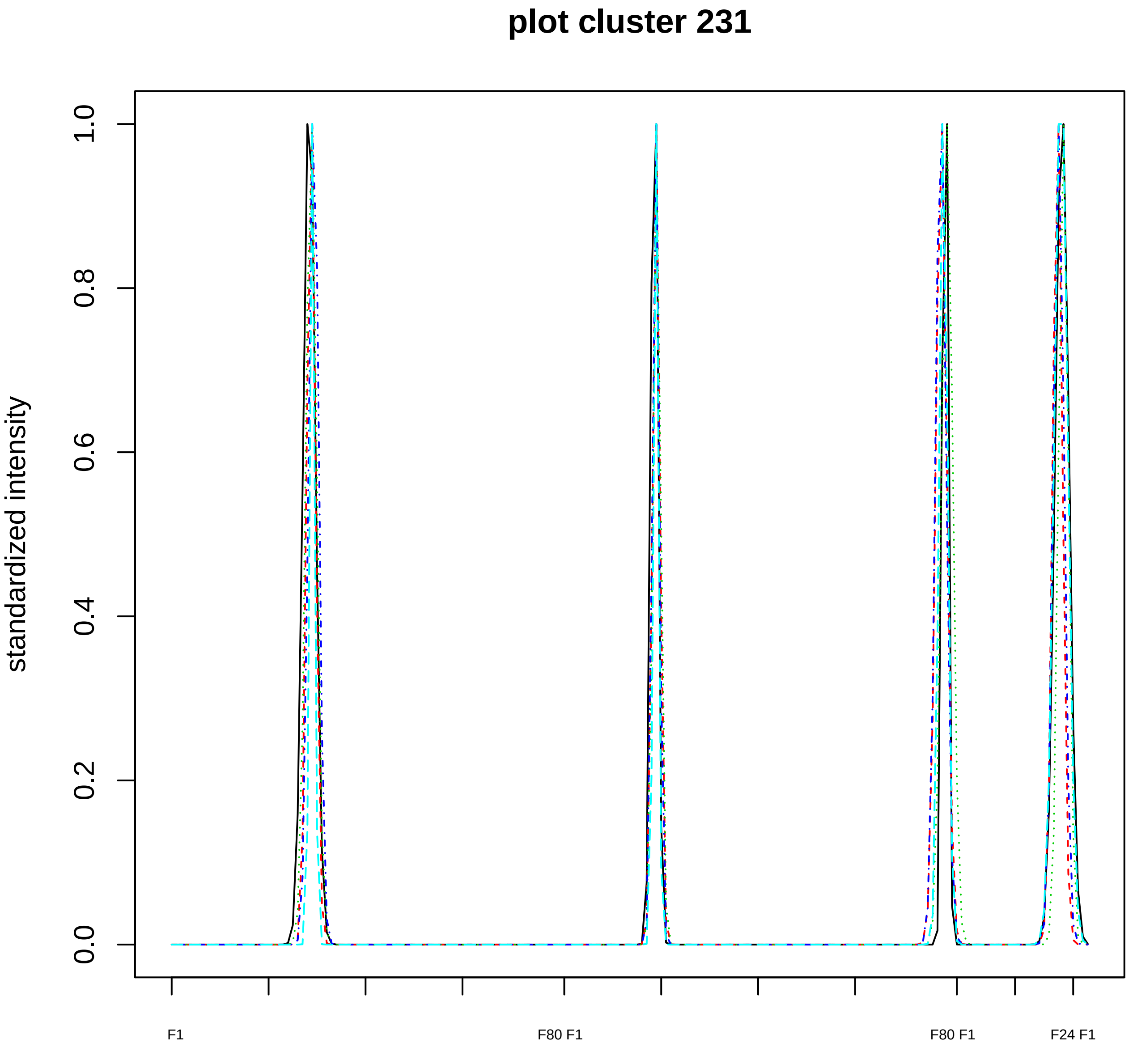
plot cluster 231
standardized intensity
0.0
0.2
0.4
0.6
0.8
1.0
F1
F80 F1
F80 F1
F24 F1

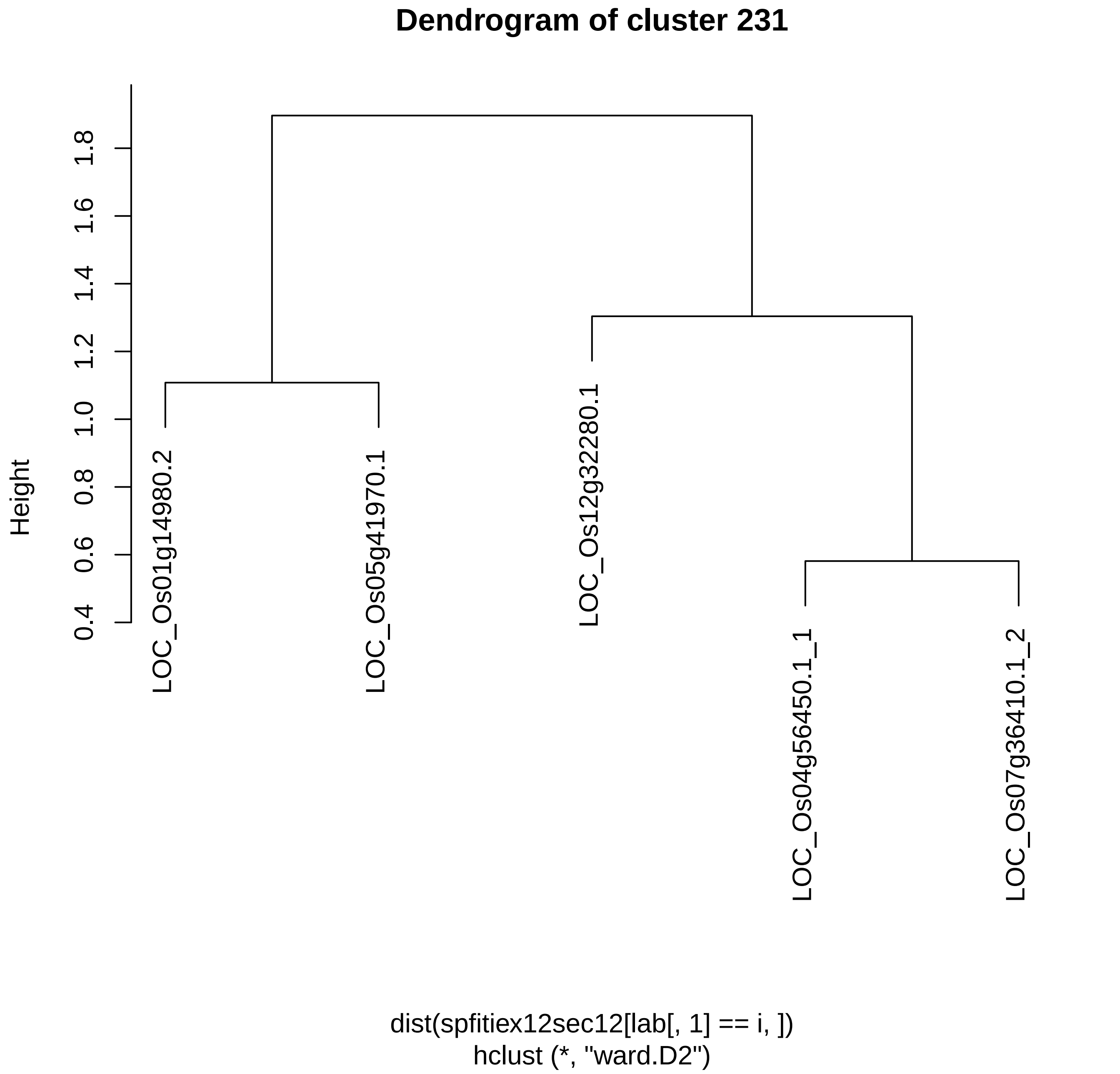

Dendrogram of cluster 231
Height
0.4
0.6
0.8
1.0
1.2
1.4
1.6
1.8
LOC_Os01g14980.2
LOC_Os05g41970.1
LOC_Os12g32280.1
LOC_Os04g56450.1_1
LOC_Os07g36410.1_2
dist(spfitiex12sec12[lab[, 1] == i, ])
hclust (*, "ward.D2")

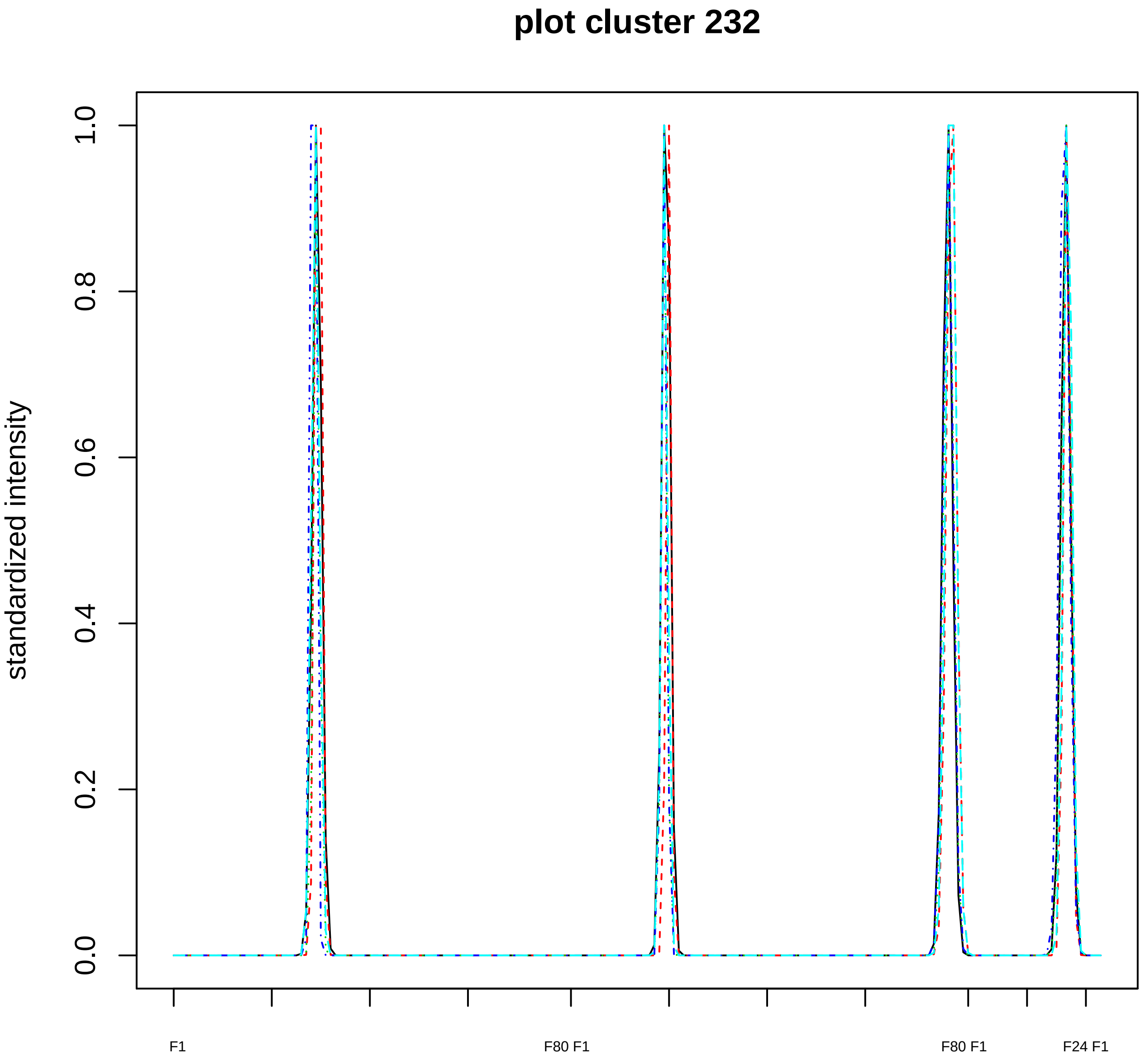

plot cluster 232
standardized intensity
1.0
0.8
0.6
0.4
0.2
0.0
F1
F80 F1
F80 F1
F24 F1

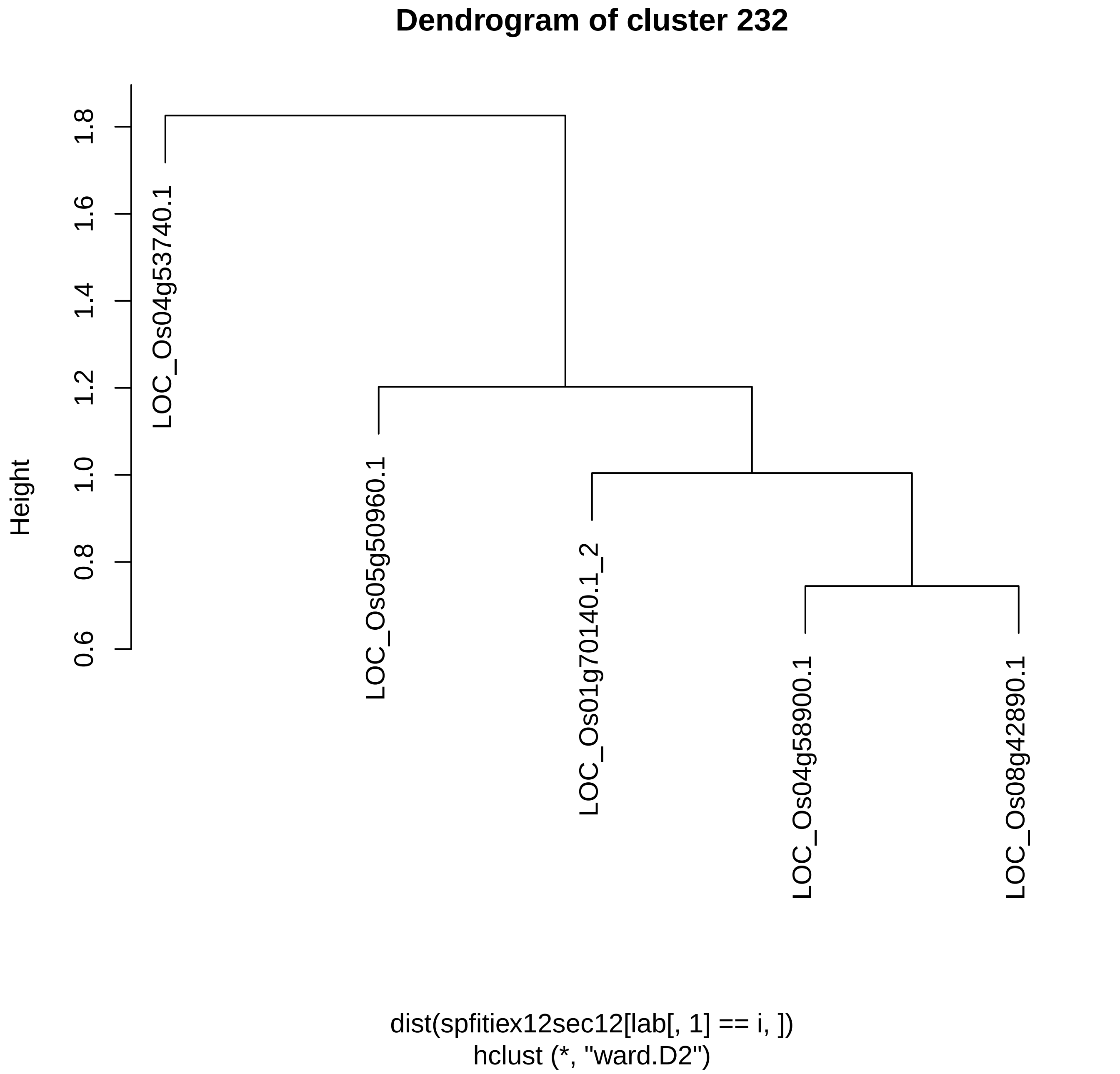

Dendrogram of cluster 232
Height
1.8
1.6
1.4
1.2
1.0
0.8
0.6
LOC_Os04g53740.1
LOC_Os05g50960.1
LOC_Os01g70140.1_2
LOC_Os04g58900.1
LOC_Os08g42890.1
dist(spfitiex12sec12[lab[, 1] == i, ])
hclust (*, "ward.D2")

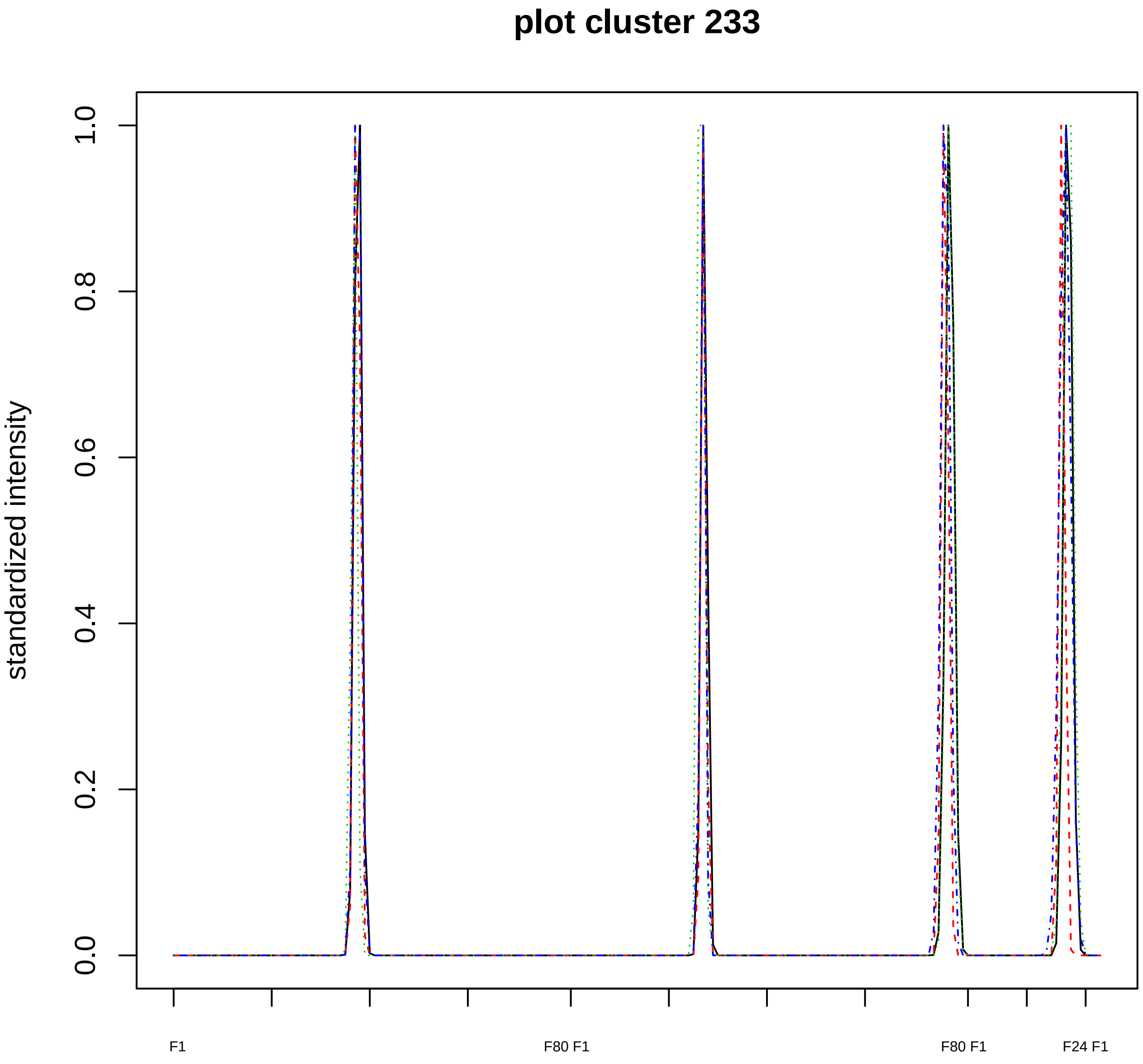
plot cluster 233
standardized intensity
0.0
0.2
0.4
0.6
0.8
1.0
F1
F80 F1
F80 F1
F24 F1

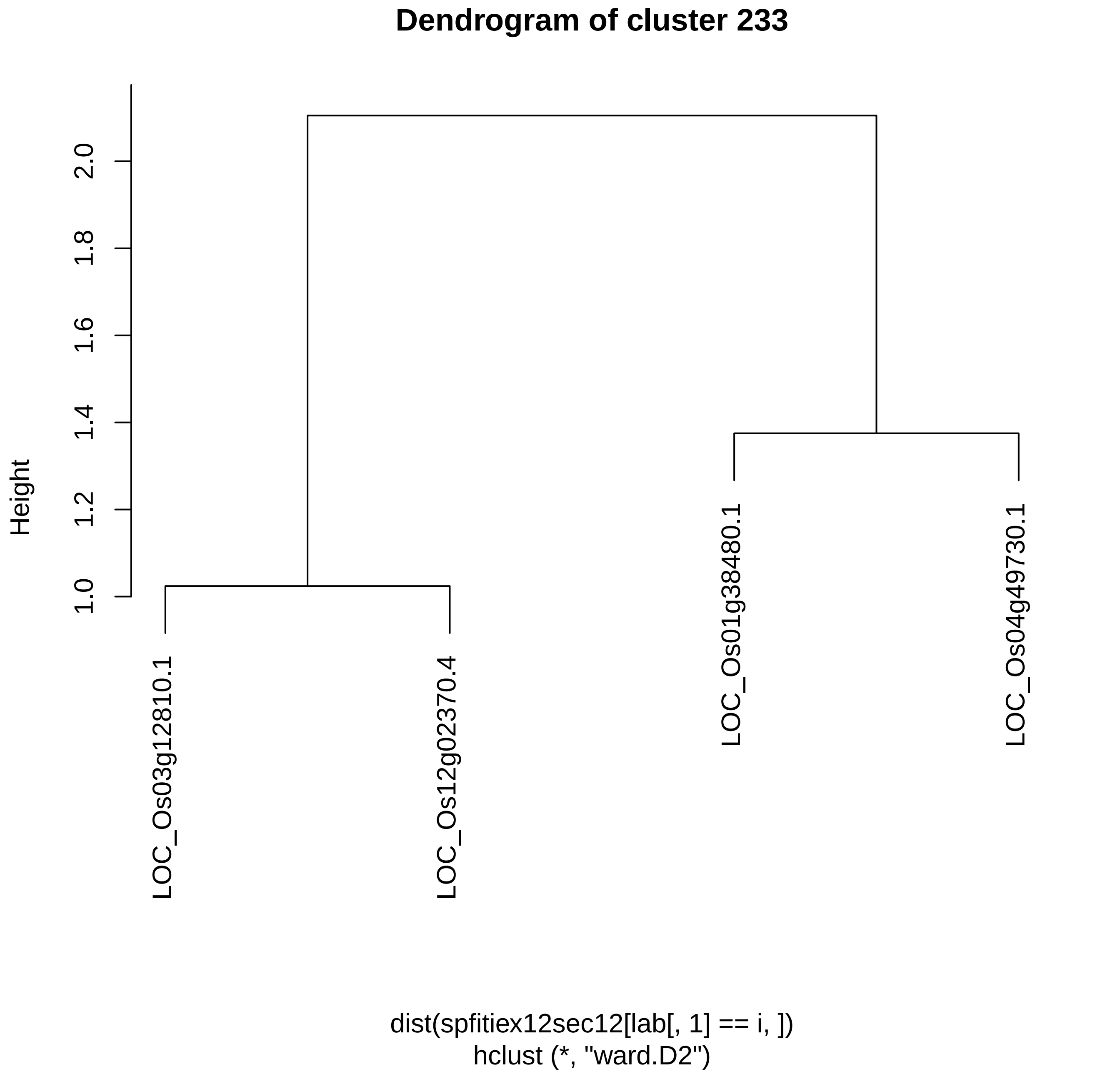
Dendrogram of cluster 233
Height
2.0
1.8
1.6
1.4
1.2
1.0
LOC_Os03g12810.1
LOC_Os12g02370.4
LOC_Os01g38480.1
LOC_Os04g49730.1
dist(spfitiex12sec12[lab[, 1] == i, ])
hclust (*, "ward.D2")

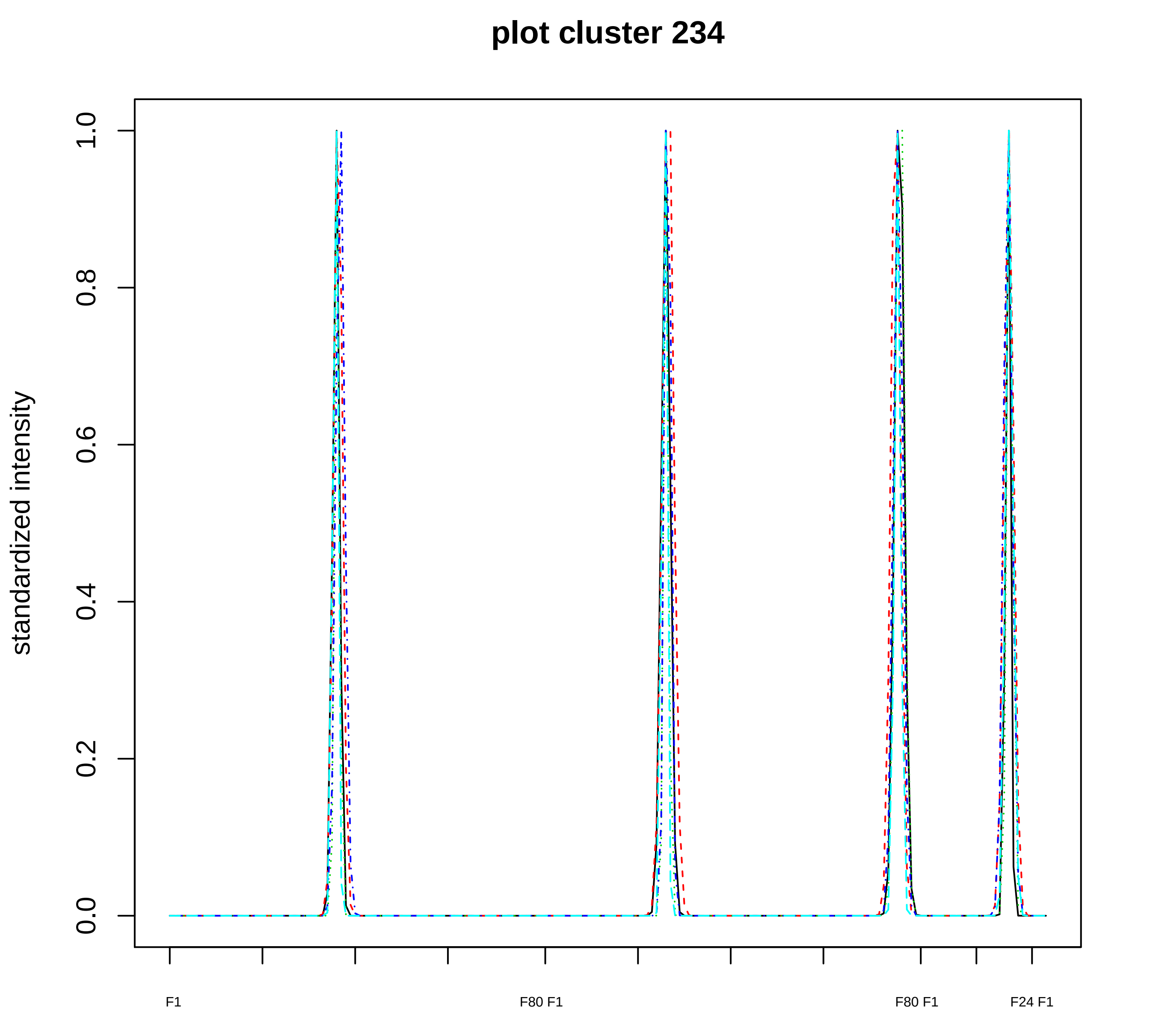
plot cluster 234
standardized intensity
0.0
0.2
0.4
0.6
0.8
1.0
F1
F80 F1
F80 F1
F24 F1

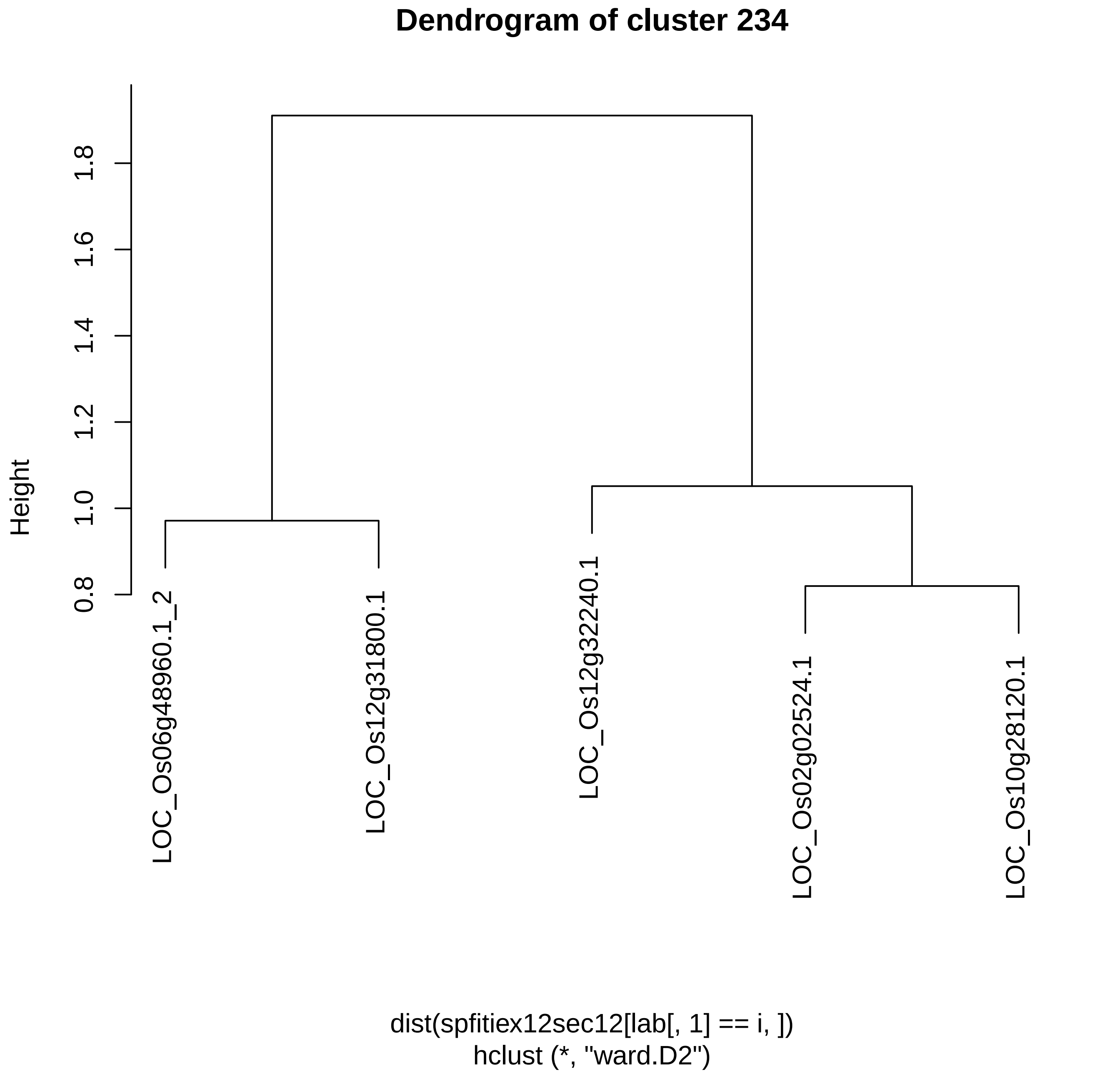

Dendrogram of cluster 234
Height
1.8
1.6
1.4
1.2
1.0
0.8
LOC_Os06g48960.1_2
LOC_Os12g31800.1
LOC_Os12g32240.1
LOC_Os02g02524.1
LOC_Os10g28120.1
dist(spfitiex12sec12[lab[, 1] == i, ])
hclust (*, "ward.D2")

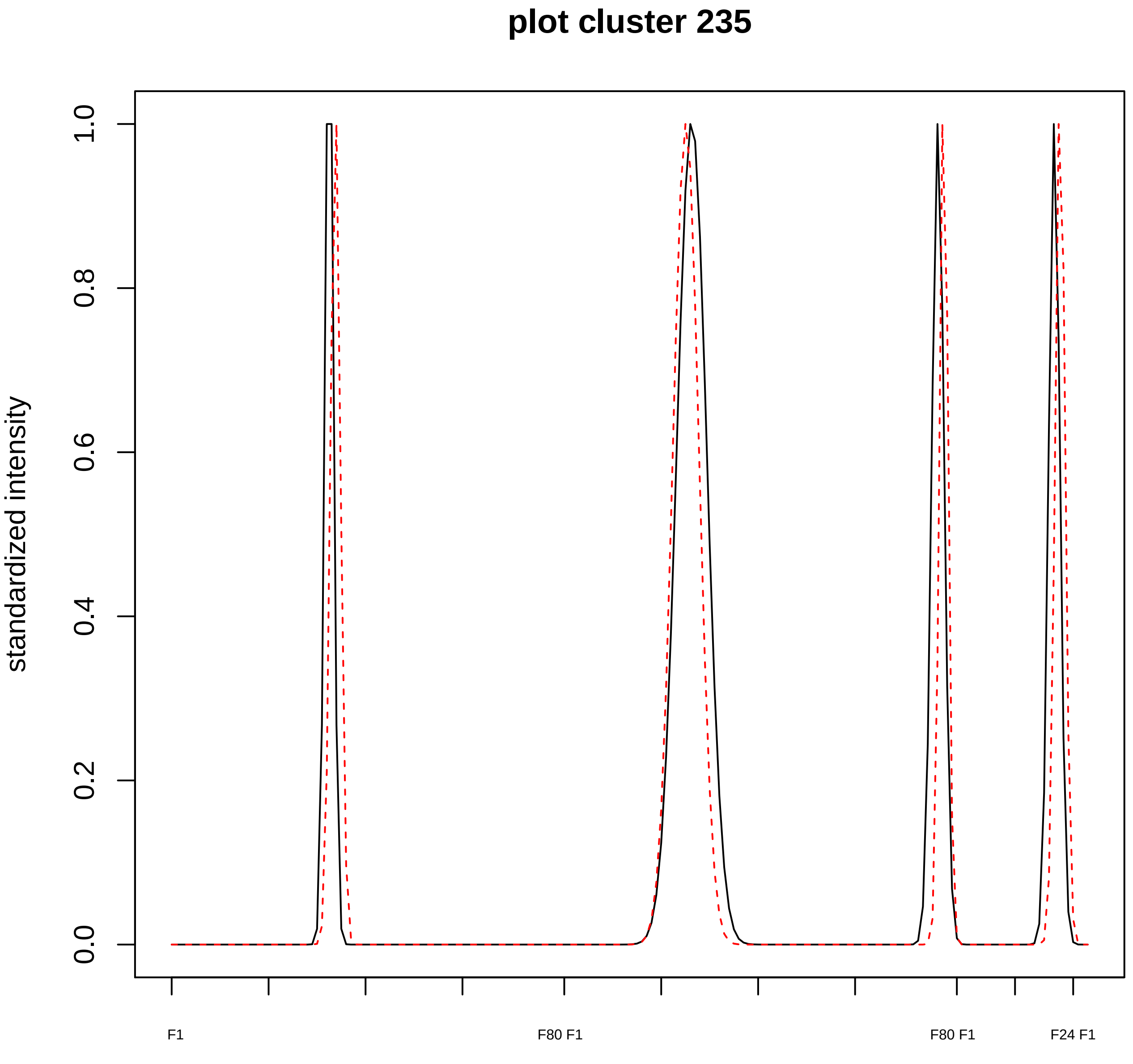
plot cluster 235
standardized intensity
0.0
0.2
0.4
0.6
0.8
1.0
F1
F80 F1
F80 F1
F24 F1

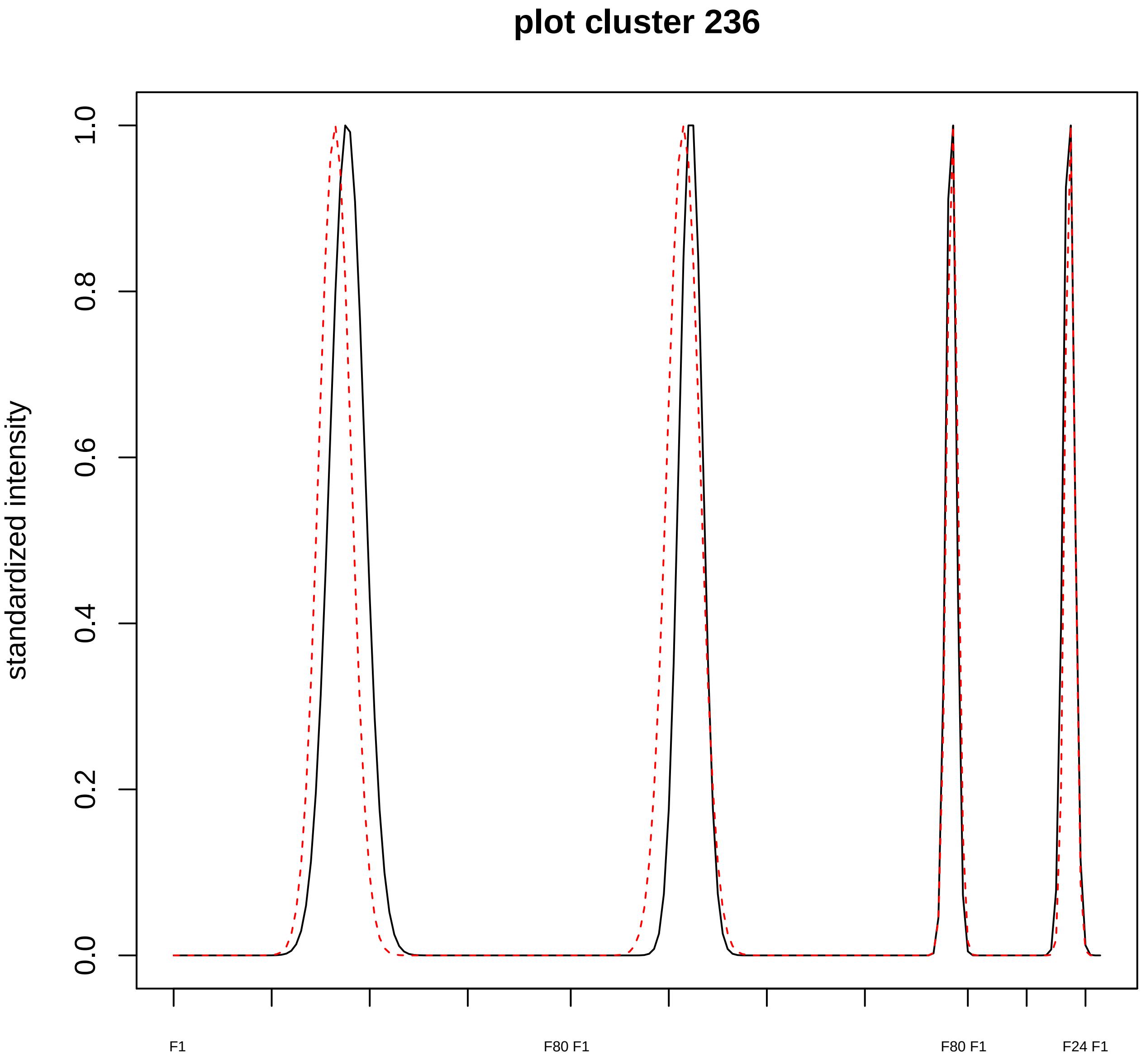

plot cluster 236
standardized intensity
0.0
0.2
0.4
0.6
0.8
1.0
F1
F80 F1
F80 F1
F24 F1

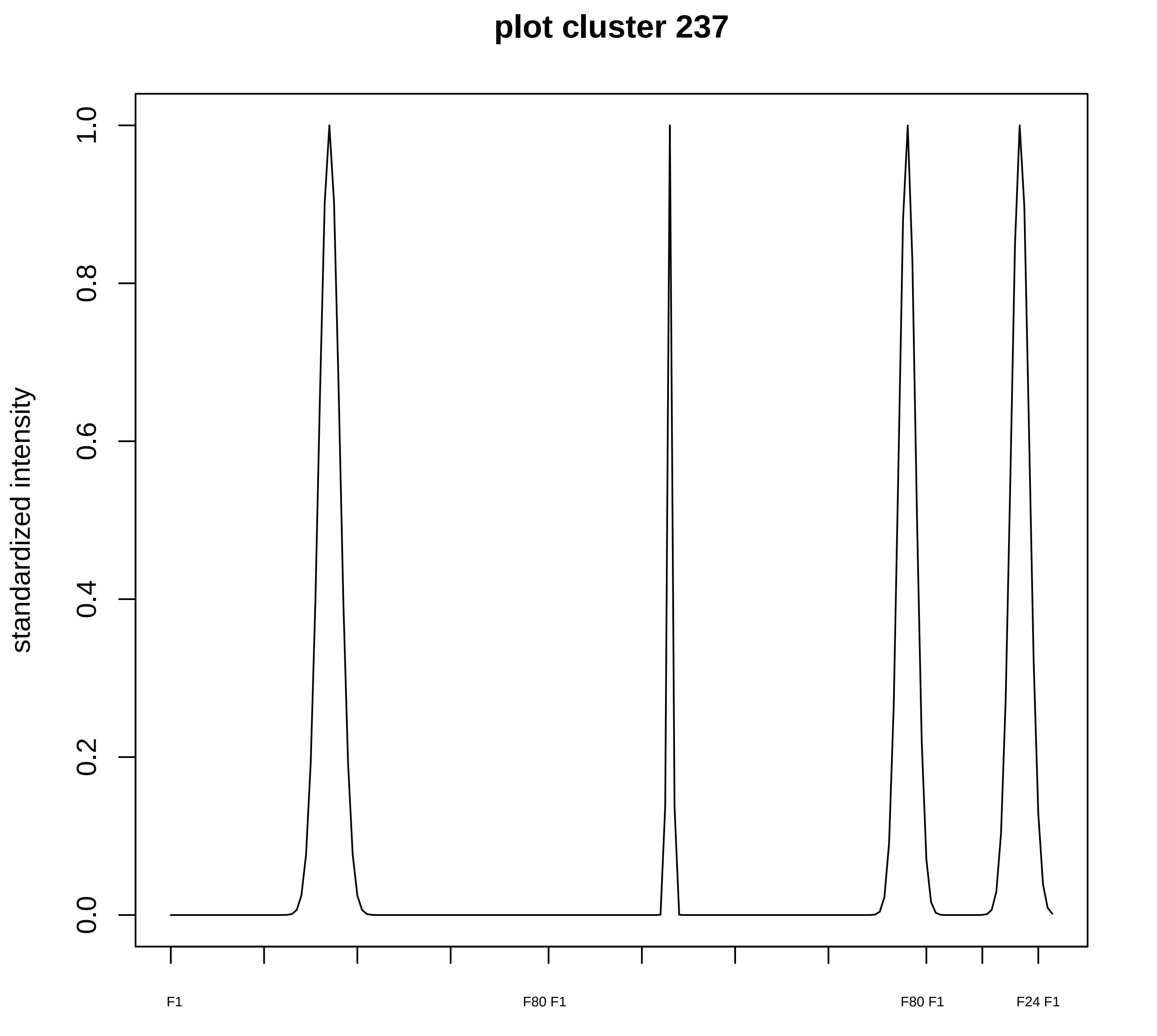
plot cluster 237
standardized intensity
0.0
0.2
0.4
0.6
0.8
1.0
F1
F80 F1
F80 F1
F24 F1

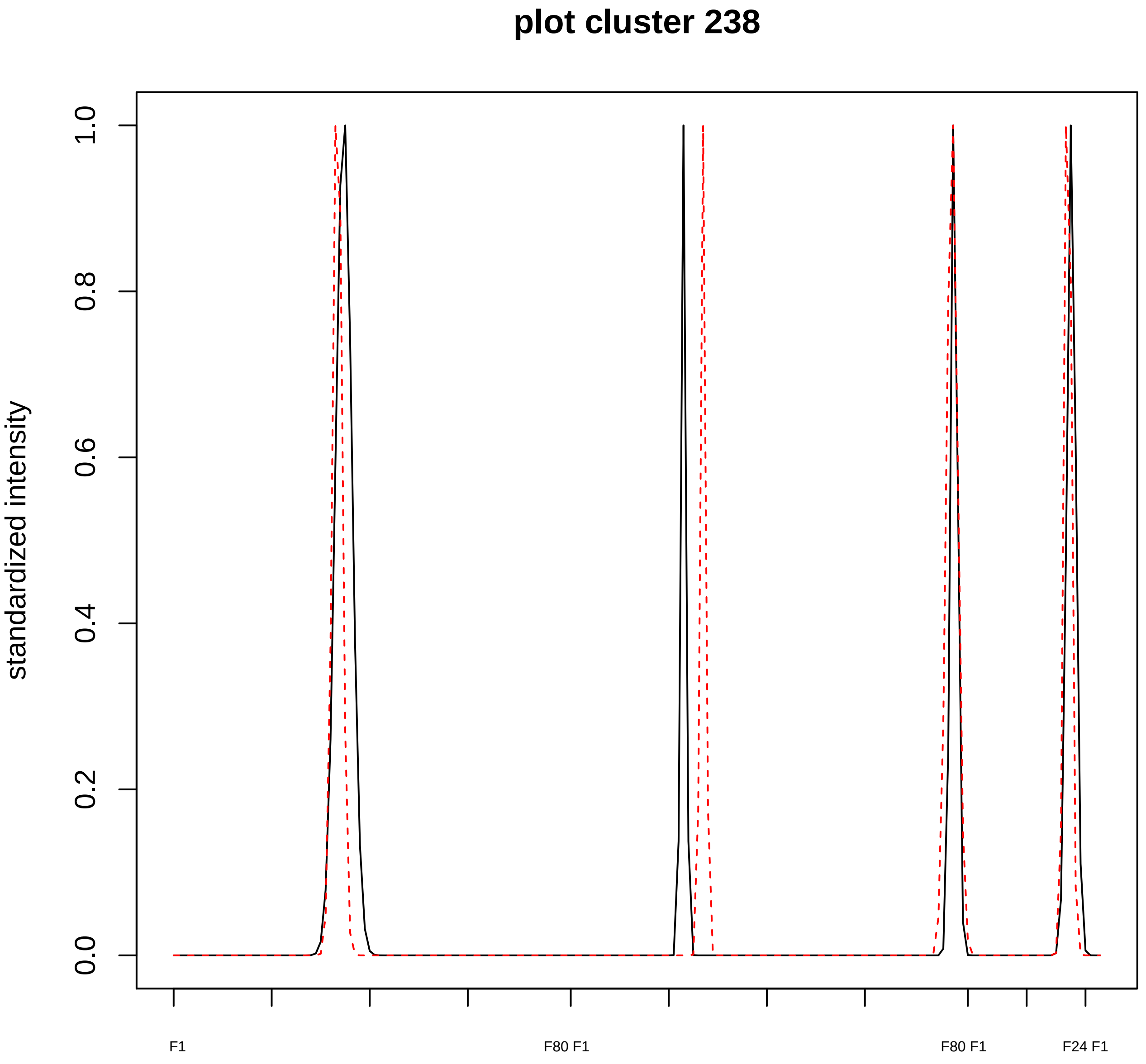
plot cluster 238
standardized intensity
0.0
0.2
0.4
0.6
0.8
1.0
F1
F80 F1
F80 F1
F24 F1

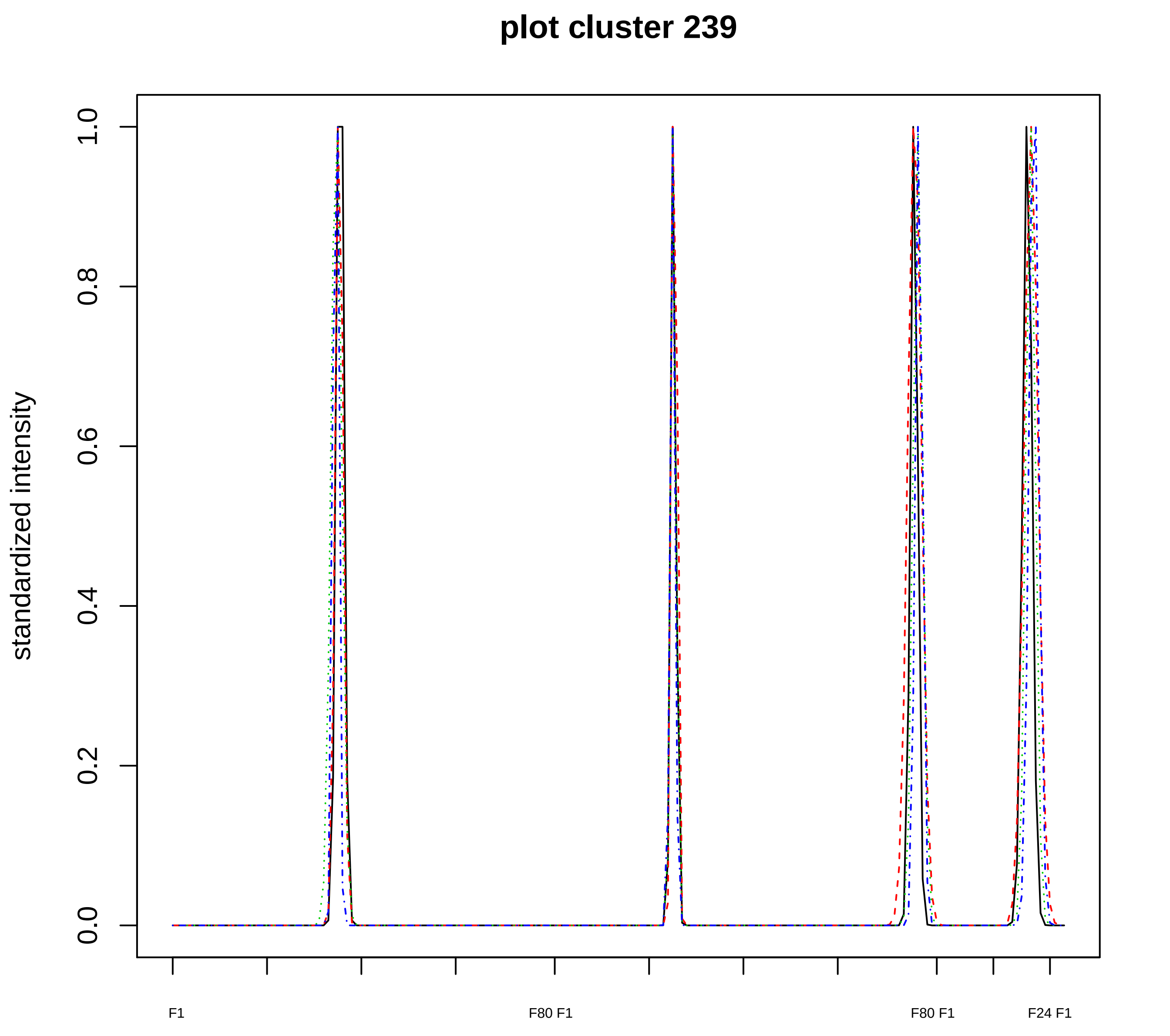

plot cluster 239
standardized intensity
0.0
0.2
0.4
0.6
0.8
1.0
F1
F80 F1
F80 F1
F24 F1

### Dendrogram of cluster 239

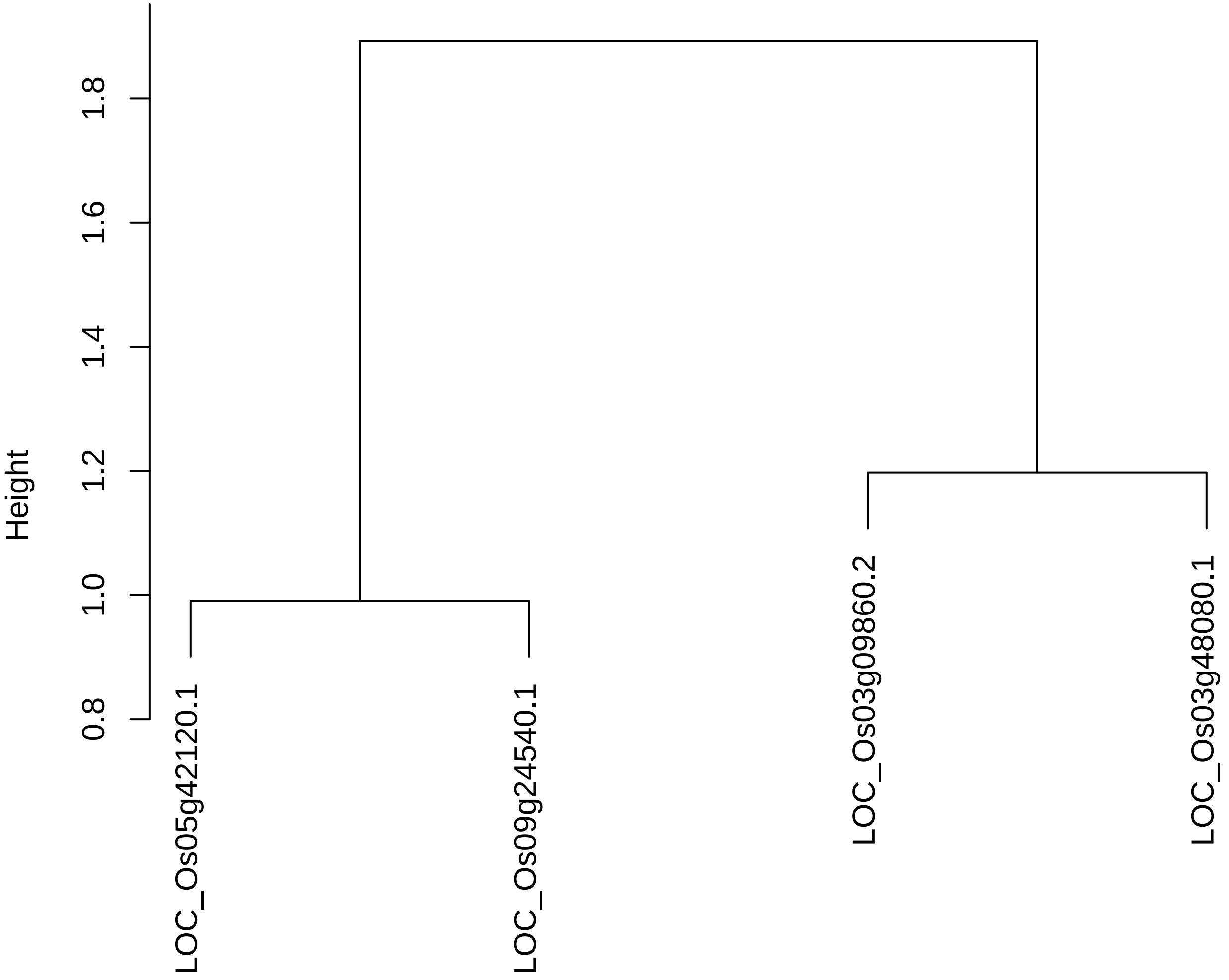


dist(spfitiex12sec12[lab[, 1] == i, ])
hclust (*, "ward.D2")

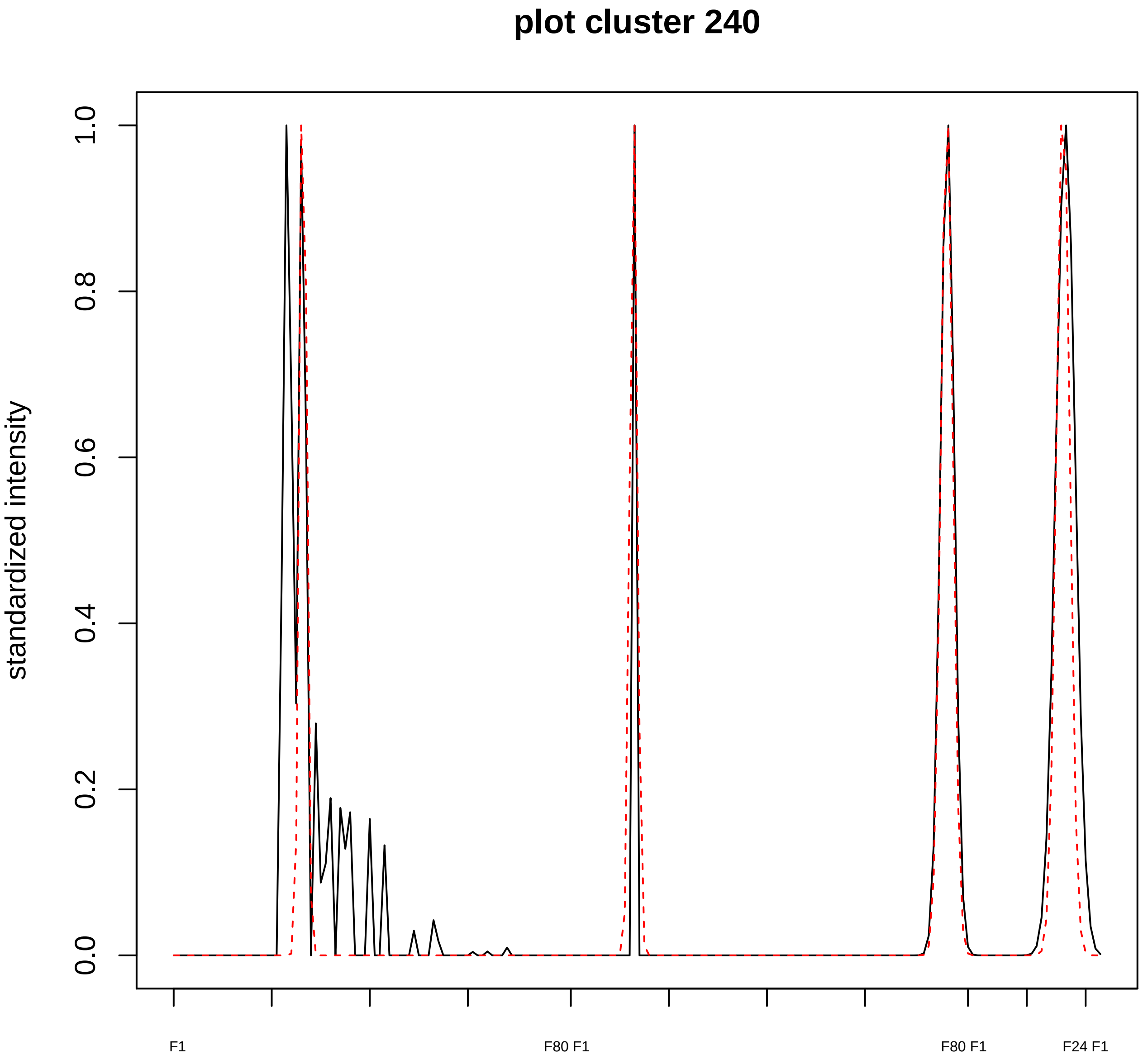

plot cluster 240
standardized intensity
0.0
0.2
0.4
0.6
0.8
1.0
F1
F80 F1
F80 F1
F24 F1

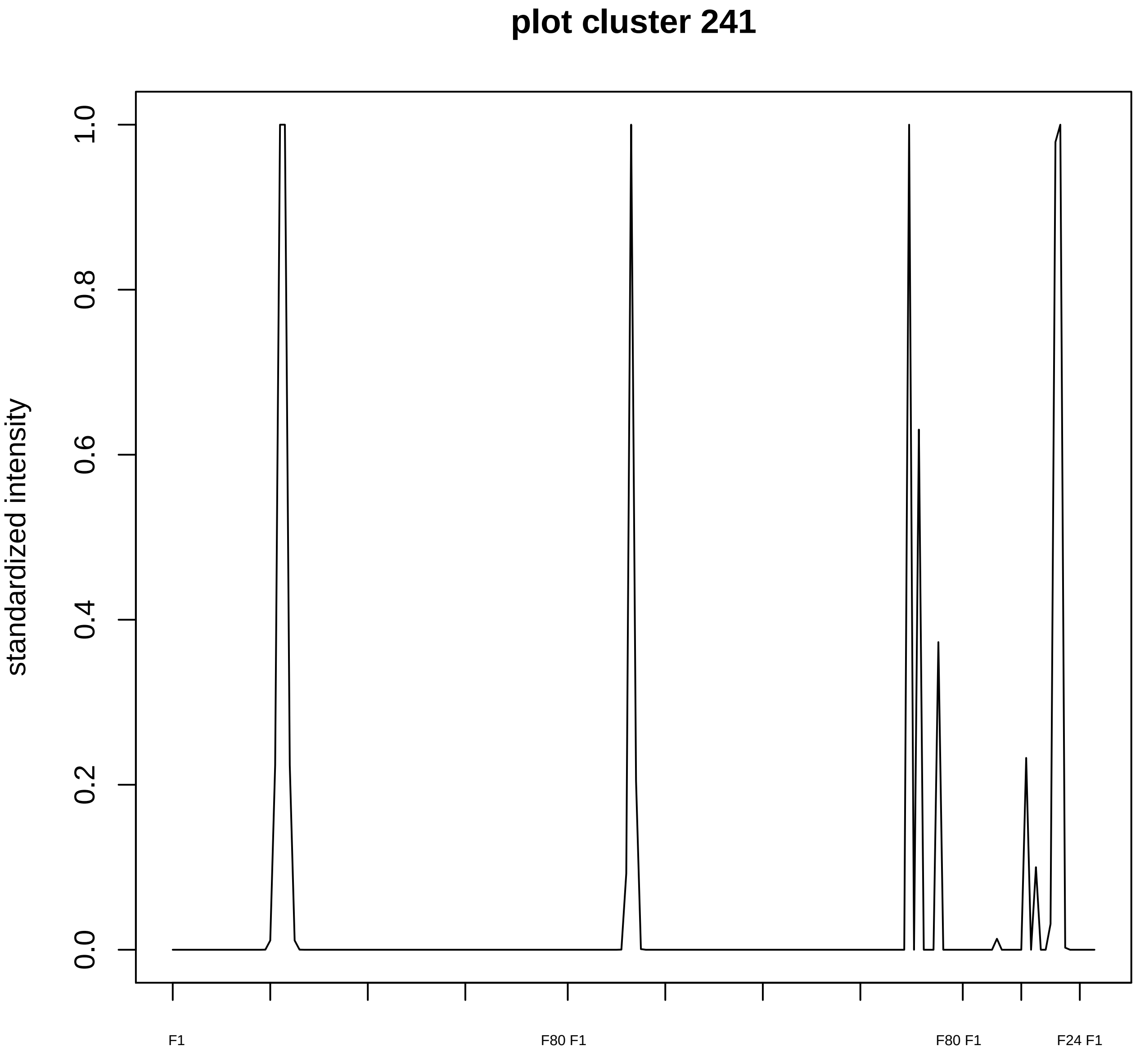
plot cluster 241
standardized intensity
0.0
0.2
0.4
0.6
0.8
1.0
F1
F80 F1
F80 F1
F24 F1

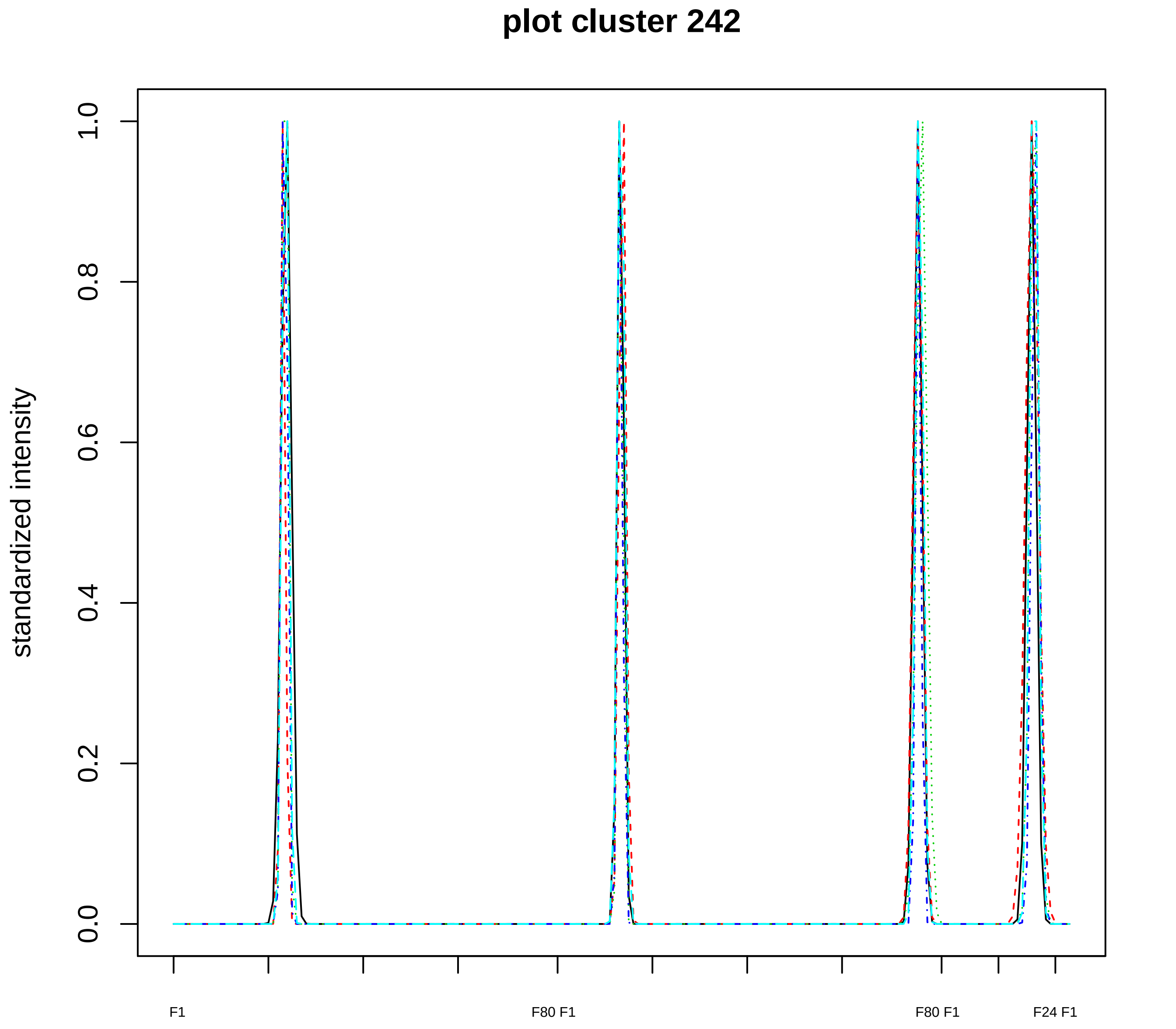
plot cluster 242
standardized intensity
0.0
0.2
0.4
0.6
0.8
1.0
F1
F80 F1
F80 F1
F24 F1

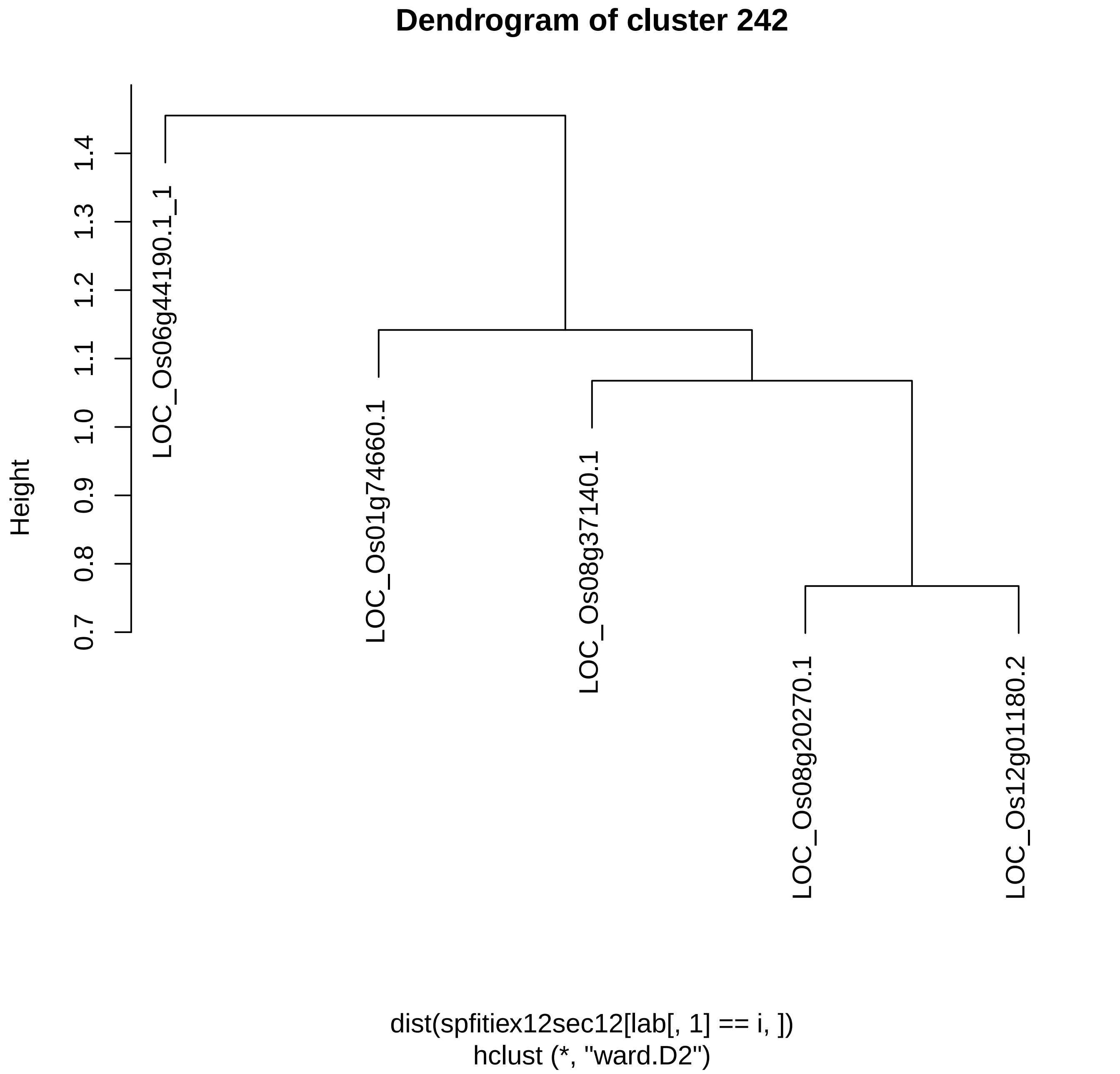
Dendrogram of cluster 242
Height
0.7
0.8
0.9
1.0
1.1
1.2
1.3
1.4
LOC_Os06g44190.1_1
LOC_Os01g74660.1
LOC_Os08g37140.1
LOC_Os08g20270.1
LOC_Os12g01180.2
dist(spfitiex12sec12[lab[, 1] == i, ])
hclust (*, "ward.D2")

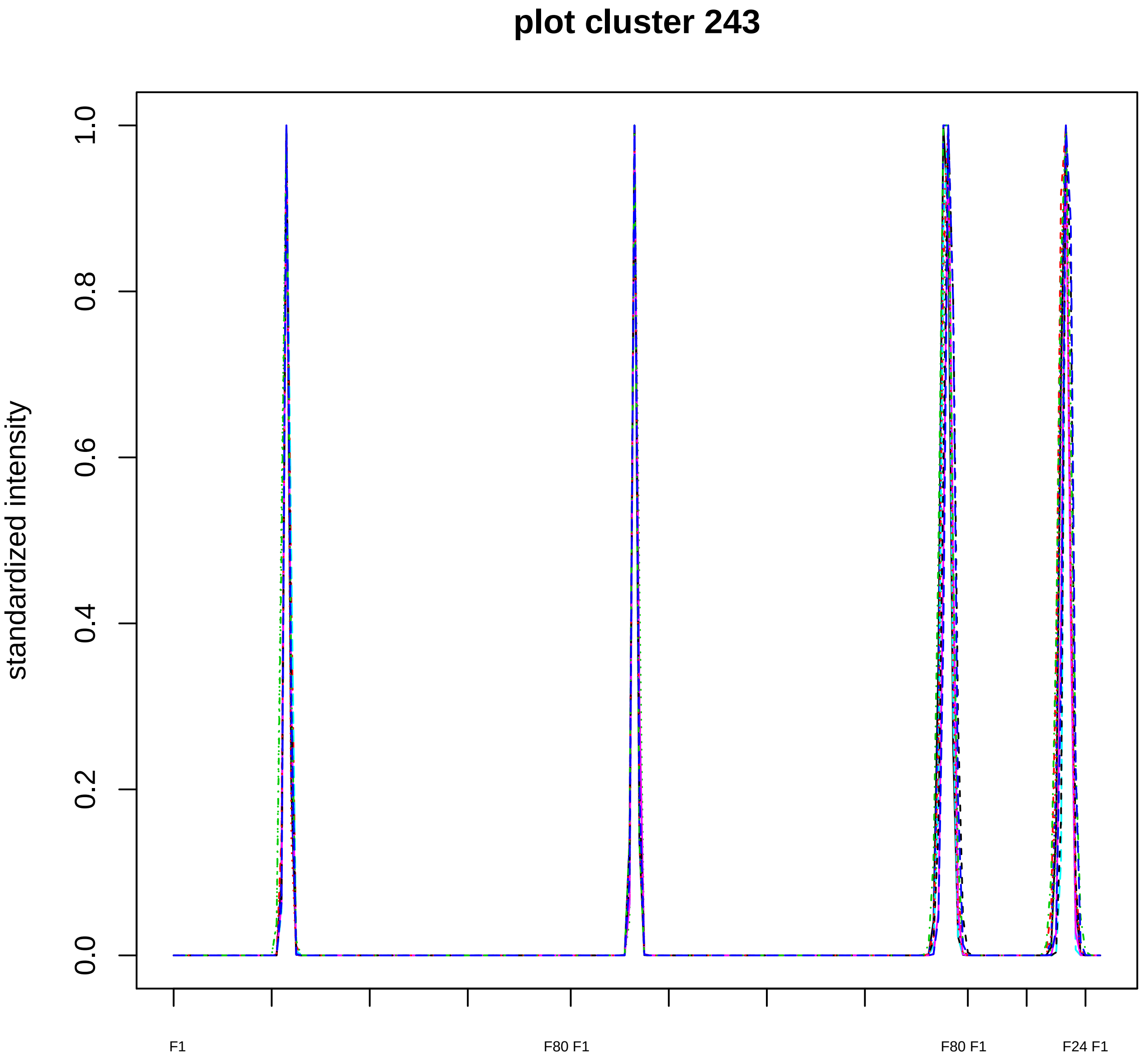
plot cluster 243
standardized intensity
0.0
0.2
0.4
0.6
0.8
1.0
F1
F80 F1
F80 F1
F24 F1

#### Dendrogram of cluster 243

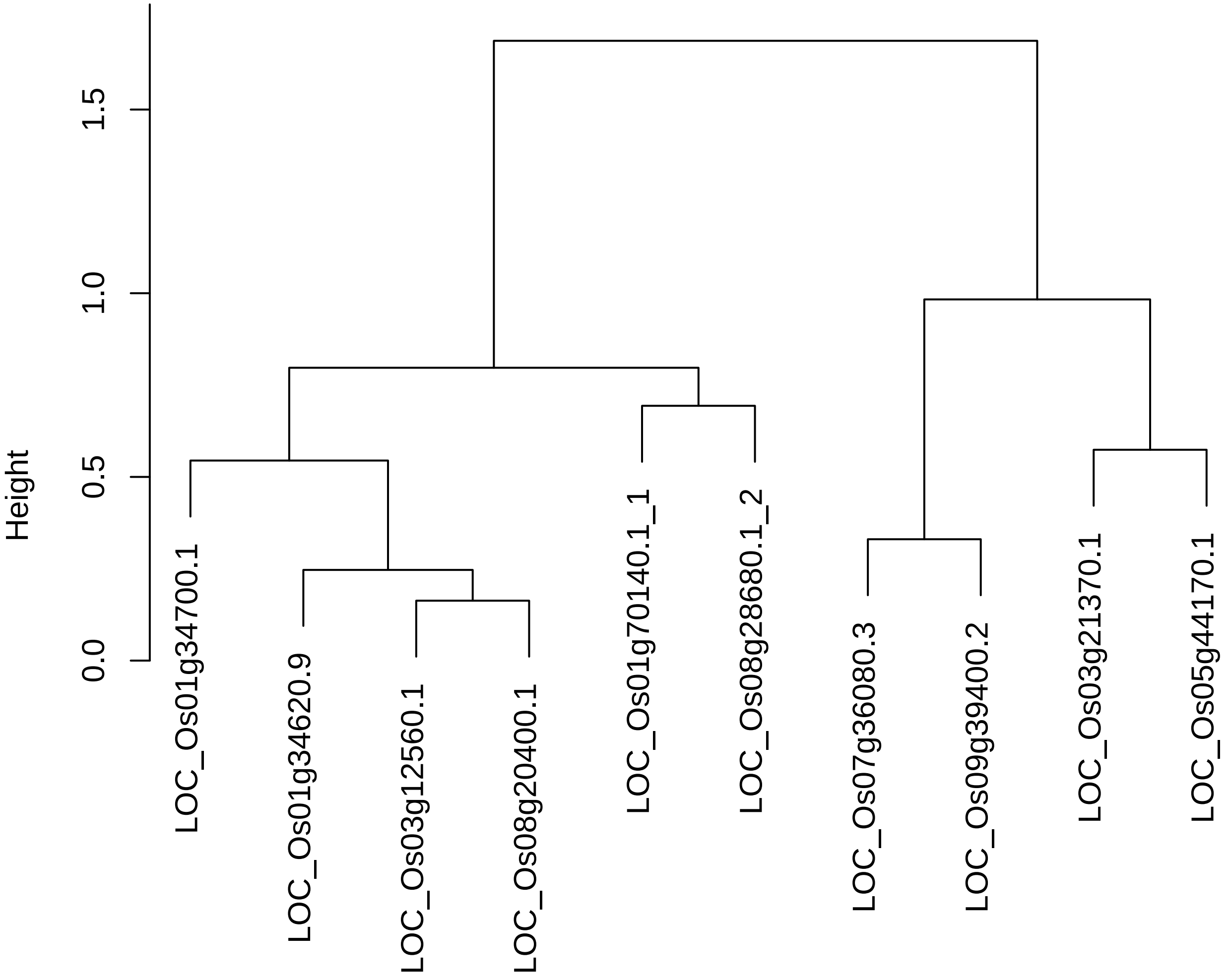


dist(spfitiex12sec12[lab[, 1] == i, ])
hclust (*, "ward.D2")

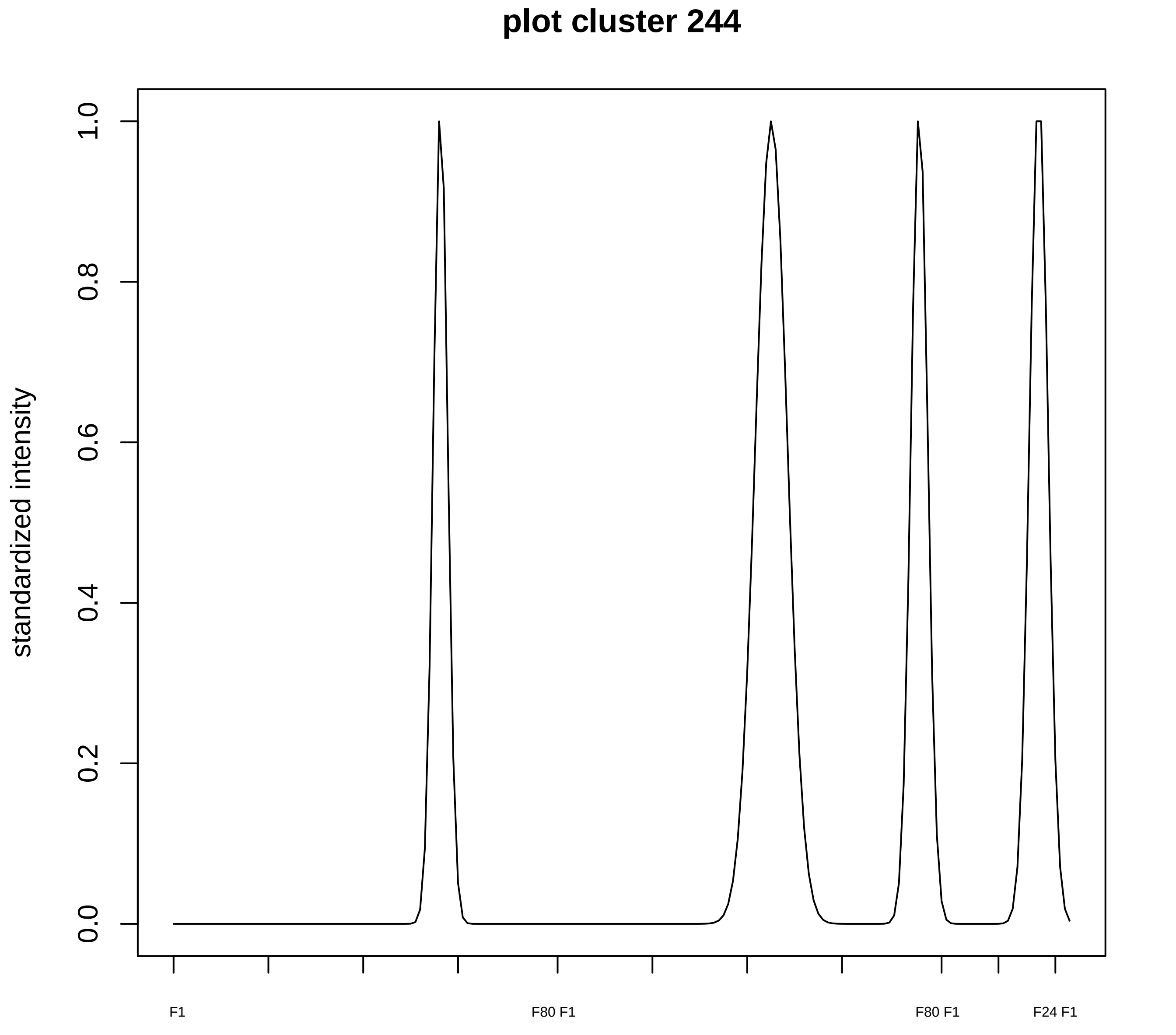
plot cluster 244
standardized intensity
0.0
0.2
0.4
0.6
0.8
1.0
F1
F80 F1
F80 F1
F24 F1

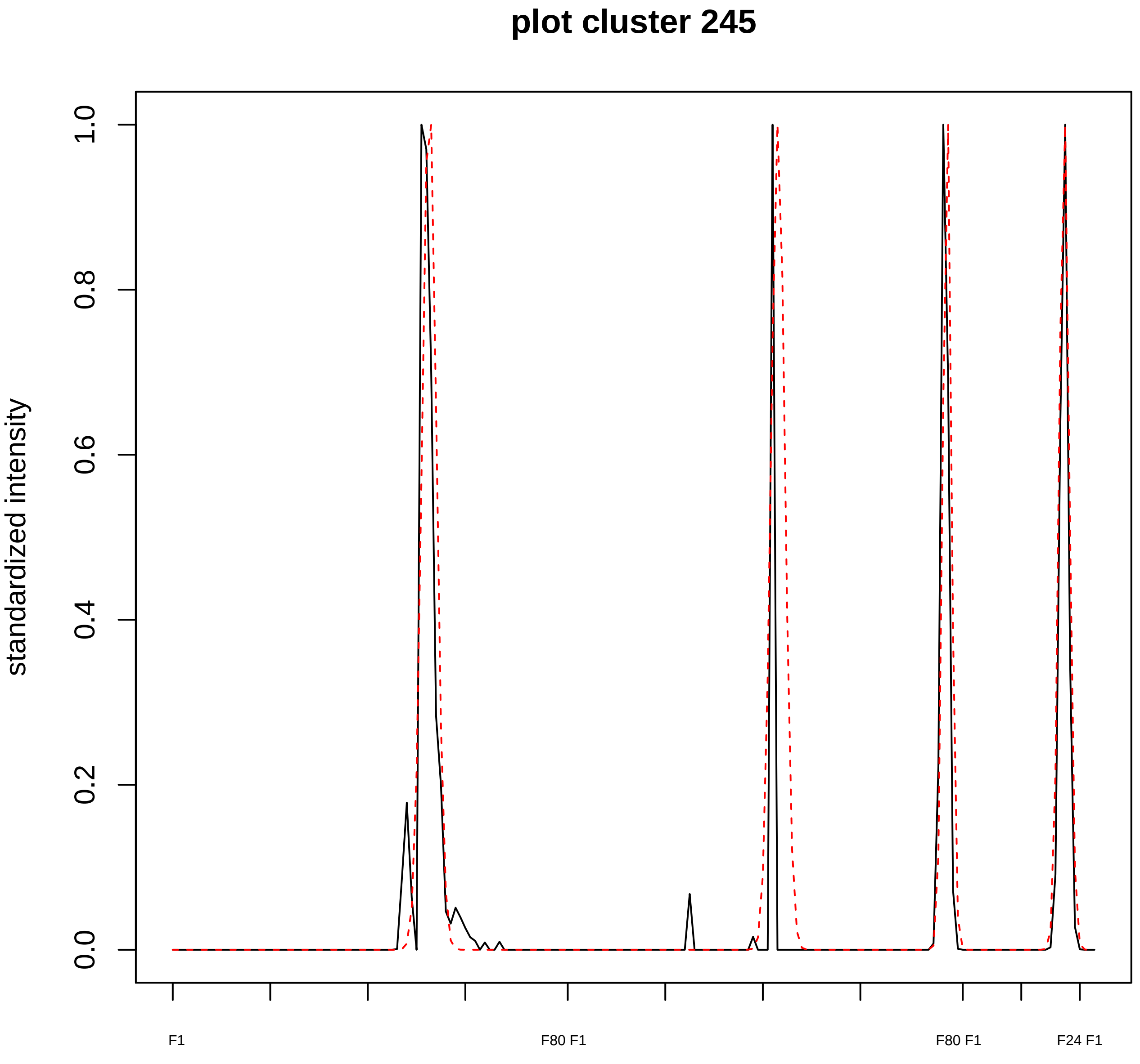

plot cluster 245
standardized intensity
0.0
0.2
0.4
0.6
0.8
1.0
F1
F80 F1
F80 F1
F24 F1

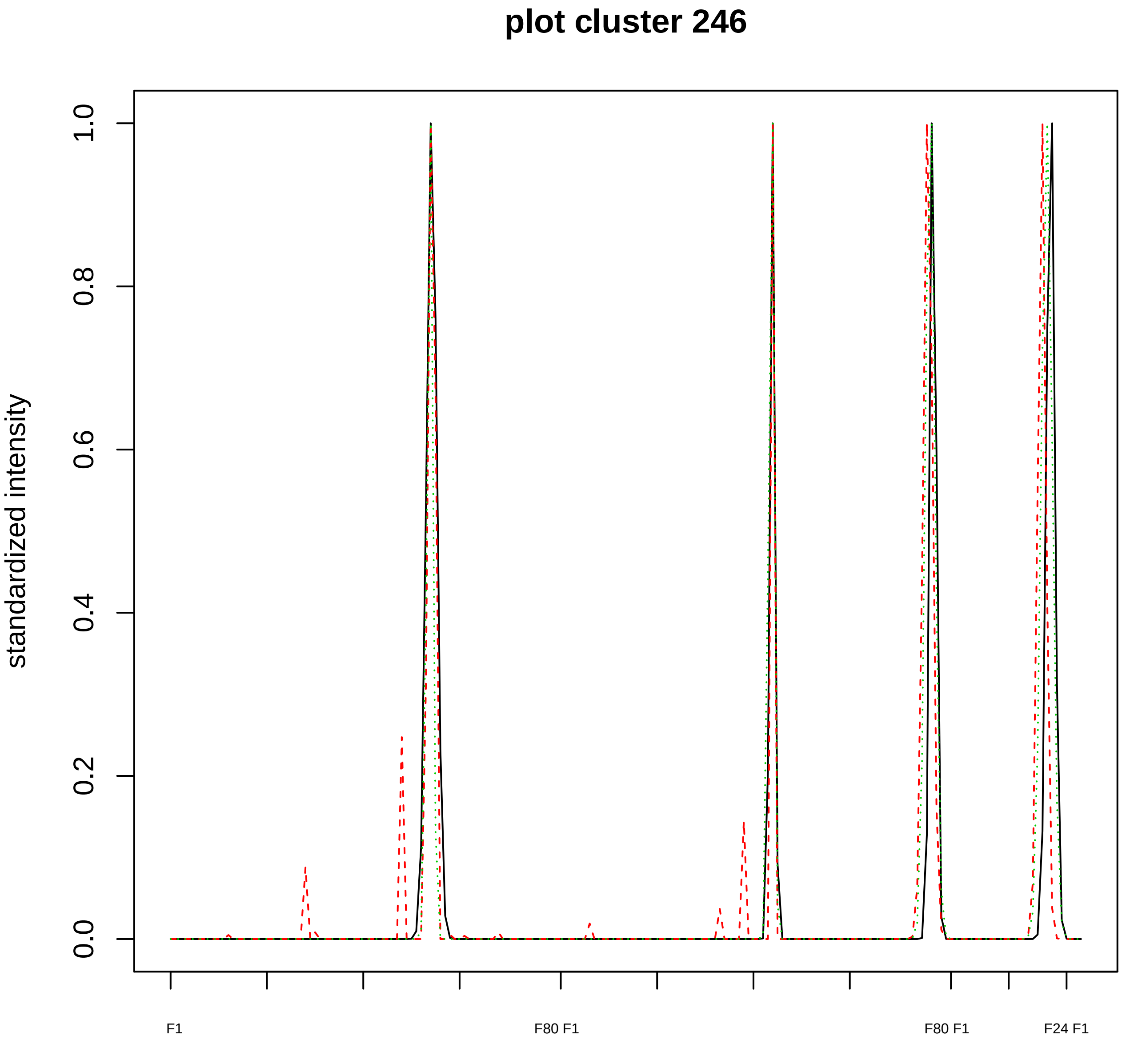
plot cluster 246
standardized intensity
0.0
0.2
0.4
0.6
0.8
1.0
F1
F80 F1
F80 F1
F24 F1

### Dendrogram of cluster 246

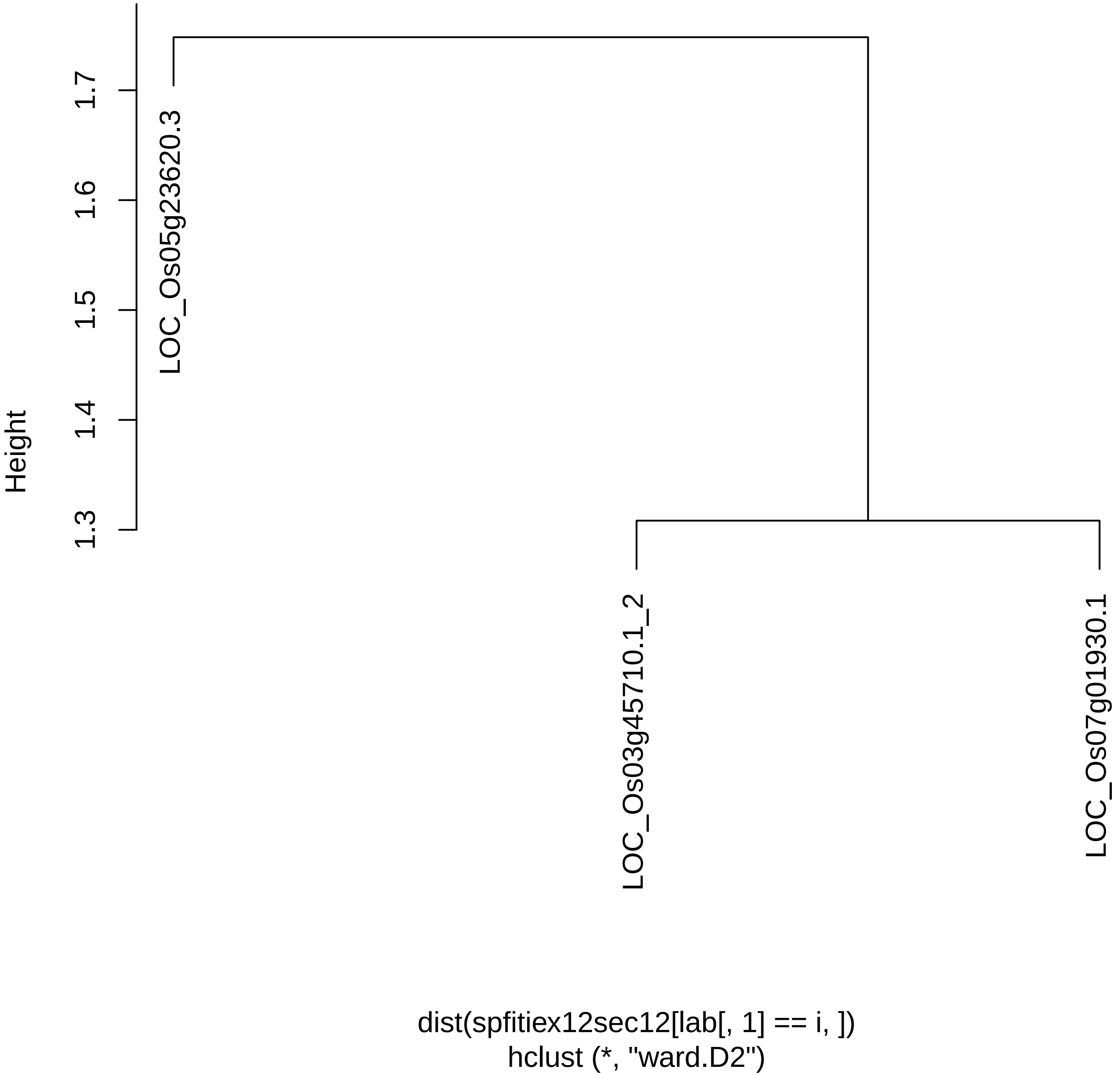

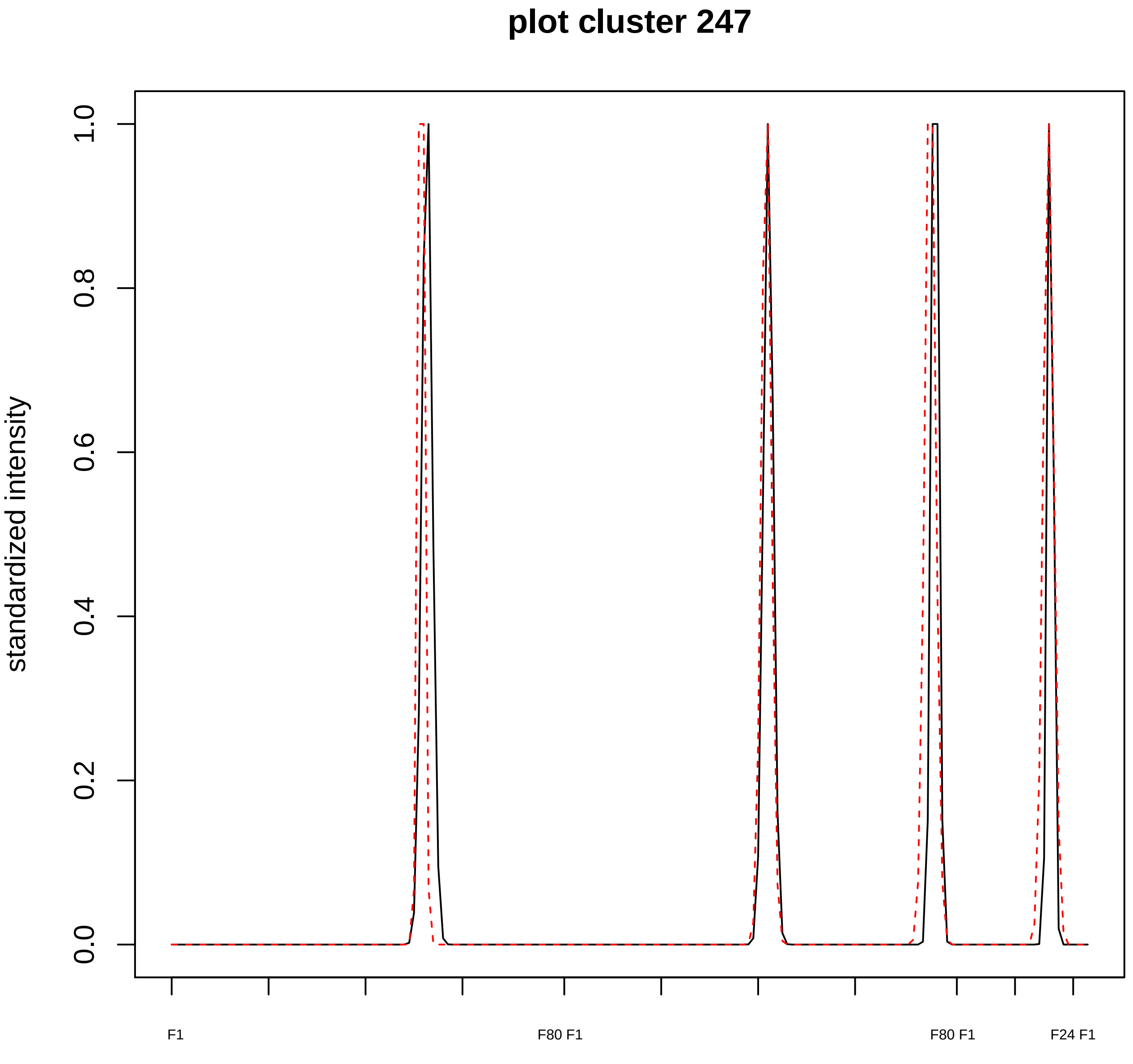

plot cluster 247
standardized intensity
0.0
0.2
0.4
0.6
0.8
1.0
F1
F80 F1
F80 F1
F24 F1

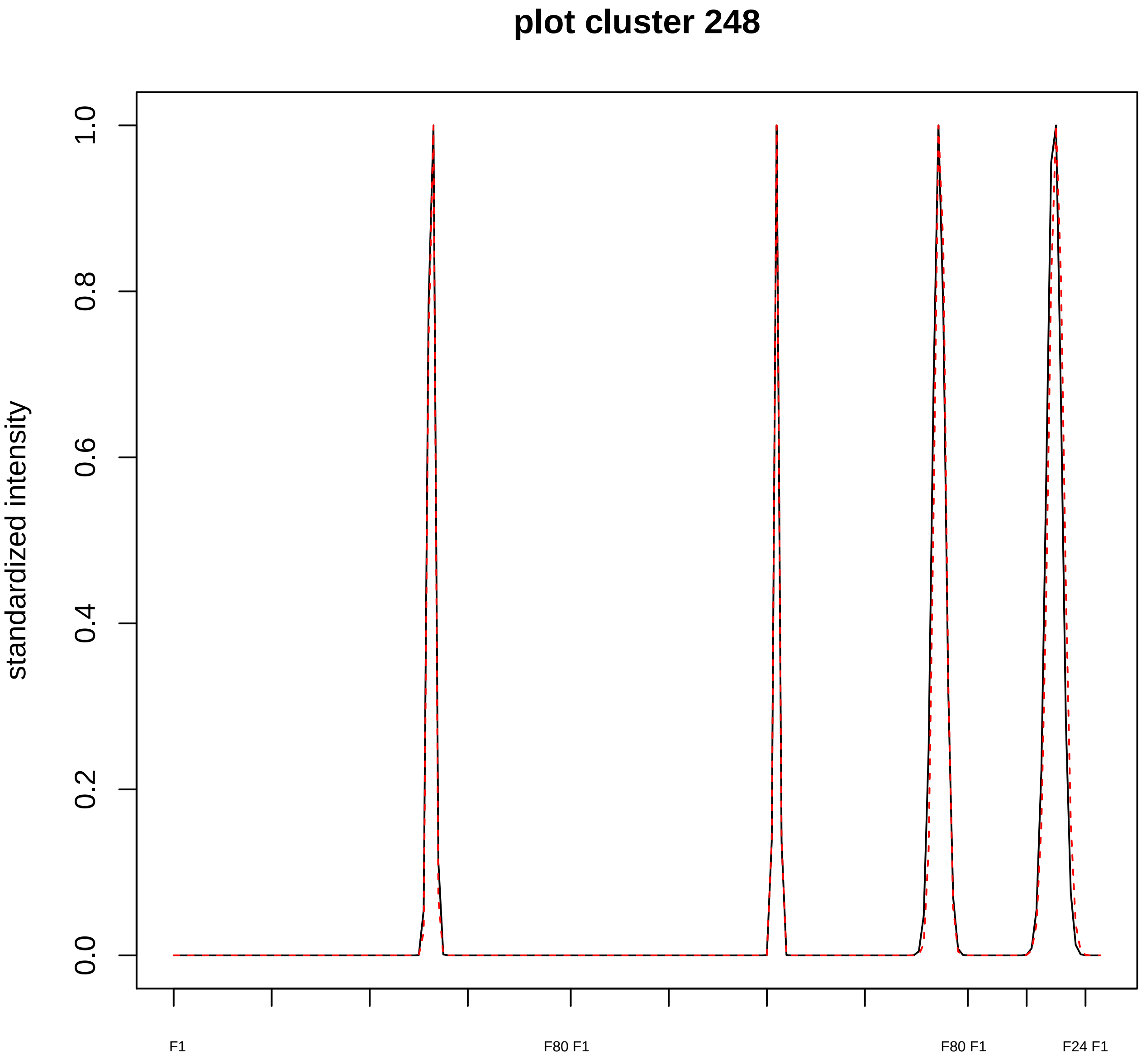
plot cluster 248
standardized intensity
0.0
0.2
0.4
0.6
0.8
1.0
F1
F80 F1
F80 F1
F24 F1

### plot cluster 249

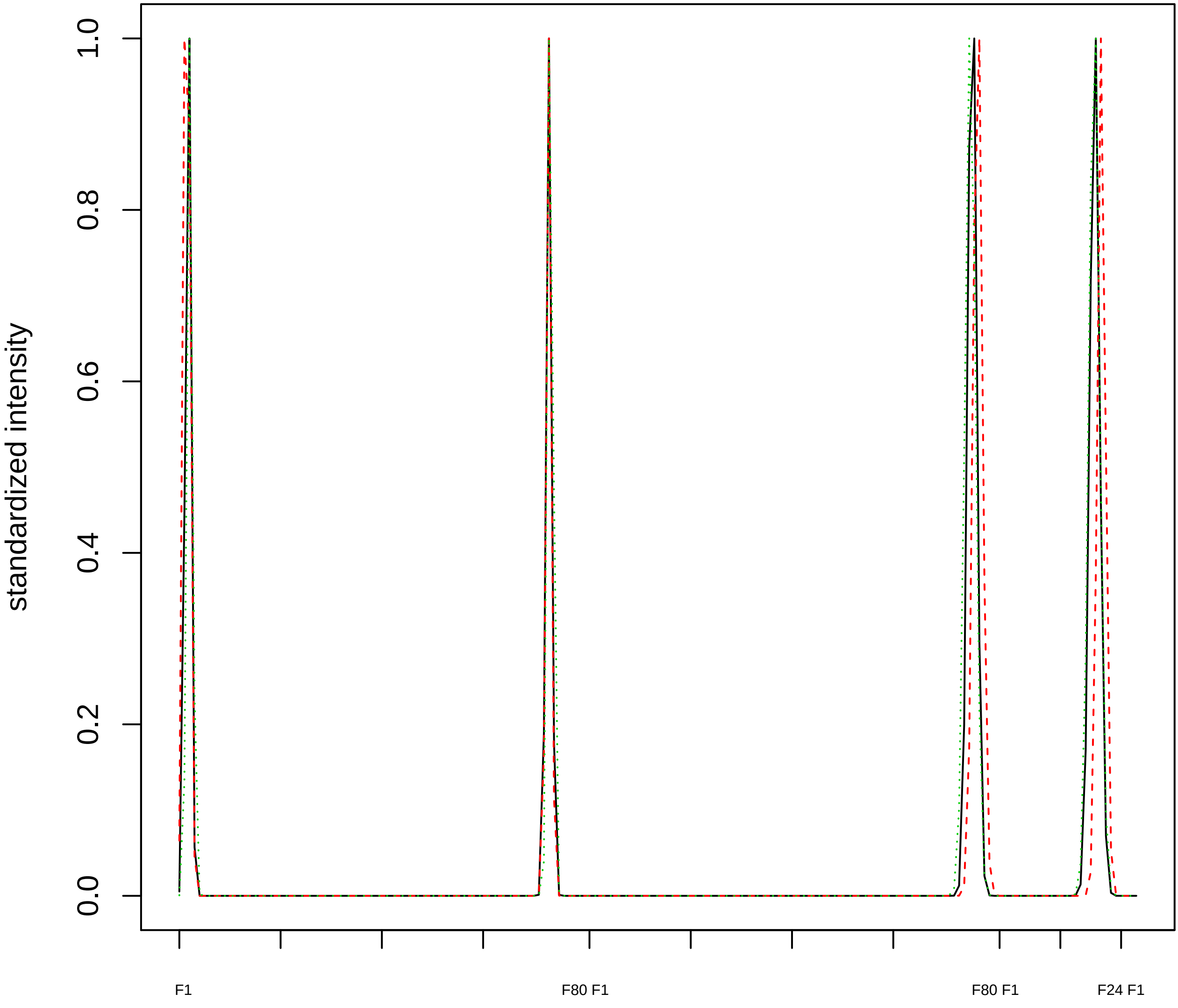

### Dendrogram of cluster 249

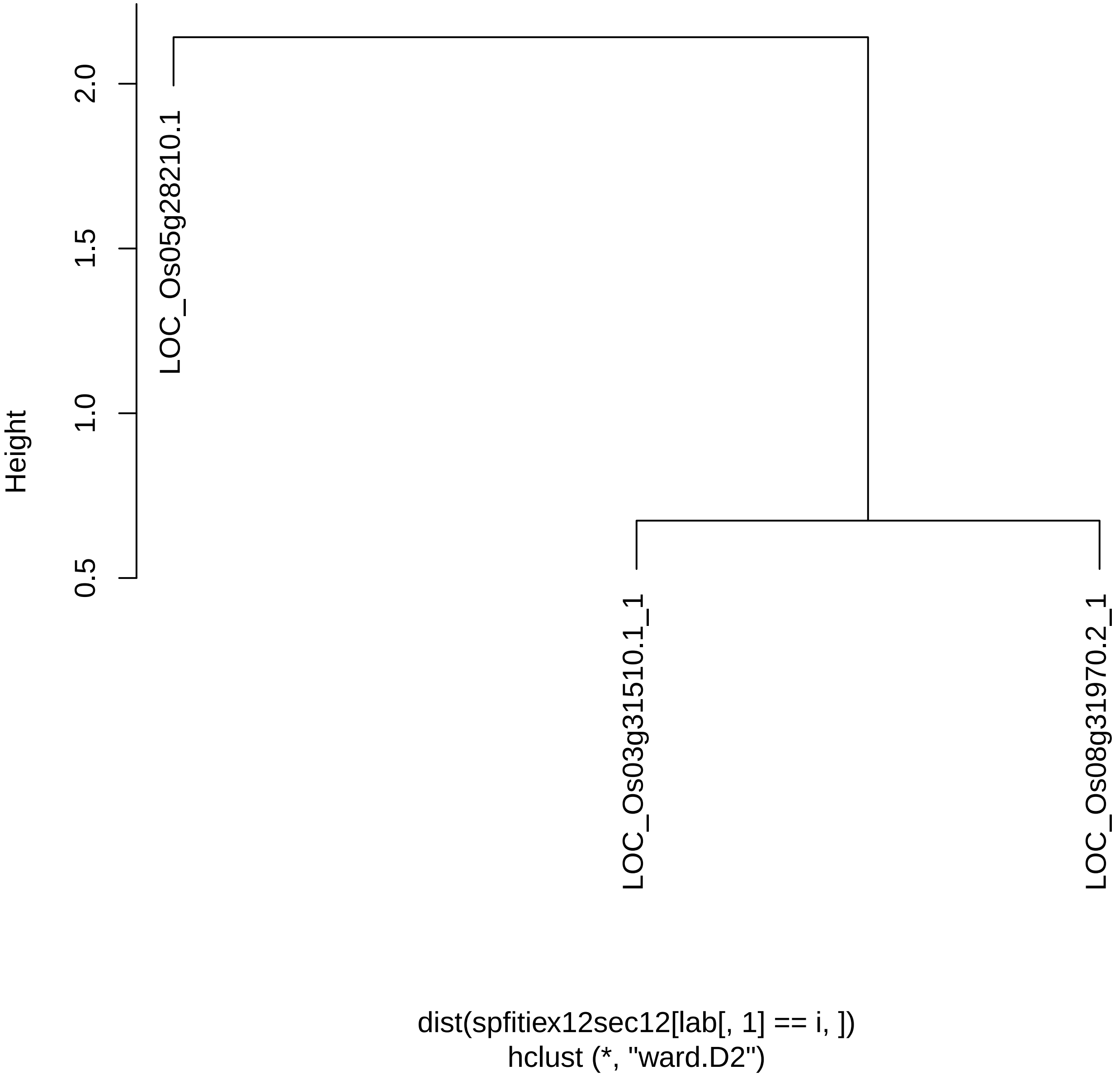

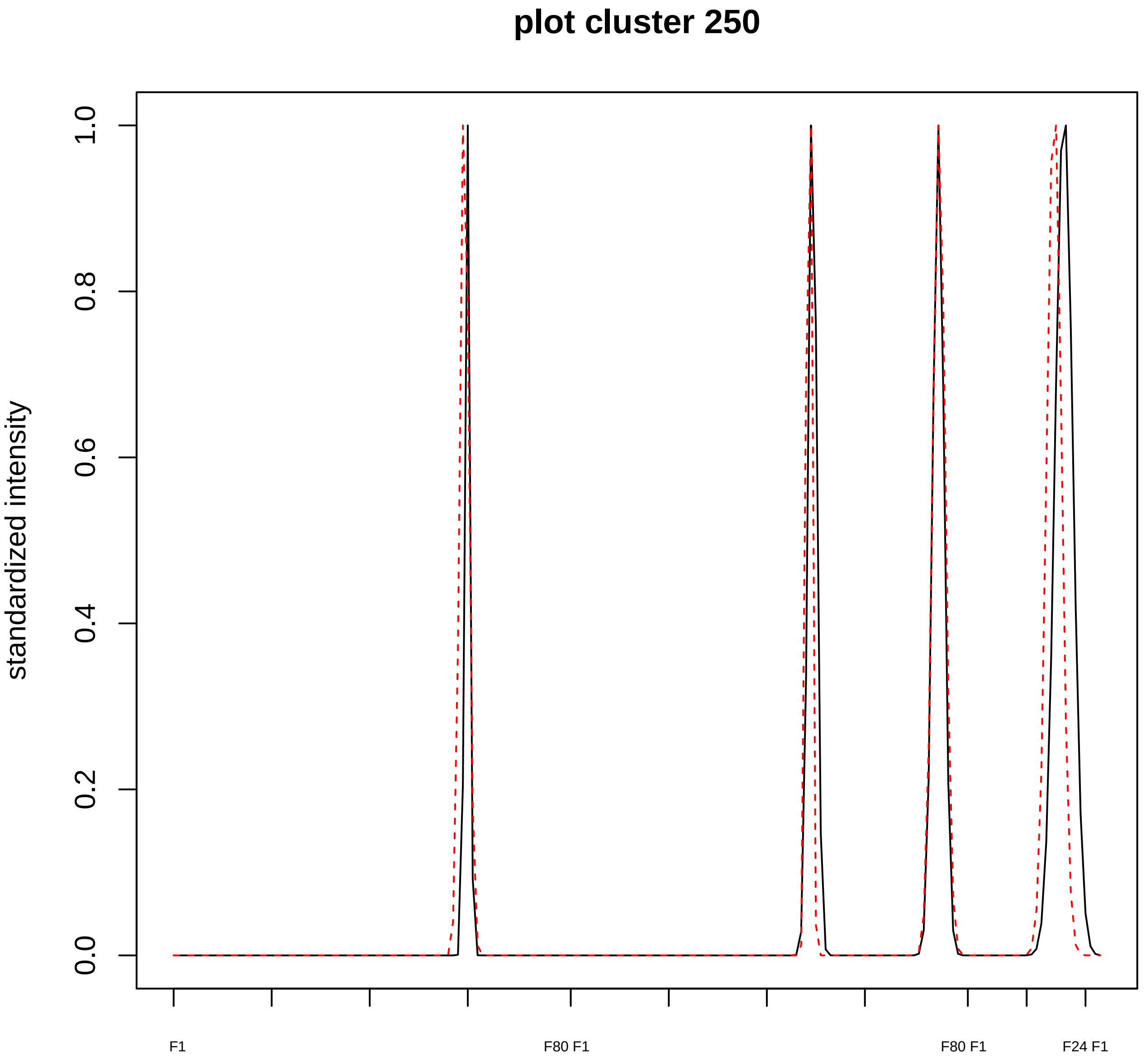

plot cluster 250
standardized intensity
0.0
0.2
0.4
0.6
0.8
1.0
F1
F80 F1
F80 F1
F24 F1

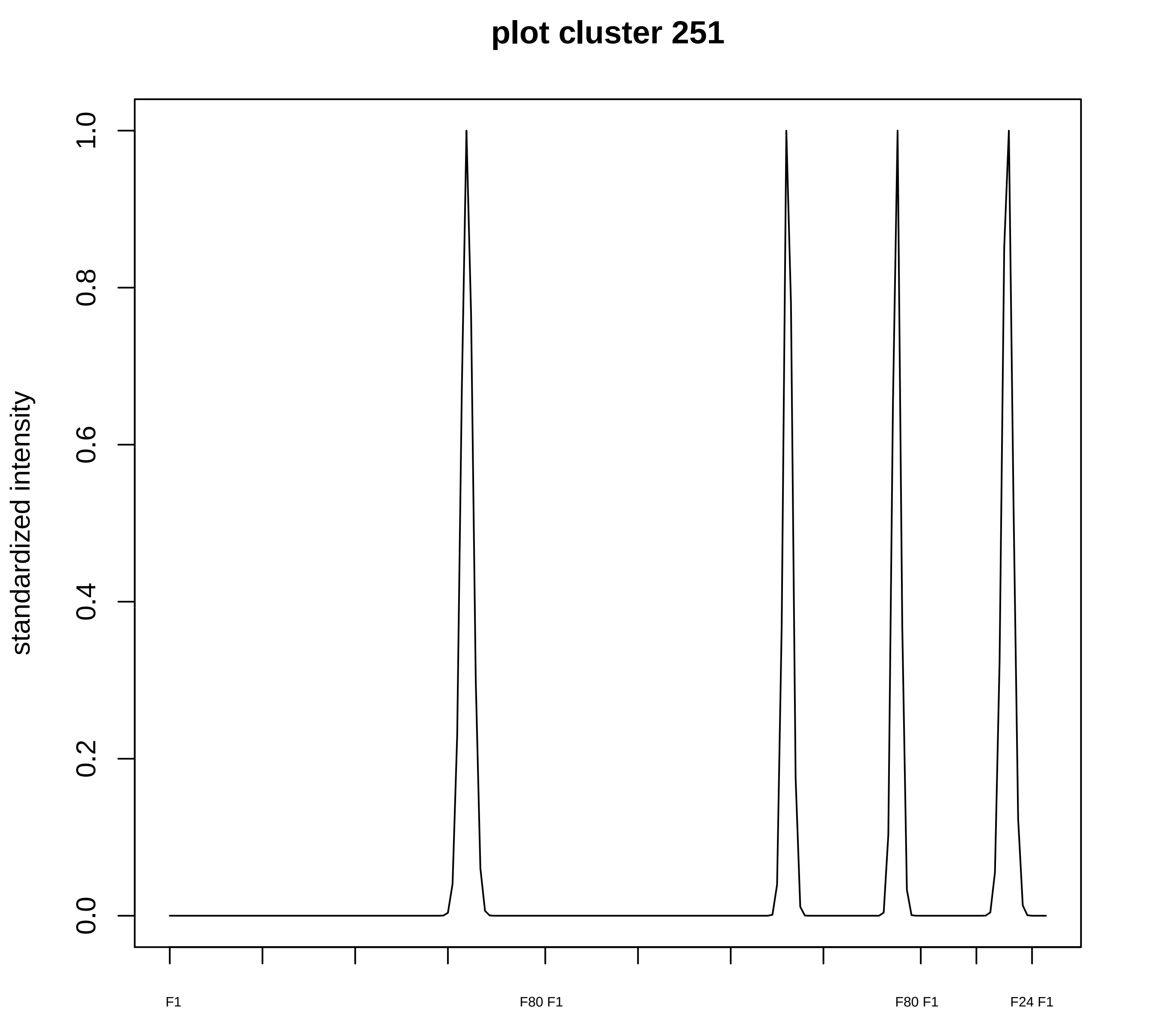
plot cluster 251
standardized intensity
0.0
0.2
0.4
0.6
0.8
1.0
F1
F80 F1
F80 F1
F24 F1

### plot cluster 252

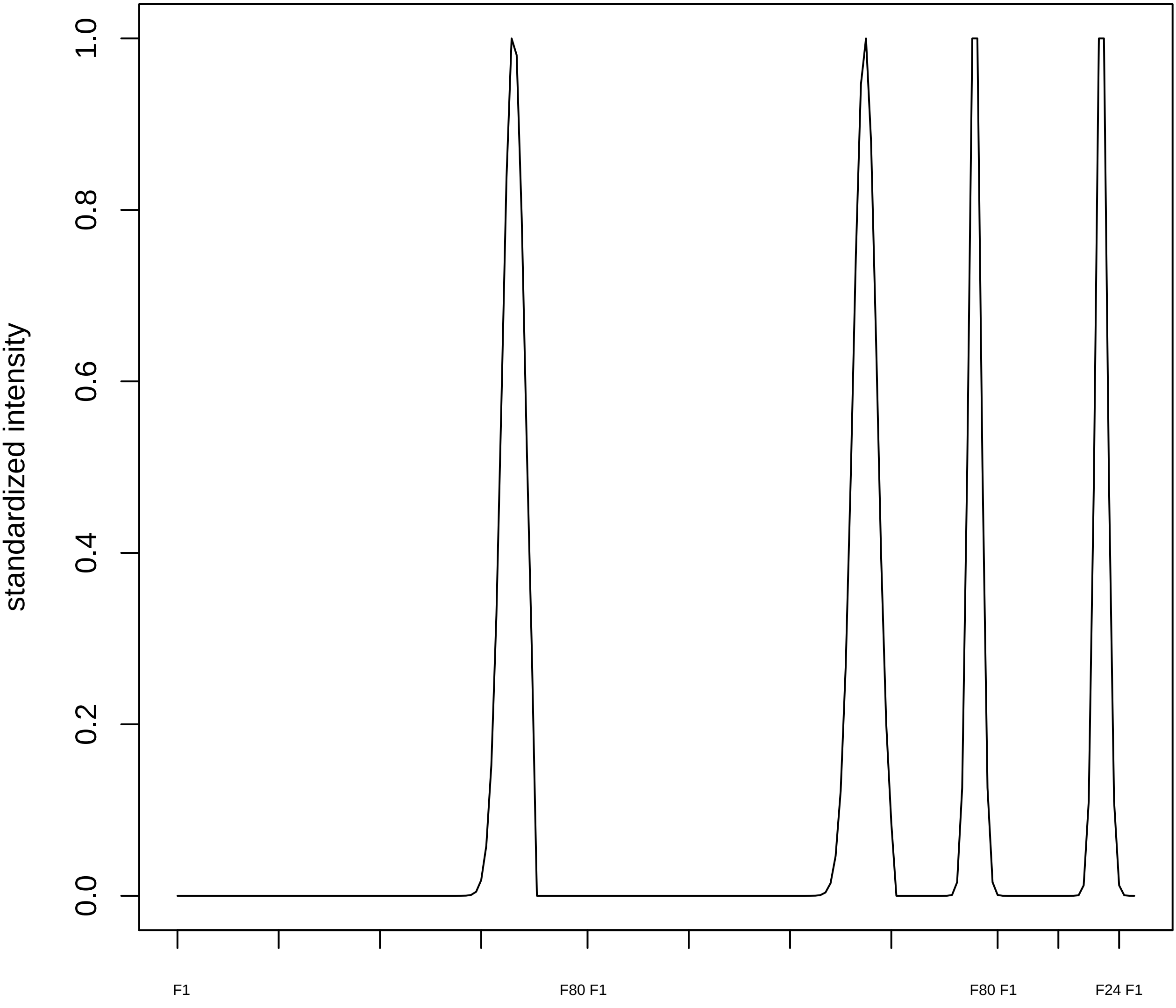

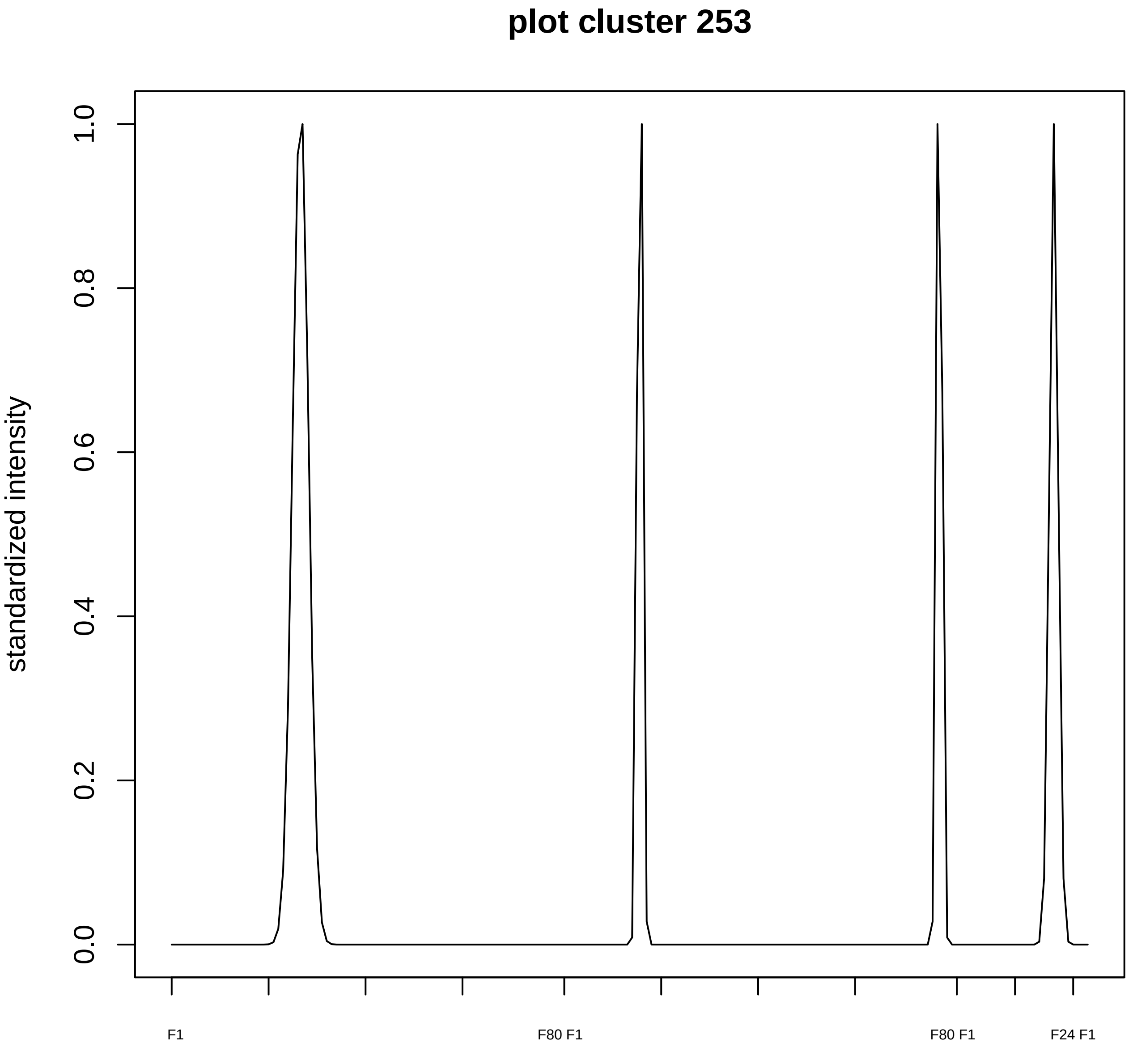
plot cluster 253
standardized intensity
0.0
0.2
0.4
0.6
0.8
1.0
F1
F80 F1
F80 F1
F24 F1

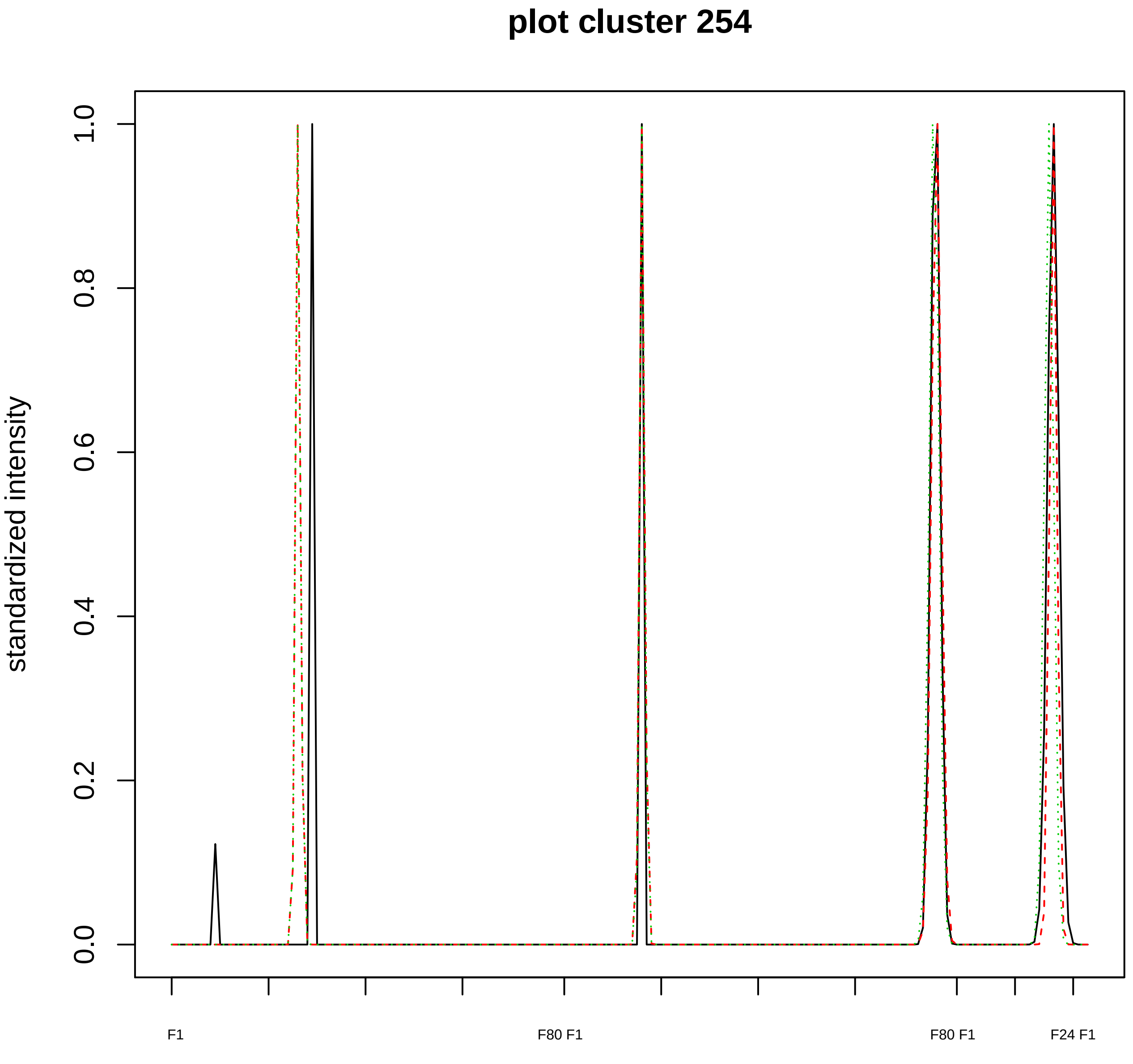

plot cluster 254
standardized intensity
1.0
0.8
0.6
0.4
0.2
0.0
F1
F80 F1
F80 F1
F24 F1

#### Dendrogram of cluster 254

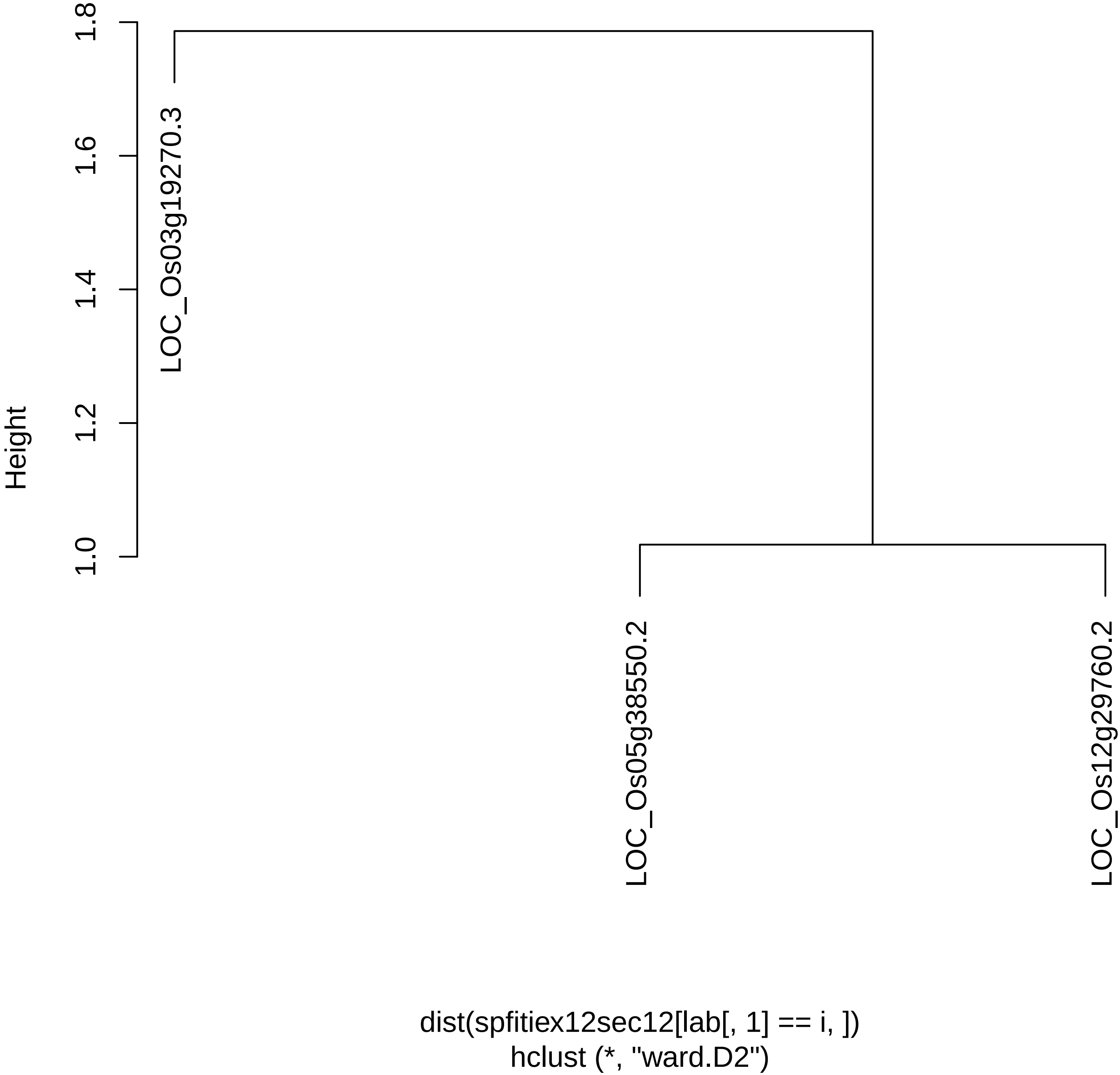

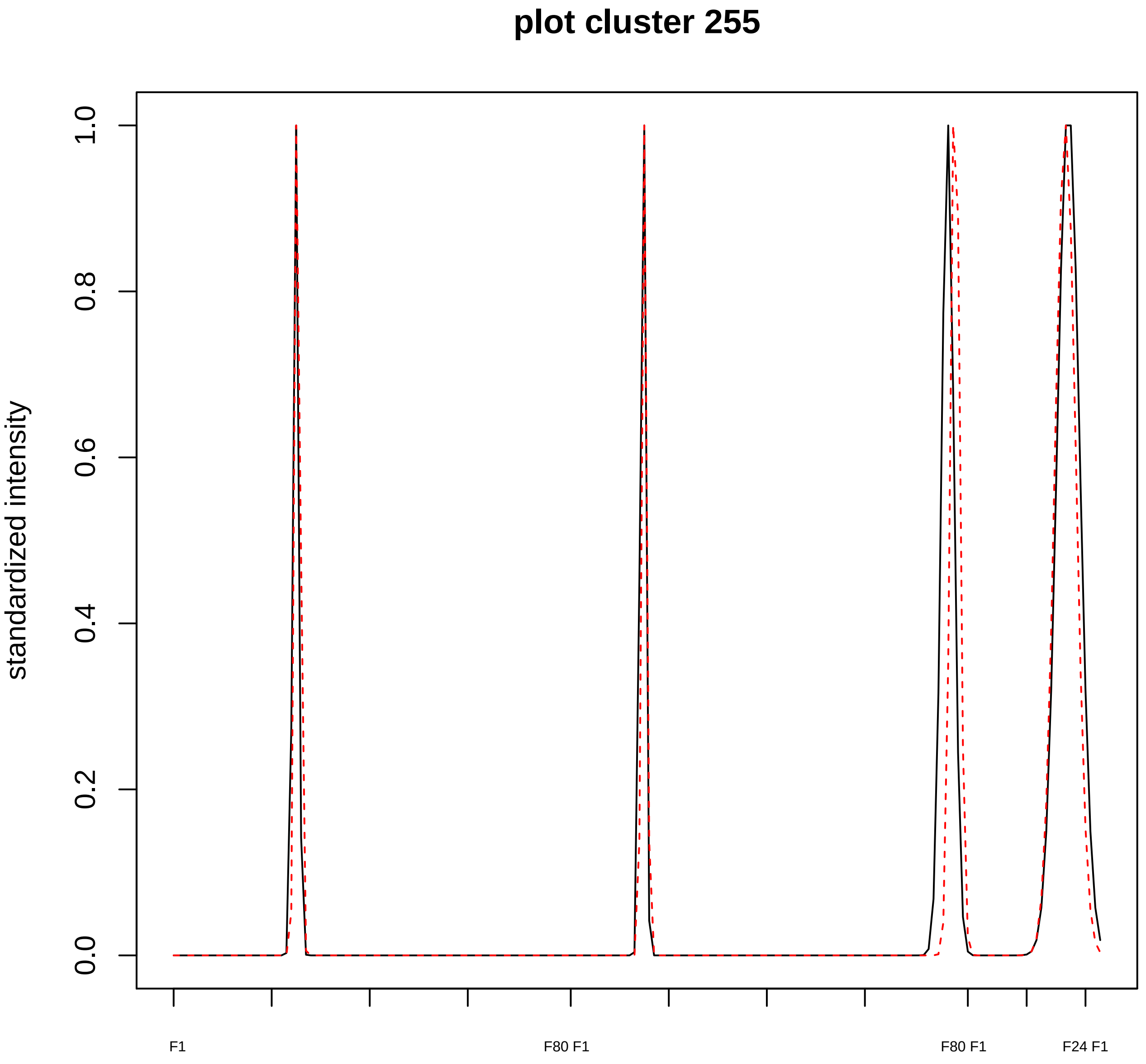
plot cluster 255
standardized intensity
0.0
0.2
0.4
0.6
0.8
1.0
F1
F80 F1
F80 F1
F24 F1

### plot cluster 256

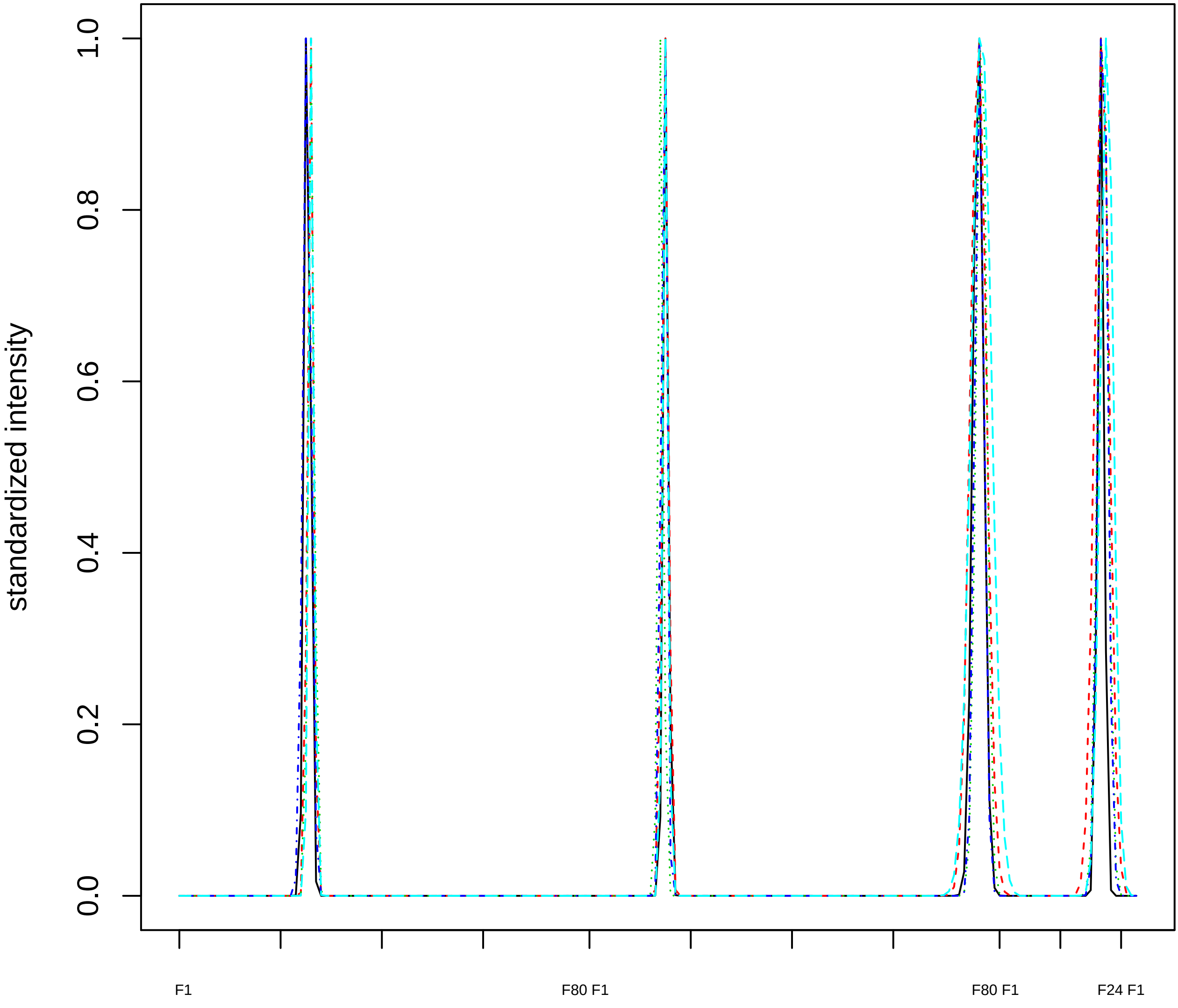

#### Dendrogram of cluster 256

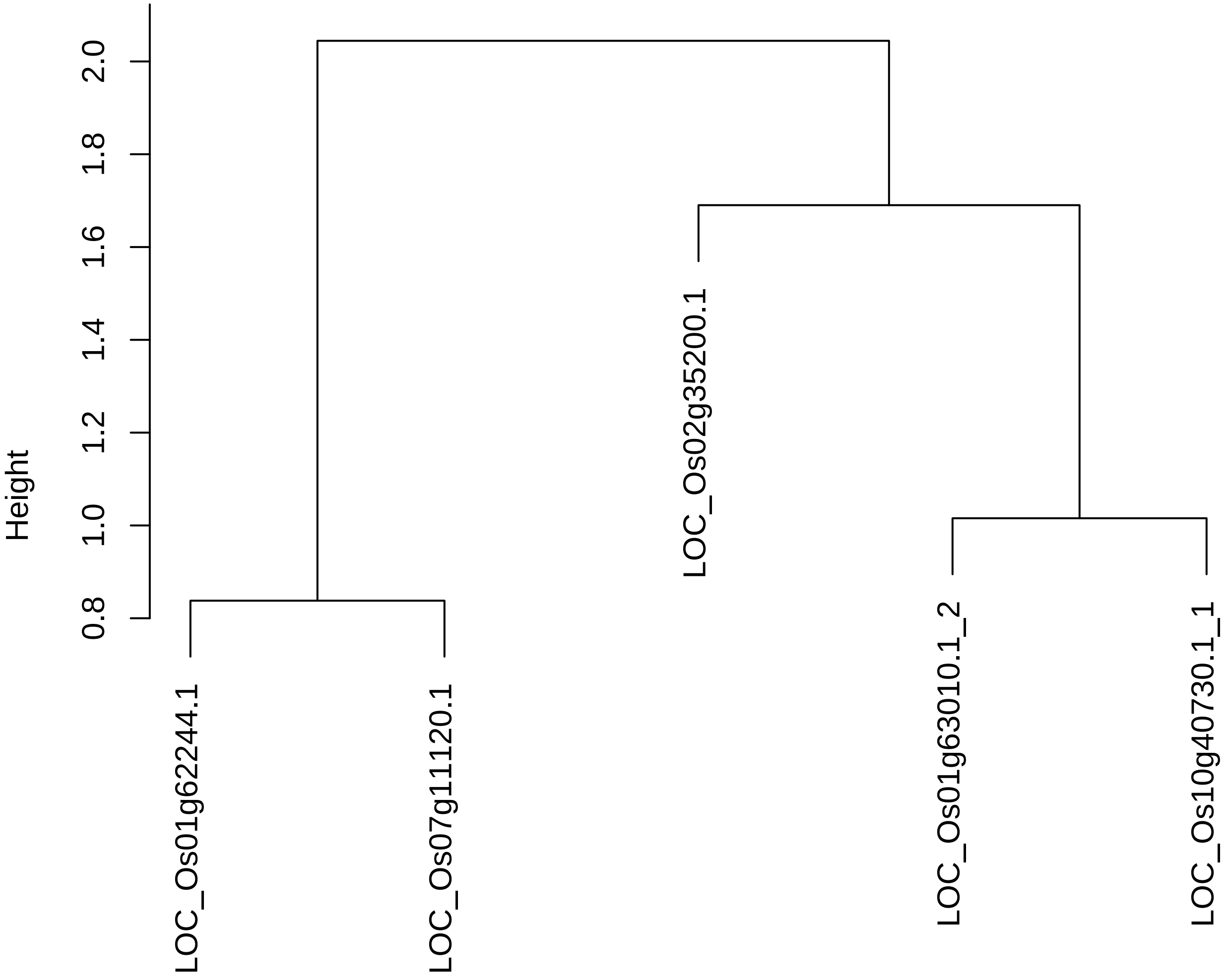


dist(spfitiex12sec12[lab[, 1] == i, ])
hclust (*, "ward.D2")

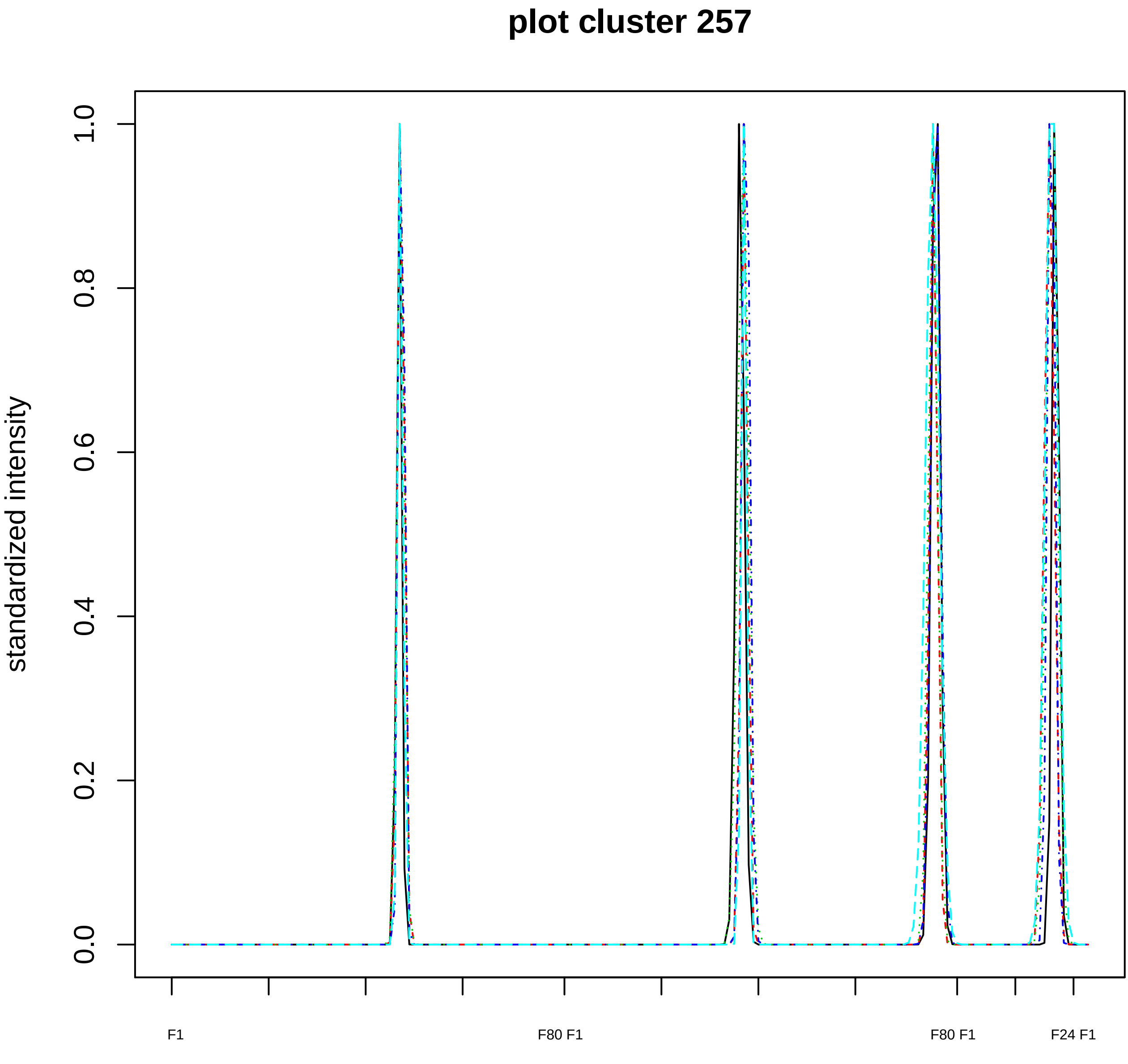

plot cluster 257
standardized intensity
0.0
0.2
0.4
0.6
0.8
1.0
F1
F80 F1
F80 F1
F24 F1

#### Dendrogram of cluster 257

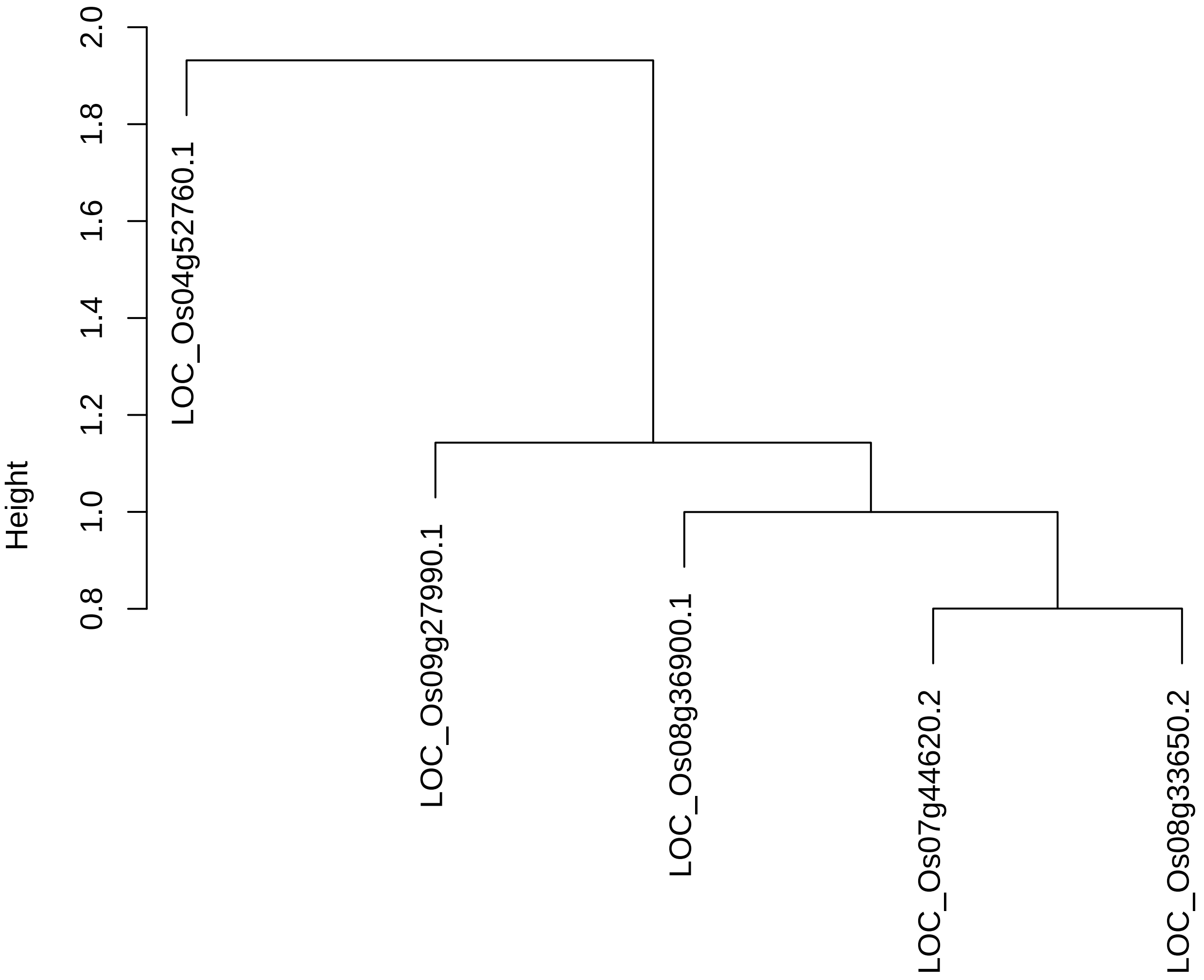


dist(spfitiex12sec12[lab[, 1] == i, ])
hclust (*, "ward.D2")

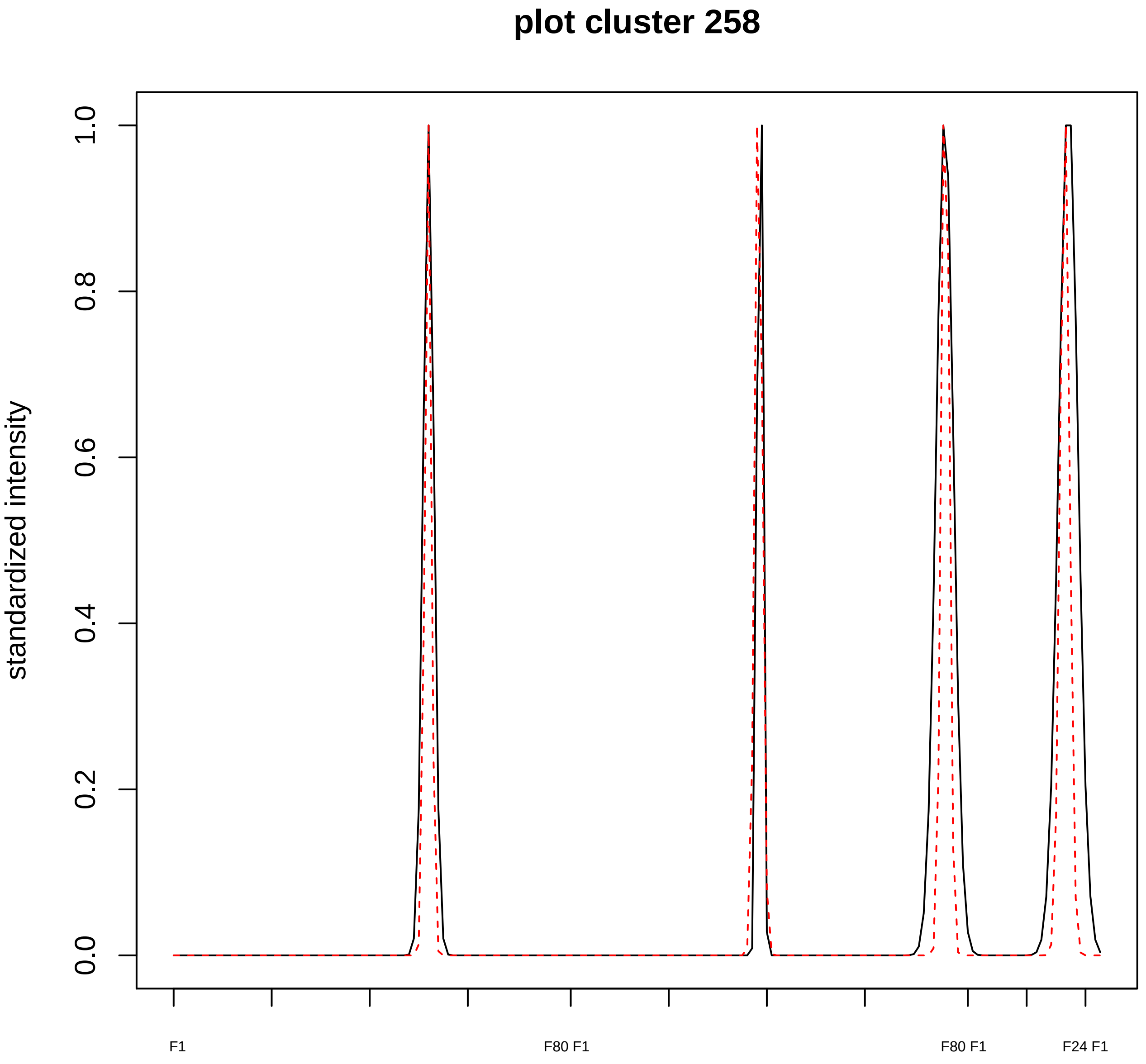

plot cluster 258
standardized intensity
0.0
0.2
0.4
0.6
0.8
1.0
F1
F80 F1
F80 F1
F24 F1

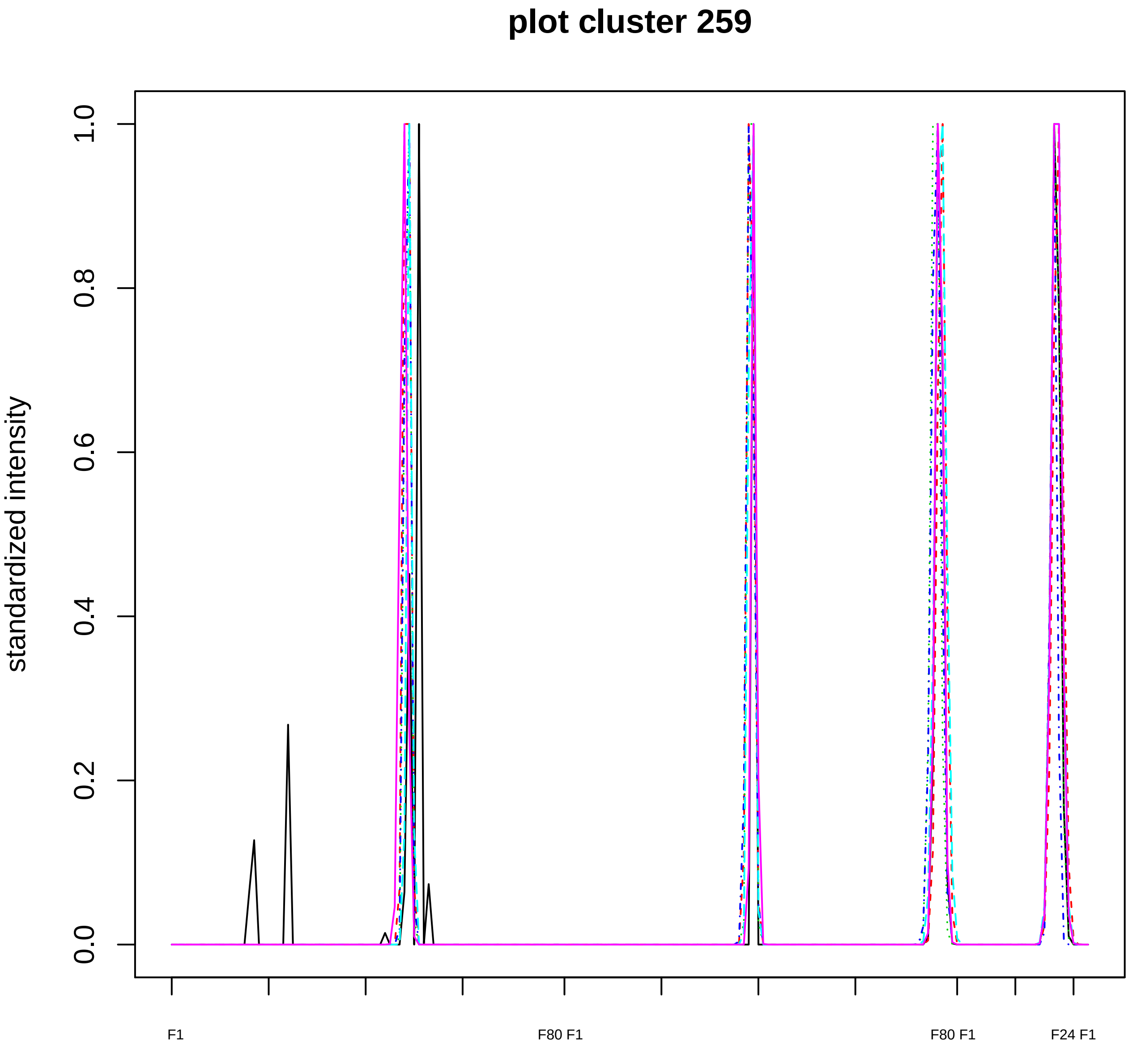

plot cluster 259
standardized intensity
0.0
0.2
0.4
0.6
0.8
1.0
F1
F80 F1
F80 F1
F24 F1

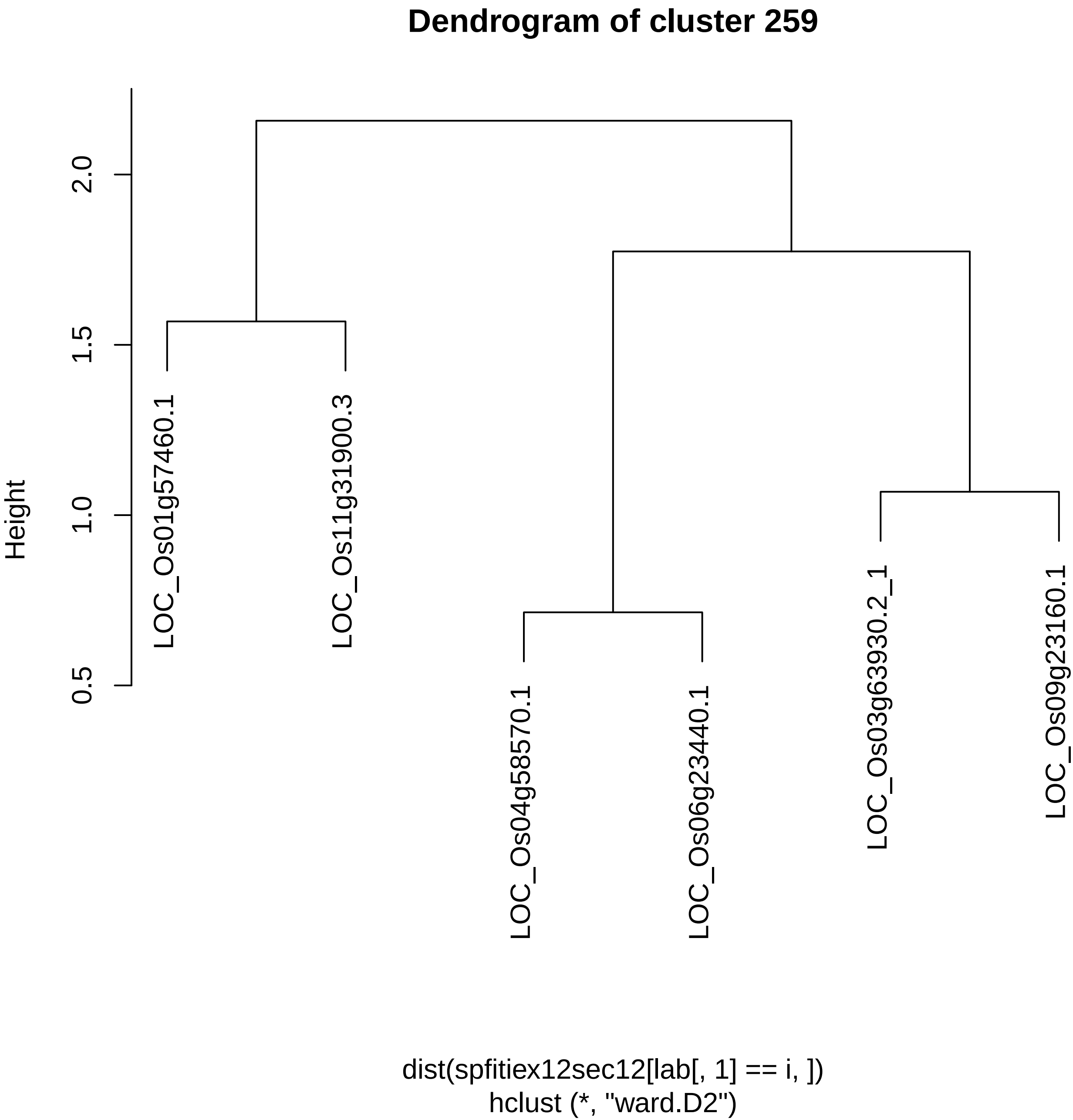

Dendrogram of cluster 259
Height
0.5
1.0
1.5
2.0
LOC_Os01g57460.1
LOC_Os11g31900.3
LOC_Os04g58570.1
LOC_Os06g23440.1
LOC_Os03g63930.2_1
LOC_Os09g23160.1
dist(spfitiex12sec12[lab[, 1] == i, ])
hclust (*, "ward.D2")

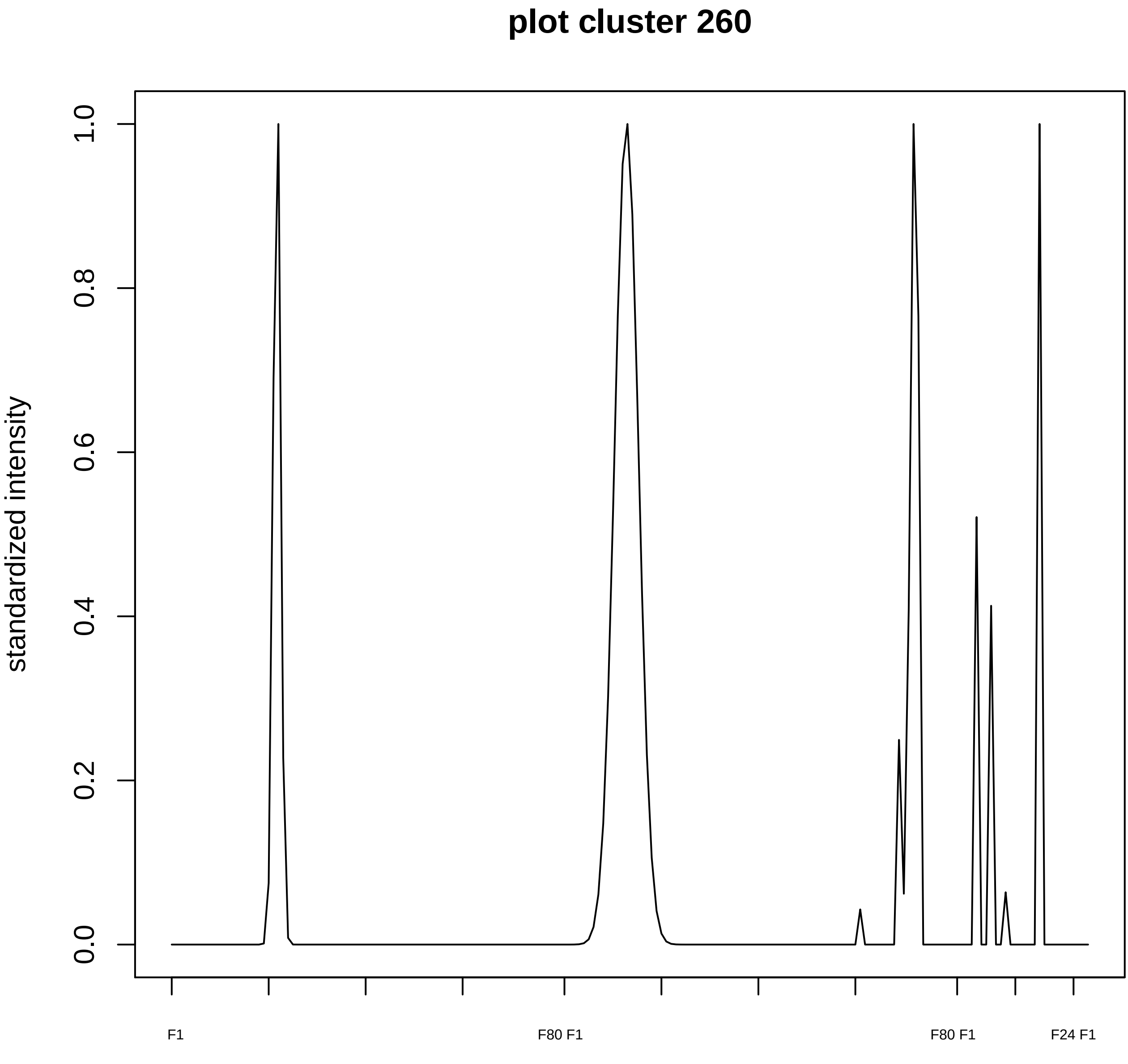
plot cluster 260
standardized intensity
0.0
0.2
0.4
0.6
0.8
1.0
F1
F80 F1
F80 F1
F24 F1

### plot cluster 261

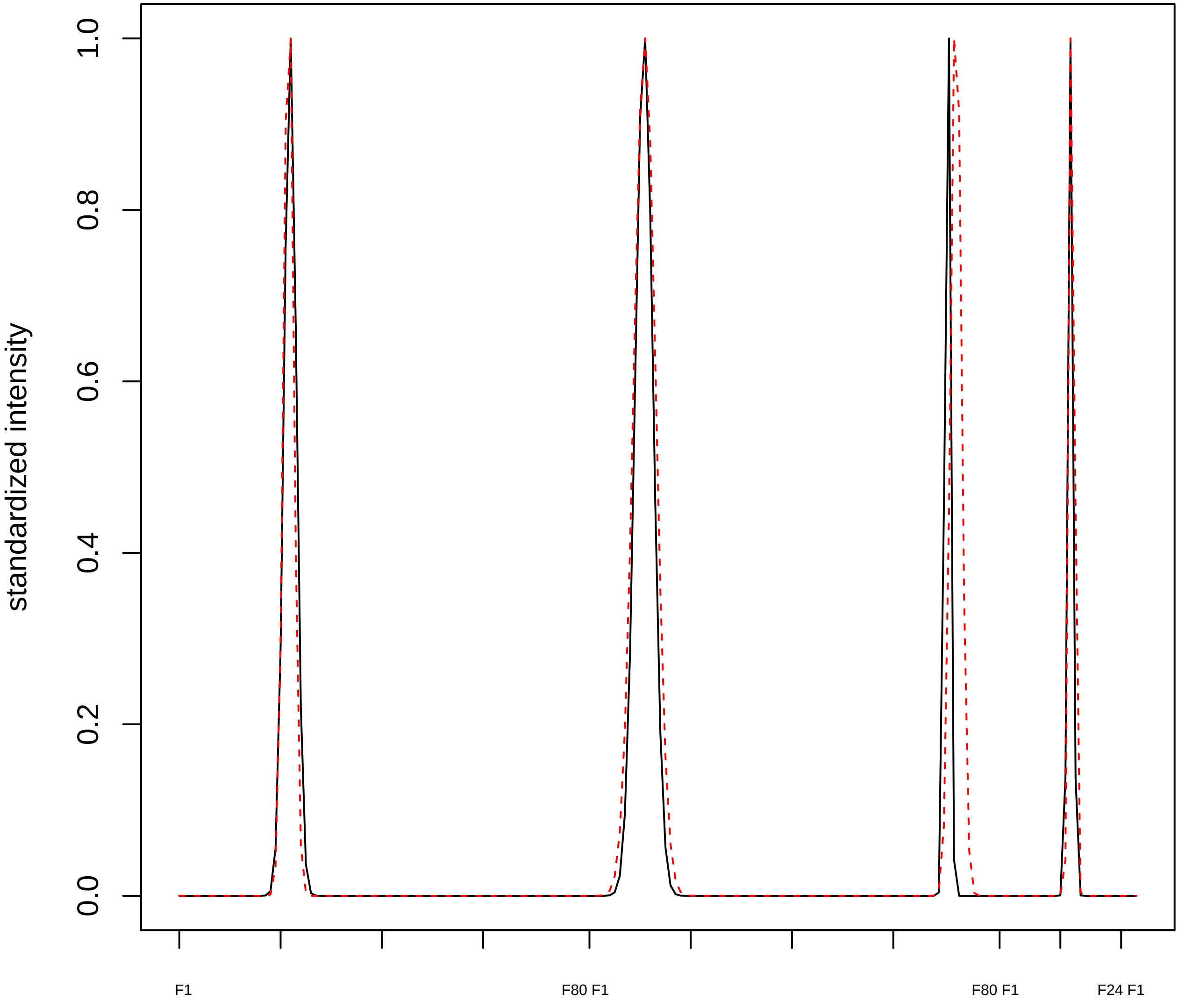

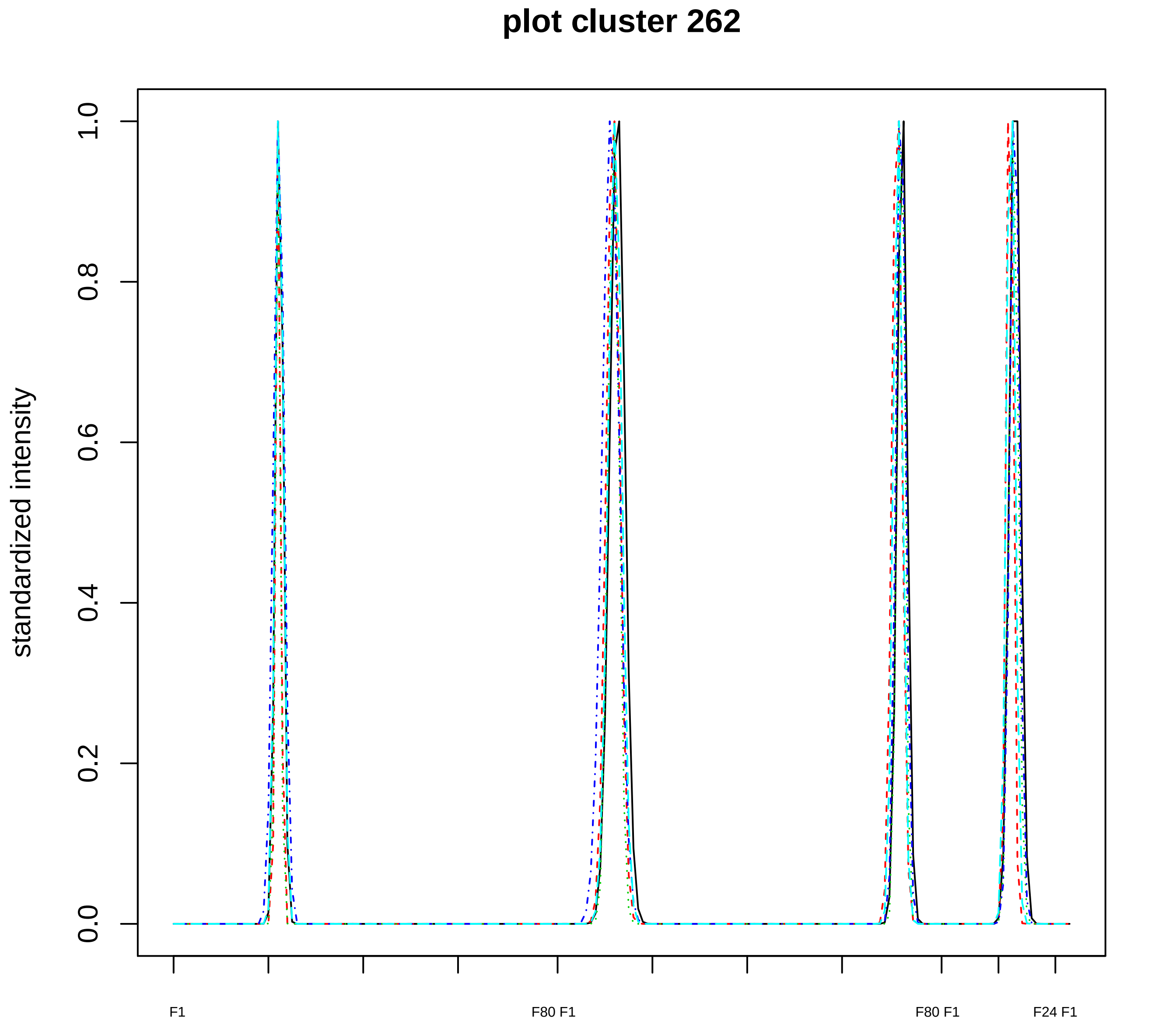

plot cluster 262
standardized intensity
0.0
0.2
0.4
0.6
0.8
1.0
F1
F80 F1
F80 F1
F24 F1

### Dendrogram of cluster 262

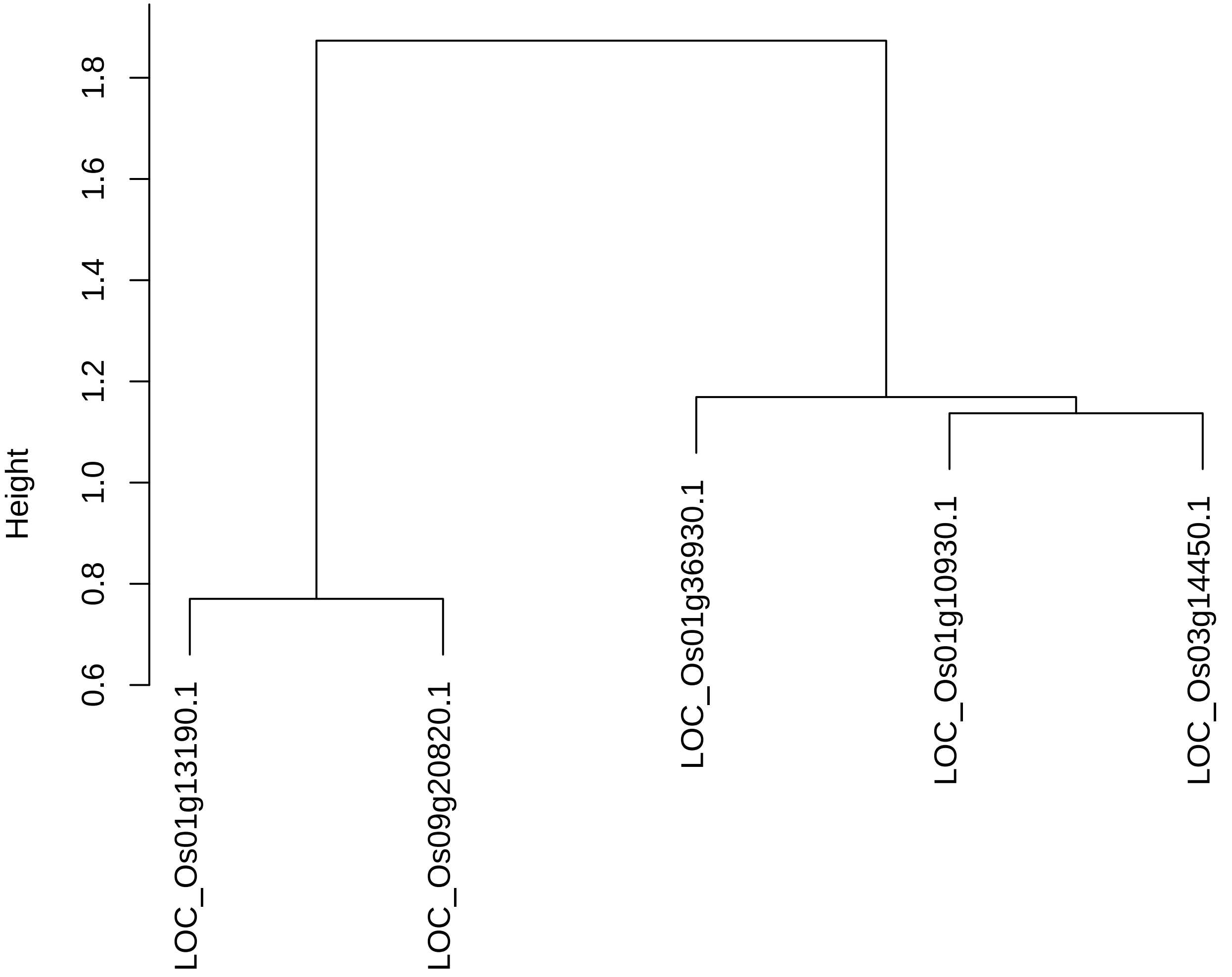


dist(spfitiex12sec12[lab[, 1] == i, ])
hclust (*, "ward.D2")

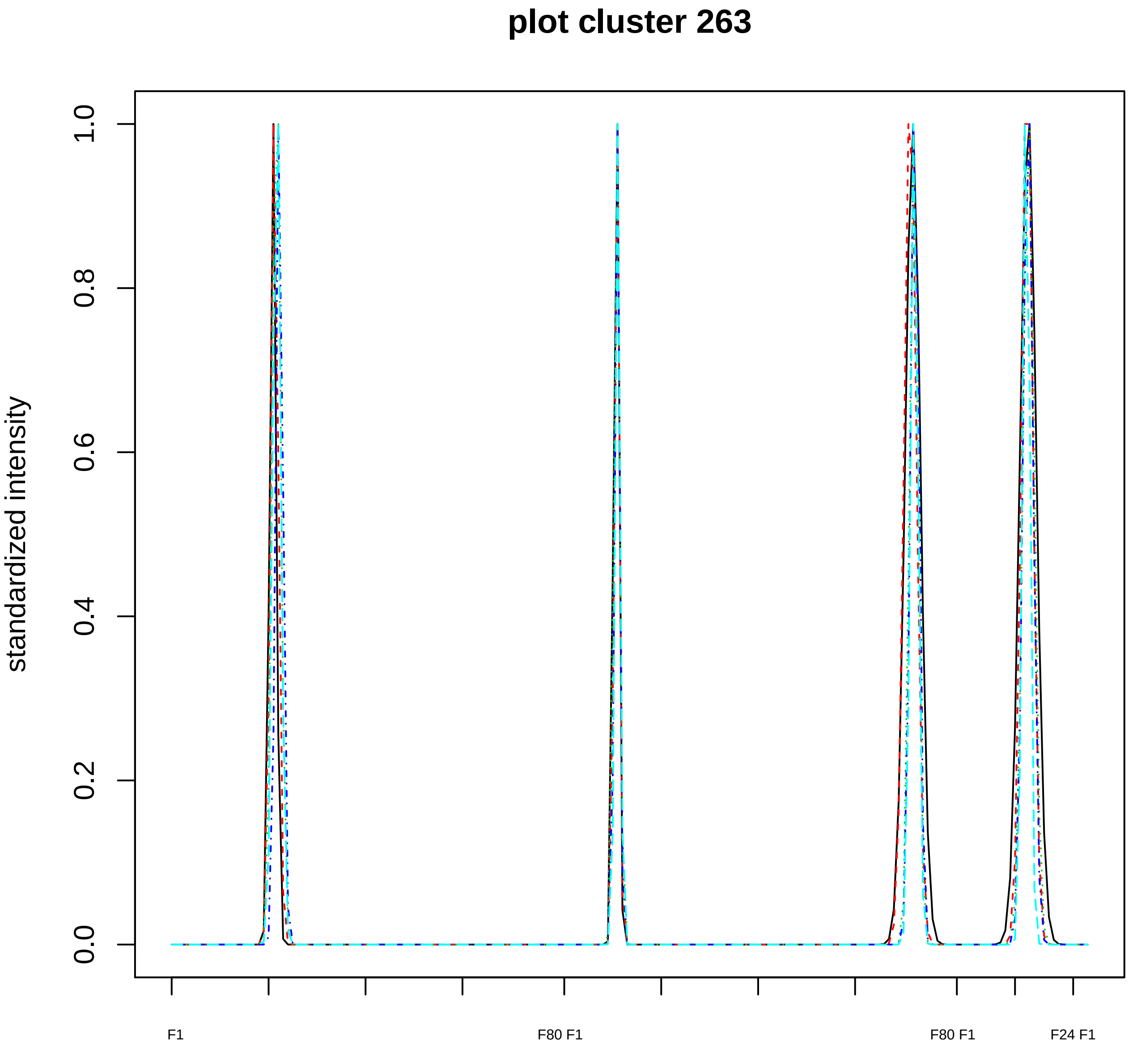
plot cluster 263
standardized intensity
0.0
0.2
0.4
0.6
0.8
1.0
F1
F80 F1
F80 F1
F24 F1

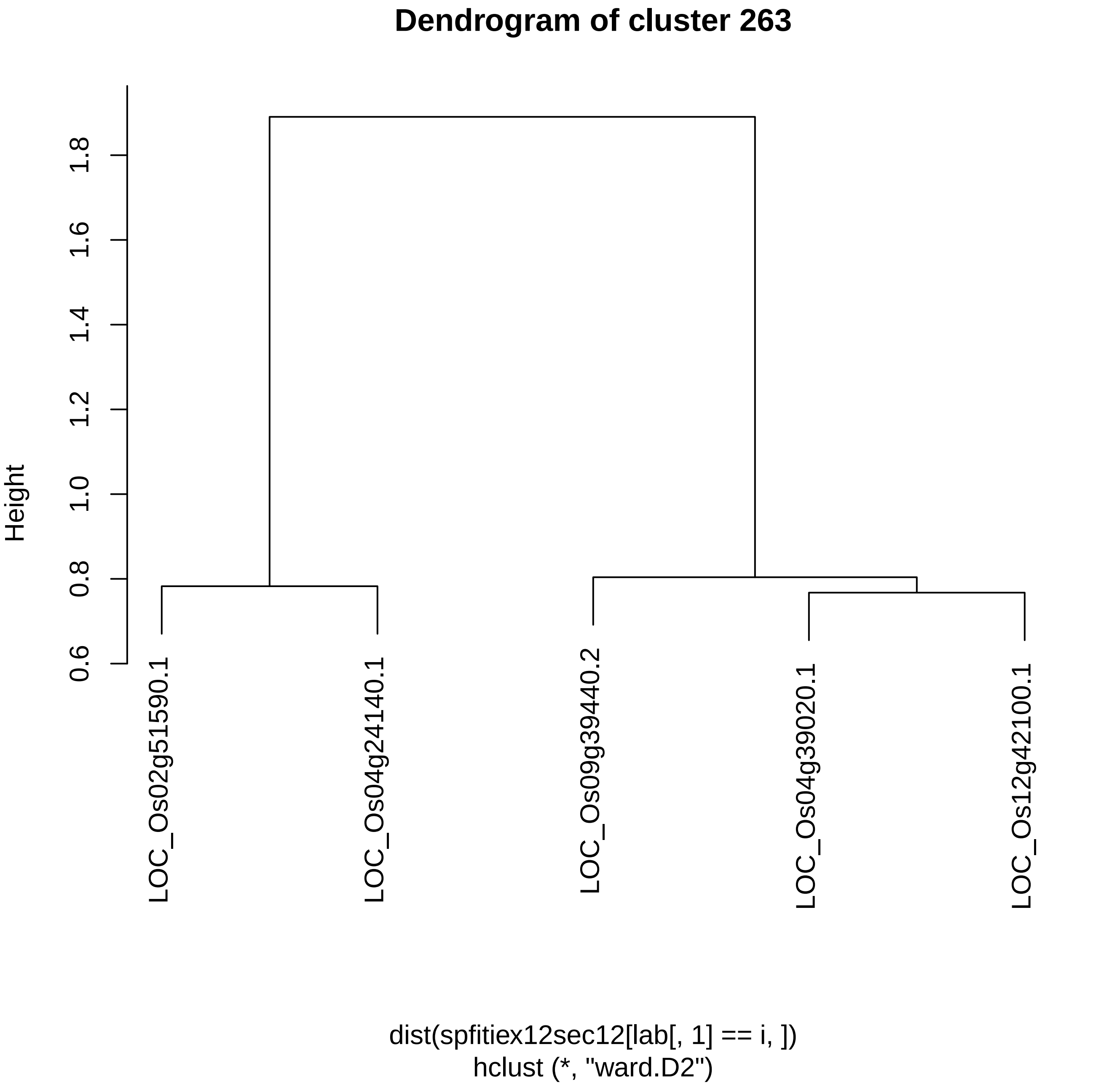
Dendrogram of cluster 263
Height
0.6
0.8
1.0
1.2
1.4
1.6
1.8
LOC_Os02g51590.1
LOC_Os04g24140.1
LOC_Os09g39440.2
LOC_Os04g39020.1
LOC_Os12g42100.1
dist(spfitiex12sec12[lab[, 1] == i, ])
hclust (*, "ward.D2")

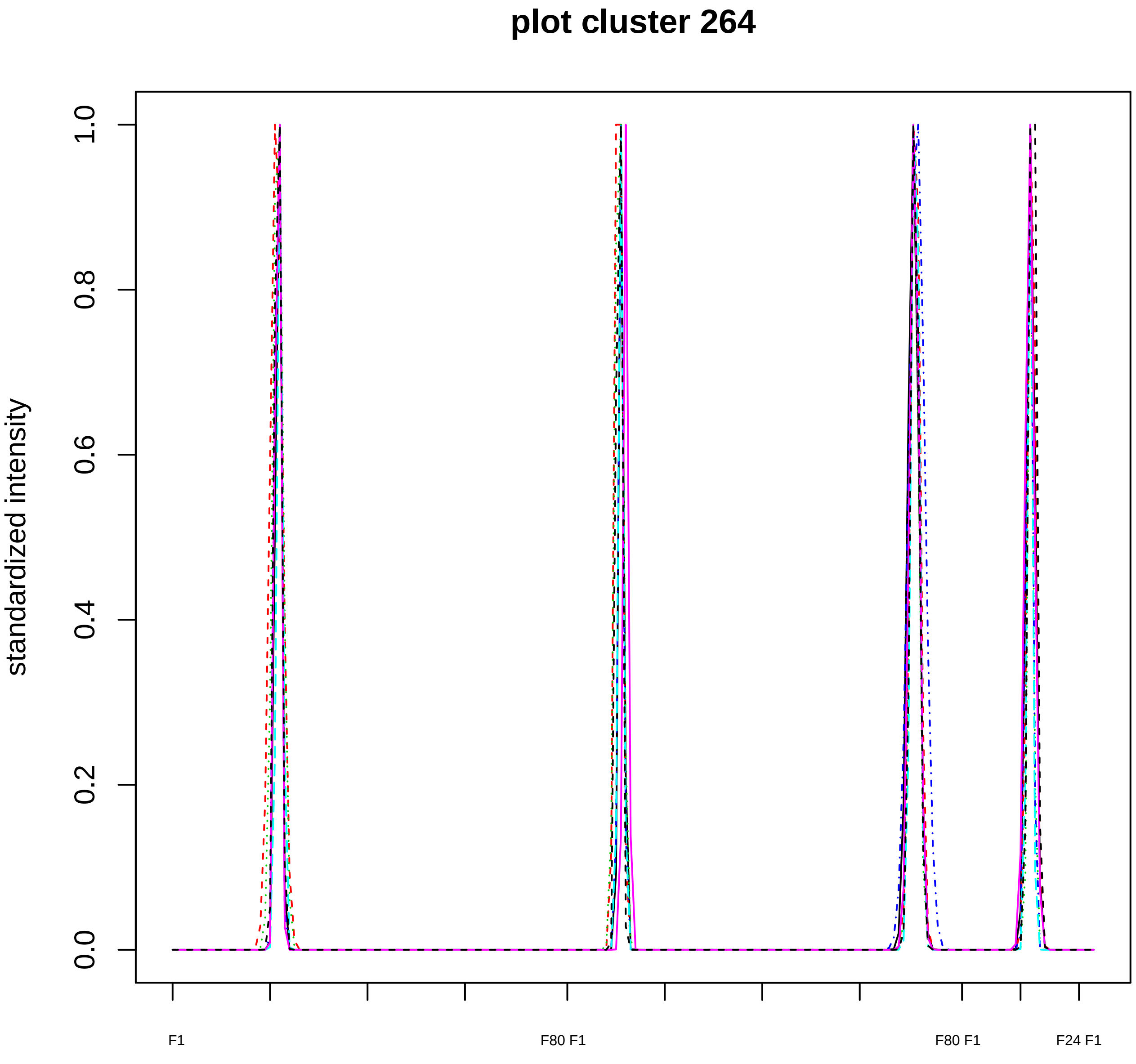

plot cluster 264
standardized intensity
0.0
0.2
0.4
0.6
0.8
1.0
F1
F80 F1
F80 F1
F24 F1

### Dendrogram of cluster 264

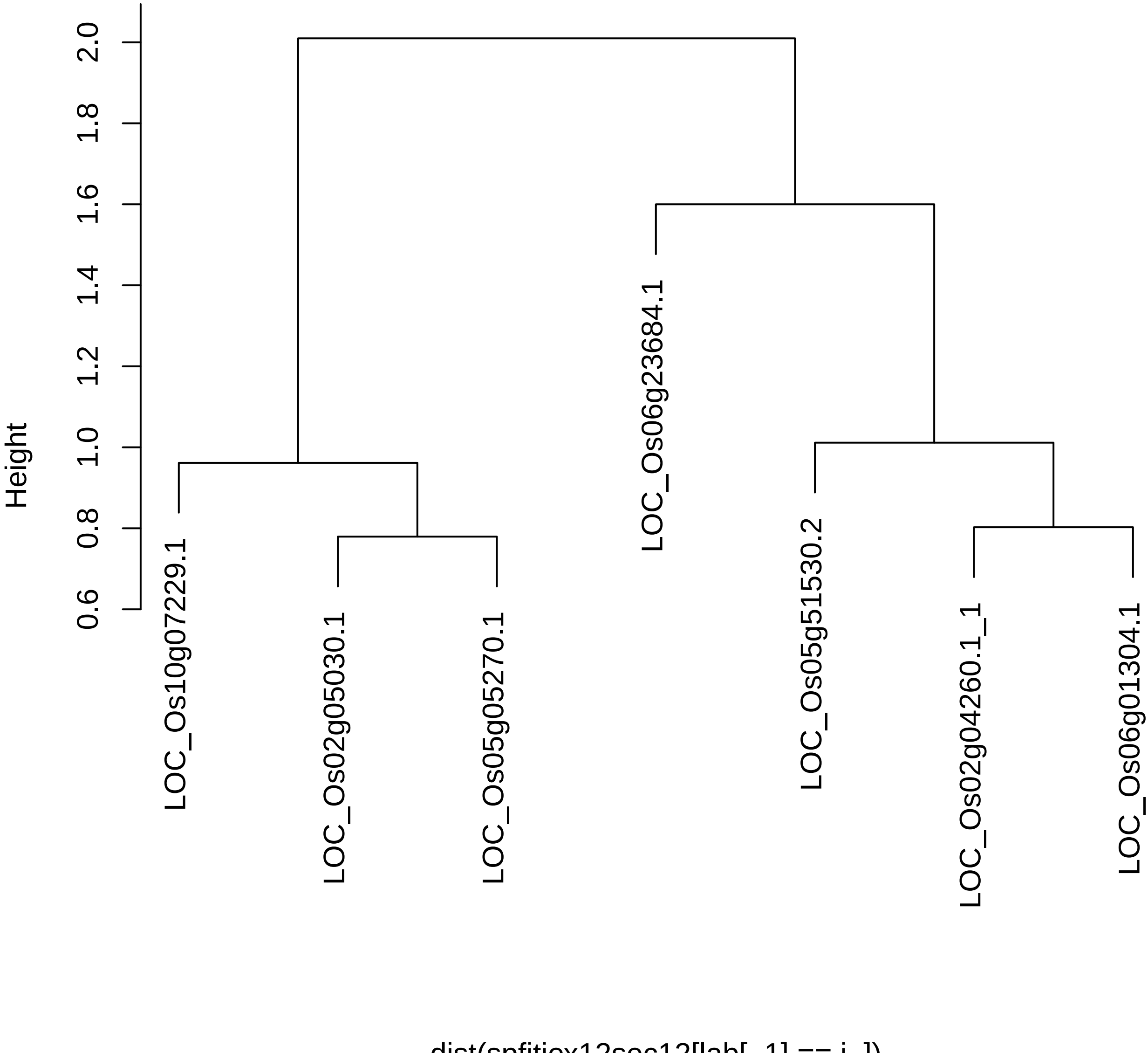


dist(spfitiex12sec12[lab[, 1] == i, ])
hclust (*, "ward.D2")

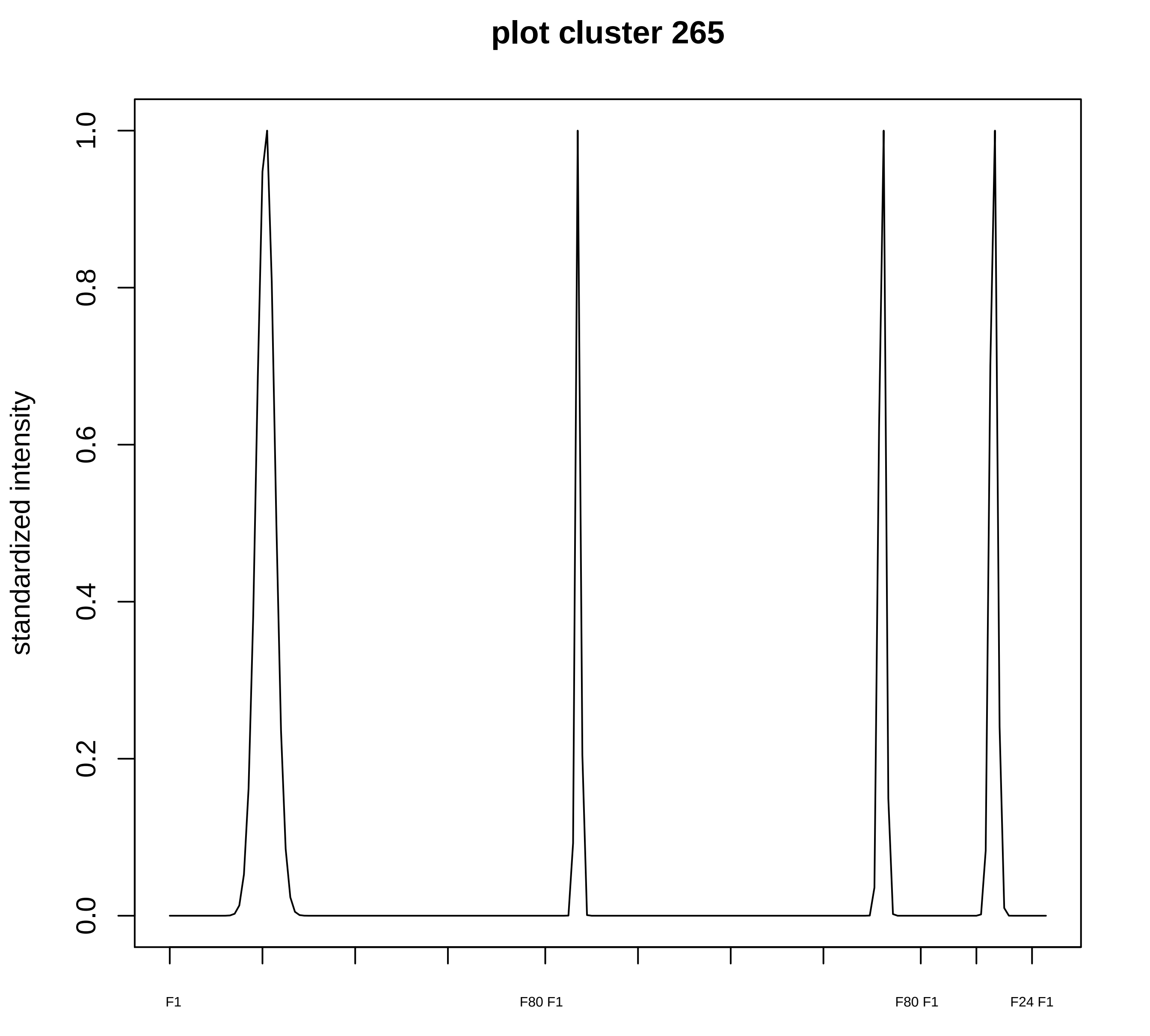
plot cluster 265
standardized intensity
0.0
0.2
0.4
0.6
0.8
1.0
F1
F80 F1
F80 F1
F24 F1

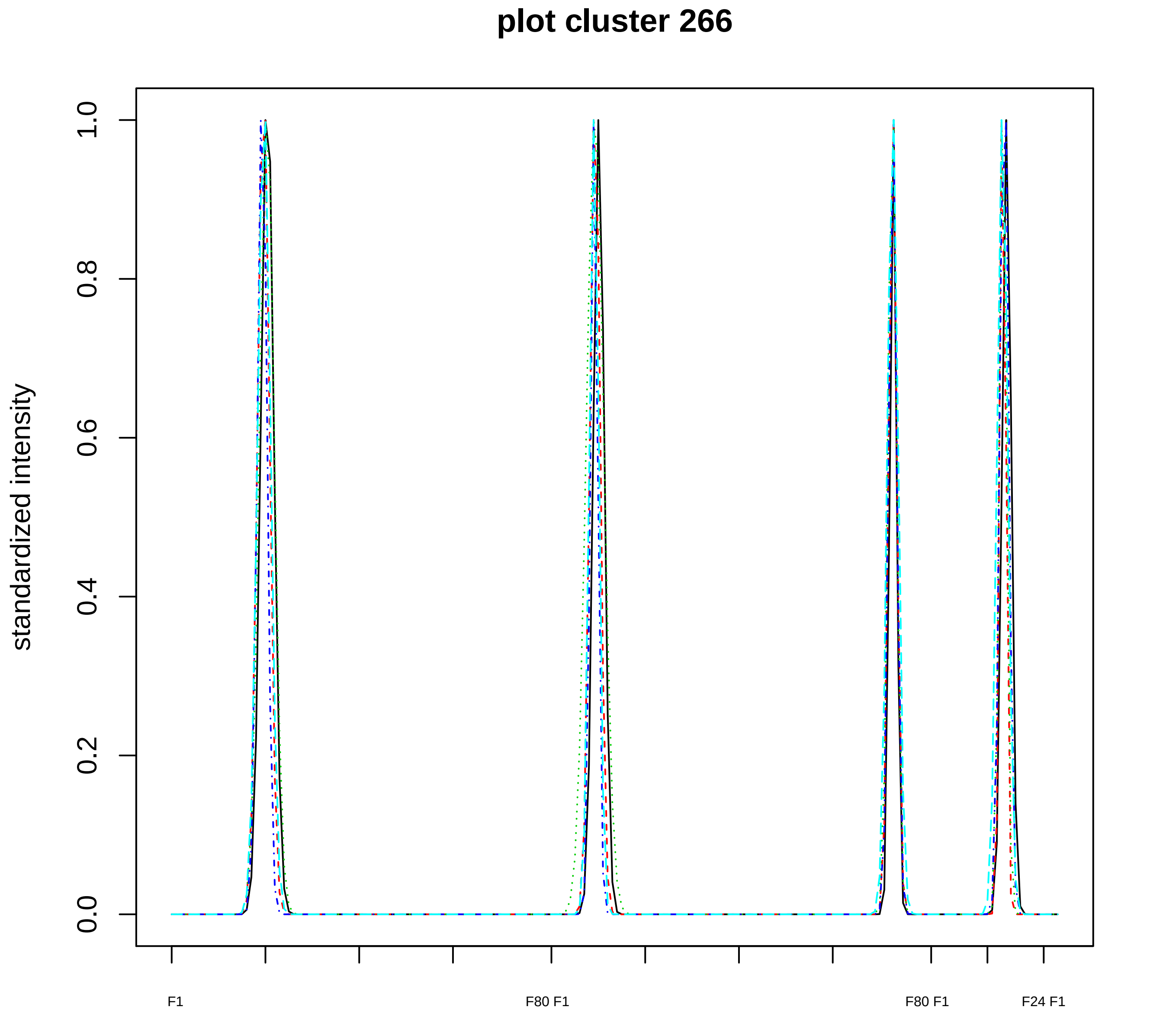
plot cluster 266
standardized intensity
0.0
0.2
0.4
0.6
0.8
1.0
F1
F80 F1
F80 F1
F24 F1

### Dendrogram of cluster 266

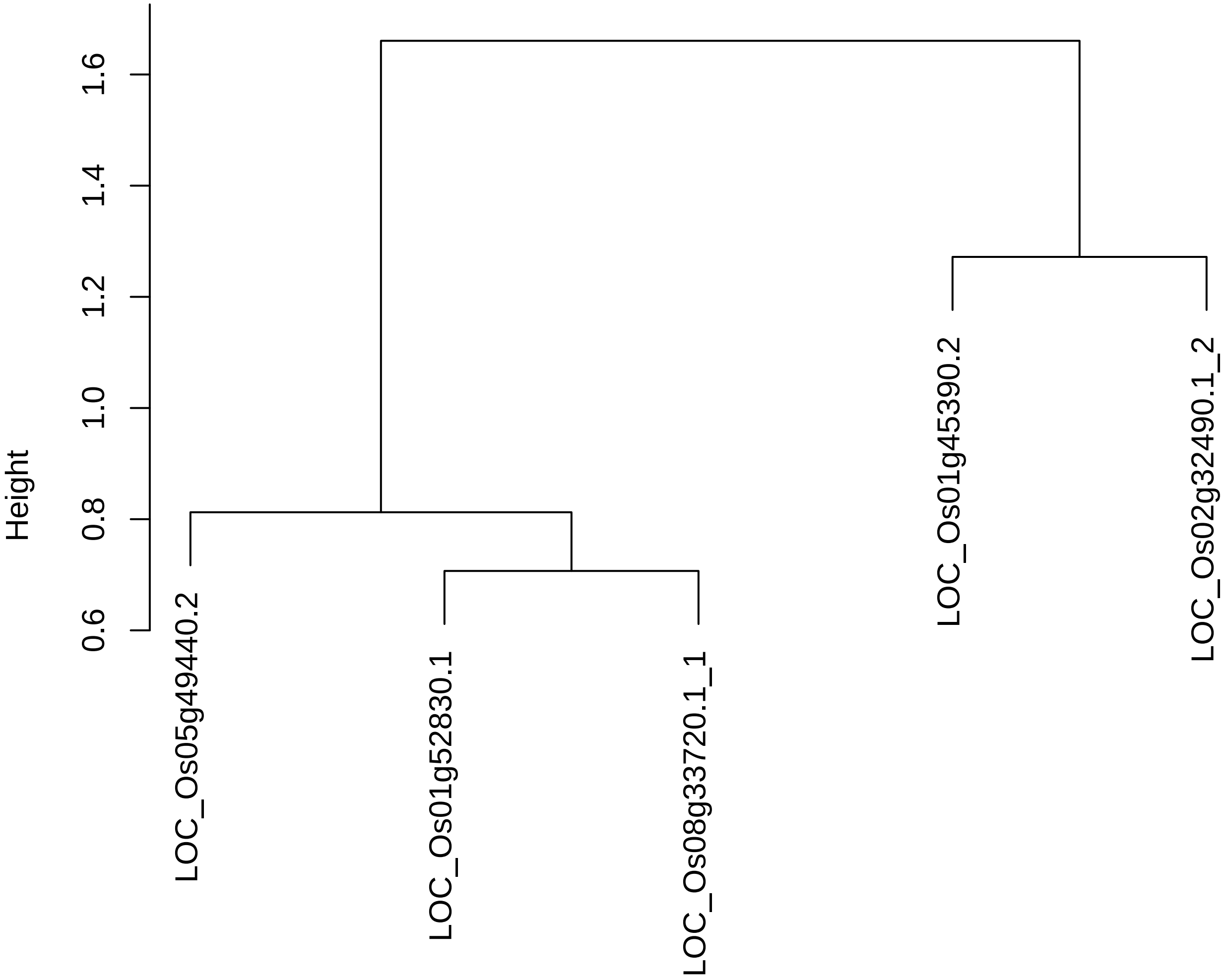


dist(spfitiex12sec12[lab[, 1] == i, ])
hclust (*, "ward.D2")

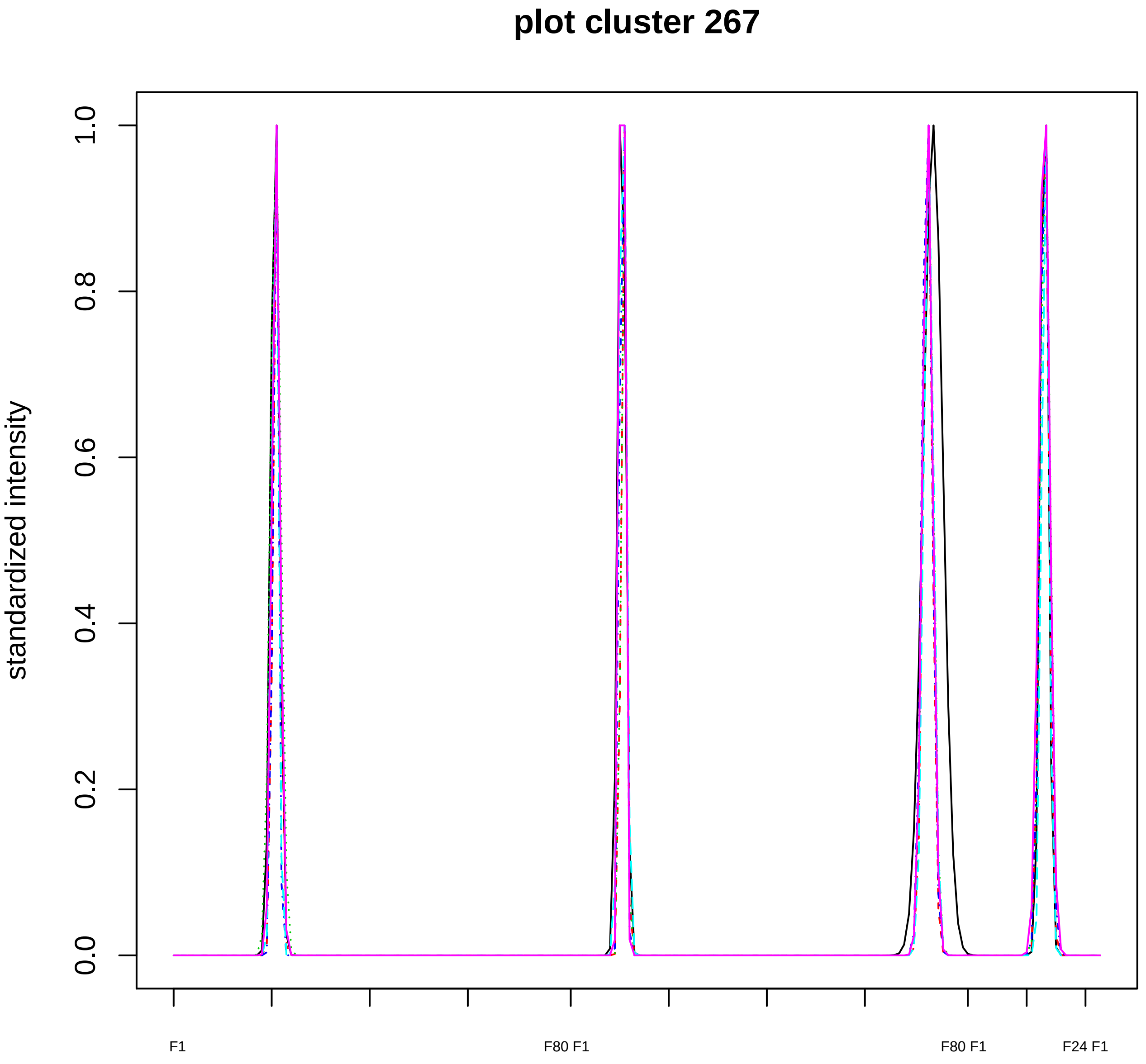
plot cluster 267
standardized intensity
0.0
0.2
0.4
0.6
0.8
1.0
F1
F80 F1
F80 F1
F24 F1

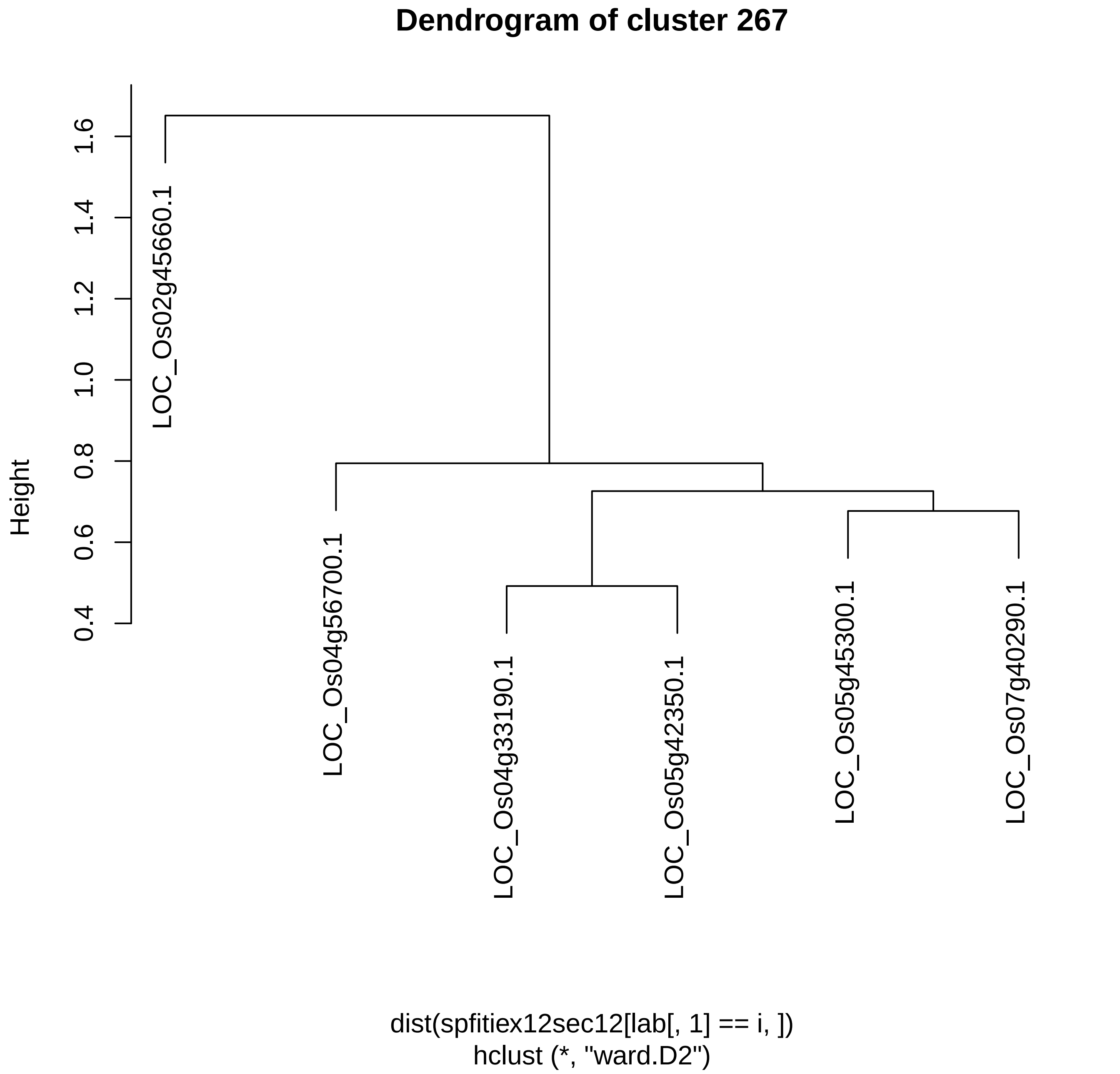
Dendrogram of cluster 267
Height
0.4
0.6
0.8
1.0
1.2
1.4
1.6
LOC_Os02g45660.1
LOC_Os04g56700.1
LOC_Os04g33190.1
LOC_Os05g42350.1
LOC_Os05g45300.1
LOC_Os07g40290.1
dist(spfitiex12sec12[lab[, 1] == i, ])
hclust (*, "ward.D2")

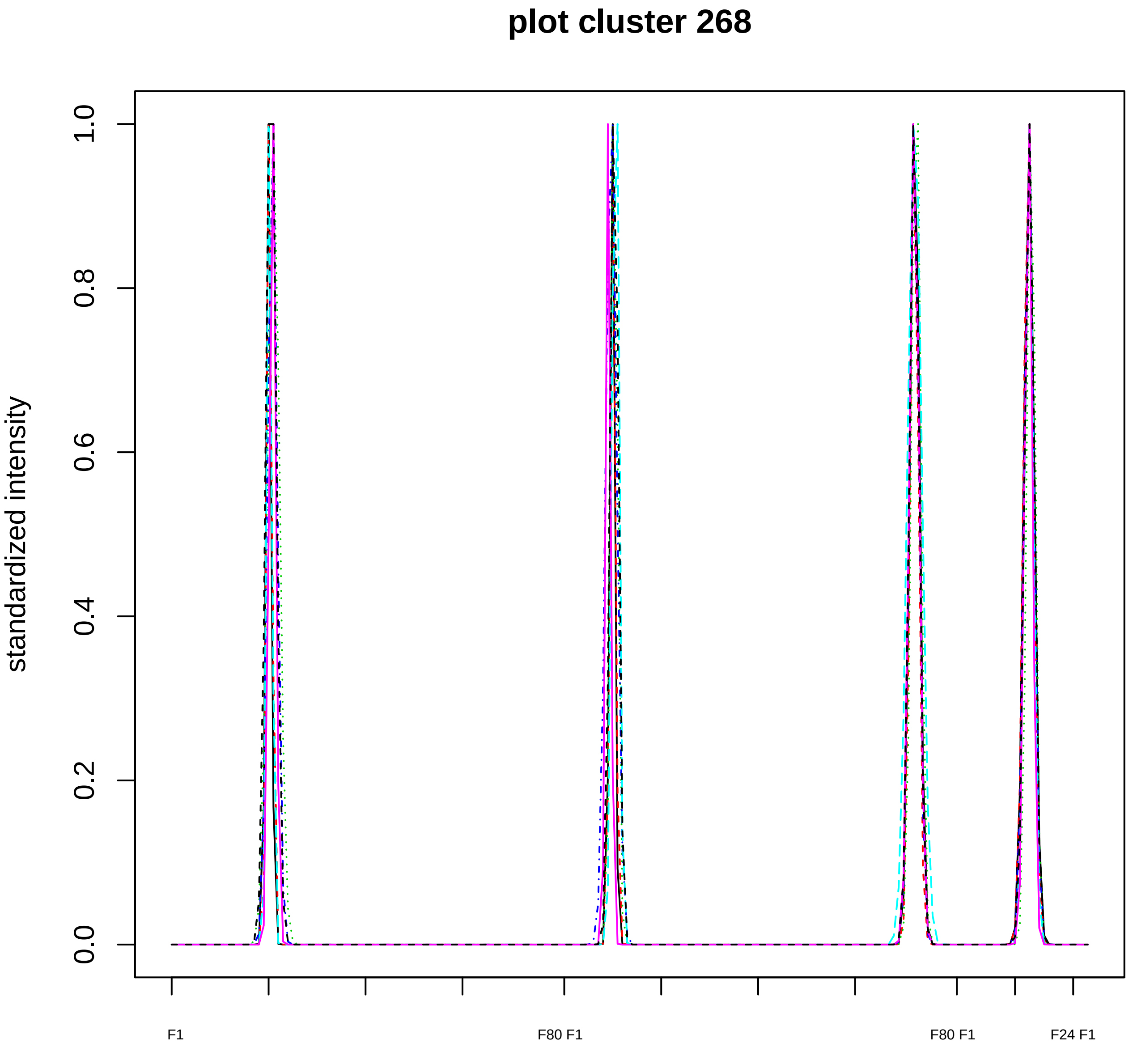
plot cluster 268
standardized intensity
0.0
0.2
0.4
0.6
0.8
1.0
F1
F80 F1
F80 F1
F24 F1

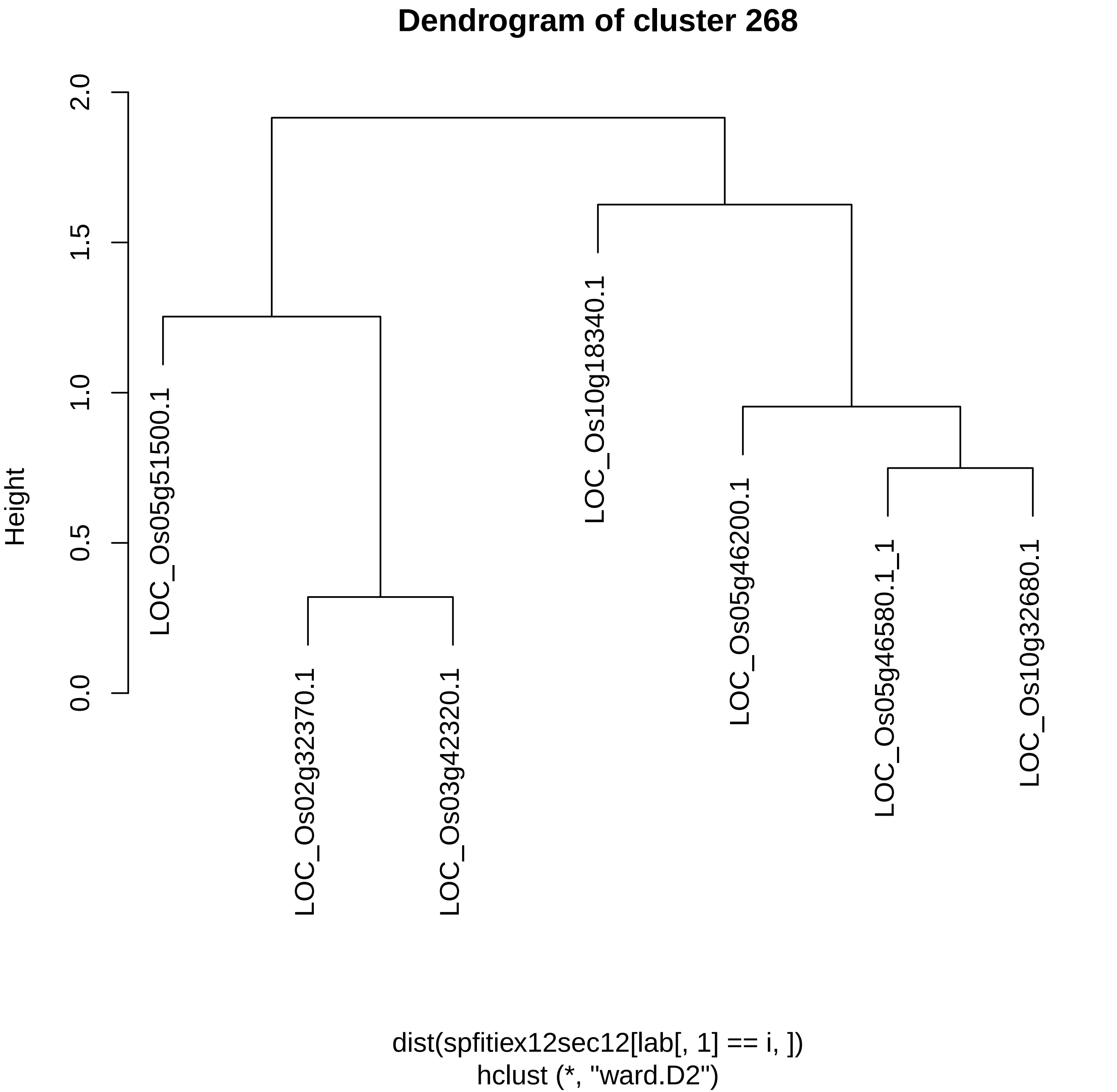
Dendrogram of cluster 268
Height
0.0
0.5
1.0
1.5
2.0
LOC_Os05g51500.1
LOC_Os02g32370.1
LOC_Os03g42320.1
LOC_Os10g18340.1
LOC_Os05g46200.1
LOC_Os05g46580.1_1
LOC_Os10g32680.1
dist(spfitiex12sec12[lab[, 1] == i, ])
hclust (*, "ward.D2")

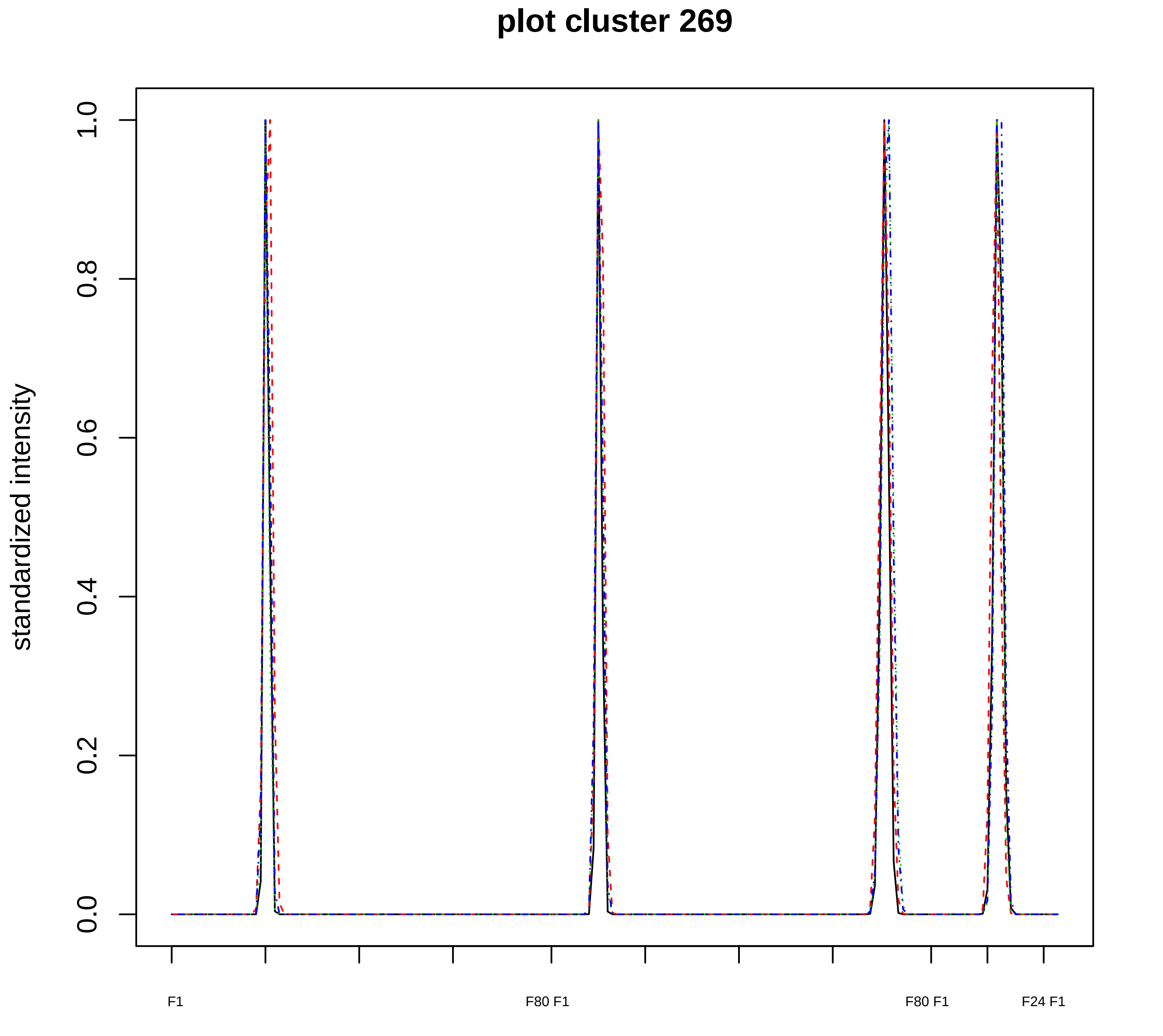

plot cluster 269
standardized intensity
0.0
0.2
0.4
0.6
0.8
1.0
F1
F80 F1
F80 F1
F24 F1

#### Dendrogram of cluster 269

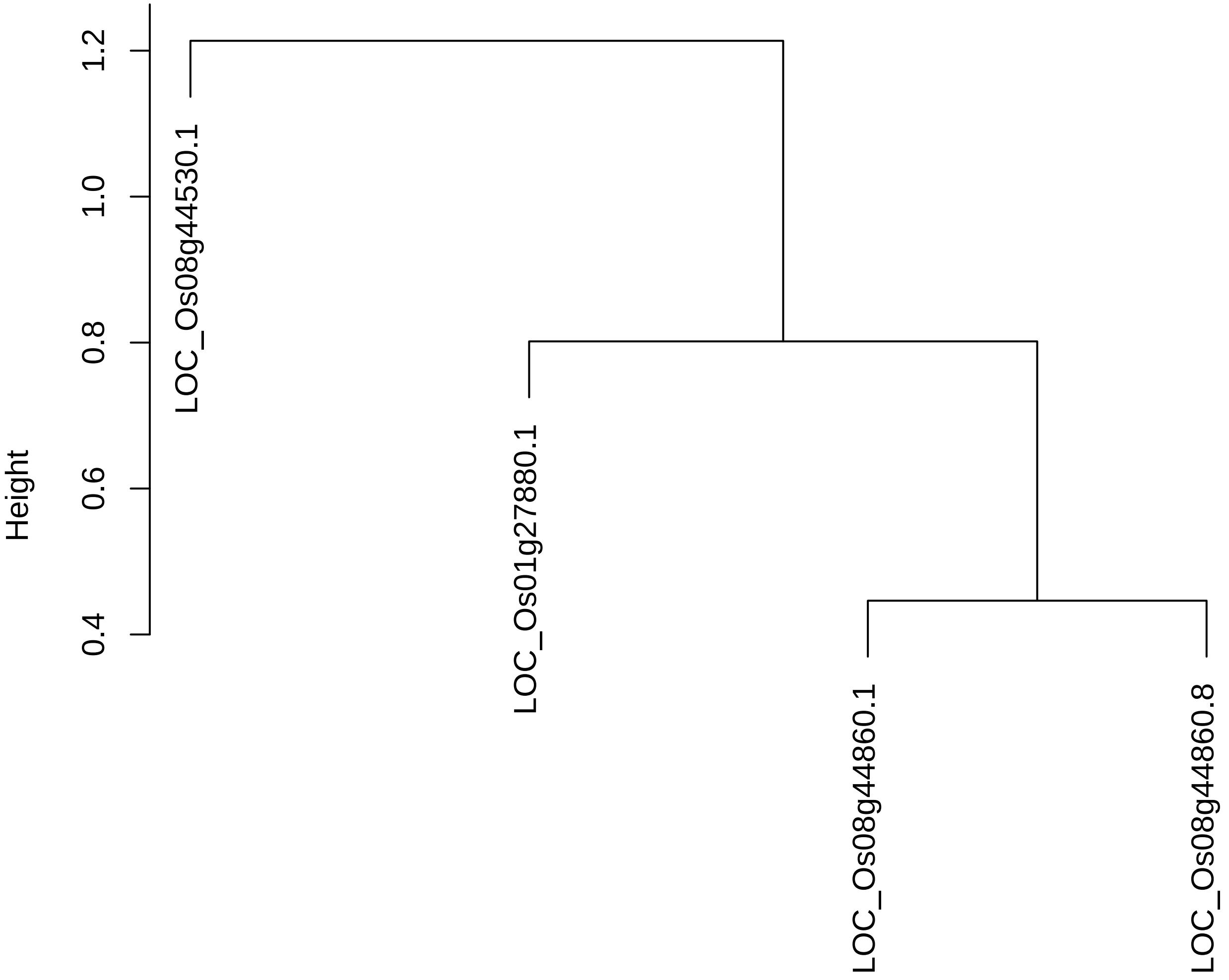


dist(spfitiex12sec12[lab[, 1] == i, ])
hclust (*, "ward.D2")

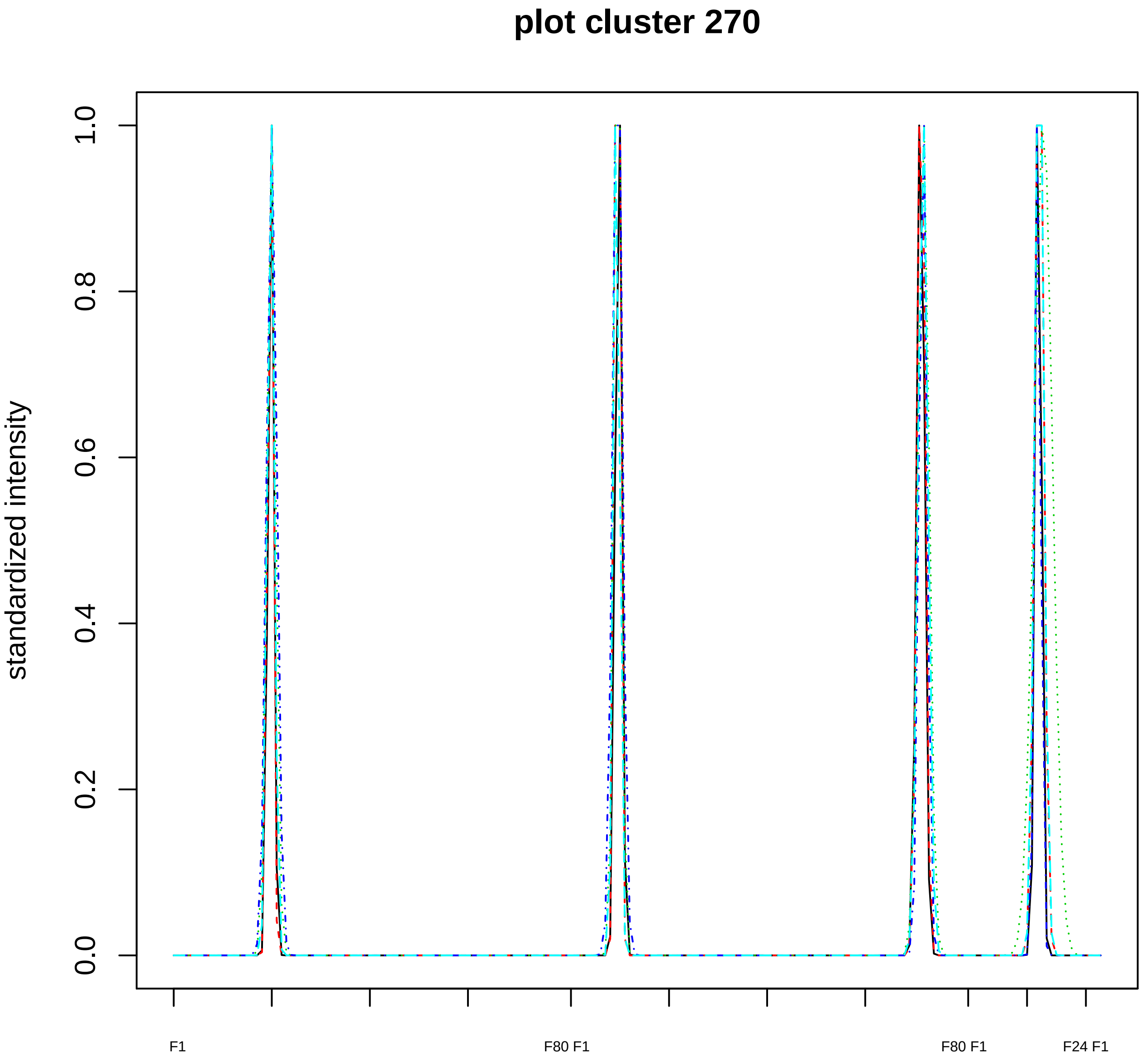

plot cluster 270
standardized intensity
0.0
0.2
0.4
0.6
0.8
1.0
F1
F80 F1
F80 F1
F24 F1

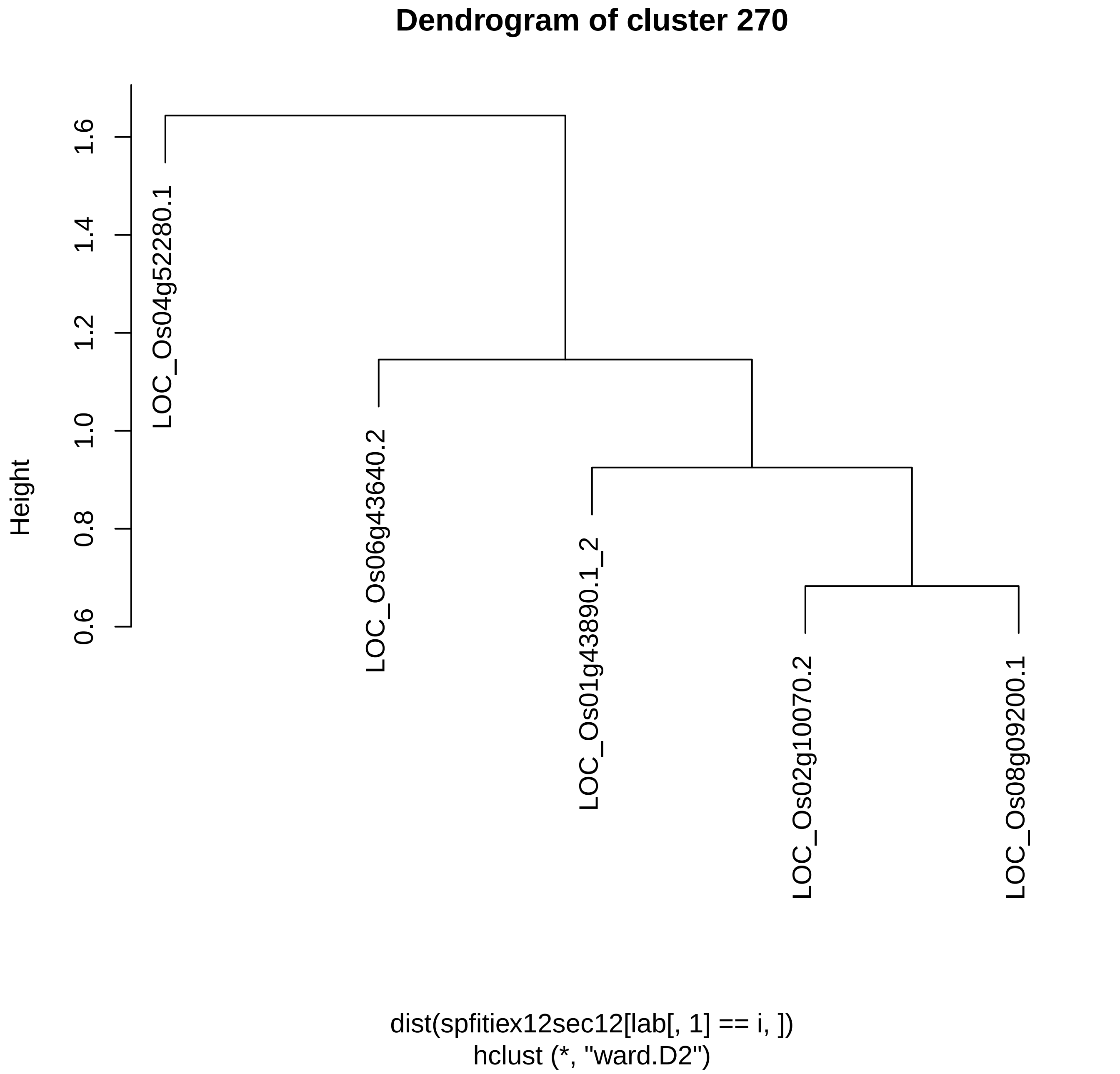
Dendrogram of cluster 270
Height
1.6
1.4
1.2
1.0
0.8
0.6
LOC_Os04g52280.1
LOC_Os06g43640.2
LOC_Os01g43890.1_2
LOC_Os02g10070.2
LOC_Os08g09200.1
dist(spfitiex12sec12[lab[, 1] == i, ])
hclust (*, "ward.D2")

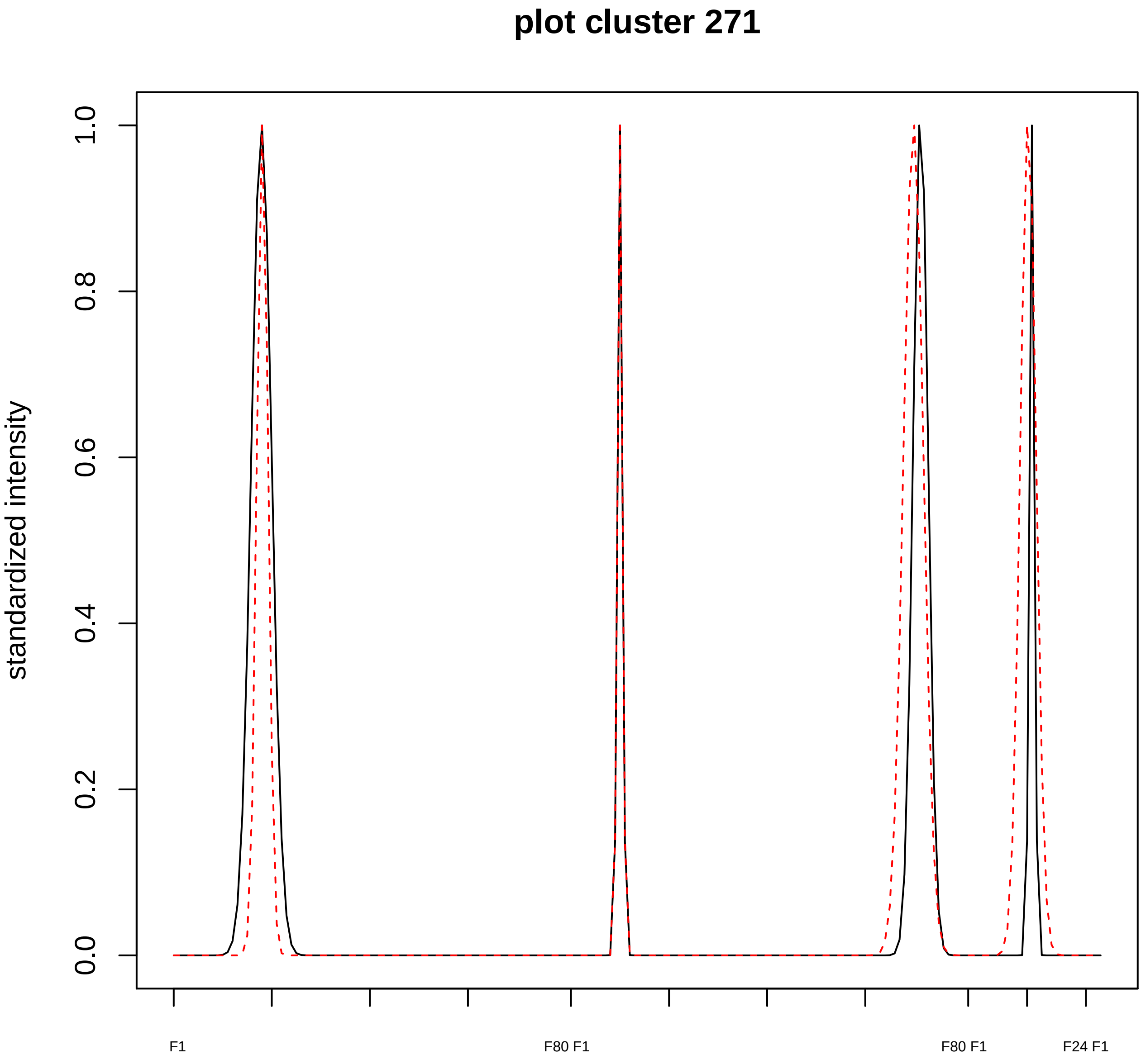

plot cluster 271
standardized intensity
0.0
0.2
0.4
0.6
0.8
1.0
F1
F80 F1
F80 F1
F24 F1

#### plot cluster 272

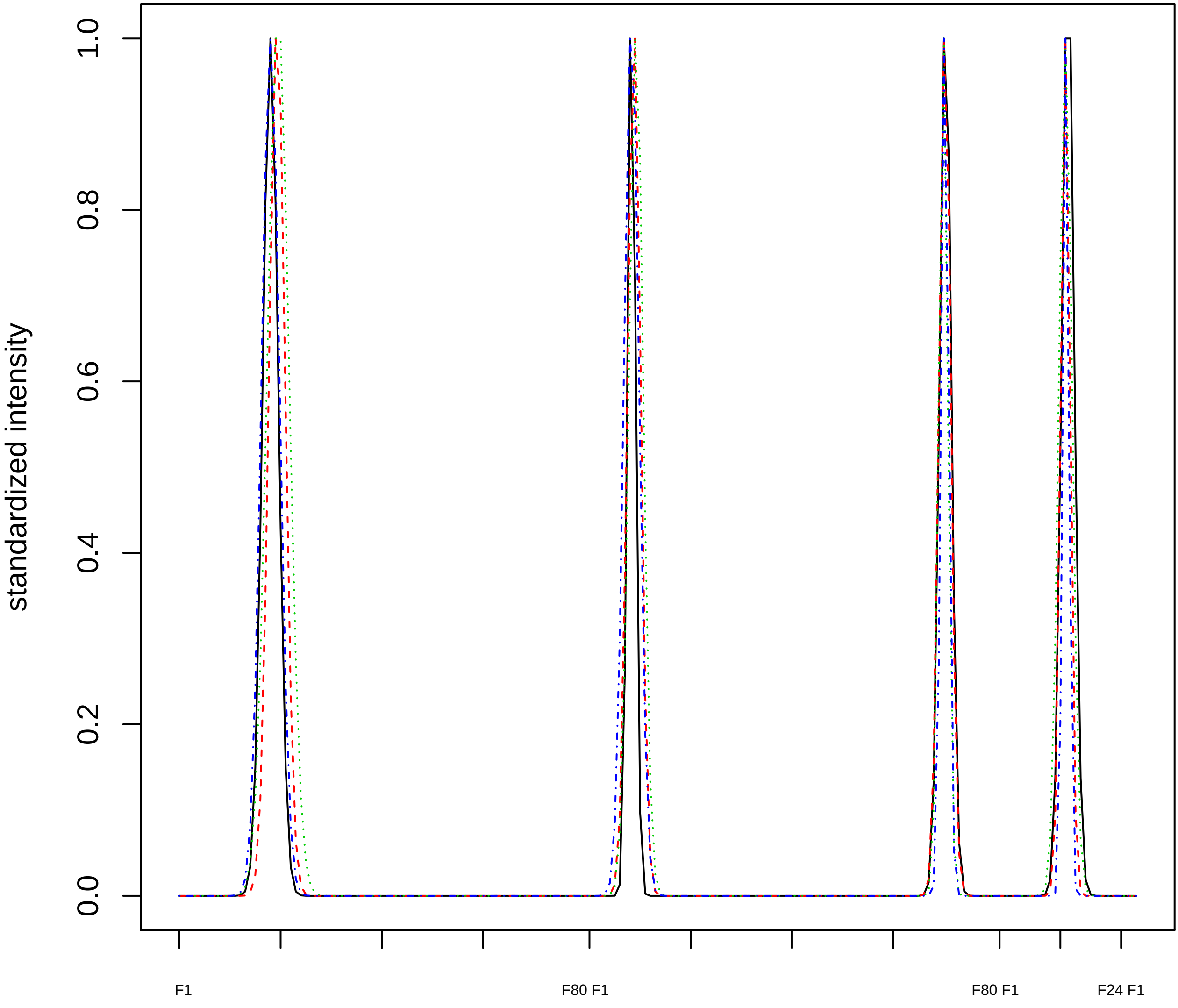

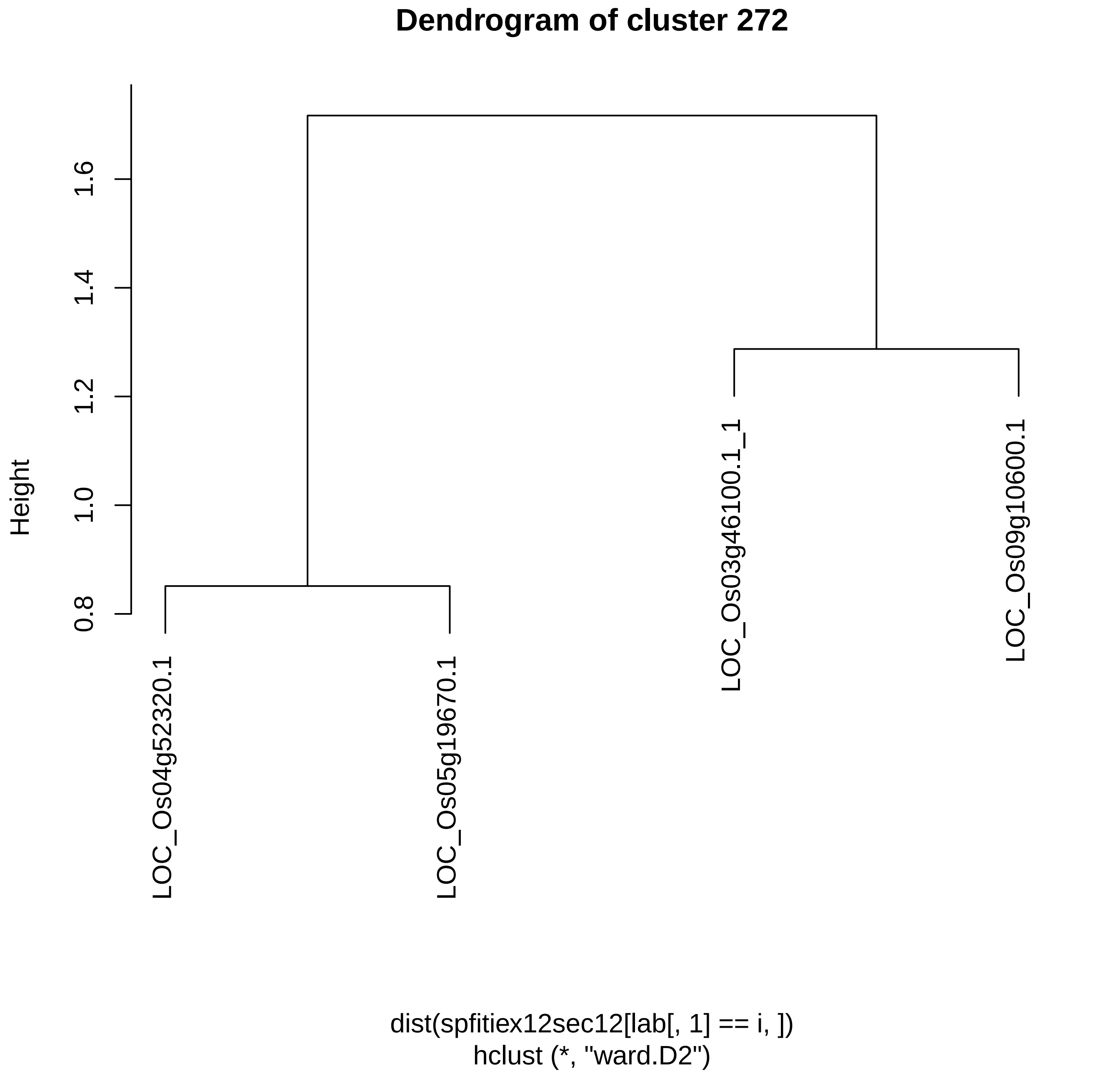
Dendrogram of cluster 272
Height
0.8
1.0
1.2
1.4
1.6
LOC_Os04g52320.1
LOC_Os05g19670.1
LOC_Os03g46100.1_1
LOC_Os09g10600.1
dist(spfitiex12sec12[lab[, 1] == i, ])
hclust (*, "ward.D2")

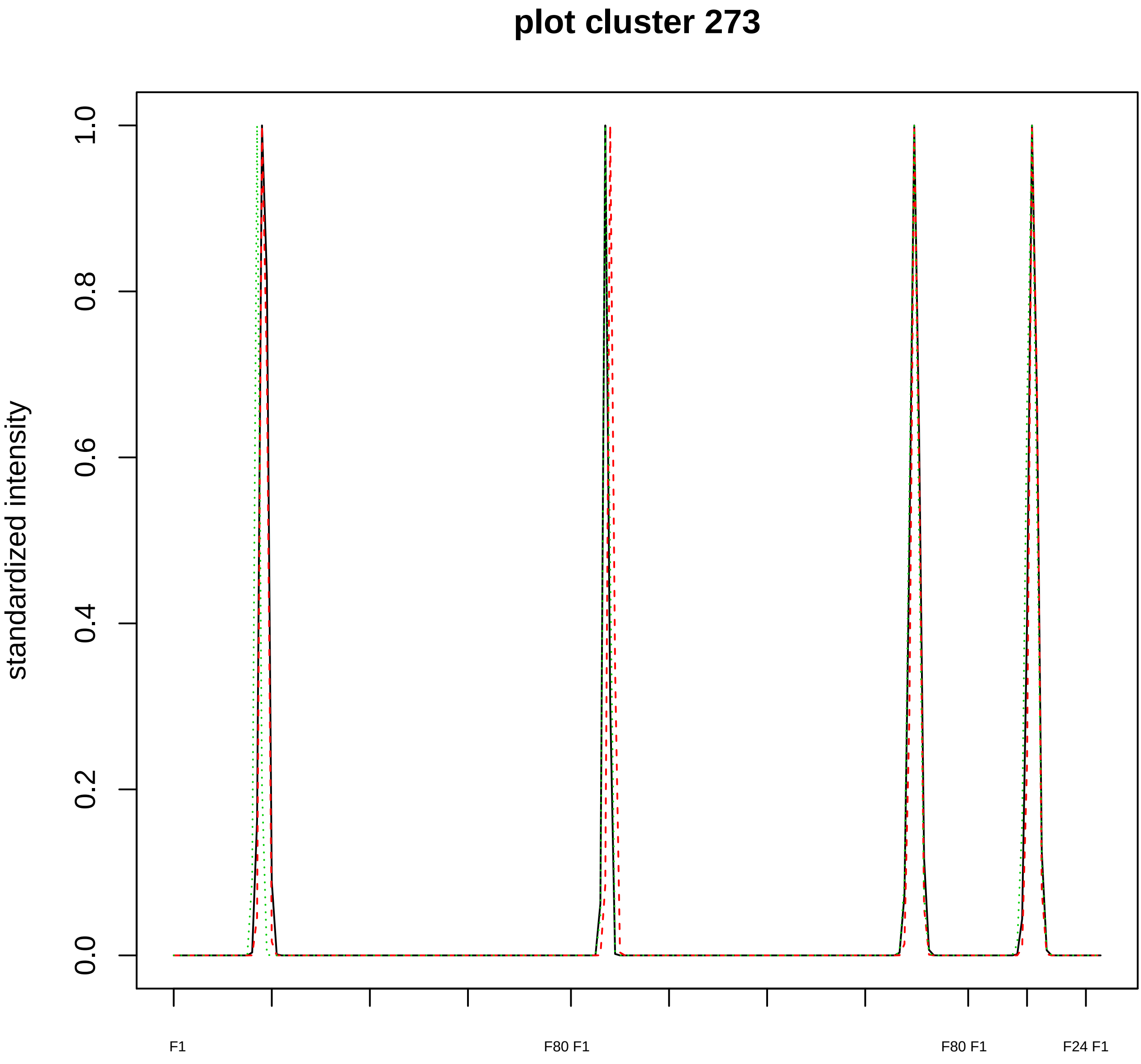

plot cluster 273
standardized intensity
0.0
0.2
0.4
0.6
0.8
1.0
F1
F80 F1
F80 F1
F24 F1

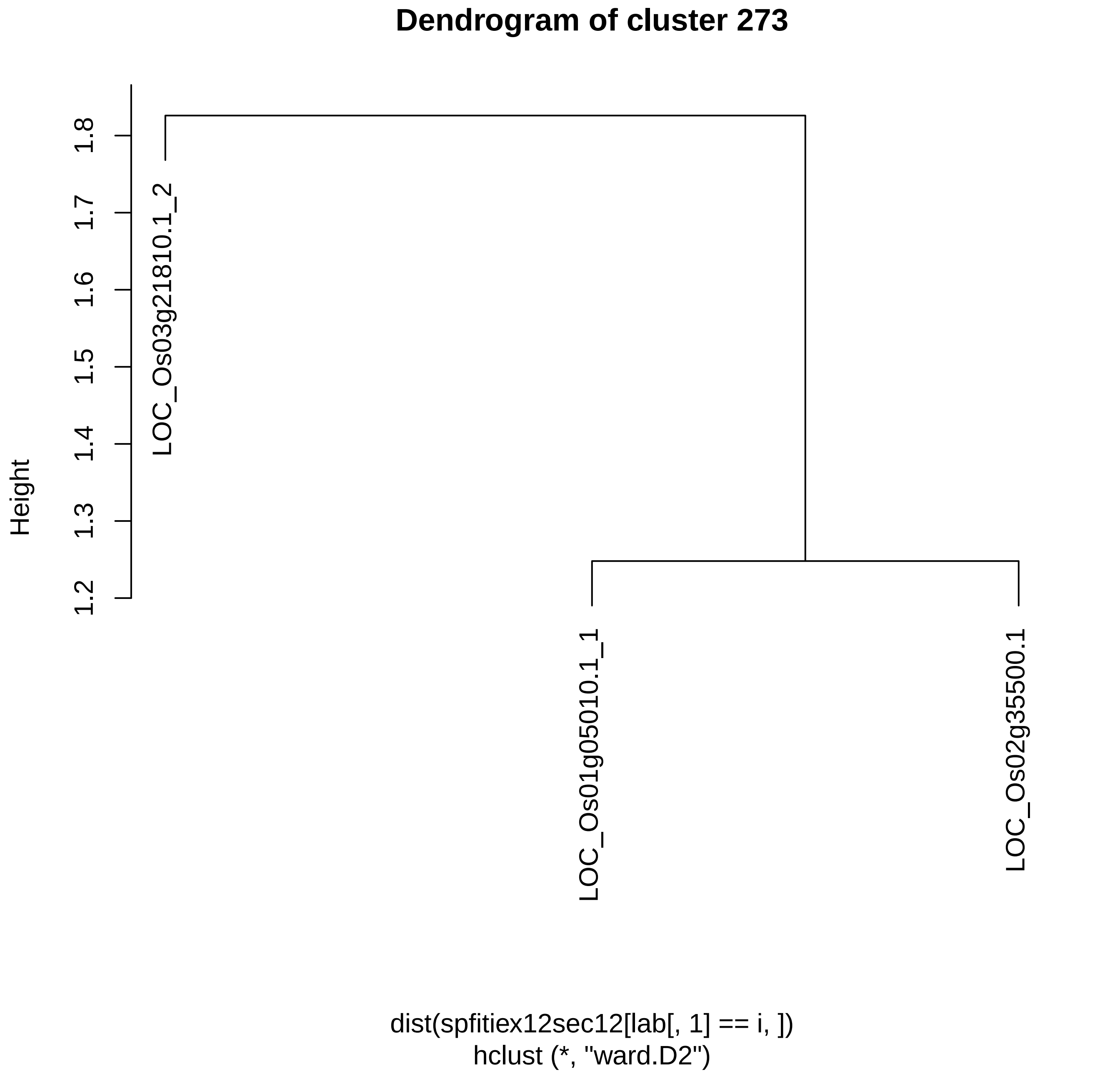
Dendrogram of cluster 273
Height
1.2
1.3
1.4
1.5
1.6
1.7
1.8
LOC_Os03g21810.1_2
LOC_Os01g05010.1_1
LOC_Os02g35500.1
dist(spfitiex12sec12[lab[, 1] == i, ])
hclust (*, "ward.D2")

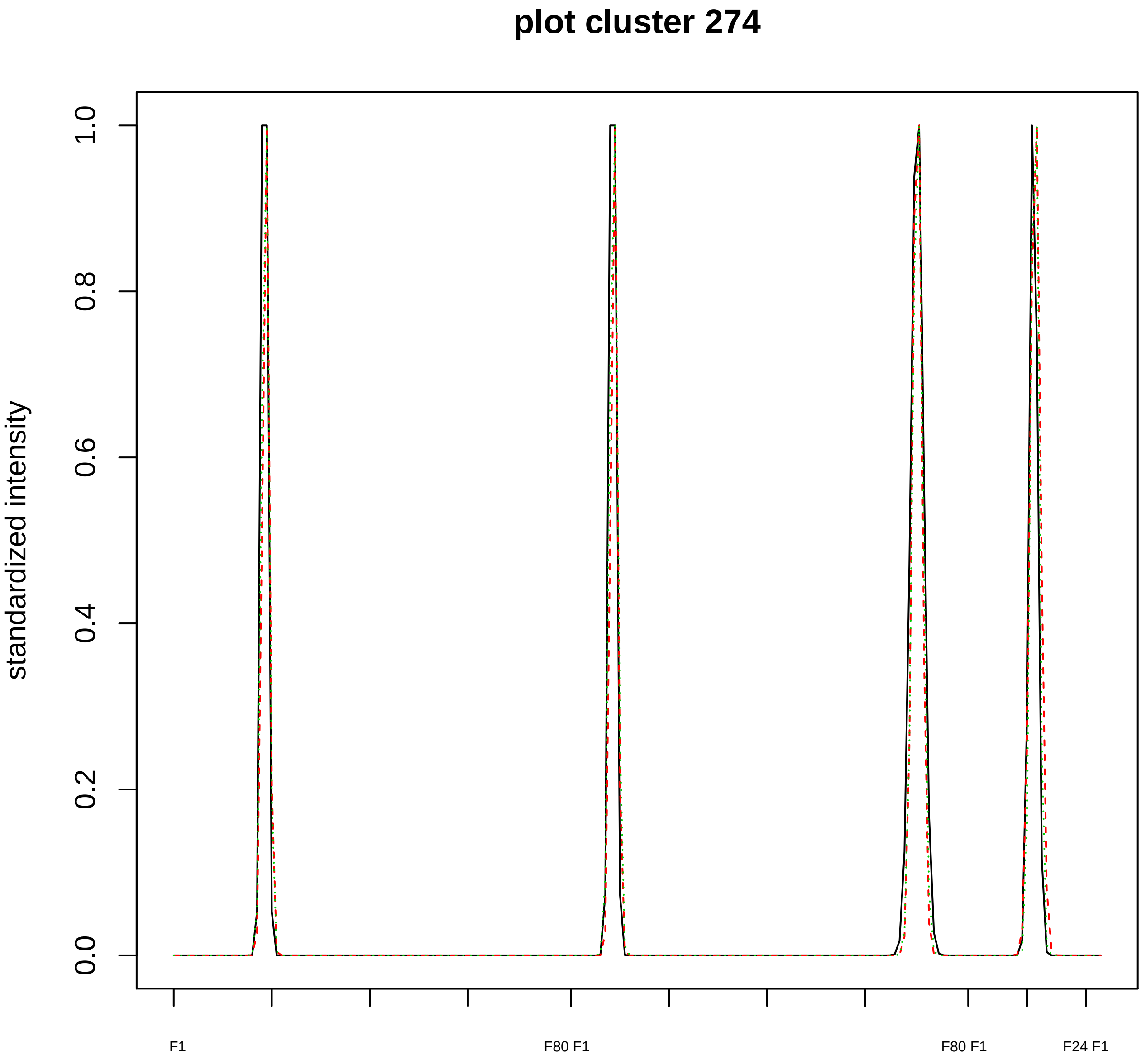
plot cluster 274
standardized intensity
0.0
0.2
0.4
0.6
0.8
1.0
F1
F80 F1
F80 F1
F24 F1

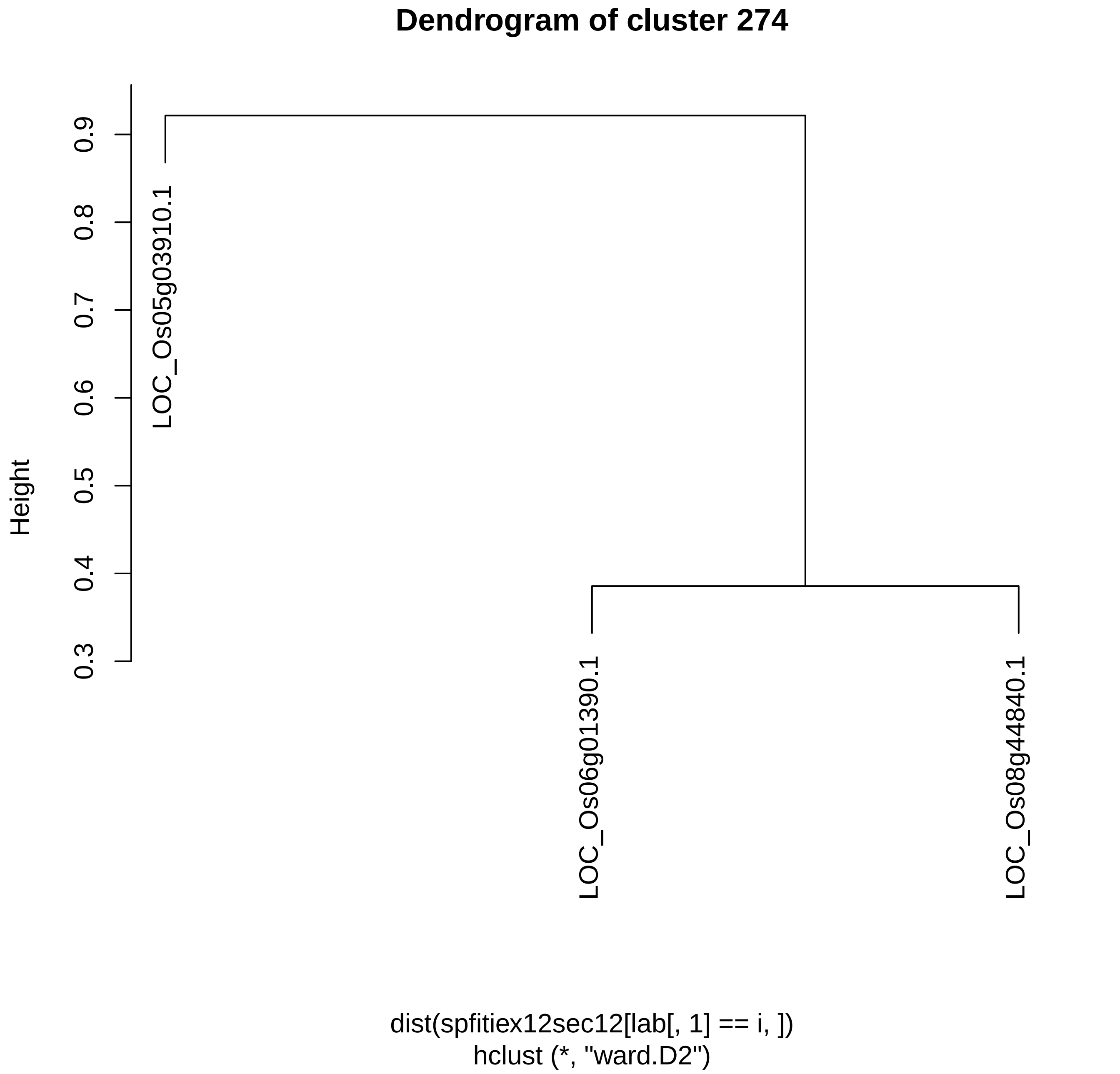

Dendrogram of cluster 274
Height
0.3
0.4
0.5
0.6
0.7
0.8
0.9
LOC_Os05g03910.1
LOC_Os06g01390.1
LOC_Os08g44840.1
dist(spfitiex12sec12[lab[, 1] == i, ])
hclust (*, "ward.D2")

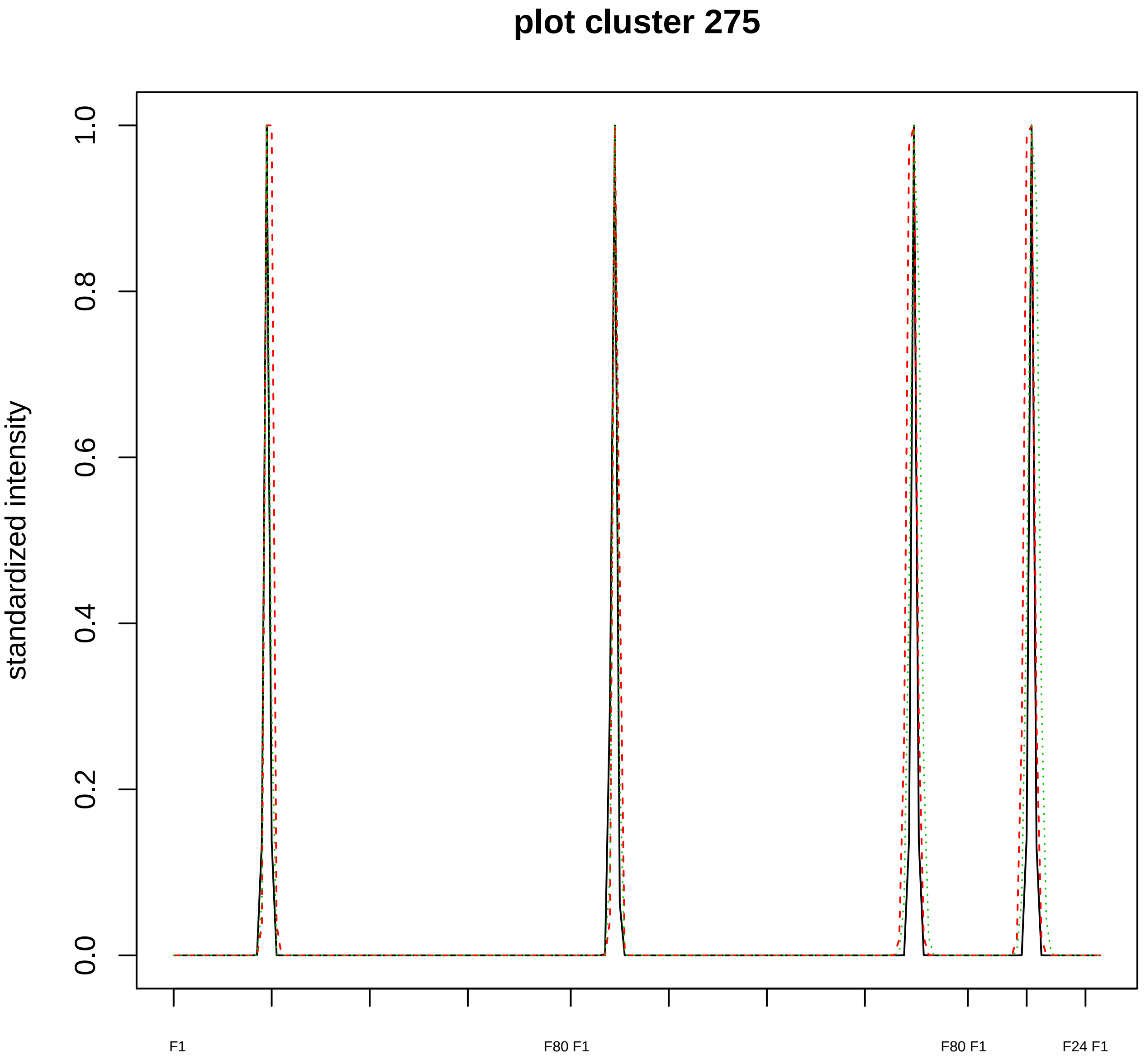
plot cluster 275
standardized intensity
0.0
0.2
0.4
0.6
0.8
1.0
F1
F80 F1
F80 F1
F24 F1

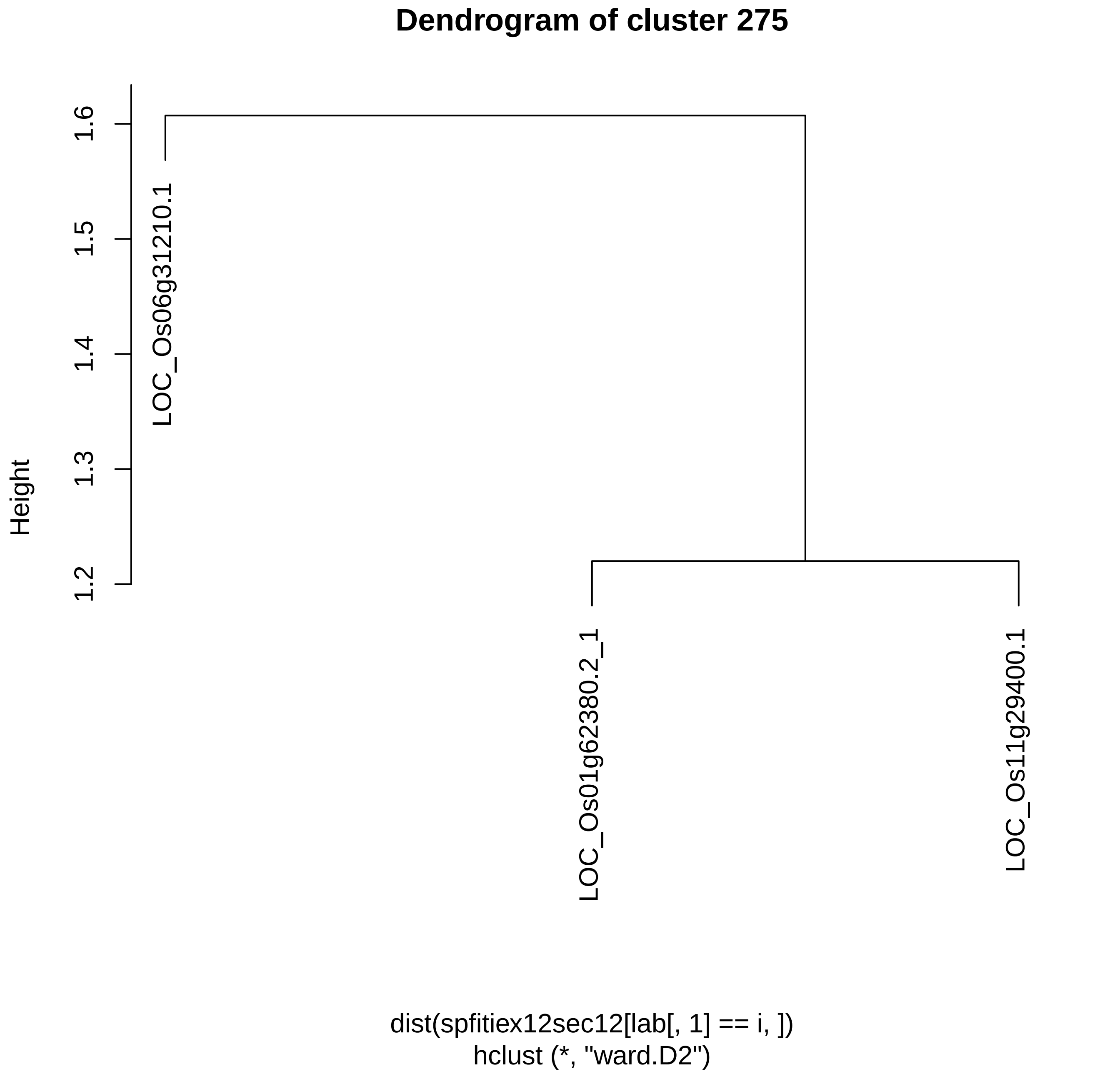
Dendrogram of cluster 275
Height
1.2
1.3
1.4
1.5
1.6
LOC_Os06g31210.1
LOC_Os01g62380.2_1
LOC_Os11g29400.1
dist(spfitiex12sec12[lab[, 1] == i, ])
hclust (*, "ward.D2")

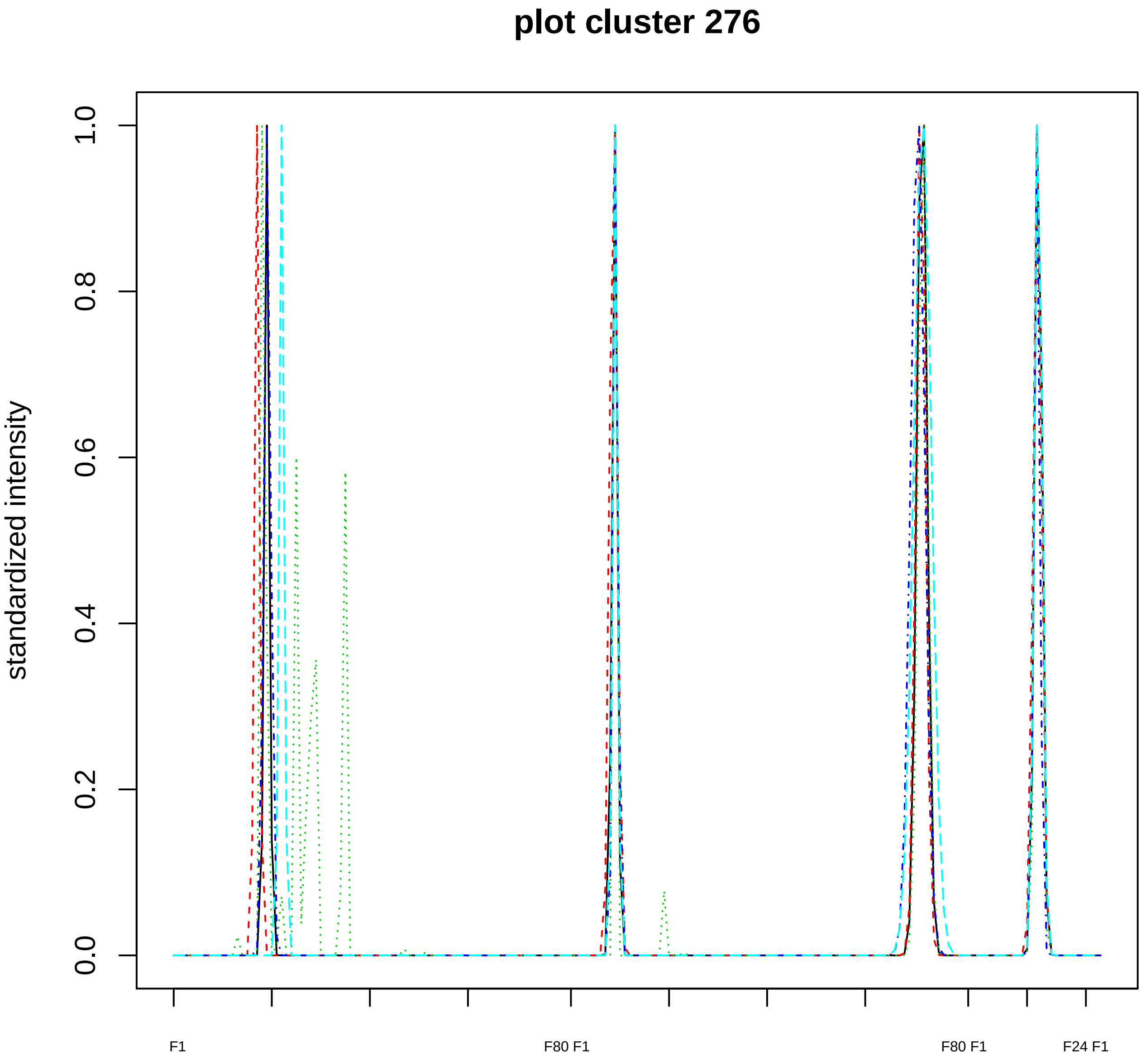
plot cluster 276
standardized intensity
0.0
0.2
0.4
0.6
0.8
1.0
F1
F80 F1
F80 F1
F24 F1

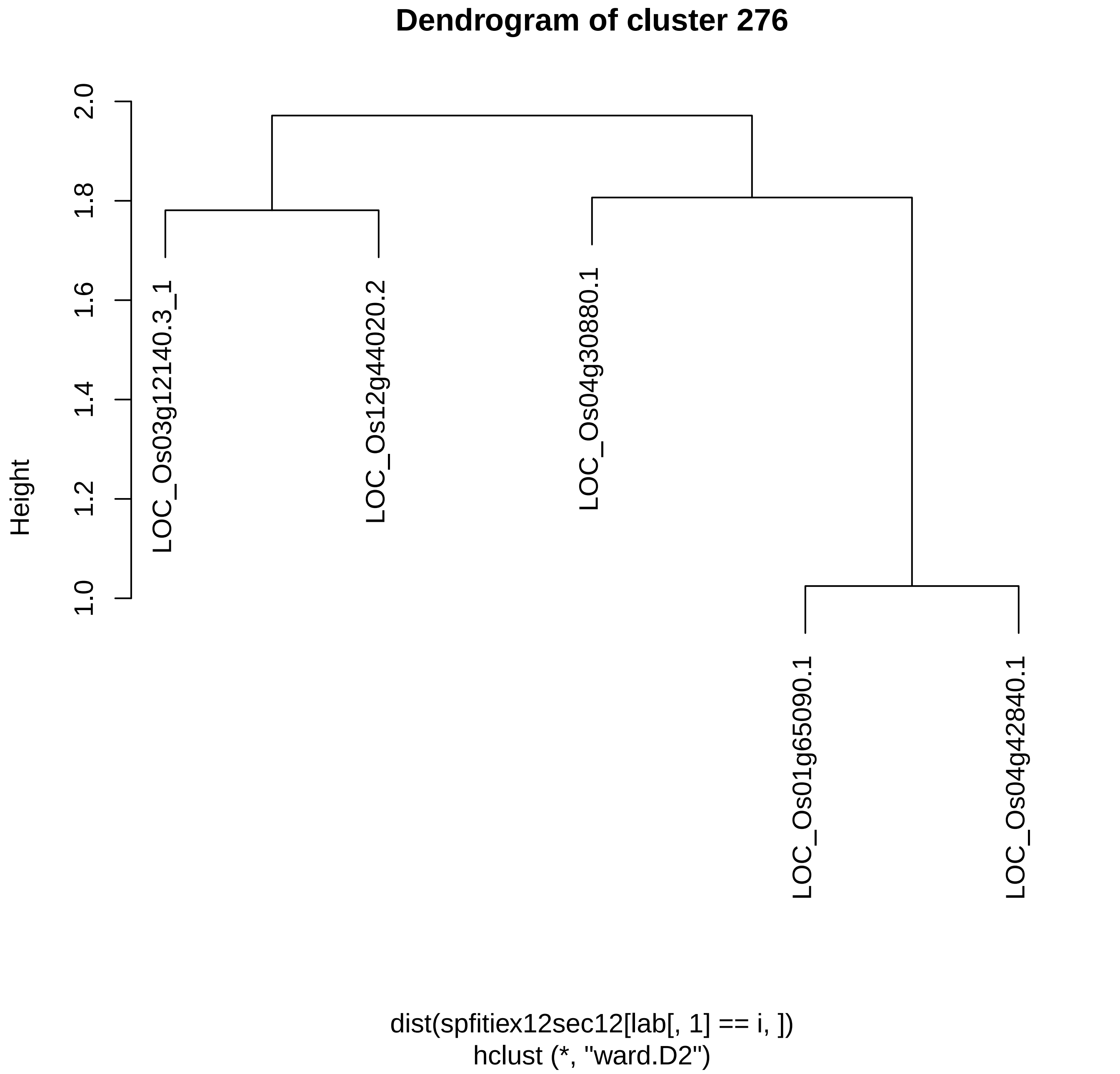
Dendrogram of cluster 276
Height
2.0
1.8
1.6
1.4
1.2
1.0
LOC_Os03g12140.3_1
LOC_Os12g44020.2
LOC_Os04g30880.1
LOC_Os01g65090.1
LOC_Os04g42840.1
dist(spfitiex12sec12[lab[, 1] == i, ])
hclust (*, "ward.D2")

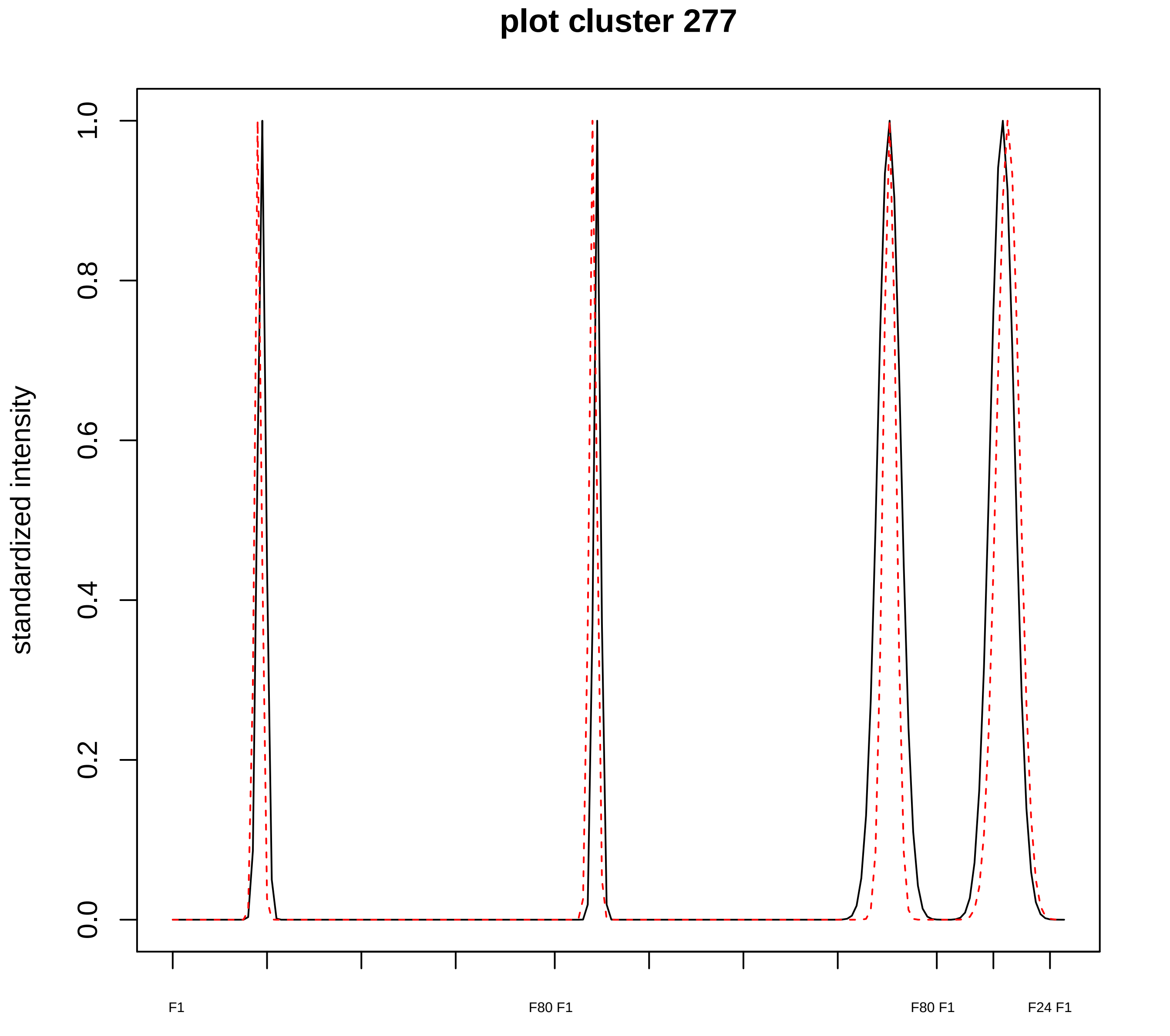
plot cluster 277
standardized intensity
0.0
0.2
0.4
0.6
0.8
1.0
F1
F80 F1
F80 F1
F24 F1

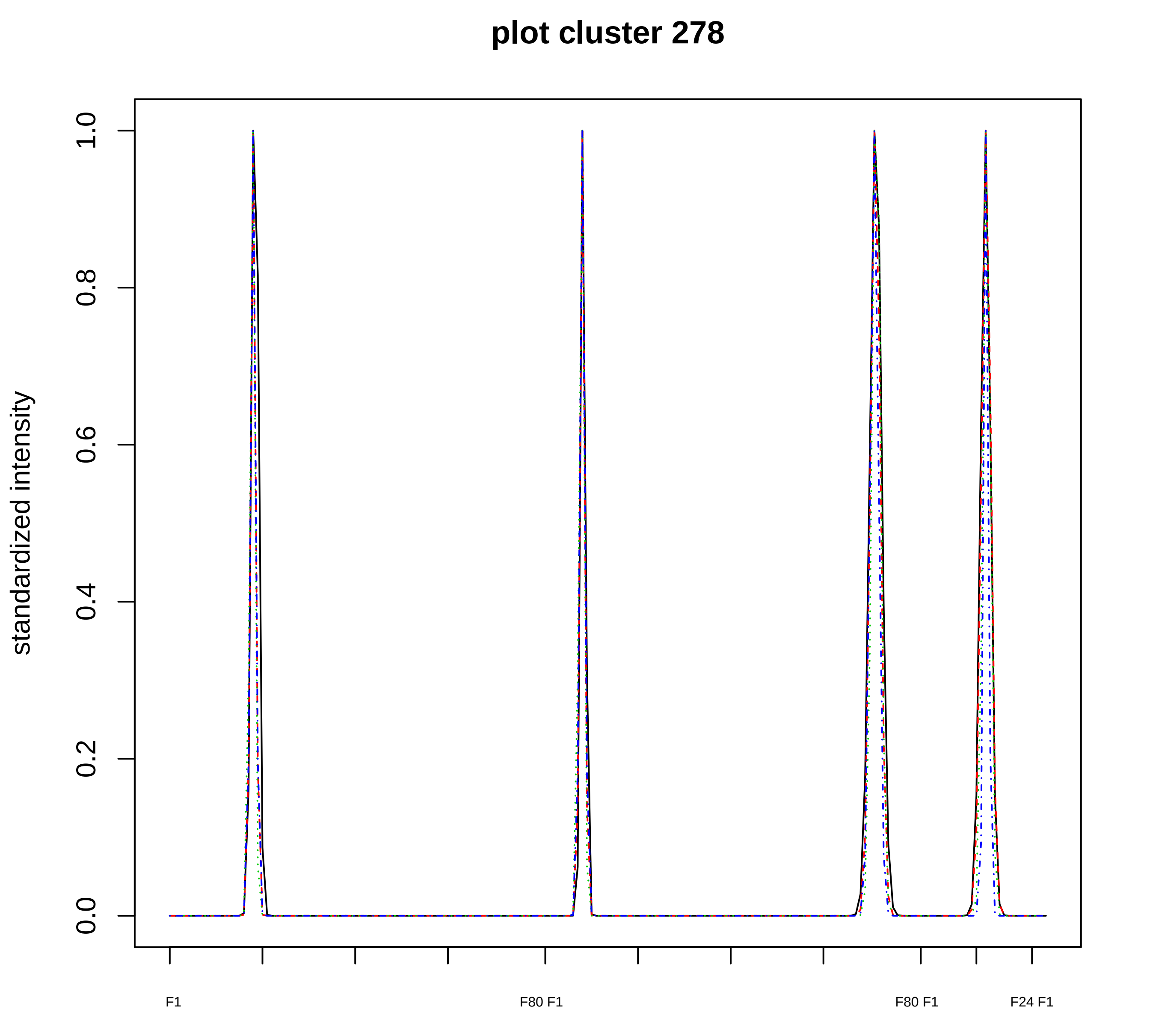
plot cluster 278
standardized intensity
0.0
0.2
0.4
0.6
0.8
1.0
F1
F80 F1
F80 F1
F24 F1

#### Dendrogram of cluster 278

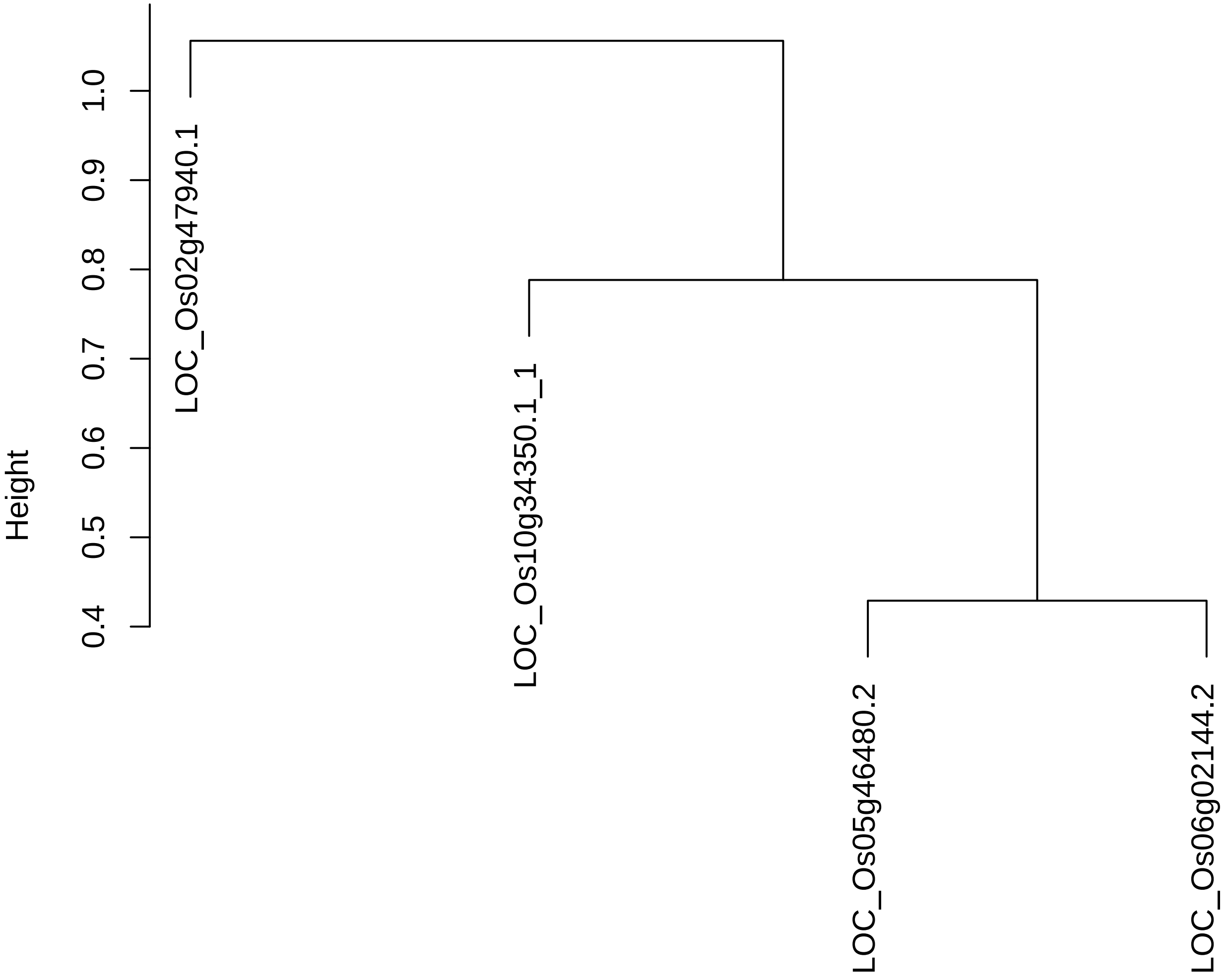


dist(spfitiex12sec12[lab[, 1] == i, ])
hclust (*, "ward.D2")

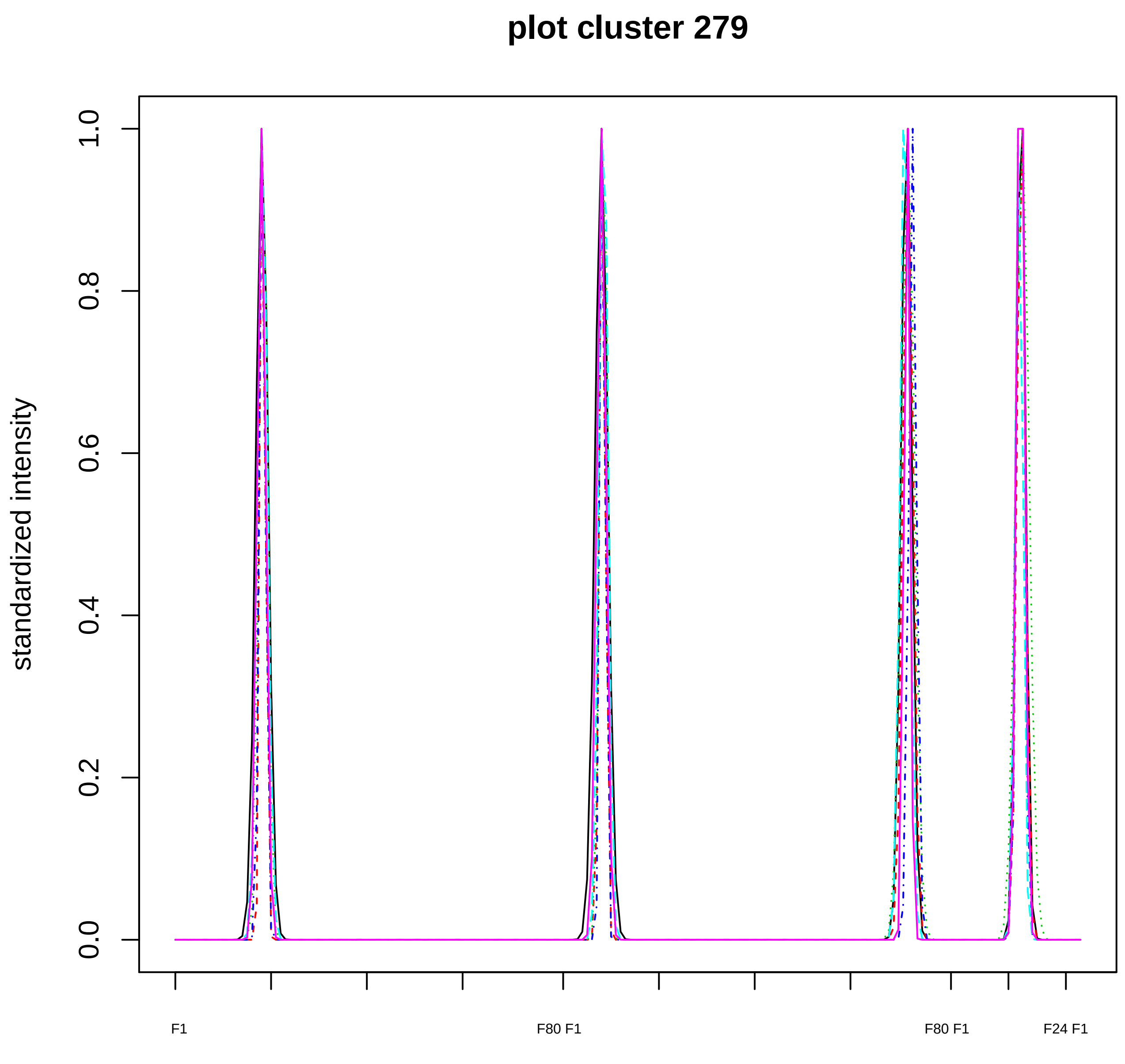
plot cluster 279
standardized intensity
0.0
0.2
0.4
0.6
0.8
1.0
F1
F80 F1
F80 F1
F24 F1

### Dendrogram of cluster 279

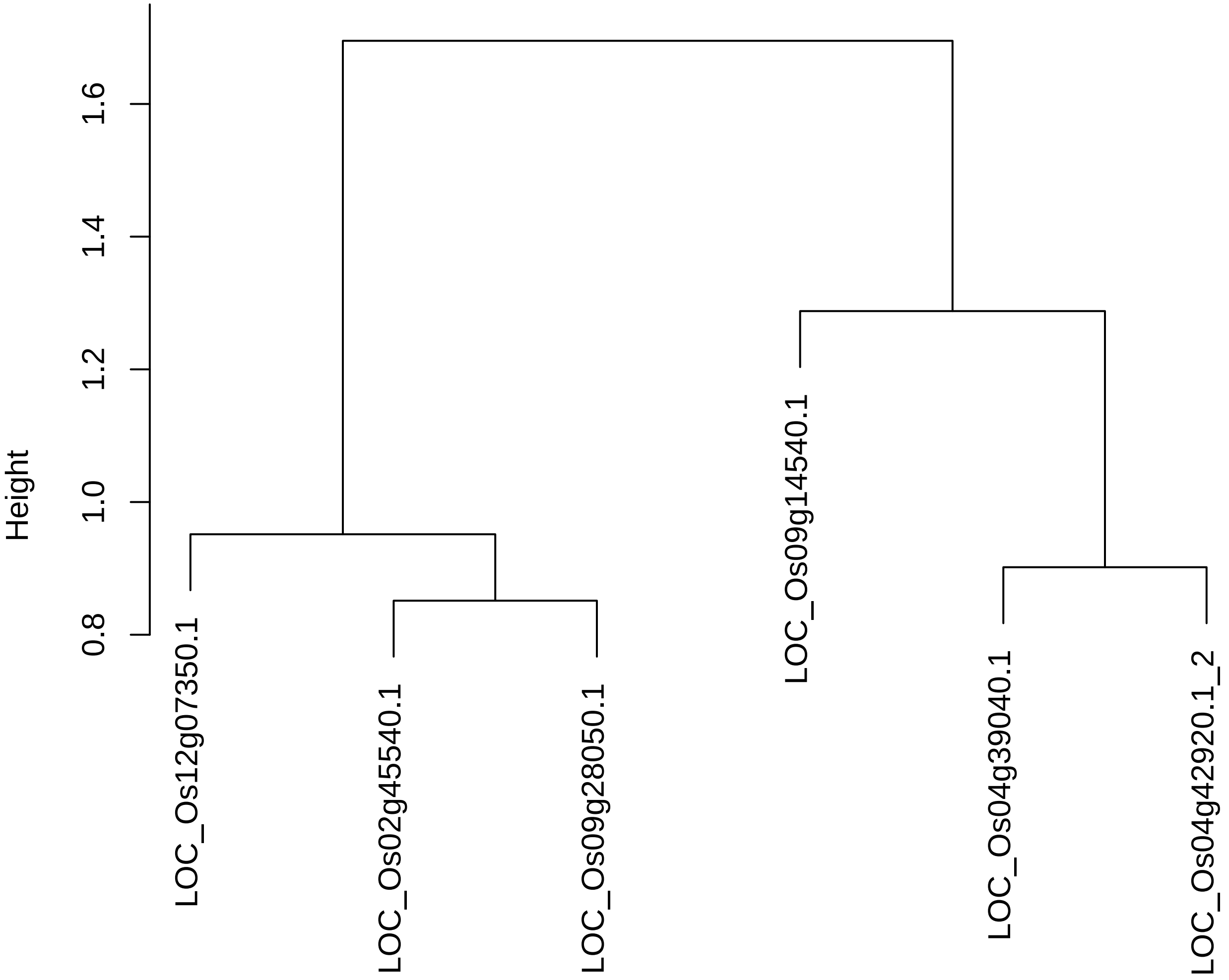


dist(spfitiex12sec12[lab[, 1] == i, ])
hclust (*, "ward.D2")

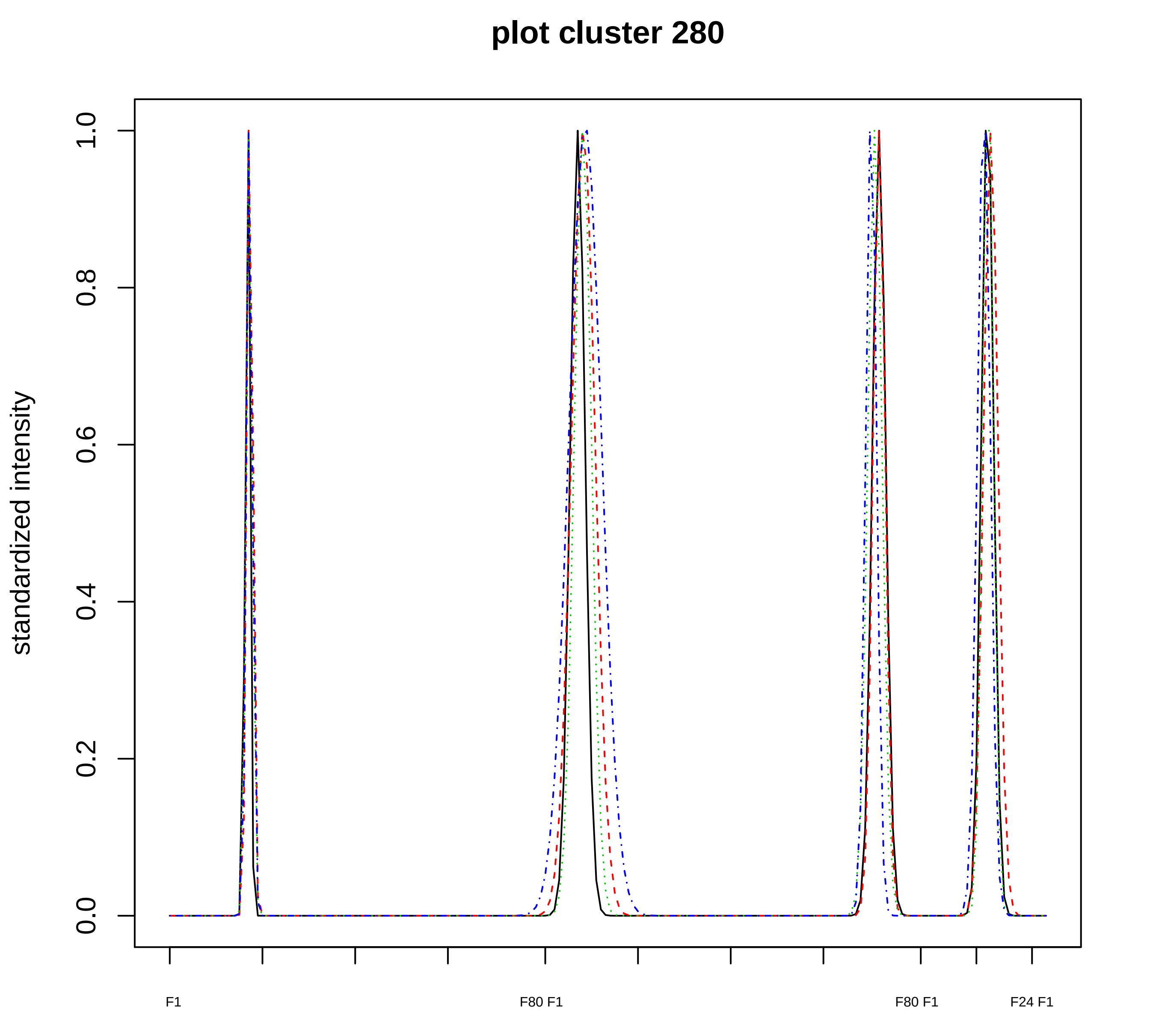
plot cluster 280
standardized intensity
0.0
0.2
0.4
0.6
0.8
1.0
F1
F80 F1
F80 F1
F24 F1

#### Dendrogram of cluster 280

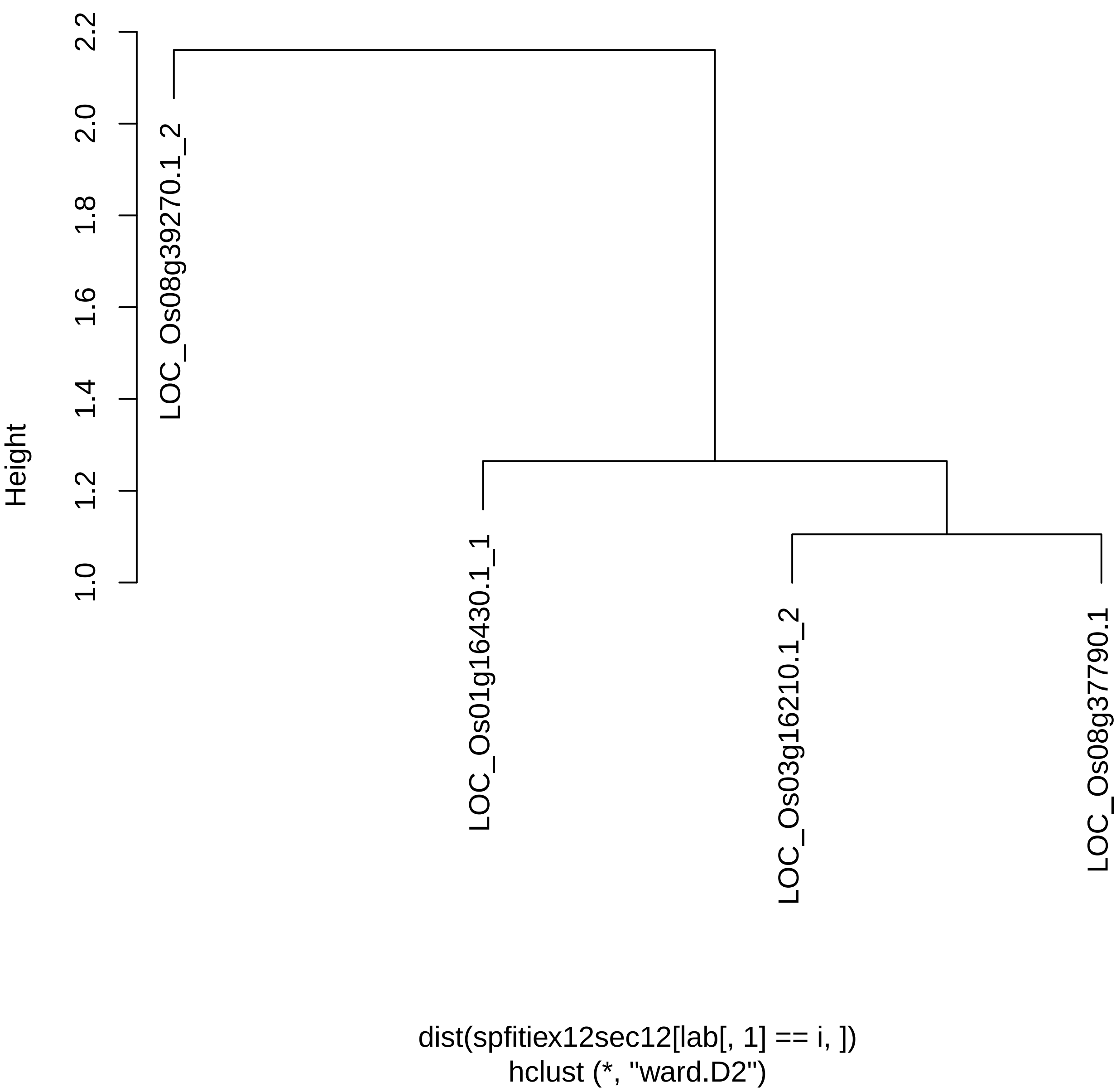

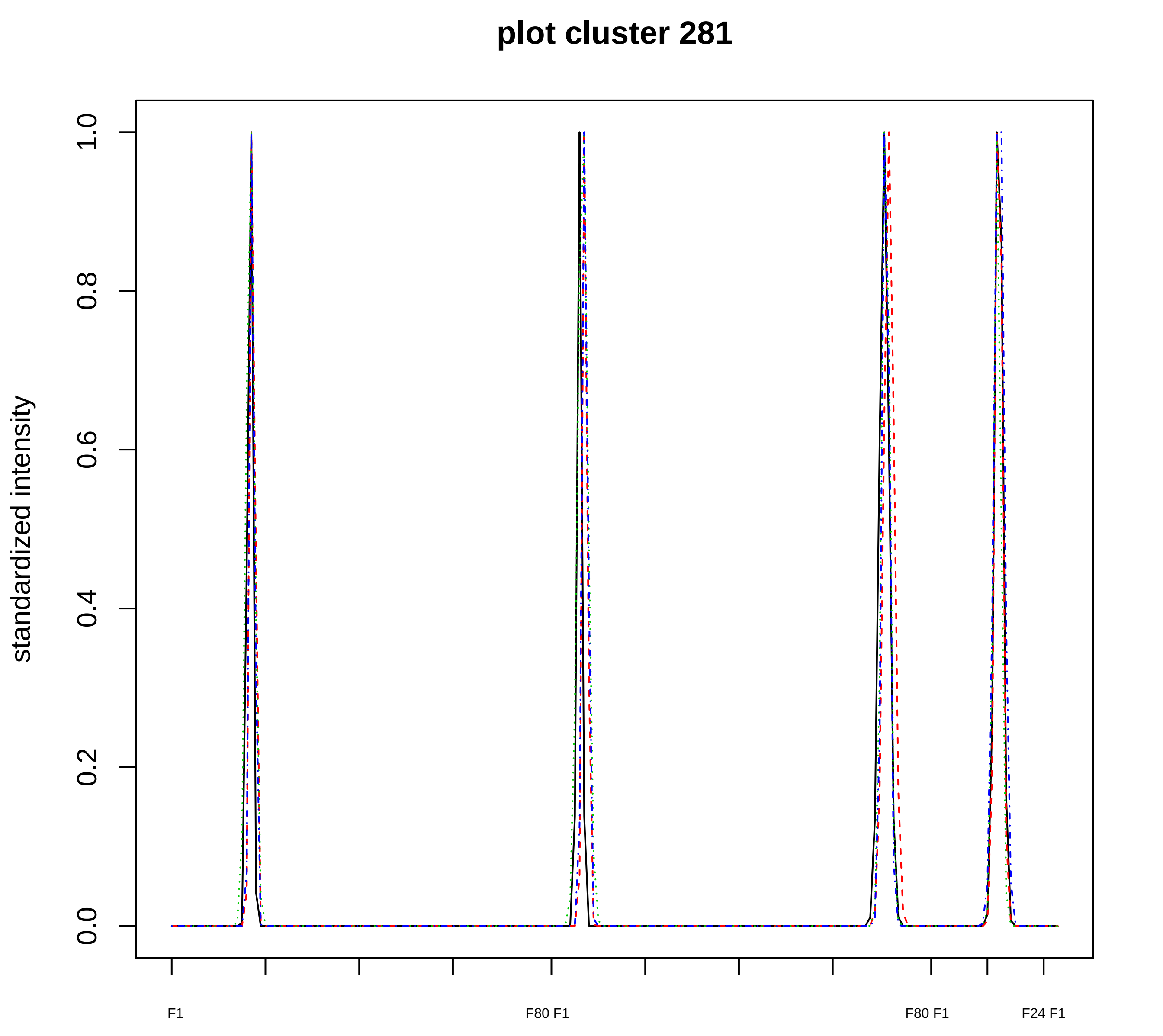
plot cluster 281
standardized intensity
0.0
0.2
0.4
0.6
0.8
1.0
F1
F80 F1
F80 F1
F24 F1

### Dendrogram of cluster 281

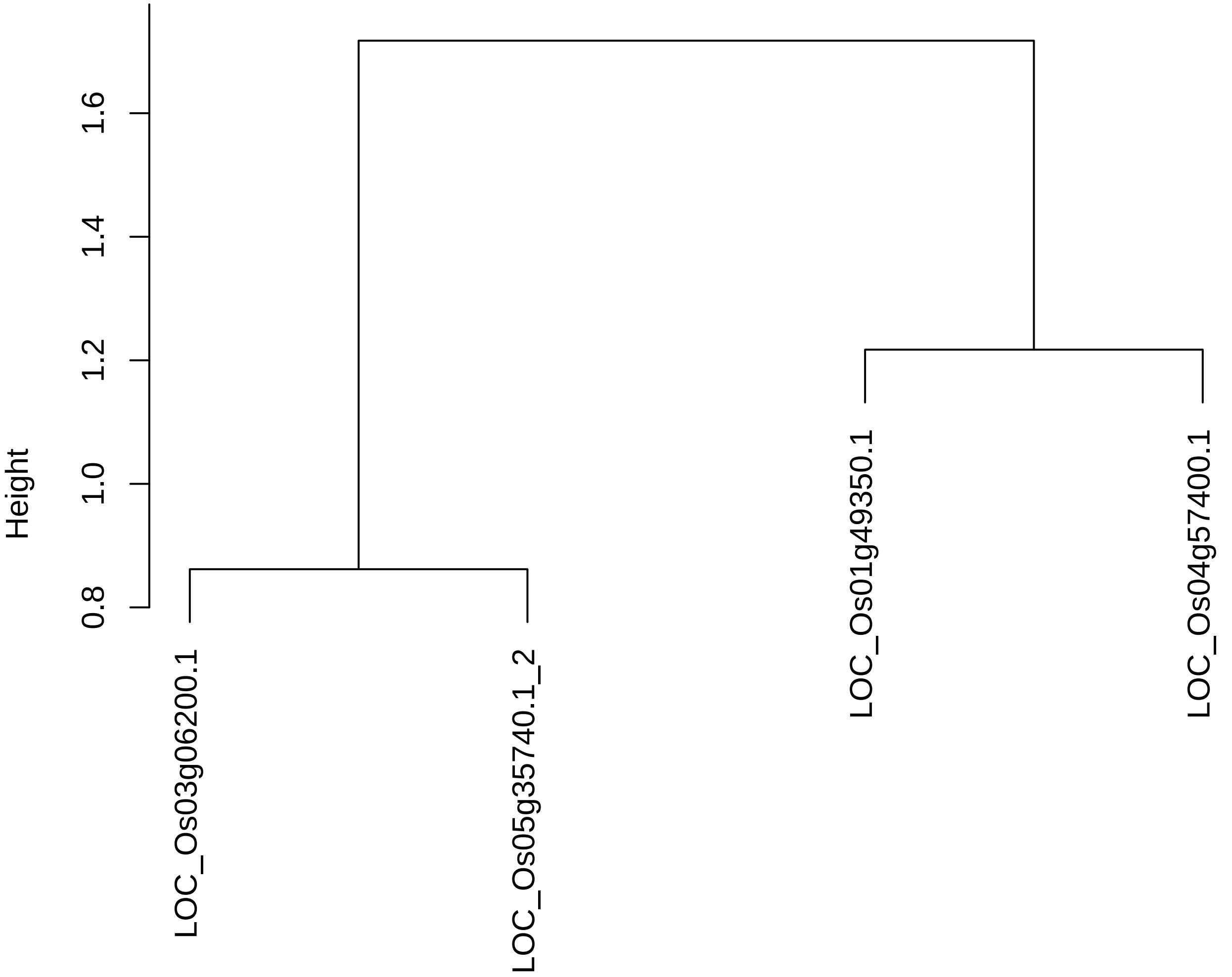


dist(spfitiex12sec12[lab[, 1] == i, ])
hclust (*, "ward.D2")

### plot cluster 282

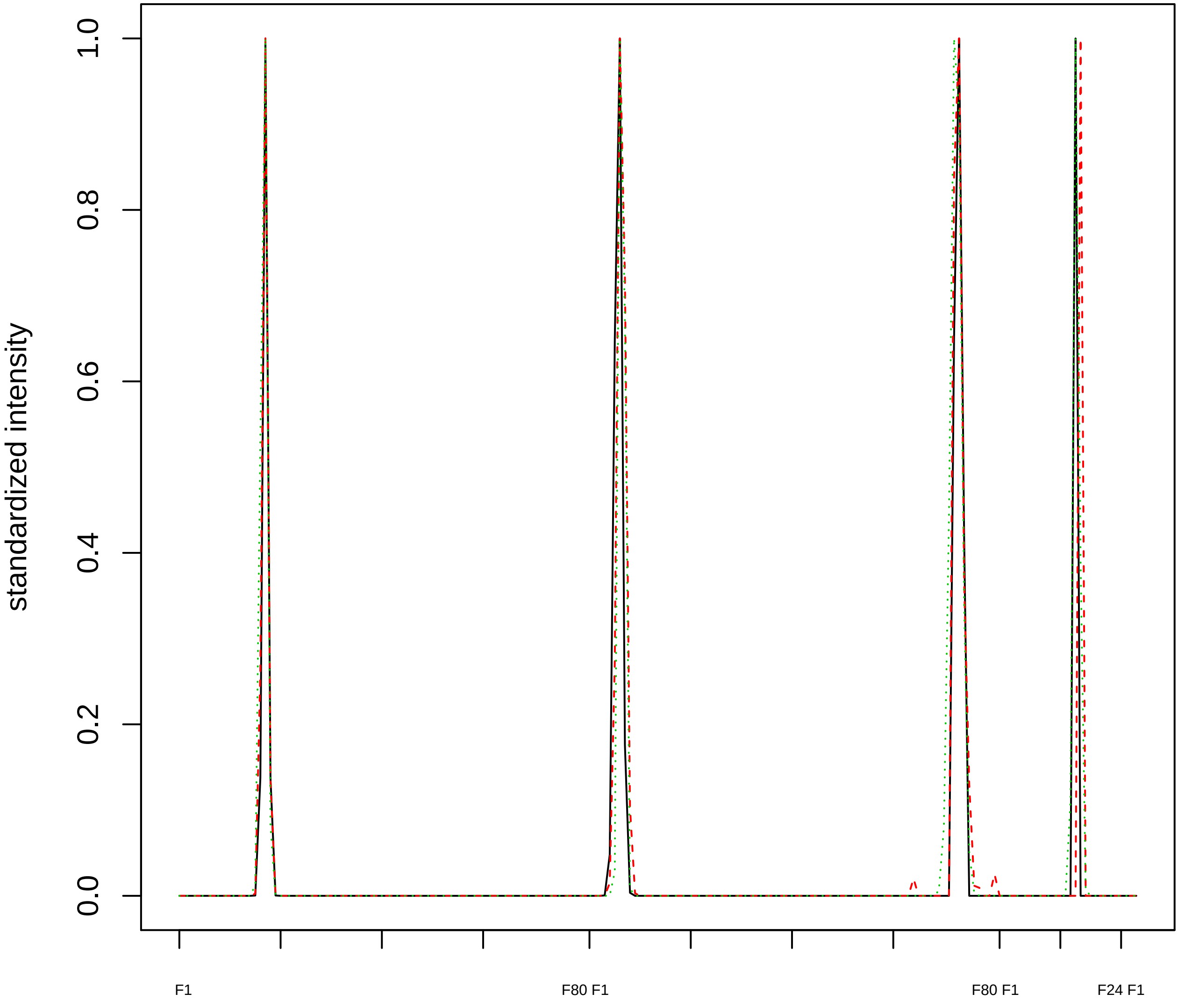

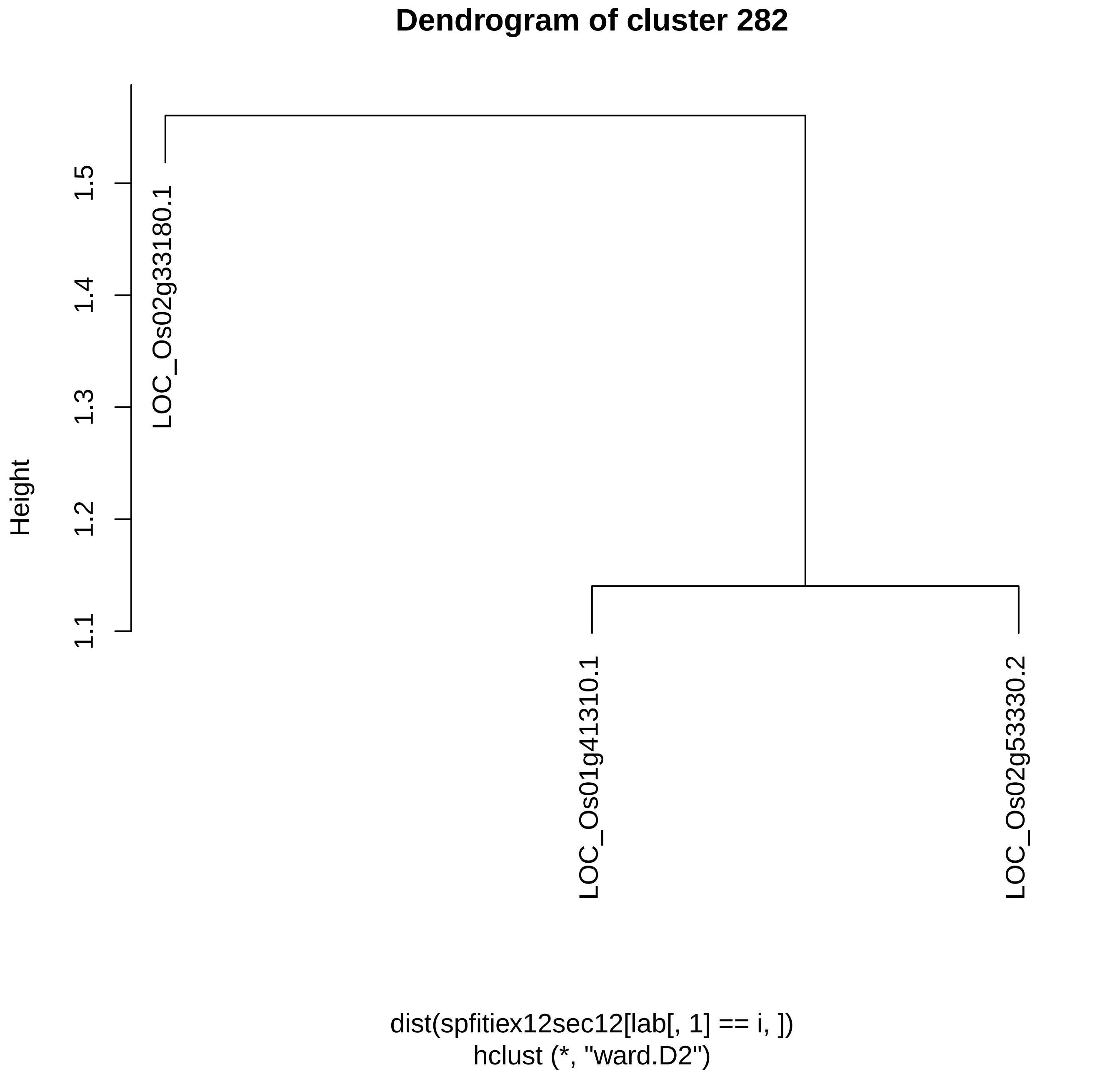
Dendrogram of cluster 282
Height
1.1
1.2
1.3
1.4
1.5
LOC_Os02g33180.1
LOC_Os01g41310.1
LOC_Os02g53330.2
dist(spfitiex12sec12[lab[, 1] == i, ])
hclust (*, "ward.D2")

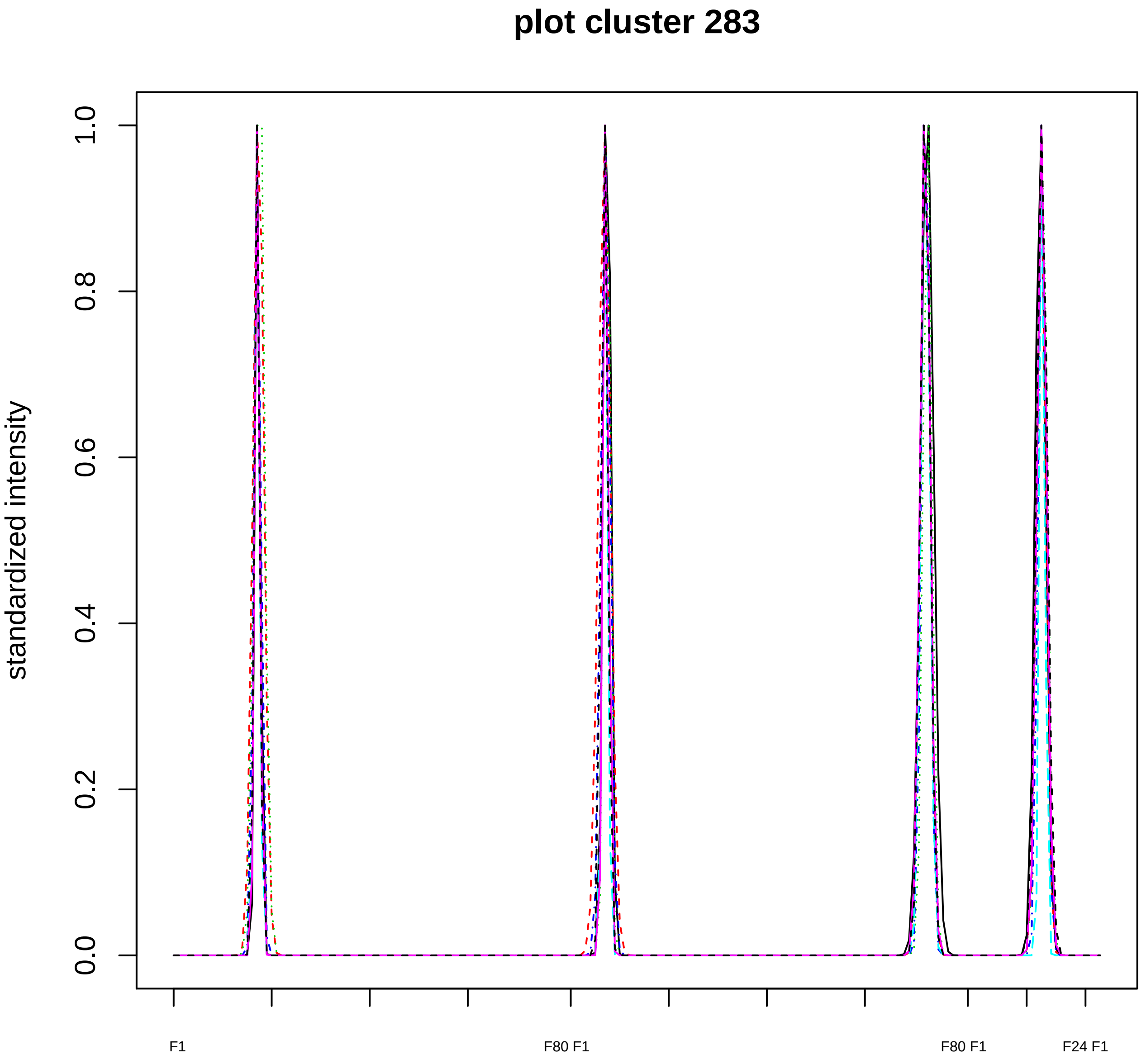

plot cluster 283
standardized intensity
0.0
0.2
0.4
0.6
0.8
1.0
F1
F80 F1
F80 F1
F24 F1

### Dendrogram of cluster 283

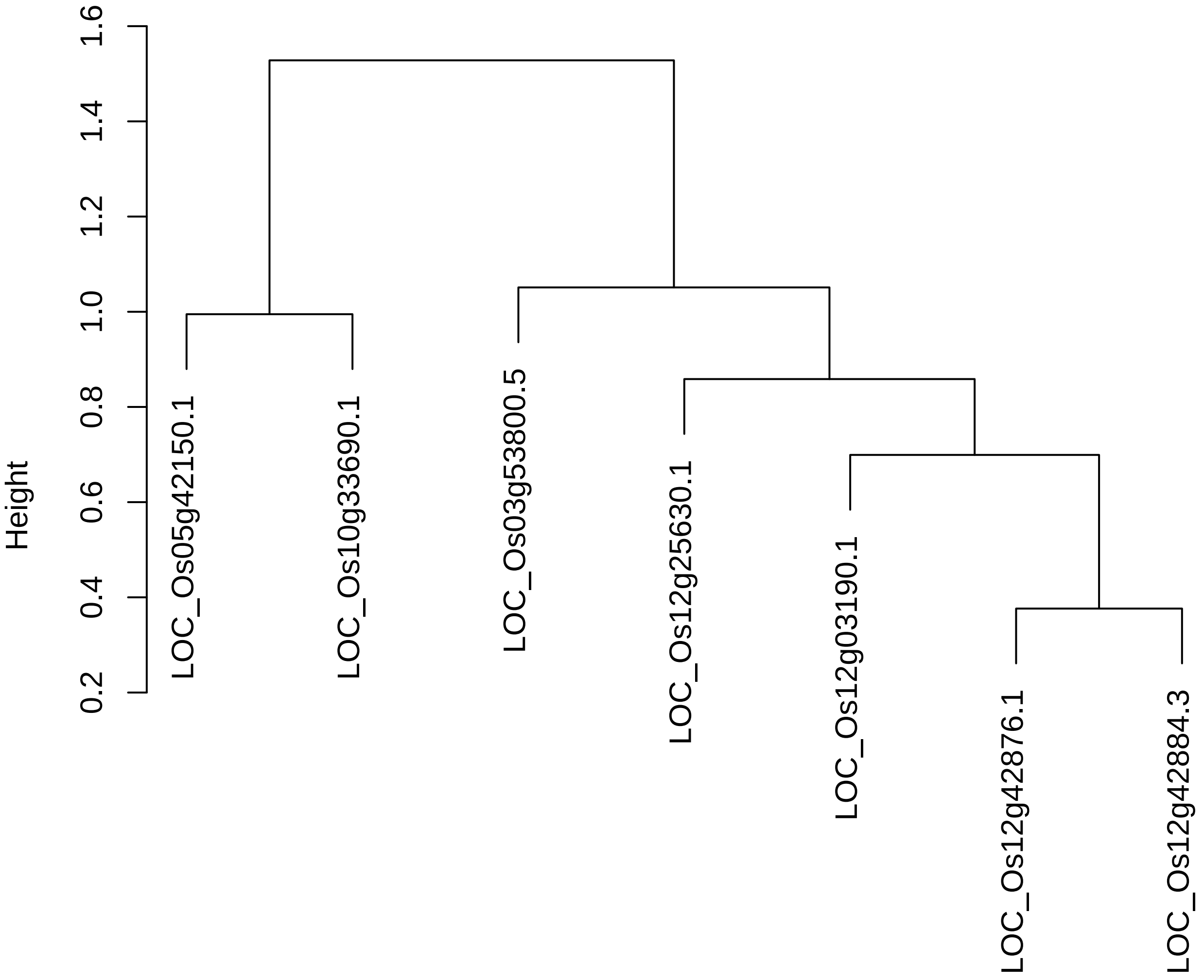


dist(spfitiex12sec12[lab[, 1] == i, ])
hclust (*, "ward.D2")

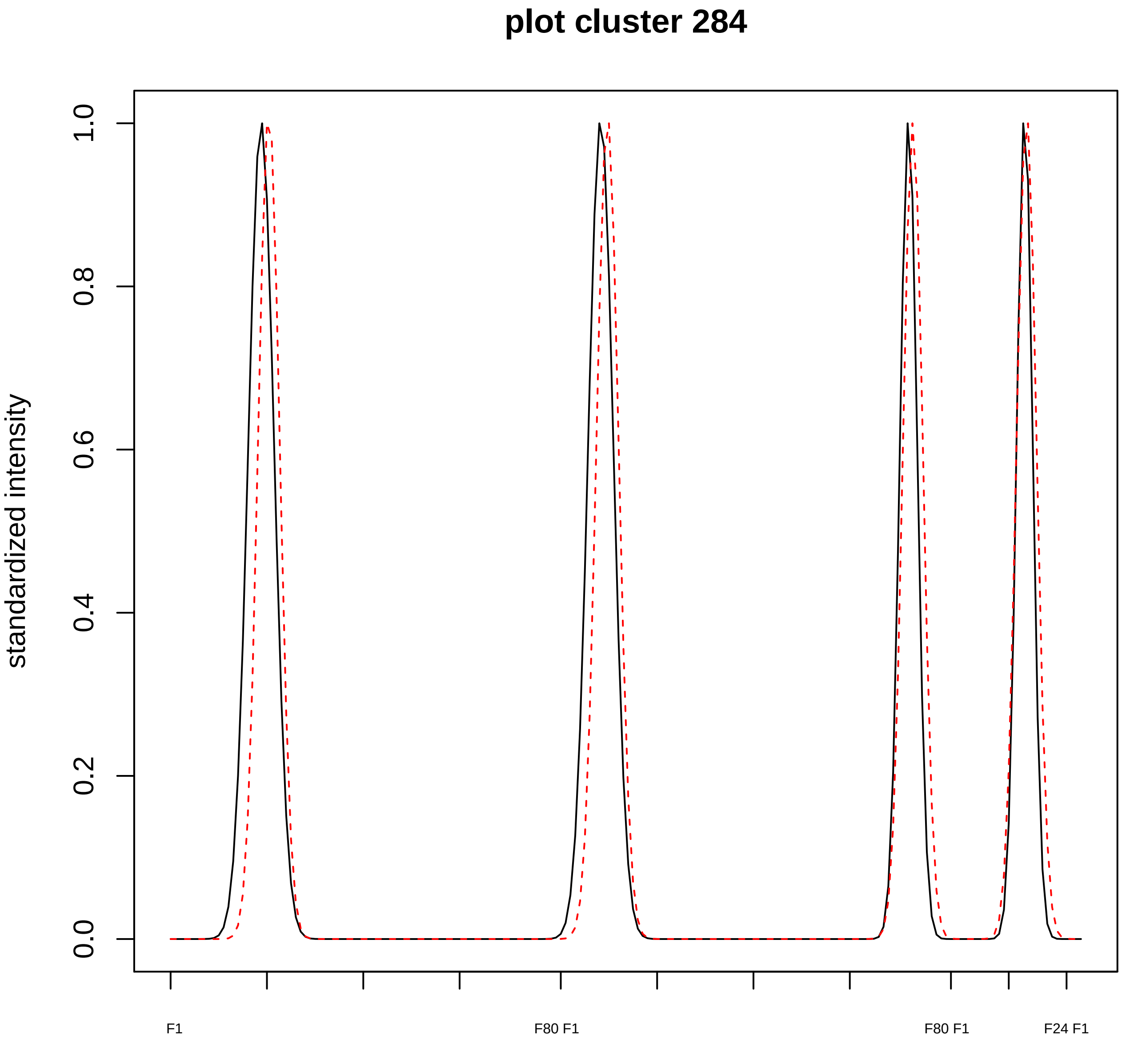
plot cluster 284
standardized intensity
0.0
0.2
0.4
0.6
0.8
1.0
F1
F80 F1
F80 F1
F24 F1

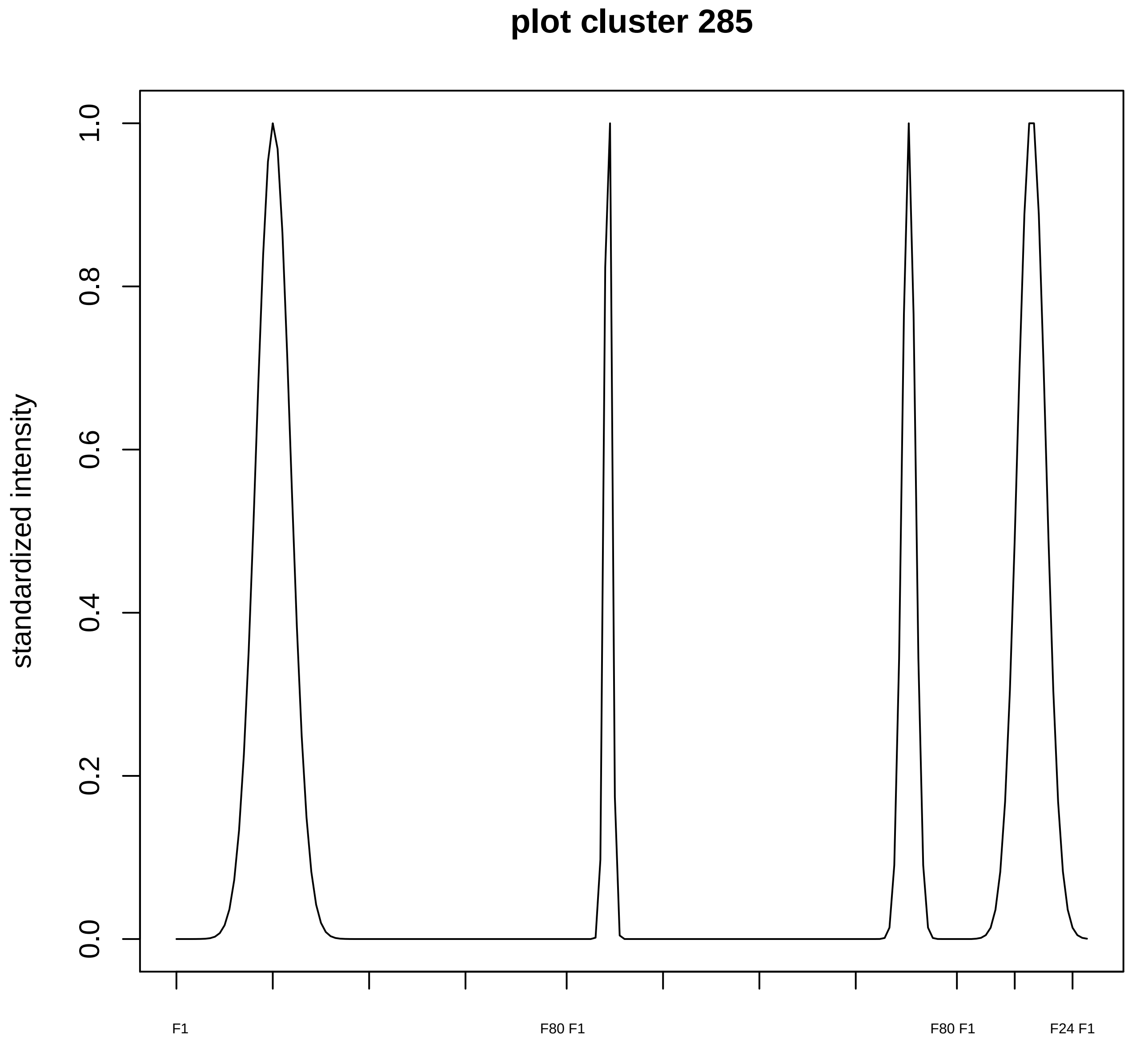

plot cluster 285
standardized intensity
0.0
0.2
0.4
0.6
0.8
1.0
F1
F80 F1
F80 F1
F24 F1

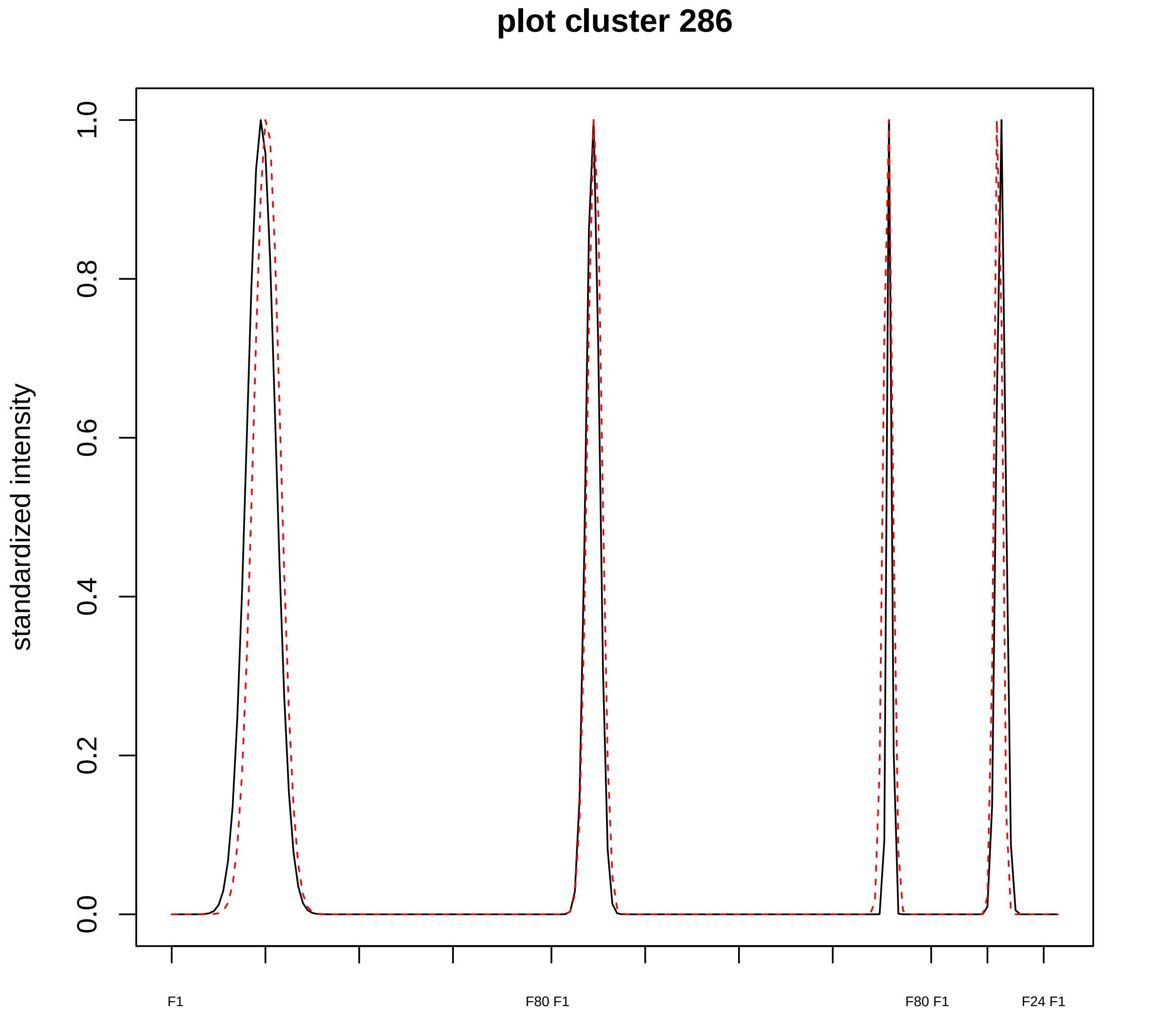
plot cluster 286
standardized intensity
0.0
0.2
0.4
0.6
0.8
1.0
F1
F80 F1
F80 F1
F24 F1

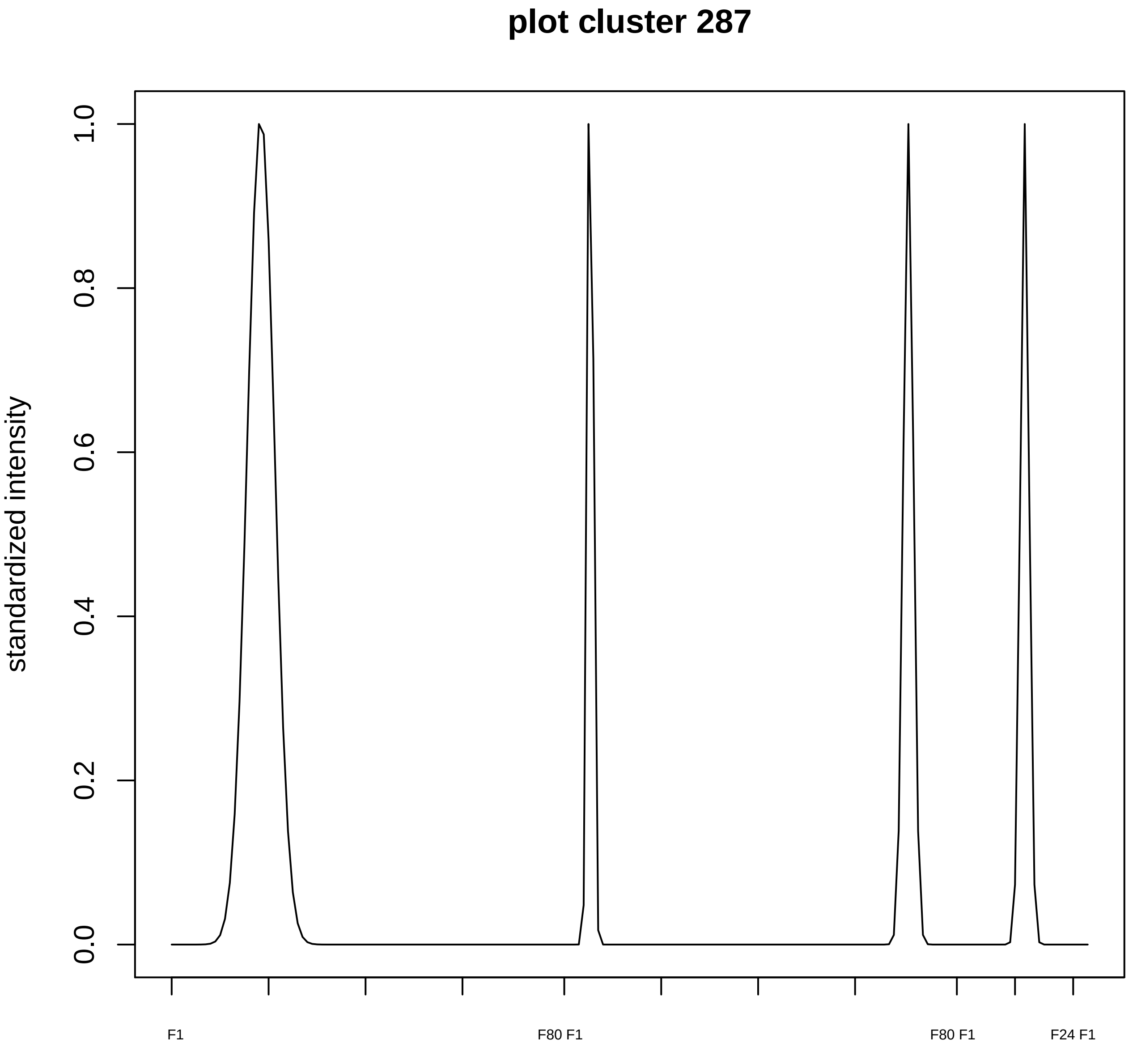

plot cluster 287
standardized intensity
0.0
0.2
0.4
0.6
0.8
1.0
F1
F80 F1
F80 F1
F24 F1

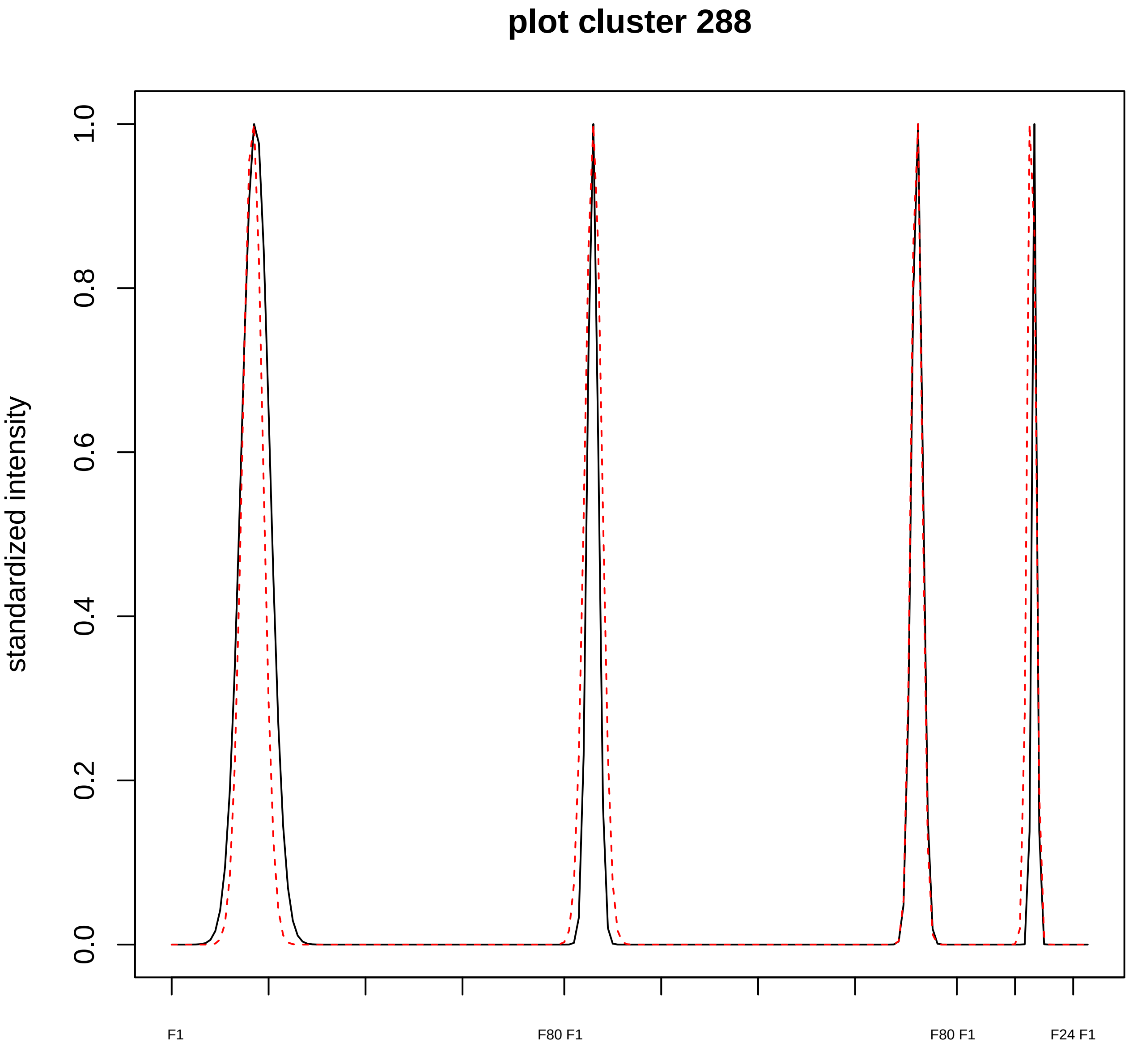
plot cluster 288
standardized intensity
0.0
0.2
0.4
0.6
0.8
1.0
F1
F80 F1
F80 F1
F24 F1

### plot cluster 289

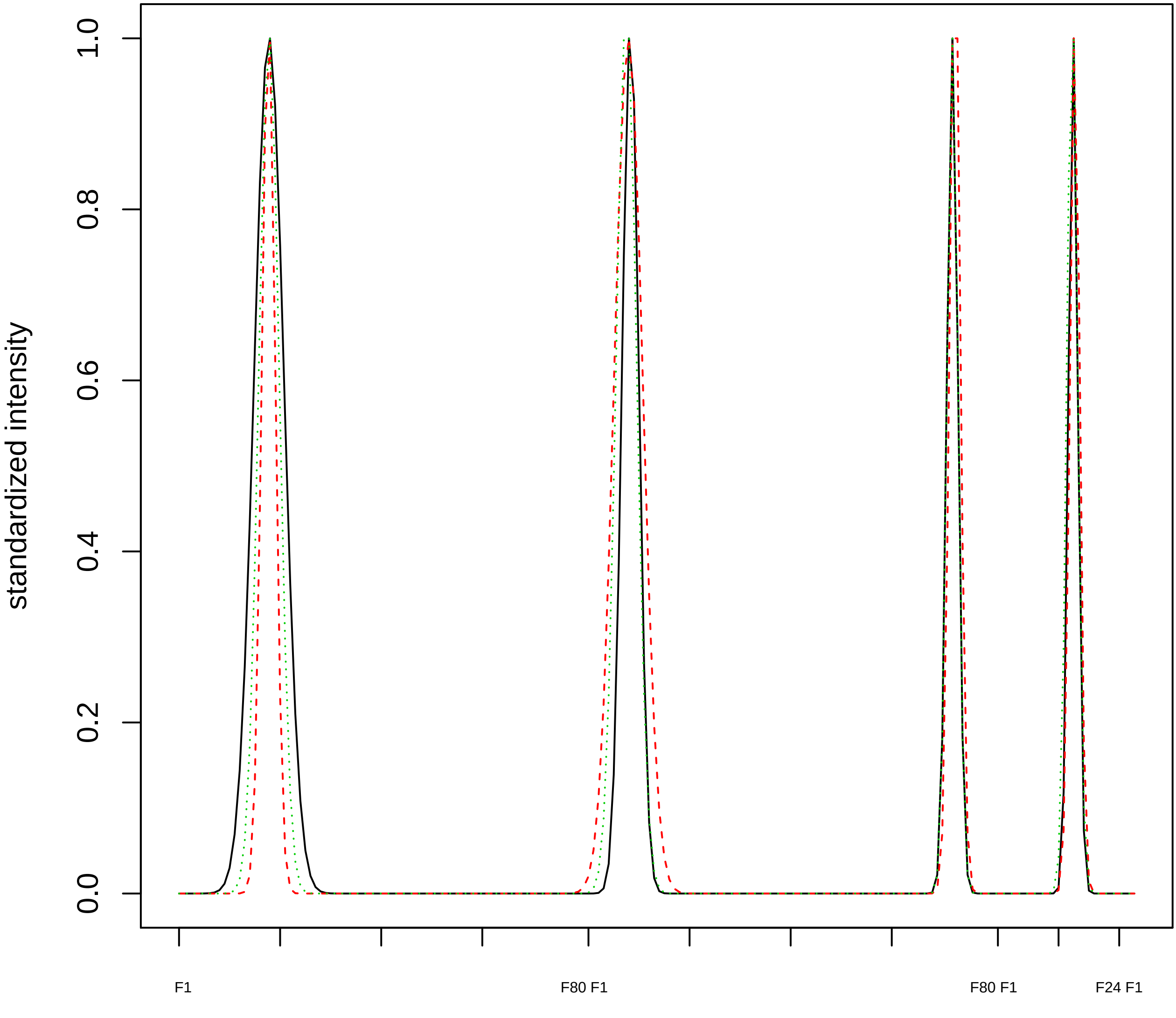

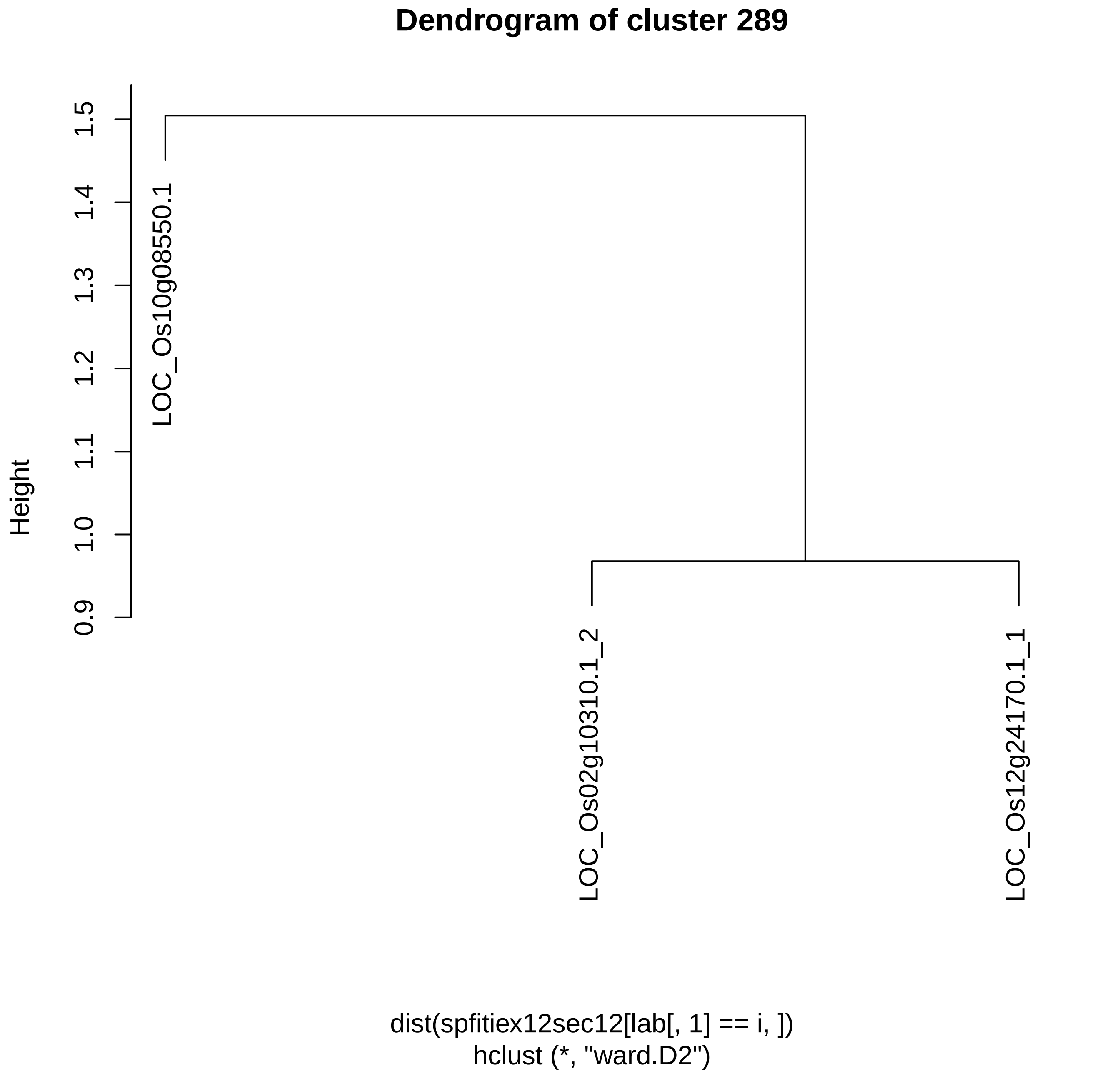
Dendrogram of cluster 289
Height
1.5
1.4
1.3
1.2
1.1
1.0
0.9
LOC_Os10g08550.1
LOC_Os02g10310.1_2
LOC_Os12g24170.1_1
dist(spfitiex12sec12[lab[, 1] == i, ])
hclust (*, "ward.D2")

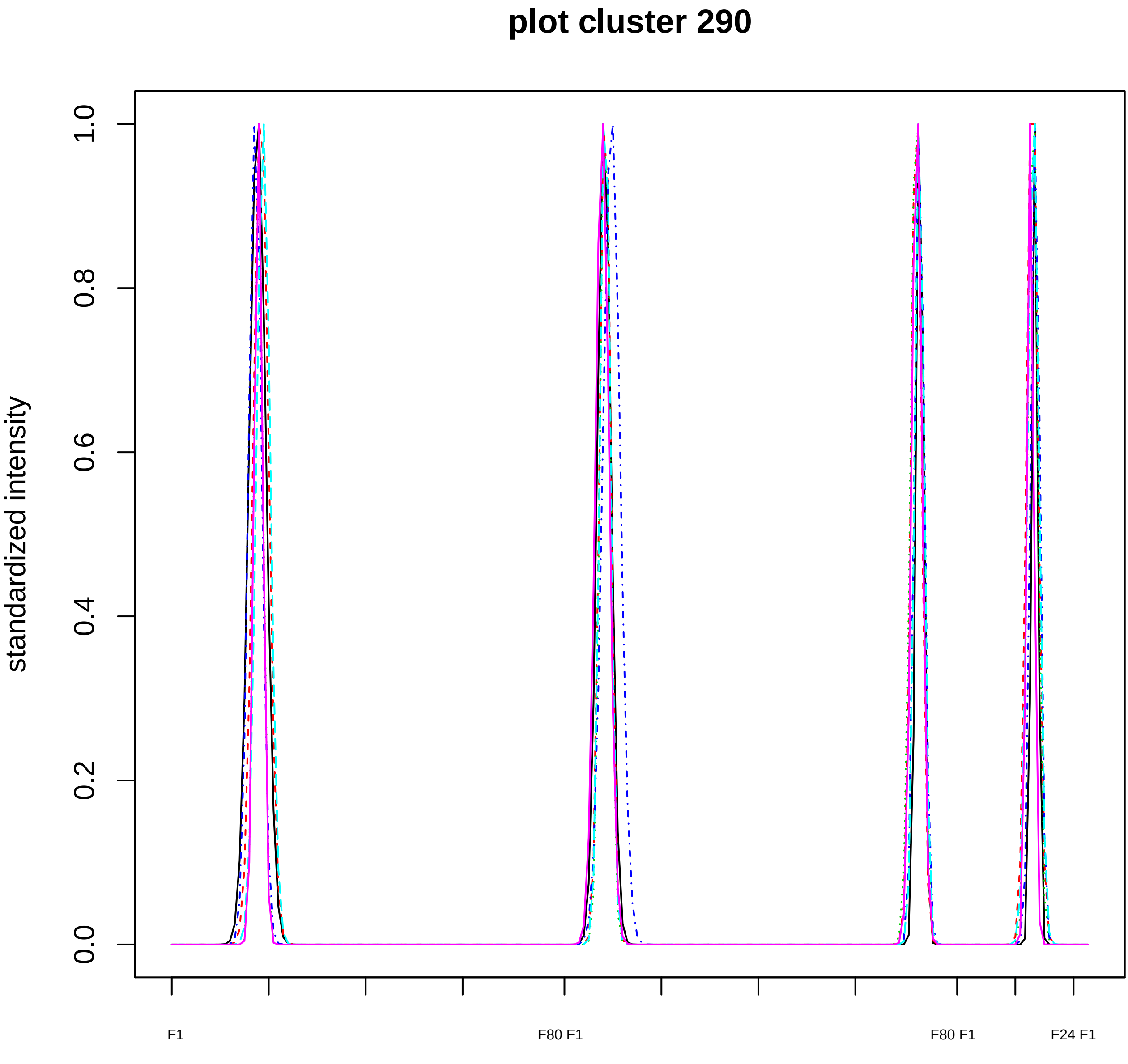
plot cluster 290
standardized intensity
0.0
0.2
0.4
0.6
0.8
1.0
F1
F80 F1
F80 F1
F24 F1

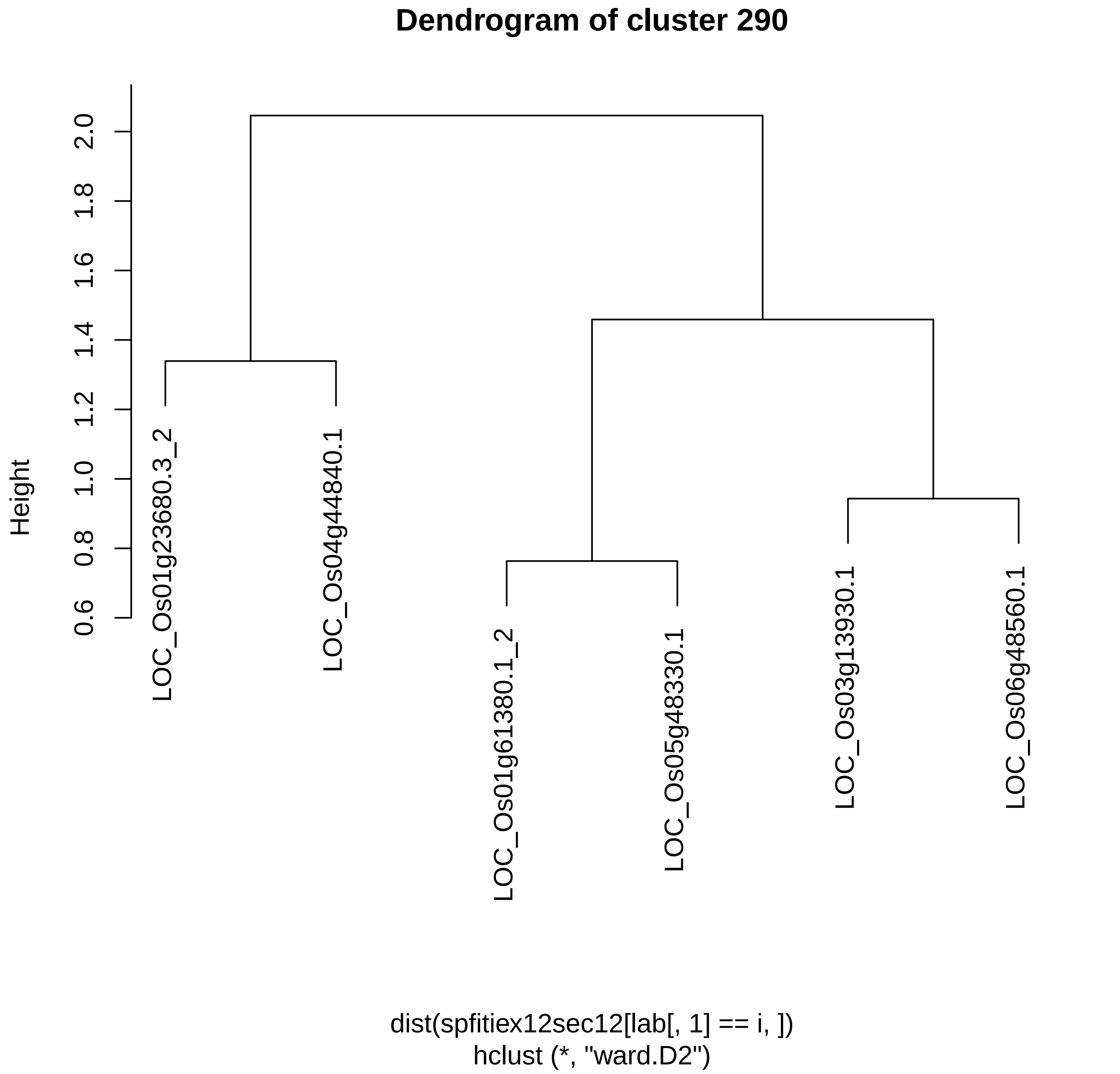

Dendrogram of cluster 290
Height
2.0
1.8
1.6
1.4
1.2
1.0
0.8
0.6
LOC_Os01g23680.3_2
LOC_Os04g44840.1
LOC_Os01g61380.1_2
LOC_Os05g48330.1
LOC_Os03g13930.1
LOC_Os06g48560.1
dist(spfitiex12sec12[lab[, 1] == i, ])
hclust (*, "ward.D2")

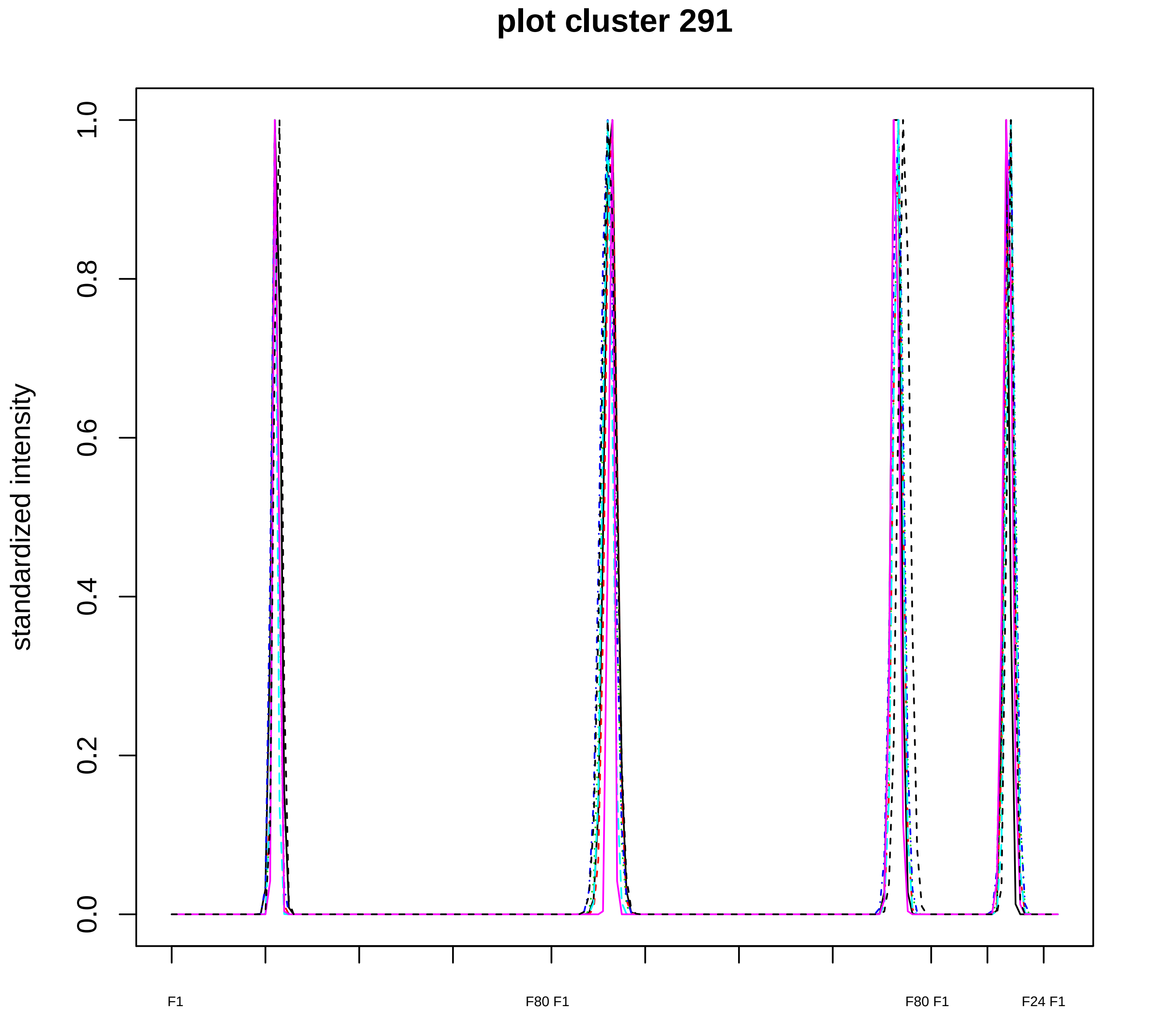
plot cluster 291
standardized intensity
0.0
0.2
0.4
0.6
0.8
1.0
F1
F80 F1
F80 F1
F24 F1

### Dendrogram of cluster 291

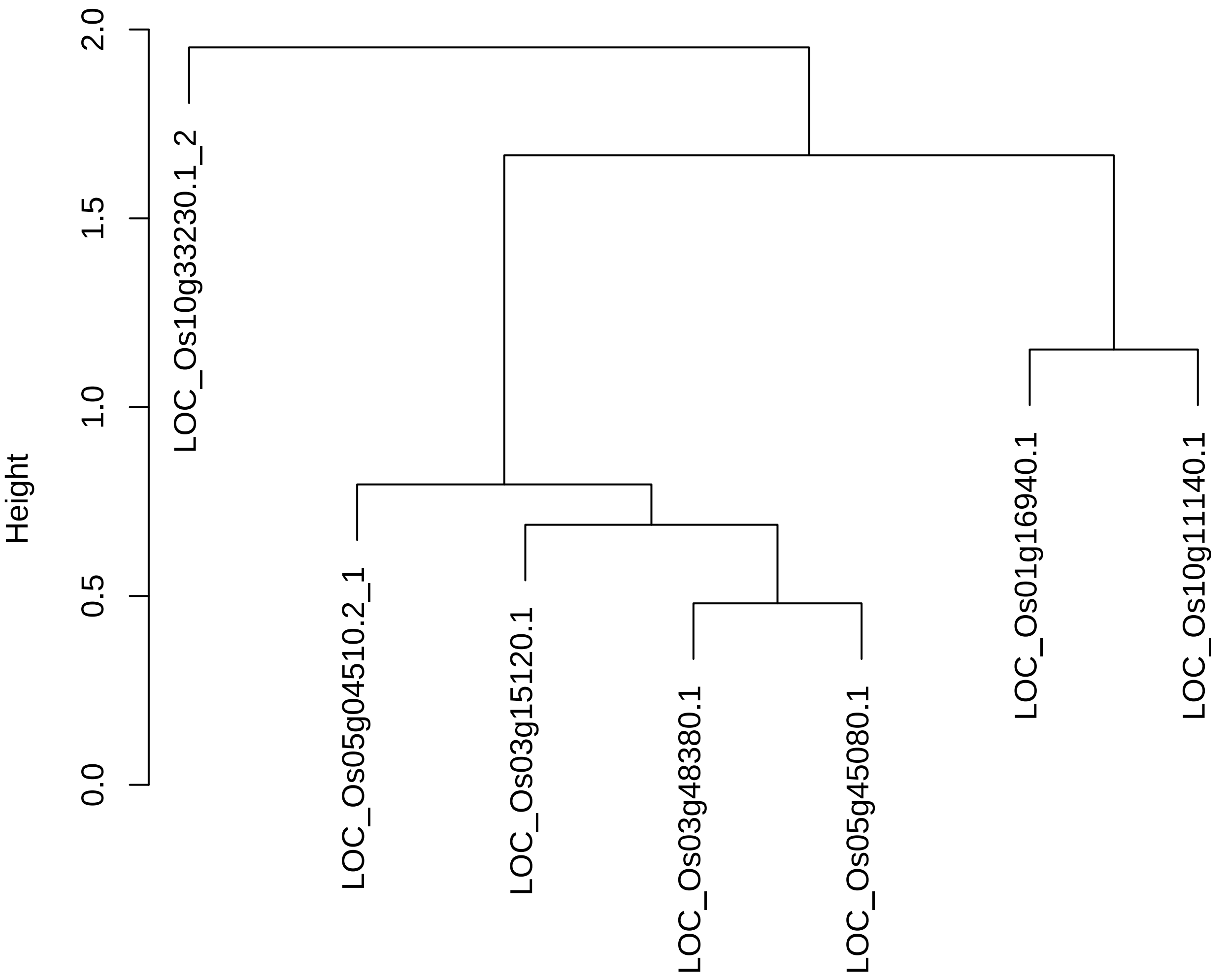


dist(spfitiex12sec12[lab[, 1] == i, ])
hclust (*, "ward.D2")

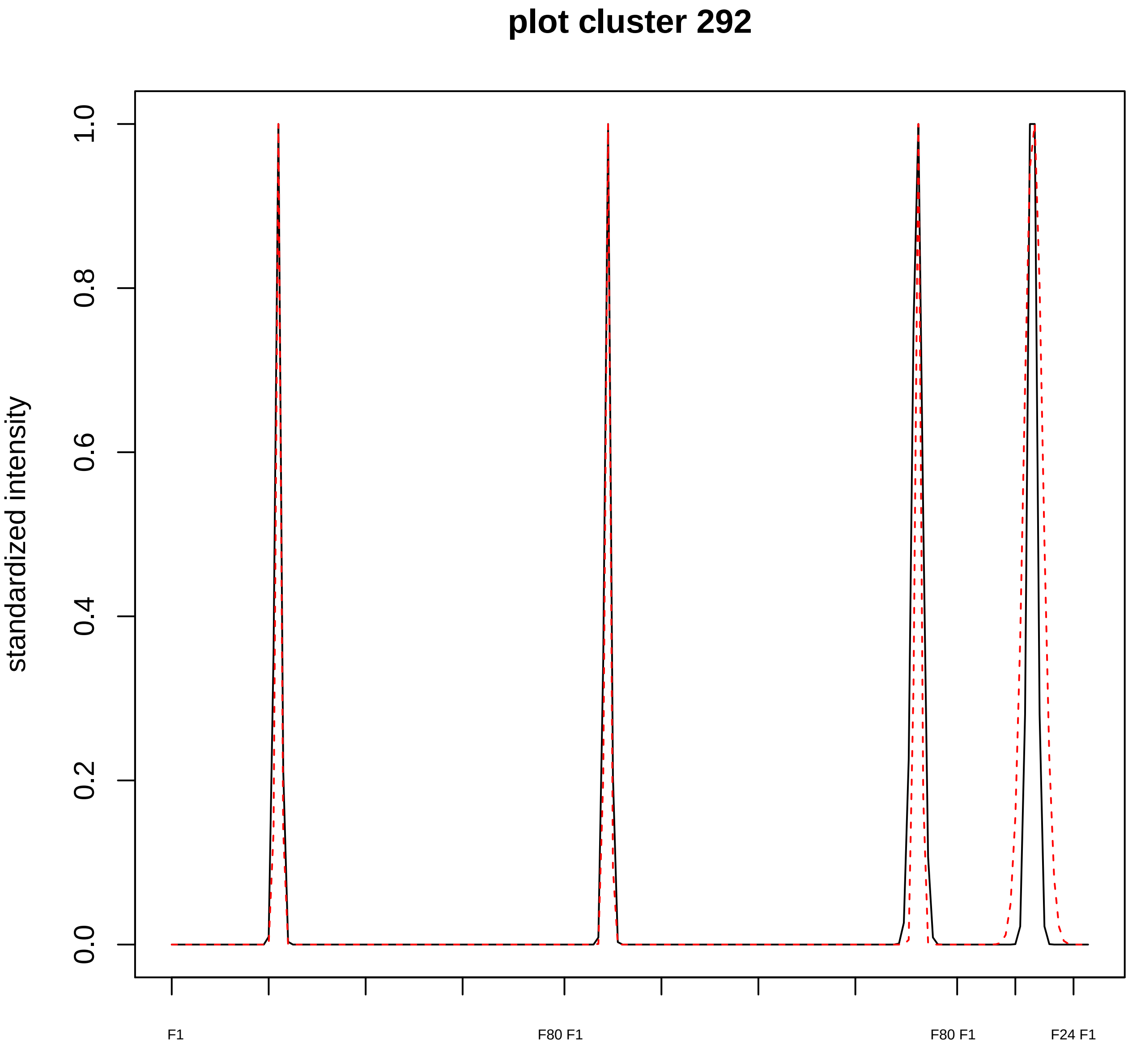

plot cluster 292
standardized intensity
0.0
0.2
0.4
0.6
0.8
1.0
F1
F80 F1
F80 F1
F24 F1

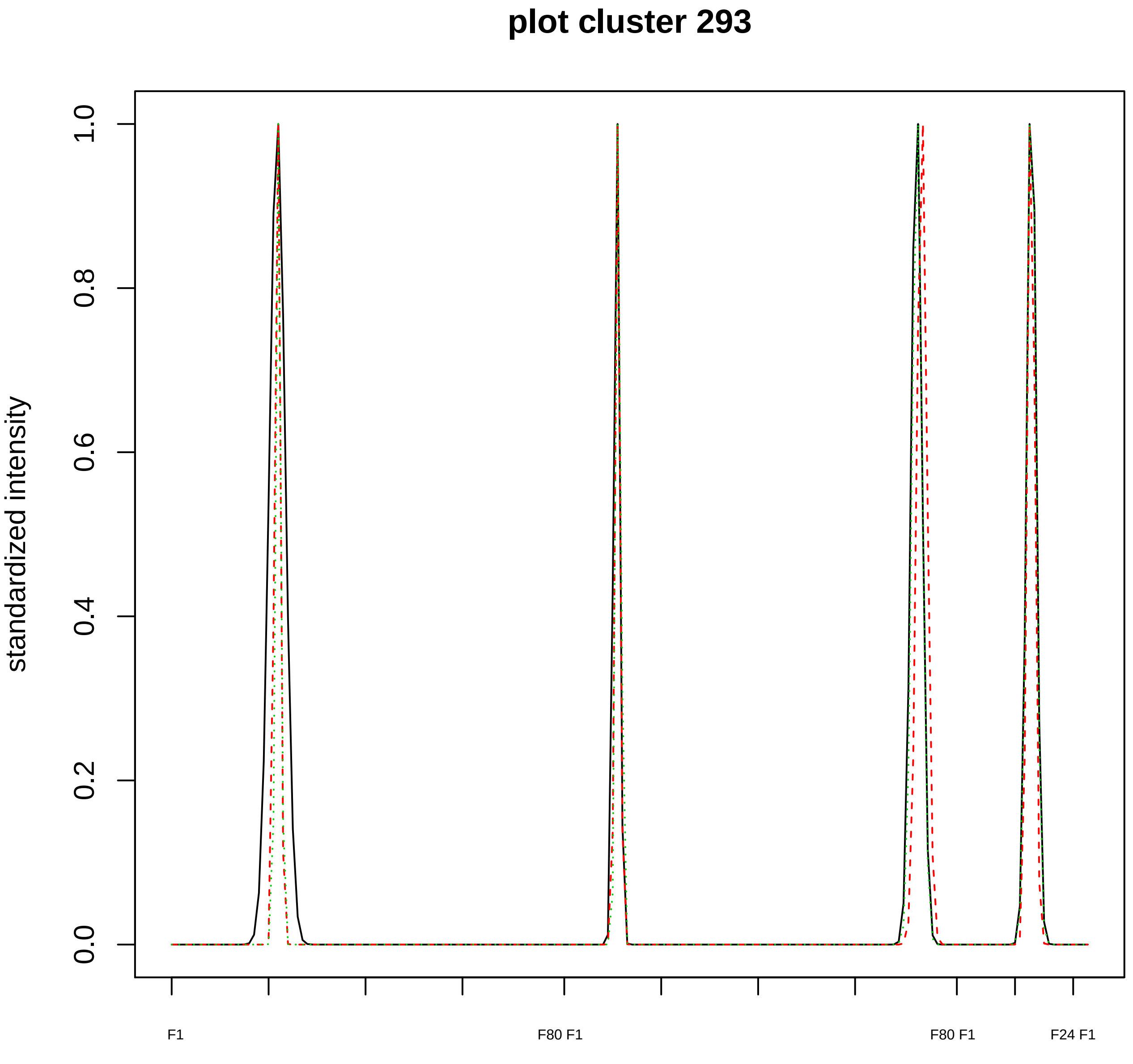
plot cluster 293
standardized intensity
0.0
0.2
0.4
0.6
0.8
1.0
F1
F80 F1
F80 F1
F24 F1

#### Dendrogram of cluster 293

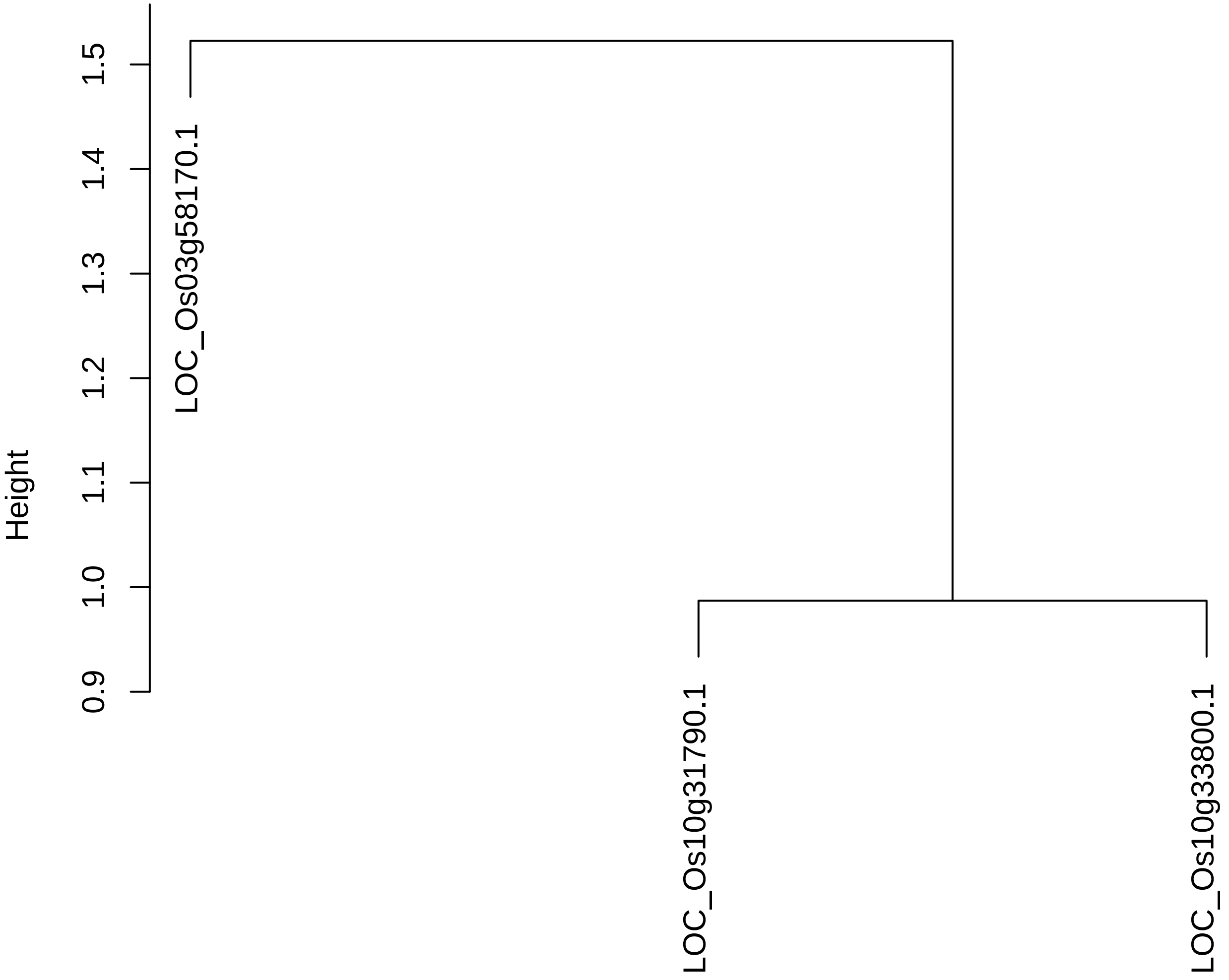


dist(spfitiex12sec12[lab[, 1] == i, ])
hclust (*, "ward.D2")

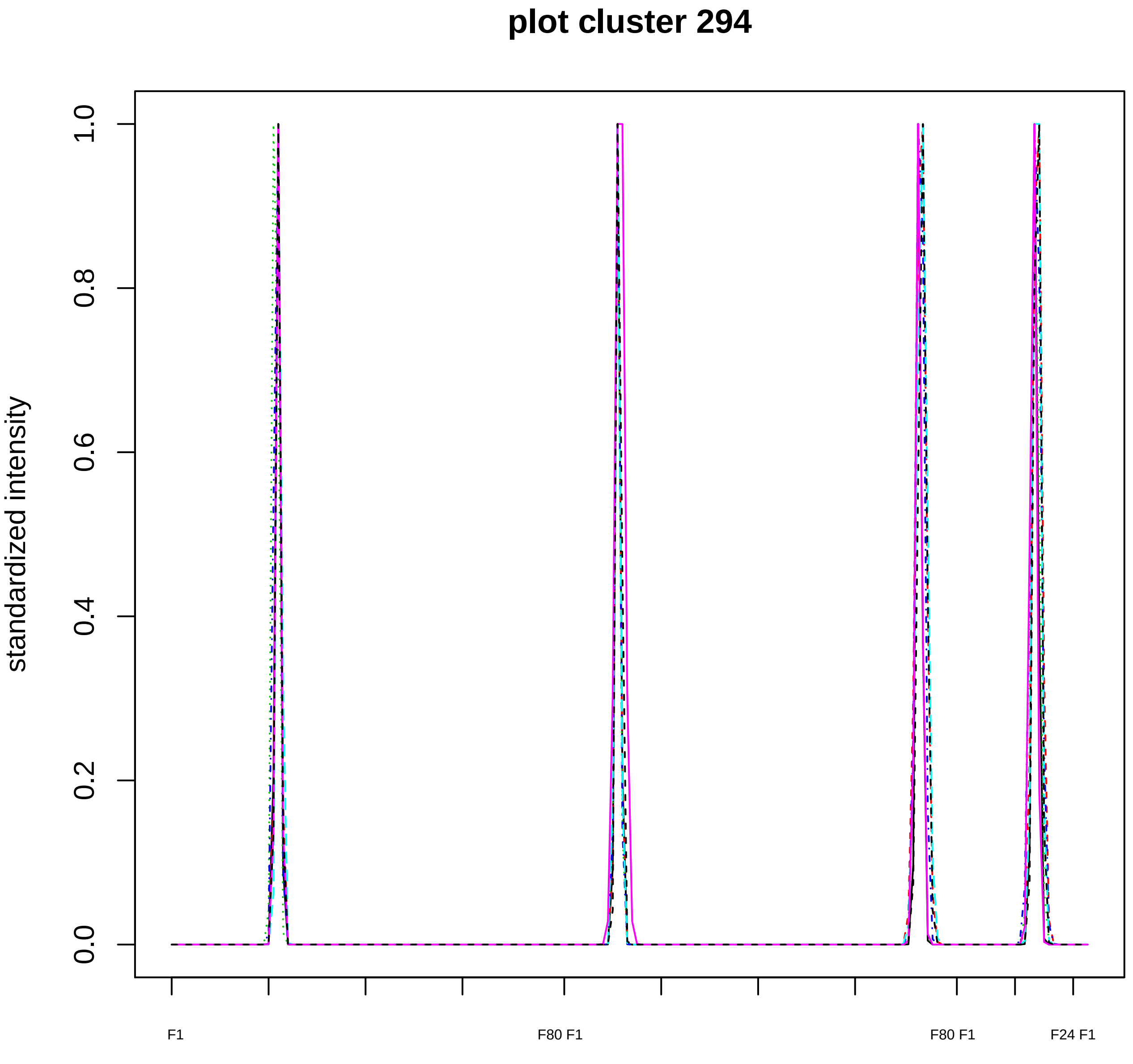
plot cluster 294
standardized intensity
0.0
0.2
0.4
0.6
0.8
1.0
F1
F80 F1
F80 F1
F24 F1

### Dendrogram of cluster 294

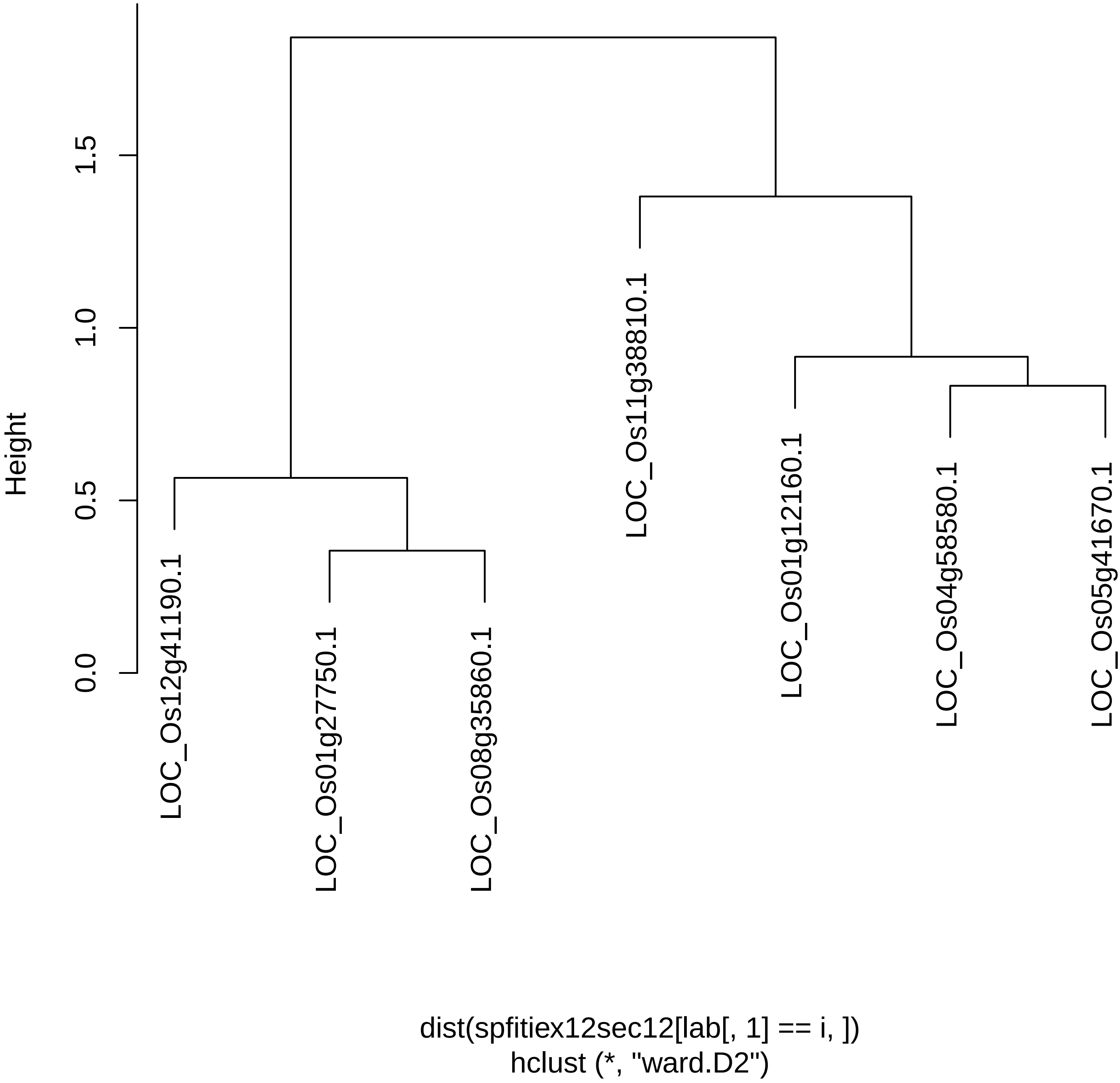

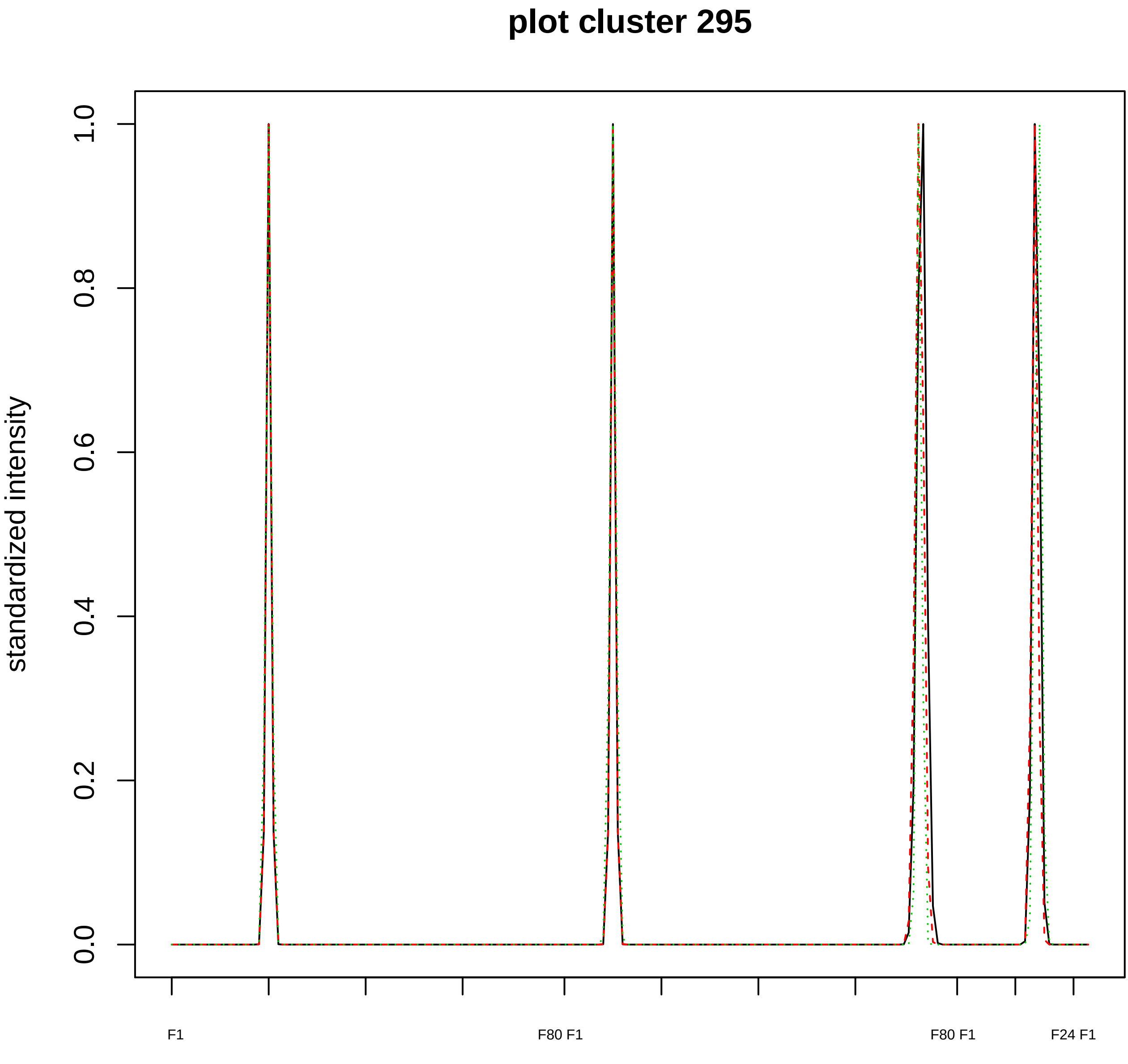
plot cluster 295
standardized intensity
1.0
0.8
0.6
0.4
0.2
0.0
F1
F80 F1
F80 F1
F24 F1

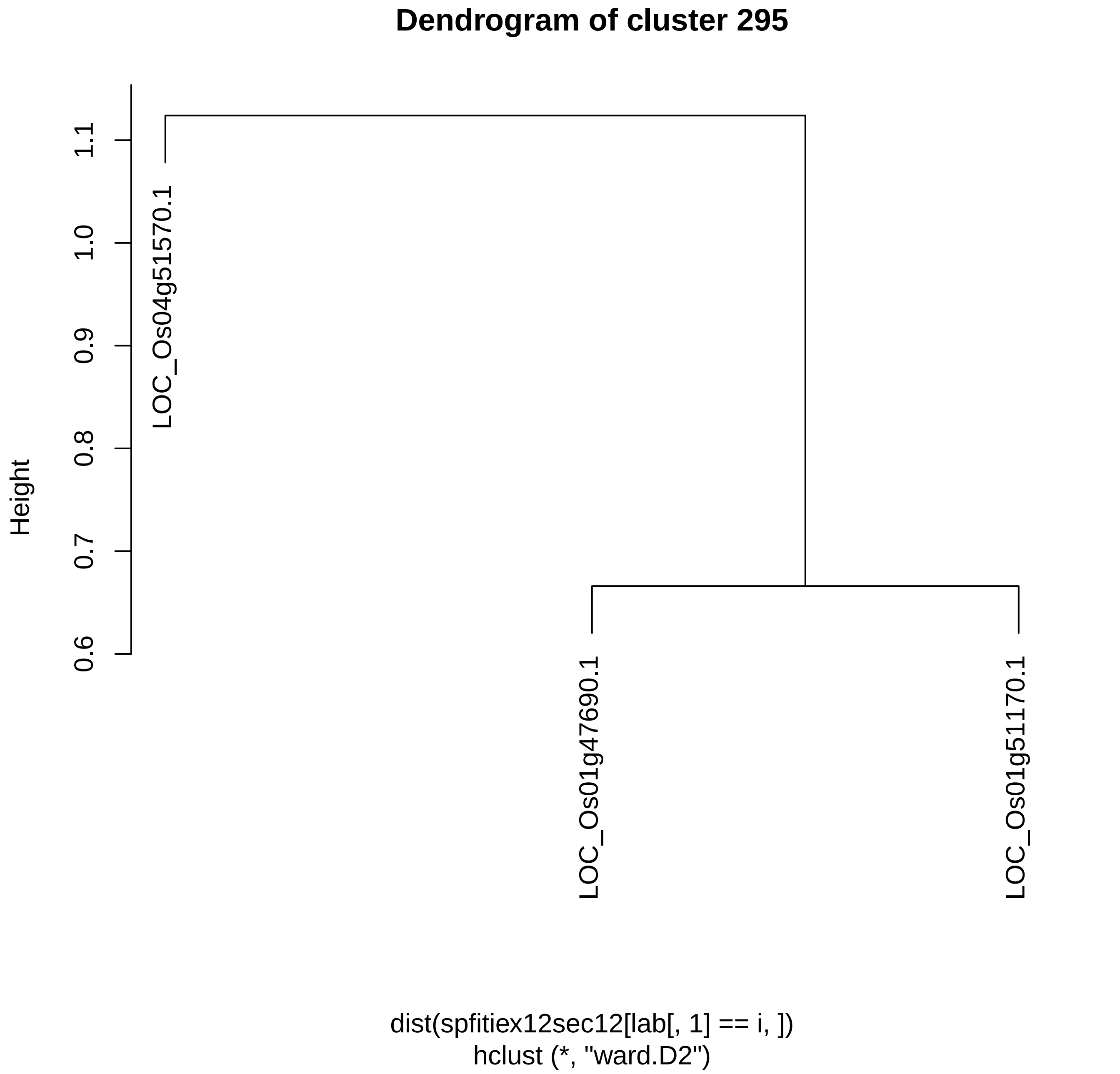
Dendrogram of cluster 295
Height
0.6
0.7
0.8
0.9
1.0
1.1
LOC_Os04g51570.1
LOC_Os01g47690.1
LOC_Os01g51170.1
dist(spfitiex12sec12[lab[, 1] == i, ])
hclust (*, "ward.D2")

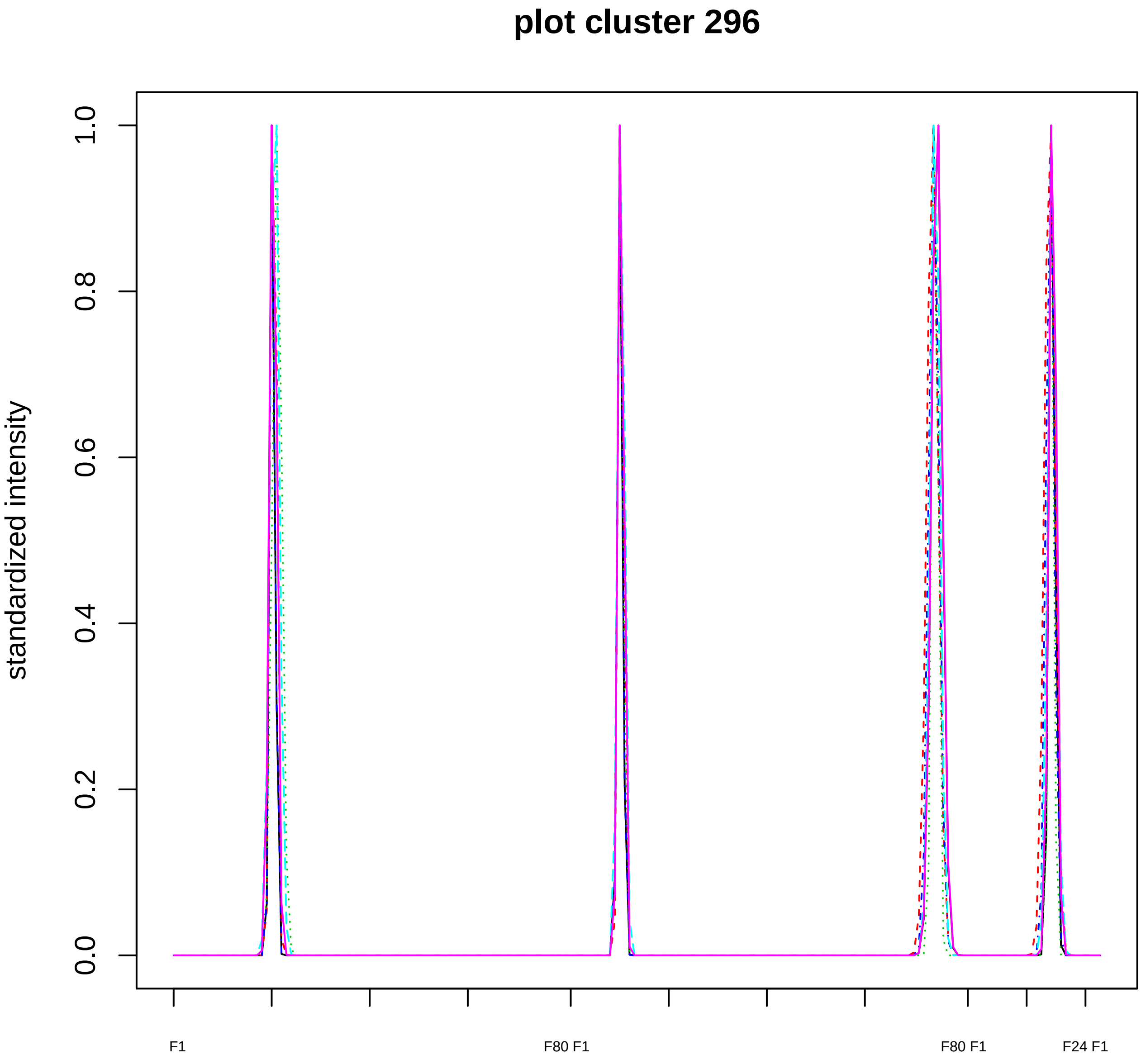

plot cluster 296
standardized intensity
0.0
0.2
0.4
0.6
0.8
1.0
F1
F80 F1
F80 F1
F24 F1

#### Dendrogram of cluster 296

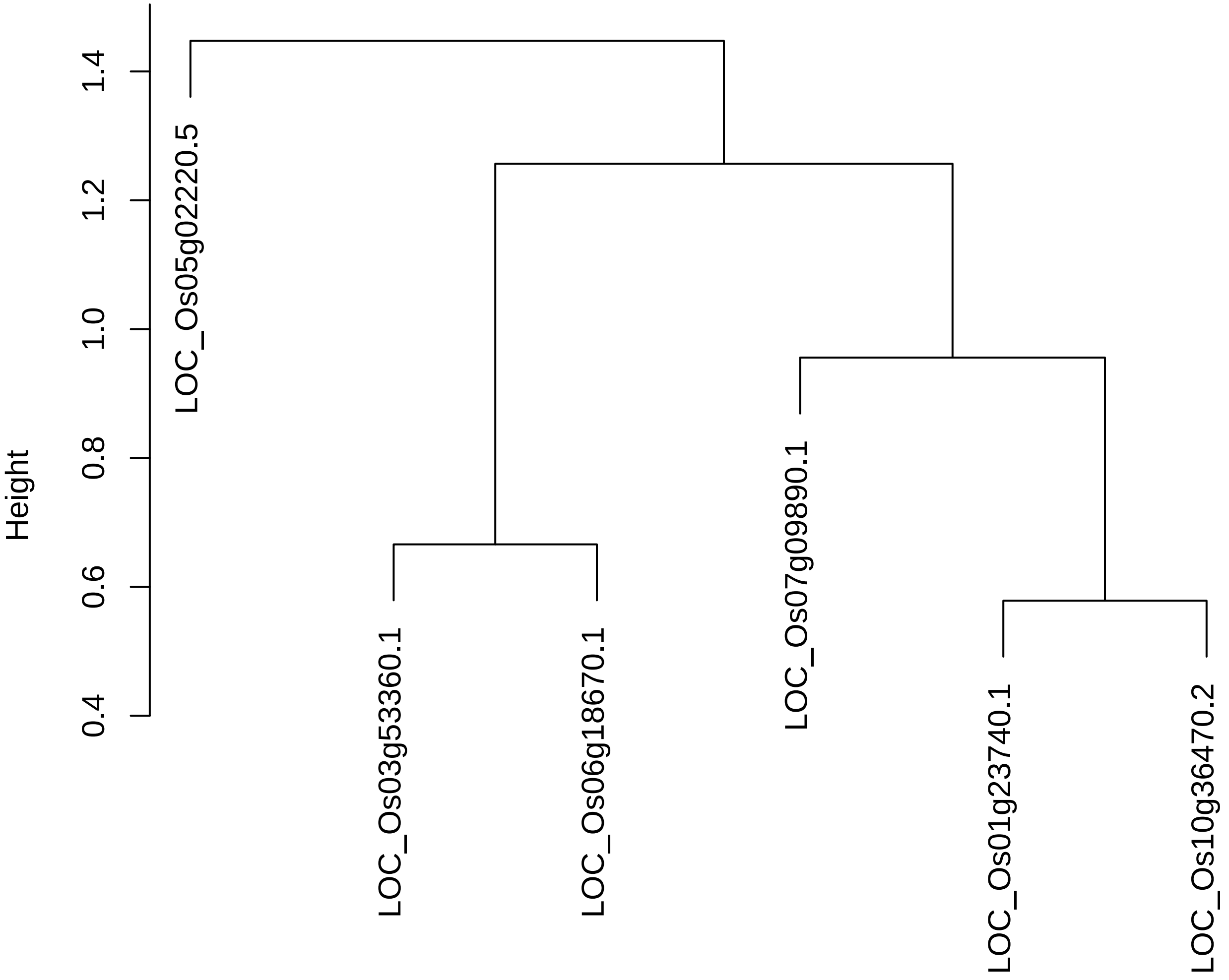


dist(spfitiex12sec12[lab[, 1] == i, ])
hclust (*, "ward.D2")

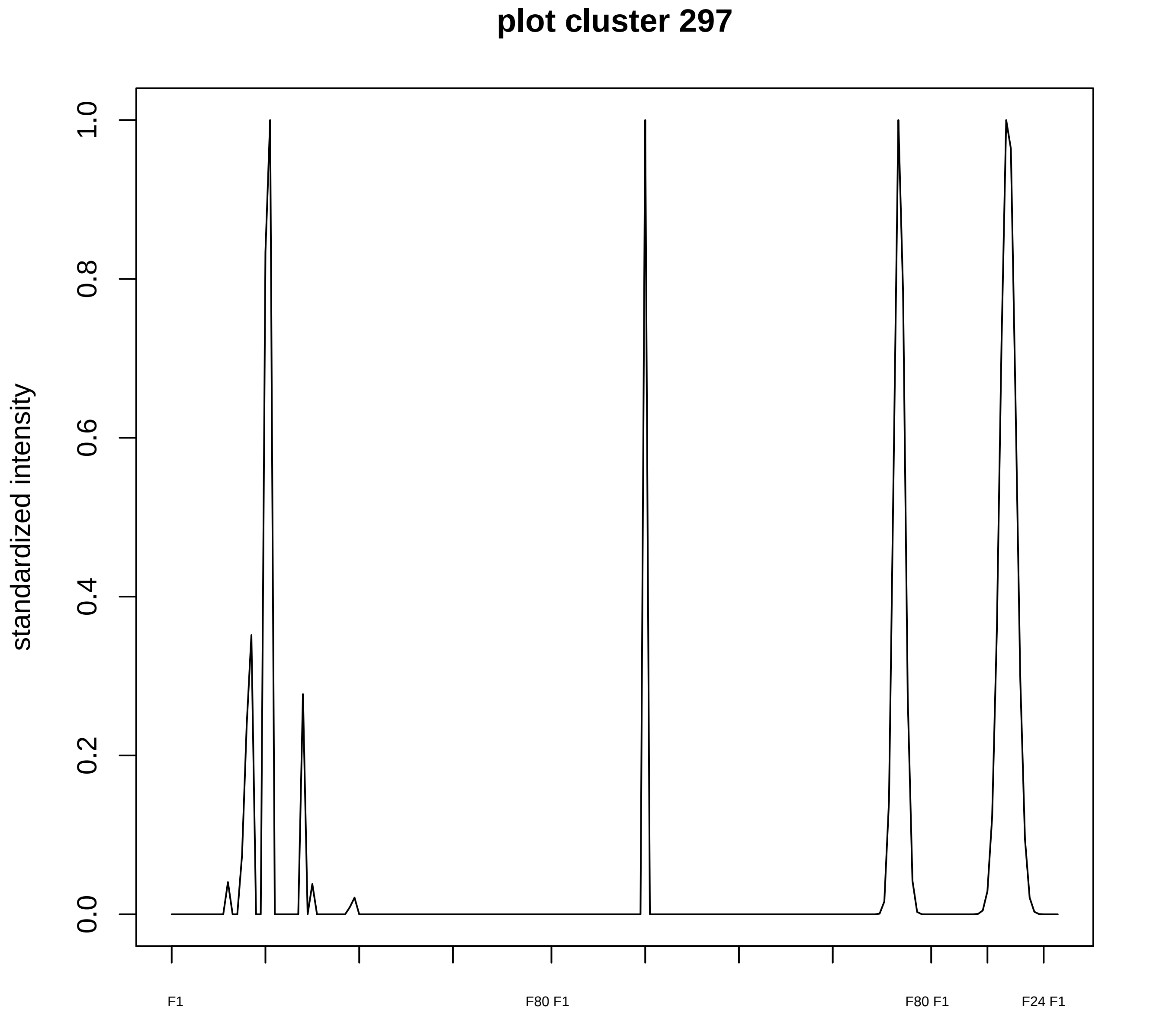

plot cluster 297
standardized intensity
0.0
0.2
0.4
0.6
0.8
1.0
F1
F80 F1
F80 F1
F24 F1

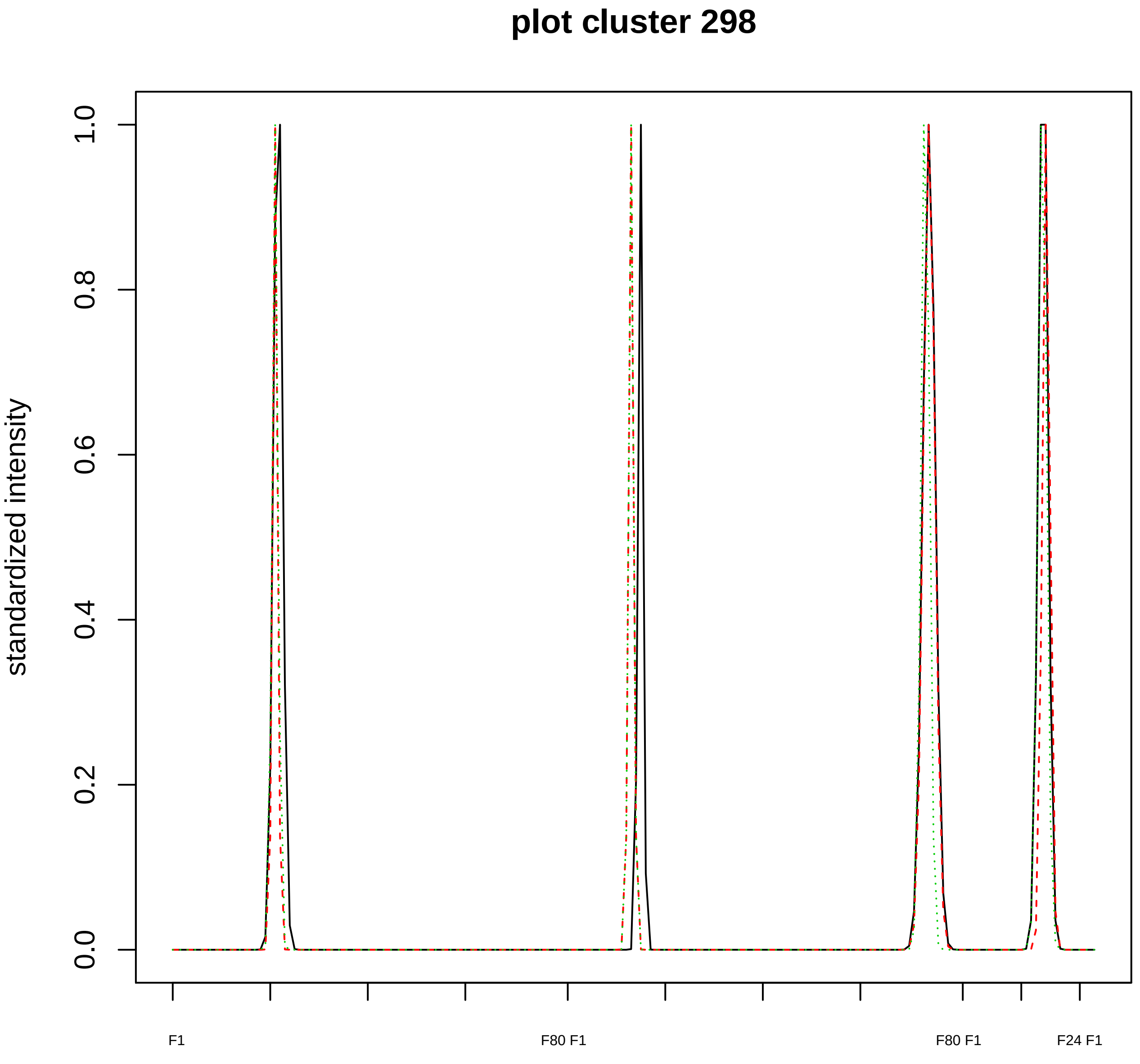

plot cluster 298
standardized intensity
0.0
0.2
0.4
0.6
0.8
1.0
F1
F80 F1
F80 F1
F24 F1

#### Dendrogram of cluster 298

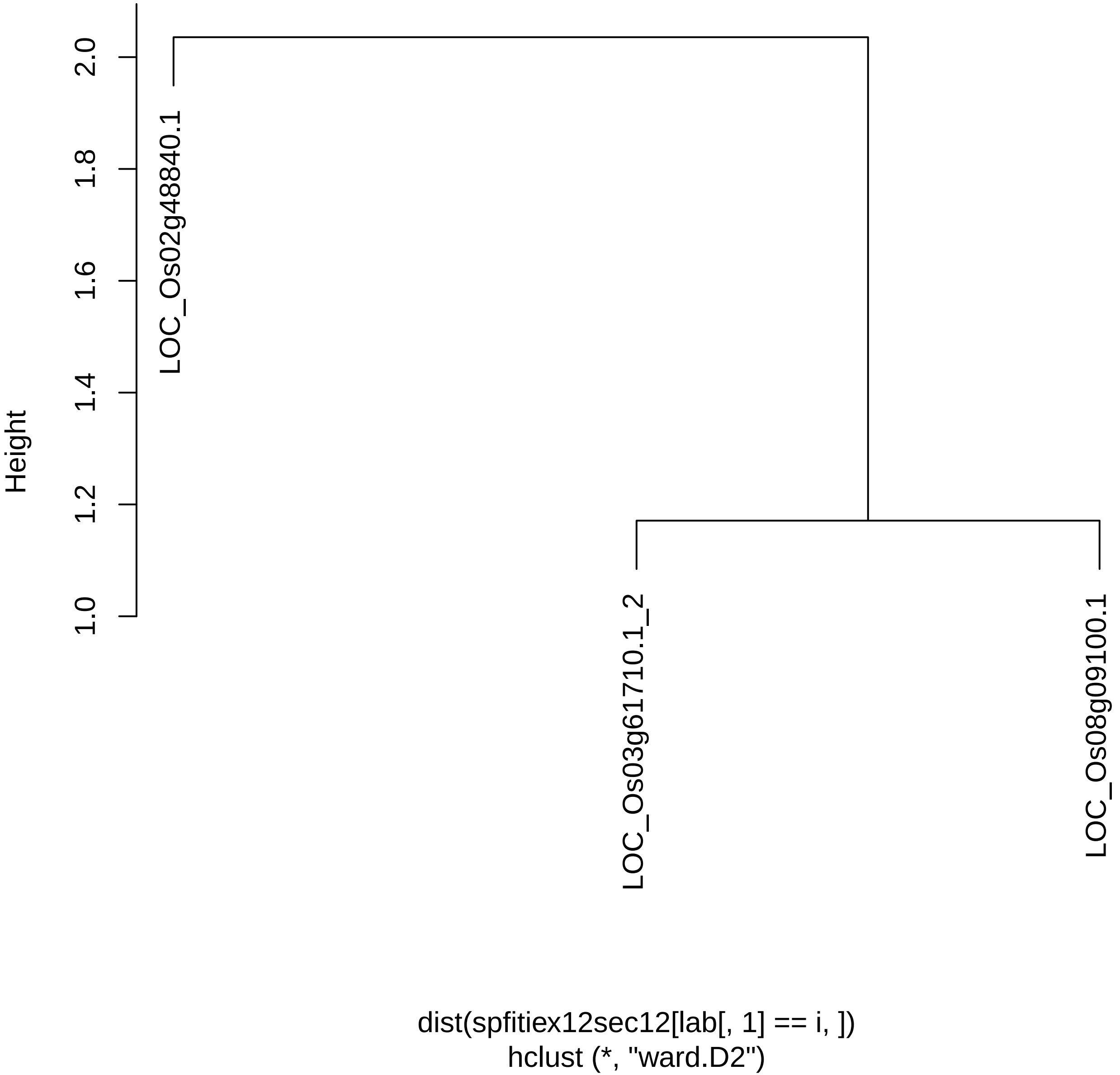

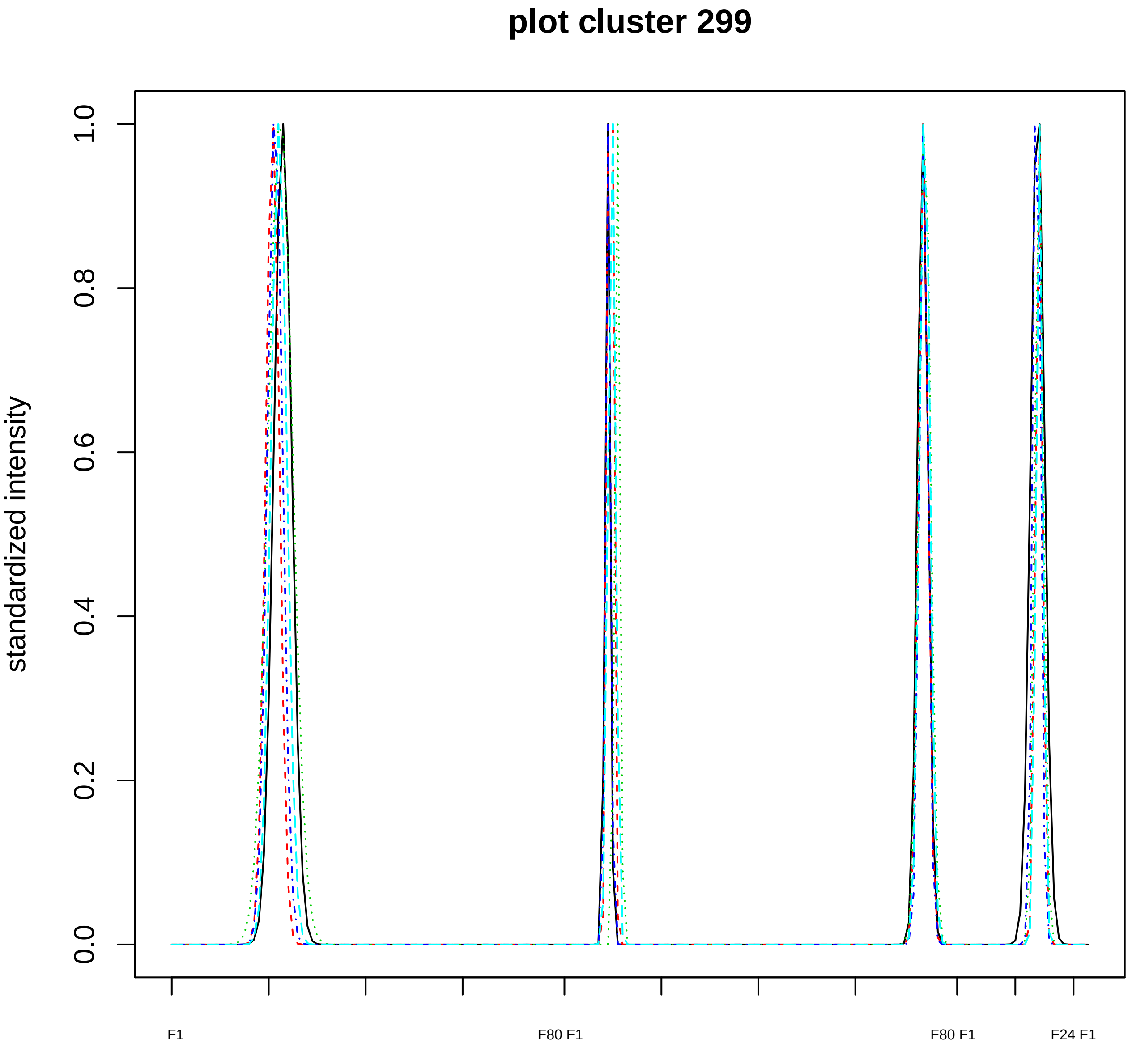
plot cluster 299
standardized intensity
0.0
0.2
0.4
0.6
0.8
1.0
F1
F80 F1
F80 F1
F24 F1

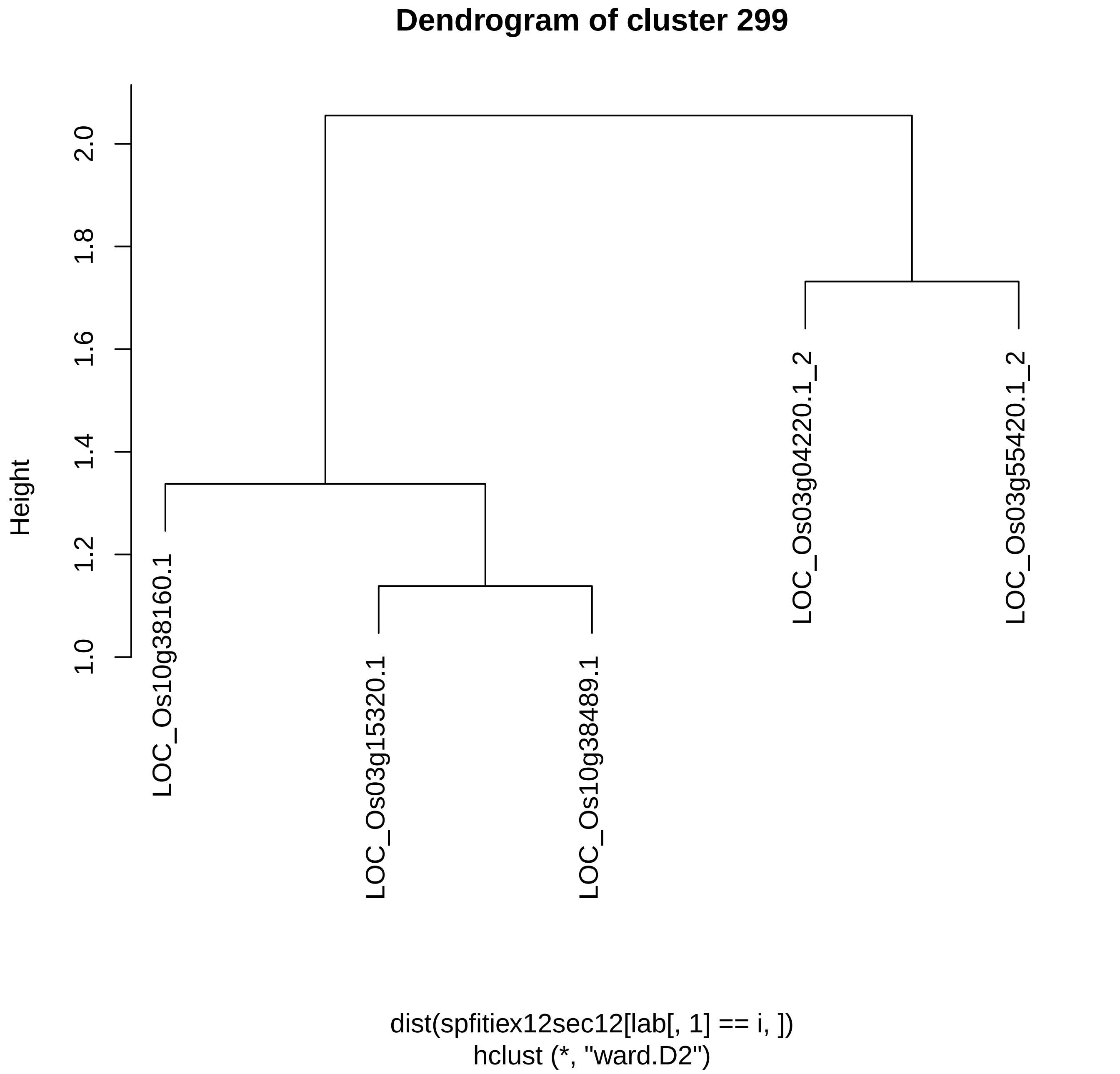
Dendrogram of cluster 299
Height
2.0
1.8
1.6
1.4
1.2
1.0
LOC_Os10g38160.1
LOC_Os03g15320.1
LOC_Os10g38489.1
LOC_Os03g04220.1_2
LOC_Os03g55420.1_2
dist(spfitiex12sec12[lab[, 1] == i, ])
hclust (*, "ward.D2")

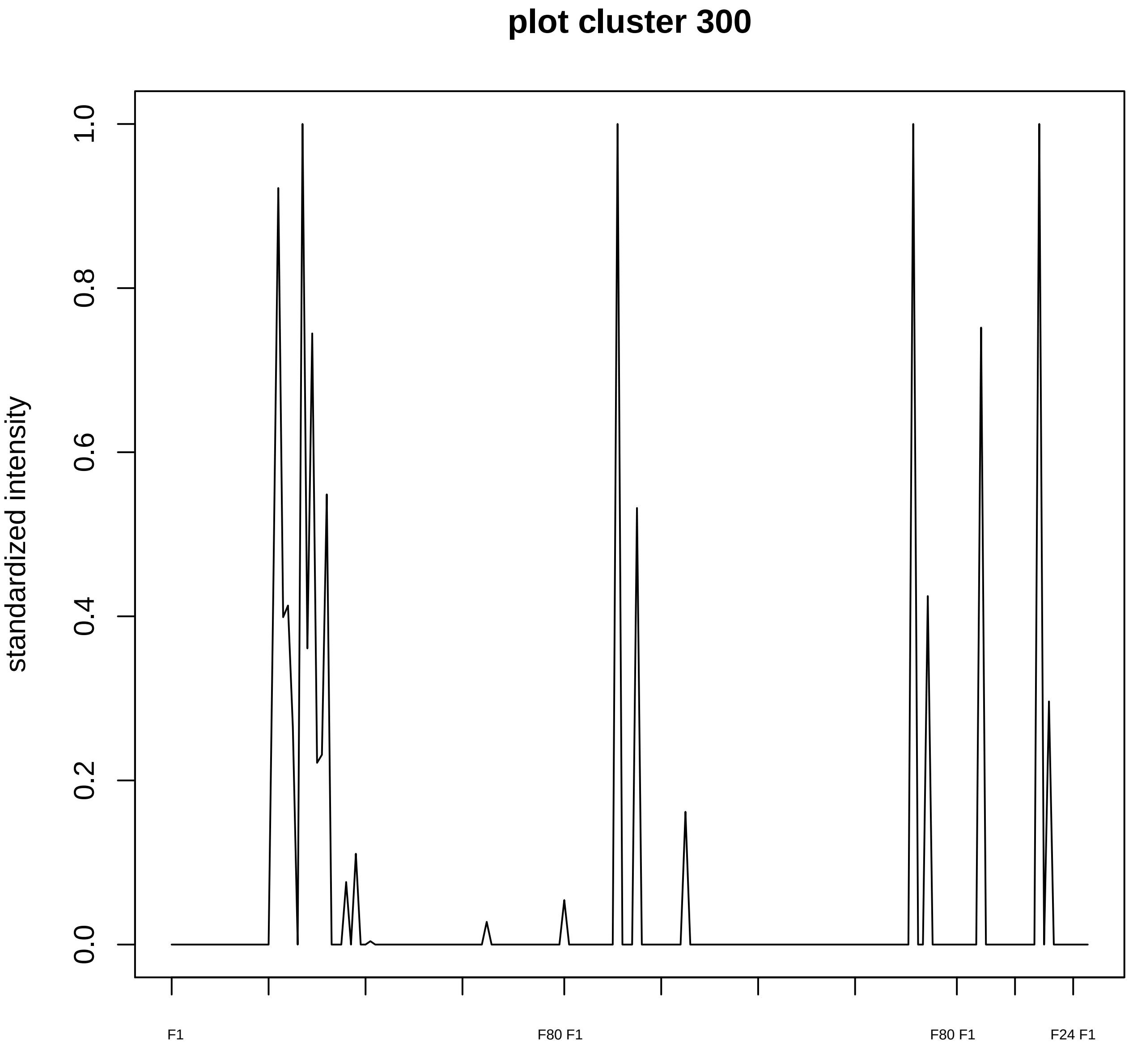
plot cluster 300
standardized intensity
0.0
0.2
0.4
0.6
0.8
1.0
F1
F80 F1
F80 F1
F24 F1

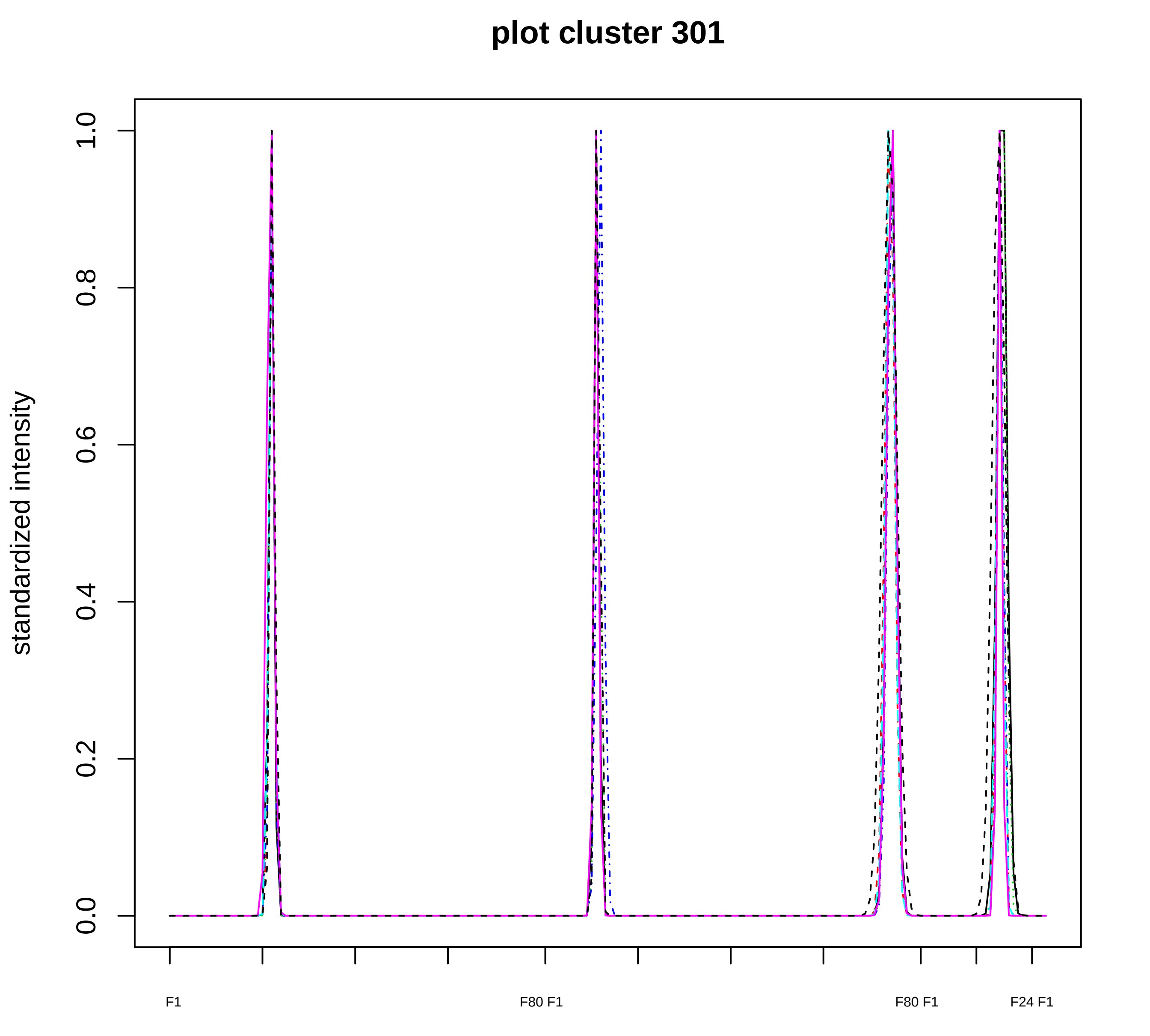

plot cluster 301
standardized intensity
0.0
0.2
0.4
0.6
0.8
1.0
F1
F80 F1
F80 F1
F24 F1

### Dendrogram of cluster 301

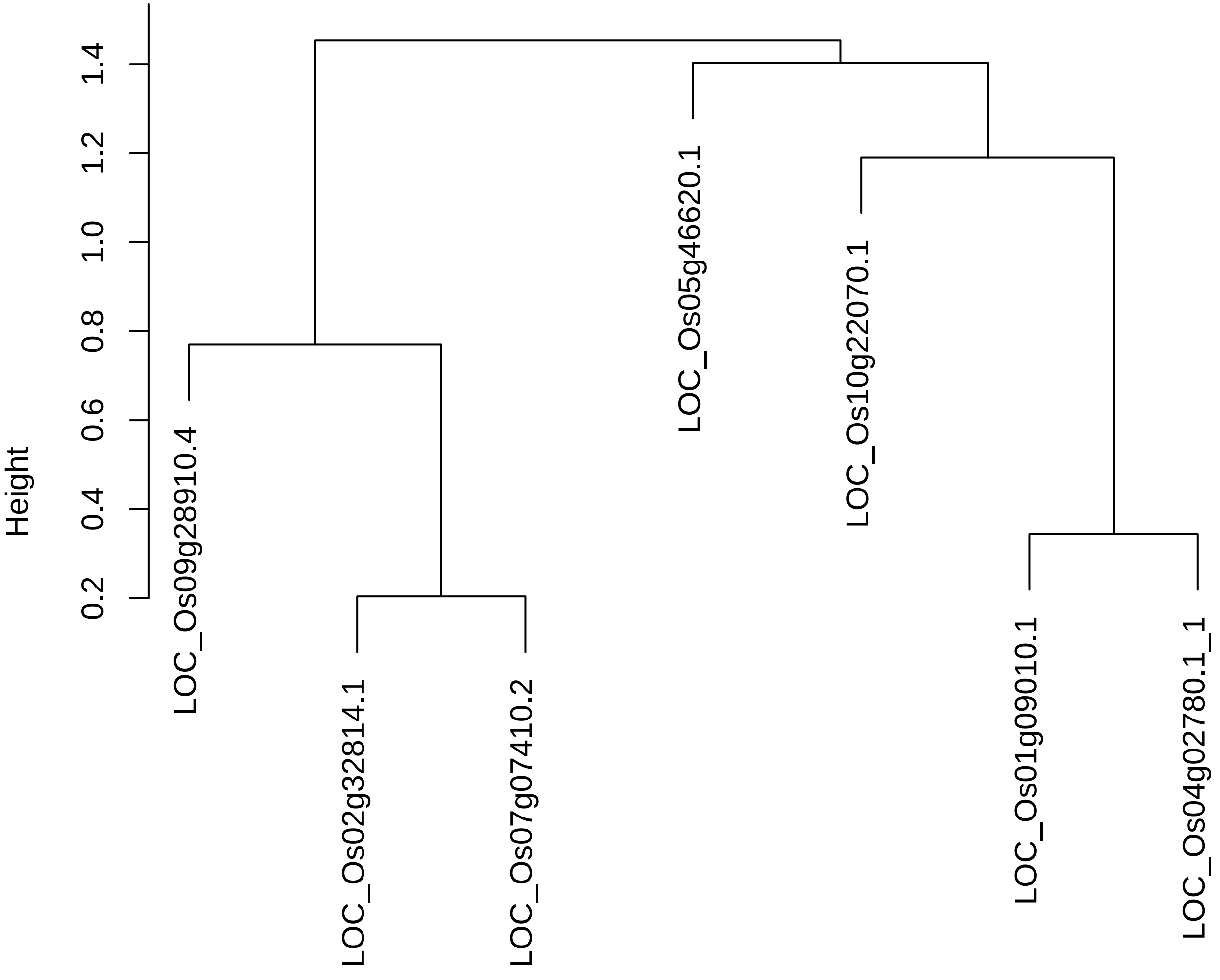


dist(spfitiex12sec12[lab[, 1] == i, ])
hclust (*, "ward.D2")

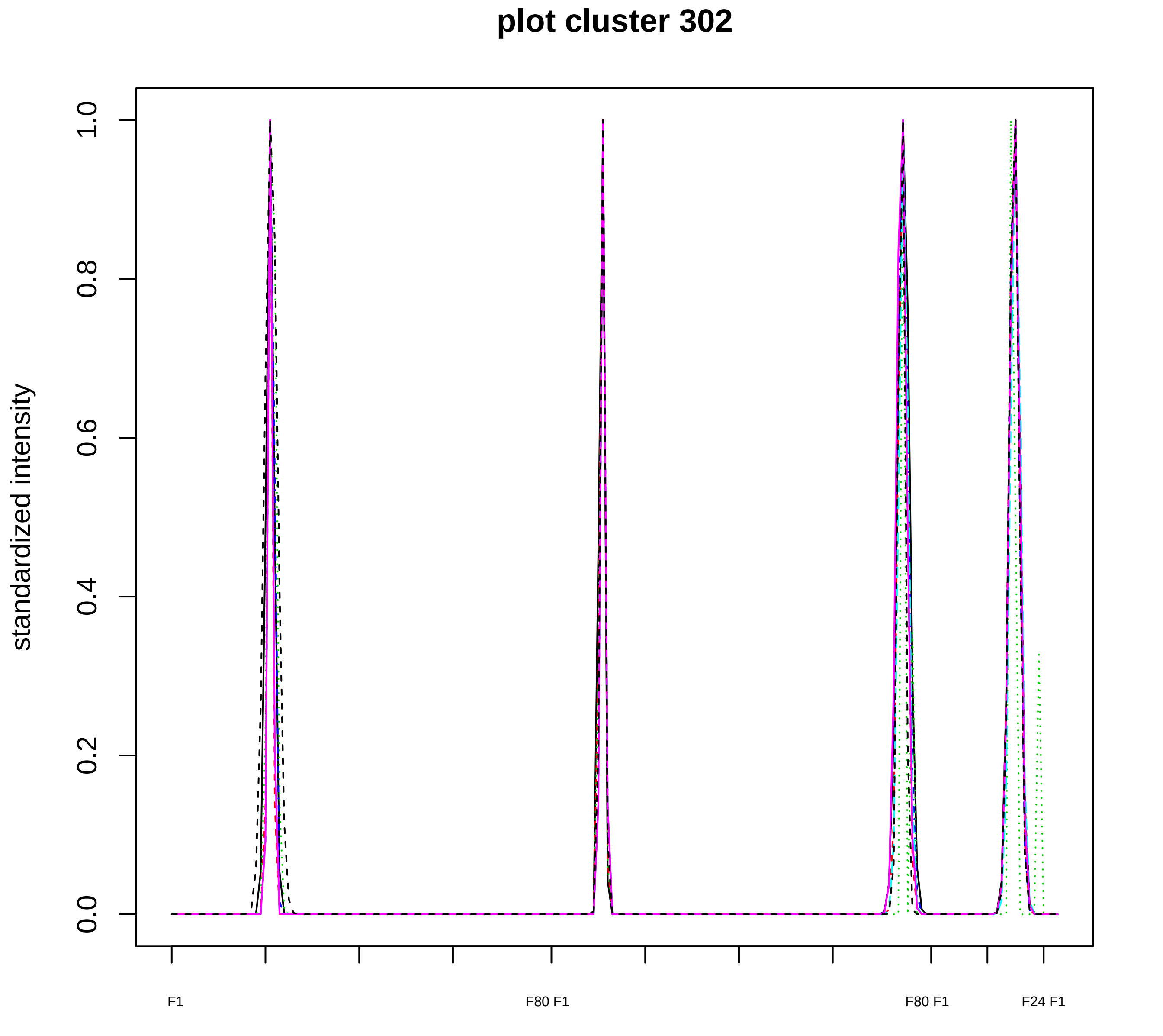
plot cluster 302
standardized intensity
0.0
0.2
0.4
0.6
0.8
1.0
F1
F80 F1
F80 F1
F24 F1

### Dendrogram of cluster 302

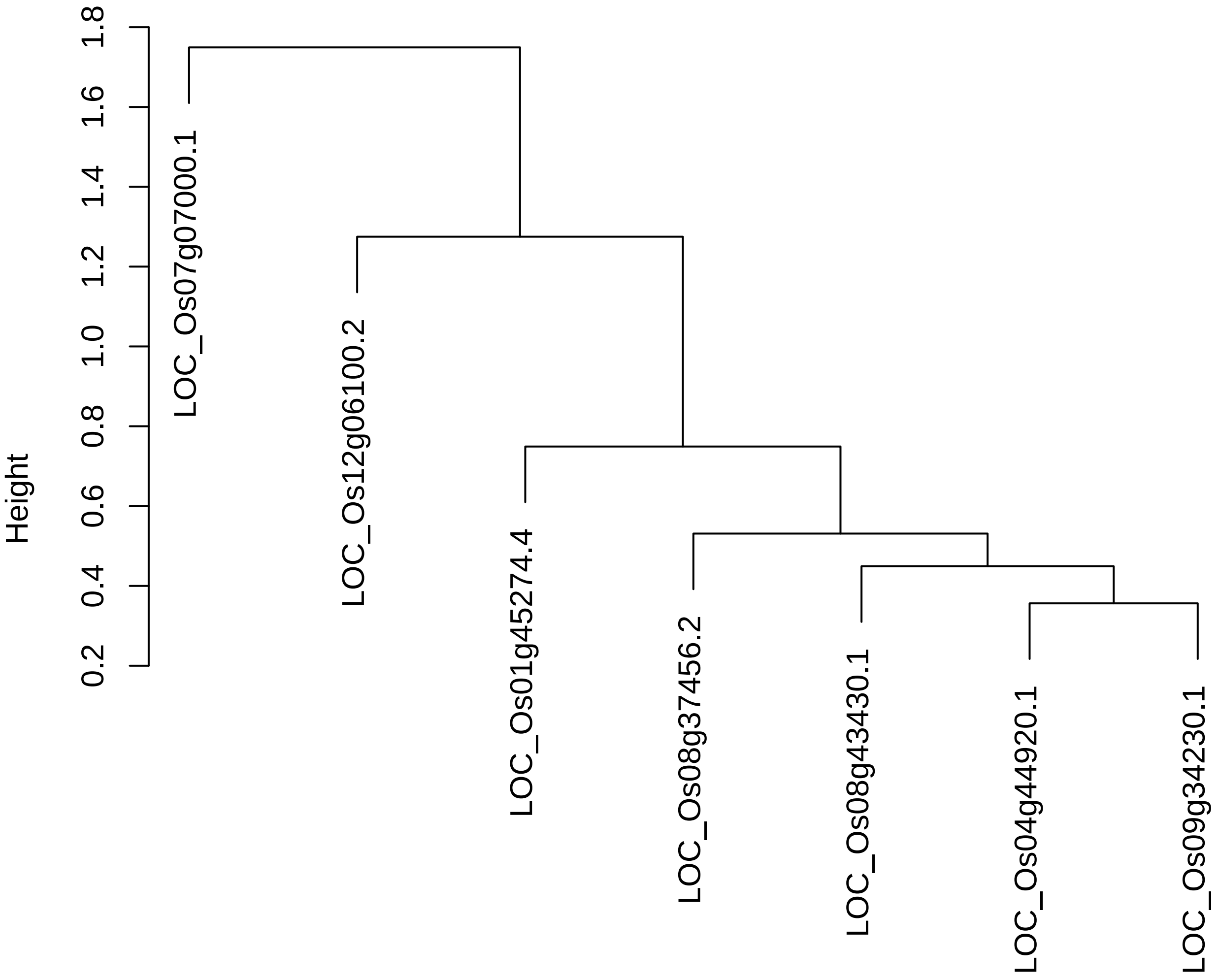


dist(spfitiex12sec12[lab[, 1] == i, ])
hclust (*, "ward.D2")

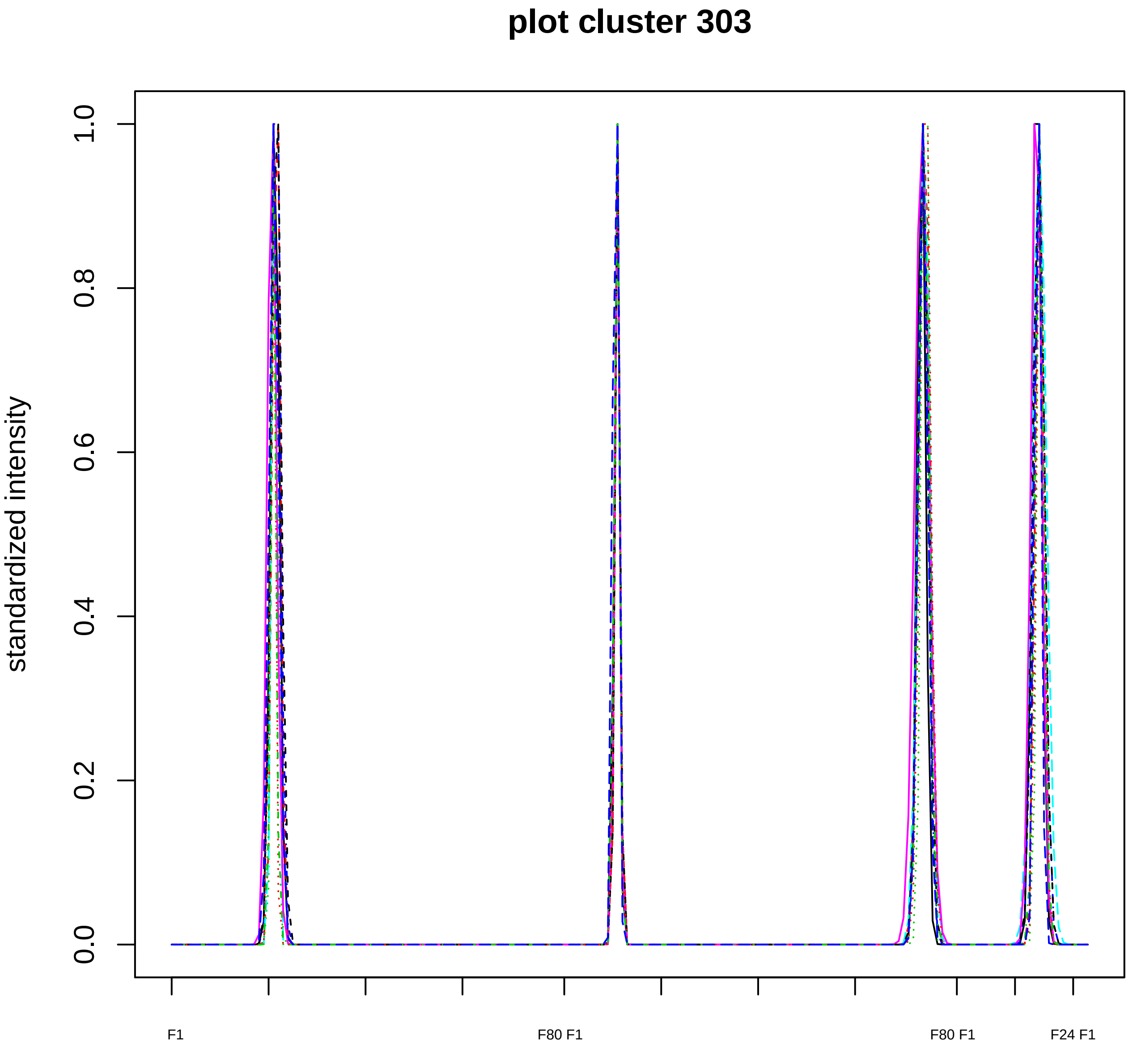
plot cluster 303
standardized intensity
1.0
0.8
0.6
0.4
0.2
0.0
F1
F80 F1
F80 F1
F24 F1

### Dendrogram of cluster 303

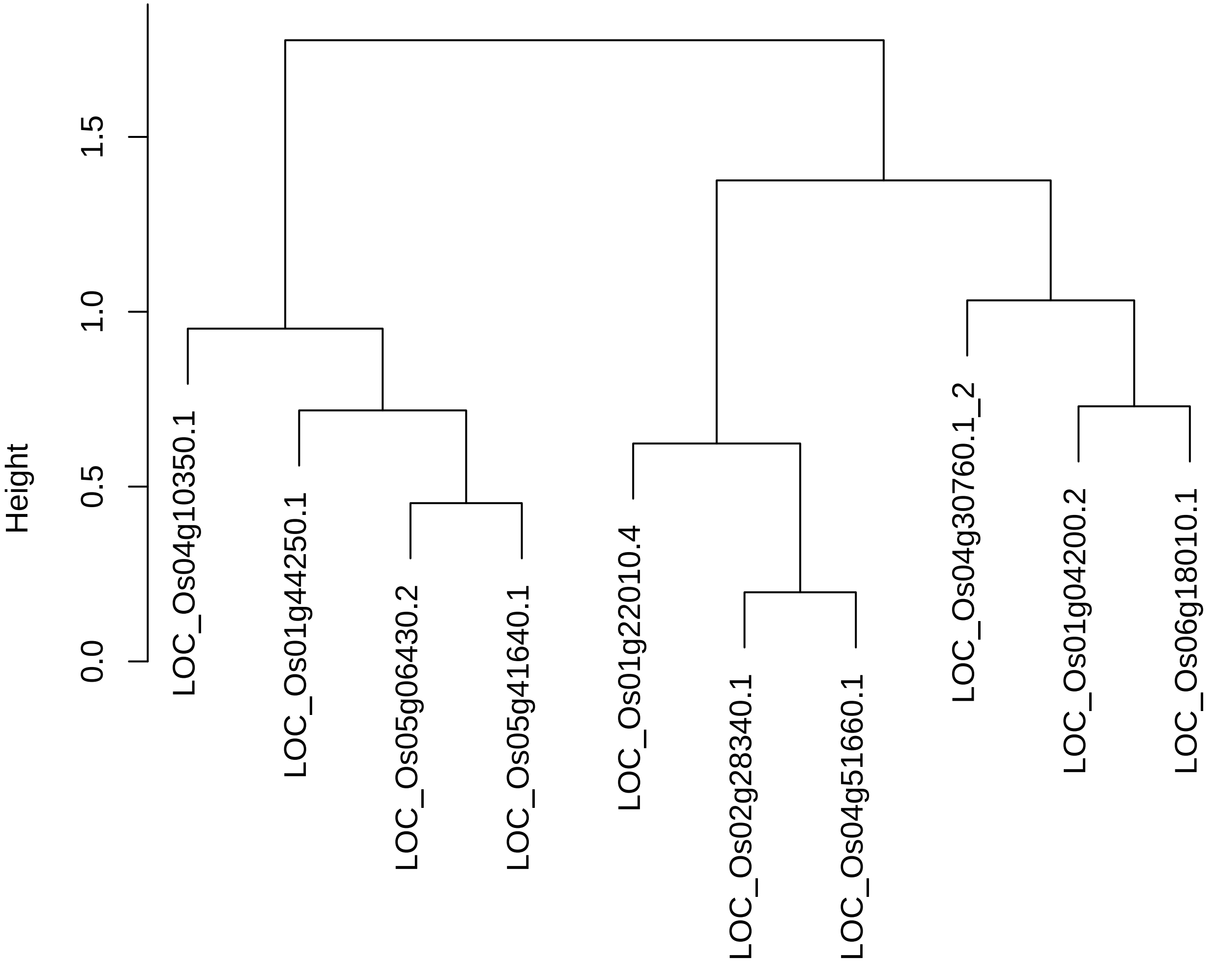


dist(spfitiex12sec12[lab[, 1] == i, ])
hclust (*, "ward.D2")

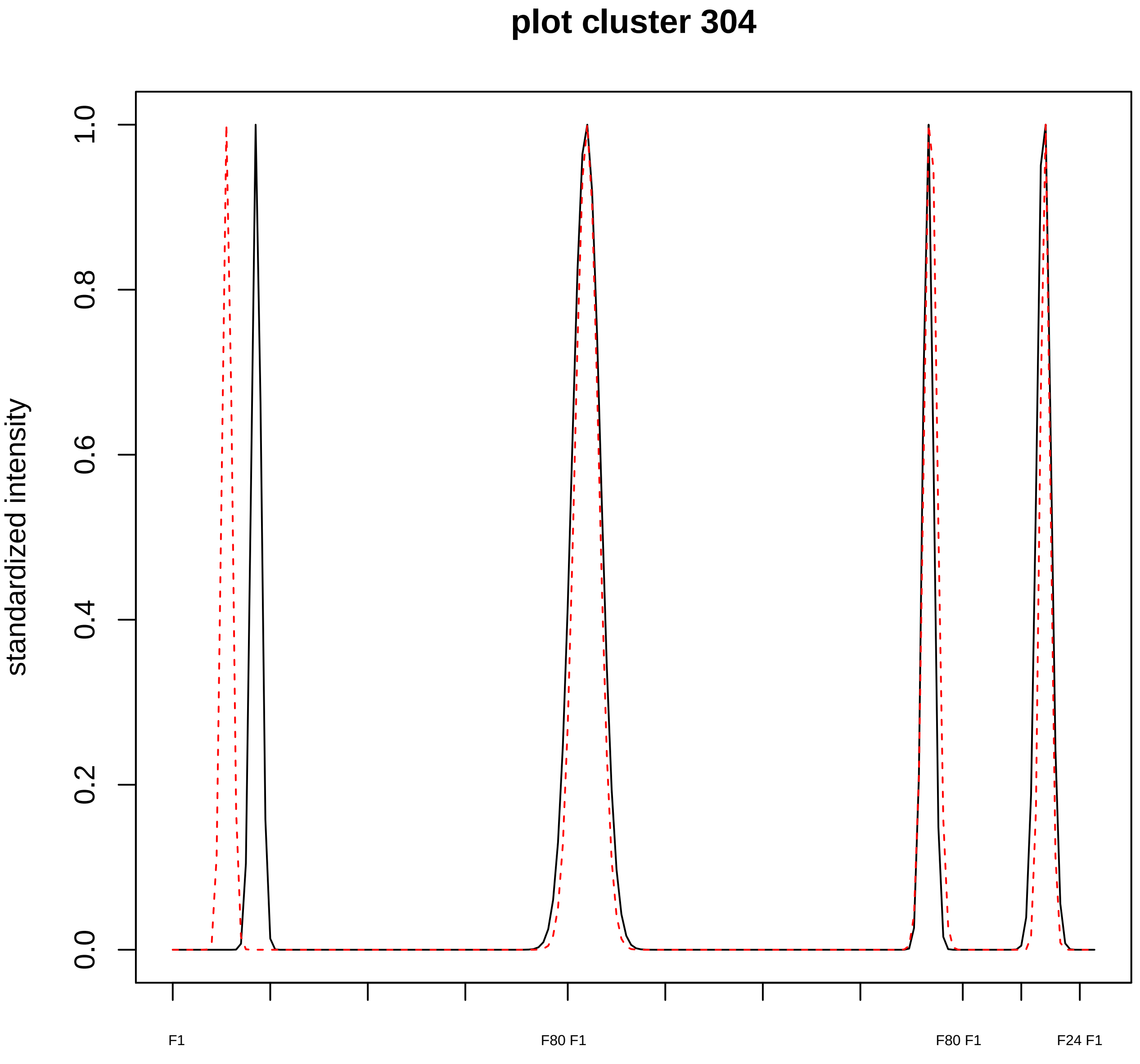

plot cluster 304
standardized intensity
0.0
0.2
0.4
0.6
0.8
1.0
F1
F80 F1
F80 F1
F24 F1

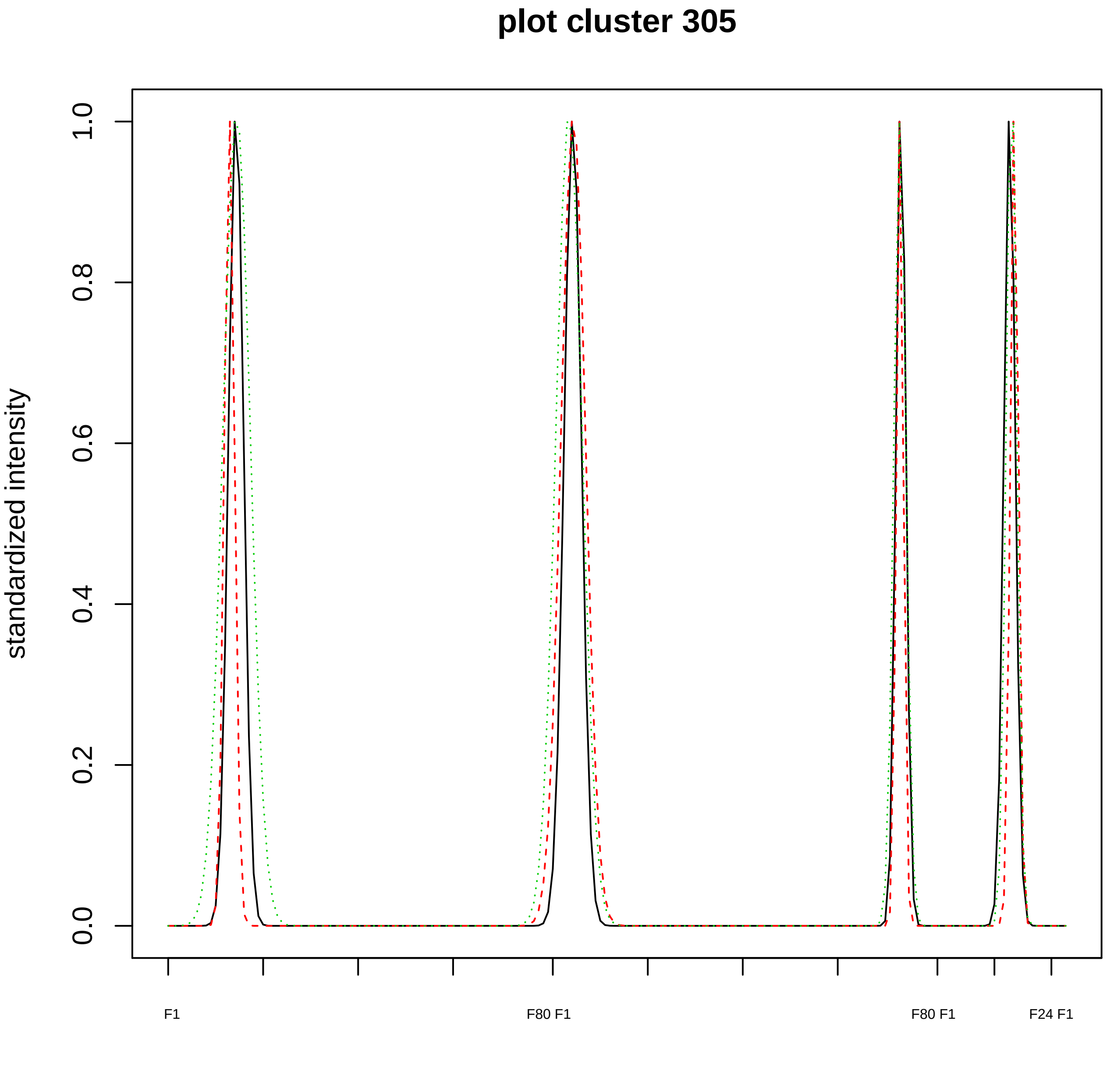
plot cluster 305
standardized intensity
0.0
0.2
0.4
0.6
0.8
1.0
F1
F80 F1
F80 F1
F24 F1

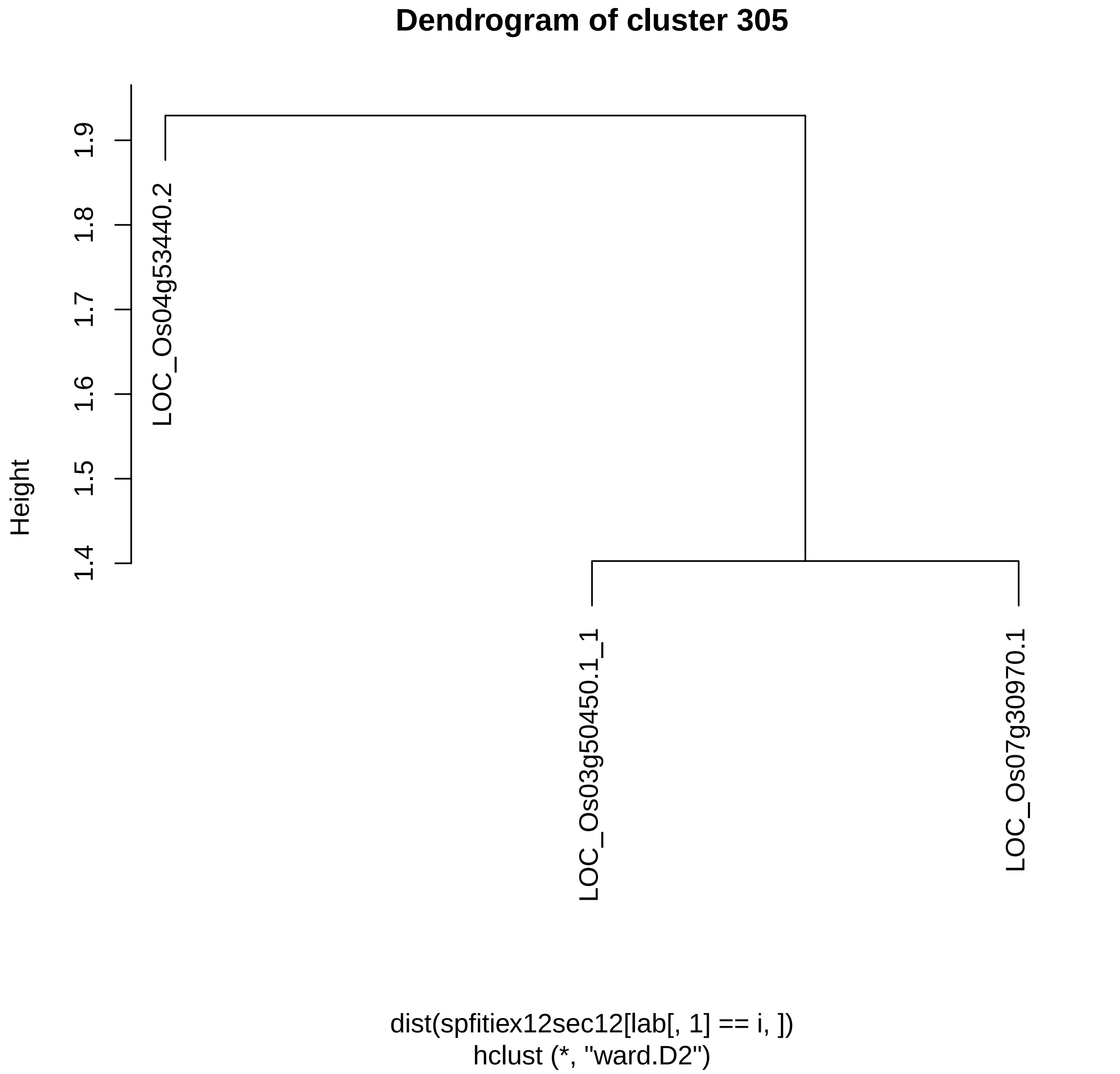
Dendrogram of cluster 305
Height
1.4
1.5
1.6
1.7
1.8
1.9
LOC_Os04g53440.2
LOC_Os03g50450.1_1
LOC_Os07g30970.1
dist(spfitiex12sec12[lab[, 1] == i, ])
hclust (*, "ward.D2")

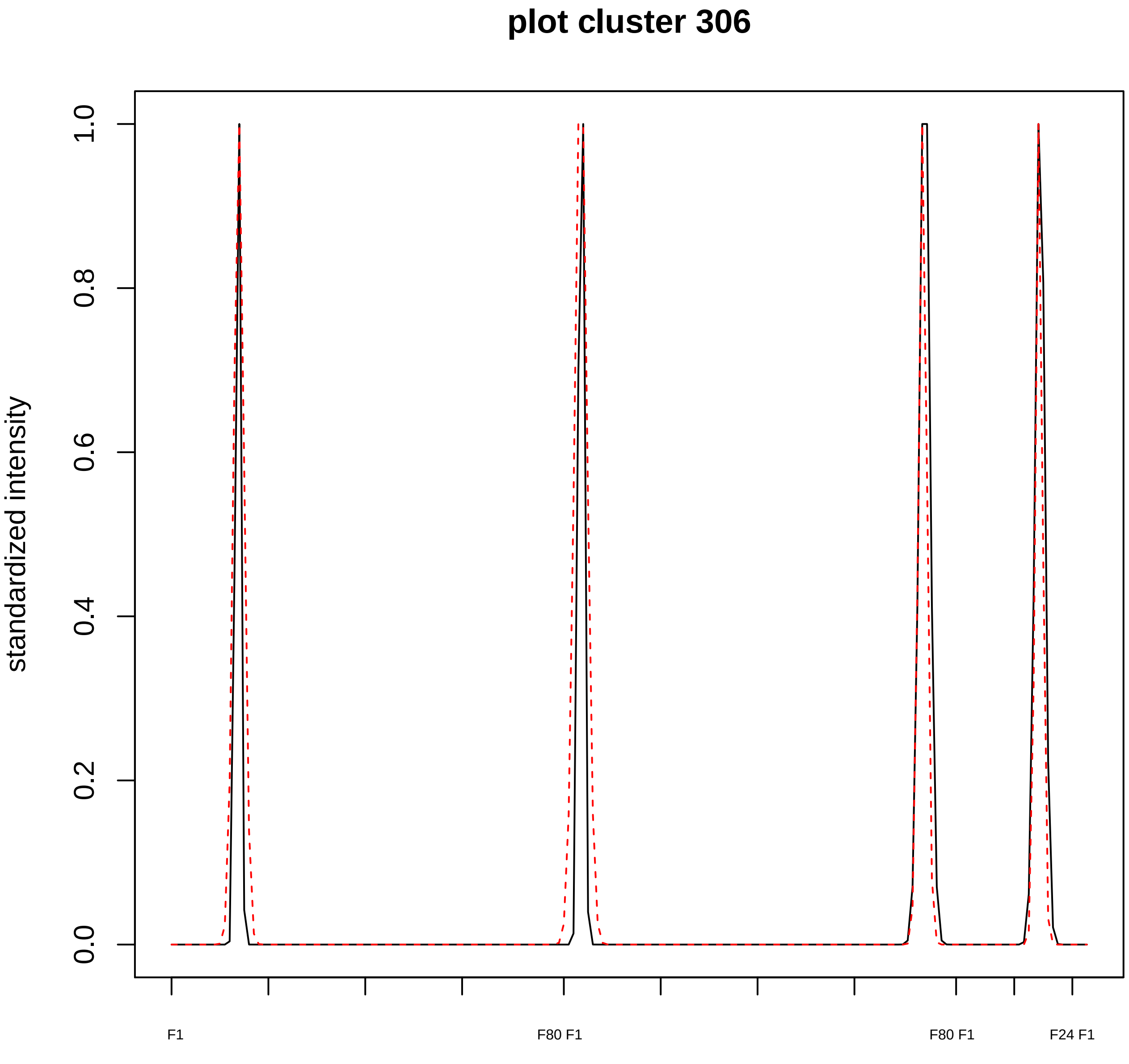

plot cluster 306
standardized intensity
0.0
0.2
0.4
0.6
0.8
1.0
F1
F80 F1
F80 F1
F24 F1

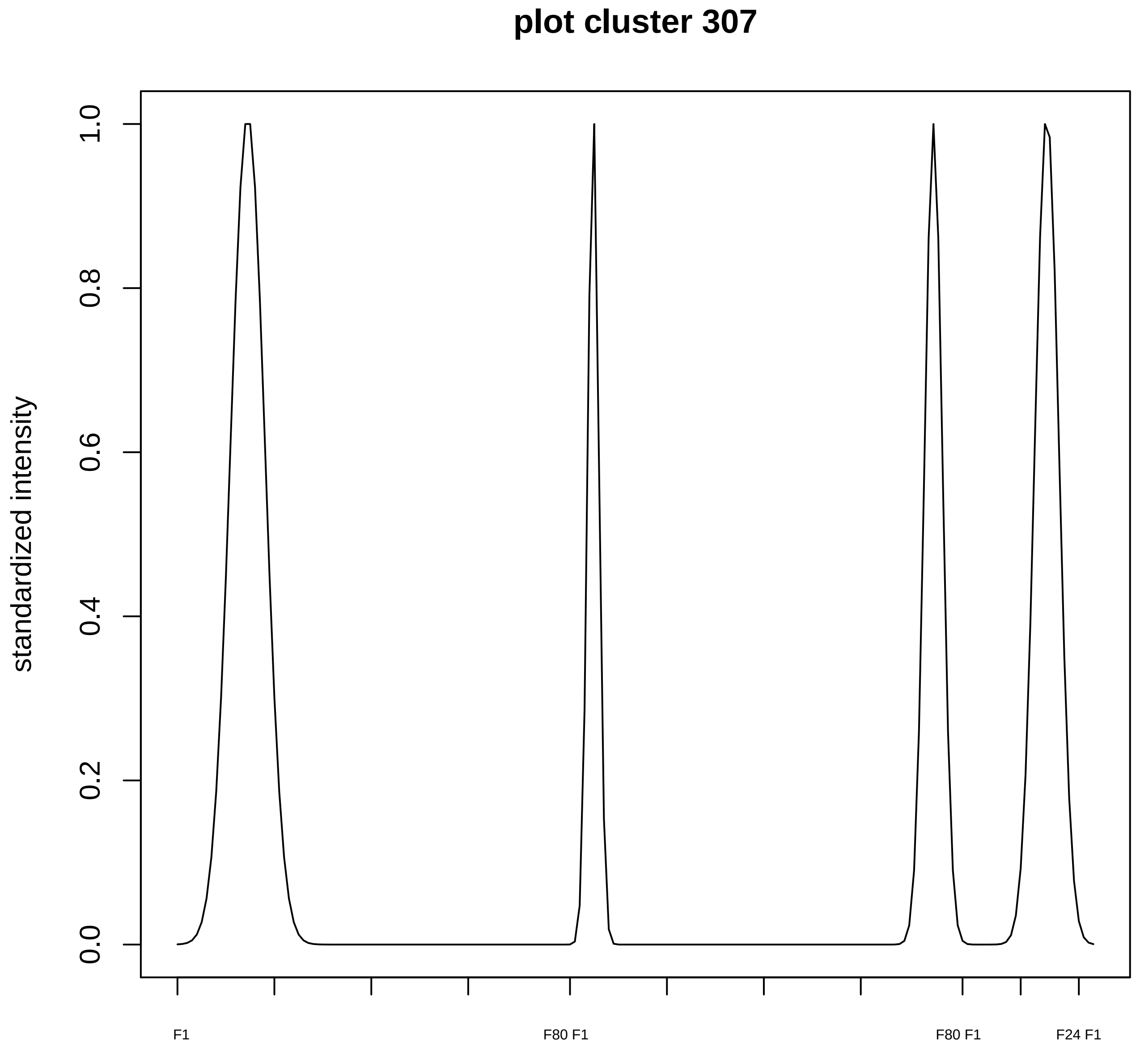
plot cluster 307
standardized intensity
0.0
0.2
0.4
0.6
0.8
1.0
F1
F80 F1
F80 F1
F24 F1

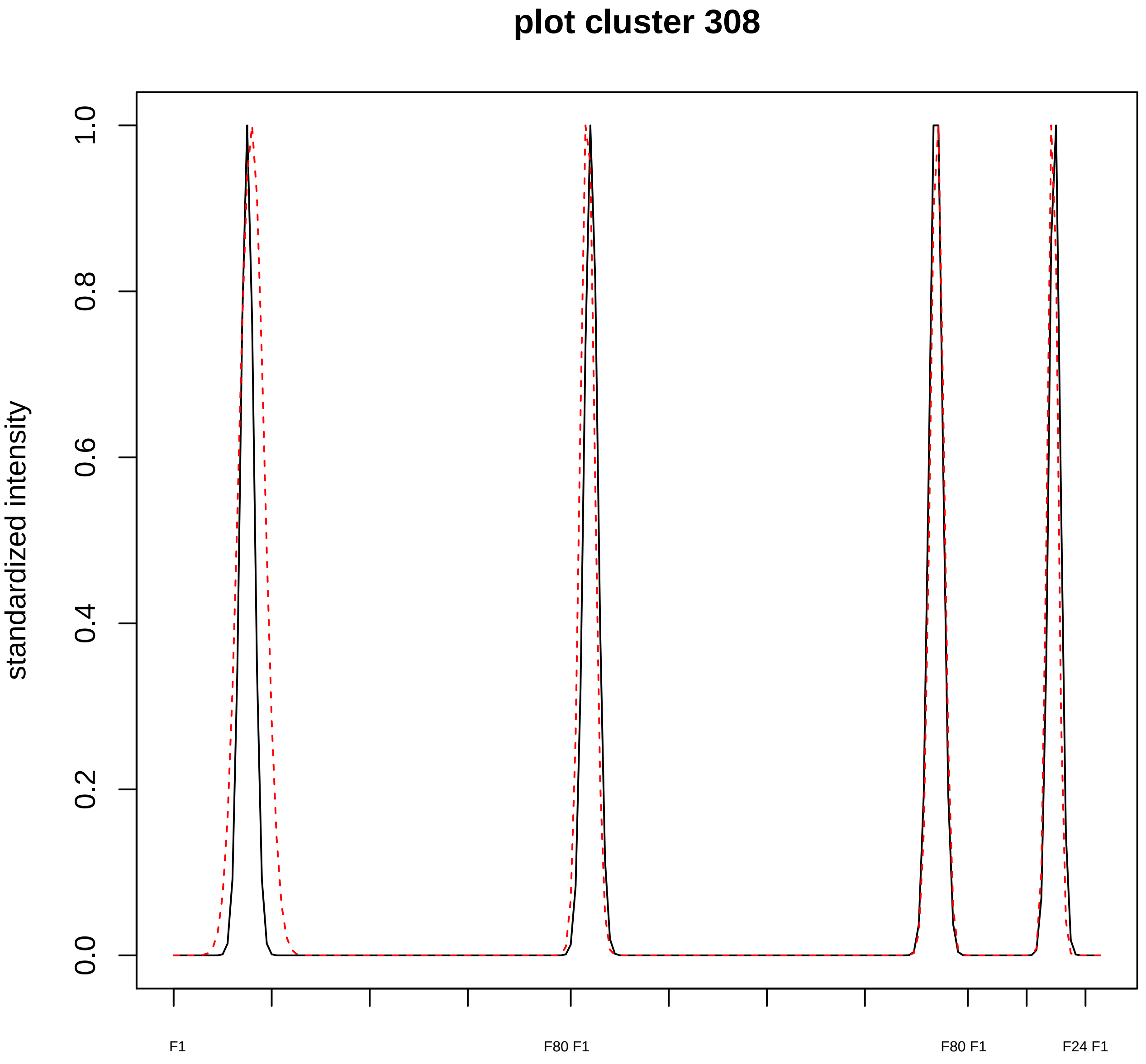
plot cluster 308
standardized intensity
0.0
0.2
0.4
0.6
0.8
1.0
F1
F80 F1
F80 F1
F24 F1

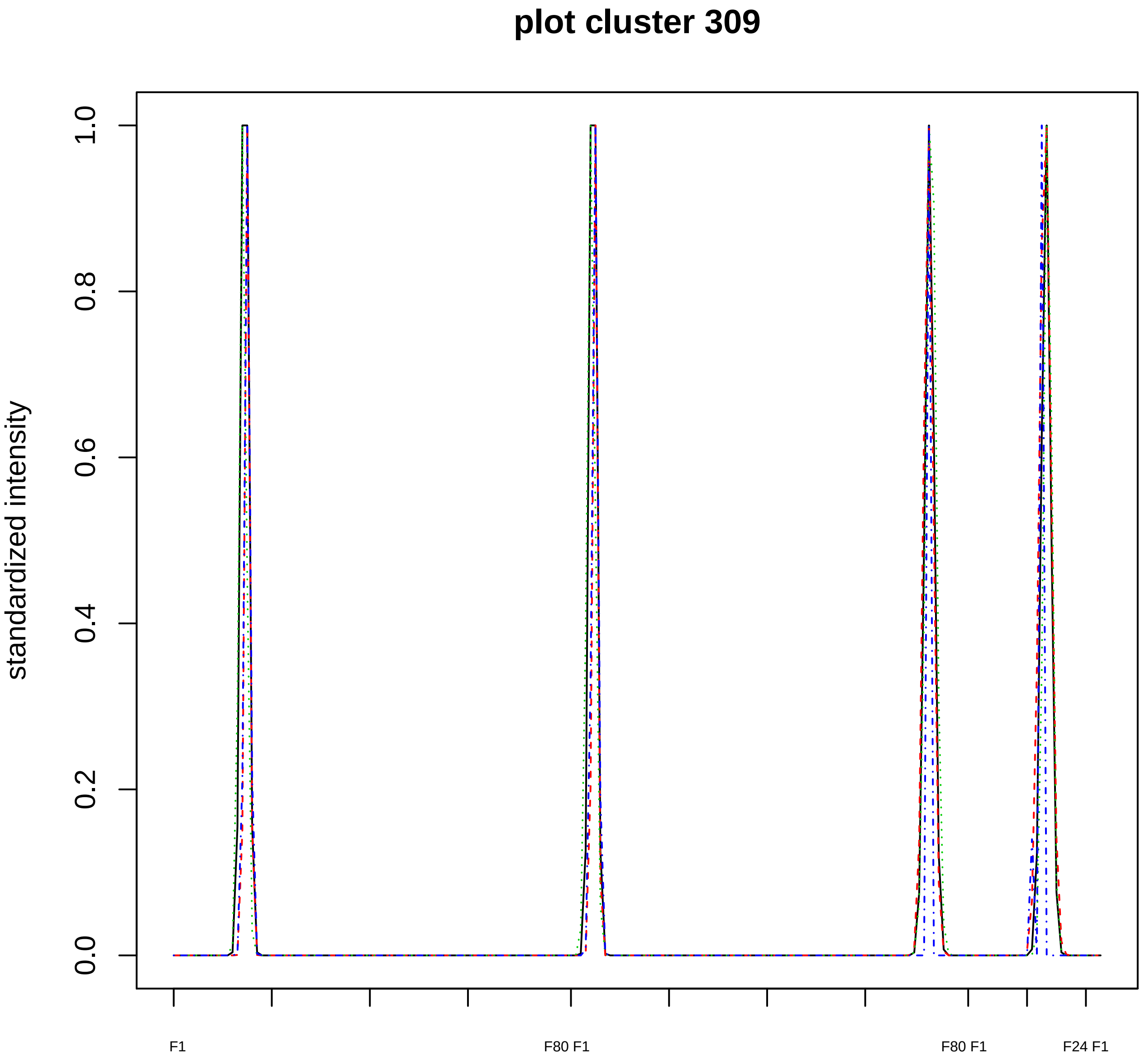
plot cluster 309
standardized intensity
1.0
0.8
0.6
0.4
0.2
0.0
F1
F80 F1
F80 F1
F24 F1

### Dendrogram of cluster 309

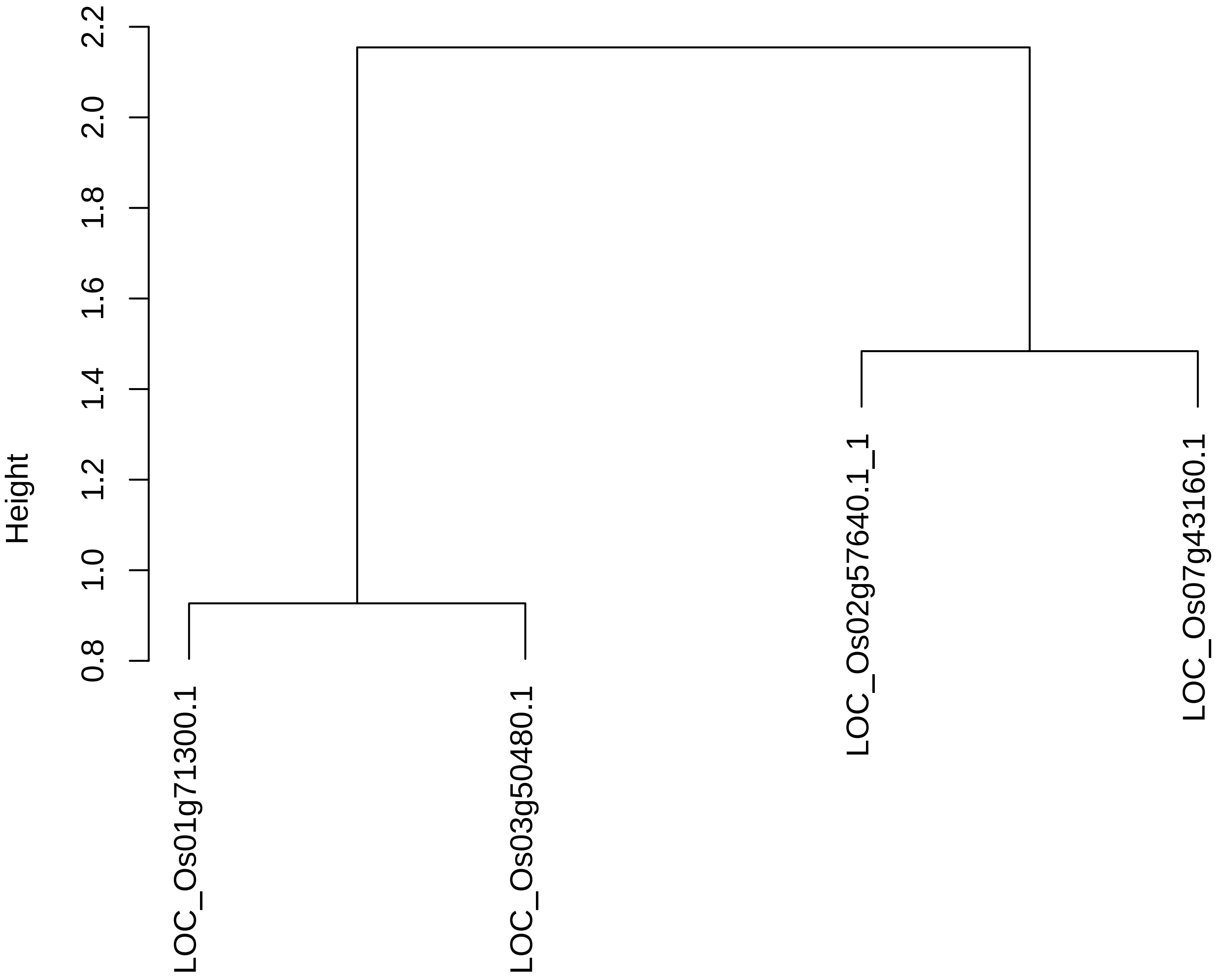


dist(spfitiex12sec12[lab[, 1] == i, ])
hclust (*, "ward.D2")

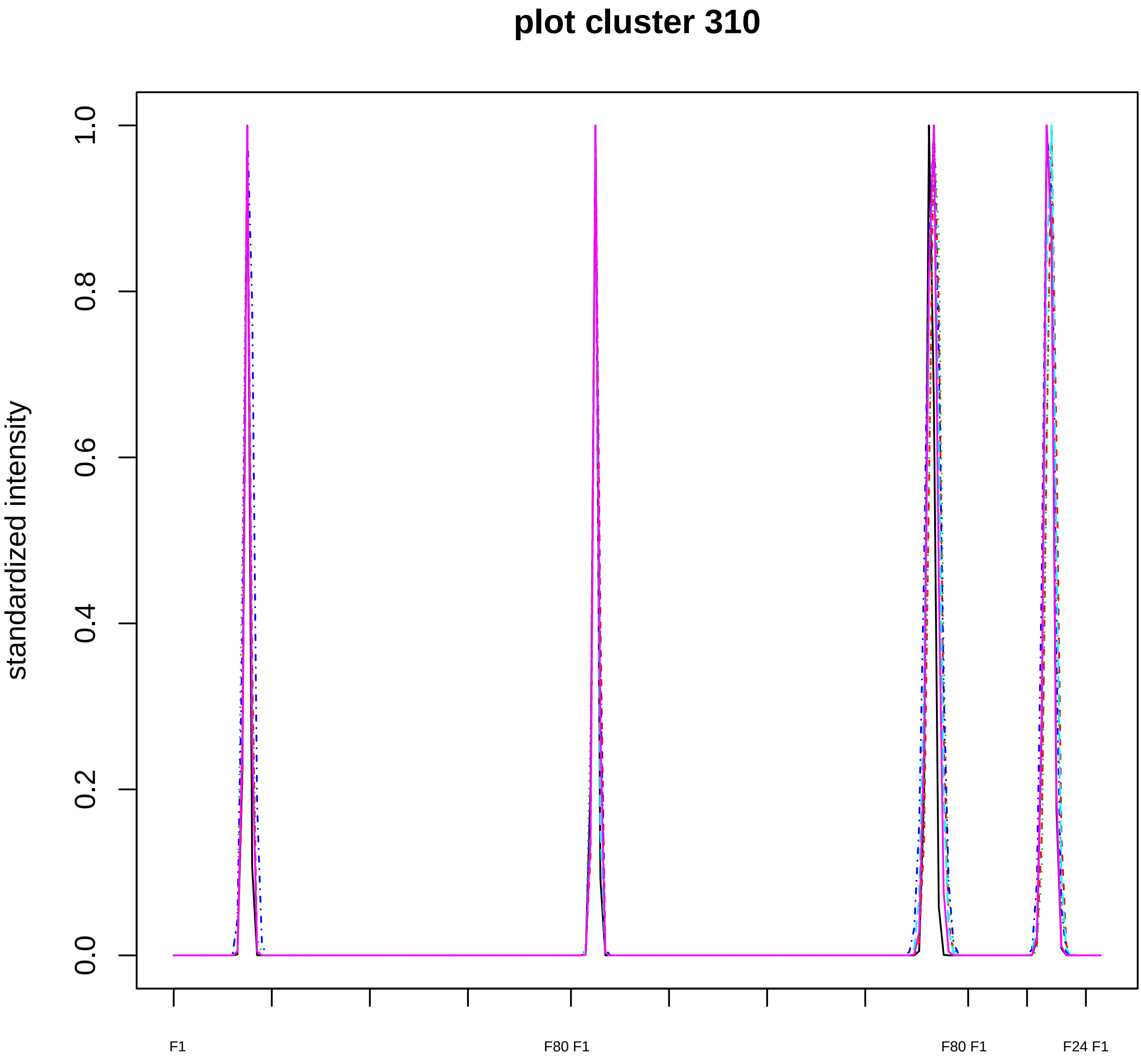
plot cluster 310
standardized intensity
0.0
0.2
0.4
0.6
0.8
1.0
F1
F80 F1
F80 F1
F24 F1

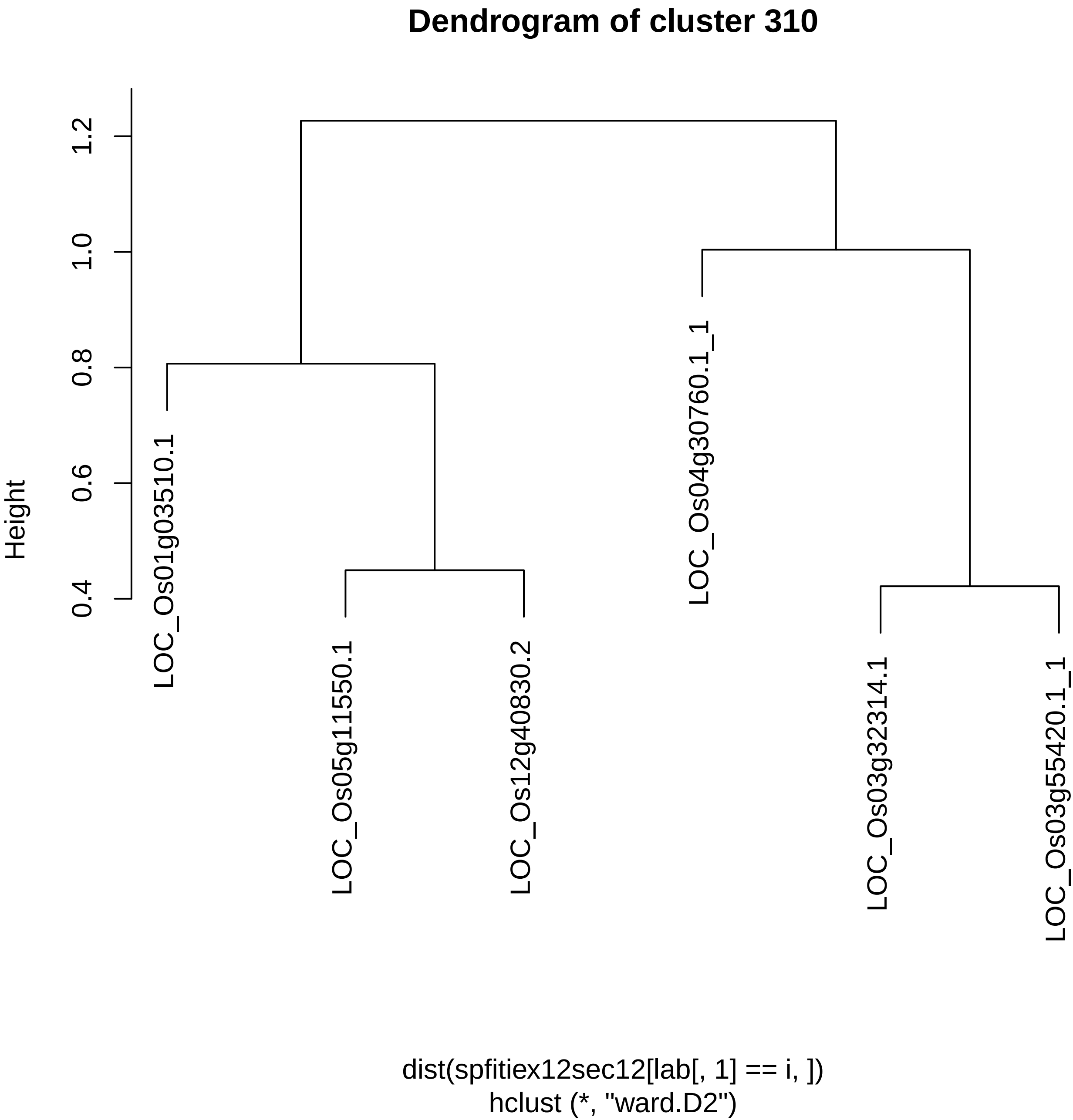

Dendrogram of cluster 310
Height
0.4
0.6
0.8
1.0
1.2
LOC_Os01g03510.1
LOC_Os05g11550.1
LOC_Os12g40830.2
LOC_Os04g30760.1_1
LOC_Os03g32314.1
LOC_Os03g55420.1_1
dist(spfitiex12sec12[lab[, 1] == i, ])
hclust (*, "ward.D2")

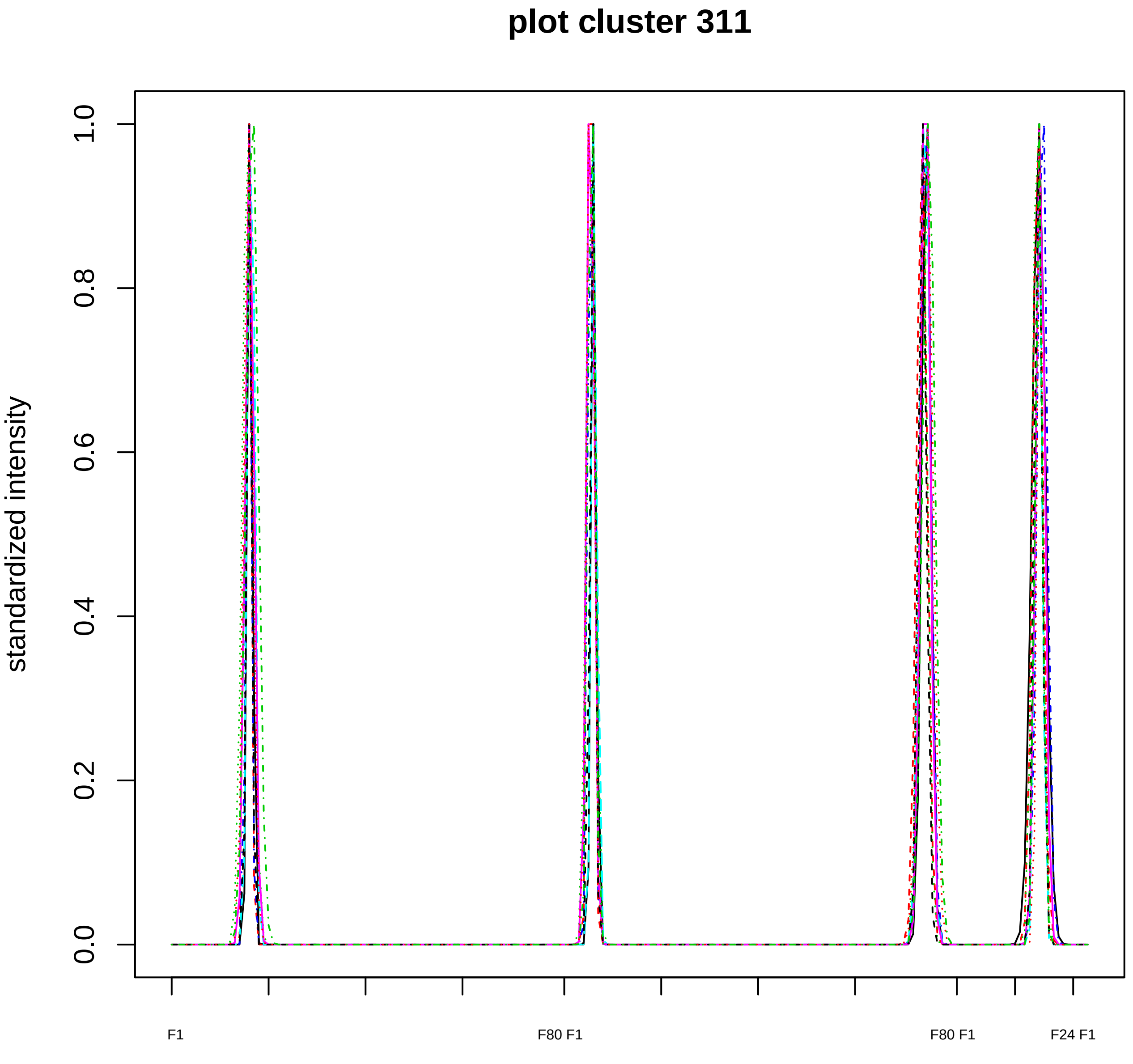
plot cluster 311
standardized intensity
0.0
0.2
0.4
0.6
0.8
1.0
F1
F80 F1
F80 F1
F24 F1

### Dendrogram of cluster 311

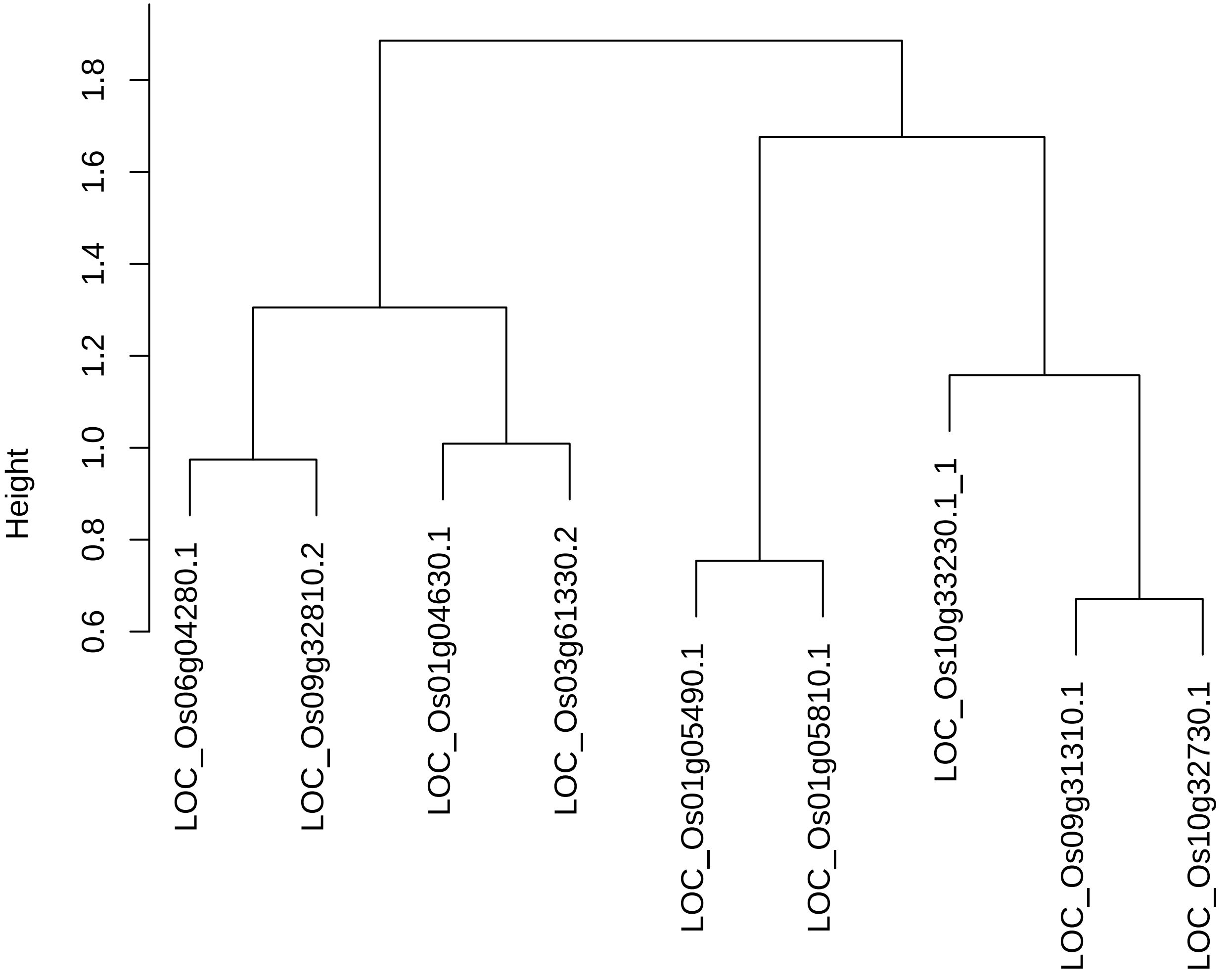


dist(spfitiex12sec12[lab[, 1] == i, ])
hclust (*, "ward.D2")

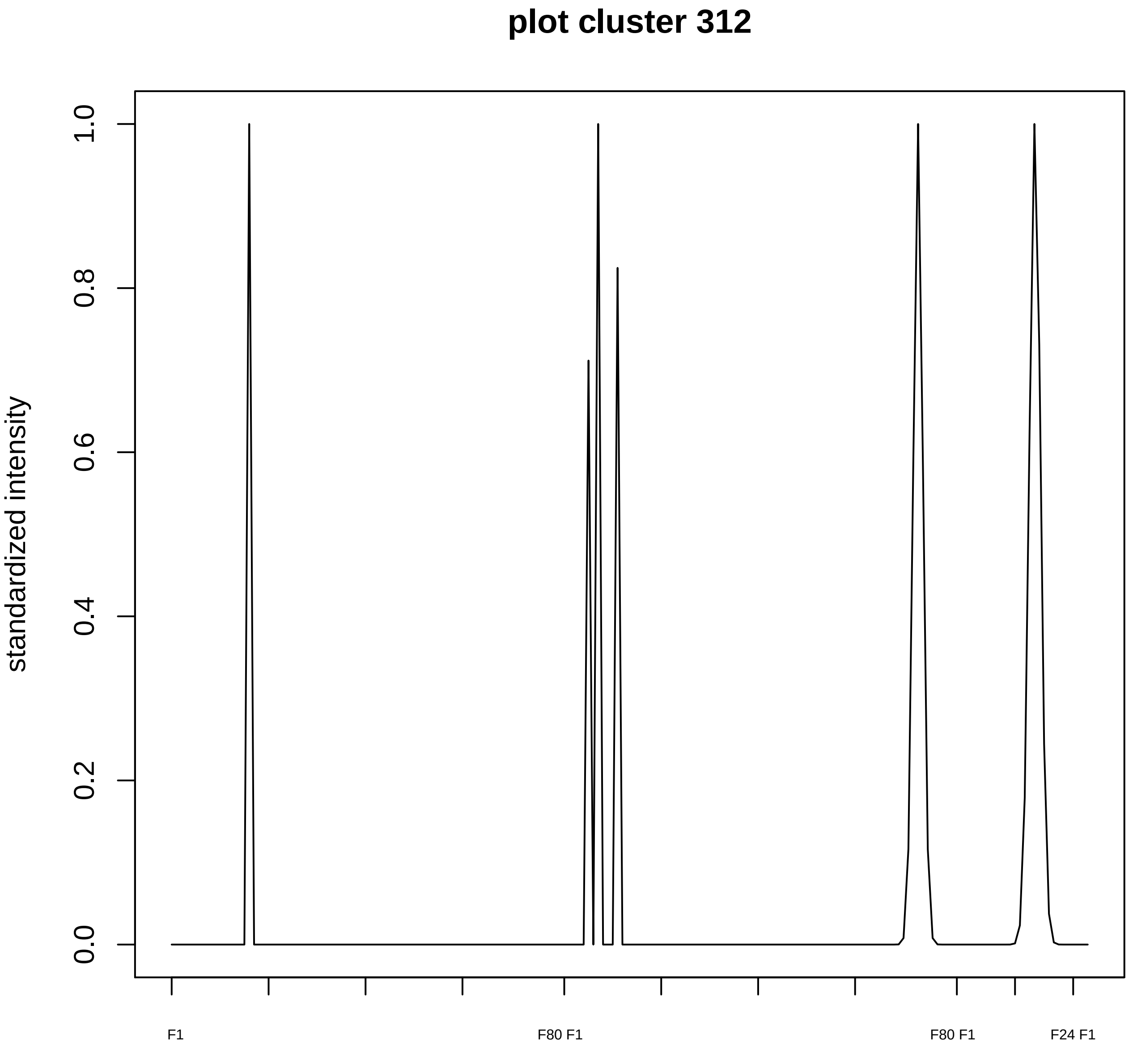
plot cluster 312
standardized intensity
0.0
0.2
0.4
0.6
0.8
1.0
F1
F80 F1
F80 F1
F24 F1

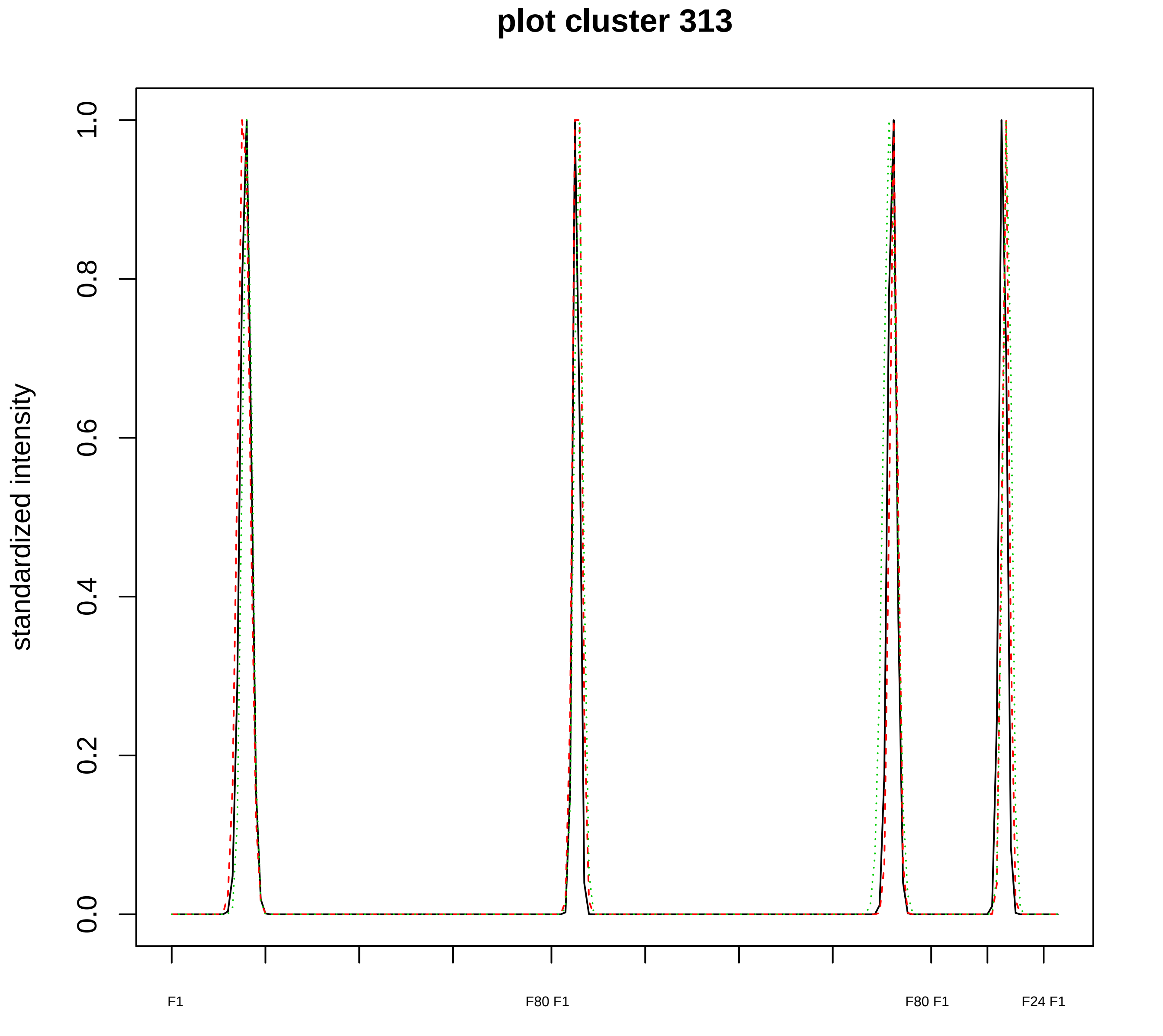
plot cluster 313
standardized intensity
0.0
0.2
0.4
0.6
0.8
1.0
F1
F80 F1
F80 F1
F24 F1

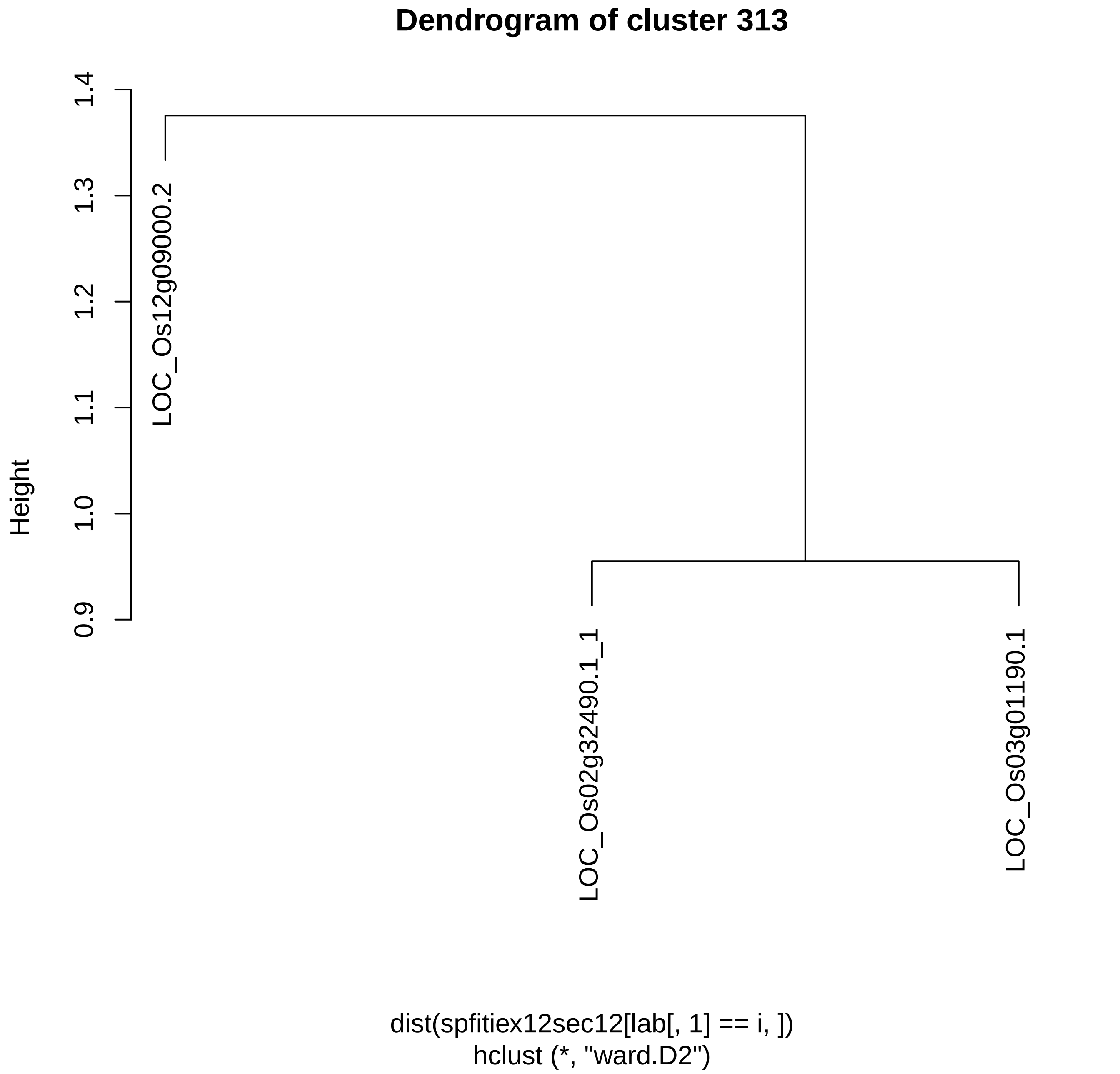
Dendrogram of cluster 313
Height
0.9
1.0
1.1
1.2
1.3
1.4
LOC_Os12g09000.2
LOC_Os02g32490.1_1
LOC_Os03g01190.1
dist(spfitiex12sec12[lab[, 1] == i, ])
hclust (*, "ward.D2")

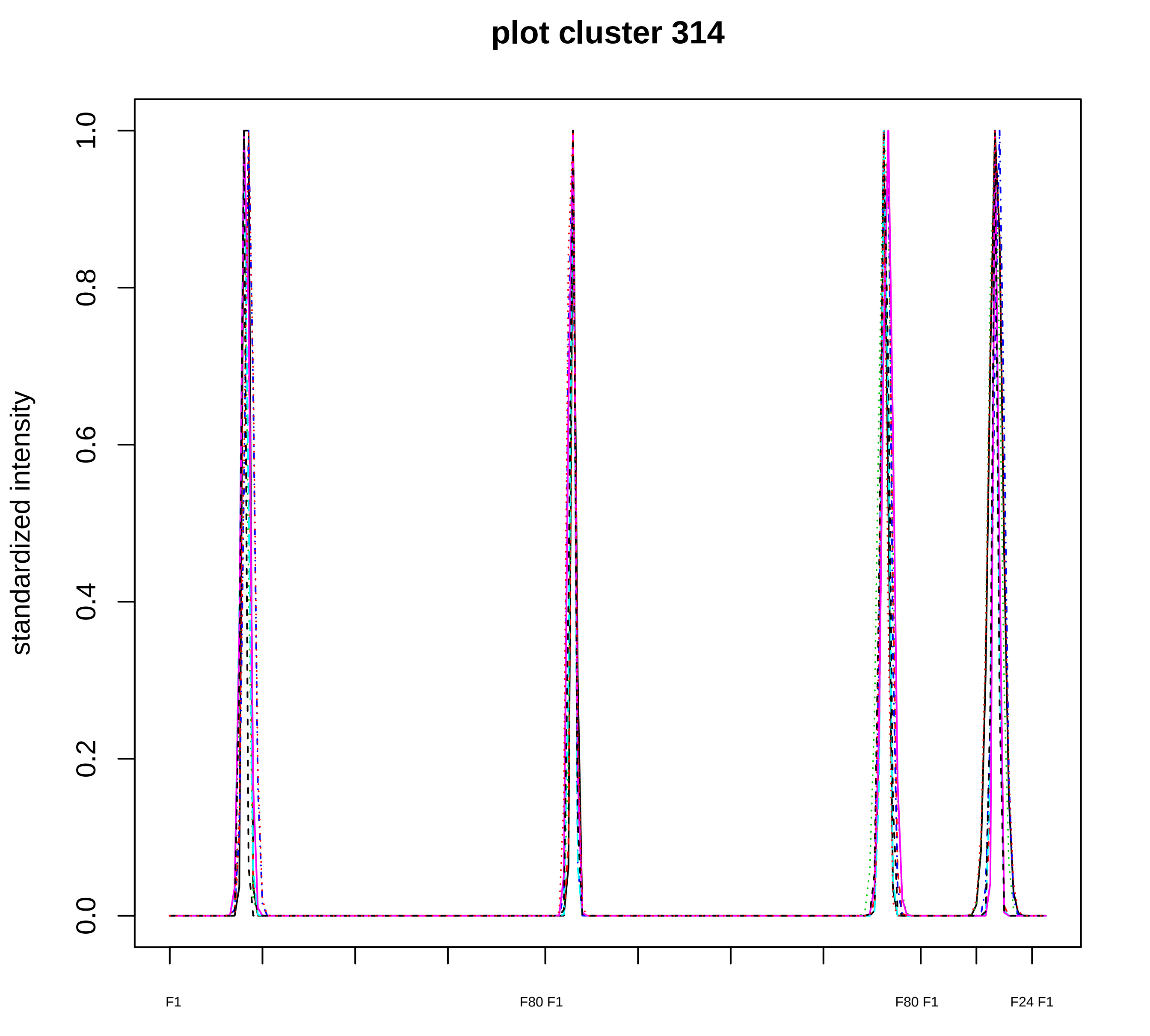
plot cluster 314
standardized intensity
0.0
0.2
0.4
0.6
0.8
1.0
F1
F80 F1
F80 F1
F24 F1

#### Dendrogram of cluster 314

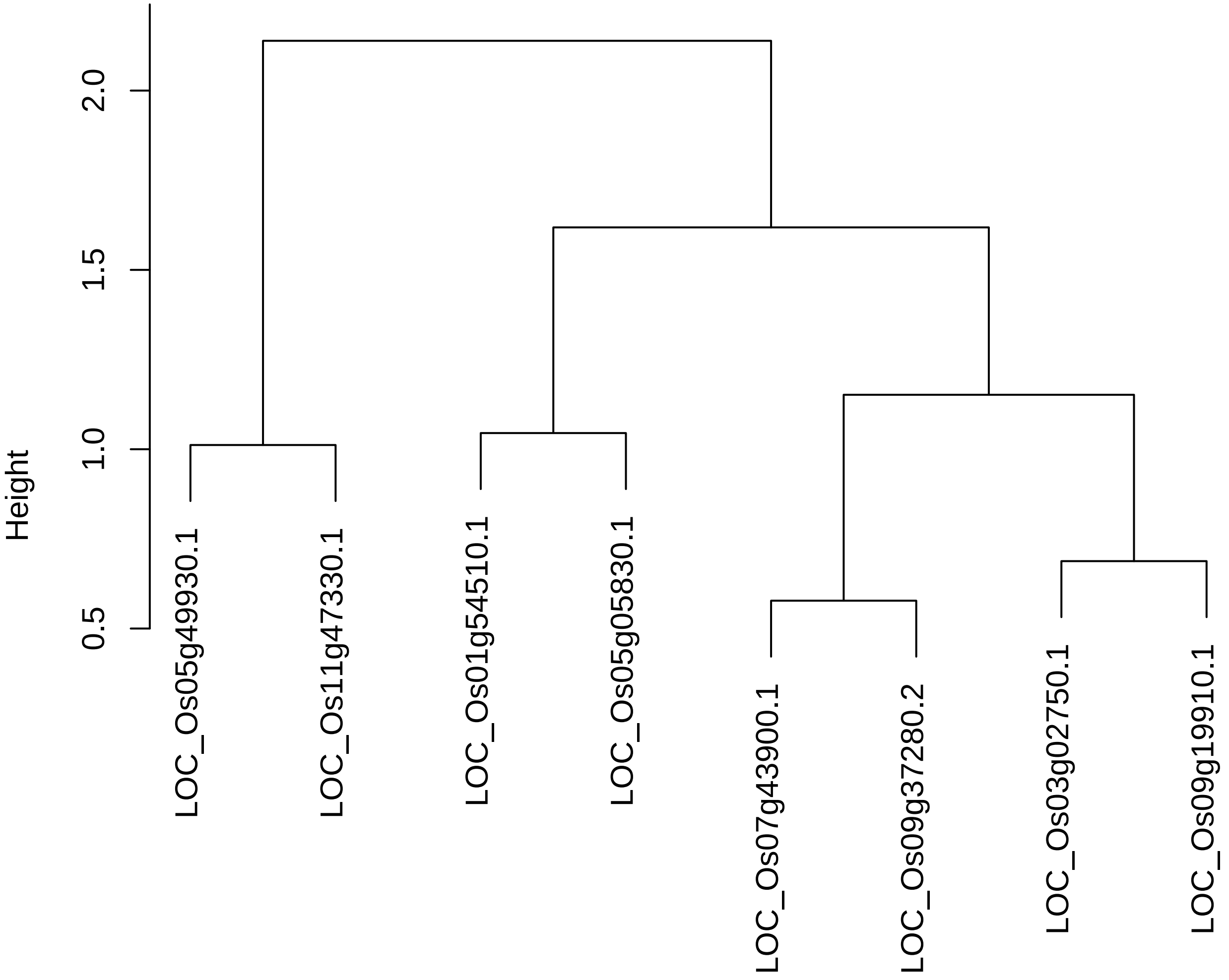


dist(spfitiex12sec12[lab[, 1] == i, ])
hclust (*, "ward.D2")

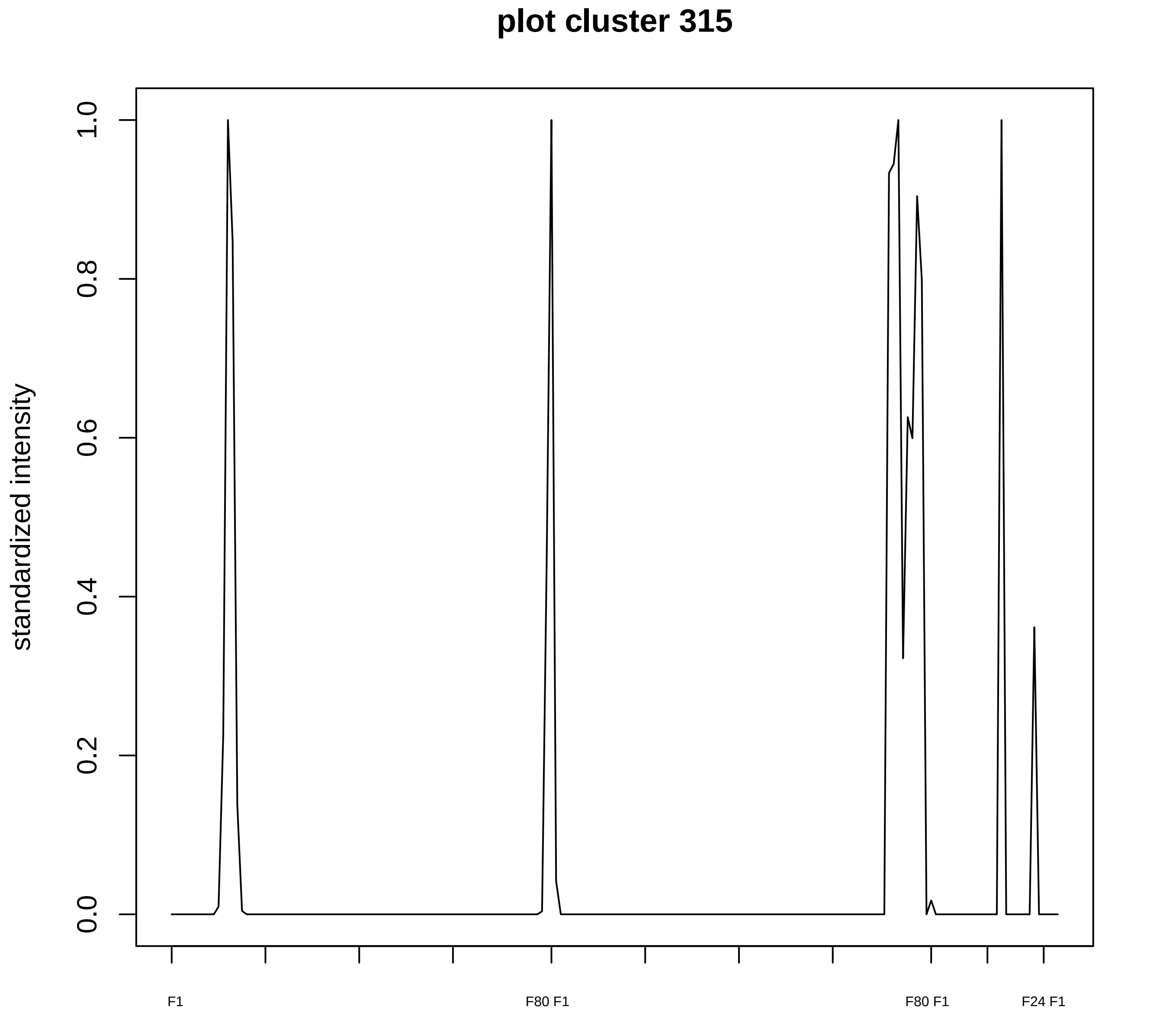
plot cluster 315
standardized intensity
0.0
0.2
0.4
0.6
0.8
1.0
F1
F80 F1
F80 F1
F24 F1

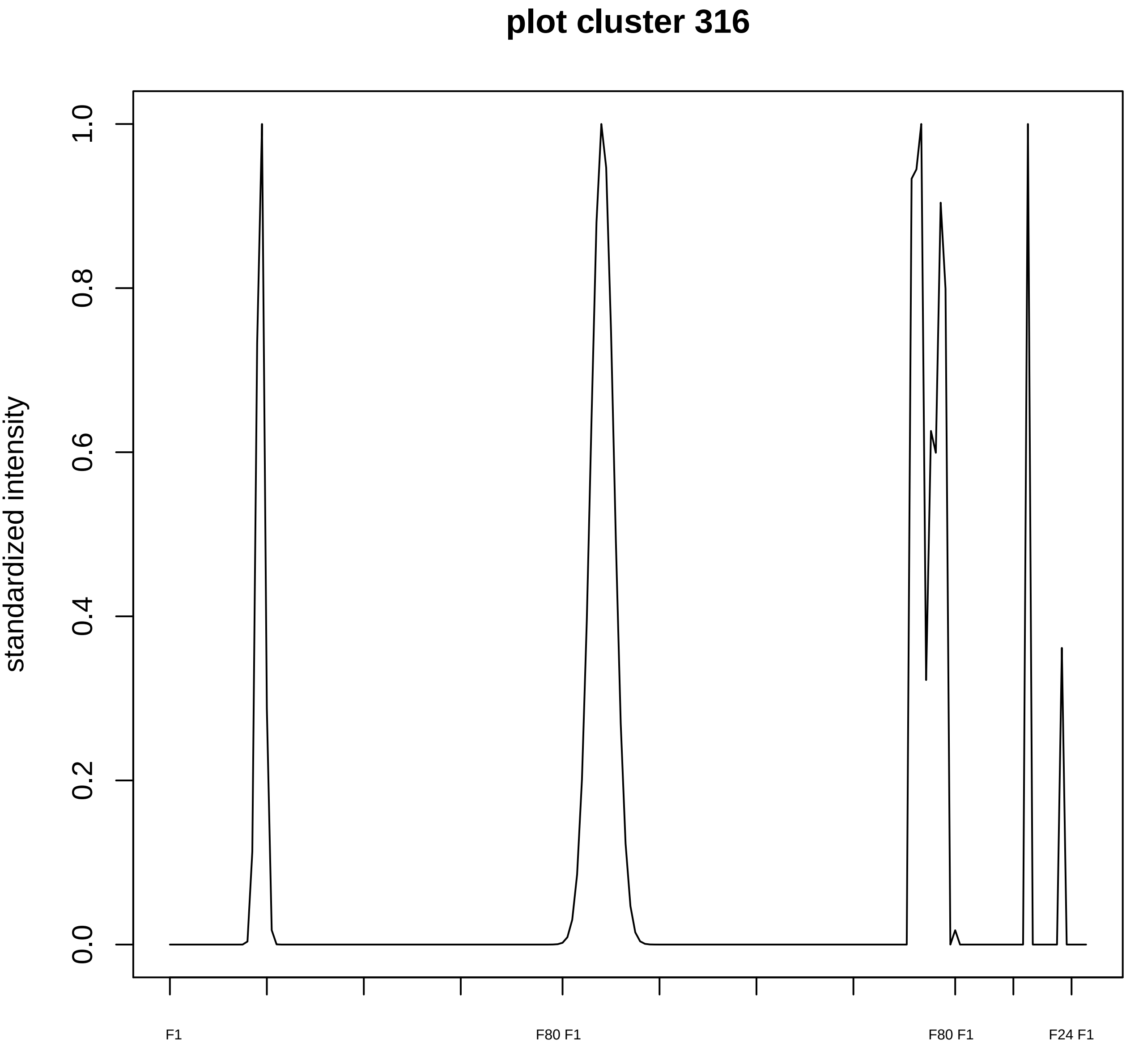
plot cluster 316
standardized intensity
0.0
0.2
0.4
0.6
0.8
1.0
F1
F80 F1
F80 F1
F24 F1

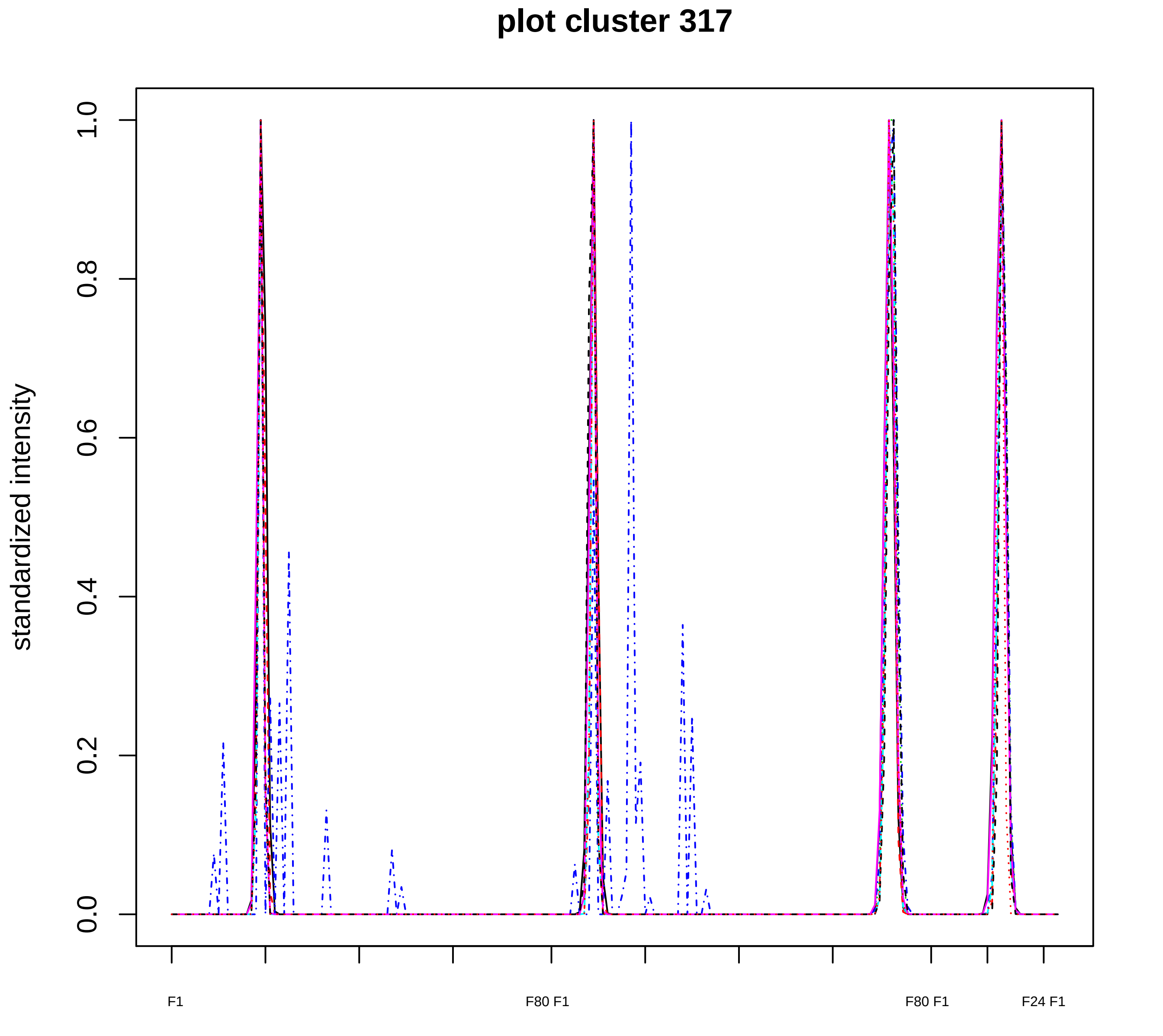
plot cluster 317
standardized intensity
0.0
0.2
0.4
0.6
0.8
1.0
F1
F80 F1
F80 F1
F24 F1

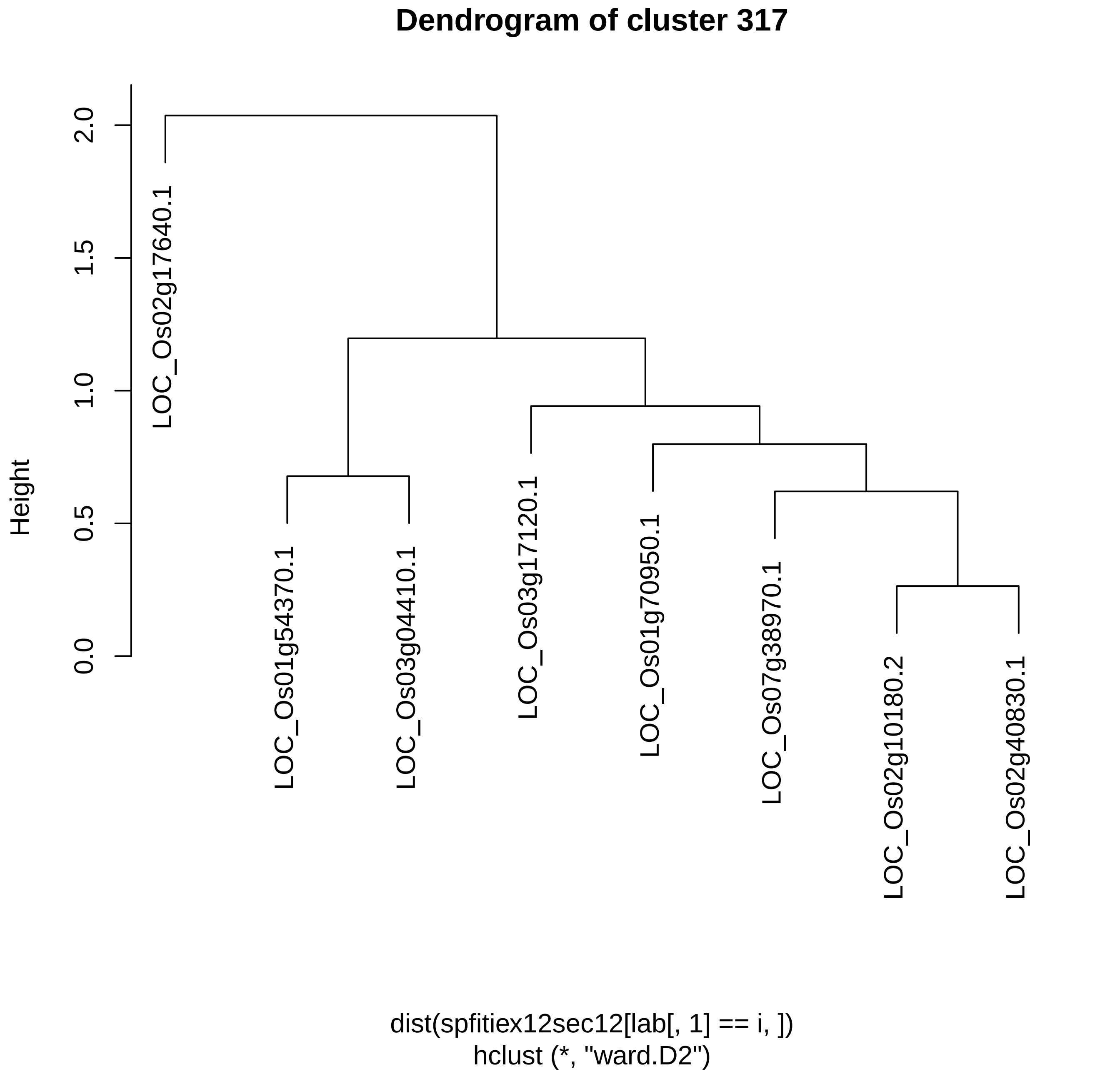

Dendrogram of cluster 317
Height
0.0
0.5
1.0
1.5
2.0
LOC_Os02g17640.1
LOC_Os01g54370.1
LOC_Os03g04410.1
LOC_Os03g17120.1
LOC_Os01g70950.1
LOC_Os07g38970.1
LOC_Os02g10180.2
LOC_Os02g40830.1
dist(spfitiex12sec12[lab[, 1] == i, ])
hclust (*, "ward.D2")

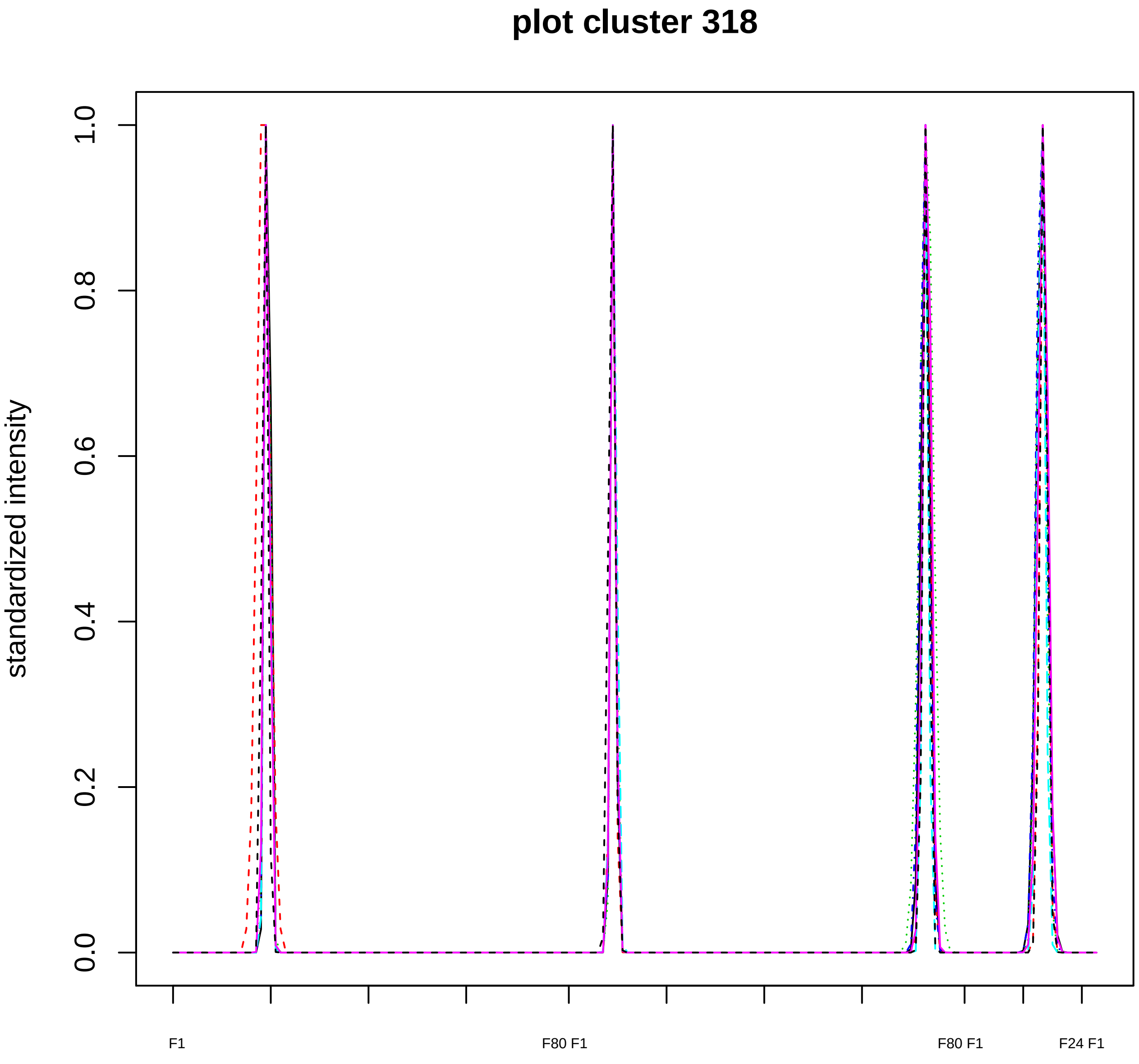
plot cluster 318
standardized intensity
0.0
0.2
0.4
0.6
0.8
1.0
F1
F80 F1
F80 F1
F24 F1

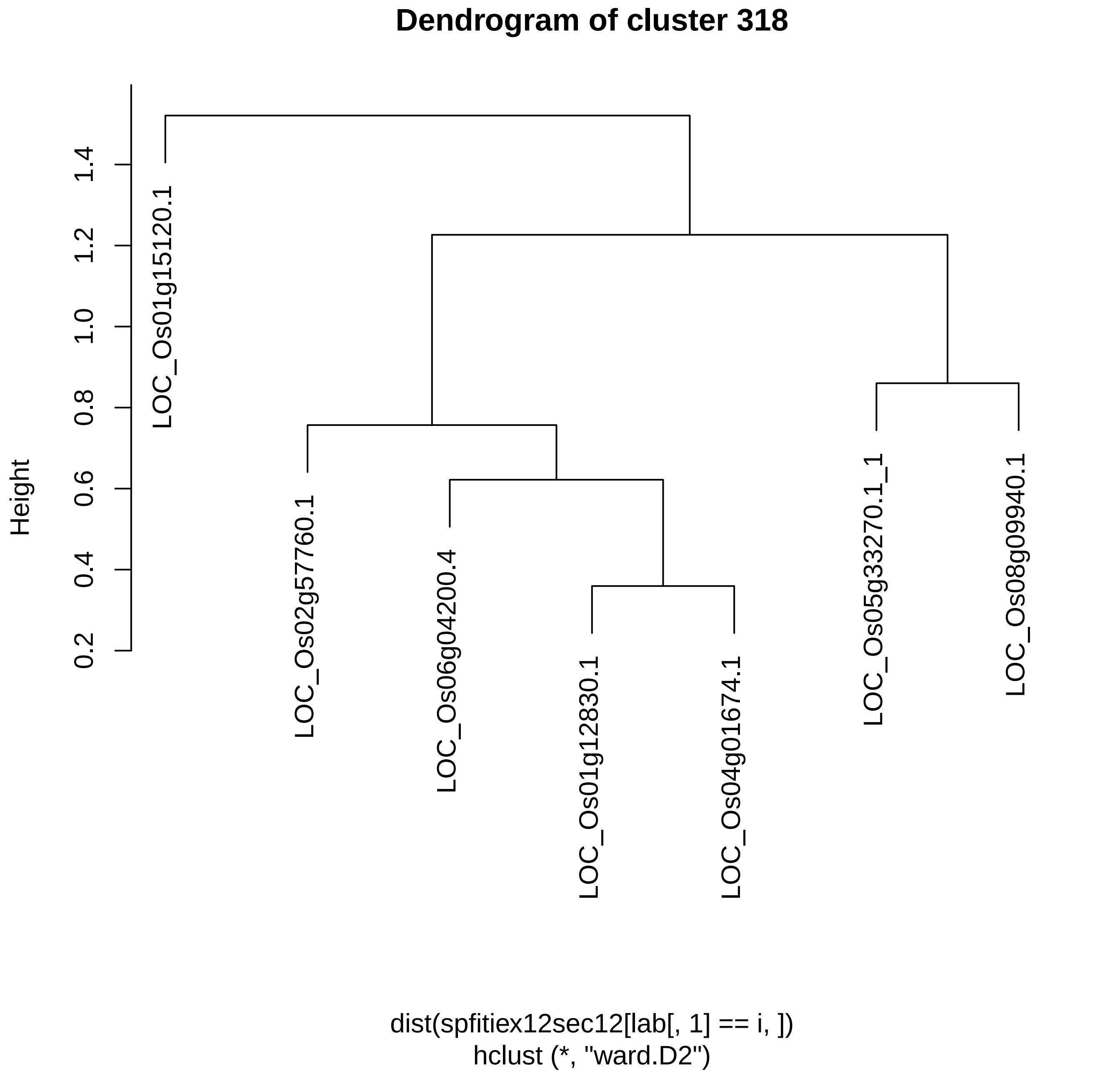

Dendrogram of cluster 318
Height
0.2
0.4
0.6
0.8
1.0
1.2
1.4
LOC_Os01g15120.1
LOC_Os02g57760.1
LOC_Os06g04200.4
LOC_Os01g12830.1
LOC_Os04g01674.1
LOC_Os05g33270.1_1
LOC_Os08g09940.1
dist(spfitiex12sec12[lab[, 1] == i, ])
hclust (*, "ward.D2")

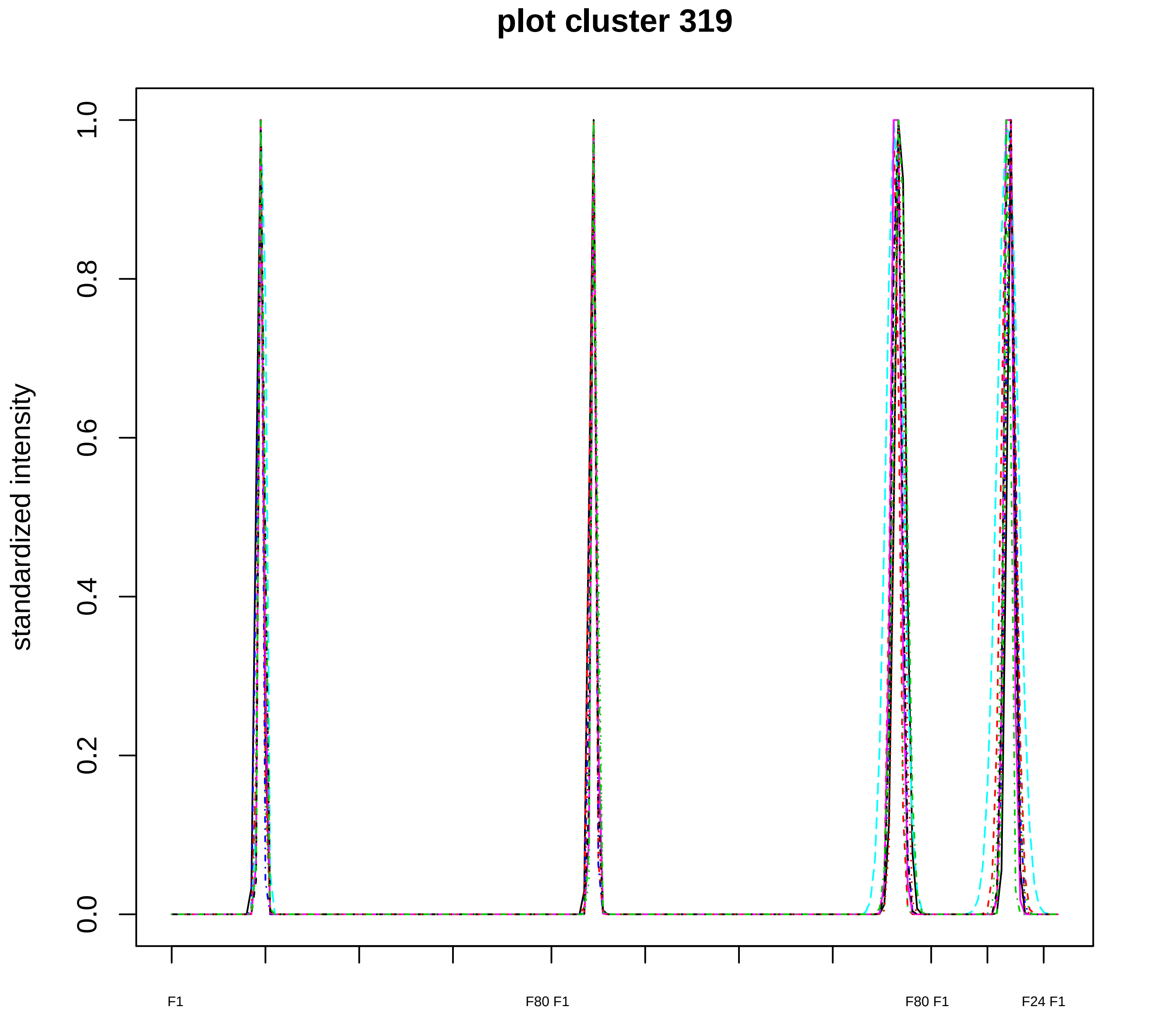

plot cluster 319
standardized intensity
0.0
0.2
0.4
0.6
0.8
1.0
F1
F80 F1
F80 F1
F24 F1

#### Dendrogram of cluster 319

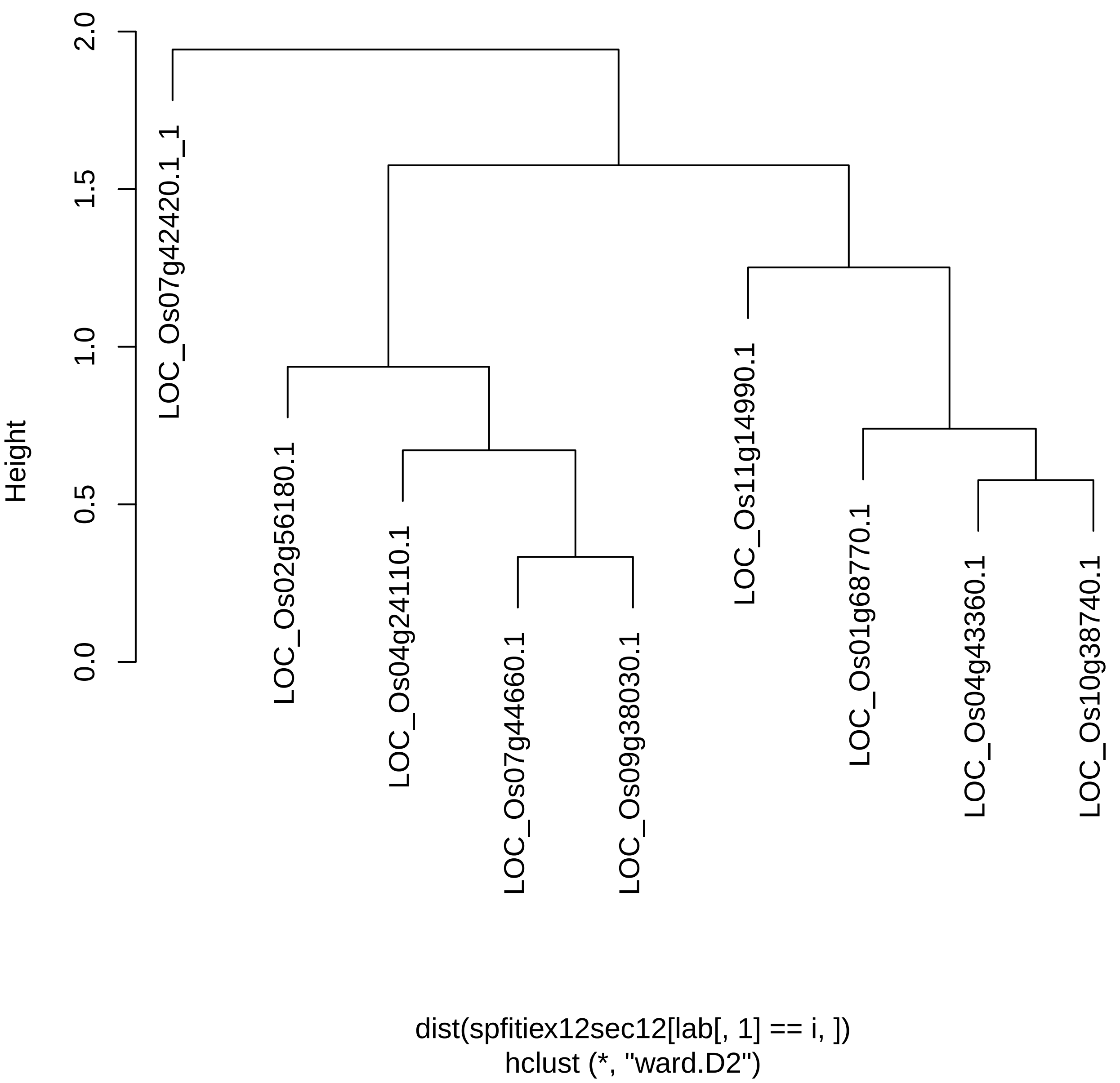

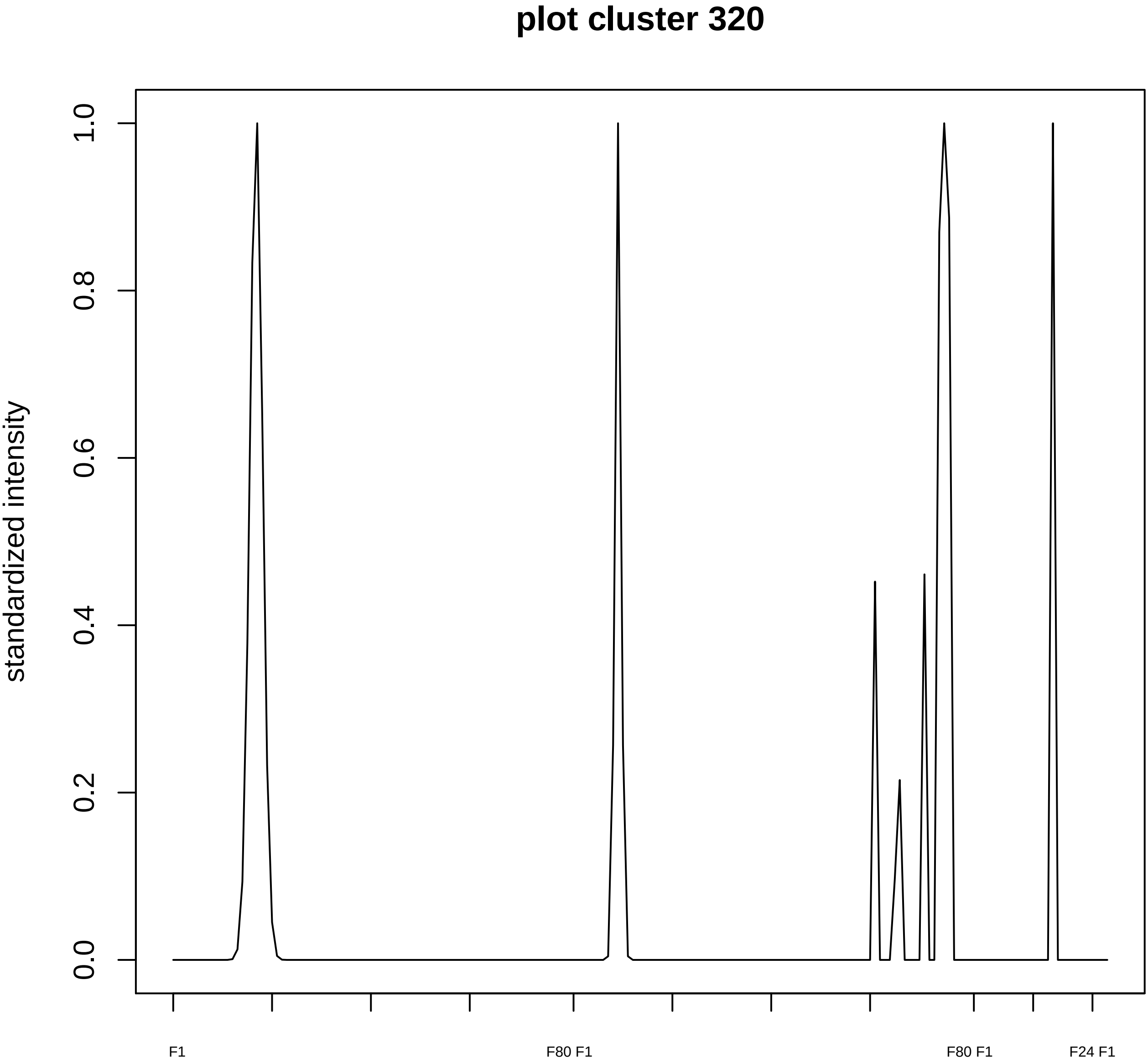
plot cluster 320
standardized intensity
0.0
0.2
0.4
0.6
0.8
1.0
F1
F80 F1
F80 F1
F24 F1

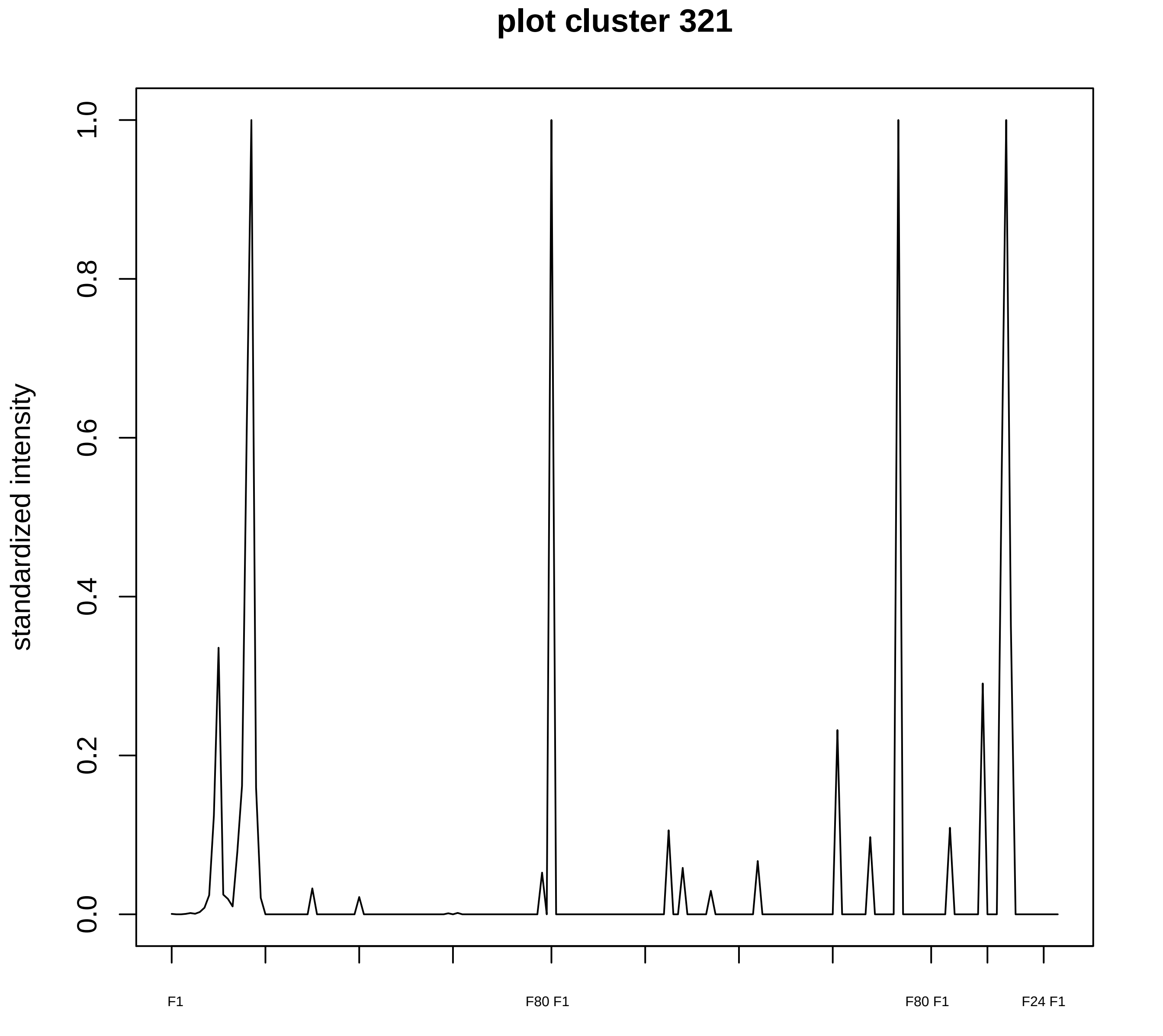
plot cluster 321
standardized intensity
1.0
0.8
0.6
0.4
0.2
0.0
F1
F80 F1
F80 F1
F24 F1

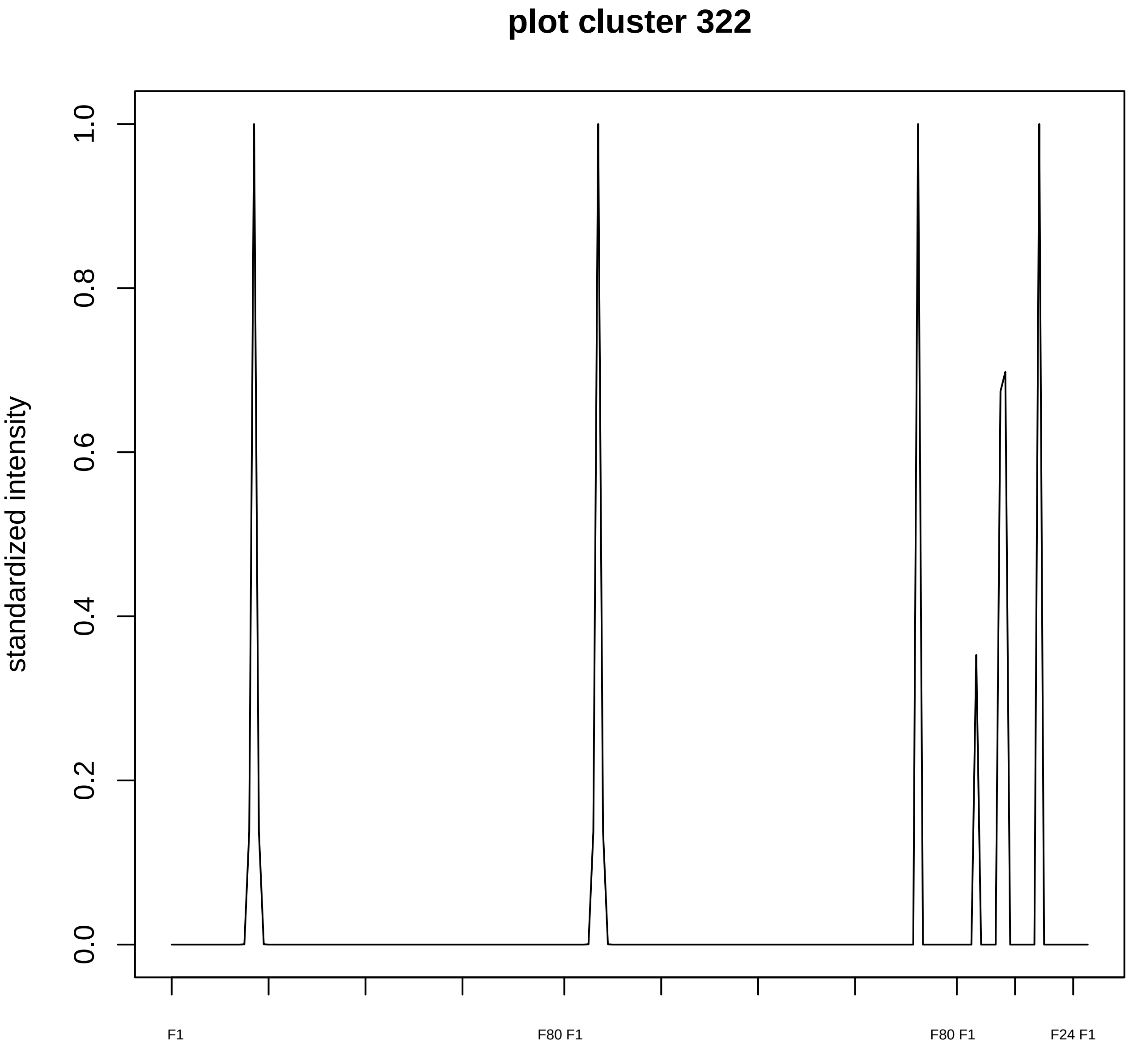

plot cluster 322
standardized intensity
0.0
0.2
0.4
0.6
0.8
1.0
F1
F80 F1
F80 F1
F24 F1

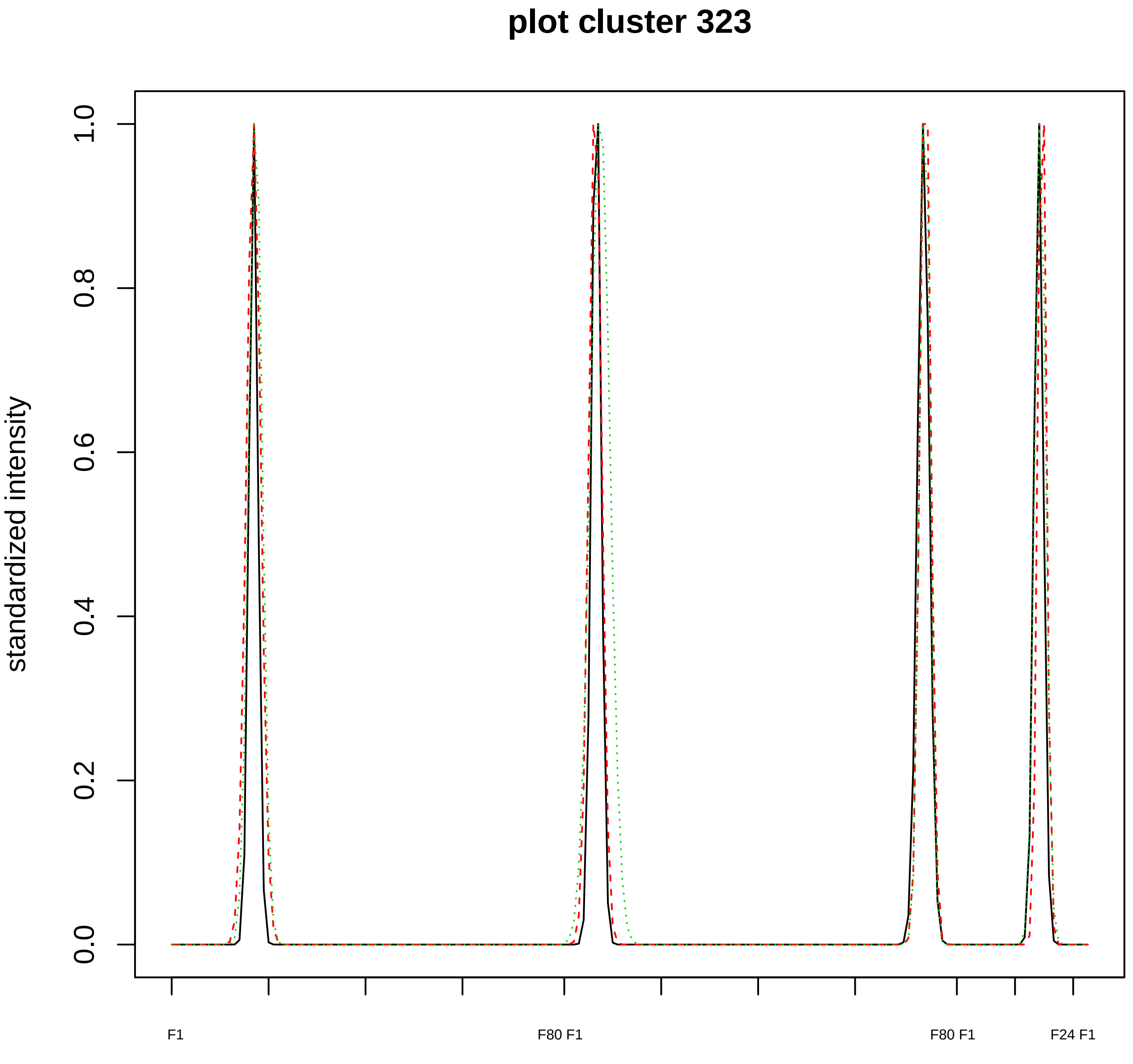
plot cluster 323
standardized intensity
1.0
0.8
0.6
0.4
0.2
0.0
F1
F80 F1
F80 F1
F24 F1

### Dendrogram of cluster 323

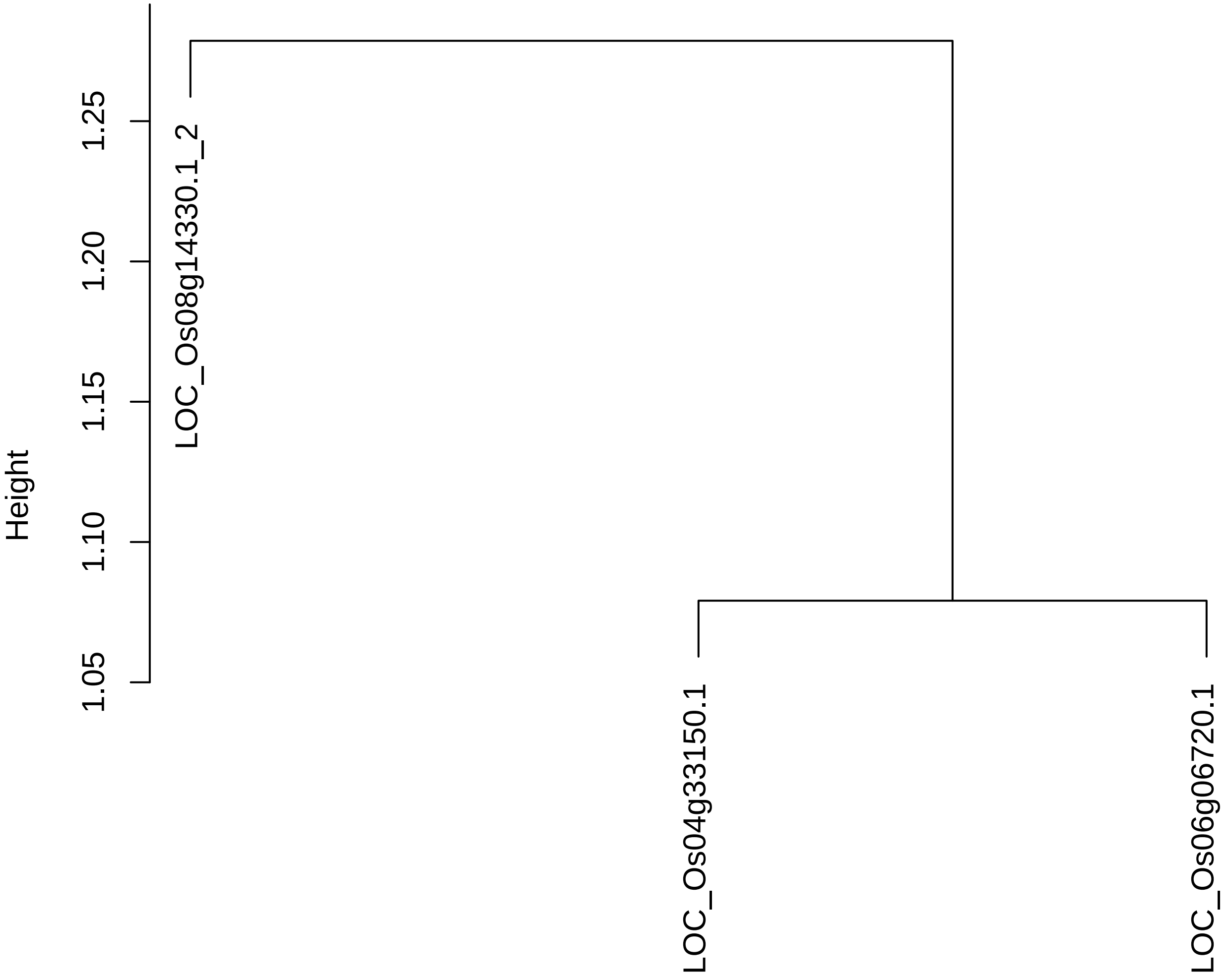


dist(spfitiex12sec12[lab[, 1] == i, ])
hclust (*, "ward.D2")

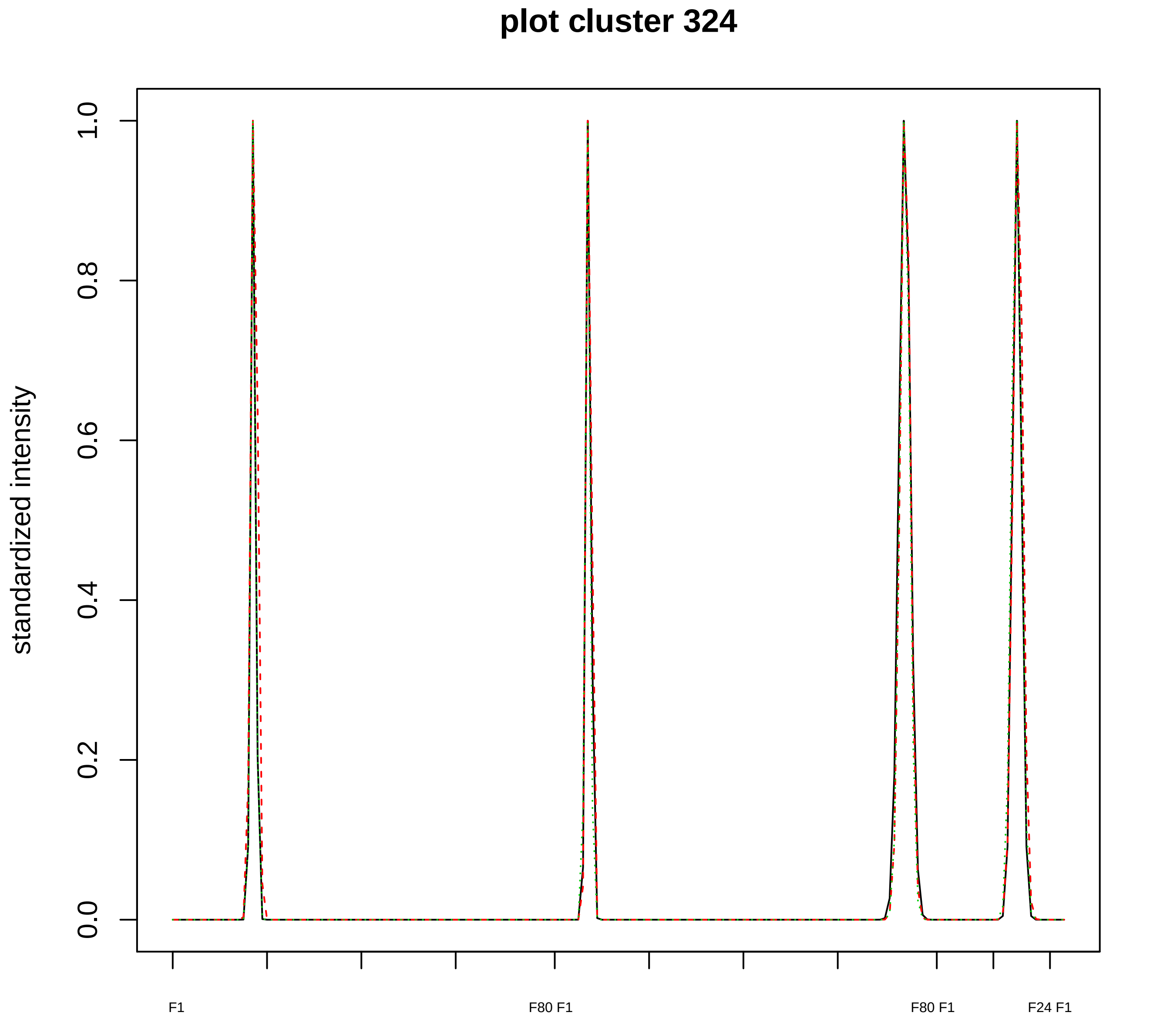
plot cluster 324
standardized intensity
0.0
0.2
0.4
0.6
0.8
1.0
F1
F80 F1
F80 F1
F24 F1

#### Dendrogram of cluster 324

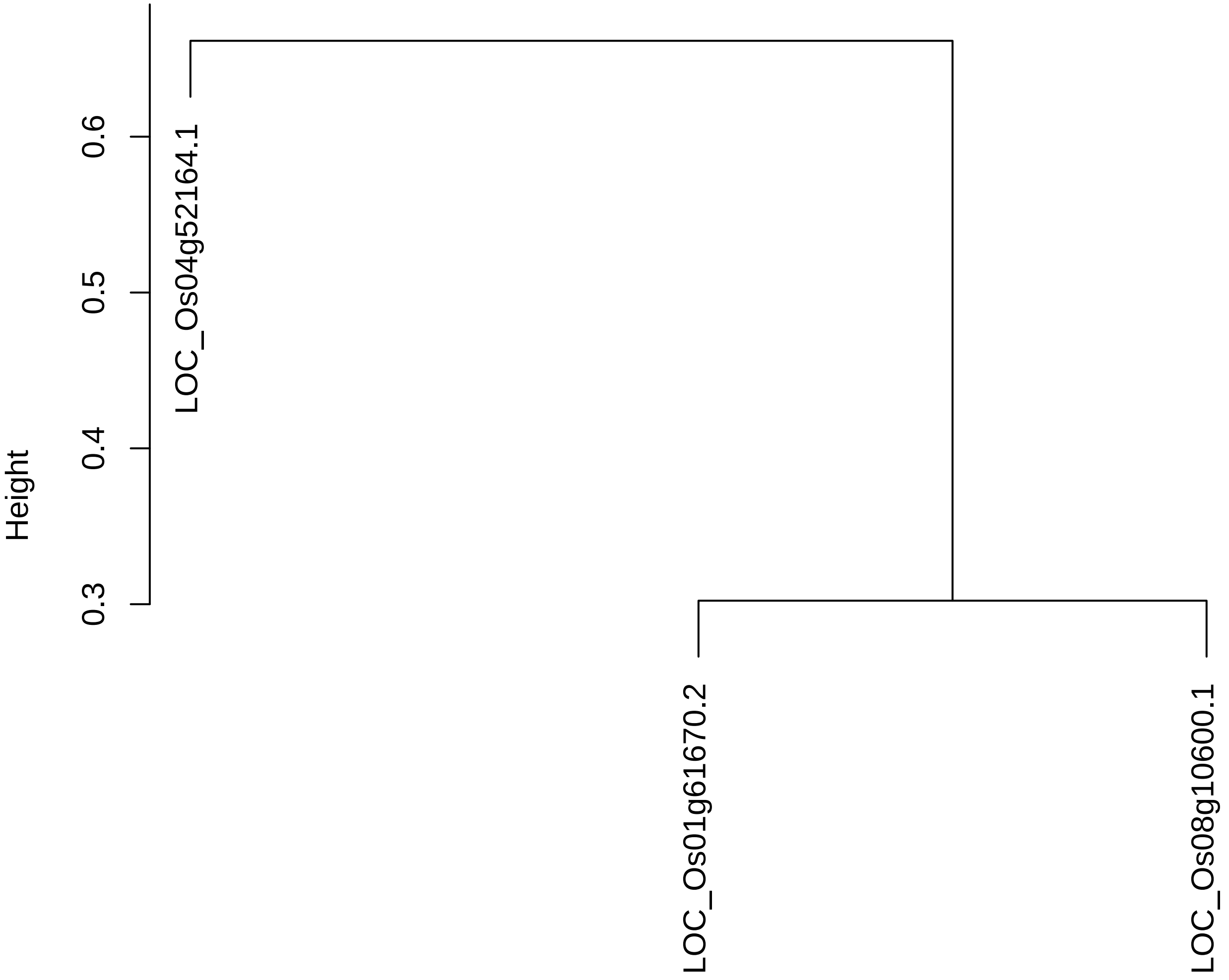


dist(spfitiex12sec12[lab[, 1] == i, ])
hclust (*, "ward.D2")

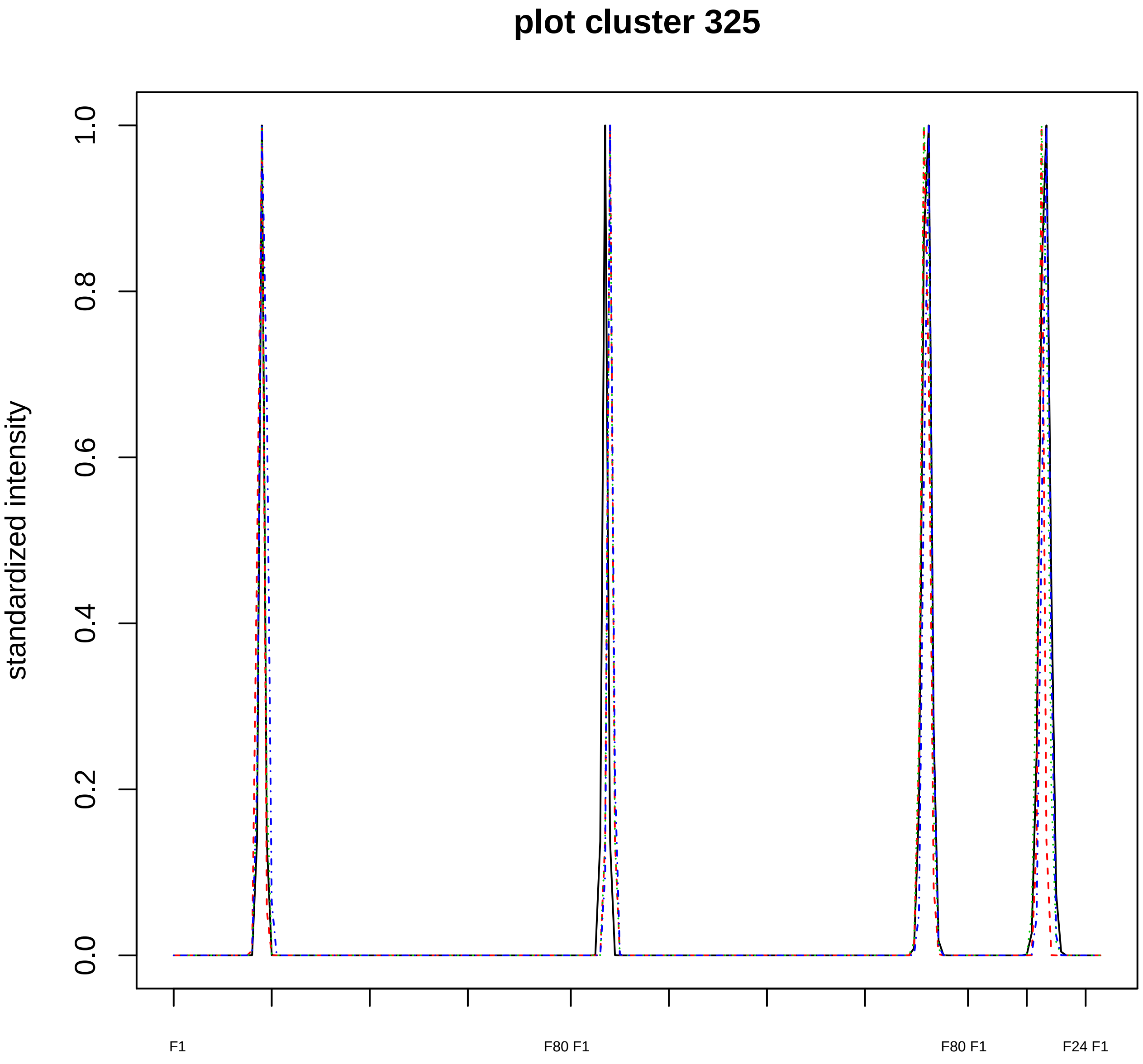
plot cluster 325
standardized intensity
0.0
0.2
0.4
0.6
0.8
1.0
F1
F80 F1
F80 F1
F24 F1

### Dendrogram of cluster 325

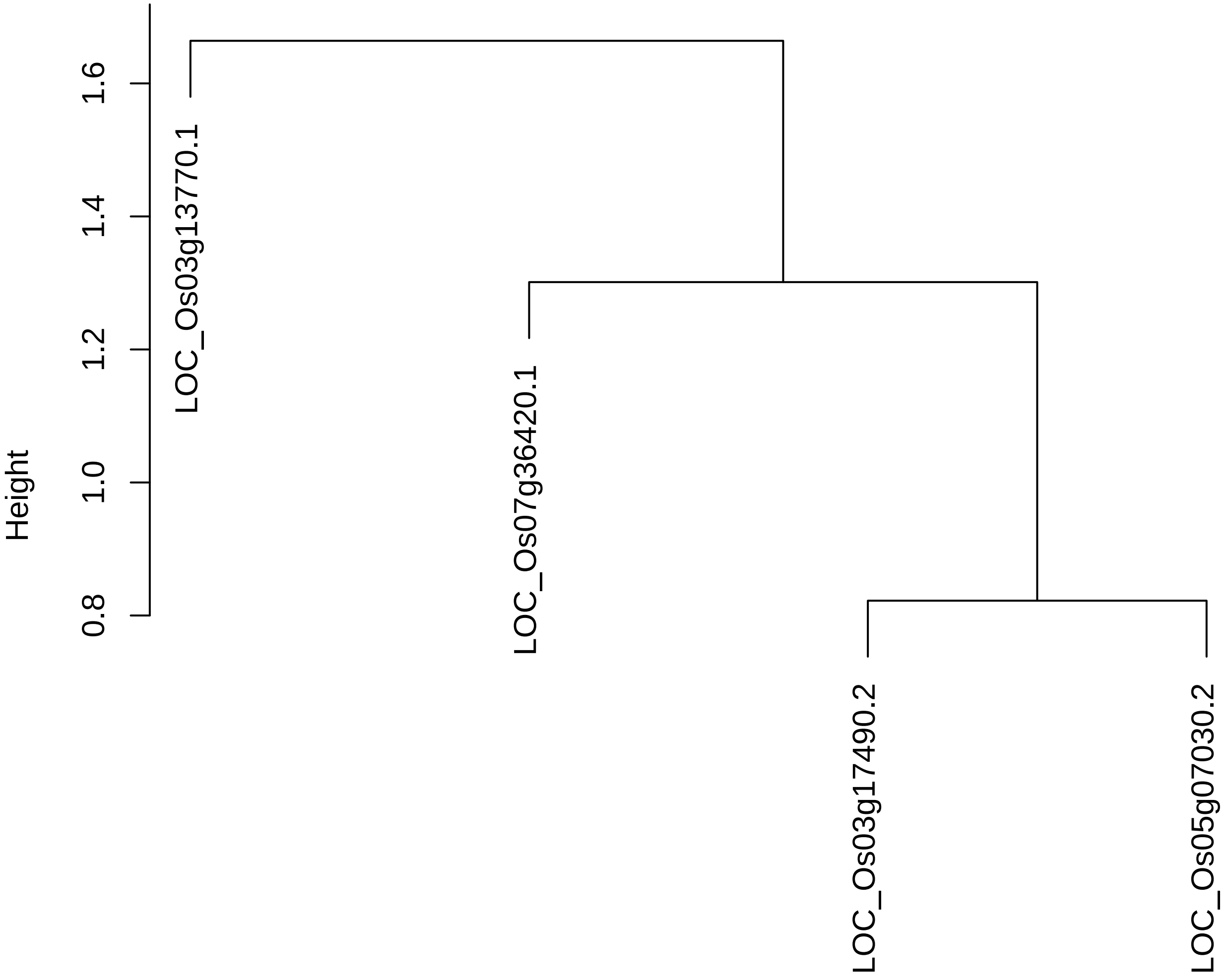


dist(spfitiex12sec12[lab[, 1] == i, ])
hclust (*, "ward.D2")

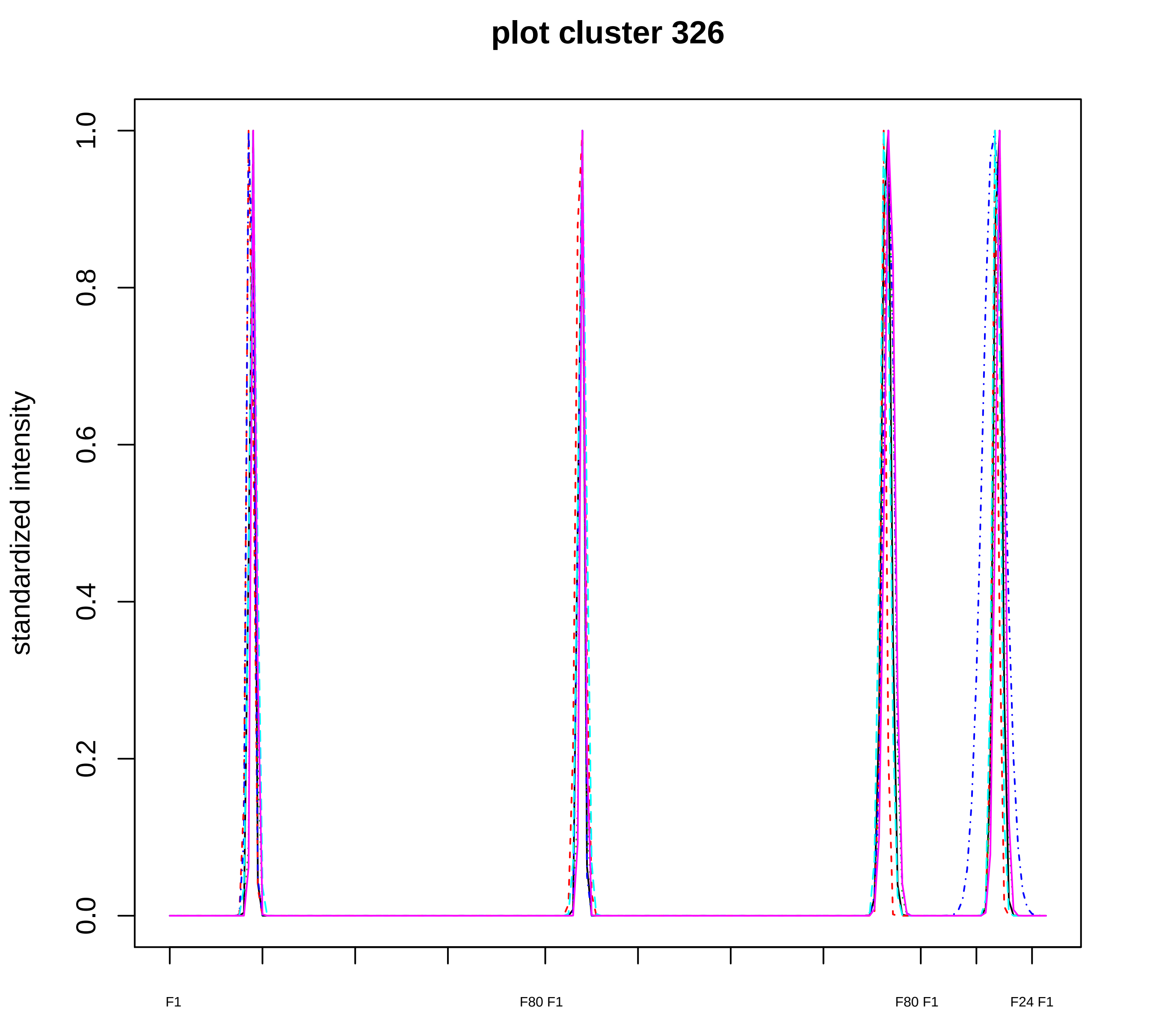

plot cluster 326
standardized intensity
0.0
0.2
0.4
0.6
0.8
1.0
F1
F80 F1
F80 F1
F24 F1

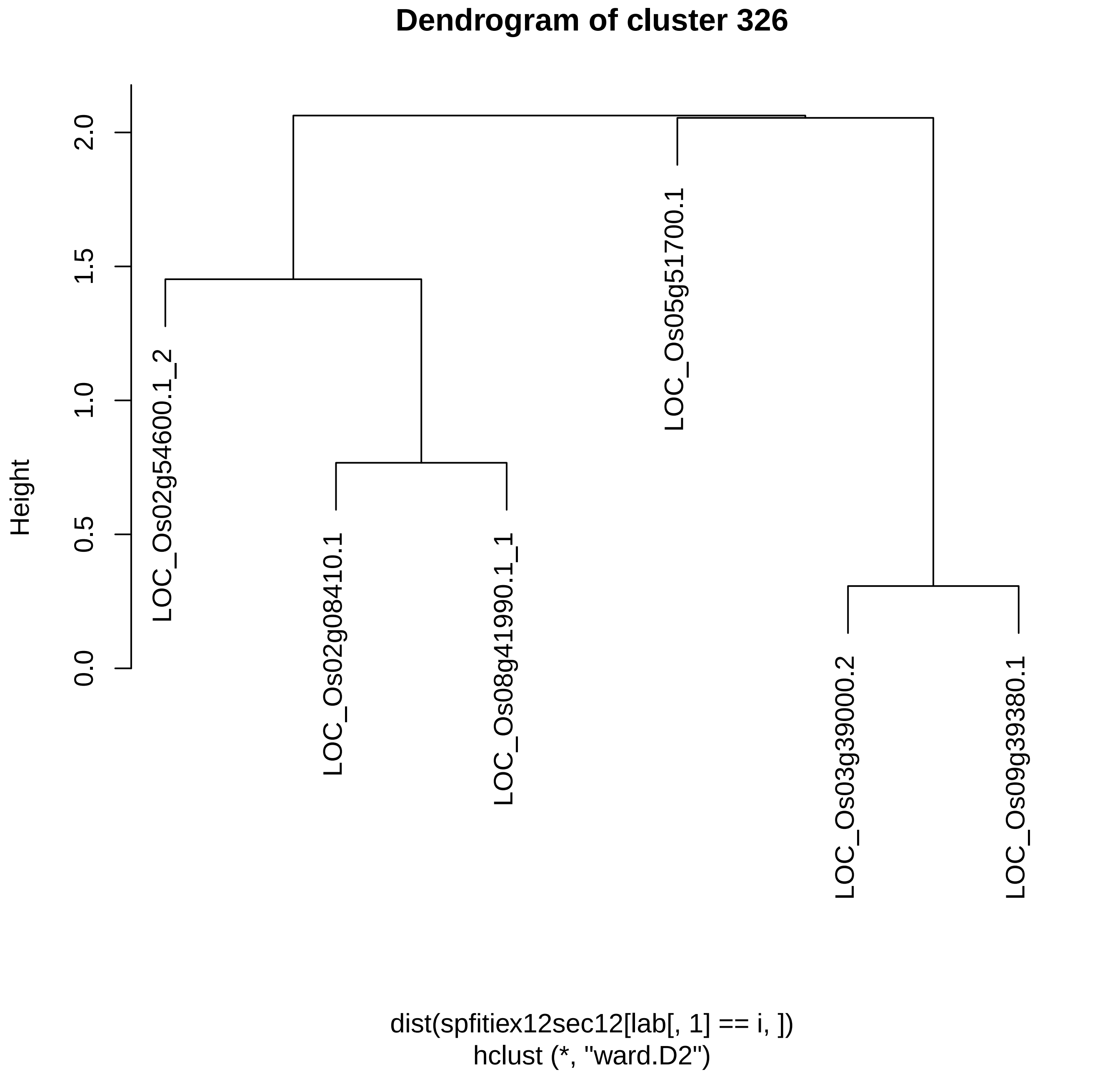

Dendrogram of cluster 326
Height
0.0
0.5
1.0
1.5
2.0
LOC_Os02g54600.1_2
LOC_Os02g08410.1
LOC_Os08g41990.1_1
LOC_Os05g51700.1
LOC_Os03g39000.2
LOC_Os09g39380.1
dist(spfitiex12sec12[lab[, 1] == i, ])
hclust (*, "ward.D2")

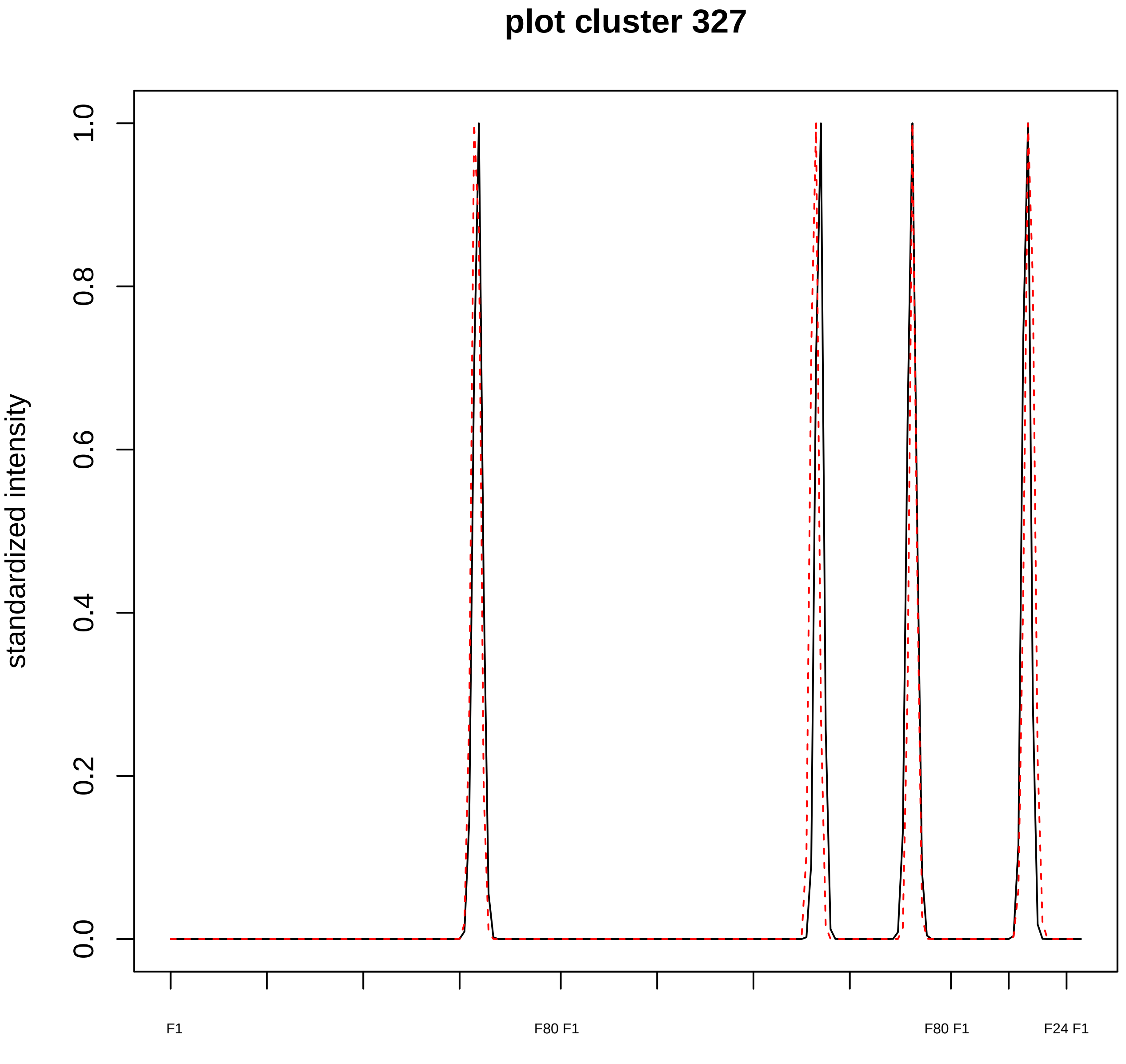
plot cluster 327
standardized intensity
0.0
0.2
0.4
0.6
0.8
1.0
F1
F80 F1
F80 F1
F24 F1

### plot cluster 328

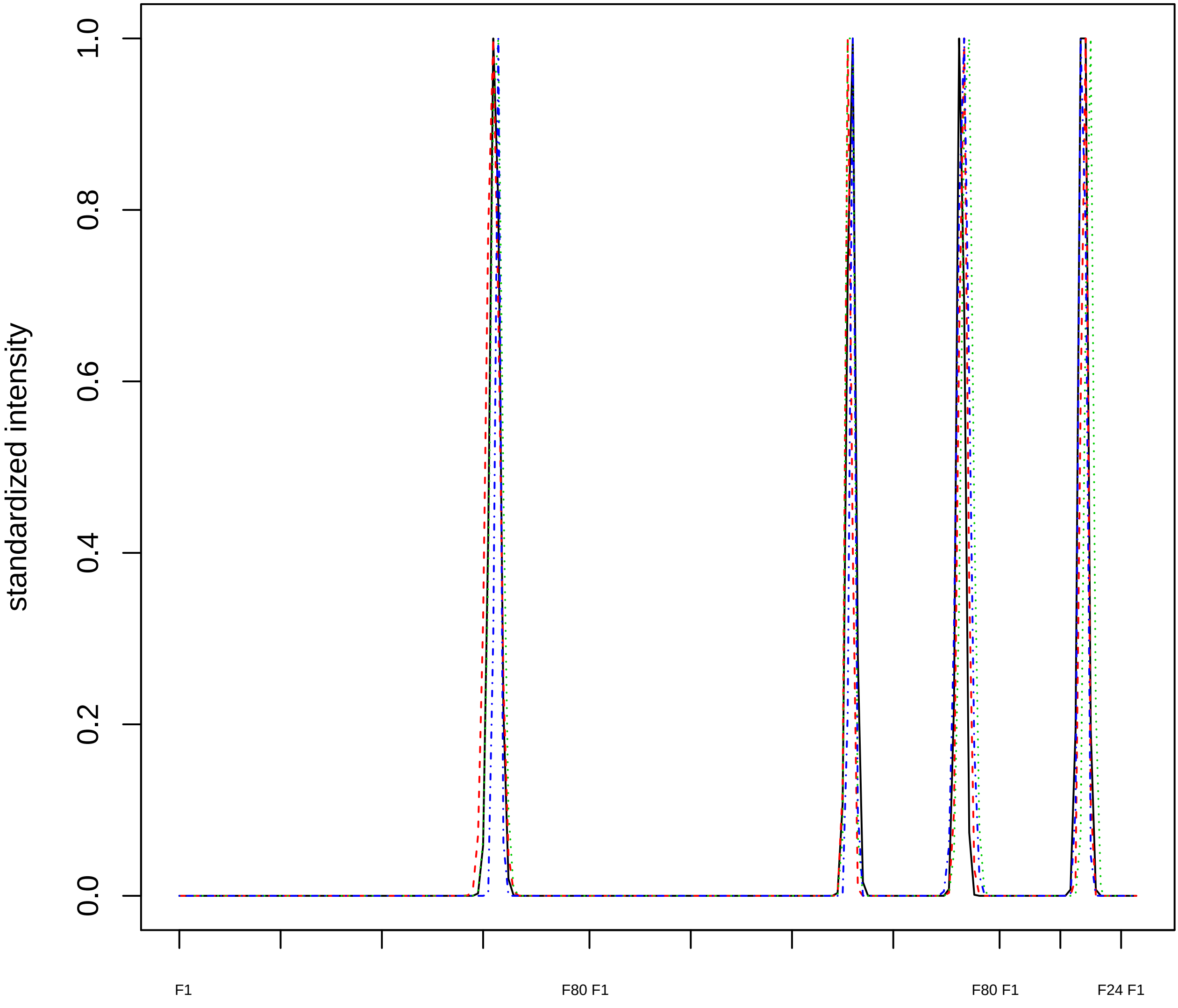

#### Dendrogram of cluster 328

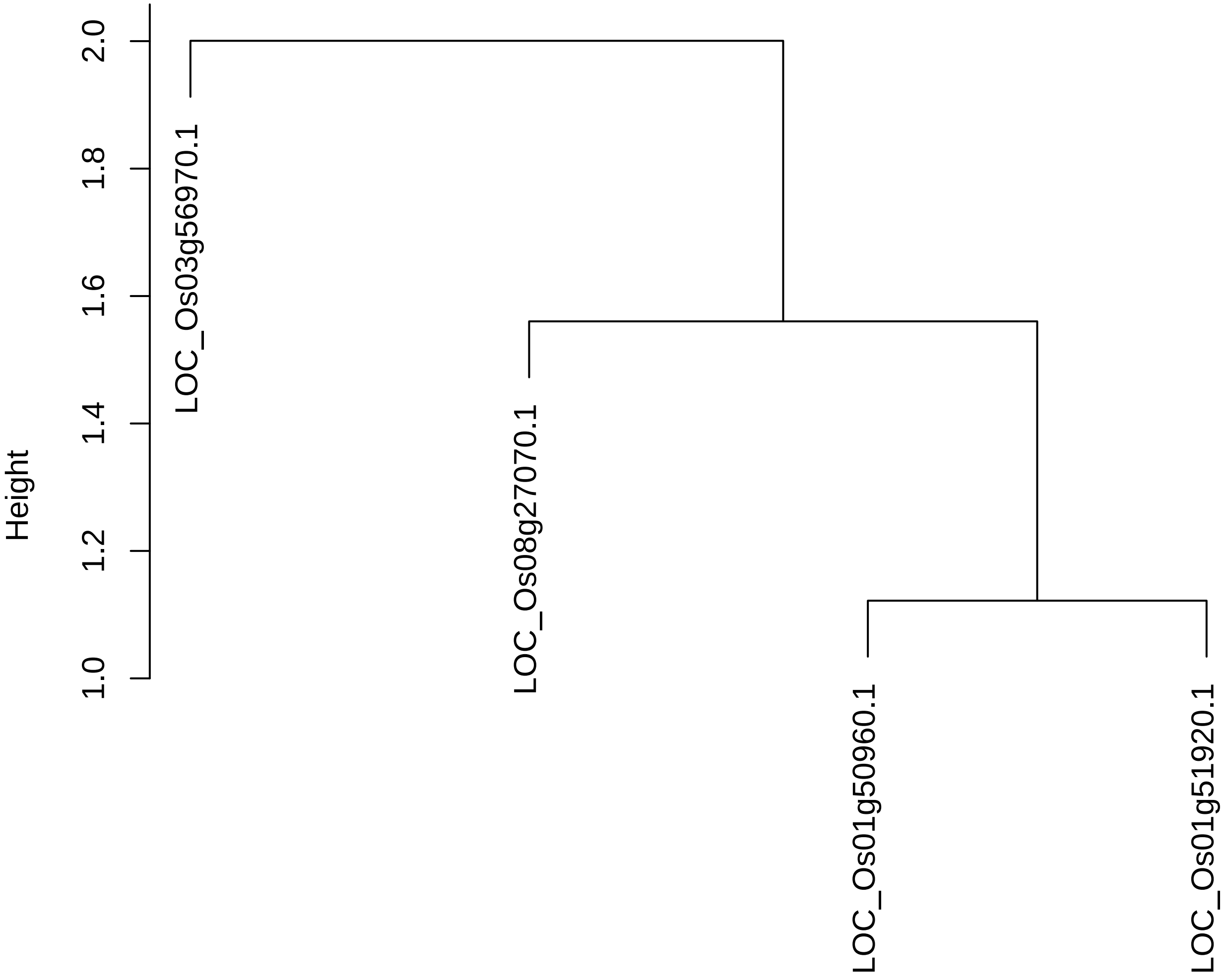


dist(spfitiex12sec12[lab[, 1] == i, ])
hclust (*, "ward.D2")

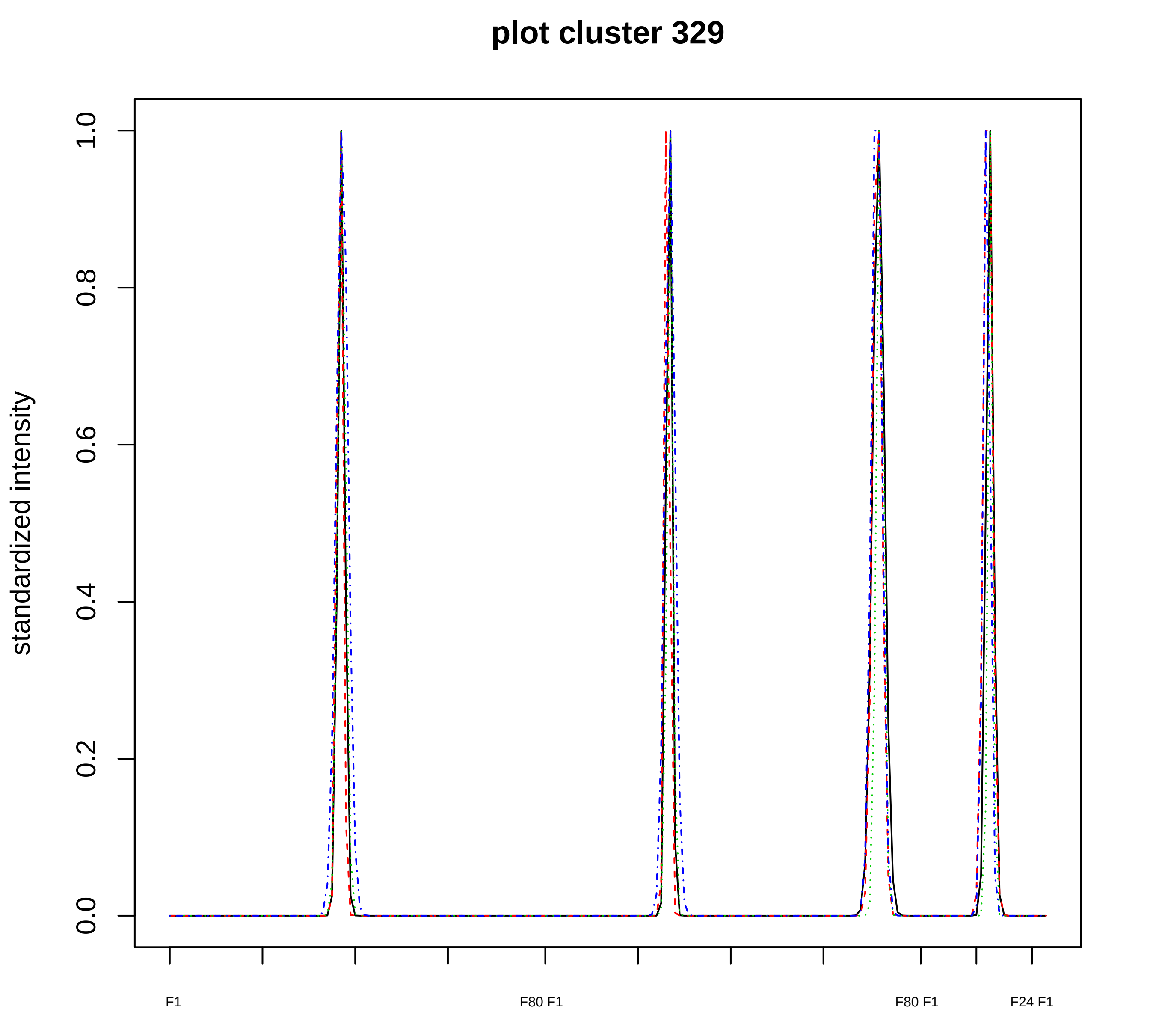
plot cluster 329
standardized intensity
0.0
0.2
0.4
0.6
0.8
1.0
F1
F80 F1
F80 F1
F24 F1

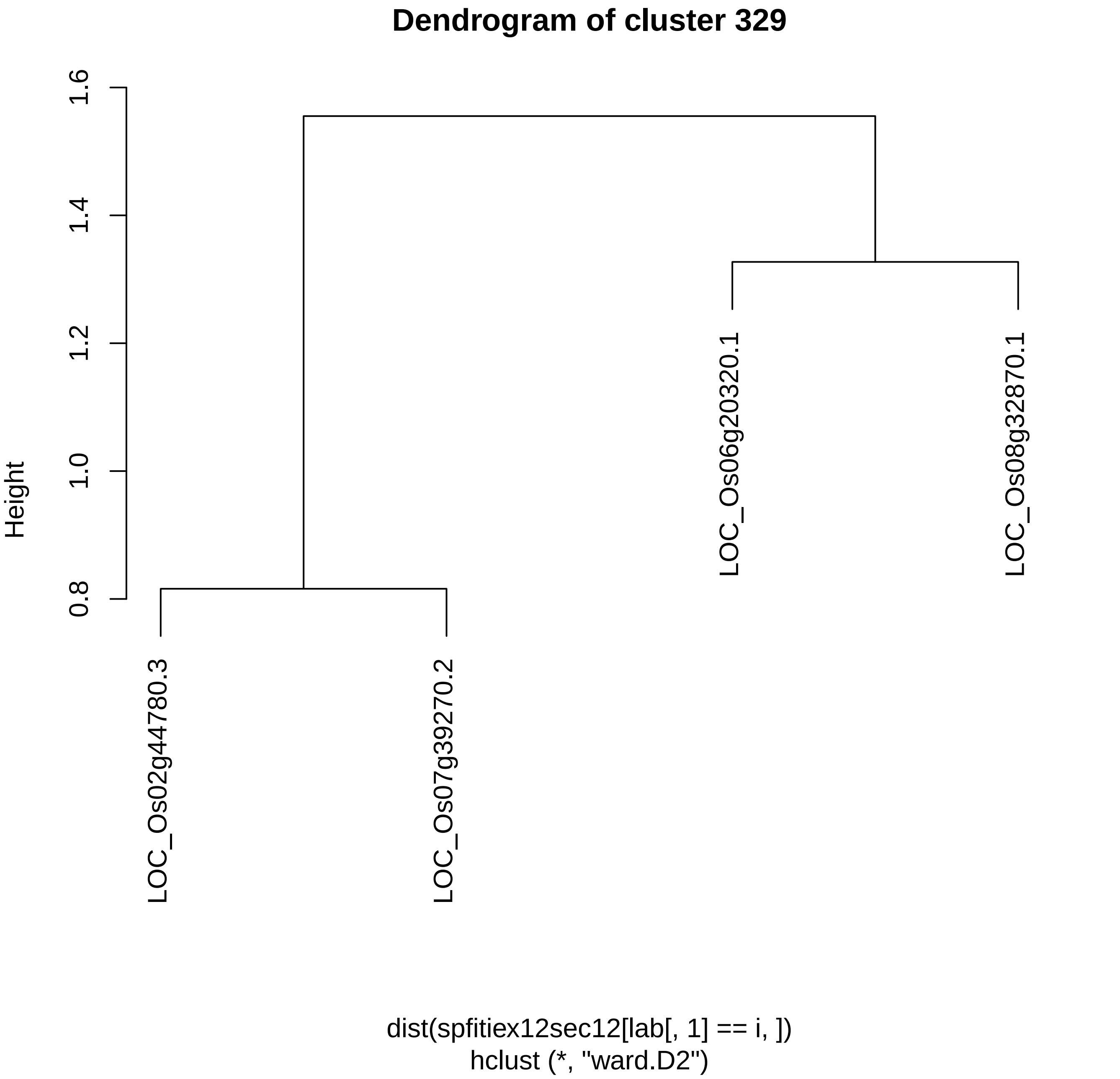
Dendrogram of cluster 329
Height
1.6
1.4
1.2
1.0
0.8
LOC_Os02g44780.3
LOC_Os07g39270.2
LOC_Os06g20320.1
LOC_Os08g32870.1
dist(spfitiex12sec12[lab[, 1] == i, ])
hclust (*, "ward.D2")

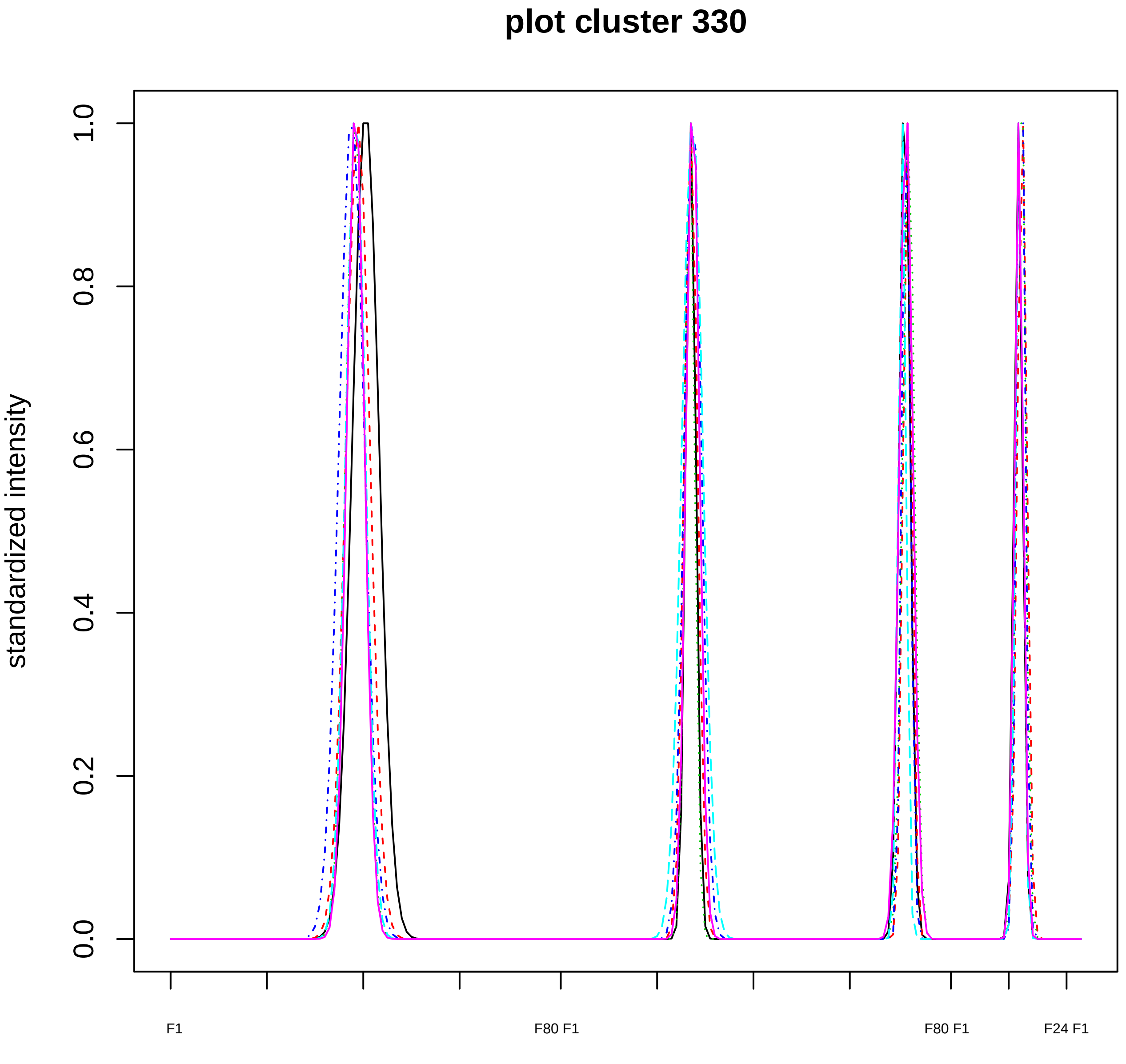
plot cluster 330
standardized intensity
0.0
0.2
0.4
0.6
0.8
1.0
F1
F80 F1
F80 F1
F24 F1

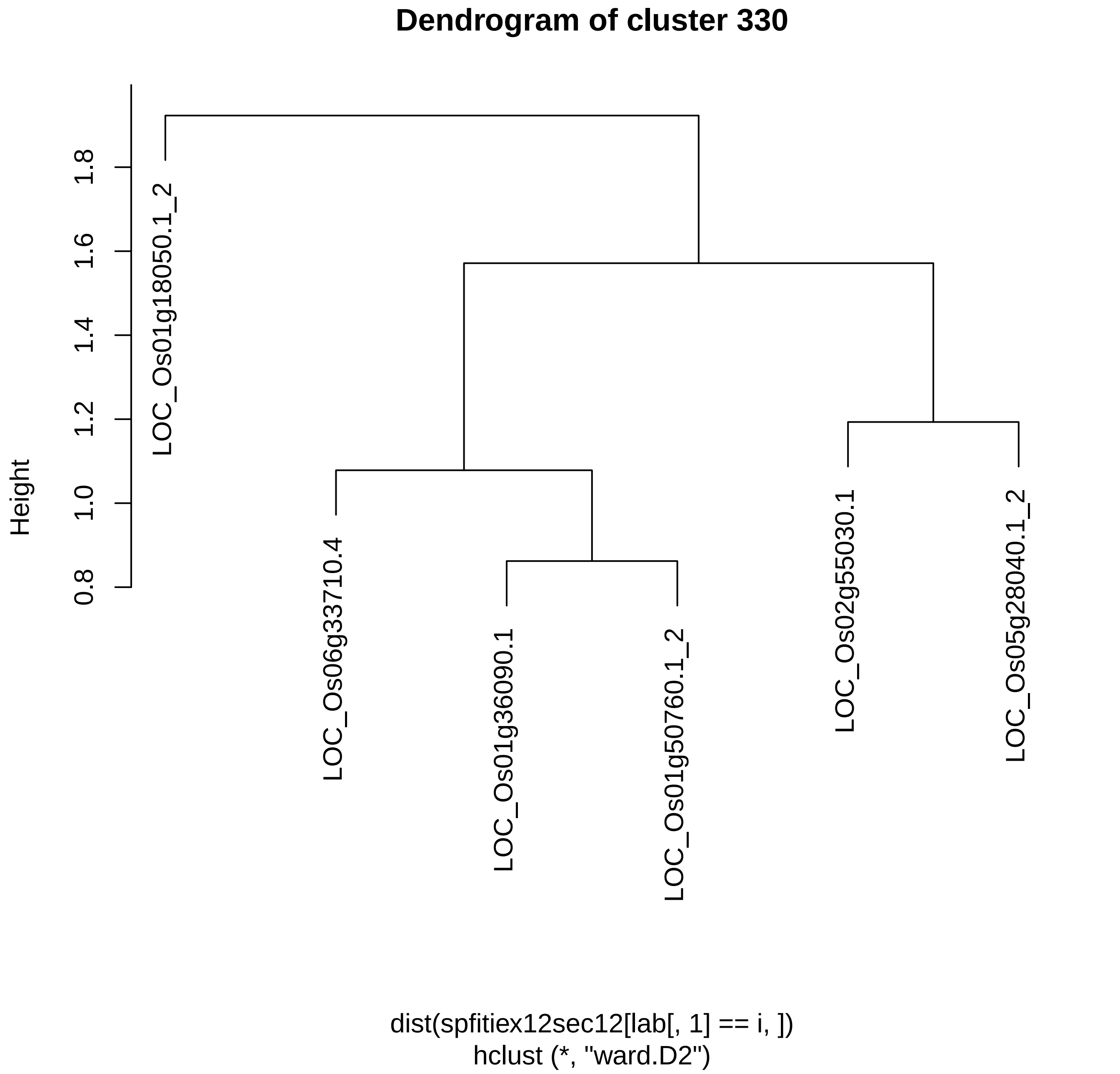
Dendrogram of cluster 330
Height
0.8
1.0
1.2
1.4
1.6
1.8
LOC_Os01g18050.1_2
LOC_Os06g33710.4
LOC_Os01g36090.1
LOC_Os01g50760.1_2
LOC_Os02g55030.1
LOC_Os05g28040.1_2
dist(spfitiex12sec12[lab[, 1] == i, ])
hclust (*, "ward.D2")

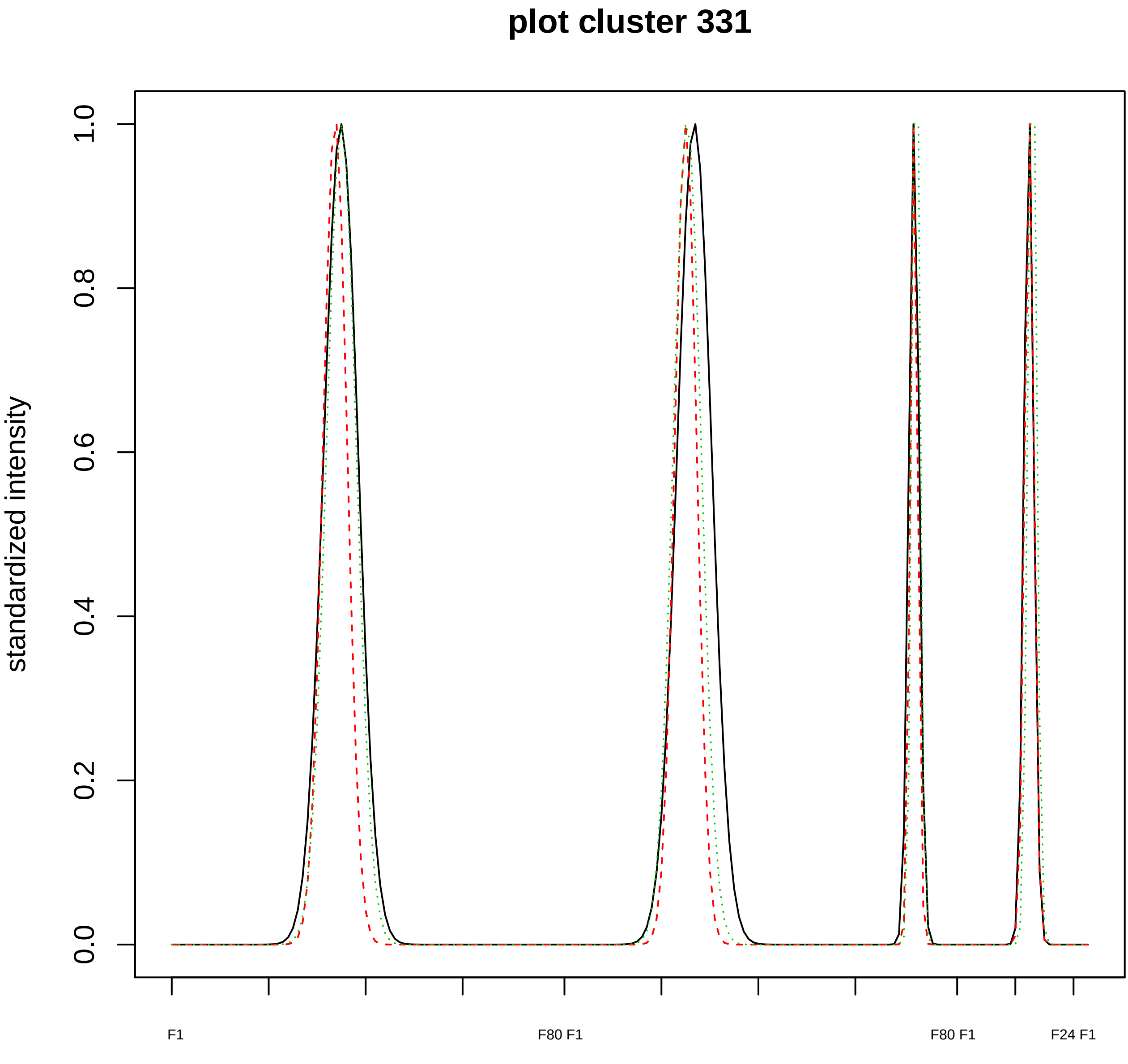
plot cluster 331
standardized intensity
0.0
0.2
0.4
0.6
0.8
1.0
F1
F80 F1
F80 F1
F24 F1

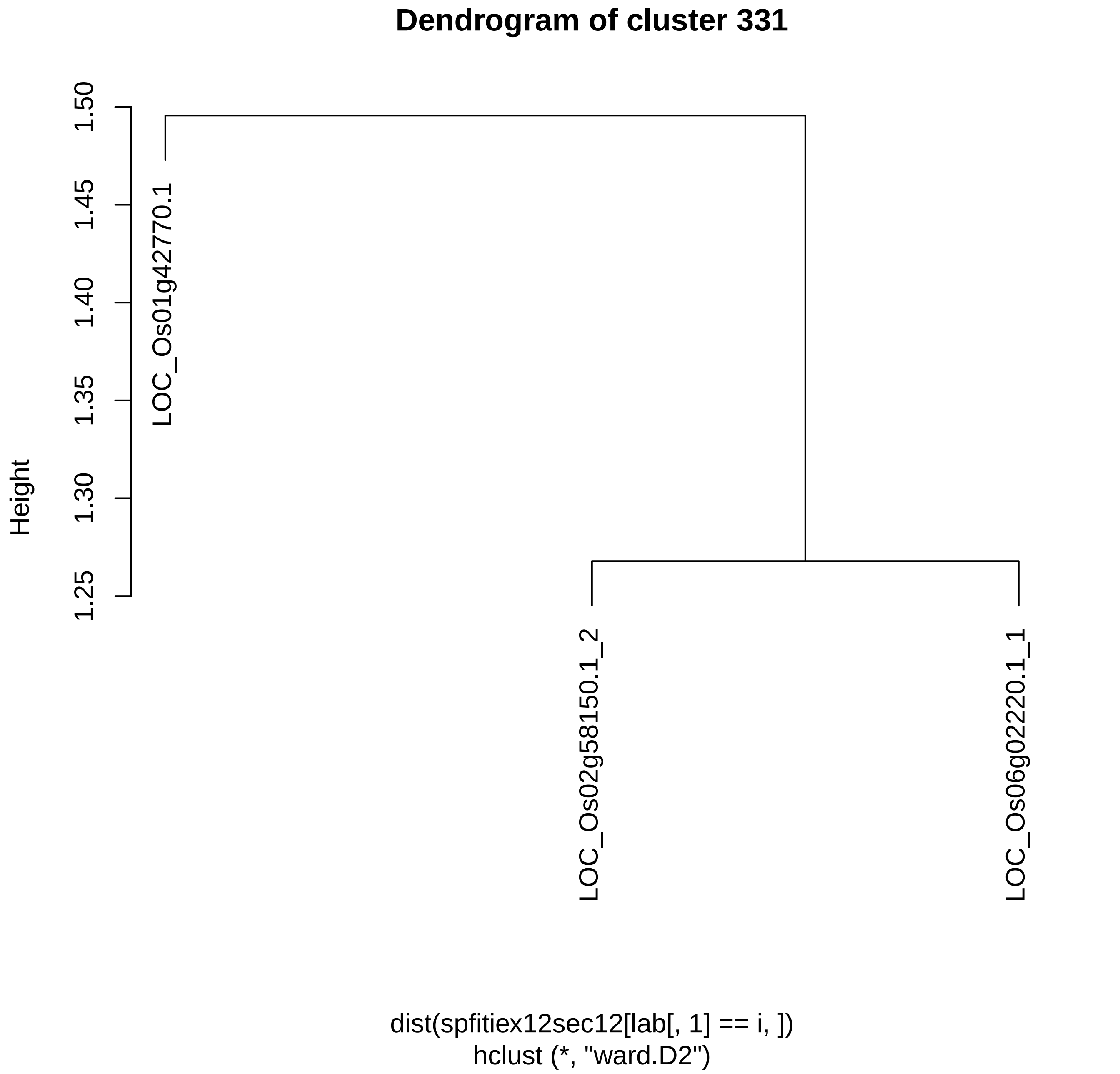
Dendrogram of cluster 331
Height
1.25
1.30
1.35
1.40
1.45
1.50
LOC_Os01g42770.1
LOC_Os02g58150.1_2
LOC_Os06g02220.1_1
dist(spfitiex12sec12[lab[, 1] == i, ])
hclust (*, "ward.D2")

#### plot cluster 332

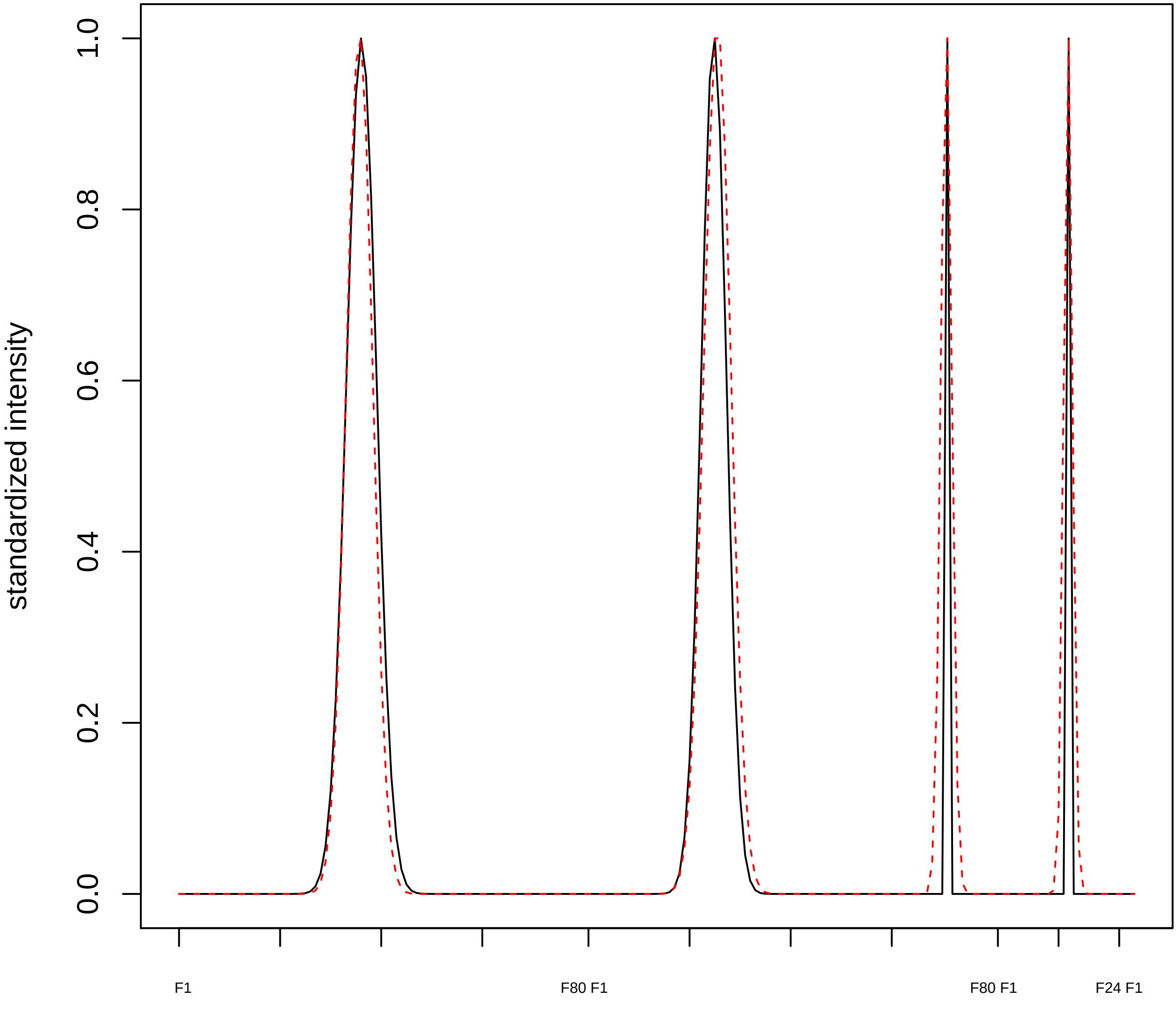

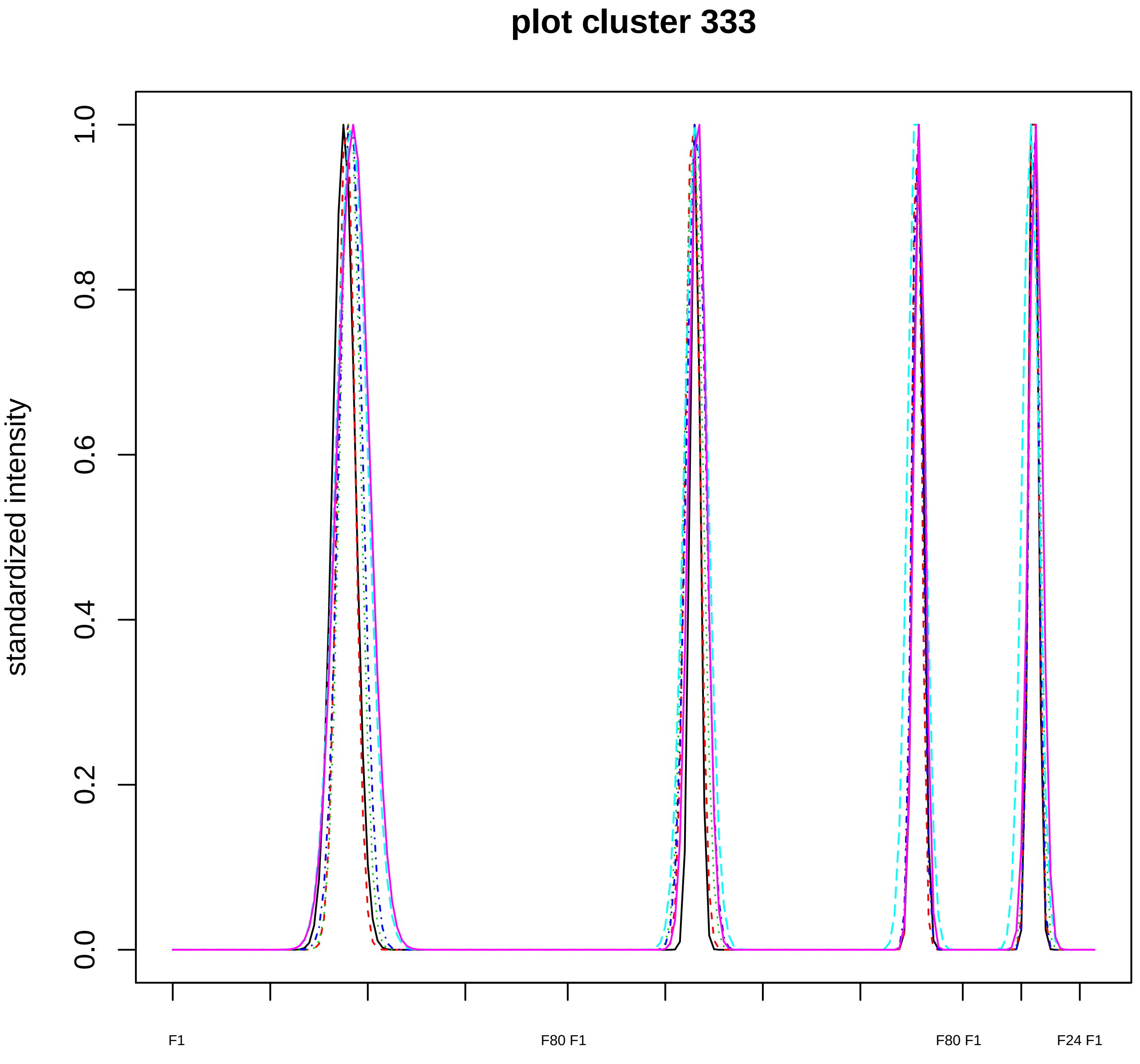
plot cluster 333
standardized intensity
0.0
0.2
0.4
0.6
0.8
1.0
F1
F80 F1
F80 F1
F24 F1

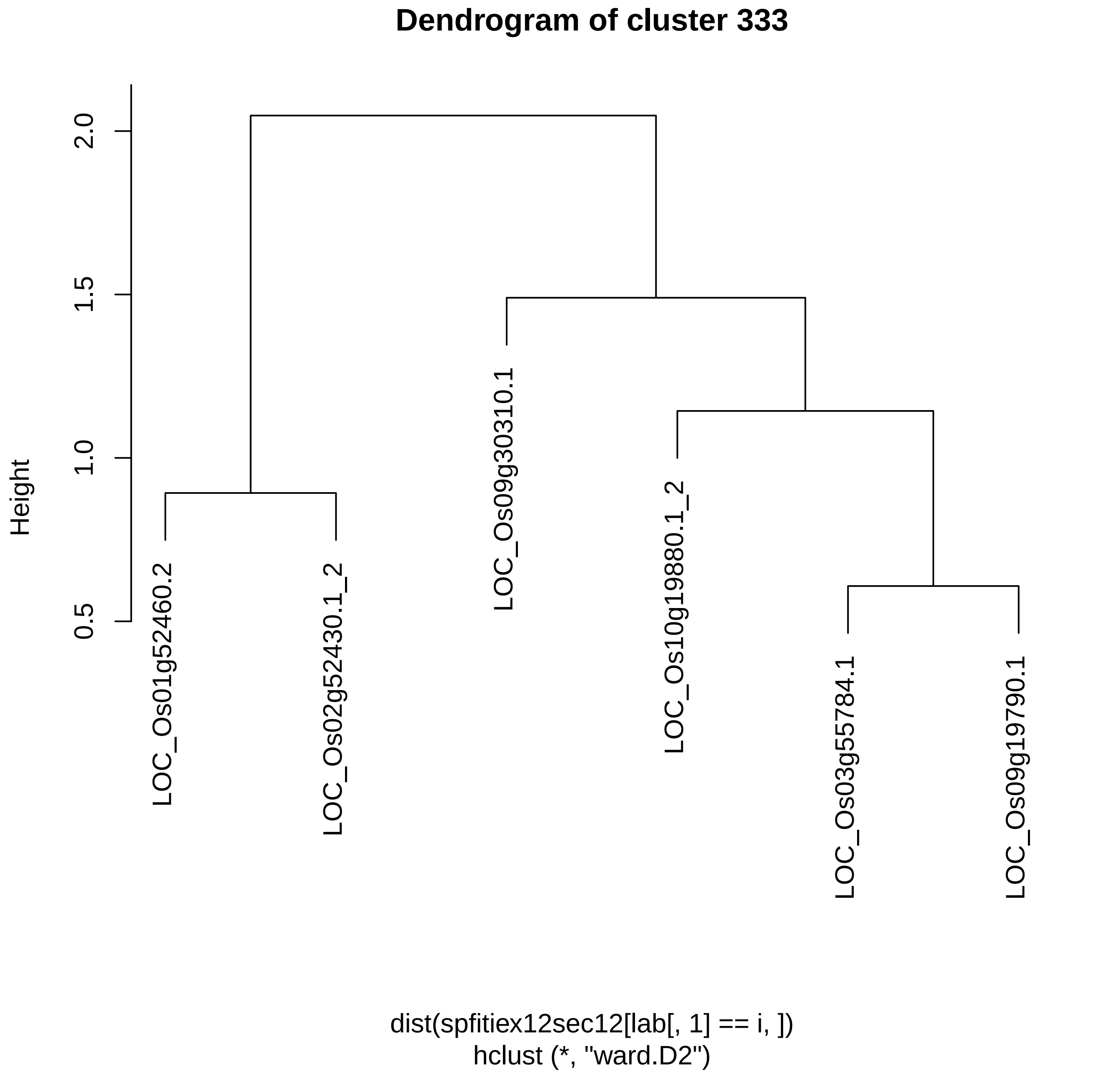
Dendrogram of cluster 333
Height
2.0
1.5
1.0
0.5
LOC_Os01g52460.2
LOC_Os02g52430.1_2
LOC_Os09g30310.1
LOC_Os10g19880.1_2
LOC_Os03g55784.1
LOC_Os09g19790.1
dist(spfitiex12sec12[lab[, 1] == i, ])
hclust (*, "ward.D2")

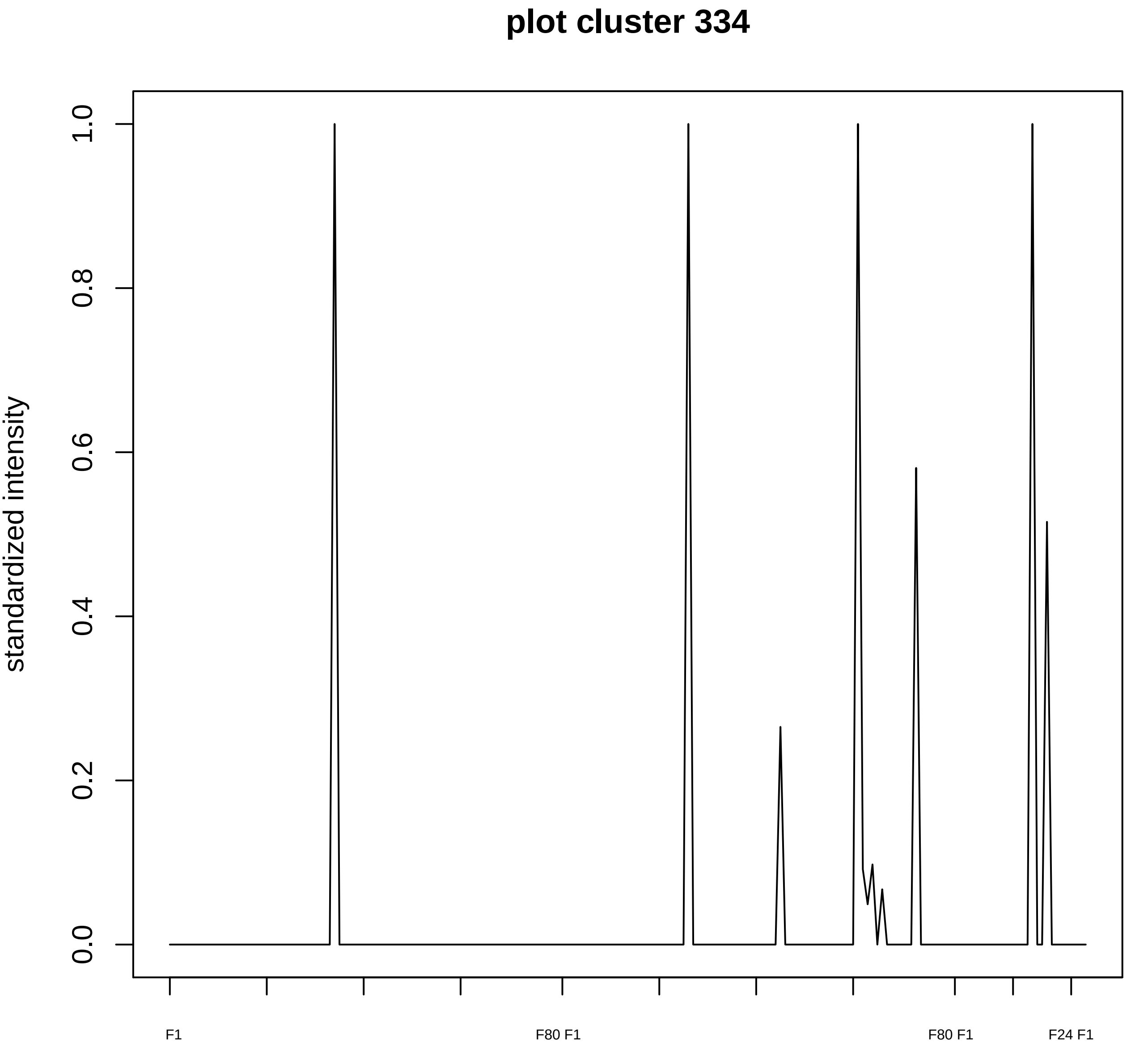
plot cluster 334
standardized intensity
0.0
0.2
0.4
0.6
0.8
1.0
F1
F80 F1
F80 F1
F24 F1

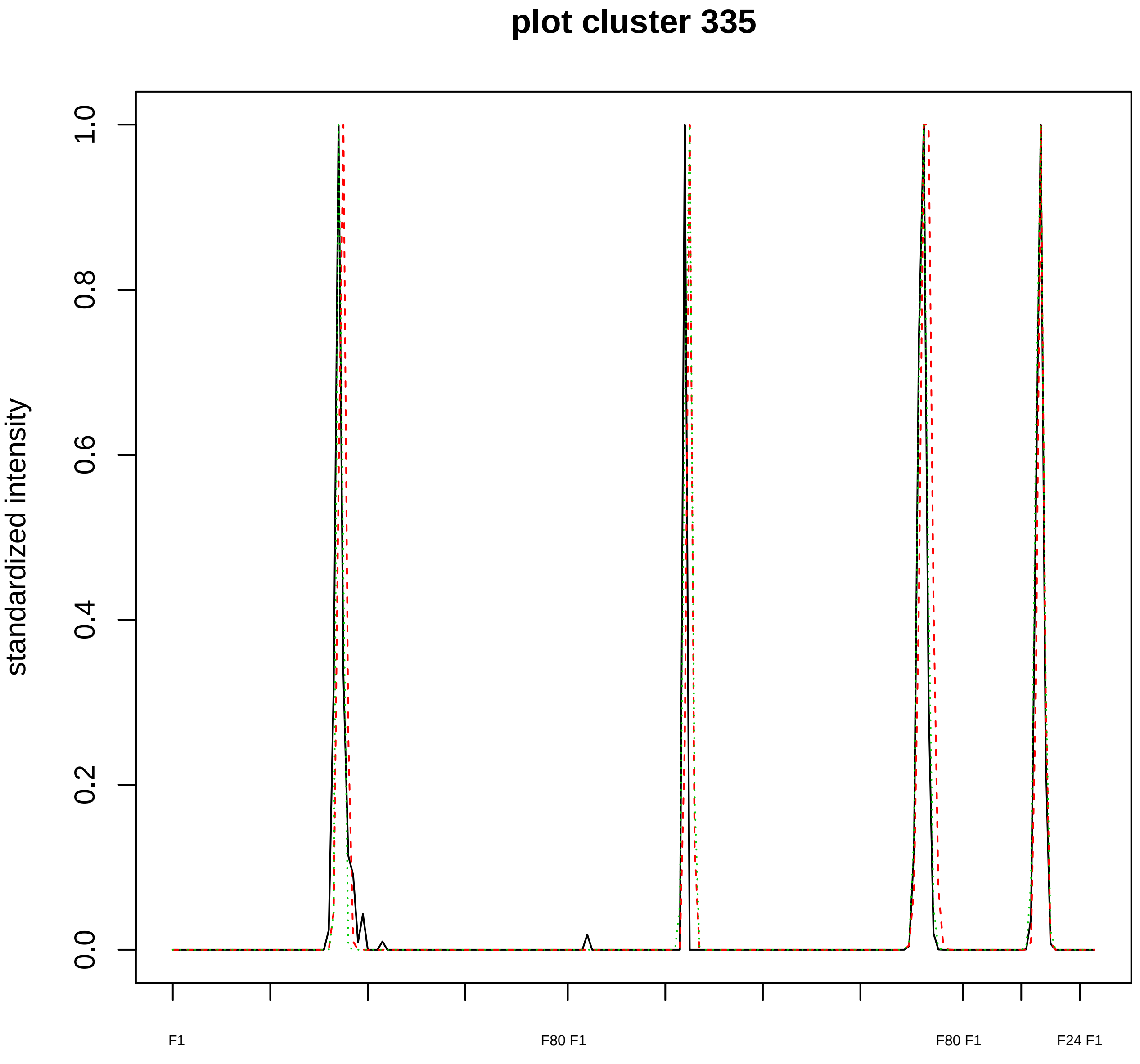
plot cluster 335
standardized intensity
0.0
0.2
0.4
0.6
0.8
1.0
F1
F80 F1
F80 F1
F24 F1

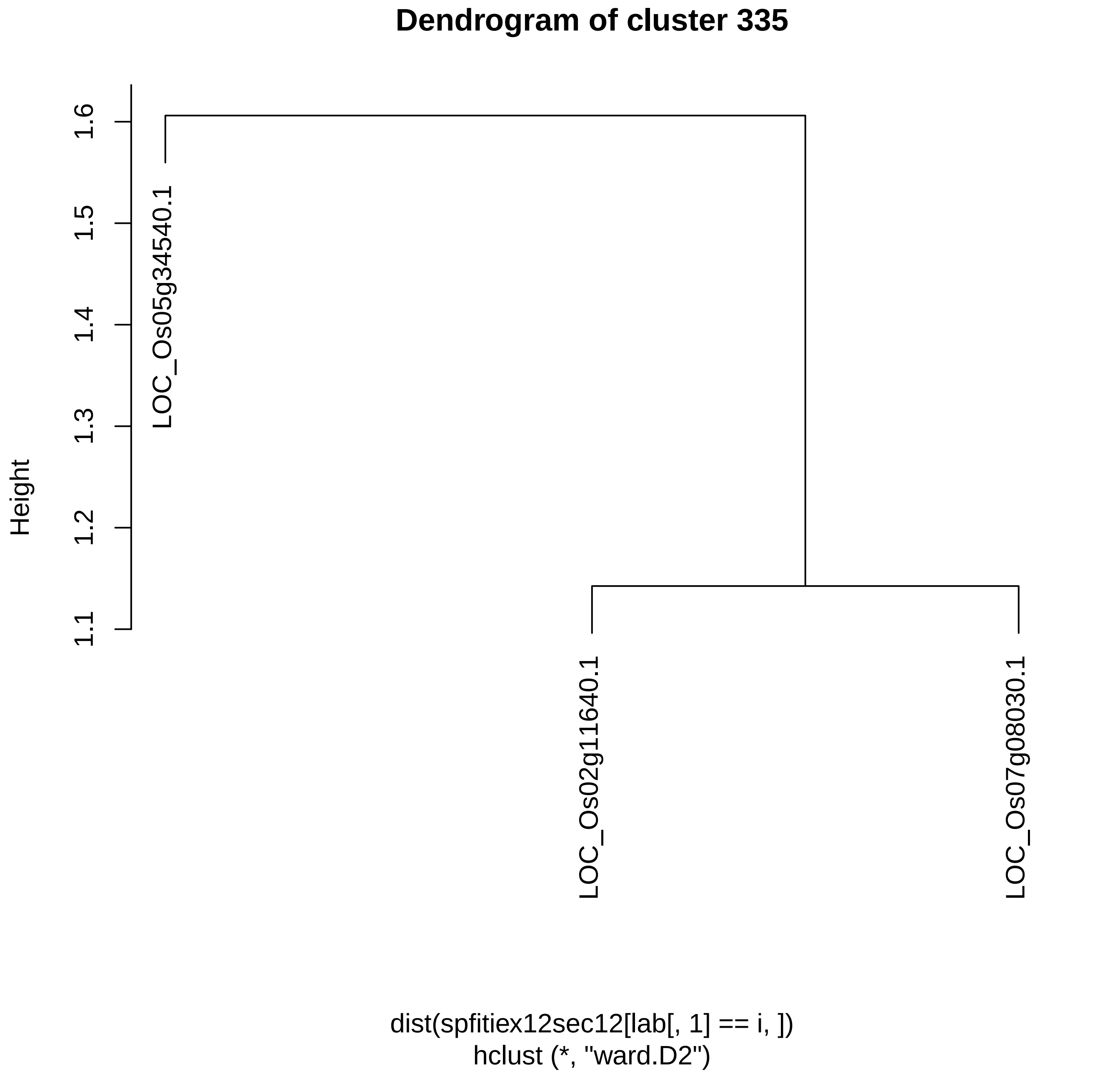

Dendrogram of cluster 335
Height
1.6
1.5
1.4
1.3
1.2
1.1
LOC_Os05g34540.1
LOC_Os02g11640.1
LOC_Os07g08030.1
dist(spfitiex12sec12[lab[, 1] == i, ])
hclust (*, "ward.D2")

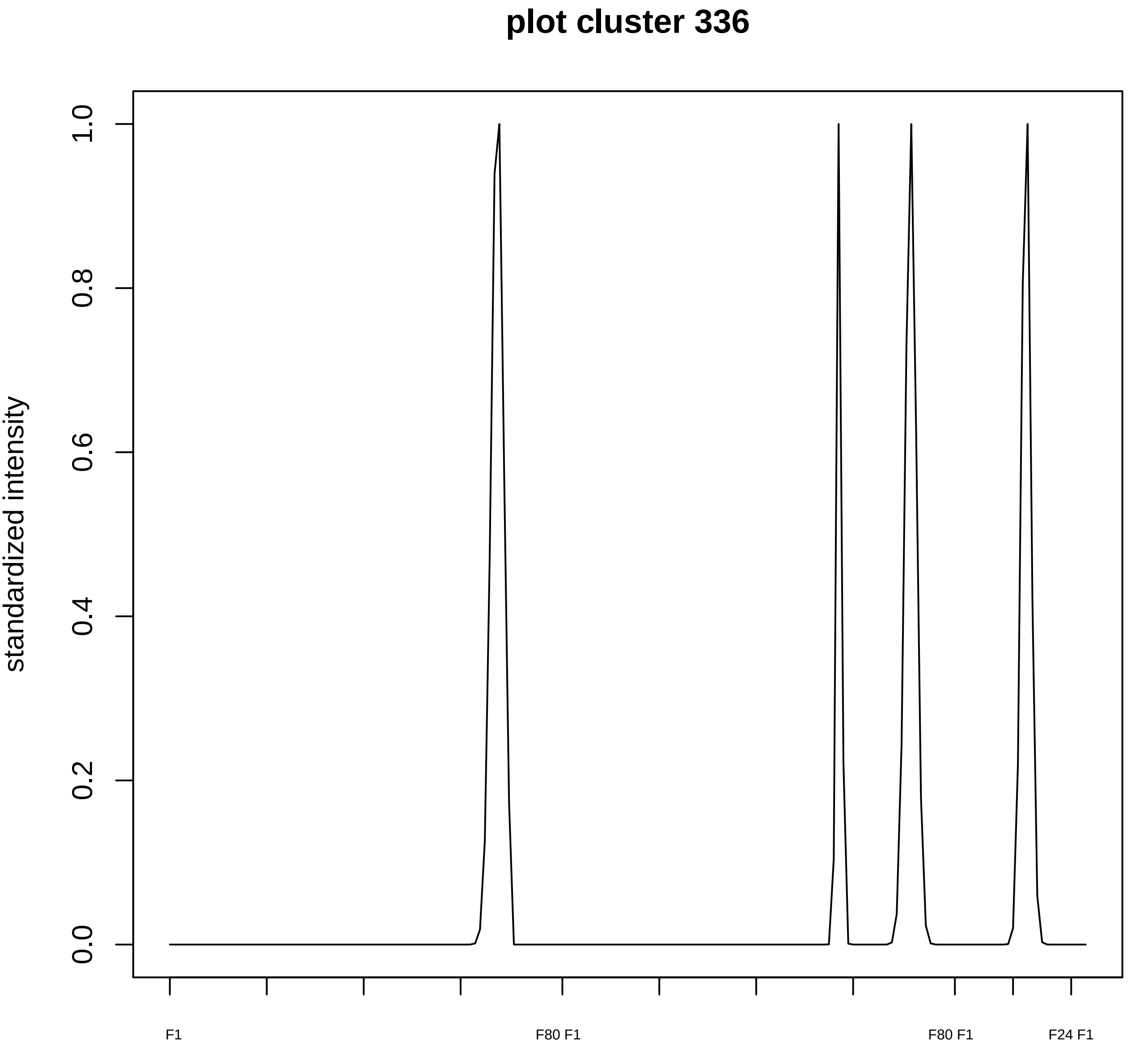

plot cluster 336
standardized intensity
0.0
0.2
0.4
0.6
0.8
1.0
F1
F80 F1
F80 F1
F24 F1

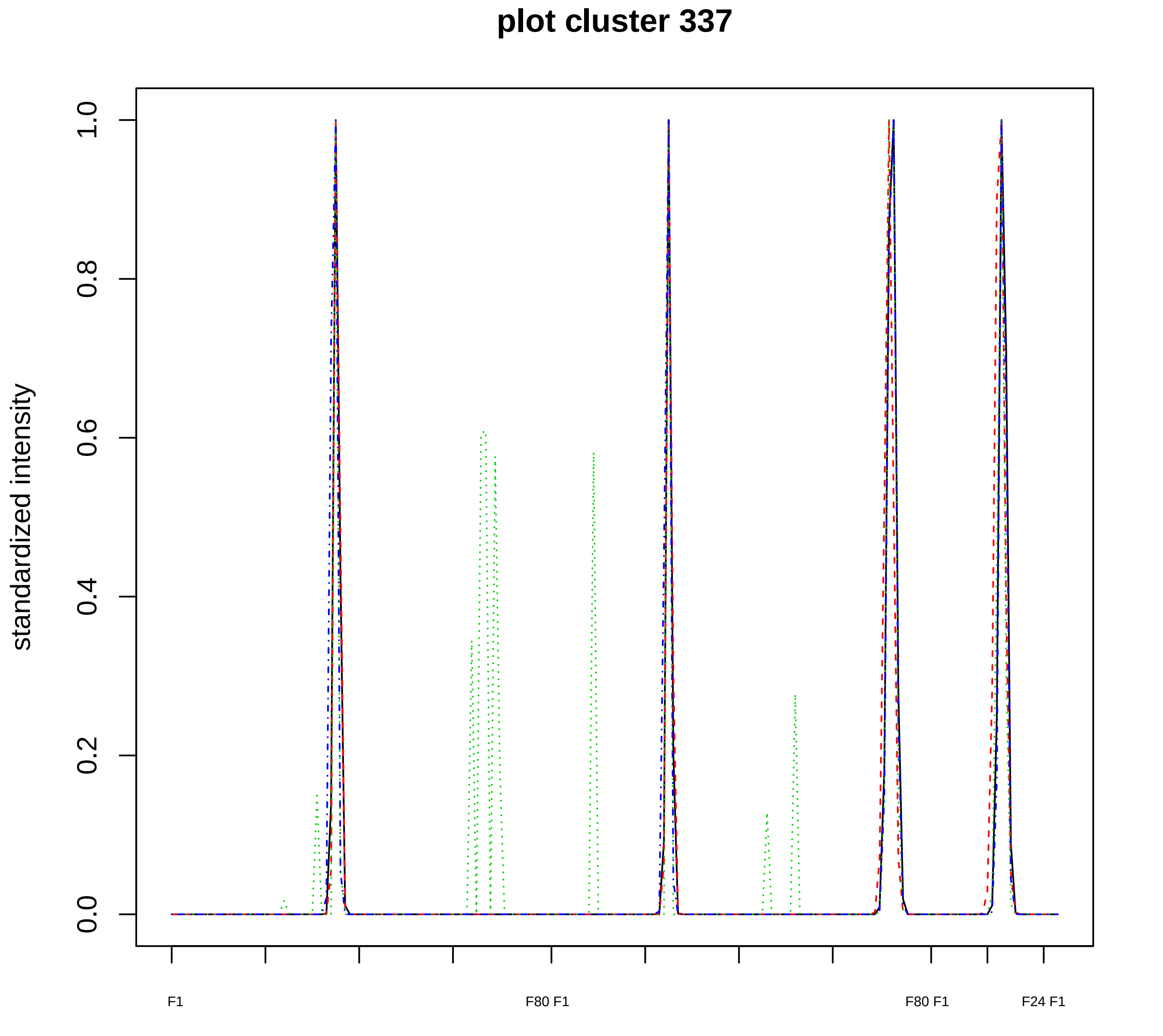
plot cluster 337
standardized intensity
0.0
0.2
0.4
0.6
0.8
1.0
F1
F80 F1
F80 F1
F24 F1

### Dendrogram of cluster 337

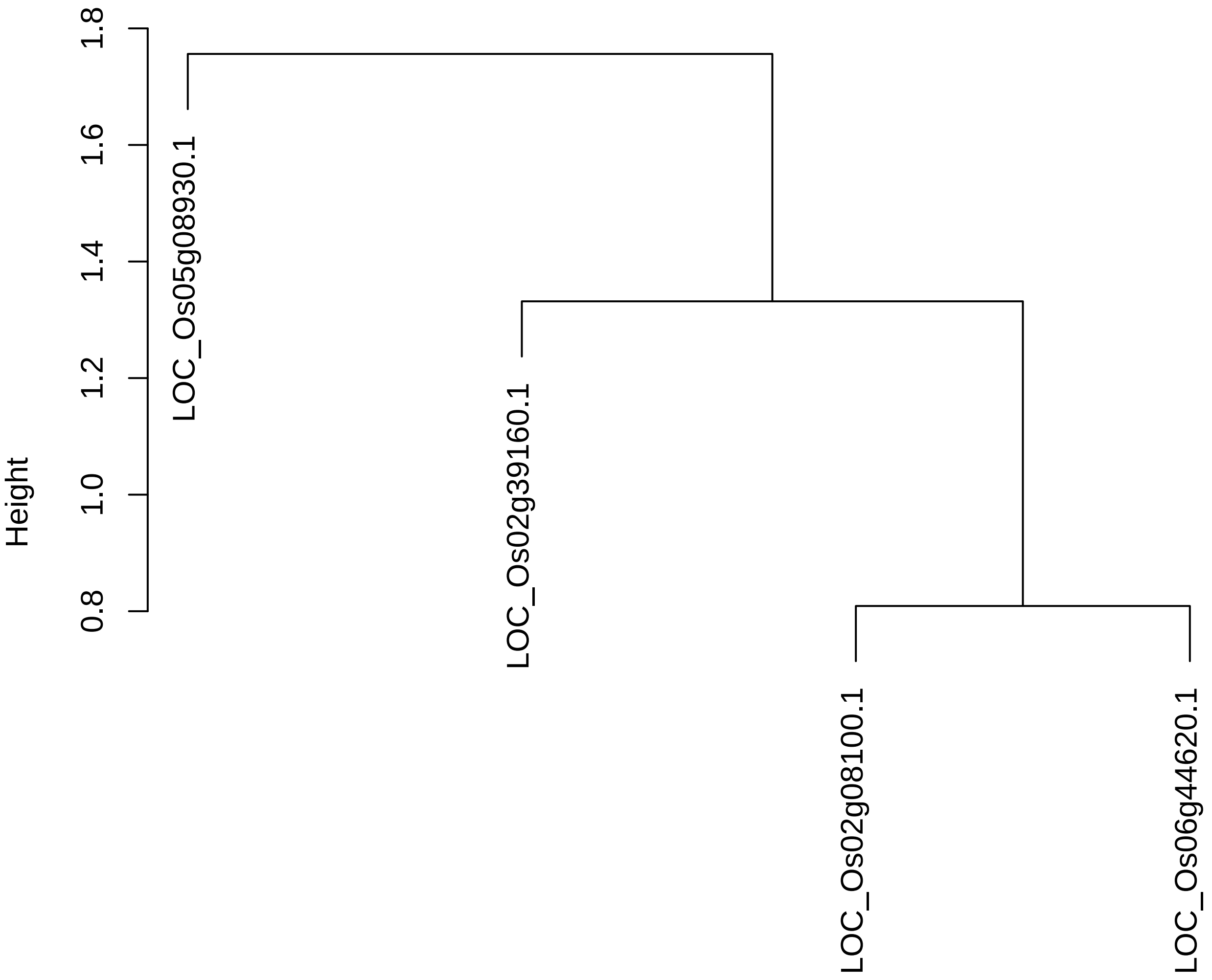


dist(spfitiex12sec12[lab[, 1] == i, ])
hclust (*, "ward.D2")

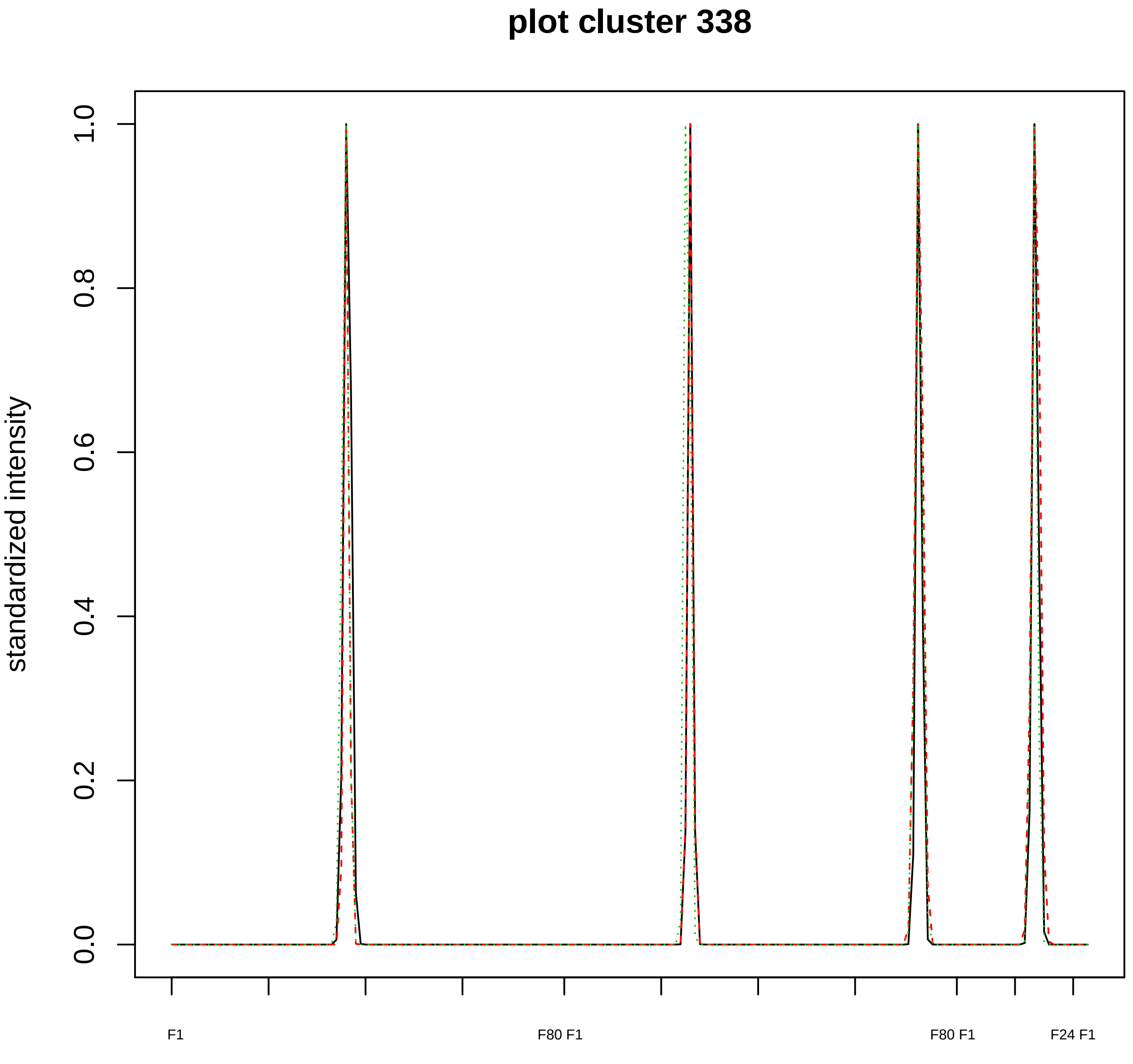
plot cluster 338
standardized intensity
0.0
0.2
0.4
0.6
0.8
1.0
F1
F80 F1
F80 F1
F24 F1

### Dendrogram of cluster 338

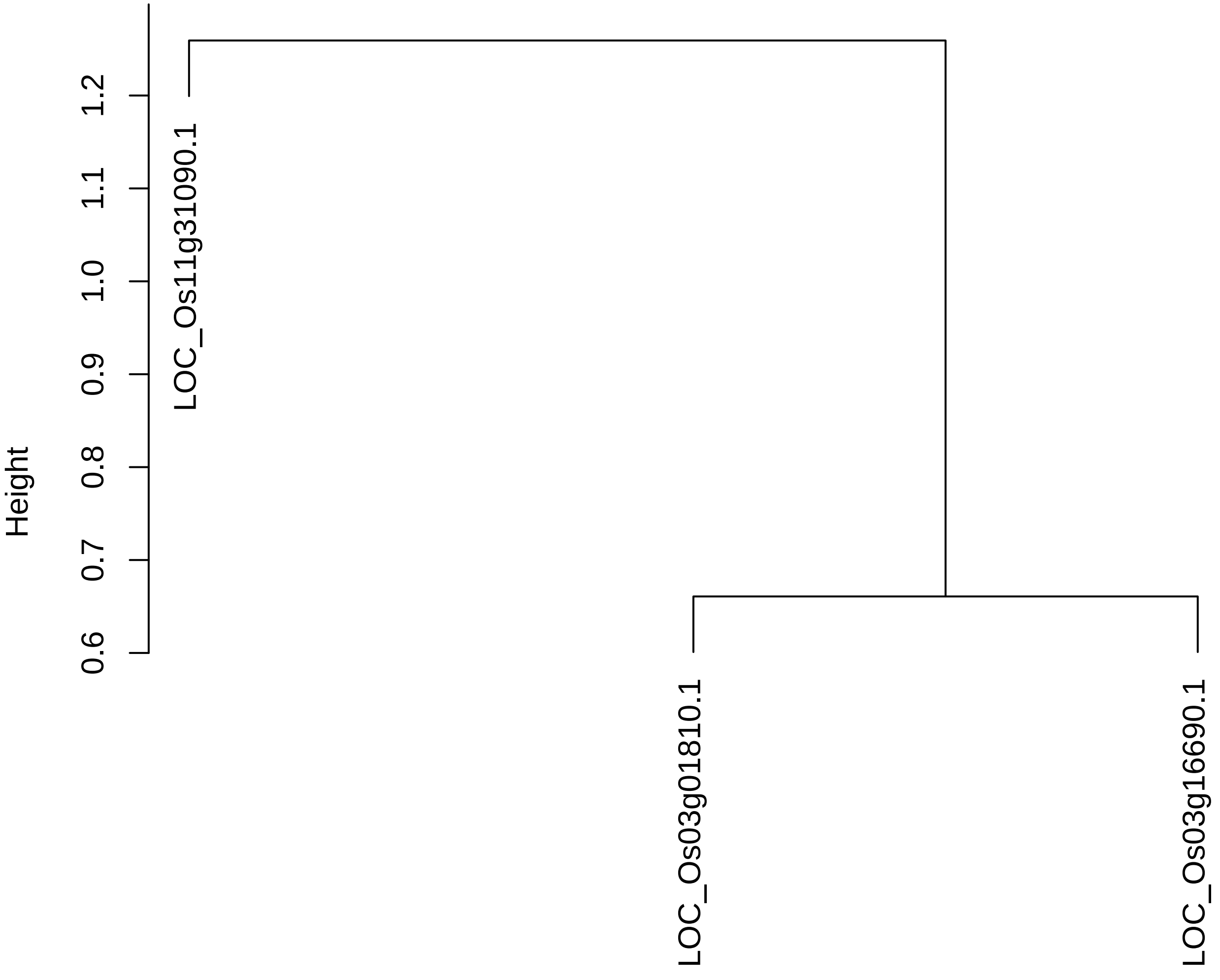


dist(spfitiex12sec12[lab[, 1] == i, ])
hclust (*, "ward.D2")

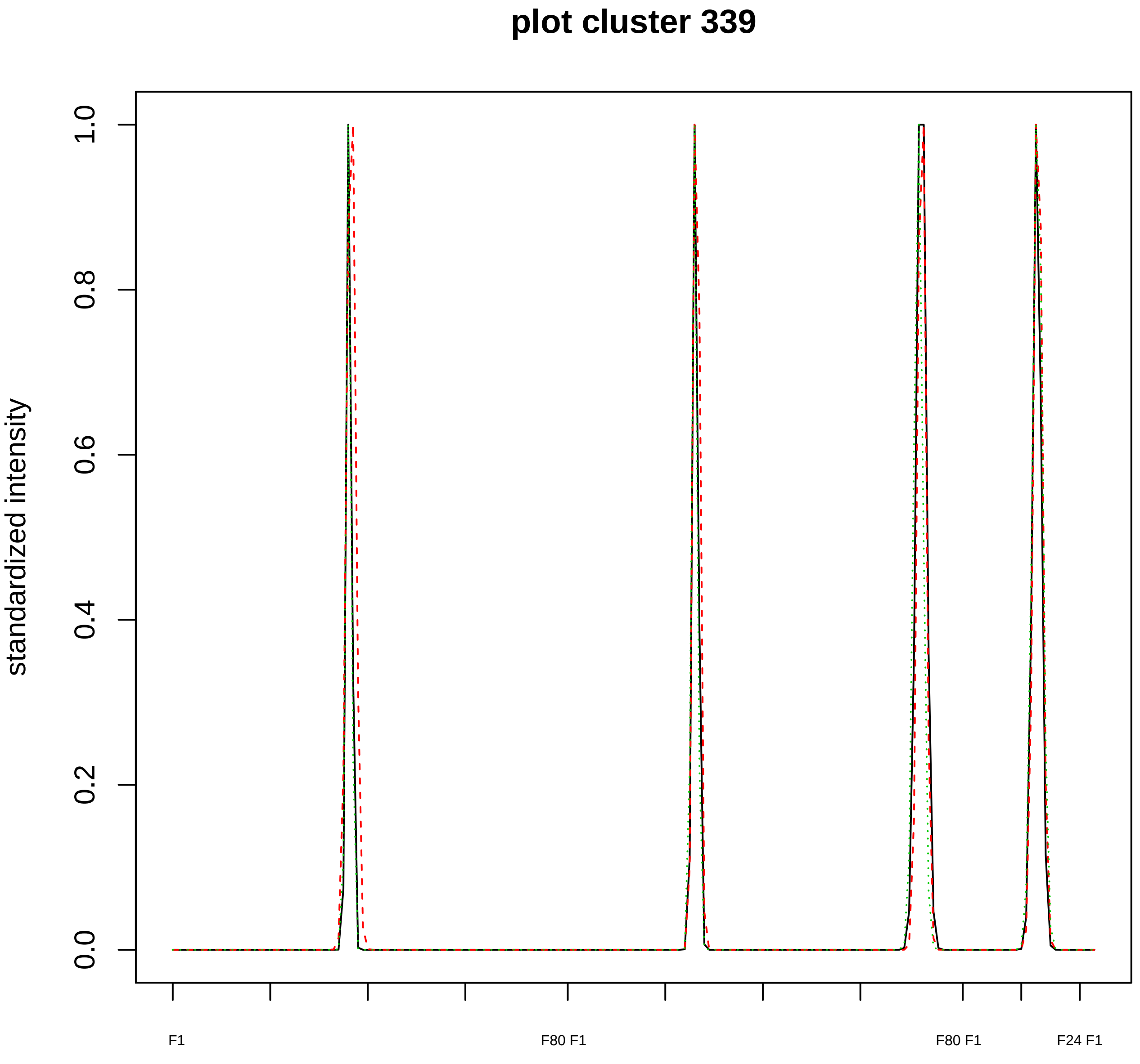
plot cluster 339
standardized intensity
0.0
0.2
0.4
0.6
0.8
1.0
F1
F80 F1
F80 F1
F24 F1

### Dendrogram of cluster 339

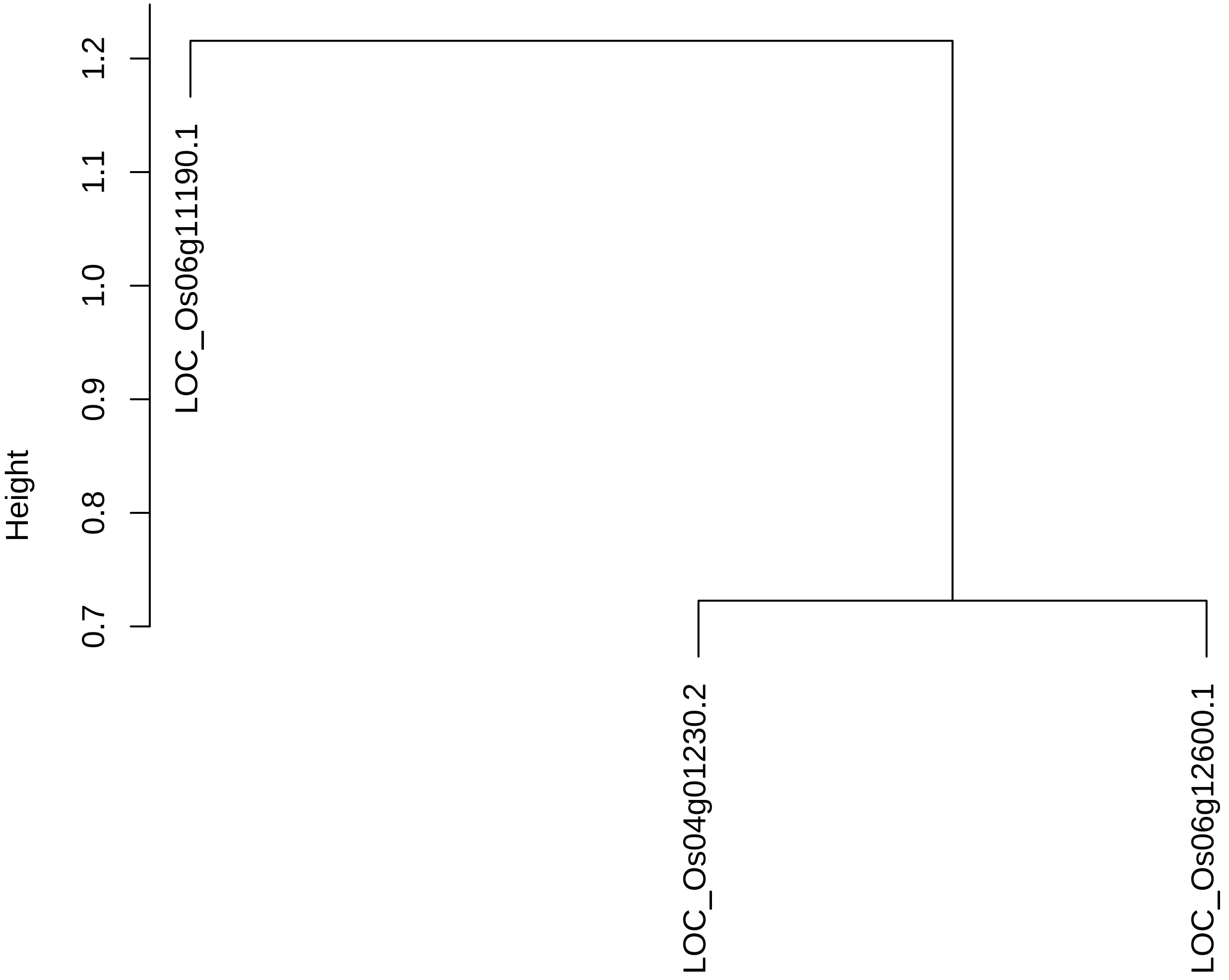


dist(spfitiex12sec12[lab[, 1] == i, ])
hclust (*, "ward.D2")

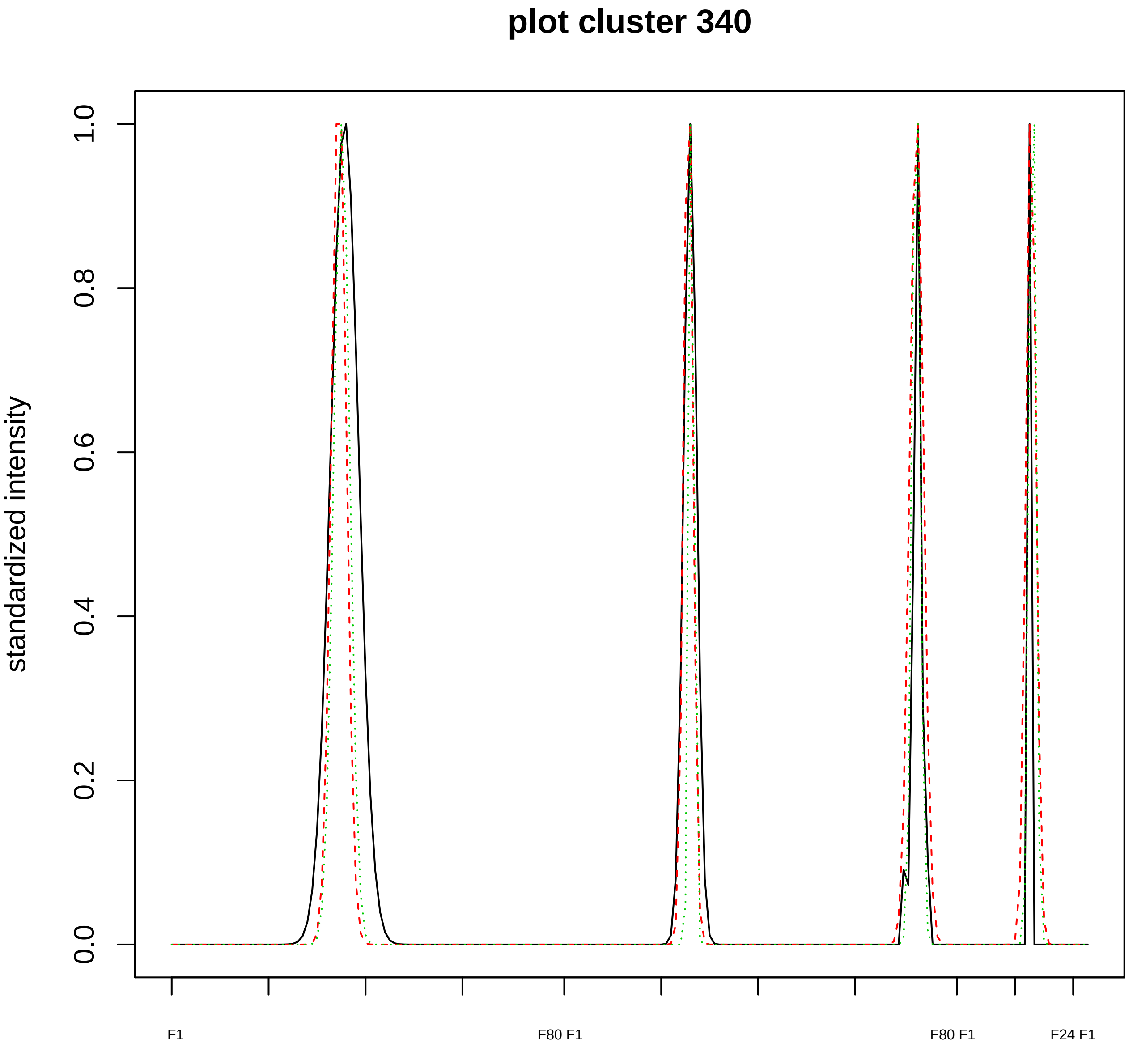
plot cluster 340
standardized intensity
0.0
0.2
0.4
0.6
0.8
1.0
F1
F80 F1
F80 F1
F24 F1

### Dendrogram of cluster 340

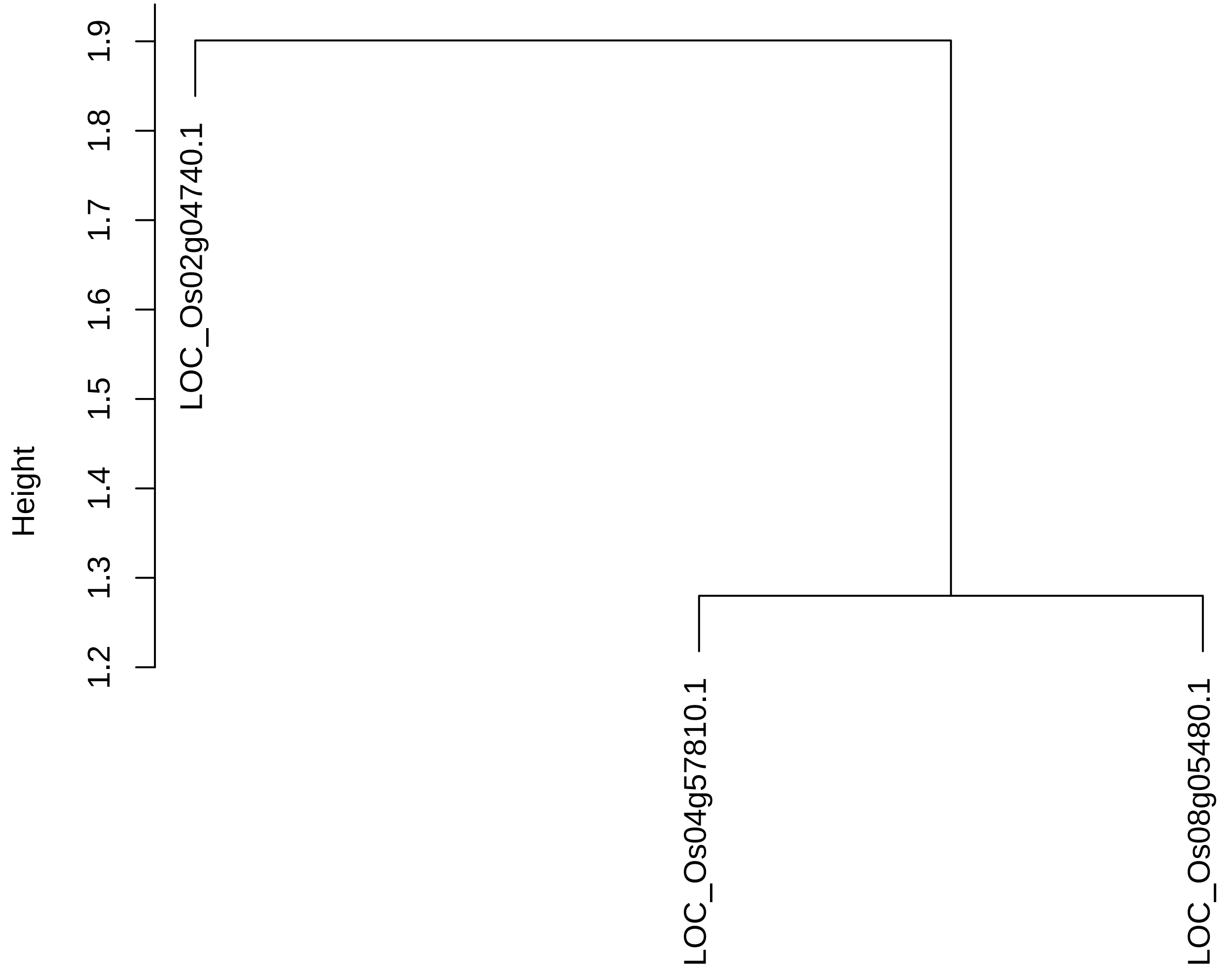


dist(spfitiex12sec12[lab[, 1] == i, ])
hclust (*, "ward.D2")

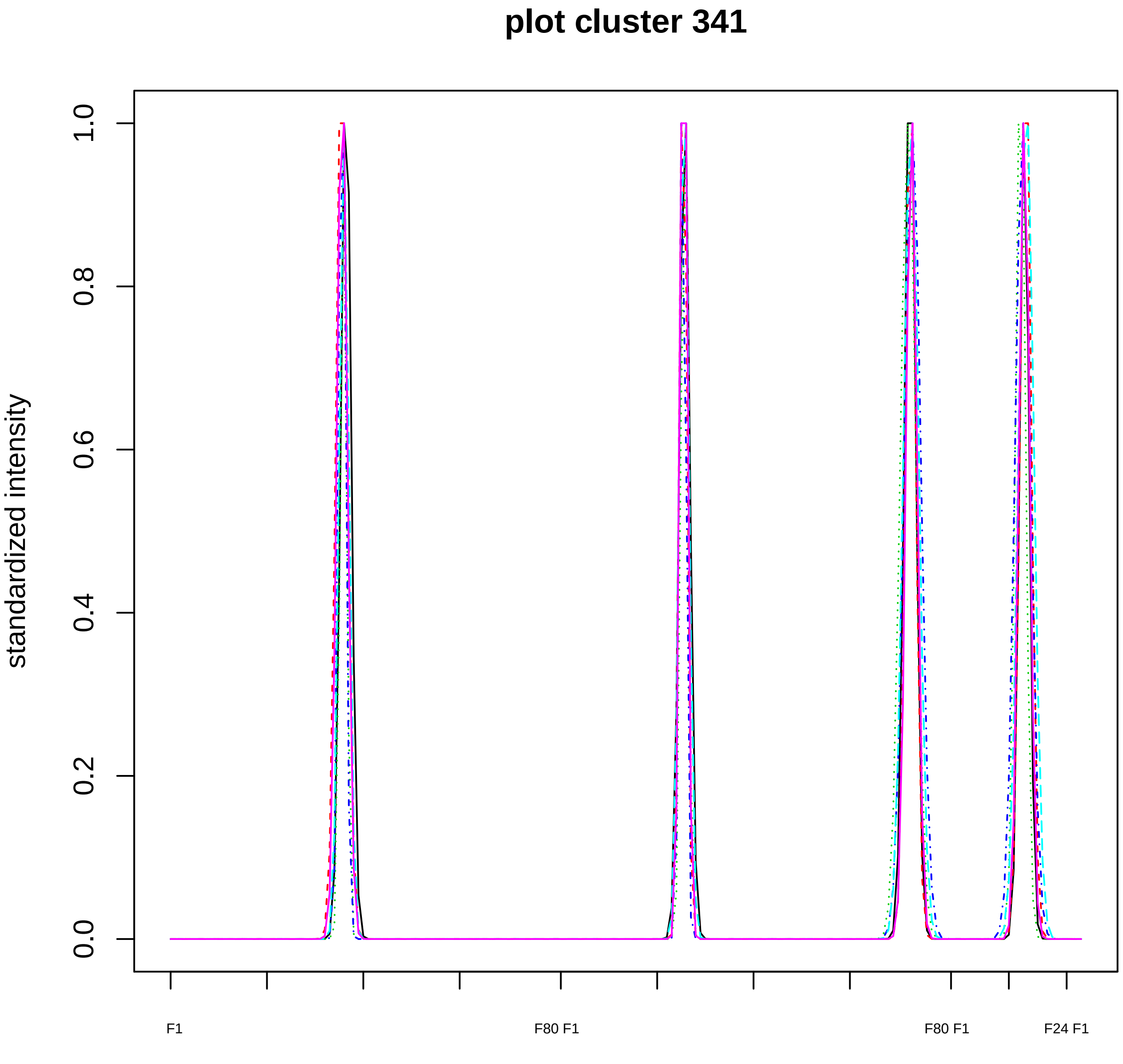
plot cluster 341
standardized intensity
0.0
0.2
0.4
0.6
0.8
1.0
F1
F80 F1
F80 F1
F24 F1

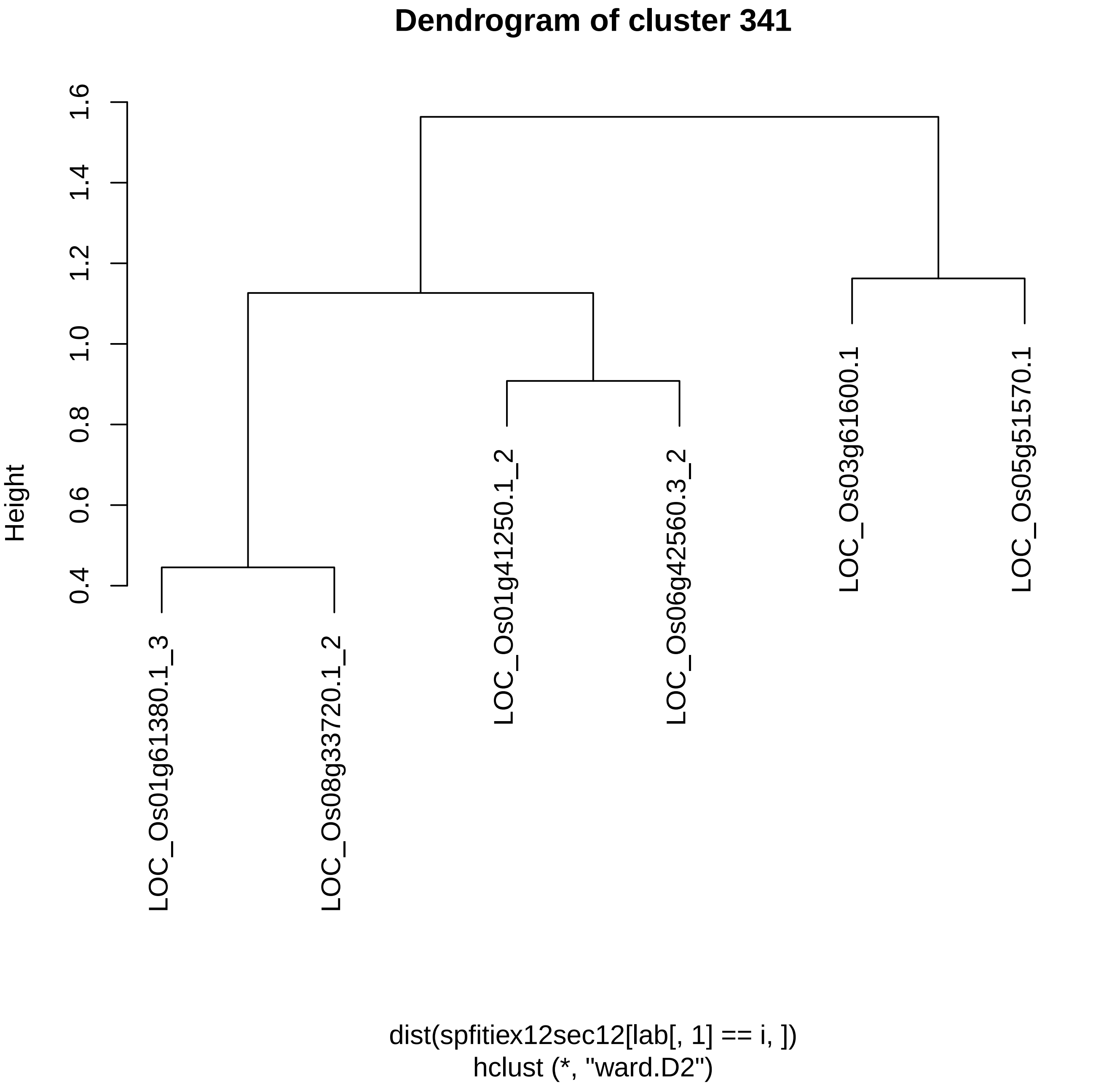
Dendrogram of cluster 341
Height
0.4
0.6
0.8
1.0
1.2
1.4
1.6
LOC_Os01g61380.1_3
LOC_Os08g33720.1_2
LOC_Os01g41250.1_2
LOC_Os06g42560.3_2
LOC_Os03g61600.1
LOC_Os05g51570.1
dist(spfitiex12sec12[lab[, 1] == i, ])
hclust (*, "ward.D2")

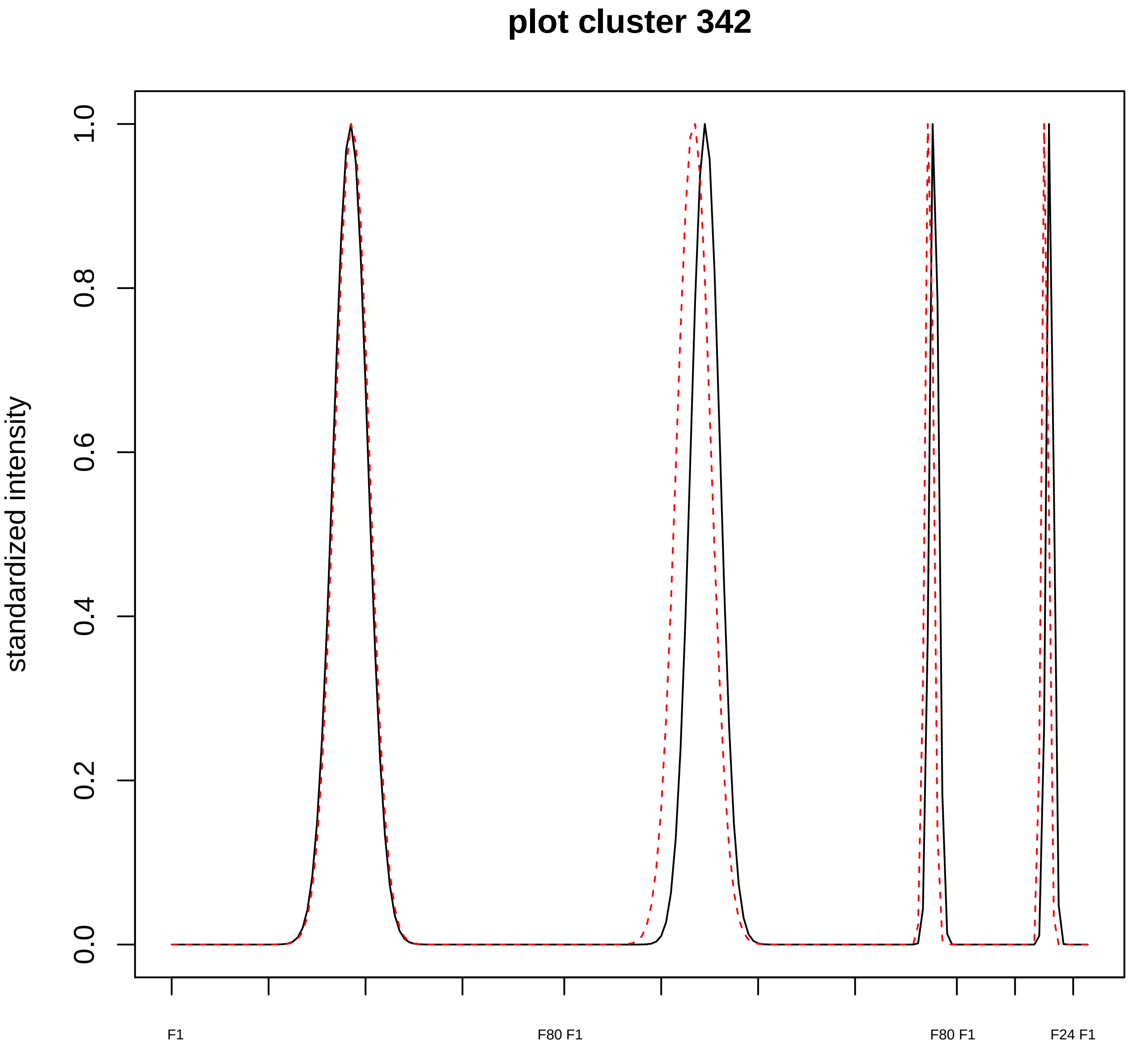

plot cluster 342
standardized intensity
0.0
0.2
0.4
0.6
0.8
1.0
F1
F80 F1
F80 F1
F24 F1

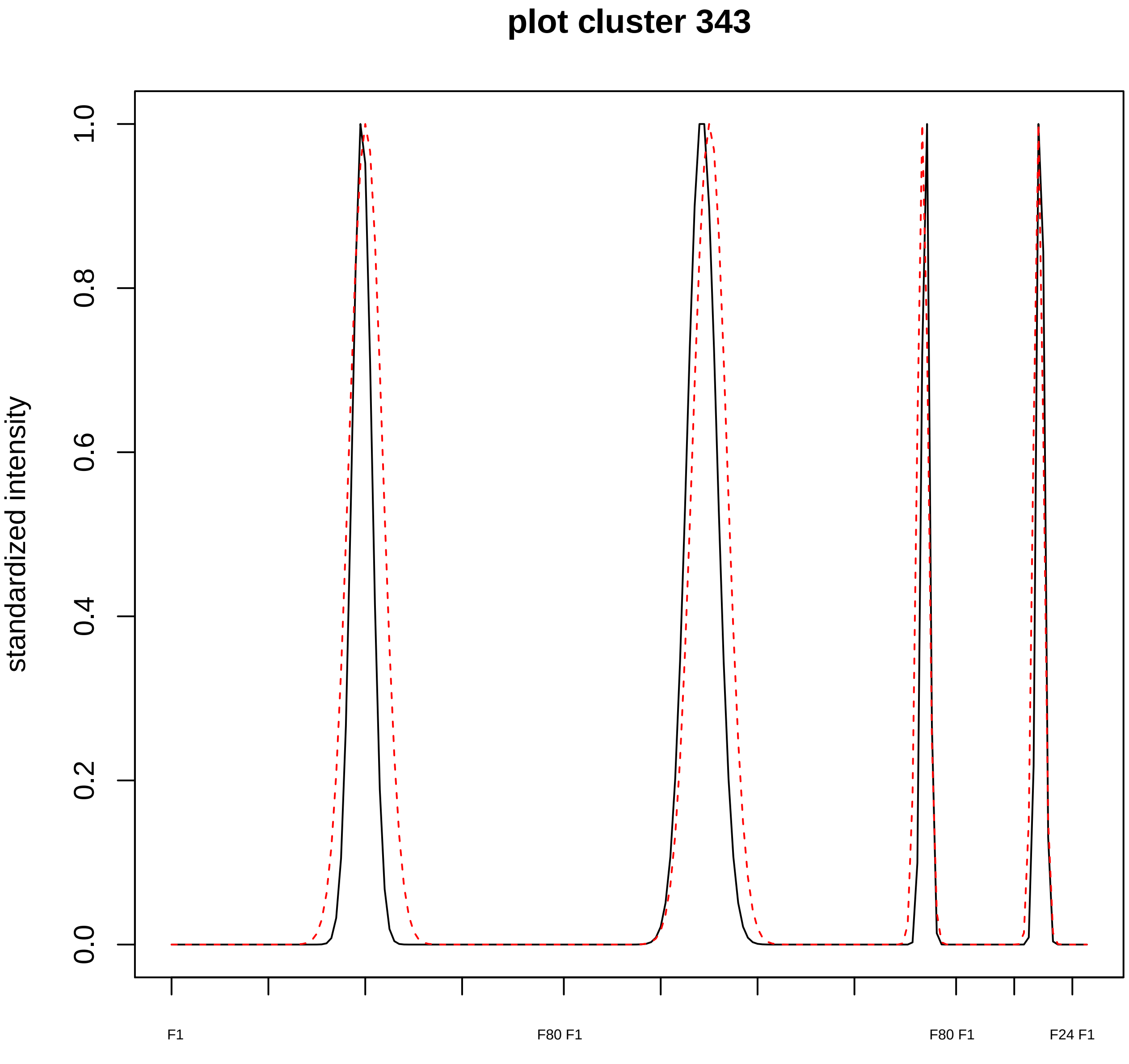
plot cluster 343
standardized intensity
0.0
0.2
0.4
0.6
0.8
1.0
F1
F80 F1
F80 F1
F24 F1

### plot cluster 344

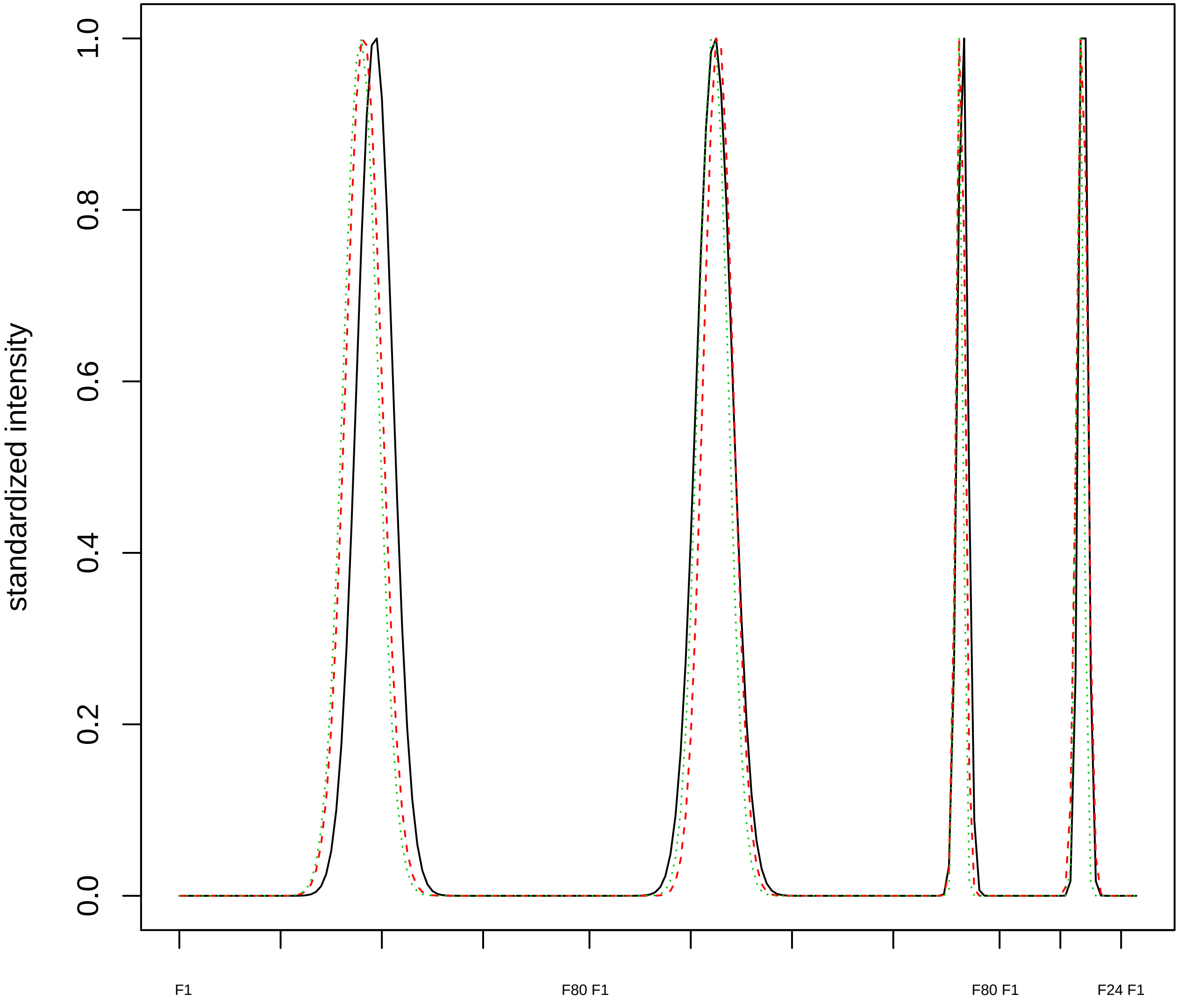

### Dendrogram of cluster 344

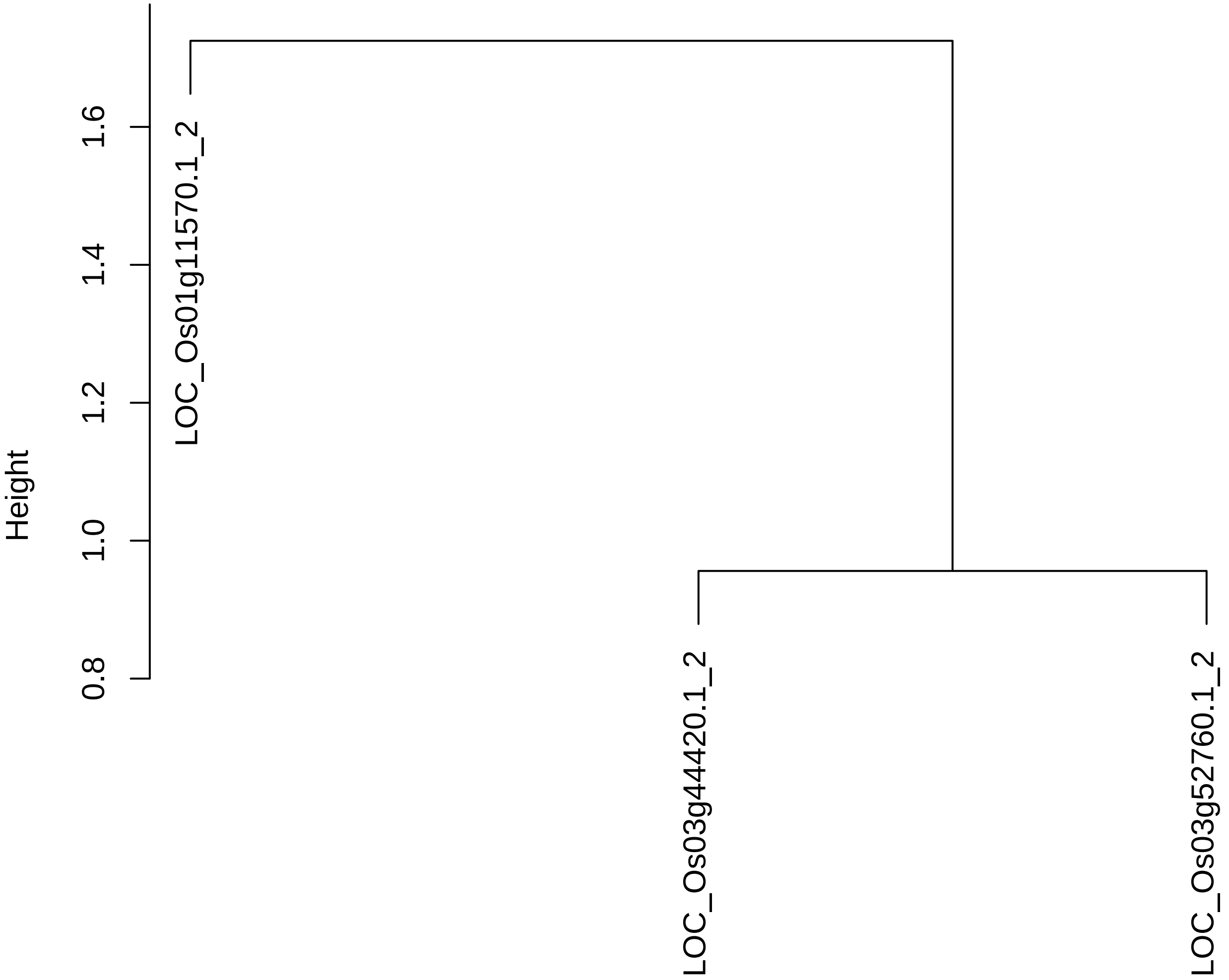


dist(spfitiex12sec12[lab[, 1] == i, ])
hclust (*, "ward.D2")

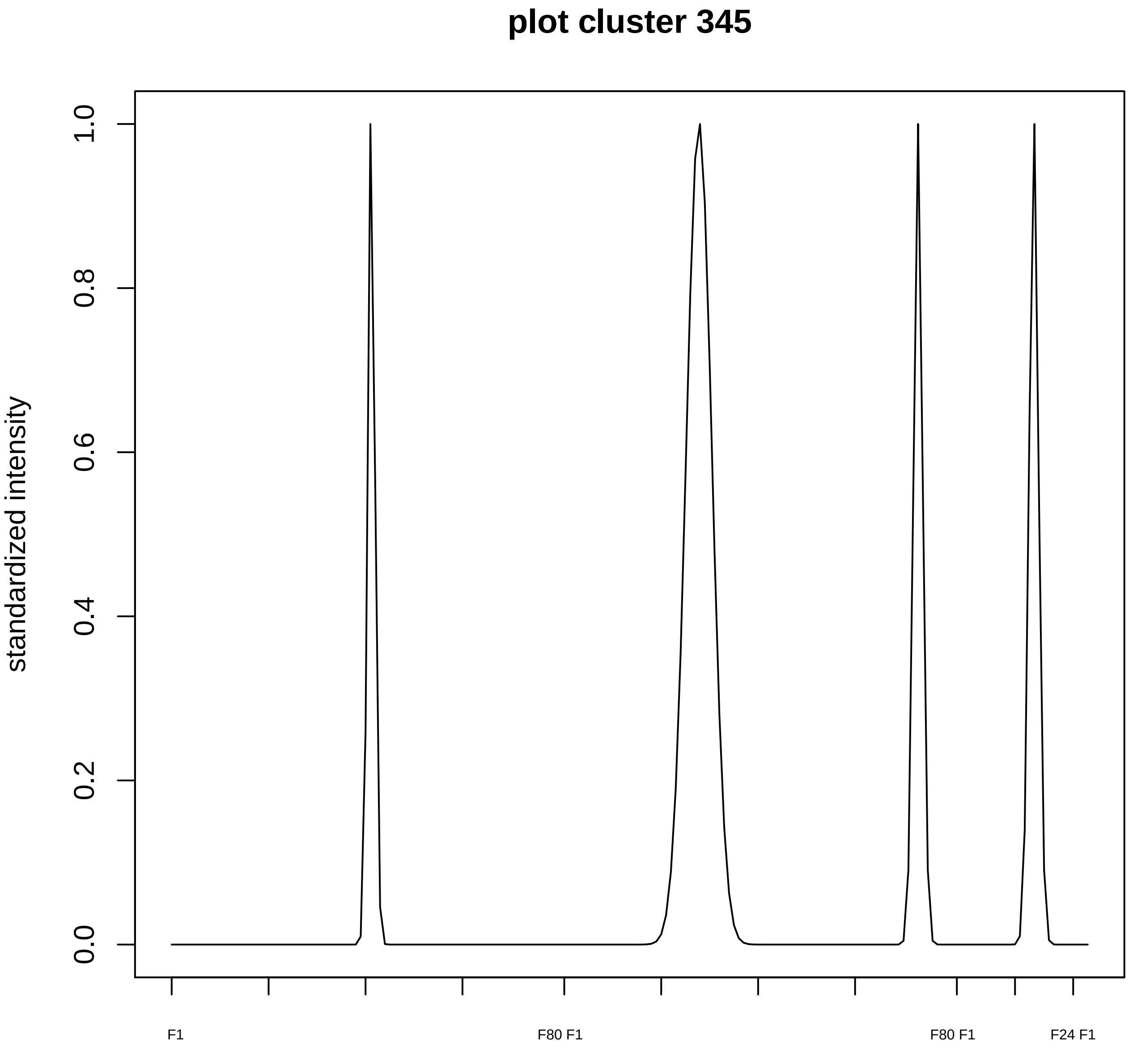
plot cluster 345
standardized intensity
0.0
0.2
0.4
0.6
0.8
1.0
F1
F80 F1
F80 F1
F24 F1

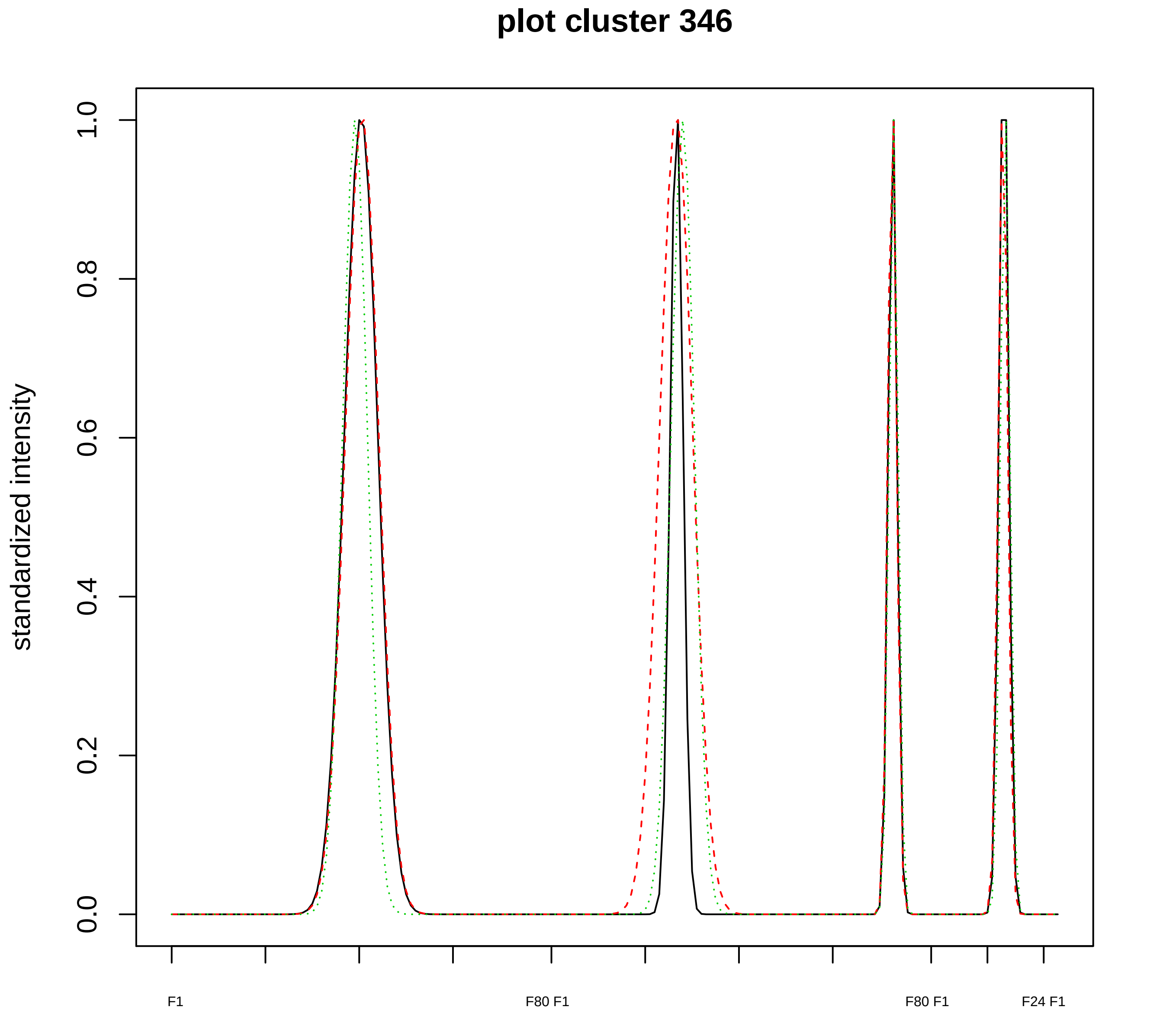

plot cluster 346
standardized intensity
0.0
0.2
0.4
0.6
0.8
1.0
F1
F80 F1
F80 F1
F24 F1

### Dendrogram of cluster 346

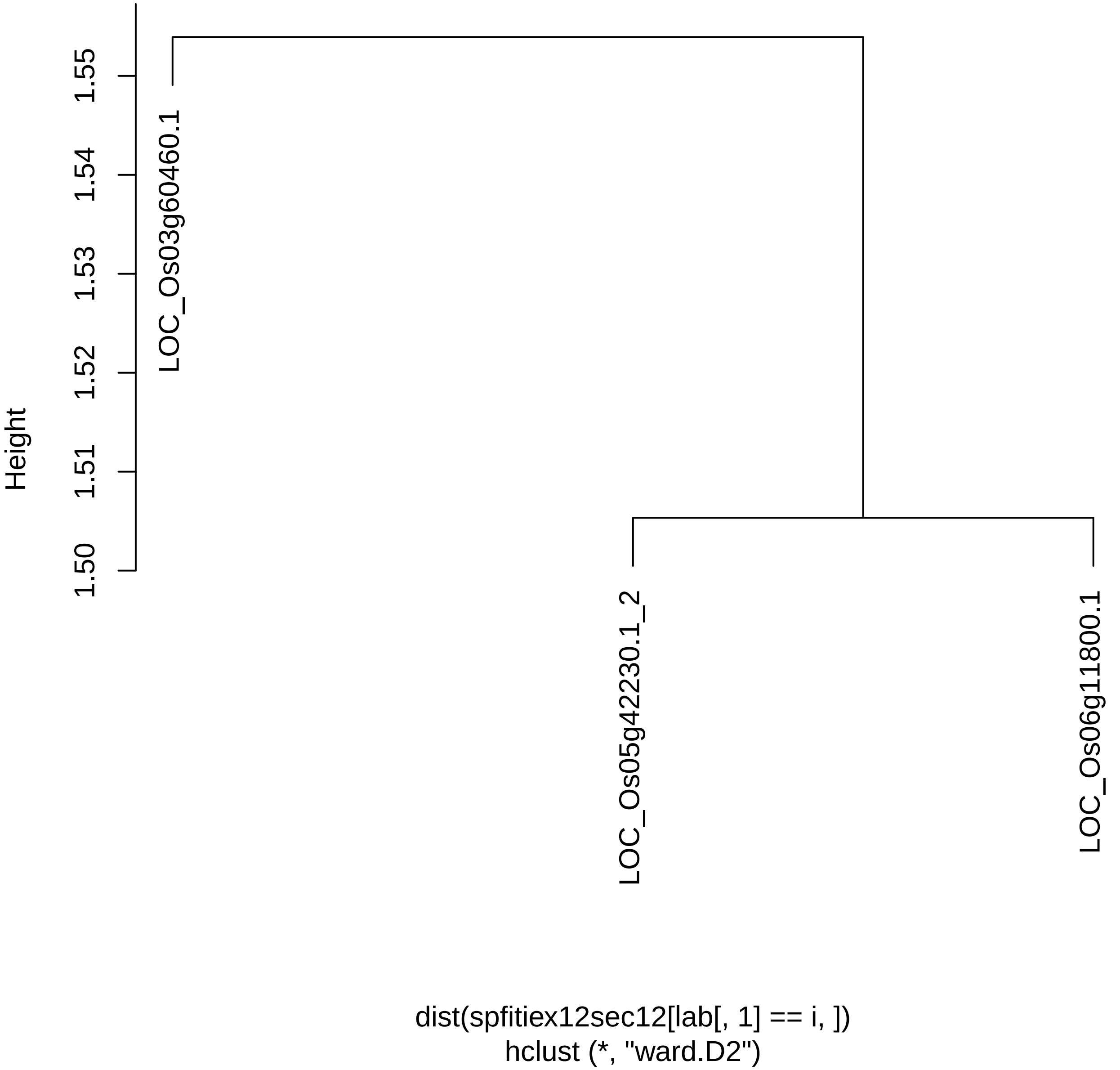

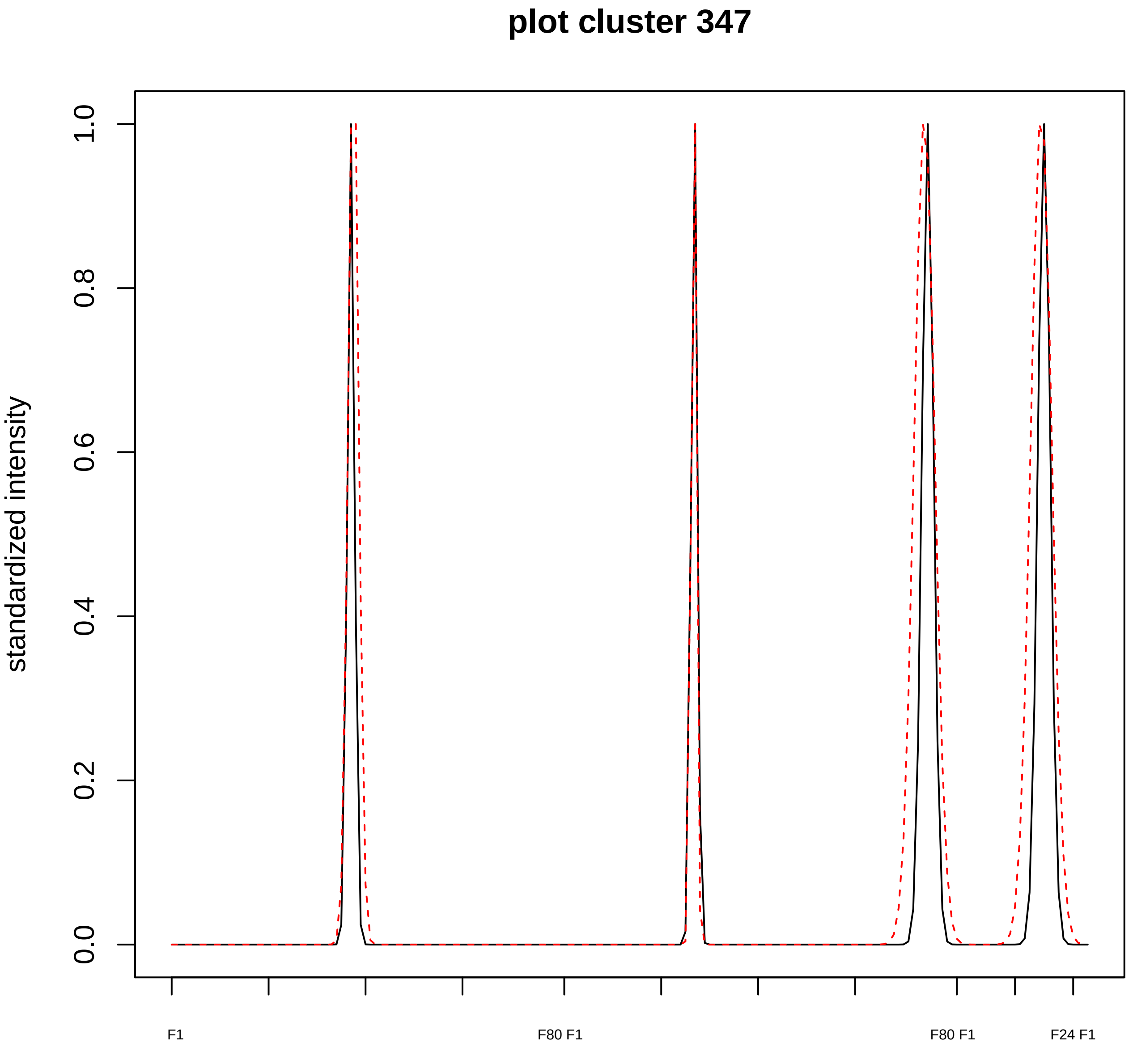

plot cluster 347
standardized intensity
0.0
0.2
0.4
0.6
0.8
1.0
F1
F80 F1
F80 F1
F24 F1

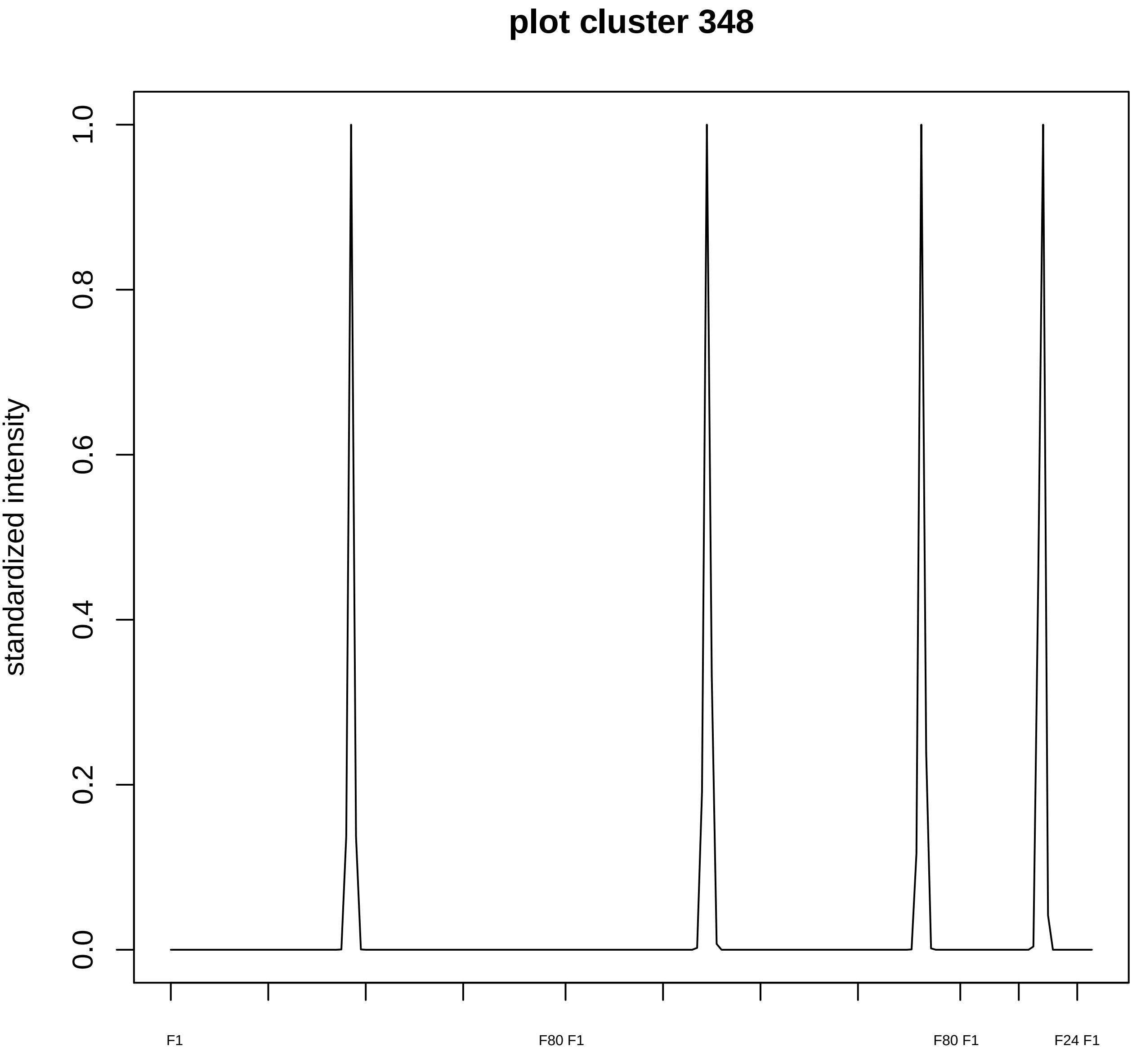
plot cluster 348
standardized intensity
0.0
0.2
0.4
0.6
0.8
1.0
F1
F80 F1
F80 F1
F24 F1

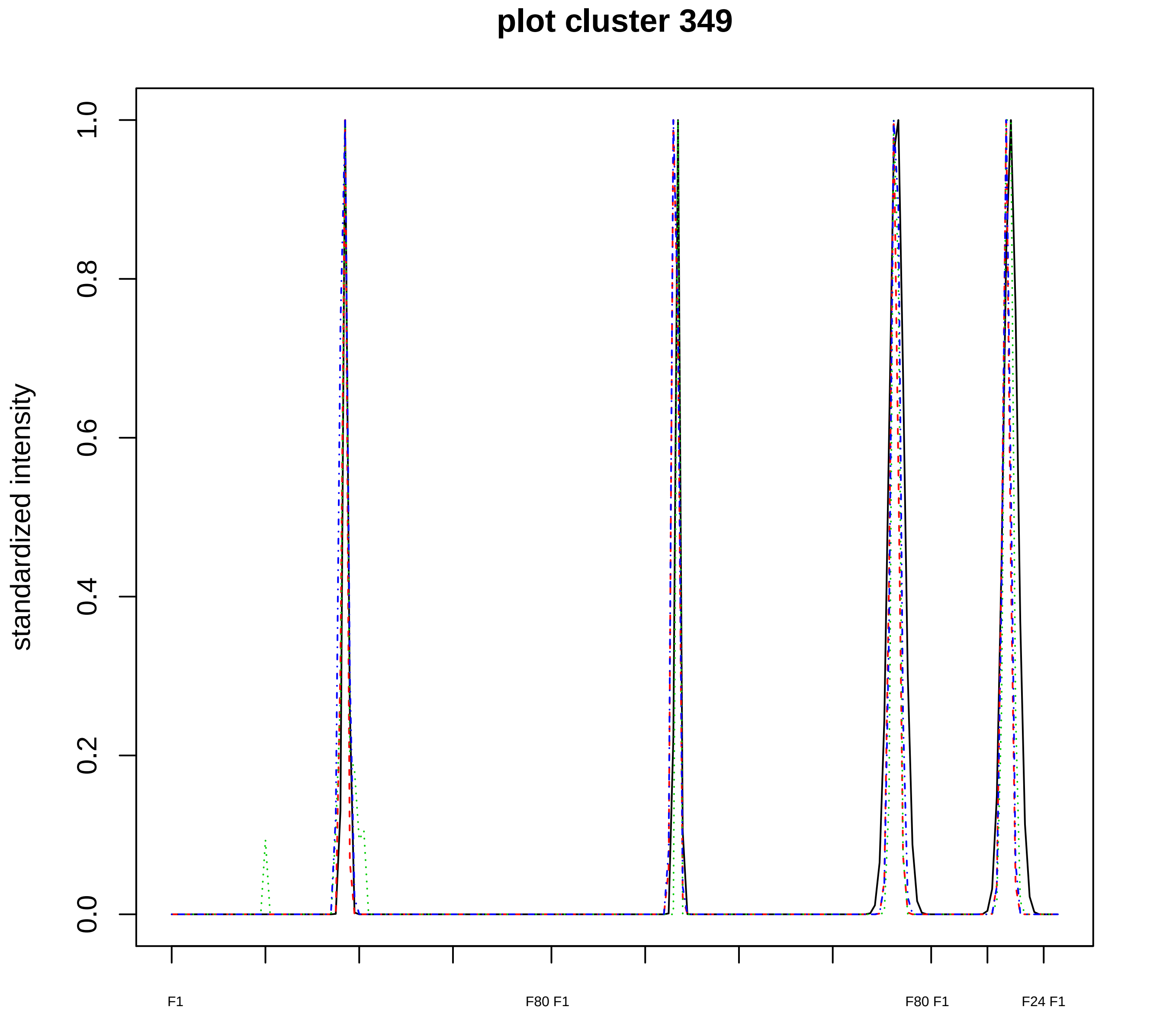
plot cluster 349
standardized intensity
0.0
0.2
0.4
0.6
0.8
1.0
F1
F80 F1
F80 F1
F24 F1

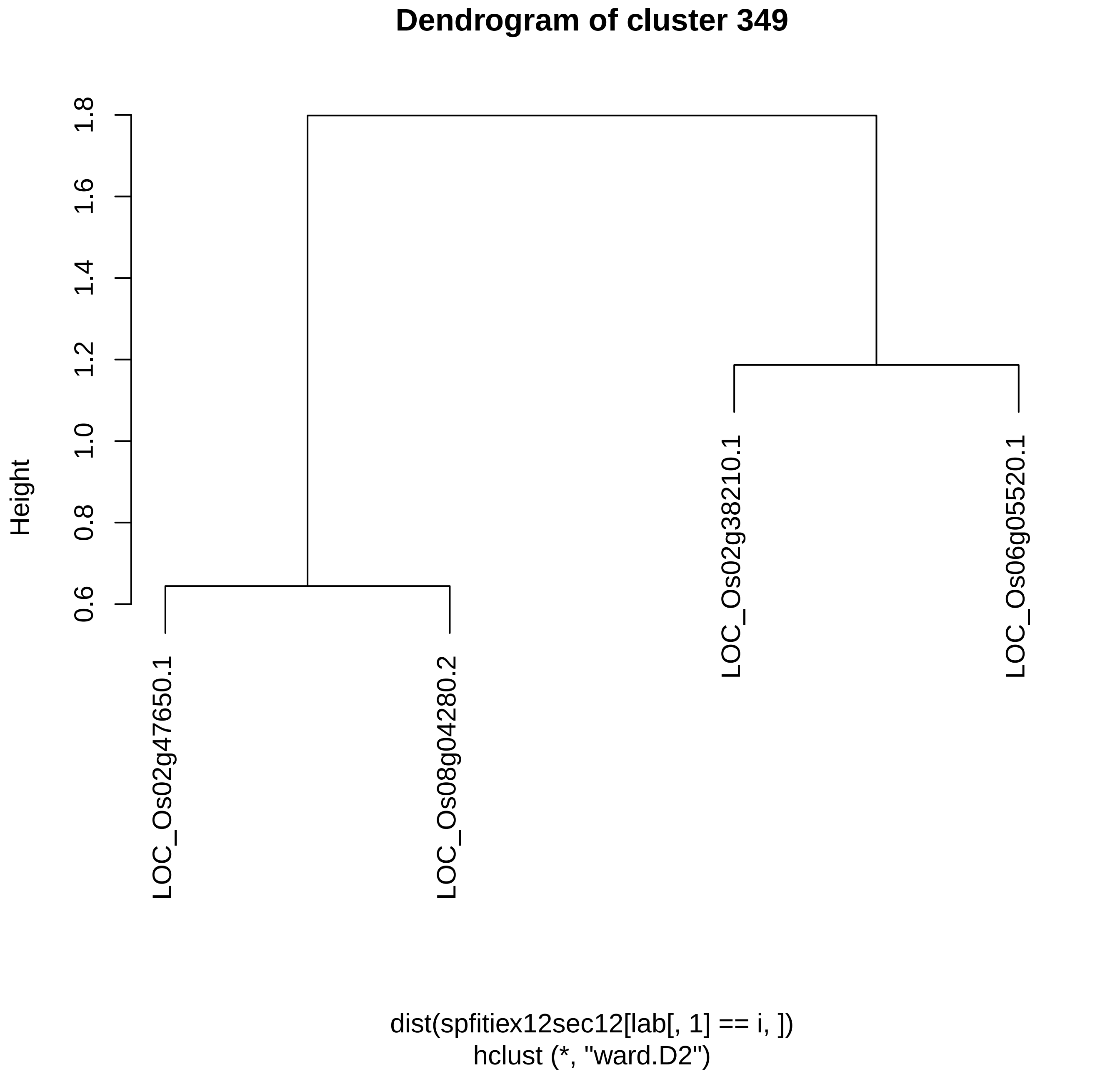
Dendrogram of cluster 349
Height
0.6
0.8
1.0
1.2
1.4
1.6
1.8
LOC_Os02g47650.1
LOC_Os08g04280.2
LOC_Os02g38210.1
LOC_Os06g05520.1
dist(spfitiex12sec12[lab[, 1] == i, ])
hclust (*, "ward.D2")

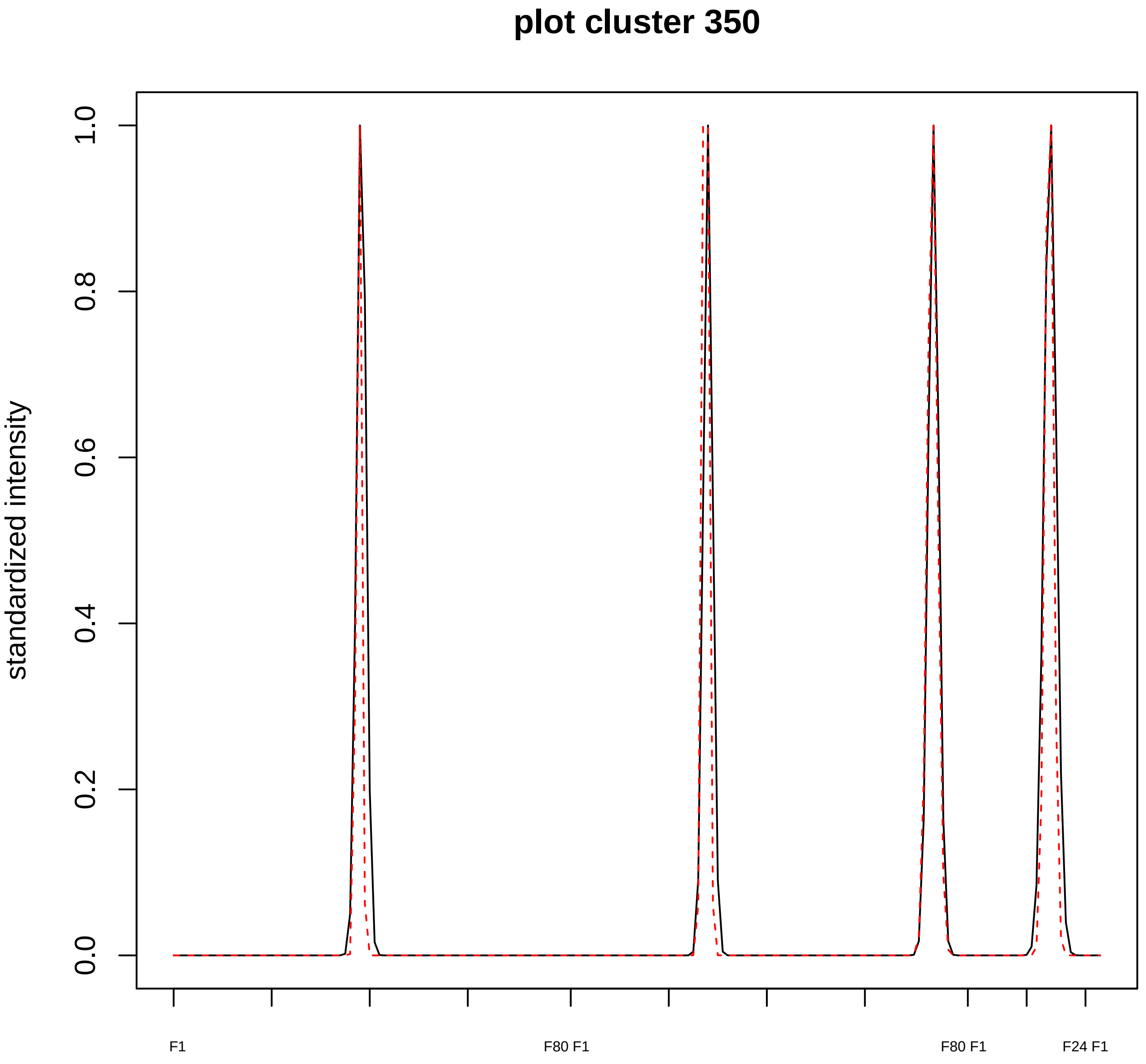
plot cluster 350
standardized intensity
0.0
0.2
0.4
0.6
0.8
1.0
F1
F80 F1
F80 F1
F24 F1

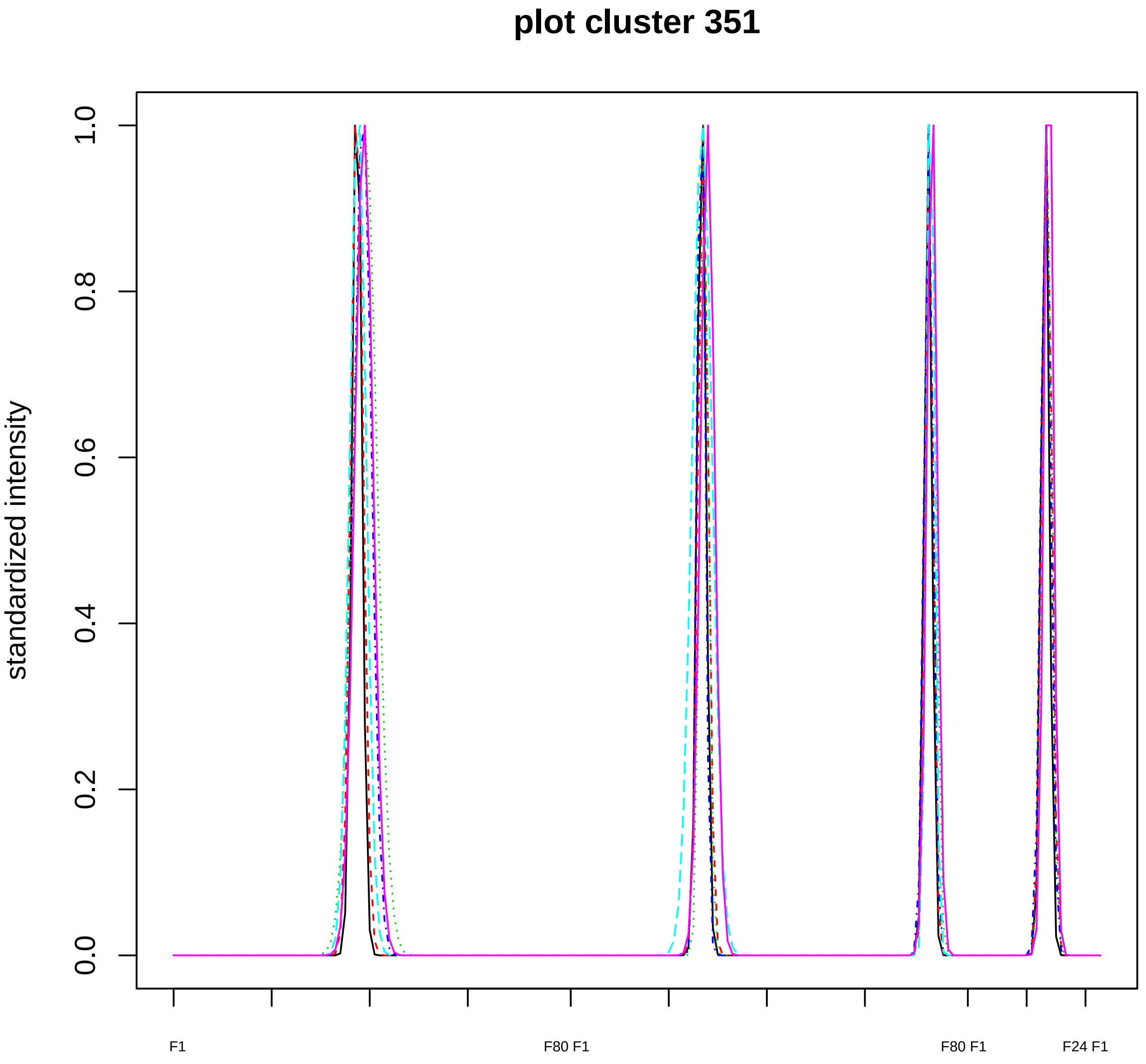
plot cluster 351
standardized intensity
0.0
0.2
0.4
0.6
0.8
1.0
F1
F80 F1
F80 F1
F24 F1

### Dendrogram of cluster 351

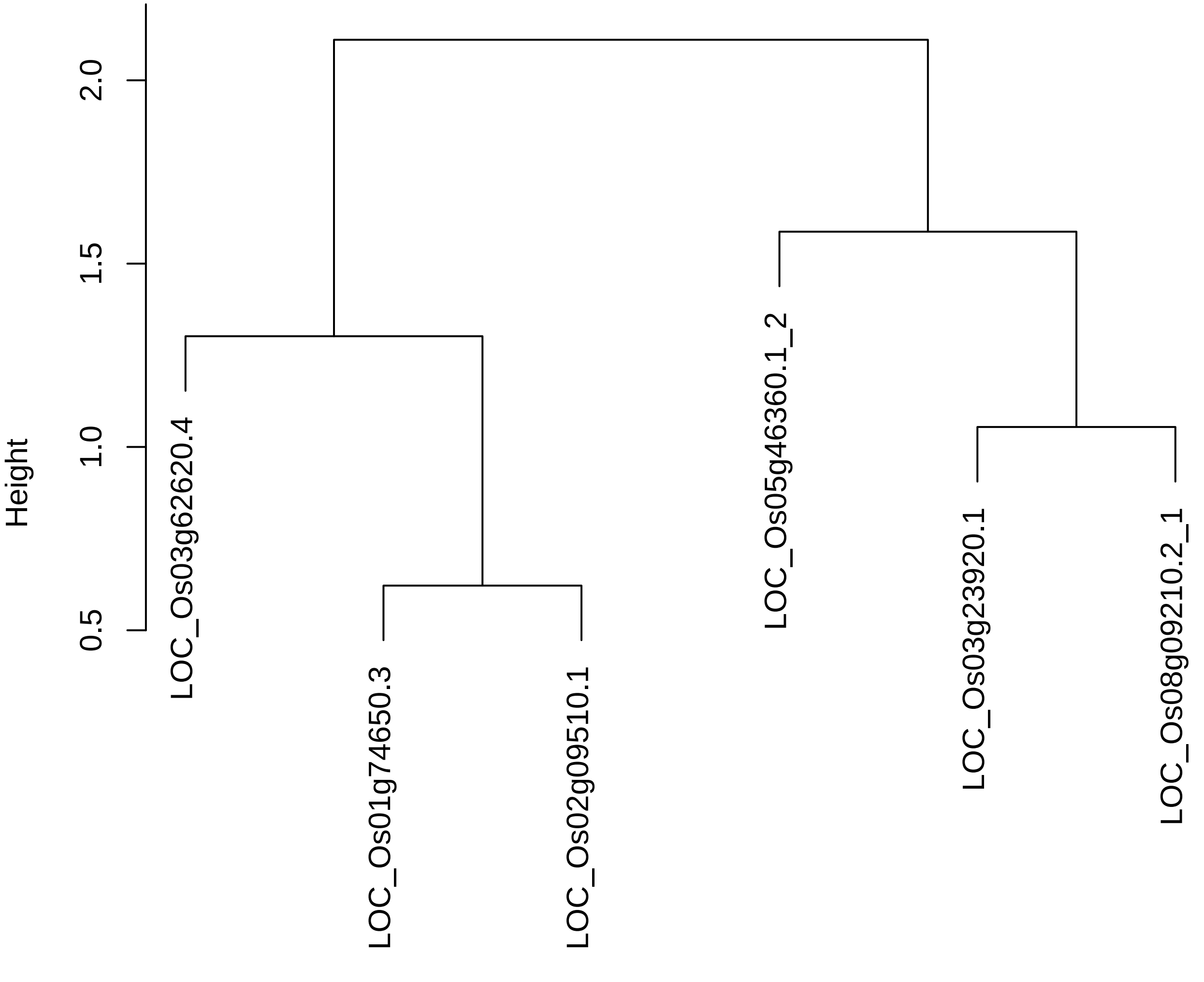


dist(spfitiex12sec12[lab[, 1] == i, ])
hclust (*, "ward.D2")

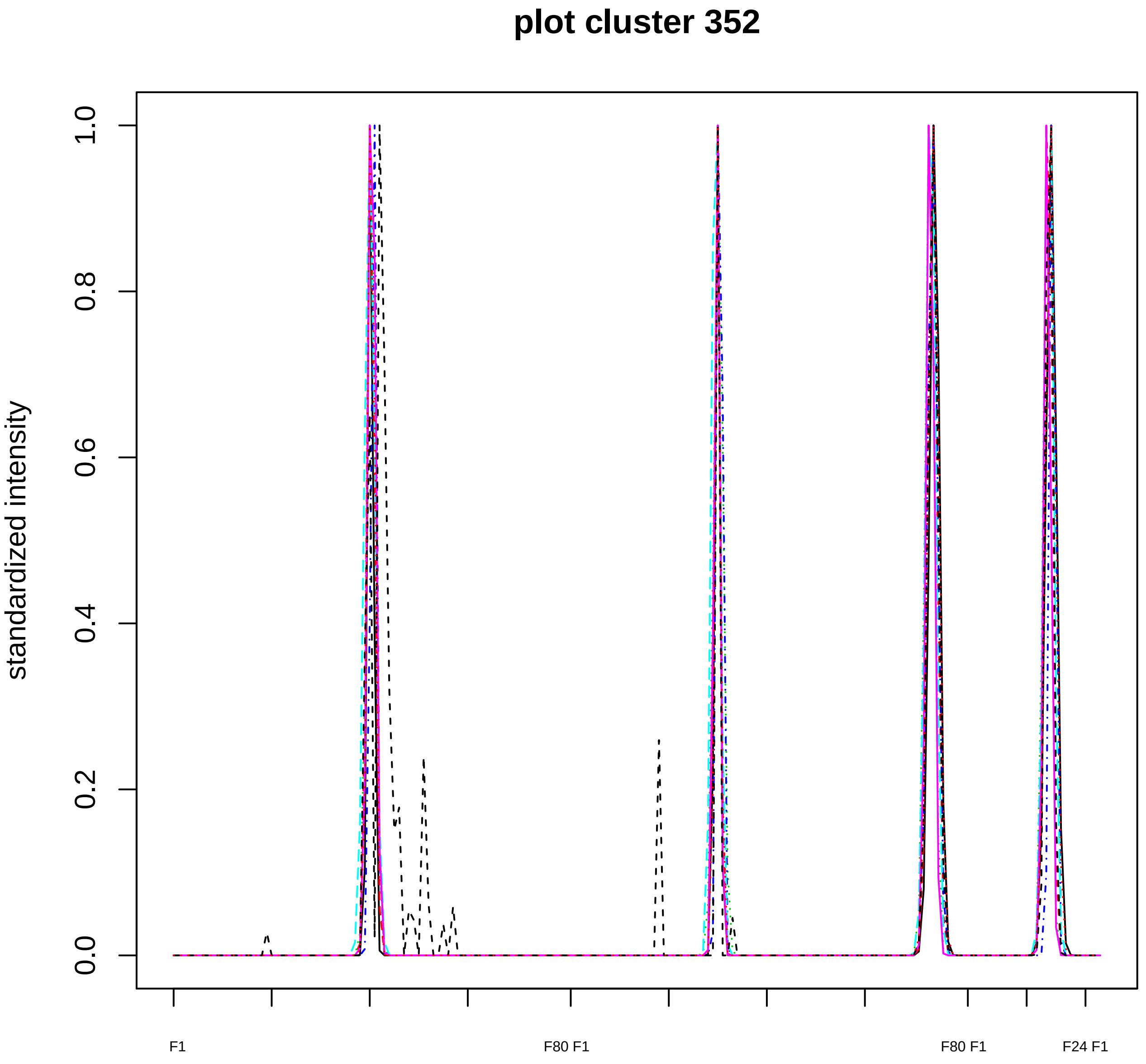
plot cluster 352
standardized intensity
0.0
0.2
0.4
0.6
0.8
1.0
F1
F80 F1
F80 F1
F24 F1

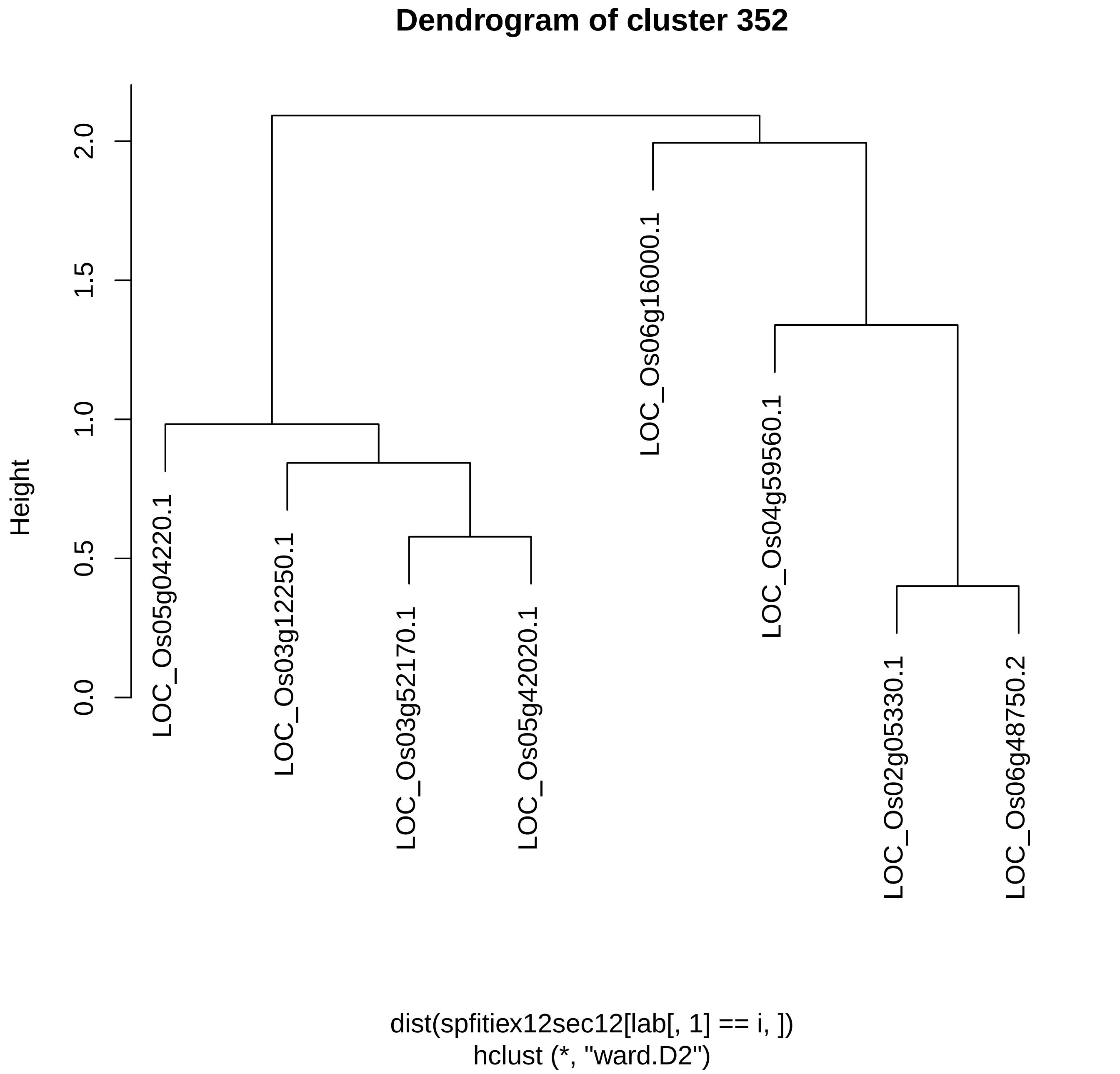

Dendrogram of cluster 352
Height
2.0
1.5
1.0
0.5
0.0
LOC_Os05g04220.1
LOC_Os03g12250.1
LOC_Os03g52170.1
LOC_Os05g42020.1
LOC_Os06g16000.1
LOC_Os04g59560.1
LOC_Os02g05330.1
LOC_Os06g48750.2
dist(spfitiex12sec12[lab[, 1] == i, ])
hclust (*, "ward.D2")

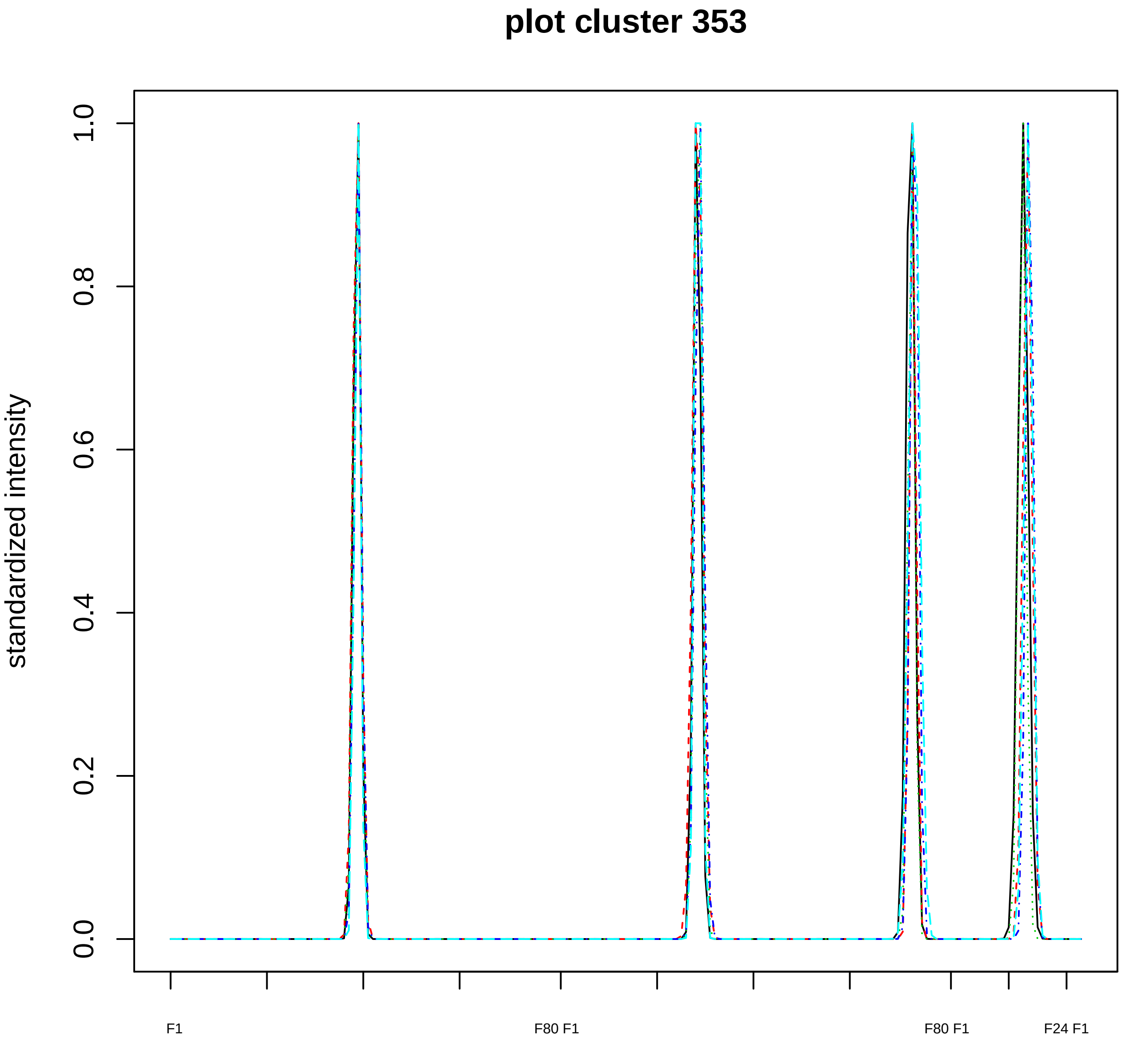

plot cluster 353
standardized intensity
0.0
0.2
0.4
0.6
0.8
1.0
F1
F80 F1
F80 F1
F24 F1

#### Dendrogram of cluster 353

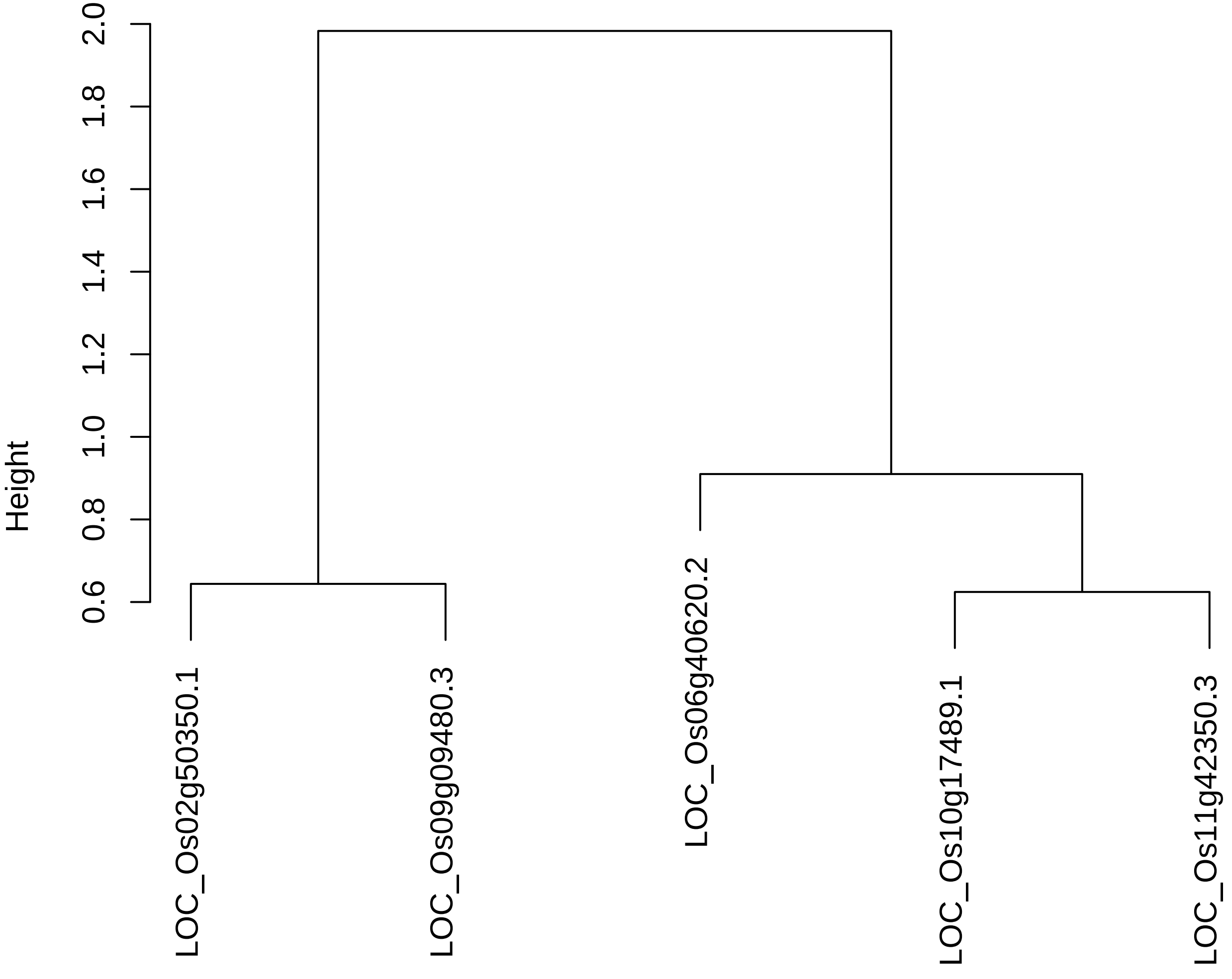


dist(spfitiex12sec12[lab[, 1] == i, ])
hclust (*, "ward.D2")

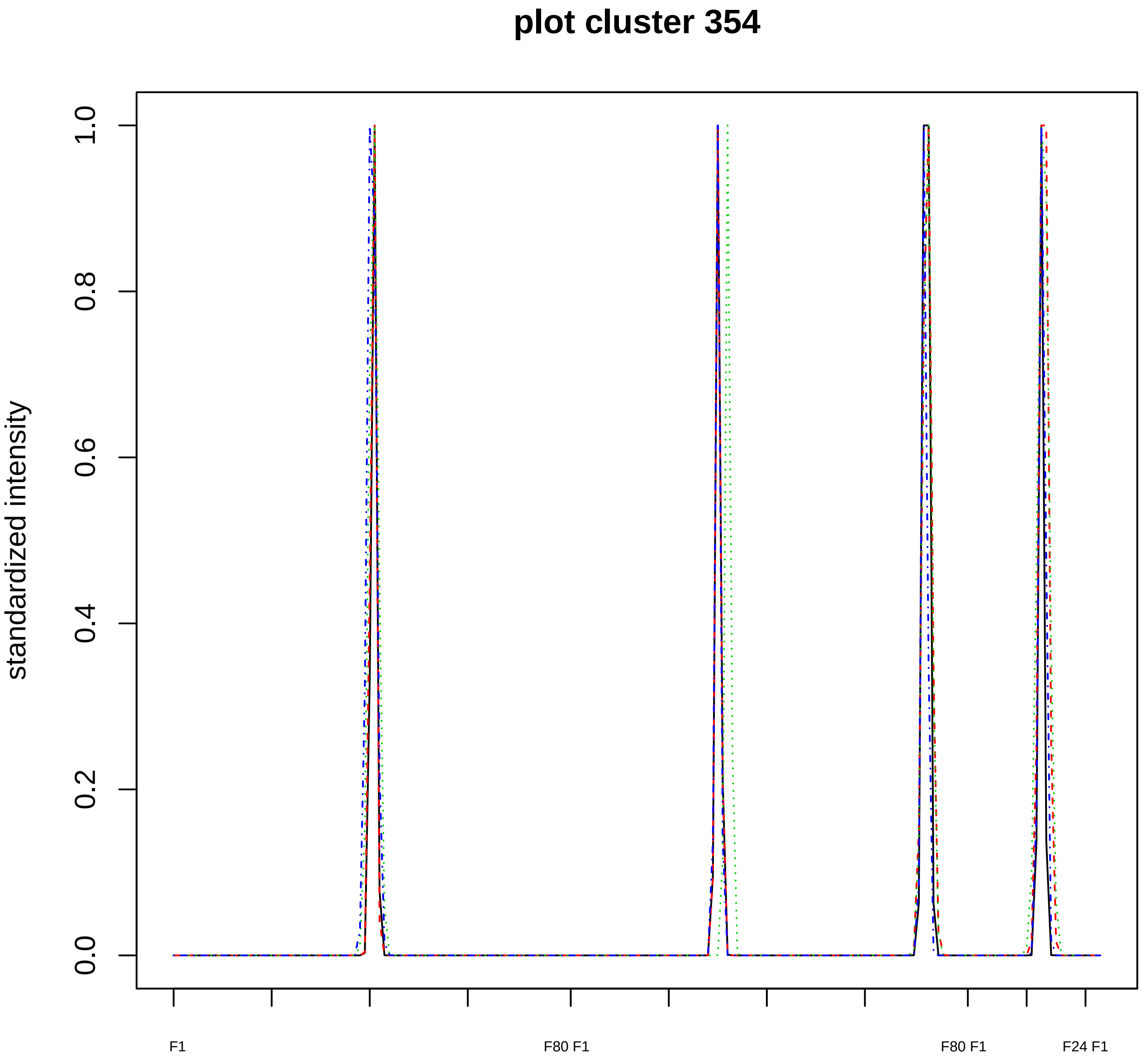
plot cluster 354
standardized intensity
1.0
0.8
0.6
0.4
0.2
0.0
F1
F80 F1
F80 F1
F24 F1

### Dendrogram of cluster 354

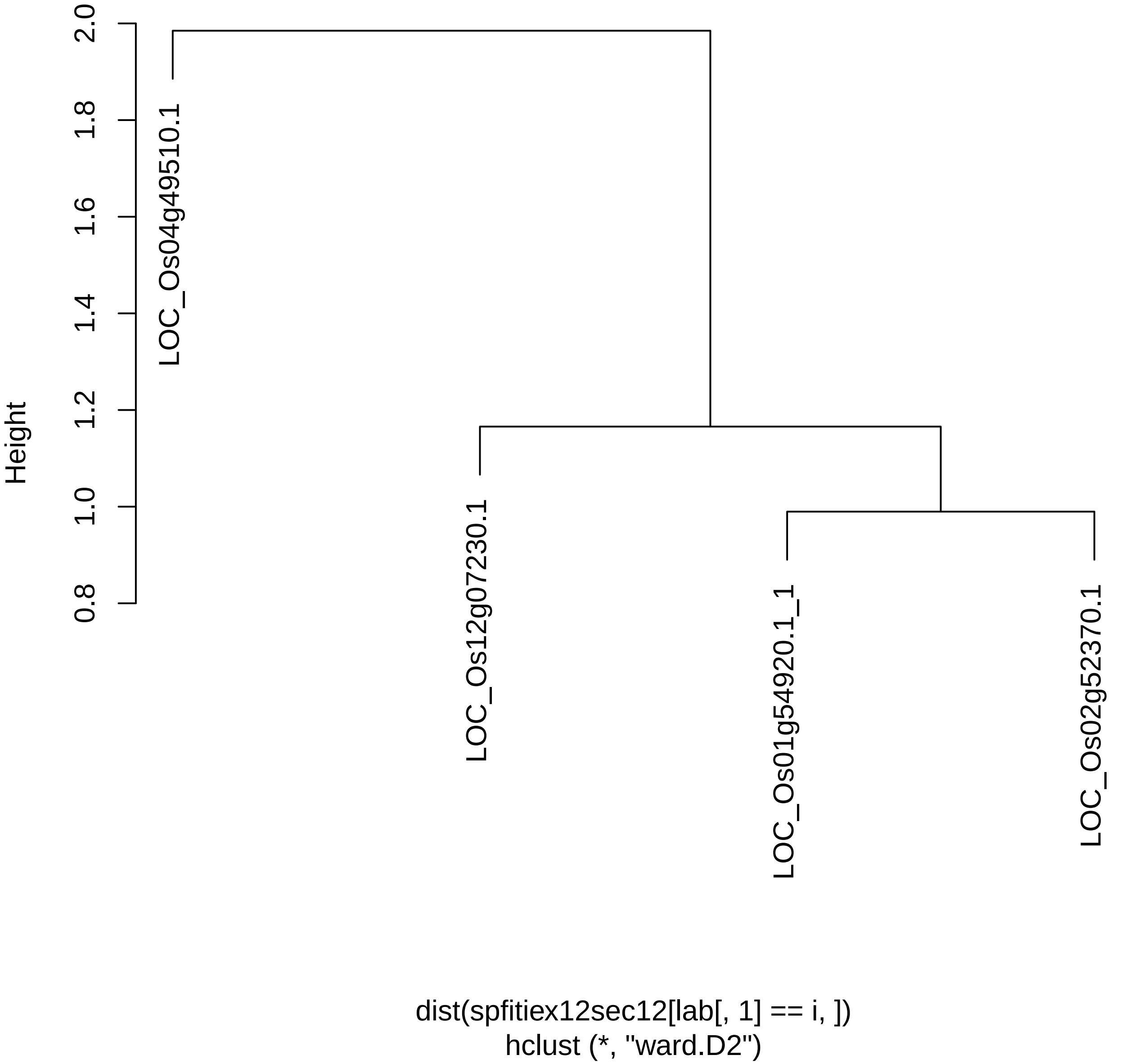

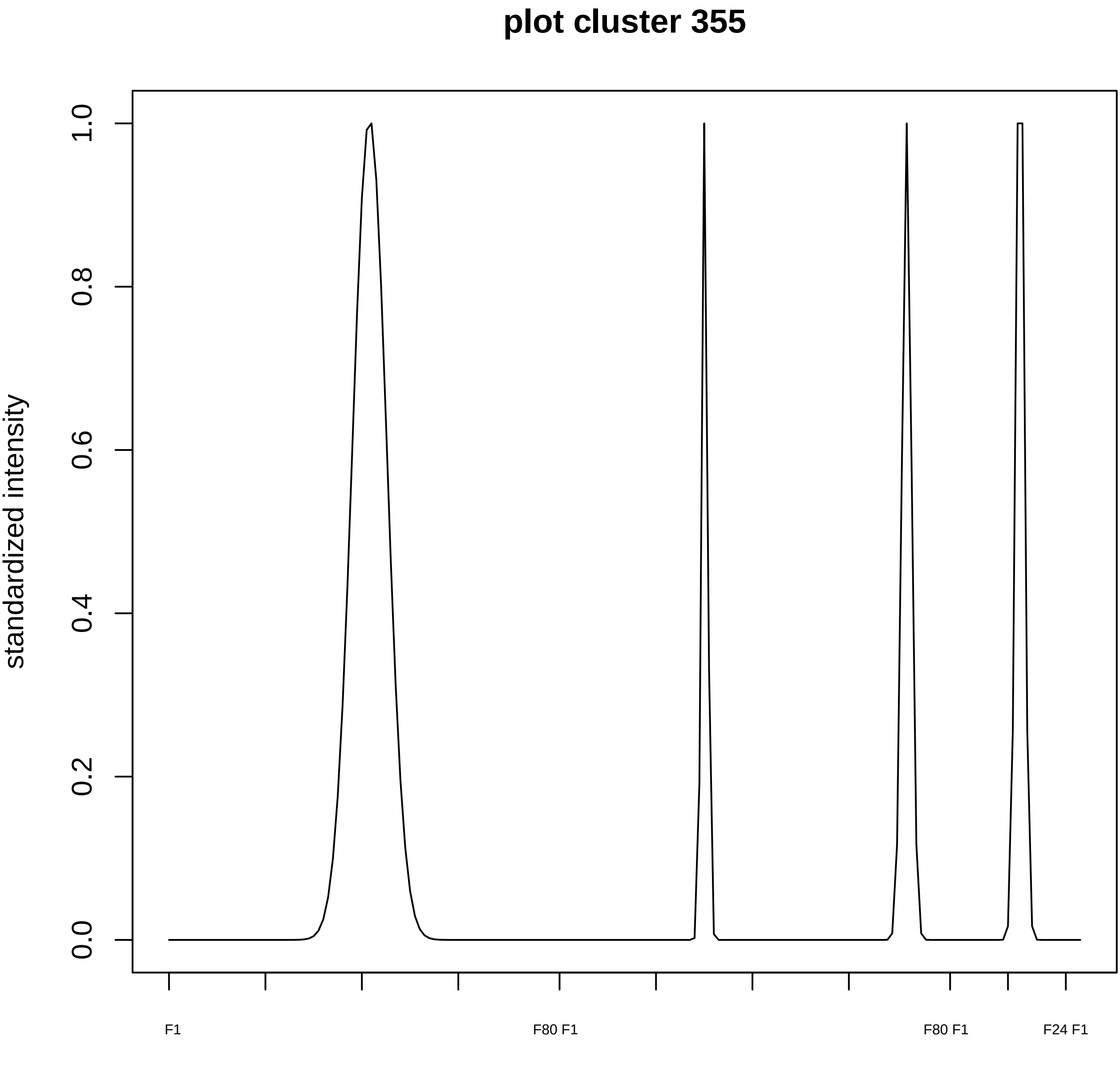

plot cluster 355
standardized intensity
0.0
0.2
0.4
0.6
0.8
1.0
F1
F80 F1
F80 F1
F24 F1

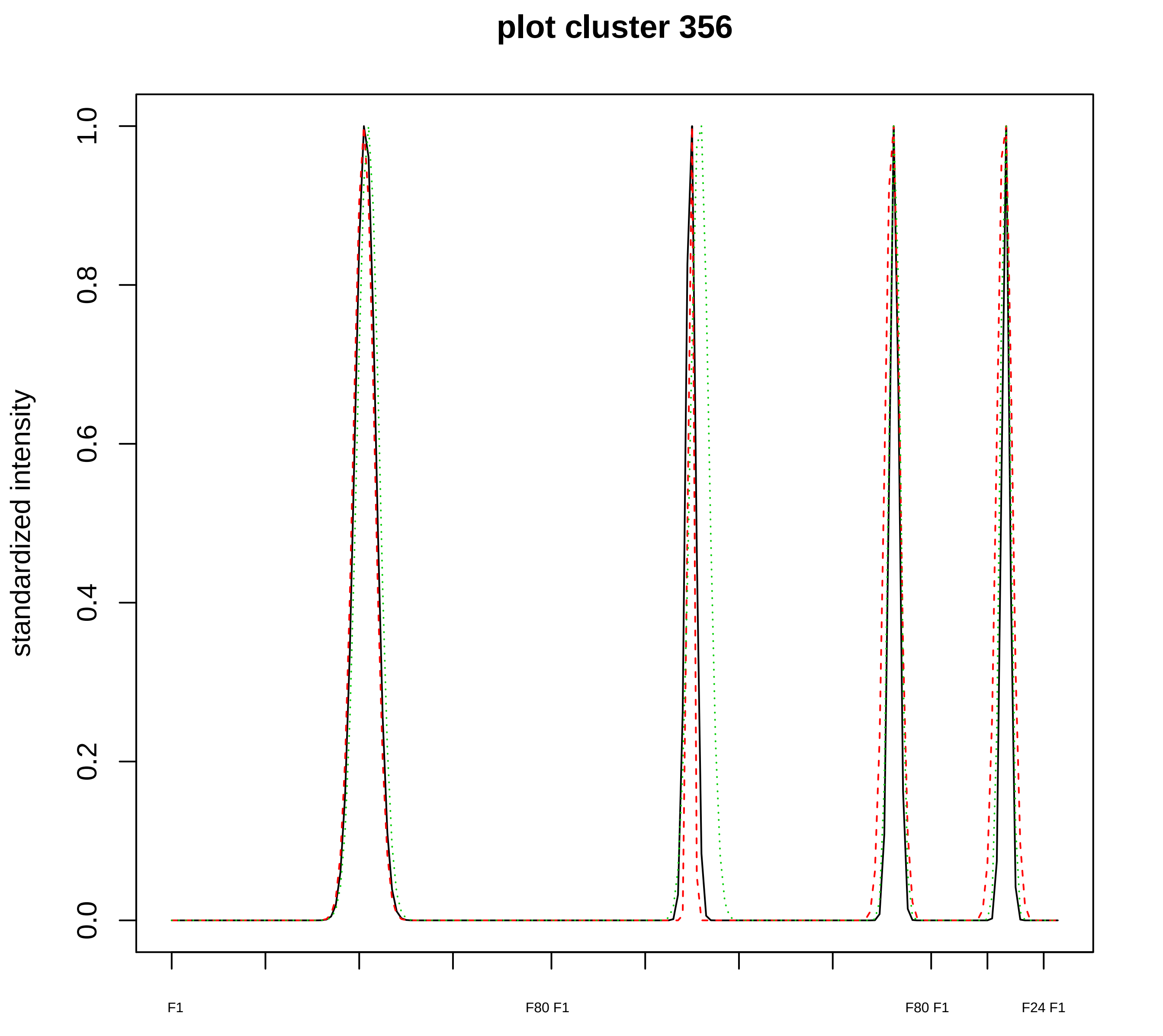

plot cluster 356
standardized intensity
0.0
0.2
0.4
0.6
0.8
1.0
F1
F80 F1
F80 F1
F24 F1

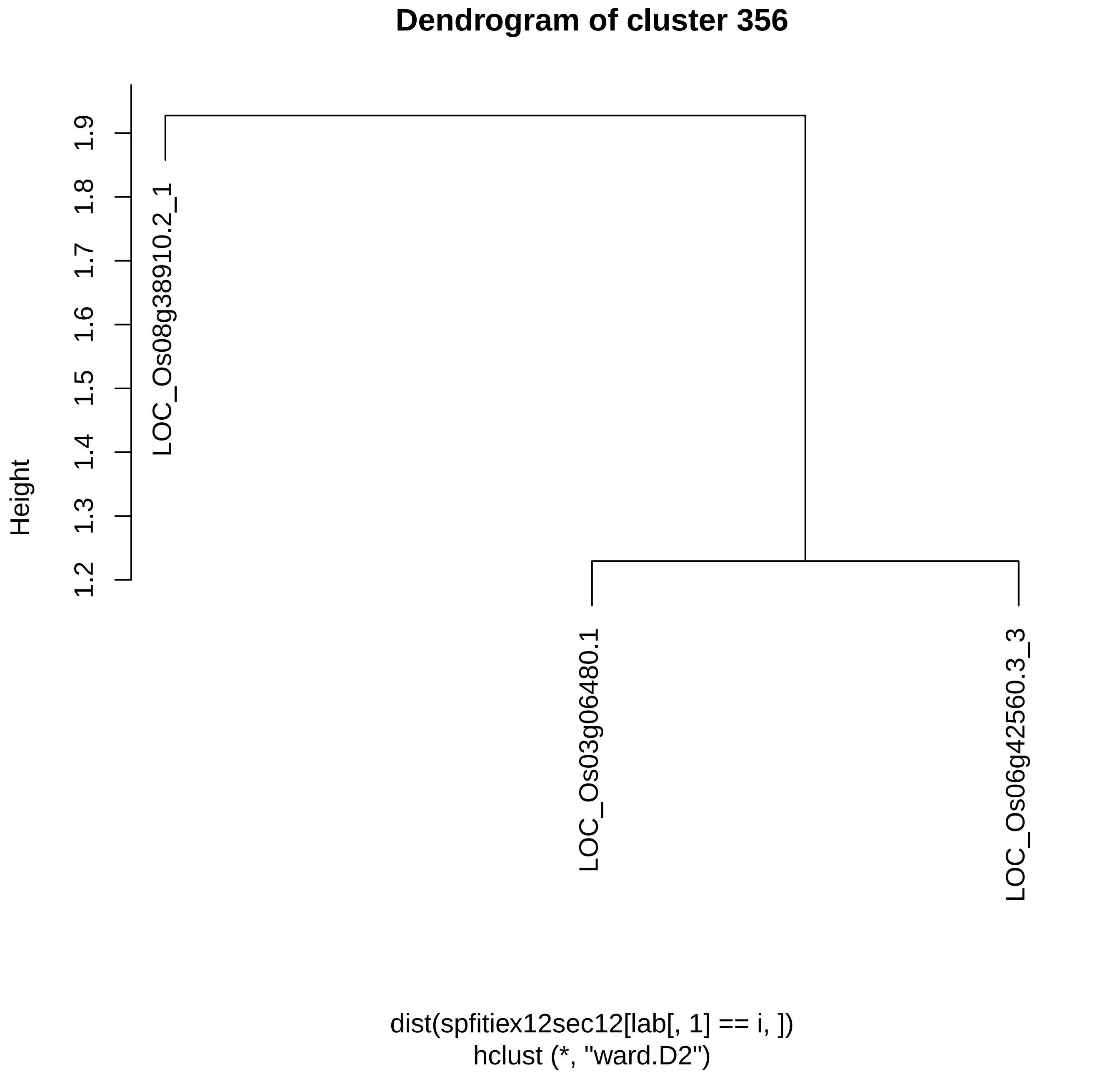
Dendrogram of cluster 356
Height
1.2
1.3
1.4
1.5
1.6
1.7
1.8
1.9
LOC_Os08g38910.2_1
LOC_Os03g06480.1
LOC_Os06g42560.3_3
dist(spfitiex12sec12[lab[, 1] == i, ])
hclust (*, "ward.D2")

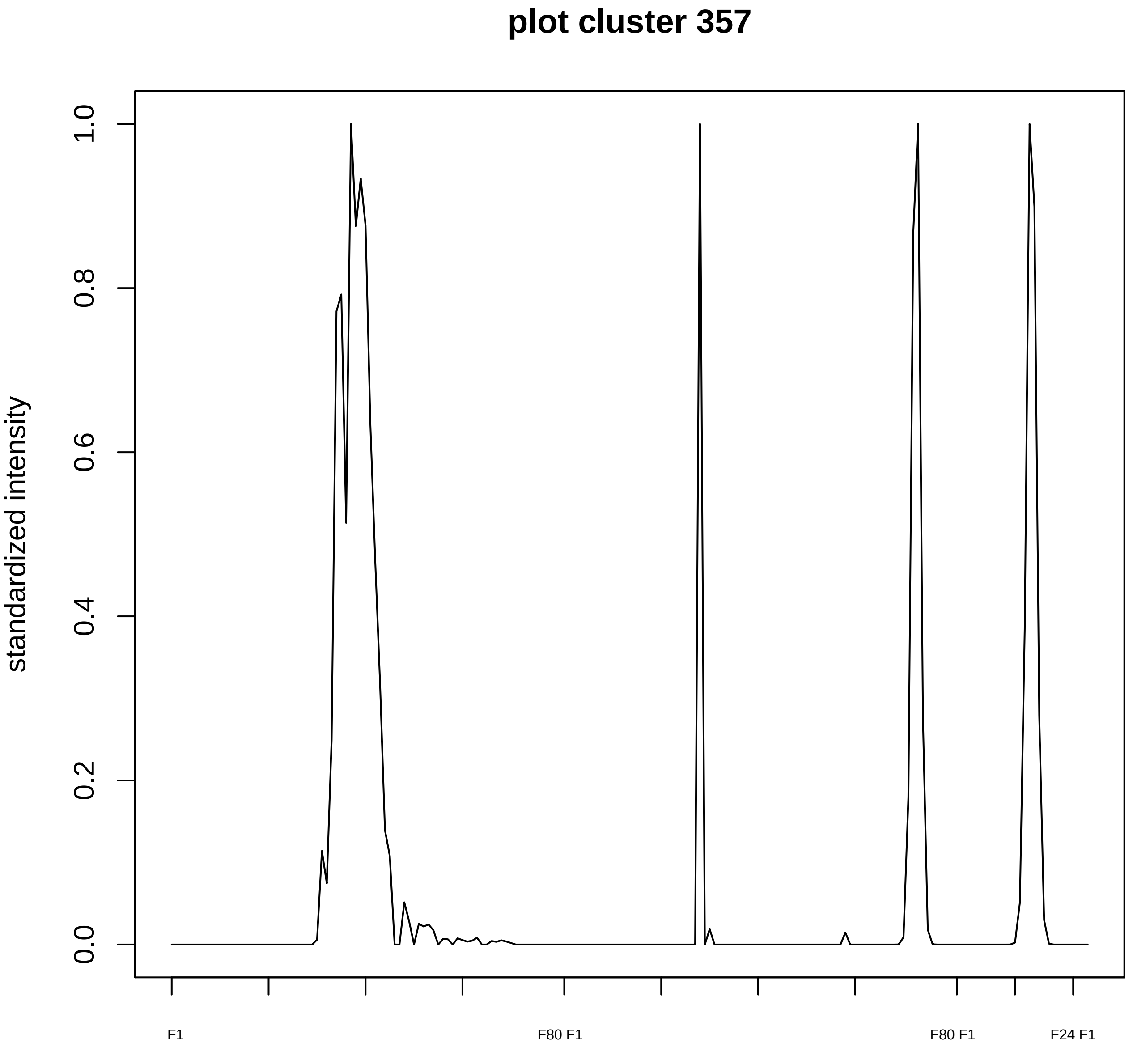

plot cluster 357
standardized intensity
0.0
0.2
0.4
0.6
0.8
1.0
F1
F80 F1
F80 F1
F24 F1

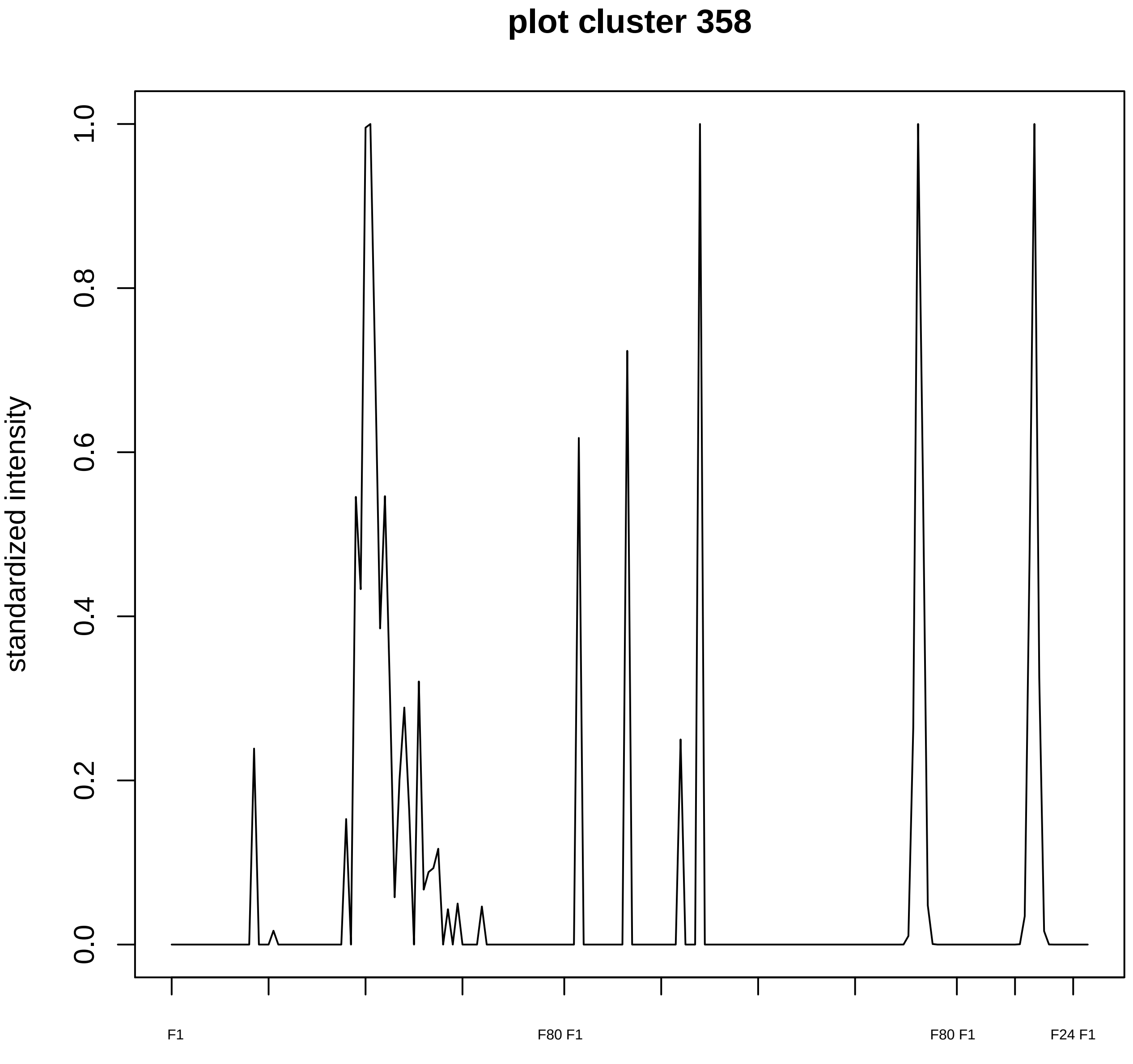

plot cluster 358
standardized intensity
0.0
0.2
0.4
0.6
0.8
1.0
F1
F80 F1
F80 F1
F24 F1

#### plot cluster 359

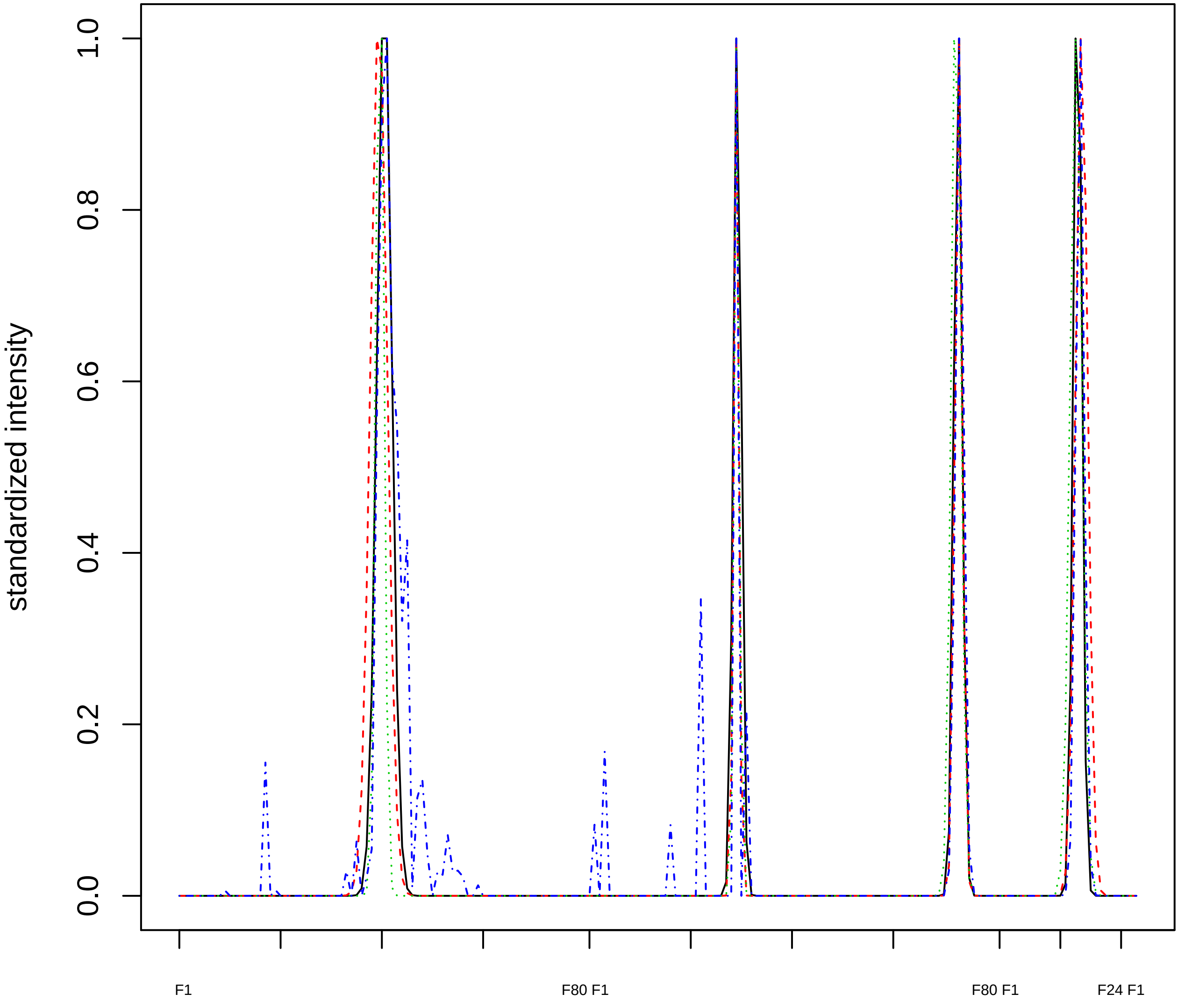

### Dendrogram of cluster 359

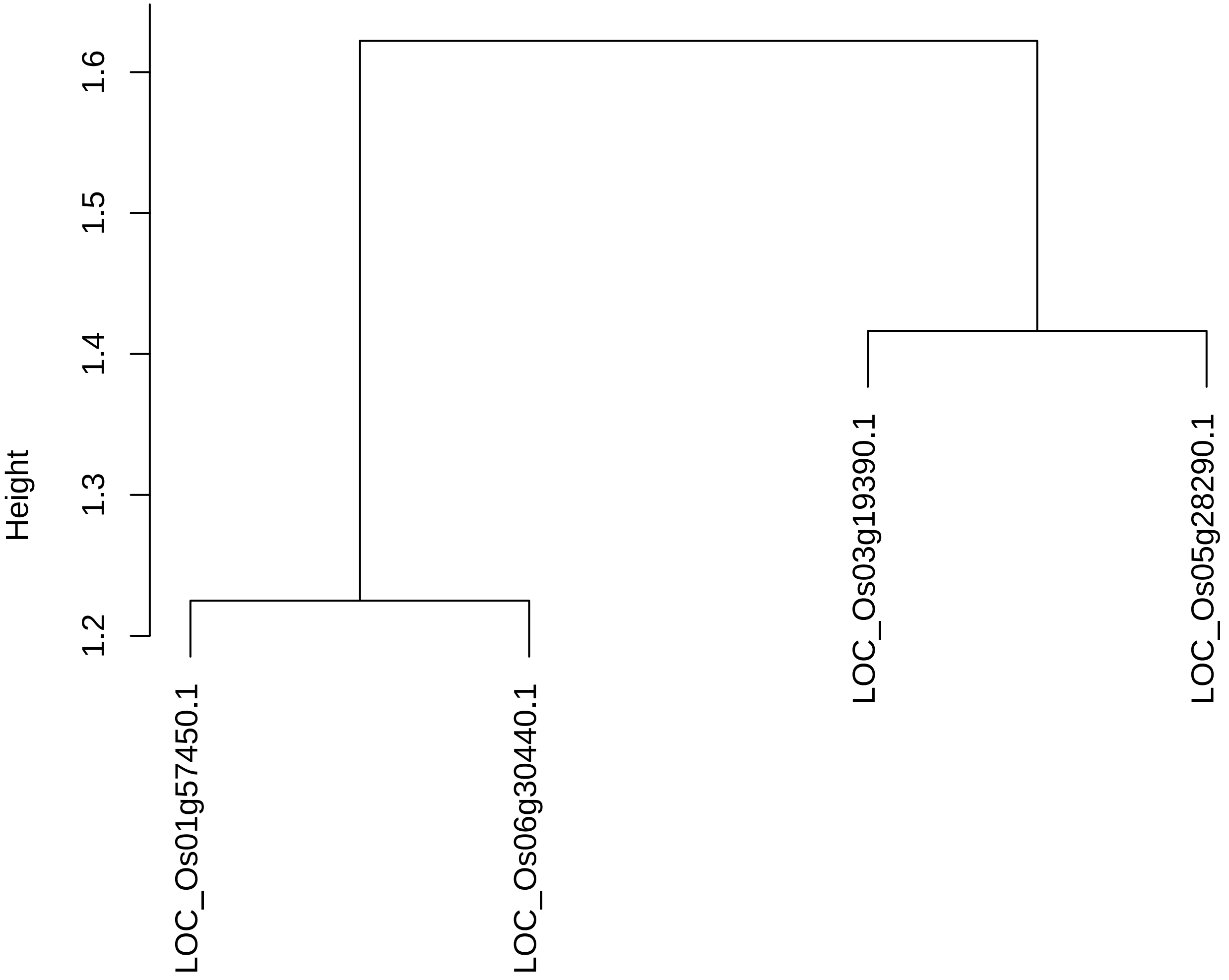


dist(spfitiex12sec12[lab[, 1] == i, ])
hclust (*, "ward.D2")

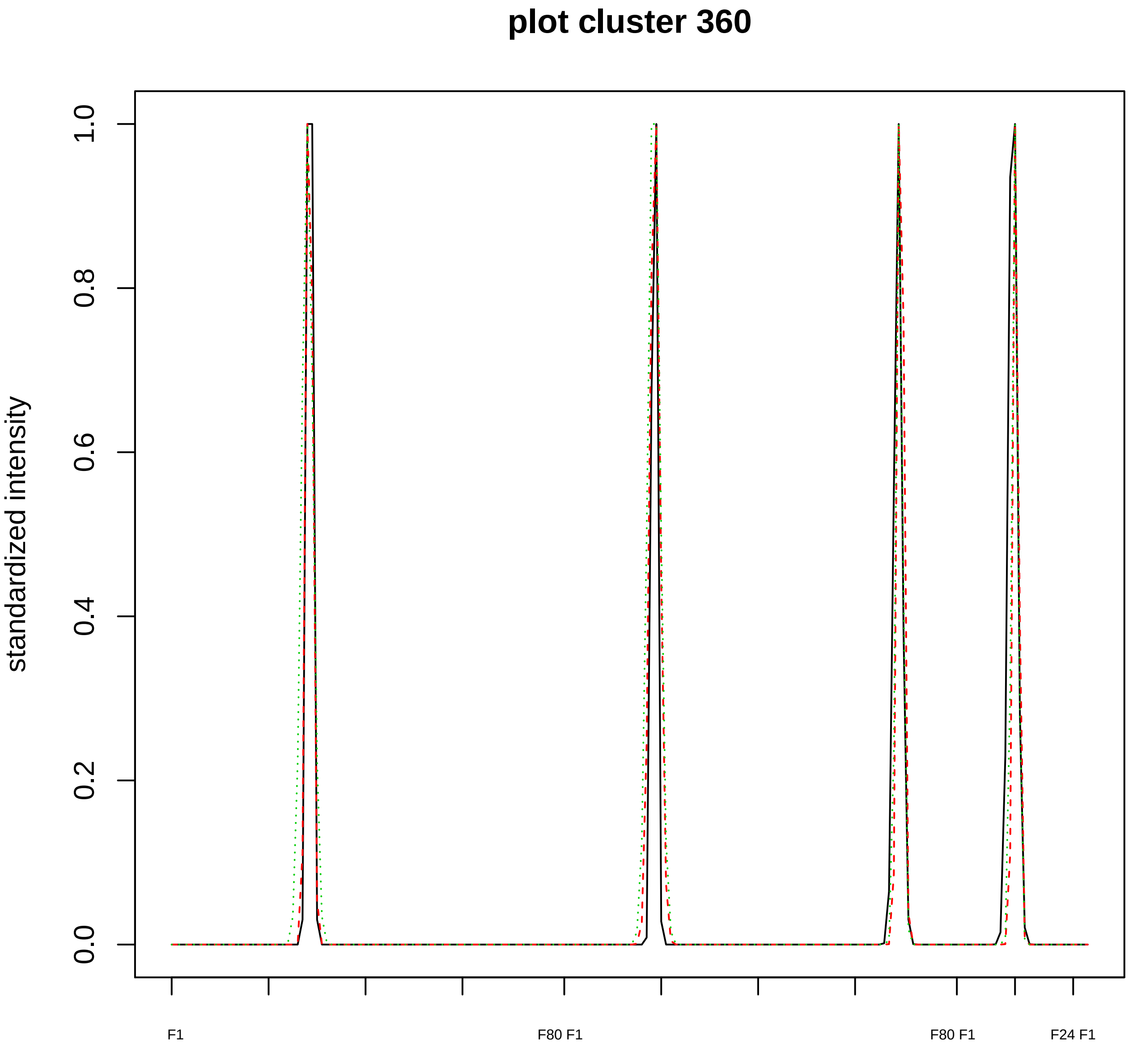
plot cluster 360
standardized intensity
0.0
0.2
0.4
0.6
0.8
1.0
F1
F80 F1
F80 F1
F24 F1

### Dendrogram of cluster 360

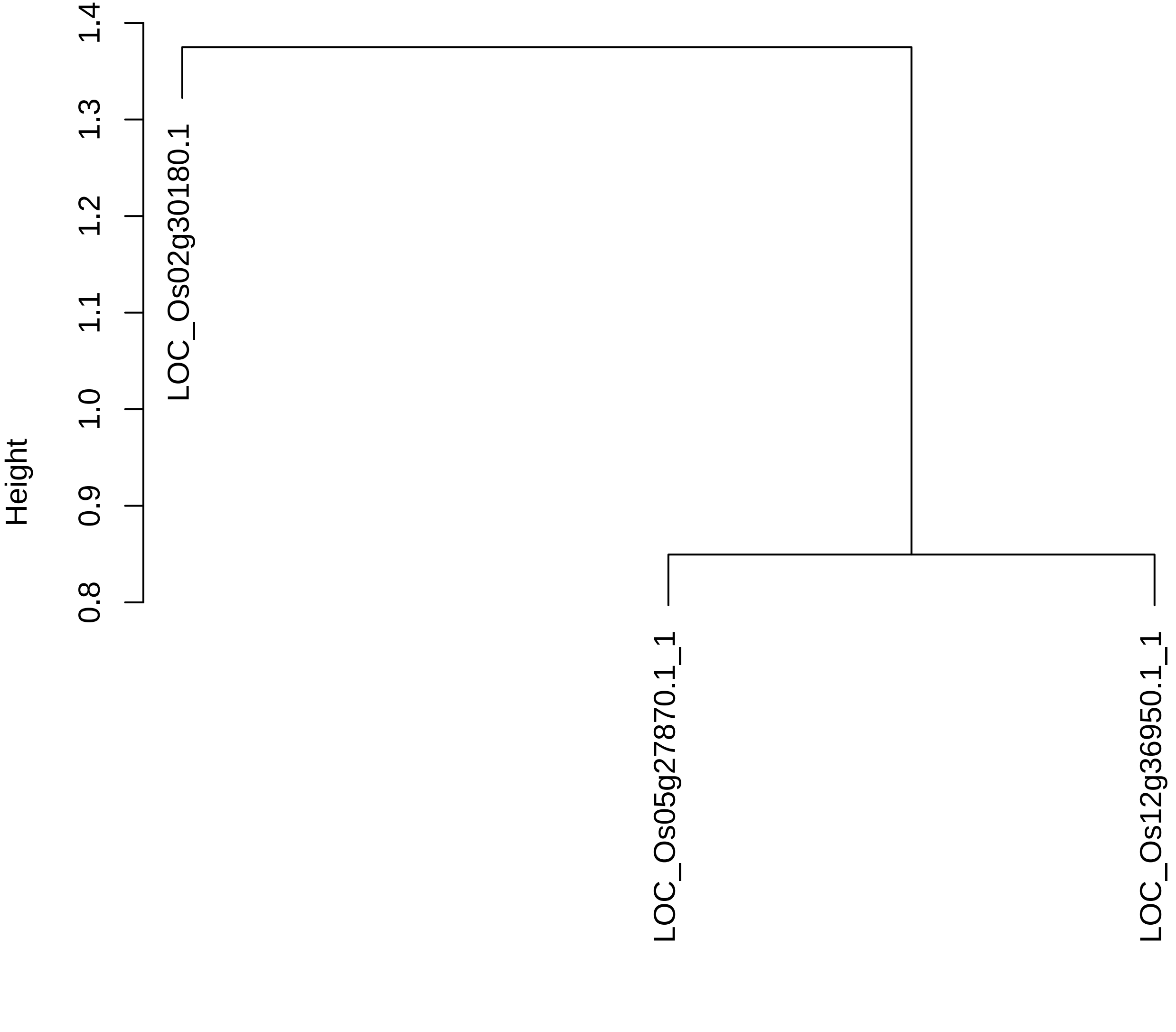


dist(spfitiex12sec12[lab[, 1] == i, ])
hclust (*, "ward.D2")

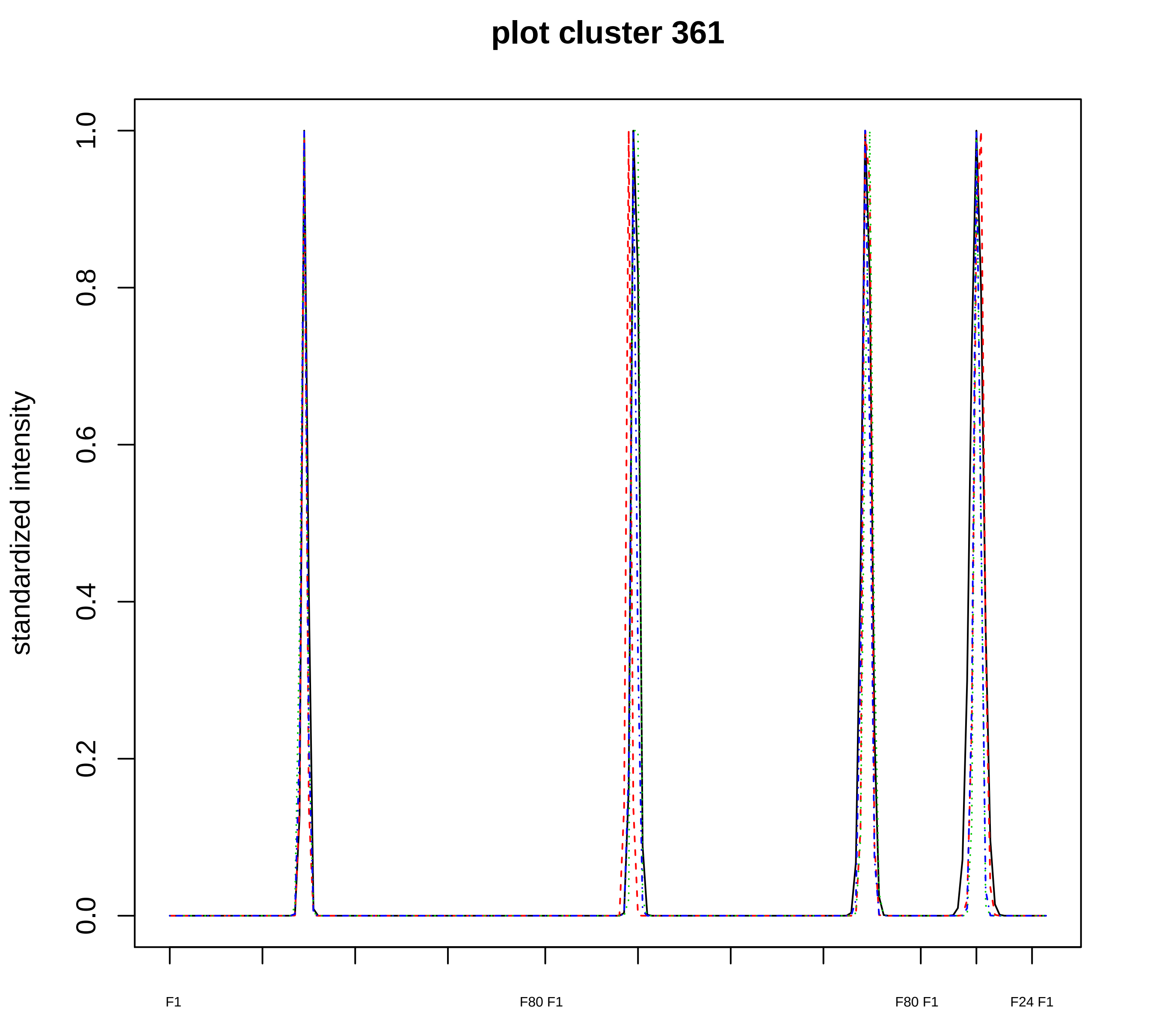

plot cluster 361
standardized intensity
0.0
0.2
0.4
0.6
0.8
1.0
F1
F80 F1
F80 F1
F24 F1

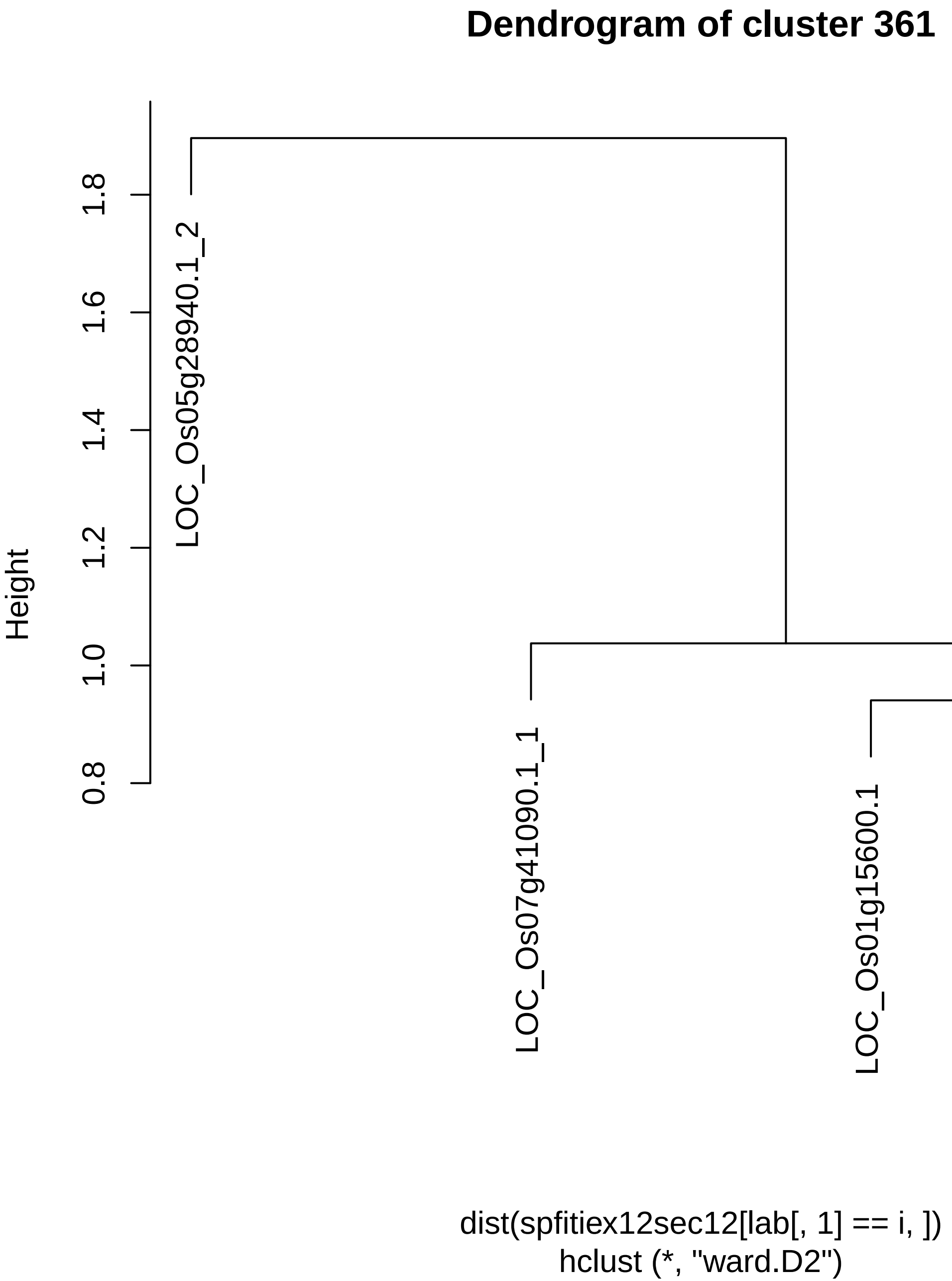
Dendrogram of cluster 361
Height
0.8
1.0
1.2
1.4
1.6
1.8
LOC_Os05g28940.1_2
LOC_Os07g41090.1_1
LOC_Os01g15600.1
LOC_Os07g46460.1
dist(spfitiex12sec12[lab[, 1] == i, ])
hclust (*, "ward.D2")

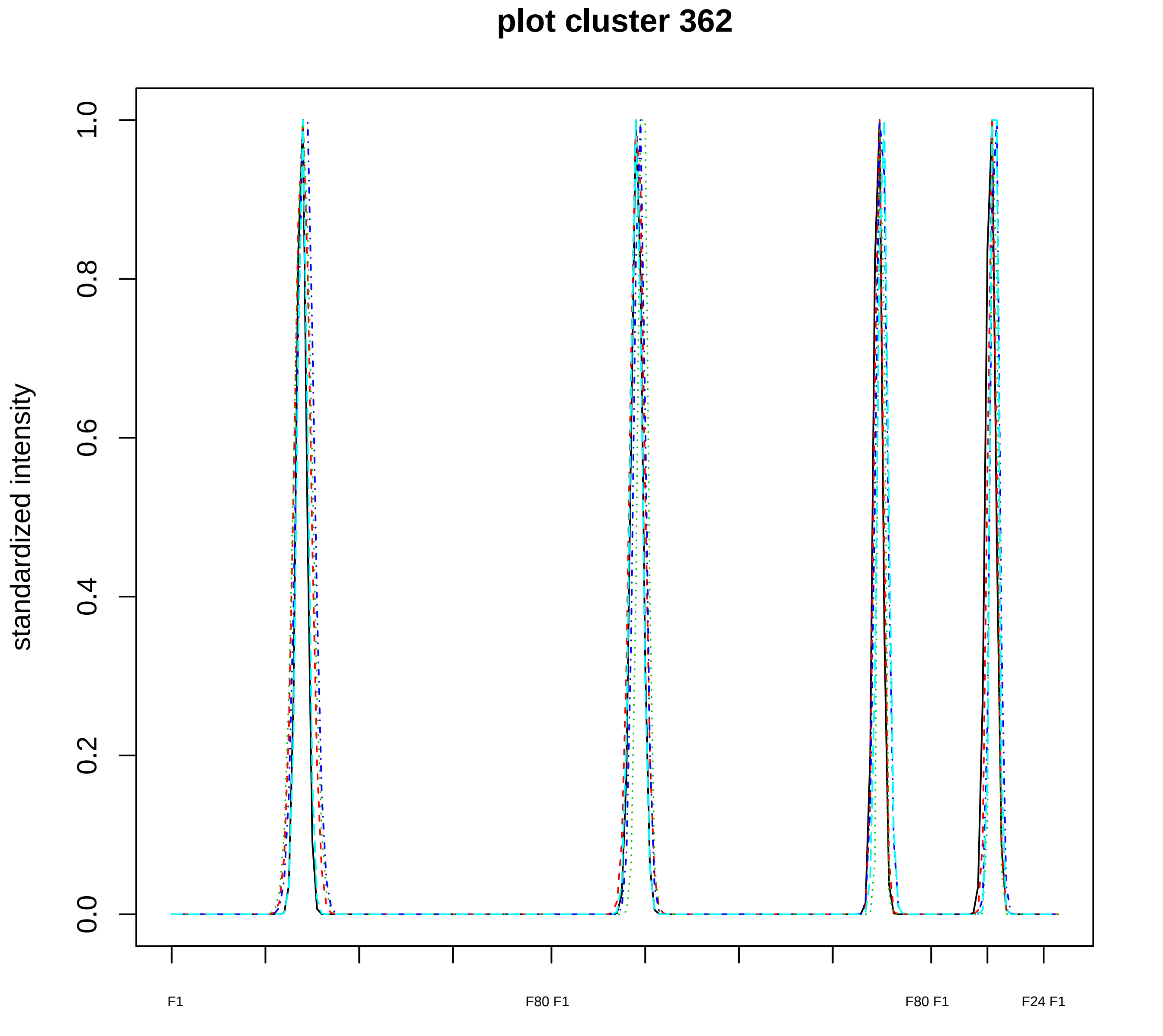

plot cluster 362
standardized intensity
0.0
0.2
0.4
0.6
0.8
1.0
F1
F80 F1
F80 F1
F24 F1

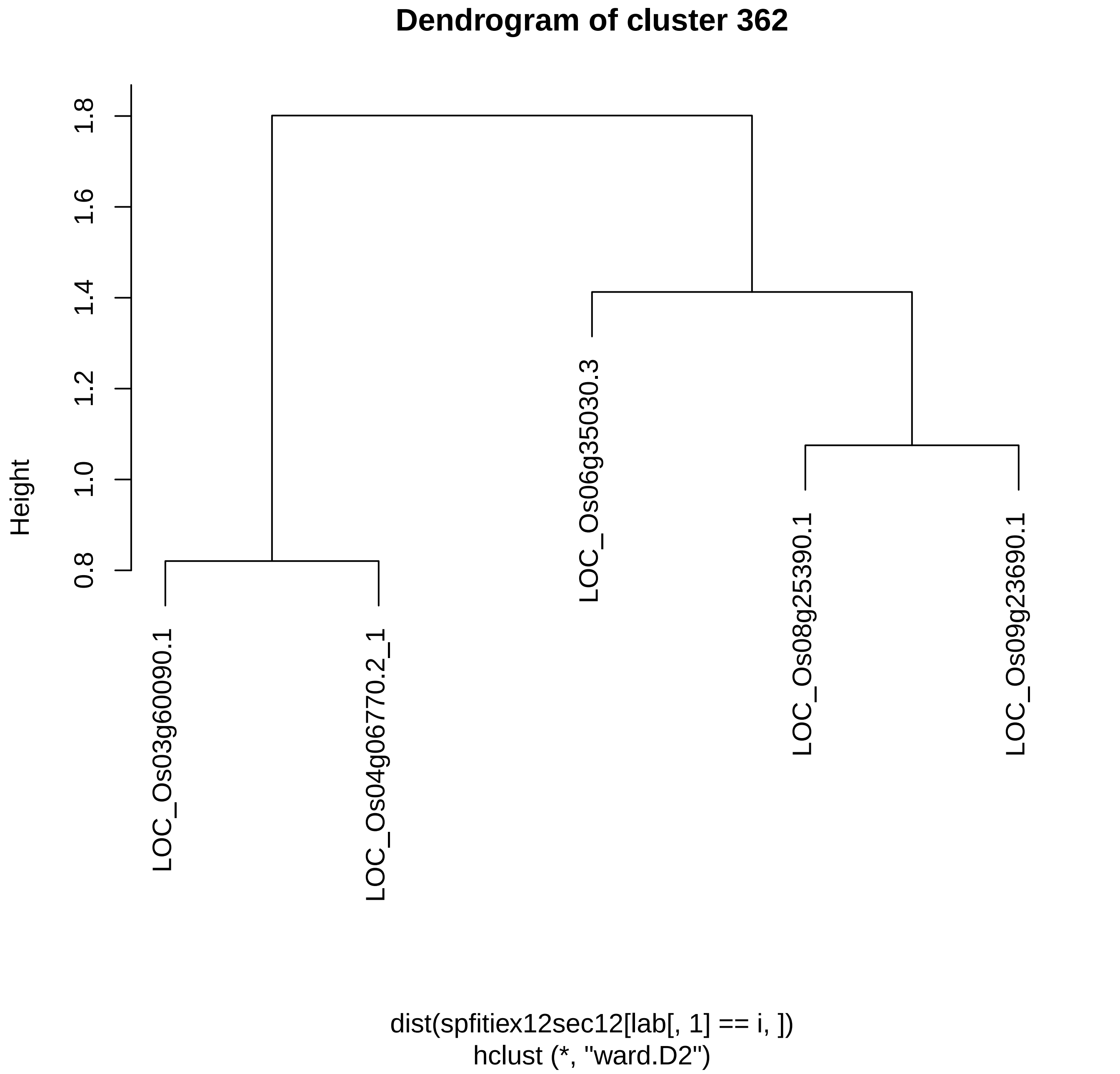
Dendrogram of cluster 362
Height
0.8
1.0
1.2
1.4
1.6
1.8
LOC_Os03g60090.1
LOC_Os04g06770.2_1
LOC_Os06g35030.3
LOC_Os08g25390.1
LOC_Os09g23690.1
dist(spfitiex12sec12[lab[, 1] == i, ])
hclust (*, "ward.D2")

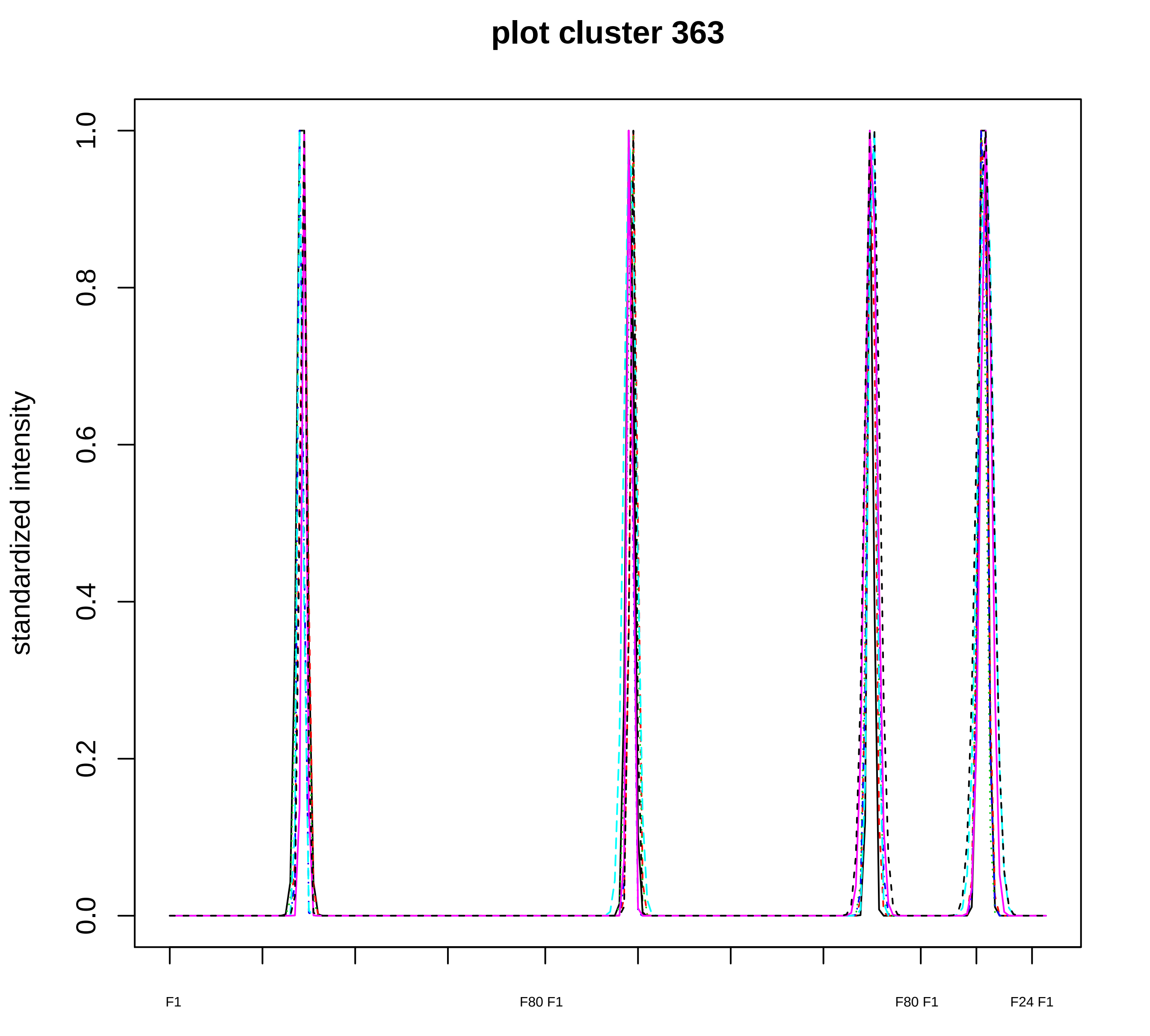

plot cluster 363
standardized intensity
1.0
0.8
0.6
0.4
0.2
0.0
F1
F80 F1
F80 F1
F24 F1

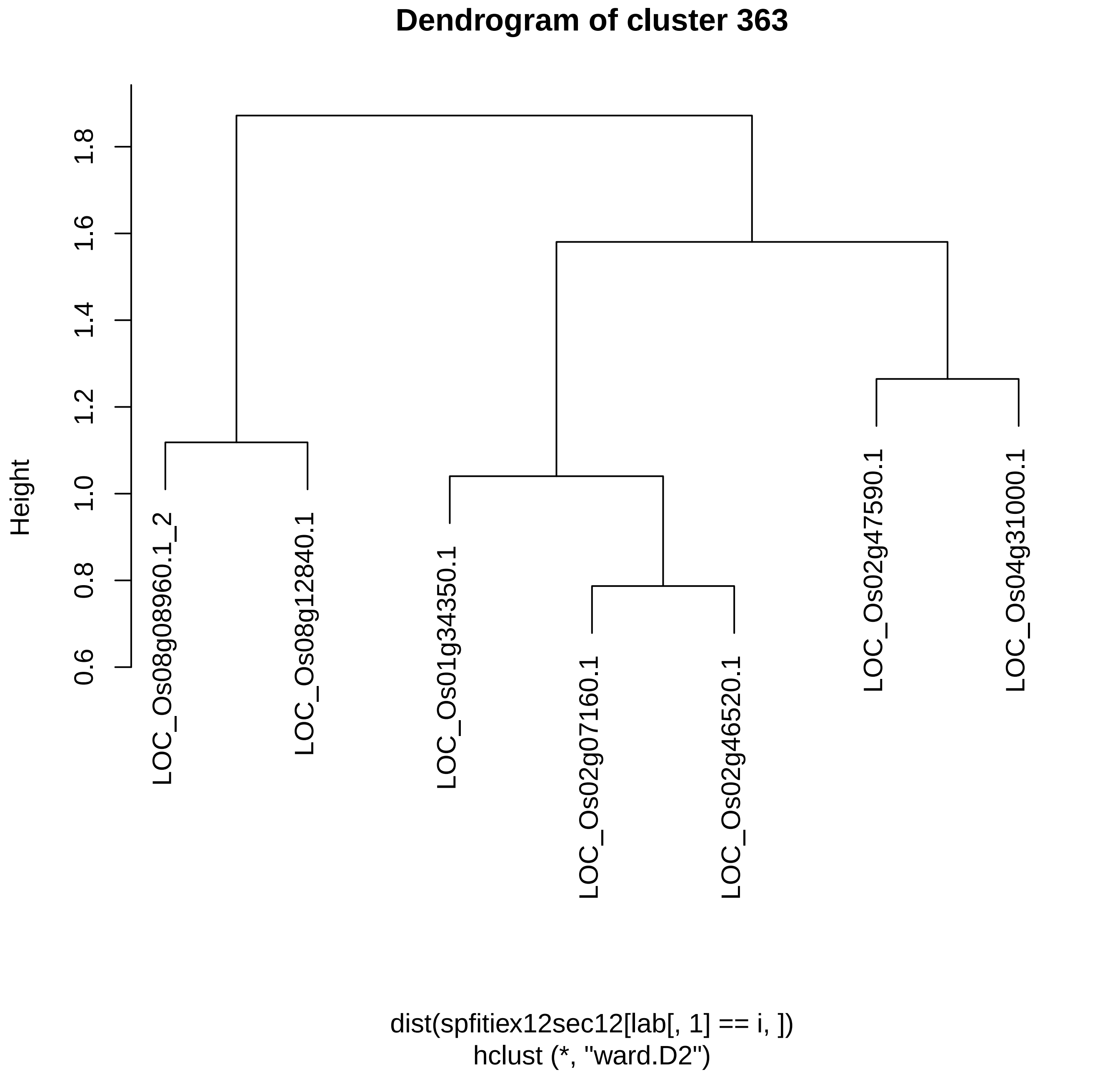
Dendrogram of cluster 363
Height
0.6
0.8
1.0
1.2
1.4
1.6
1.8
LOC_Os08g08960.1_2
LOC_Os08g12840.1
LOC_Os01g34350.1
LOC_Os02g07160.1
LOC_Os02g46520.1
LOC_Os02g47590.1
LOC_Os04g31000.1
dist(spfitiex12sec12[lab[, 1] == i, ])
hclust (*, "ward.D2")

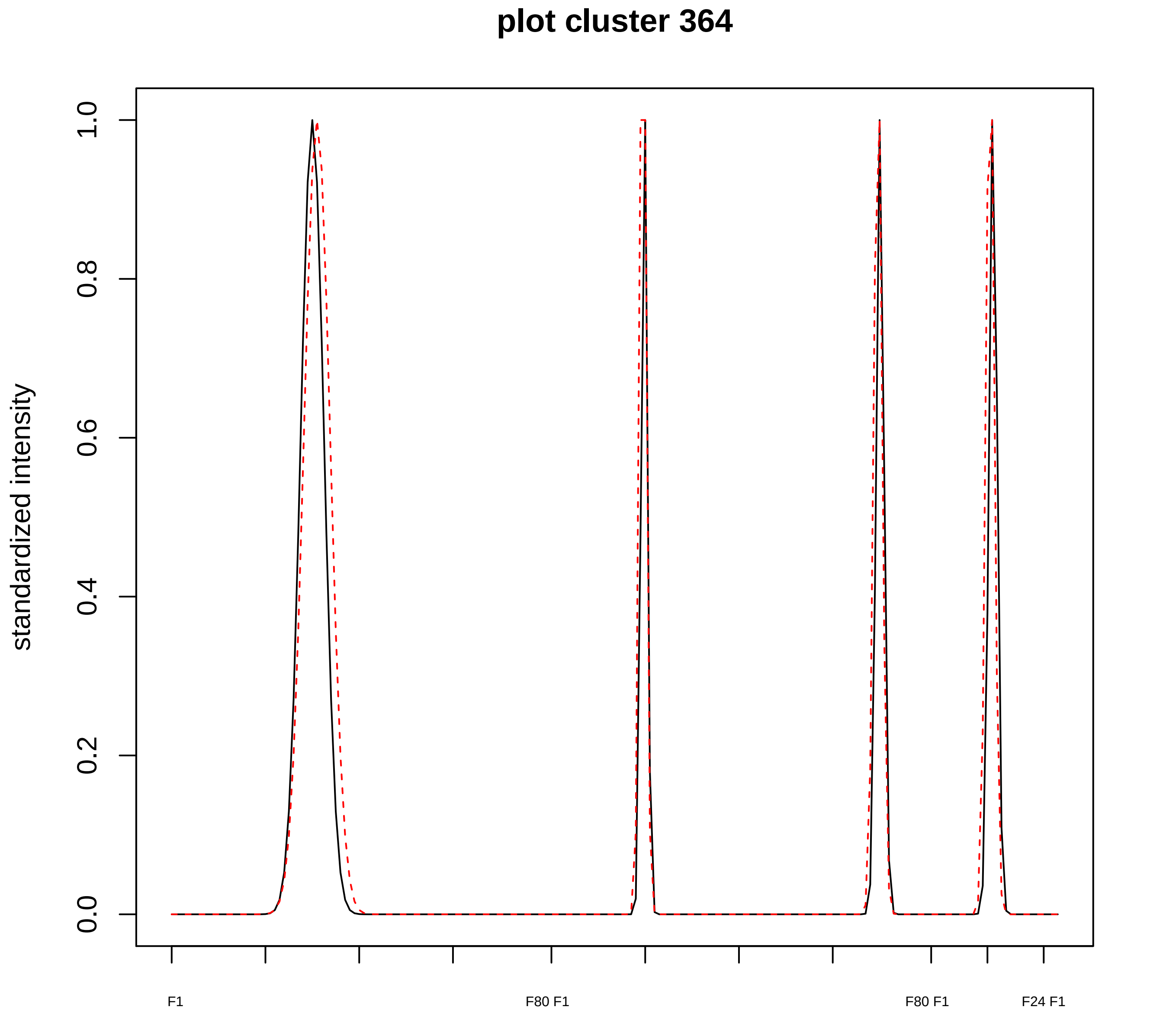
plot cluster 364
standardized intensity
0.0
0.2
0.4
0.6
0.8
1.0
F1
F80 F1
F80 F1
F24 F1

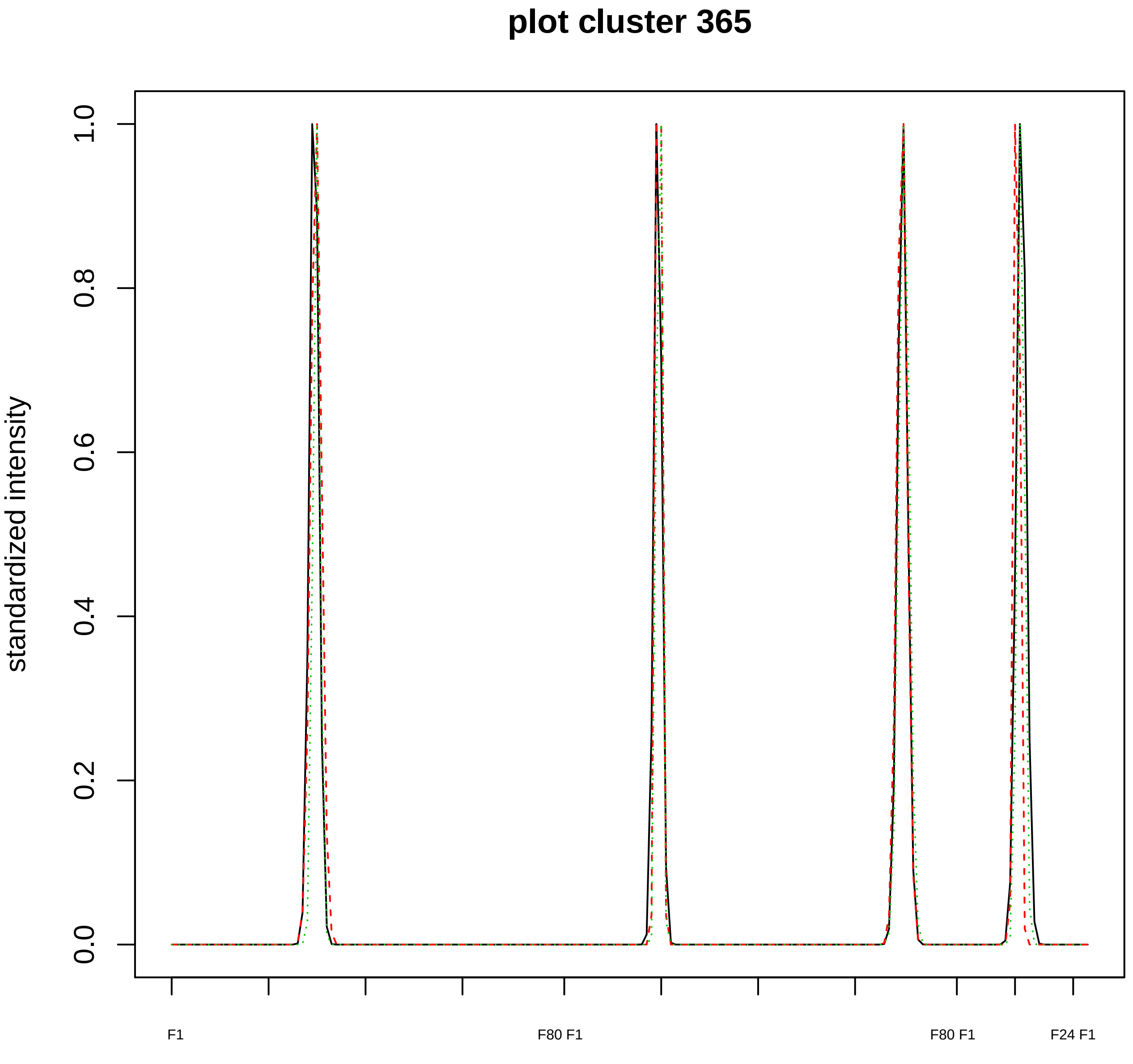
plot cluster 365
standardized intensity
0.0
0.2
0.4
0.6
0.8
1.0
F1
F80 F1
F80 F1
F24 F1

### Dendrogram of cluster 365

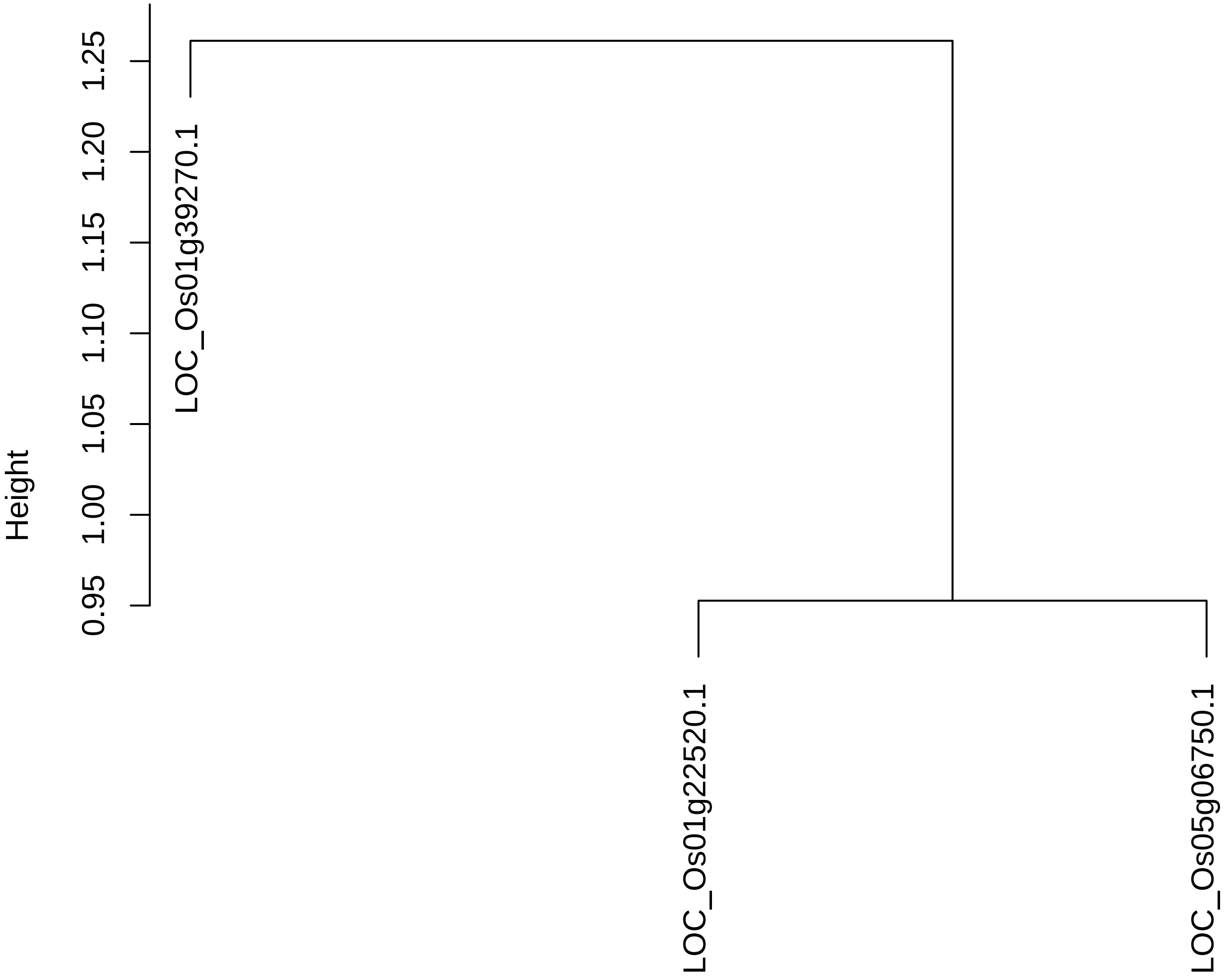


dist(spfitiex12sec12[lab[, 1] == i, ])
hclust (*, "ward.D2")

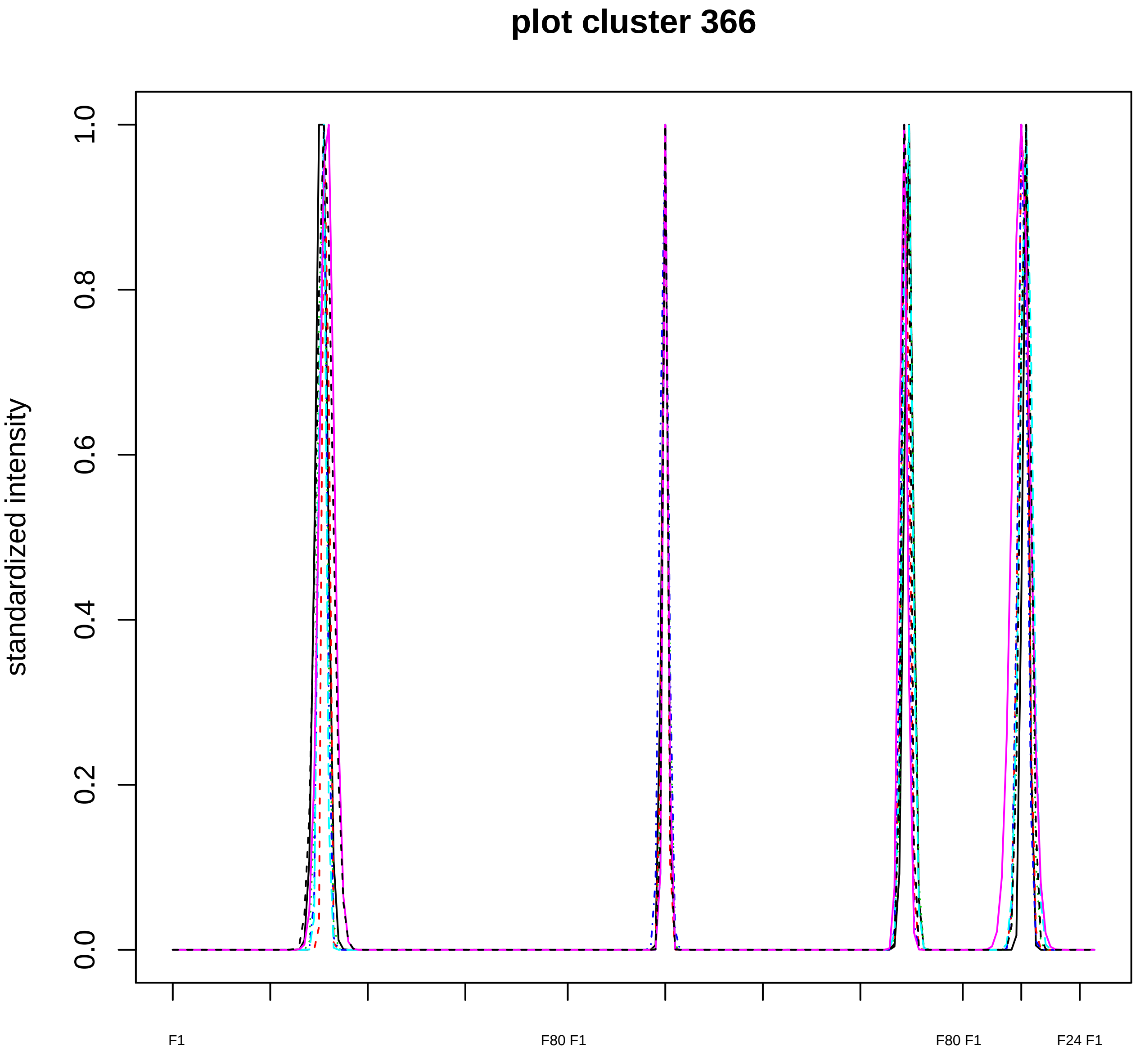
plot cluster 366
standardized intensity
0.0
0.2
0.4
0.6
0.8
1.0
F1
F80 F1
F80 F1
F24 F1

### Dendrogram of cluster 366

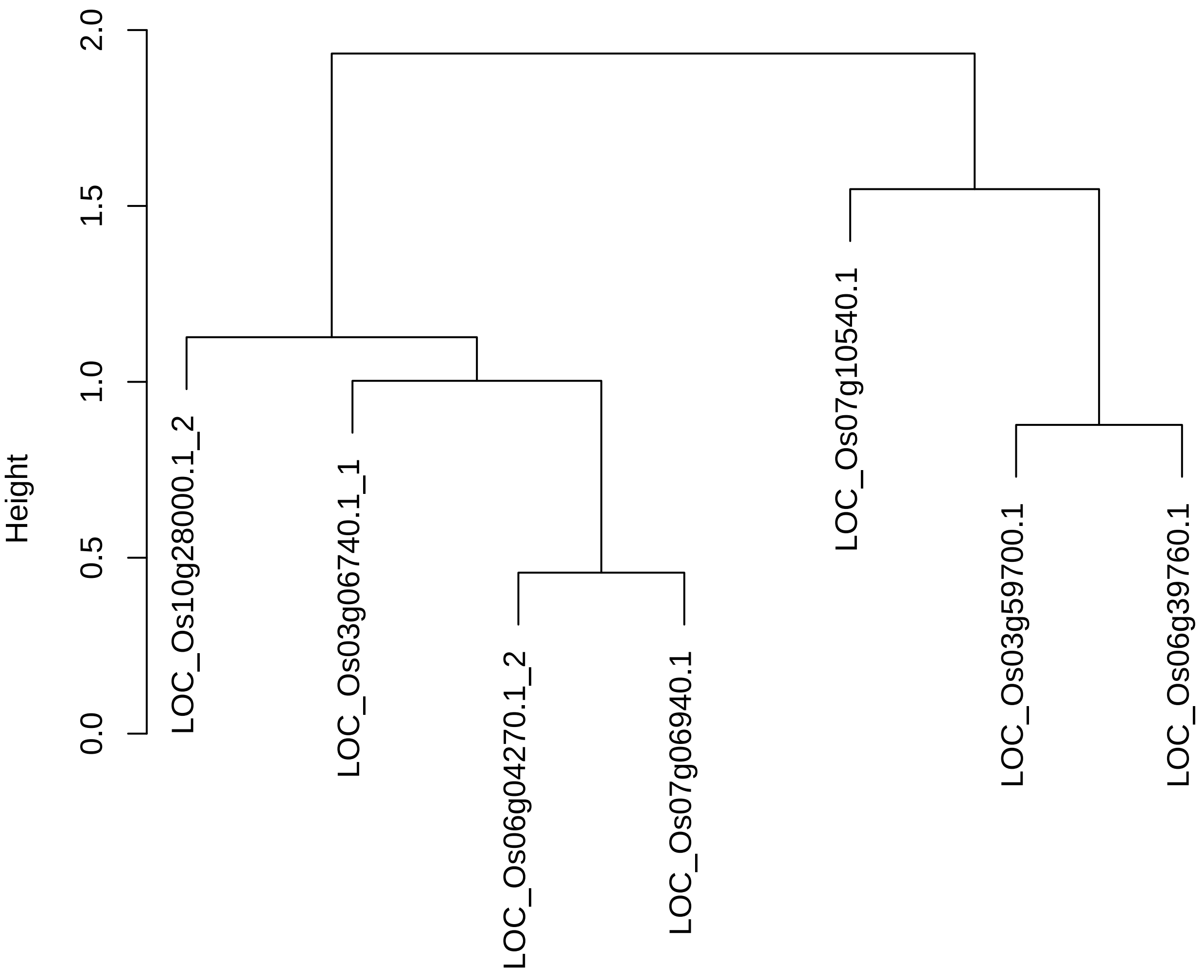


dist(spfitiex12sec12[lab[, 1] == i, ])
hclust (*, "ward.D2")

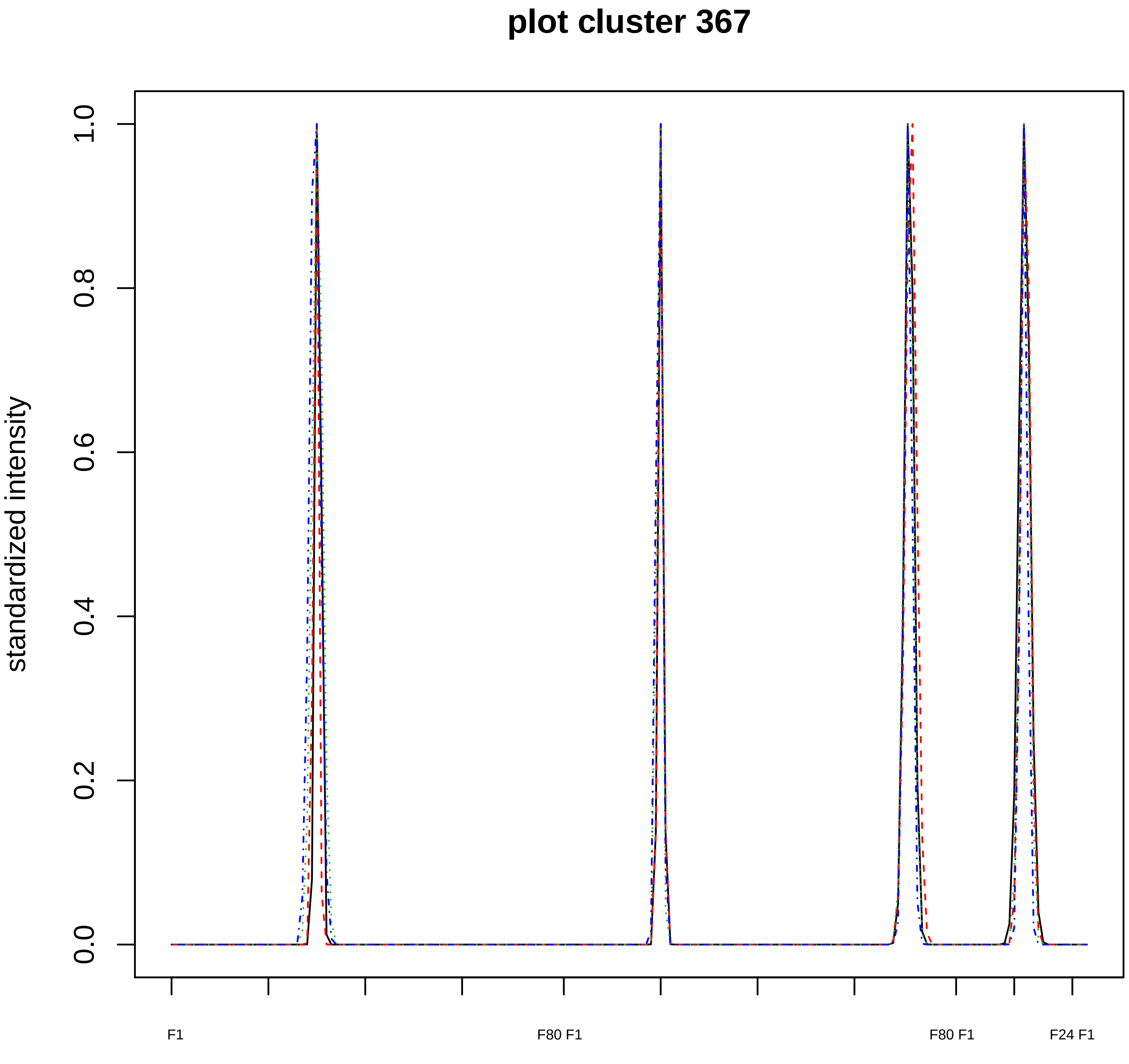
plot cluster 367
standardized intensity
0.0
0.2
0.4
0.6
0.8
1.0
F1
F80 F1
F80 F1
F24 F1

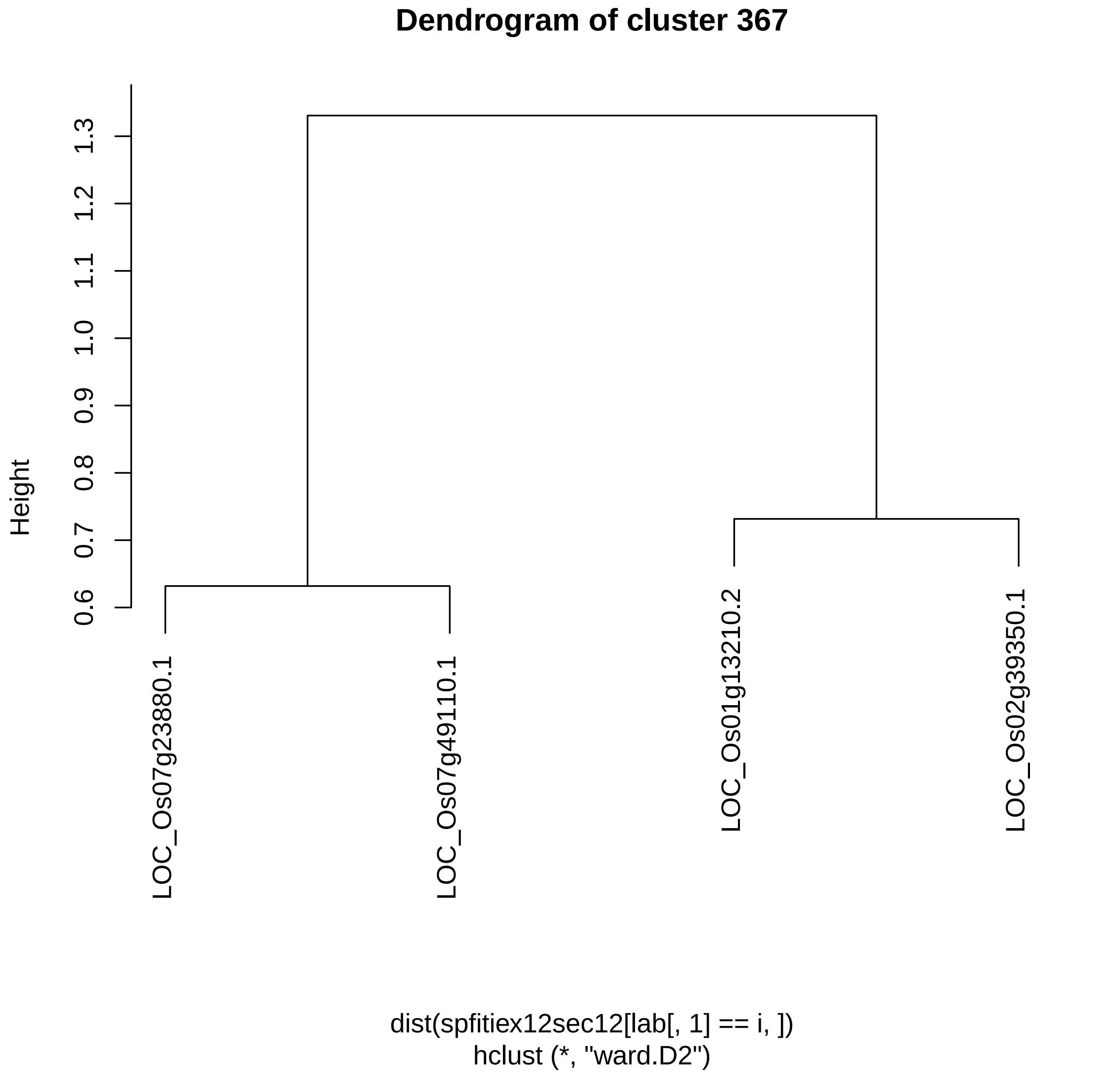
Dendrogram of cluster 367
Height
0.6
0.7
0.8
0.9
1.0
1.1
1.2
1.3
LOC_Os07g23880.1
LOC_Os07g49110.1
LOC_Os01g13210.2
LOC_Os02g39350.1
dist(spfitiex12sec12[lab[, 1] == i, ])
hclust (*, "ward.D2")

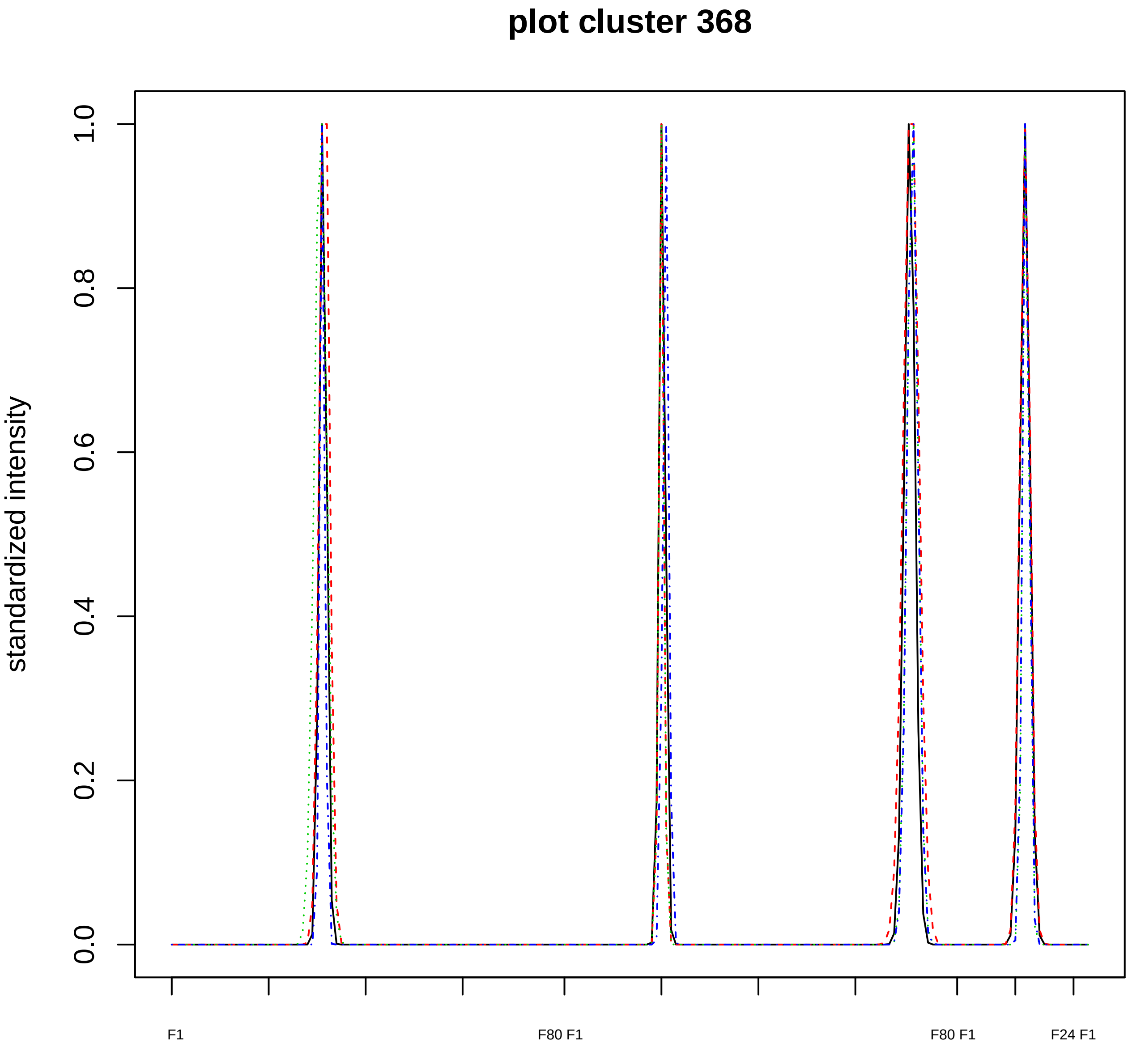
plot cluster 368
standardized intensity
0.0
0.2
0.4
0.6
0.8
1.0
F1
F80 F1
F80 F1
F24 F1

### Dendrogram of cluster 368

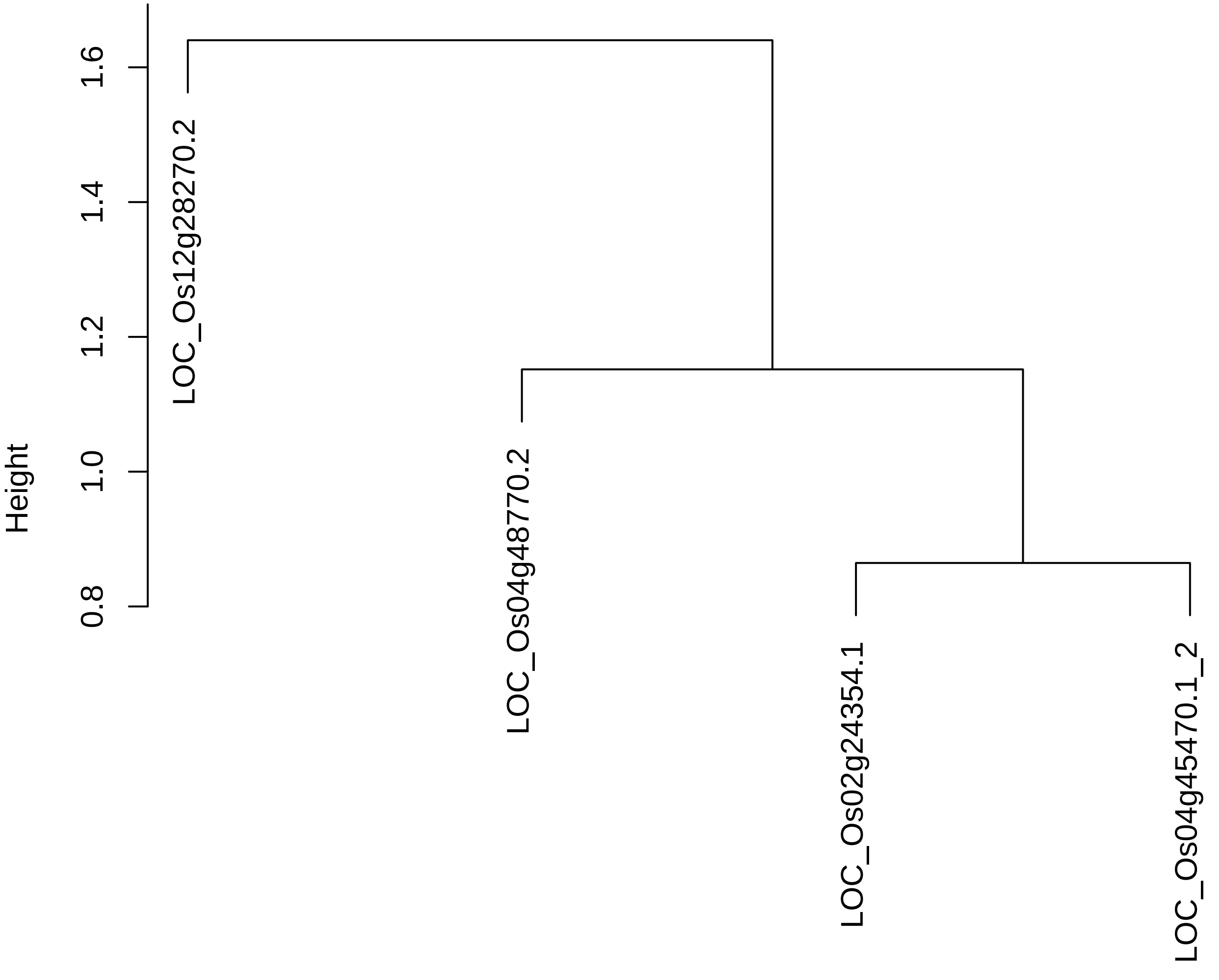


dist(spfitiex12sec12[lab[, 1] == i, ])
hclust (*, "ward.D2")

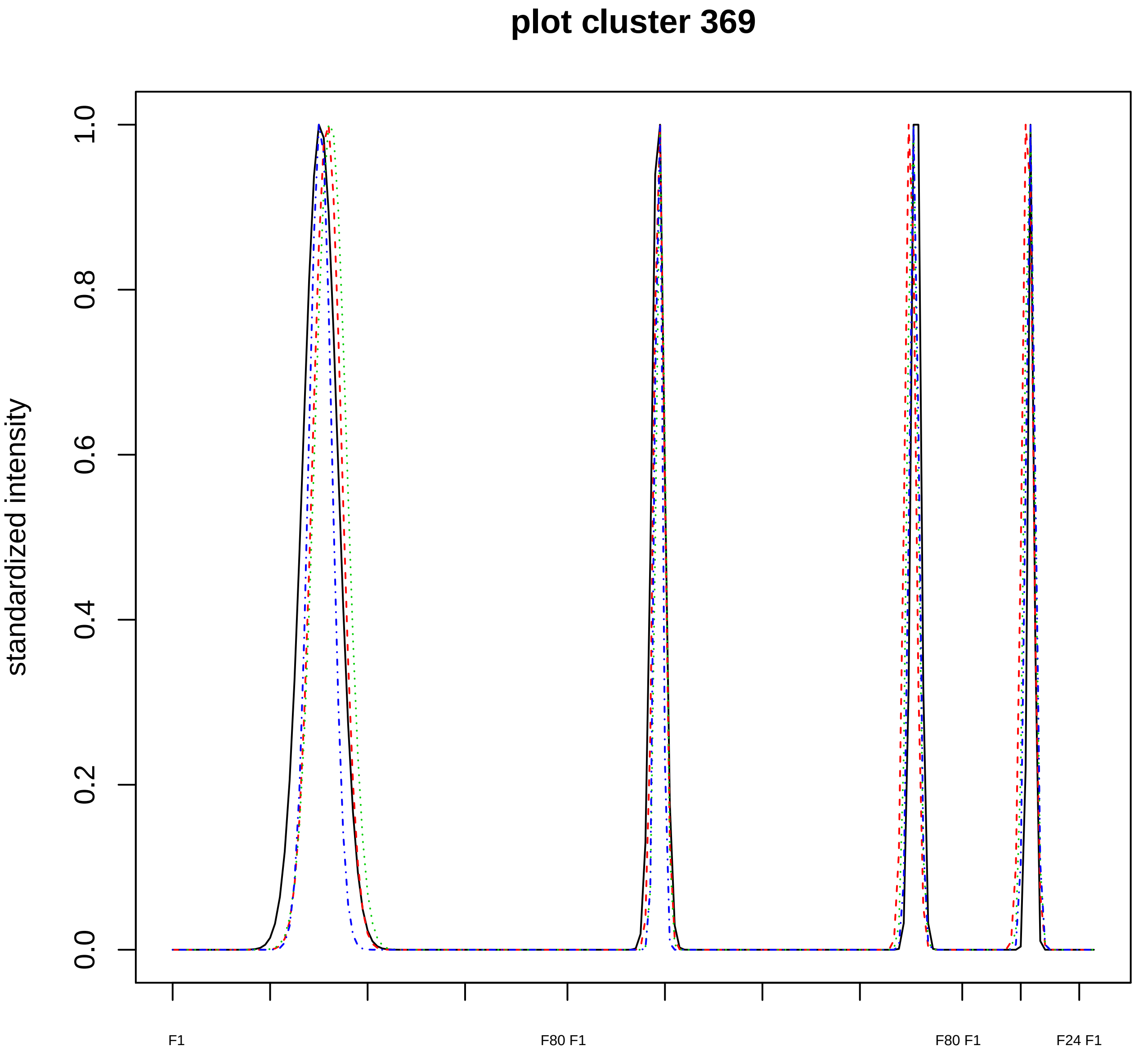

plot cluster 369
standardized intensity
0.0
0.2
0.4
0.6
0.8
1.0
F1
F80 F1
F80 F1
F24 F1

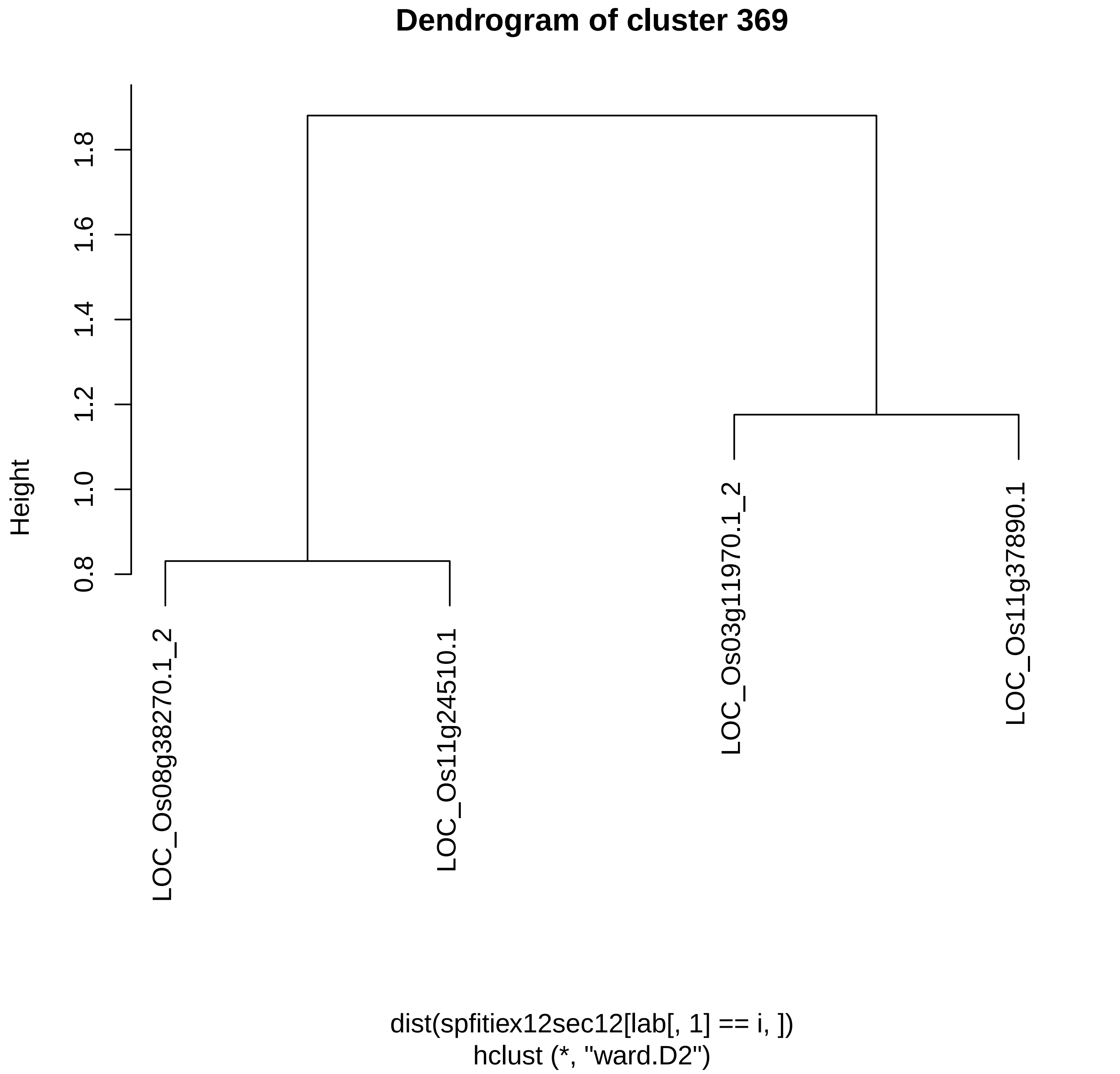
Dendrogram of cluster 369
Height
0.8
1.0
1.2
1.4
1.6
1.8
LOC_Os08g38270.1_2
LOC_Os11g24510.1
LOC_Os03g11970.1_2
LOC_Os11g37890.1
dist(spfitiex12sec12[lab[, 1] == i, ])
hclust (*, "ward.D2")

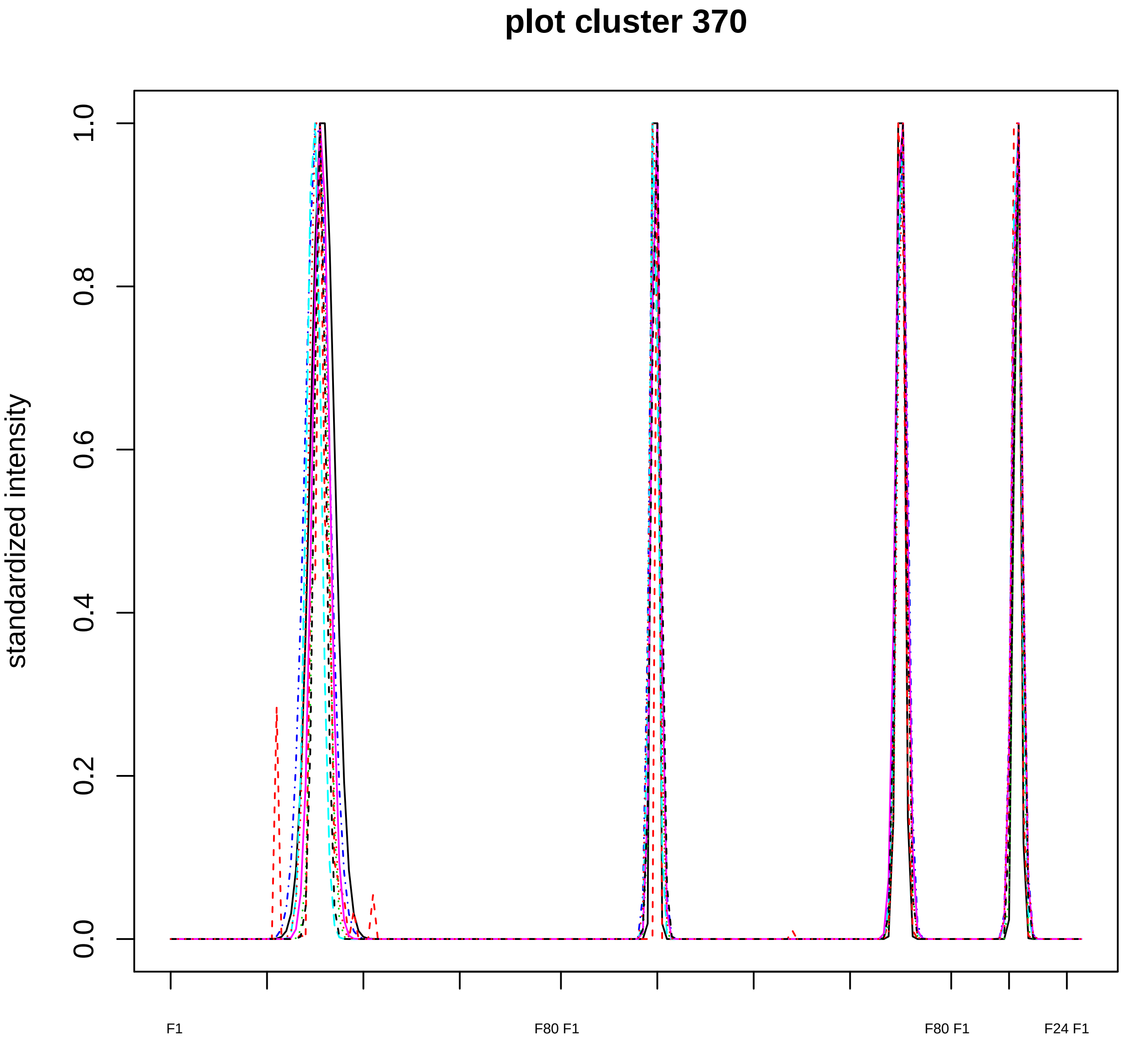

plot cluster 370
standardized intensity
0.0
0.2
0.4
0.6
0.8
1.0
F1
F80 F1
F80 F1
F24 F1

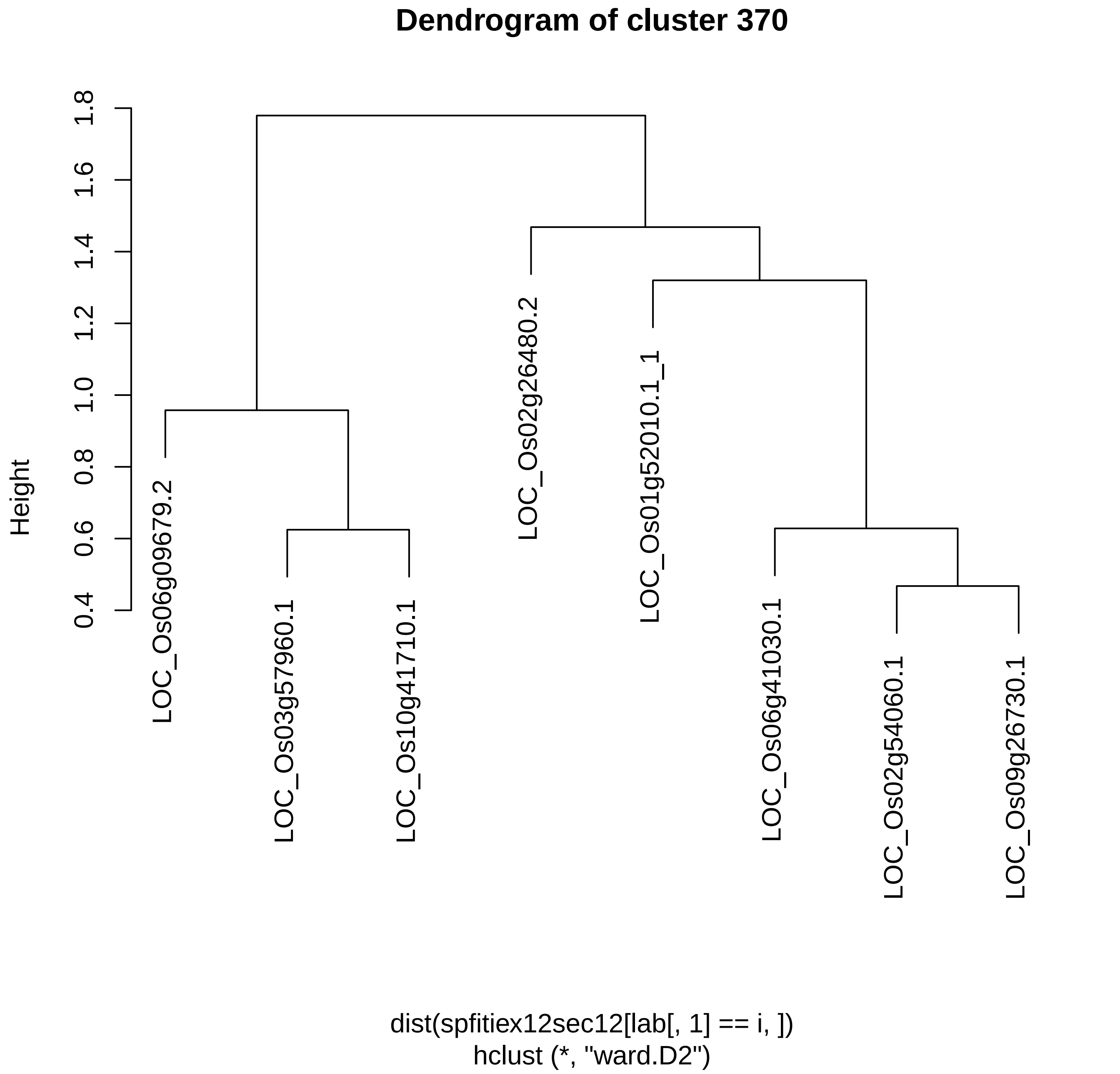

Dendrogram of cluster 370
Height
0.4
0.6
0.8
1.0
1.2
1.4
1.6
1.8
LOC_Os06g09679.2
LOC_Os03g57960.1
LOC_Os10g41710.1
LOC_Os02g26480.2
LOC_Os01g52010.1_1
LOC_Os06g41030.1
LOC_Os02g54060.1
LOC_Os09g26730.1
dist(spfitiex12sec12[lab[, 1] == i, ])
hclust (*, "ward.D2")

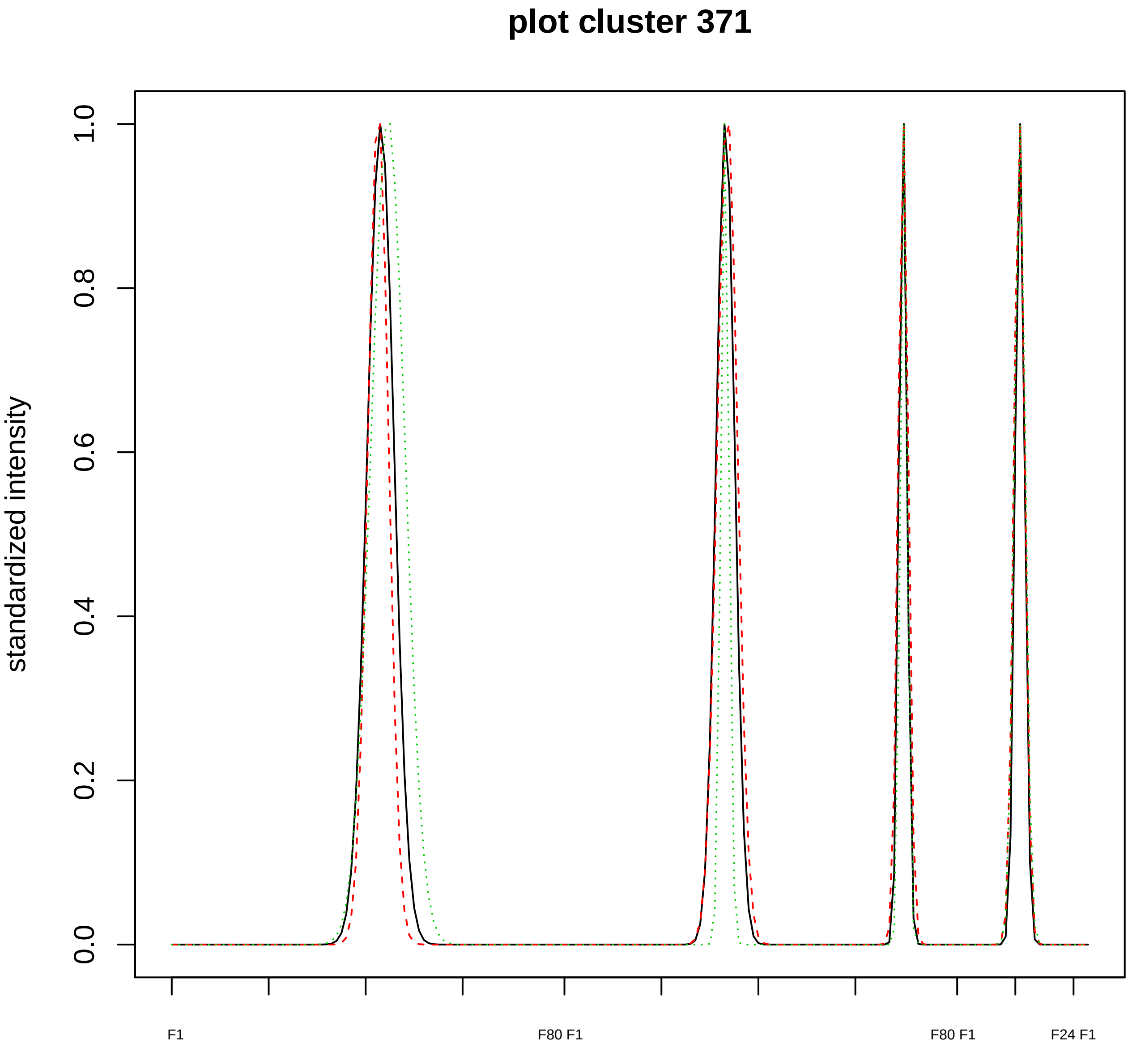
plot cluster 371
standardized intensity
0.0
0.2
0.4
0.6
0.8
1.0
F1
F80 F1
F80 F1
F24 F1

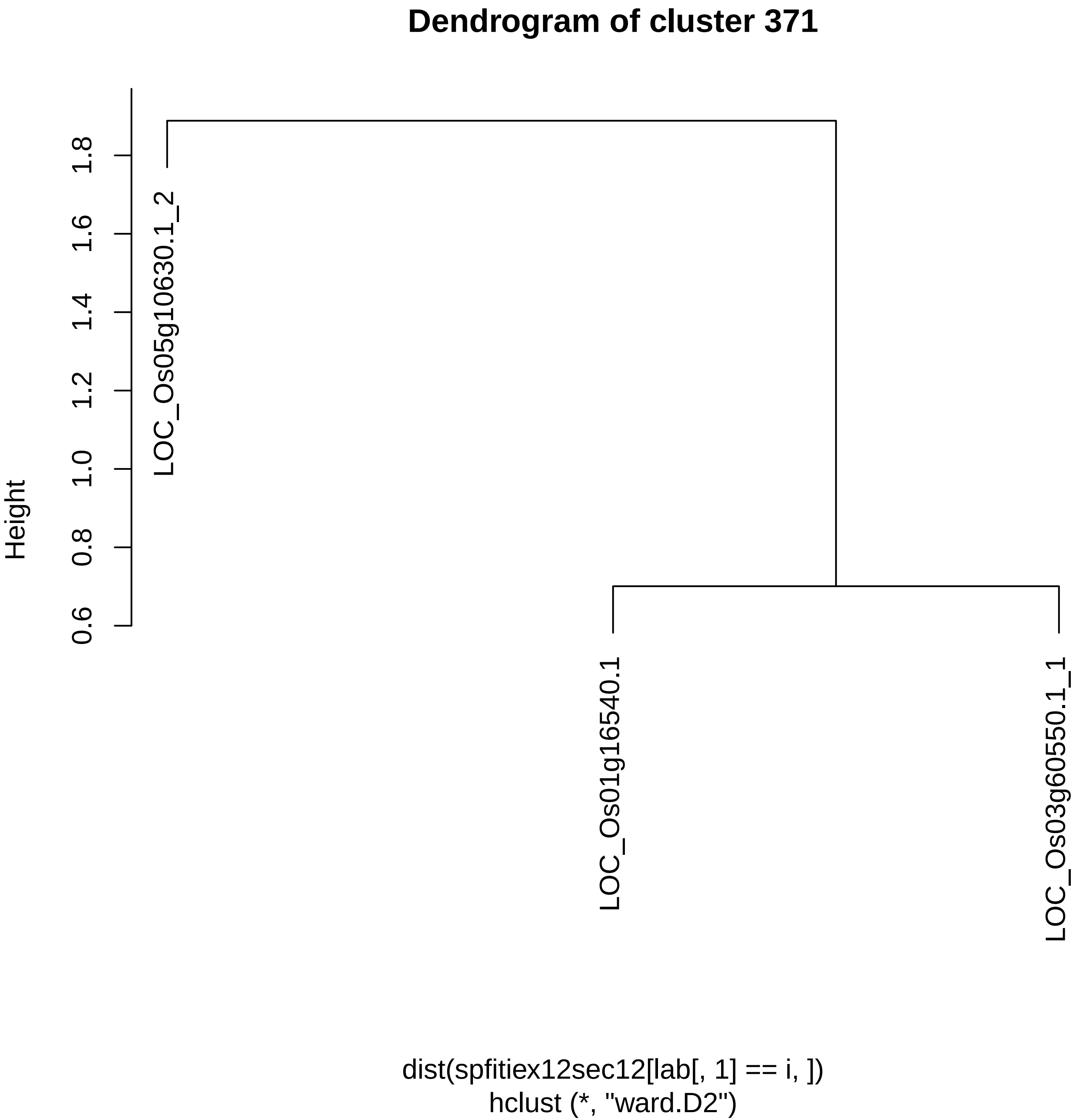
Dendrogram of cluster 371
Height
0.6
0.8
1.0
1.2
1.4
1.6
1.8
LOC_Os05g10630.1_2
LOC_Os01g16540.1
LOC_Os03g60550.1_1
dist(spfitiex12sec12[lab[, 1] == i, ])
hclust (*, "ward.D2")

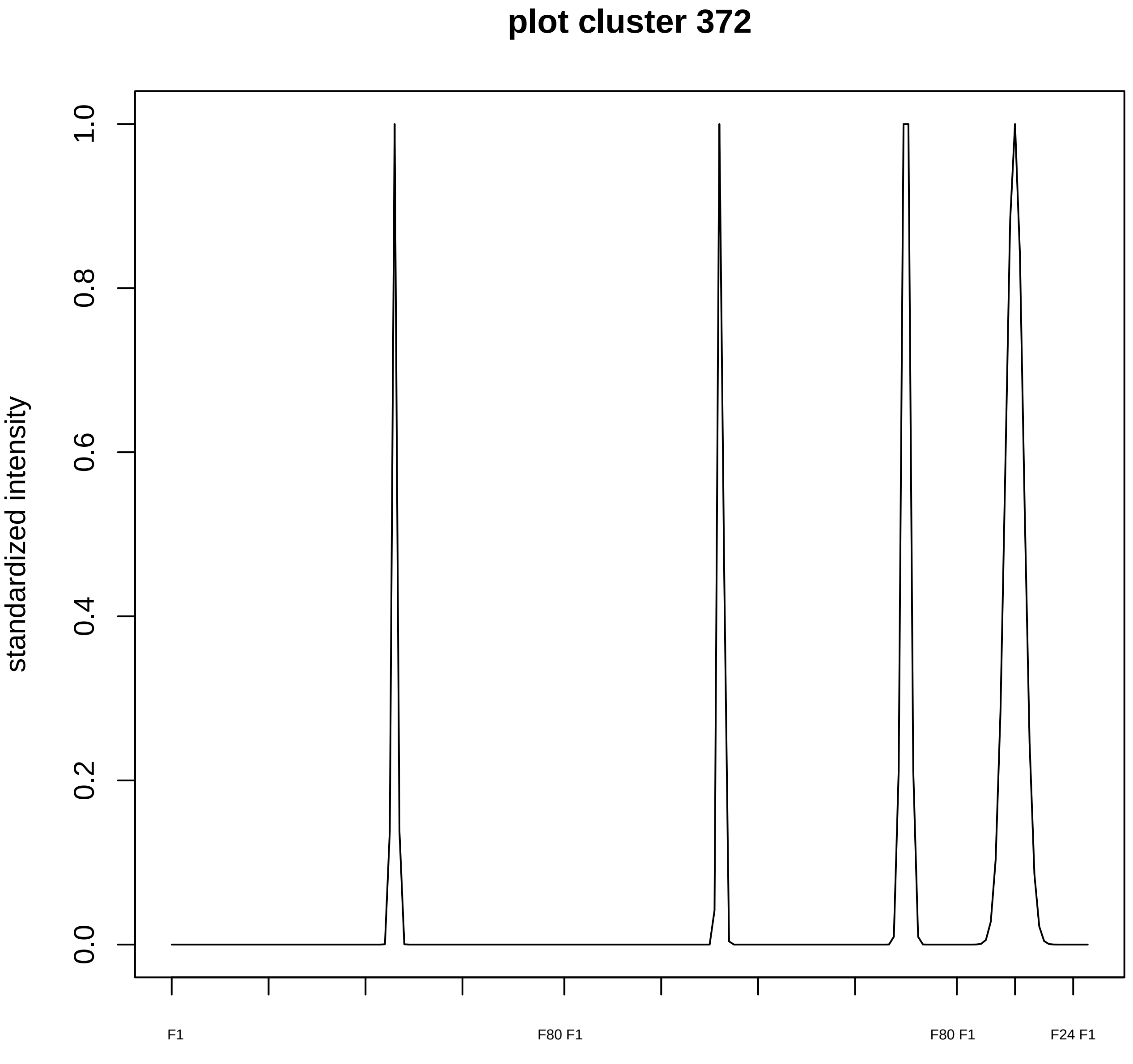

plot cluster 372
standardized intensity
0.0
0.2
0.4
0.6
0.8
1.0
F1
F80 F1
F80 F1
F24 F1

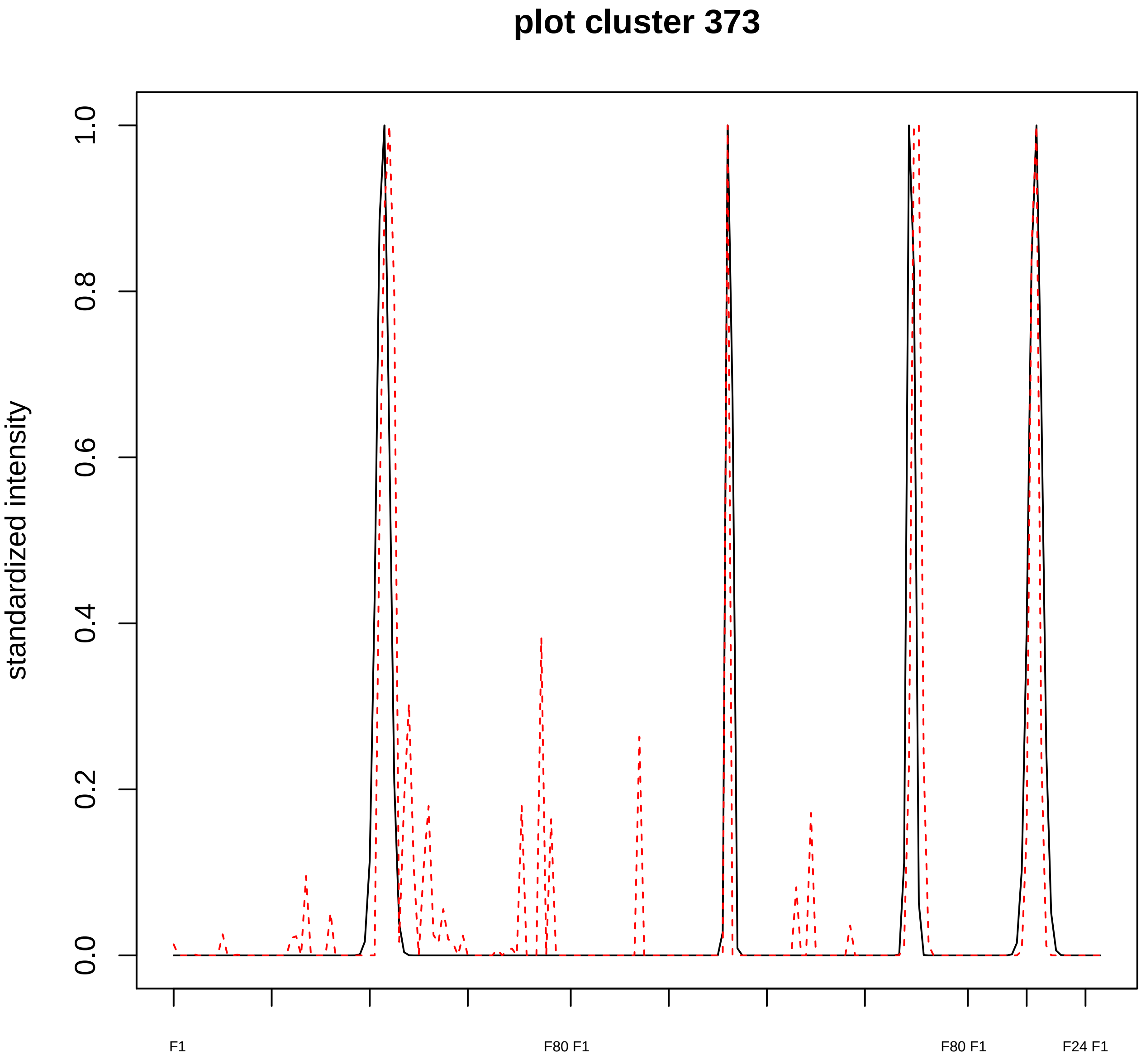

plot cluster 373
standardized intensity
0.0
0.2
0.4
0.6
0.8
1.0
F1
F80 F1
F80 F1
F24 F1

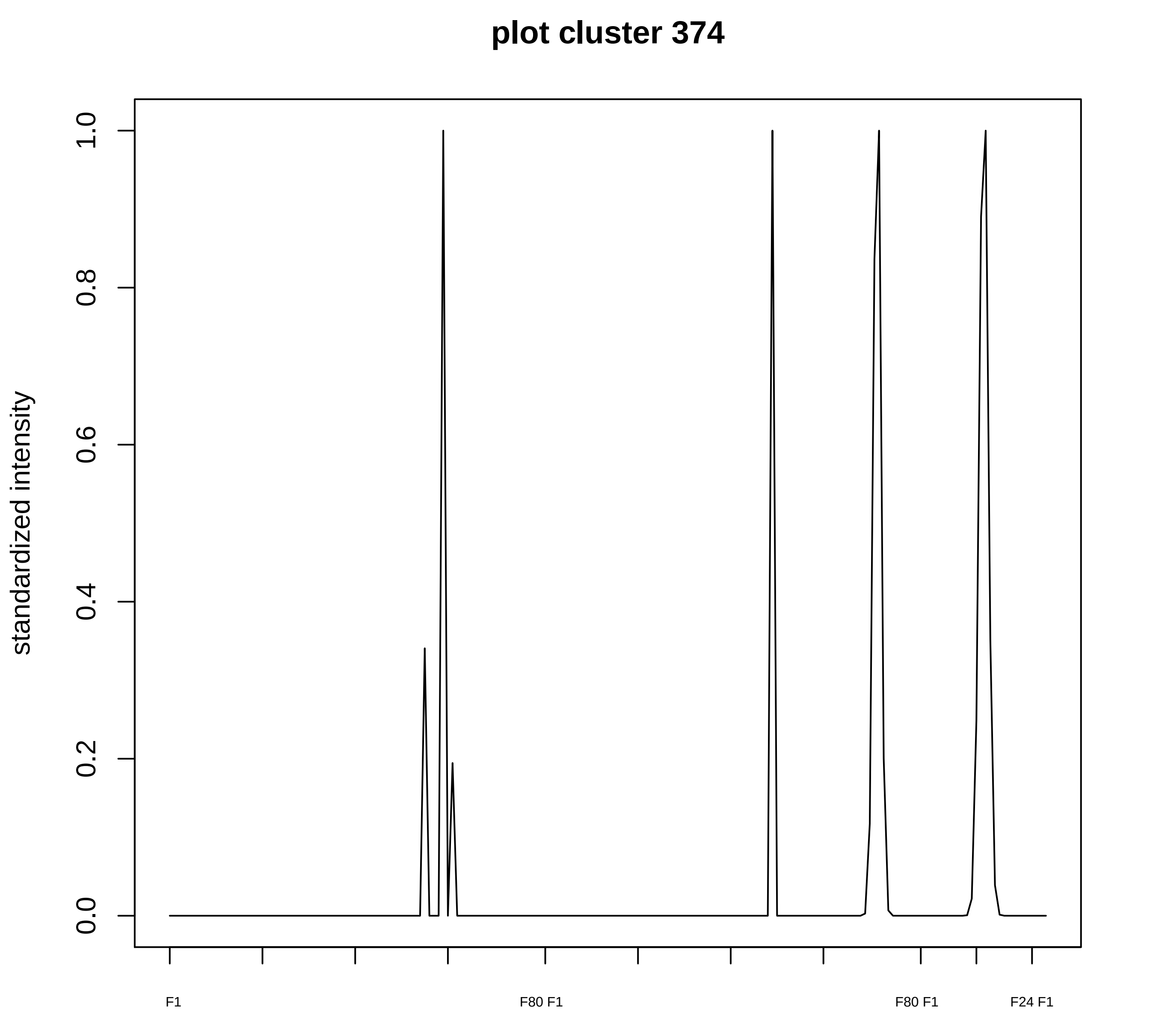

plot cluster 374
standardized intensity
0.0
0.2
0.4
0.6
0.8
1.0
F1
F80 F1
F80 F1
F24 F1

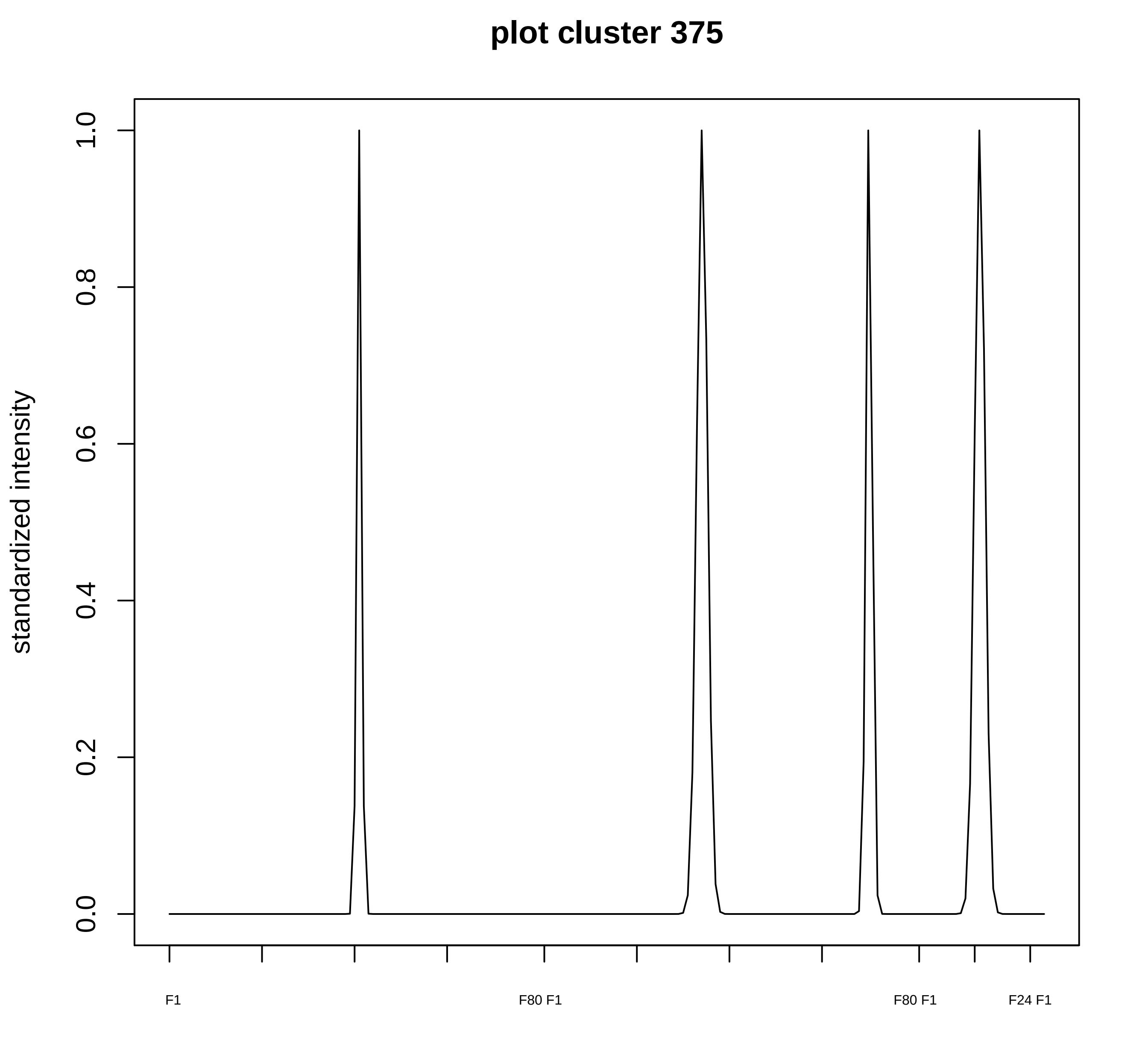
plot cluster 375
standardized intensity
0.0
0.2
0.4
0.6
0.8
1.0
F1
F80 F1
F80 F1
F24 F1

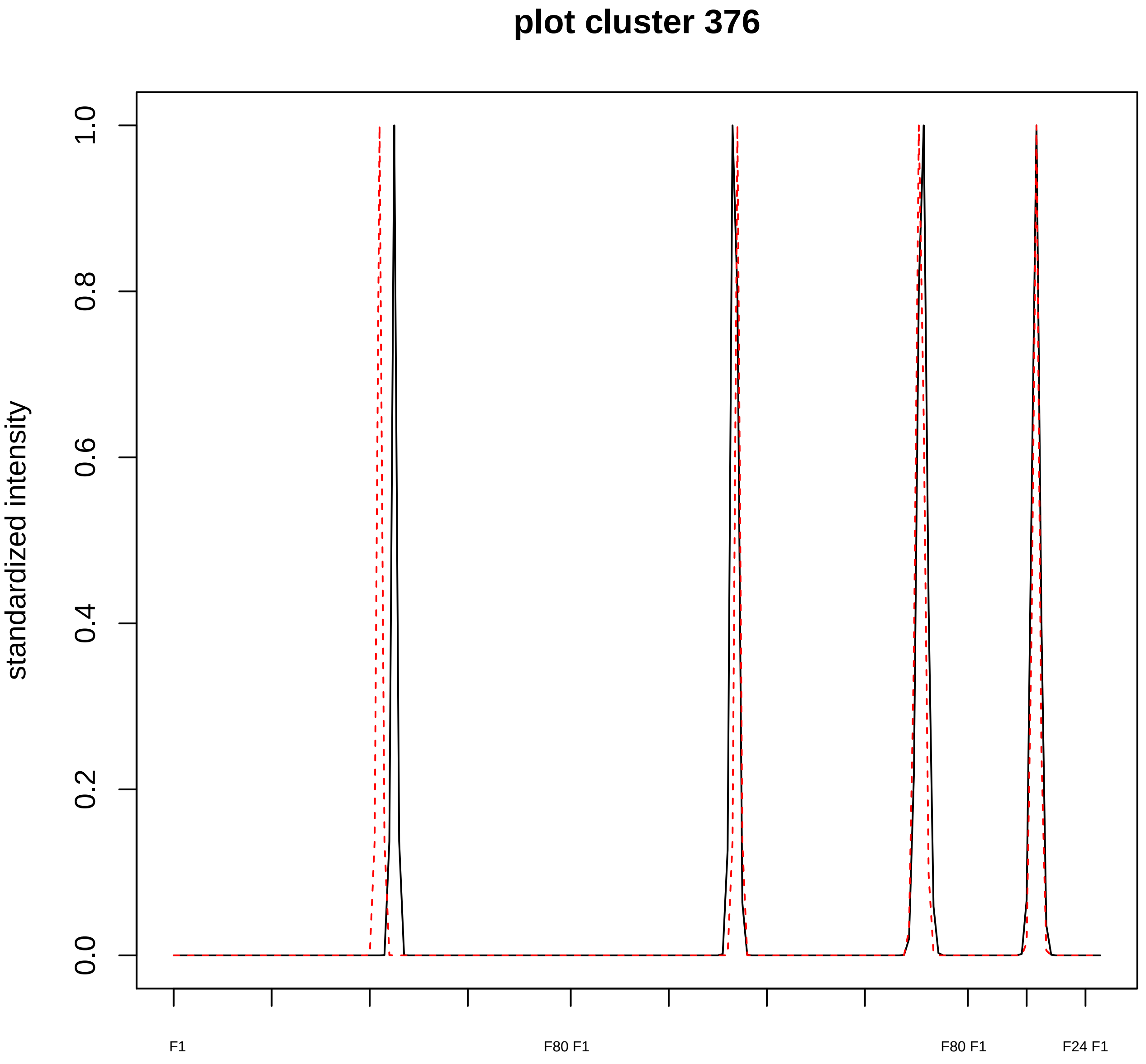
plot cluster 376
standardized intensity
0.0
0.2
0.4
0.6
0.8
1.0
F1
F80 F1
F80 F1
F24 F1

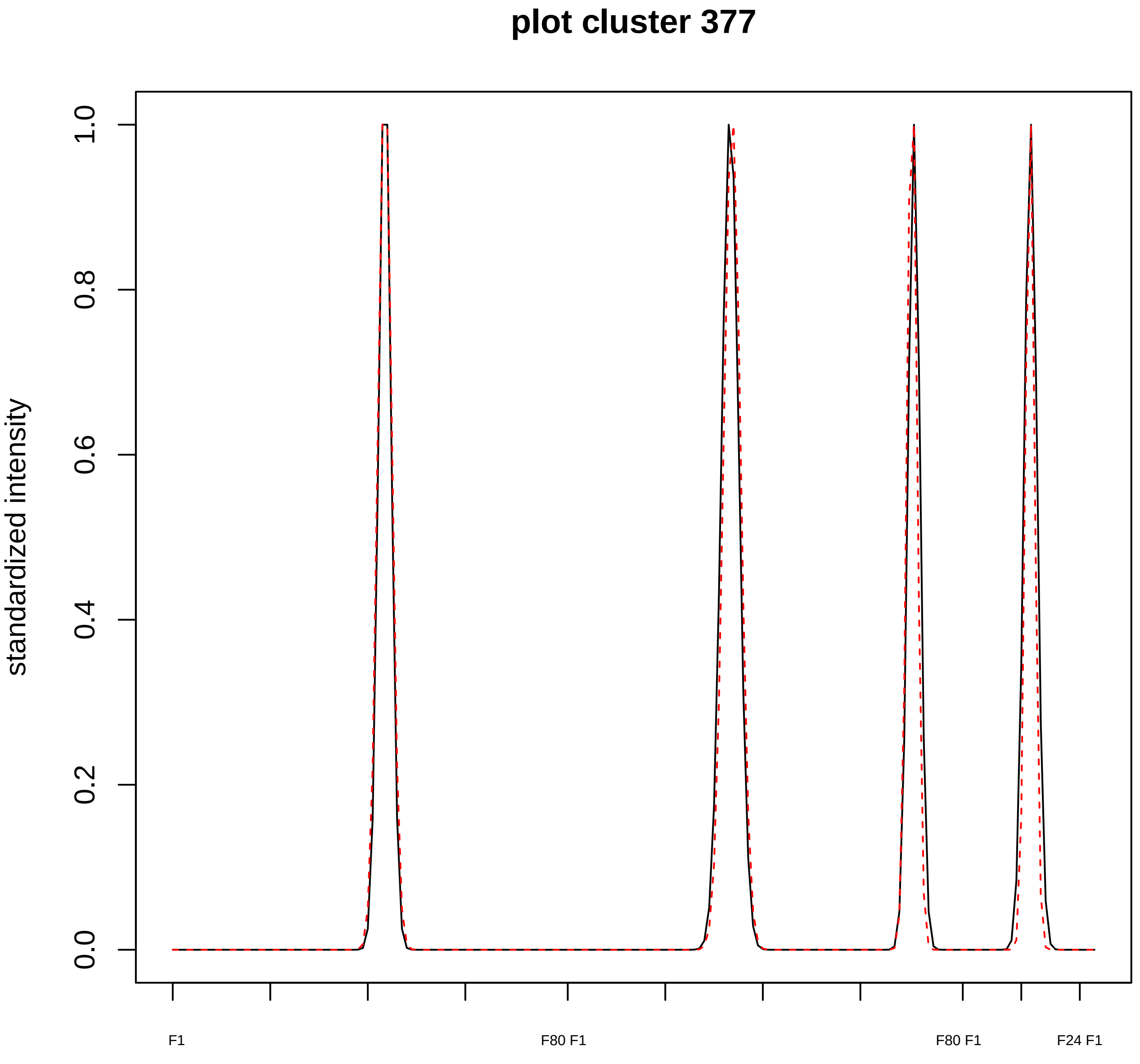

plot cluster 377
standardized intensity
0.0
0.2
0.4
0.6
0.8
1.0
F1
F80 F1
F80 F1
F24 F1

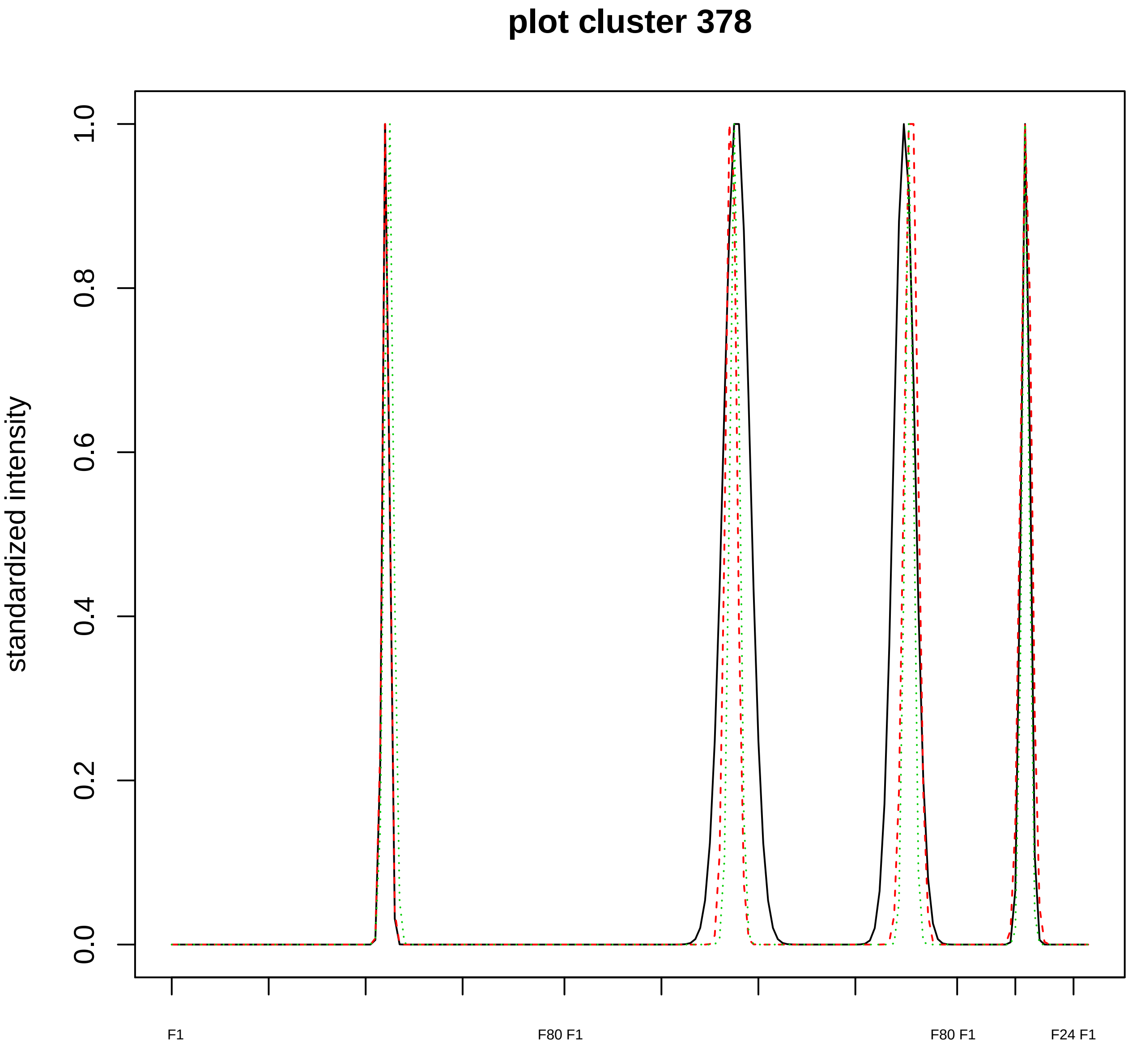
plot cluster 378
standardized intensity
0.0
0.2
0.4
0.6
0.8
1.0
F1
F80 F1
F80 F1
F24 F1

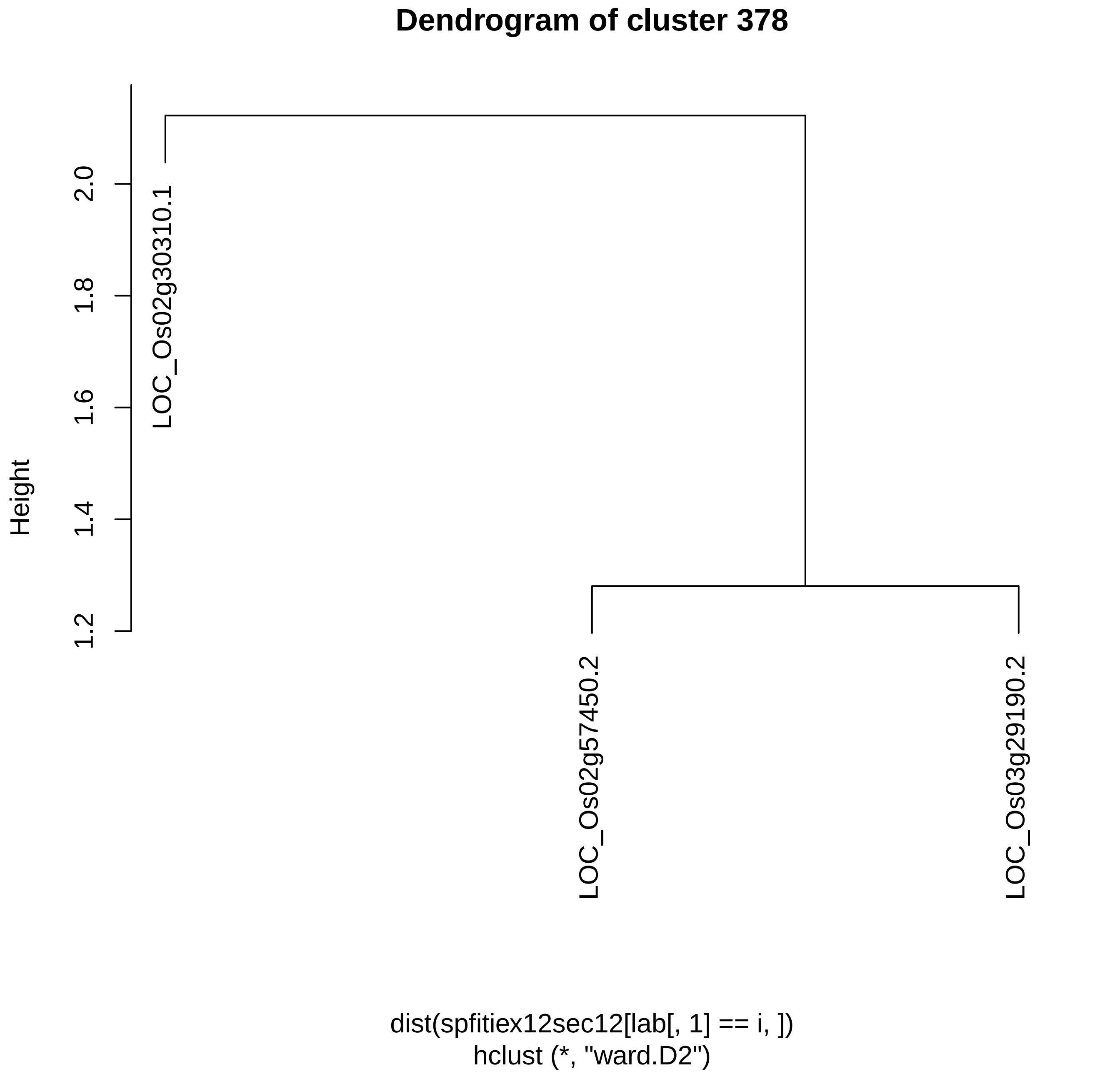
Dendrogram of cluster 378
Height
2.0
1.8
1.6
1.4
1.2
LOC_Os02g30310.1
LOC_Os02g57450.2
LOC_Os03g29190.2
dist(spfitiex12sec12[lab[, 1] == i, ])
hclust (*, "ward.D2")

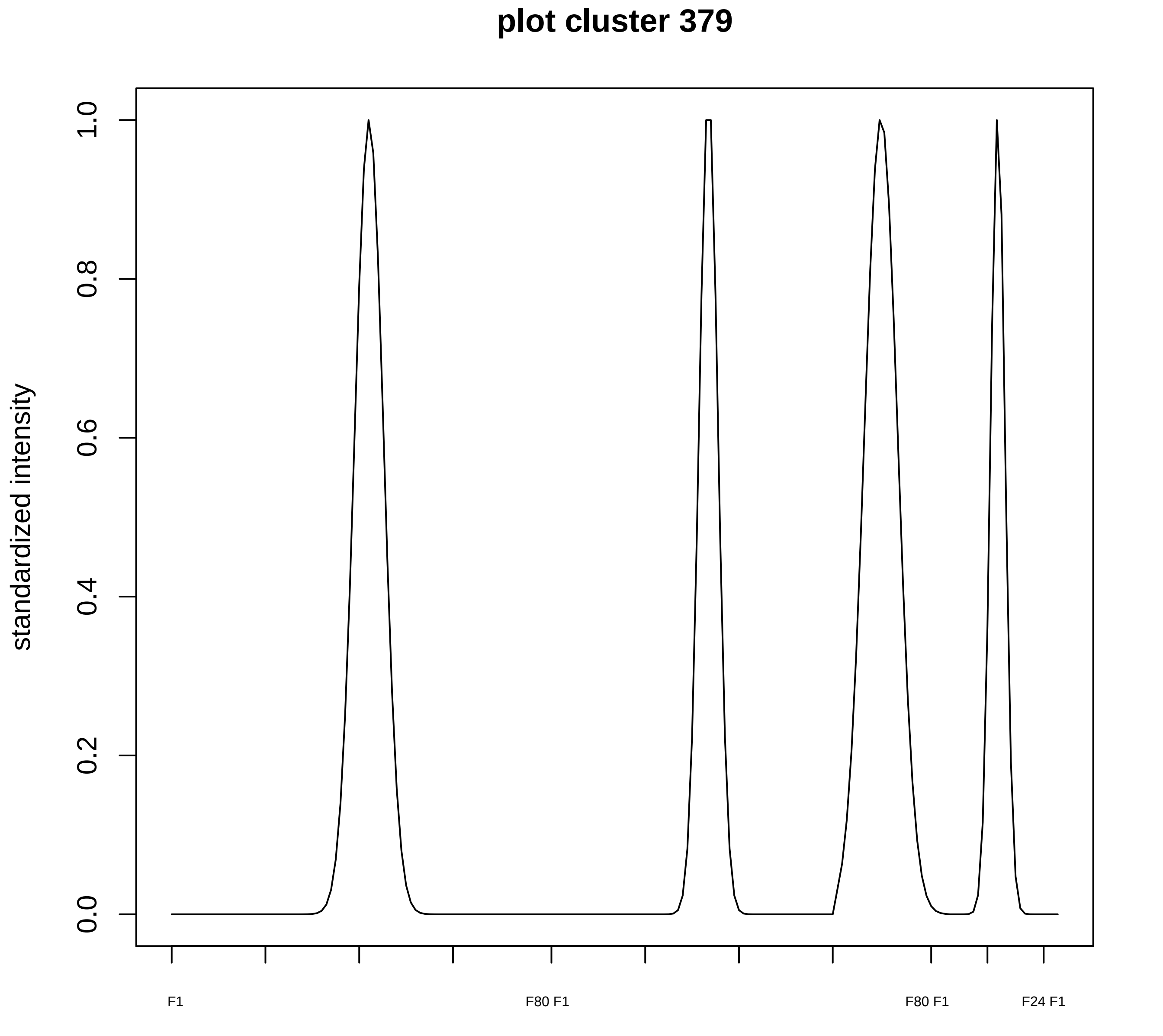
plot cluster 379
standardized intensity
0.0
0.2
0.4
0.6
0.8
1.0
F1
F80 F1
F80 F1
F24 F1

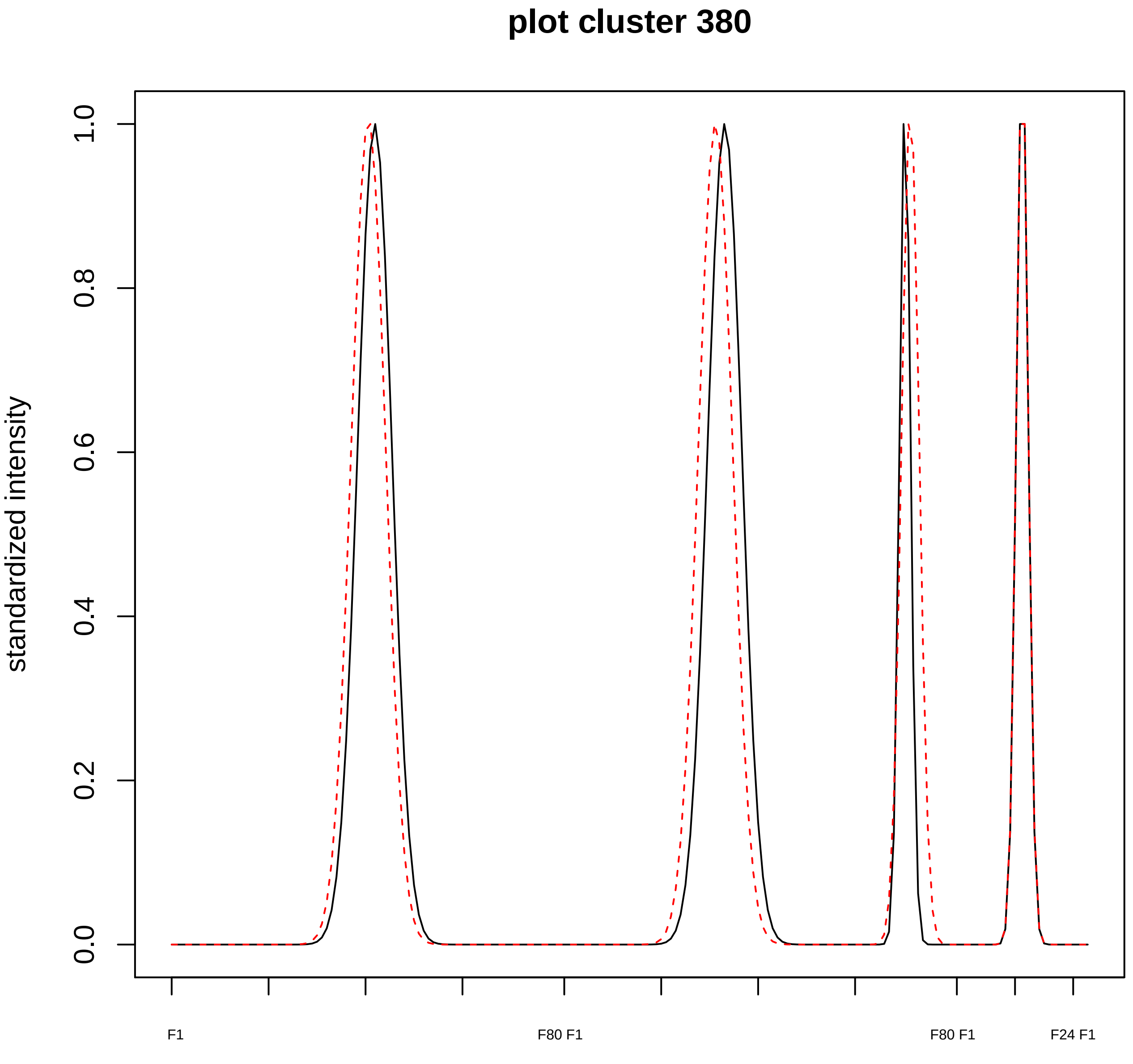

plot cluster 380
standardized intensity
0.0
0.2
0.4
0.6
0.8
1.0
F1
F80 F1
F80 F1
F24 F1

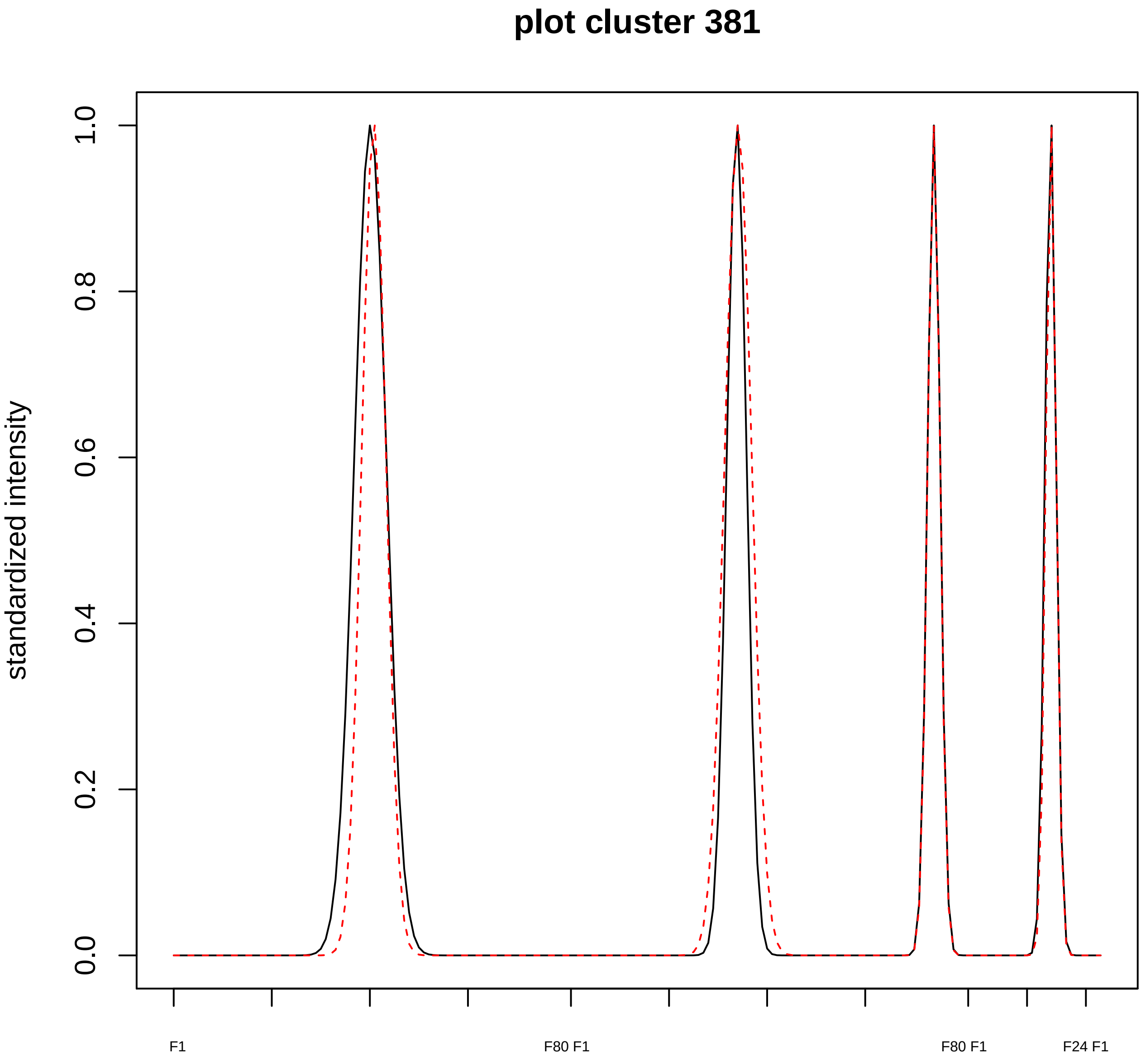
plot cluster 381
standardized intensity
0.0
0.2
0.4
0.6
0.8
1.0
F1
F80 F1
F80 F1
F24 F1

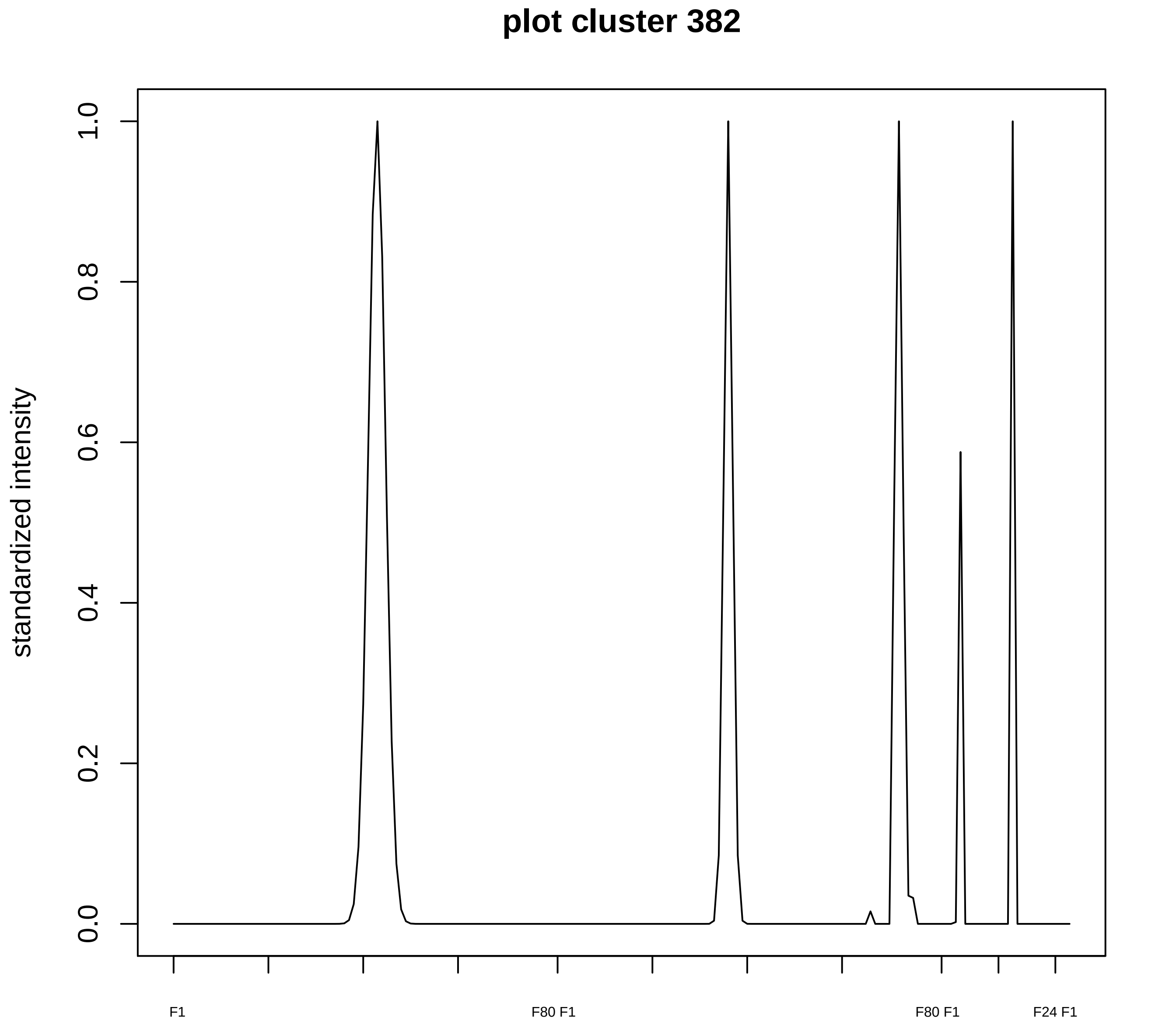
plot cluster 382
standardized intensity
0.0
0.2
0.4
0.6
0.8
1.0
F1
F80 F1
F80 F1
F24 F1

### plot cluster 383

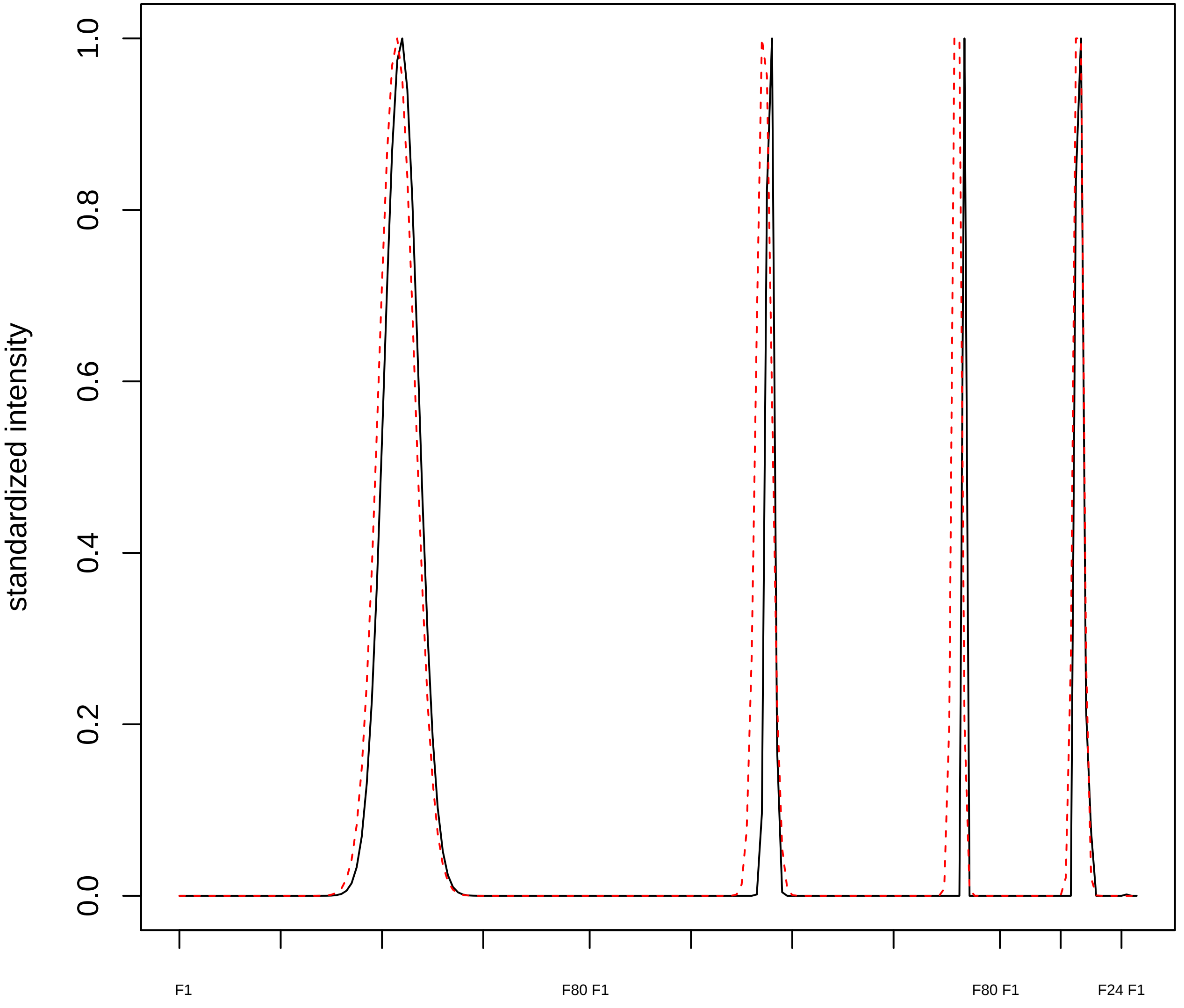

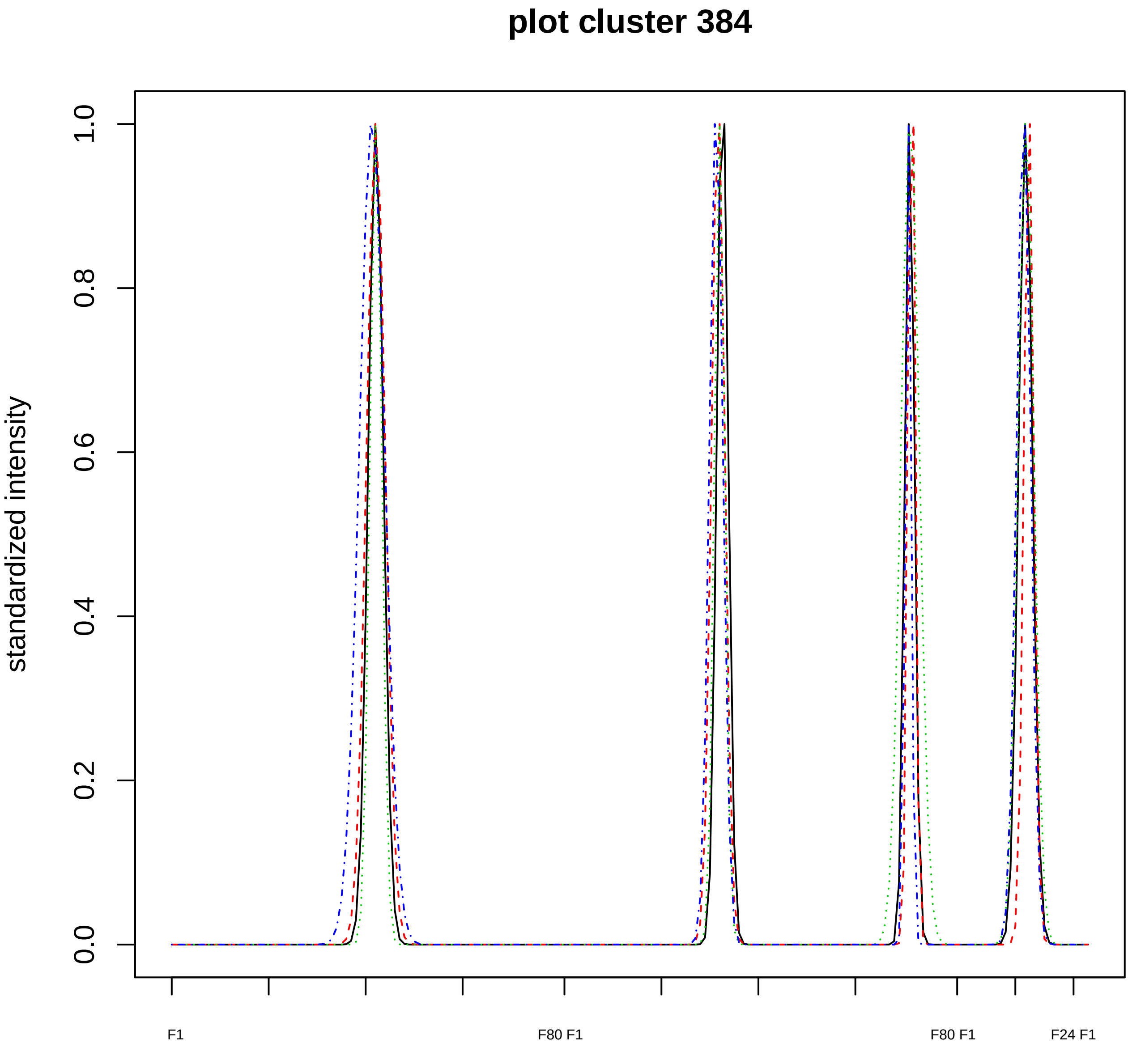
plot cluster 384
standardized intensity
0.0
0.2
0.4
0.6
0.8
1.0
F1
F80 F1
F80 F1
F24 F1

#### Dendrogram of cluster 384

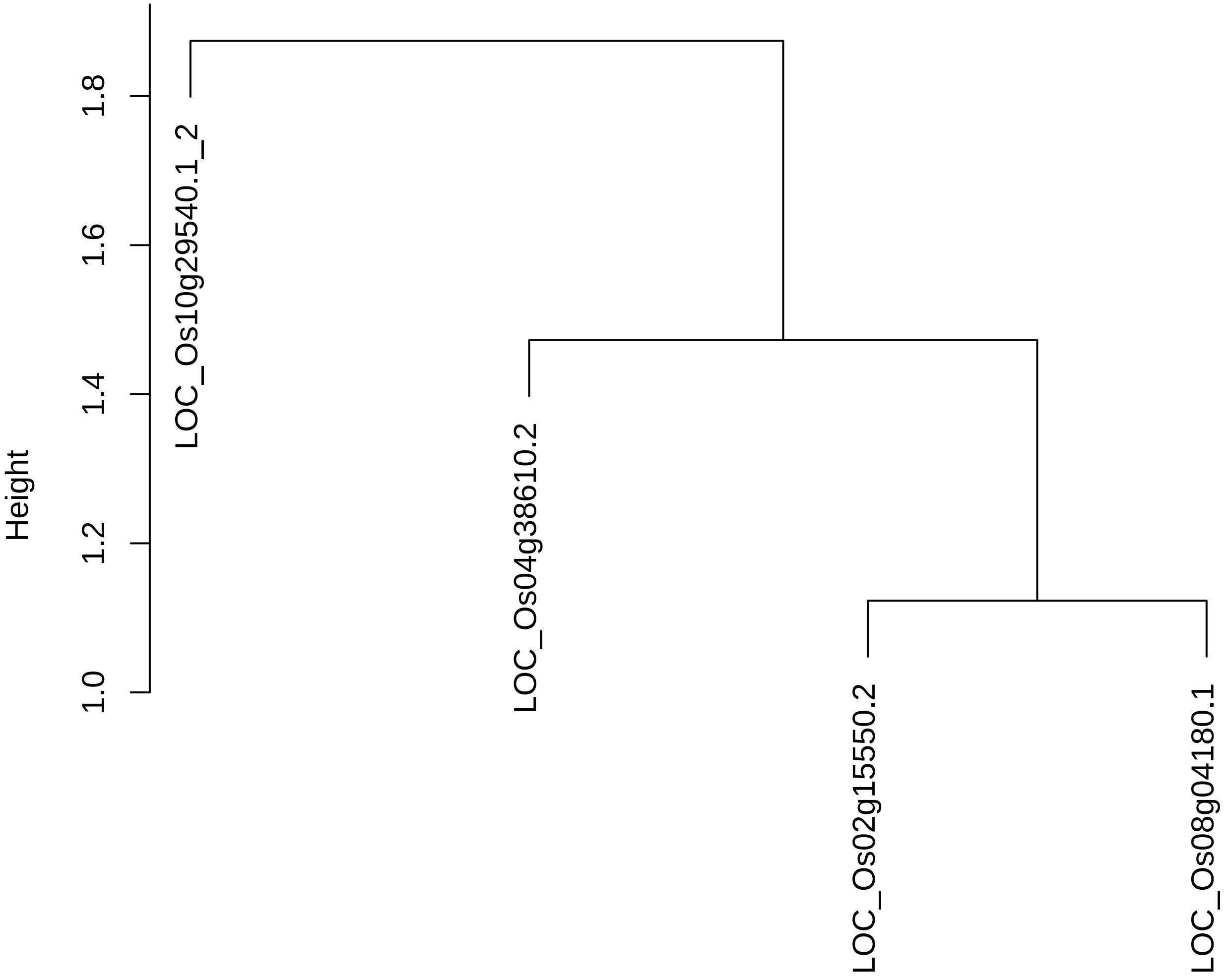


dist(spfitiex12sec12[lab[, 1] == i, ])
hclust (*, "ward.D2")

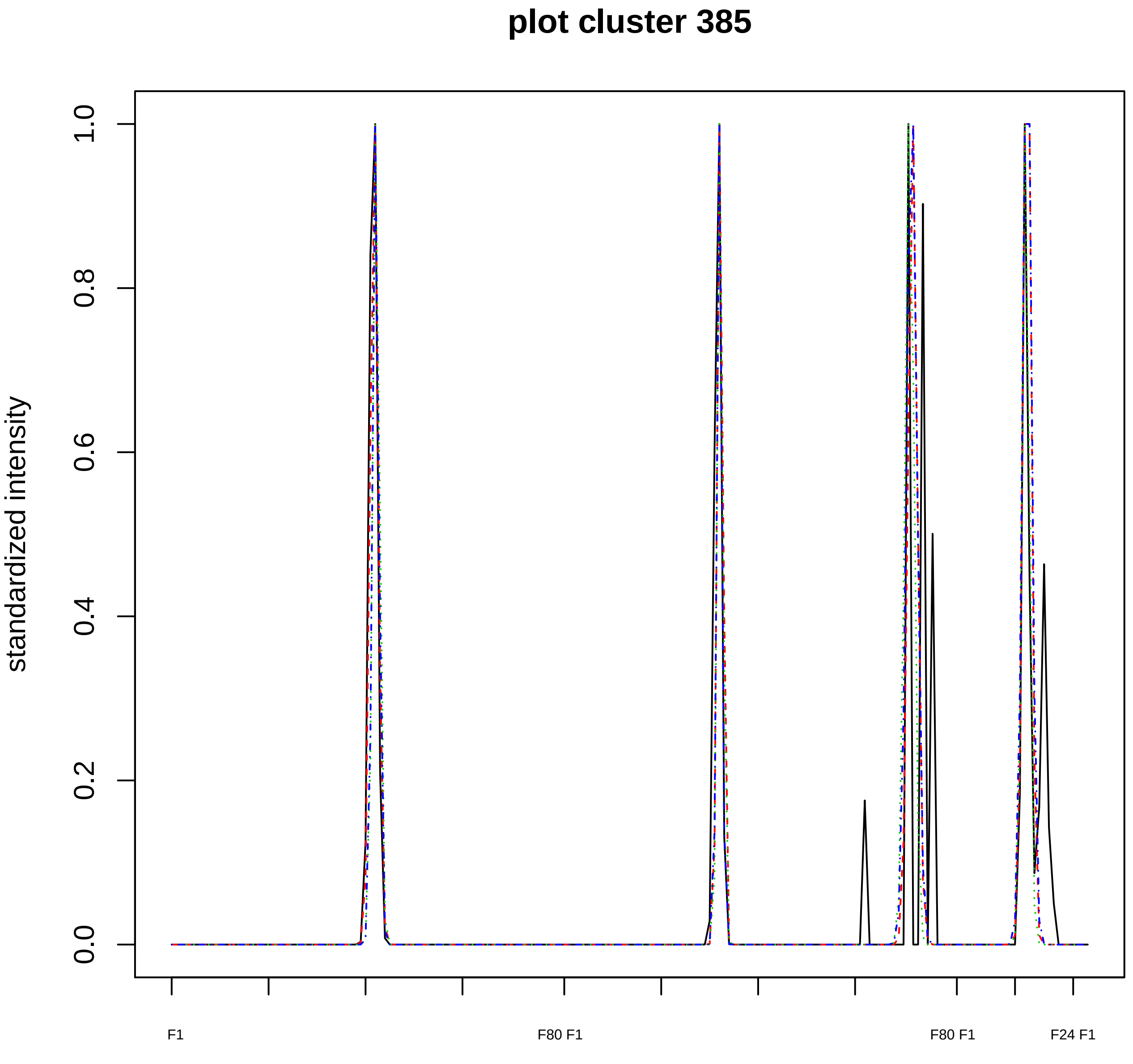
plot cluster 385
standardized intensity
0.0
0.2
0.4
0.6
0.8
1.0
F1
F80 F1
F80 F1
F24 F1

### Dendrogram of cluster 385

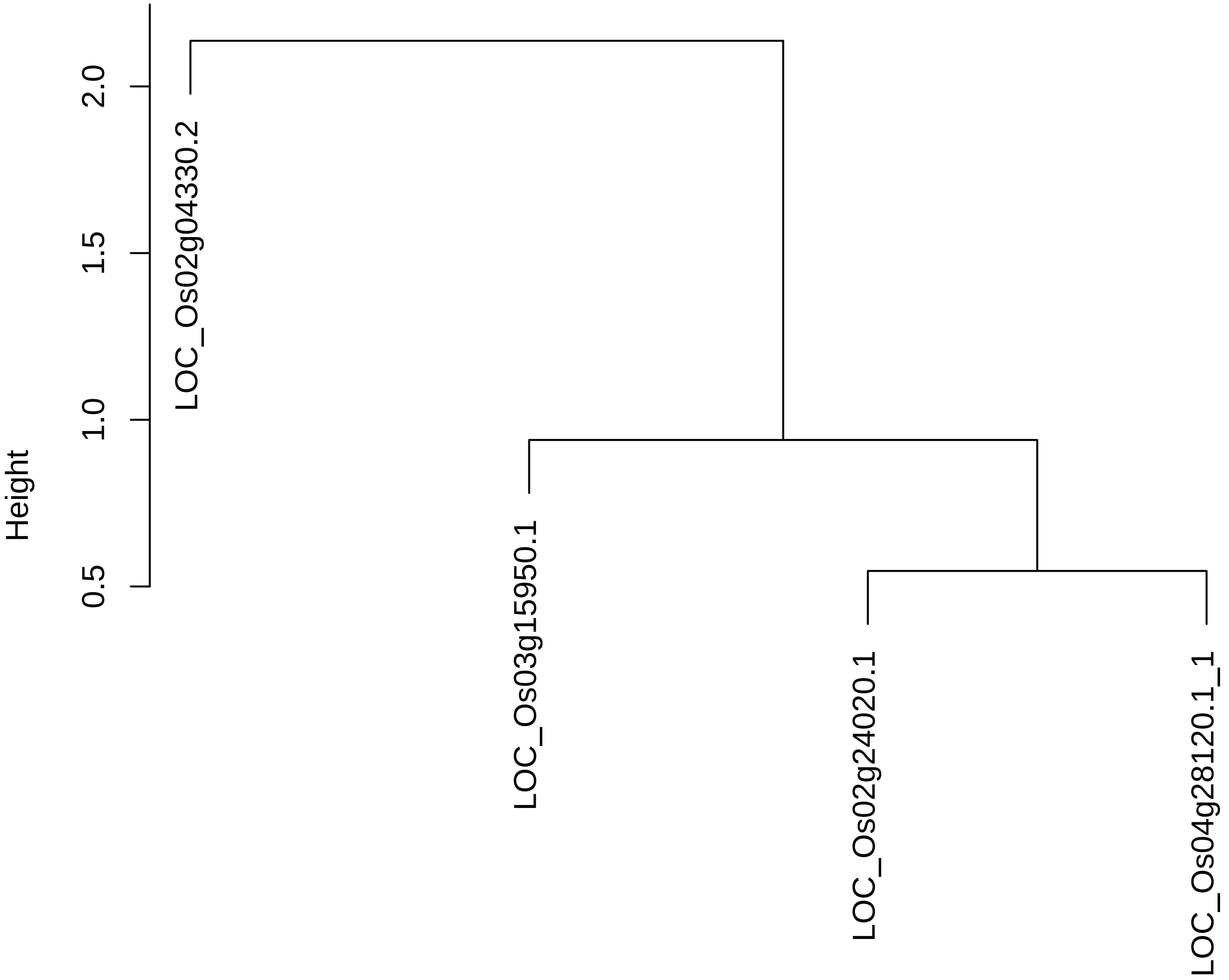


dist(spfitiex12sec12[lab[, 1] == i, ])
hclust (*, "ward.D2")

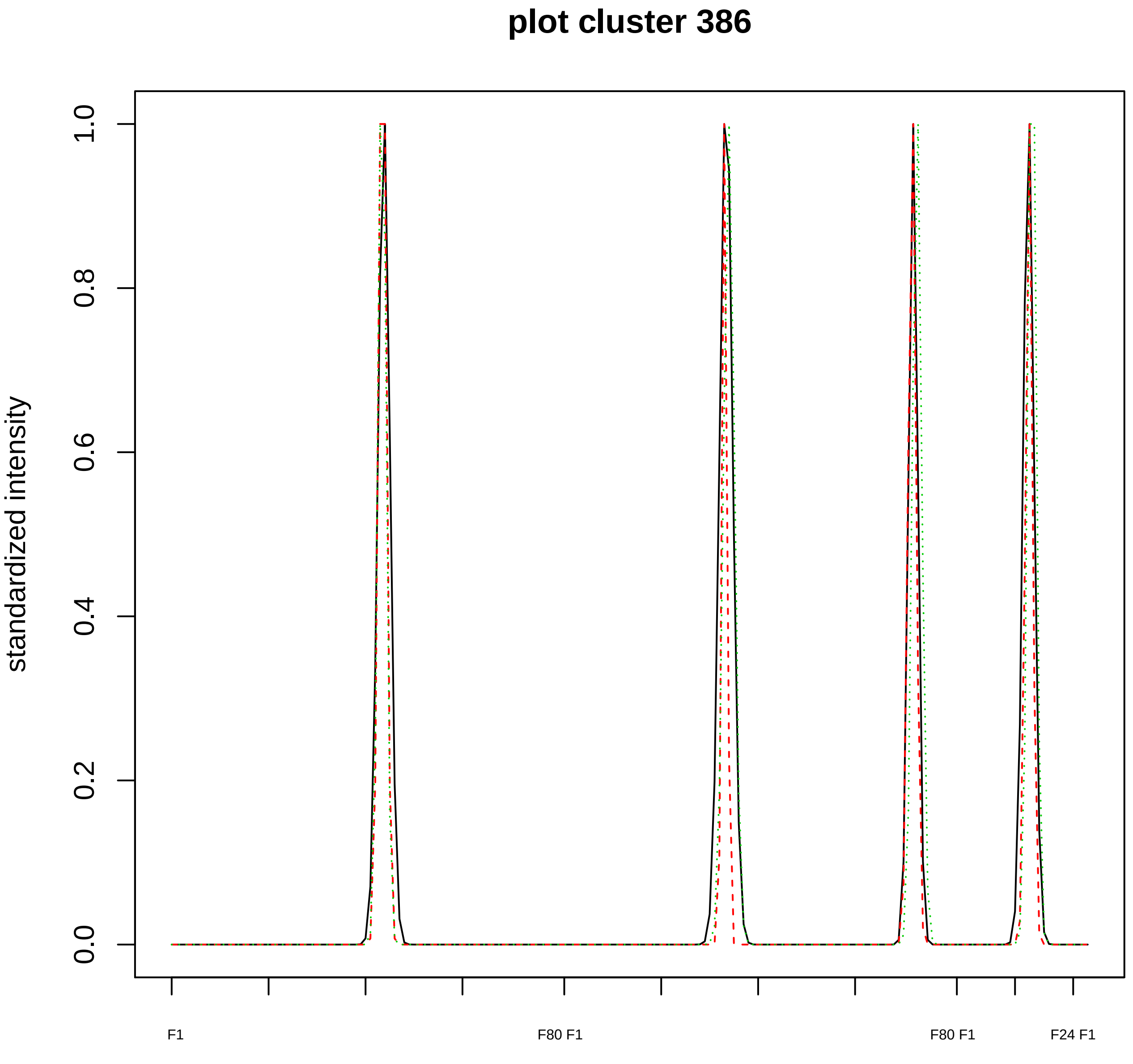
plot cluster 386
standardized intensity
0.0
0.2
0.4
0.6
0.8
1.0
F1
F80 F1
F80 F1
F24 F1

#### Dendrogram of cluster 386

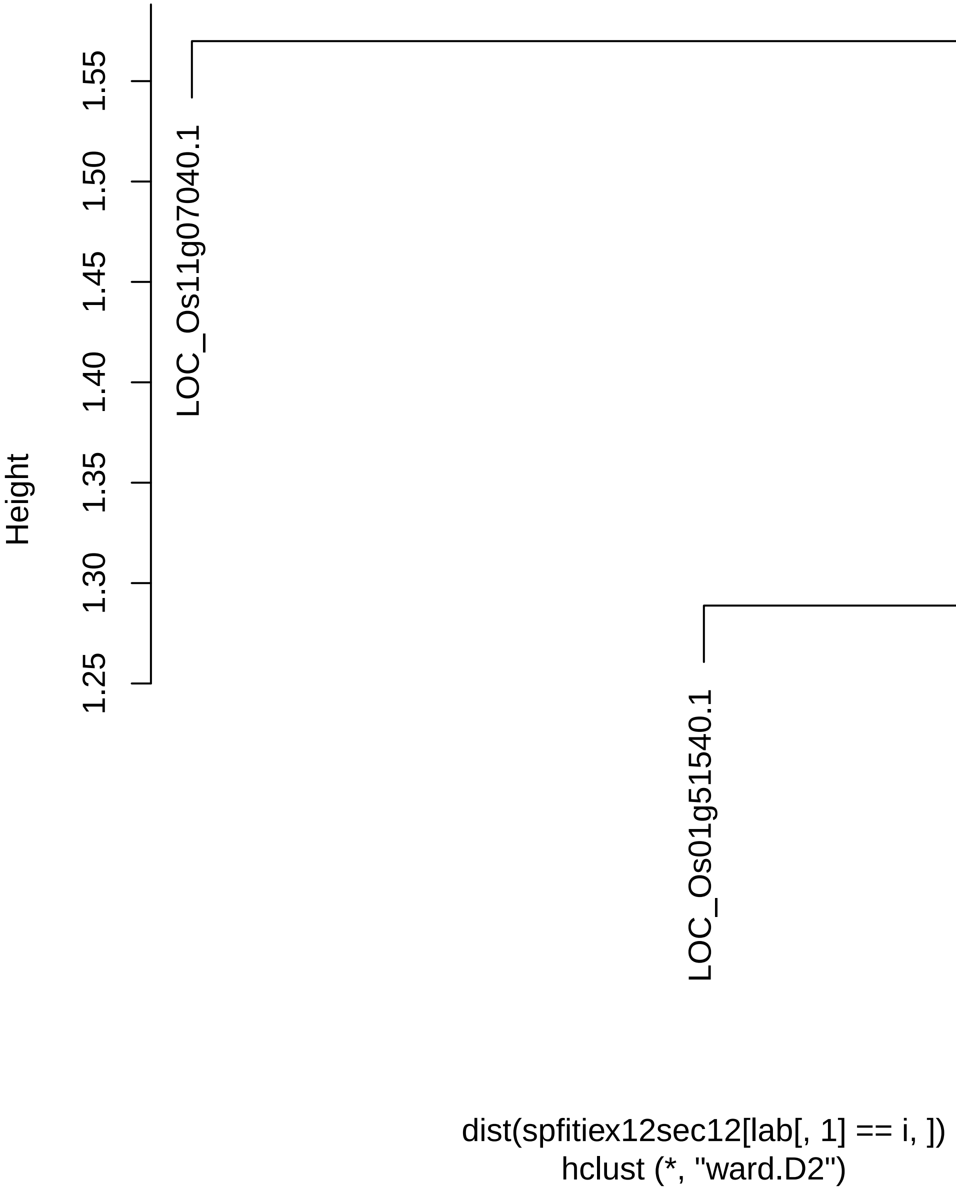

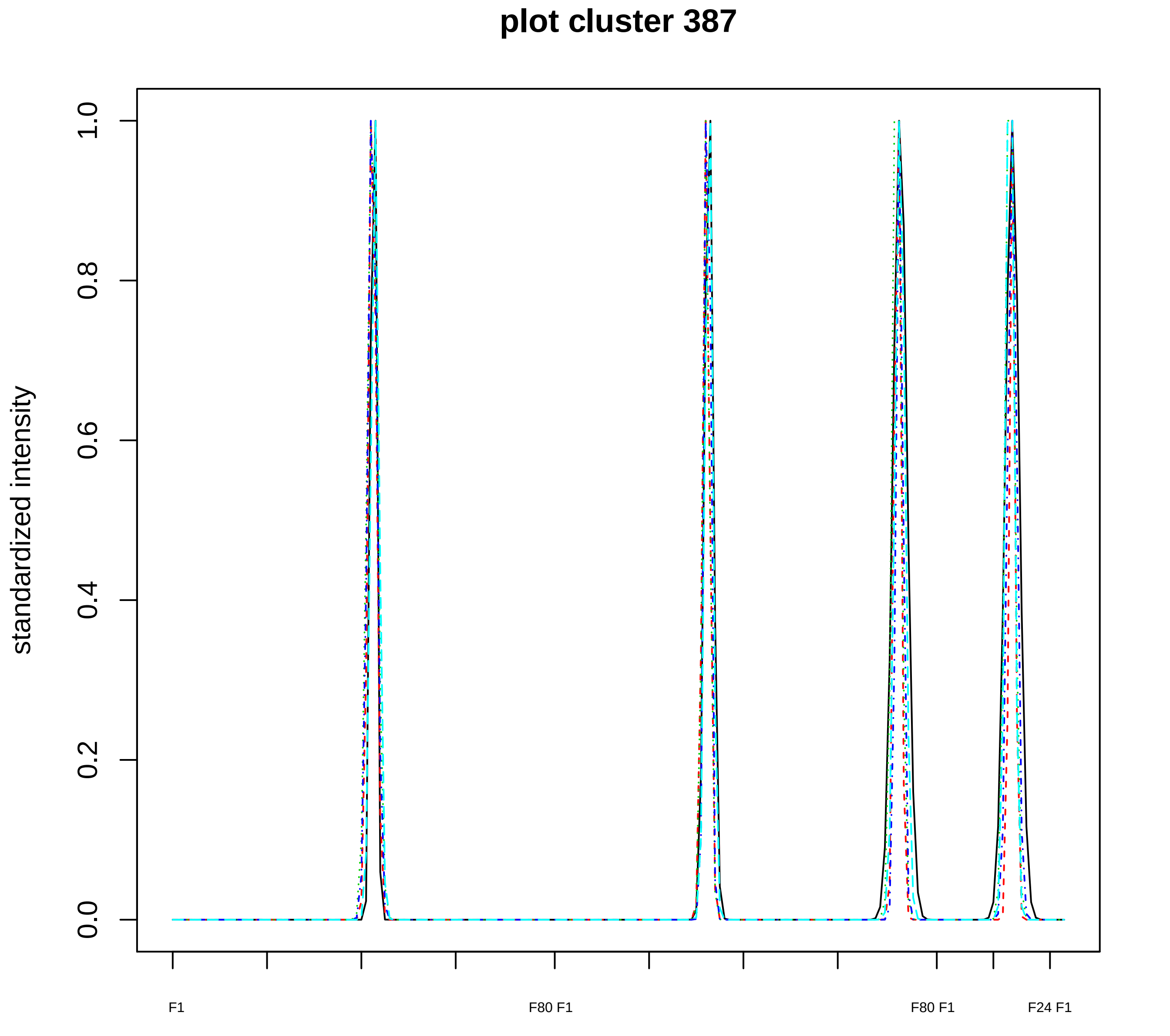
plot cluster 387
standardized intensity
1.0
0.8
0.6
0.4
0.2
0.0
F1
F80 F1
F80 F1
F24 F1

#### Dendrogram of cluster 387

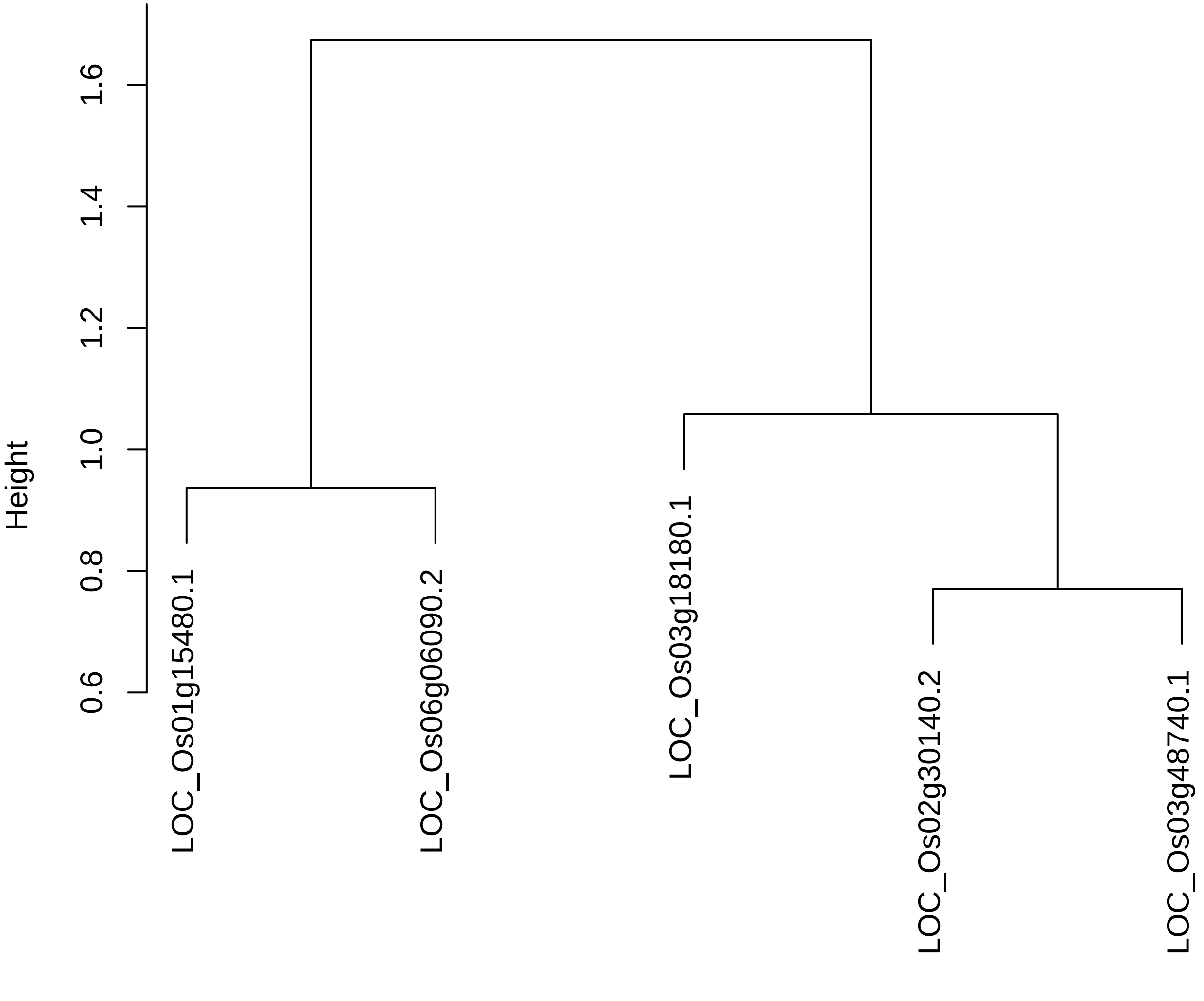


dist(spfitiex12sec12[lab[, 1] == i, ])
hclust (*, "ward.D2")

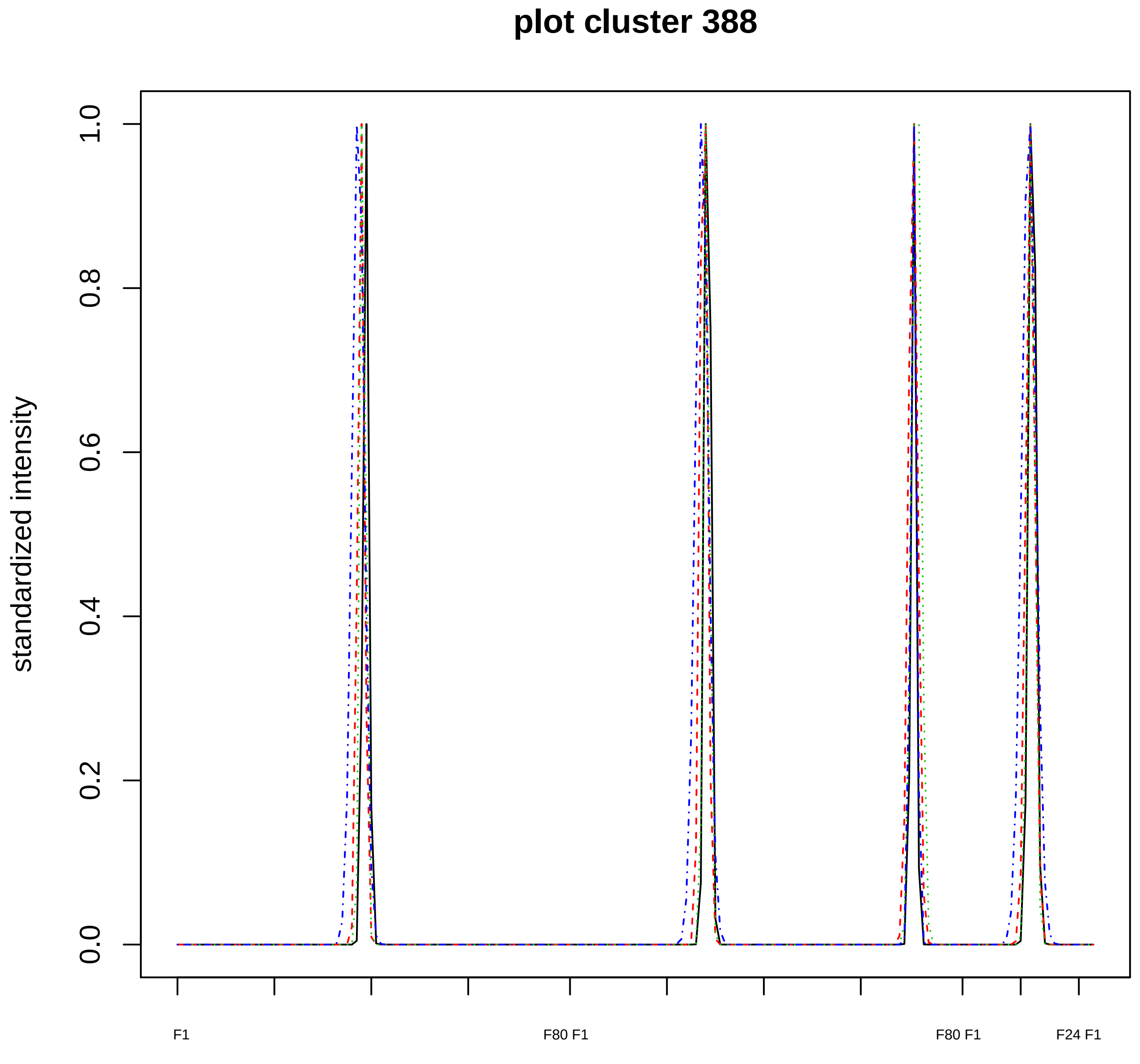
plot cluster 388
standardized intensity
0.0
0.2
0.4
0.6
0.8
1.0
F1
F80 F1
F80 F1
F24 F1

### Dendrogram of cluster 388

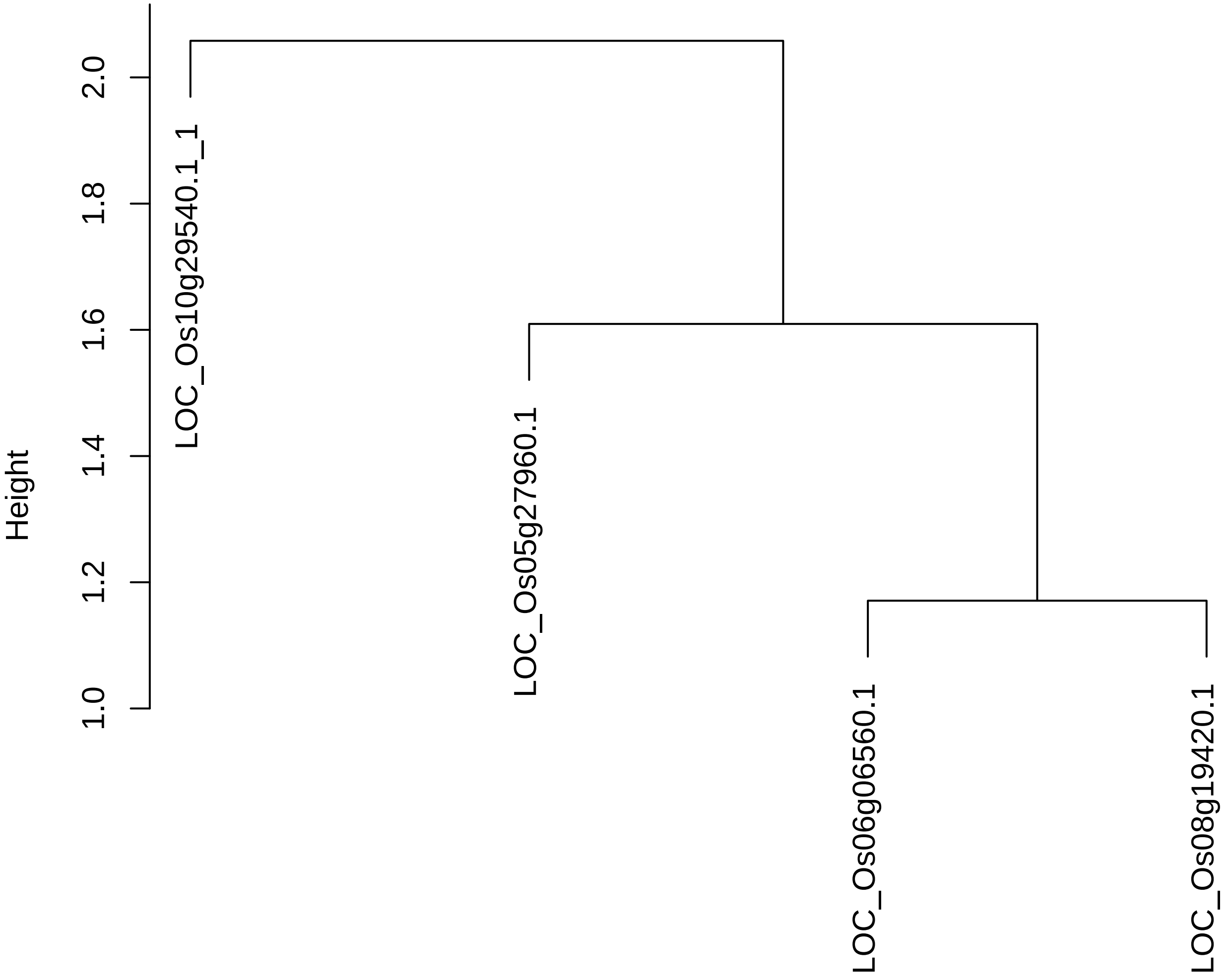


dist(spfitiex12sec12[lab[, 1] == i, ])
hclust (*, "ward.D2")

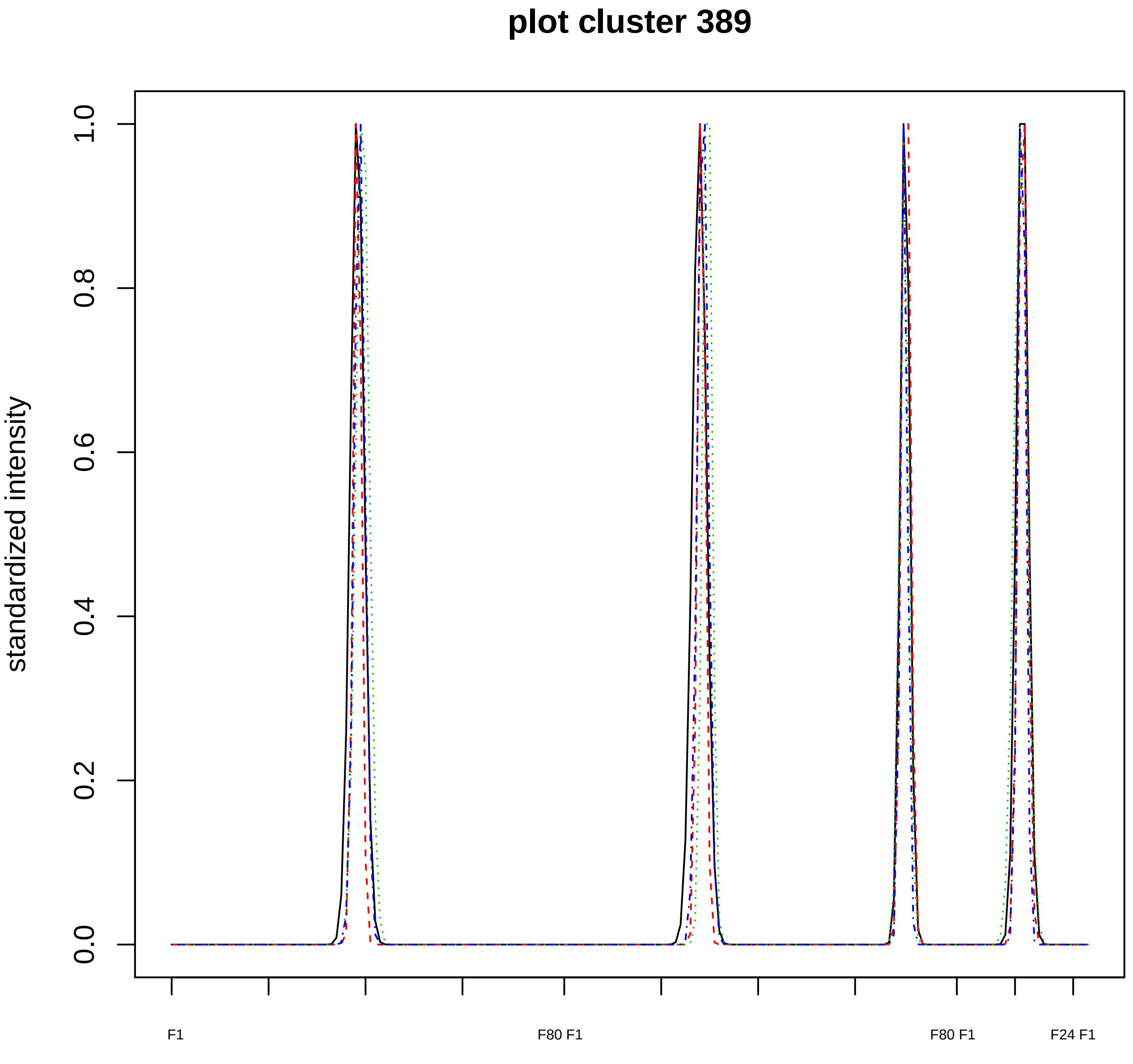
plot cluster 389
standardized intensity
0.0
0.2
0.4
0.6
0.8
1.0
F1
F80 F1
F80 F1
F24 F1

### Dendrogram of cluster 389

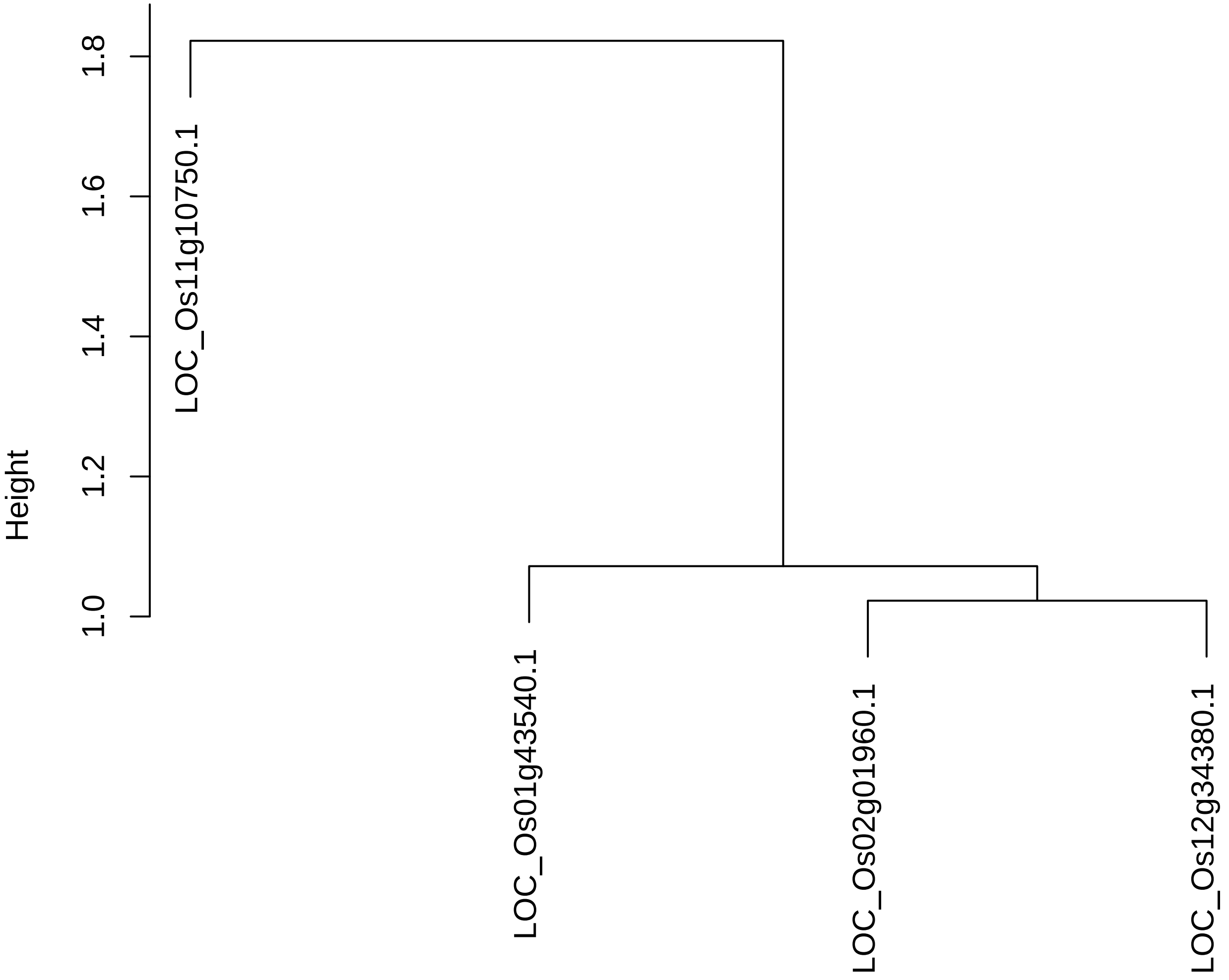


dist(spfitiex12sec12[lab[, 1] == i, ])
hclust (*, "ward.D2")

### plot cluster 390

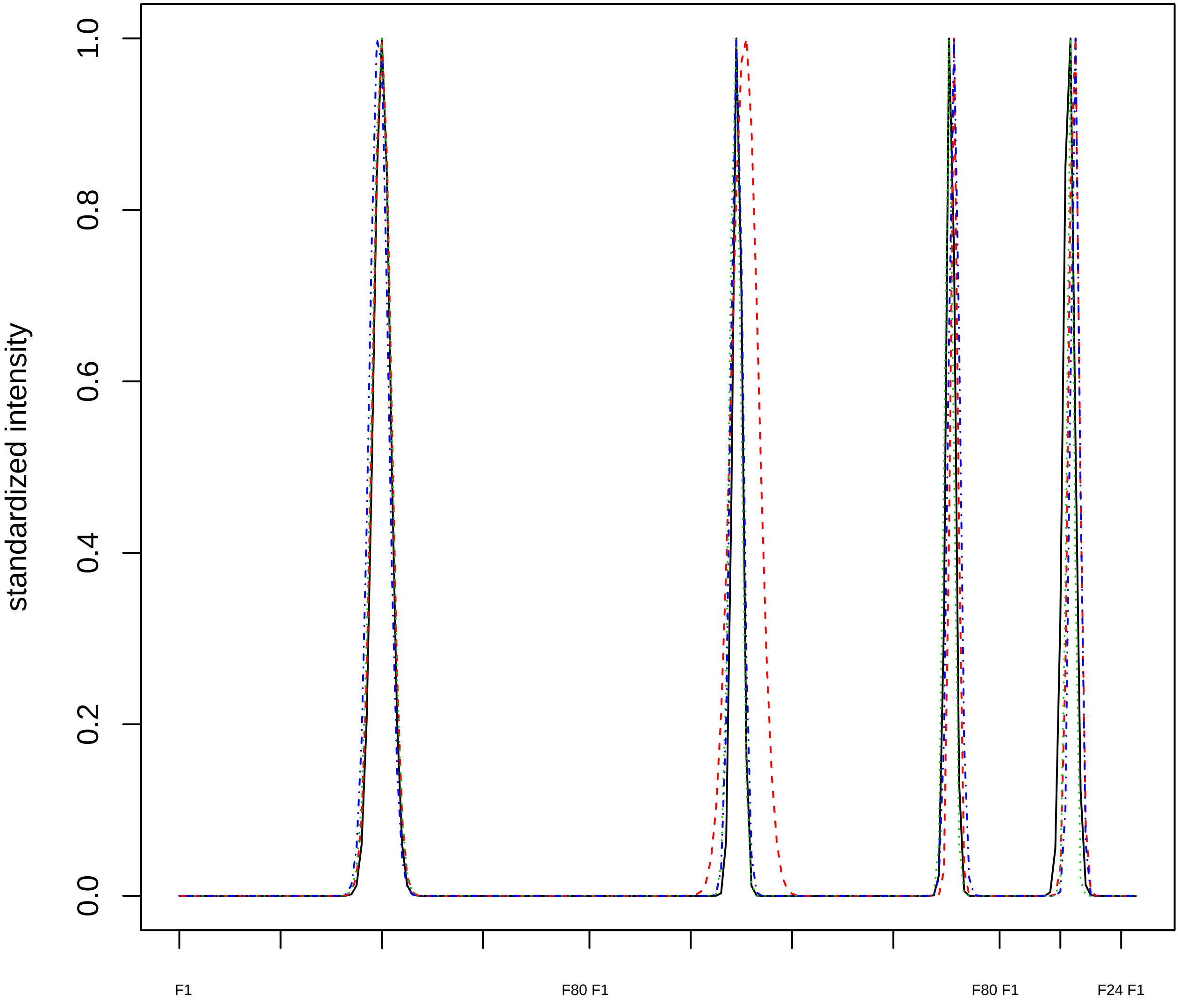

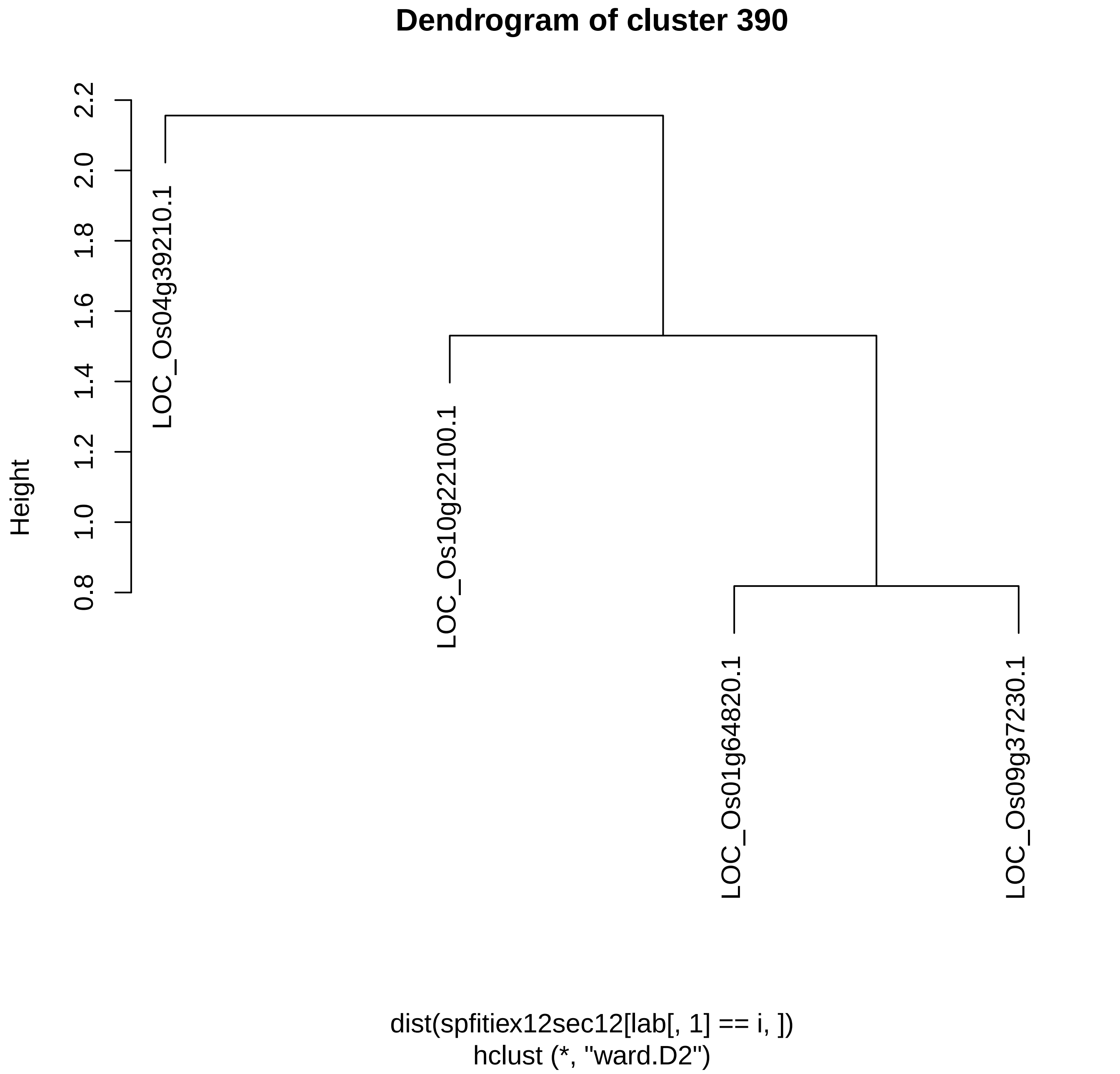
Dendrogram of cluster 390
Height
2.2
2.0
1.8
1.6
1.4
1.2
1.0
0.8
LOC_Os04g39210.1
LOC_Os10g22100.1
LOC_Os01g64820.1
LOC_Os09g37230.1
dist(spfitiex12sec12[lab[, 1] == i, ])
hclust (*, "ward.D2")

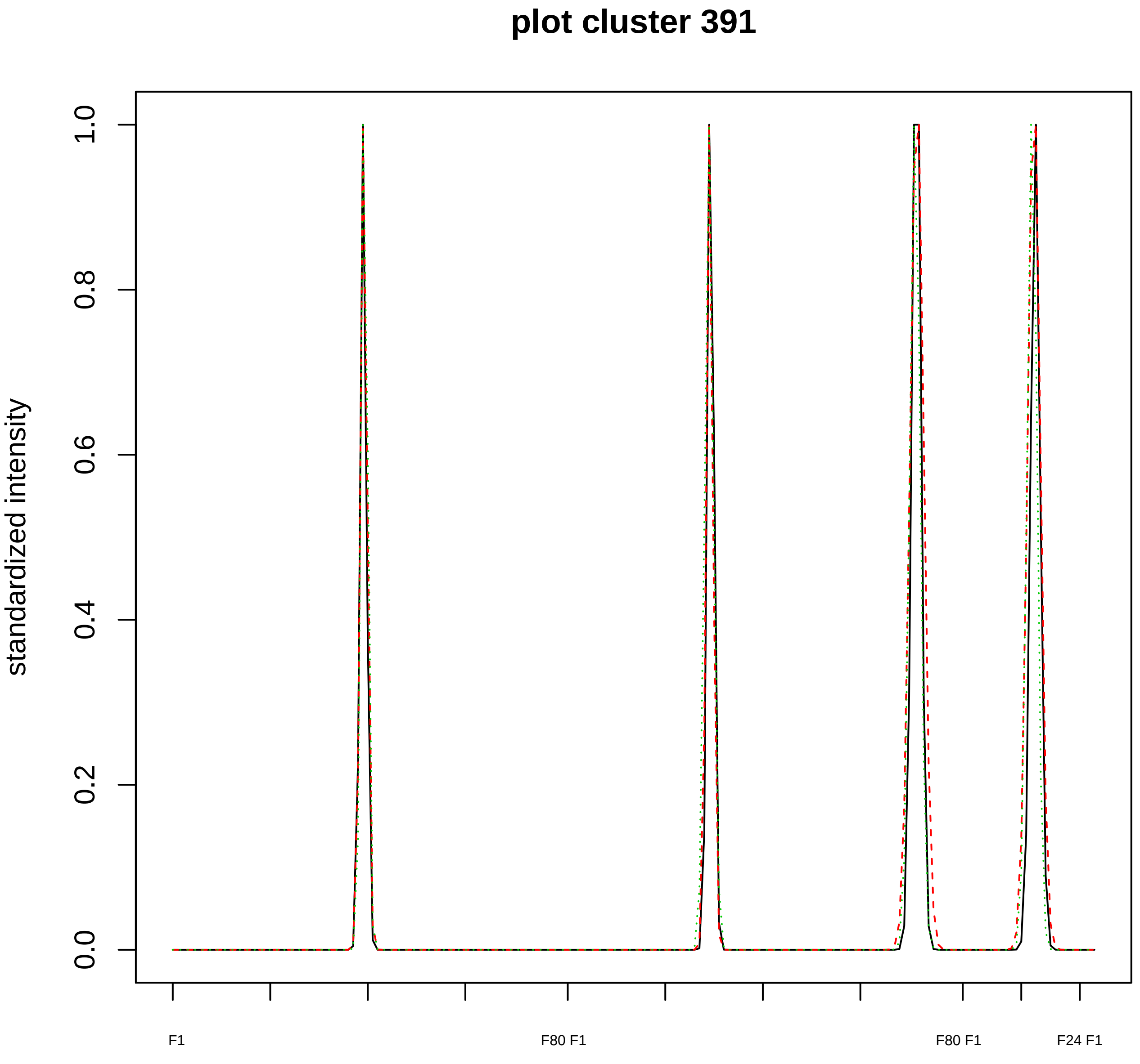
plot cluster 391
standardized intensity
0.0
0.2
0.4
0.6
0.8
1.0
F1
F80 F1
F80 F1
F24 F1

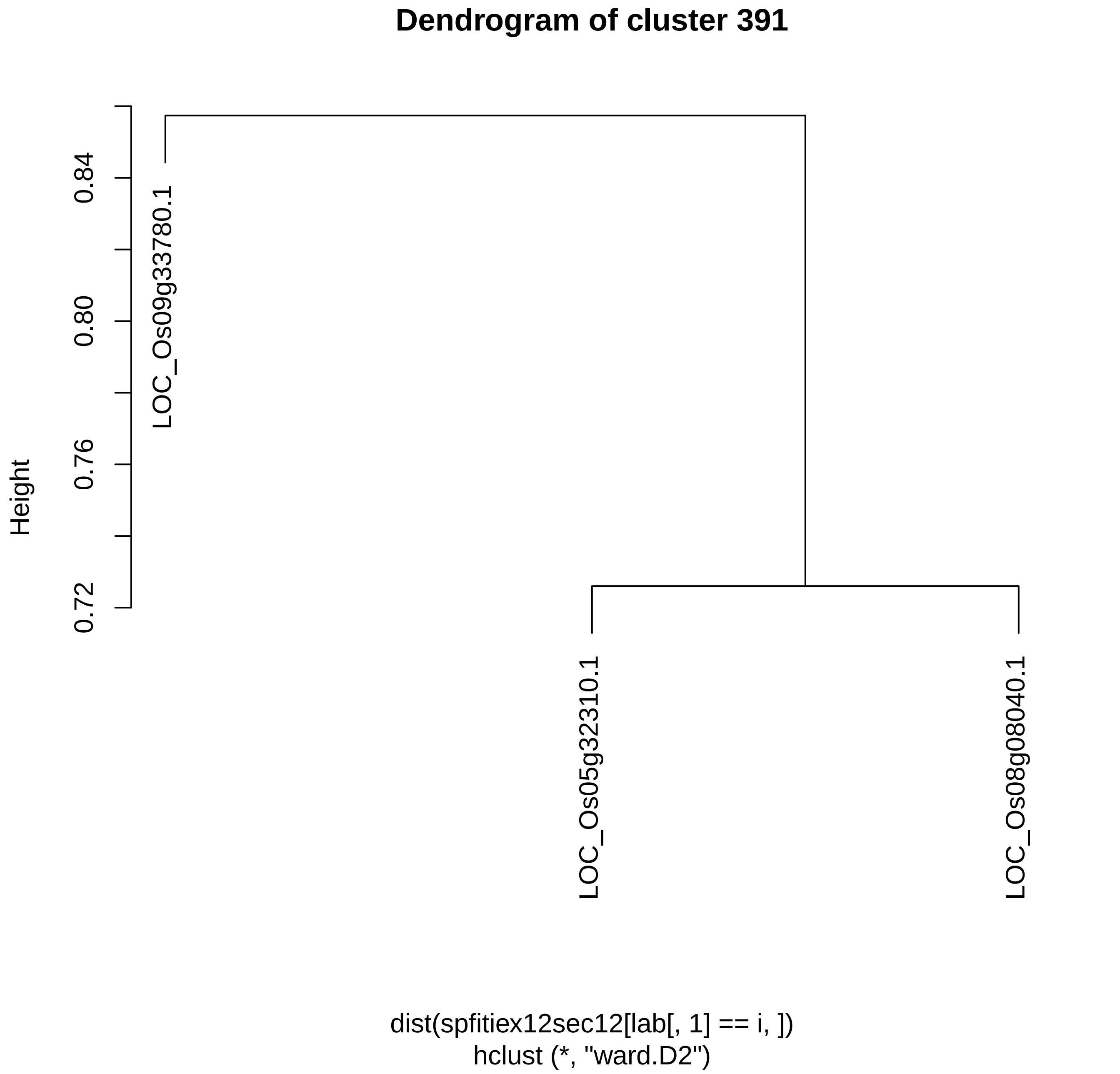
Dendrogram of cluster 391
Height
0.72
0.76
0.80
0.84
LOC_Os09g33780.1
LOC_Os05g32310.1
LOC_Os08g08040.1
dist(spfitiex12sec12[lab[, 1] == i, ])
hclust (*, "ward.D2")

### plot cluster 392

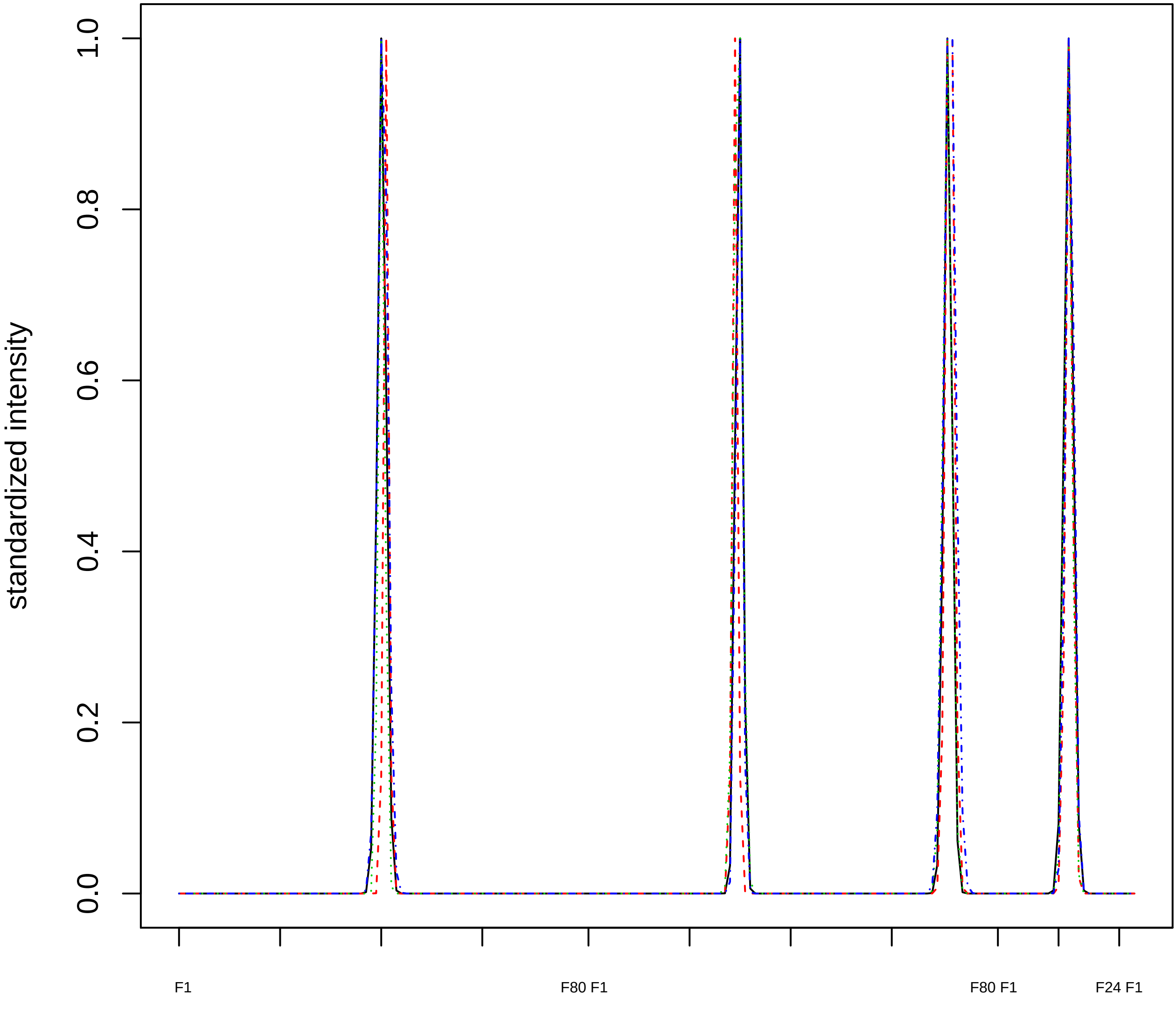

### Dendrogram of cluster 392

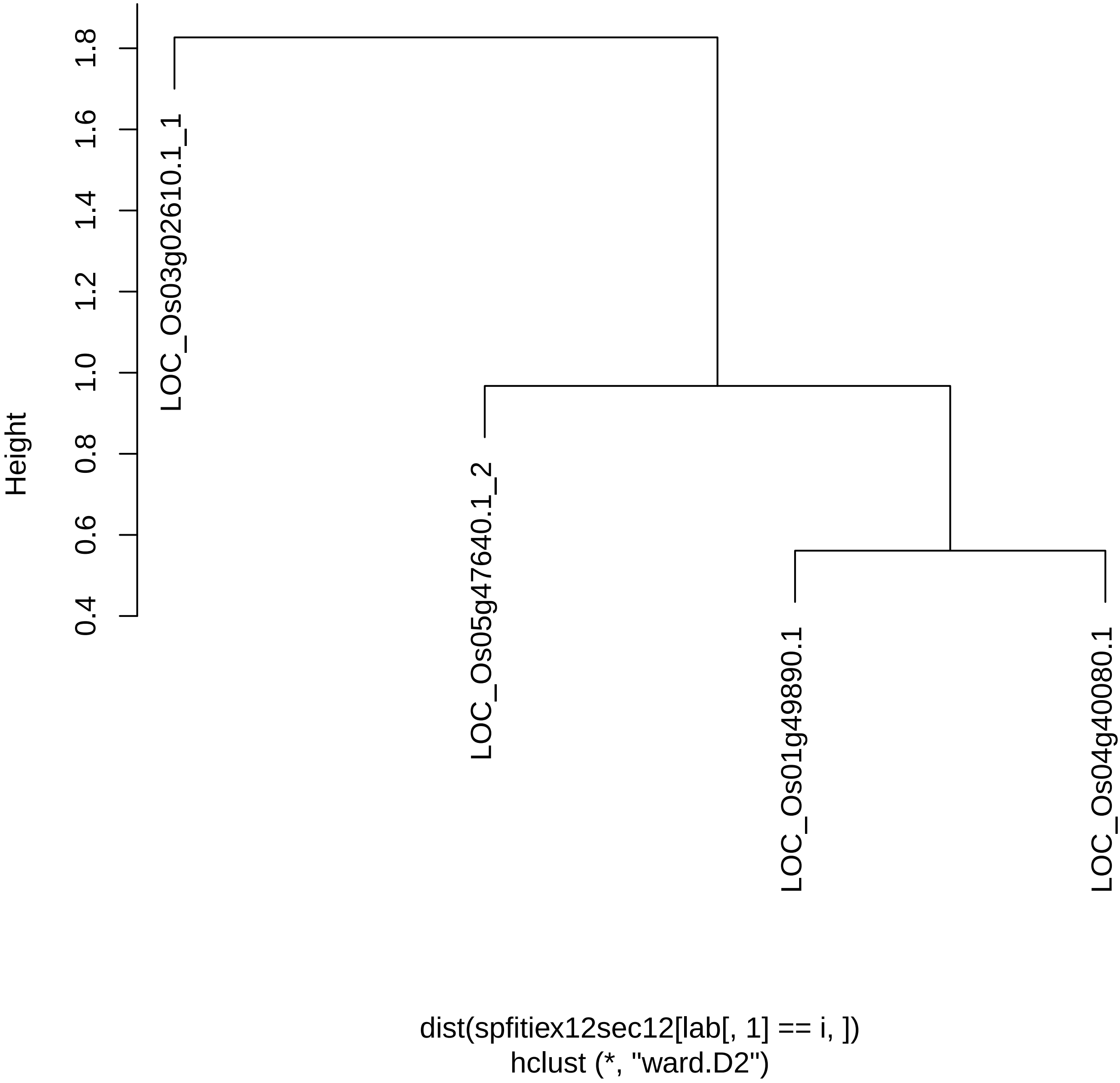

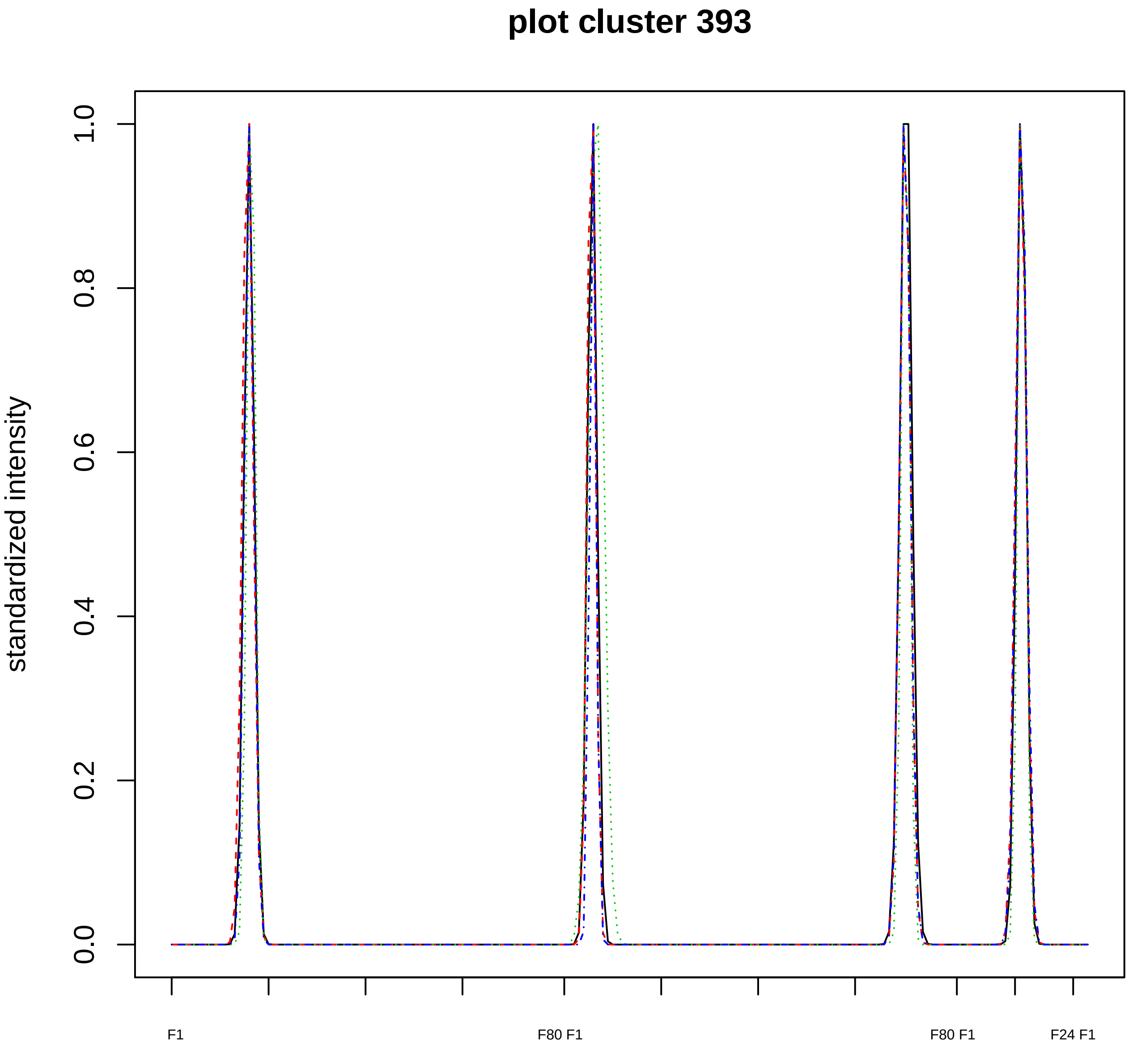

plot cluster 393
standardized intensity
0.0
0.2
0.4
0.6
0.8
1.0
F1
F80 F1
F80 F1
F24 F1

#### Dendrogram of cluster 393

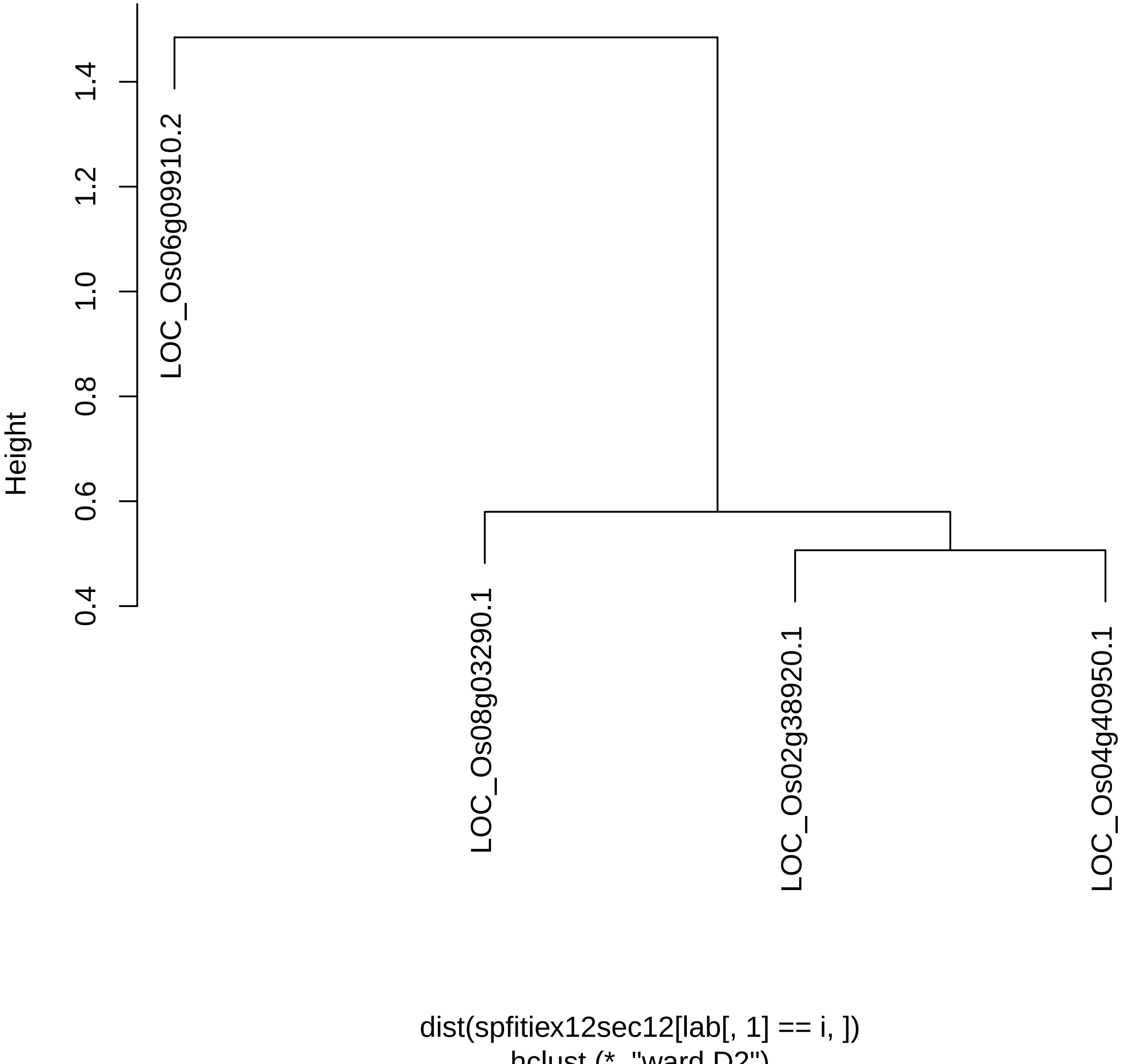

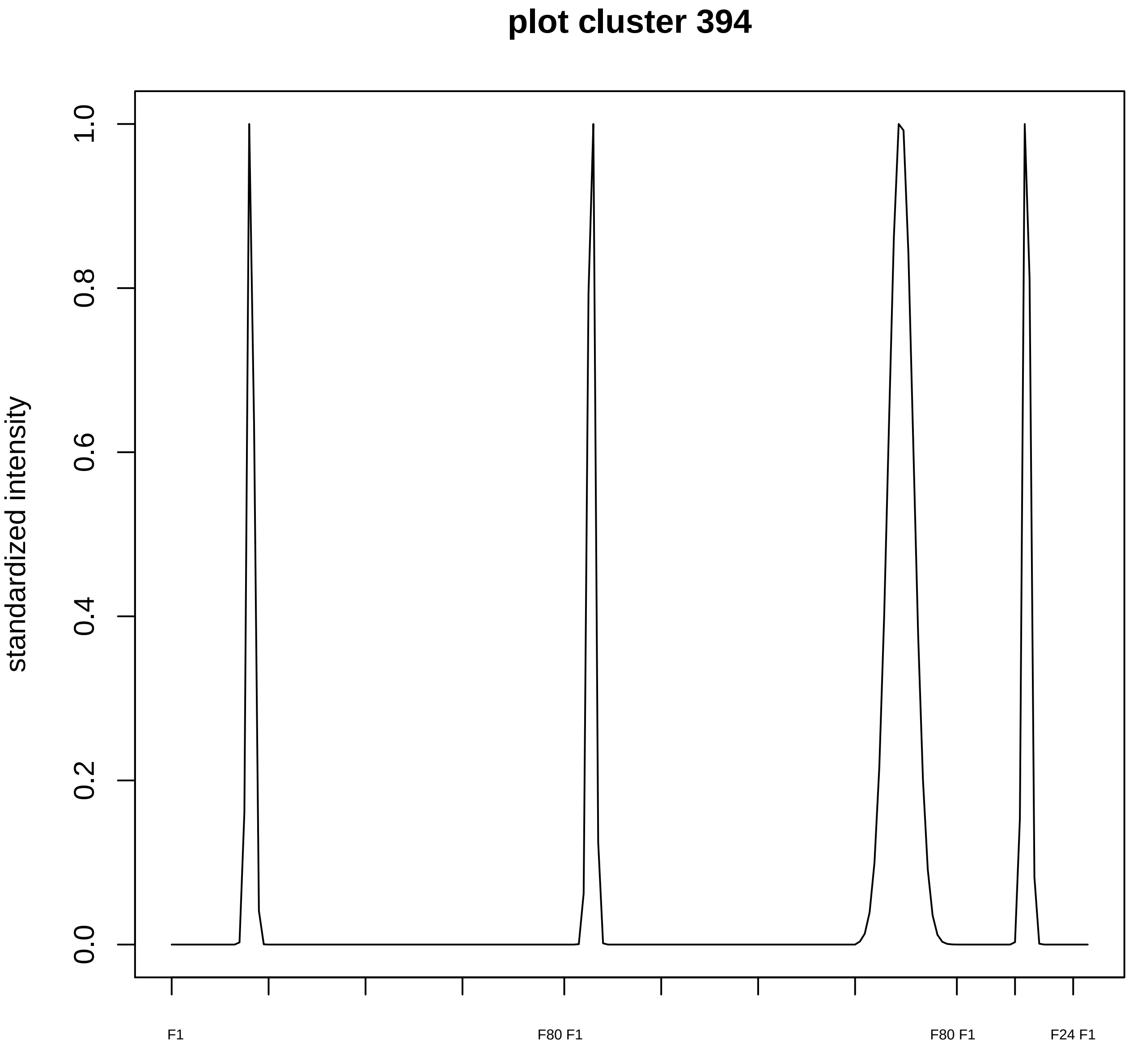

plot cluster 394
standardized intensity
0.0
0.2
0.4
0.6
0.8
1.0
F1
F80 F1
F80 F1
F24 F1

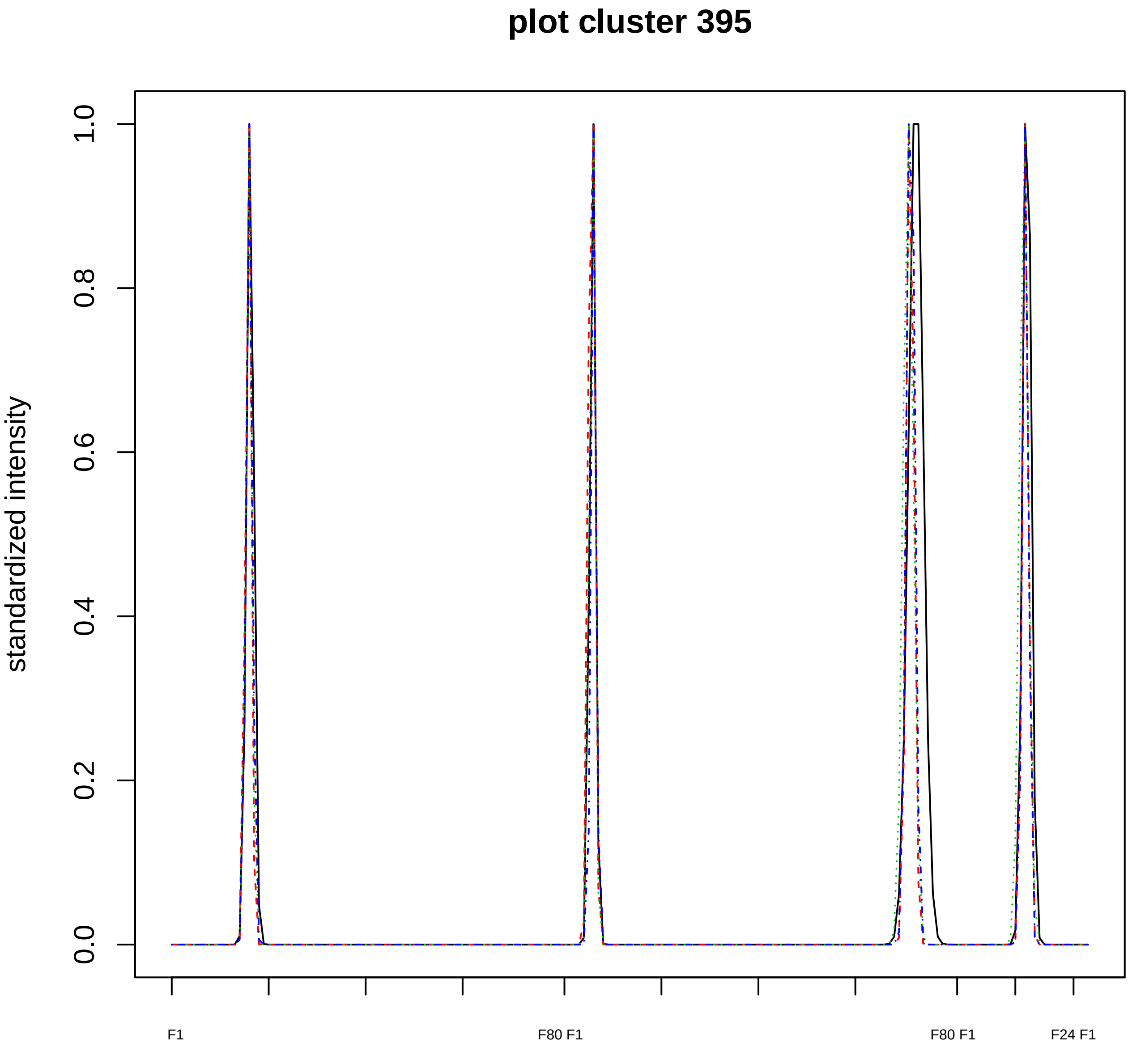

plot cluster 395
standardized intensity
0.0
0.2
0.4
0.6
0.8
1.0
F1
F80 F1
F80 F1
F24 F1

### Dendrogram of cluster 395

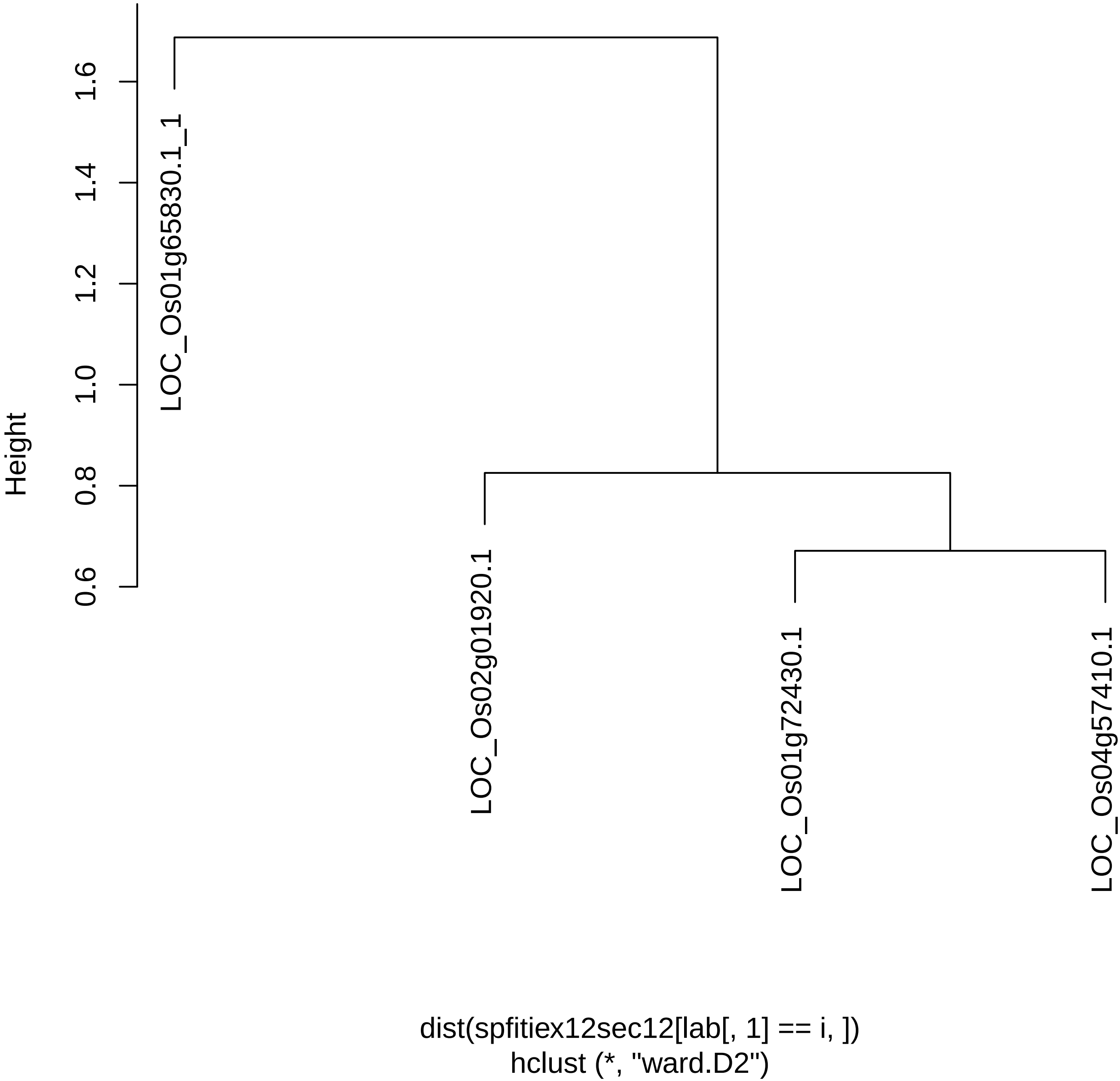

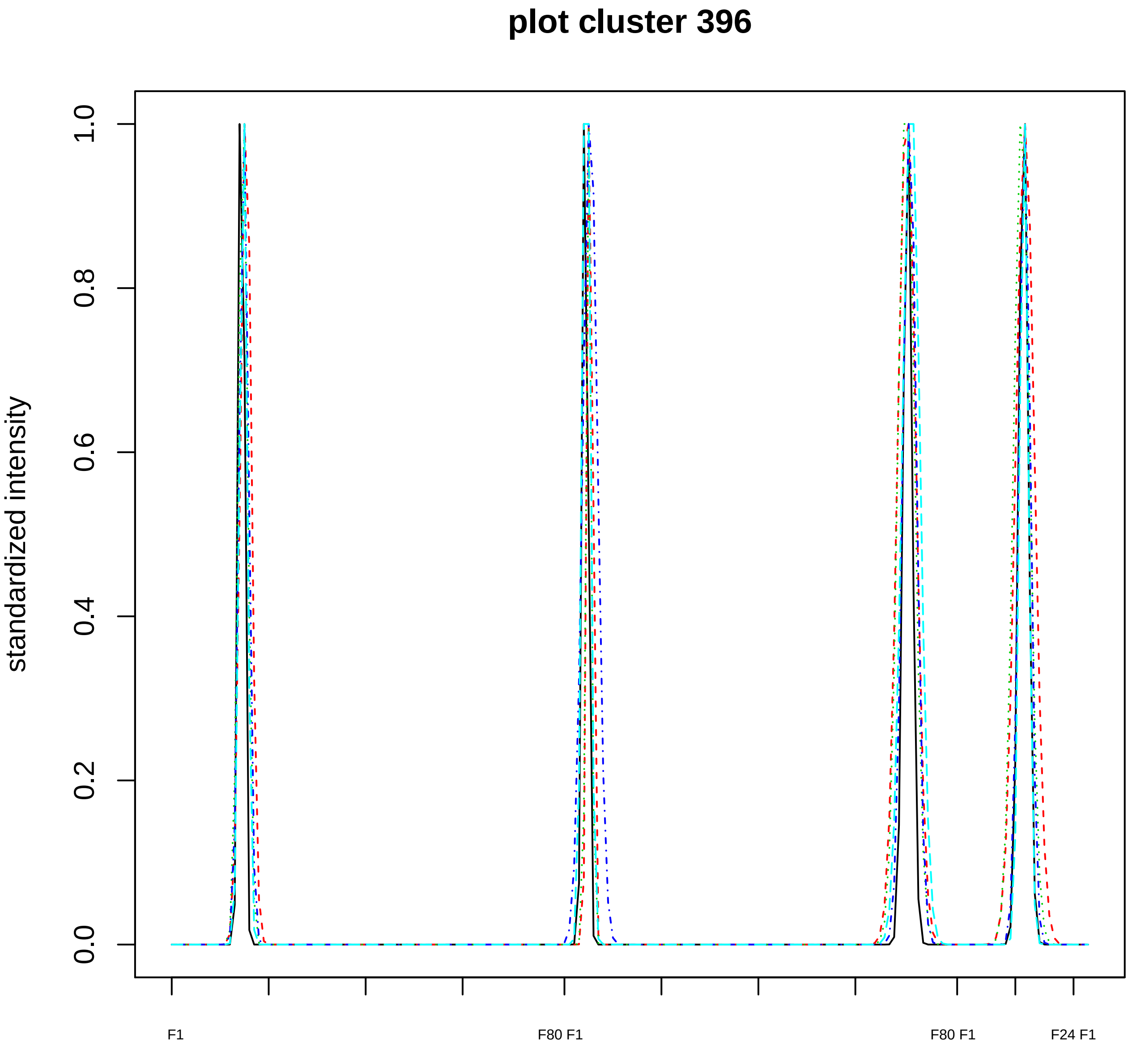
plot cluster 396
standardized intensity
0.0
0.2
0.4
0.6
0.8
1.0
F1
F80 F1
F80 F1
F24 F1

### Dendrogram of cluster 396

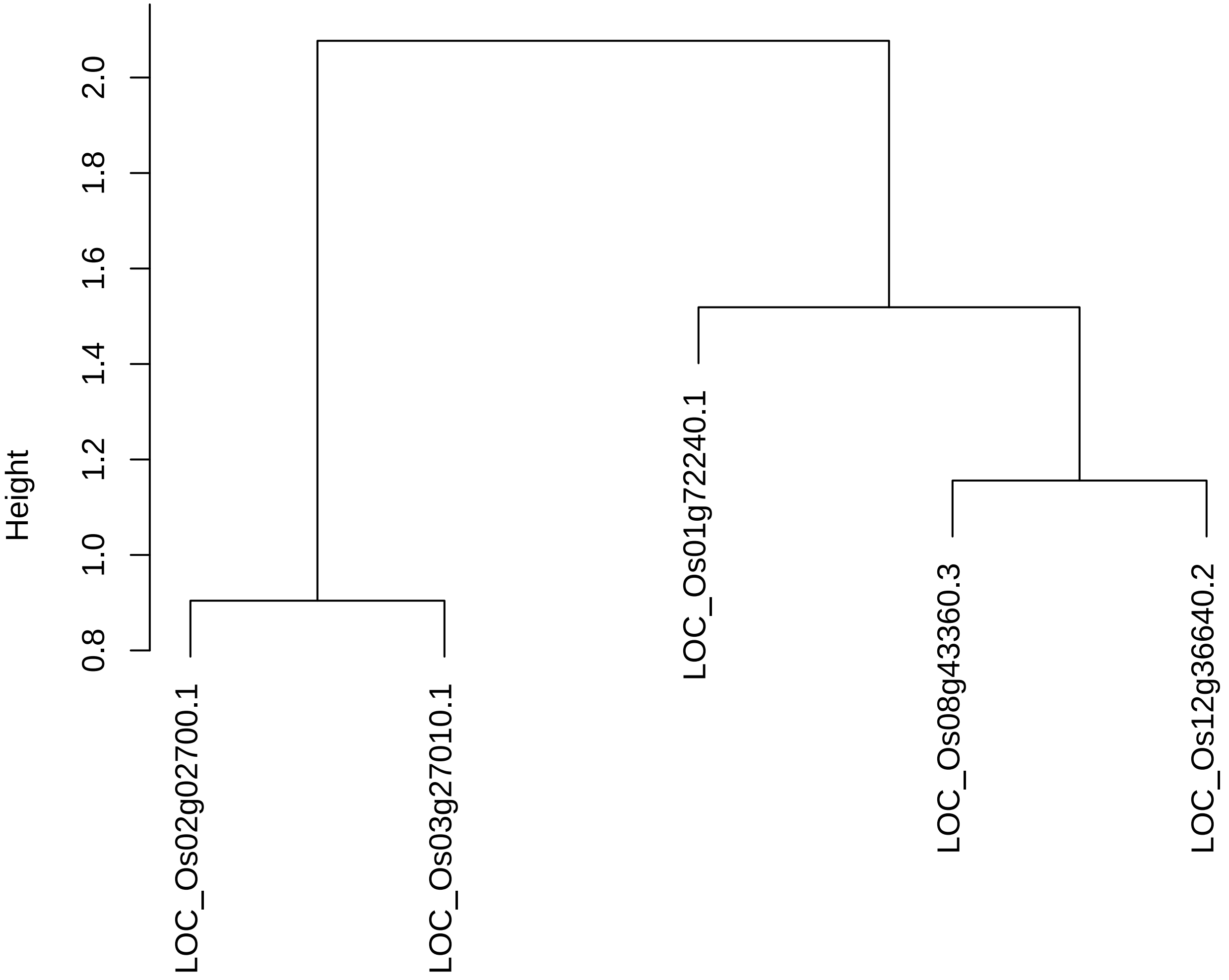


dist(spfitiex12sec12[lab[, 1] == i, ])
hclust (*, "ward.D2")

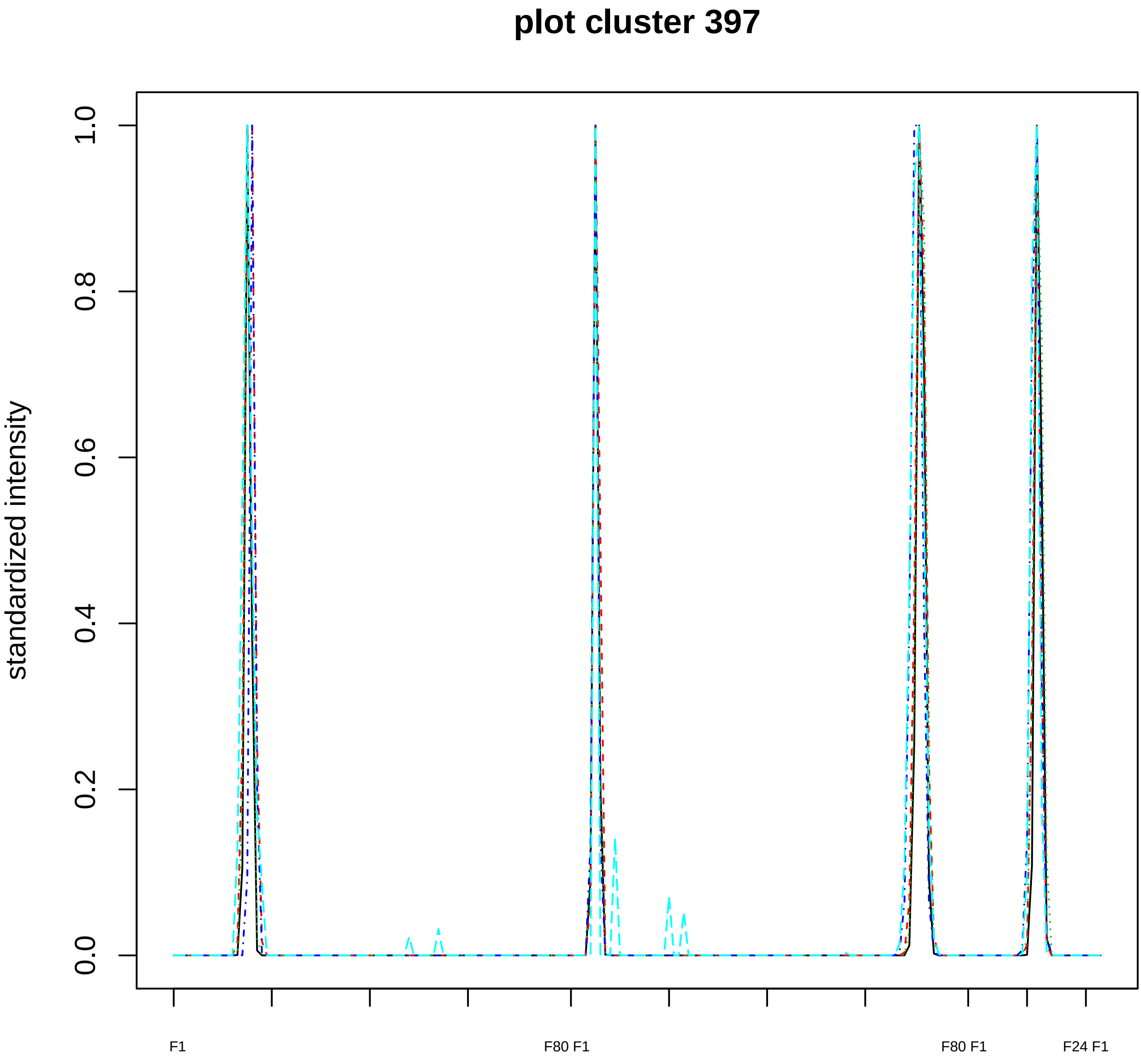

plot cluster 397
standardized intensity
0.0
0.2
0.4
0.6
0.8
1.0
F1
F80 F1
F80 F1
F24 F1

### Dendrogram of cluster 397

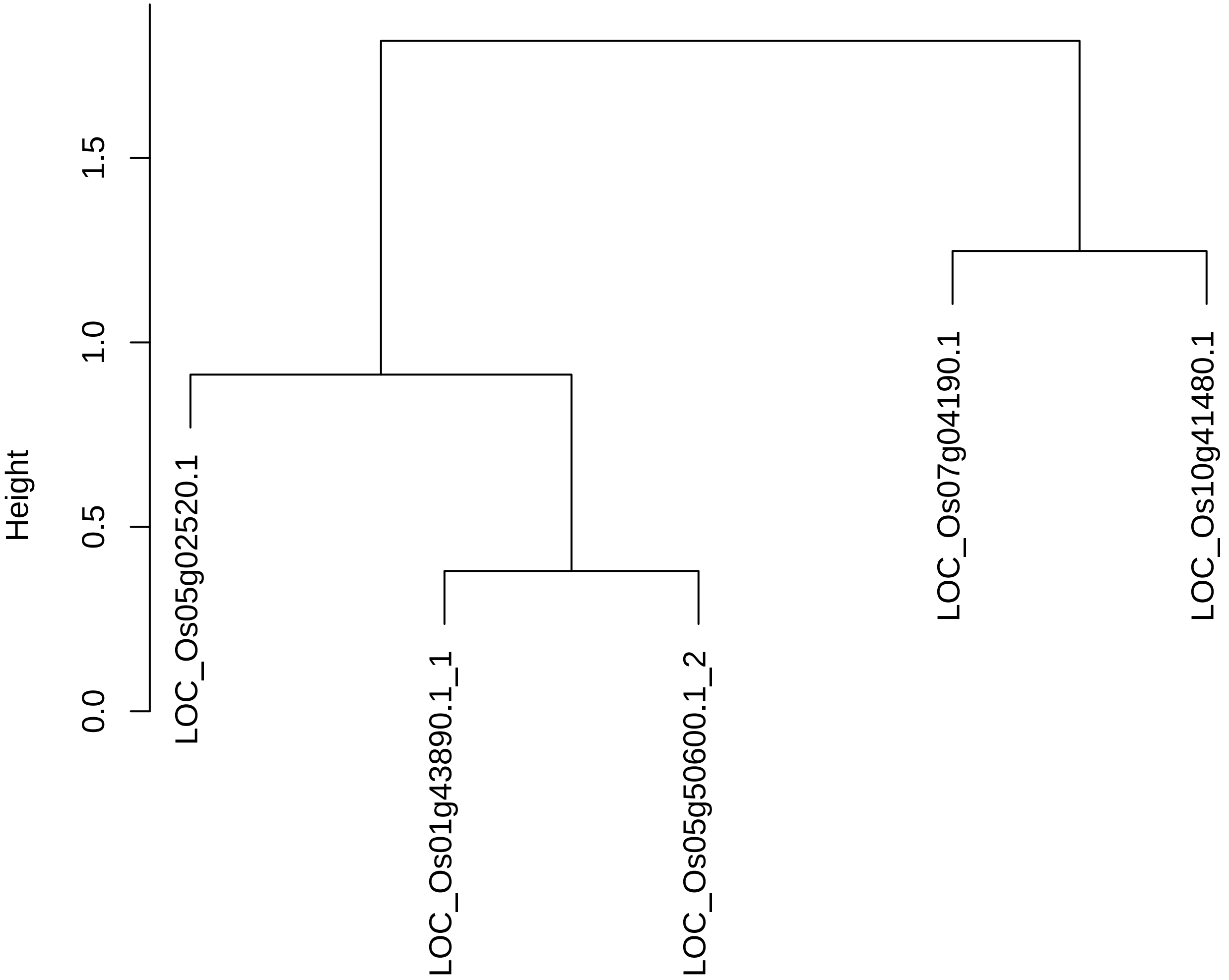


dist(spfitiex12sec12[lab[, 1] == i, ])
hclust (*, "ward.D2")

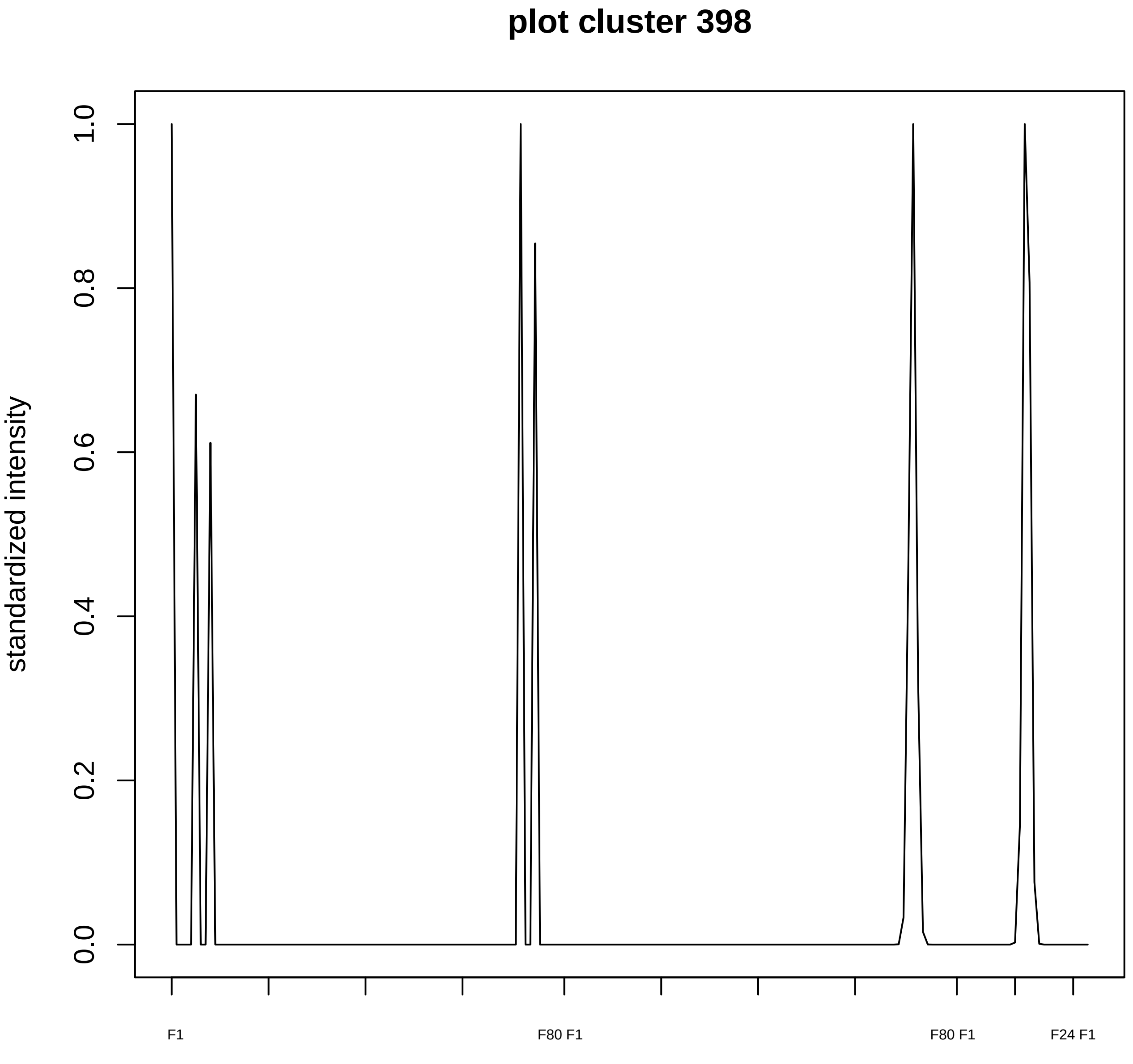
plot cluster 398
standardized intensity
0.0
0.2
0.4
0.6
0.8
1.0
F1
F80 F1
F80 F1
F24 F1

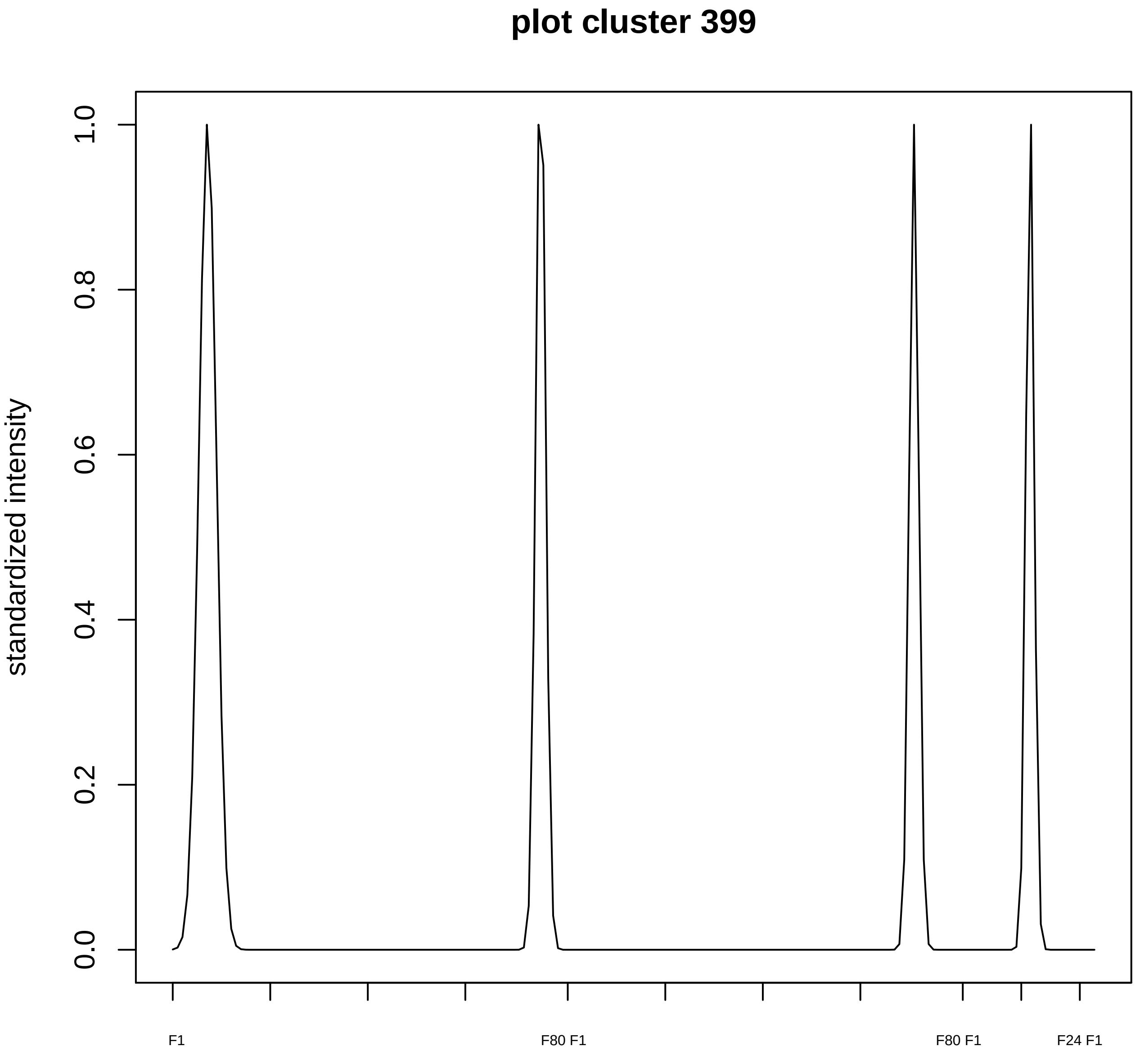
plot cluster 399
standardized intensity
0.0
0.2
0.4
0.6
0.8
1.0
F1
F80 F1
F80 F1
F24 F1

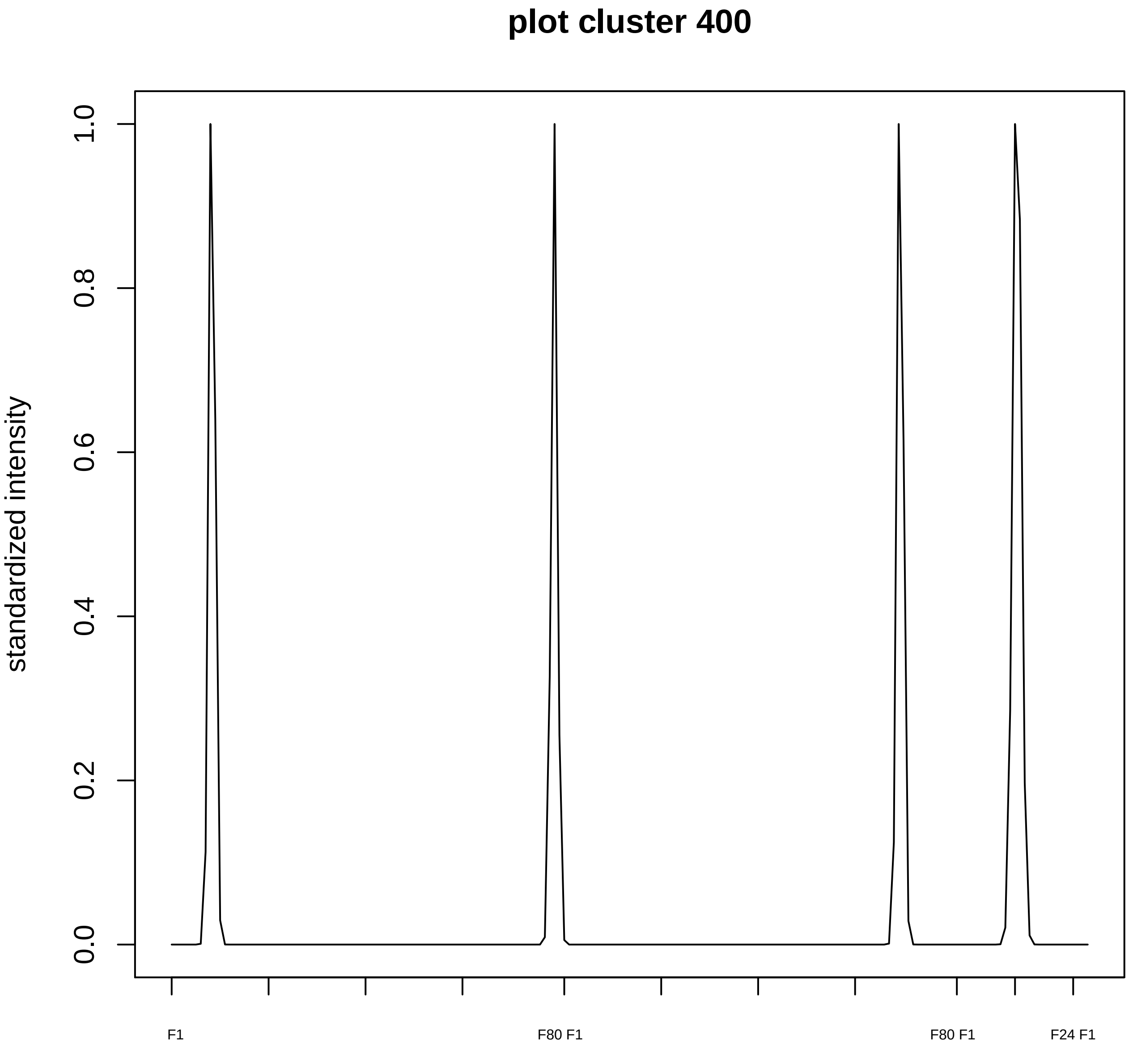
plot cluster 400
standardized intensity
0.0
0.2
0.4
0.6
0.8
1.0
F1
F80 F1
F80 F1
F24 F1

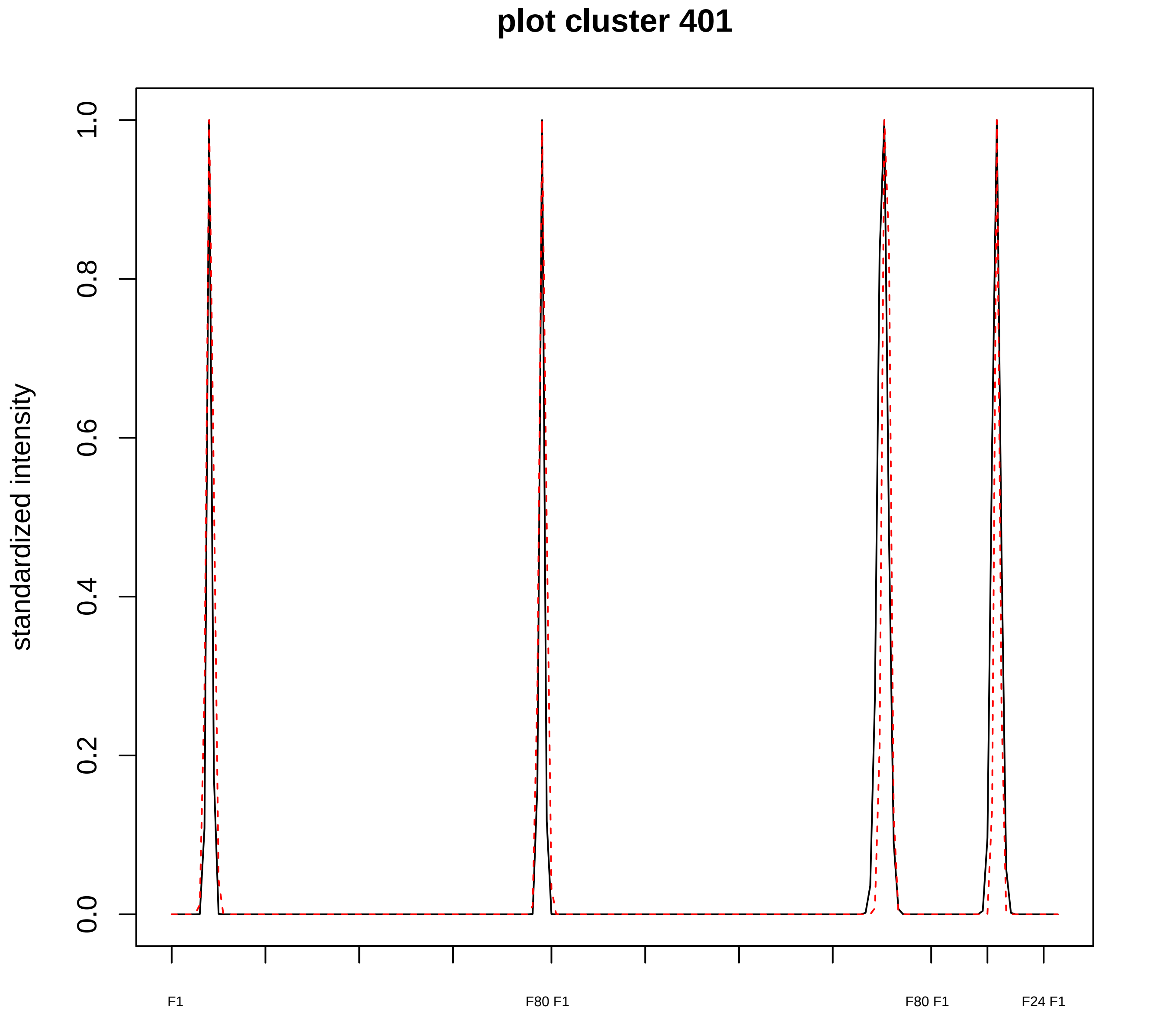
plot cluster 401
standardized intensity
0.0
0.2
0.4
0.6
0.8
1.0
F1
F80 F1
F80 F1
F24 F1

#### plot cluster 402

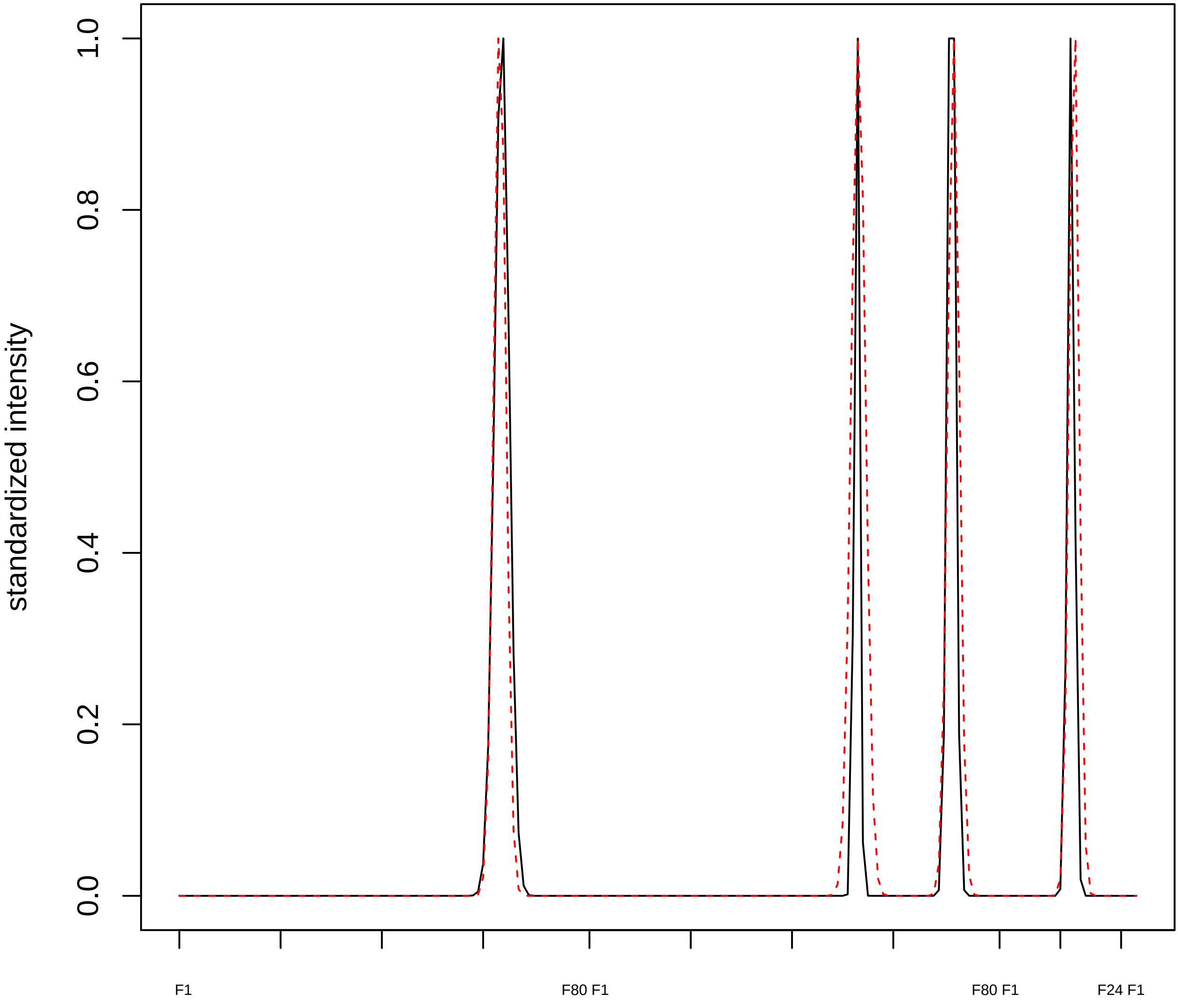

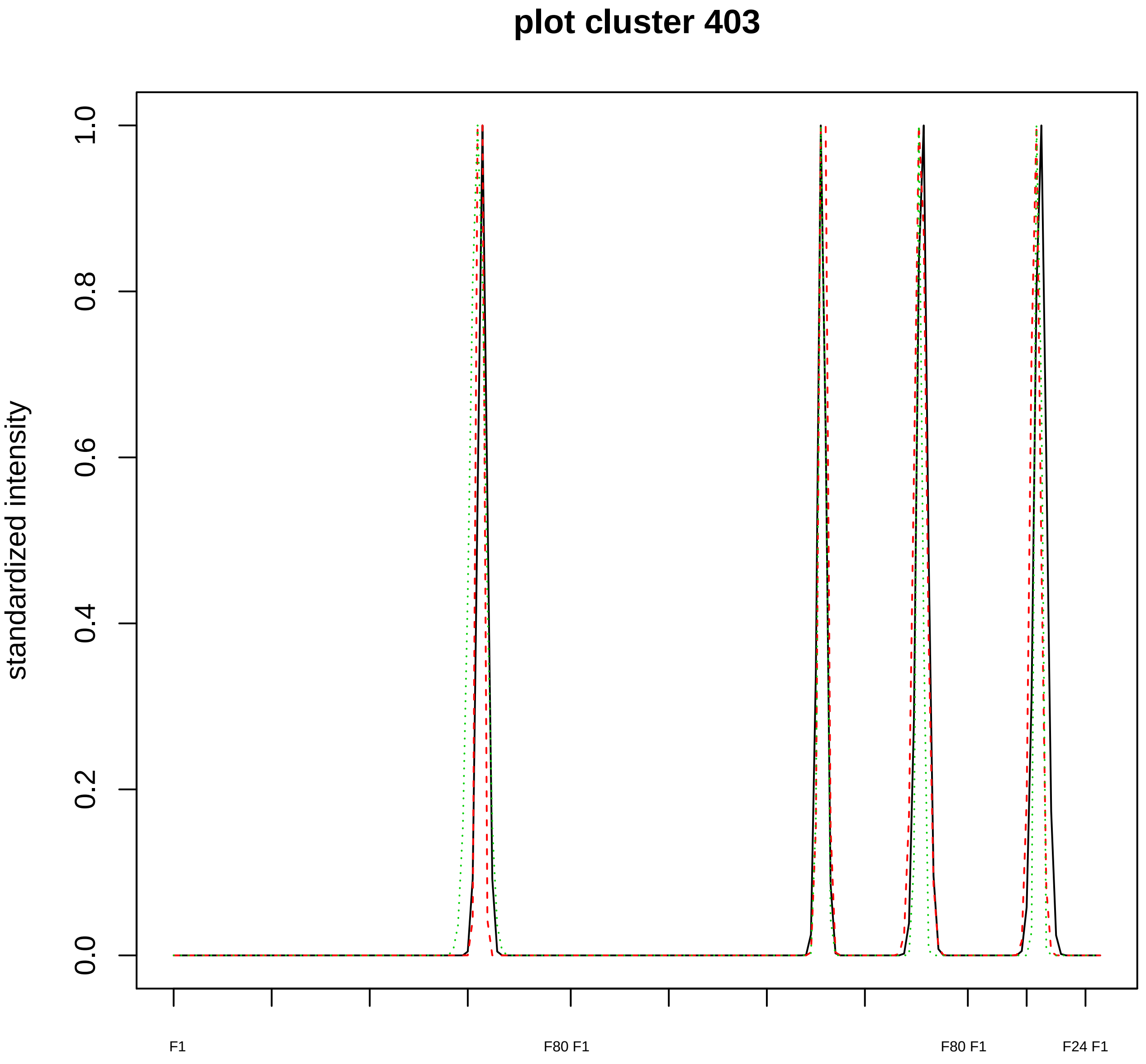

plot cluster 403
standardized intensity
0.0
0.2
0.4
0.6
0.8
1.0
F1
F80 F1
F80 F1
F24 F1

#### Dendrogram of cluster 403

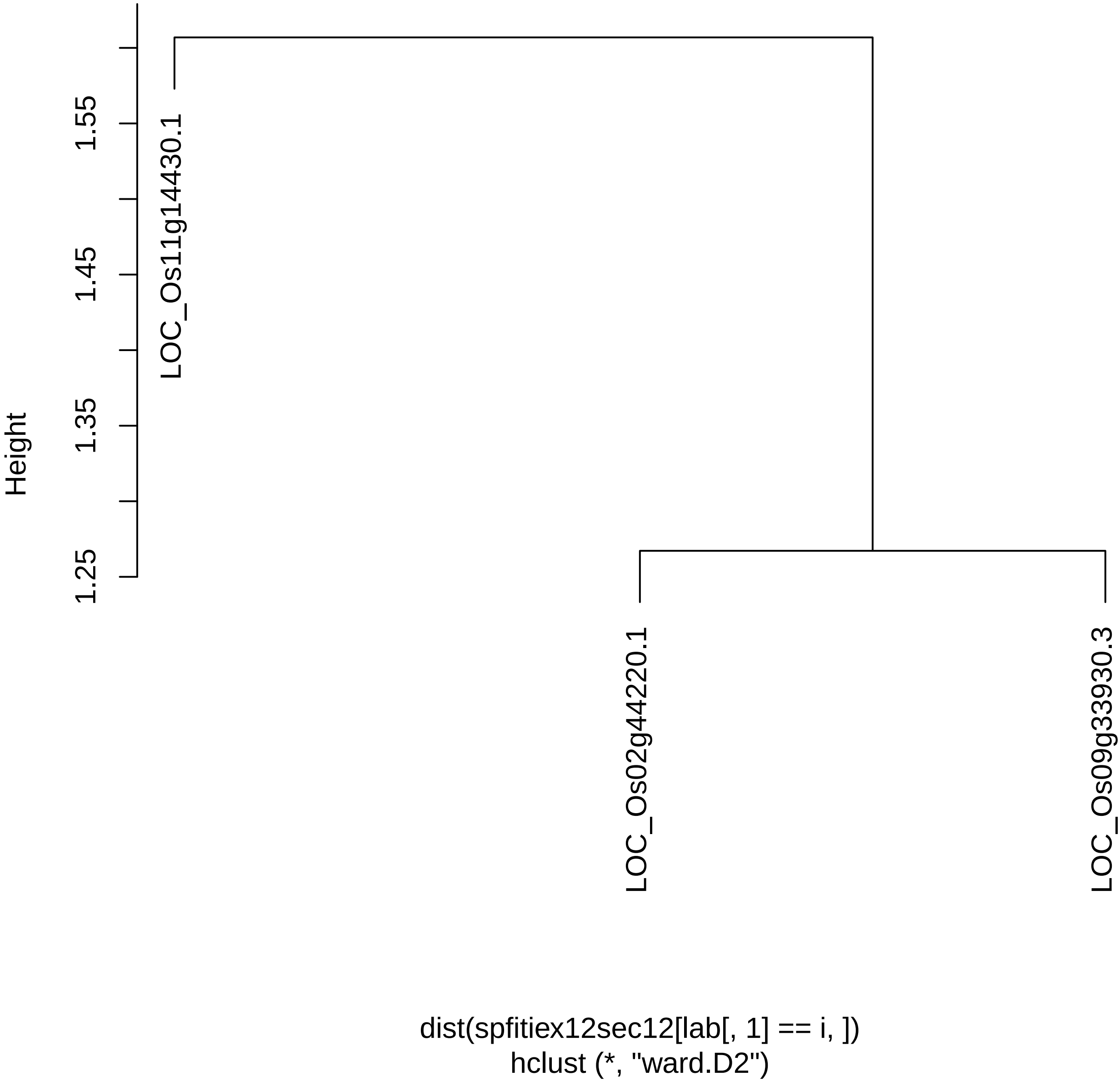

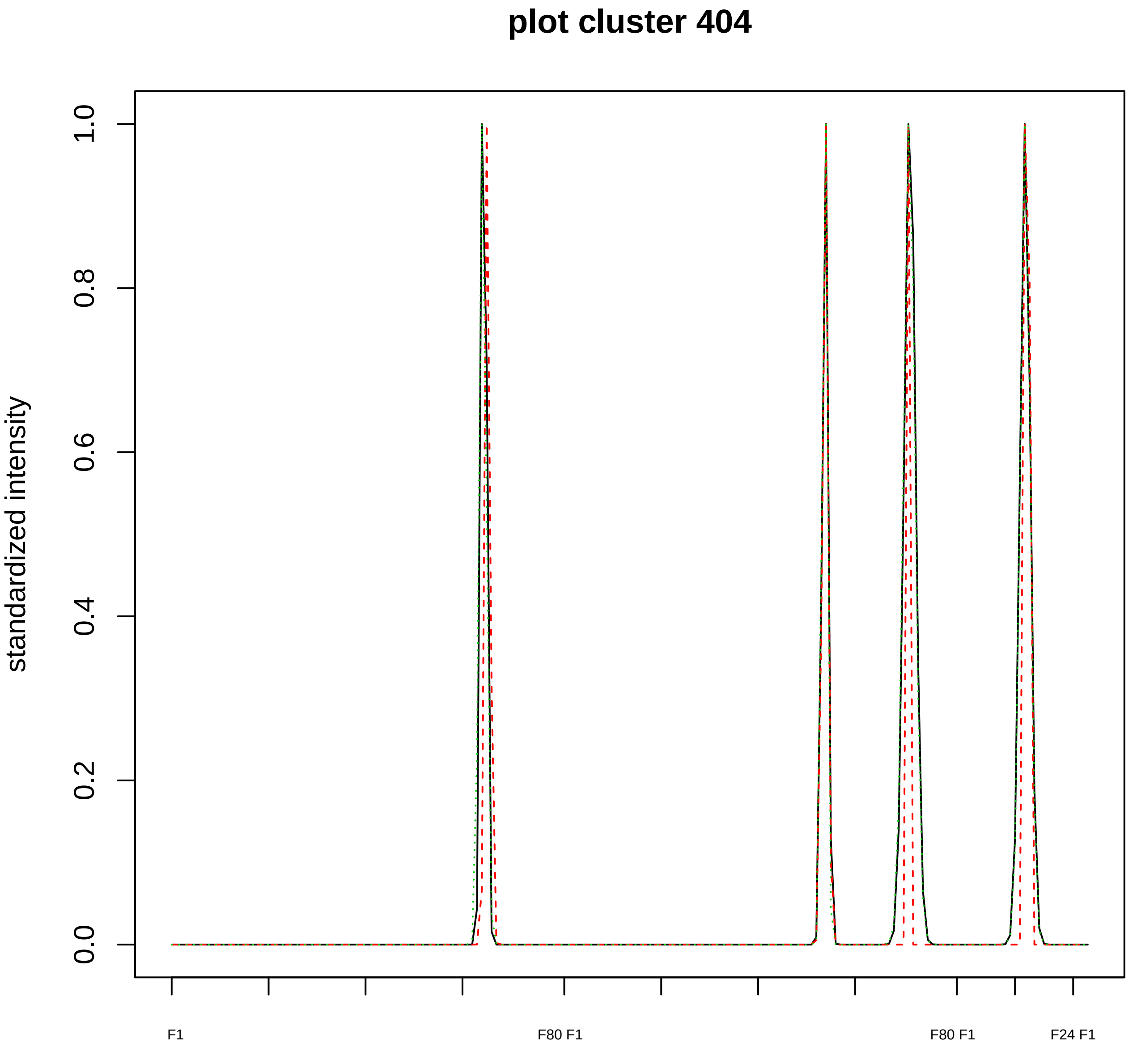

plot cluster 404
standardized intensity
0.0
0.2
0.4
0.6
0.8
1.0
F1
F80 F1
F80 F1
F24 F1

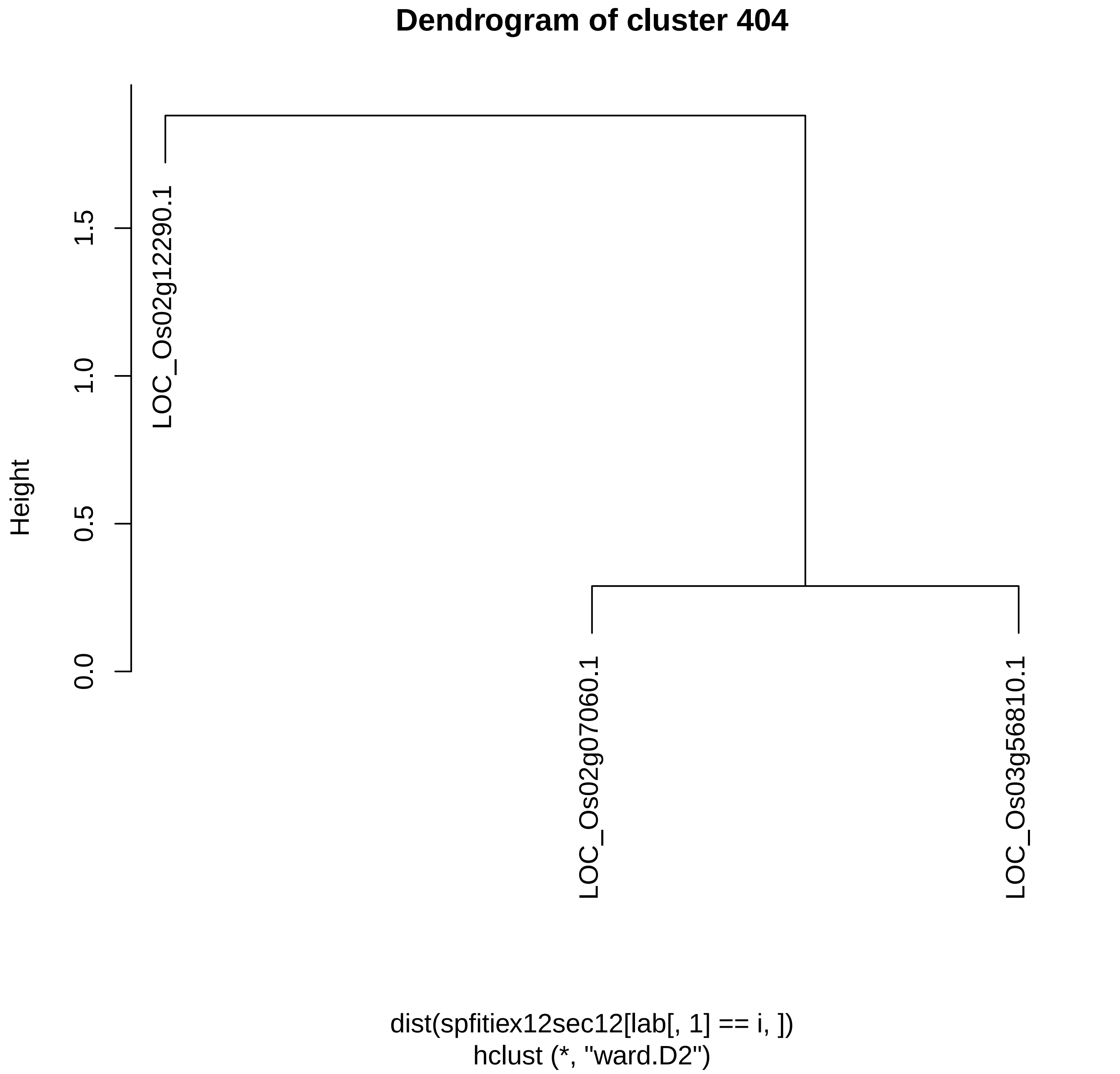

Dendrogram of cluster 404
Height
0.0
0.5
1.0
1.5
LOC_Os02g12290.1
LOC_Os02g07060.1
LOC_Os03g56810.1
dist(spfitiex12sec12[lab[, 1] == i, ])
hclust (*, "ward.D2")

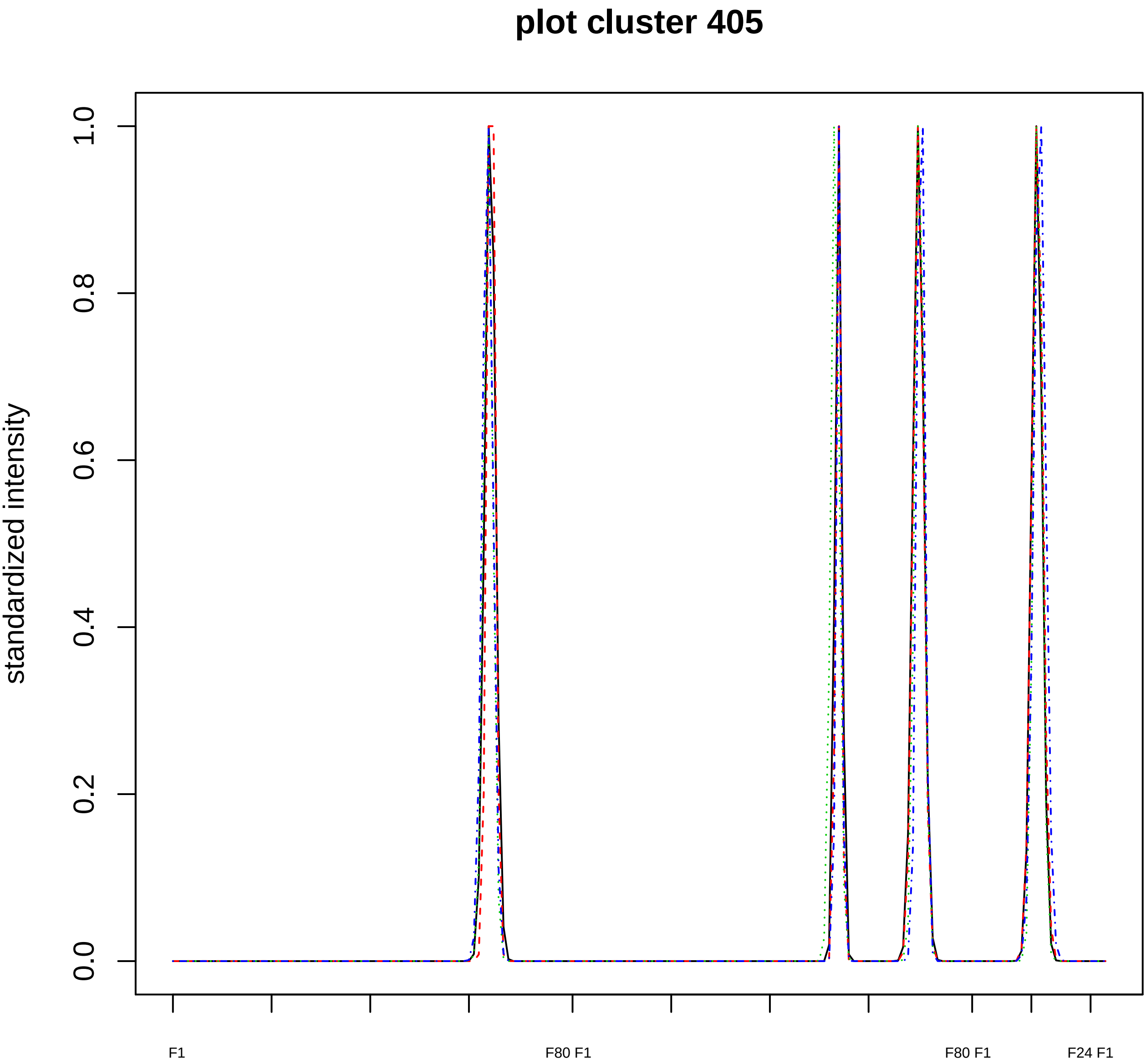
plot cluster 405
standardized intensity
0.0
0.2
0.4
0.6
0.8
1.0
F1
F80 F1
F80 F1
F24 F1

### Dendrogram of cluster 405

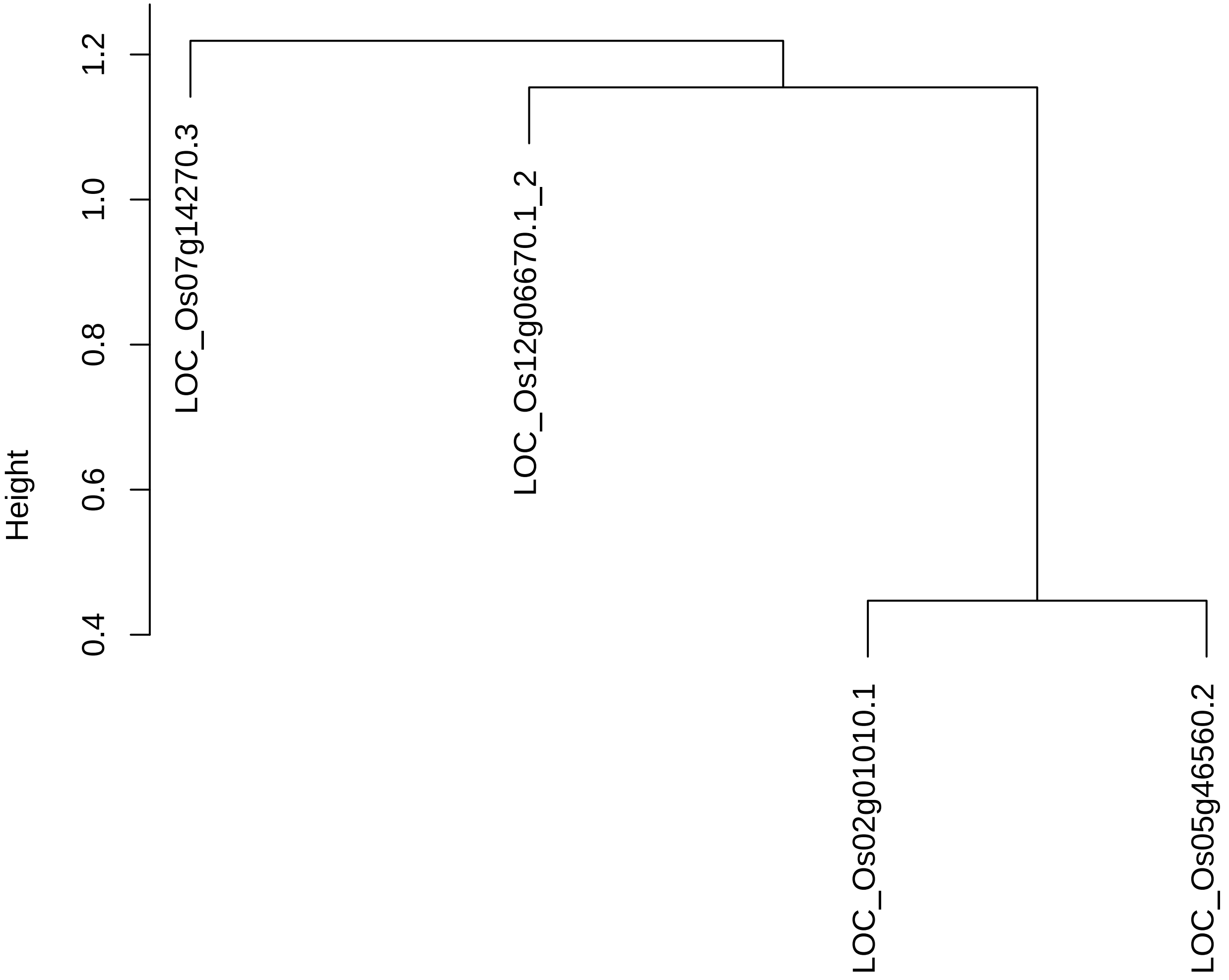


dist(spfitiex12sec12[lab[, 1] == i, ])
hclust (*, "ward.D2")

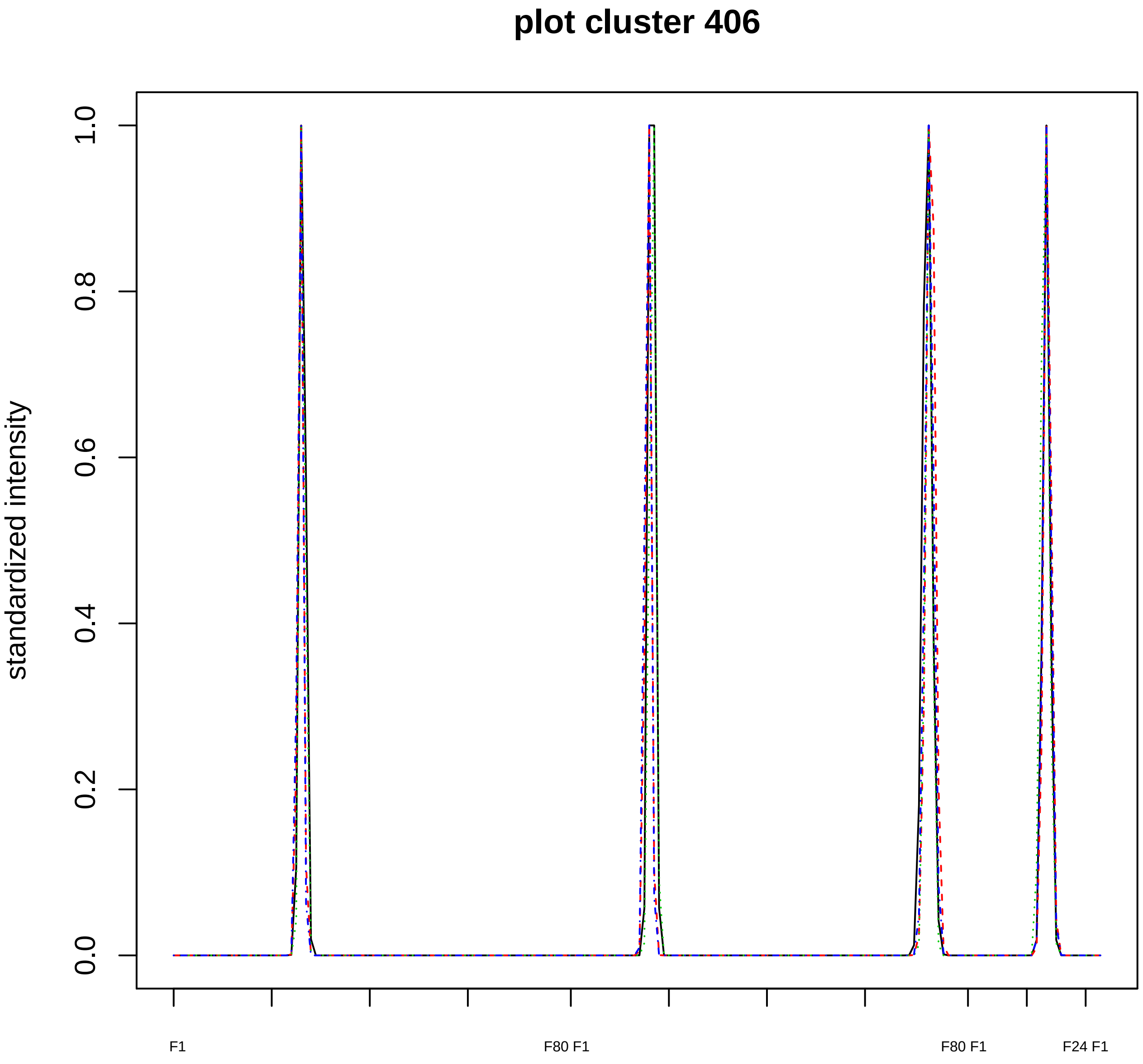
plot cluster 406
standardized intensity
0.0
0.2
0.4
0.6
0.8
1.0
F1
F80 F1
F80 F1
F24 F1

### Dendrogram of cluster 406

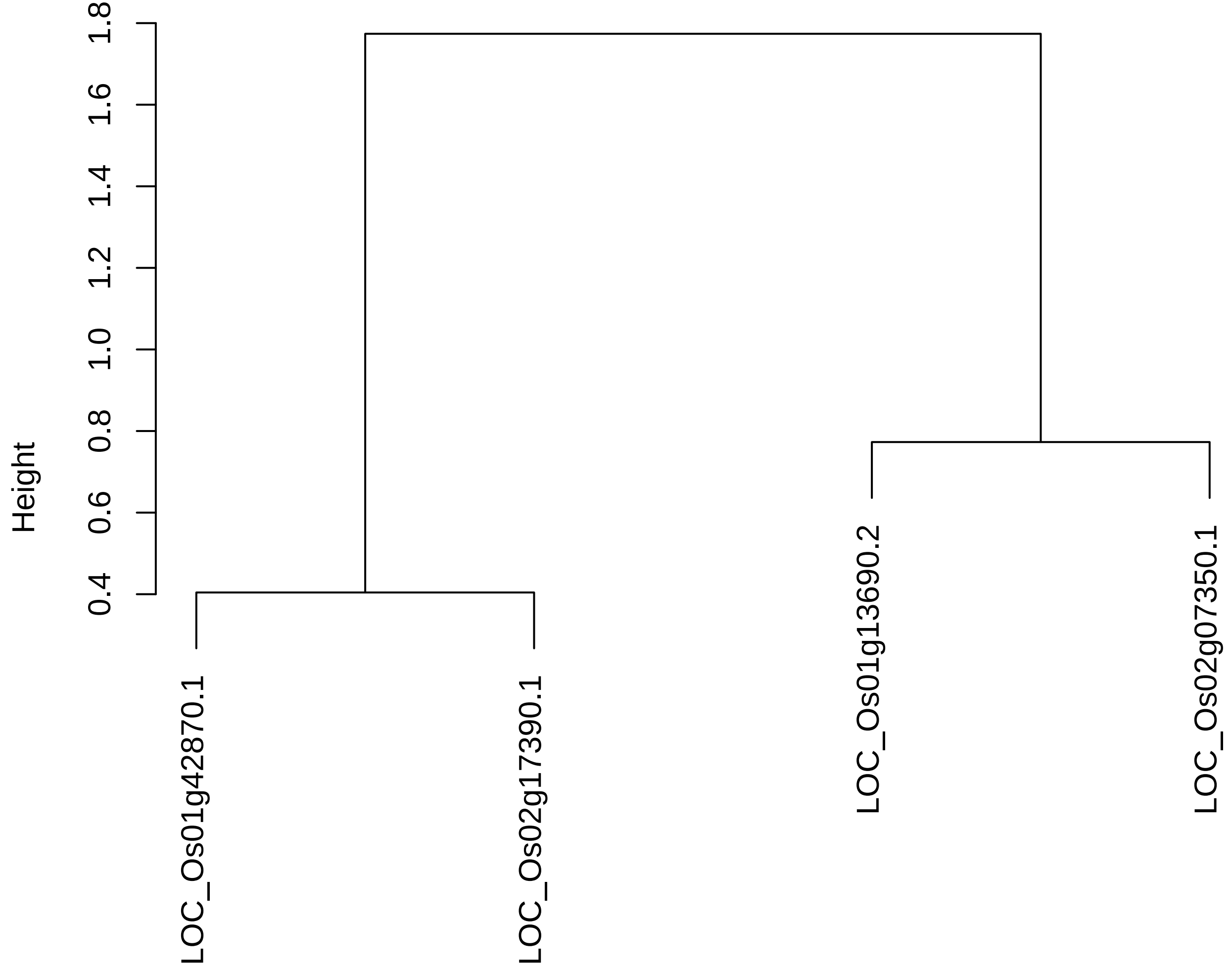


dist(spfitiex12sec12[lab[, 1] == i, ])
hclust (*, "ward.D2")

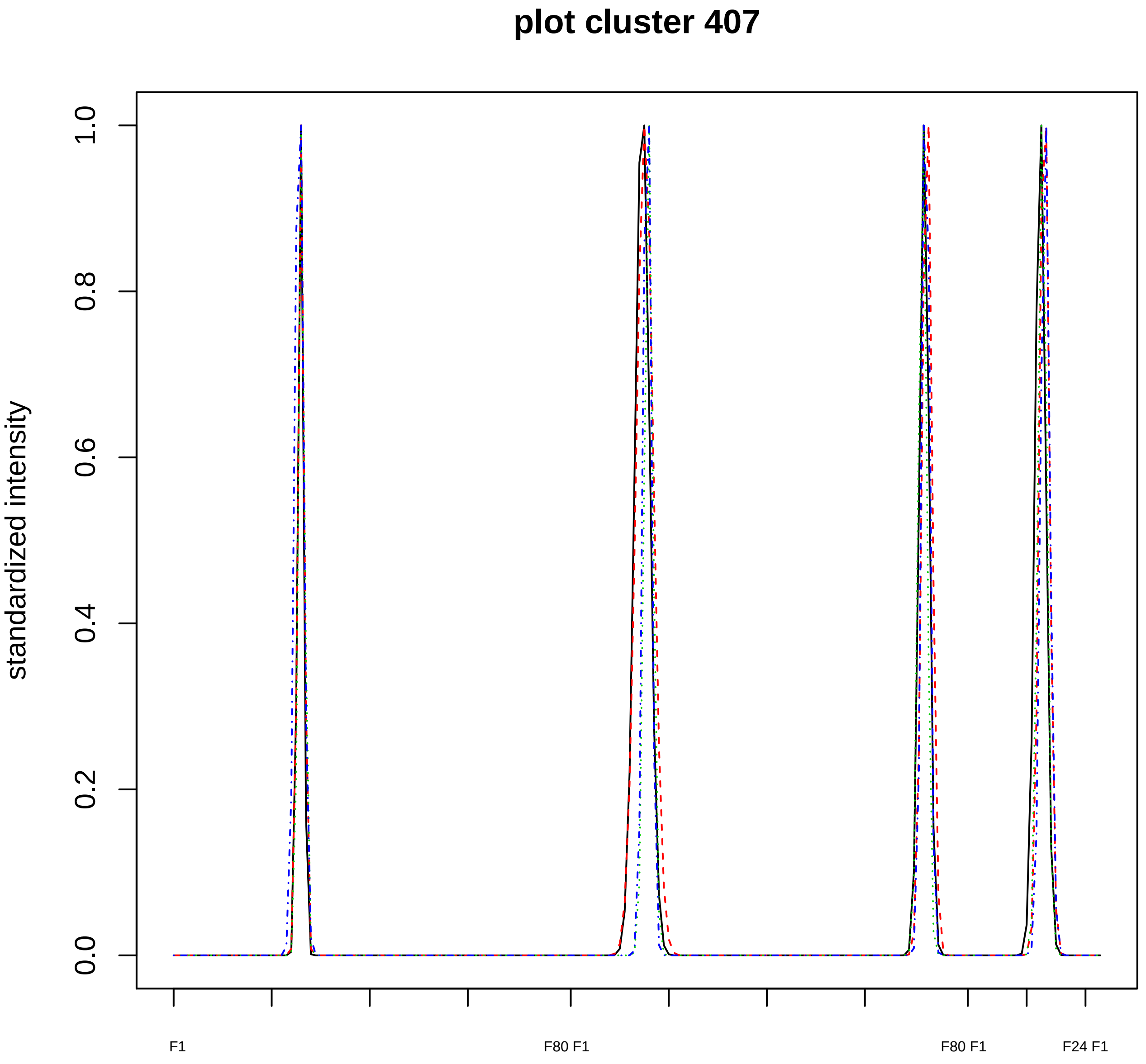

plot cluster 407
standardized intensity
0.0
0.2
0.4
0.6
0.8
1.0
F1
F80 F1
F80 F1
F24 F1

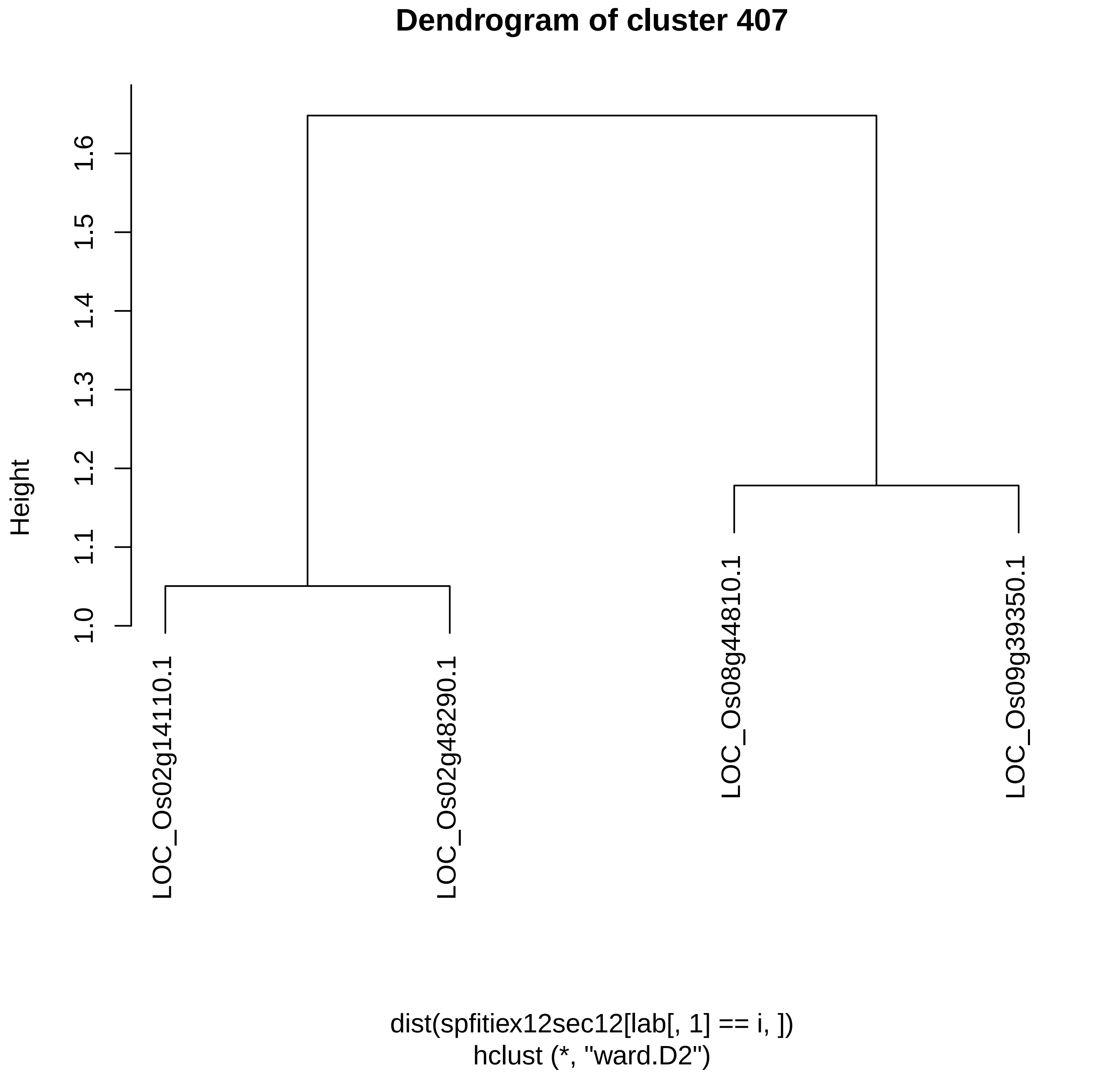
Dendrogram of cluster 407
Height
1.0
1.1
1.2
1.3
1.4
1.5
1.6
LOC_Os02g14110.1
LOC_Os02g48290.1
LOC_Os08g44810.1
LOC_Os09g39350.1
dist(spfitiex12sec12[lab[, 1] == i, ])
hclust (*, "ward.D2")

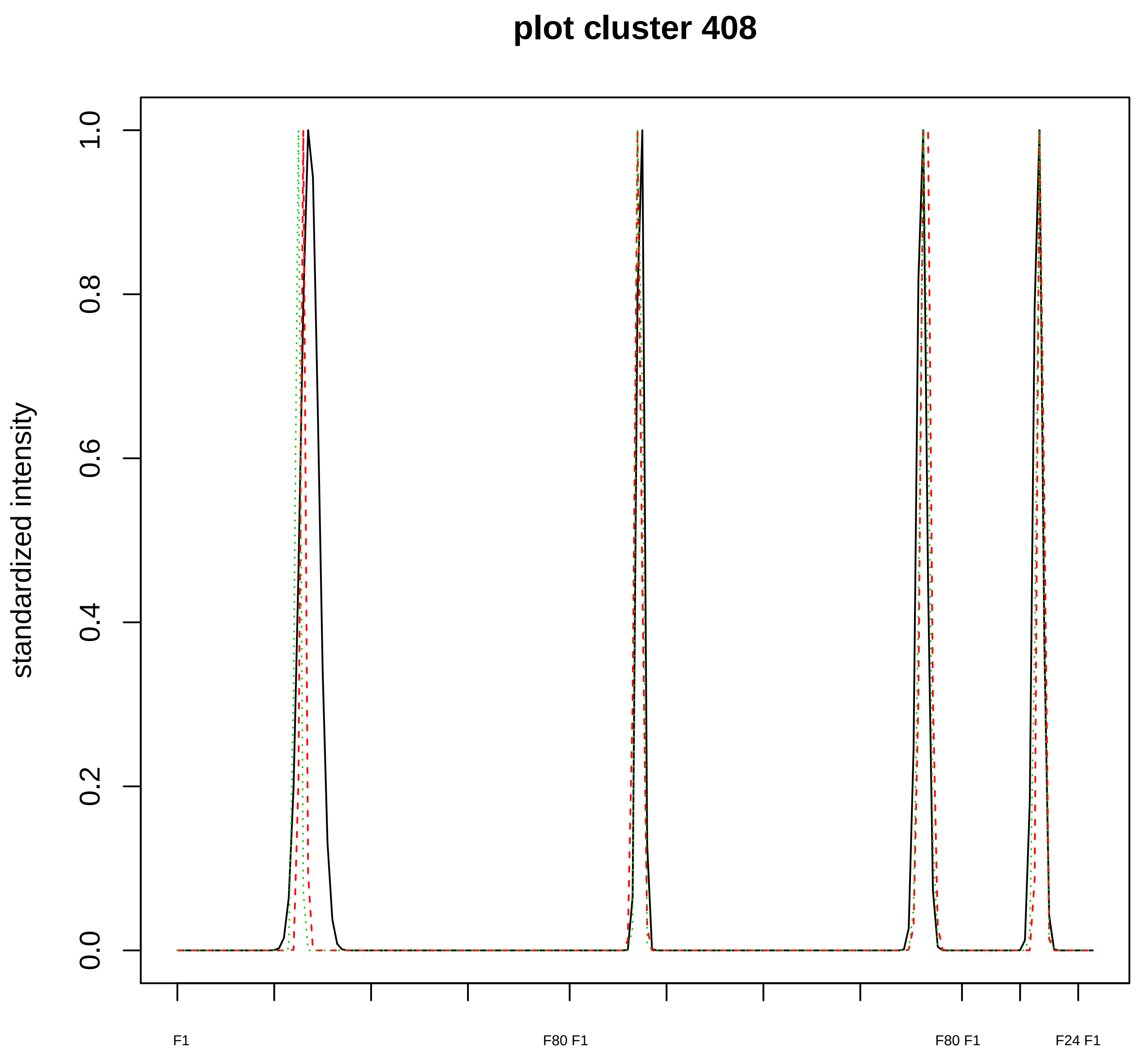
plot cluster 408
standardized intensity
0.0
0.2
0.4
0.6
0.8
1.0
F1
F80 F1
F80 F1
F24 F1

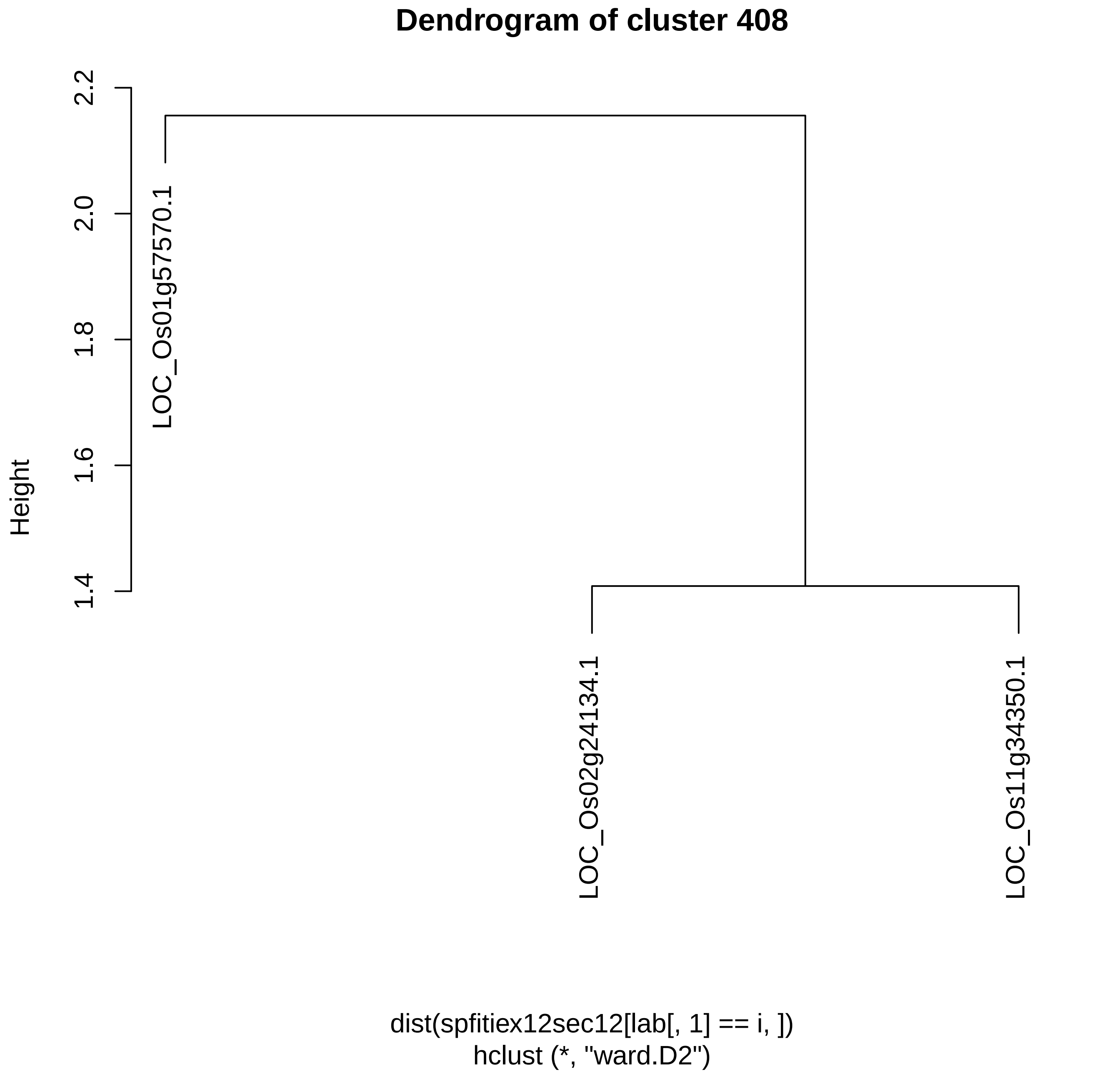
Dendrogram of cluster 408
Height
2.2
2.0
1.8
1.6
1.4
LOC_Os01g57570.1
LOC_Os02g24134.1
LOC_Os11g34350.1
dist(spfitiex12sec12[lab[, 1] == i, ])
hclust (*, "ward.D2")

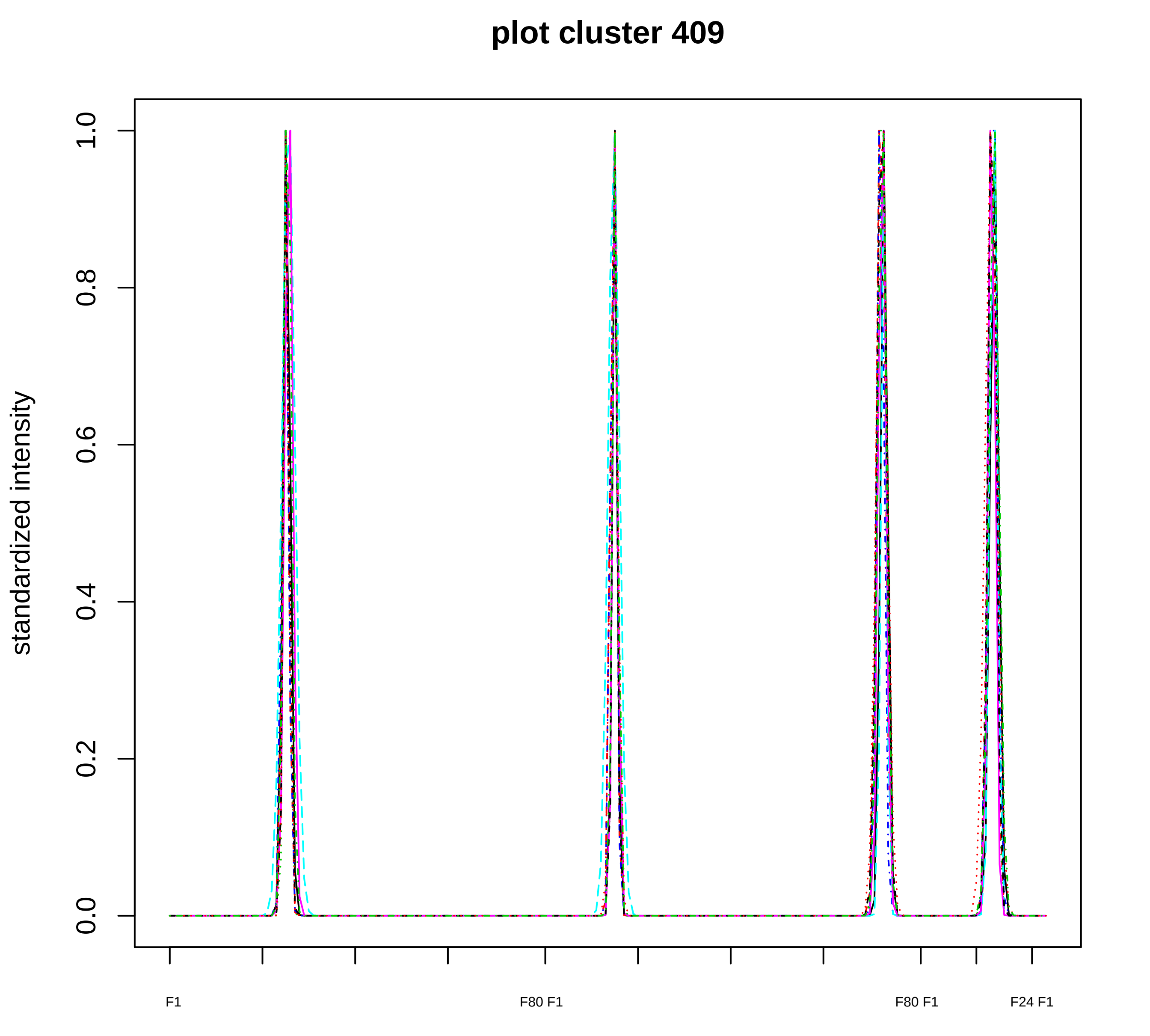
plot cluster 409
standardized intensity
0.0
0.2
0.4
0.6
0.8
1.0
F1
F80 F1
F80 F1
F24 F1

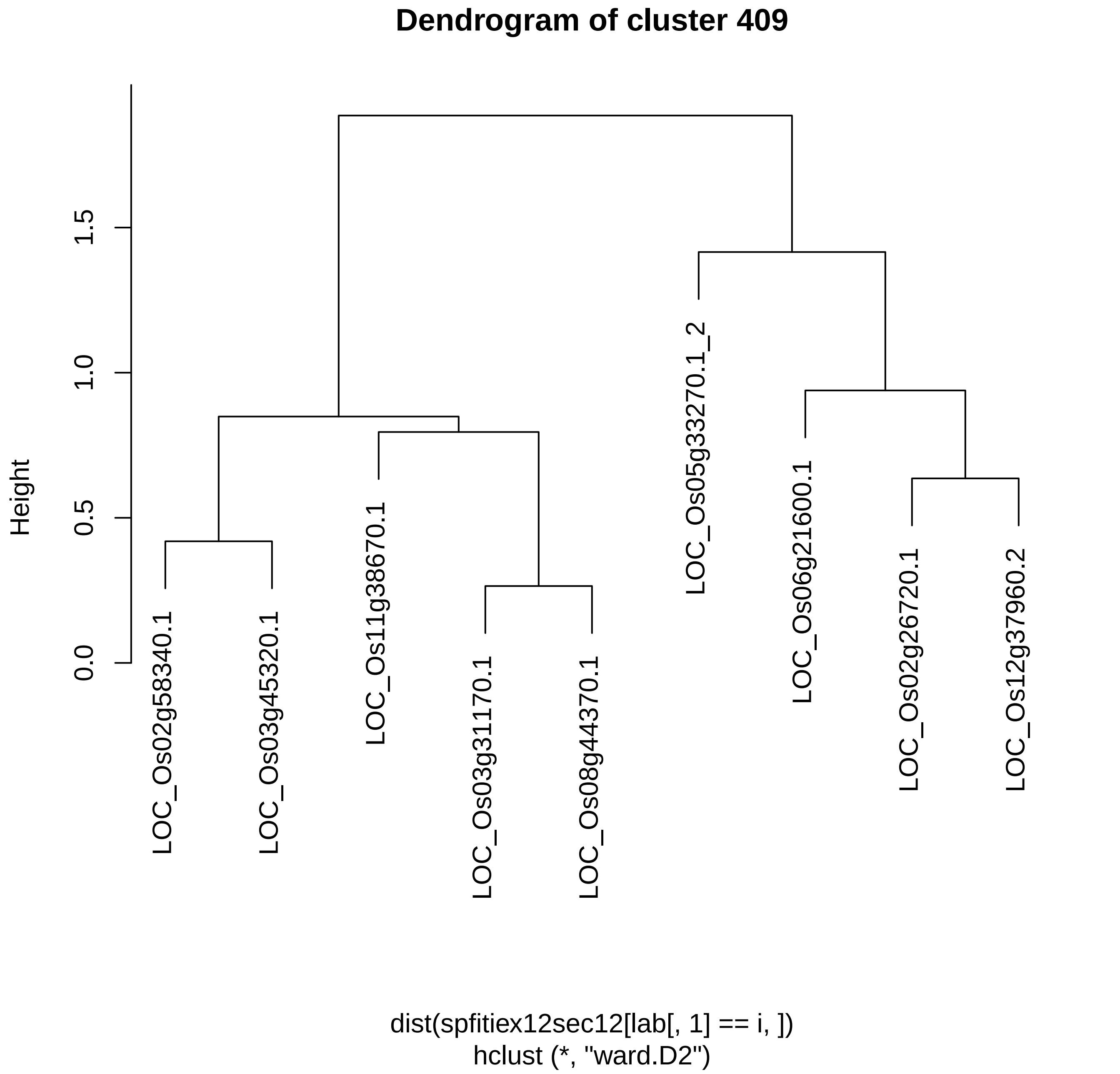
Dendrogram of cluster 409
Height
0.0
0.5
1.0
1.5
LOC_Os02g58340.1
LOC_Os03g45320.1
LOC_Os11g38670.1
LOC_Os03g31170.1
LOC_Os08g44370.1
LOC_Os05g33270.1_2
LOC_Os06g21600.1
LOC_Os02g26720.1
LOC_Os12g37960.2
dist(spfitiex12sec12[lab[, 1] == i, ])
hclust (*, "ward.D2")

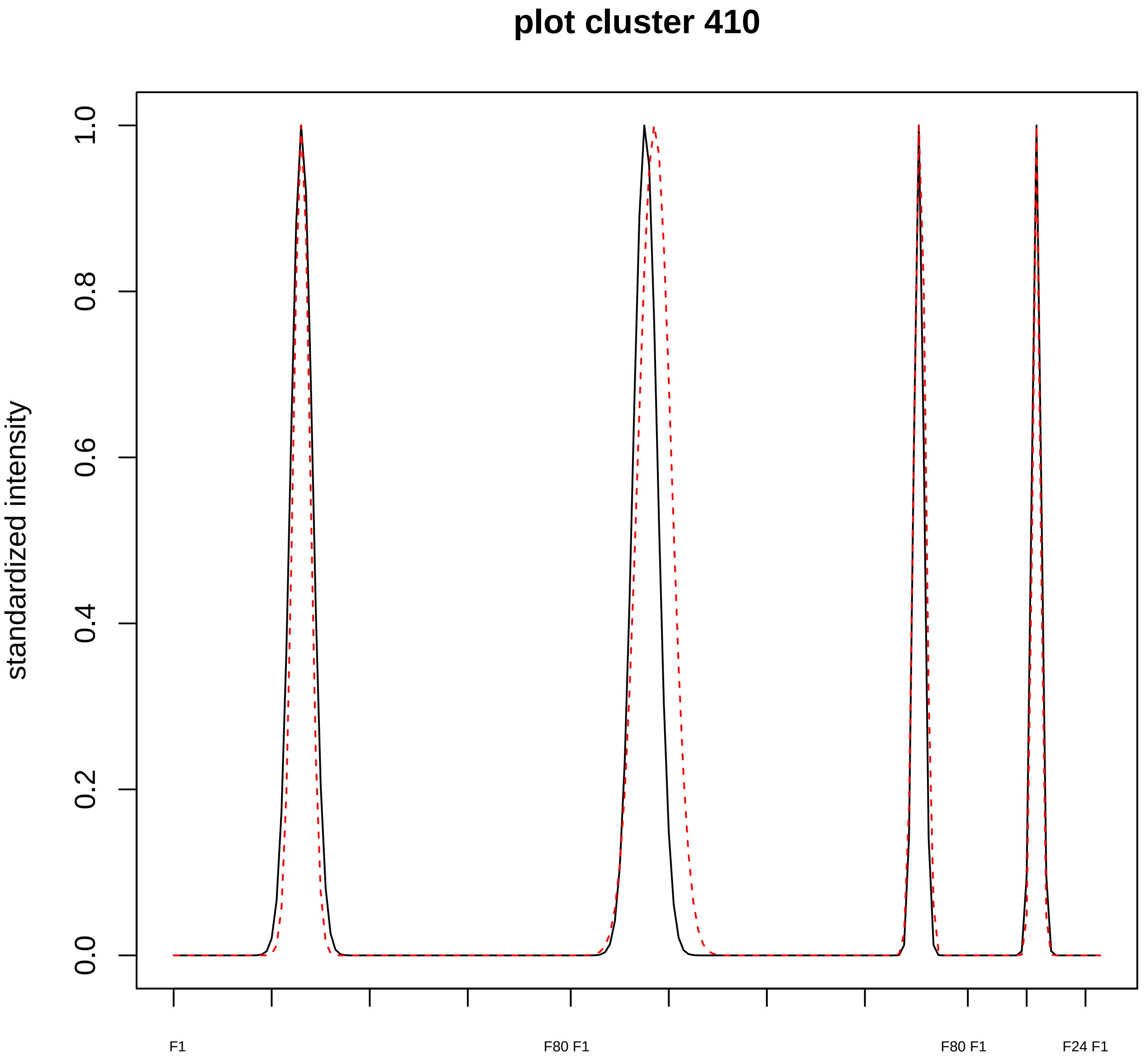
plot cluster 410
standardized intensity
0.0
0.2
0.4
0.6
0.8
1.0
F1
F80 F1
F80 F1
F24 F1

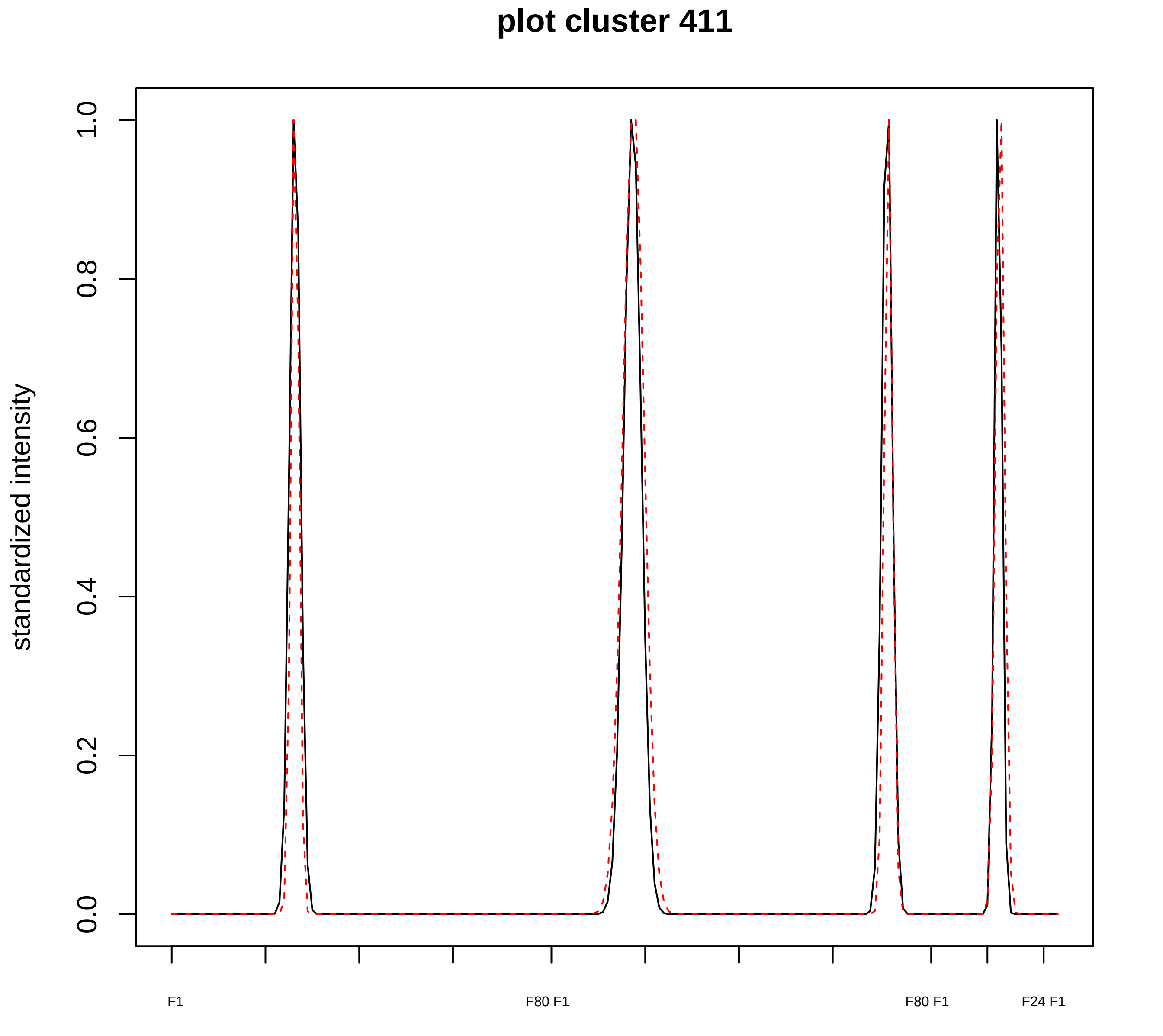

plot cluster 411
standardized intensity
0.0
0.2
0.4
0.6
0.8
1.0
F1
F80 F1
F80 F1
F24 F1

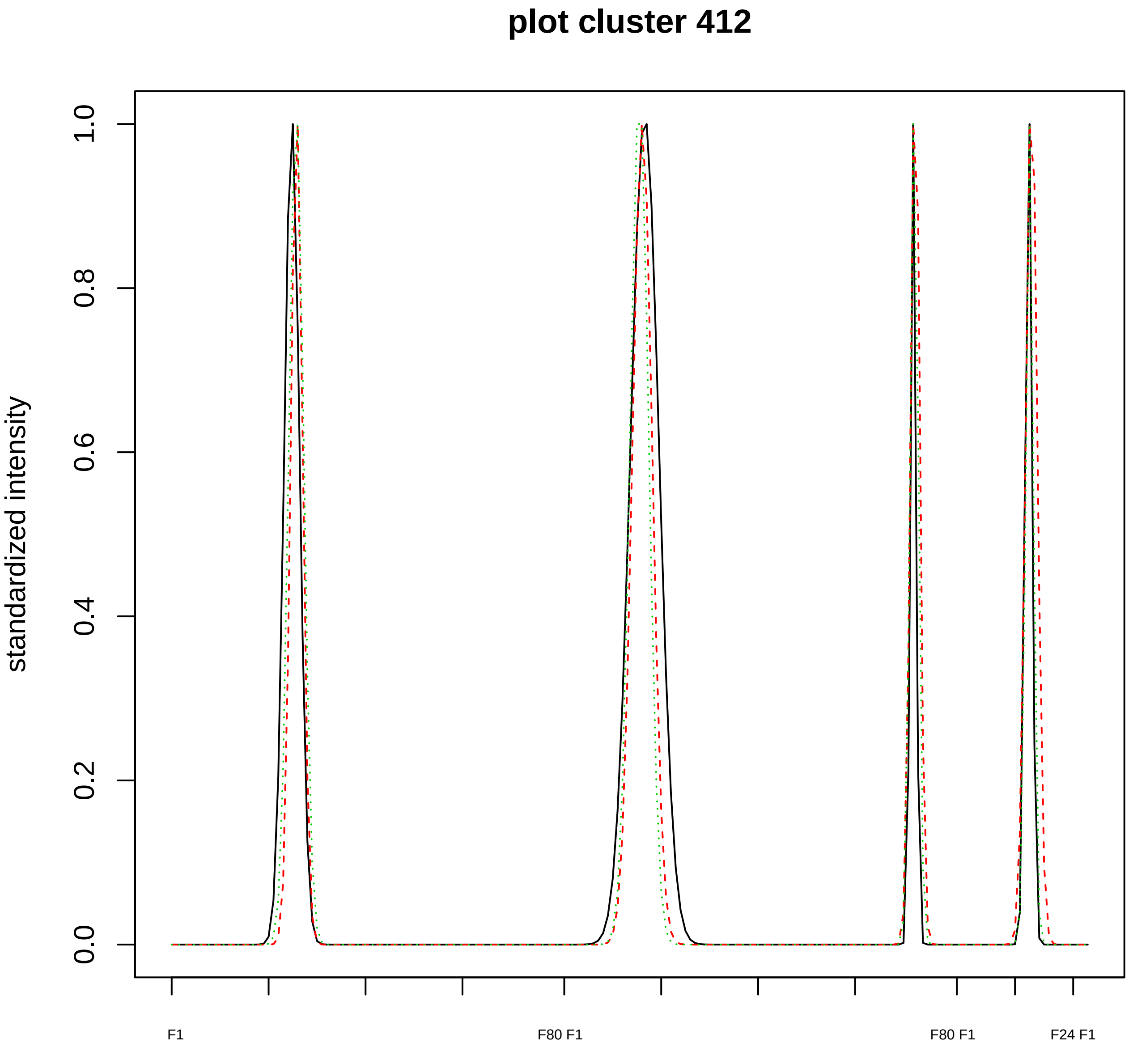
plot cluster 412
standardized intensity
0.0
0.2
0.4
0.6
0.8
1.0
F1
F80 F1
F80 F1
F24 F1

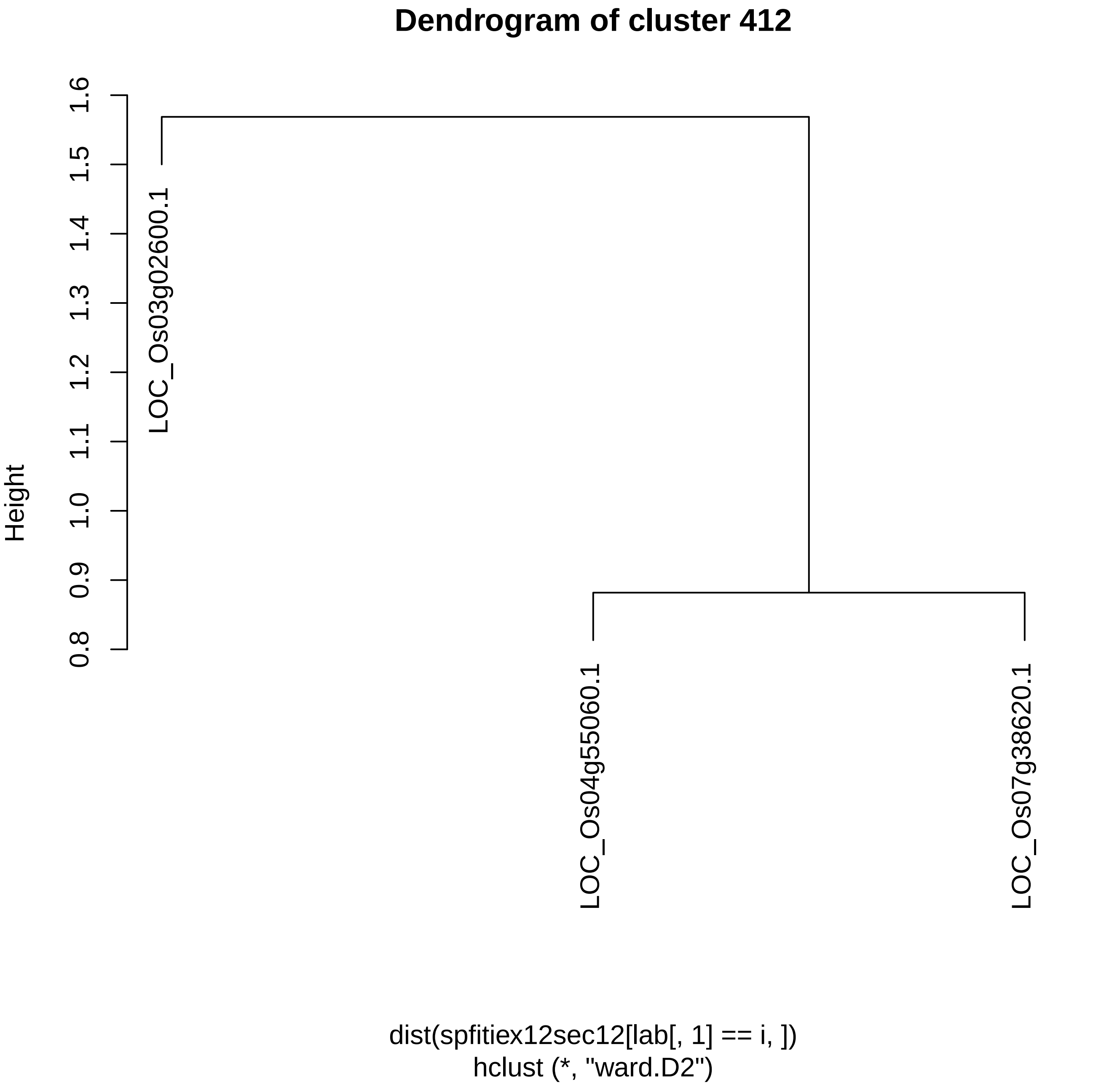
Dendrogram of cluster 412
Height
0.8
0.9
1.0
1.1
1.2
1.3
1.4
1.5
1.6
LOC_Os03g02600.1
LOC_Os04g55060.1
LOC_Os07g38620.1
dist(spfitiex12sec12[lab[, 1] == i, ])
hclust (*, "ward.D2")

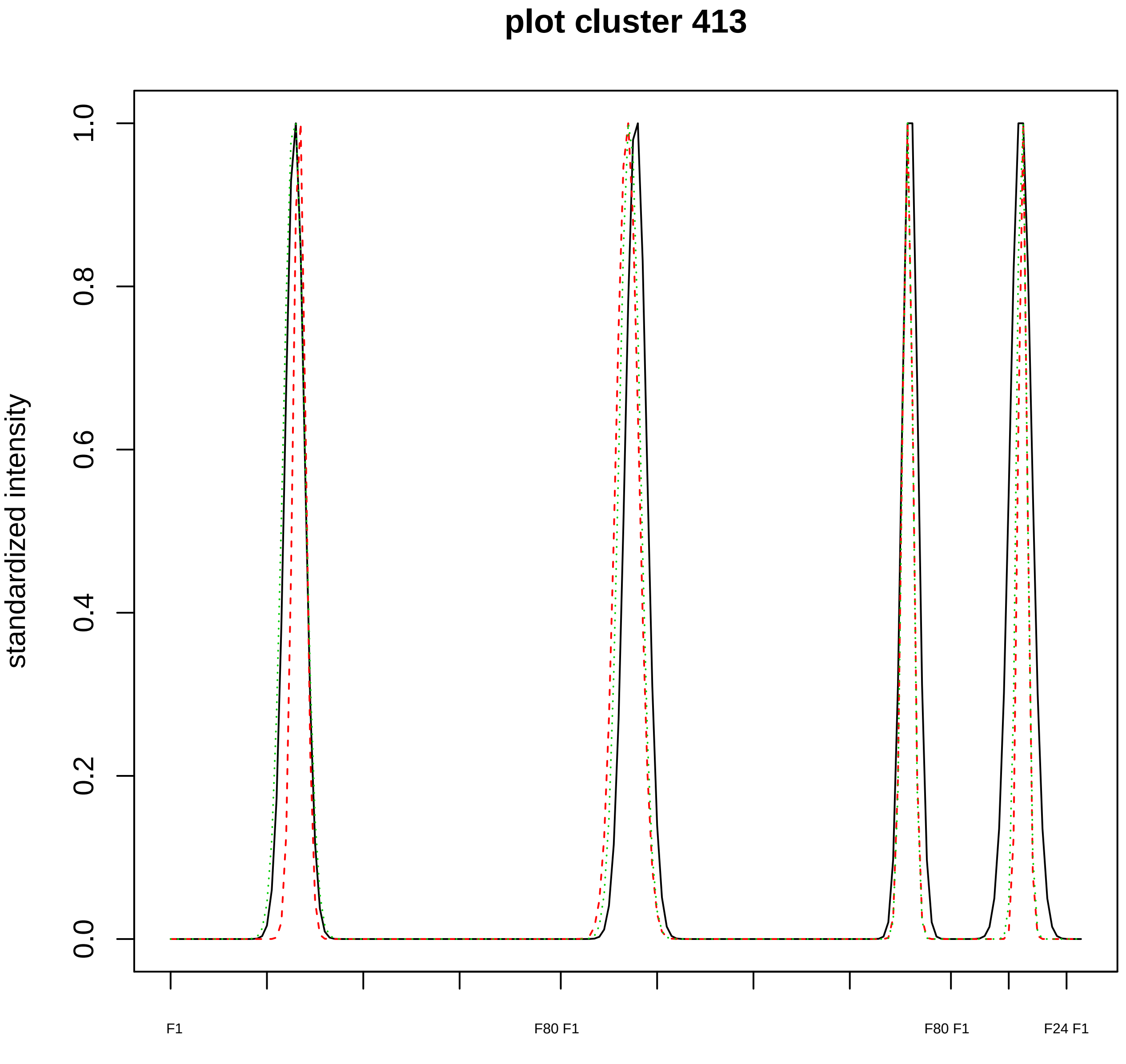

plot cluster 413
standardized intensity
0.0
0.2
0.4
0.6
0.8
1.0
F1
F80 F1
F80 F1
F24 F1

### Dendrogram of cluster 413

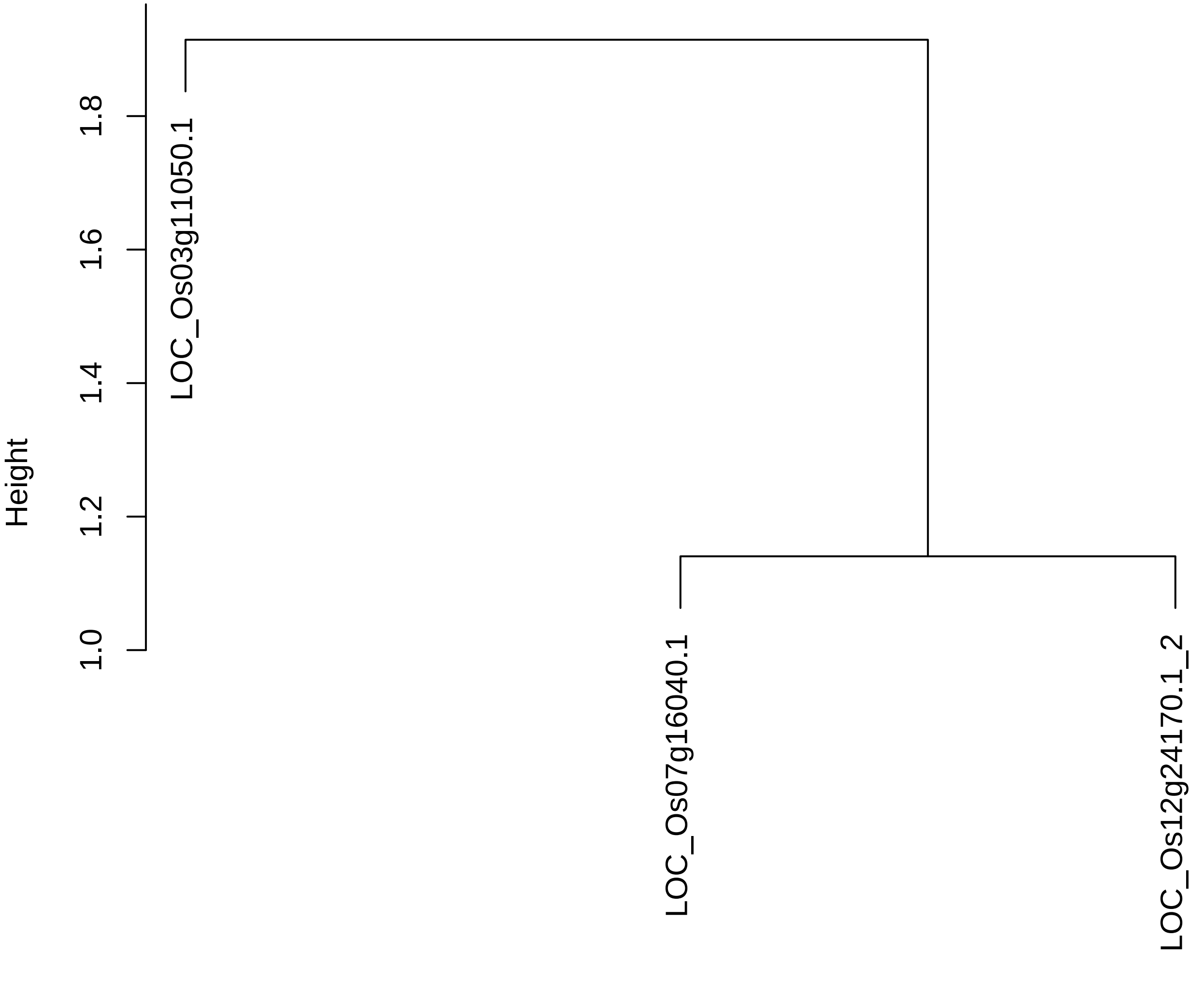


dist(spfitiex12sec12[lab[, 1] == i, ])
hclust (*, "ward.D2")

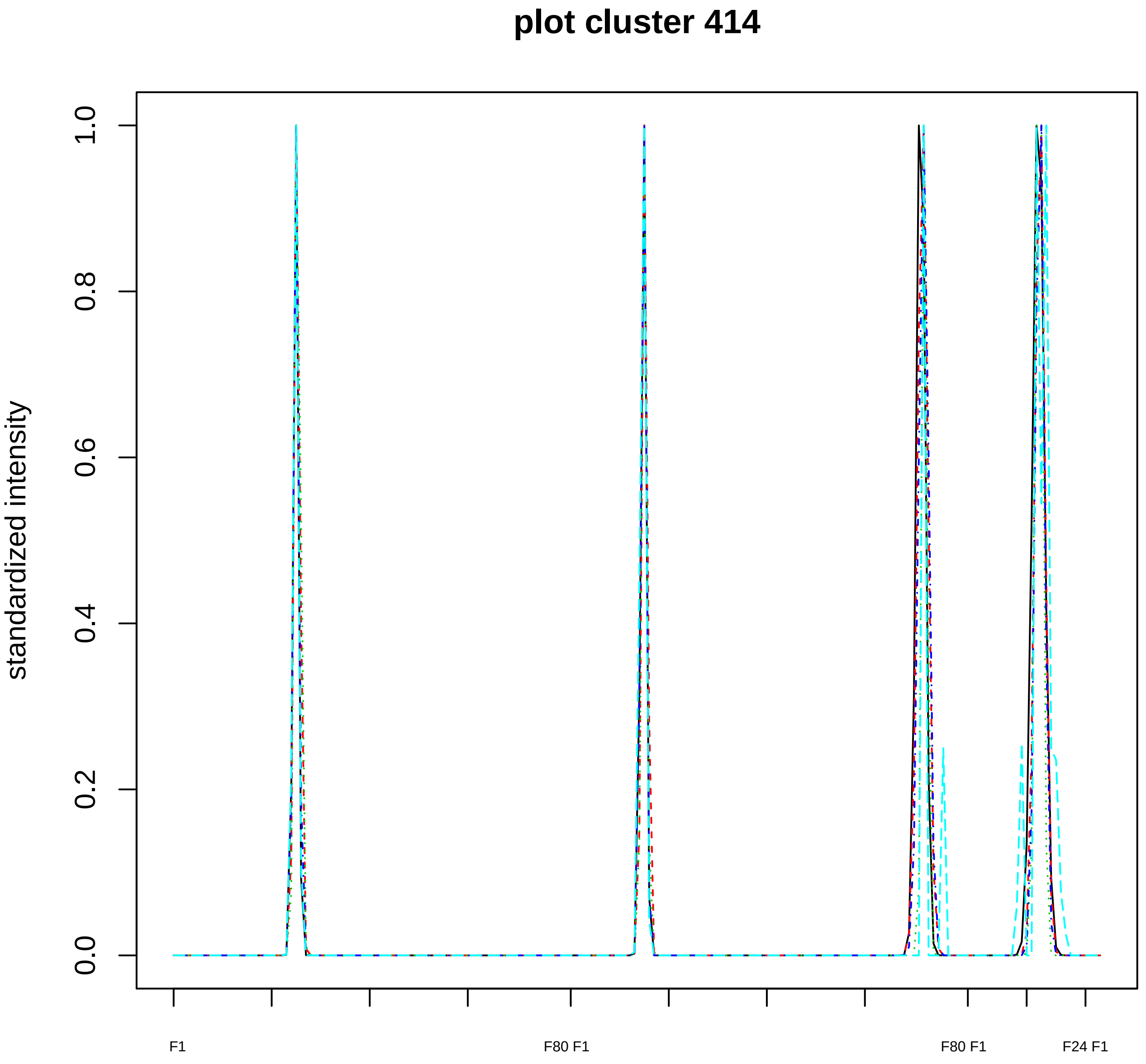
plot cluster 414
standardized intensity
0.0
0.2
0.4
0.6
0.8
1.0
F1
F80 F1
F80 F1
F24 F1

### Dendrogram of cluster 414

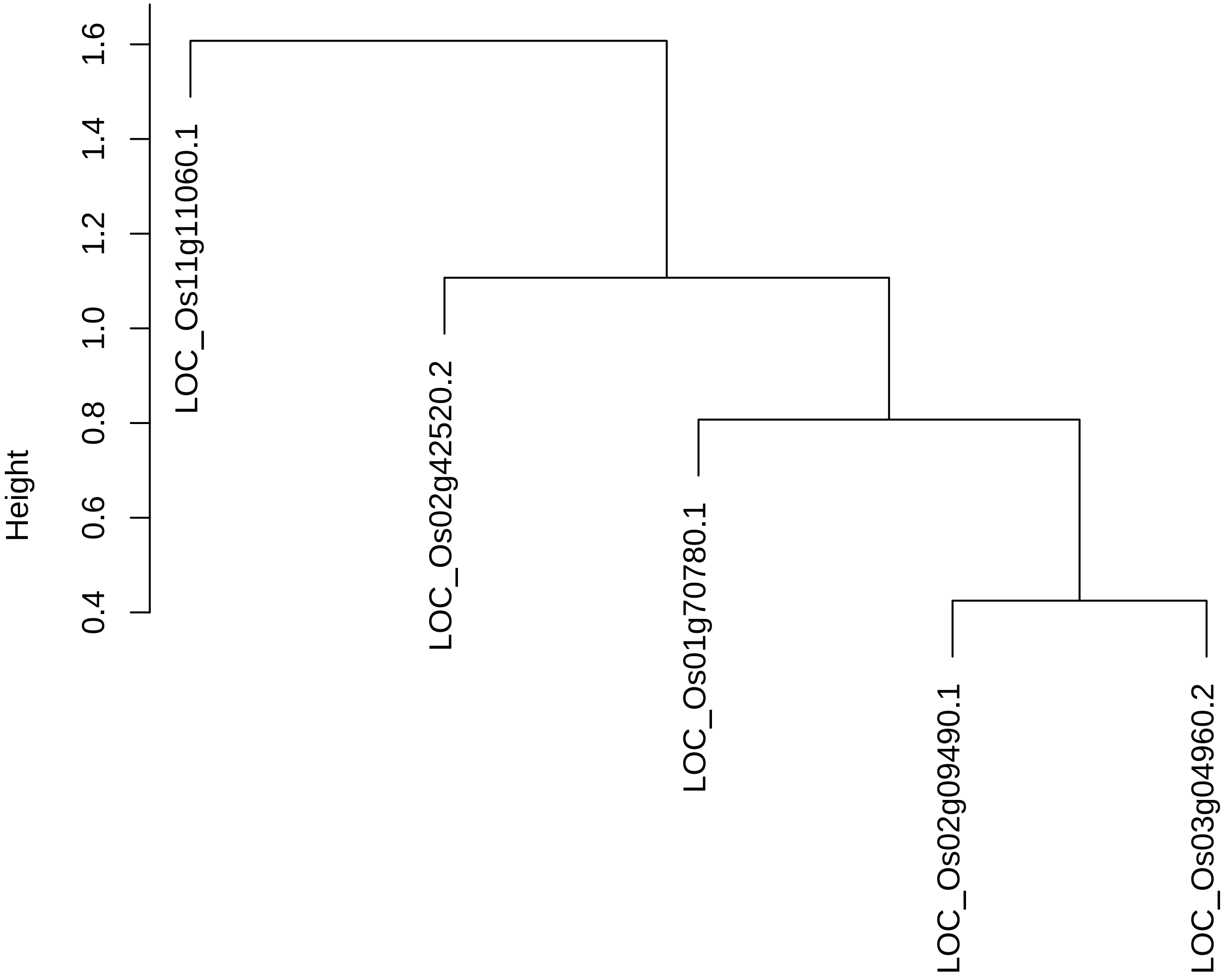


dist(spfitiex12sec12[lab[, 1] == i, ])
hclust (*, "ward.D2")

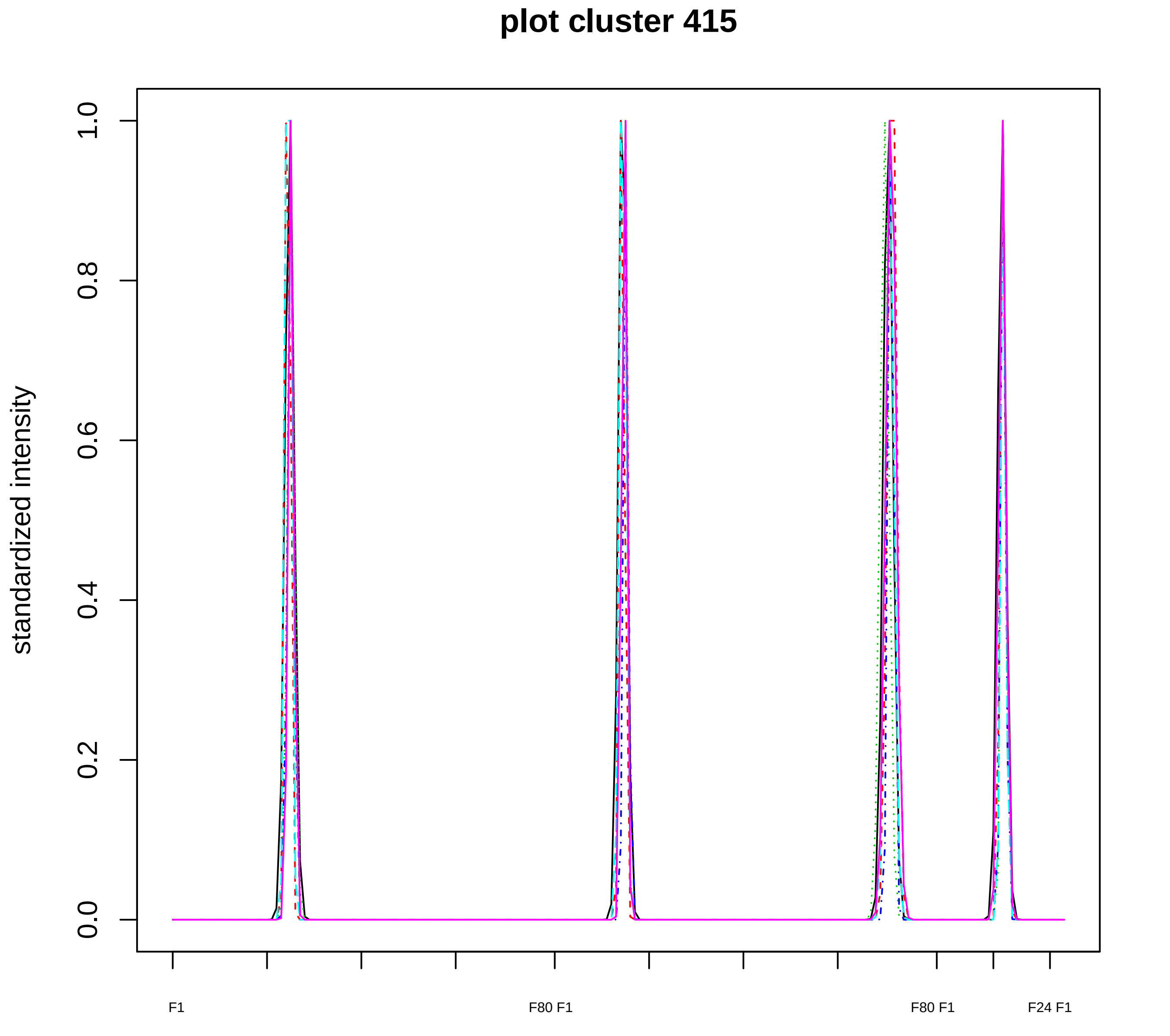

plot cluster 415
standardized intensity
0.0
0.2
0.4
0.6
0.8
1.0
F1
F80 F1
F80 F1
F24 F1

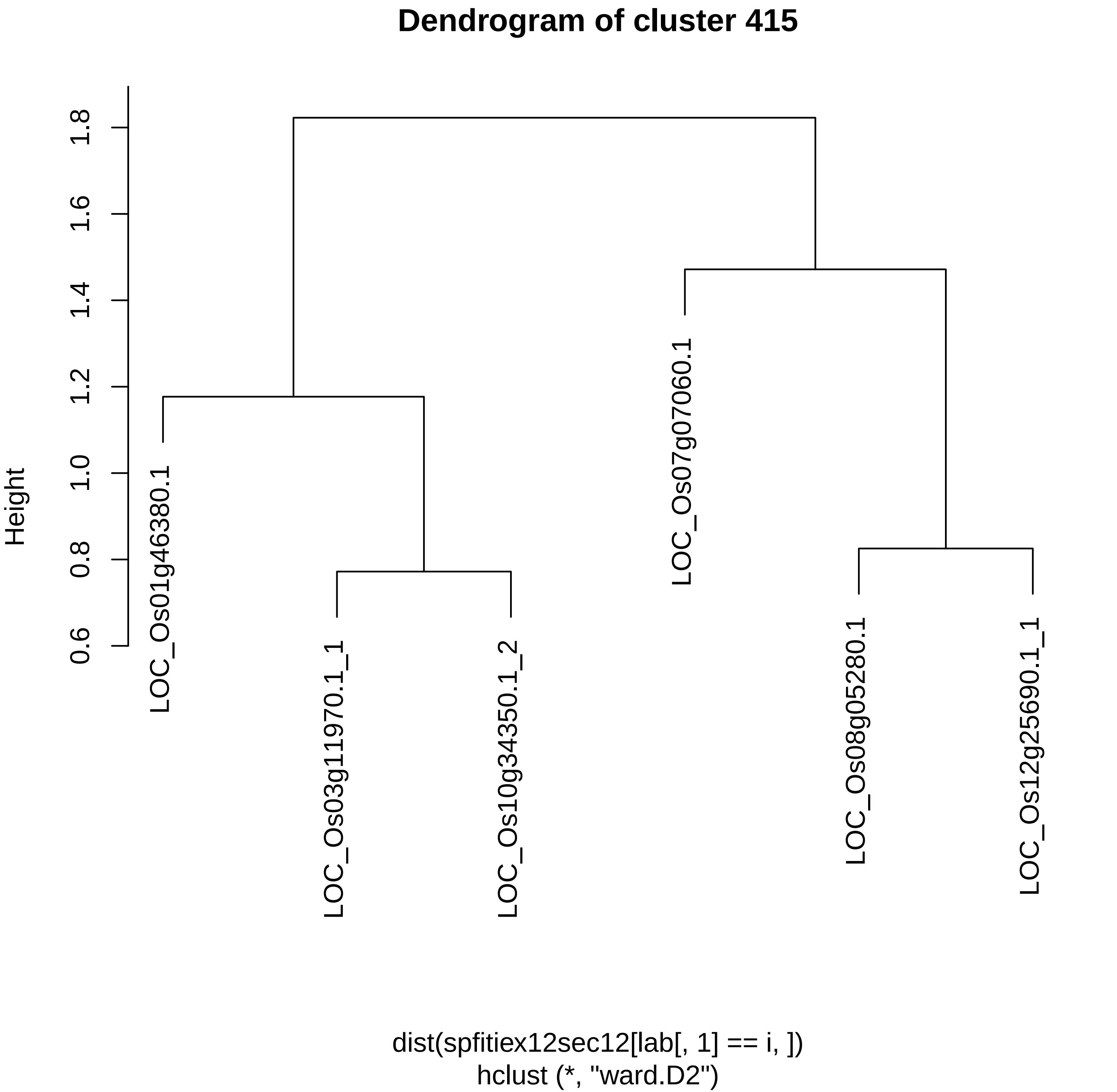
Dendrogram of cluster 415
Height
0.6
0.8
1.0
1.2
1.4
1.6
1.8
LOC_Os01g46380.1
LOC_Os03g11970.1_1
LOC_Os10g34350.1_2
LOC_Os07g07060.1
LOC_Os08g05280.1
LOC_Os12g25690.1_1
dist(spfitiex12sec12[lab[, 1] == i, ])
hclust (*, "ward.D2")

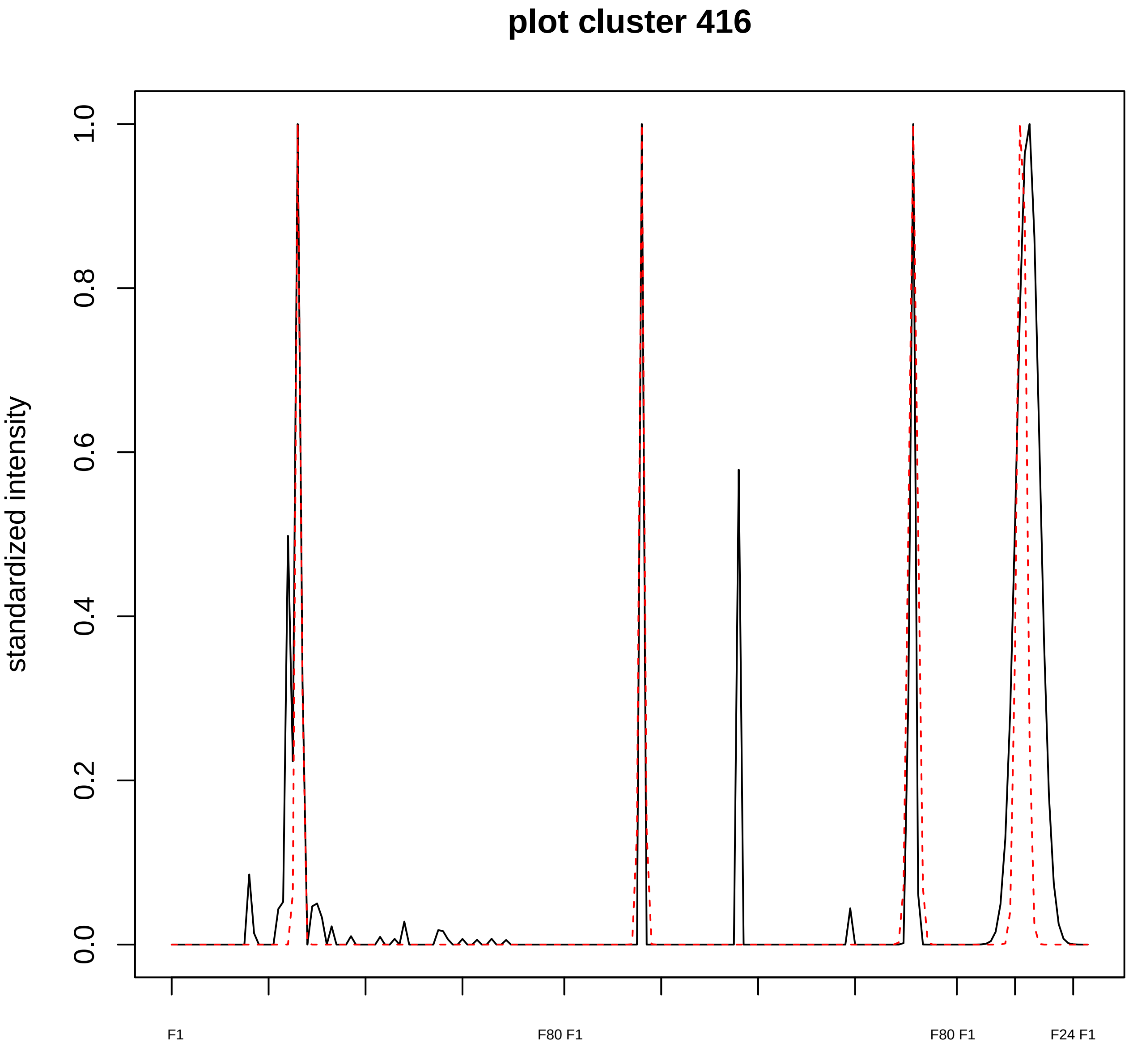
plot cluster 416
standardized intensity
0.0
0.2
0.4
0.6
0.8
1.0
F1
F80 F1
F80 F1
F24 F1

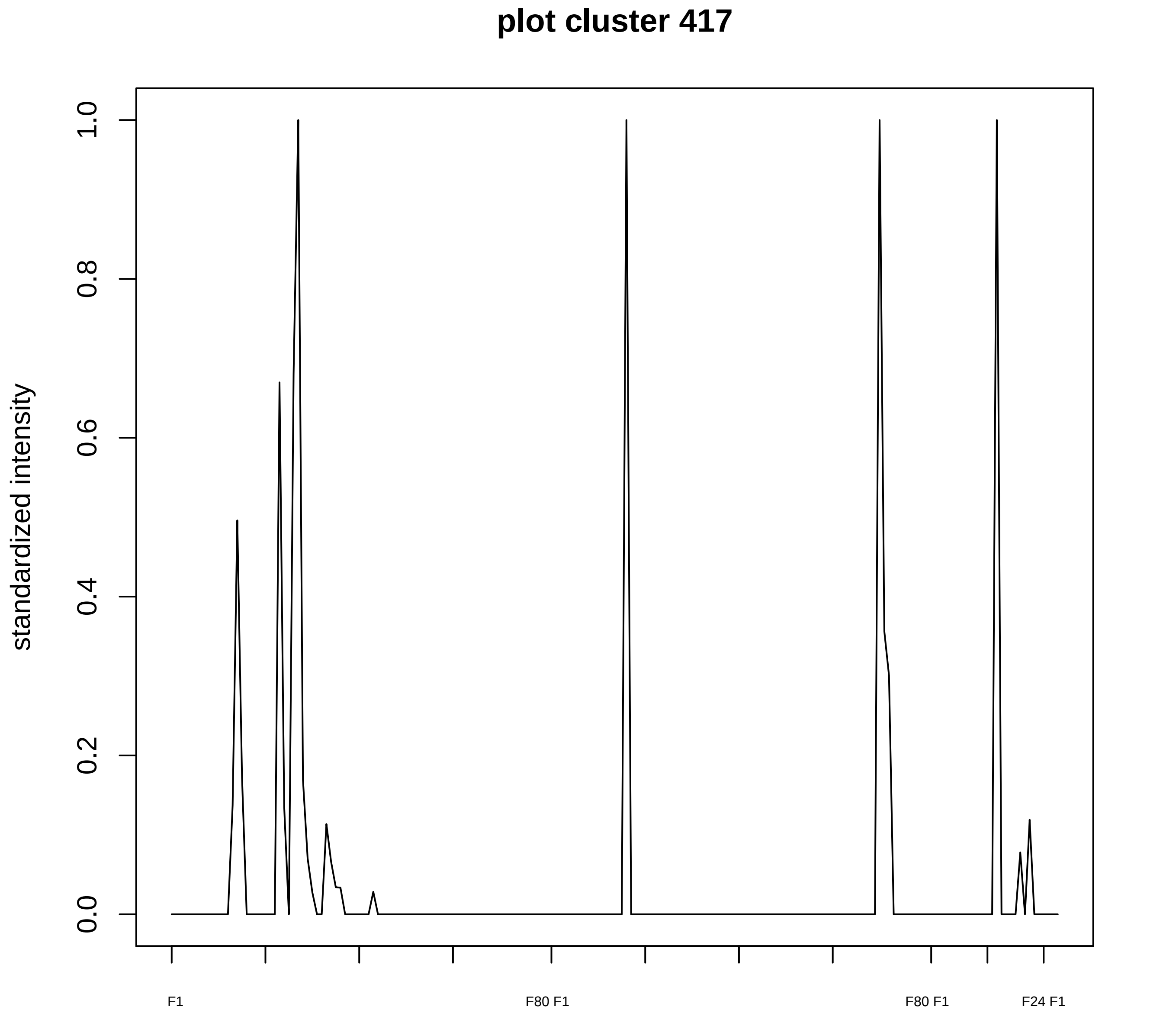

plot cluster 417
standardized intensity
0.0
0.2
0.4
0.6
0.8
1.0
F1
F80 F1
F80 F1
F24 F1

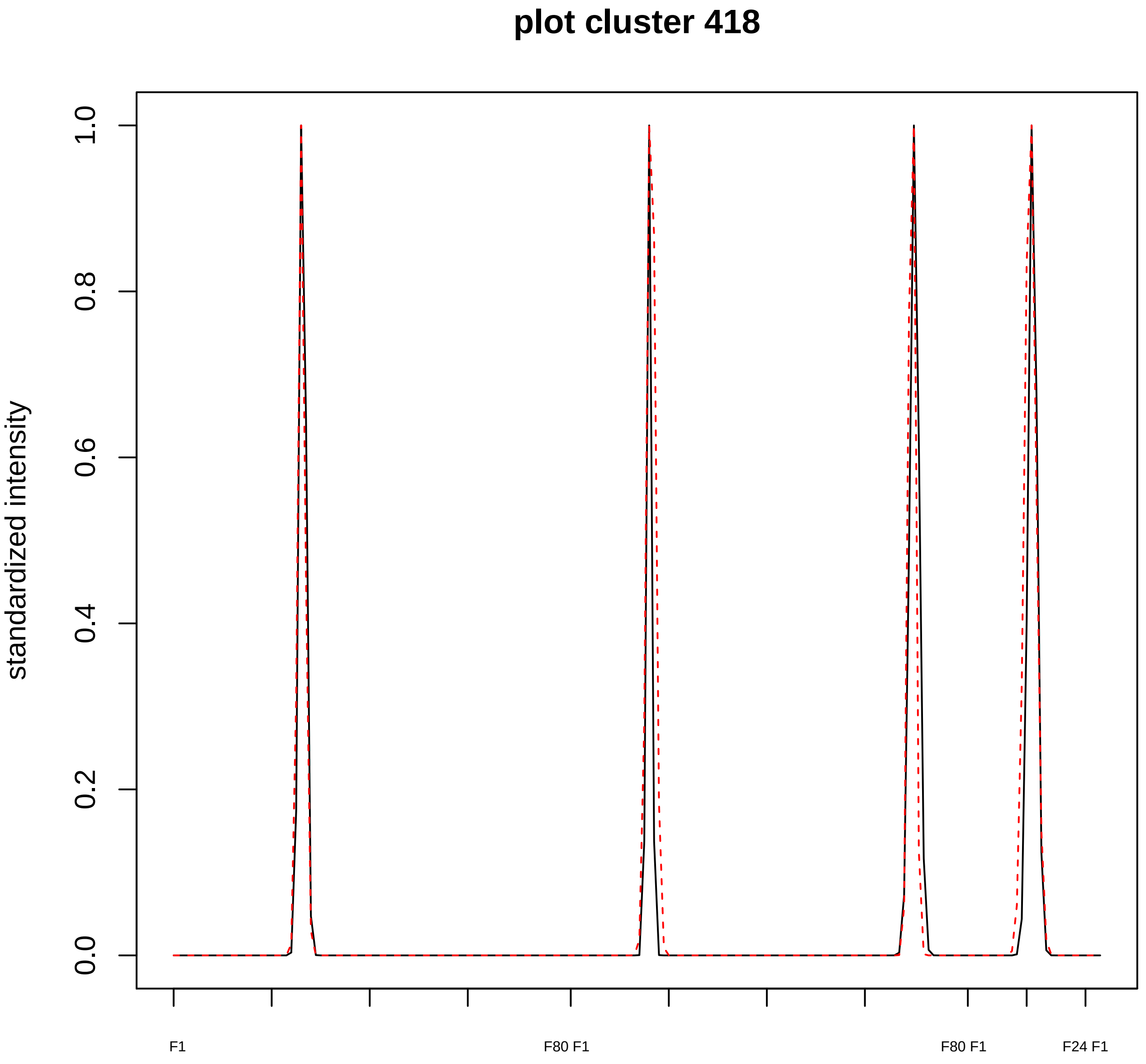

plot cluster 418
standardized intensity
0.0
0.2
0.4
0.6
0.8
1.0
F1
F80 F1
F80 F1
F24 F1

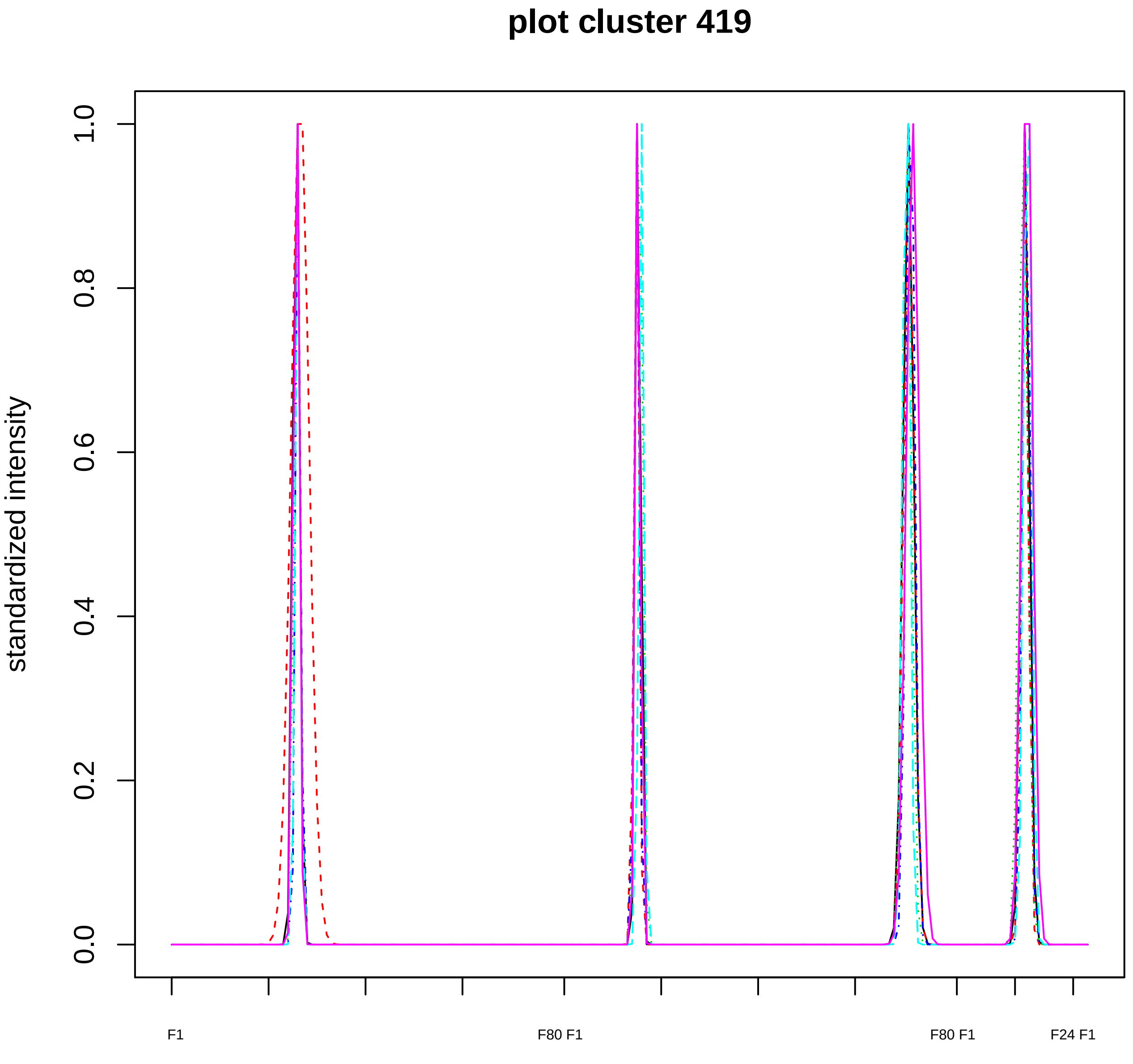

plot cluster 419
standardized intensity
0.0
0.2
0.4
0.6
0.8
1.0
F1
F80 F1
F80 F1
F24 F1

### Dendrogram of cluster 419

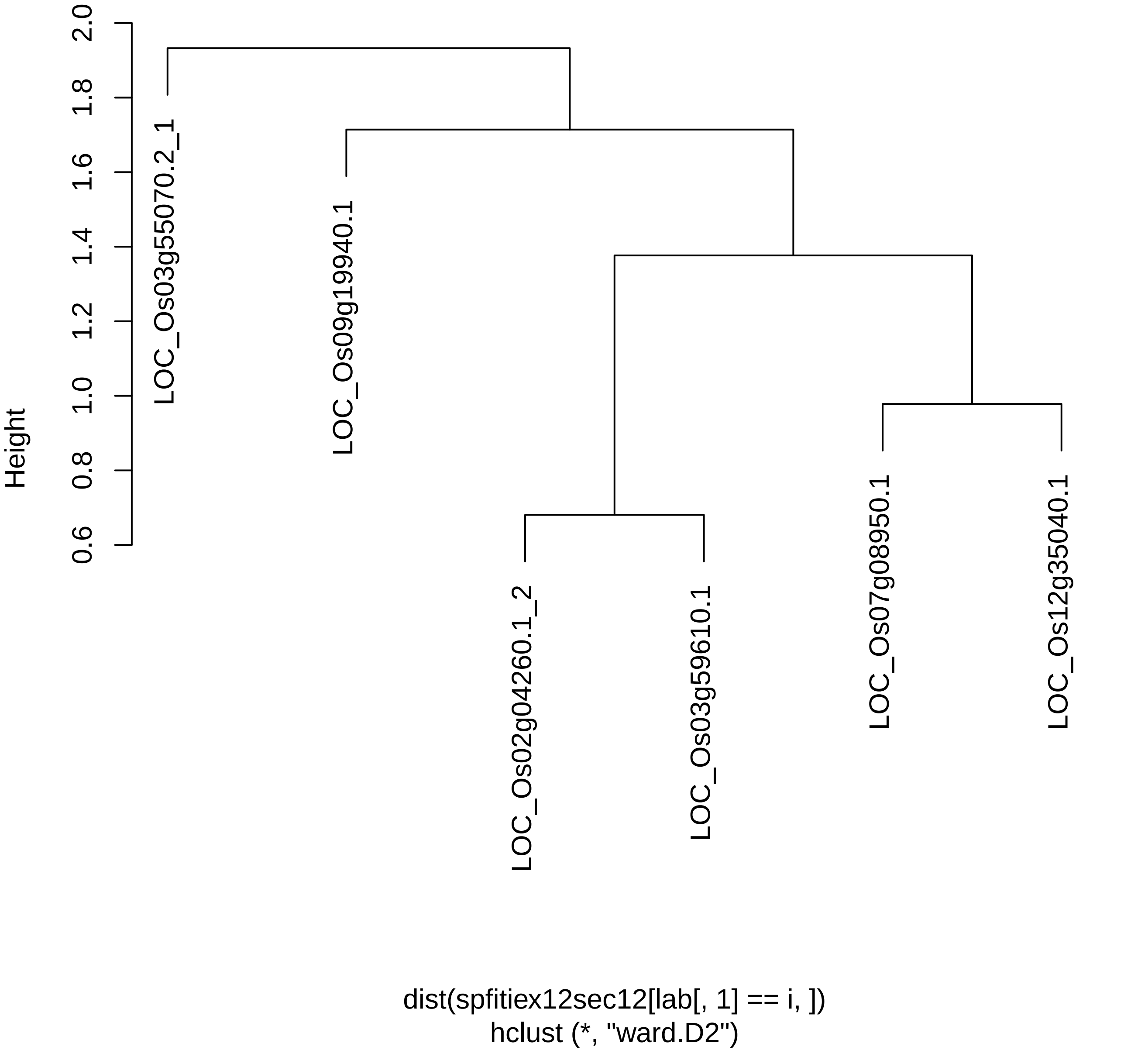

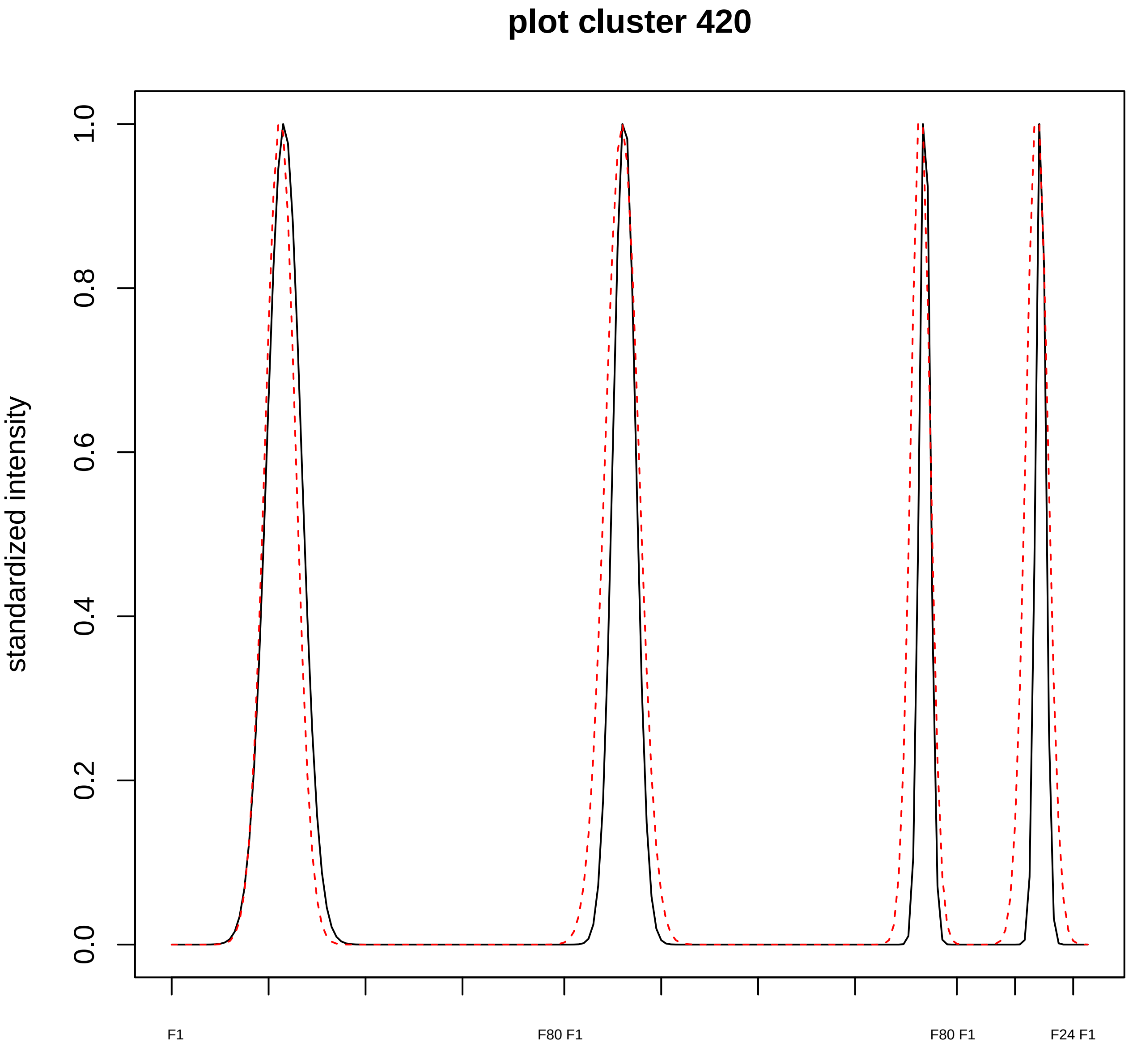
plot cluster 420
standardized intensity
0.0
0.2
0.4
0.6
0.8
1.0
F1
F80 F1
F80 F1
F24 F1

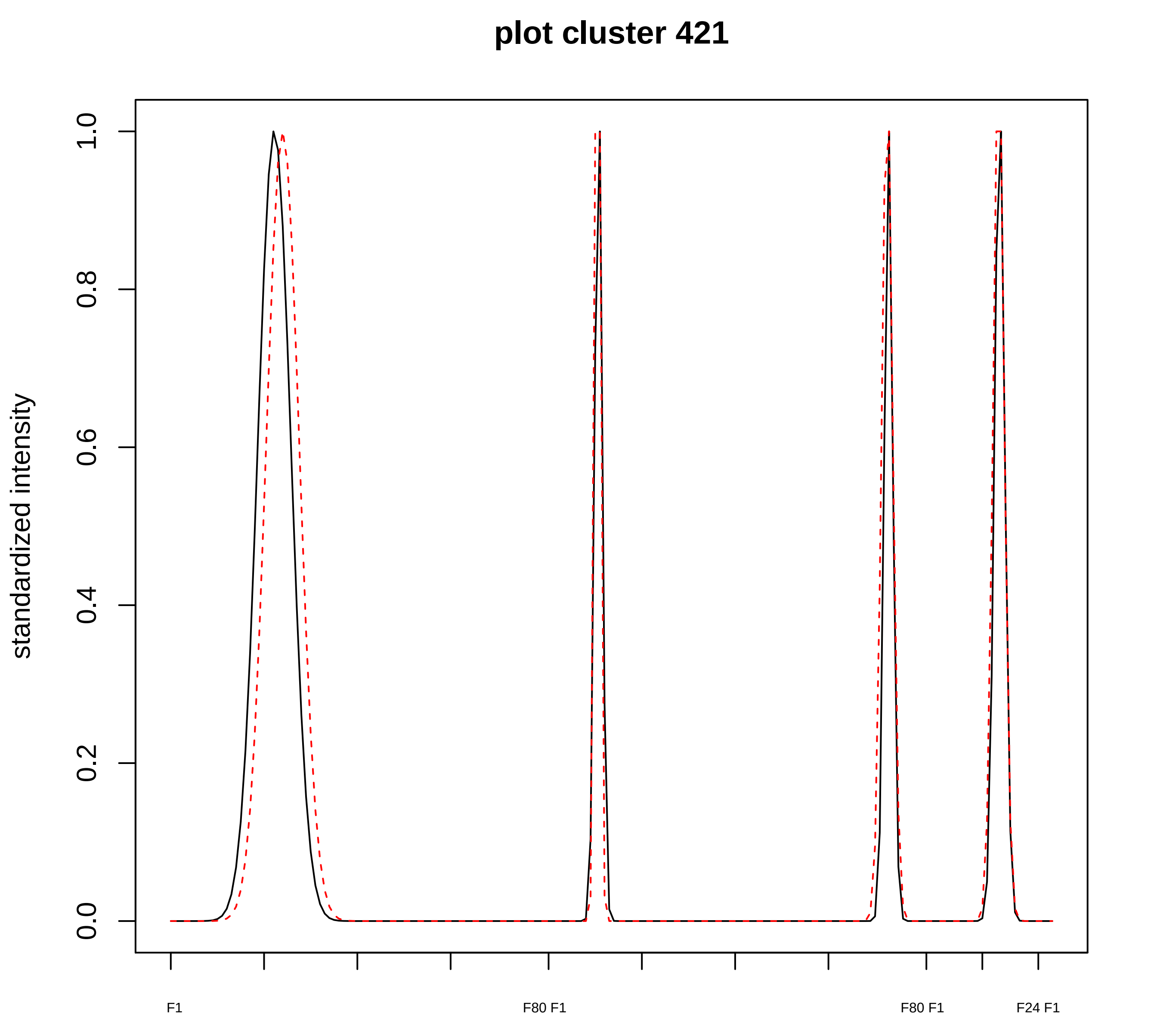

plot cluster 421
standardized intensity
1.0
0.8
0.6
0.4
0.2
0.0
F1
F80 F1
F80 F1
F24 F1

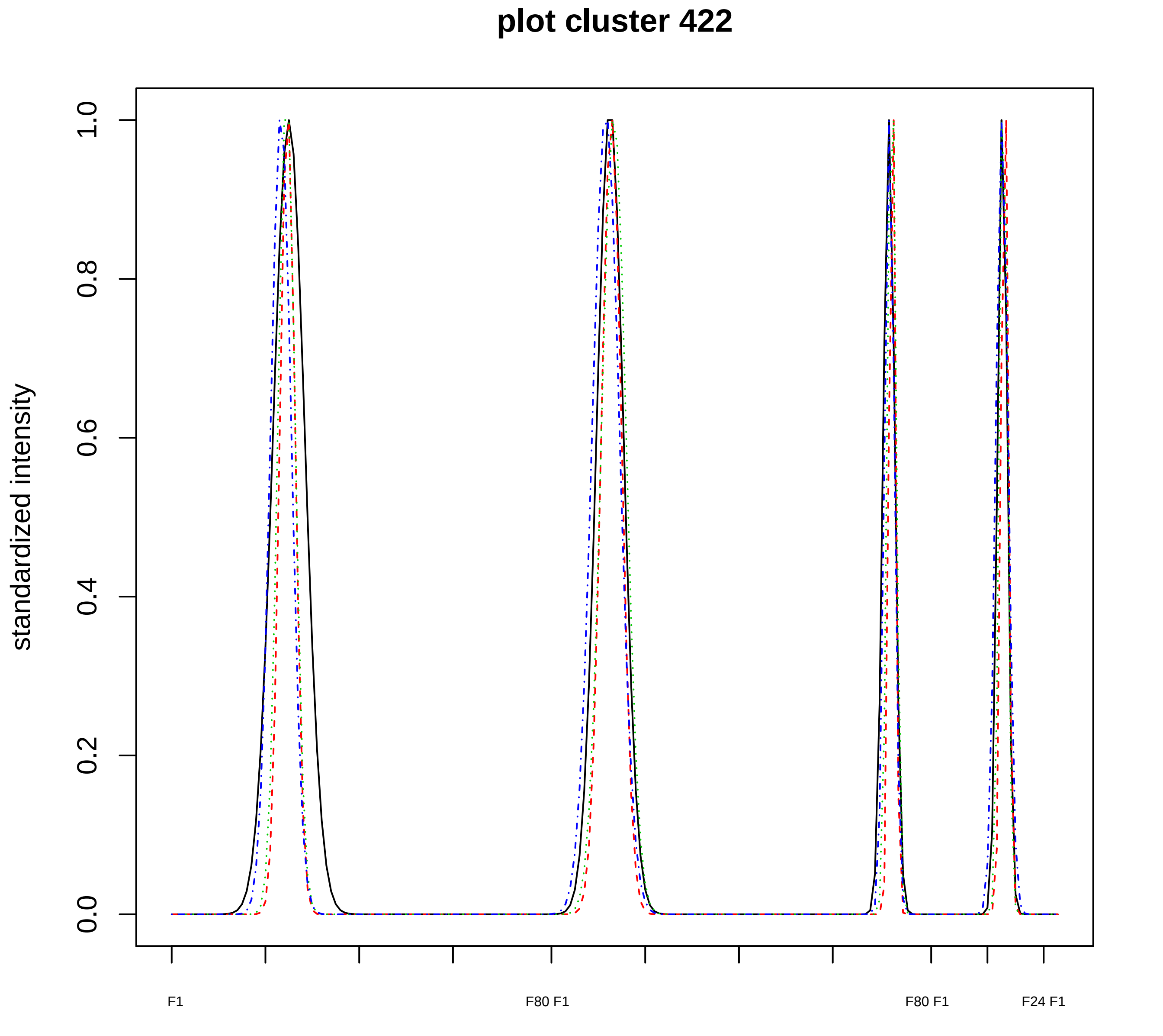
plot cluster 422
standardized intensity
0.0
0.2
0.4
0.6
0.8
1.0
F1
F80 F1
F80 F1
F24 F1

### Dendrogram of cluster 422

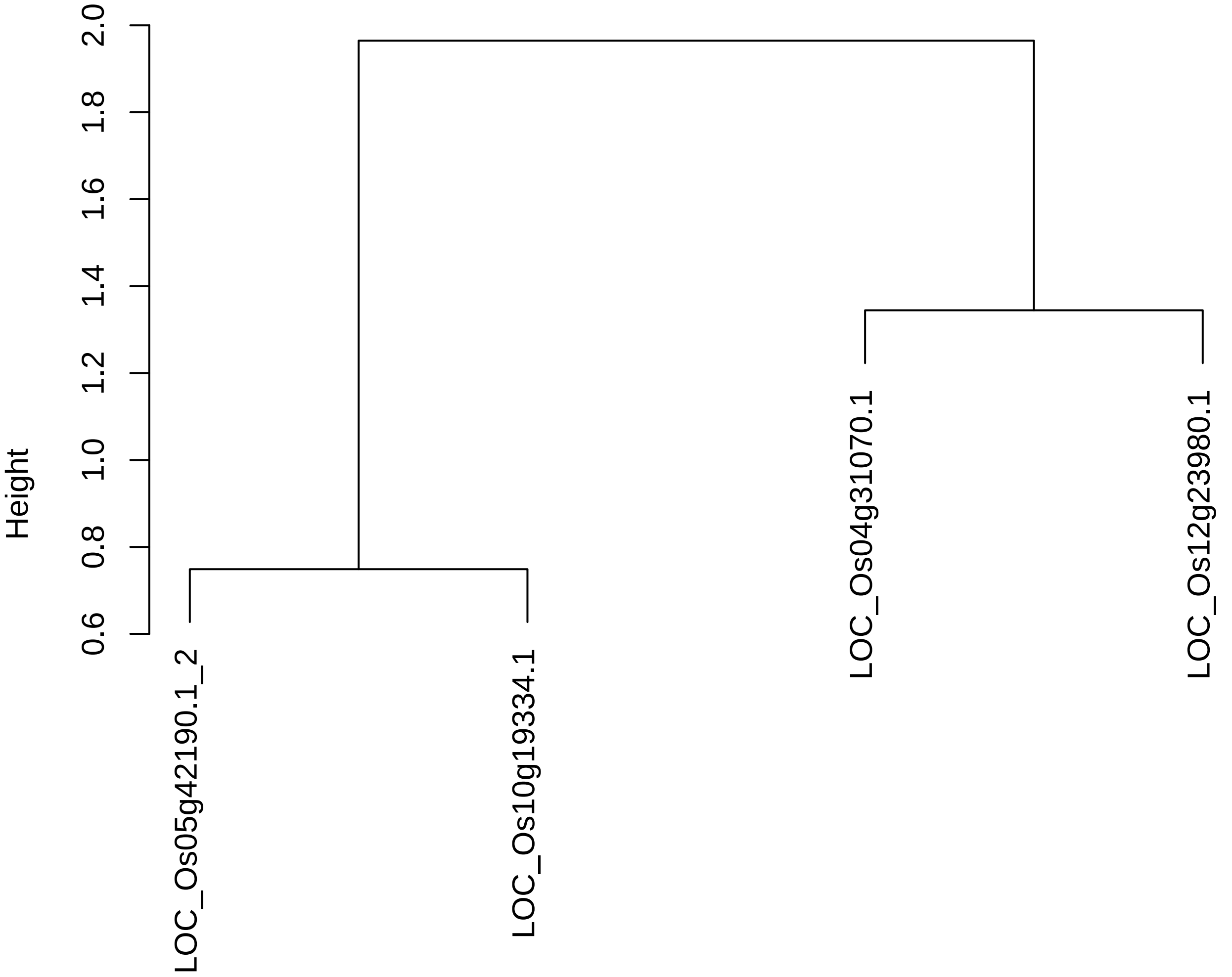


dist(spfitiex12sec12[lab[, 1] == i, ])
hclust (*, "ward.D2")

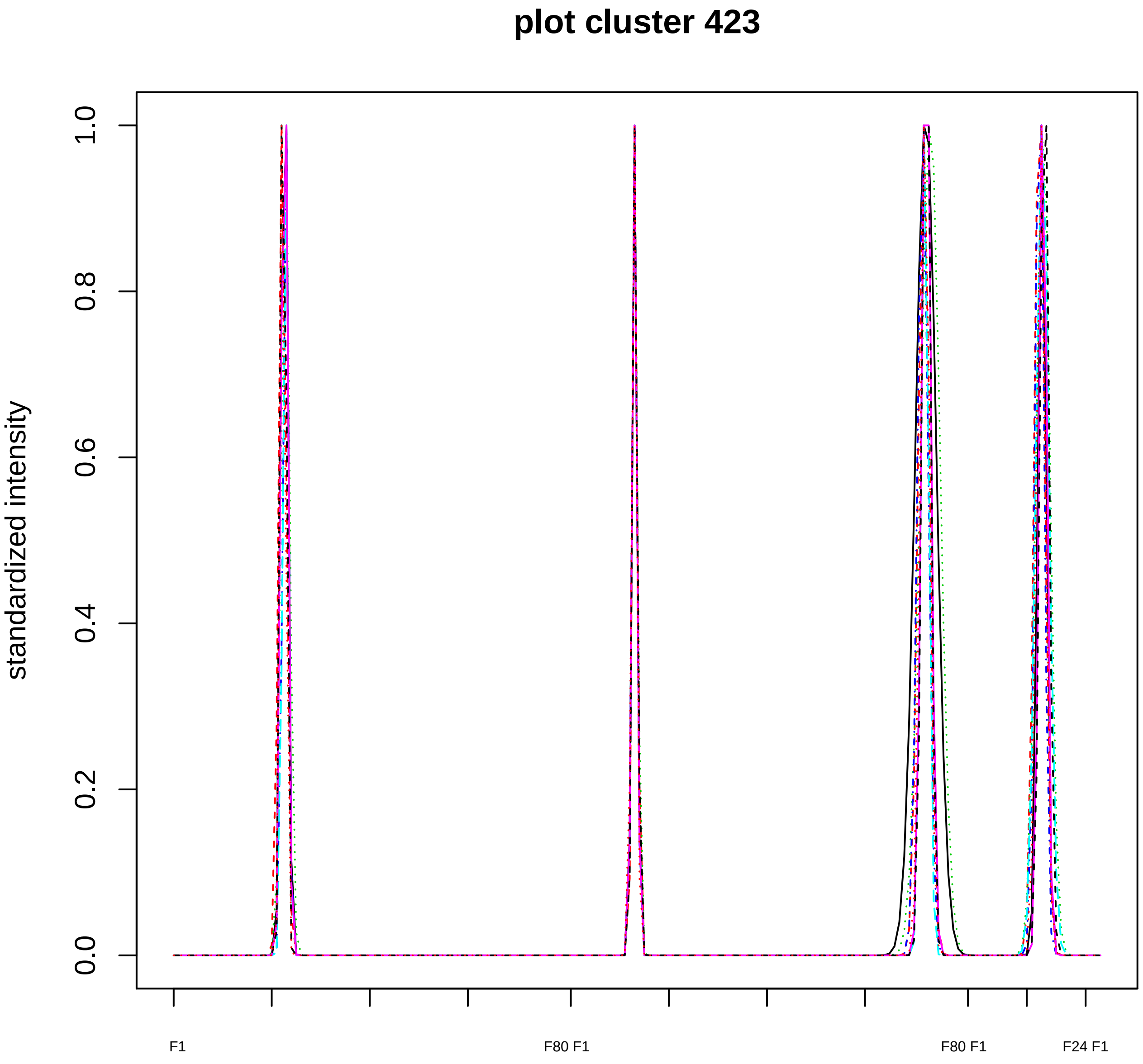
plot cluster 423
standardized intensity
0.0
0.2
0.4
0.6
0.8
1.0
F1
F80 F1
F80 F1
F24 F1

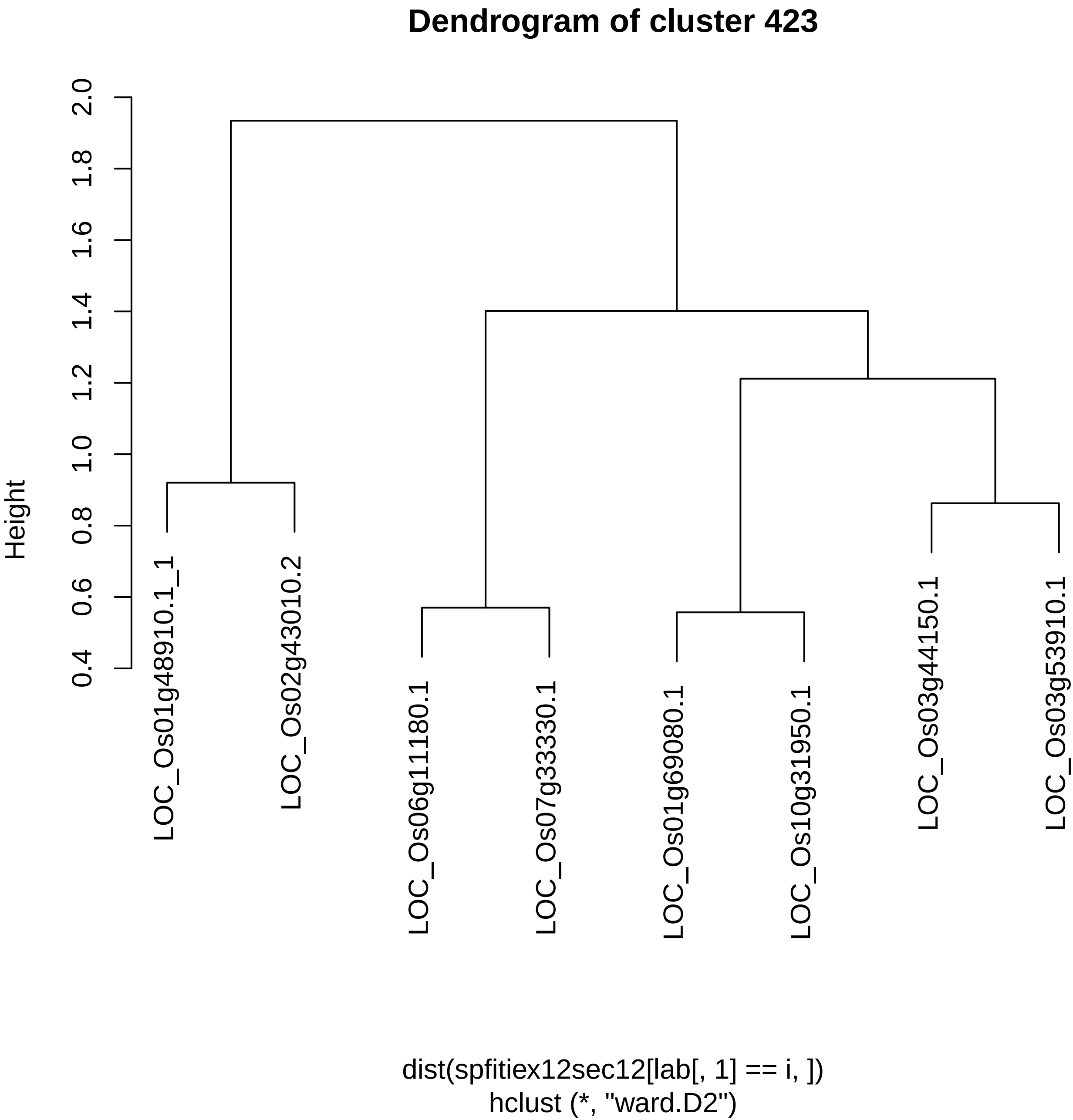
Dendrogram of cluster 423
Height
2.0
1.8
1.6
1.4
1.2
1.0
0.8
0.6
0.4
LOC_Os01g48910.1_1
LOC_Os02g43010.2
LOC_Os06g11180.1
LOC_Os07g33330.1
LOC_Os01g69080.1
LOC_Os10g31950.1
LOC_Os03g44150.1
LOC_Os03g53910.1
dist(spfitiex12sec12[lab[, 1] == i, ])
hclust (*, "ward.D2")

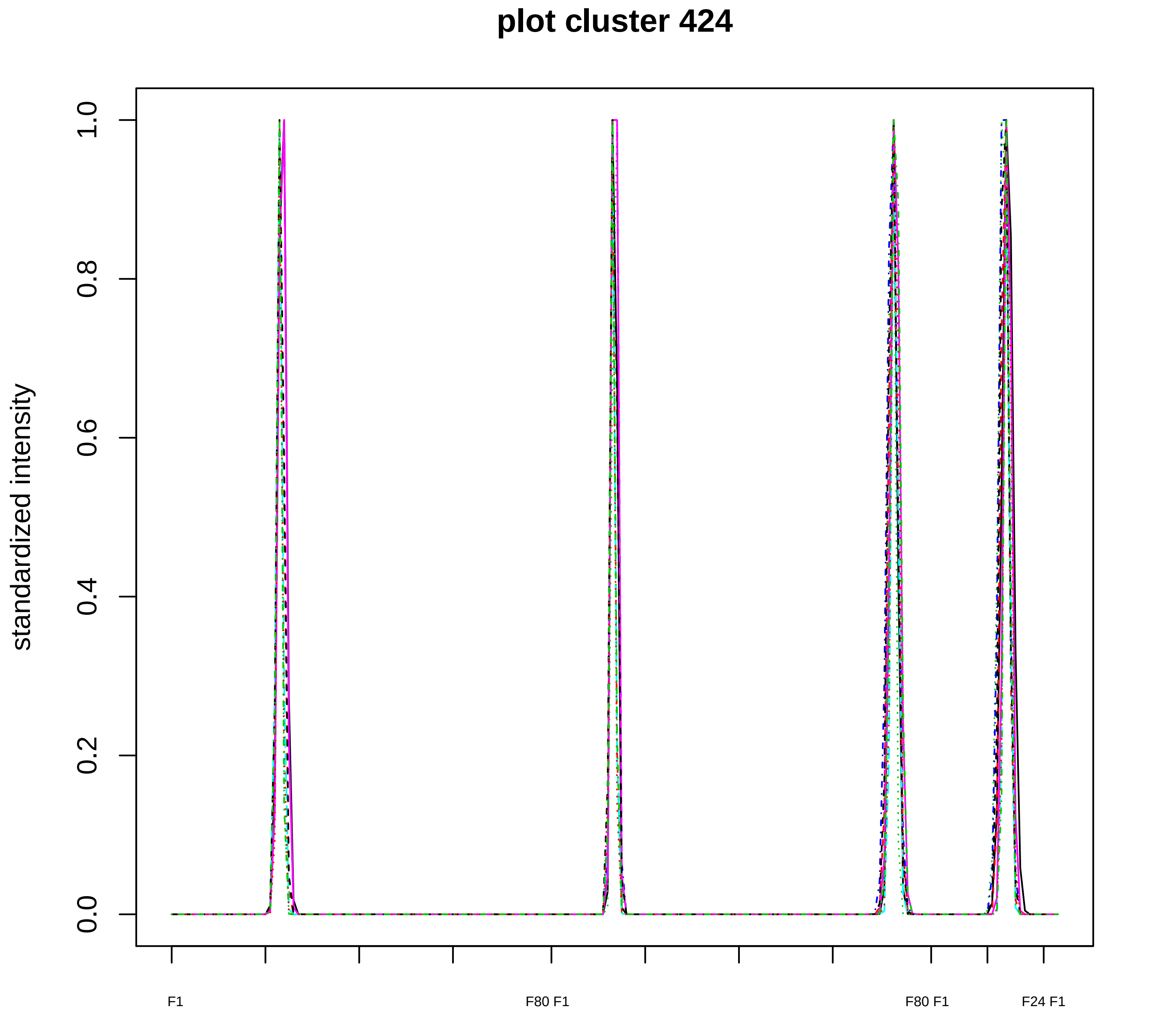

plot cluster 424
standardized intensity
0.0
0.2
0.4
0.6
0.8
1.0
F1
F80 F1
F80 F1
F24 F1

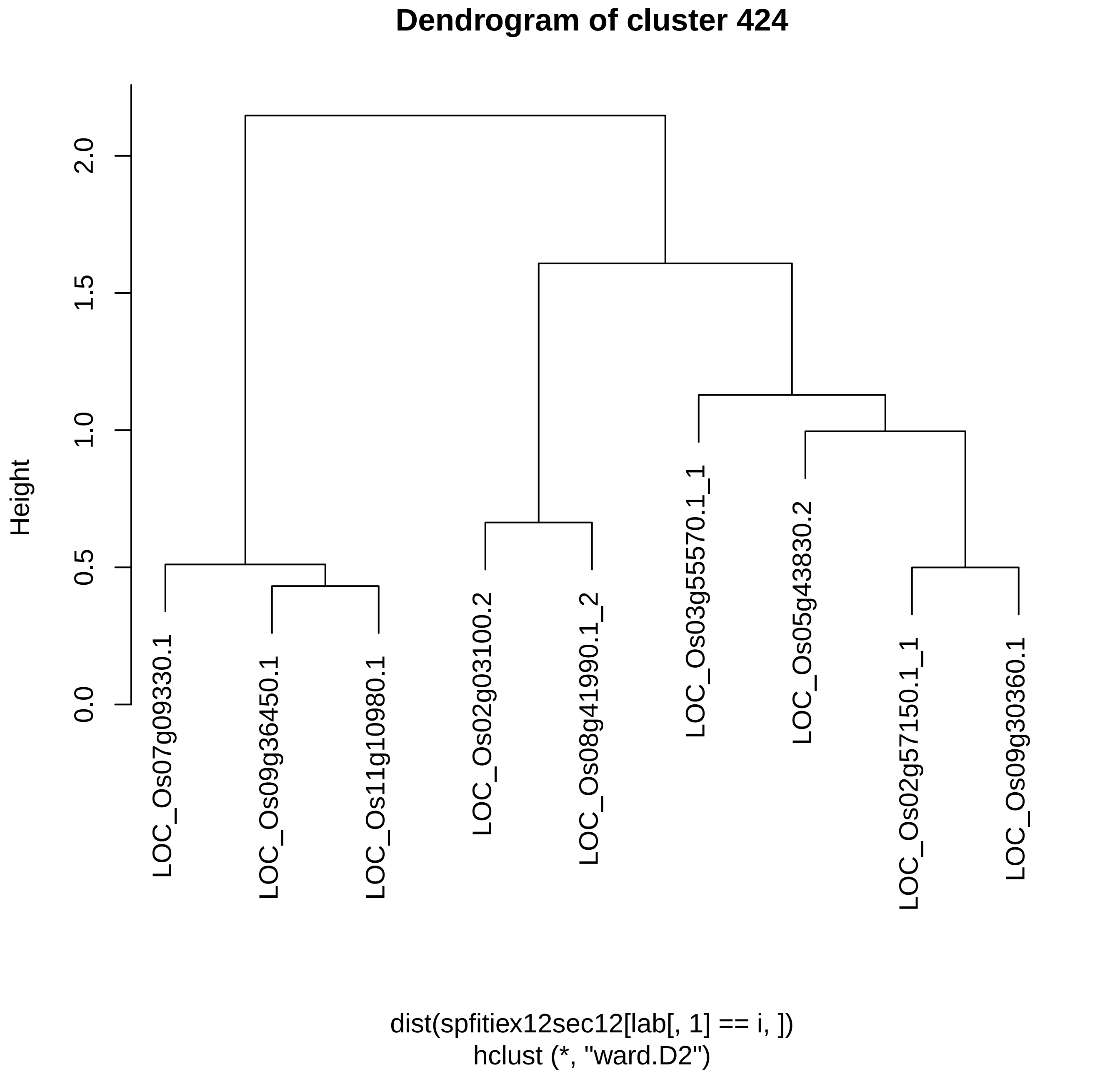
Dendrogram of cluster 424
Height
0.0
0.5
1.0
1.5
2.0
LOC_Os07g09330.1
LOC_Os09g36450.1
LOC_Os11g10980.1
LOC_Os02g03100.2
LOC_Os08g41990.1_2
LOC_Os03g55570.1_1
LOC_Os05g43830.2
LOC_Os02g57150.1_1
LOC_Os09g30360.1
dist(spfitiex12sec12[lab[, 1] == i, ])
hclust (*, "ward.D2")

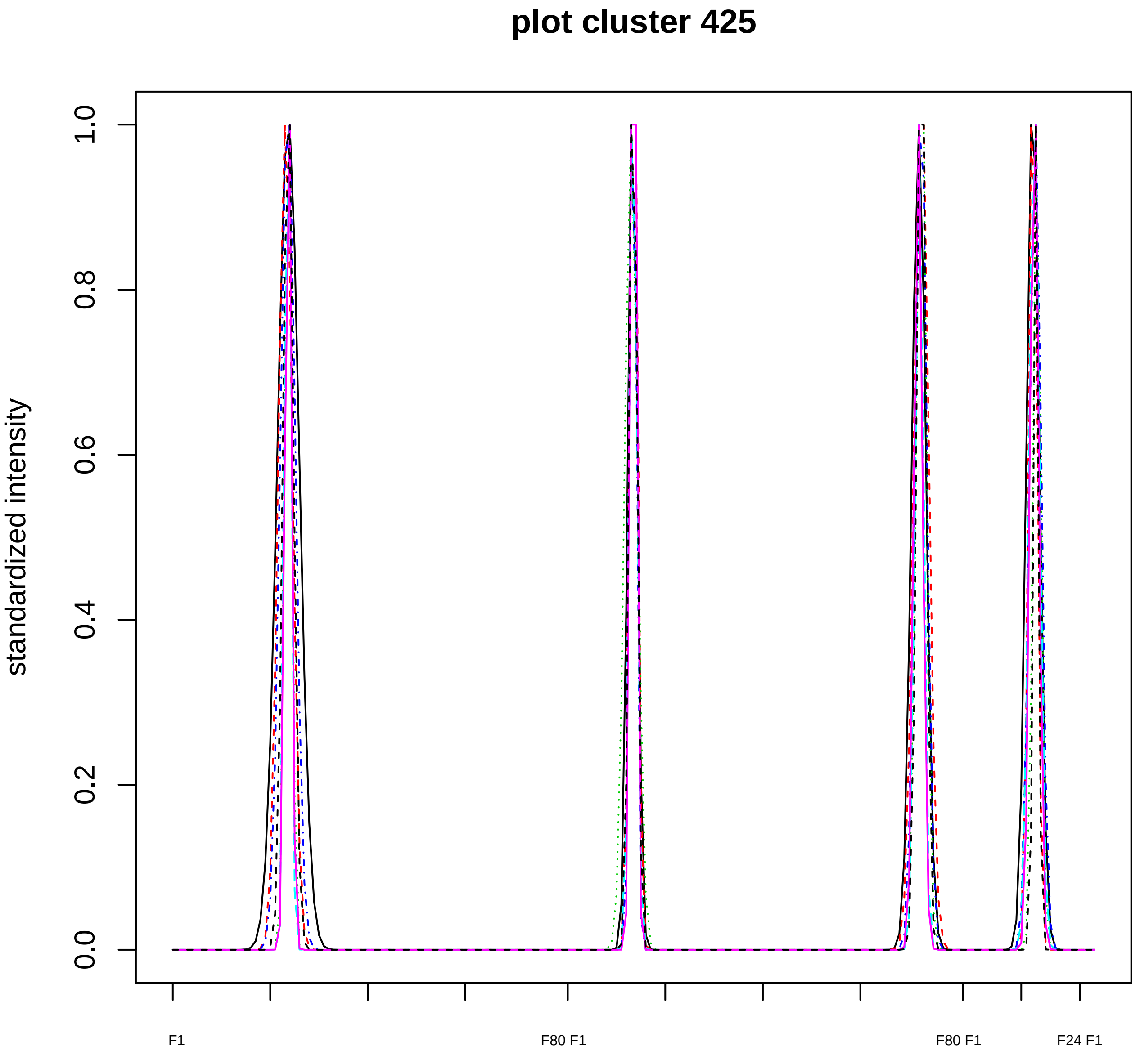
plot cluster 425
standardized intensity
0.0
0.2
0.4
0.6
0.8
1.0
F1
F80 F1
F80 F1
F24 F1

### Dendrogram of cluster 425

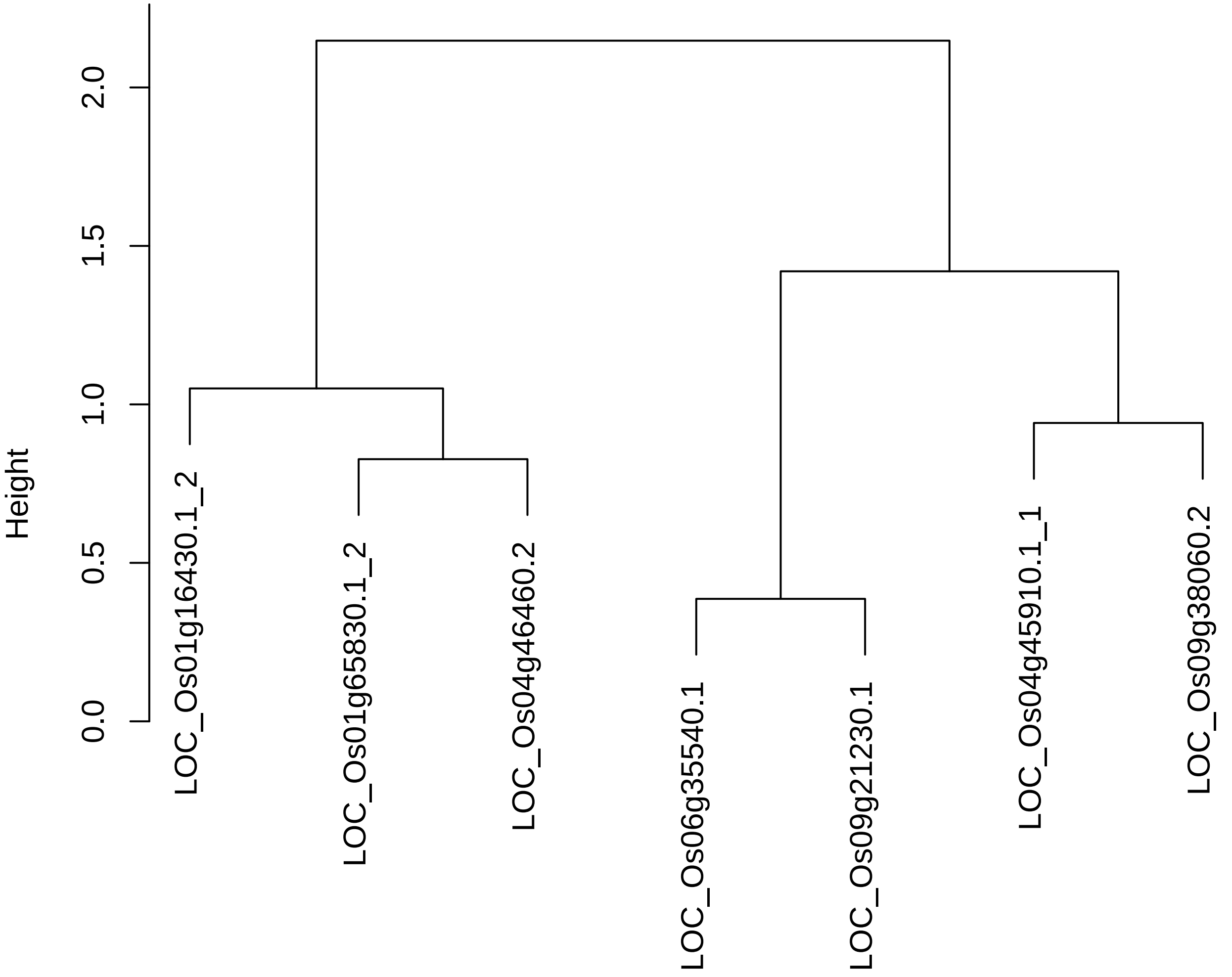


dist(spfitiex12sec12[lab[, 1] == i, ])
hclust (*, "ward.D2")

#### plot cluster 426

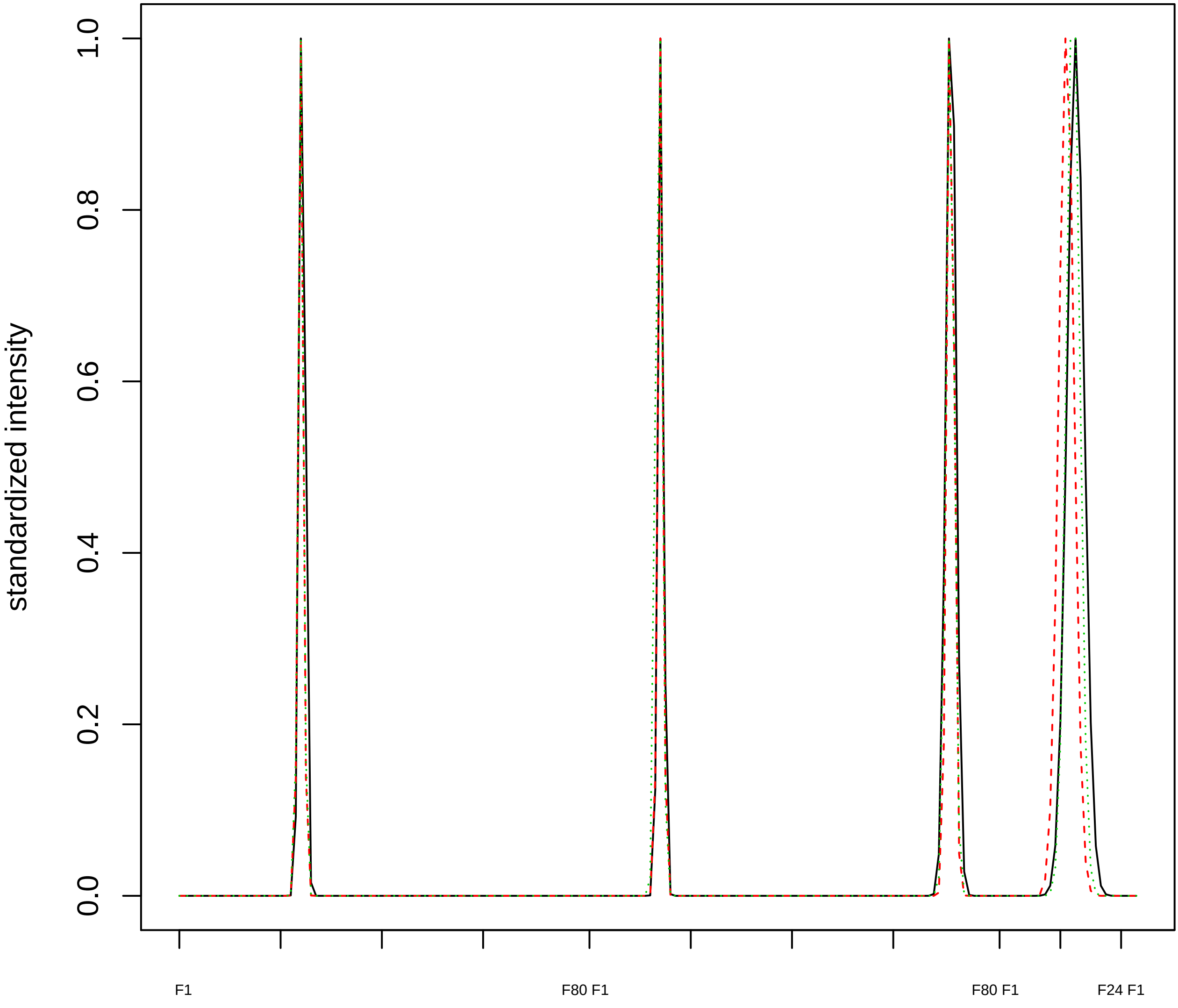

#### Dendrogram of cluster 426

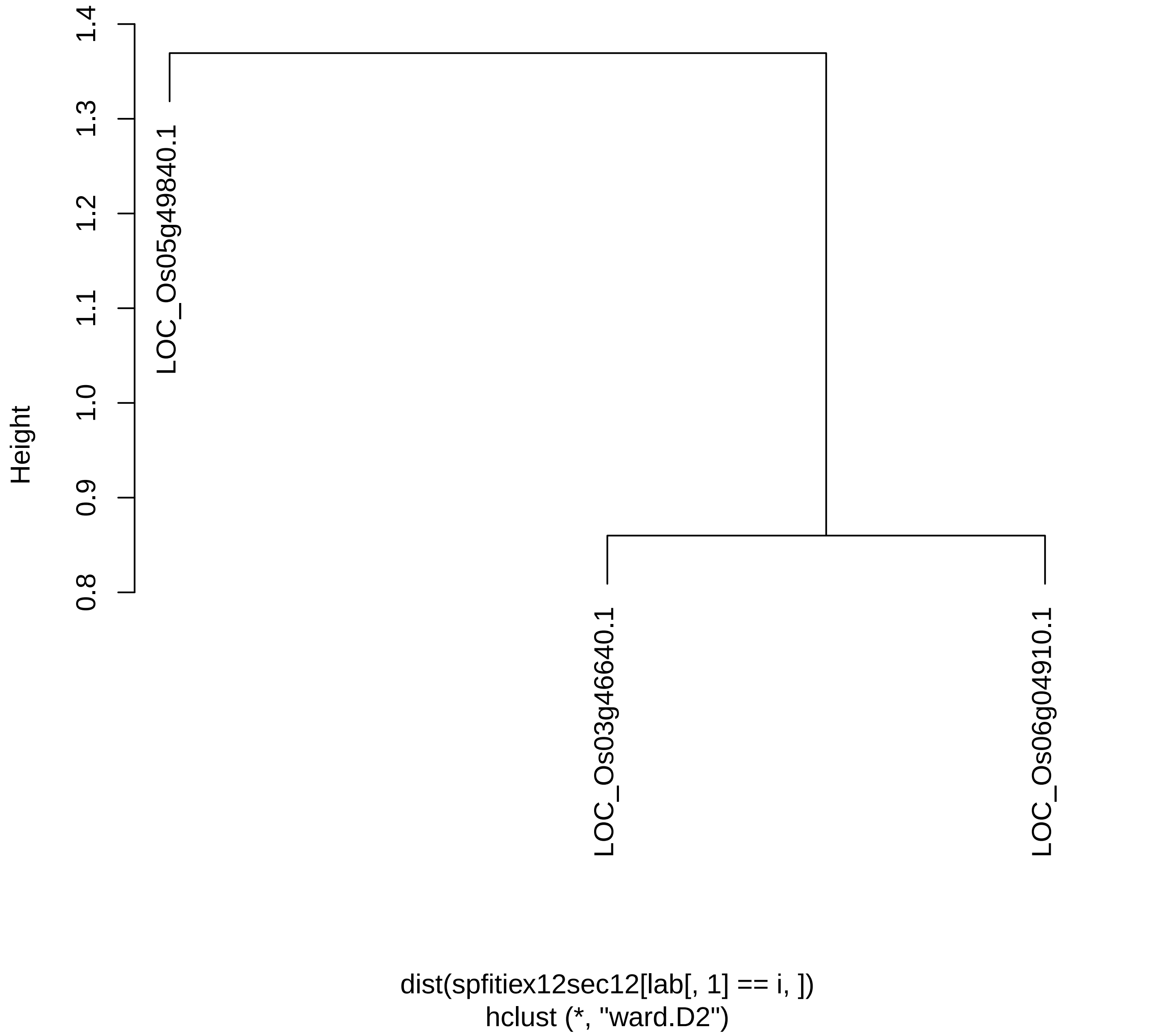

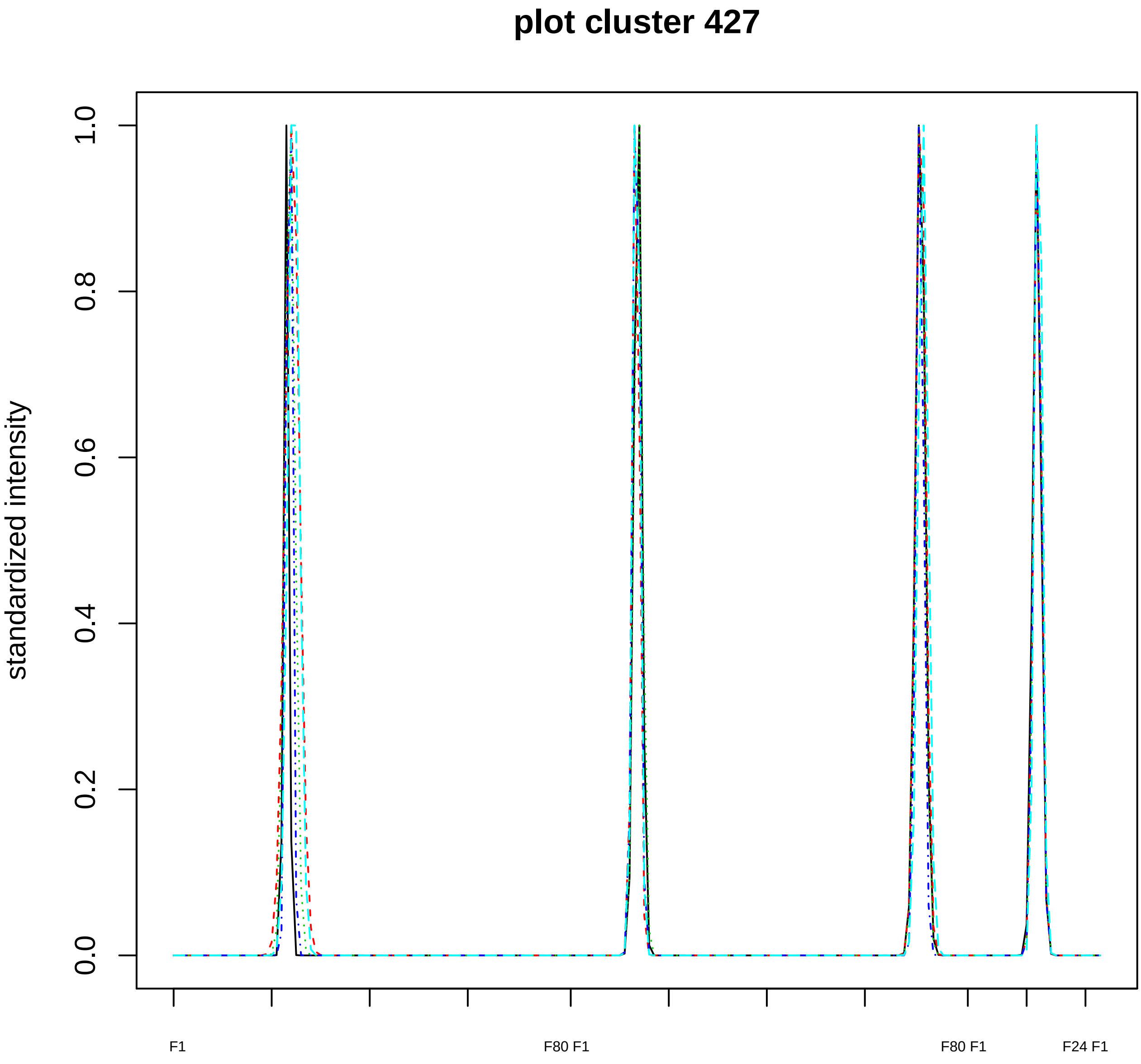
plot cluster 427
standardized intensity
0.0
0.2
0.4
0.6
0.8
1.0
F1
F80 F1
F80 F1
F24 F1

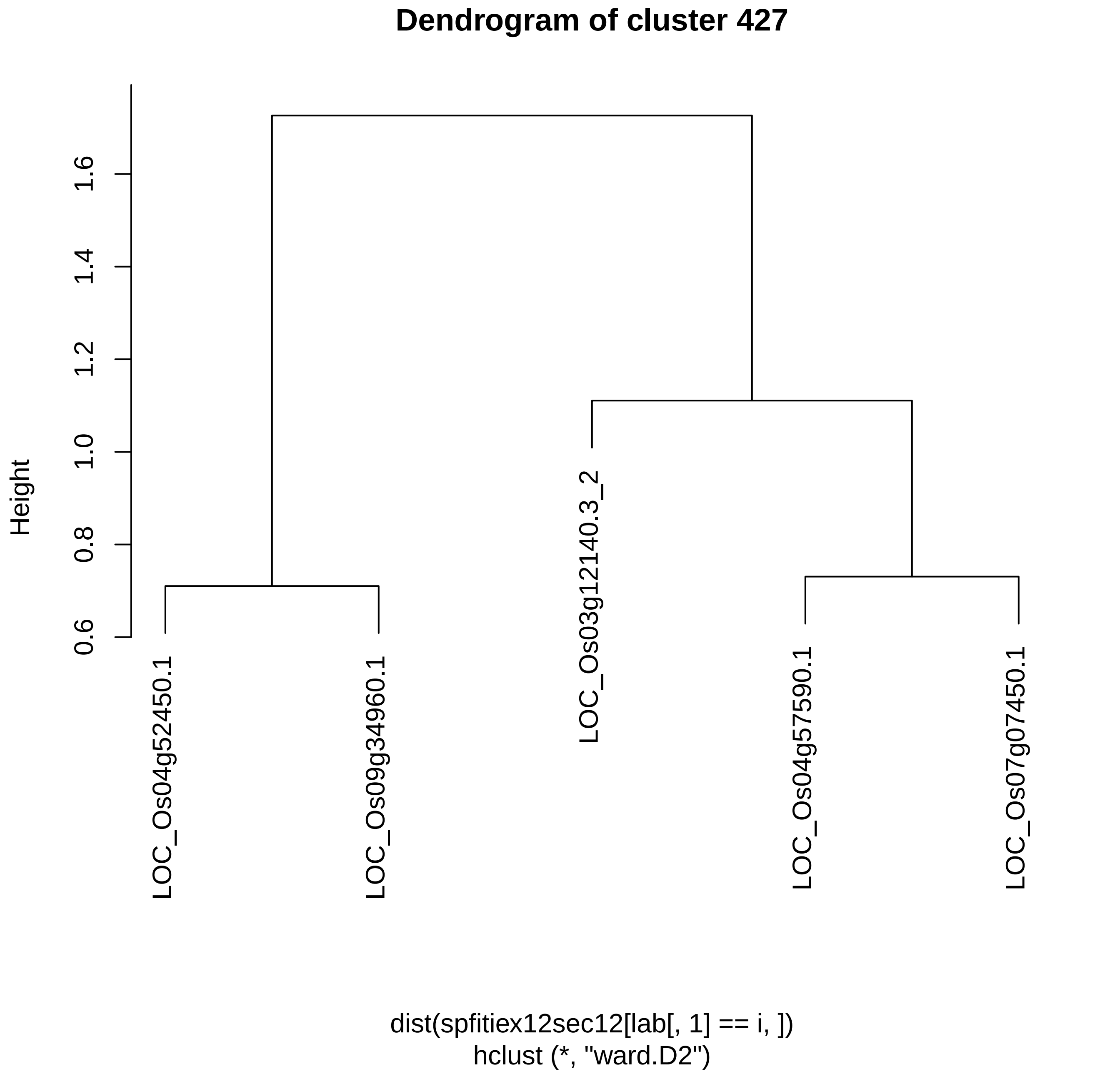
Dendrogram of cluster 427
Height
0.6
0.8
1.0
1.2
1.4
1.6
LOC_Os04g52450.1
LOC_Os09g34960.1
LOC_Os03g12140.3_2
LOC_Os04g57590.1
LOC_Os07g07450.1
dist(spfitiex12sec12[lab[, 1] == i, ])
hclust (*, "ward.D2")

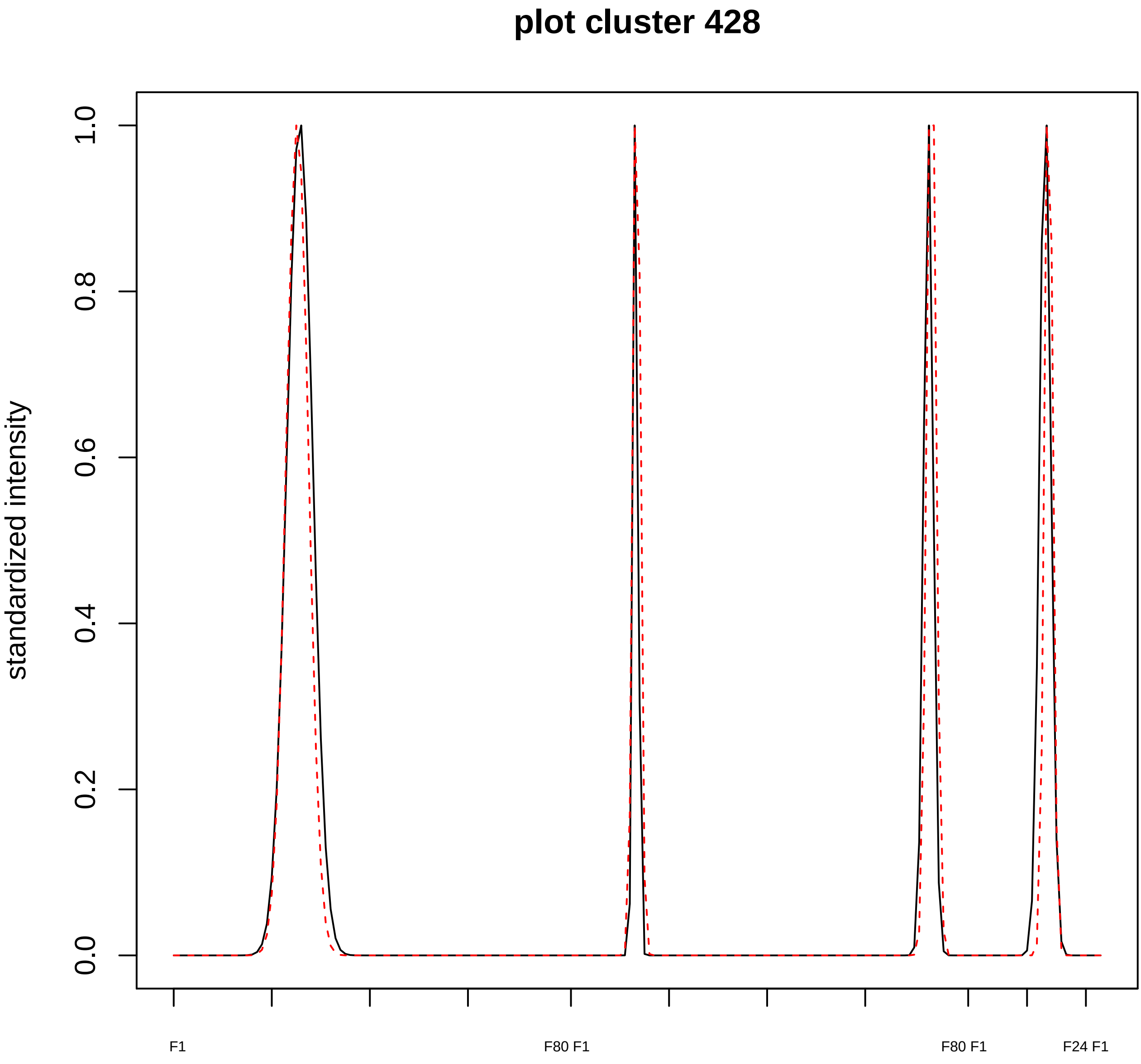
plot cluster 428
standardized intensity
0.0
0.2
0.4
0.6
0.8
1.0
F1
F80 F1
F80 F1
F24 F1

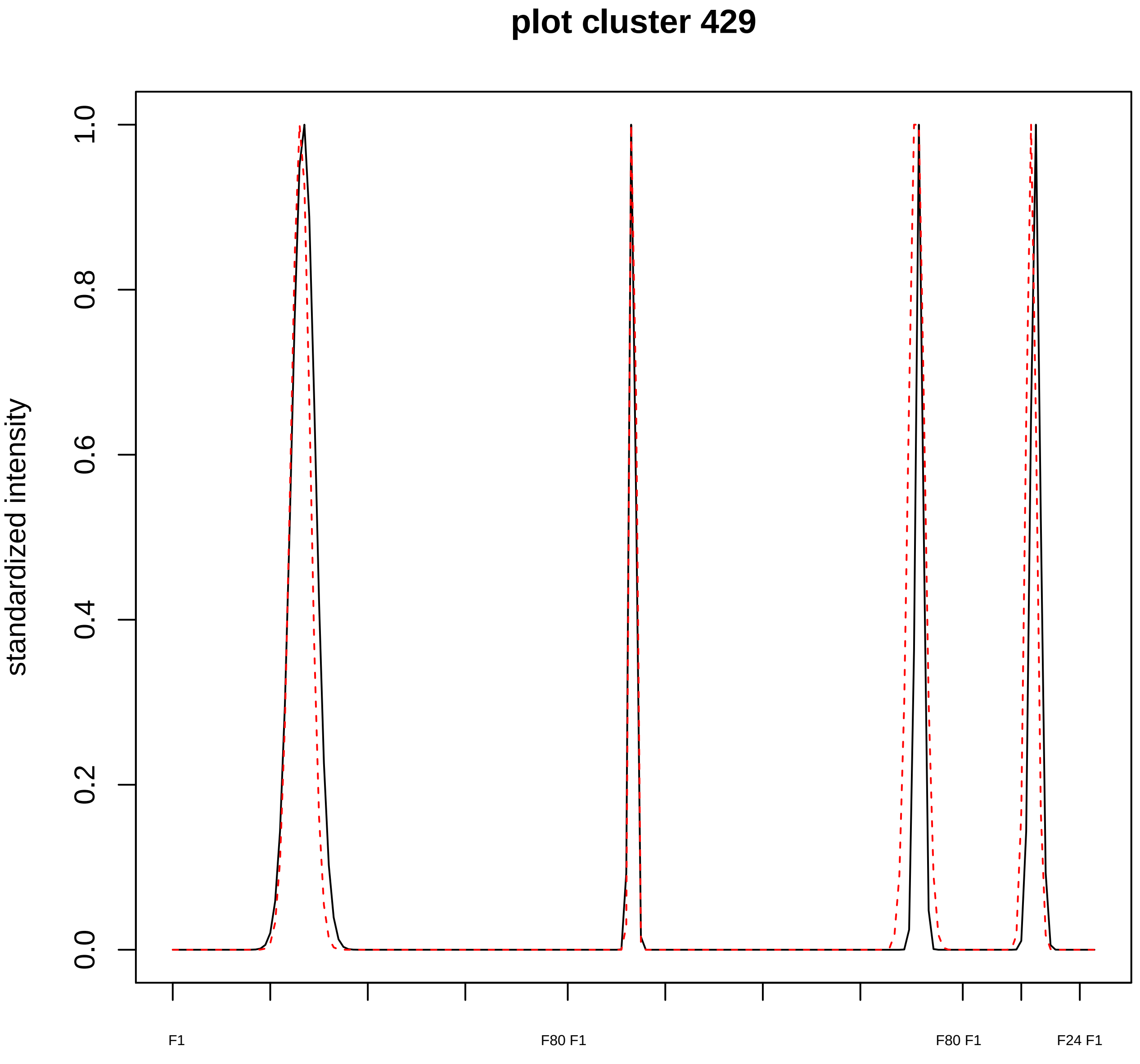
plot cluster 429
standardized intensity
0.0
0.2
0.4
0.6
0.8
1.0
F1
F80 F1
F80 F1
F24 F1

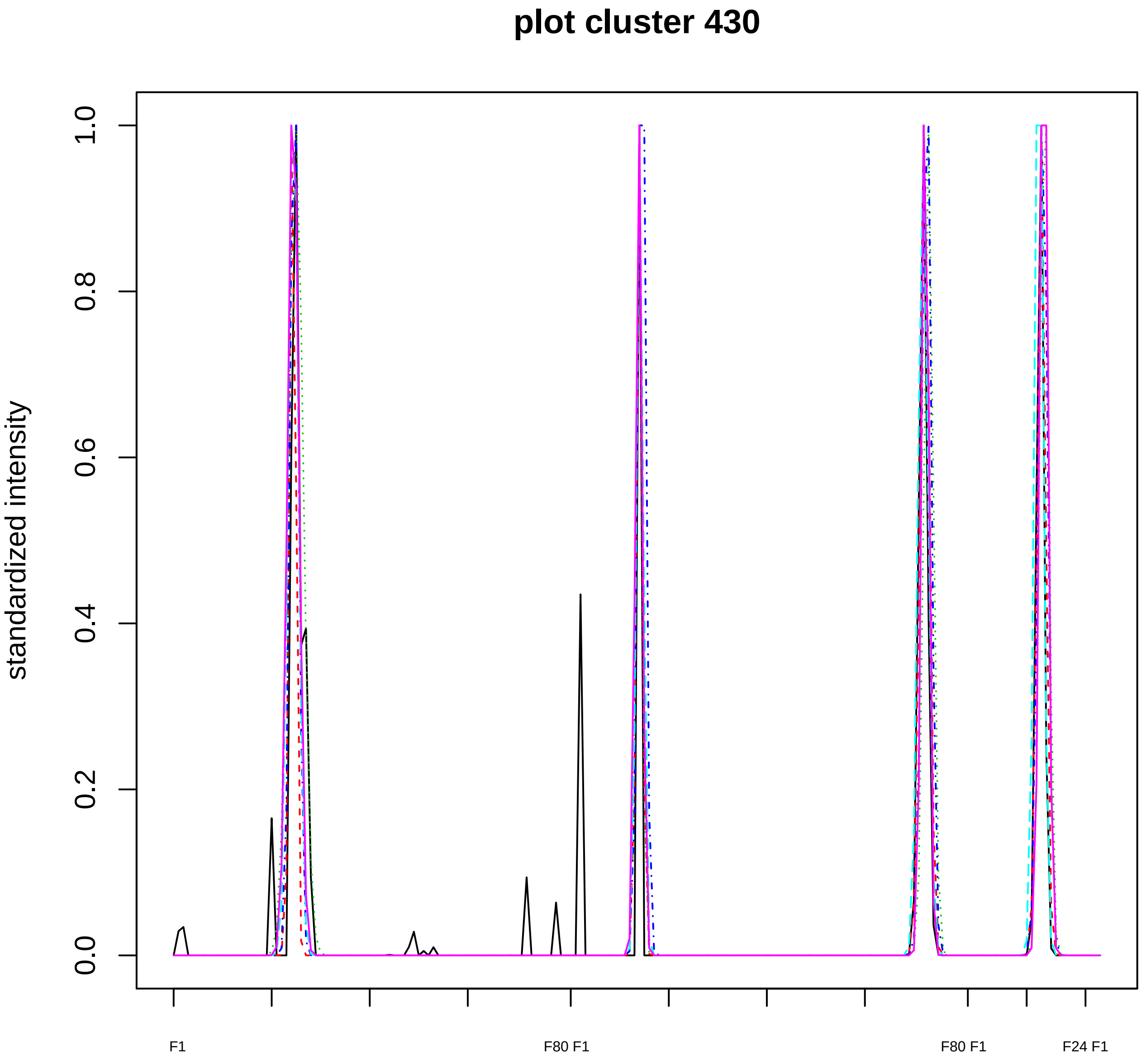

plot cluster 430
standardized intensity
0.0
0.2
0.4
0.6
0.8
1.0
F1
F80 F1
F80 F1
F24 F1

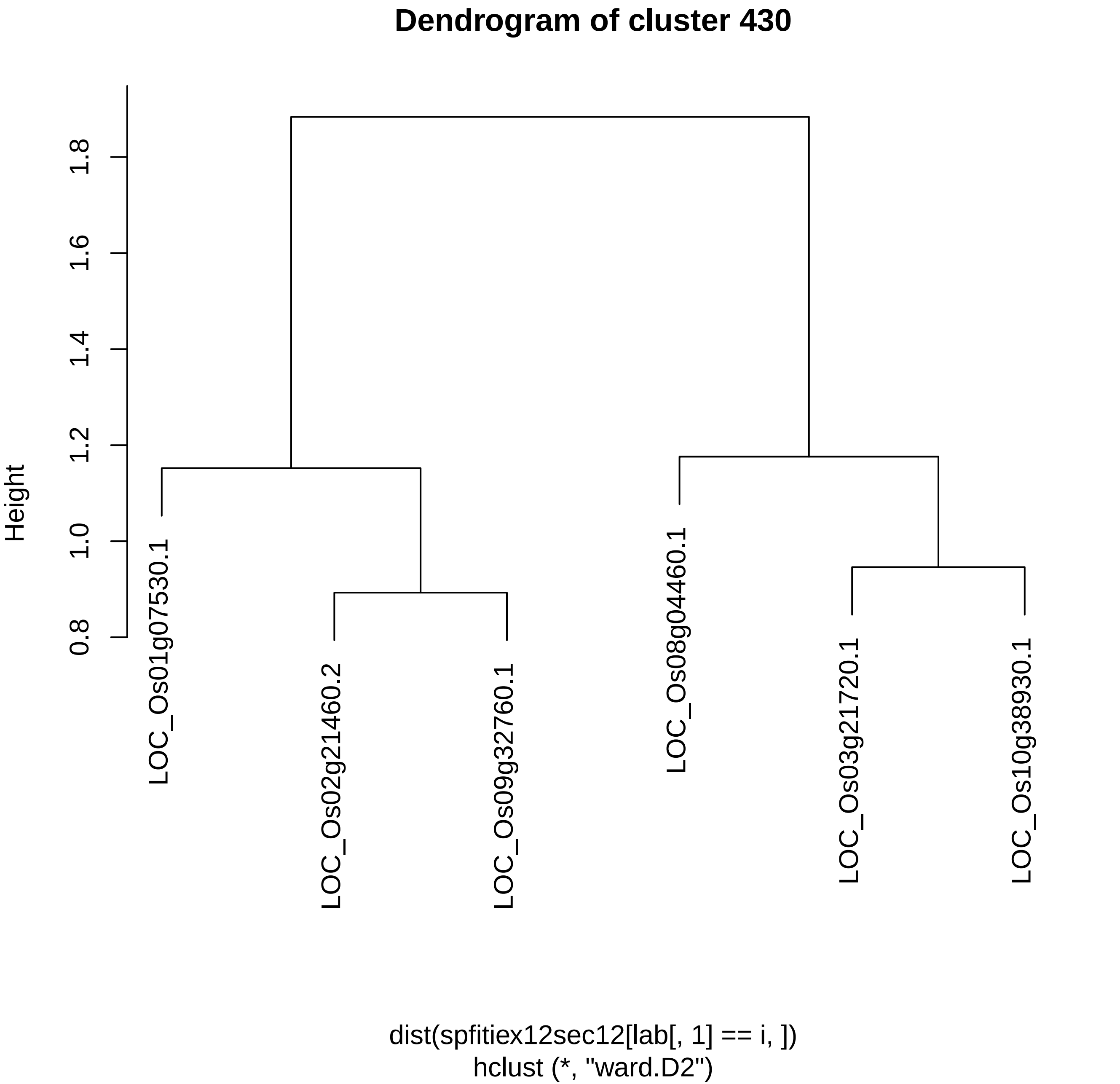
Dendrogram of cluster 430
Height
1.8
1.6
1.4
1.2
1.0
0.8
LOC_Os01g07530.1
LOC_Os02g21460.2
LOC_Os09g32760.1
LOC_Os08g04460.1
LOC_Os03g21720.1
LOC_Os10g38930.1
dist(spfitiex12sec12[lab[, 1] == i, ])
hclust (*, "ward.D2")

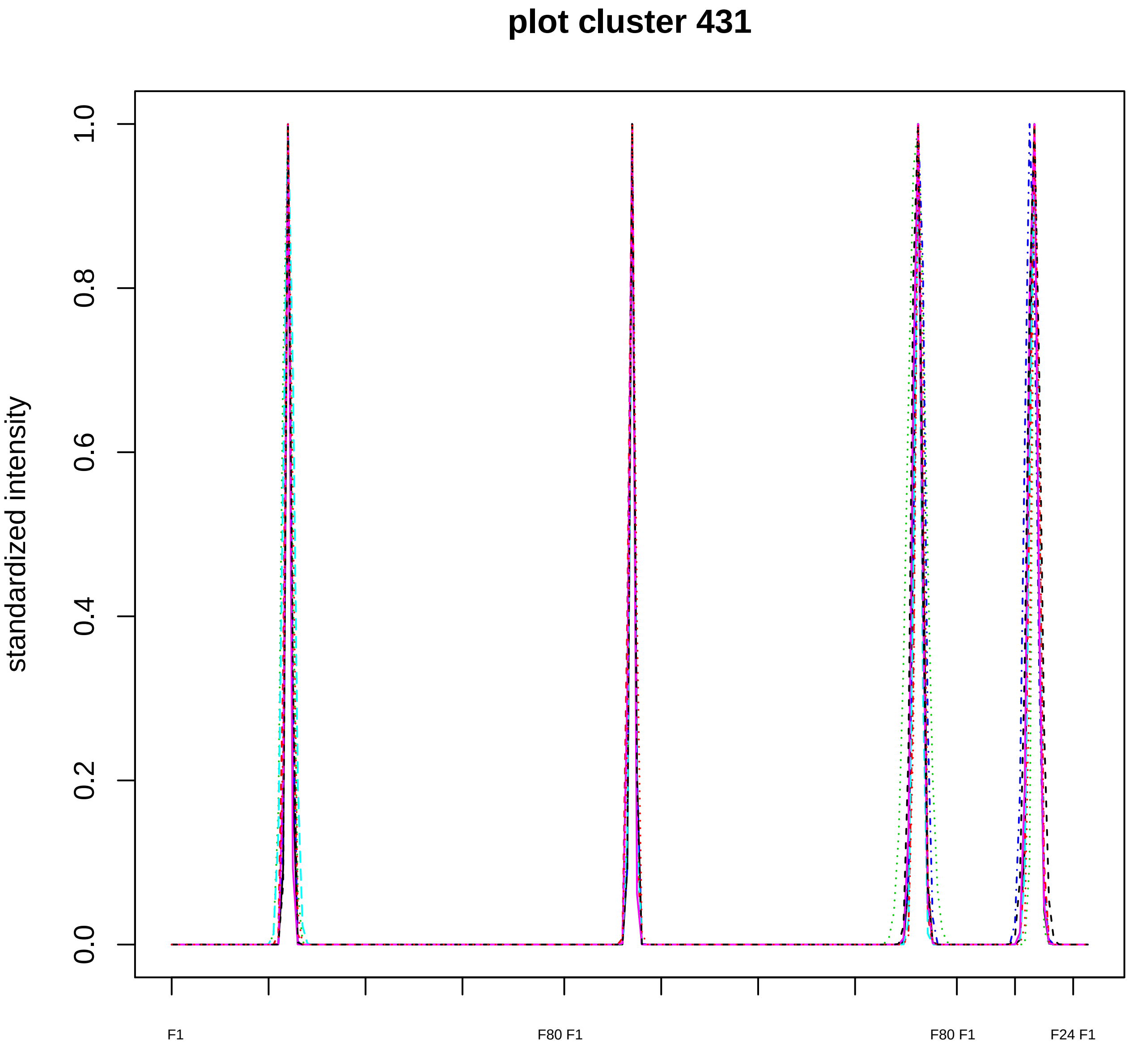

plot cluster 431
standardized intensity
0.0
0.2
0.4
0.6
0.8
1.0
F1
F80 F1
F80 F1
F24 F1

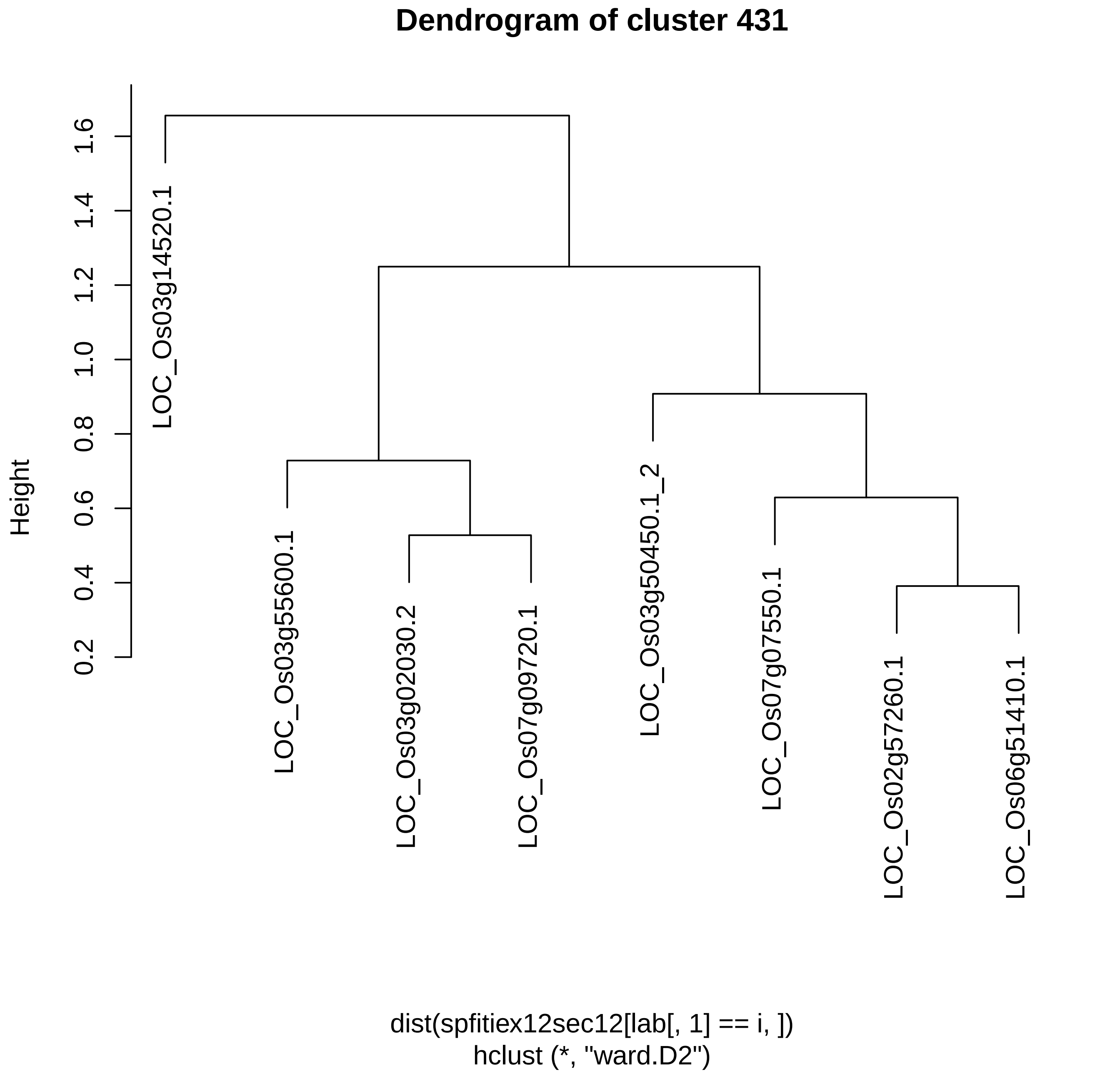
Dendrogram of cluster 431
Height
0.2
0.4
0.6
0.8
1.0
1.2
1.4
1.6
LOC_Os03g14520.1
LOC_Os03g55600.1
LOC_Os03g02030.2
LOC_Os07g09720.1
LOC_Os03g50450.1_2
LOC_Os07g07550.1
LOC_Os02g57260.1
LOC_Os06g51410.1
dist(spfitiex12sec12[lab[, 1] == i, ])
hclust (*, "ward.D2")

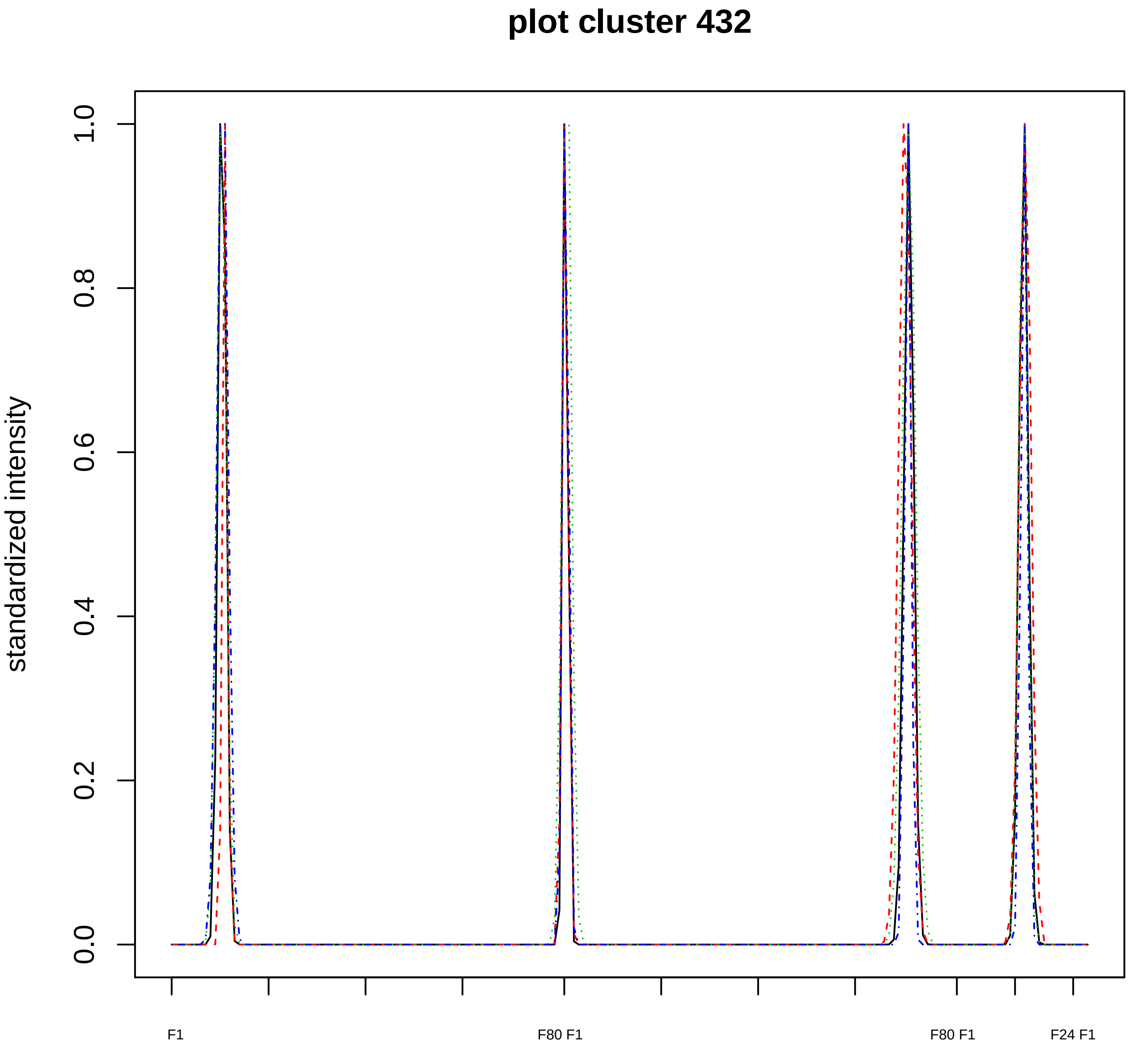
plot cluster 432
standardized intensity
0.0
0.2
0.4
0.6
0.8
1.0
F1
F80 F1
F80 F1
F24 F1

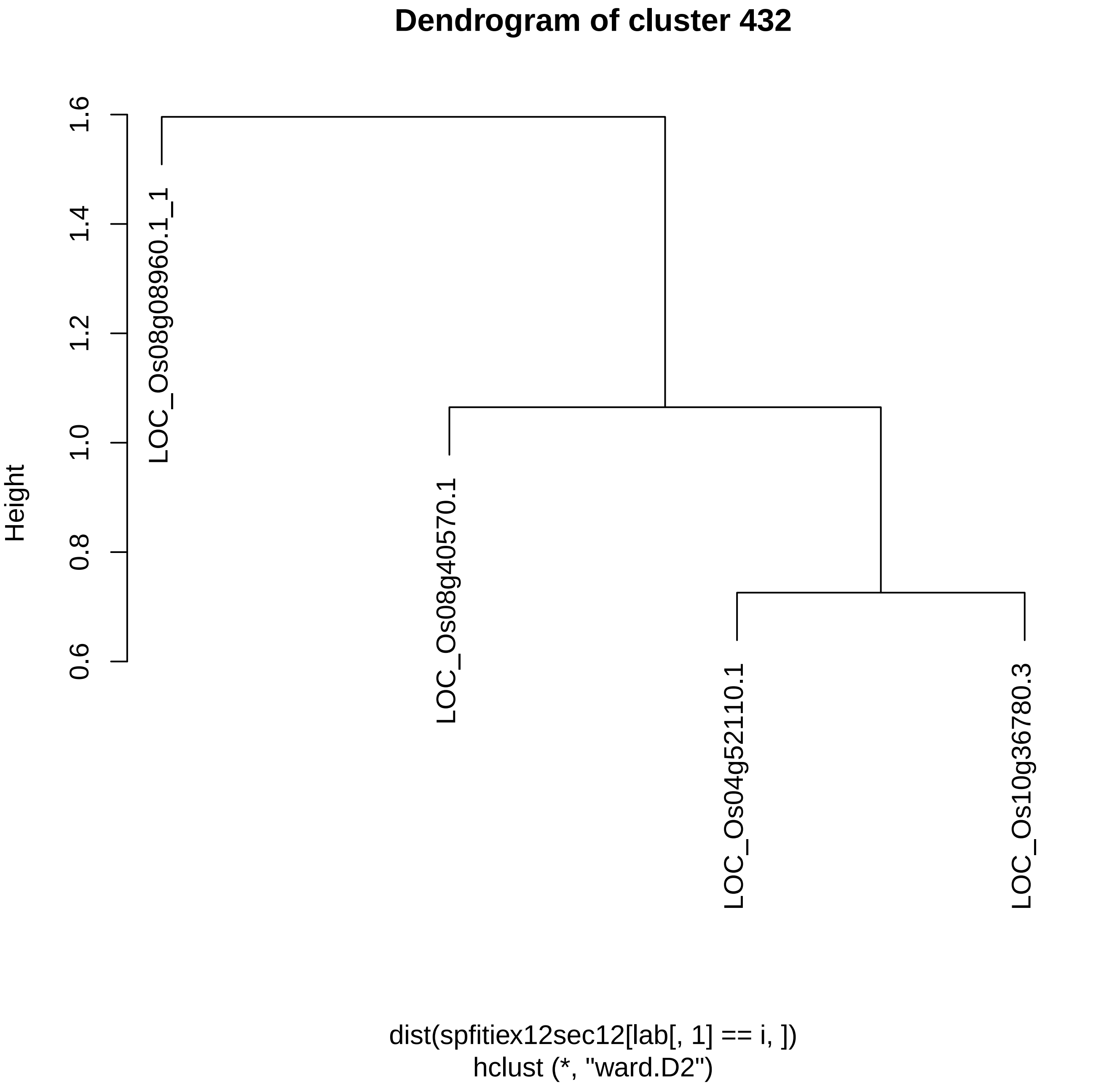
Dendrogram of cluster 432
Height
1.6
1.4
1.2
1.0
0.8
0.6
LOC_Os08g08960.1_1
LOC_Os08g40570.1
LOC_Os04g52110.1
LOC_Os10g36780.3
dist(spfitiex12sec12[lab[, 1] == i, ])
hclust (*, "ward.D2")

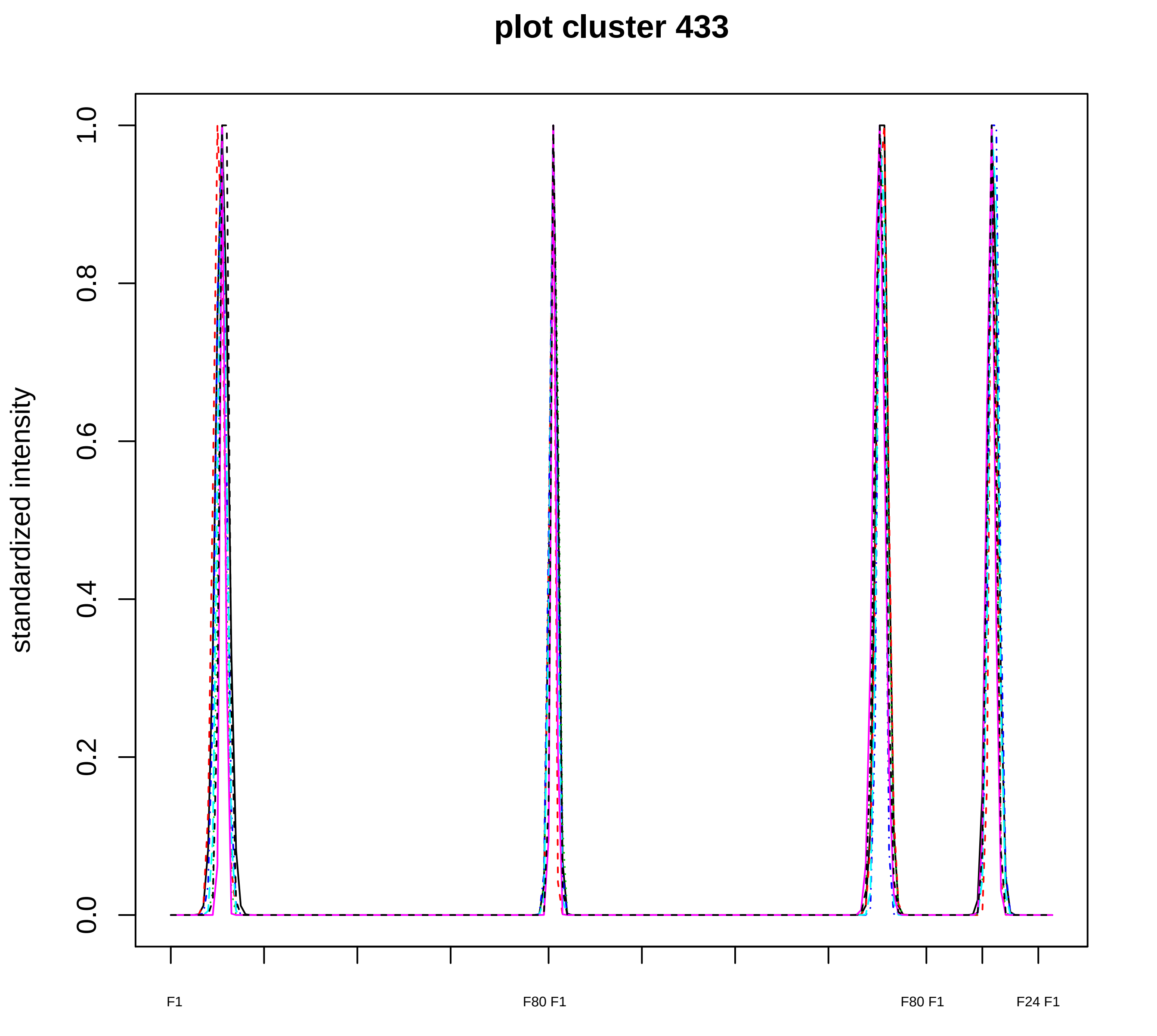
plot cluster 433
standardized intensity
0.0
0.2
0.4
0.6
0.8
1.0
F1
F80 F1
F80 F1
F24 F1

### Dendrogram of cluster 433

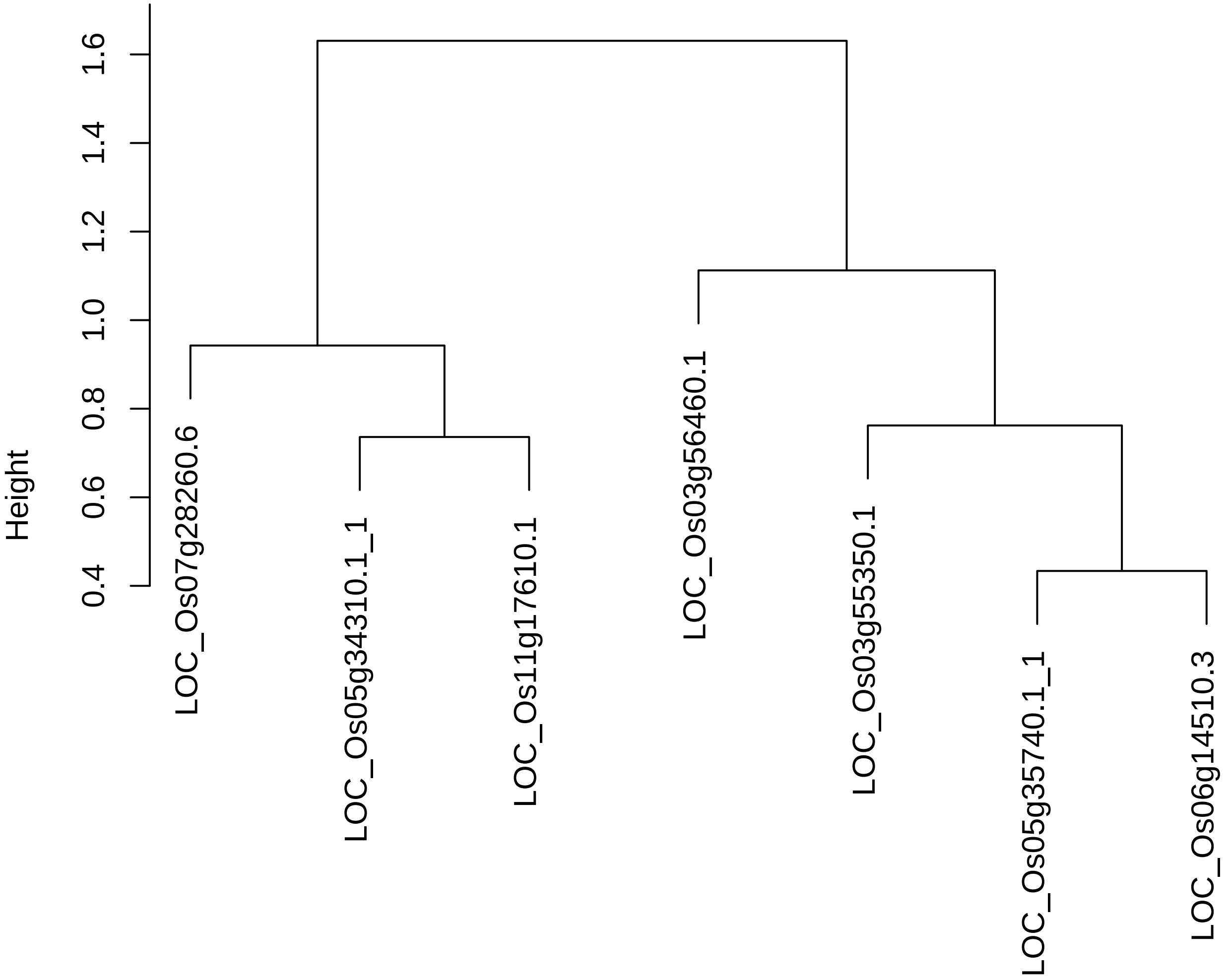


dist(spfitiex12sec12[lab[, 1] == i, ])
hclust (*, "ward.D2")

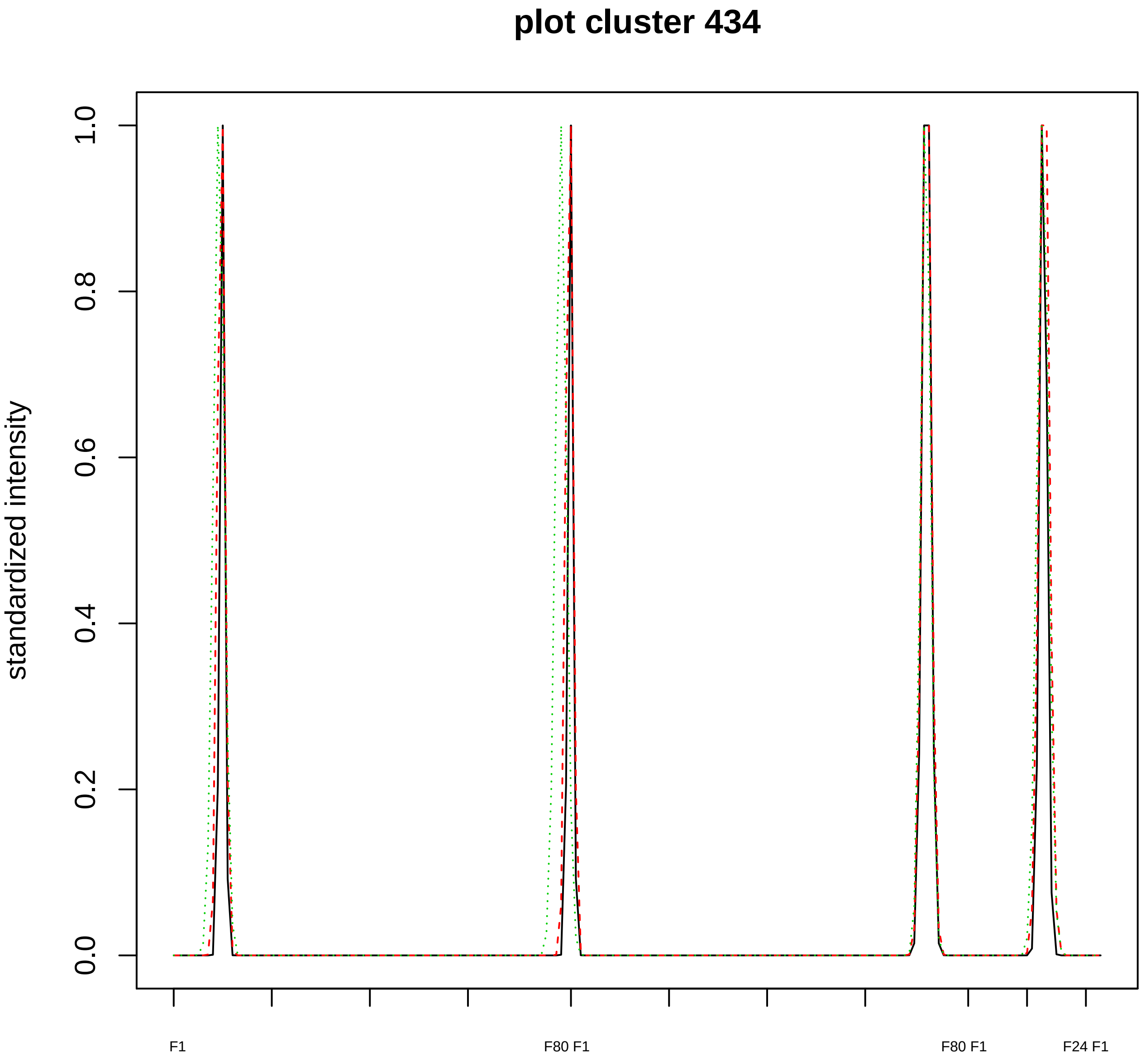
plot cluster 434
standardized intensity
0.0
0.2
0.4
0.6
0.8
1.0
F1
F80 F1
F80 F1
F24 F1

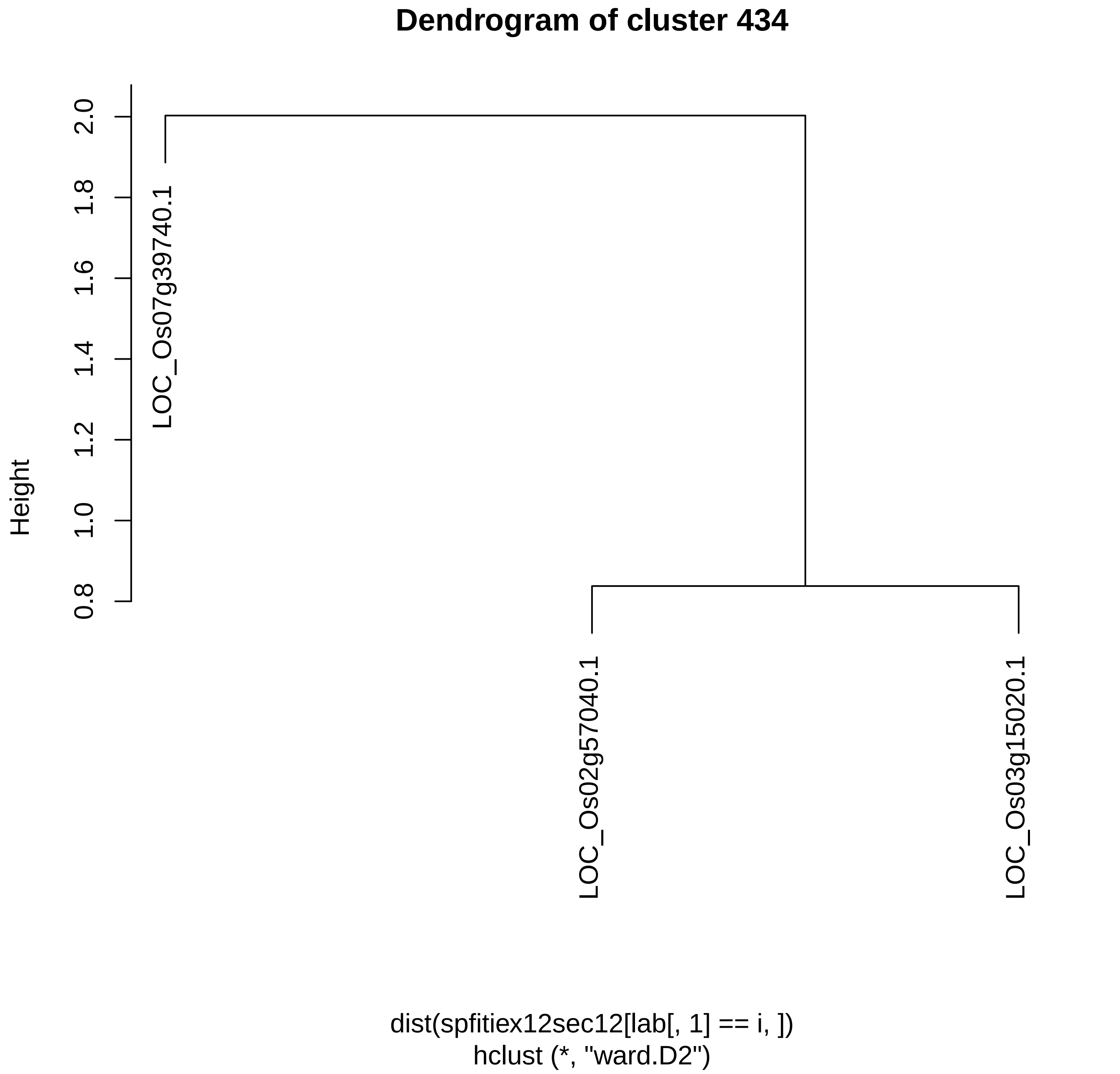
Dendrogram of cluster 434
Height
2.0
1.8
1.6
1.4
1.2
1.0
0.8
LOC_Os07g39740.1
LOC_Os02g57040.1
LOC_Os03g15020.1
dist(spfitiex12sec12[lab[, 1] == i, ])
hclust (*, "ward.D2")

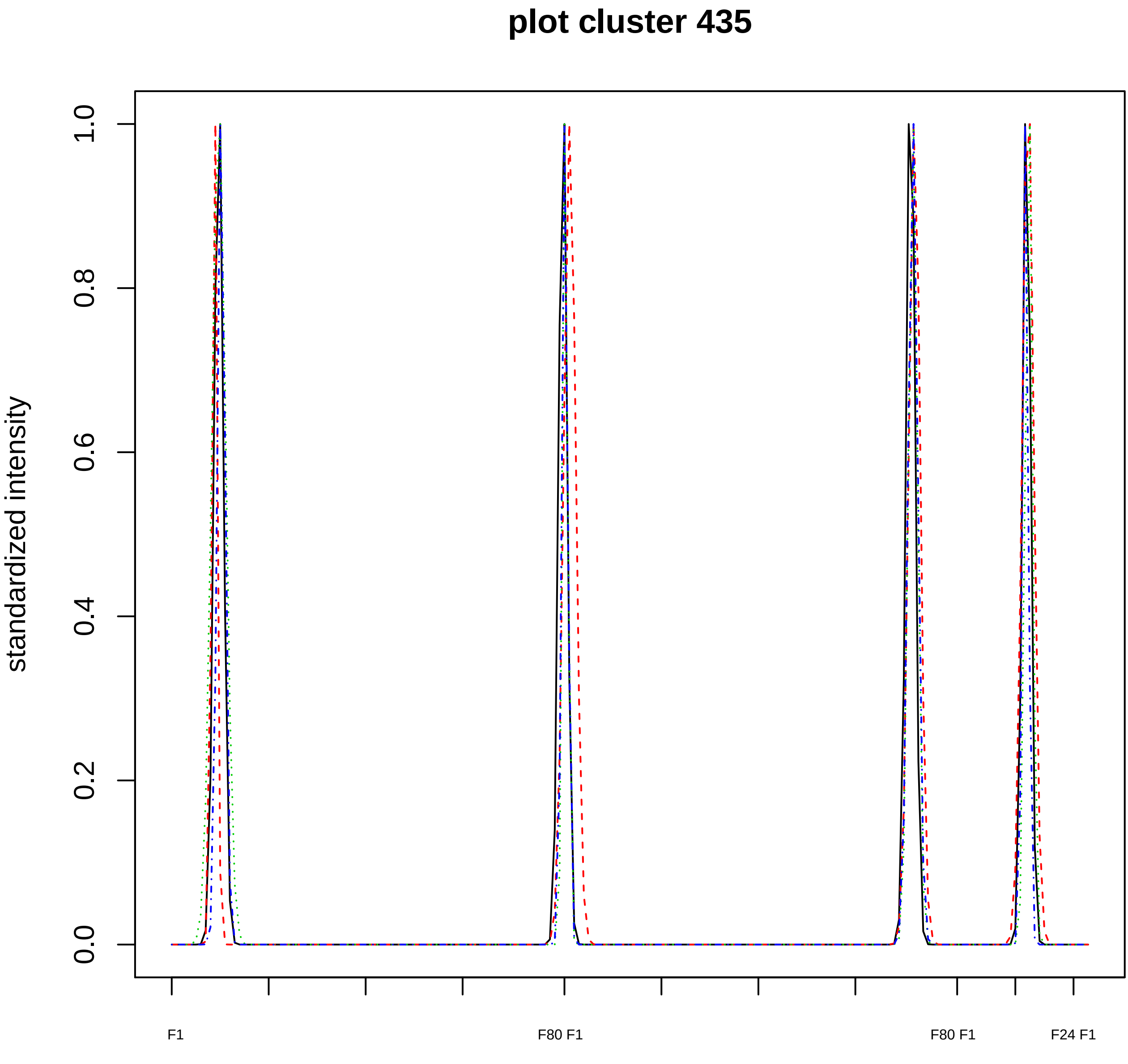

plot cluster 435
standardized intensity
0.0
0.2
0.4
0.6
0.8
1.0
F1
F80 F1
F80 F1
F24 F1

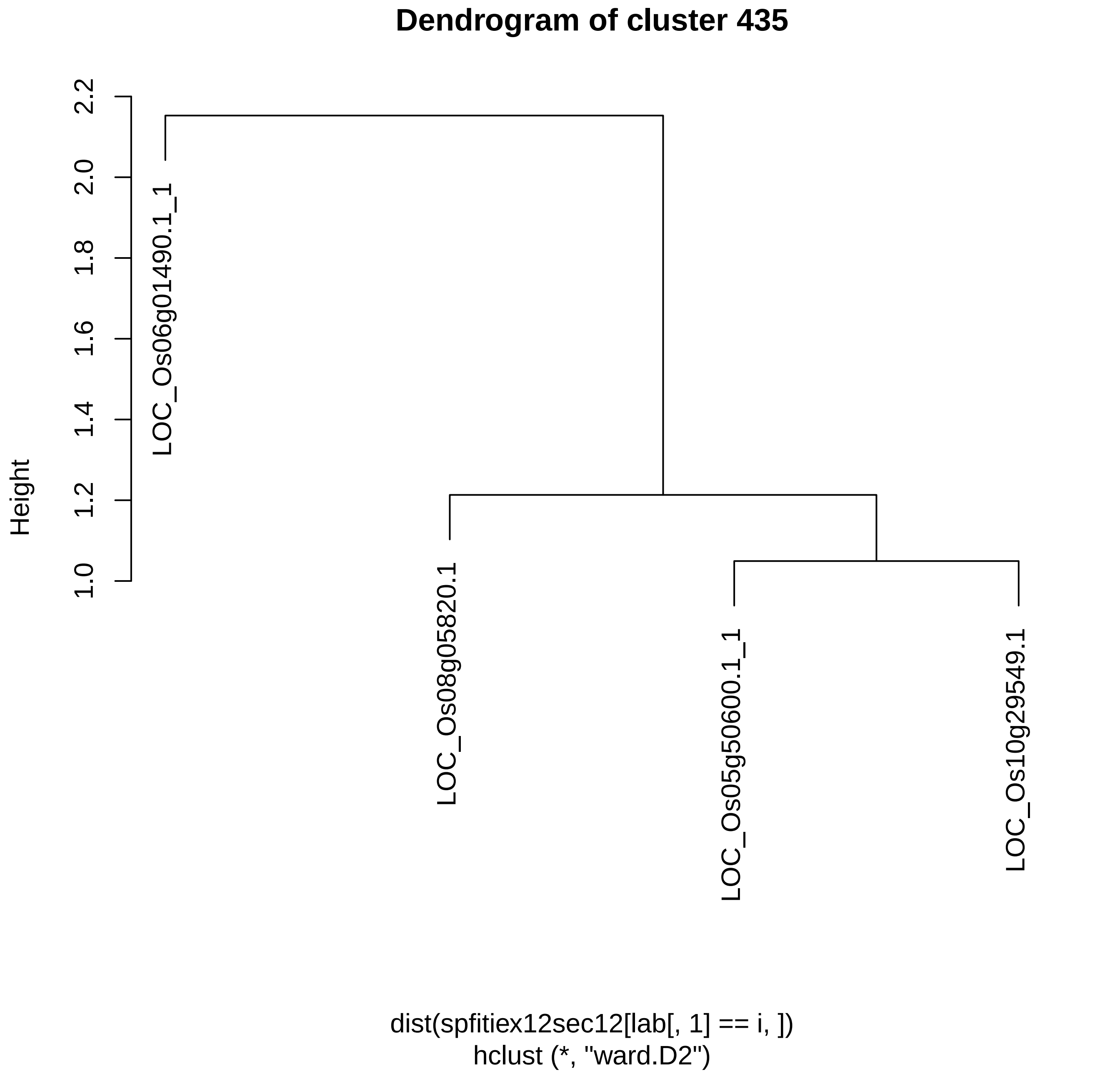
Dendrogram of cluster 435
Height
2.2
2.0
1.8
1.6
1.4
1.2
1.0
LOC_Os06g01490.1_1
LOC_Os08g05820.1
LOC_Os05g50600.1_1
LOC_Os10g29549.1
dist(spfitiex12sec12[lab[, 1] == i, ])
hclust (*, "ward.D2")

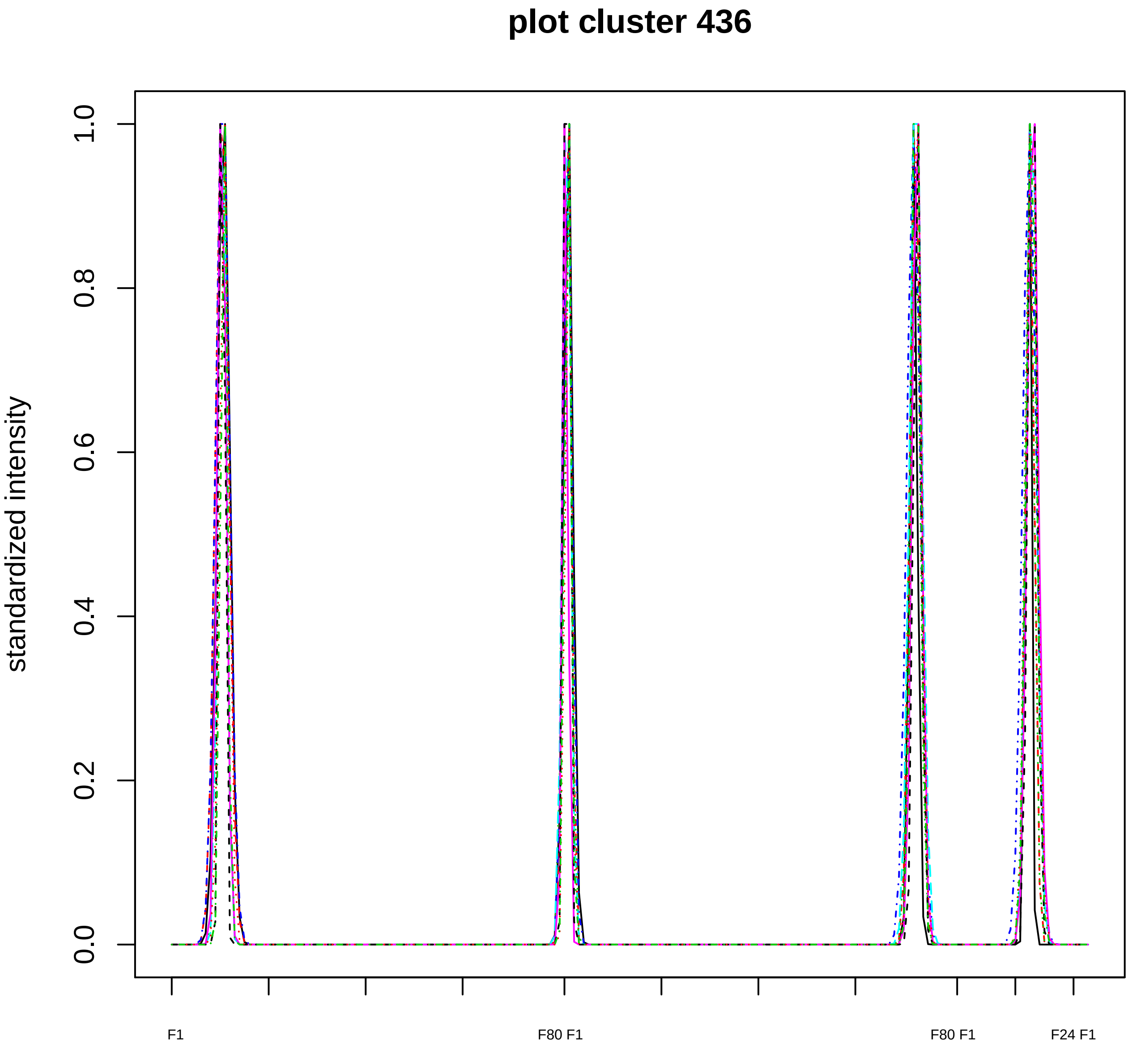

plot cluster 436
standardized intensity
0.0
0.2
0.4
0.6
0.8
1.0
F1
F80 F1
F80 F1
F24 F1

### Dendrogram of cluster 436

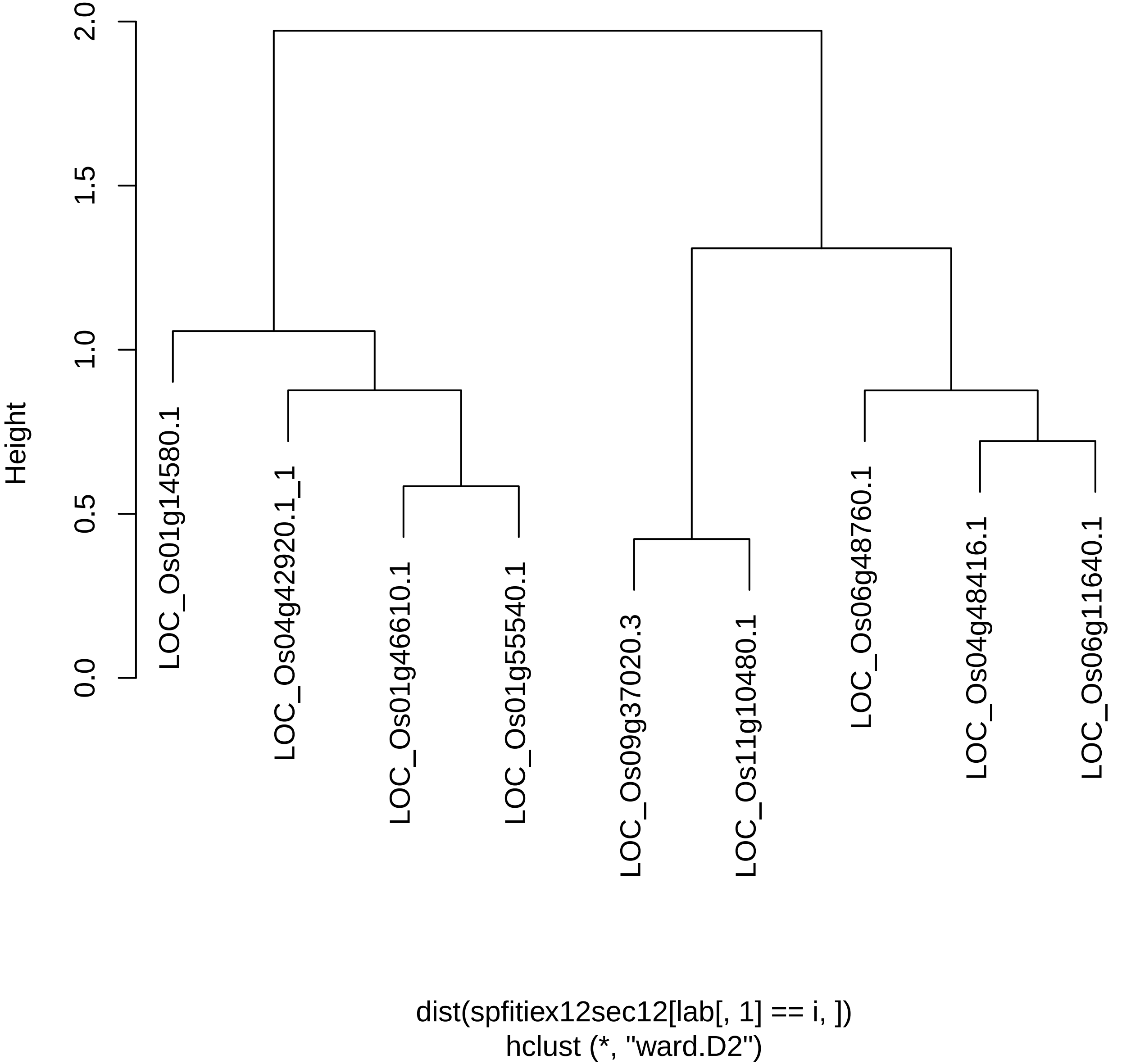

### plot cluster 437

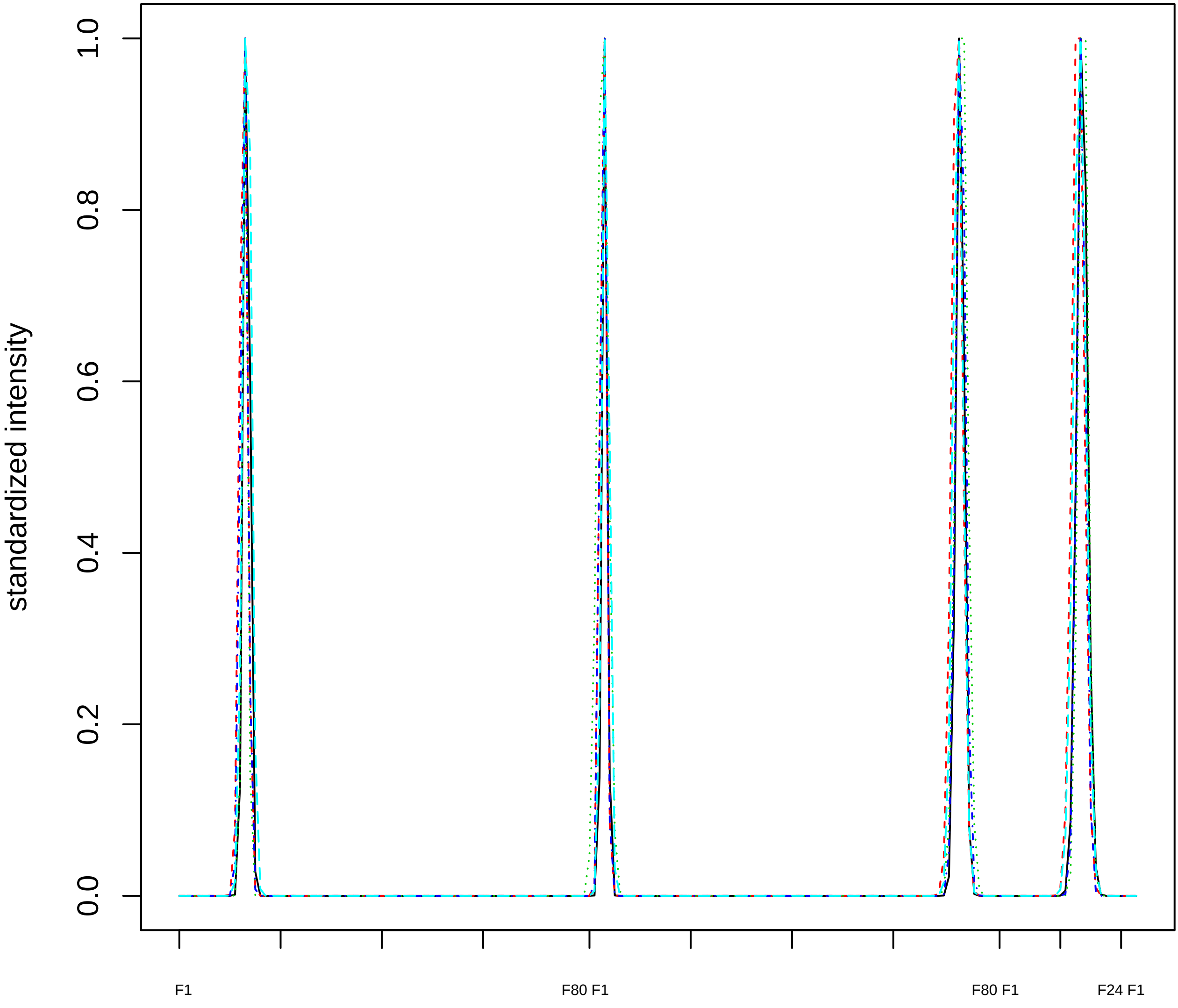

#### Dendrogram of cluster 437

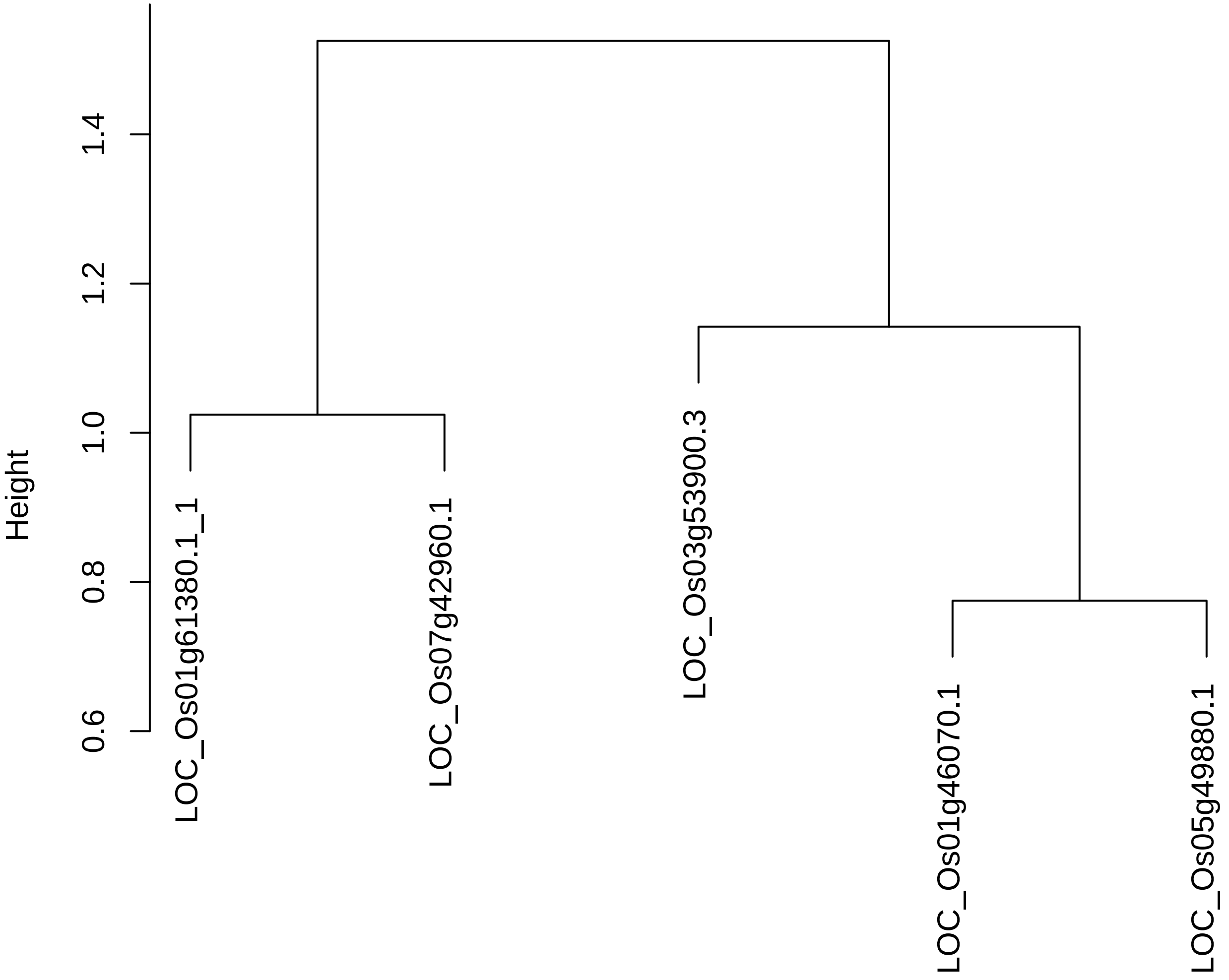


dist(spfitiex12sec12[lab[, 1] == i, ])
hclust (*, "ward.D2")

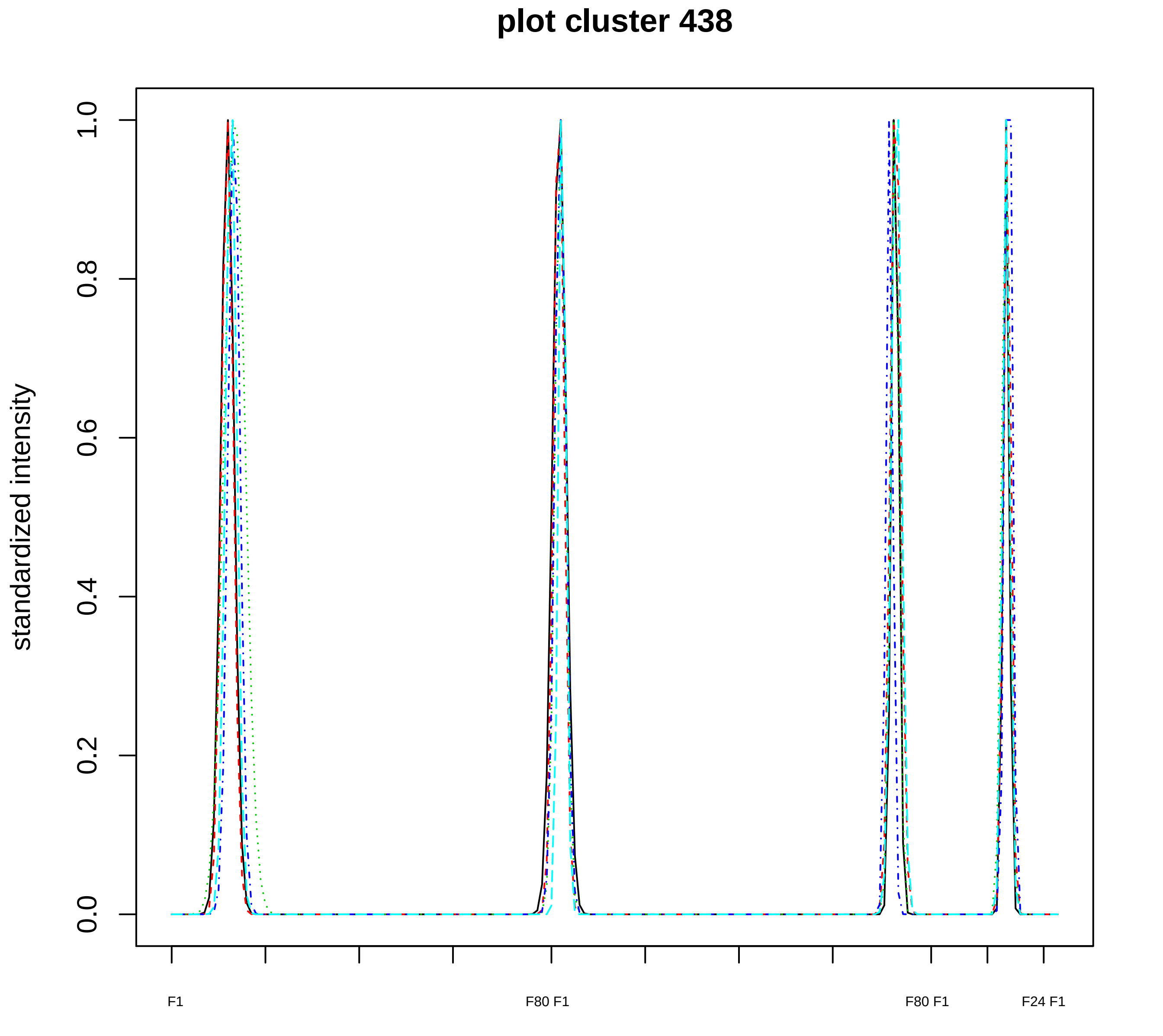
plot cluster 438
standardized intensity
0.0
0.2
0.4
0.6
0.8
1.0
F1
F80 F1
F80 F1
F24 F1

### Dendrogram of cluster 438

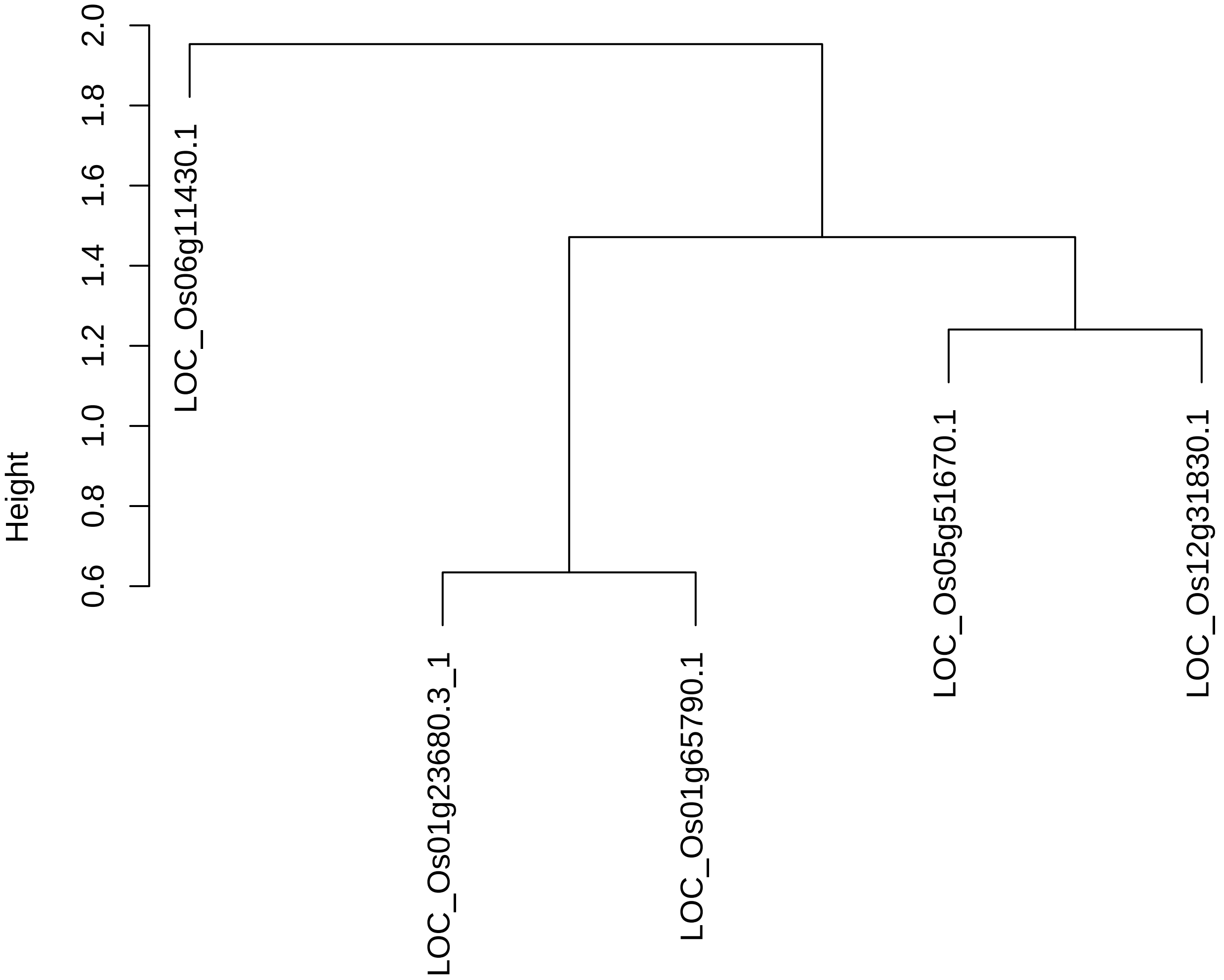


dist(spfitiex12sec12[lab[, 1] == i, ])
hclust (*, "ward.D2")

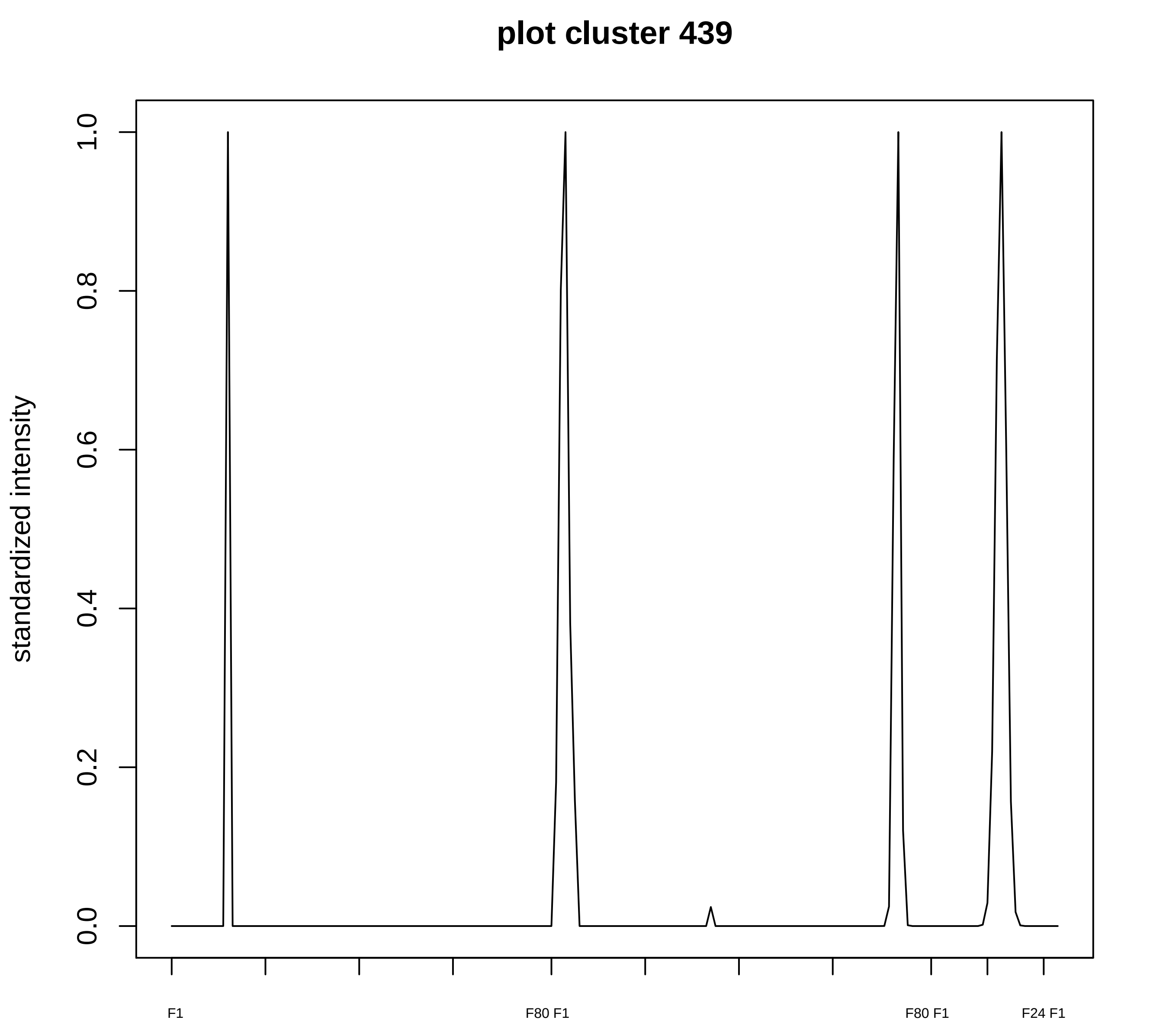

plot cluster 439
standardized intensity
0.0
0.2
0.4
0.6
0.8
1.0
F1
F80 F1
F80 F1
F24 F1

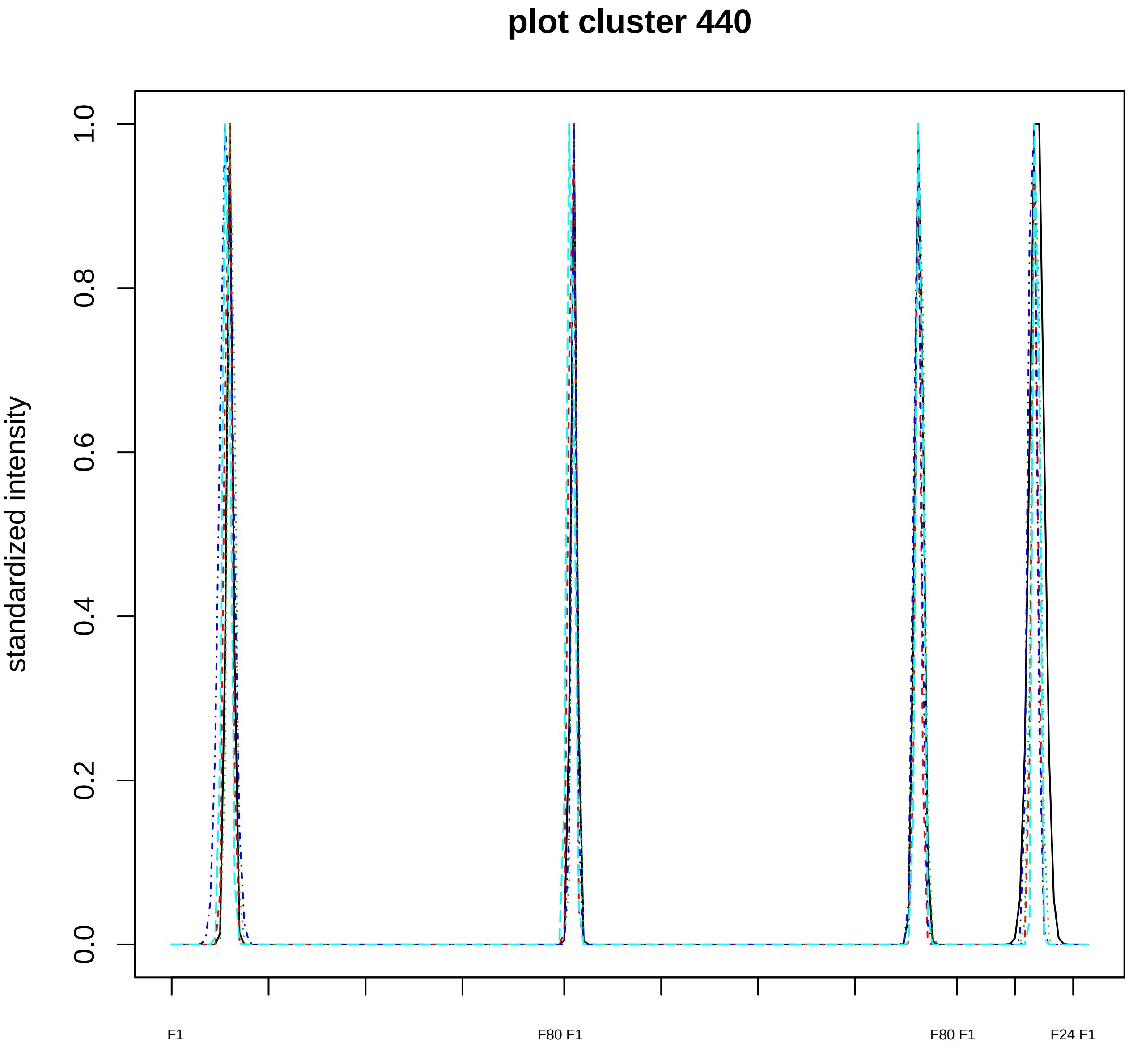
plot cluster 440
standardized intensity
0.0
0.2
0.4
0.6
0.8
1.0
F1
F80 F1
F80 F1
F24 F1

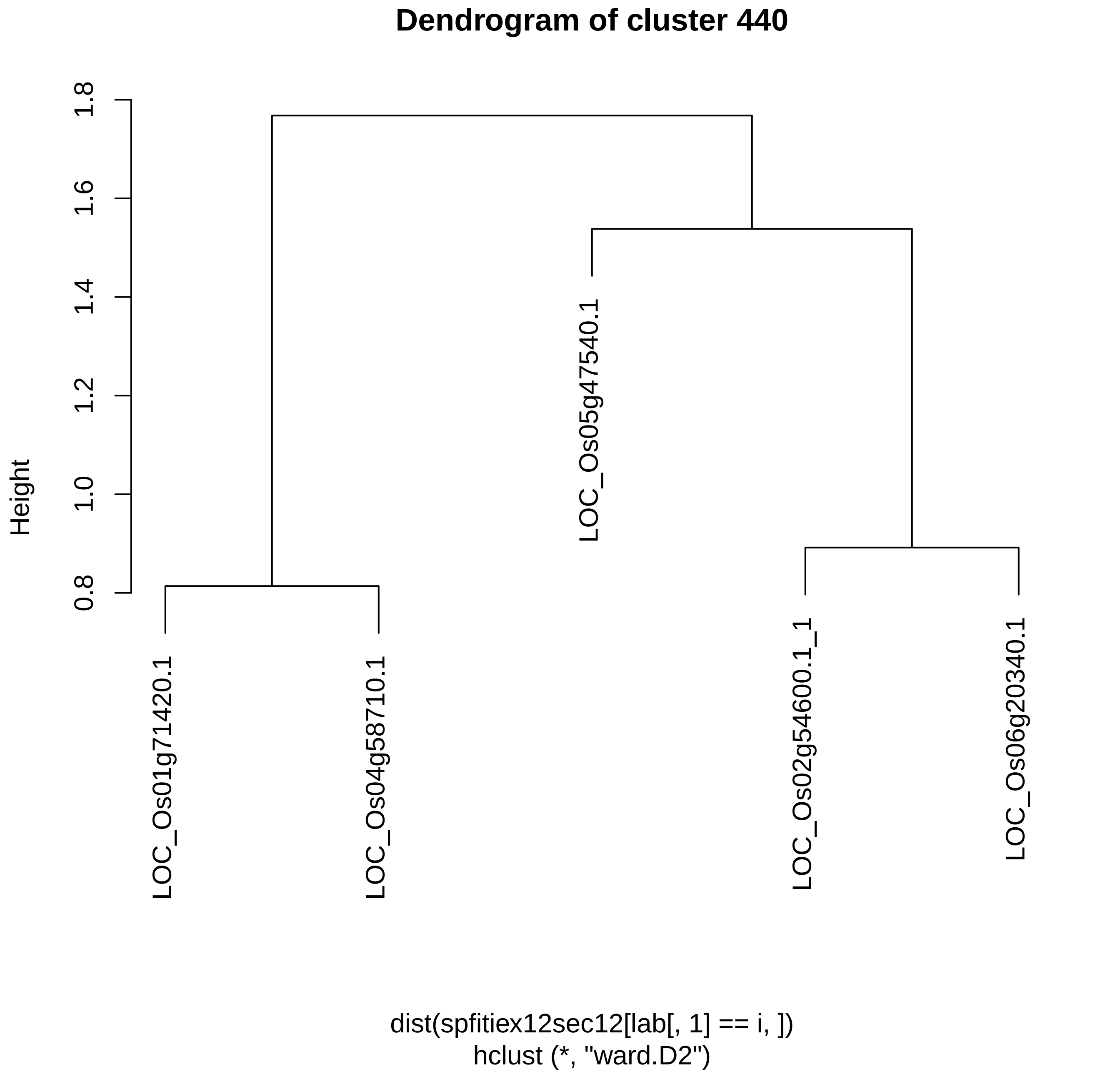
Dendrogram of cluster 440
Height
0.8
1.0
1.2
1.4
1.6
1.8
LOC_Os01g71420.1
LOC_Os04g58710.1
LOC_Os05g47540.1
LOC_Os02g54600.1_1
LOC_Os06g20340.1
dist(spfitiex12sec12[lab[, 1] == i, ])
hclust (*, "ward.D2")

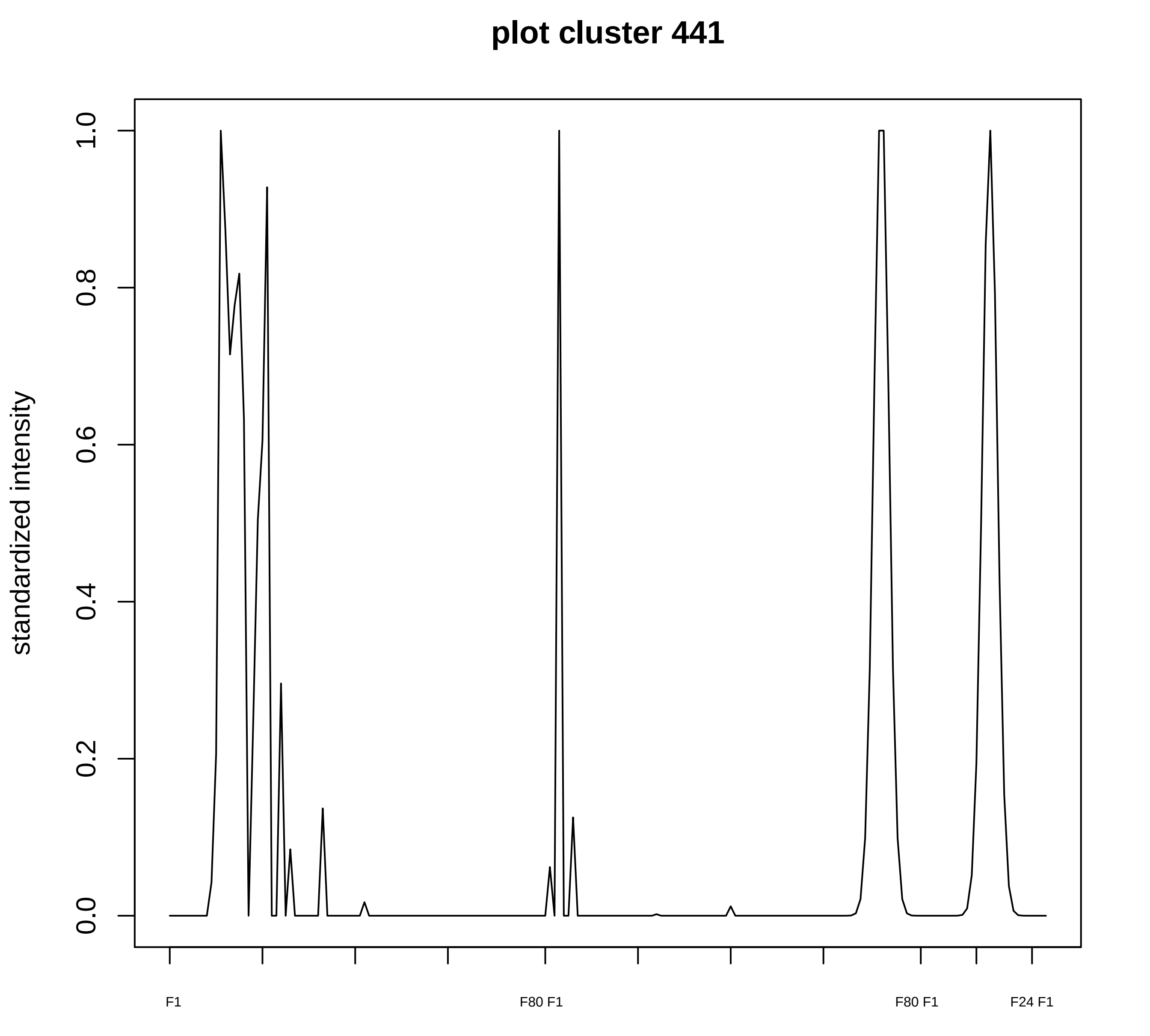
plot cluster 441
standardized intensity
0.0
0.2
0.4
0.6
0.8
1.0
F1
F80 F1
F80 F1
F24 F1

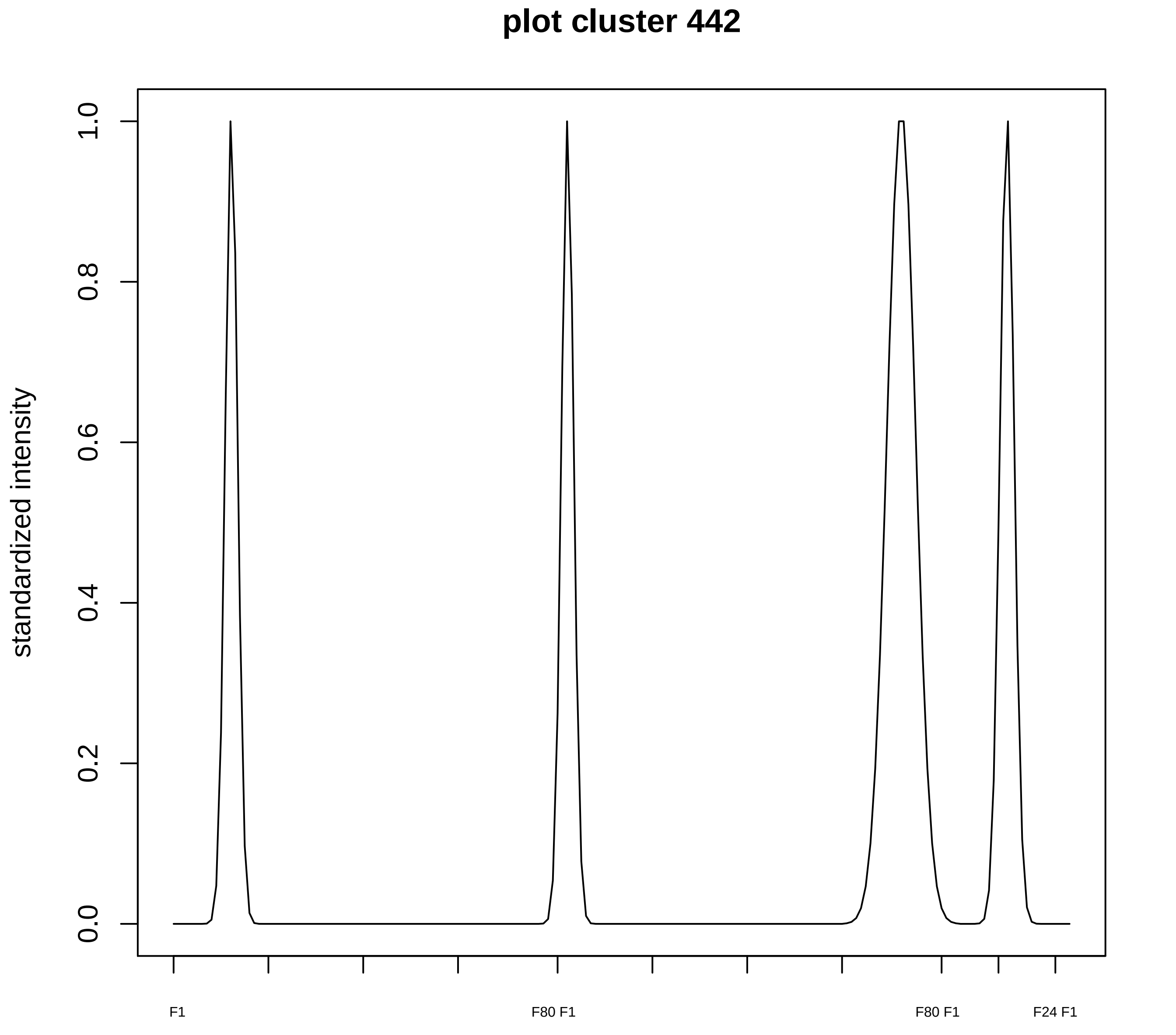
plot cluster 442
standardized intensity
0.0
0.2
0.4
0.6
0.8
1.0
F1
F80 F1
F80 F1
F24 F1

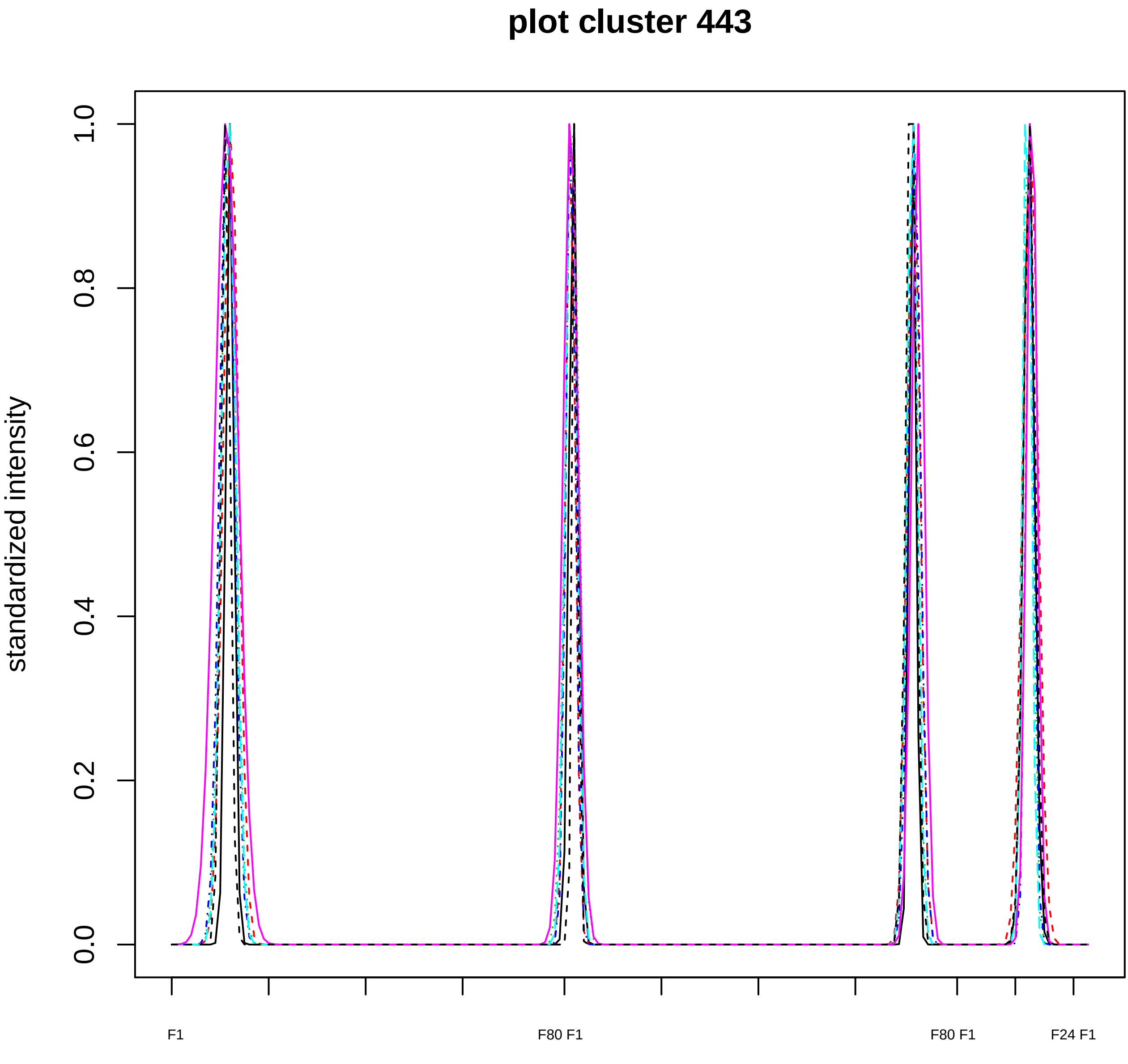
plot cluster 443
standardized intensity
0.0
0.2
0.4
0.6
0.8
1.0
F1
F80 F1
F80 F1
F24 F1

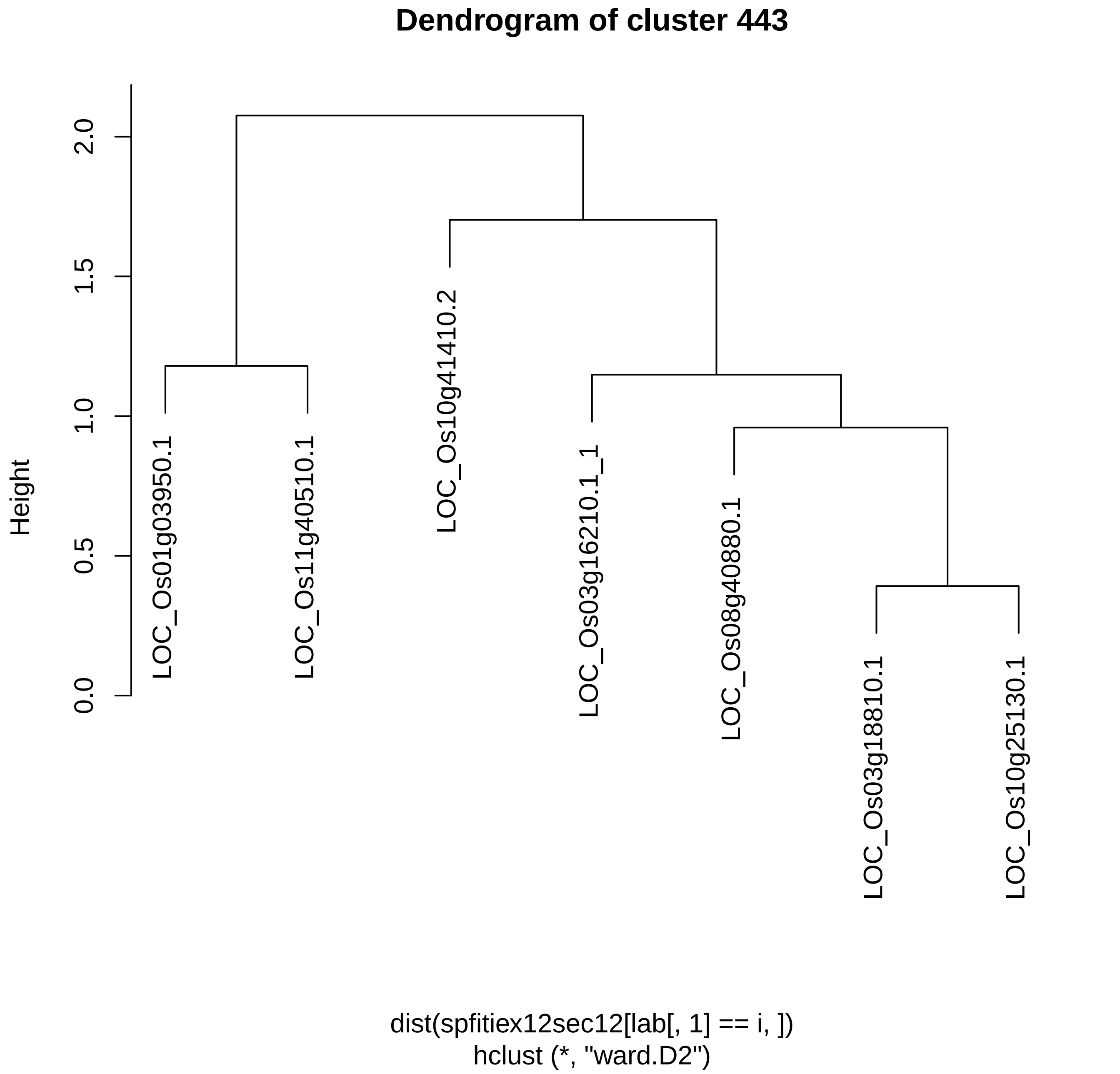
Dendrogram of cluster 443
Height
0.0
0.5
1.0
1.5
2.0
LOC_Os01g03950.1
LOC_Os11g40510.1
LOC_Os10g41410.2
LOC_Os03g16210.1_1
LOC_Os08g40880.1
LOC_Os03g18810.1
LOC_Os10g25130.1
dist(spfitiex12sec12[lab[, 1] == i, ])
hclust (*, "ward.D2")

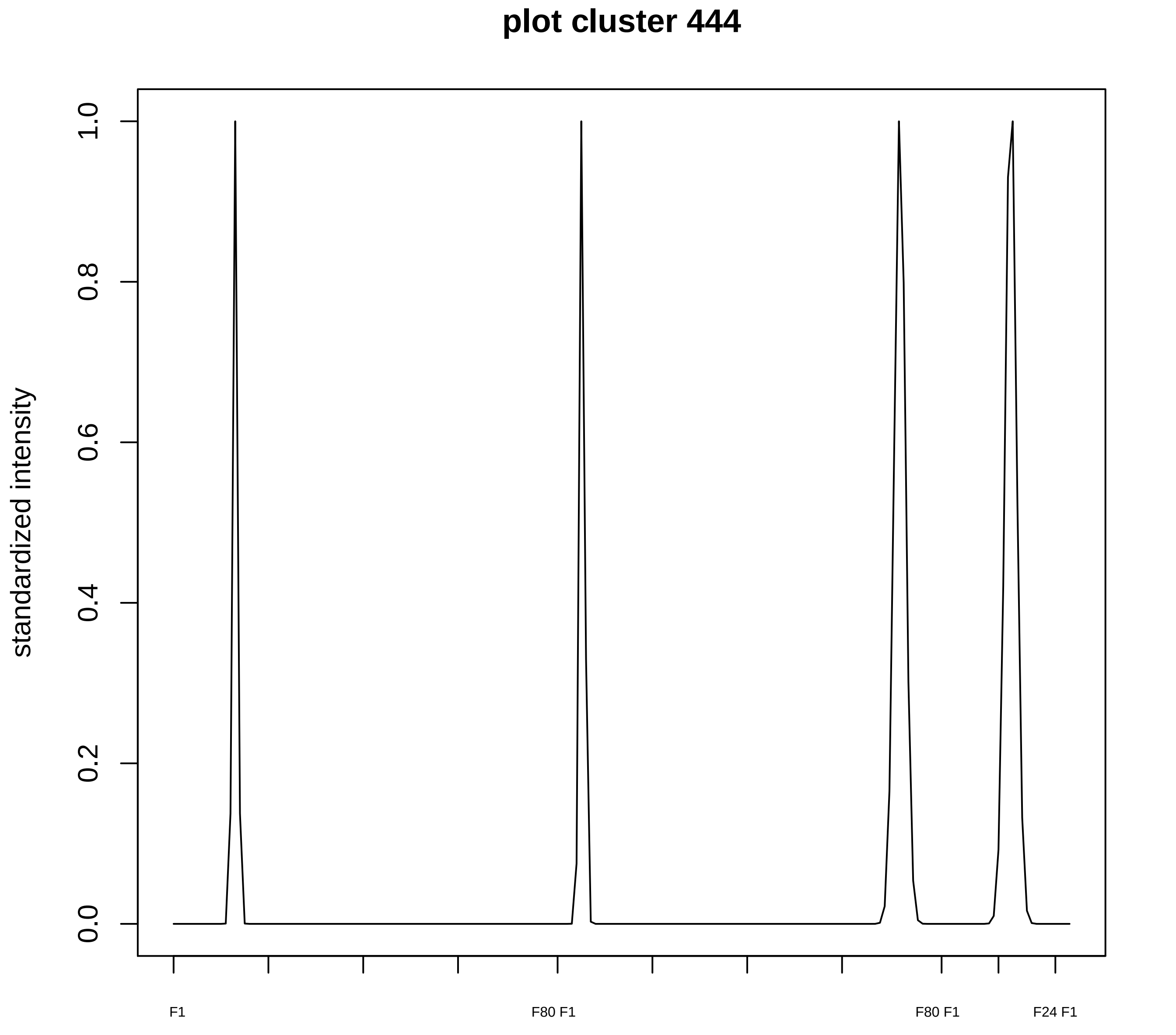
plot cluster 444
standardized intensity
0.0
0.2
0.4
0.6
0.8
1.0
F1
F80 F1
F80 F1
F24 F1

### plot cluster 445

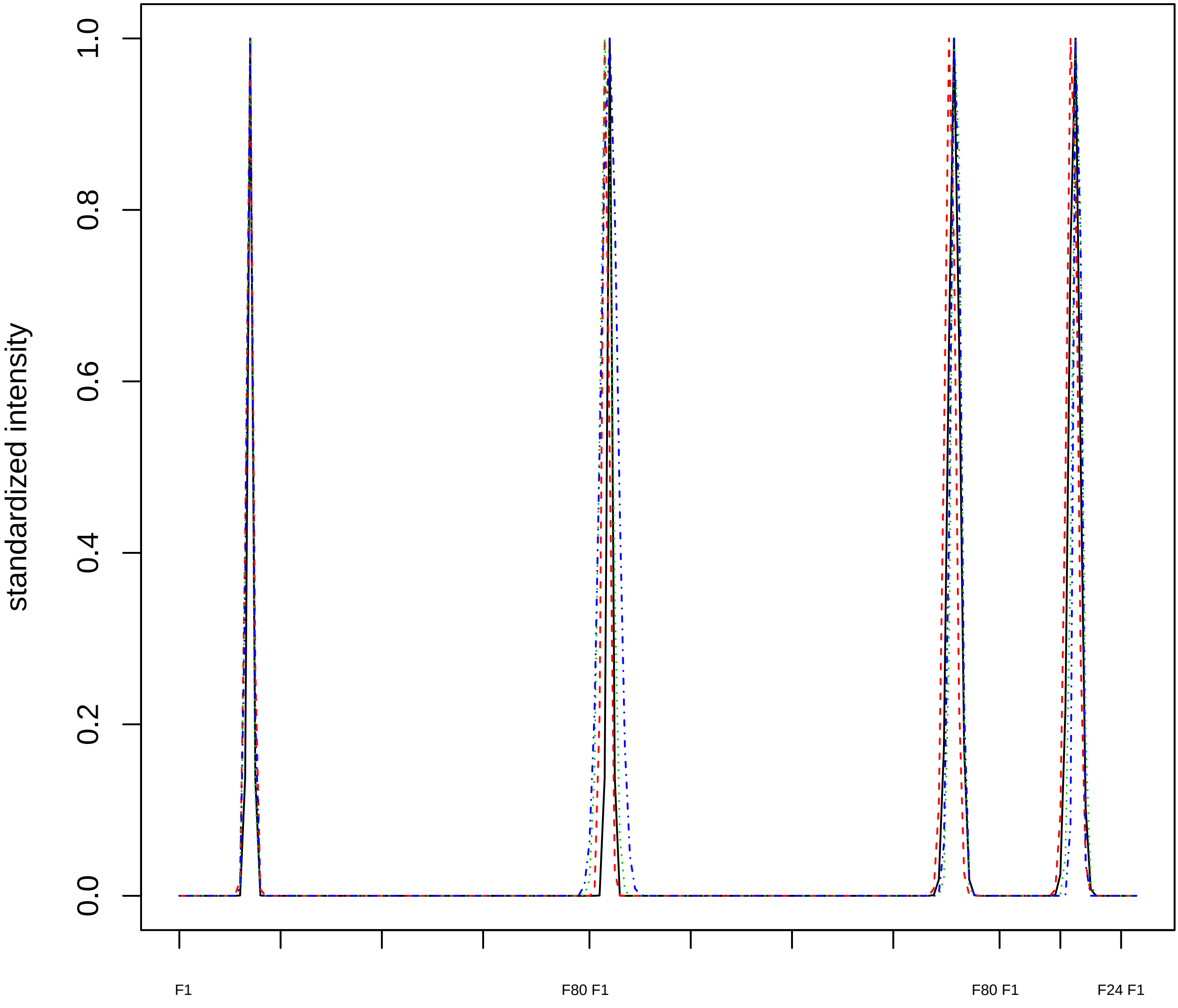

### Dendrogram of cluster 445

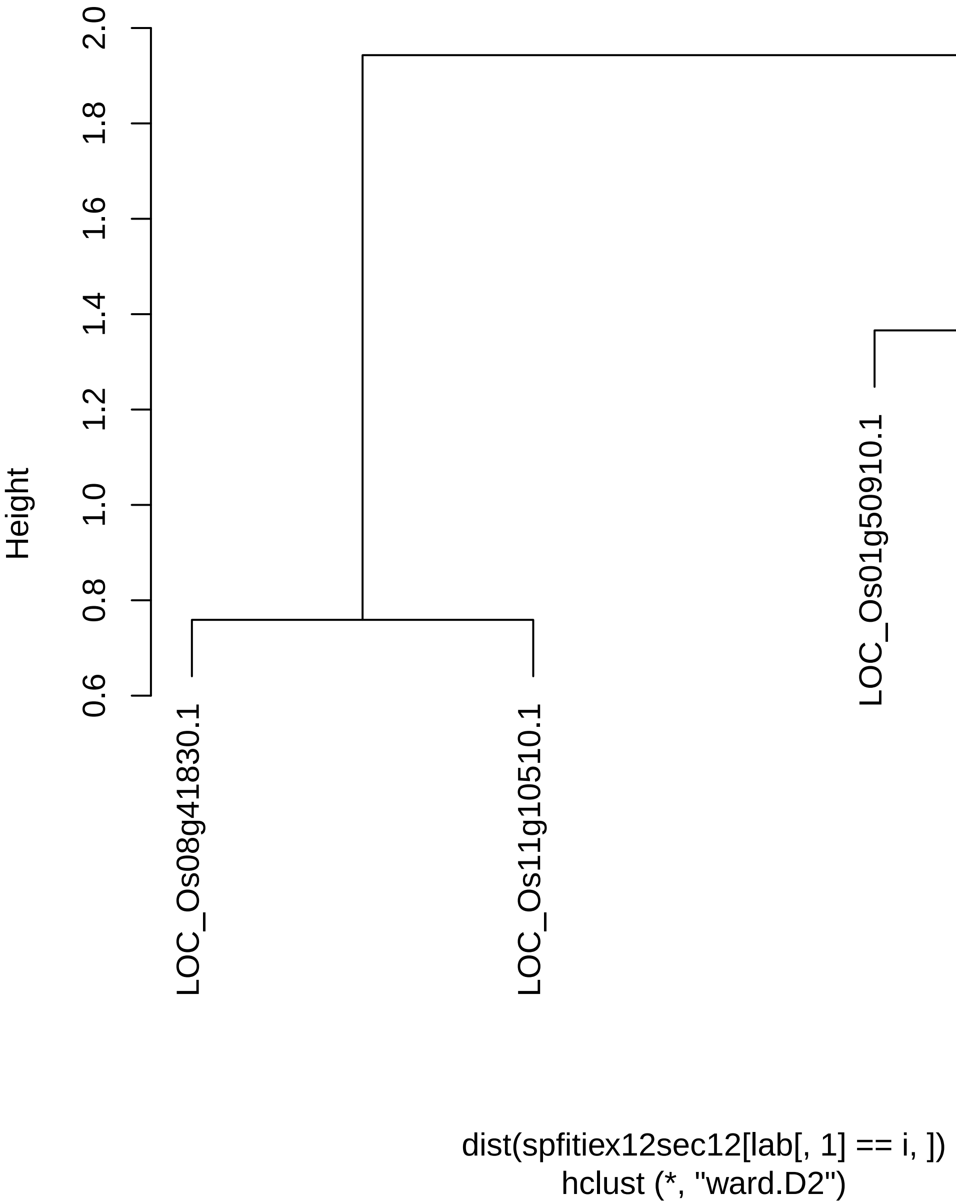

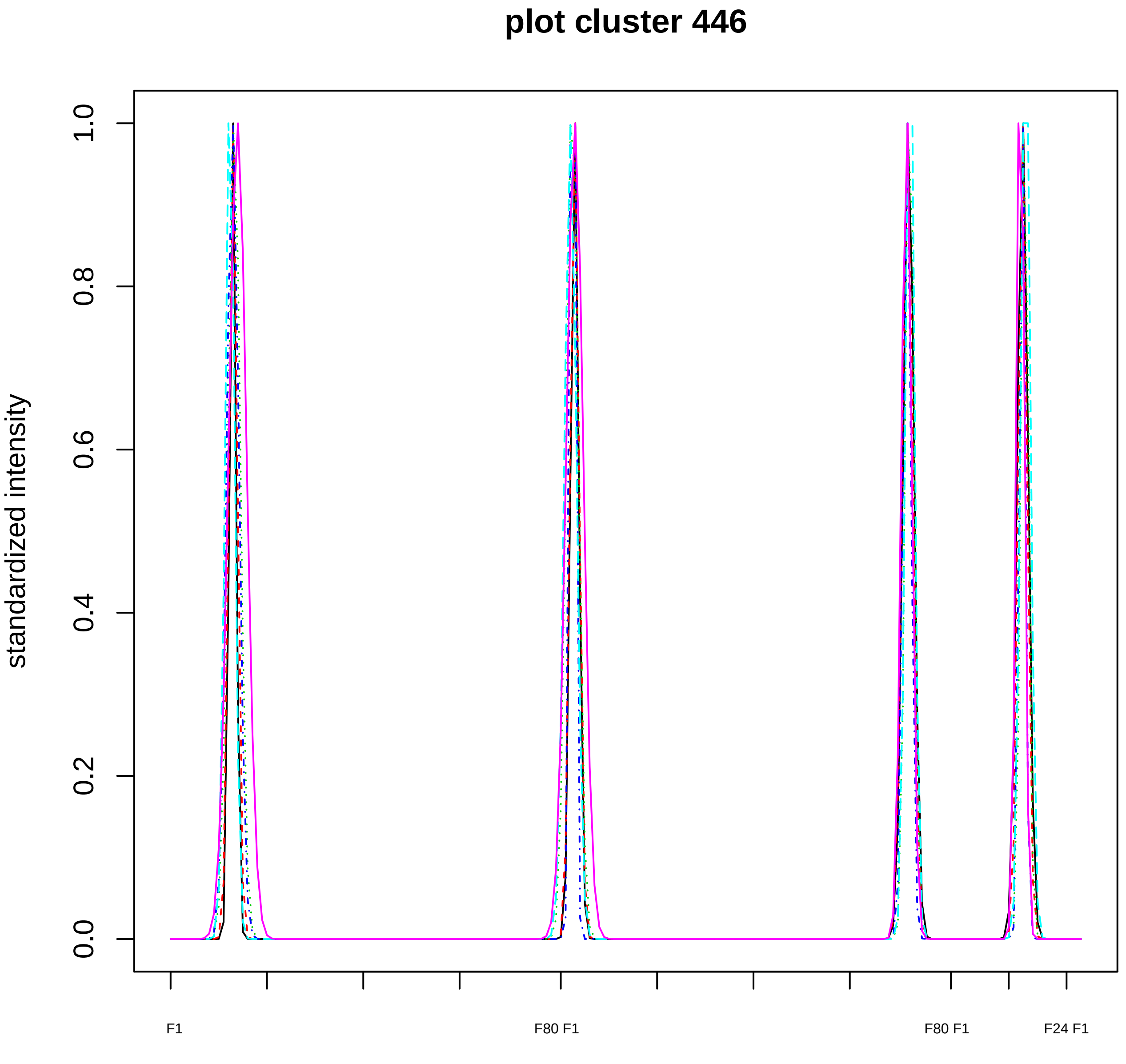
plot cluster 446
standardized intensity
0.0
0.2
0.4
0.6
0.8
1.0
F1
F80 F1
F80 F1
F24 F1

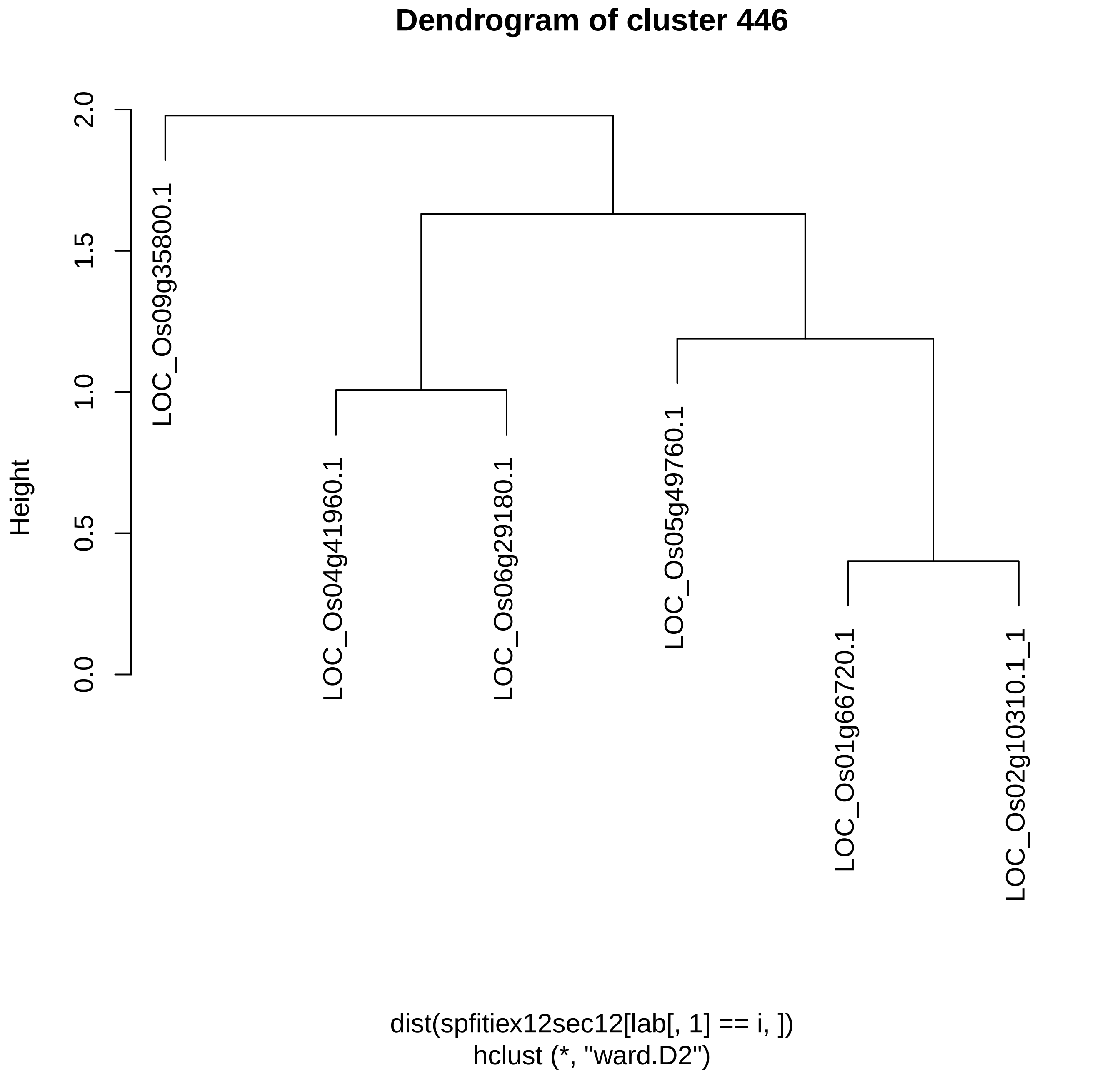
Dendrogram of cluster 446
Height
2.0
1.5
1.0
0.5
0.0
LOC_Os09g35800.1
LOC_Os04g41960.1
LOC_Os06g29180.1
LOC_Os05g49760.1
LOC_Os01g66720.1
LOC_Os02g10310.1_1
dist(spfitiex12sec12[lab[, 1] == i, ])
hclust (*, "ward.D2")

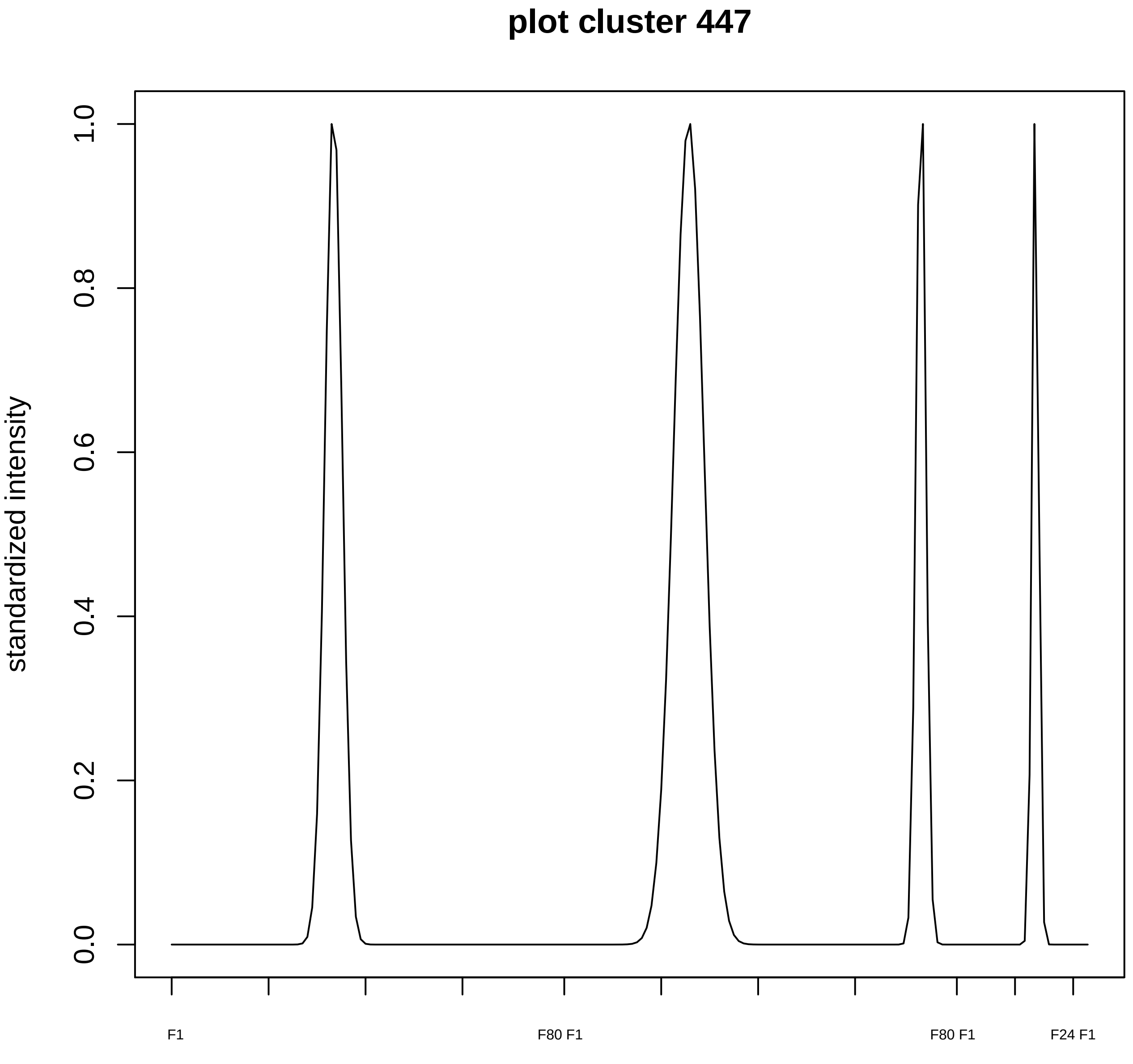
plot cluster 447
standardized intensity
0.0
0.2
0.4
0.6
0.8
1.0
F1
F80 F1
F80 F1
F24 F1

### plot cluster 448

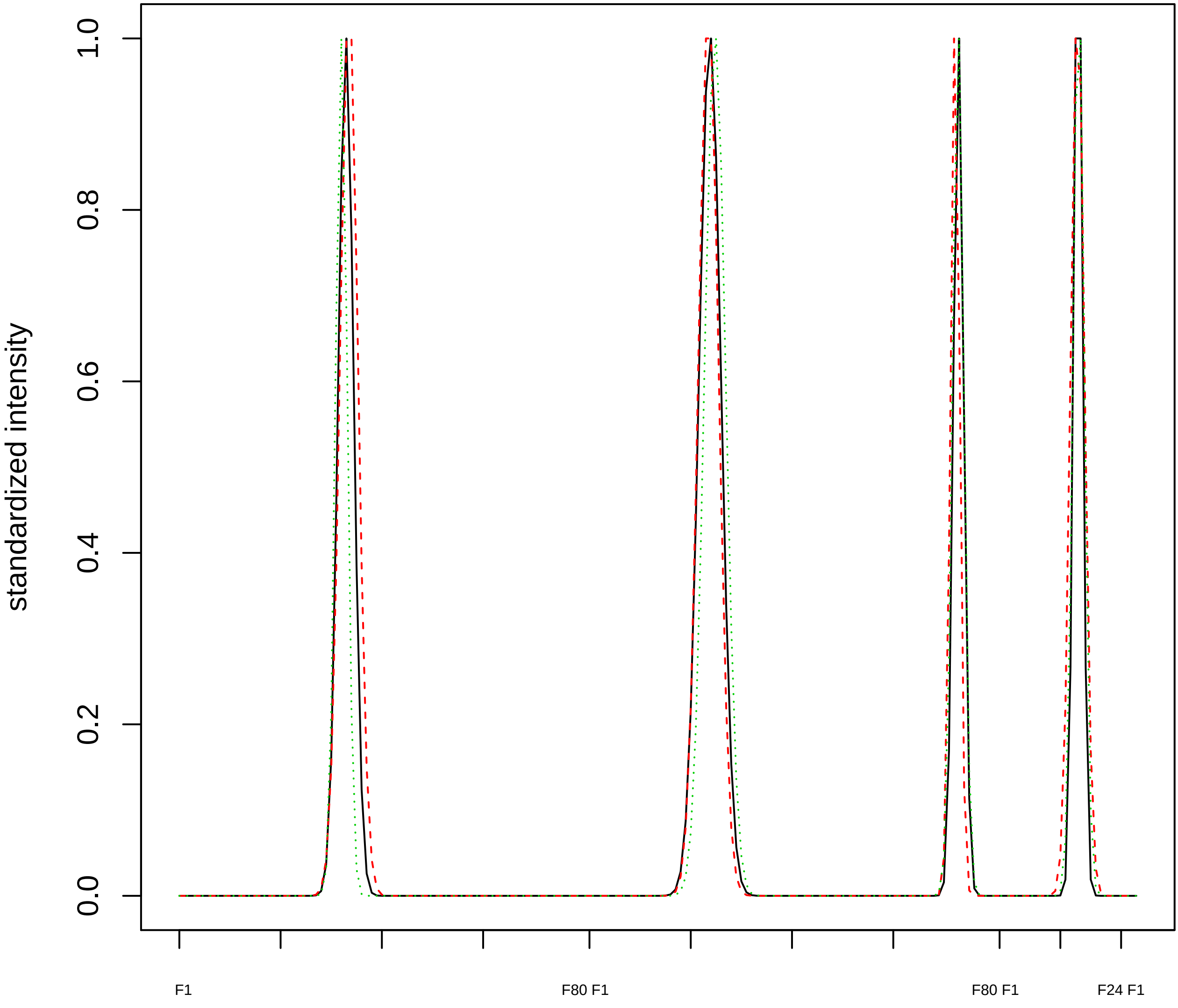

### Dendrogram of cluster 448

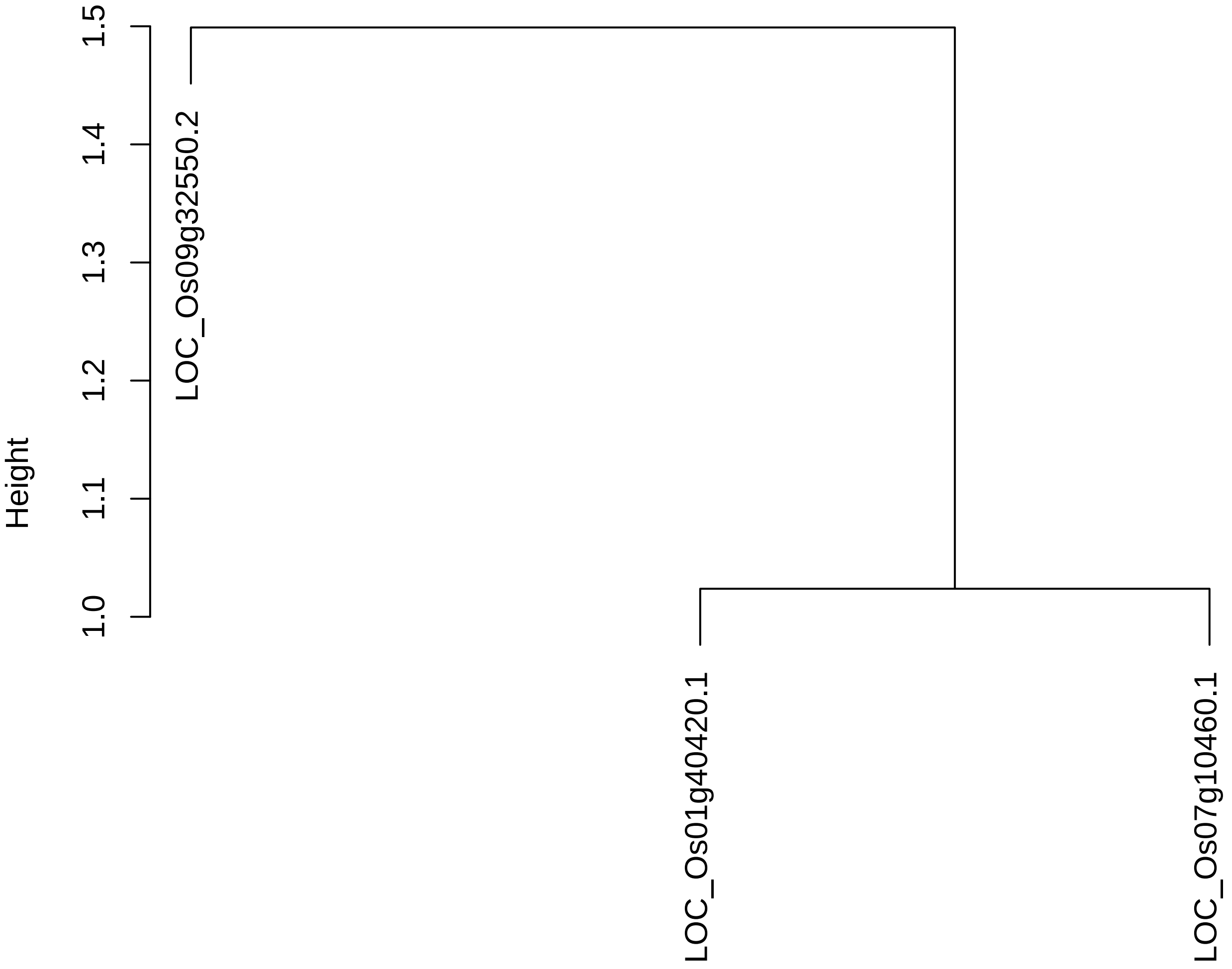


dist(spfitiex12sec12[lab[, 1] == i, ])
hclust (*, "ward.D2")

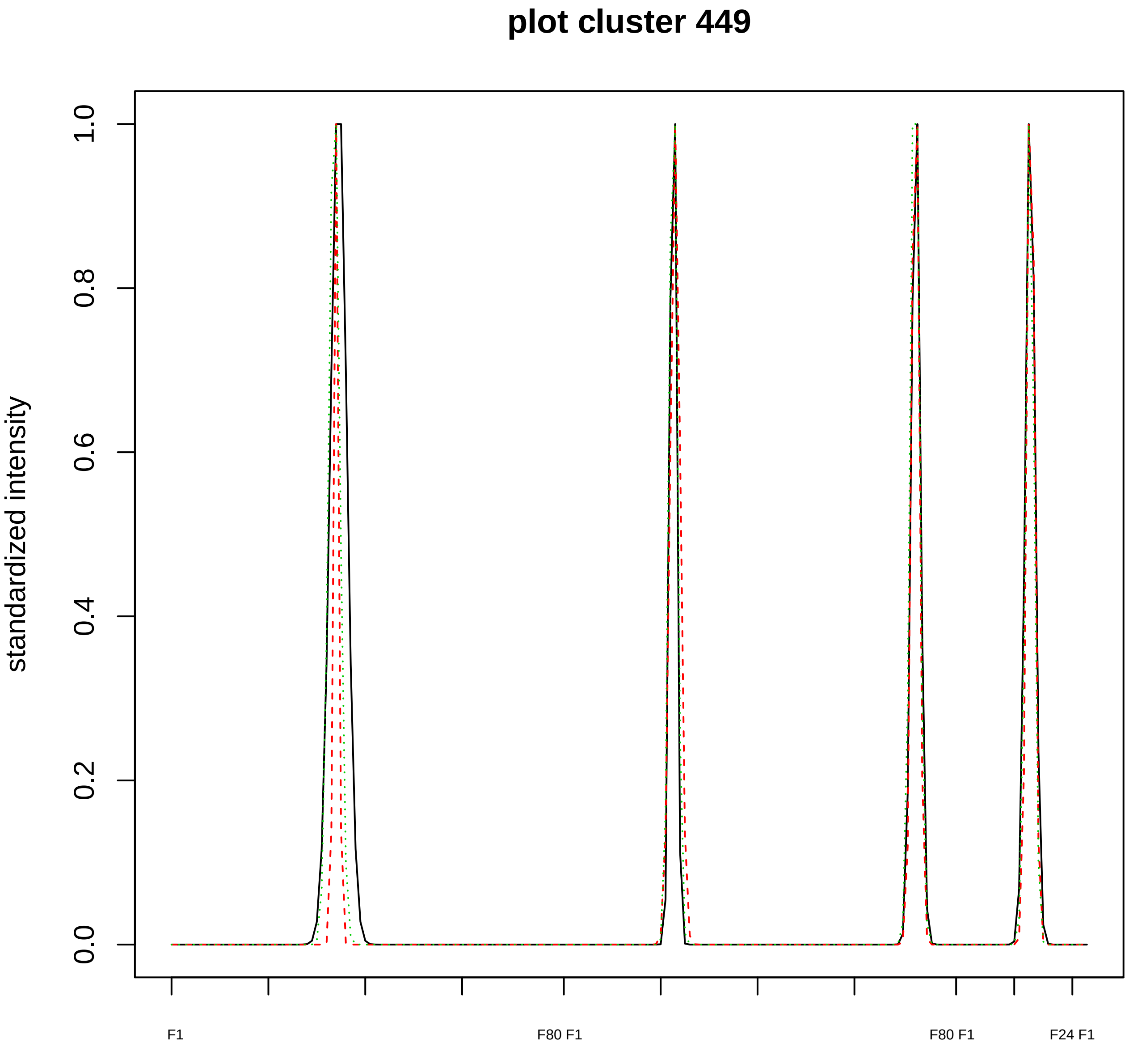
plot cluster 449
standardized intensity
0.0
0.2
0.4
0.6
0.8
1.0
F1
F80 F1
F80 F1
F24 F1

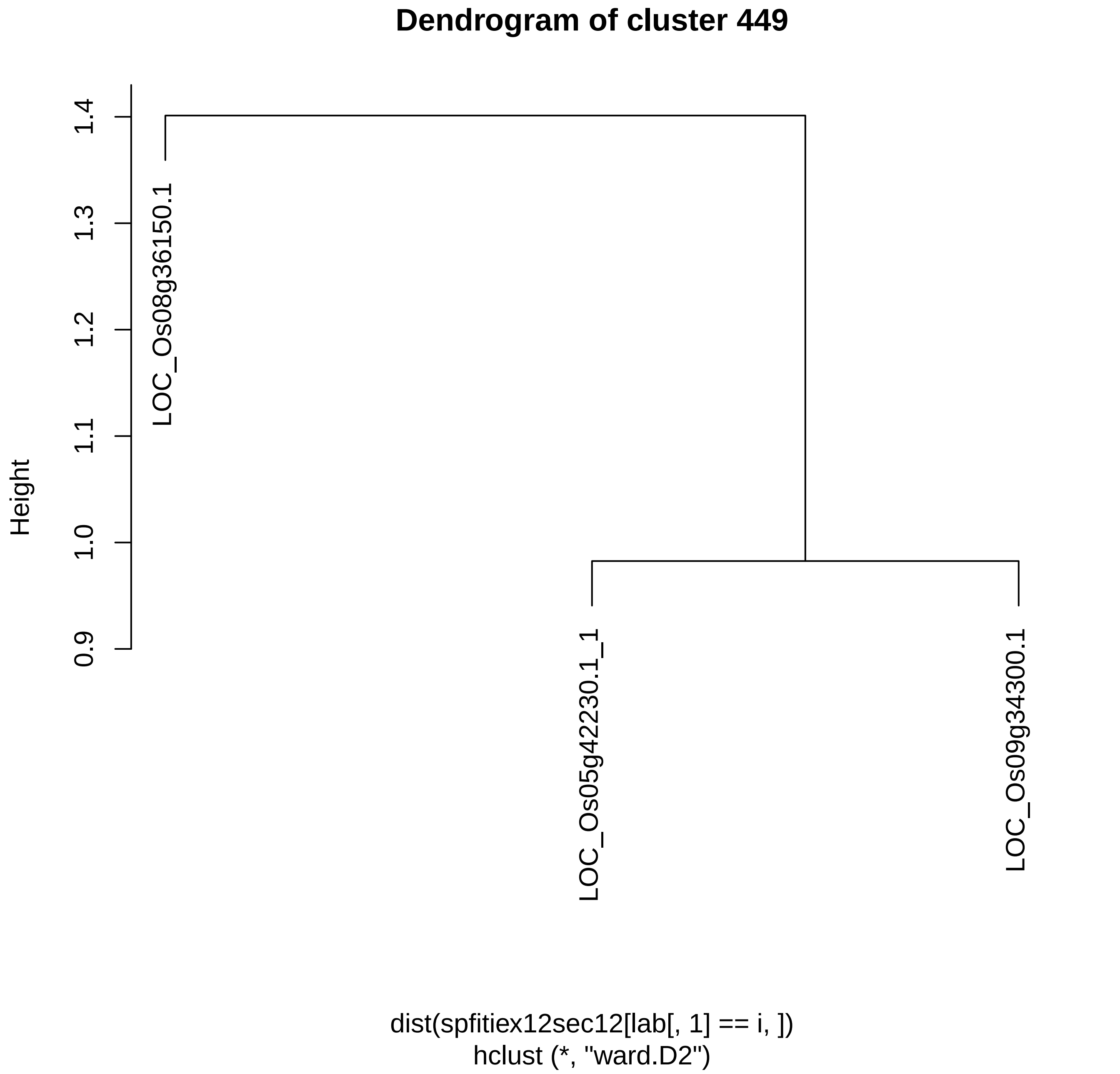

Dendrogram of cluster 449
Height
0.9
1.0
1.1
1.2
1.3
1.4
LOC_Os08g36150.1
LOC_Os05g42230.1_1
LOC_Os09g34300.1
dist(spfitiex12sec12[lab[, 1] == i, ])
hclust (*, "ward.D2")

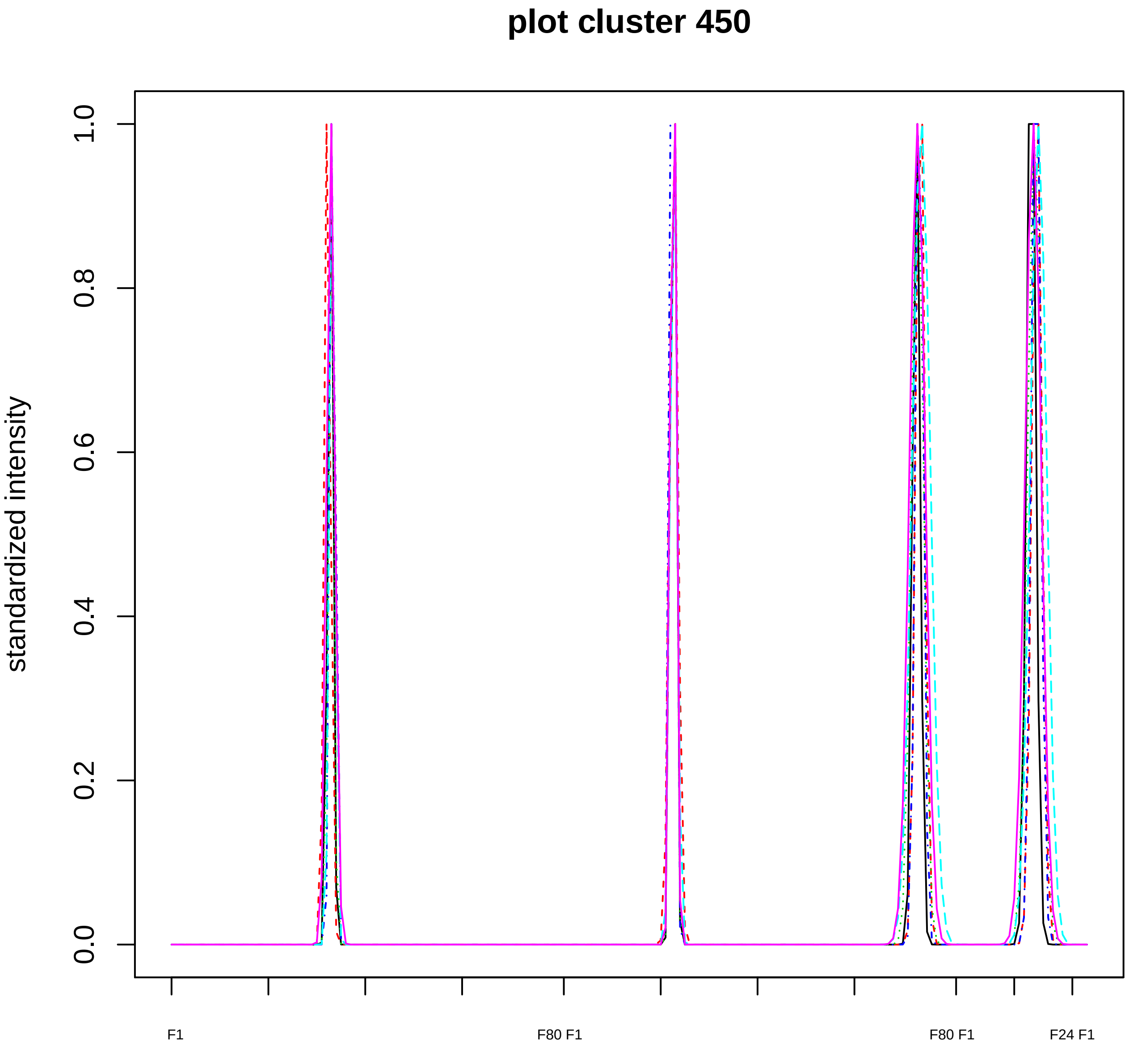

plot cluster 450
standardized intensity
0.0
0.2
0.4
0.6
0.8
1.0
F1
F80 F1
F80 F1
F24 F1

### Dendrogram of cluster 450

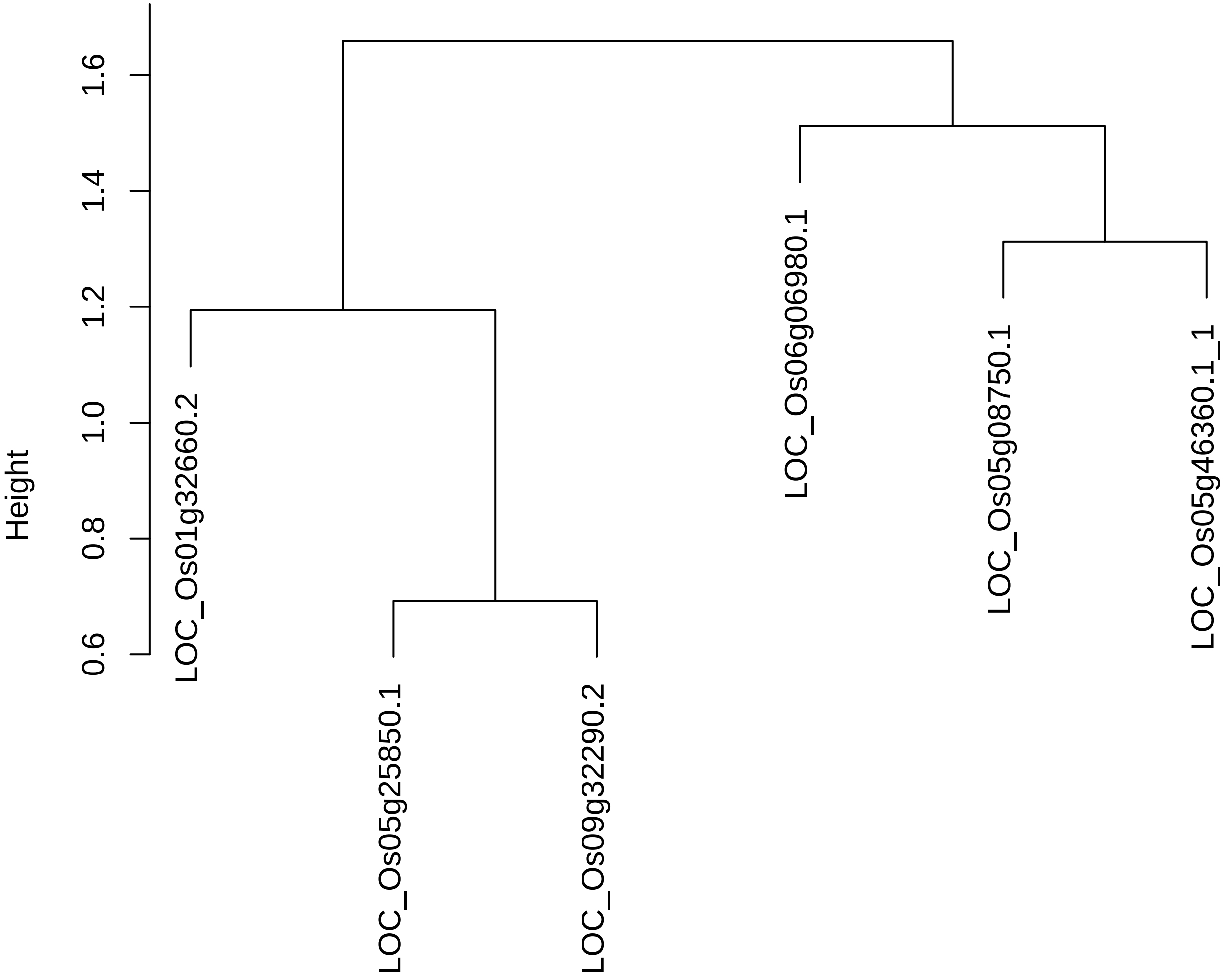


dist(spfitiex12sec12[lab[, 1] == i, ])
hclust (*, "ward.D2")

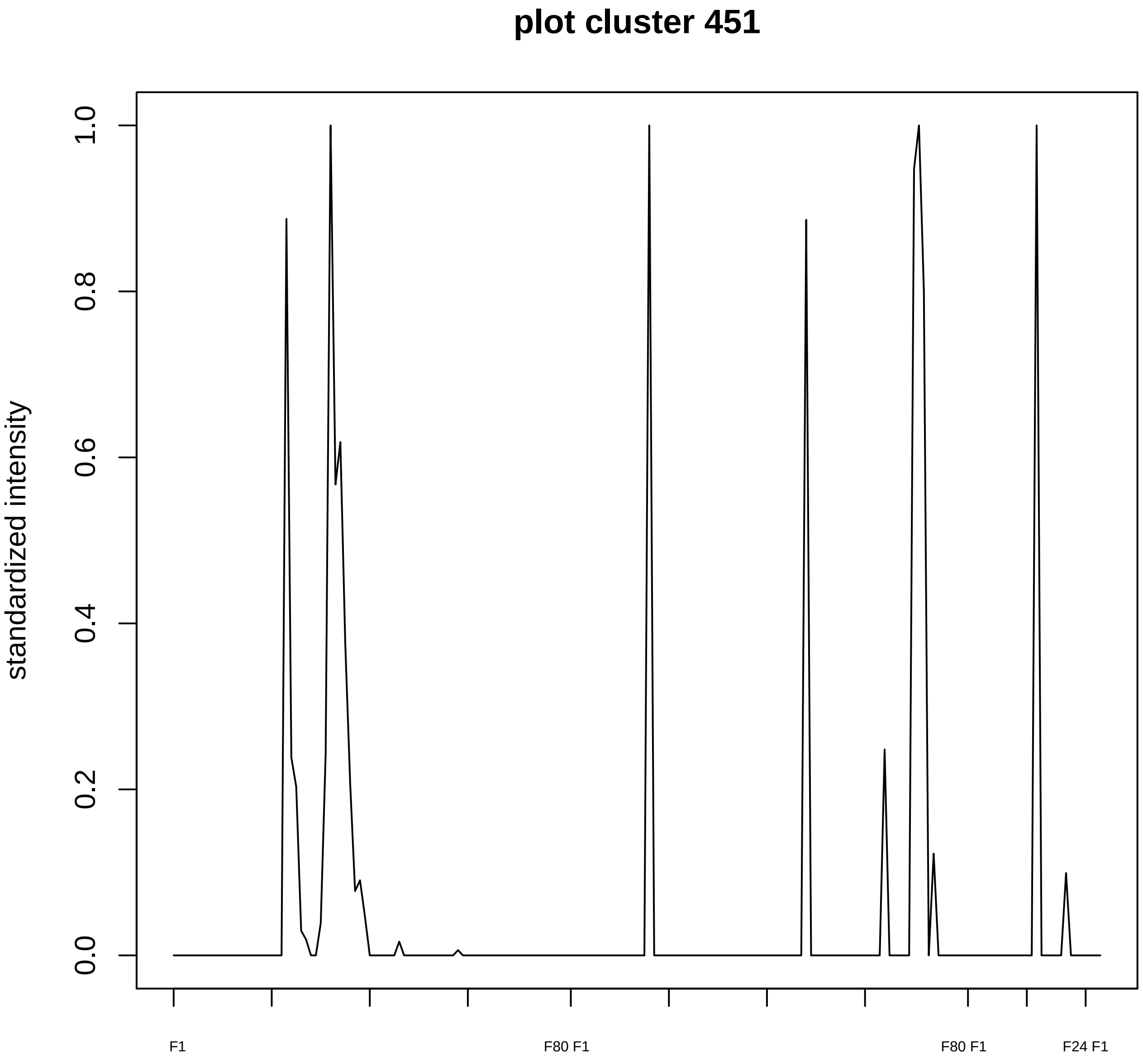

plot cluster 451
standardized intensity
0.0
0.2
0.4
0.6
0.8
1.0
F1
F80 F1
F80 F1
F24 F1

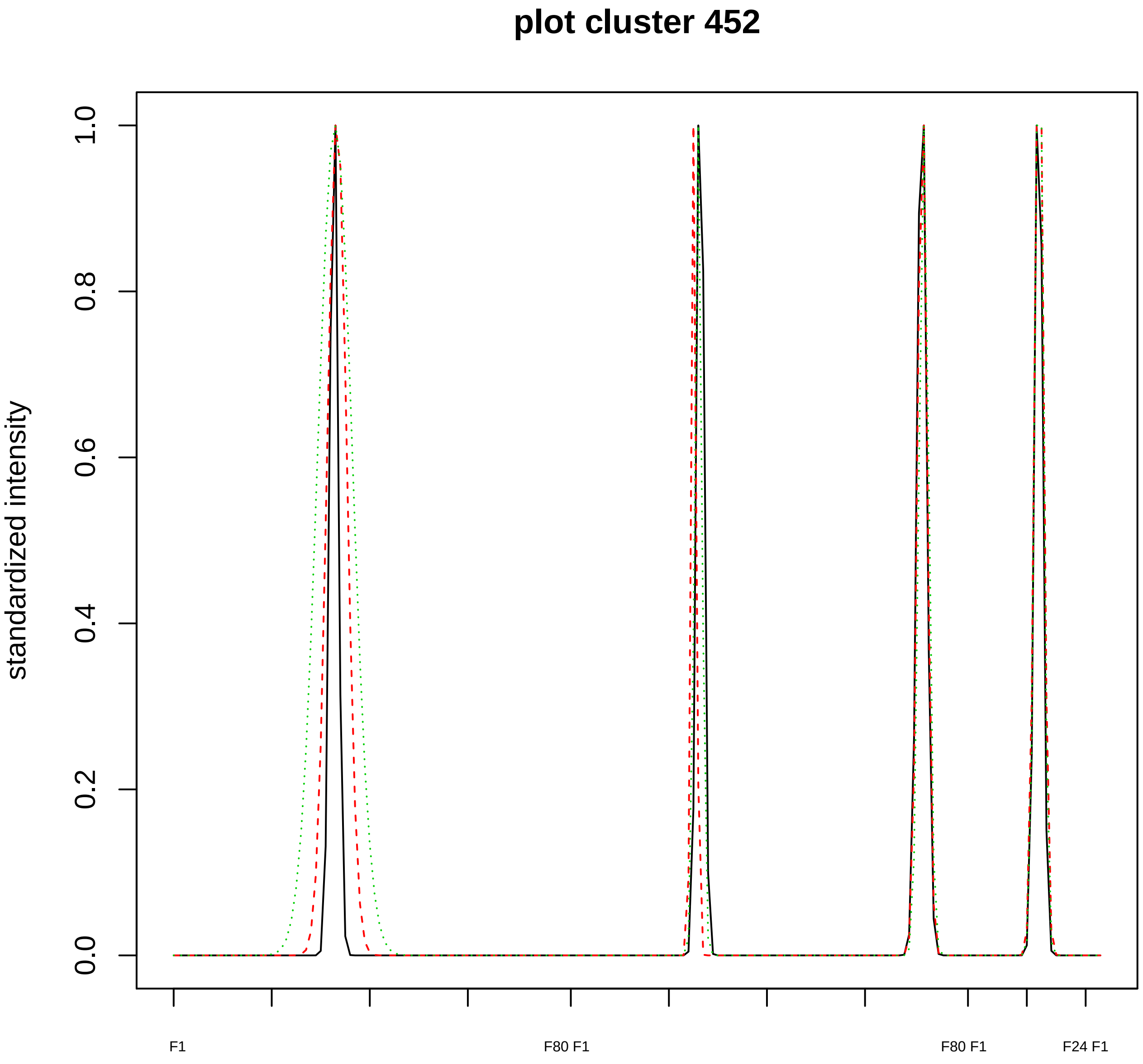
plot cluster 452
standardized intensity
0.0
0.2
0.4
0.6
0.8
1.0
F1
F80 F1
F80 F1
F24 F1

### Dendrogram of cluster 452

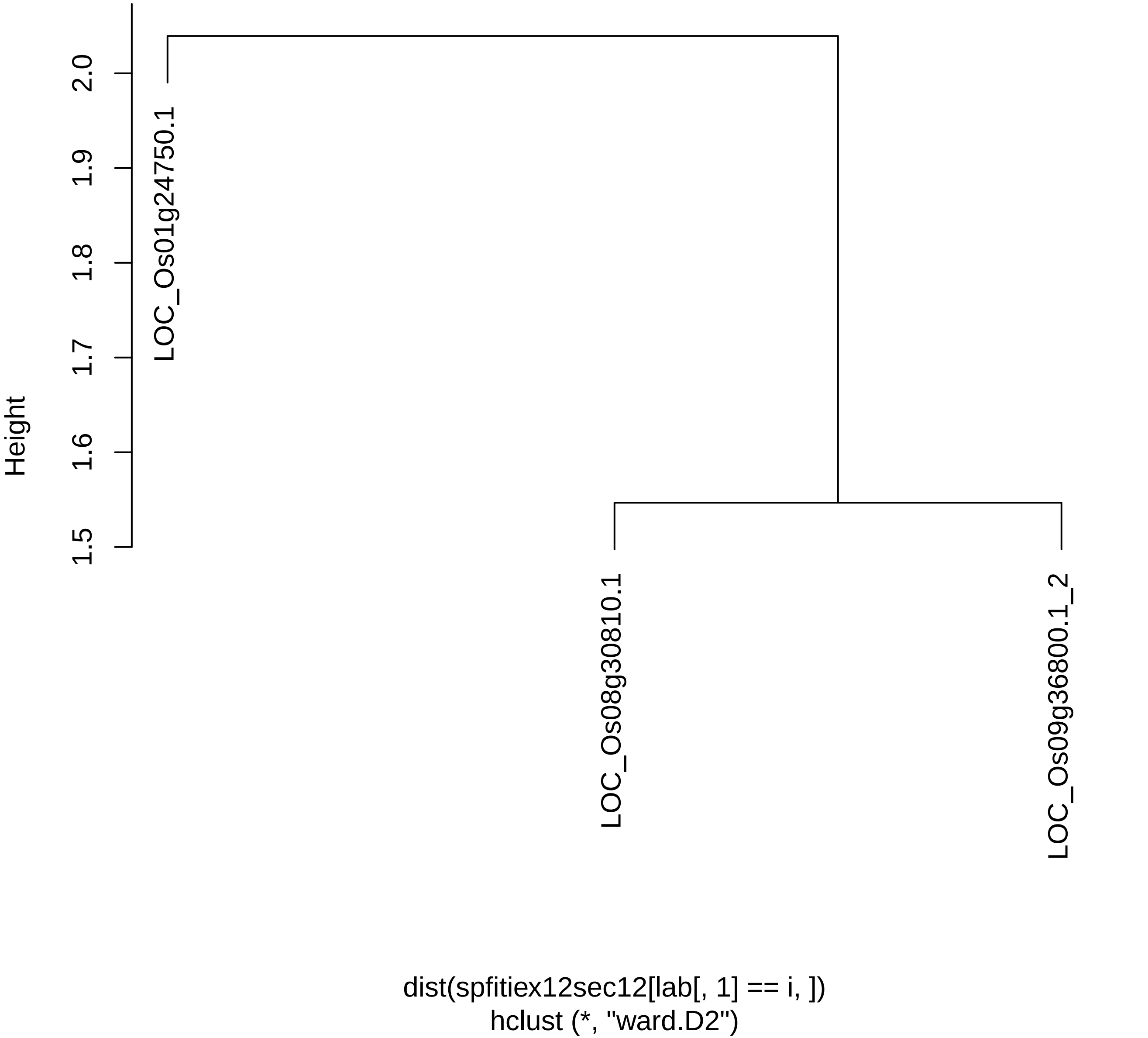

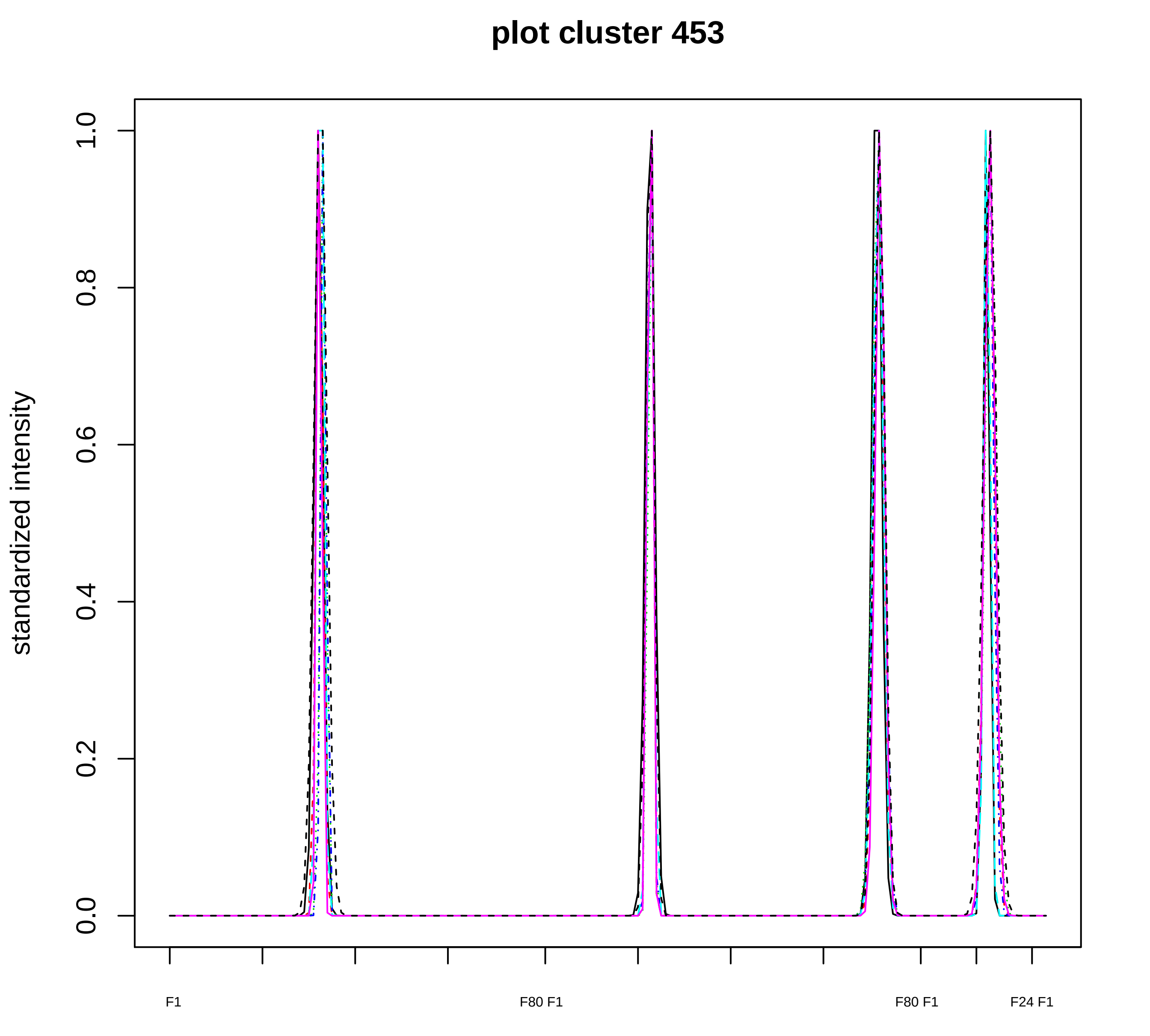

plot cluster 453
standardized intensity
0.0
0.2
0.4
0.6
0.8
1.0
F1
F80 F1
F80 F1
F24 F1

### Dendrogram of cluster 453

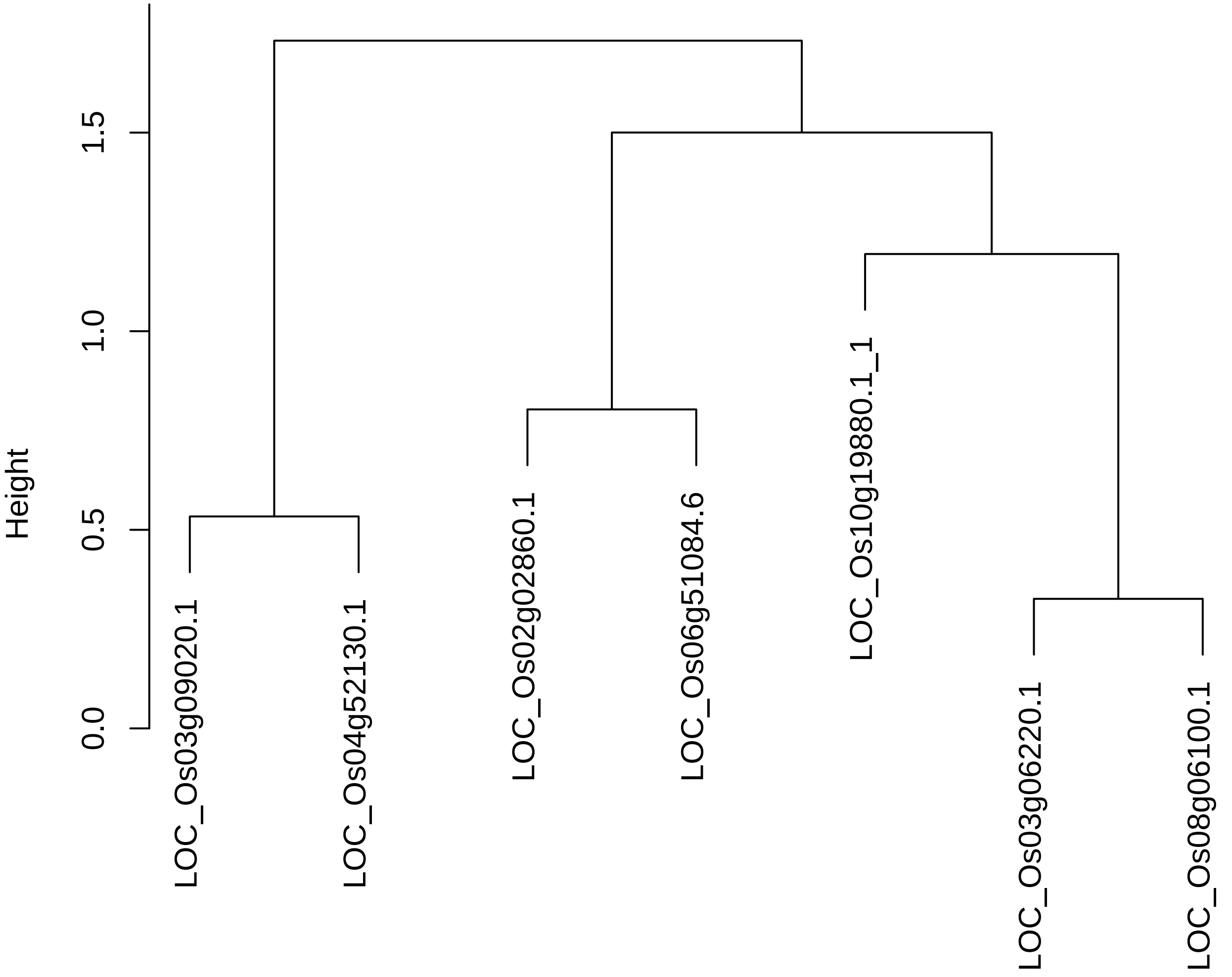


dist(spfitiex12sec12[lab[, 1] == i, ])
hclust (*, "ward.D2")

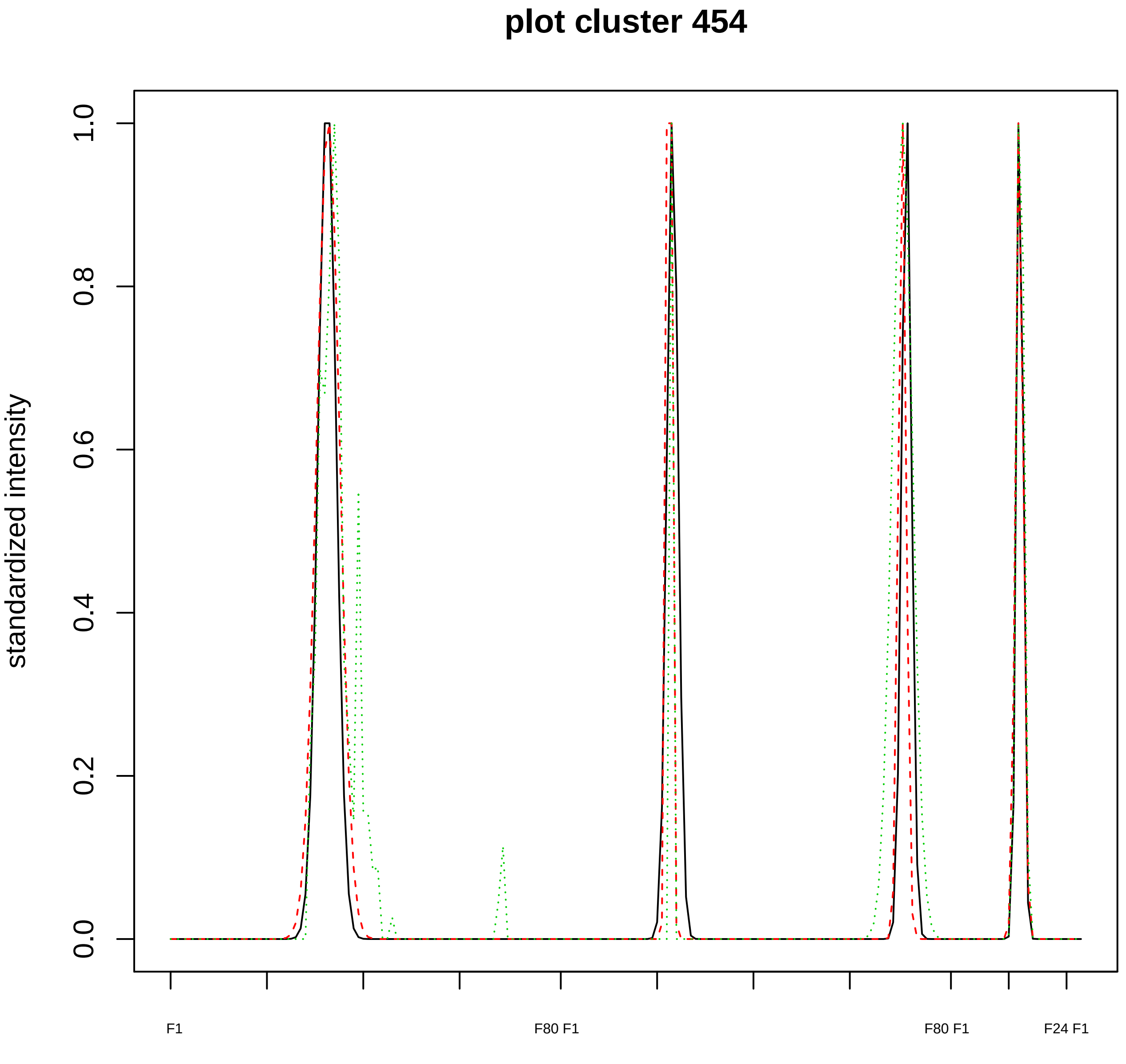

plot cluster 454
standardized intensity
1.0
0.8
0.6
0.4
0.2
0.0
F1
F80 F1
F80 F1
F24 F1

### Dendrogram of cluster 454

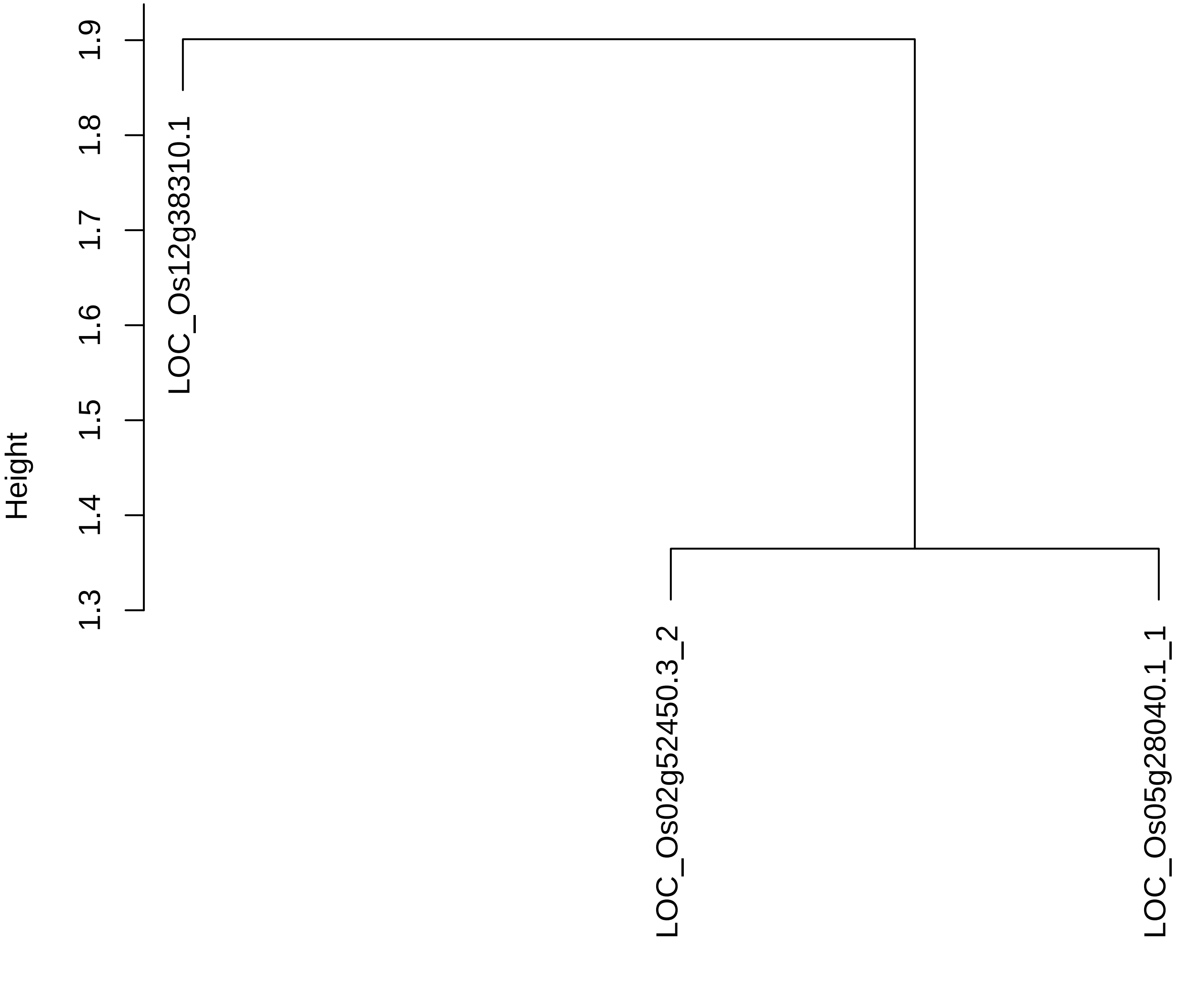


dist(spfitiex12sec12[lab[, 1] == i, ])
hclust (*, "ward.D2")

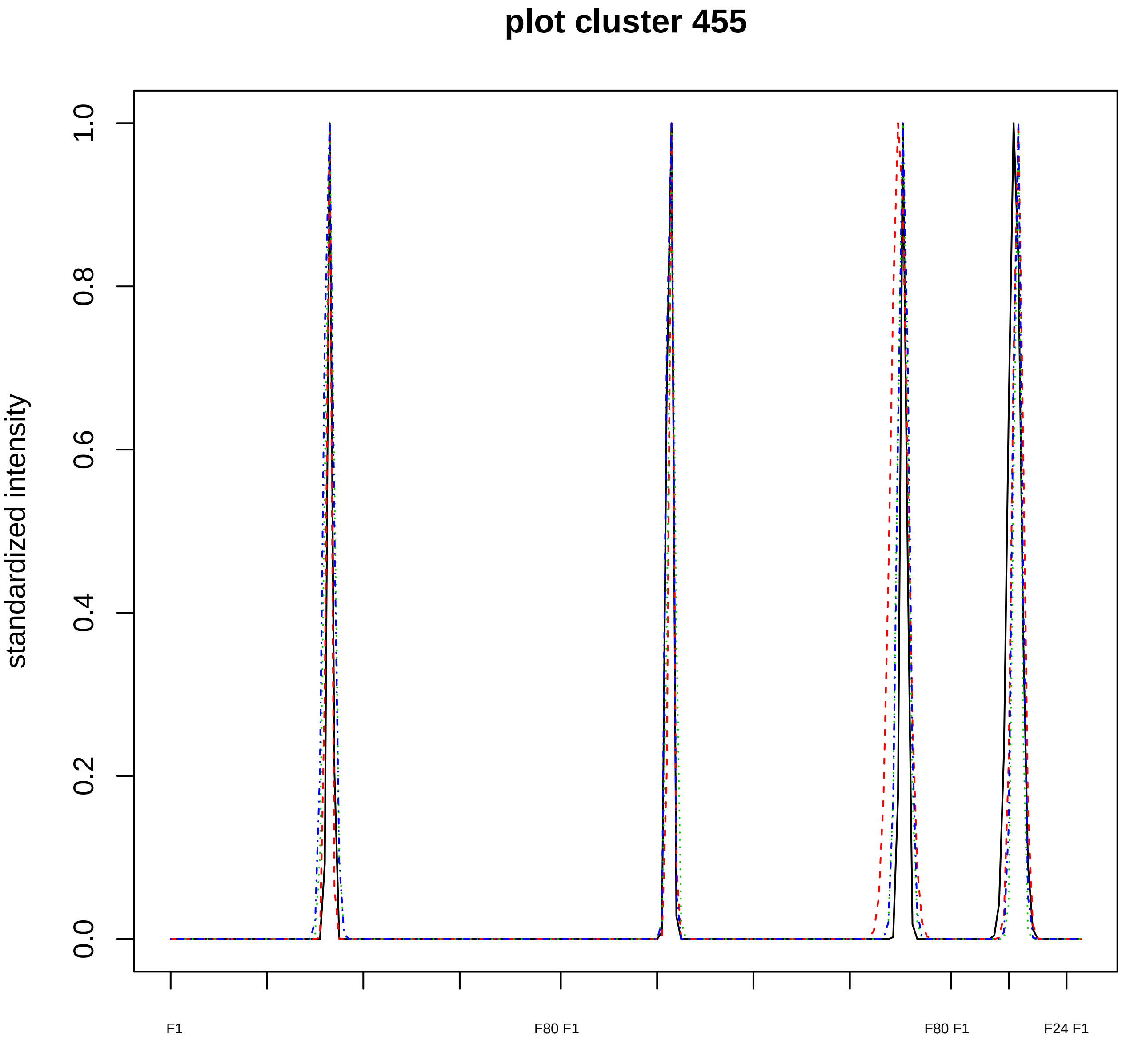

plot cluster 455
standardized intensity
0.0
0.2
0.4
0.6
0.8
1.0
F1
F80 F1
F80 F1
F24 F1

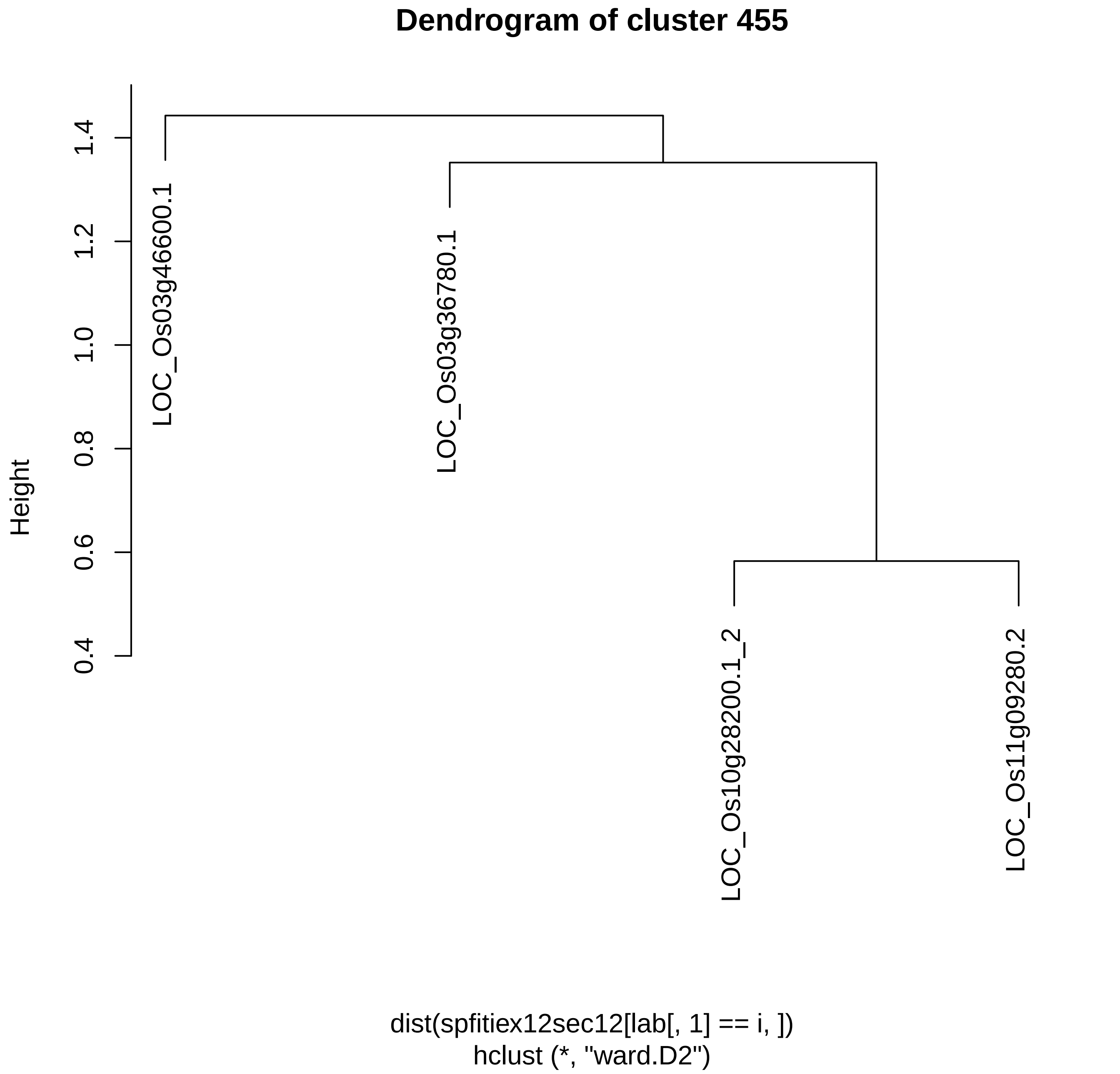
Dendrogram of cluster 455
Height
1.4
1.2
1.0
0.8
0.6
0.4
LOC_Os03g46600.1
LOC_Os03g36780.1
LOC_Os10g28200.1_2
LOC_Os11g09280.2
dist(spfitiex12sec12[lab[, 1] == i, ])
hclust (*, "ward.D2")

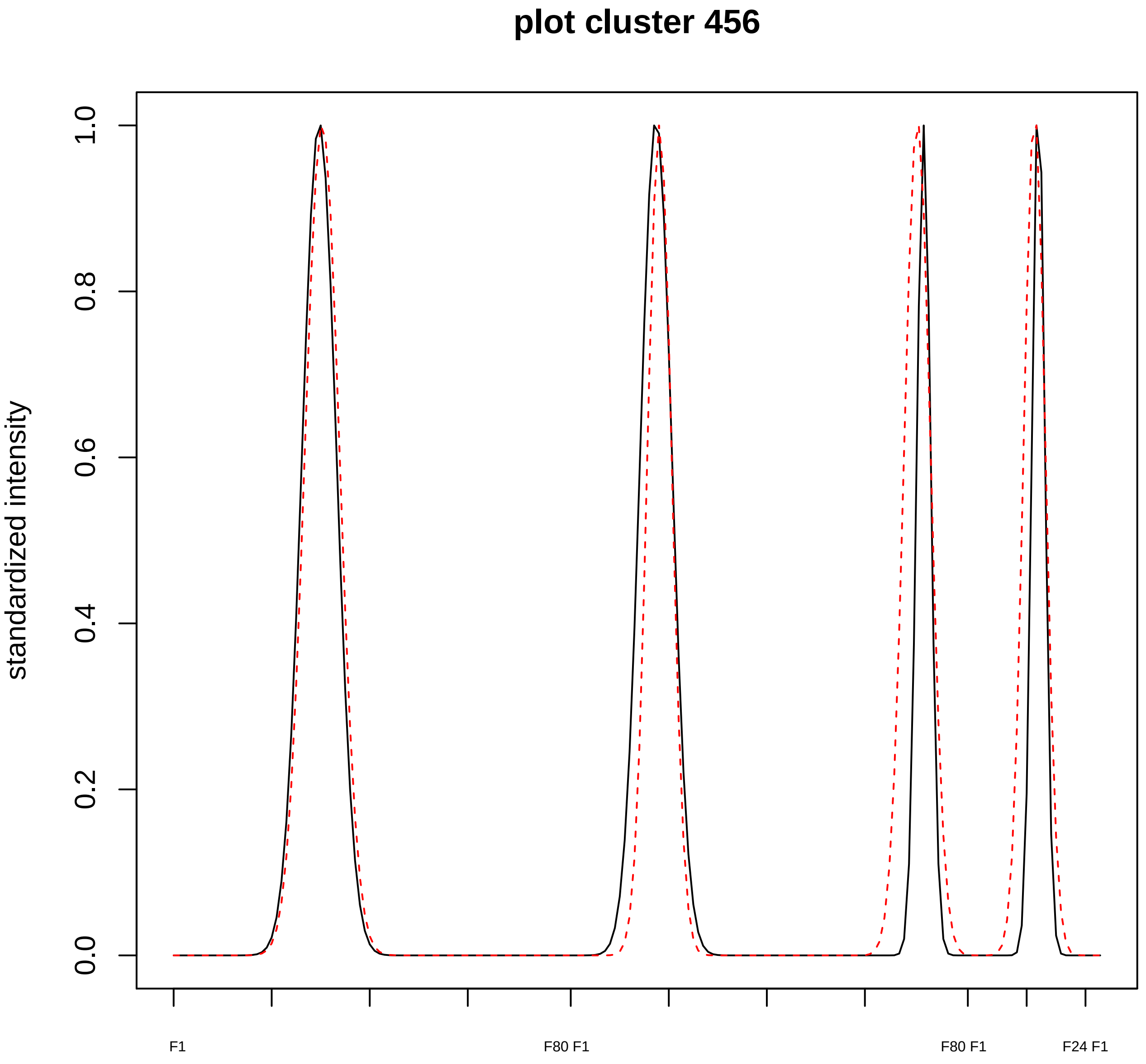
plot cluster 456
standardized intensity
0.0
0.2
0.4
0.6
0.8
1.0
F1
F80 F1
F80 F1
F24 F1

plot cluster 457

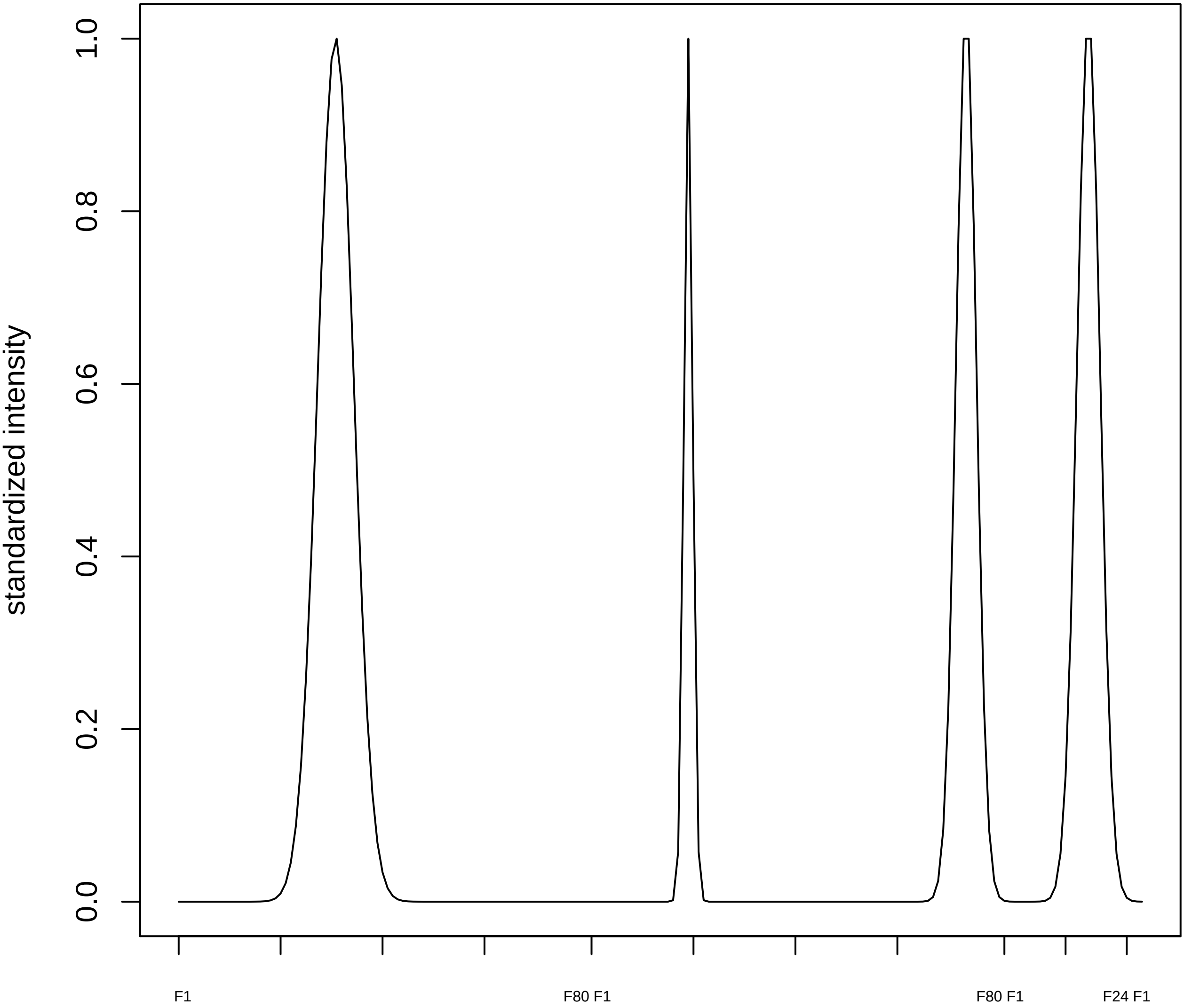

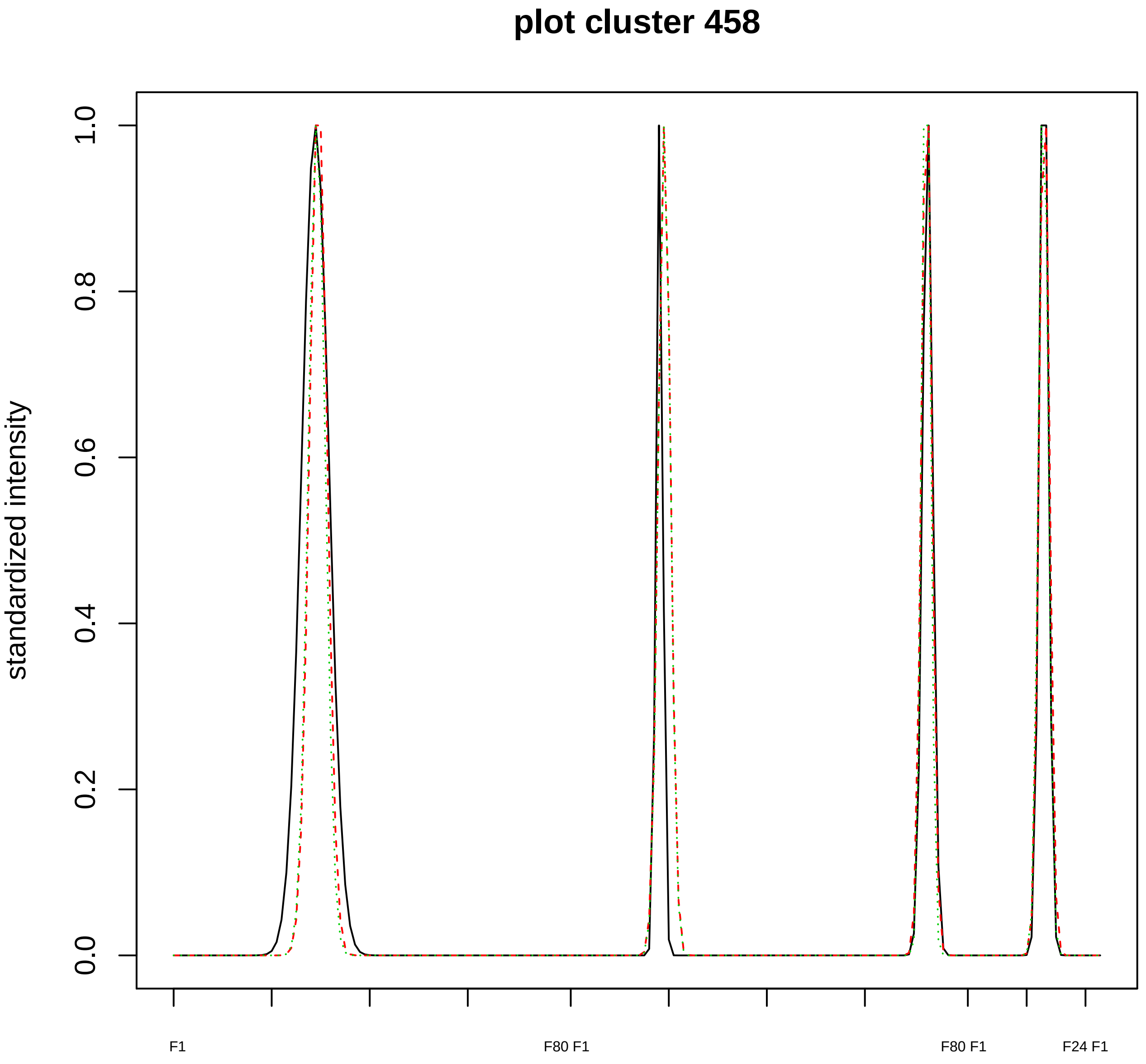
plot cluster 458
standardized intensity
0.0
0.2
0.4
0.6
0.8
1.0
F1
F80 F1
F80 F1
F24 F1

#### Dendrogram of cluster 458

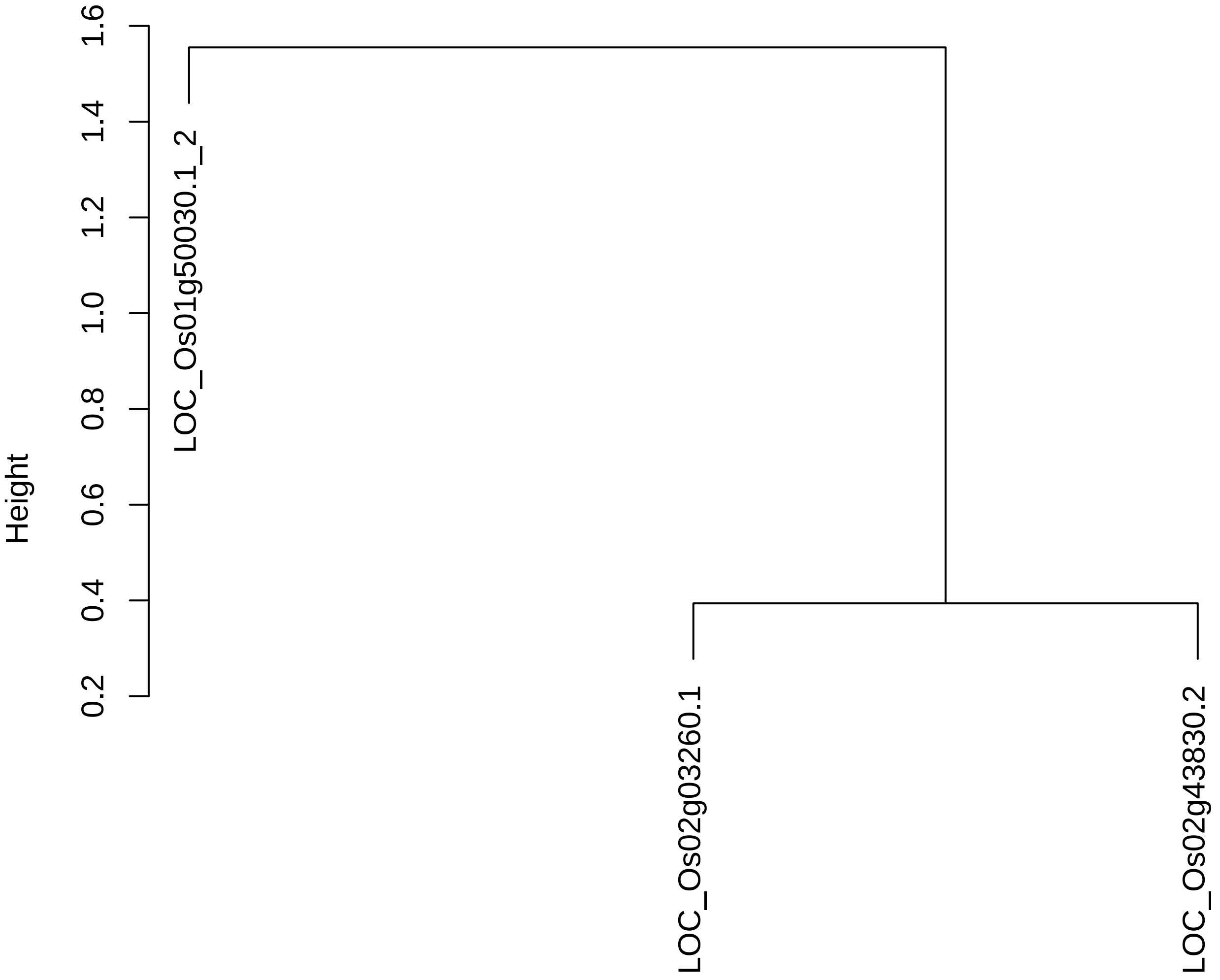


dist(spfitiex12sec12[lab[, 1] == i, ])
hclust (*, "ward.D2")

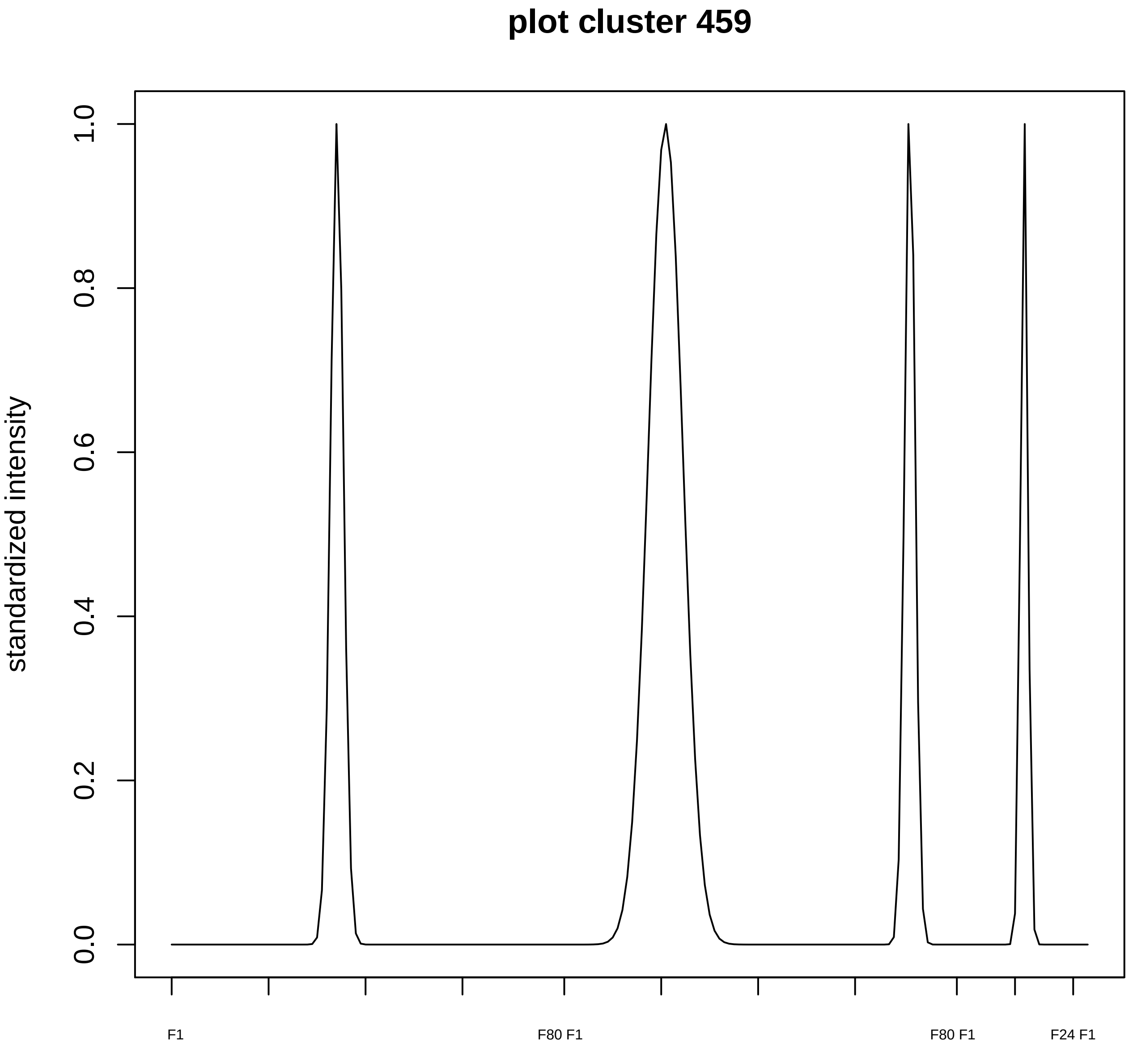
plot cluster 459
standardized intensity
0.0
0.2
0.4
0.6
0.8
1.0
F1
F80 F1
F80 F1
F24 F1

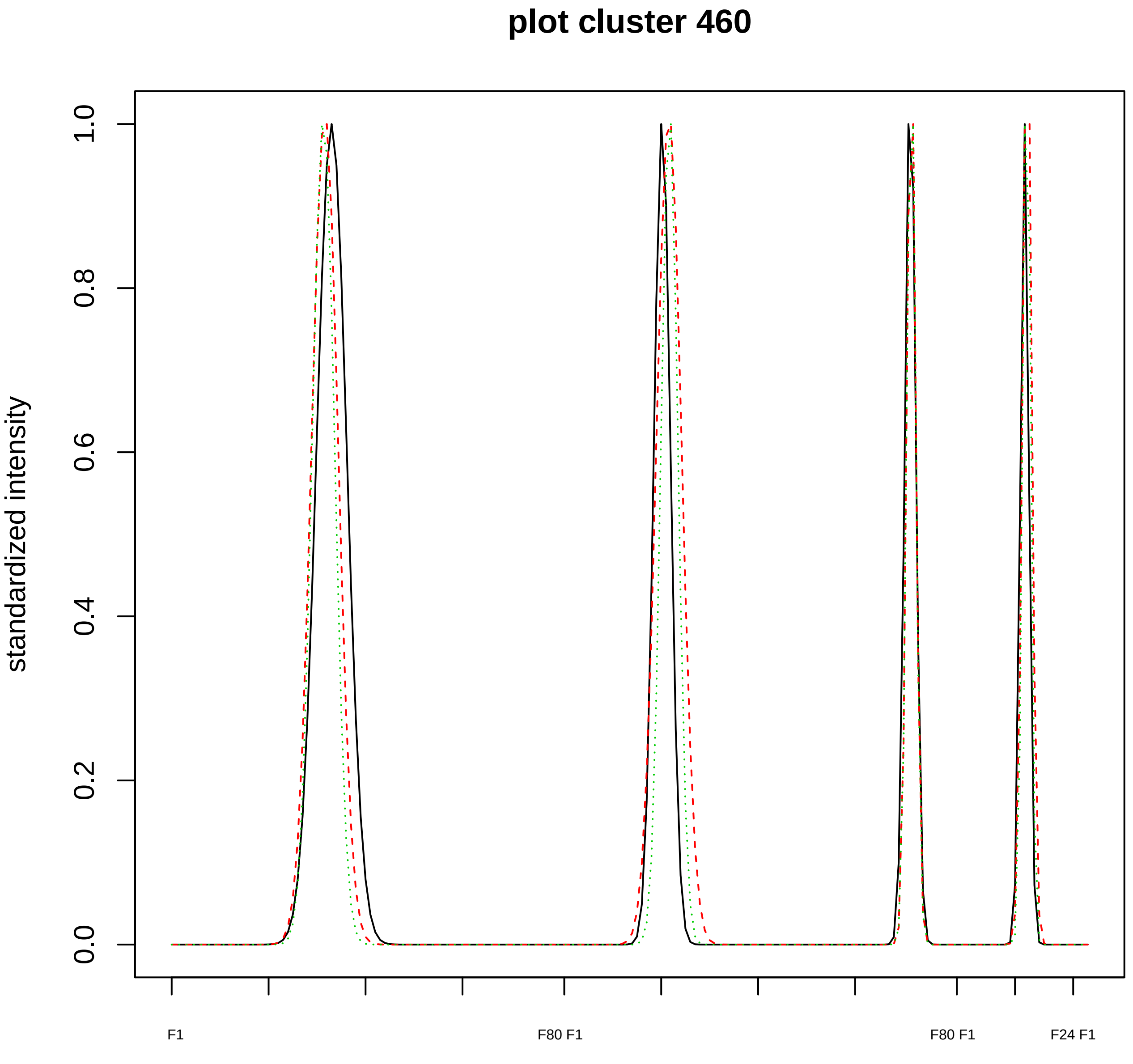
plot cluster 460
standardized intensity
0.0
0.2
0.4
0.6
0.8
1.0
F1
F80 F1
F80 F1
F24 F1

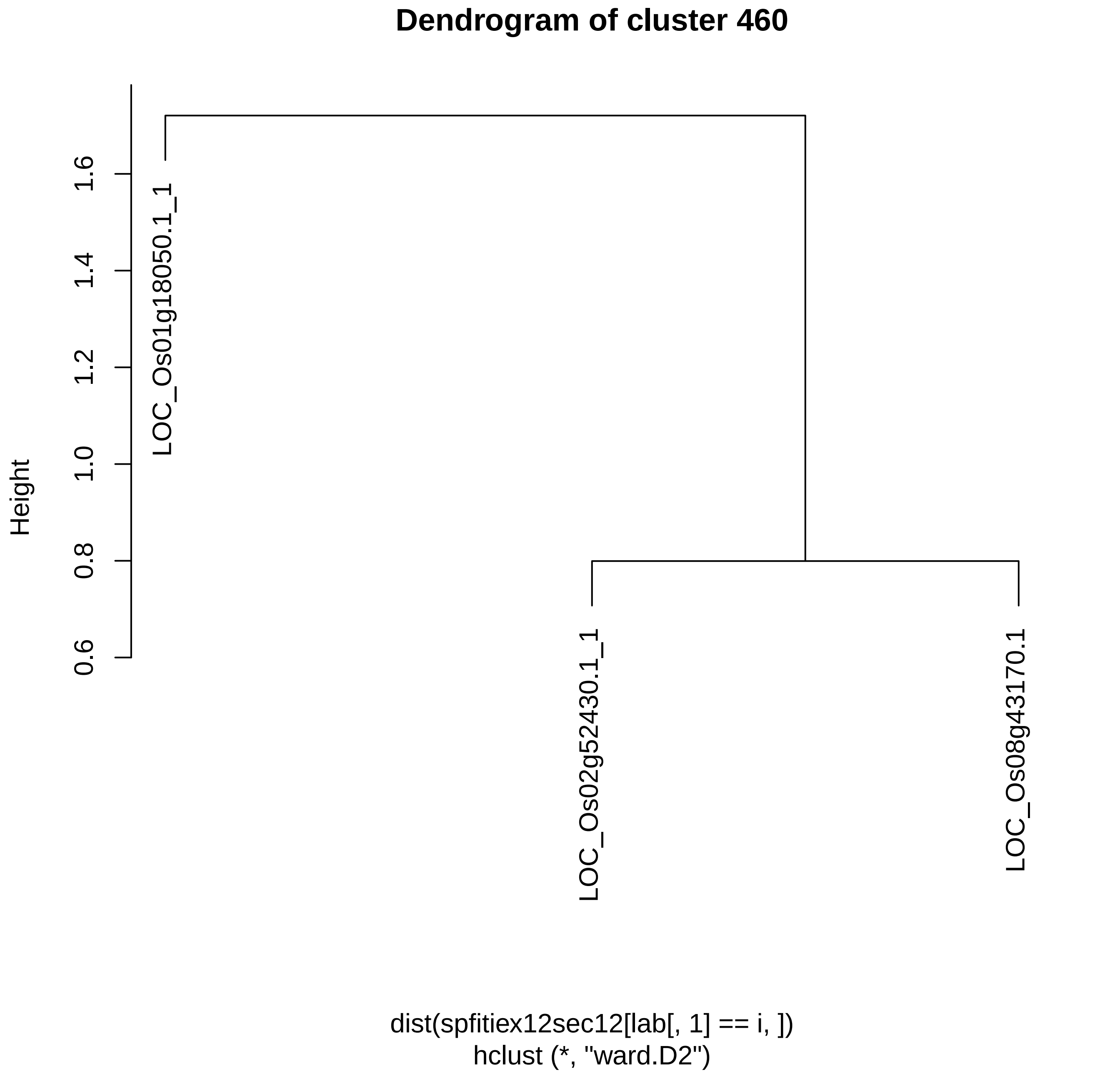
Dendrogram of cluster 460
Height
0.6
0.8
1.0
1.2
1.4
1.6
LOC_Os01g18050.1_1
LOC_Os02g52430.1_1
LOC_Os08g43170.1
dist(spfitiex12sec12[lab[, 1] == i, ])
hclust (*, "ward.D2")

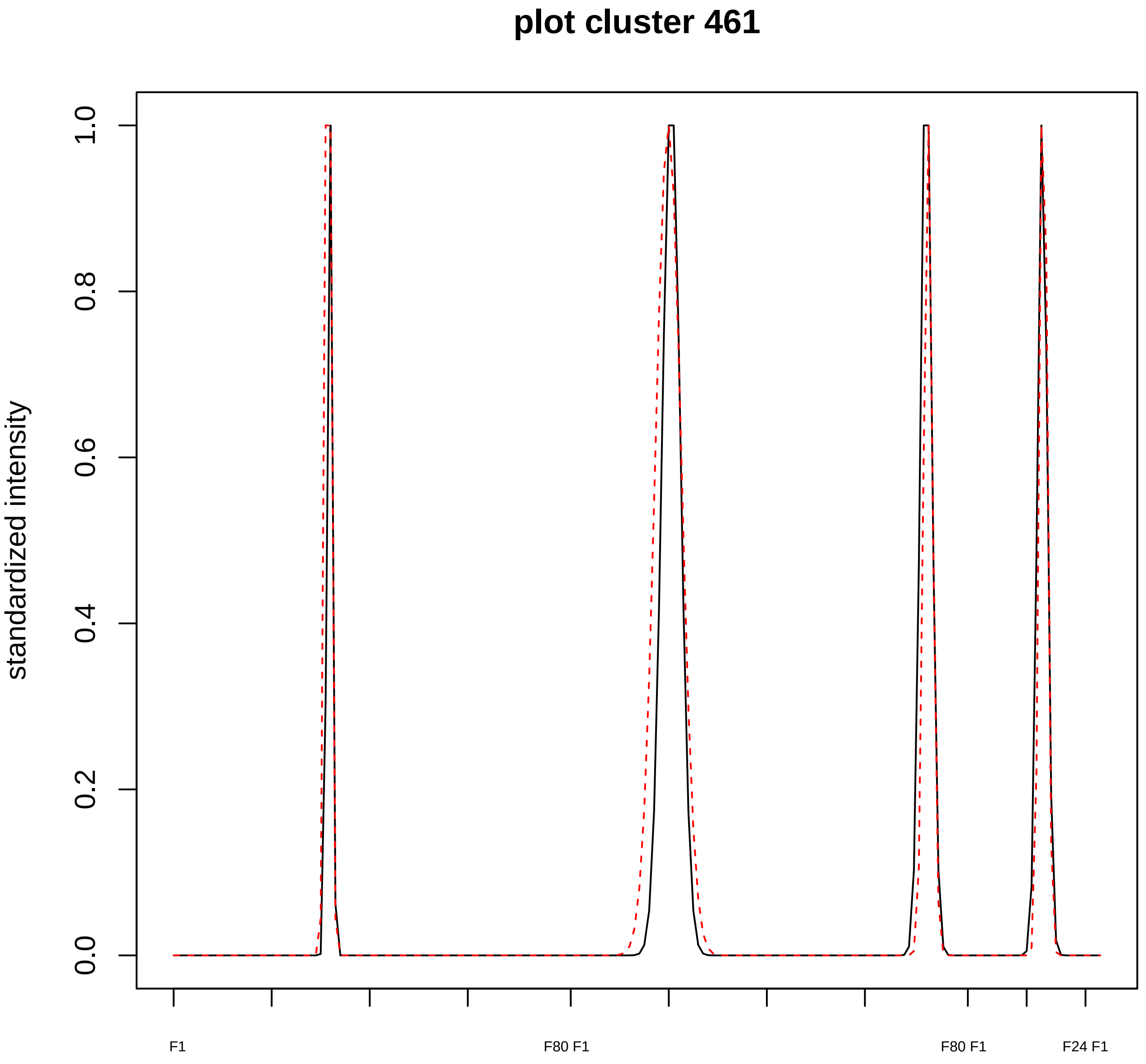
plot cluster 461
standardized intensity
0.0
0.2
0.4
0.6
0.8
1.0
F1
F80 F1
F80 F1
F24 F1

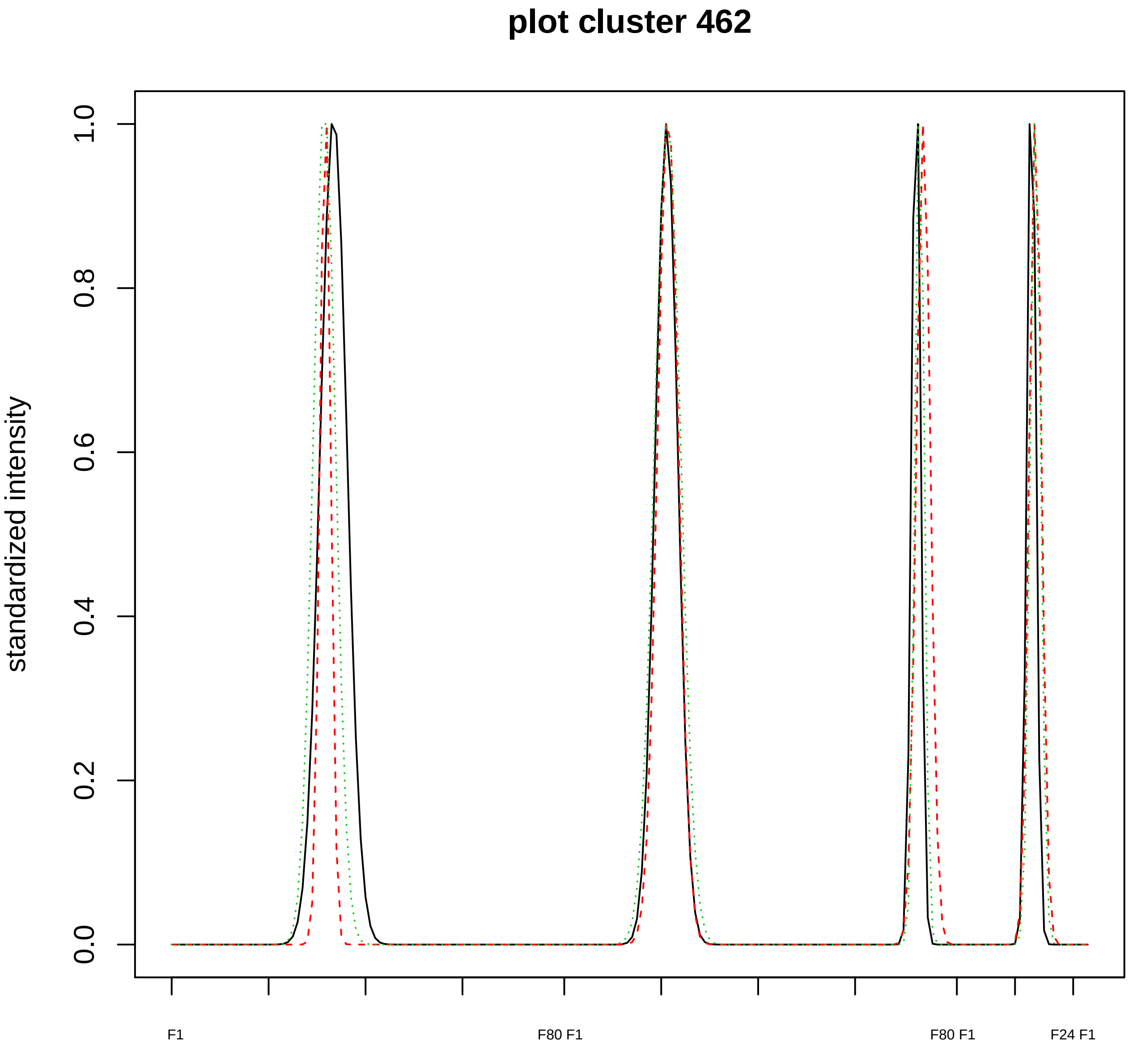

plot cluster 462
standardized intensity
0.0
0.2
0.4
0.6
0.8
1.0
F1
F80 F1
F80 F1
F24 F1

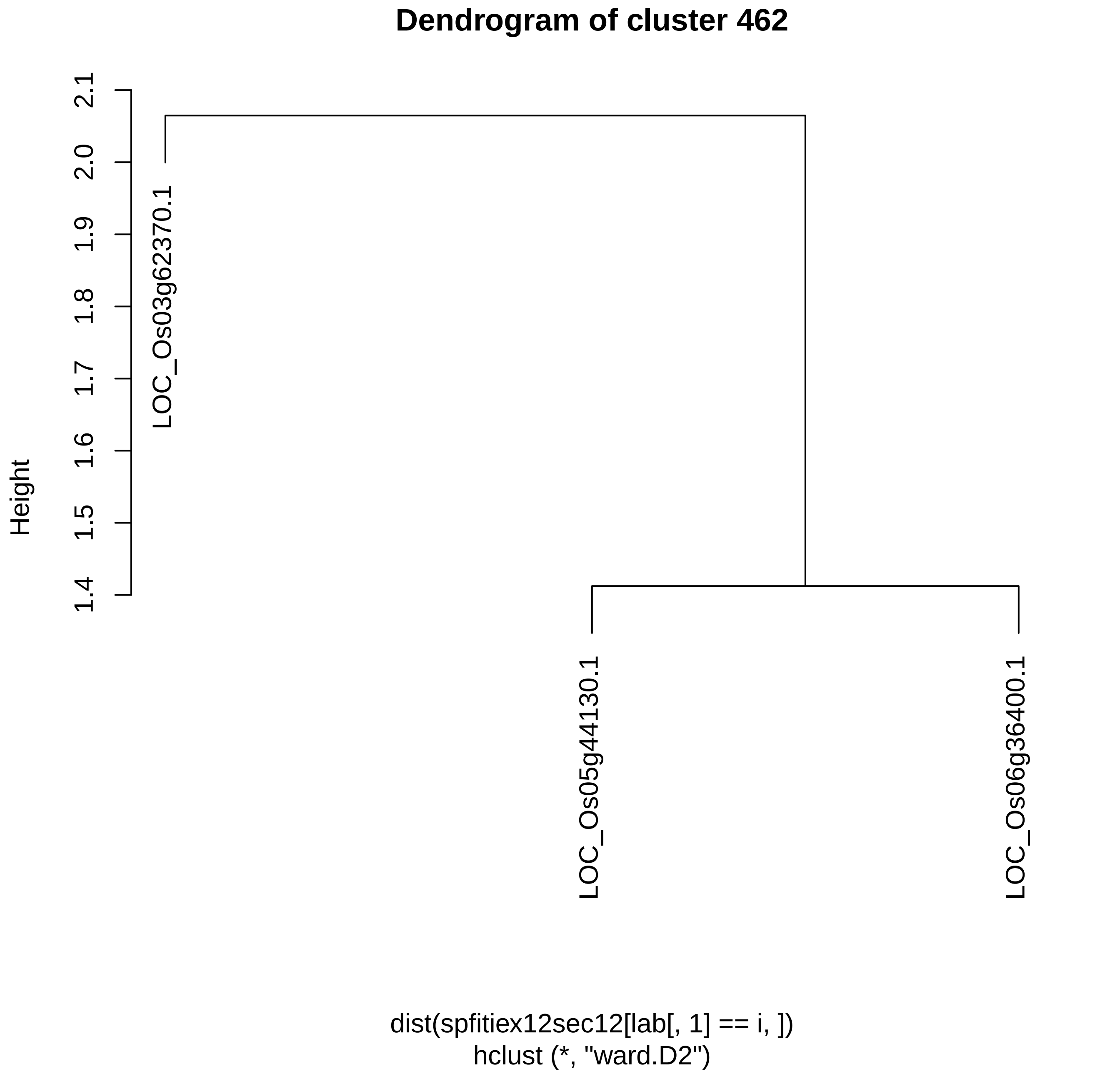
Dendrogram of cluster 462
Height
2.1
2.0
1.9
1.8
1.7
1.6
1.5
1.4
LOC_Os03g62370.1
LOC_Os05g44130.1
LOC_Os06g36400.1
dist(spfitiex12sec12[lab[, 1] == i, ])
hclust (*, "ward.D2")

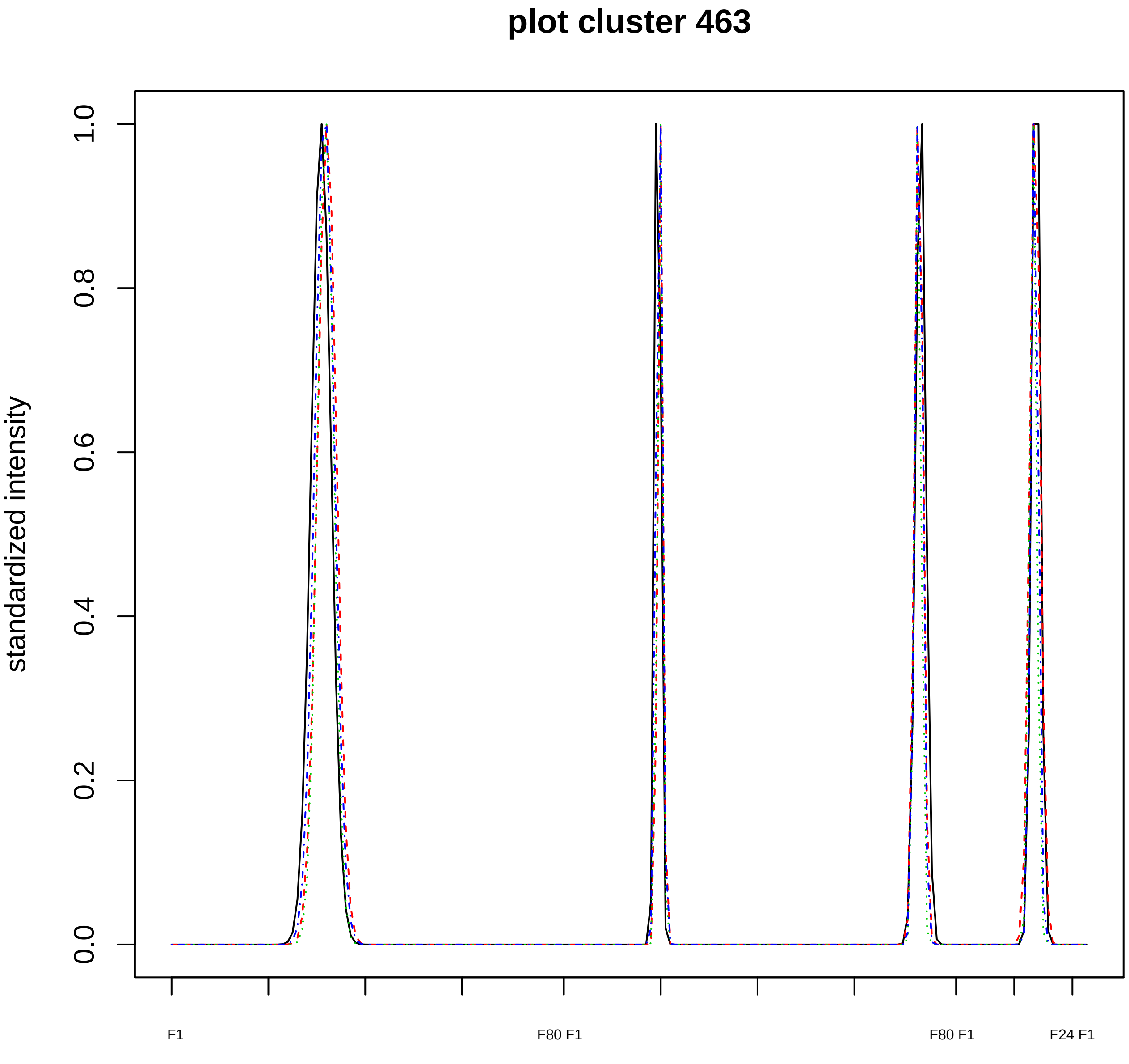

plot cluster 463
standardized intensity
0.0
0.2
0.4
0.6
0.8
1.0
F1
F80 F1
F80 F1
F24 F1

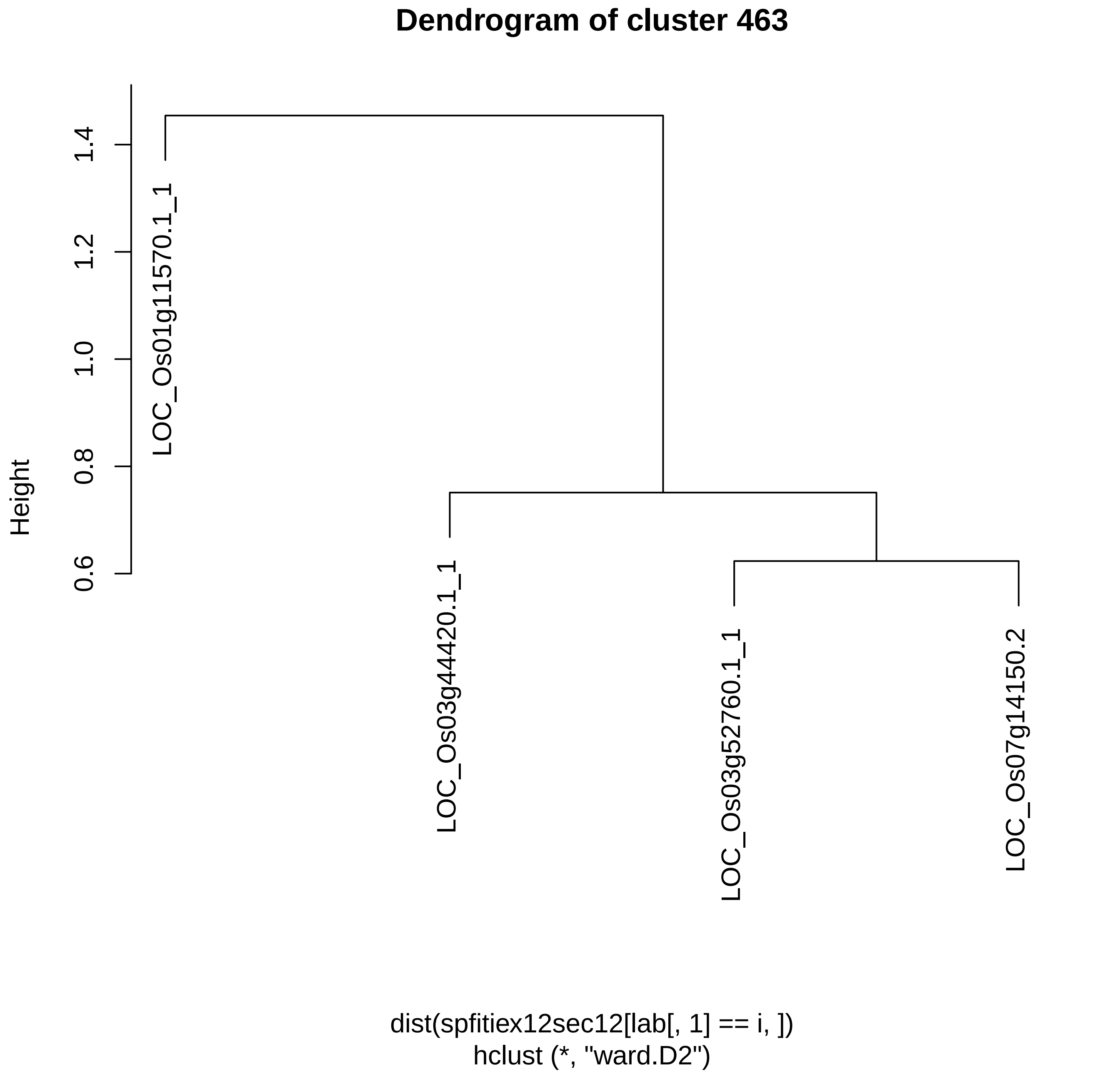
Dendrogram of cluster 463
Height
1.4
1.2
1.0
0.8
0.6
LOC_Os01g11570.1_1
LOC_Os03g44420.1_1
LOC_Os03g52760.1_1
LOC_Os07g14150.2
dist(spfitiex12sec12[lab[, 1] == i, ])
hclust (*, "ward.D2")

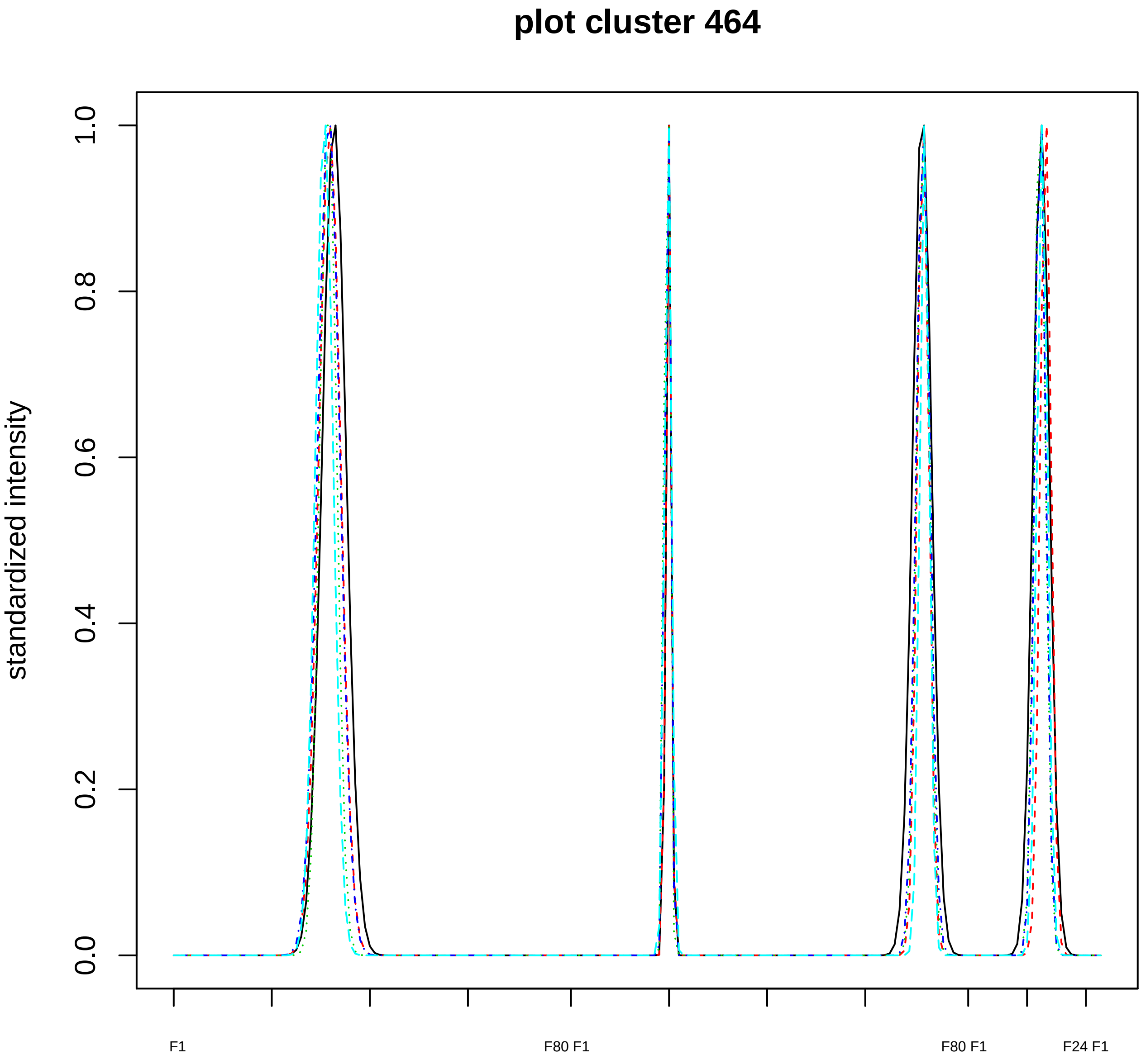

plot cluster 464
standardized intensity
0.0
0.2
0.4
0.6
0.8
1.0
F1
F80 F1
F80 F1
F24 F1

### Dendrogram of cluster 464

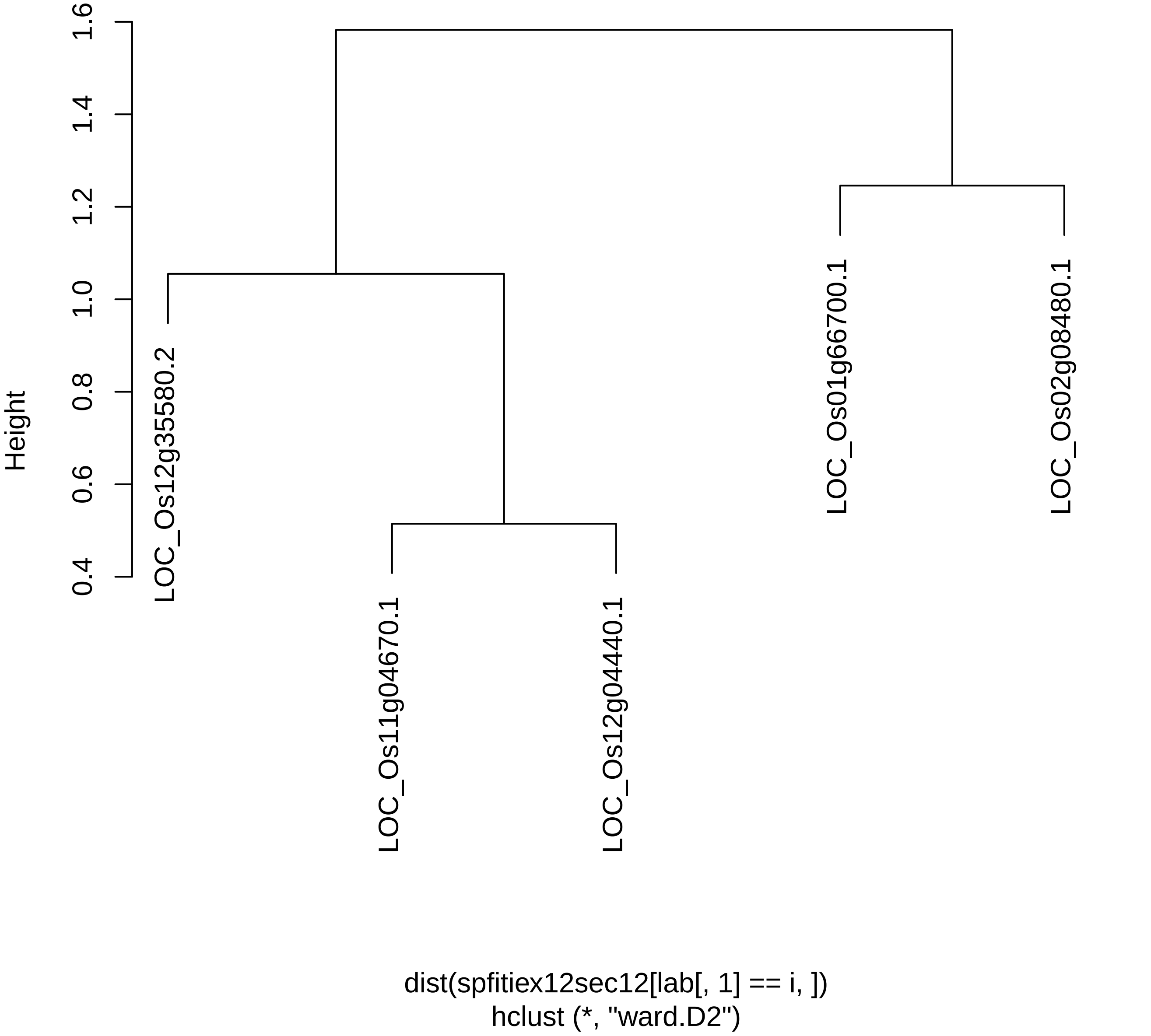

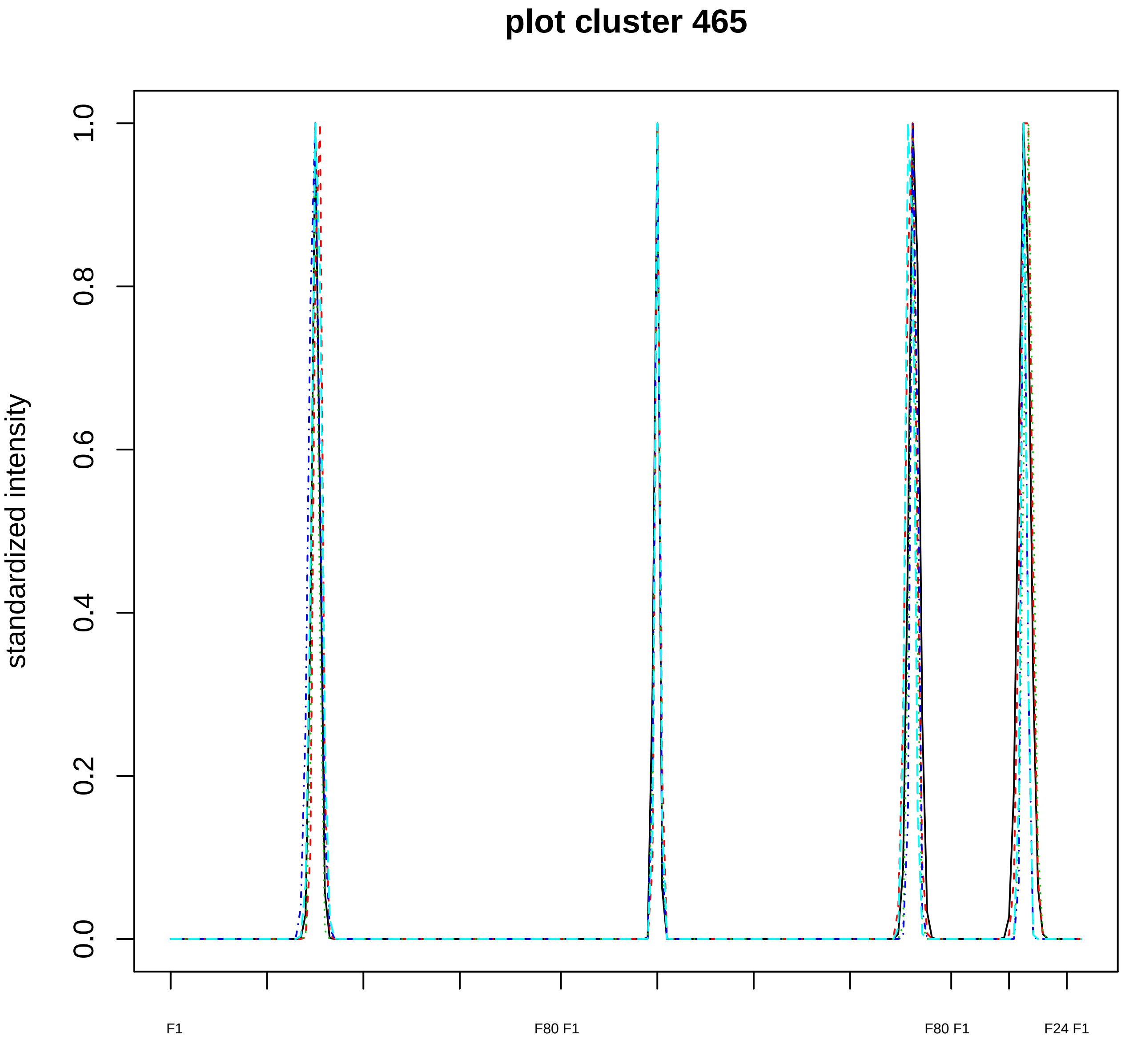
plot cluster 465
standardized intensity
0.0
0.2
0.4
0.6
0.8
1.0
F1
F80 F1
F80 F1
F24 F1

### Dendrogram of cluster 465

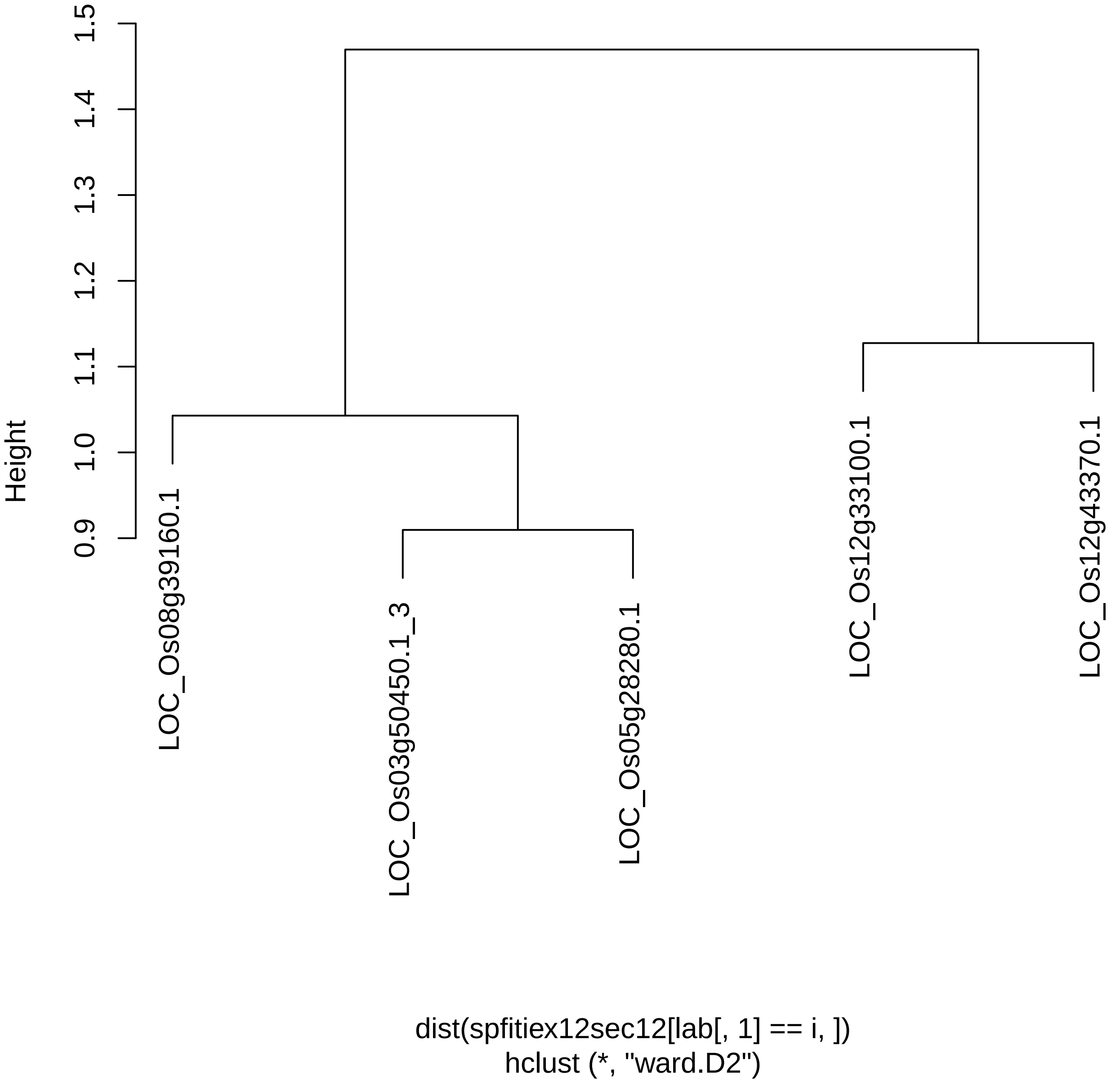

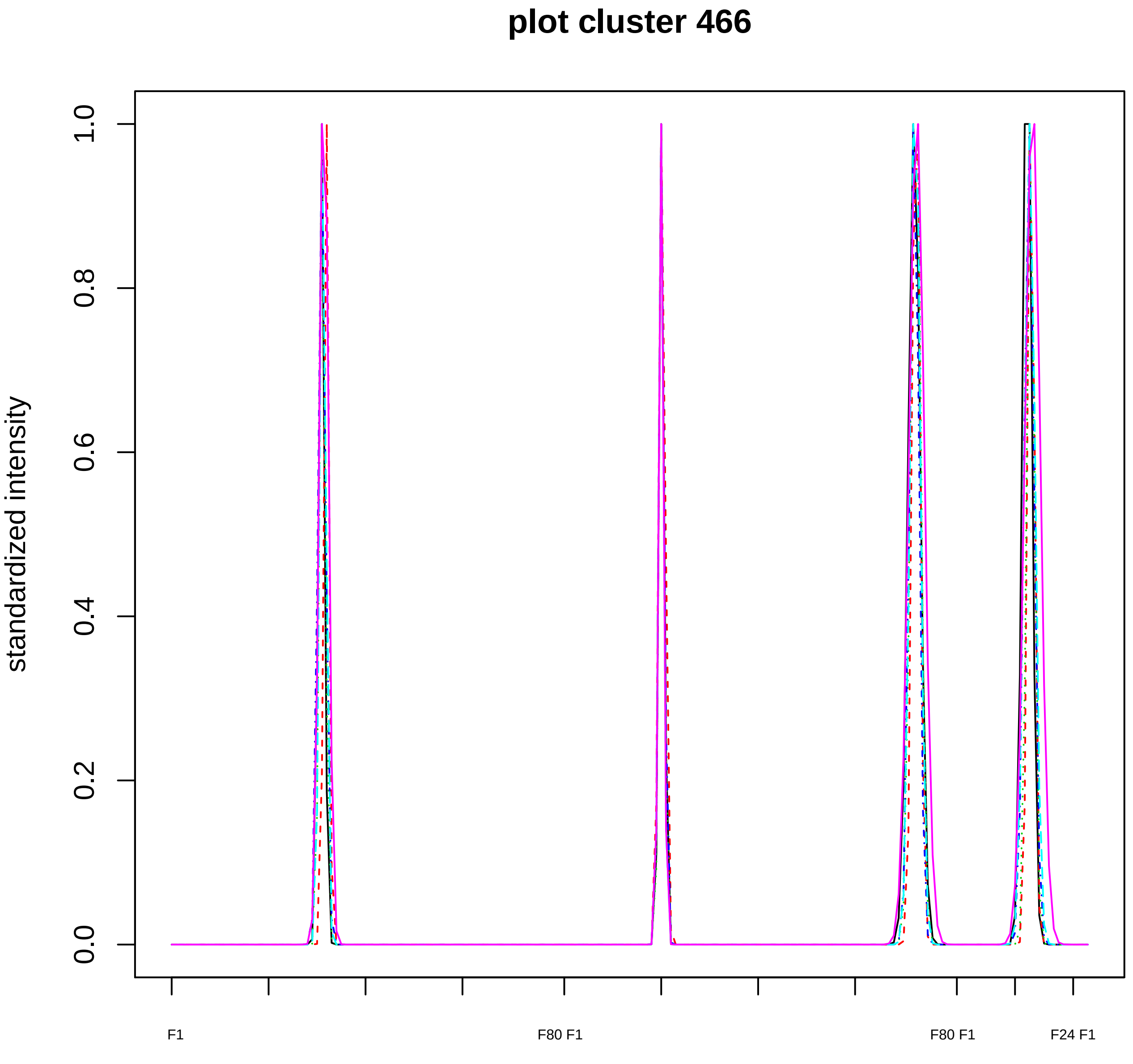

plot cluster 466
standardized intensity
0.0
0.2
0.4
0.6
0.8
1.0
F1
F80 F1
F80 F1
F24 F1

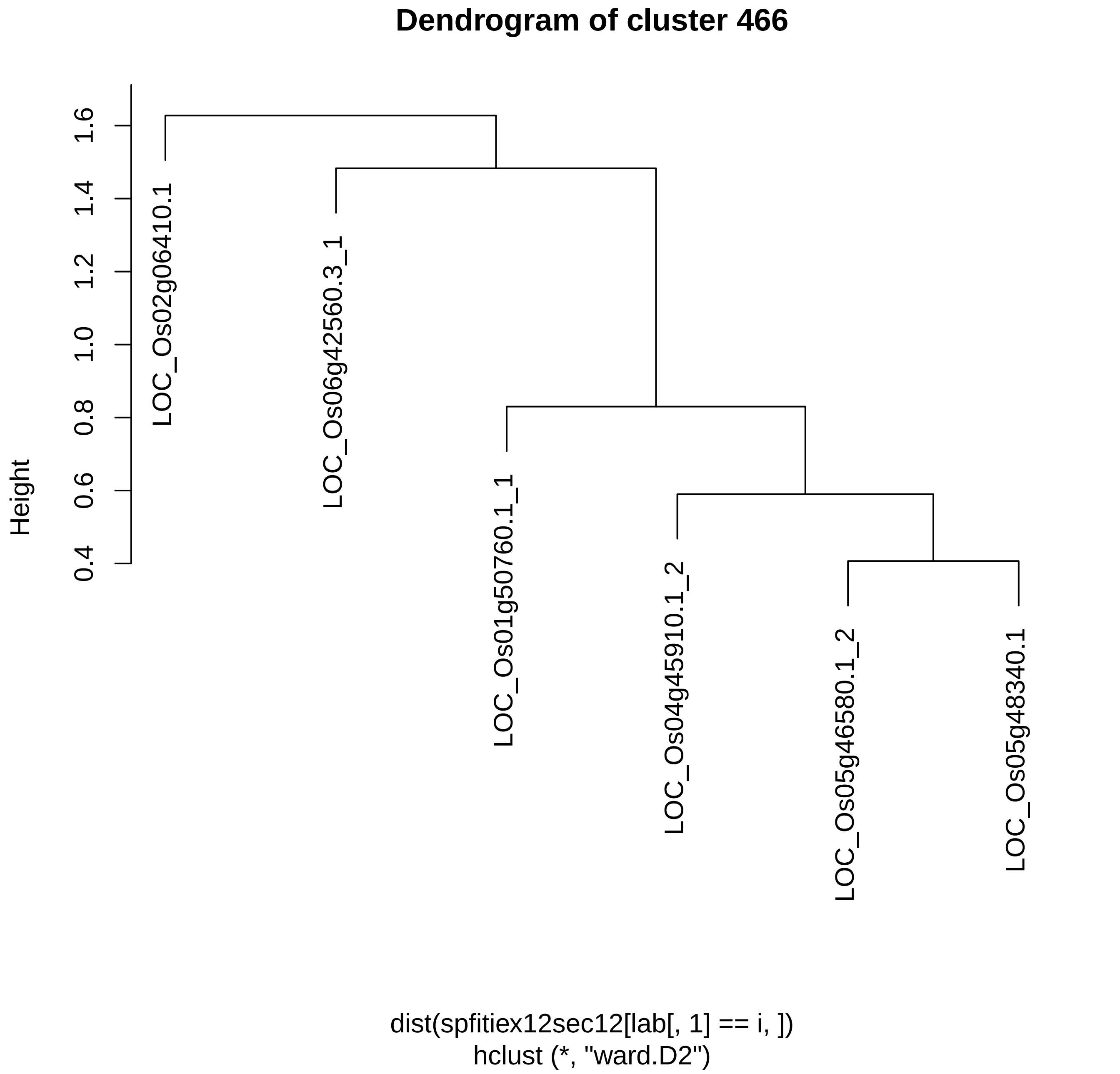

Dendrogram of cluster 466
Height
1.6
1.4
1.2
1.0
0.8
0.6
0.4
LOC_Os02g06410.1
LOC_Os06g42560.3_1
LOC_Os01g50760.1_1
LOC_Os04g45910.1_2
LOC_Os05g46580.1_2
LOC_Os05g48340.1
dist(spfitiex12sec12[lab[, 1] == i, ])
hclust (*, "ward.D2")

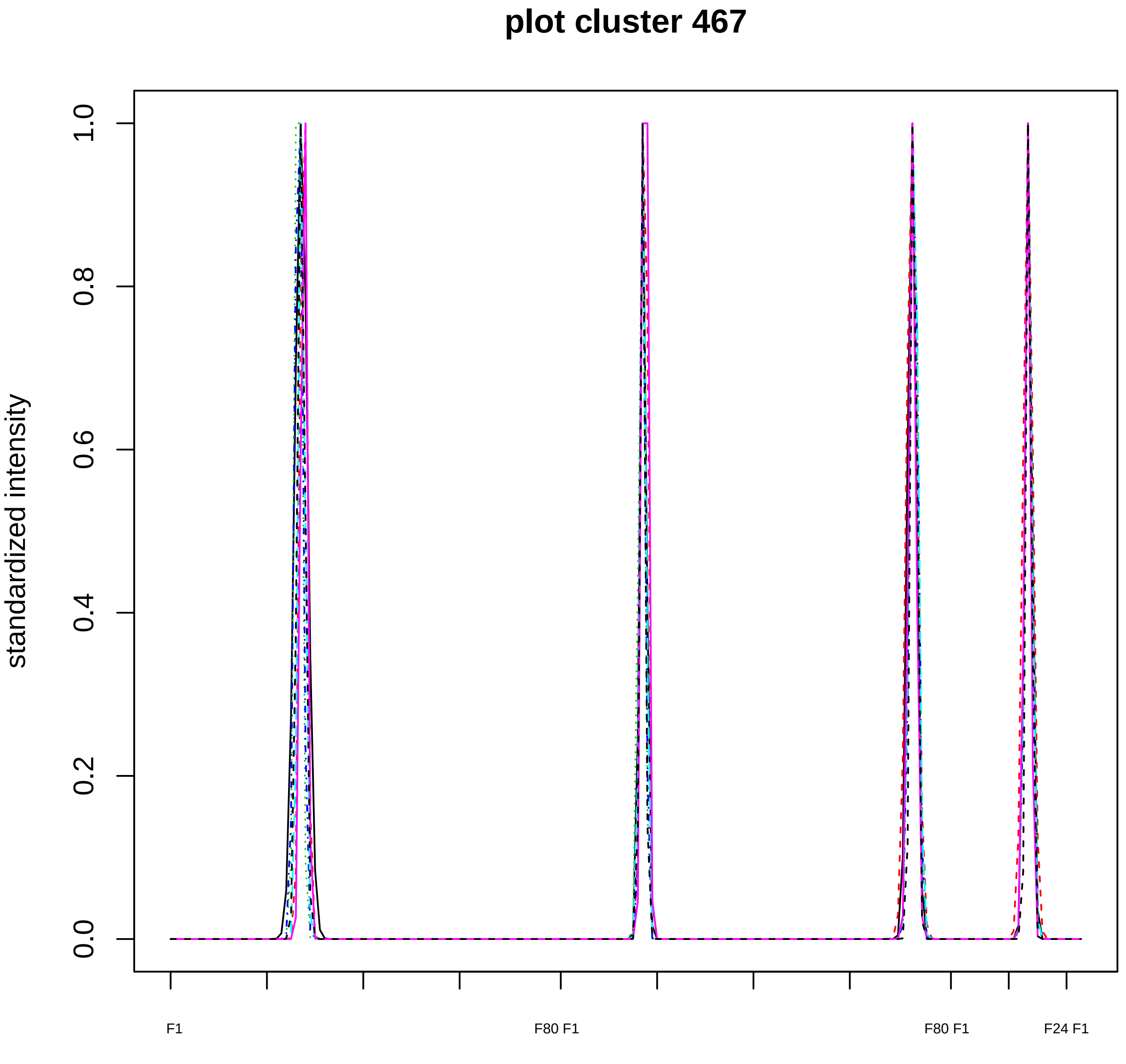
plot cluster 467
standardized intensity
0.0
0.2
0.4
0.6
0.8
1.0
F1
F80 F1
F80 F1
F24 F1

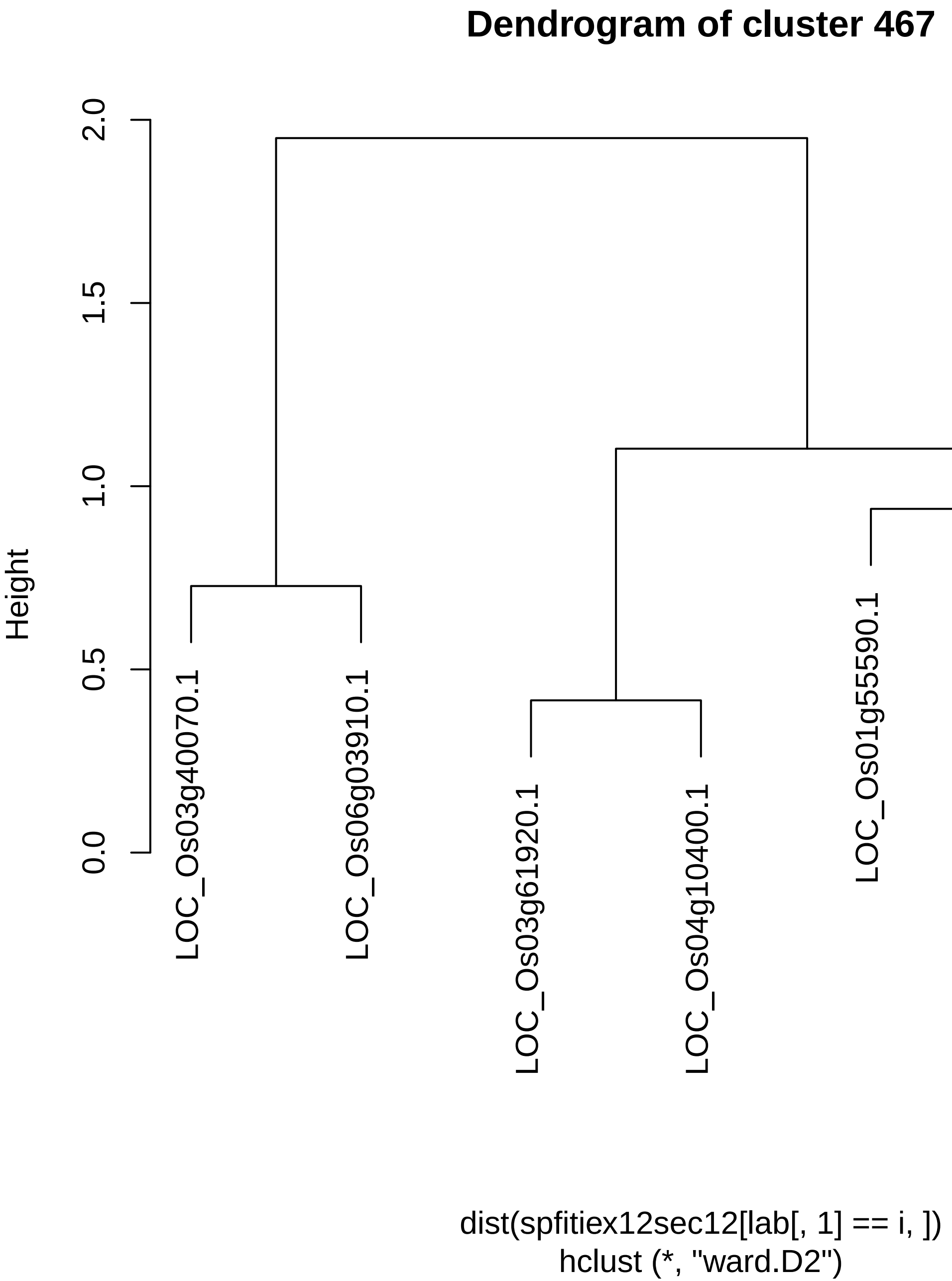
Dendrogram of cluster 467
Height
2.0
1.5
1.0
0.5
0.0
LOC_Os03g40070.1
LOC_Os06g03910.1
LOC_Os03g61920.1
LOC_Os04g10400.1
LOC_Os01g55590.1
LOC_Os05g40420.1
LOC_Os07g40320.1
dist(spfitiex12sec12[lab[, 1] == i, ])
hclust (*, "ward.D2")

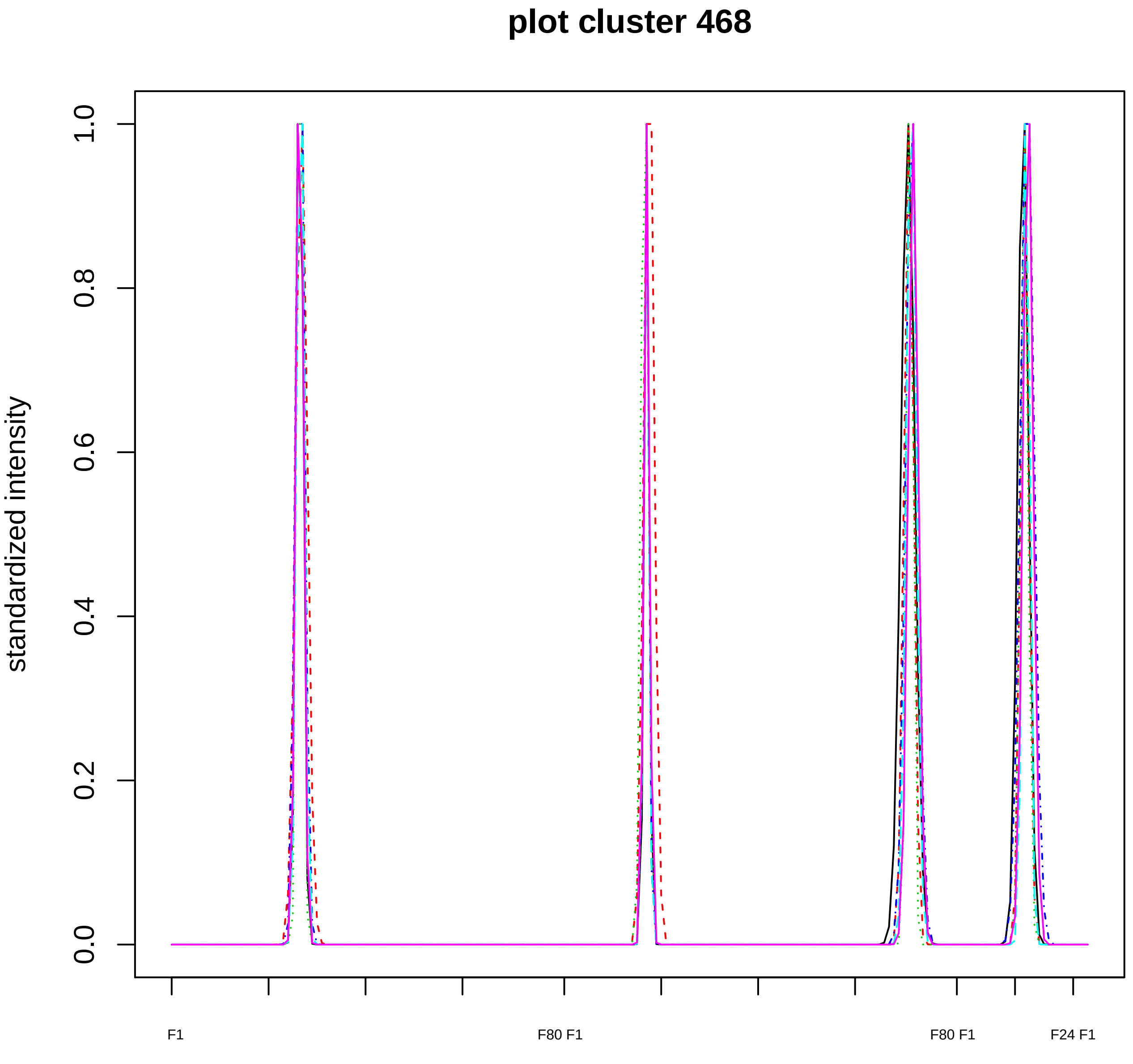

plot cluster 468
standardized intensity
0.0
0.2
0.4
0.6
0.8
1.0
F1
F80 F1
F80 F1
F24 F1

### Dendrogram of cluster 468

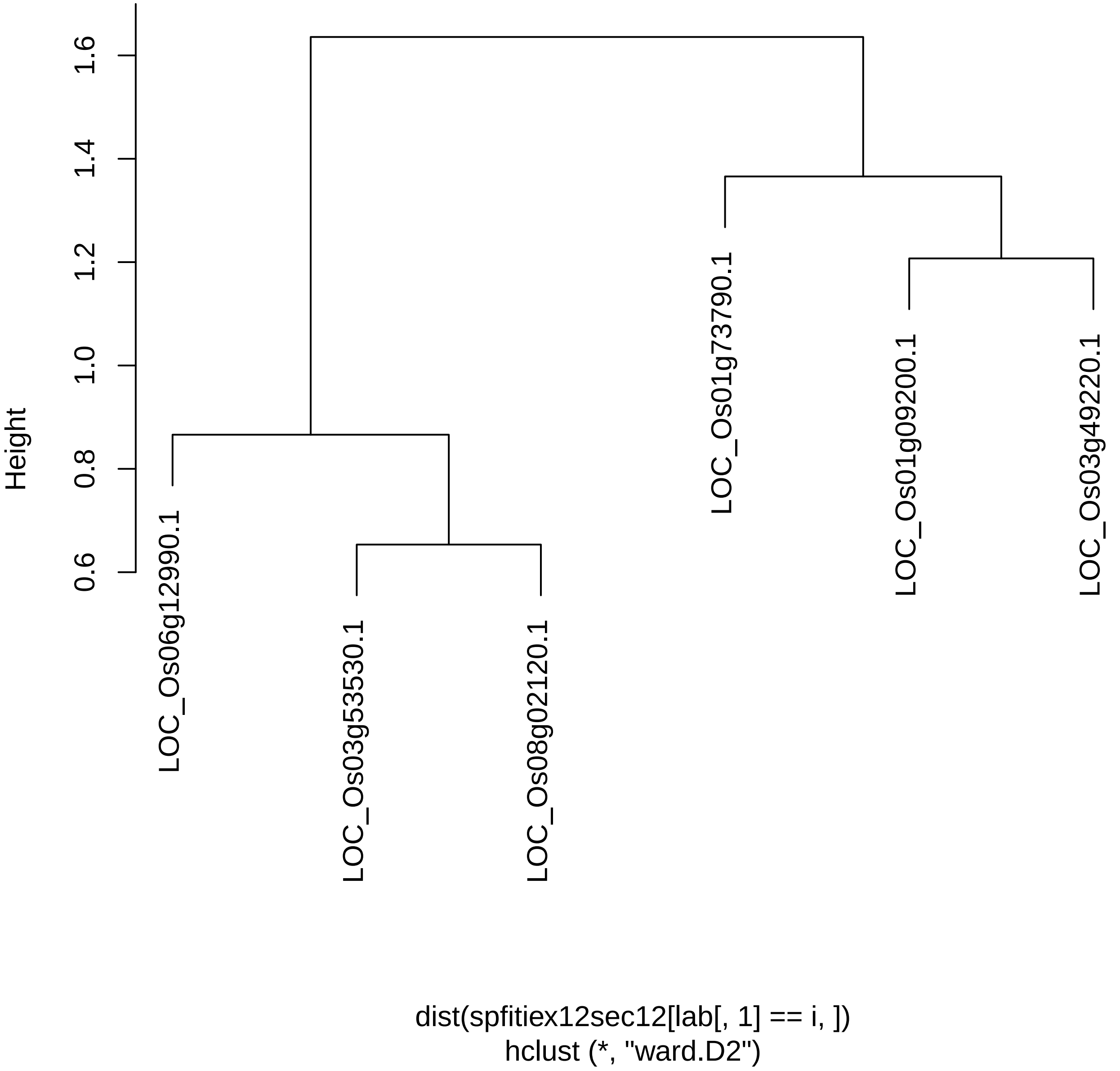

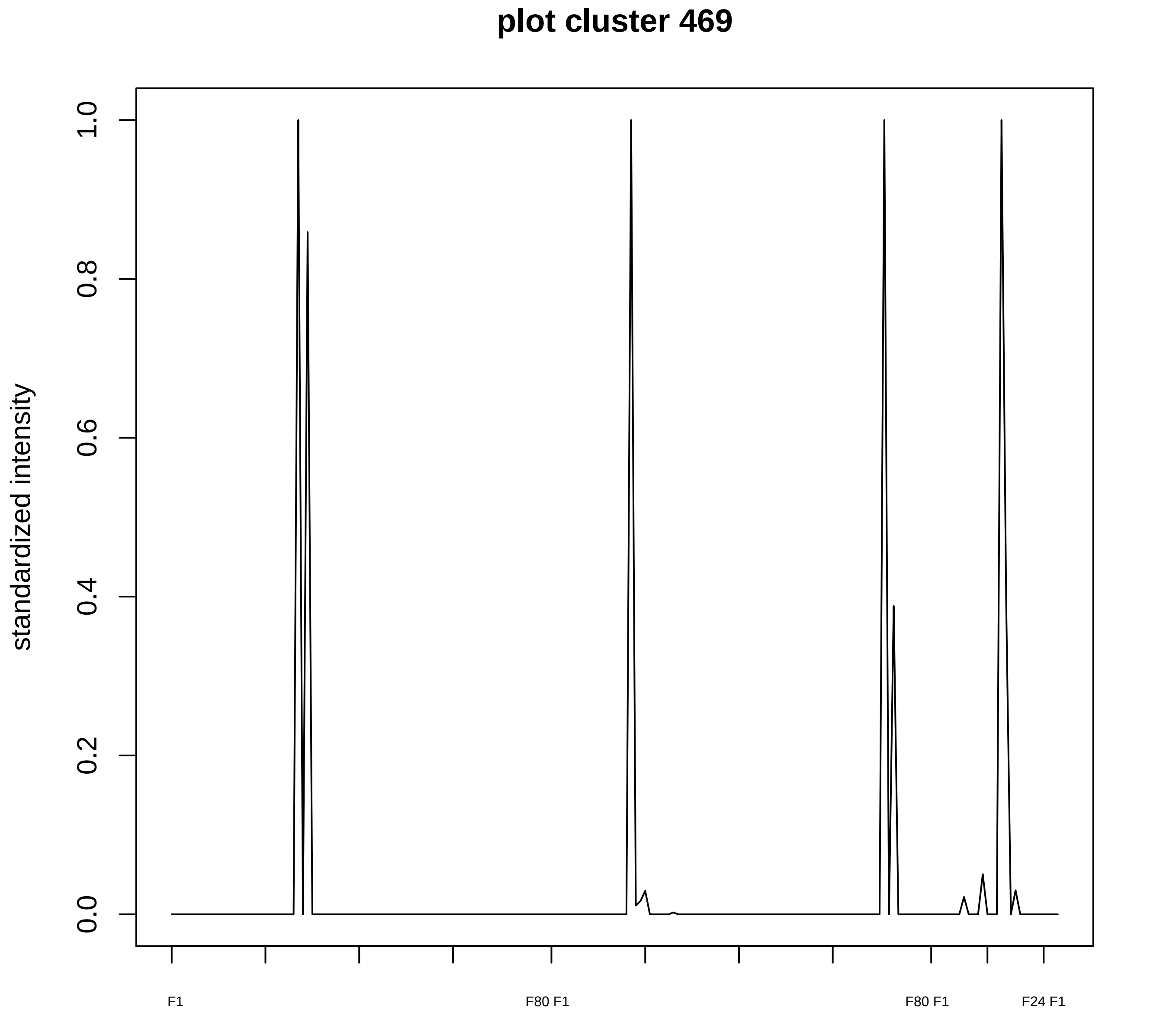
plot cluster 469
standardized intensity
0.0
0.2
0.4
0.6
0.8
1.0
F1
F80 F1
F80 F1
F24 F1

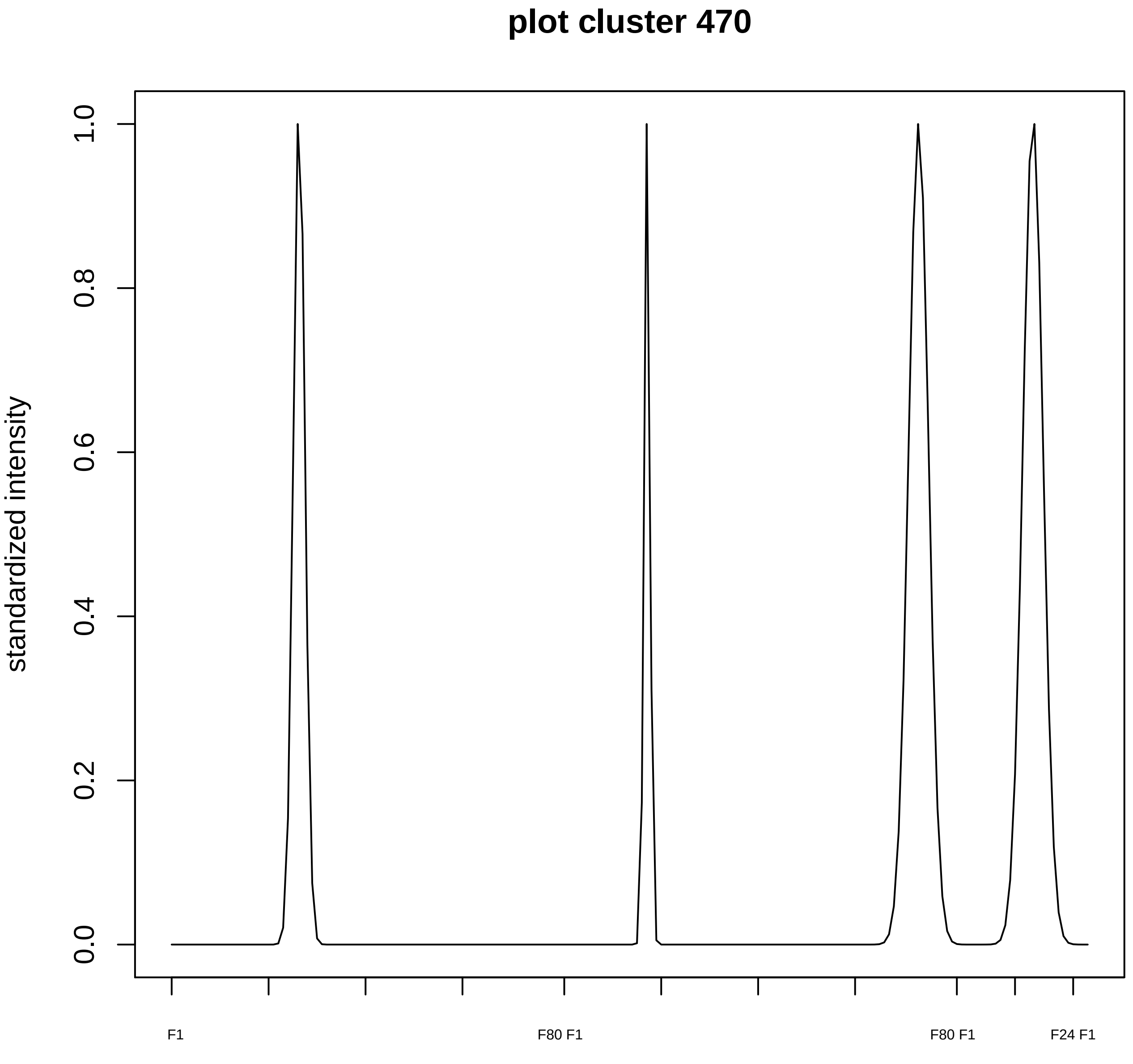
plot cluster 470
standardized intensity
0.0
0.2
0.4
0.6
0.8
1.0
F1
F80 F1
F80 F1
F24 F1

### plot cluster 471

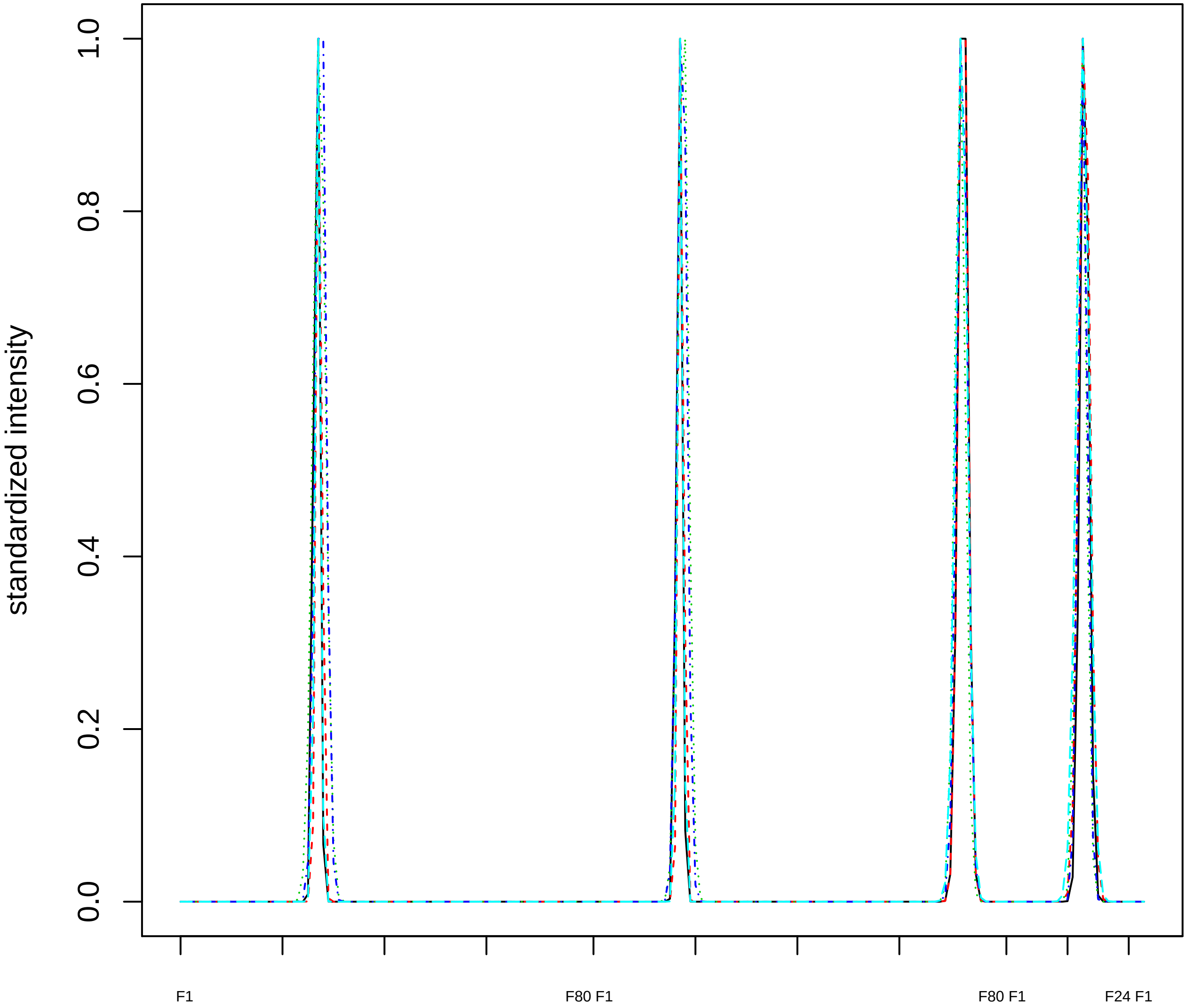

#### Dendrogram of cluster 471

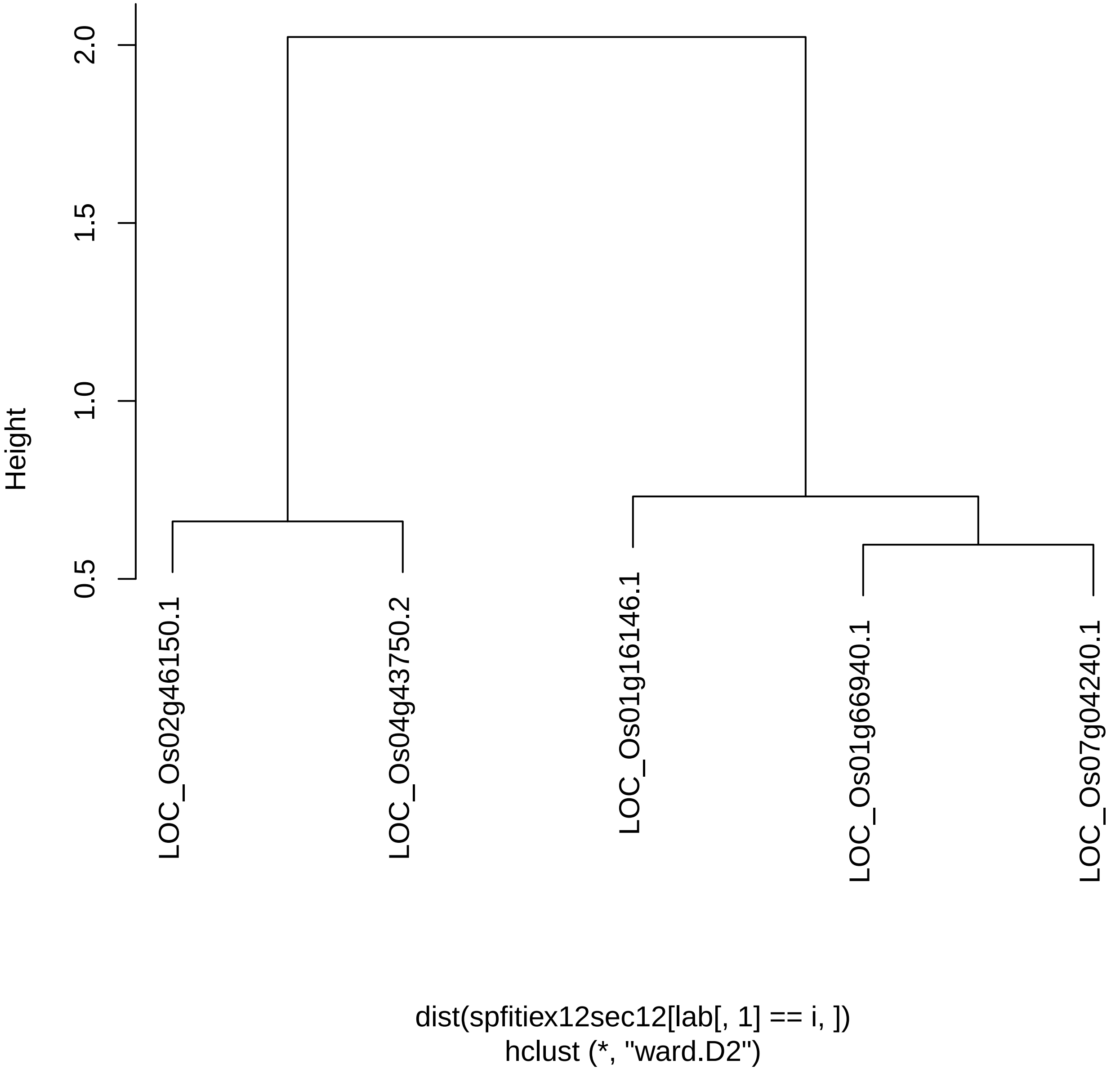

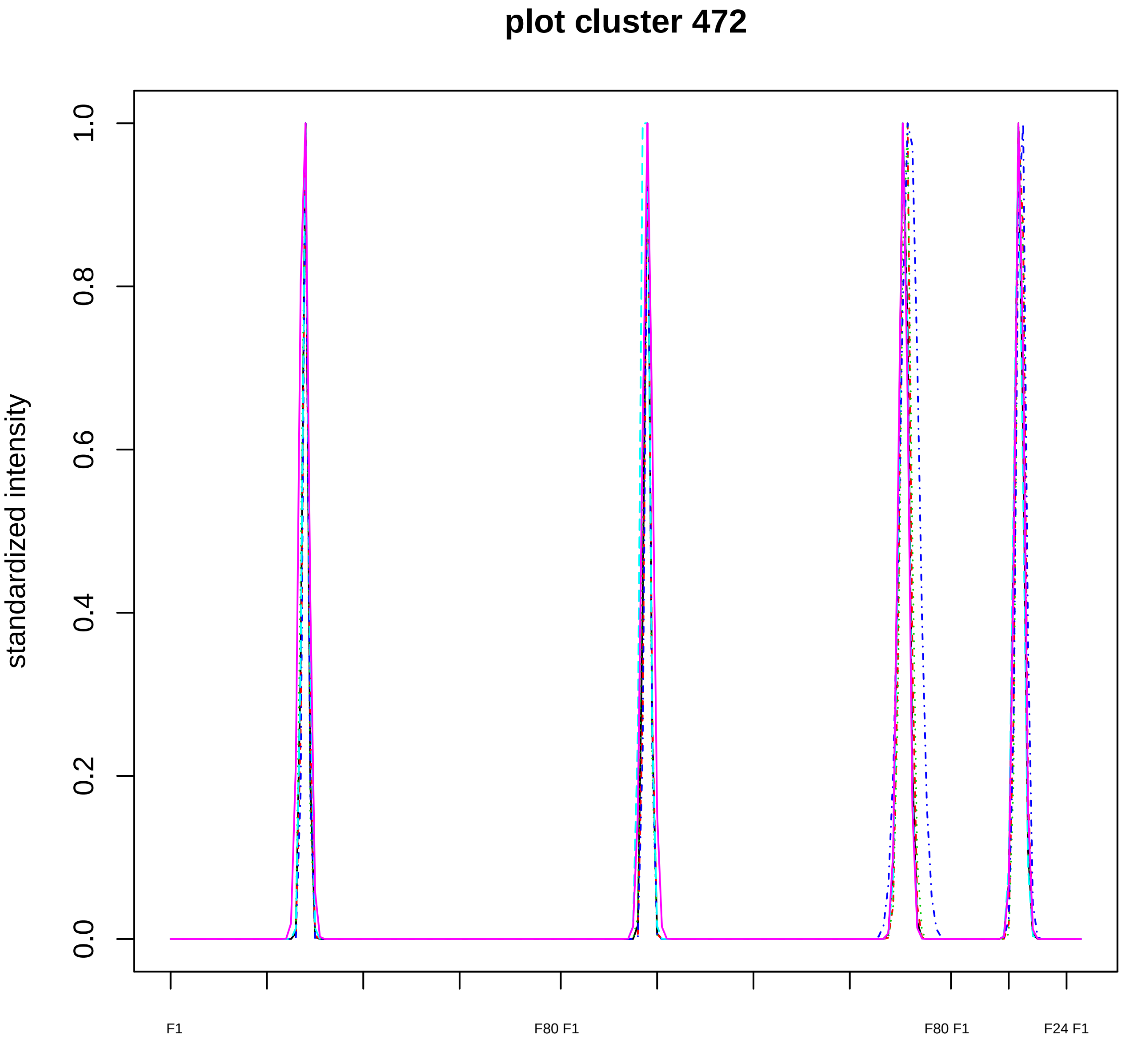
plot cluster 472
standardized intensity
0.0
0.2
0.4
0.6
0.8
1.0
F1
F80 F1
F80 F1
F24 F1

### Dendrogram of cluster 472

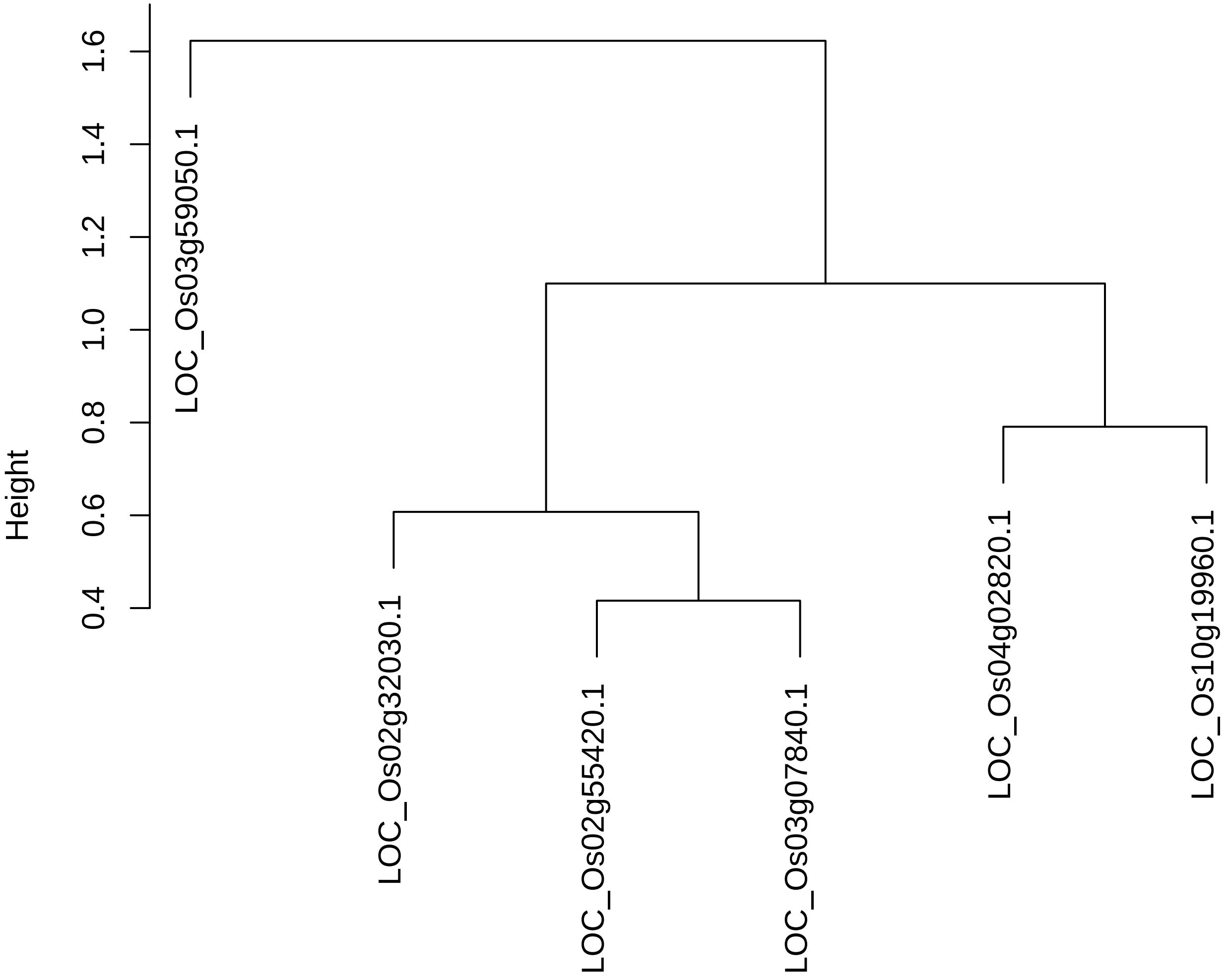


dist(spfitiex12sec12[lab[, 1] == i, ])
hclust (*, "ward.D2")

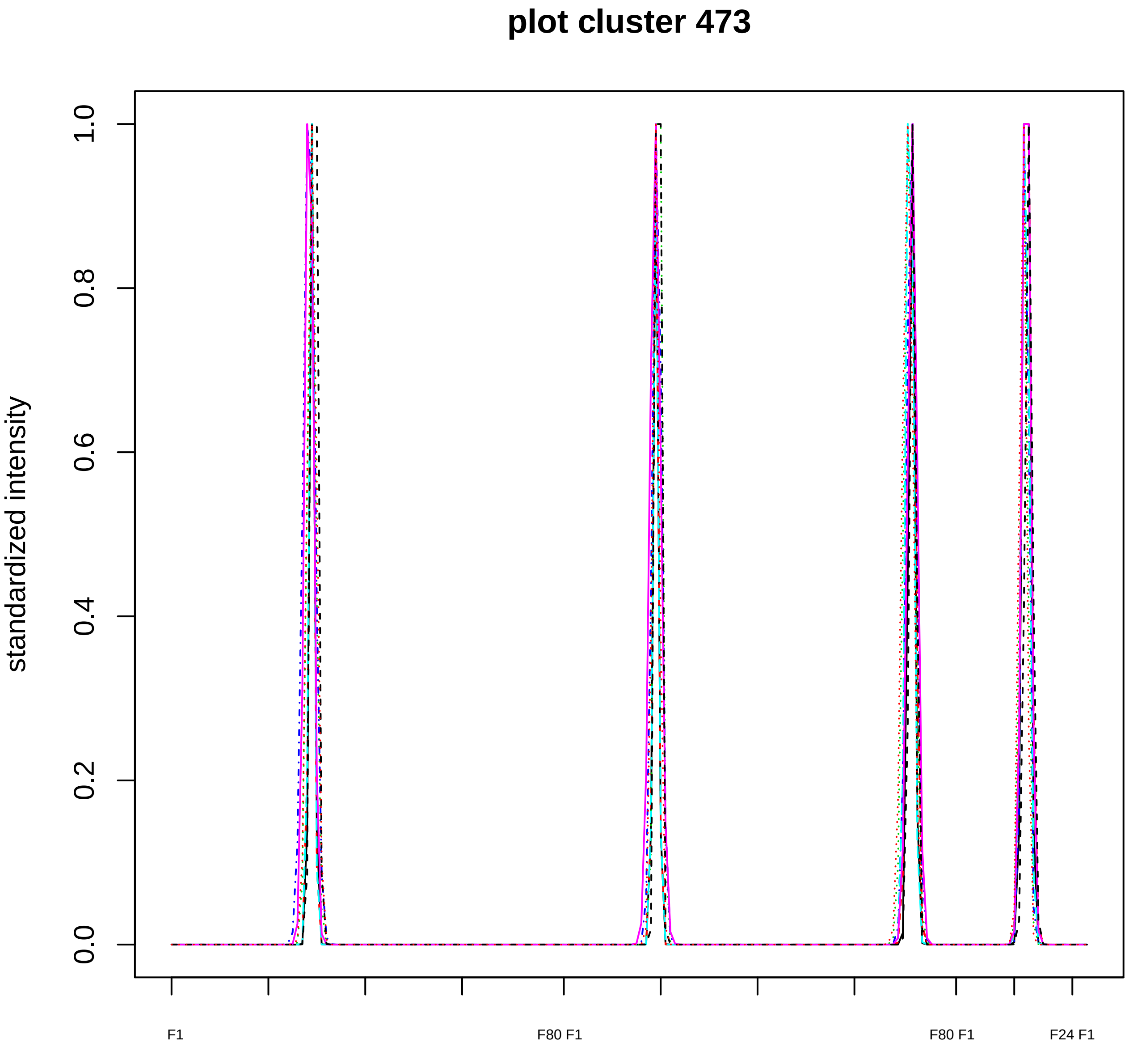

plot cluster 473
standardized intensity
0.0
0.2
0.4
0.6
0.8
1.0
F1
F80 F1
F80 F1
F24 F1

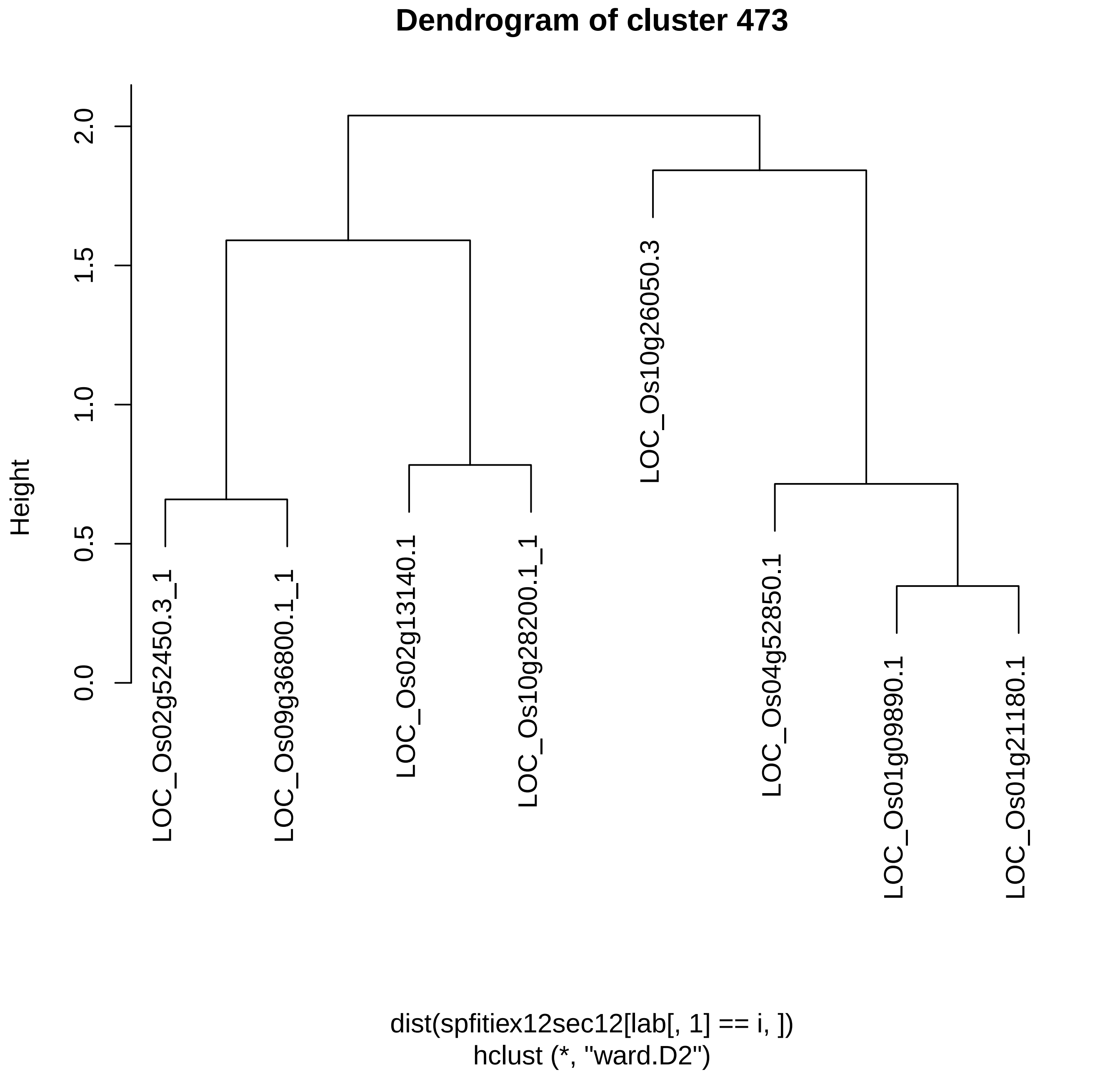

Dendrogram of cluster 473
Height
0.0
0.5
1.0
1.5
2.0
LOC_Os02g52450.3_1
LOC_Os09g36800.1_1
LOC_Os02g13140.1
LOC_Os10g28200.1_1
LOC_Os10g26050.3
LOC_Os04g52850.1
LOC_Os01g09890.1
LOC_Os01g21180.1
dist(spfitiex12sec12[lab[, 1] == i, ])
hclust (*, "ward.D2")

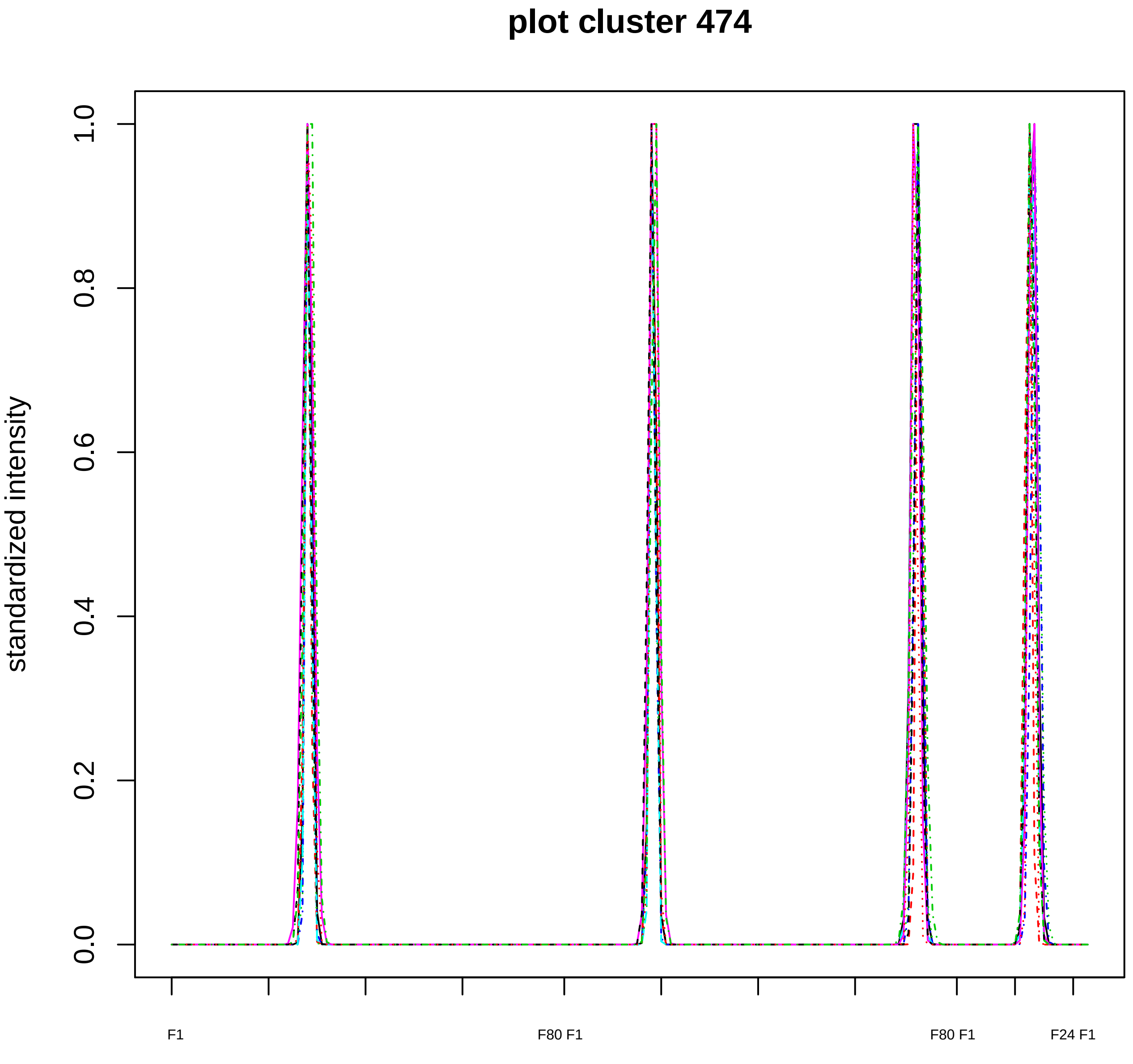
plot cluster 474
standardized intensity
0.0
0.2
0.4
0.6
0.8
1.0
F1
F80 F1
F80 F1
F24 F1

### Dendrogram of cluster 474

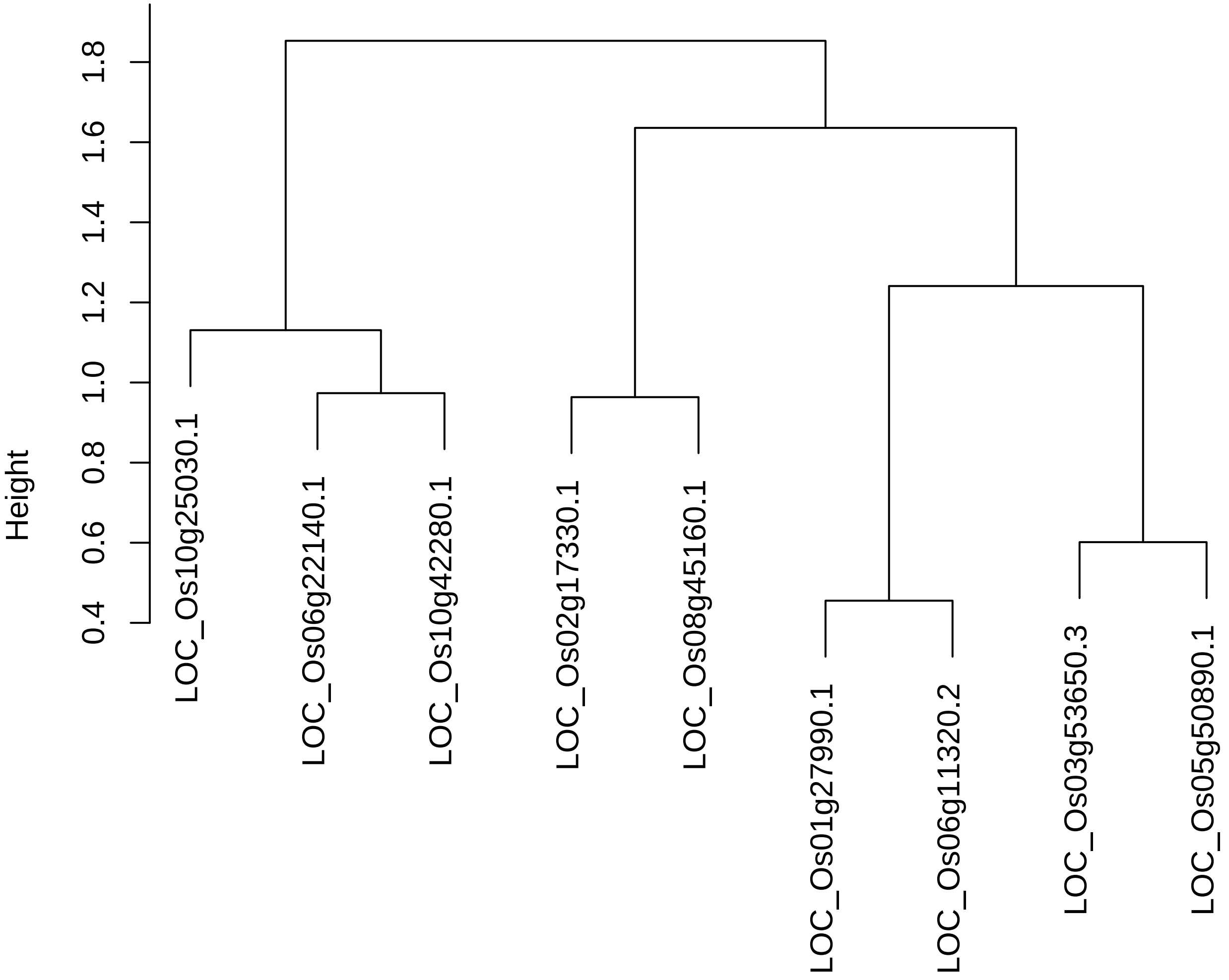


dist(spfitiex12sec12[lab[, 1] == i, ])
hclust (*, "ward.D2")

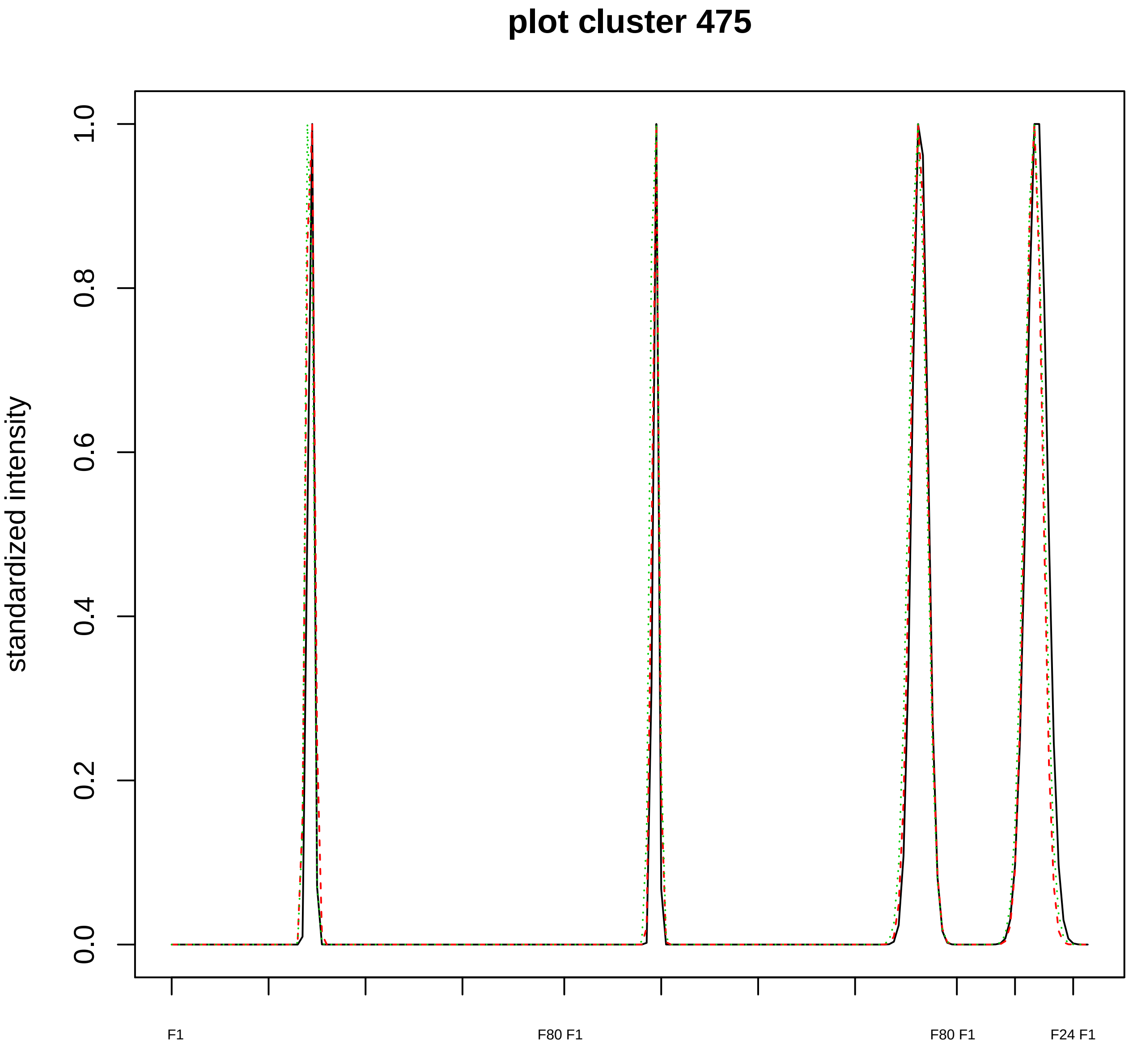

plot cluster 475
standardized intensity
0.0
0.2
0.4
0.6
0.8
1.0
F1
F80 F1
F80 F1
F24 F1

#### Dendrogram of cluster 475

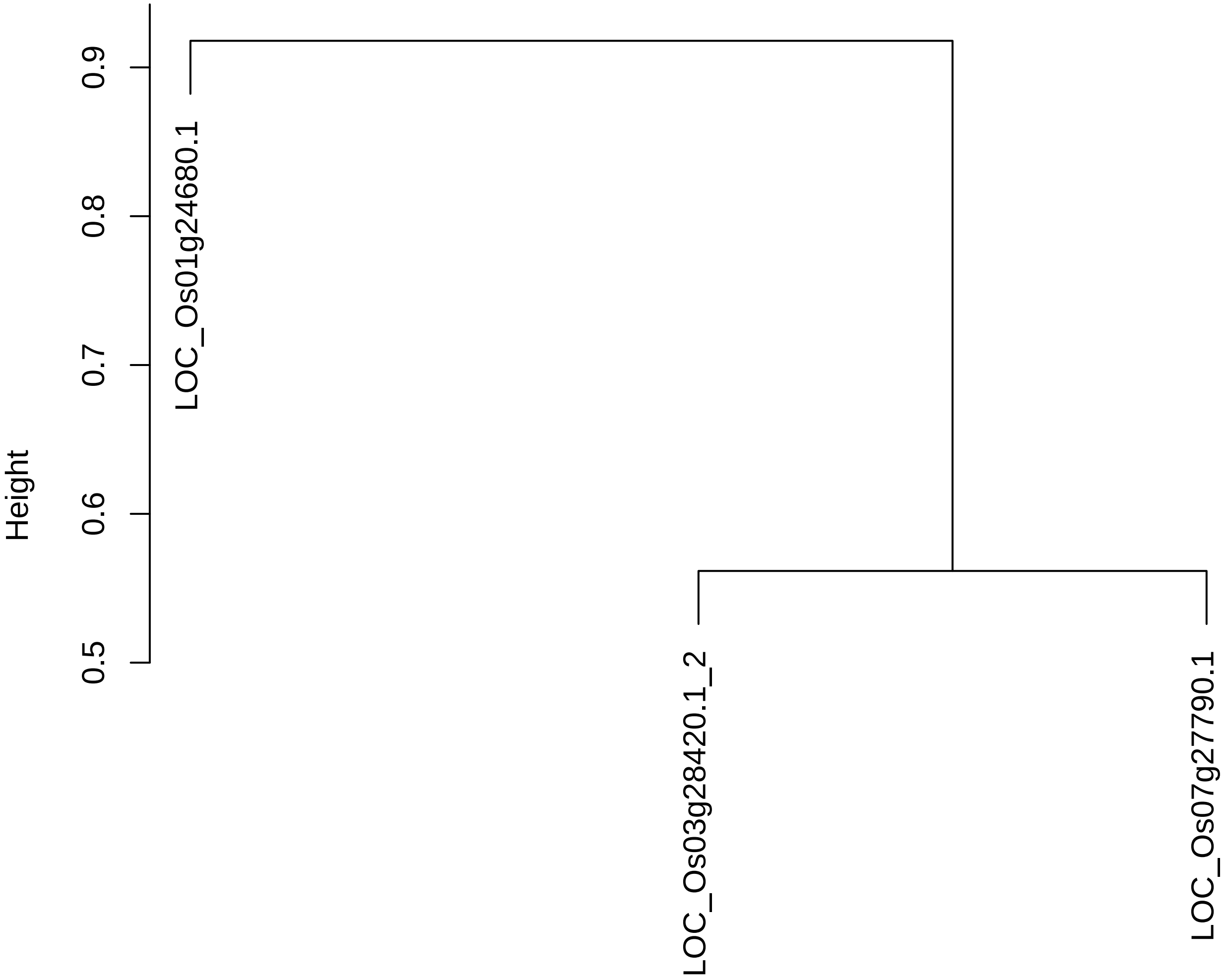


dist(spfitiex12sec12[lab[, 1] == i, ])
hclust (*, "ward.D2")

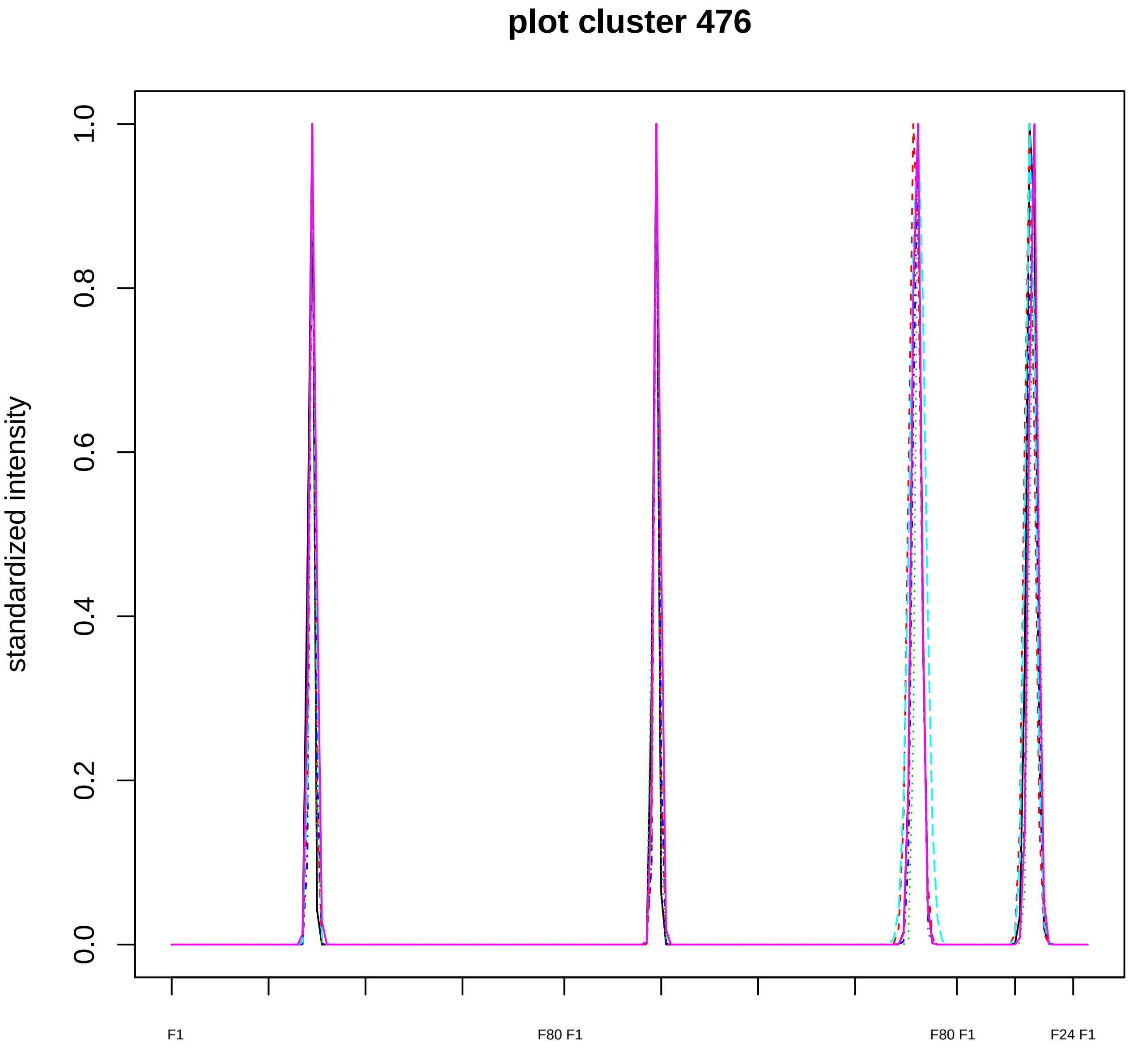
plot cluster 476
standardized intensity
0.0
0.2
0.4
0.6
0.8
1.0
F1
F80 F1
F80 F1
F24 F1

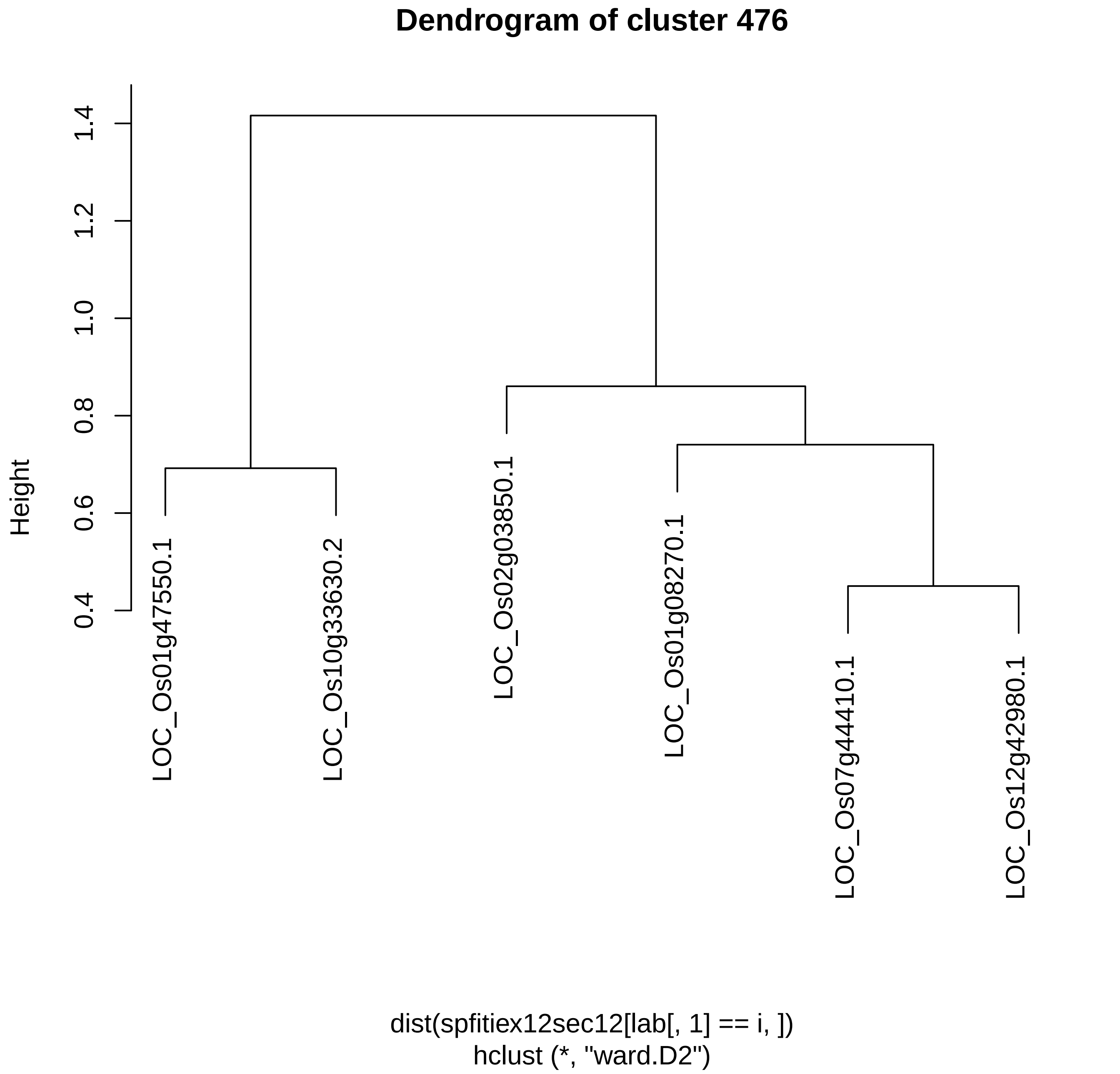
Dendrogram of cluster 476
Height
0.4
0.6
0.8
1.0
1.2
1.4
LOC_Os01g47550.1
LOC_Os10g33630.2
LOC_Os02g03850.1
LOC_Os01g08270.1
LOC_Os07g44410.1
LOC_Os12g42980.1
dist(spfitiex12sec12[lab[, 1] == i, ])
hclust (*, "ward.D2")

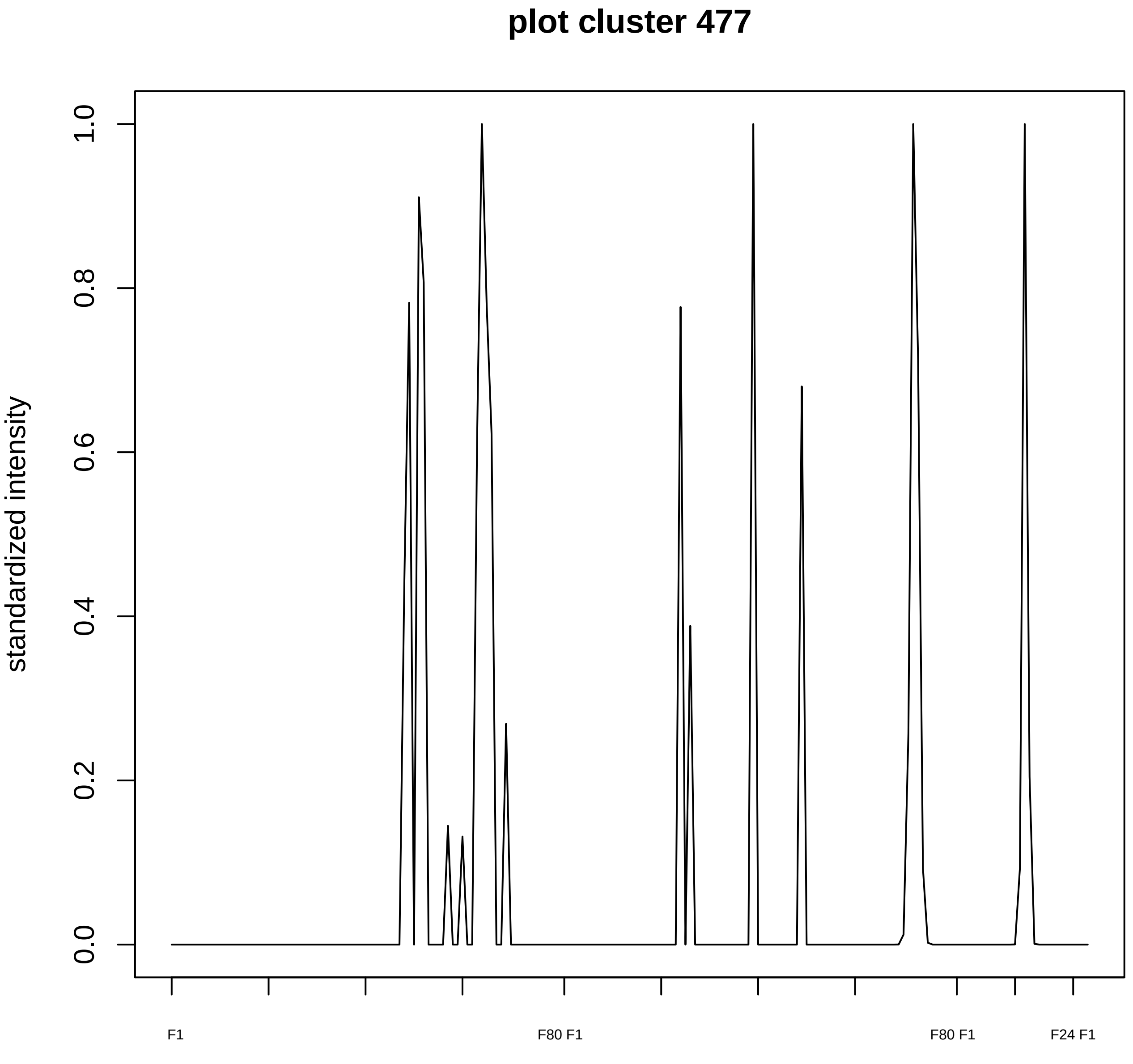

plot cluster 477
standardized intensity
0.0
0.2
0.4
0.6
0.8
1.0
F1
F80 F1
F80 F1
F24 F1

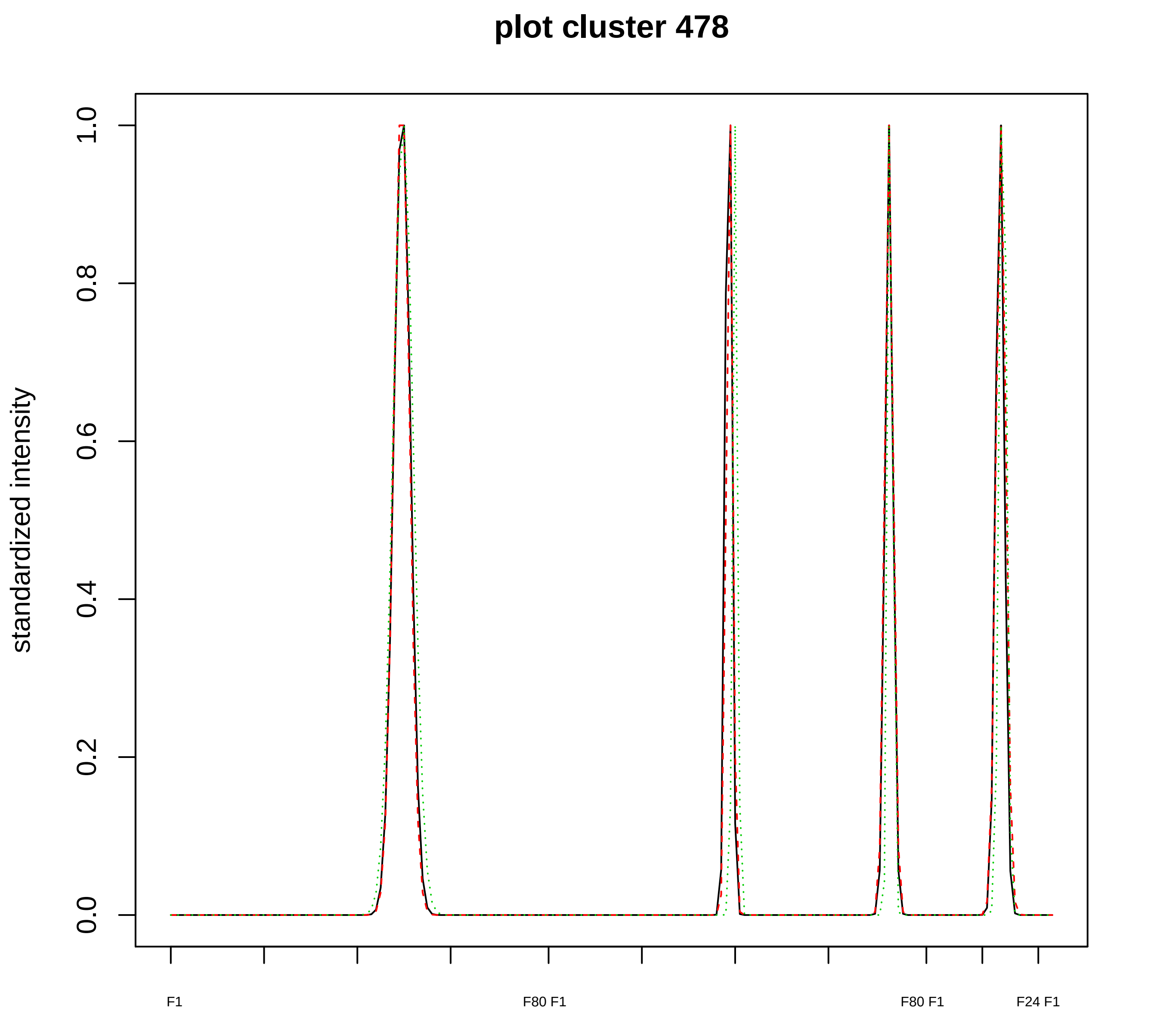

plot cluster 478
standardized intensity
0.0
0.2
0.4
0.6
0.8
1.0
F1
F80 F1
F80 F1
F24 F1

### Dendrogram of cluster 478

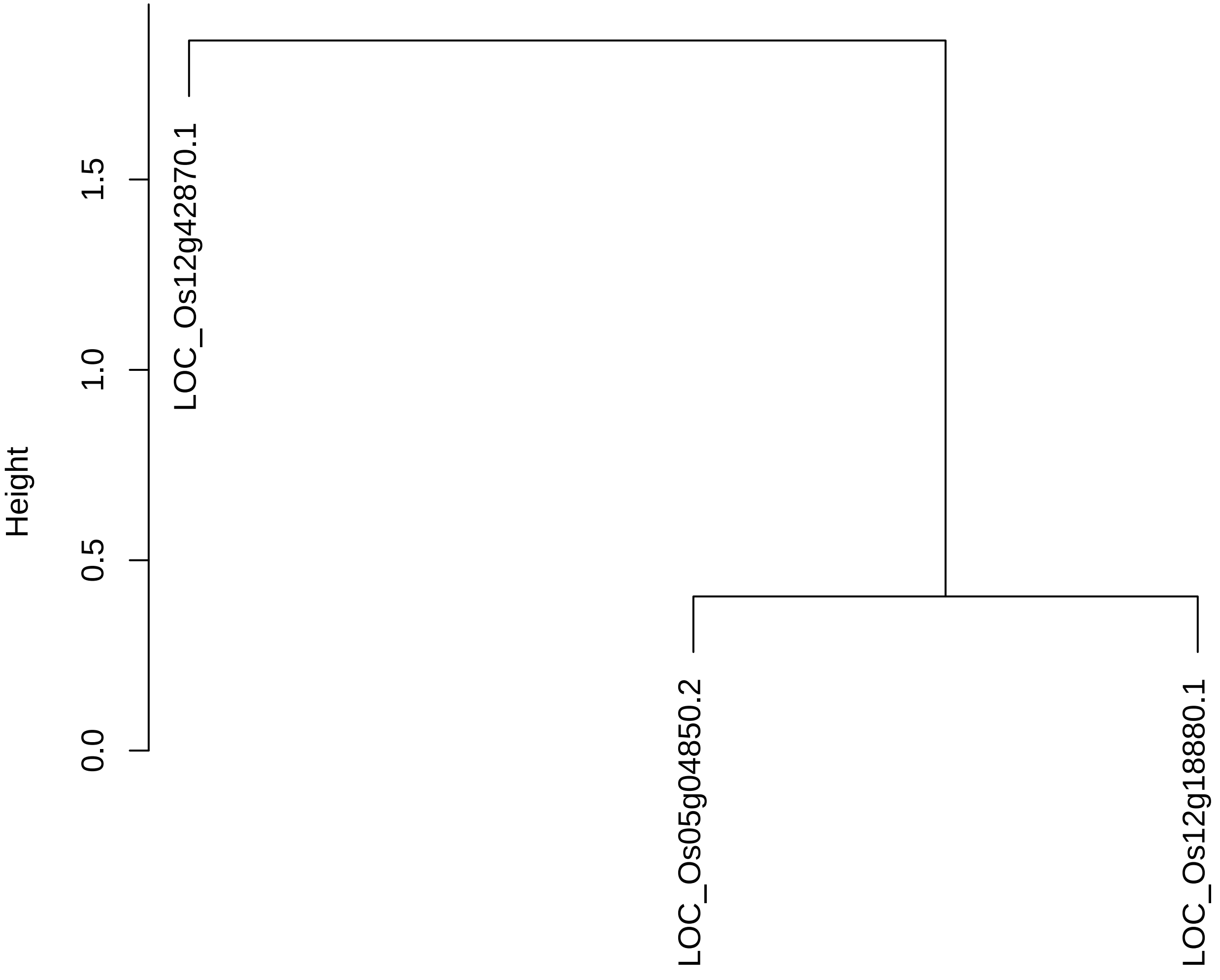


dist(spfitiex12sec12[lab[, 1] == i, ])
hclust (*, "ward.D2")

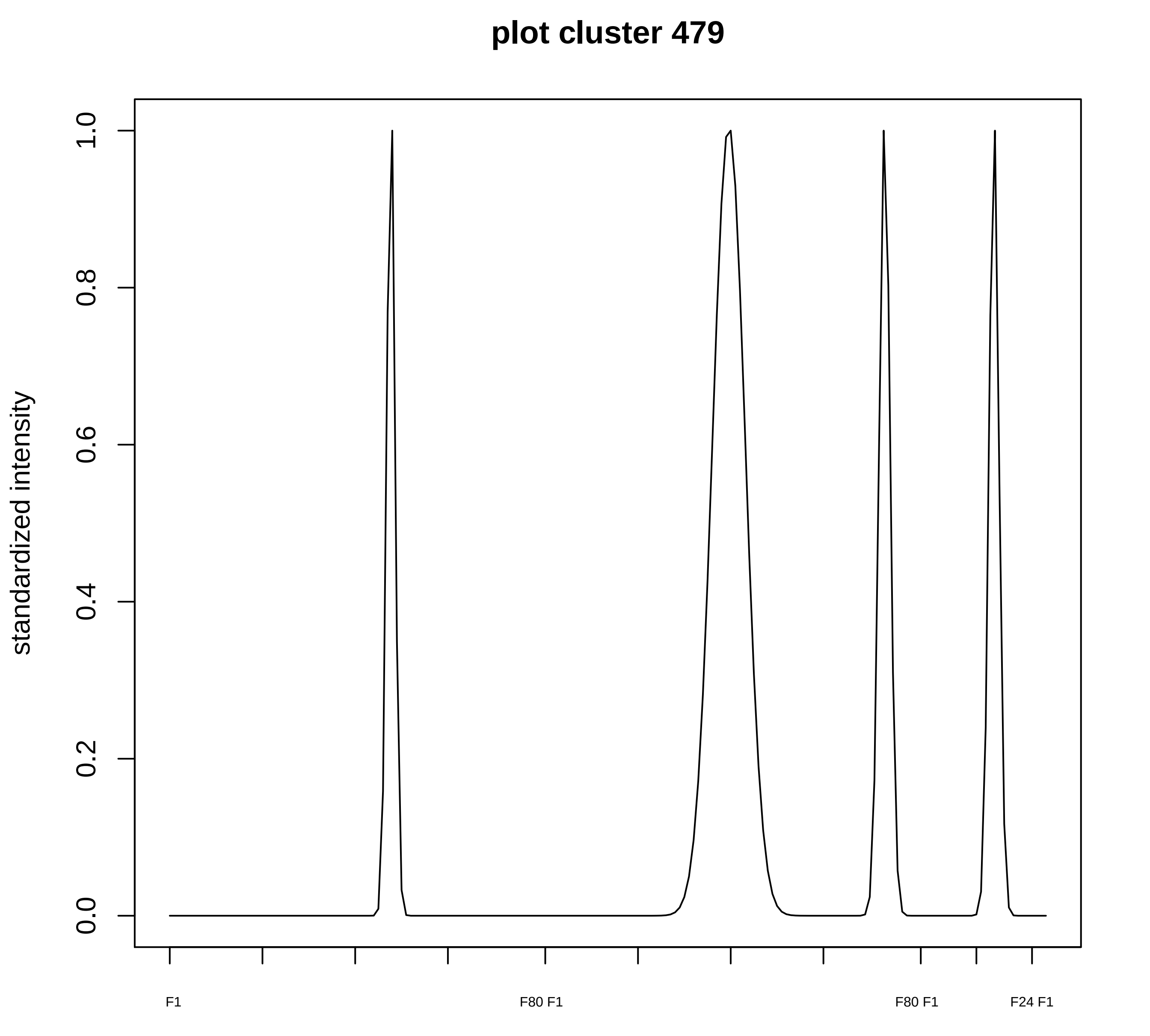
plot cluster 479
standardized intensity
1.0
0.8
0.6
0.4
0.2
0.0
F1
F80 F1
F80 F1
F24 F1

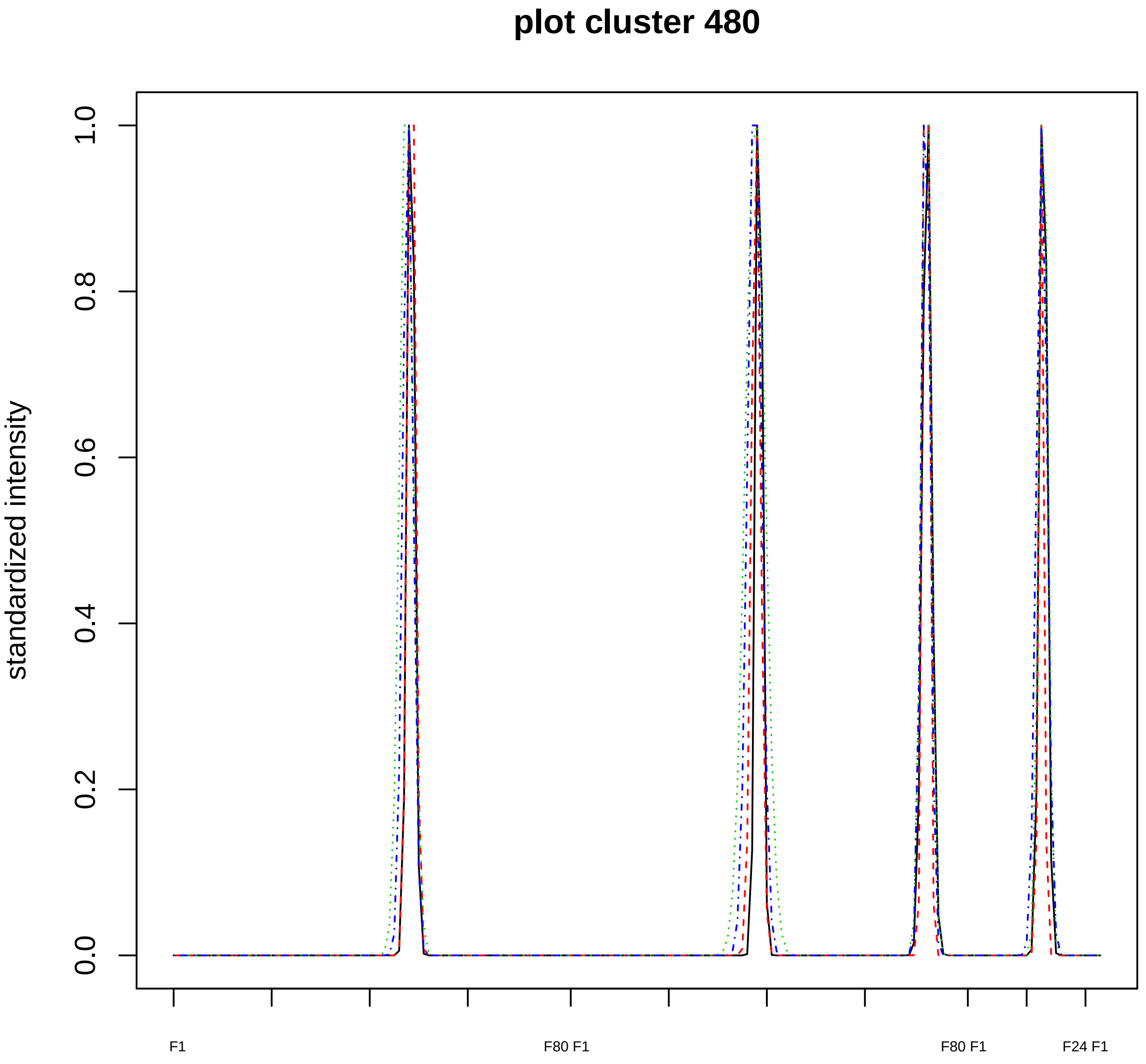
plot cluster 480
standardized intensity
0.0
0.2
0.4
0.6
0.8
1.0
F1
F80 F1
F80 F1
F24 F1

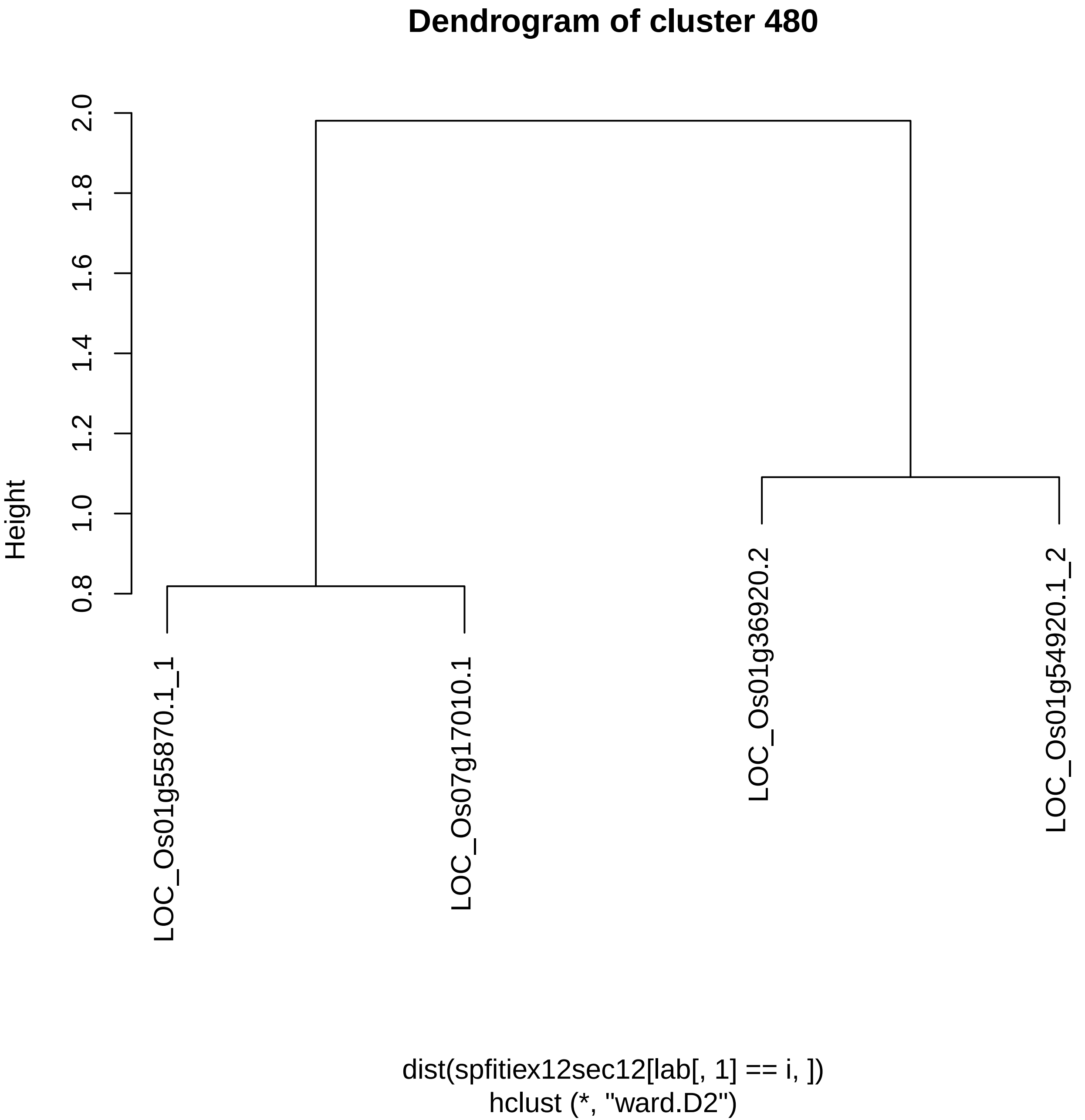
Dendrogram of cluster 480
Height
2.0
1.8
1.6
1.4
1.2
1.0
0.8
LOC_Os01g55870.1_1
LOC_Os07g17010.1
LOC_Os01g36920.2
LOC_Os01g54920.1_2
dist(spfitiex12sec12[lab[, 1] == i, ])
hclust (*, "ward.D2")

#### plot cluster 481

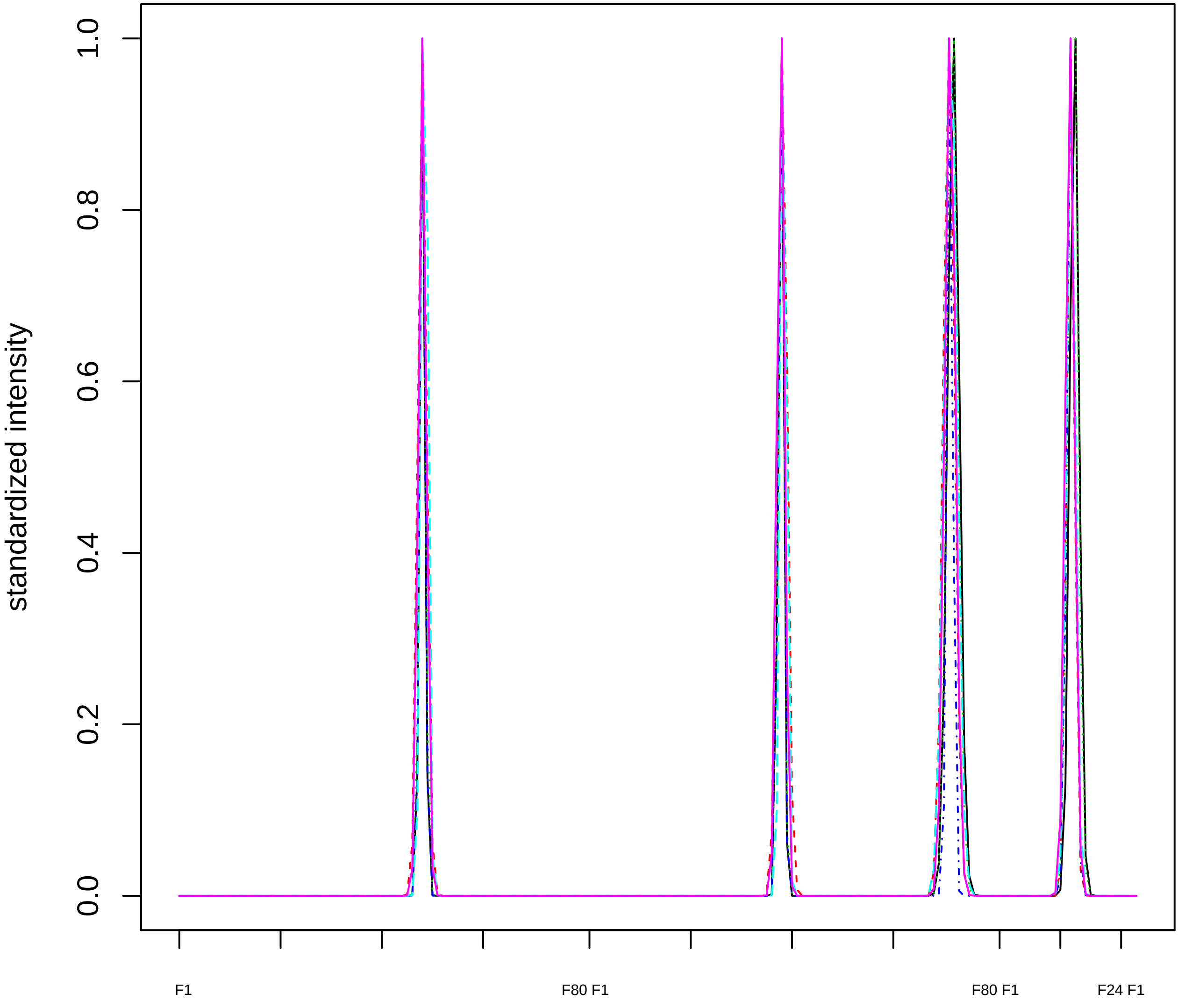

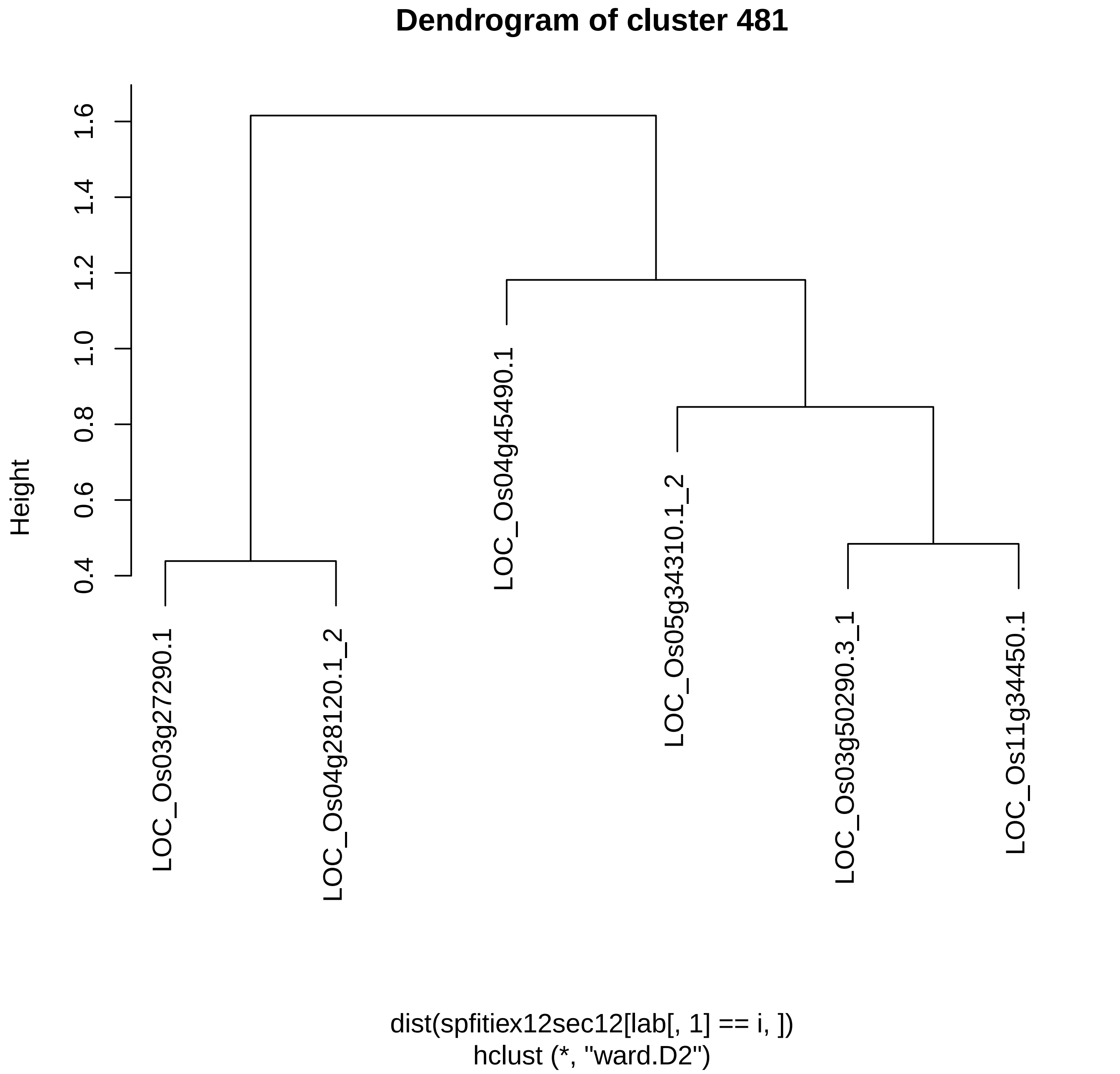
Dendrogram of cluster 481
Height
0.4
0.6
0.8
1.0
1.2
1.4
1.6
LOC_Os03g27290.1
LOC_Os04g28120.1_2
LOC_Os04g45490.1
LOC_Os05g34310.1_2
LOC_Os03g50290.3_1
LOC_Os11g34450.1
dist(spfitiex12sec12[lab[, 1] == i, ])
hclust (*, "ward.D2")

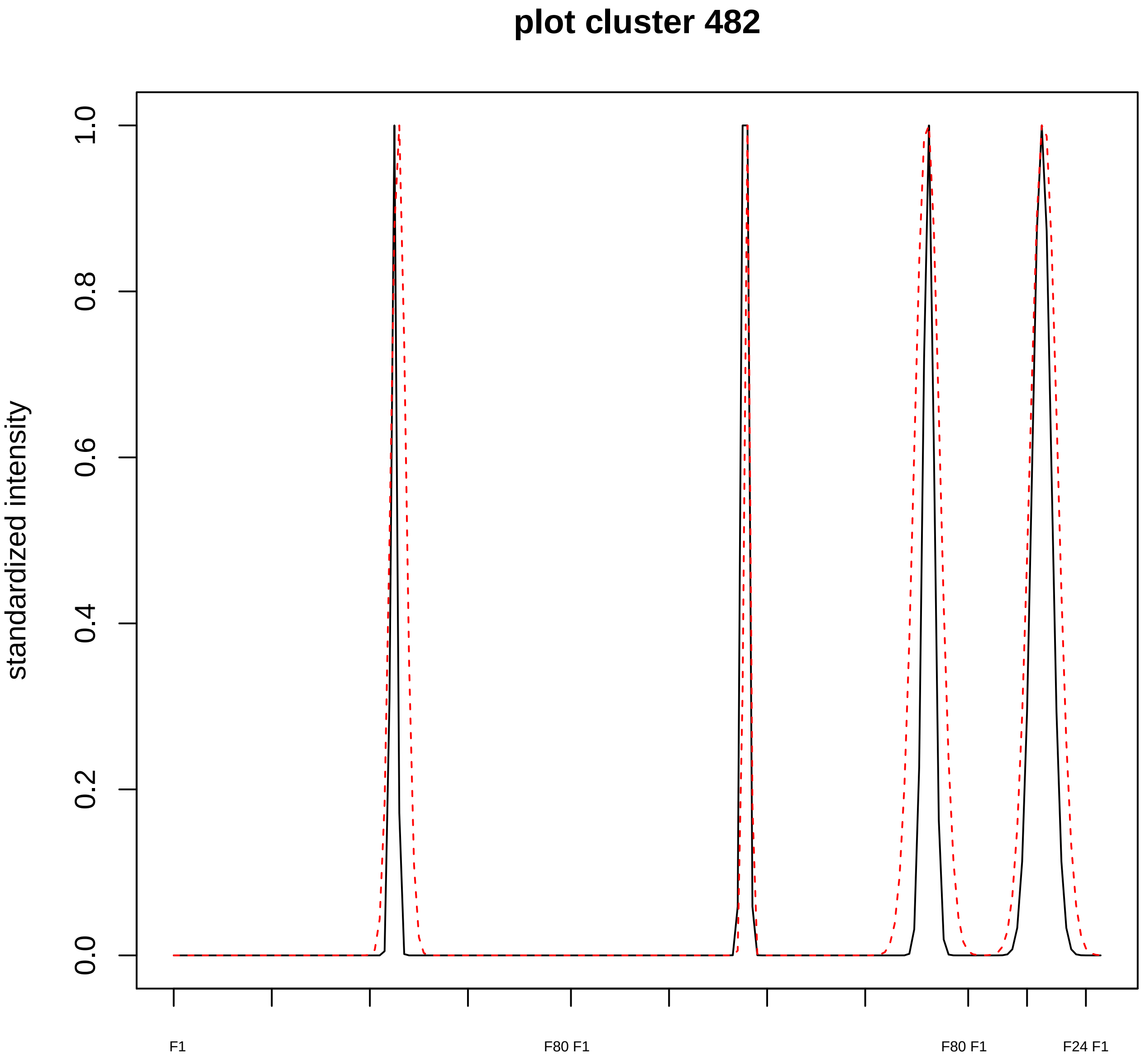
plot cluster 482
standardized intensity
0.0
0.2
0.4
0.6
0.8
1.0
F1
F80 F1
F80 F1
F24 F1

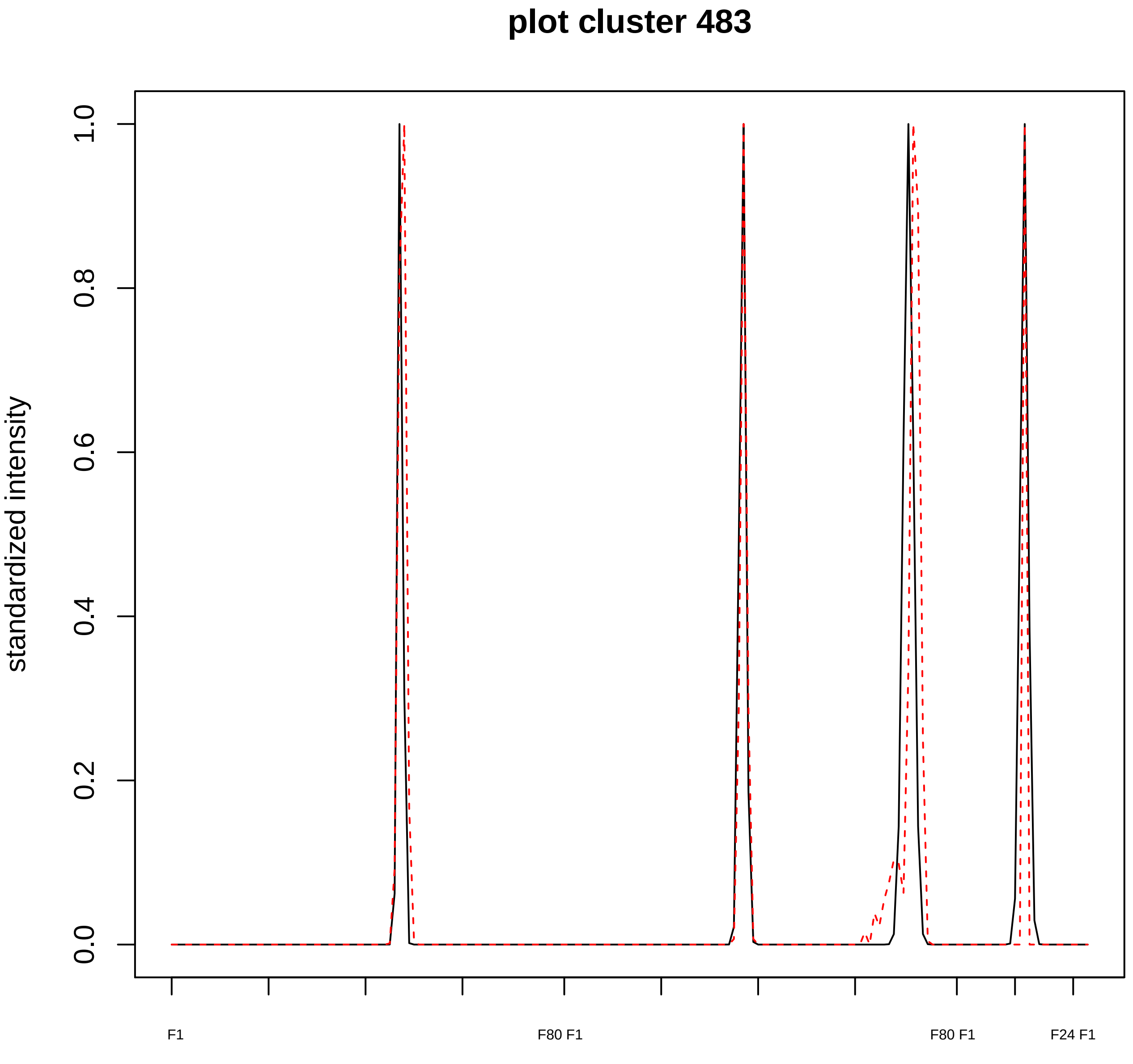

plot cluster 483
standardized intensity
0.0
0.2
0.4
0.6
0.8
1.0
F1
F80 F1
F80 F1
F24 F1

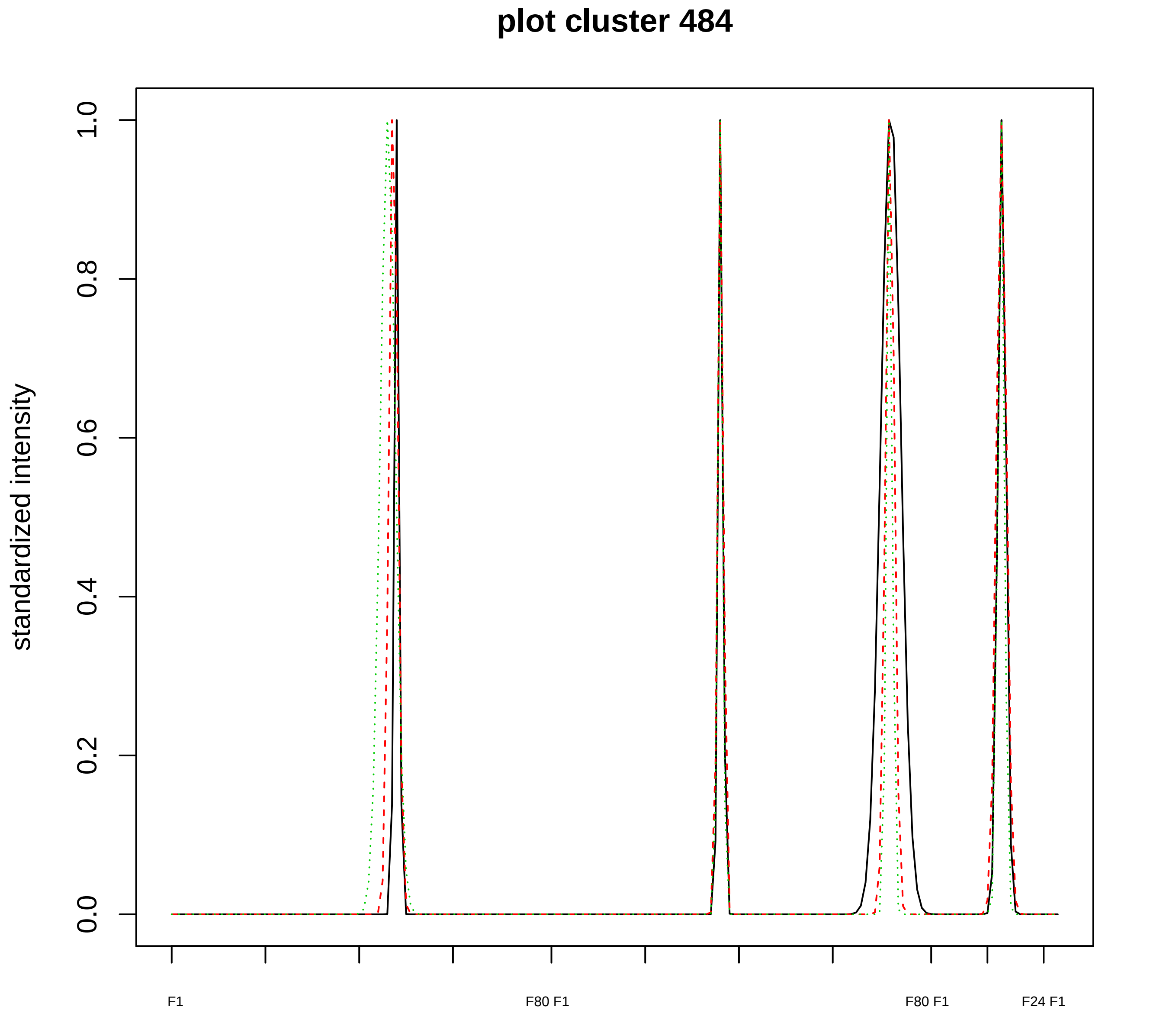

plot cluster 484
standardized intensity
0.0
0.2
0.4
0.6
0.8
1.0
F1
F80 F1
F80 F1
F24 F1

#### Dendrogram of cluster 484

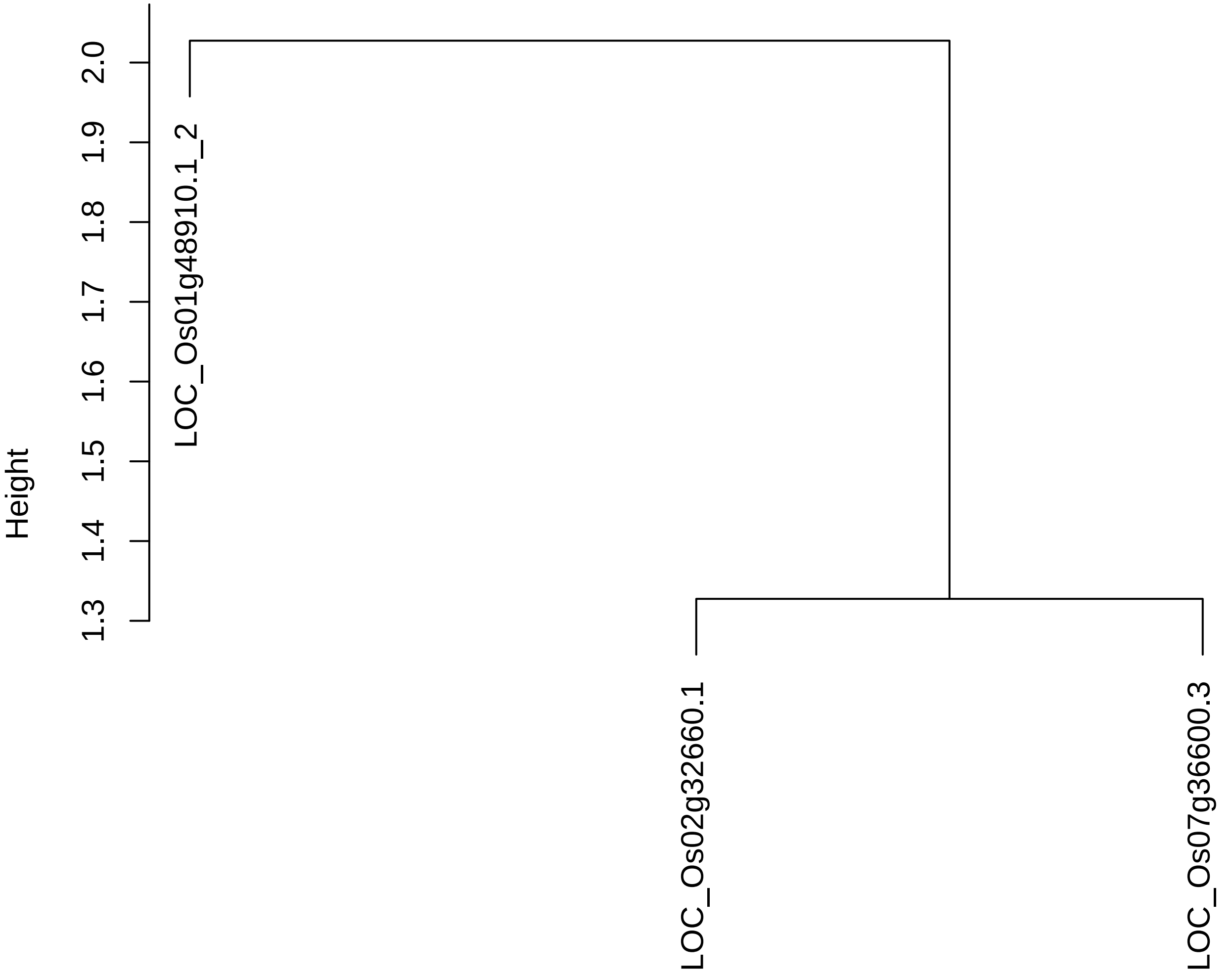


dist(spfitiex12sec12[lab[, 1] == i, ])
hclust (*, "ward.D2")

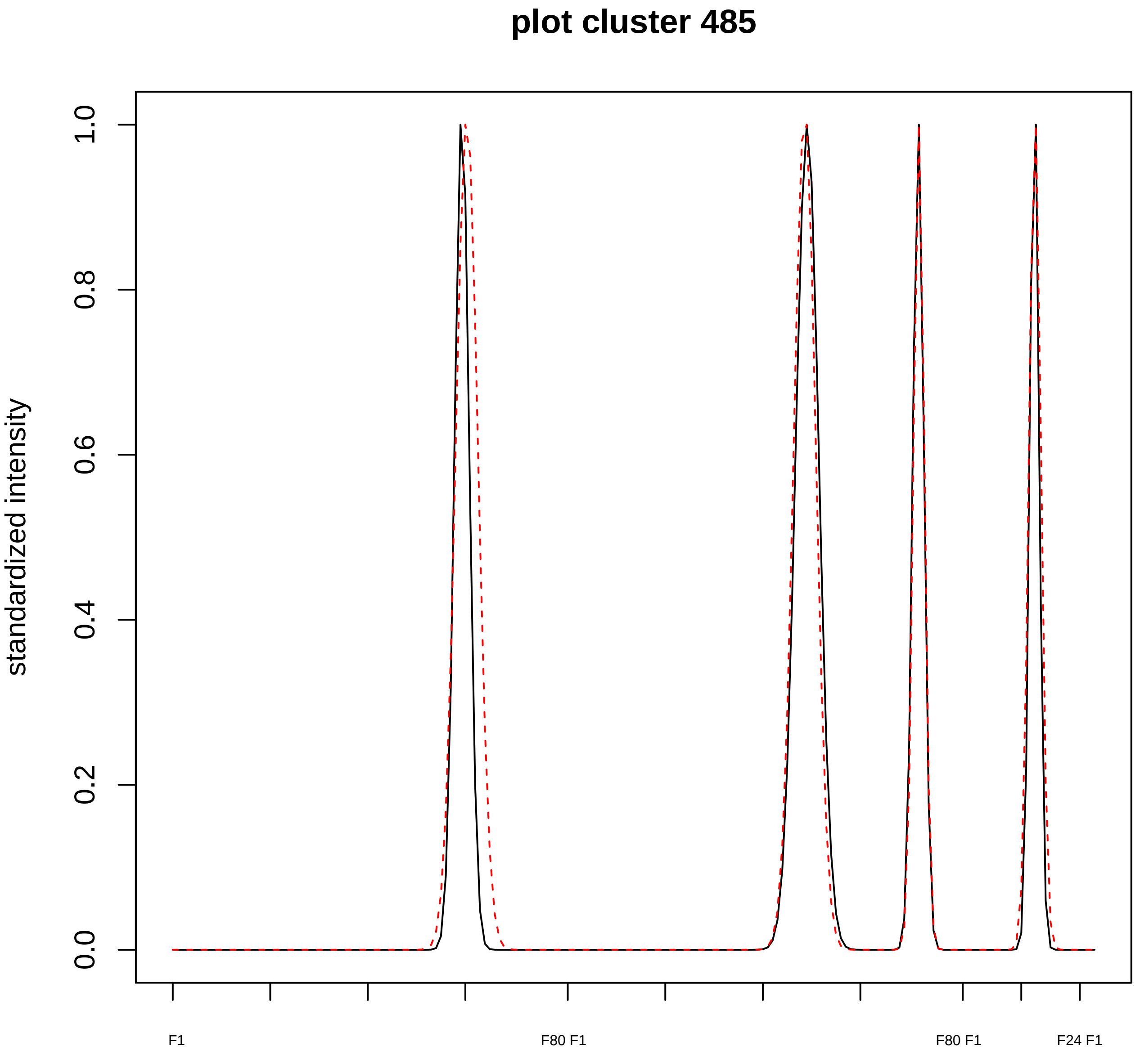

plot cluster 485
standardized intensity
0.0
0.2
0.4
0.6
0.8
1.0
F1
F80 F1
F80 F1
F24 F1

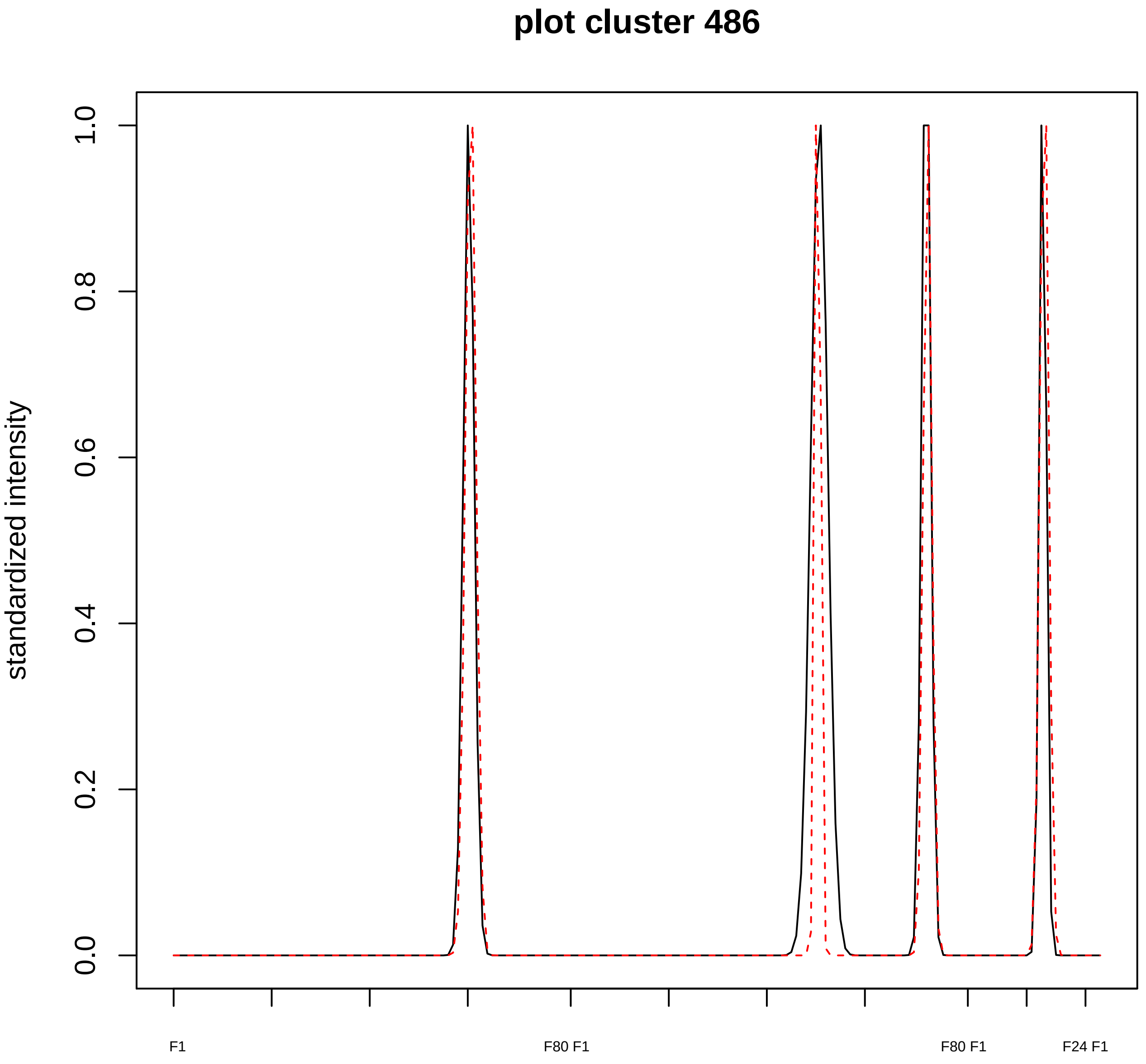
plot cluster 486
standardized intensity
0.0
0.2
0.4
0.6
0.8
1.0
F1
F80 F1
F80 F1
F24 F1

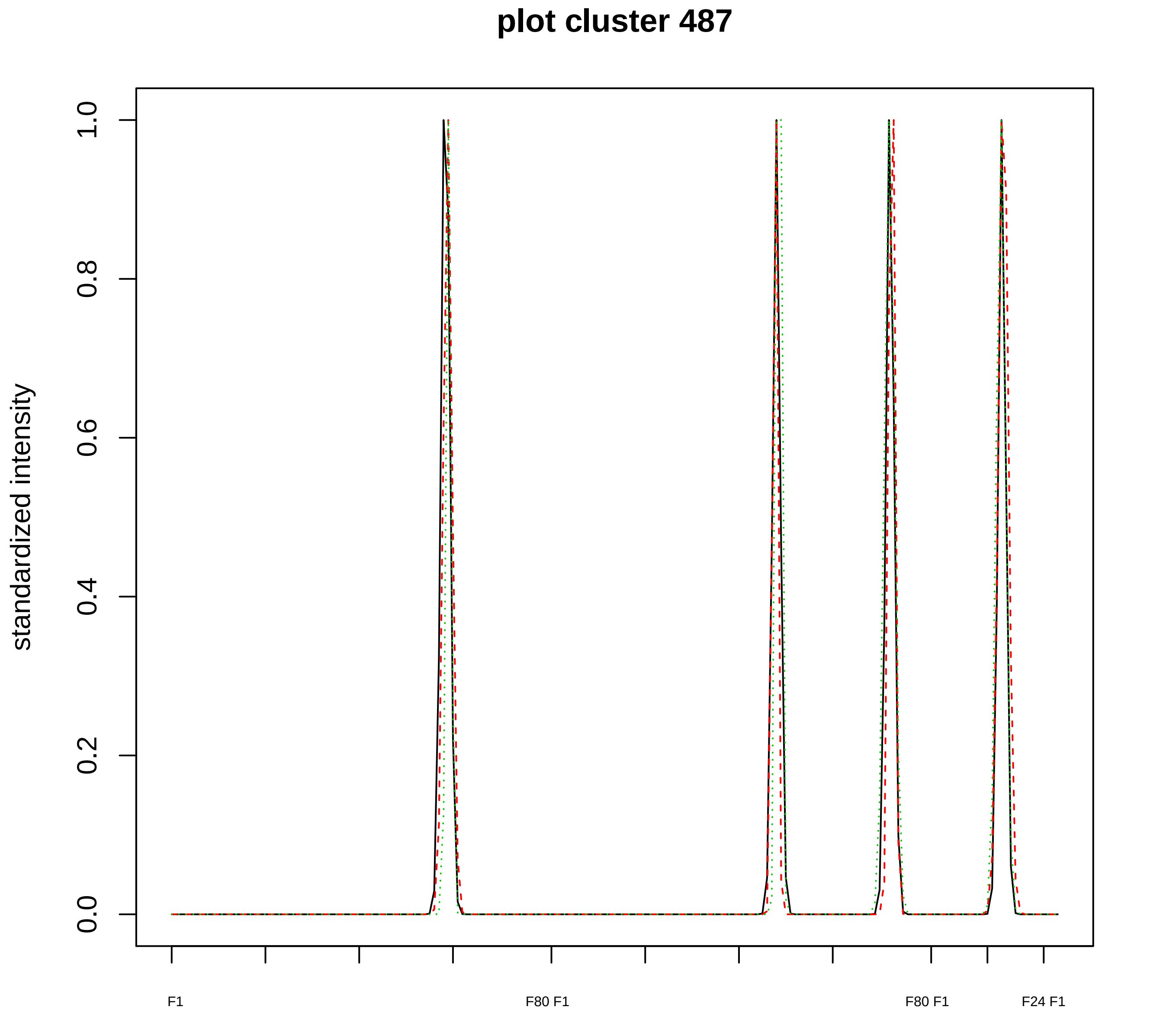

plot cluster 487
standardized intensity
0.0
0.2
0.4
0.6
0.8
1.0
F1
F80 F1
F80 F1
F24 F1

#### Dendrogram of cluster 487

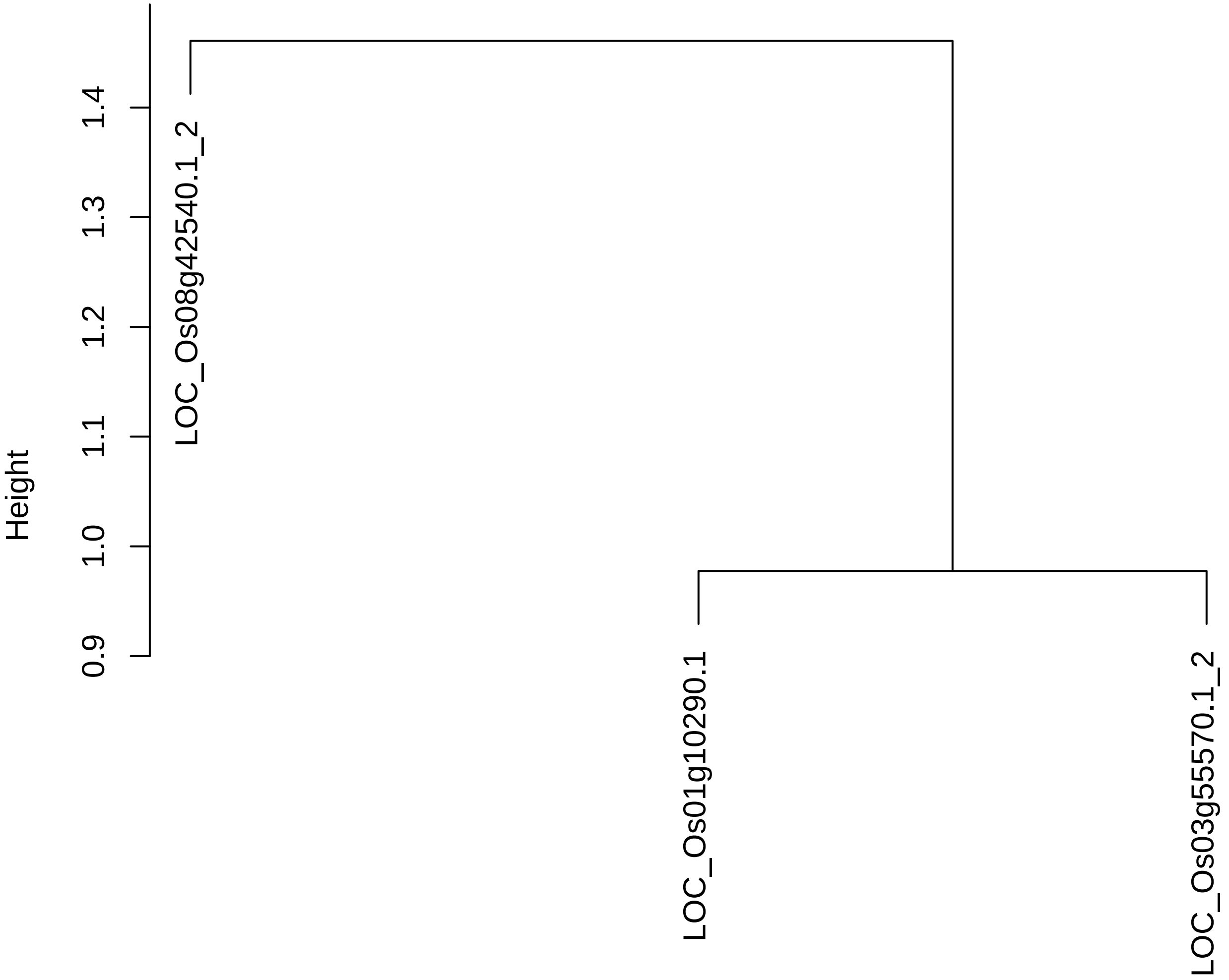


dist(spfitiex12sec12[lab[, 1] == i, ])
hclust (*, "ward.D2")

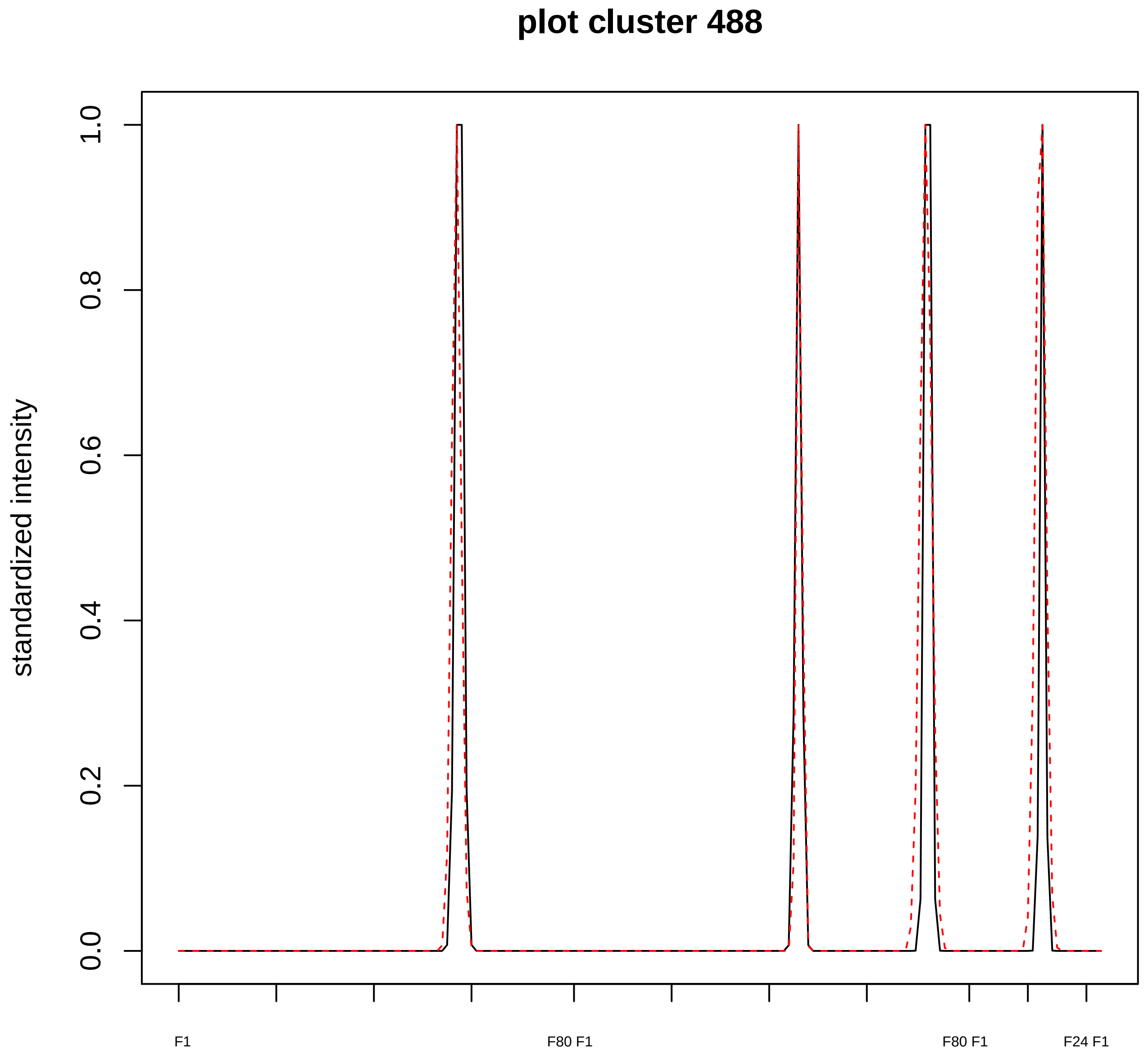

plot cluster 488
standardized intensity
0.0
0.2
0.4
0.6
0.8
1.0
F1
F80 F1
F80 F1
F24 F1

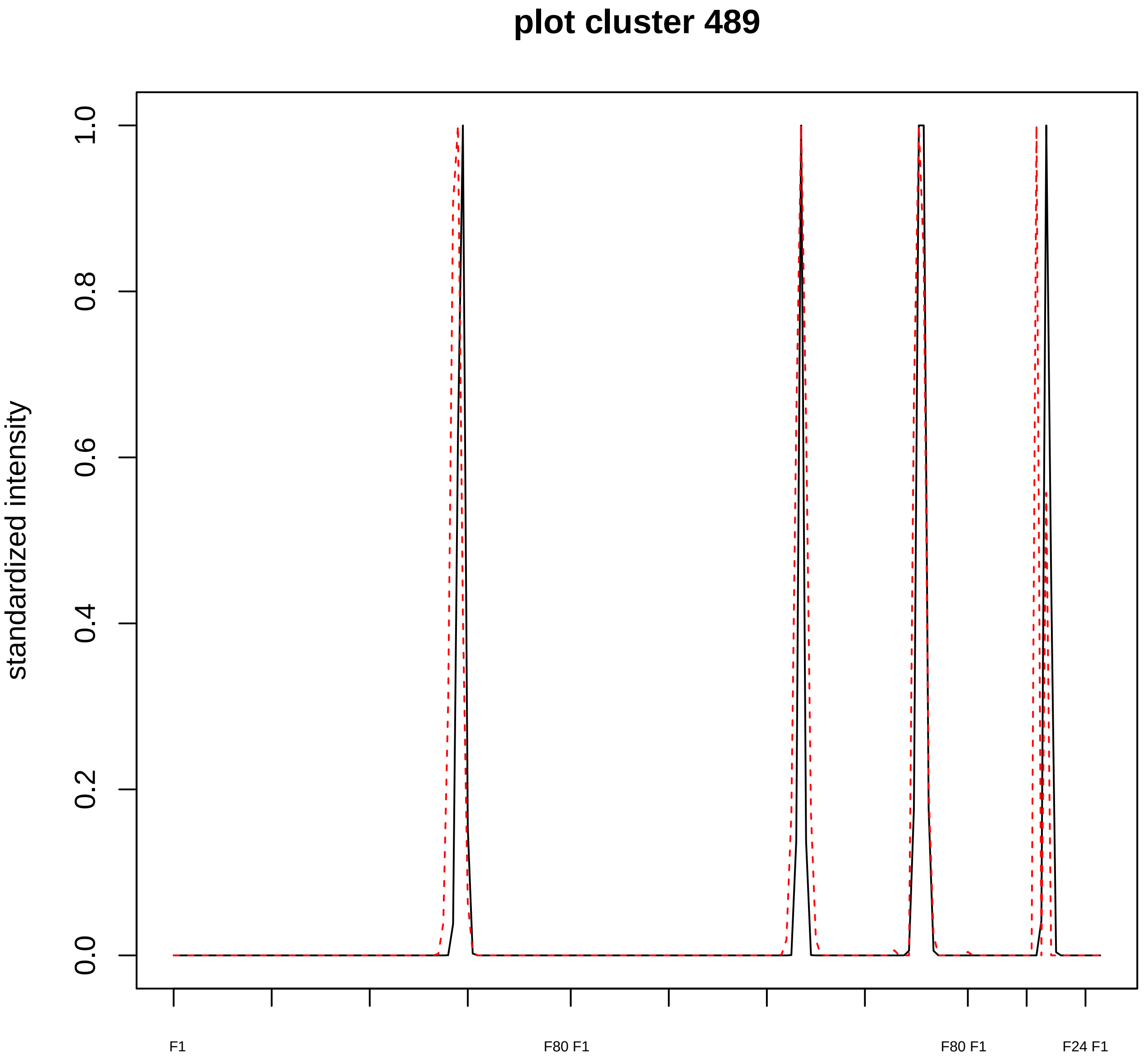

plot cluster 489
standardized intensity
0.0
0.2
0.4
0.6
0.8
1.0
F1
F80 F1
F80 F1
F24 F1

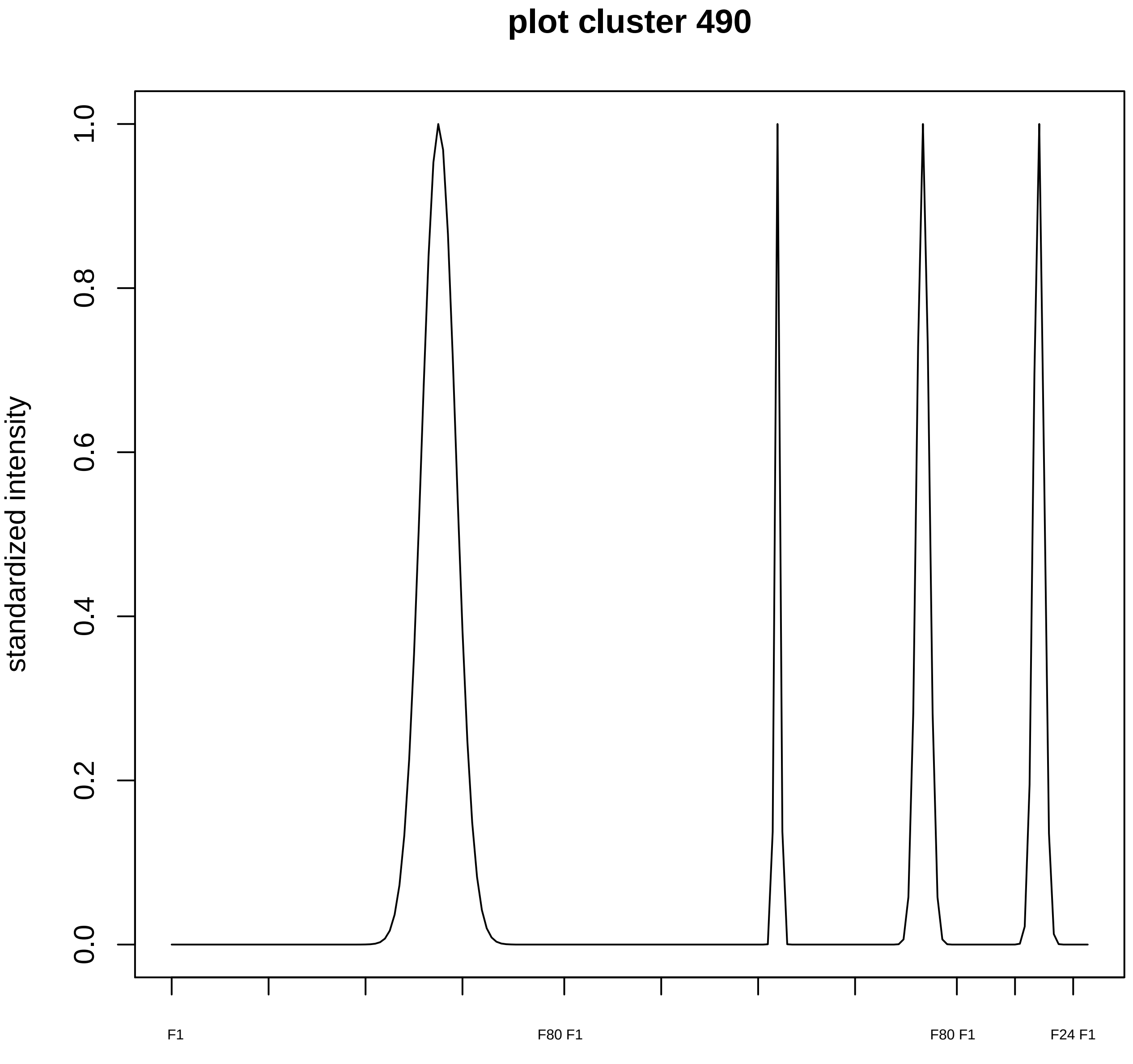
plot cluster 490
standardized intensity
0.0
0.2
0.4
0.6
0.8
1.0
F1
F80 F1
F80 F1
F24 F1

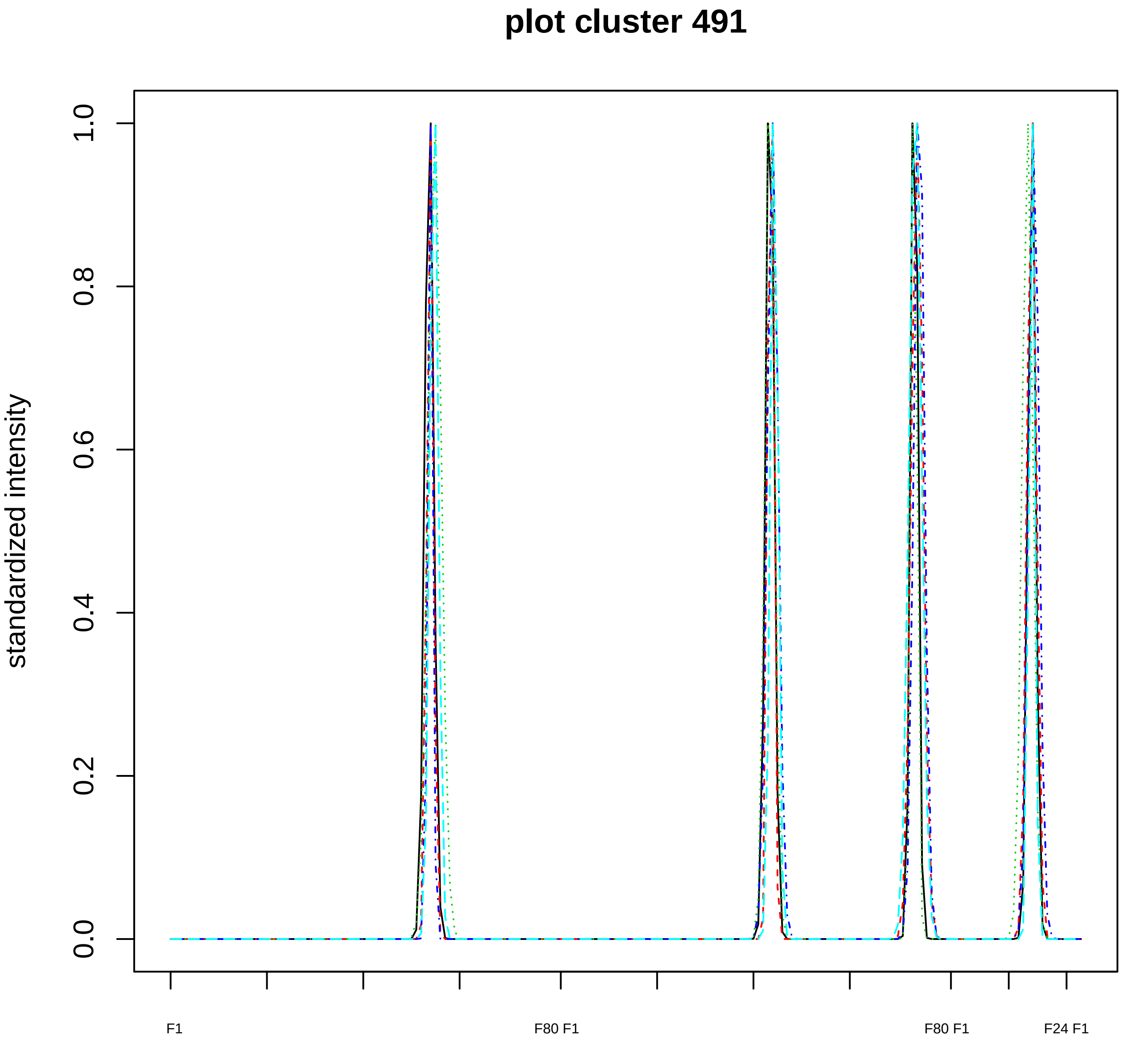

plot cluster 491
standardized intensity
0.0
0.2
0.4
0.6
0.8
1.0
F1
F80 F1
F80 F1
F24 F1

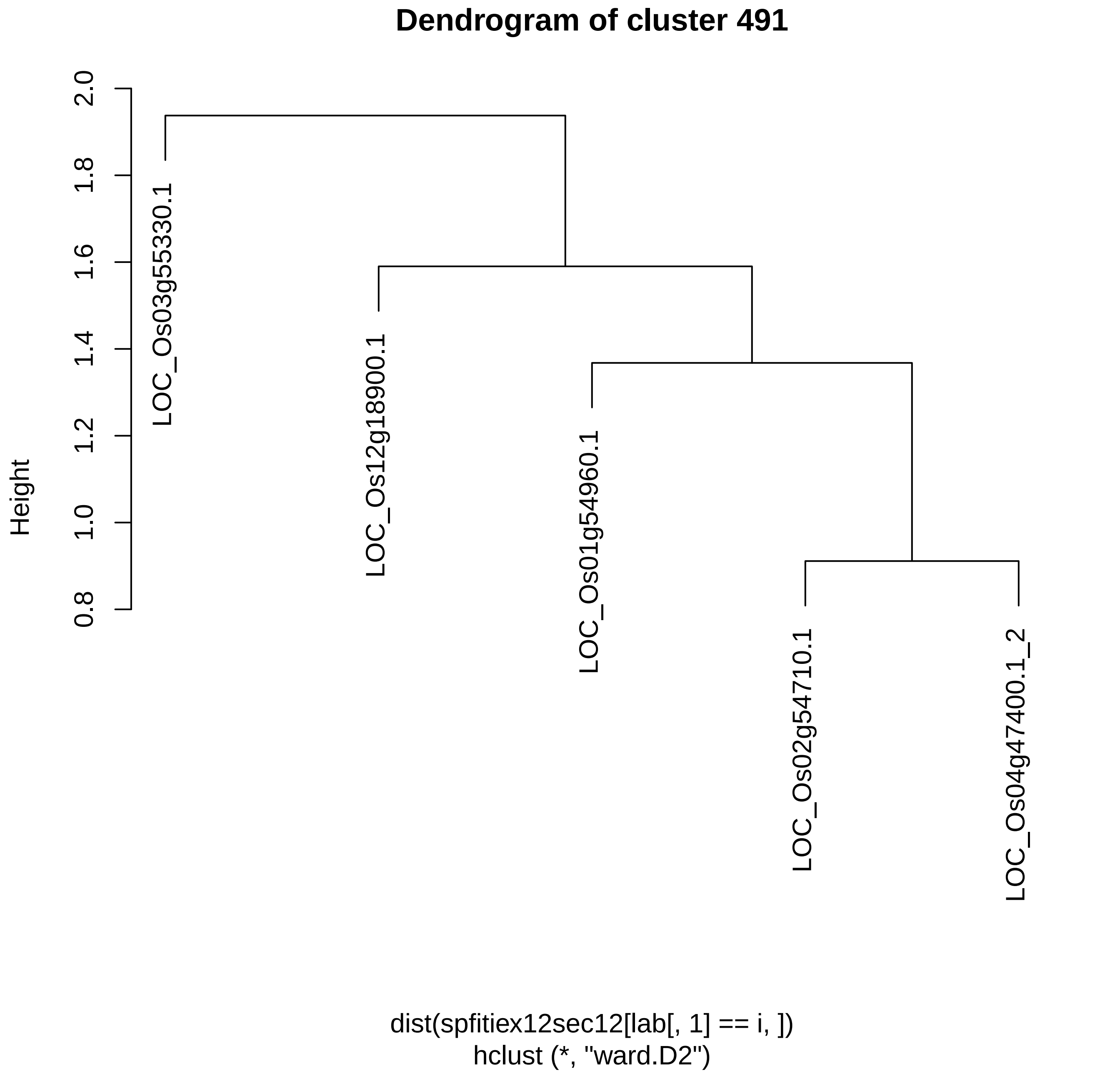
Dendrogram of cluster 491
Height
2.0
1.8
1.6
1.4
1.2
1.0
0.8
LOC_Os03g55330.1
LOC_Os12g18900.1
LOC_Os01g54960.1
LOC_Os02g54710.1
LOC_Os04g47400.1_2
dist(spfitiex12sec12[lab[, 1] == i, ])
hclust (*, "ward.D2")

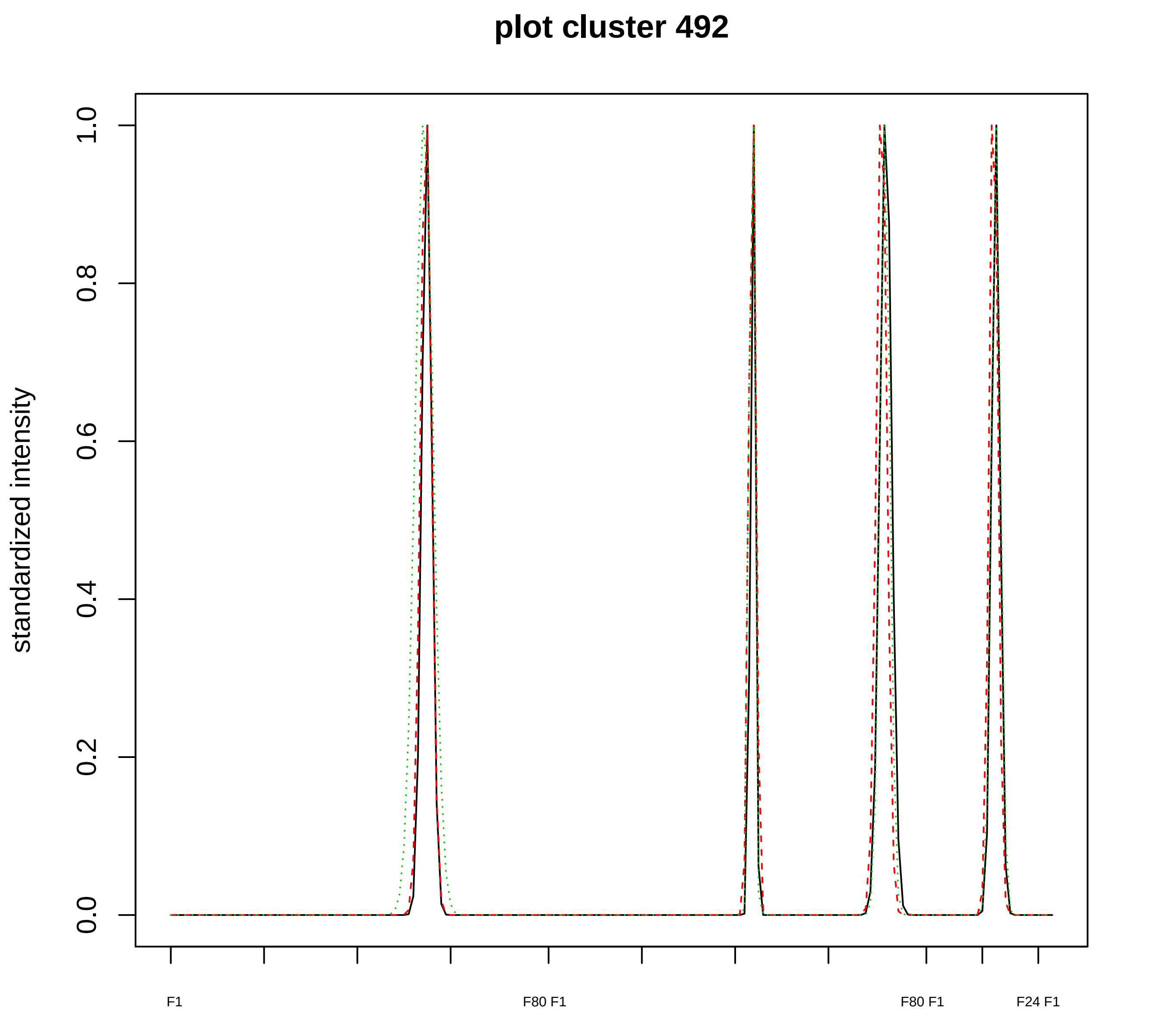
plot cluster 492
standardized intensity
0.0
0.2
0.4
0.6
0.8
1.0
F1
F80 F1
F80 F1
F24 F1

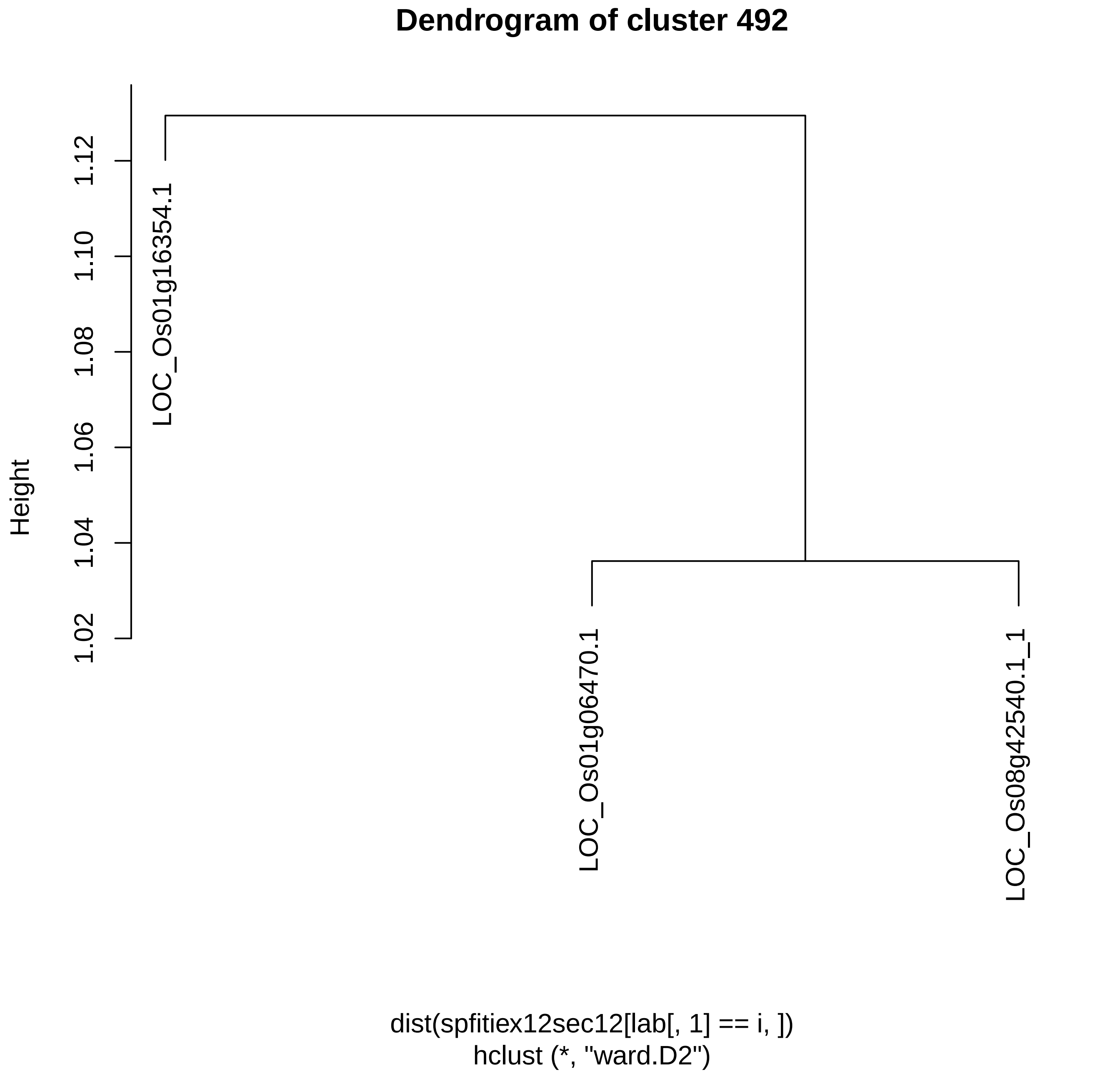
Dendrogram of cluster 492
Height
1.02
1.04
1.06
1.08
1.10
1.12
LOC_Os01g16354.1
LOC_Os01g06470.1
LOC_Os08g42540.1_1
dist(spfitiex12sec12[lab[, 1] == i, ])
hclust (*, "ward.D2")

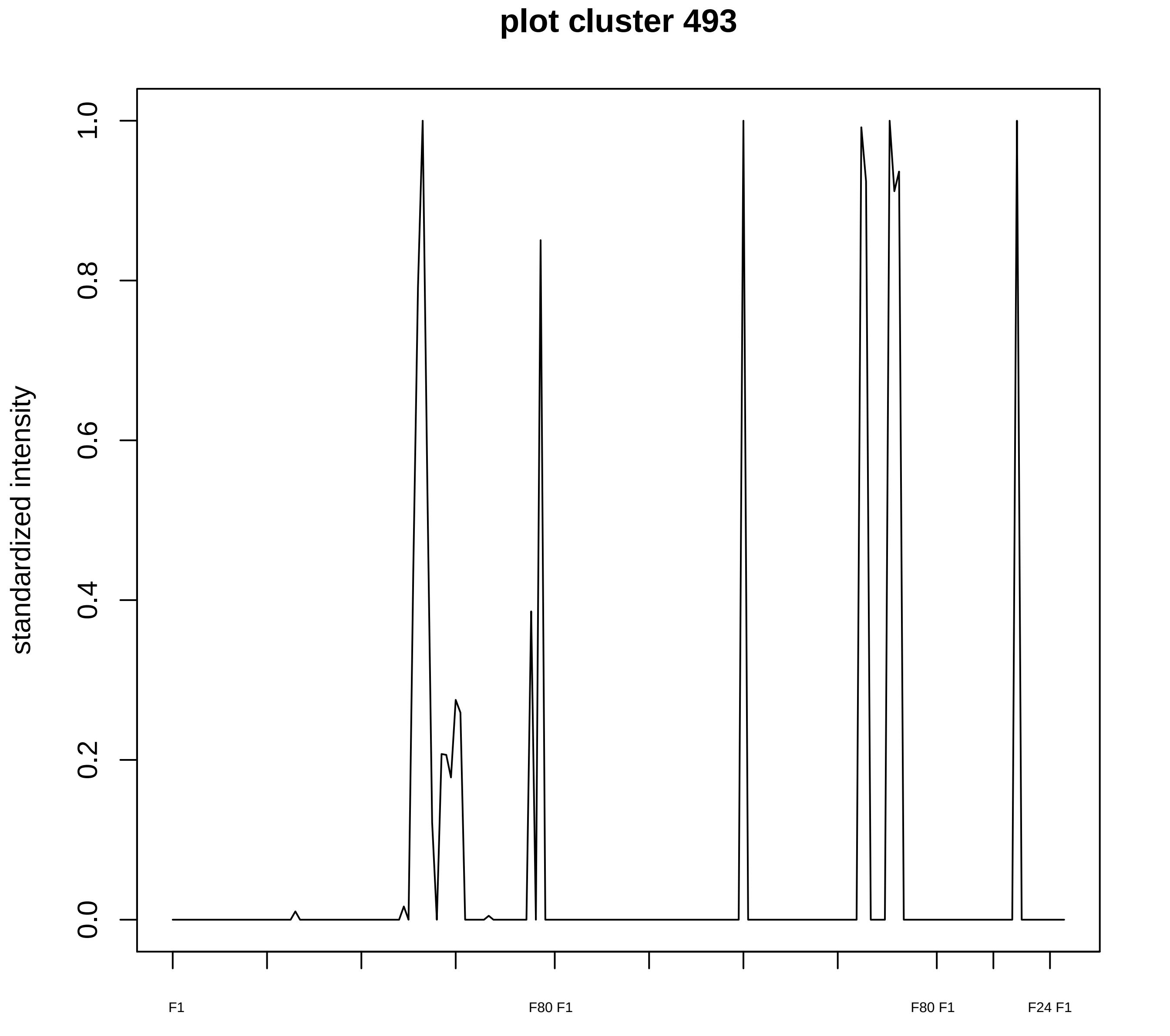

plot cluster 493
standardized intensity
0.0
0.2
0.4
0.6
0.8
1.0
F1
F80 F1
F80 F1
F24 F1

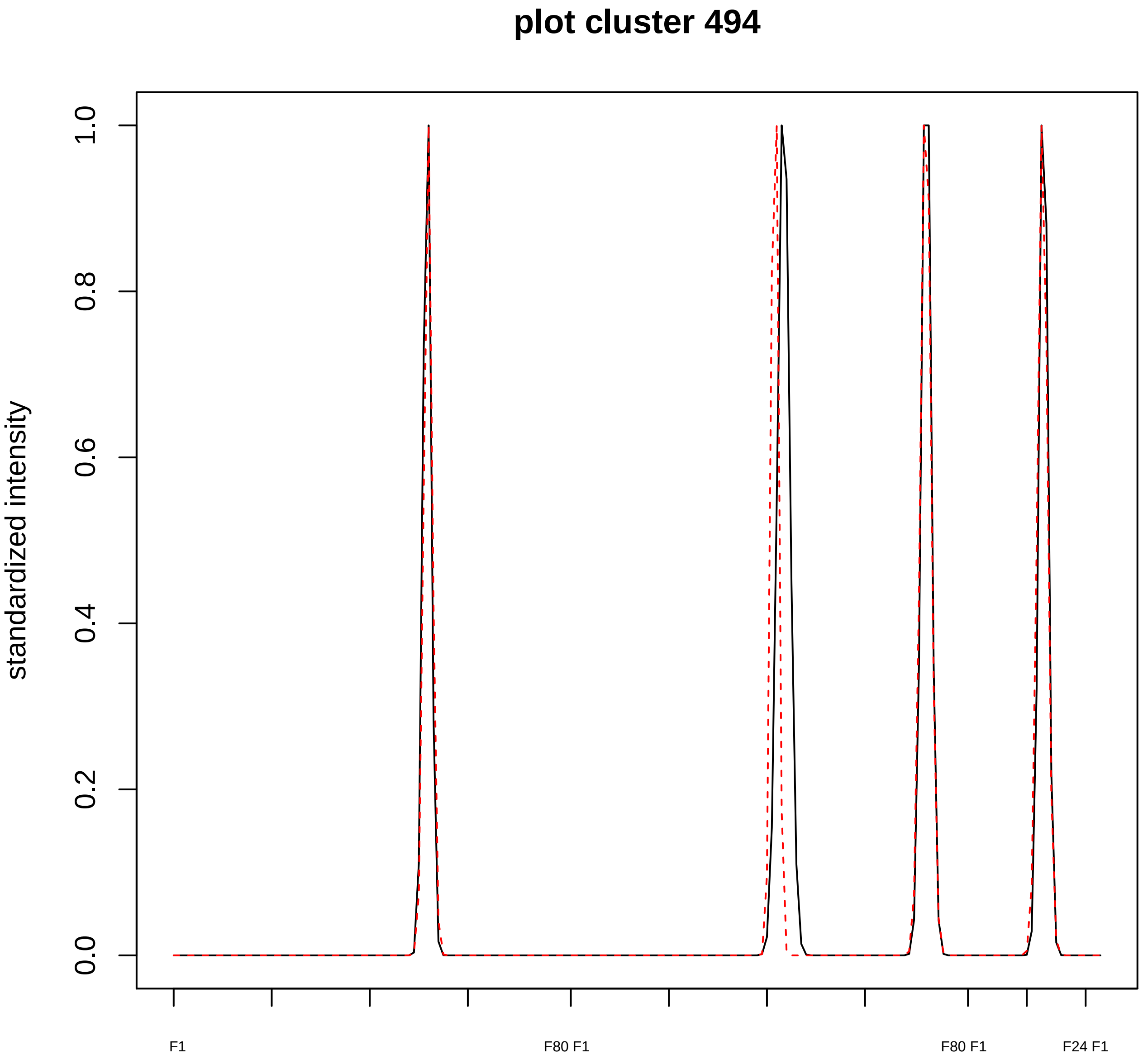

plot cluster 494
standardized intensity
0.0
0.2
0.4
0.6
0.8
1.0
F1
F80 F1
F80 F1
F24 F1

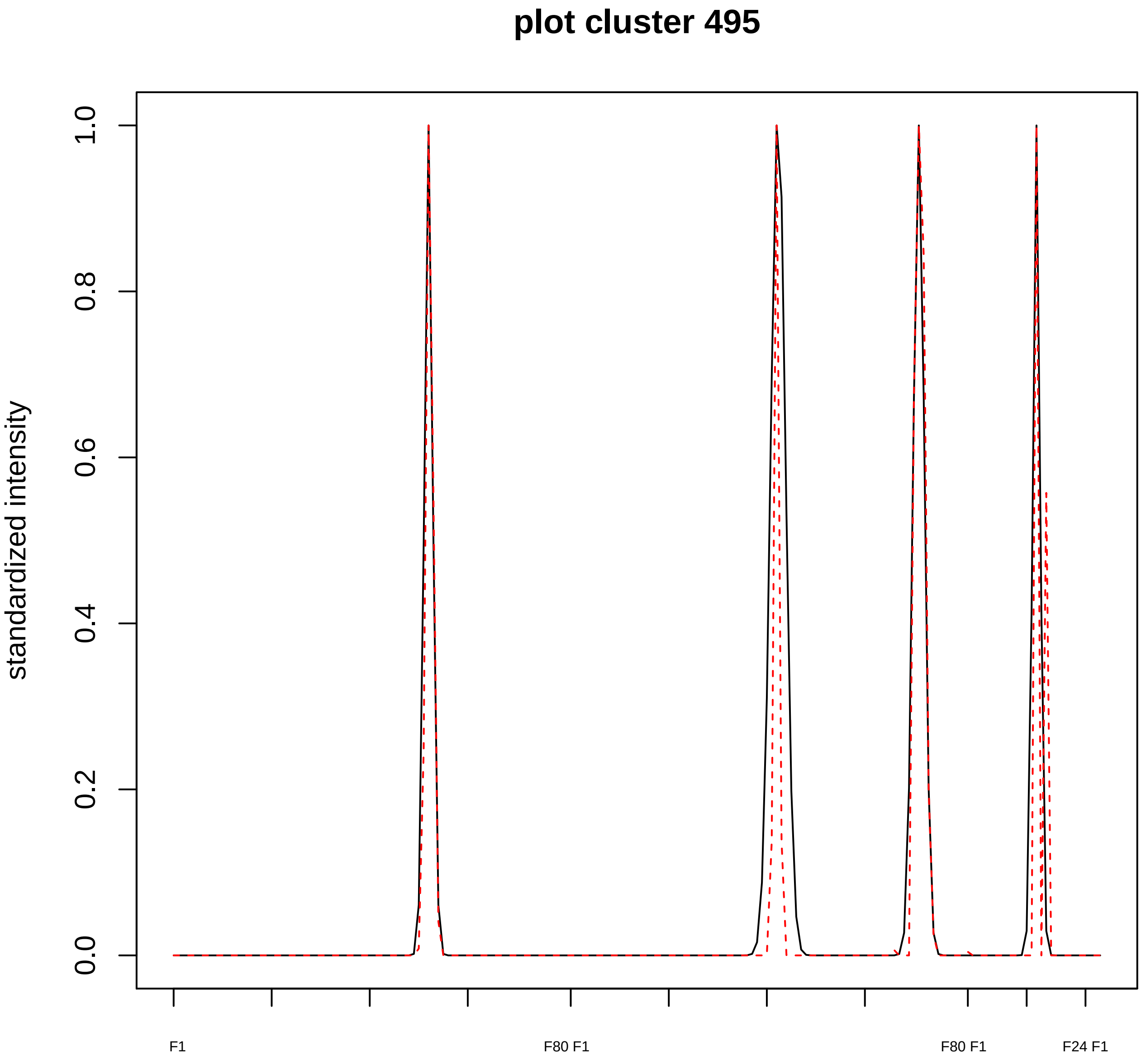

plot cluster 495
standardized intensity
0.0
0.2
0.4
0.6
0.8
1.0
F1
F80 F1
F80 F1
F24 F1

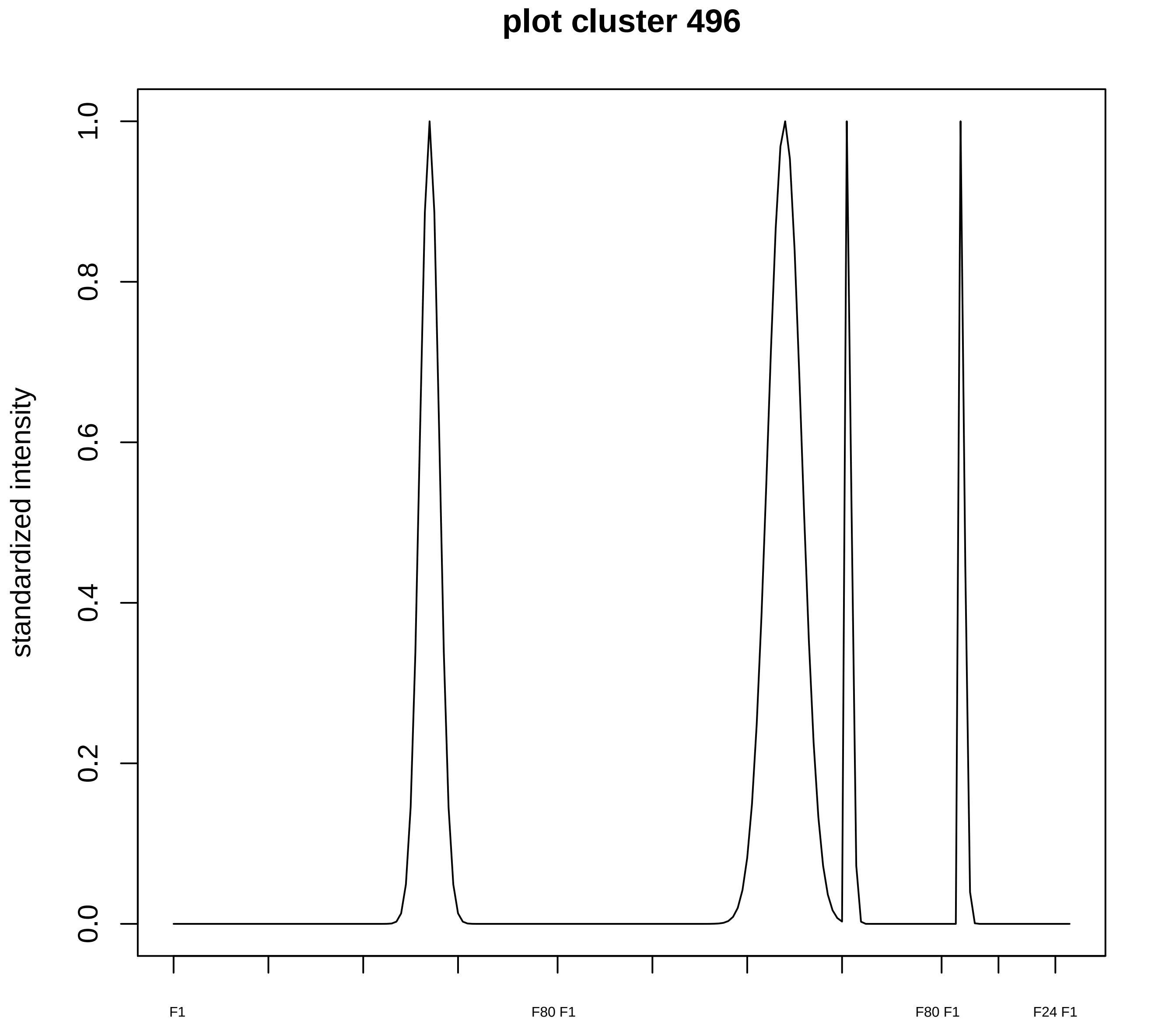
plot cluster 496
standardized intensity
0.0
0.2
0.4
0.6
0.8
1.0
F1
F80 F1
F80 F1
F24 F1

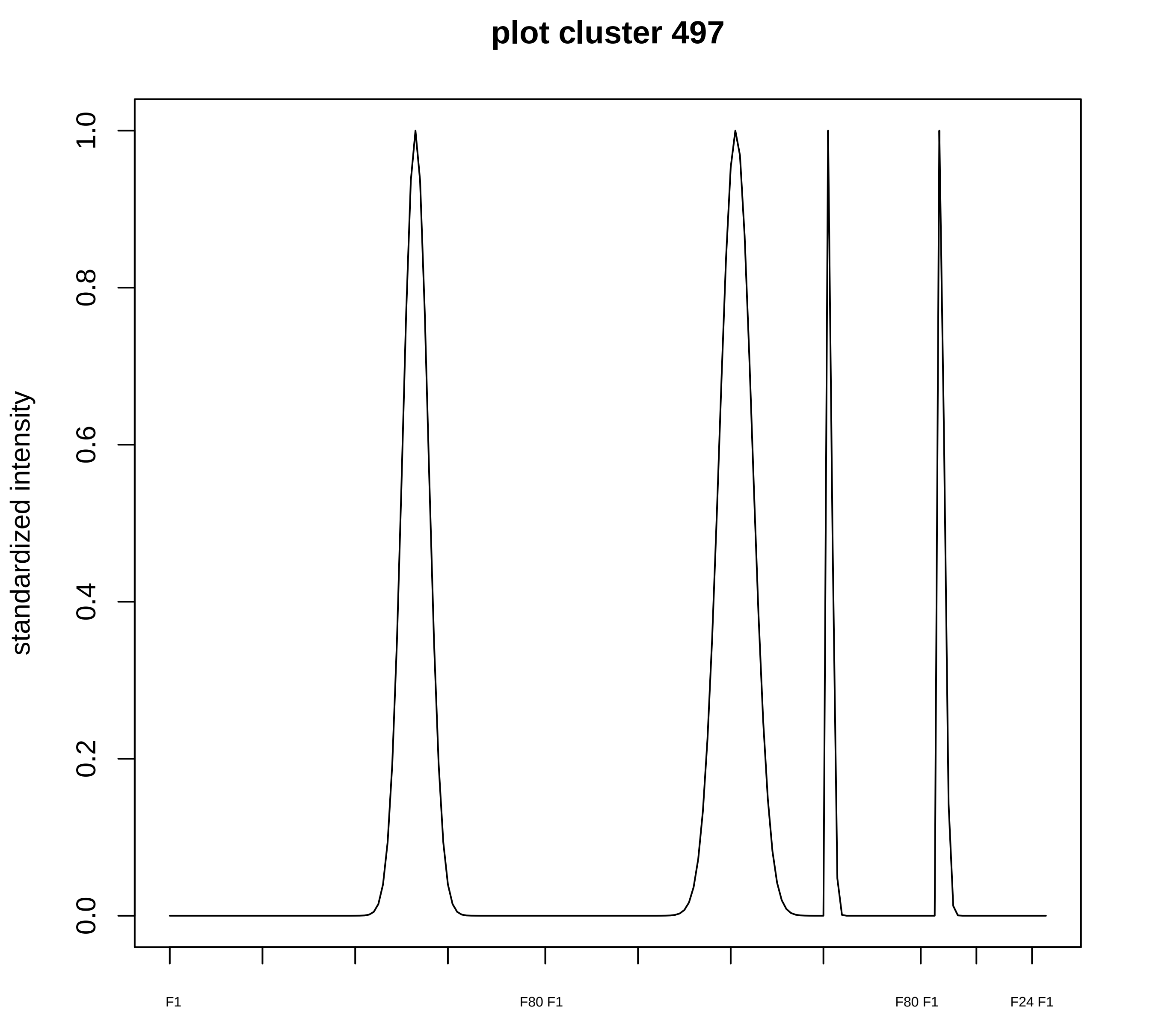

plot cluster 497
standardized intensity
0.0
0.2
0.4
0.6
0.8
1.0
F1
F80 F1
F80 F1
F24 F1

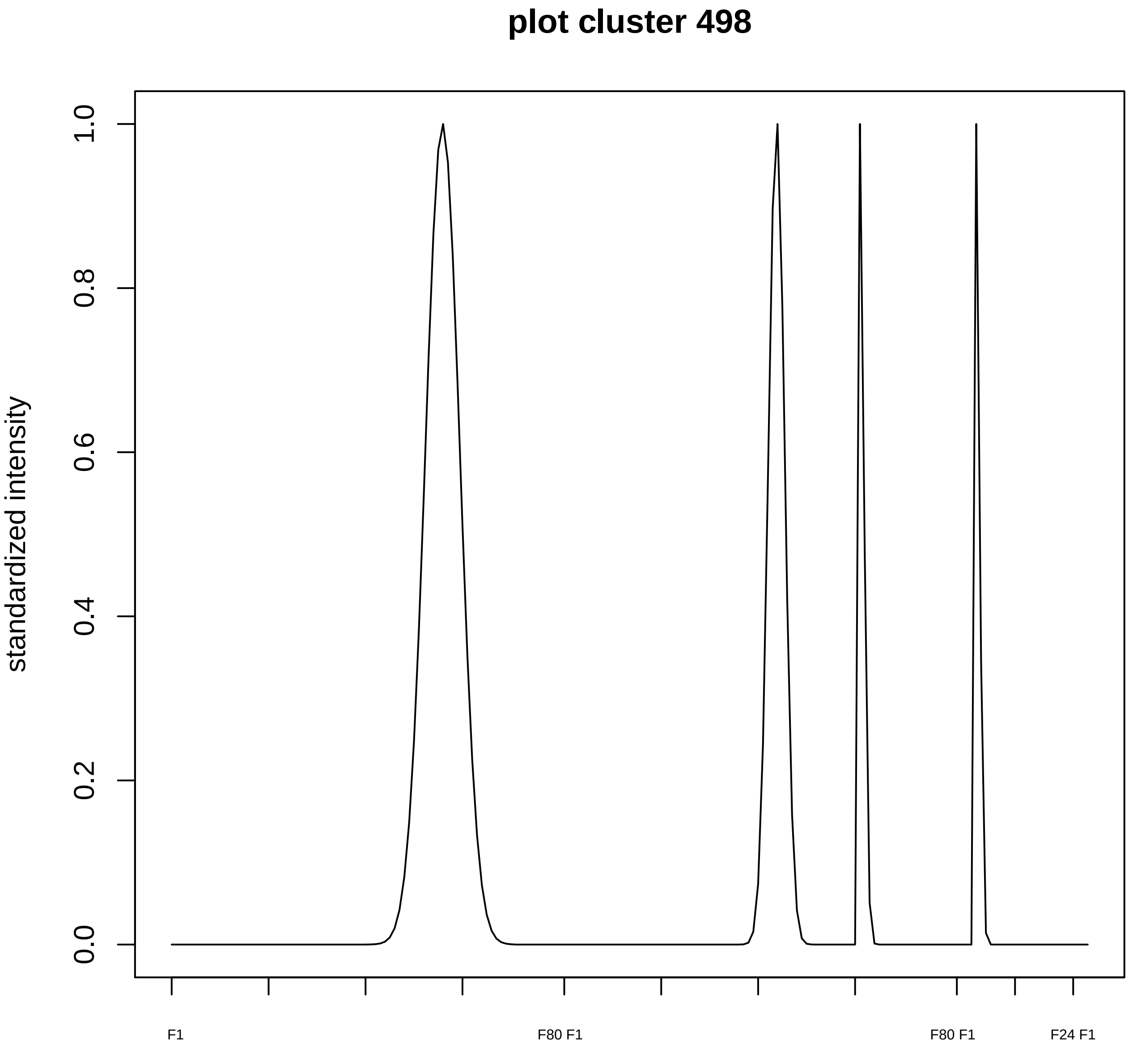

plot cluster 498
standardized intensity
0.0
0.2
0.4
0.6
0.8
1.0
F1
F80 F1
F80 F1
F24 F1

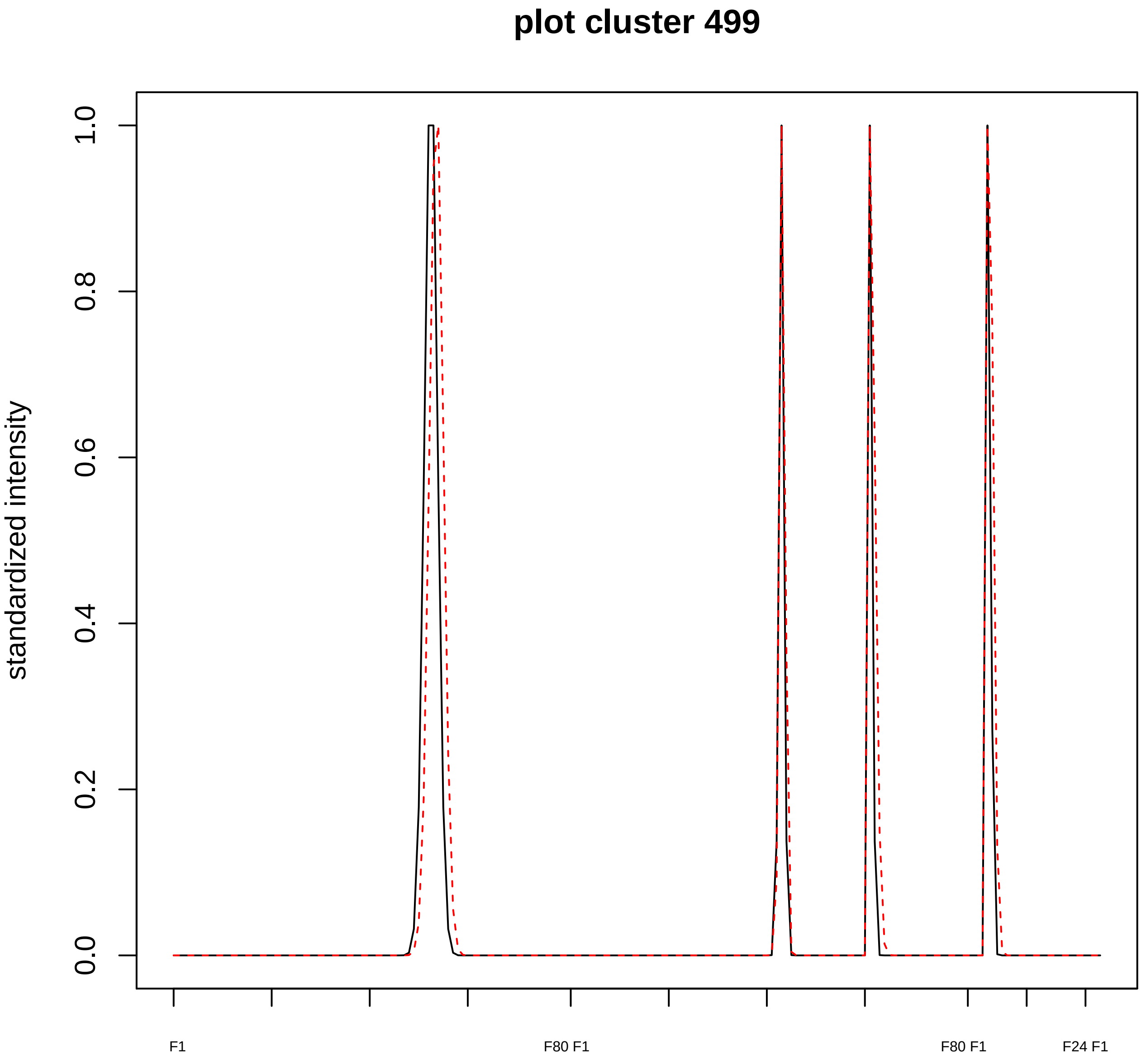

plot cluster 499
standardized intensity
0.0
0.2
0.4
0.6
0.8
1.0
F1
F80 F1
F80 F1
F24 F1

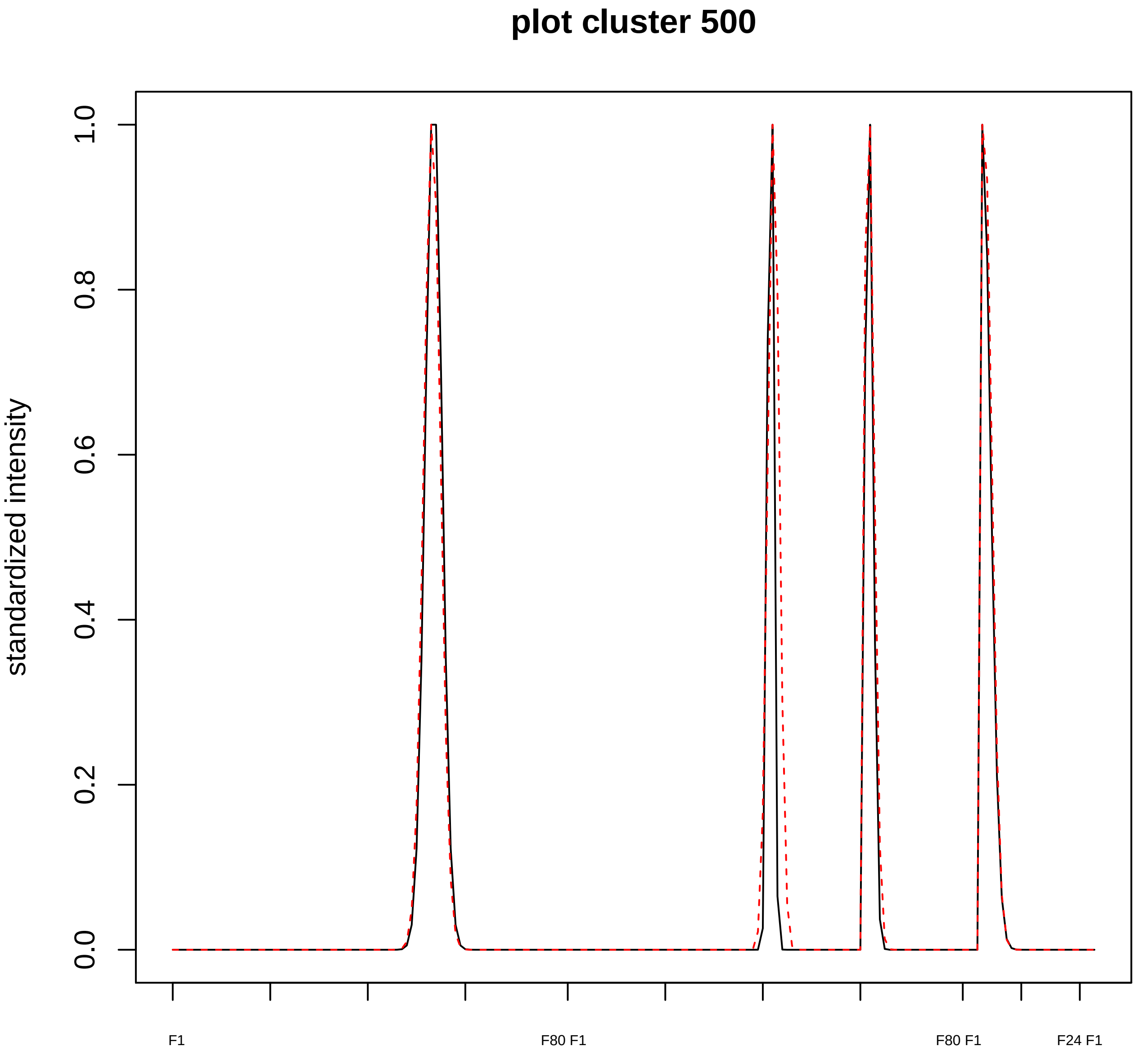
plot cluster 500
standardized intensity
0.0
0.2
0.4
0.6
0.8
1.0
F1
F80 F1
F80 F1
F24 F1

### plot cluster 501

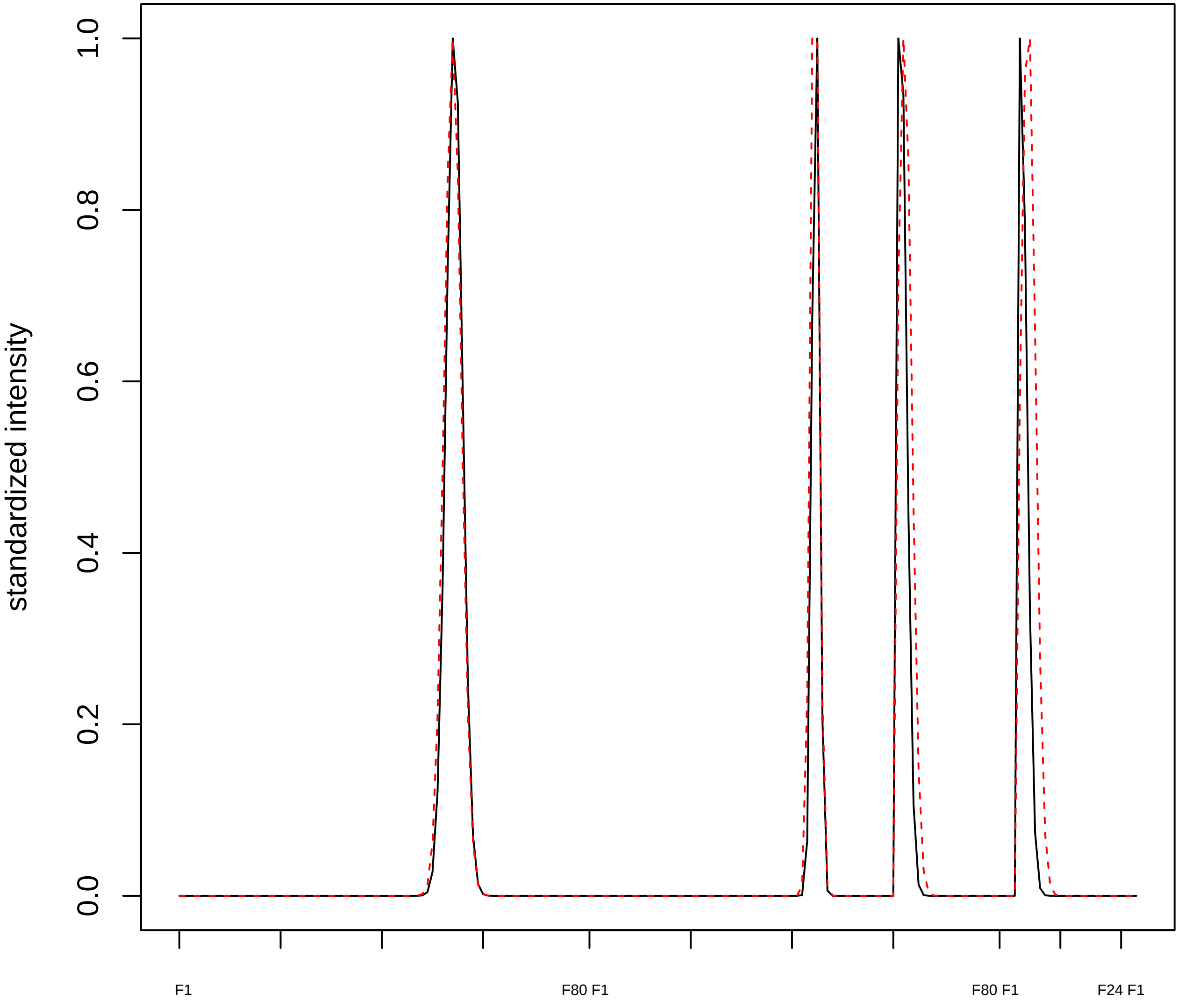

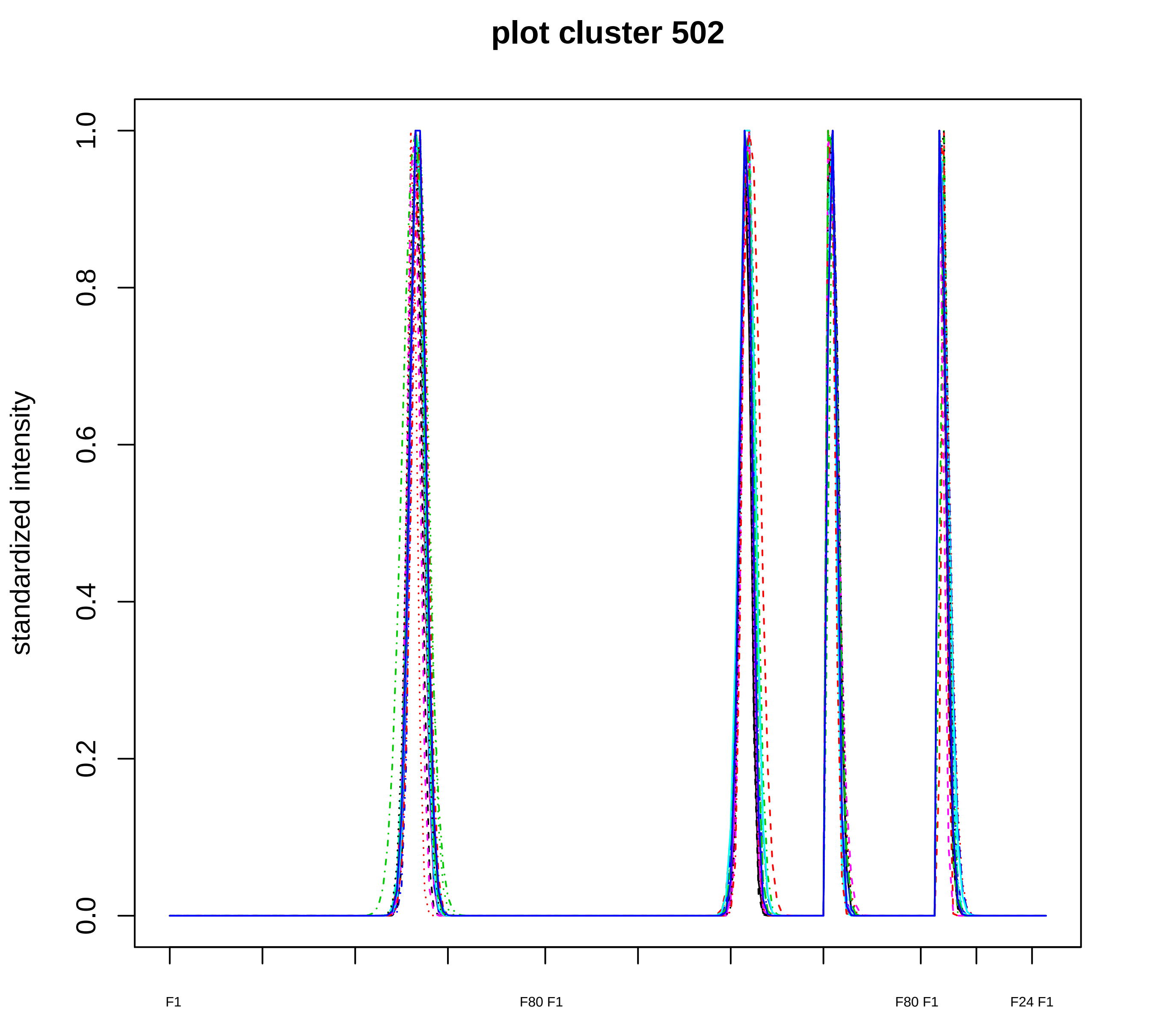
plot cluster 502
standardized intensity
0.0
0.2
0.4
0.6
0.8
1.0
F1
F80 F1
F80 F1
F24 F1

#### Dendrogram of cluster 502

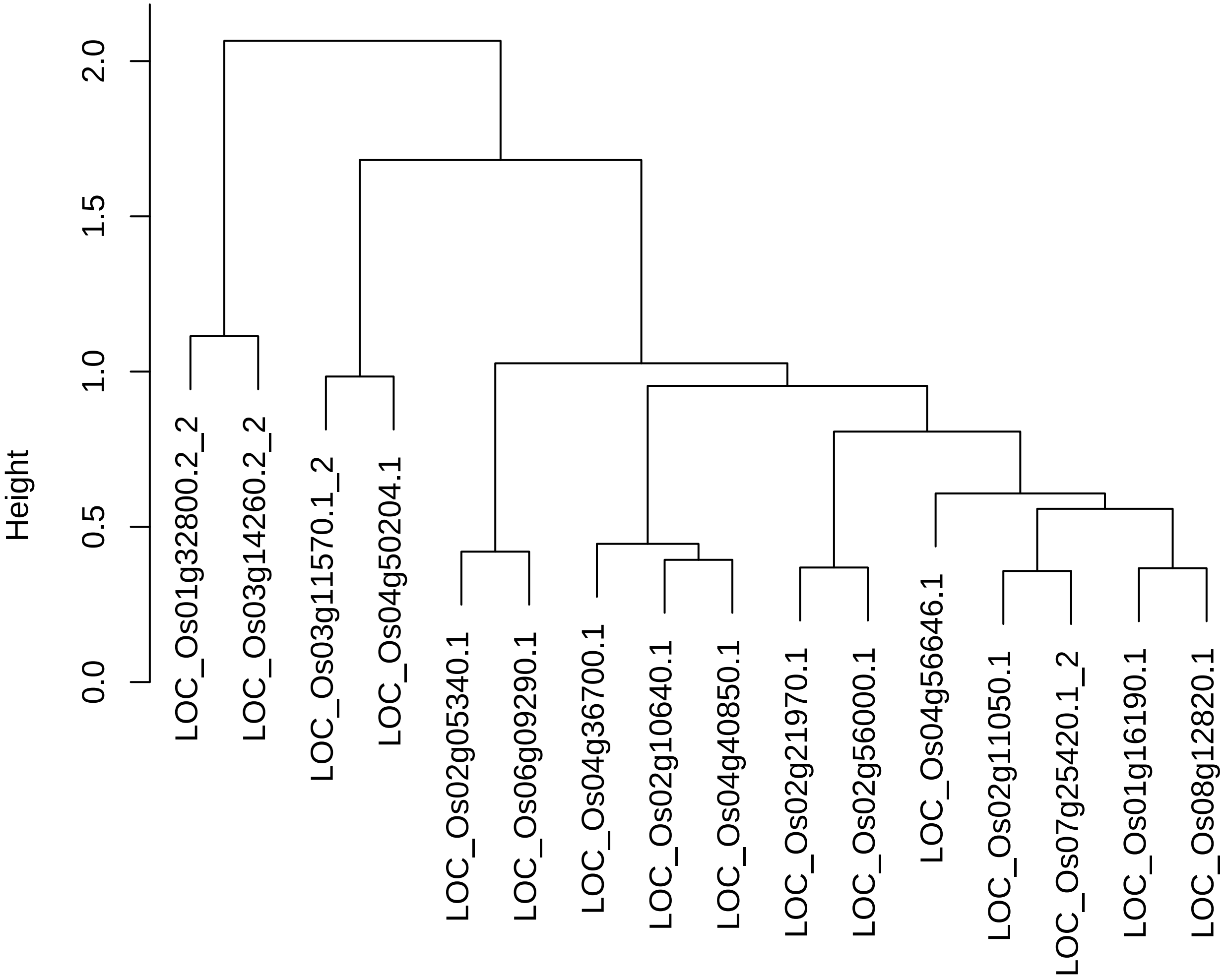


dist(spfitiex12sec12[lab[, 1] == i, ])
hclust (*, "ward.D2")

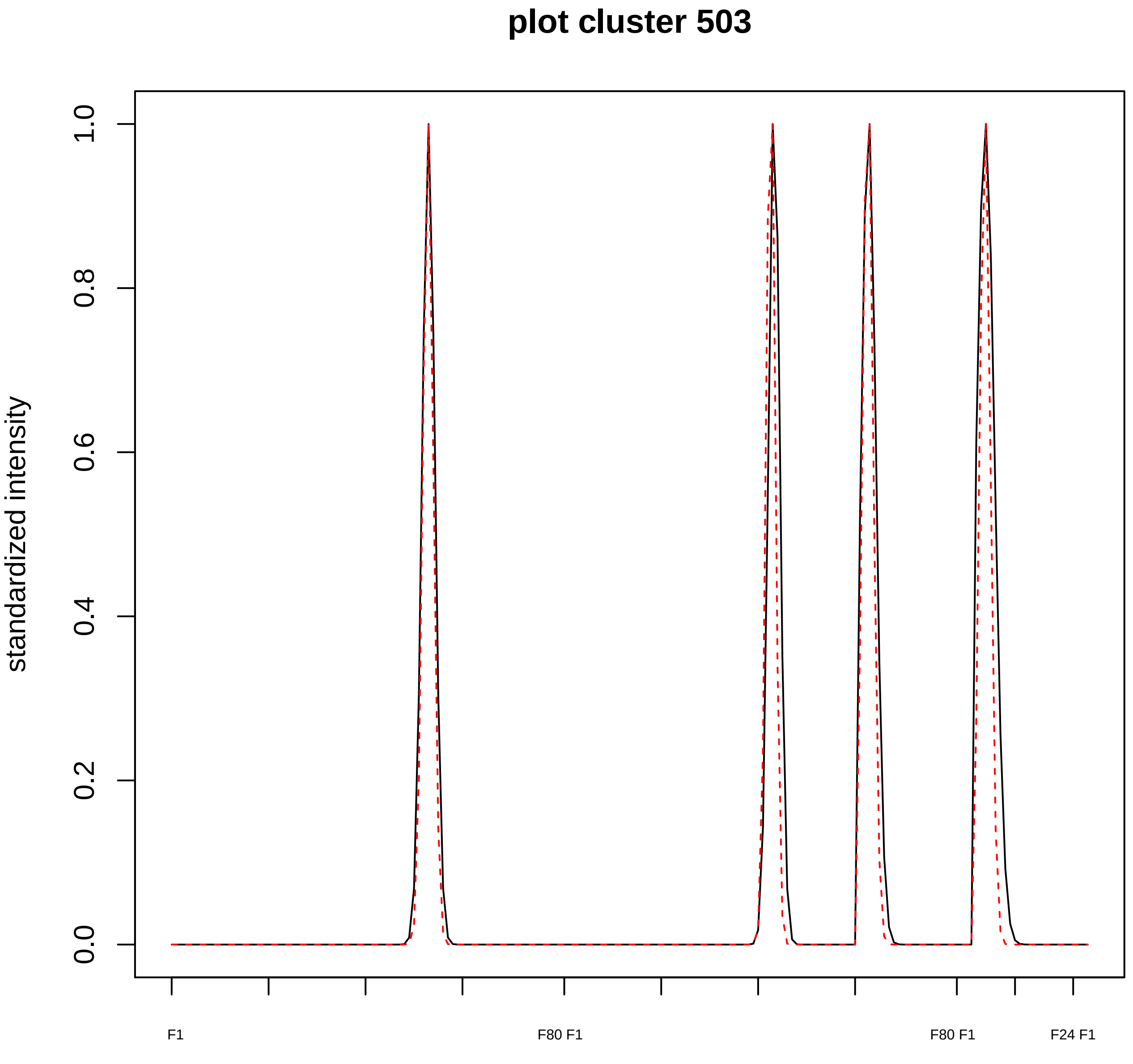
plot cluster 503
standardized intensity
0.0
0.2
0.4
0.6
0.8
1.0
F1
F80 F1
F80 F1
F24 F1

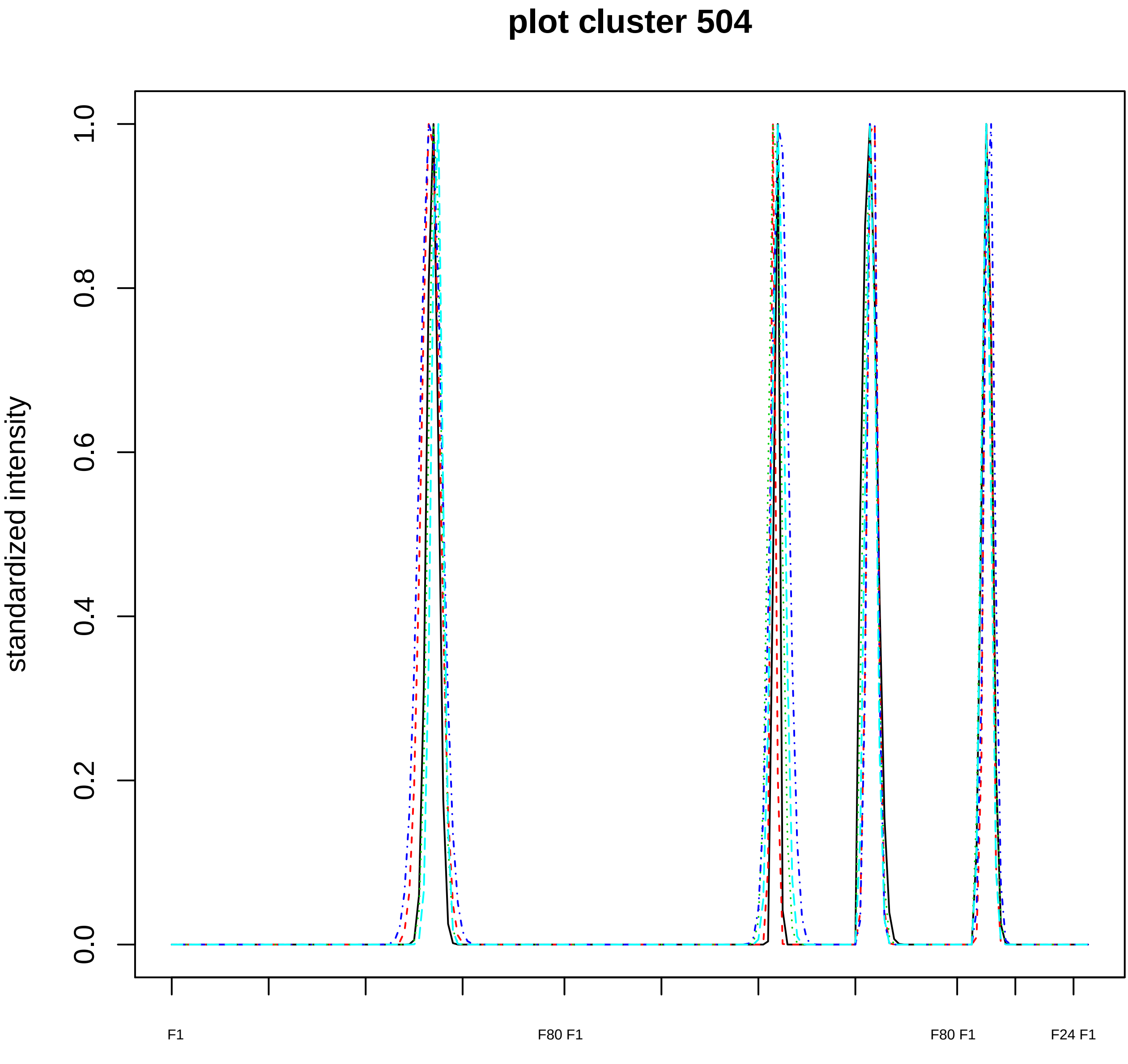
plot cluster 504
standardized intensity
0.0
0.2
0.4
0.6
0.8
1.0
F1
F80 F1
F80 F1
F24 F1

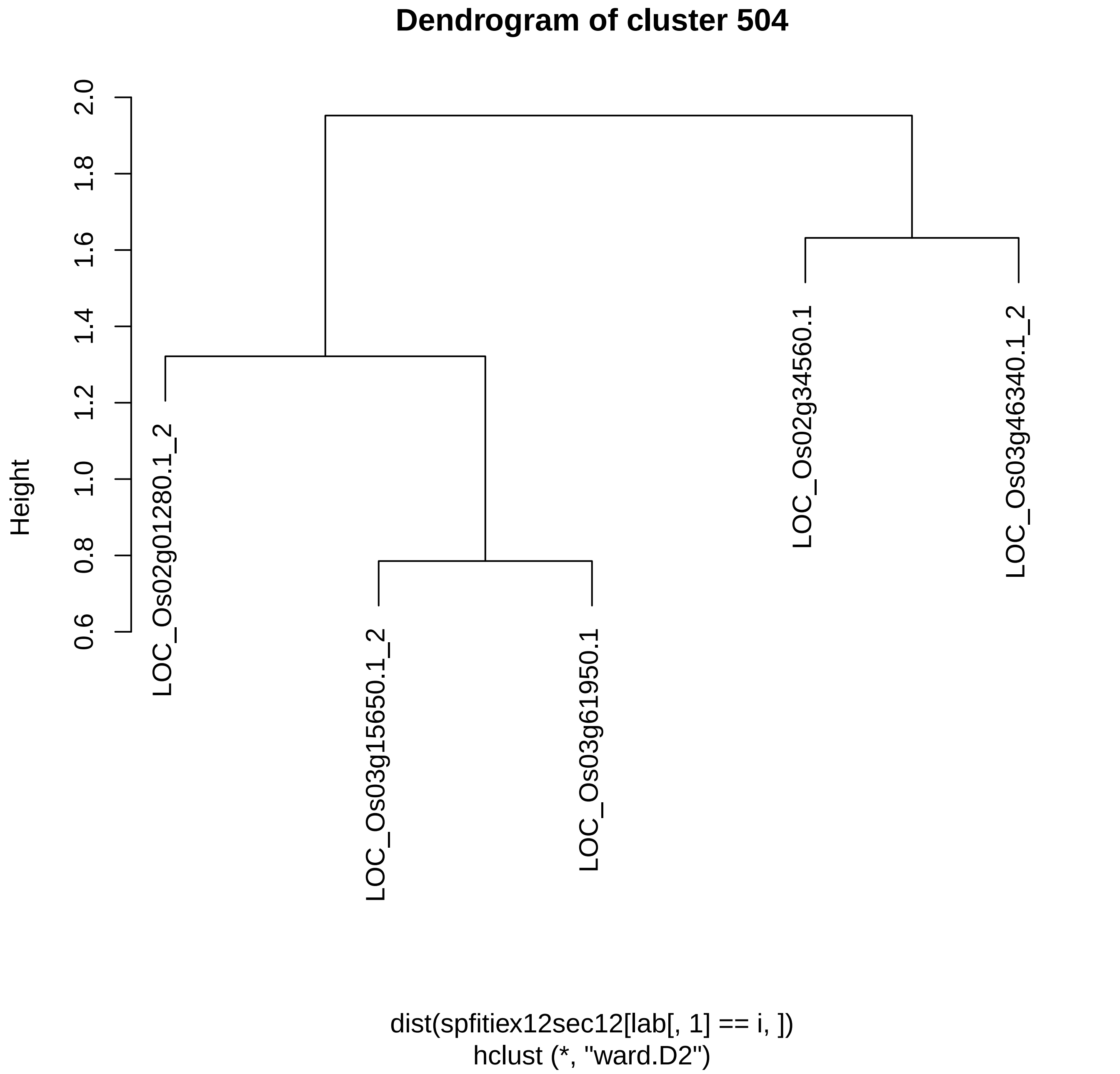
Dendrogram of cluster 504
Height
2.0
1.8
1.6
1.4
1.2
1.0
0.8
0.6
LOC_Os02g01280.1_2
LOC_Os03g15650.1_2
LOC_Os03g61950.1
LOC_Os02g34560.1
LOC_Os03g46340.1_2
dist(spfitiex12sec12[lab[, 1] == i, ])
hclust (*, "ward.D2")

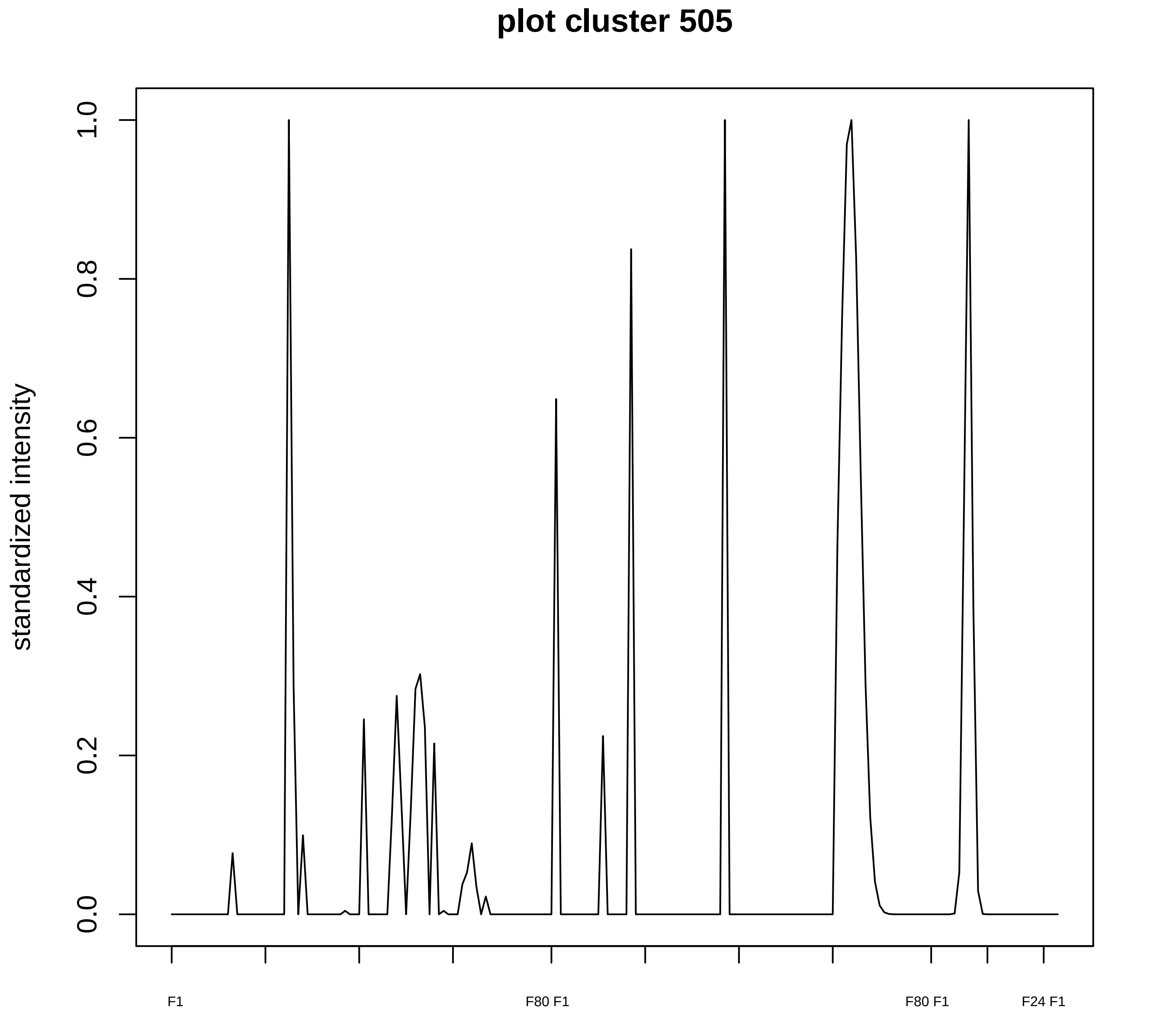
plot cluster 505
standardized intensity
0.0
0.2
0.4
0.6
0.8
1.0
F1
F80 F1
F80 F1
F24 F1

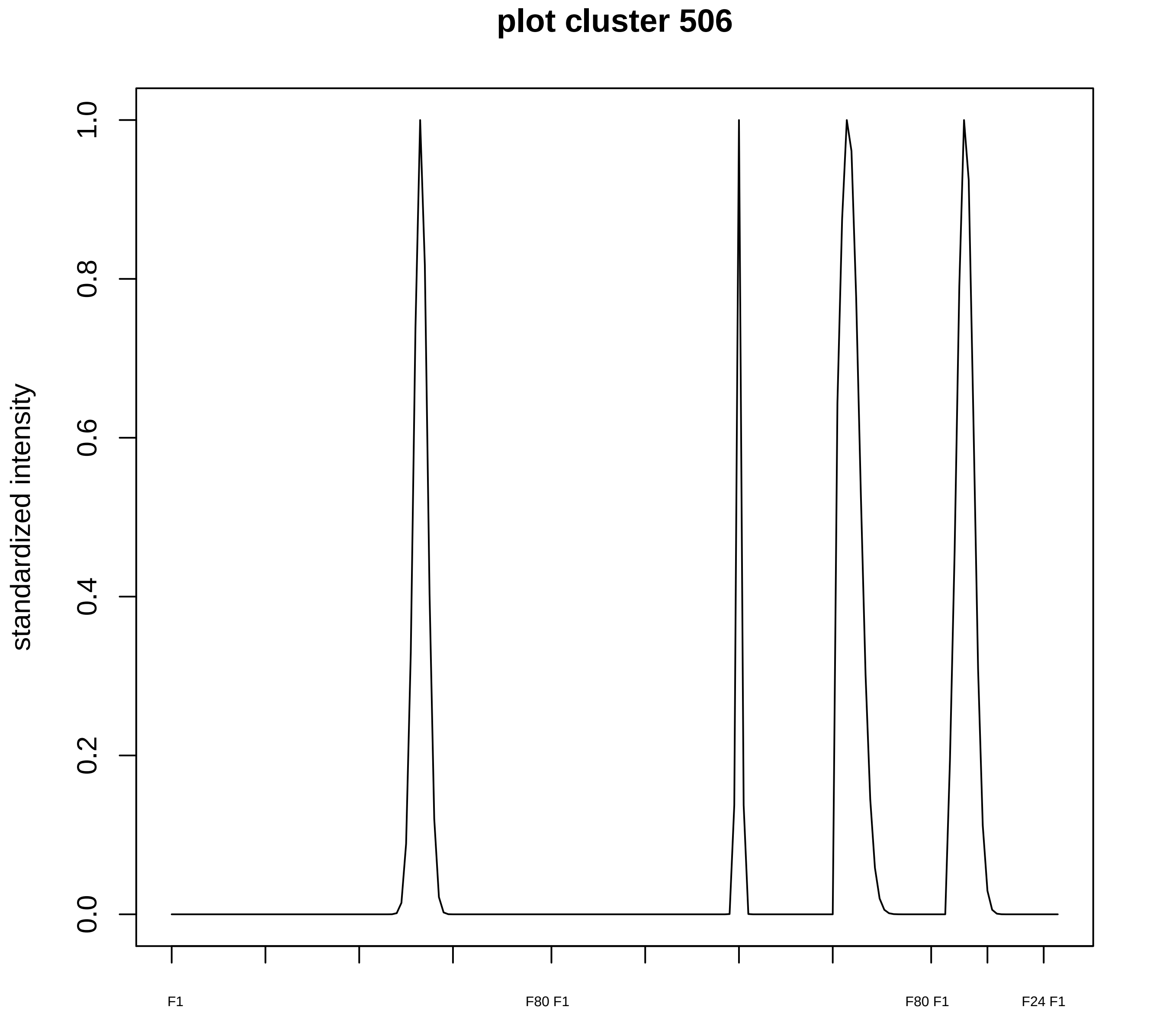
plot cluster 506
standardized intensity
0.0
0.2
0.4
0.6
0.8
1.0
F1
F80 F1
F80 F1
F24 F1

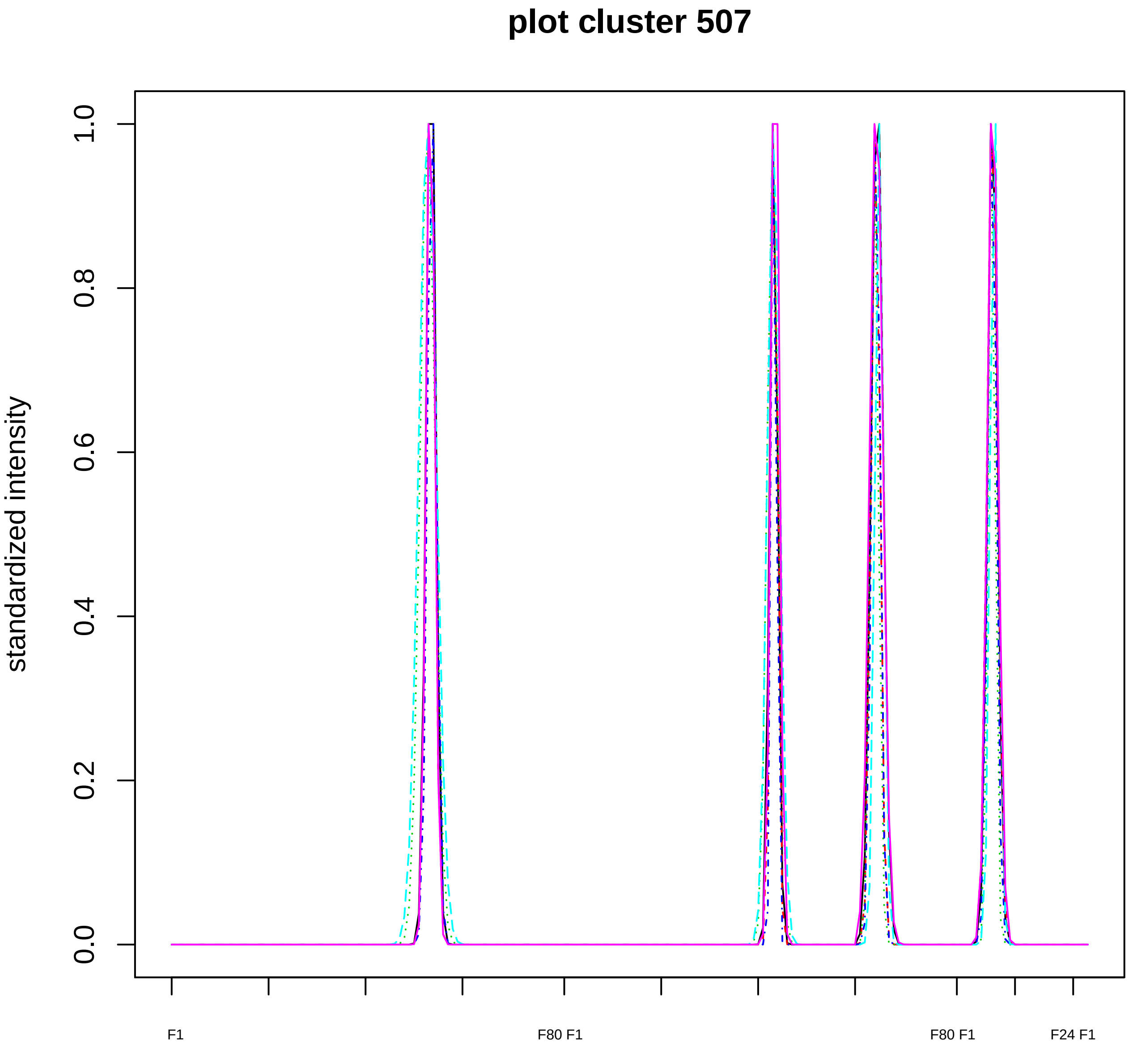
plot cluster 507
standardized intensity
0.0
0.2
0.4
0.6
0.8
1.0
F1
F80 F1
F80 F1
F24 F1

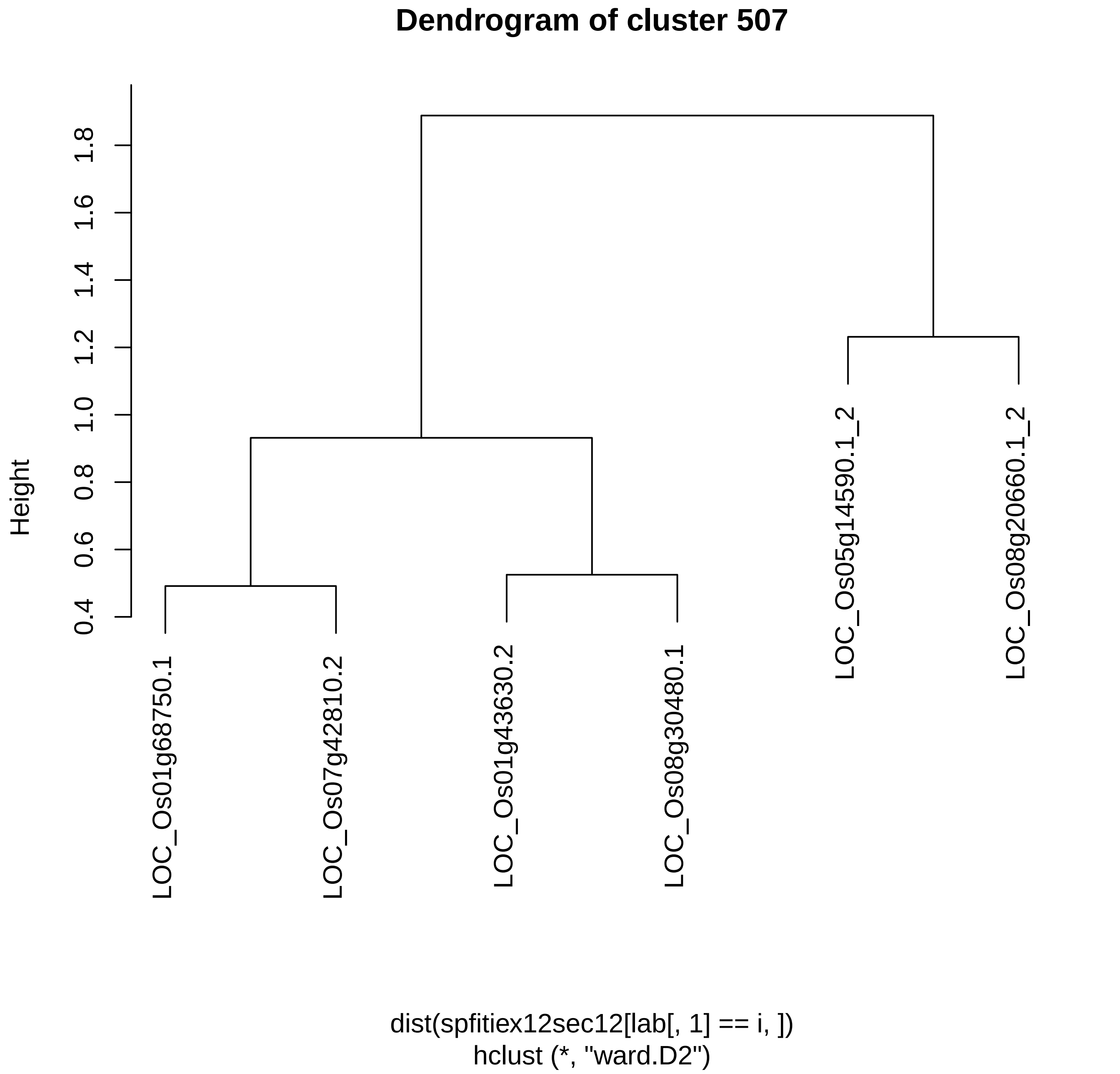
Dendrogram of cluster 507
Height
0.4
0.6
0.8
1.0
1.2
1.4
1.6
1.8
LOC_Os01g68750.1
LOC_Os07g42810.2
LOC_Os01g43630.2
LOC_Os08g30480.1
LOC_Os05g14590.1_2
LOC_Os08g20660.1_2
dist(spfitiex12sec12[lab[, 1] == i, ])
hclust (*, "ward.D2")

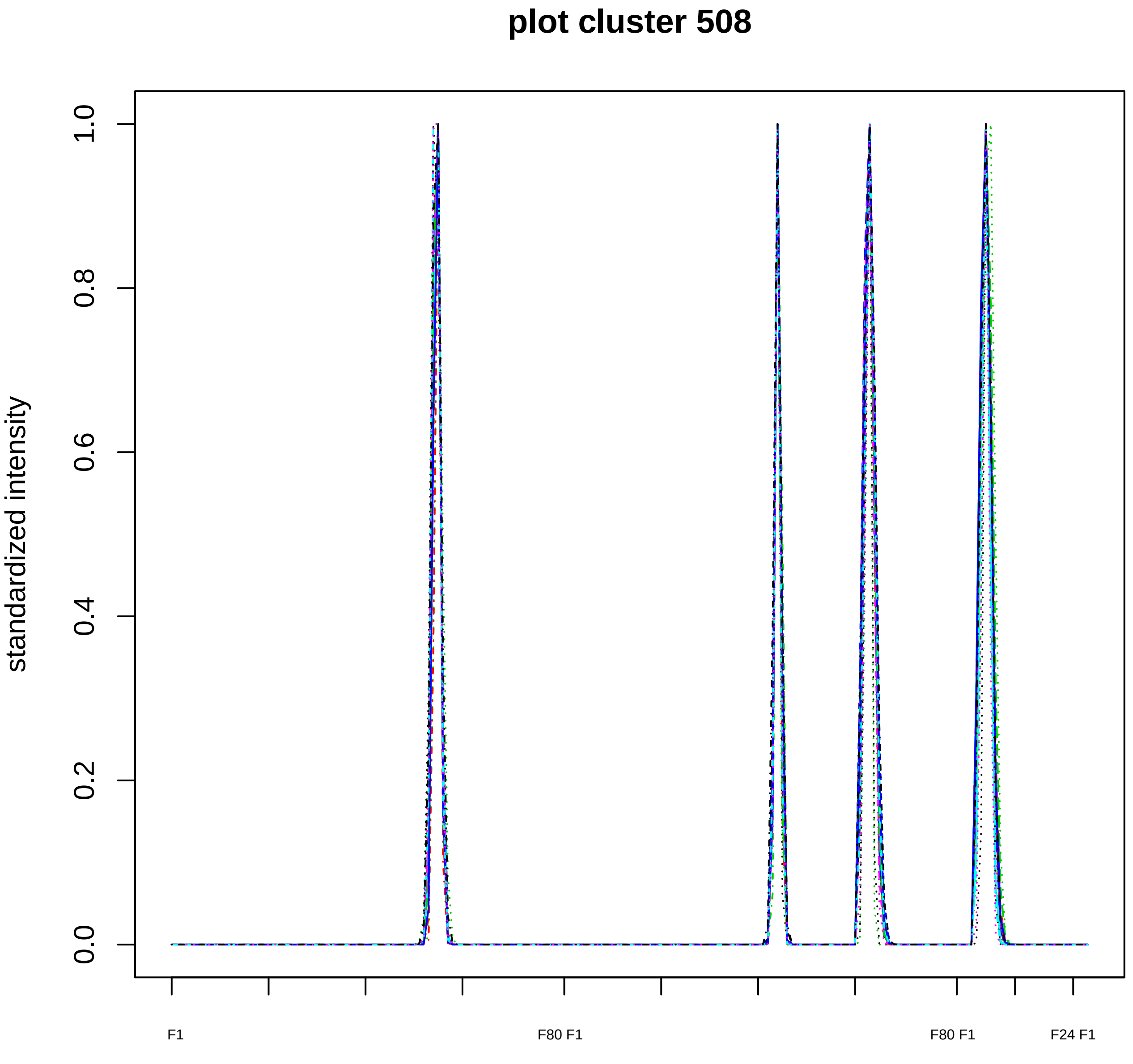

plot cluster 508
standardized intensity
0.0
0.2
0.4
0.6
0.8
1.0
F1
F80 F1
F80 F1
F24 F1

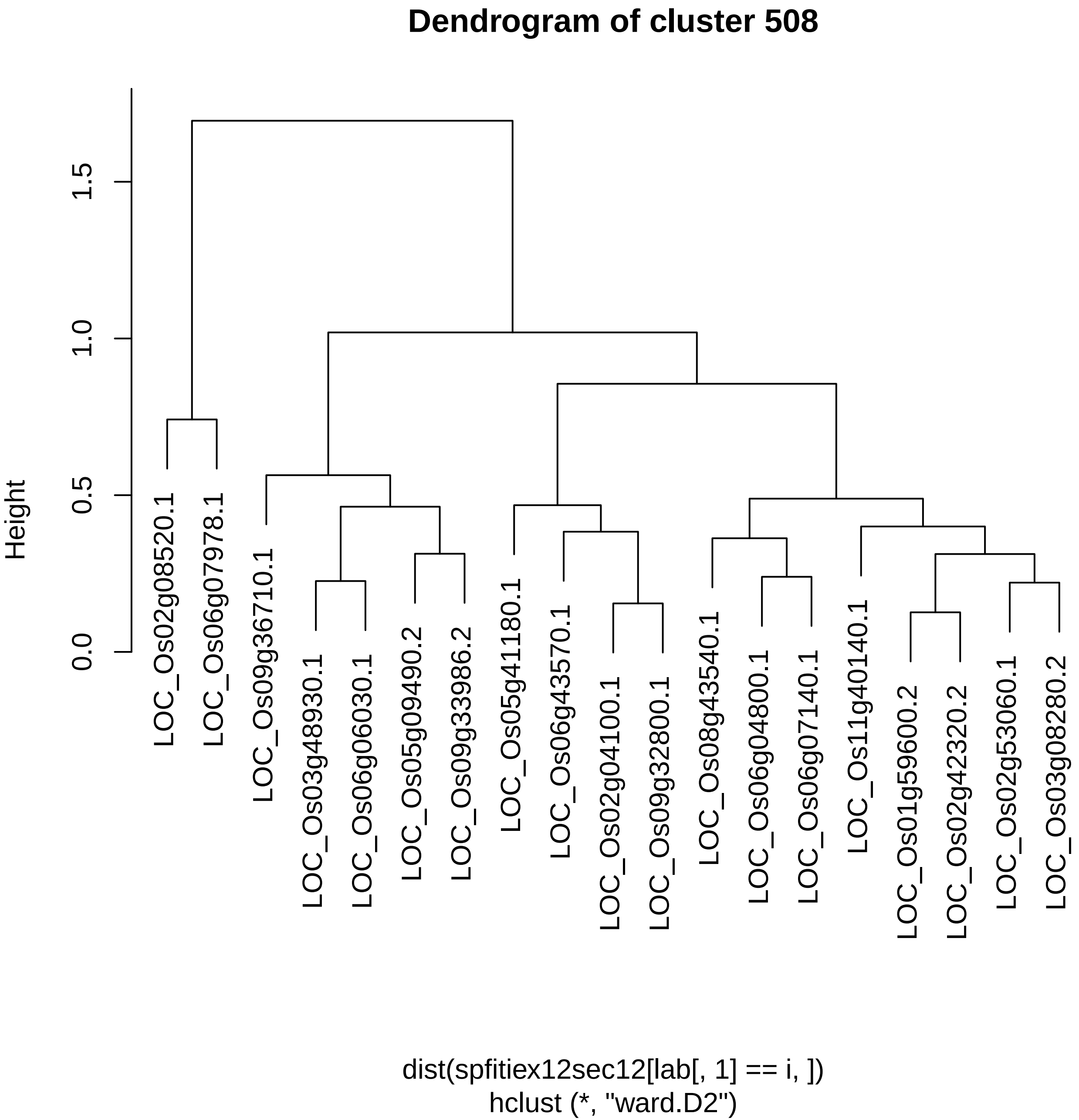
Dendrogram of cluster 508
Height
0.0
0.5
1.0
1.5
LOC_Os02g08520.1
LOC_Os06g07978.1
LOC_Os09g36710.1
LOC_Os03g48930.1
LOC_Os06g06030.1
LOC_Os05g09490.2
LOC_Os09g33986.2
LOC_Os05g41180.1
LOC_Os06g43570.1
LOC_Os02g04100.1
LOC_Os09g32800.1
LOC_Os08g43540.1
LOC_Os06g04800.1
LOC_Os06g07140.1
LOC_Os11g40140.1
LOC_Os01g59600.2
LOC_Os02g42320.2
LOC_Os02g53060.1
LOC_Os03g08280.2
dist(spfitiex12sec12[lab[, 1] == i, ])
hclust (*, "ward.D2")

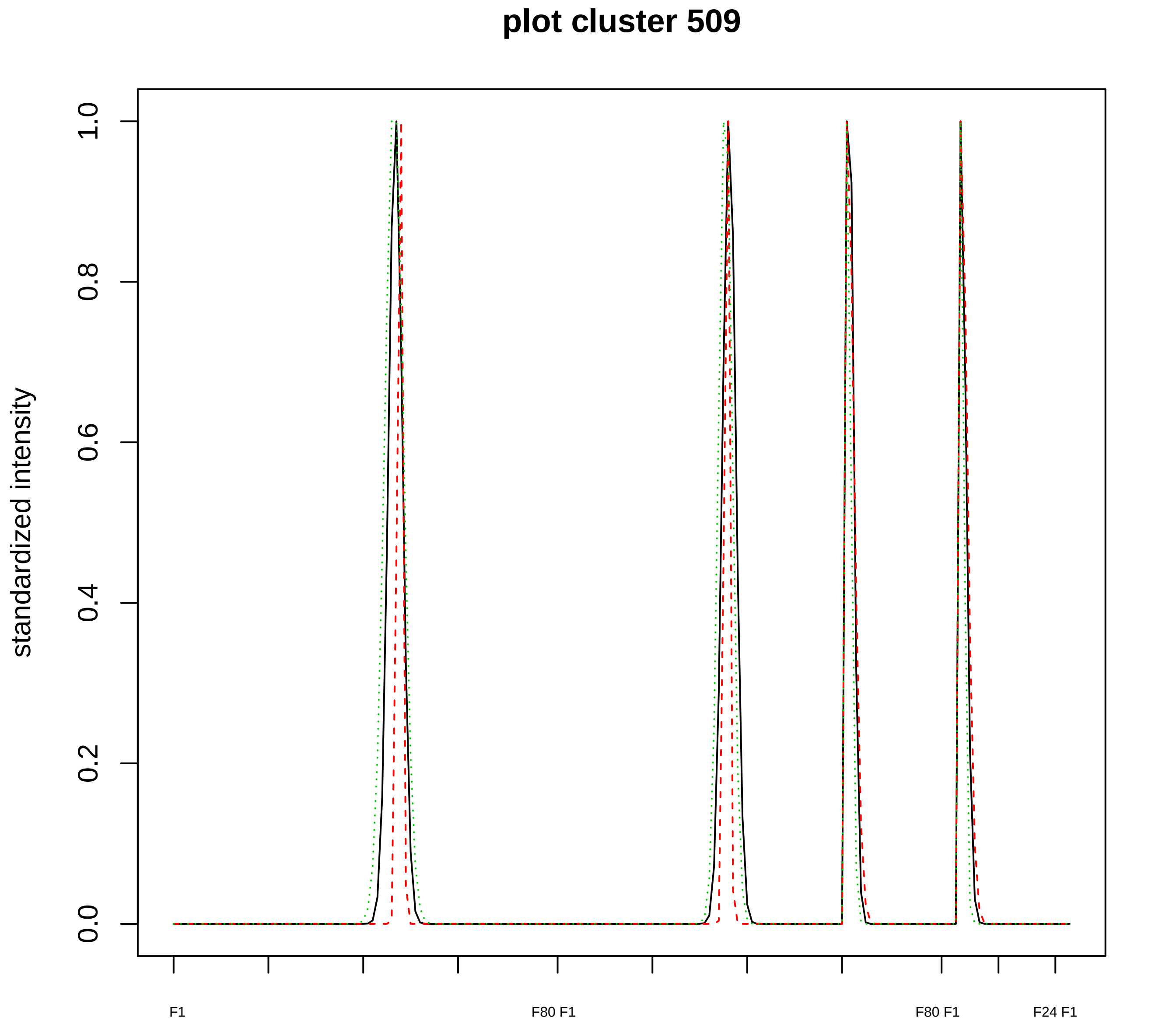

plot cluster 509
standardized intensity
0.0
0.2
0.4
0.6
0.8
1.0
F1
F80 F1
F80 F1
F24 F1

#### Dendrogram of cluster 509

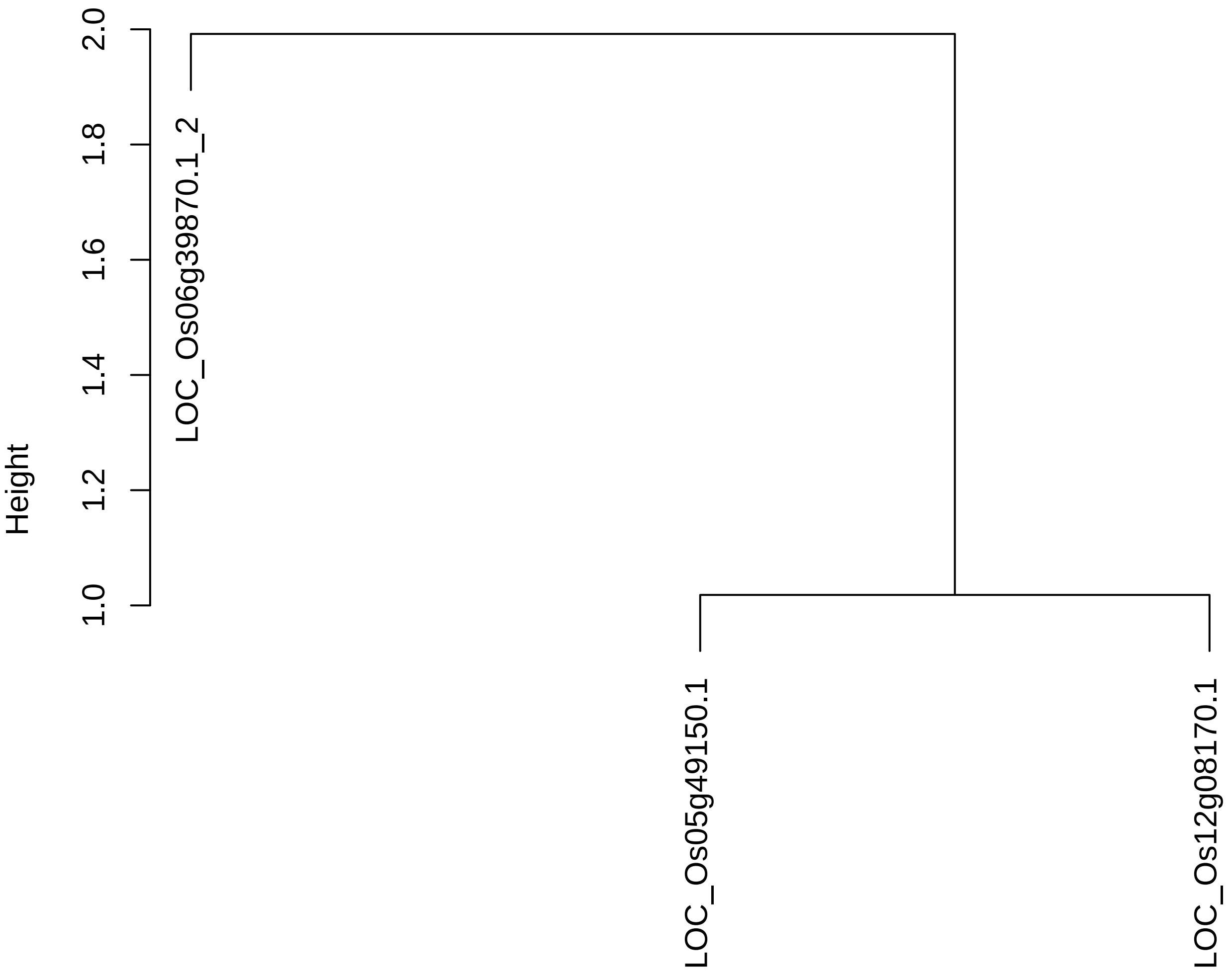


dist(spfitiex12sec12[lab[, 1] == i, ])
hclust (*, "ward.D2")

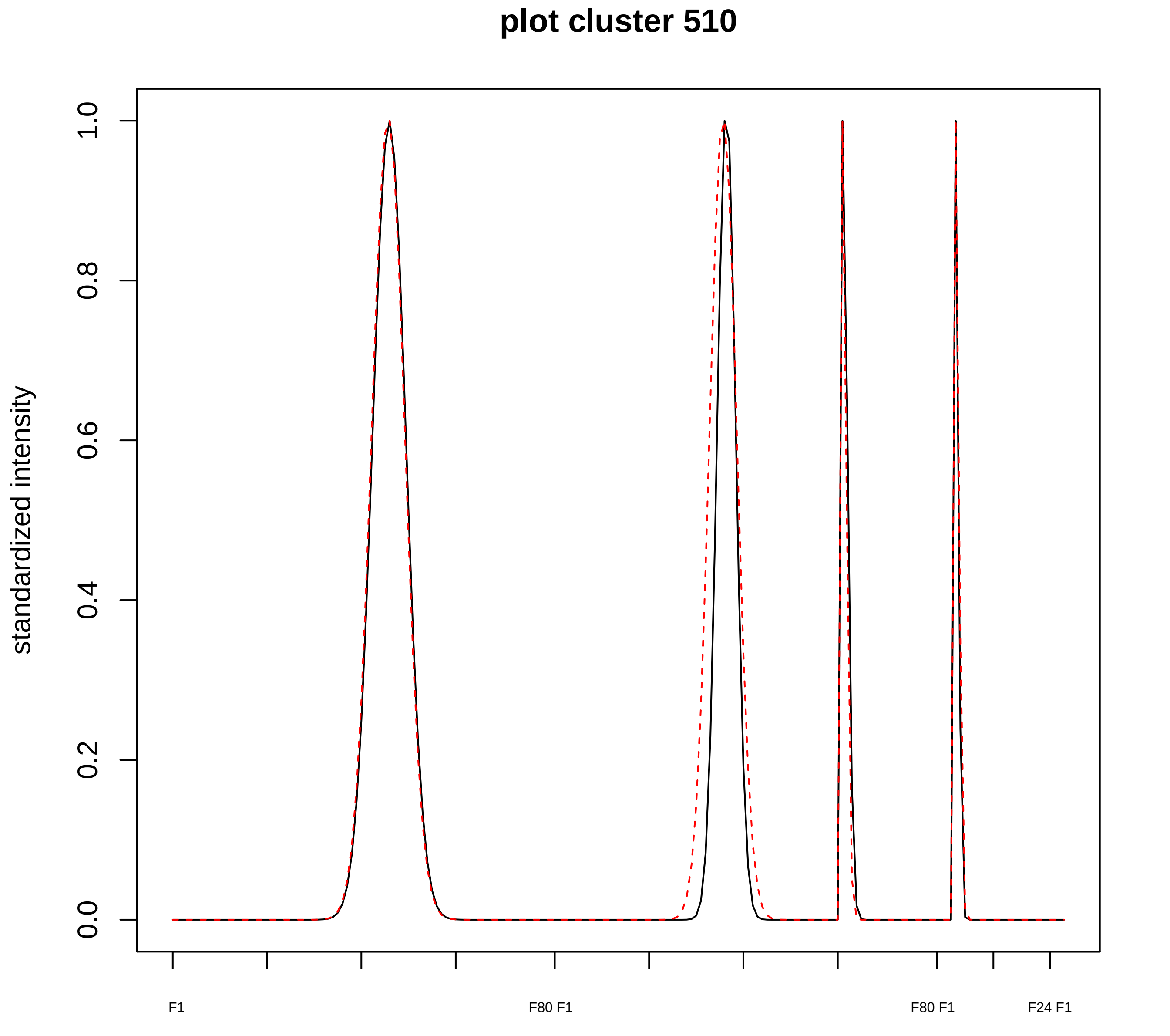
plot cluster 510
standardized intensity
0.0
0.2
0.4
0.6
0.8
1.0
F1
F80 F1
F80 F1
F24 F1

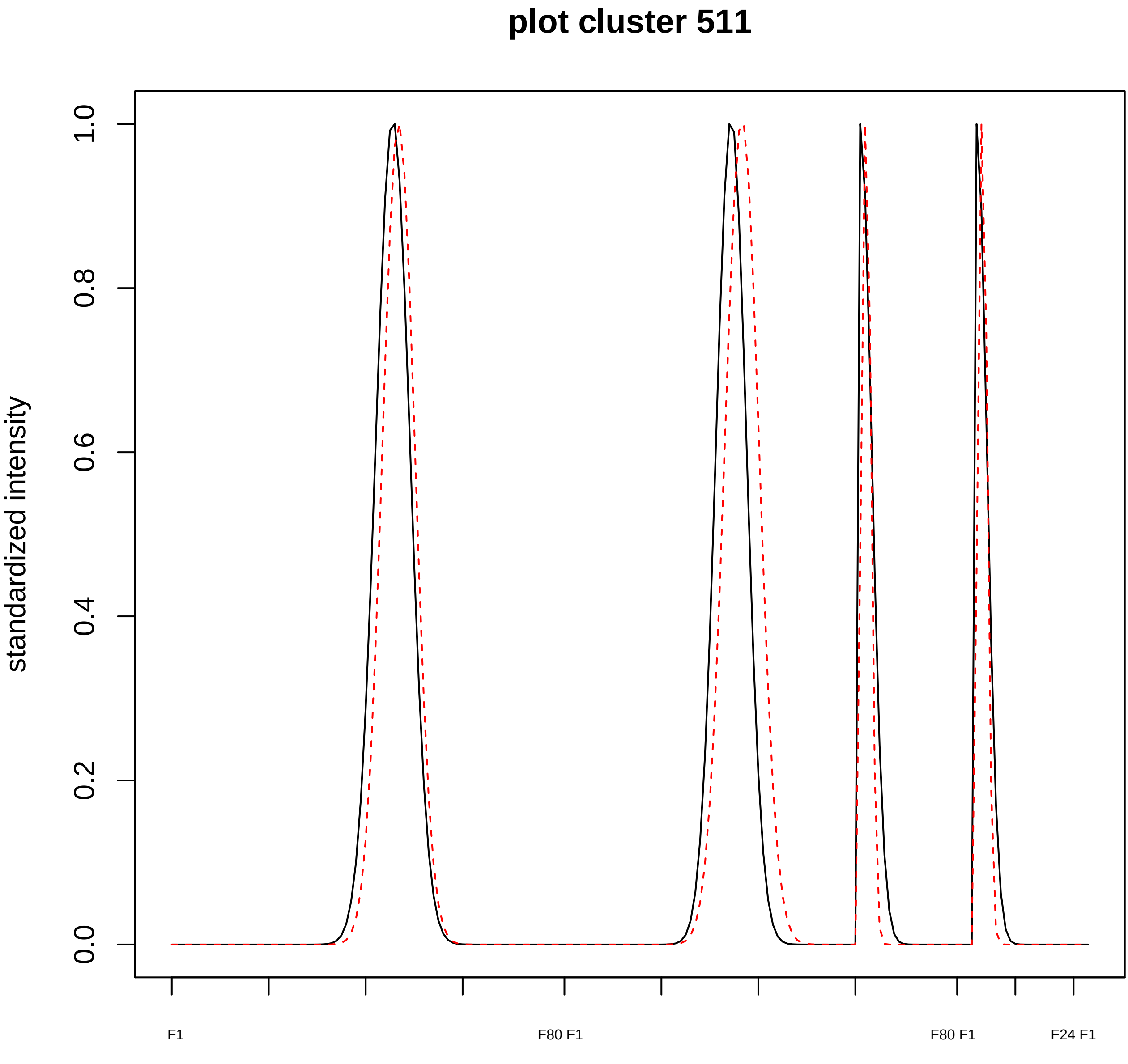

plot cluster 511
standardized intensity
0.0
0.2
0.4
0.6
0.8
1.0
F1
F80 F1
F80 F1
F24 F1

### plot cluster 512

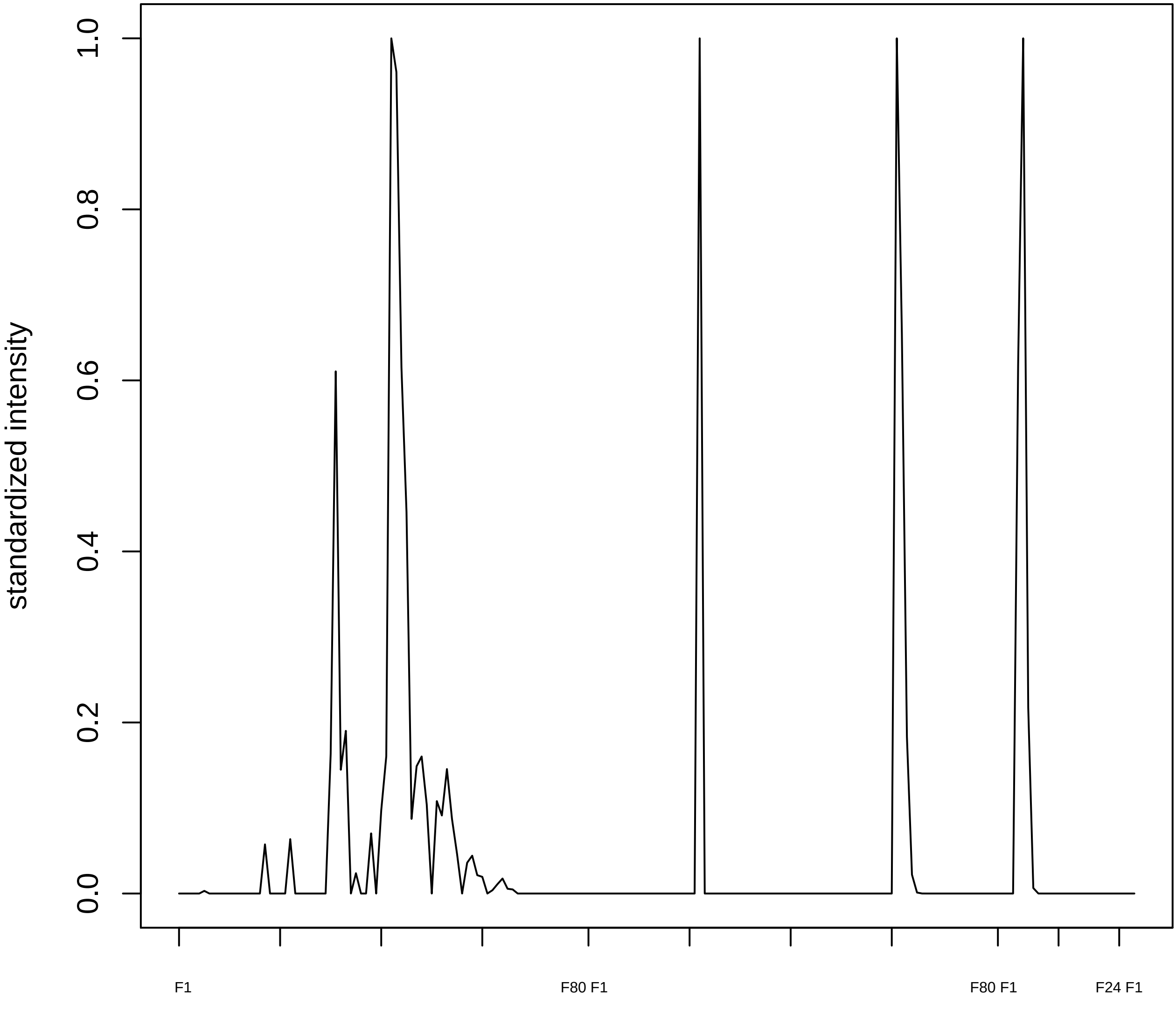

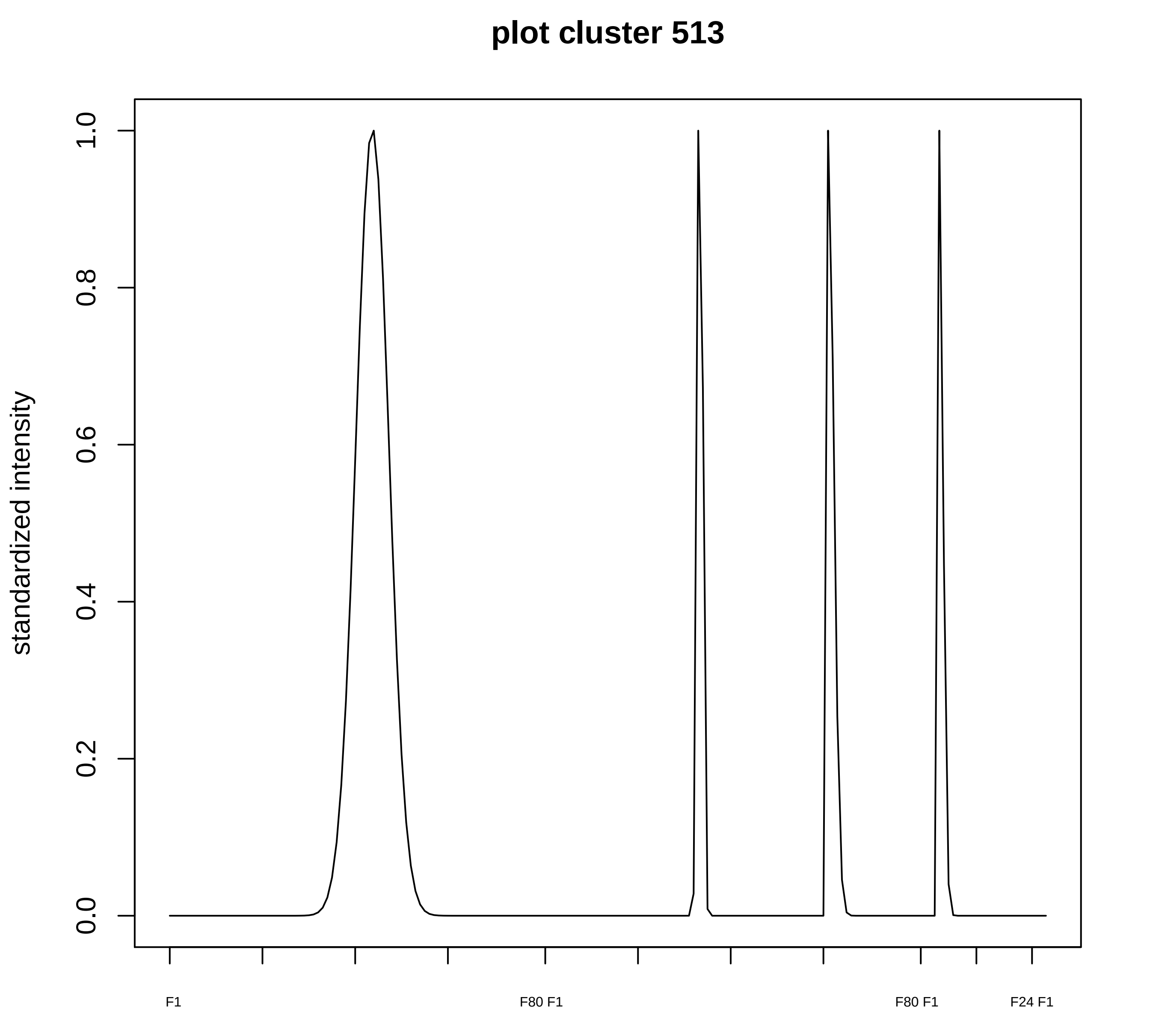
plot cluster 513
standardized intensity
0.0
0.2
0.4
0.6
0.8
1.0
F1
F80 F1
F80 F1
F24 F1

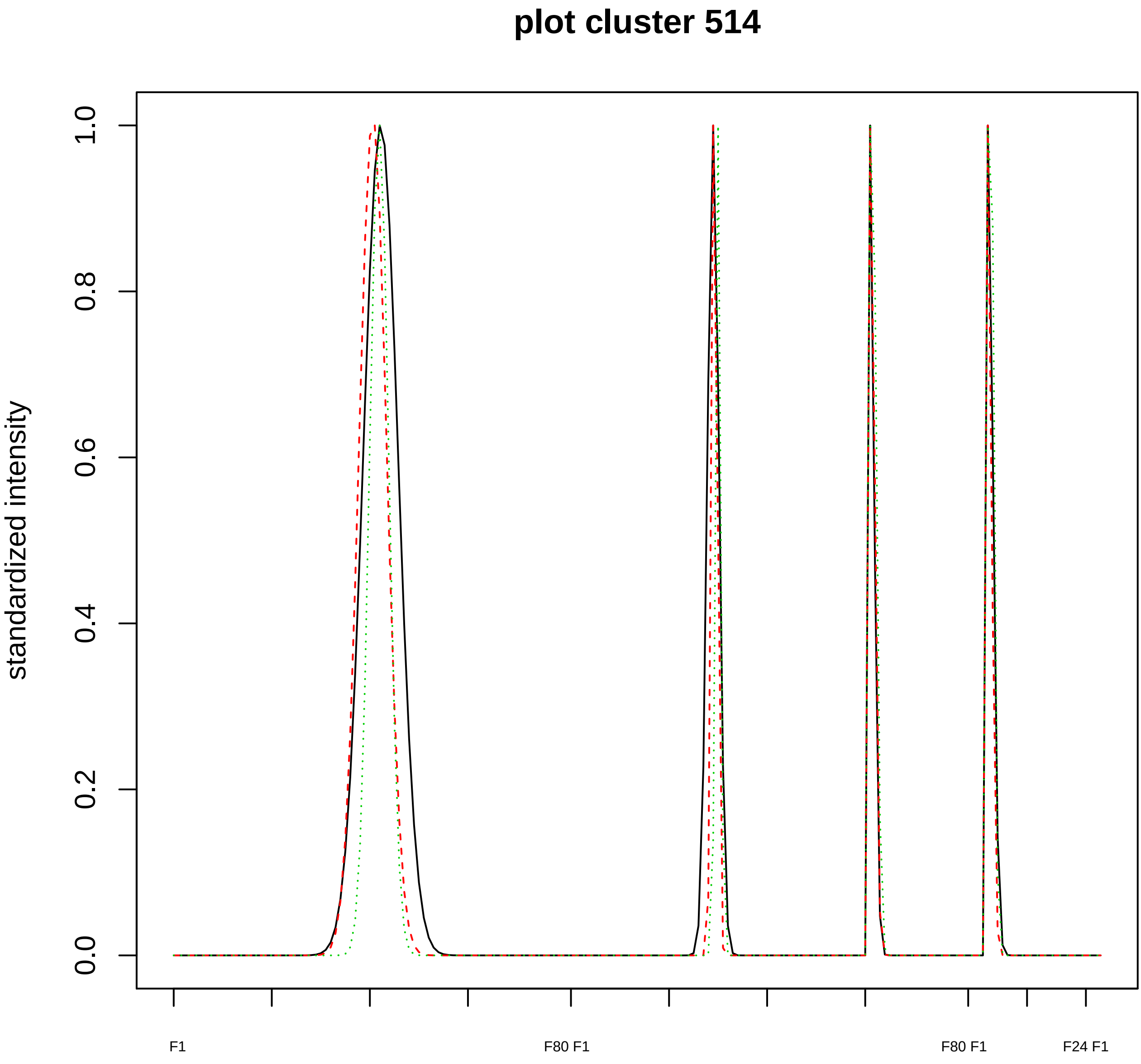
plot cluster 514
standardized intensity
0.0
0.2
0.4
0.6
0.8
1.0
F1
F80 F1
F80 F1
F24 F1

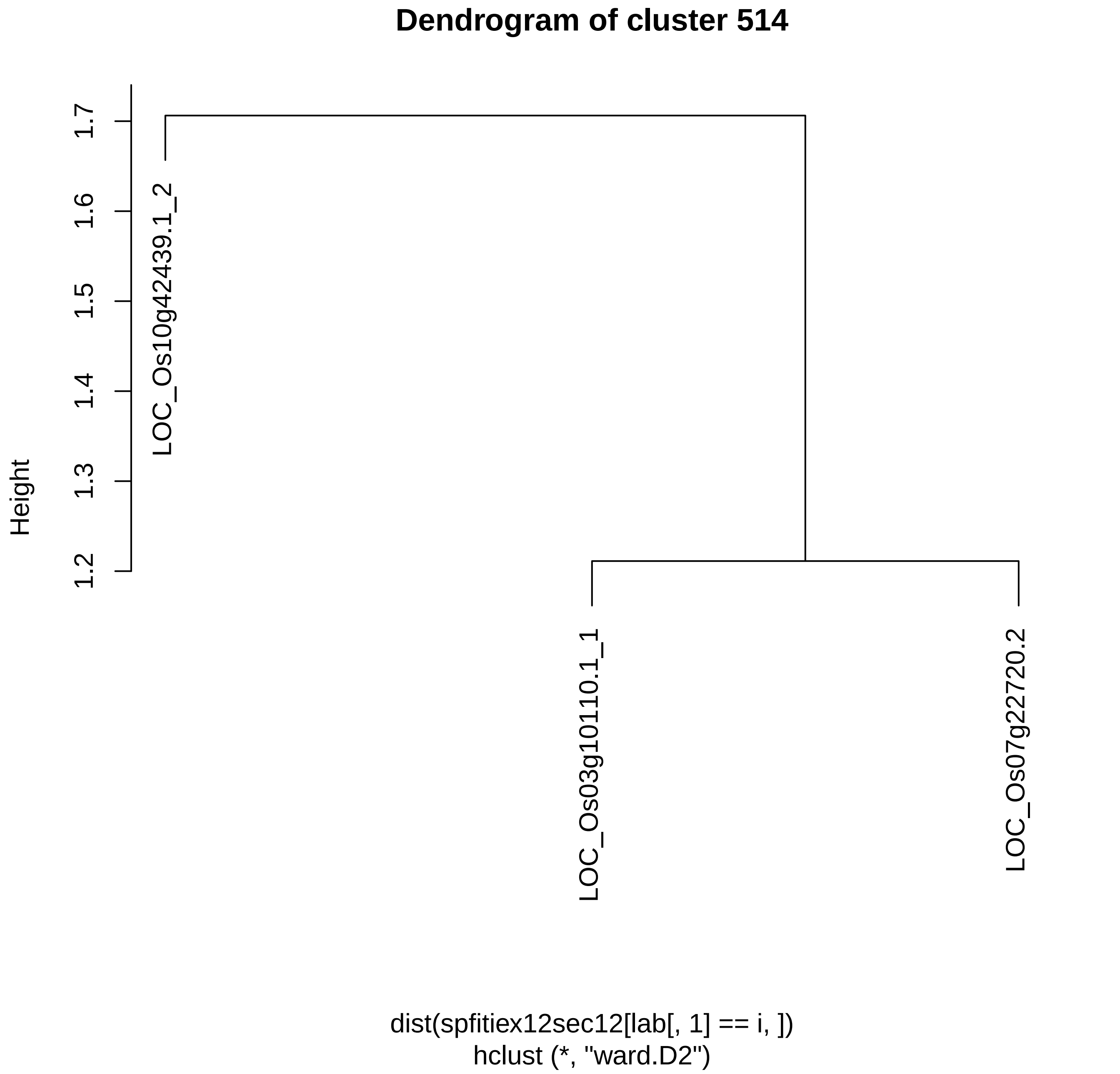
Dendrogram of cluster 514
Height
1.2
1.3
1.4
1.5
1.6
1.7
LOC_Os10g42439.1_2
LOC_Os03g10110.1_1
LOC_Os07g22720.2
dist(spfitiex12sec12[lab[, 1] == i, ])
hclust (*, "ward.D2")

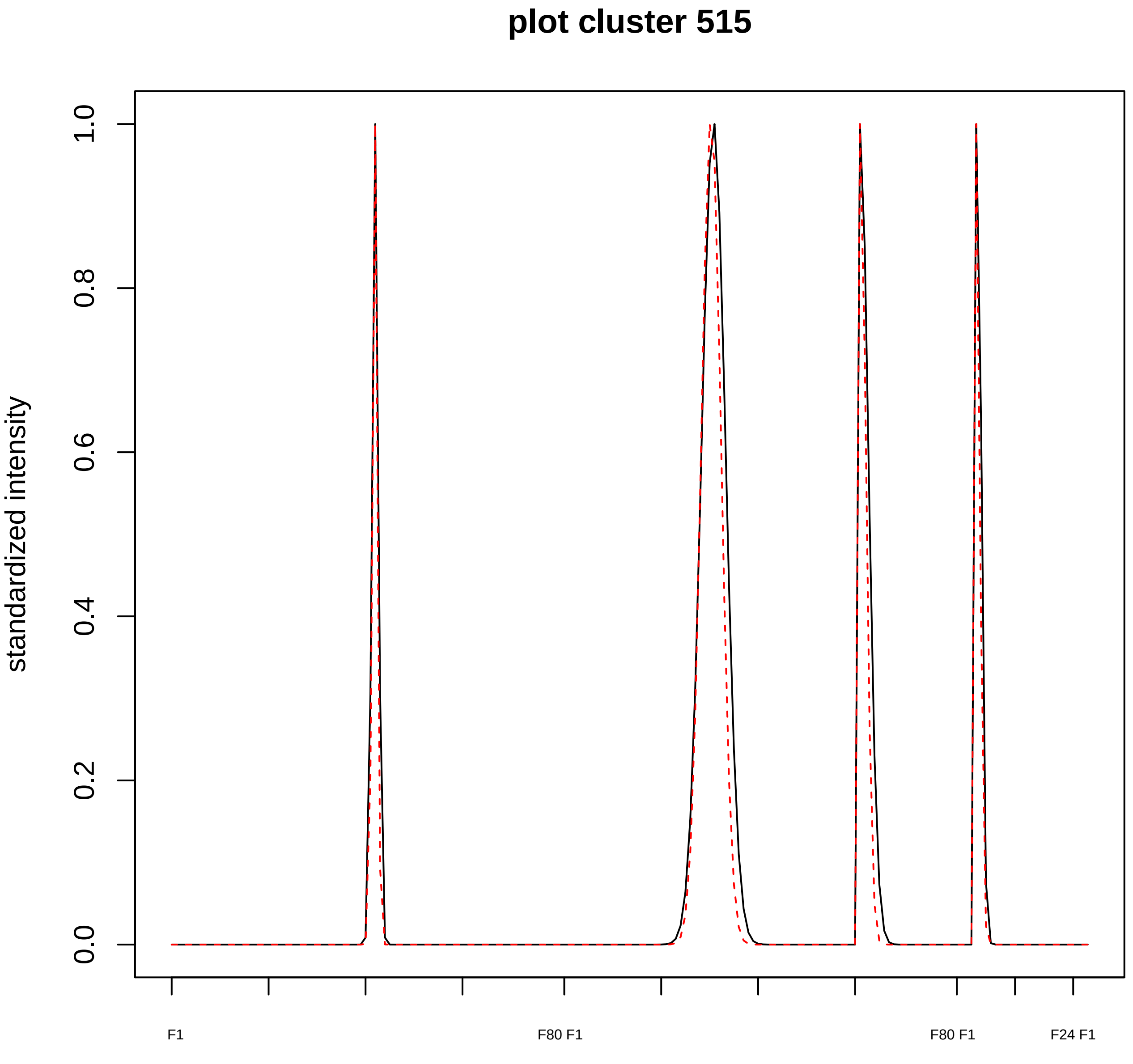

plot cluster 515
standardized intensity
0.0
0.2
0.4
0.6
0.8
1.0
F1
F80 F1
F80 F1
F24 F1

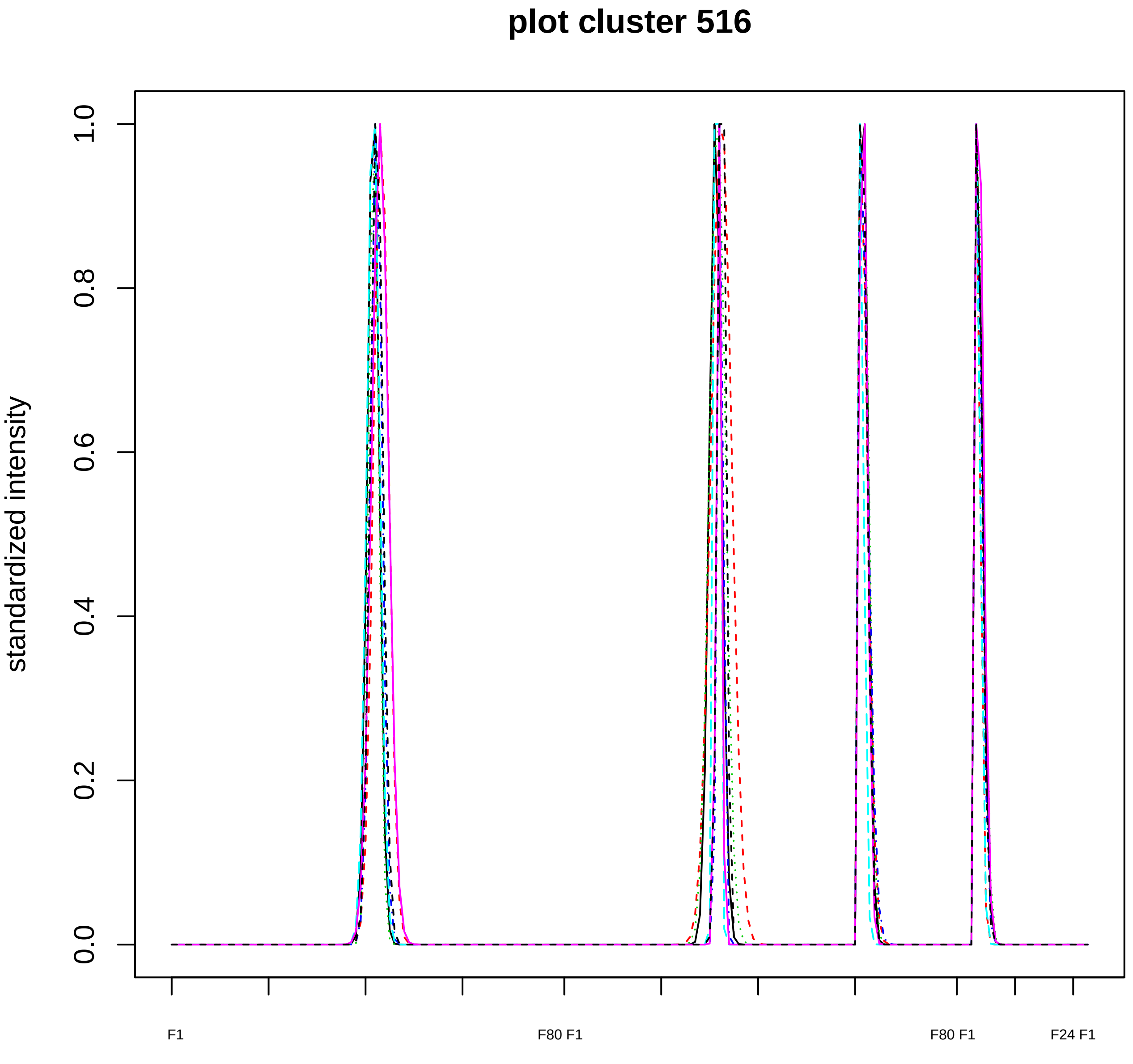

plot cluster 516
standardized intensity
0.0
0.2
0.4
0.6
0.8
1.0
F1
F80 F1
F80 F1
F24 F1

### Dendrogram of cluster 516

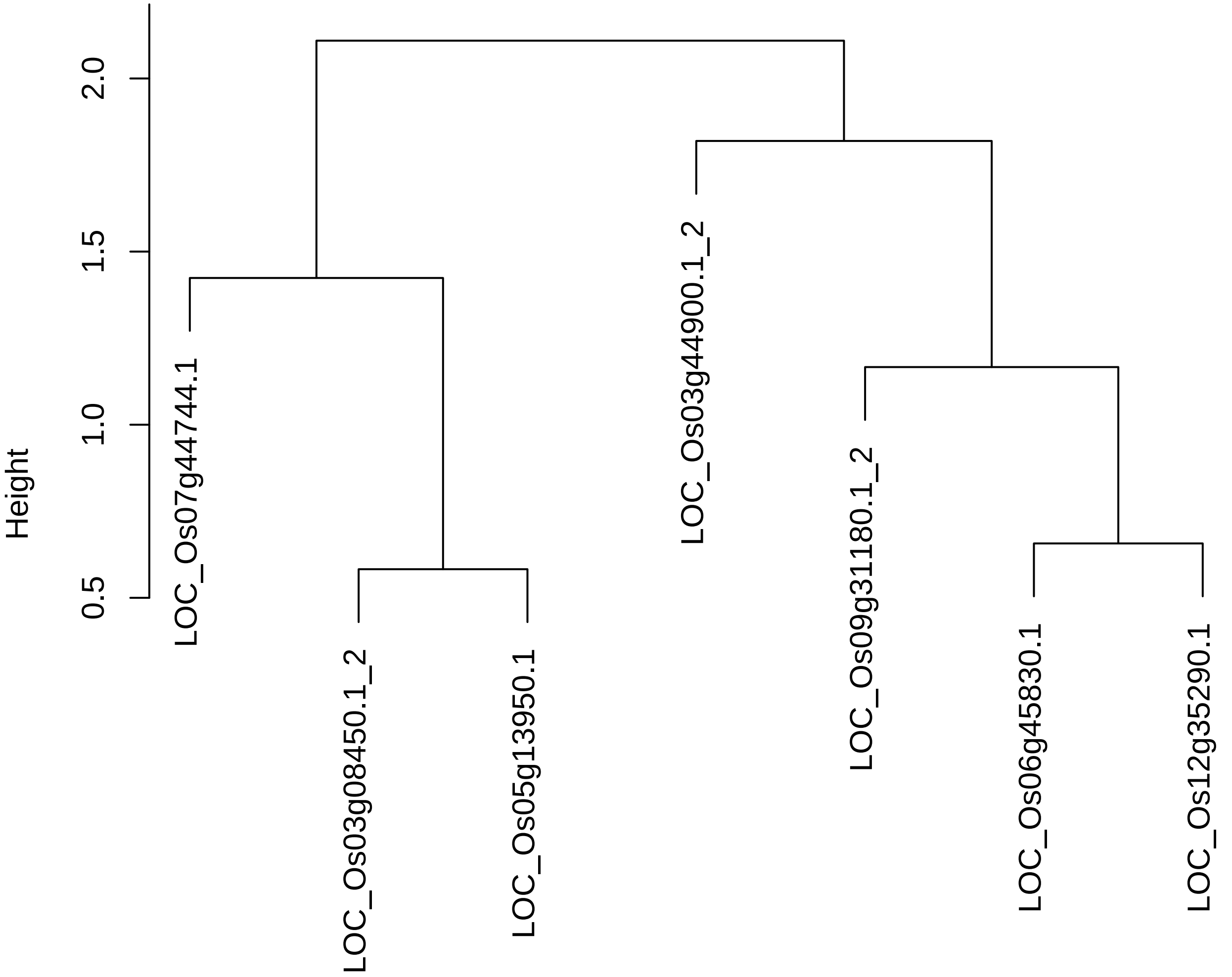


dist(spfitiex12sec12[lab[, 1] == i, ])
hclust (*, "ward.D2")

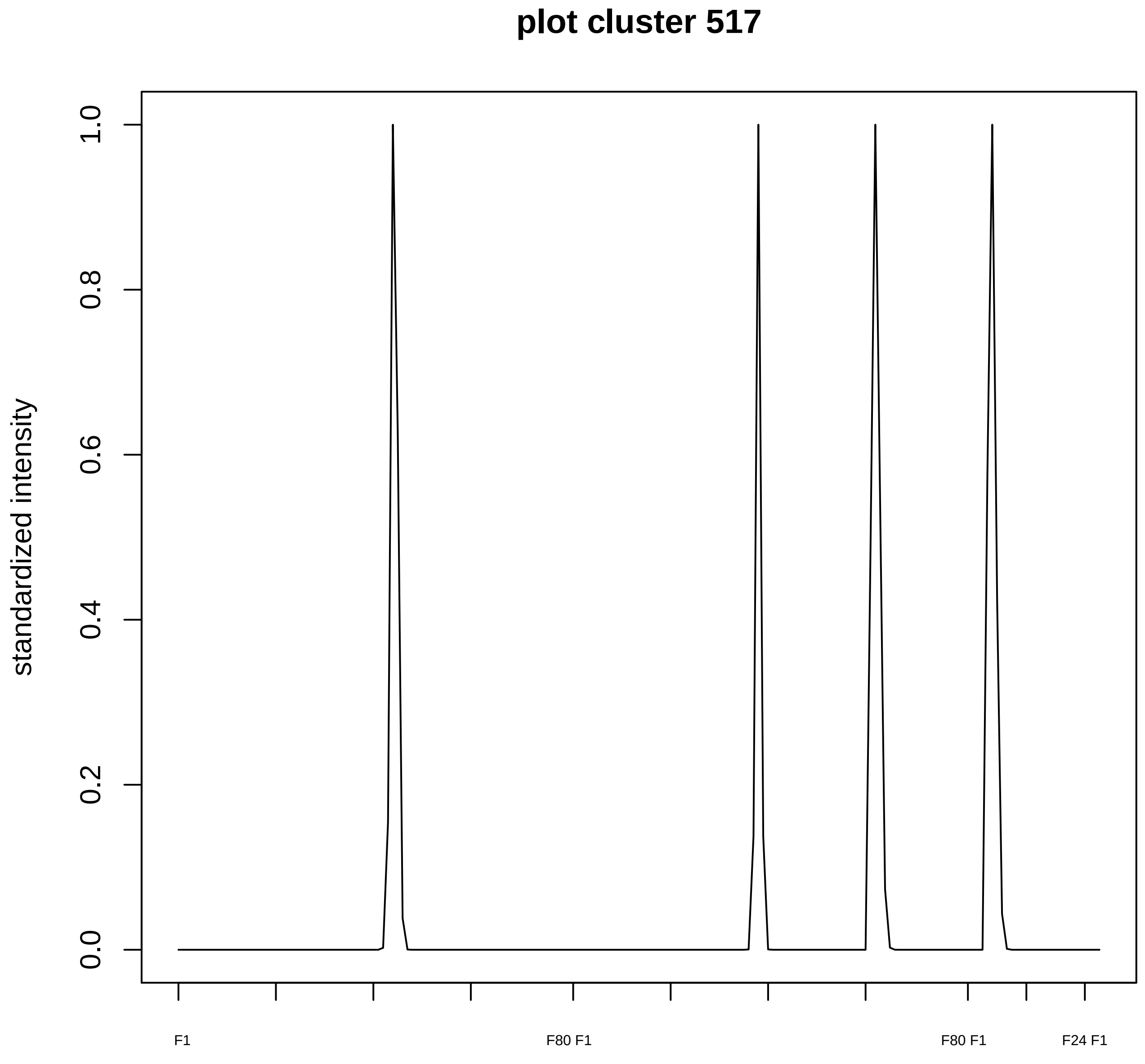
plot cluster 517
standardized intensity
0.0
0.2
0.4
0.6
0.8
1.0
F1
F80 F1
F80 F1
F24 F1

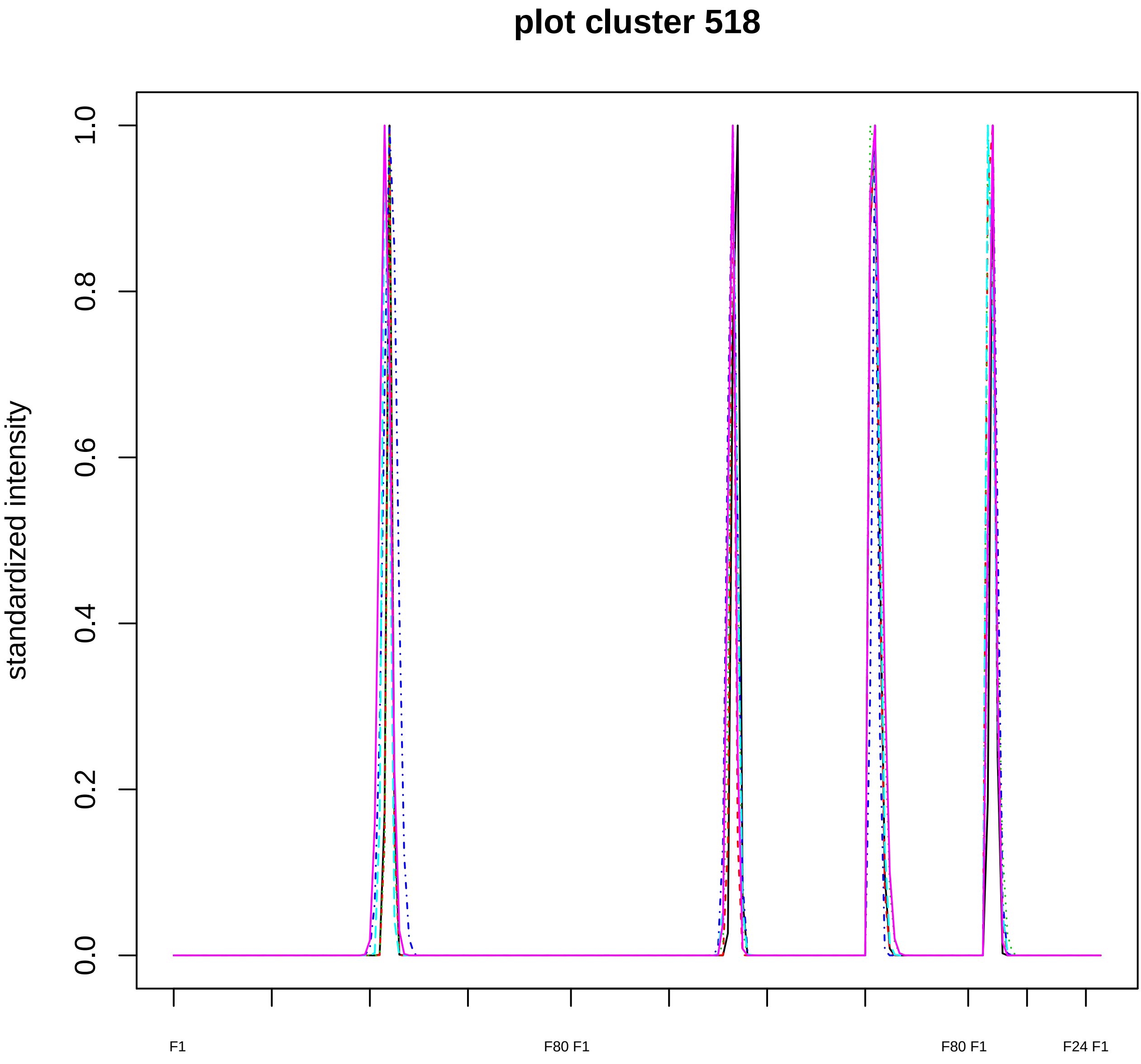
plot cluster 518
standardized intensity
0.0
0.2
0.4
0.6
0.8
1.0
F1
F80 F1
F80 F1
F24 F1

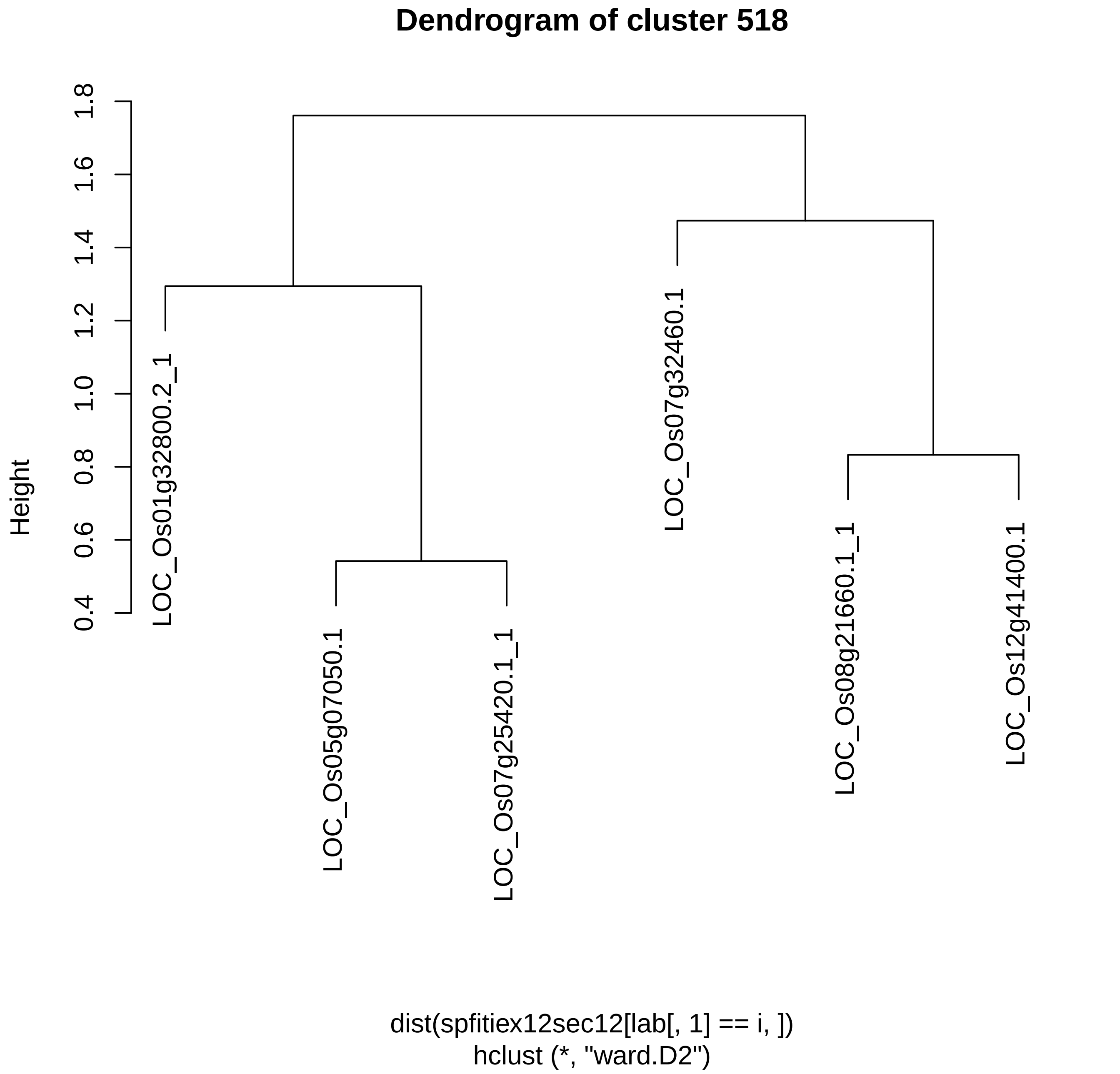
Dendrogram of cluster 518
Height
0.4
0.6
0.8
1.0
1.2
1.4
1.6
1.8
LOC_Os01g32800.2_1
LOC_Os05g07050.1
LOC_Os07g25420.1_1
LOC_Os07g32460.1
LOC_Os08g21660.1_1
LOC_Os12g41400.1
dist(spfitiex12sec12[lab[, 1] == i, ])
hclust (*, "ward.D2")

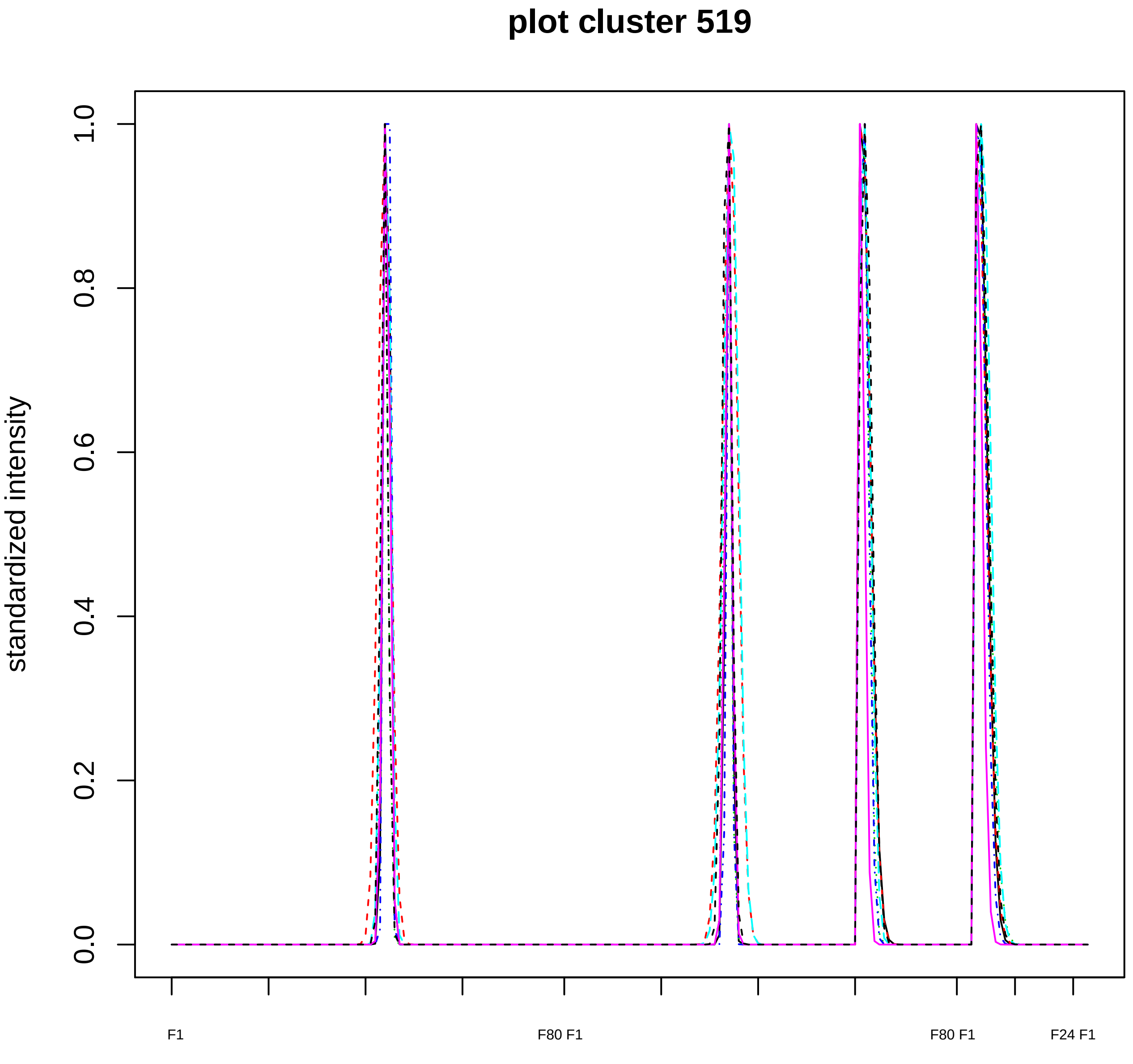

plot cluster 519
standardized intensity
1.0
0.8
0.6
0.4
0.2
0.0
F1
F80 F1
F80 F1
F24 F1

#### Dendrogram of cluster 519

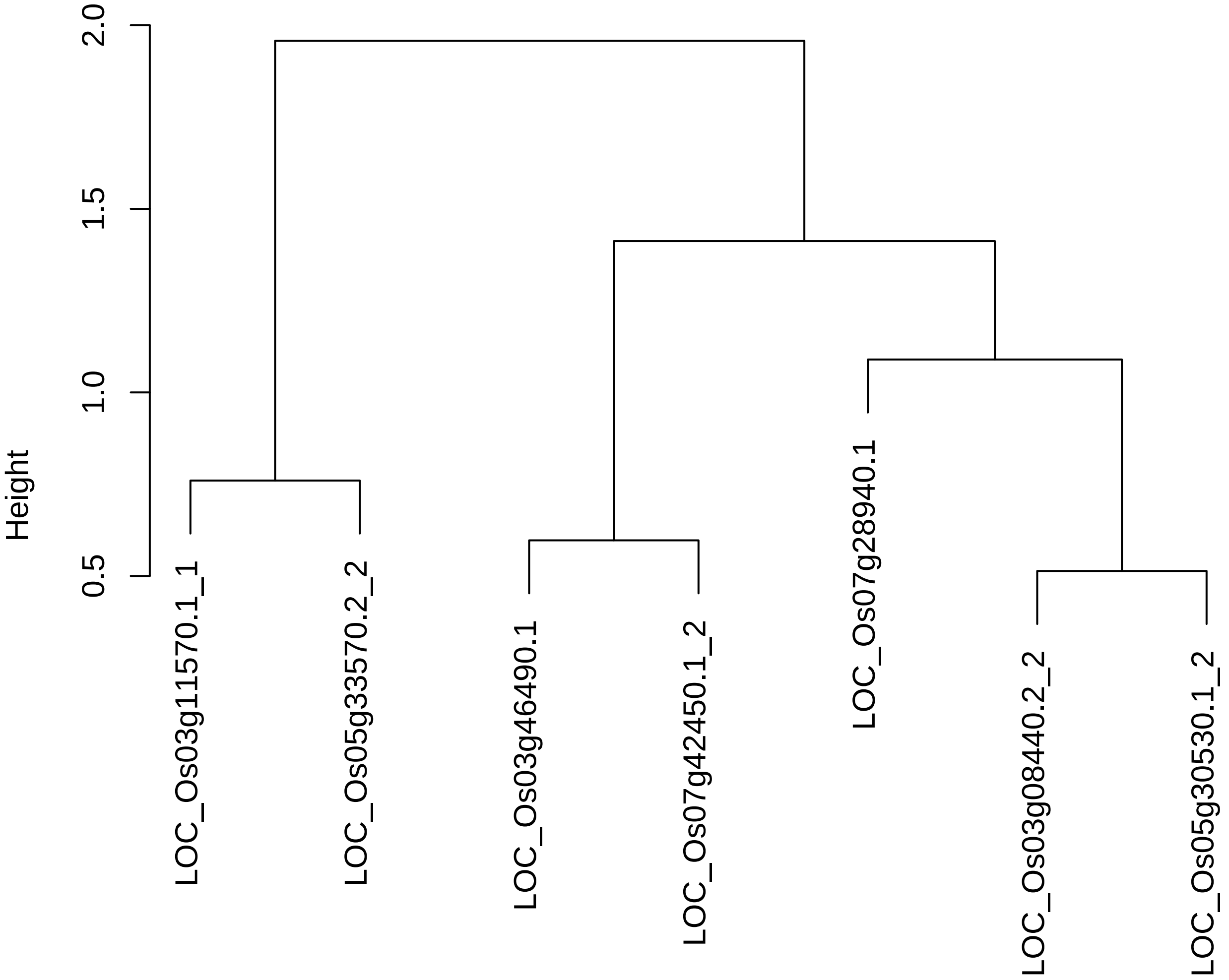


dist(spfitiex12sec12[lab[, 1] == i, ])
hclust (*, "ward.D2")

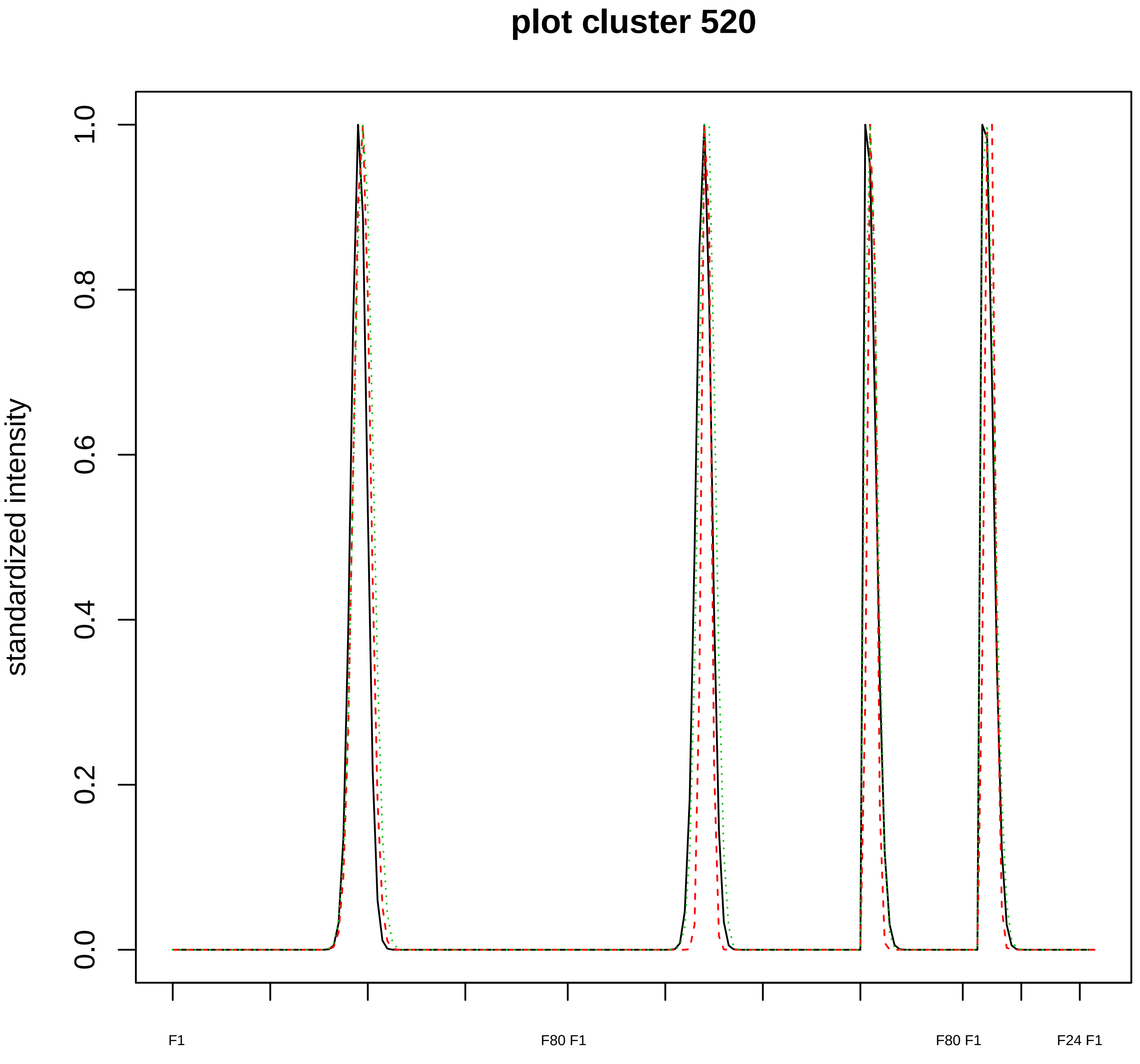
plot cluster 520
standardized intensity
0.0
0.2
0.4
0.6
0.8
1.0
F1
F80 F1
F80 F1
F24 F1

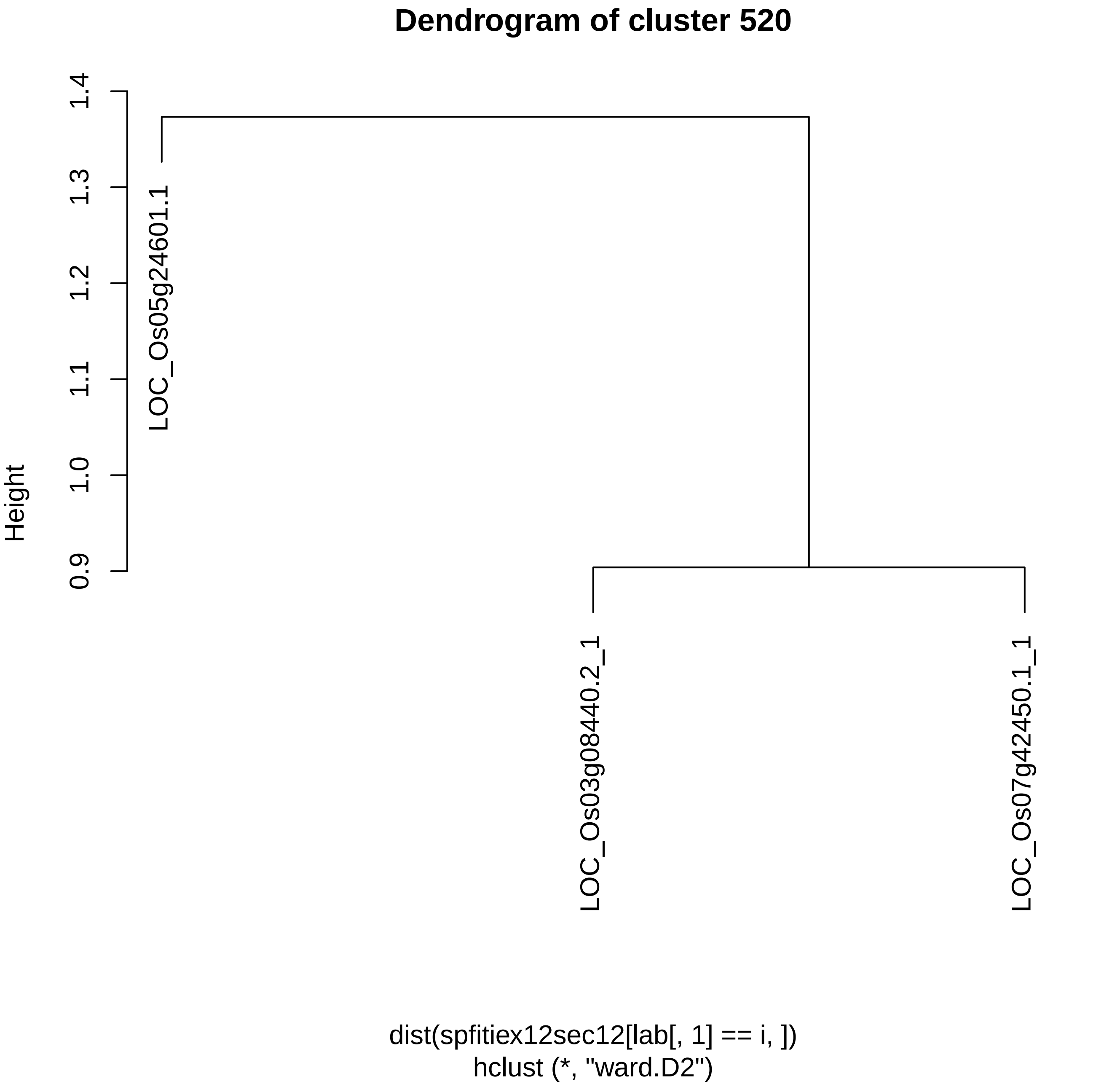
Dendrogram of cluster 520
Height
1.4
1.3
1.2
1.1
1.0
0.9
LOC_Os05g24601.1
LOC_Os03g08440.2_1
LOC_Os07g42450.1_1
dist(spfitiex12sec12[lab[, 1] == i, ])
hclust (*, "ward.D2")

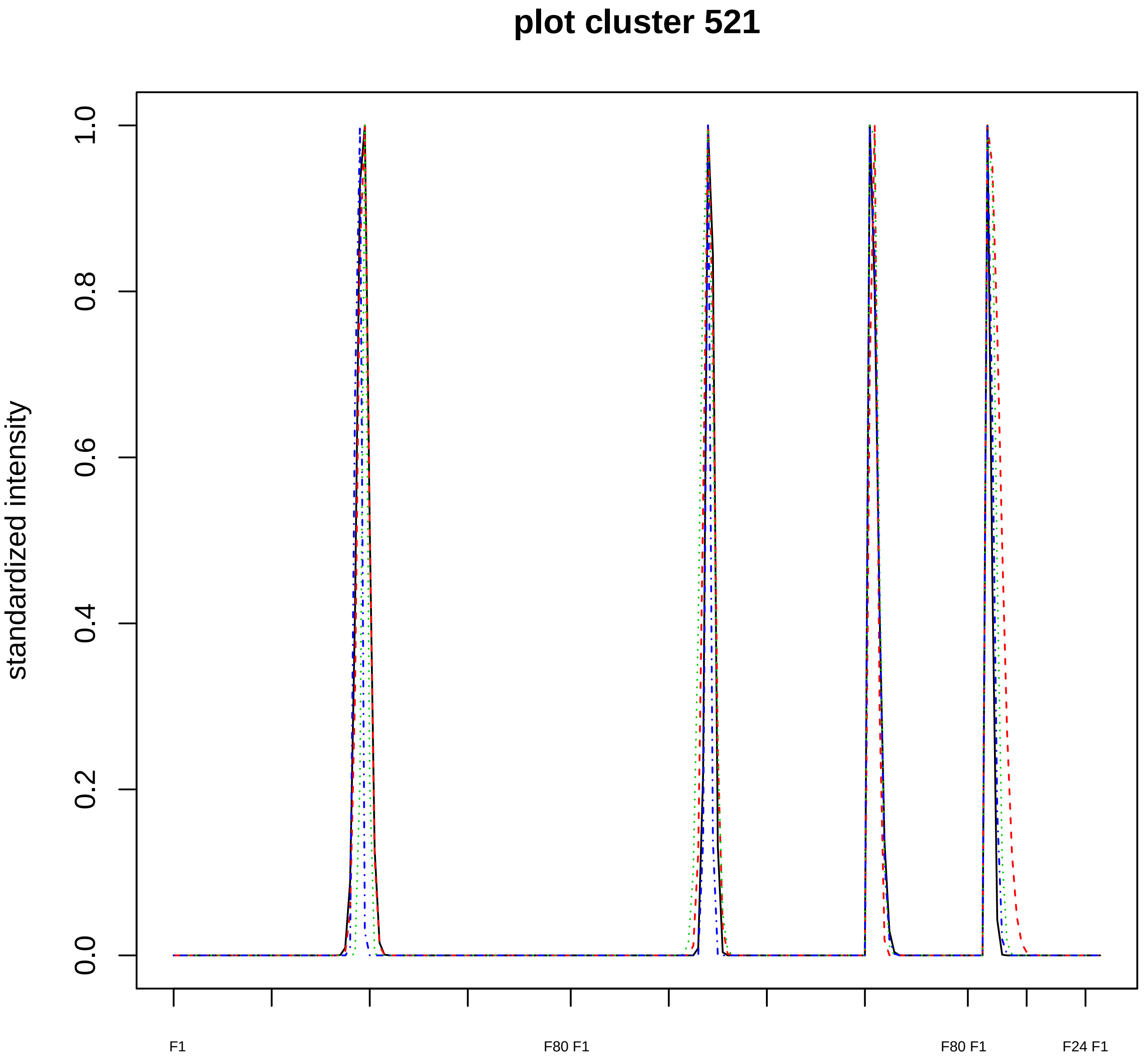
plot cluster 521
standardized intensity
0.0
0.2
0.4
0.6
0.8
1.0
F1
F80 F1
F80 F1
F24 F1

#### Dendrogram of cluster 521

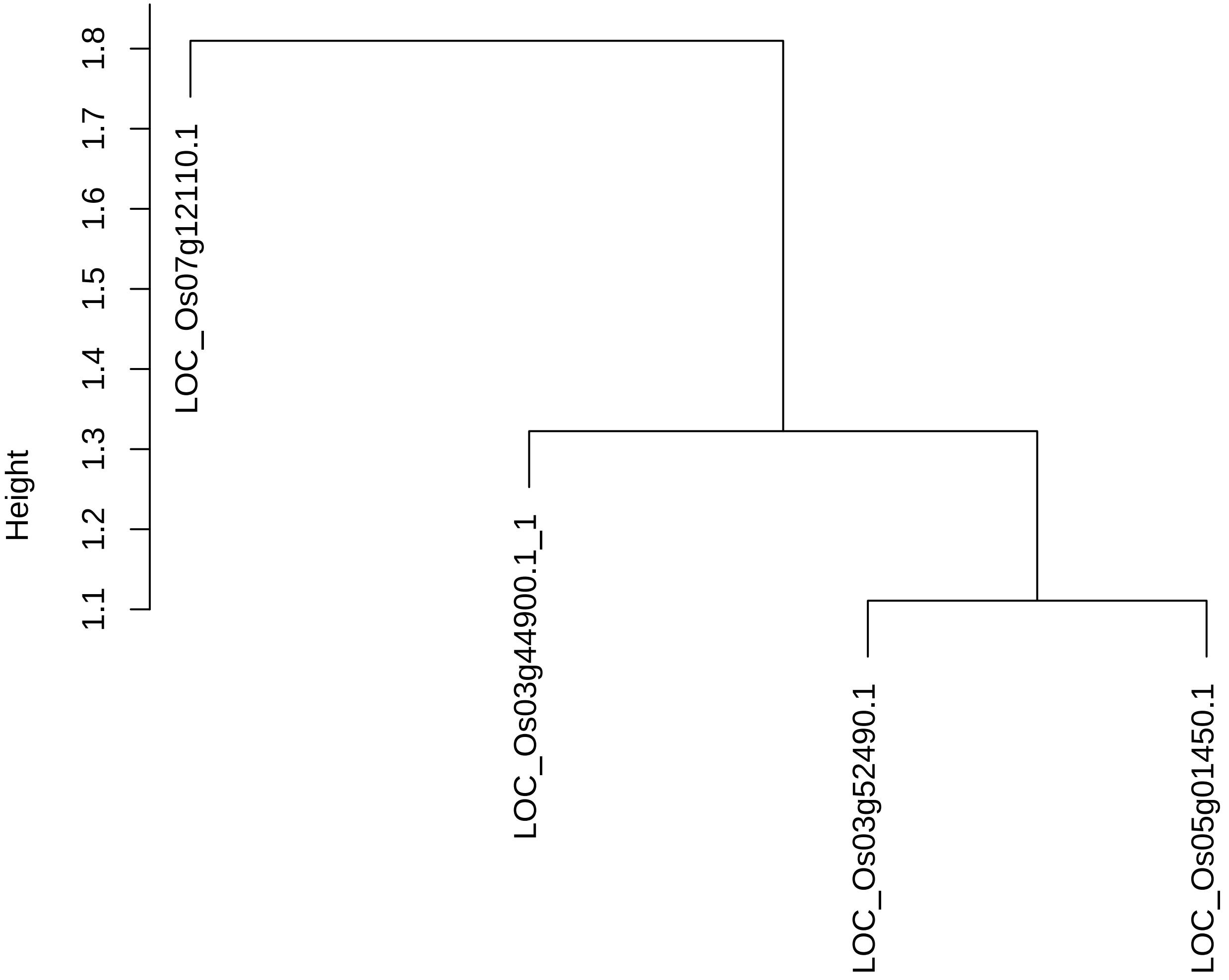


dist(spfitiex12sec12[lab[, 1] == i, ])
hclust (*, "ward.D2")

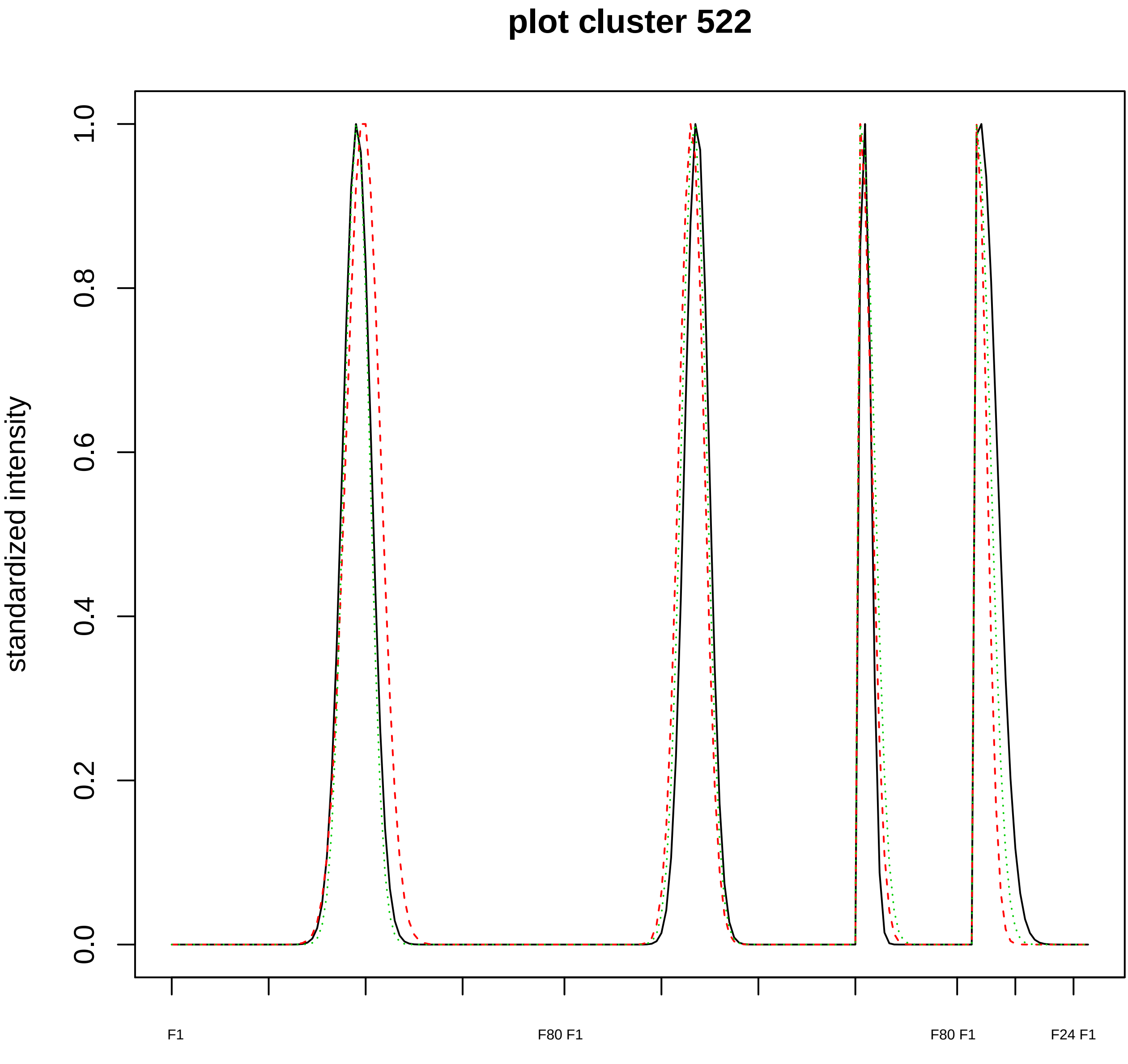
plot cluster 522
standardized intensity
0.0
0.2
0.4
0.6
0.8
1.0
F1
F80 F1
F80 F1
F24 F1

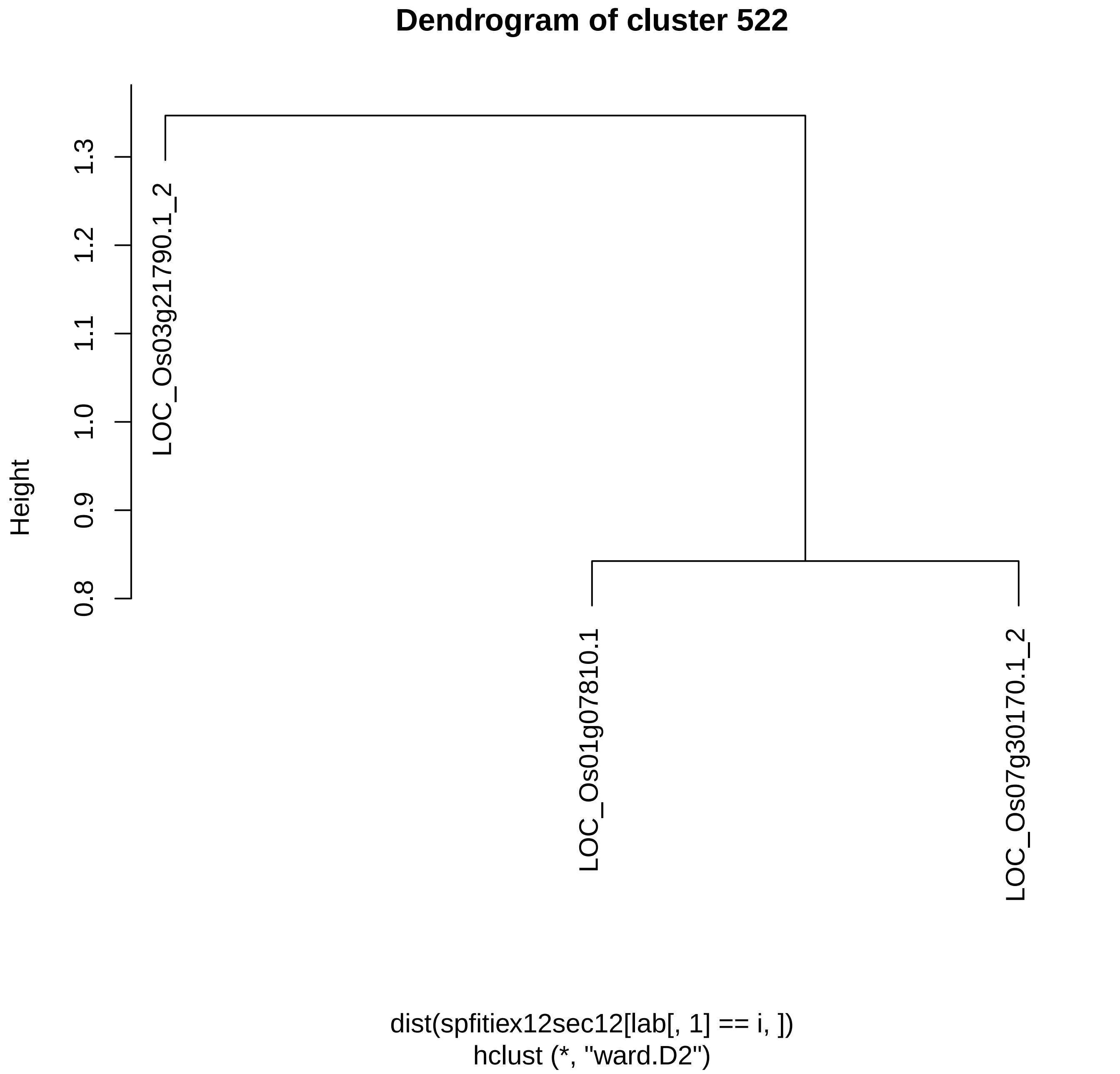
Dendrogram of cluster 522
Height
0.8
0.9
1.0
1.1
1.2
1.3
LOC_Os03g21790.1_2
LOC_Os01g07810.1
LOC_Os07g30170.1_2
dist(spfitiex12sec12[lab[, 1] == i, ])
hclust (*, "ward.D2")

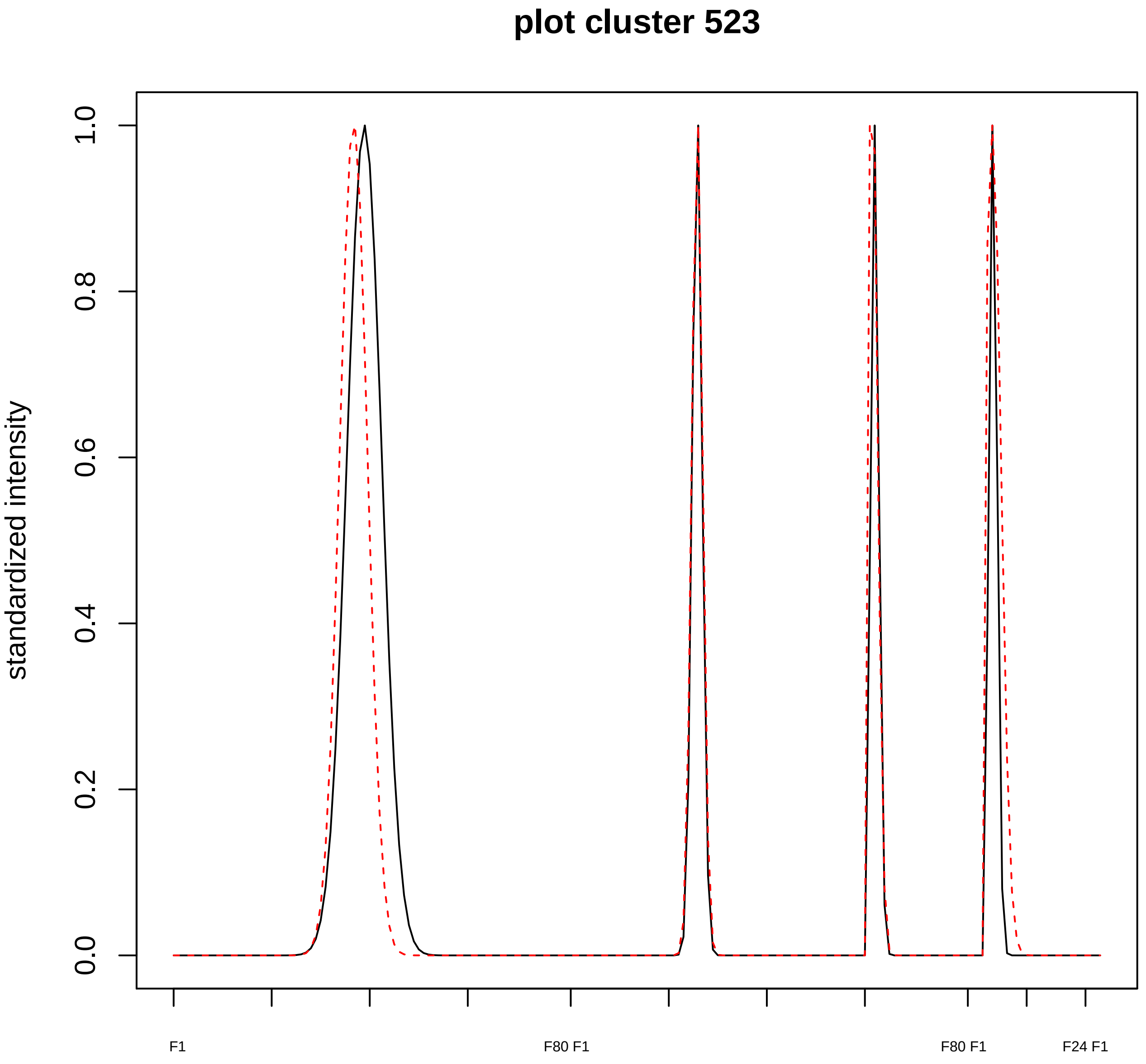
plot cluster 523
standardized intensity
0.0
0.2
0.4
0.6
0.8
1.0
F1
F80 F1
F80 F1
F24 F1

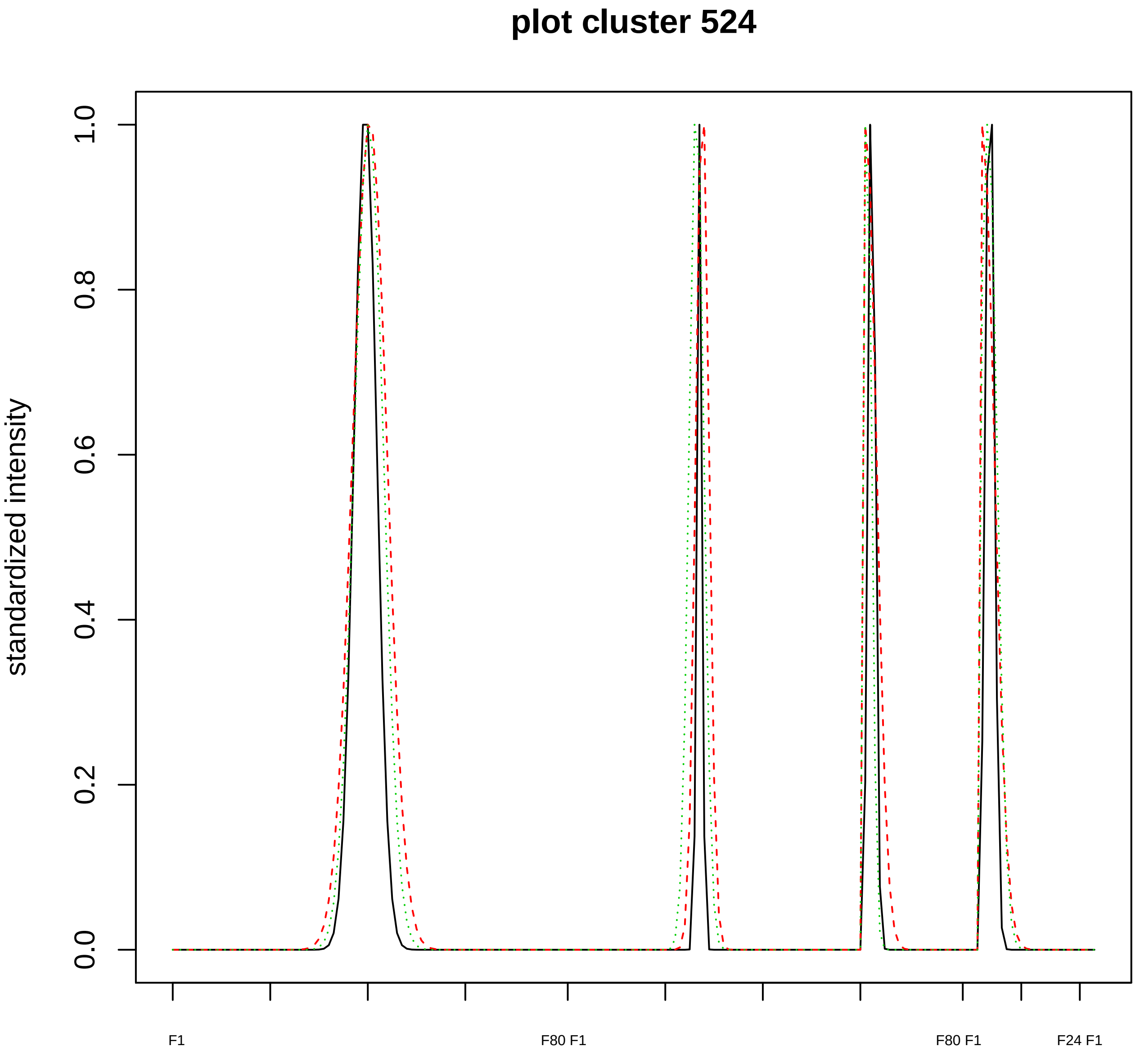
plot cluster 524
standardized intensity
0.0
0.2
0.4
0.6
0.8
1.0
F1
F80 F1
F80 F1
F24 F1

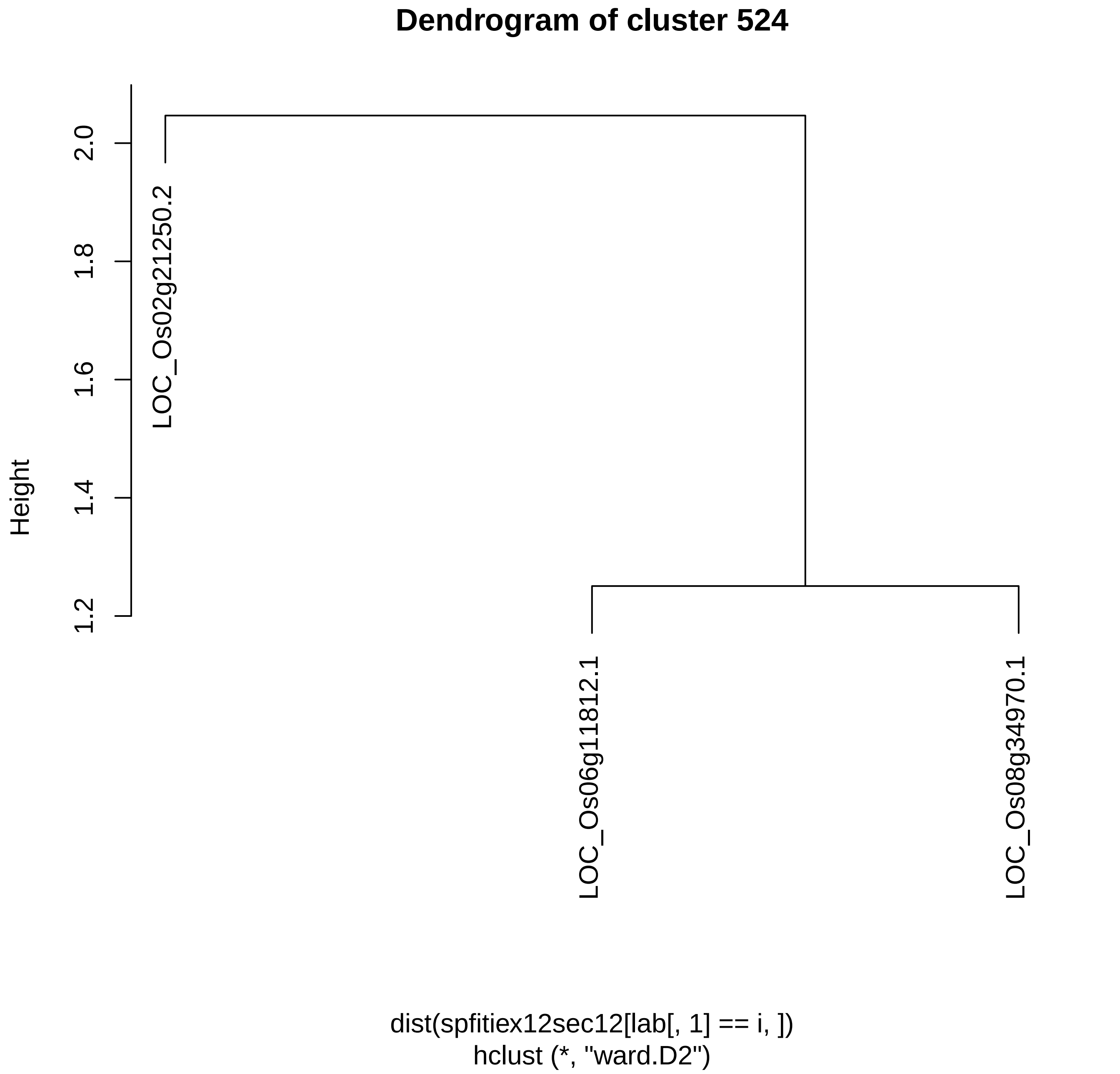
Dendrogram of cluster 524
Height
2.0
1.8
1.6
1.4
1.2
LOC_Os02g21250.2
LOC_Os06g11812.1
LOC_Os08g34970.1
dist(spfitiex12sec12[lab[, 1] == i, ])
hclust (*, "ward.D2")

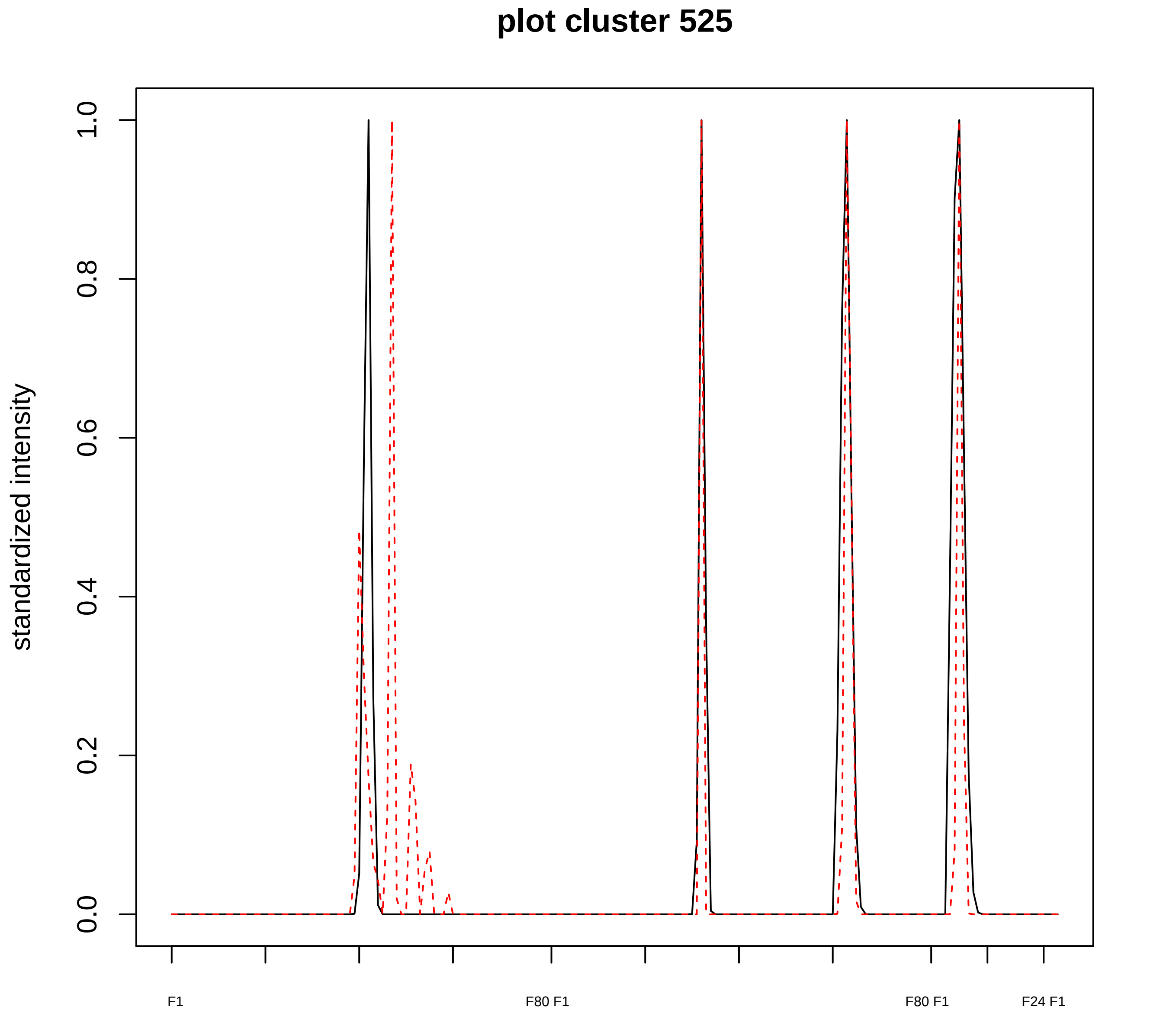

plot cluster 525
standardized intensity
0.0
0.2
0.4
0.6
0.8
1.0
F1
F80 F1
F80 F1
F24 F1

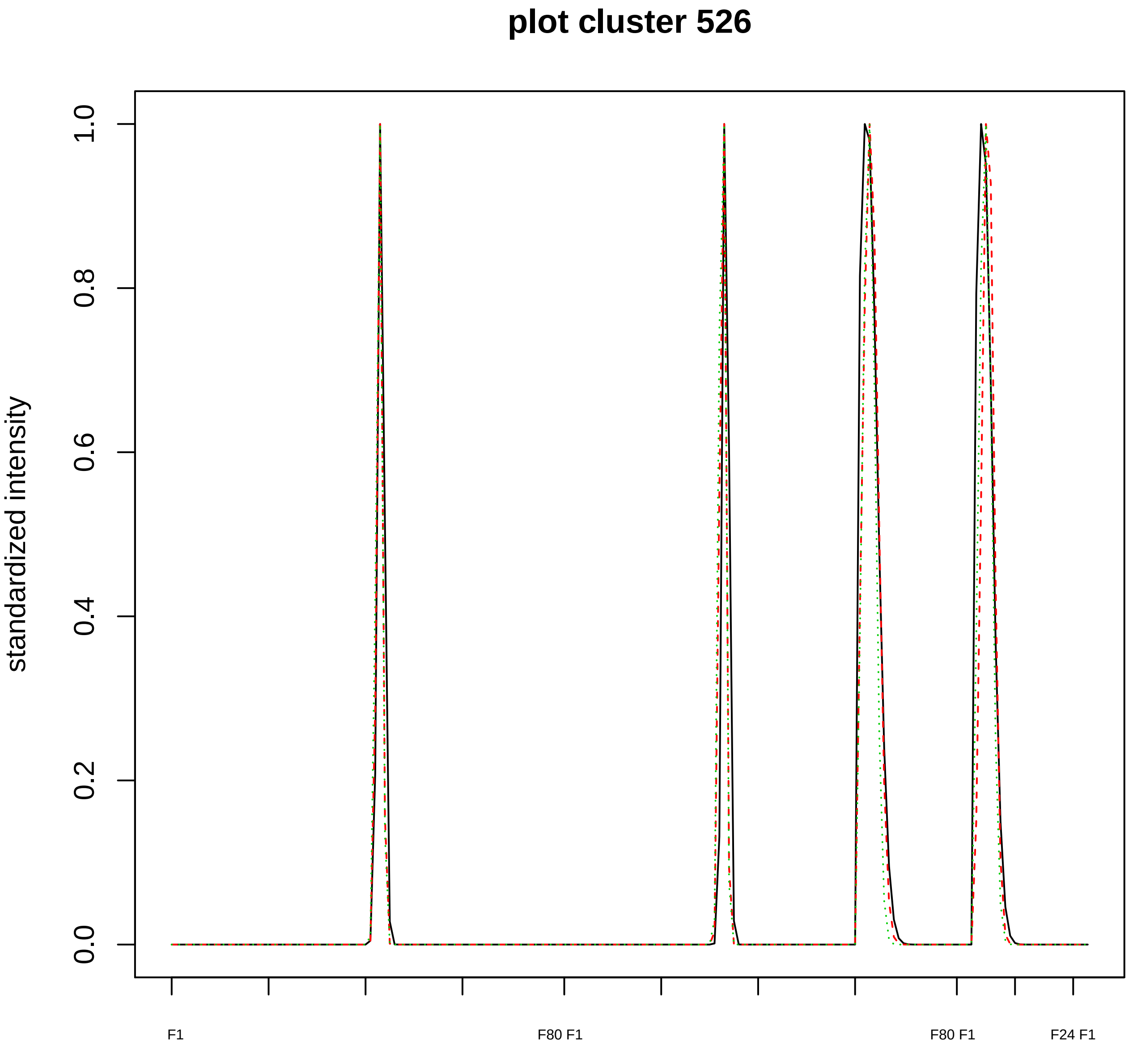

plot cluster 526
standardized intensity
1.0
0.8
0.6
0.4
0.2
0.0
F1
F80 F1
F80 F1
F24 F1

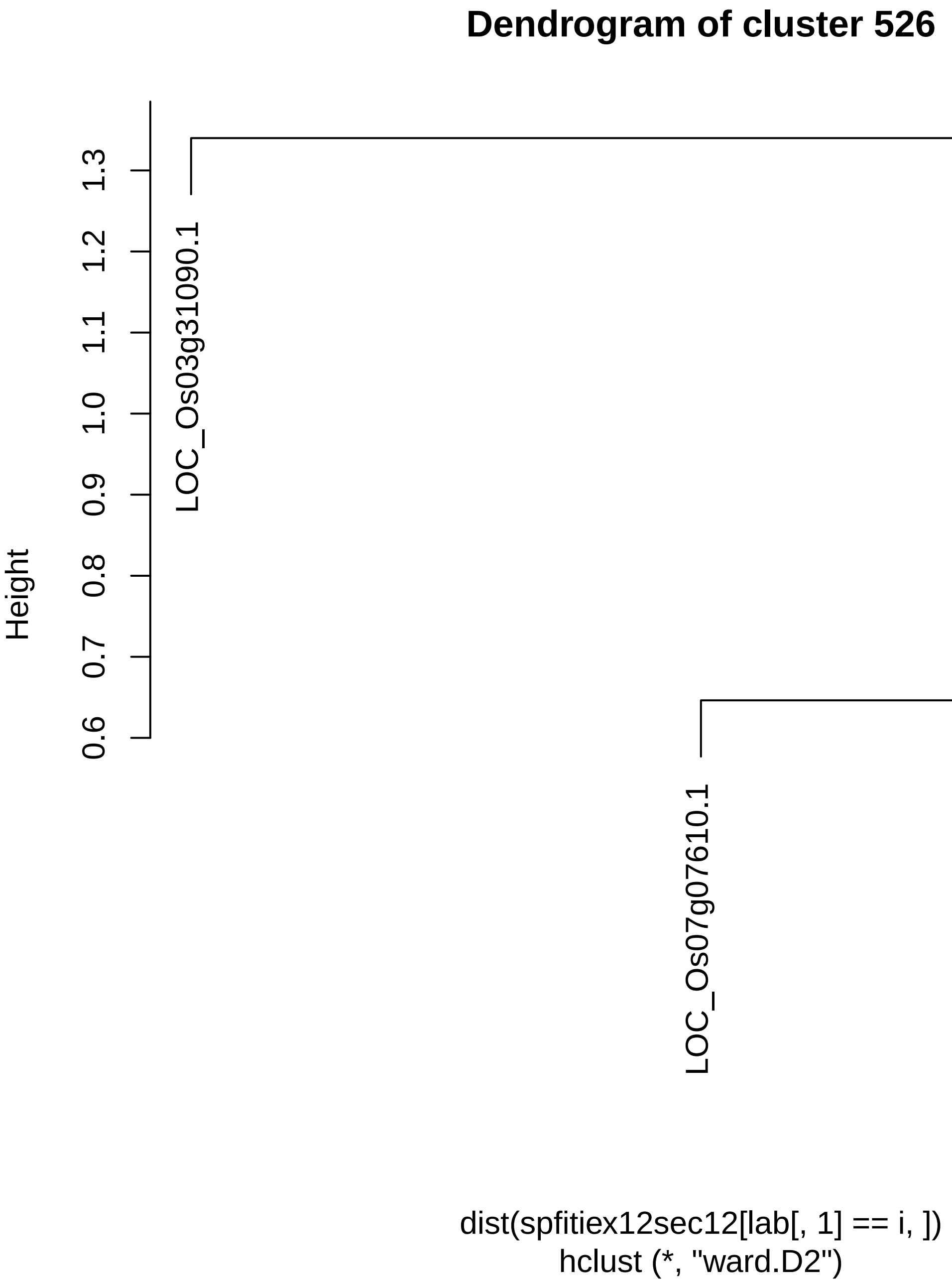
Dendrogram of cluster 526
Height
0.6
0.7
0.8
0.9
1.0
1.1
1.2
1.3
LOC_Os03g31090.1
LOC_Os07g07610.1
LOC_Os09g02284.1
dist(spfitiex12sec12[lab[, 1] == i, ])
hclust (*, "ward.D2")

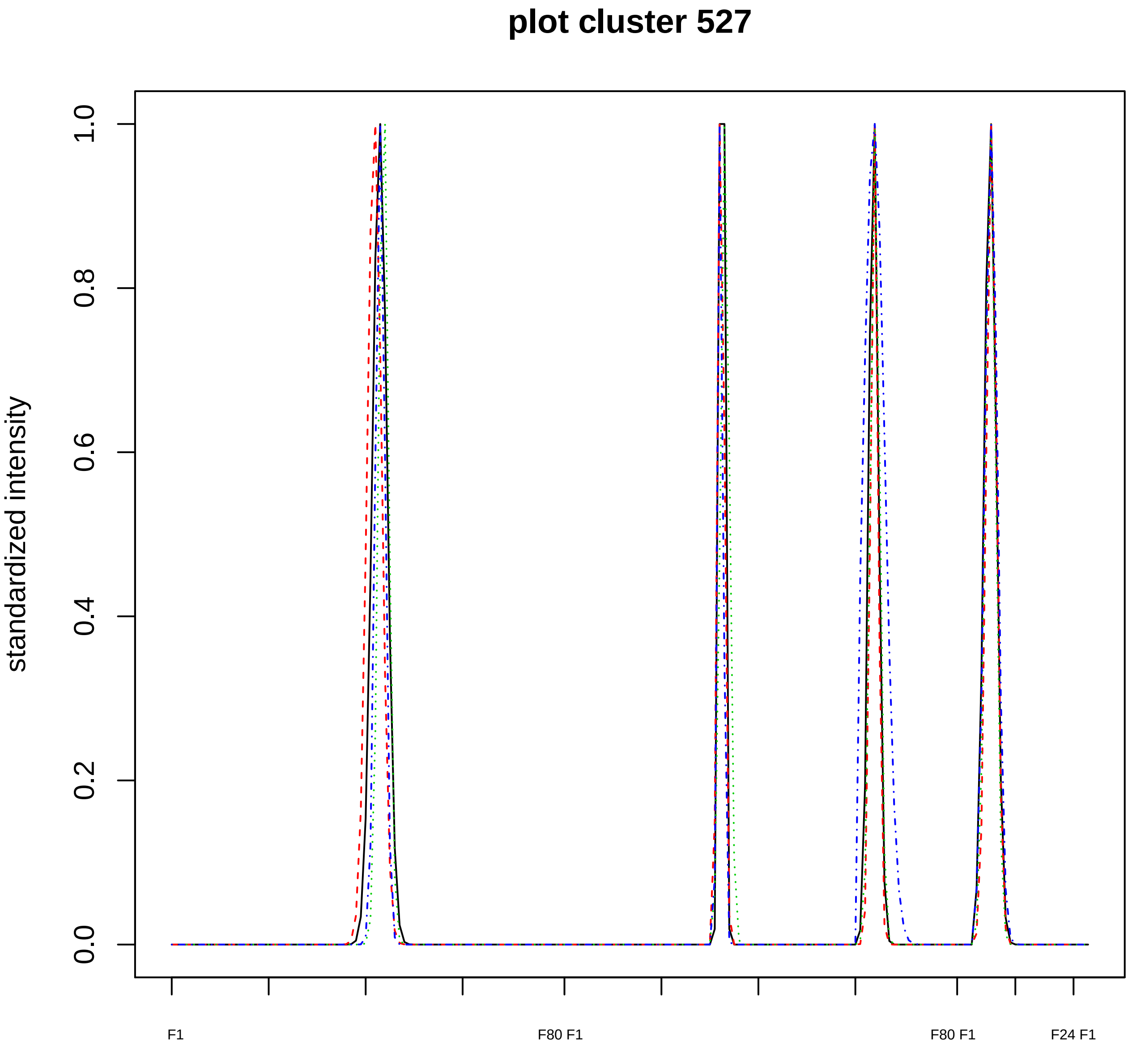
plot cluster 527
standardized intensity
0.0
0.2
0.4
0.6
0.8
1.0
F1
F80 F1
F80 F1
F24 F1

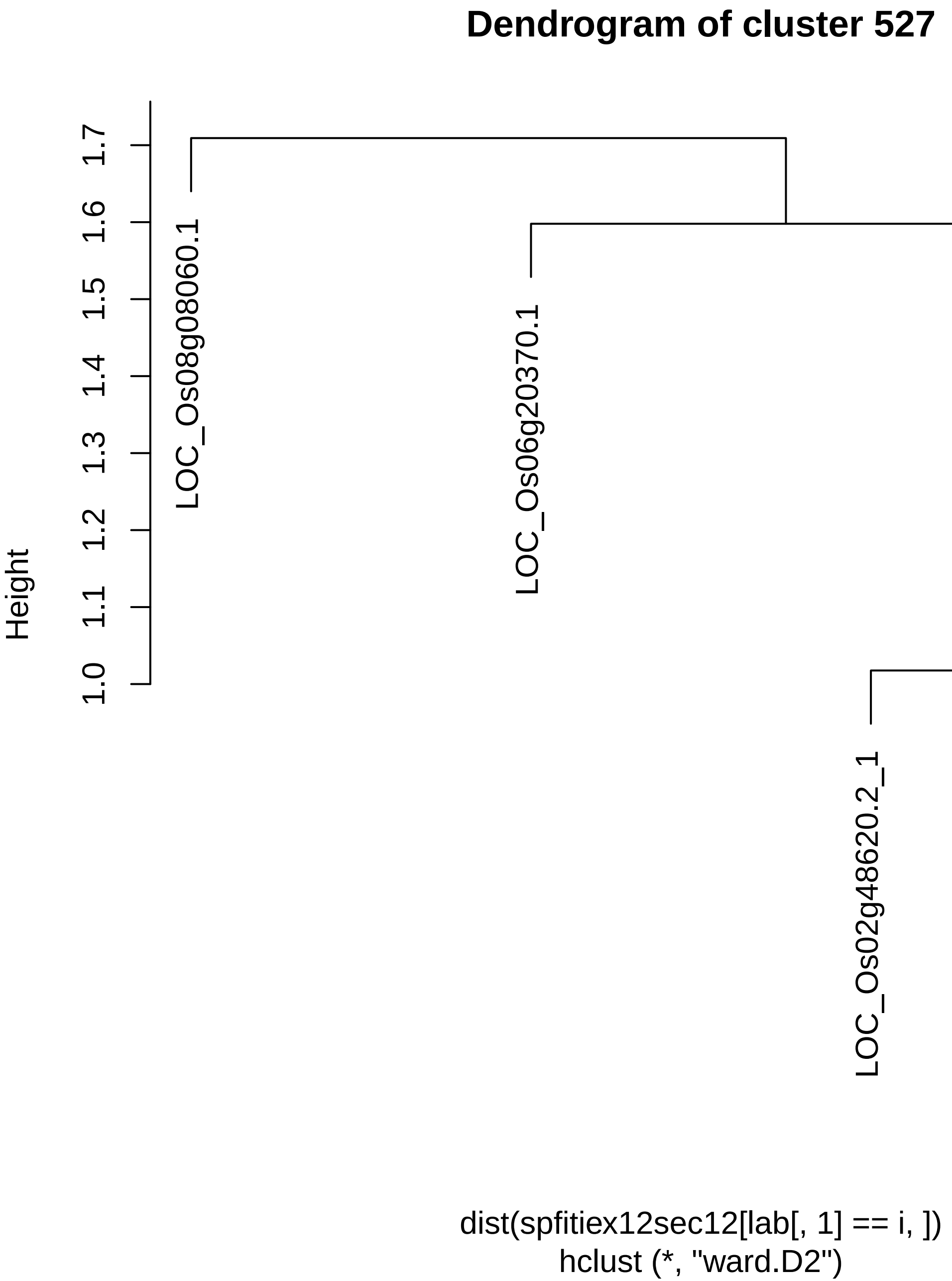
Dendrogram of cluster 527
Height
1.0
1.1
1.2
1.3
1.4
1.5
1.6
1.7
LOC_Os08g08060.1
LOC_Os06g20370.1
LOC_Os02g48620.2_1
LOC_Os02g48830.1
dist(spfitiex12sec12[lab[, 1] == i, ])
hclust (*, "ward.D2")

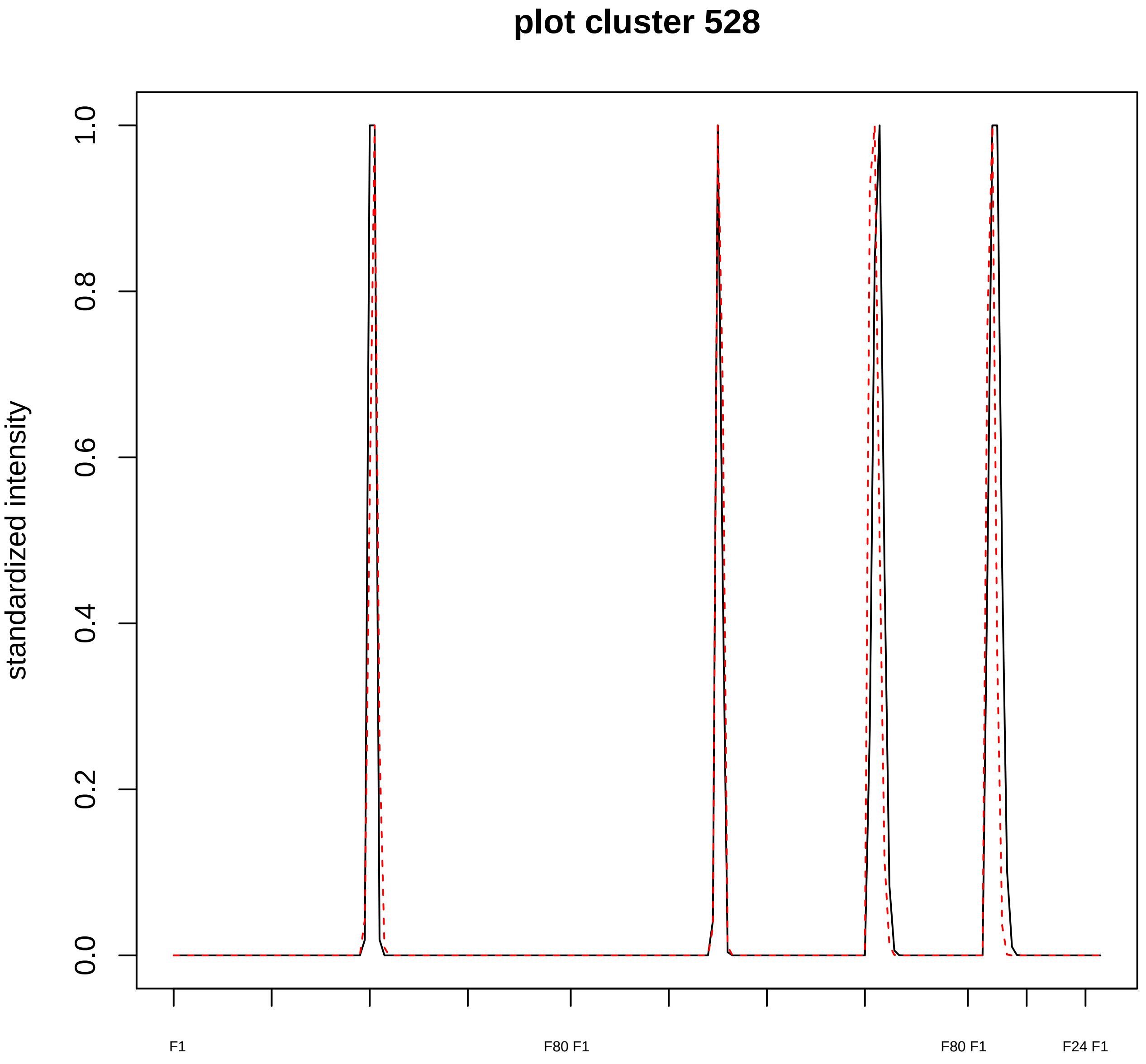
plot cluster 528
standardized intensity
0.0
0.2
0.4
0.6
0.8
1.0
F1
F80 F1
F80 F1
F24 F1

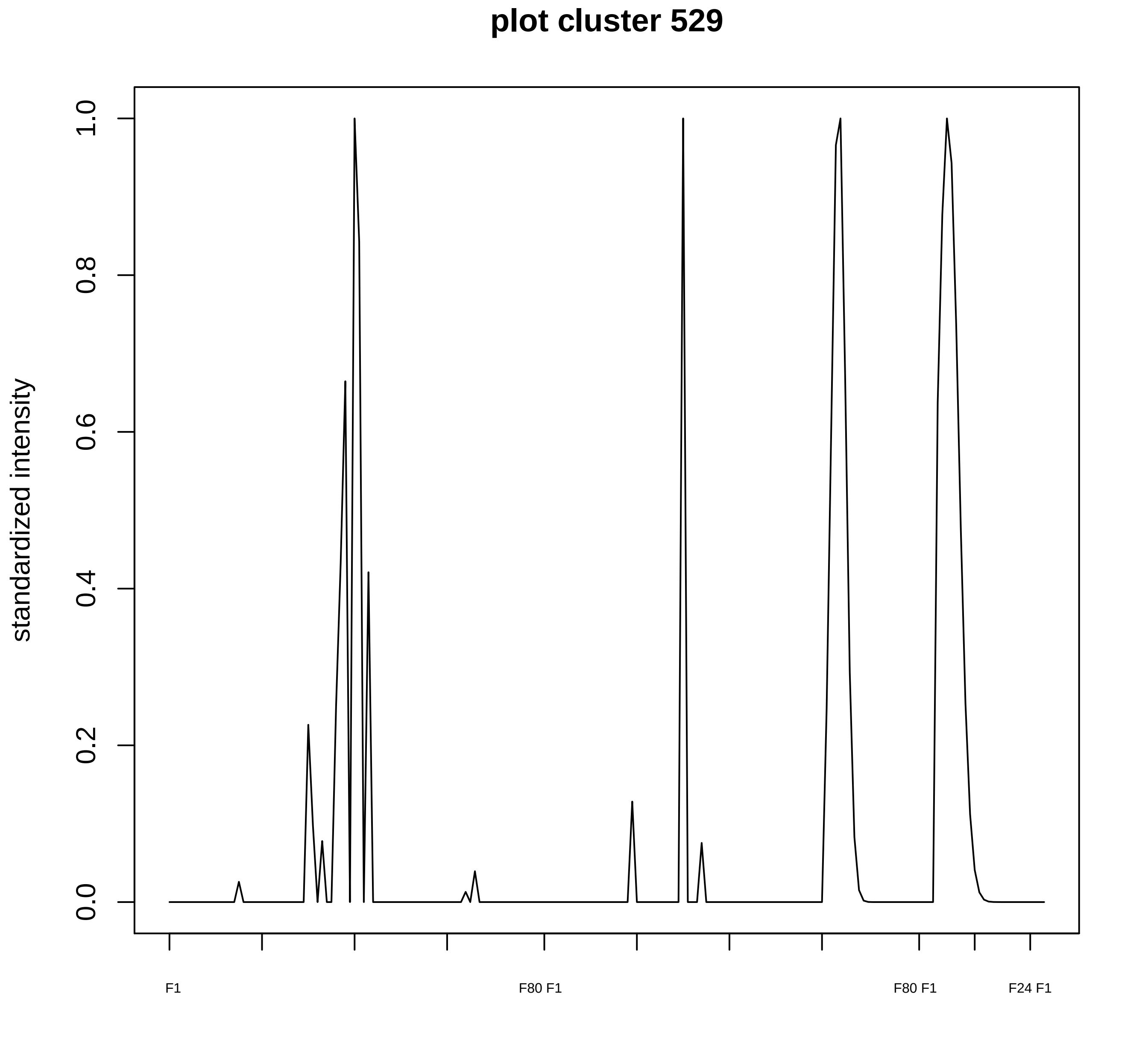

plot cluster 529
standardized intensity
0.0
0.2
0.4
0.6
0.8
1.0
F1
F80 F1
F80 F1
F24 F1

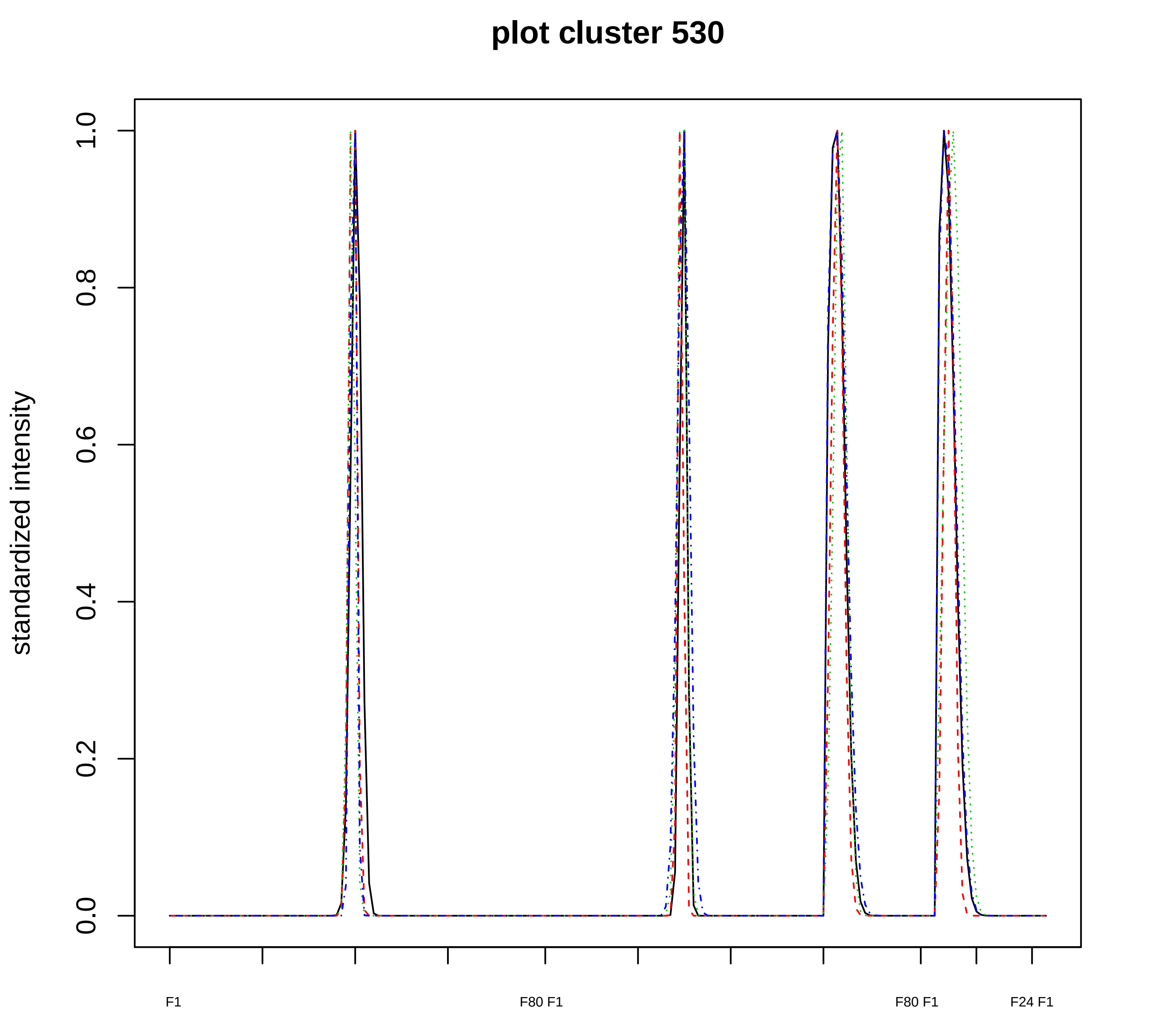

plot cluster 530
standardized intensity
0.0
0.2
0.4
0.6
0.8
1.0
F1
F80 F1
F80 F1
F24 F1

### Dendrogram of cluster 530

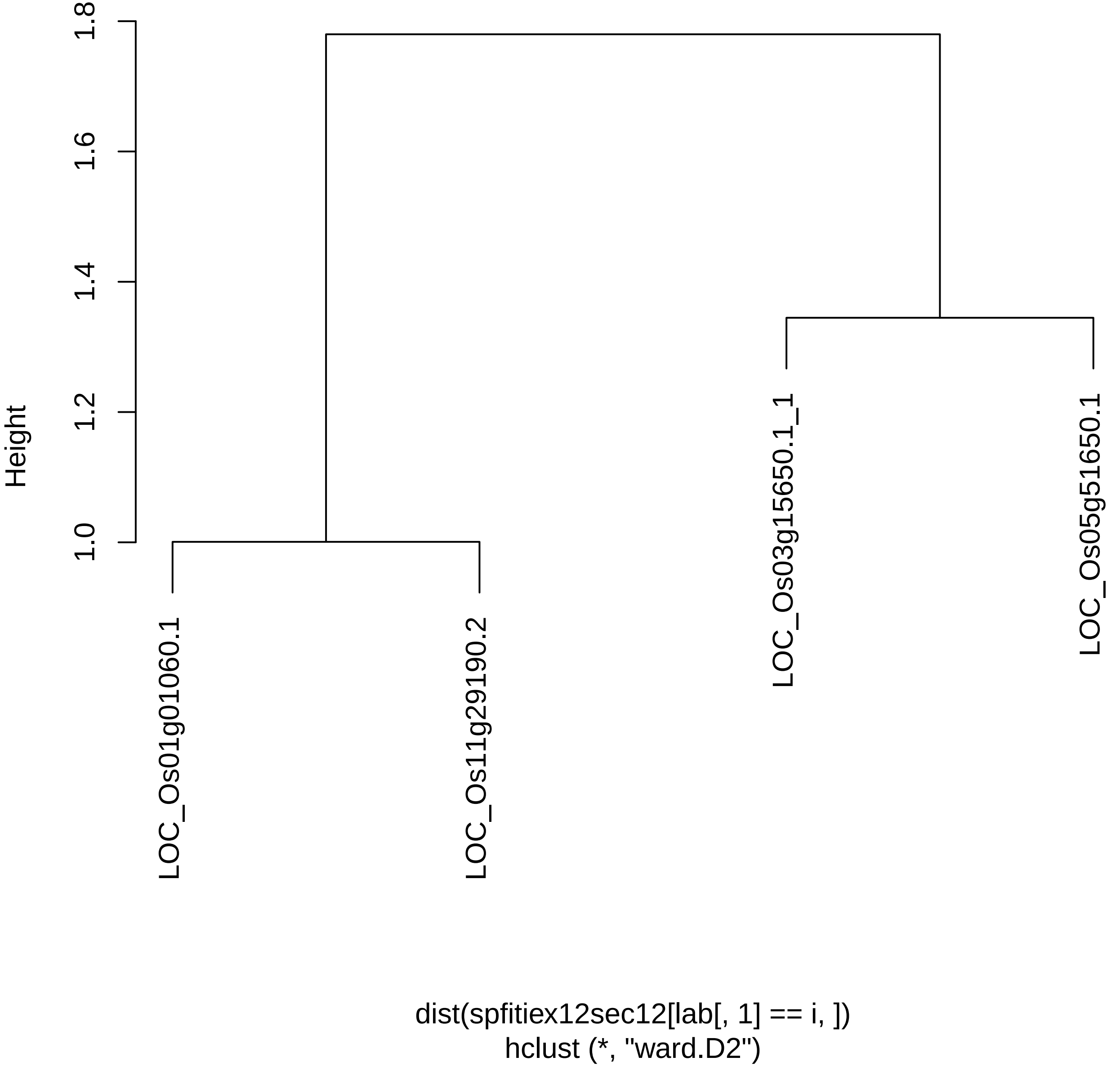

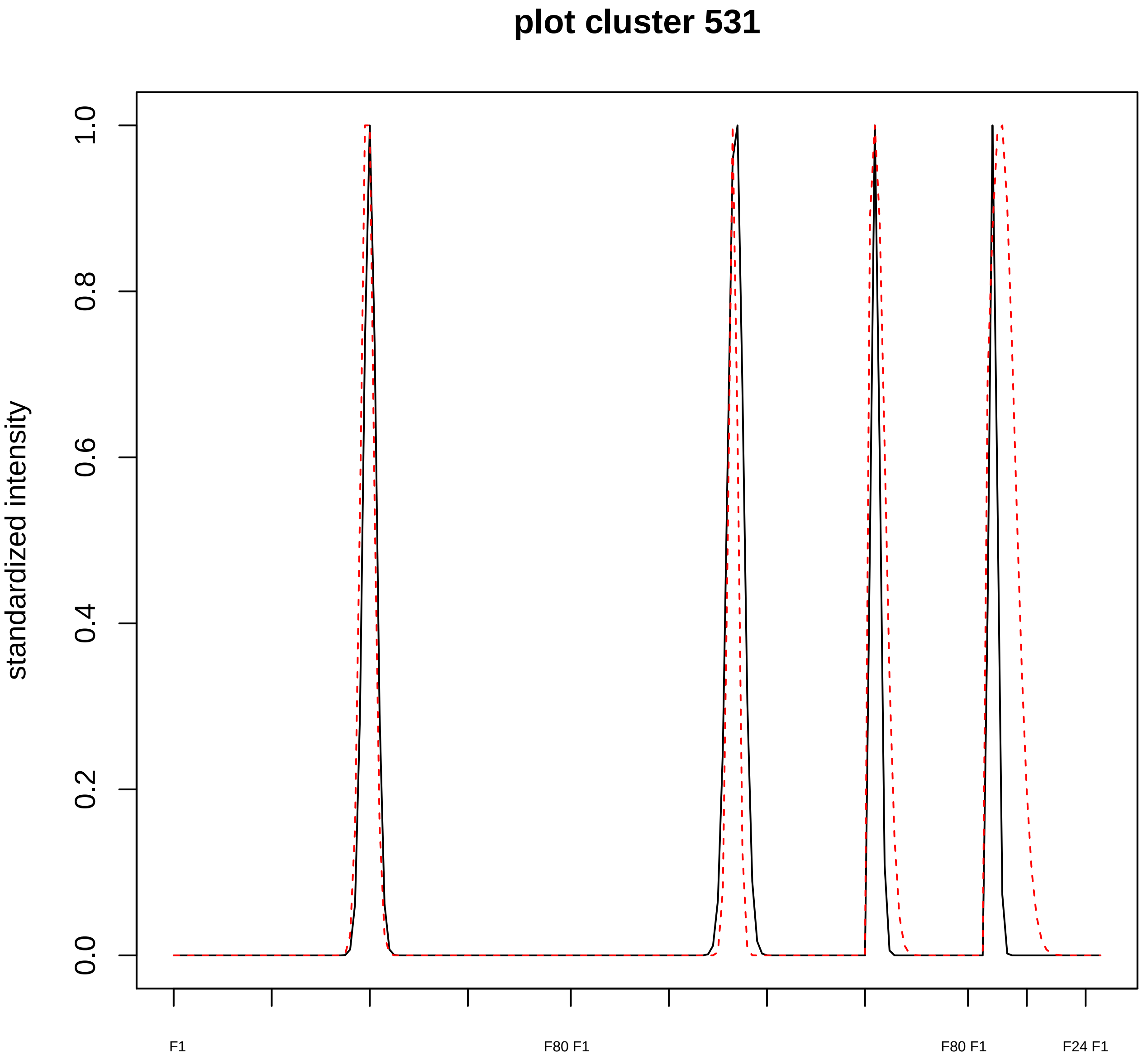
plot cluster 531
standardized intensity
0.0
0.2
0.4
0.6
0.8
1.0
F1
F80 F1
F80 F1
F24 F1

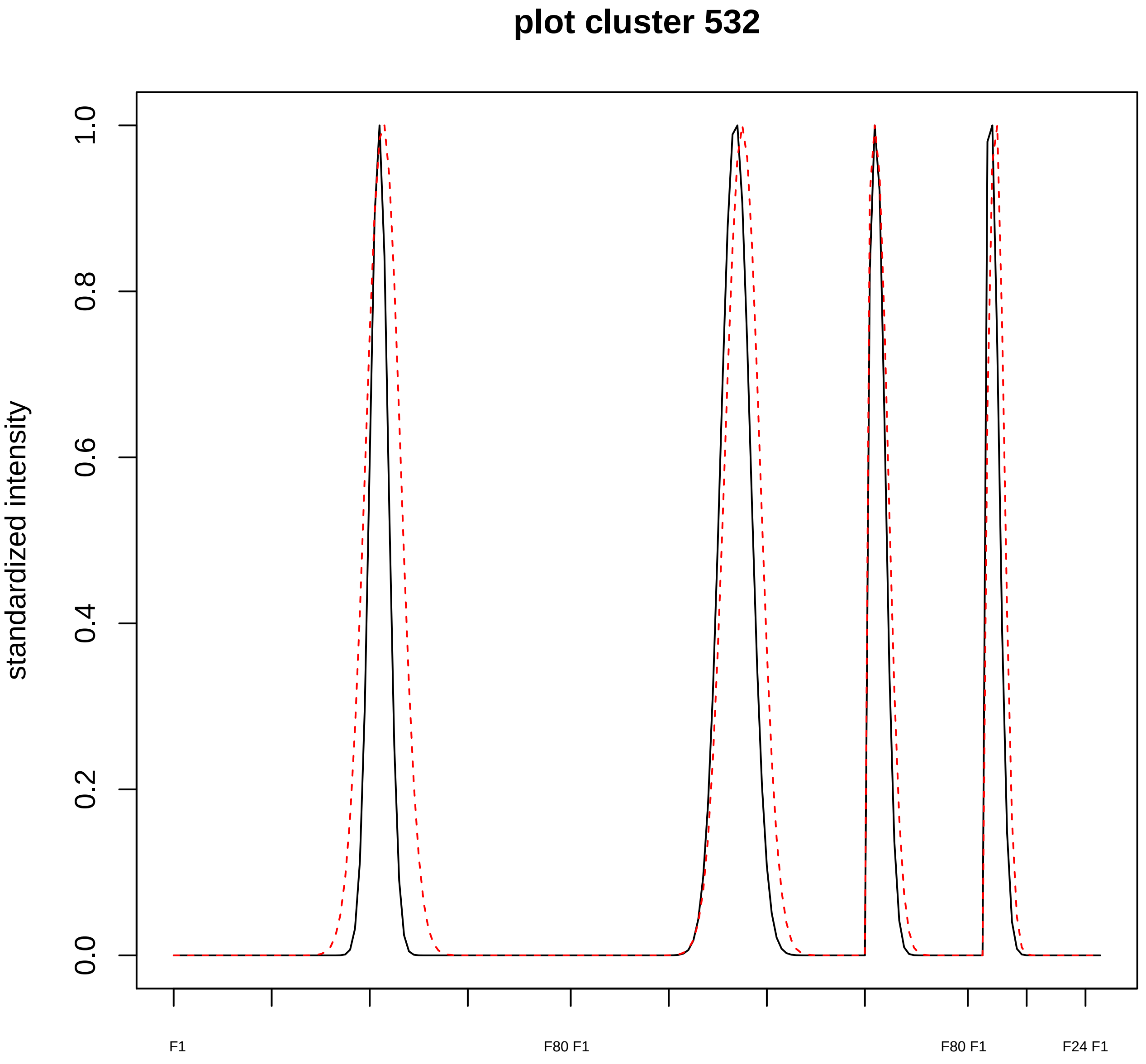

plot cluster 532
standardized intensity
0.0
0.2
0.4
0.6
0.8
1.0
F1
F80 F1
F80 F1
F24 F1

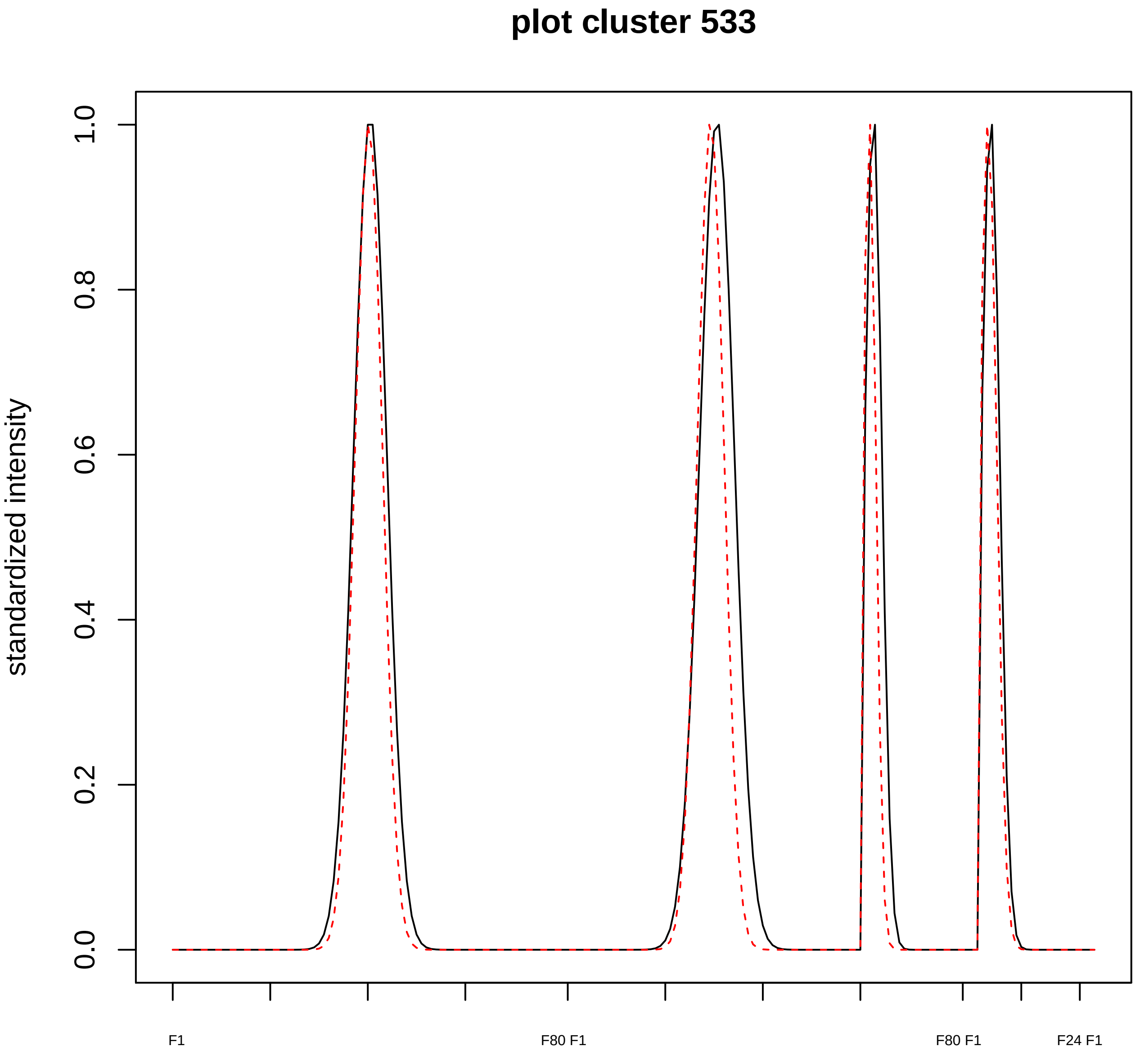

plot cluster 533
standardized intensity
0.0
0.2
0.4
0.6
0.8
1.0
F1
F80 F1
F80 F1
F24 F1

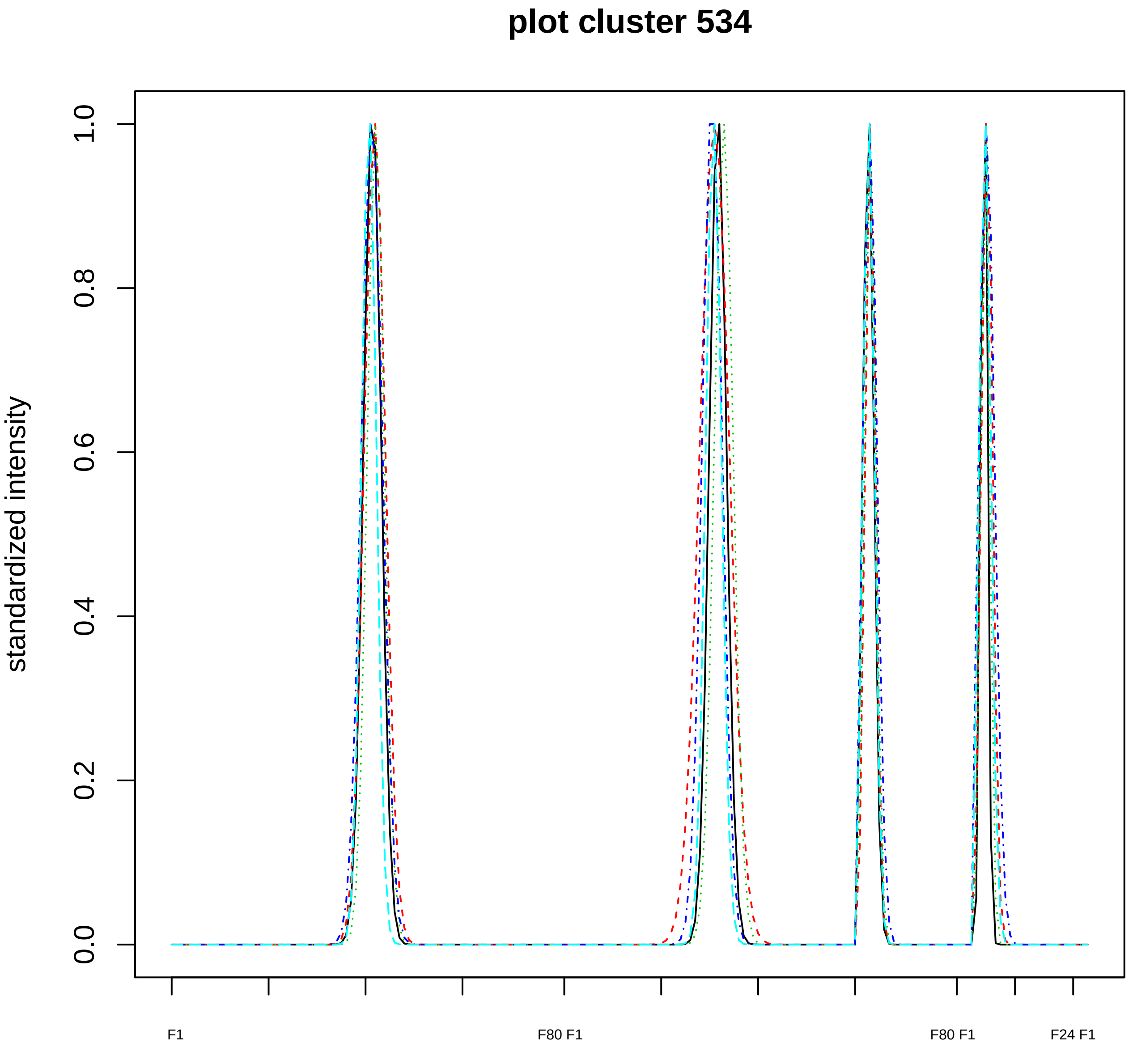
plot cluster 534
standardized intensity
0.0
0.2
0.4
0.6
0.8
1.0
F1
F80 F1
F80 F1
F24 F1

### Dendrogram of cluster 534

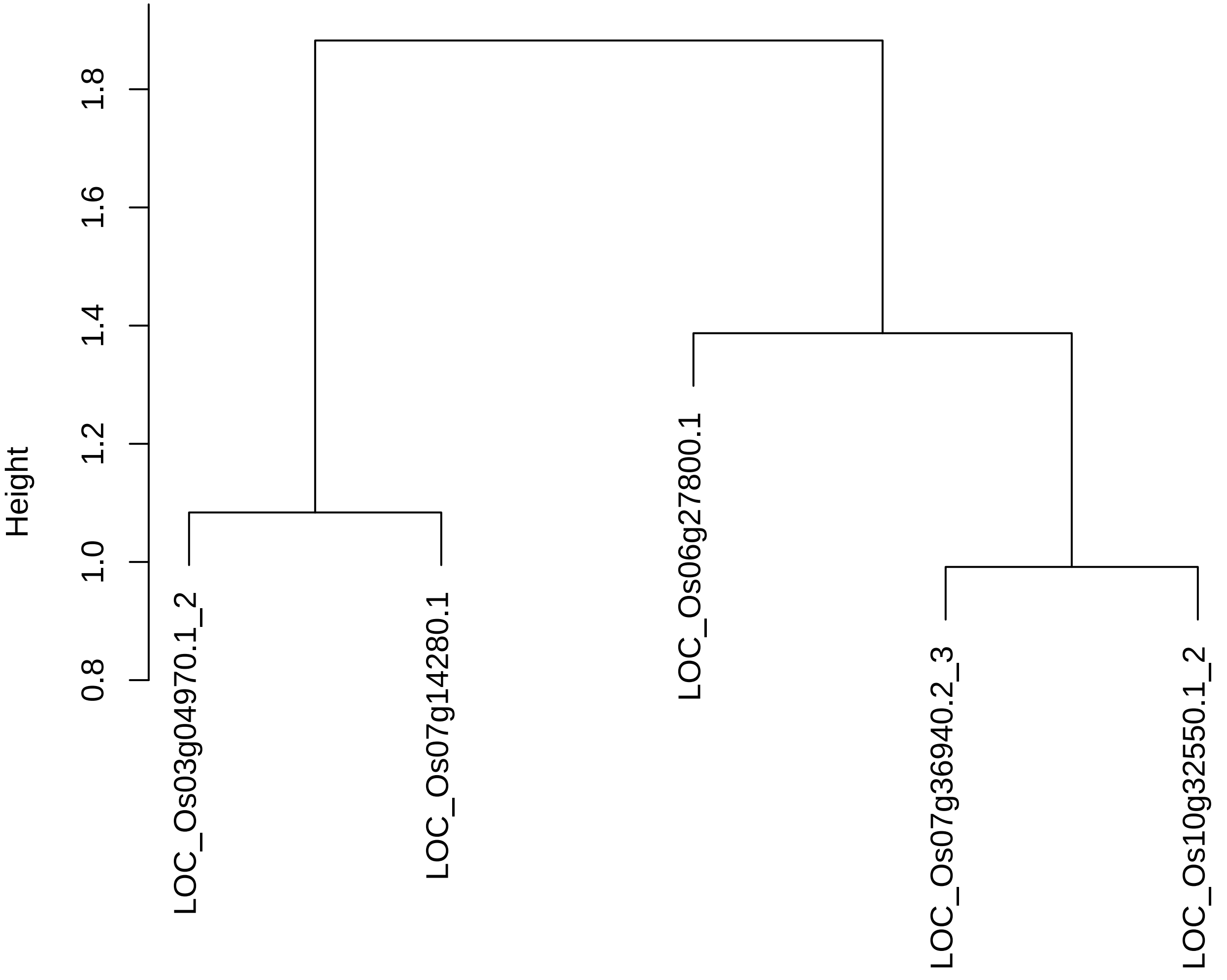


dist(spfitiex12sec12[lab[, 1] == i, ])
hclust (*, "ward.D2")

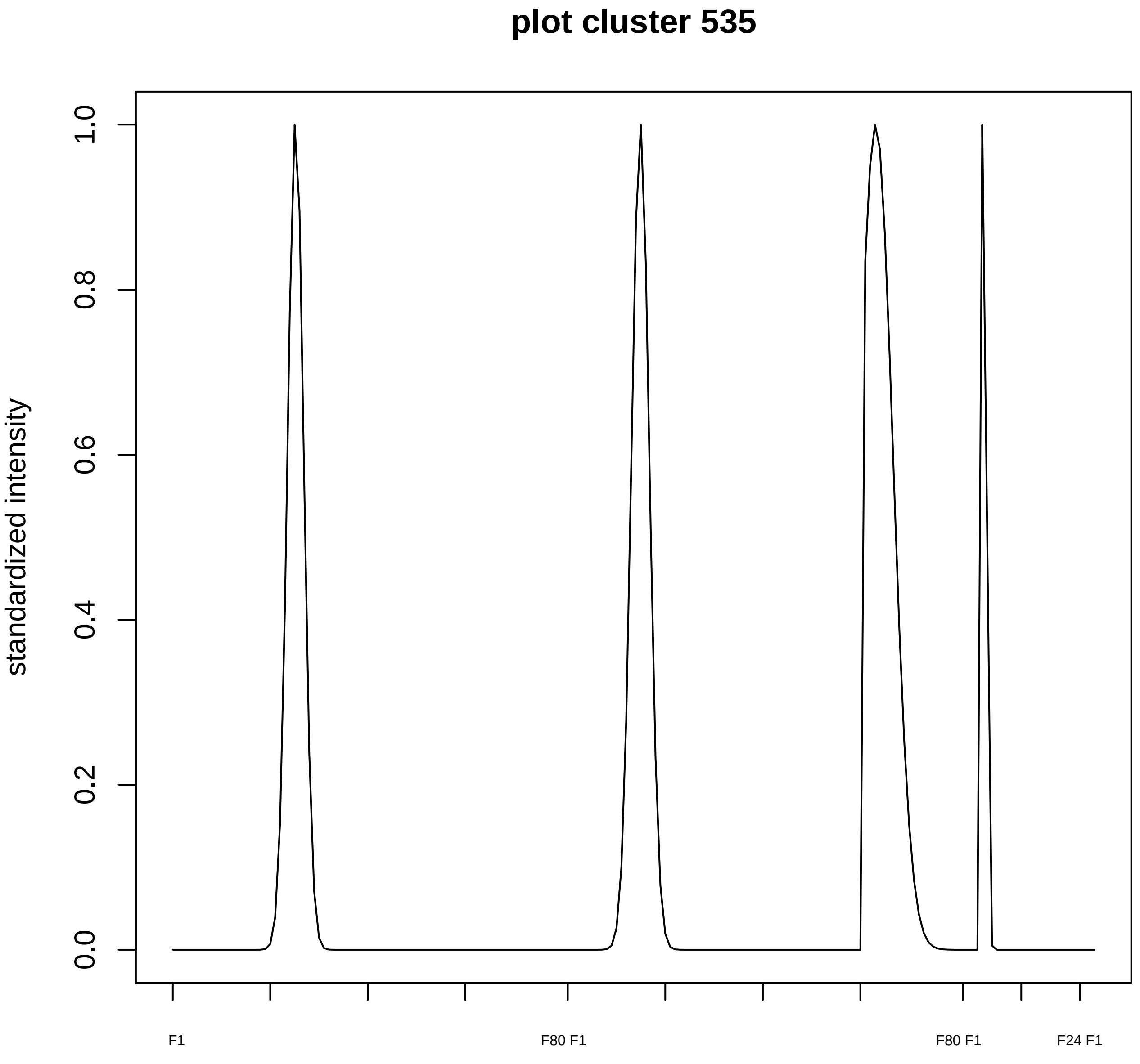
plot cluster 535
standardized intensity
0.0
0.2
0.4
0.6
0.8
1.0
F1
F80 F1
F80 F1
F24 F1

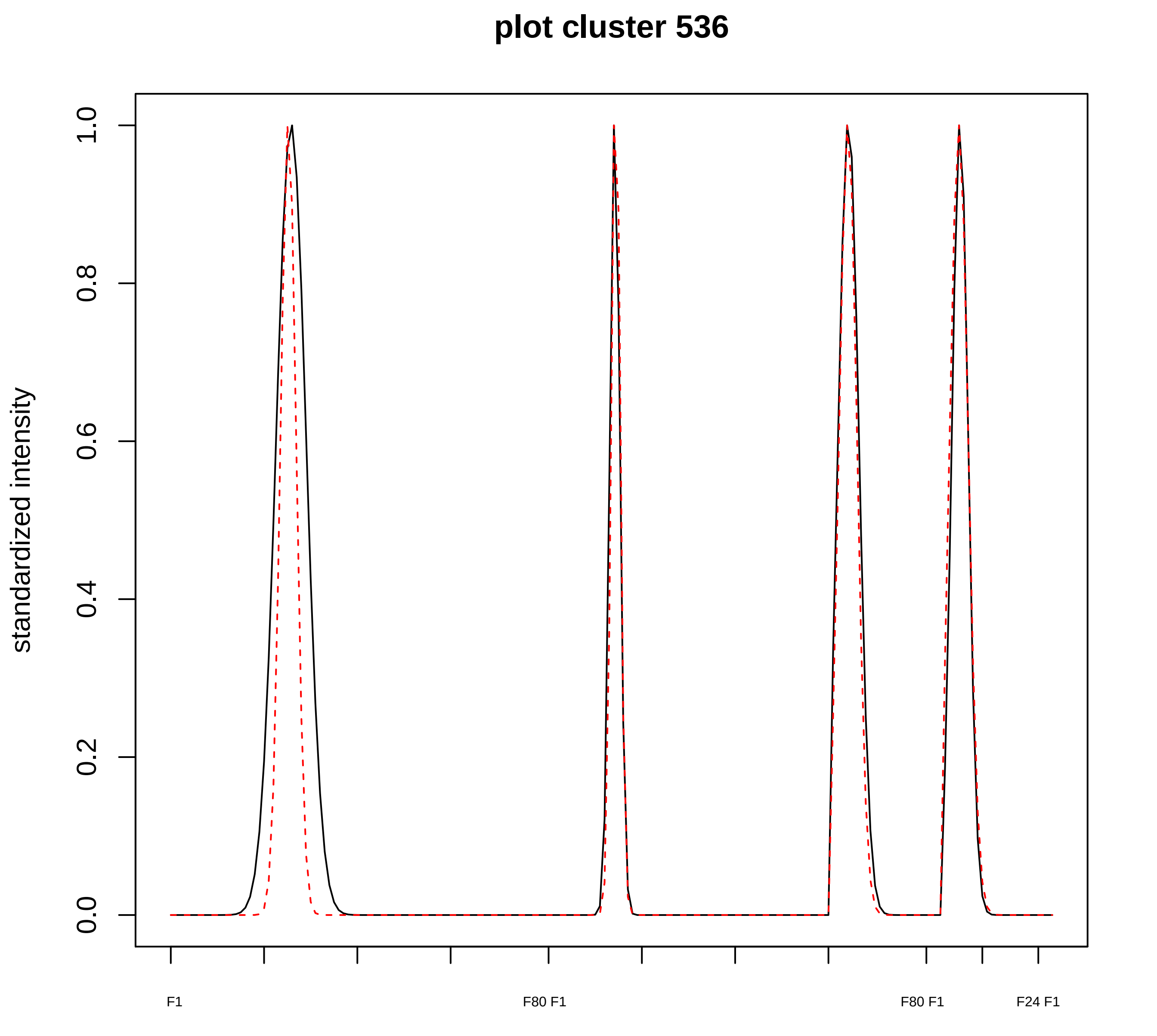
plot cluster 536
standardized intensity
0.0
0.2
0.4
0.6
0.8
1.0
F1
F80 F1
F80 F1
F24 F1

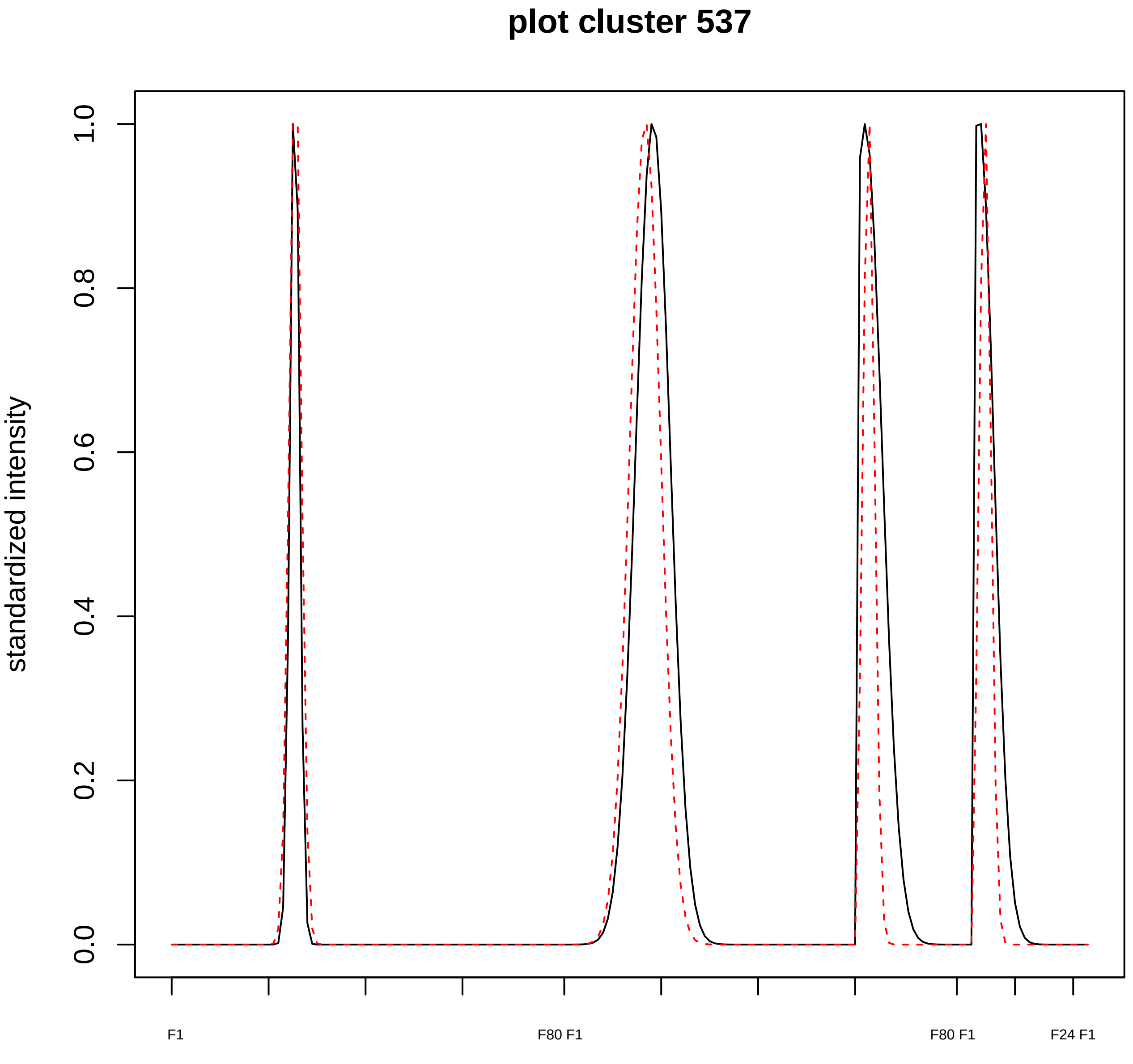

plot cluster 537
standardized intensity
0.0
0.2
0.4
0.6
0.8
1.0
F1
F80 F1
F80 F1
F24 F1

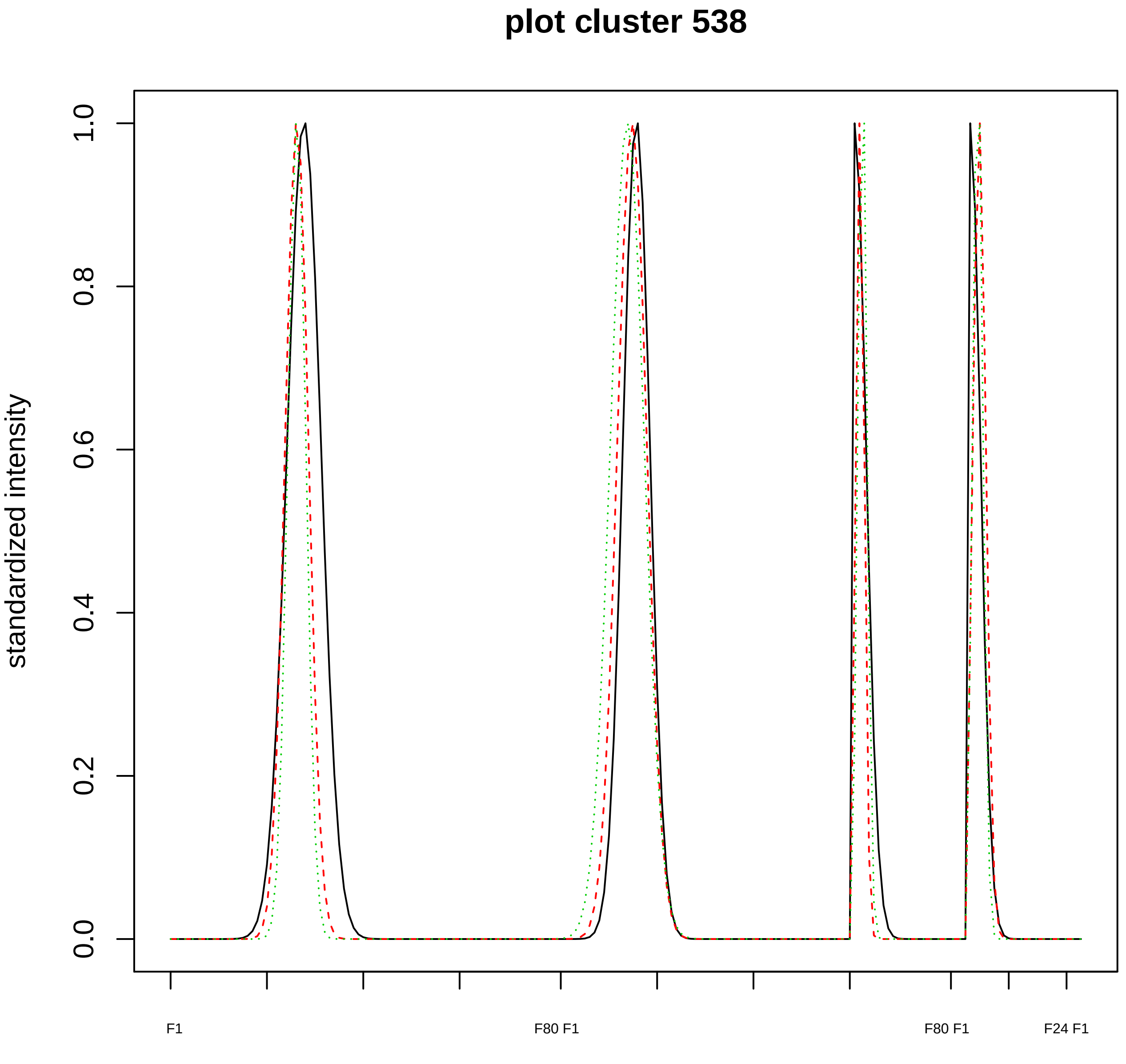

plot cluster 538
standardized intensity
0.0
0.2
0.4
0.6
0.8
1.0
F1
F80 F1
F80 F1
F24 F1

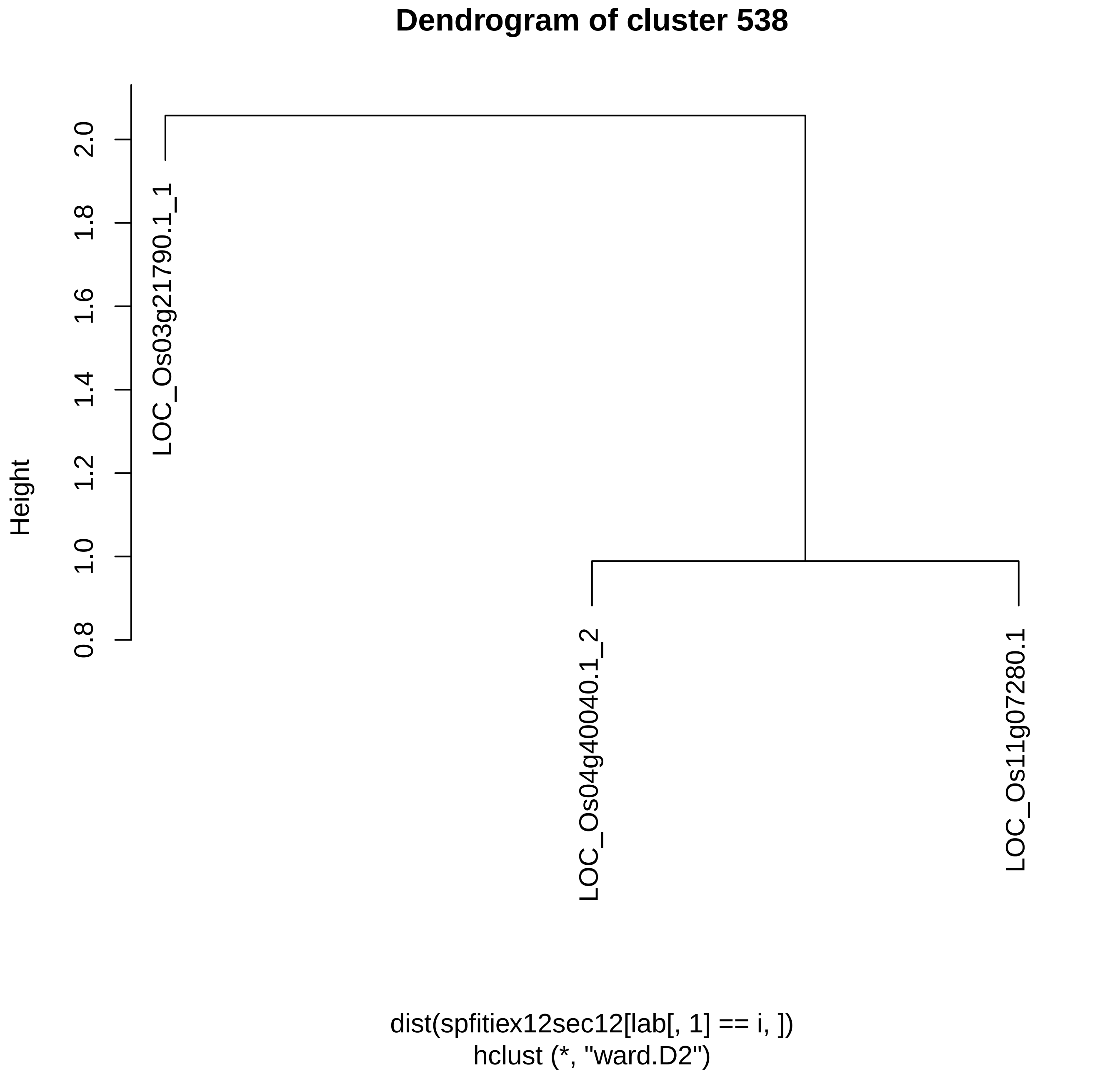
Dendrogram of cluster 538
Height
2.0
1.8
1.6
1.4
1.2
1.0
0.8
LOC_Os03g21790.1_1
LOC_Os04g40040.1_2
LOC_Os11g07280.1
dist(spfitiex12sec12[lab[, 1] == i, ])
hclust (*, "ward.D2")

### plot cluster 539

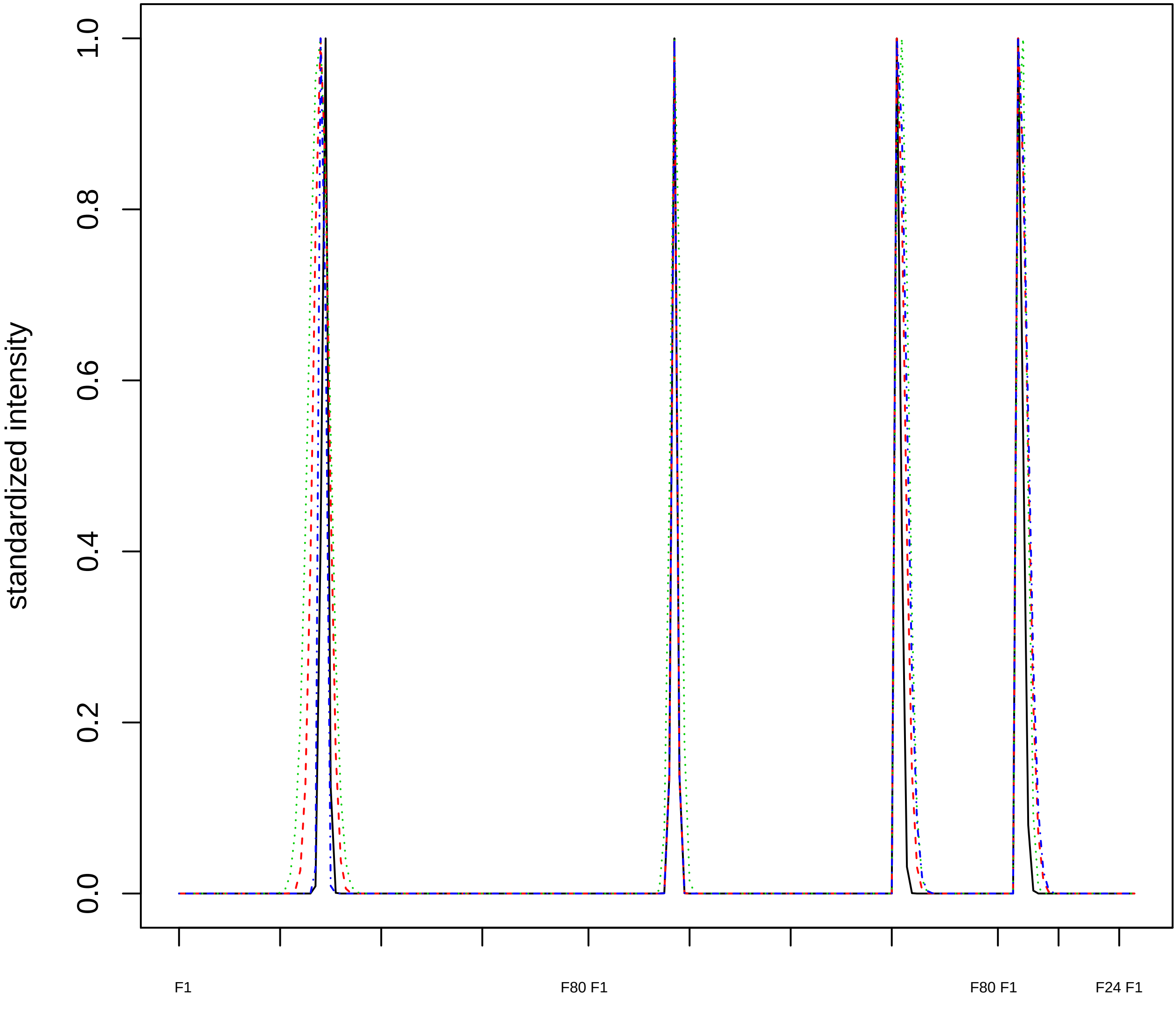

#### Dendrogram of cluster 539

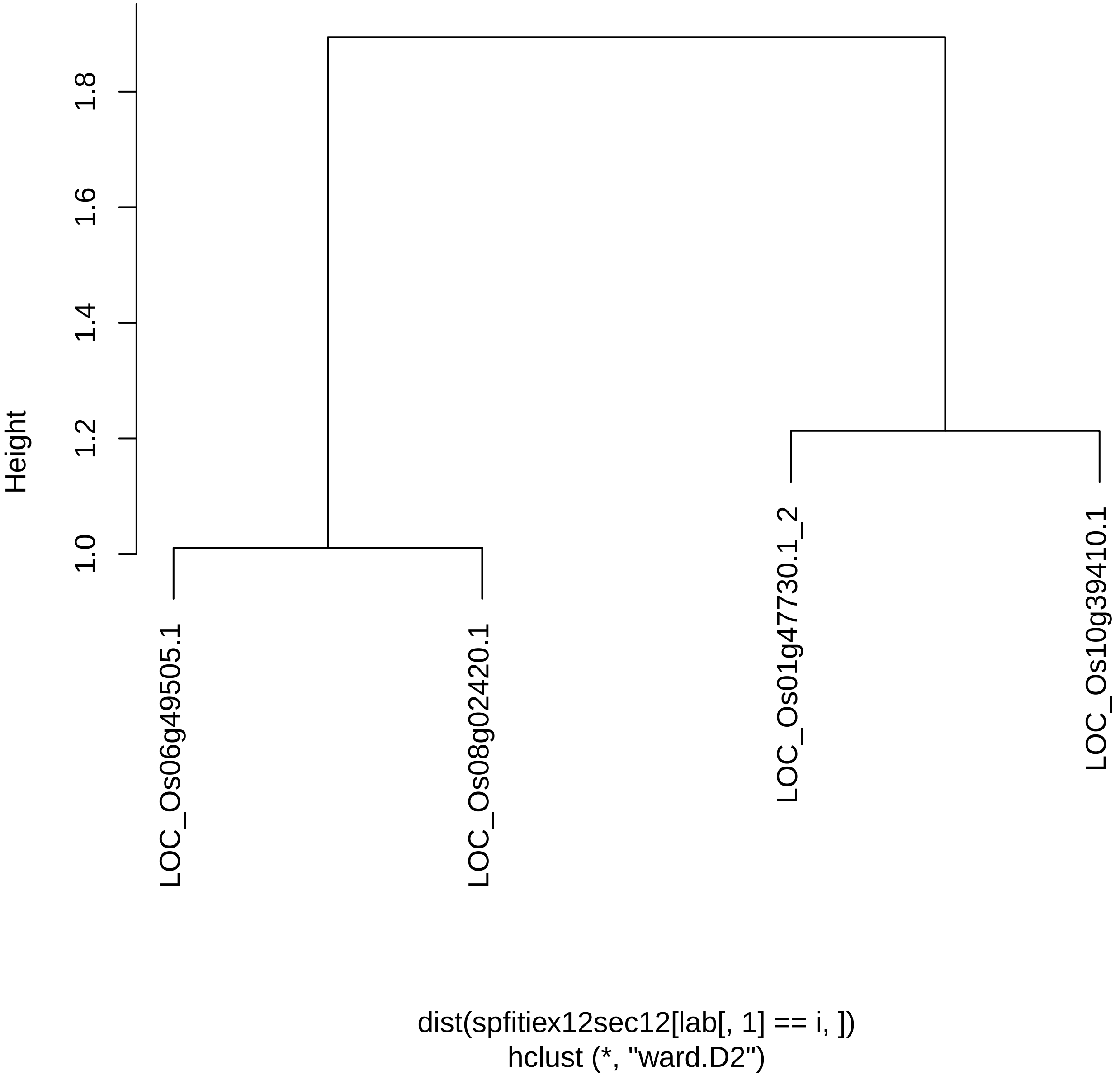

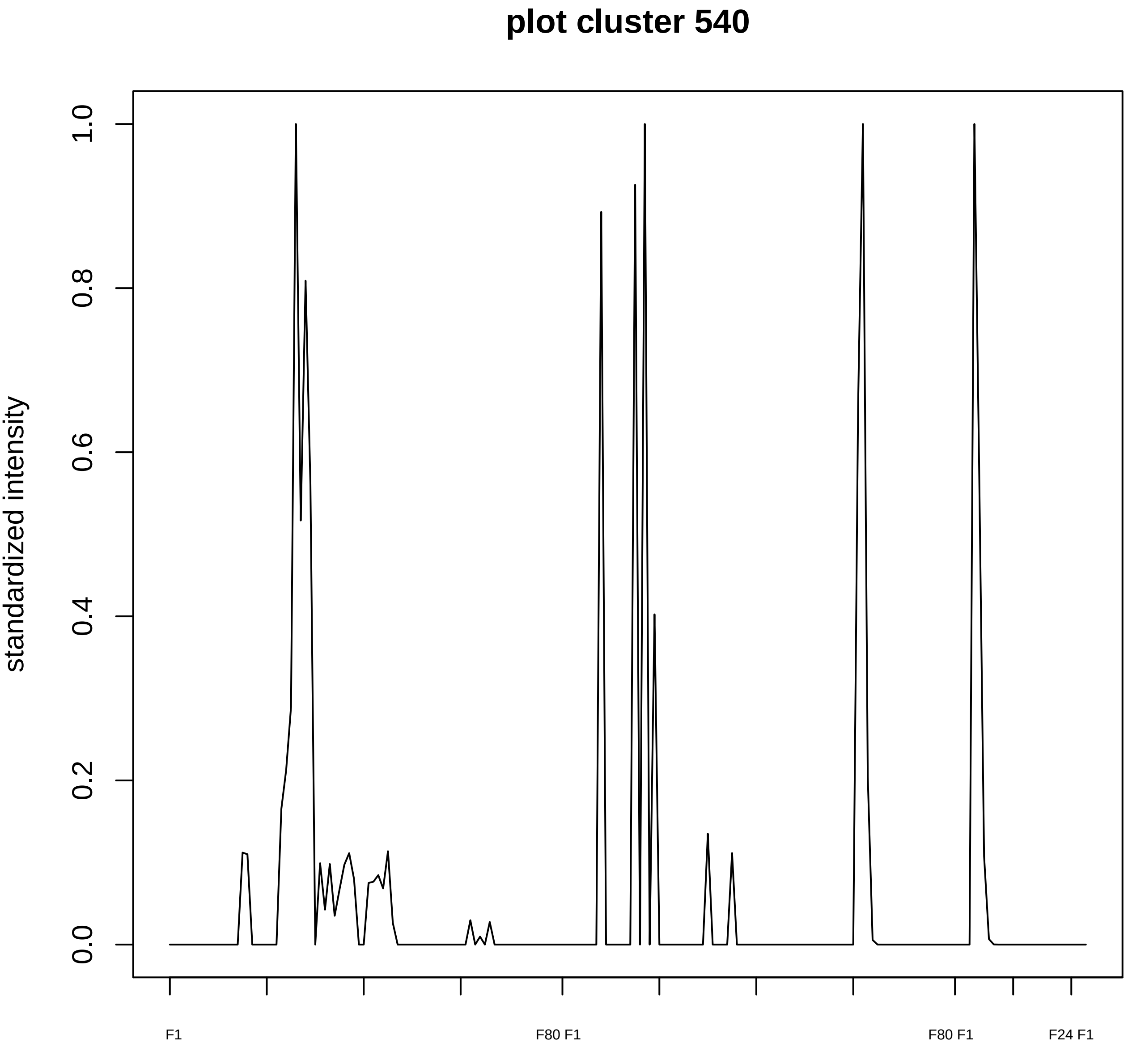
plot cluster 540
standardized intensity
0.0
0.2
0.4
0.6
0.8
1.0
F1
F80 F1
F80 F1
F24 F1

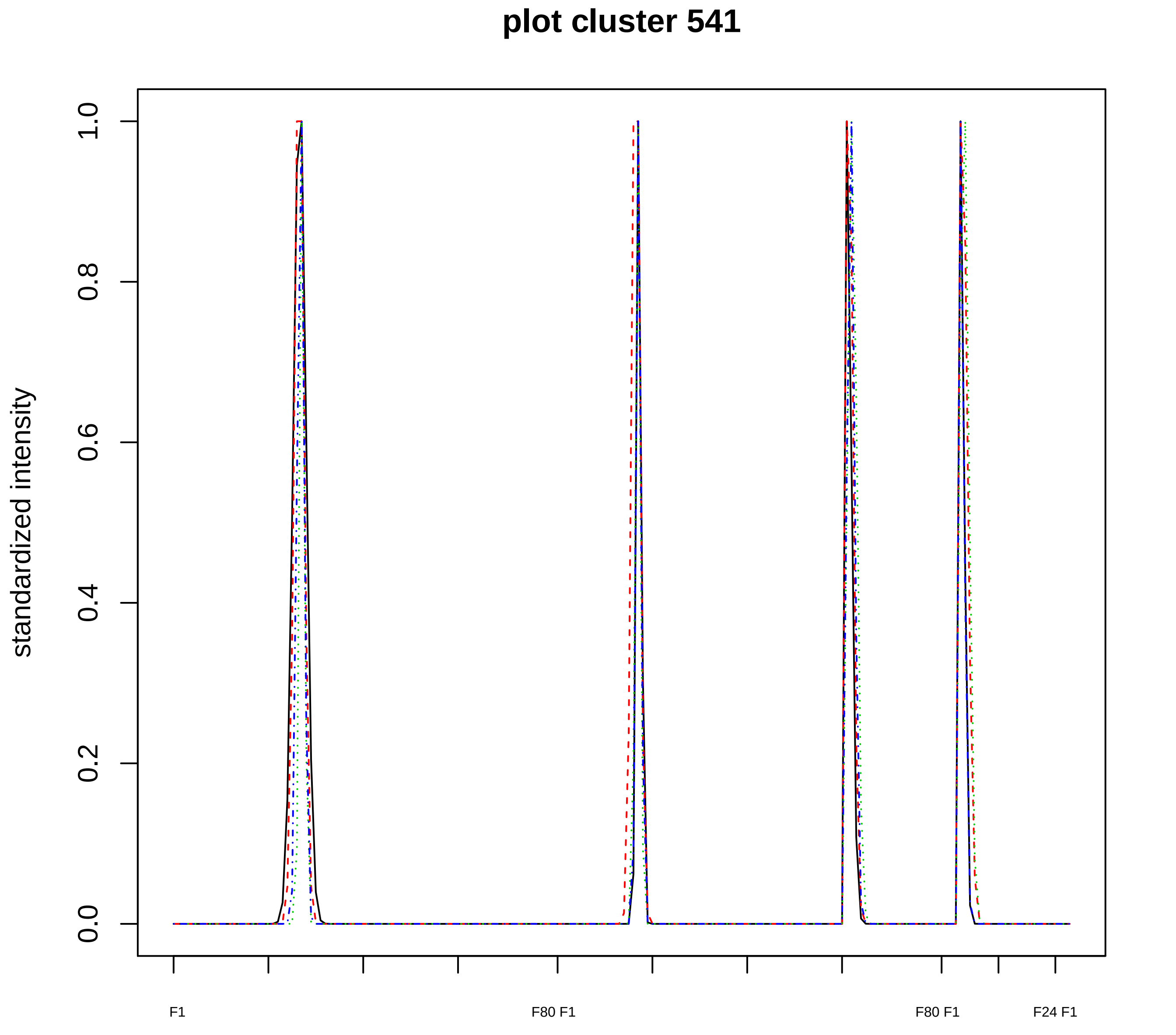
plot cluster 541
standardized intensity
0.0
0.2
0.4
0.6
0.8
1.0
F1
F80 F1
F80 F1
F24 F1

### Dendrogram of cluster 541

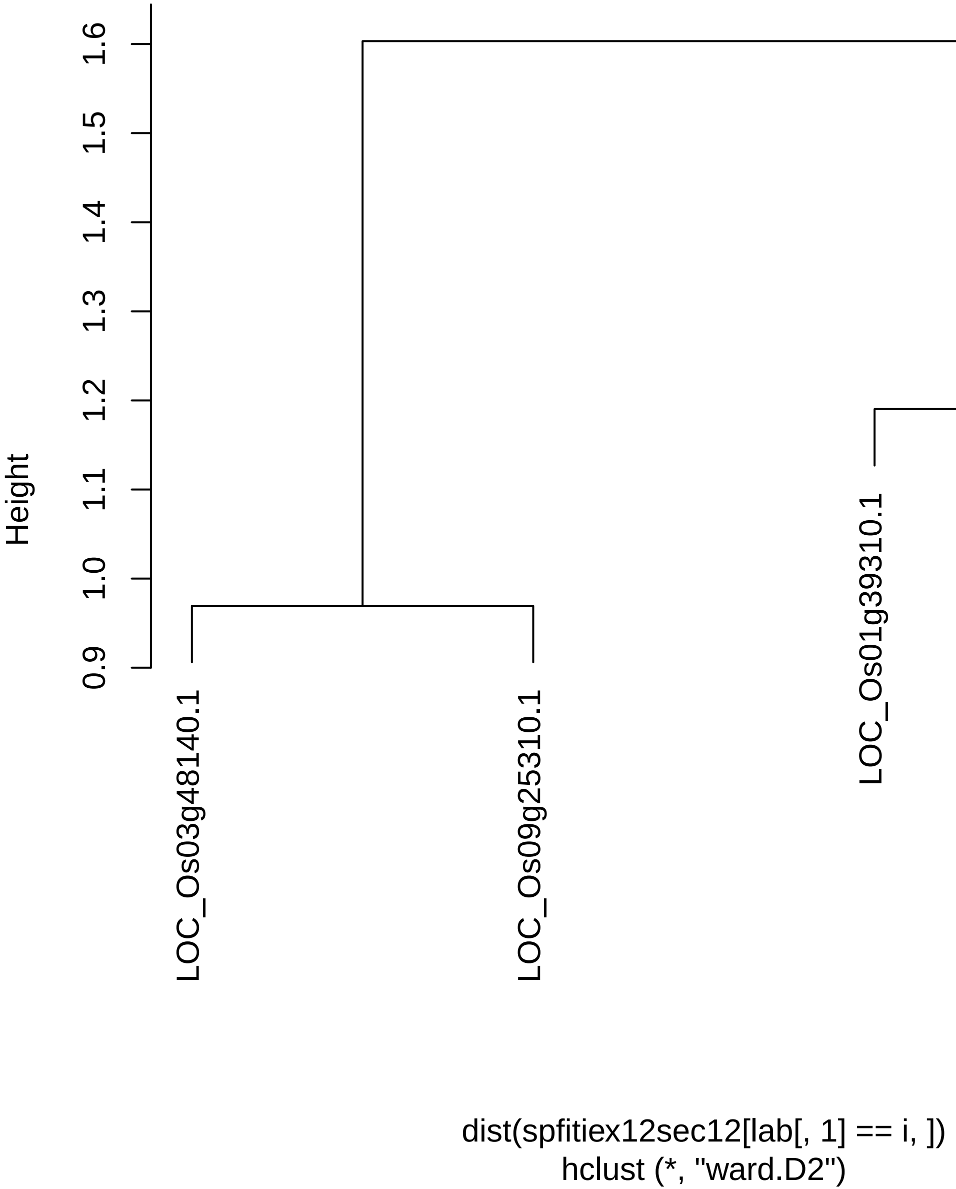

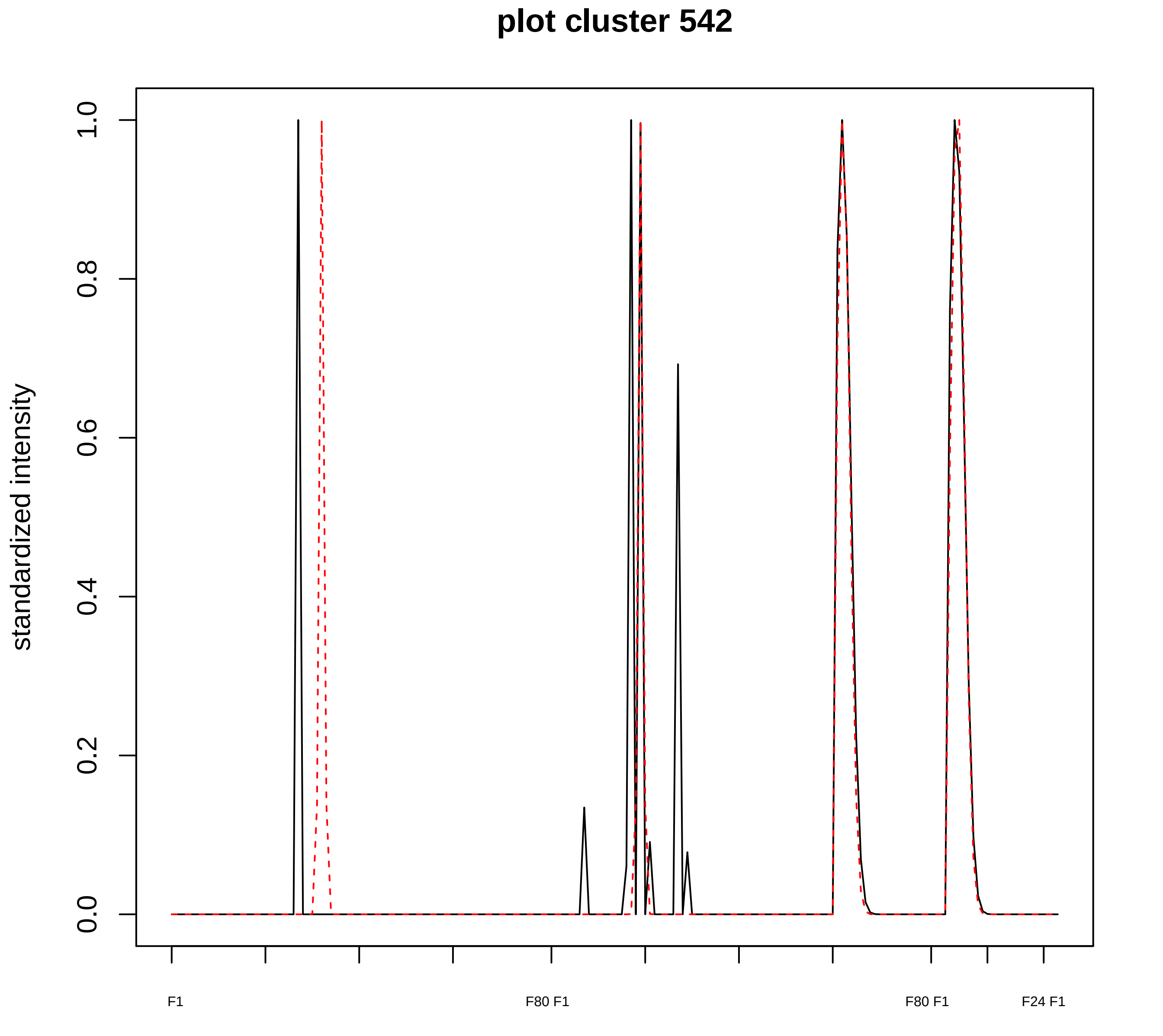
plot cluster 542
standardized intensity
0.0
0.2
0.4
0.6
0.8
1.0
F1
F80 F1
F80 F1
F24 F1

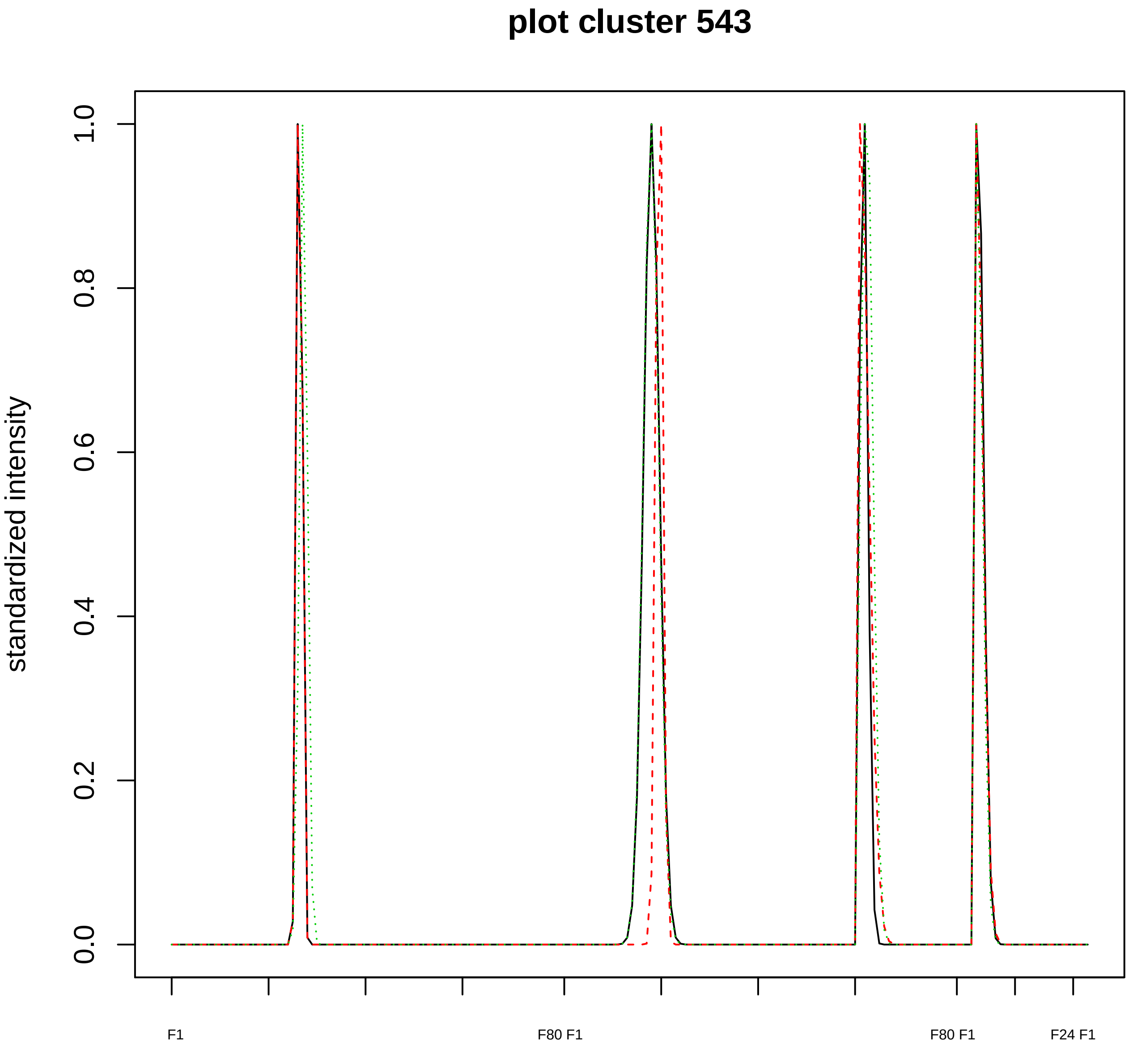
plot cluster 543
standardized intensity
0.0
0.2
0.4
0.6
0.8
1.0
F1
F80 F1
F80 F1
F24 F1

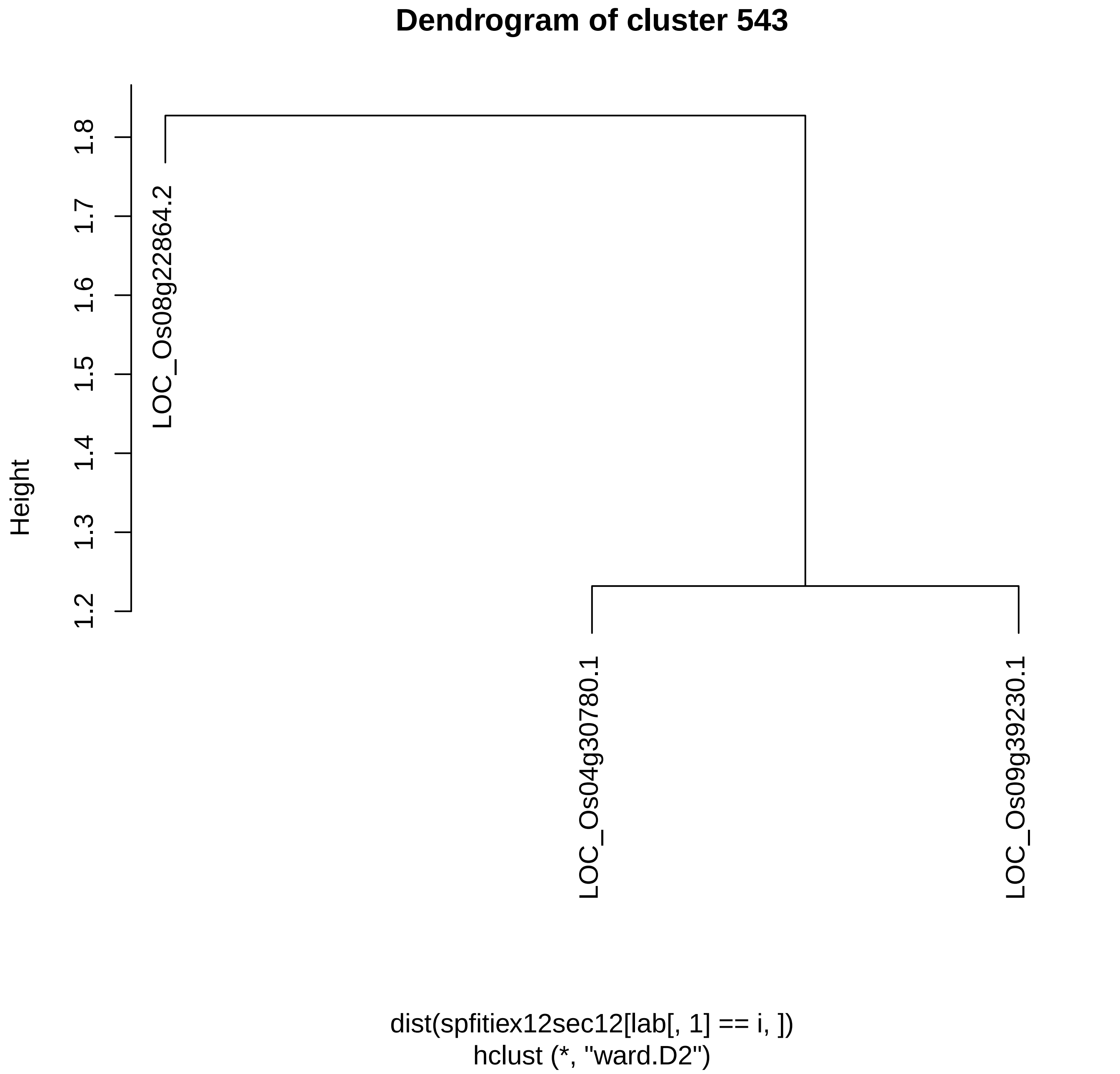
Dendrogram of cluster 543
Height
1.2
1.3
1.4
1.5
1.6
1.7
1.8
LOC_Os08g22864.2
LOC_Os04g30780.1
LOC_Os09g39230.1
dist(spfitiex12sec12[lab[, 1] == i, ])
hclust (*, "ward.D2")

### plot cluster 544

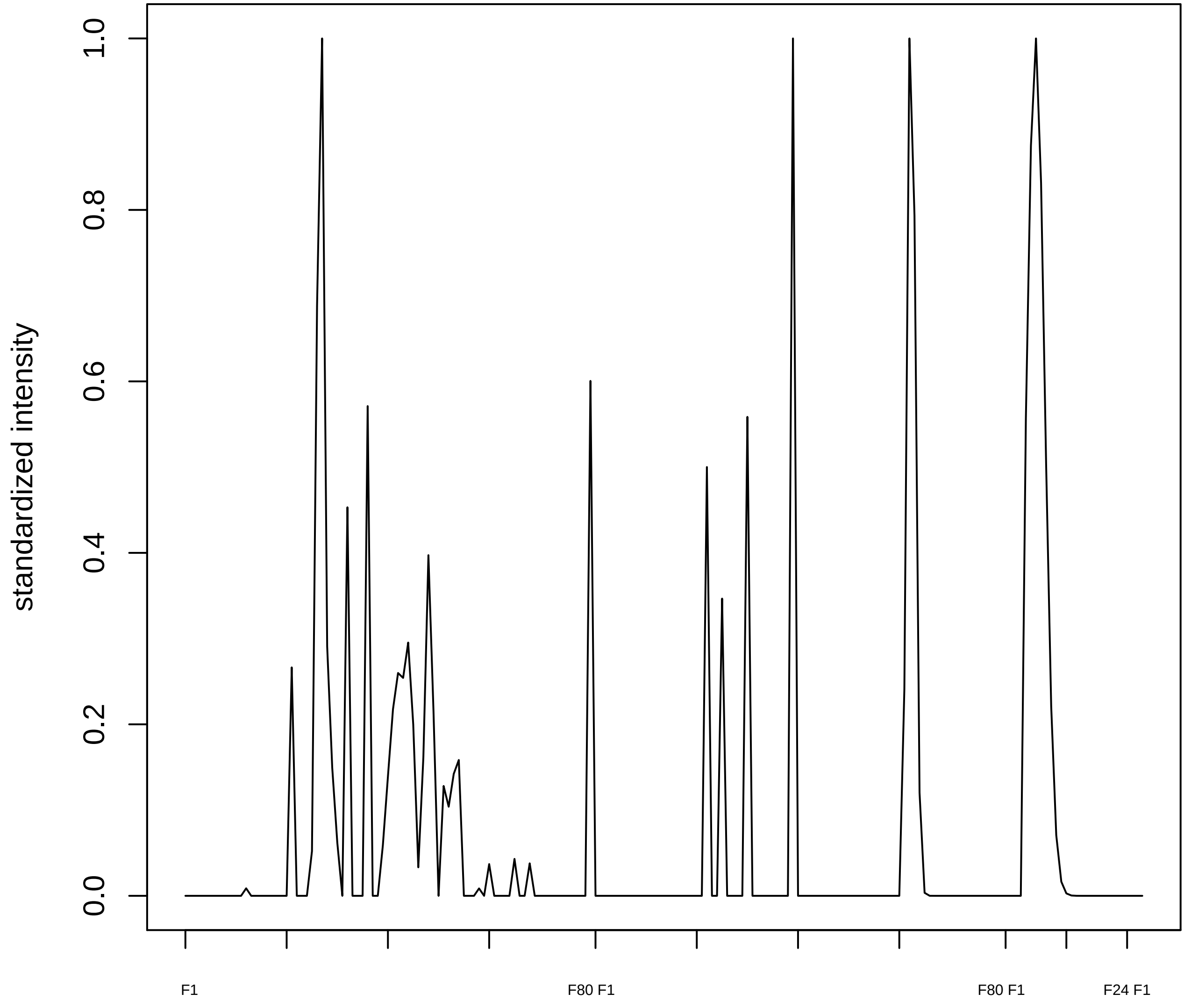

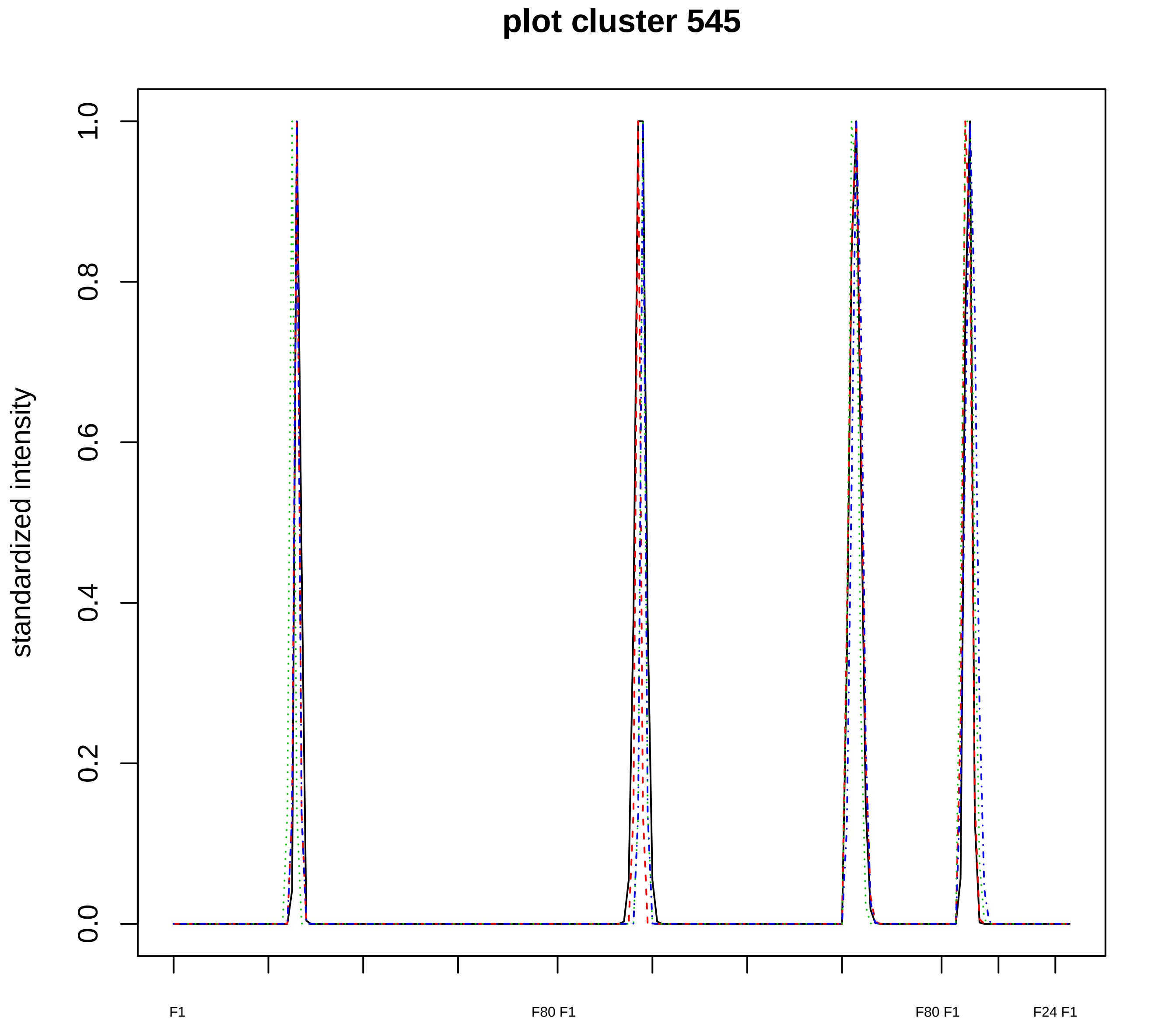
plot cluster 545
standardized intensity
0.0
0.2
0.4
0.6
0.8
1.0
F1
F80 F1
F80 F1
F24 F1

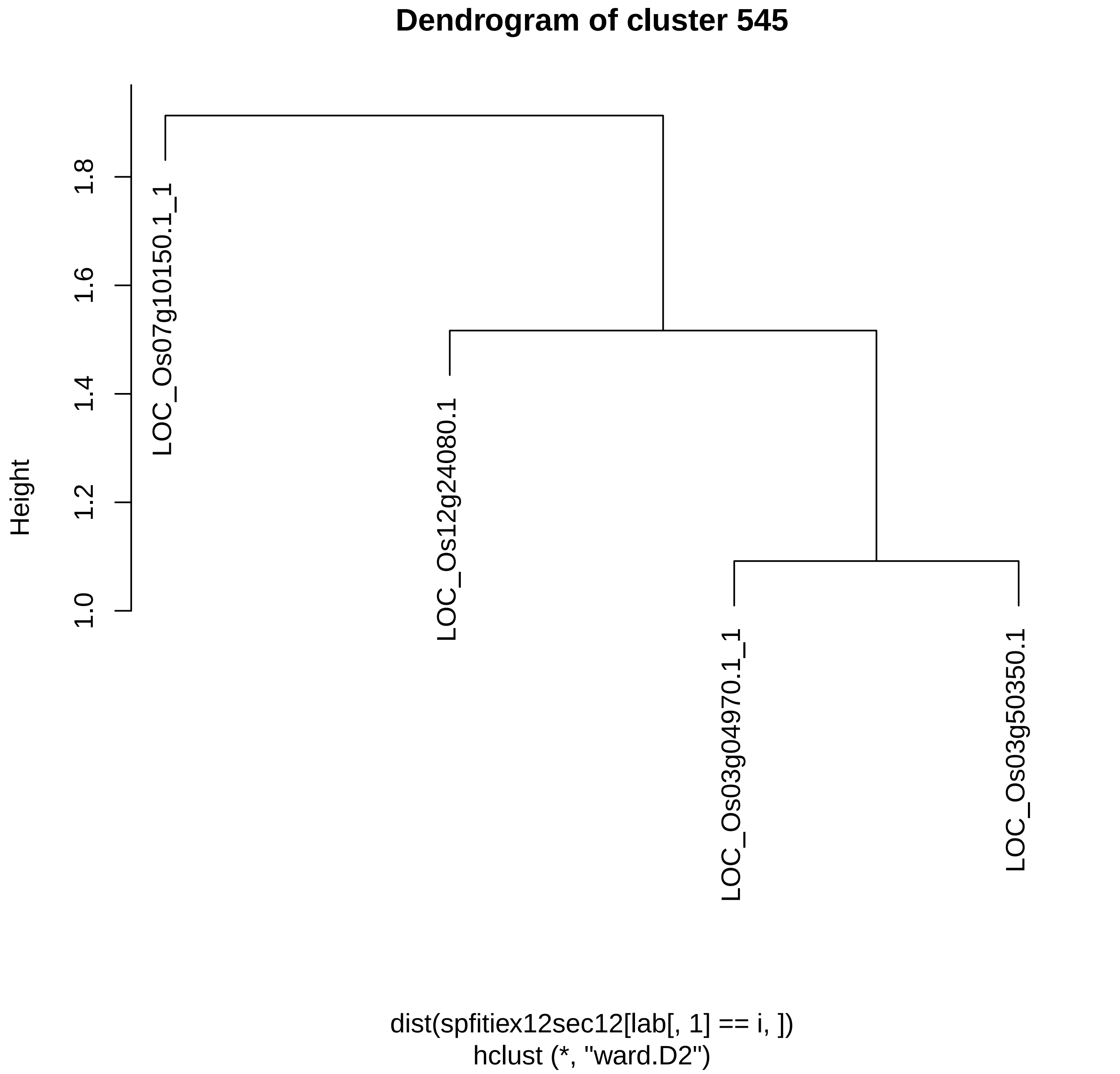

Dendrogram of cluster 545
Height
1.0
1.2
1.4
1.6
1.8
LOC_Os07g10150.1_1
LOC_Os12g24080.1
LOC_Os03g04970.1_1
LOC_Os03g50350.1
dist(spfitiex12sec12[lab[, 1] == i, ])
hclust (*, "ward.D2")

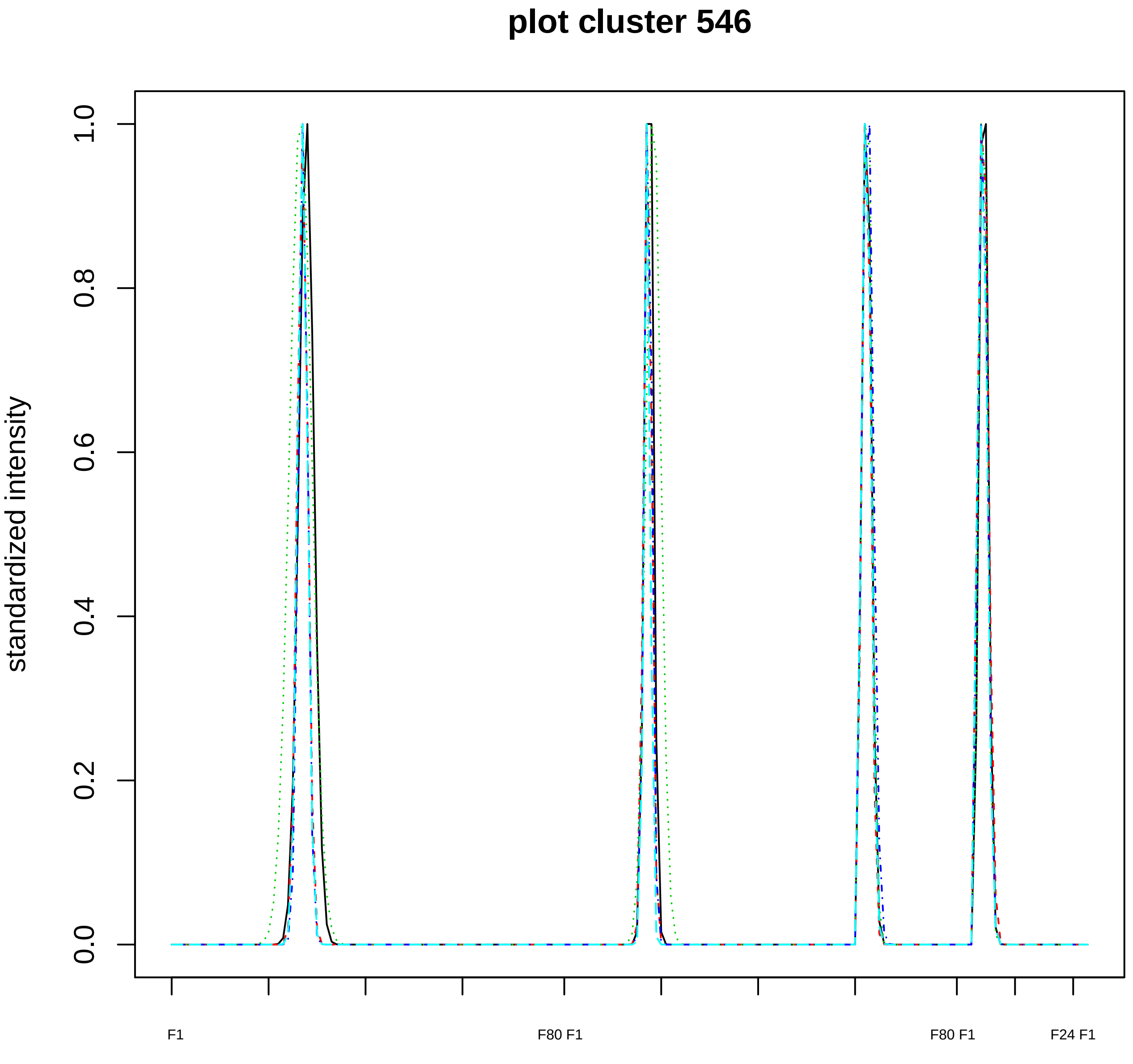
plot cluster 546
standardized intensity
0.0
0.2
0.4
0.6
0.8
1.0
F1
F80 F1
F80 F1
F24 F1

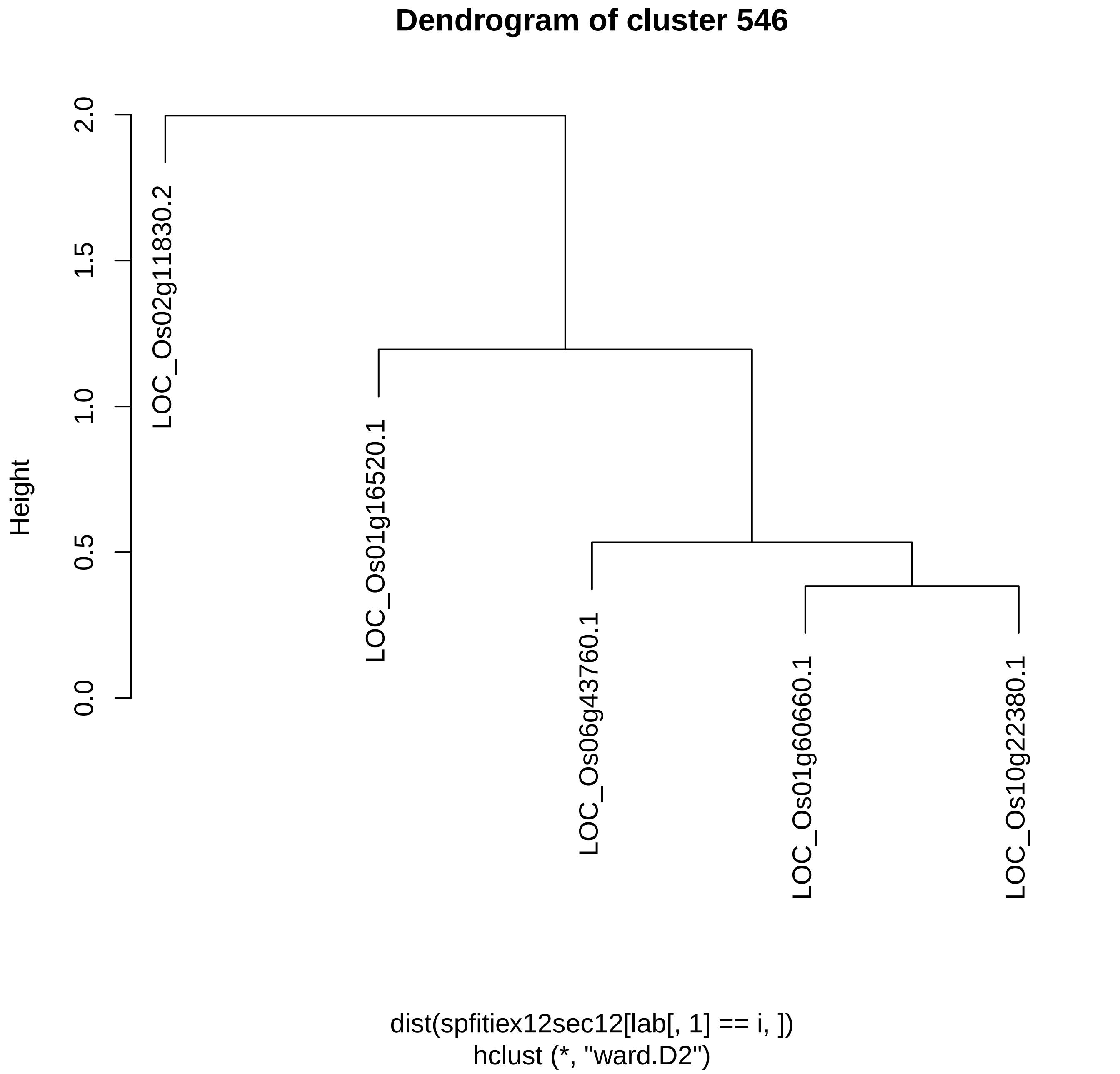
Dendrogram of cluster 546
Height
0.0
0.5
1.0
1.5
2.0
LOC_Os02g11830.2
LOC_Os01g16520.1
LOC_Os06g43760.1
LOC_Os01g60660.1
LOC_Os10g22380.1
dist(spfitiex12sec12[lab[, 1] == i, ])
hclust (*, "ward.D2")

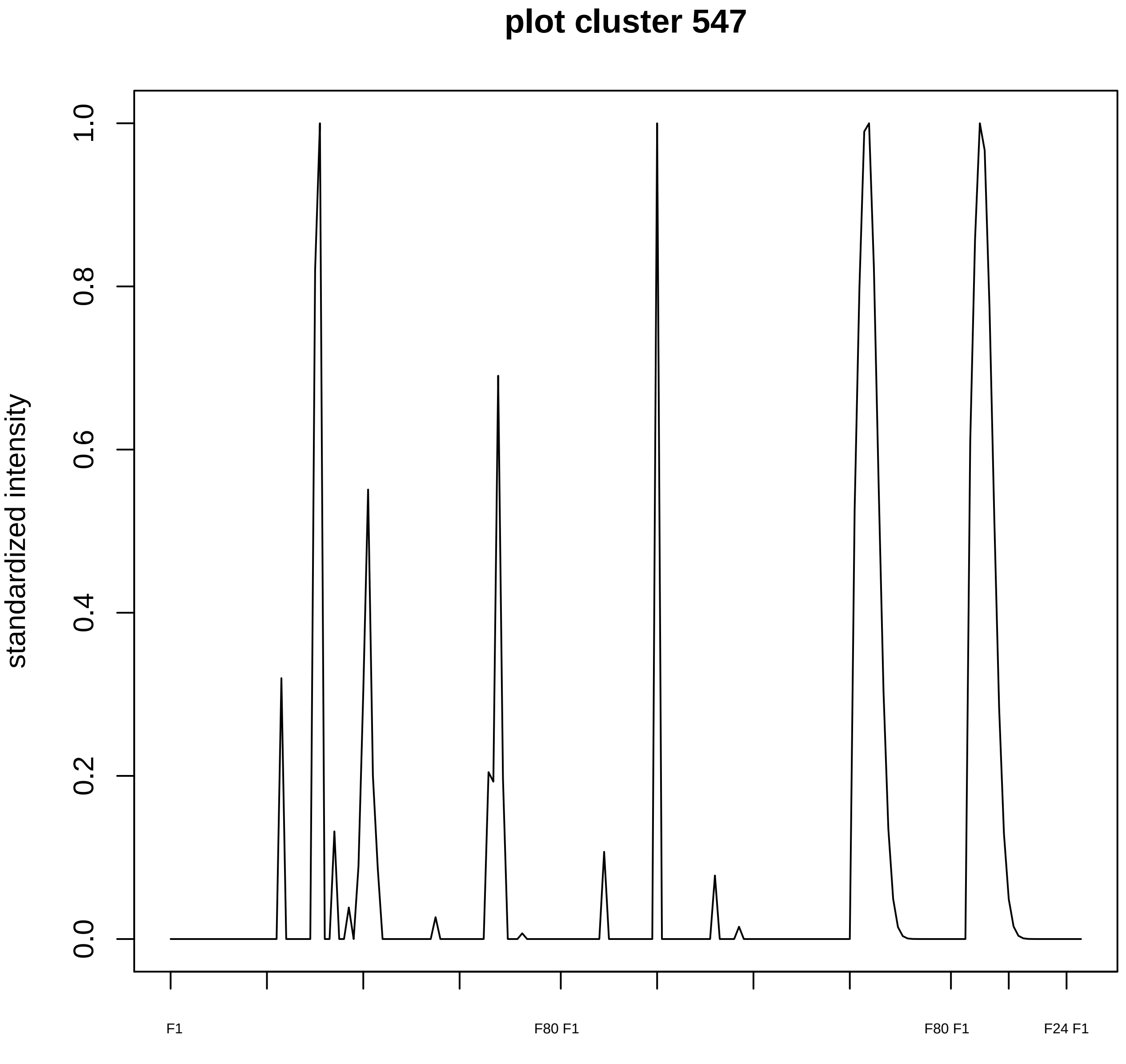
plot cluster 547
standardized intensity
0.0
0.2
0.4
0.6
0.8
1.0
F1
F80 F1
F80 F1
F24 F1

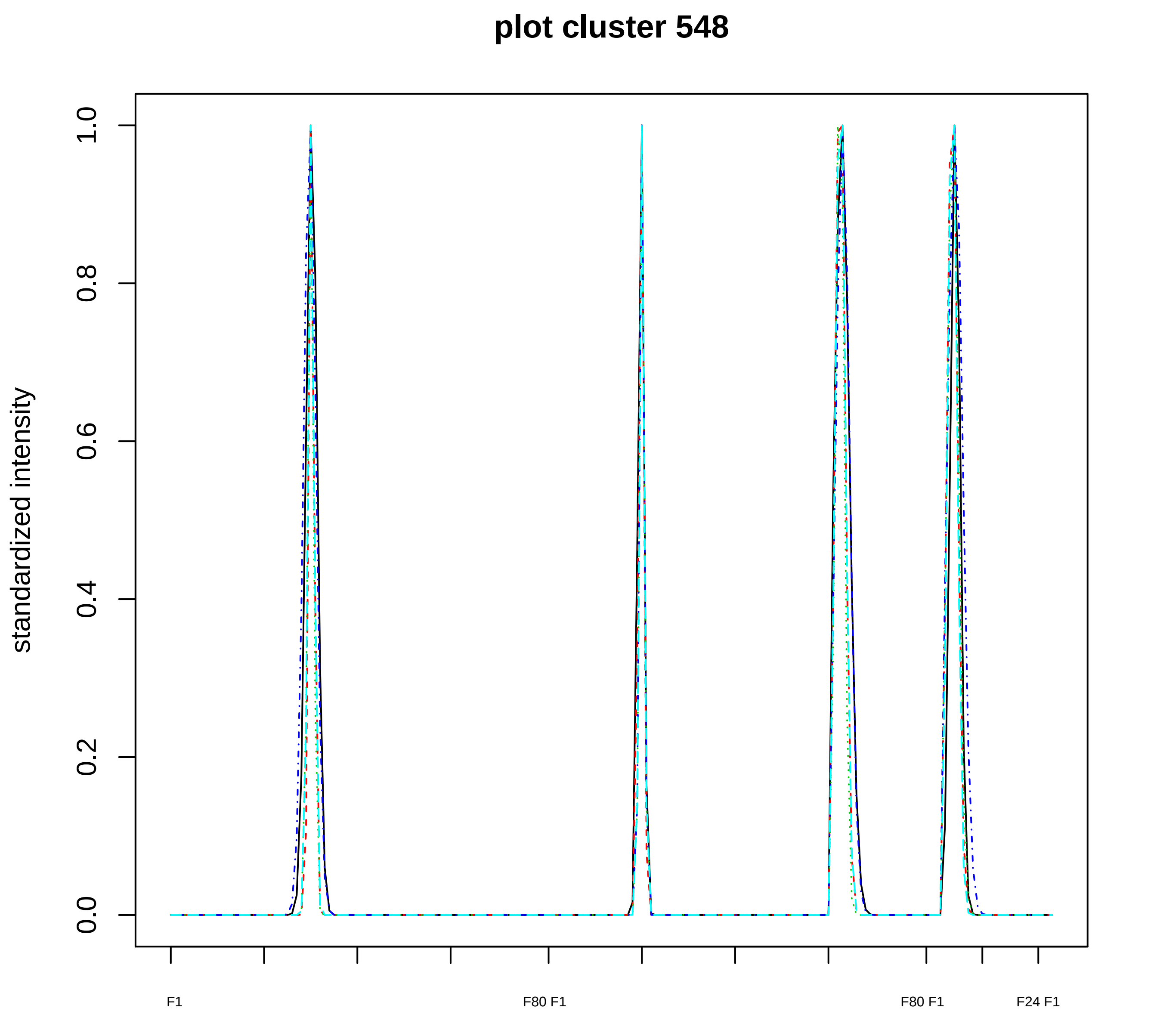
plot cluster 548
standardized intensity
0.0
0.2
0.4
0.6
0.8
1.0
F1
F80 F1
F80 F1
F24 F1

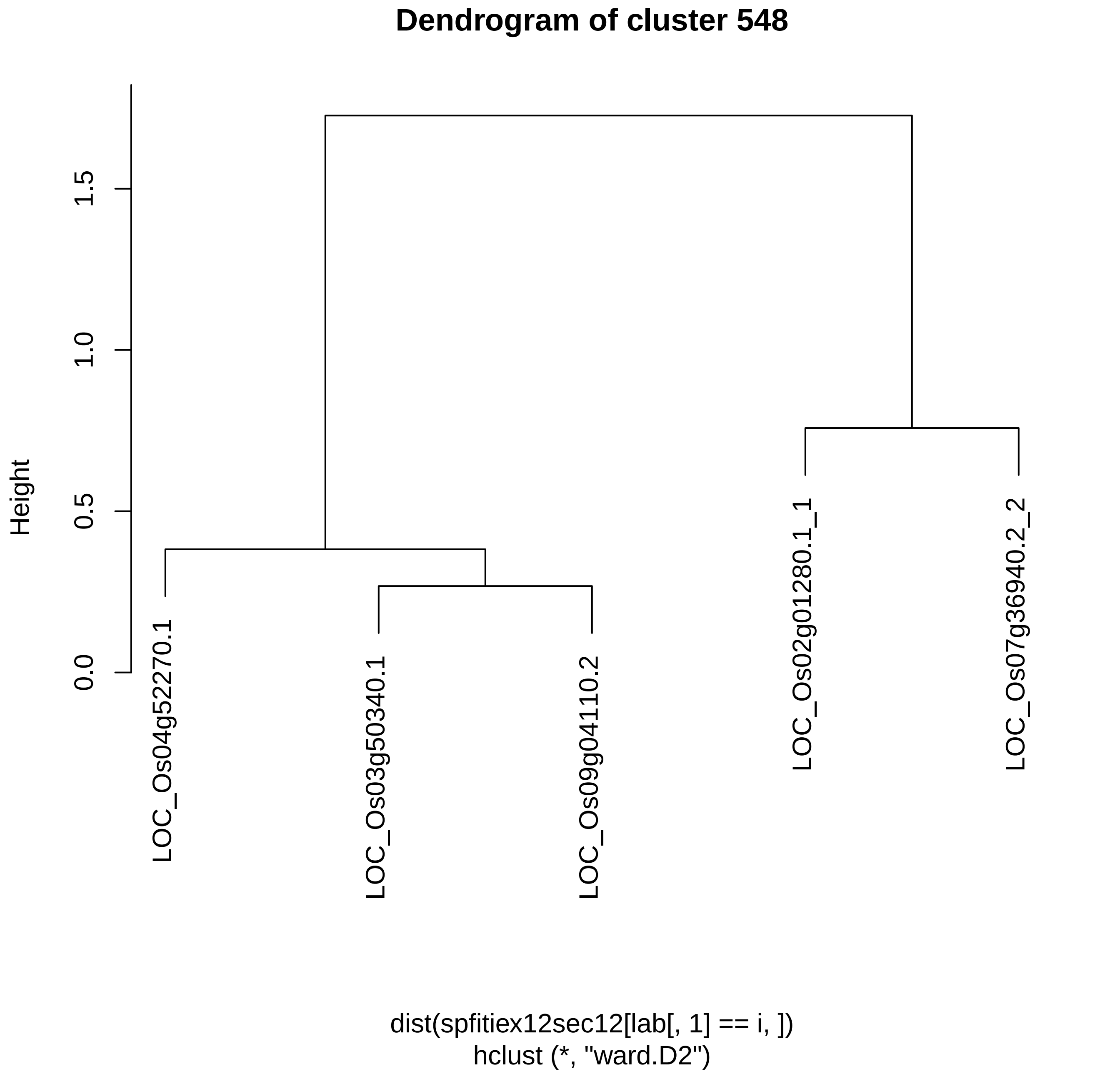
Dendrogram of cluster 548
Height
0.0
0.5
1.0
1.5
LOC_Os04g52270.1
LOC_Os03g50340.1
LOC_Os09g04110.2
LOC_Os02g01280.1_1
LOC_Os07g36940.2_2
dist(spfitiex12sec12[lab[, 1] == i, ])
hclust (*, "ward.D2")

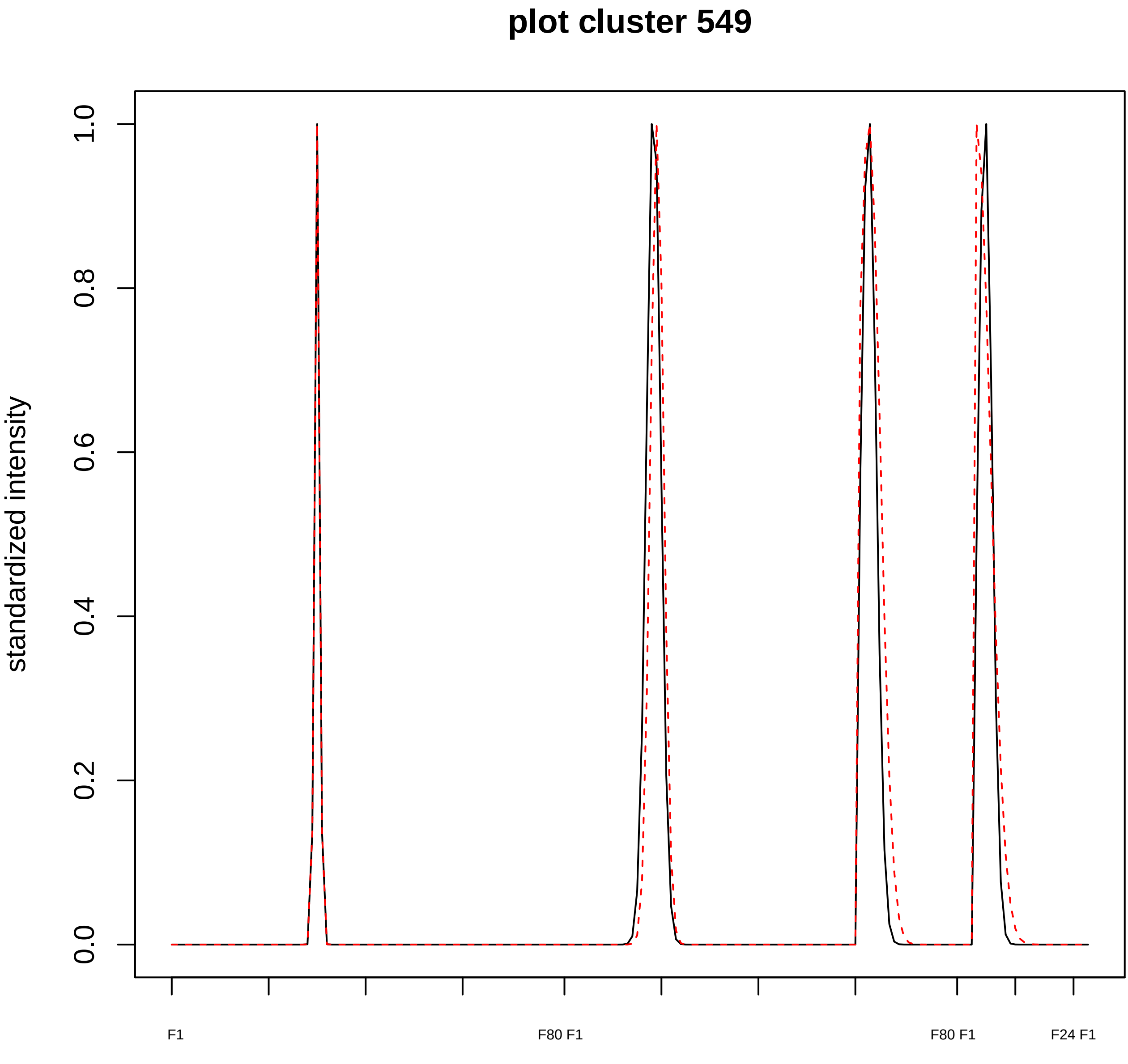
plot cluster 549
standardized intensity
0.0
0.2
0.4
0.6
0.8
1.0
F1
F80 F1
F80 F1
F24 F1

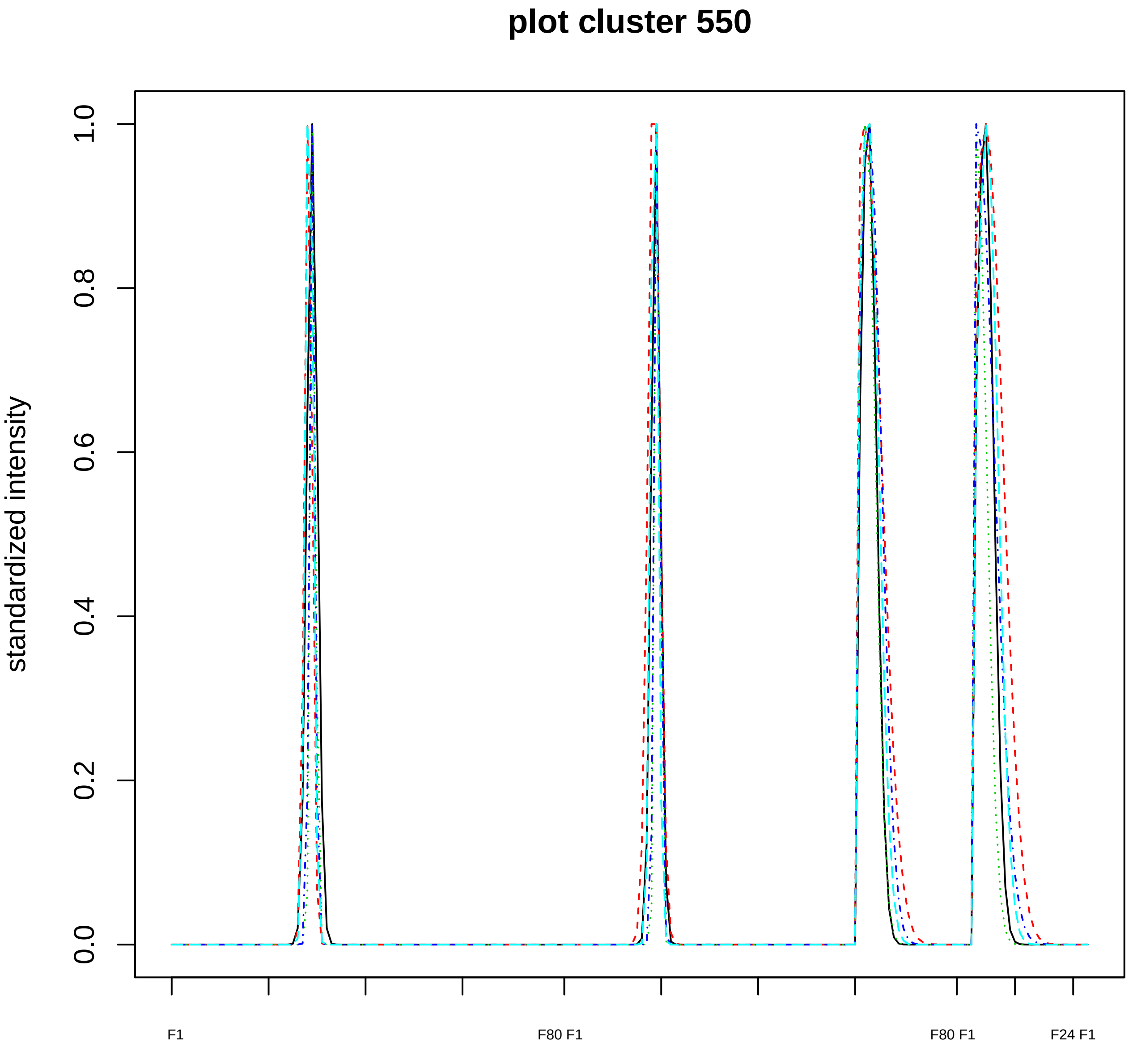
plot cluster 550
standardized intensity
0.0
0.2
0.4
0.6
0.8
1.0
F1
F80 F1
F80 F1
F24 F1

#### Dendrogram of cluster 550

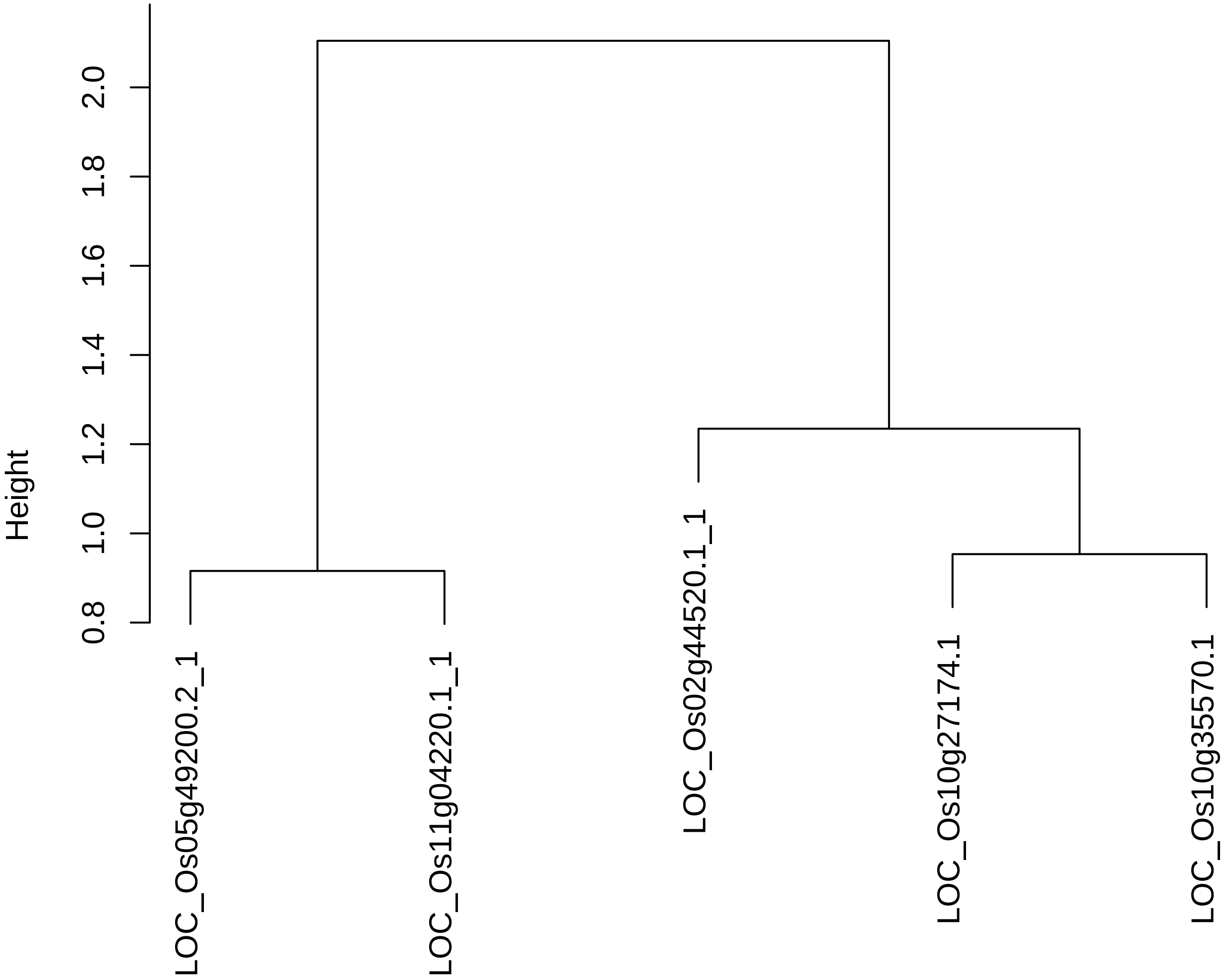


dist(spfitiex12sec12[lab[, 1] == i, ])
hclust (*, "ward.D2")

### plot cluster 551

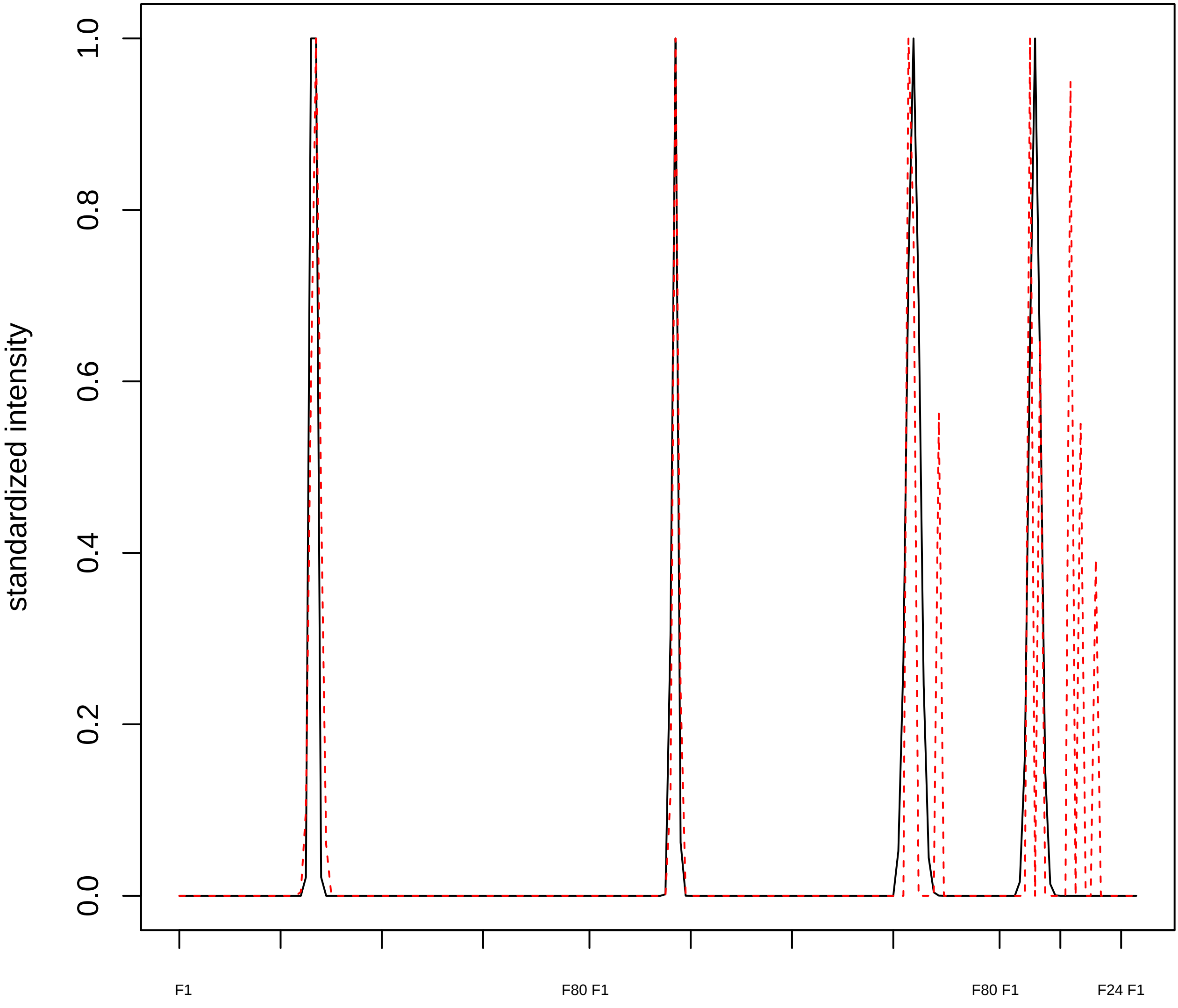

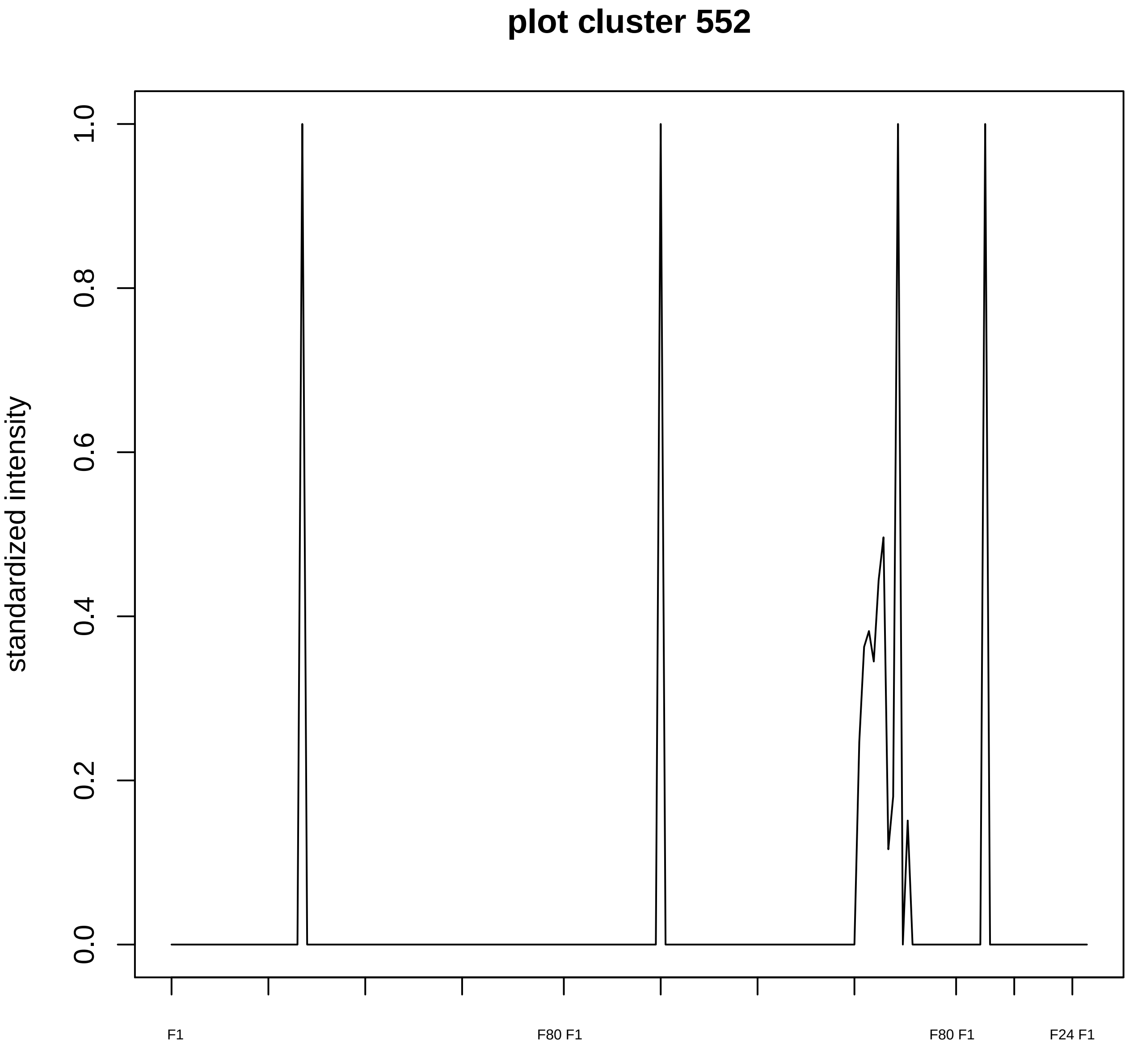
plot cluster 552
standardized intensity
0.0
0.2
0.4
0.6
0.8
1.0
F1
F80 F1
F80 F1
F24 F1

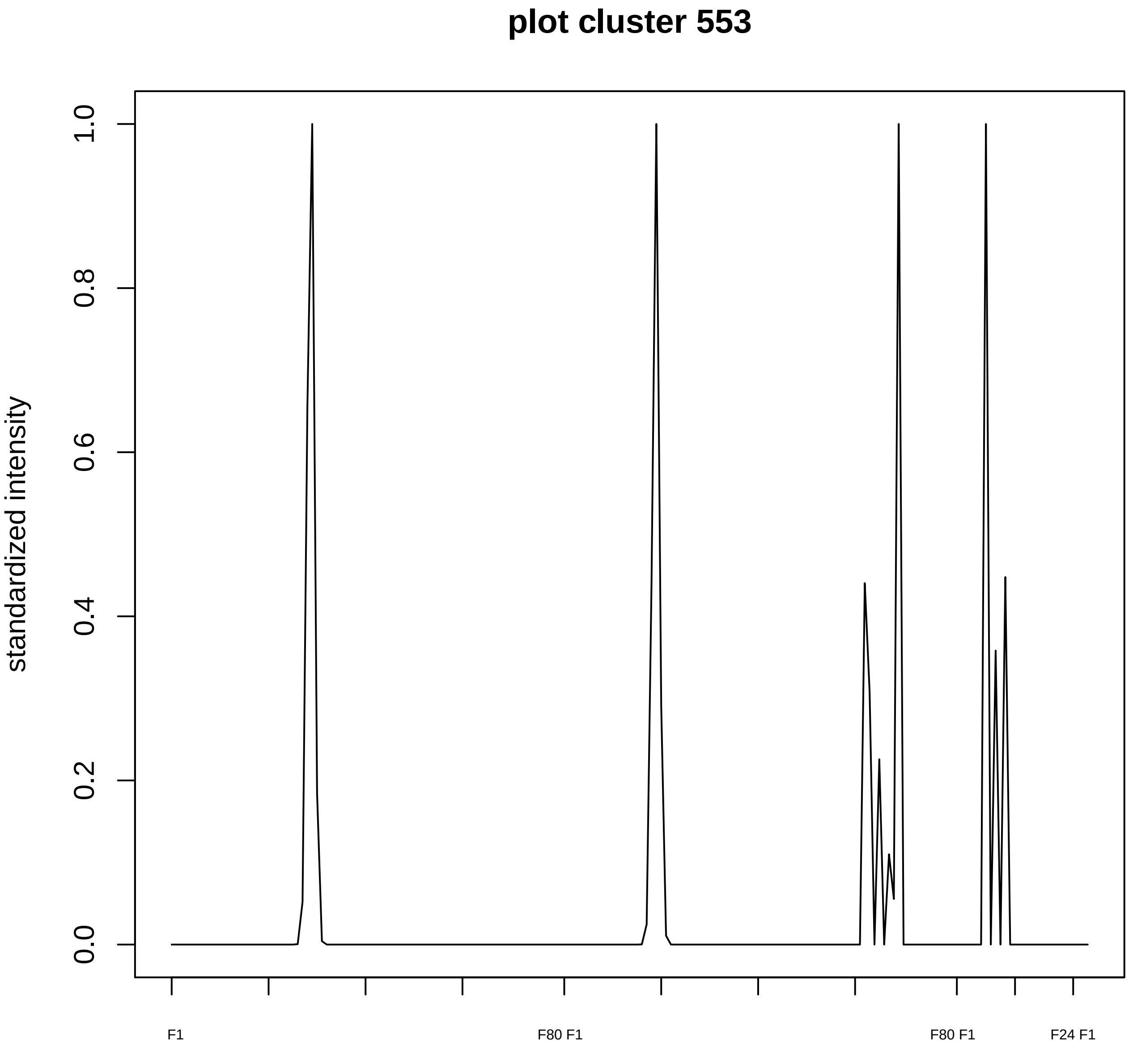
plot cluster 553
standardized intensity
0.0
0.2
0.4
0.6
0.8
1.0
F1
F80 F1
F80 F1
F24 F1

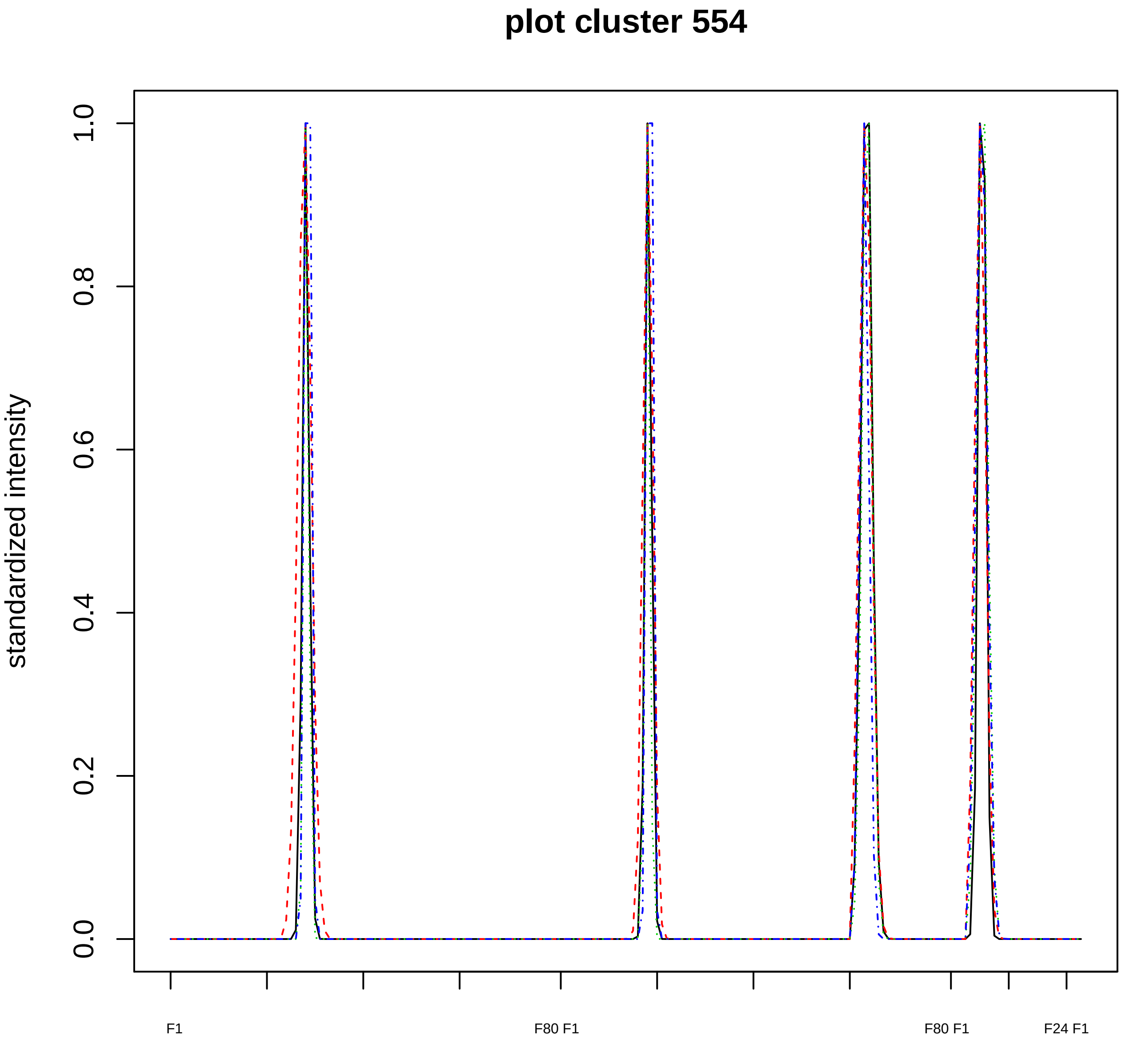
plot cluster 554
standardized intensity
0.0
0.2
0.4
0.6
0.8
1.0
F1
F80 F1
F80 F1
F24 F1

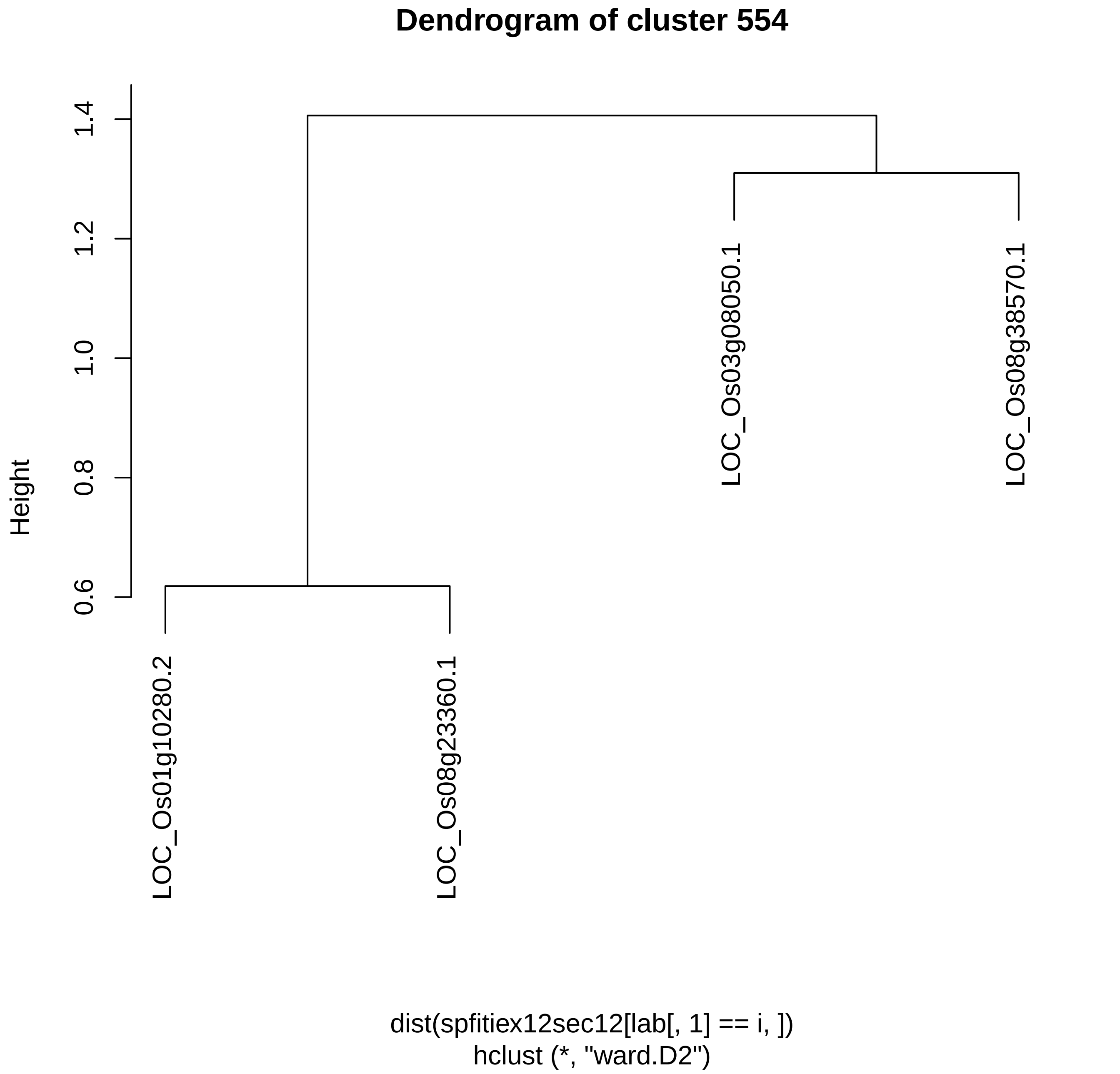
Dendrogram of cluster 554
Height
0.6
0.8
1.0
1.2
1.4
LOC_Os01g10280.2
LOC_Os08g23360.1
LOC_Os03g08050.1
LOC_Os08g38570.1
dist(spfitiex12sec12[lab[, 1] == i, ])
hclust (*, "ward.D2")

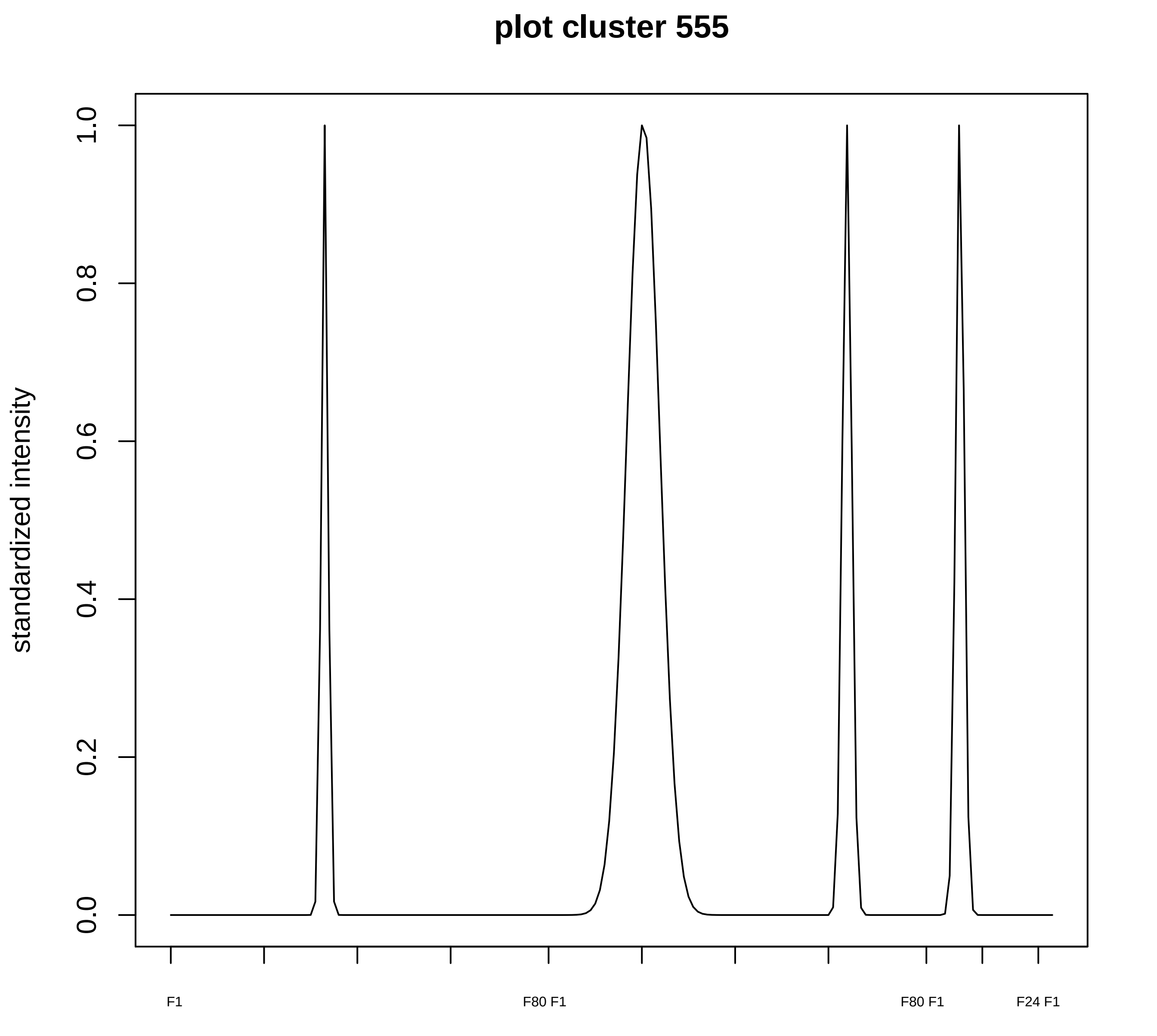

plot cluster 555
standardized intensity
0.0
0.2
0.4
0.6
0.8
1.0
F1
F80 F1
F80 F1
F24 F1

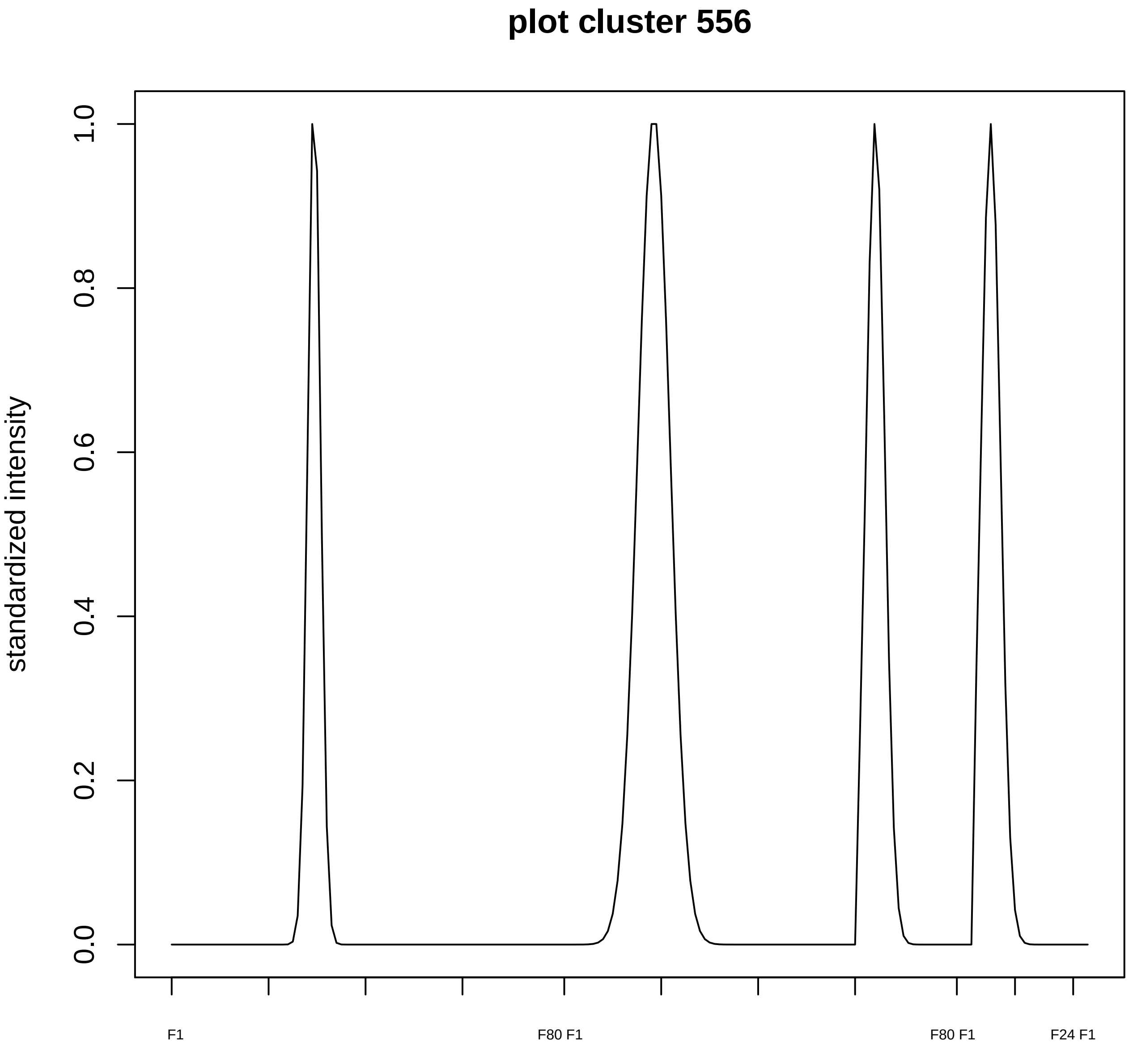
plot cluster 556
standardized intensity
0.0
0.2
0.4
0.6
0.8
1.0
F1
F80 F1
F80 F1
F24 F1

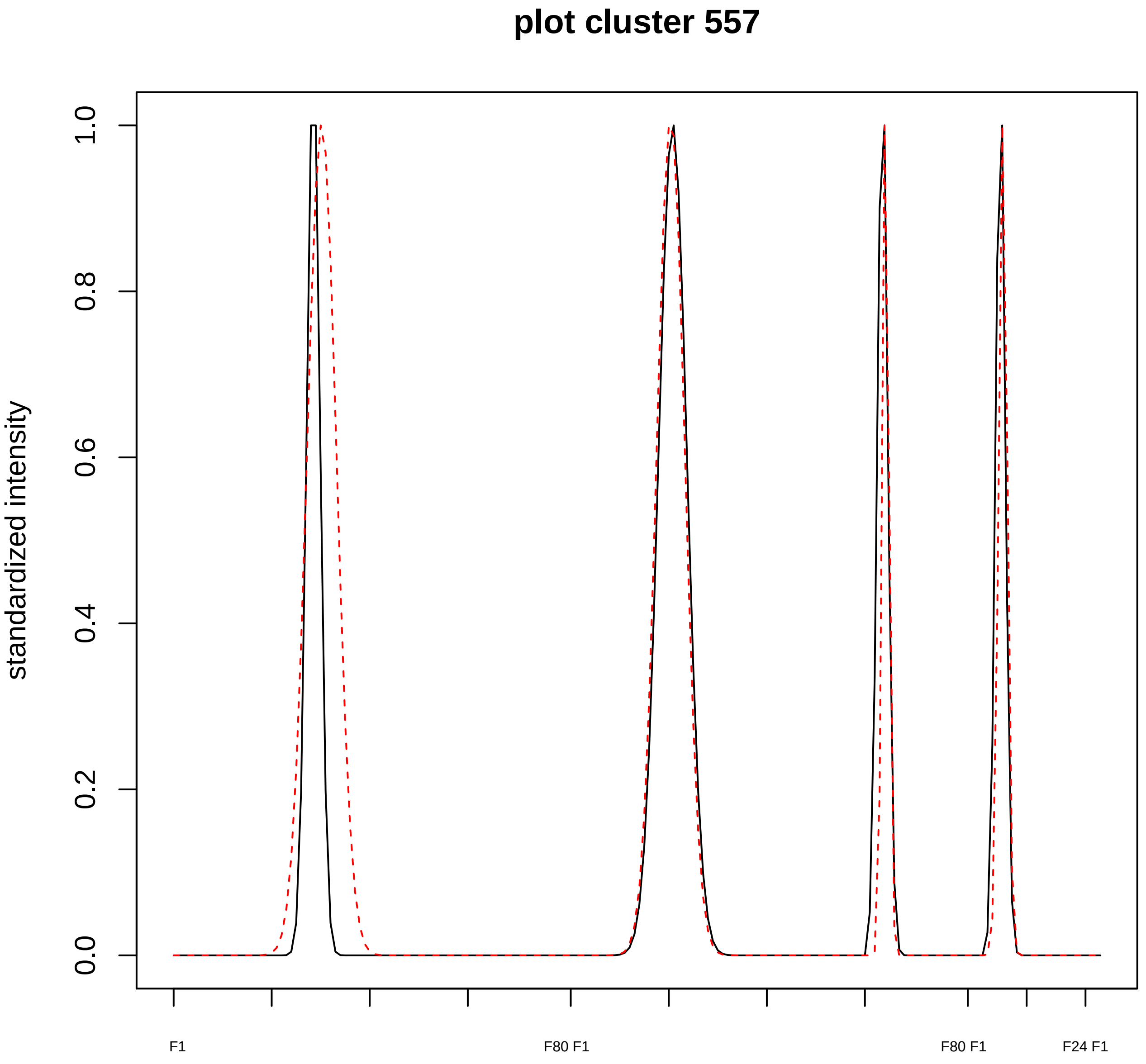

plot cluster 557
standardized intensity
0.0
0.2
0.4
0.6
0.8
1.0
F1
F80 F1
F80 F1
F24 F1

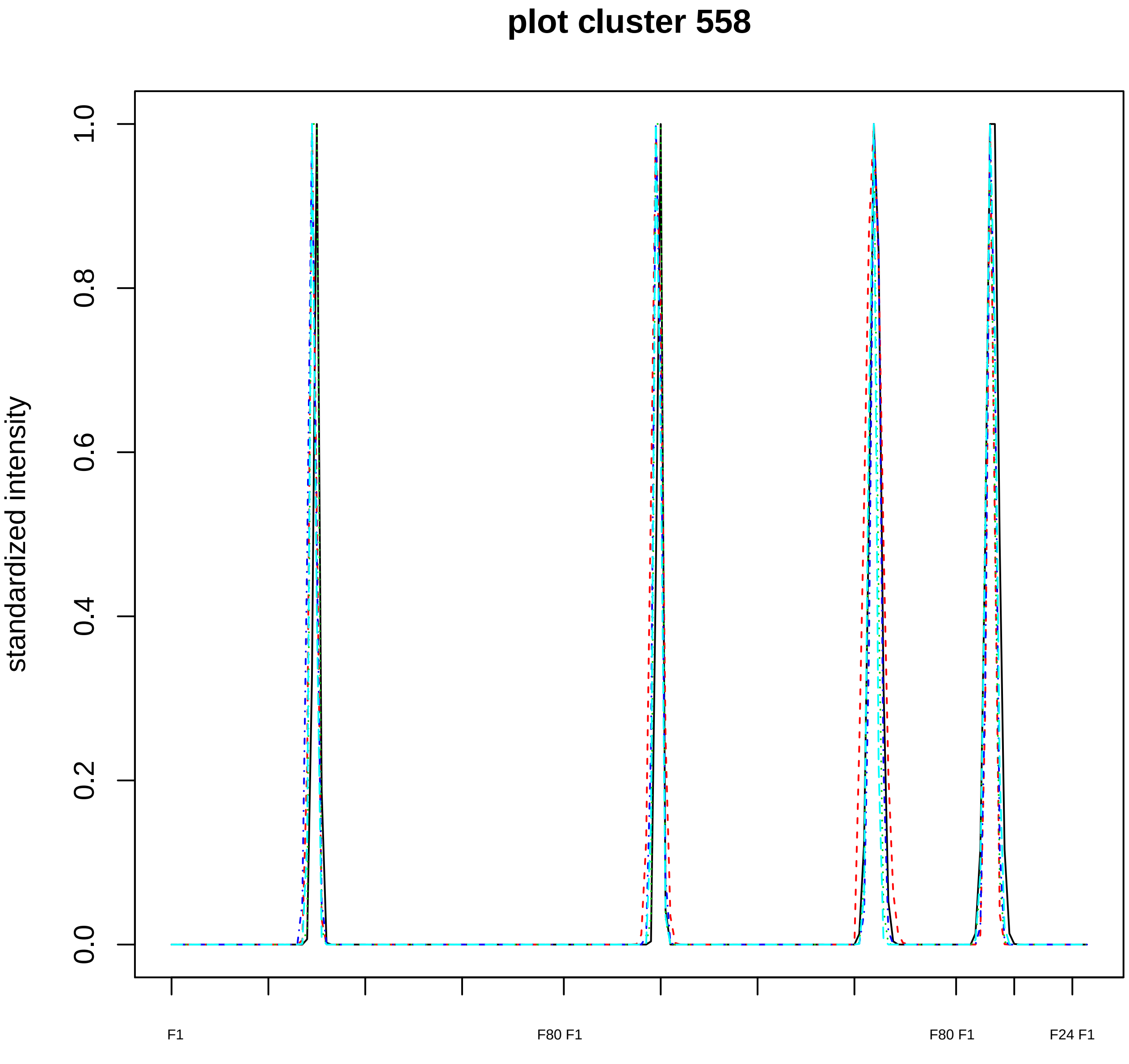

plot cluster 558
standardized intensity
0.0
0.2
0.4
0.6
0.8
1.0
F1
F80 F1
F80 F1
F24 F1

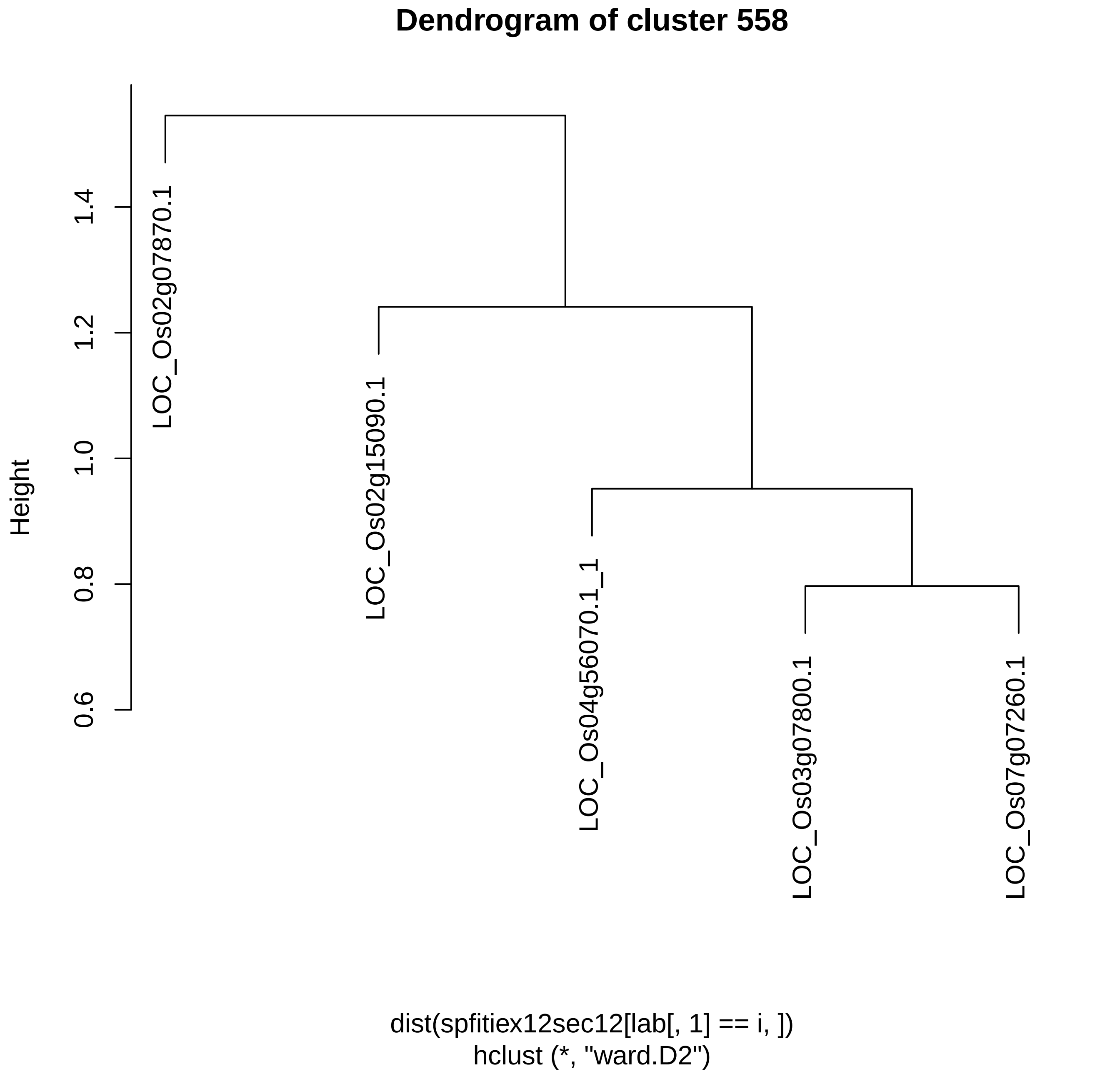

Dendrogram of cluster 558
Height
0.6
0.8
1.0
1.2
1.4
LOC_Os02g07870.1
LOC_Os02g15090.1
LOC_Os04g56070.1_1
LOC_Os03g07800.1
LOC_Os07g07260.1
dist(spfitiex12sec12[lab[, 1] == i, ])
hclust (*, "ward.D2")

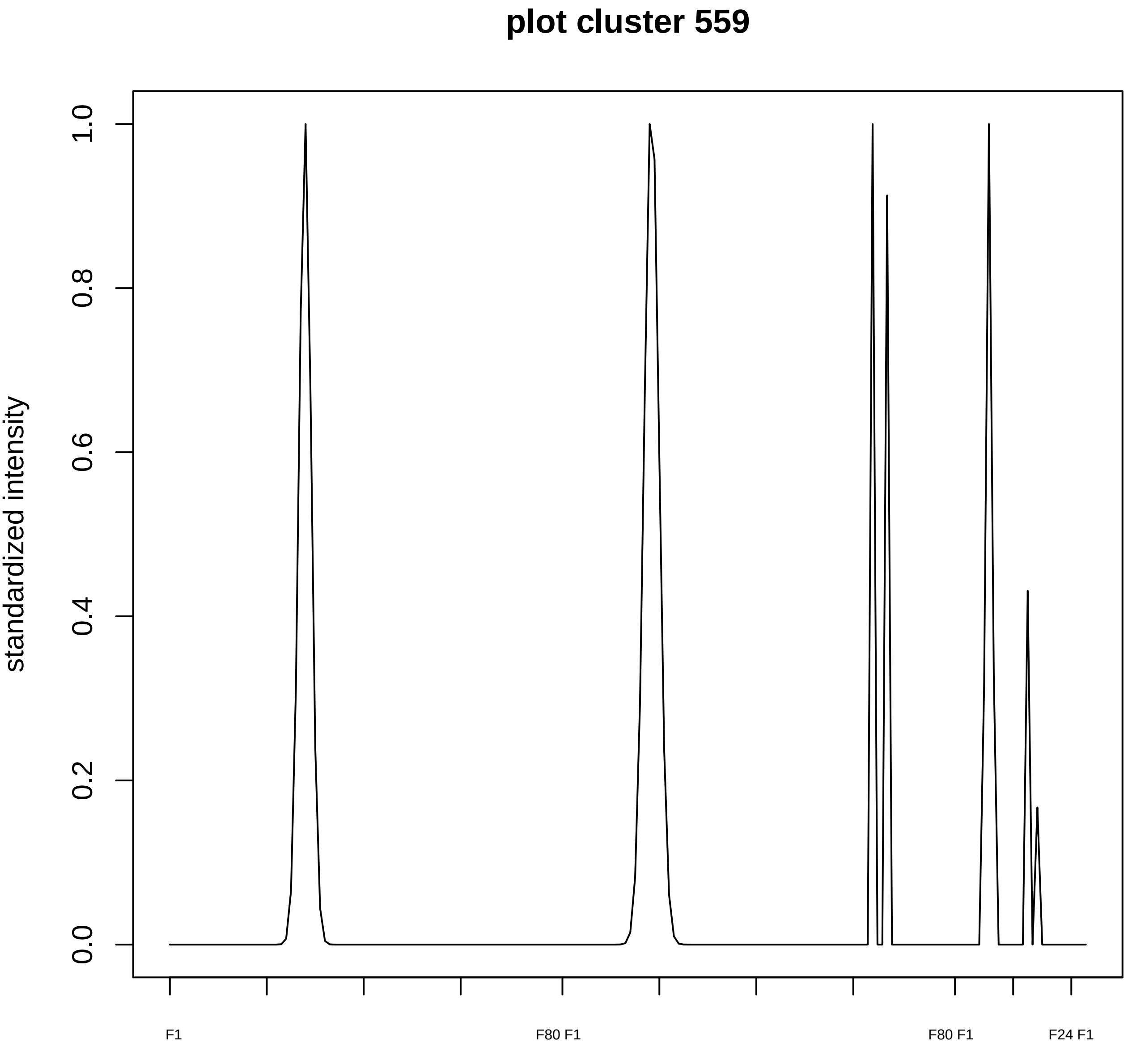
plot cluster 559
standardized intensity
1.0
0.8
0.6
0.4
0.2
0.0
F1
F80 F1
F80 F1
F24 F1

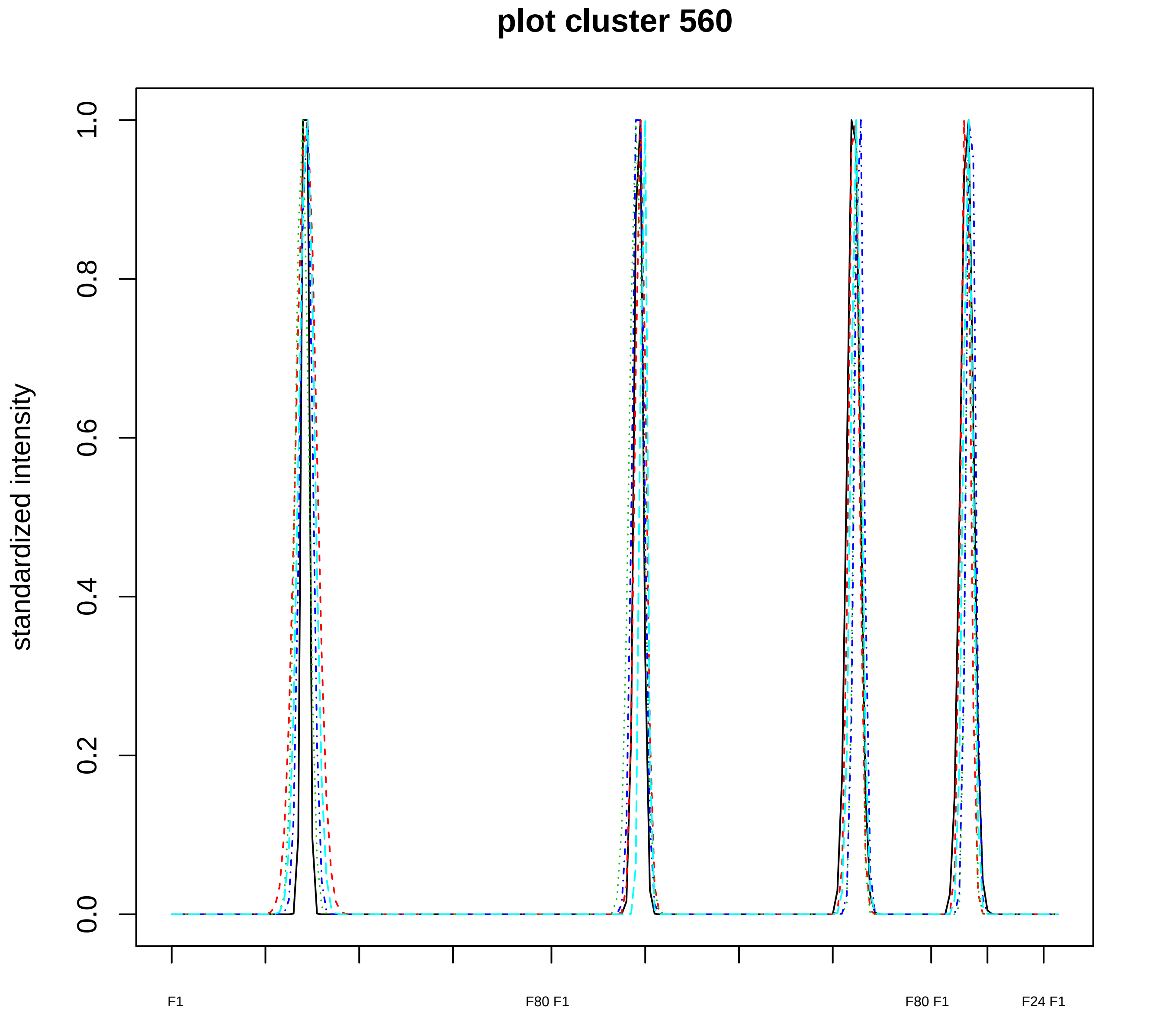
plot cluster 560
standardized intensity
0.0
0.2
0.4
0.6
0.8
1.0
F1
F80 F1
F80 F1
F24 F1

#### Dendrogram of cluster 560

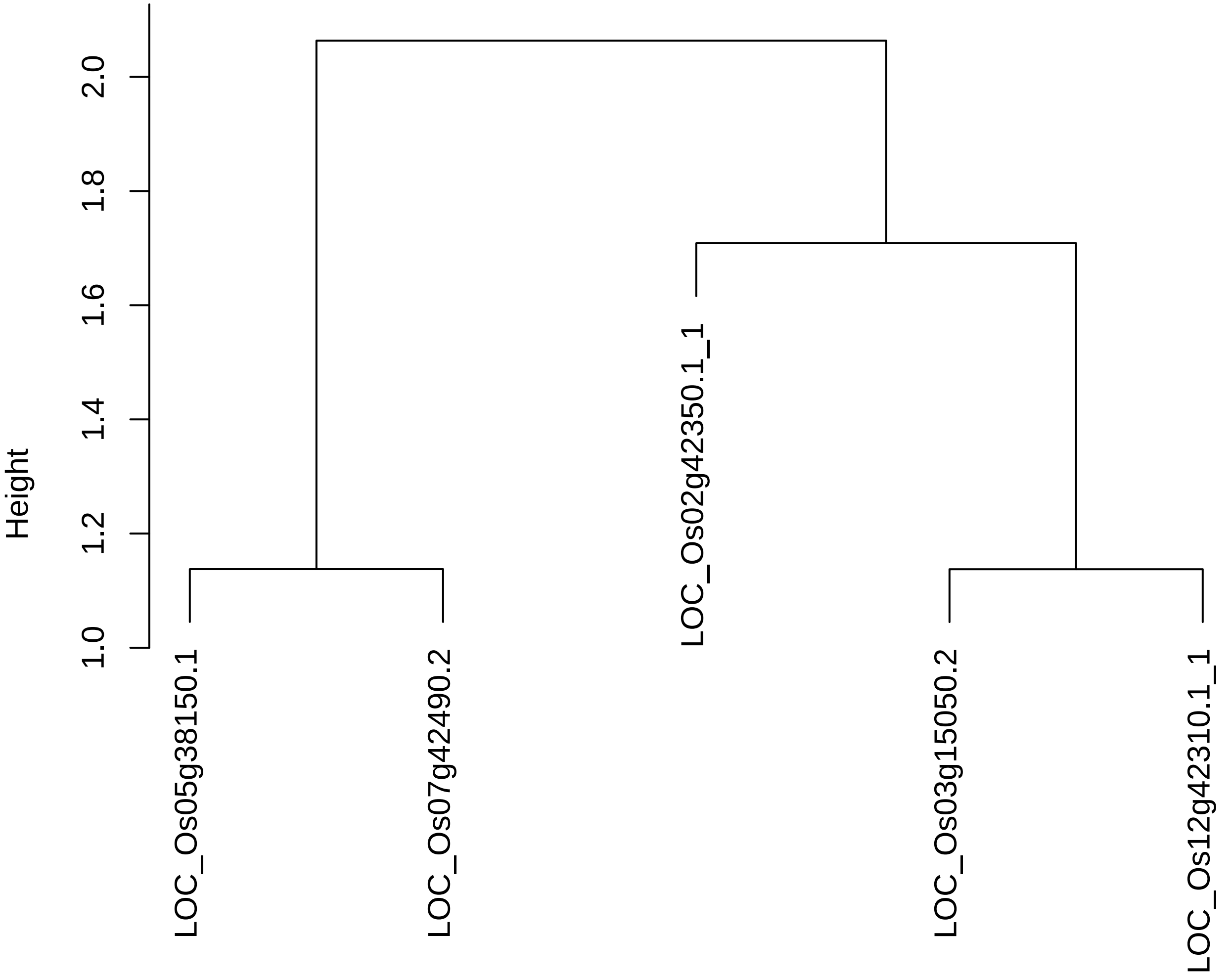


dist(spfitiex12sec12[lab[, 1] == i, ])
hclust (*, "ward.D2")

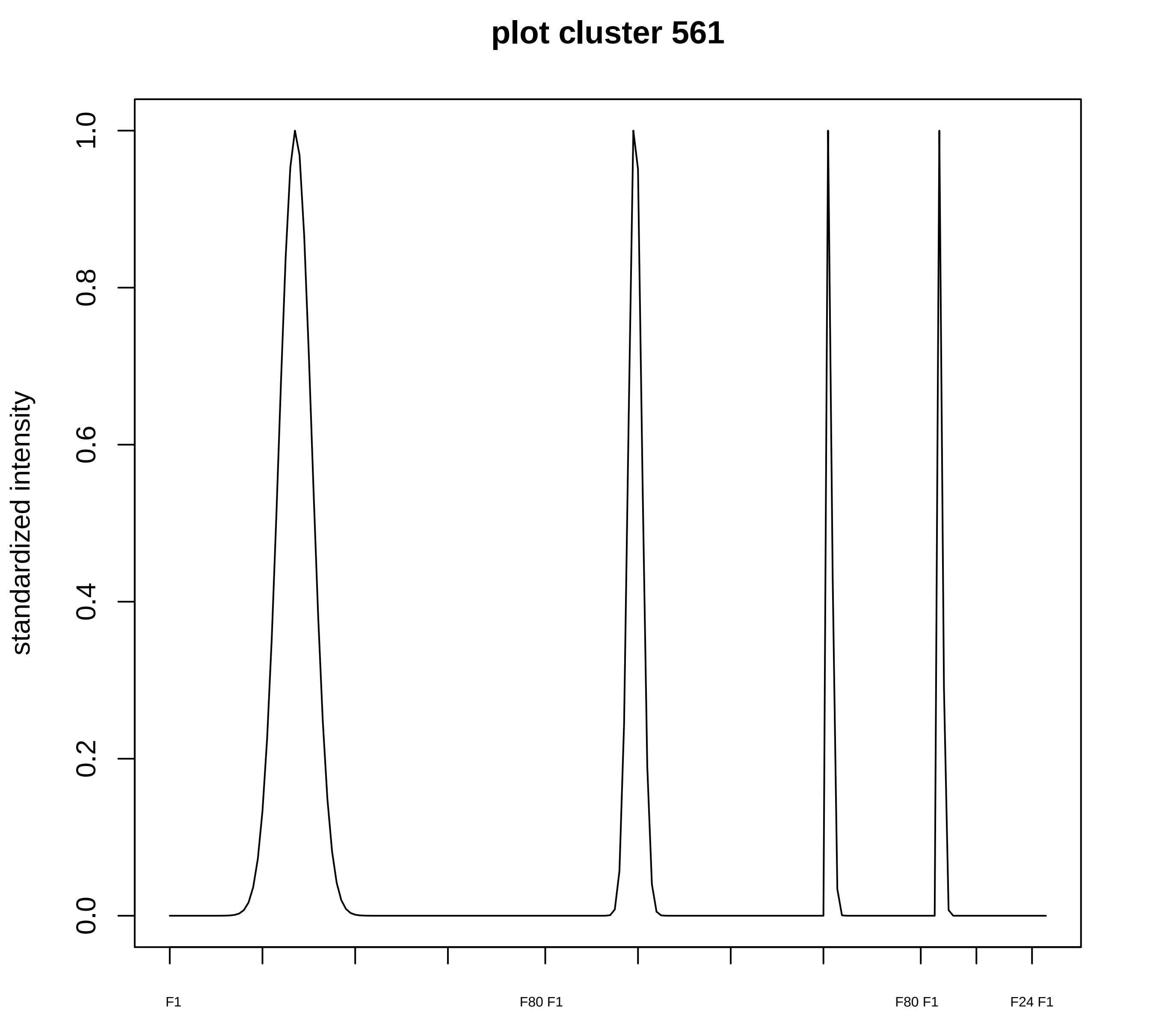

plot cluster 561
standardized intensity
0.0
0.2
0.4
0.6
0.8
1.0
F1
F80 F1
F80 F1
F24 F1

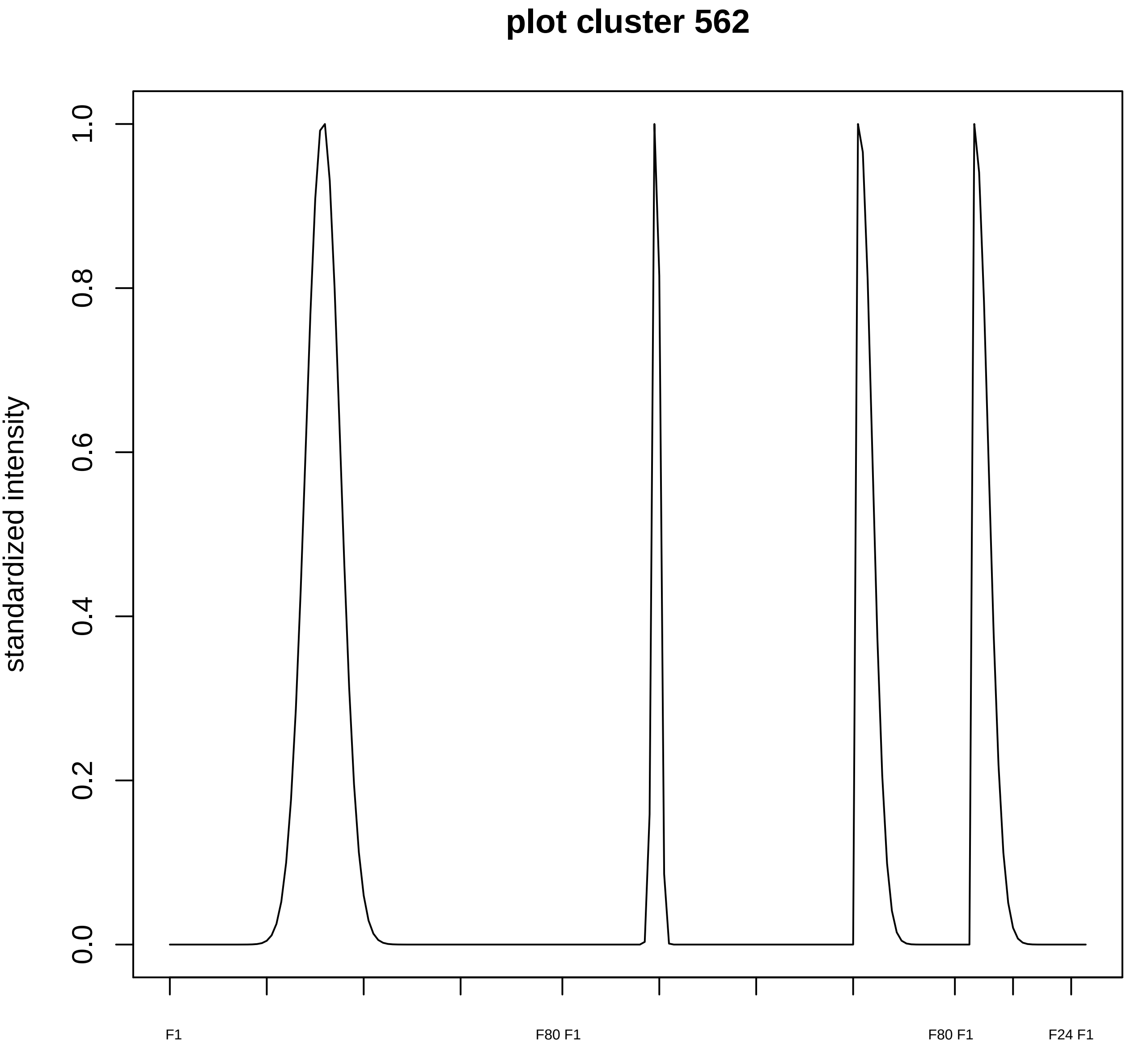
plot cluster 562
standardized intensity
0.0
0.2
0.4
0.6
0.8
1.0
F1
F80 F1
F80 F1
F24 F1

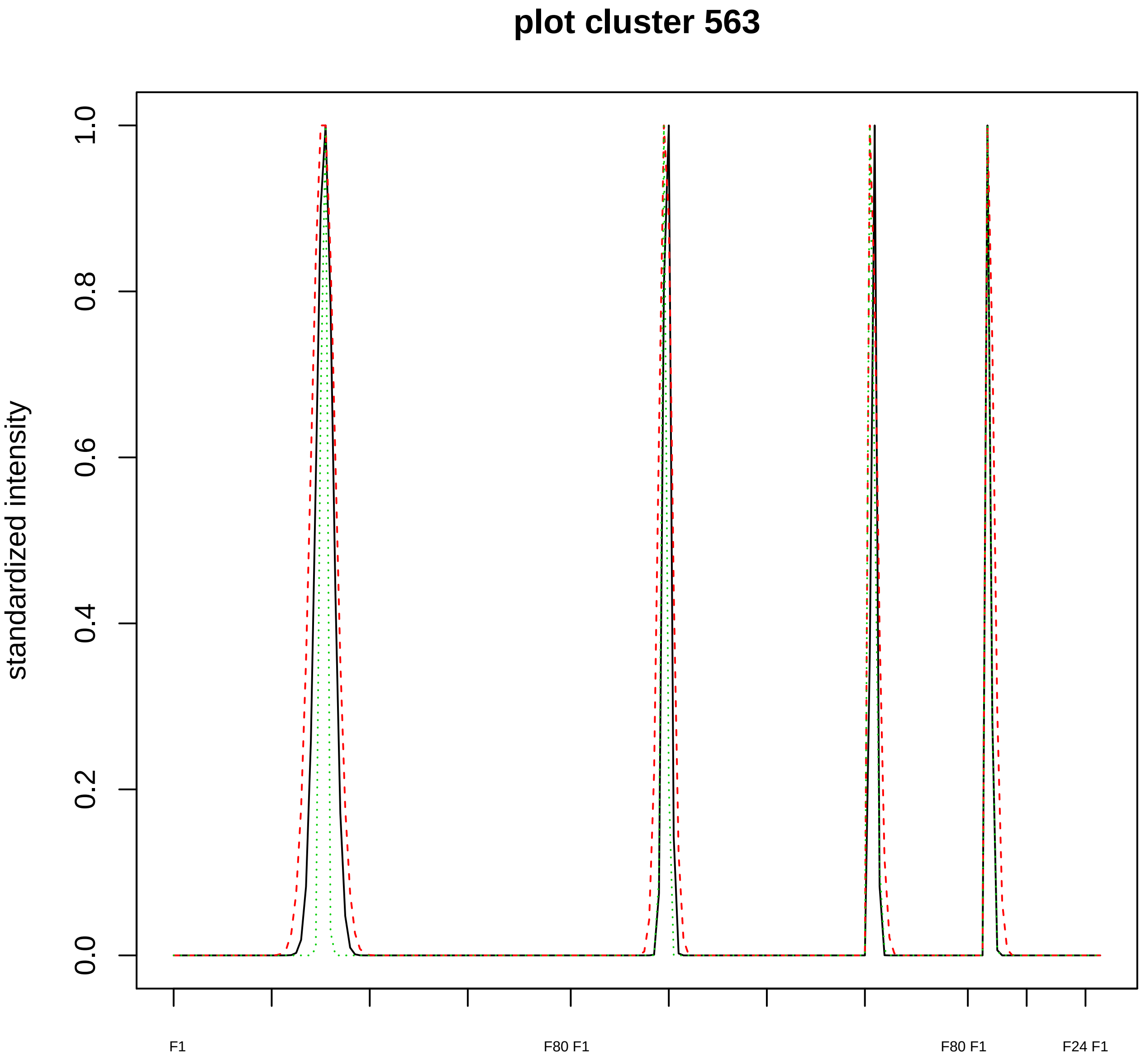

plot cluster 563
standardized intensity
0.0
0.2
0.4
0.6
0.8
1.0
F1
F80 F1
F80 F1
F24 F1

### Dendrogram of cluster 563

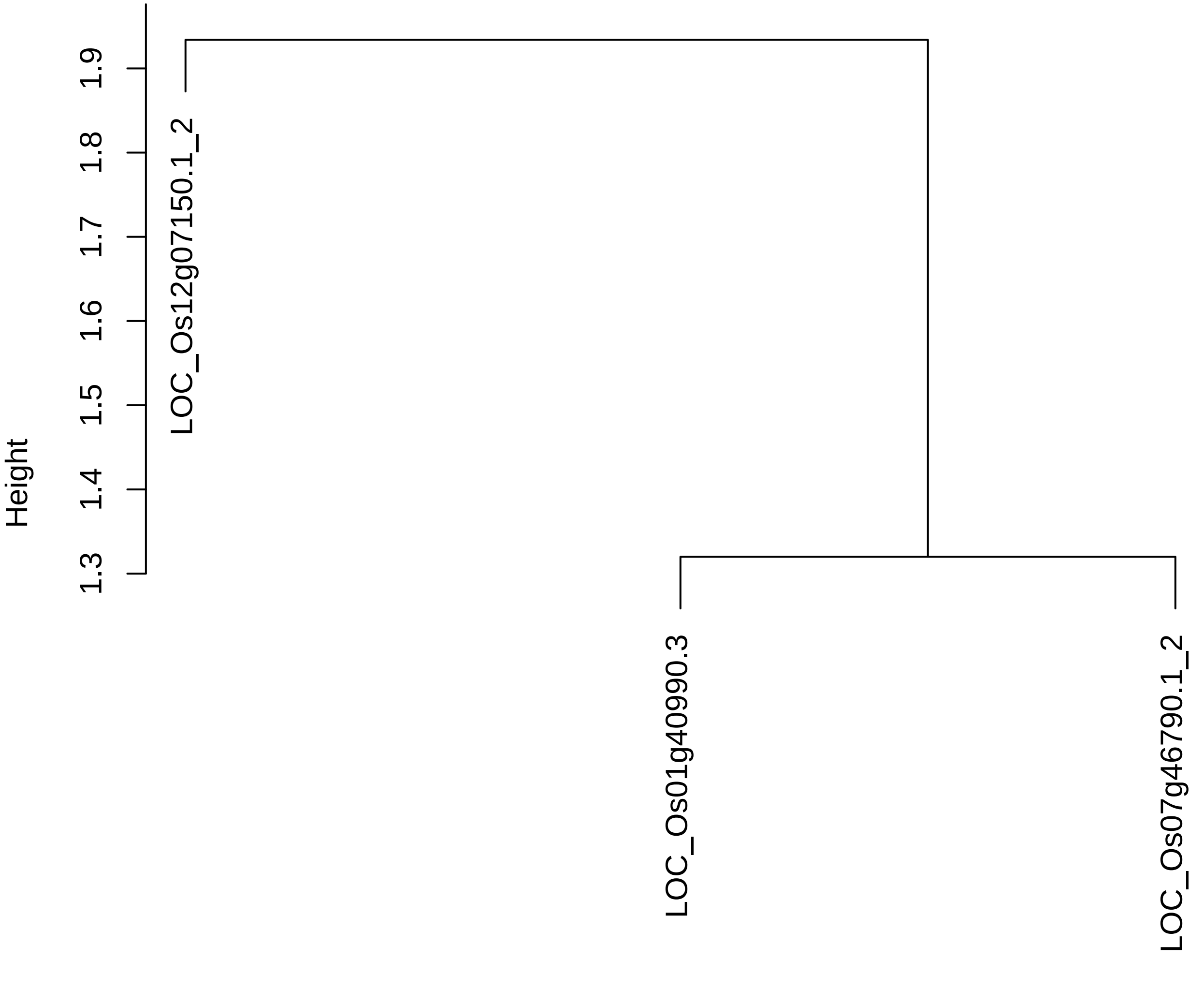


dist(spfitiex12sec12[lab[, 1] == i, ])
hclust (*, "ward.D2")

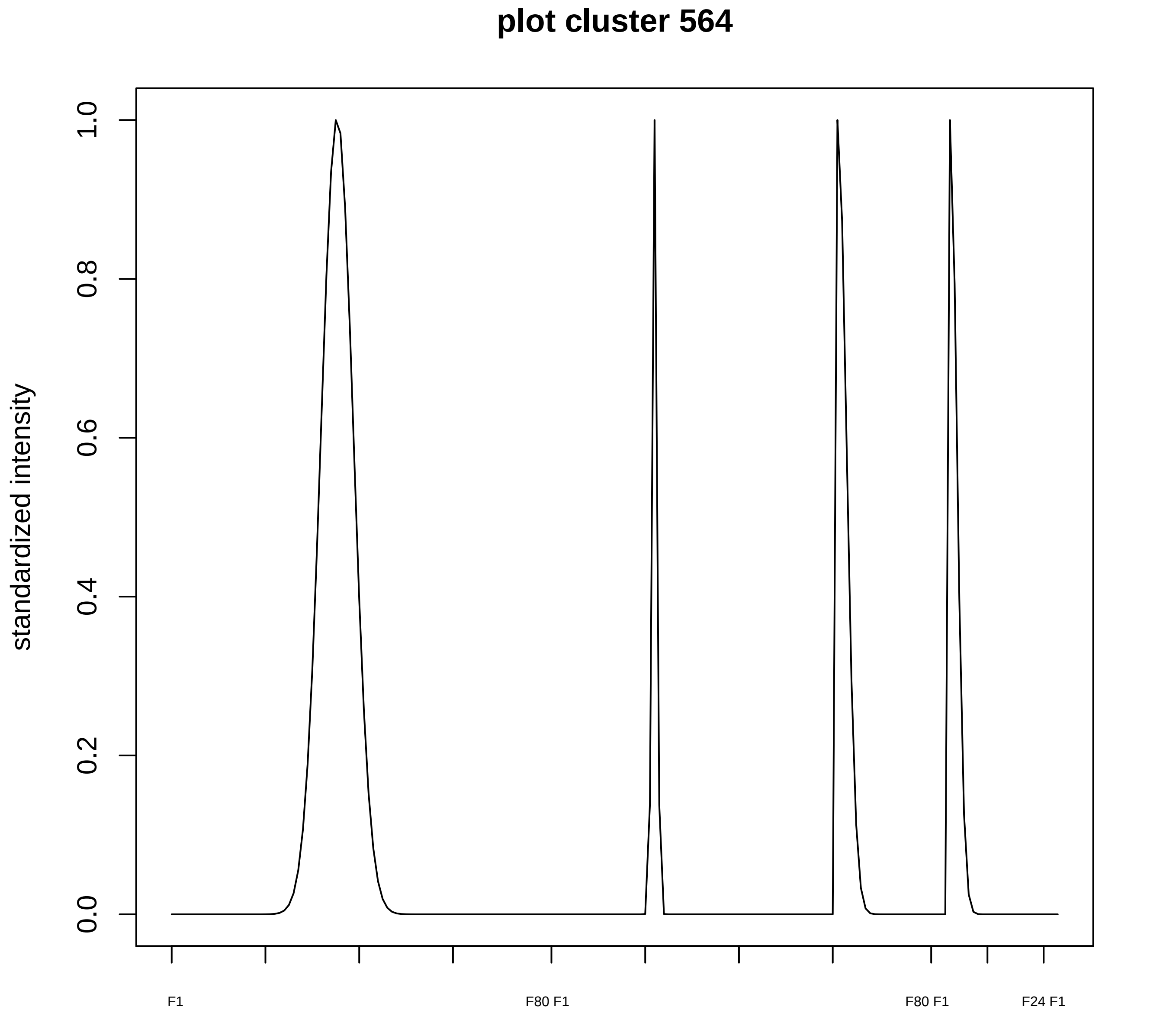
plot cluster 564
standardized intensity
0.0
0.2
0.4
0.6
0.8
1.0
F1
F80 F1
F80 F1
F24 F1

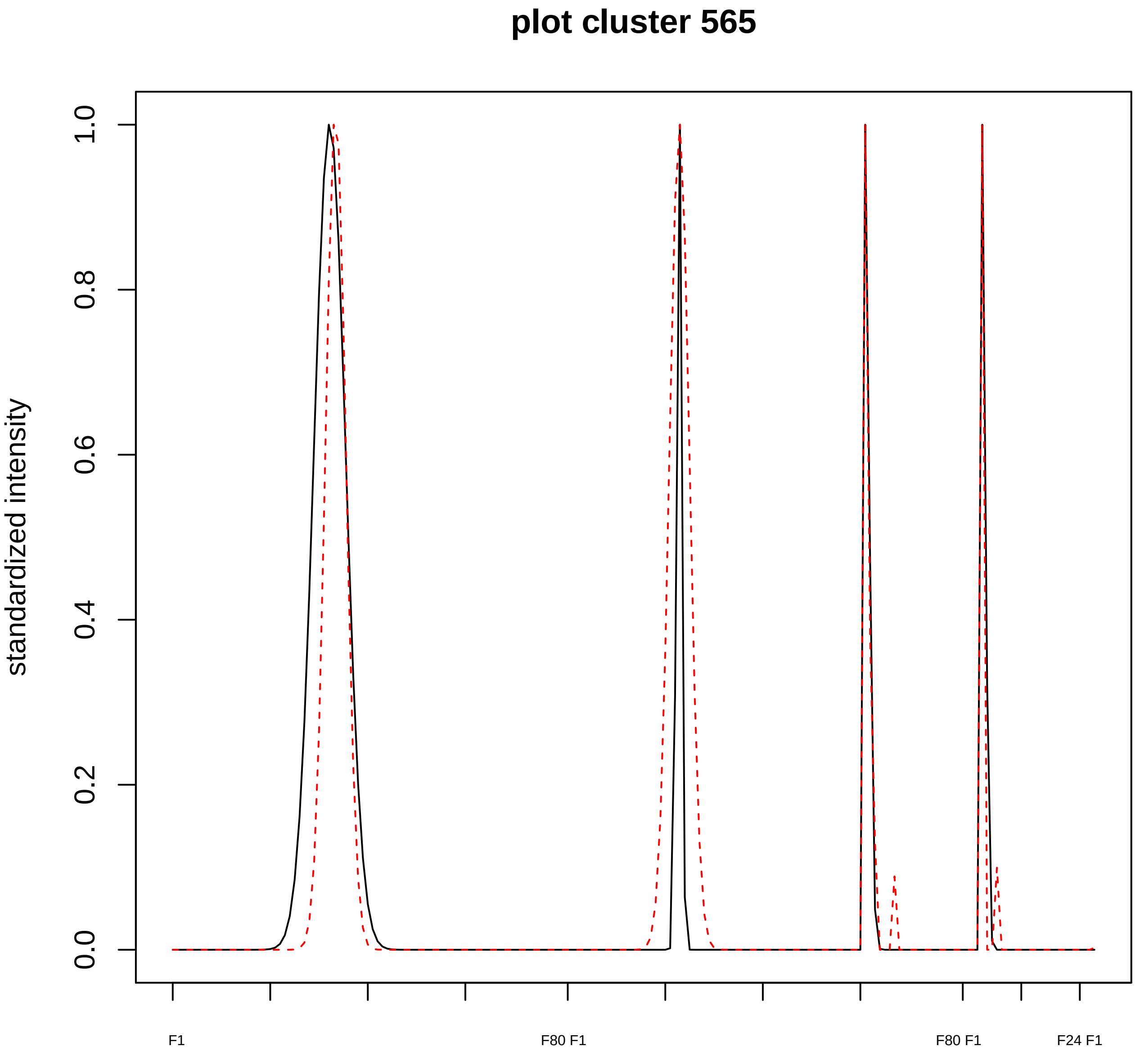

plot cluster 565
standardized intensity
0.0
0.2
0.4
0.6
0.8
1.0
F1
F80 F1
F80 F1
F24 F1

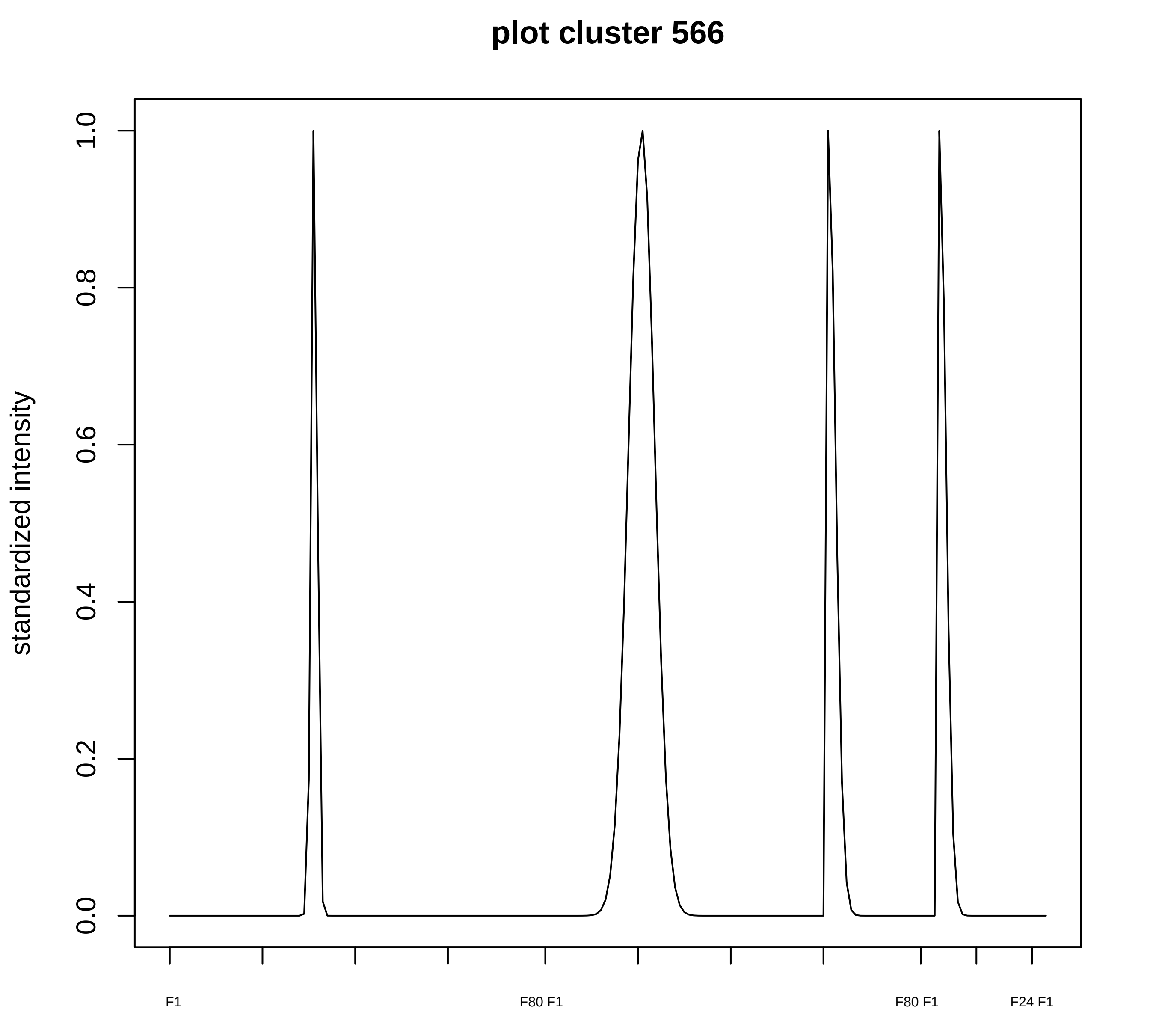
plot cluster 566
standardized intensity
0.0
0.2
0.4
0.6
0.8
1.0
F1
F80 F1
F80 F1
F24 F1

### plot cluster 567

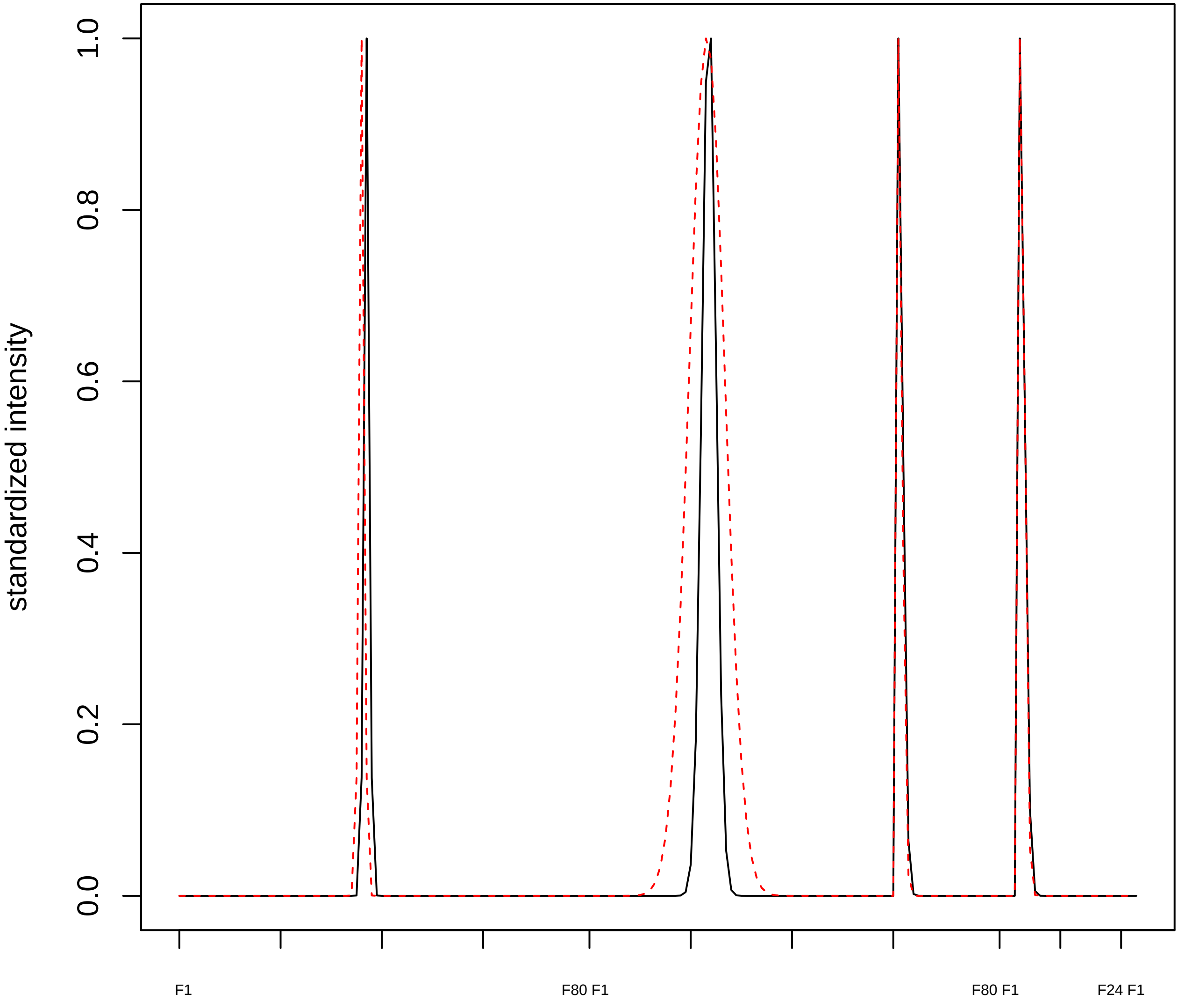

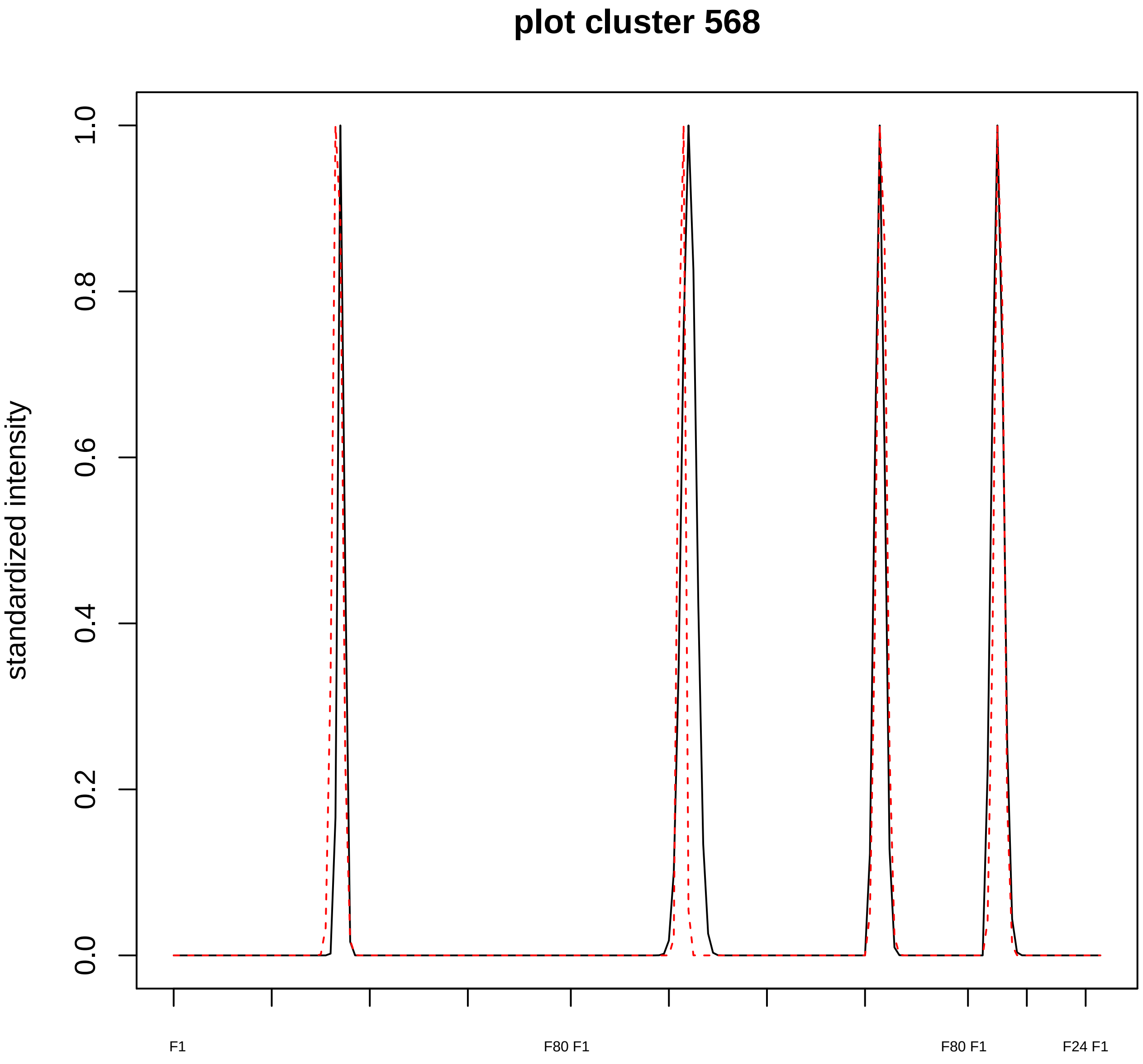
plot cluster 568
standardized intensity
0.0
0.2
0.4
0.6
0.8
1.0
F1
F80 F1
F80 F1
F24 F1

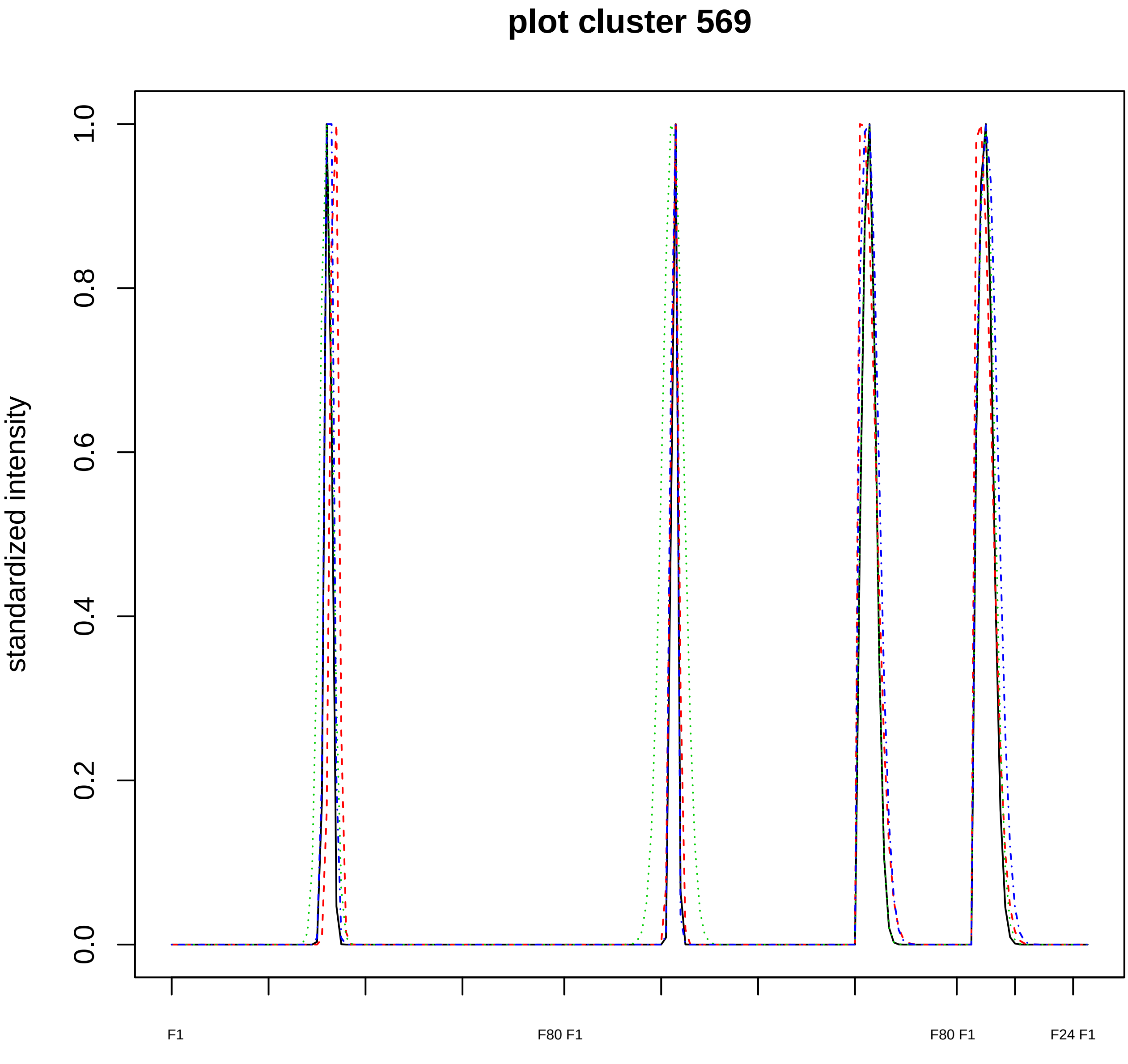
plot cluster 569
standardized intensity
0.0
0.2
0.4
0.6
0.8
1.0
F1
F80 F1
F80 F1
F24 F1

### Dendrogram of cluster 569

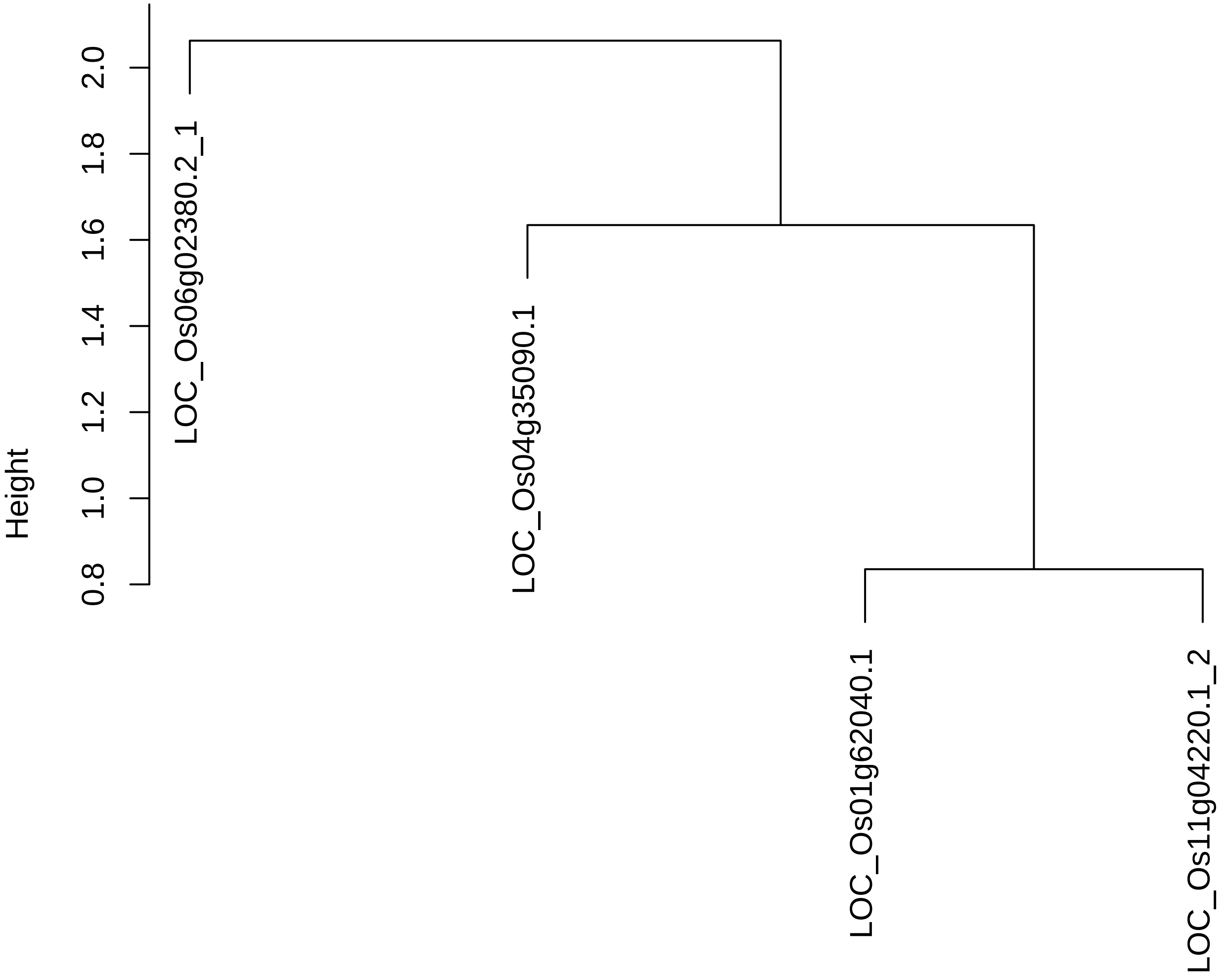


dist(spfitiex12sec12[lab[, 1] == i, ])
hclust (*, "ward.D2")

### plot cluster 570

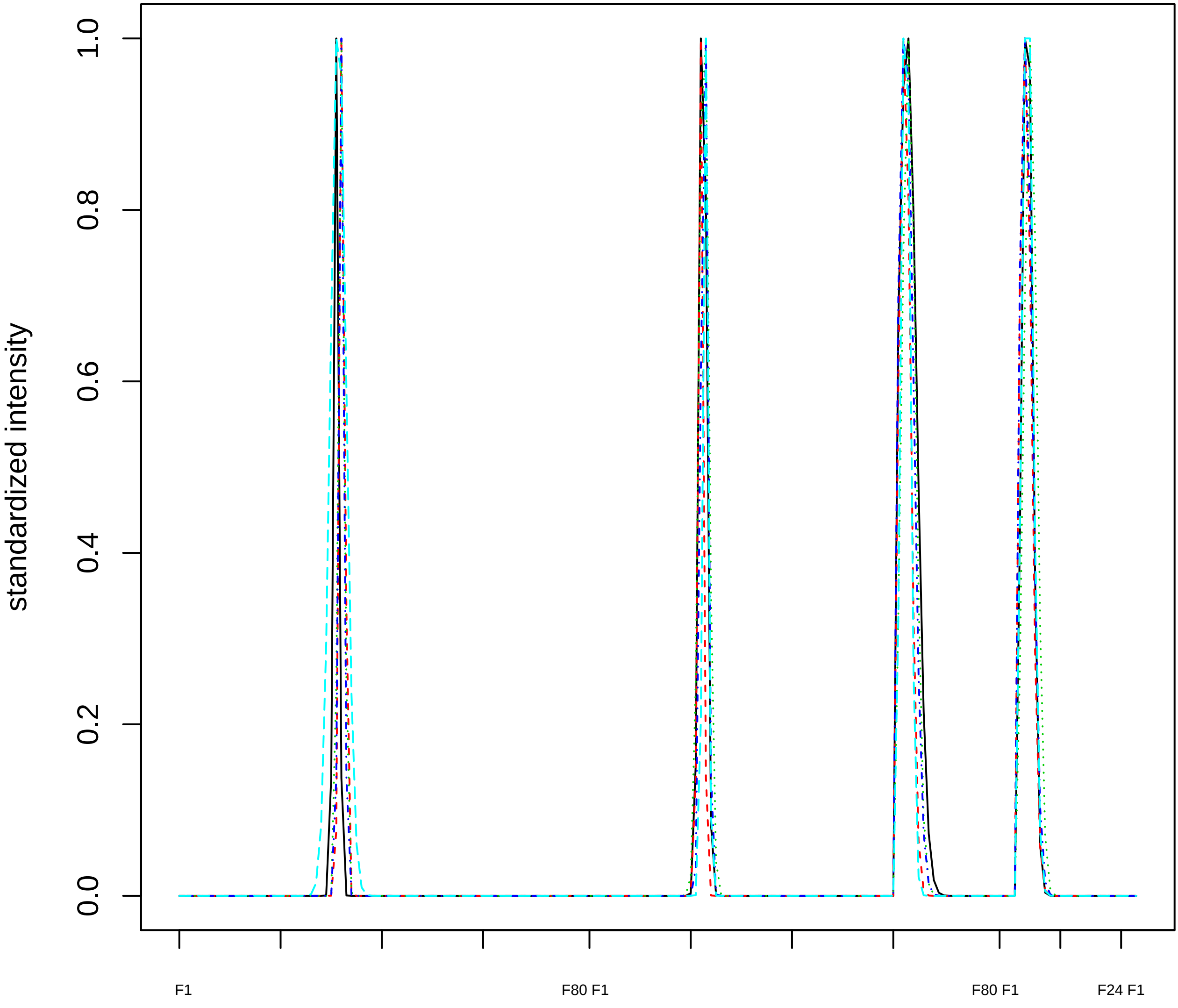

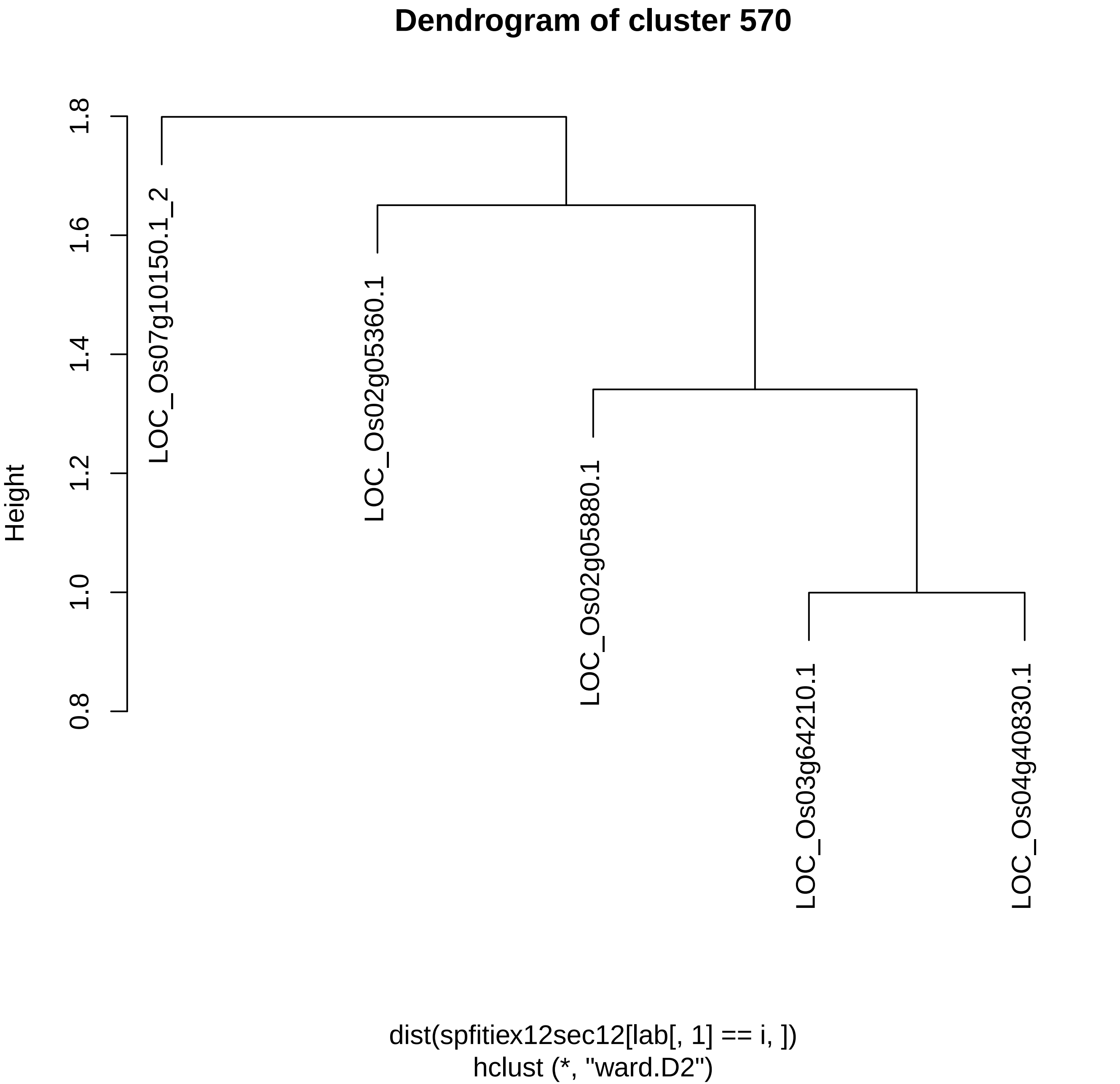

Dendrogram of cluster 570
Height
1.8
1.6
1.4
1.2
1.0
0.8
LOC_Os07g10150.1_2
LOC_Os02g05360.1
LOC_Os02g05880.1
LOC_Os03g64210.1
LOC_Os04g40830.1
dist(spfitiex12sec12[lab[, 1] == i, ])
hclust (*, "ward.D2")

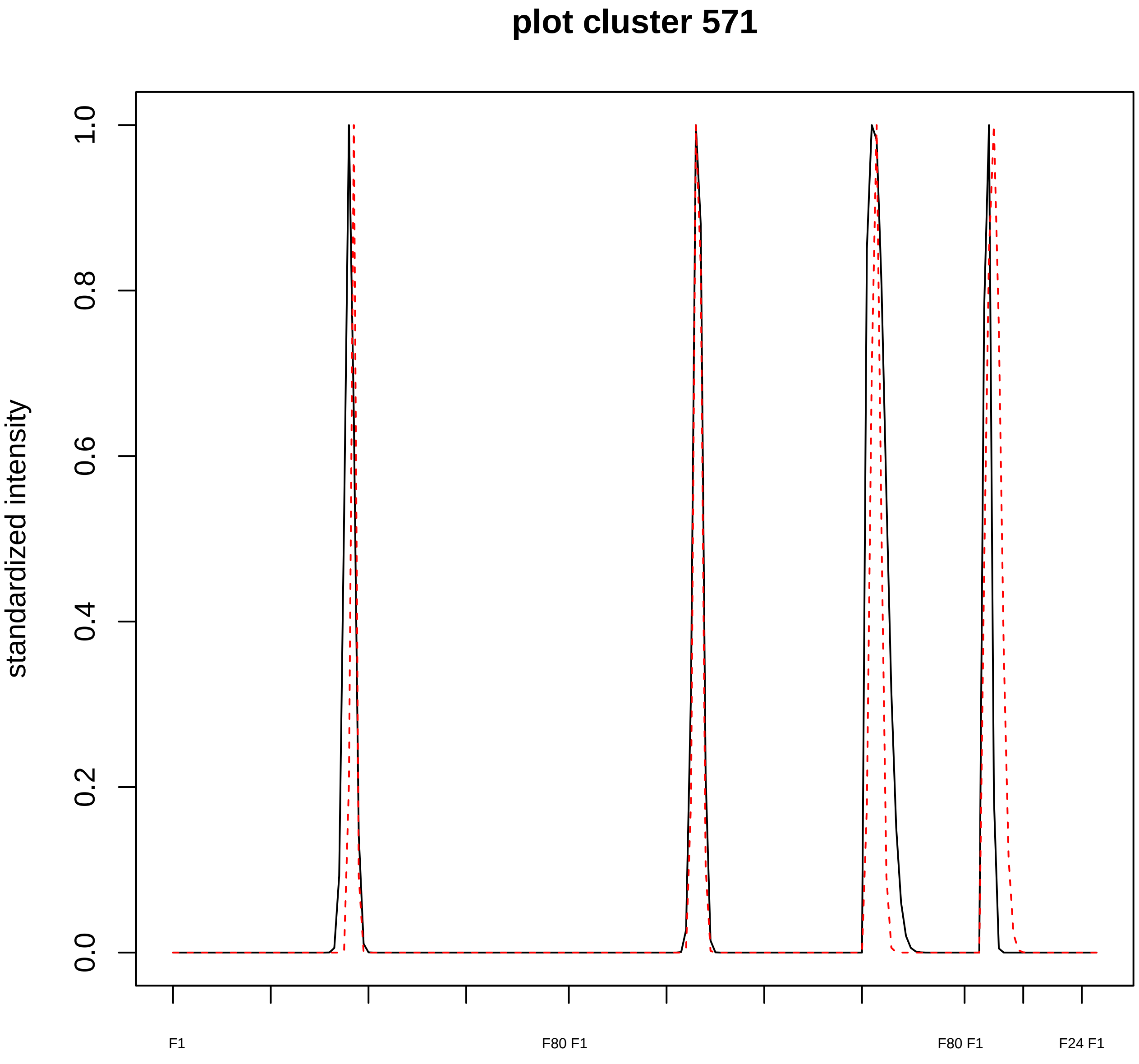
plot cluster 571
standardized intensity
0.0
0.2
0.4
0.6
0.8
1.0
F1
F80 F1
F80 F1
F24 F1

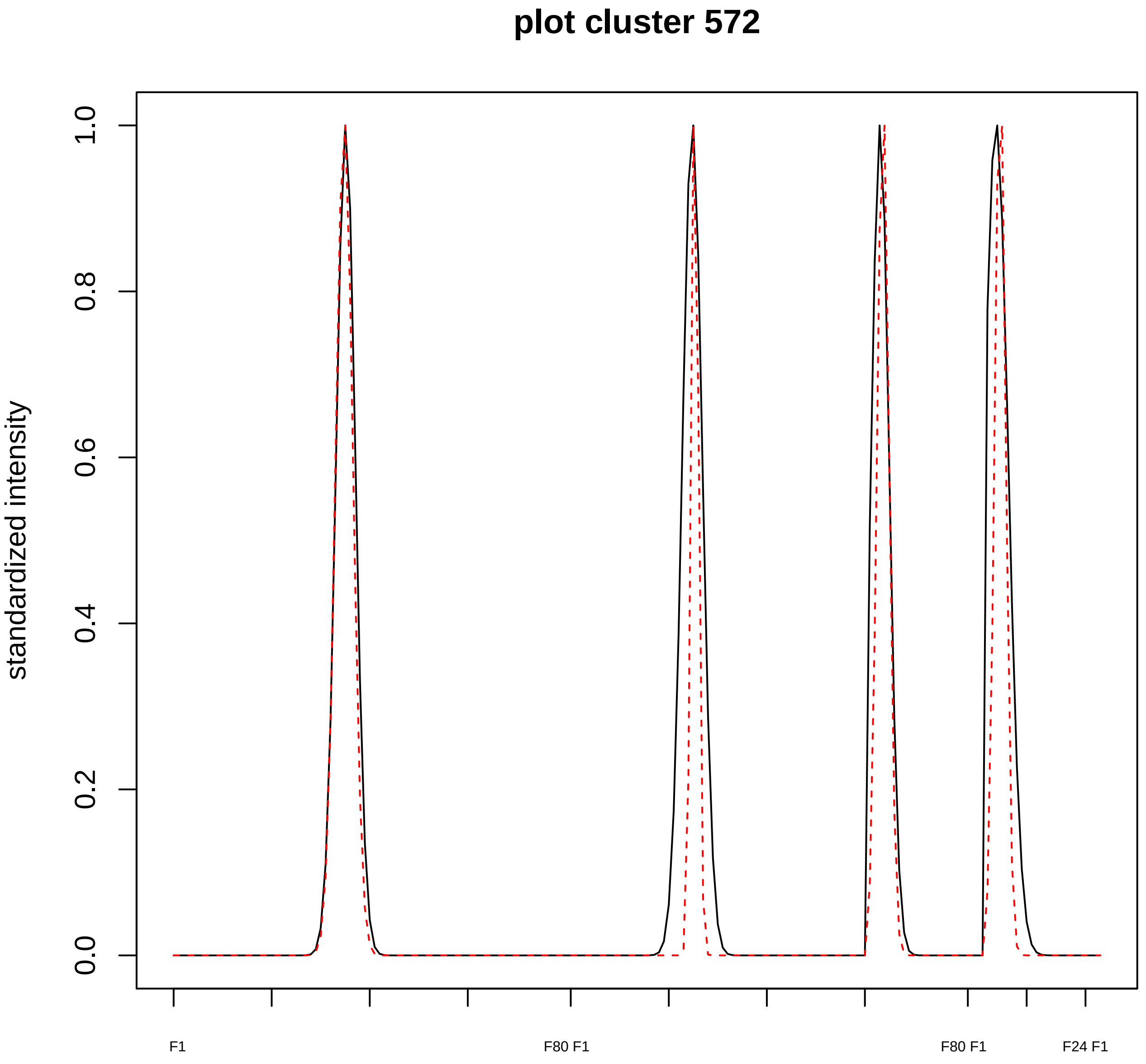
plot cluster 572
standardized intensity
0.0
0.2
0.4
0.6
0.8
1.0
F1
F80 F1
F80 F1
F24 F1

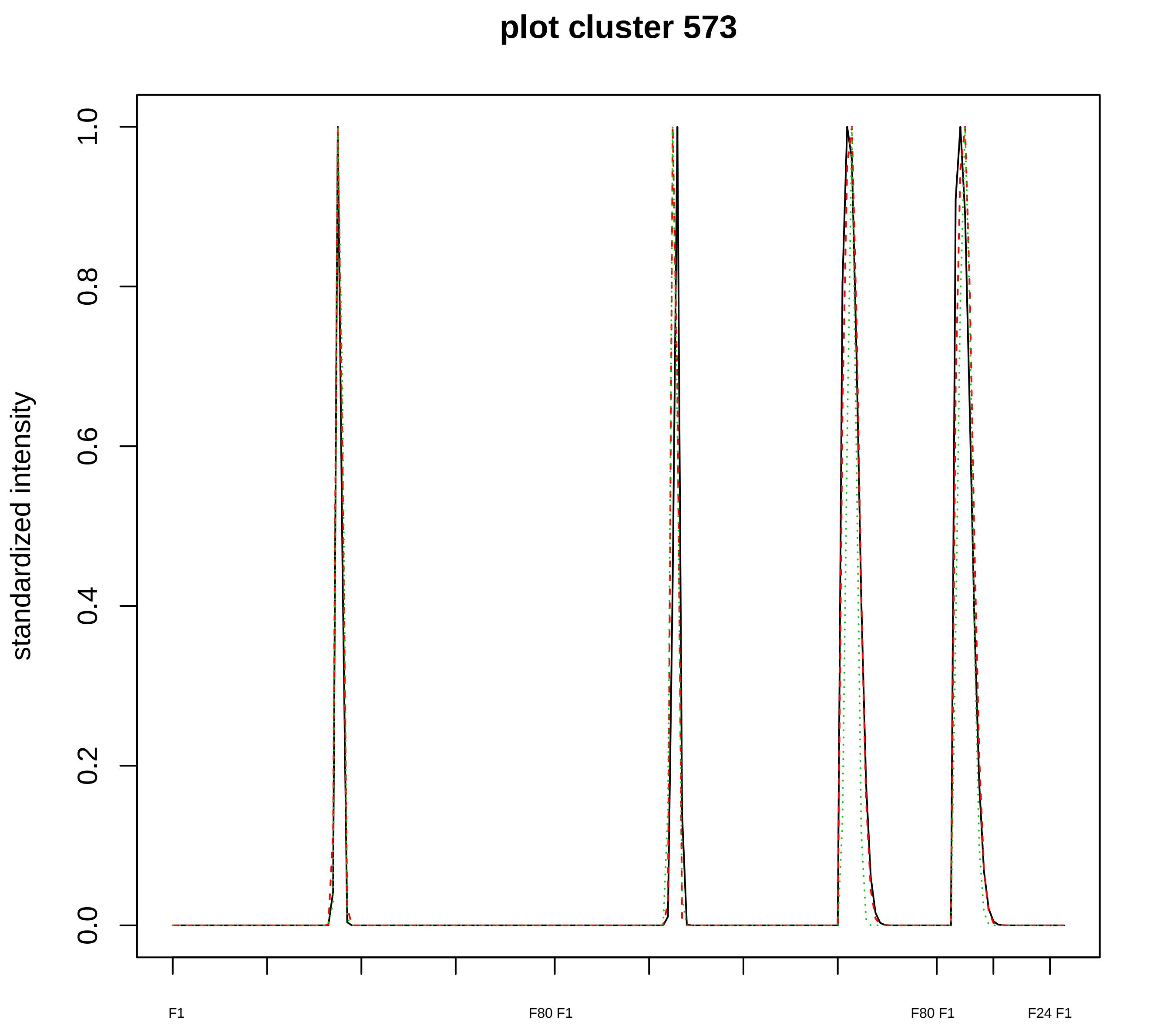

plot cluster 573
standardized intensity
0.0
0.2
0.4
0.6
0.8
1.0
F1
F80 F1
F80 F1
F24 F1

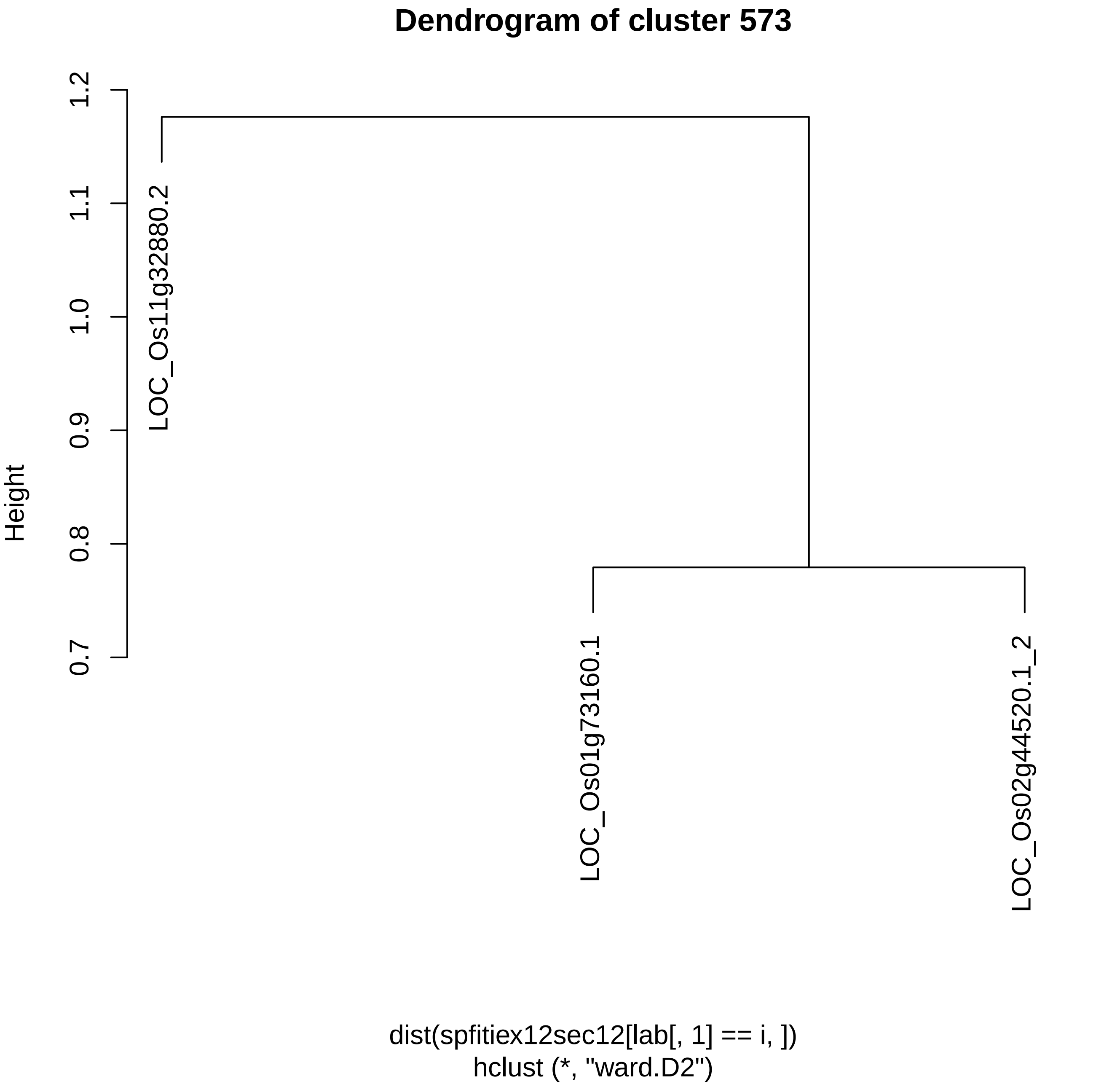
Dendrogram of cluster 573
Height
1.2
1.1
1.0
0.9
0.8
0.7
LOC_Os11g32880.2
LOC_Os01g73160.1
LOC_Os02g44520.1_2
dist(spfitiex12sec12[lab[, 1] == i, ])
hclust (*, "ward.D2")

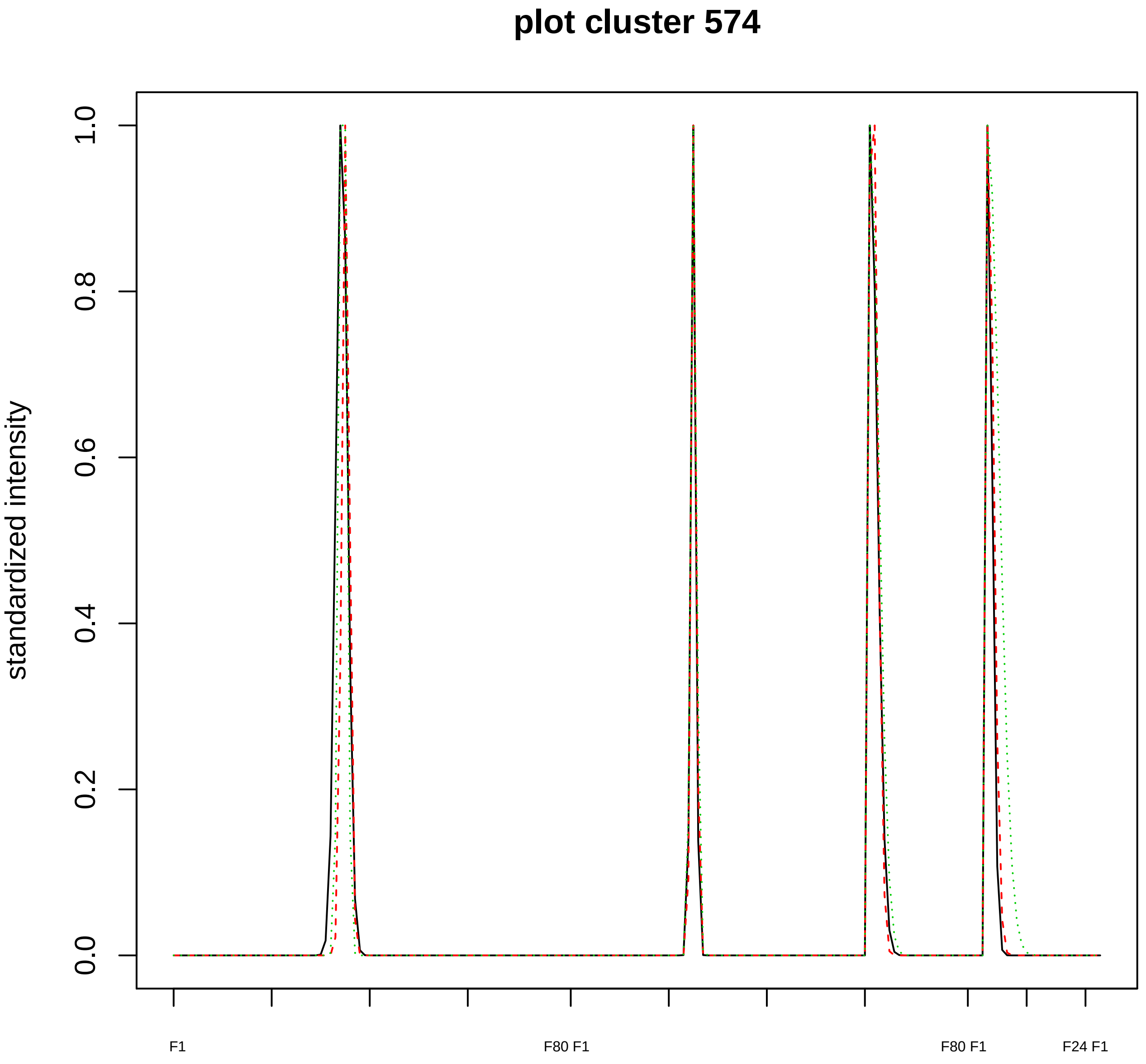
plot cluster 574
standardized intensity
1.0
0.8
0.6
0.4
0.2
0.0
F1
F80 F1
F80 F1
F24 F1

### Dendrogram of cluster 574

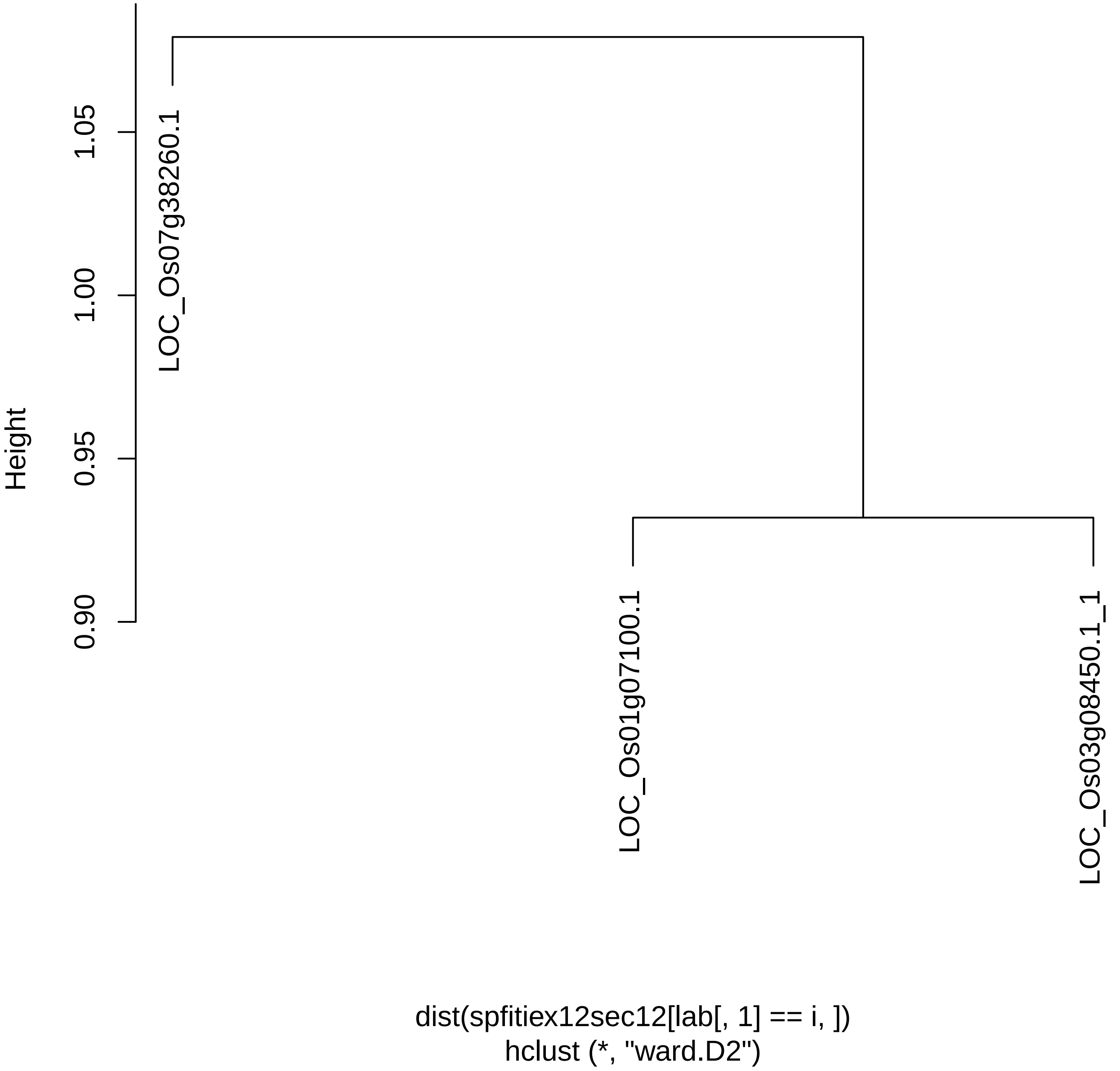

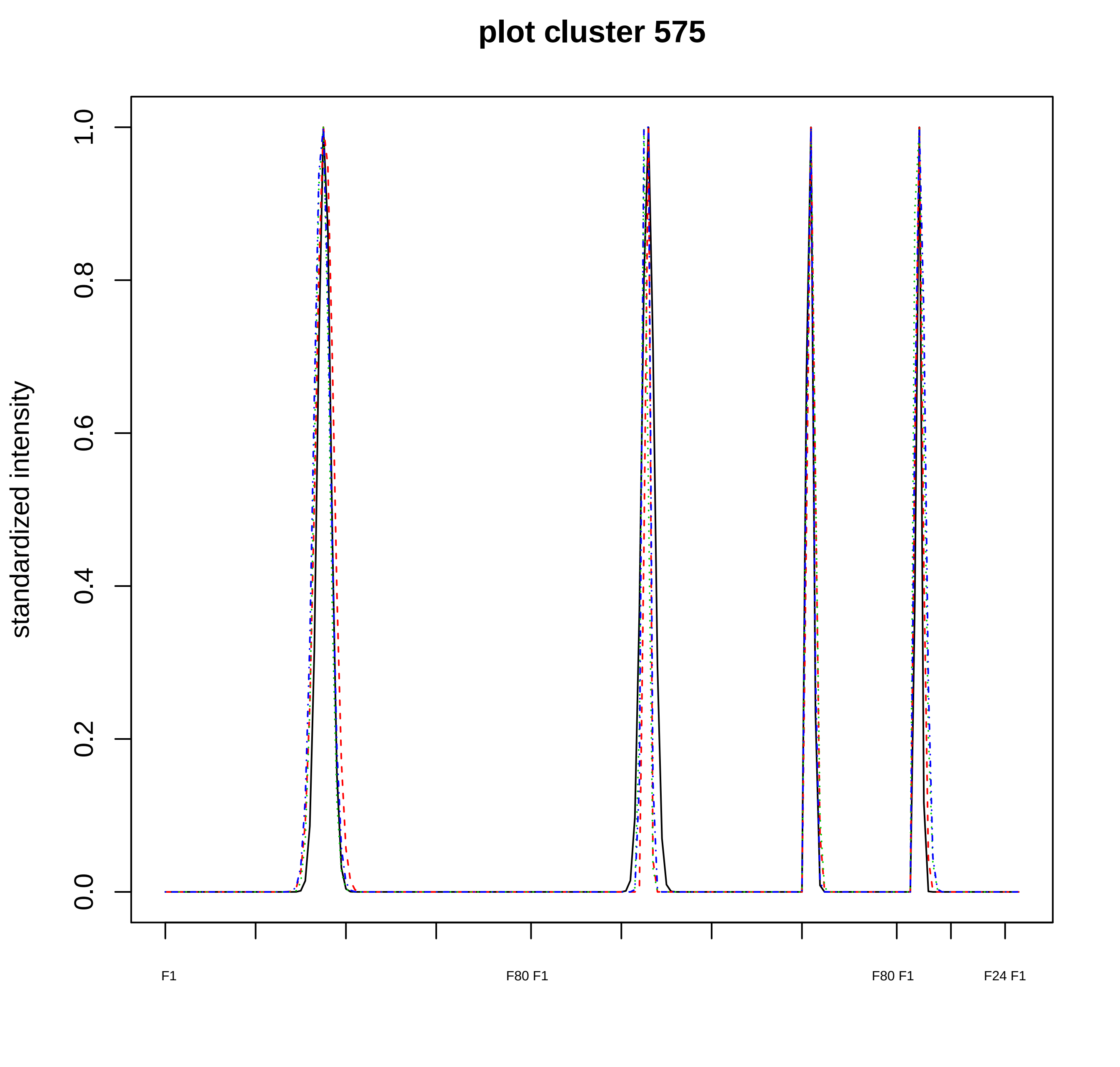
plot cluster 575
standardized intensity
0.0
0.2
0.4
0.6
0.8
1.0
F1
F80 F1
F80 F1
F24 F1

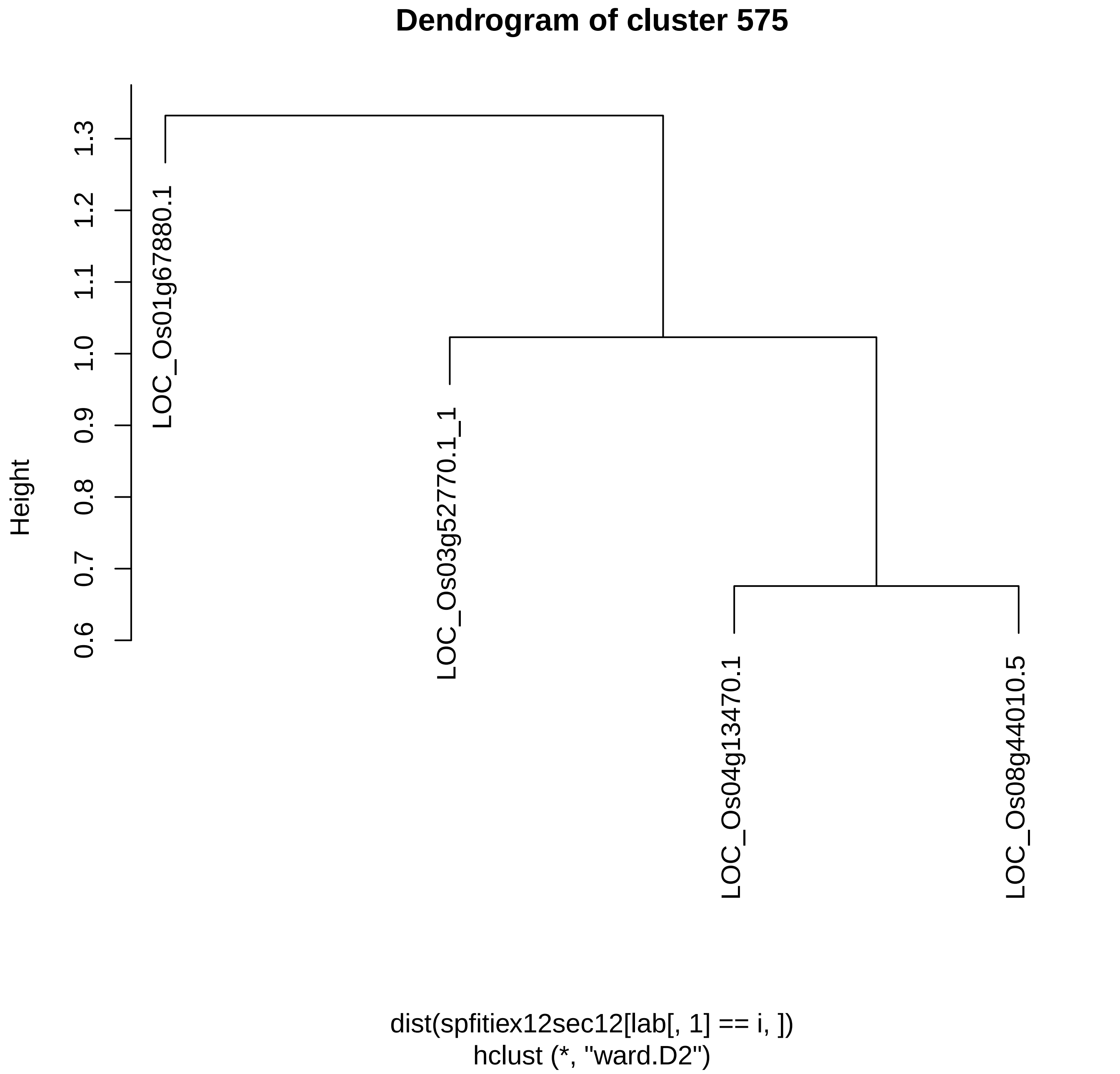
Dendrogram of cluster 575
Height
0.6
0.7
0.8
0.9
1.0
1.1
1.2
1.3
LOC_Os01g67880.1
LOC_Os03g52770.1_1
LOC_Os04g13470.1
LOC_Os08g44010.5
dist(spfitiex12sec12[lab[, 1] == i, ])
hclust (*, "ward.D2")

### plot cluster 576

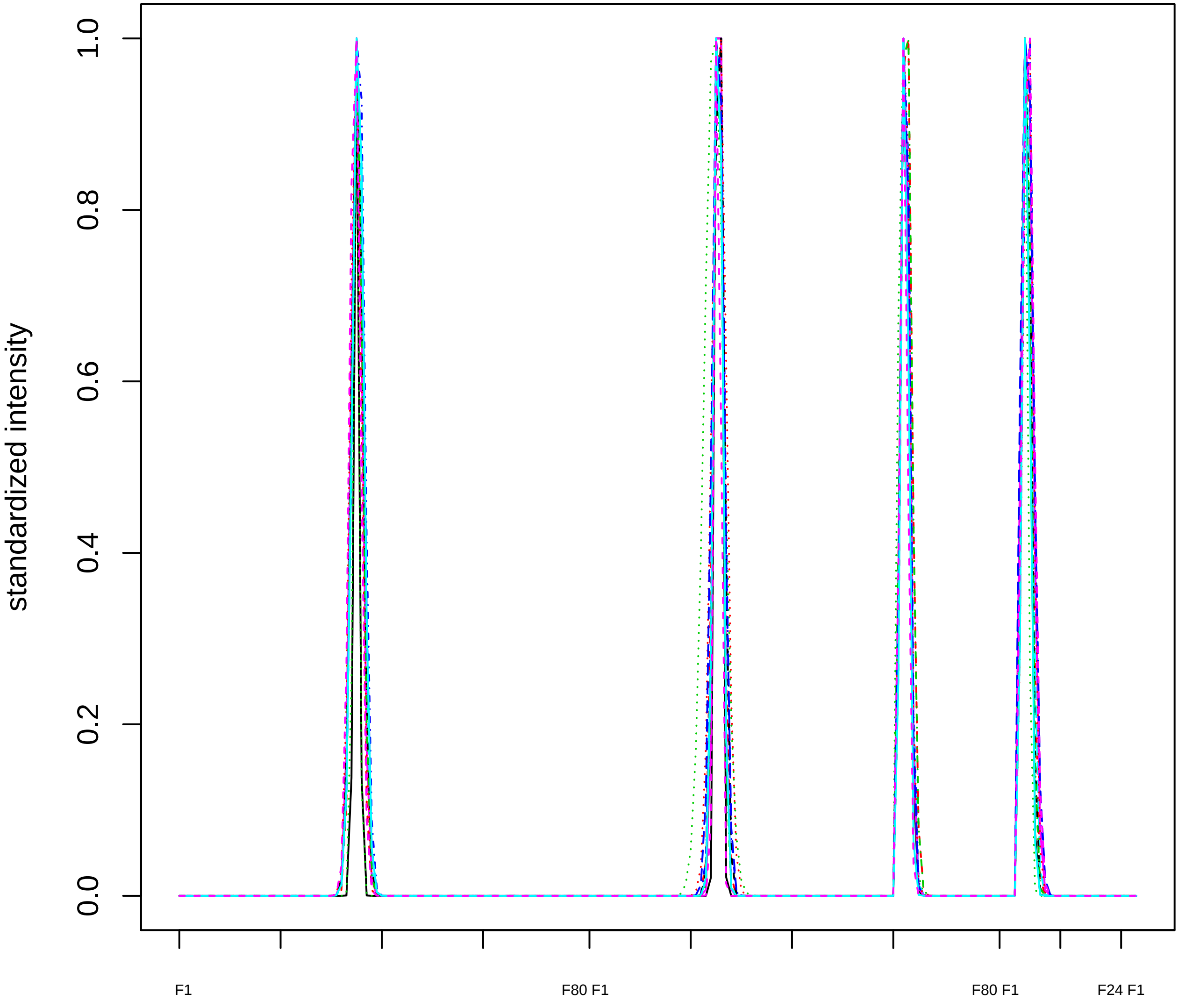

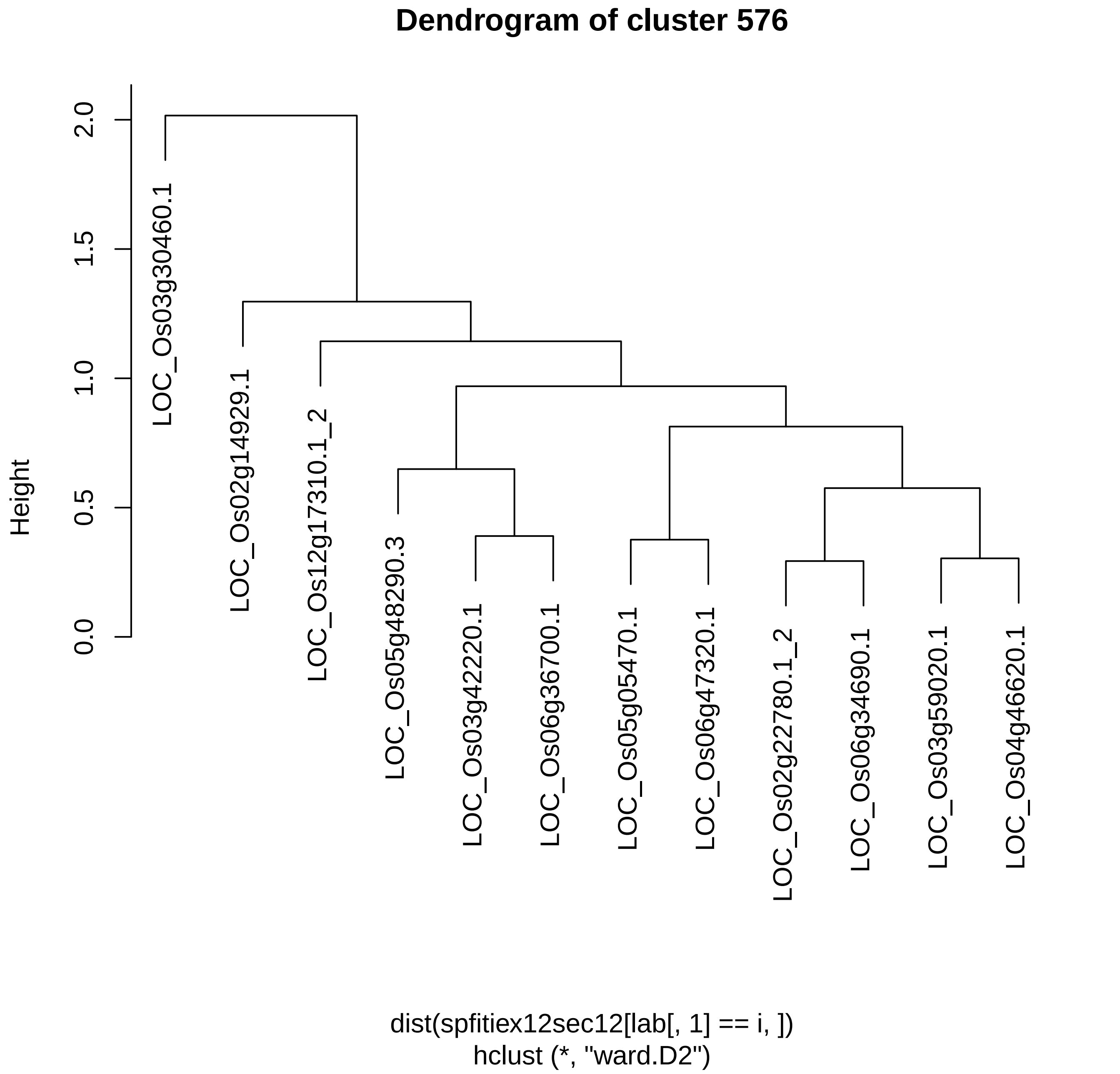
Dendrogram of cluster 576
Height
0.0
0.5
1.0
1.5
2.0
LOC_Os03g30460.1
LOC_Os02g14929.1
LOC_Os12g17310.1_2
LOC_Os05g48290.3
LOC_Os03g42220.1
LOC_Os06g36700.1
LOC_Os05g05470.1
LOC_Os06g47320.1
LOC_Os02g22780.1_2
LOC_Os06g34690.1
LOC_Os03g59020.1
LOC_Os04g46620.1
dist(spfitiex12sec12[lab[, 1] == i, ])
hclust (*, "ward.D2")

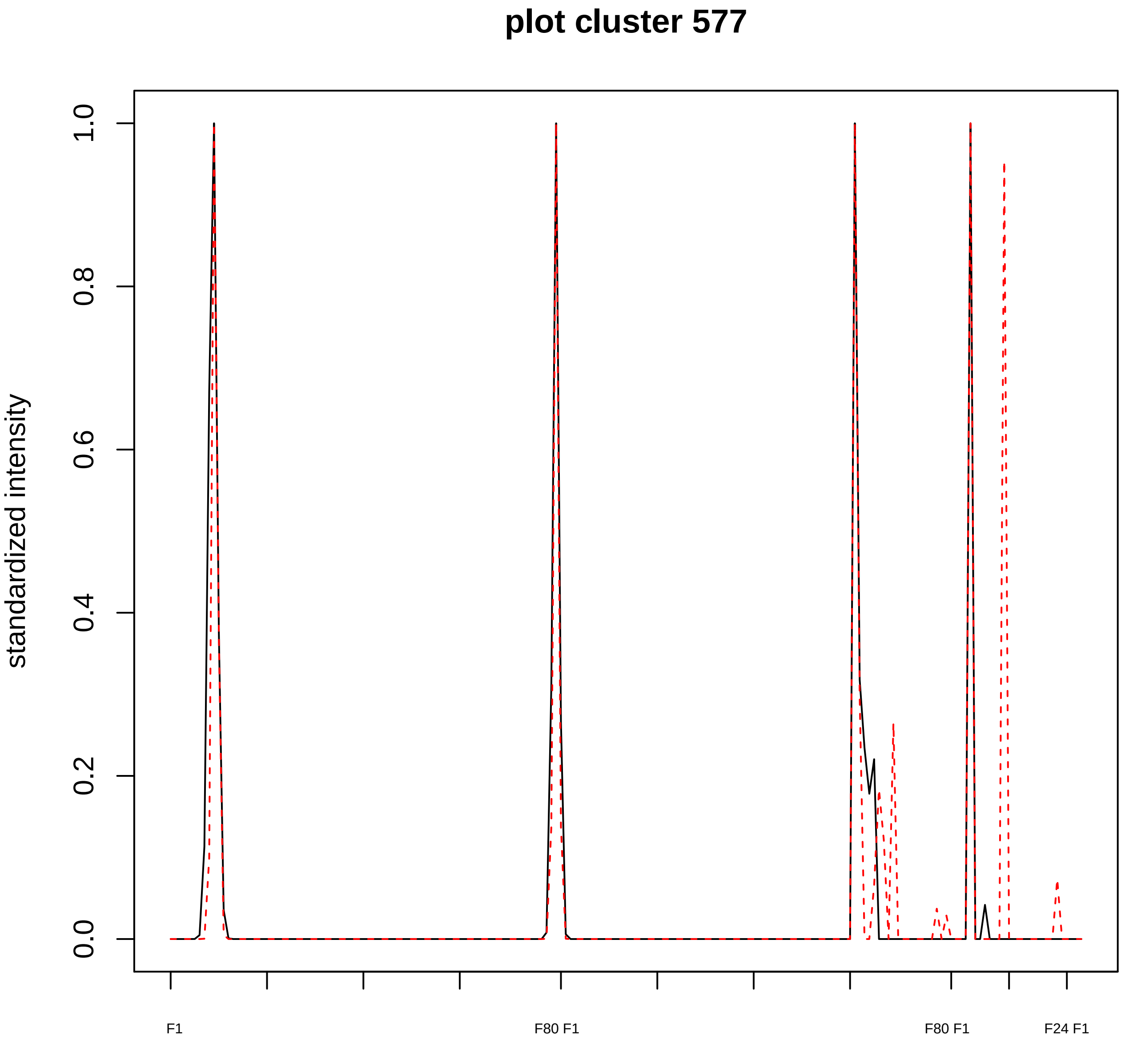
plot cluster 577
standardized intensity
0.0
0.2
0.4
0.6
0.8
1.0
F1
F80 F1
F80 F1
F24 F1

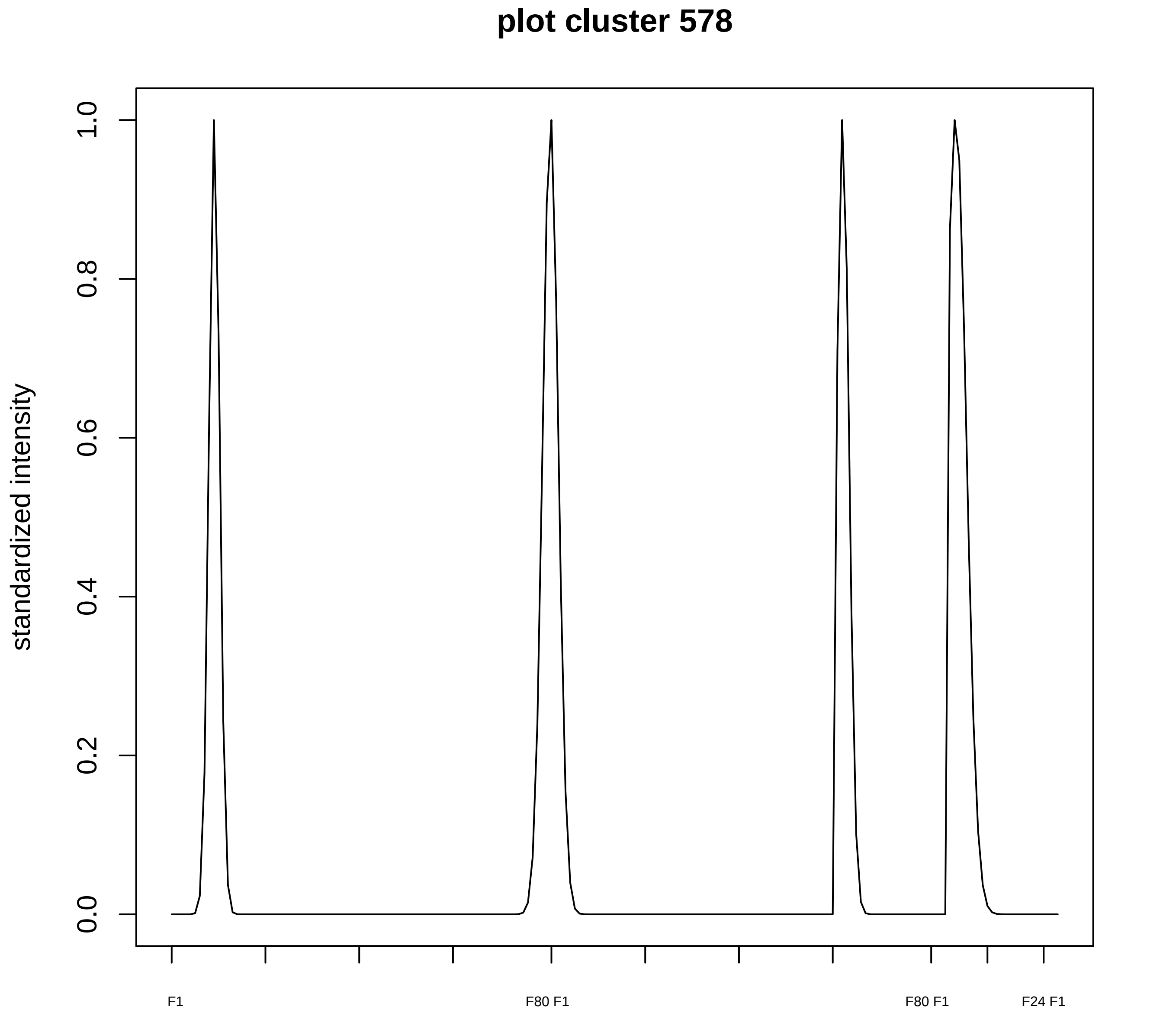
plot cluster 578
standardized intensity
0.0
0.2
0.4
0.6
0.8
1.0
F1
F80 F1
F80 F1
F24 F1

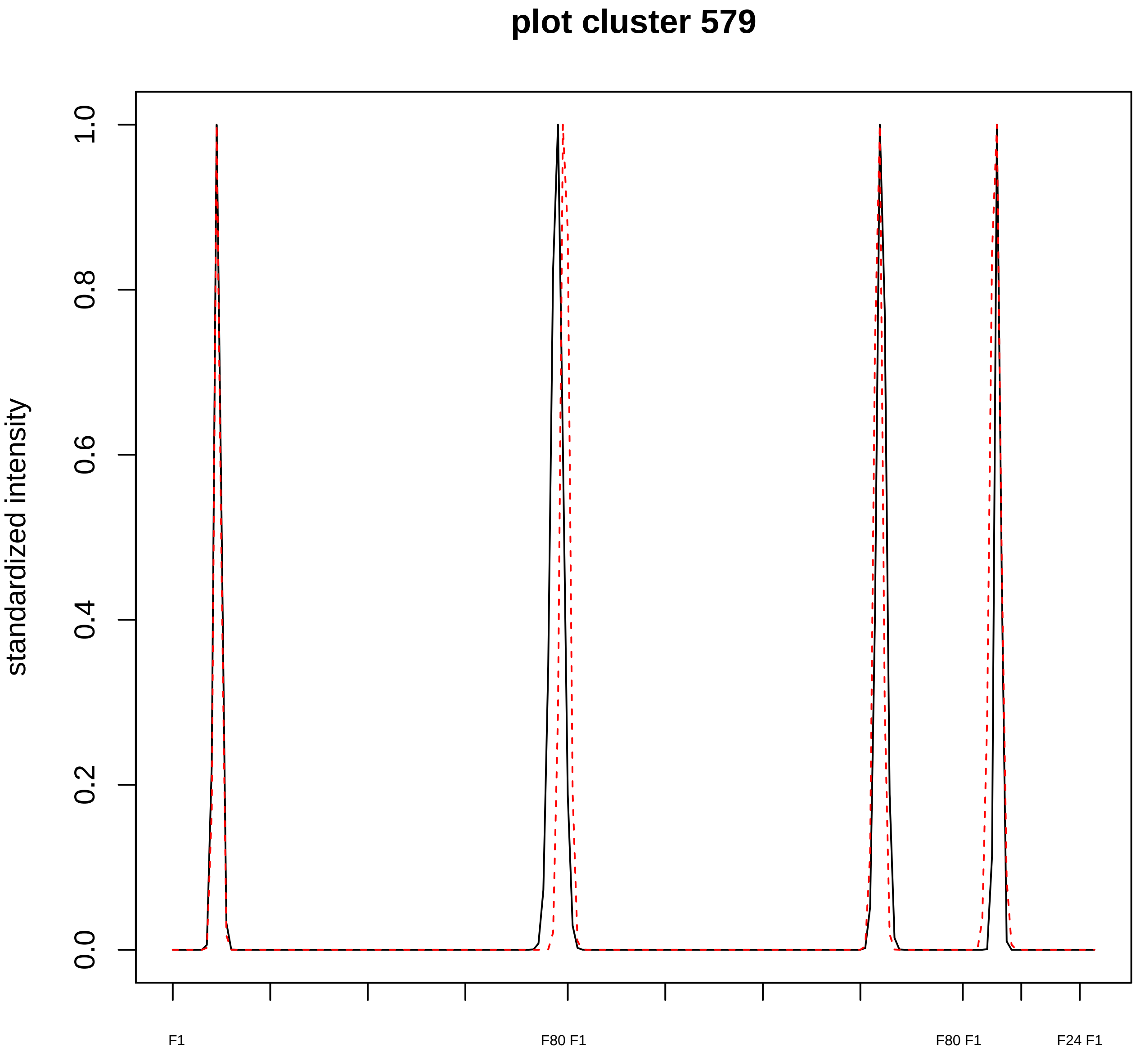

plot cluster 579
standardized intensity
0.0
0.2
0.4
0.6
0.8
1.0
F1
F80 F1
F80 F1
F24 F1

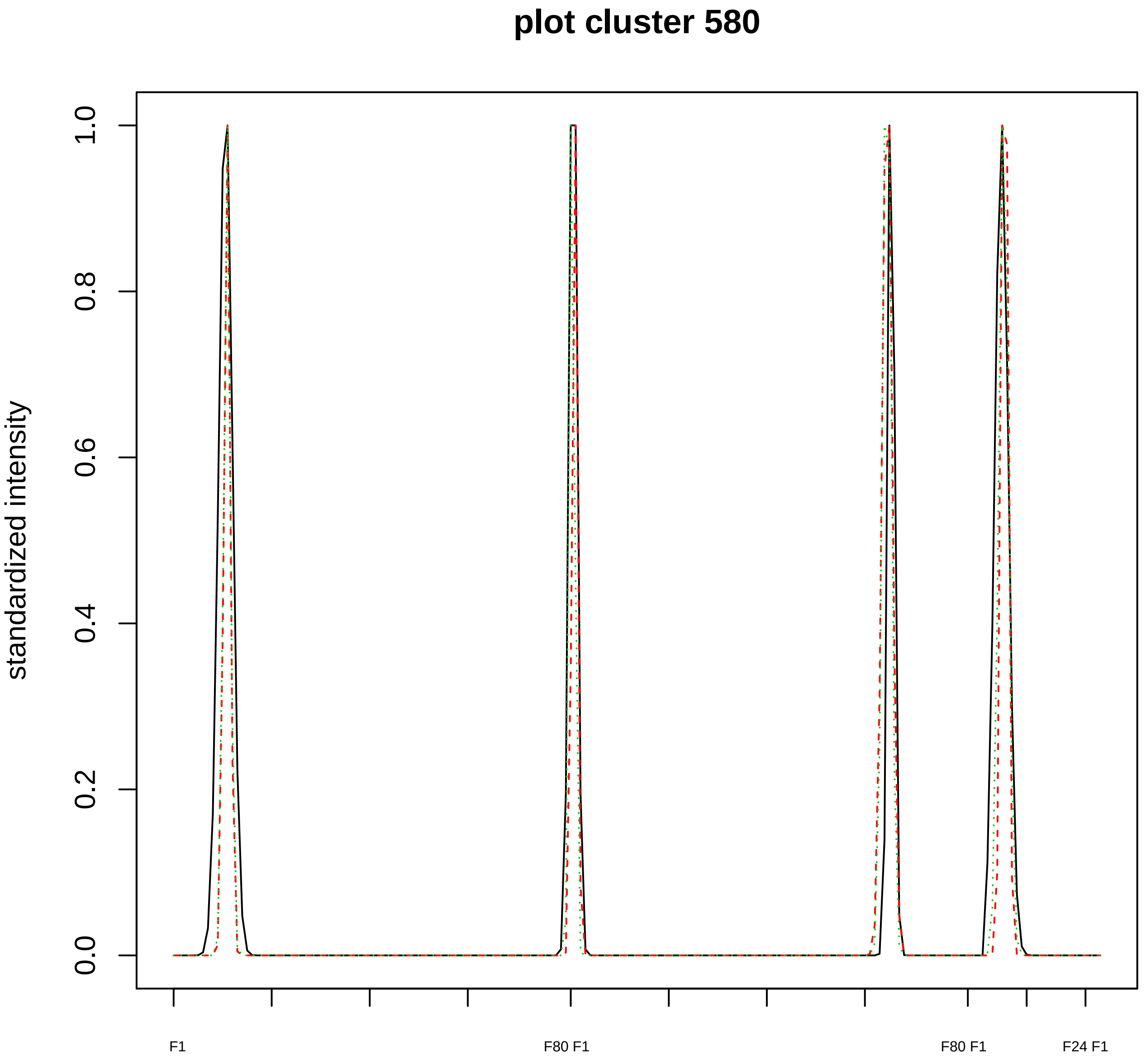

plot cluster 580
standardized intensity
0.0
0.2
0.4
0.6
0.8
1.0
F1
F80 F1
F80 F1
F24 F1

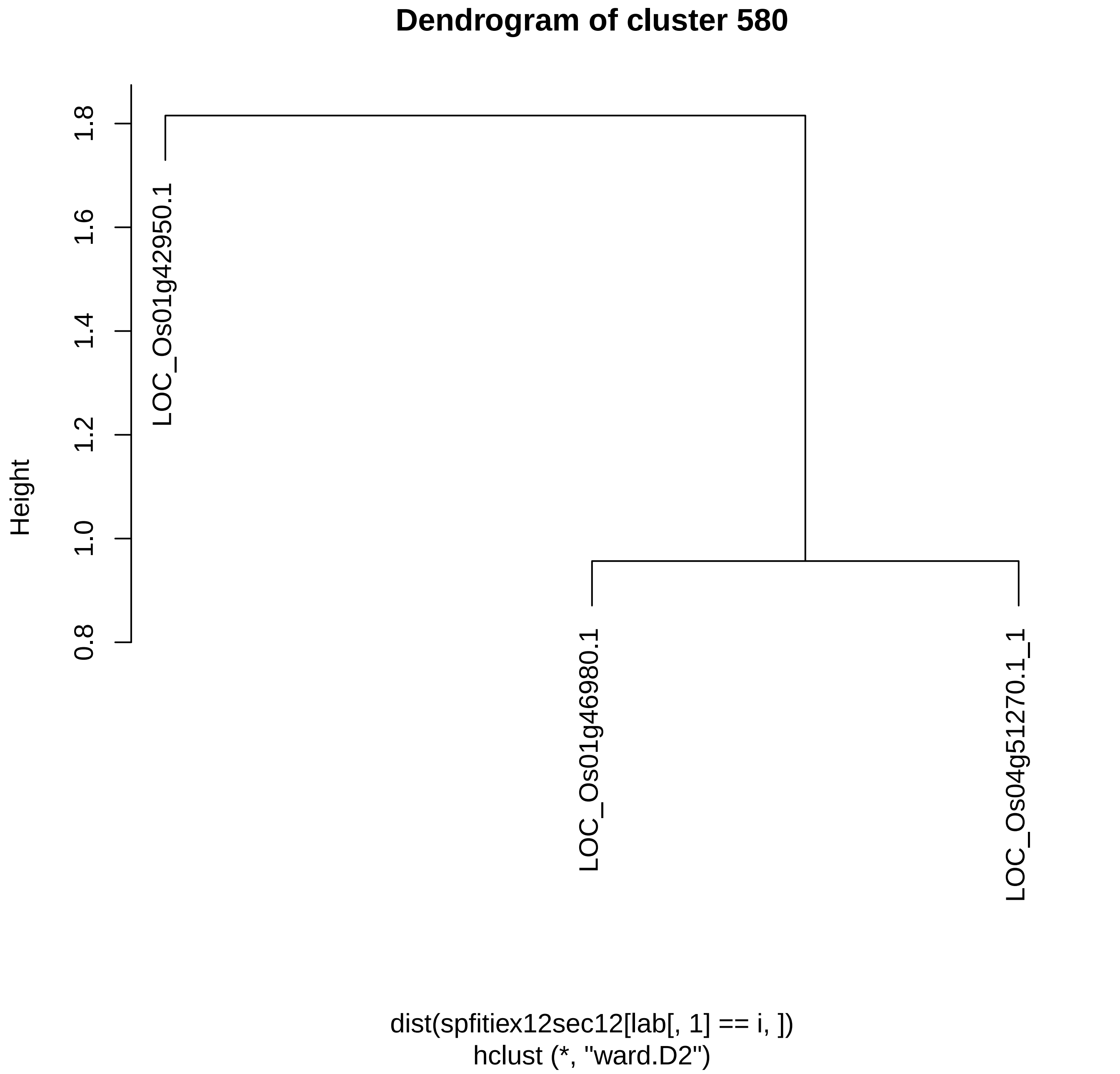
Dendrogram of cluster 580
Height
0.8
1.0
1.2
1.4
1.6
1.8
LOC_Os01g42950.1
LOC_Os01g46980.1
LOC_Os04g51270.1_1
dist(spfitiex12sec12[lab[, 1] == i, ])
hclust (*, "ward.D2")

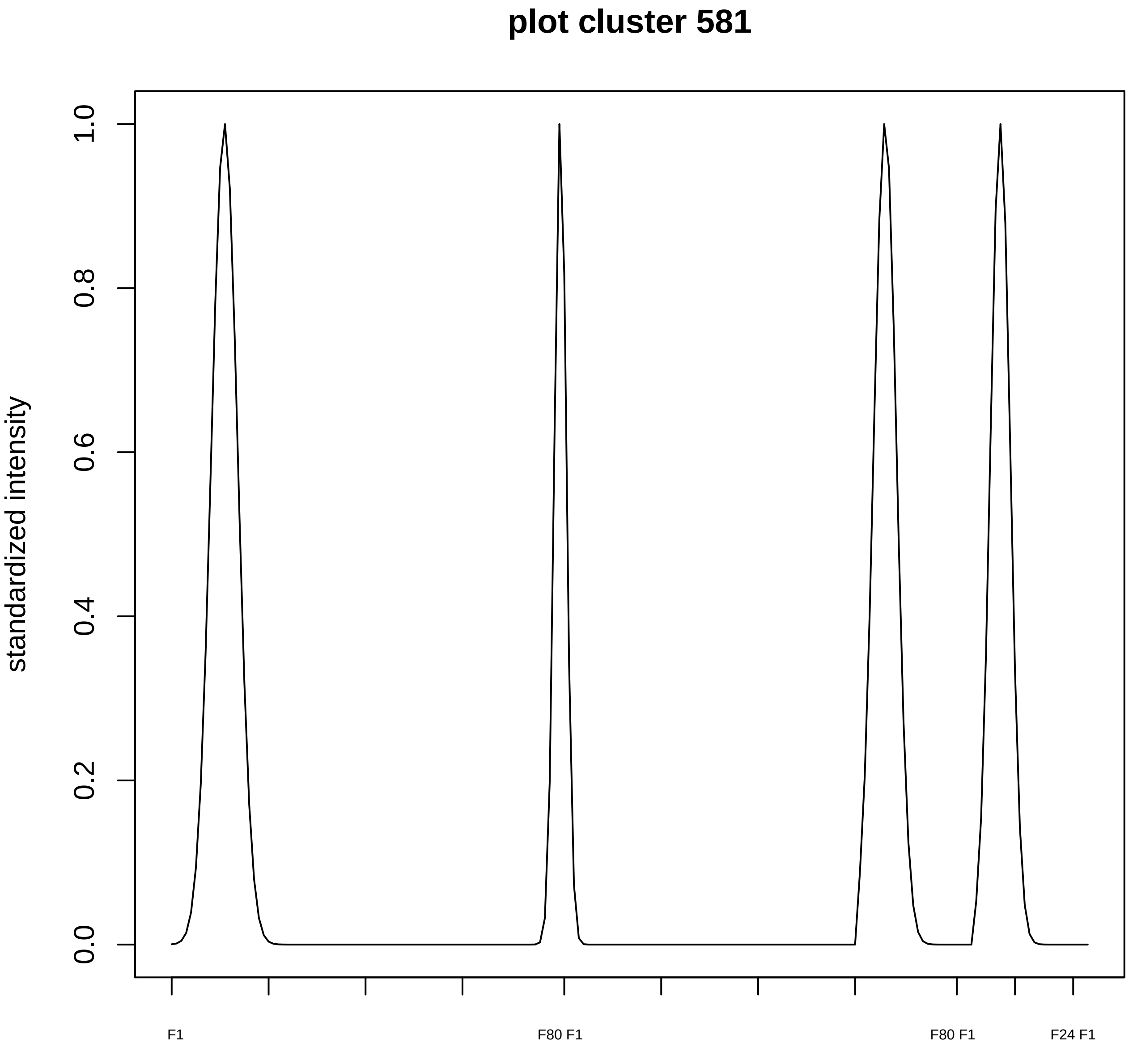

plot cluster 581
standardized intensity
0.0
0.2
0.4
0.6
0.8
1.0
F1
F80 F1
F80 F1
F24 F1

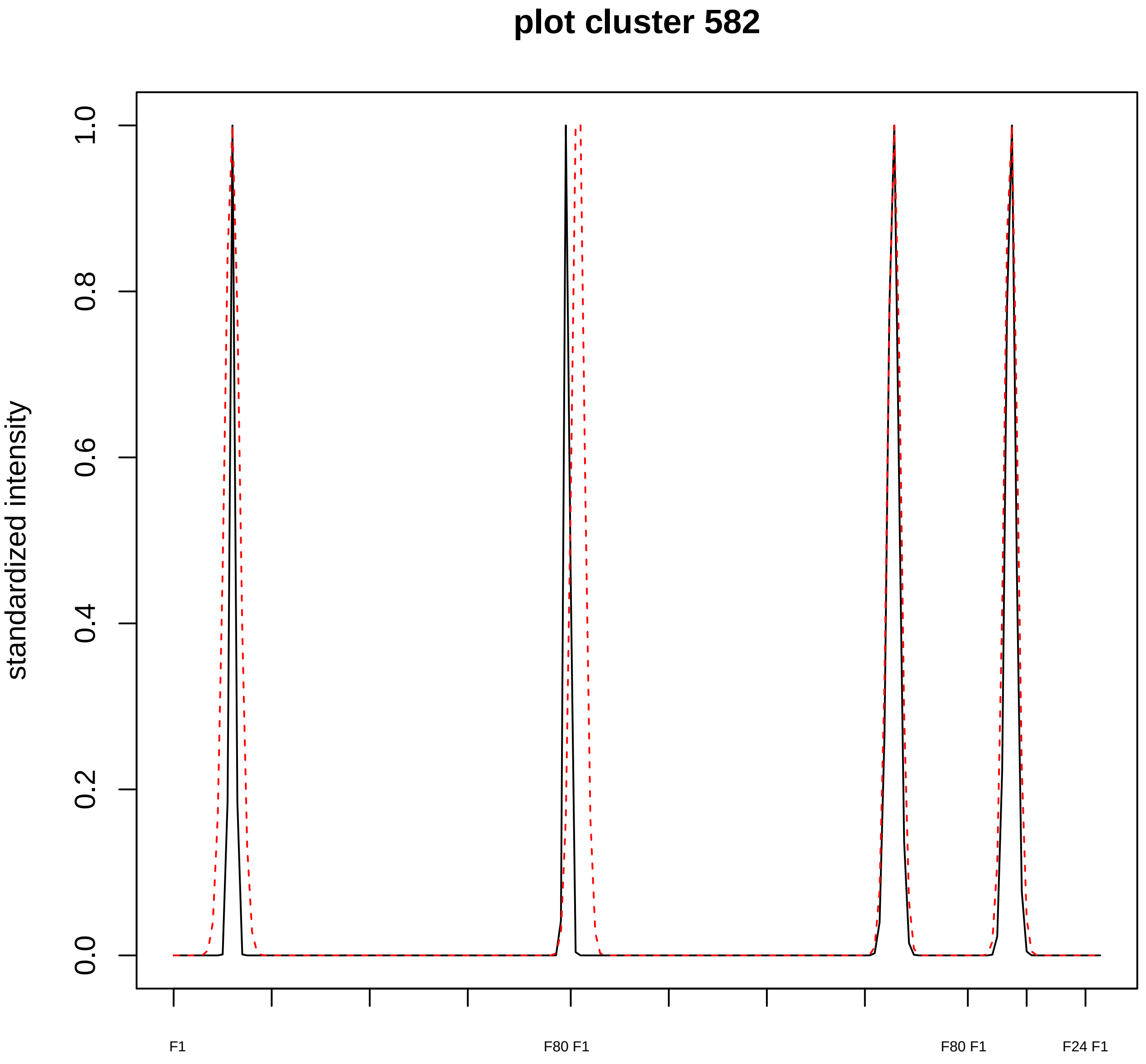
plot cluster 582
standardized intensity
0.0
0.2
0.4
0.6
0.8
1.0
F1
F80 F1
F80 F1
F24 F1

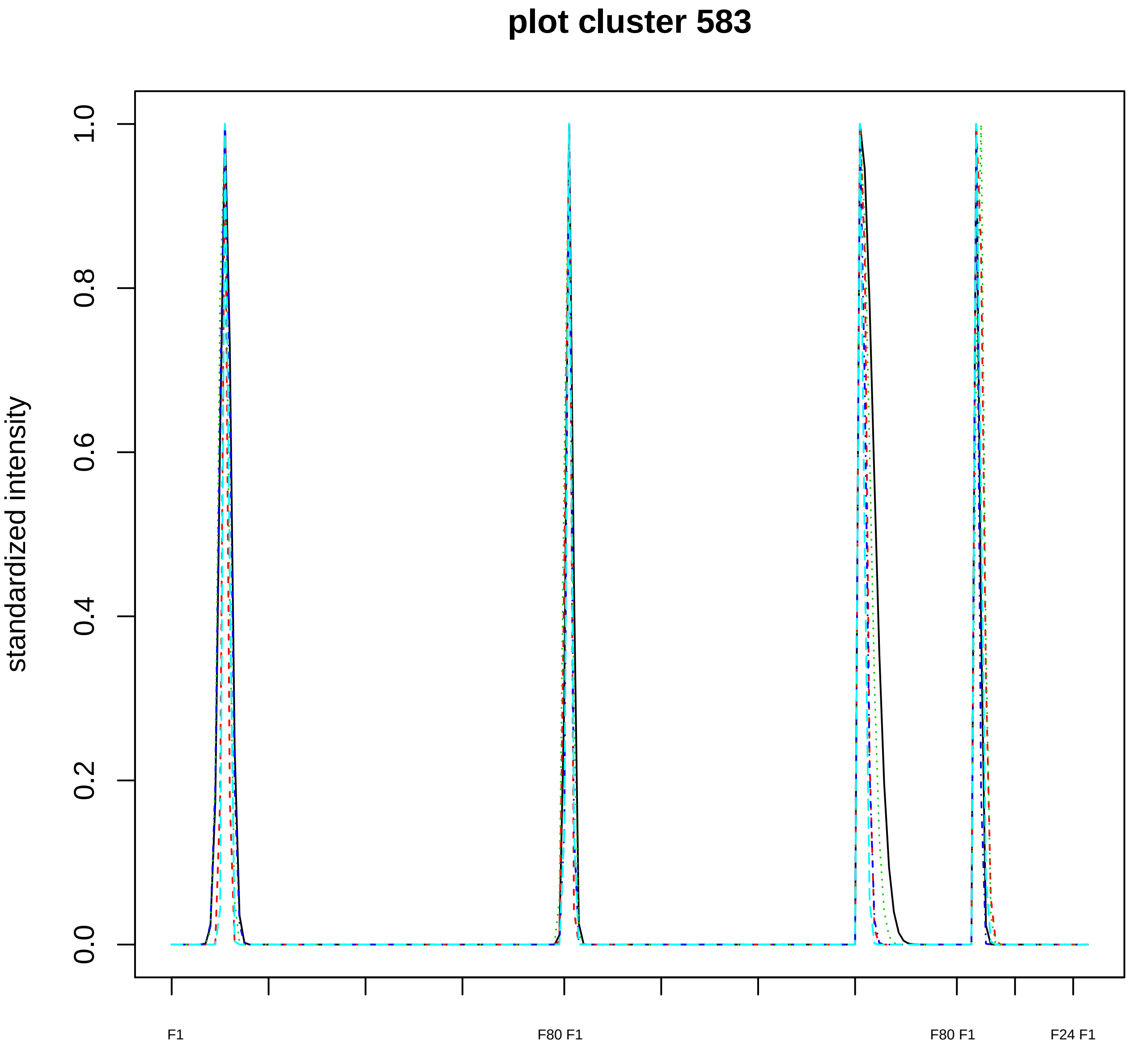
plot cluster 583
standardized intensity
0.0
0.2
0.4
0.6
0.8
1.0
F1
F80 F1
F80 F1
F24 F1

### Dendrogram of cluster 583

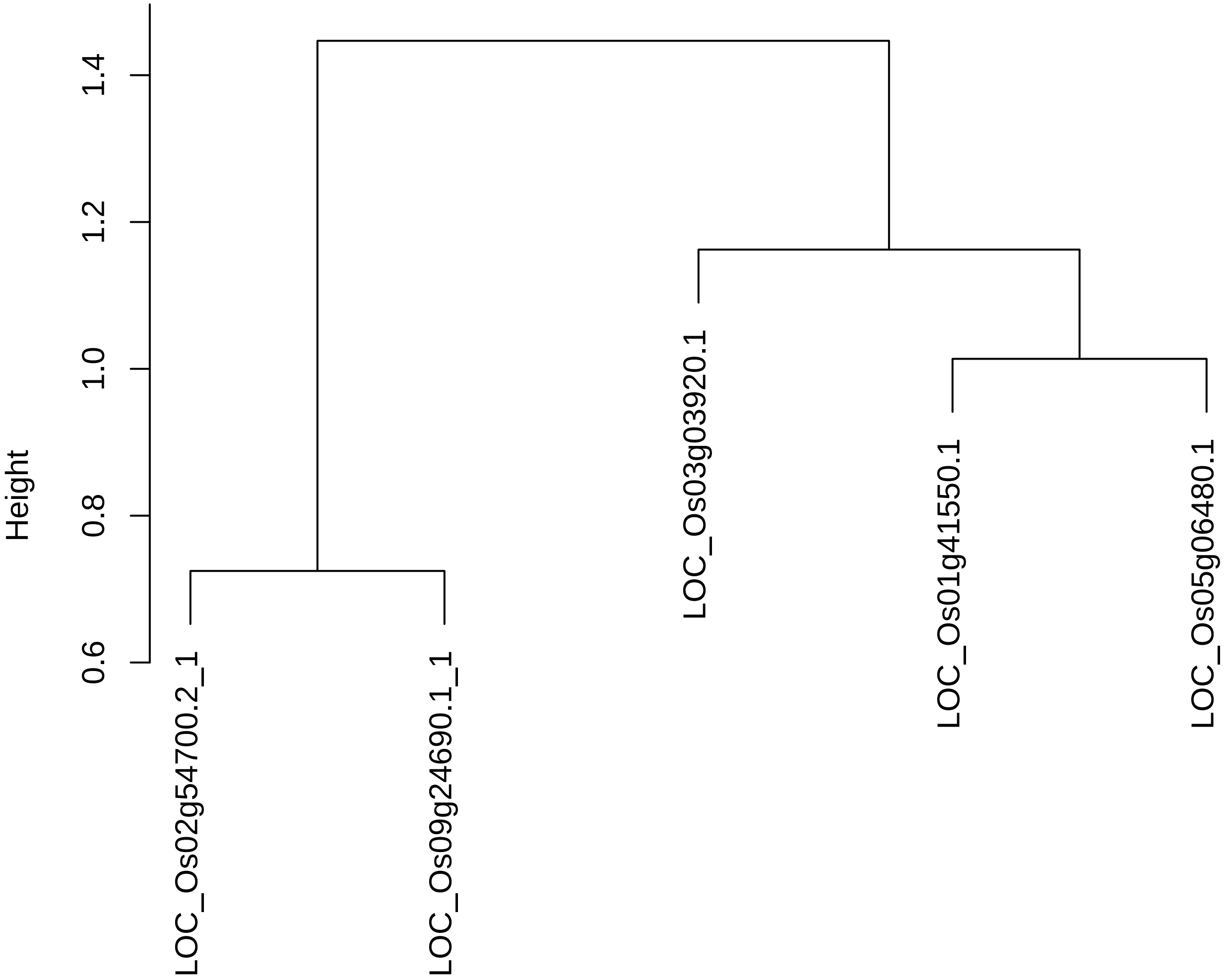


dist(spfitiex12sec12[lab[, 1] == i, ])
hclust (*, "ward.D2")

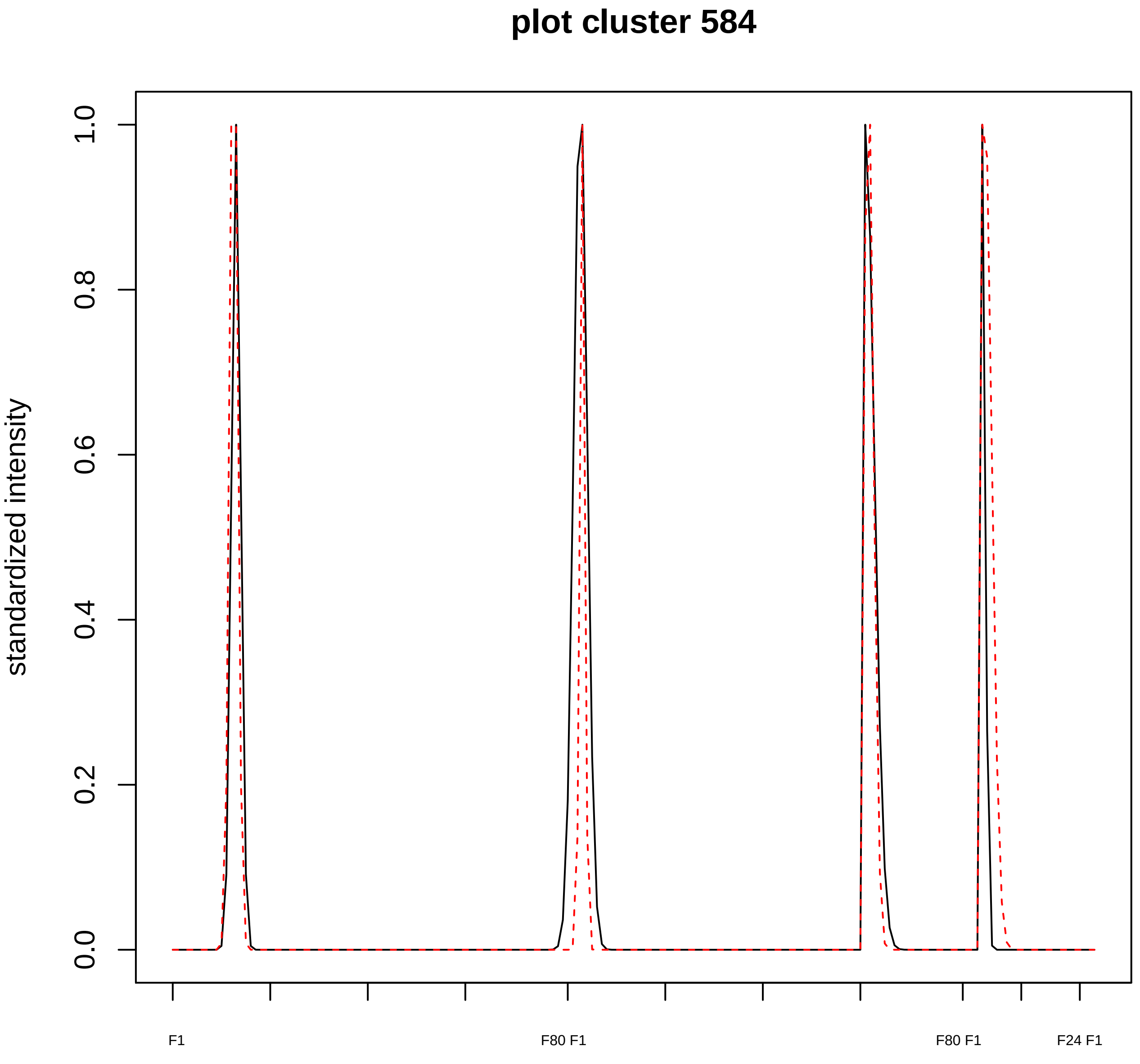

plot cluster 584
standardized intensity
0.0
0.2
0.4
0.6
0.8
1.0
F1
F80 F1
F80 F1
F24 F1

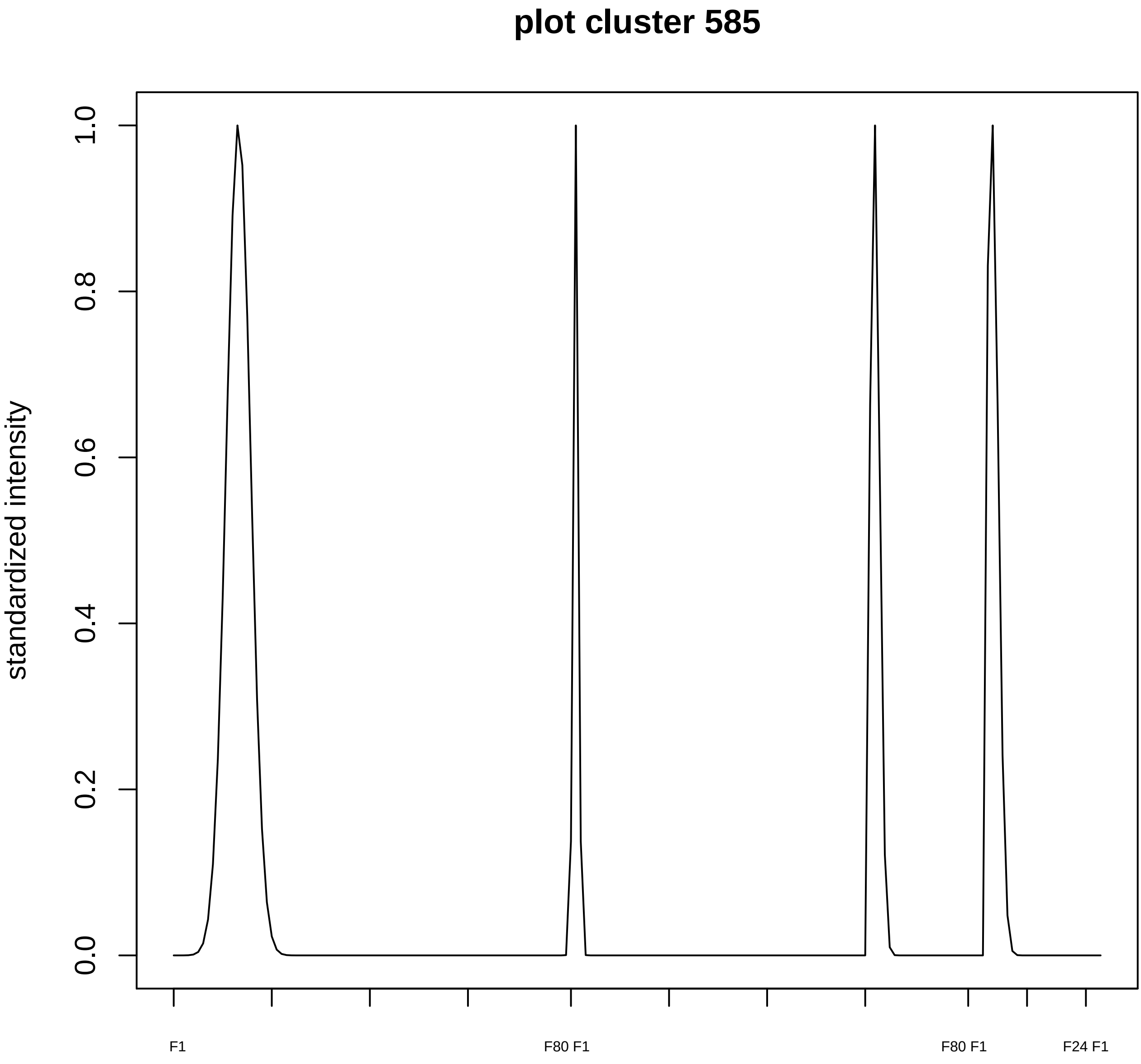

plot cluster 585
standardized intensity
0.0
0.2
0.4
0.6
0.8
1.0
F1
F80 F1
F80 F1
F24 F1

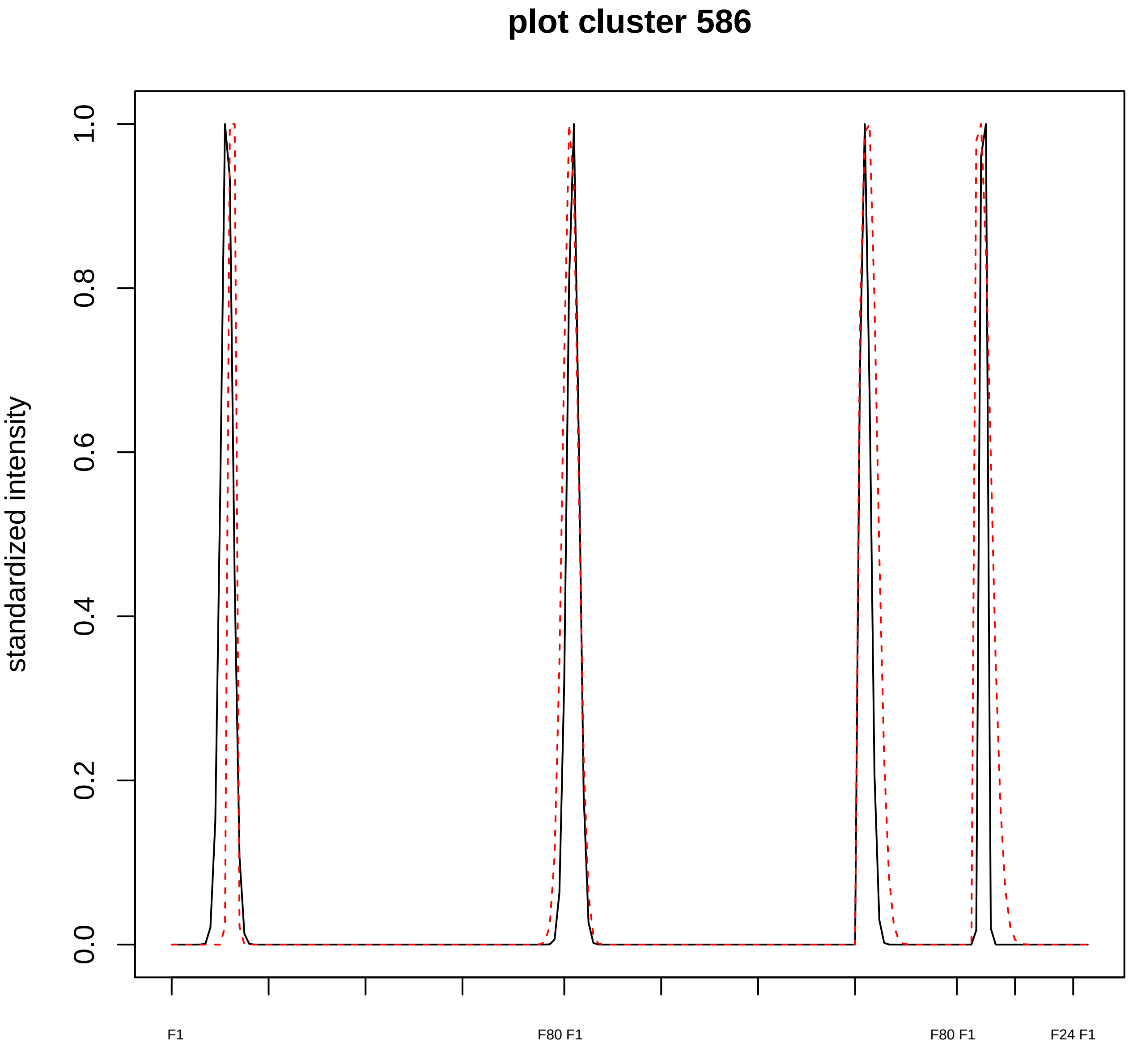
plot cluster 586
standardized intensity
0.0
0.2
0.4
0.6
0.8
1.0
F1
F80 F1
F80 F1
F24 F1

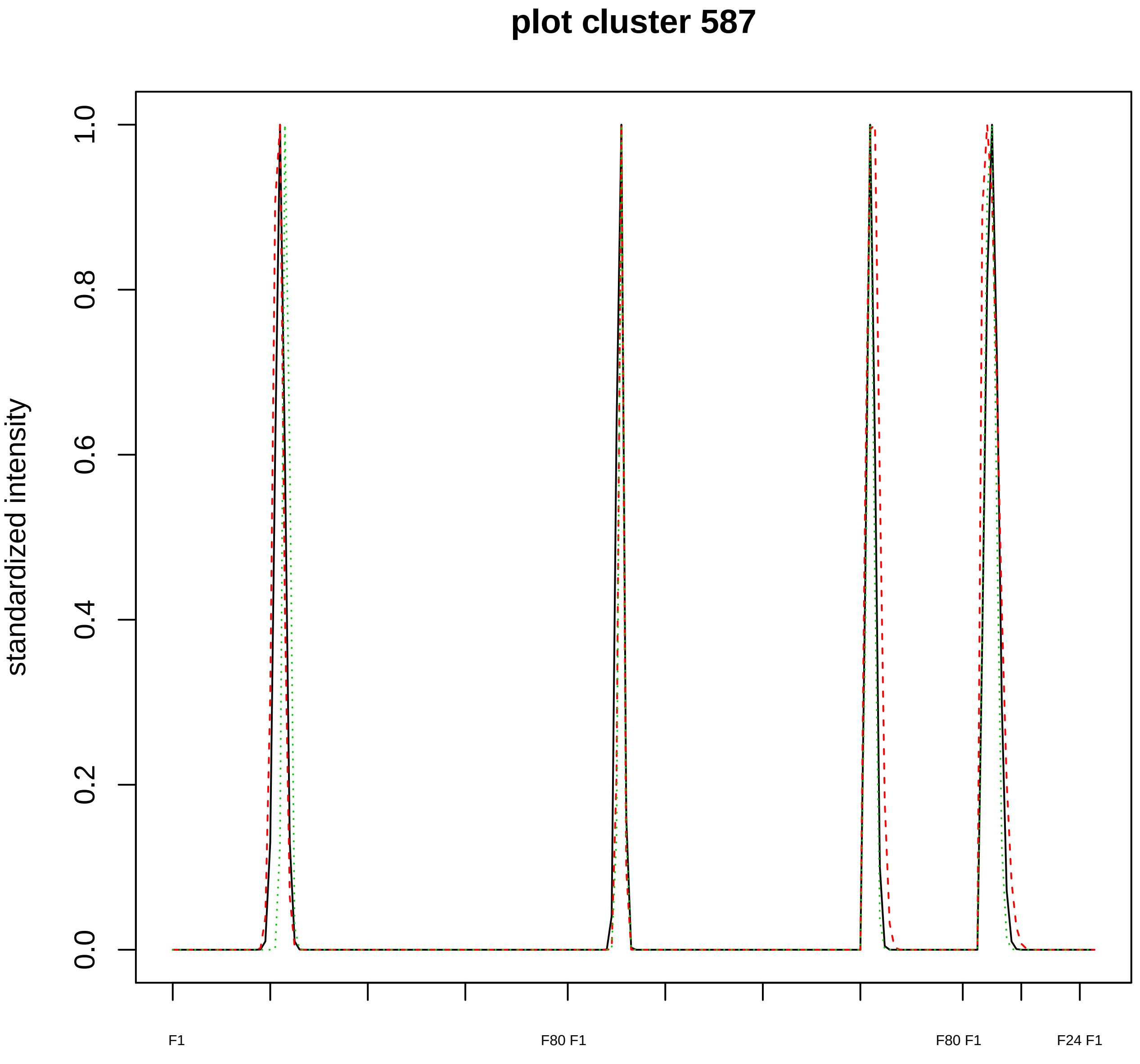
plot cluster 587
standardized intensity
0.0
0.2
0.4
0.6
0.8
1.0
F1
F80 F1
F80 F1
F24 F1

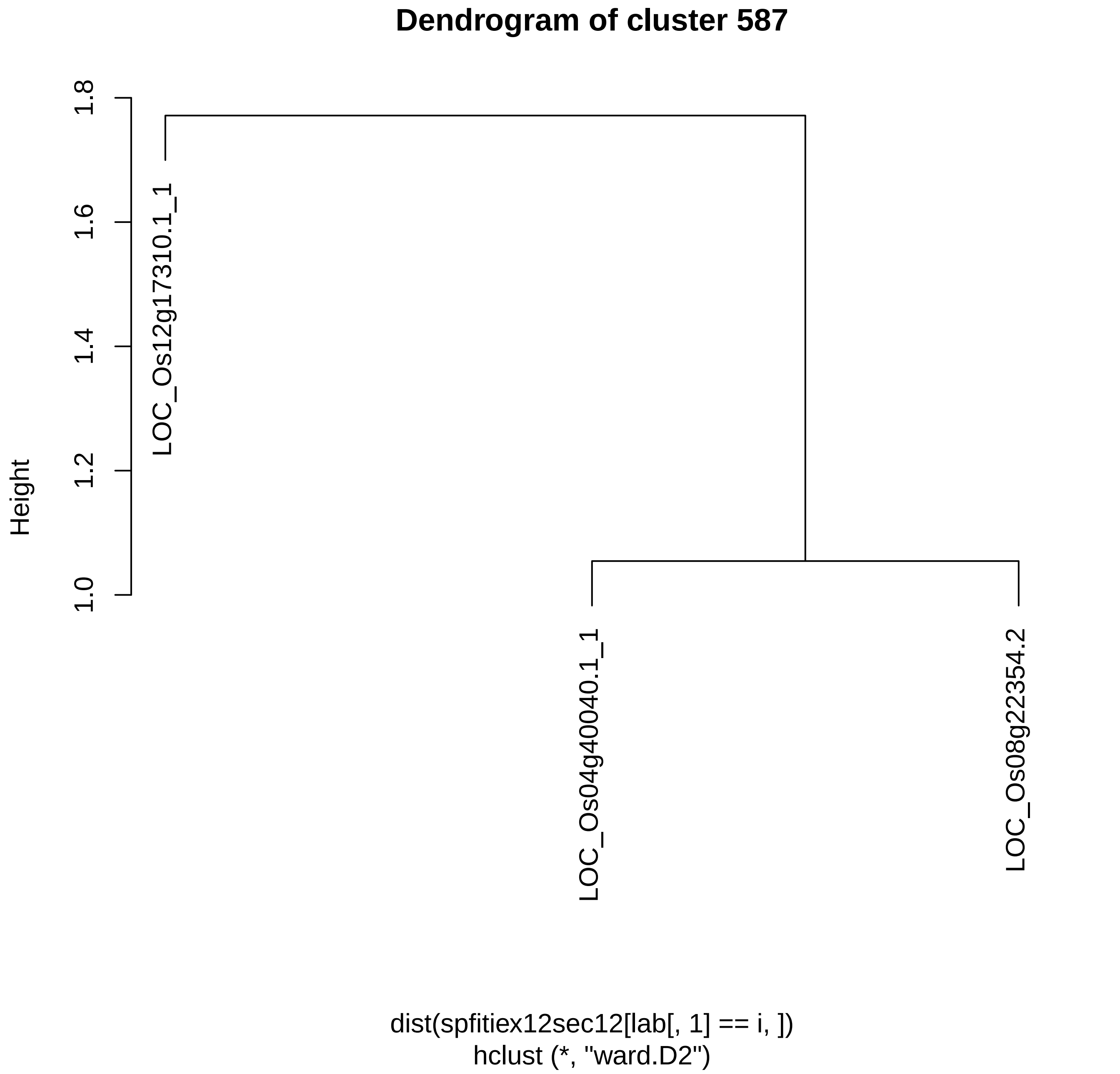
Dendrogram of cluster 587
Height
1.0
1.2
1.4
1.6
1.8
LOC_Os12g17310.1_1
LOC_Os04g40040.1_1
LOC_Os08g22354.2
dist(spfitiex12sec12[lab[, 1] == i, ])
hclust (*, "ward.D2")

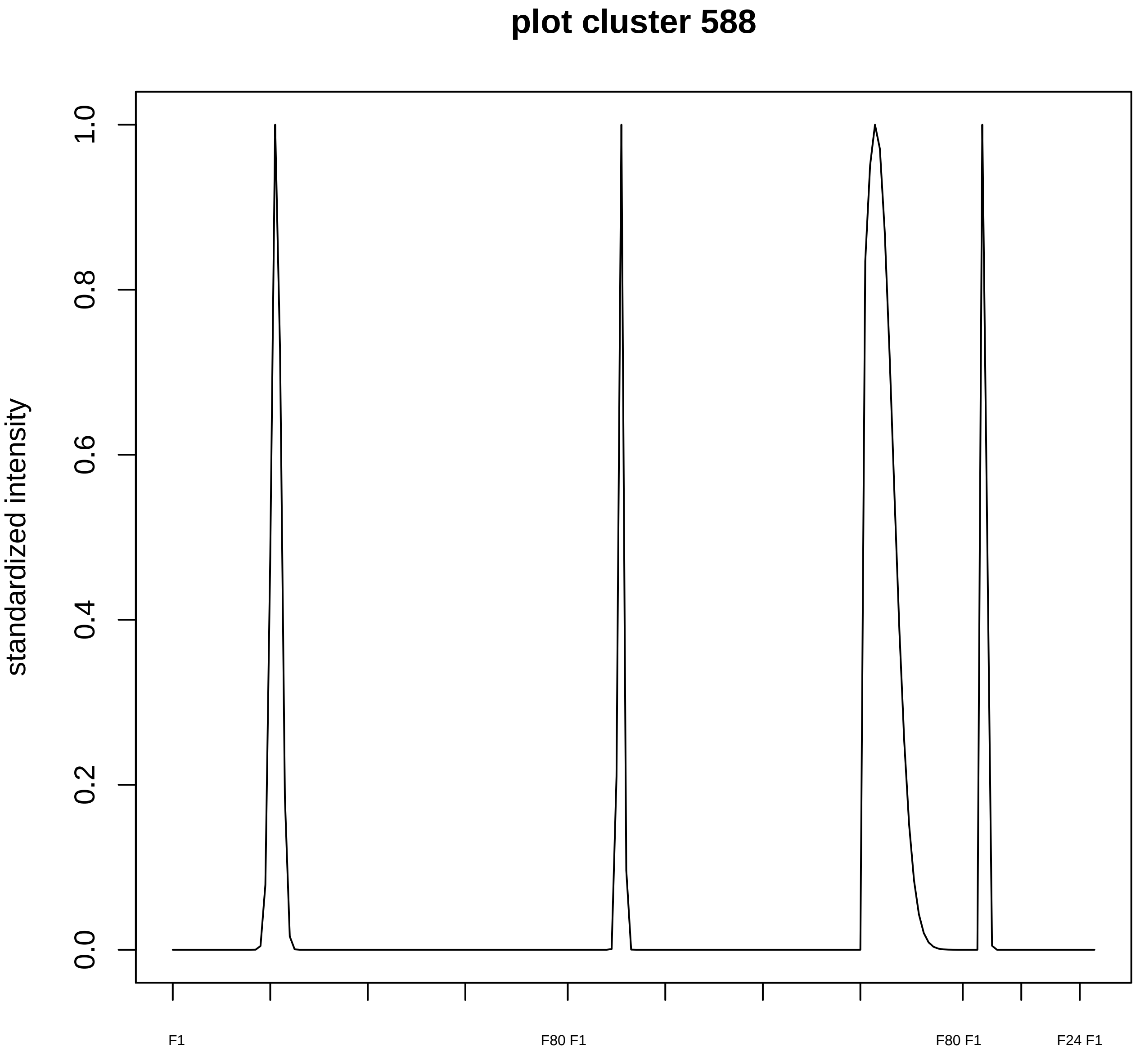
plot cluster 588
standardized intensity
0.0
0.2
0.4
0.6
0.8
1.0
F1
F80 F1
F80 F1
F24 F1

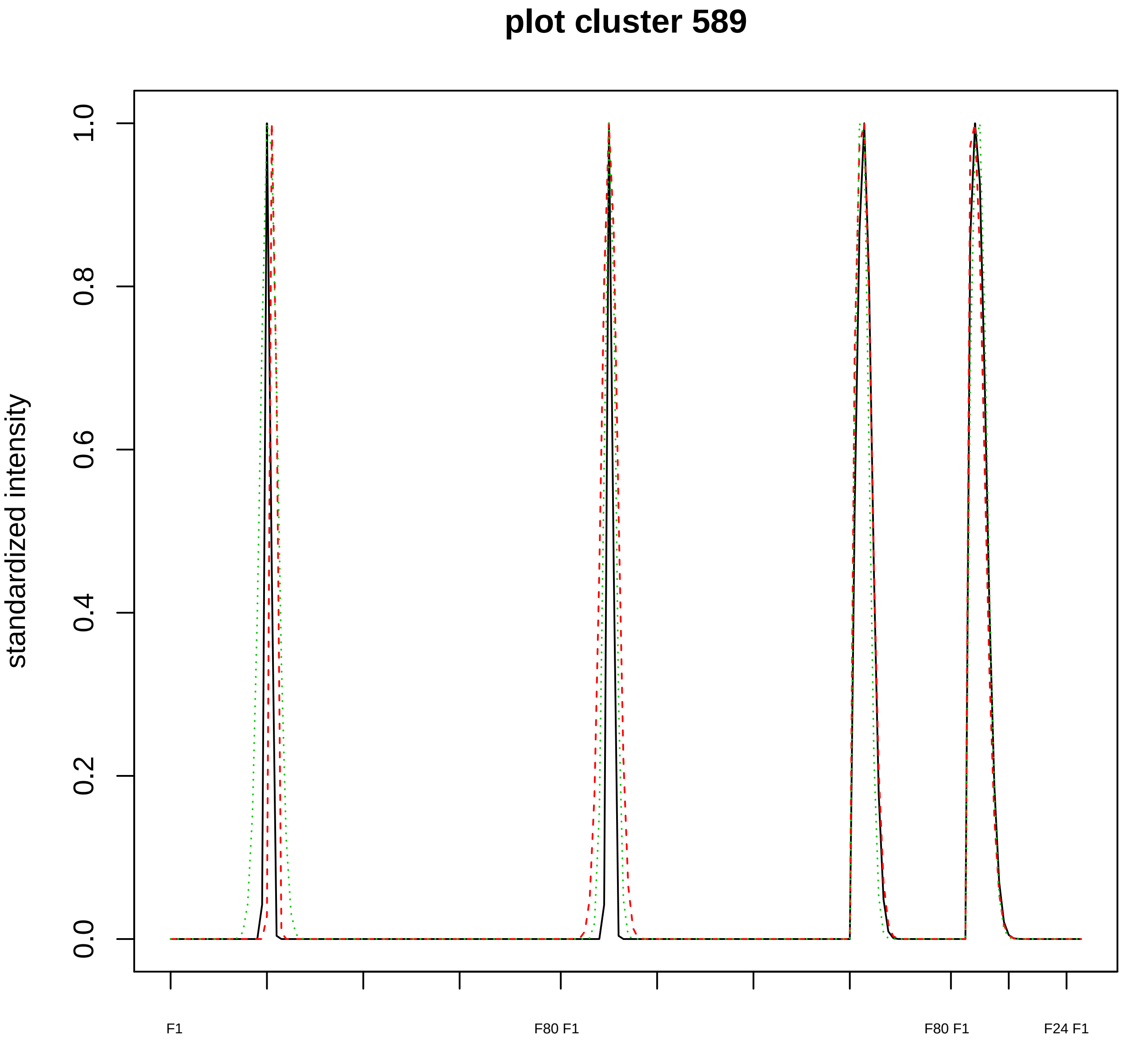

plot cluster 589
standardized intensity
0.0
0.2
0.4
0.6
0.8
1.0
F1
F80 F1
F80 F1
F24 F1

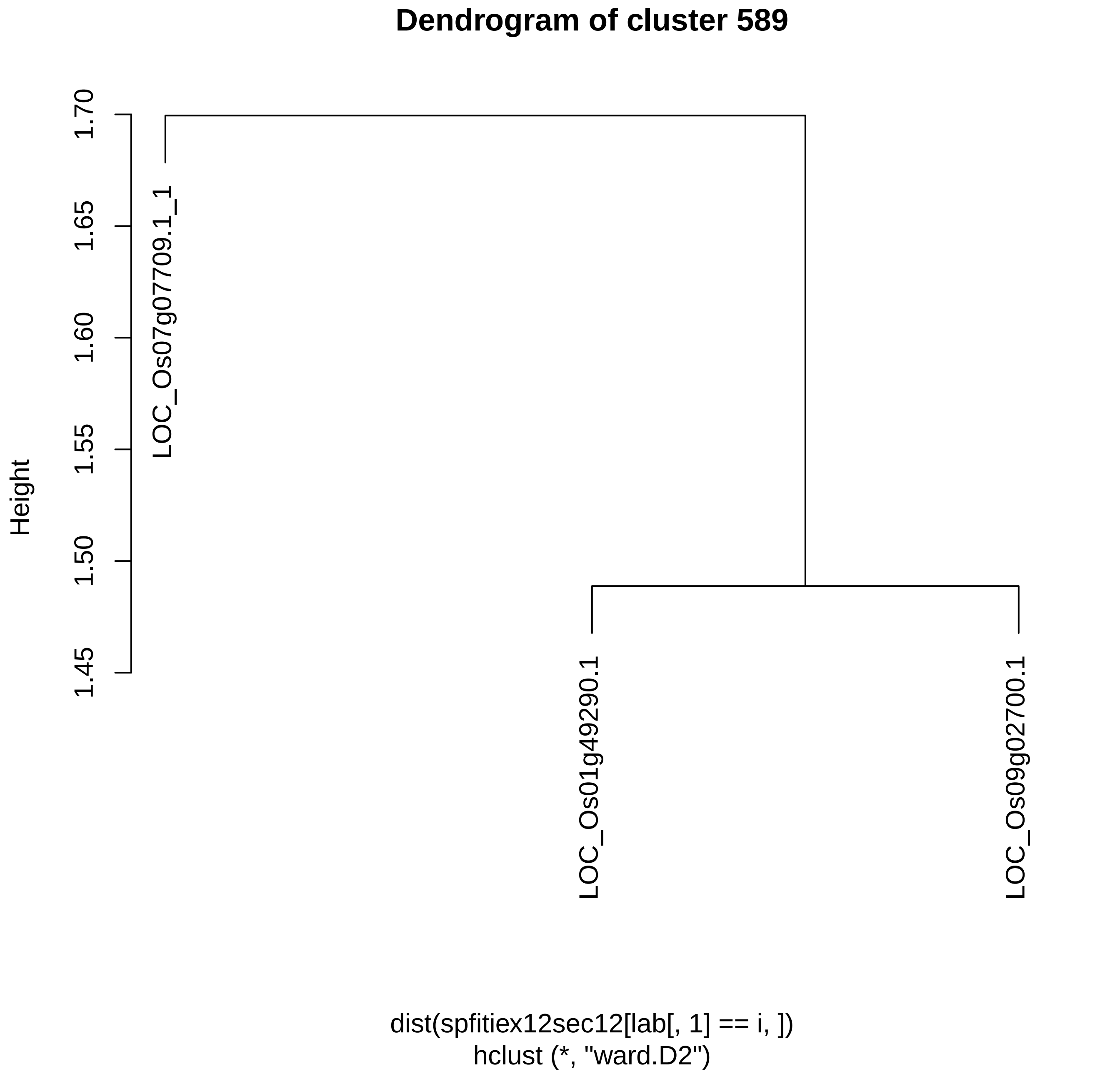
Dendrogram of cluster 589
Height
1.70
1.65
1.60
1.55
1.50
1.45
LOC_Os07g07709.1_1
LOC_Os01g49290.1
LOC_Os09g02700.1
dist(spfitiex12sec12[lab[, 1] == i, ])
hclust (*, "ward.D2")

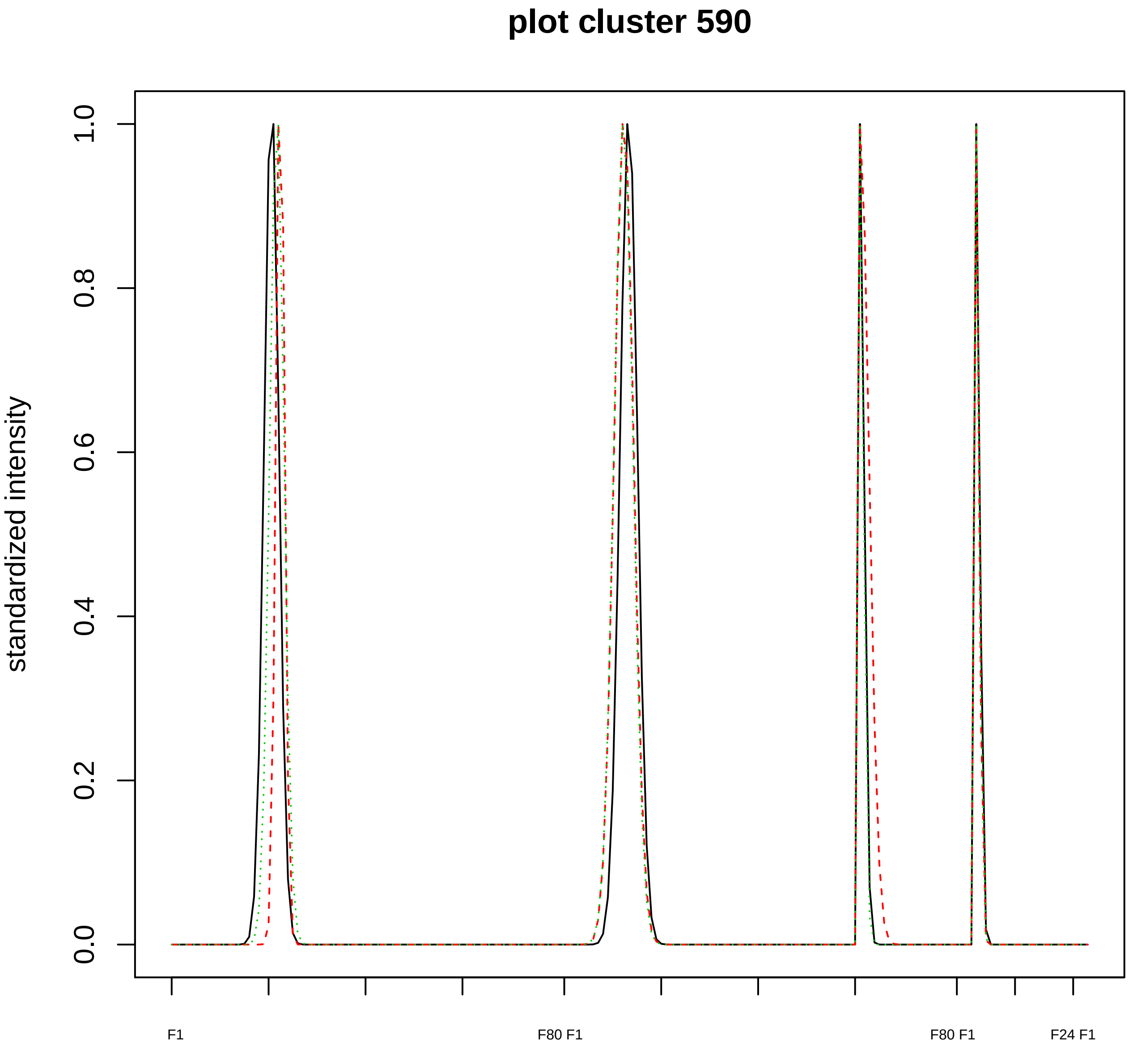

plot cluster 590
standardized intensity
0.0
0.2
0.4
0.6
0.8
1.0
F1
F80 F1
F80 F1
F24 F1

#### Dendrogram of cluster 590

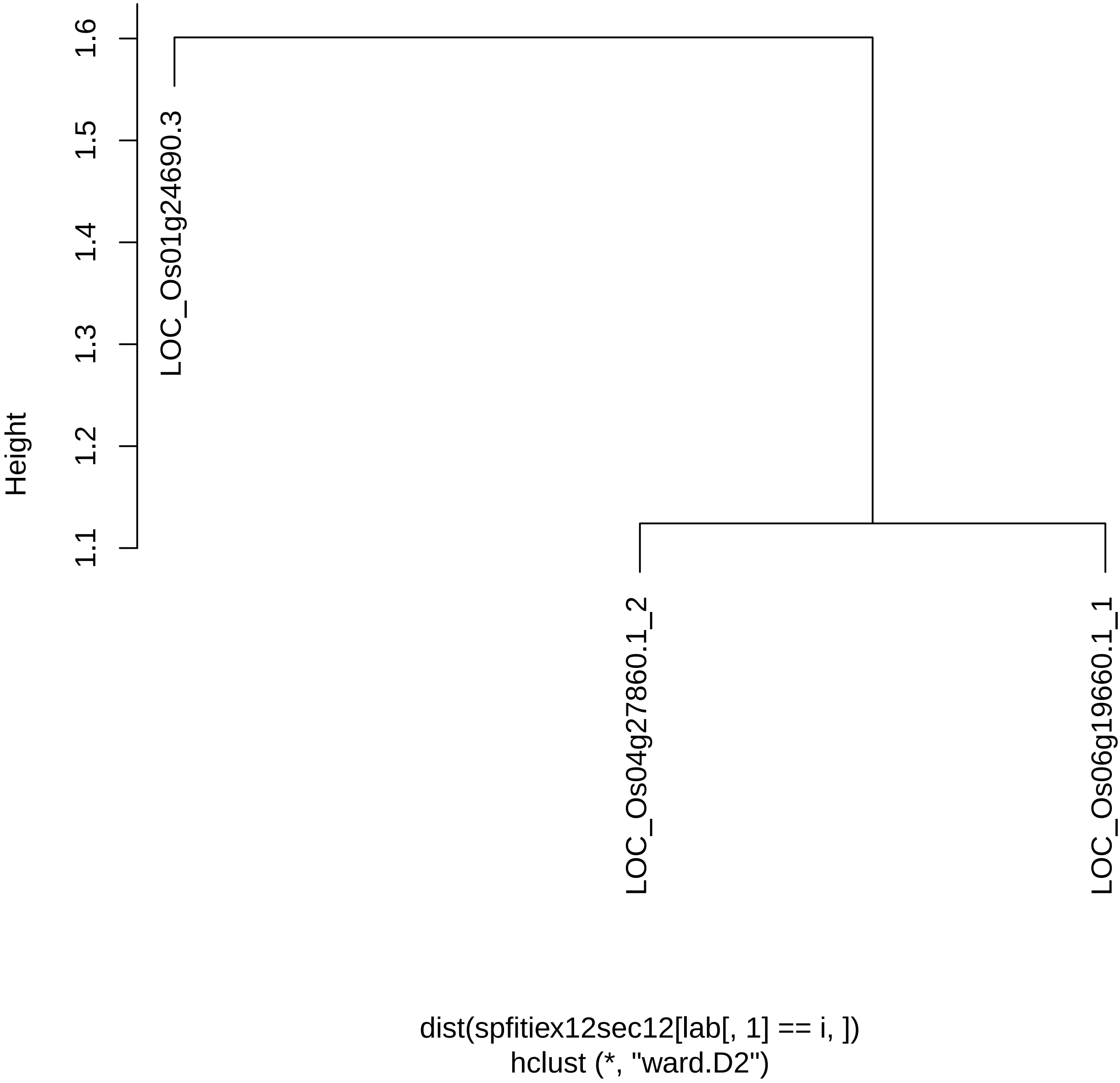

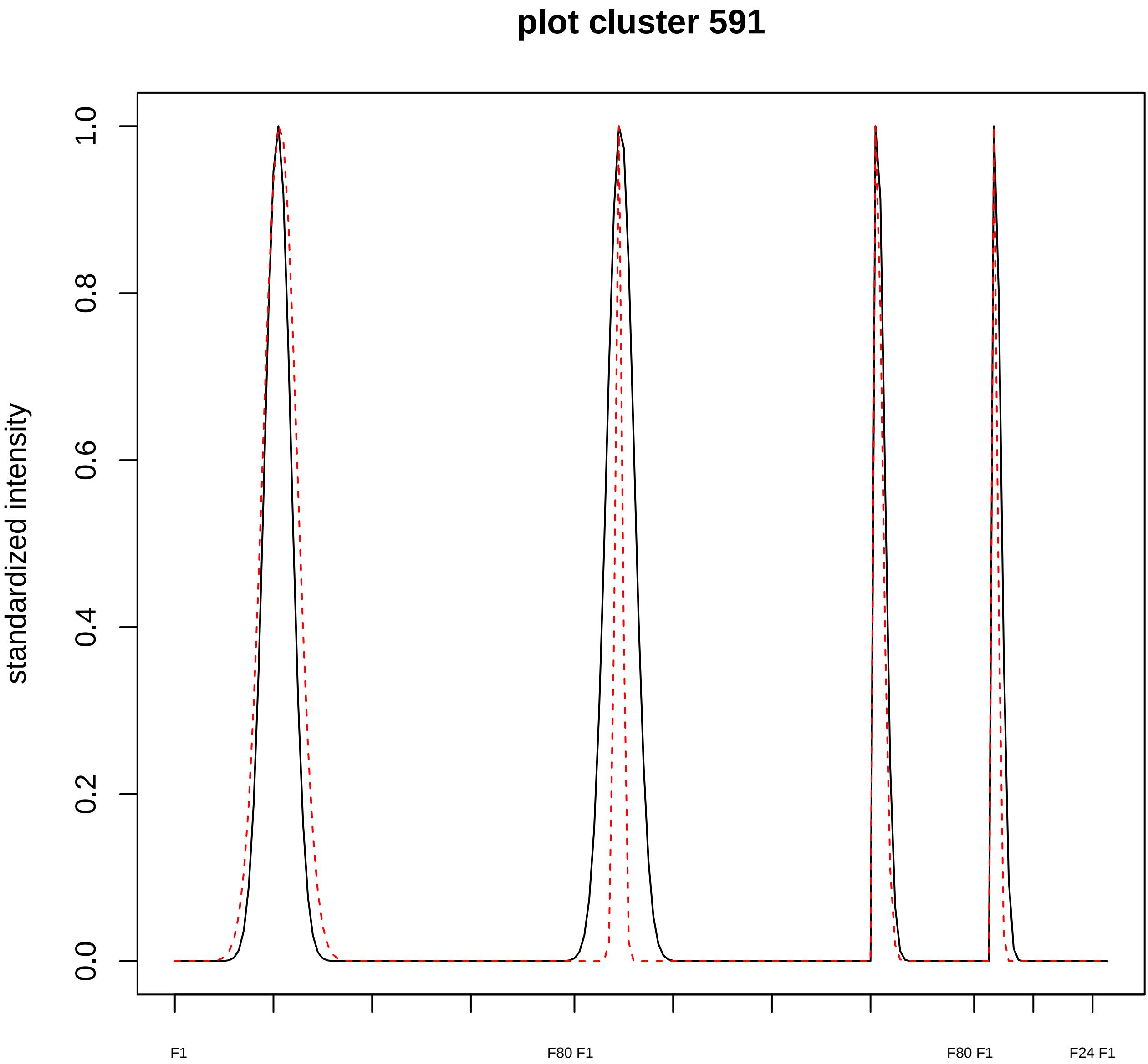

plot cluster 591
standardized intensity
0.0
0.2
0.4
0.6
0.8
1.0
F1
F80 F1
F80 F1
F24 F1

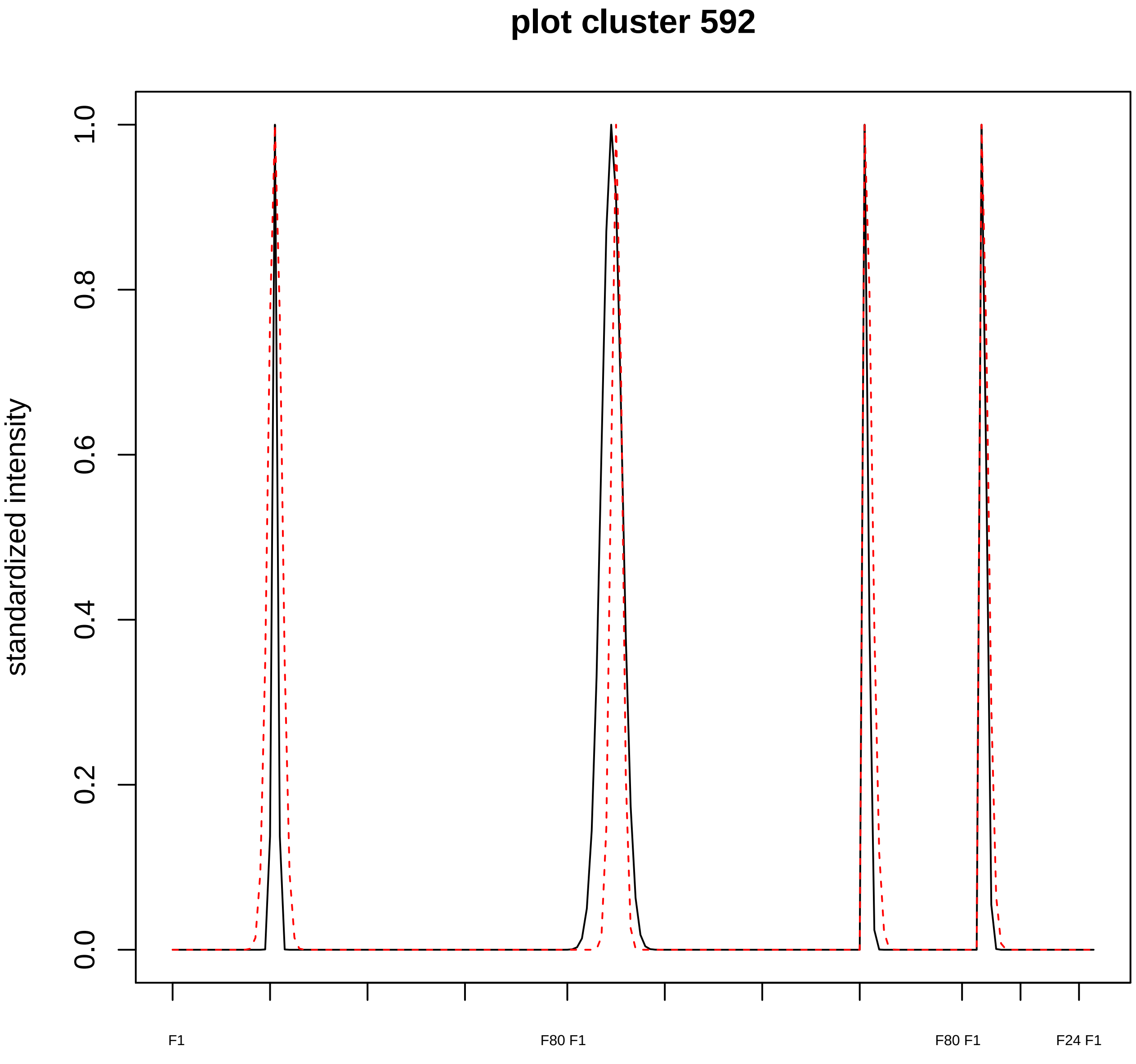
plot cluster 592
standardized intensity
0.0
0.2
0.4
0.6
0.8
1.0
F1
F80 F1
F80 F1
F24 F1

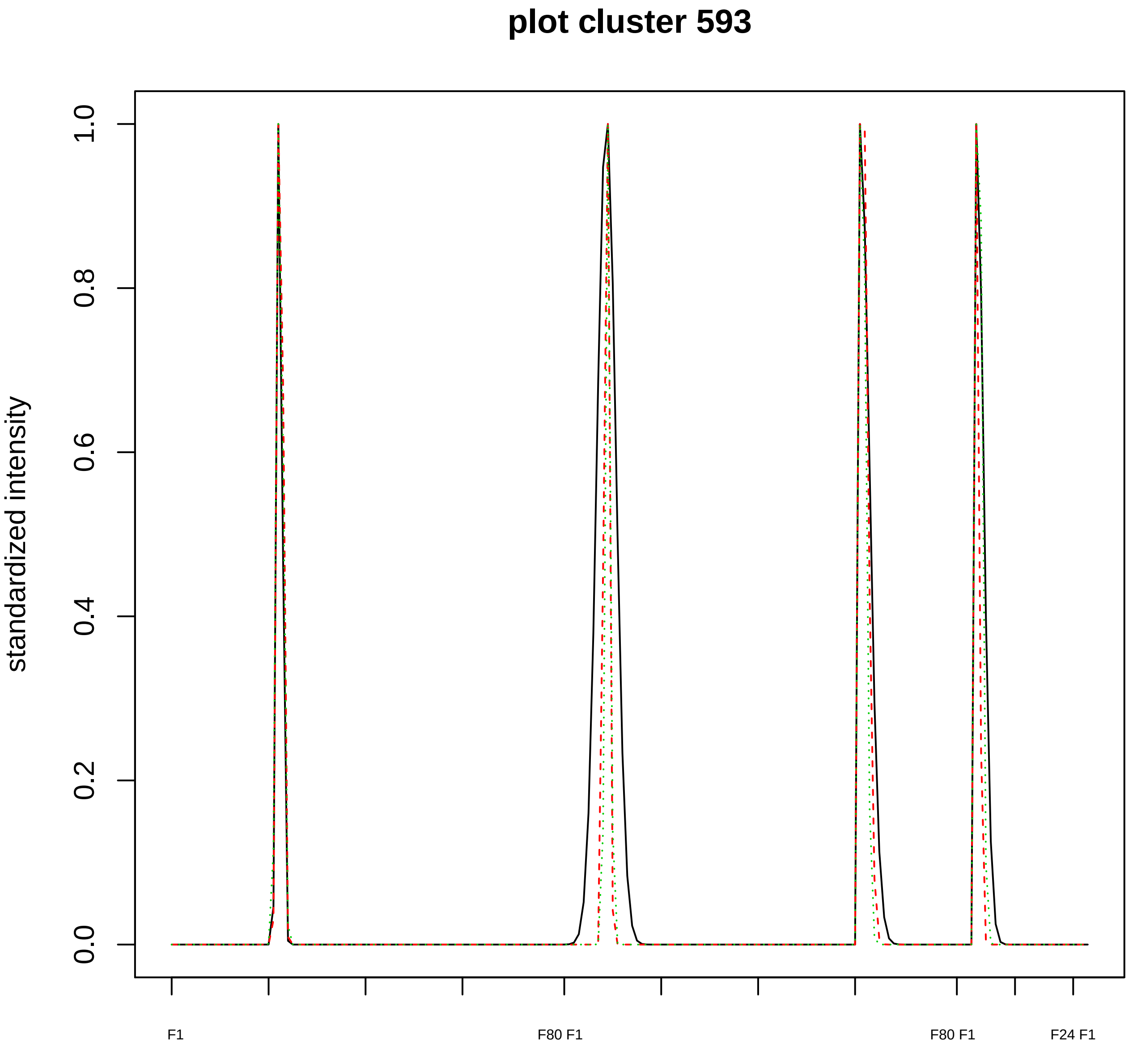

plot cluster 593
standardized intensity
0.0
0.2
0.4
0.6
0.8
1.0
F1
F80 F1
F80 F1
F24 F1

### Dendrogram of cluster 593

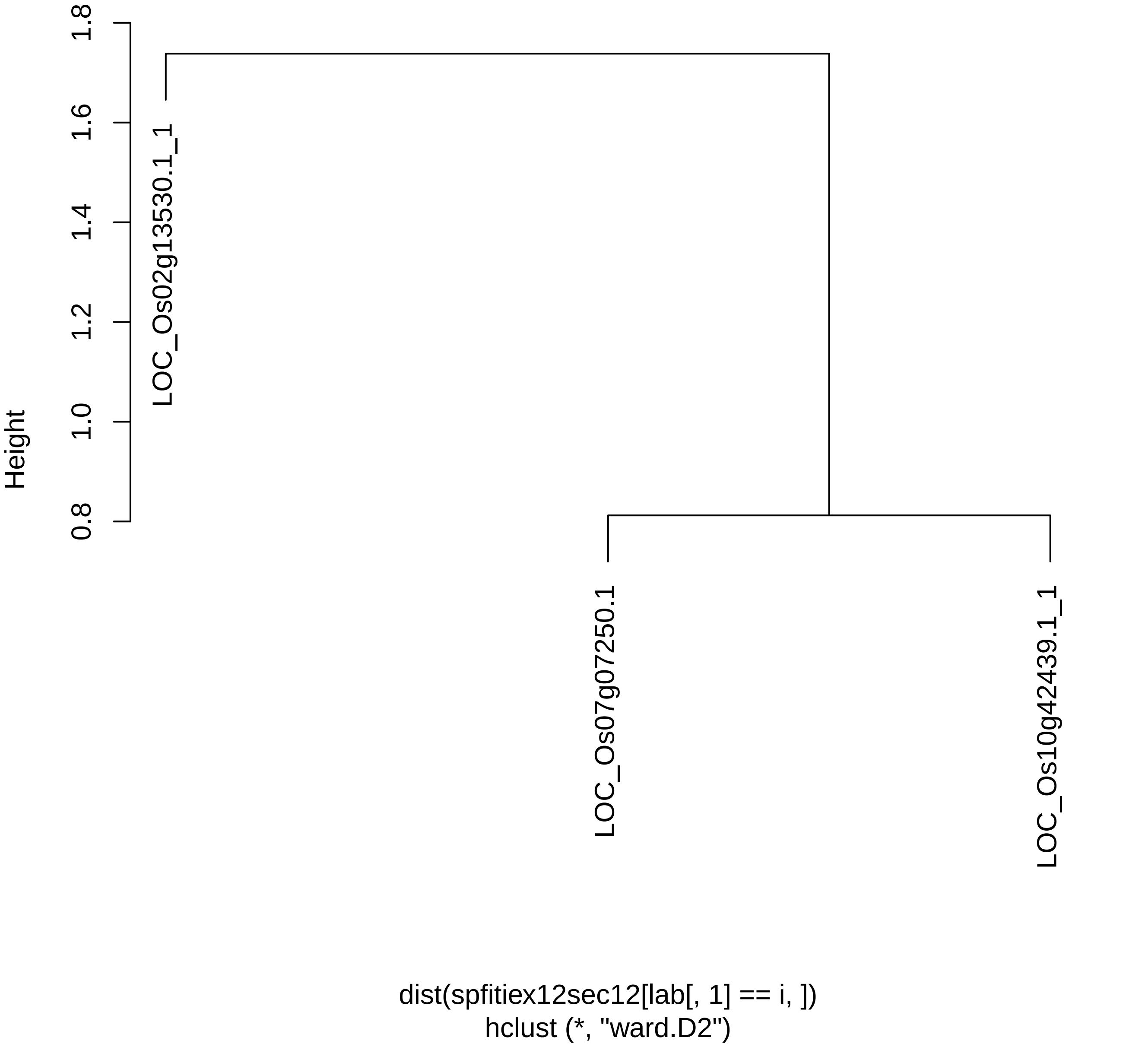

### plot cluster 594

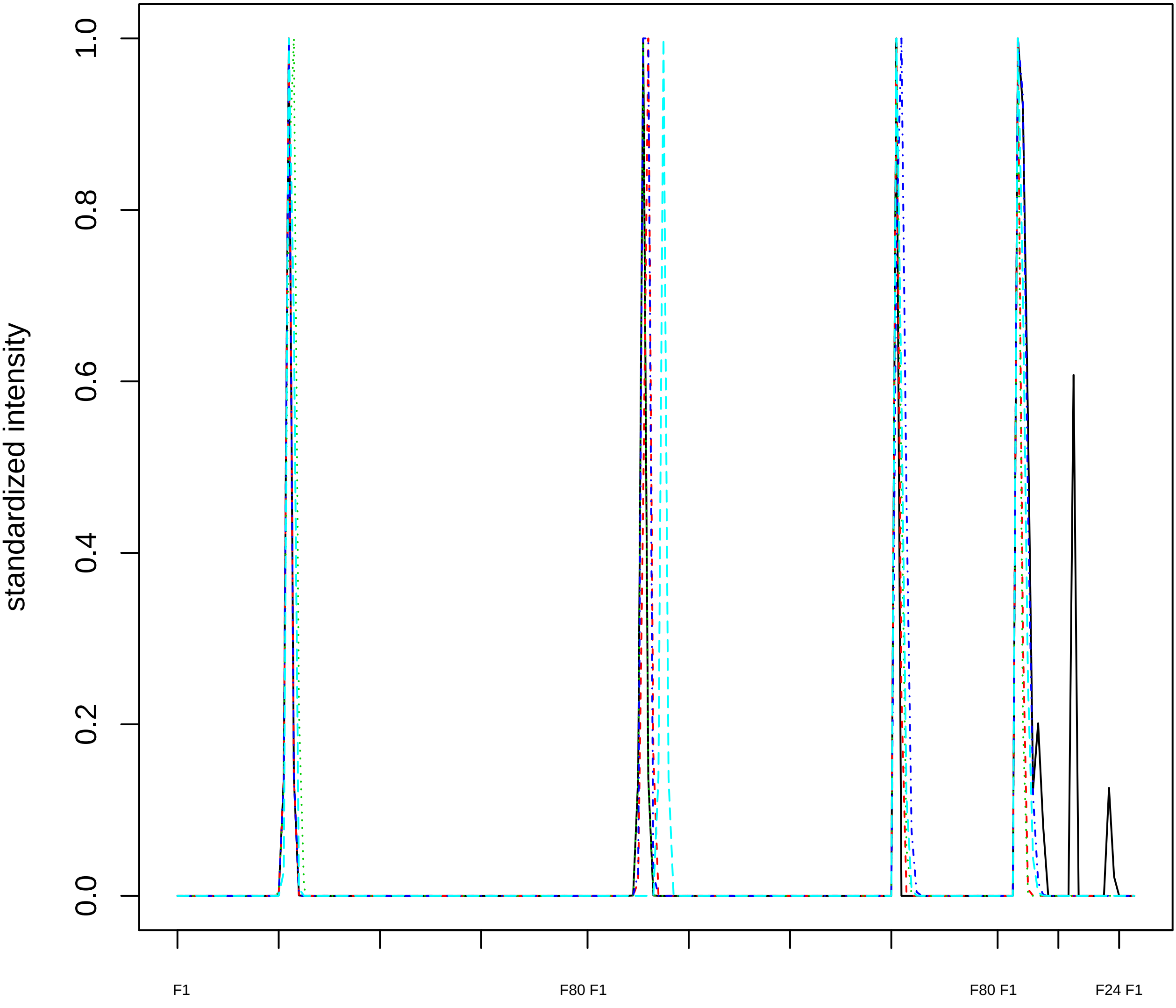

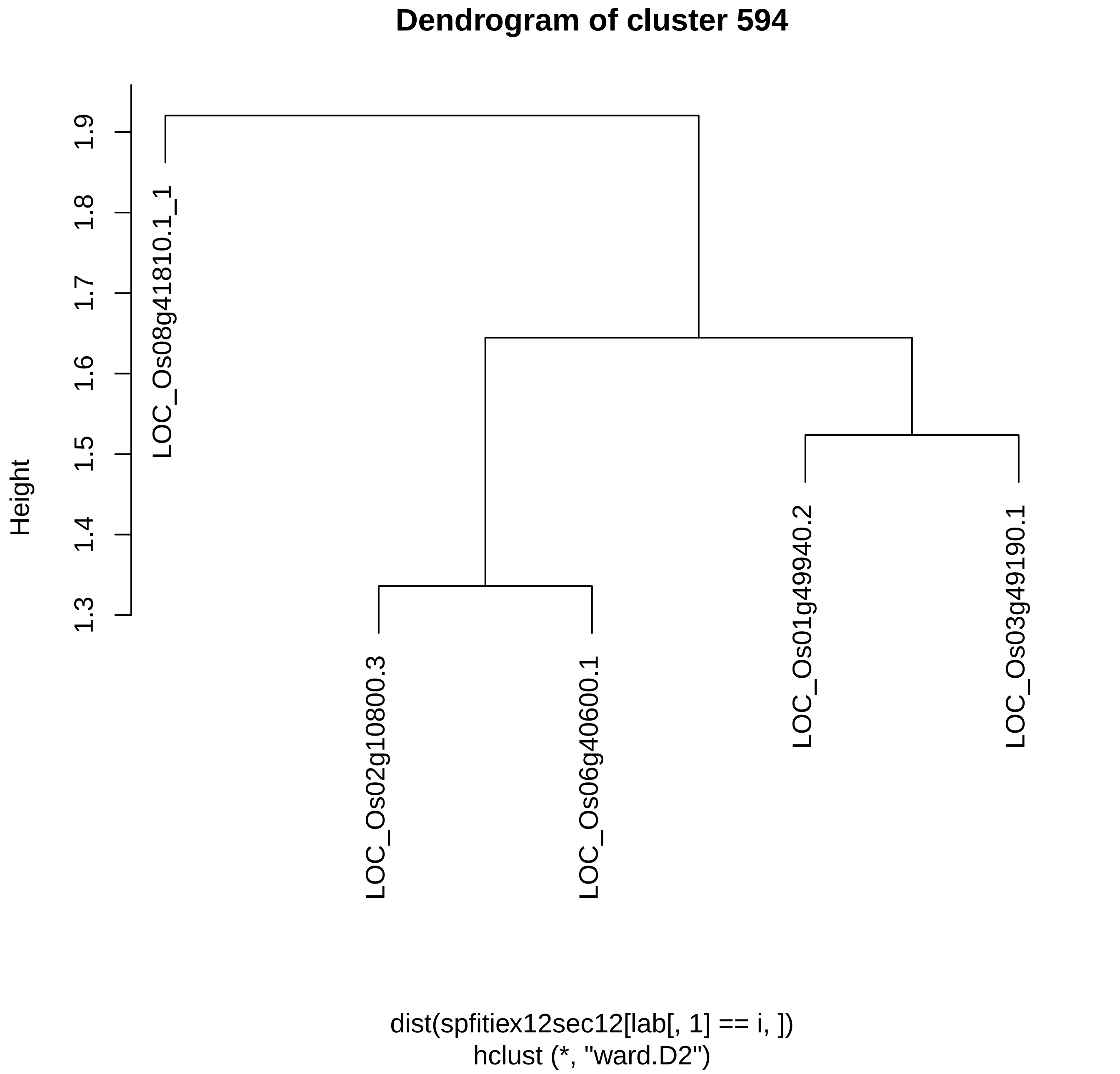

Dendrogram of cluster 594
Height
1.3
1.4
1.5
1.6
1.7
1.8
1.9
LOC_Os08g41810.1_1
LOC_Os02g10800.3
LOC_Os06g40600.1
LOC_Os01g49940.2
LOC_Os03g49190.1
dist(spfitiex12sec12[lab[, 1] == i, ])
hclust (*, "ward.D2")

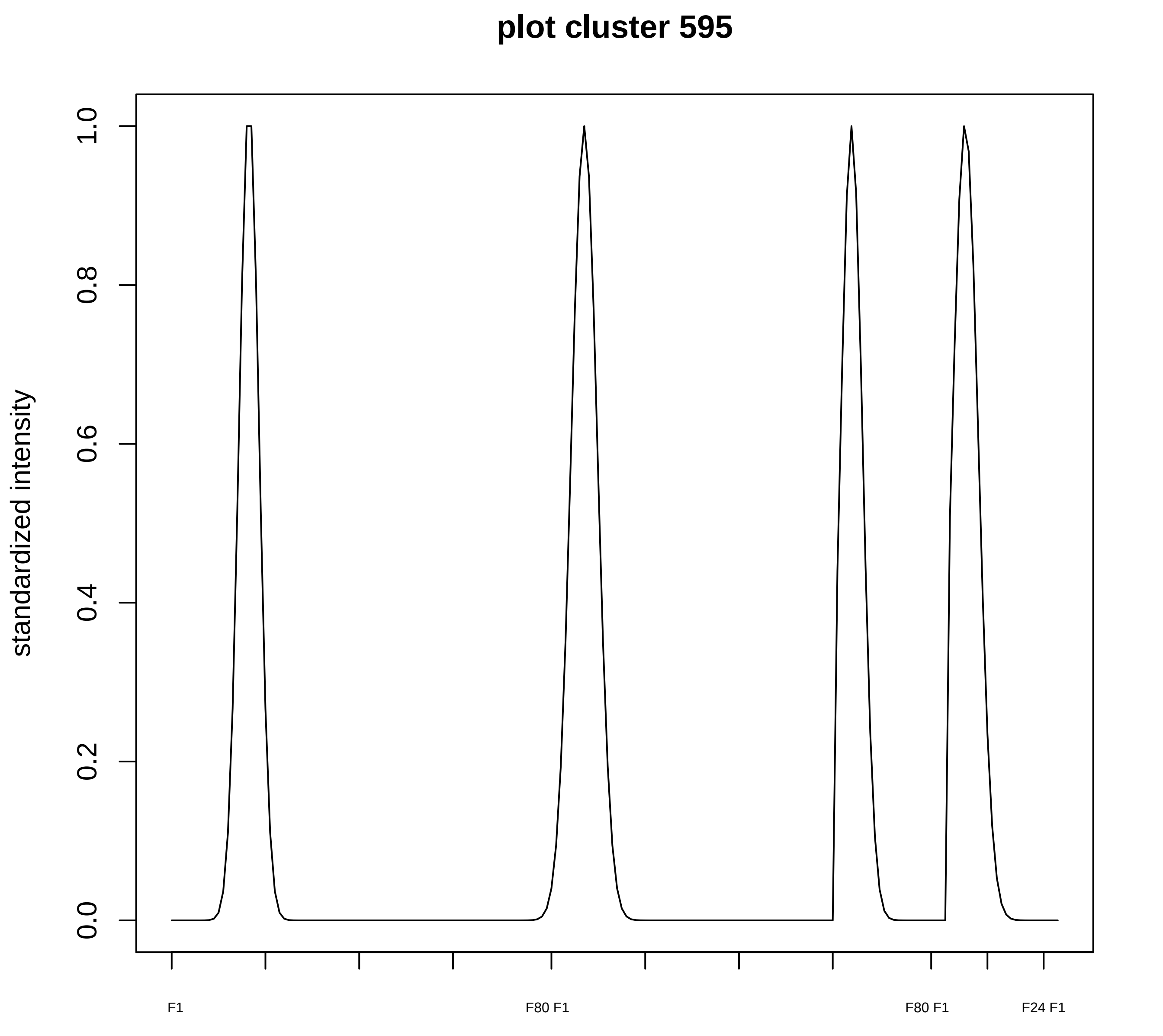
plot cluster 595
standardized intensity
0.0
0.2
0.4
0.6
0.8
1.0
F1
F80 F1
F80 F1
F24 F1

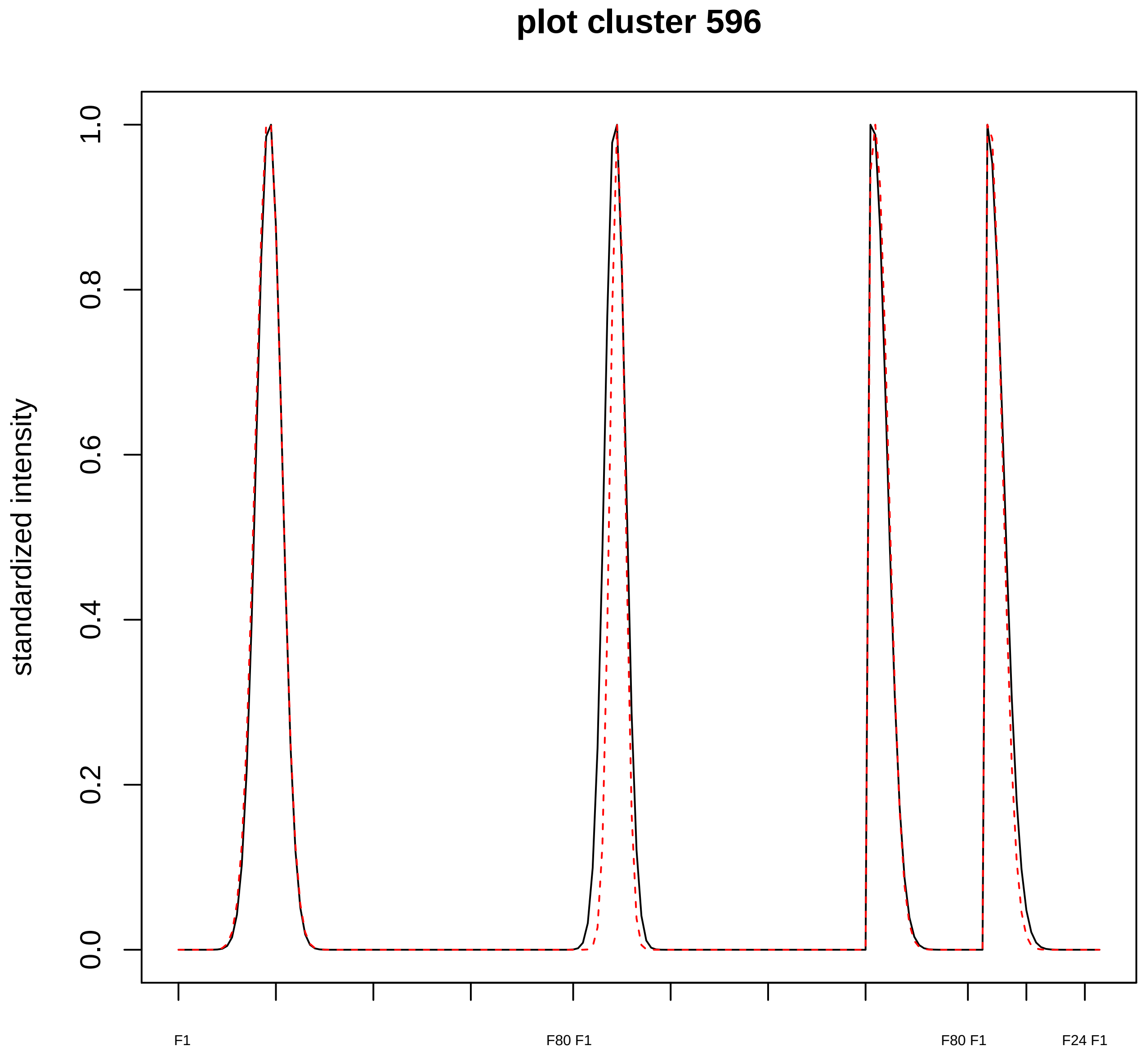
plot cluster 596
standardized intensity
0.0
0.2
0.4
0.6
0.8
1.0
F1
F80 F1
F80 F1
F24 F1

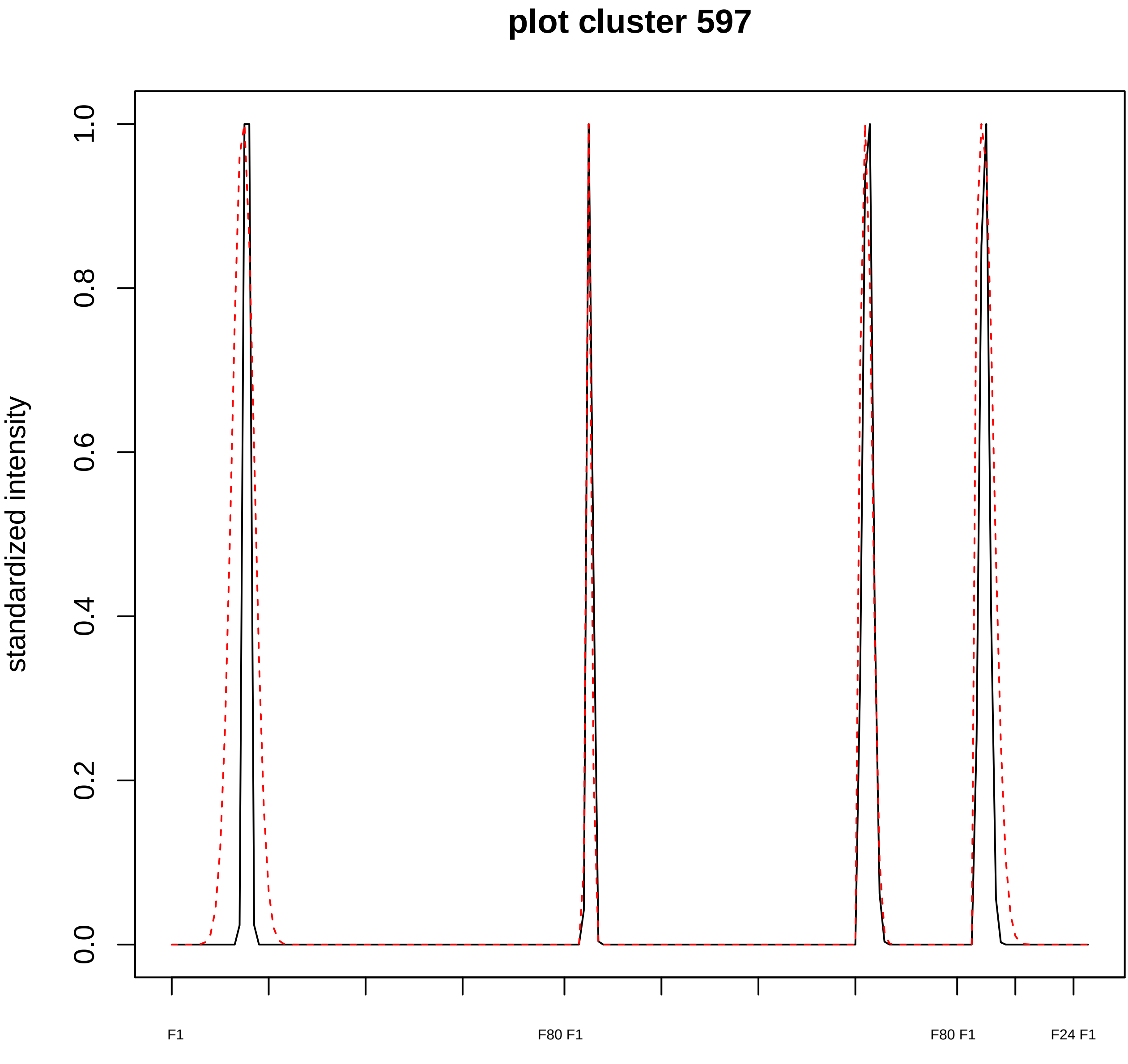
plot cluster 597
standardized intensity
0.0
0.2
0.4
0.6
0.8
1.0
F1
F80 F1
F80 F1
F24 F1

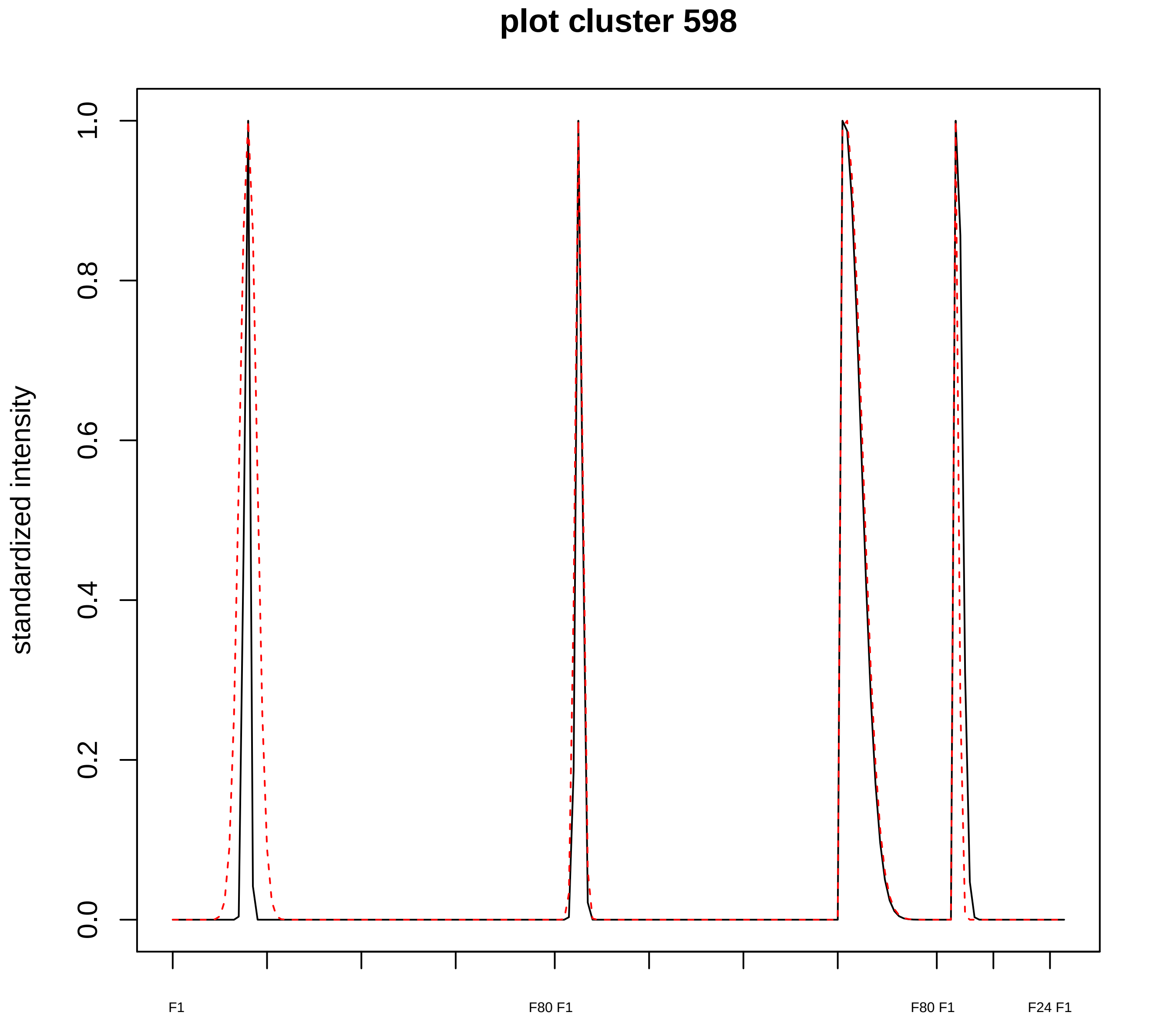

plot cluster 598
standardized intensity
0.0
0.2
0.4
0.6
0.8
1.0
F1
F80 F1
F80 F1
F24 F1

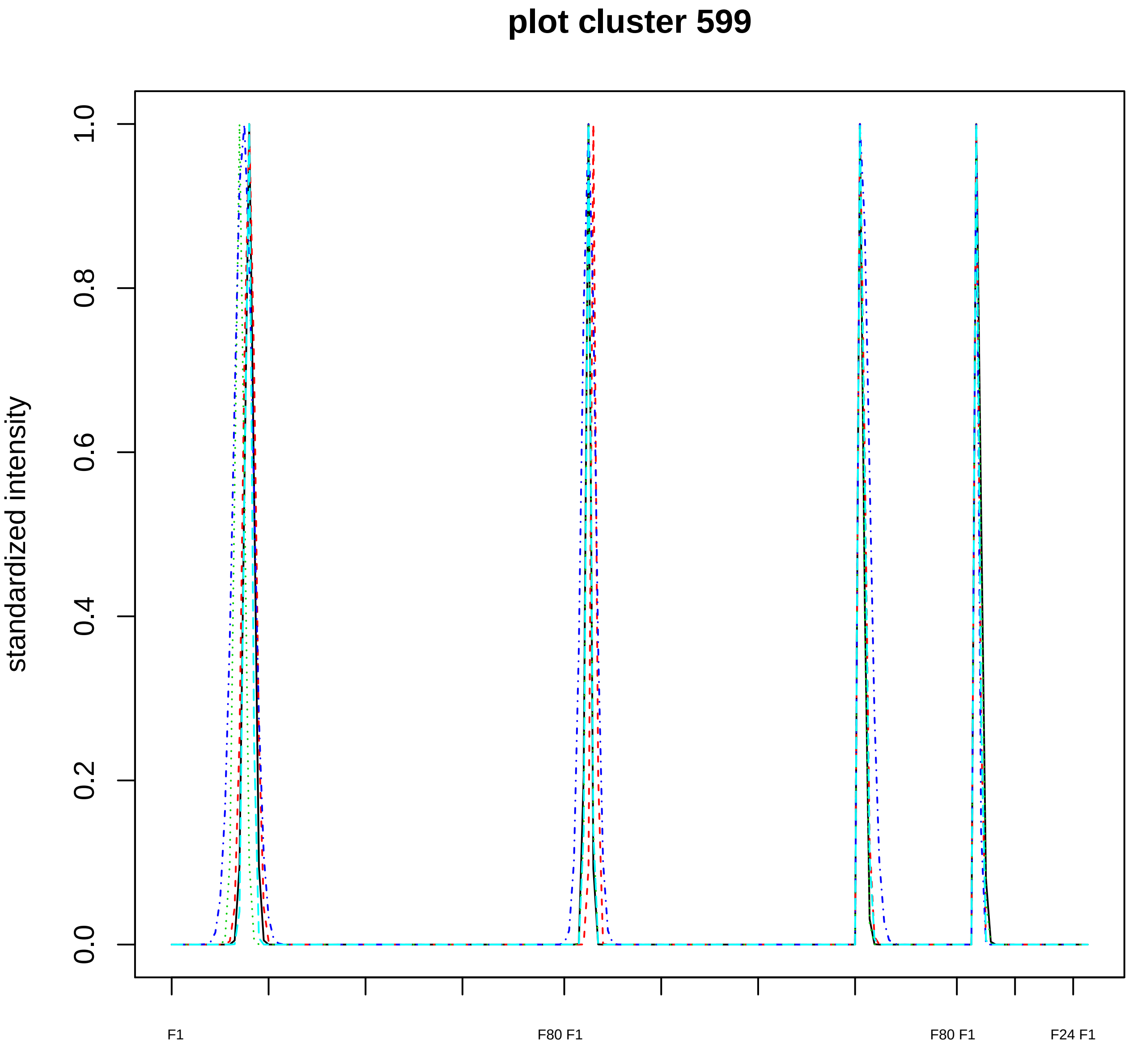

plot cluster 599
standardized intensity
0.0
0.2
0.4
0.6
0.8
1.0
F1
F80 F1
F80 F1
F24 F1

#### Dendrogram of cluster 599

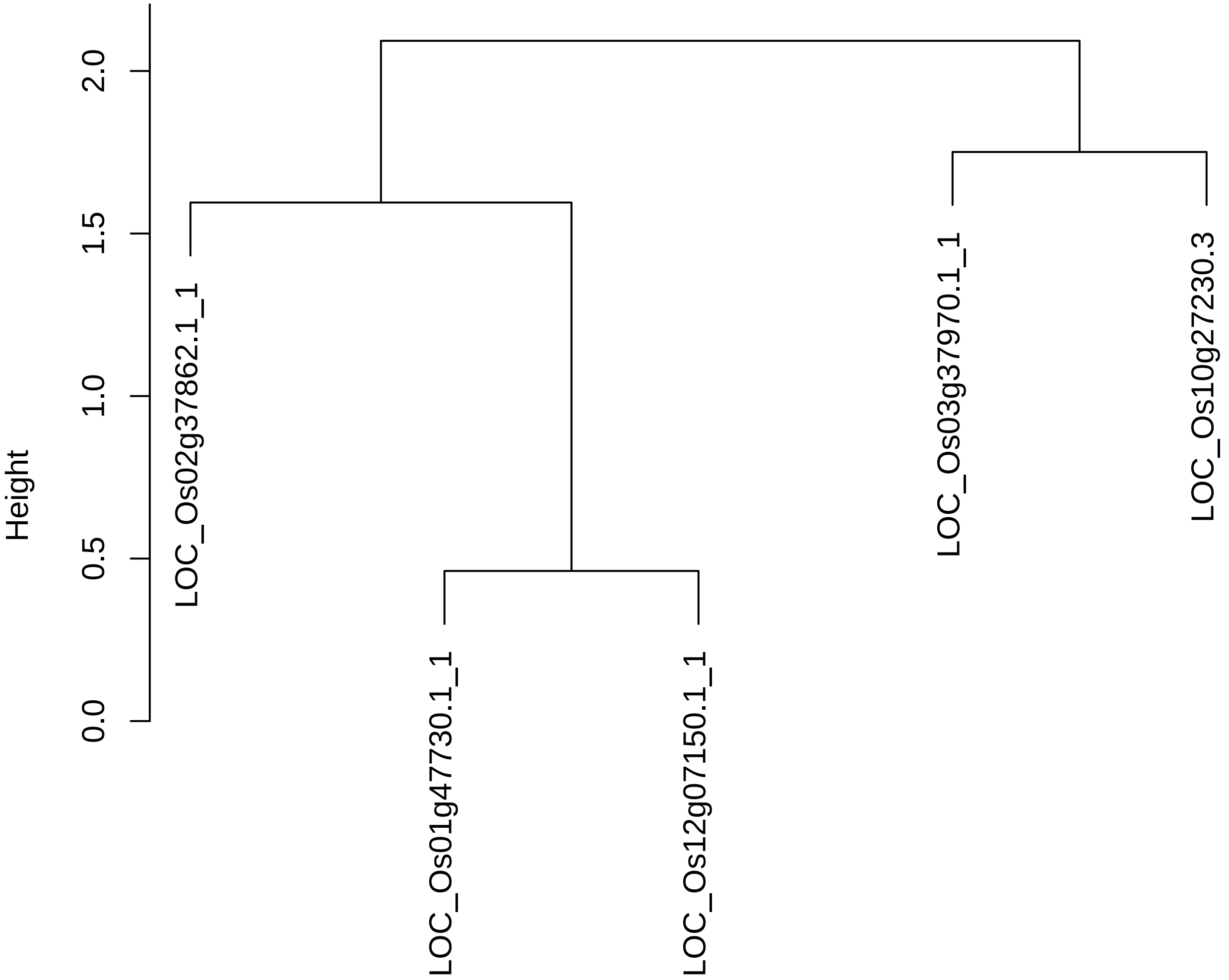


dist(spfitiex12sec12[lab[, 1] == i, ])
hclust (*, "ward.D2")

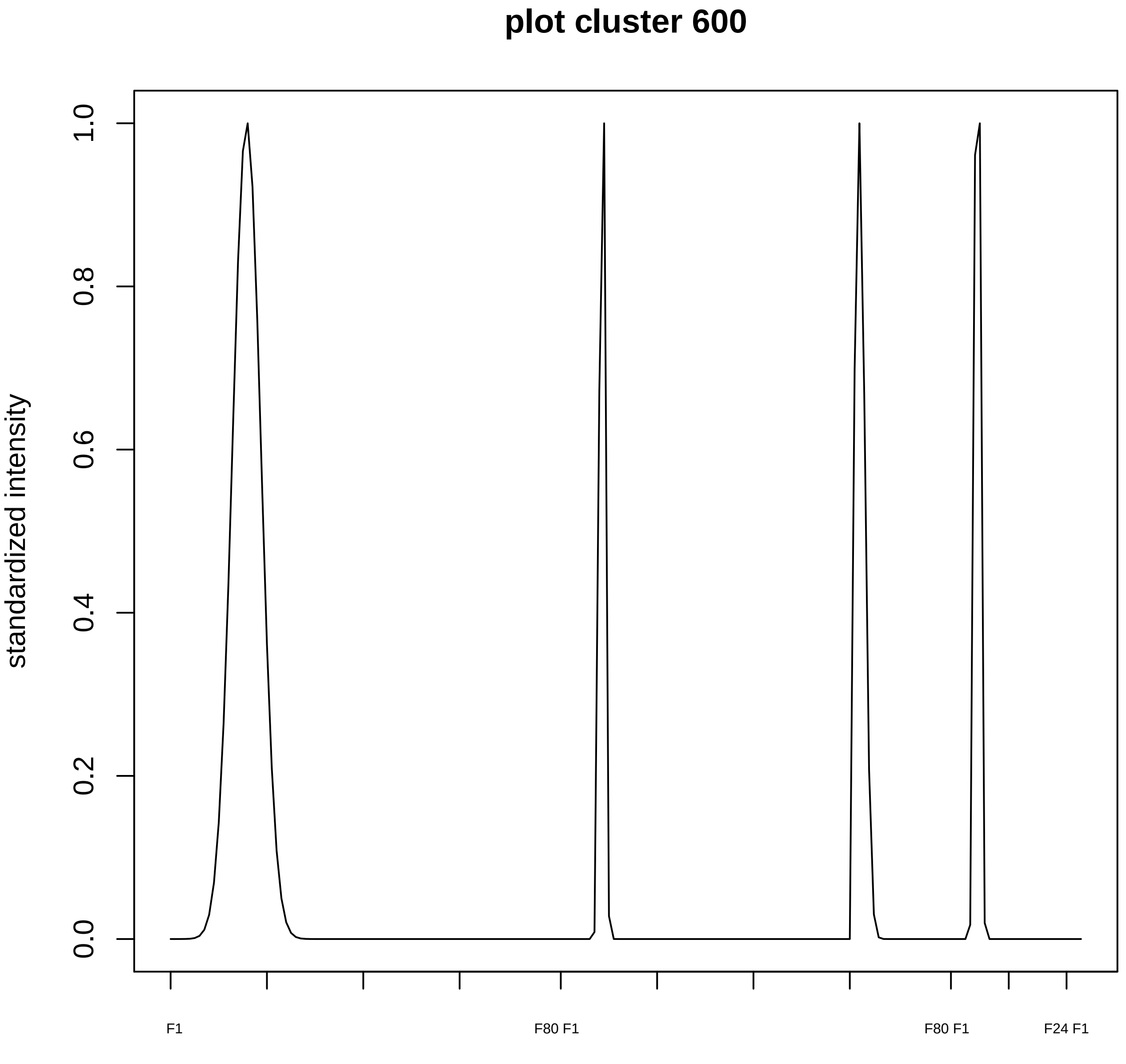
plot cluster 600
standardized intensity
0.0
0.2
0.4
0.6
0.8
1.0
F1
F80 F1
F80 F1
F24 F1

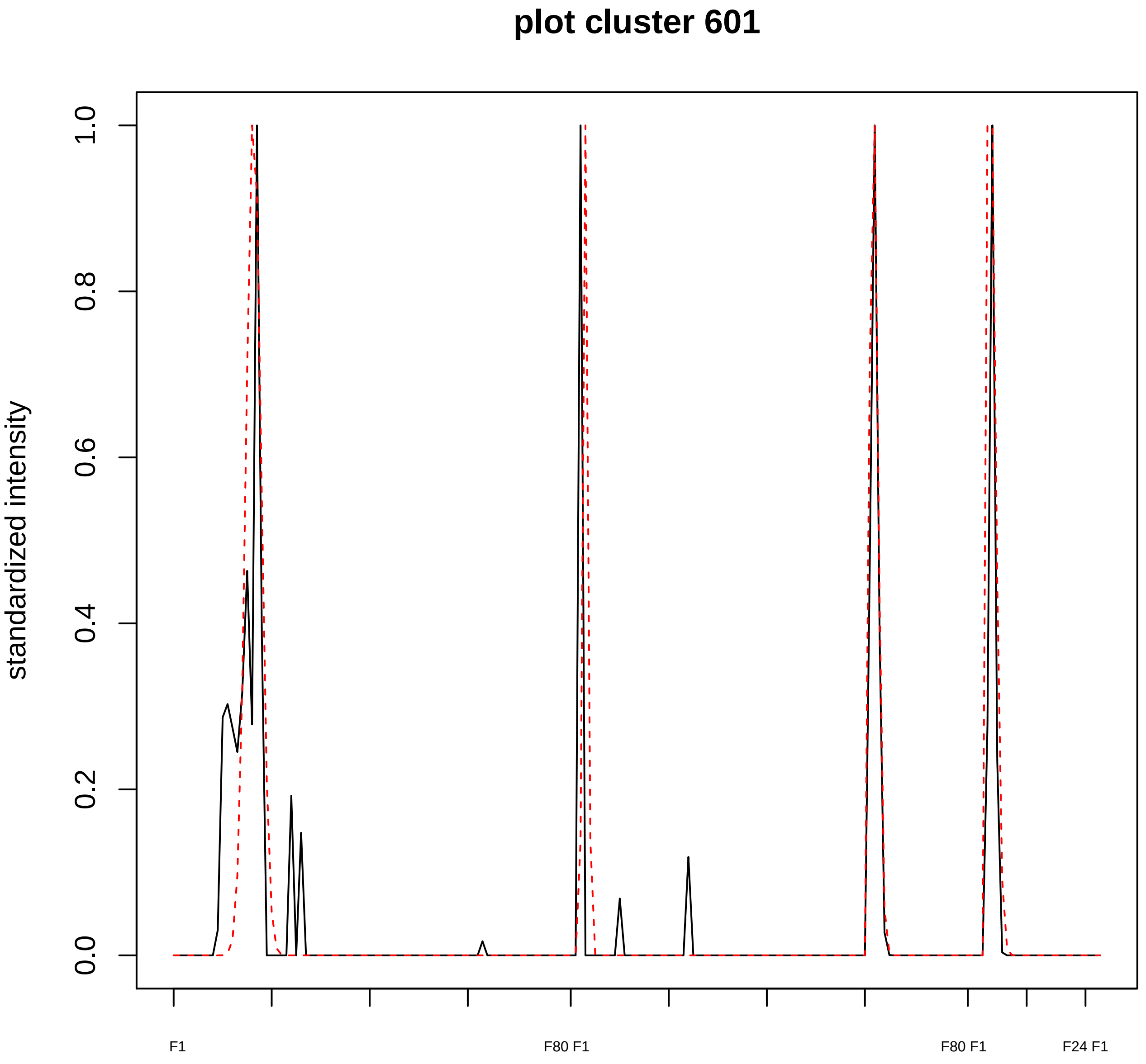

plot cluster 601
standardized intensity
0.0
0.2
0.4
0.6
0.8
1.0
F1
F80 F1
F80 F1
F24 F1

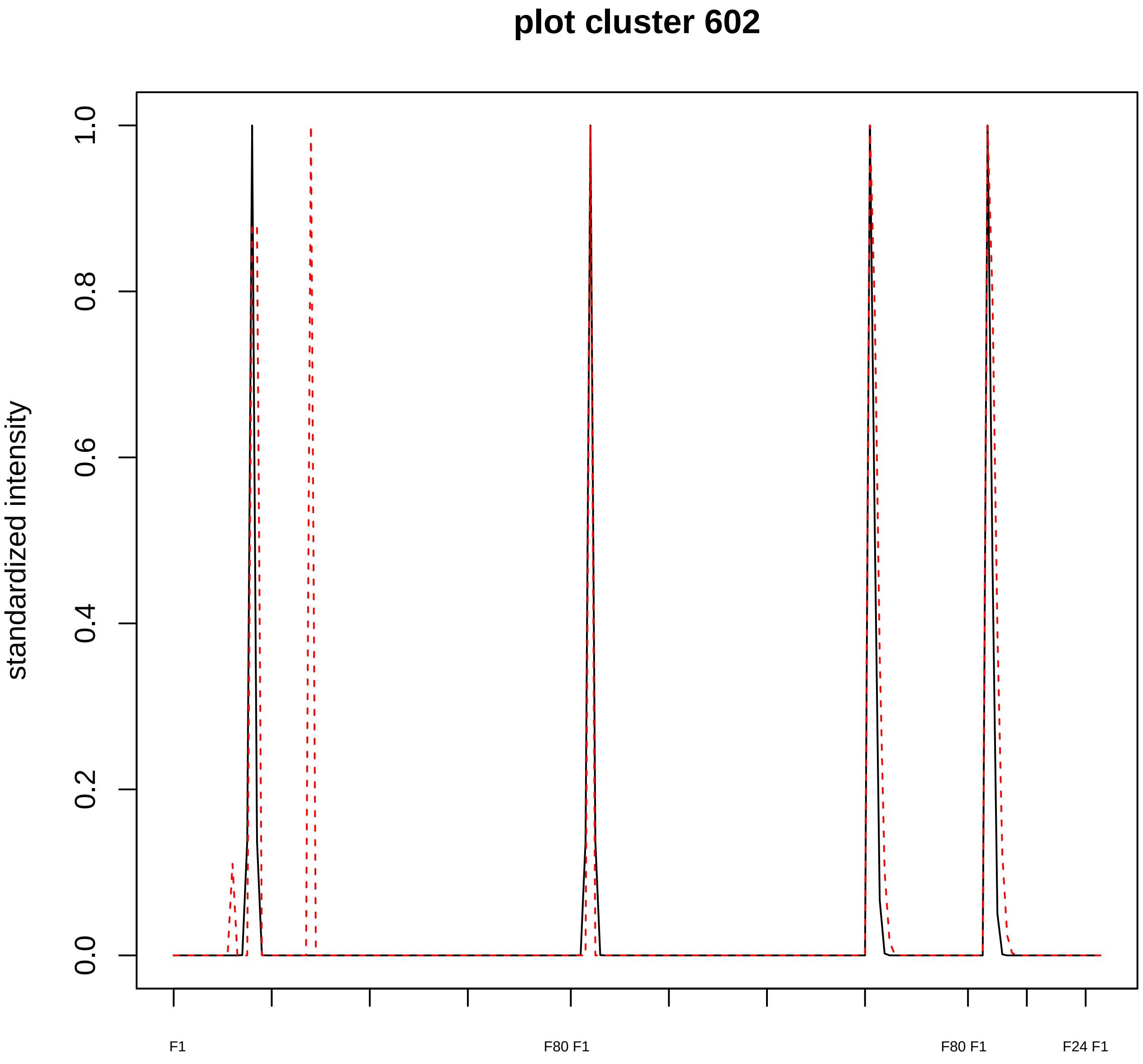
plot cluster 602
standardized intensity
1.0
0.8
0.6
0.4
0.2
0.0
F1
F80 F1
F80 F1
F24 F1

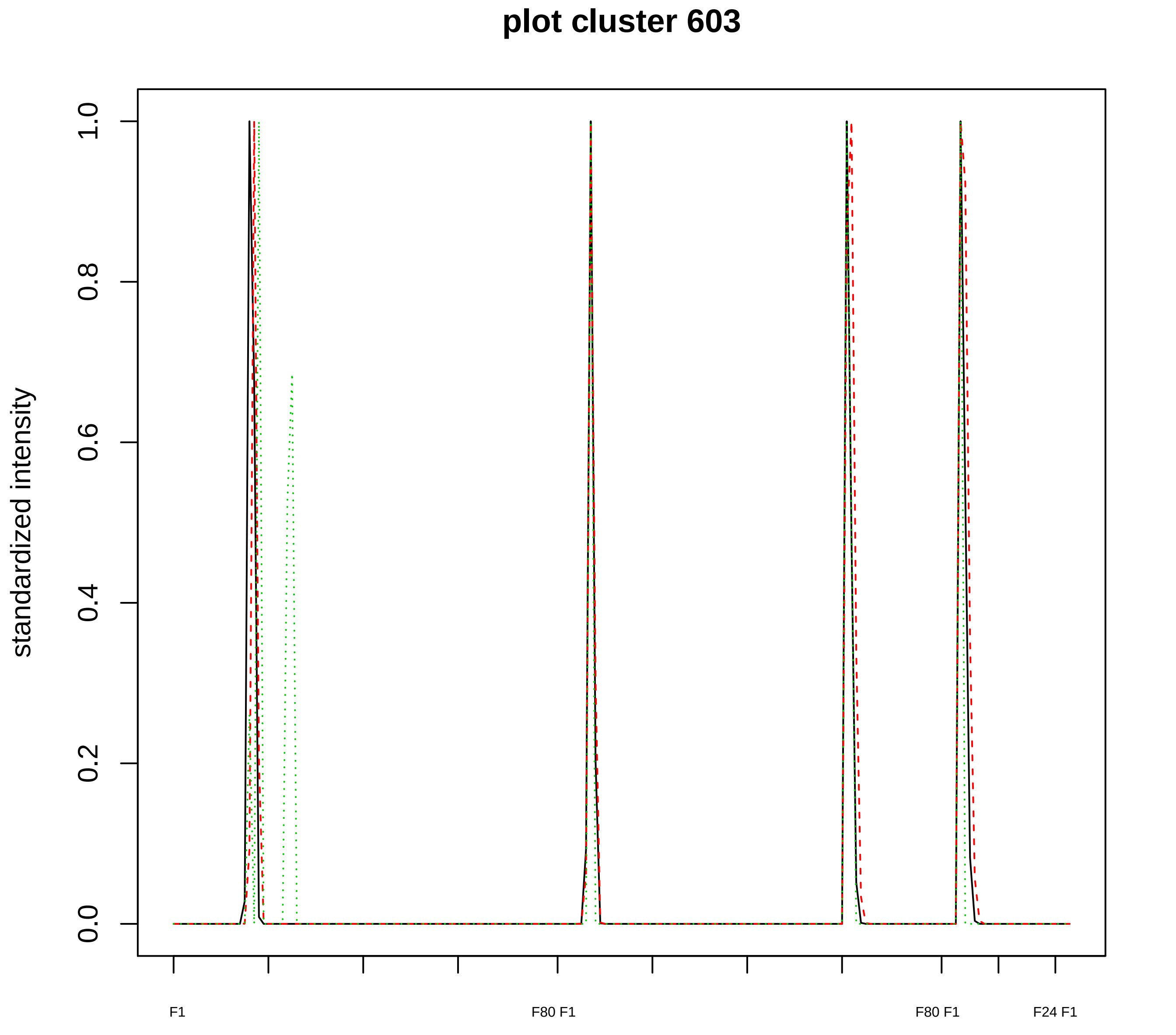

plot cluster 603
standardized intensity
0.0
0.2
0.4
0.6
0.8
1.0
F1
F80 F1
F80 F1
F24 F1

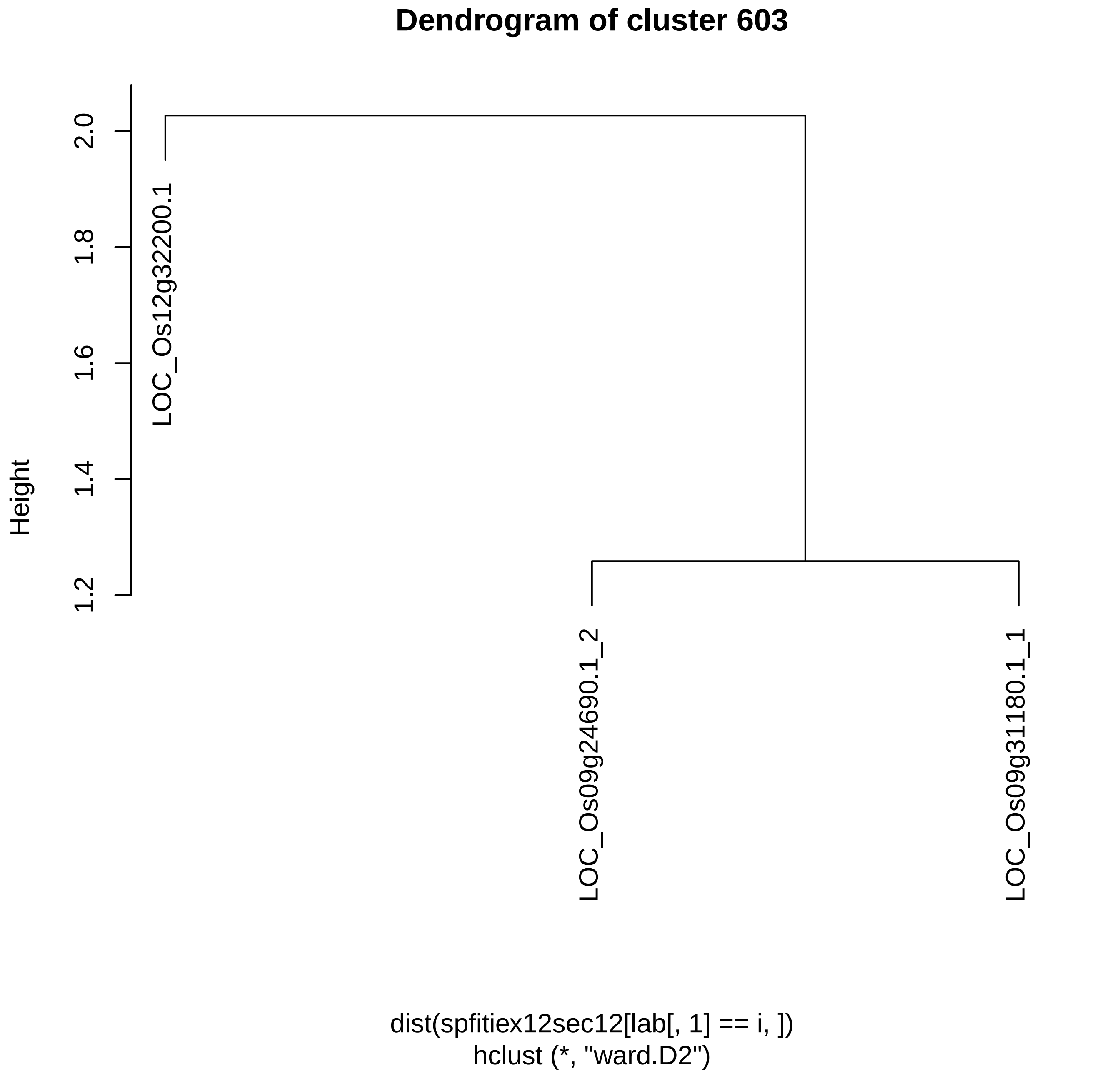

Dendrogram of cluster 603
Height
2.0
1.8
1.6
1.4
1.2
LOC_Os12g32200.1
LOC_Os09g24690.1_2
LOC_Os09g31180.1_1
dist(spfitiex12sec12[lab[, 1] == i, ])
hclust (*, "ward.D2")

#### plot cluster 604

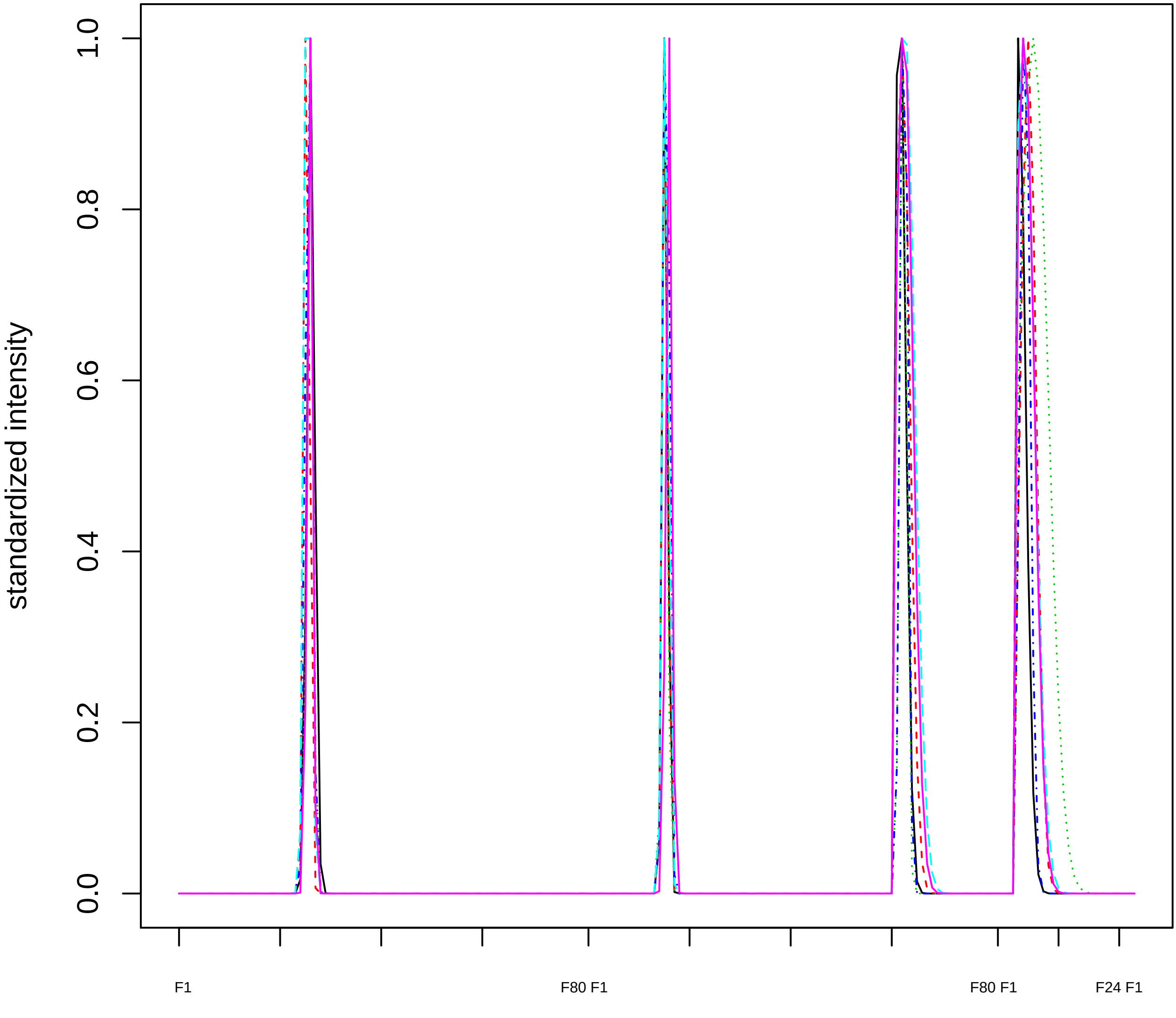

#### Dendrogram of cluster 604

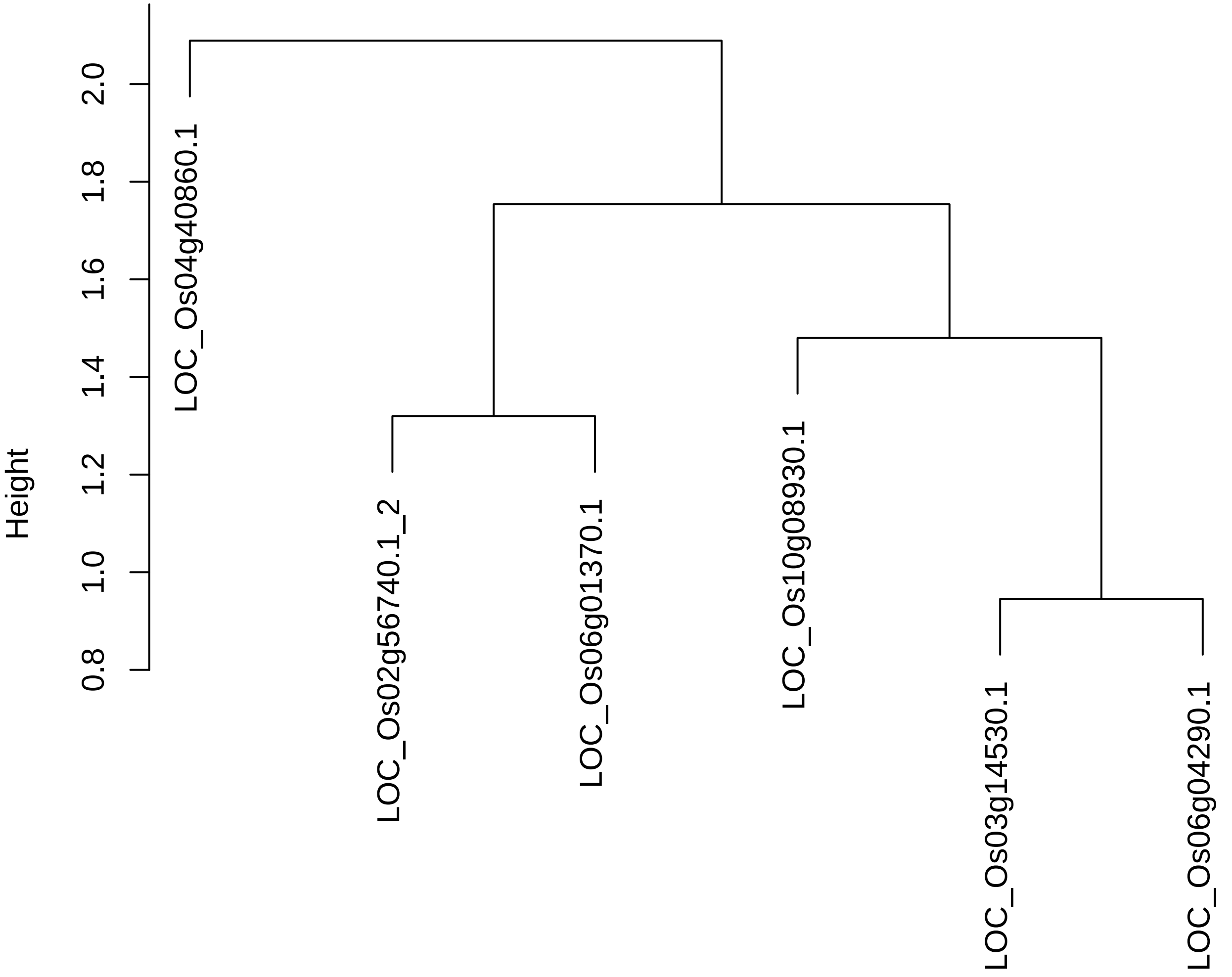


dist(spfitiex12sec12[lab[, 1] == i, ])
hclust (*, "ward.D2")

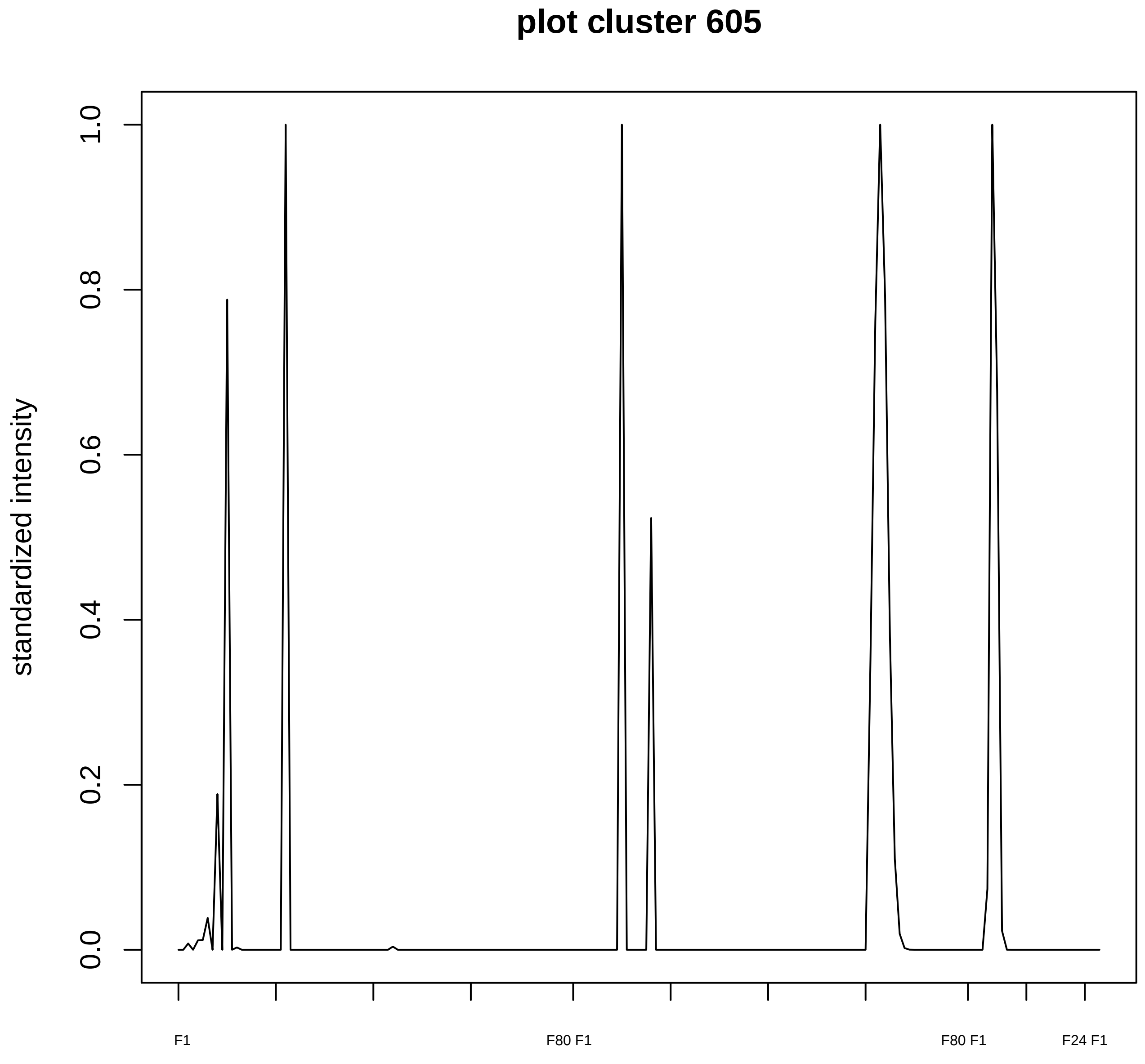
plot cluster 605
standardized intensity
0.0
0.2
0.4
0.6
0.8
1.0
F1
F80 F1
F80 F1
F24 F1

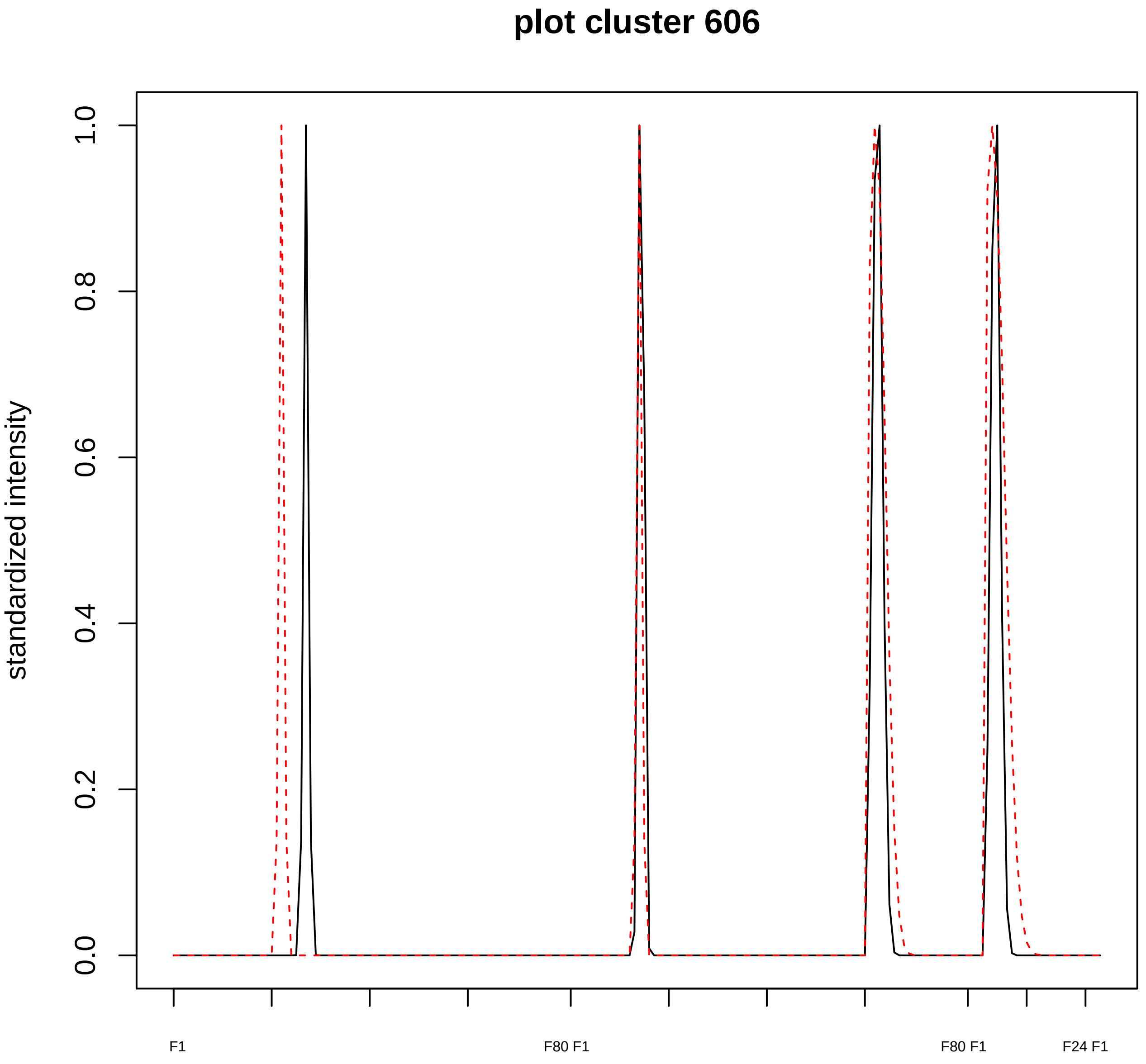

plot cluster 606
standardized intensity
0.0
0.2
0.4
0.6
0.8
1.0
F1
F80 F1
F80 F1
F24 F1

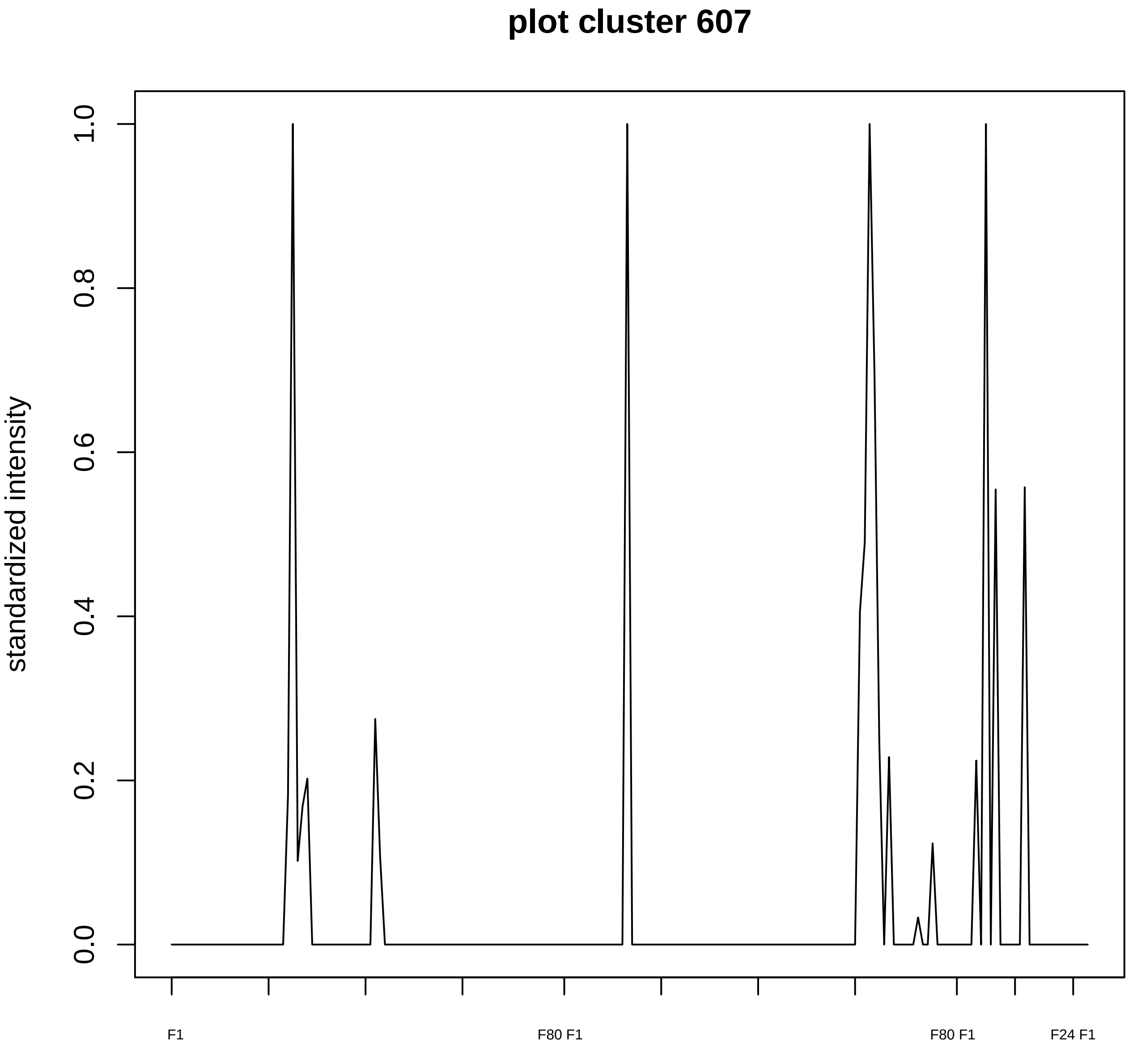

plot cluster 607
standardized intensity
0.0
0.2
0.4
0.6
0.8
1.0
F1
F80 F1
F80 F1
F24 F1

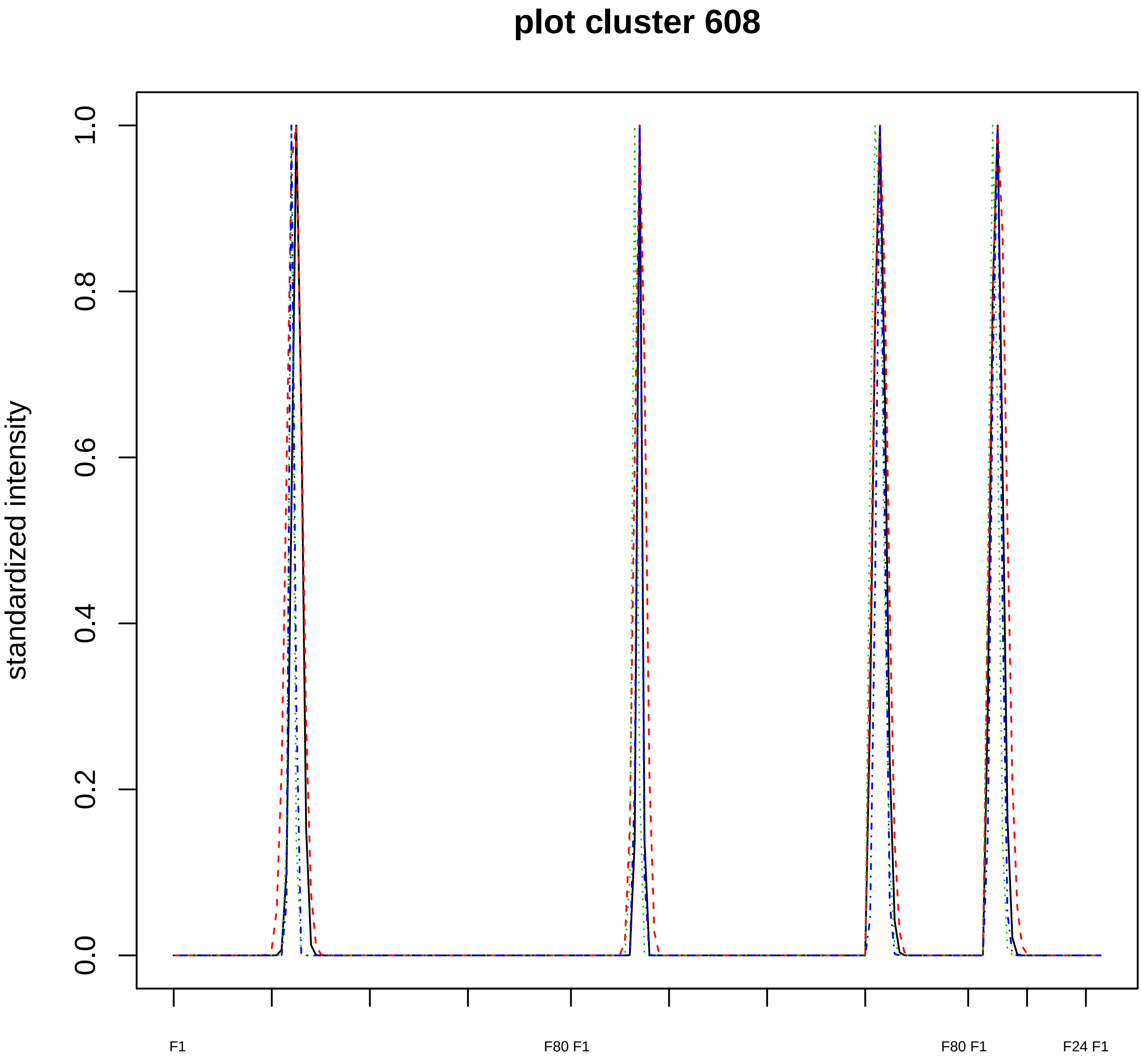
plot cluster 608
standardized intensity
0.0
0.2
0.4
0.6
0.8
1.0
F1
F80 F1
F80 F1
F24 F1

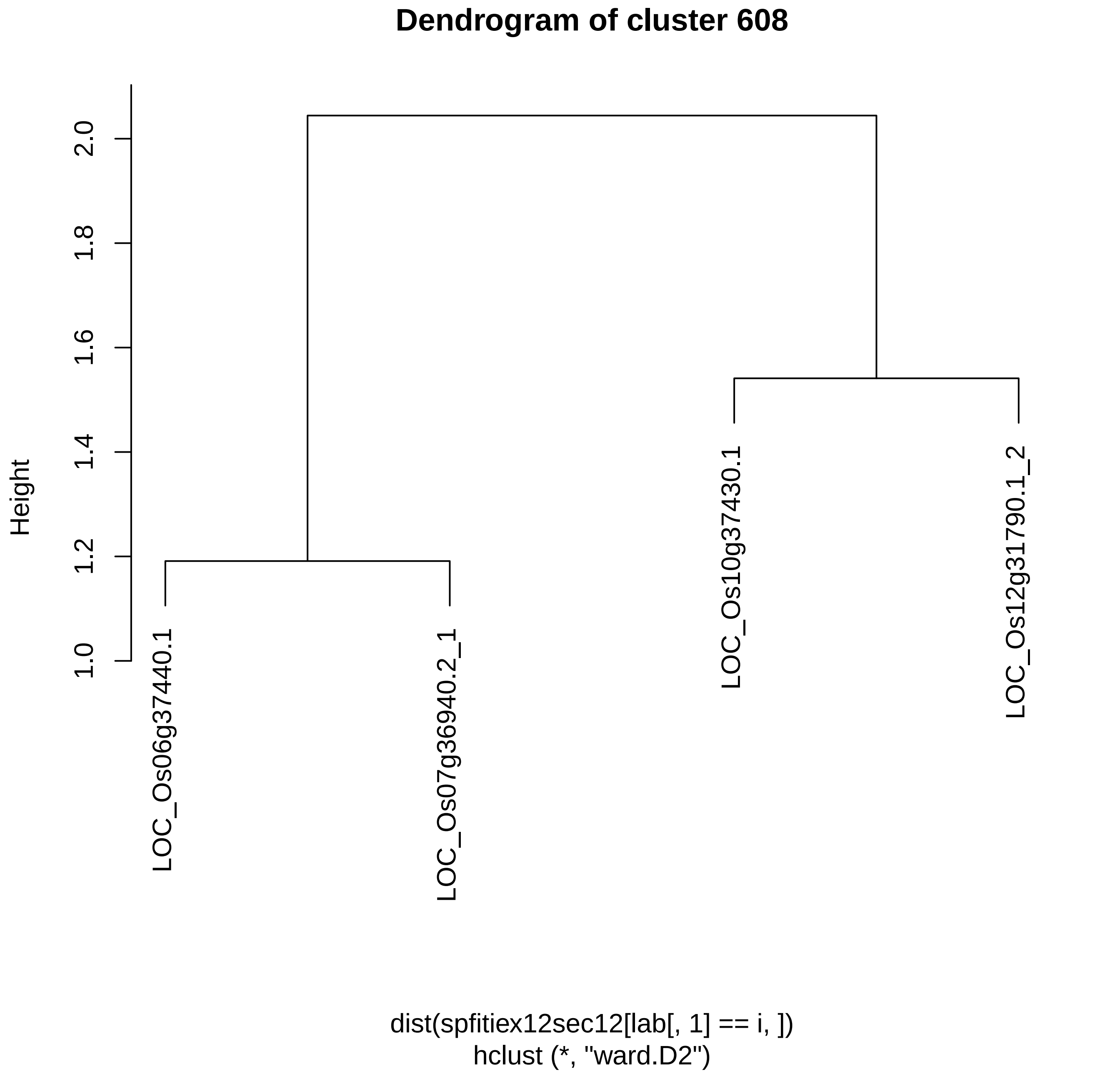
Dendrogram of cluster 608
Height
2.0
1.8
1.6
1.4
1.2
1.0
LOC_Os06g37440.1
LOC_Os07g36940.2_1
LOC_Os10g37430.1
LOC_Os12g31790.1_2
dist(spfitiex12sec12[lab[, 1] == i, ])
hclust (*, "ward.D2")

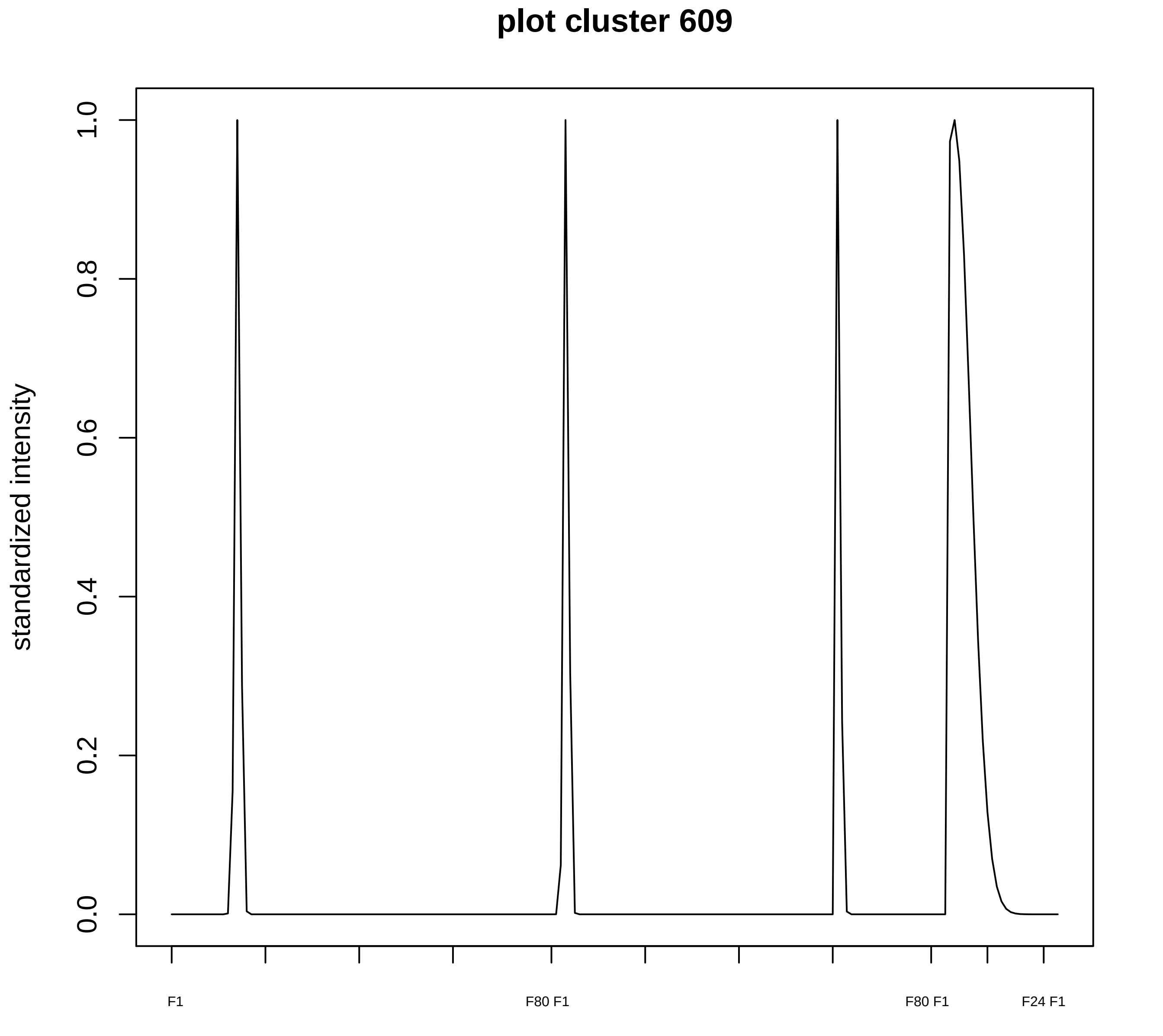
plot cluster 609
standardized intensity
0.0
0.2
0.4
0.6
0.8
1.0
F1
F80 F1
F80 F1
F24 F1

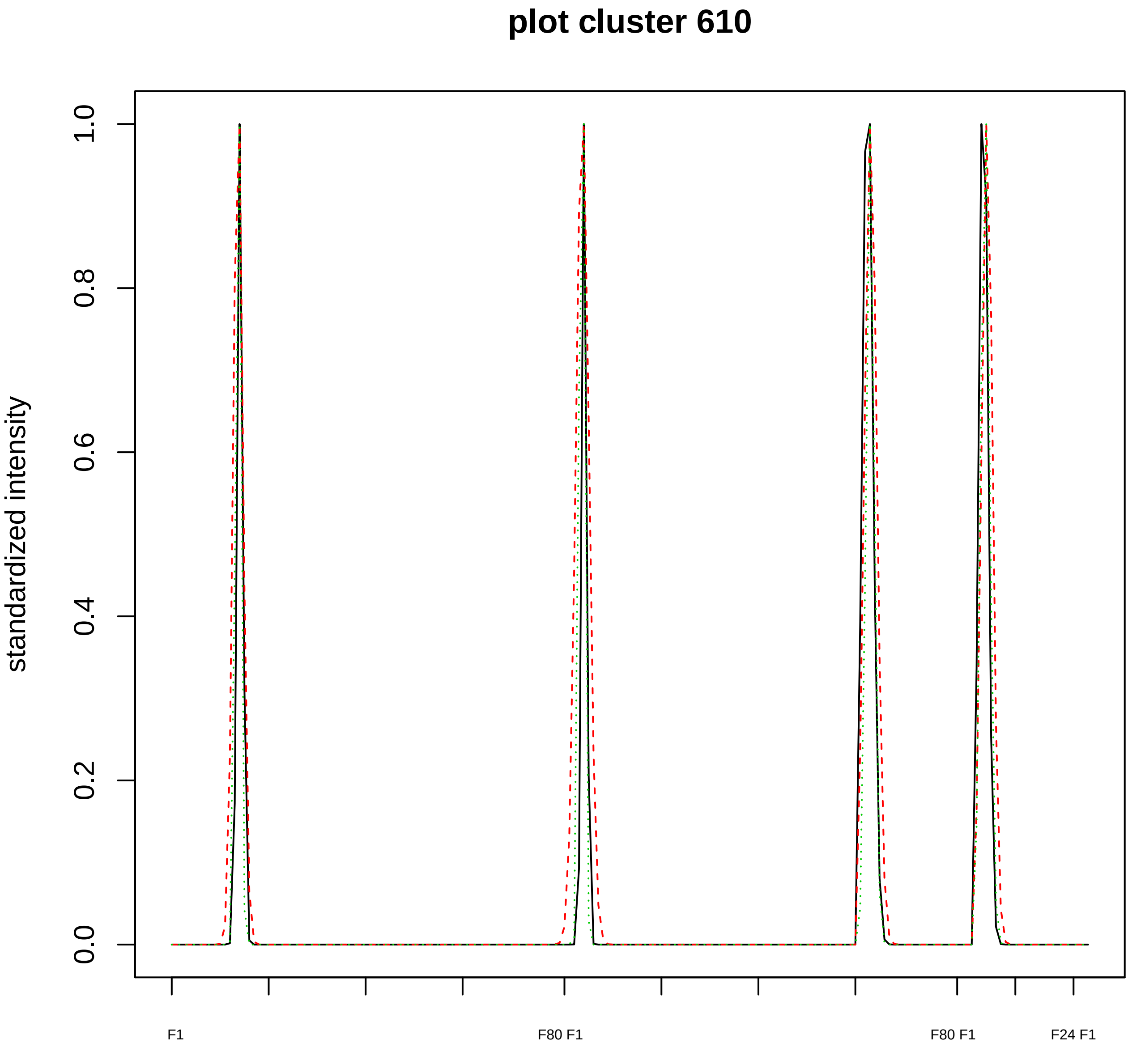
plot cluster 610
standardized intensity
0.0
0.2
0.4
0.6
0.8
1.0
F1
F80 F1
F80 F1
F24 F1

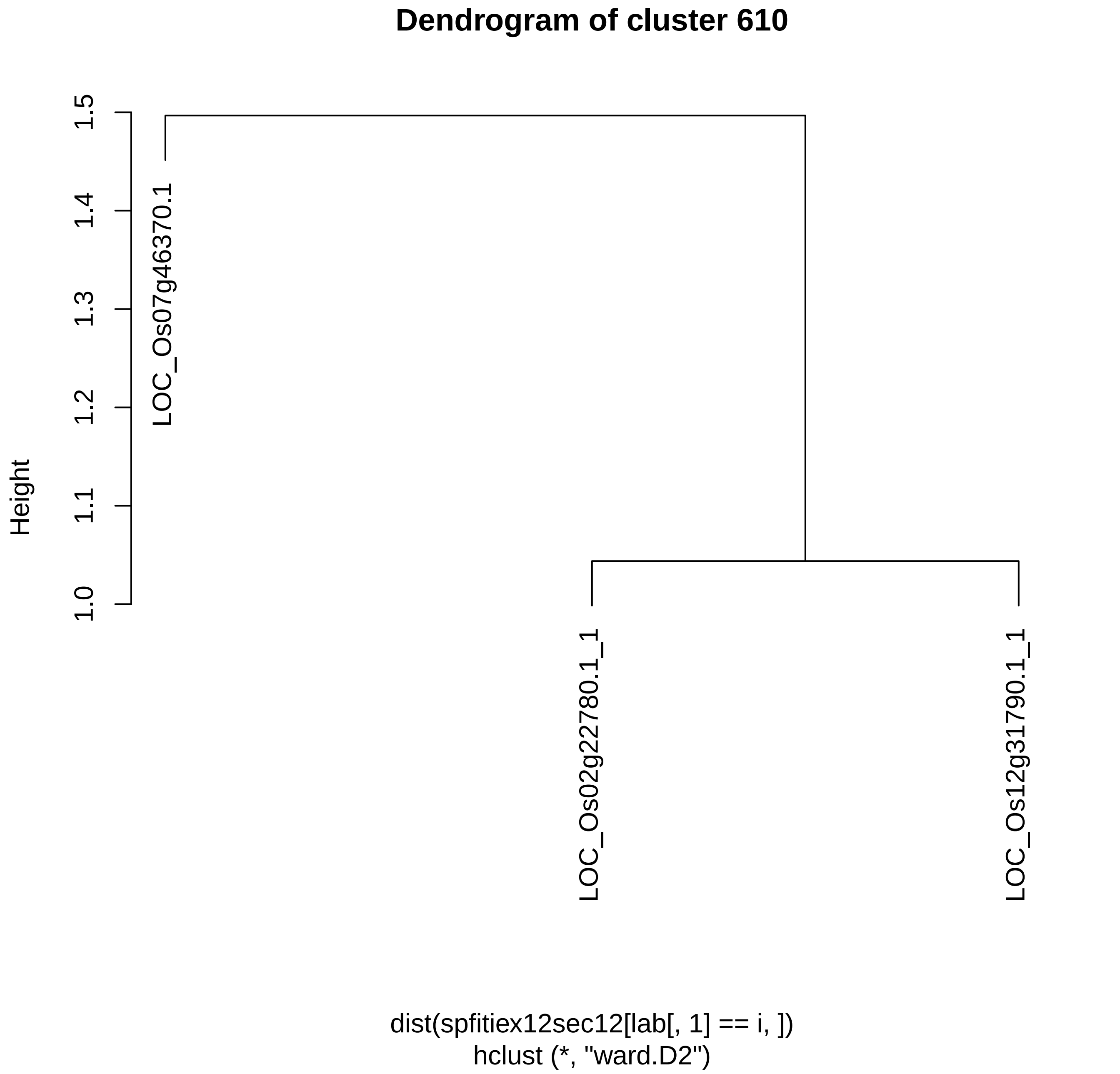
Dendrogram of cluster 610
Height
1.0
1.1
1.2
1.3
1.4
1.5
LOC_Os07g46370.1
LOC_Os02g22780.1_1
LOC_Os12g31790.1_1
dist(spfitiex12sec12[lab[, 1] == i, ])
hclust (*, "ward.D2")

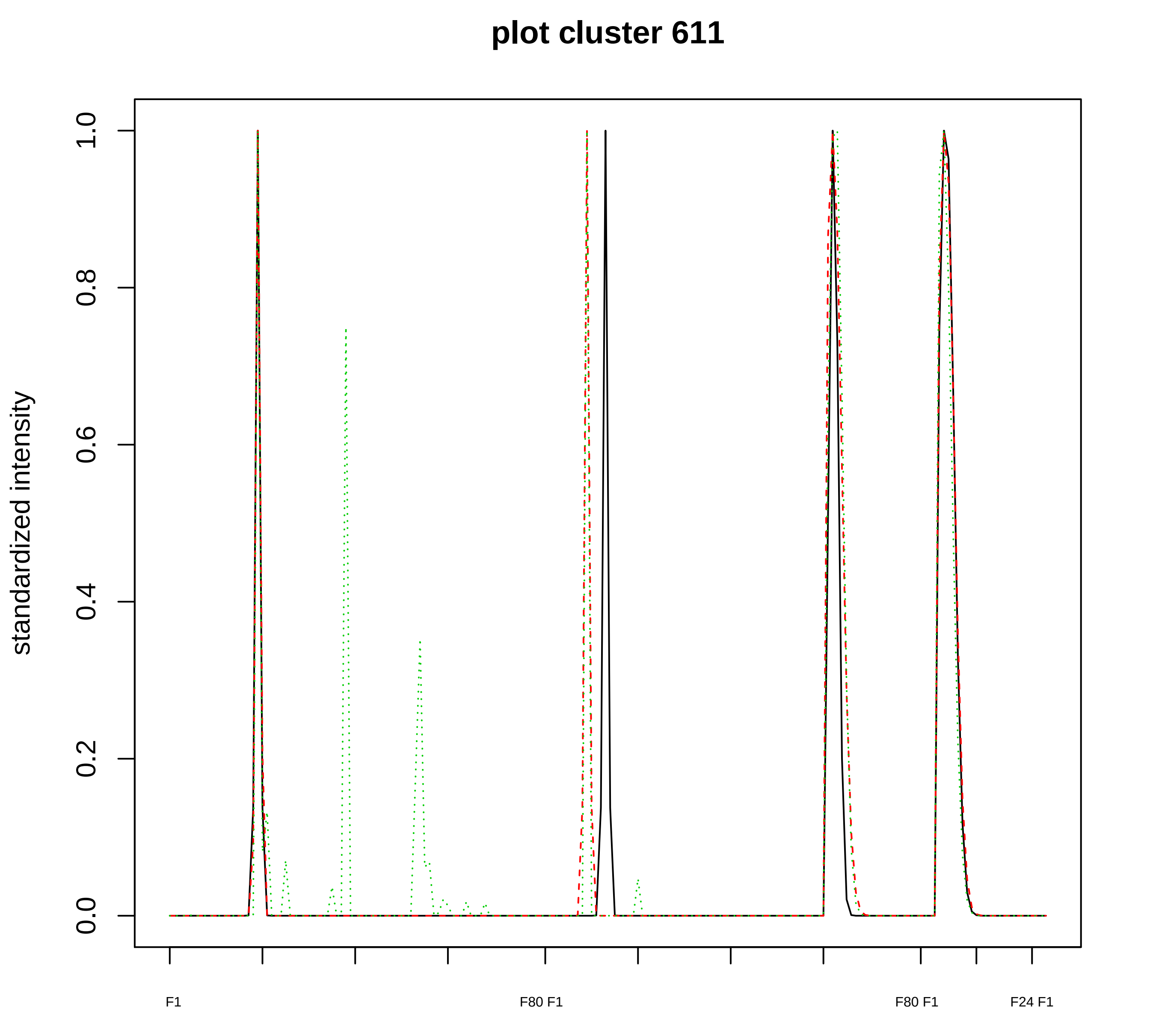
plot cluster 611
standardized intensity
0.0
0.2
0.4
0.6
0.8
1.0
F1
F80 F1
F80 F1
F24 F1

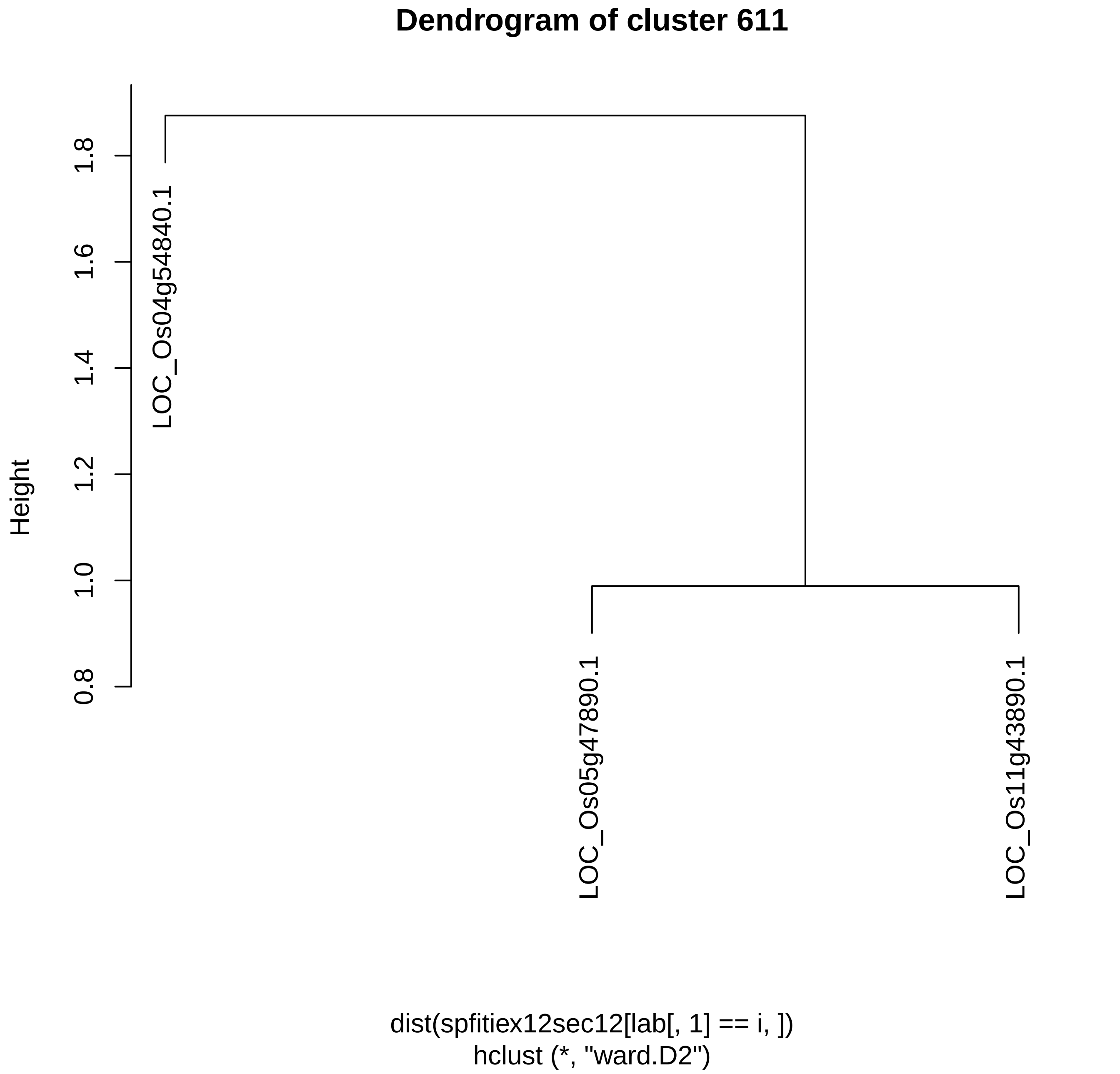
Dendrogram of cluster 611
Height
0.8
1.0
1.2
1.4
1.6
1.8
LOC_Os04g54840.1
LOC_Os05g47890.1
LOC_Os11g43890.1
dist(spfitiex12sec12[lab[, 1] == i, ])
hclust (*, "ward.D2")

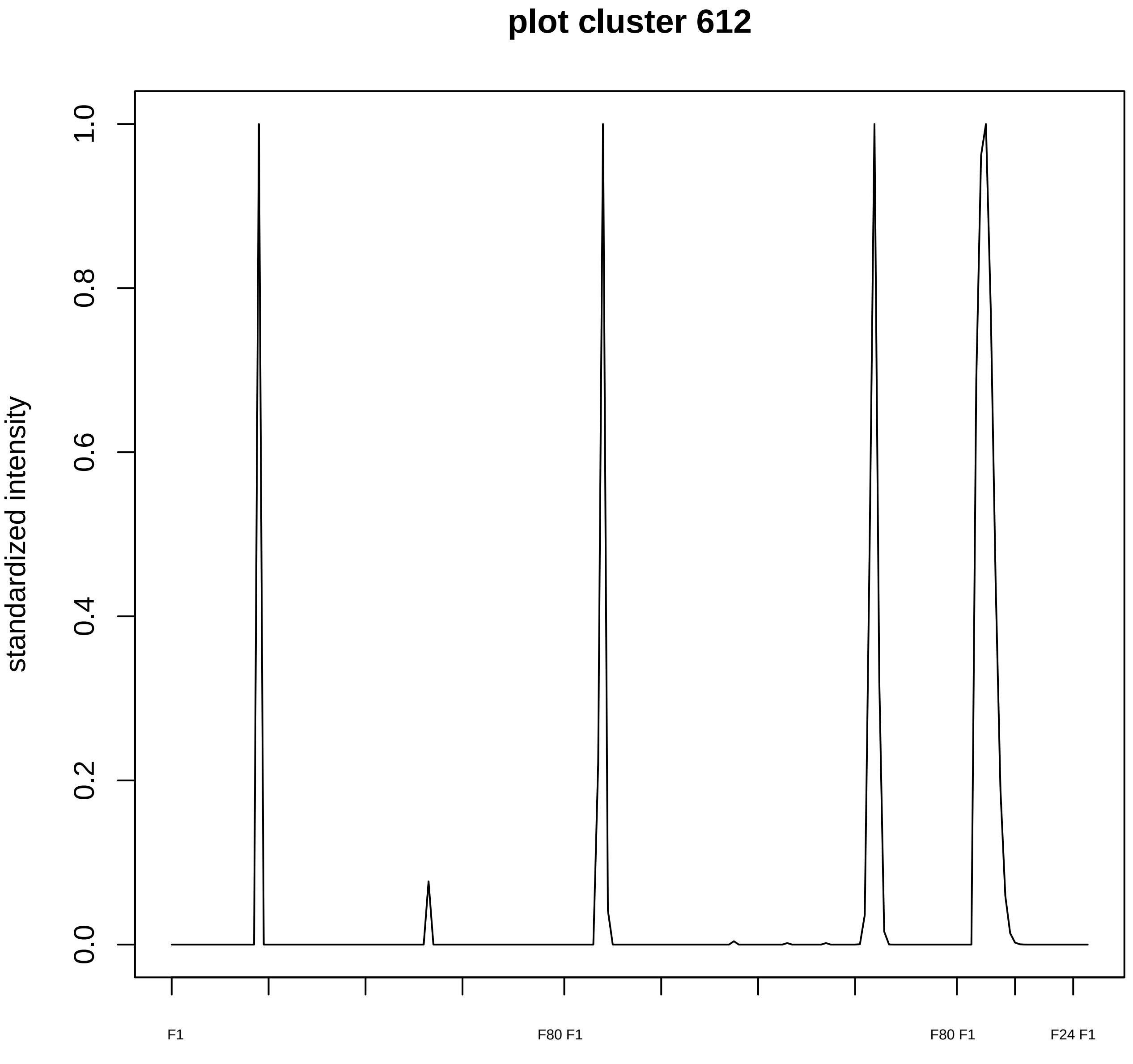

plot cluster 612
standardized intensity
0.0
0.2
0.4
0.6
0.8
1.0
F1
F80 F1
F80 F1
F24 F1

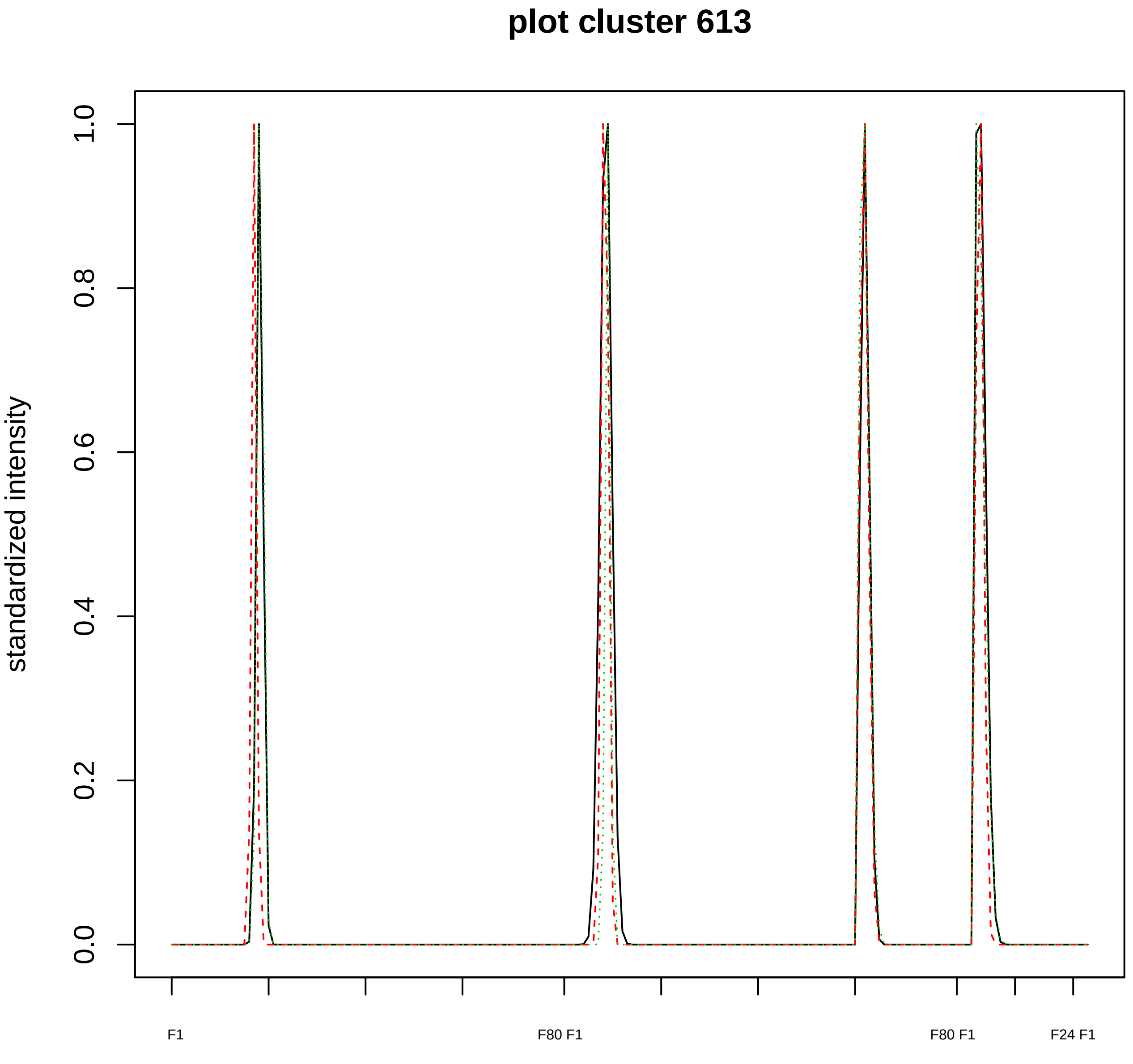

plot cluster 613
standardized intensity
0.0
0.2
0.4
0.6
0.8
1.0
F1
F80 F1
F80 F1
F24 F1

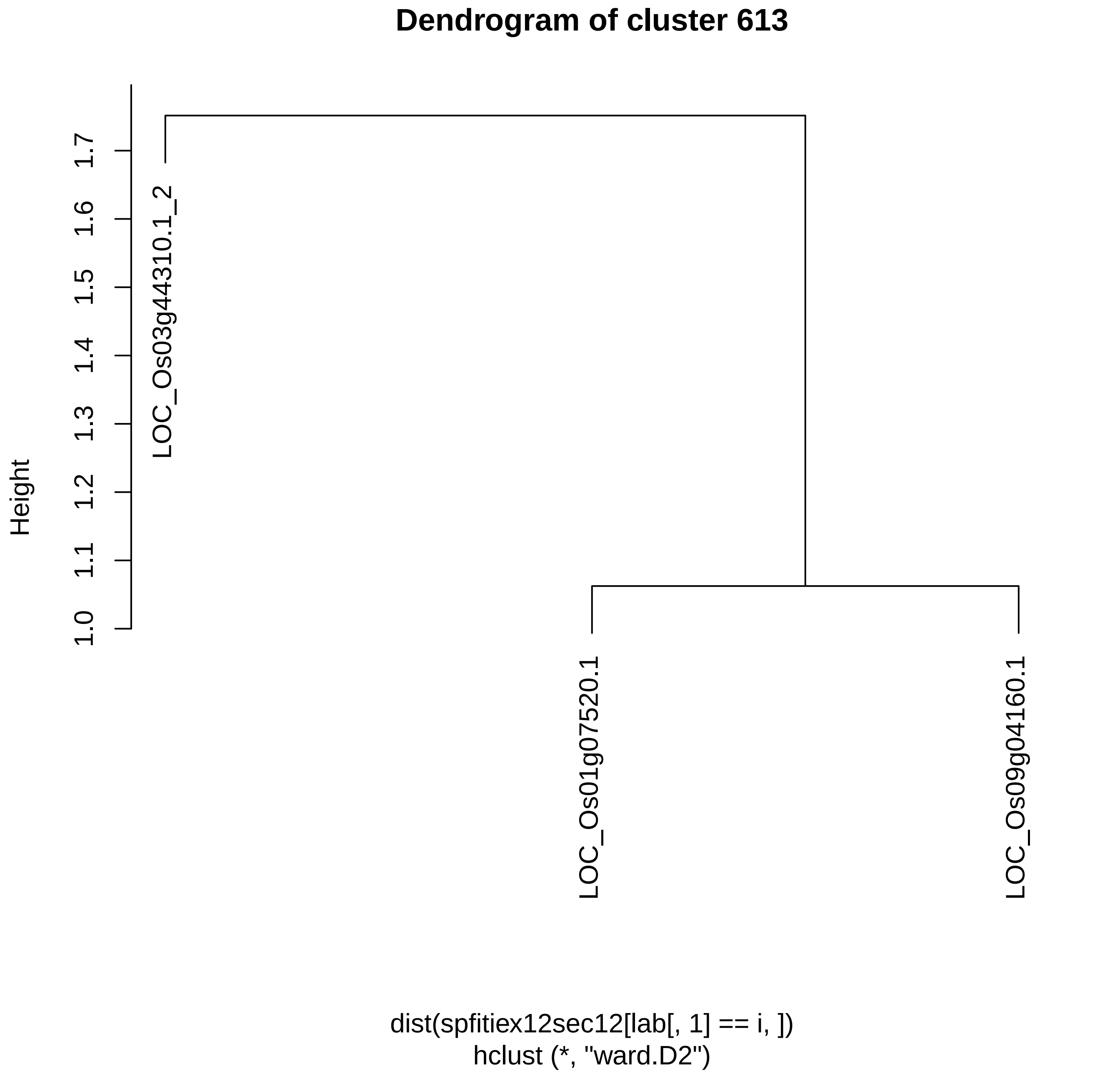
Dendrogram of cluster 613
Height
1.0
1.1
1.2
1.3
1.4
1.5
1.6
1.7
LOC_Os03g44310.1_2
LOC_Os01g07520.1
LOC_Os09g04160.1
dist(spfitiex12sec12[lab[, 1] == i, ])
hclust (*, "ward.D2")

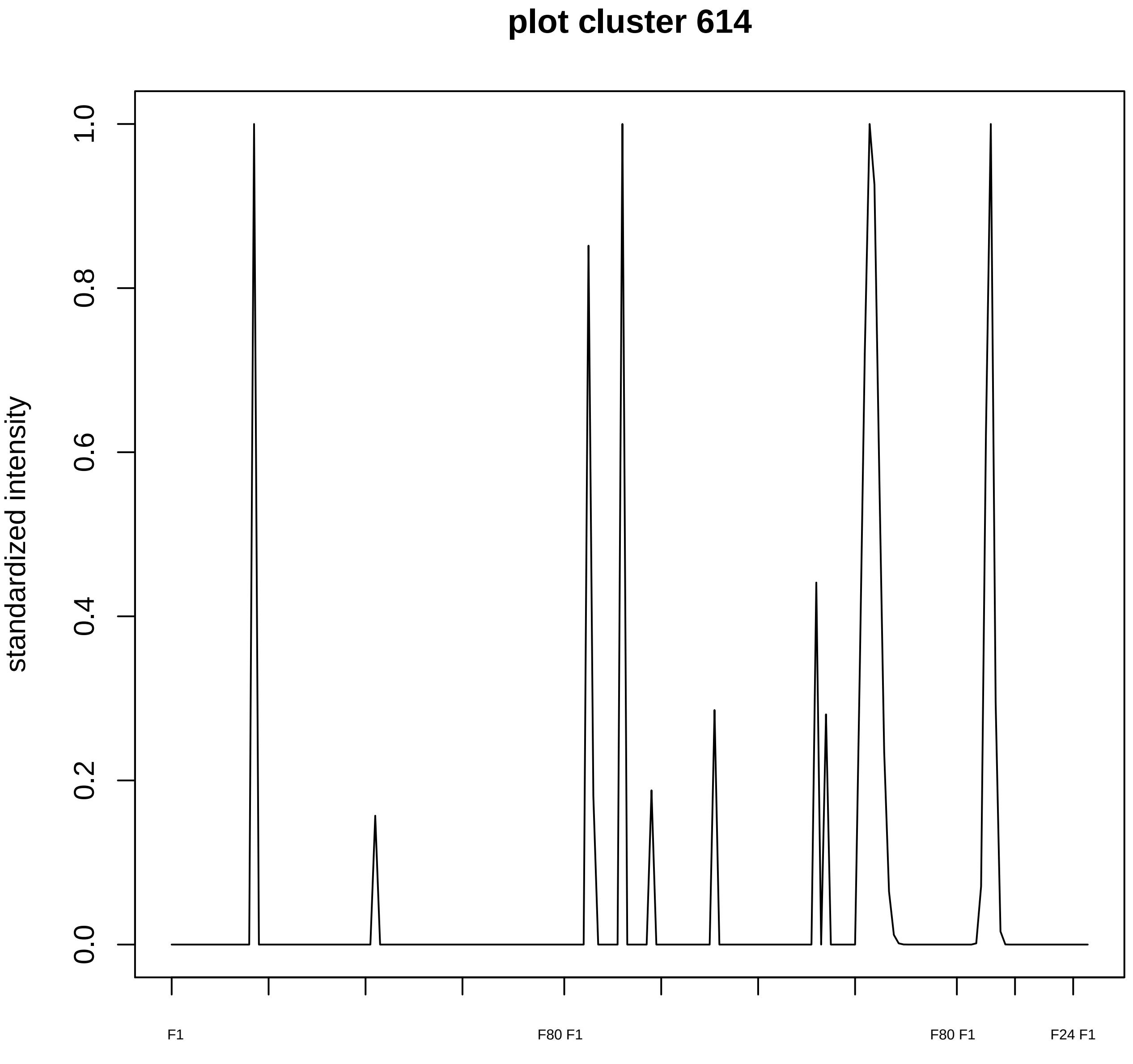

plot cluster 614
standardized intensity
0.0
0.2
0.4
0.6
0.8
1.0
F1
F80 F1
F80 F1
F24 F1

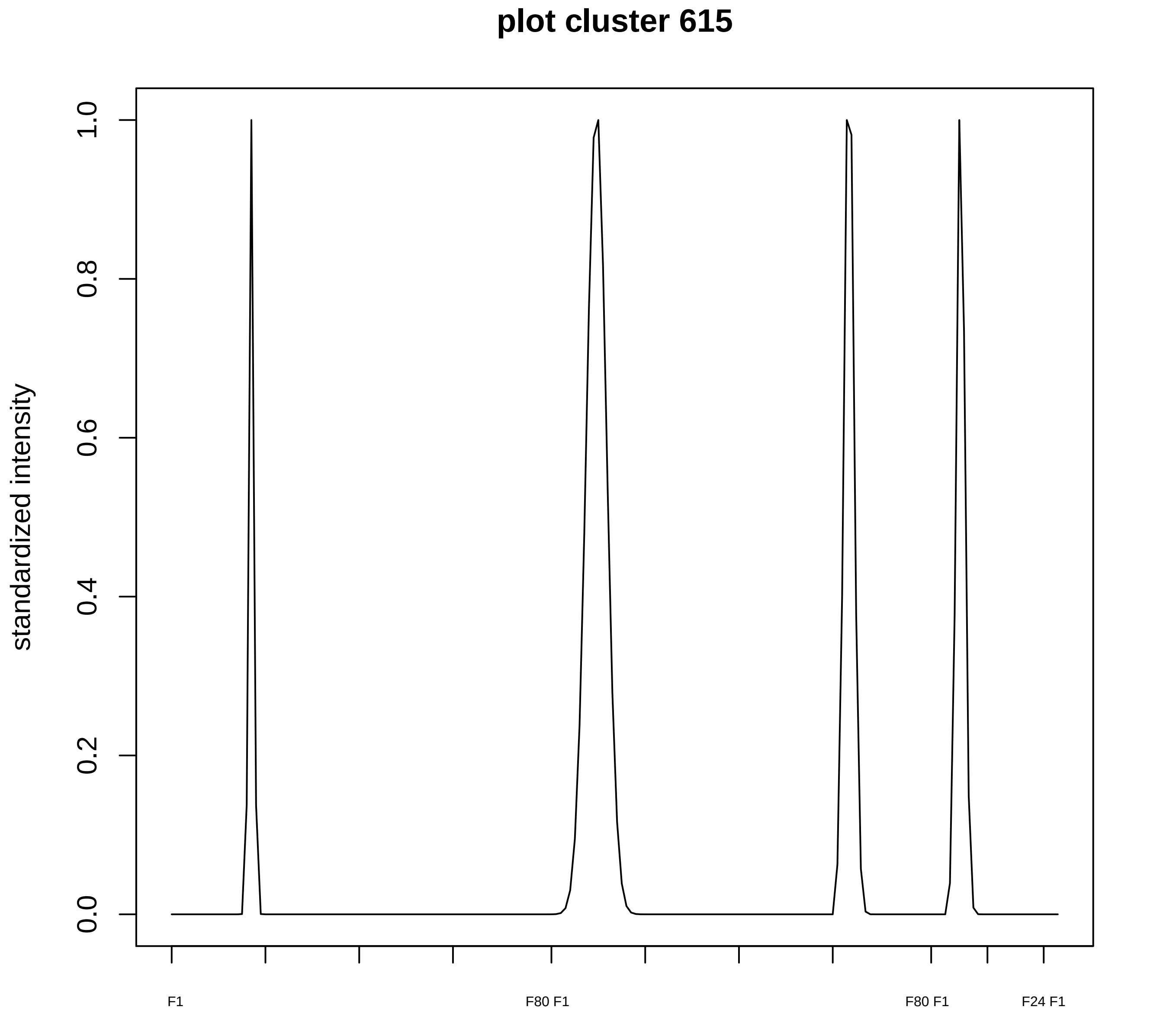
plot cluster 615
standardized intensity
0.0
0.2
0.4
0.6
0.8
1.0
F1
F80 F1
F80 F1
F24 F1

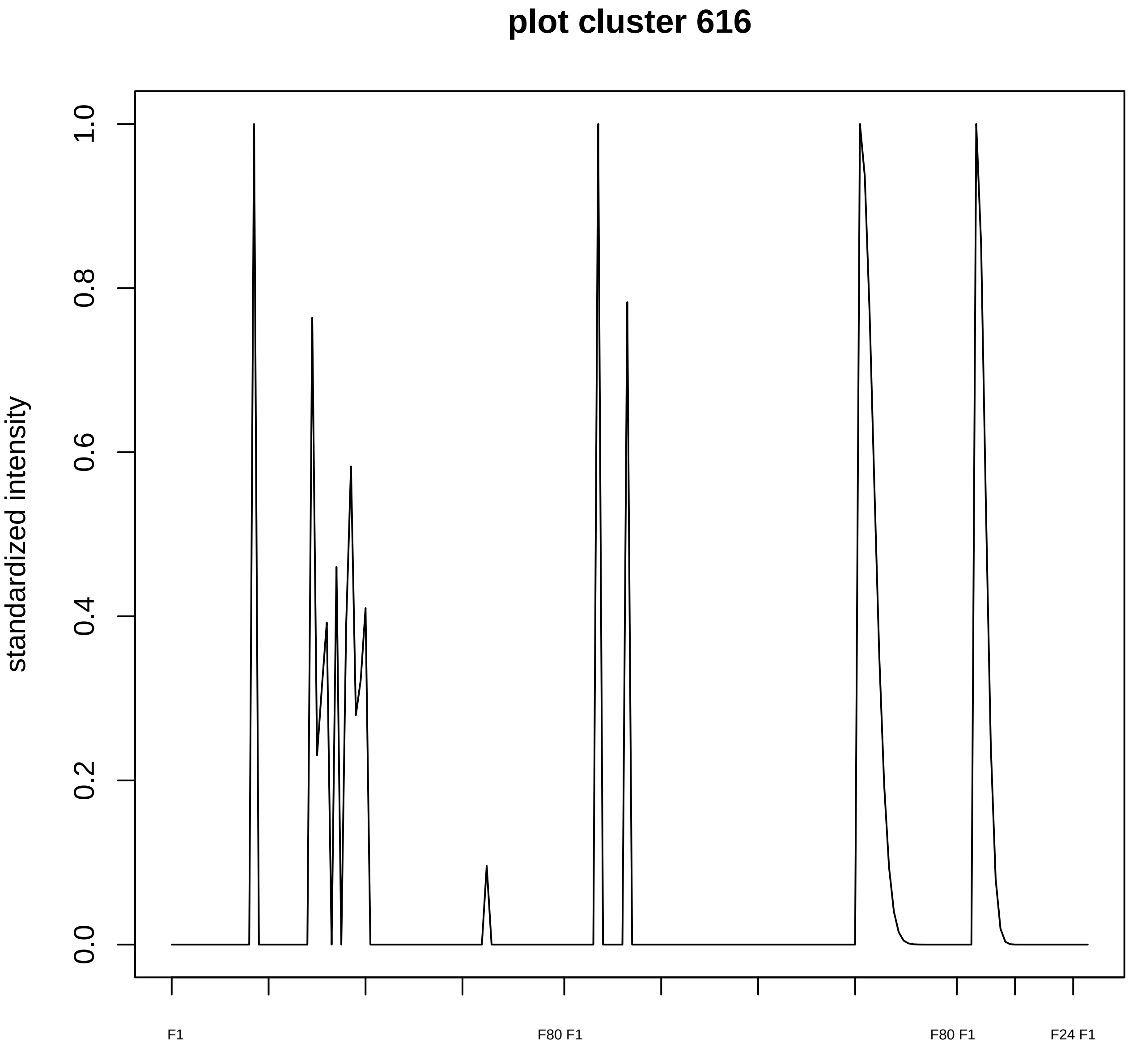
plot cluster 616
standardized intensity
0.0
0.2
0.4
0.6
0.8
1.0
F1
F80 F1
F80 F1
F24 F1

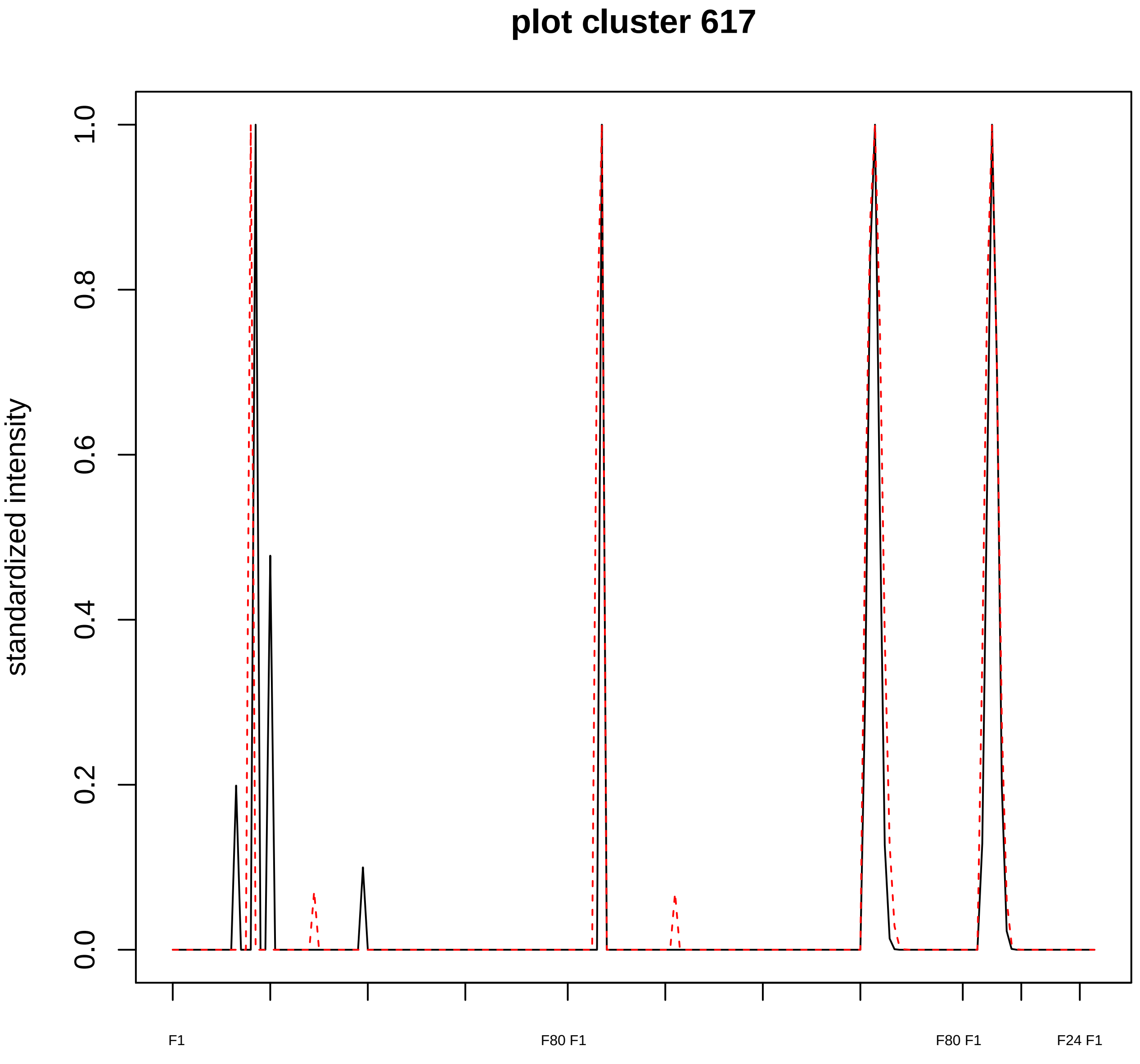
plot cluster 617
standardized intensity
0.0
0.2
0.4
0.6
0.8
1.0
F1
F80 F1
F80 F1
F24 F1

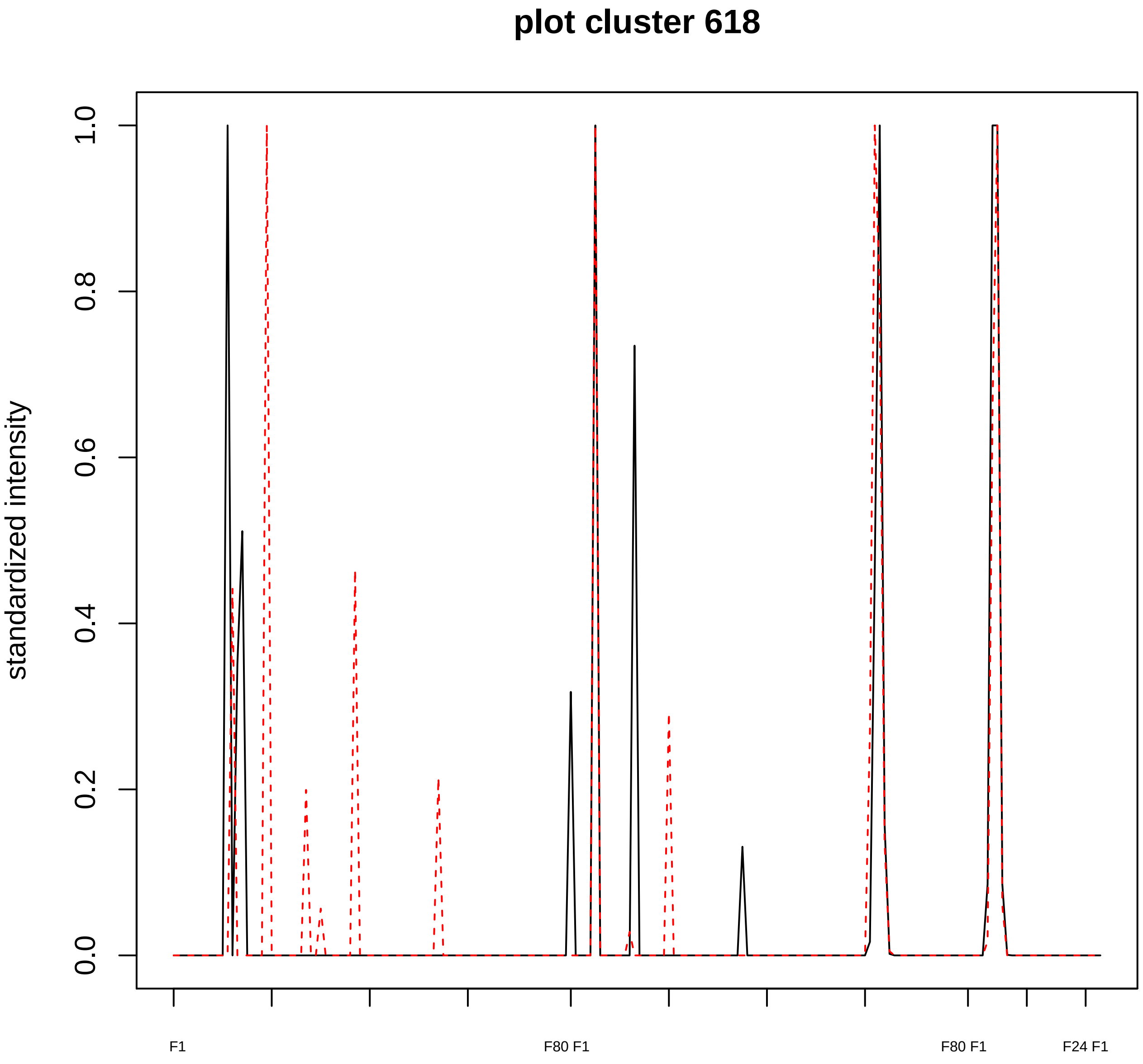
plot cluster 618
standardized intensity
0.0
0.2
0.4
0.6
0.8
1.0
F1
F80 F1
F80 F1
F24 F1

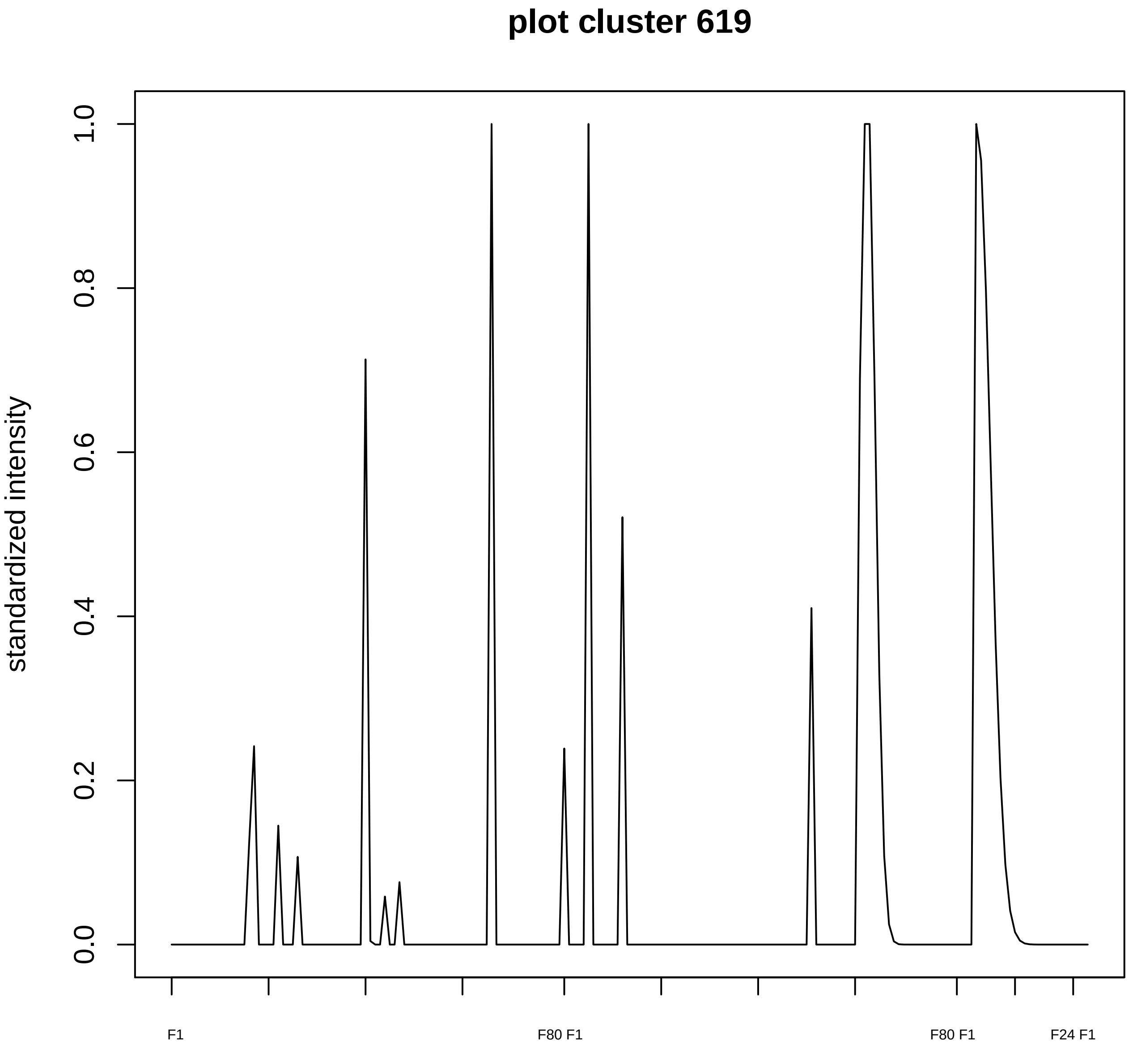
plot cluster 619
standardized intensity
0.0
0.2
0.4
0.6
0.8
1.0
F1
F80 F1
F80 F1
F24 F1

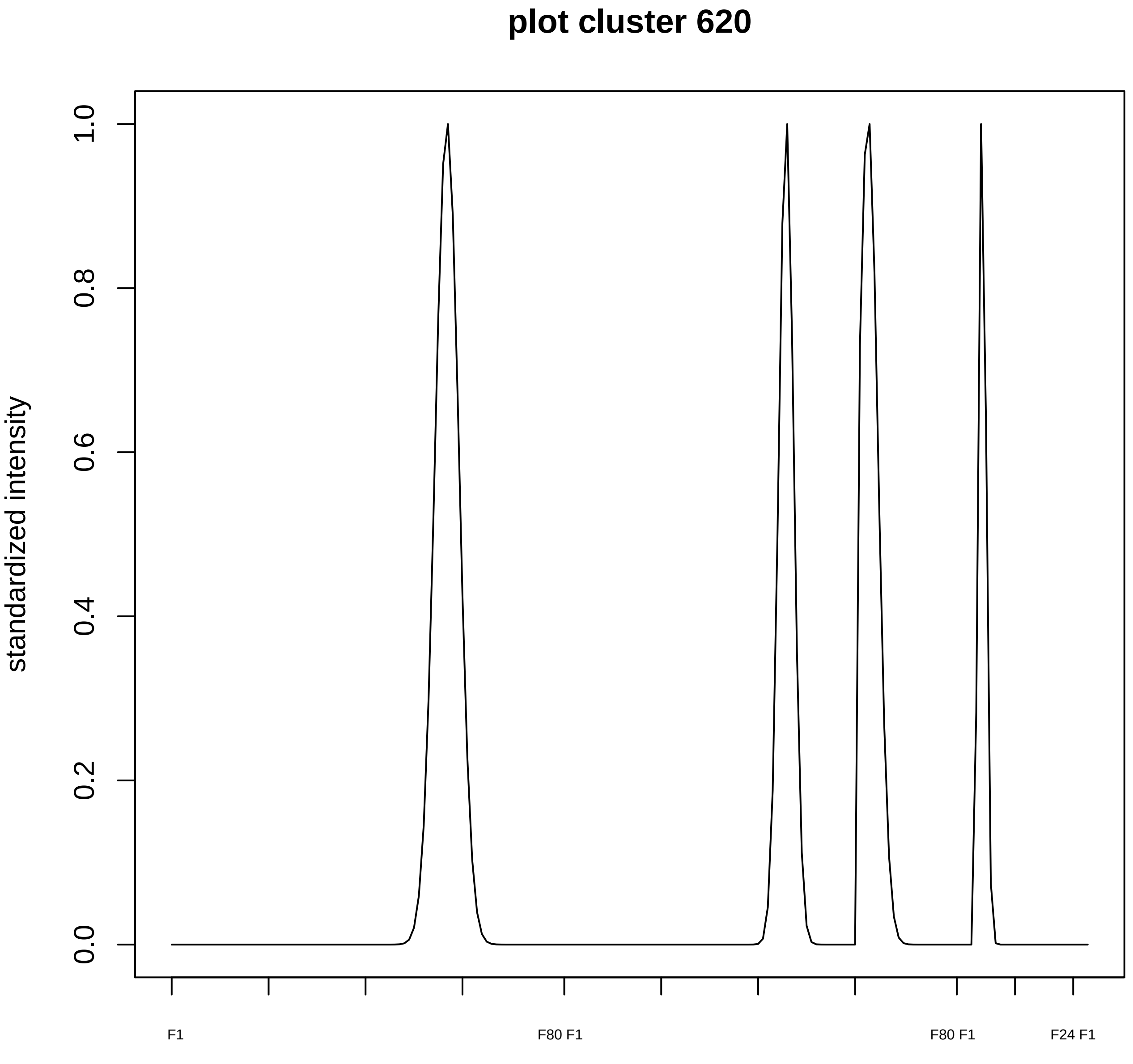

plot cluster 620
standardized intensity
0.0
0.2
0.4
0.6
0.8
1.0
F1
F80 F1
F80 F1
F24 F1

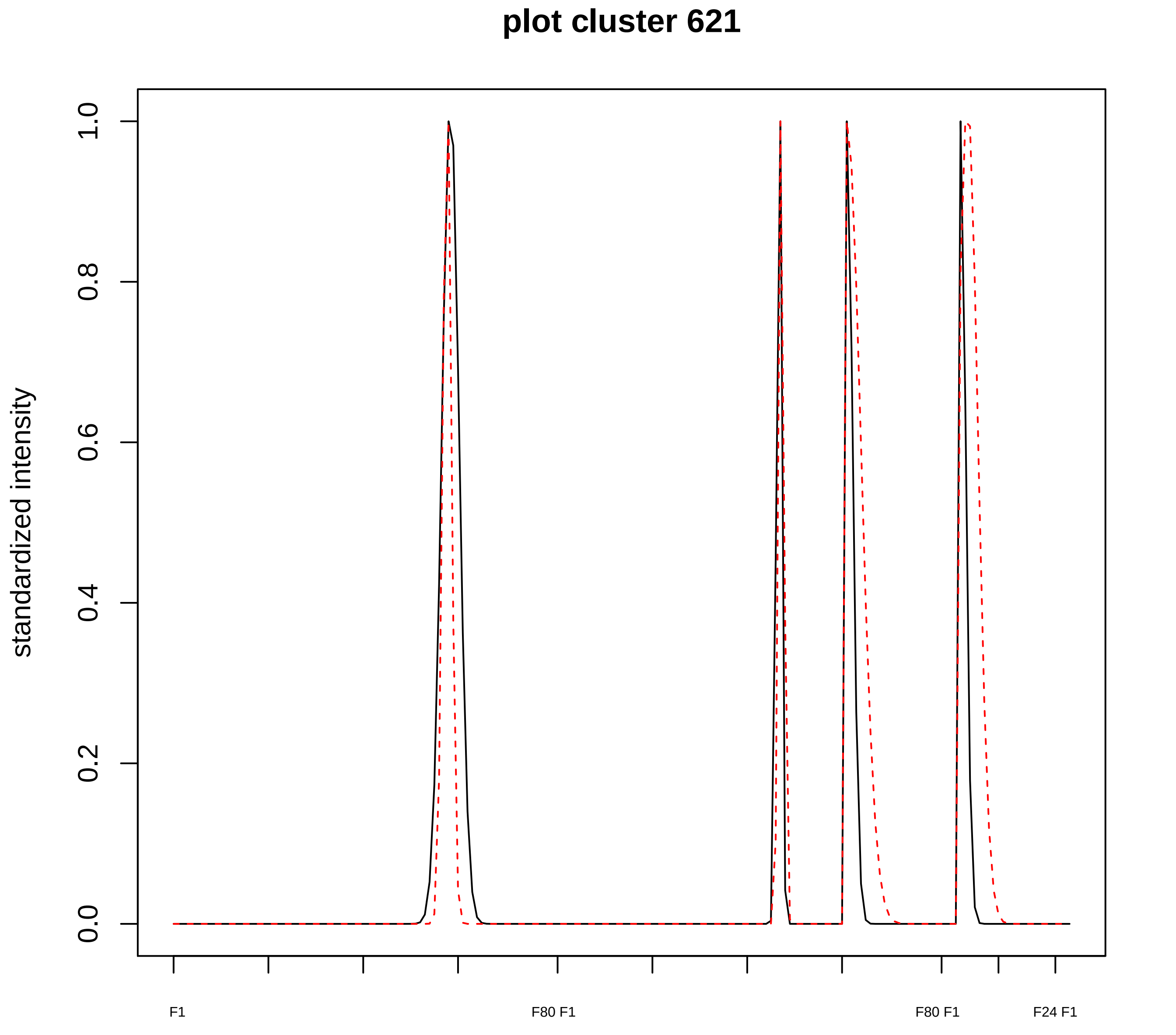
plot cluster 621
standardized intensity
0.0
0.2
0.4
0.6
0.8
1.0
F1
F80 F1
F80 F1
F24 F1

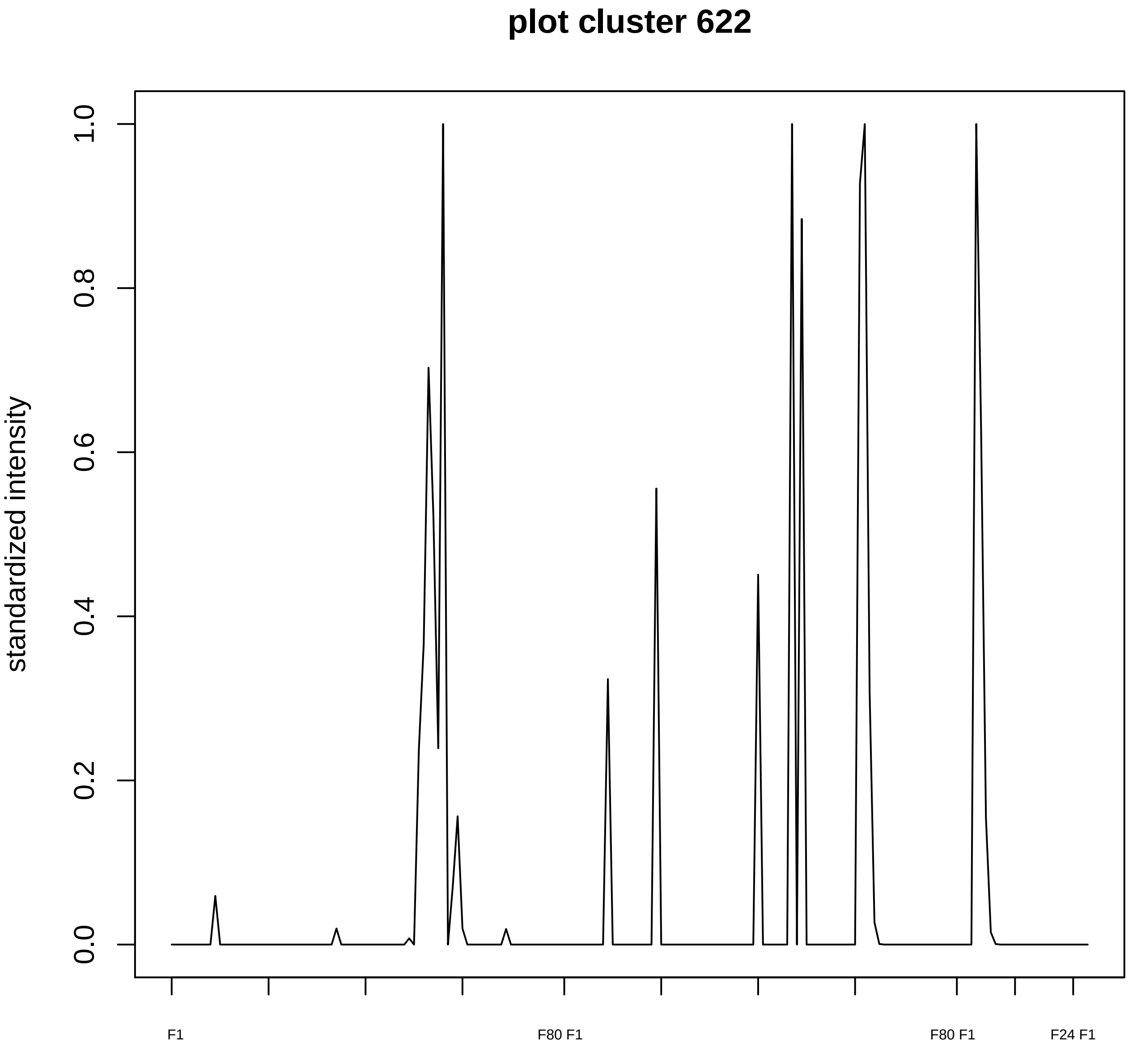
plot cluster 622
standardized intensity
0.0
0.2
0.4
0.6
0.8
1.0
F1
F80 F1
F80 F1
F24 F1

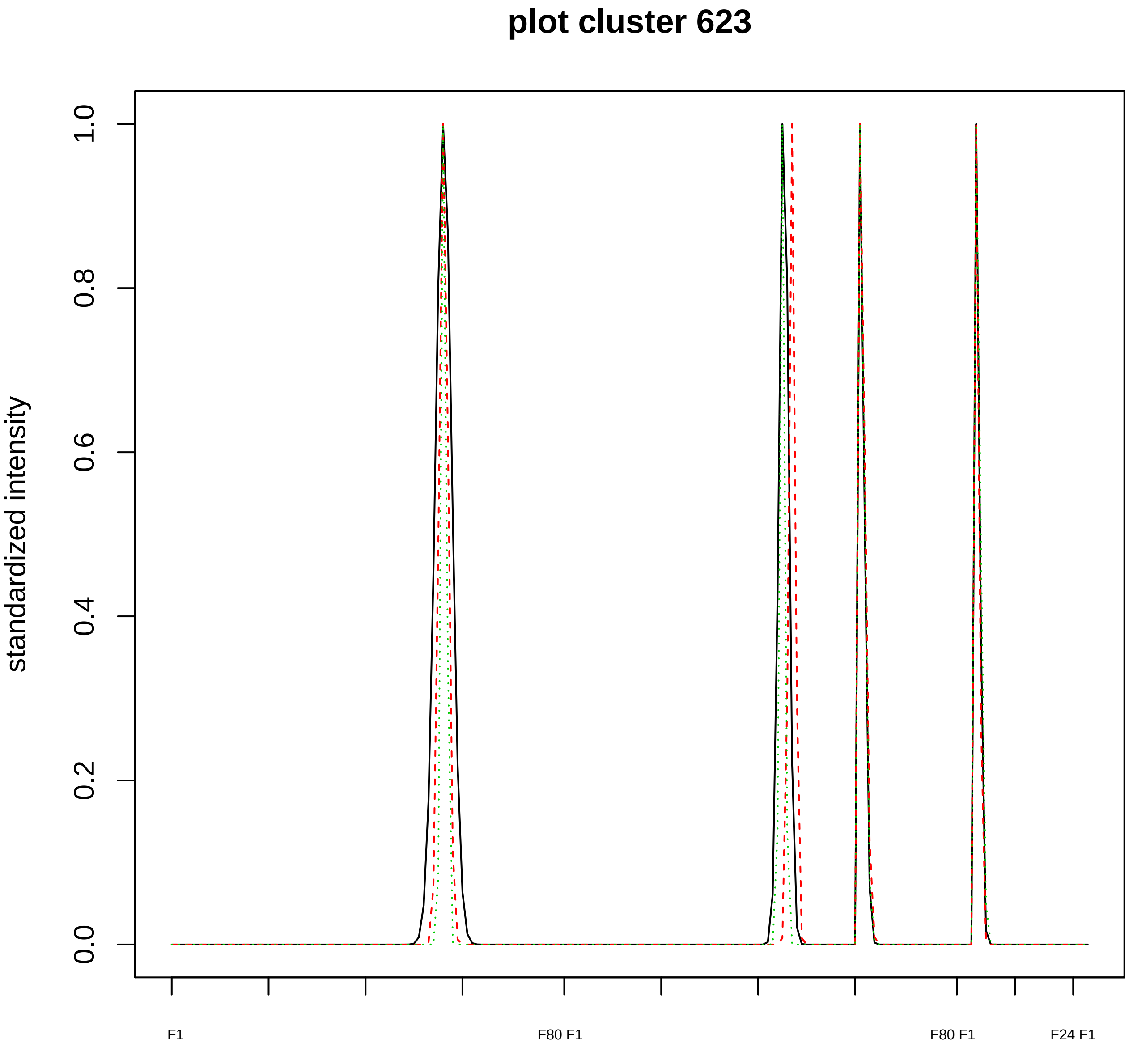

plot cluster 623
standardized intensity
0.0
0.2
0.4
0.6
0.8
1.0
F1
F80 F1
F80 F1
F24 F1

### Dendrogram of cluster 623

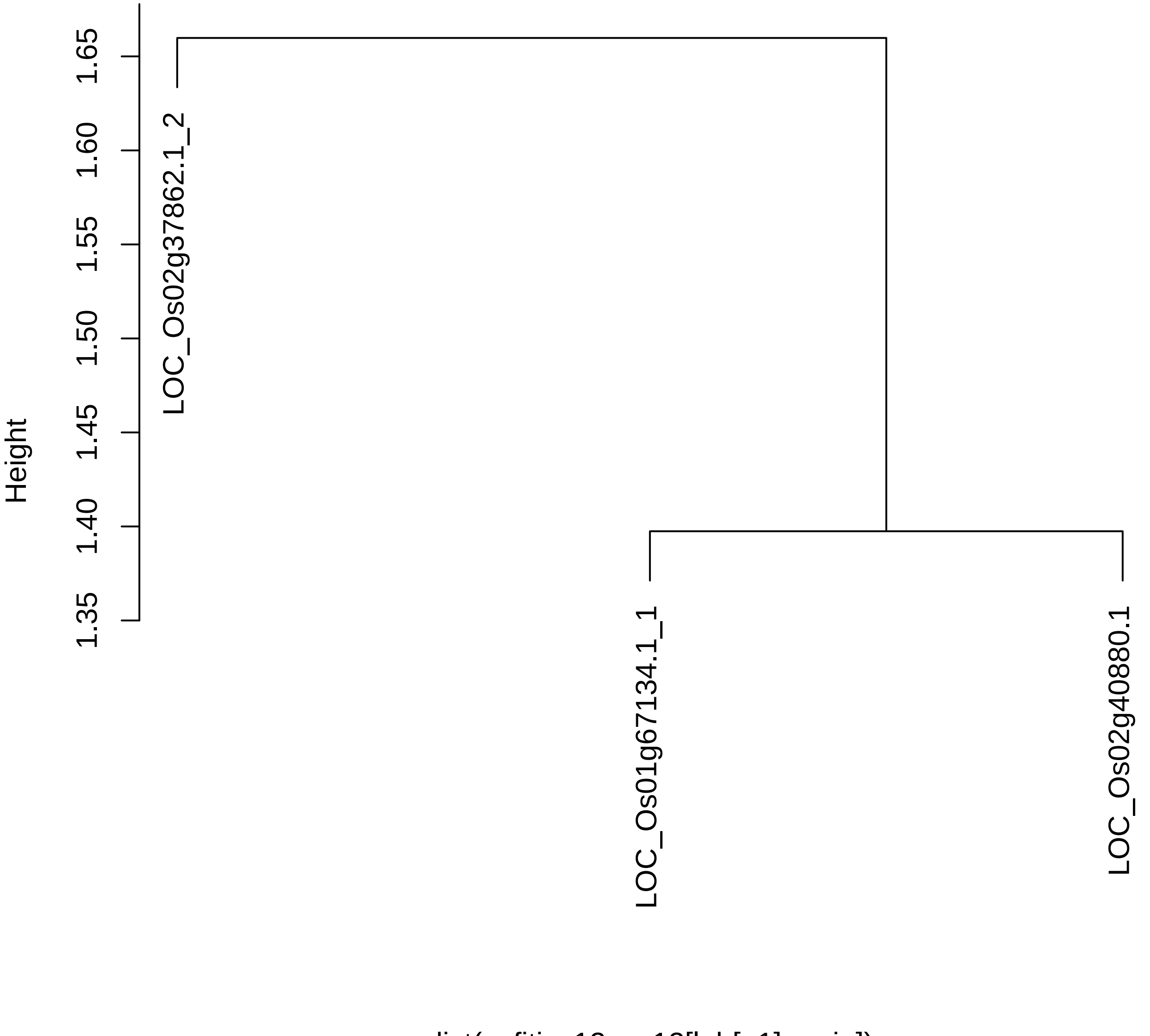


dist(spfitiex12sec12[lab[, 1] == i, ])
hclust (*, "ward.D2")

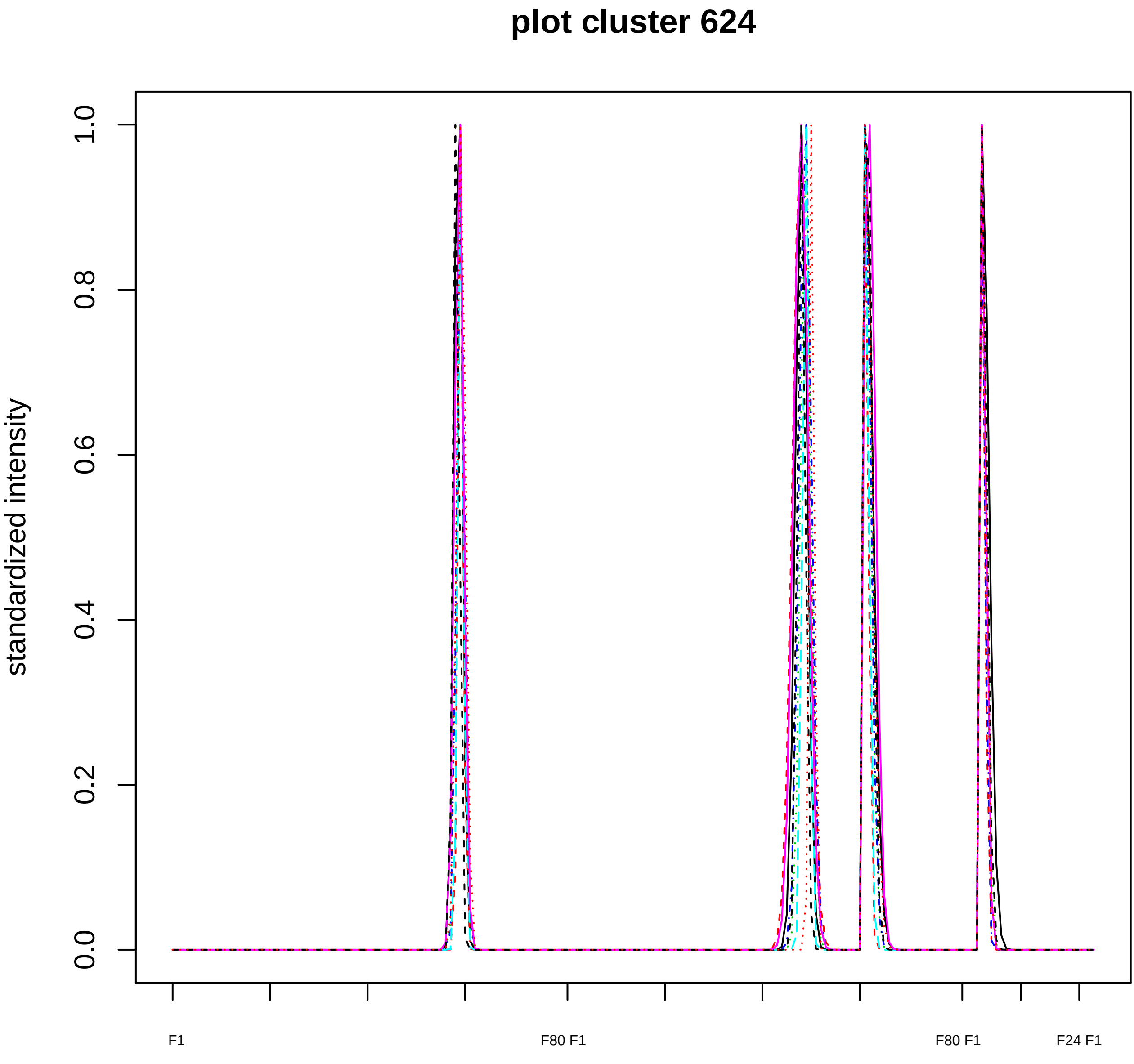

plot cluster 624
standardized intensity
1.0
0.8
0.6
0.4
0.2
0.0
F1
F80 F1
F80 F1
F24 F1

### Dendrogram of cluster 624

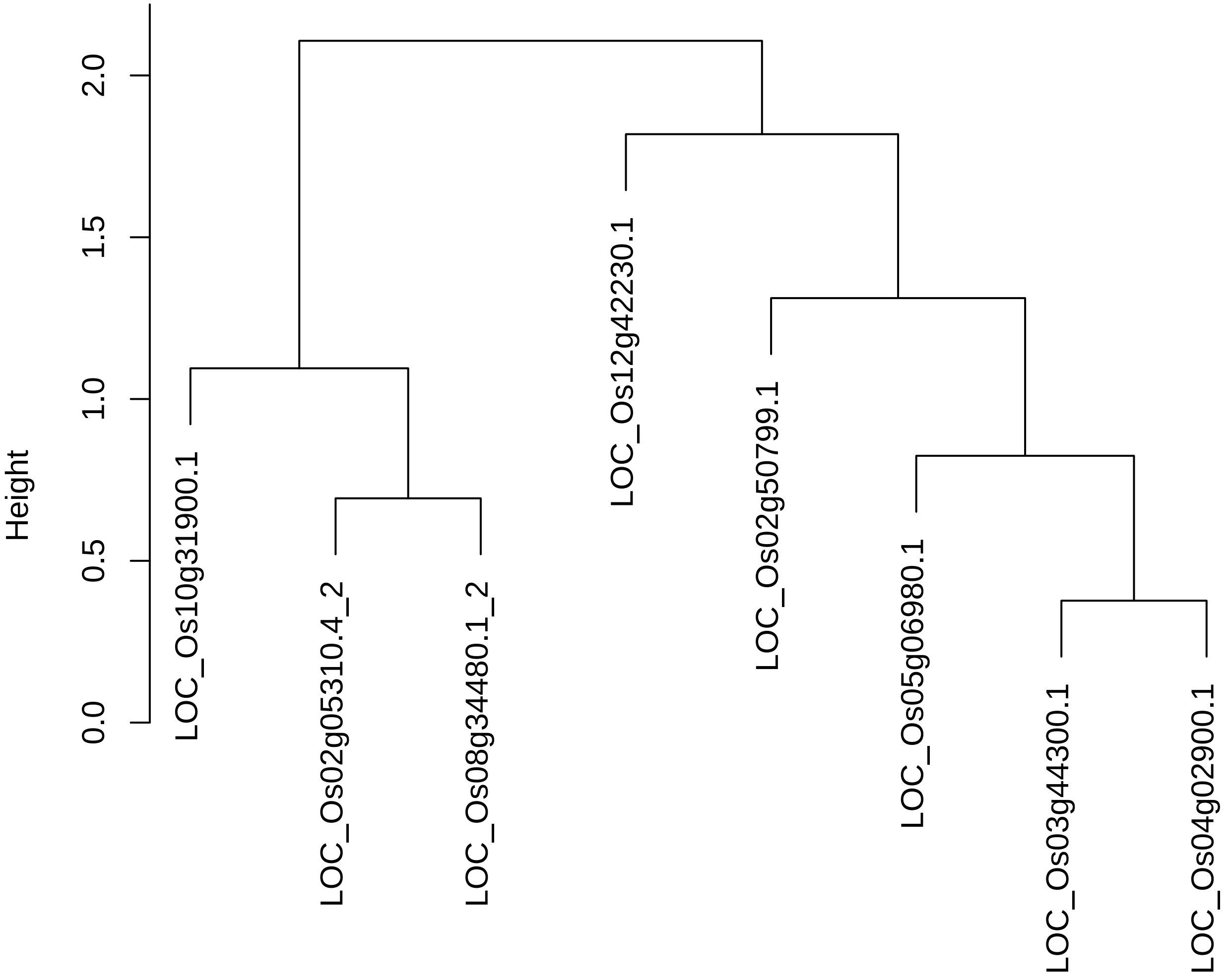


dist(spfitiex12sec12[lab[, 1] == i, ])
hclust (*, "ward.D2")

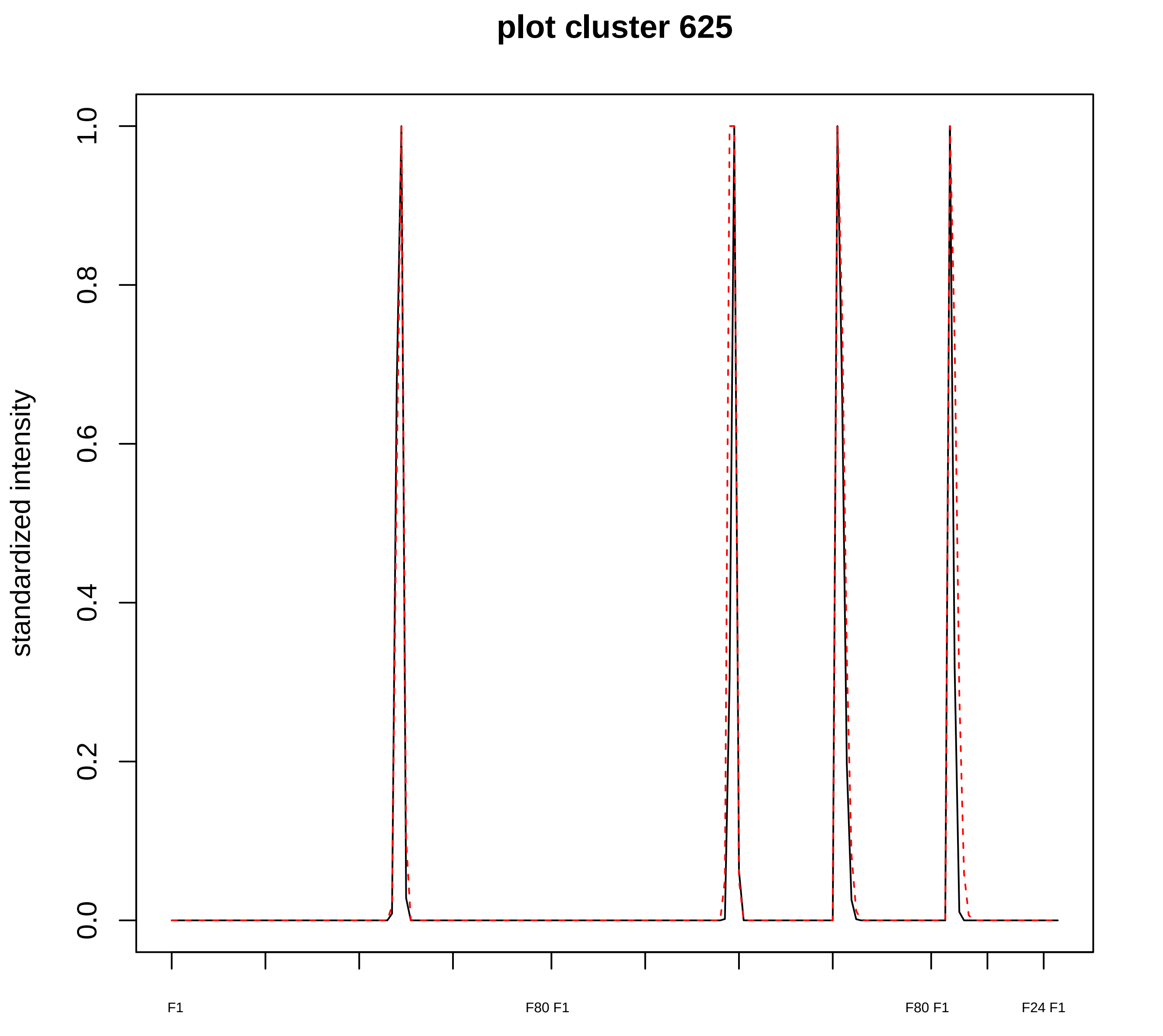
plot cluster 625
standardized intensity
0.0
0.2
0.4
0.6
0.8
1.0
F1
F80 F1
F80 F1
F24 F1

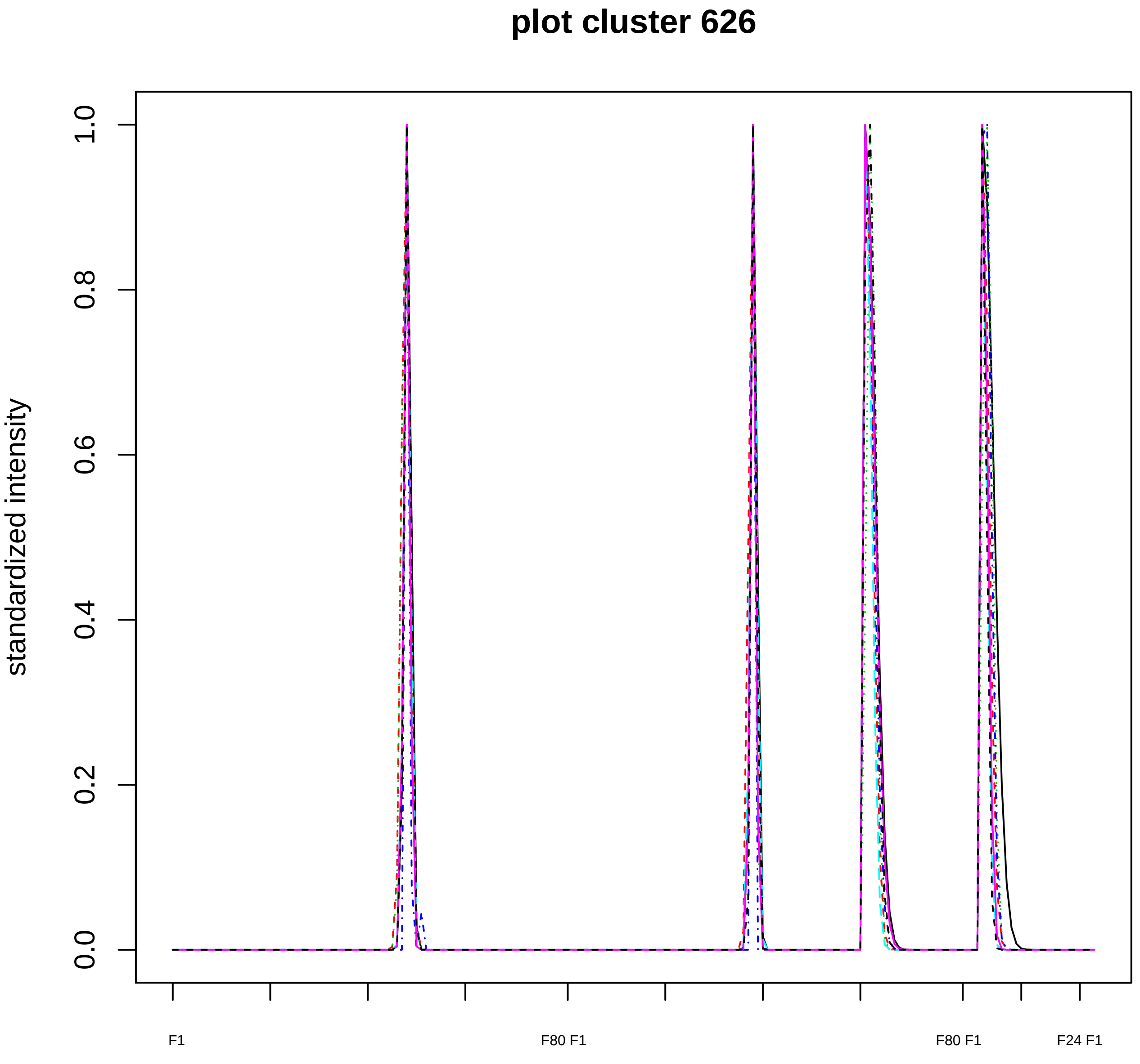
plot cluster 626
standardized intensity
0.0
0.2
0.4
0.6
0.8
1.0
F1
F80 F1
F80 F1
F24 F1

### Dendrogram of cluster 626

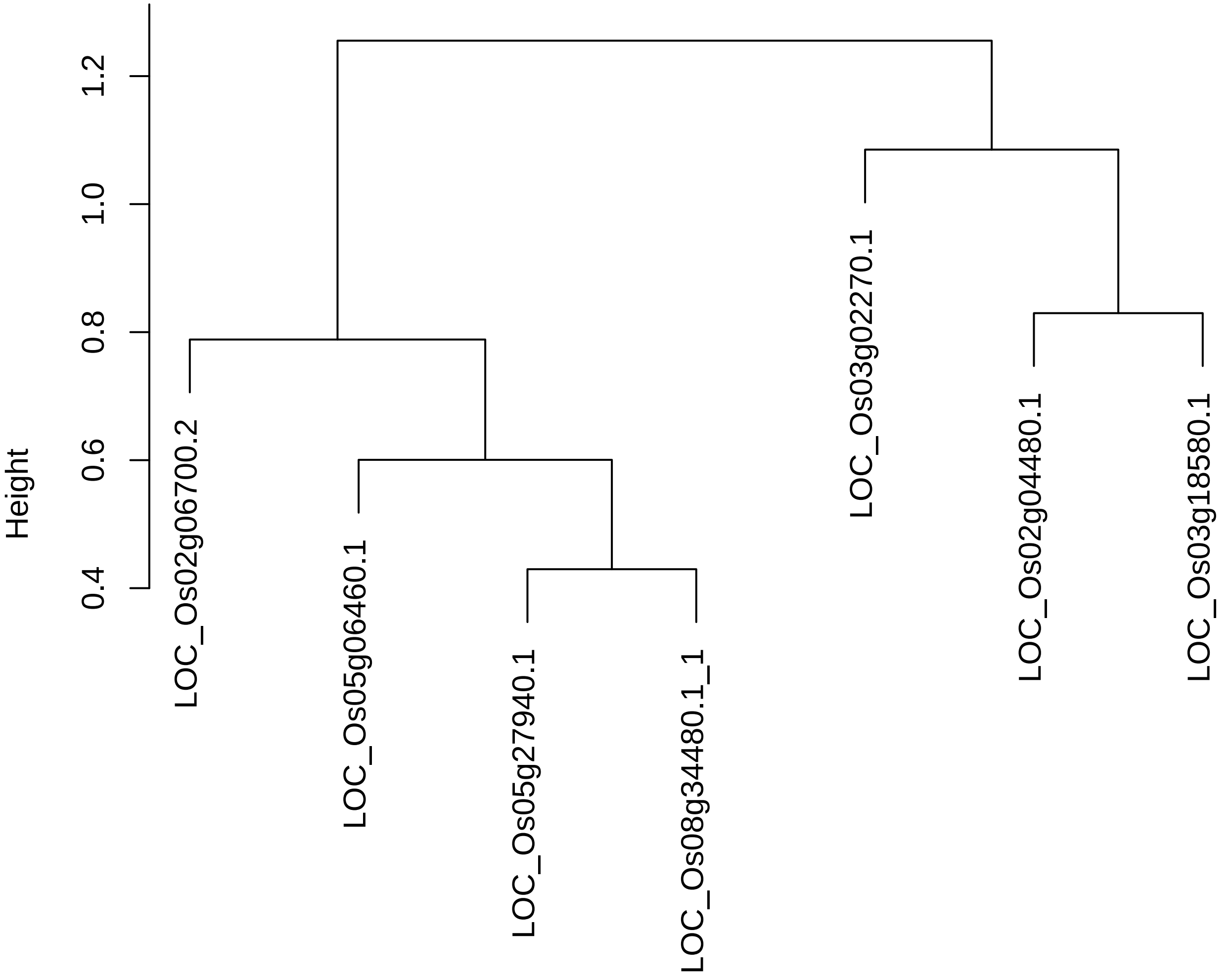


dist(spfitiex12sec12[lab[, 1] == i, ])
hclust (*, "ward.D2")

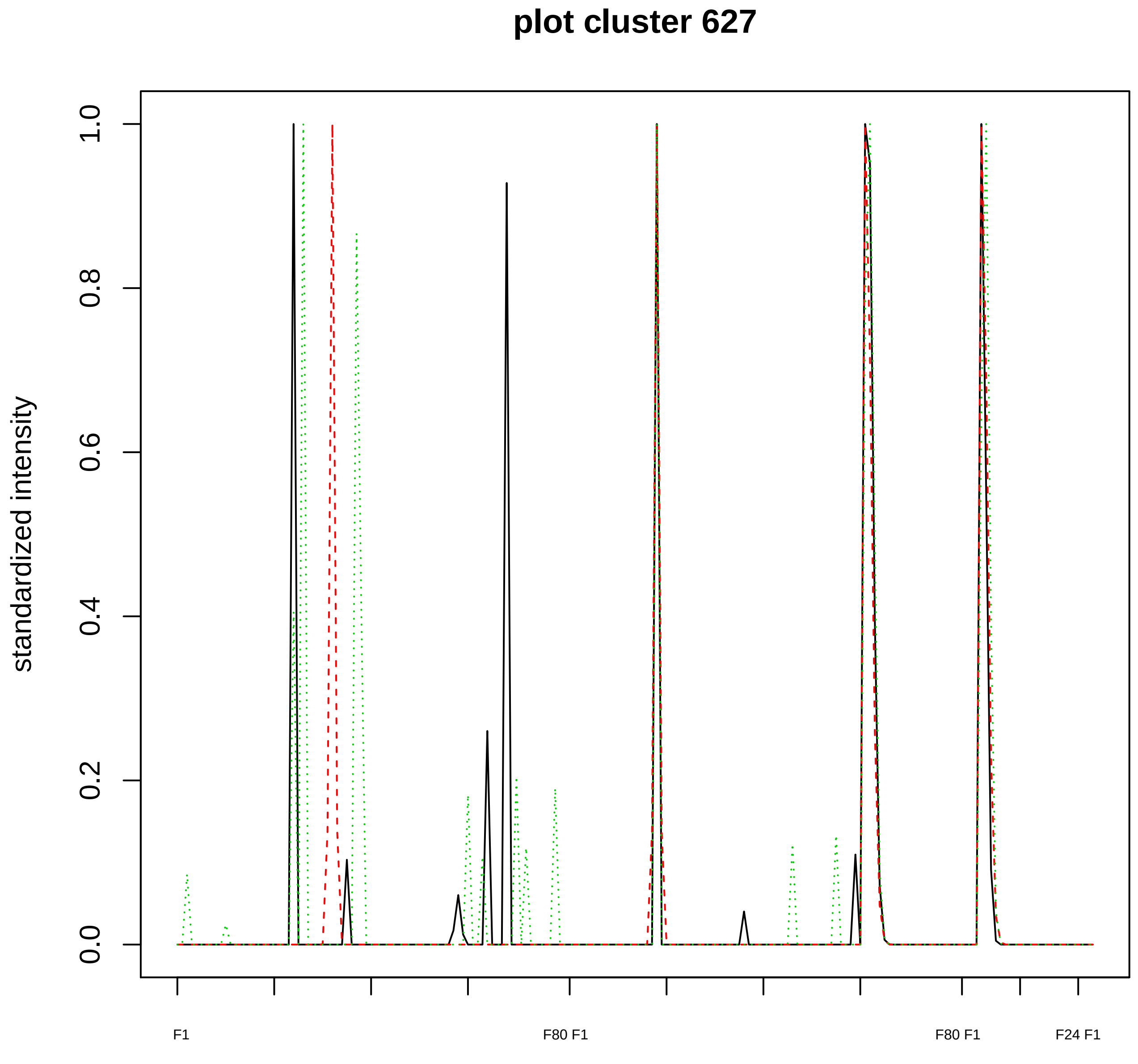
plot cluster 627
standardized intensity
0.0
0.2
0.4
0.6
0.8
1.0
F1
F80 F1
F80 F1
F24 F1

### Dendrogram of cluster 627

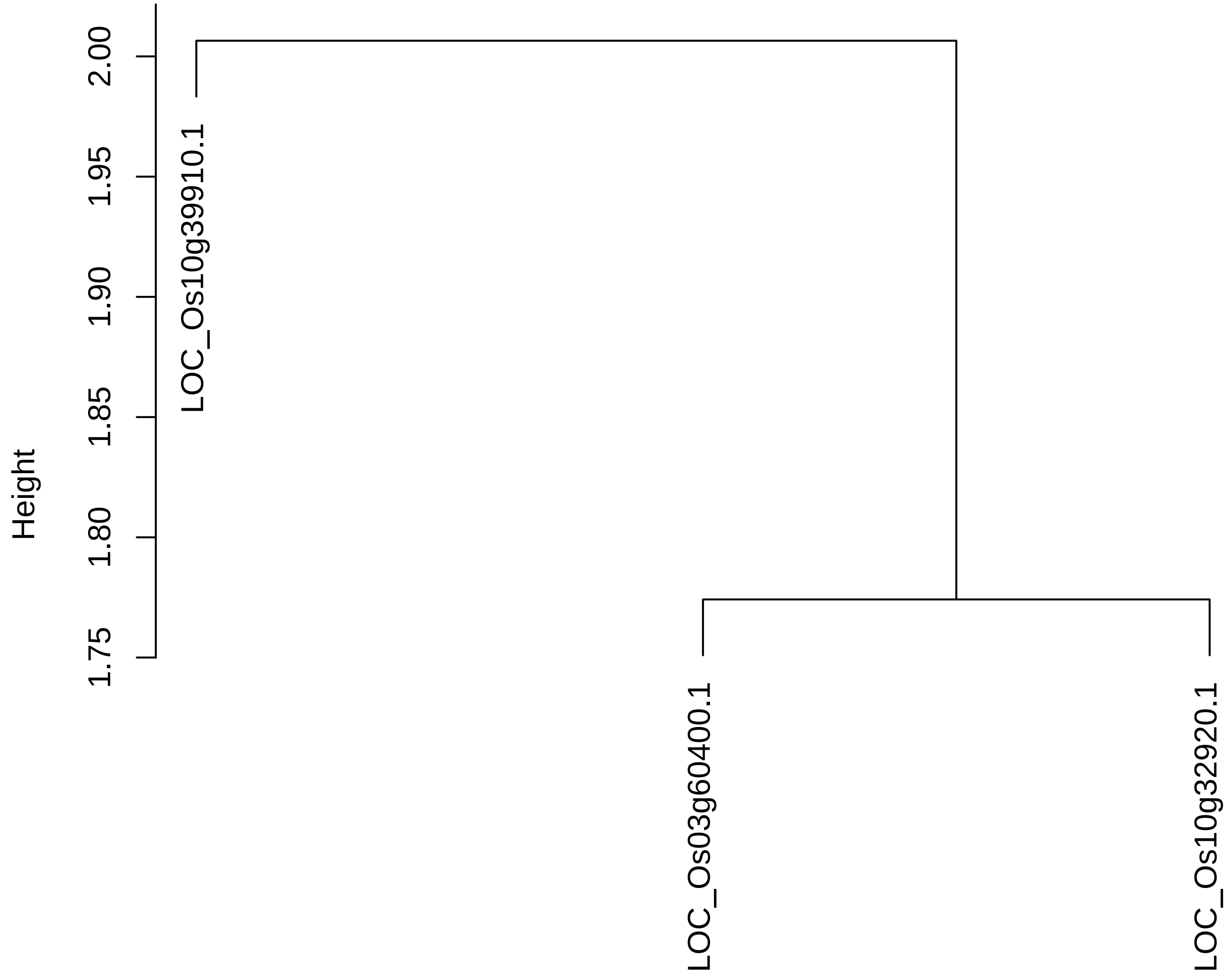


dist(spfitiex12sec12[lab[, 1] == i, ])
hclust (*, "ward.D2")

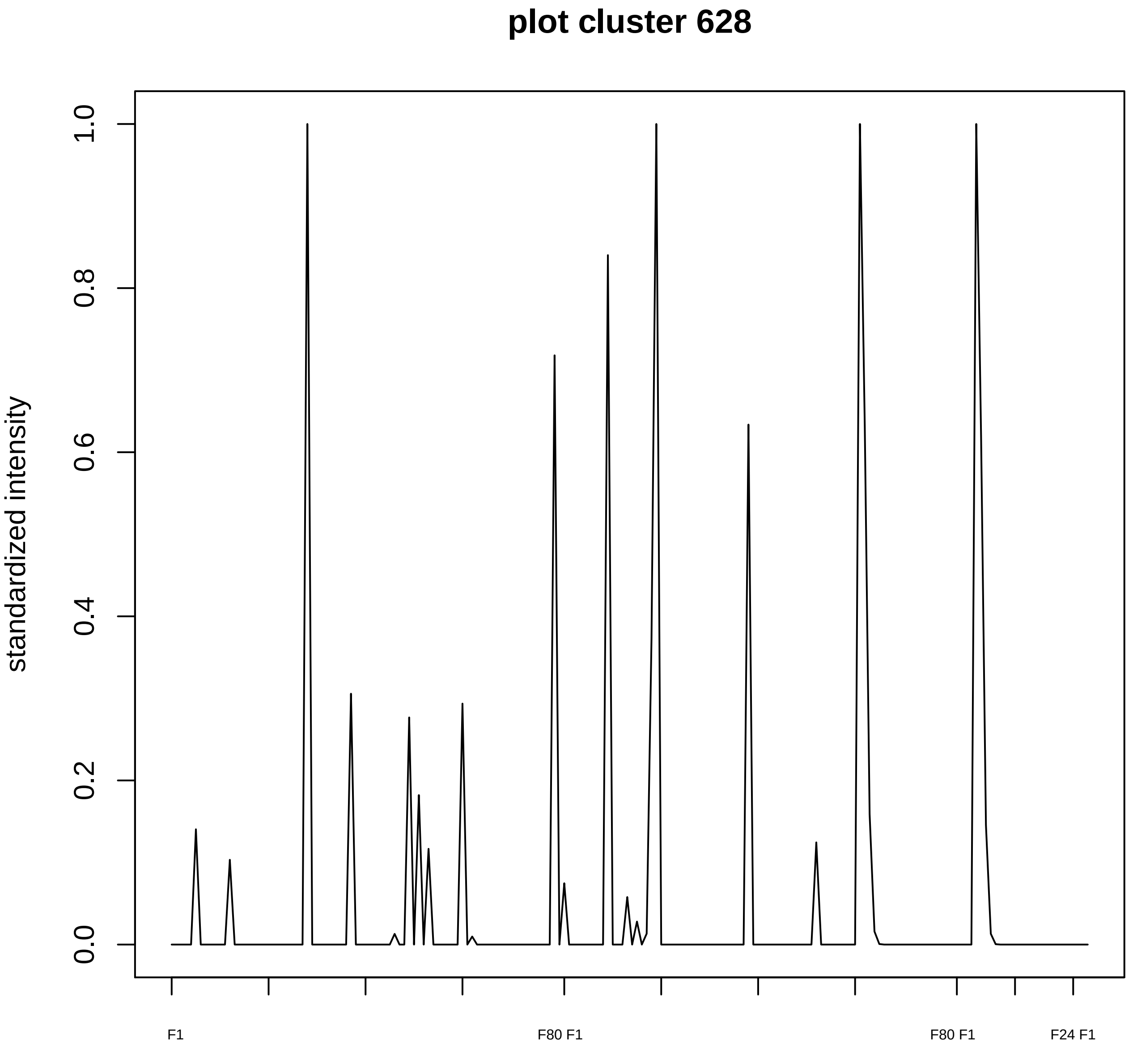
plot cluster 628
standardized intensity
0.0
0.2
0.4
0.6
0.8
1.0
F1
F80 F1
F80 F1
F24 F1

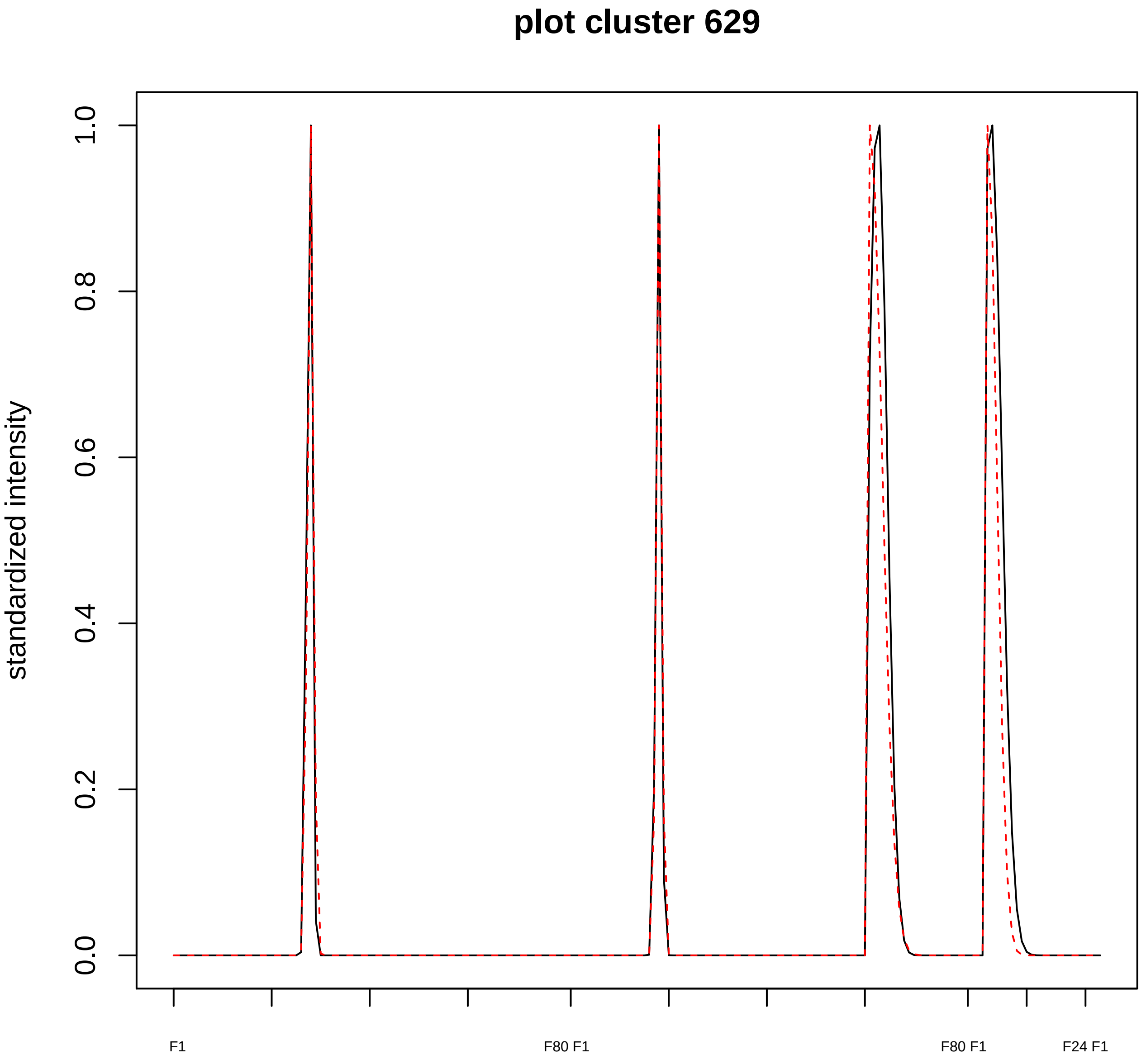

plot cluster 629
standardized intensity
0.0
0.2
0.4
0.6
0.8
1.0
F1
F80 F1
F80 F1
F24 F1

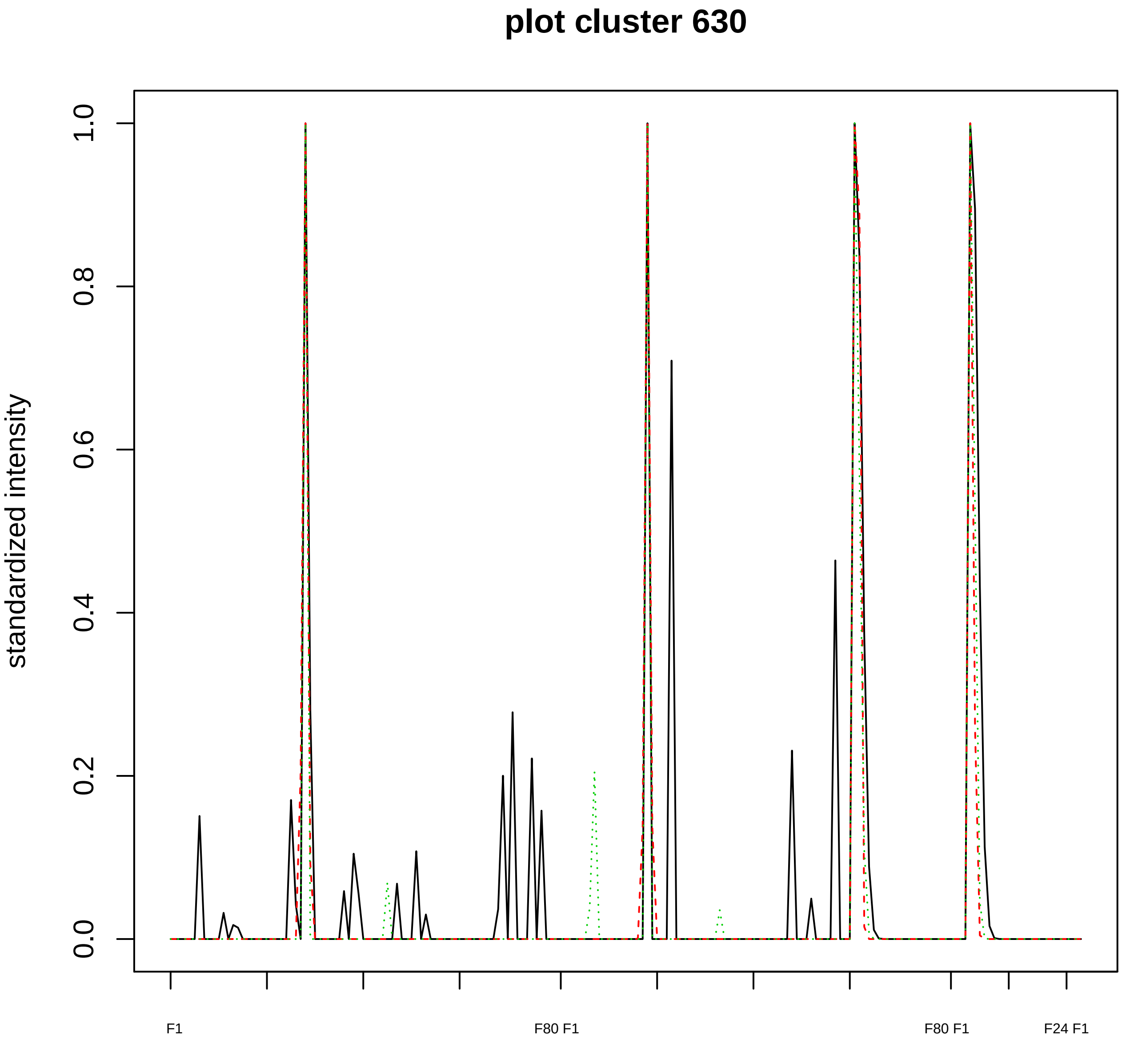

plot cluster 630
standardized intensity
0.0
0.2
0.4
0.6
0.8
1.0
F1
F80 F1
F80 F1
F24 F1

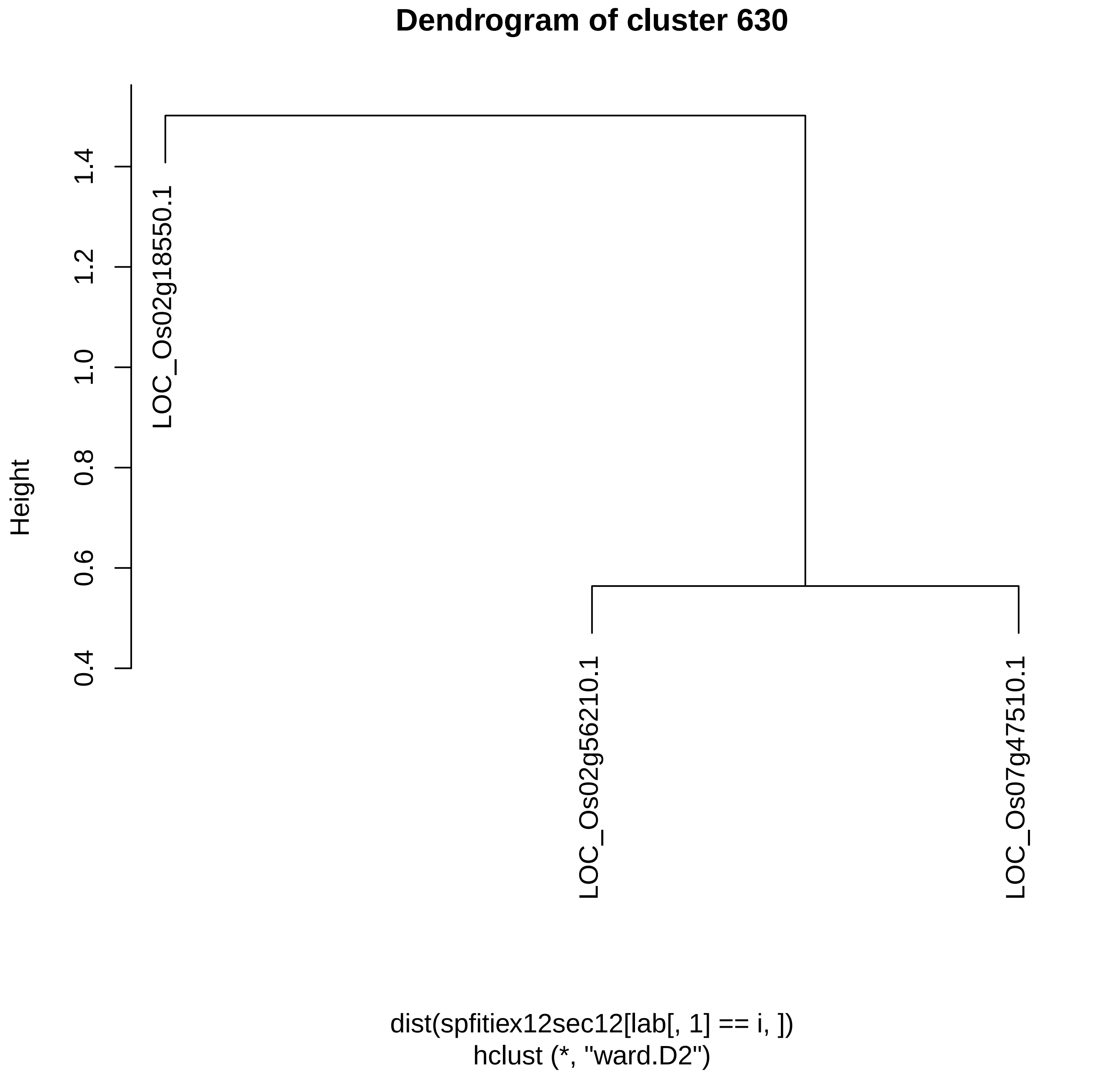
Dendrogram of cluster 630
Height
0.4
0.6
0.8
1.0
1.2
1.4
LOC_Os02g18550.1
LOC_Os02g56210.1
LOC_Os07g47510.1
dist(spfitiex12sec12[lab[, 1] == i, ])
hclust (*, "ward.D2")

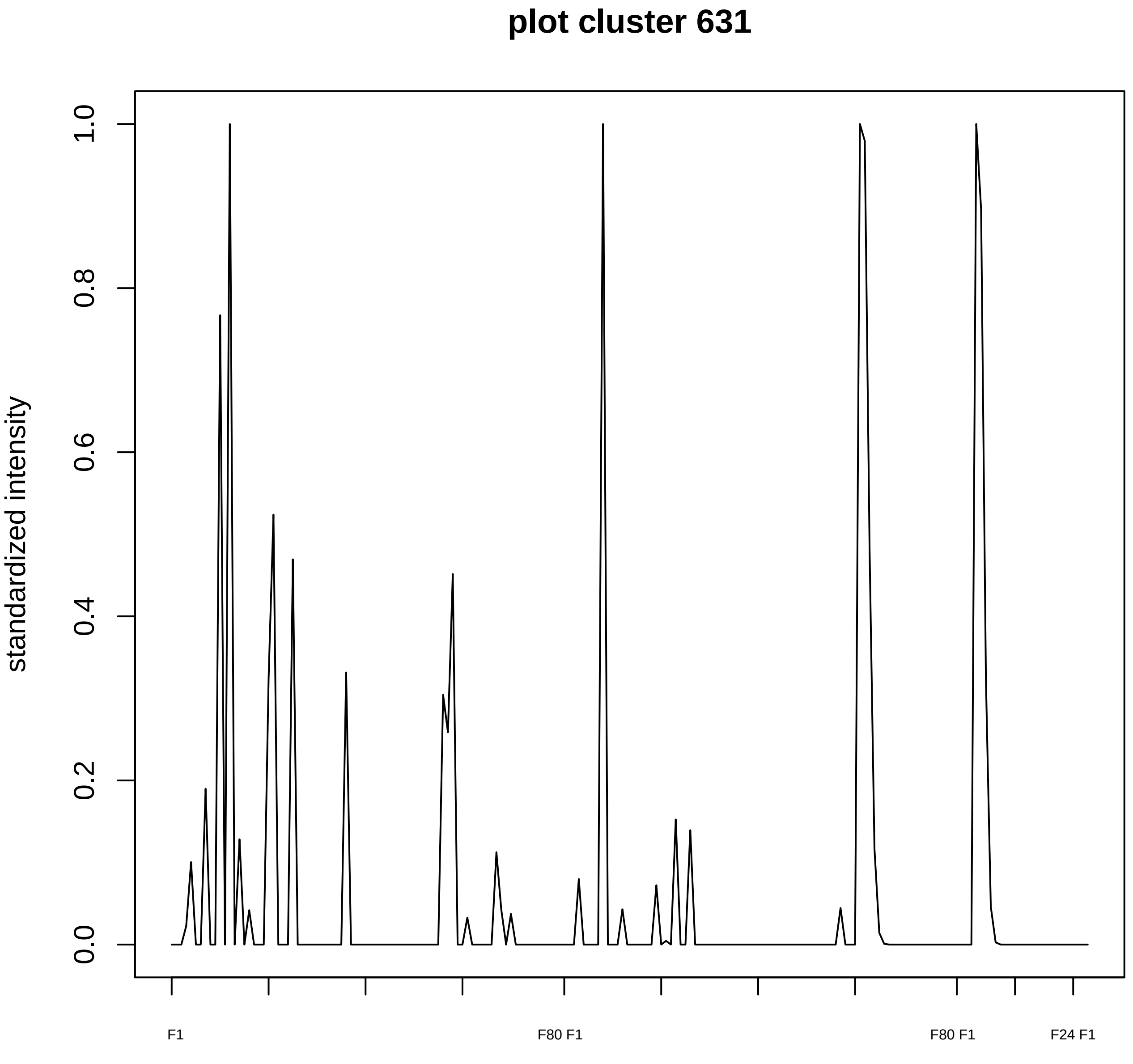

plot cluster 631
standardized intensity
0.0
0.2
0.4
0.6
0.8
1.0
F1
F80 F1
F80 F1
F24 F1

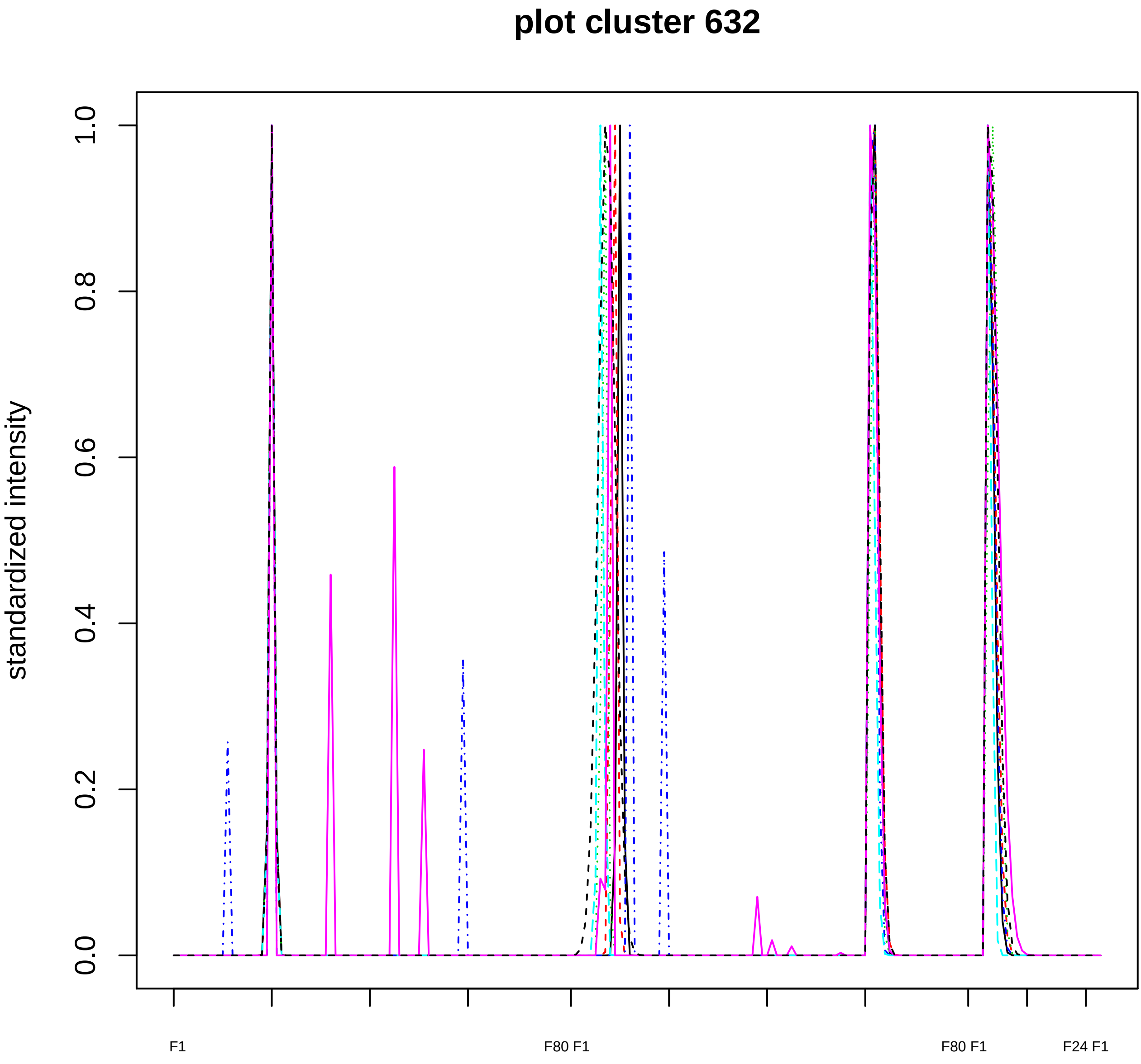

plot cluster 632
standardized intensity
0.0
0.2
0.4
0.6
0.8
1.0
F1
F80 F1
F80 F1
F24 F1

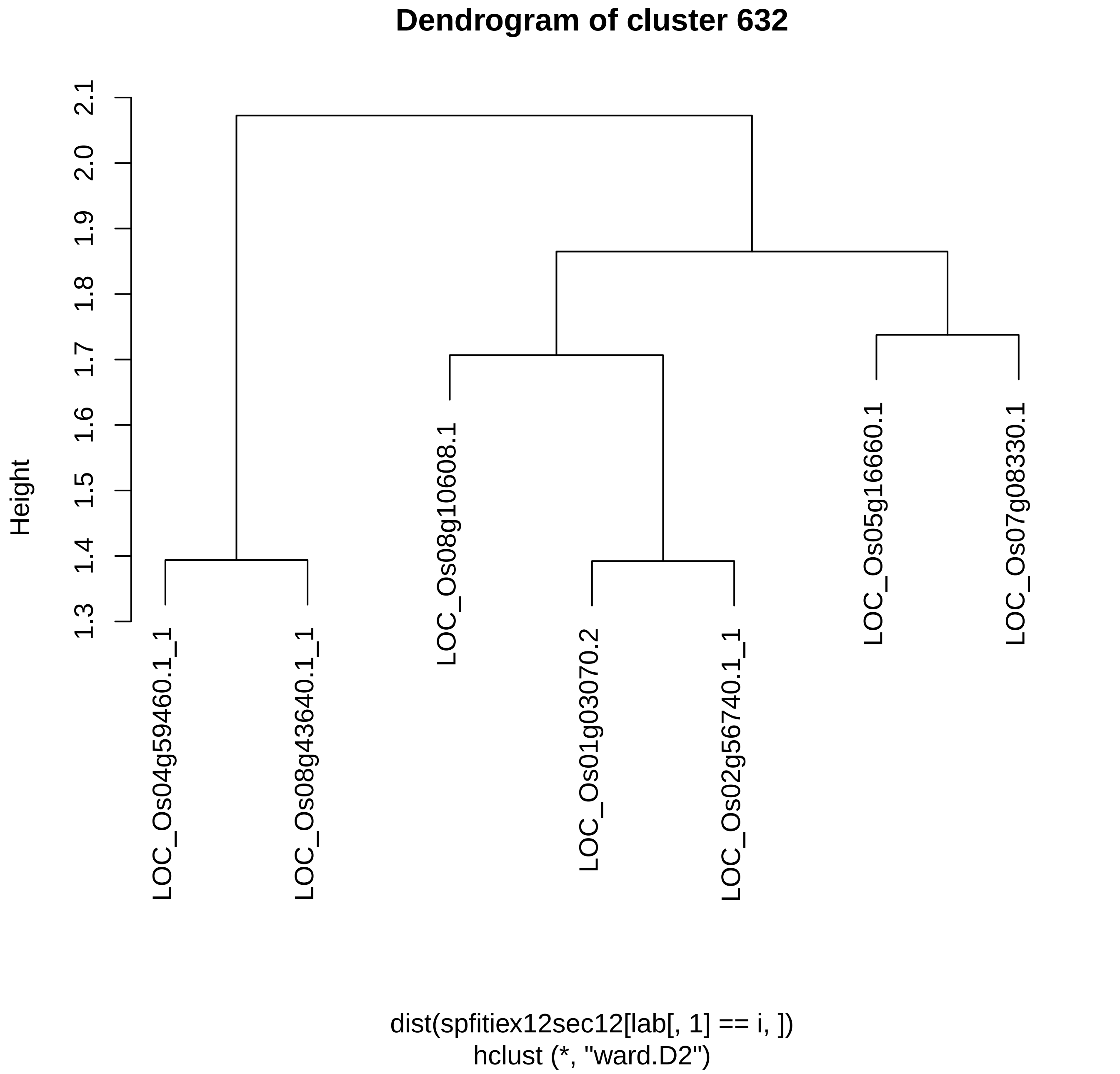

Dendrogram of cluster 632
Height
1.3
1.4
1.5
1.6
1.7
1.8
1.9
2.0
2.1
LOC_Os04g59460.1_1
LOC_Os08g43640.1_1
LOC_Os08g10608.1
LOC_Os01g03070.2
LOC_Os02g56740.1_1
LOC_Os05g16660.1
LOC_Os07g08330.1
dist(spfitiex12sec12[lab[, 1] == i, ])
hclust (*, "ward.D2")

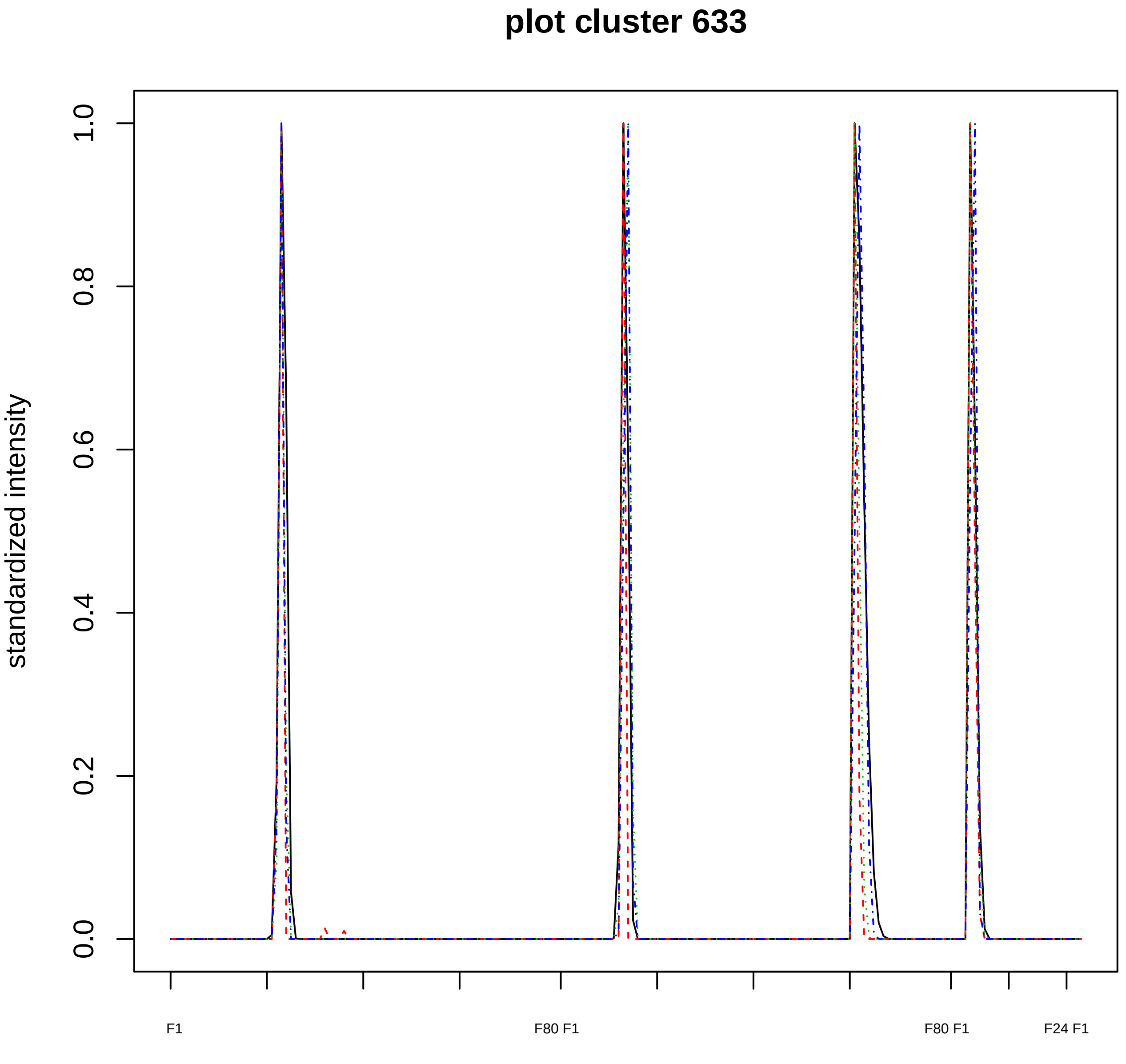
plot cluster 633
standardized intensity
1.0
0.8
0.6
0.4
0.2
0.0
F1
F80 F1
F80 F1
F24 F1

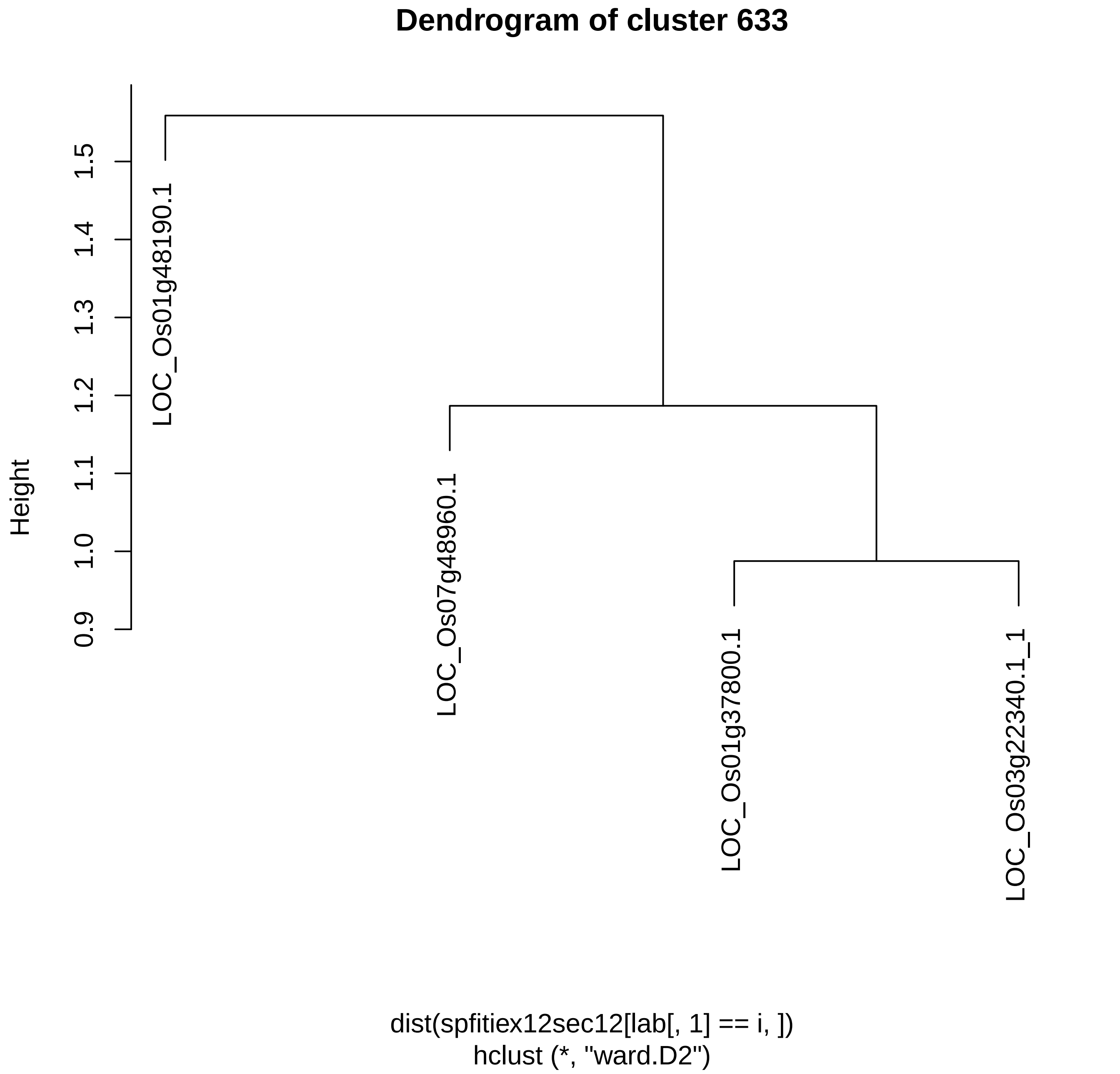
Dendrogram of cluster 633
Height
0.9
1.0
1.1
1.2
1.3
1.4
1.5
LOC_Os01g48190.1
LOC_Os07g48960.1
LOC_Os01g37800.1
LOC_Os03g22340.1_1
dist(spfitiex12sec12[lab[, 1] == i, ])
hclust (*, "ward.D2")

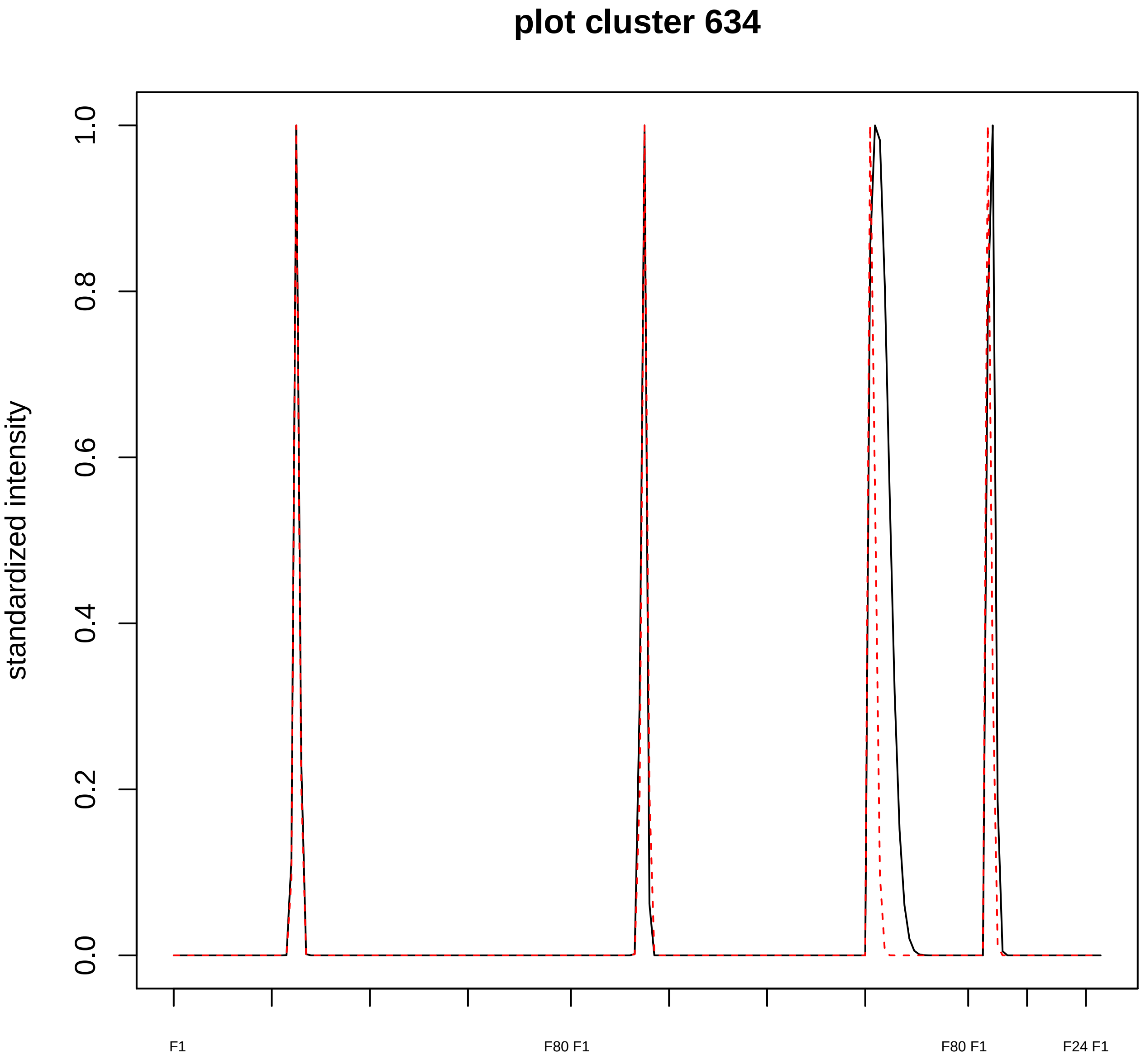
plot cluster 634
standardized intensity
0.0
0.2
0.4
0.6
0.8
1.0
F1
F80 F1
F80 F1
F24 F1

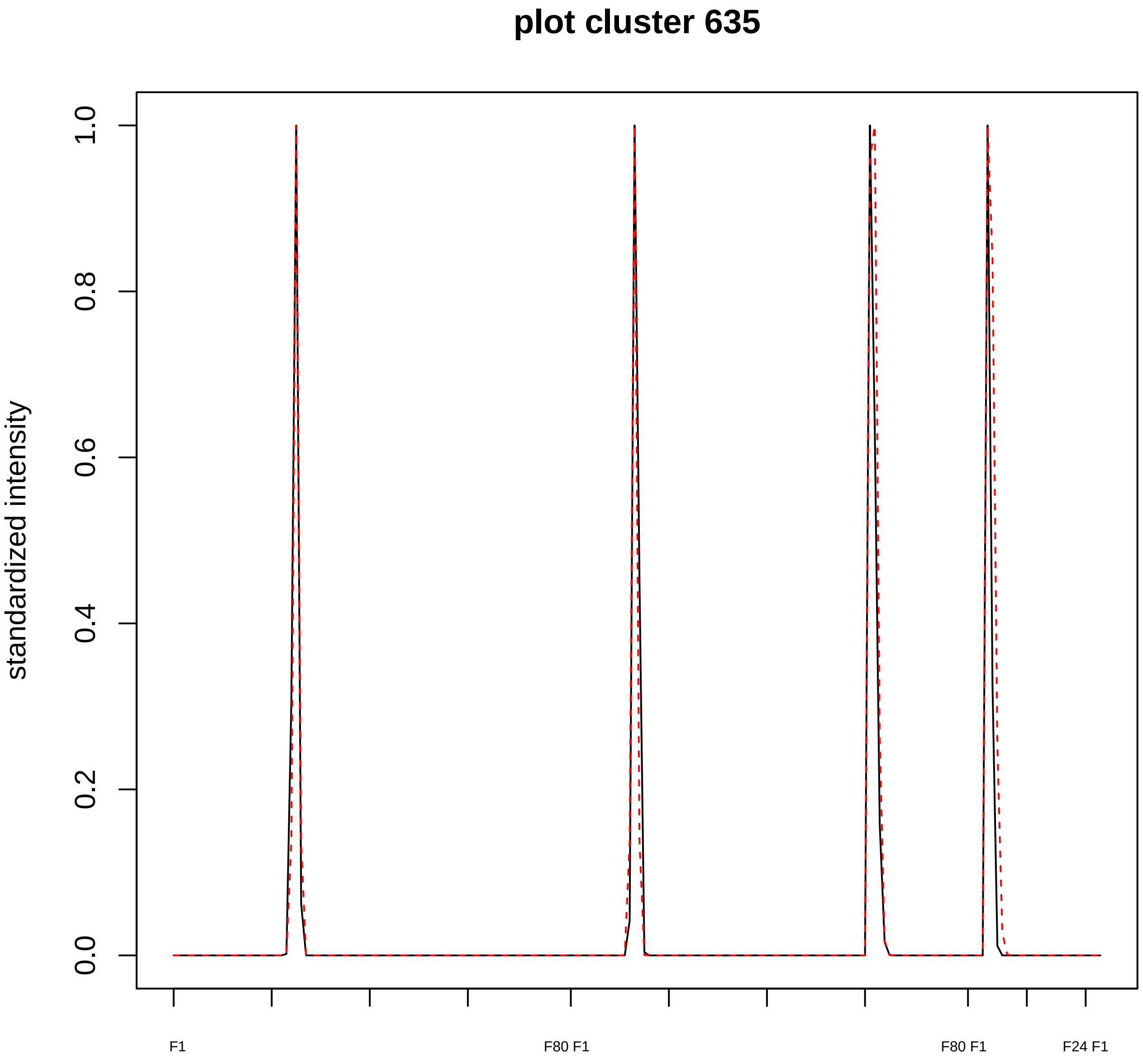

plot cluster 635
standardized intensity
0.0
0.2
0.4
0.6
0.8
1.0
F1
F80 F1
F80 F1
F24 F1

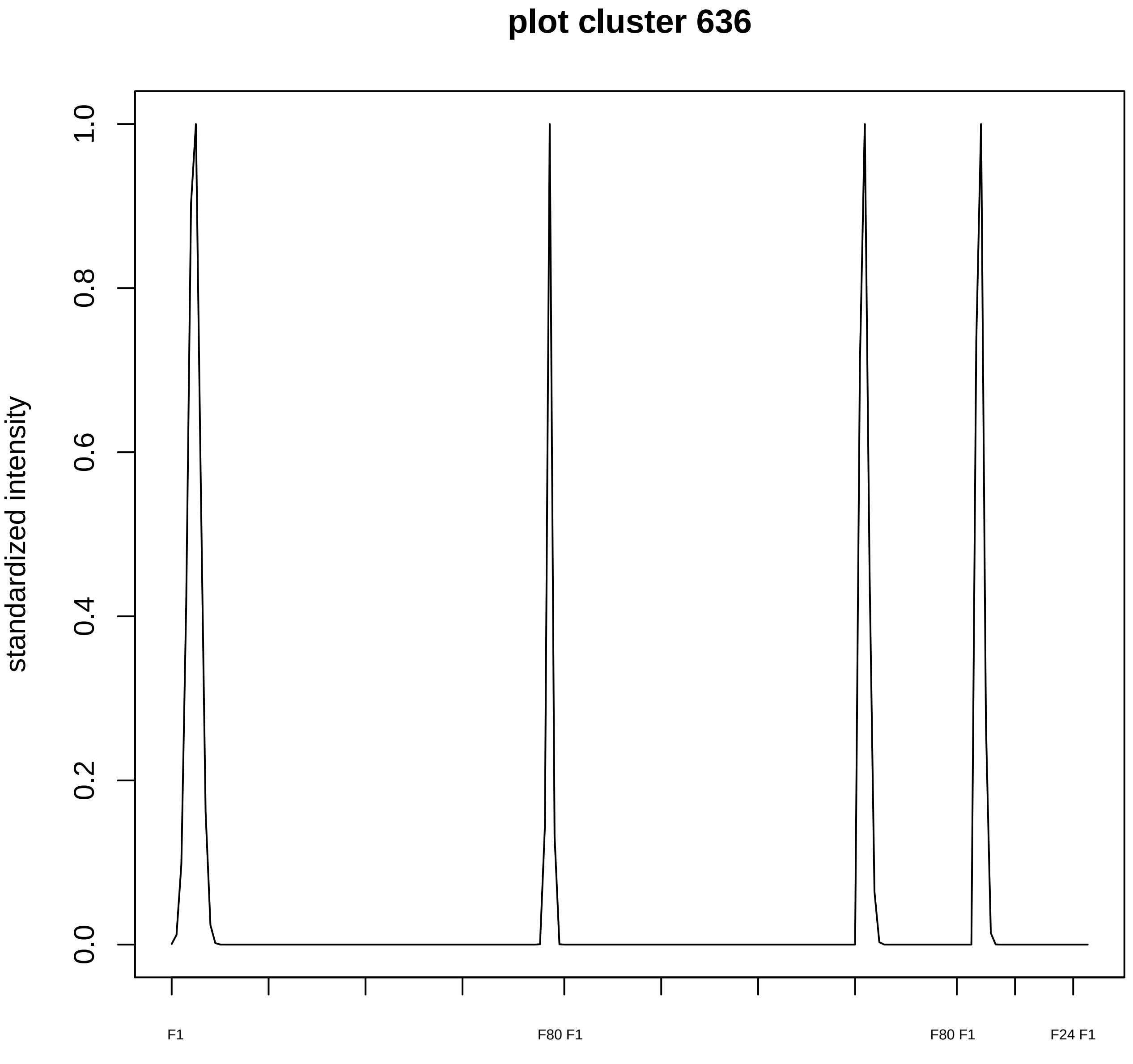

plot cluster 636
standardized intensity
0.0
0.2
0.4
0.6
0.8
1.0
F1
F80 F1
F80 F1
F24 F1

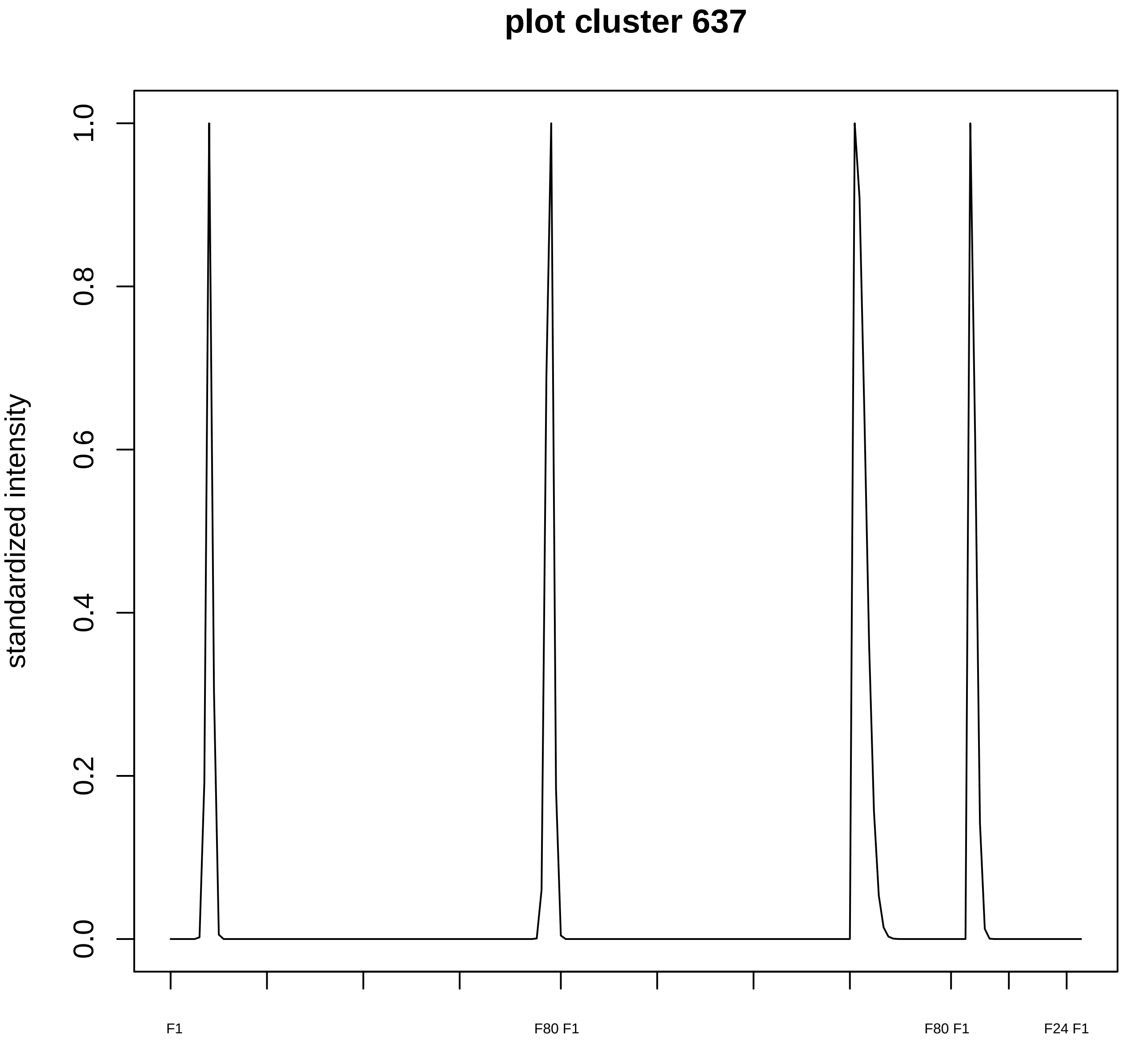

plot cluster 637
standardized intensity
0.0
0.2
0.4
0.6
0.8
1.0
F1
F80 F1
F80 F1
F24 F1

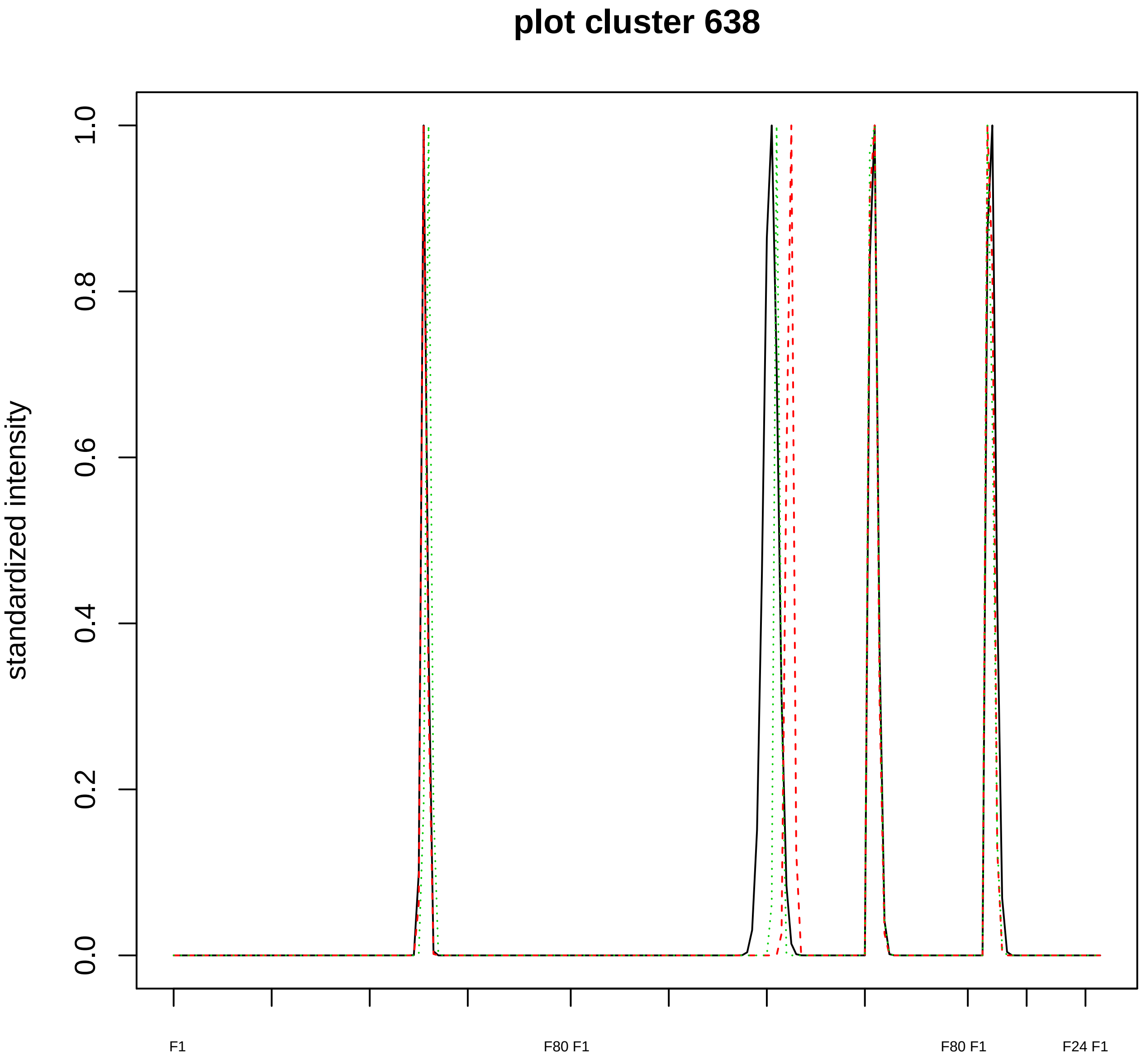
plot cluster 638
standardized intensity
0.0
0.2
0.4
0.6
0.8
1.0
F1
F80 F1
F80 F1
F24 F1

### Dendrogram of cluster 638

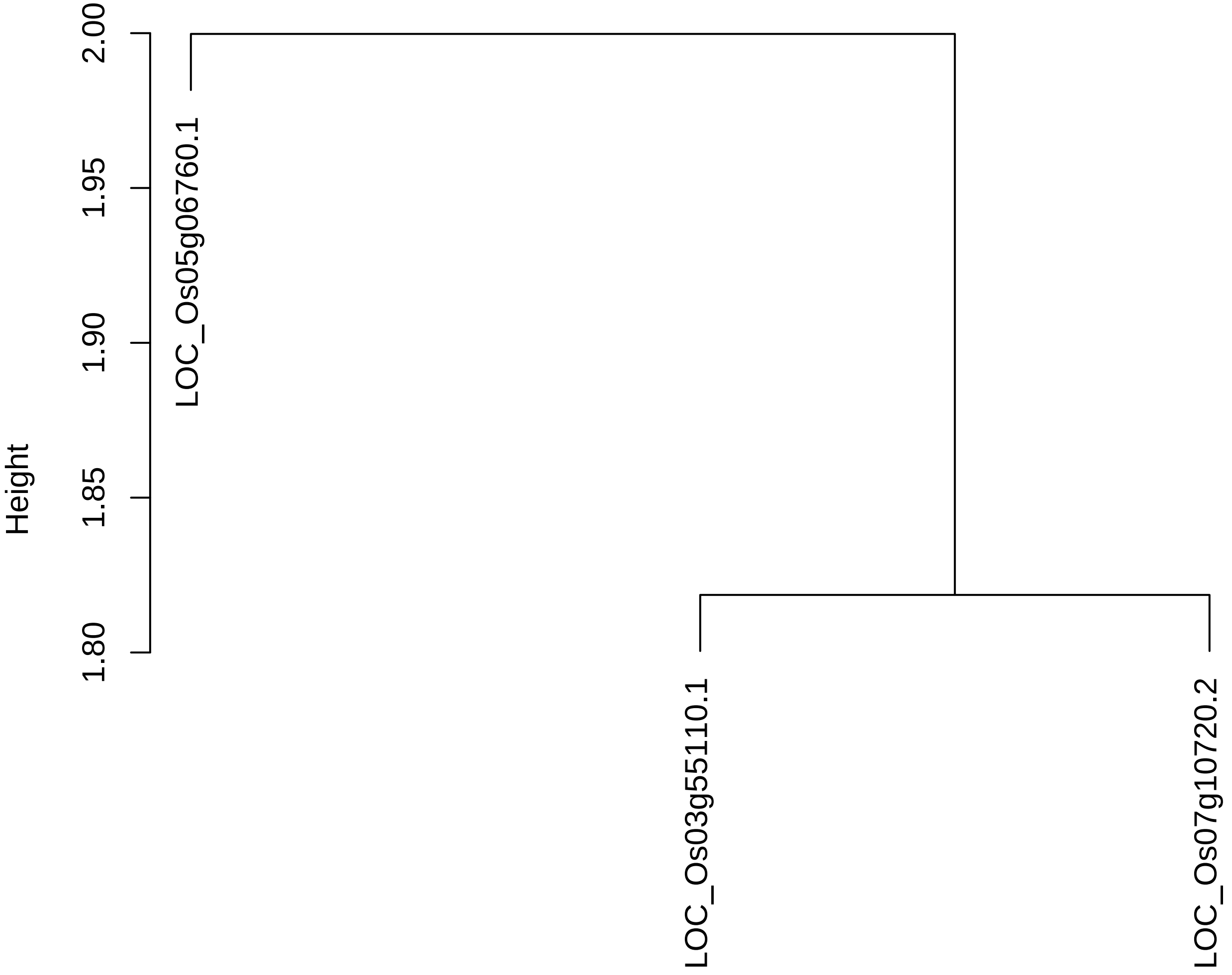


dist(spfitiex12sec12[lab[, 1] == i, ])
hclust (*, "ward.D2")

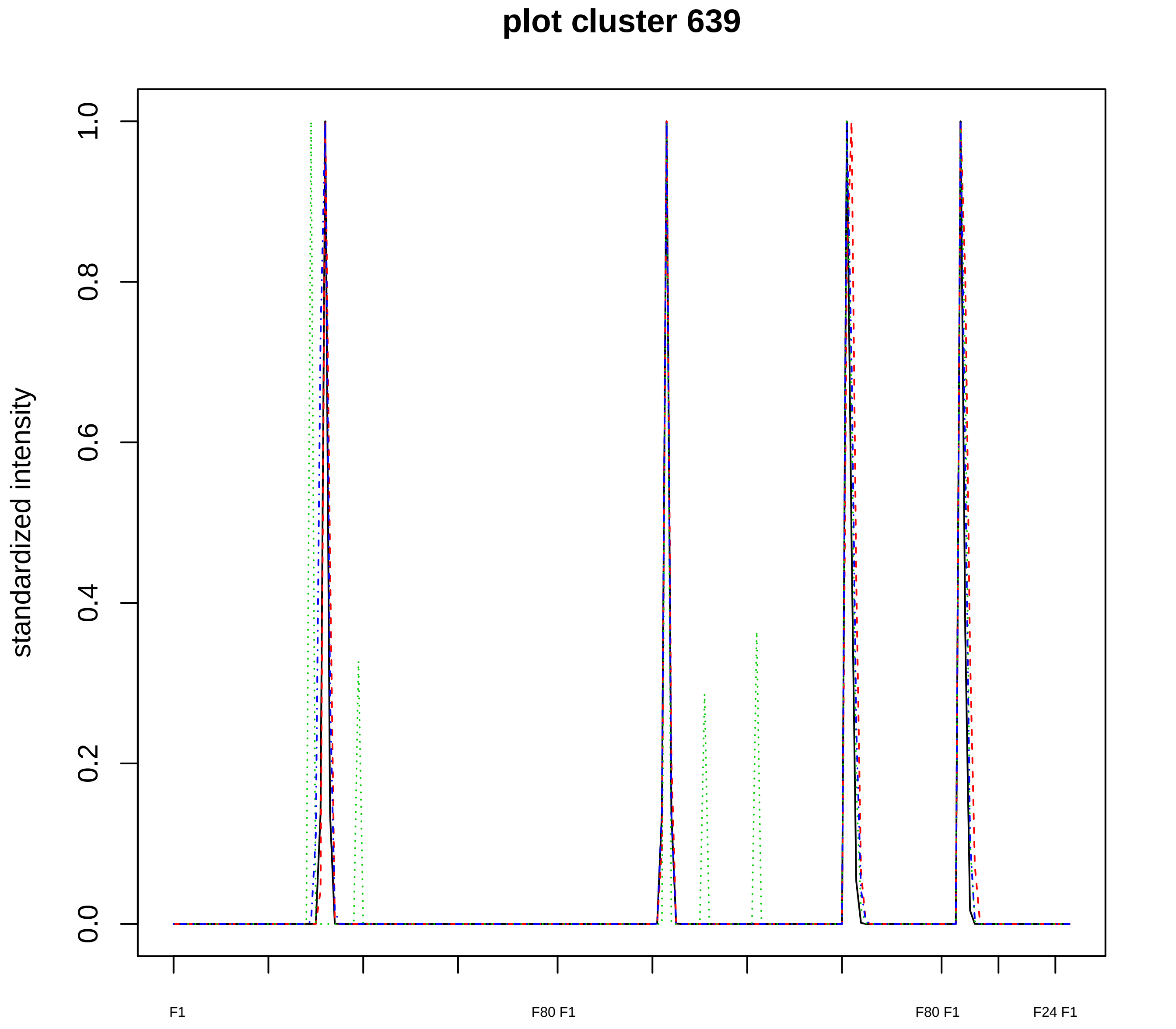
plot cluster 639
standardized intensity
0.0
0.2
0.4
0.6
0.8
1.0
F1
F80 F1
F80 F1
F24 F1

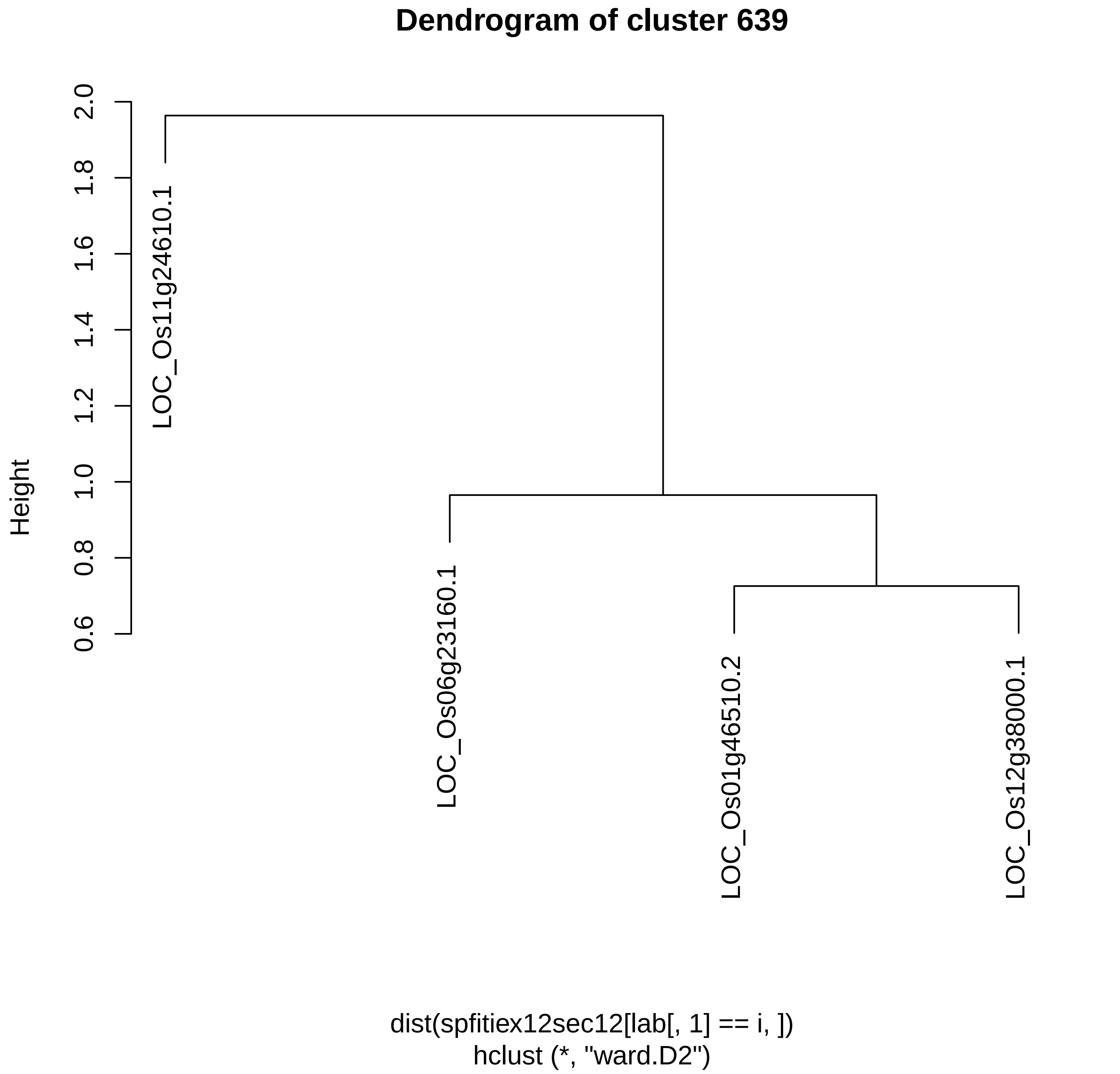
Dendrogram of cluster 639
Height
2.0
1.8
1.6
1.4
1.2
1.0
0.8
0.6
LOC_Os11g24610.1
LOC_Os06g23160.1
LOC_Os01g46510.2
LOC_Os12g38000.1
dist(spfitiex12sec12[lab[, 1] == i, ])
hclust (*, "ward.D2")

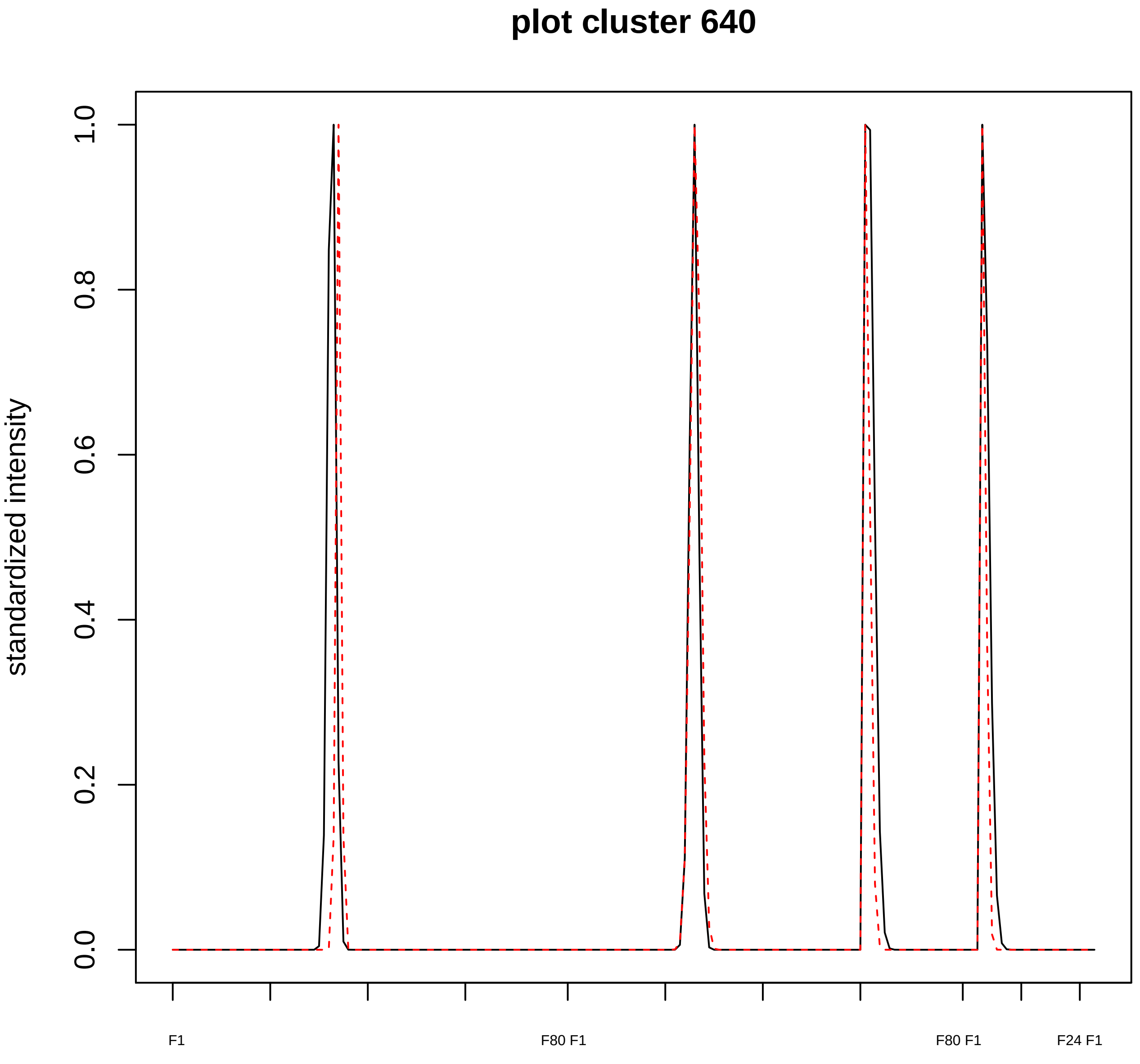
plot cluster 640
standardized intensity
0.0
0.2
0.4
0.6
0.8
1.0
F1
F80 F1
F80 F1
F24 F1

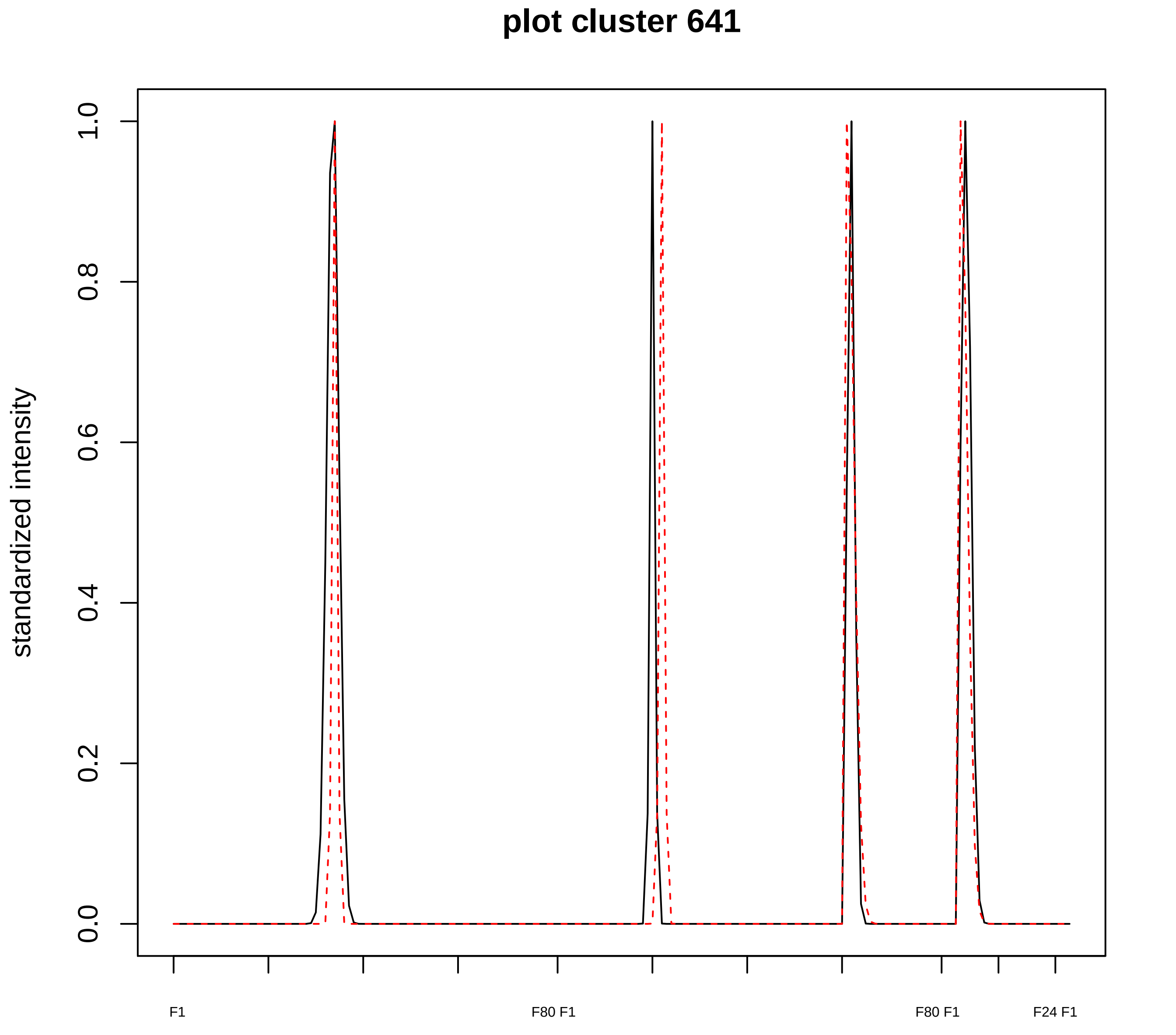

plot cluster 641
standardized intensity
0.0
0.2
0.4
0.6
0.8
1.0
F1
F80 F1
F80 F1
F24 F1

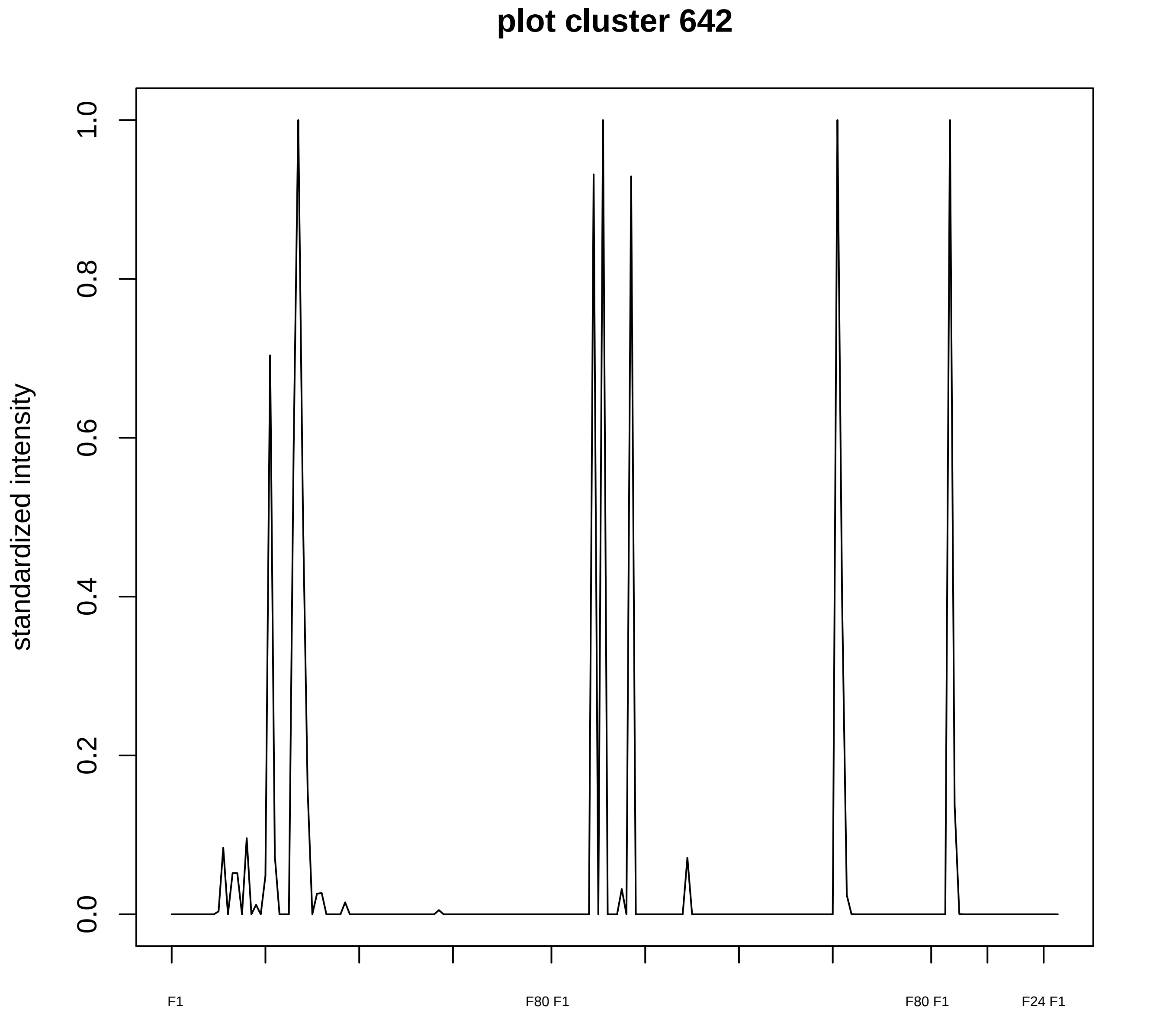
plot cluster 642
standardized intensity
0.0
0.2
0.4
0.6
0.8
1.0
F1
F80 F1
F80 F1
F24 F1

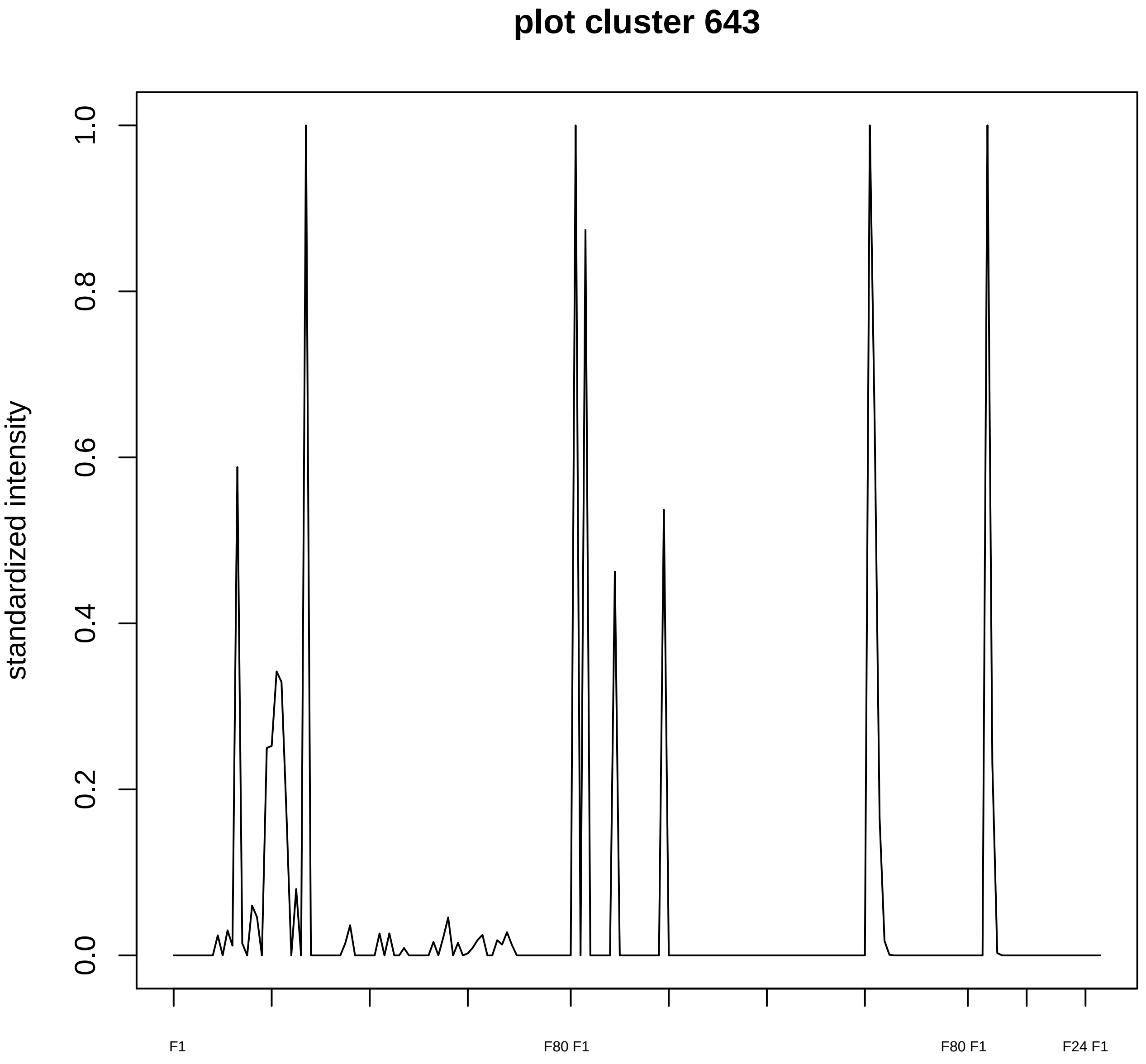
plot cluster 643
standardized intensity
0.0
0.2
0.4
0.6
0.8
1.0
F1
F80 F1
F80 F1
F24 F1

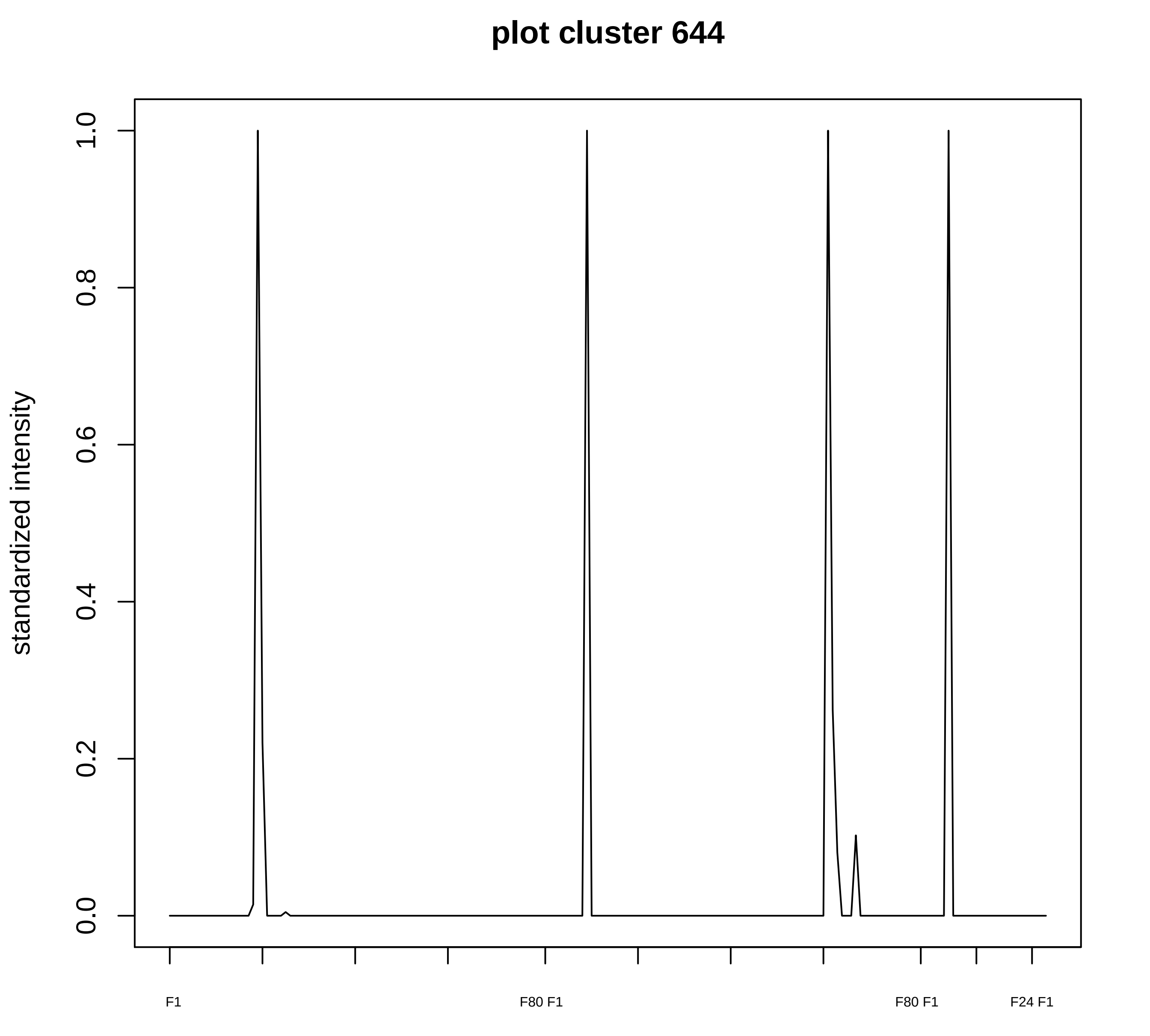

plot cluster 644
standardized intensity
0.0
0.2
0.4
0.6
0.8
1.0
F1
F80 F1
F80 F1
F24 F1

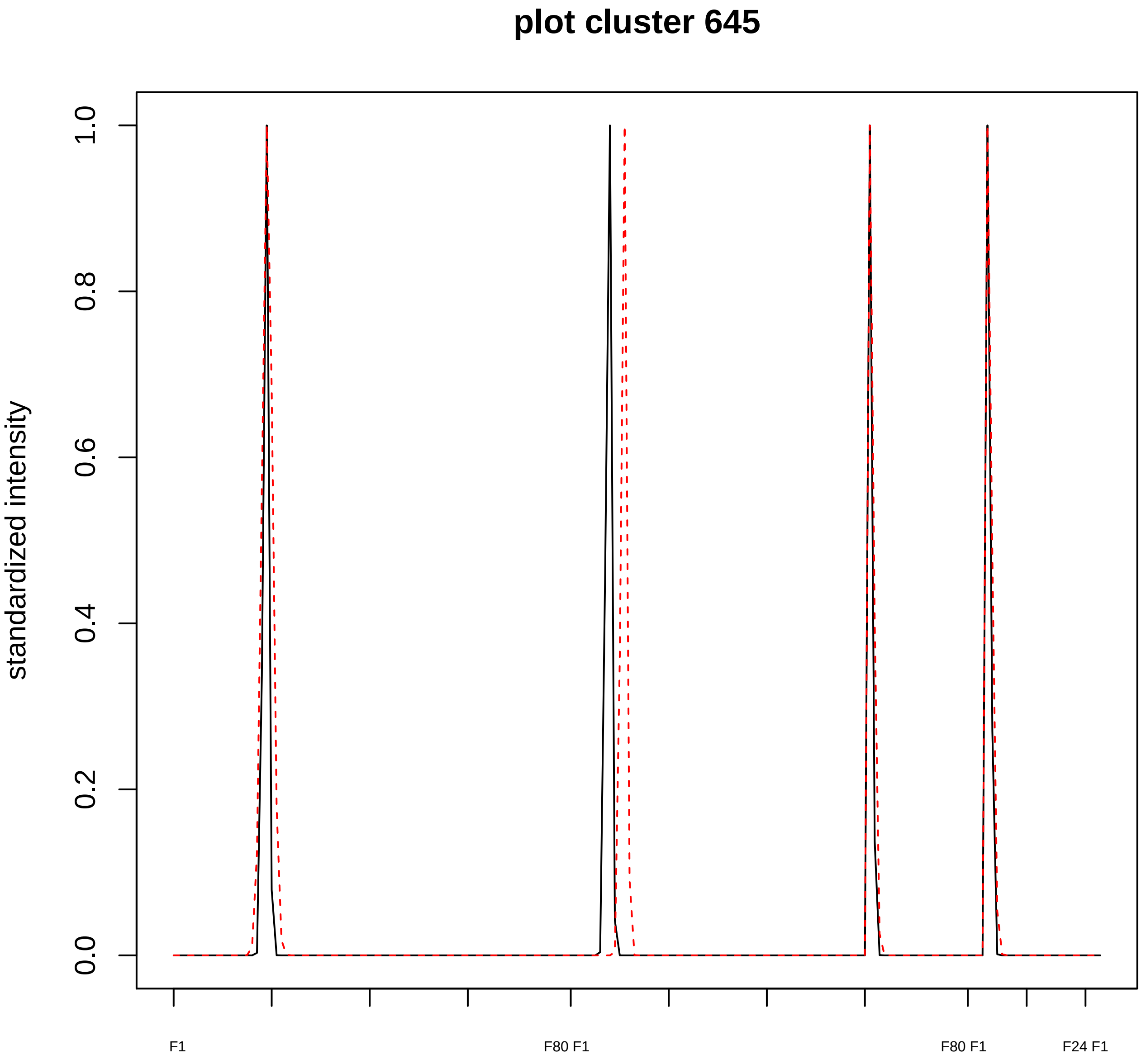
plot cluster 645
standardized intensity
0.0
0.2
0.4
0.6
0.8
1.0
F1
F80 F1
F80 F1
F24 F1

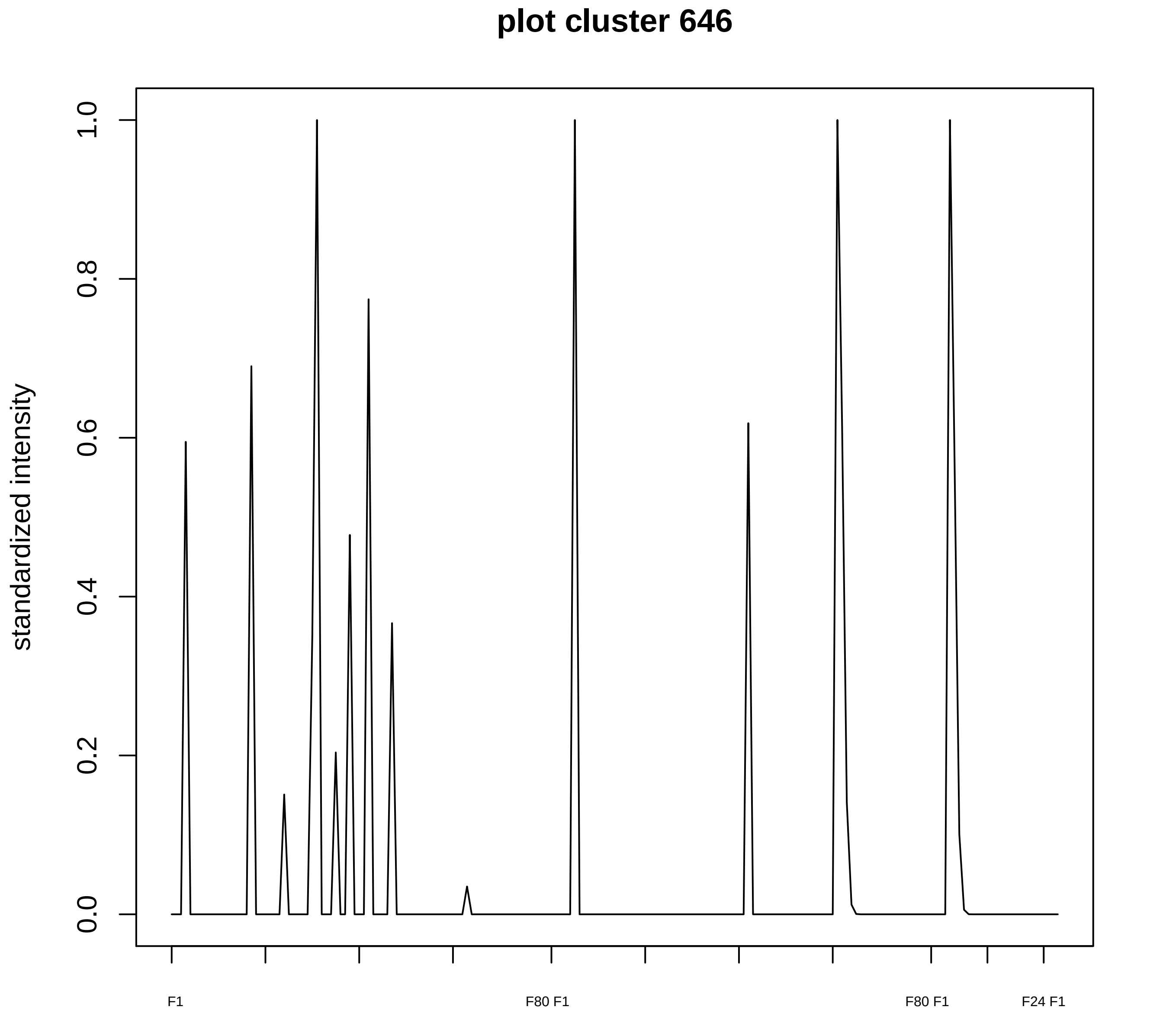

plot cluster 646
standardized intensity
0.0
0.2
0.4
0.6
0.8
1.0
F1
F80 F1
F80 F1
F24 F1

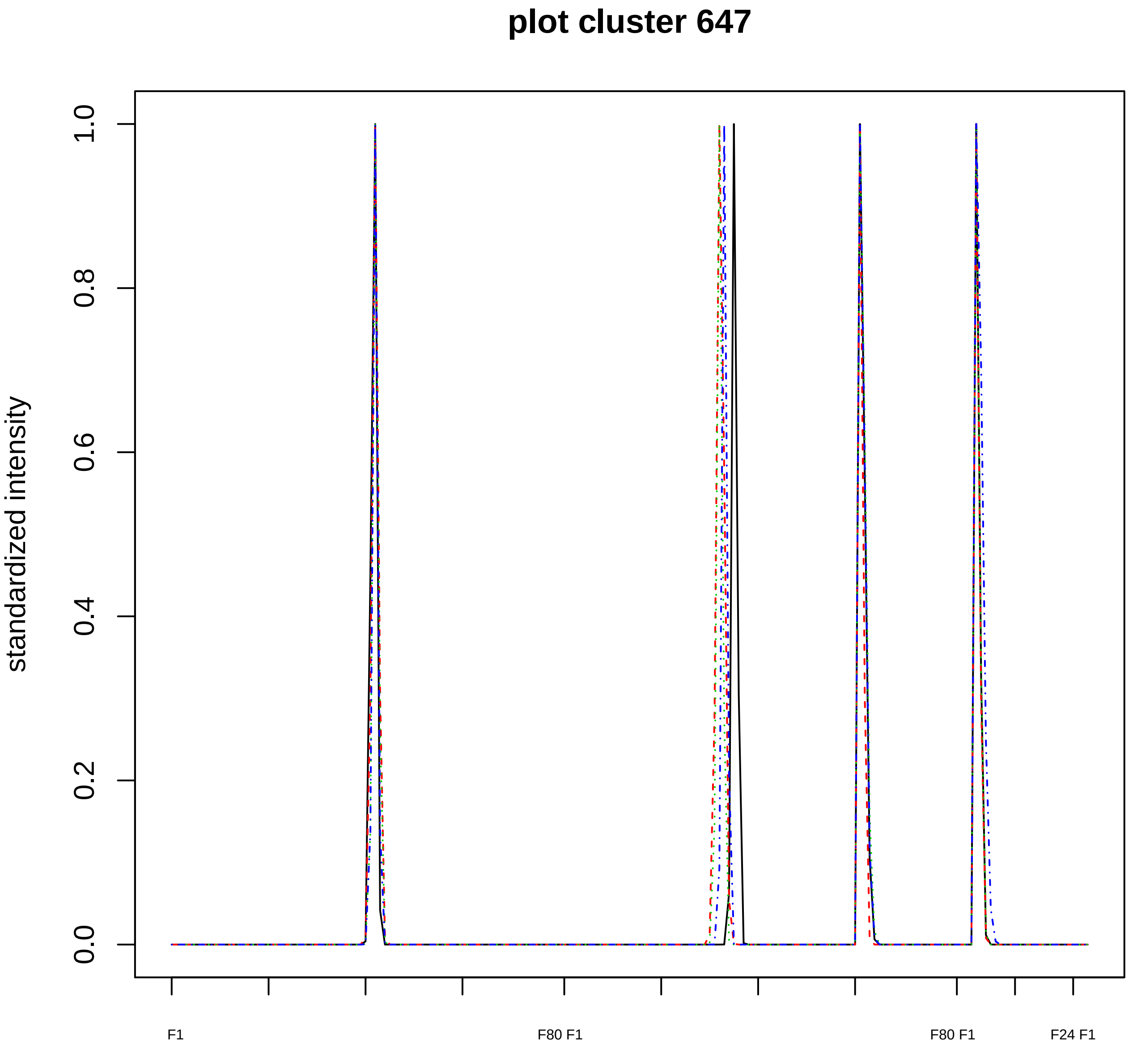
plot cluster 647
standardized intensity
0.0
0.2
0.4
0.6
0.8
1.0
F1
F80 F1
F80 F1
F24 F1

#### Dendrogram of cluster 647

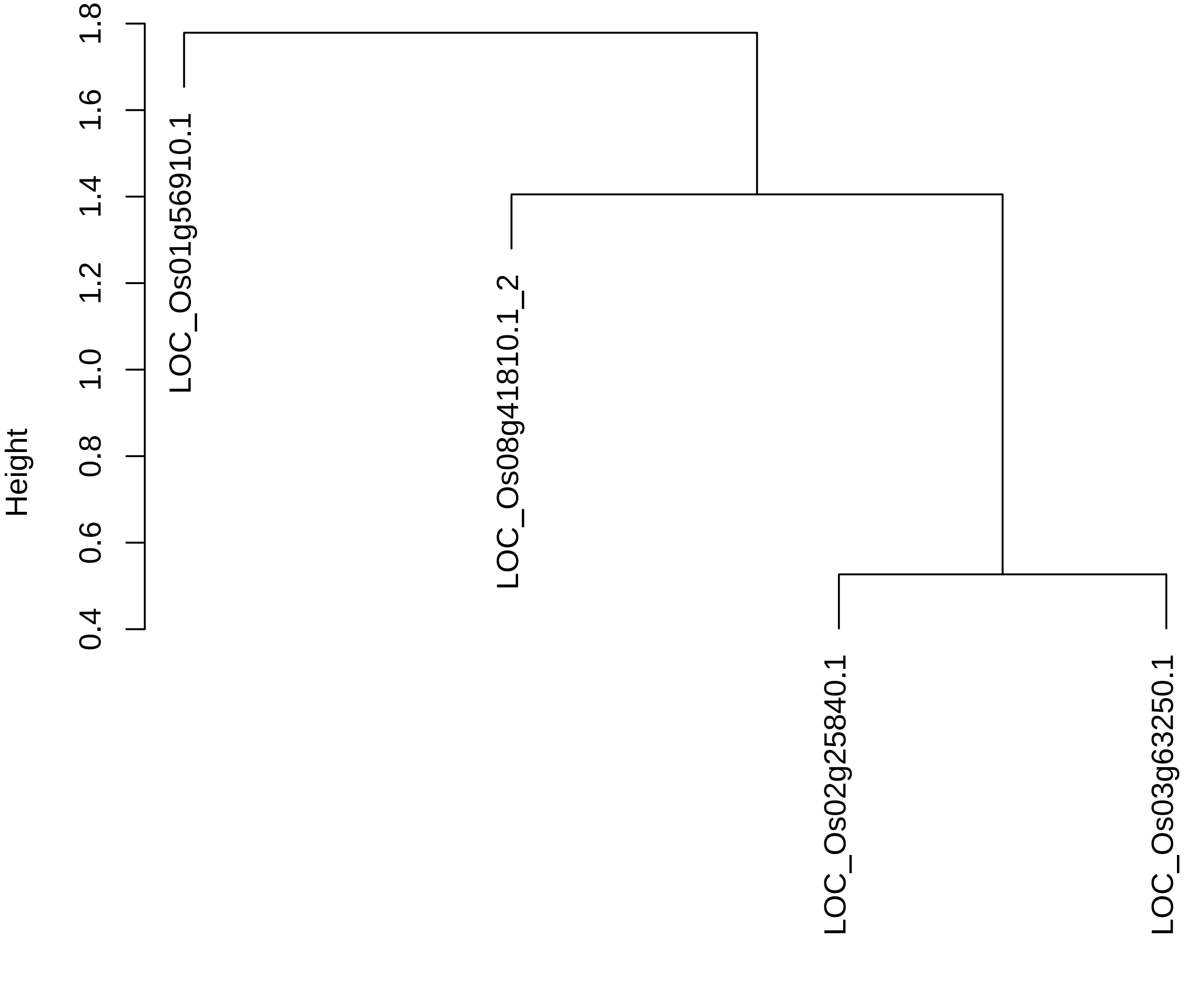


dist(spfitiex12sec12[lab[, 1] == i, ])
hclust (*, "ward.D2")

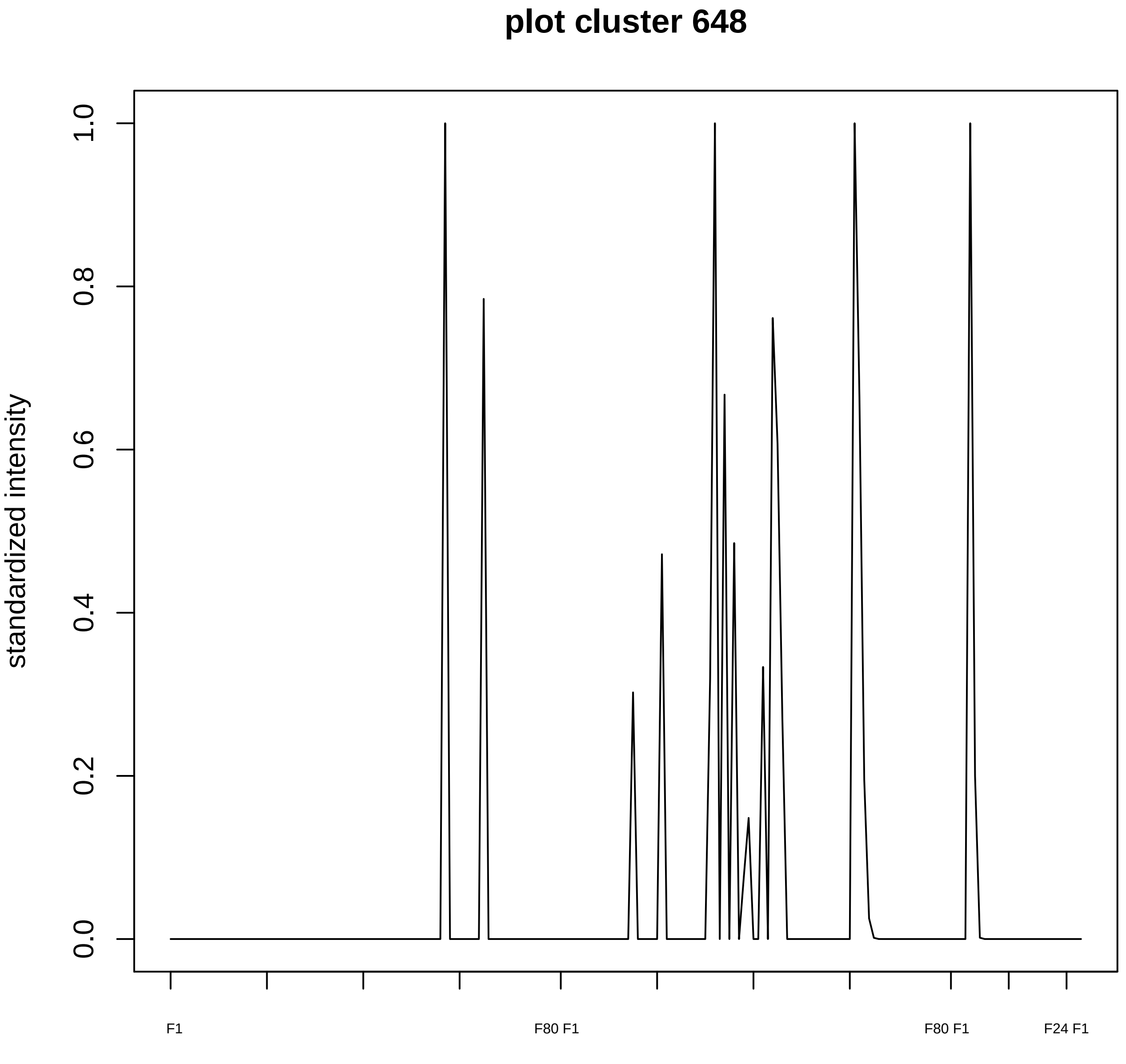
plot cluster 648
standardized intensity
0.0
0.2
0.4
0.6
0.8
1.0
F1
F80 F1
F80 F1
F24 F1

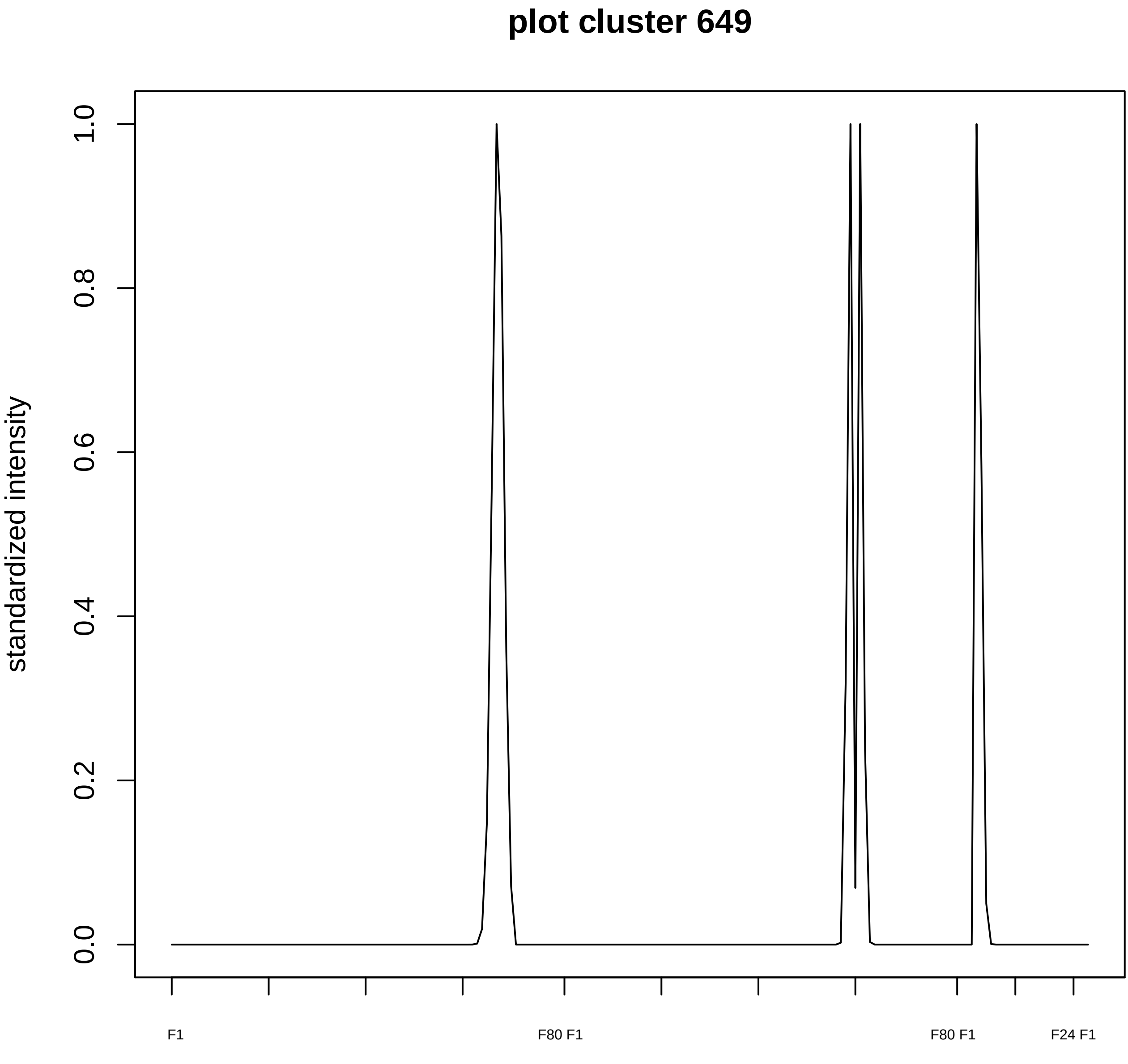
plot cluster 649
standardized intensity
0.0
0.2
0.4
0.6
0.8
1.0
F1
F80 F1
F80 F1
F24 F1

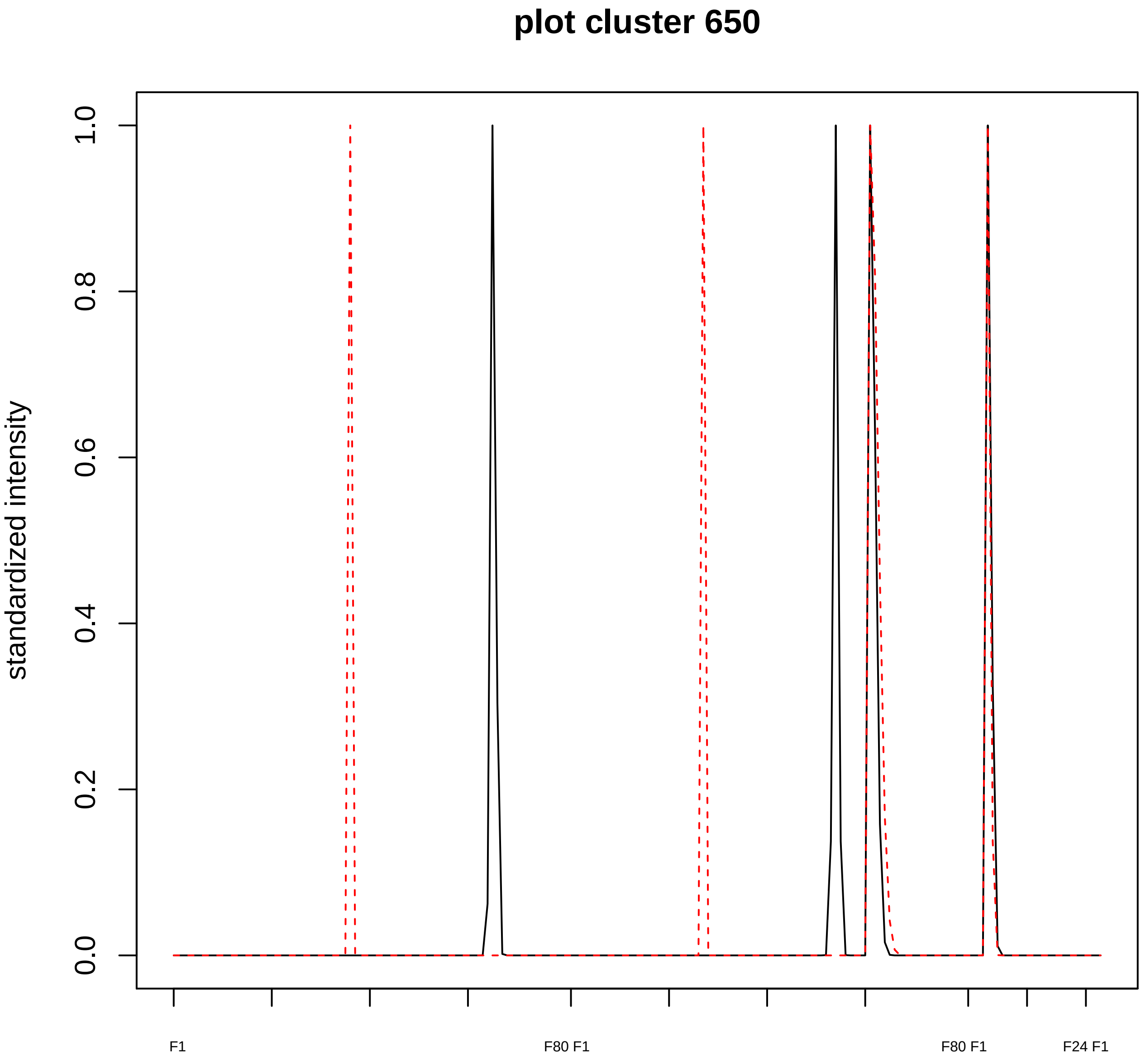
plot cluster 650
standardized intensity
0.0
0.2
0.4
0.6
0.8
1.0
F1
F80 F1
F80 F1
F24 F1

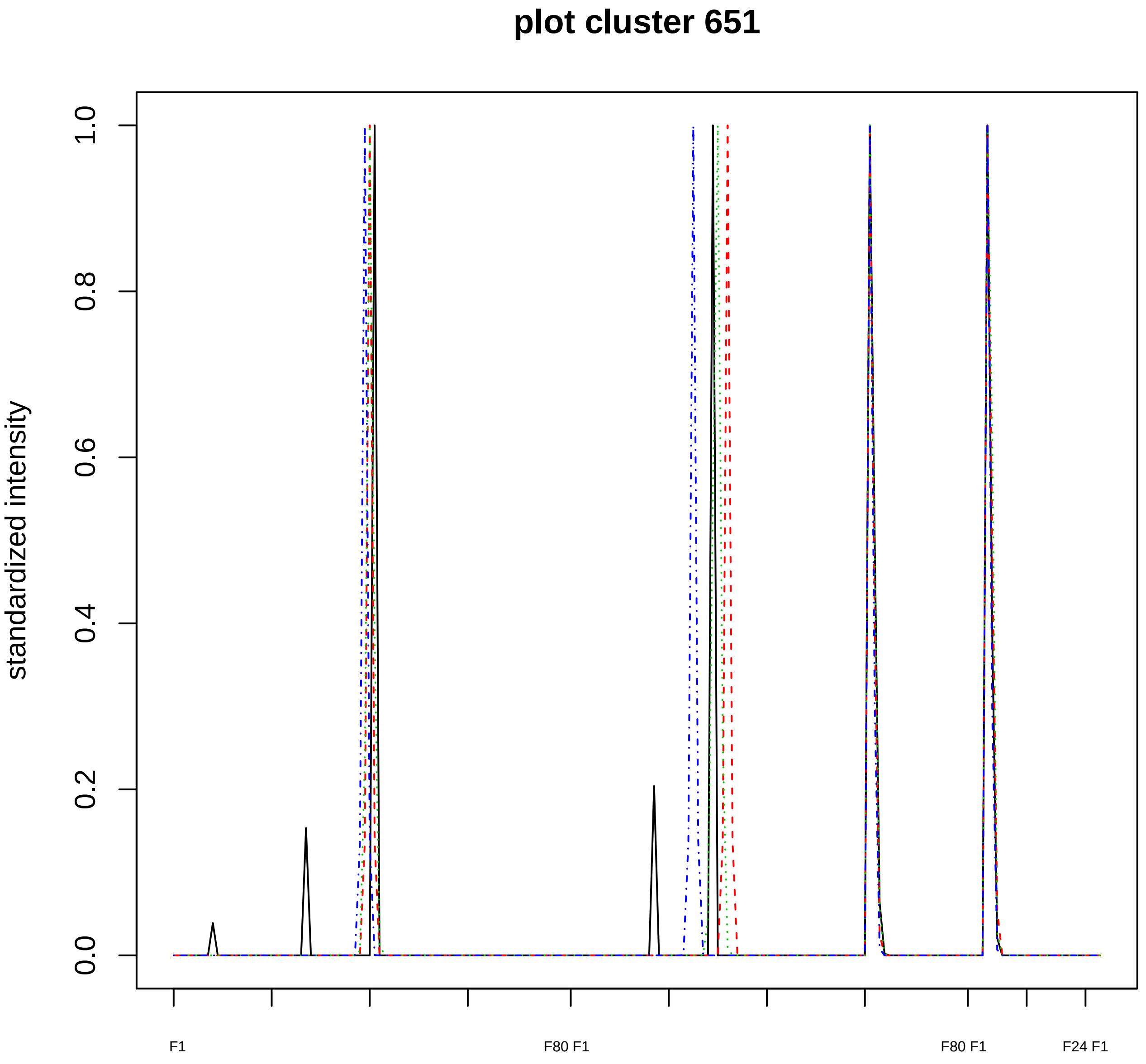
plot cluster 651
standardized intensity
0.0
0.2
0.4
0.6
0.8
1.0
F1
F80 F1
F80 F1
F24 F1

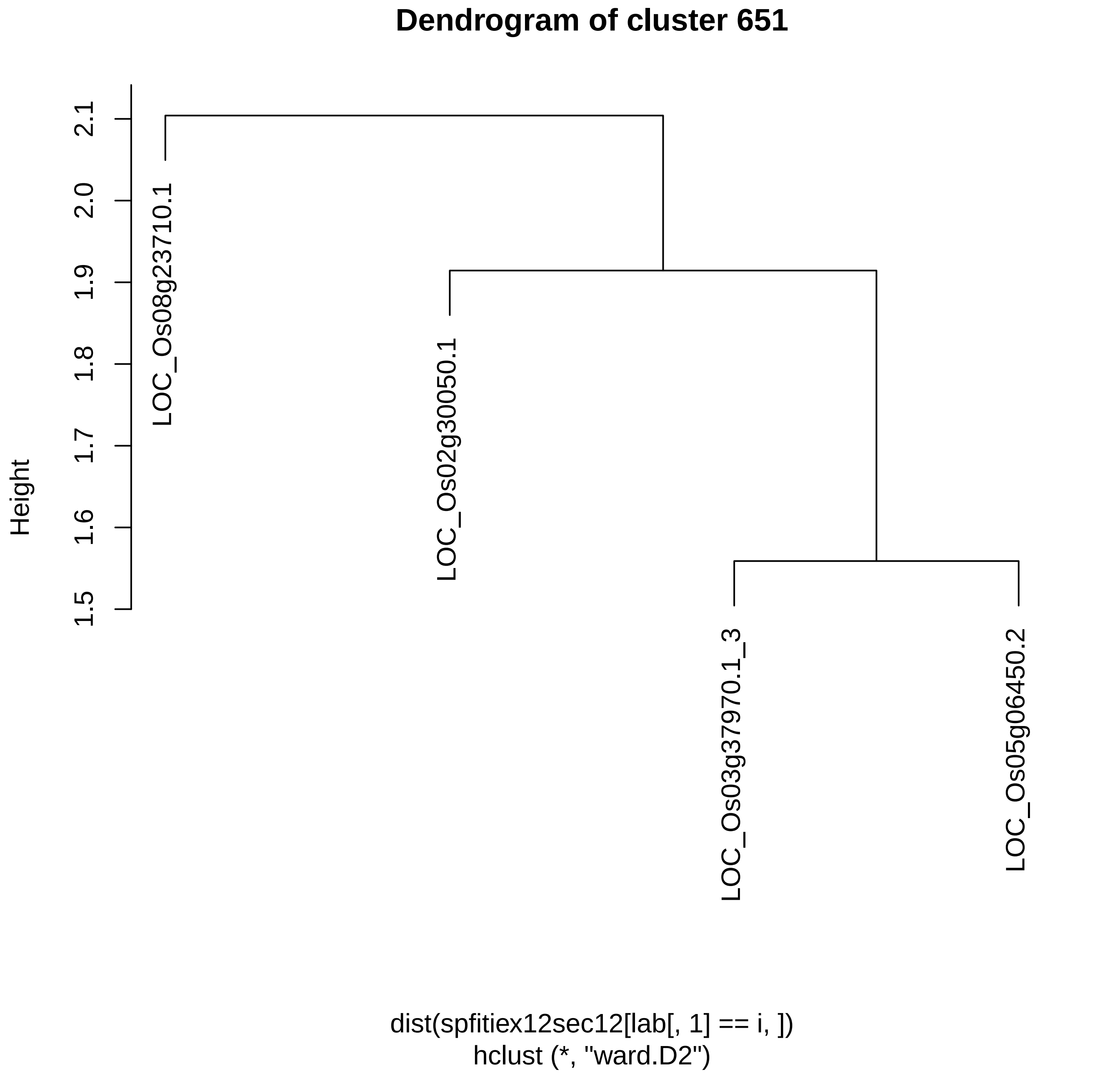
Dendrogram of cluster 651
Height
2.1
2.0
1.9
1.8
1.7
1.6
1.5
LOC_Os08g23710.1
LOC_Os02g30050.1
LOC_Os03g37970.1_3
LOC_Os05g06450.2
dist(spfitiex12sec12[lab[, 1] == i, ])
hclust (*, "ward.D2")

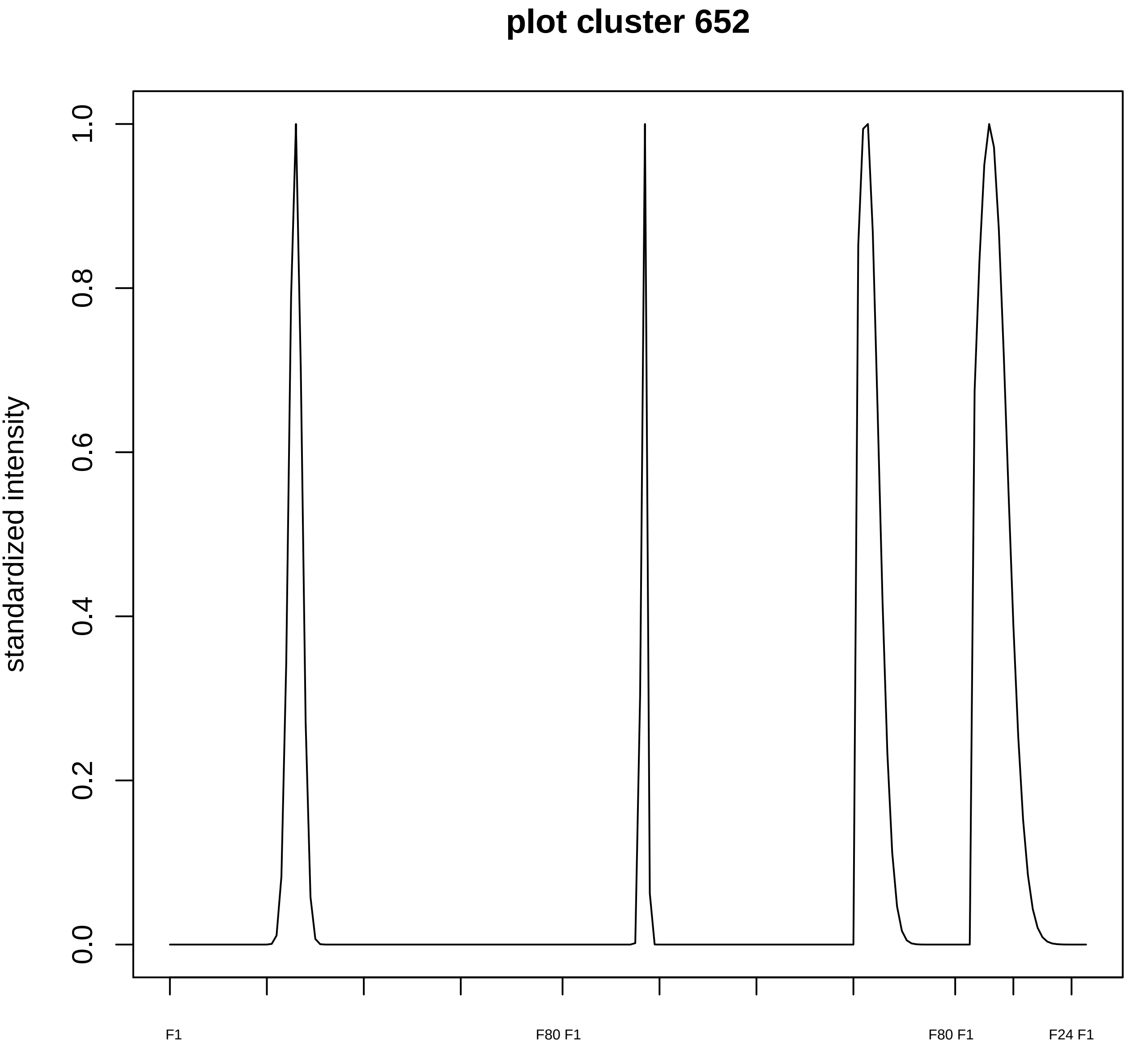

plot cluster 652
standardized intensity
0.0
0.2
0.4
0.6
0.8
1.0
F1
F80 F1
F80 F1
F24 F1

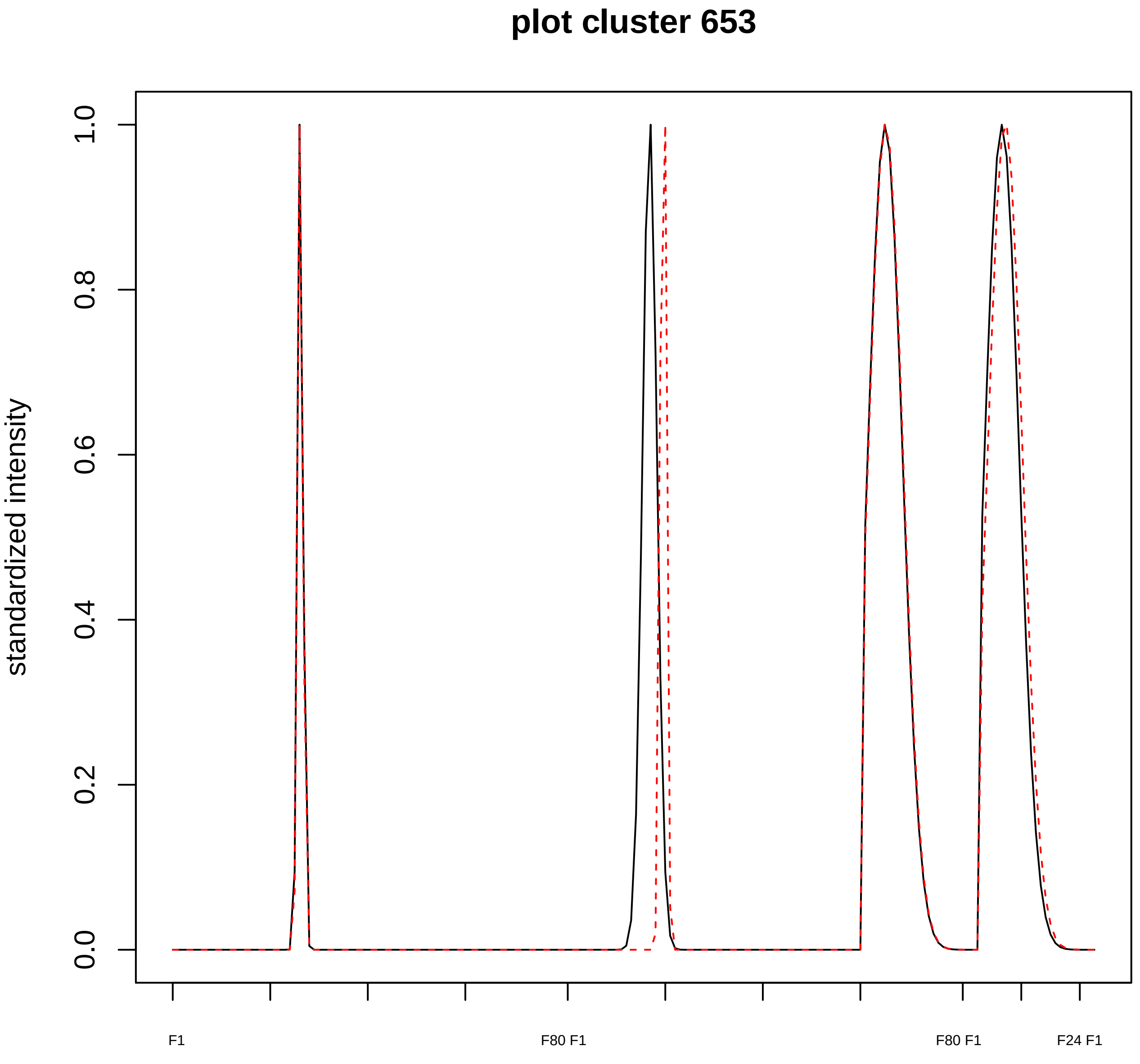

plot cluster 653
standardized intensity
0.0
0.2
0.4
0.6
0.8
1.0
F1
F80 F1
F80 F1
F24 F1

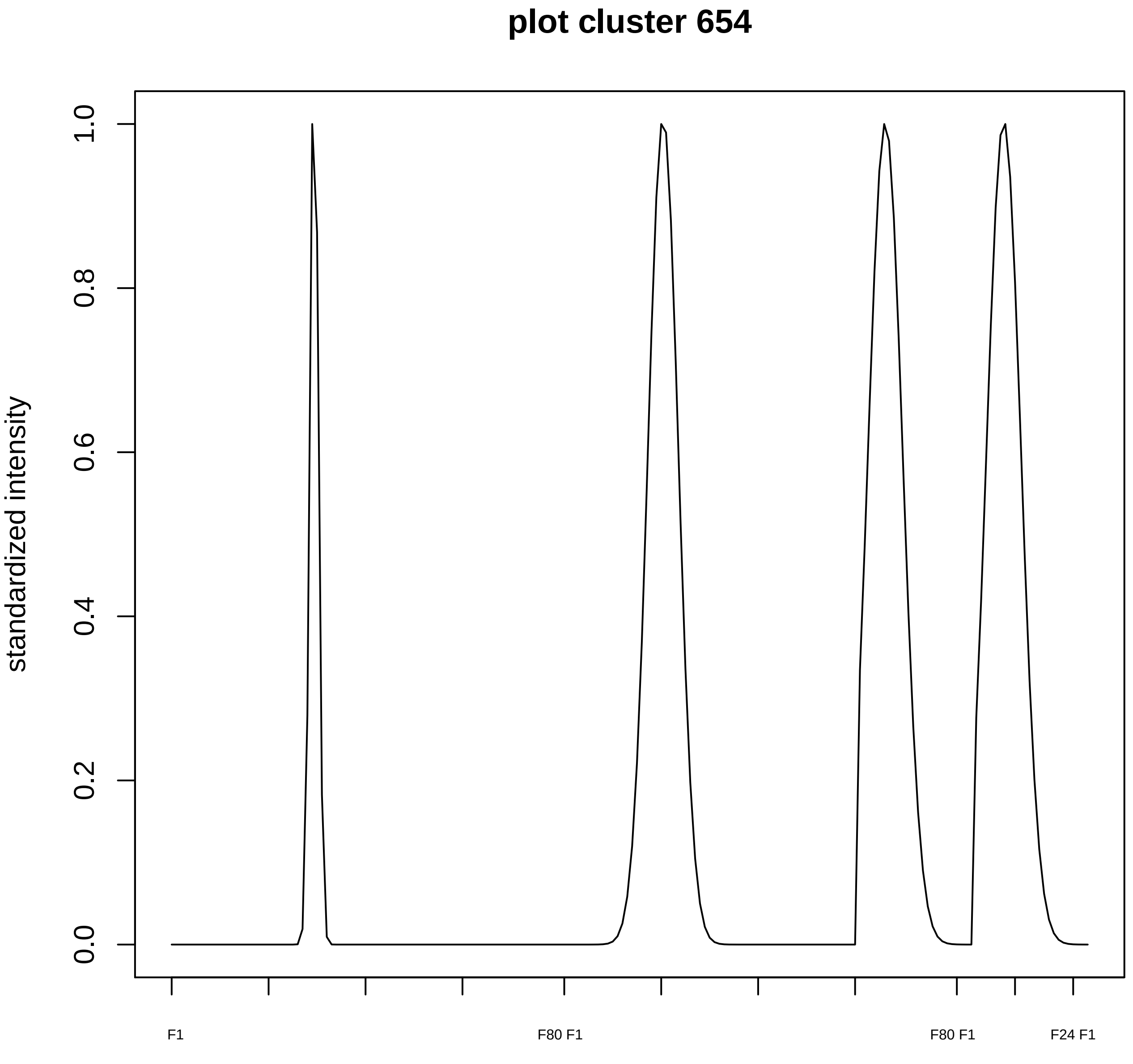

plot cluster 654
standardized intensity
1.0
0.8
0.6
0.4
0.2
0.0
F1
F80 F1
F80 F1
F24 F1

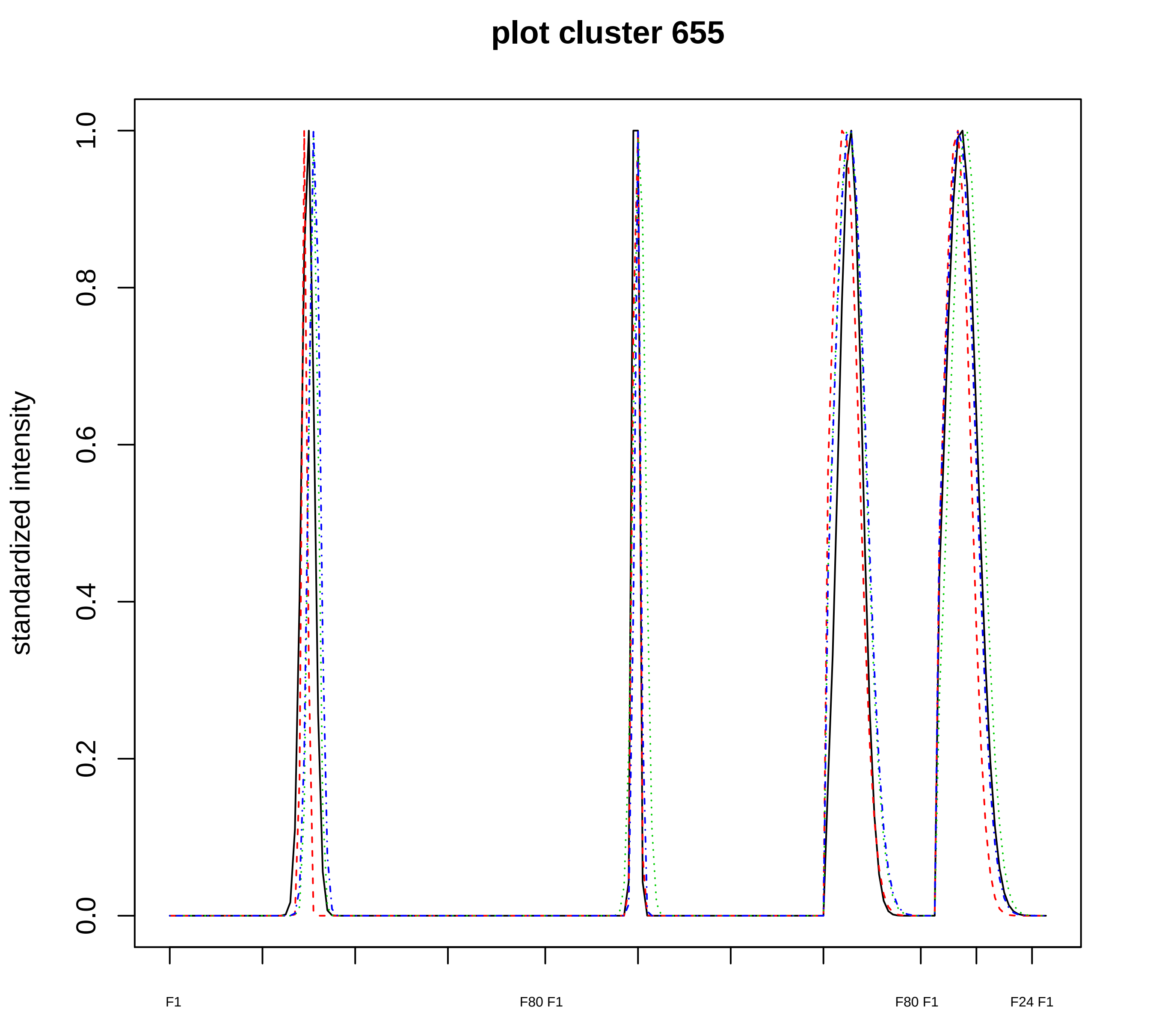
plot cluster 655
standardized intensity
0.0
0.2
0.4
0.6
0.8
1.0
F1
F80 F1
F80 F1
F24 F1

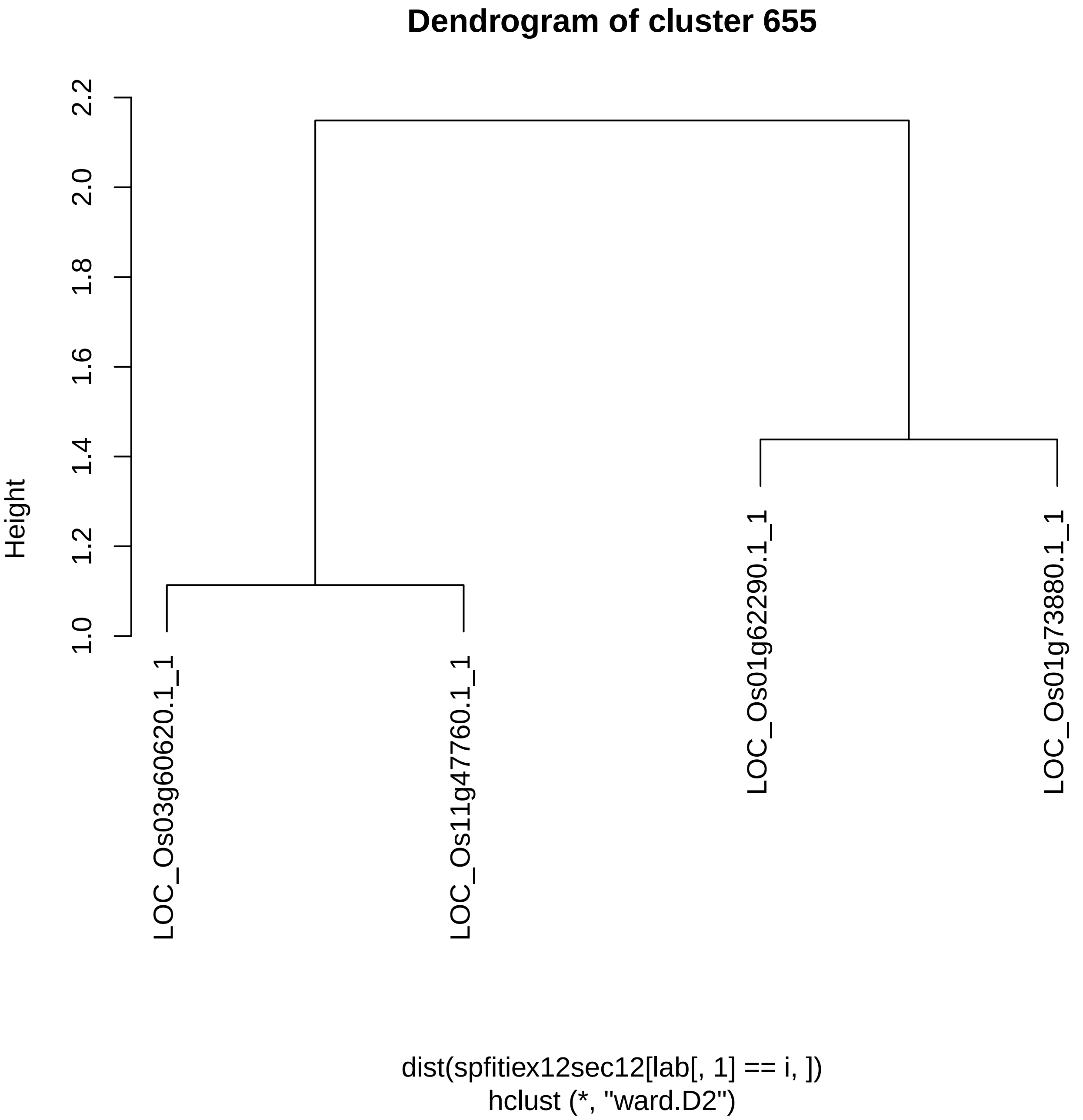
Dendrogram of cluster 655
Height
2.2
2.0
1.8
1.6
1.4
1.2
1.0
LOC_Os03g60620.1_1_
LOC_Os11g47760.1_1_
LOC_Os01g62290.1_1_
LOC_Os01g73880.1_1_
dist(spfitiex12sec12[lab[, 1] == i, ])
hclust (*, "ward.D2")

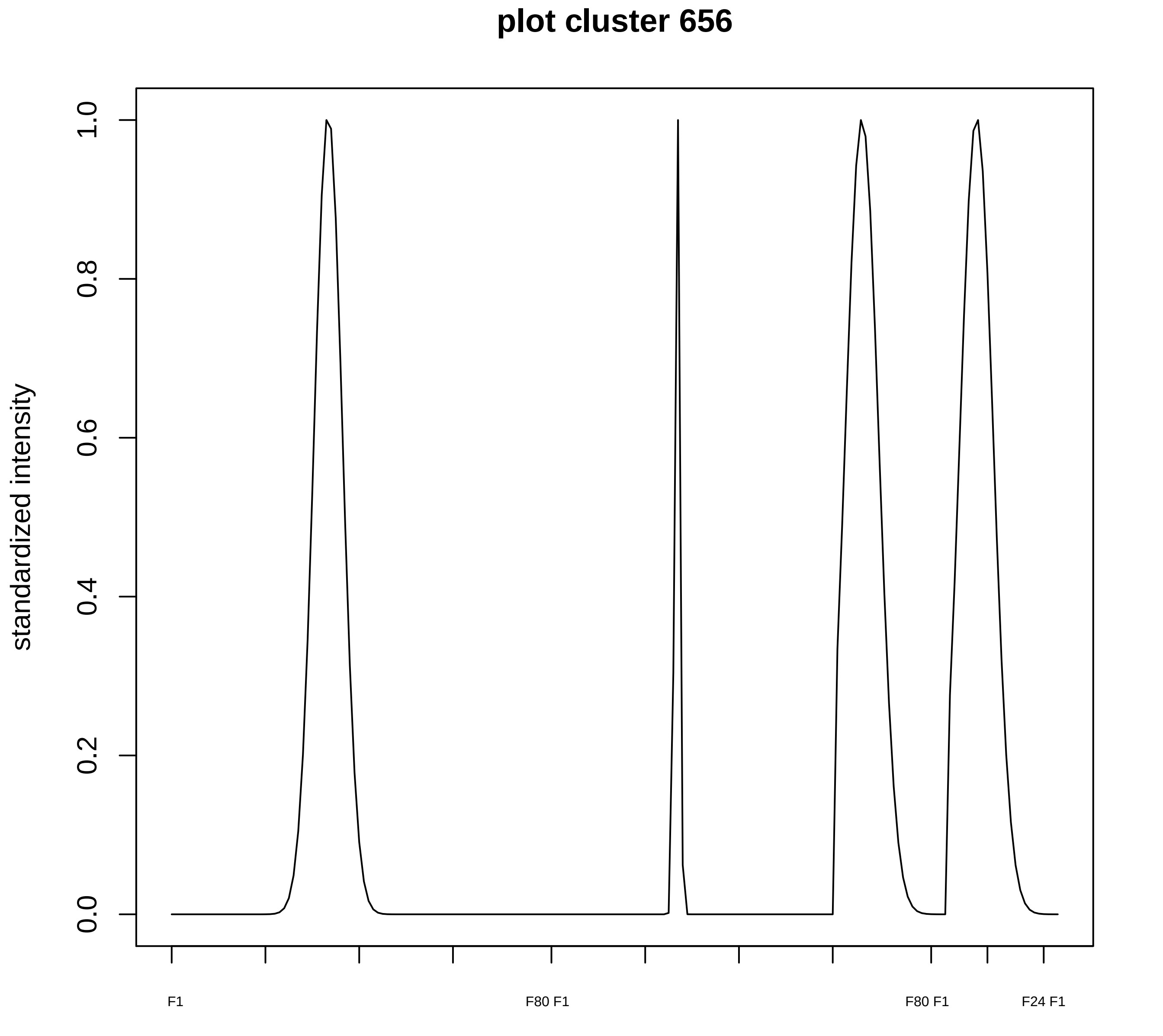

plot cluster 656
standardized intensity
0.0
0.2
0.4
0.6
0.8
1.0
F1
F80 F1
F80 F1
F24 F1

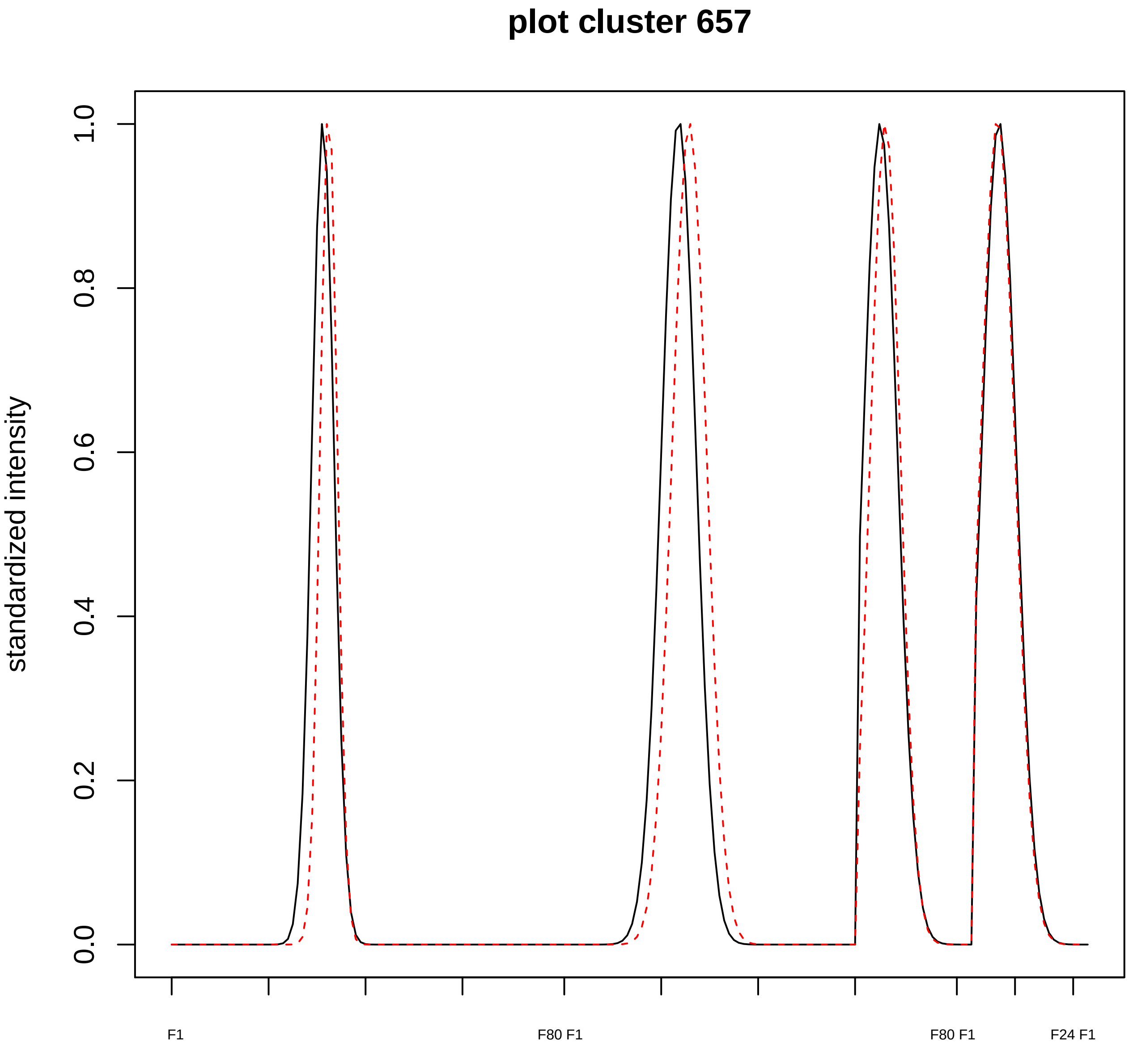
plot cluster 657
standardized intensity
0.0
0.2
0.4
0.6
0.8
1.0
F1
F80 F1
F80 F1
F24 F1

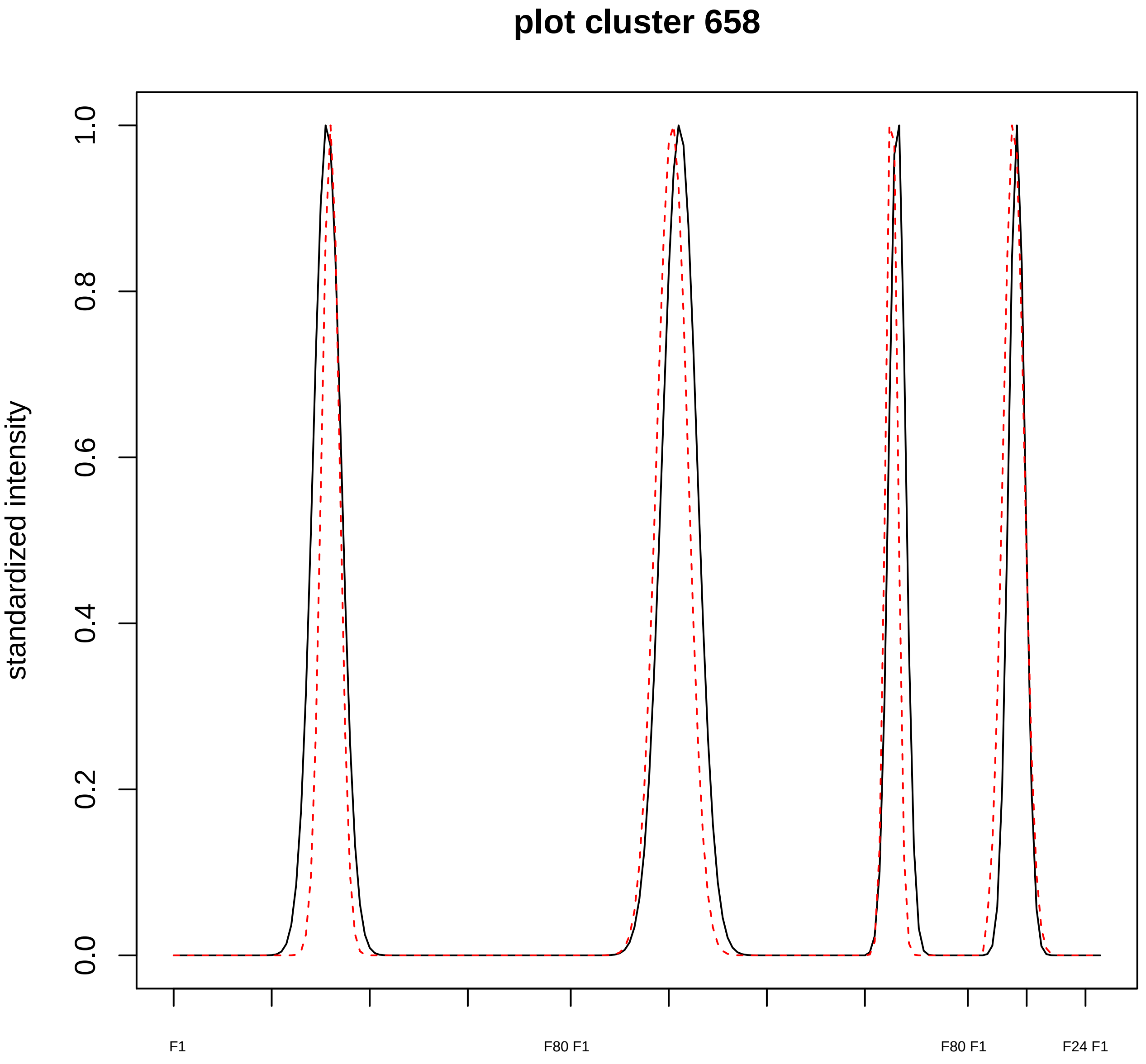

plot cluster 658
standardized intensity
0.0
0.2
0.4
0.6
0.8
1.0
F1
F80 F1
F80 F1
F24 F1

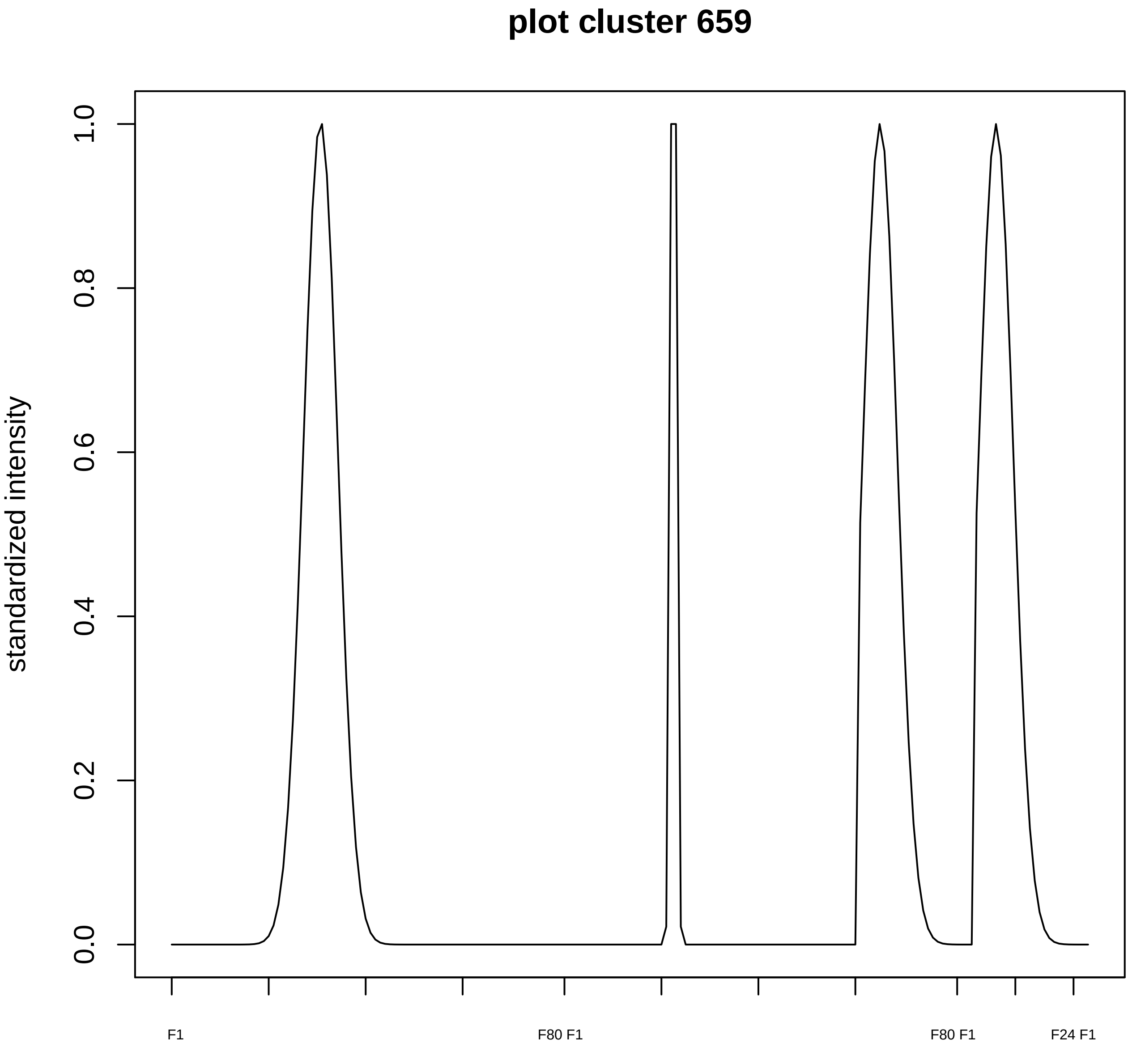
plot cluster 659
standardized intensity
0.0
0.2
0.4
0.6
0.8
1.0
F1
F80 F1
F80 F1
F24 F1

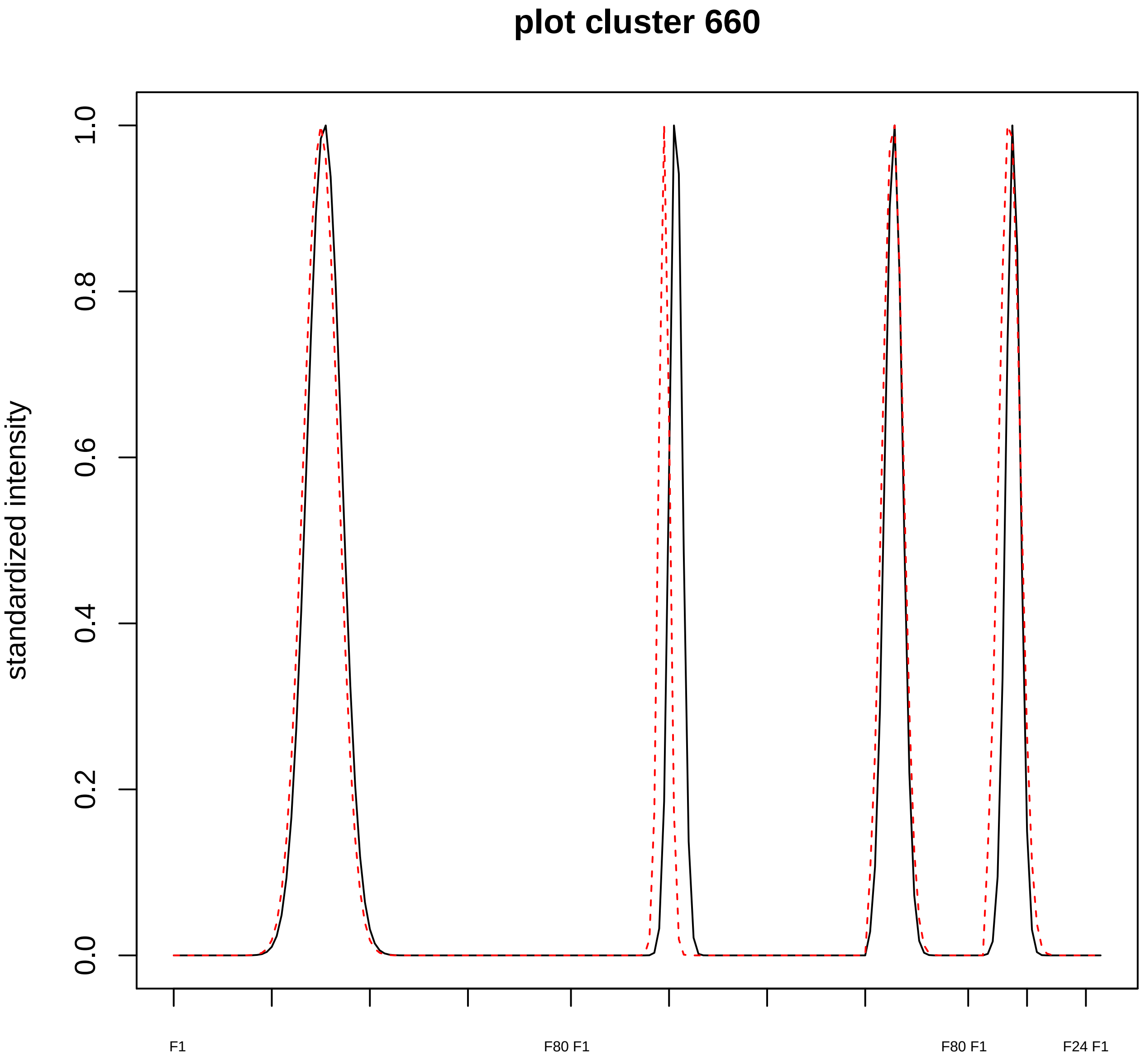
plot cluster 660
standardized intensity
0.0
0.2
0.4
0.6
0.8
1.0
F1
F80 F1
F80 F1
F24 F1

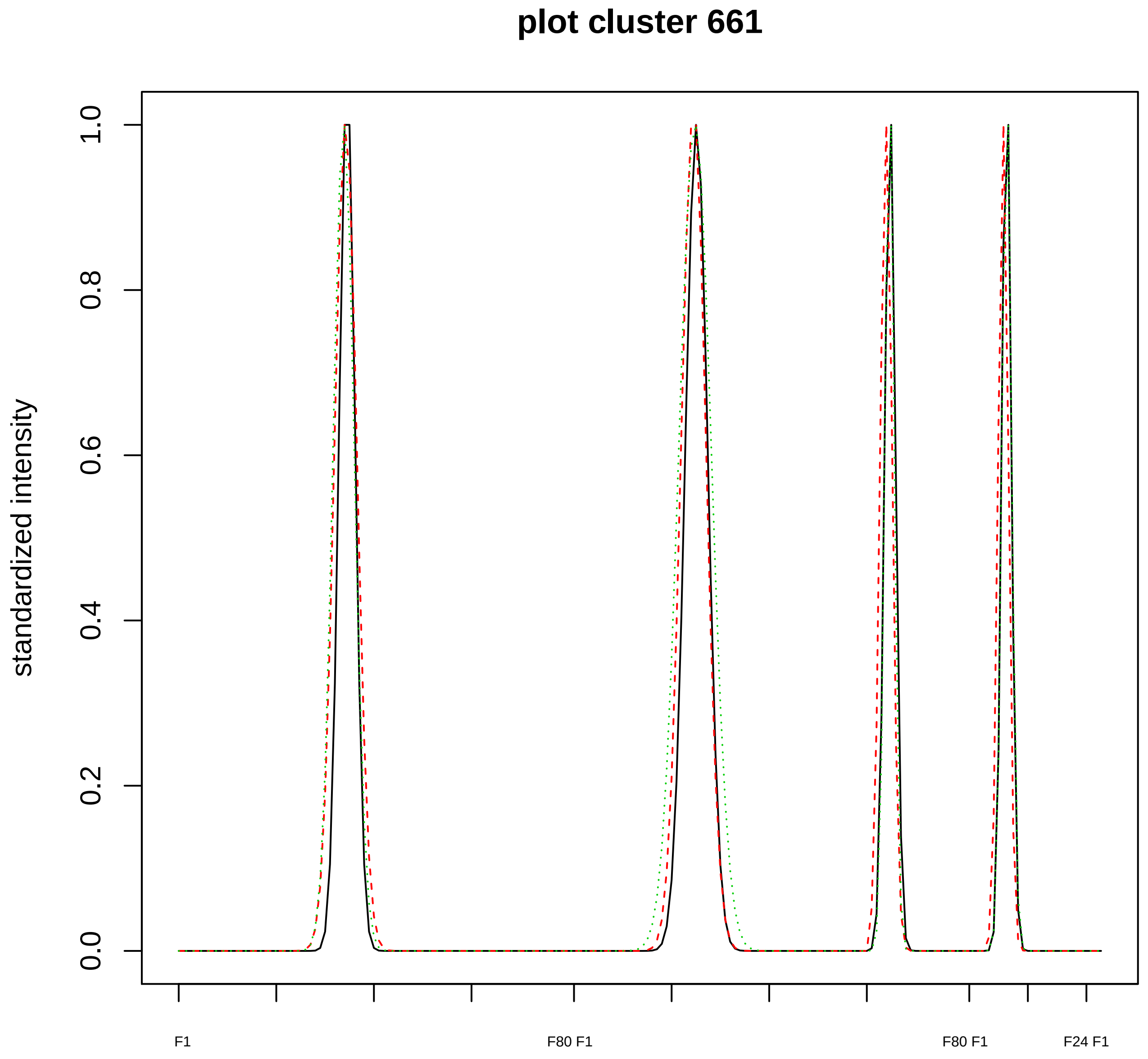
plot cluster 661
standardized intensity
0.0
0.2
0.4
0.6
0.8
1.0
F1
F80 F1
F80 F1
F24 F1

#### Dendrogram of cluster 661

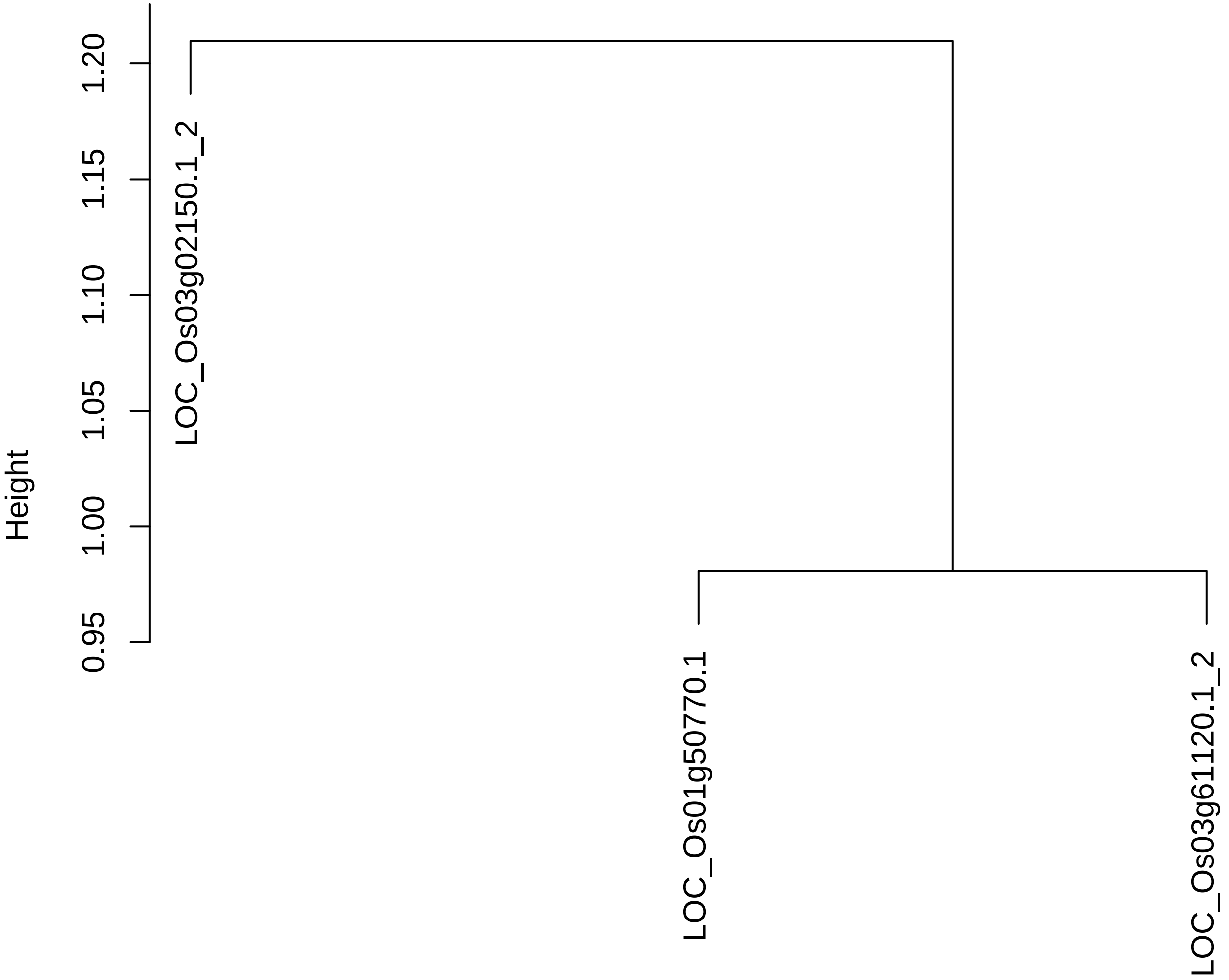


dist(spfitiex12sec12[lab[, 1] == i, ])
hclust (*, "ward.D2")

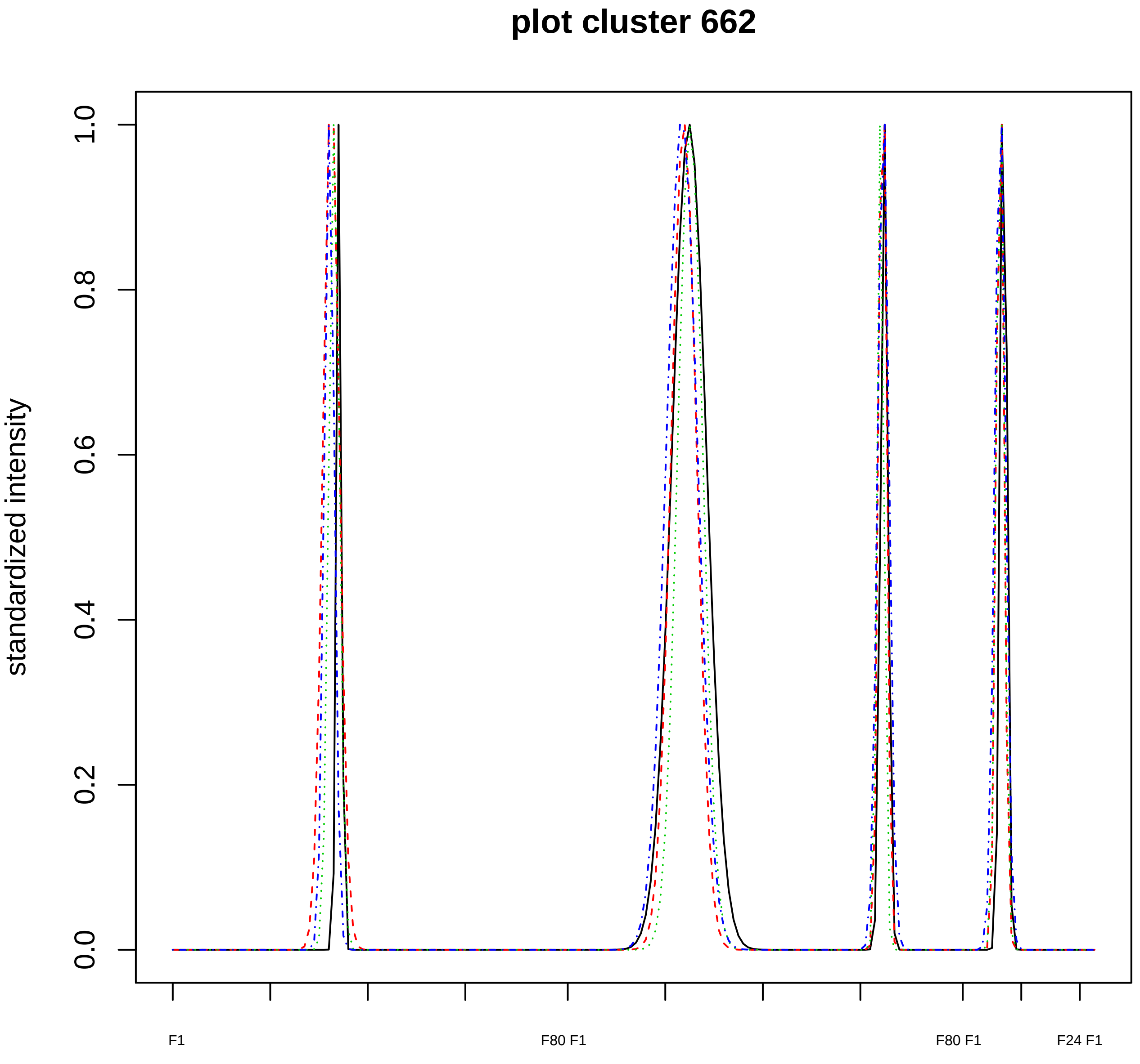

plot cluster 662
standardized intensity
0.0
0.2
0.4
0.6
0.8
1.0
F1
F80 F1
F80 F1
F24 F1

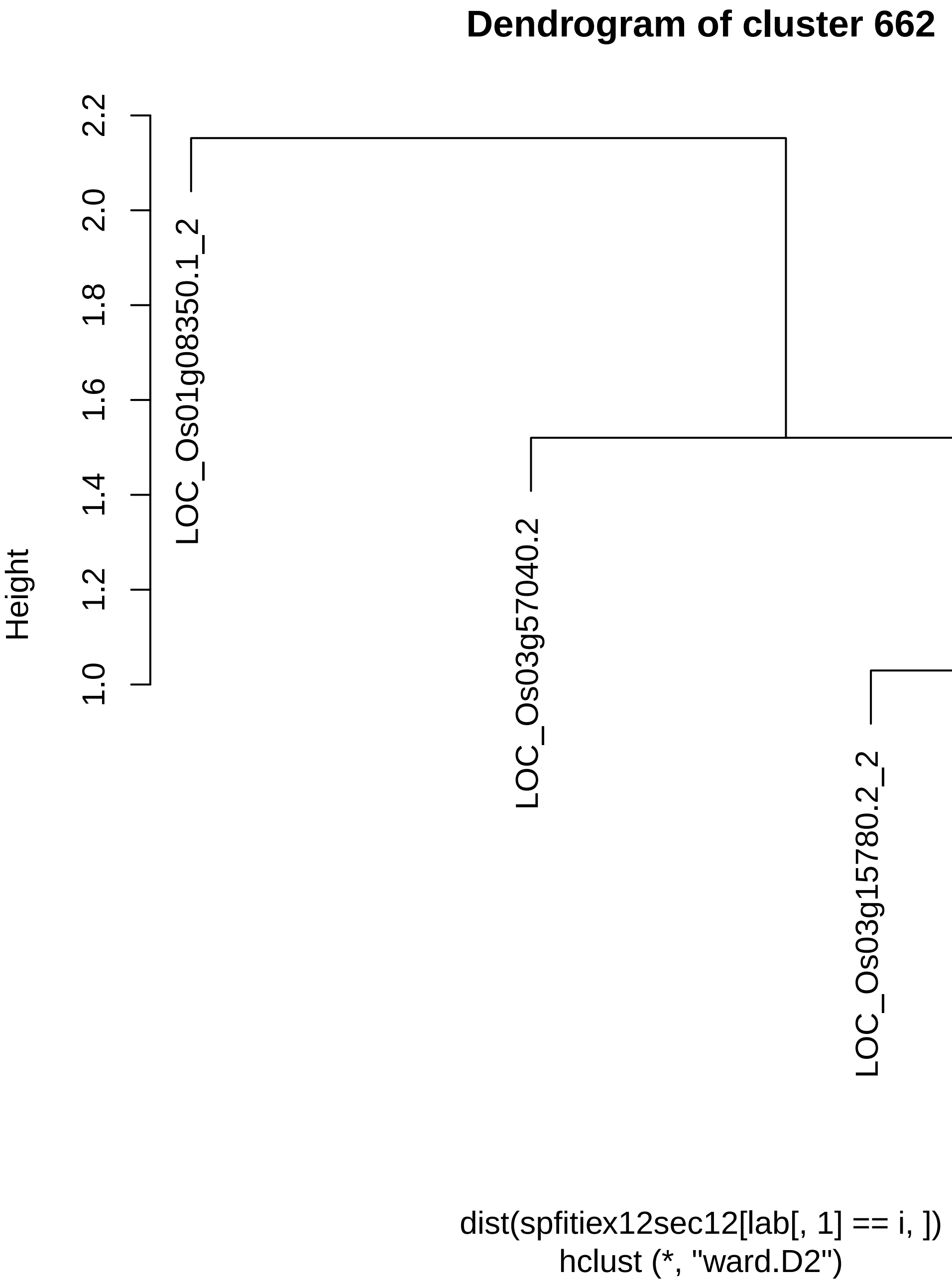
Dendrogram of cluster 662
Height
2.2
2.0
1.8
1.6
1.4
1.2
1.0
LOC_Os01g08350.1_2
LOC_Os03g57040.2
LOC_Os03g15780.2_2
LOC_Os07g01020.1
dist(spfitiex12sec12[lab[, 1] == i, ])
hclust (*, "ward.D2")

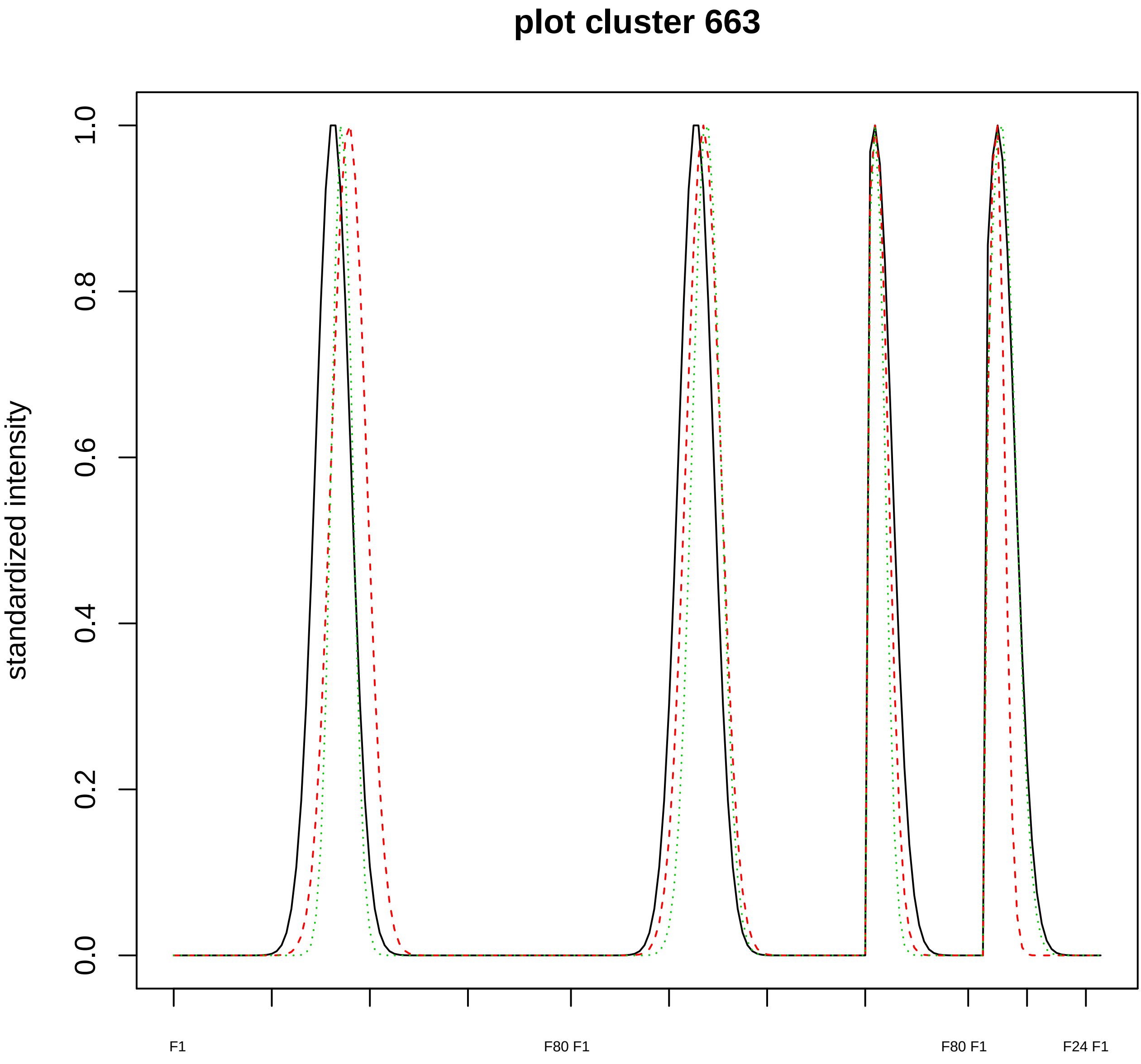
plot cluster 663
standardized intensity
0.0
0.2
0.4
0.6
0.8
1.0
F1
F80 F1
F80 F1
F24 F1

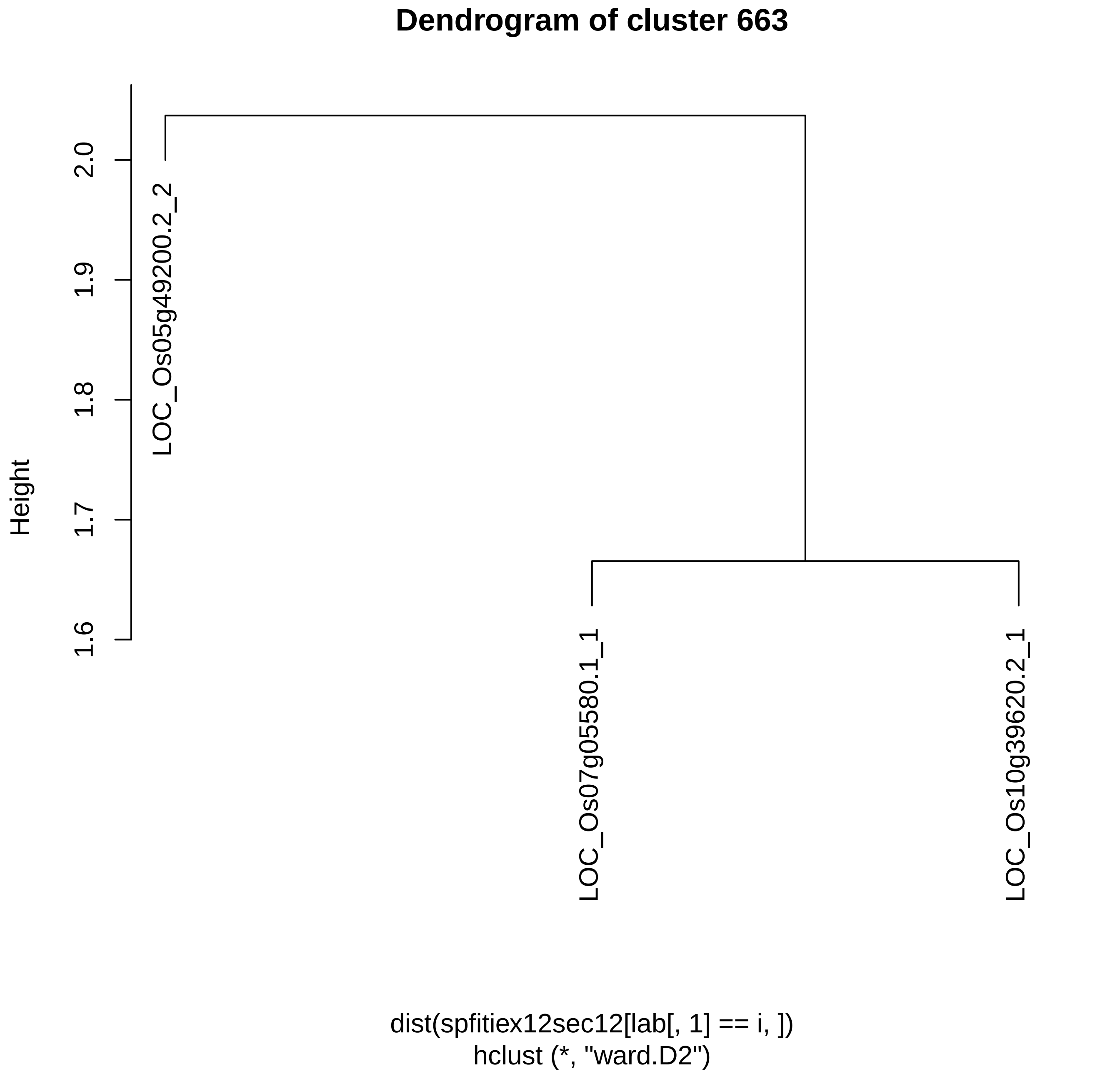

Dendrogram of cluster 663
Height
2.0
1.9
1.8
1.7
1.6
LOC_Os05g49200.2_2
LOC_Os07g05580.1_1
LOC_Os10g39620.2_1
dist(spfitiex12sec12[lab[, 1] == i, ])
hclust (*, "ward.D2")

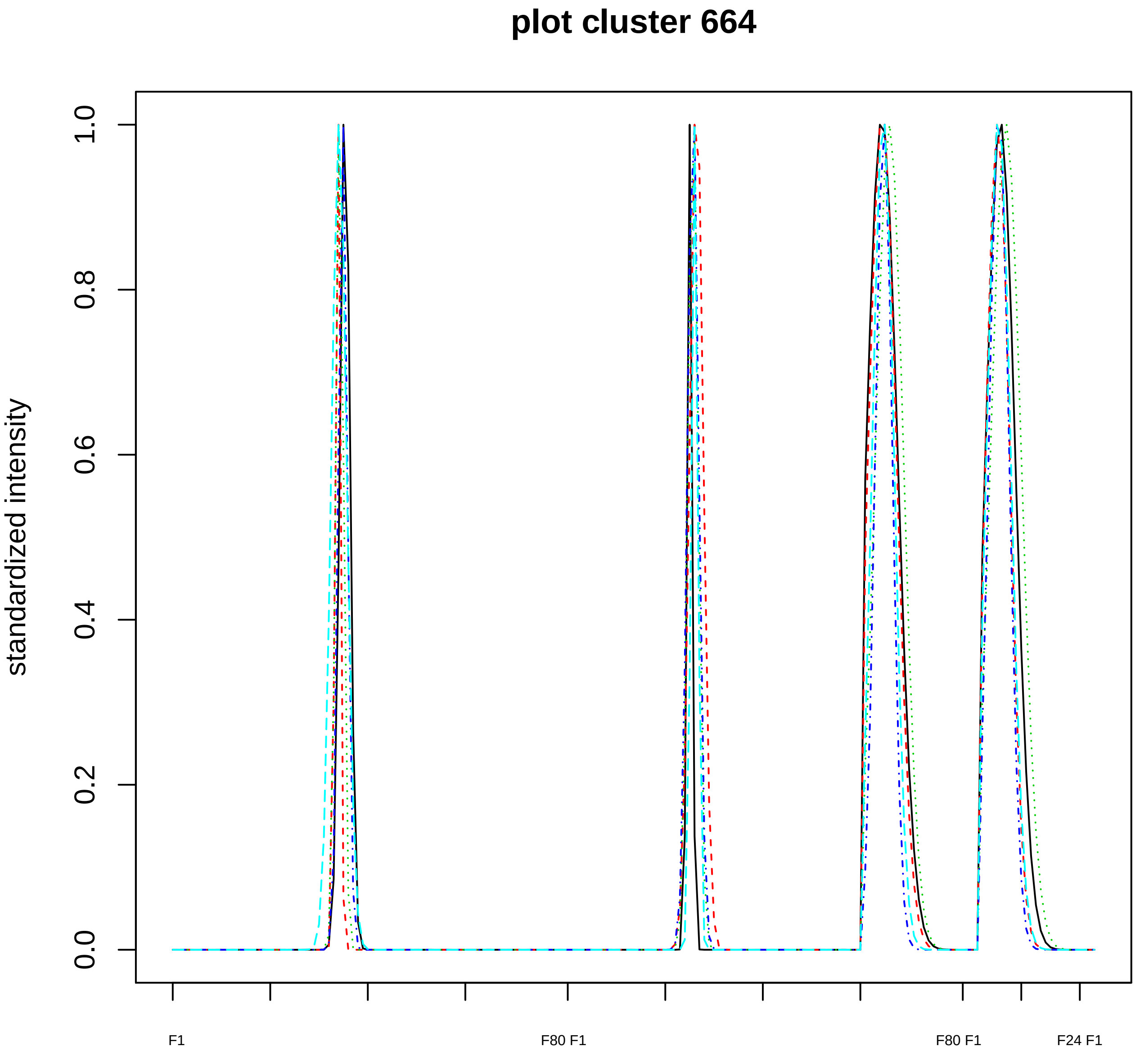

plot cluster 664
standardized intensity
0.0
0.2
0.4
0.6
0.8
1.0
F1
F80 F1
F80 F1
F24 F1

### Dendrogram of cluster 664

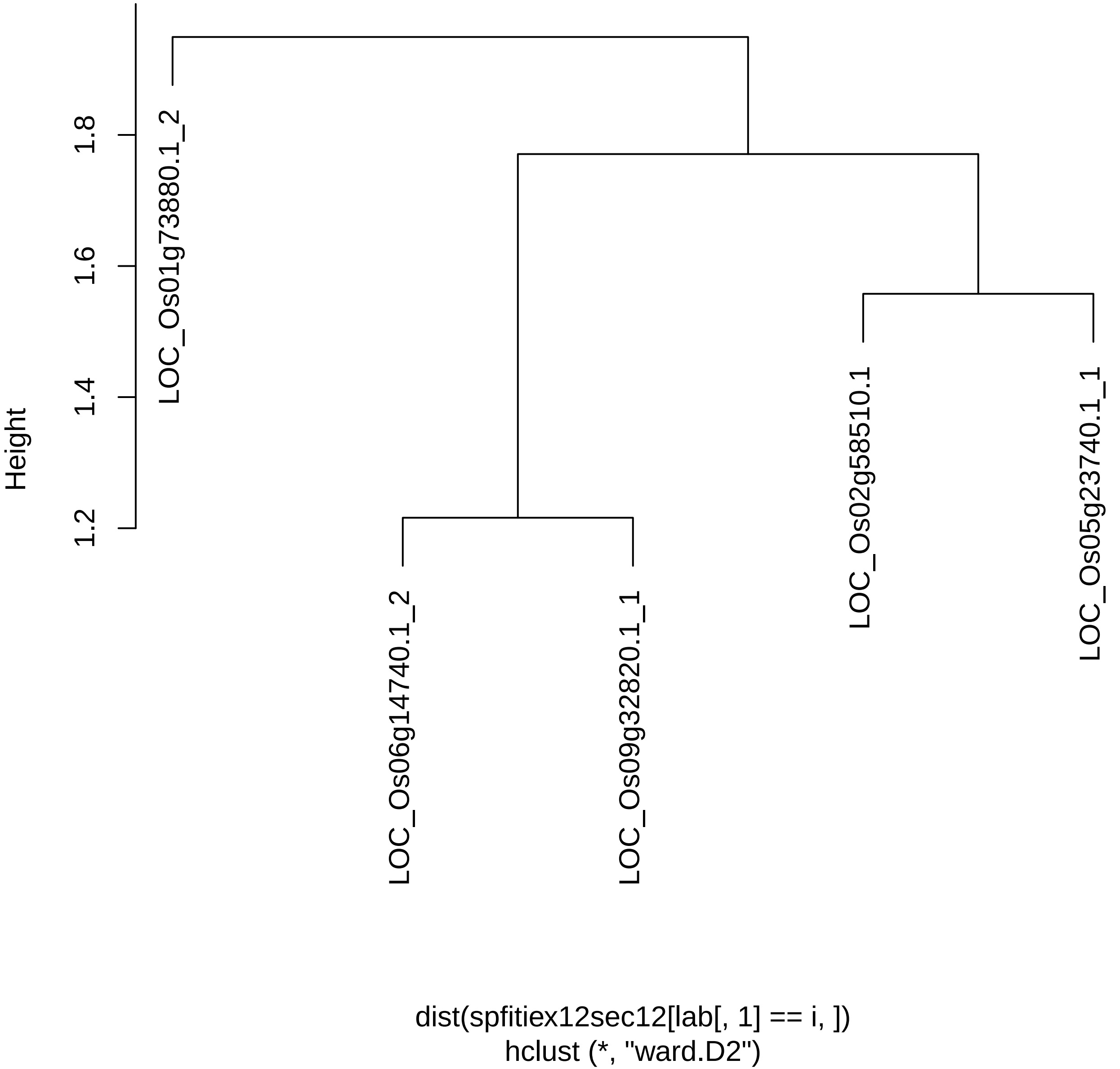

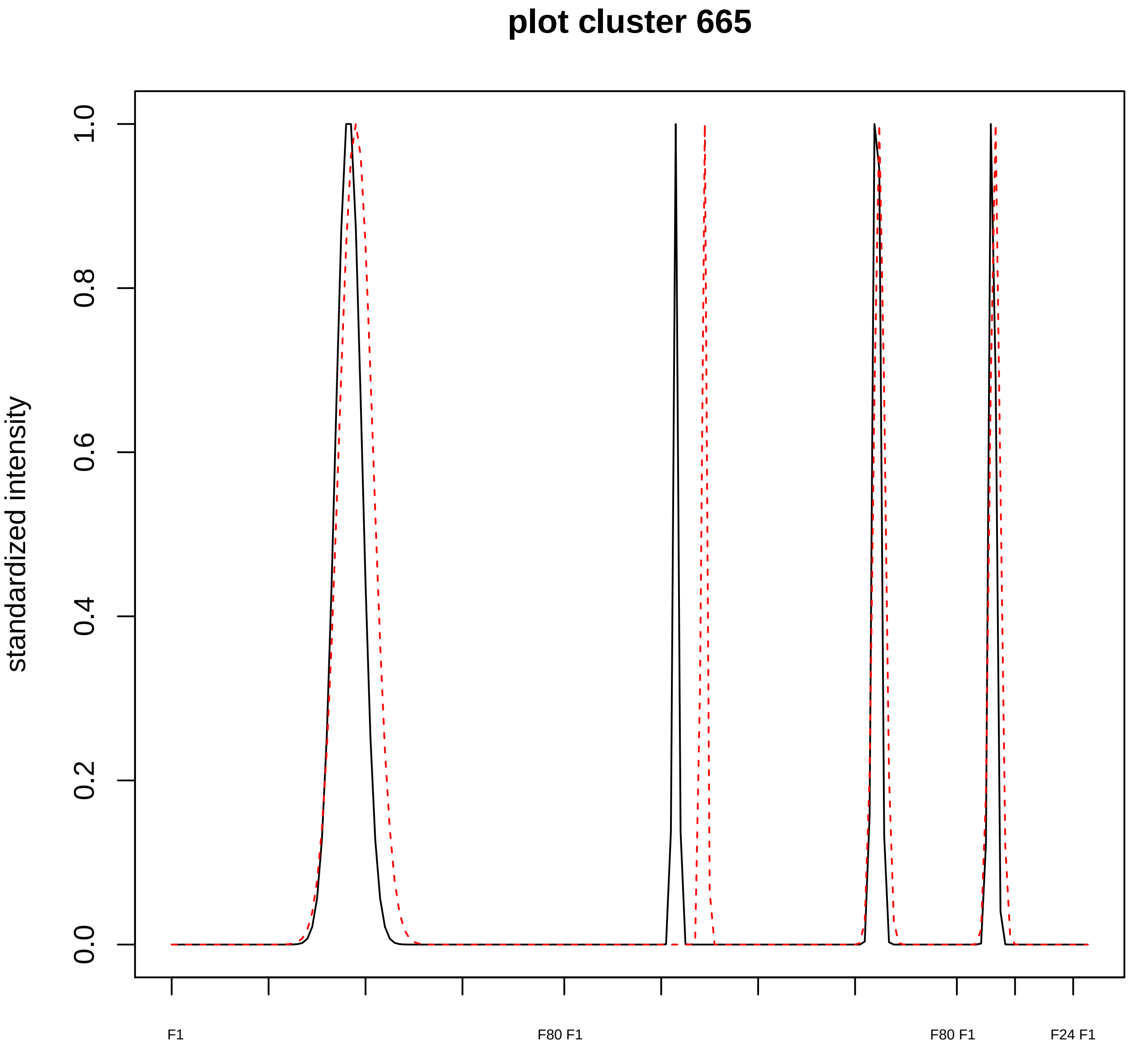
plot cluster 665
standardized intensity
1.0
0.8
0.6
0.4
0.2
0.0
F1
F80 F1
F80 F1
F24 F1

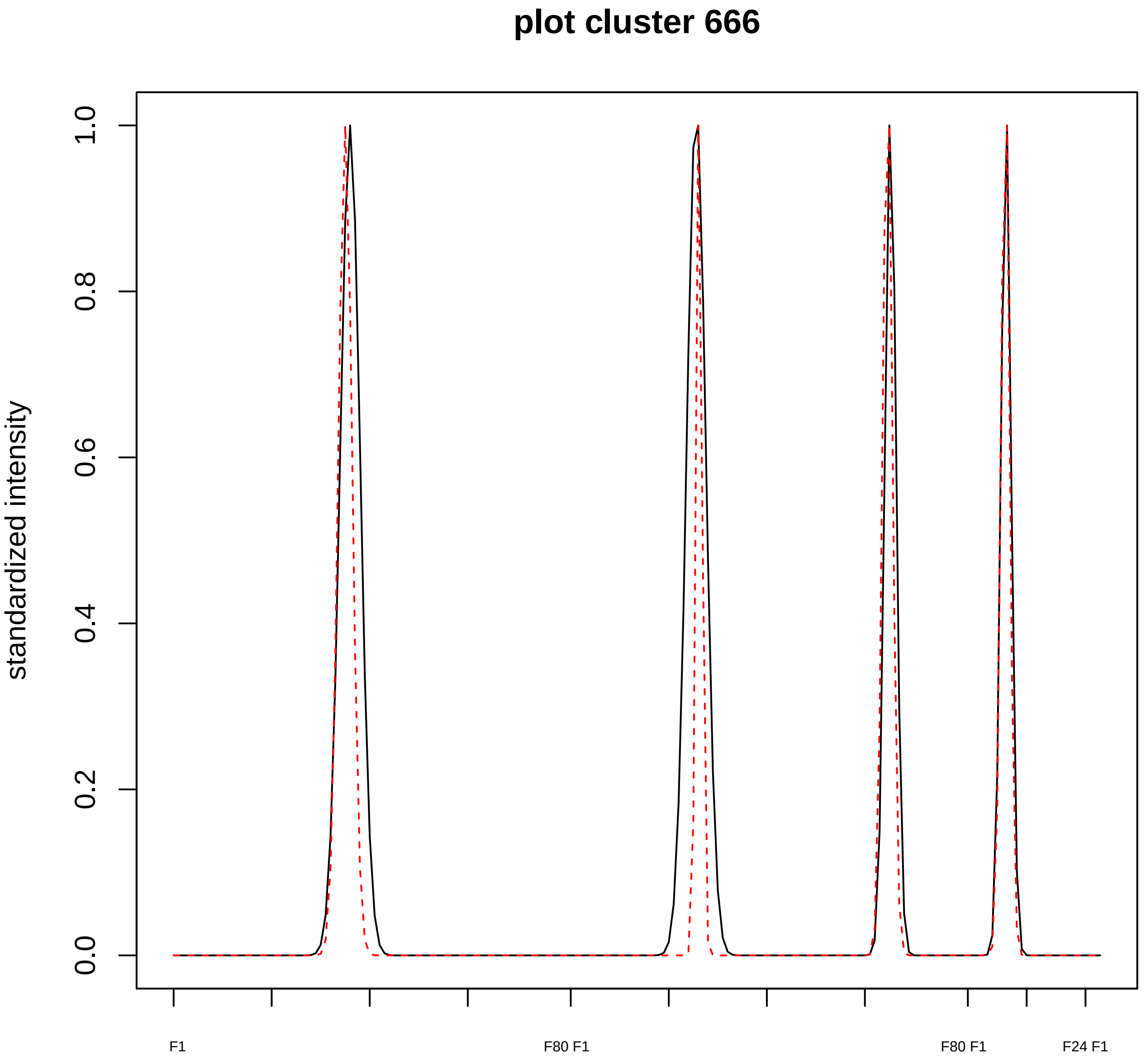

plot cluster 666
standardized intensity
0.0
0.2
0.4
0.6
0.8
1.0
F1
F80 F1
F80 F1
F24 F1

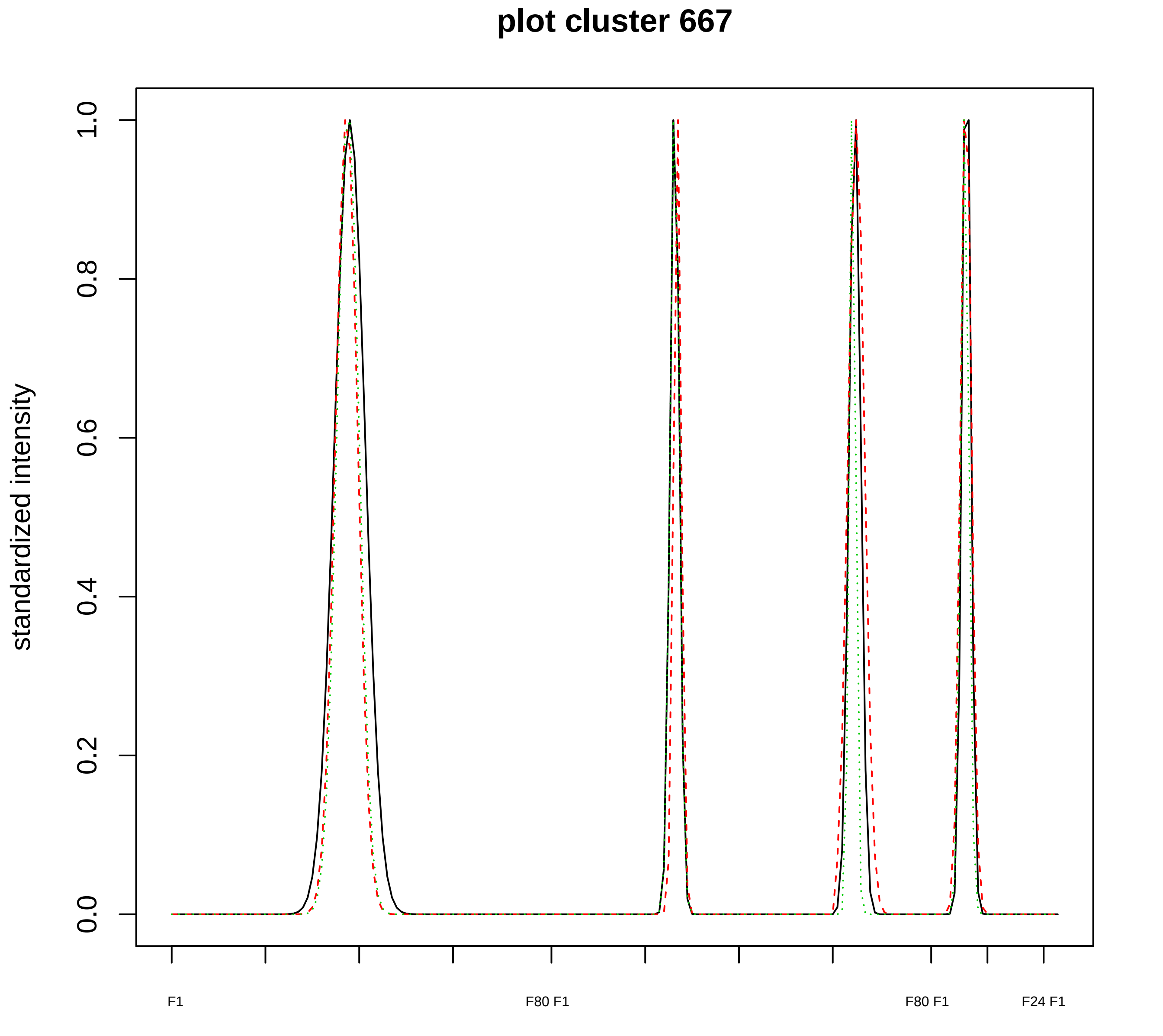
plot cluster 667
standardized intensity
0.0
0.2
0.4
0.6
0.8
1.0
F1
F80 F1
F80 F1
F24 F1

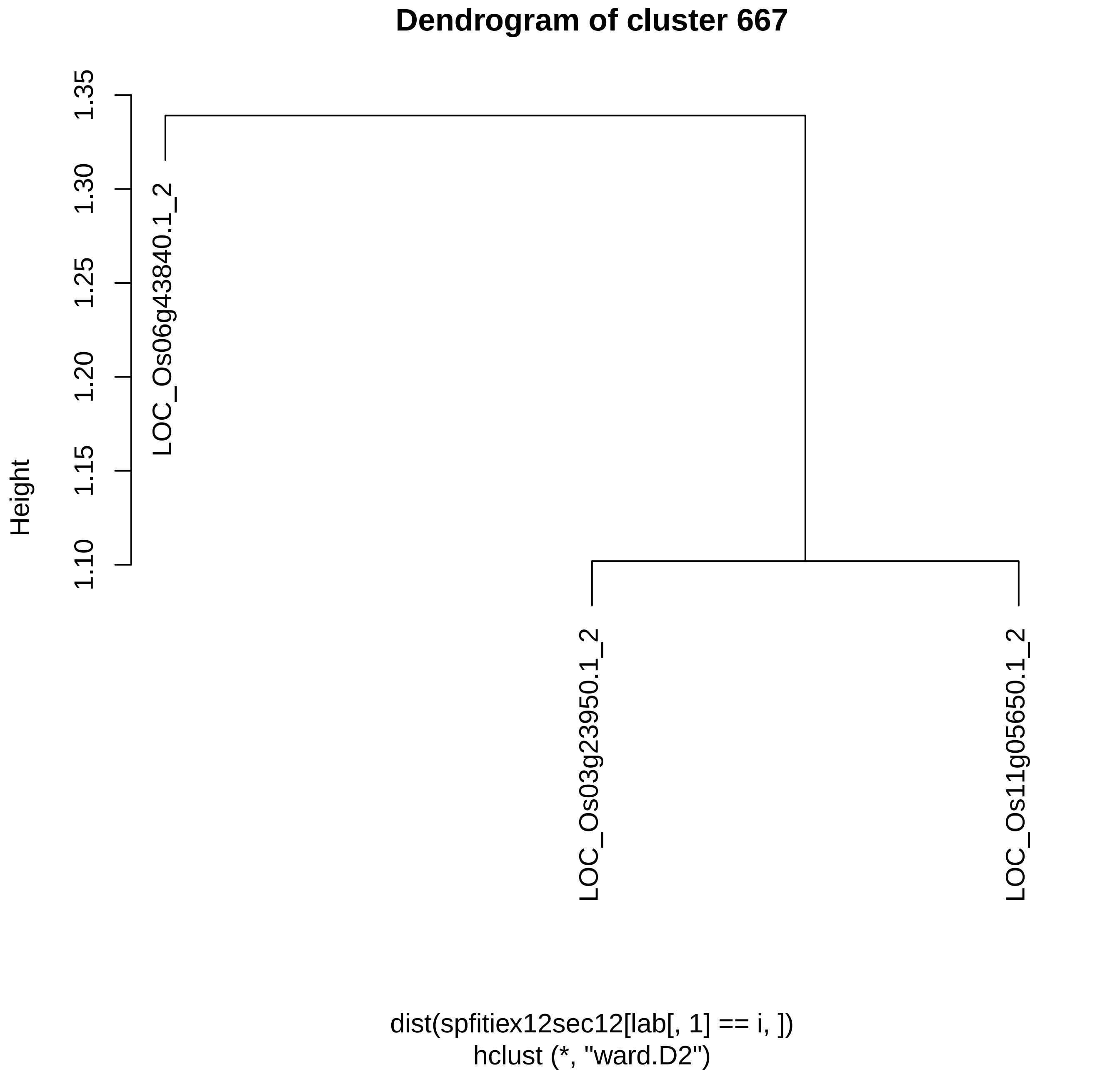
Dendrogram of cluster 667
Height
1.10
1.15
1.20
1.25
1.30
1.35
LOC_Os06g43840.1_2
LOC_Os03g23950.1_2
LOC_Os11g05650.1_2
dist(spfitiex12sec12[lab[, 1] == i, ])
hclust (*, "ward.D2")

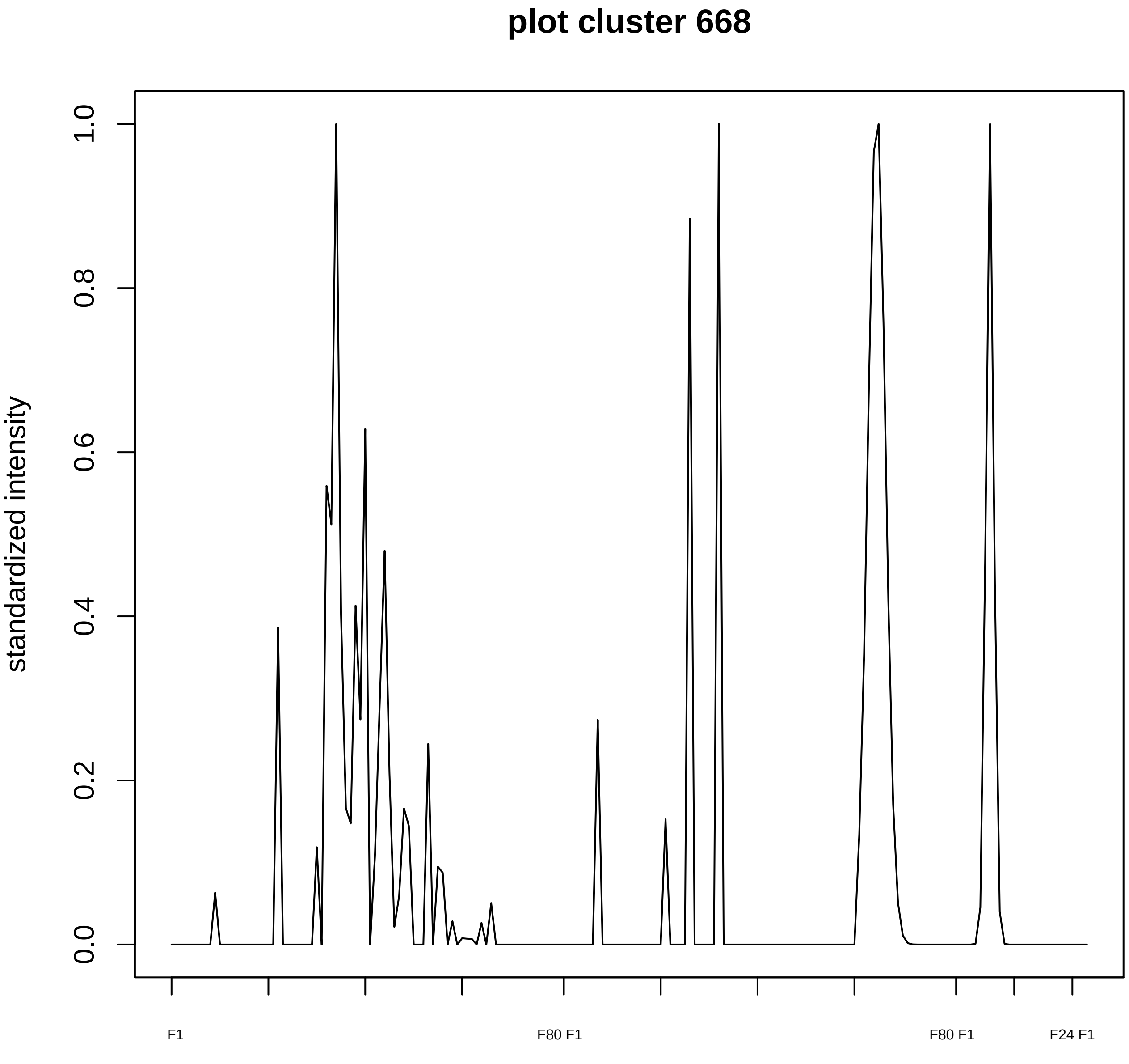
plot cluster 668
standardized intensity
0.0
0.2
0.4
0.6
0.8
1.0
F1
F80 F1
F80 F1
F24 F1

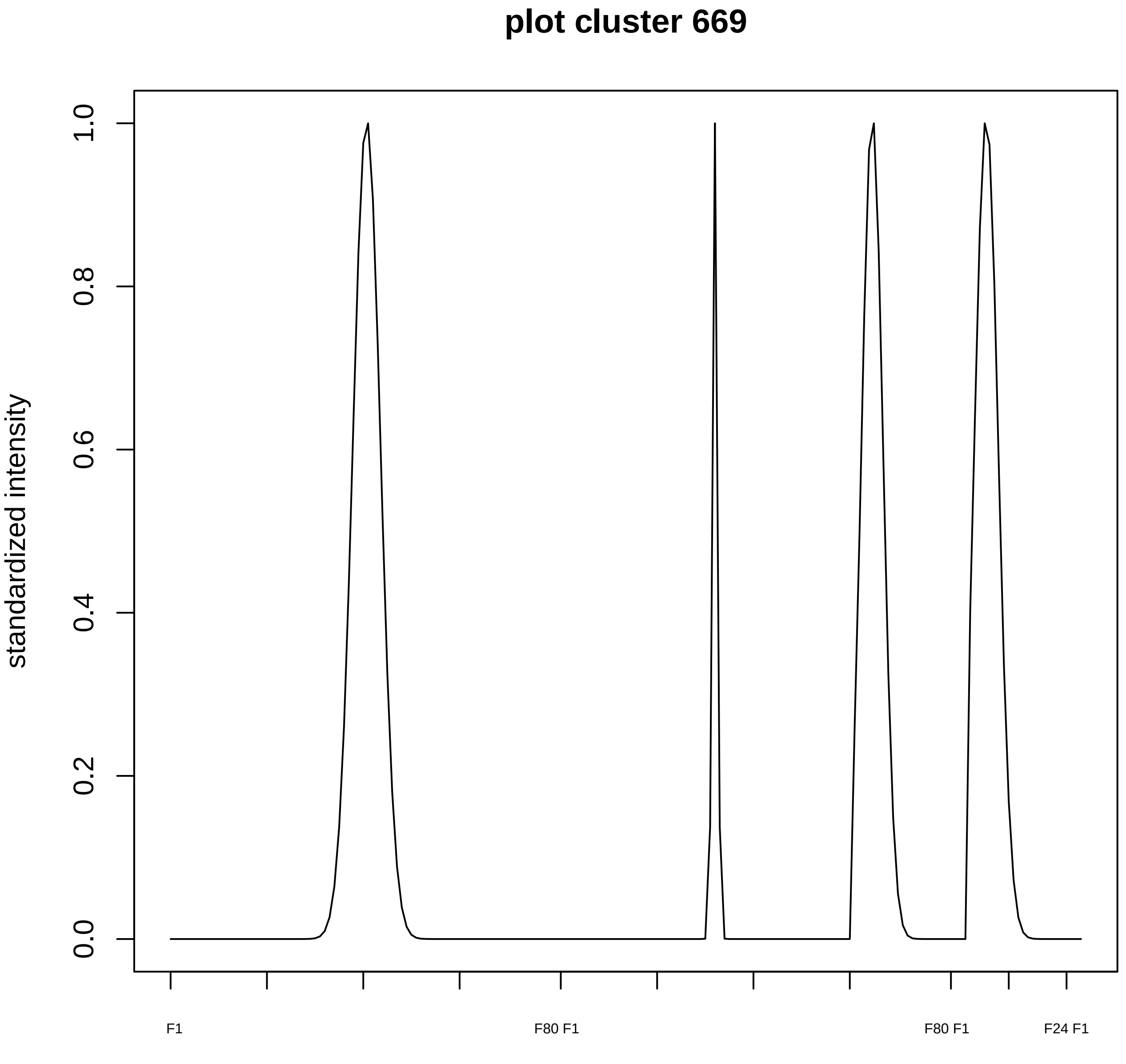

plot cluster 669
standardized intensity
0.0
0.2
0.4
0.6
0.8
1.0
F1
F80 F1
F80 F1
F24 F1

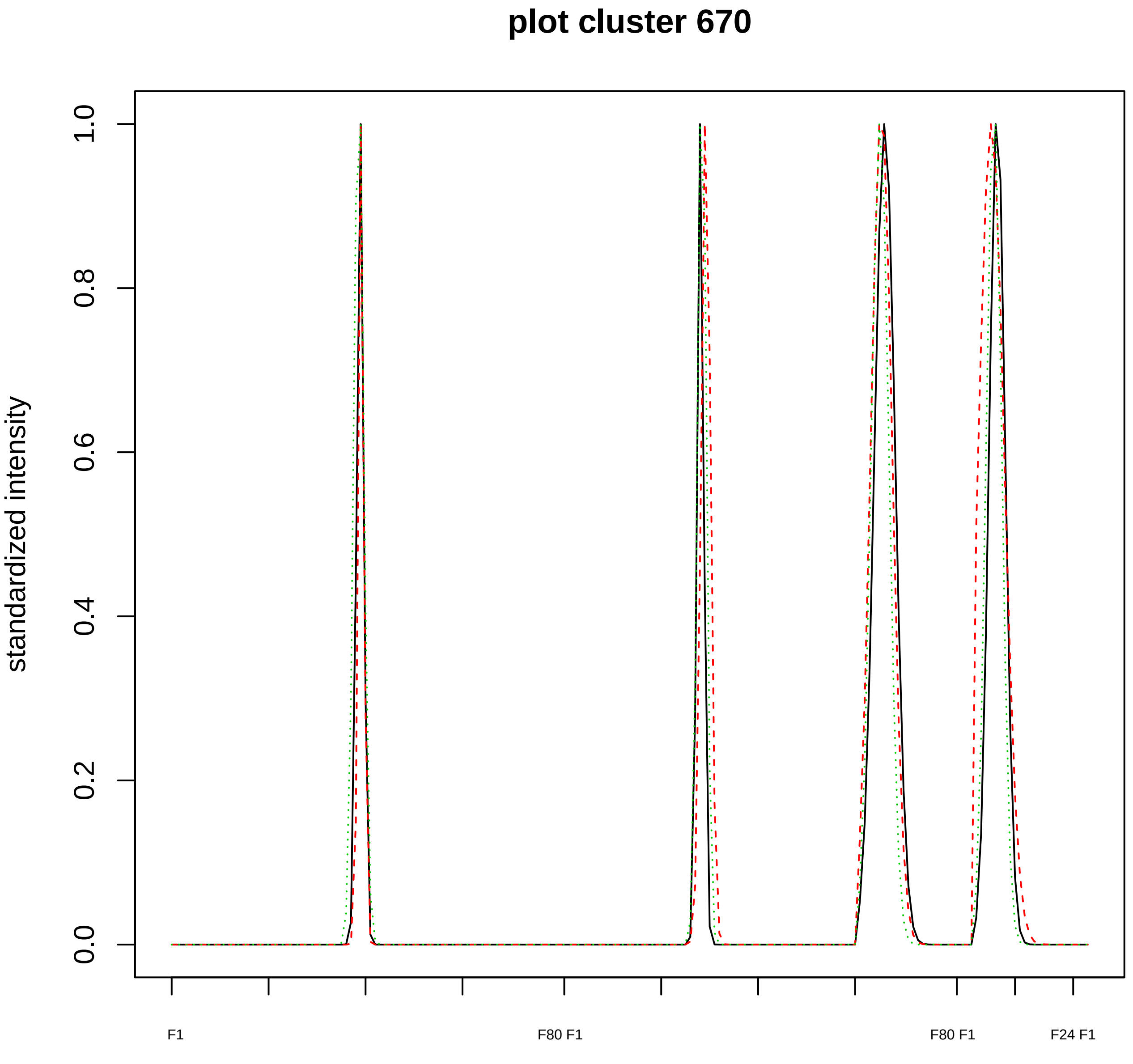

plot cluster 670
standardized intensity
0.0
0.2
0.4
0.6
0.8
1.0
F1
F80 F1
F80 F1
F24 F1

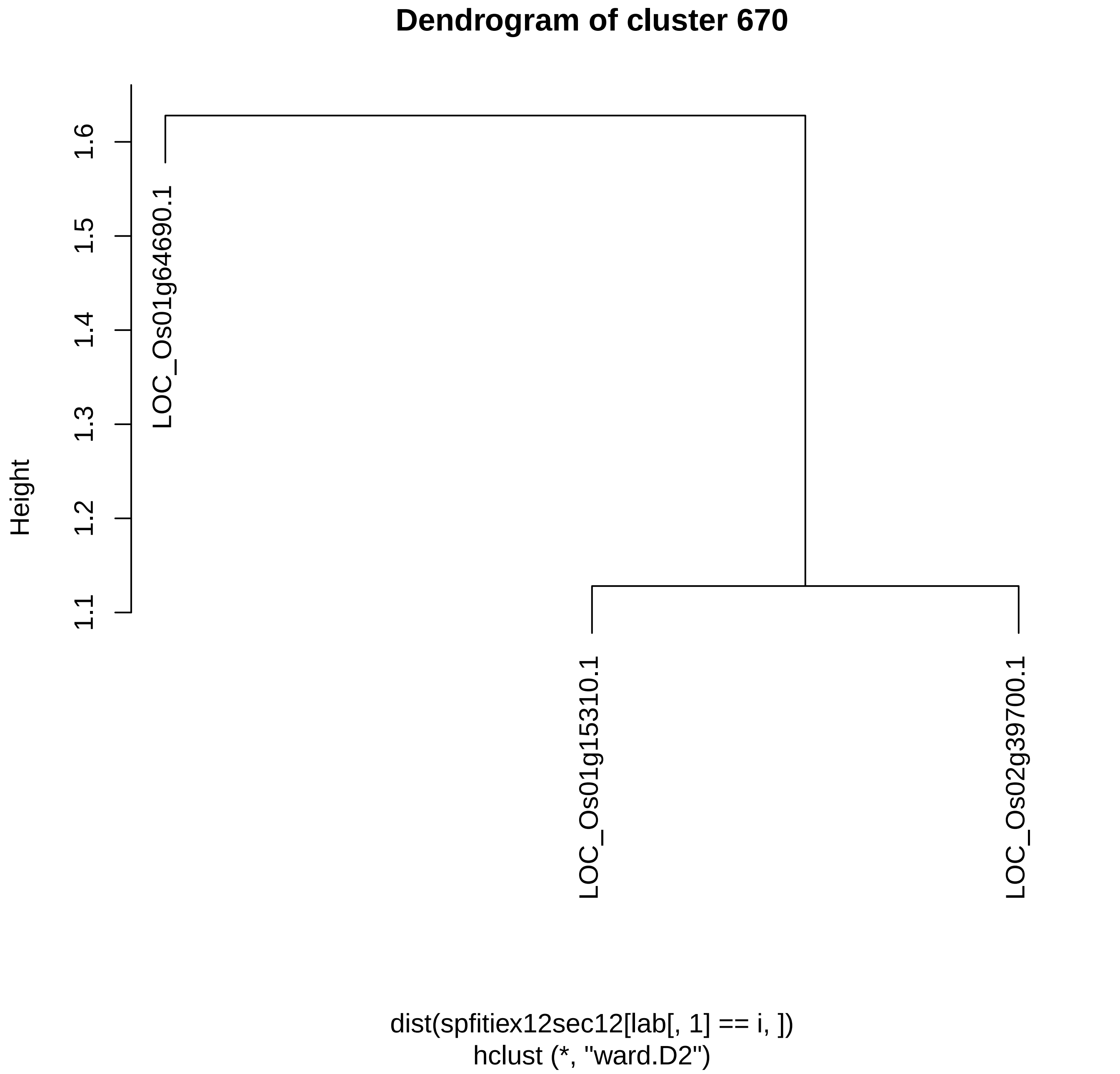
Dendrogram of cluster 670
Height
1.1
1.2
1.3
1.4
1.5
1.6
LOC_Os01g64690.1
LOC_Os01g15310.1
LOC_Os02g39700.1
dist(spfitiex12sec12[lab[, 1] == i, ])
hclust (*, "ward.D2")

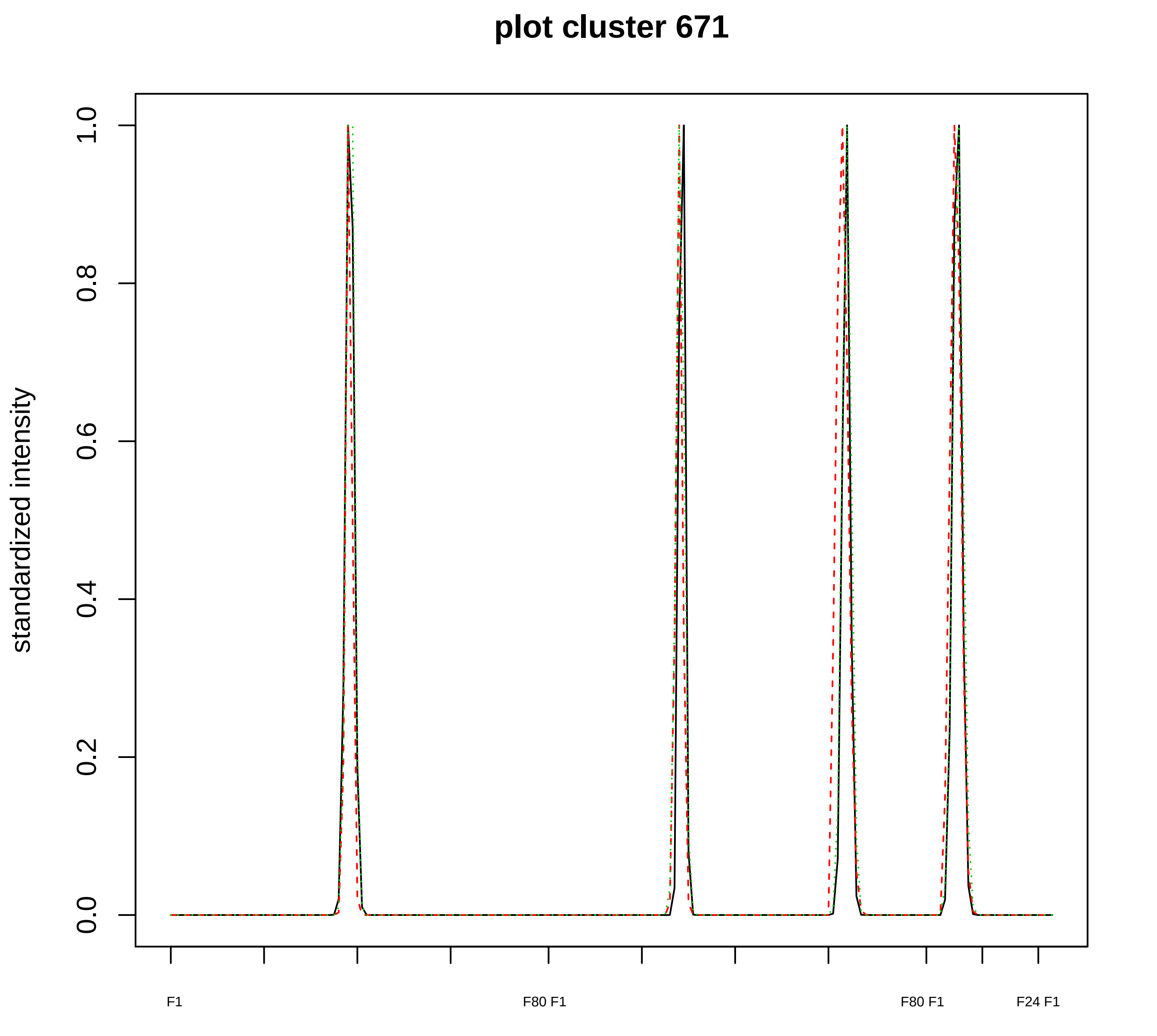
plot cluster 671
standardized intensity
0.0
0.2
0.4
0.6
0.8
1.0
F1
F80 F1
F80 F1
F24 F1

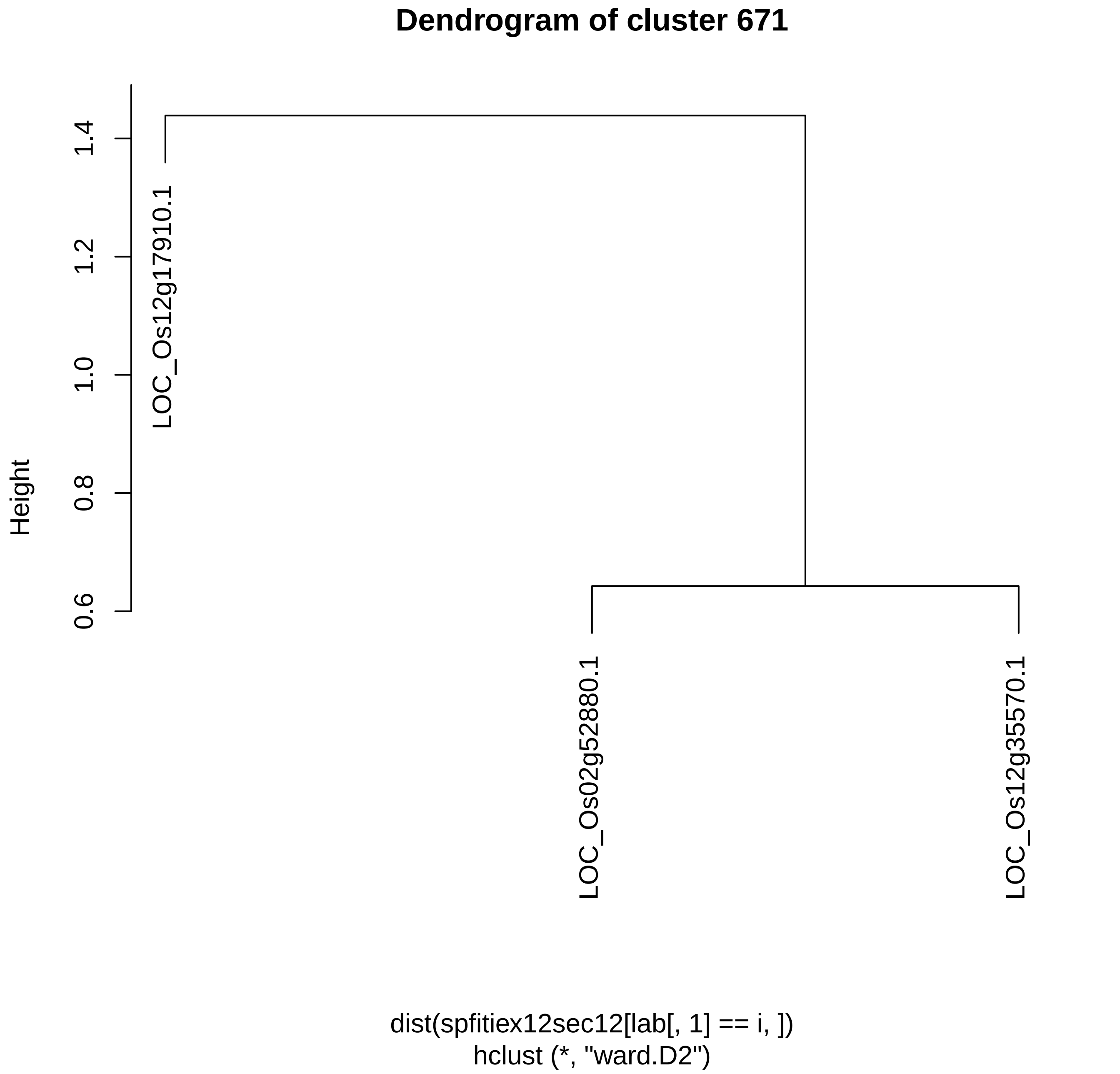

Dendrogram of cluster 671
Height
1.4
1.2
1.0
0.8
0.6
LOC_Os12g17910.1
LOC_Os02g52880.1
LOC_Os12g35570.1
dist(spfitiex12sec12[lab[, 1] == i, ])
hclust (*, "ward.D2")

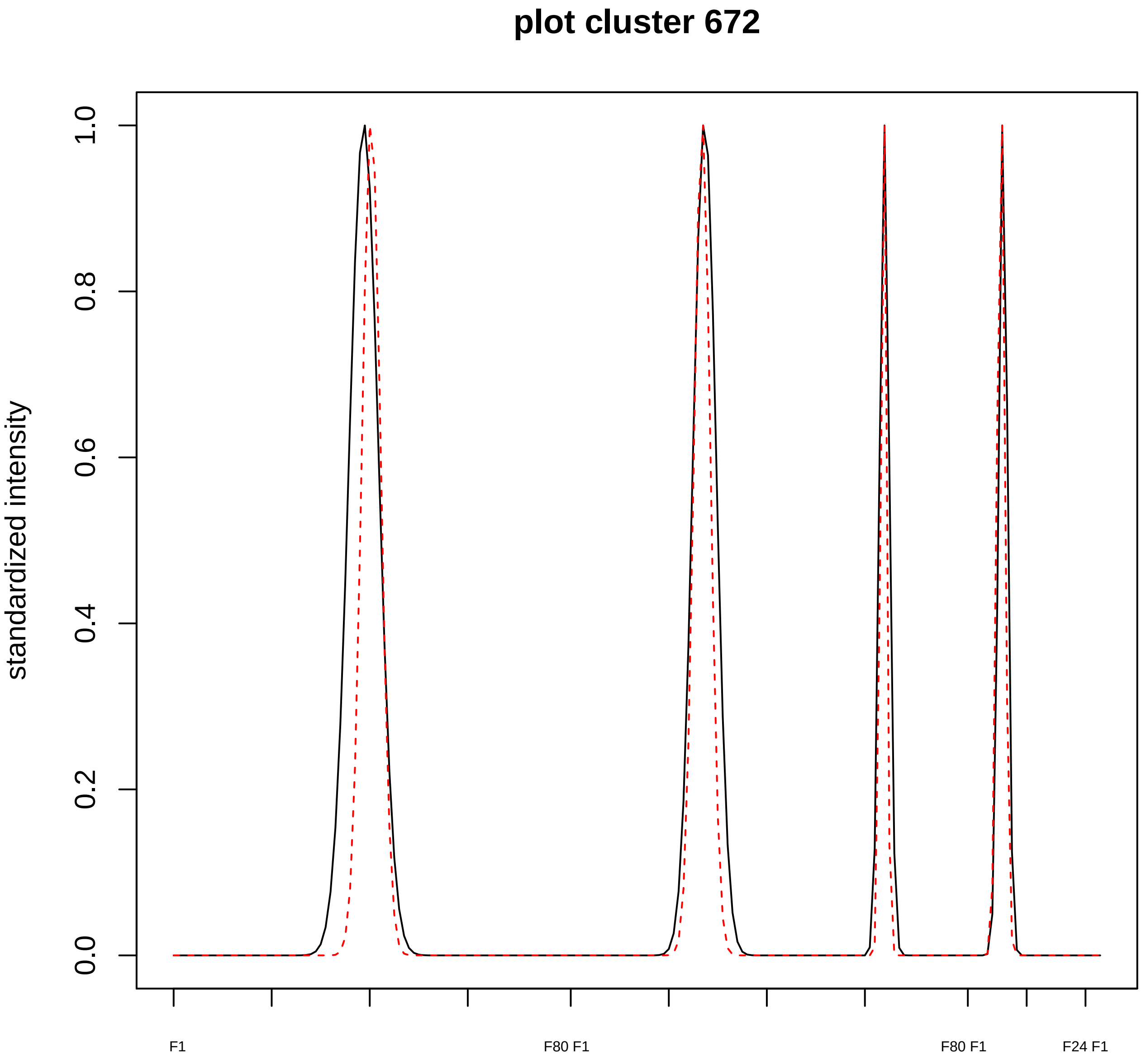

plot cluster 672
standardized intensity
0.0
0.2
0.4
0.6
0.8
1.0
F1
F80 F1
F80 F1
F24 F1

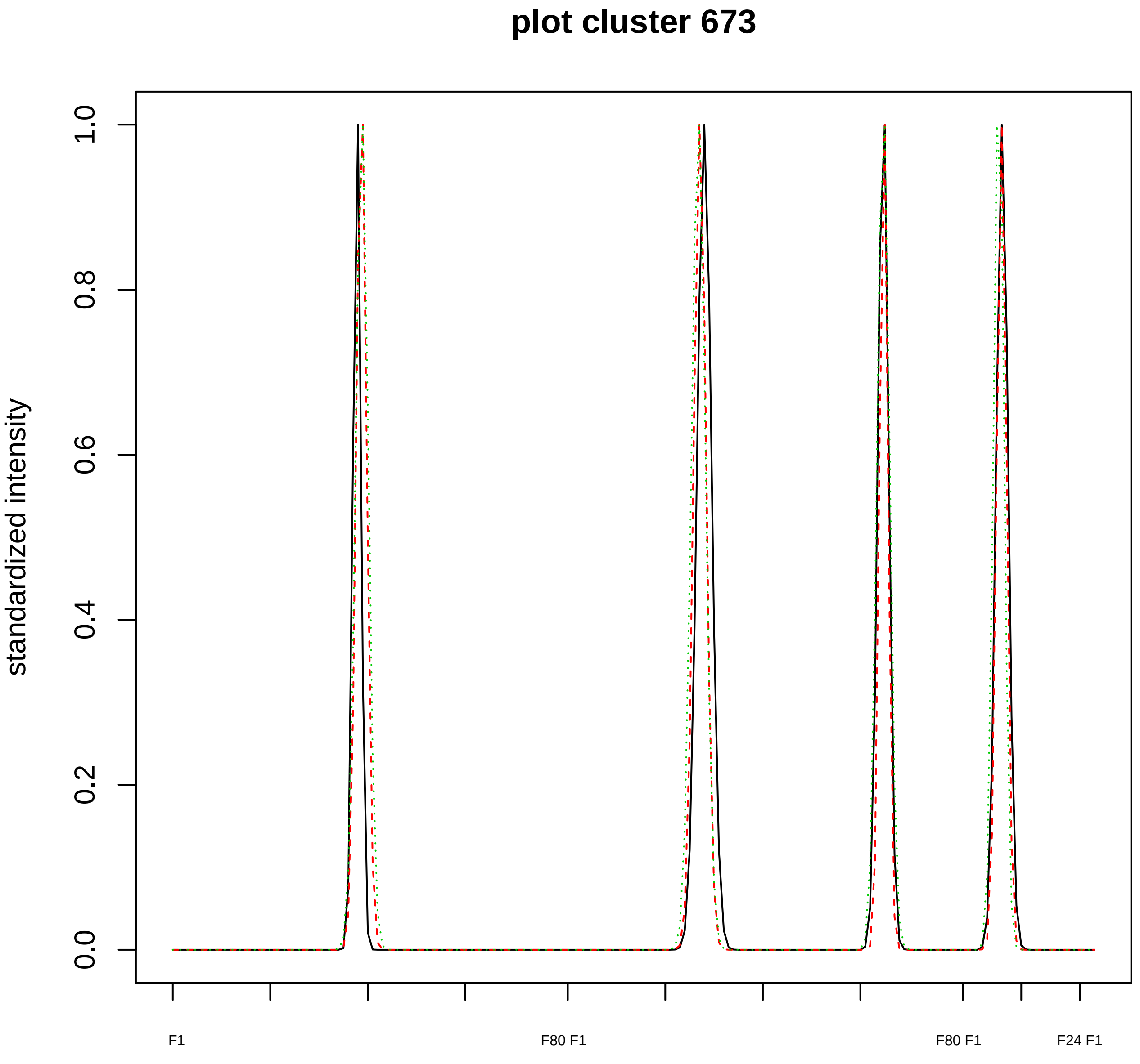
plot cluster 673
standardized intensity
1.0
0.8
0.6
0.4
0.2
0.0
F1
F80 F1
F80 F1
F24 F1

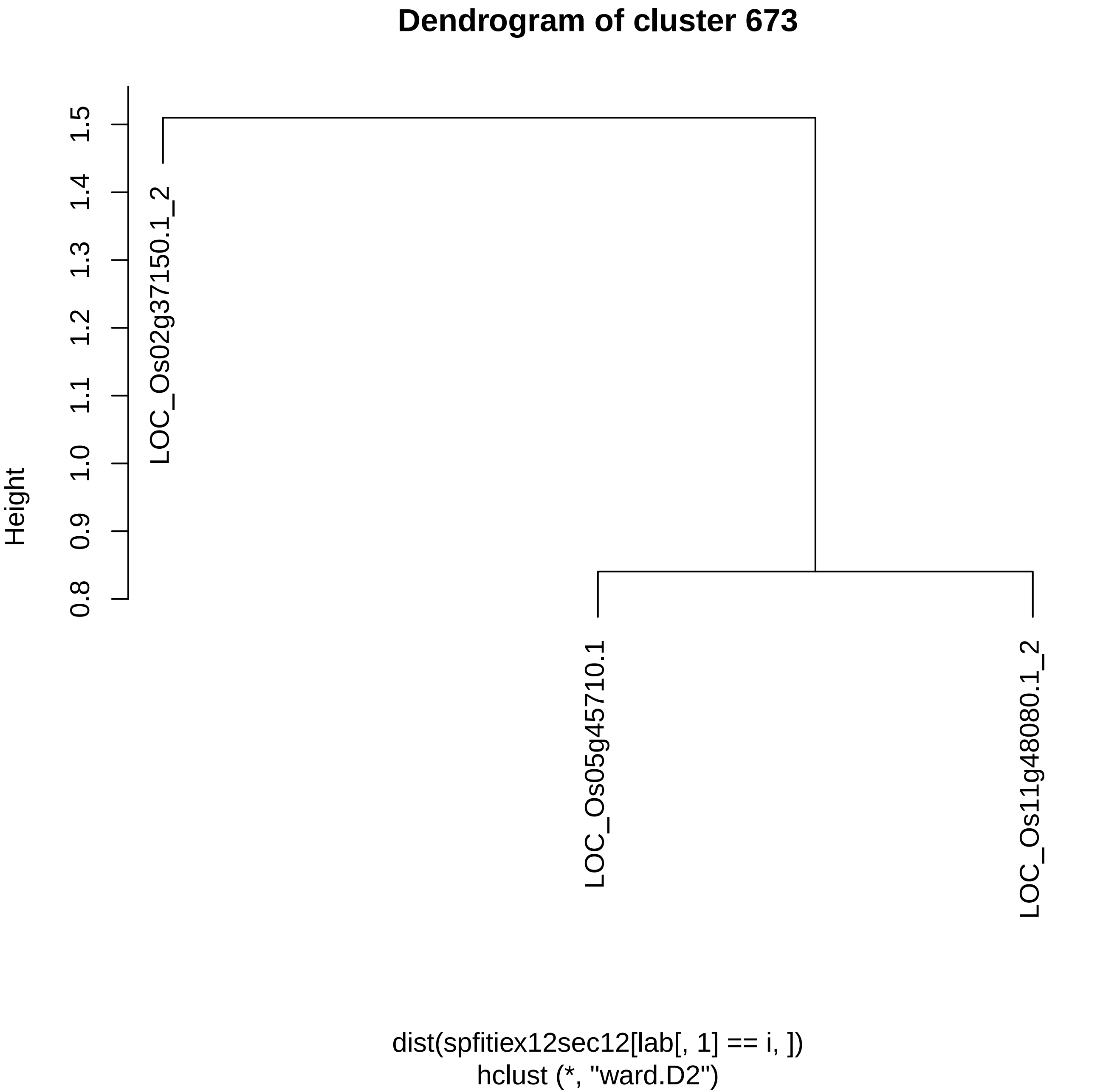
Dendrogram of cluster 673
Height
0.8
0.9
1.0
1.1
1.2
1.3
1.4
1.5
LOC_Os02g37150.1_2
LOC_Os05g45710.1
LOC_Os11g48080.1_2
dist(spfitiex12sec12[lab[, 1] == i, ])
hclust (*, "ward.D2")

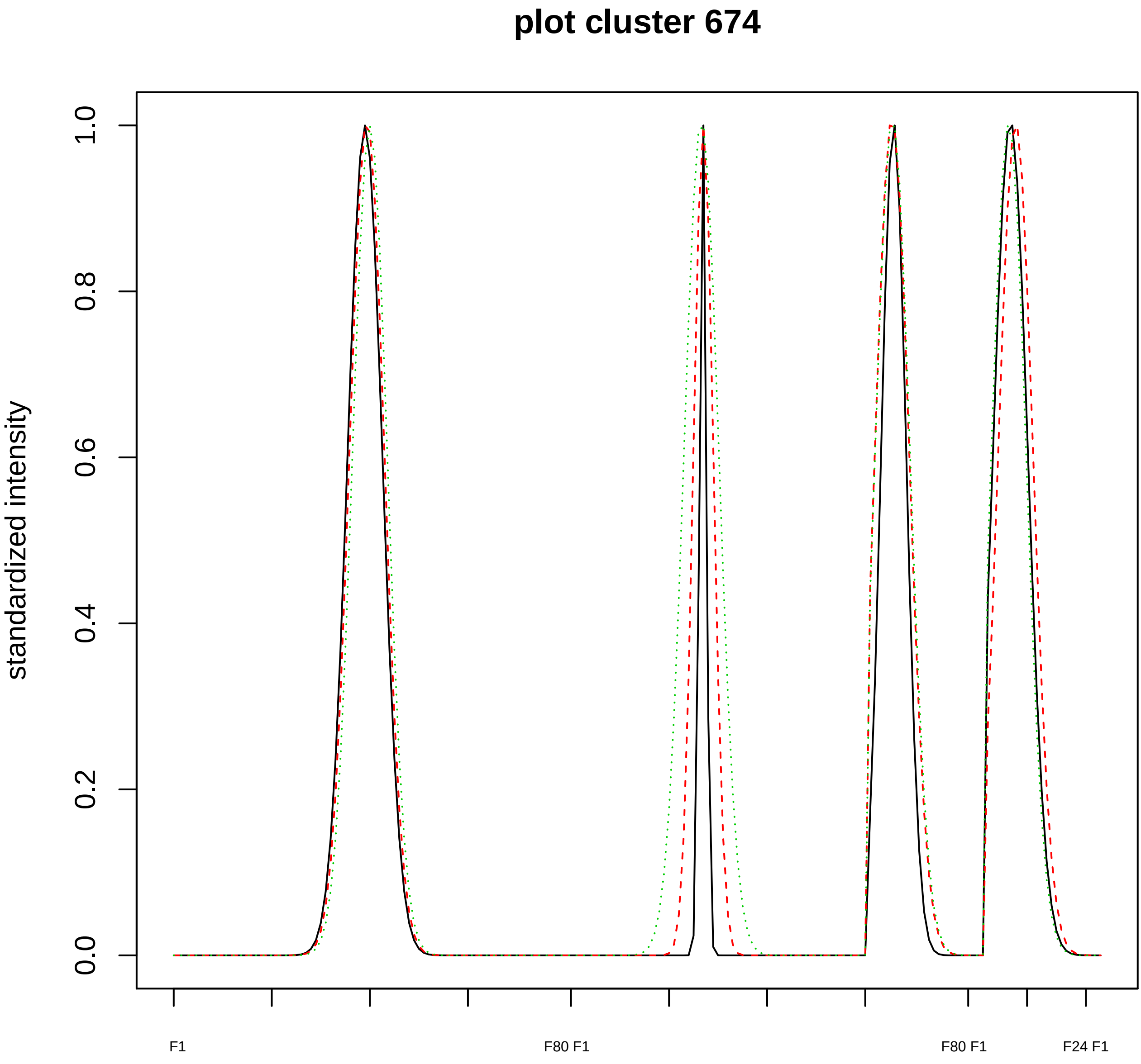

plot cluster 674
standardized intensity
0.0
0.2
0.4
0.6
0.8
1.0
F1
F80 F1
F80 F1
F24 F1

### Dendrogram of cluster 674

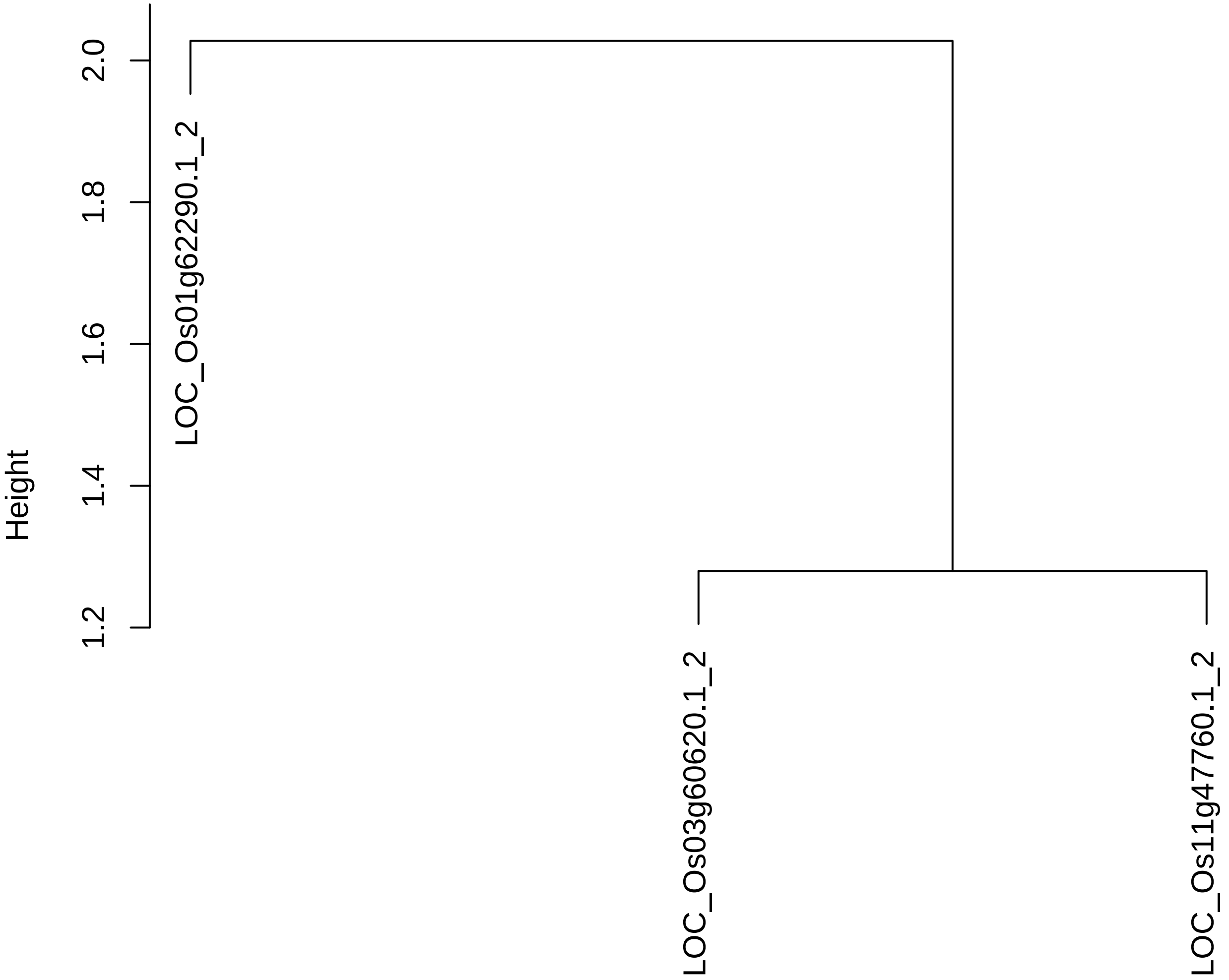


dist(spfitiex12sec12[lab[, 1] == i, ])
hclust (*, "ward.D2")

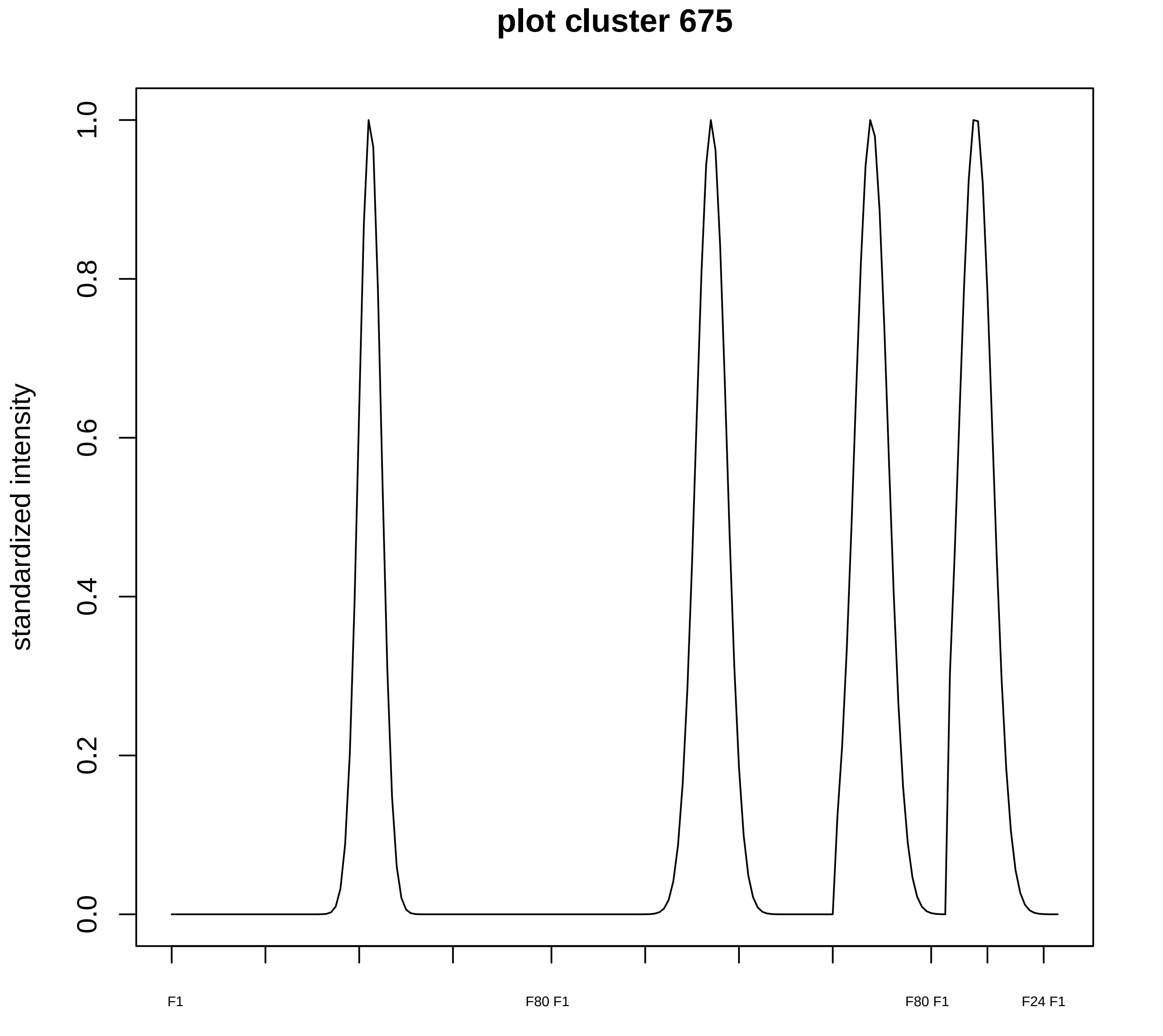

plot cluster 675
standardized intensity
0.0
0.2
0.4
0.6
0.8
1.0
F1
F80 F1
F80 F1
F24 F1

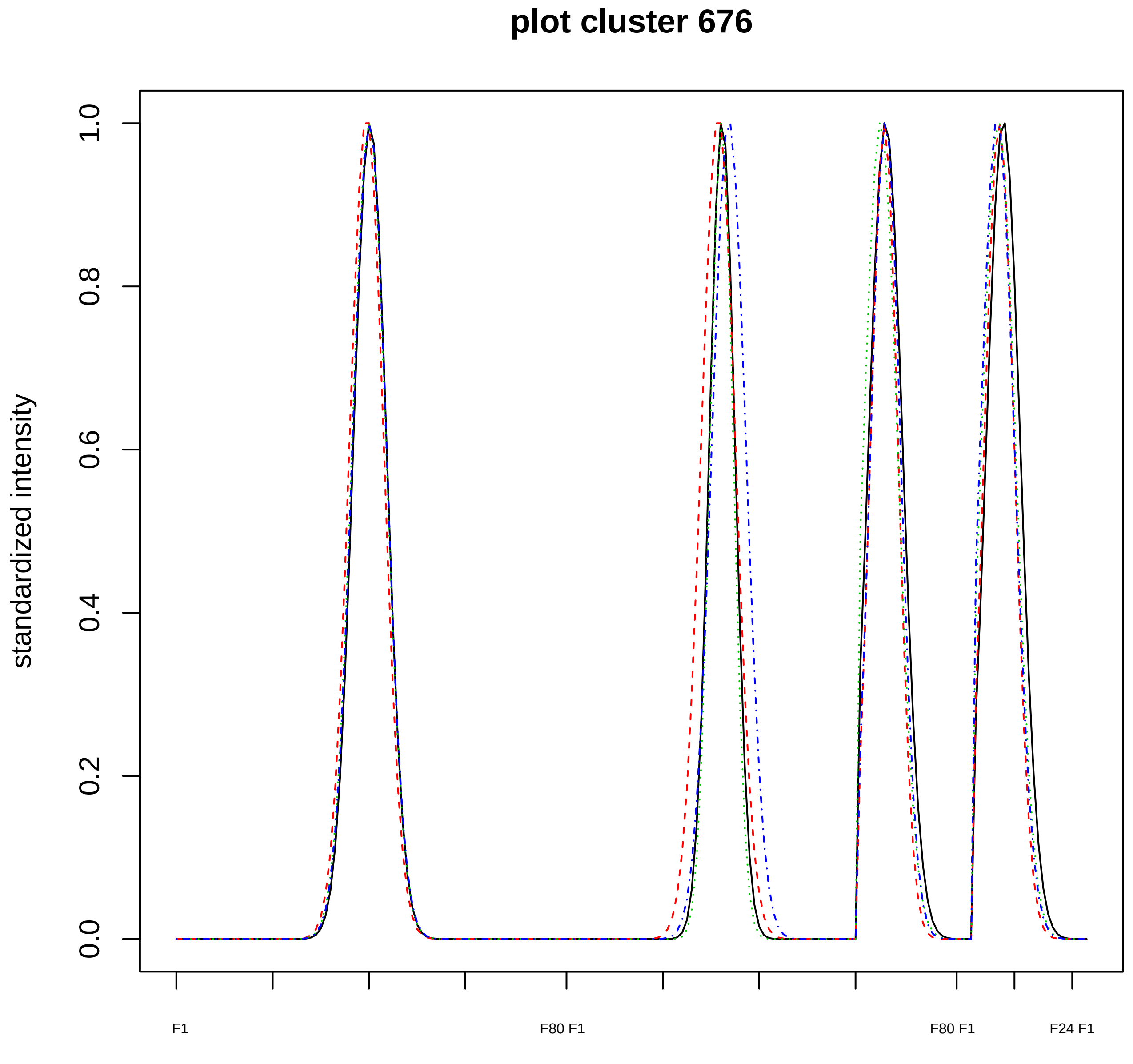
plot cluster 676
standardized intensity
0.0
0.2
0.4
0.6
0.8
1.0
F1
F80 F1
F80 F1
F24 F1

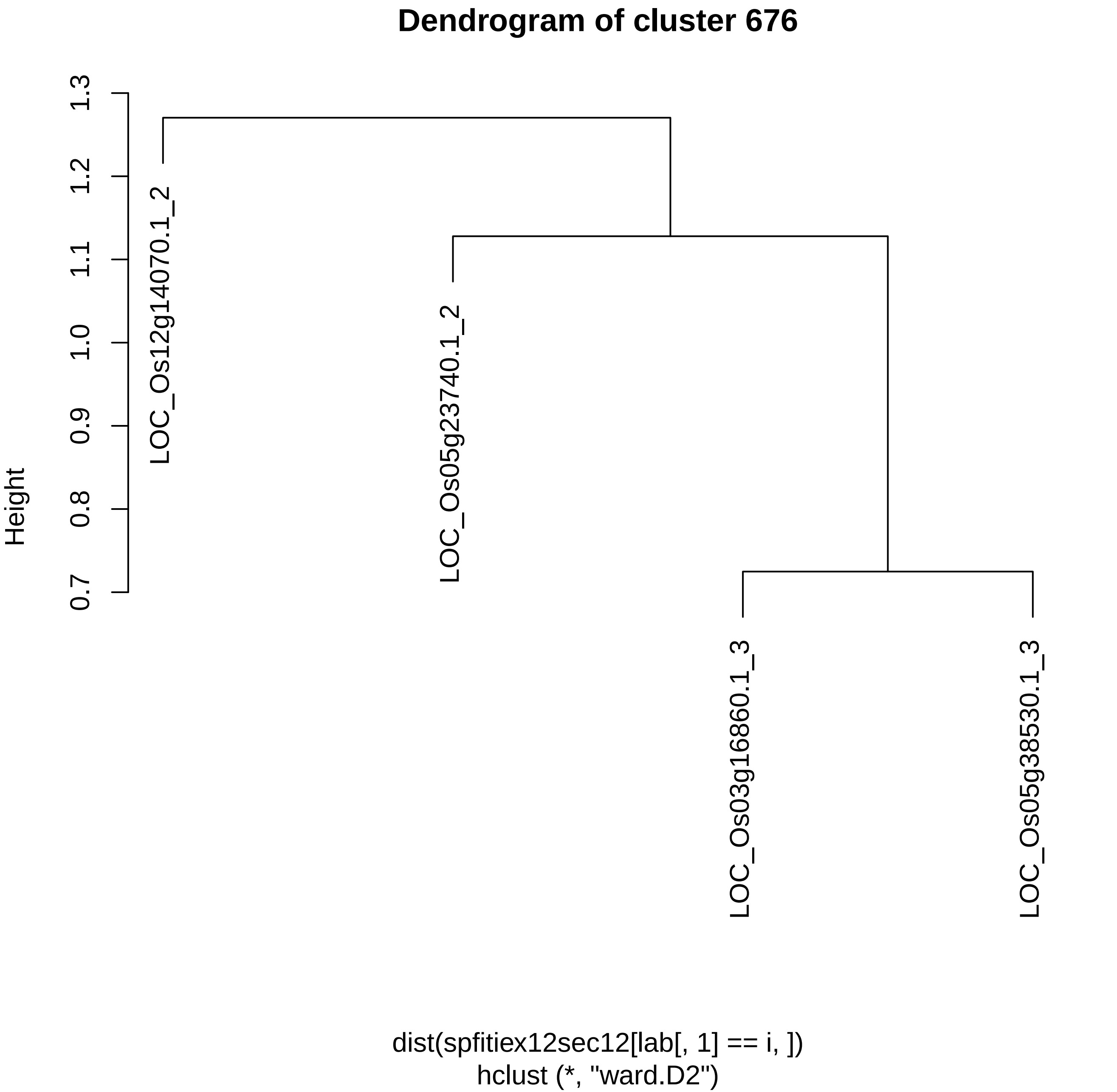

Dendrogram of cluster 676
Height
1.3
1.2
1.1
1.0
0.9
0.8
0.7
LOC_Os12g14070.1_2
LOC_Os05g23740.1_2
LOC_Os03g16860.1_3
LOC_Os05g38530.1_3
dist(spfitiex12sec12[lab[, 1] == i, ])
hclust (*, "ward.D2")

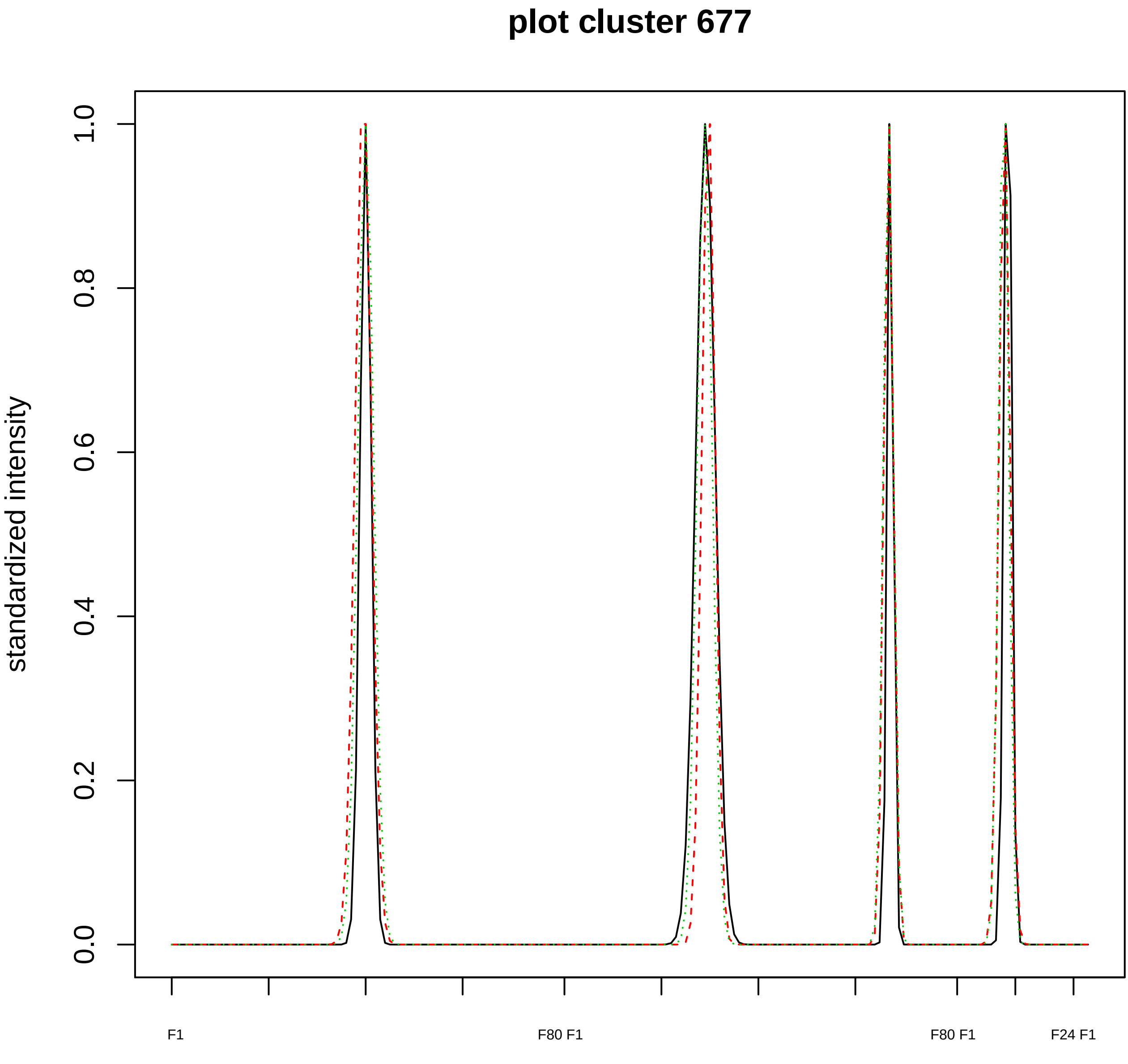
plot cluster 677
standardized intensity
1.0
0.8
0.6
0.4
0.2
0.0
F1
F80 F1
F80 F1
F24 F1

### Dendrogram of cluster 677

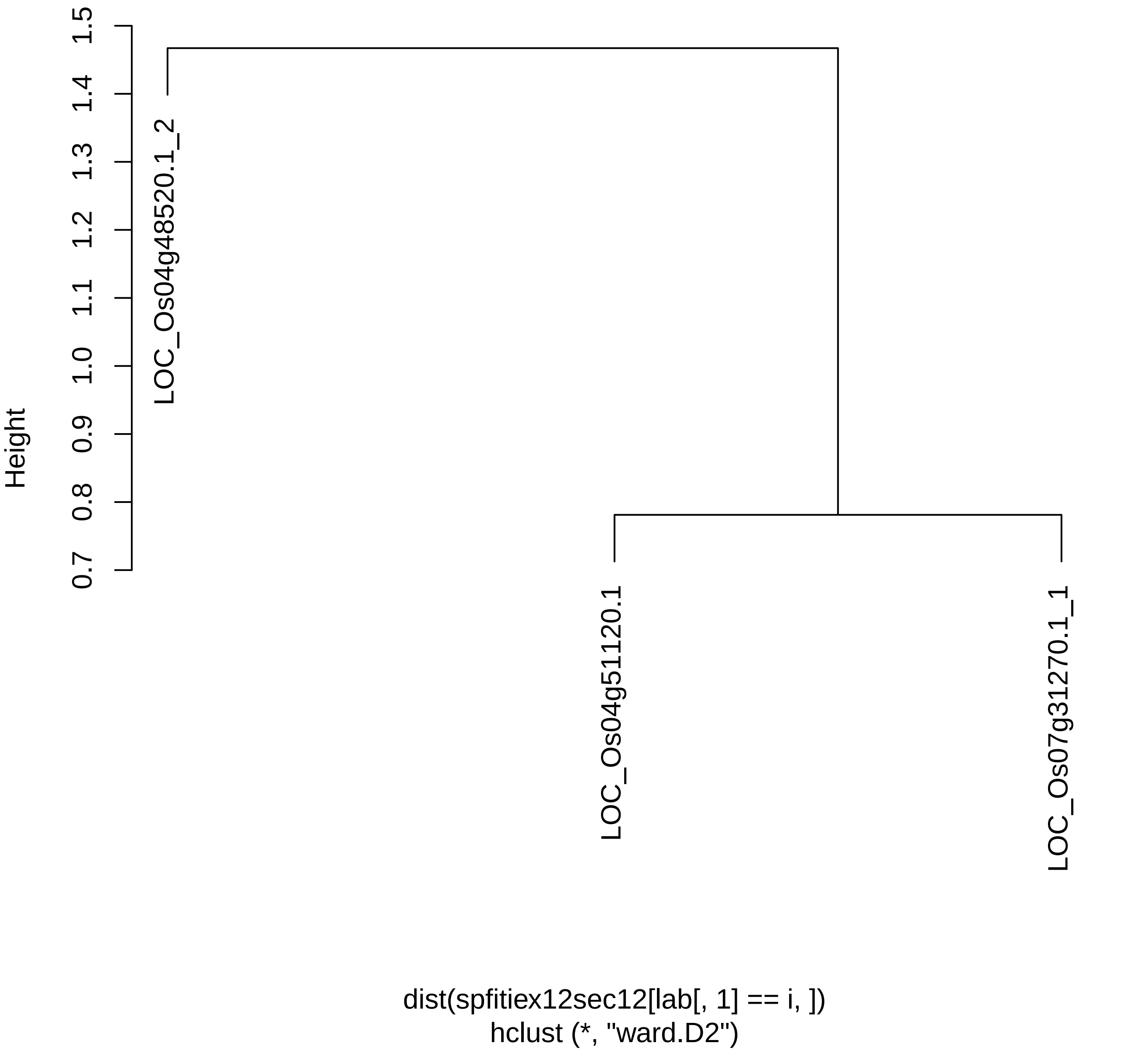

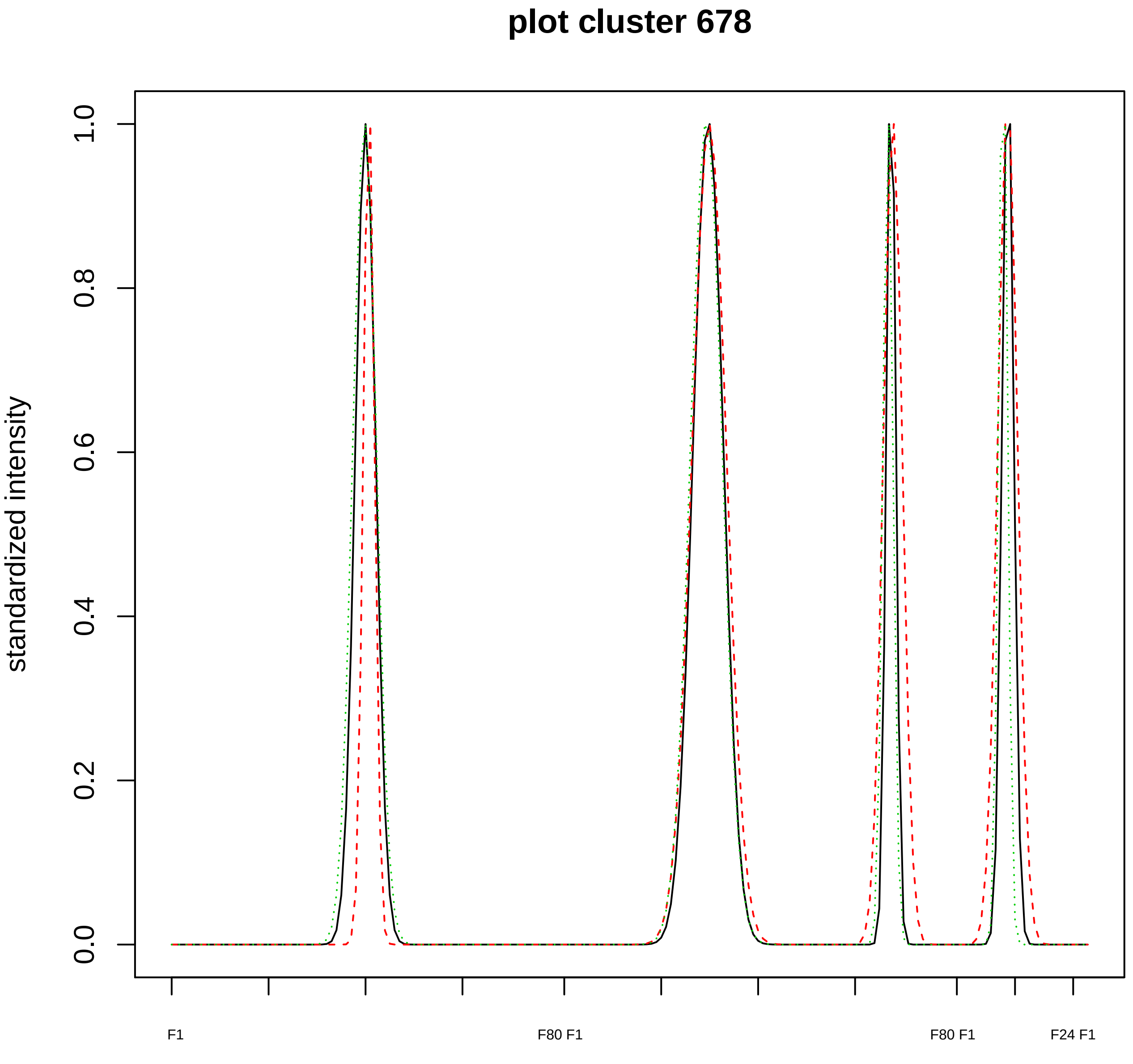

plot cluster 678
standardized intensity
0.0
0.2
0.4
0.6
0.8
1.0
F1
F80 F1
F80 F1
F24 F1

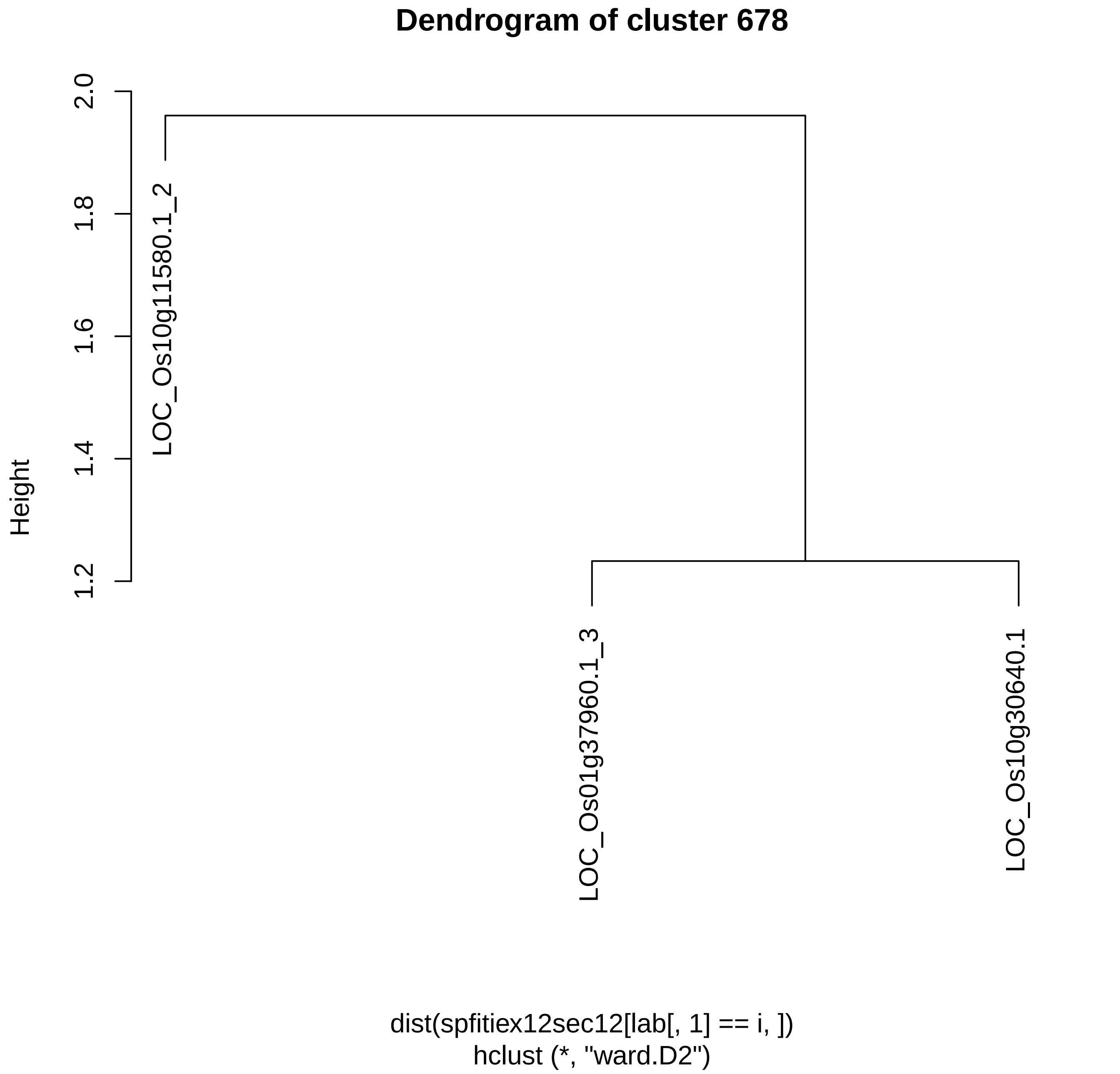

Dendrogram of cluster 678
Height
2.0
1.8
1.6
1.4
1.2
LOC_Os10g11580.1_2
LOC_Os01g37960.1_3
LOC_Os10g30640.1
dist(spfitiex12sec12[lab[, 1] == i, ])
hclust (*, "ward.D2")

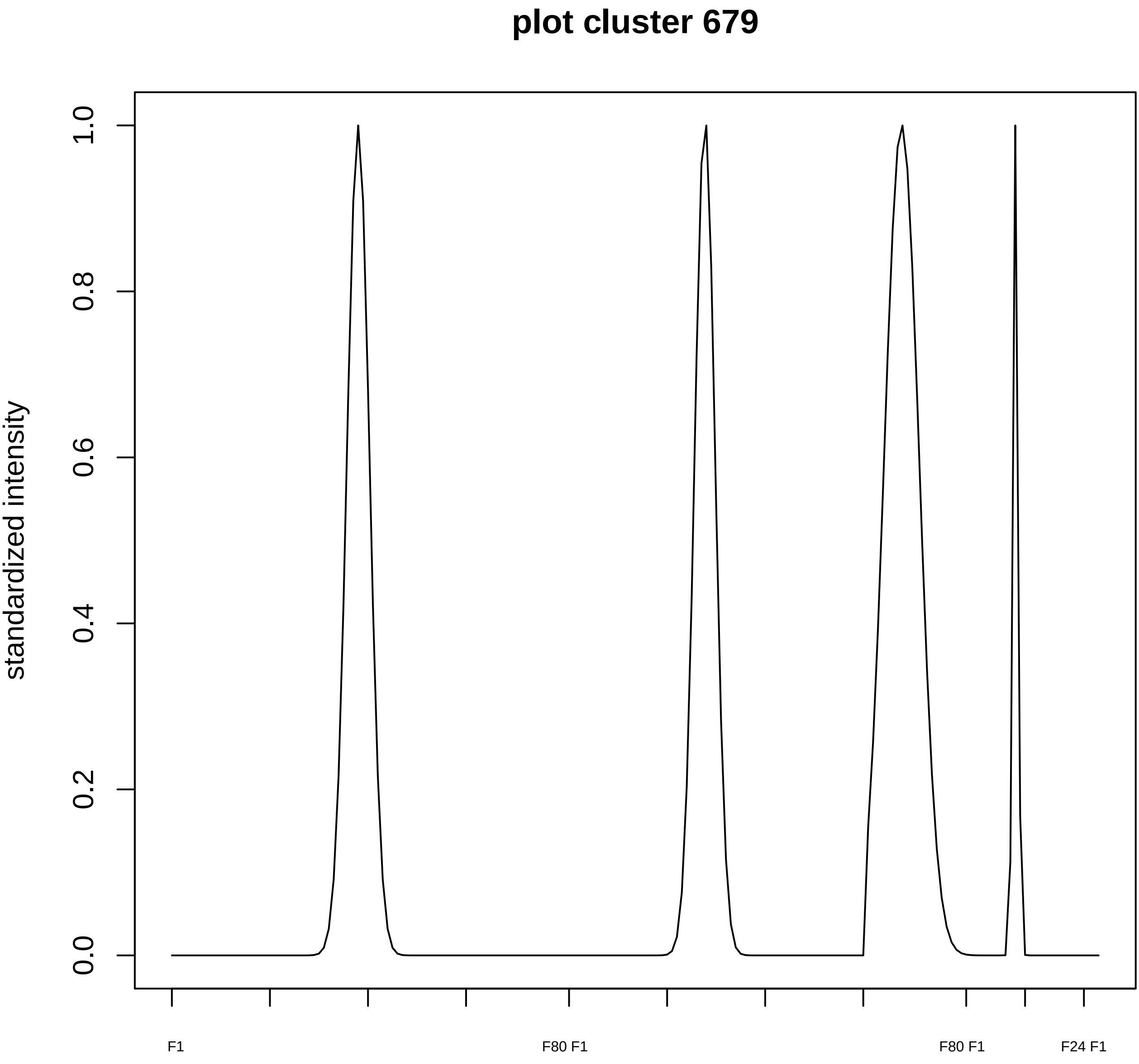
plot cluster 679
standardized intensity
0.0
0.2
0.4
0.6
0.8
1.0
F1
F80 F1
F80 F1
F24 F1

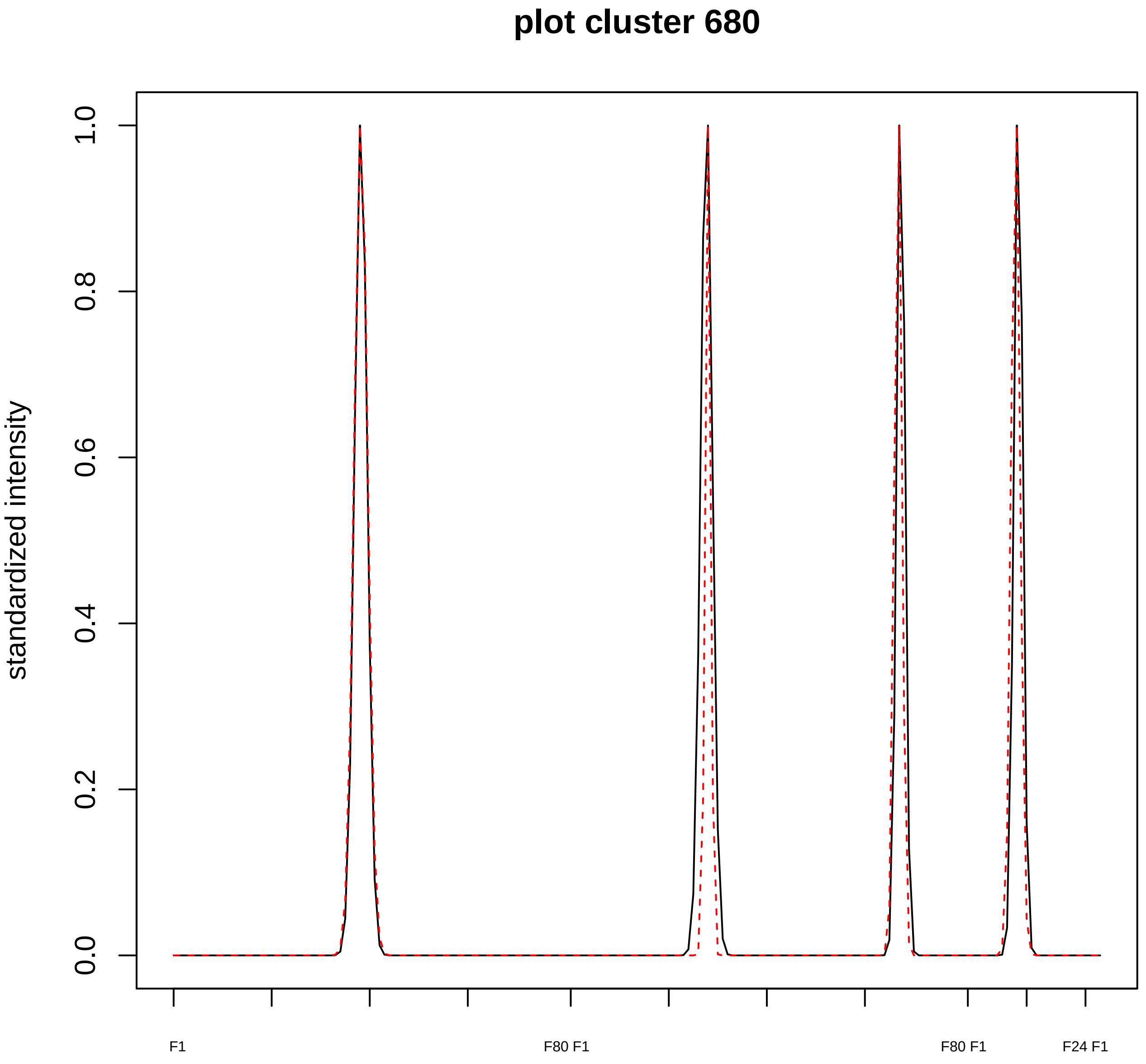
plot cluster 680
standardized intensity
1.0
0.8
0.6
0.4
0.2
0.0
F1
F80 F1
F80 F1
F24 F1

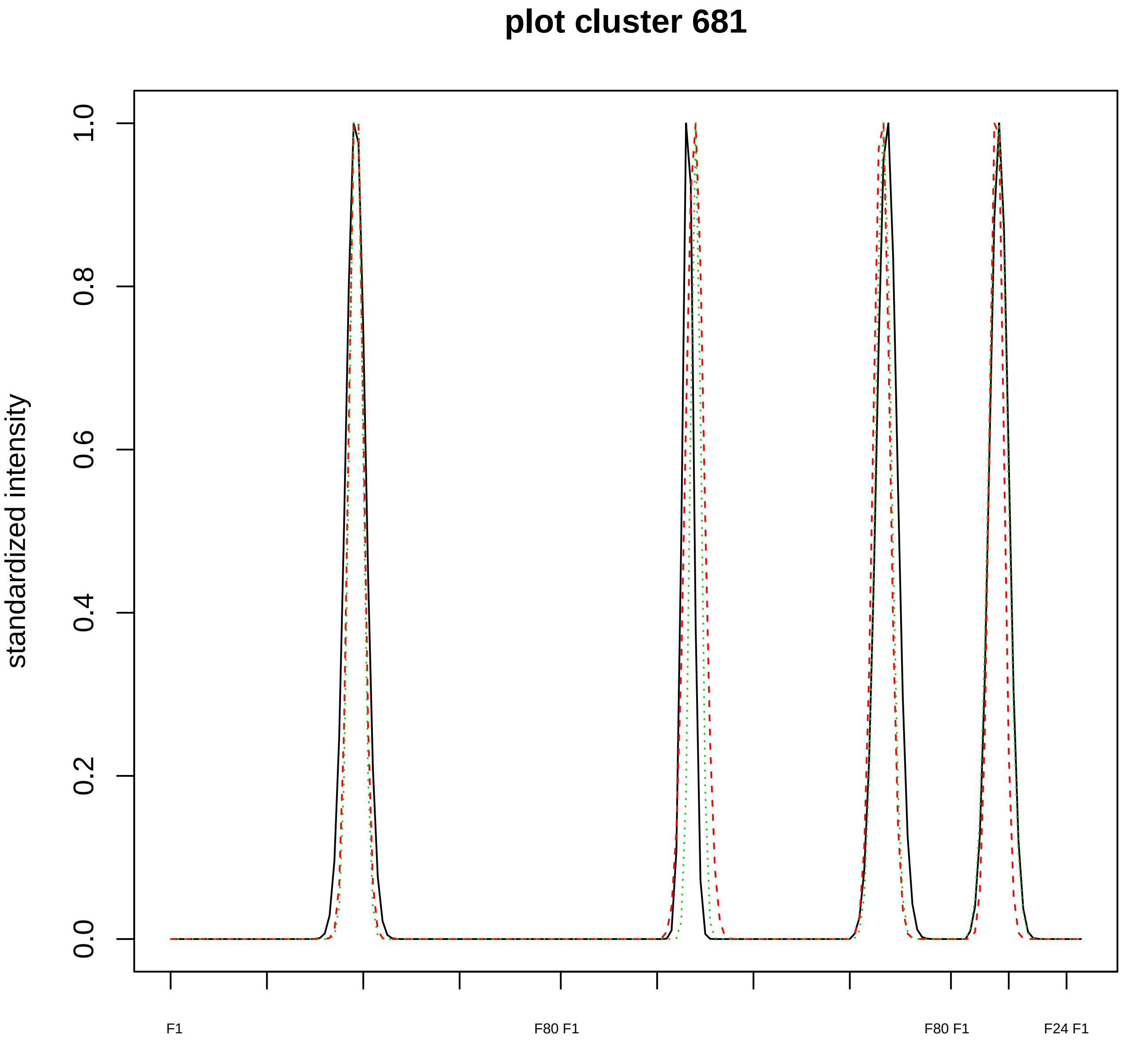

plot cluster 681
standardized intensity
0.0
0.2
0.4
0.6
0.8
1.0
F1
F80 F1
F80 F1
F24 F1

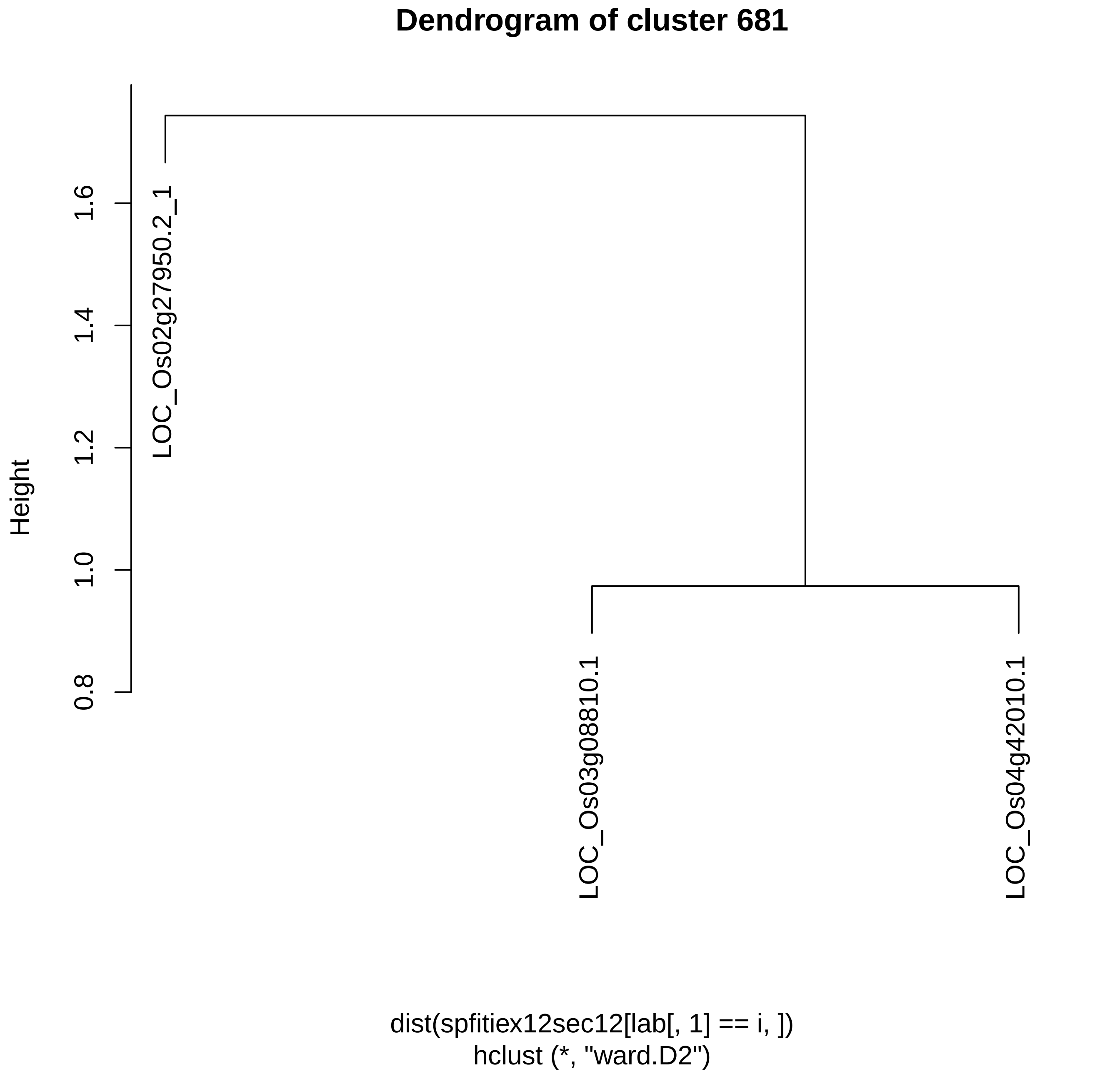
Dendrogram of cluster 681
Height
0.8
1.0
1.2
1.4
1.6
LOC_Os02g27950.2_1
LOC_Os03g08810.1
LOC_Os04g42010.1
dist(spfitiex12sec12[lab[, 1] == i, ])
hclust (*, "ward.D2")

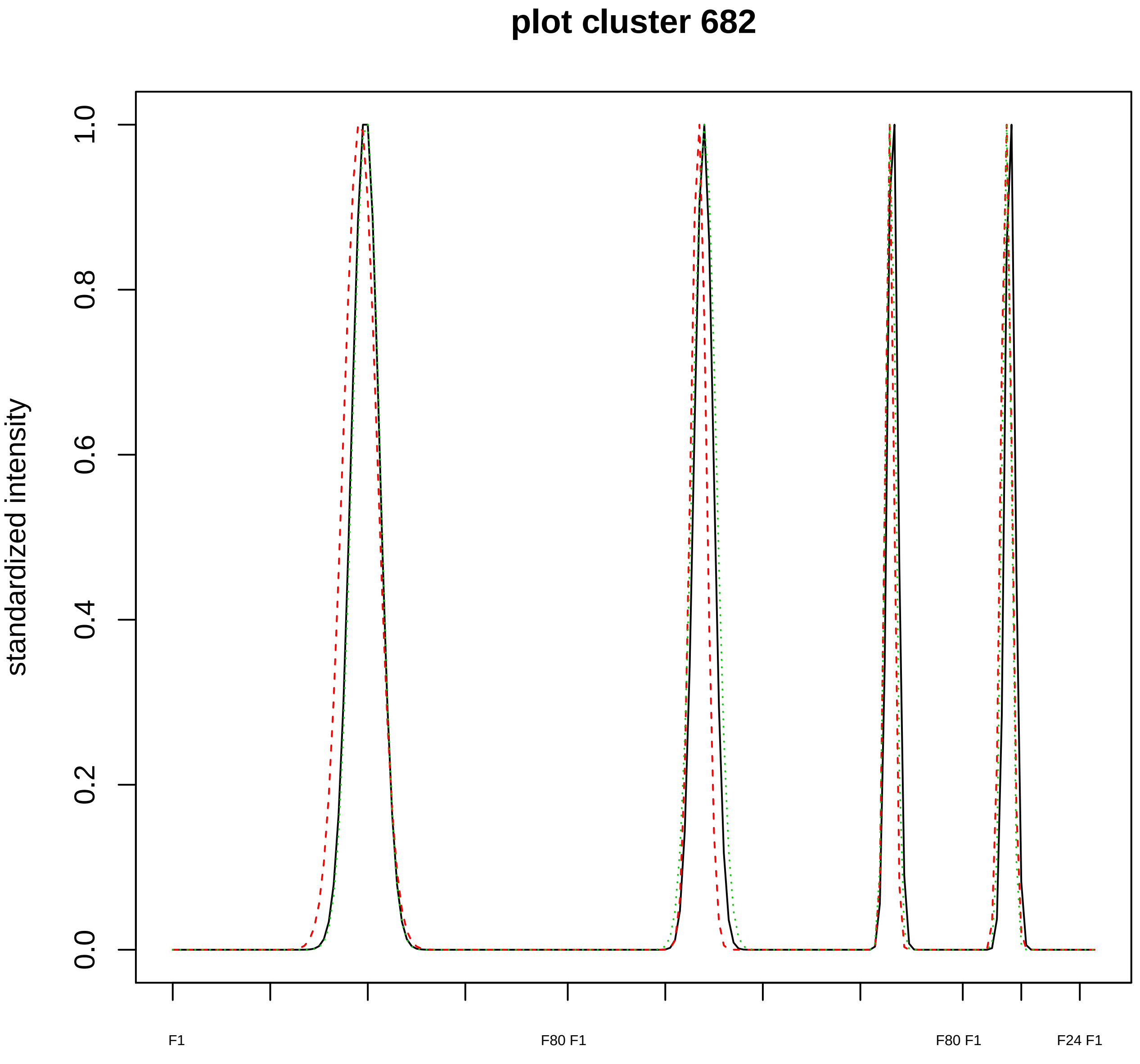

plot cluster 682
standardized intensity
0.0
0.2
0.4
0.6
0.8
1.0
F1
F80 F1
F80 F1
F24 F1

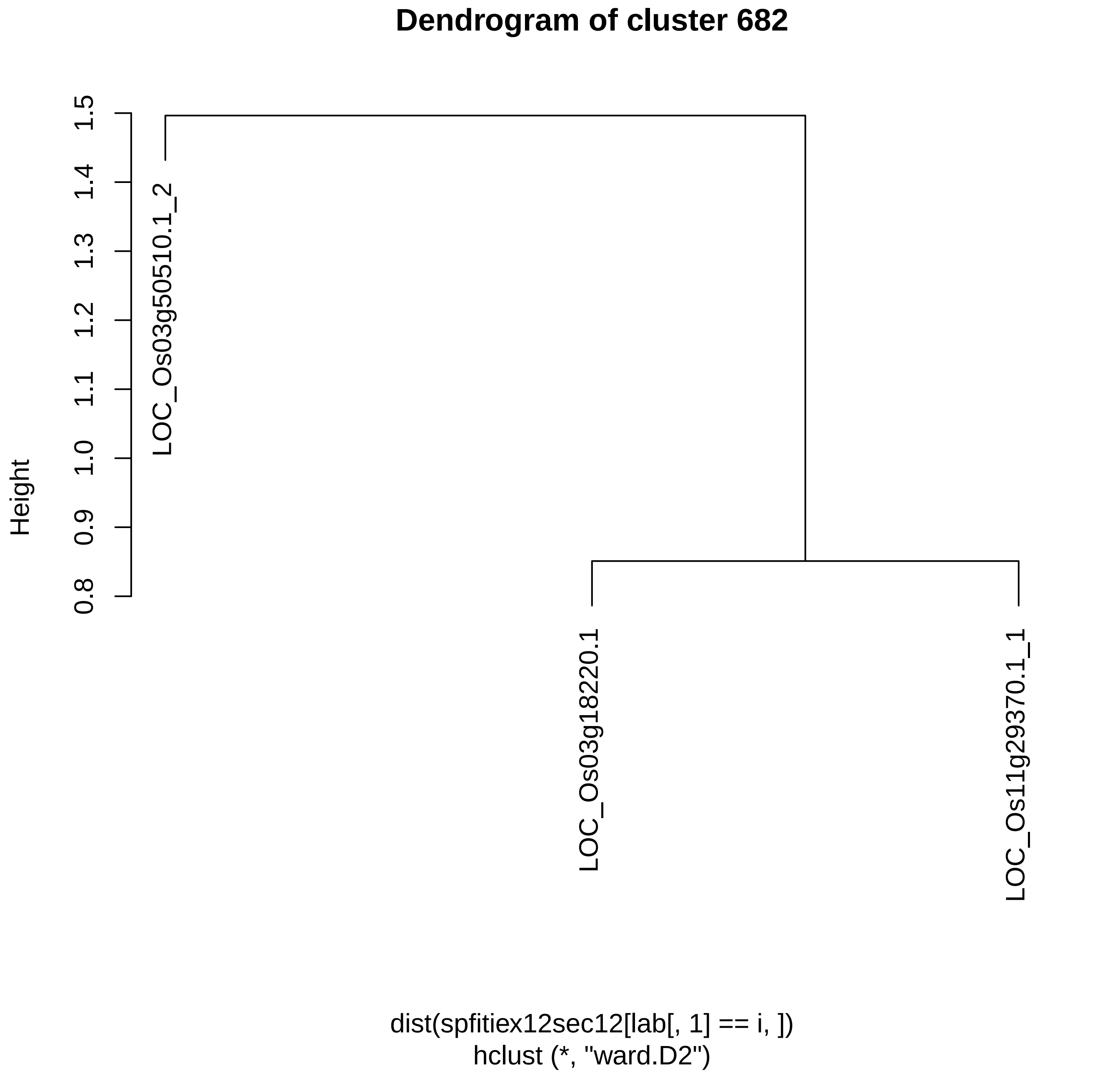
Dendrogram of cluster 682
Height
0.8
0.9
1.0
1.1
1.2
1.3
1.4
1.5
LOC_Os03g50510.1_2
LOC_Os03g18220.1
LOC_Os11g29370.1_1
dist(spfitiex12sec12[lab[, 1] == i, ])
hclust (*, "ward.D2")

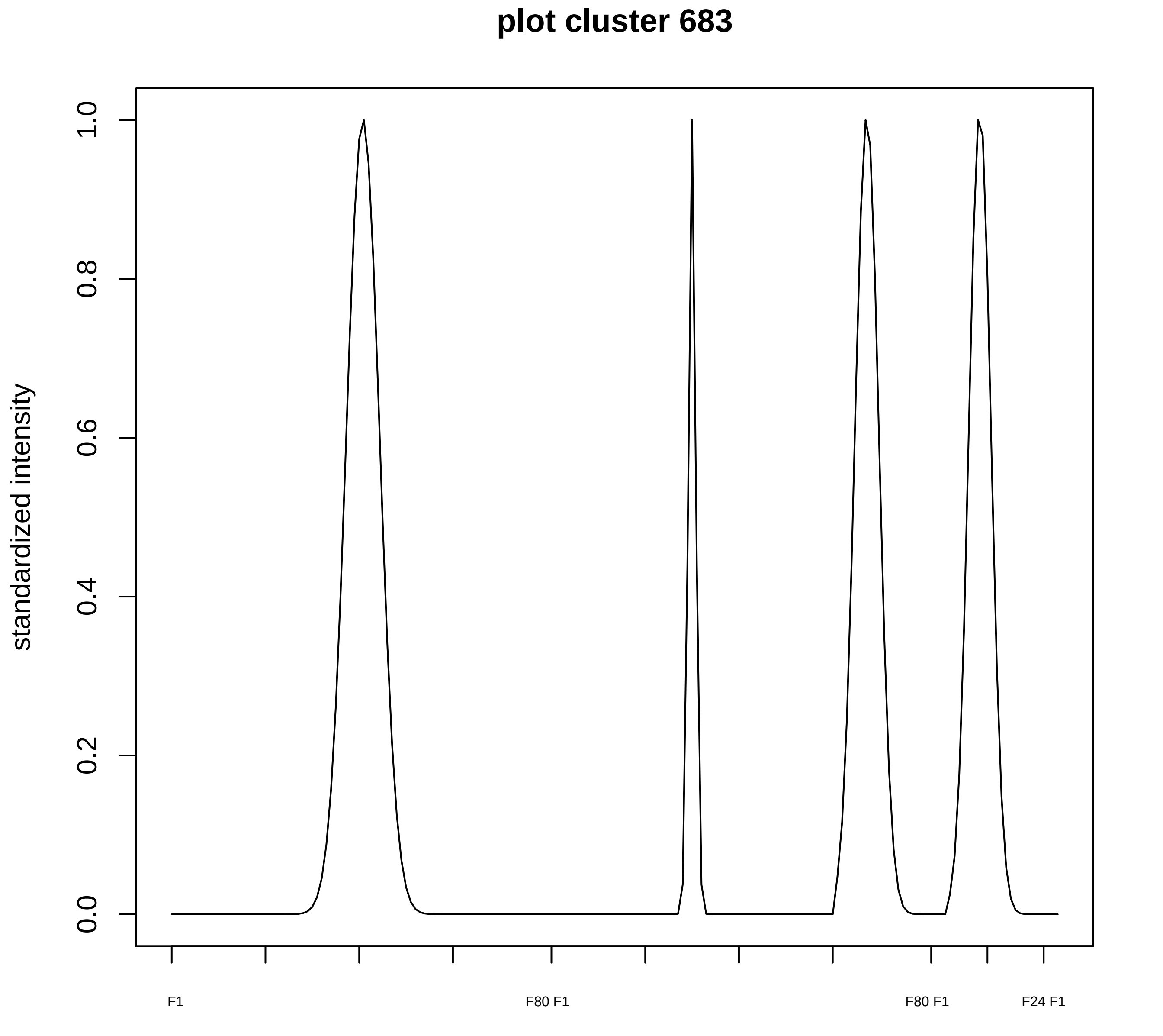

plot cluster 683
standardized intensity
0.0
0.2
0.4
0.6
0.8
1.0
F1
F80 F1
F80 F1
F24 F1

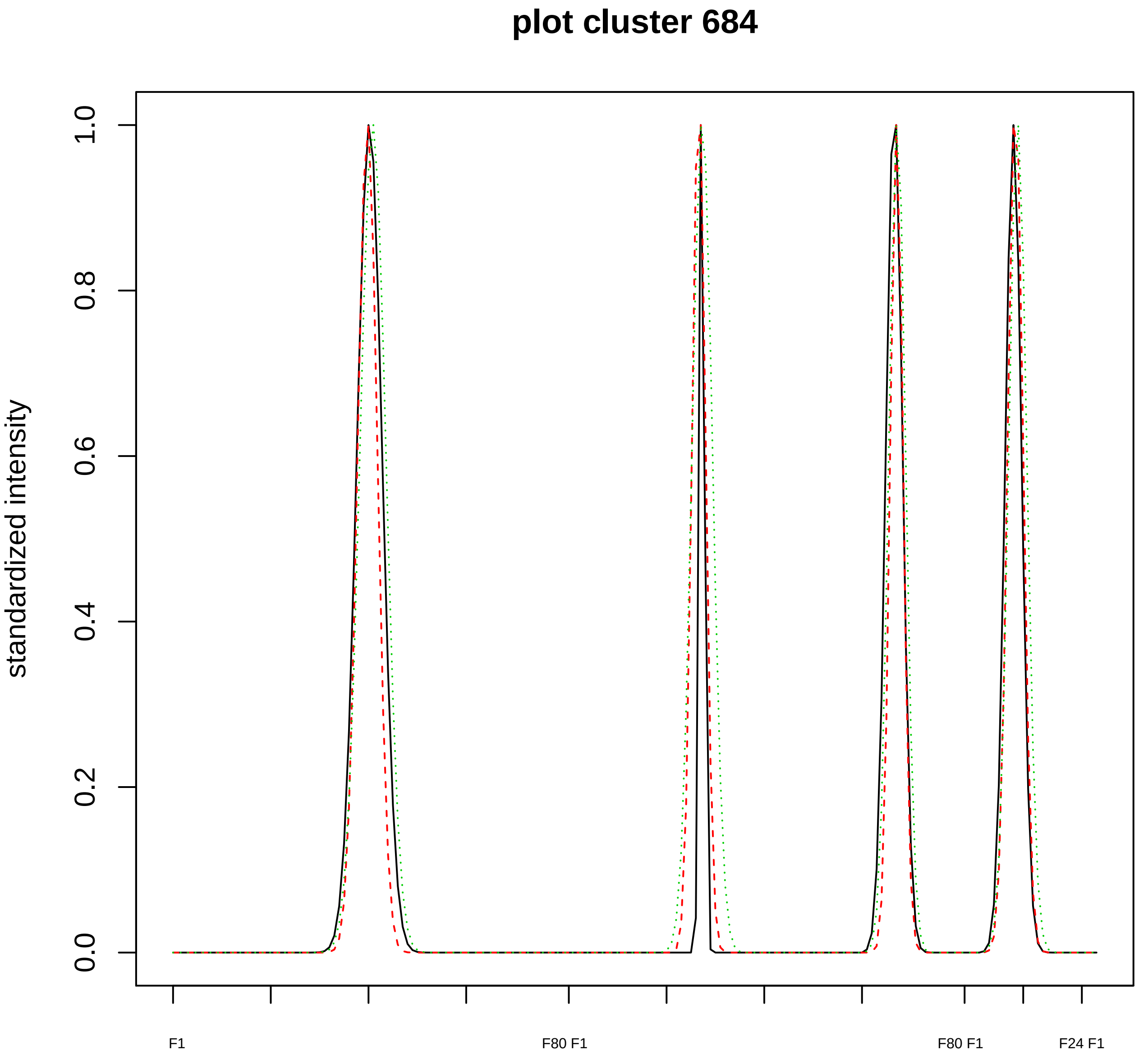
plot cluster 684
standardized intensity
0.0
0.2
0.4
0.6
0.8
1.0
F1
F80 F1
F80 F1
F24 F1

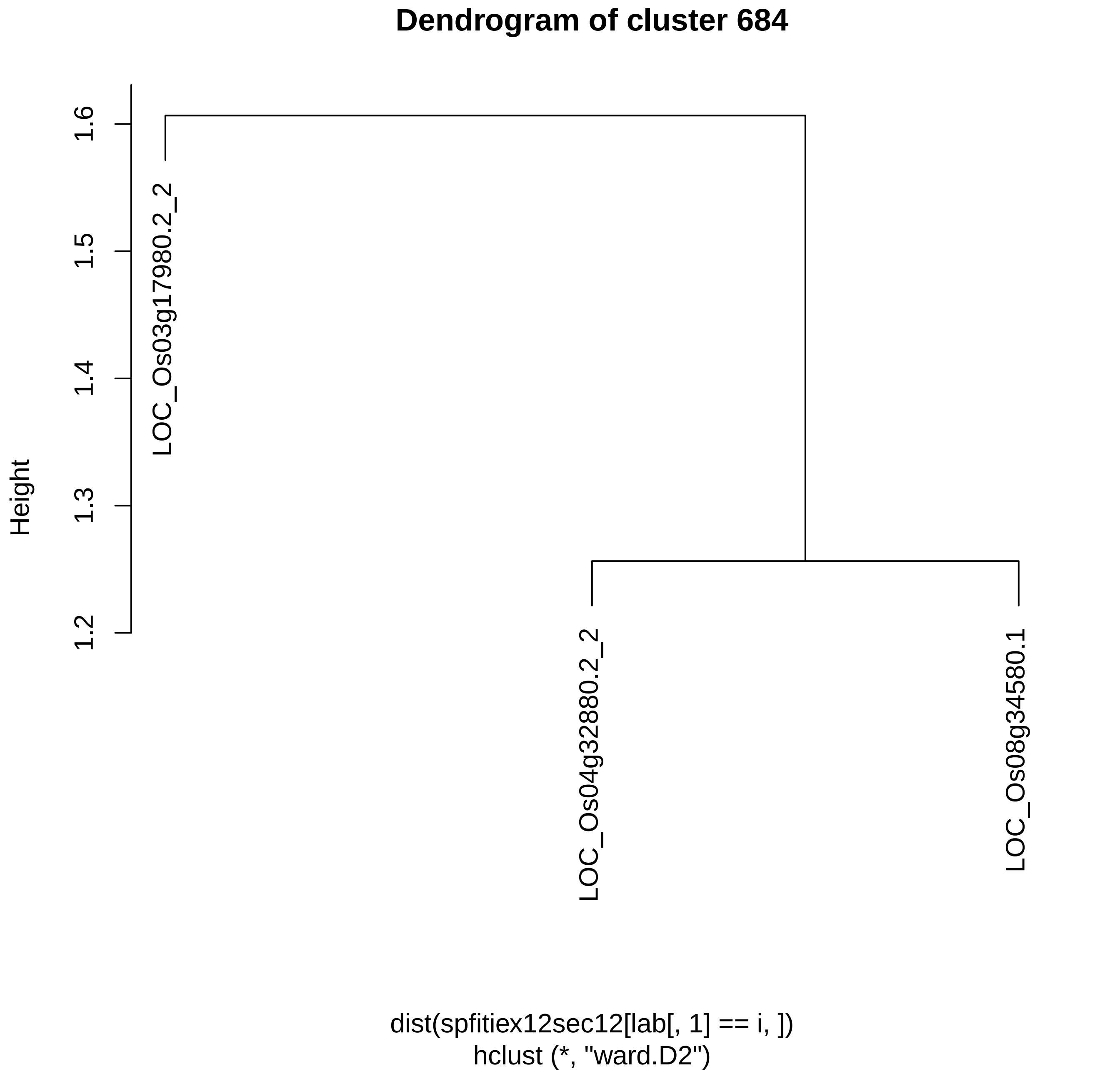
Dendrogram of cluster 684
Height
1.2
1.3
1.4
1.5
1.6
LOC_Os03g17980.2_2
LOC_Os04g32880.2_2
LOC_Os08g34580.1
dist(spfitiex12sec12[lab[, 1] == i, ])
hclust (*, "ward.D2")

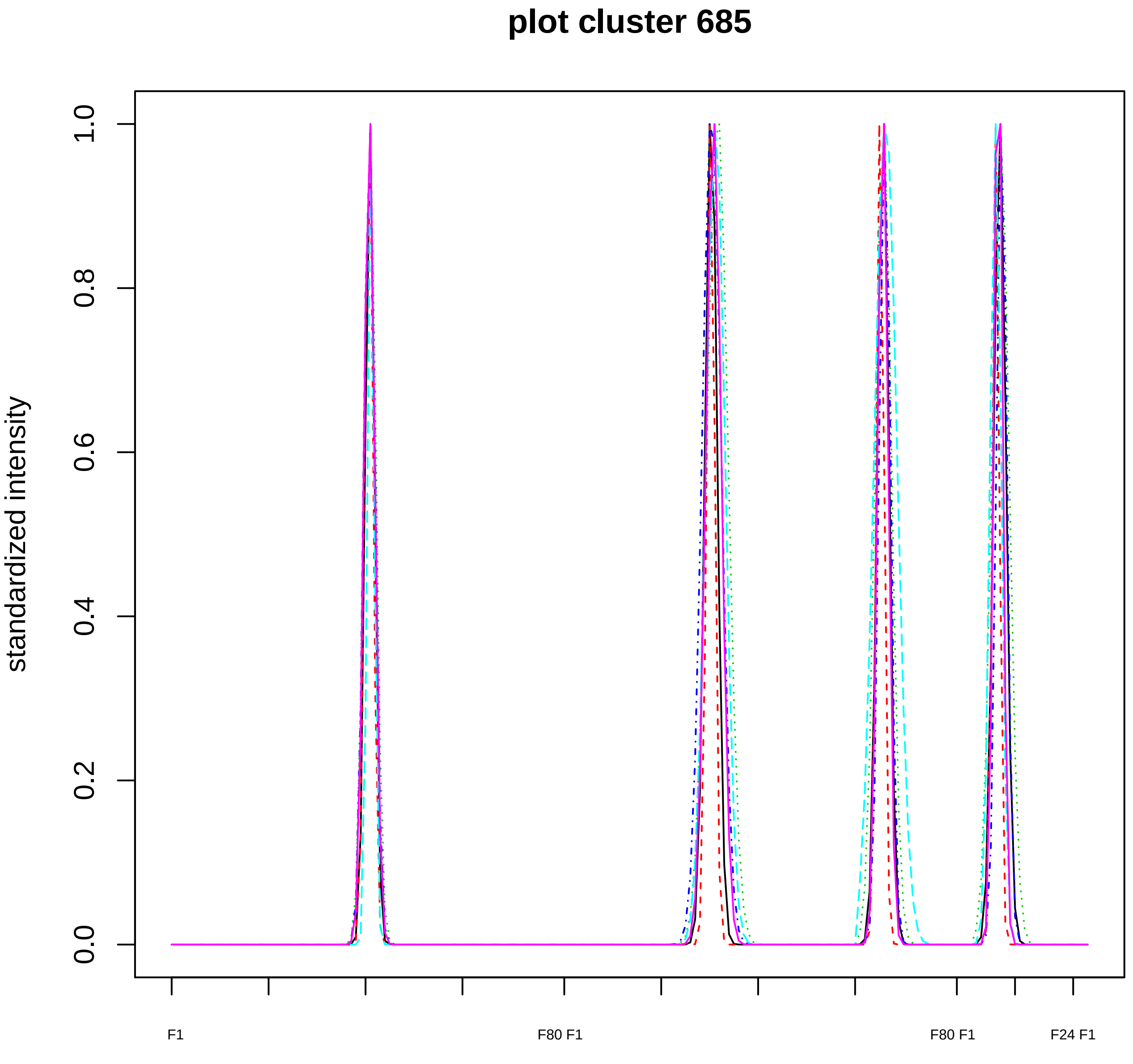

plot cluster 685
standardized intensity
0.0
0.2
0.4
0.6
0.8
1.0
F1
F80 F1
F80 F1
F24 F1

#### Dendrogram of cluster 685

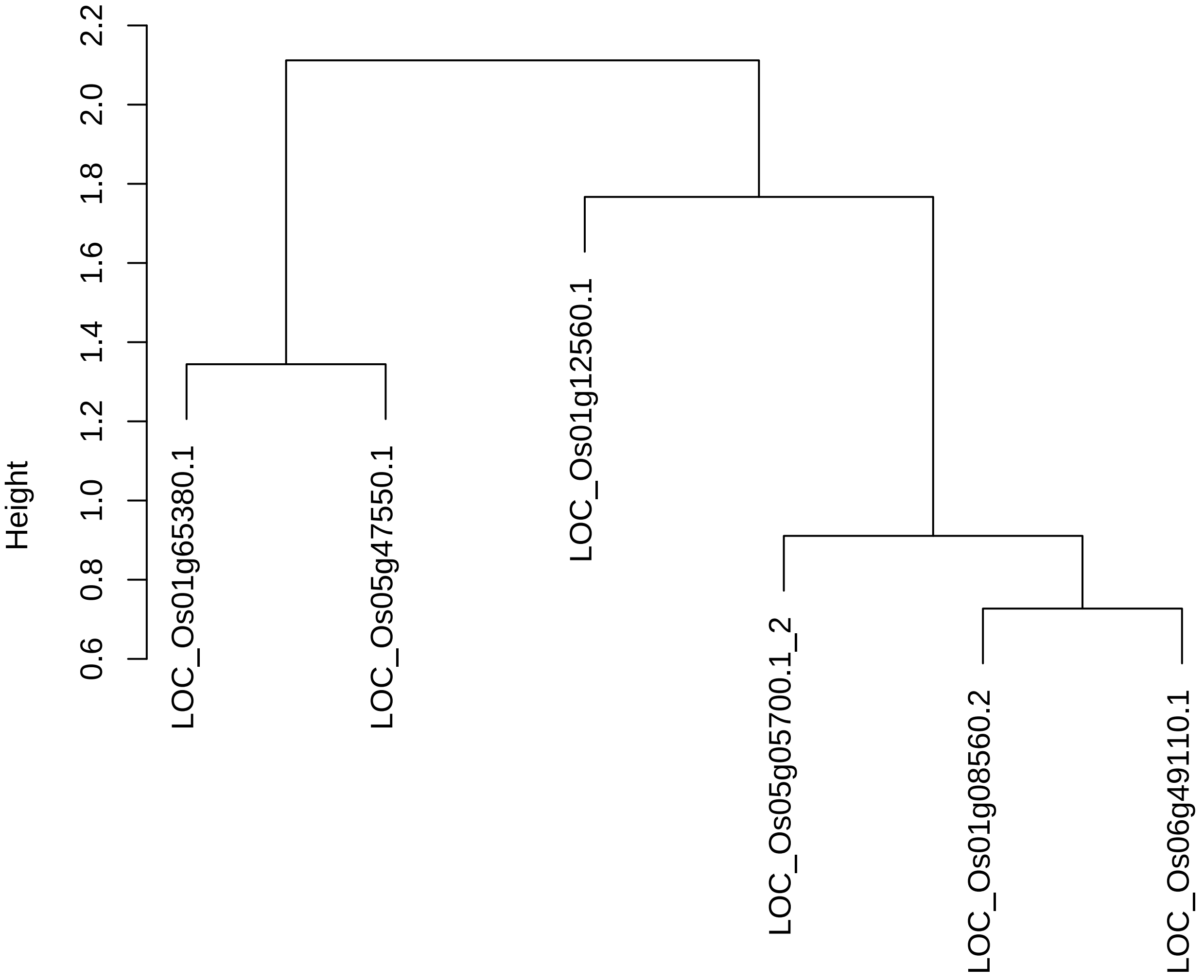


dist(spfitiex12sec12[lab[, 1] == i, ])
hclust (*, "ward.D2")

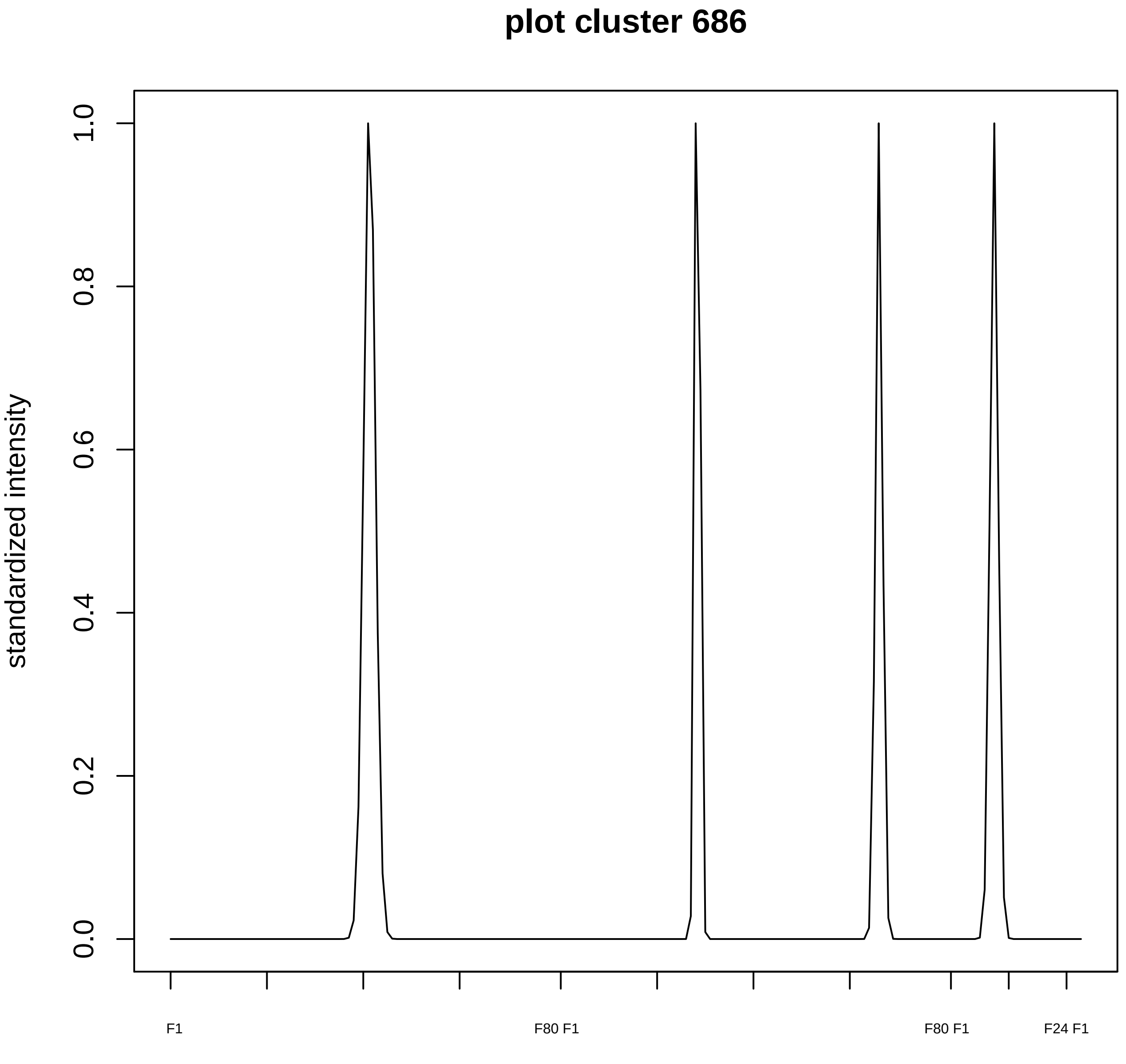
plot cluster 686
standardized intensity
0.0
0.2
0.4
0.6
0.8
1.0
F1
F80 F1
F80 F1
F24 F1

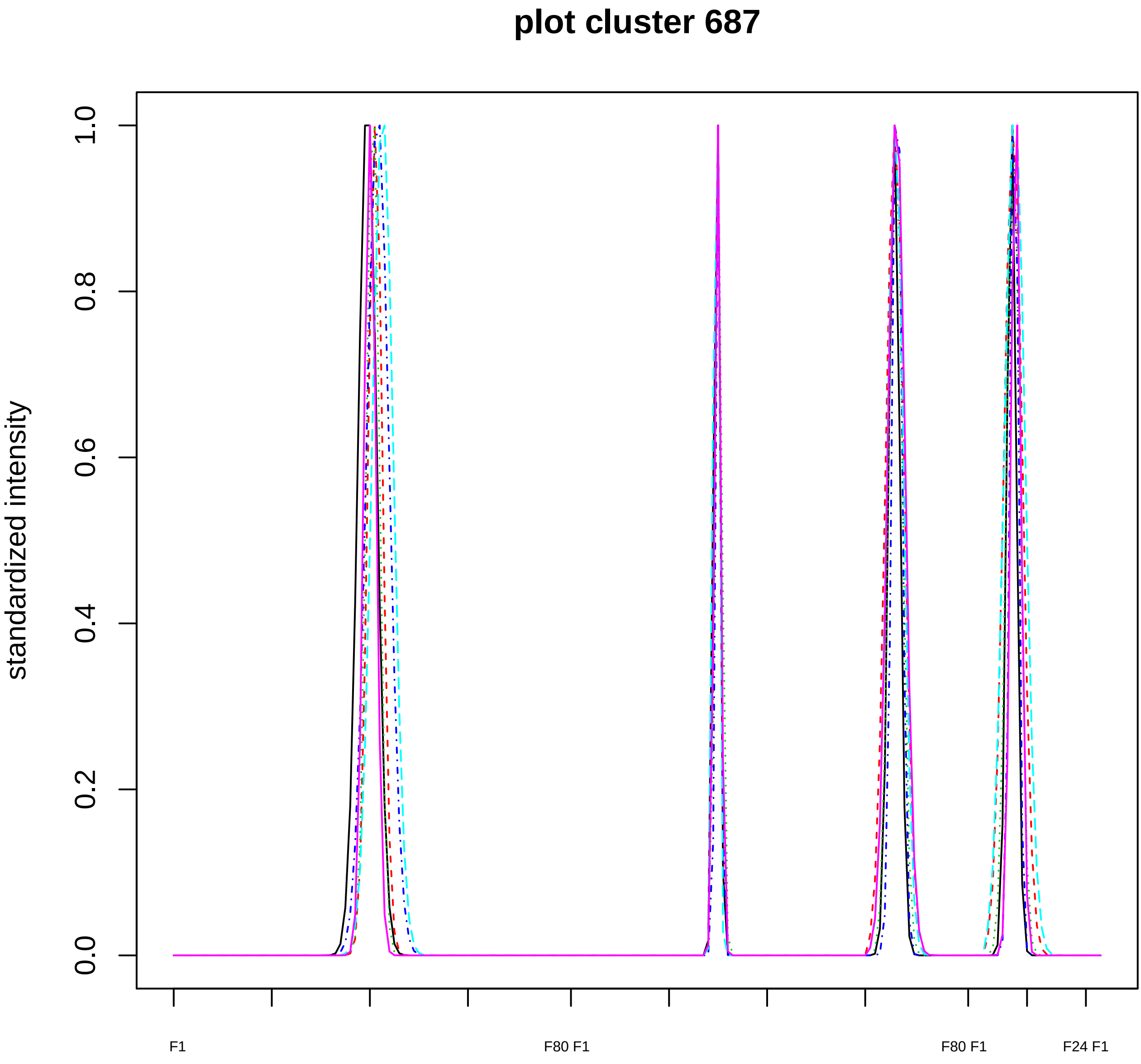
plot cluster 687
standardized intensity
0.0
0.2
0.4
0.6
0.8
1.0
F1
F80 F1
F80 F1
F24 F1

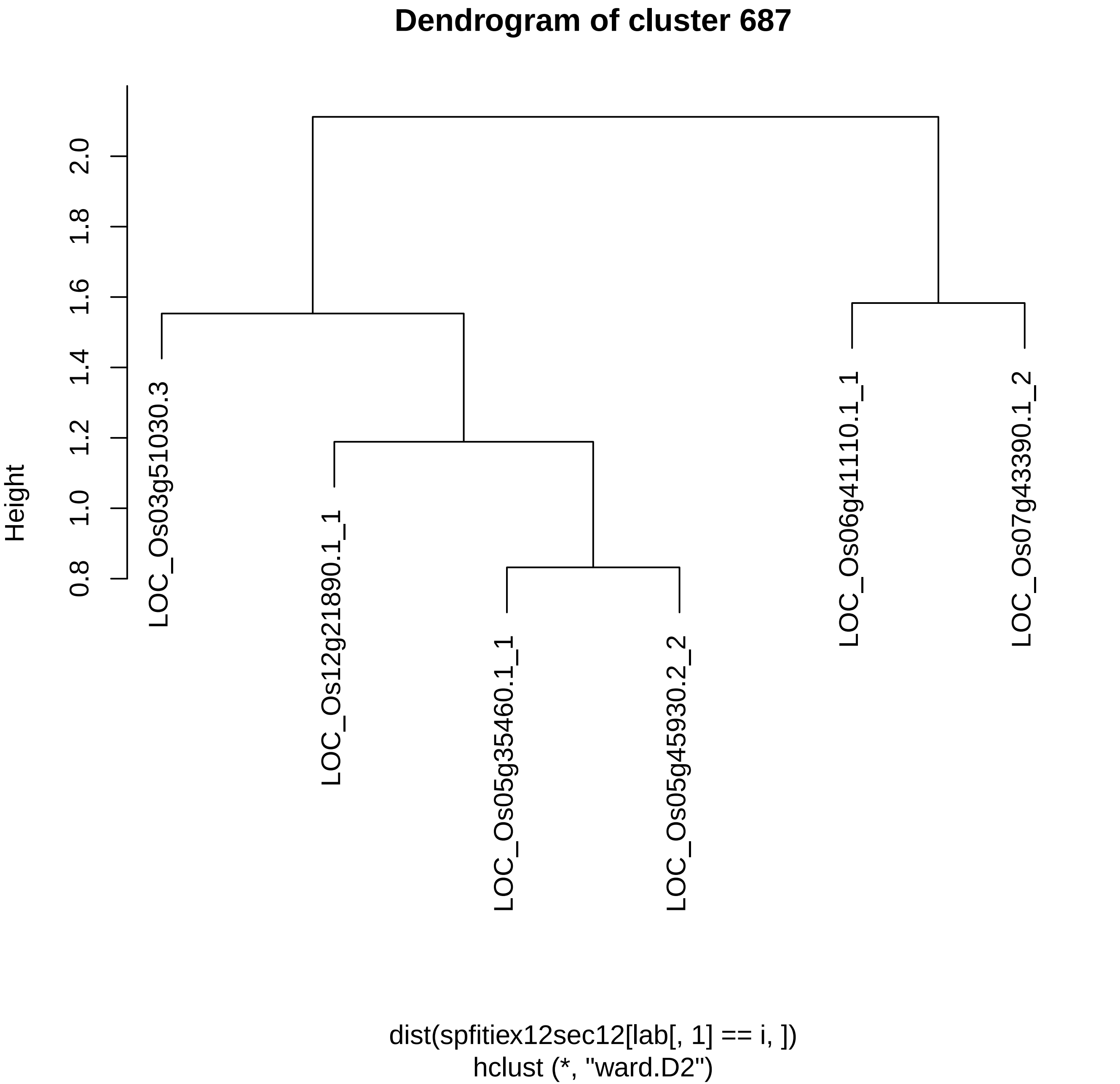
Dendrogram of cluster 687
Height
0.8
1.0
1.2
1.4
1.6
1.8
2.0
LOC_Os03g51030.3
LOC_Os12g21890.1_1
LOC_Os05g35460.1_1
LOC_Os05g45930.2_2
LOC_Os06g41110.1_1
LOC_Os07g43390.1_2
dist(spfitiex12sec12[lab[, 1] == i, ])
hclust (*, "ward.D2")

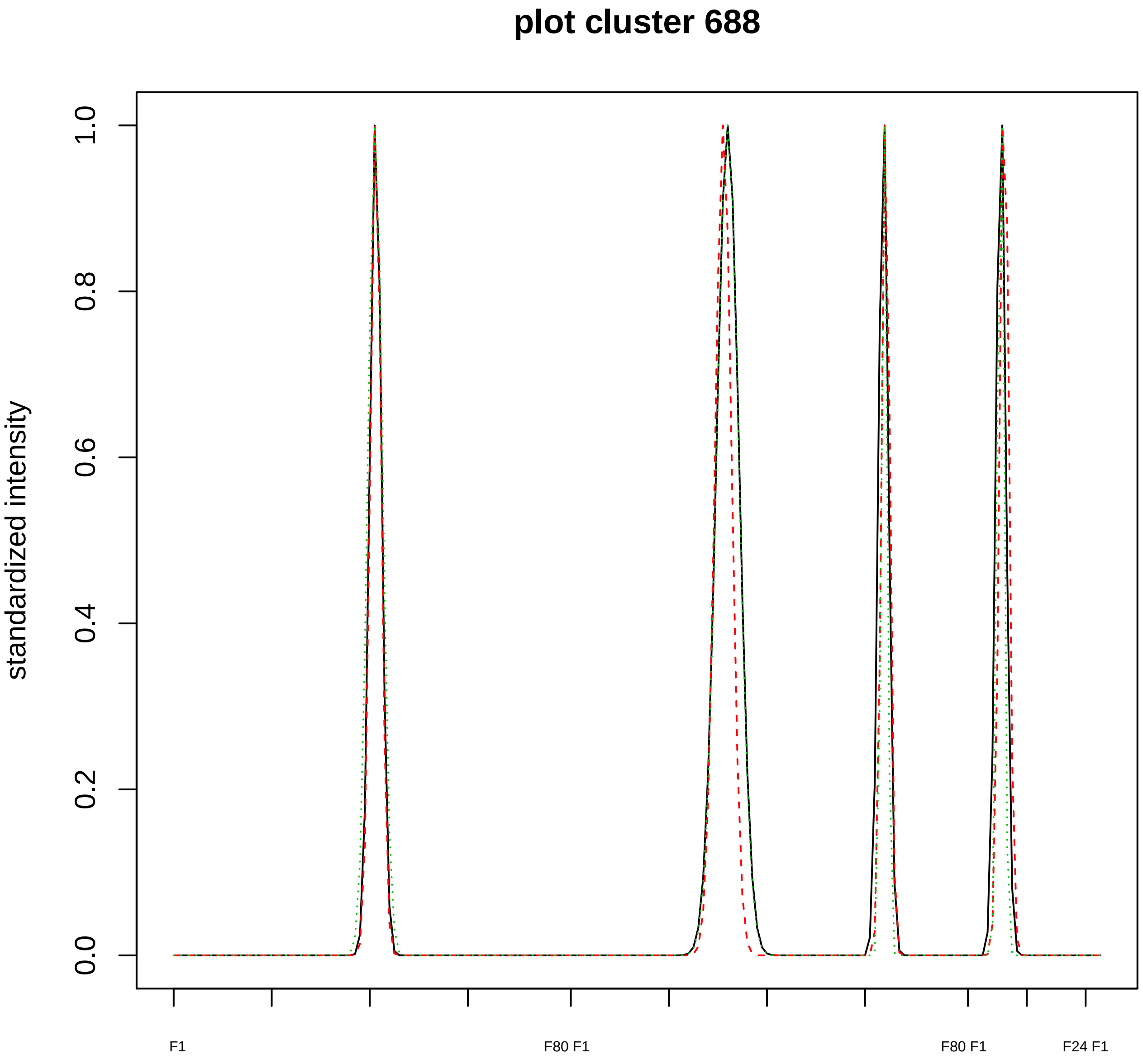

plot cluster 688
standardized intensity
0.0
0.2
0.4
0.6
0.8
1.0
F1
F80 F1
F80 F1
F24 F1

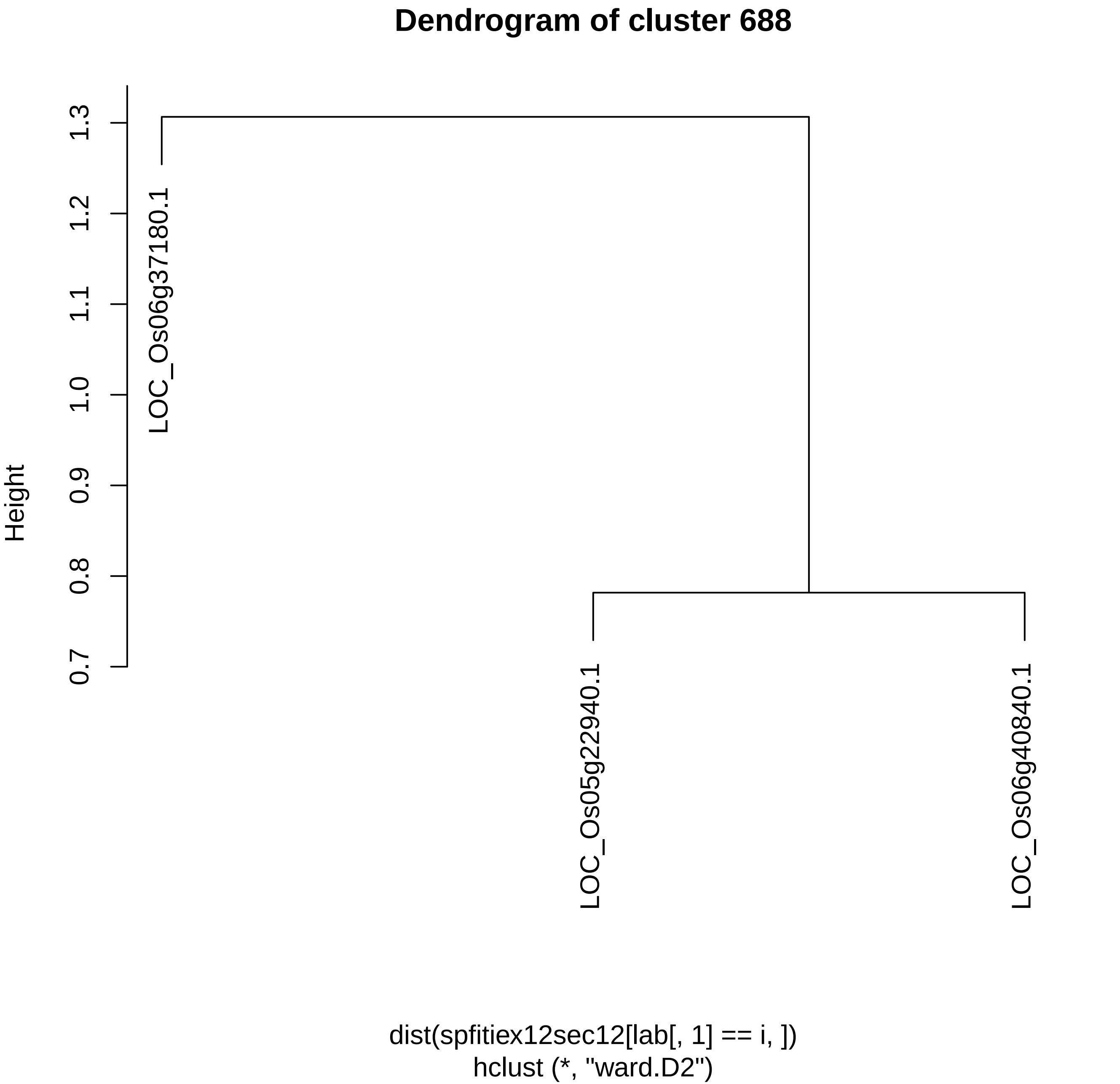
Dendrogram of cluster 688
Height
0.7
0.8
0.9
1.0
1.1
1.2
1.3
LOC_Os06g37180.1
LOC_Os05g22940.1
LOC_Os06g40840.1
dist(spfitiex12sec12[lab[, 1] == i, ])
hclust (*, "ward.D2")

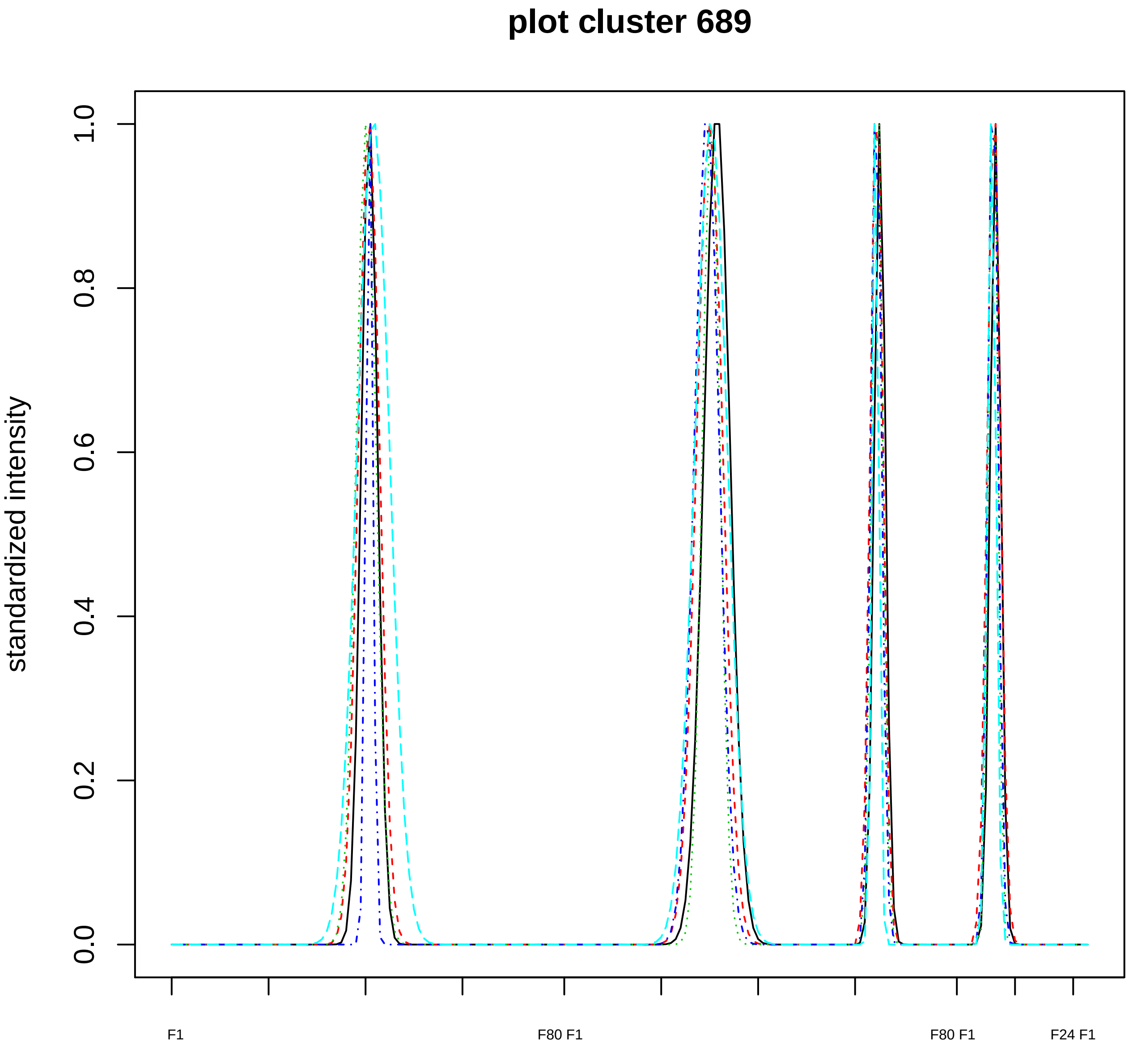

plot cluster 689
standardized intensity
0.0
0.2
0.4
0.6
0.8
1.0
F1
F80 F1
F80 F1
F24 F1

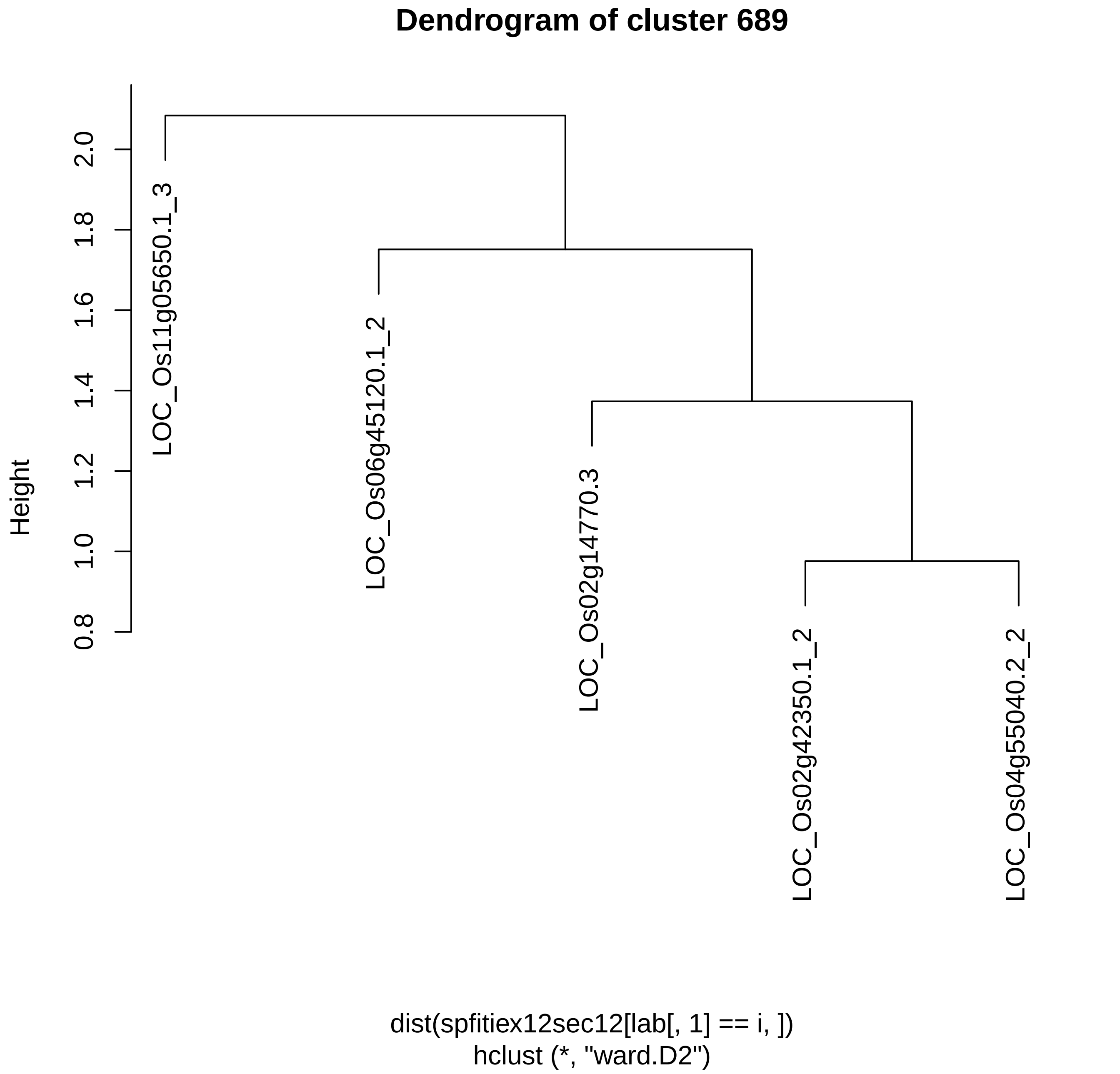
Dendrogram of cluster 689
Height
0.8
1.0
1.2
1.4
1.6
1.8
2.0
LOC_Os11g05650.1_3
LOC_Os06g45120.1_2
LOC_Os02g14770.3
LOC_Os02g42350.1_2
LOC_Os04g55040.2_2
dist(spfitiex12sec12[lab[, 1] == i, ])
hclust (*, "ward.D2")

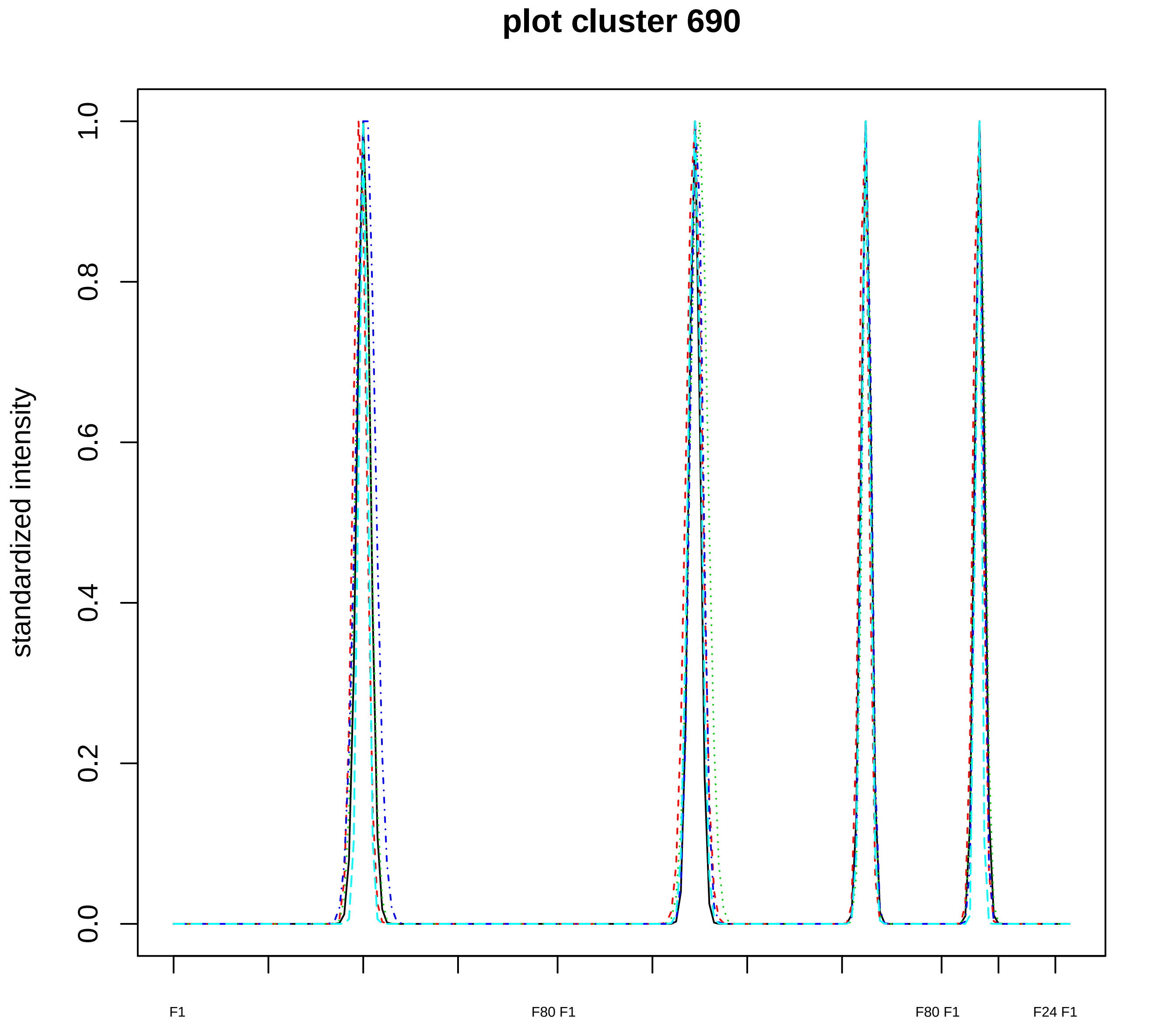
plot cluster 690
standardized intensity
0.0
0.2
0.4
0.6
0.8
1.0
F1
F80 F1
F80 F1
F24 F1

### Dendrogram of cluster 690

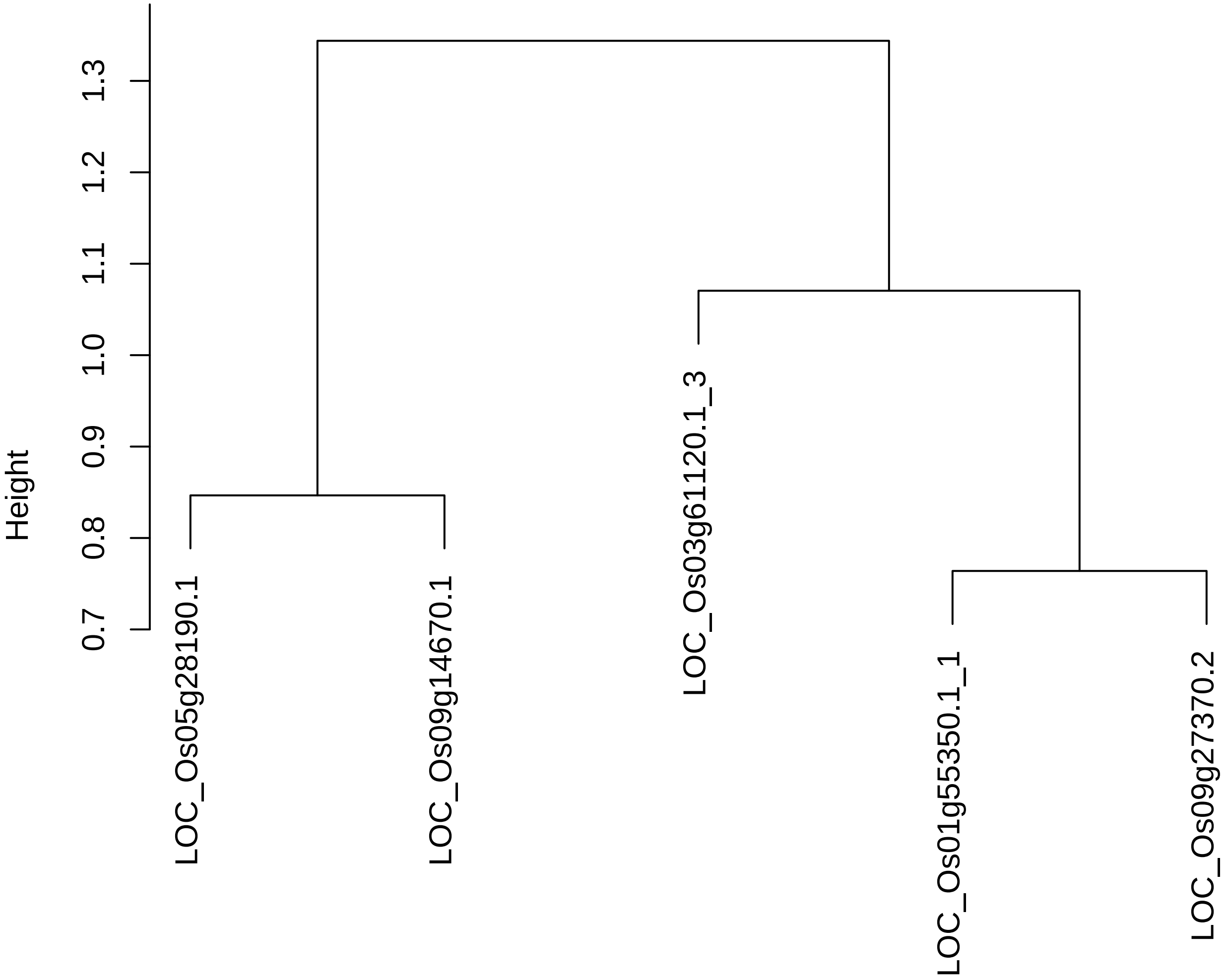


dist(spfitiex12sec12[lab[, 1] == i, ])
hclust (*, "ward.D2")

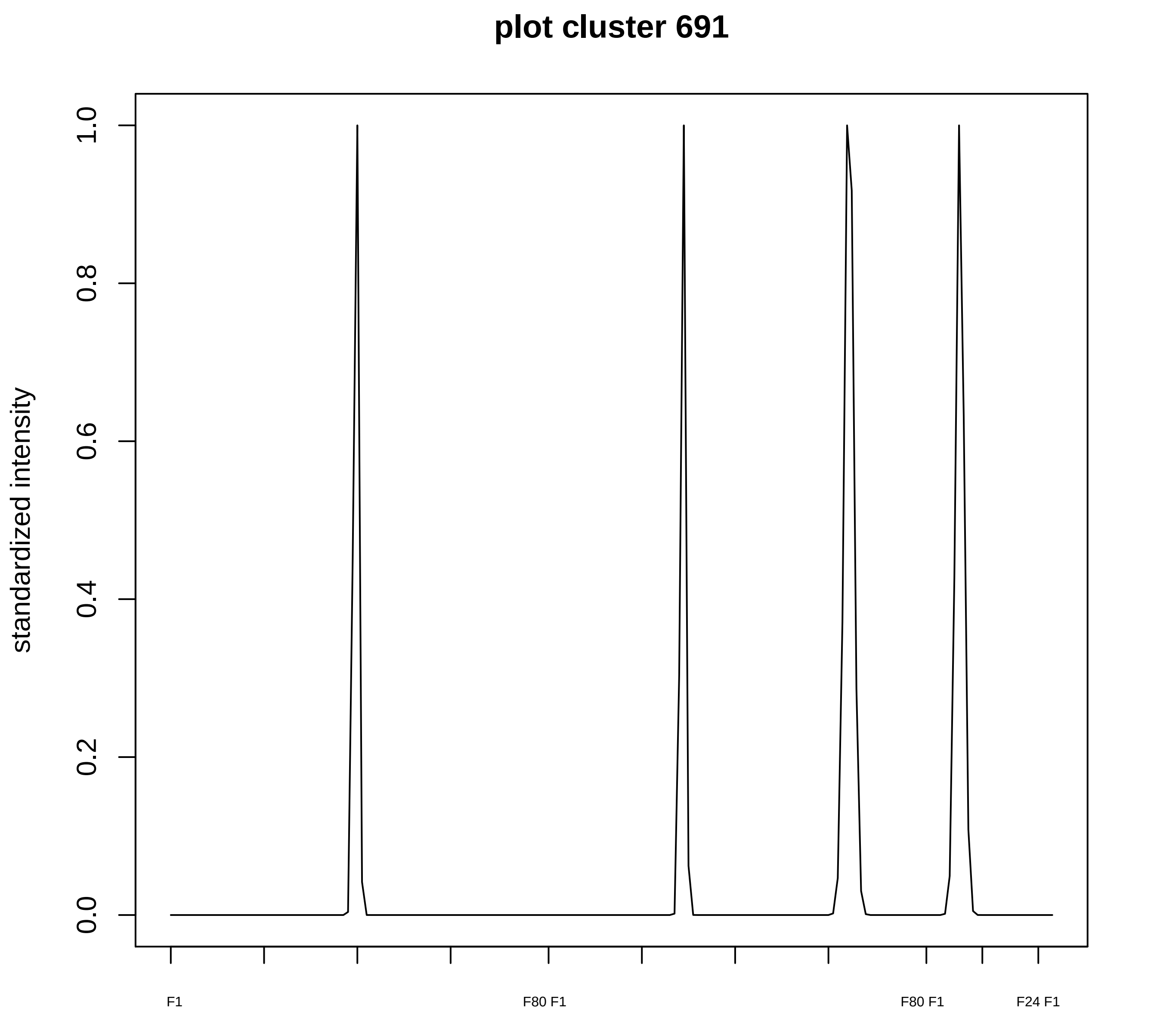
plot cluster 691
standardized intensity
0.0
0.2
0.4
0.6
0.8
1.0
F1
F80 F1
F80 F1
F24 F1

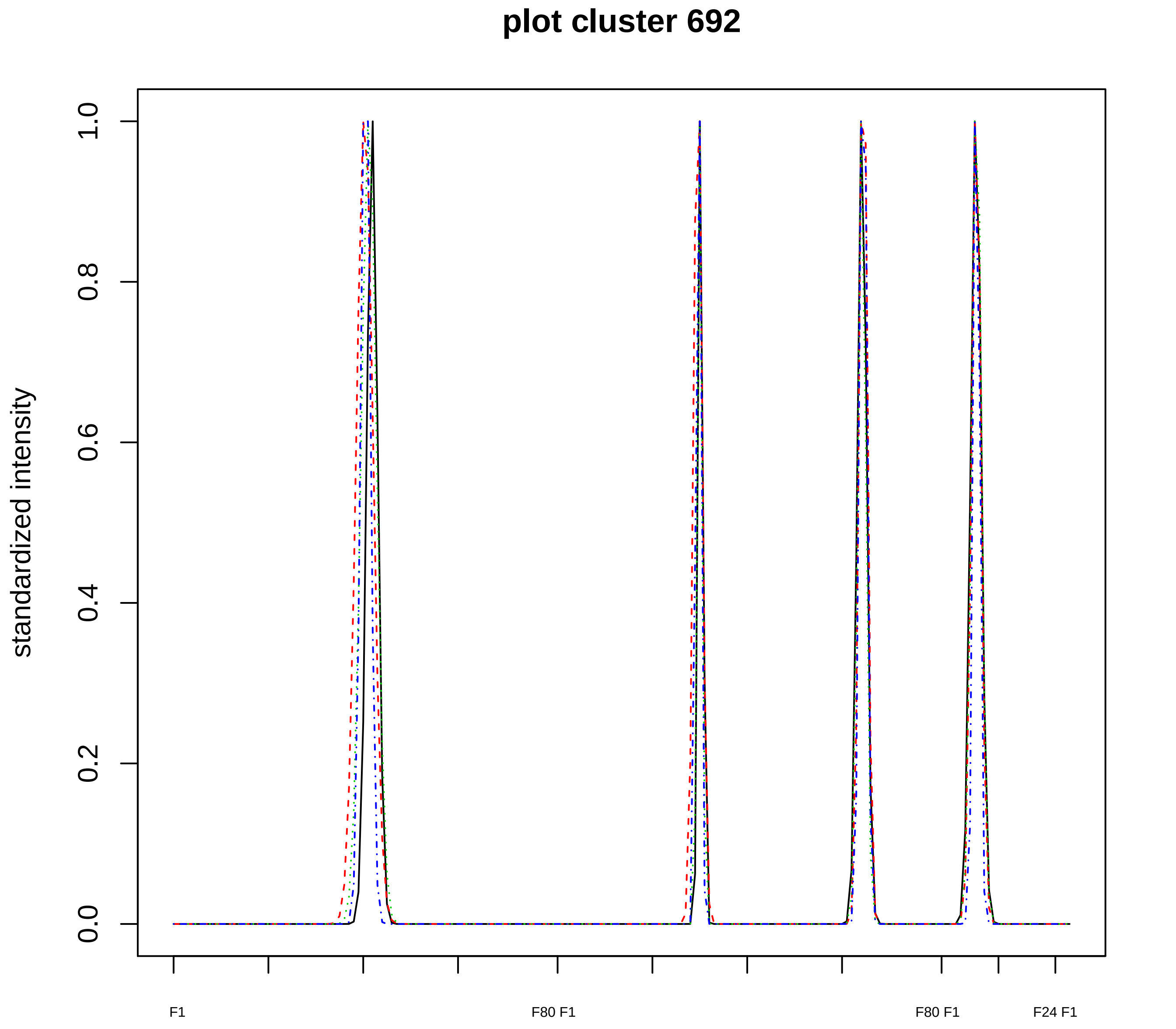
plot cluster 692
standardized intensity
0.0
0.2
0.4
0.6
0.8
1.0
F1
F80 F1
F80 F1
F24 F1

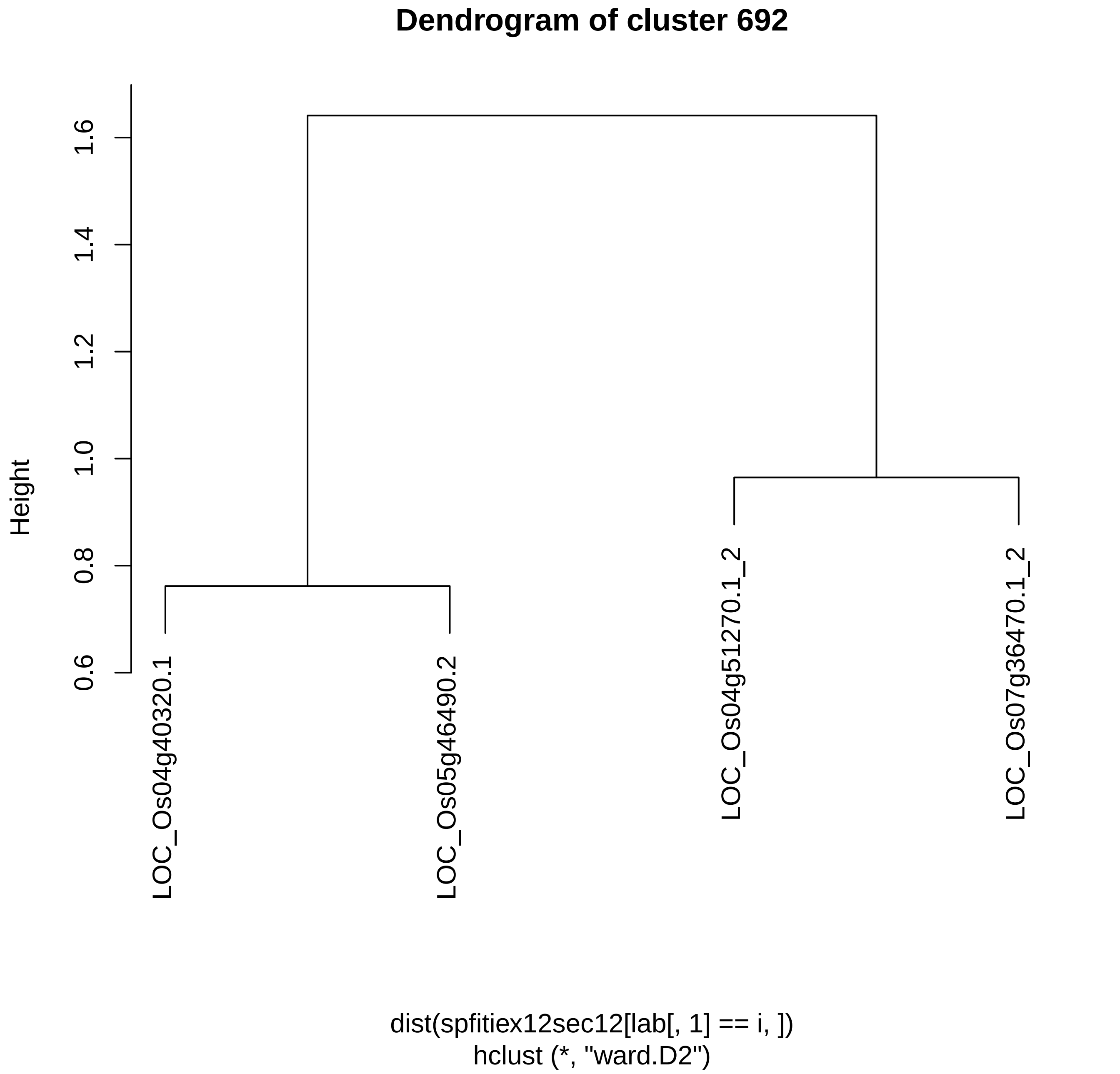
Dendrogram of cluster 692
Height
0.6
0.8
1.0
1.2
1.4
1.6
LOC_Os04g40320.1
LOC_Os05g46490.2
LOC_Os04g51270.1_2
LOC_Os07g36470.1_2
dist(spfitiex12sec12[lab[, 1] == i, ])
hclust (*, "ward.D2")

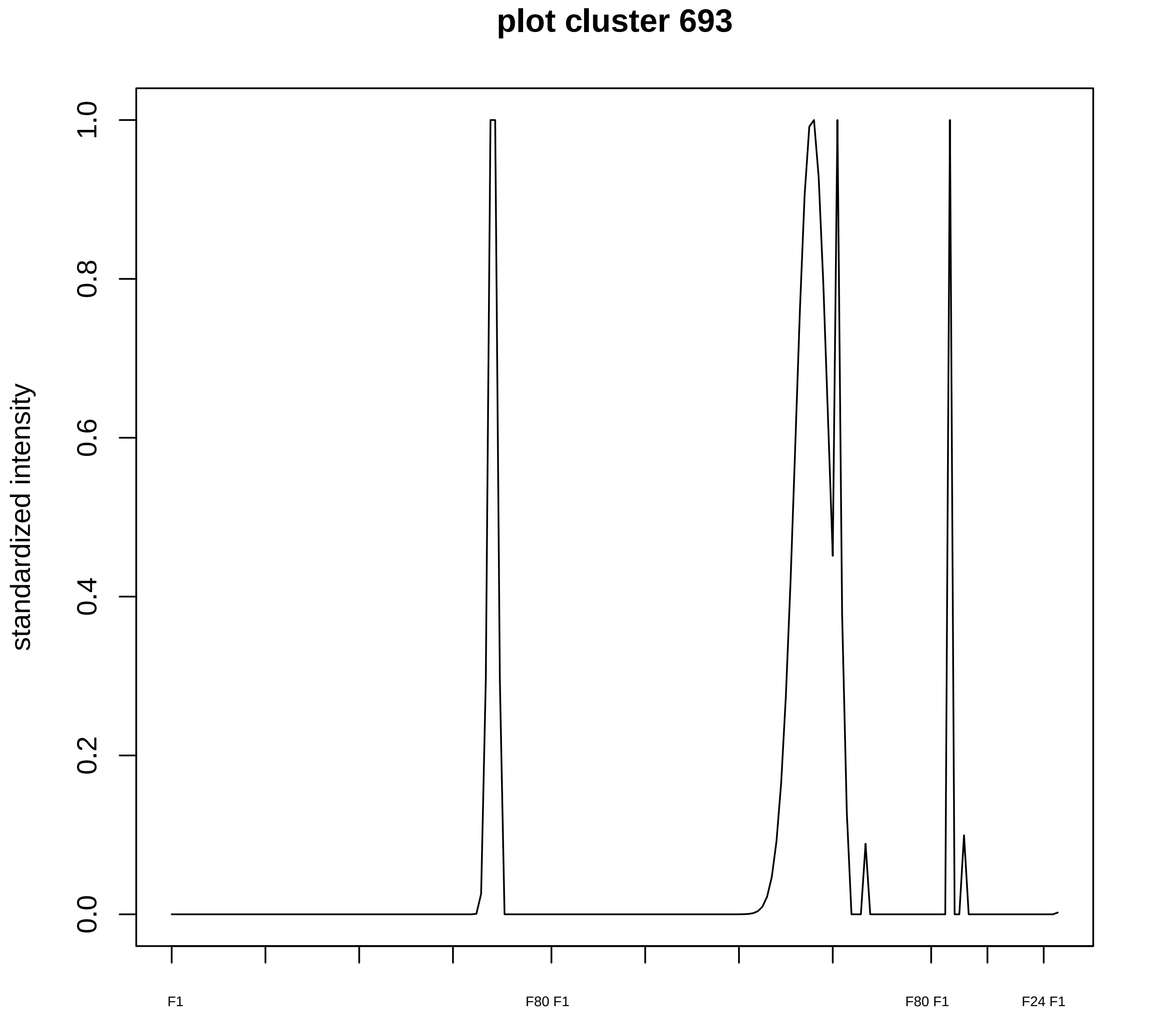
plot cluster 693
standardized intensity
0.0
0.2
0.4
0.6
0.8
1.0
F1
F80 F1
F80 F1
F24 F1

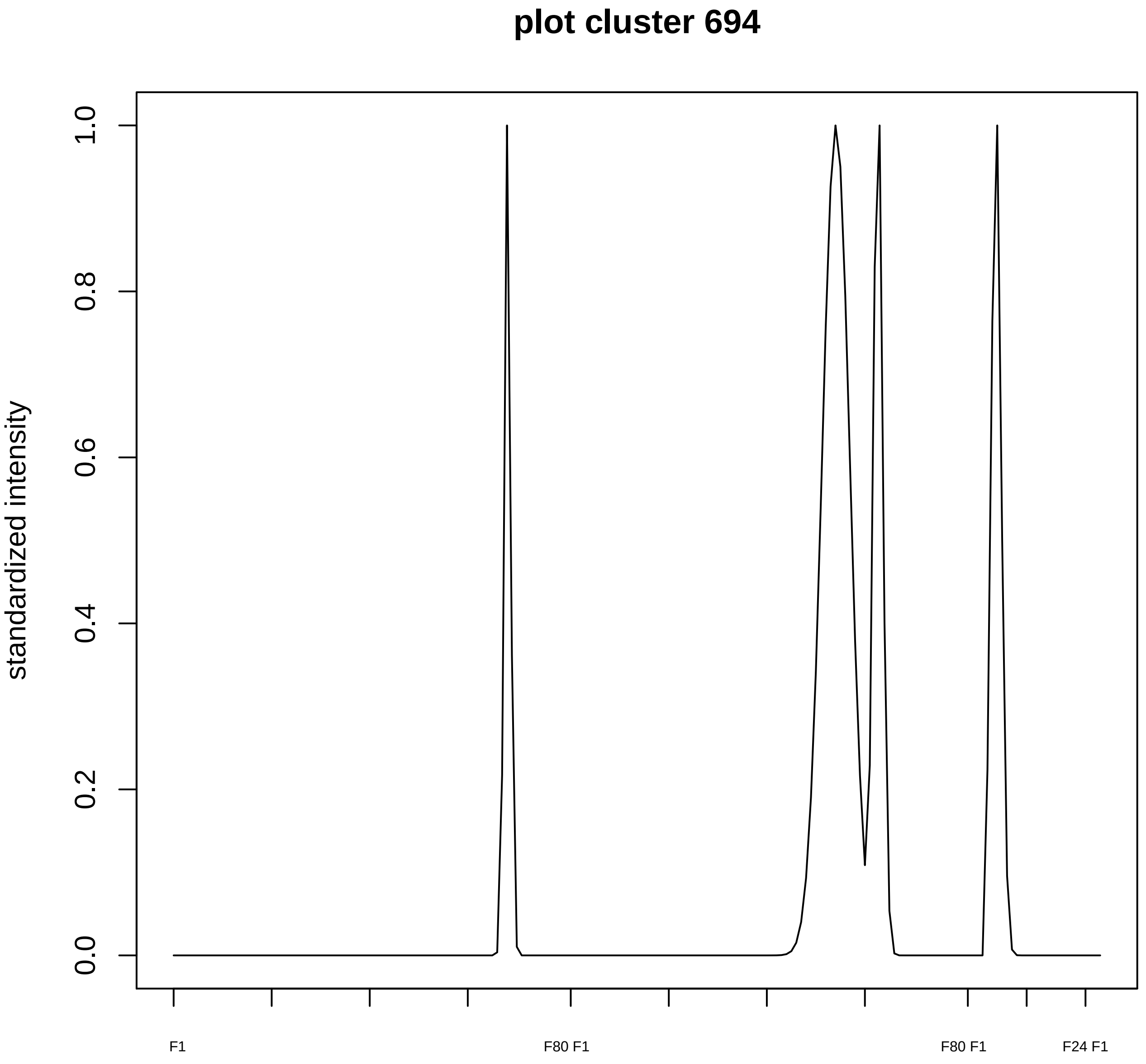

plot cluster 694
standardized intensity
0.0
0.2
0.4
0.6
0.8
1.0
F1
F80 F1
F80 F1
F24 F1

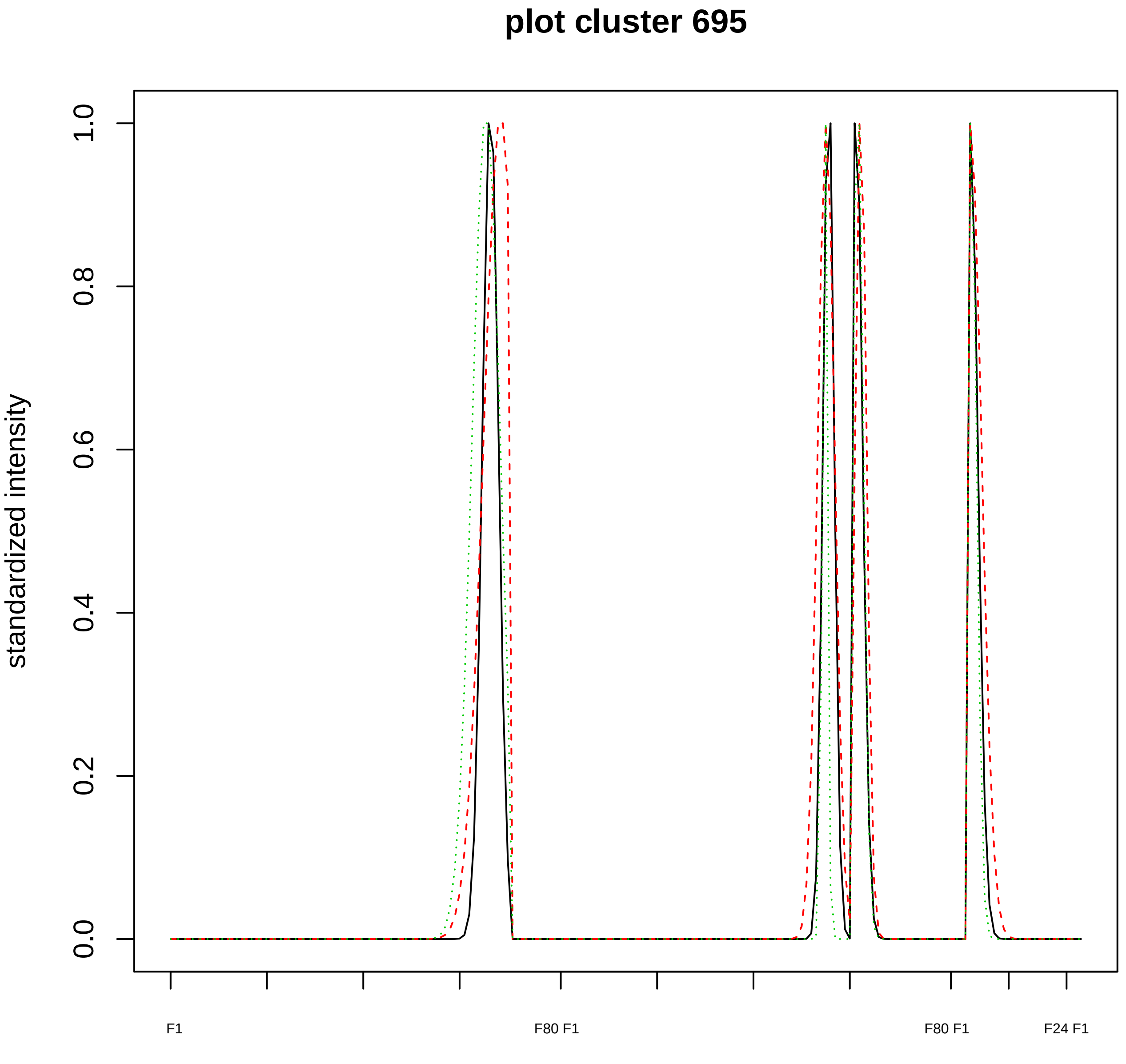
plot cluster 695
standardized intensity
0.0
0.2
0.4
0.6
0.8
1.0
F1
F80 F1
F80 F1
F24 F1

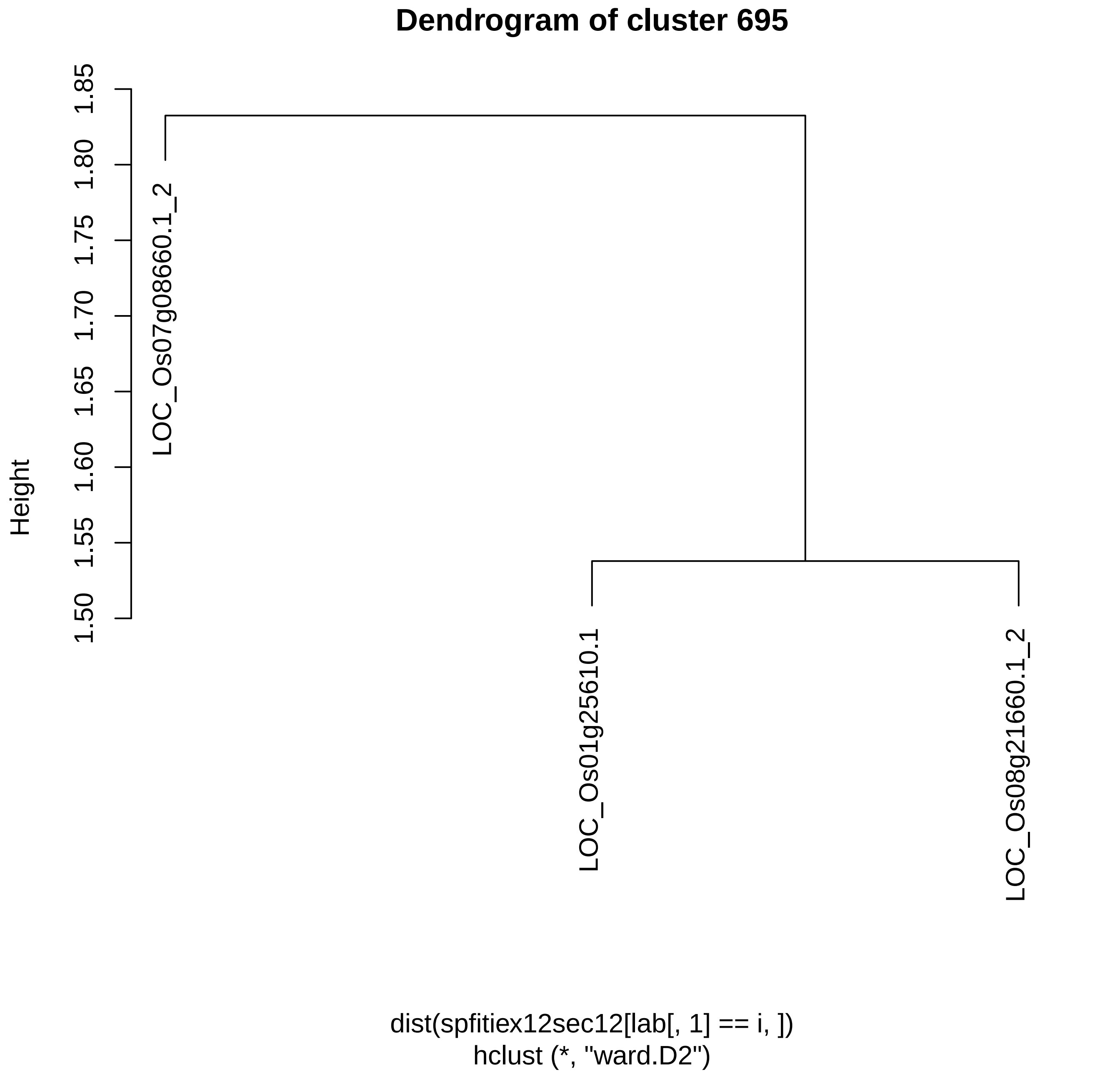
Dendrogram of cluster 695
Height
1.50
1.55
1.60
1.65
1.70
1.75
1.80
1.85
LOC_Os07g08660.1_2
LOC_Os01g25610.1
LOC_Os08g21660.1_2
dist(spfitiex12sec12[lab[, 1] == i, ])
hclust (*, "ward.D2")

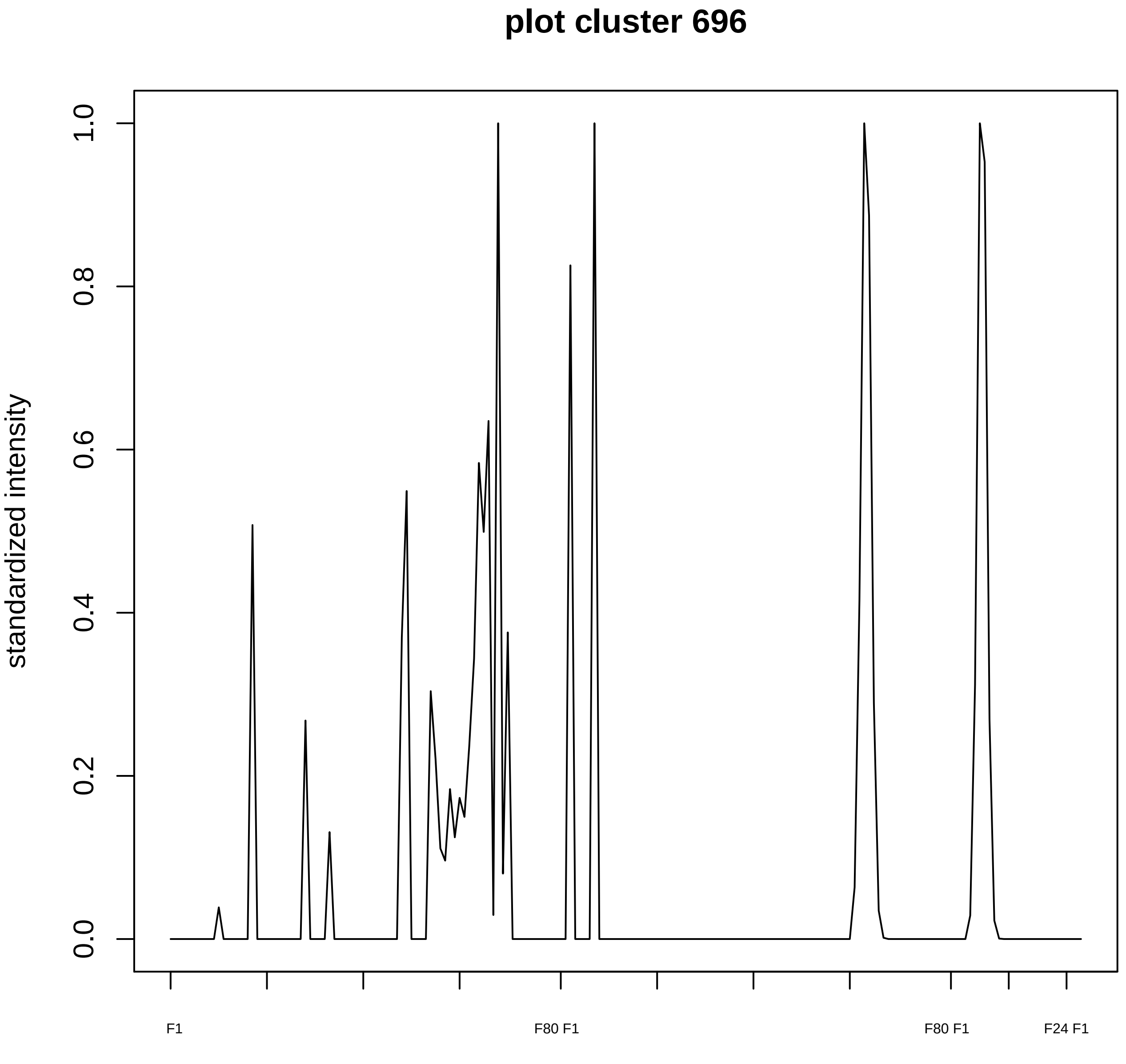
plot cluster 696
standardized intensity
0.0
0.2
0.4
0.6
0.8
1.0
F1
F80 F1
F80 F1
F24 F1

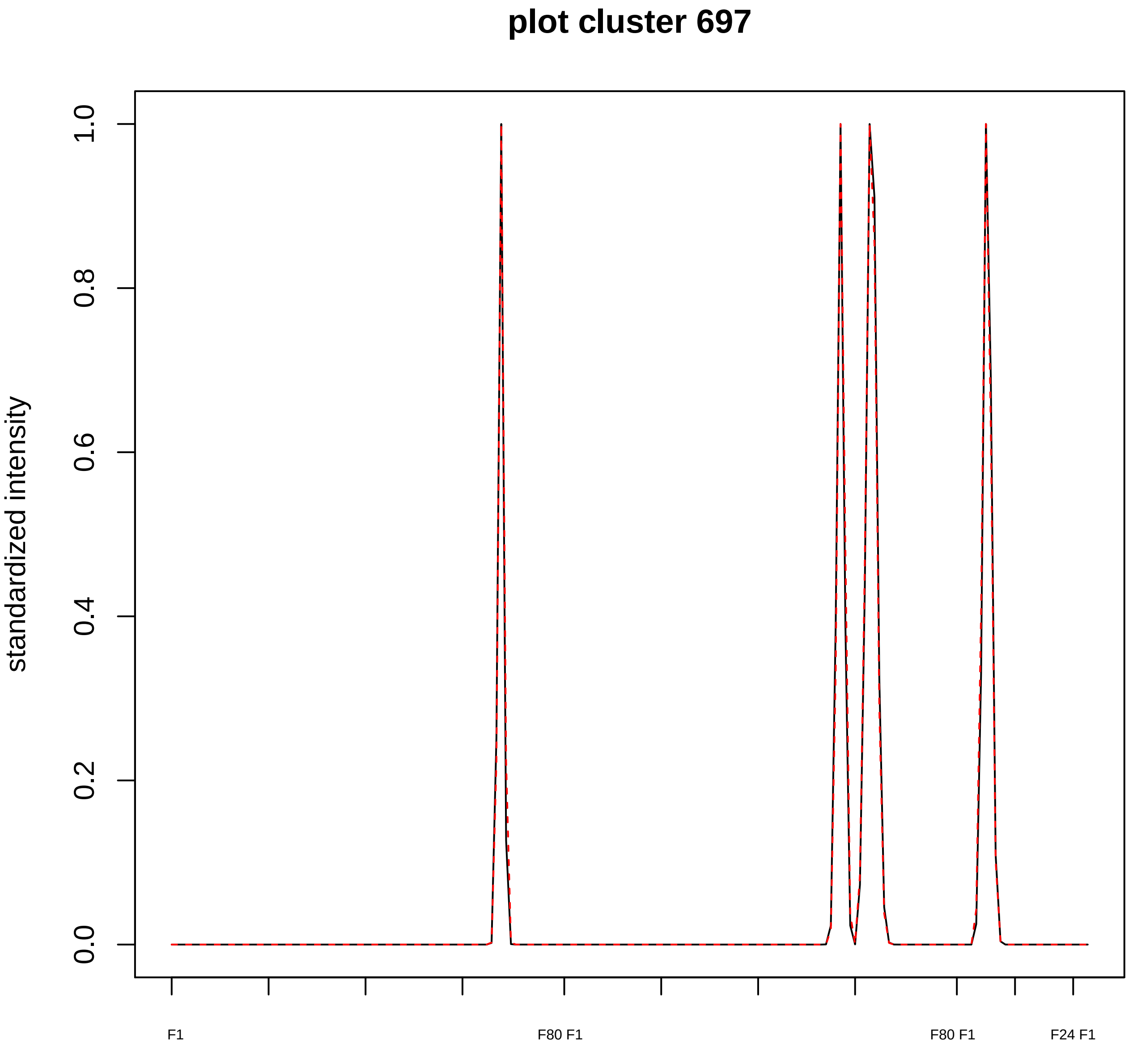
plot cluster 697
standardized intensity
0.0
0.2
0.4
0.6
0.8
1.0
F1
F80 F1
F80 F1
F24 F1

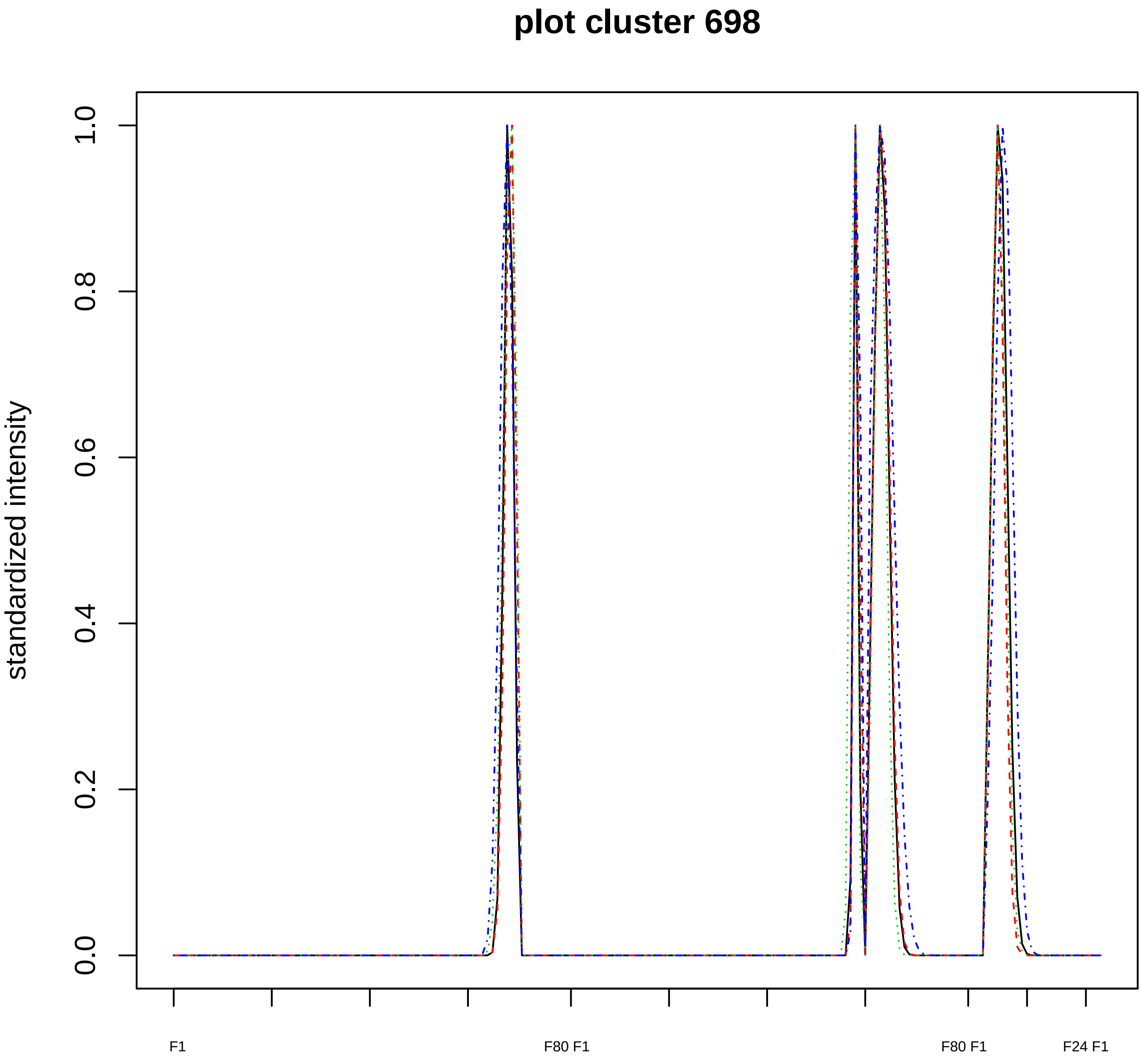
plot cluster 698
standardized intensity
0.0
0.2
0.4
0.6
0.8
1.0
F1
F80 F1
F80 F1
F24 F1

Dendrogram of cluster 698

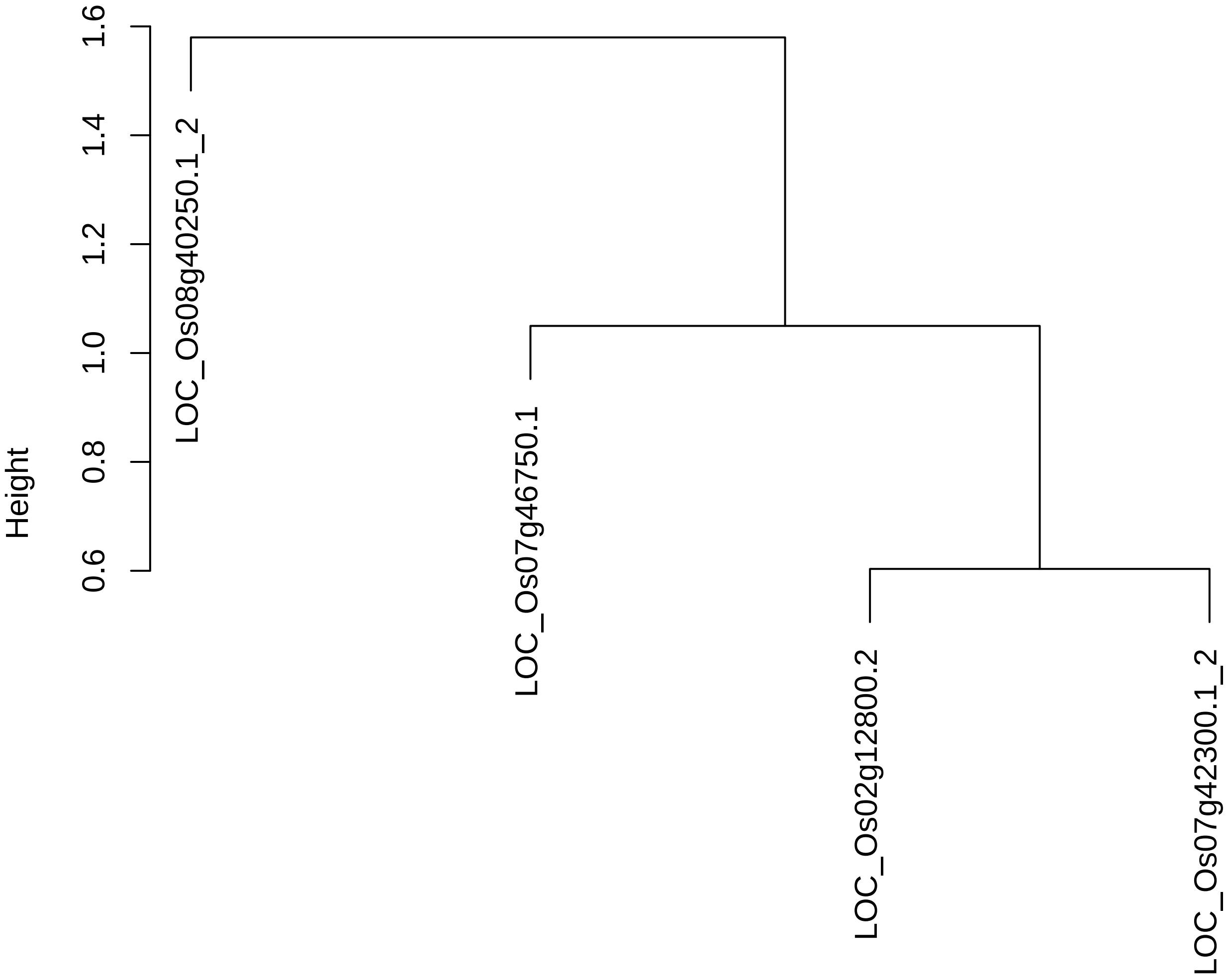


dist(spfitiex12sec12[lab[, 1] == i, ])
hclust (*, "ward.D2")

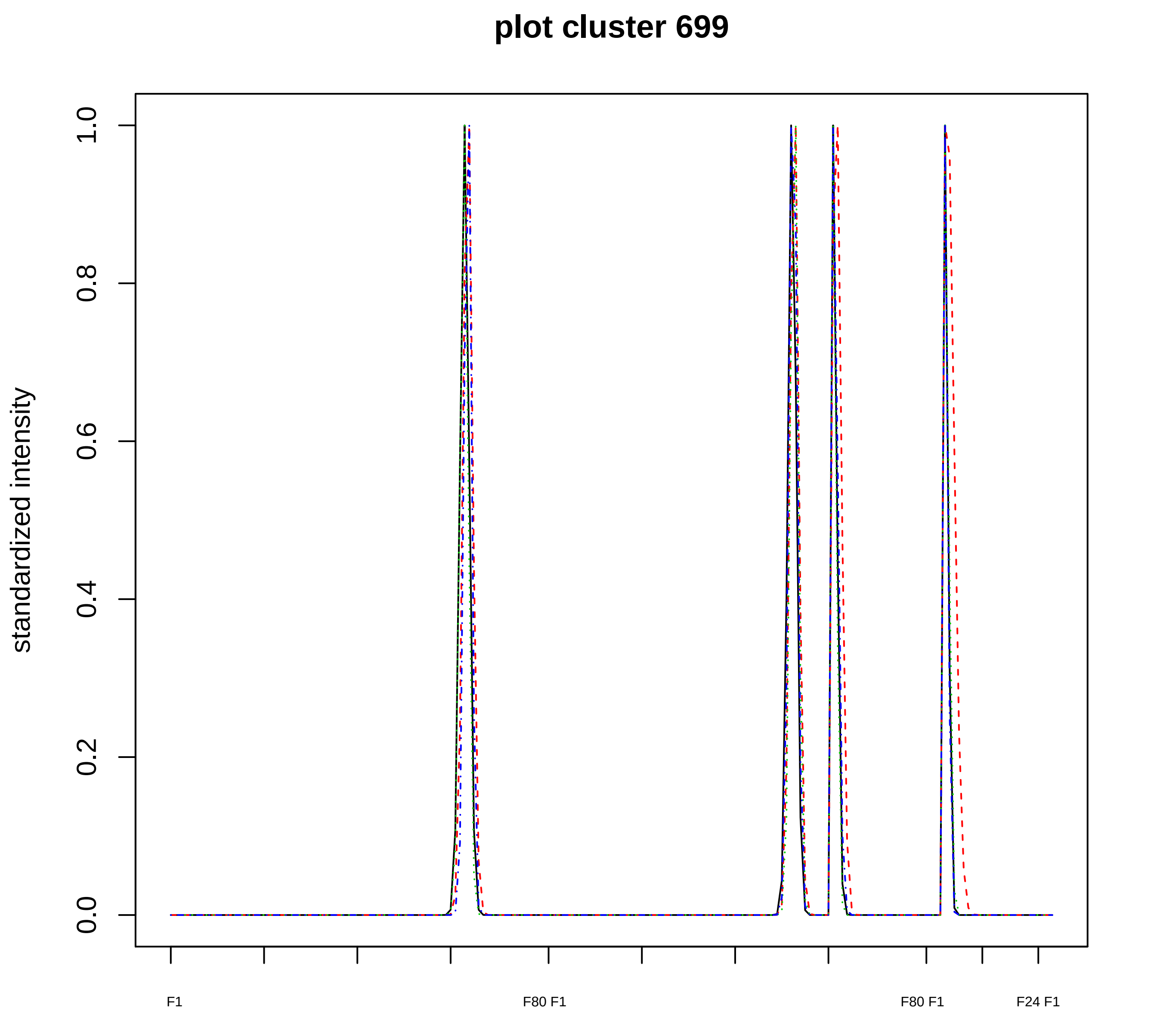

plot cluster 699
standardized intensity
0.0
0.2
0.4
0.6
0.8
1.0
F1
F80 F1
F80 F1
F24 F1

### Dendrogram of cluster 699

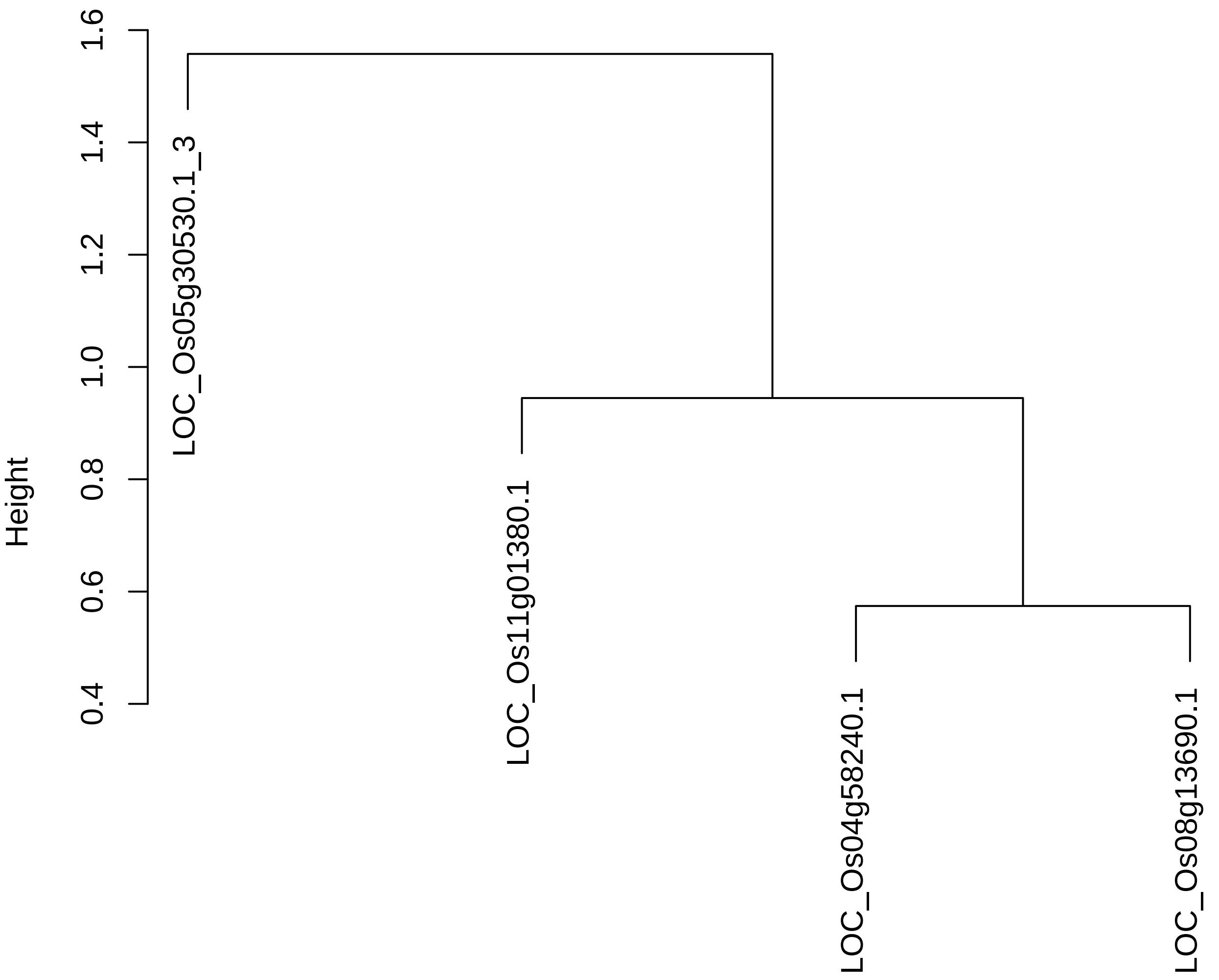


dist(spfitiex12sec12[lab[, 1] == i, ])
hclust (*, "ward.D2")

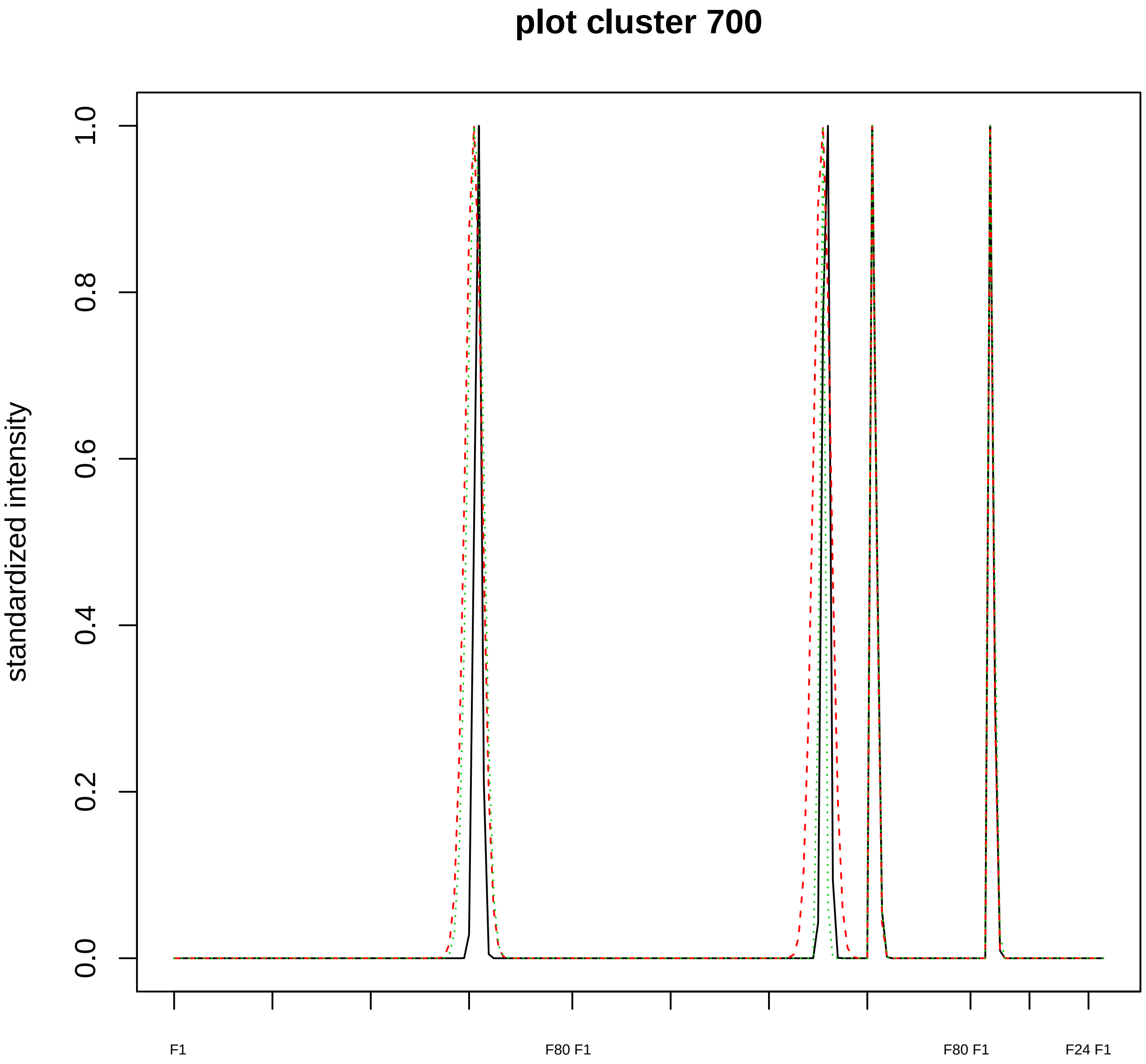
plot cluster 700
standardized intensity
0.0
0.2
0.4
0.6
0.8
1.0
F1
F80 F1
F80 F1
F24 F1

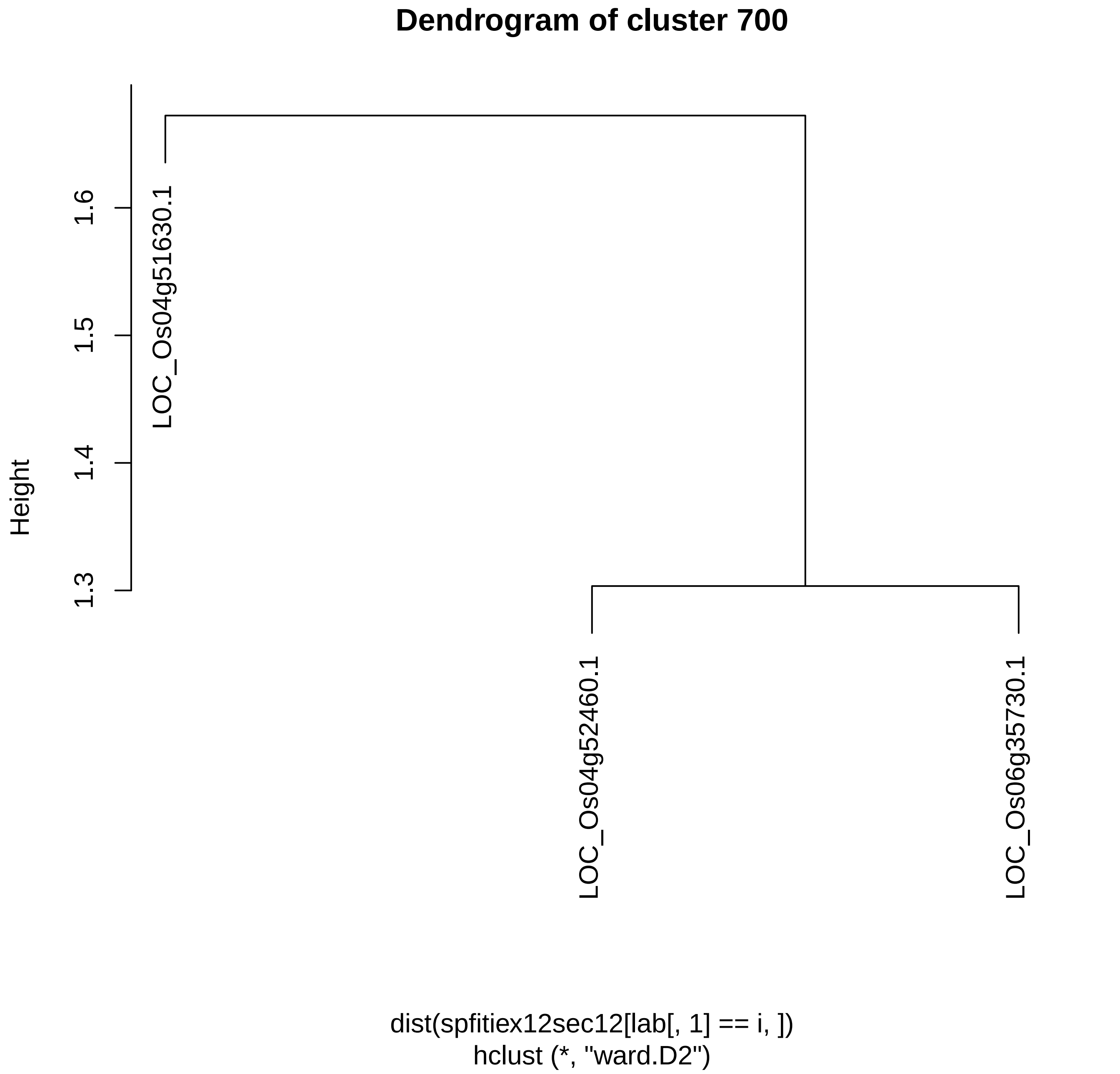

Dendrogram of cluster 700
Height
1.6
1.5
1.4
1.3
LOC_Os04g51630.1
LOC_Os04g52460.1
LOC_Os06g35730.1
dist(spfitiex12sec12[lab[, 1] == i, ])
hclust (*, "ward.D2")

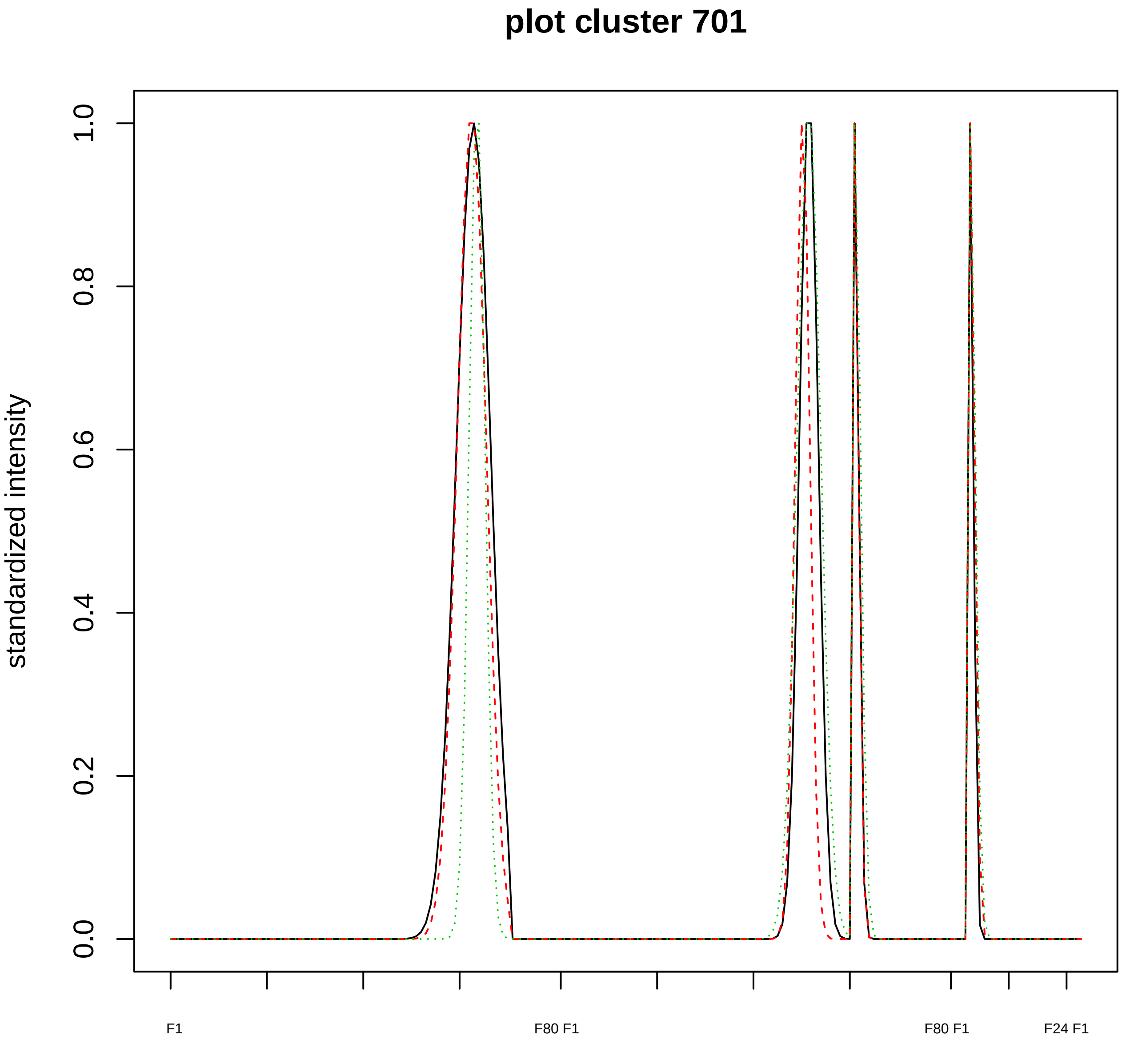

plot cluster 701
standardized intensity
0.0
0.2
0.4
0.6
0.8
1.0
F1
F80 F1
F80 F1
F24 F1

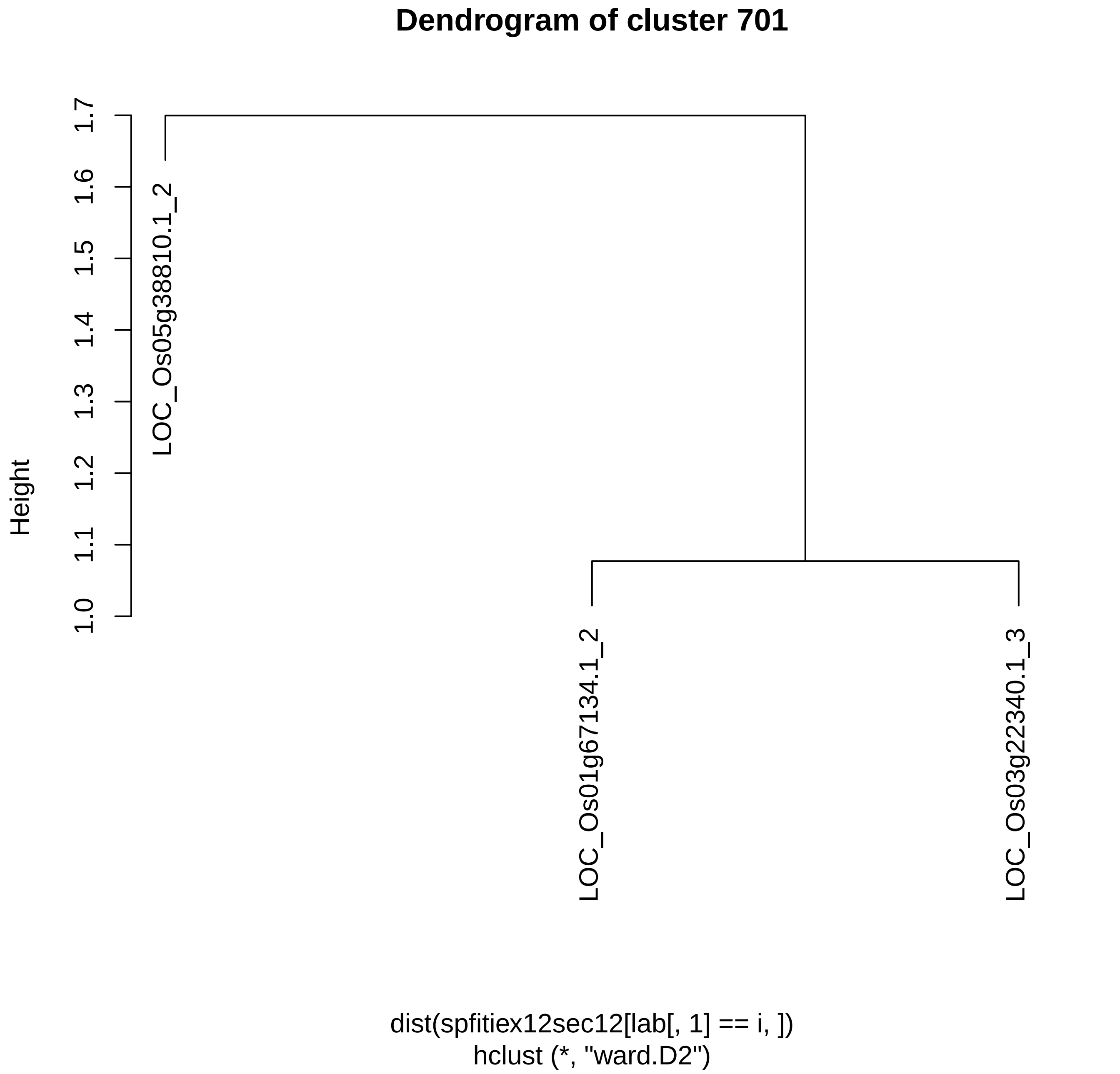

Dendrogram of cluster 701
Height
1.0
1.1
1.2
1.3
1.4
1.5
1.6
1.7
LOC_Os05g38810.1_2
LOC_Os01g67134.1_2
LOC_Os03g22340.1_3
dist(spfitiex12sec12[lab[, 1] == i, ])
hclust (*, "ward.D2")

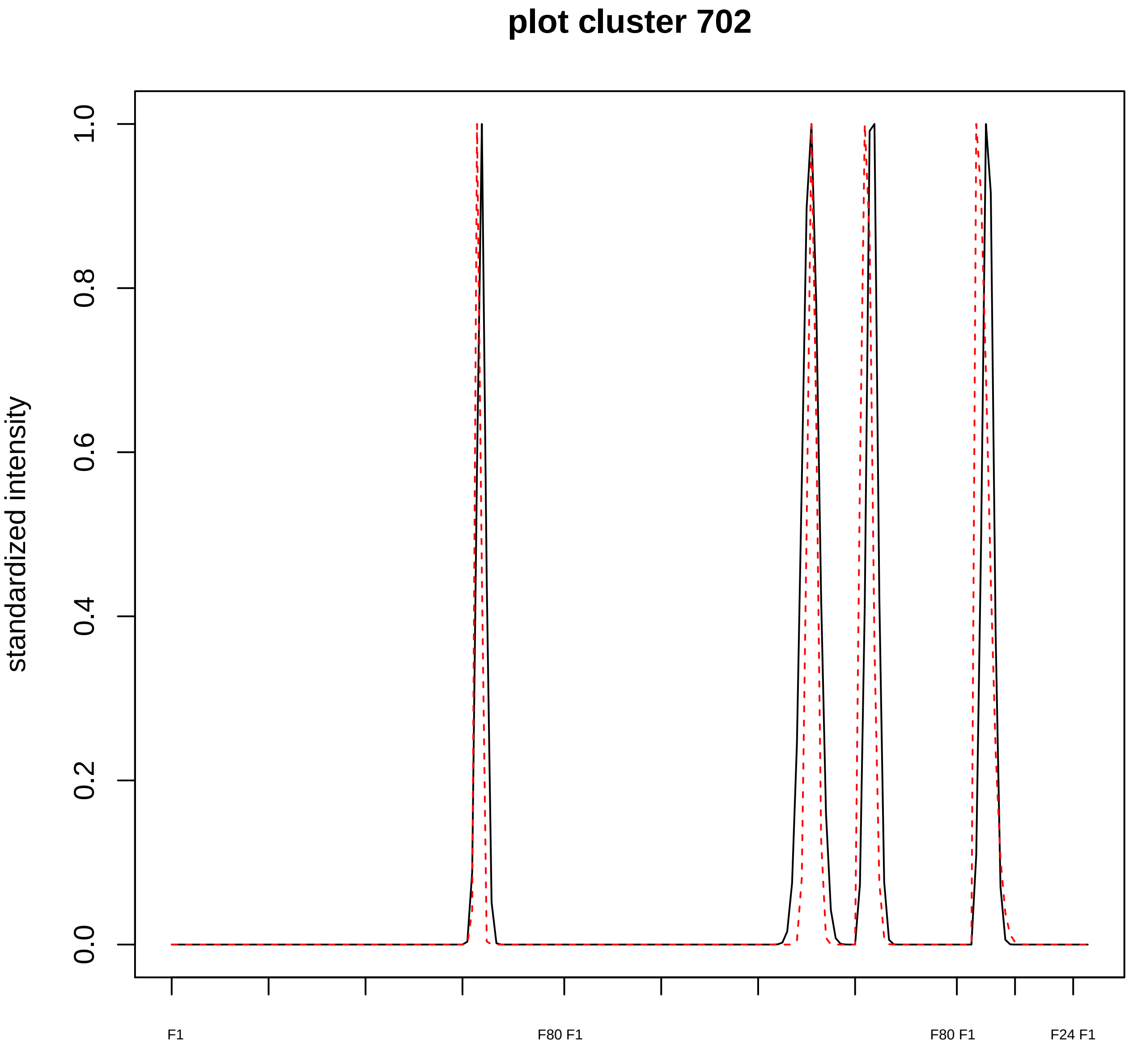
plot cluster 702
standardized intensity
0.0
0.2
0.4
0.6
0.8
1.0
F1
F80 F1
F80 F1
F24 F1

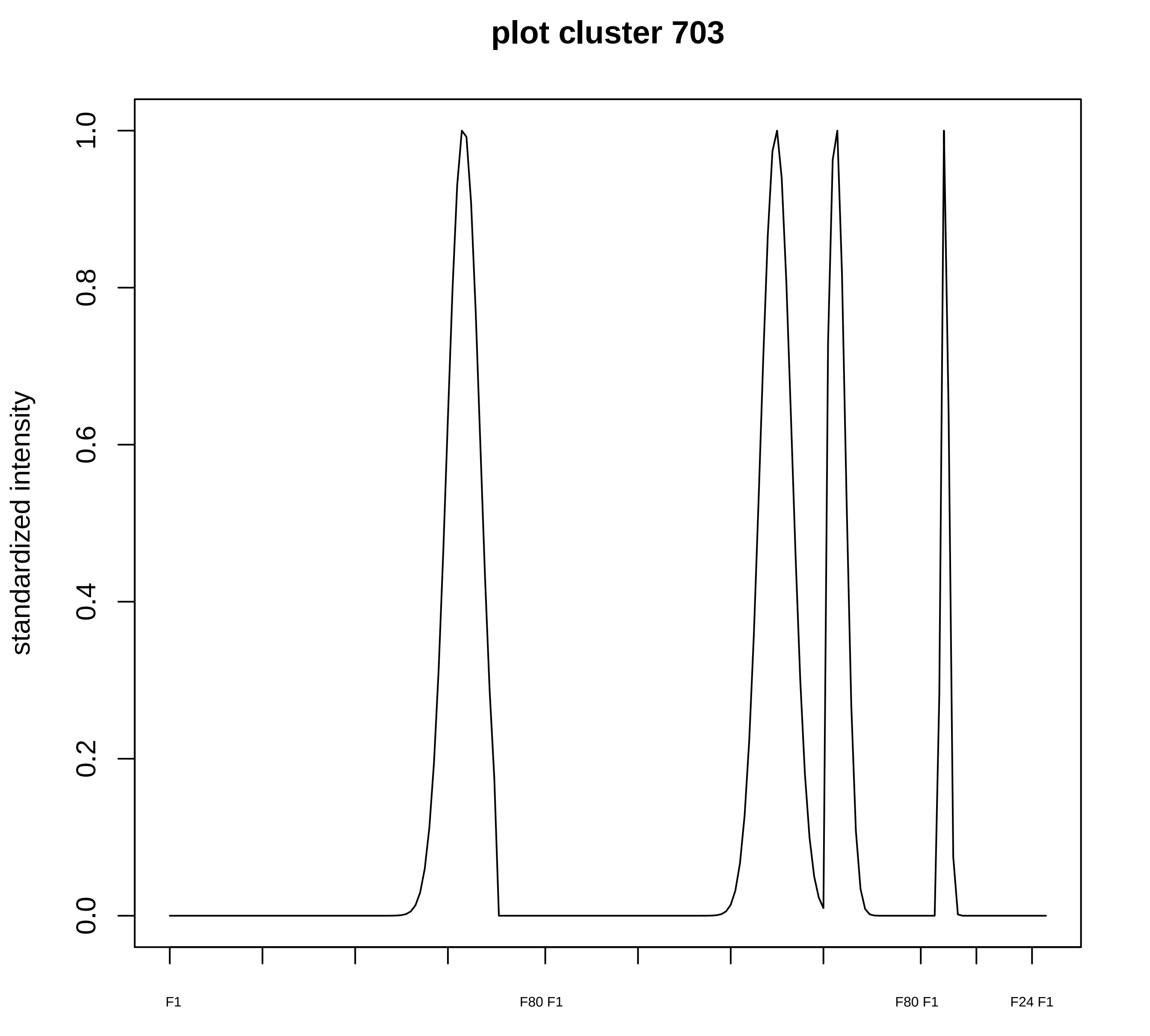

plot cluster 703
standardized intensity
0.0
0.2
0.4
0.6
0.8
1.0
F1
F80 F1
F80 F1
F24 F1

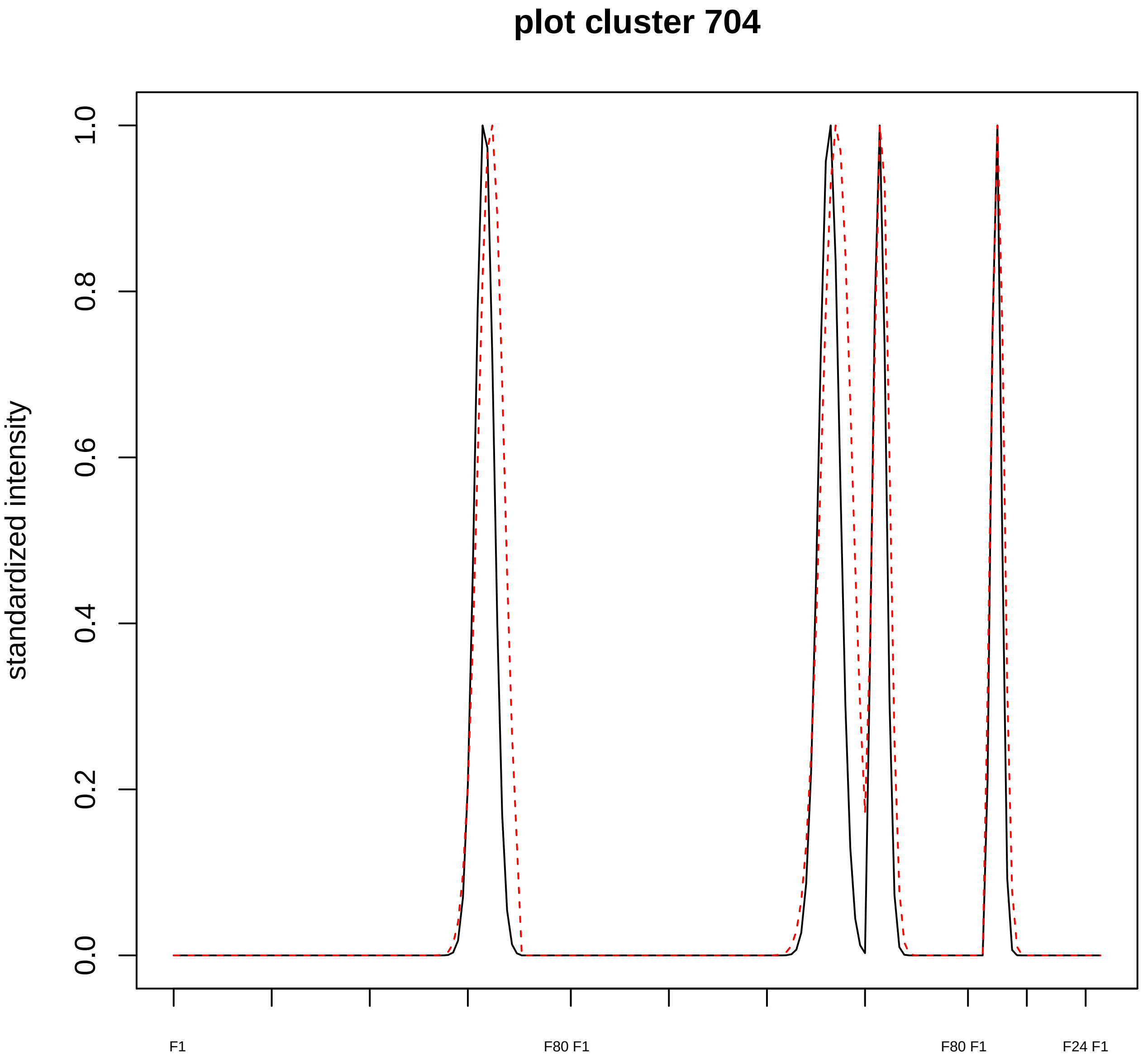

plot cluster 704
standardized intensity
0.0
0.2
0.4
0.6
0.8
1.0
F1
F80 F1
F80 F1
F24 F1

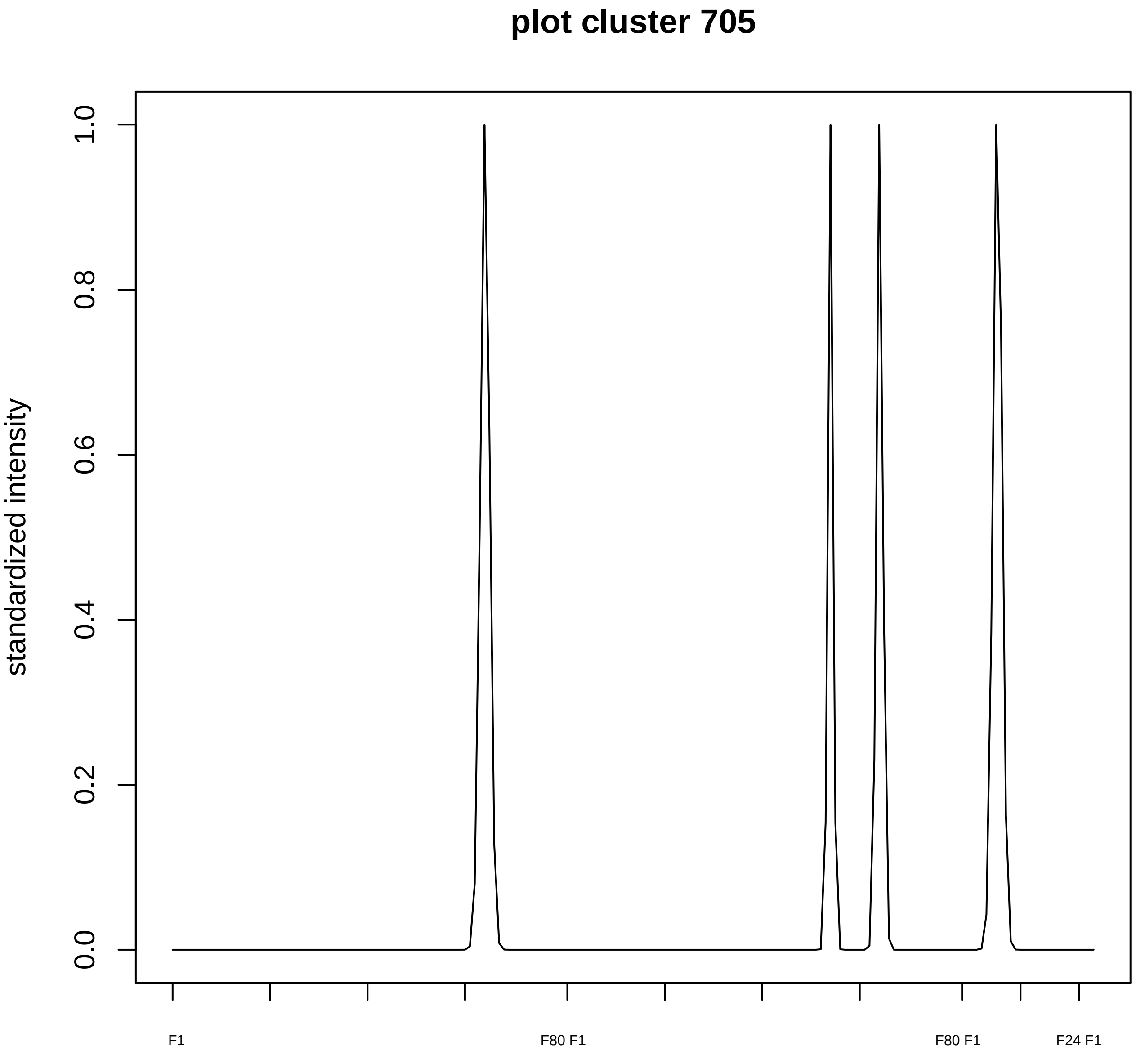

plot cluster 705
standardized intensity
0.0
0.2
0.4
0.6
0.8
1.0
F1
F80 F1
F80 F1
F24 F1

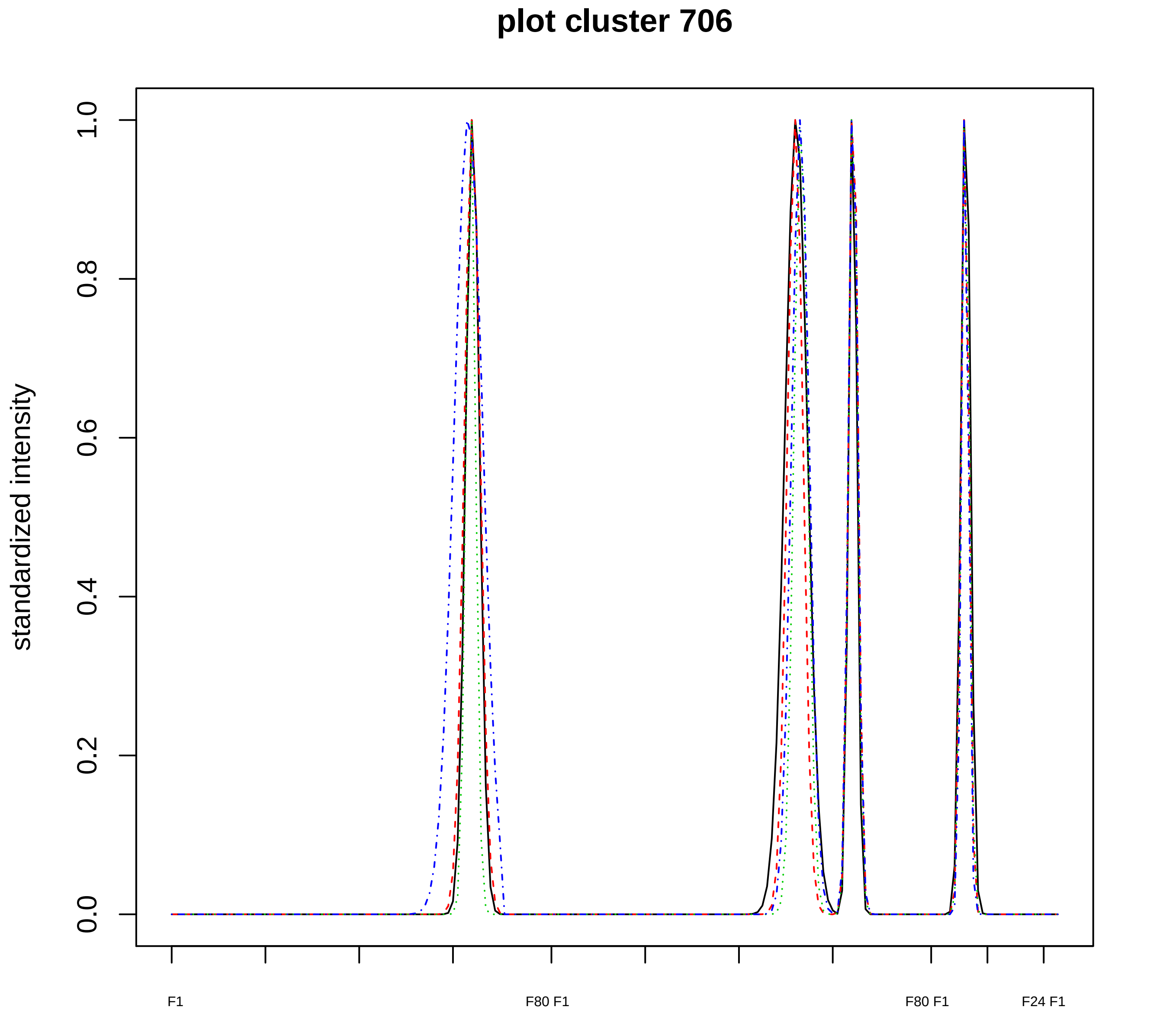
plot cluster 706
standardized intensity
0.0
0.2
0.4
0.6
0.8
1.0
F1
F80 F1
F80 F1
F24 F1

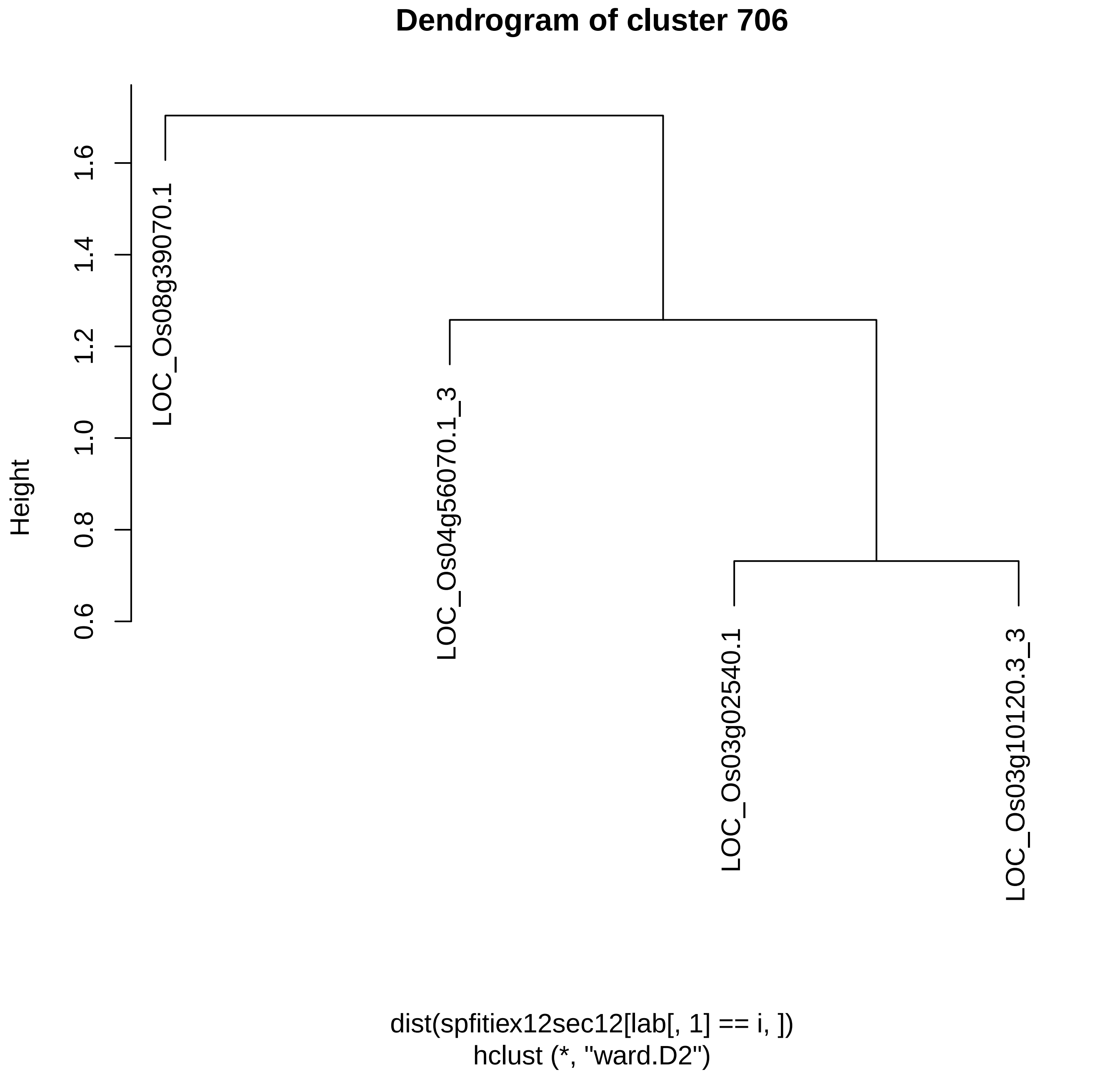
Dendrogram of cluster 706
Height
0.6
0.8
1.0
1.2
1.4
1.6
LOC_Os08g39070.1
LOC_Os04g56070.1_3
LOC_Os03g02540.1
LOC_Os03g10120.3_3
dist(spfitiex12sec12[lab[, 1] == i, ])
hclust (*, "ward.D2")

### plot cluster 707

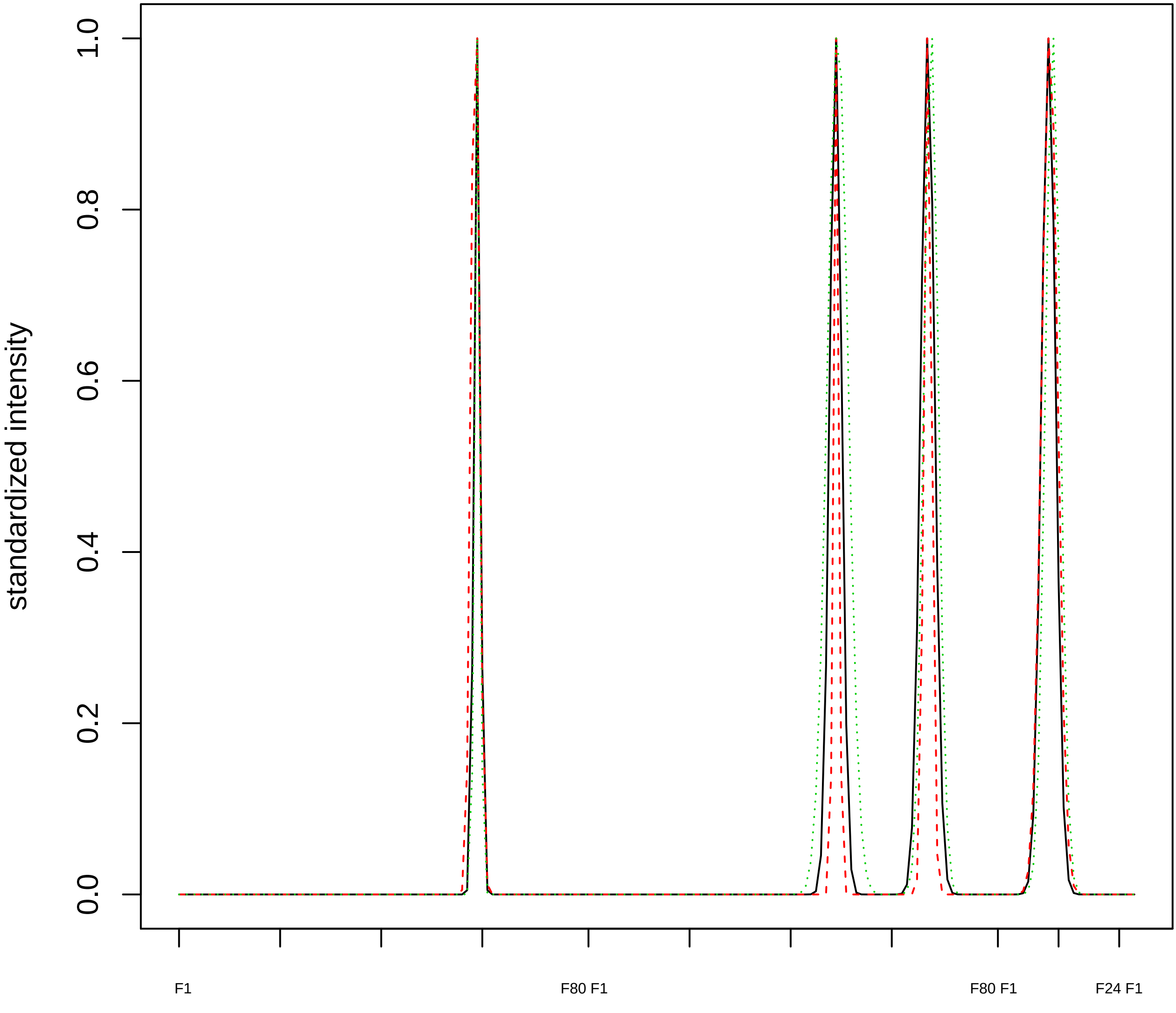

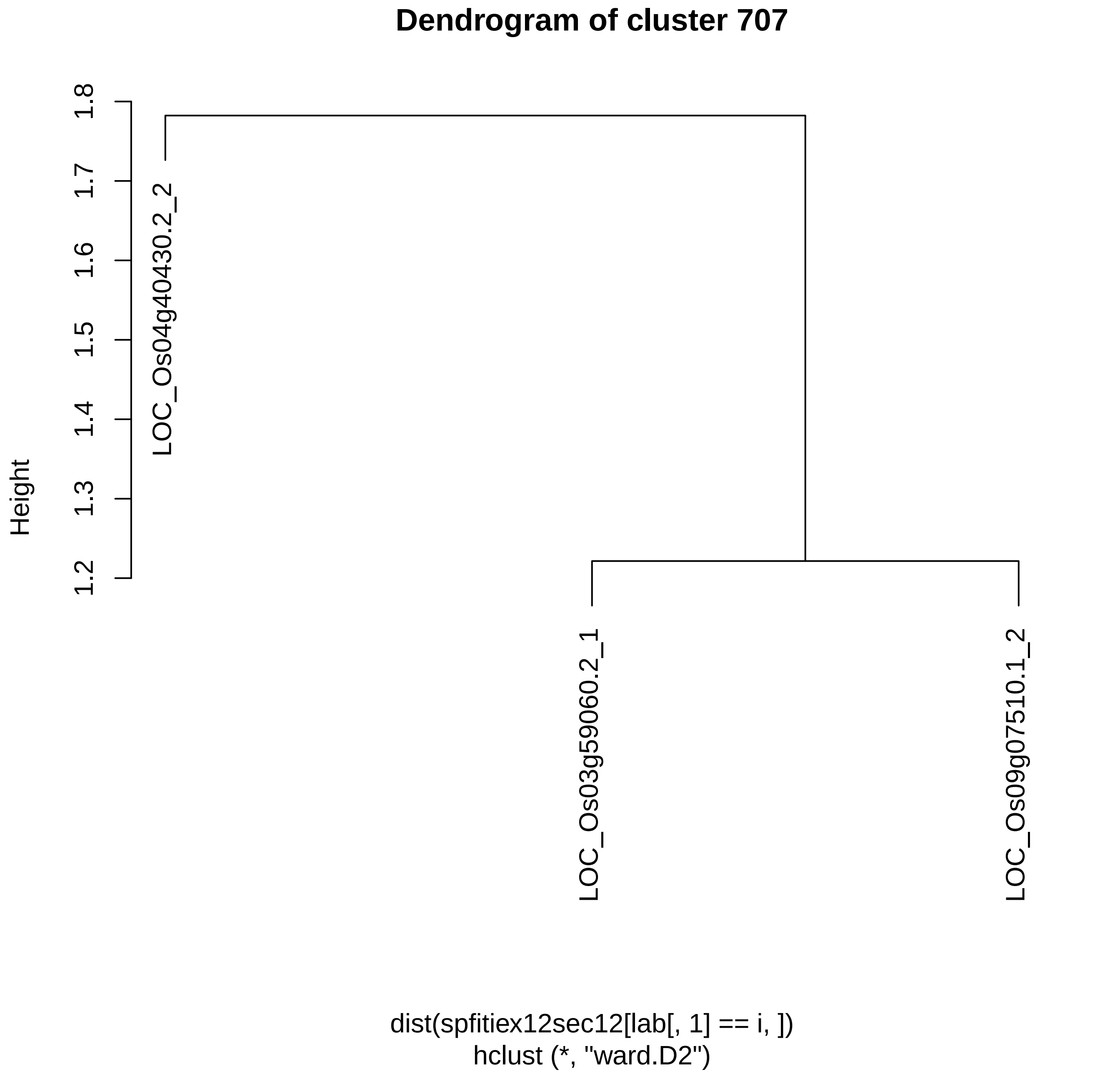
Dendrogram of cluster 707
Height
1.2
1.3
1.4
1.5
1.6
1.7
1.8
LOC_Os04g40430.2_2
LOC_Os03g59060.2_1
LOC_Os09g07510.1_2
dist(spfitiex12sec12[lab[, 1] == i, ])
hclust (*, "ward.D2")

#### plot cluster 708

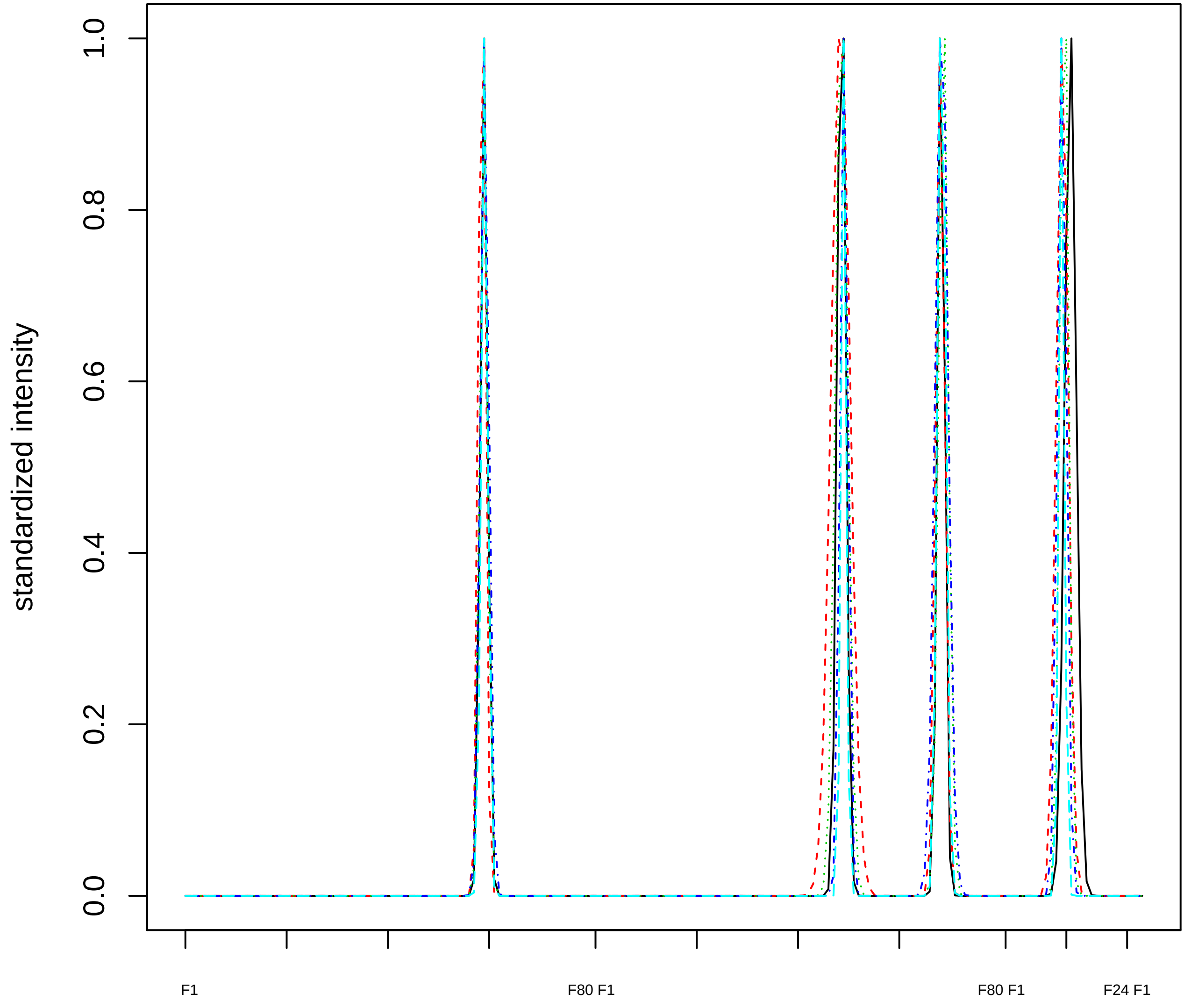

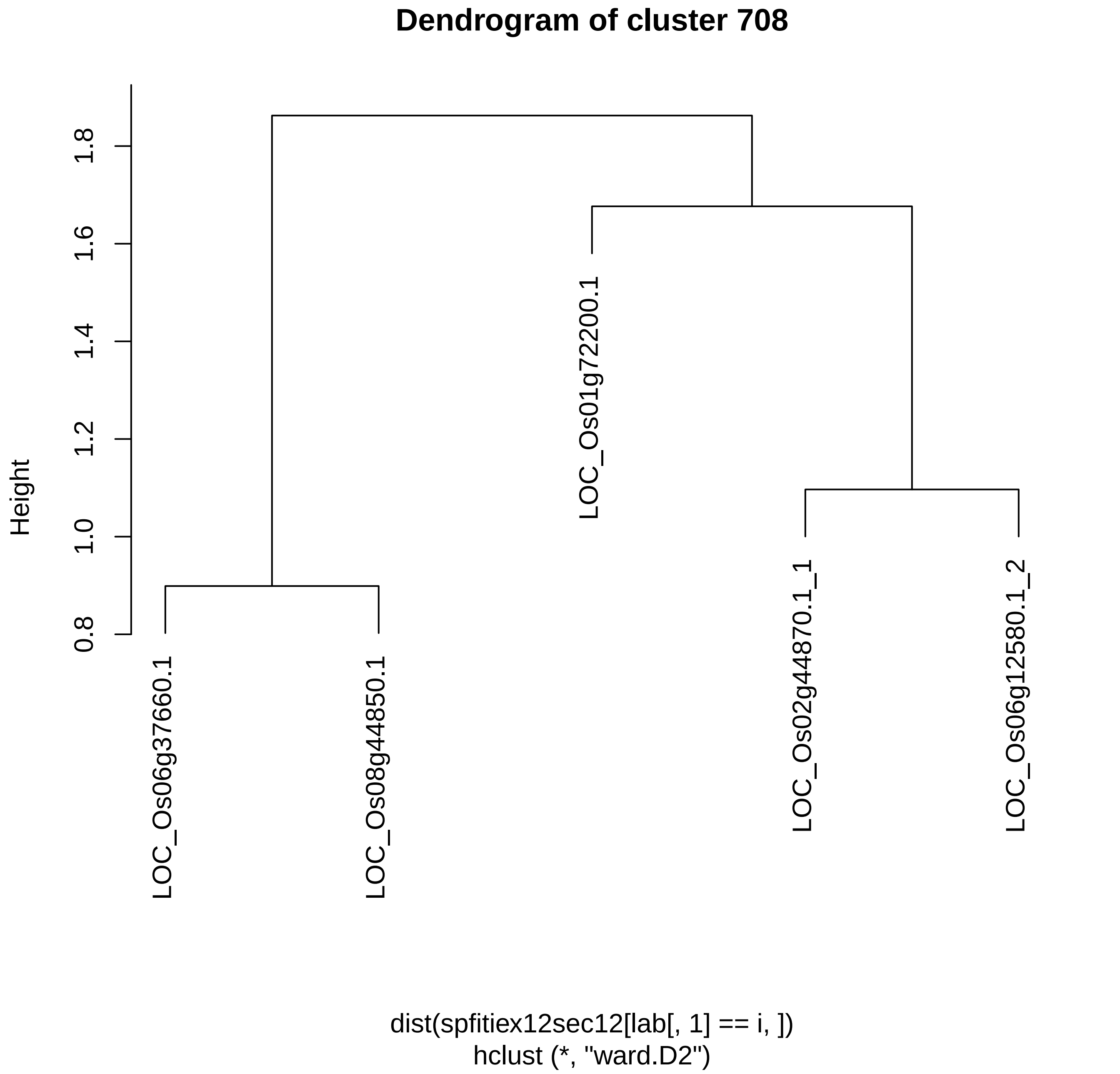

Dendrogram of cluster 708
Height
0.8
1.0
1.2
1.4
1.6
1.8
LOC_Os06g37660.1
LOC_Os08g44850.1
LOC_Os01g72200.1
LOC_Os02g44870.1_1
LOC_Os06g12580.1_2
dist(spfitiex12sec12[lab[, 1] == i, ])
hclust (*, "ward.D2")

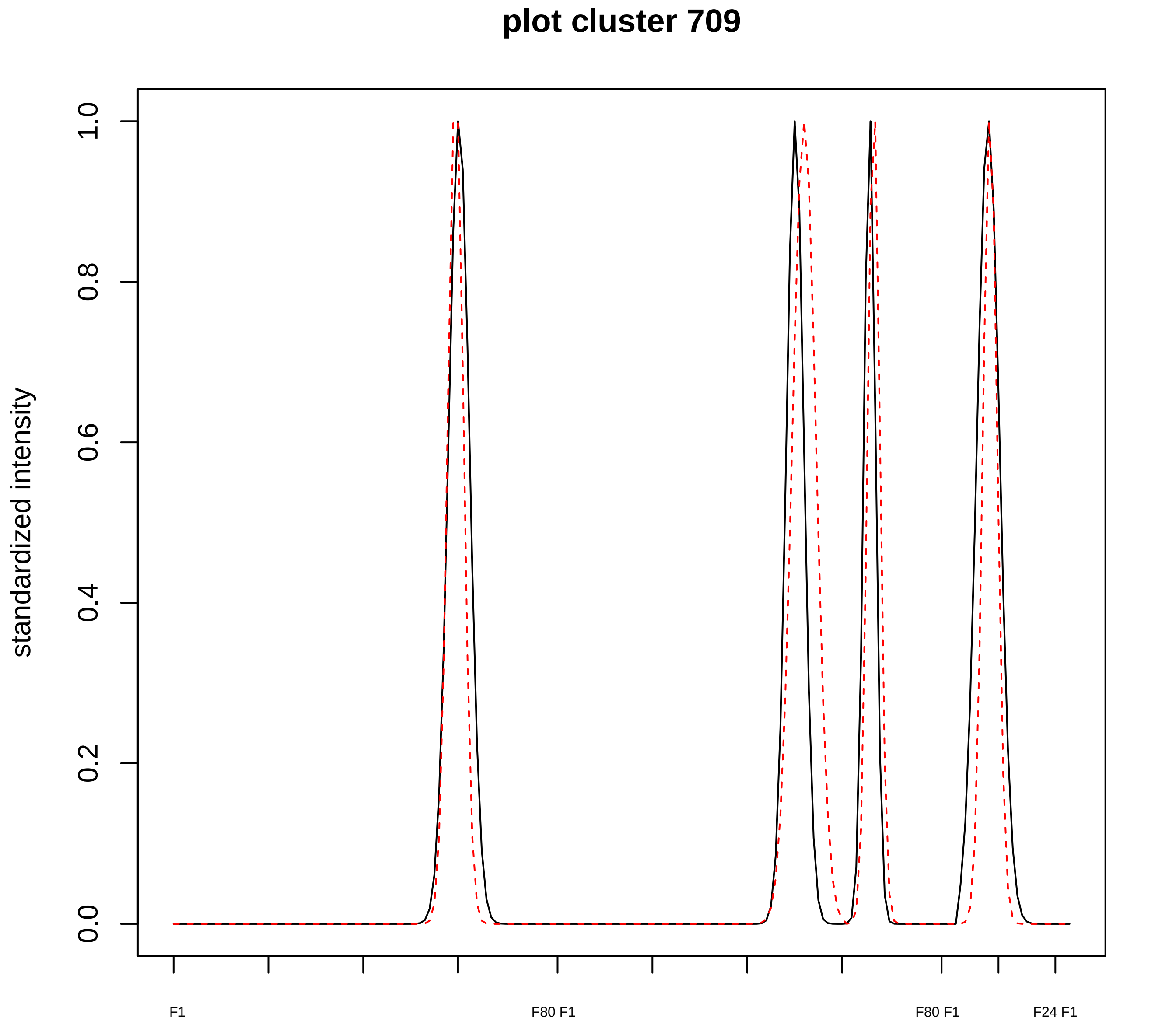

plot cluster 709
standardized intensity
0.0
0.2
0.4
0.6
0.8
1.0
F1
F80 F1
F80 F1
F24 F1

### plot cluster 710

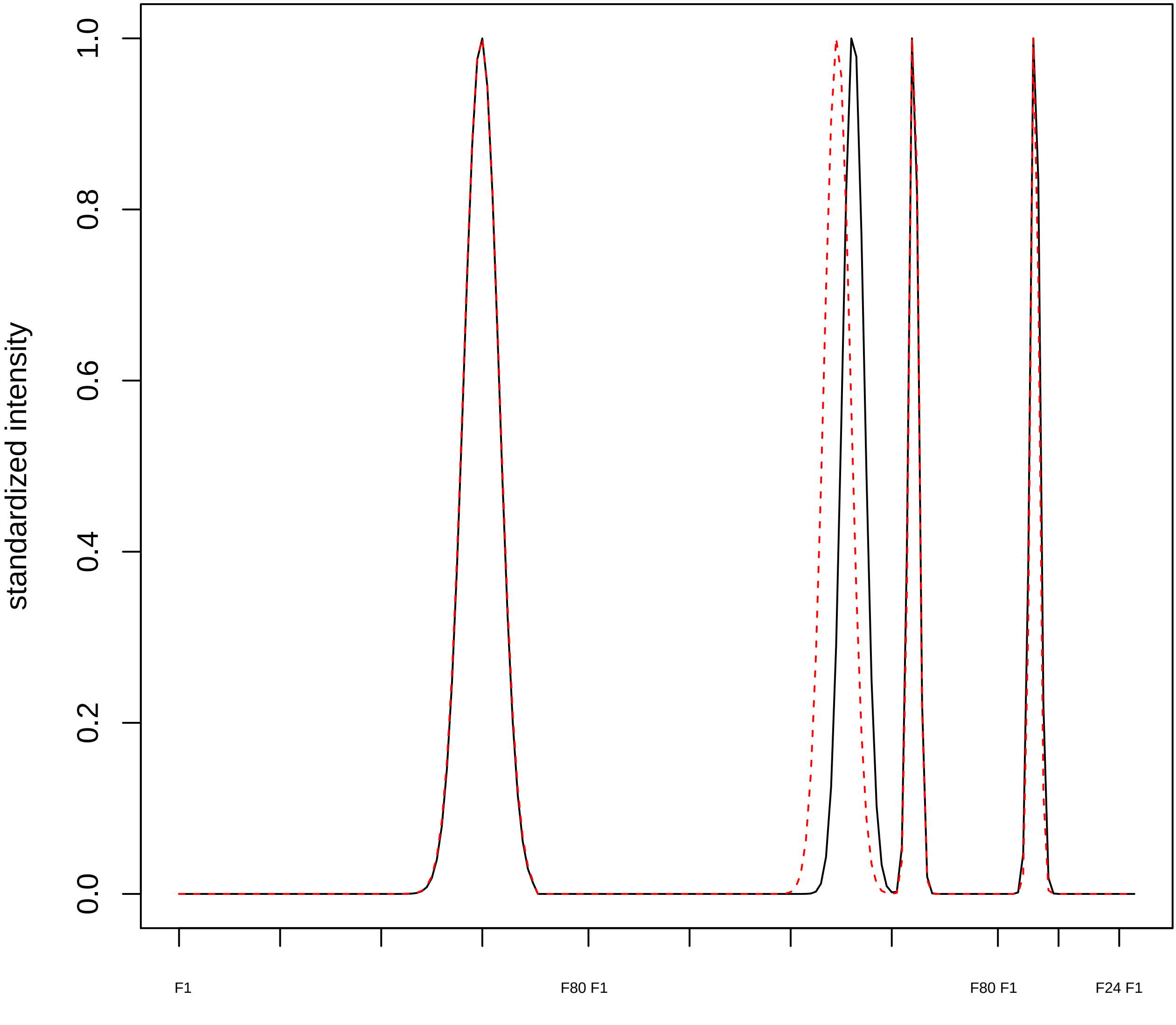

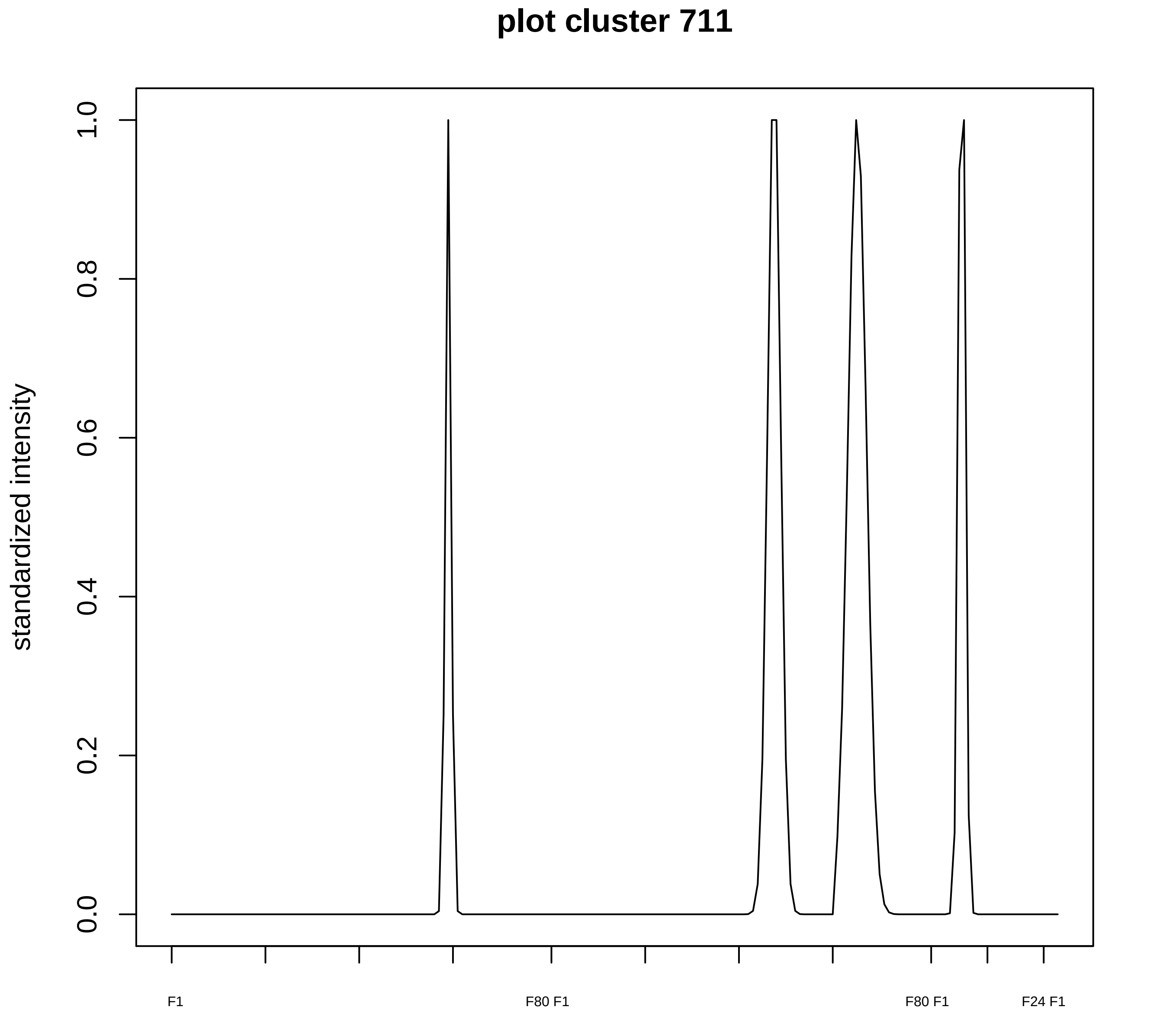
plot cluster 711
standardized intensity
0.0
0.2
0.4
0.6
0.8
1.0
F1
F80 F1
F80 F1
F24 F1

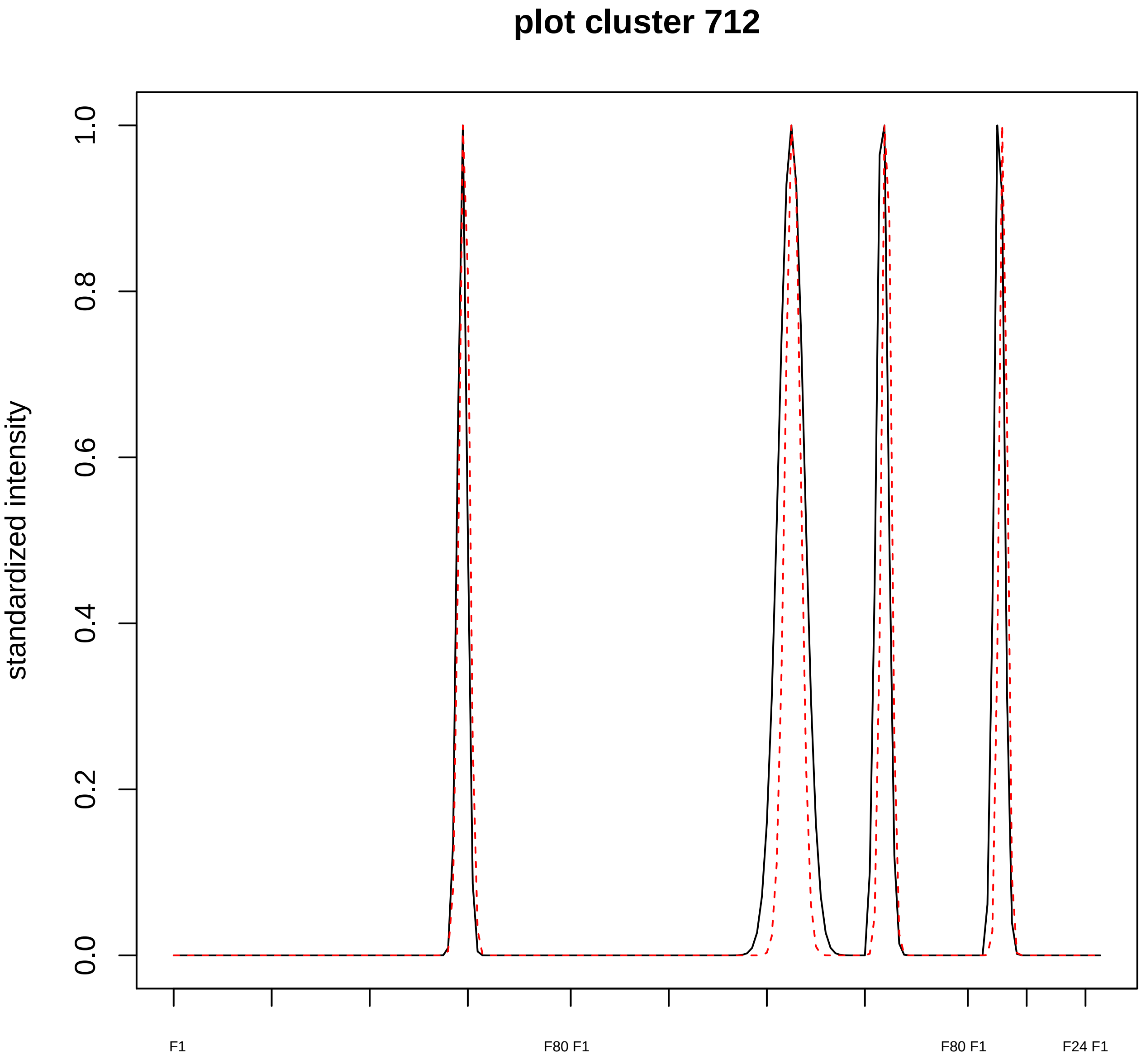

plot cluster 712
standardized intensity
0.0
0.2
0.4
0.6
0.8
1.0
F1
F80 F1
F80 F1
F24 F1

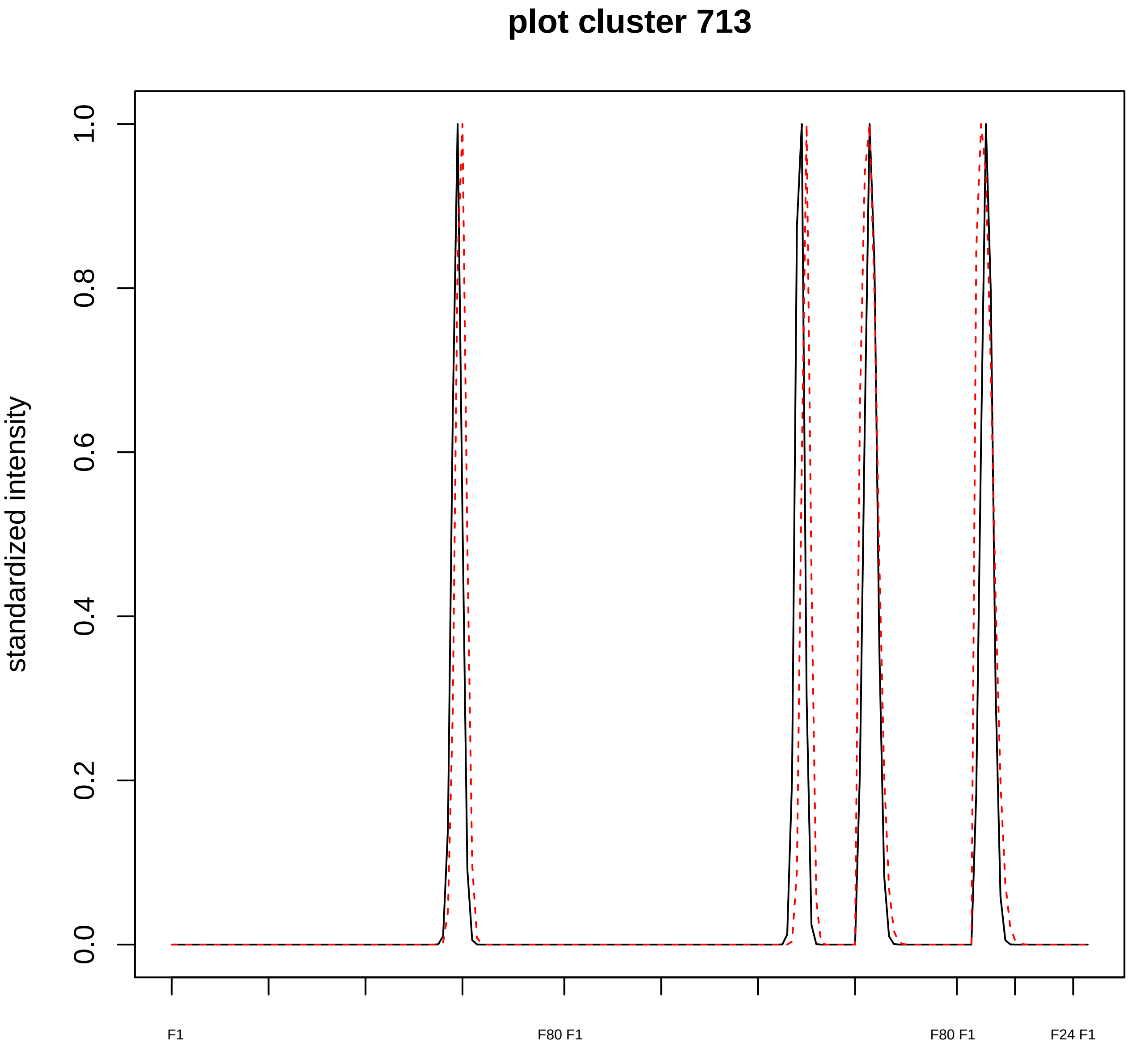
plot cluster 713
standardized intensity
0.0
0.2
0.4
0.6
0.8
1.0
F1
F80 F1
F80 F1
F24 F1

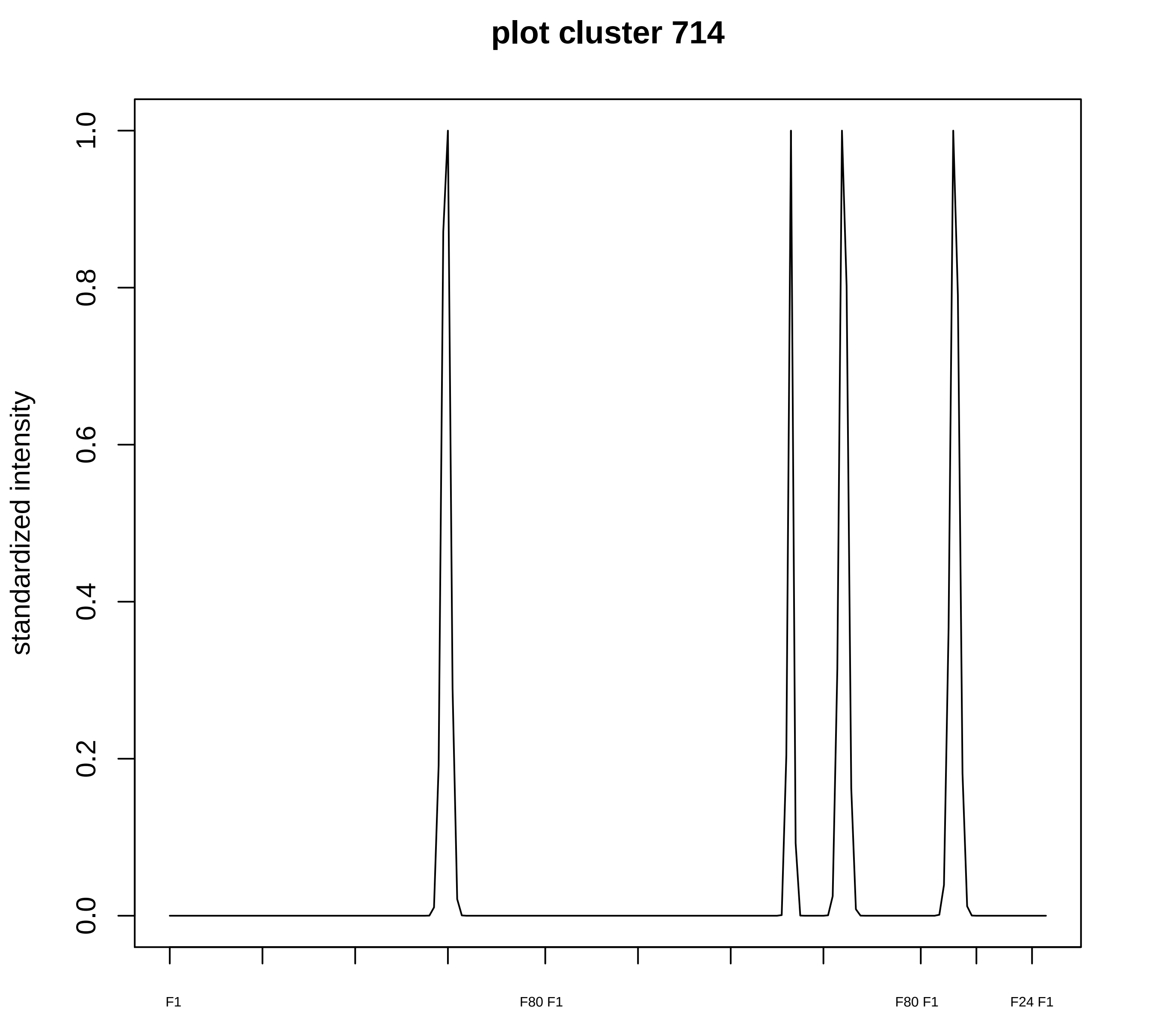
plot cluster 714
standardized intensity
0.0
0.2
0.4
0.6
0.8
1.0
F1
F80 F1
F80 F1
F24 F1

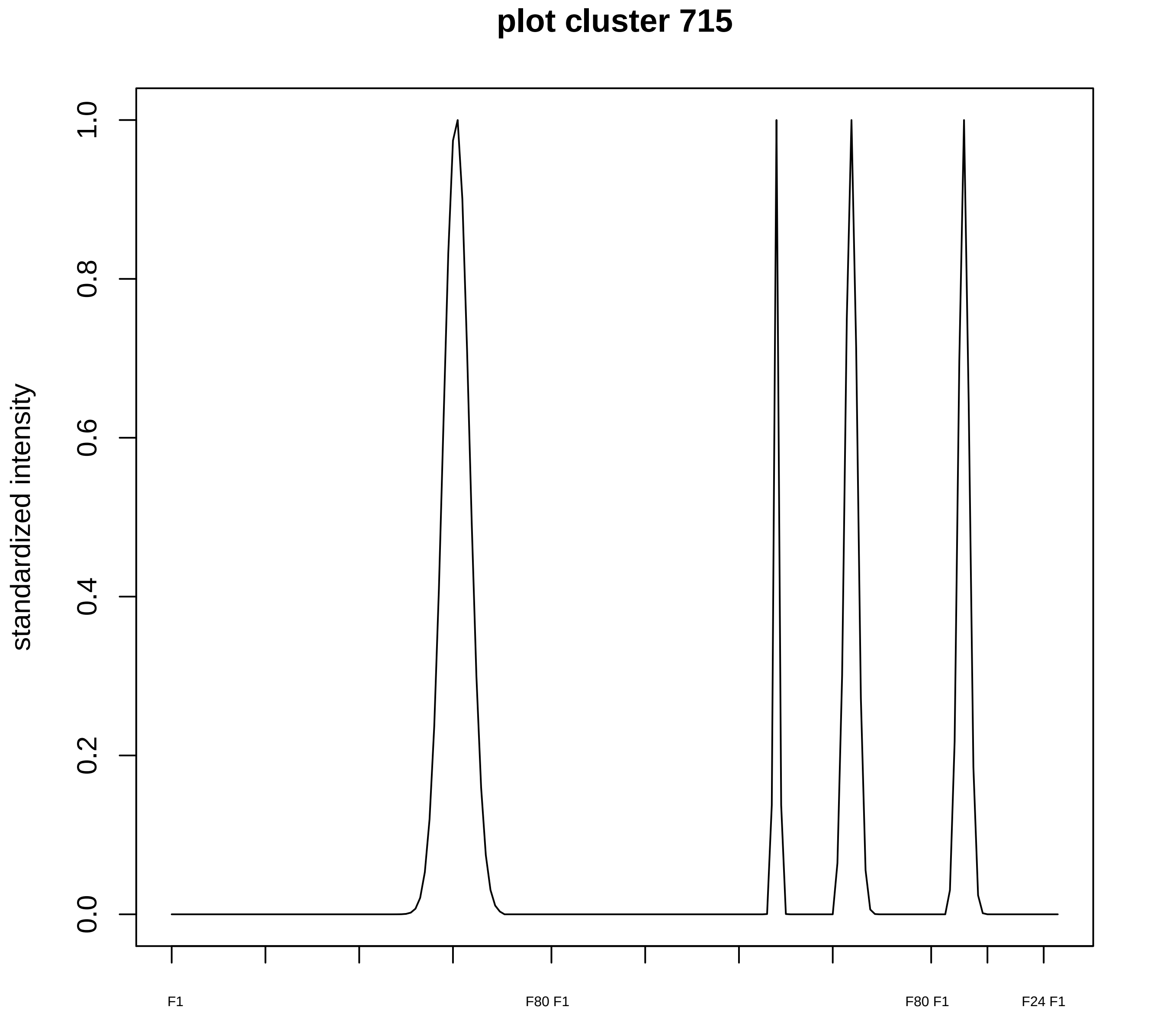

plot cluster 715
standardized intensity
0.0
0.2
0.4
0.6
0.8
1.0
F1
F80 F1
F80 F1
F24 F1

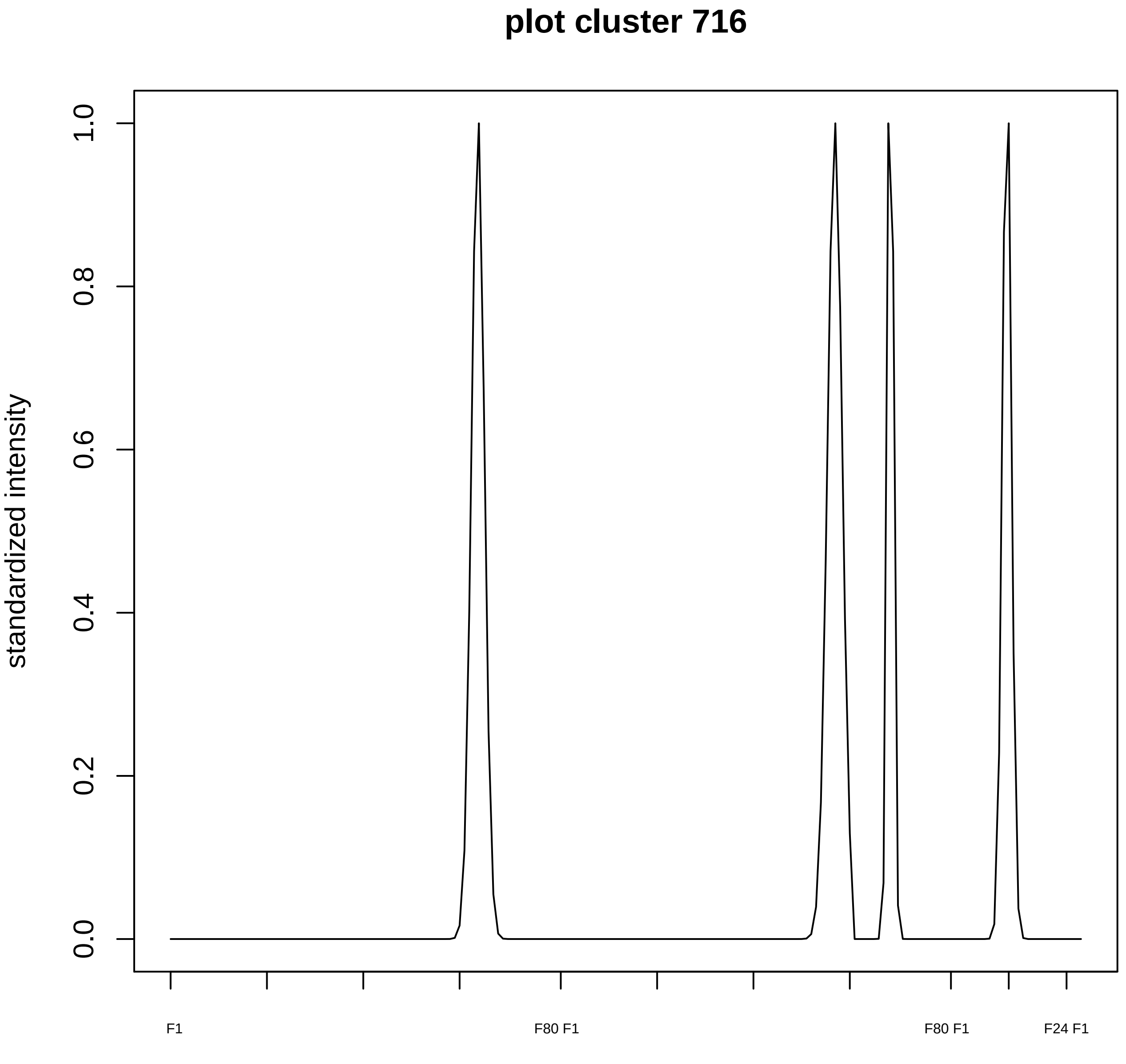

plot cluster 716
standardized intensity
0.0
0.2
0.4
0.6
0.8
1.0
F1
F80 F1
F80 F1
F24 F1

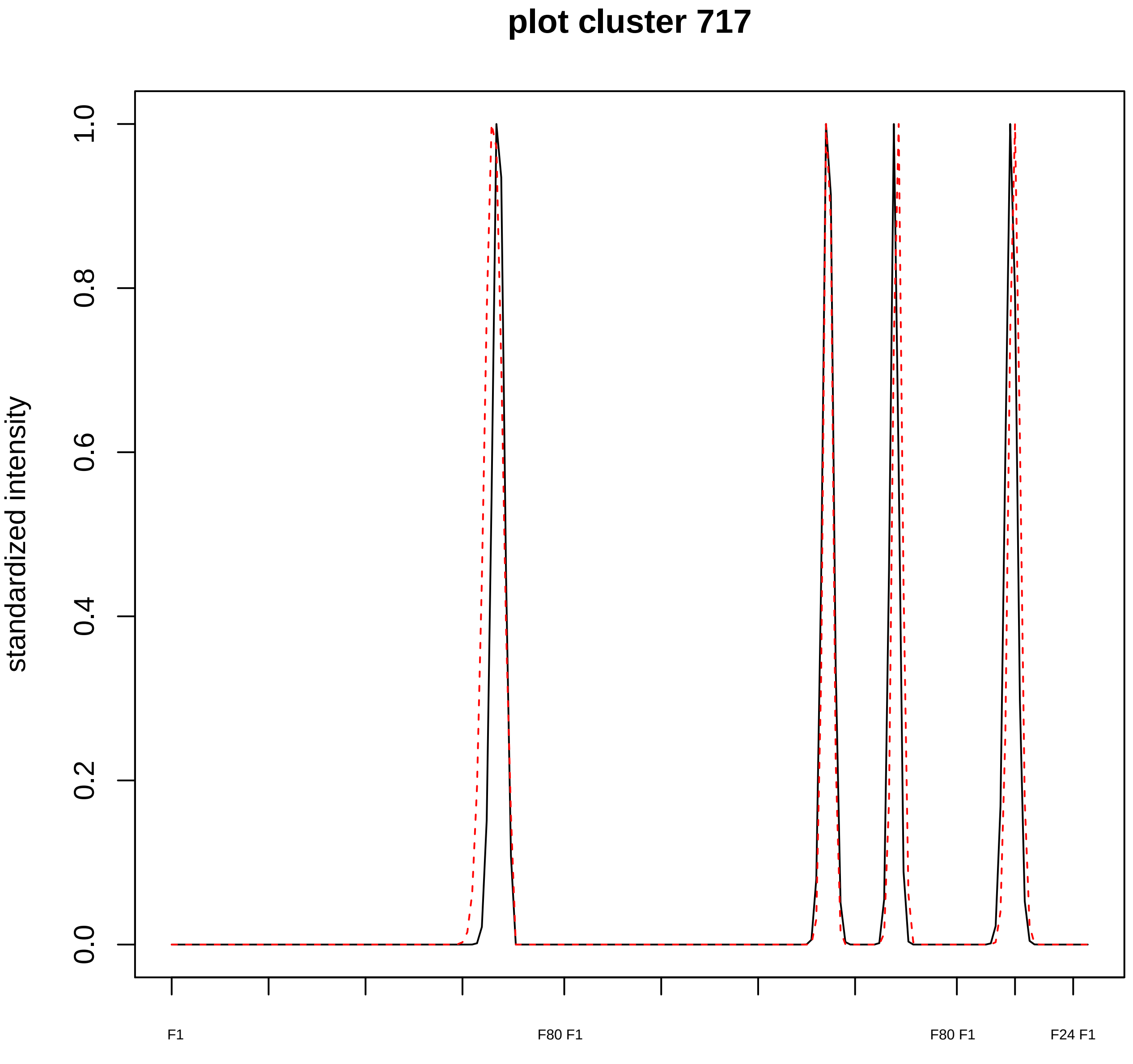
plot cluster 717
standardized intensity
0.0
0.2
0.4
0.6
0.8
1.0
F1
F80 F1
F80 F1
F24 F1

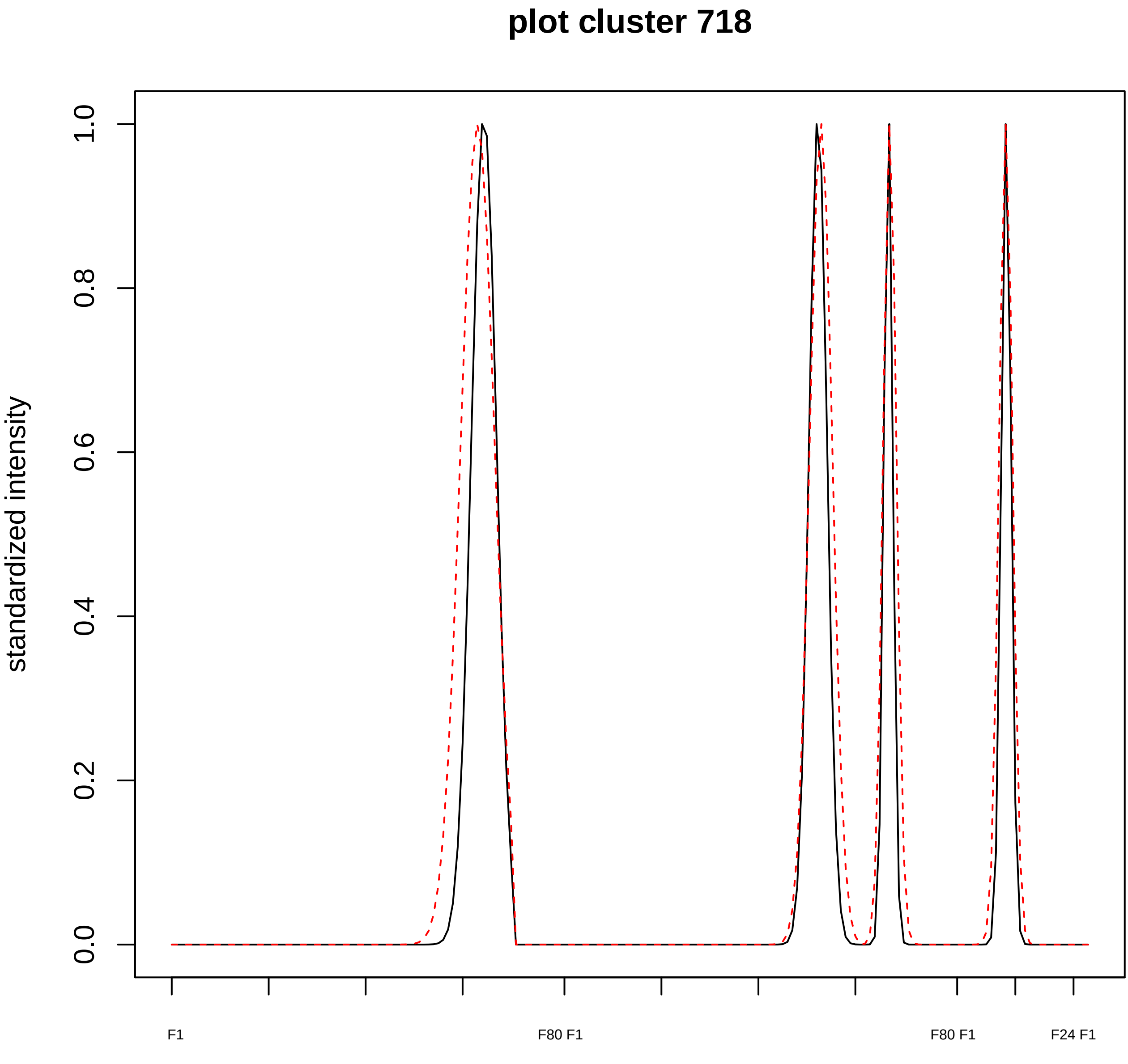

plot cluster 718
standardized intensity
0.0
0.2
0.4
0.6
0.8
1.0
F1
F80 F1
F80 F1
F24 F1

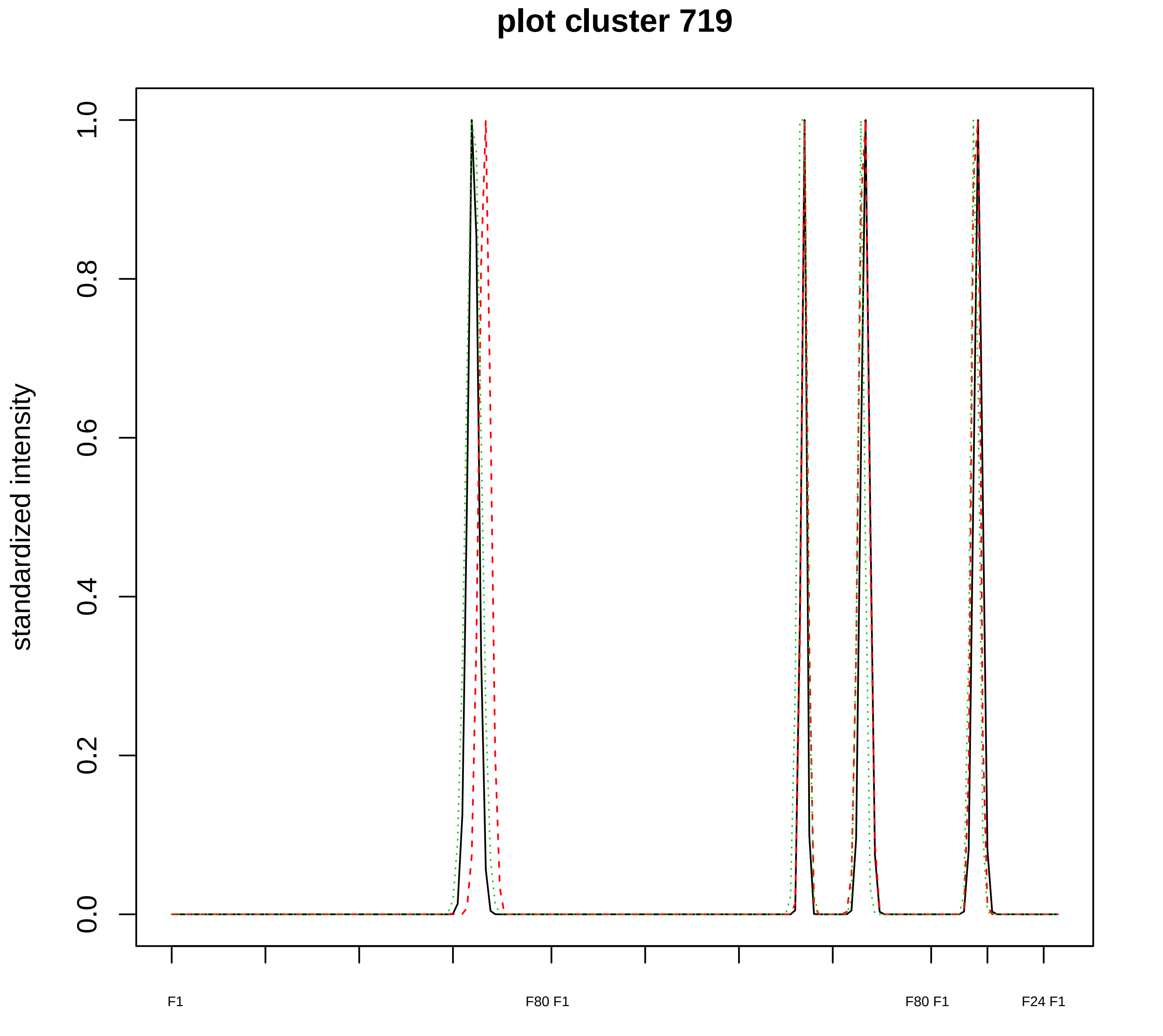
plot cluster 719
standardized intensity
0.0
0.2
0.4
0.6
0.8
1.0
F1
F80 F1
F80 F1
F24 F1

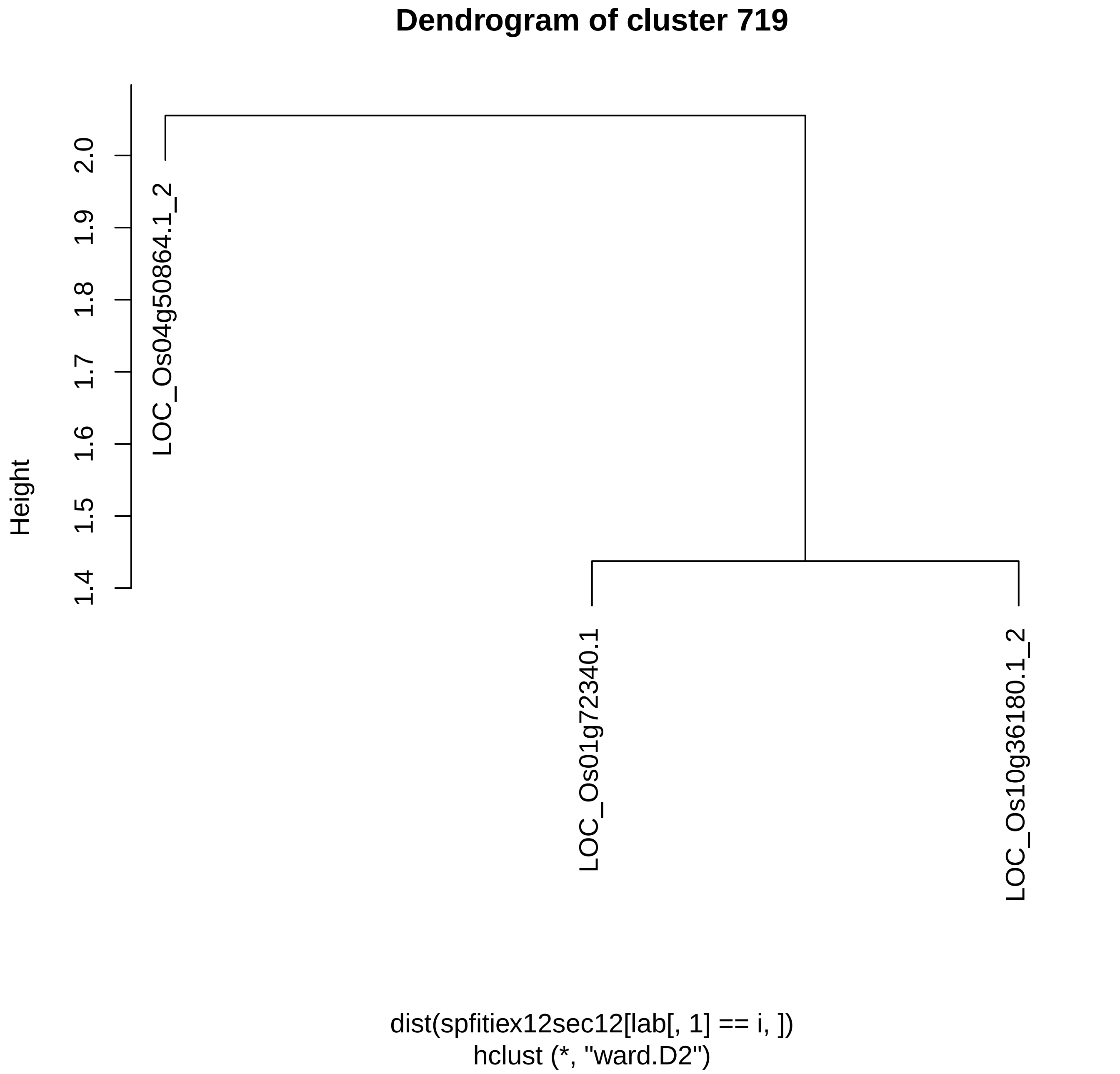
Dendrogram of cluster 719
Height
2.0
1.9
1.8
1.7
1.6
1.5
1.4
LOC_Os04g50864.1_2
LOC_Os01g72340.1
LOC_Os10g36180.1_2
dist(spfitiex12sec12[lab[, 1] == i, ])
hclust (*, "ward.D2")

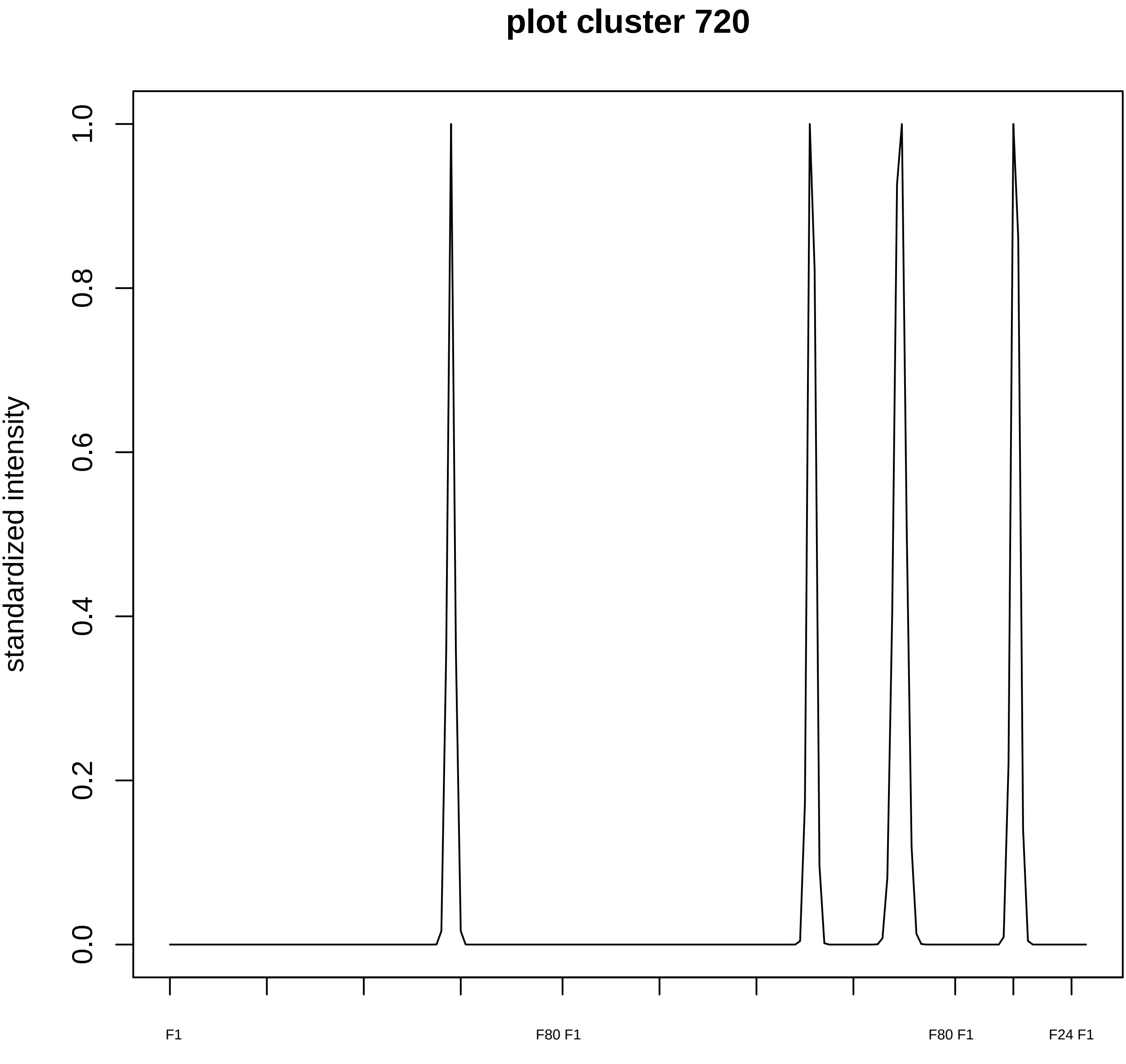

plot cluster 720
standardized intensity
0.0
0.2
0.4
0.6
0.8
1.0
F1
F80 F1
F80 F1
F24 F1

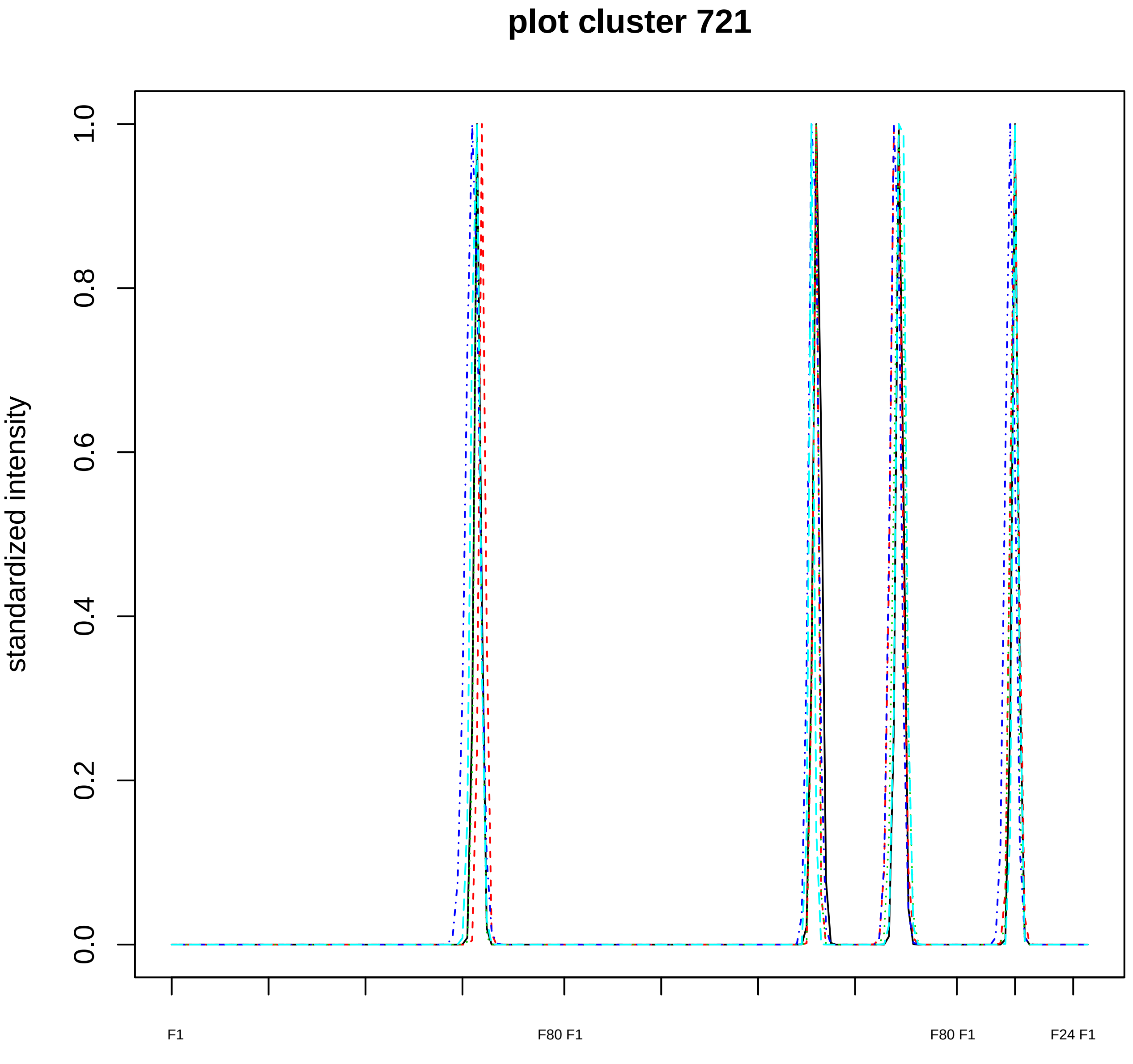
plot cluster 721
standardized intensity
0.0
0.2
0.4
0.6
0.8
1.0
F1
F80 F1
F80 F1
F24 F1

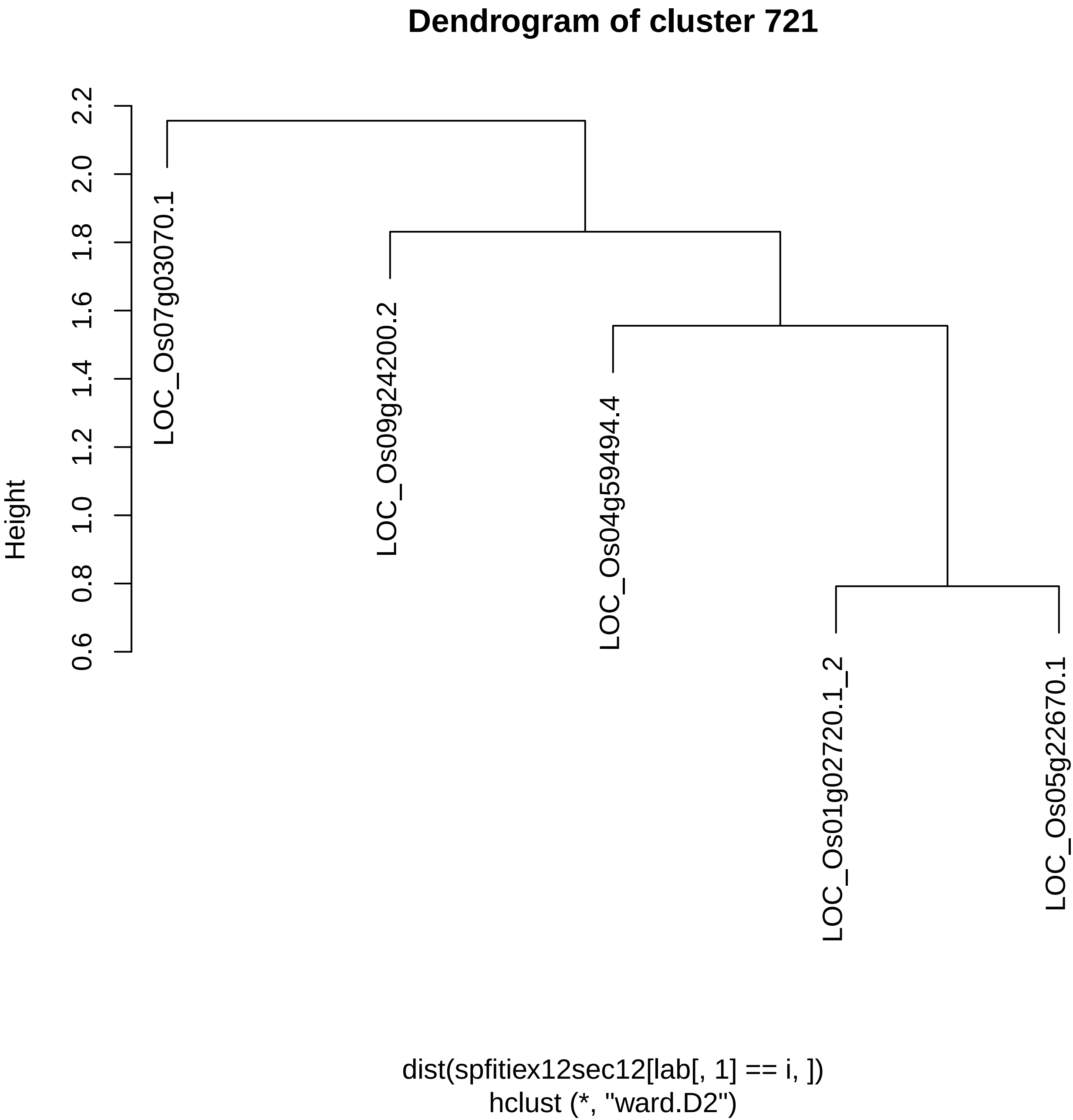

Dendrogram of cluster 721
Height
0.6
0.8
1.0
1.2
1.4
1.6
1.8
2.0
2.2
LOC_Os07g03070.1
LOC_Os09g24200.2
LOC_Os04g59494.4
LOC_Os01g02720.1_2
LOC_Os05g22670.1
dist(spfitiex12sec12[lab[, 1] == i, ])
hclust (*, "ward.D2")

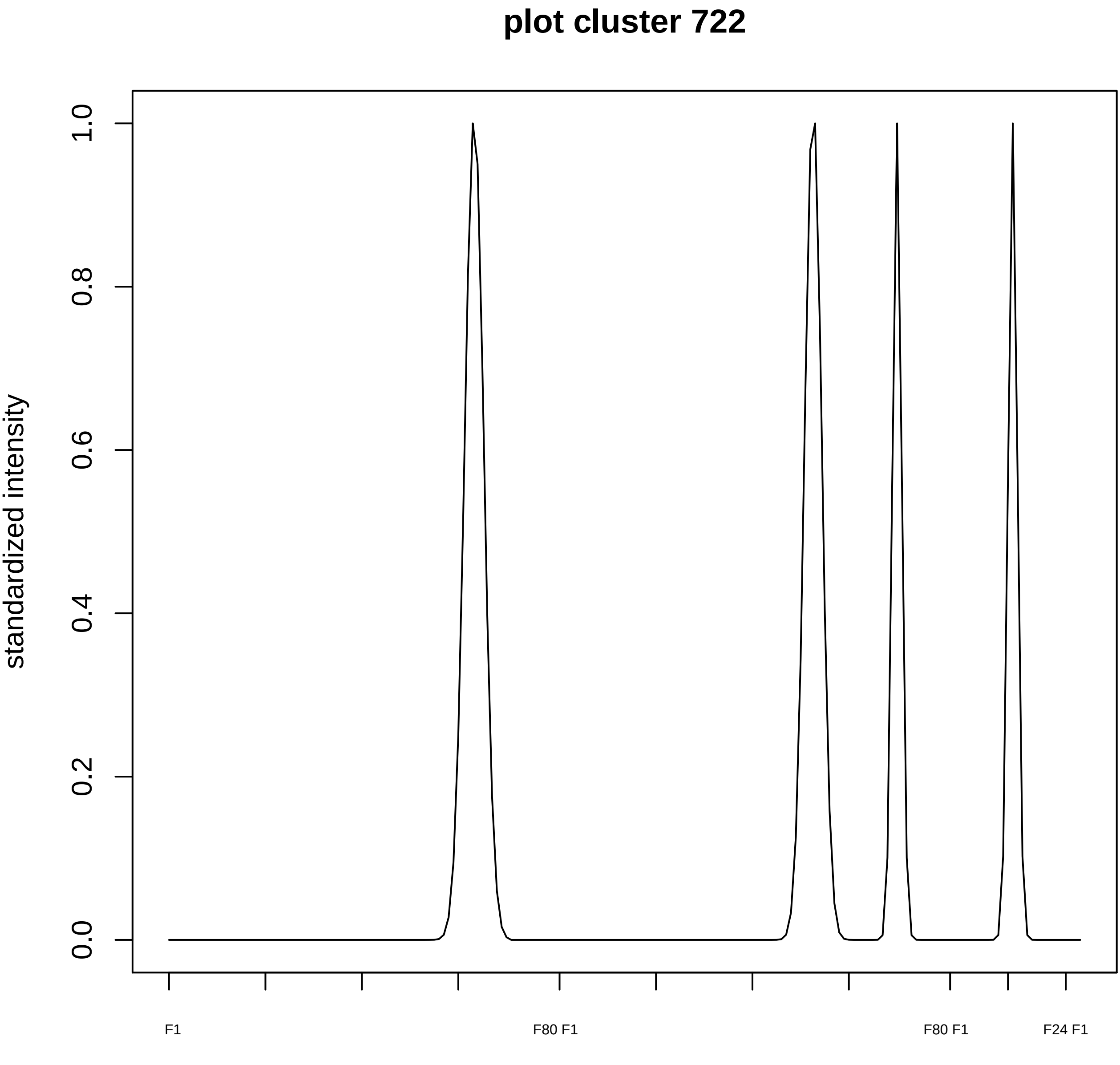

plot cluster 722
standardized intensity
0.0
0.2
0.4
0.6
0.8
1.0
F1
F80 F1
F80 F1
F24 F1

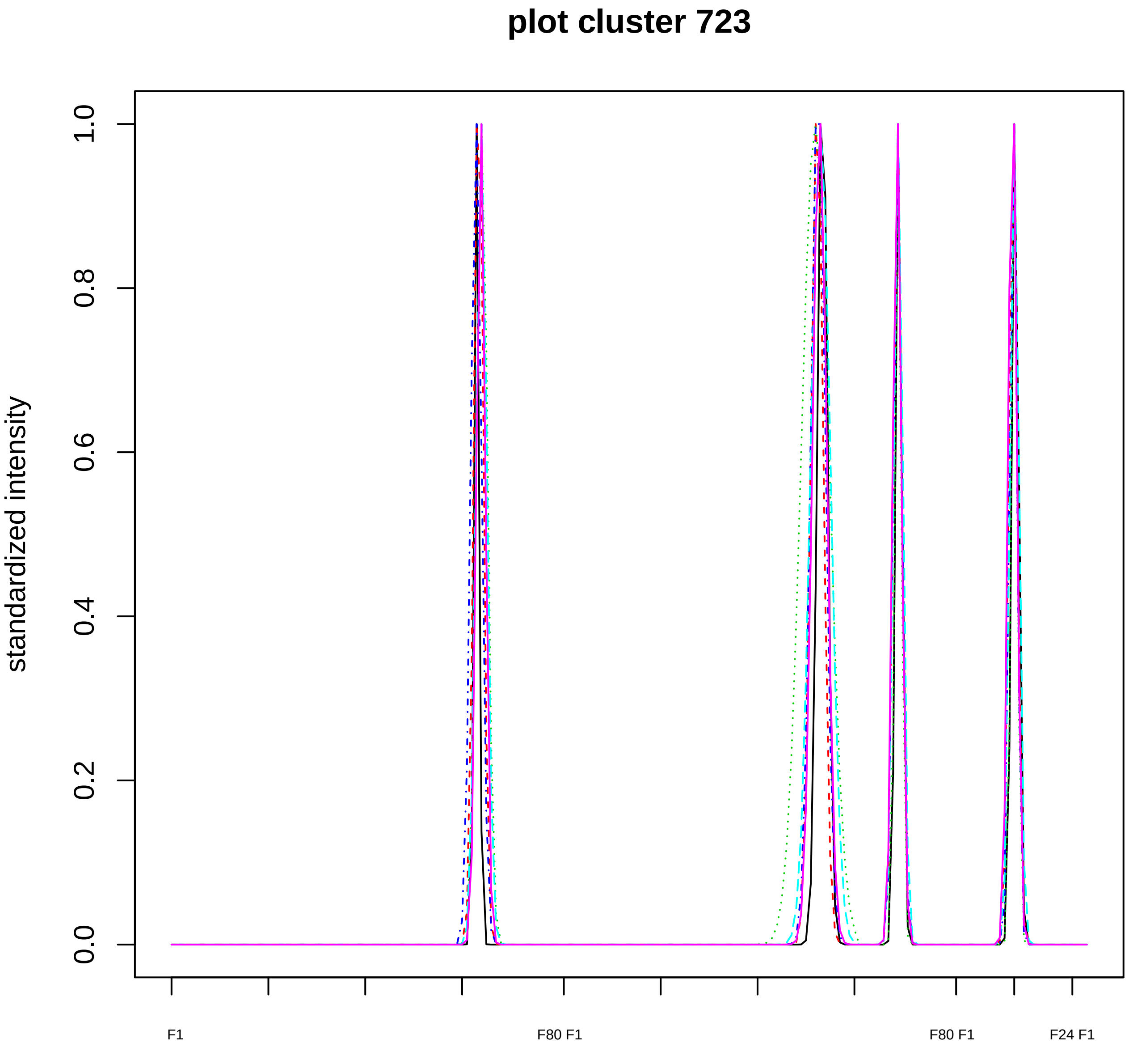
plot cluster 723
standardized intensity
1.0
0.8
0.6
0.4
0.2
0.0
F1
F80 F1
F80 F1
F24 F1

### Dendrogram of cluster 723

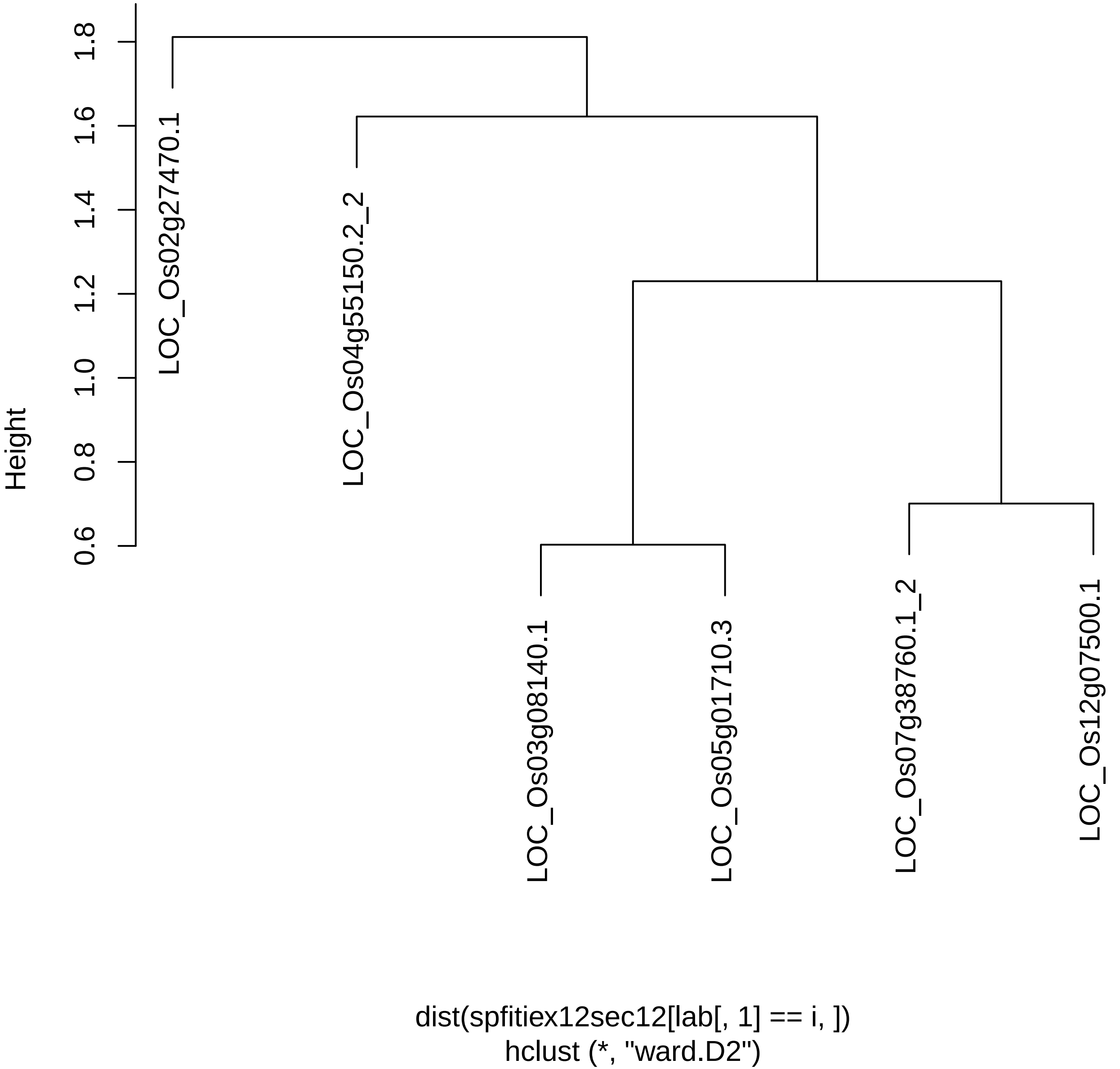

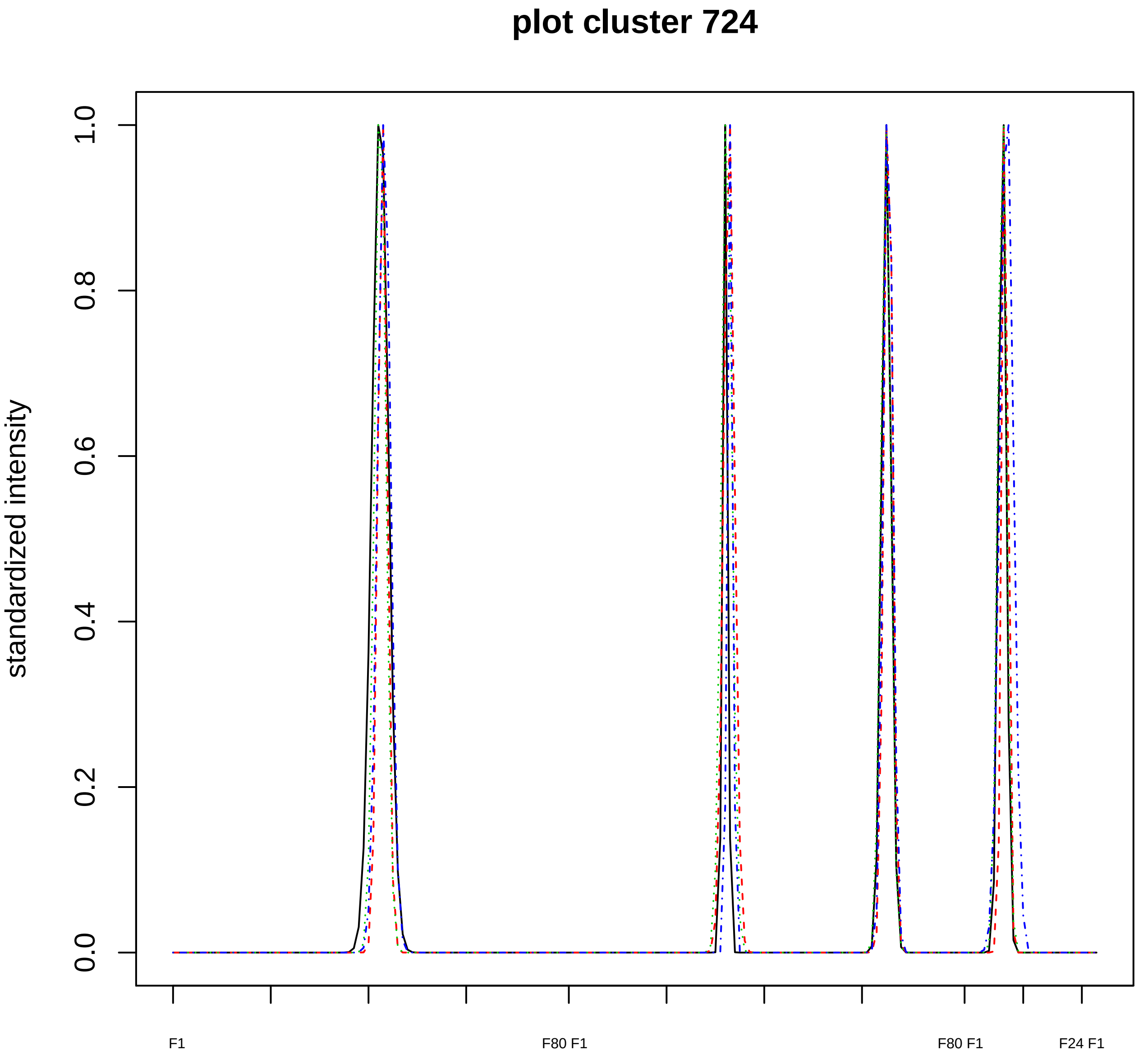

plot cluster 724
standardized intensity
0.0
0.2
0.4
0.6
0.8
1.0
F1
F80 F1
F80 F1
F24 F1

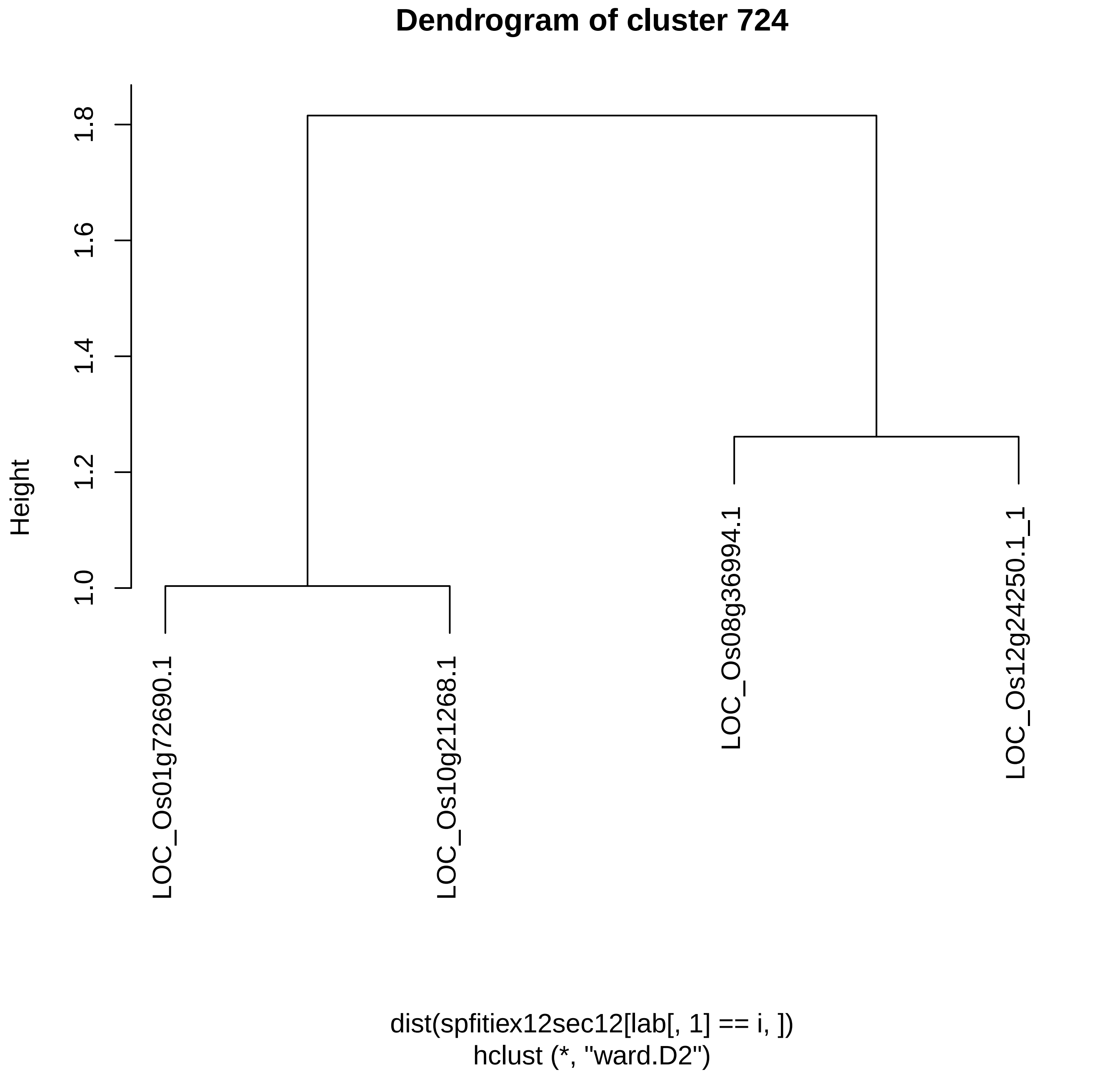
Dendrogram of cluster 724
Height
1.0
1.2
1.4
1.6
1.8
LOC_Os01g72690.1
LOC_Os10g21268.1
LOC_Os08g36994.1
LOC_Os12g24250.1_1
dist(spfitiex12sec12[lab[, 1] == i, ])
hclust (*, "ward.D2")

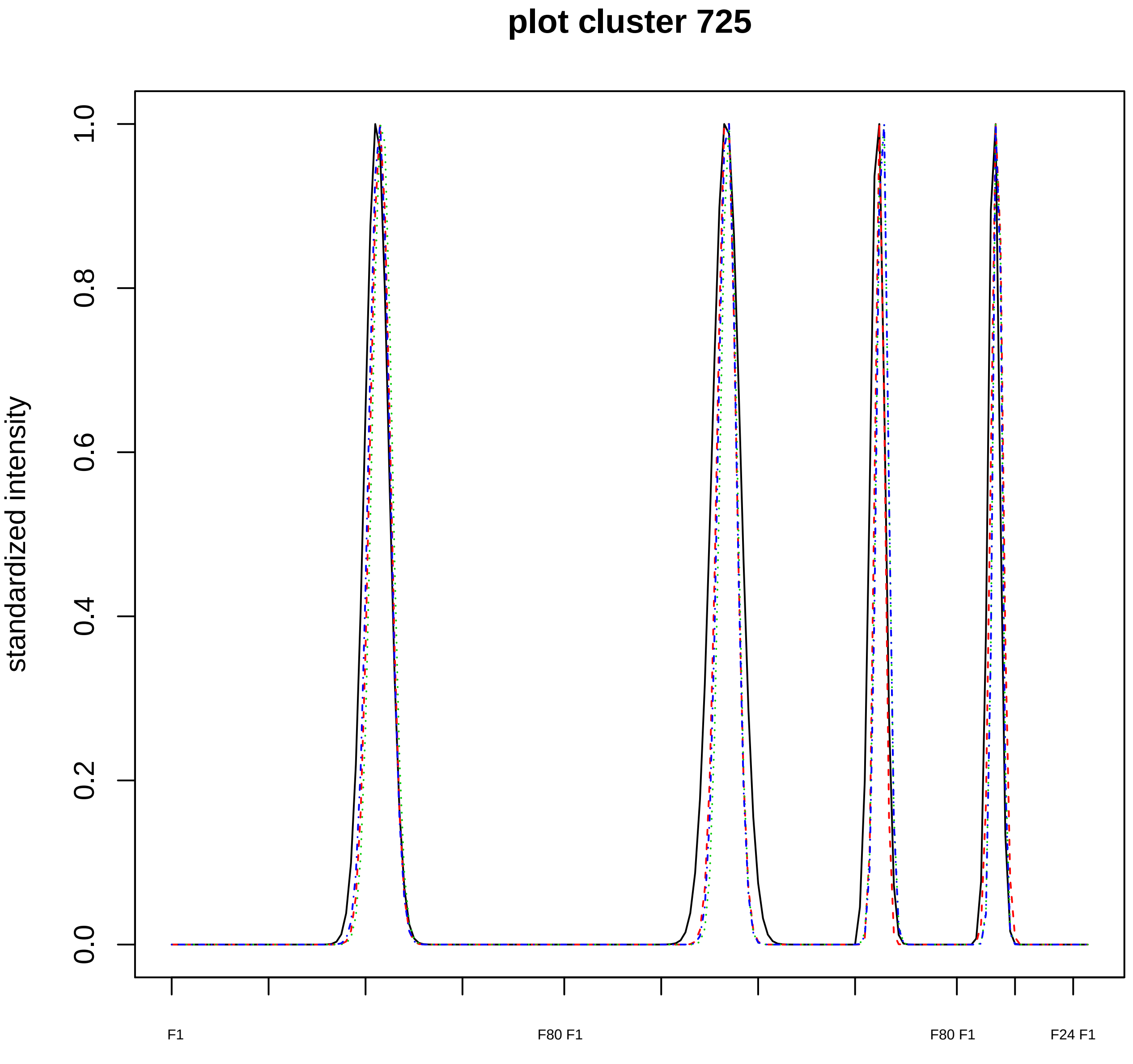
plot cluster 725
standardized intensity
0.0
0.2
0.4
0.6
0.8
1.0
F1
F80 F1
F80 F1
F24 F1

#### Dendrogram of cluster 725

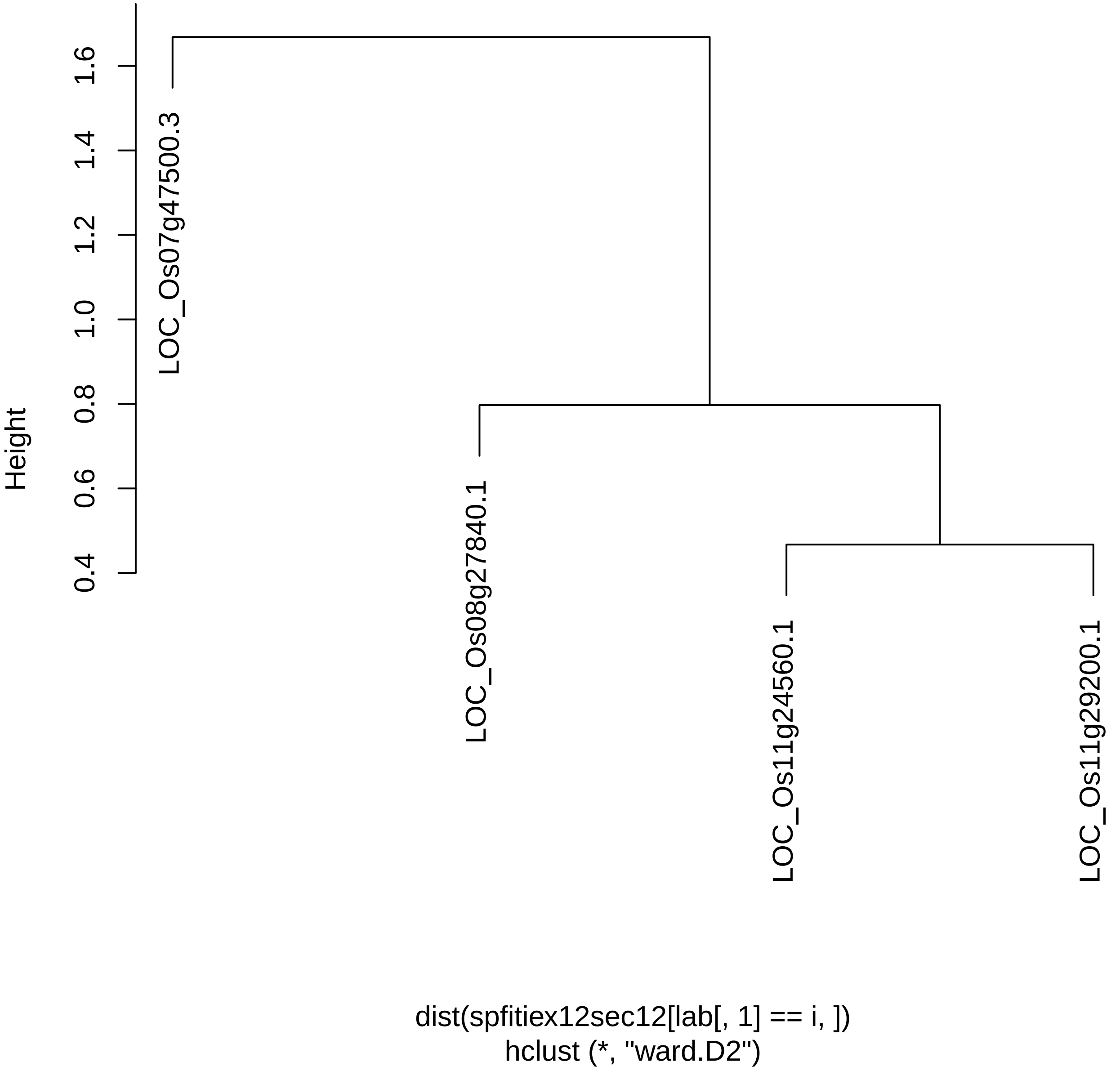

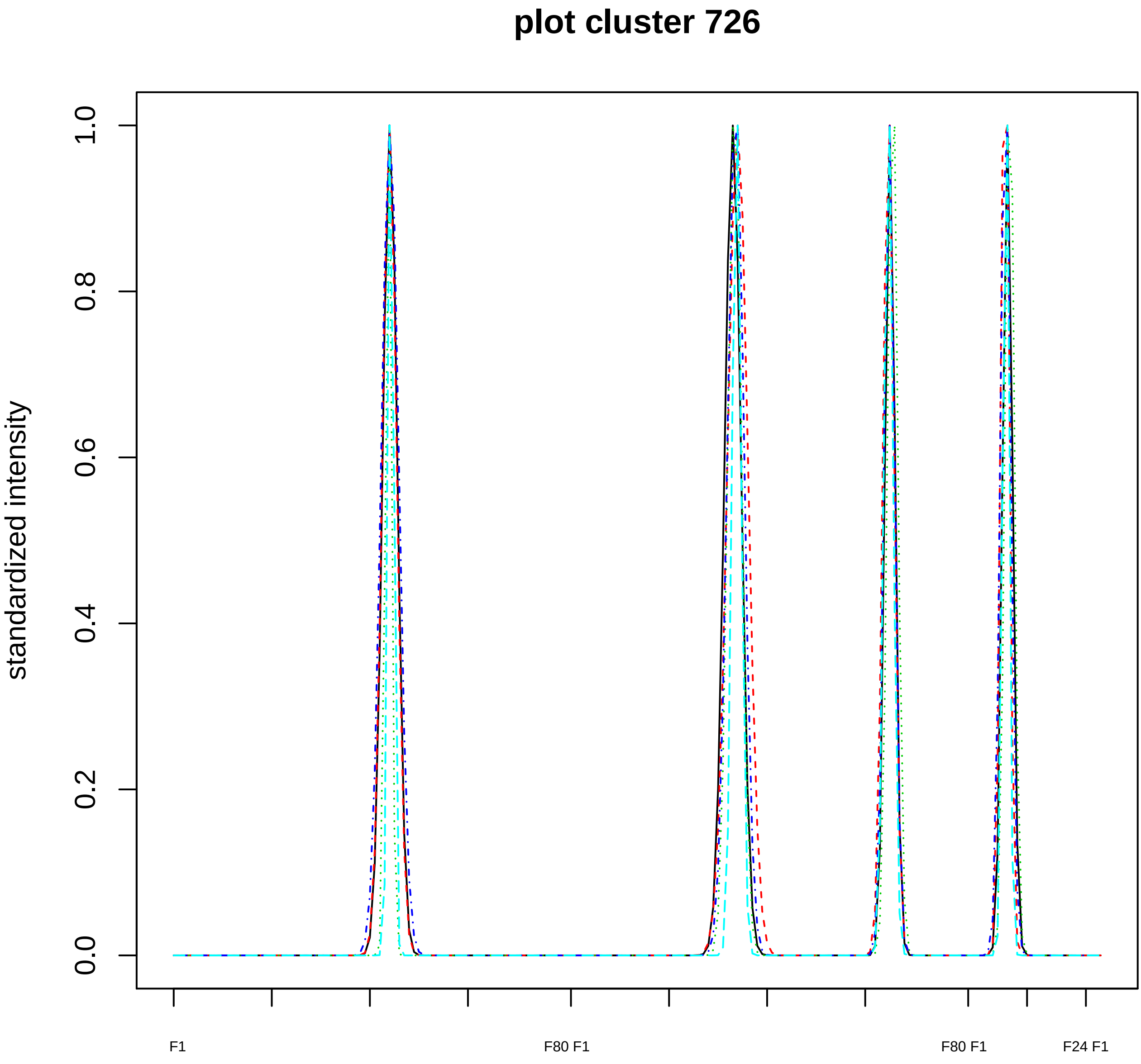
plot cluster 726
standardized intensity
0.0
0.2
0.4
0.6
0.8
1.0
F1
F80 F1
F80 F1
F24 F1

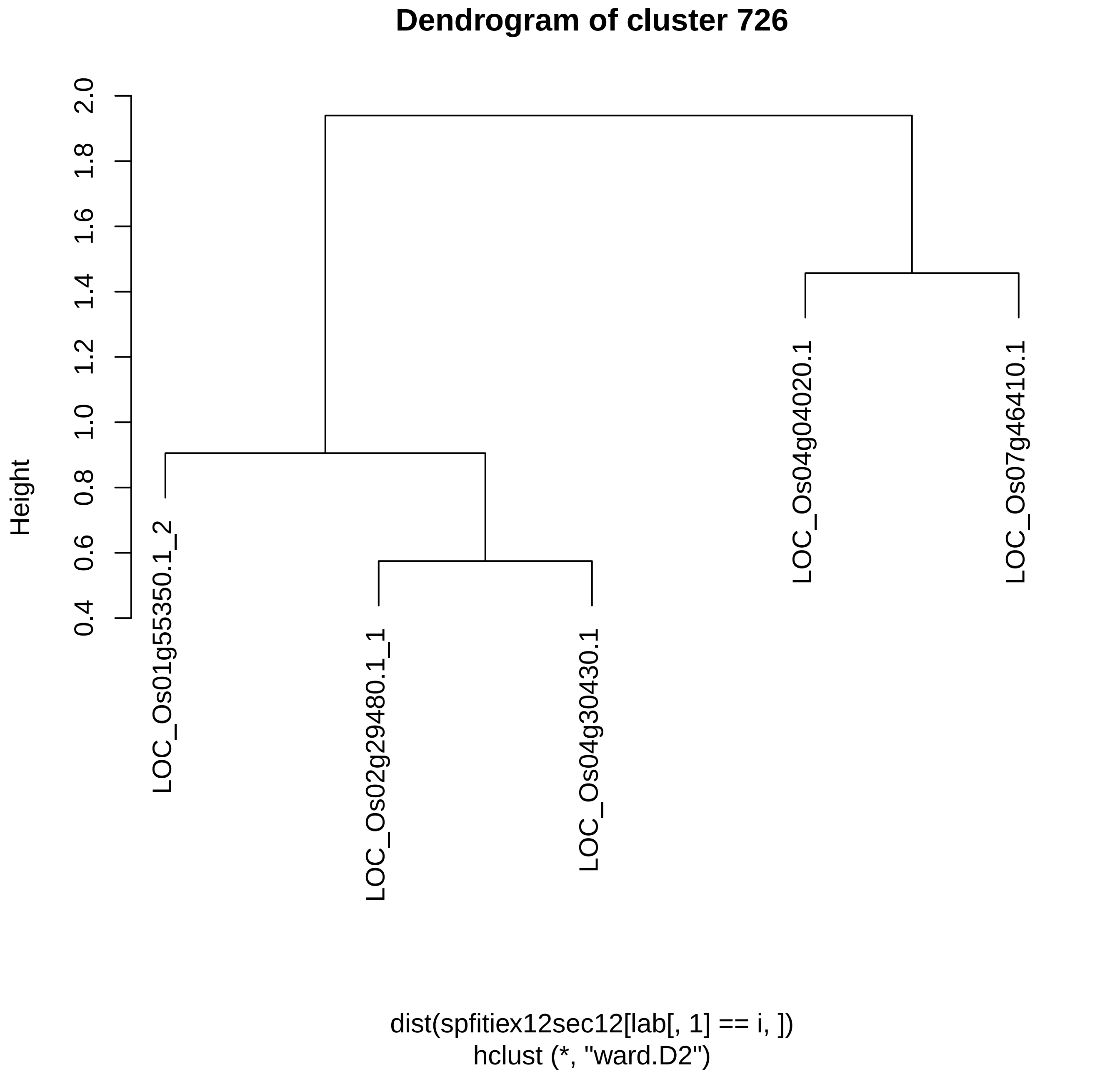
Dendrogram of cluster 726
Height
0.4
0.6
0.8
1.0
1.2
1.4
1.6
1.8
2.0
LOC_Os01g55350.1_2
LOC_Os02g29480.1_1
LOC_Os04g30430.1
LOC_Os04g04020.1
LOC_Os07g46410.1
dist(spfitiex12sec12[lab[, 1] == i, ])
hclust (*, "ward.D2")

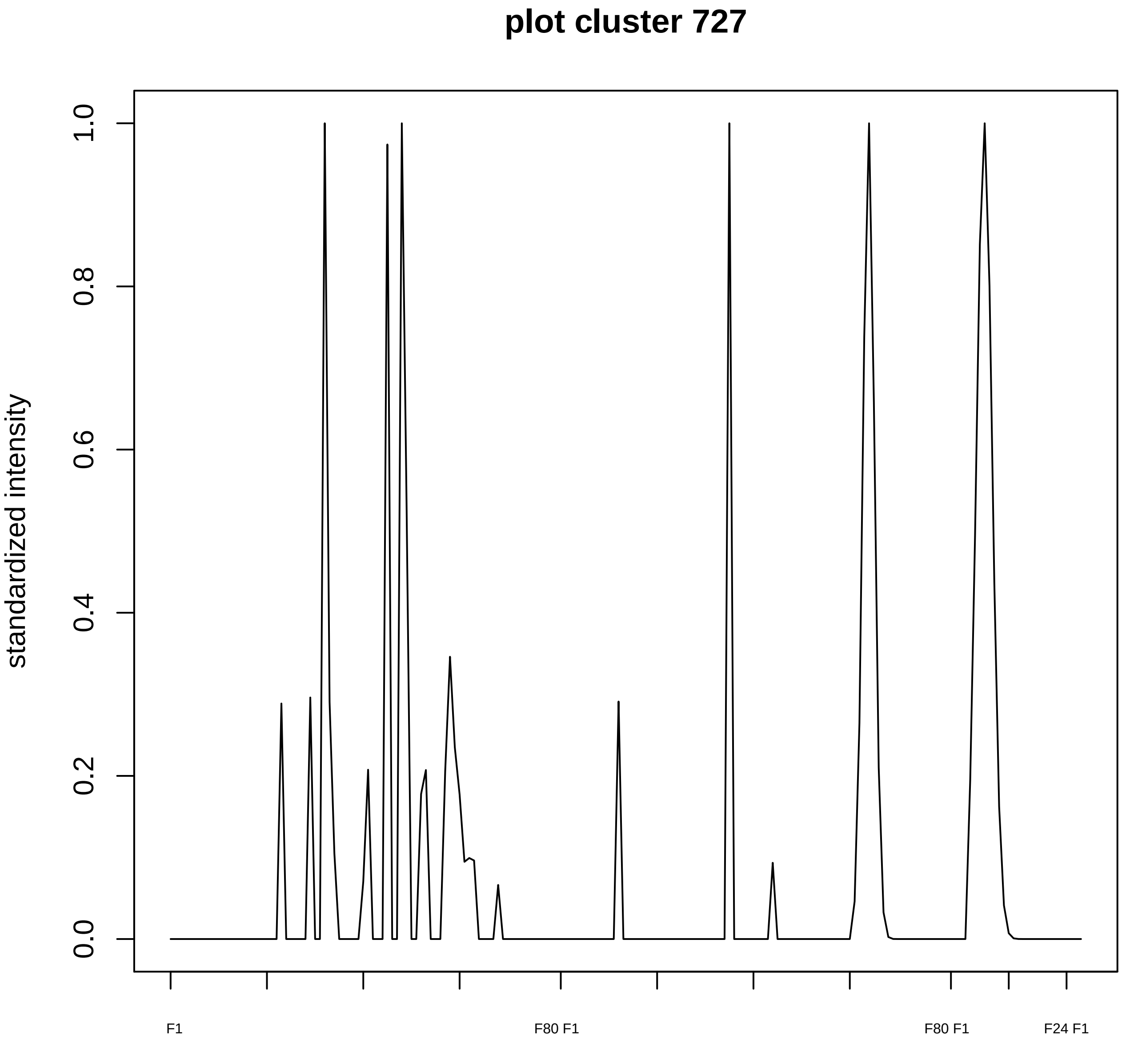

plot cluster 727
standardized intensity
0.0
0.2
0.4
0.6
0.8
1.0
F1
F80 F1
F80 F1
F24 F1

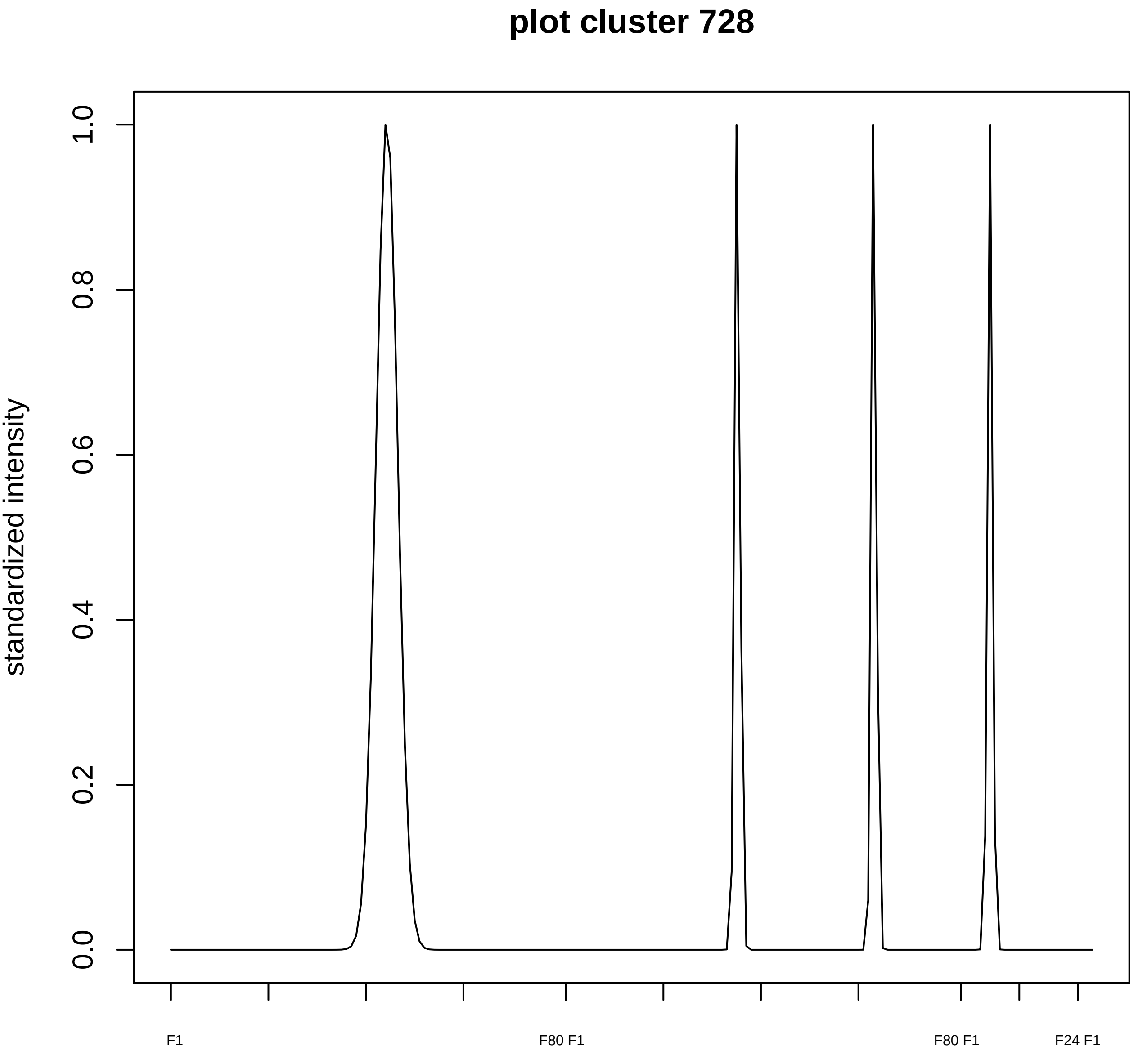

plot cluster 728
standardized intensity
0.0
0.2
0.4
0.6
0.8
1.0
F1
F80 F1
F80 F1
F24 F1

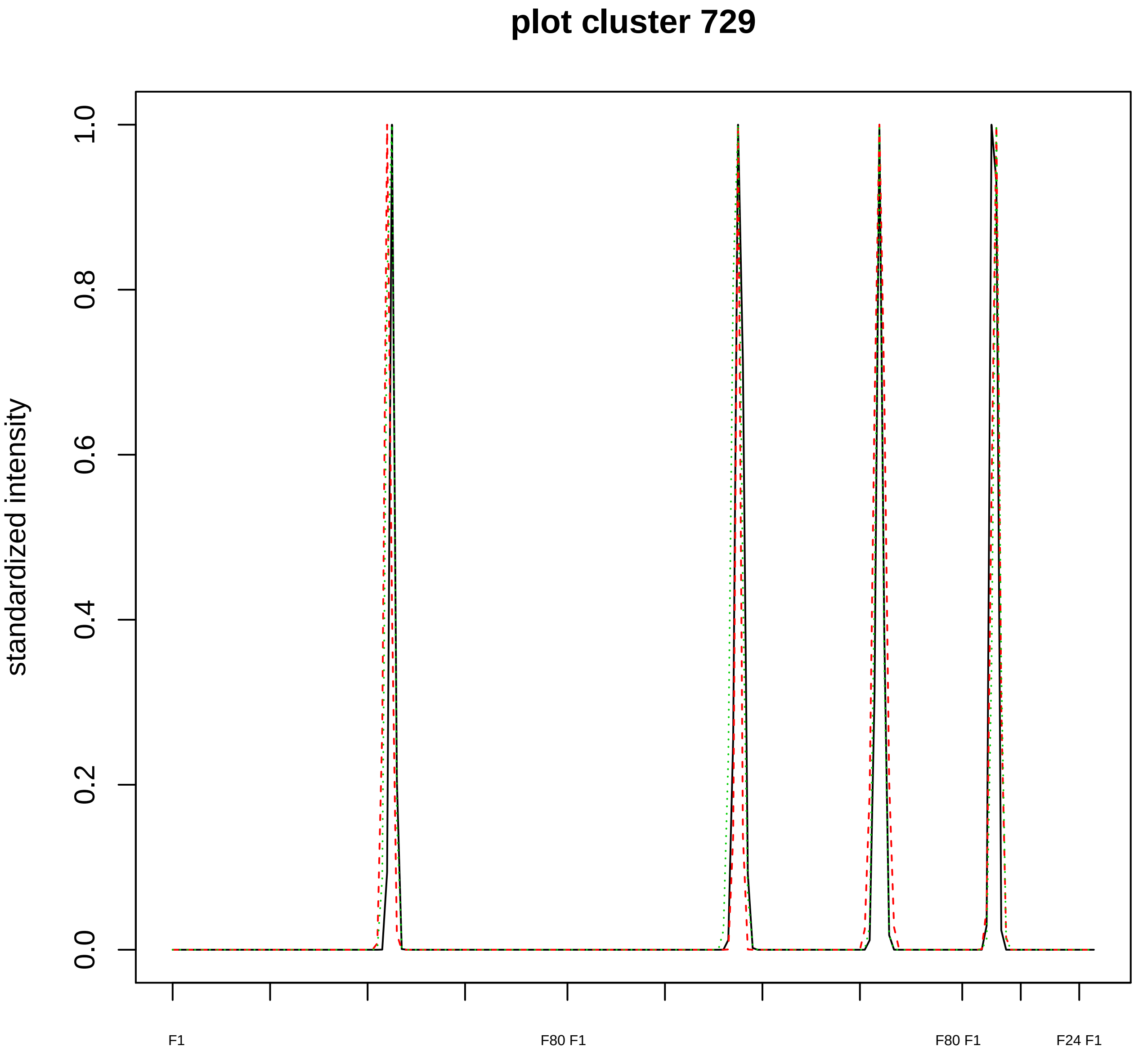
plot cluster 729
standardized intensity
0.0
0.2
0.4
0.6
0.8
1.0
F1
F80 F1
F80 F1
F24 F1

### Dendrogram of cluster 729

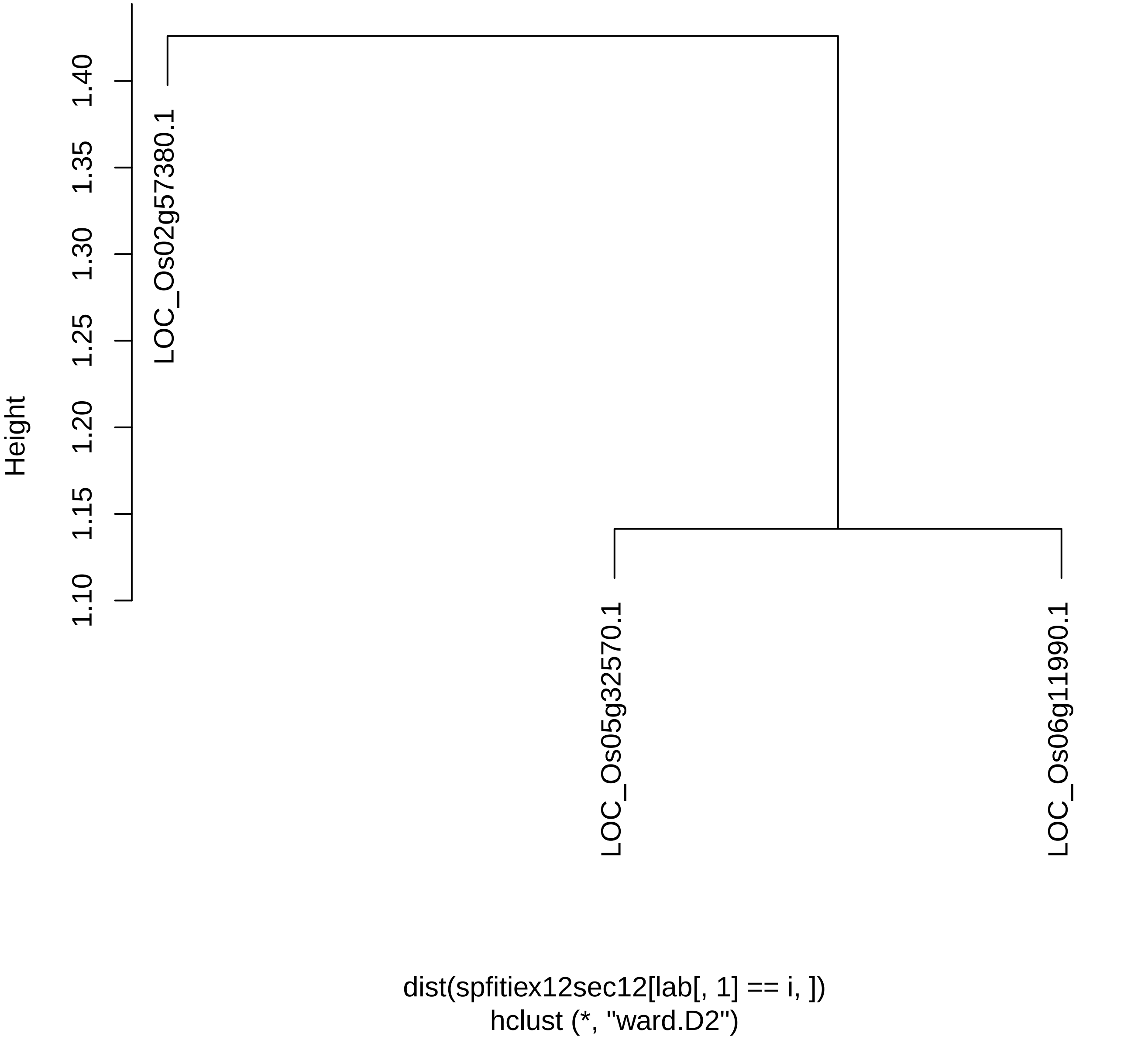

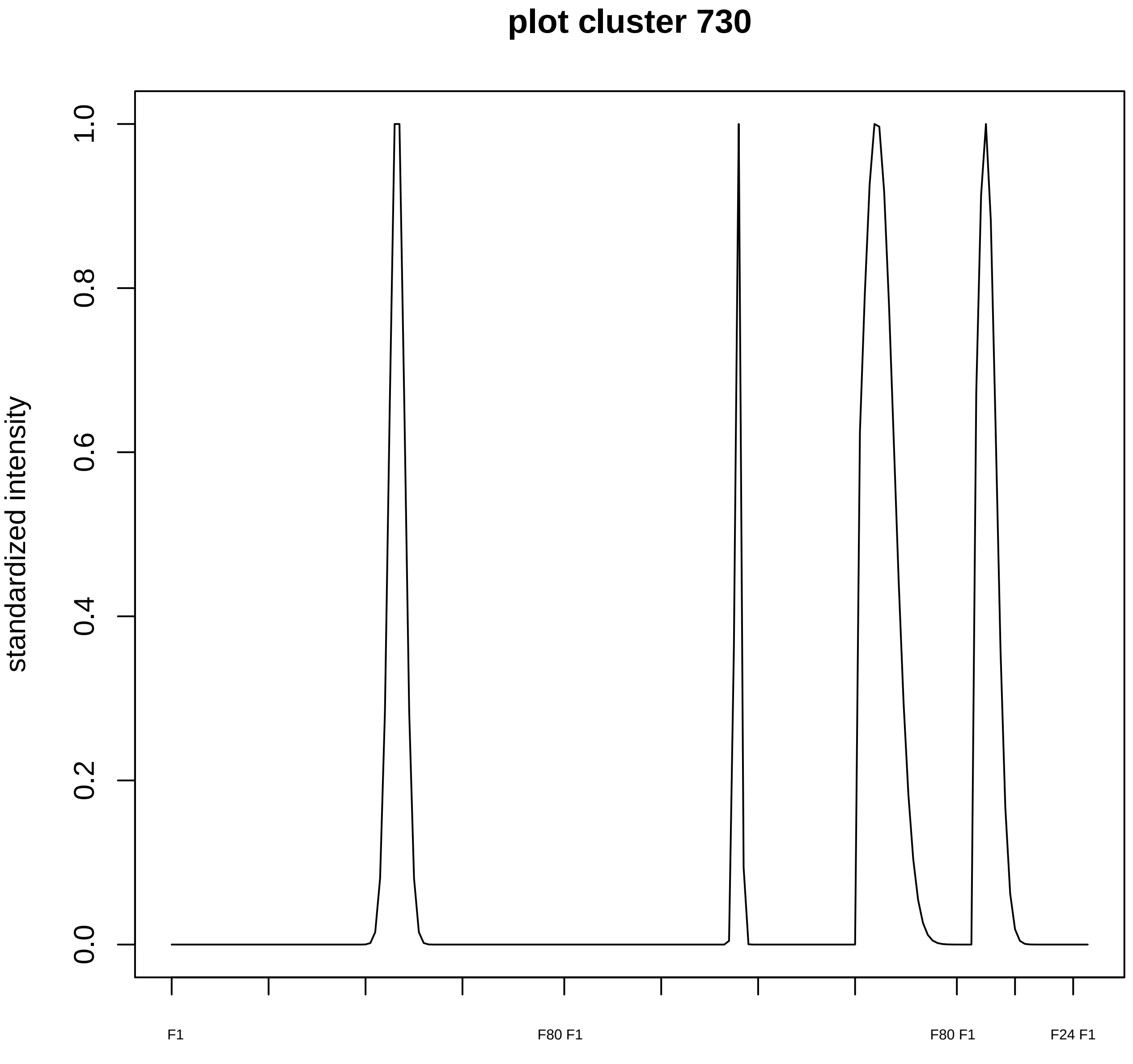

plot cluster 730
standardized intensity
0.0
0.2
0.4
0.6
0.8
1.0
F1
F80 F1
F80 F1
F24 F1

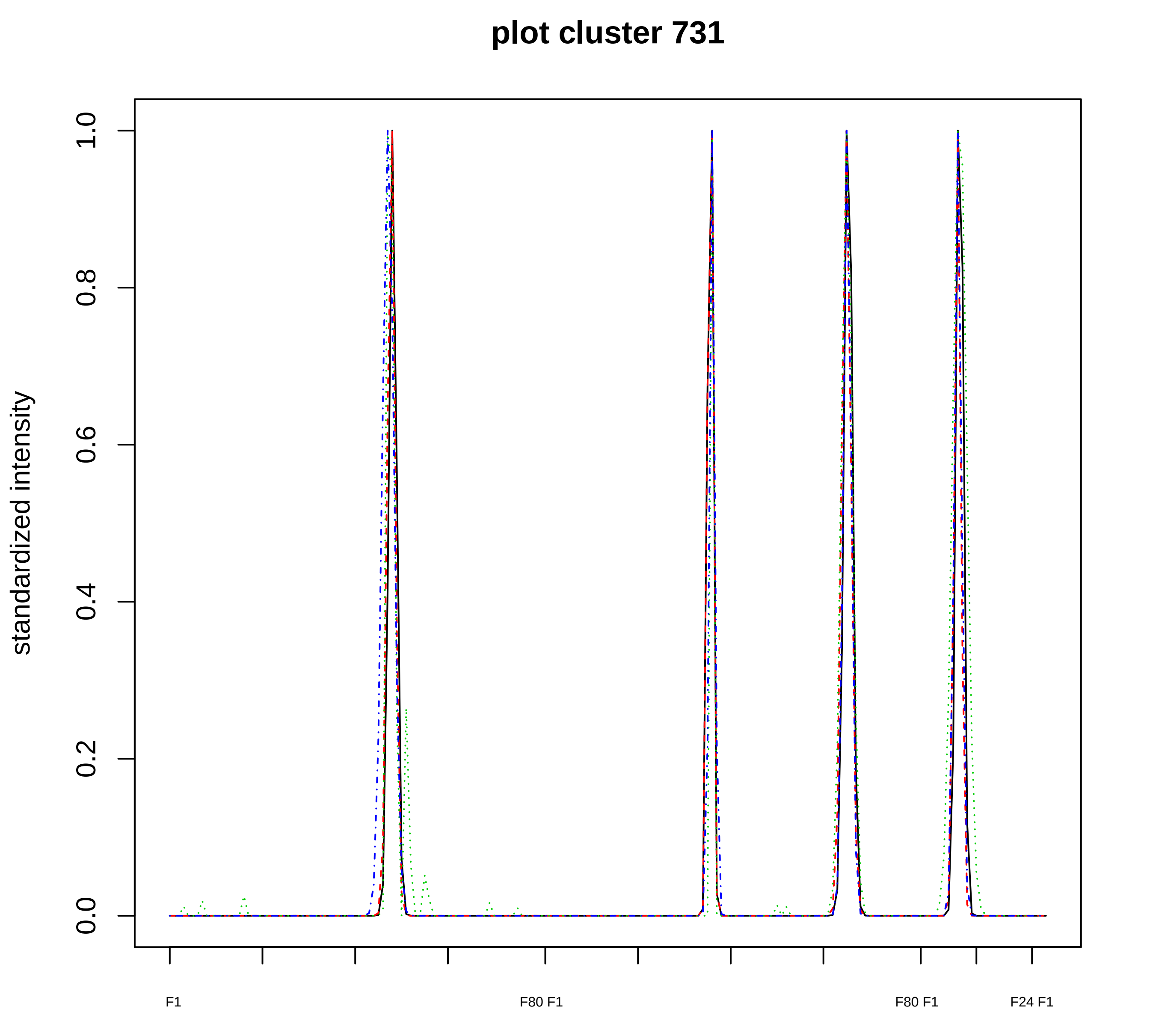

plot cluster 731
standardized intensity
0.0
0.2
0.4
0.6
0.8
1.0
F1
F80 F1
F80 F1
F24 F1

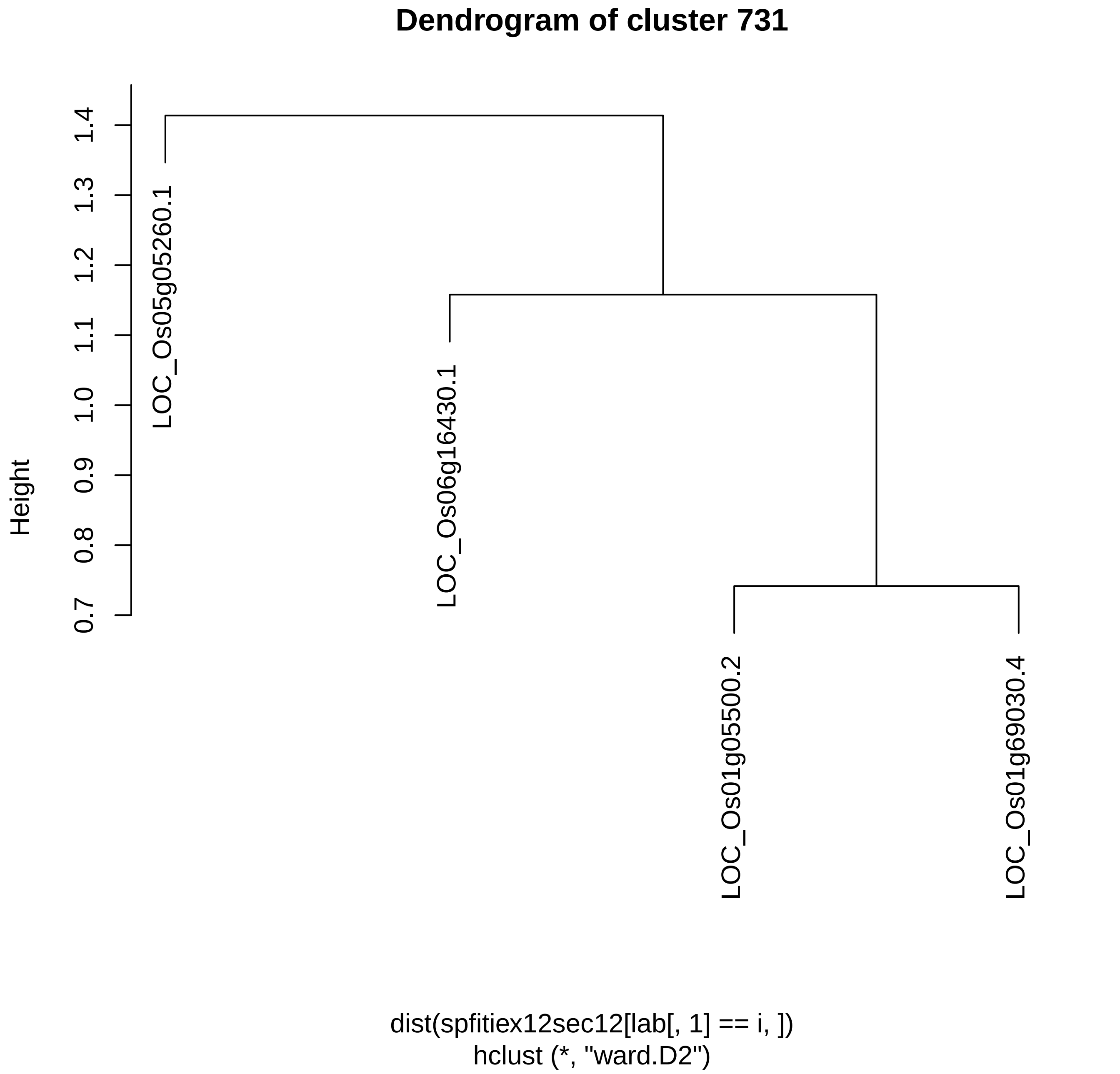
Dendrogram of cluster 731
Height
0.7
0.8
0.9
1.0
1.1
1.2
1.3
1.4
LOC_Os05g05260.1
LOC_Os06g16430.1
LOC_Os01g05500.2
LOC_Os01g69030.4
dist(spfitiex12sec12[lab[, 1] == i, ])
hclust (*, "ward.D2")

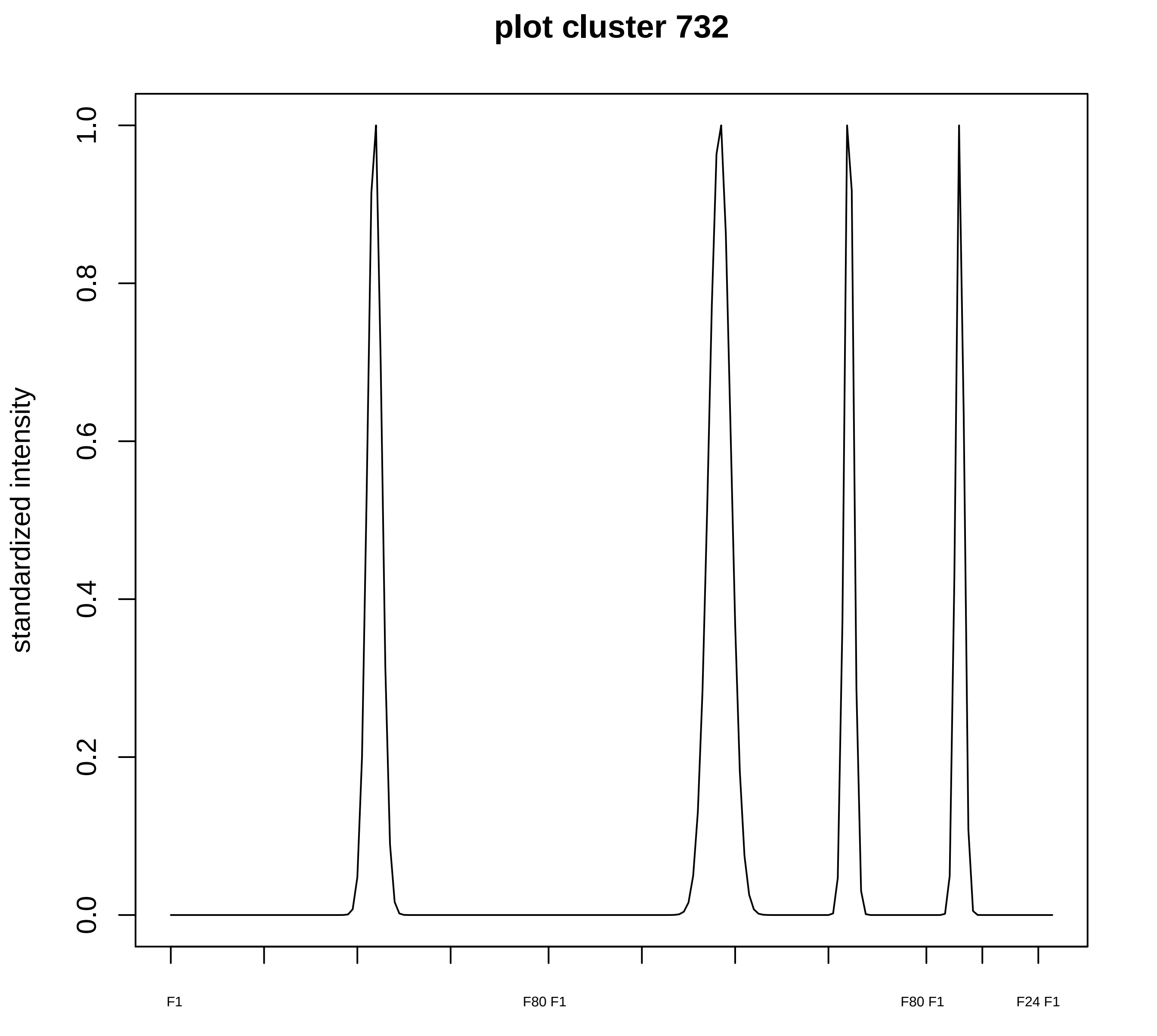
plot cluster 732
standardized intensity
0.0
0.2
0.4
0.6
0.8
1.0
F1
F80 F1
F80 F1
F24 F1

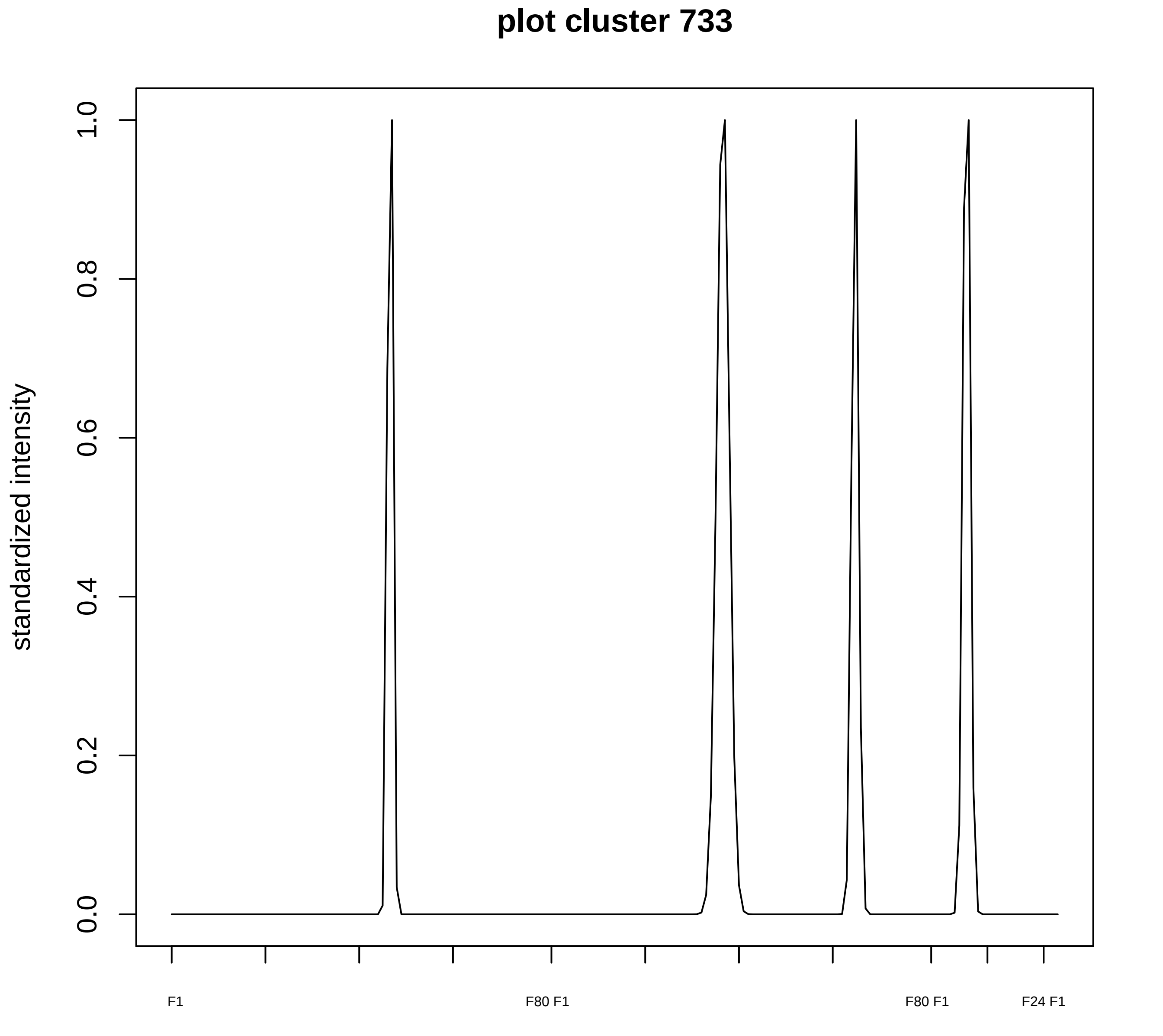

plot cluster 733
standardized intensity
0.0
0.2
0.4
0.6
0.8
1.0
F1
F80 F1
F80 F1
F24 F1

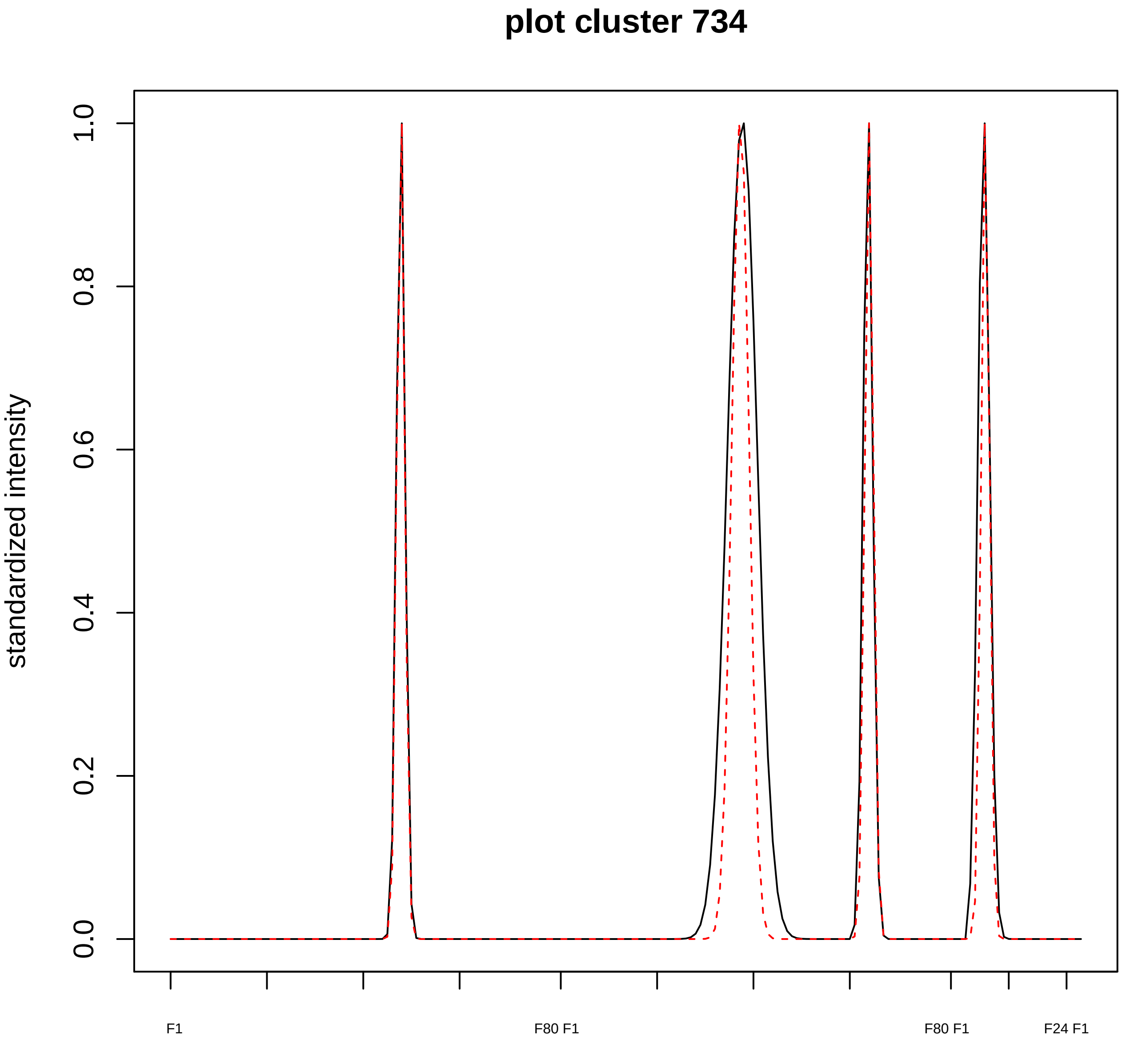

plot cluster 734
standardized intensity
0.0
0.2
0.4
0.6
0.8
1.0
F1
F80 F1
F80 F1
F24 F1

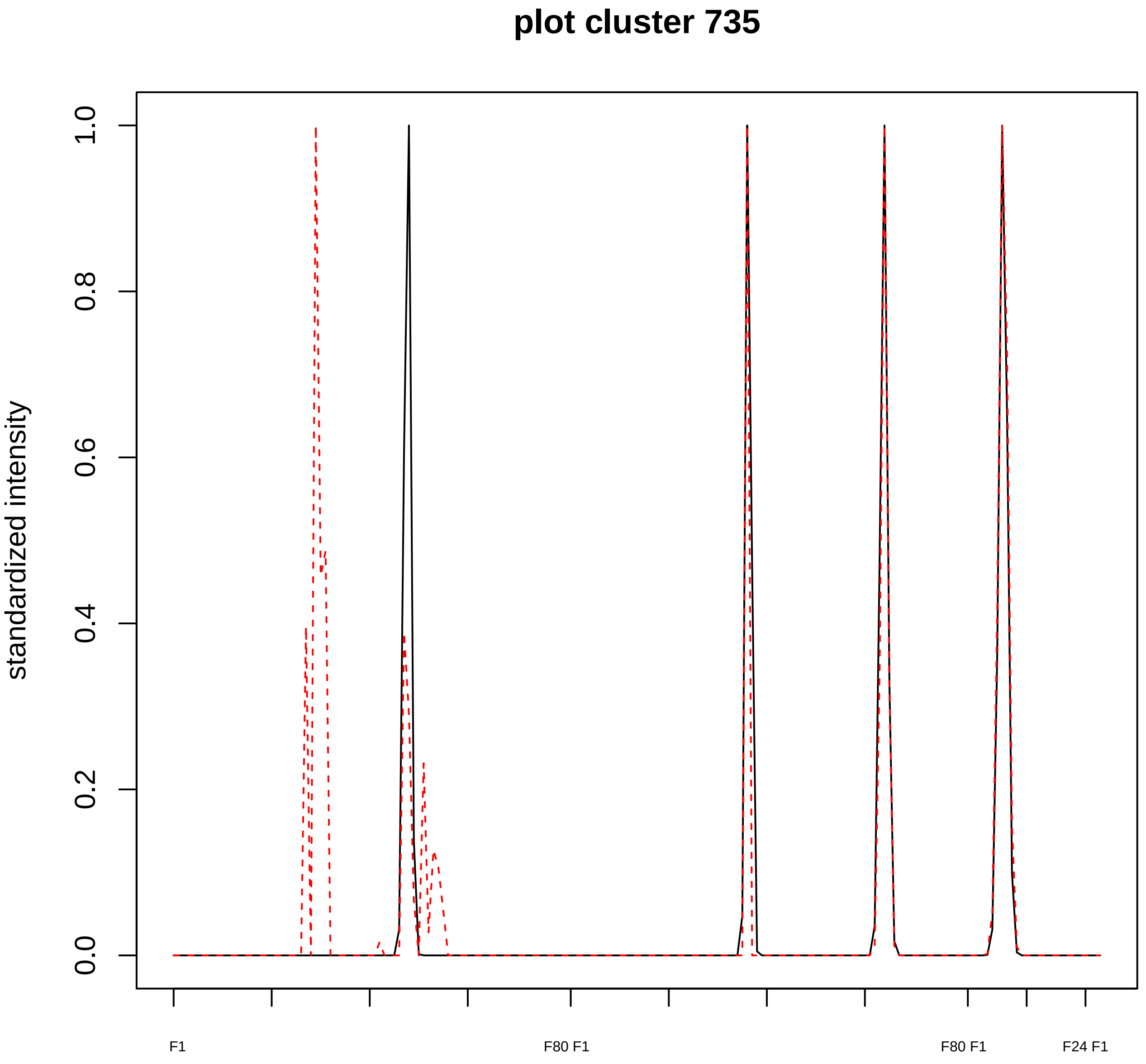
plot cluster 735
standardized intensity
0.0
0.2
0.4
0.6
0.8
1.0
F1
F80 F1
F80 F1
F24 F1

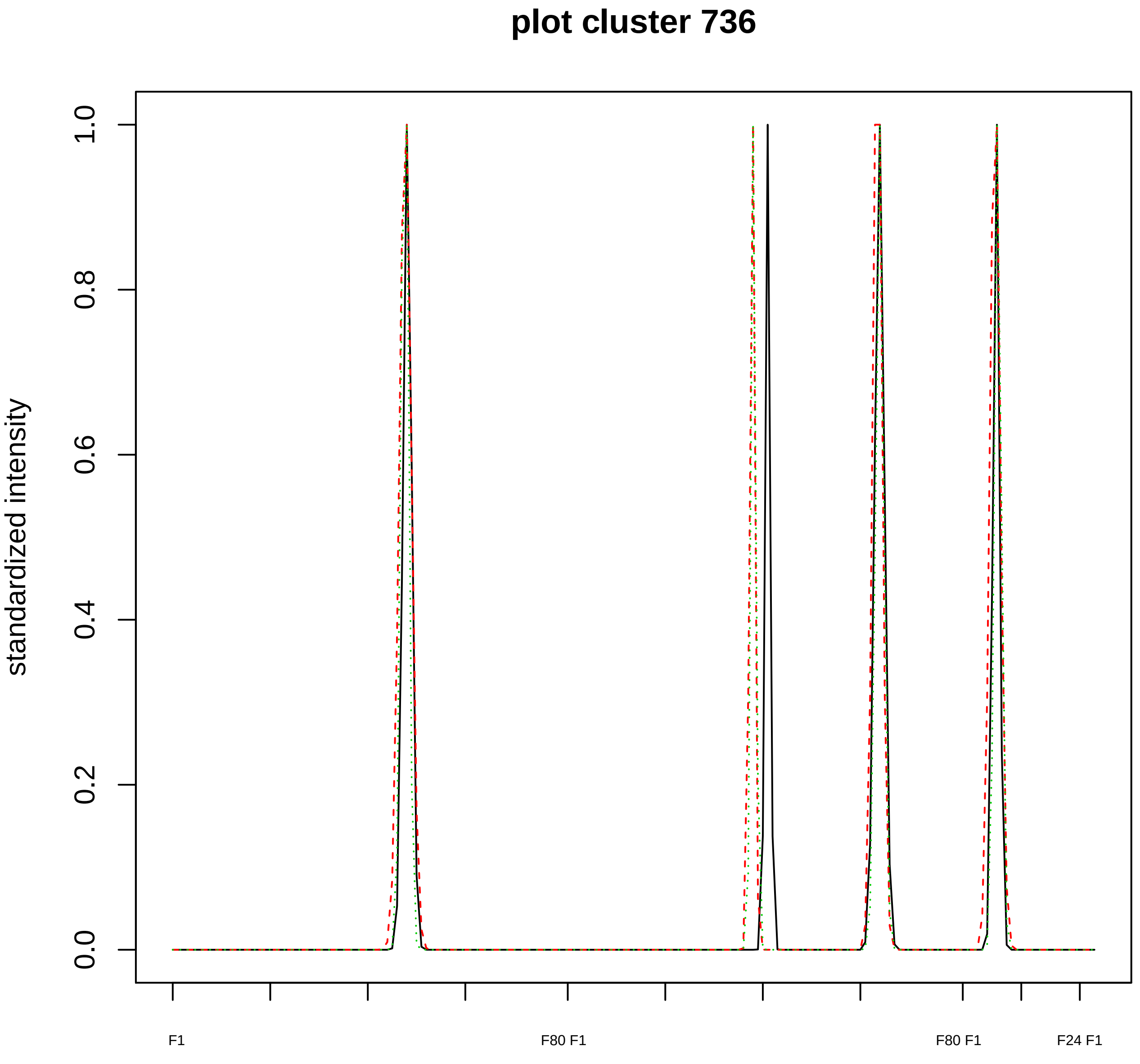
plot cluster 736
standardized intensity
0.0
0.2
0.4
0.6
0.8
1.0
F1
F80 F1
F80 F1
F24 F1

### Dendrogram of cluster 736

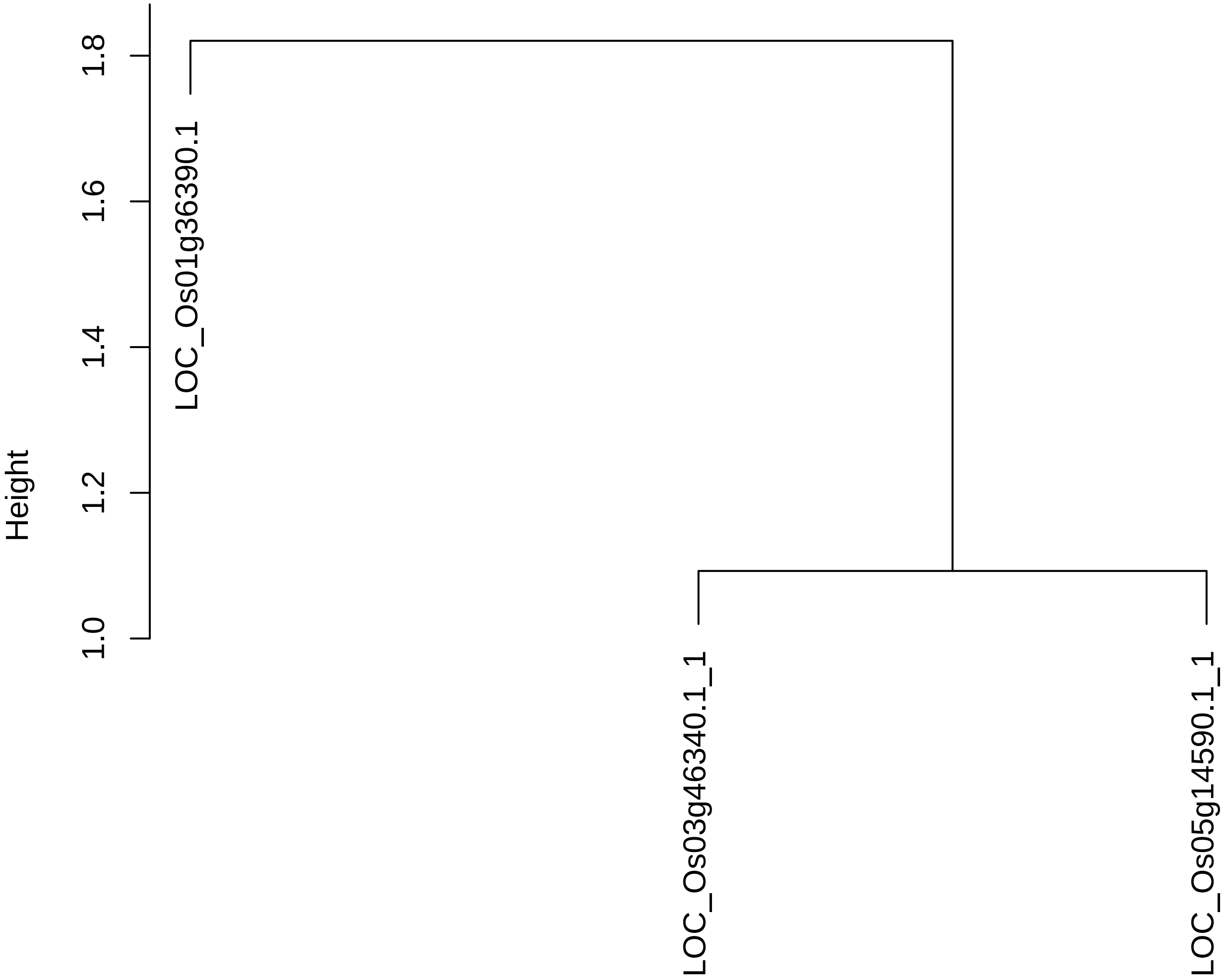


dist(spfitiex12sec12[lab[, 1] == i, ])
hclust (*, "ward.D2")

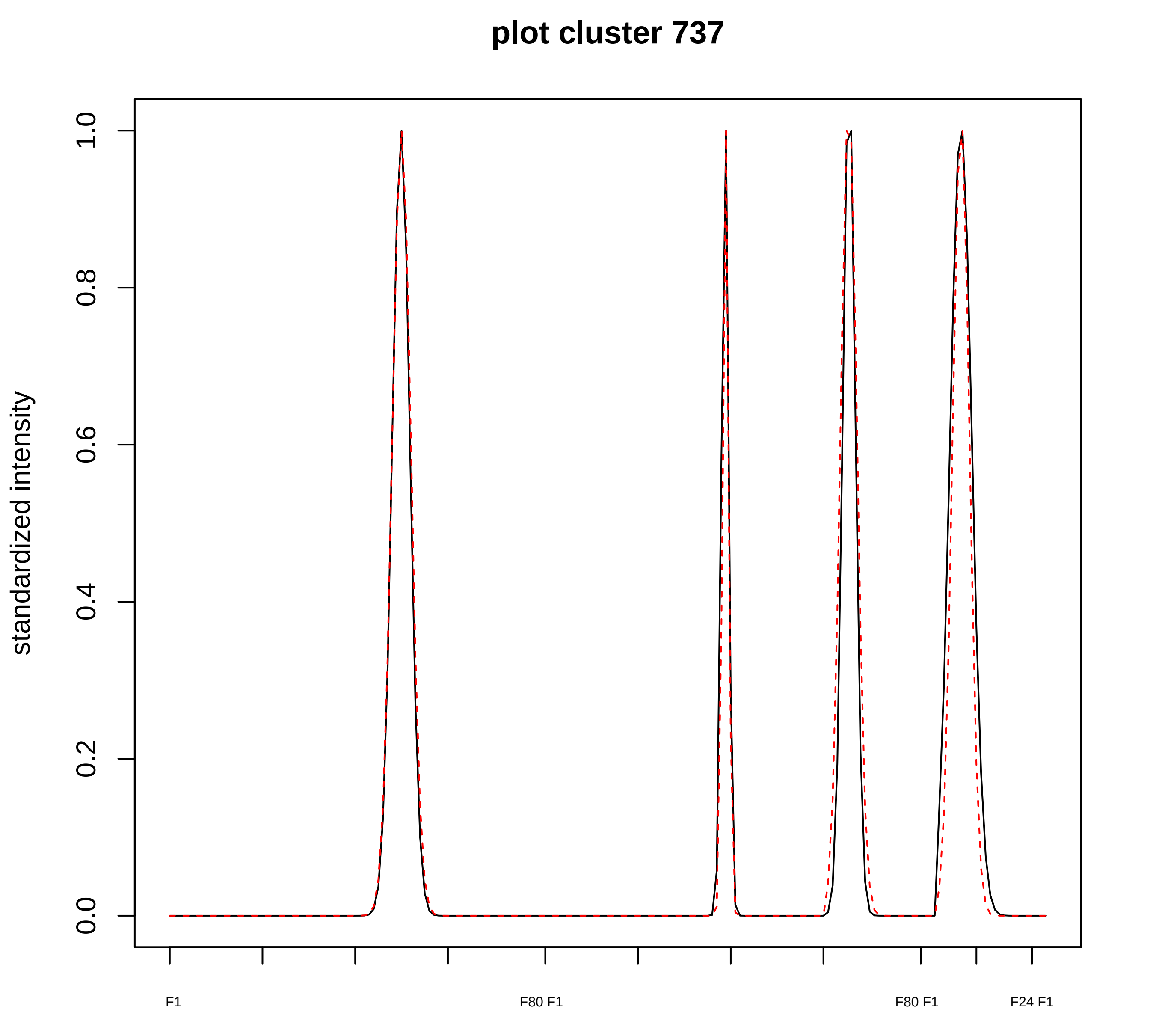
plot cluster 737
standardized intensity
0.0
0.2
0.4
0.6
0.8
1.0
F1
F80 F1
F80 F1
F24 F1

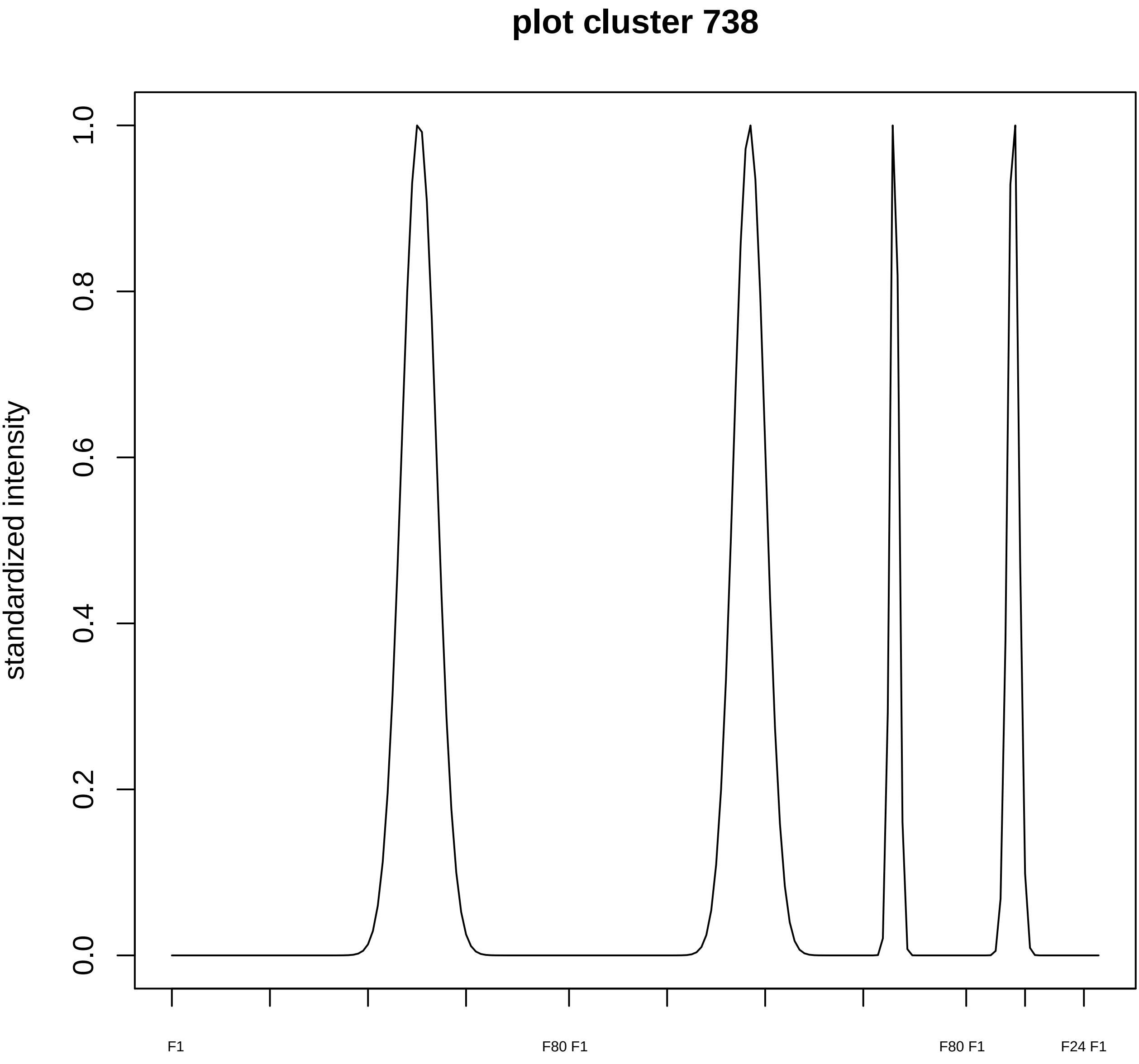
plot cluster 738
standardized intensity
0.0
0.2
0.4
0.6
0.8
1.0
F1
F80 F1
F80 F1
F24 F1

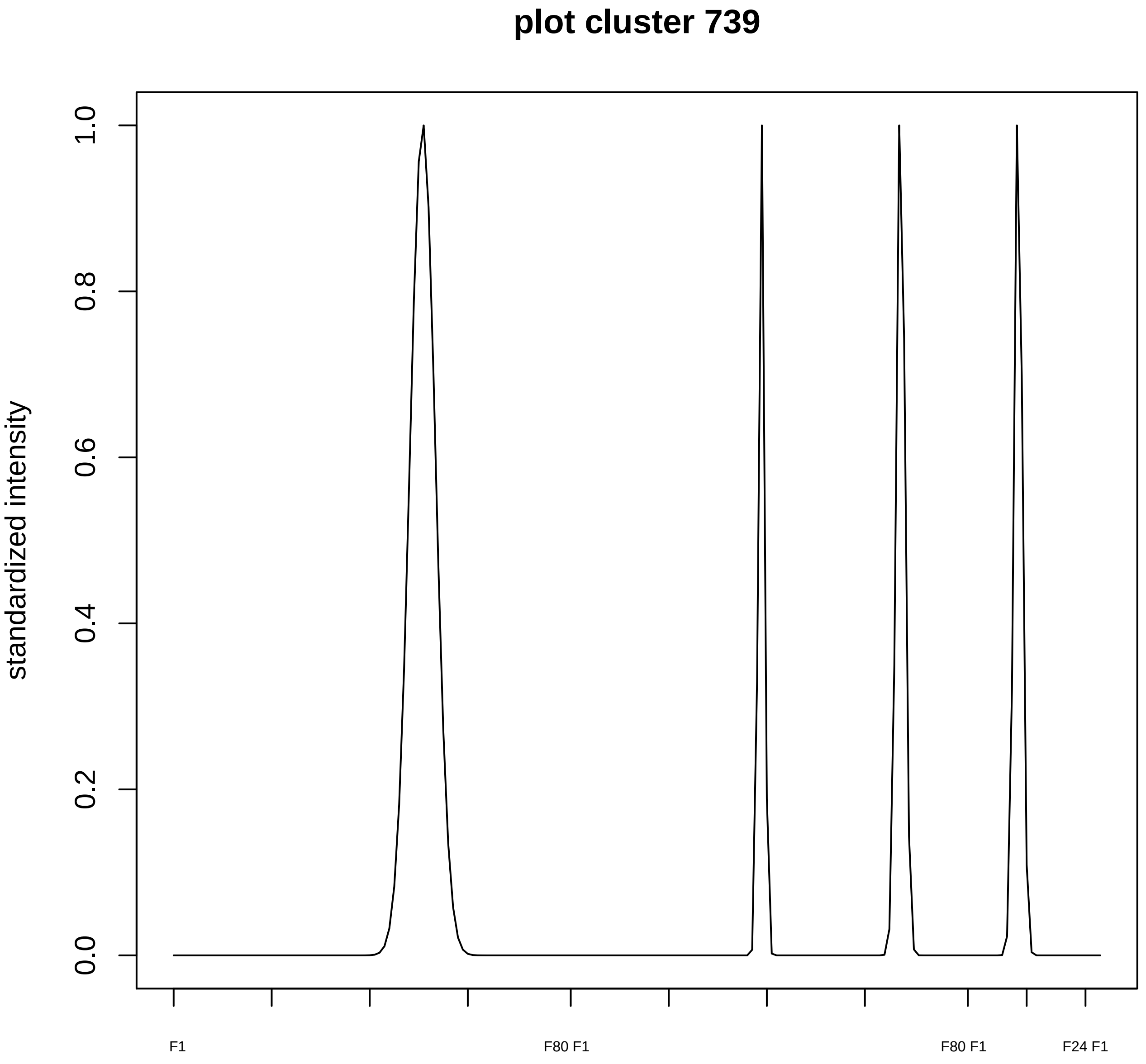

plot cluster 739
standardized intensity
0.0
0.2
0.4
0.6
0.8
1.0
F1
F80 F1
F80 F1
F24 F1

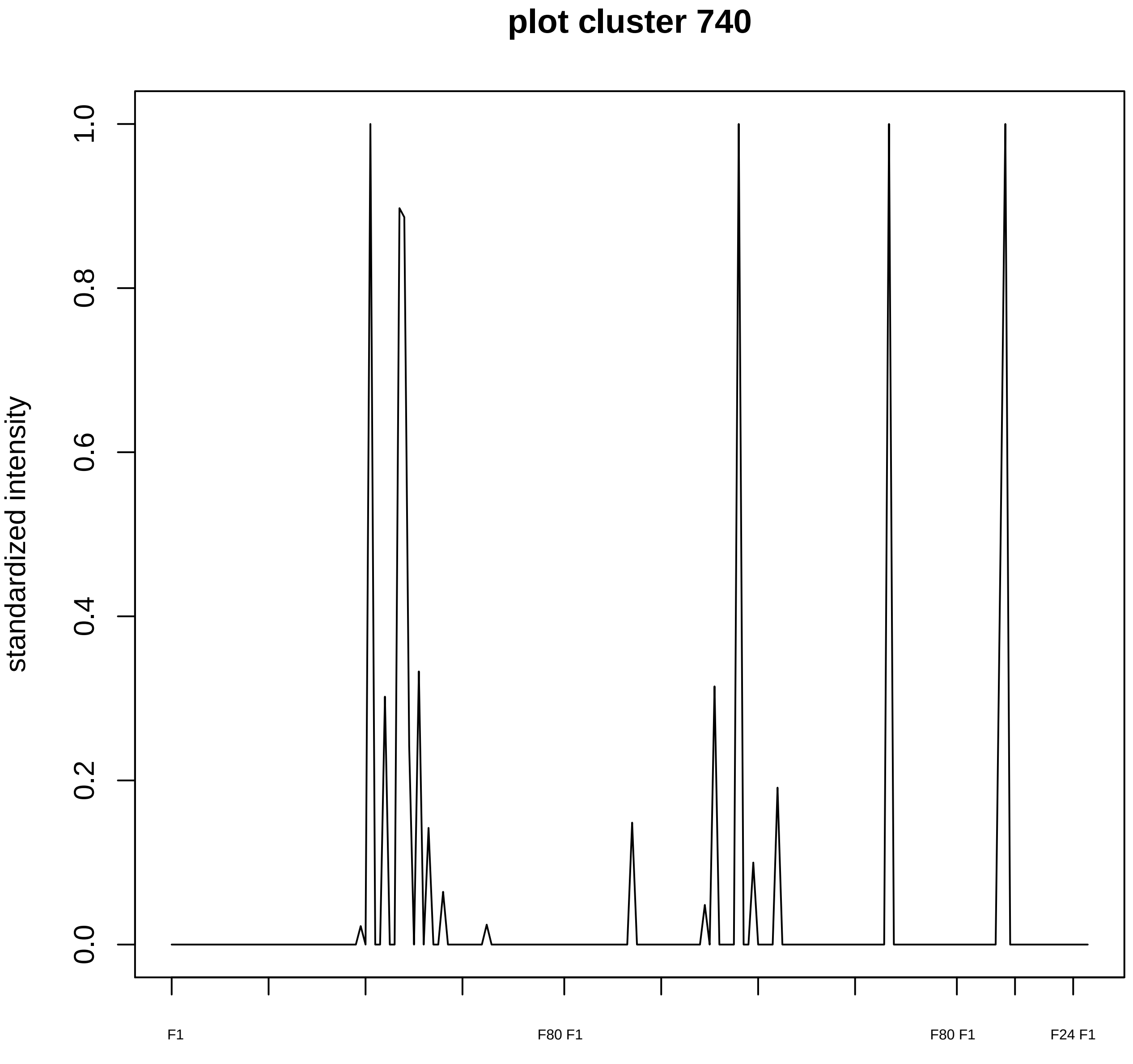
plot cluster 740
standardized intensity
0.0
0.2
0.4
0.6
0.8
1.0
F1
F80 F1
F80 F1
F24 F1

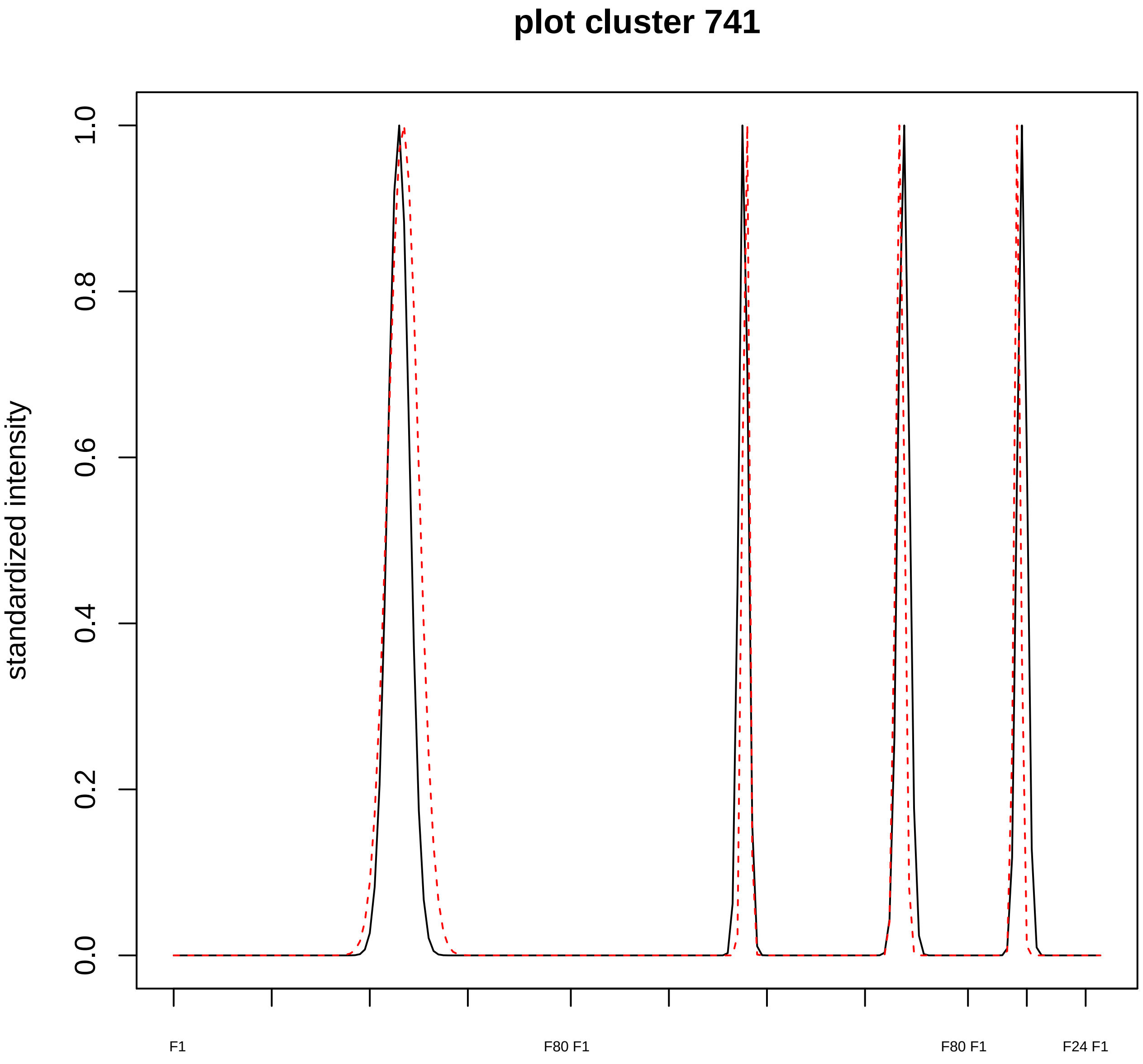
plot cluster 741
standardized intensity
1.0
0.8
0.6
0.4
0.2
0.0
F1
F80 F1
F80 F1
F24 F1

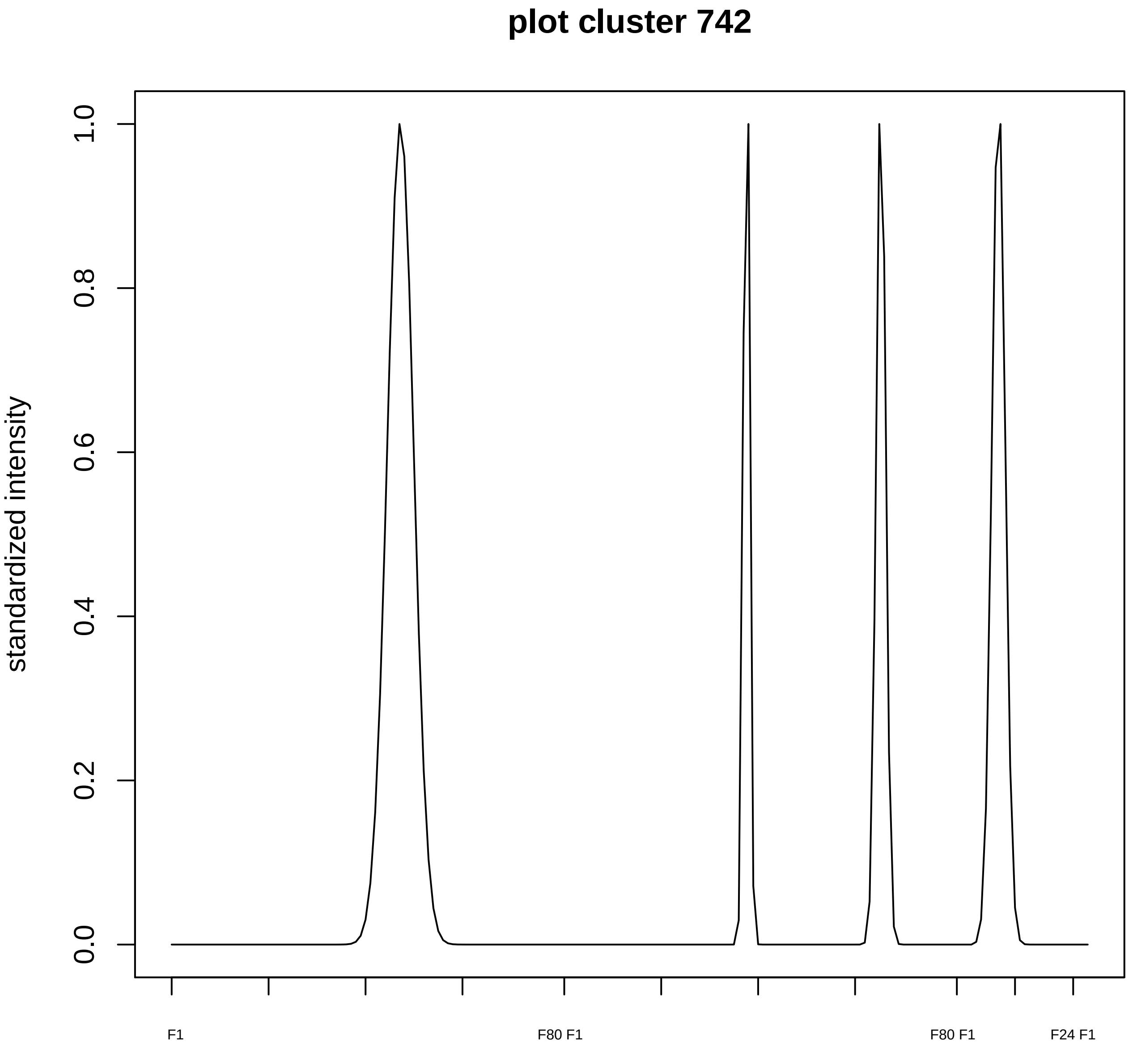

plot cluster 742
standardized intensity
0.0
0.2
0.4
0.6
0.8
1.0
F1
F80 F1
F80 F1
F24 F1

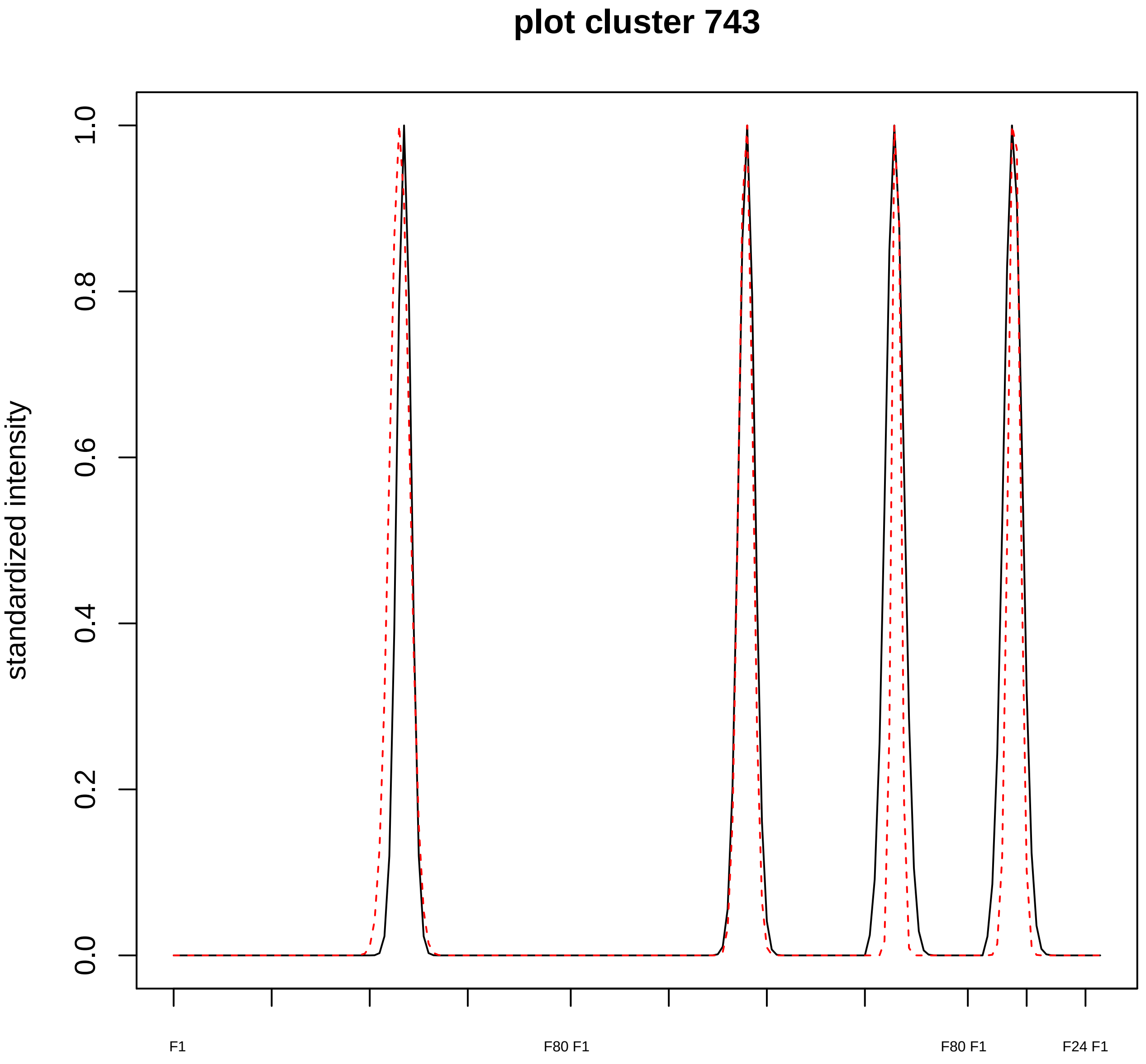

plot cluster 743
standardized intensity
1.0
0.8
0.6
0.4
0.2
0.0
F1
F80 F1
F80 F1
F24 F1

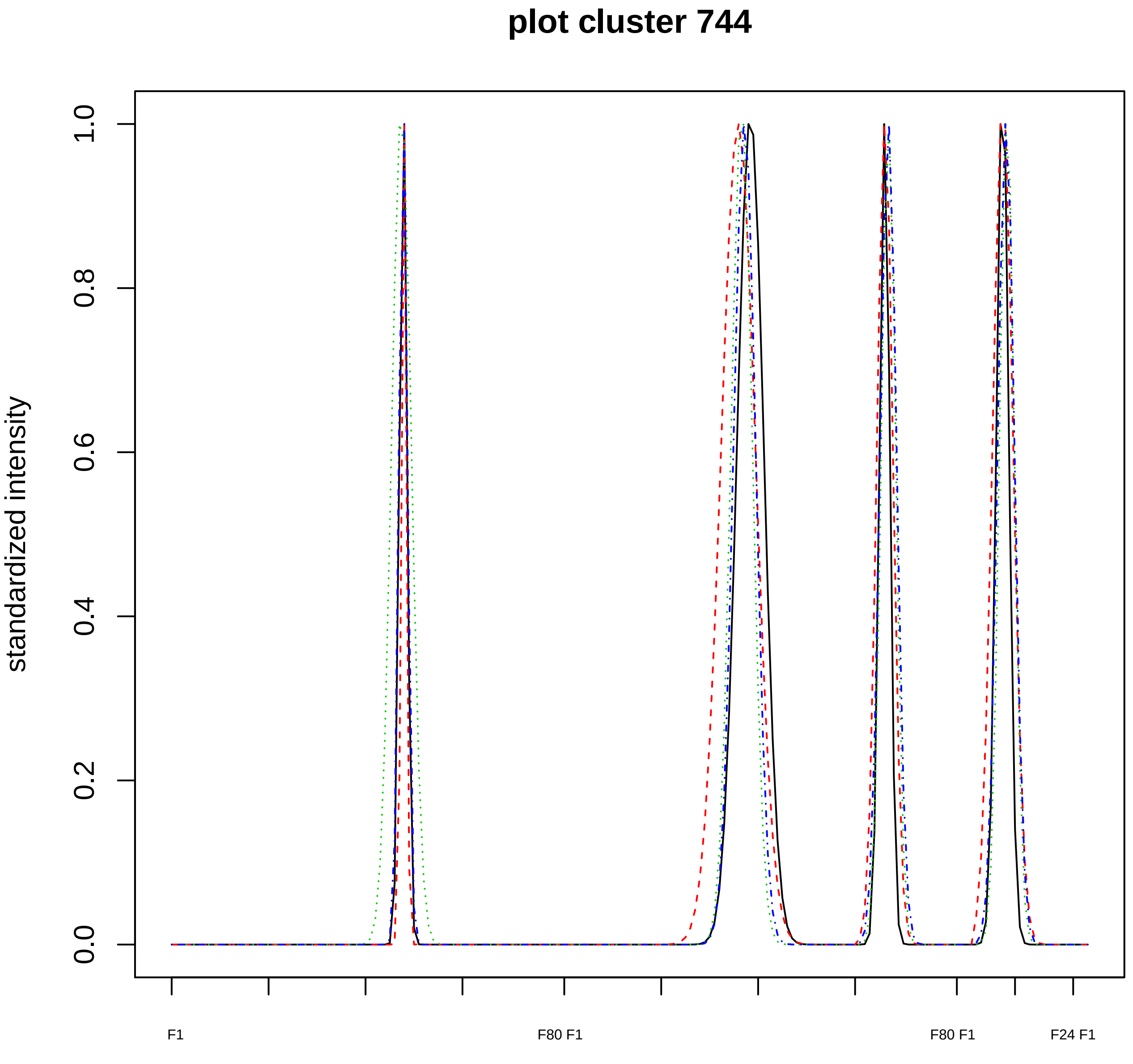

plot cluster 744
standardized intensity
0.0
0.2
0.4
0.6
0.8
1.0
F1
F80 F1
F80 F1
F24 F1

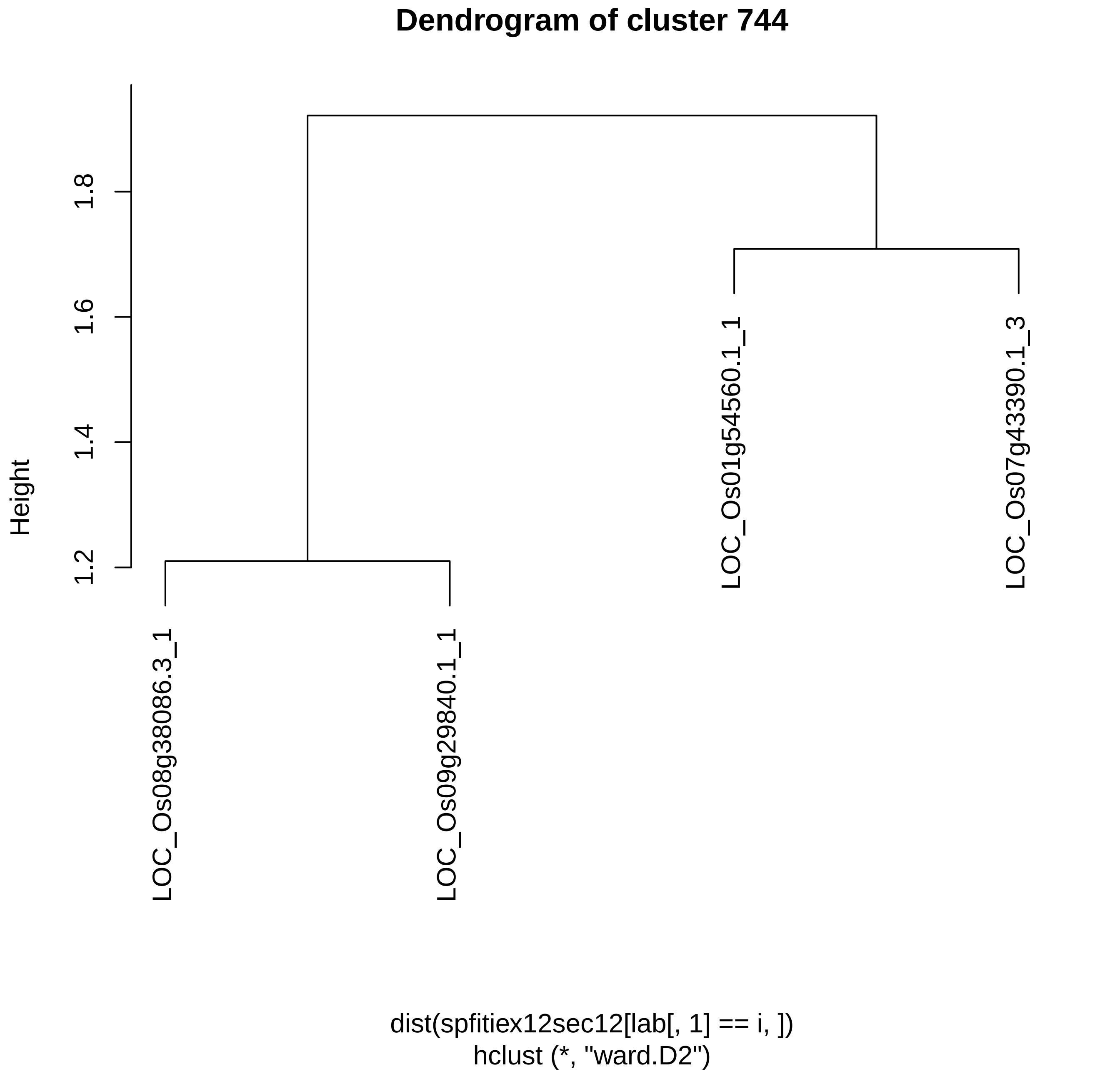

Dendrogram of cluster 744
Height
1.2
1.4
1.6
1.8
LOC_Os08g38086.3_1
LOC_Os09g29840.1_1
LOC_Os01g54560.1_1
LOC_Os07g43390.1_3
dist(spfitiex12sec12[lab[, 1] == i, ])
hclust (*, "ward.D2")

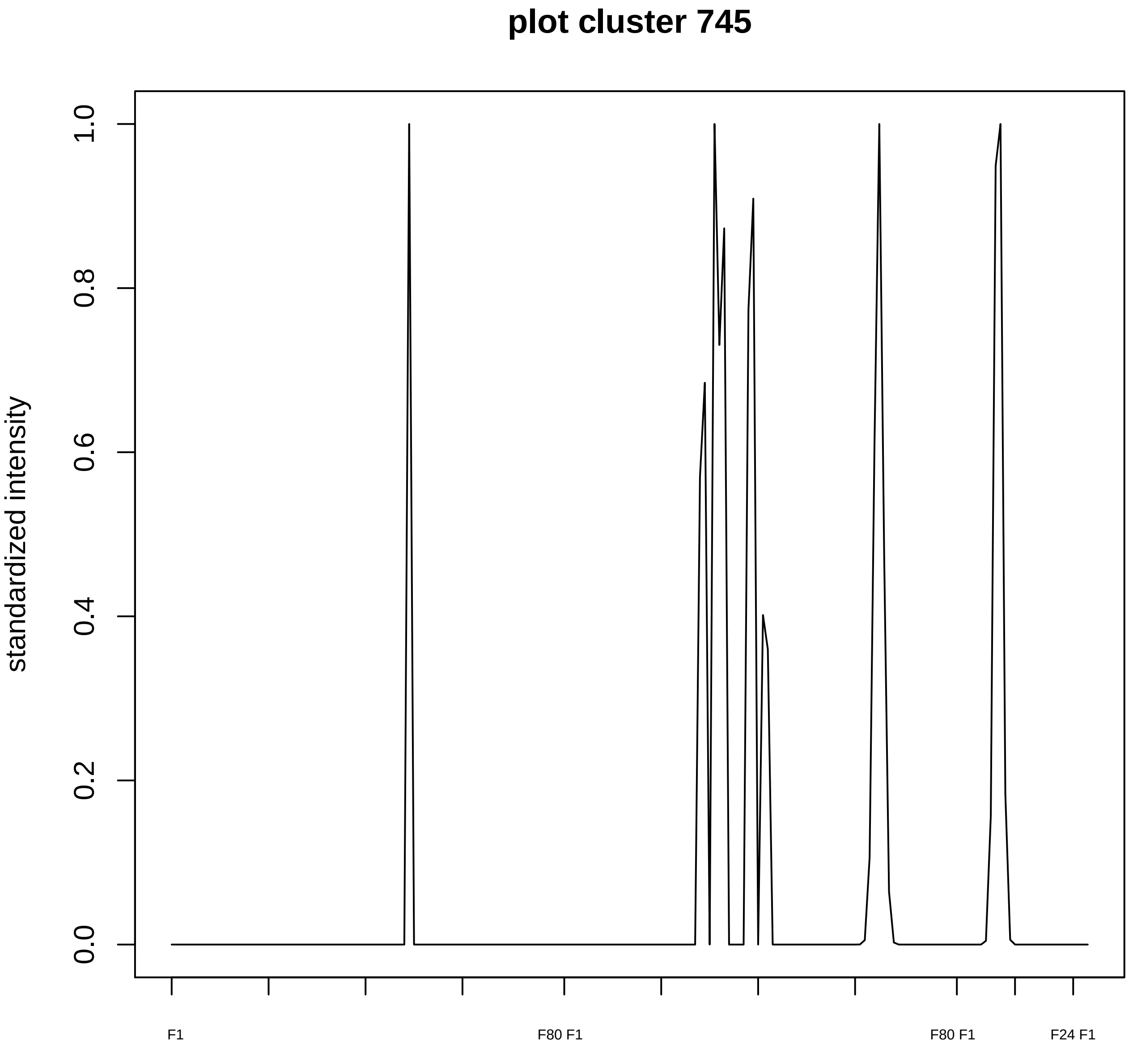

plot cluster 745
standardized intensity
0.0
0.2
0.4
0.6
0.8
1.0
F1
F80 F1
F80 F1
F24 F1

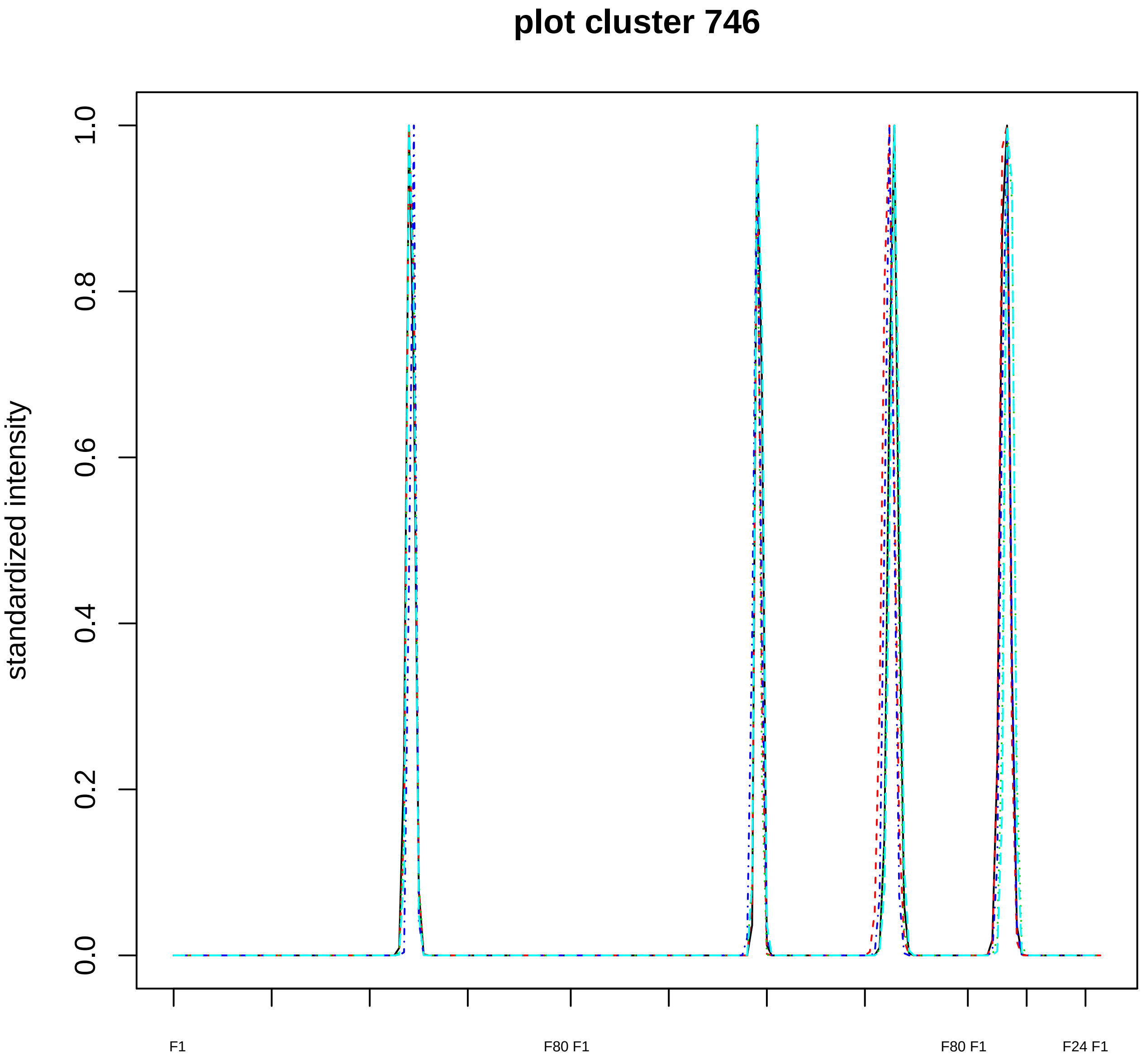

plot cluster 746
standardized intensity
0.0
0.2
0.4
0.6
0.8
1.0
F1
F80 F1
F80 F1
F24 F1

### Dendrogram of cluster 746

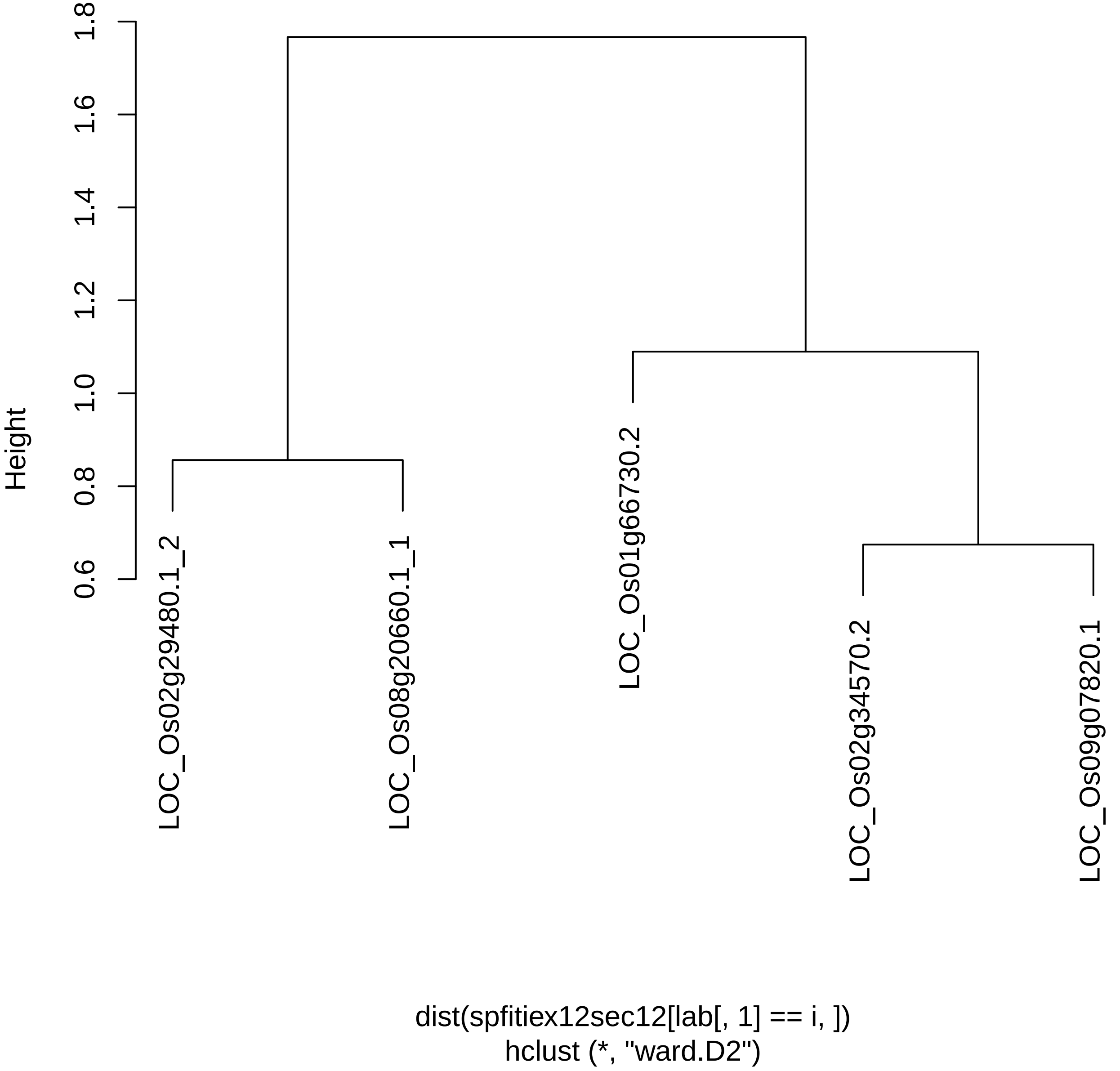

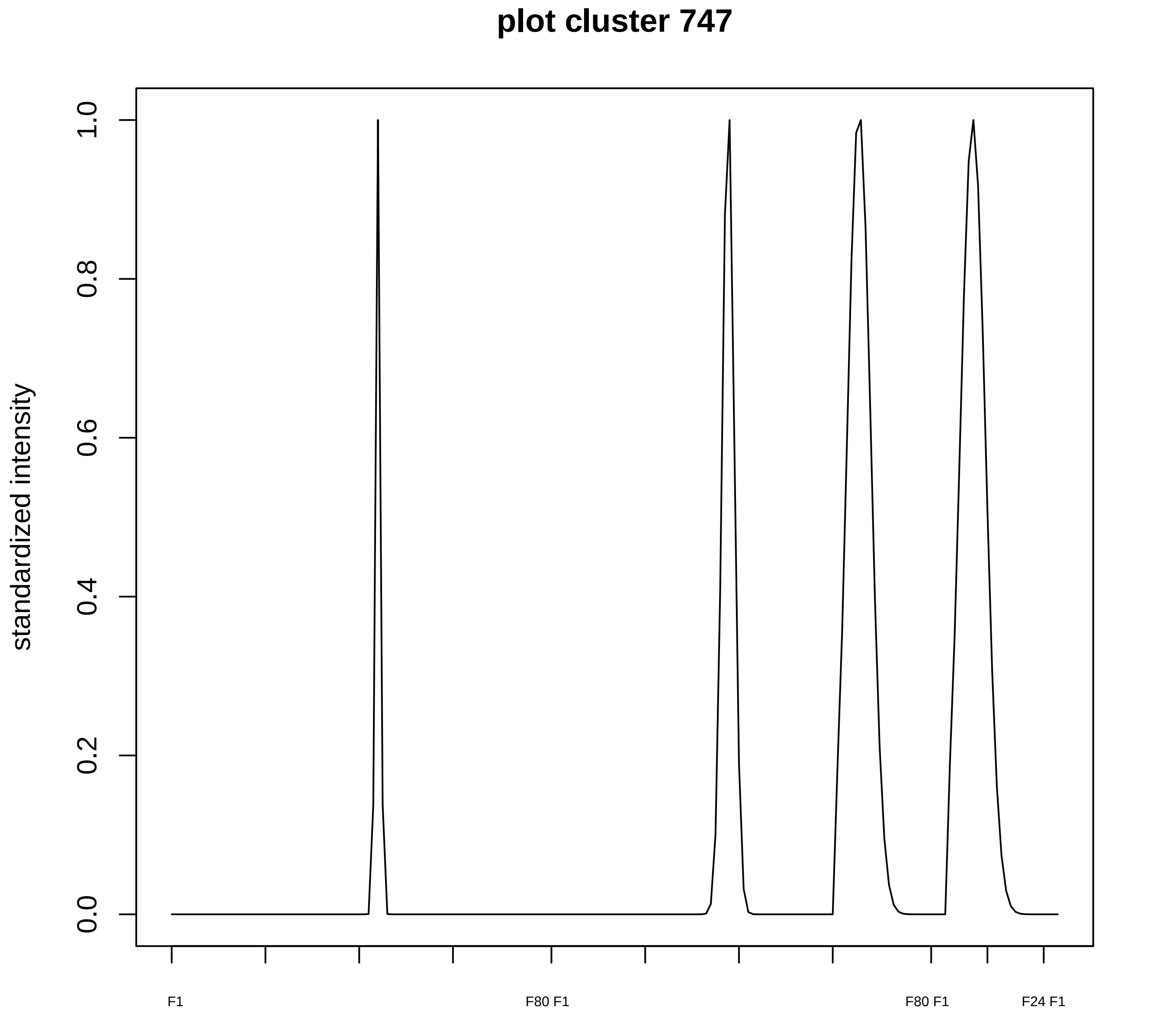
plot cluster 747
standardized intensity
0.0
0.2
0.4
0.6
0.8
1.0
F1
F80 F1
F80 F1
F24 F1

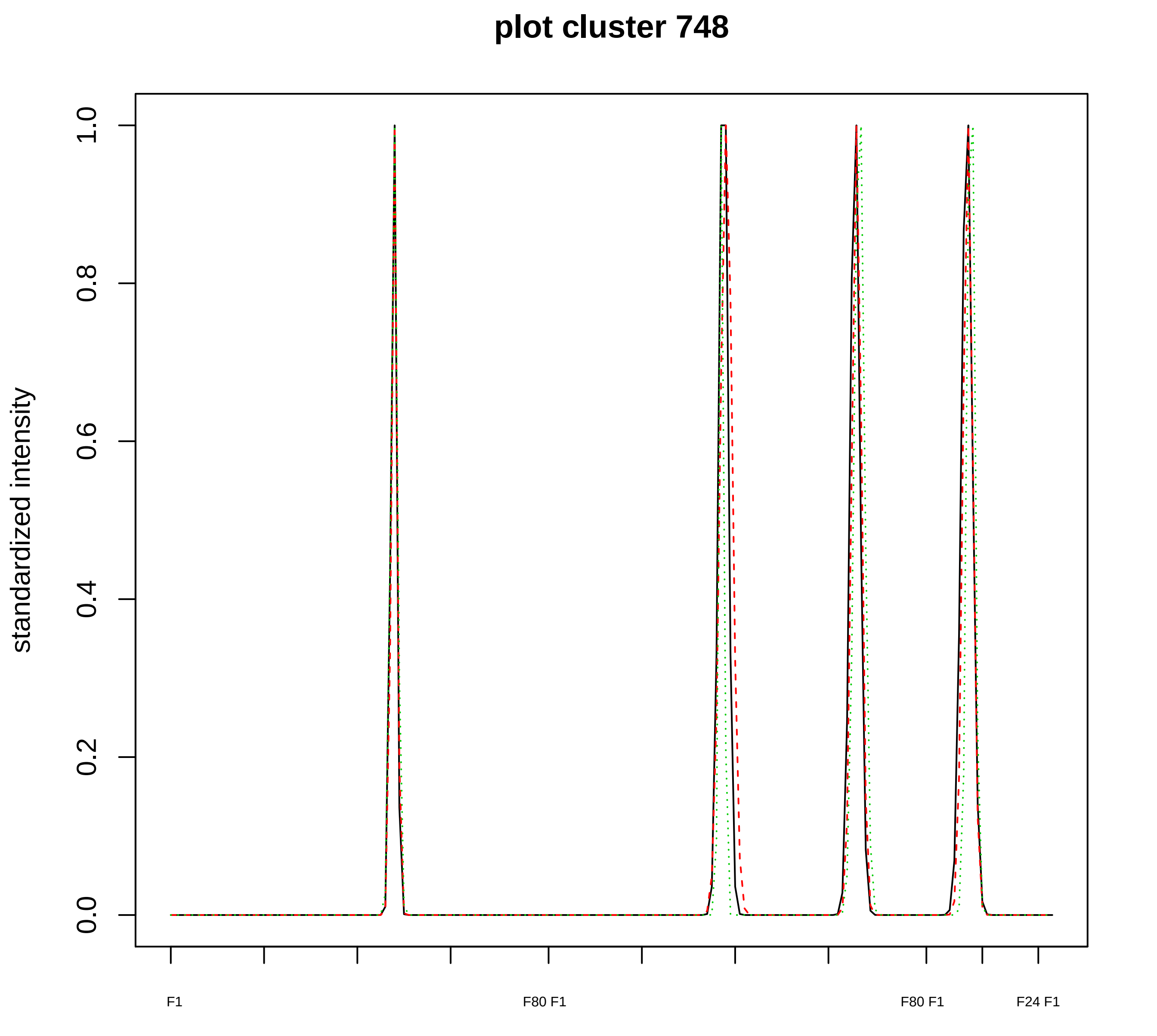

plot cluster 748
standardized intensity
0.0
0.2
0.4
0.6
0.8
1.0
F1
F80 F1
F80 F1
F24 F1

#### Dendrogram of cluster 748

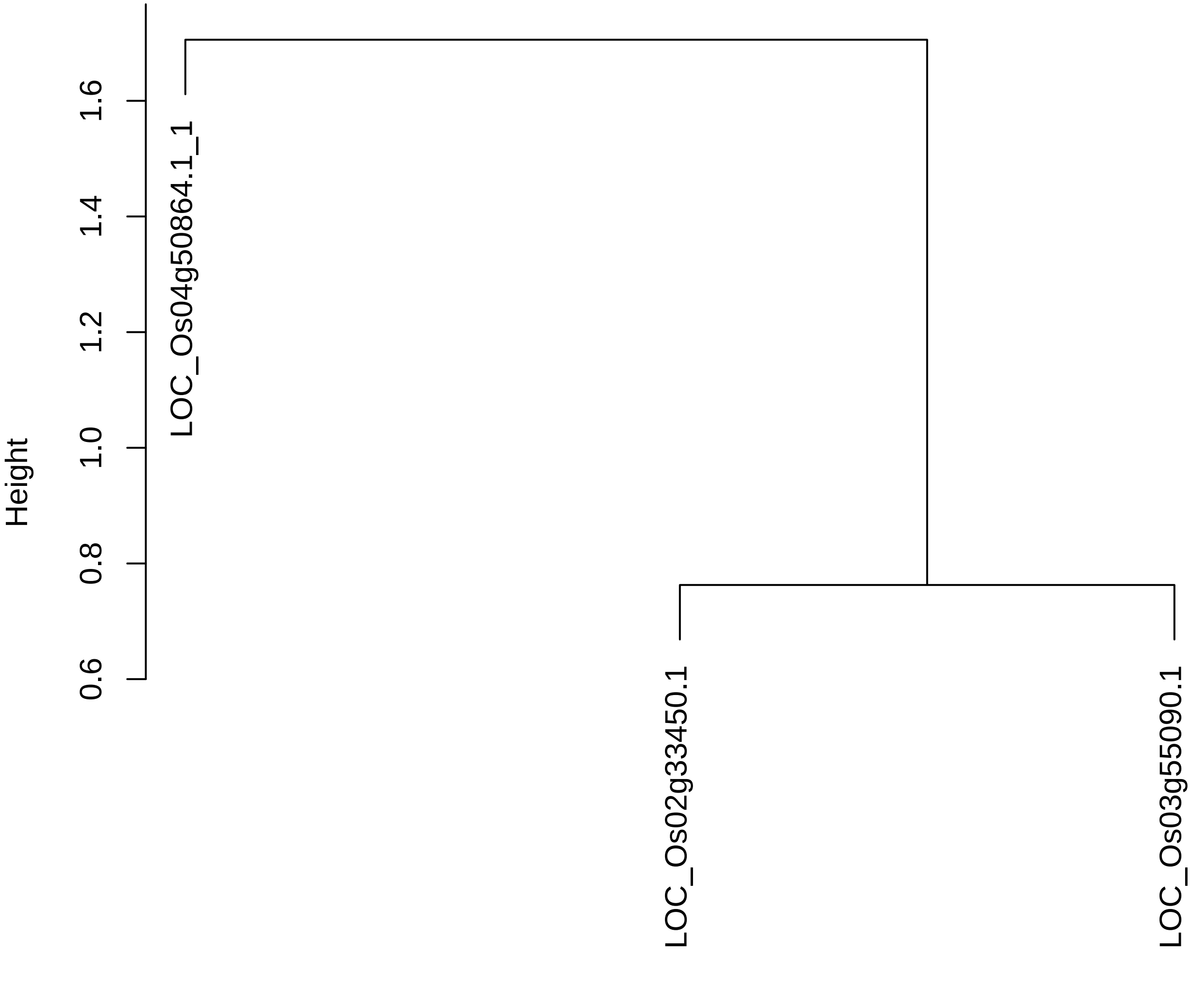


dist(spfitiex12sec12[lab[, 1] == i, ])
hclust (*, "ward.D2")

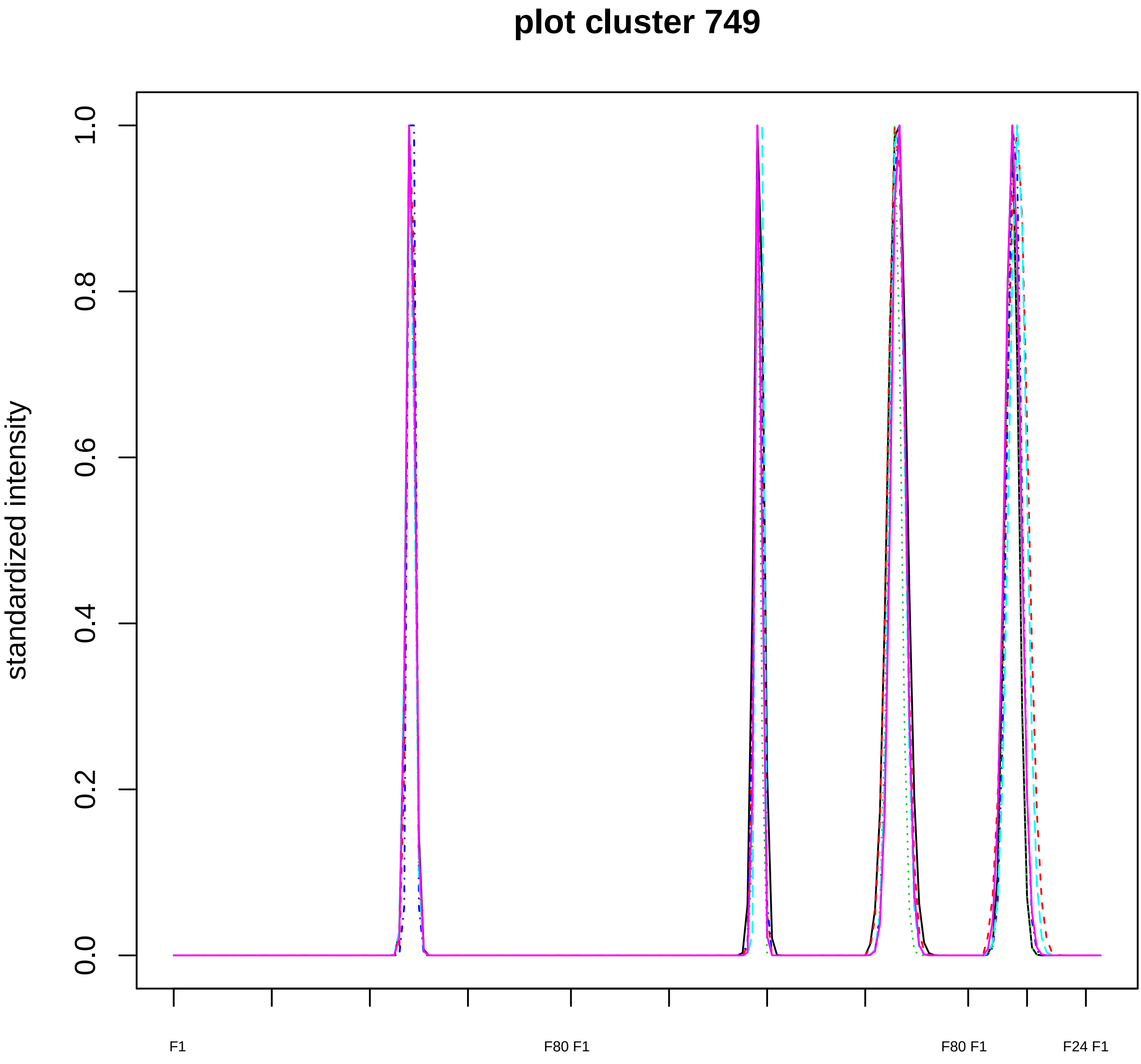

plot cluster 749
standardized intensity
0.0
0.2
0.4
0.6
0.8
1.0
F1
F80 F1
F80 F1
F24 F1

#### Dendrogram of cluster 749

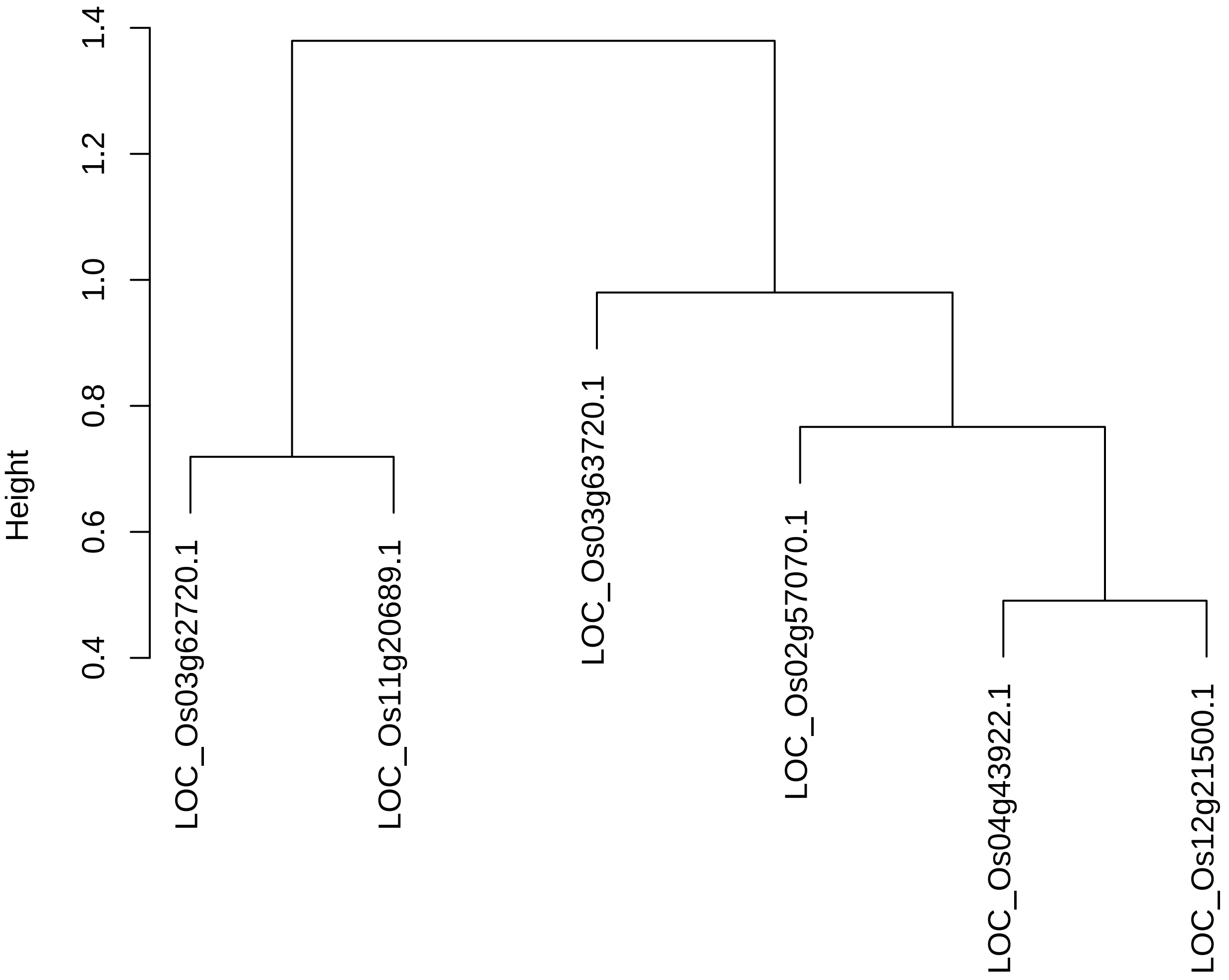


dist(spfitiex12sec12[lab[, 1] == i, ])
hclust (*, "ward.D2")

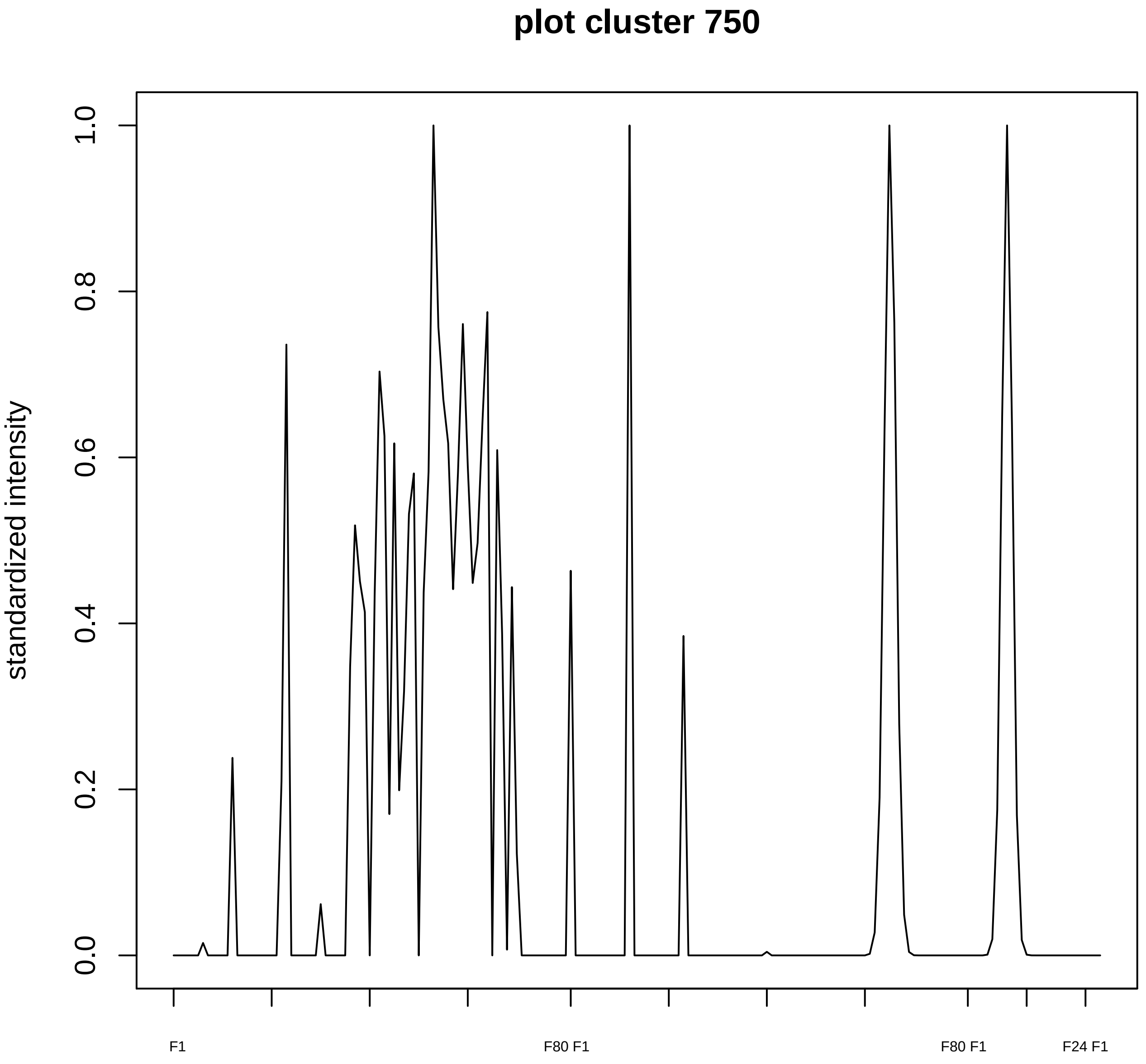

plot cluster 750
standardized intensity
0.0
0.2
0.4
0.6
0.8
1.0
F1
F80 F1
F80 F1
F24 F1

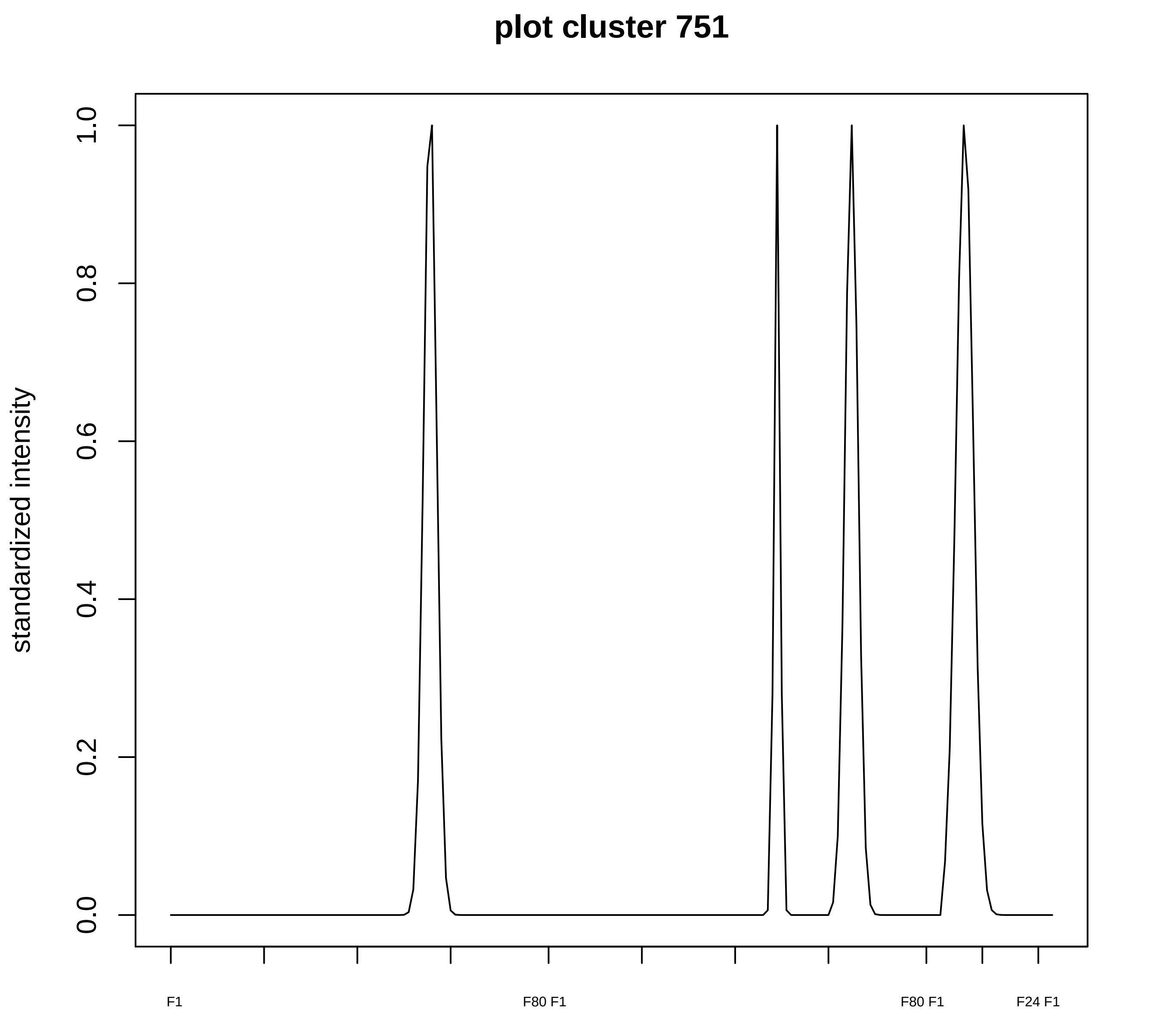

plot cluster 751
standardized intensity
0.0
0.2
0.4
0.6
0.8
1.0
F1
F80 F1
F80 F1
F24 F1

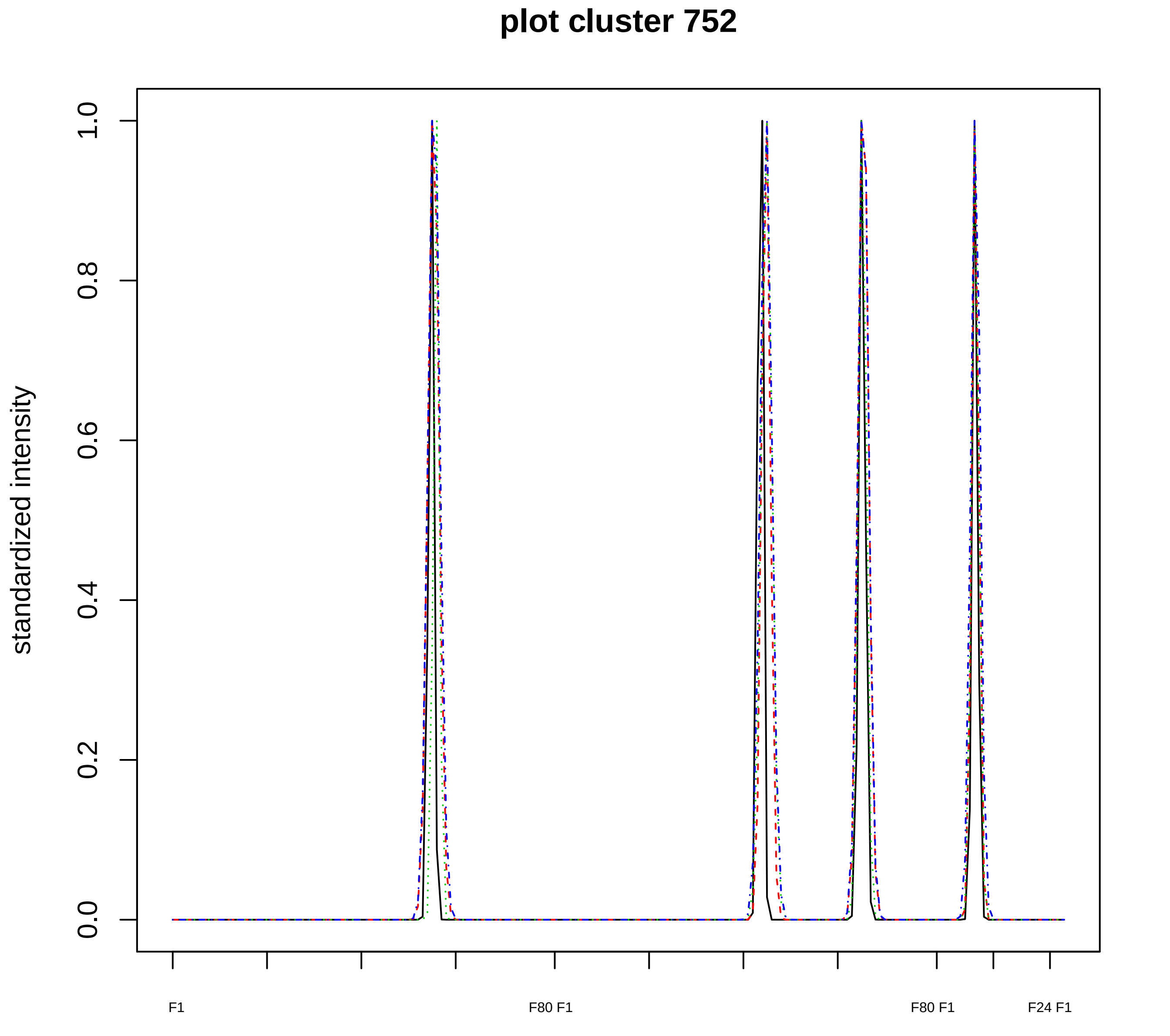
plot cluster 752
standardized intensity
0.0
0.2
0.4
0.6
0.8
1.0
F1
F80 F1
F80 F1
F24 F1

#### Dendrogram of cluster 752

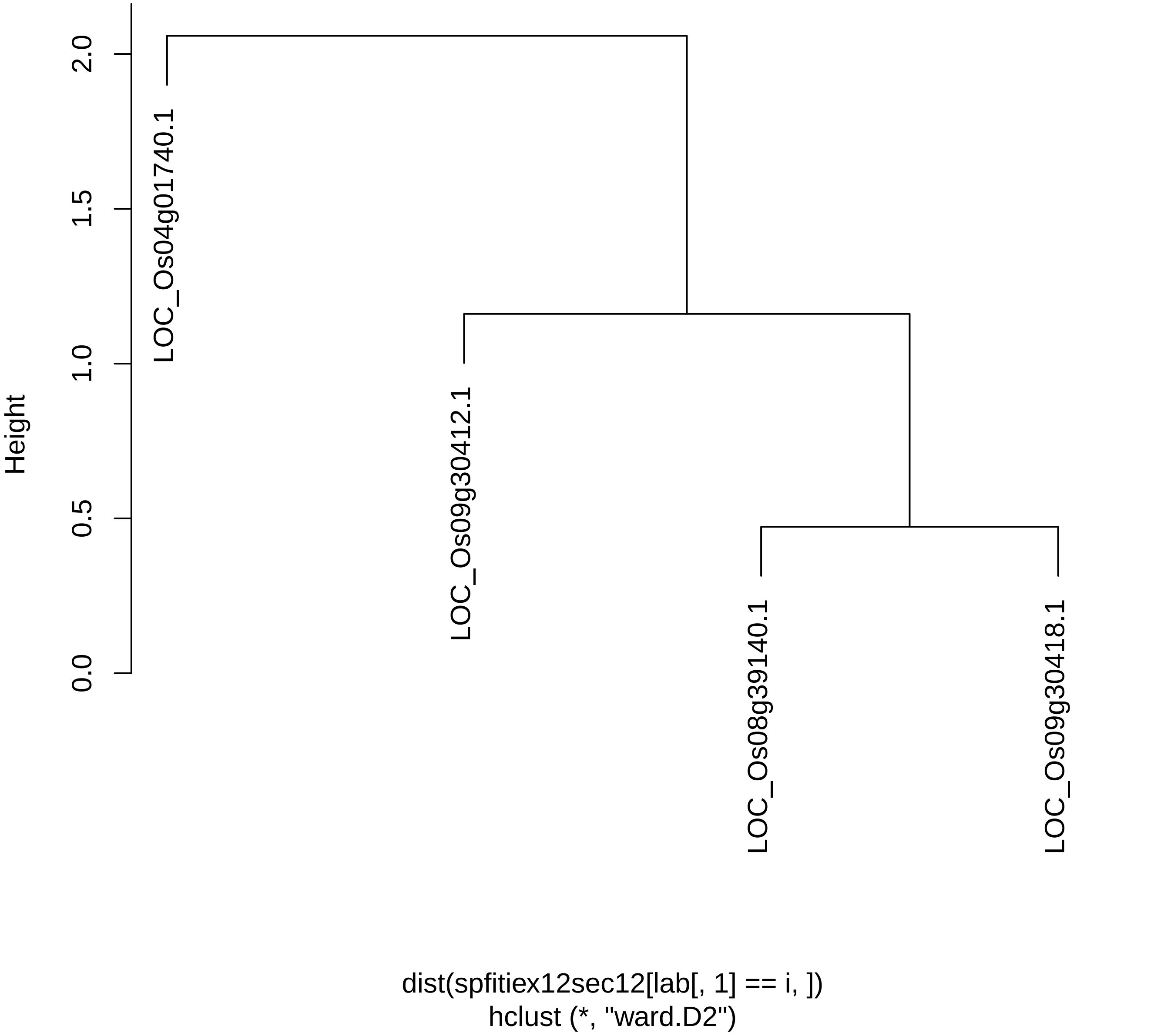

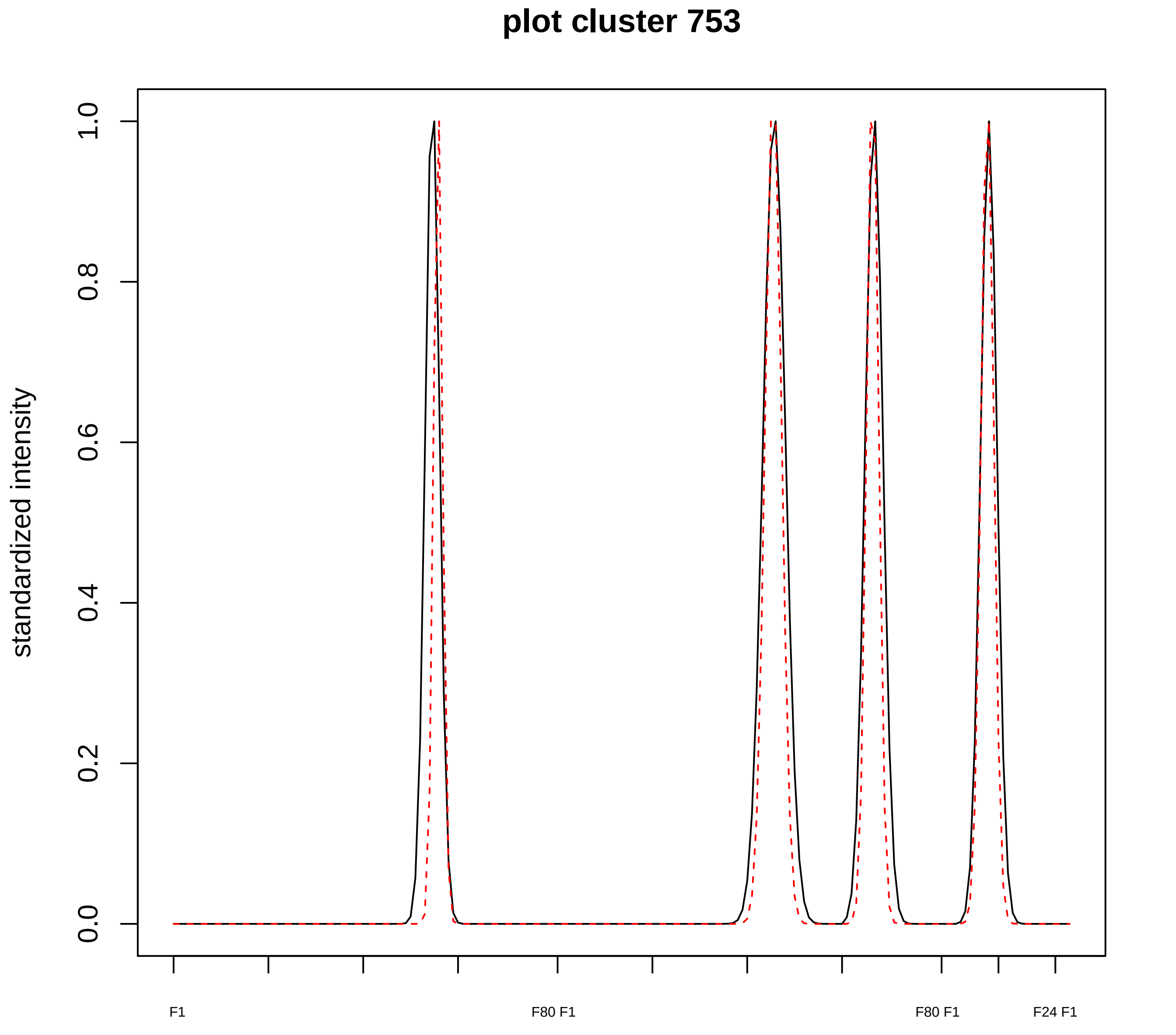
plot cluster 753
standardized intensity
0.0
0.2
0.4
0.6
0.8
1.0
F1
F80 F1
F80 F1
F24 F1

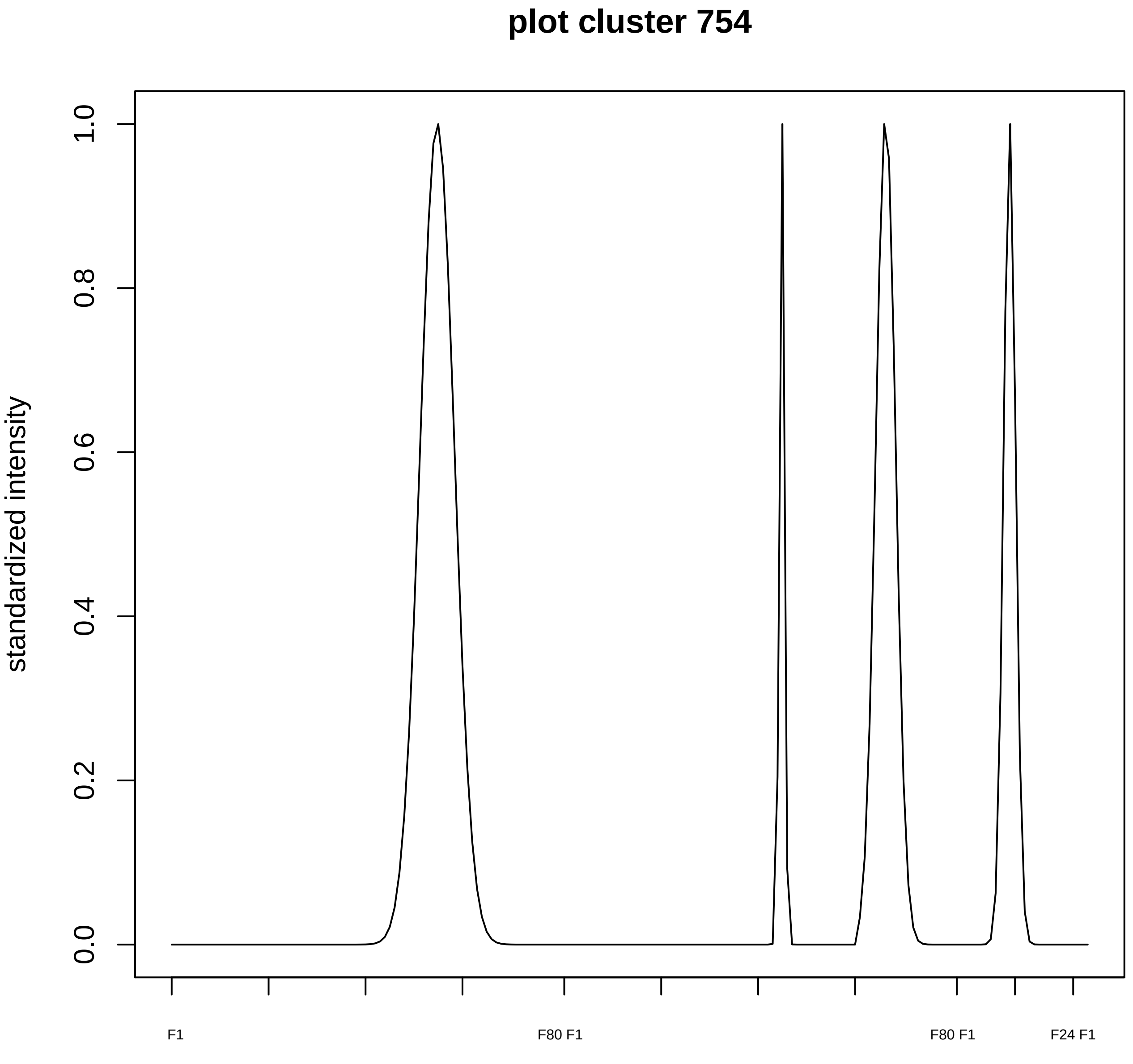
plot cluster 754
standardized intensity
0.0
0.2
0.4
0.6
0.8
1.0
F1
F80 F1
F80 F1
F24 F1

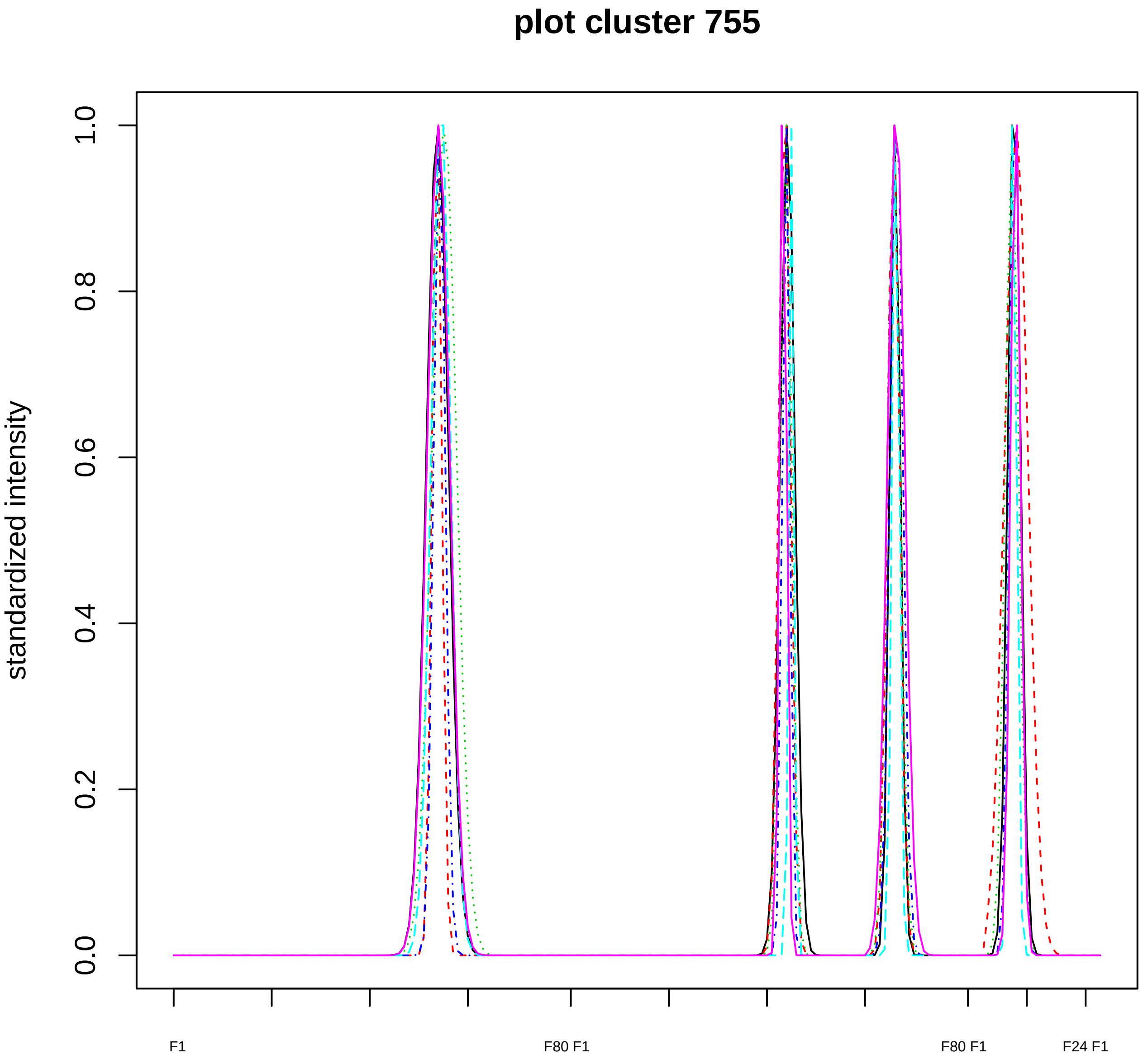

plot cluster 755
standardized intensity
0.0
0.2
0.4
0.6
0.8
1.0
F1
F80 F1
F80 F1
F24 F1

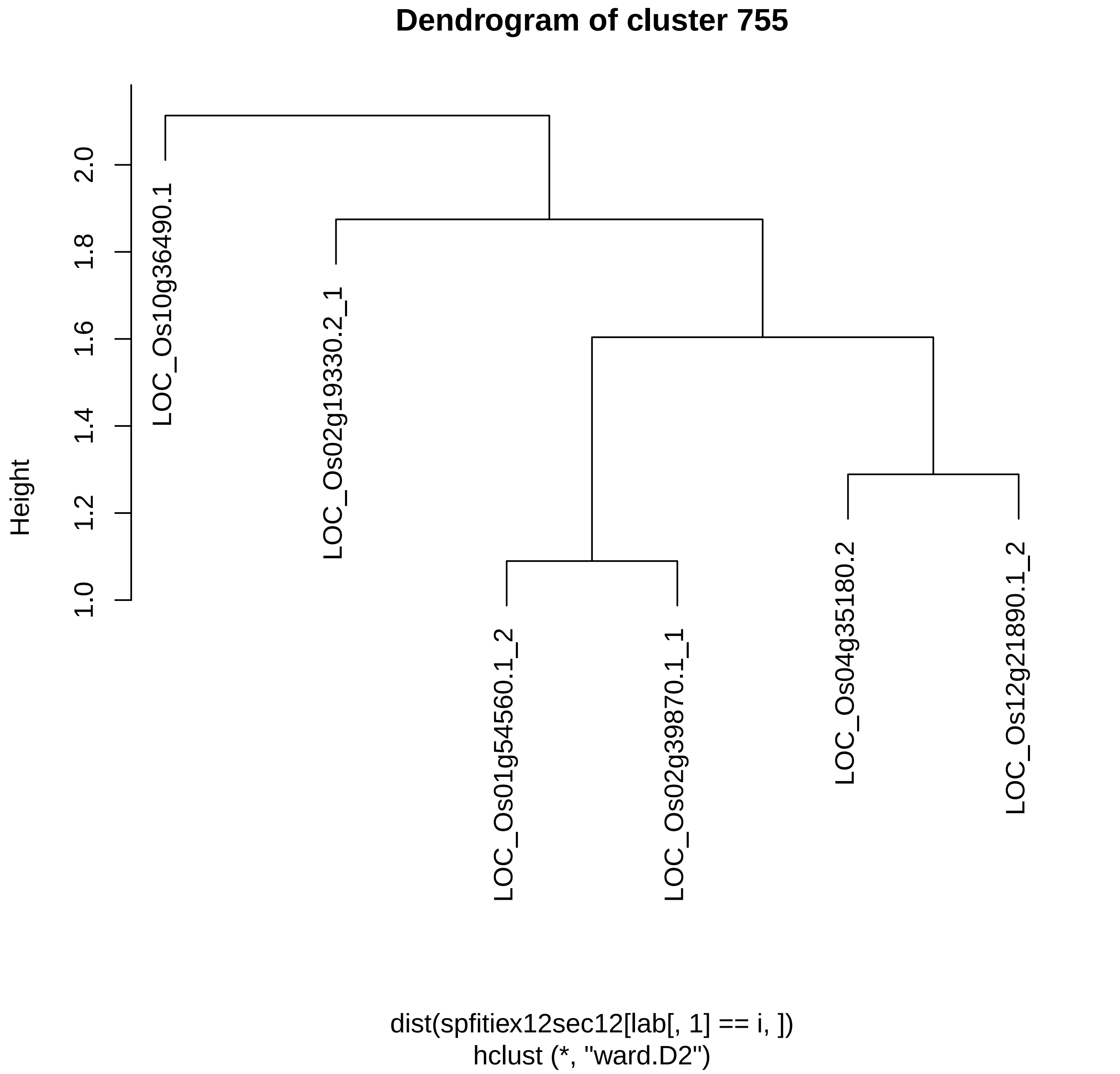

Dendrogram of cluster 755
Height
2.0
1.8
1.6
1.4
1.2
1.0
LOC_Os10g36490.1
LOC_Os02g19330.2_1
LOC_Os01g54560.1_2
LOC_Os02g39870.1_1
LOC_Os04g35180.2
LOC_Os12g21890.1_2
dist(spfitiex12sec12[lab[, 1] == i, ])
hclust (*, "ward.D2")

### plot cluster 756

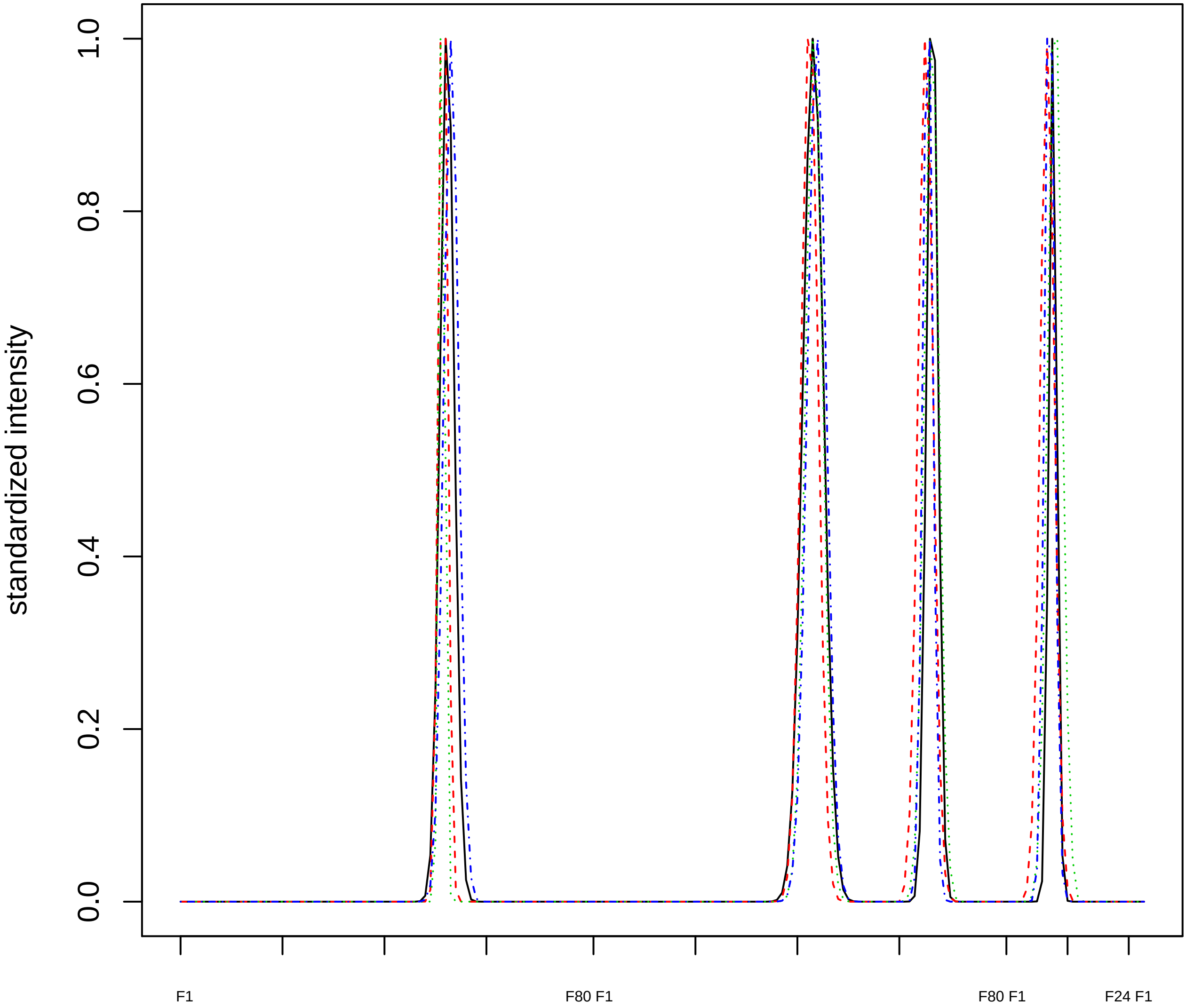

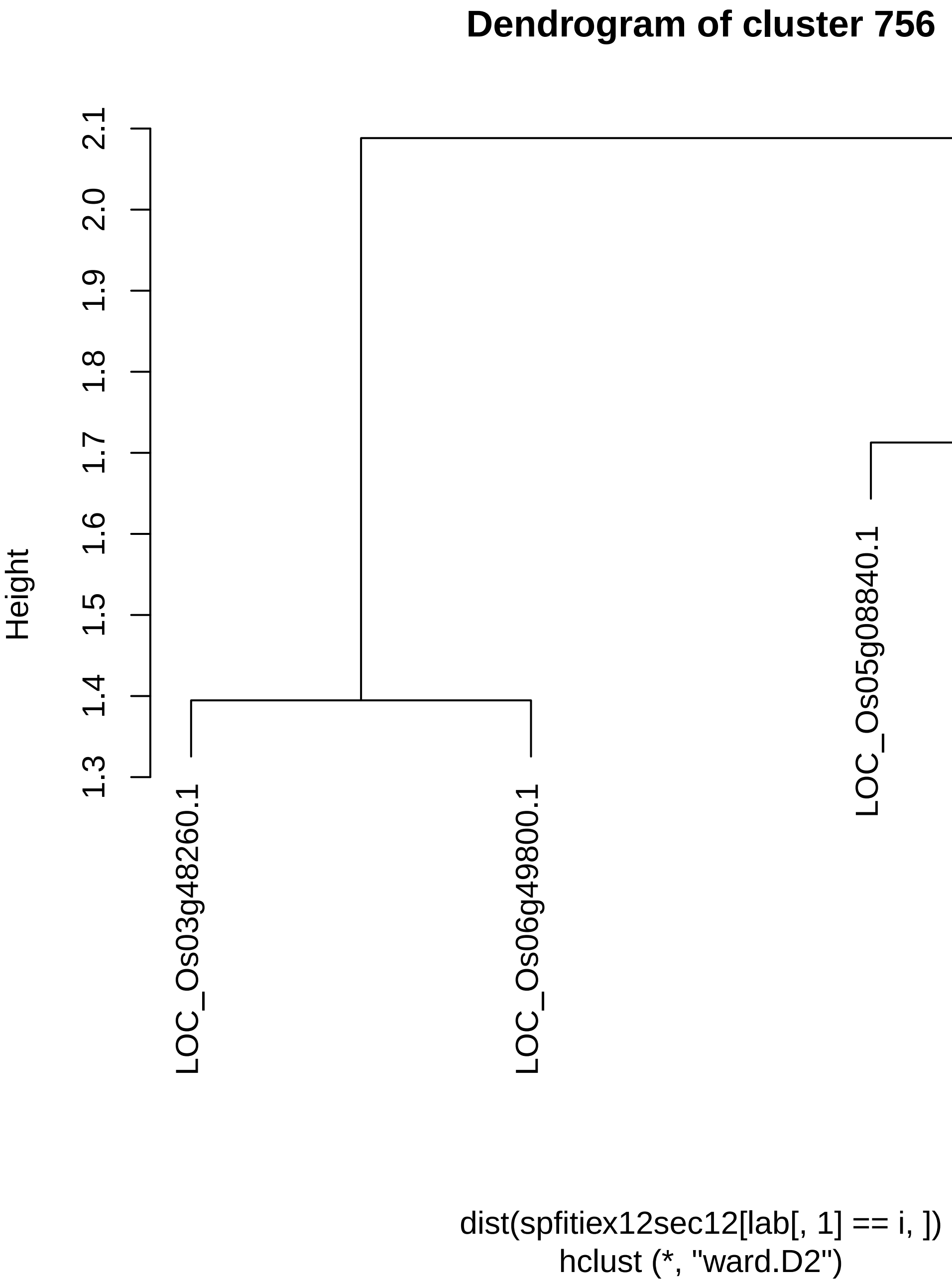
Dendrogram of cluster 756
Height
2.1
2.0
1.9
1.8
1.7
1.6
1.5
1.4
1.3
LOC_Os03g48260.1
LOC_Os06g49800.1
LOC_Os05g08840.1
LOC_Os05g44100.1
dist(spfitiex12sec12[lab[, 1] == i, ])
hclust (*, "ward.D2")

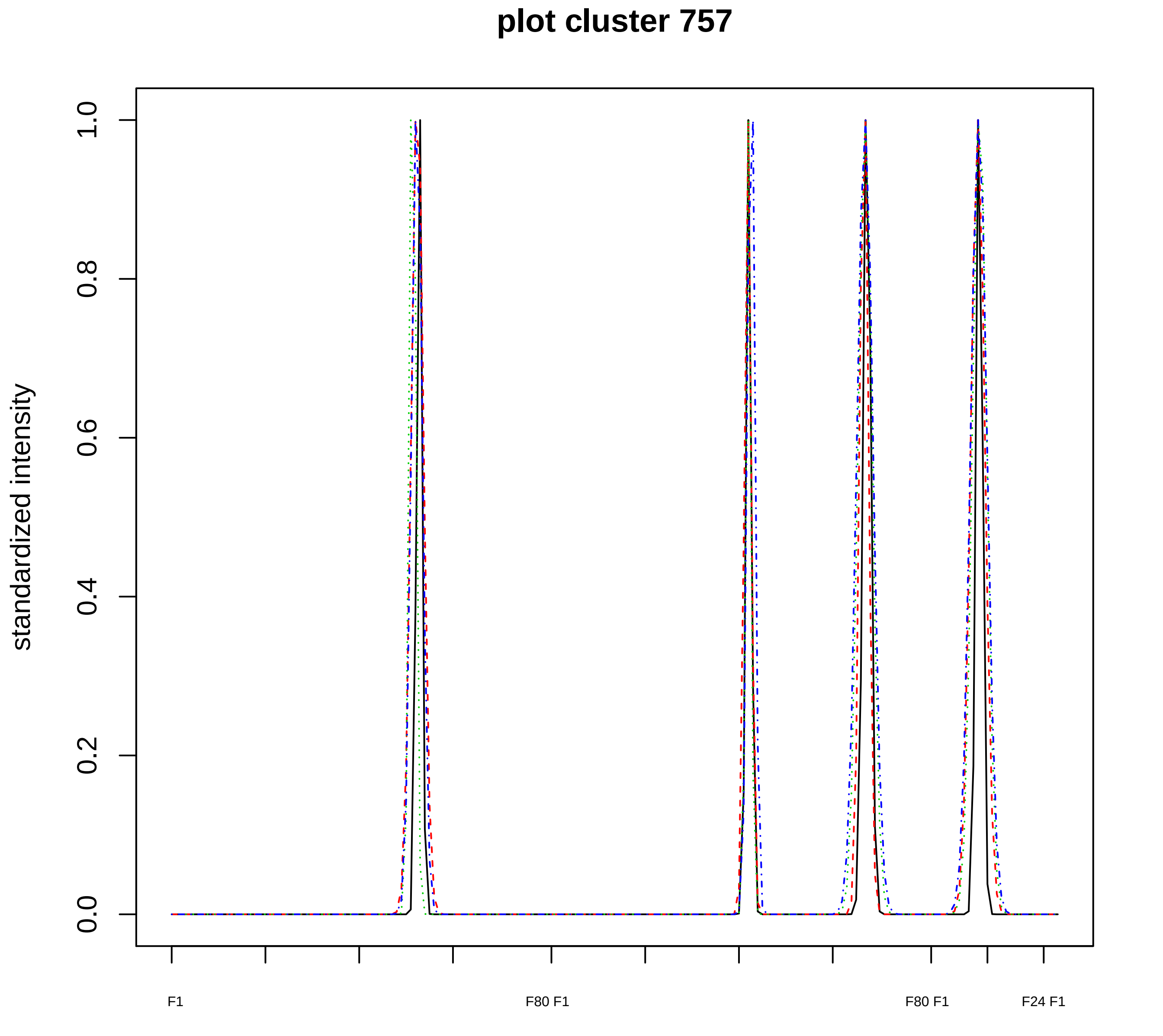

plot cluster 757
standardized intensity
0.0
0.2
0.4
0.6
0.8
1.0
F1
F80 F1
F80 F1
F24 F1

### Dendrogram of cluster 757

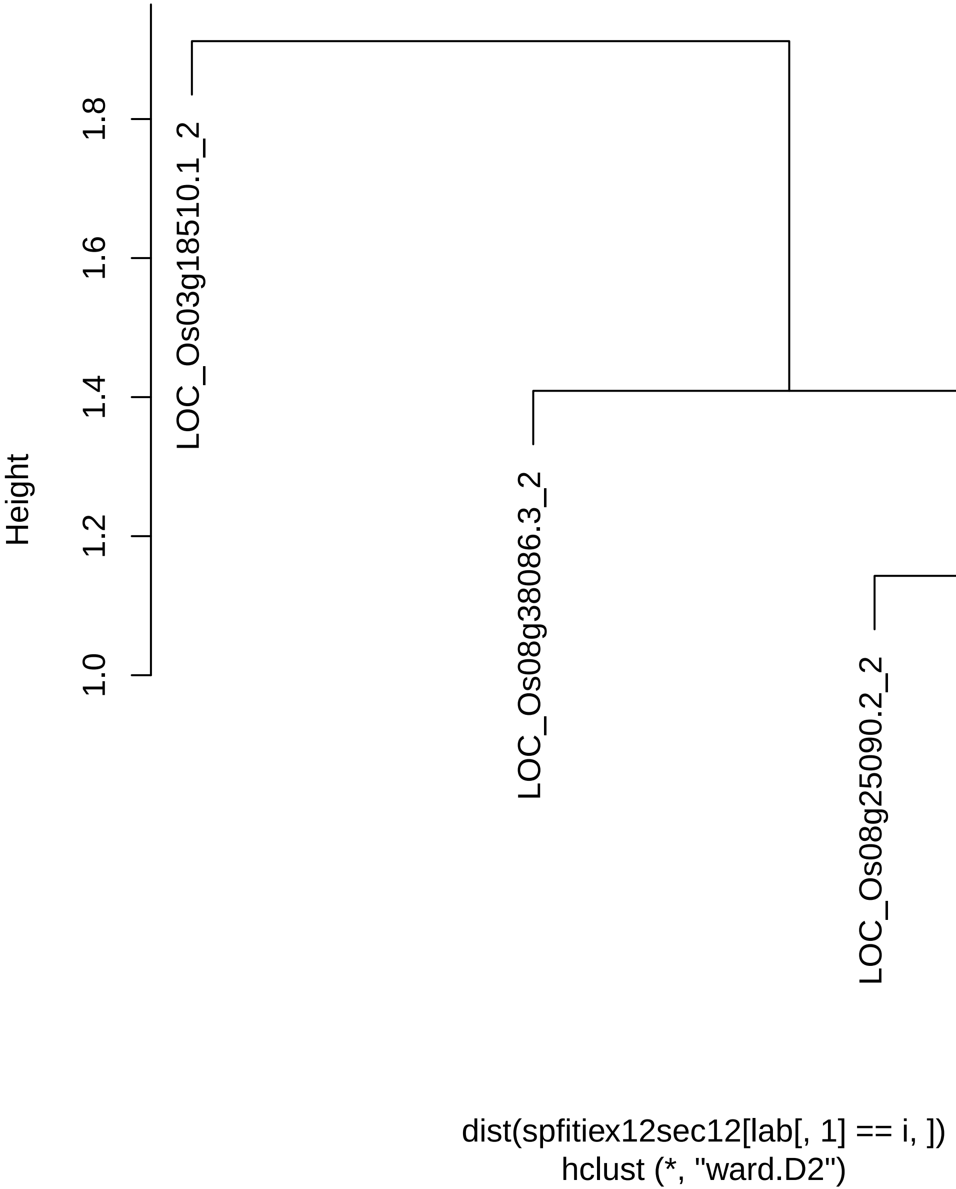

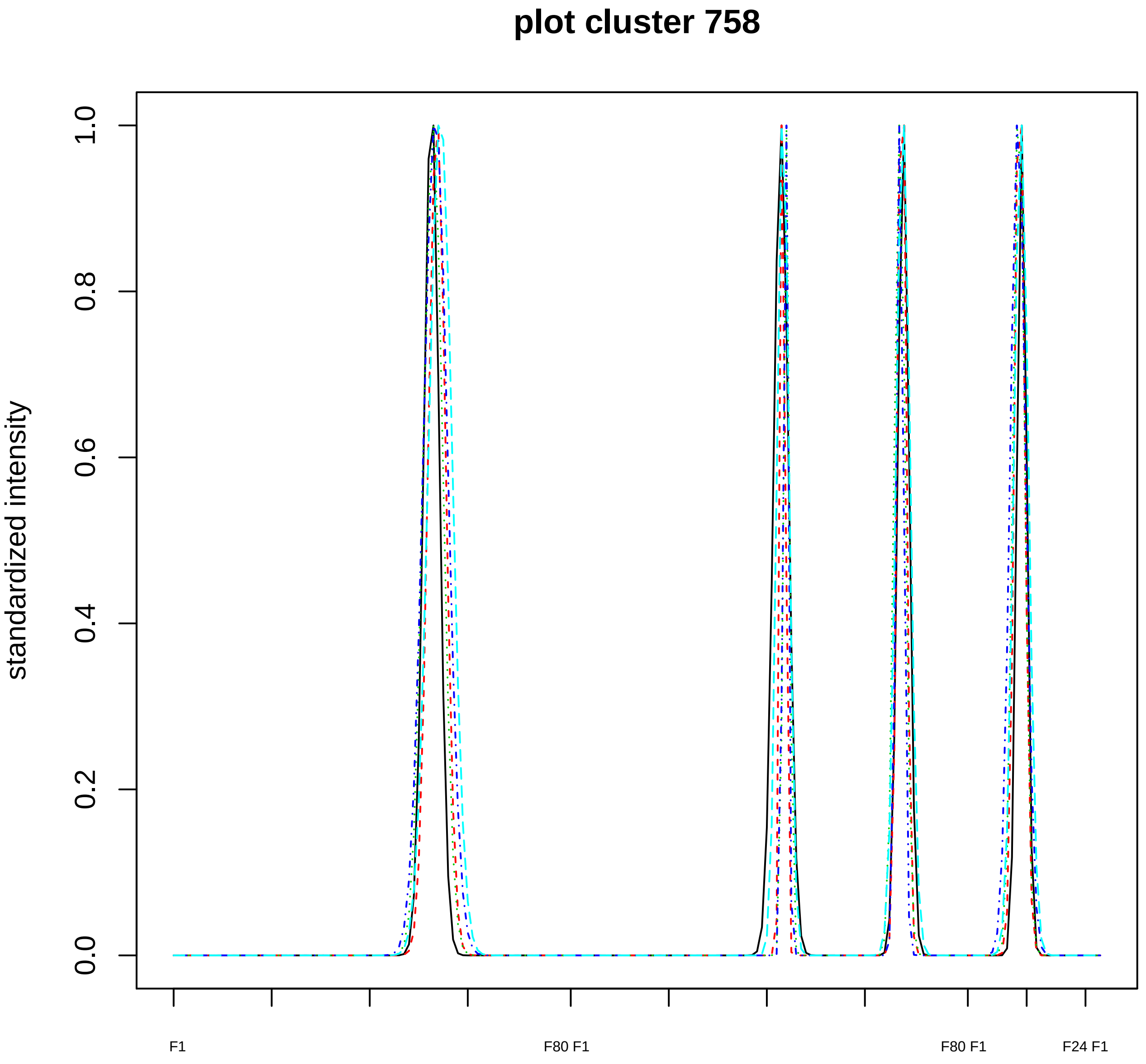

plot cluster 758
standardized intensity
0.0
0.2
0.4
0.6
0.8
1.0
F1
F80 F1
F80 F1
F24 F1

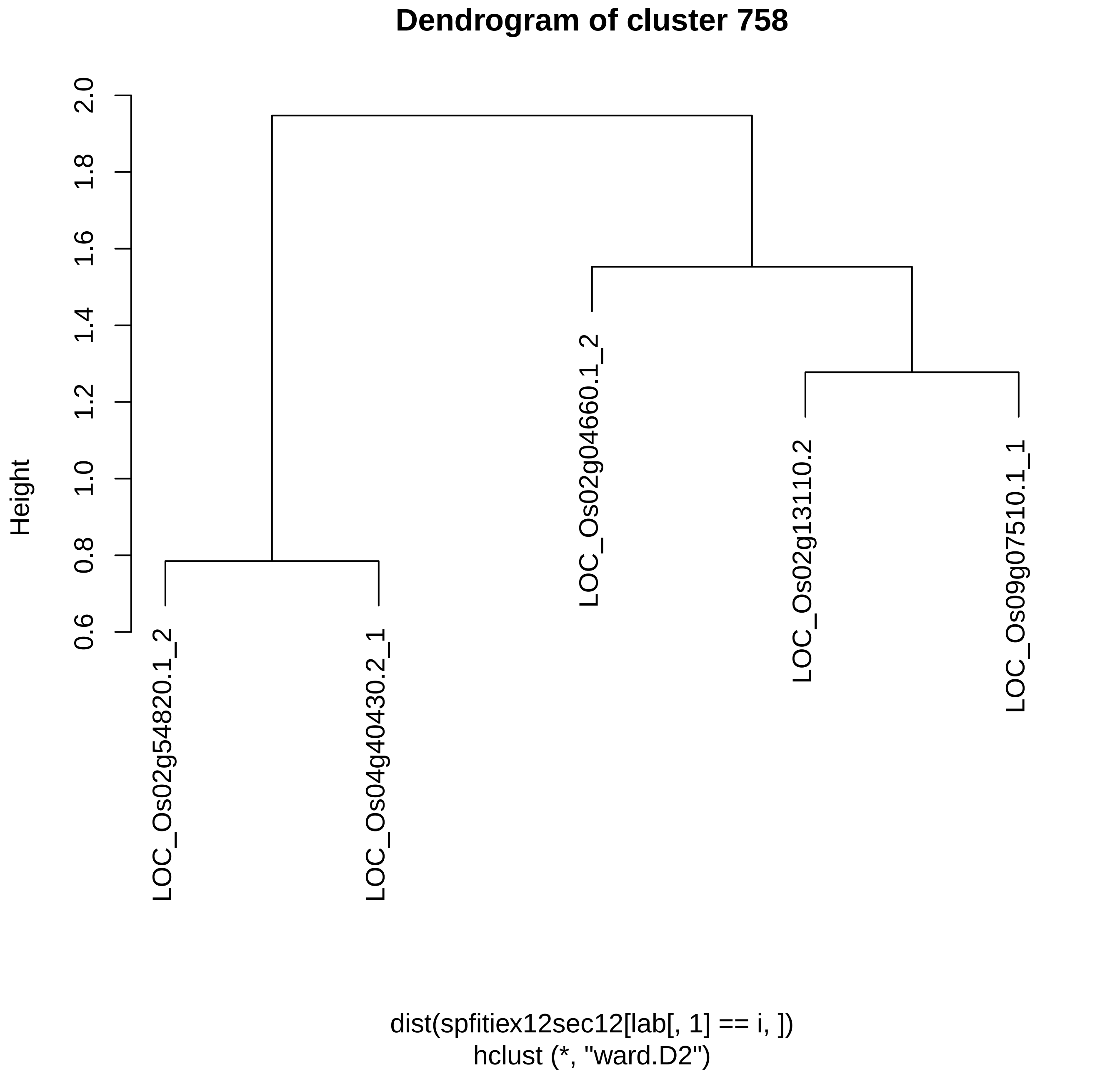
Dendrogram of cluster 758
Height
0.6
0.8
1.0
1.2
1.4
1.6
1.8
2.0
LOC_Os02g54820.1_2
LOC_Os04g40430.2_1
LOC_Os02g04660.1_2
LOC_Os02g13110.2
LOC_Os09g07510.1_1
dist(spfitiex12sec12[lab[, 1] == i, ])
hclust (*, "ward.D2")

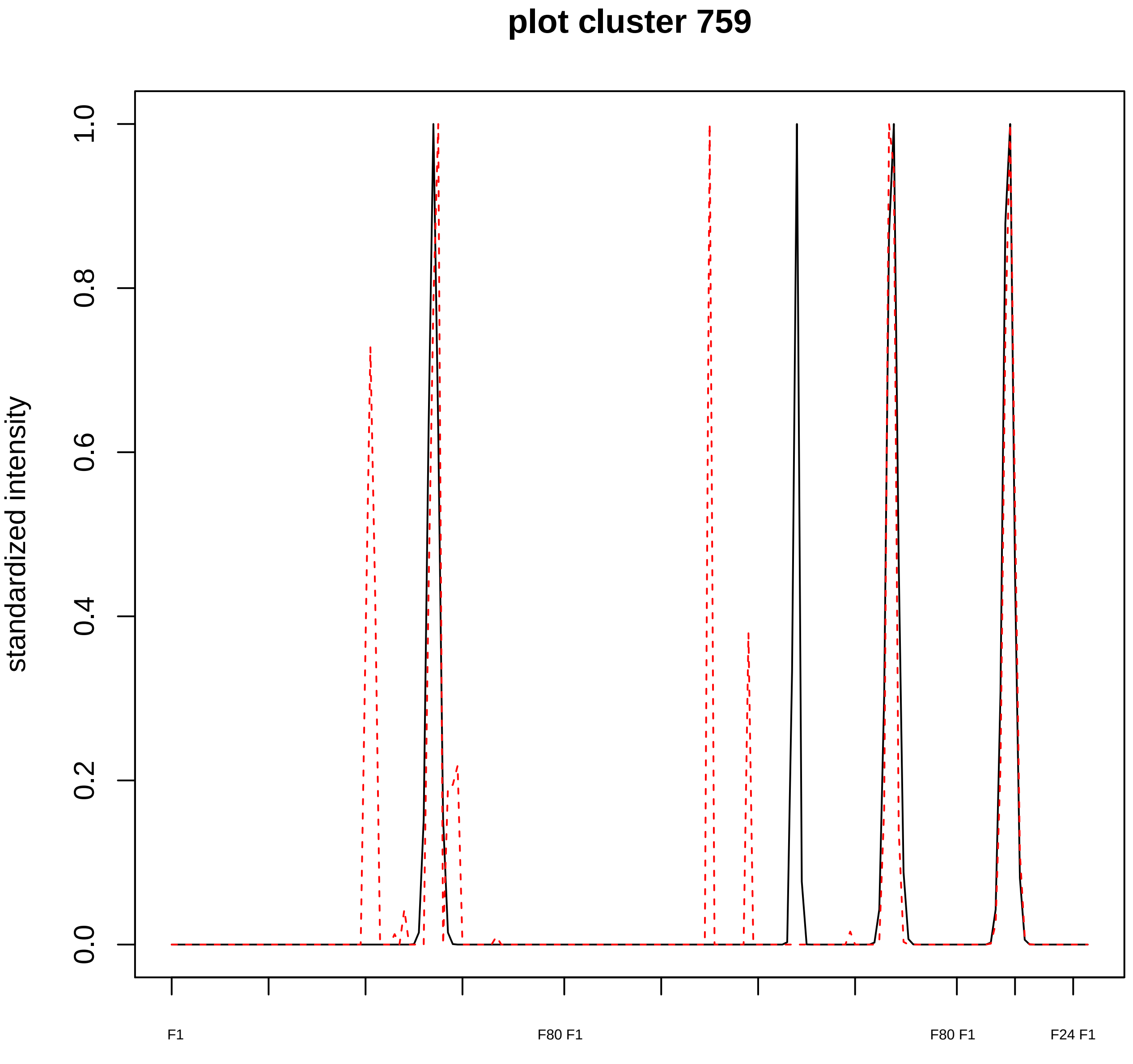
plot cluster 759
standardized intensity
0.0
0.2
0.4
0.6
0.8
1.0
F1
F80 F1
F80 F1
F24 F1

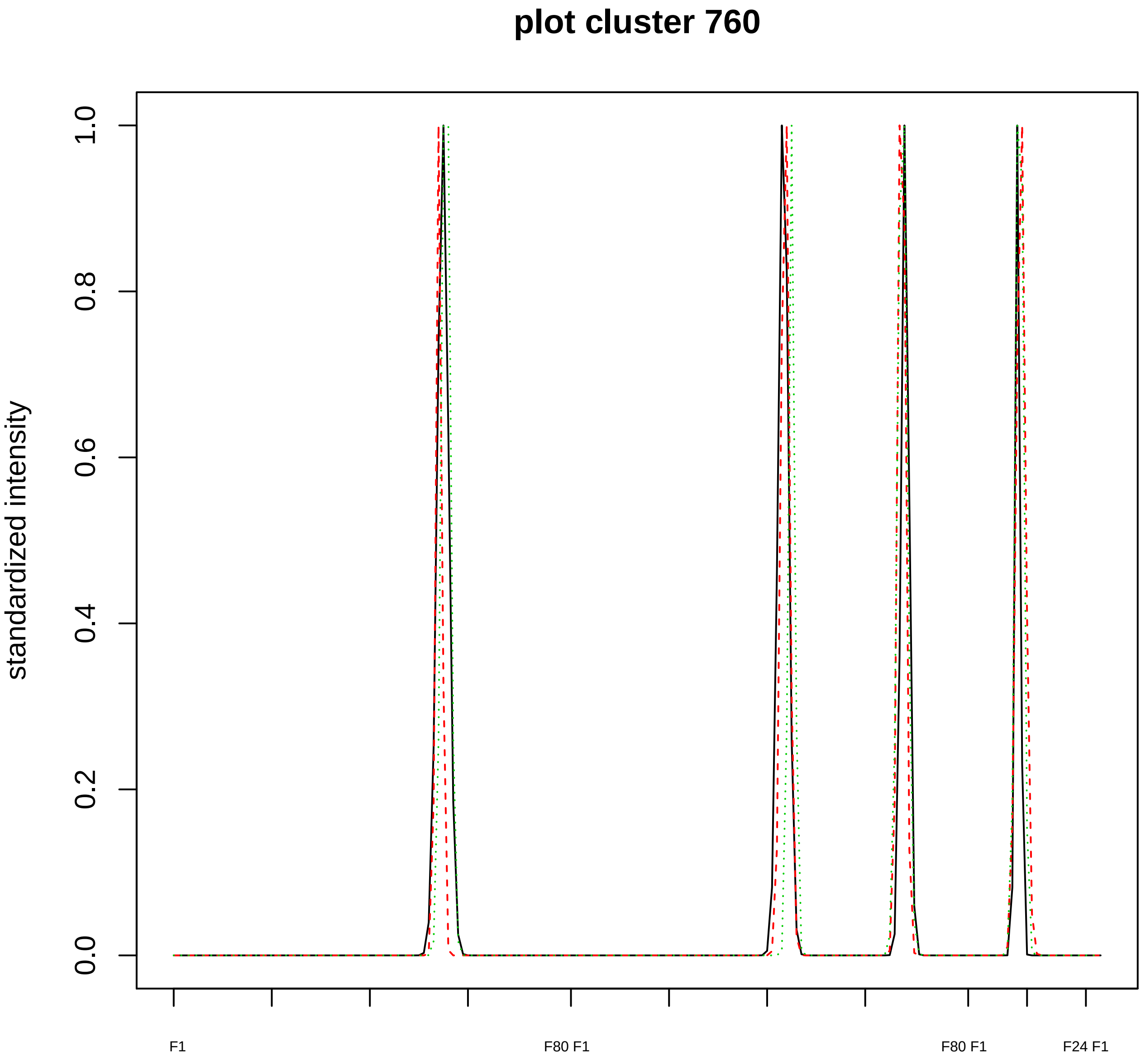

plot cluster 760
standardized intensity
0.0
0.2
0.4
0.6
0.8
1.0
F1
F80 F1
F80 F1
F24 F1

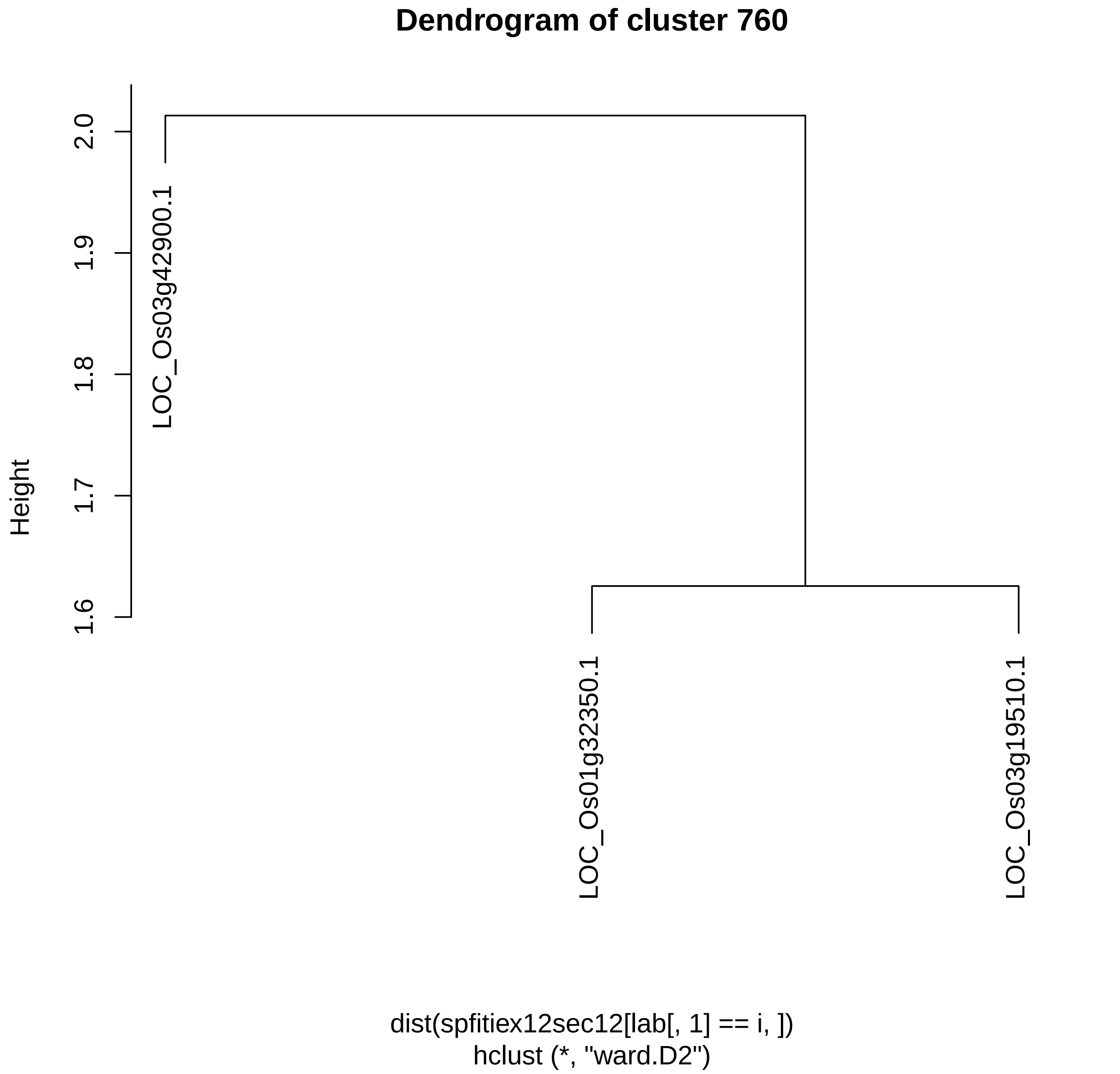
Dendrogram of cluster 760
Height
2.0
1.9
1.8
1.7
1.6
LOC_Os03g42900.1
LOC_Os01g32350.1
LOC_Os03g19510.1
dist(spfitiex12sec12[lab[, 1] == i, ])
hclust (*, "ward.D2")

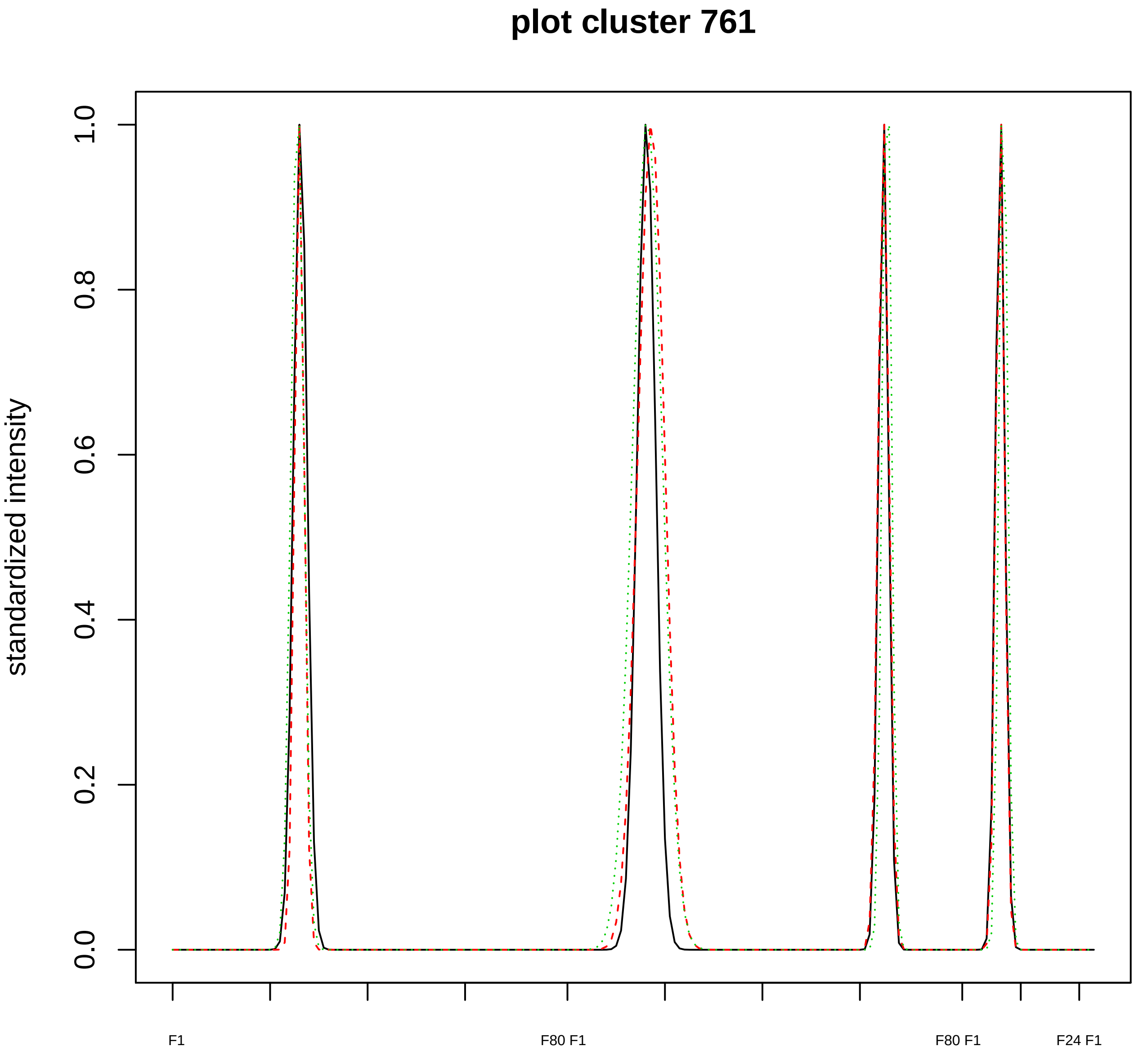

plot cluster 761
standardized intensity
0.0
0.2
0.4
0.6
0.8
1.0
F1
F80 F1
F80 F1
F24 F1

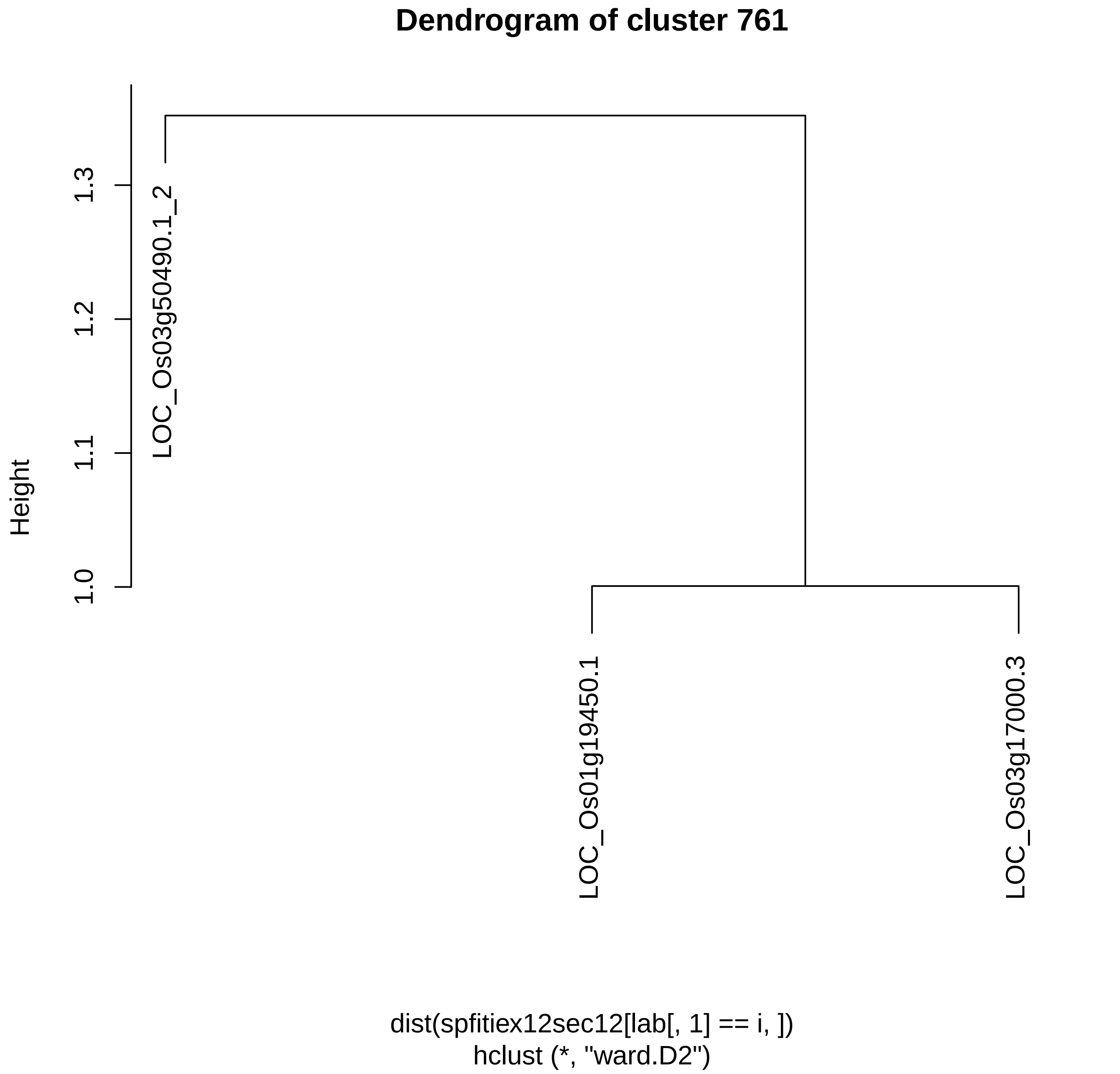
Dendrogram of cluster 761
Height
1.0
1.1
1.2
1.3
LOC_Os03g50490.1_2
LOC_Os01g19450.1
LOC_Os03g17000.3
dist(spfitiex12sec12[lab[, 1] == i, ])
hclust (*, "ward.D2")

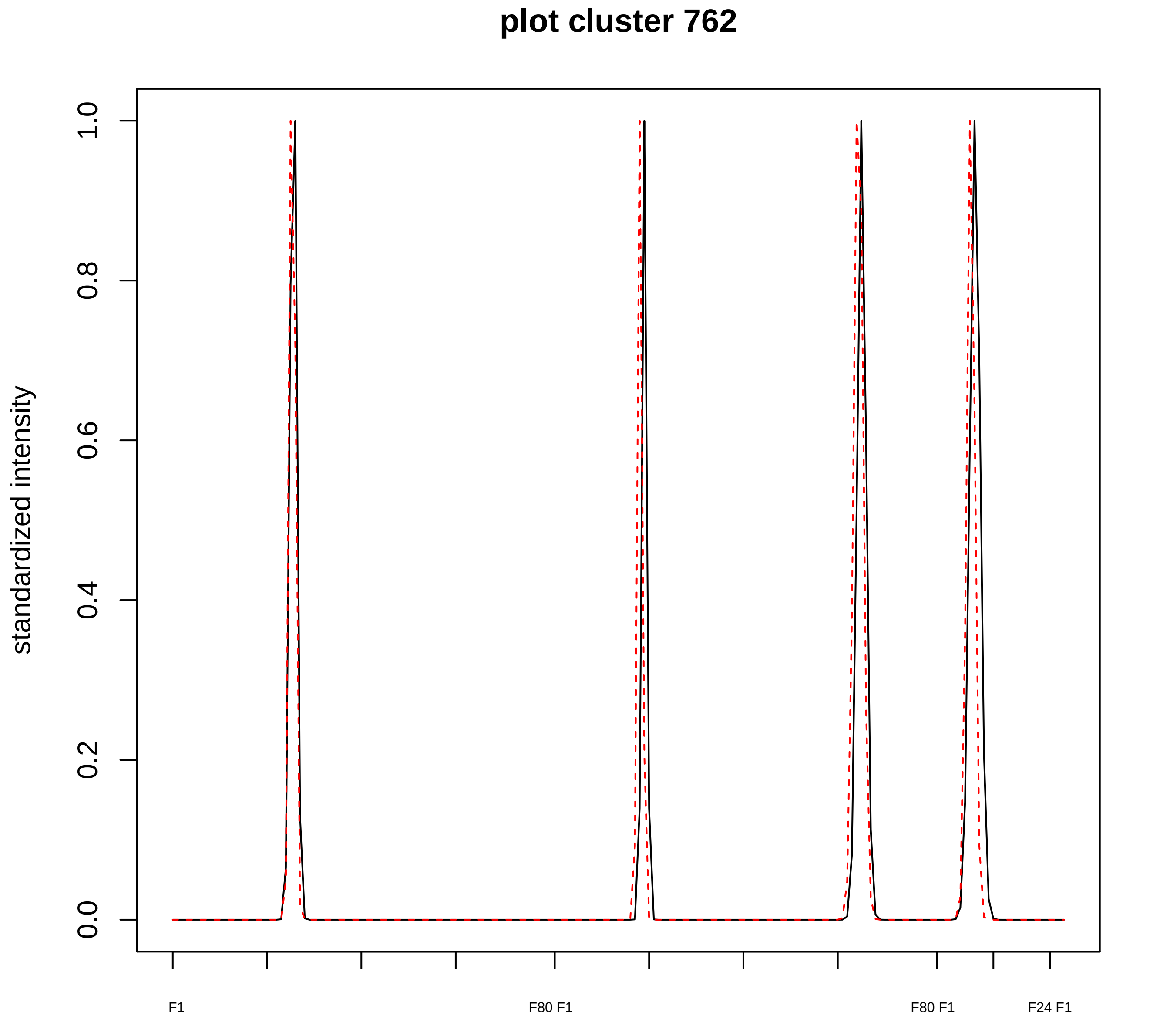

plot cluster 762
standardized intensity
1.0
0.8
0.6
0.4
0.2
0.0
F1
F80 F1
F80 F1
F24 F1

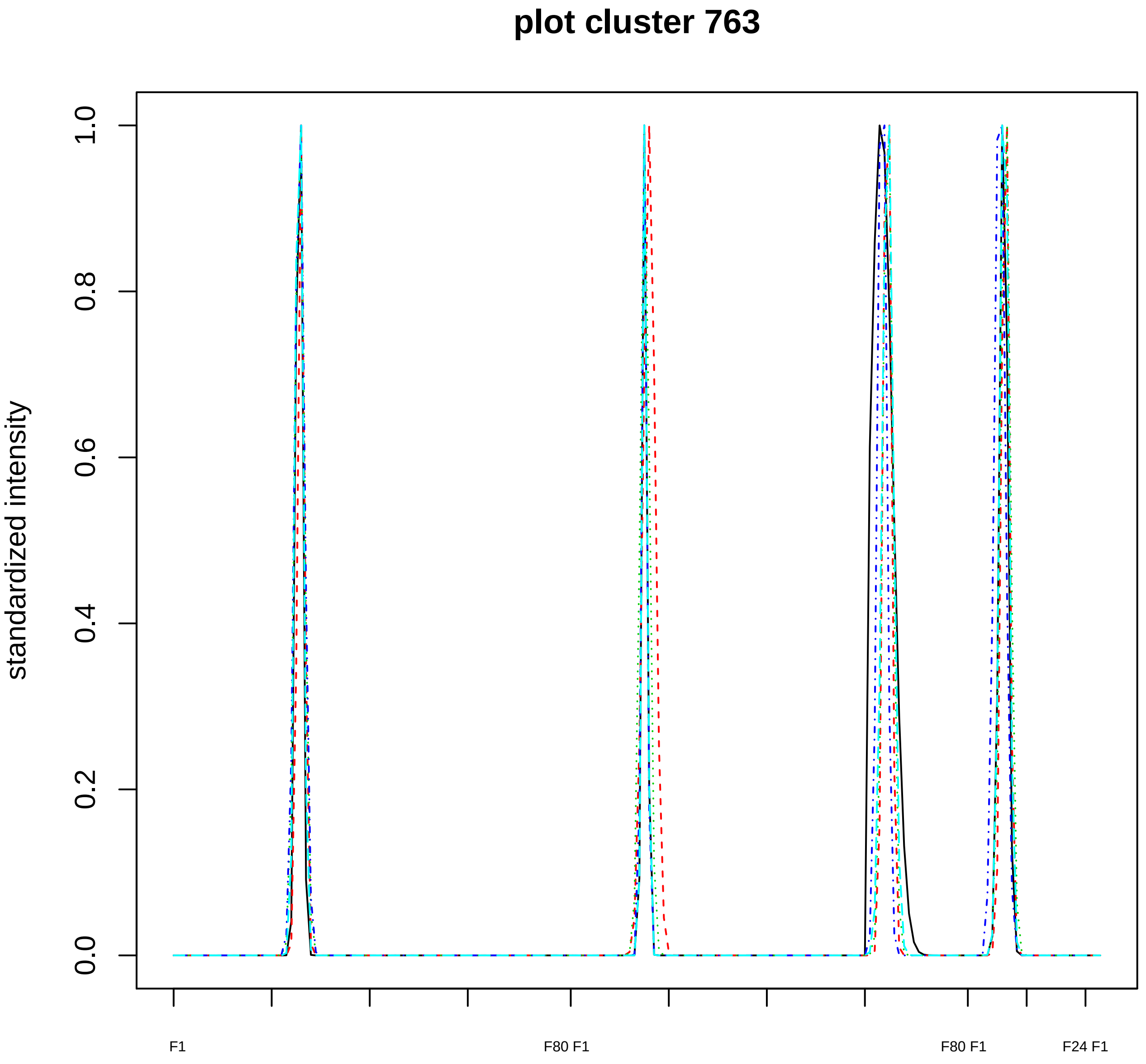

plot cluster 763
standardized intensity
0.0
0.2
0.4
0.6
0.8
1.0
F1
F80 F1
F80 F1
F24 F1

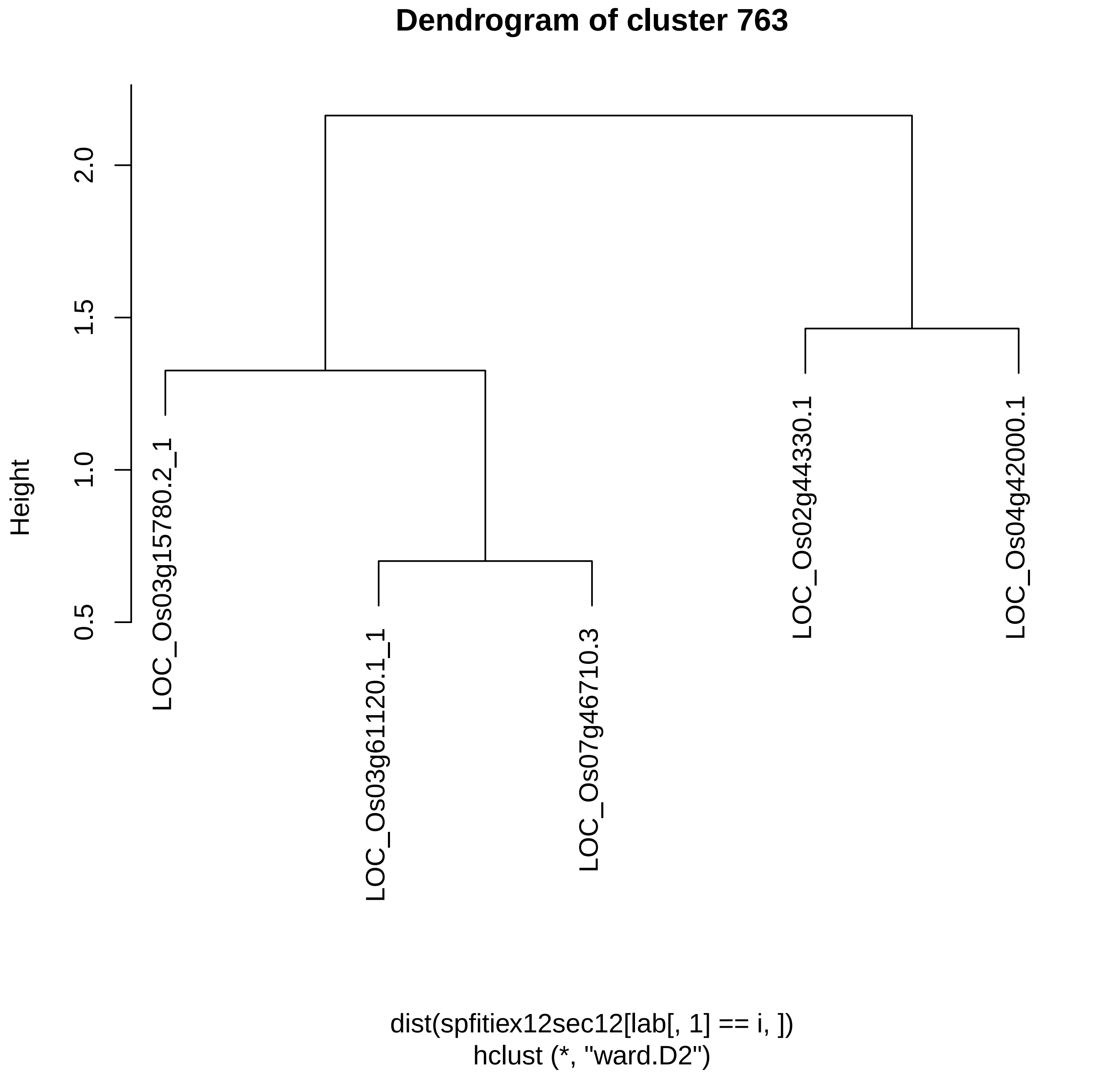

Dendrogram of cluster 763
Height
0.5
1.0
1.5
2.0
LOC_Os03g15780.2_1
LOC_Os03g61120.1_1
LOC_Os07g46710.3
LOC_Os02g44330.1
LOC_Os04g42000.1
dist(spfitiex12sec12[lab[, 1] == i, ])
hclust (*, "ward.D2")

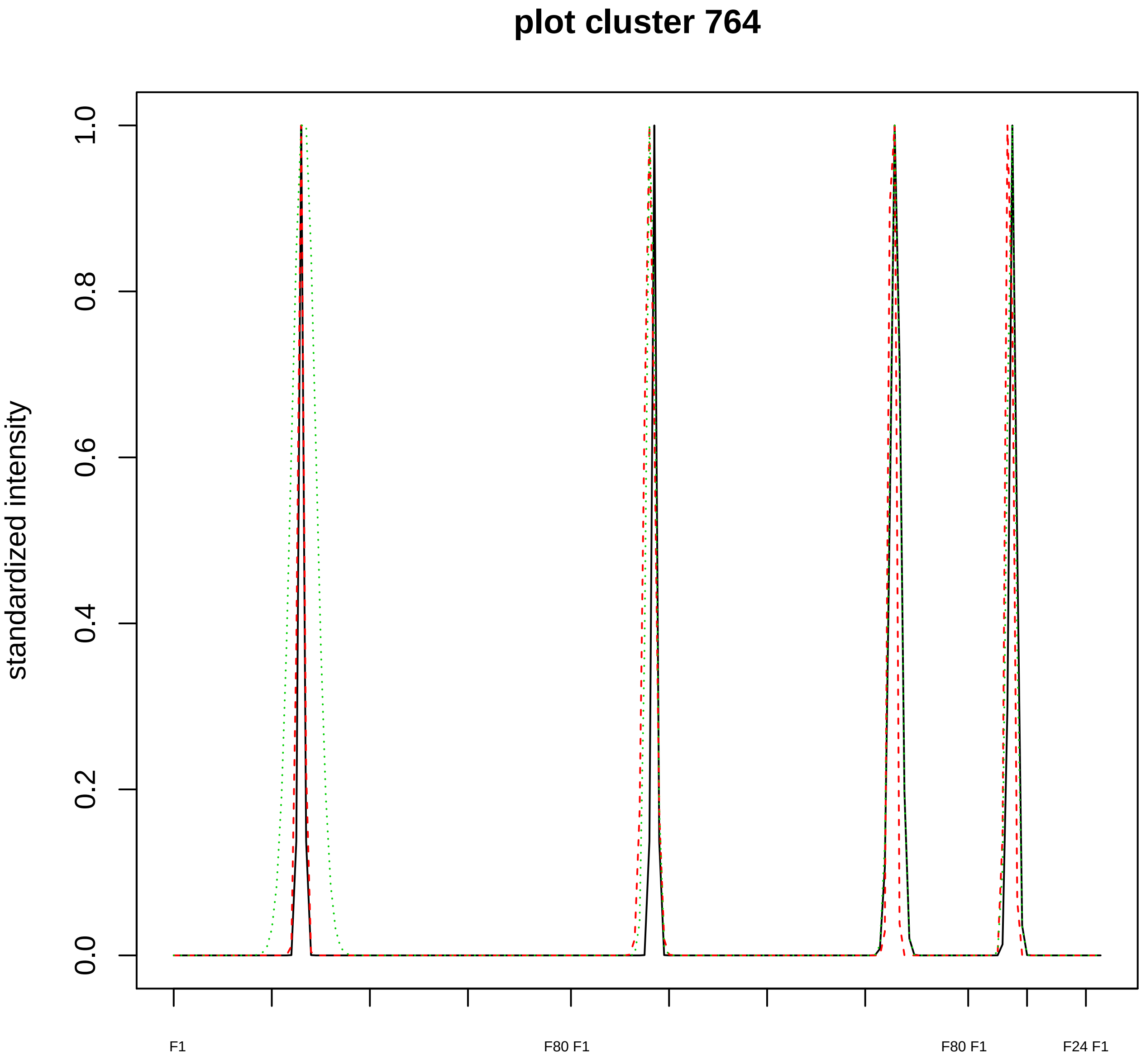

plot cluster 764
standardized intensity
0.0
0.2
0.4
0.6
0.8
1.0
F1
F80 F1
F80 F1
F24 F1

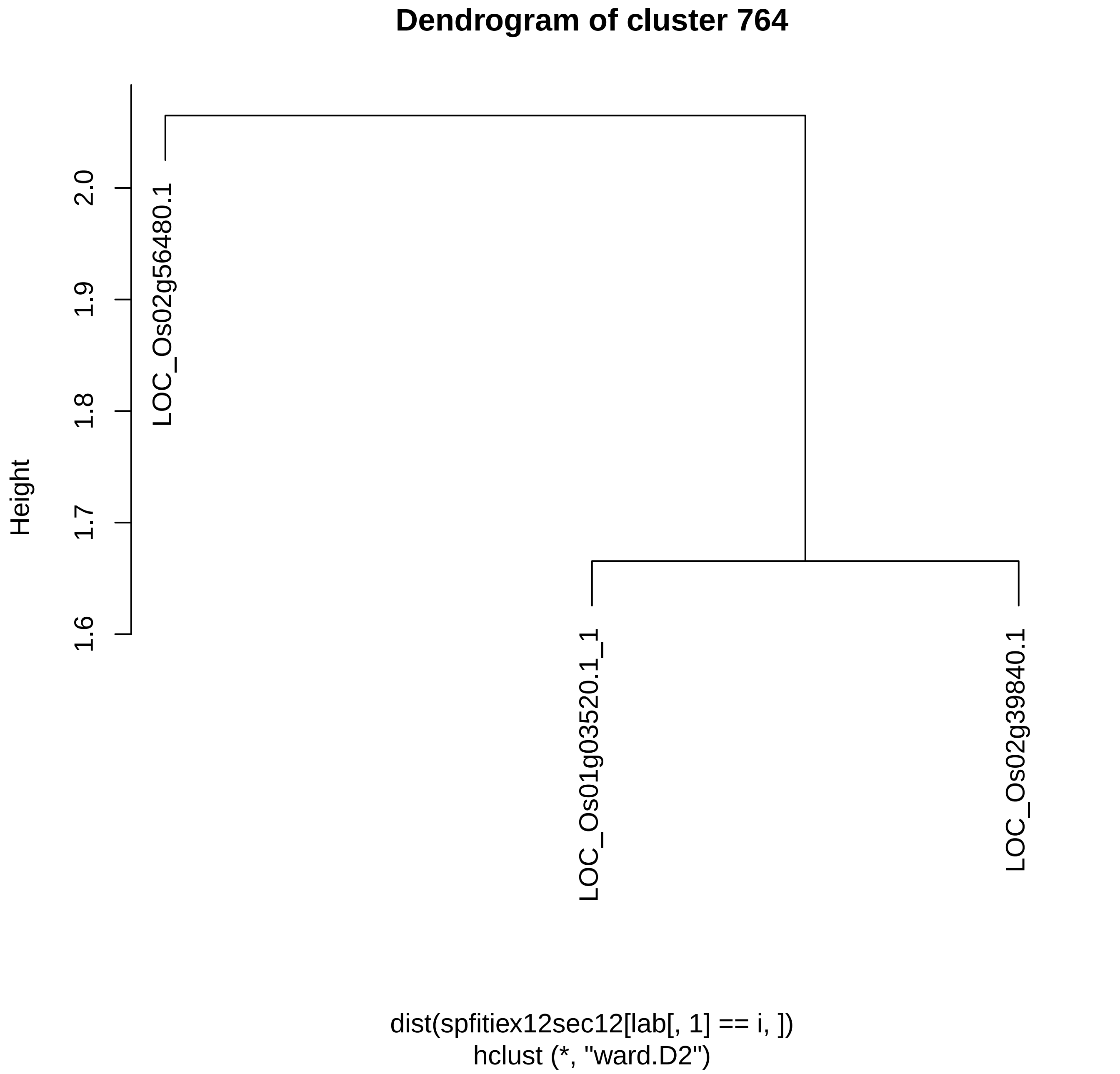
Dendrogram of cluster 764
Height
2.0
1.9
1.8
1.7
1.6
LOC_Os02g56480.1
LOC_Os01g03520.1_1
LOC_Os02g39840.1
dist(spfitiex12sec12[lab[, 1] == i, ])
hclust (*, "ward.D2")

### plot cluster 765

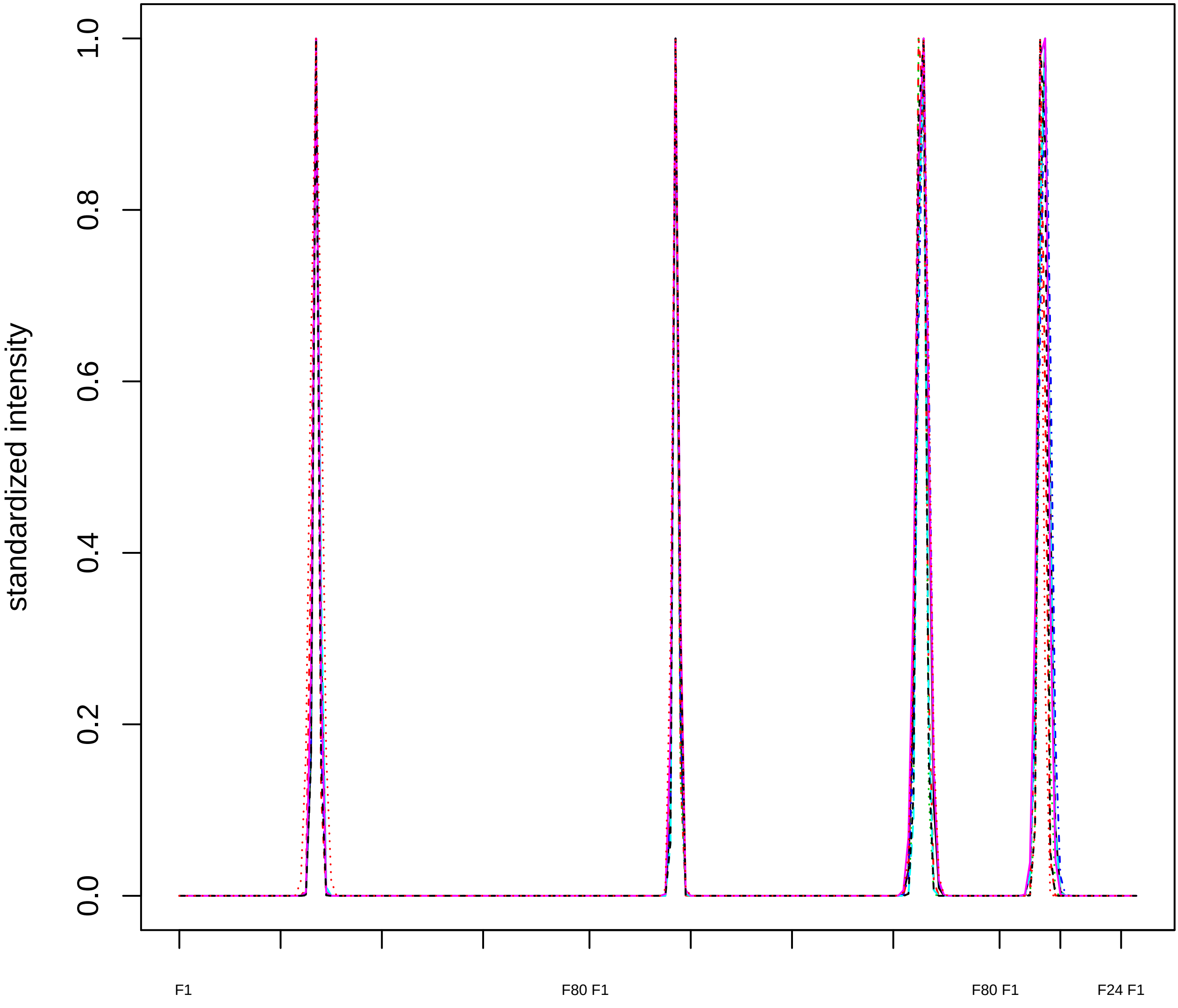

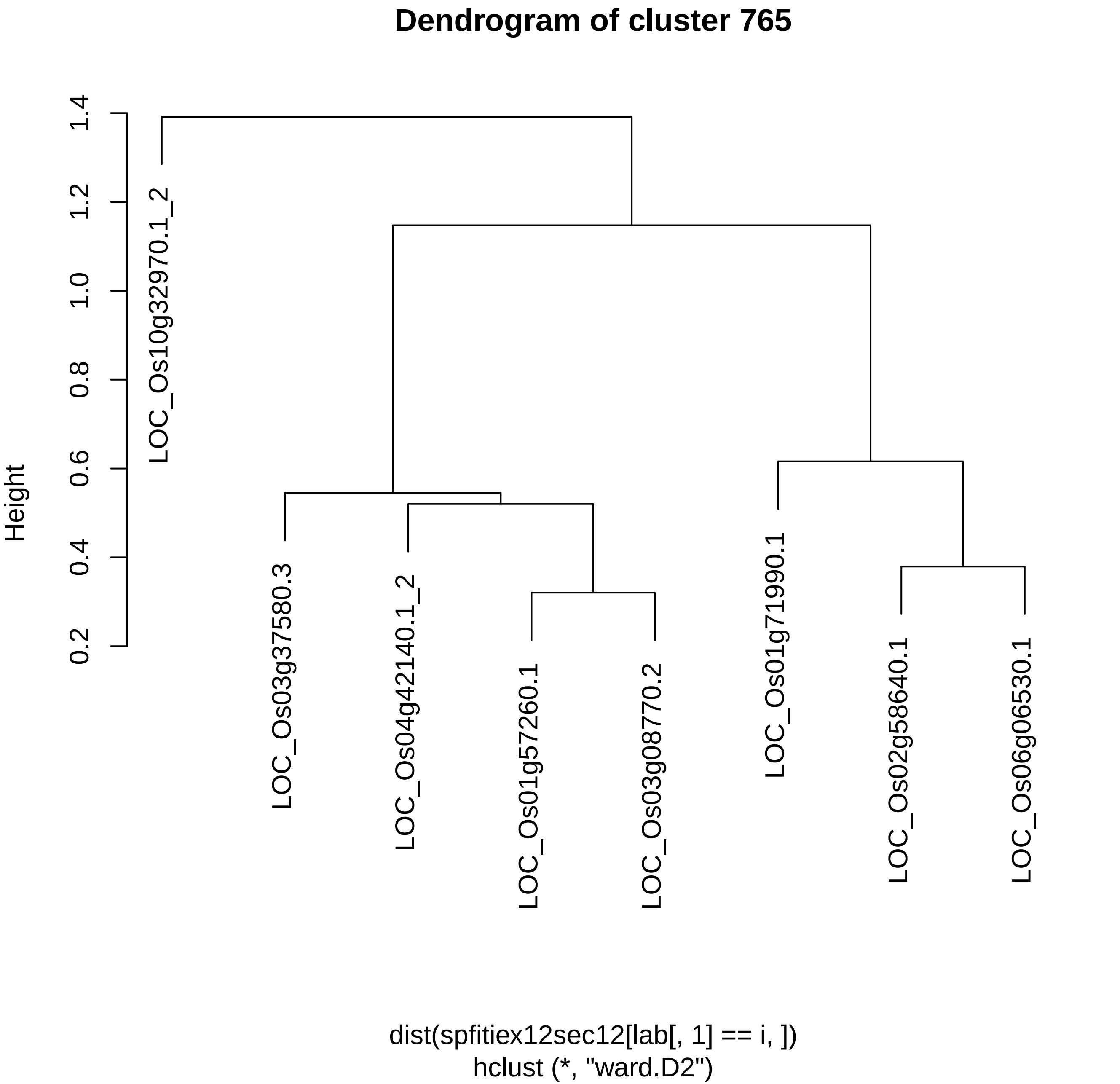
Dendrogram of cluster 765
Height
0.2
0.4
0.6
0.8
1.0
1.2
1.4
LOC_Os10g32970.1_2
LOC_Os03g37580.3
LOC_Os04g42140.1_2
LOC_Os01g57260.1
LOC_Os03g08770.2
LOC_Os01g71990.1
LOC_Os02g58640.1
LOC_Os06g06530.1
dist(spfitiex12sec12[lab[, 1] == i, ])
hclust (*, "ward.D2")

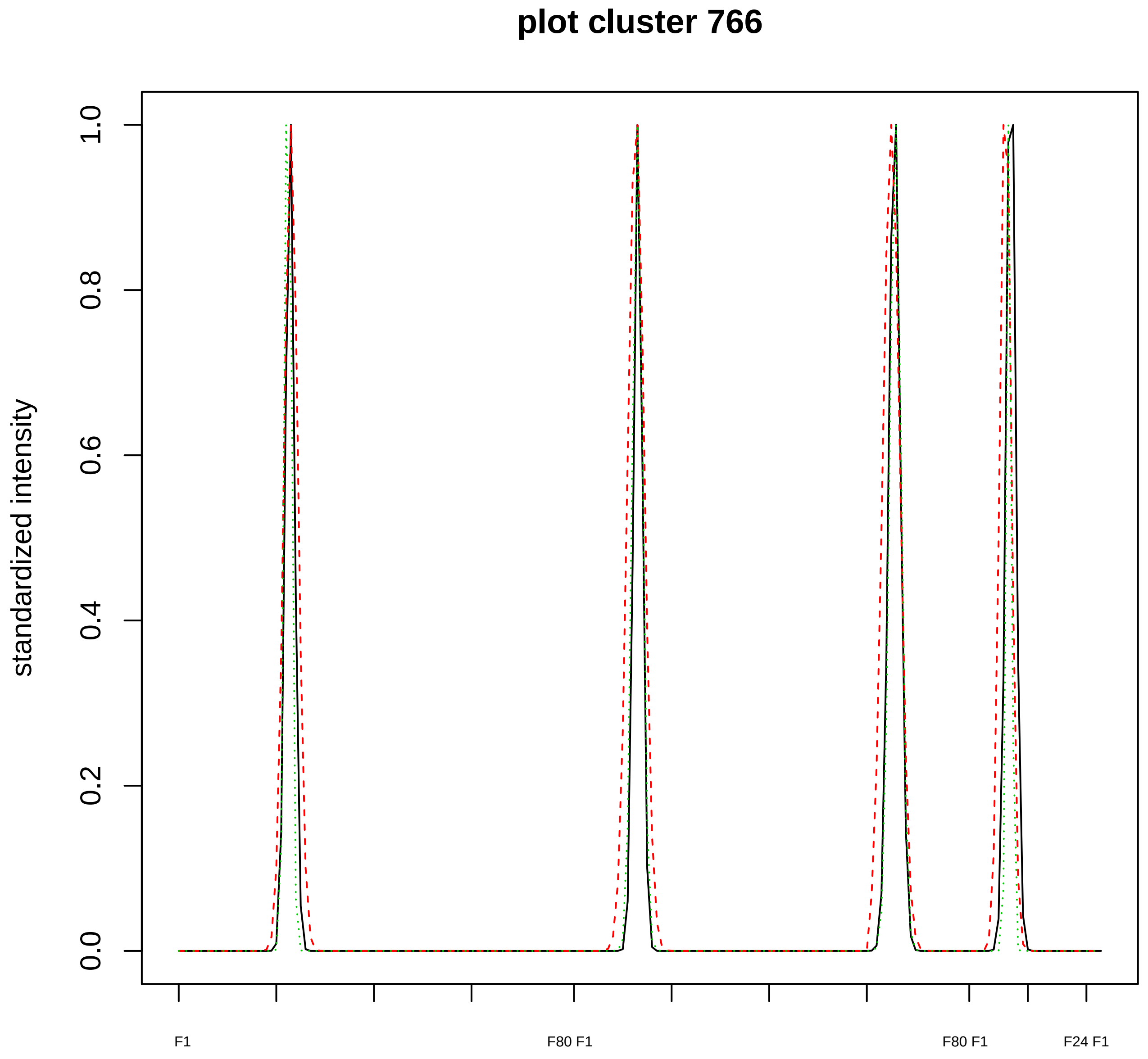
plot cluster 766
standardized intensity
0.0
0.2
0.4
0.6
0.8
1.0
F1
F80 F1
F80 F1
F24 F1

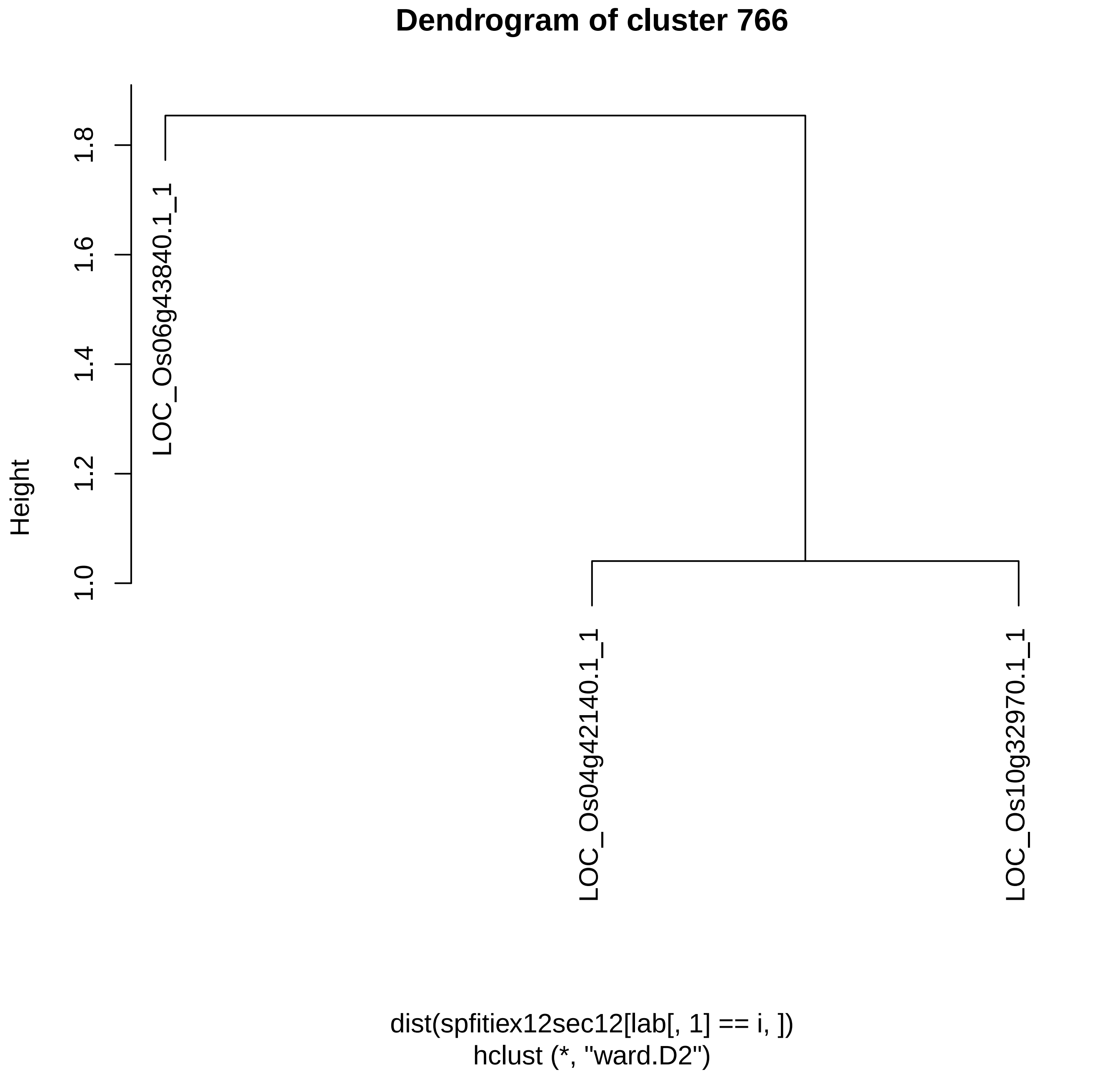
Dendrogram of cluster 766
Height
1.0
1.2
1.4
1.6
1.8
LOC_Os06g43840.1_1
LOC_Os04g42140.1_1
LOC_Os10g32970.1_1
dist(spfitiex12sec12[lab[, 1] == i, ])
hclust (*, "ward.D2")

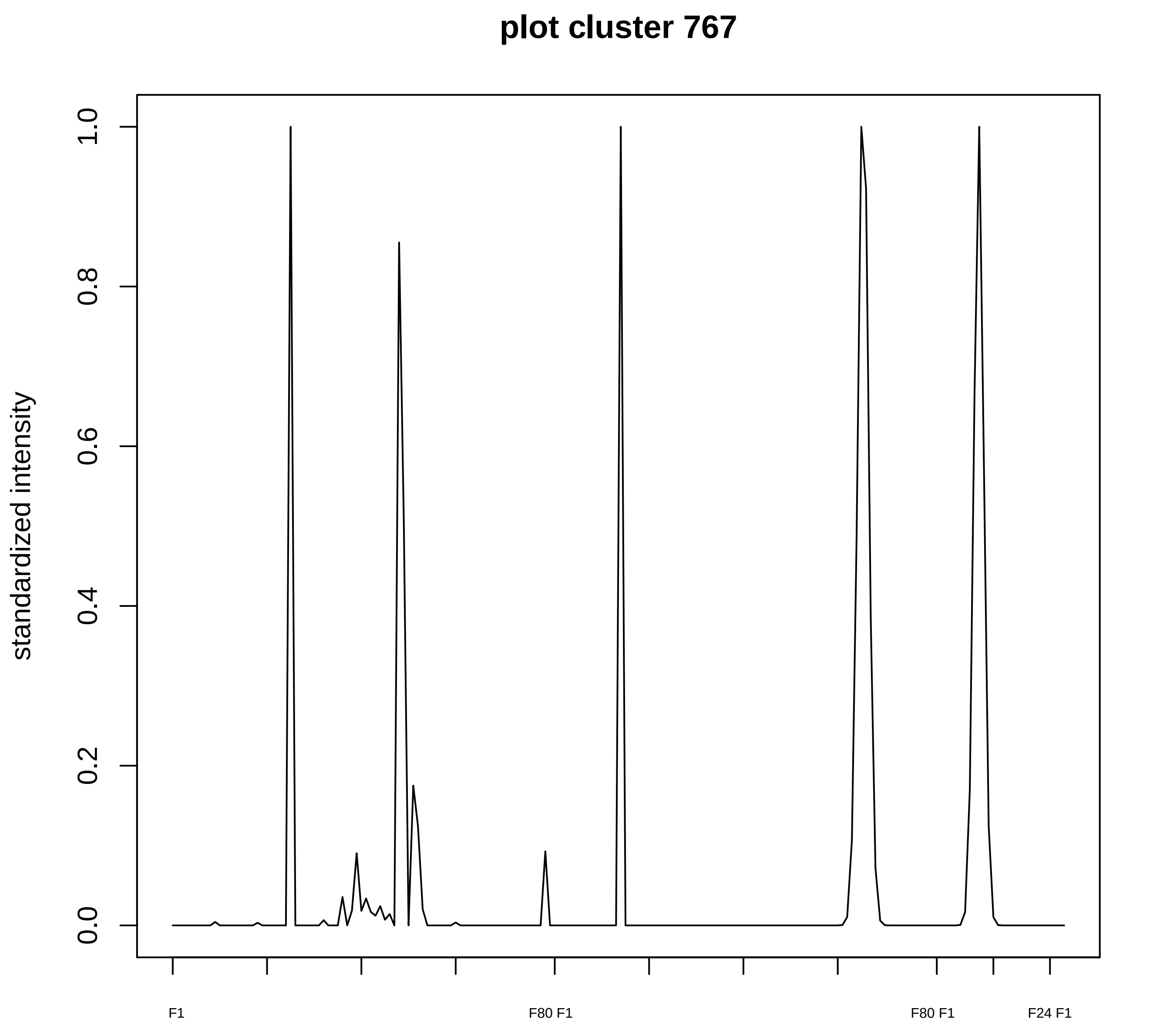
plot cluster 767
standardized intensity
0.0
0.2
0.4
0.6
0.8
1.0
F1
F80 F1
F80 F1
F24 F1

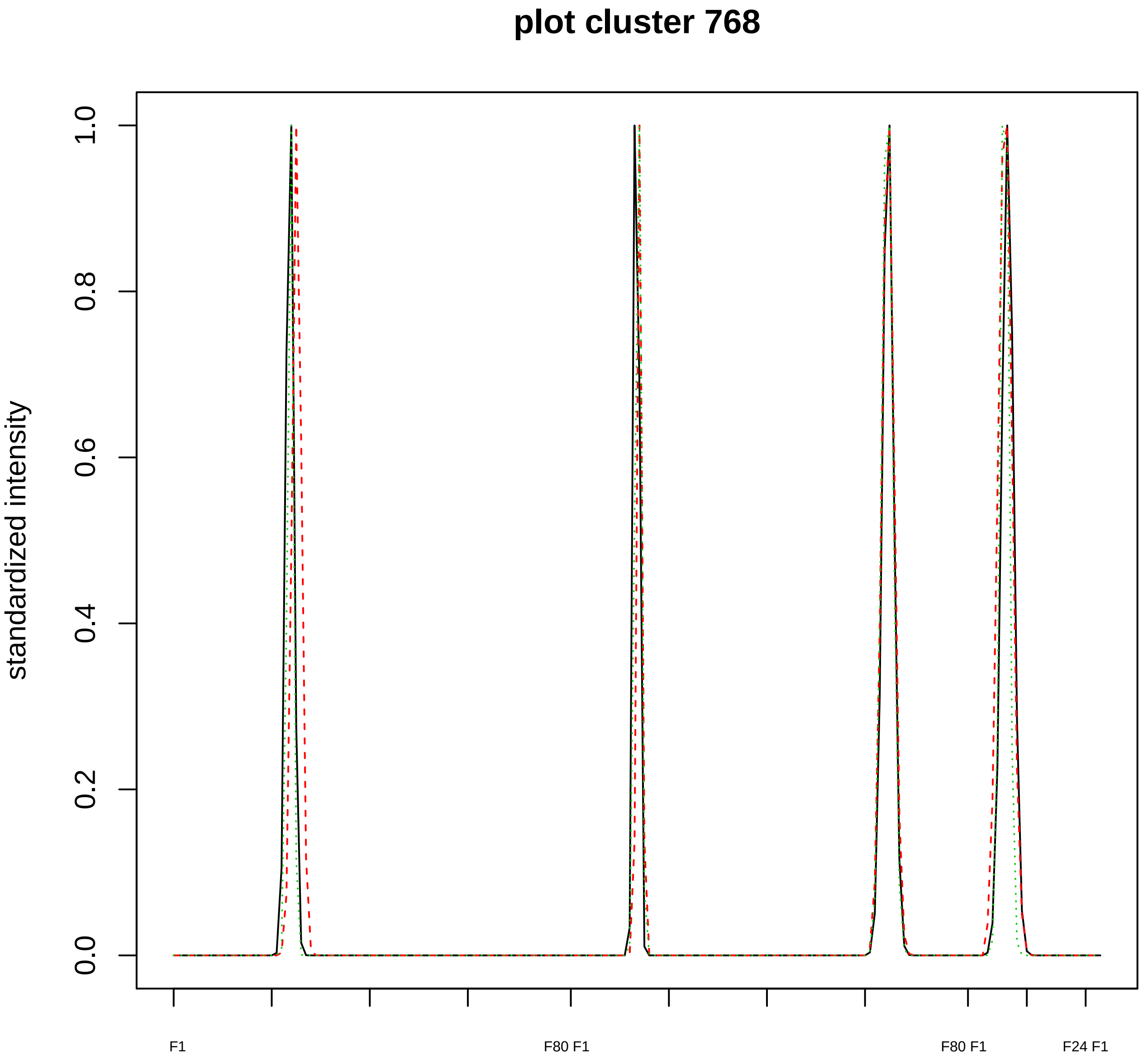
plot cluster 768
standardized intensity
0.0
0.2
0.4
0.6
0.8
1.0
F1
F80 F1
F80 F1
F24 F1

### Dendrogram of cluster 768

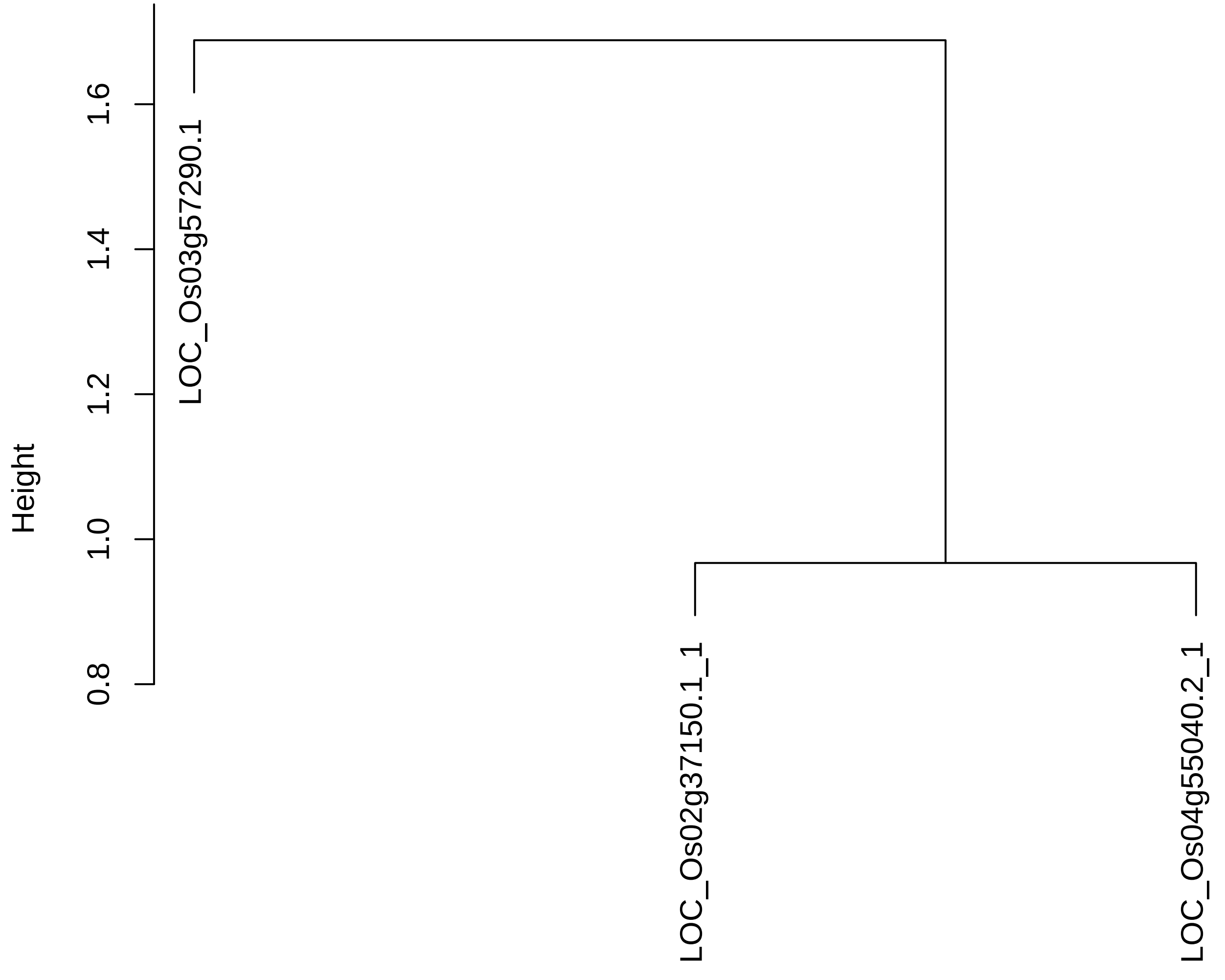


dist(spfitiex12sec12[lab[, 1] == i, ])
hclust (*, "ward.D2")

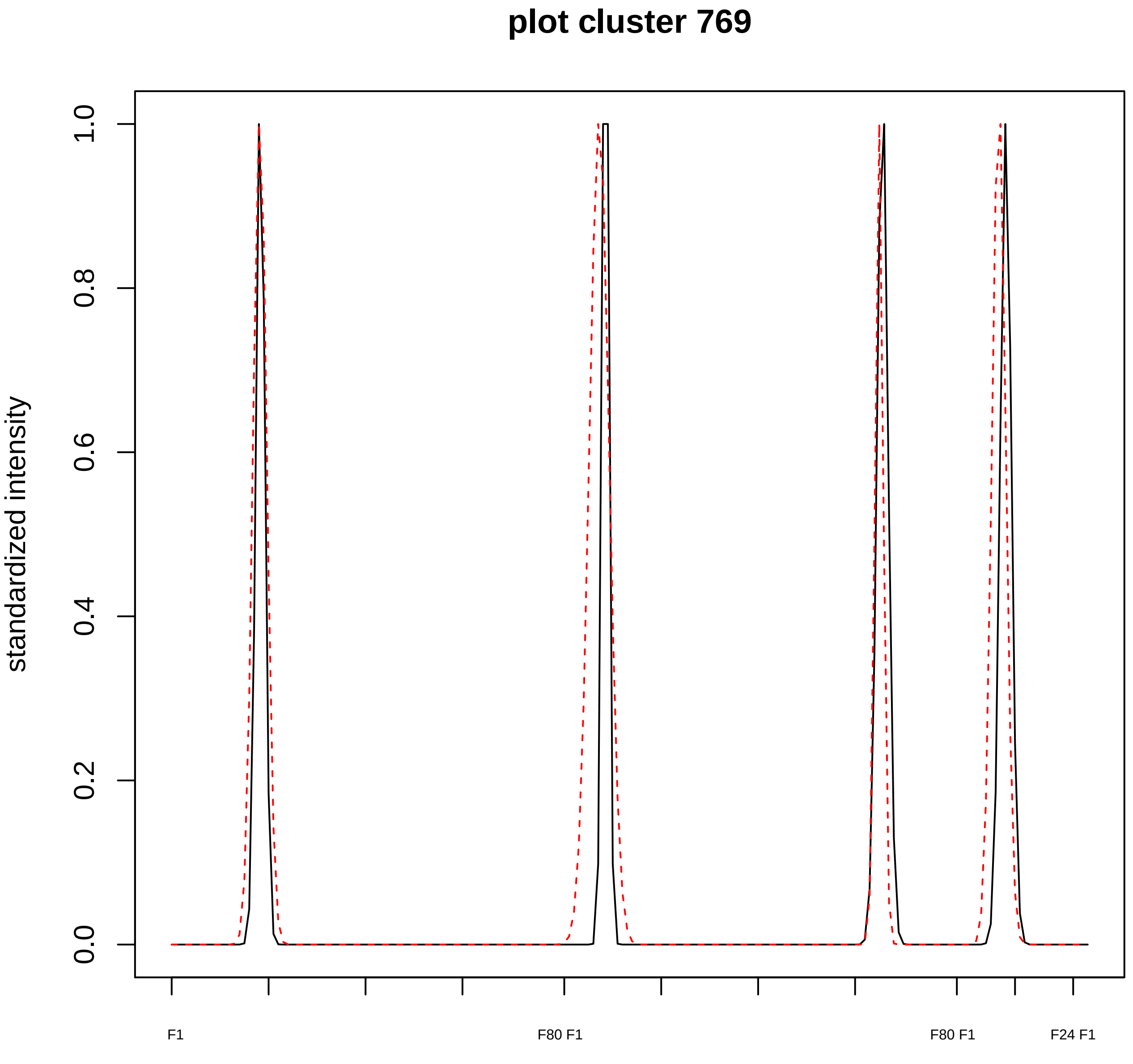

plot cluster 769
standardized intensity
0.0
0.2
0.4
0.6
0.8
1.0
F1
F80 F1
F80 F1
F24 F1

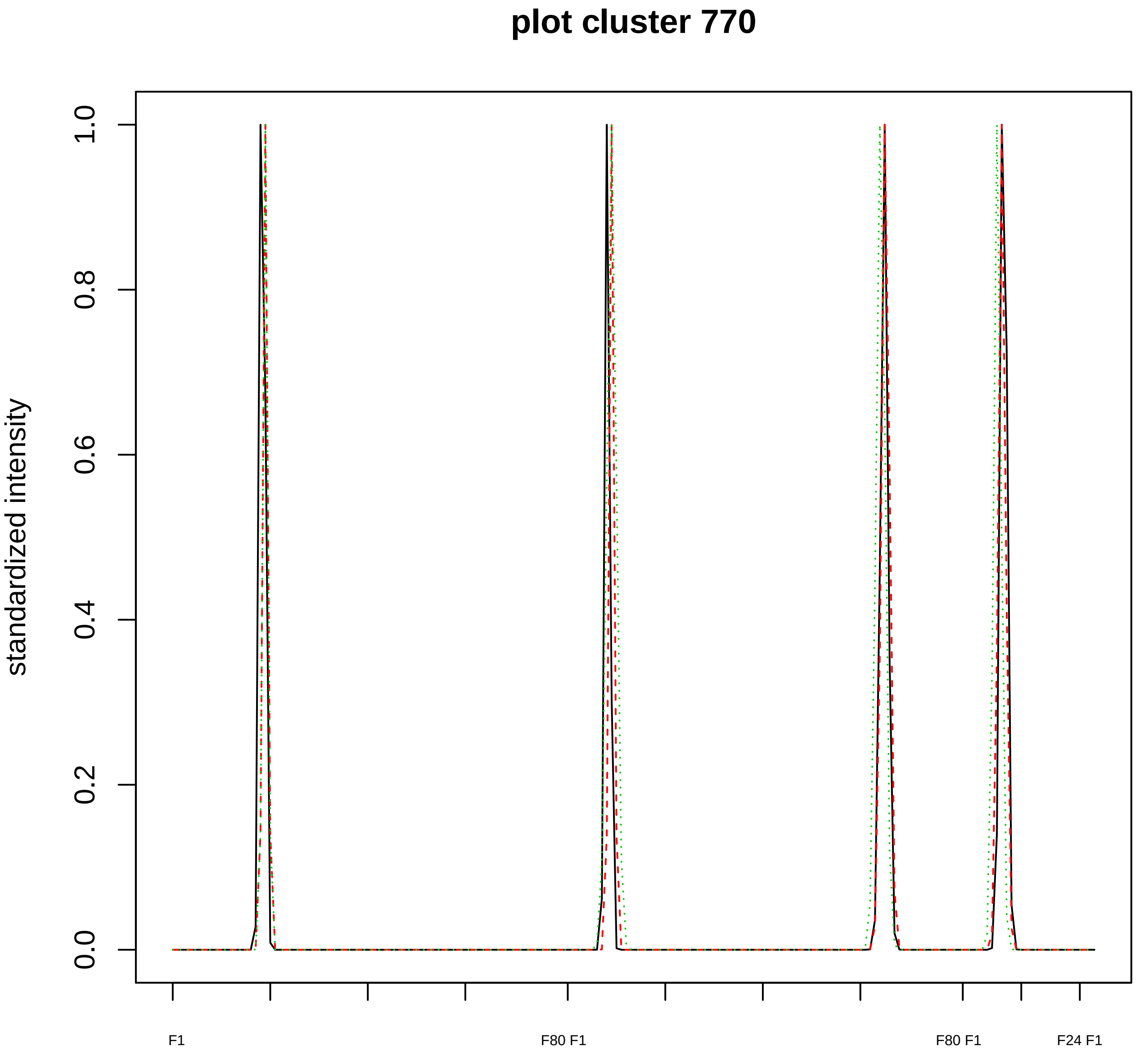
plot cluster 770
standardized intensity
0.0
0.2
0.4
0.6
0.8
1.0
F1
F80 F1
F80 F1
F24 F1

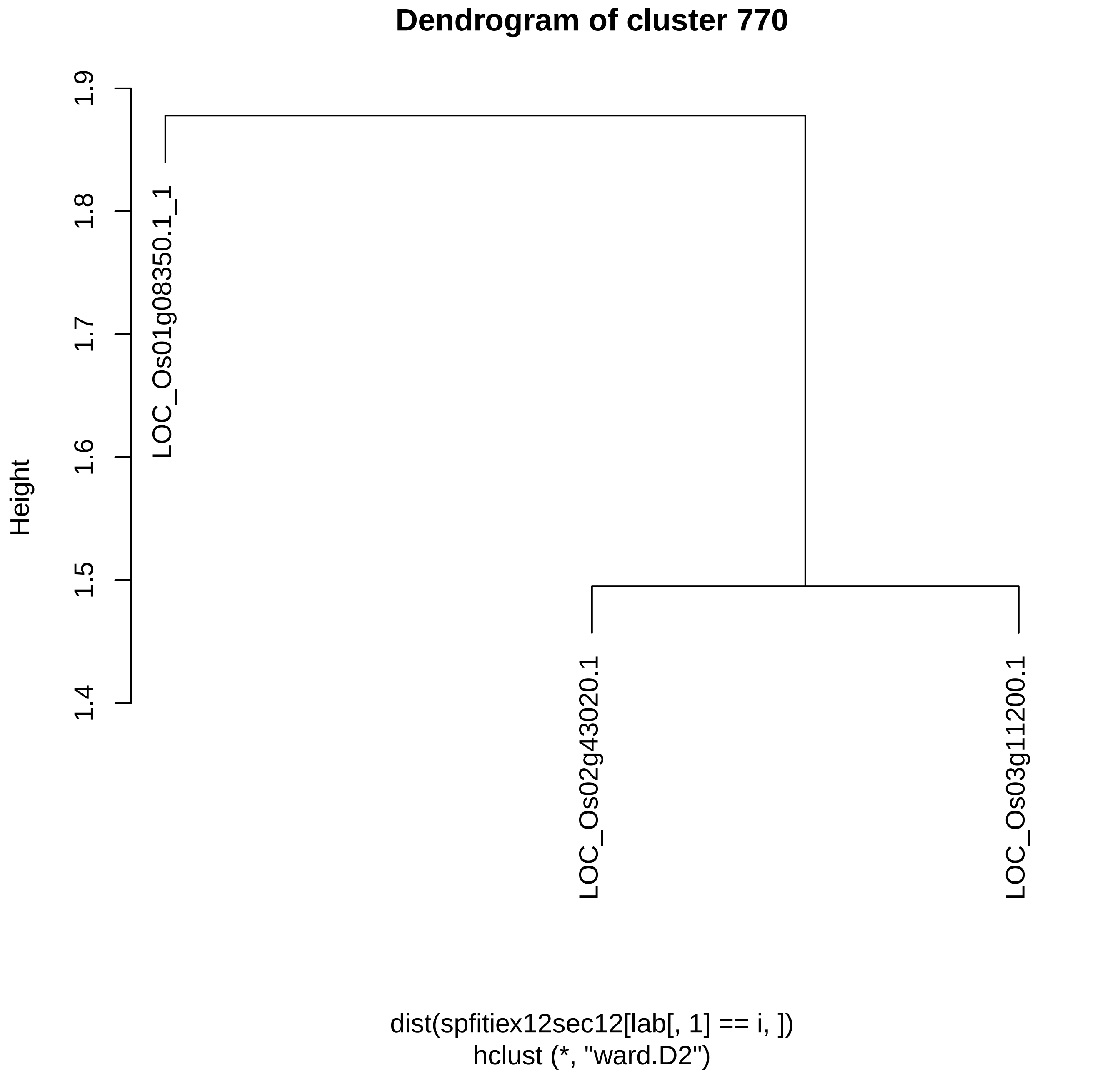
Dendrogram of cluster 770
Height
1.9
1.8
1.7
1.6
1.5
1.4
LOC_Os01g08350.1_1
LOC_Os02g43020.1
LOC_Os03g11200.1
dist(spfitiex12sec12[lab[, 1] == i, ])
hclust (*, "ward.D2")

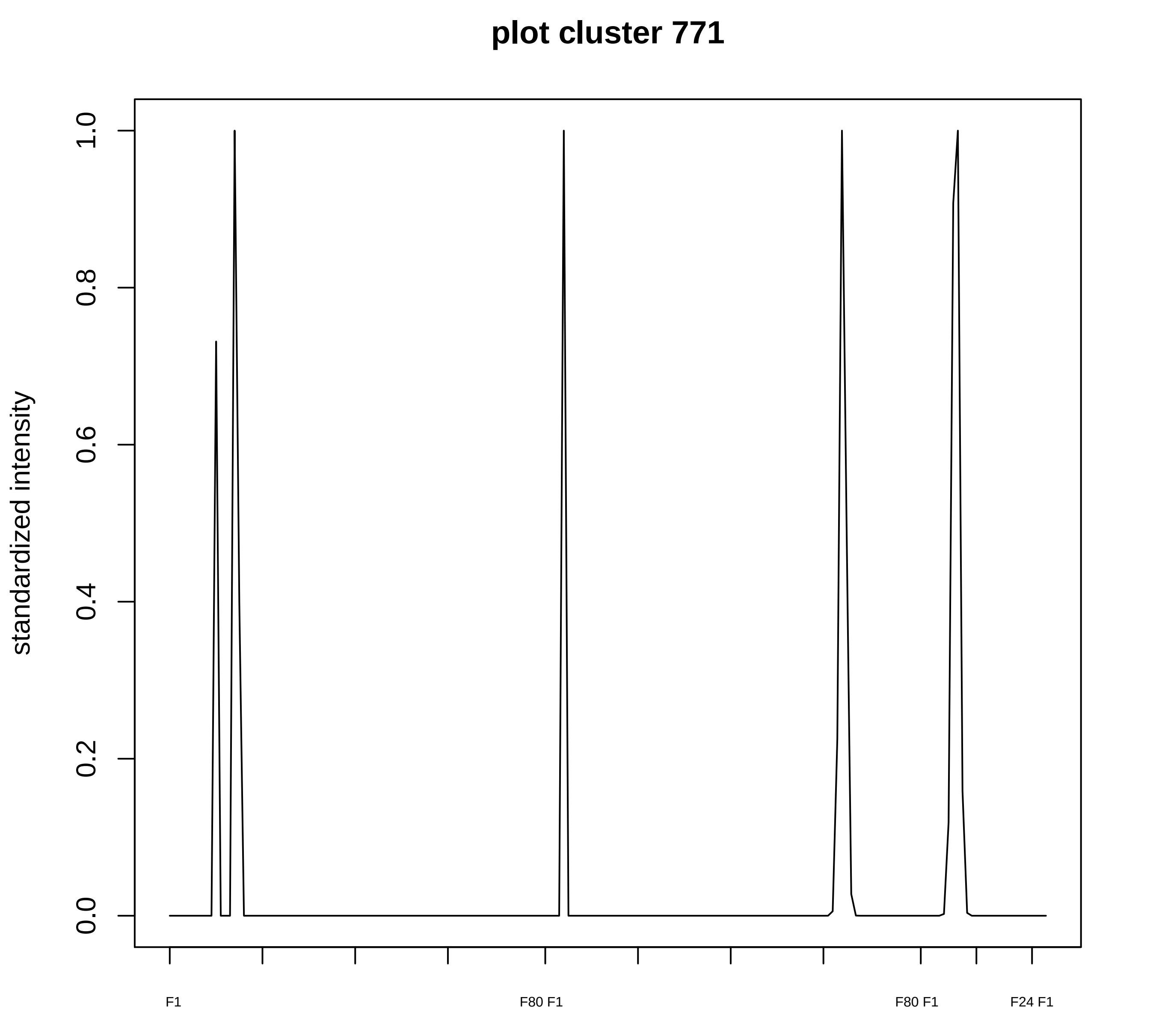
plot cluster 771
standardized intensity
0.0
0.2
0.4
0.6
0.8
1.0
F1
F80 F1
F80 F1
F24 F1

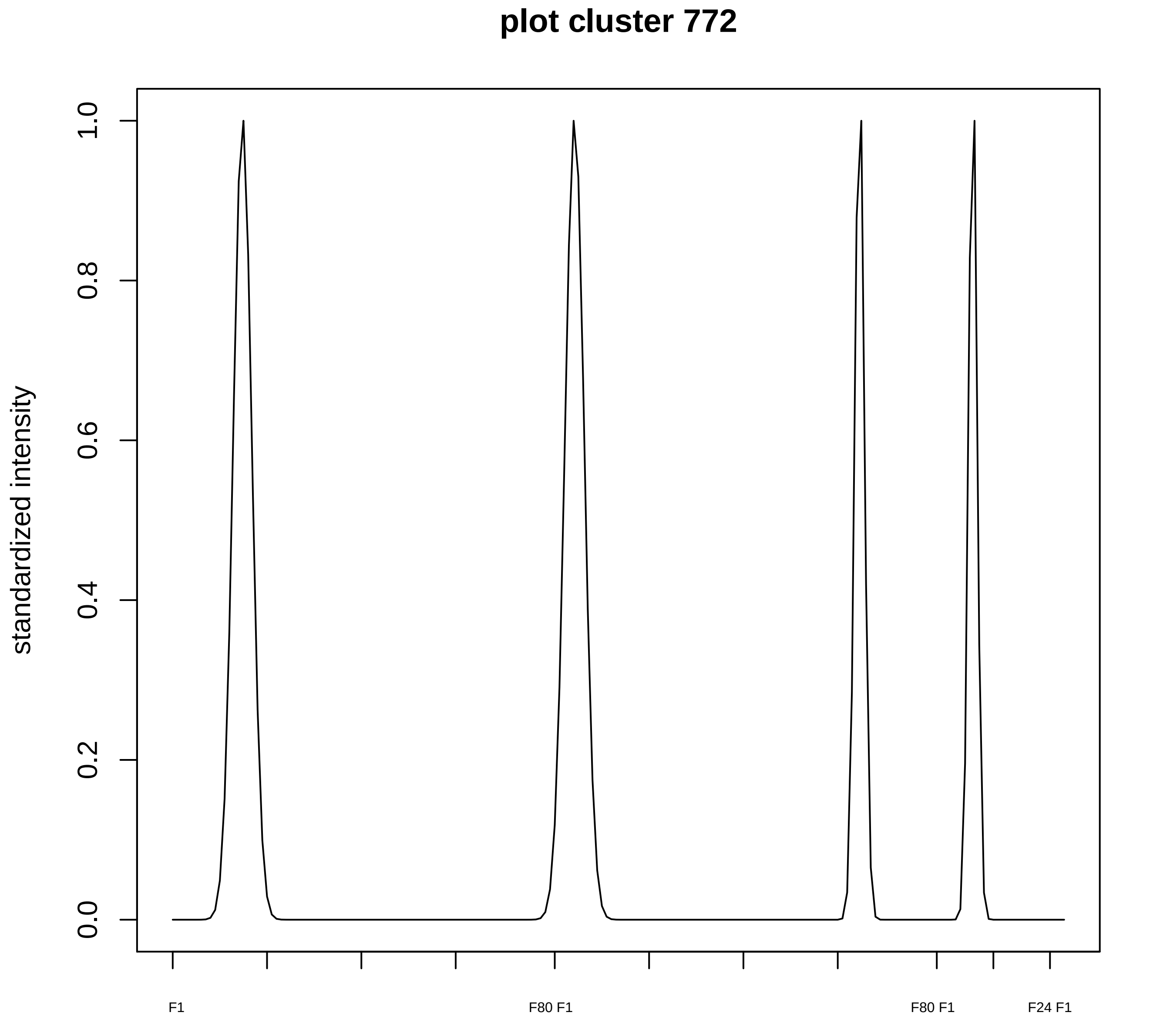

plot cluster 772
standardized intensity
0.0
0.2
0.4
0.6
0.8
1.0
F1
F80 F1
F80 F1
F24 F1

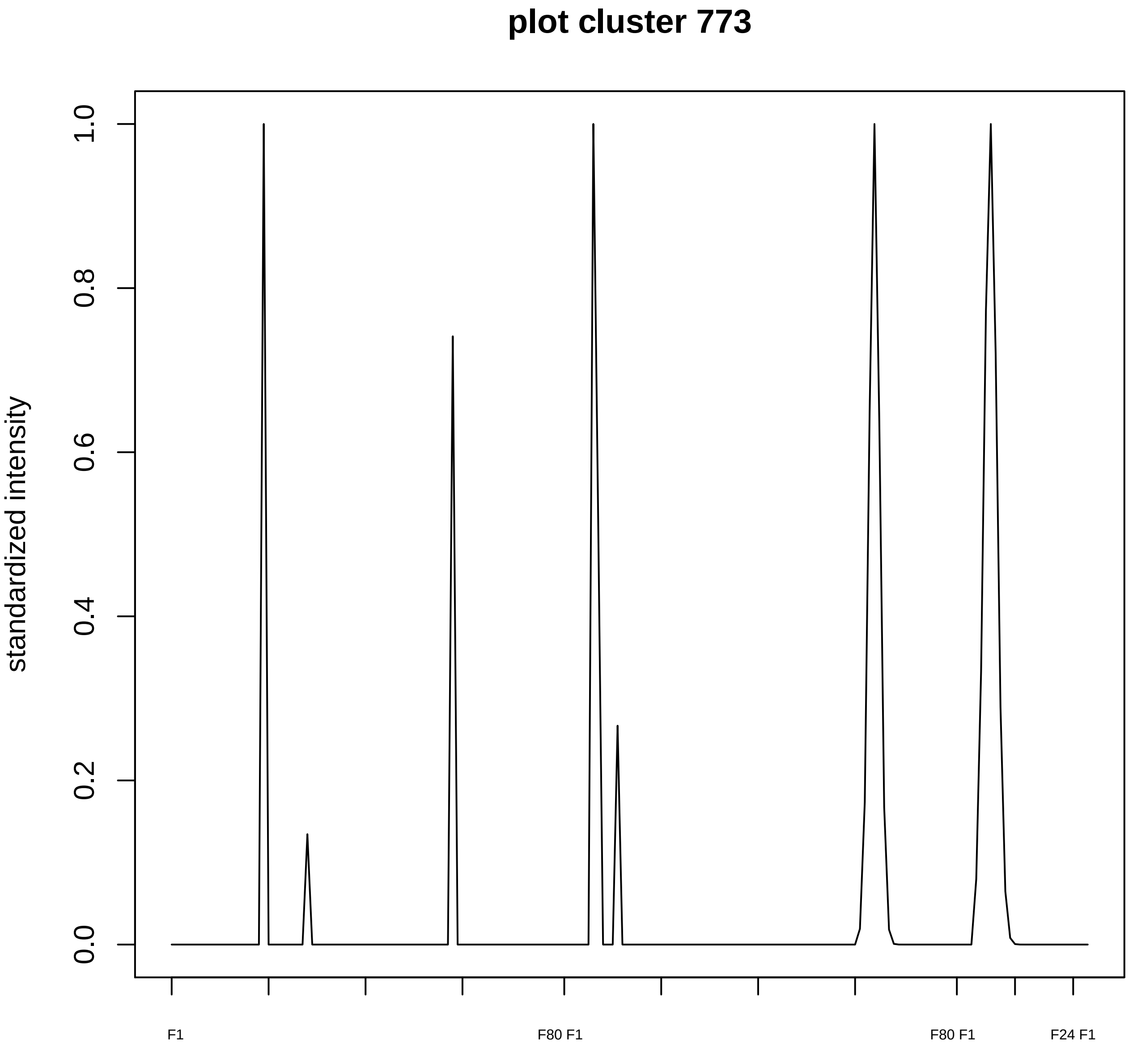
plot cluster 773
standardized intensity
0.0
0.2
0.4
0.6
0.8
1.0
F1
F80 F1
F80 F1
F24 F1

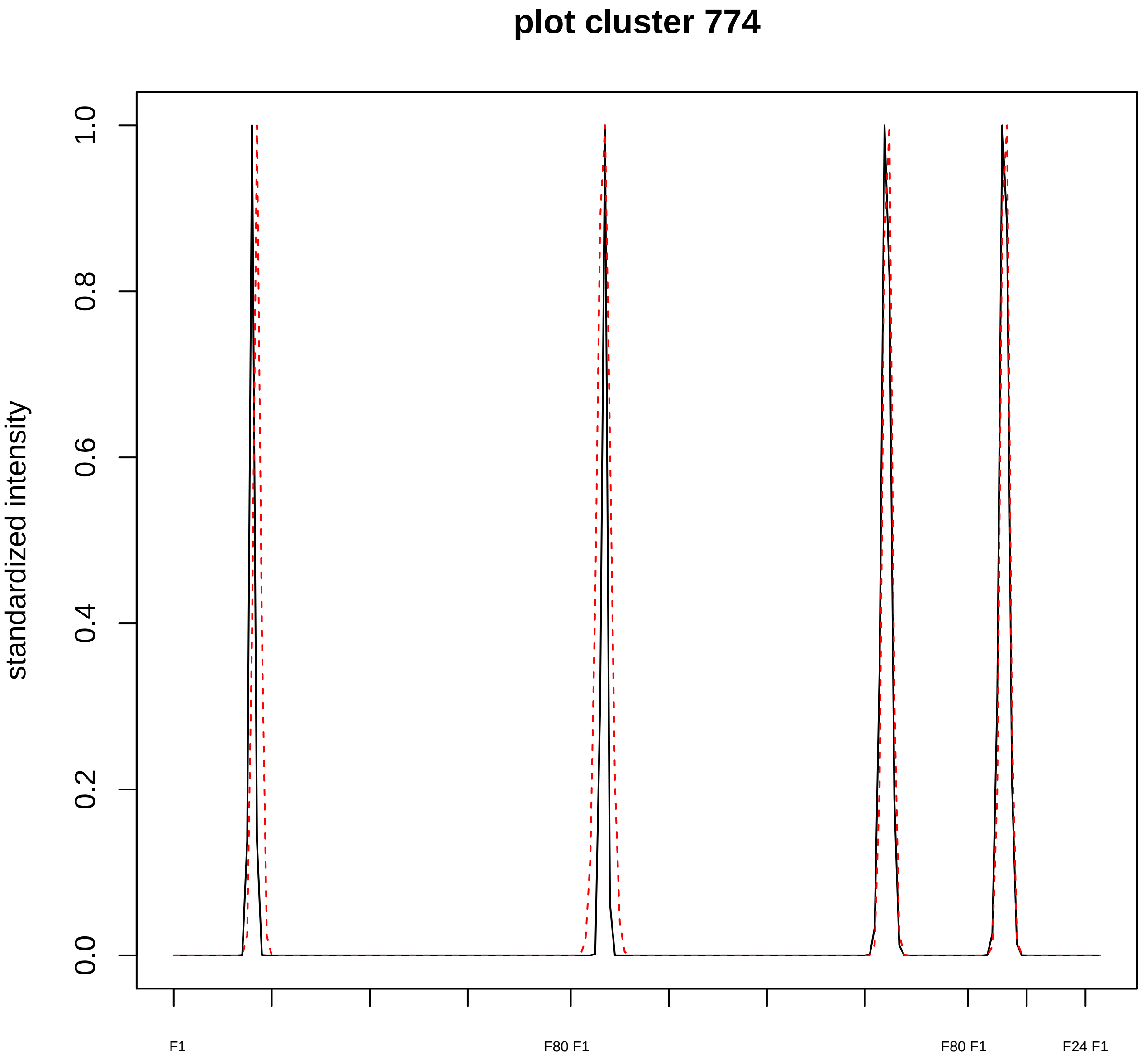

plot cluster 774
standardized intensity
0.0
0.2
0.4
0.6
0.8
1.0
F1
F80 F1
F80 F1
F24 F1

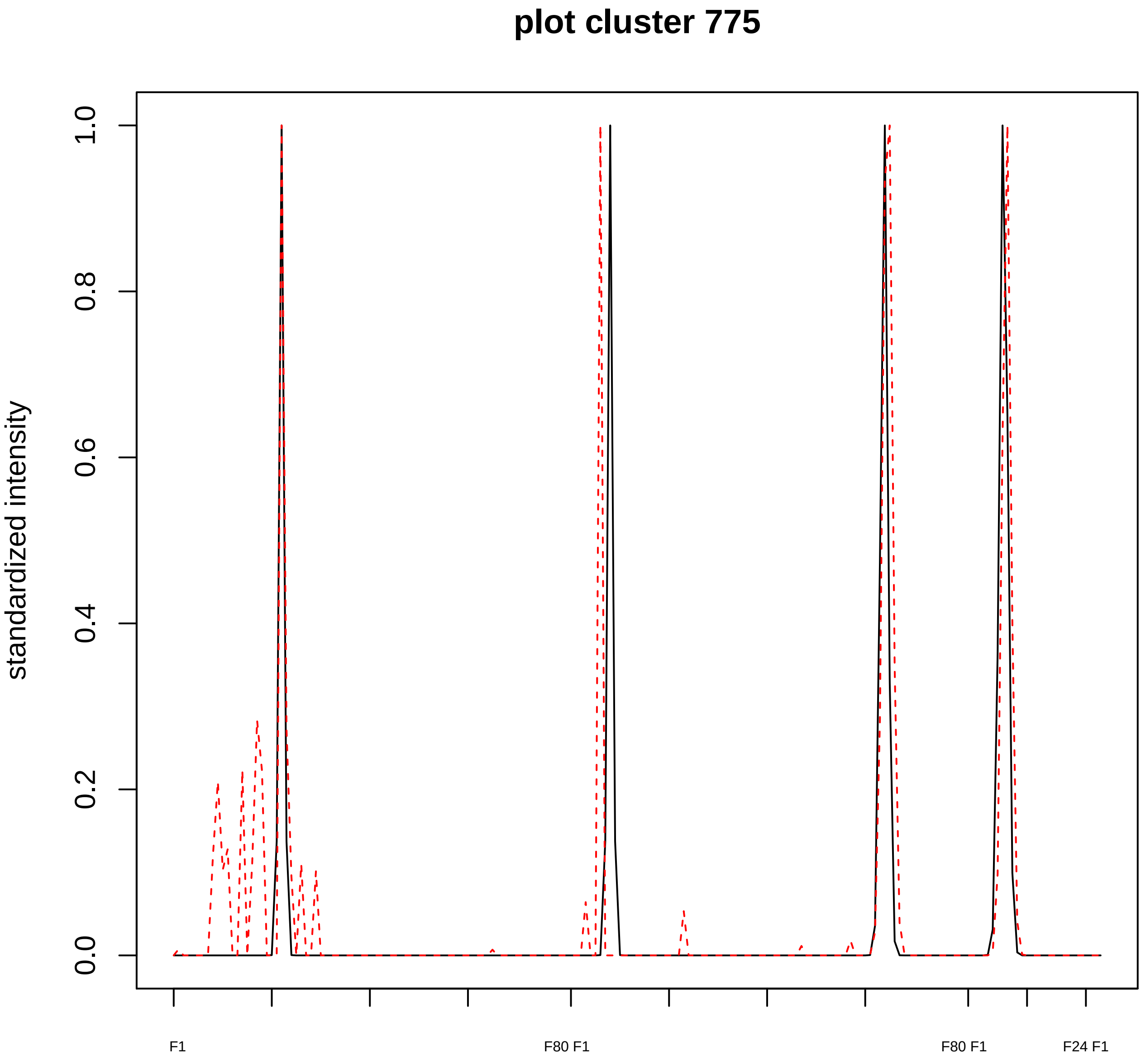

plot cluster 775
standardized intensity
0.0
0.2
0.4
0.6
0.8
1.0
F1
F80 F1
F80 F1
F24 F1

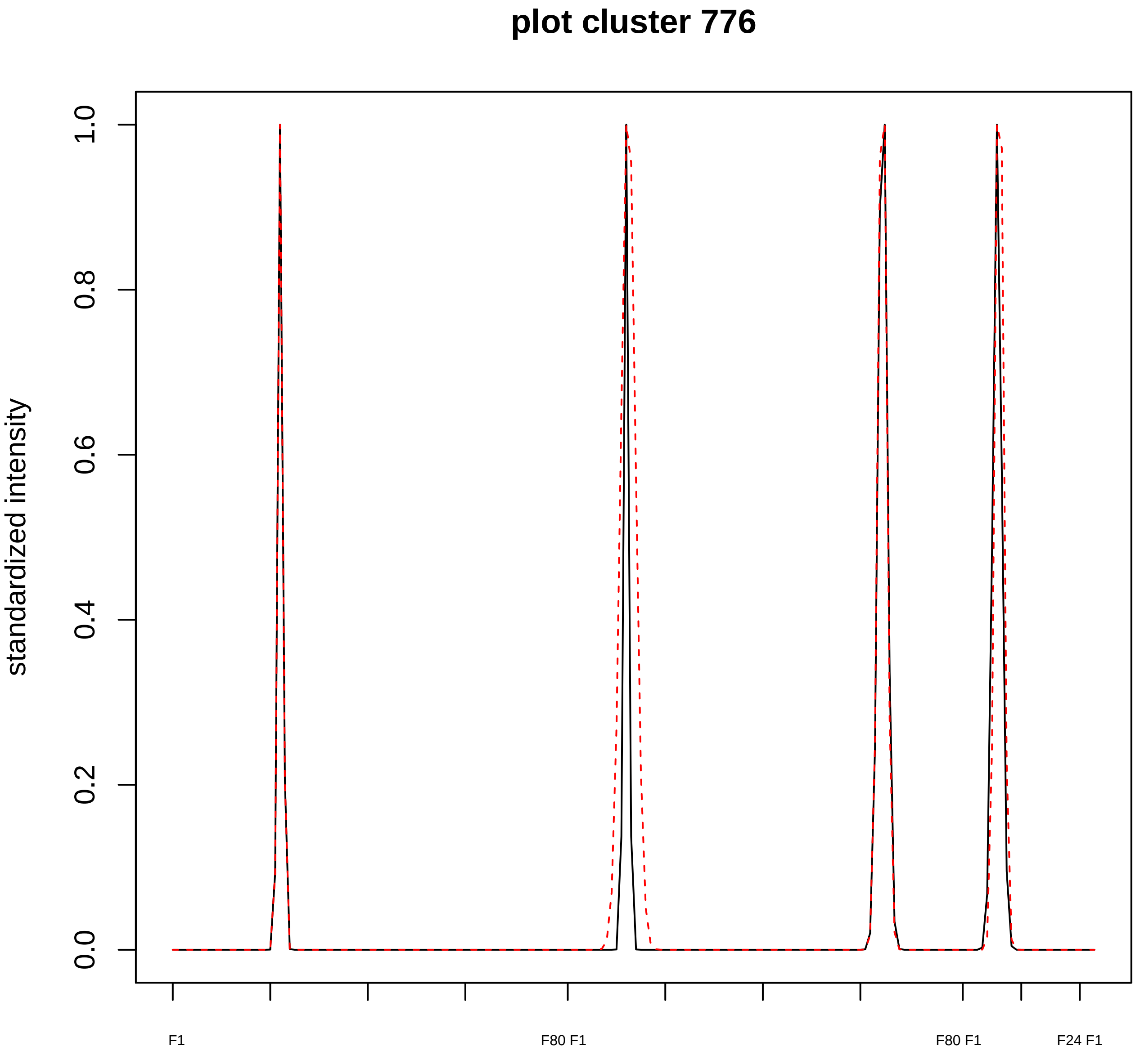

plot cluster 776
standardized intensity
0.0
0.2
0.4
0.6
0.8
1.0
F1
F80 F1
F80 F1
F24 F1

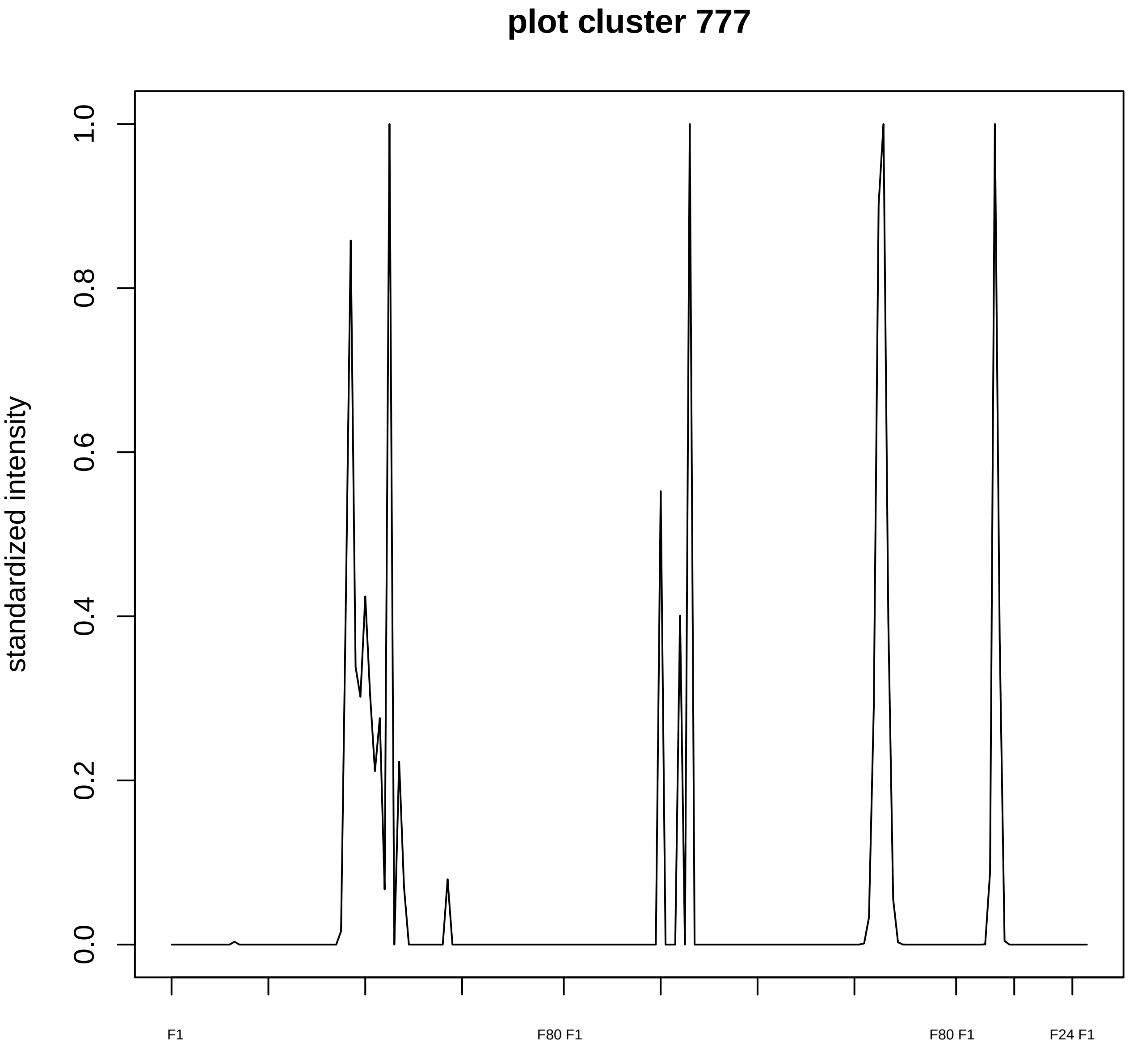

plot cluster 777
standardized intensity
0.0
0.2
0.4
0.6
0.8
1.0
F1
F80 F1
F80 F1
F24 F1

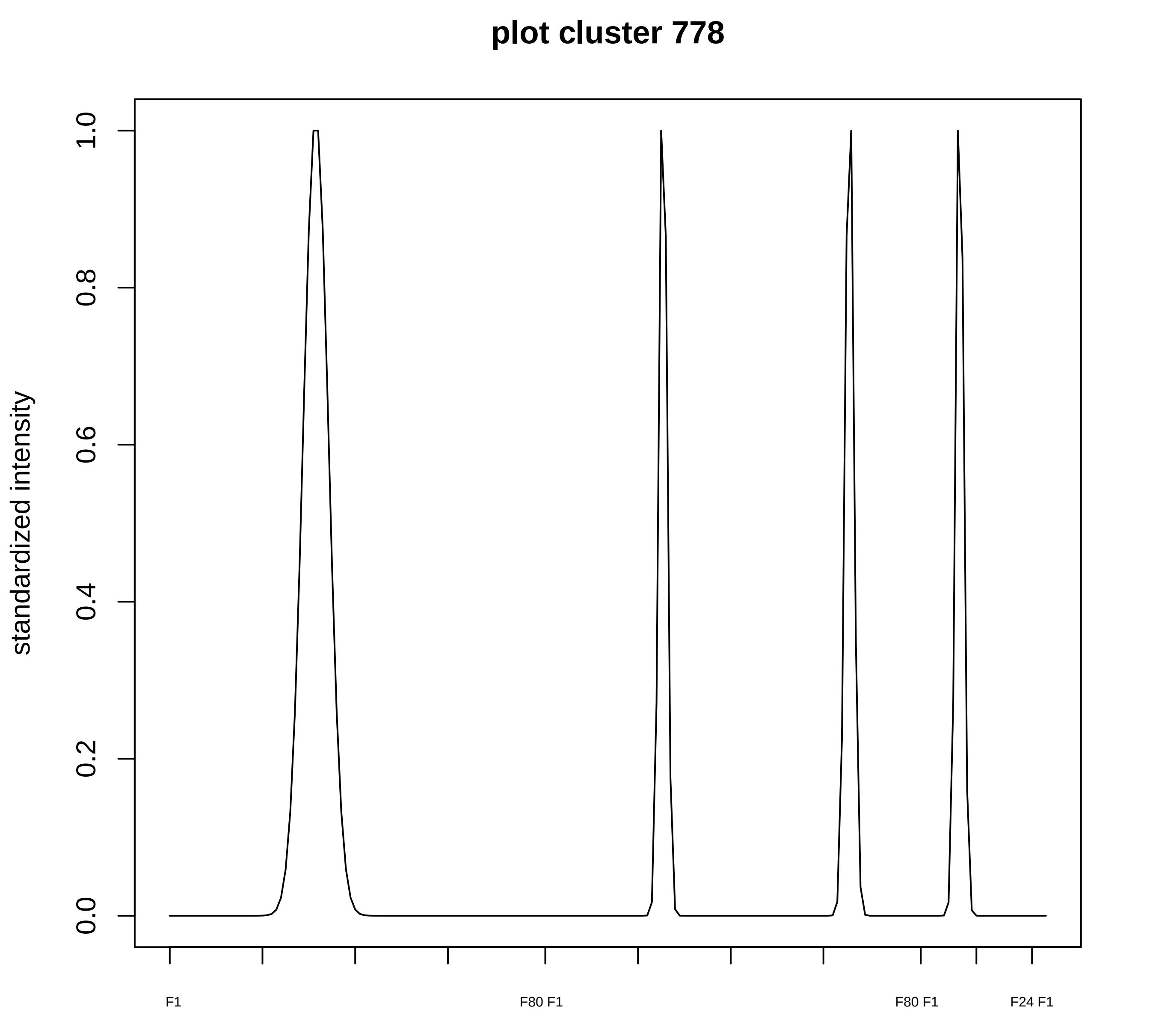

plot cluster 778
standardized intensity
0.0
0.2
0.4
0.6
0.8
1.0
F1
F80 F1
F80 F1
F24 F1

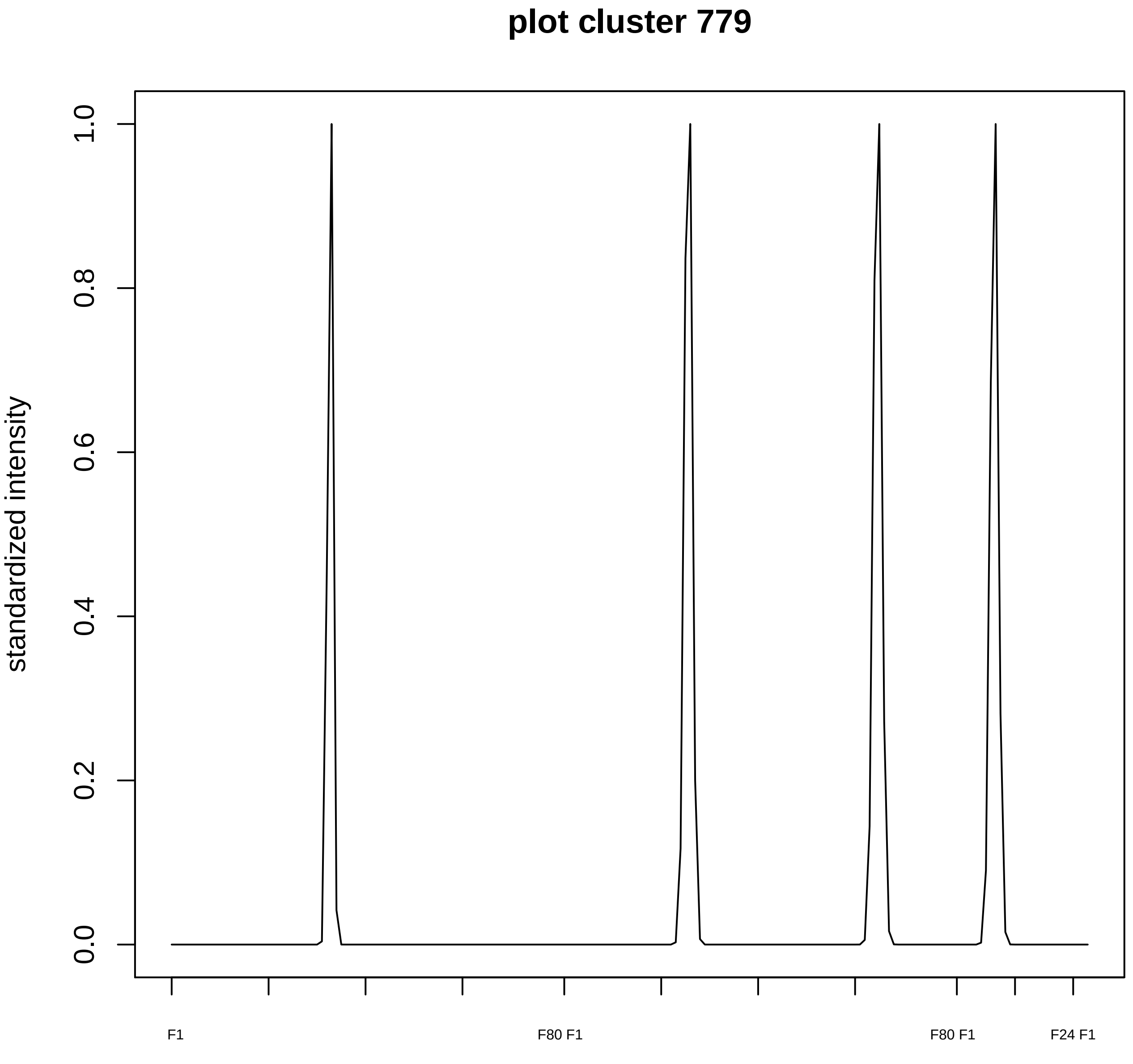
plot cluster 779
standardized intensity
0.0
0.2
0.4
0.6
0.8
1.0
F1
F80 F1
F80 F1
F24 F1

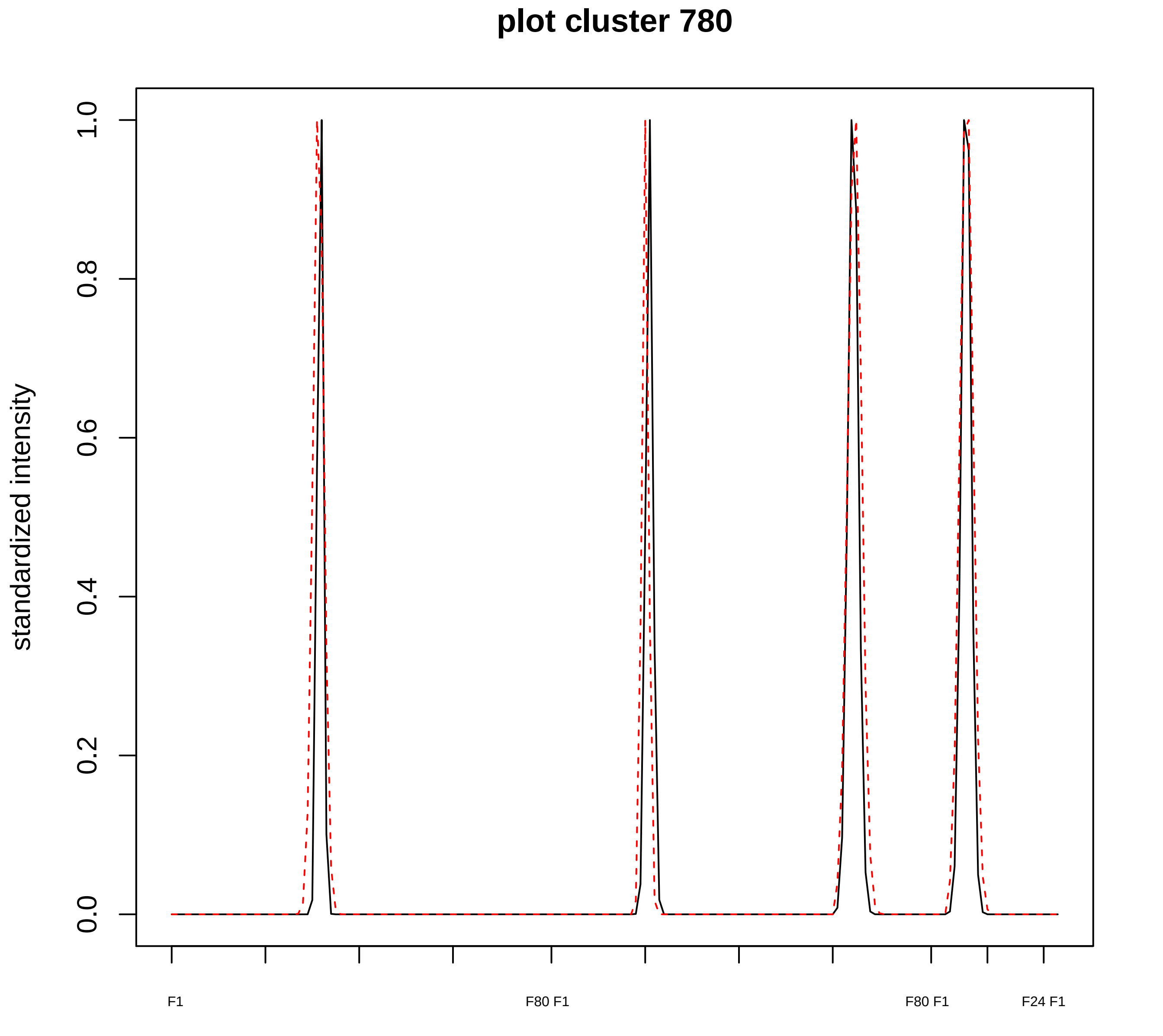
plot cluster 780
standardized intensity
0.0
0.2
0.4
0.6
0.8
1.0
F1
F80 F1
F80 F1
F24 F1

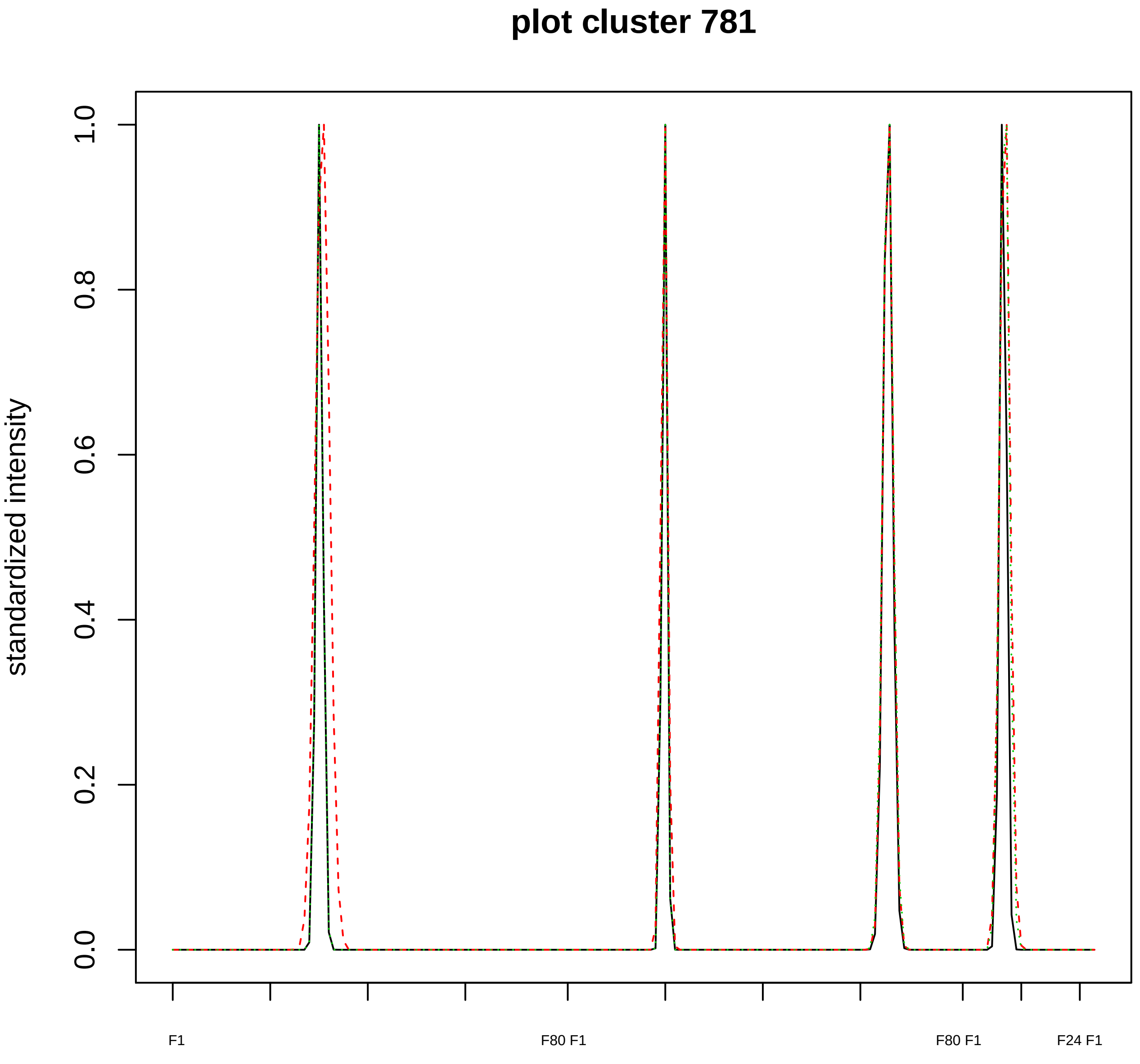
plot cluster 781
standardized intensity
0.0
0.2
0.4
0.6
0.8
1.0
F1
F80 F1
F80 F1
F24 F1

#### Dendrogram of cluster 781

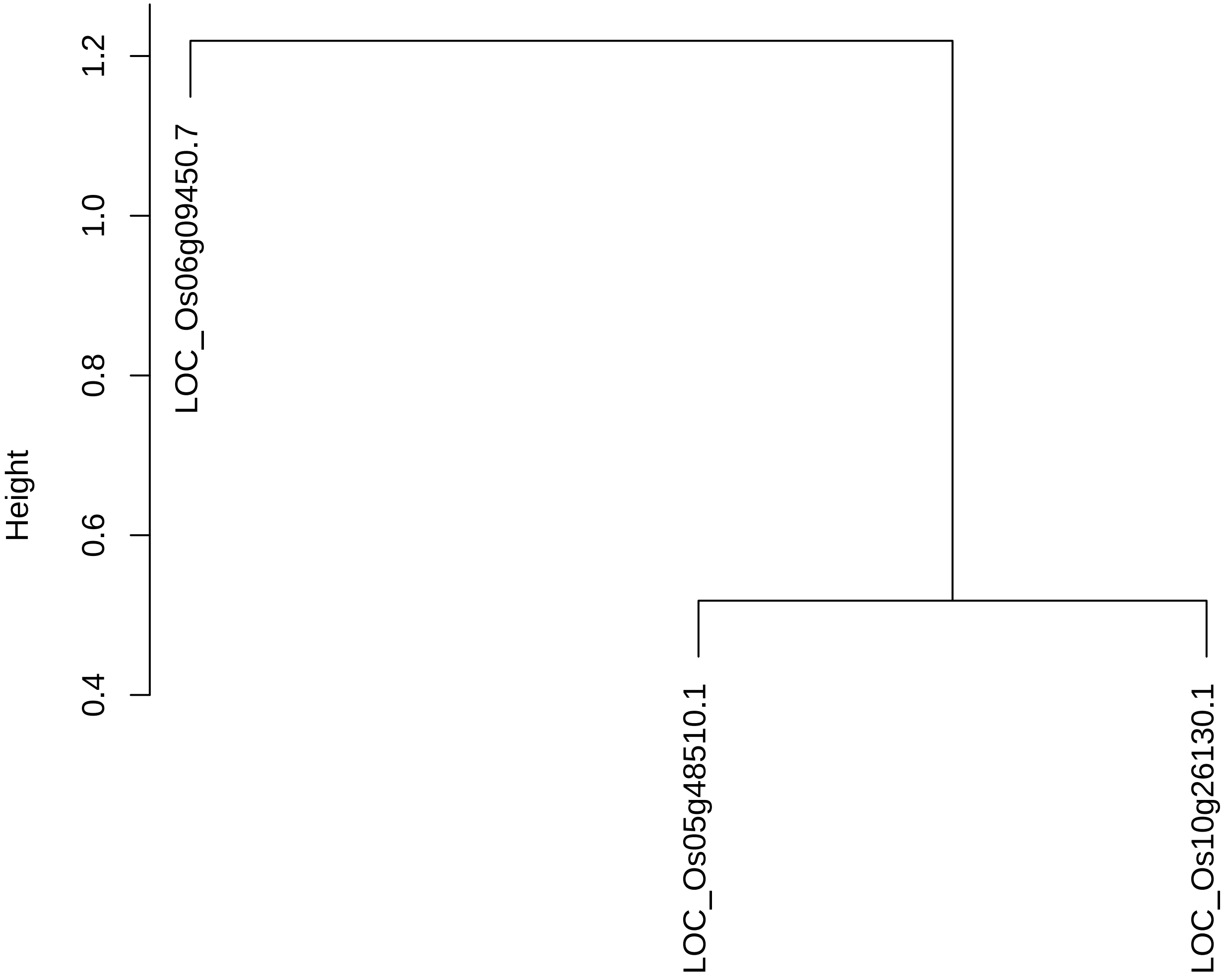


dist(spfitiex12sec12[lab[, 1] == i, ])
hclust (*, "ward.D2")

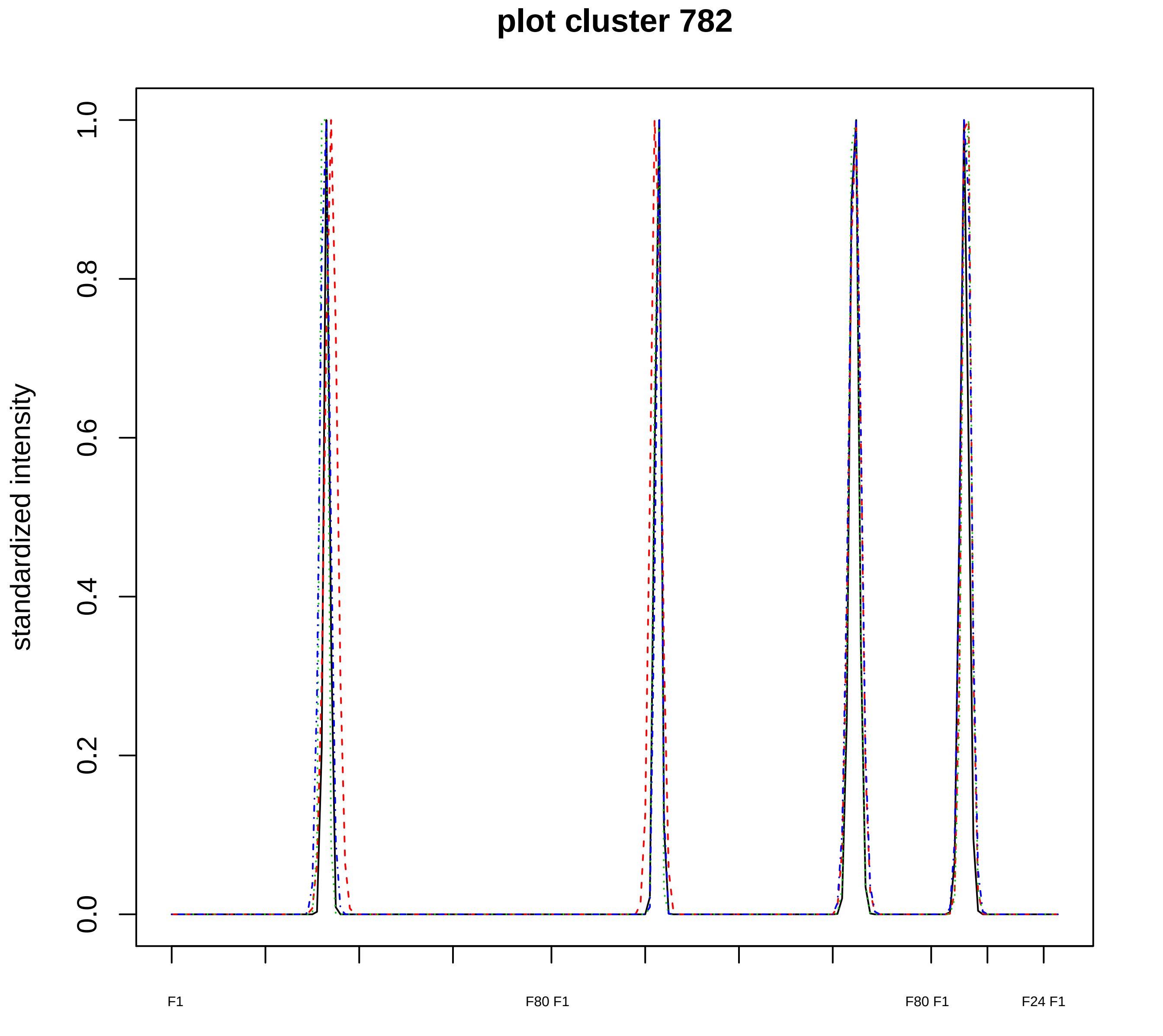

plot cluster 782
standardized intensity
0.0
0.2
0.4
0.6
0.8
1.0
F1
F80 F1
F80 F1
F24 F1

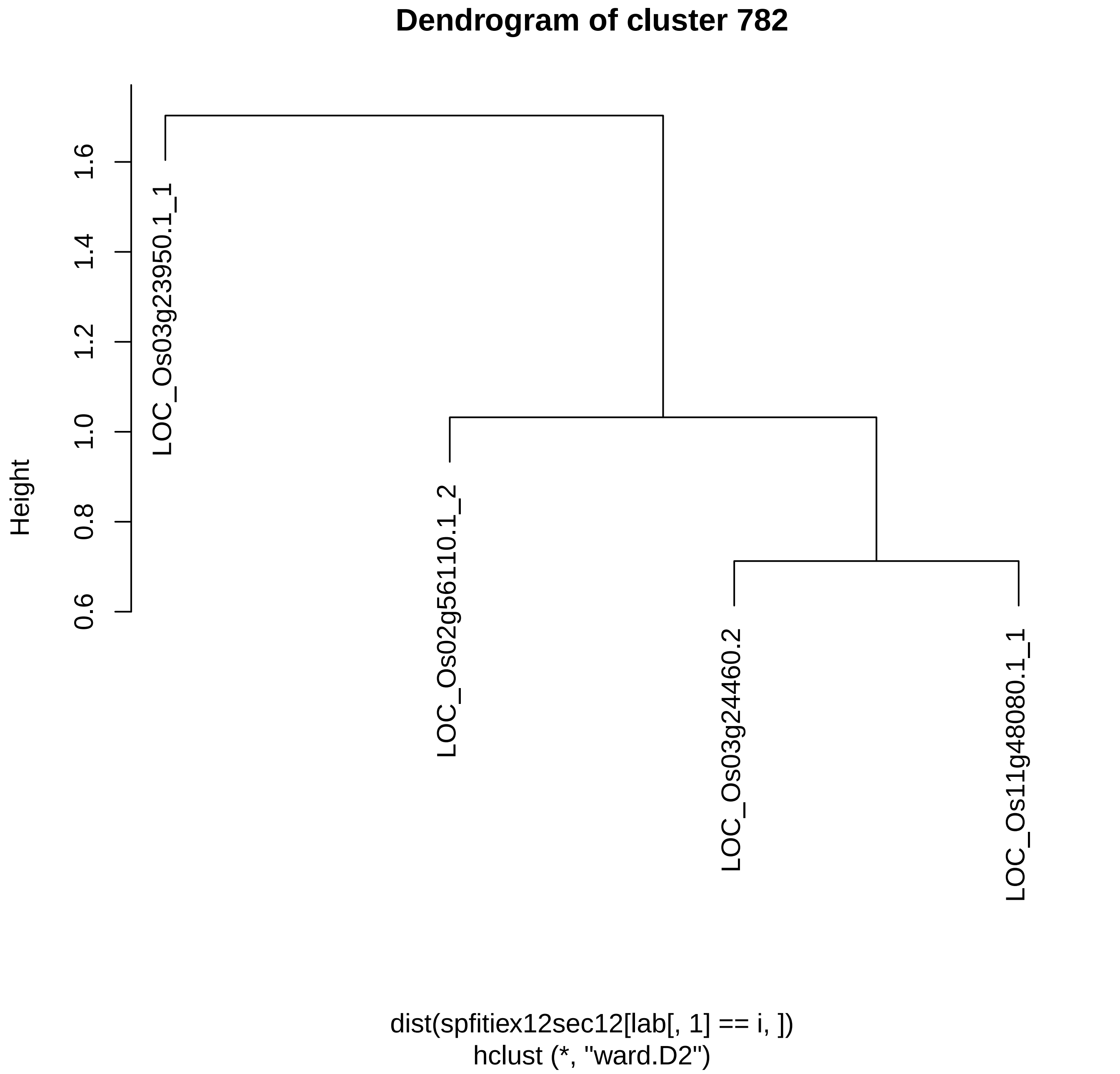
Dendrogram of cluster 782
Height
0.6
0.8
1.0
1.2
1.4
1.6
LOC_Os03g23950.1_1
LOC_Os02g56110.1_2
LOC_Os03g24460.2
LOC_Os11g48080.1_1
dist(spfitiex12sec12[lab[, 1] == i, ])
hclust (*, "ward.D2")

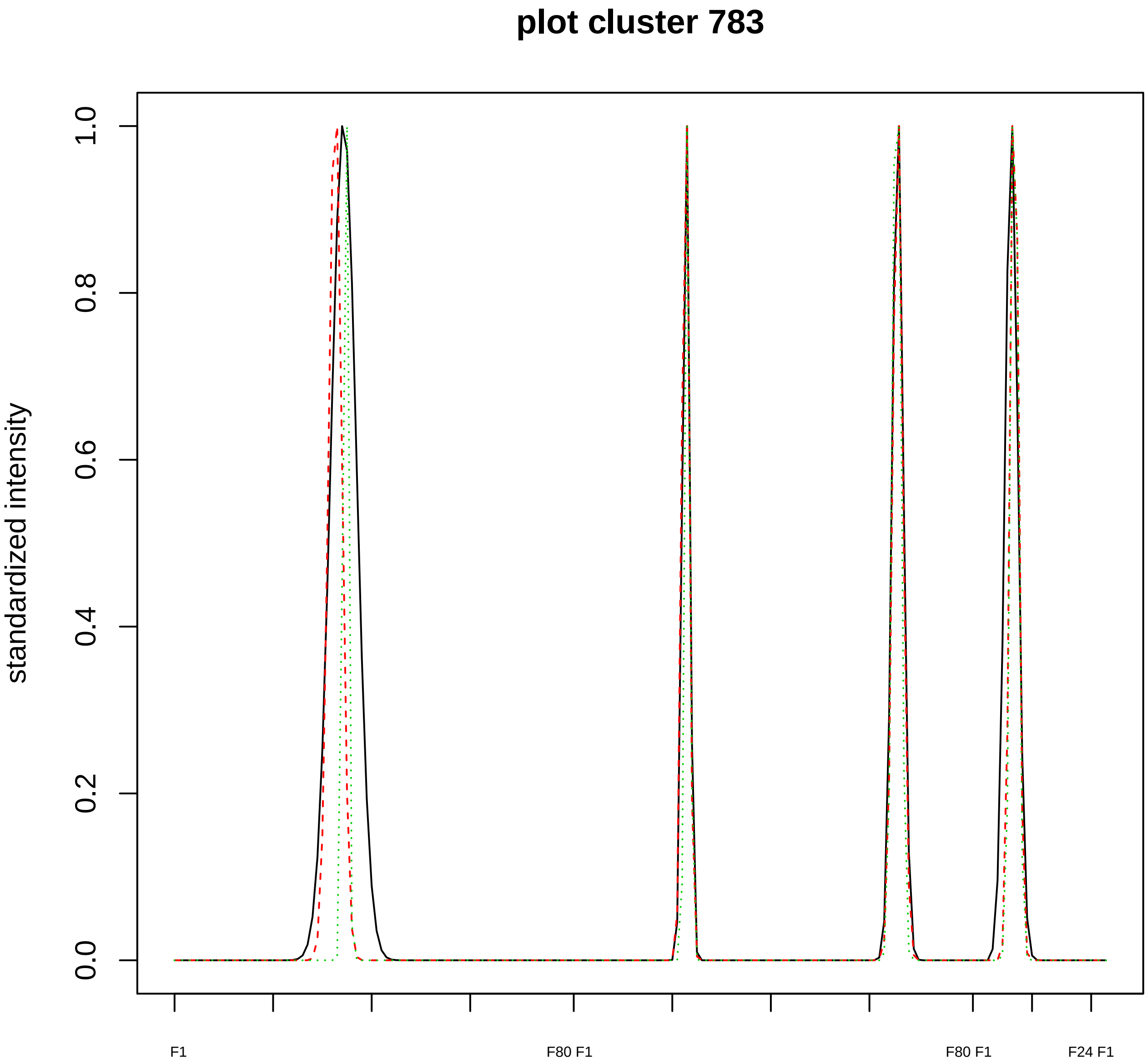

plot cluster 783
standardized intensity
0.0
0.2
0.4
0.6
0.8
1.0
F1
F80 F1
F80 F1
F24 F1

### Dendrogram of cluster 783

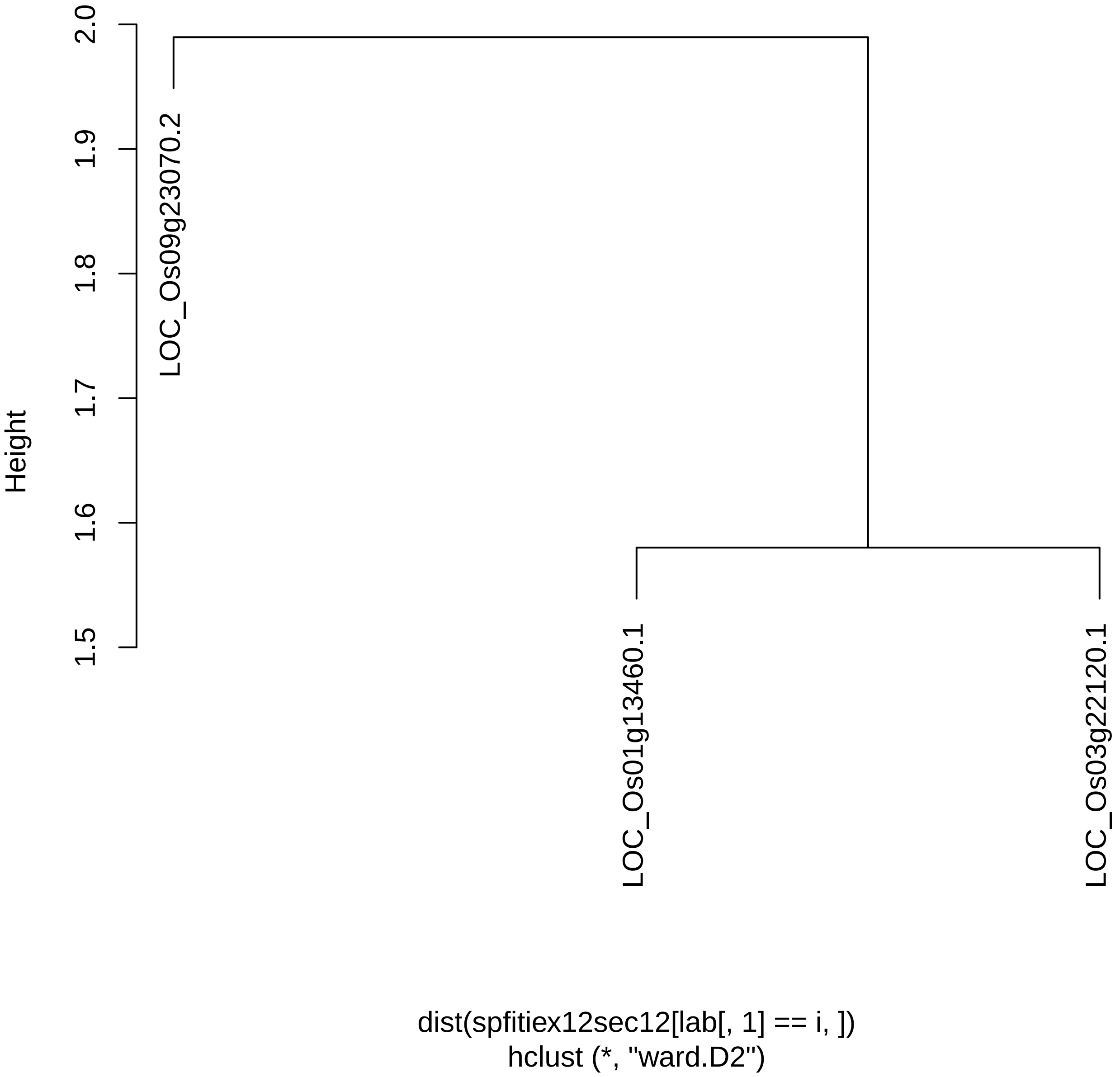

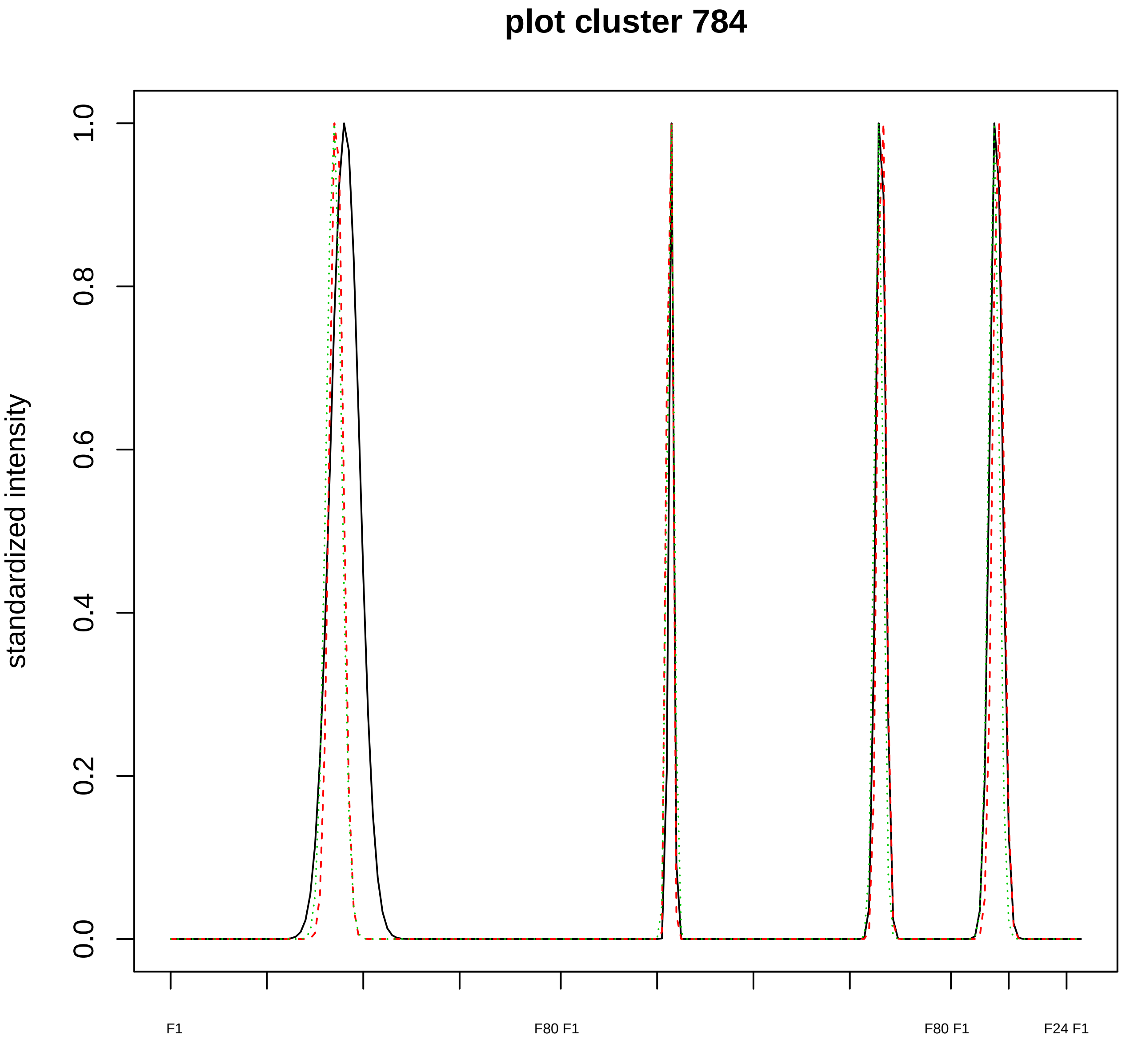
plot cluster 784
standardized intensity
0.0
0.2
0.4
0.6
0.8
1.0
F1
F80 F1
F80 F1
F24 F1

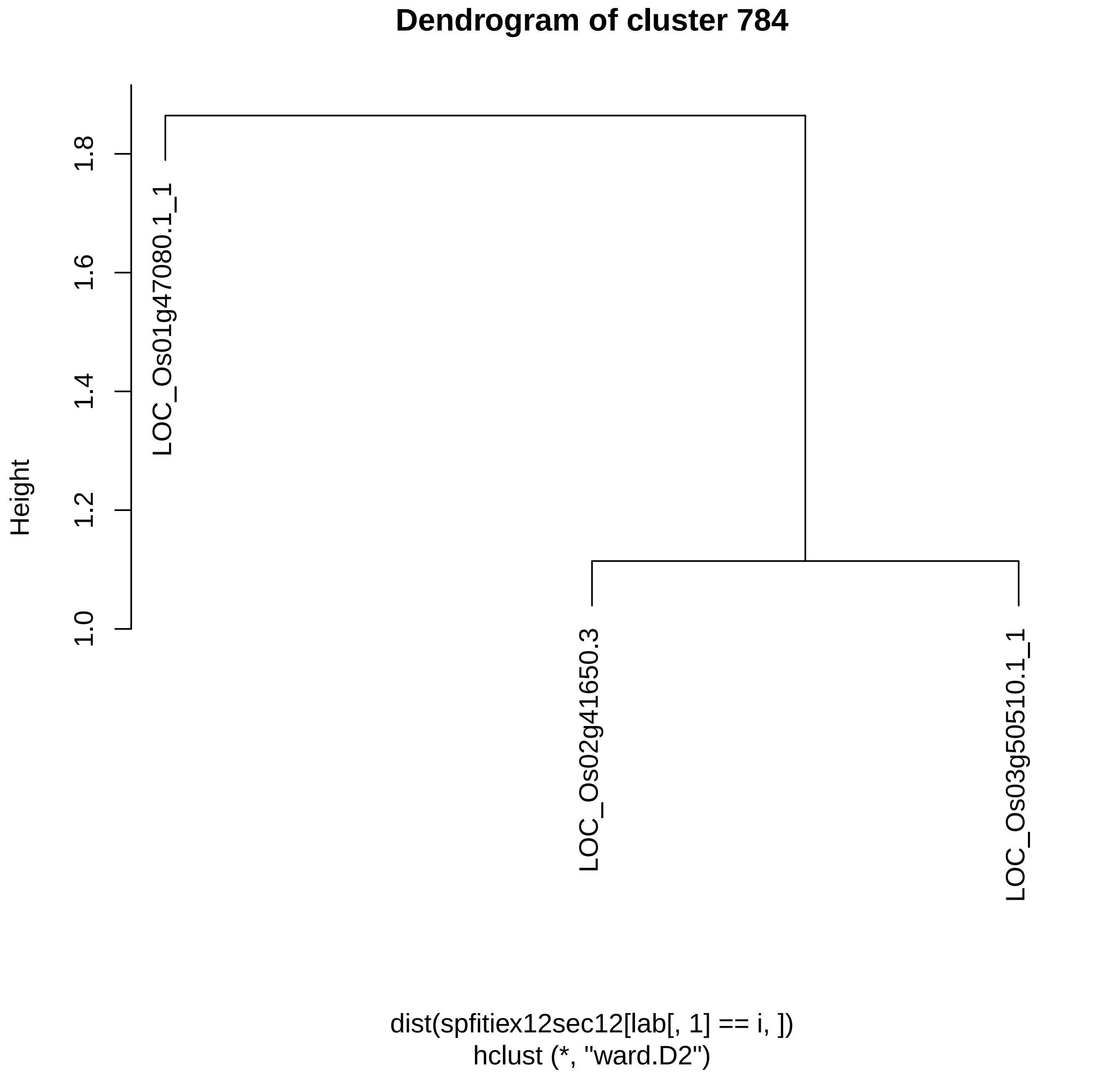
Dendrogram of cluster 784
Height
1.0
1.2
1.4
1.6
1.8
LOC_Os01g47080.1_1
LOC_Os02g41650.3
LOC_Os03g50510.1_1
dist(spfitiex12sec12[lab[, 1] == i, ])
hclust (*, "ward.D2")

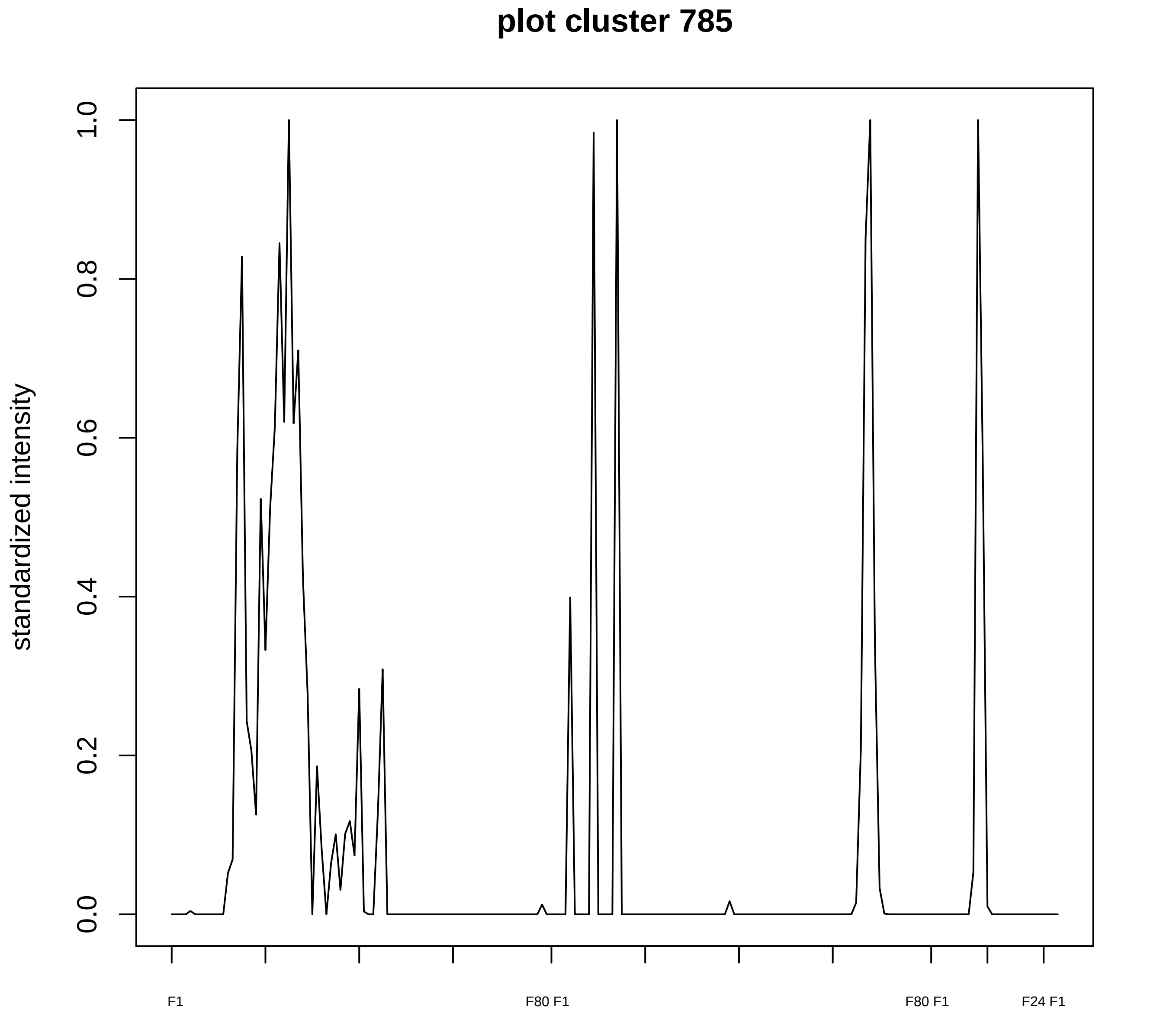
plot cluster 785
standardized intensity
0.0
0.2
0.4
0.6
0.8
1.0
F1
F80 F1
F80 F1
F24 F1

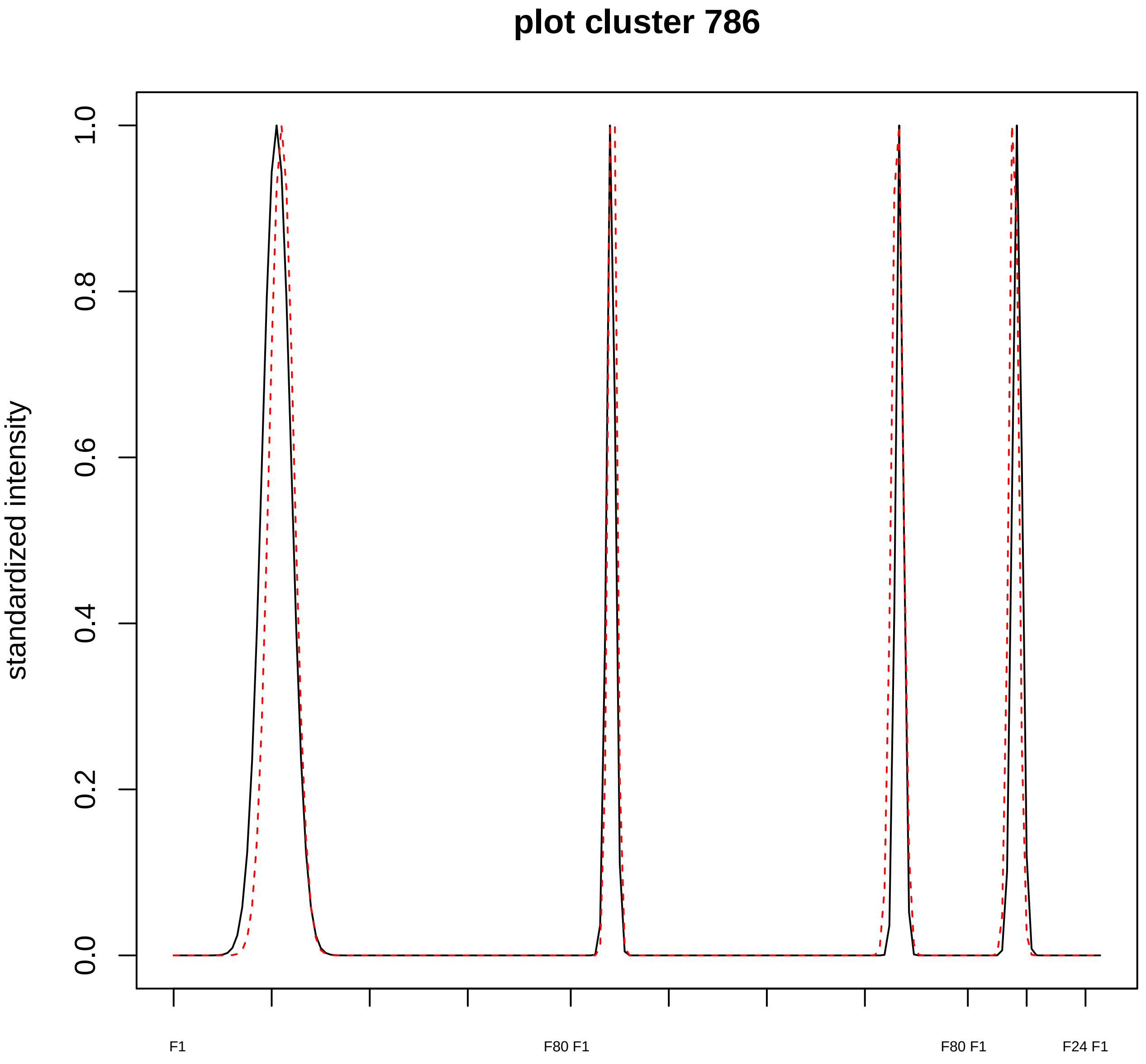

plot cluster 786
standardized intensity
0.0
0.2
0.4
0.6
0.8
1.0
F1
F80 F1
F80 F1
F24 F1

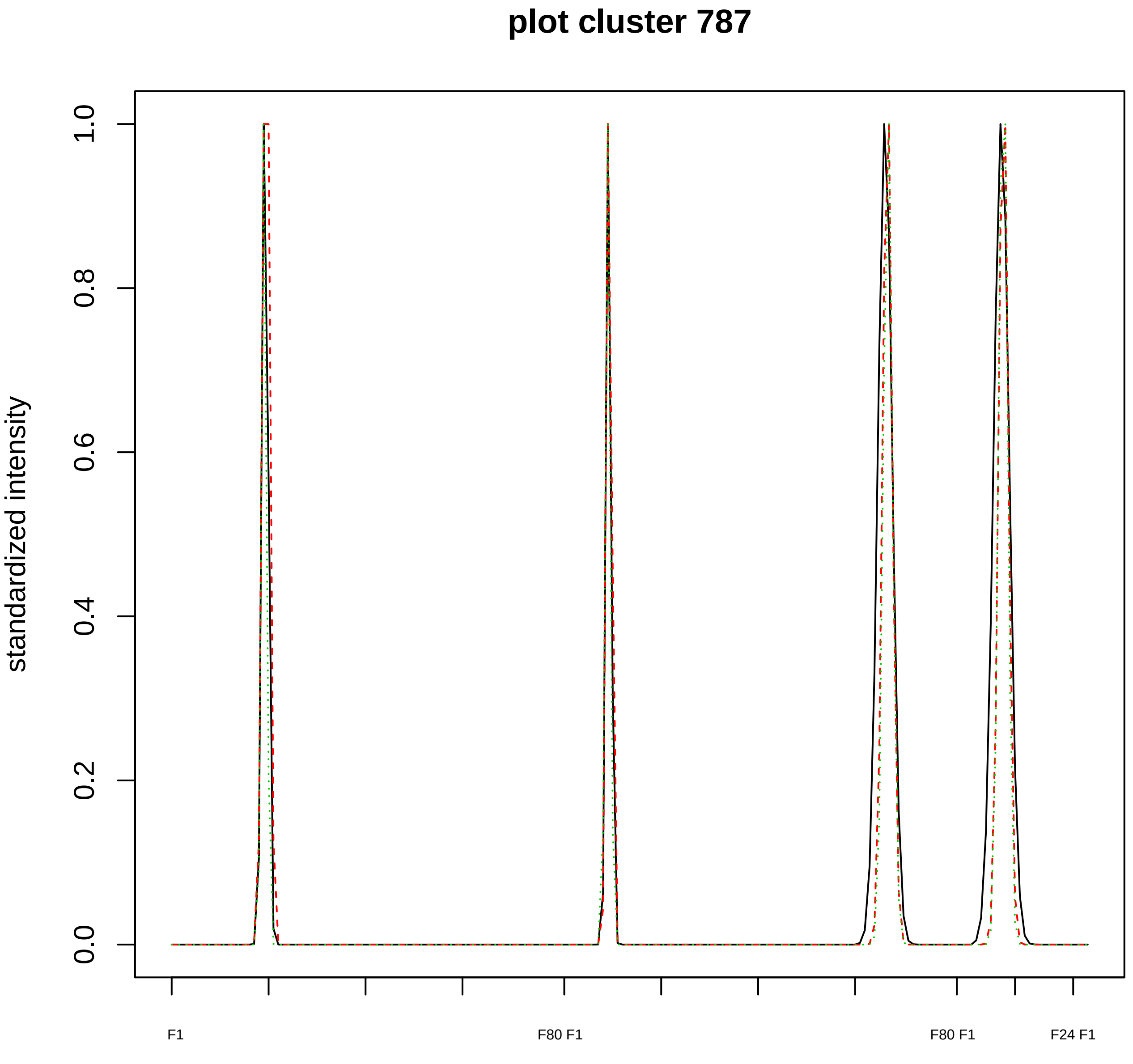

plot cluster 787
standardized intensity
0.0
0.2
0.4
0.6
0.8
1.0
F1
F80 F1
F80 F1
F24 F1

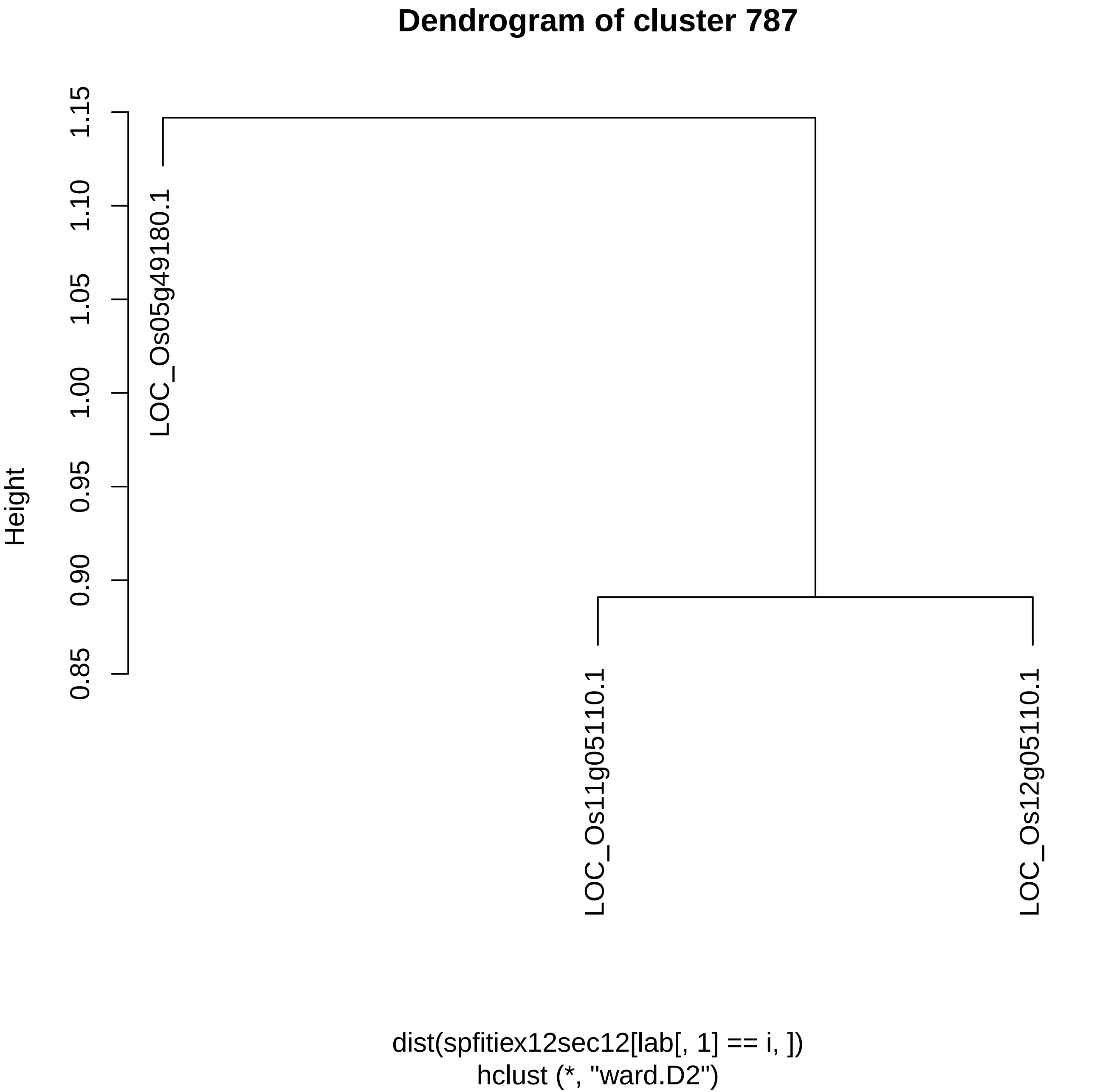
Dendrogram of cluster 787
Height
0.85
0.90
0.95
1.00
1.05
1.10
1.15
LOC_Os05g49180.1
LOC_Os11g05110.1
LOC_Os12g05110.1
dist(spfitiex12sec12[lab[, 1] == i, ])
hclust (*, "ward.D2")

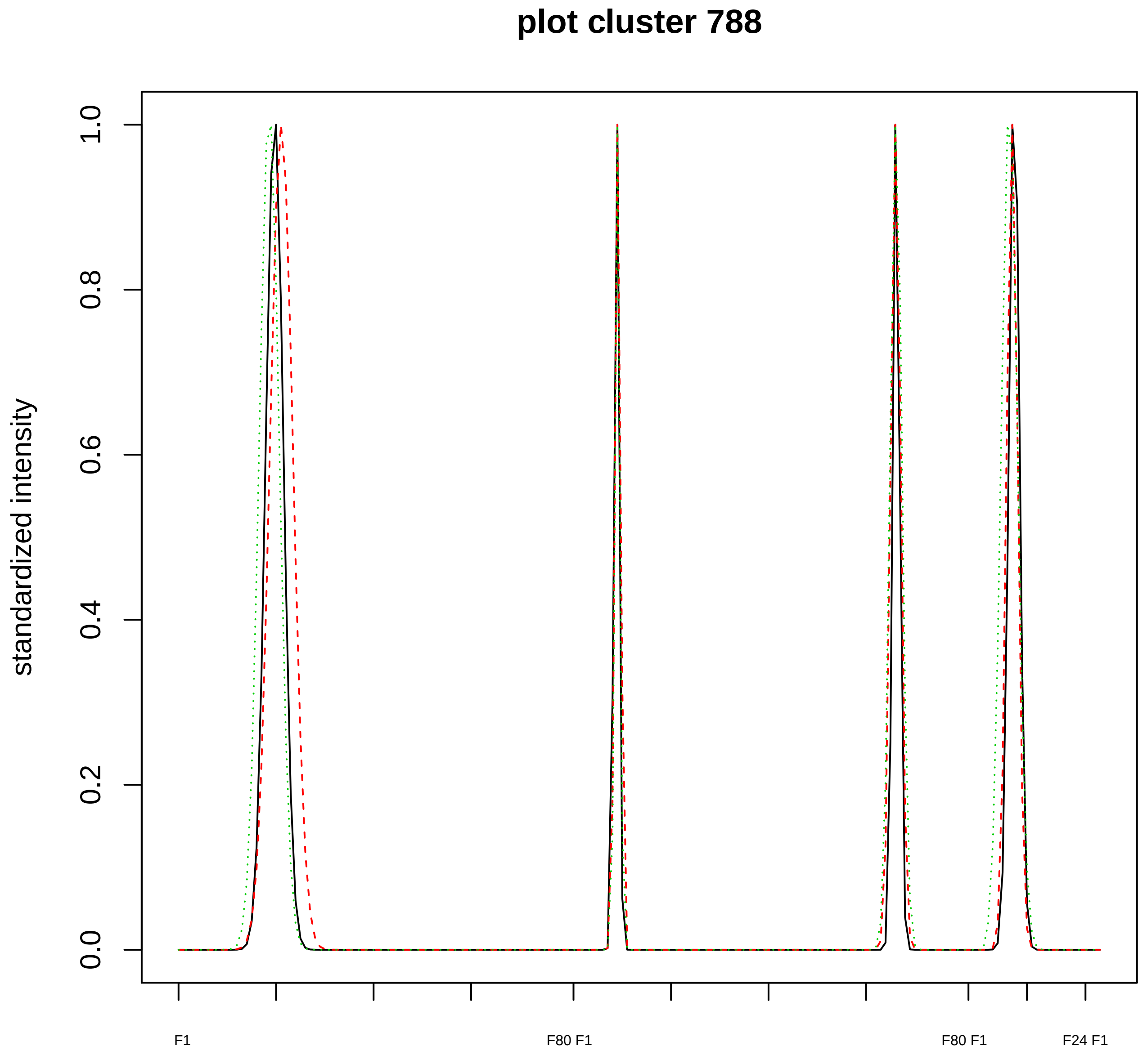

plot cluster 788
standardized intensity
0.0
0.2
0.4
0.6
0.8
1.0
F1
F80 F1
F80 F1
F24 F1

#### Dendrogram of cluster 788

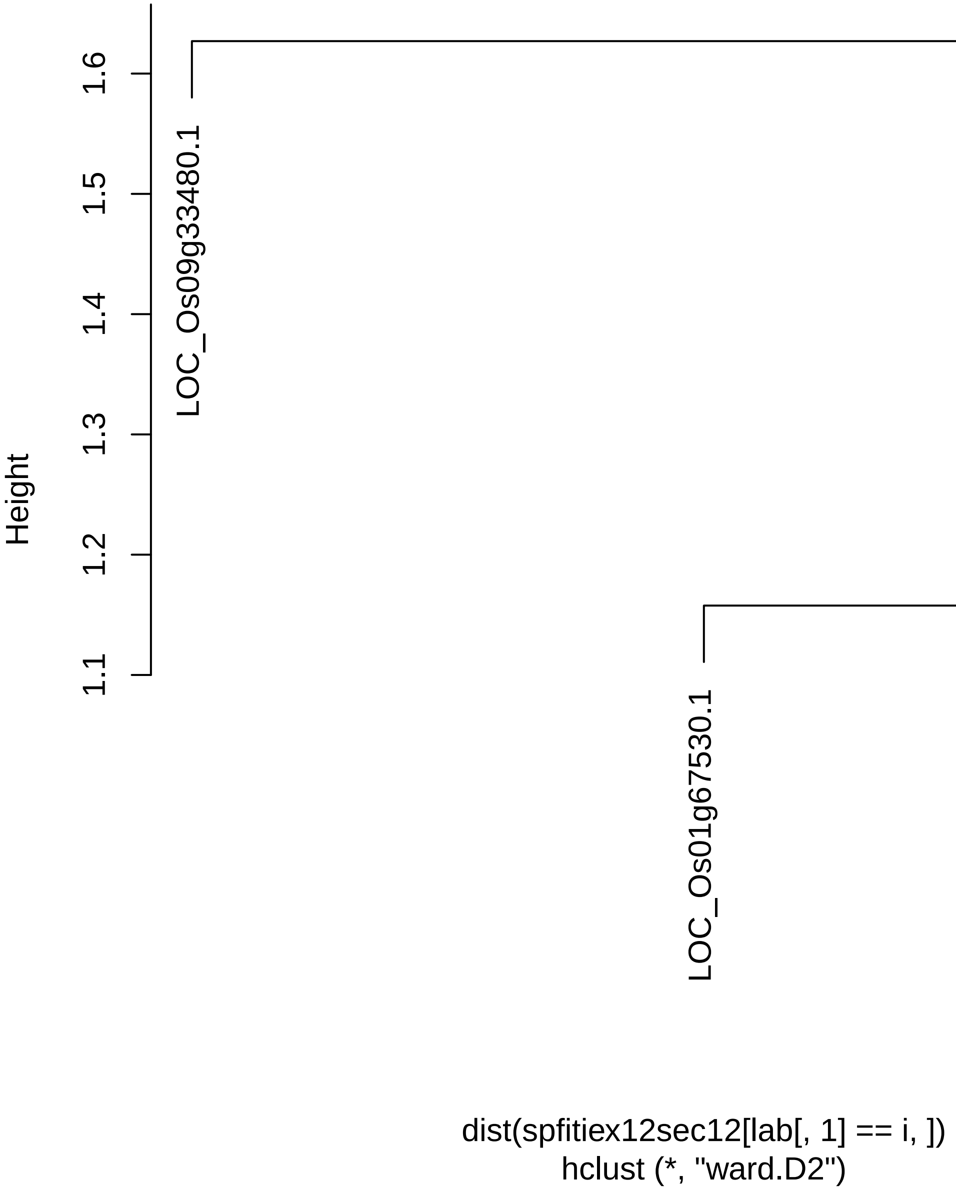

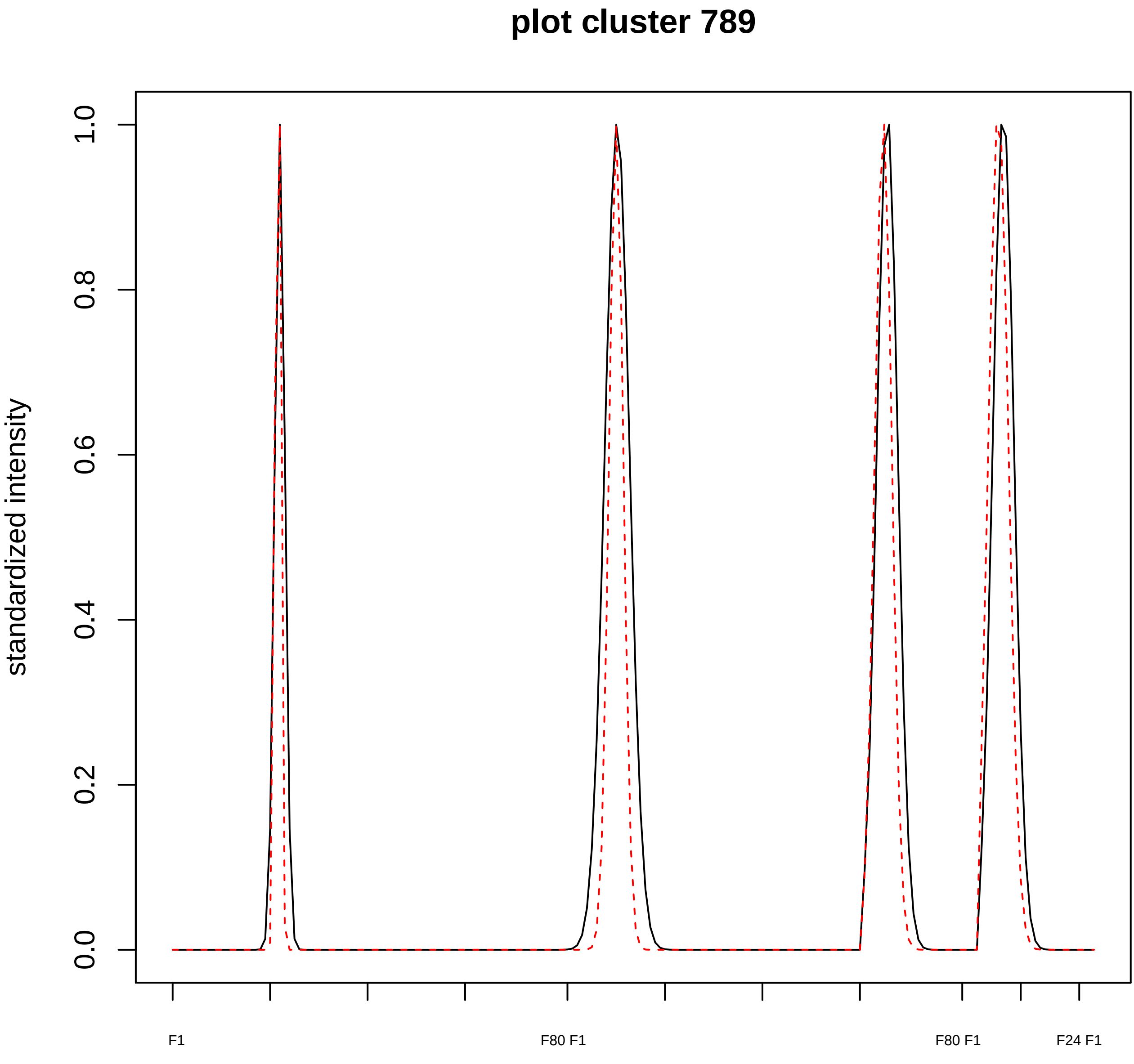
plot cluster 789
standardized intensity
0.0
0.2
0.4
0.6
0.8
1.0
F1
F80 F1
F80 F1
F24 F1

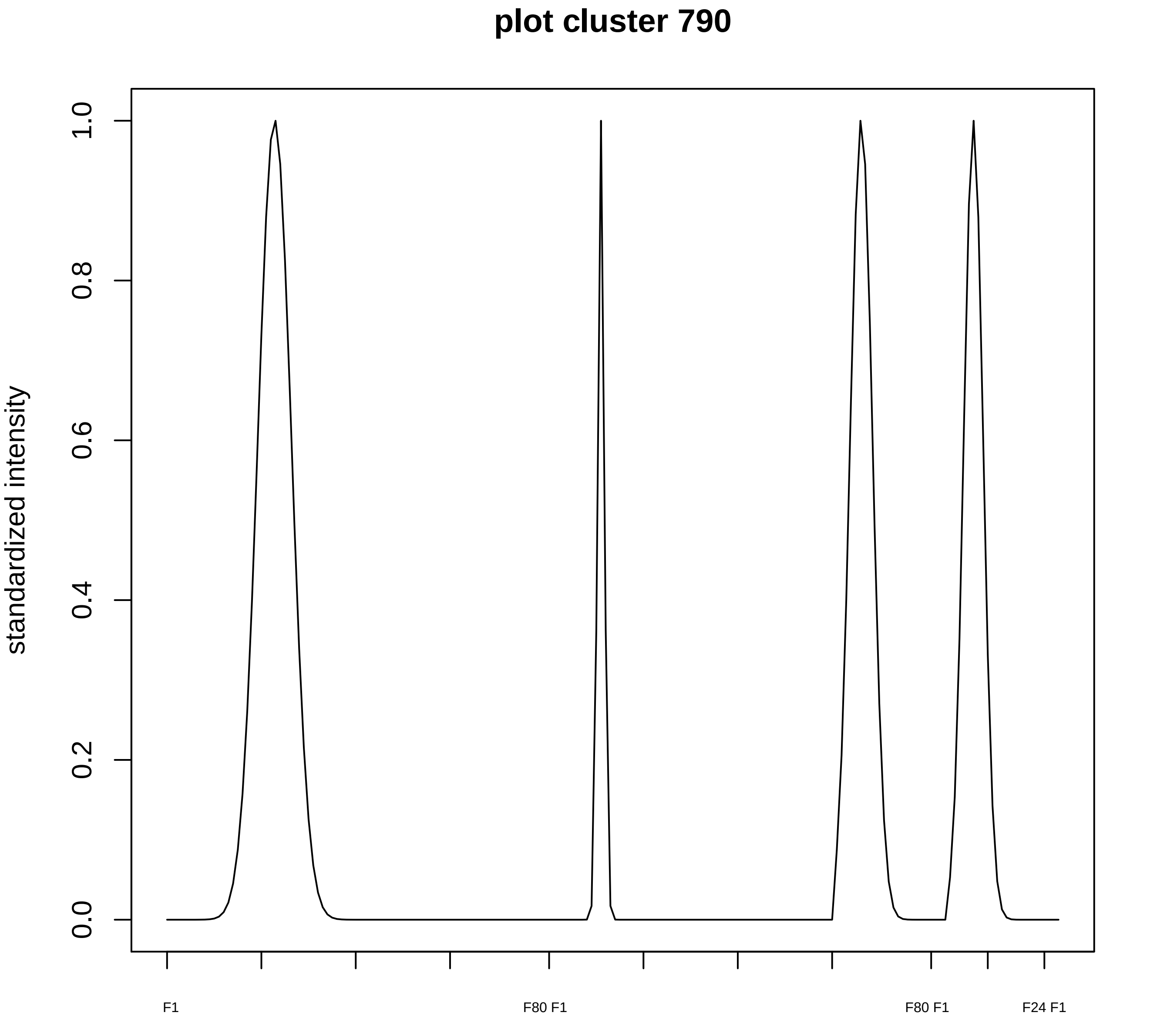

plot cluster 790
standardized intensity
0.0
0.2
0.4
0.6
0.8
1.0
F1
F80 F1
F80 F1
F24 F1

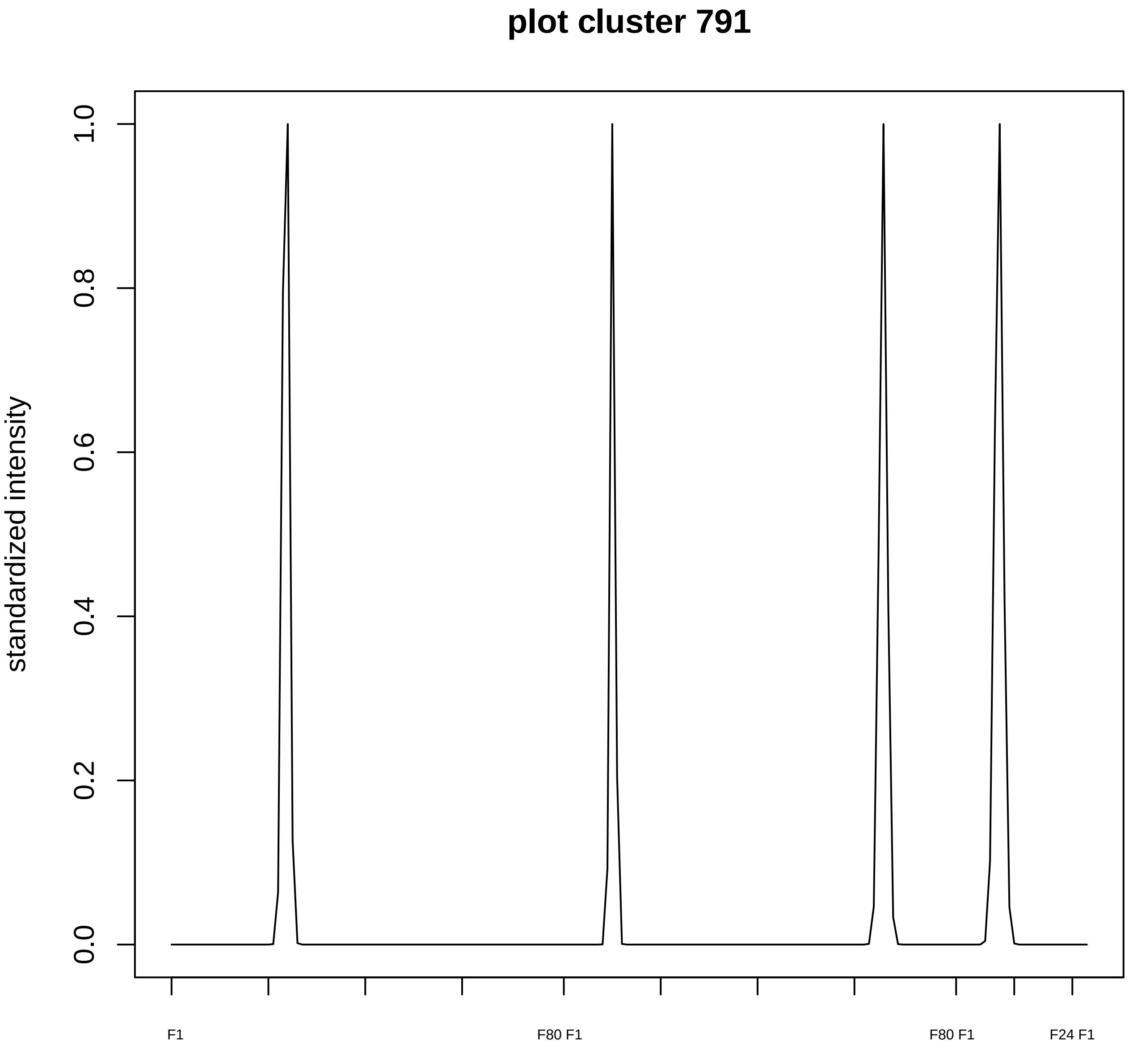

plot cluster 791
standardized intensity
0.0
0.2
0.4
0.6
0.8
1.0
F1
F80 F1
F80 F1
F24 F1

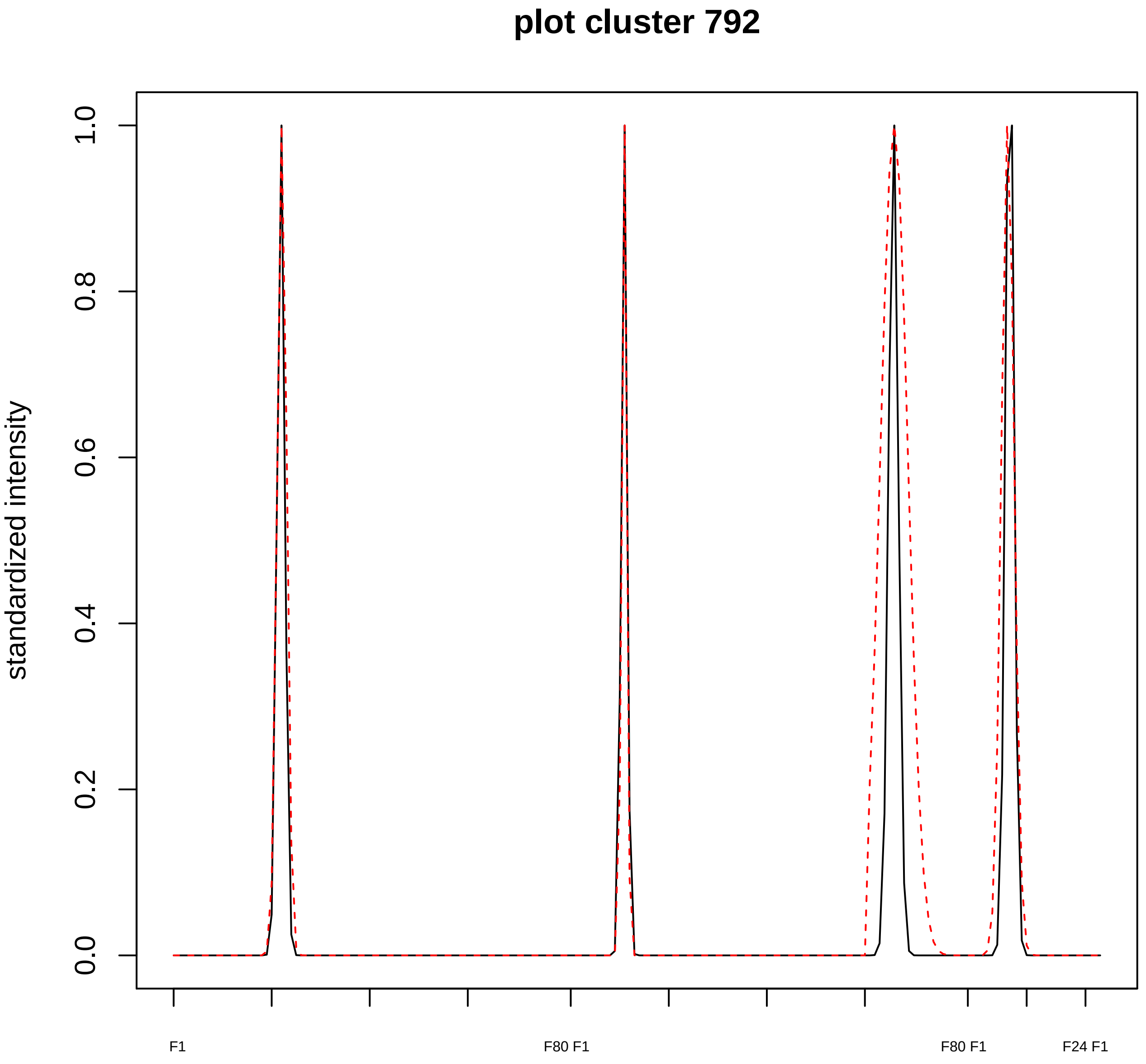
plot cluster 792
standardized intensity
0.0
0.2
0.4
0.6
0.8
1.0
F1
F80 F1
F80 F1
F24 F1

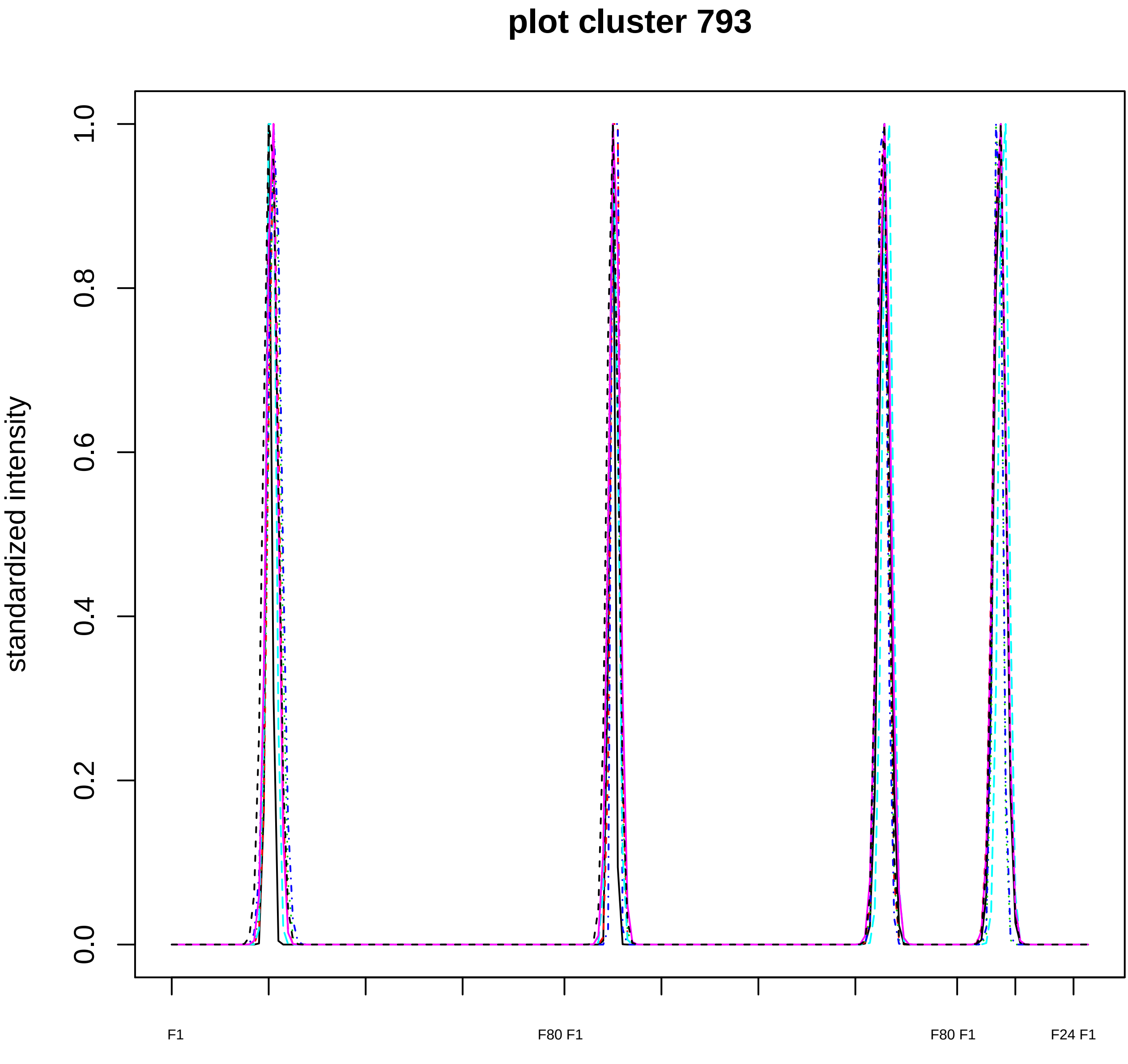

plot cluster 793
standardized intensity
0.0
0.2
0.4
0.6
0.8
1.0
F1
F80 F1
F80 F1
F24 F1

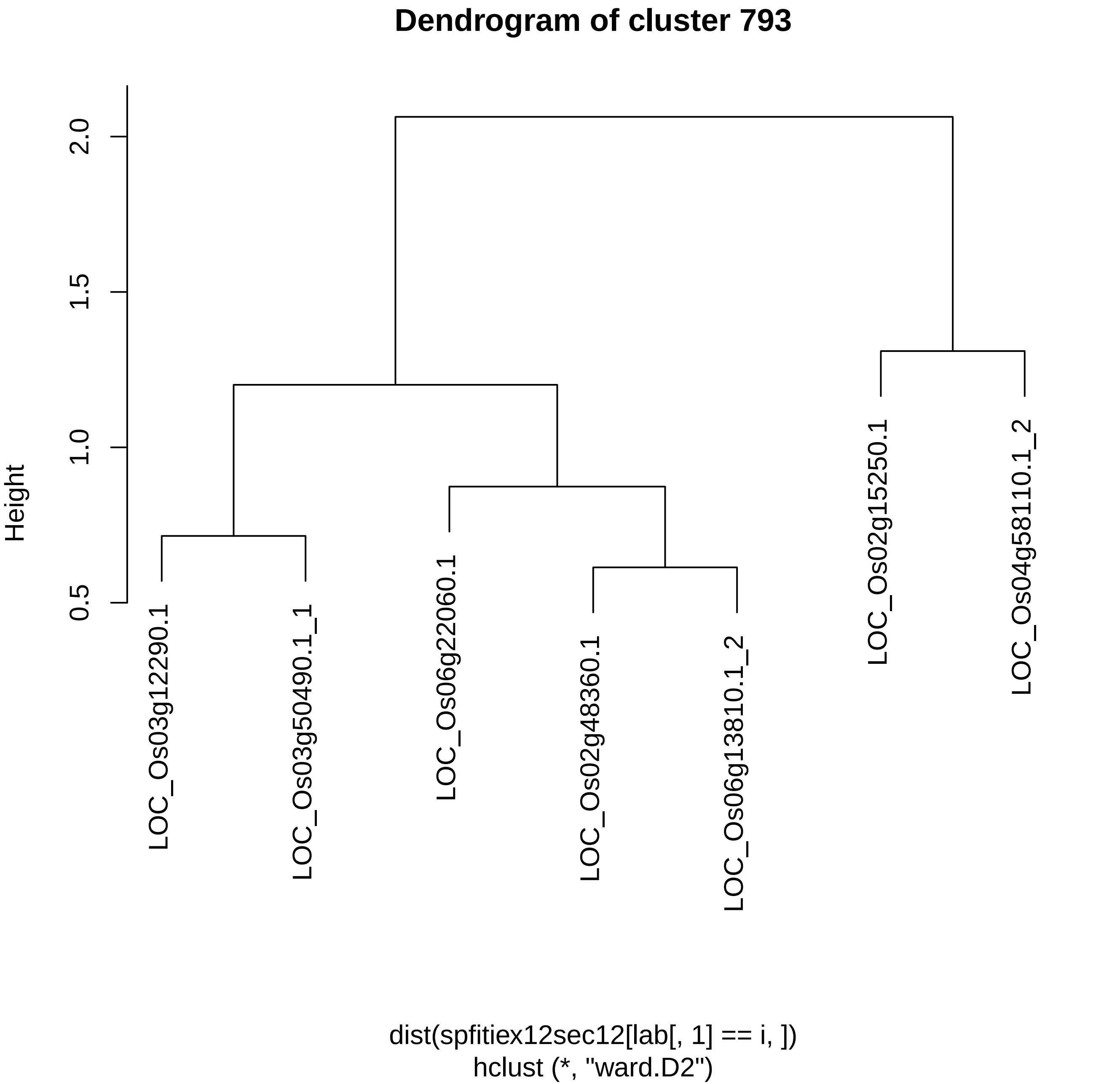
Dendrogram of cluster 793
Height
0.5
1.0
1.5
2.0
LOC_Os03g12290.1
LOC_Os03g50490.1_1
LOC_Os06g22060.1
LOC_Os02g48360.1
LOC_Os06g13810.1_2
LOC_Os02g15250.1
LOC_Os04g58110.1_2
dist(spfitiex12sec12[lab[, 1] == i, ])
hclust (*, "ward.D2")

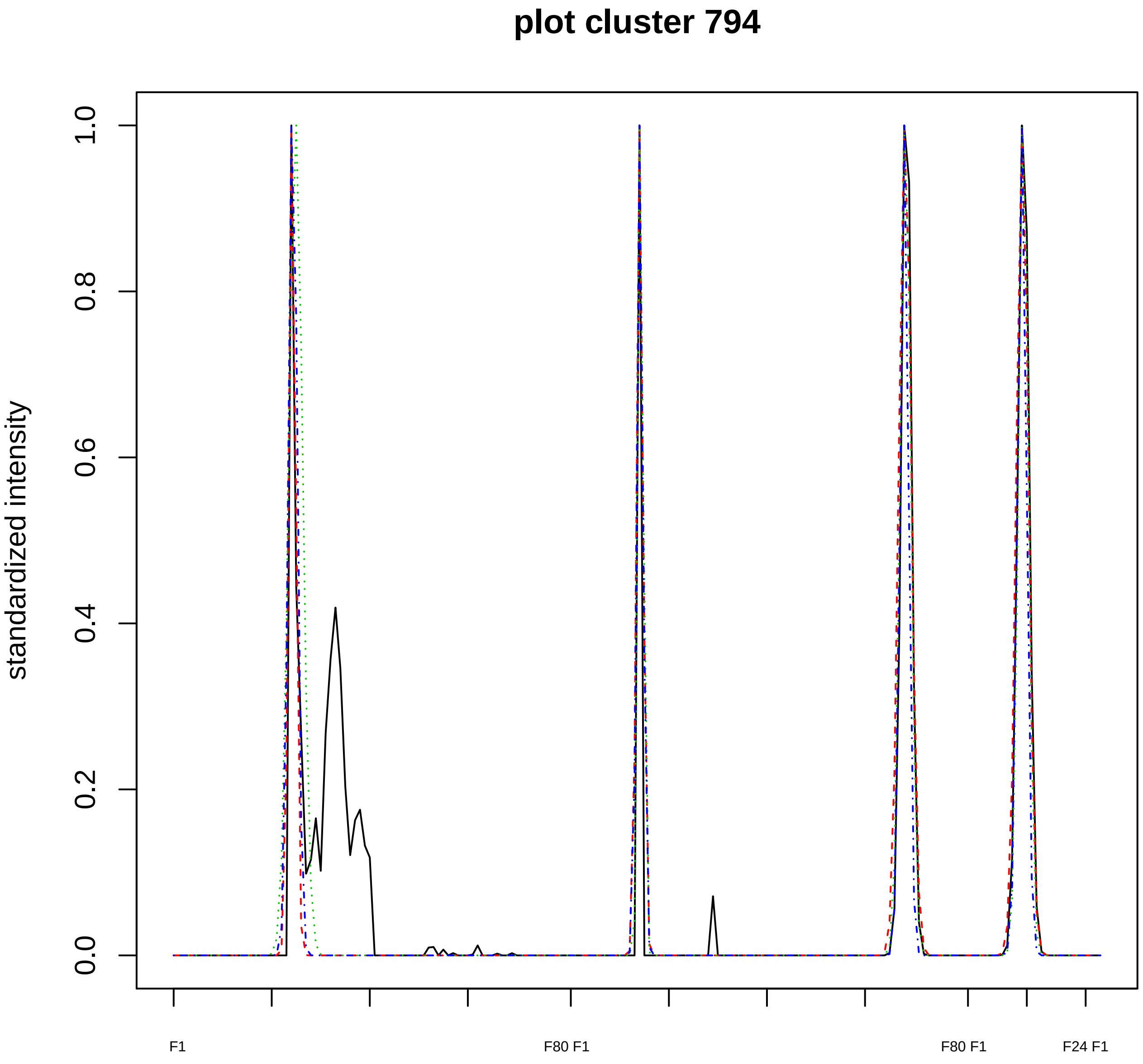

plot cluster 794
standardized intensity
1.0
0.8
0.6
0.4
0.2
0.0
F1
F80 F1
F80 F1
F24 F1

#### Dendrogram of cluster 794

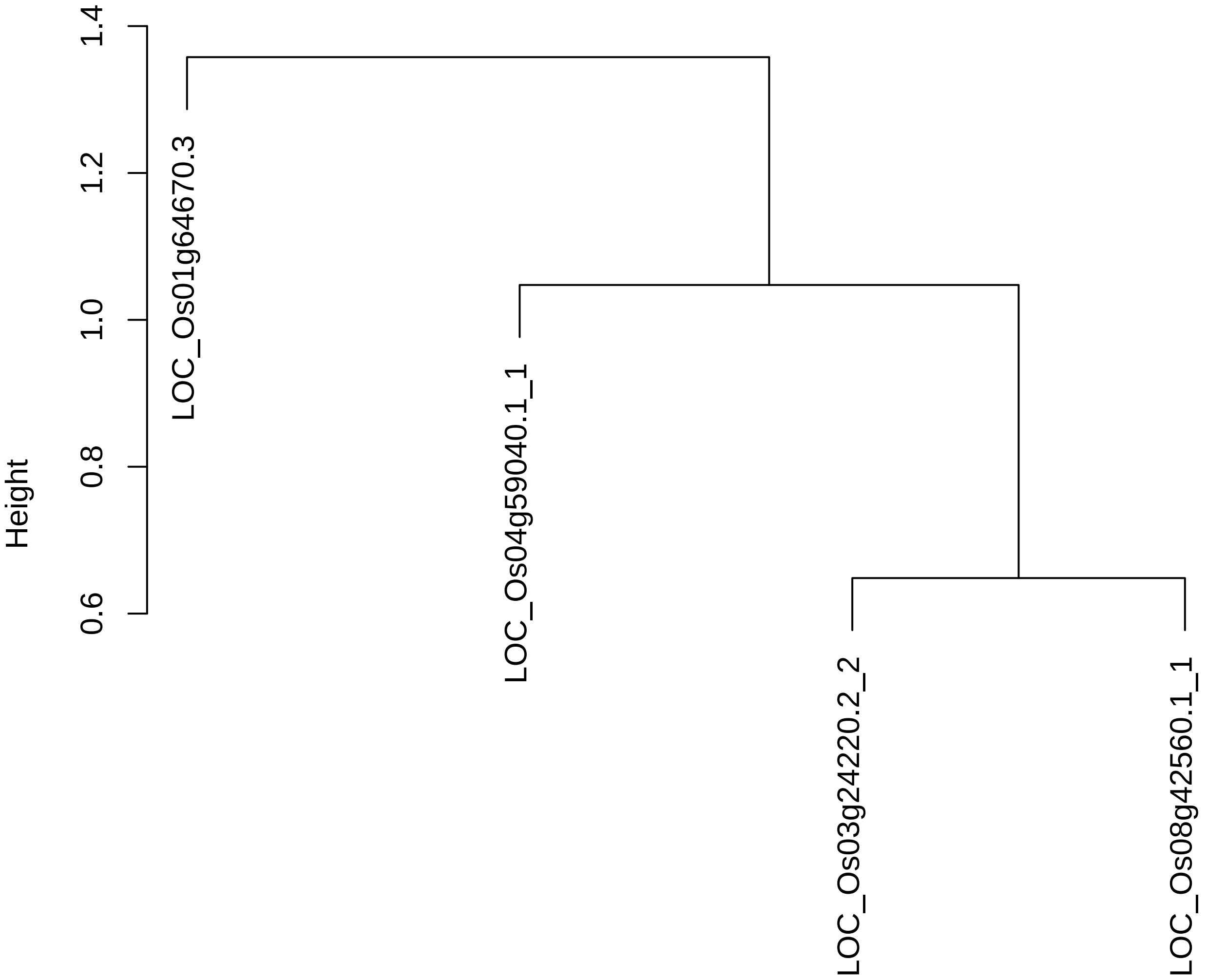


dist(spfitiex12sec12[lab[, 1] == i, ])
hclust (*, "ward.D2")

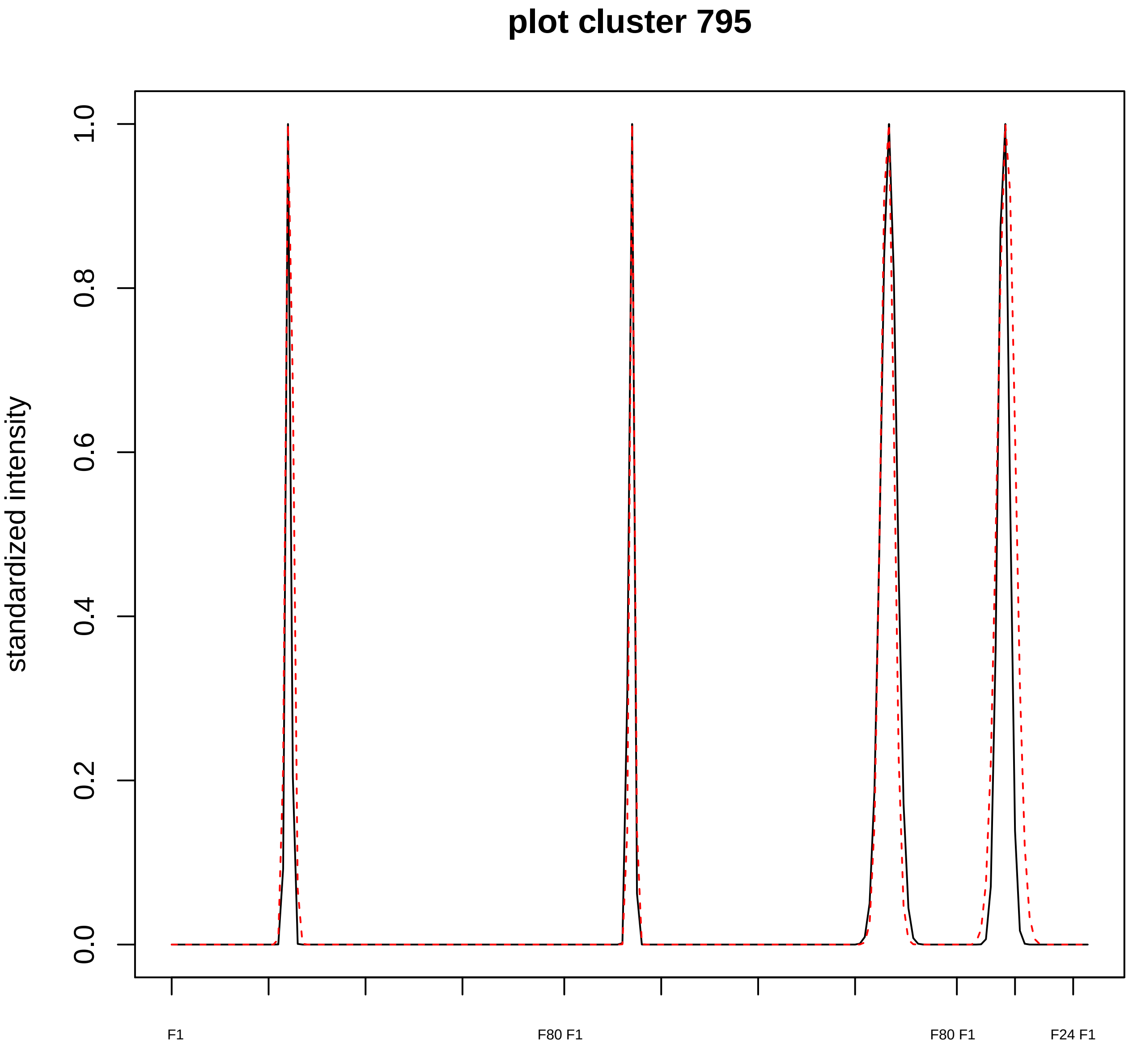
plot cluster 795
standardized intensity
0.0
0.2
0.4
0.6
0.8
1.0
F1
F80 F1
F80 F1
F24 F1

#### plot cluster 796

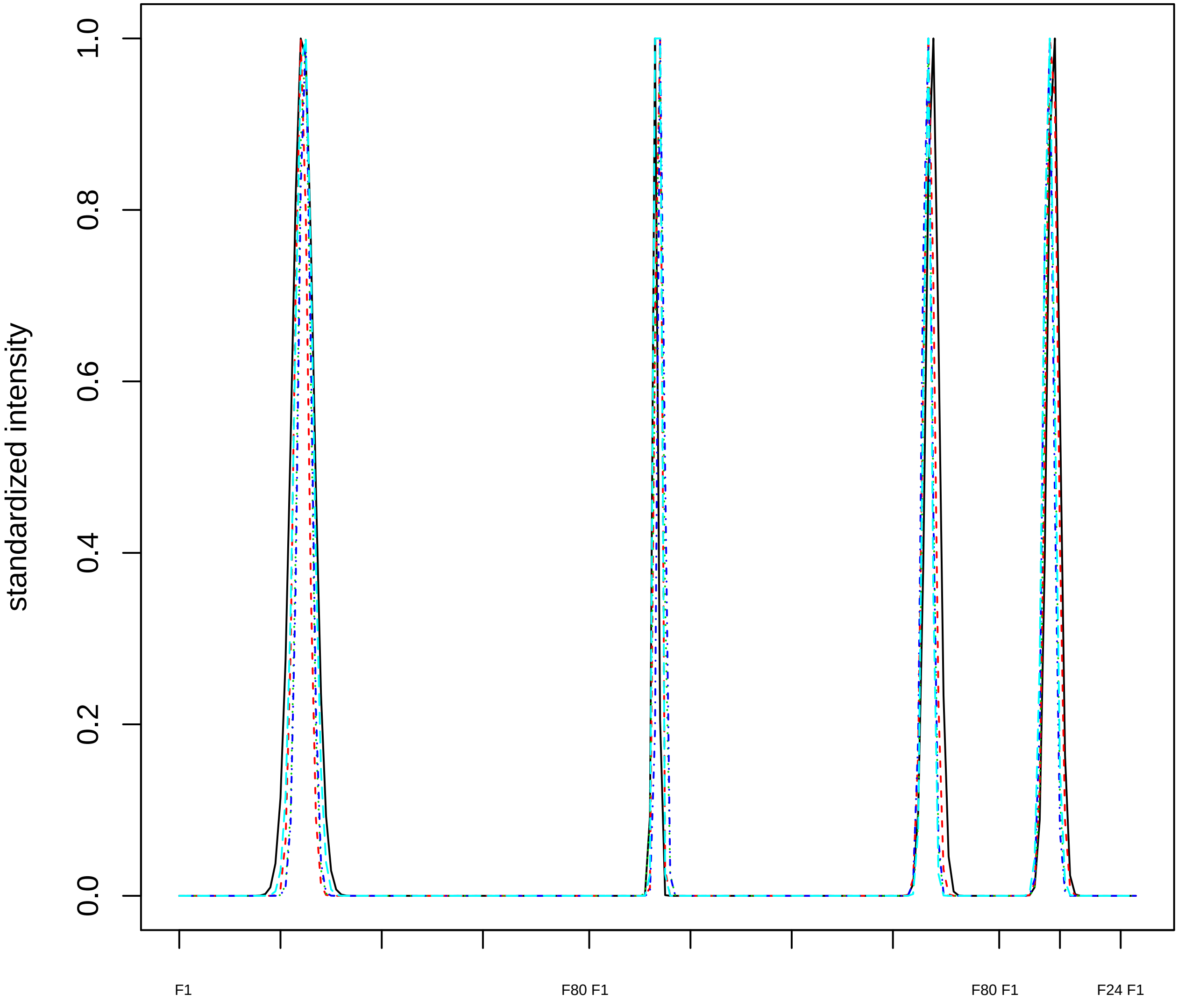

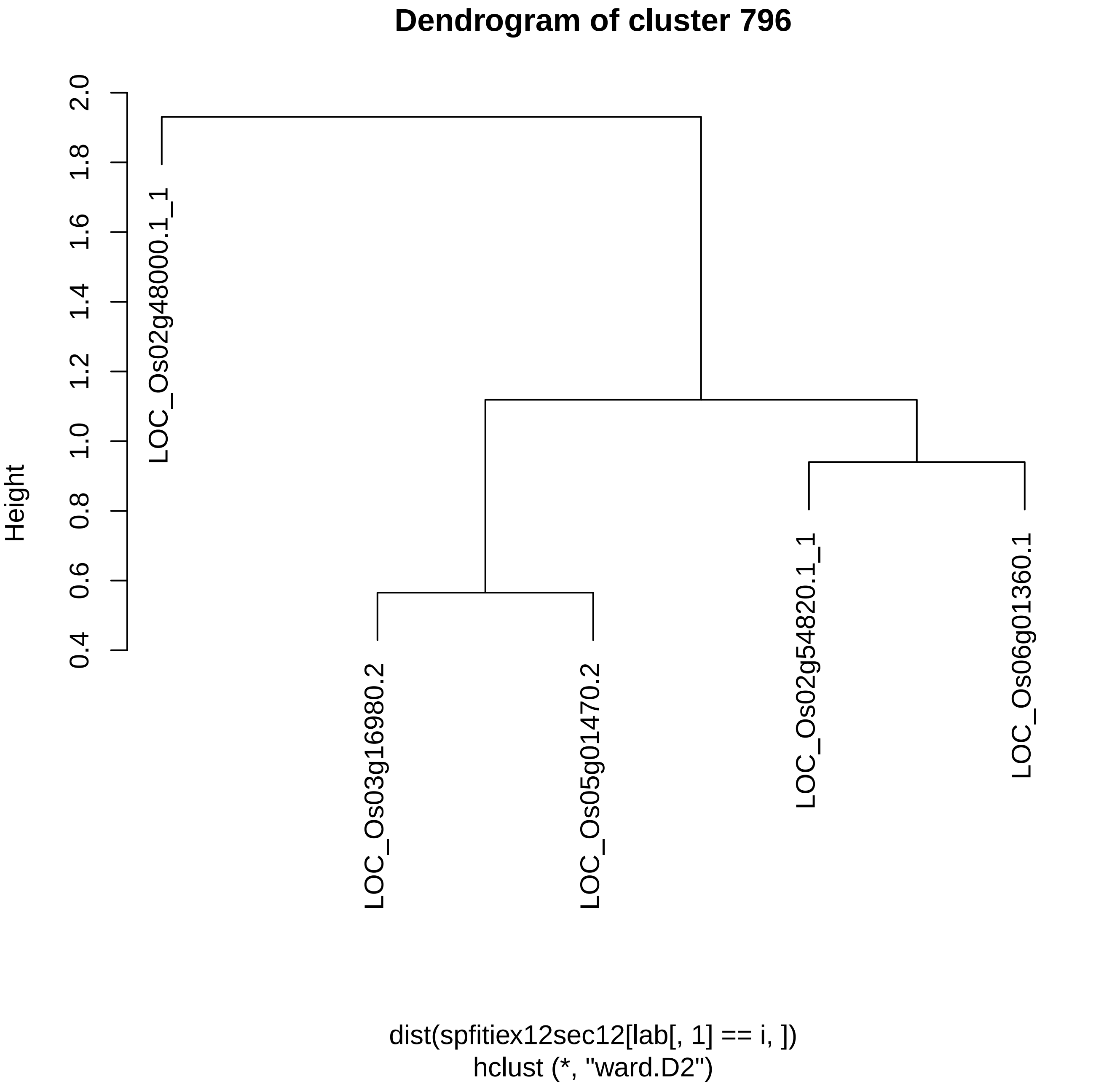
Dendrogram of cluster 796
Height
0.4
0.6
0.8
1.0
1.2
1.4
1.6
1.8
2.0
LOC_Os02g48000.1_1
LOC_Os03g16980.2
LOC_Os05g01470.2
LOC_Os02g54820.1_1
LOC_Os06g01360.1
dist(spfitiex12sec12[lab[, 1] == i, ])
hclust (*, "ward.D2")

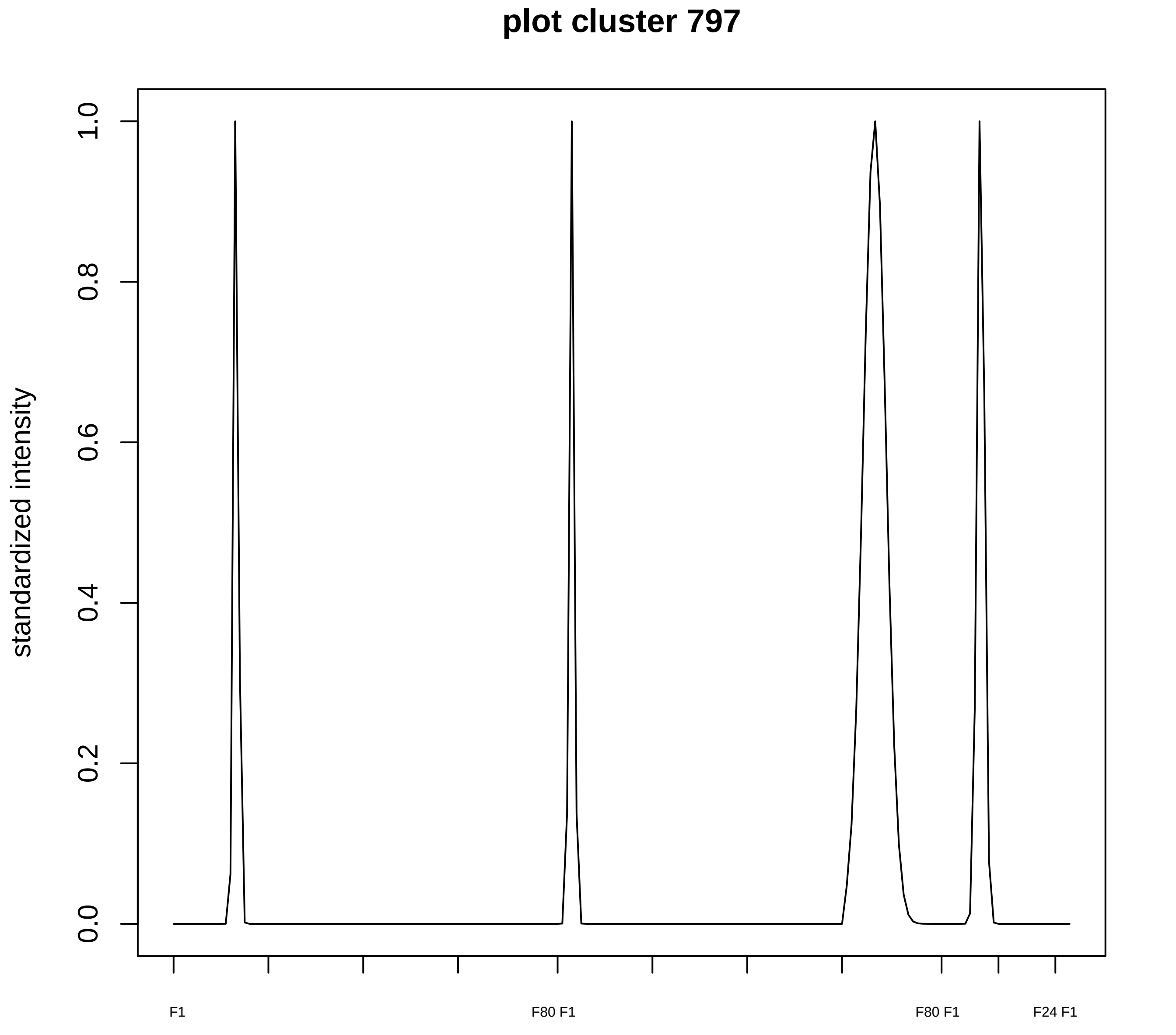

plot cluster 797
standardized intensity
0.0
0.2
0.4
0.6
0.8
1.0
F1
F80 F1
F80 F1
F24 F1

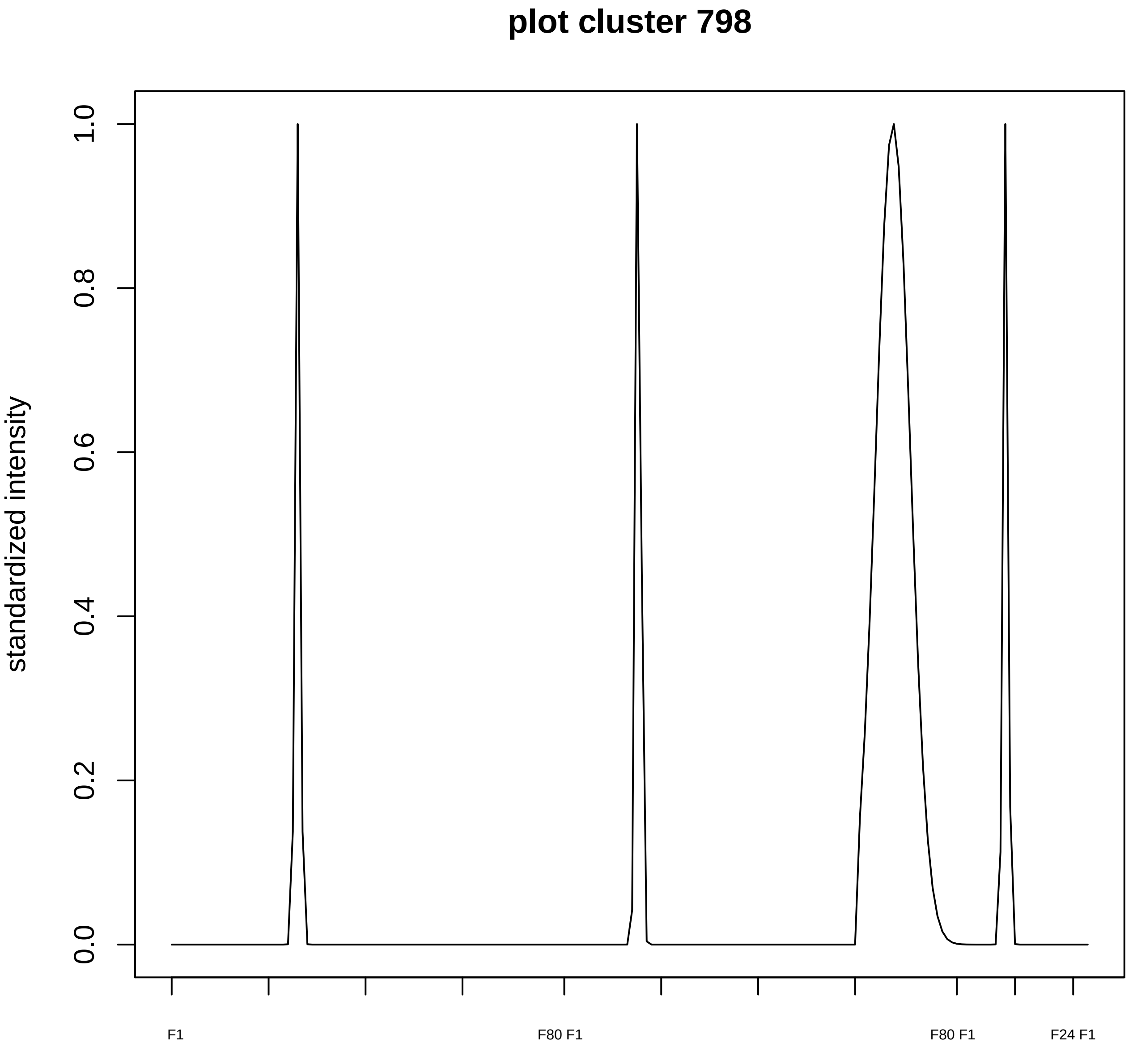
plot cluster 798
standardized intensity
0.0
0.2
0.4
0.6
0.8
1.0
F1
F80 F1
F80 F1
F24 F1

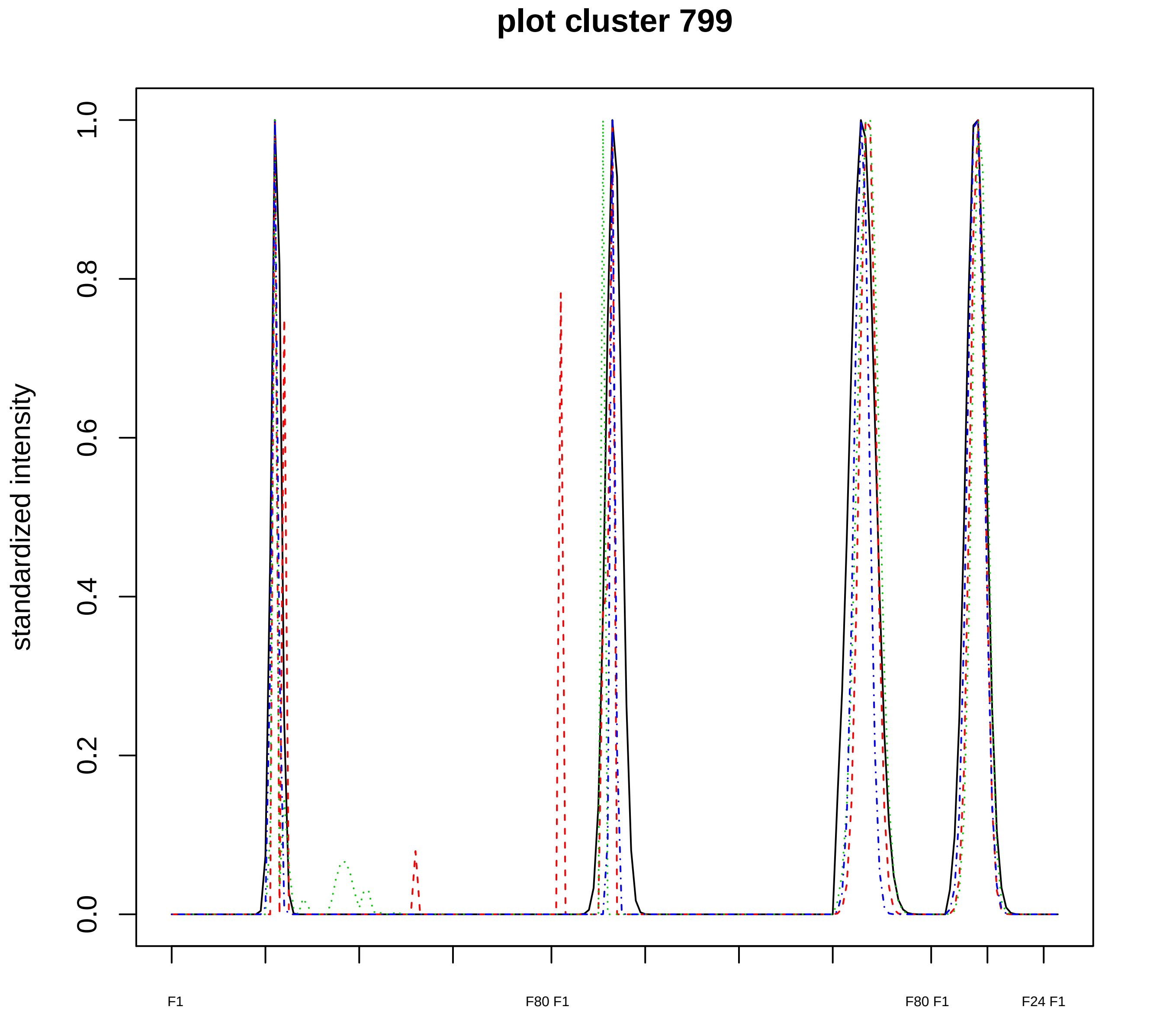

plot cluster 799
standardized intensity
0.0
0.2
0.4
0.6
0.8
1.0
F1
F80 F1
F80 F1
F24 F1

### Dendrogram of cluster 799

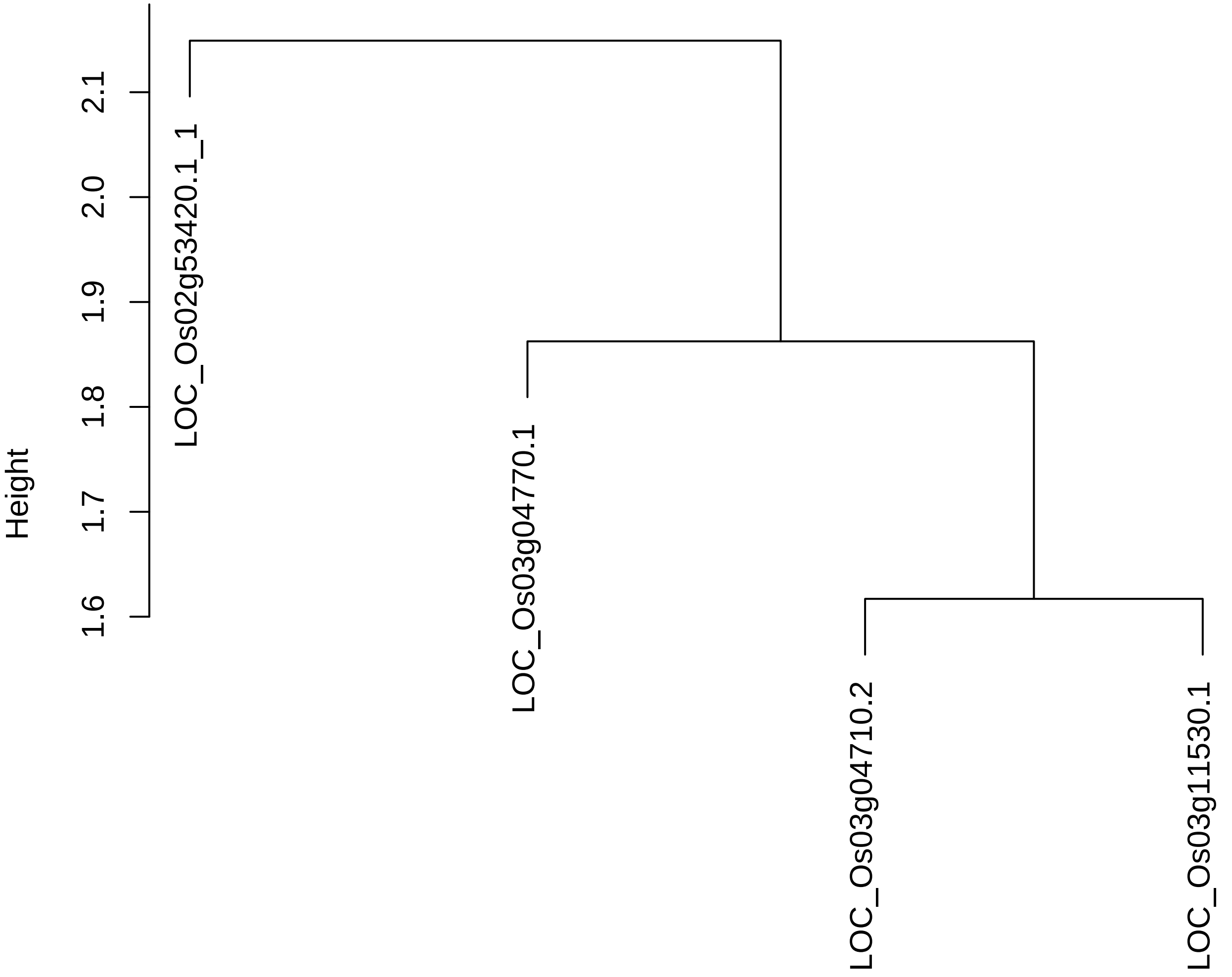


dist(spfitiex12sec12[lab[, 1] == i, ])
hclust (*, "ward.D2")

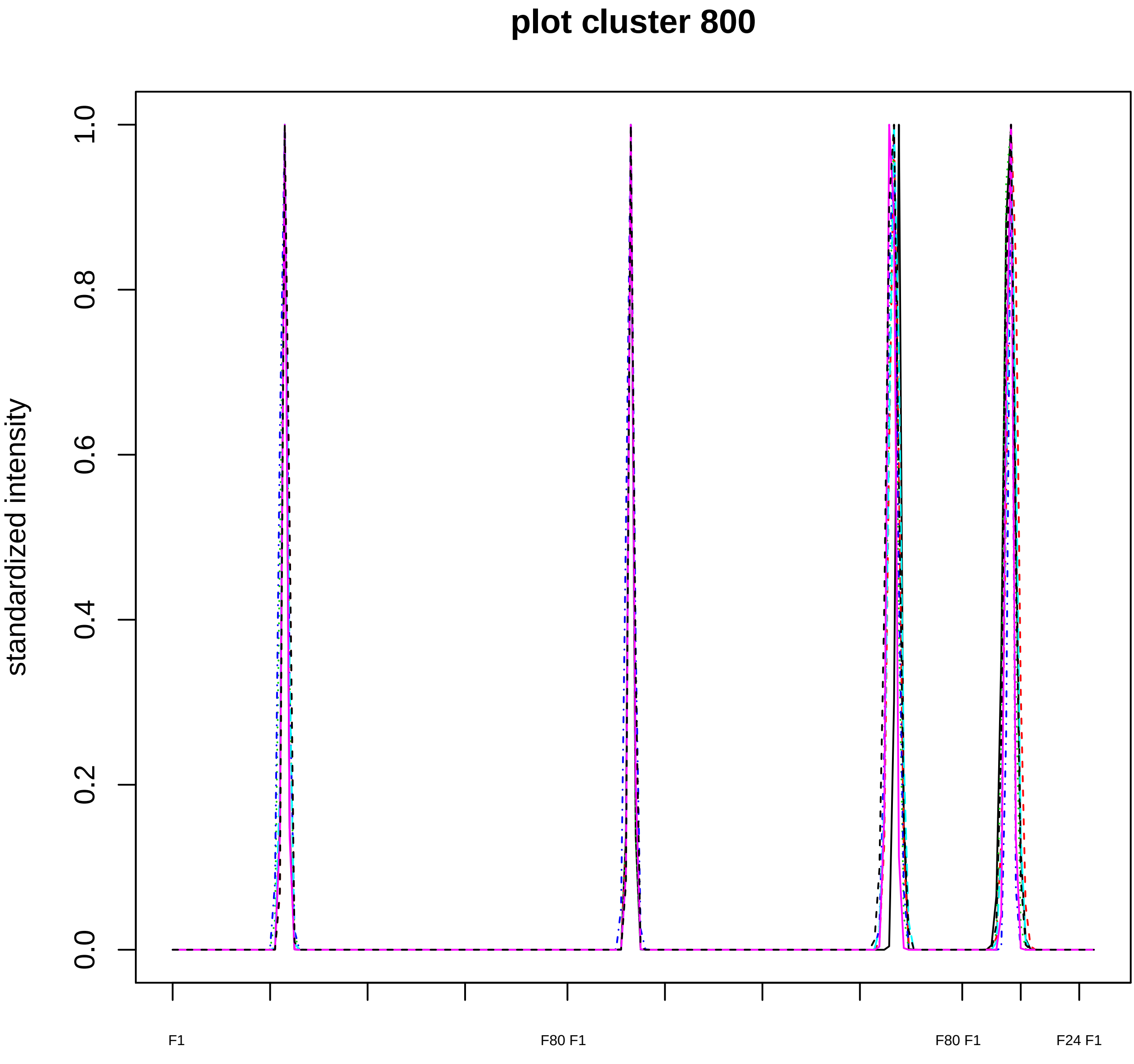

plot cluster 800
standardized intensity
0.0
0.2
0.4
0.6
0.8
1.0
F1
F80 F1
F80 F1
F24 F1

### Dendrogram of cluster 800

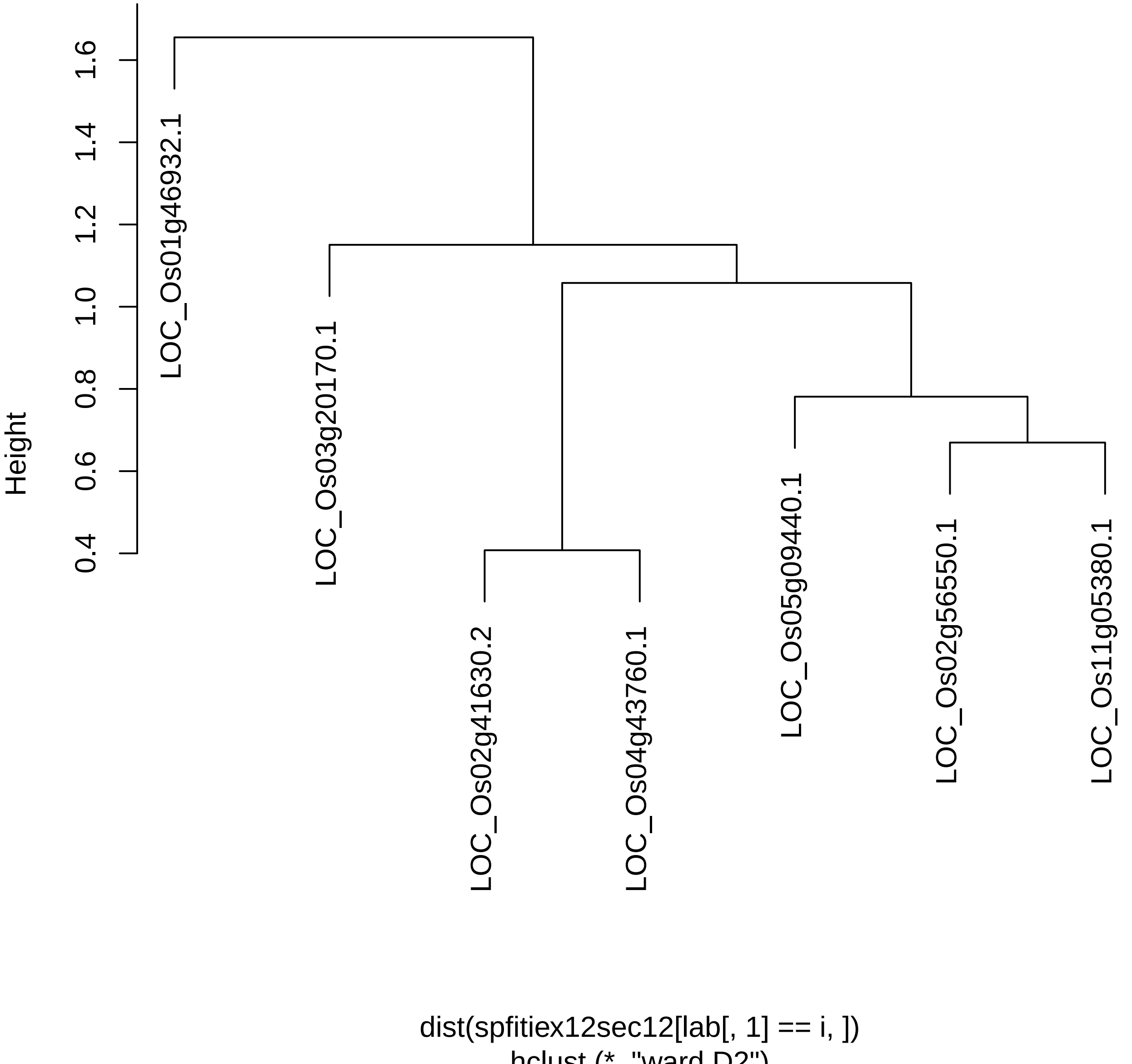

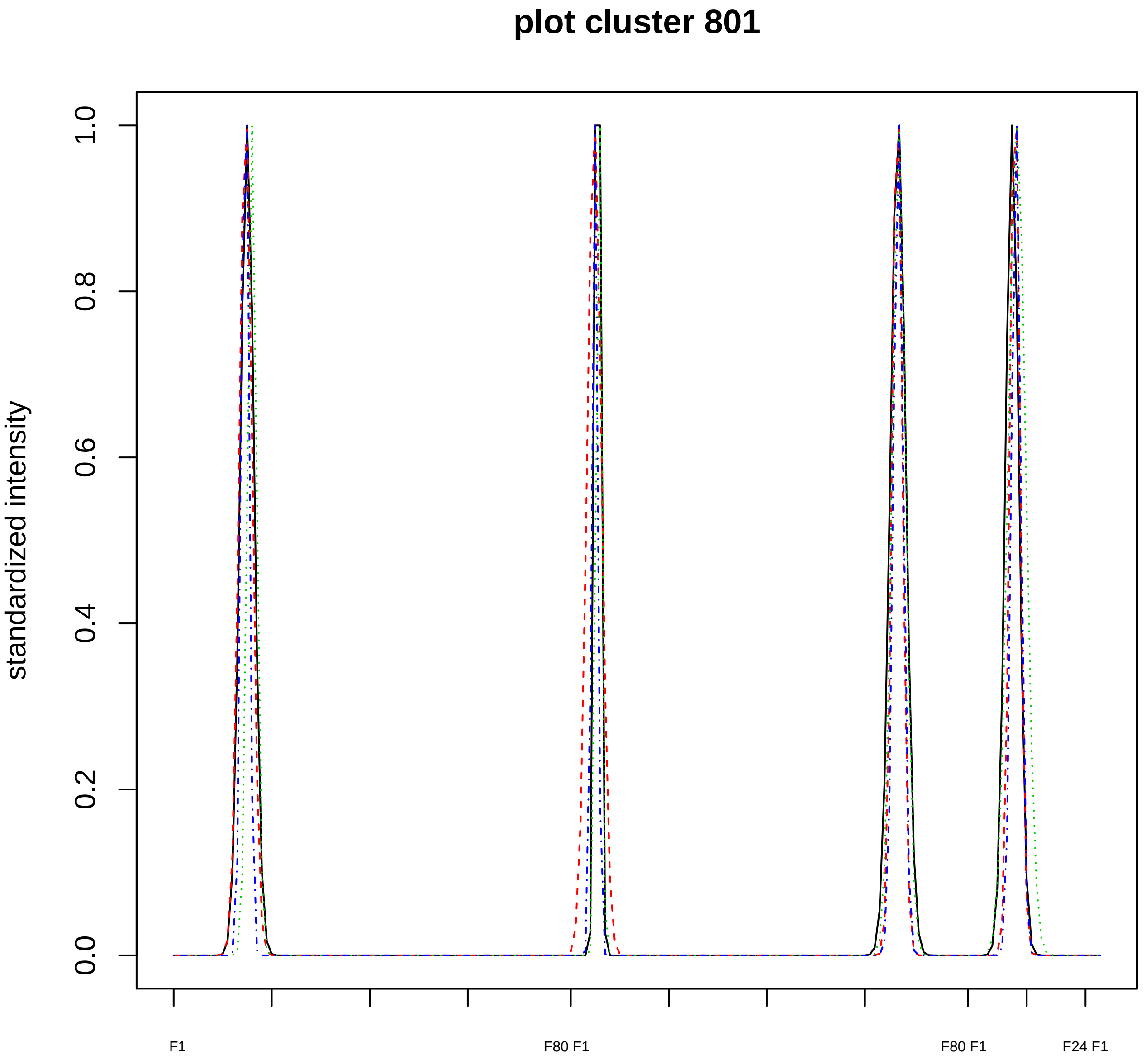
plot cluster 801
standardized intensity
0.0
0.2
0.4
0.6
0.8
1.0
F1
F80 F1
F80 F1
F24 F1

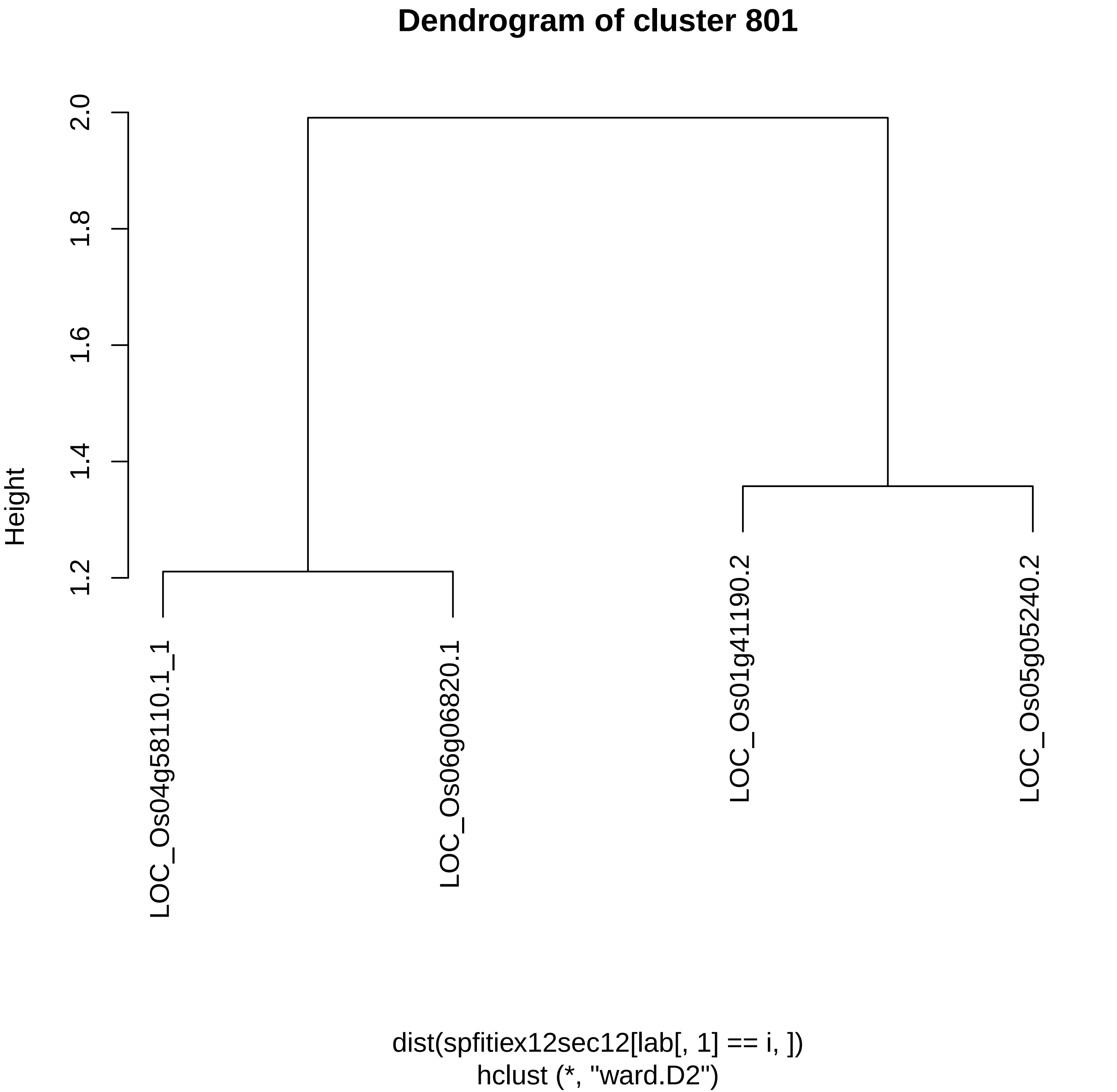
Dendrogram of cluster 801
Height
2.0
1.8
1.6
1.4
1.2
LOC_Os04g58110.1_1
LOC_Os06g06820.1
LOC_Os01g41190.2
LOC_Os05g05240.2
dist(spfitiex12sec12[lab[, 1] == i, ])
hclust (*, "ward.D2")

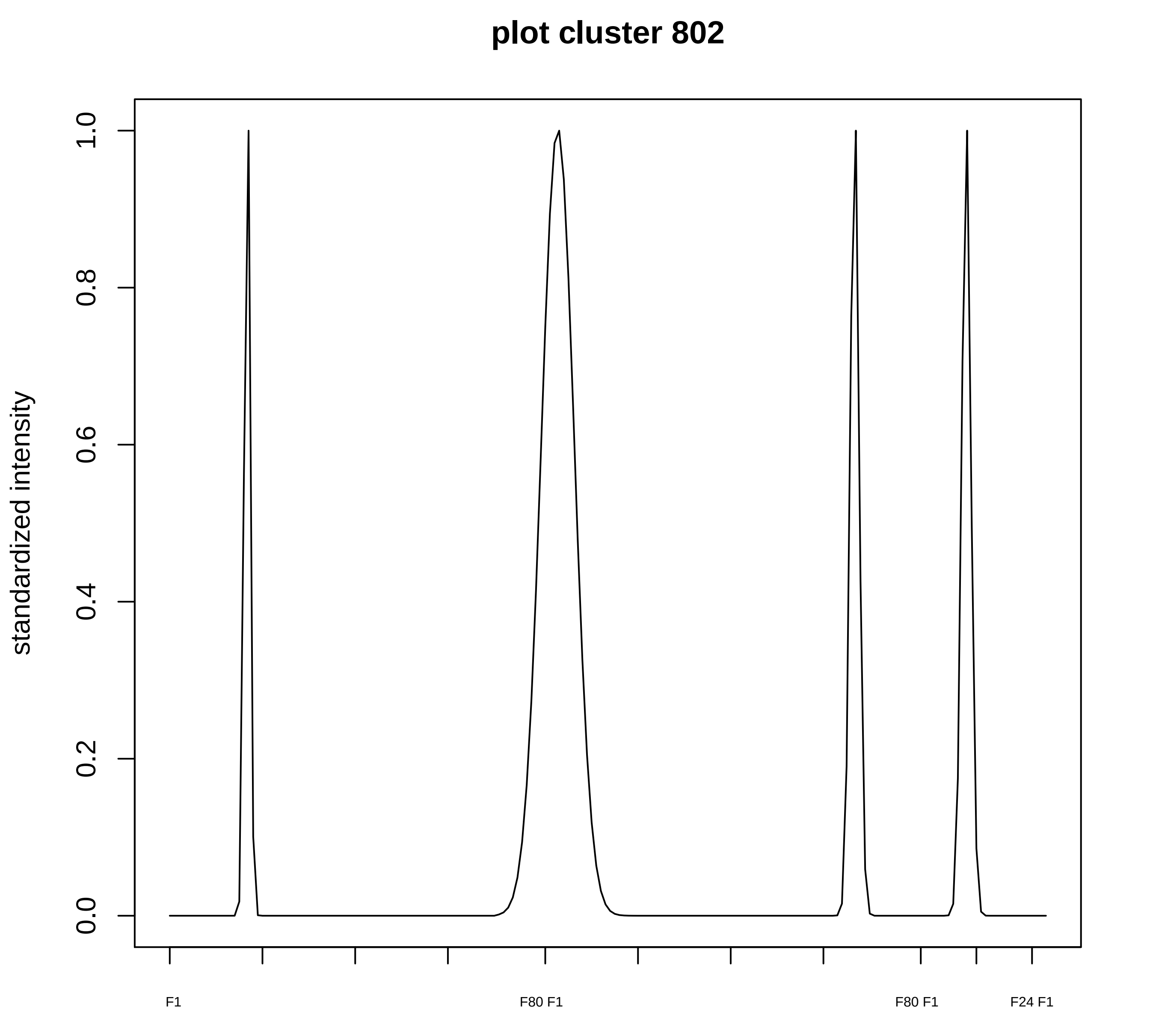

plot cluster 802
standardized intensity
0.0
0.2
0.4
0.6
0.8
1.0
F1
F80 F1
F80 F1
F24 F1

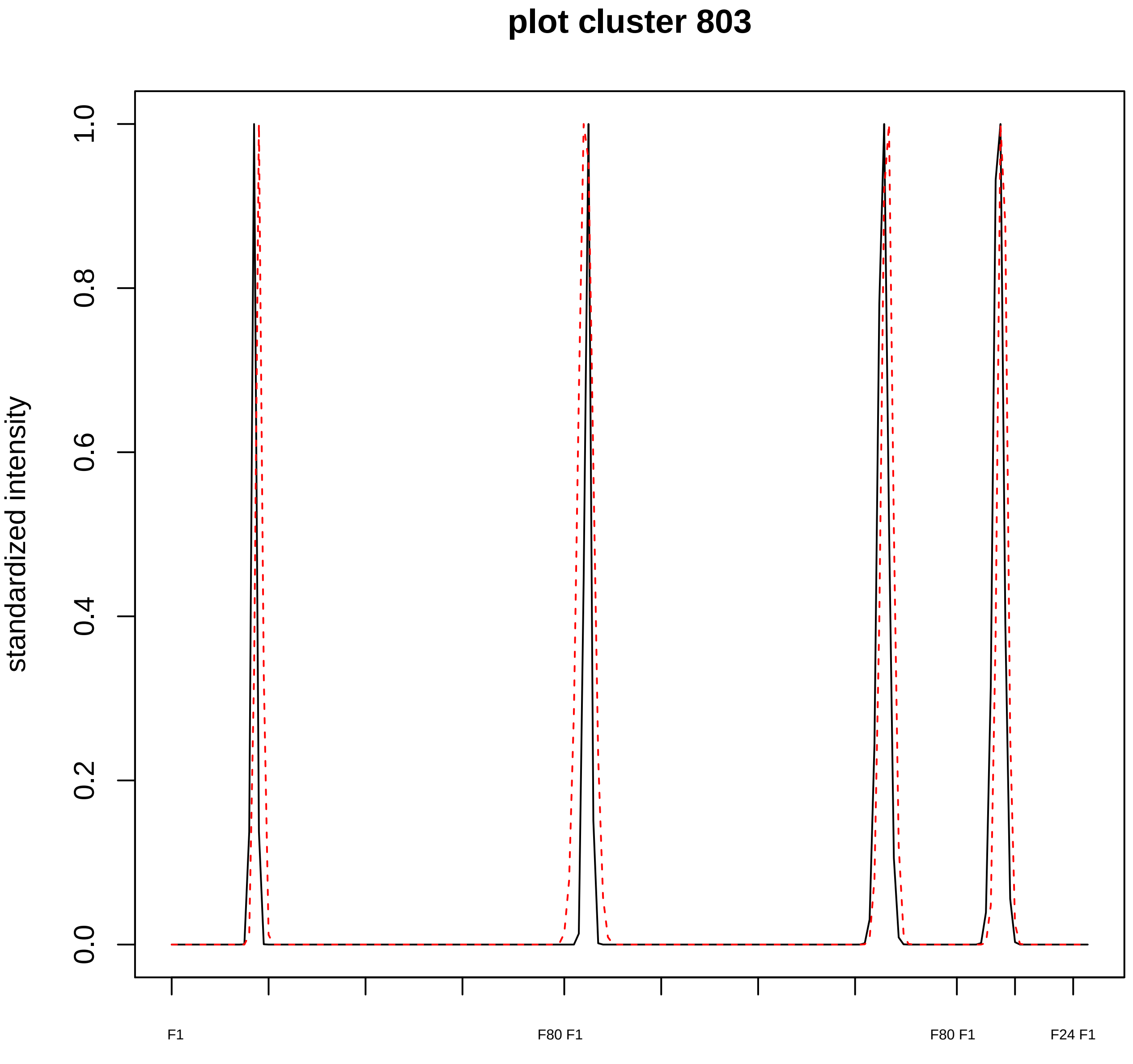

plot cluster 803
standardized intensity
0.0
0.2
0.4
0.6
0.8
1.0
F1
F80 F1
F80 F1
F24 F1

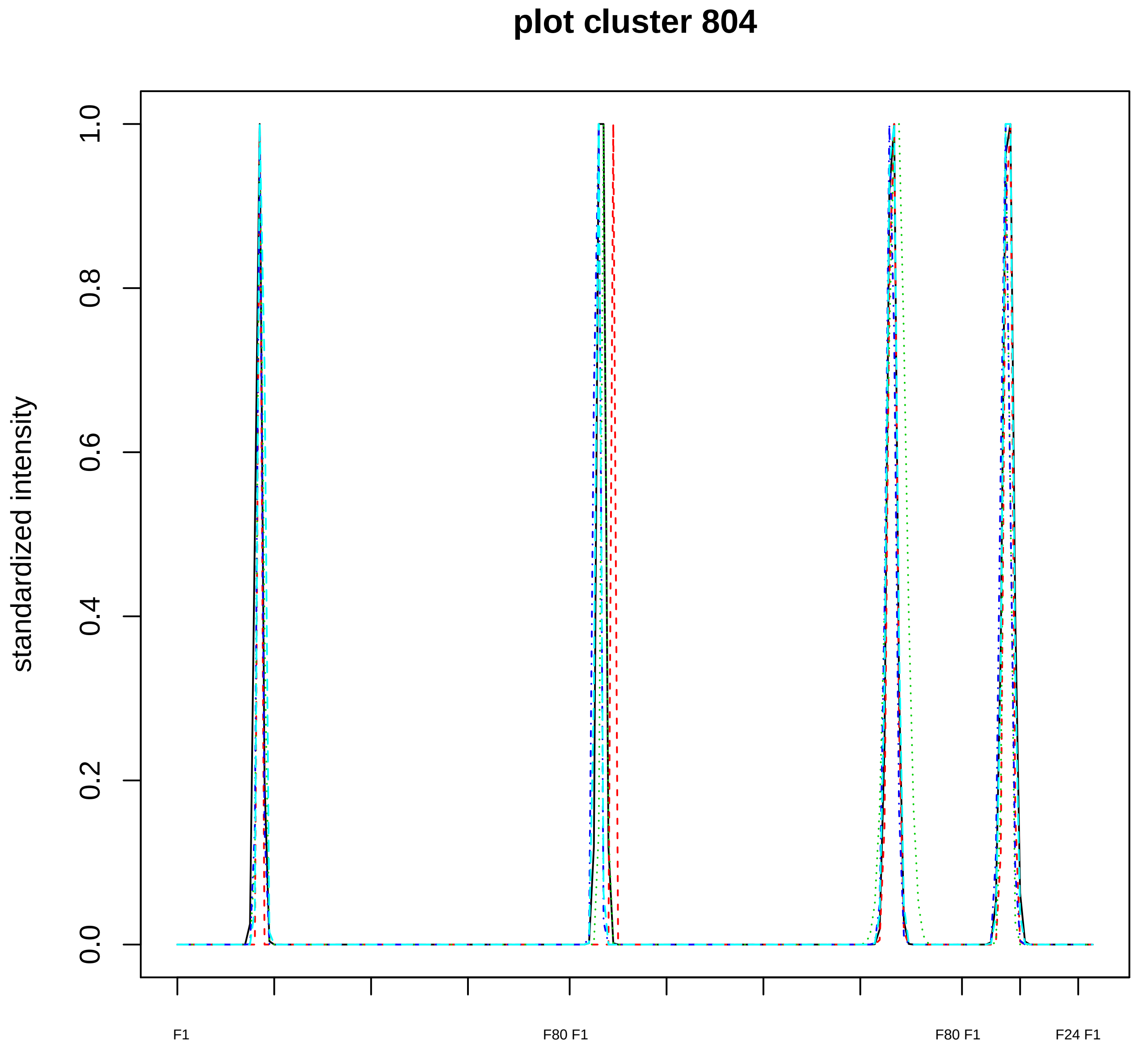
plot cluster 804
standardized intensity
0.0
0.2
0.4
0.6
0.8
1.0
F1
F80 F1
F80 F1
F24 F1

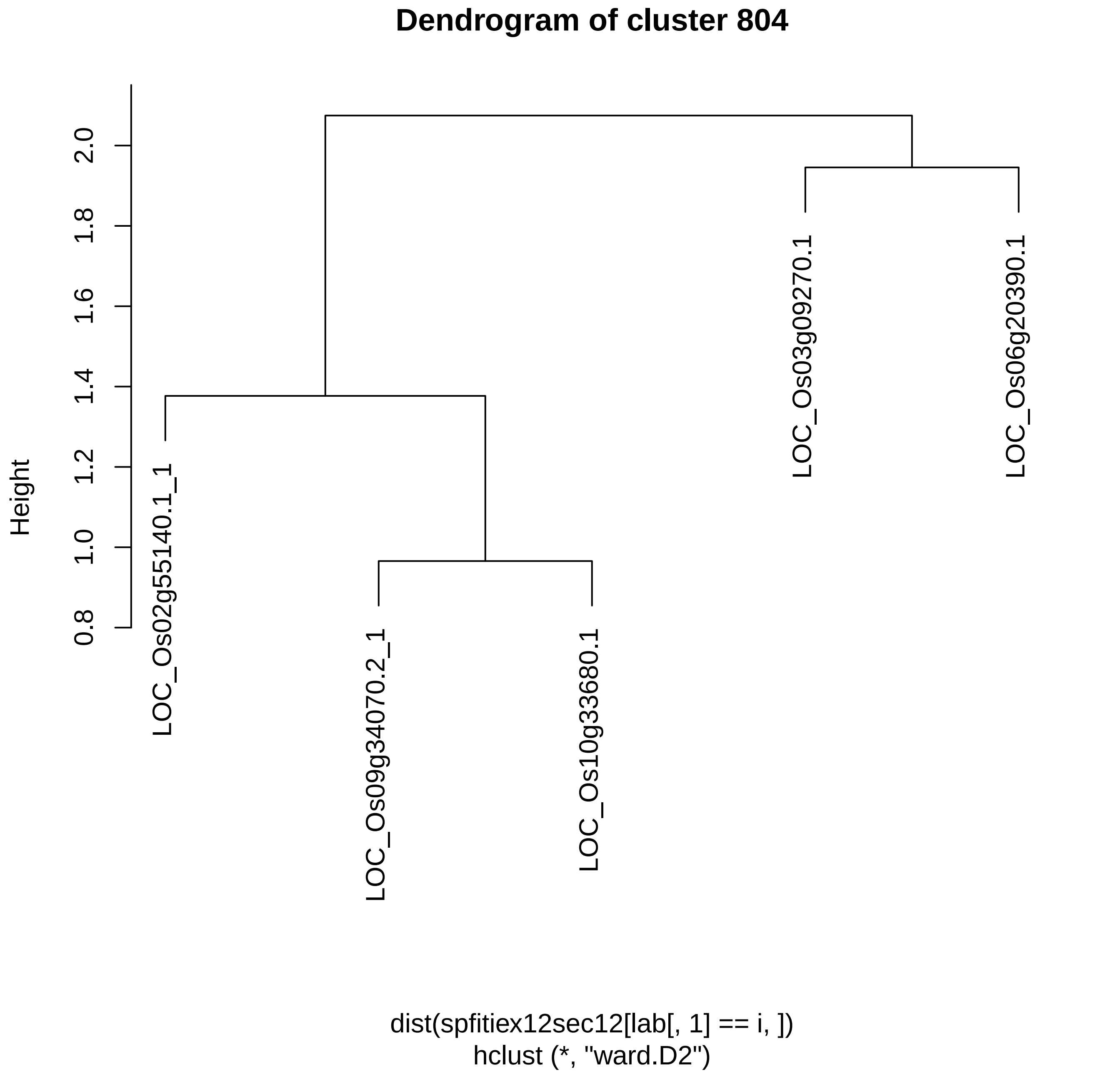
Dendrogram of cluster 804
Height
0.8
1.0
1.2
1.4
1.6
1.8
2.0
LOC_Os02g55140.1_1
LOC_Os09g34070.2_1
LOC_Os10g33680.1
LOC_Os03g09270.1
LOC_Os06g20390.1
dist(spfitiex12sec12[lab[, 1] == i, ])
hclust (*, "ward.D2")

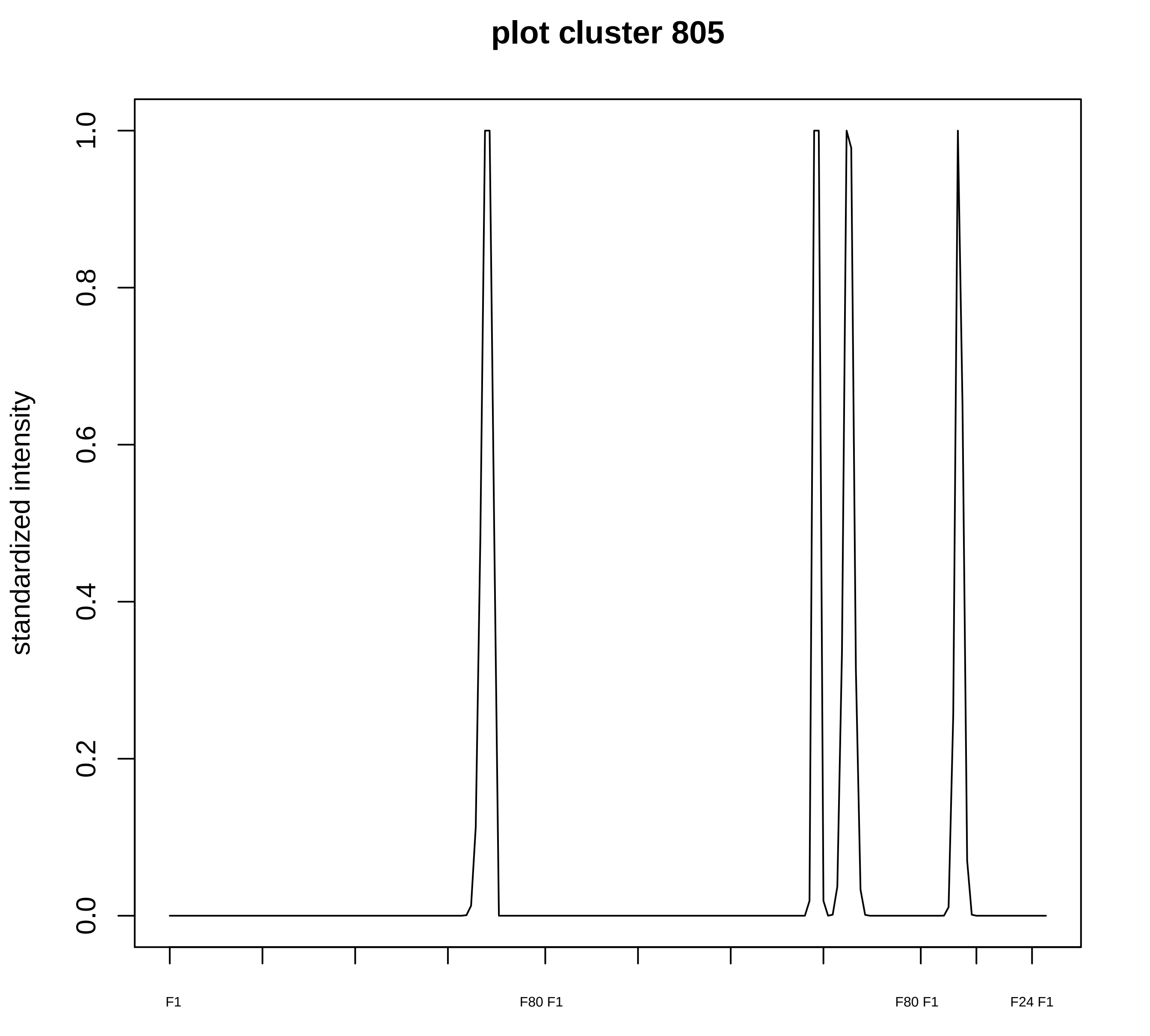
plot cluster 805
standardized intensity
0.0
0.2
0.4
0.6
0.8
1.0
F1
F80 F1
F80 F1
F24 F1

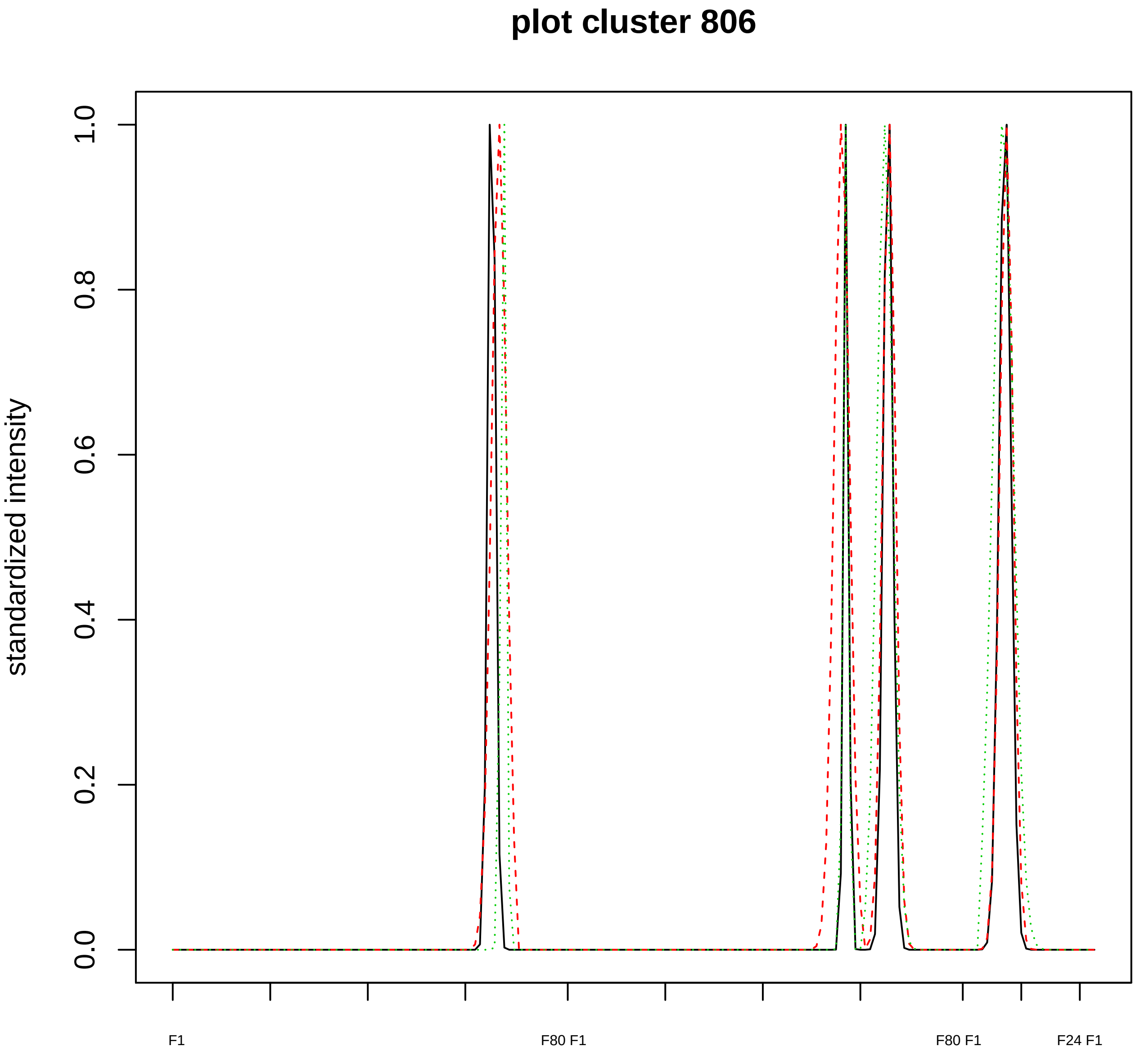

plot cluster 806
standardized intensity
1.0
0.8
0.6
0.4
0.2
0.0
F1
F80 F1
F80 F1
F24 F1

#### Dendrogram of cluster 806

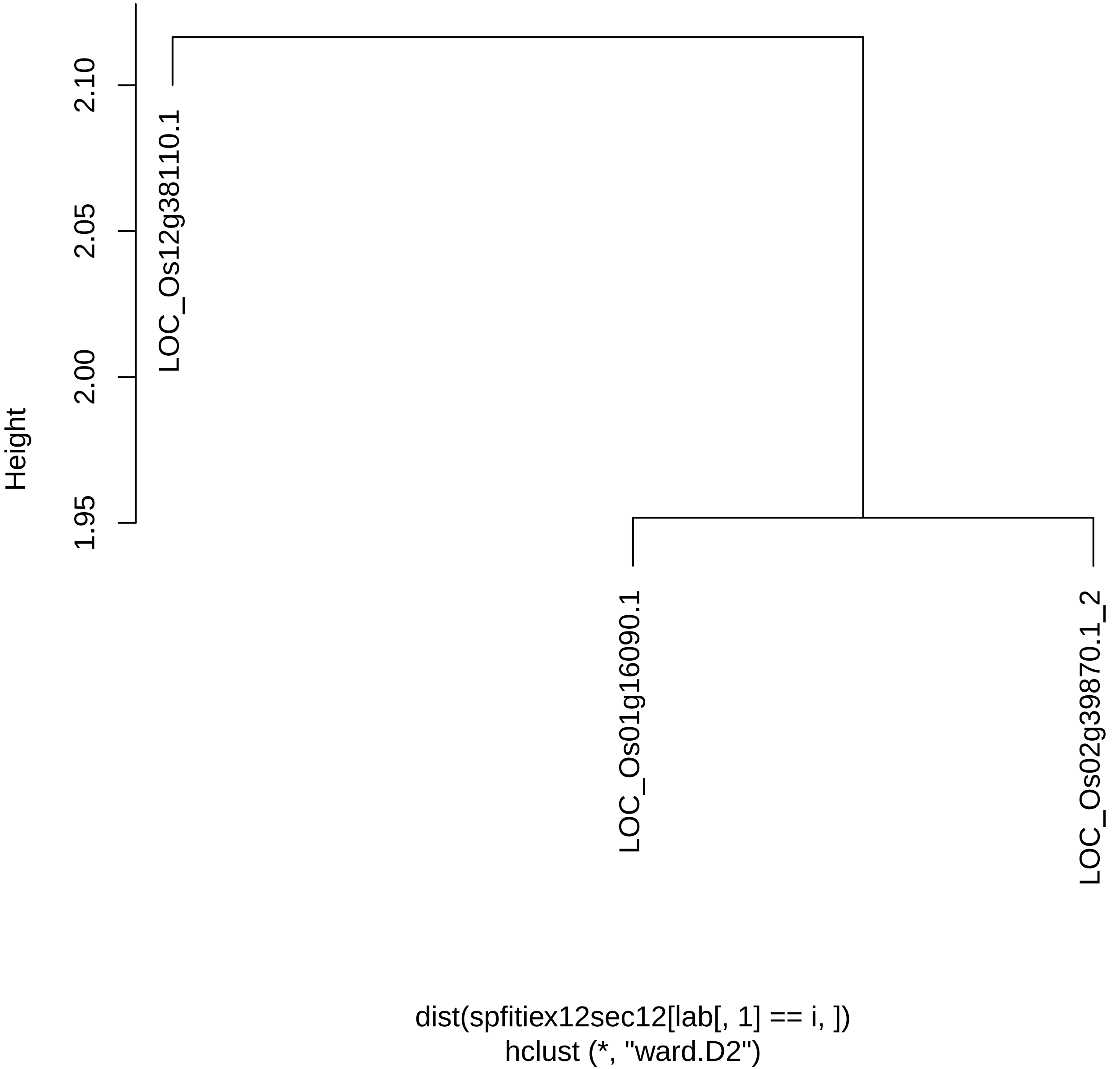

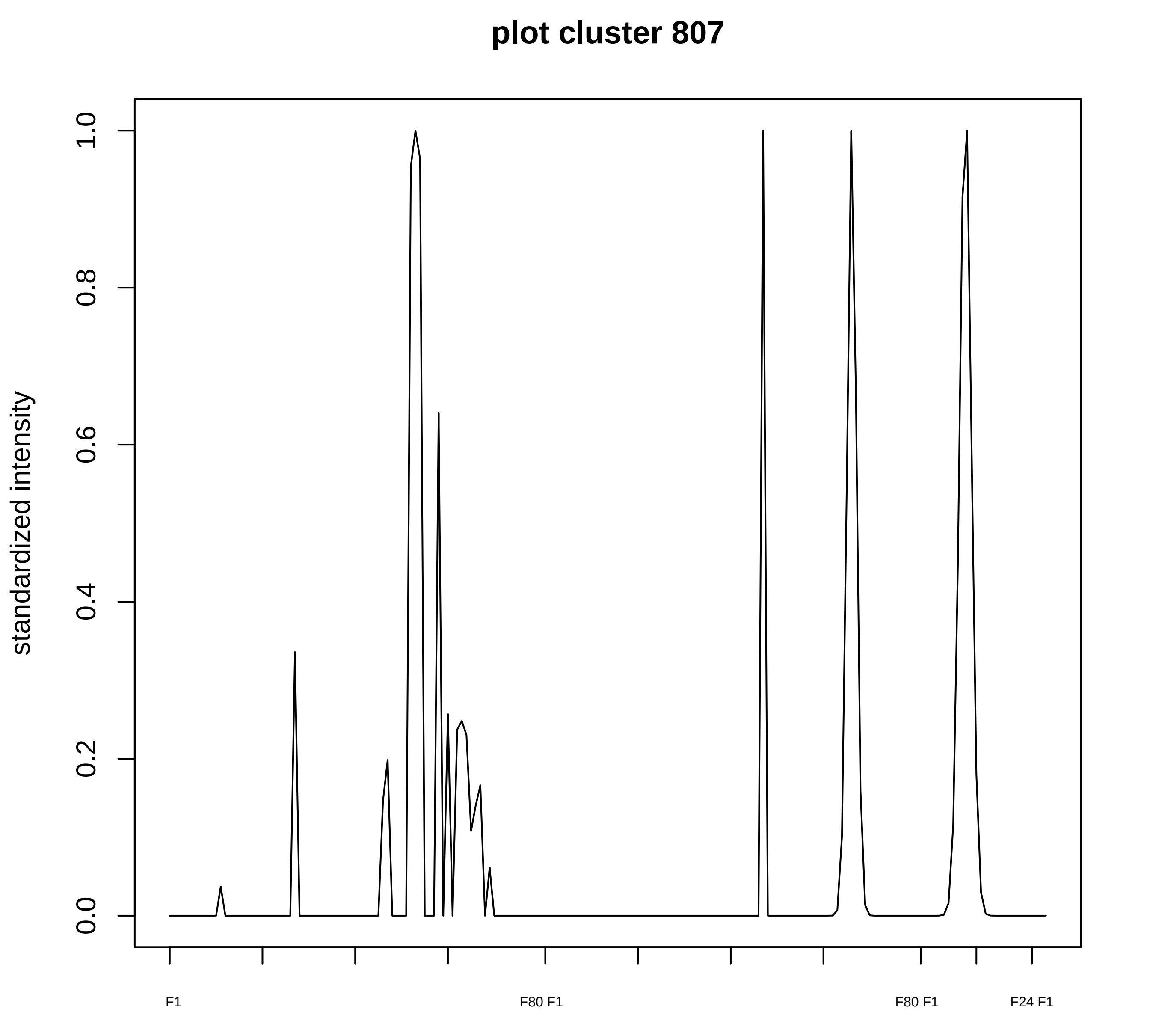

plot cluster 807
standardized intensity
0.0
0.2
0.4
0.6
0.8
1.0
F1
F80 F1
F80 F1
F24 F1

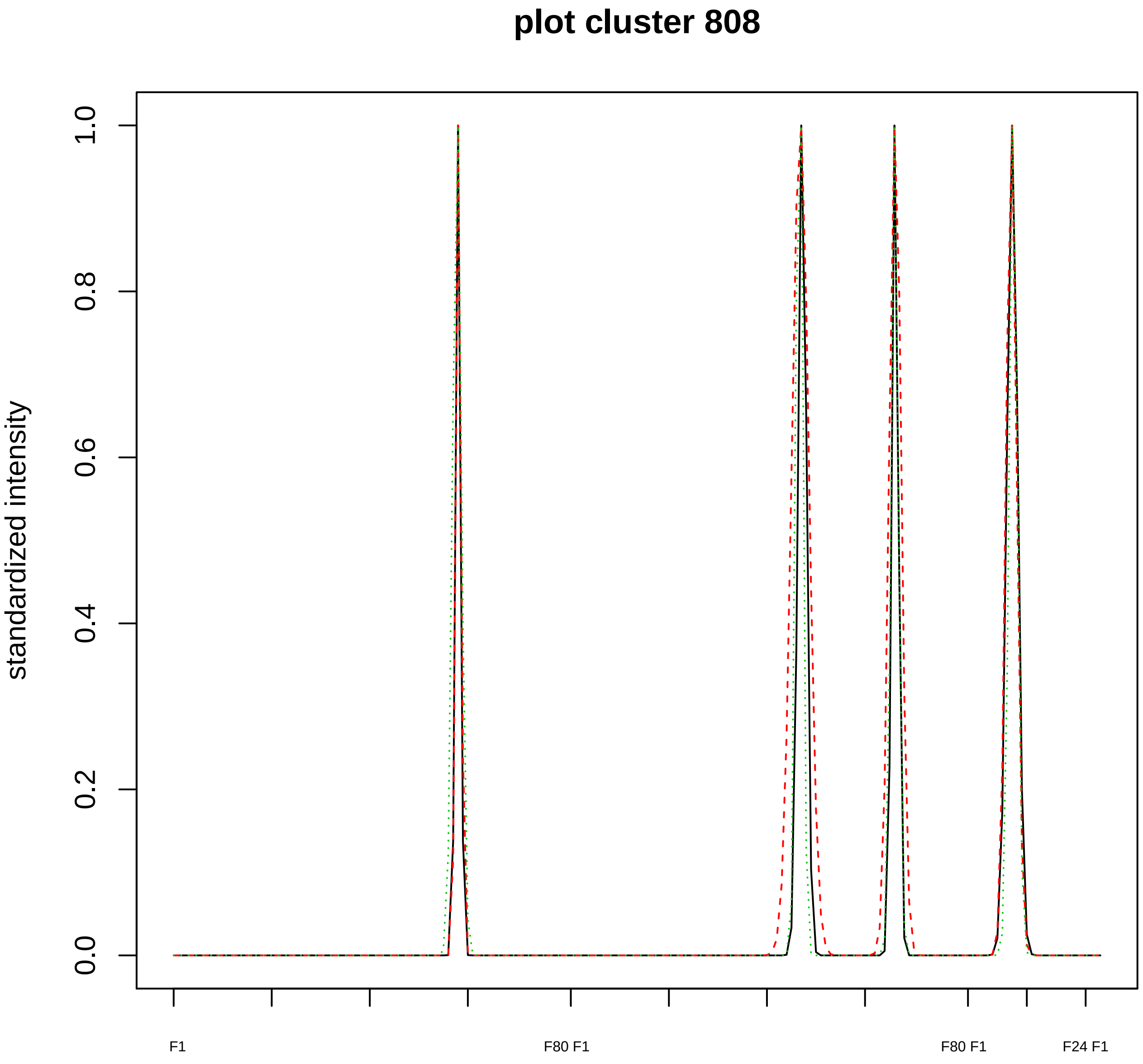
plot cluster 808
standardized intensity
0.0
0.2
0.4
0.6
0.8
1.0
F1
F80 F1
F80 F1
F24 F1

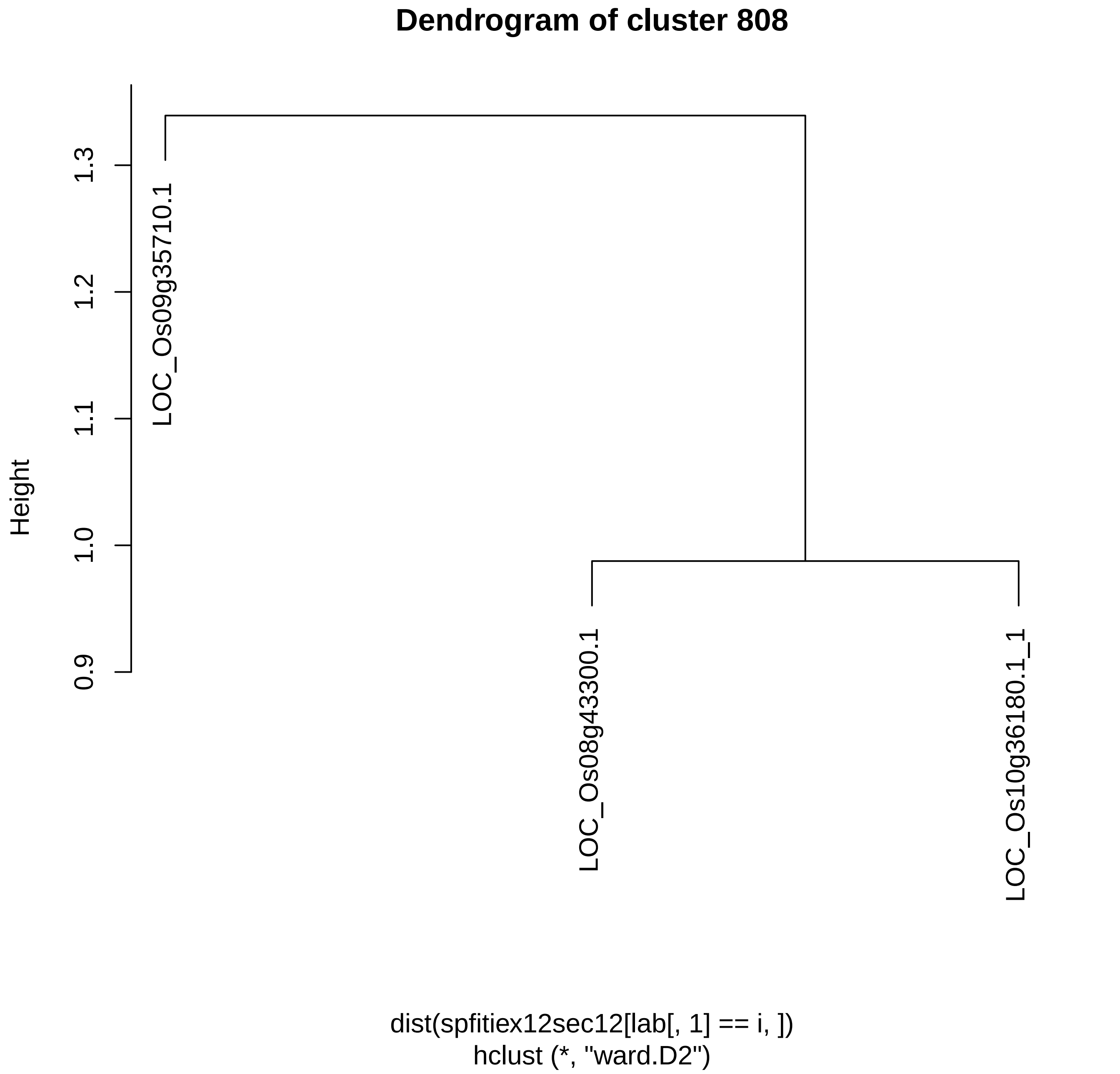
Dendrogram of cluster 808
Height
0.9
1.0
1.1
1.2
1.3
LOC_Os09g35710.1
LOC_Os08g43300.1
LOC_Os10g36180.1_1
dist(spfitiex12sec12[lab[, 1] == i, ])
hclust (*, "ward.D2")

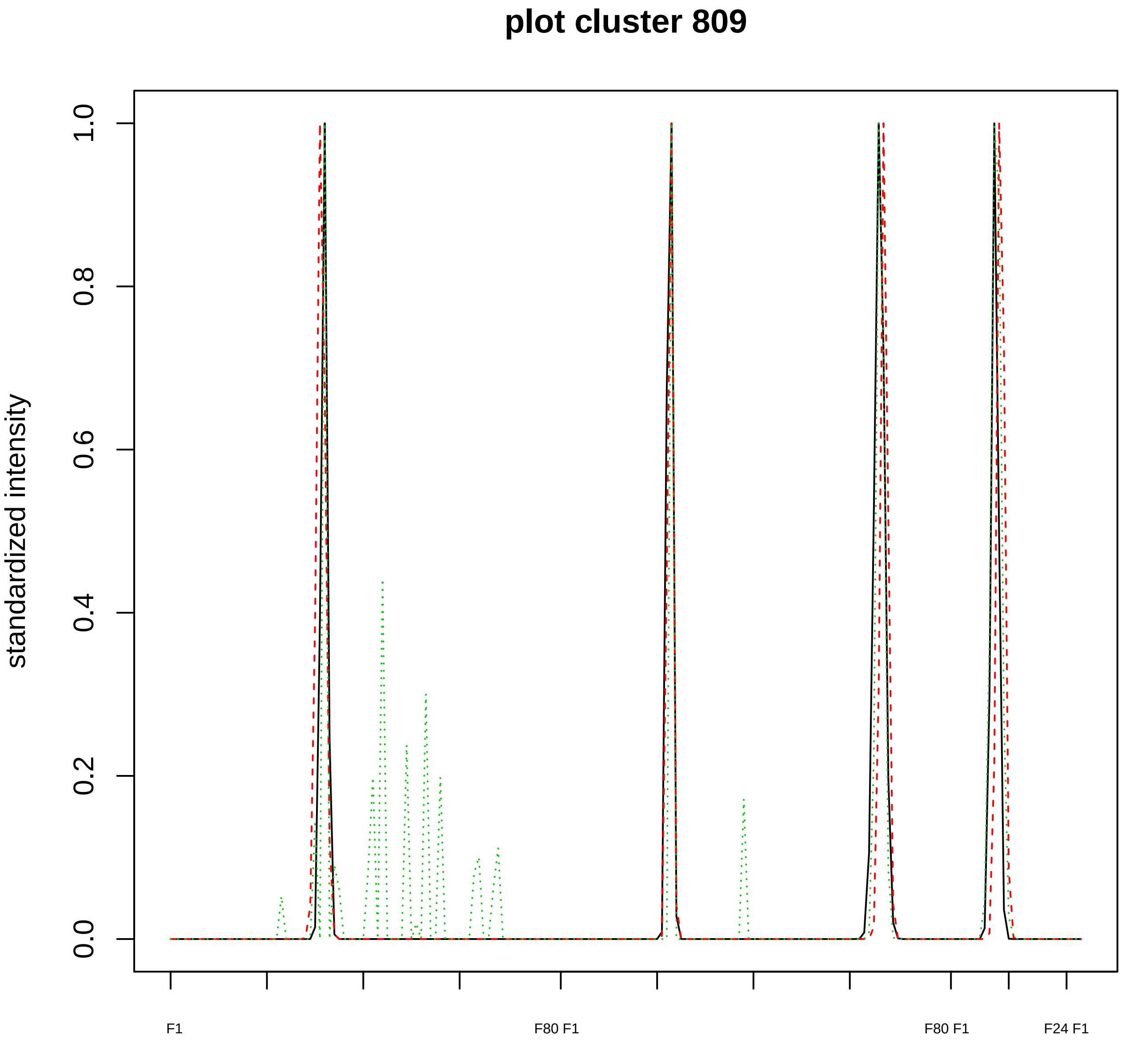

plot cluster 809
standardized intensity
0.0
0.2
0.4
0.6
0.8
1.0
F1
F80 F1
F80 F1
F24 F1

#### Dendrogram of cluster 809

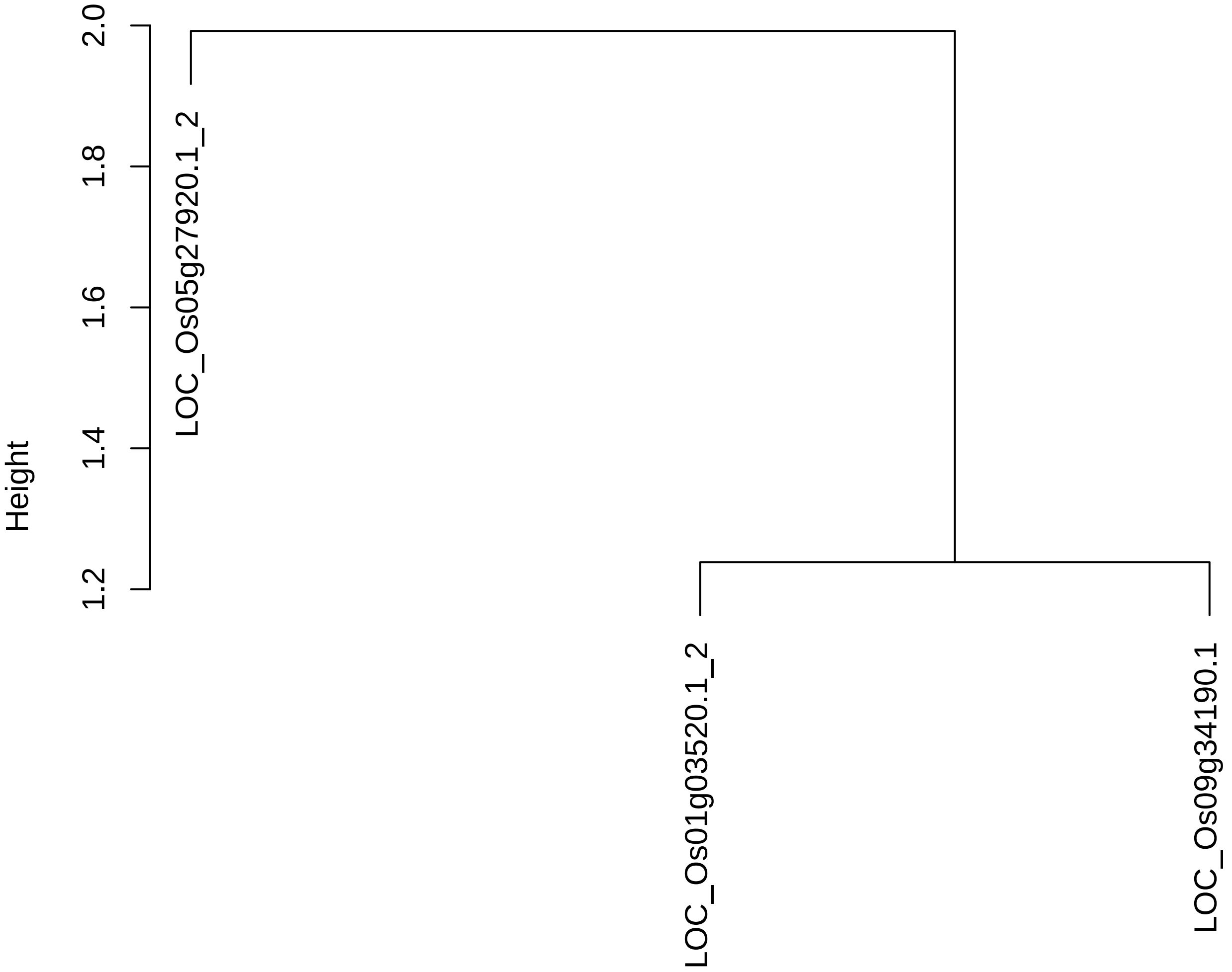


dist(spfitiex12sec12[lab[, 1] == i, ])
hclust (*, "ward.D2")

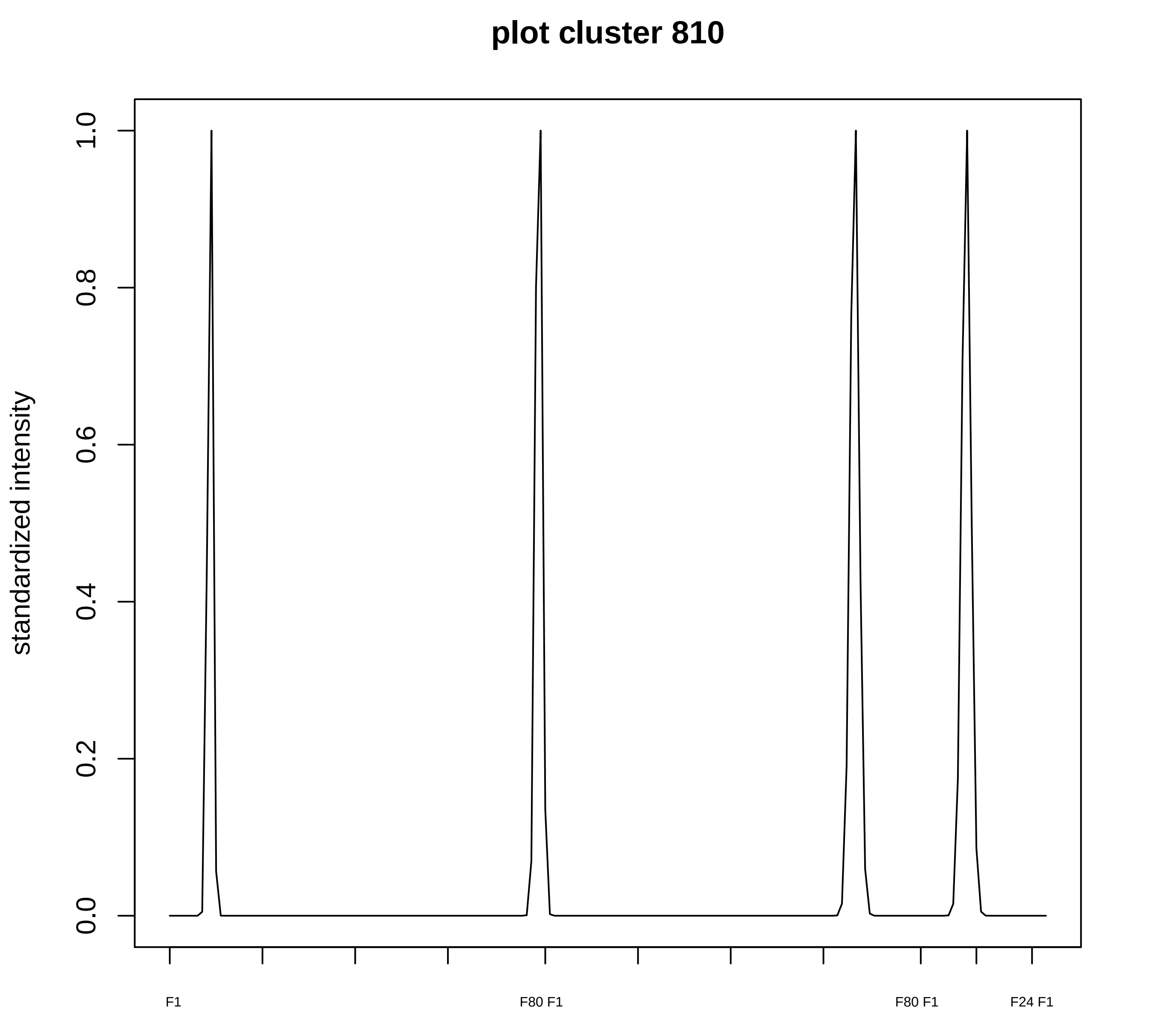

plot cluster 810
standardized intensity
0.0
0.2
0.4
0.6
0.8
1.0
F1
F80 F1
F80 F1
F24 F1

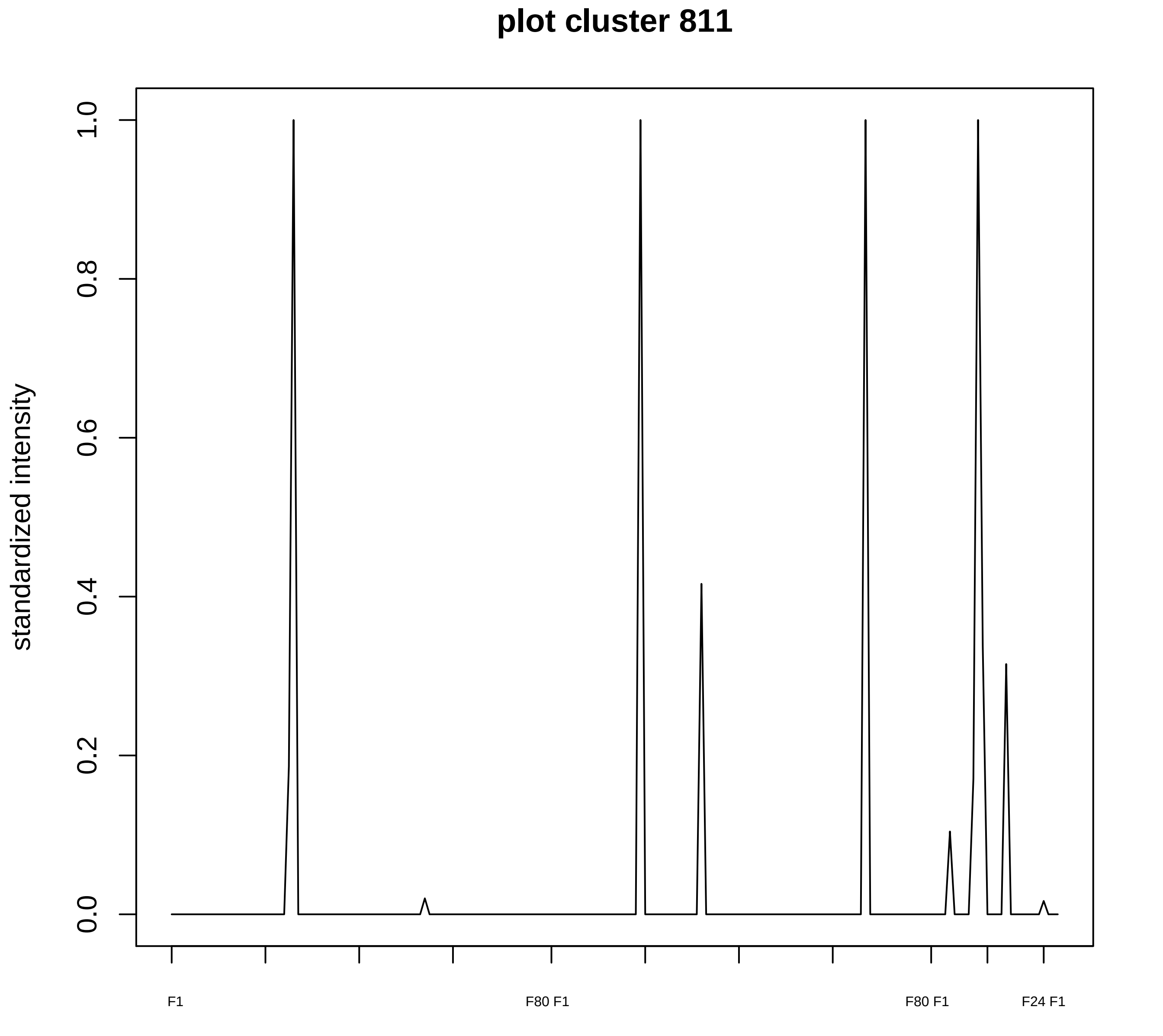

plot cluster 811
standardized intensity
0.0
0.2
0.4
0.6
0.8
1.0
F1
F80 F1
F80 F1
F24 F1

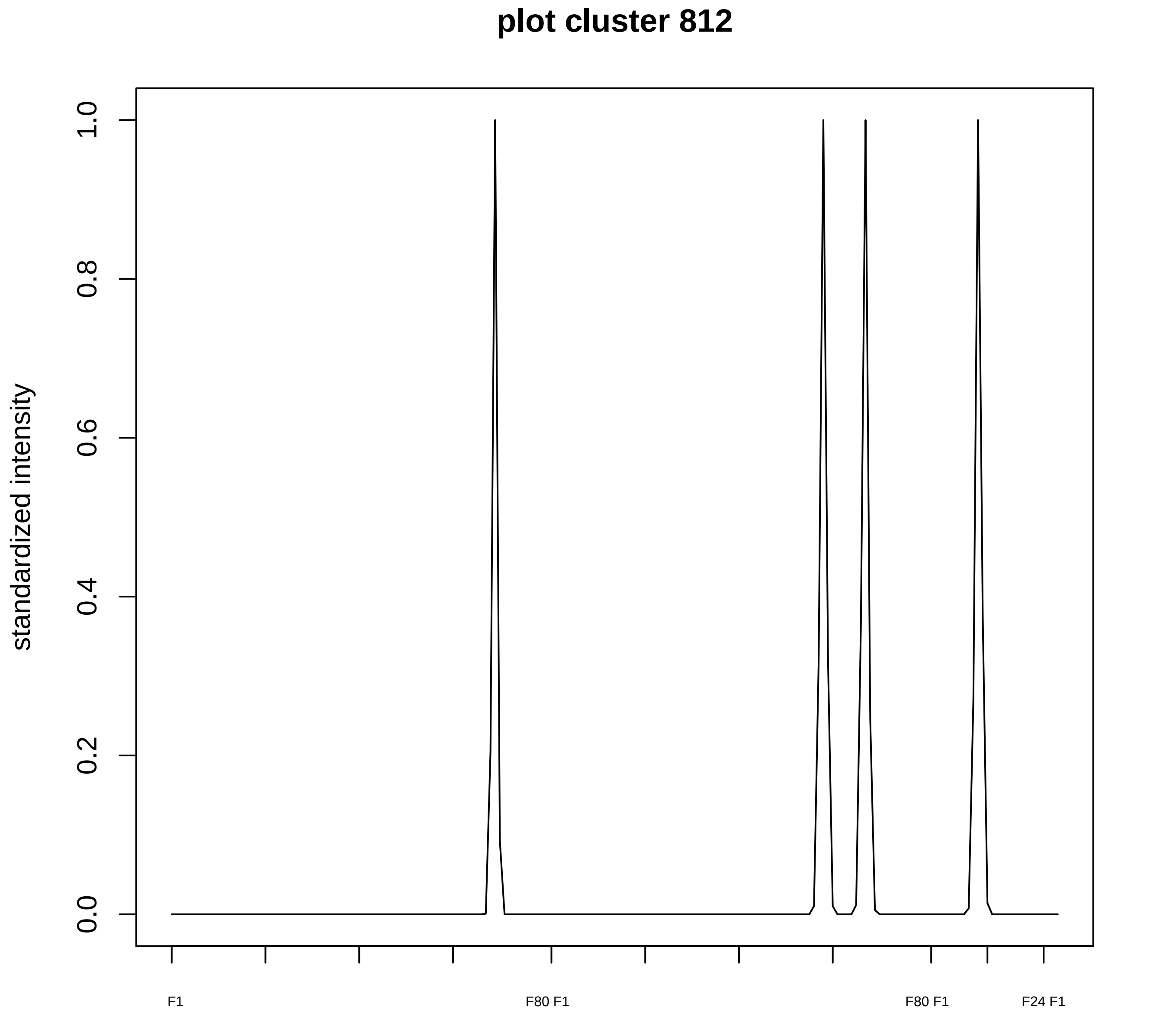

plot cluster 812
standardized intensity
1.0
0.8
0.6
0.4
0.2
0.0
F1
F80 F1
F80 F1
F24 F1

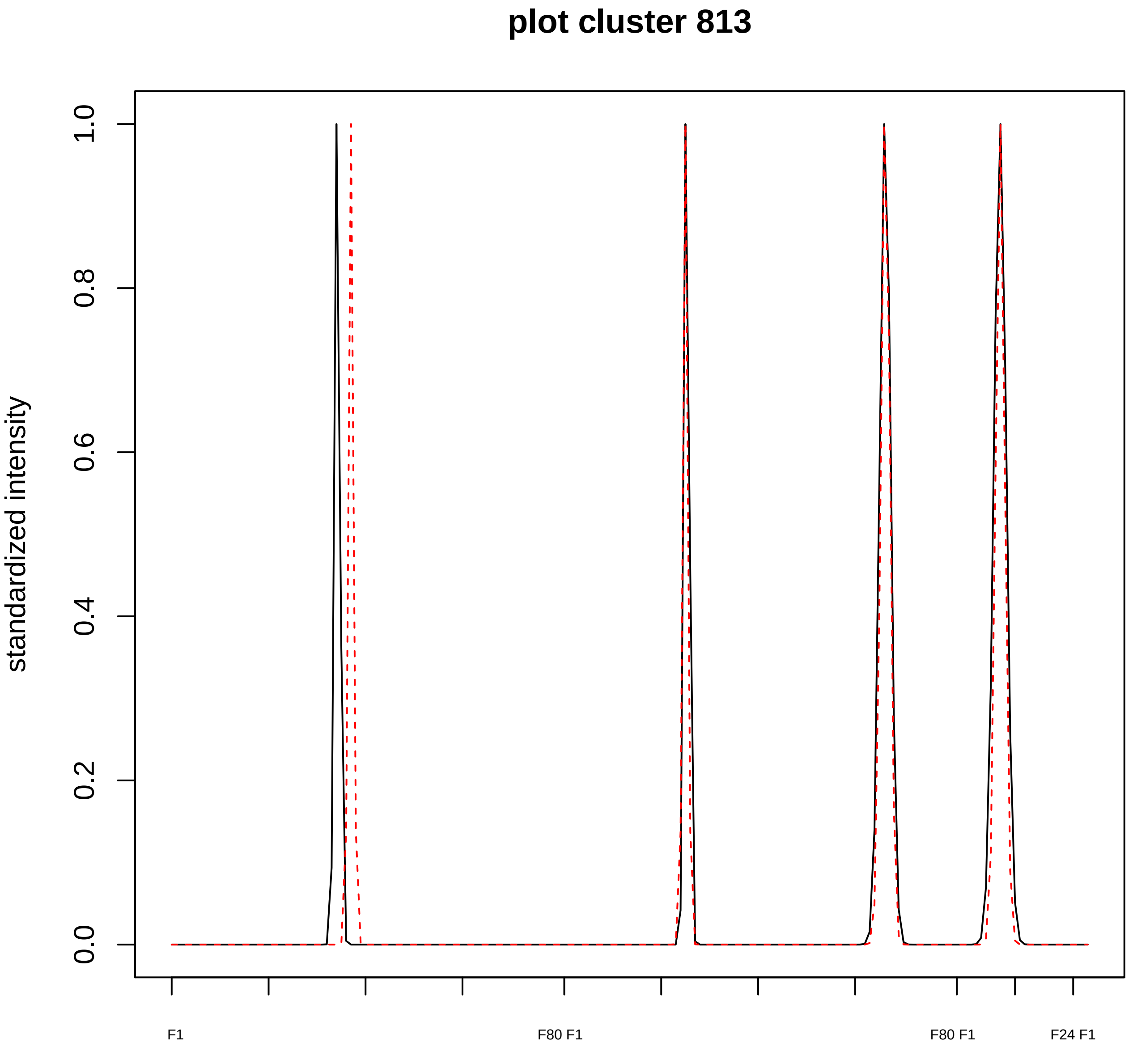
plot cluster 813
standardized intensity
1.0
0.8
0.6
0.4
0.2
0.0
F1
F80 F1
F80 F1
F24 F1

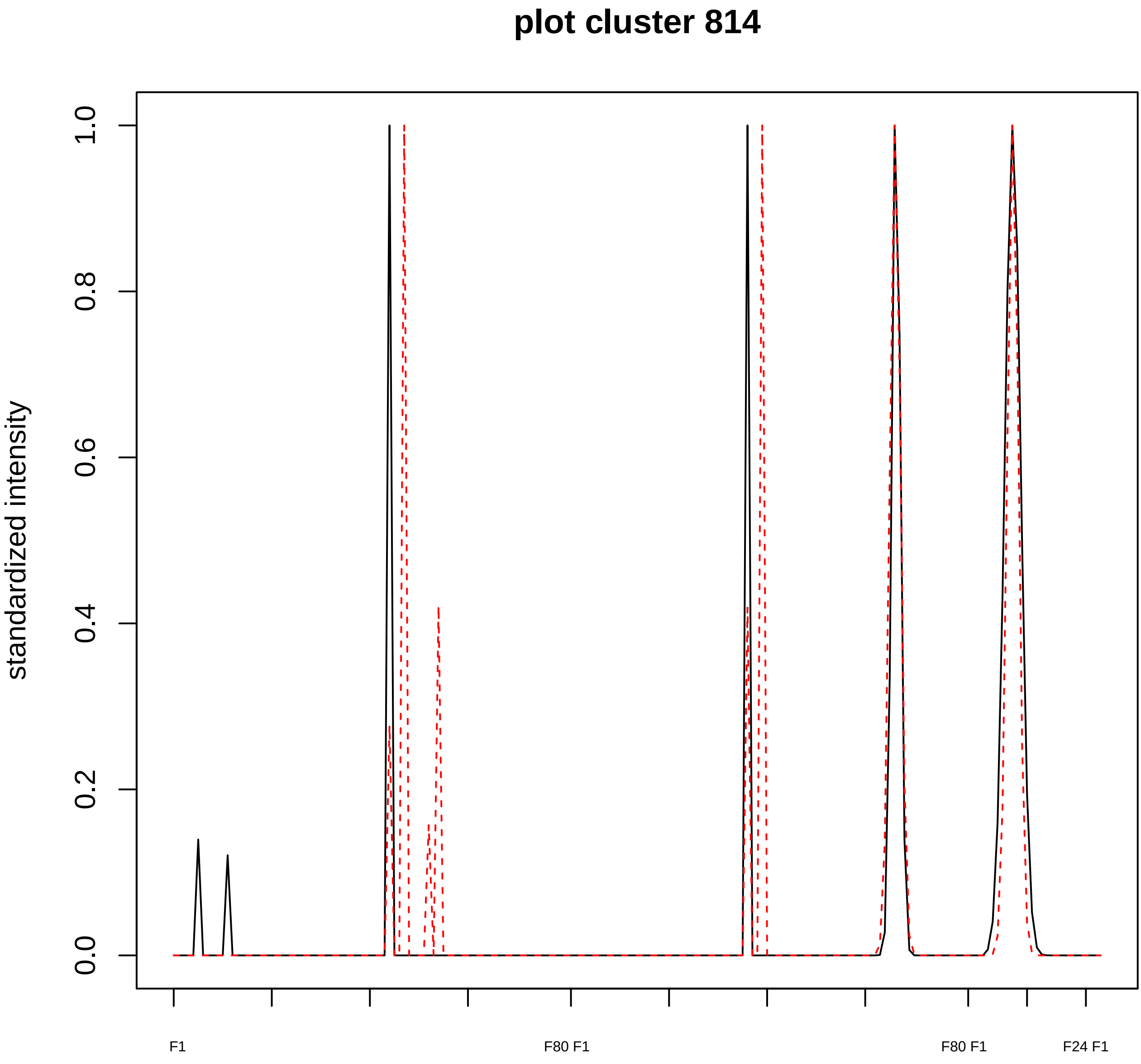
plot cluster 814
standardized intensity
0.0
0.2
0.4
0.6
0.8
1.0
F1
F80 F1
F80 F1
F24 F1

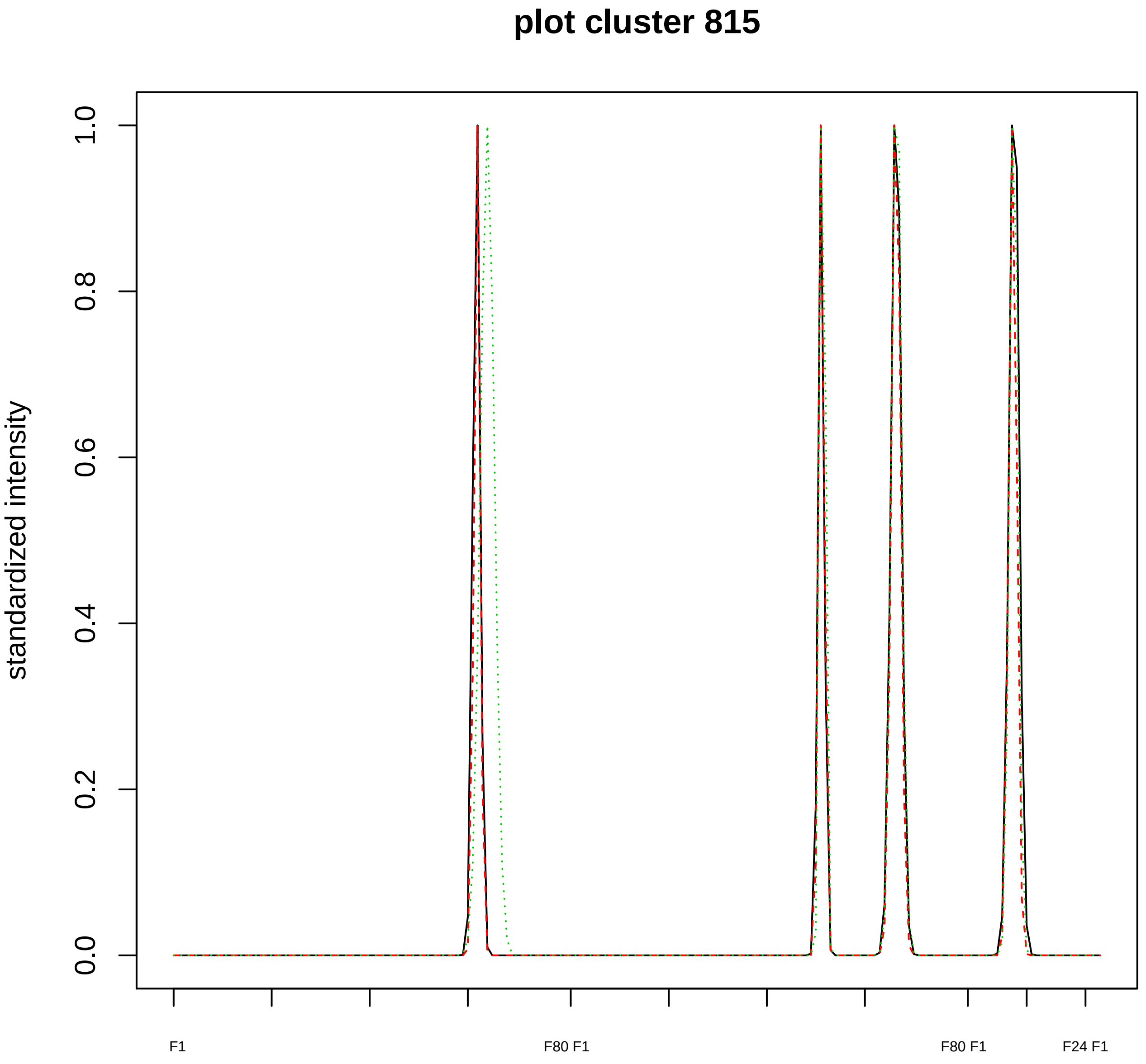

plot cluster 815
standardized intensity
0.0
0.2
0.4
0.6
0.8
1.0
F1
F80 F1
F80 F1
F24 F1

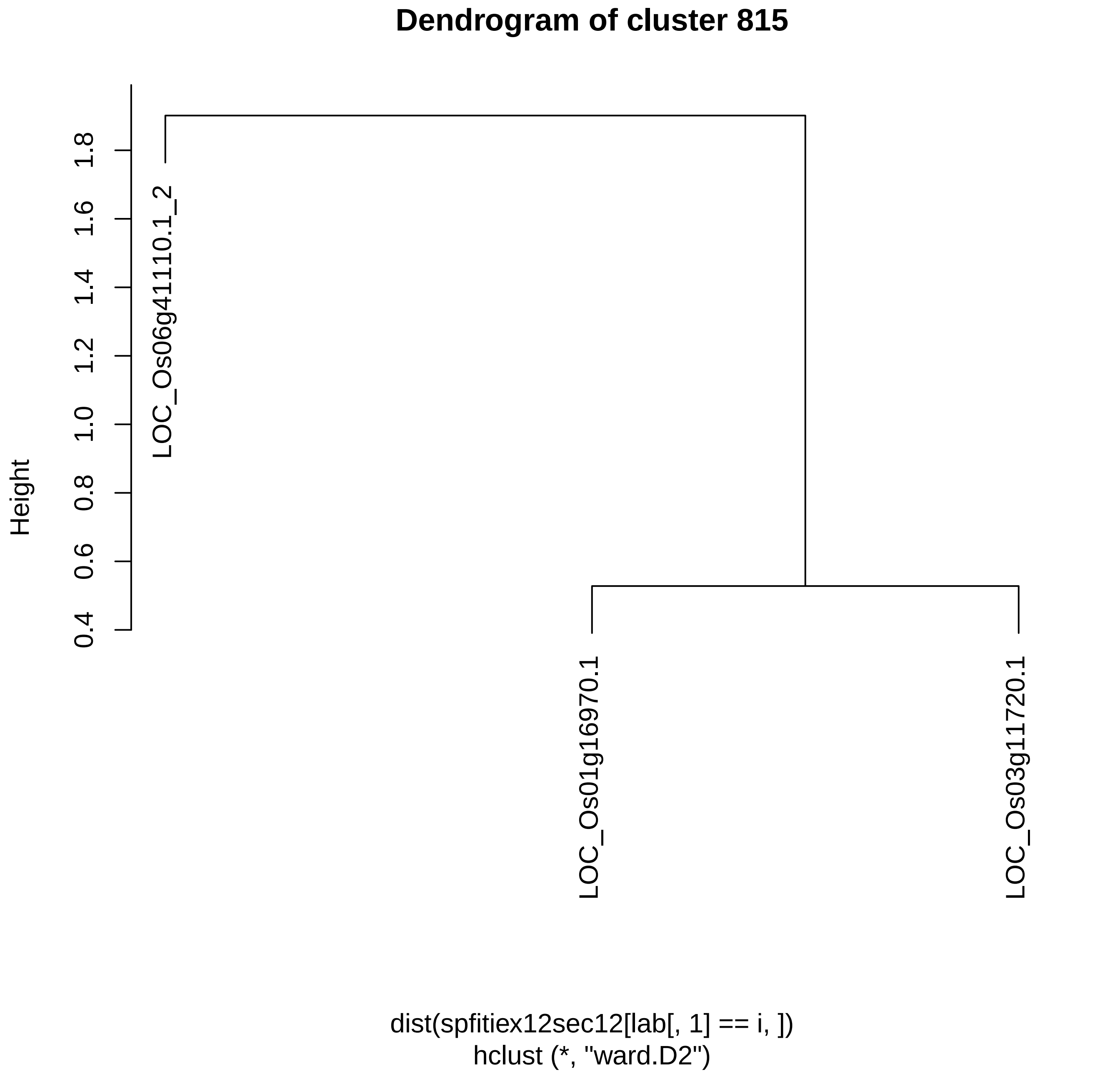
Dendrogram of cluster 815
Height
0.4
0.6
0.8
1.0
1.2
1.4
1.6
1.8
LOC_Os06g41110.1_2
LOC_Os01g16970.1
LOC_Os03g11720.1
dist(spfitiex12sec12[lab[, 1] == i, ])
hclust (*, "ward.D2")

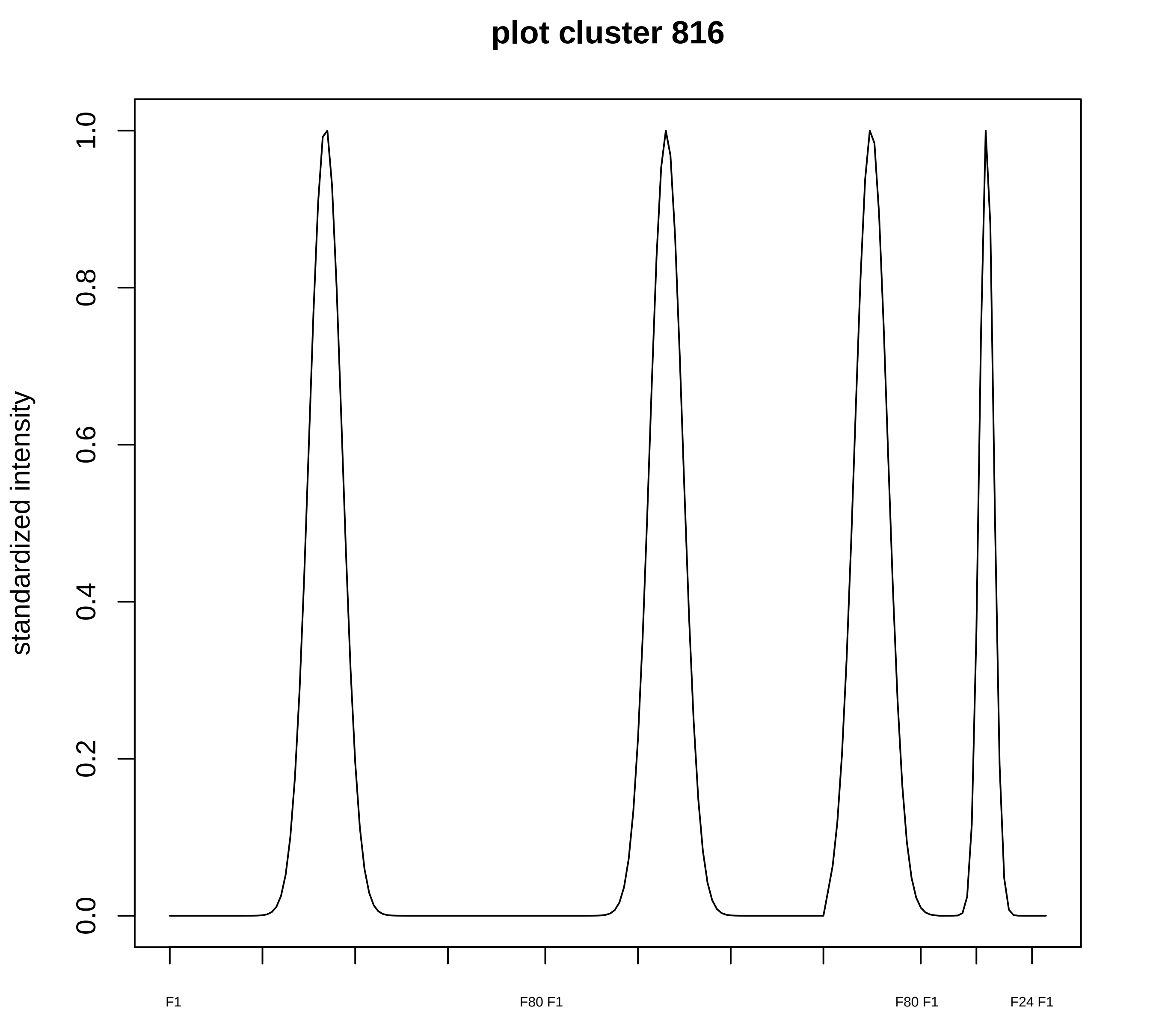

plot cluster 816
standardized intensity
0.0
0.2
0.4
0.6
0.8
1.0
F1
F80 F1
F80 F1
F24 F1

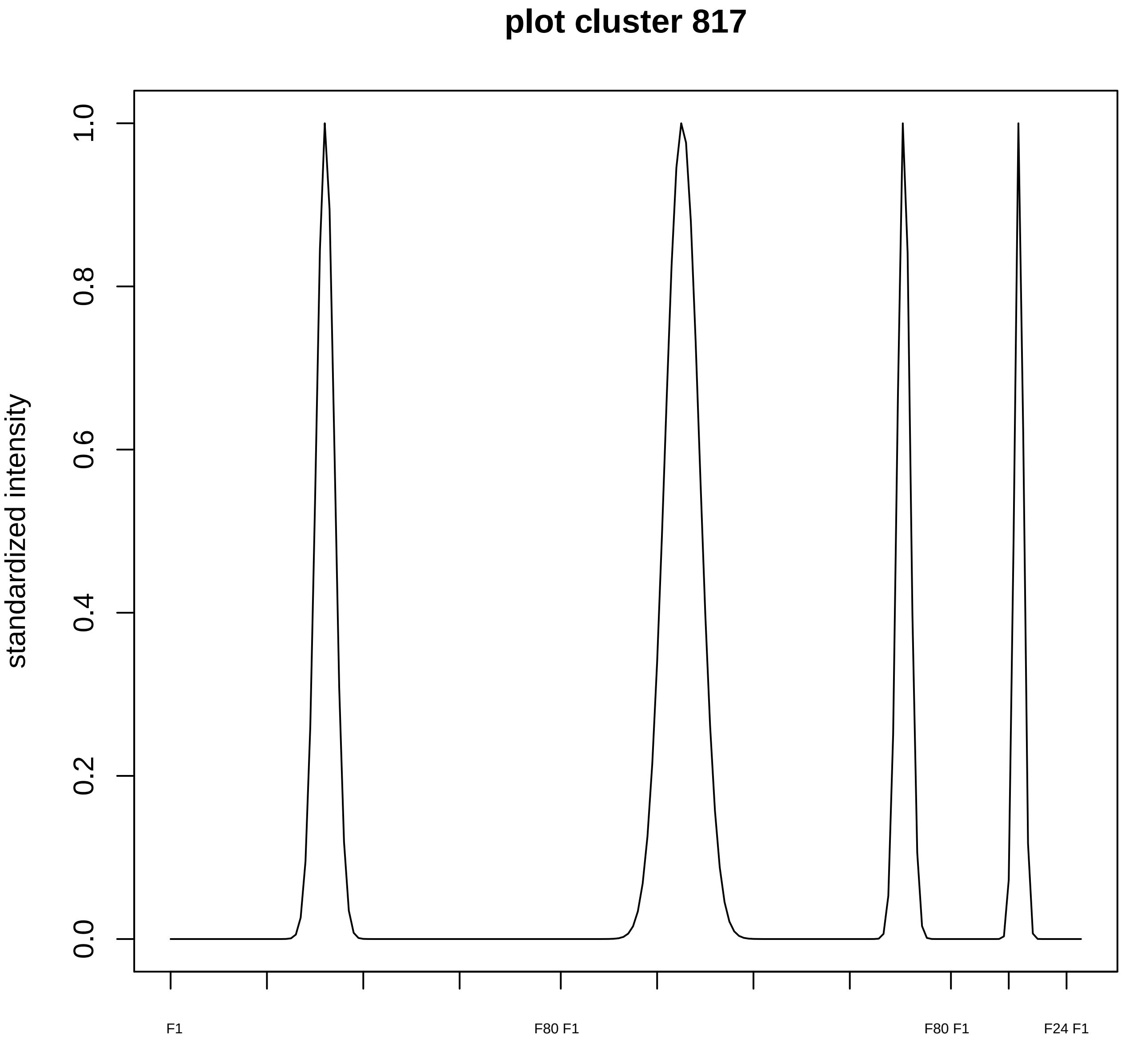
plot cluster 817
standardized intensity
0.0
0.2
0.4
0.6
0.8
1.0
F1
F80 F1
F80 F1
F24 F1

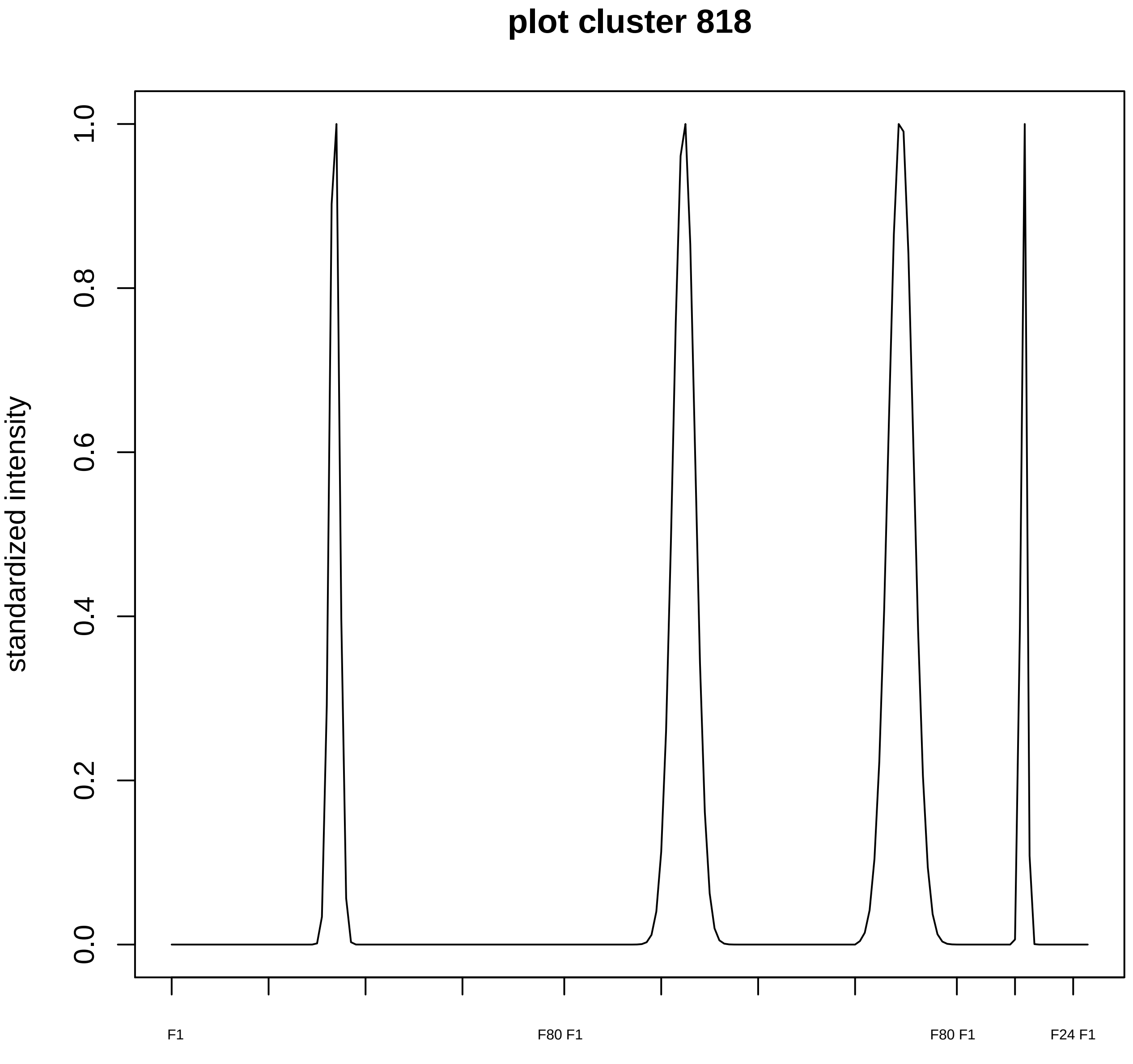
plot cluster 818
standardized intensity
0.0
0.2
0.4
0.6
0.8
1.0
F1
F80 F1
F80 F1
F24 F1

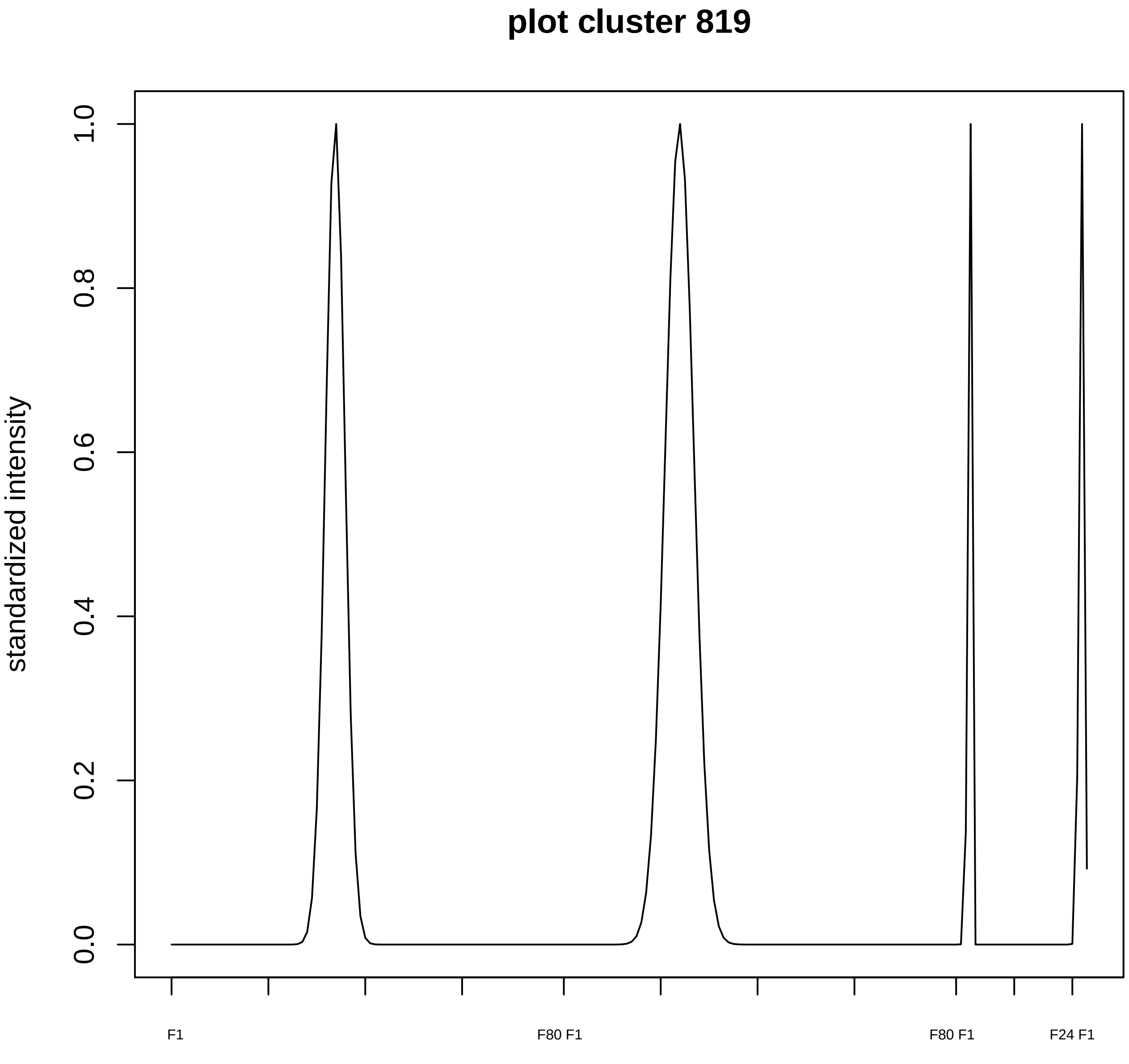

plot cluster 819
standardized intensity
0.0
0.2
0.4
0.6
0.8
1.0
F1
F80 F1
F80 F1
F24 F1

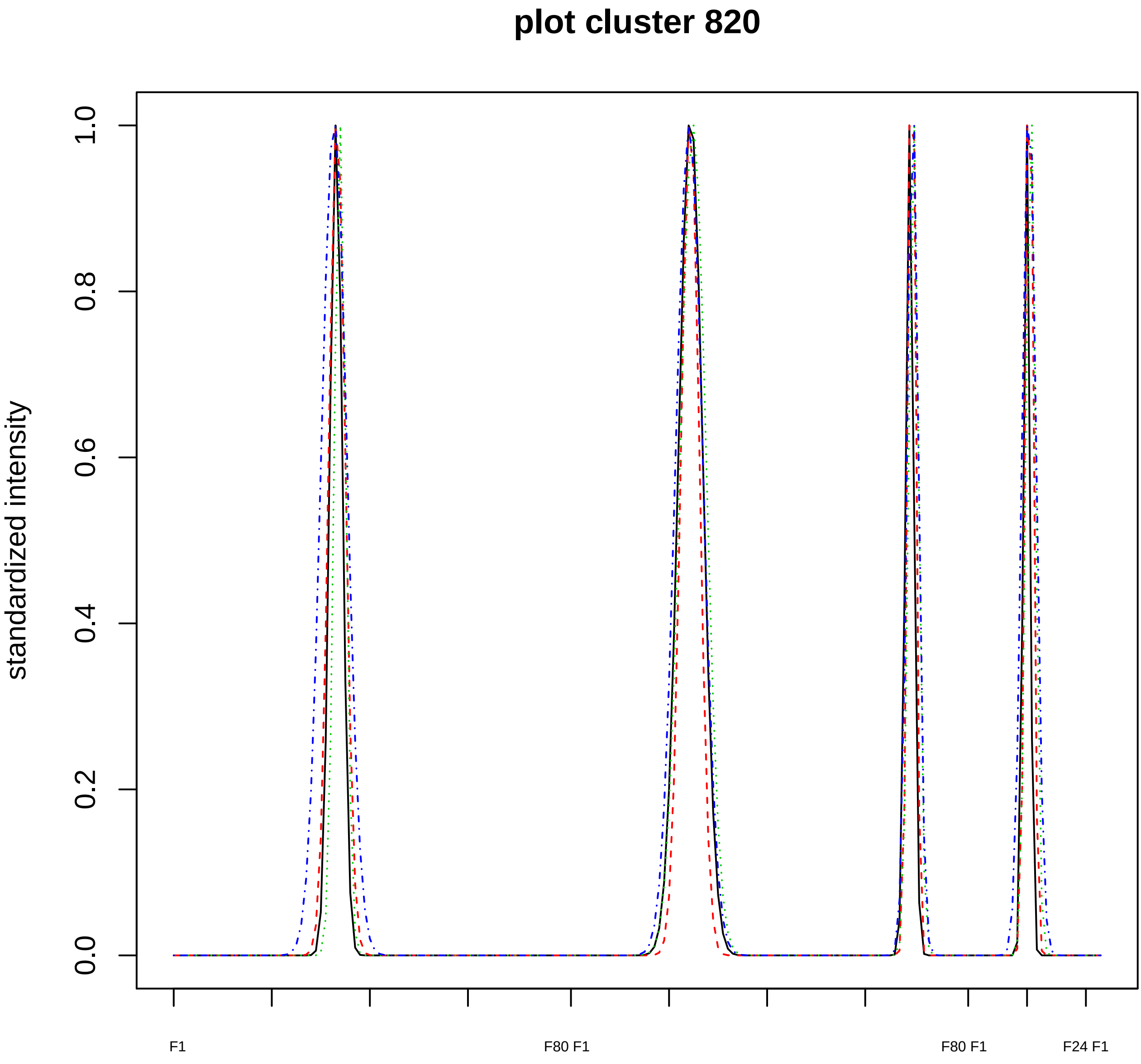

plot cluster 820
standardized intensity
0.0
0.2
0.4
0.6
0.8
1.0
F1
F80 F1
F80 F1
F24 F1

#### Dendrogram of cluster 820

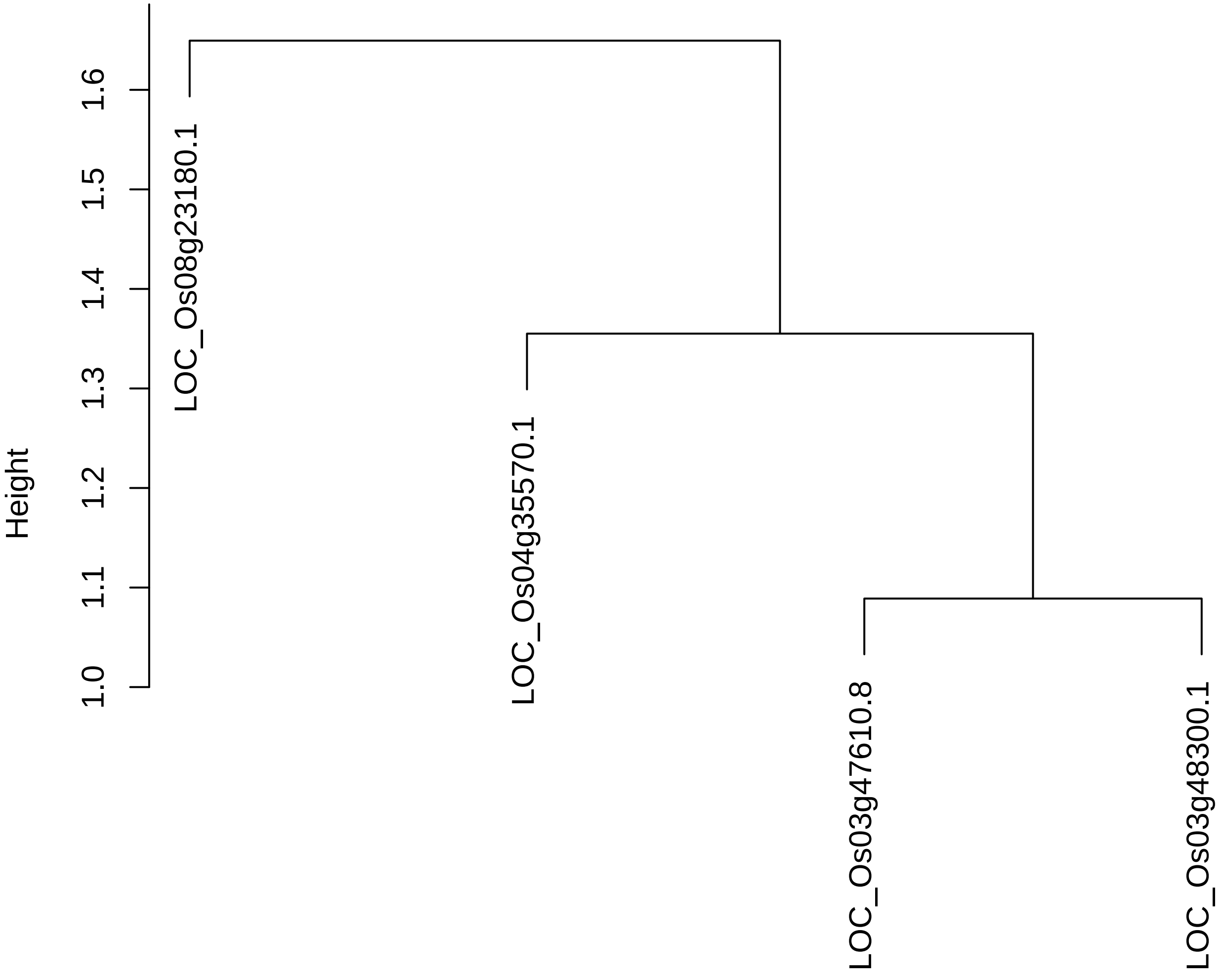


dist(spfitiex12sec12[lab[, 1] == i, ])
hclust (*, "ward.D2")

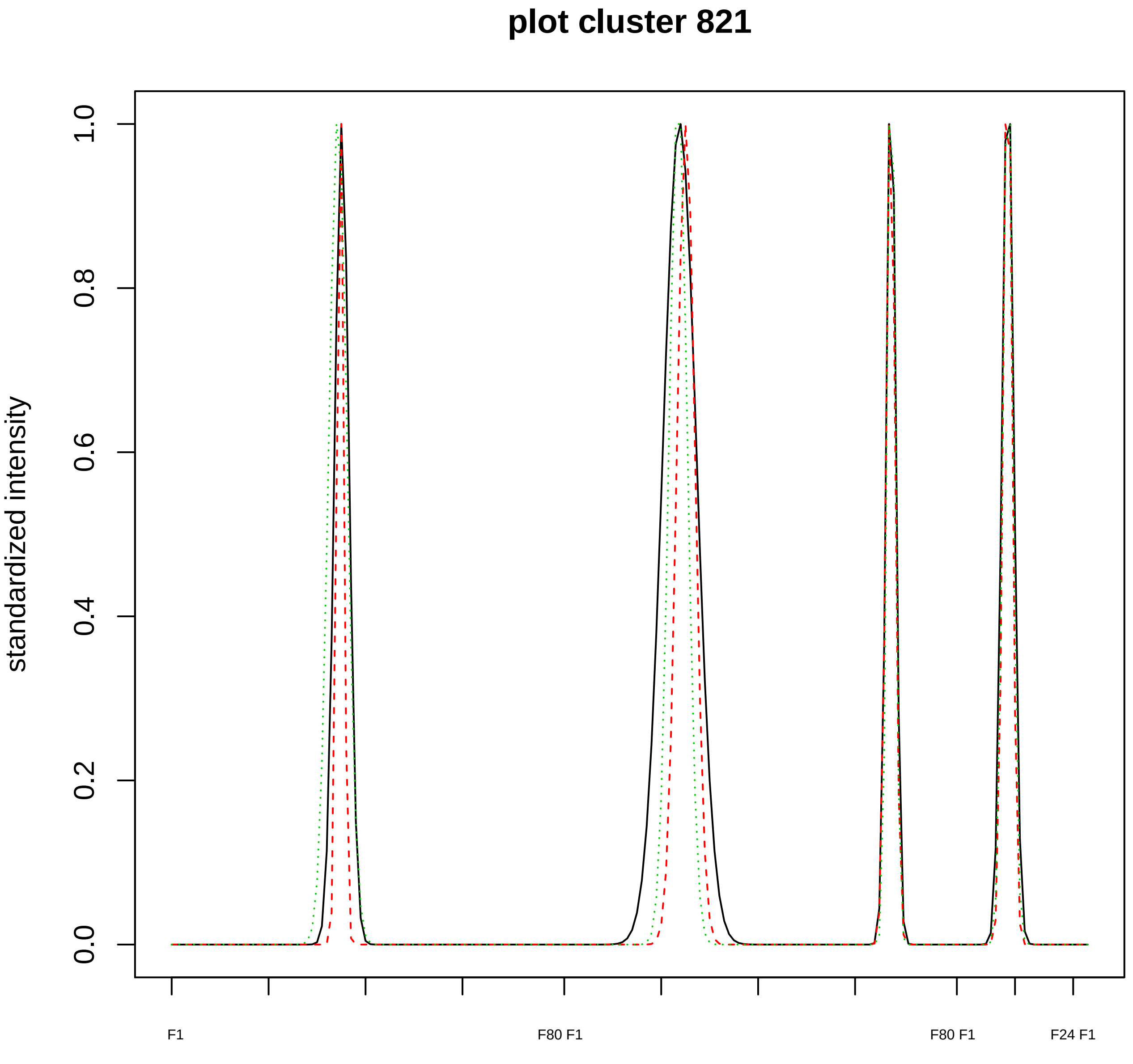

plot cluster 821
standardized intensity
0.0
0.2
0.4
0.6
0.8
1.0
F1
F80 F1
F80 F1
F24 F1

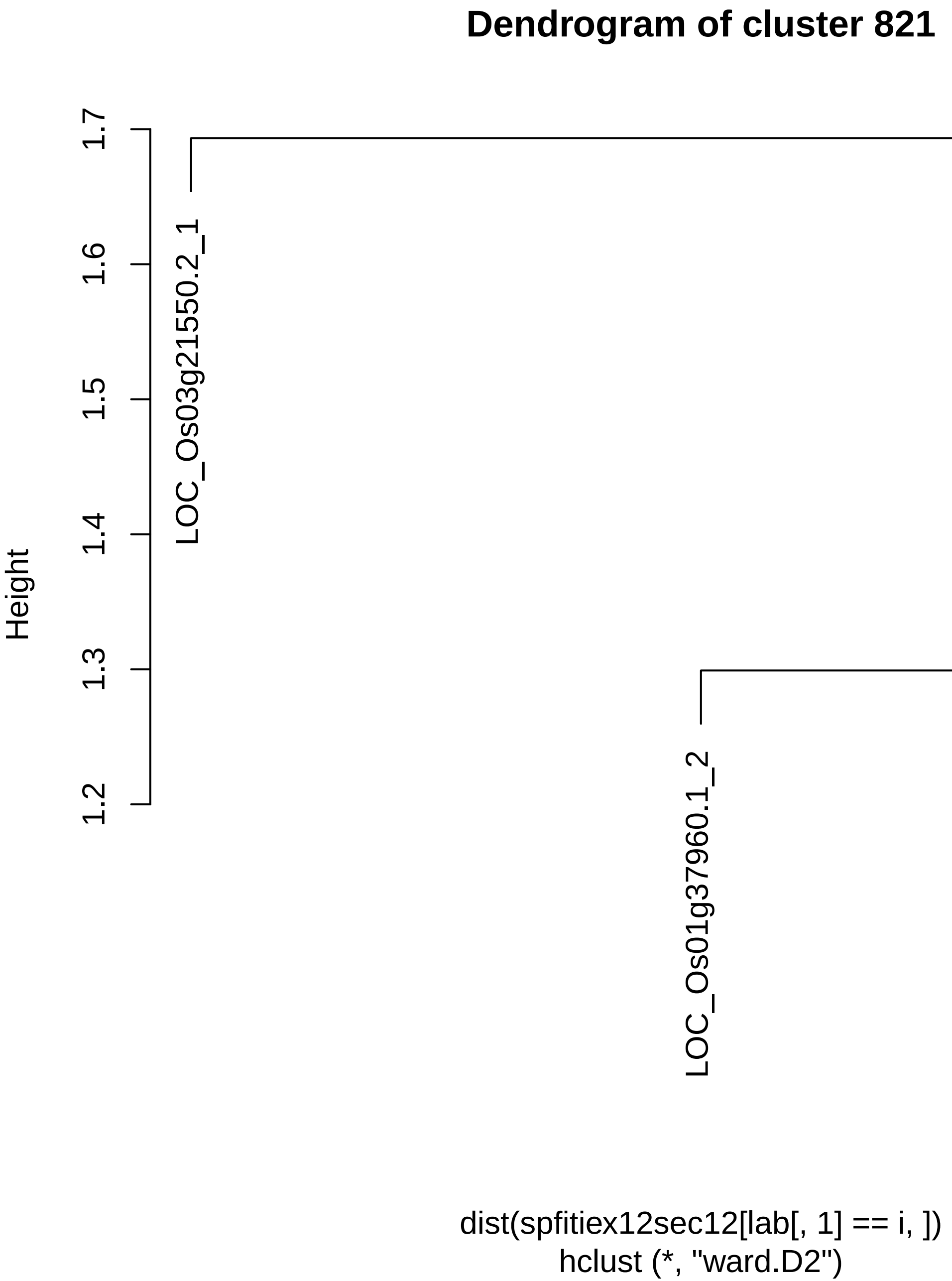
Dendrogram of cluster 821
Height
1.2
1.3
1.4
1.5
1.6
1.7
LOC_Os03g21550.2_1
LOC_Os01g37960.1_2
LOC_Os11g09329.1_1
dist(spfitiex12sec12[lab[, 1] == i, ])
hclust (*, "ward.D2")

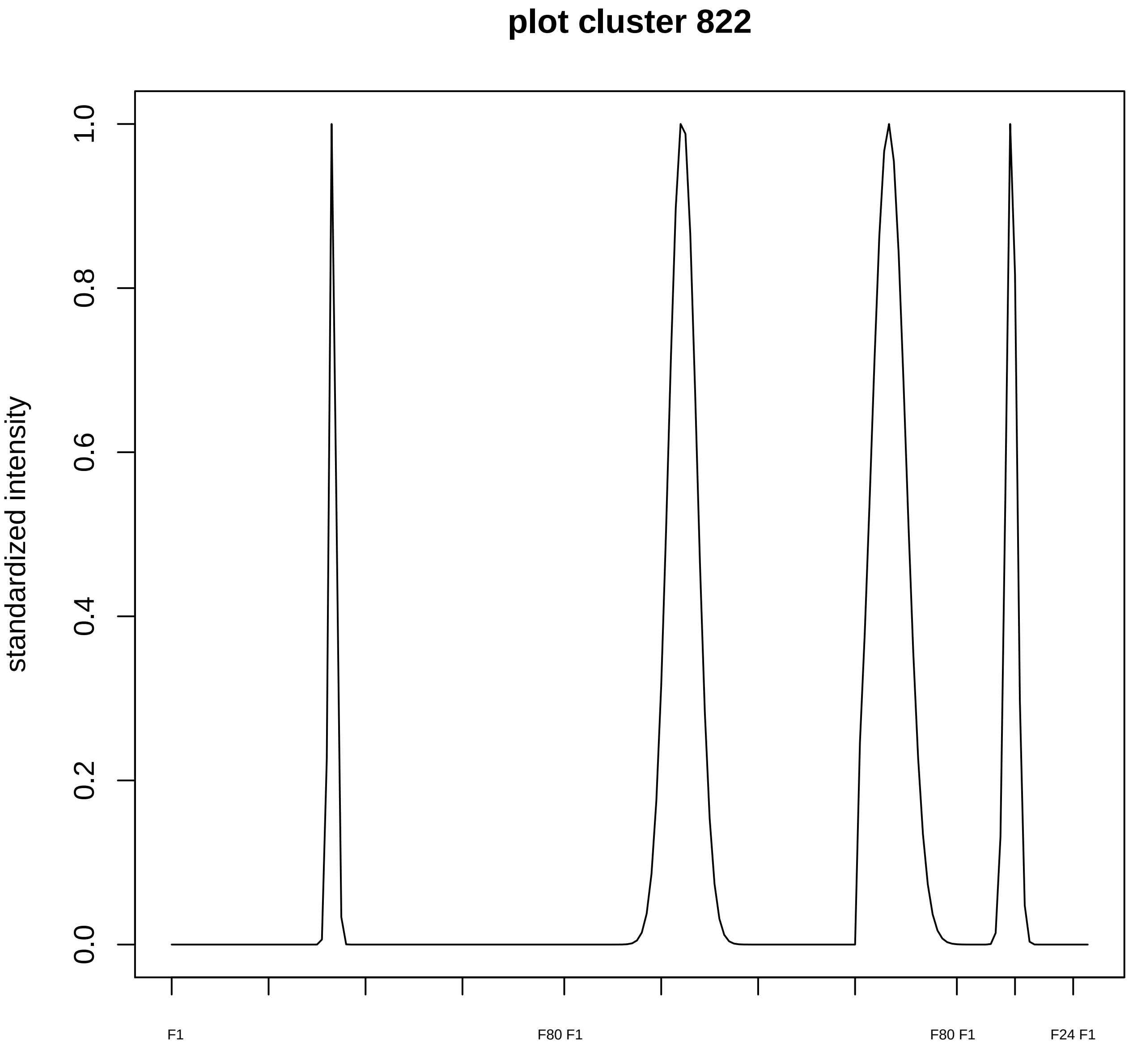

plot cluster 822
standardized intensity
0.0
0.2
0.4
0.6
0.8
1.0
F1
F80 F1
F80 F1
F24 F1

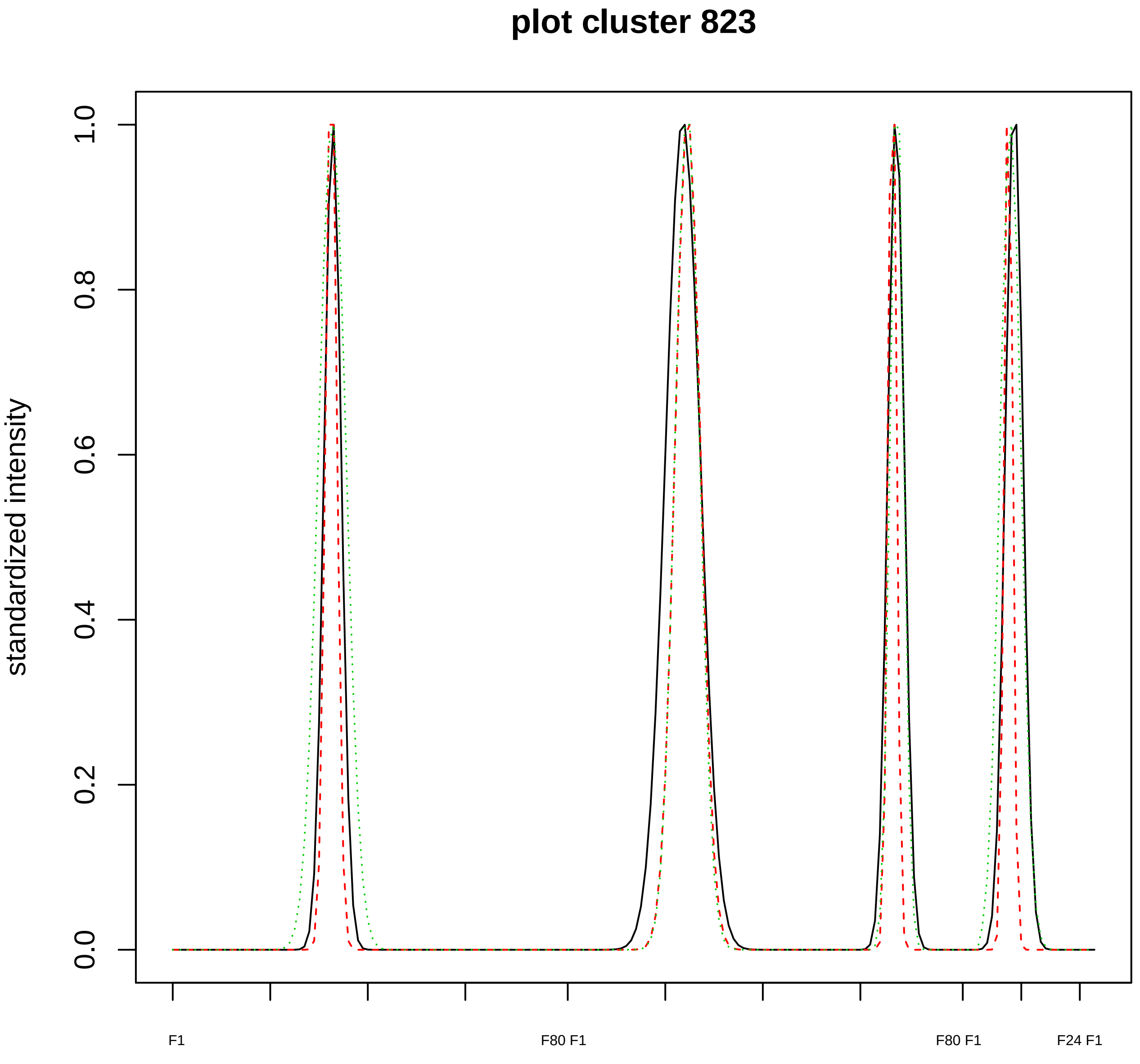
plot cluster 823
standardized intensity
0.0
0.2
0.4
0.6
0.8
1.0
F1
F80 F1
F80 F1
F24 F1

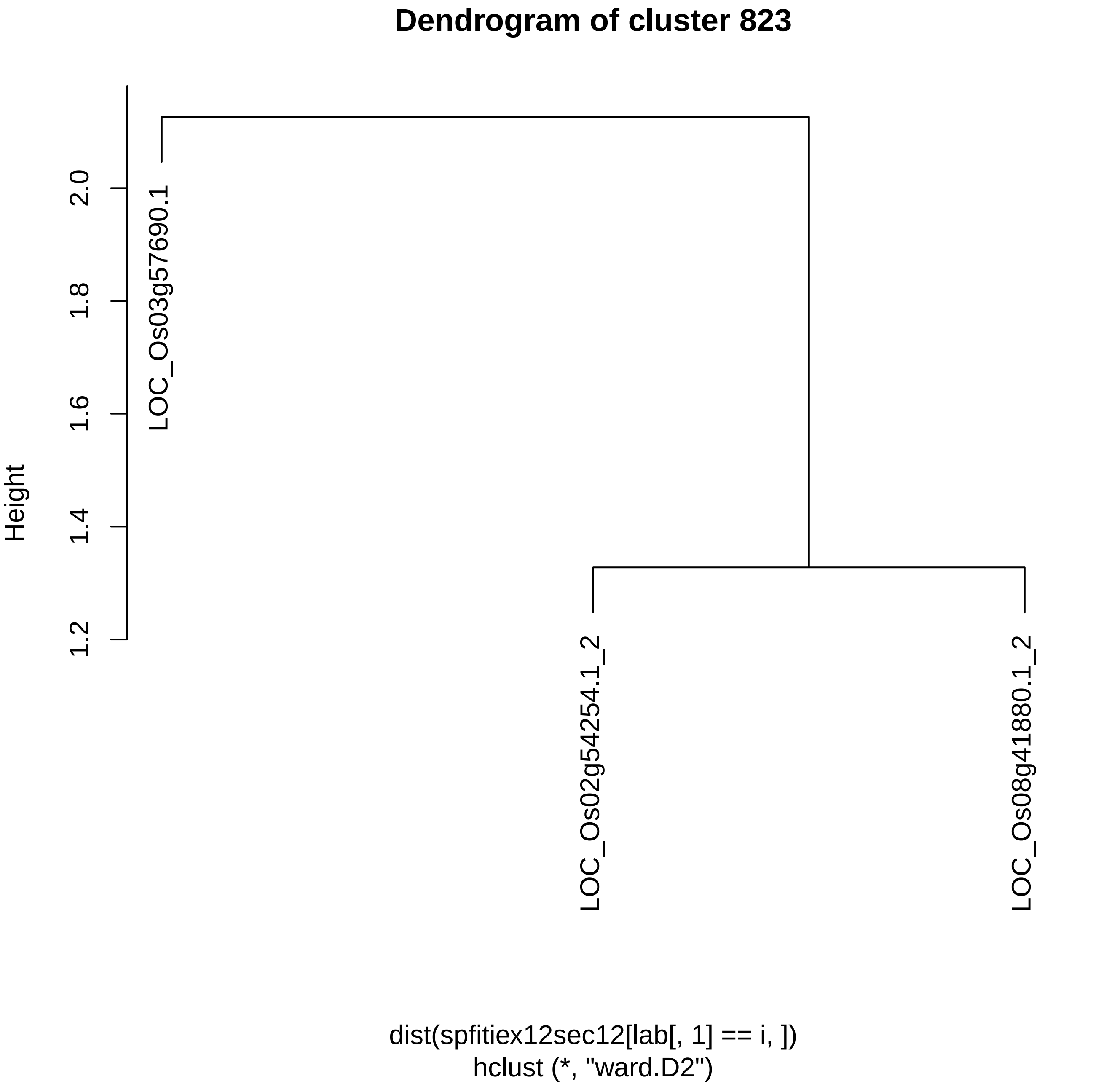
Dendrogram of cluster 823
Height
1.2
1.4
1.6
1.8
2.0
LOC_Os03g57690.1
LOC_Os02g54254.1_2
LOC_Os08g41880.1_2
dist(spfitiex12sec12[lab[, 1] == i, ])
hclust (*, "ward.D2")

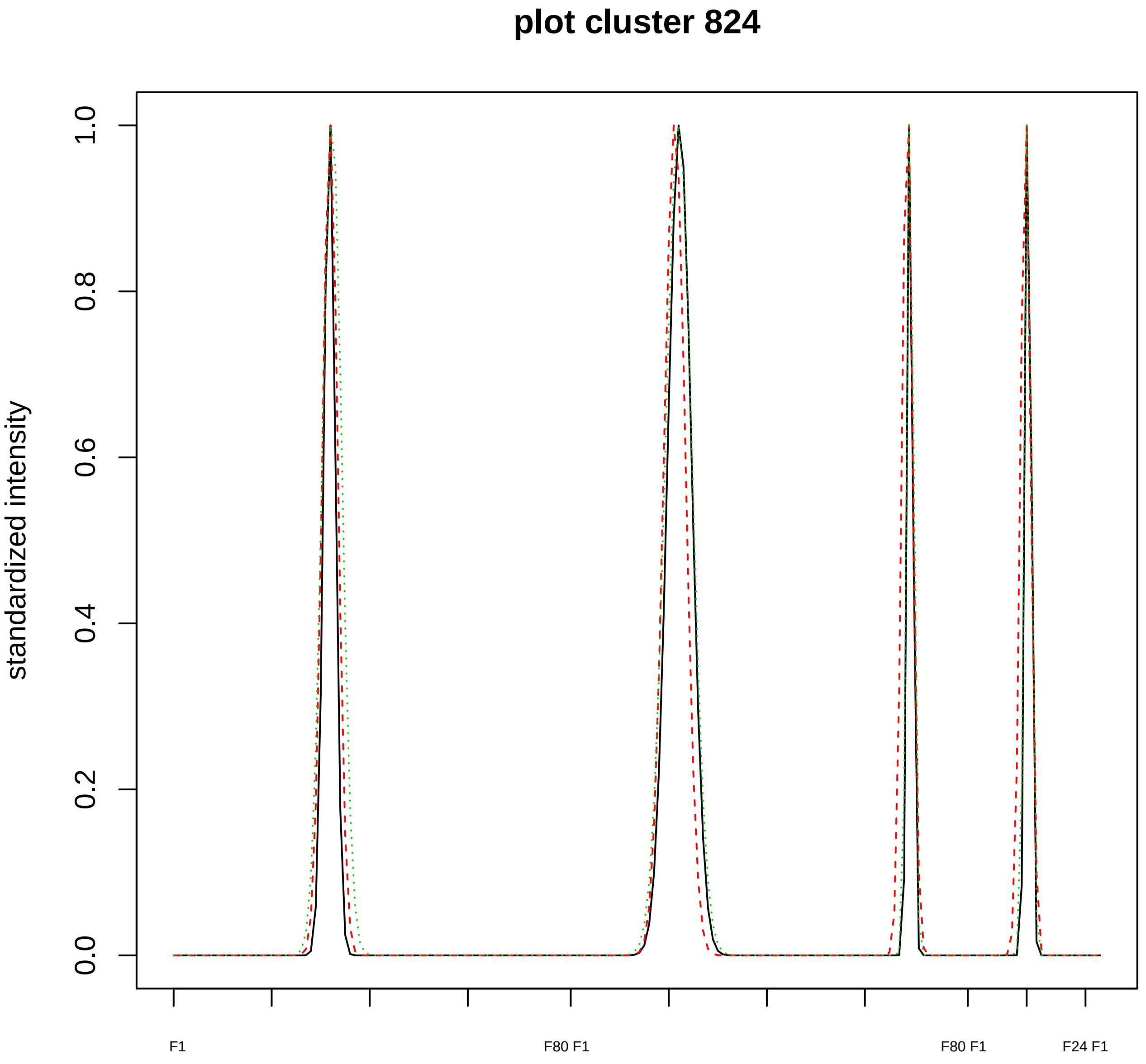
plot cluster 824
standardized intensity
0.0
0.2
0.4
0.6
0.8
1.0
F1
F80 F1
F80 F1
F24 F1

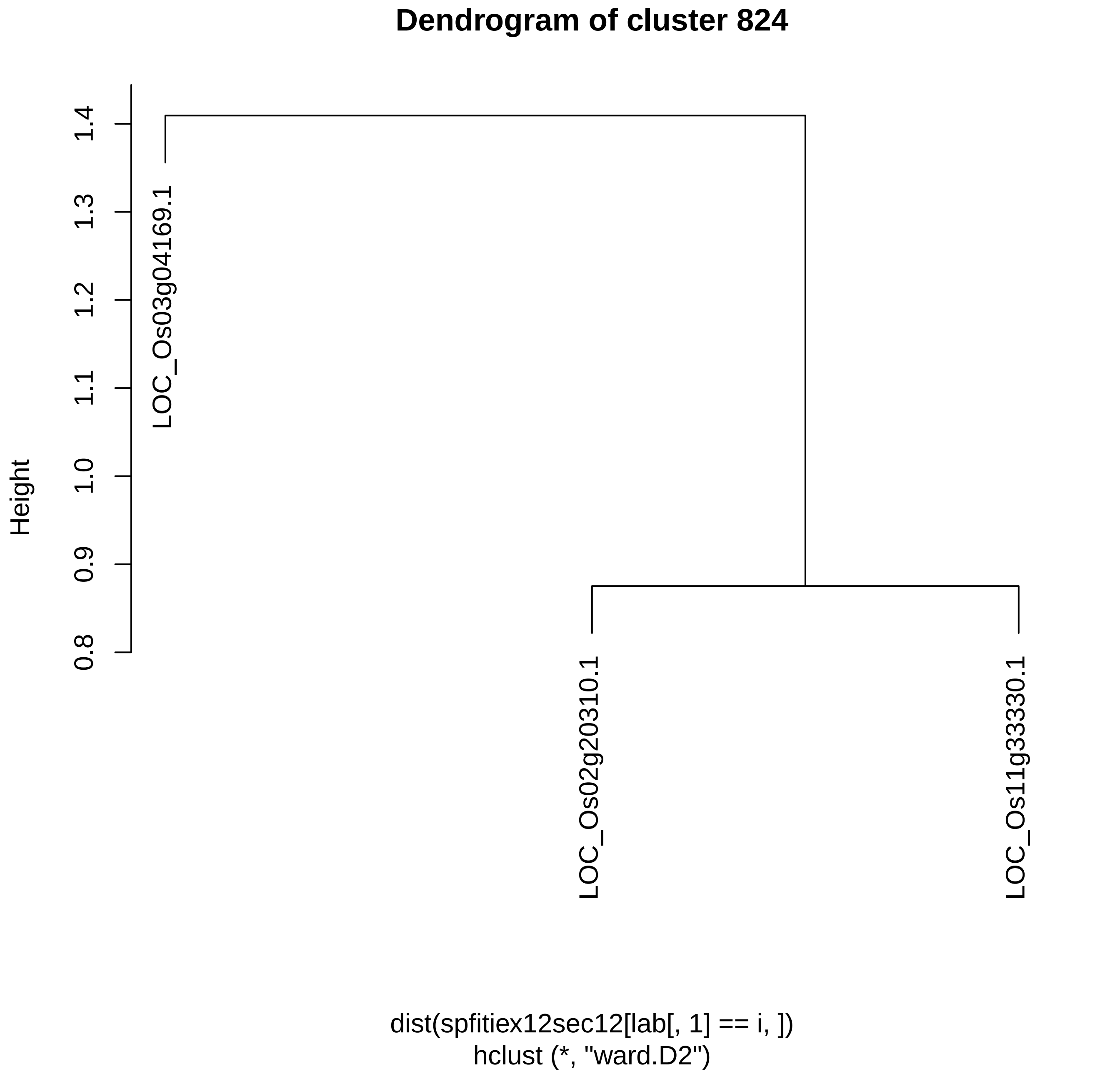
Dendrogram of cluster 824
Height
1.4
1.3
1.2
1.1
1.0
0.9
0.8
LOC_Os03g04169.1
LOC_Os02g20310.1
LOC_Os11g33330.1
dist(spfitiex12sec12[lab[, 1] == i, ])
hclust (*, "ward.D2")

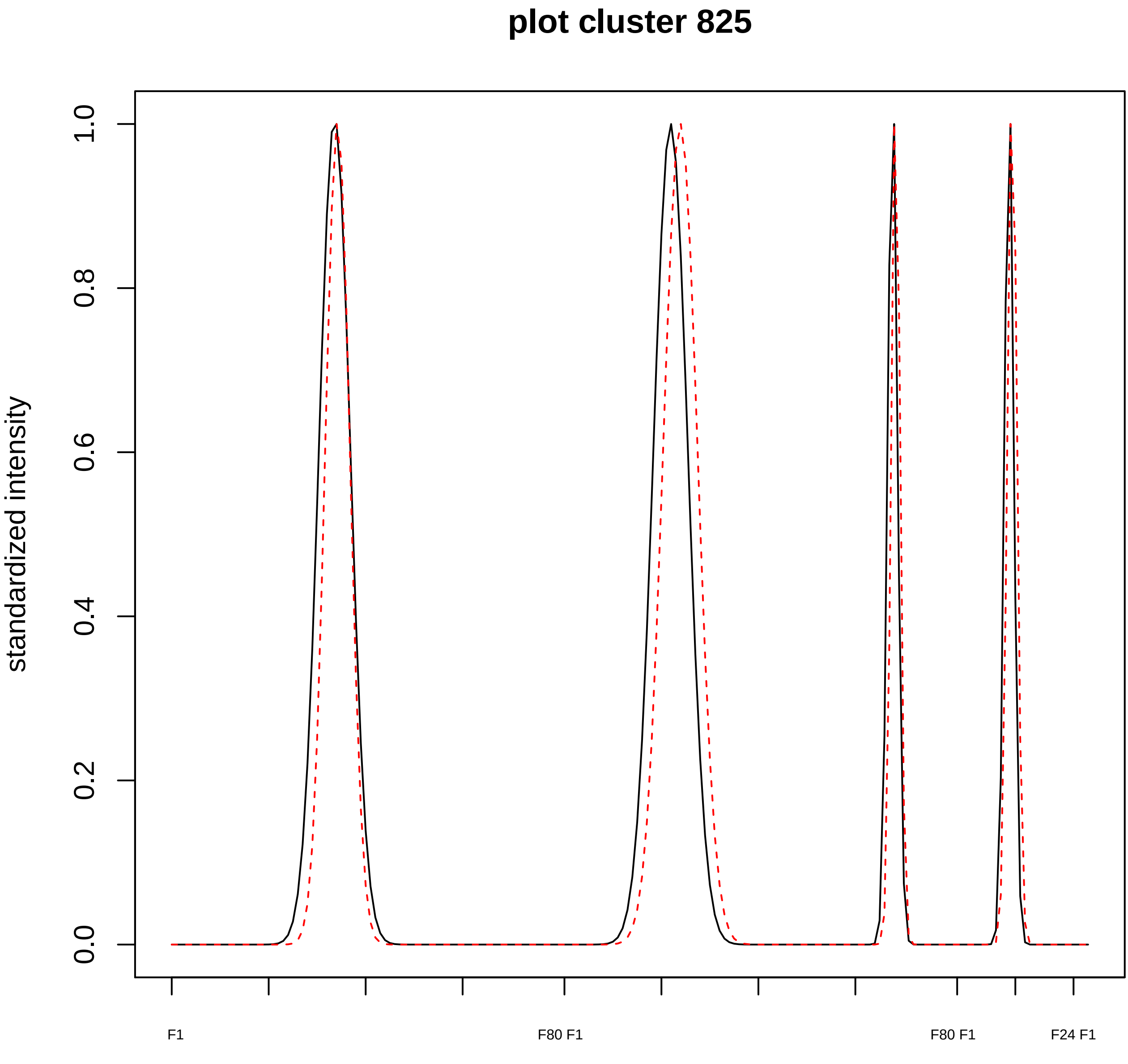

plot cluster 825
standardized intensity
0.0
0.2
0.4
0.6
0.8
1.0
F1
F80 F1
F80 F1
F24 F1

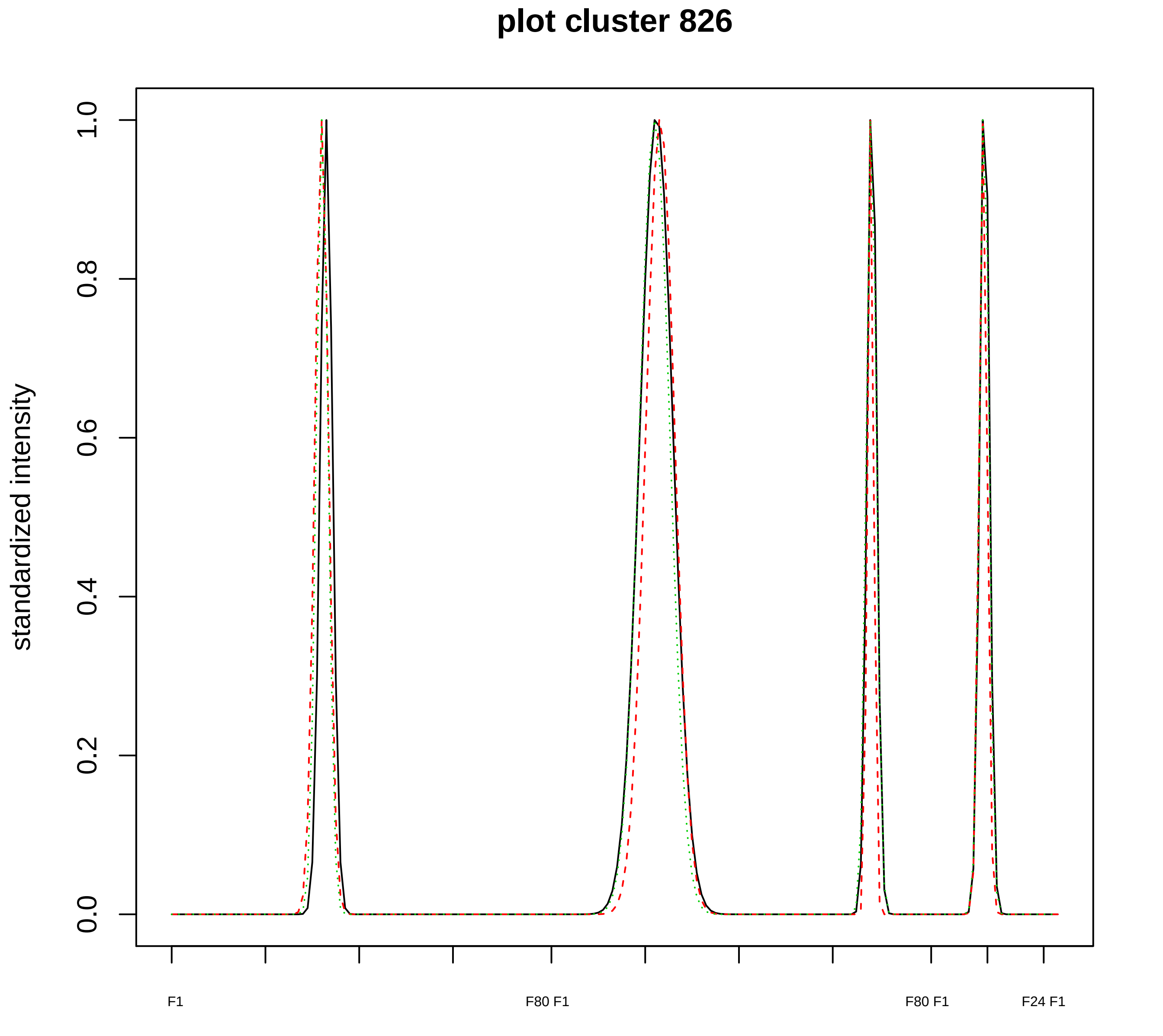

plot cluster 826
standardized intensity
0.0
0.2
0.4
0.6
0.8
1.0
F1
F80 F1
F80 F1
F24 F1

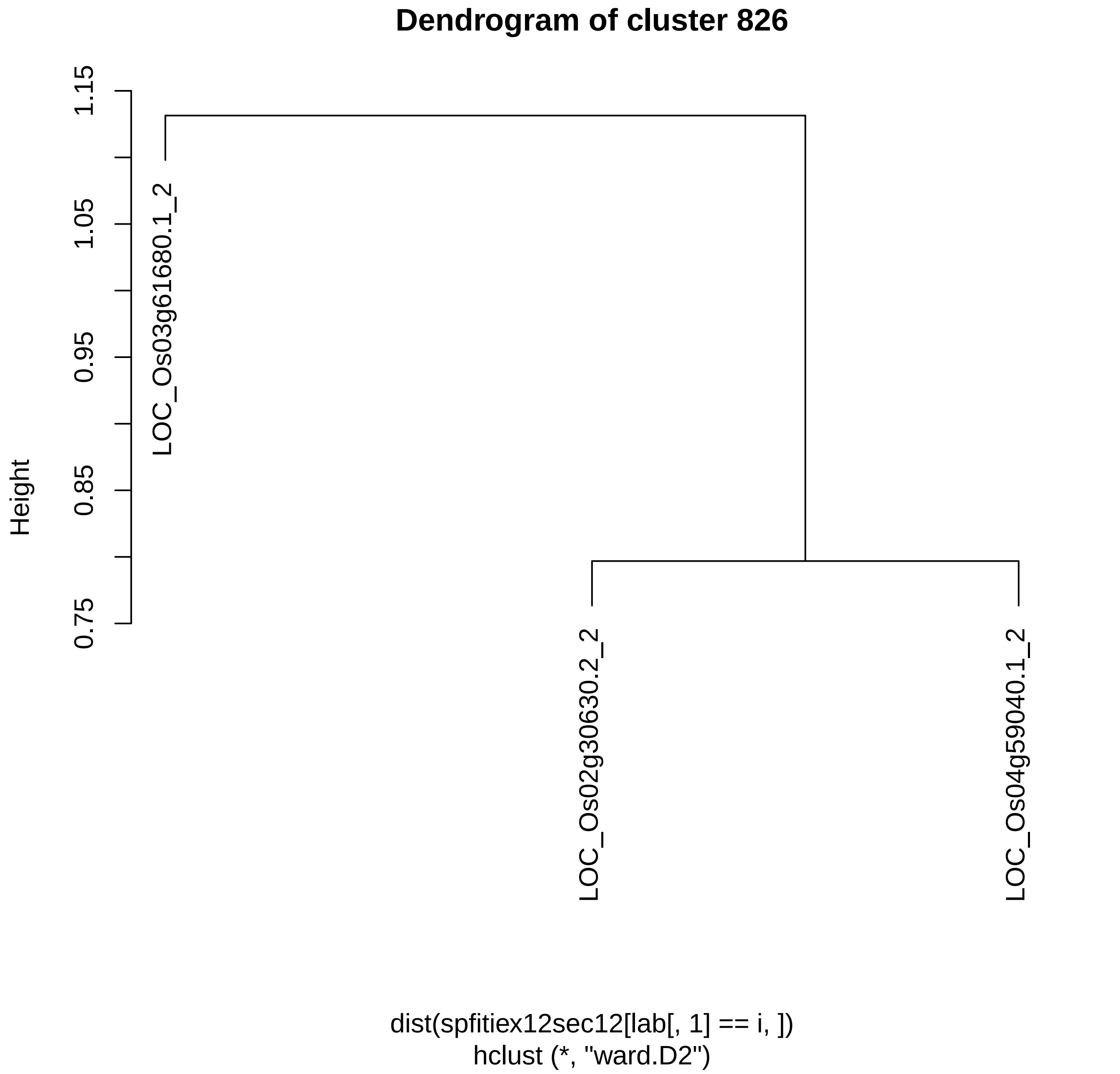

Dendrogram of cluster 826
Height
1.15
1.05
0.95
0.85
0.75
LOC_Os03g61680.1_2
LOC_Os02g30630.2_2
LOC_Os04g59040.1_2
dist(spfitiex12sec12[lab[, 1] == i, ])
hclust (*, "ward.D2")

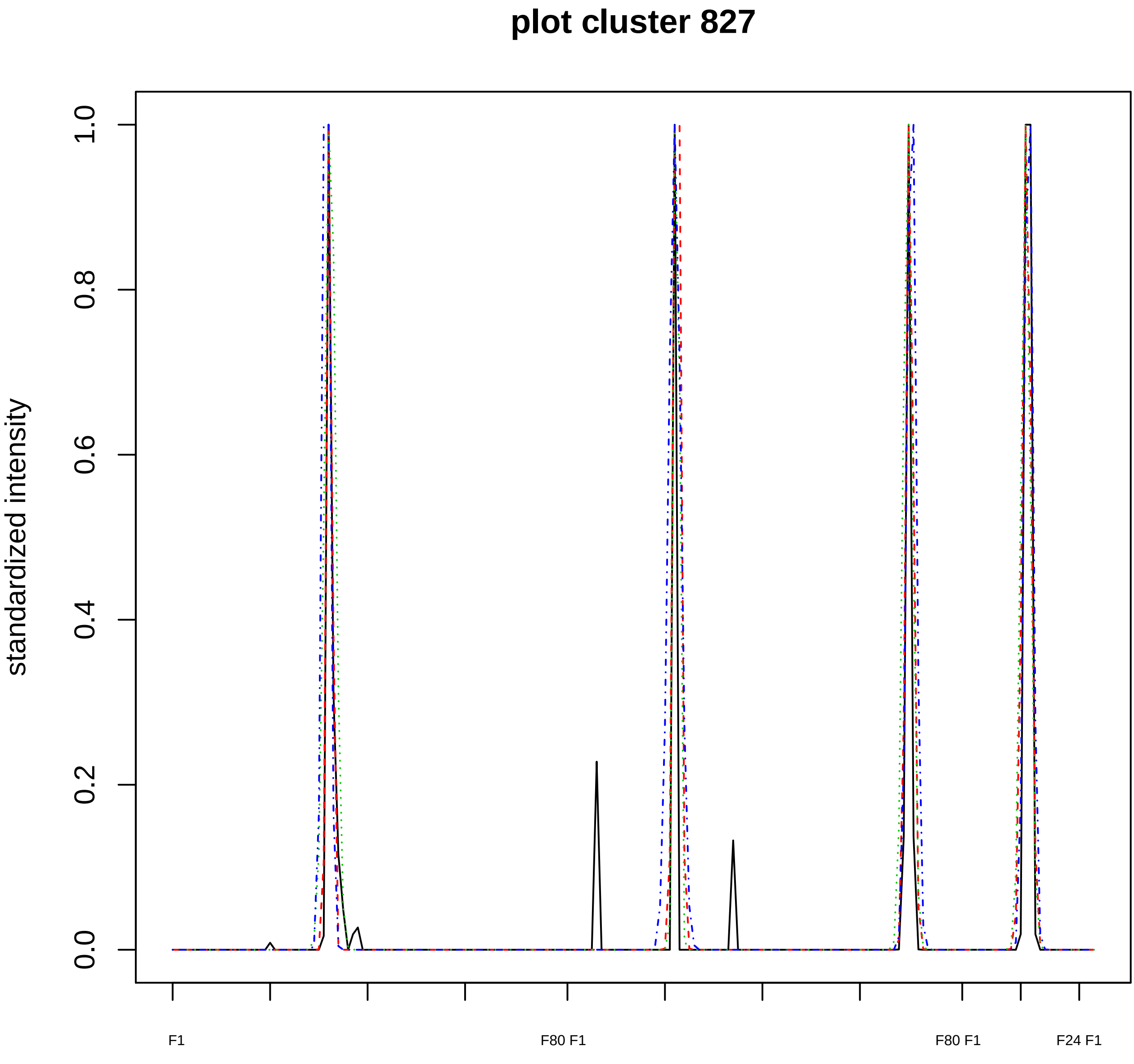

plot cluster 827
standardized intensity
0.0
0.2
0.4
0.6
0.8
1.0
F1
F80 F1
F80 F1
F24 F1

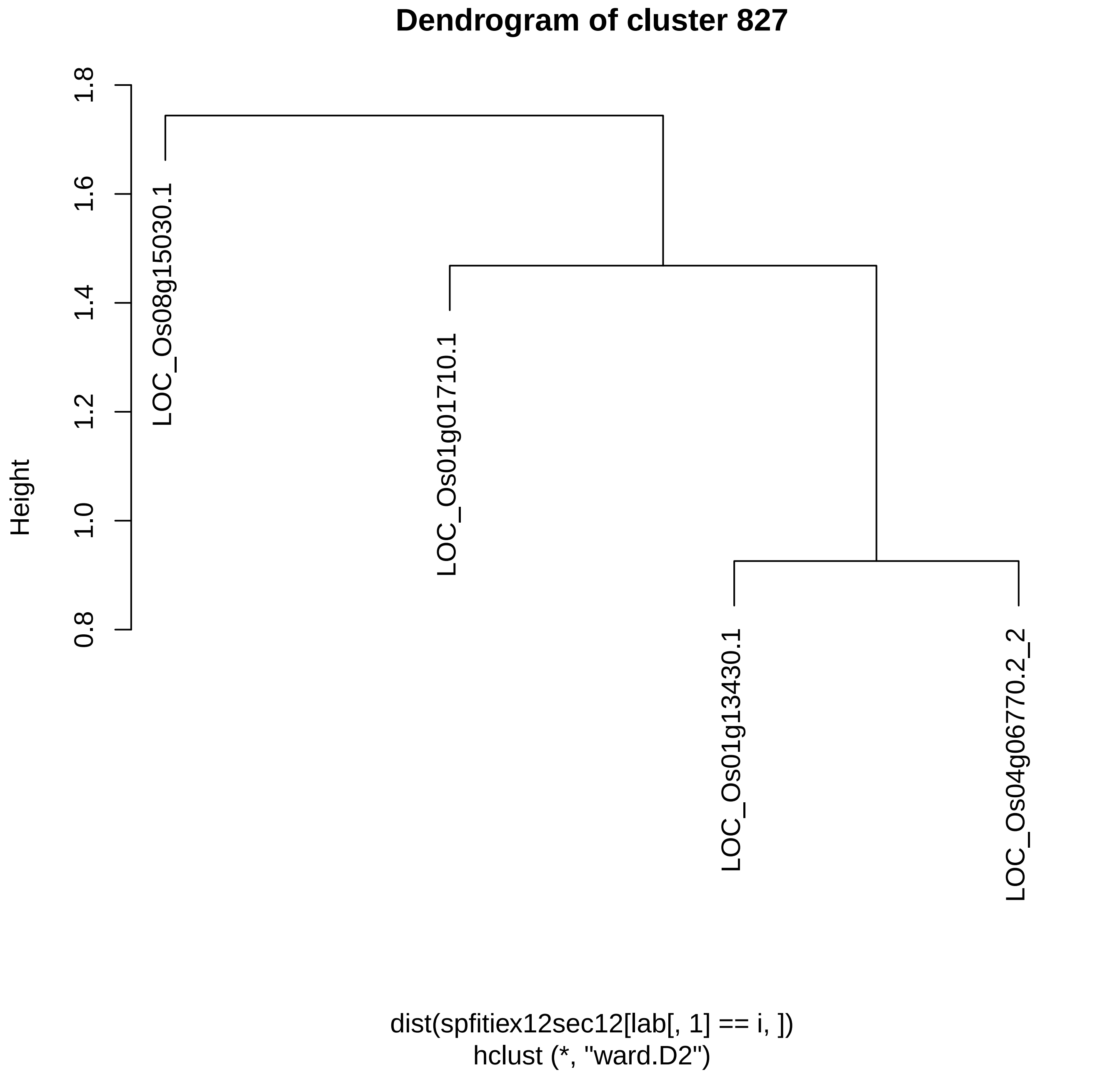
Dendrogram of cluster 827
Height
0.8
1.0
1.2
1.4
1.6
1.8
LOC_Os08g15030.1
LOC_Os01g01710.1
LOC_Os01g13430.1
LOC_Os04g06770.2_2
dist(spfitiex12sec12[lab[, 1] == i, ])
hclust (*, "ward.D2")

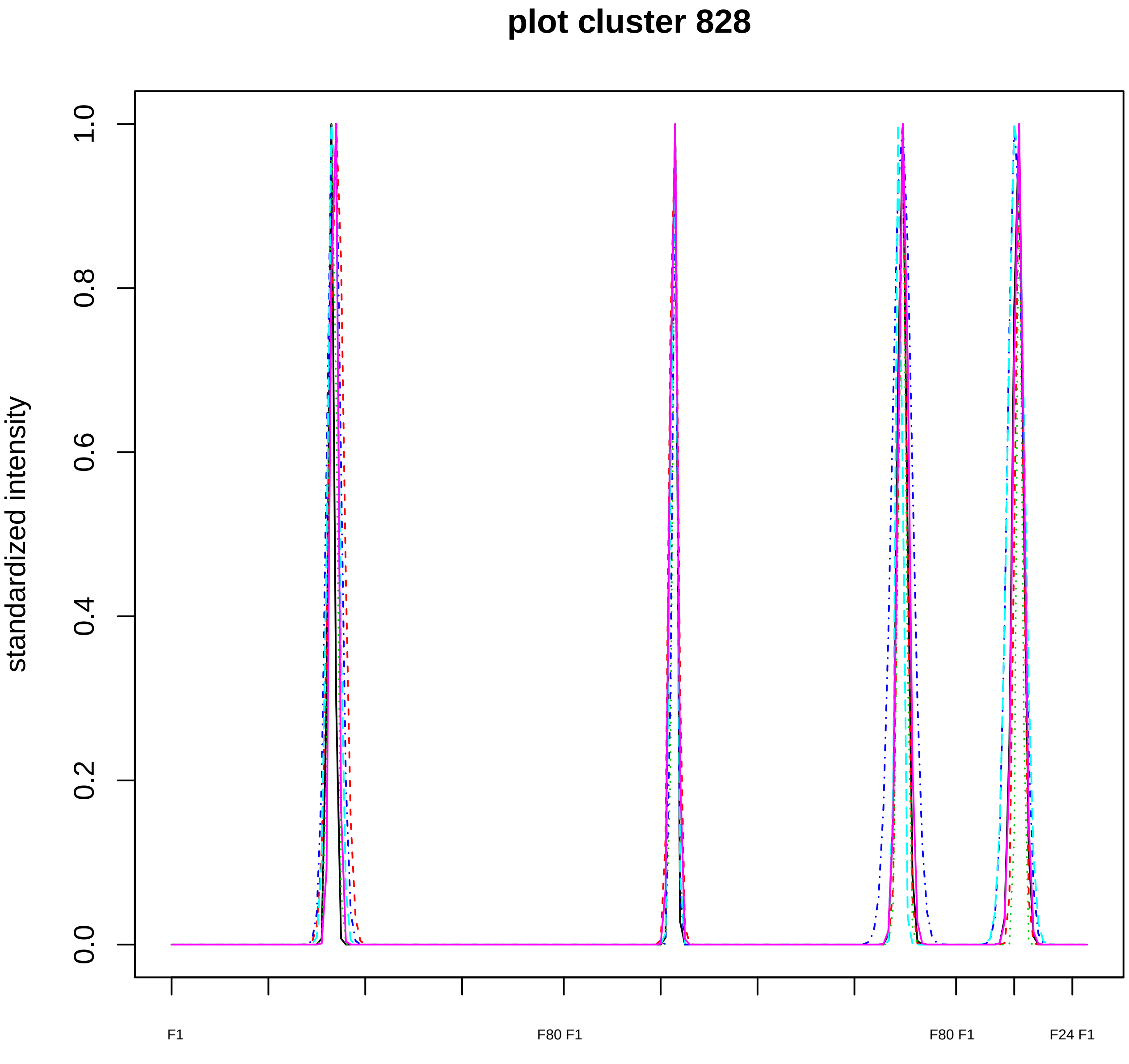
plot cluster 828
standardized intensity
0.0
0.2
0.4
0.6
0.8
1.0
F1
F80 F1
F80 F1
F24 F1

### Dendrogram of cluster 828

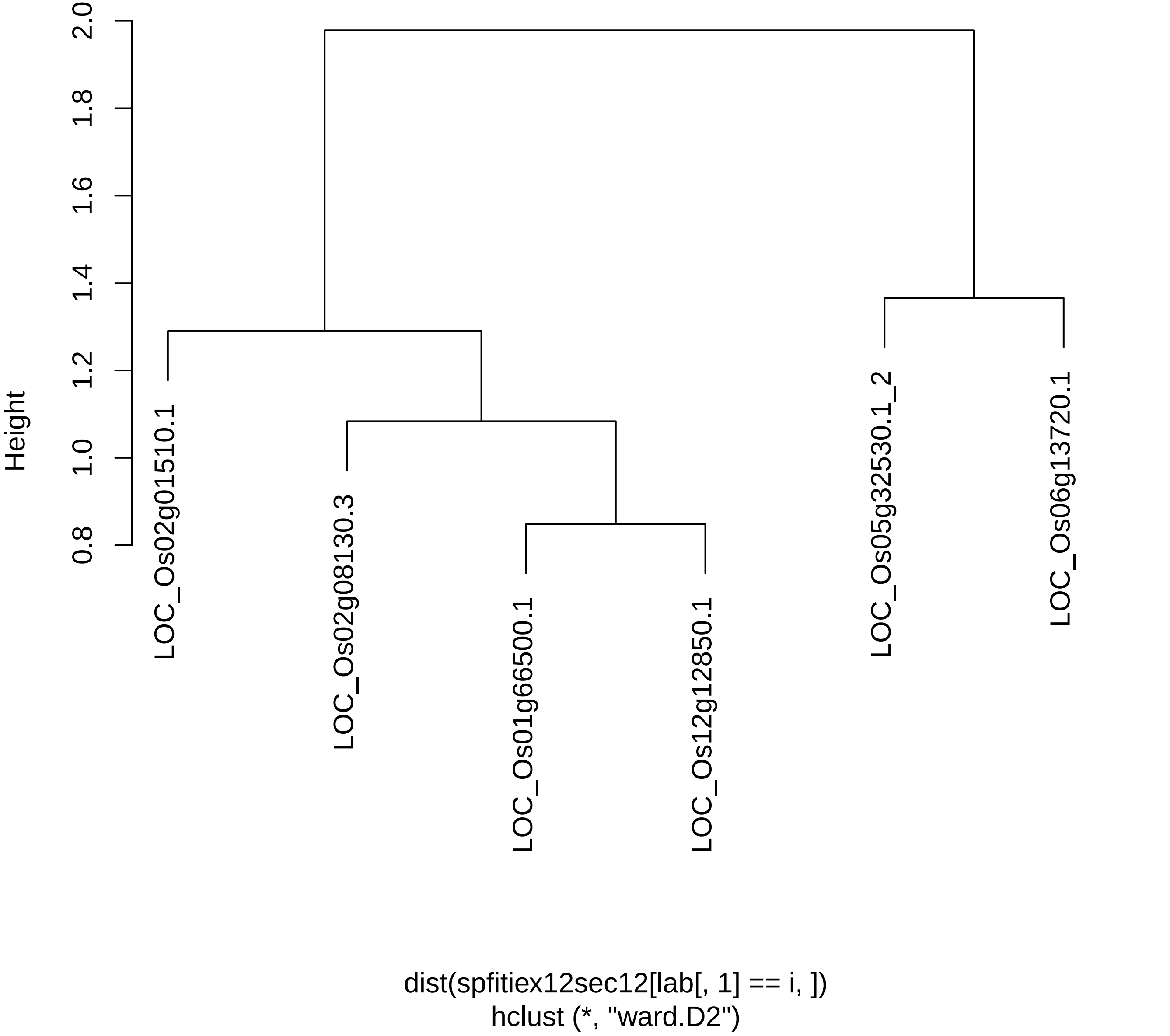

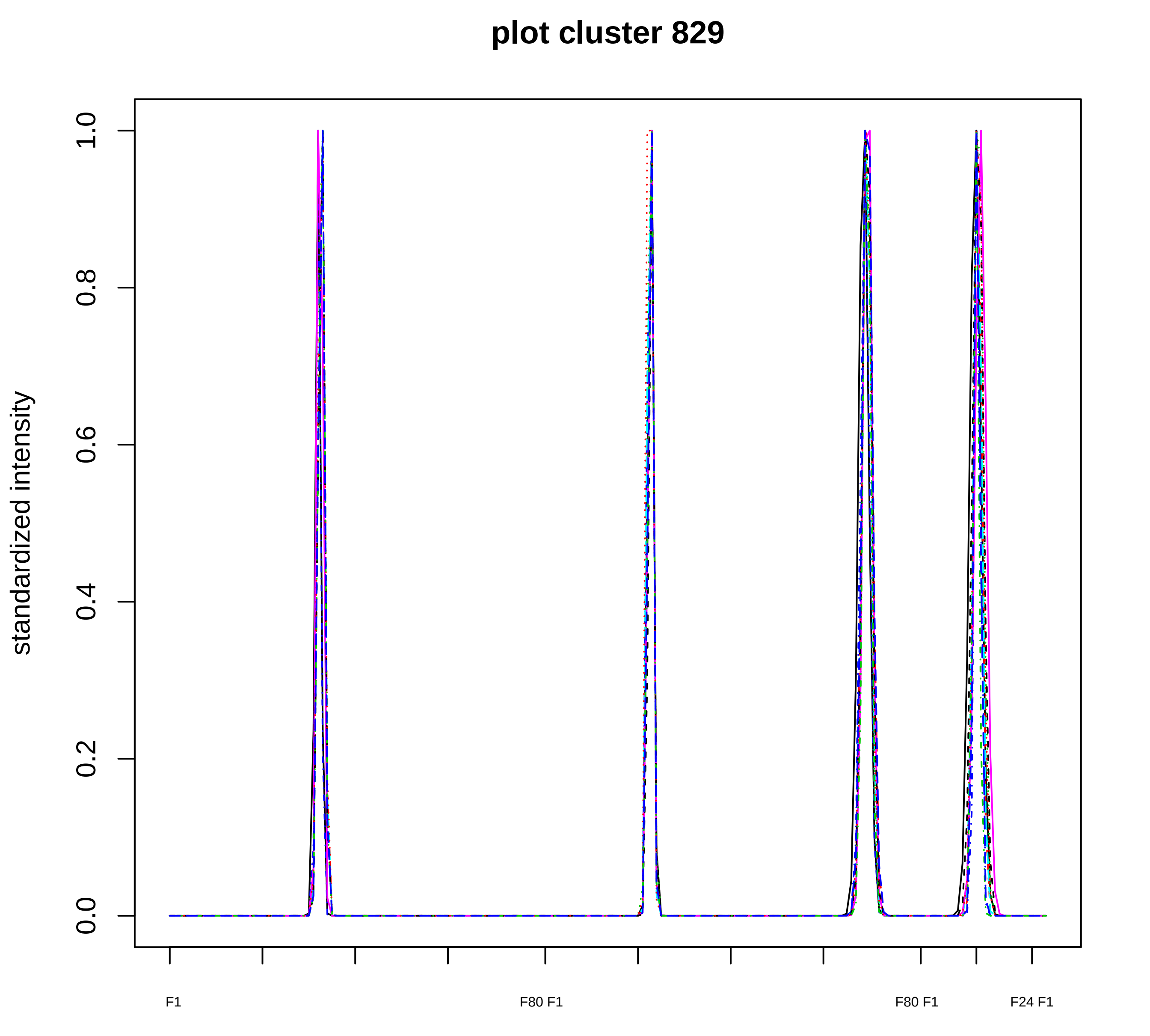

plot cluster 829
standardized intensity
0.0
0.2
0.4
0.6
0.8
1.0
F1
F80 F1
F80 F1
F24 F1

### Dendrogram of cluster 829

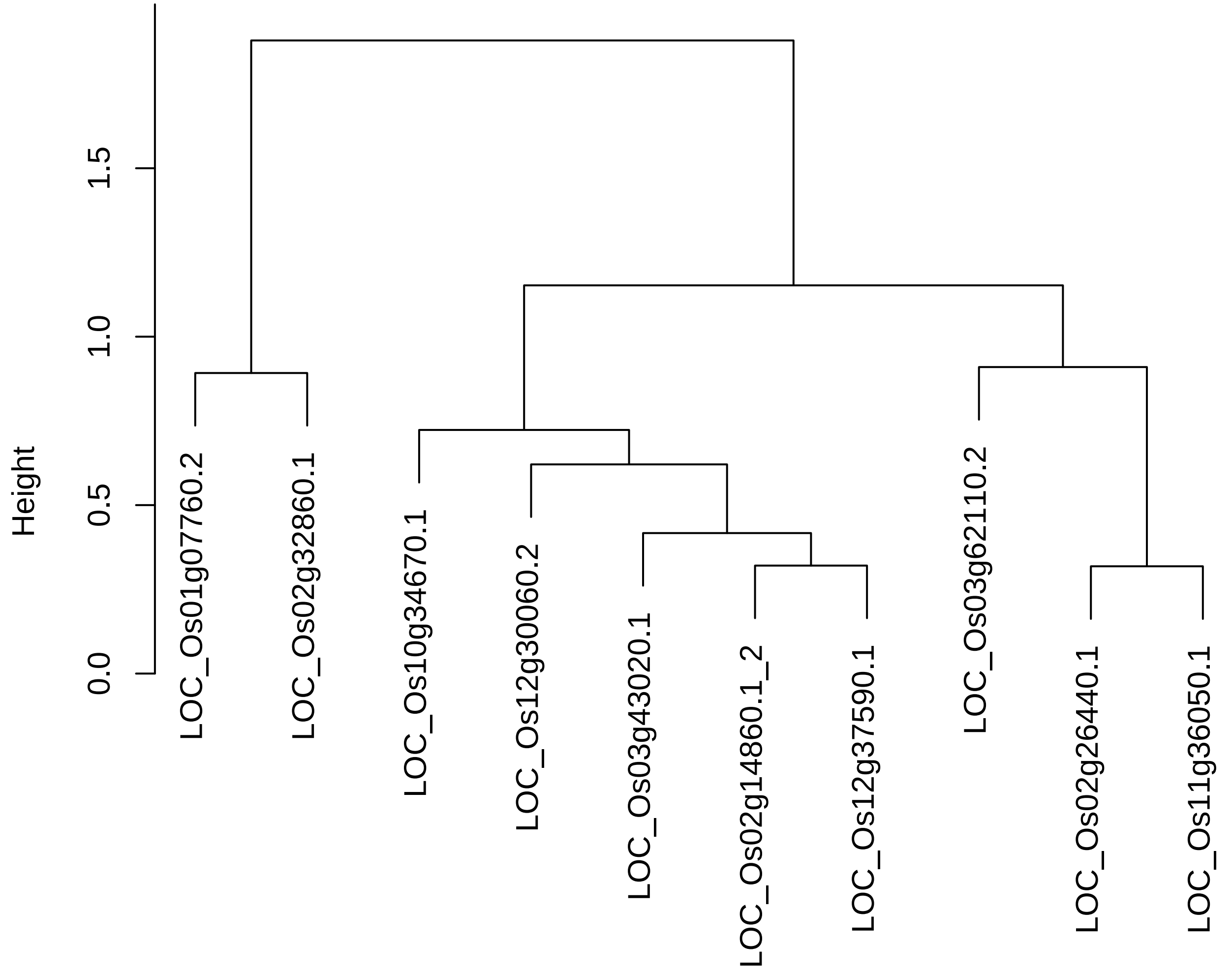


dist(spfitiex12sec12[lab[, 1] == i, ])
hclust (*, "ward.D2")

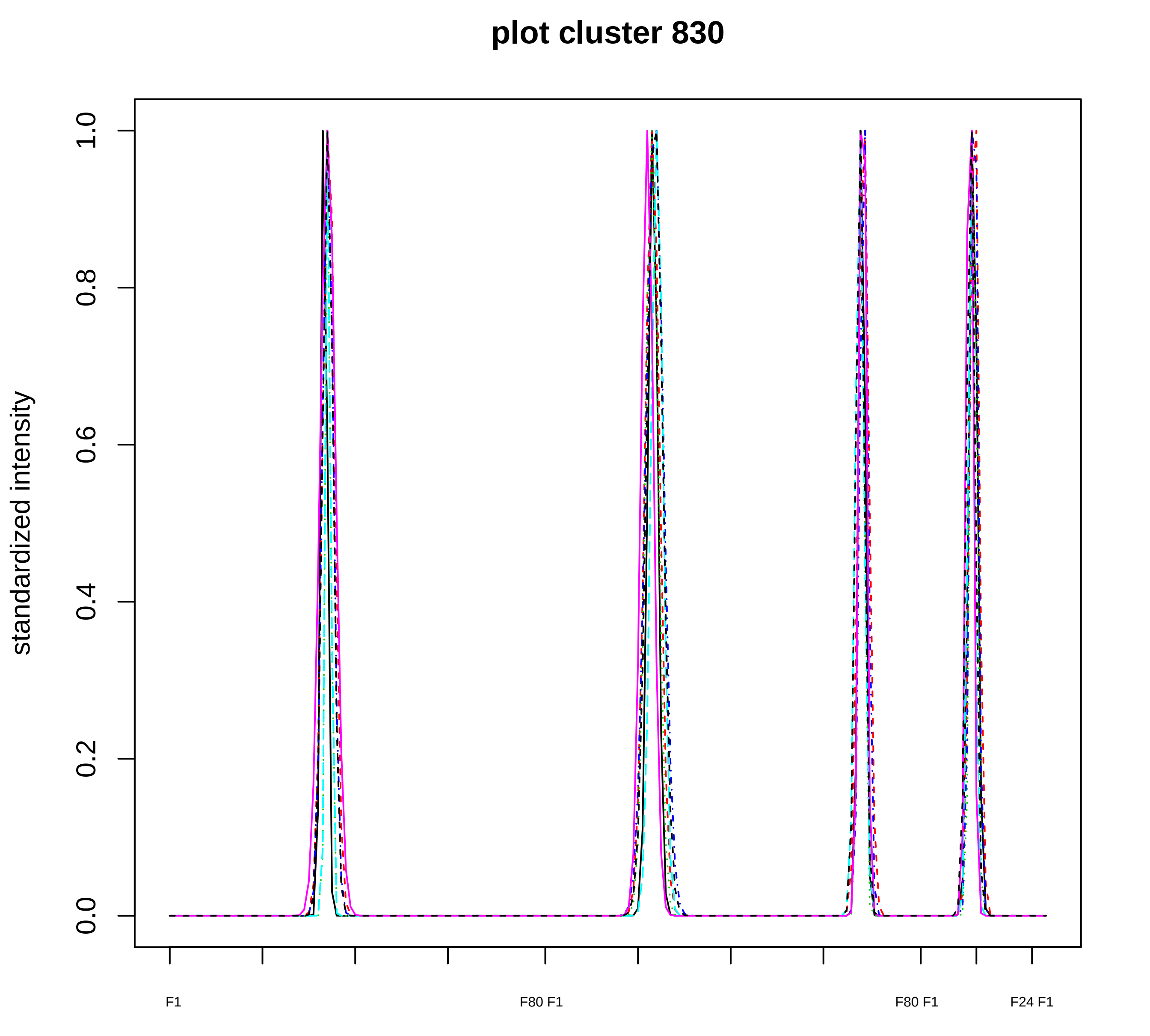

plot cluster 830
standardized intensity
0.0
0.2
0.4
0.6
0.8
1.0
F1
F80 F1
F80 F1
F24 F1

### Dendrogram of cluster 830

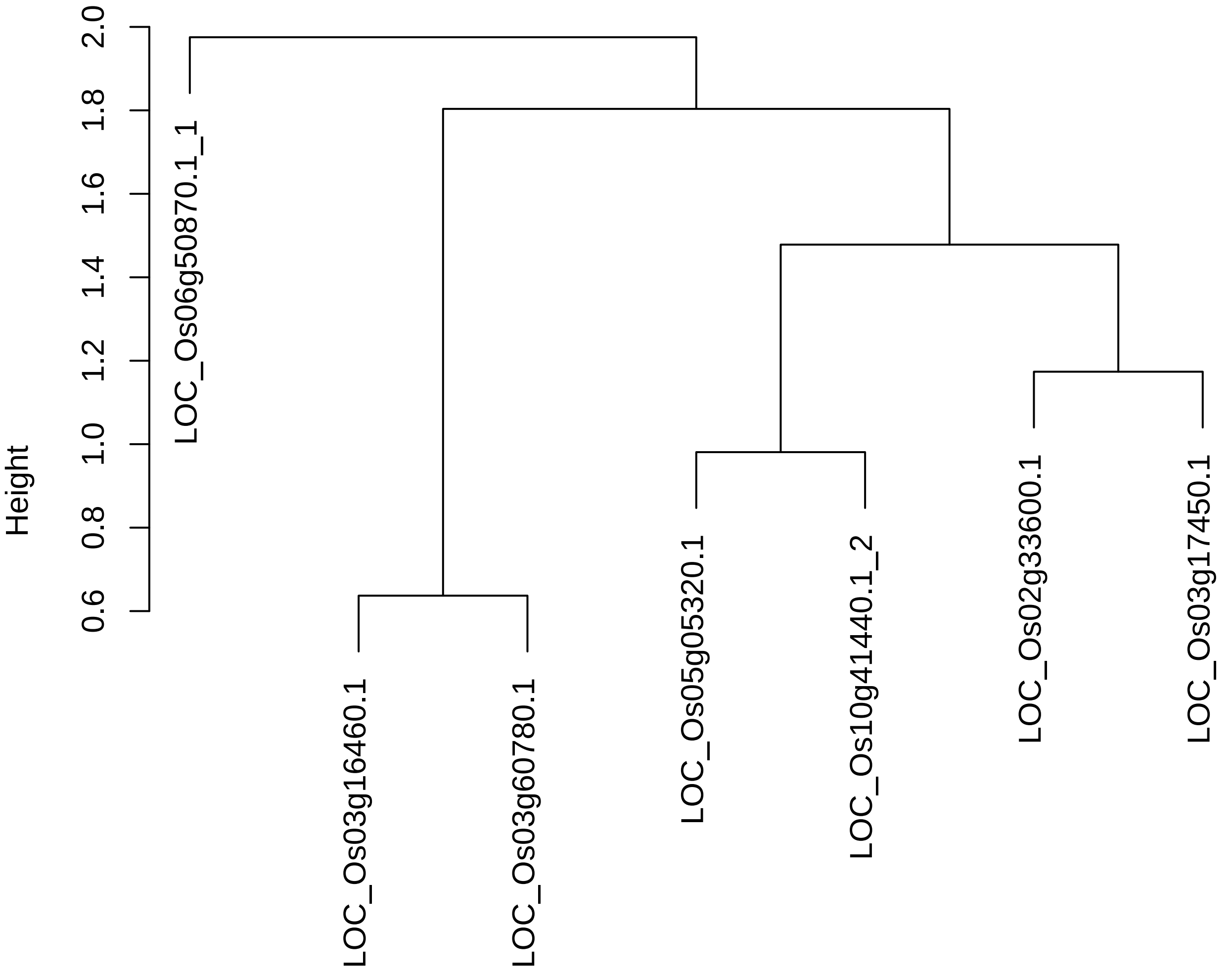


dist(spfitiex12sec12[lab[, 1] == i, ])
hclust (*, "ward.D2")

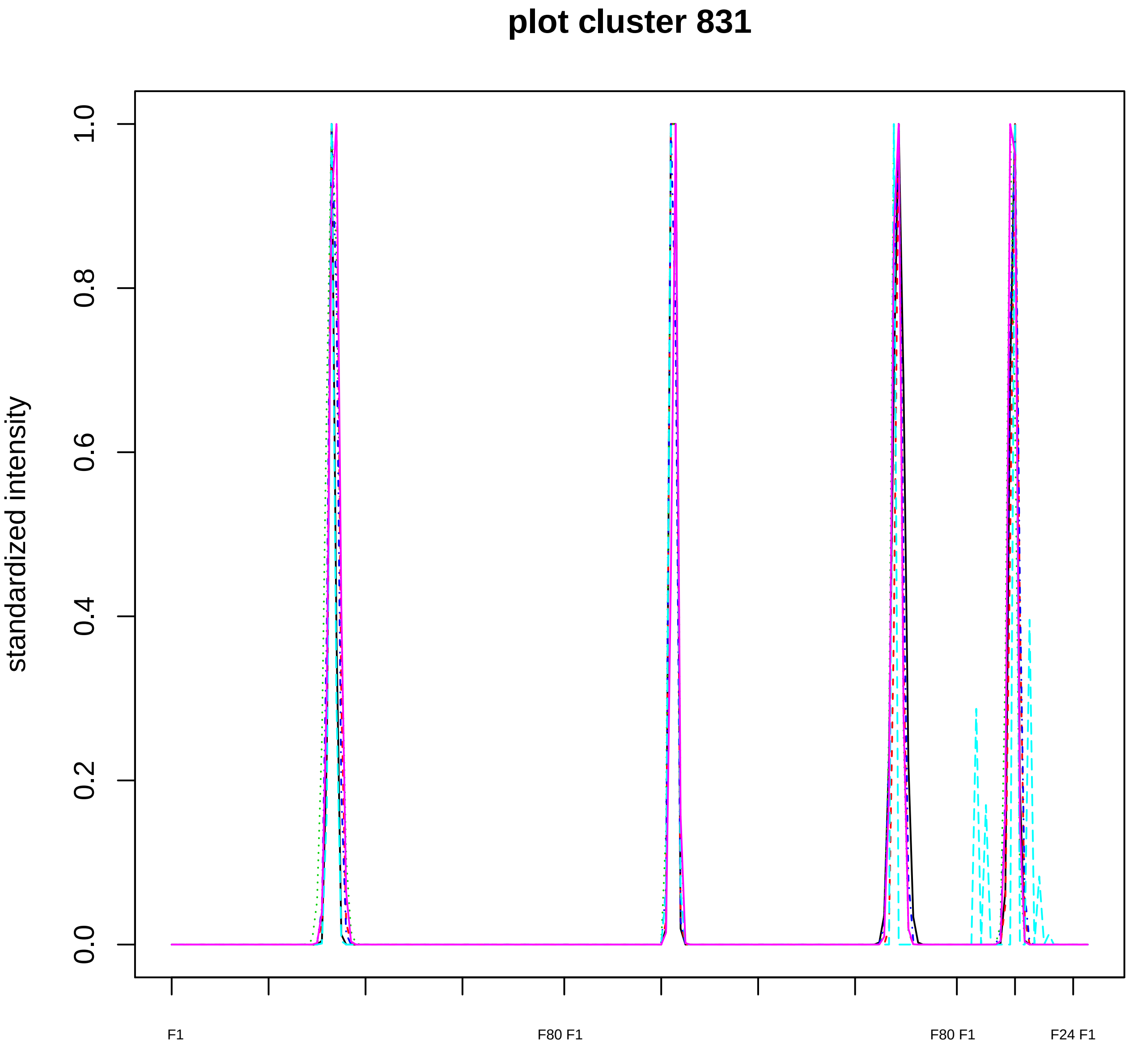
plot cluster 831
standardized intensity
0.0
0.2
0.4
0.6
0.8
1.0
F1
F80 F1
F80 F1
F24 F1

### Dendrogram of cluster 831

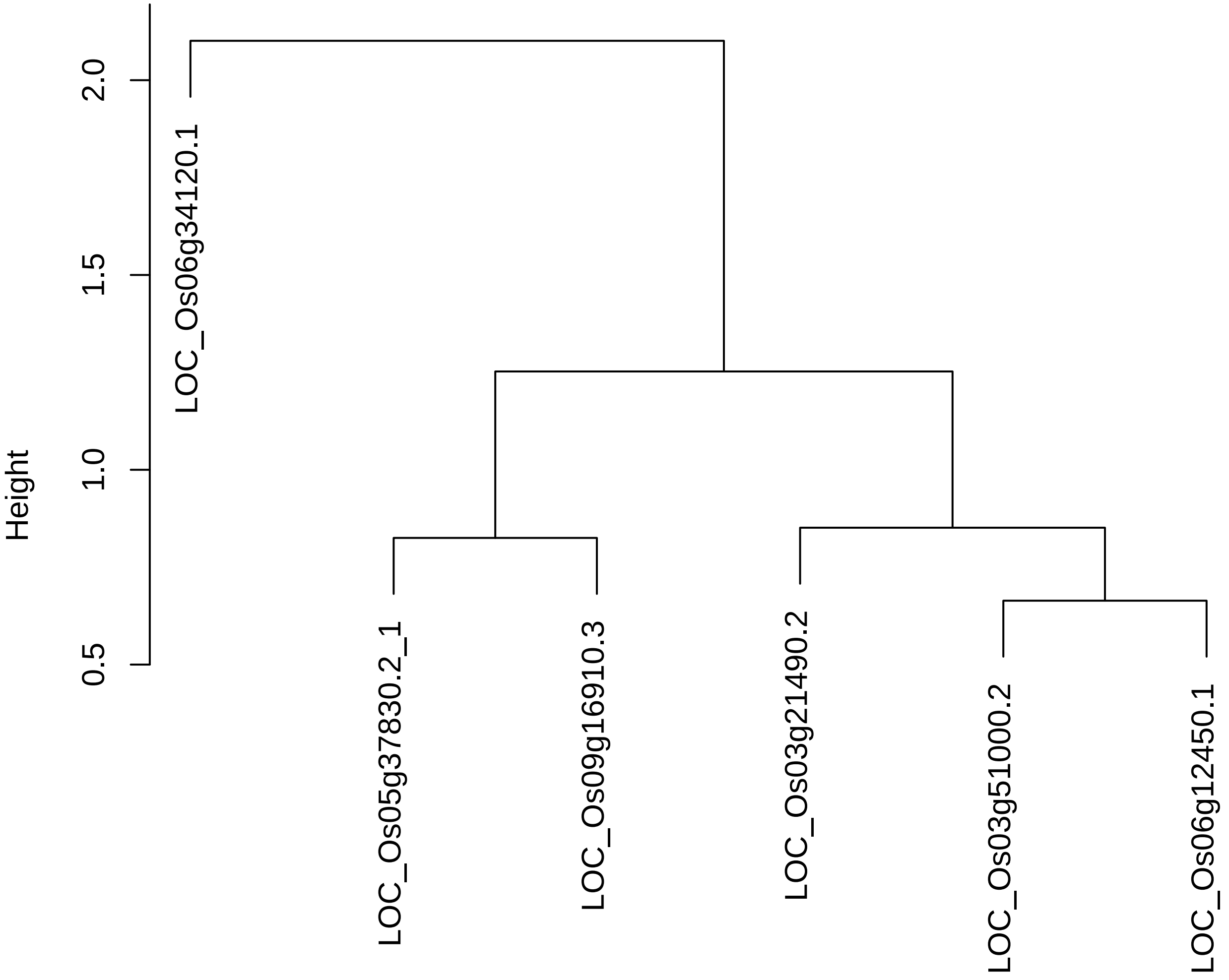


dist(spfitiex12sec12[lab[, 1] == i, ])
hclust (*, "ward.D2")

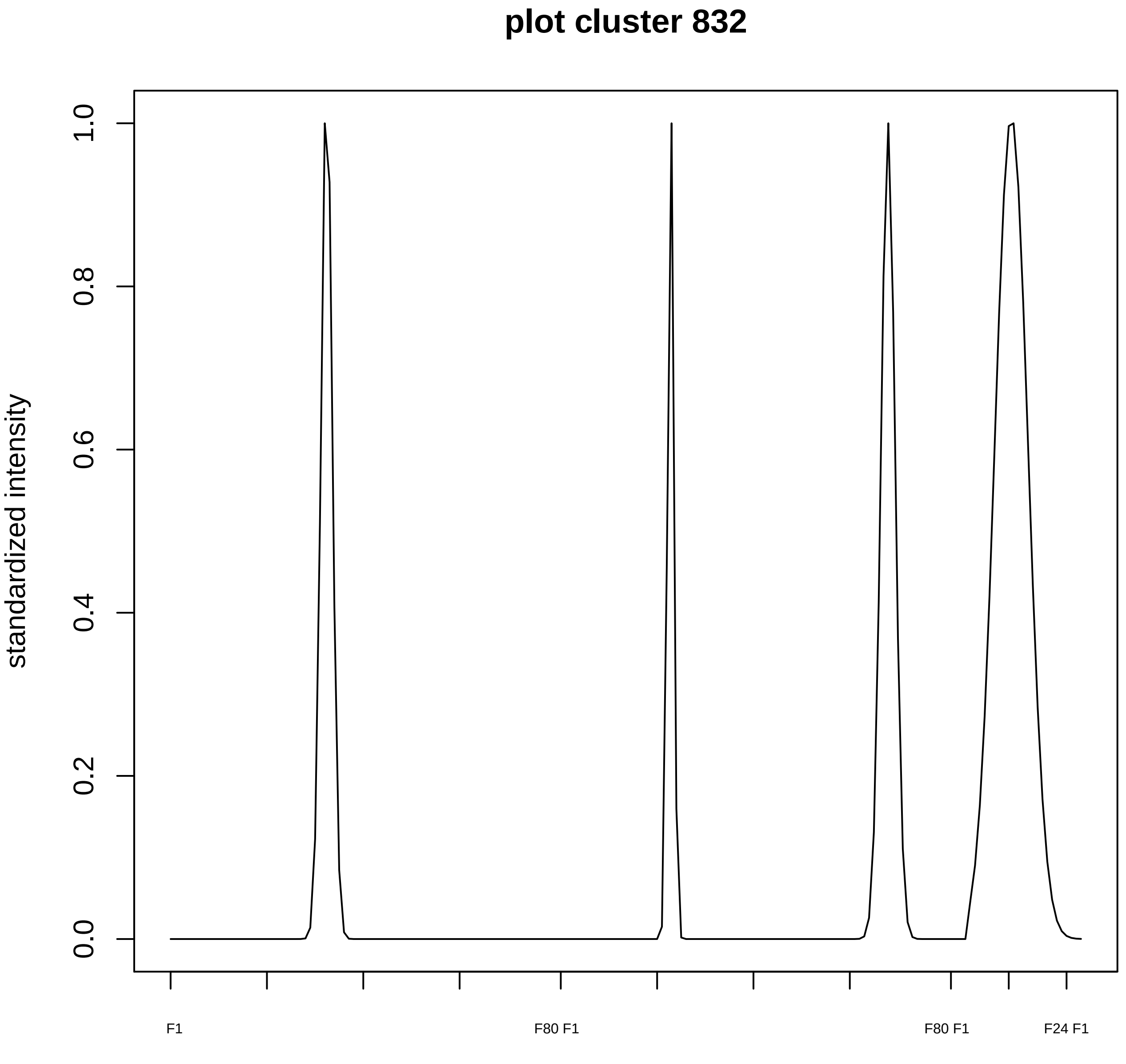
plot cluster 832
standardized intensity
0.0
0.2
0.4
0.6
0.8
1.0
F1
F80 F1
F80 F1
F24 F1

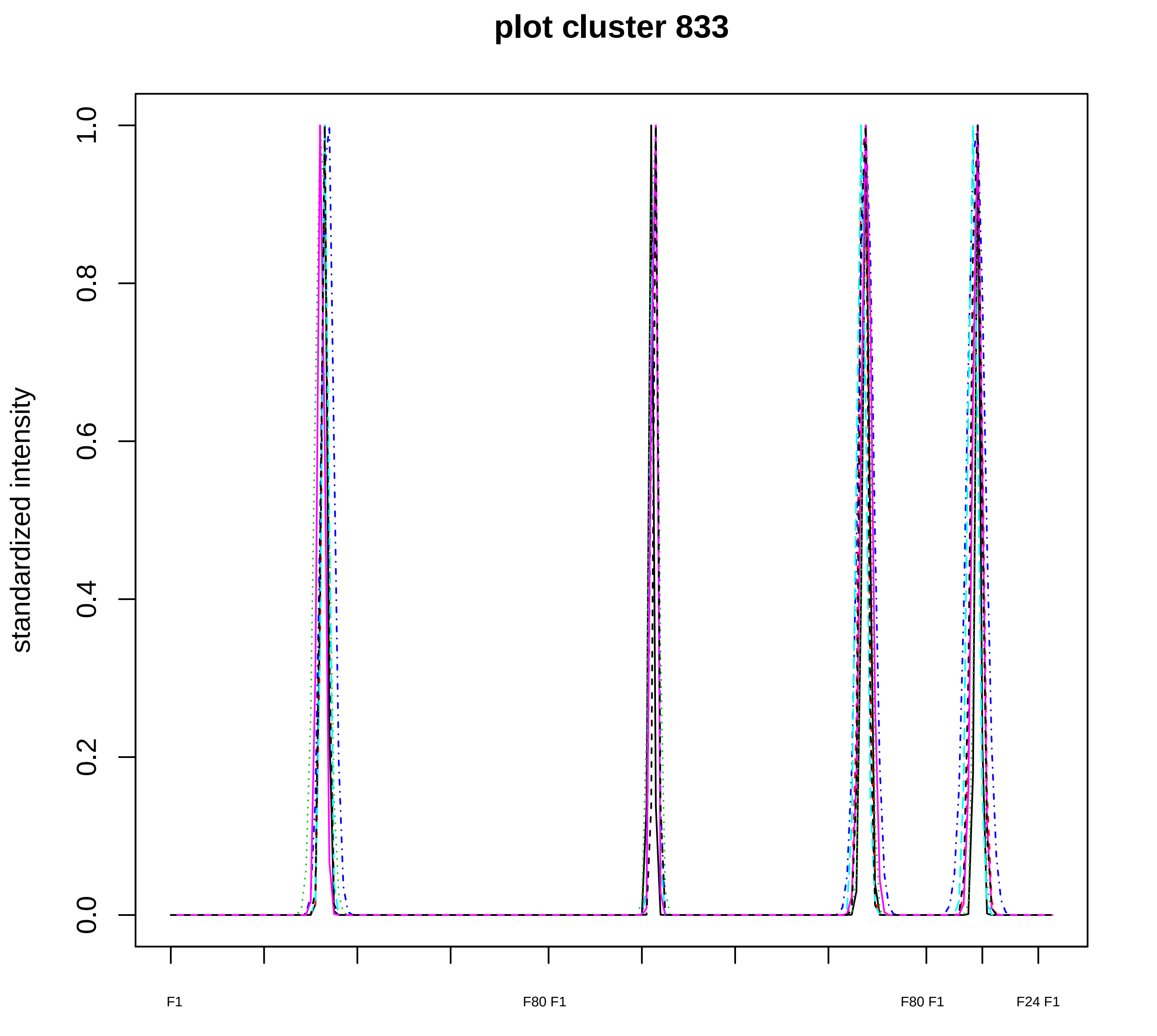
plot cluster 833
standardized intensity
0.0
0.2
0.4
0.6
0.8
1.0
F1
F80 F1
F80 F1
F24 F1

### Dendrogram of cluster 833

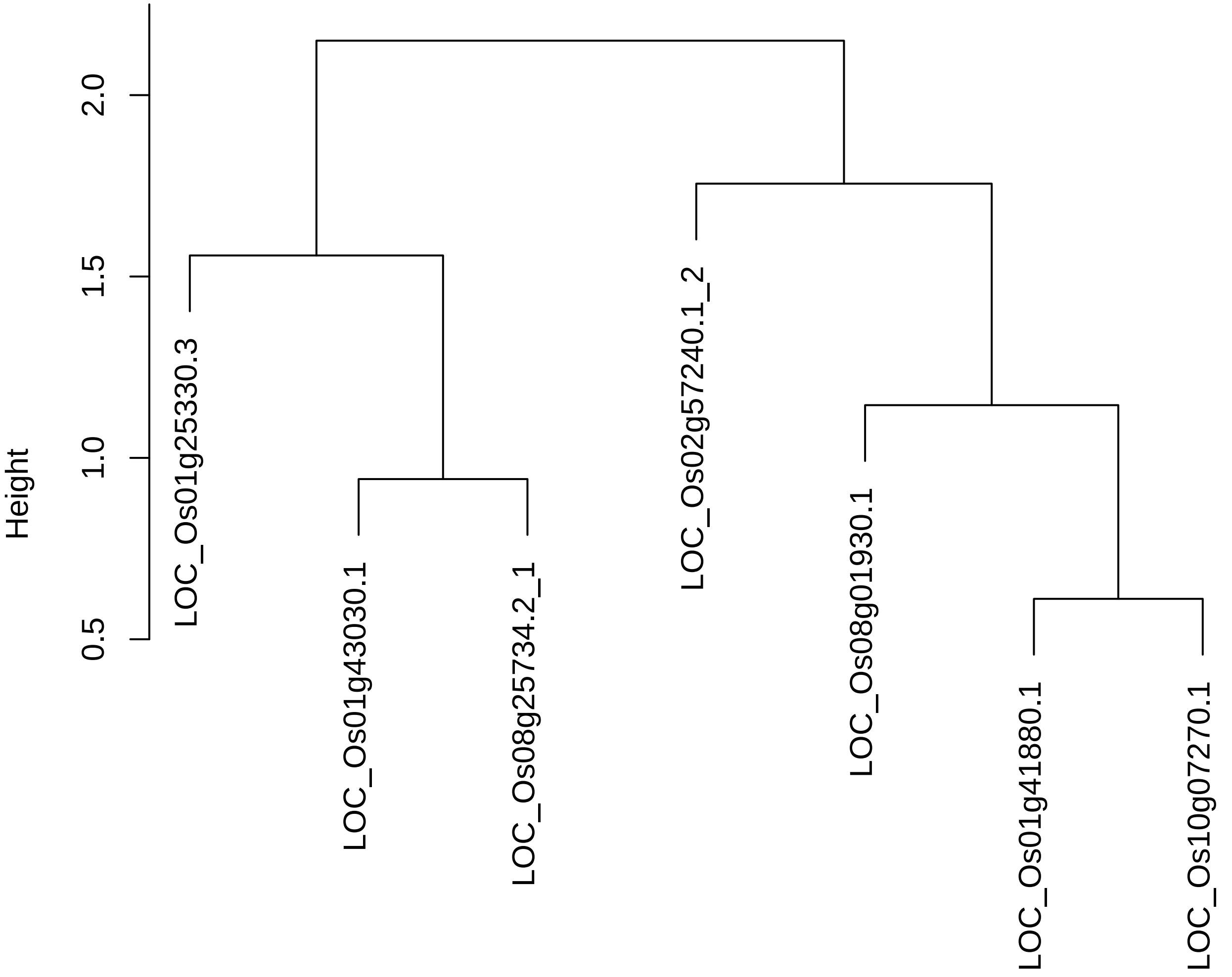


dist(spfitiex12sec12[lab[, 1] == i, ])
hclust (*, "ward.D2")

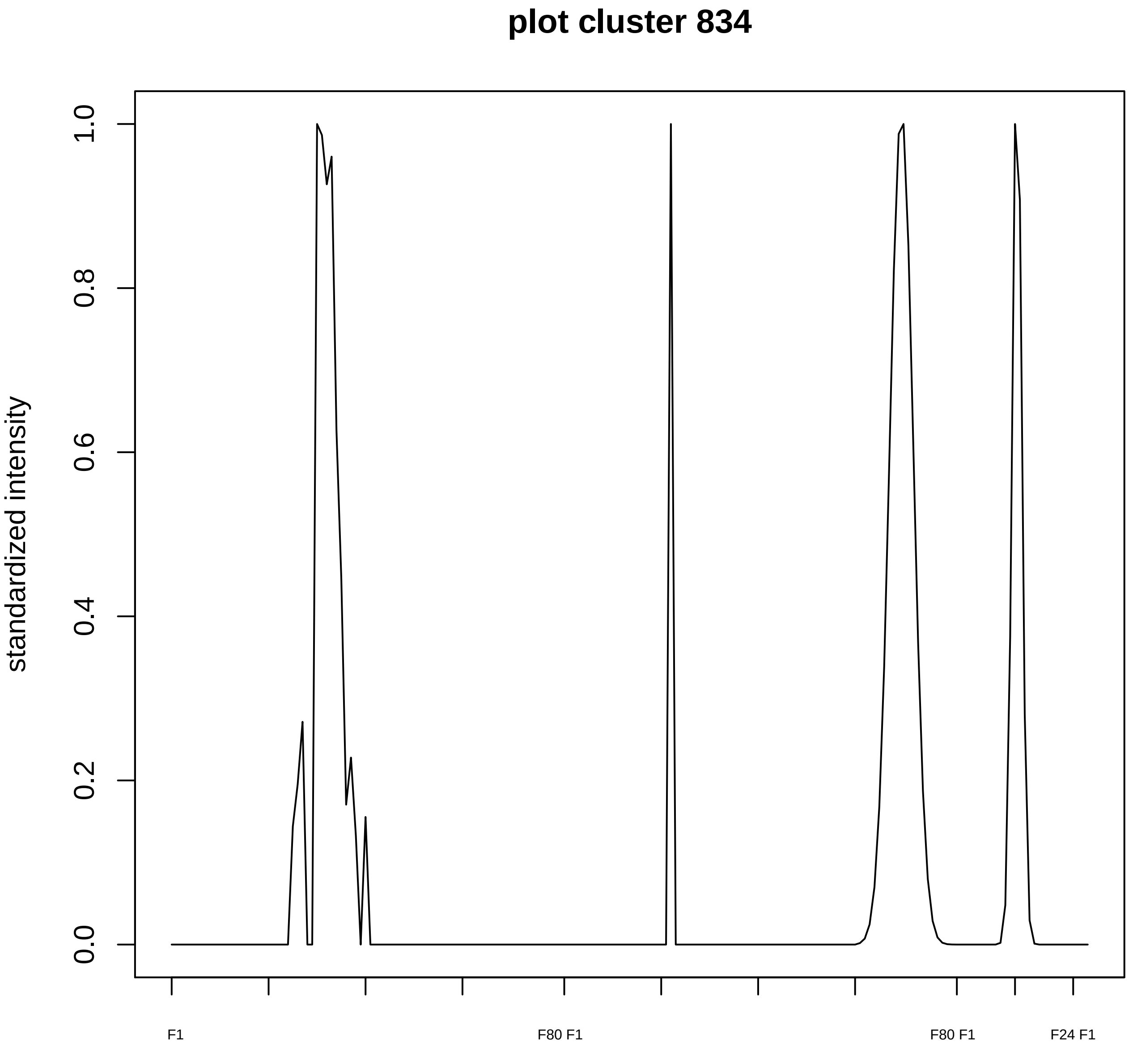
plot cluster 834
standardized intensity
0.0
0.2
0.4
0.6
0.8
1.0
F1
F80 F1
F80 F1
F24 F1

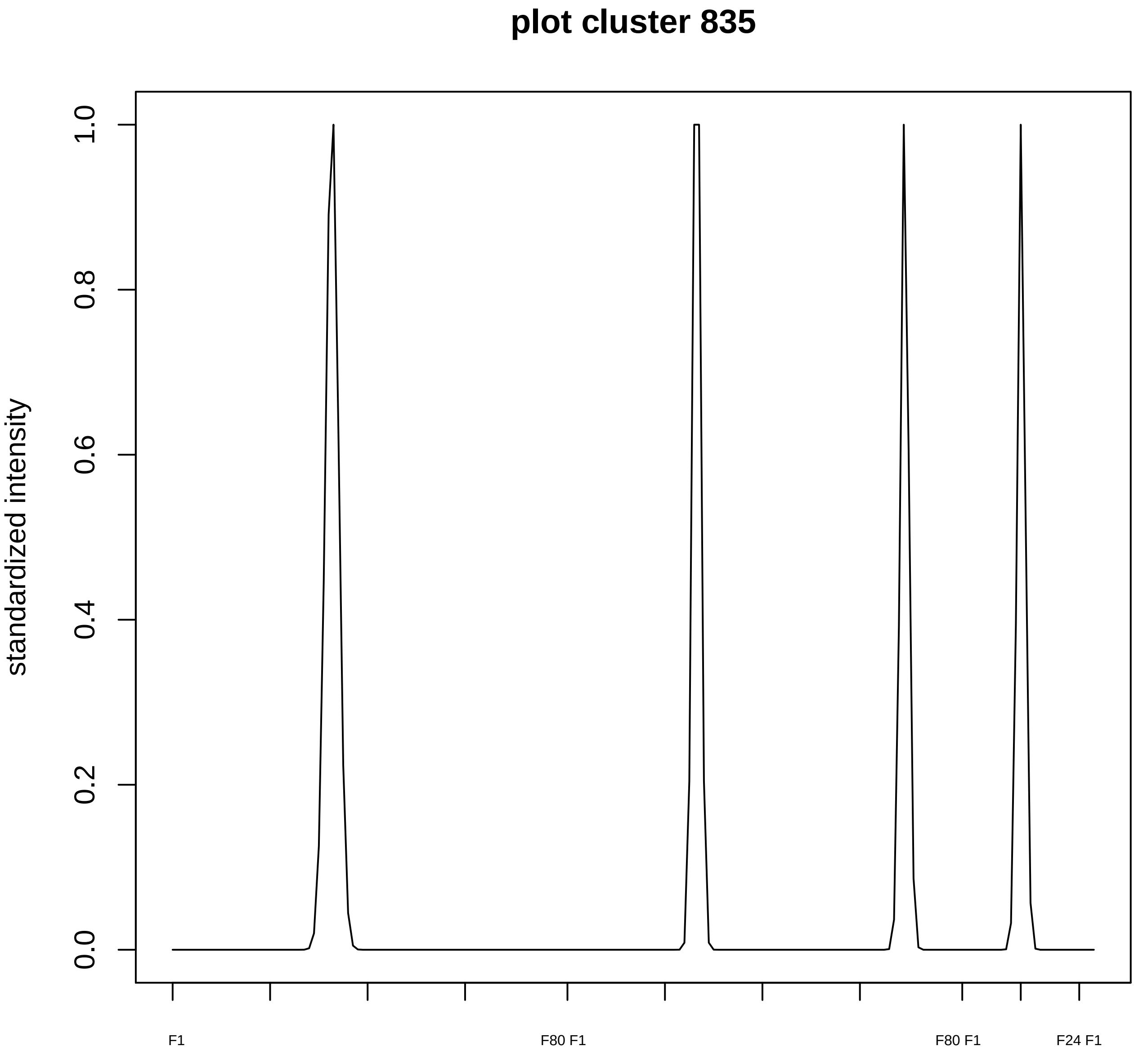

plot cluster 835
standardized intensity
0.0
0.2
0.4
0.6
0.8
1.0
F1
F80 F1
F80 F1
F24 F1

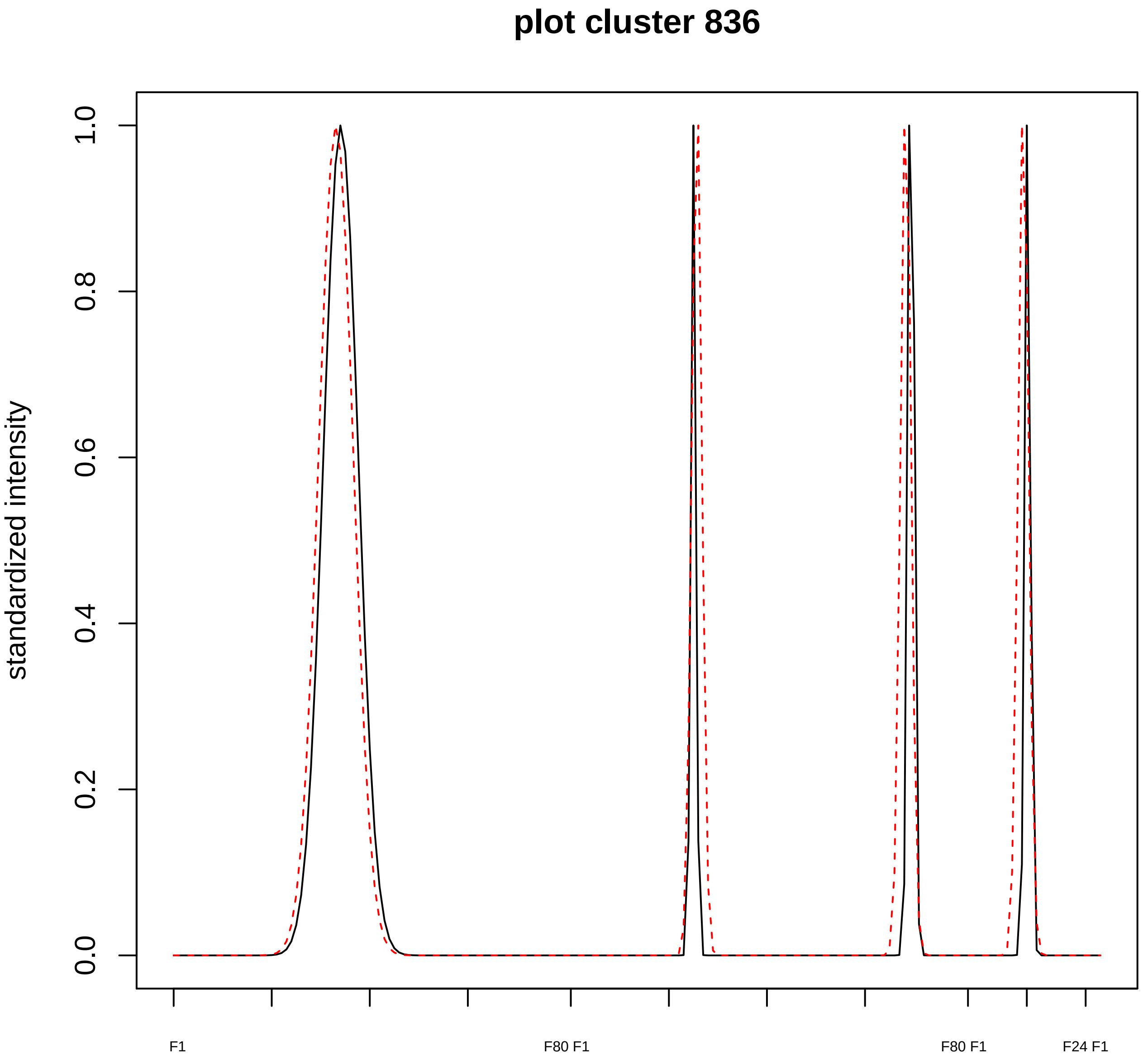
plot cluster 836
standardized intensity
0.0
0.2
0.4
0.6
0.8
1.0
F1
F80 F1
F80 F1
F24 F1

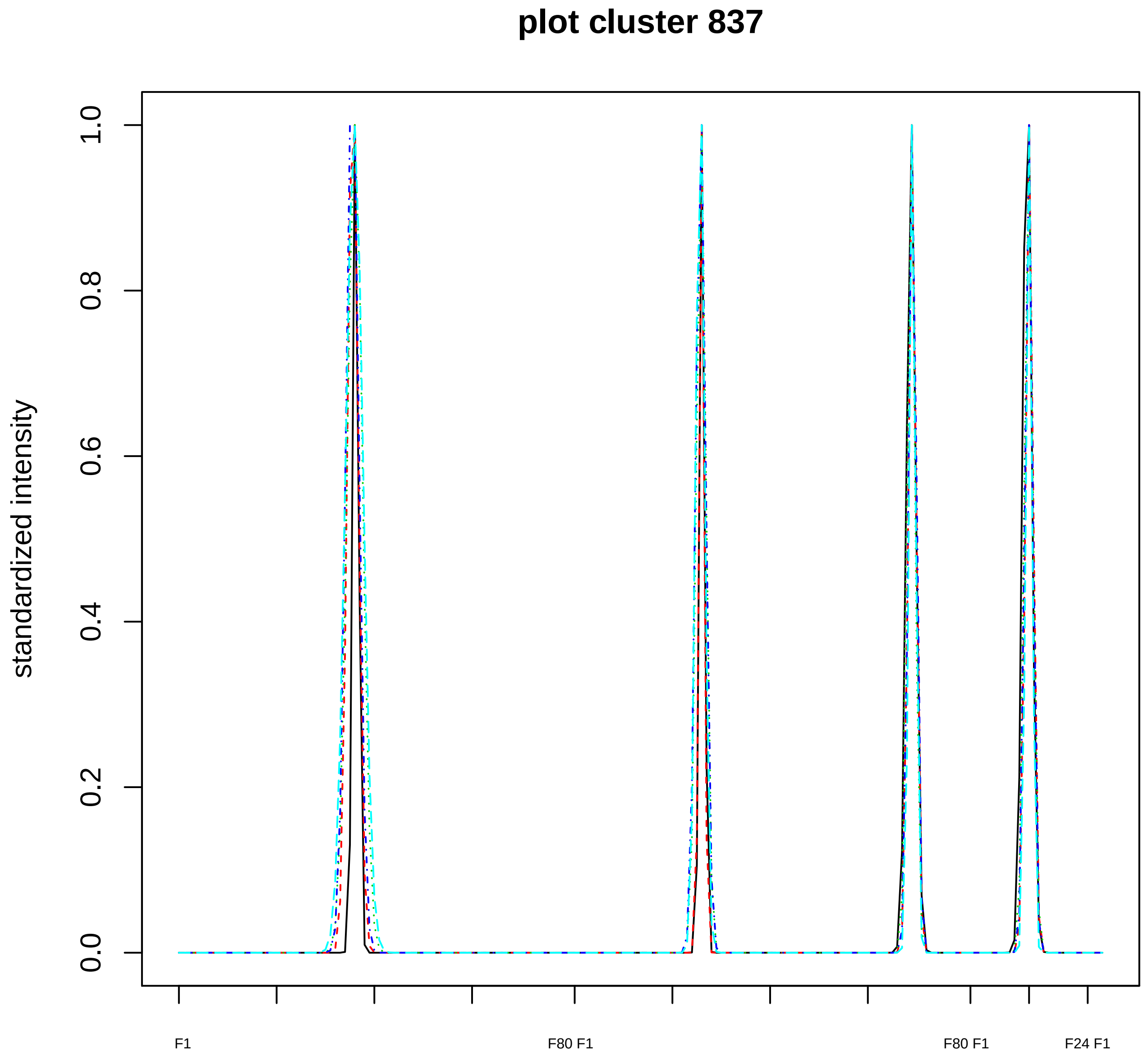

plot cluster 837
standardized intensity
0.0
0.2
0.4
0.6
0.8
1.0
F1
F80 F1
F80 F1
F24 F1

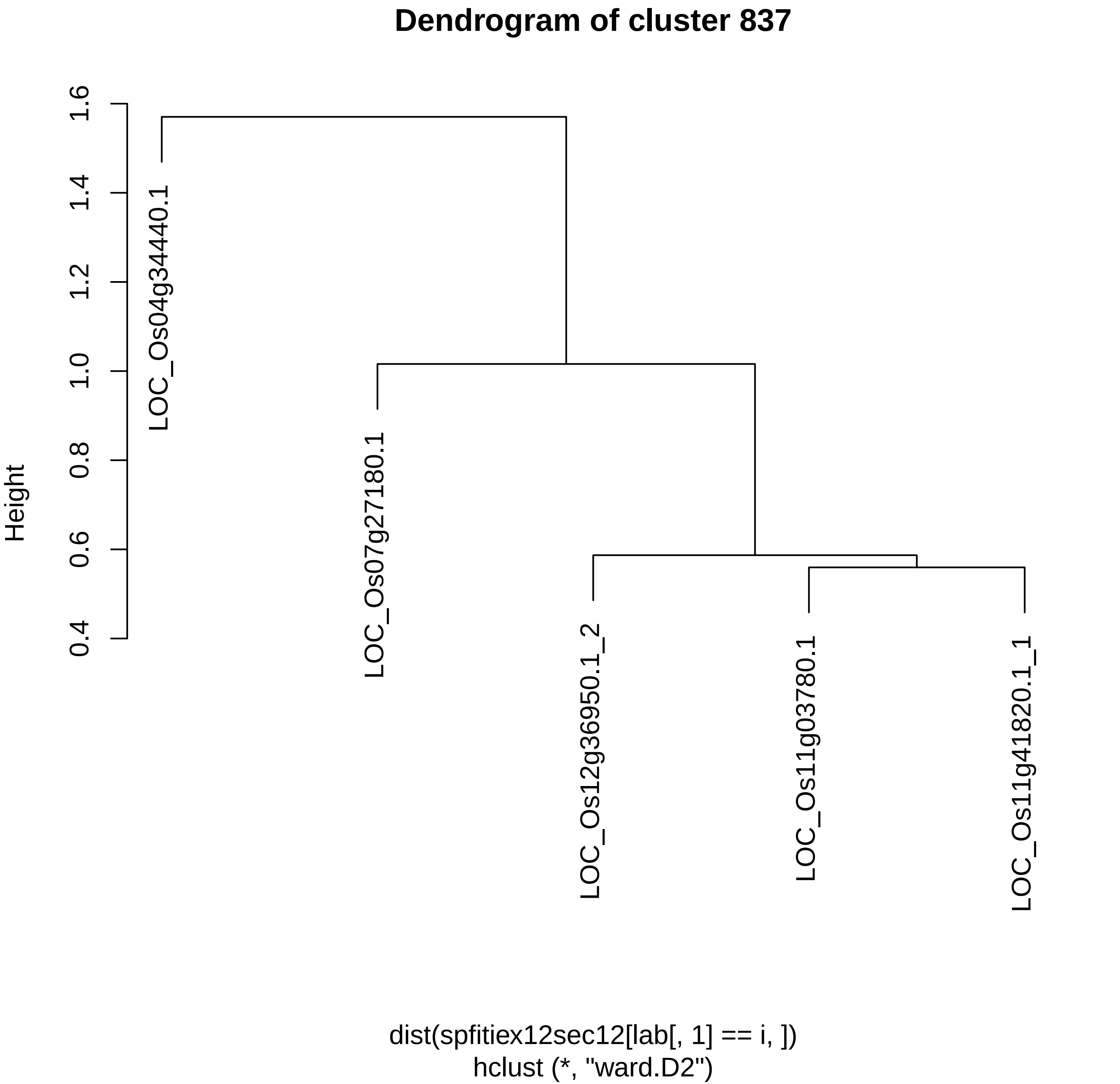
Dendrogram of cluster 837
Height
1.6
1.4
1.2
1.0
0.8
0.6
0.4
LOC_Os04g34440.1
LOC_Os07g27180.1
LOC_Os12g36950.1_2
LOC_Os11g03780.1
LOC_Os11g41820.1_1
dist(spfitiex12sec12[lab[, 1] == i, ])
hclust (*, "ward.D2")

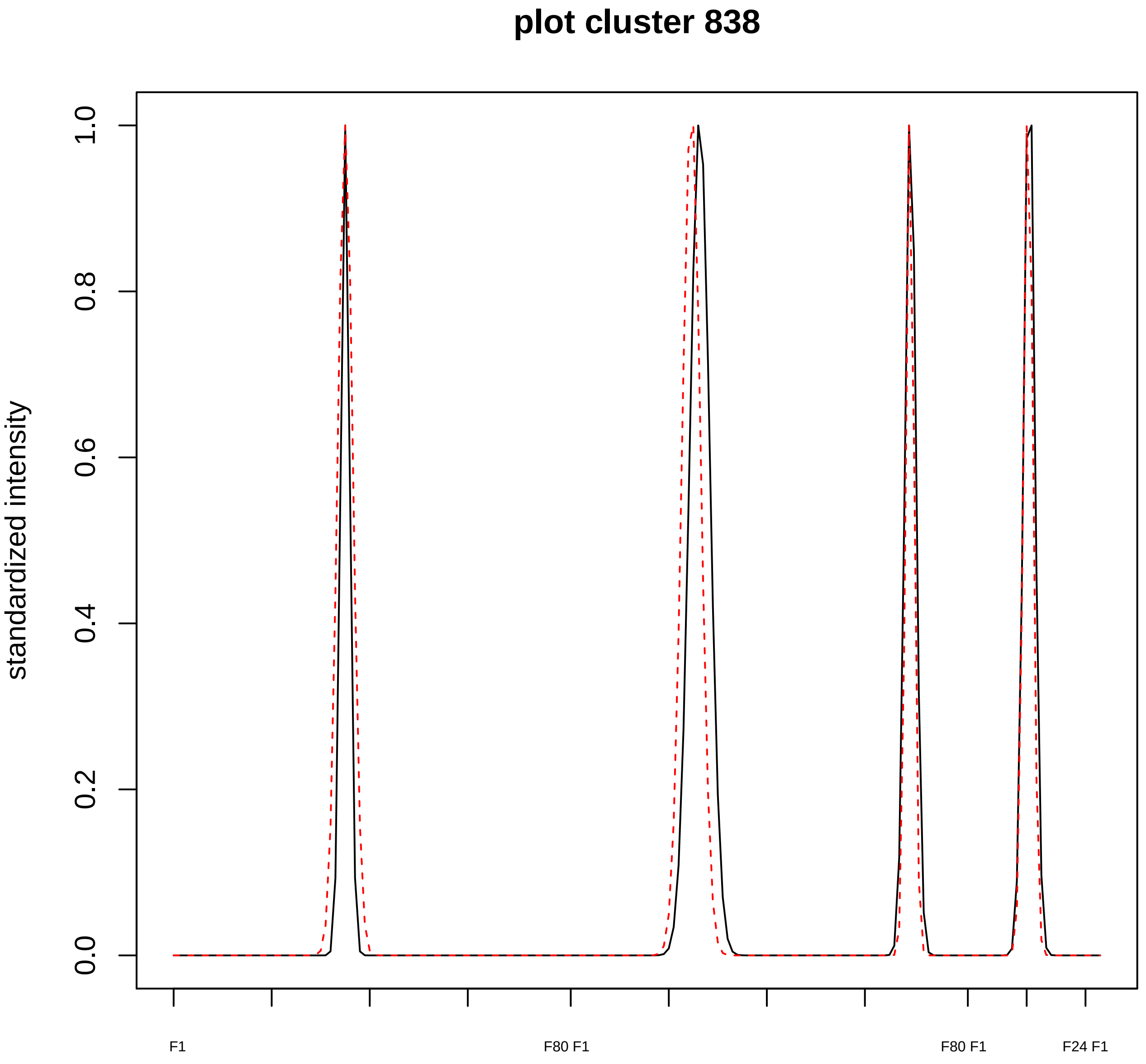

plot cluster 838
standardized intensity
0.0
0.2
0.4
0.6
0.8
1.0
F1
F80 F1
F80 F1
F24 F1

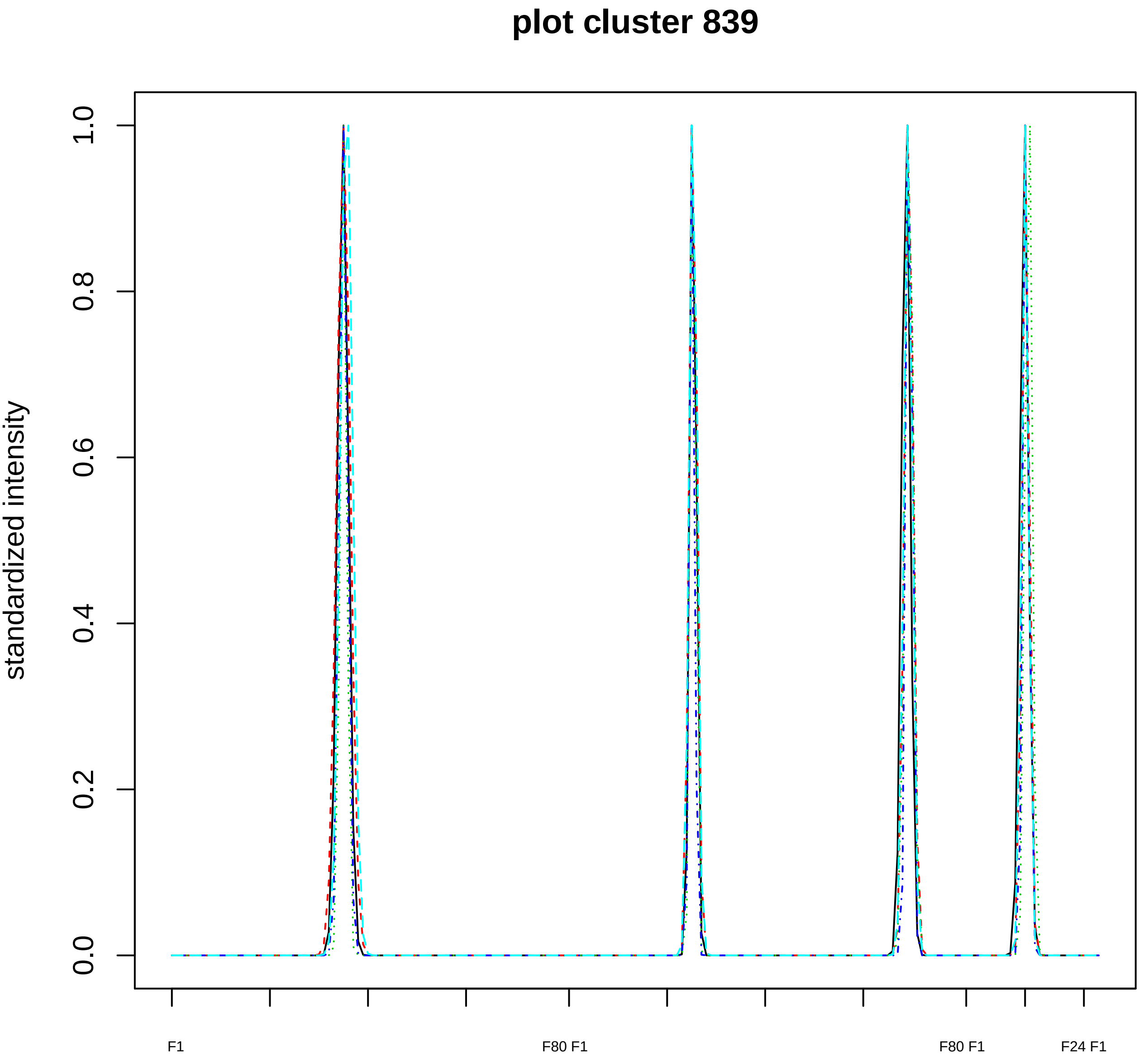
plot cluster 839
standardized intensity
1.0
0.8
0.6
0.4
0.2
0.0
F1
F80 F1
F80 F1
F24 F1

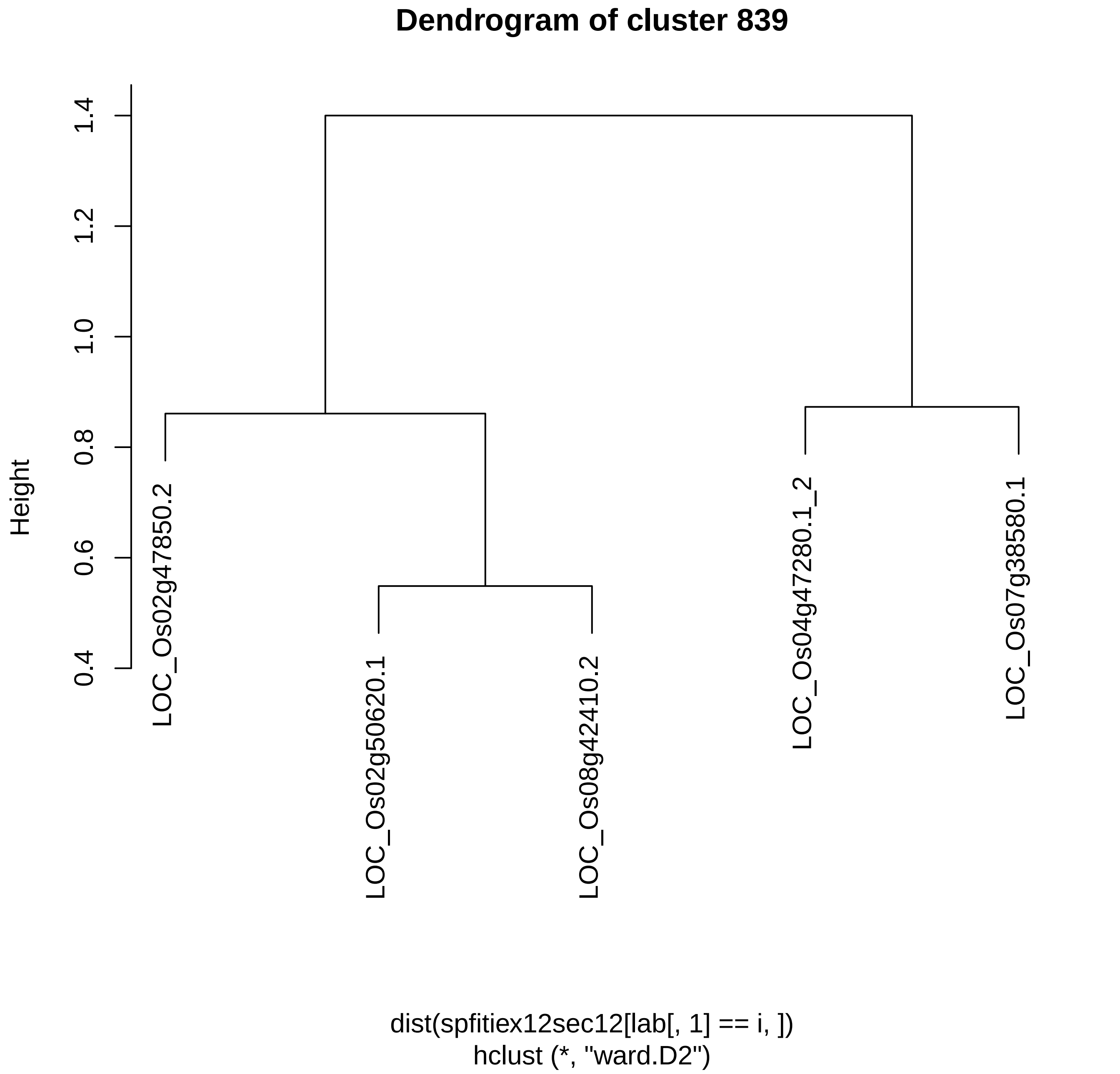
Dendrogram of cluster 839
Height
0.4
0.6
0.8
1.0
1.2
1.4
LOC_Os02g47850.2
LOC_Os02g50620.1
LOC_Os08g42410.2
LOC_Os04g47280.1_2
LOC_Os07g38580.1
dist(spfitiex12sec12[lab[, 1] == i, ])
hclust (*, "ward.D2")

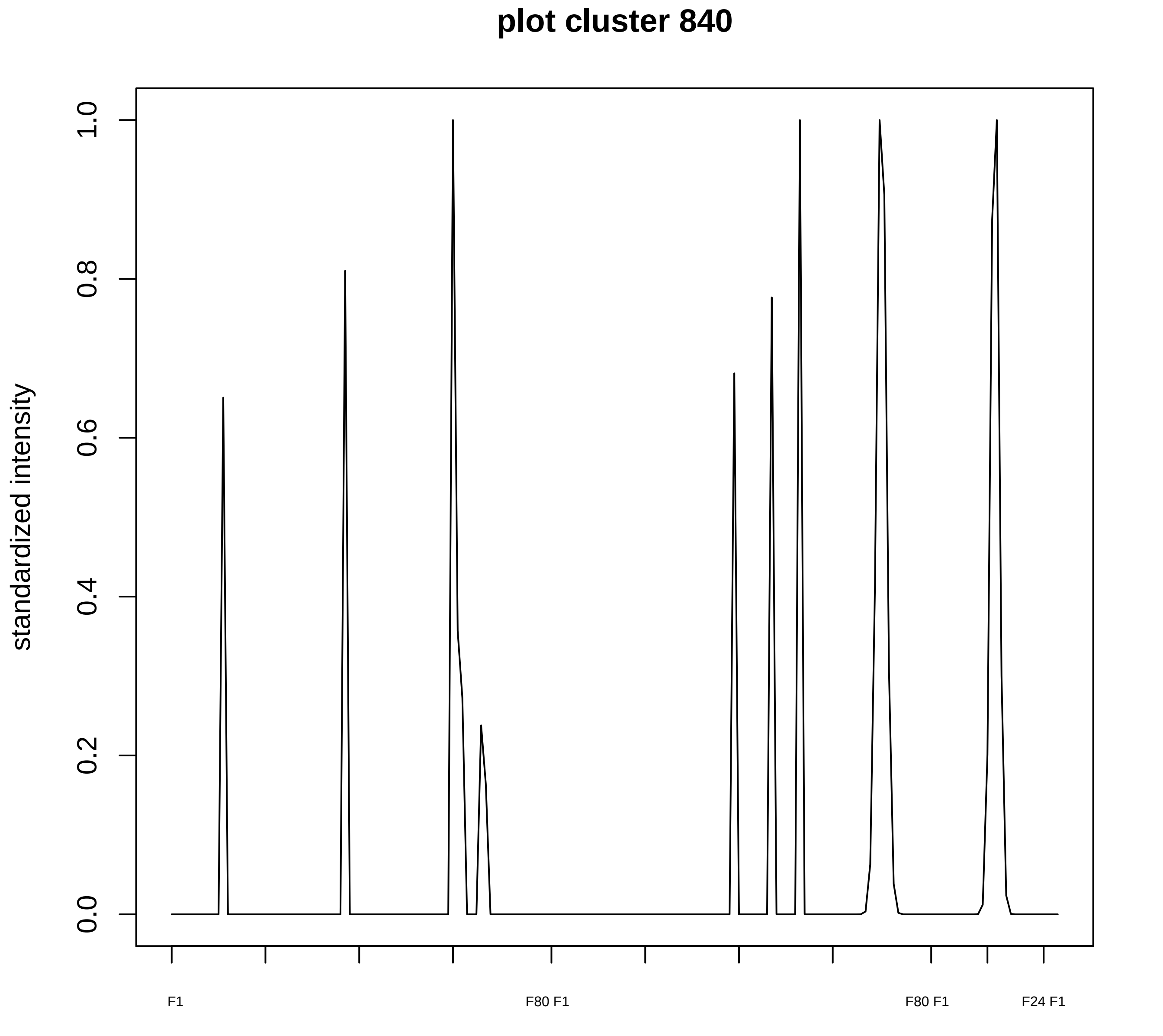

plot cluster 840
standardized intensity
0.0
0.2
0.4
0.6
0.8
1.0
F1
F80 F1
F80 F1
F24 F1

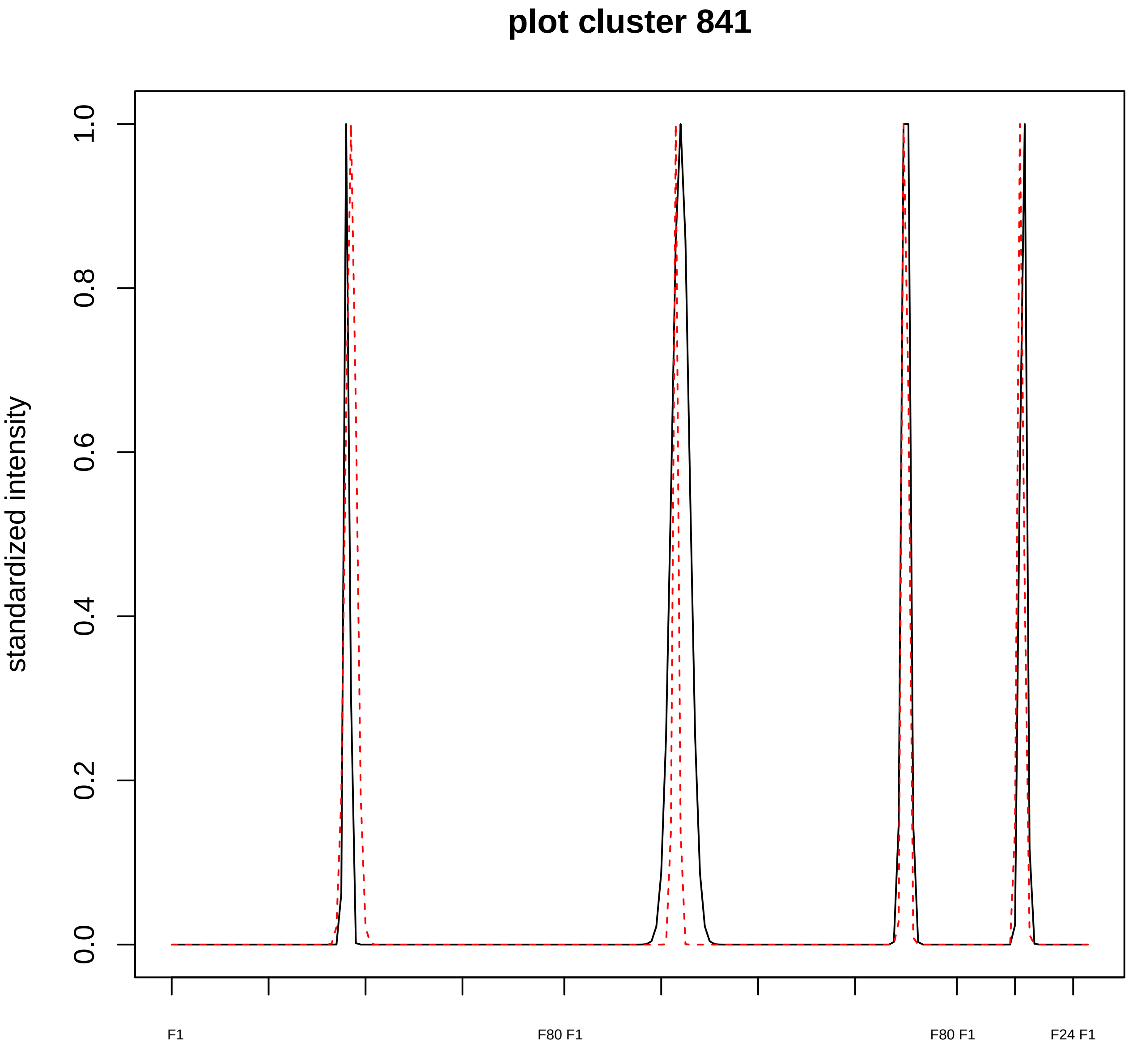

plot cluster 841
standardized intensity
0.0
0.2
0.4
0.6
0.8
1.0
F1
F80 F1
F80 F1
F24 F1

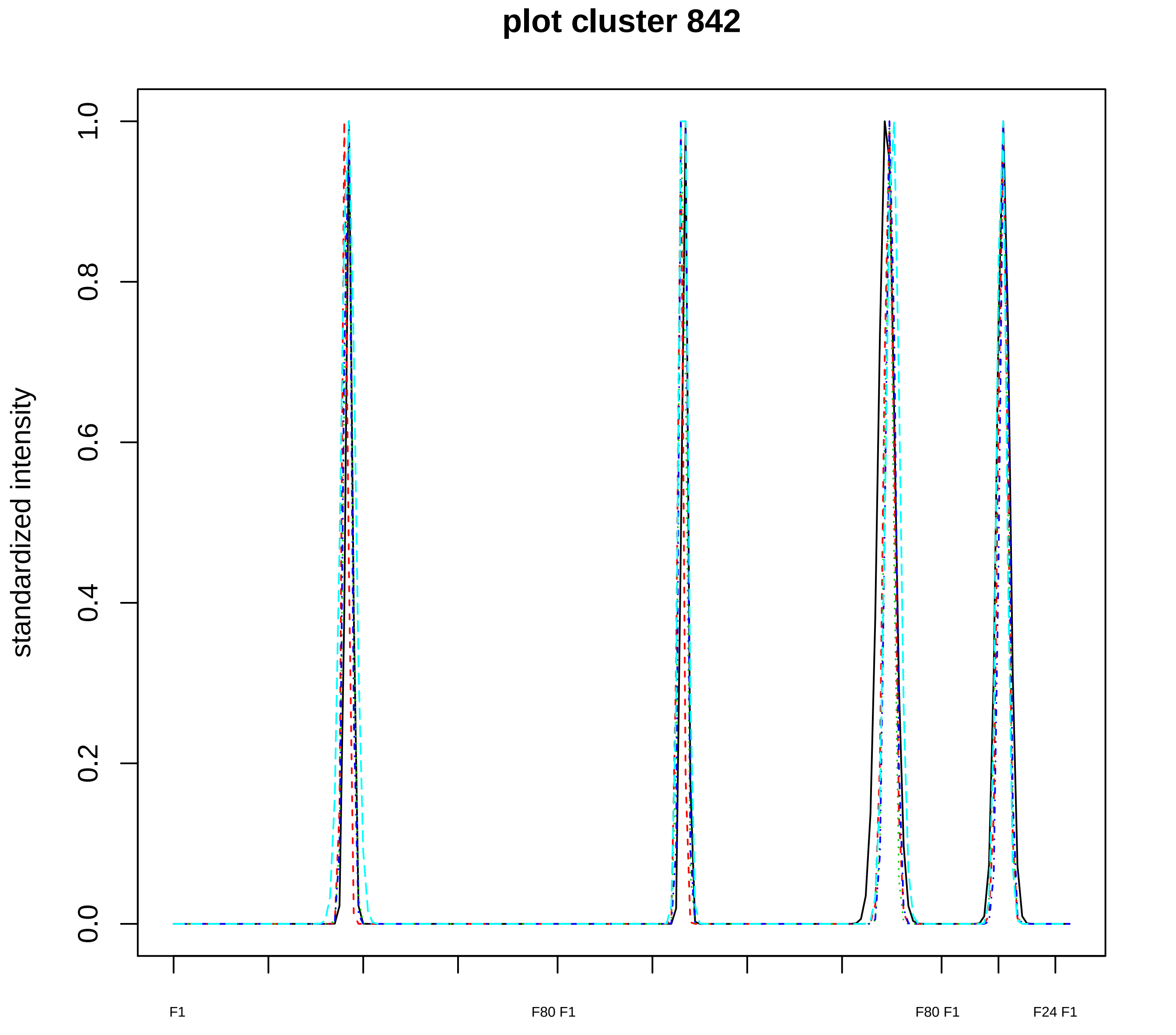
plot cluster 842
standardized intensity
0.0
0.2
0.4
0.6
0.8
1.0
F1
F80 F1
F80 F1
F24 F1

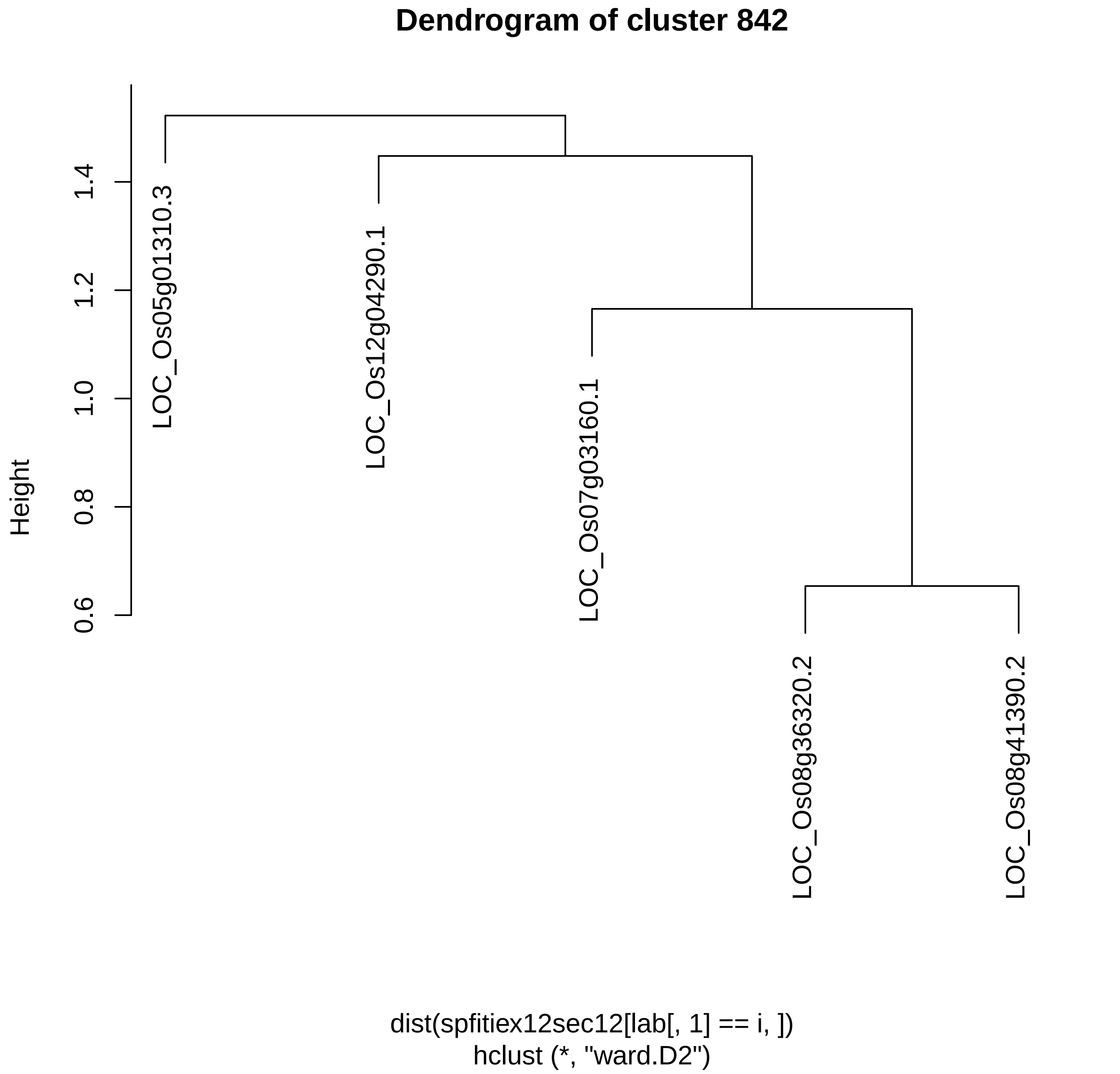
Dendrogram of cluster 842
Height
0.6
0.8
1.0
1.2
1.4
LOC_Os05g01310.3
LOC_Os12g04290.1
LOC_Os07g03160.1
LOC_Os08g36320.2
LOC_Os08g41390.2
dist(spfitiex12sec12[lab[, 1] == i, ])
hclust (*, "ward.D2")

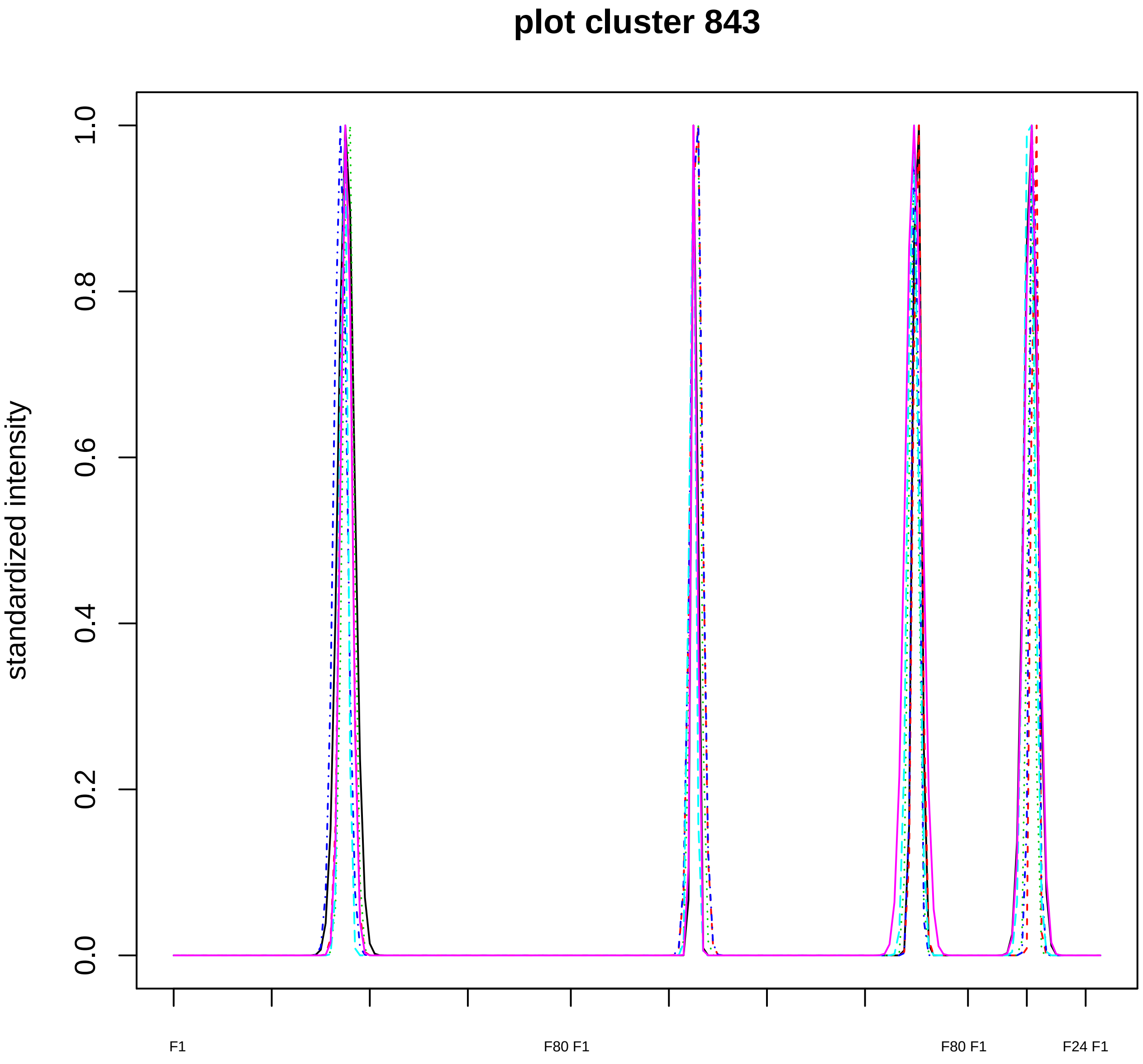

plot cluster 843
standardized intensity
0.0
0.2
0.4
0.6
0.8
1.0
F1
F80 F1
F80 F1
F24 F1

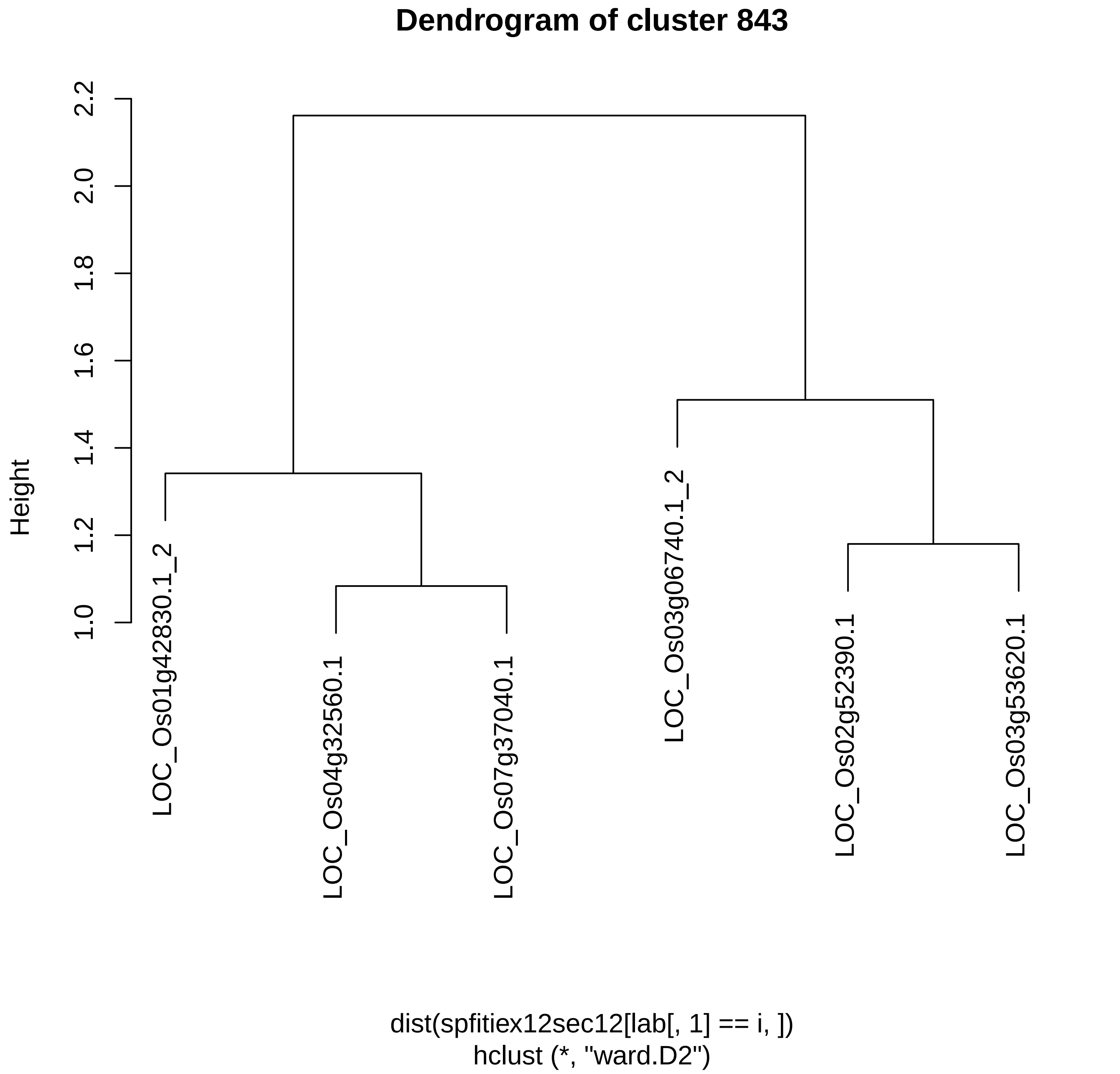
Dendrogram of cluster 843
Height
1.0
1.2
1.4
1.6
1.8
2.0
2.2
LOC_Os01g42830.1_2
LOC_Os04g32560.1
LOC_Os07g37040.1
LOC_Os03g06740.1_2
LOC_Os02g52390.1
LOC_Os03g53620.1
dist(spfitiex12sec12[lab[, 1] == i, ])
hclust (*, "ward.D2")

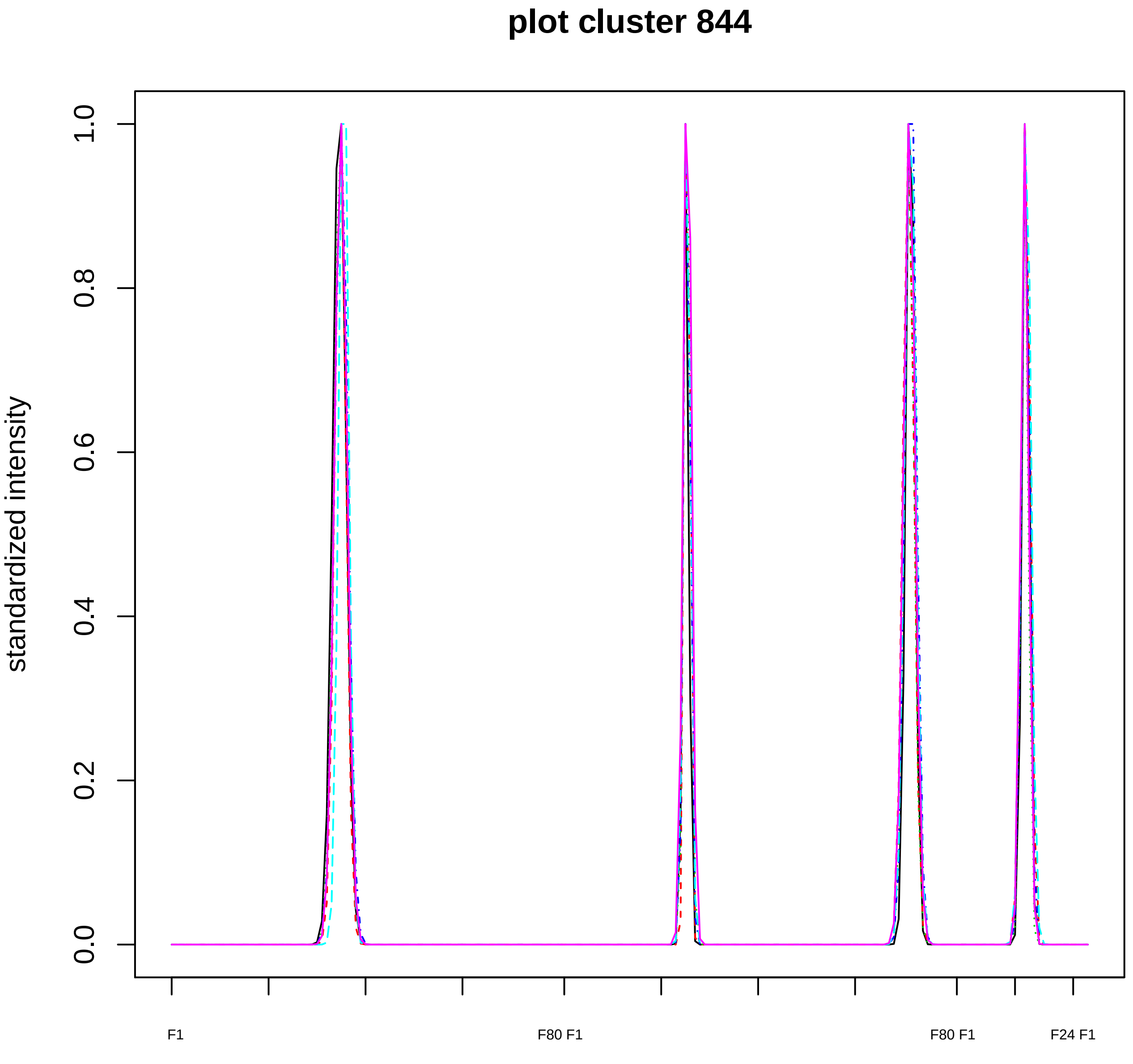

plot cluster 844
standardized intensity
1.0
0.8
0.6
0.4
0.2
0.0
F1
F80 F1
F80 F1
F24 F1

### Dendrogram of cluster 844

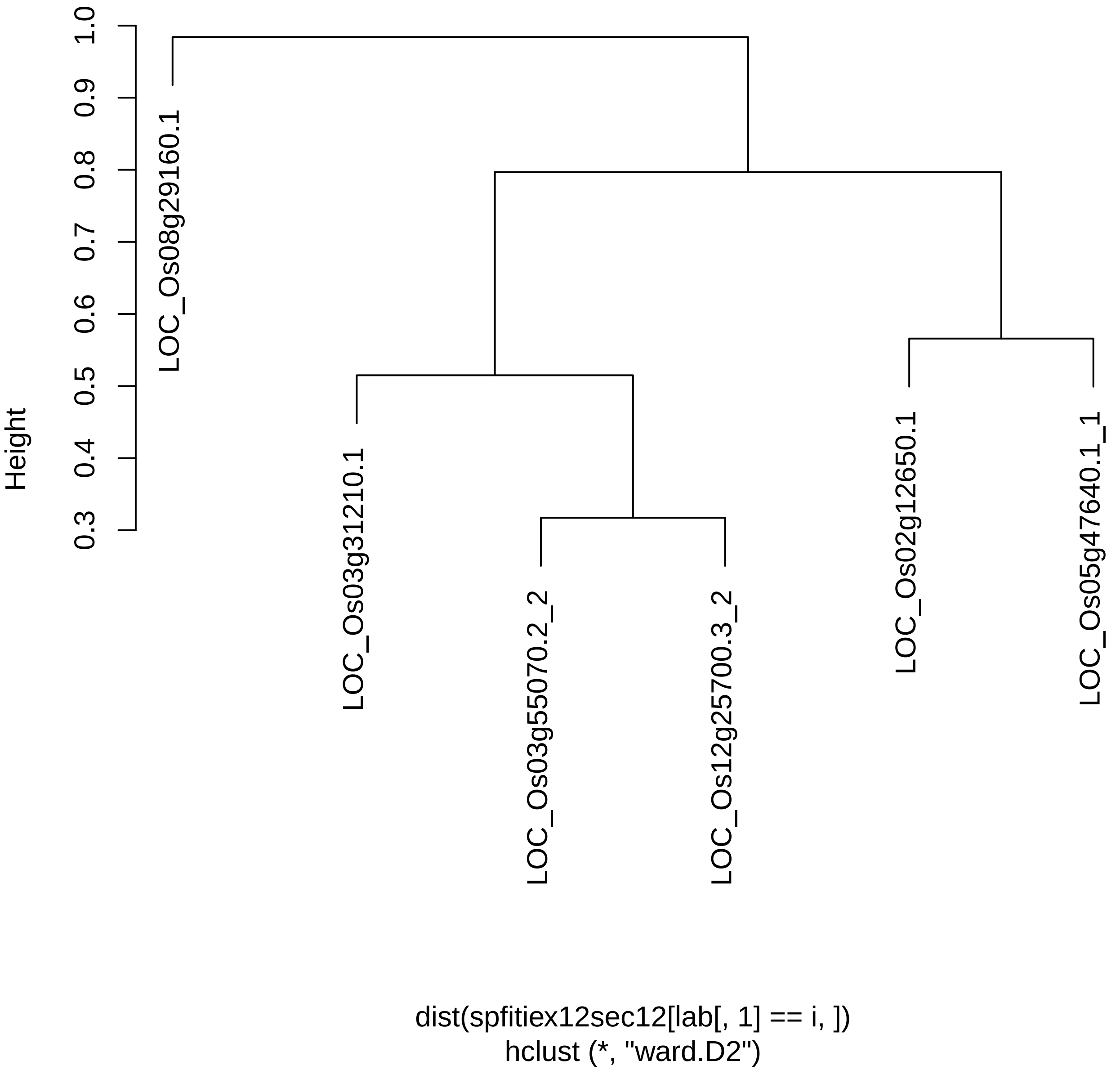

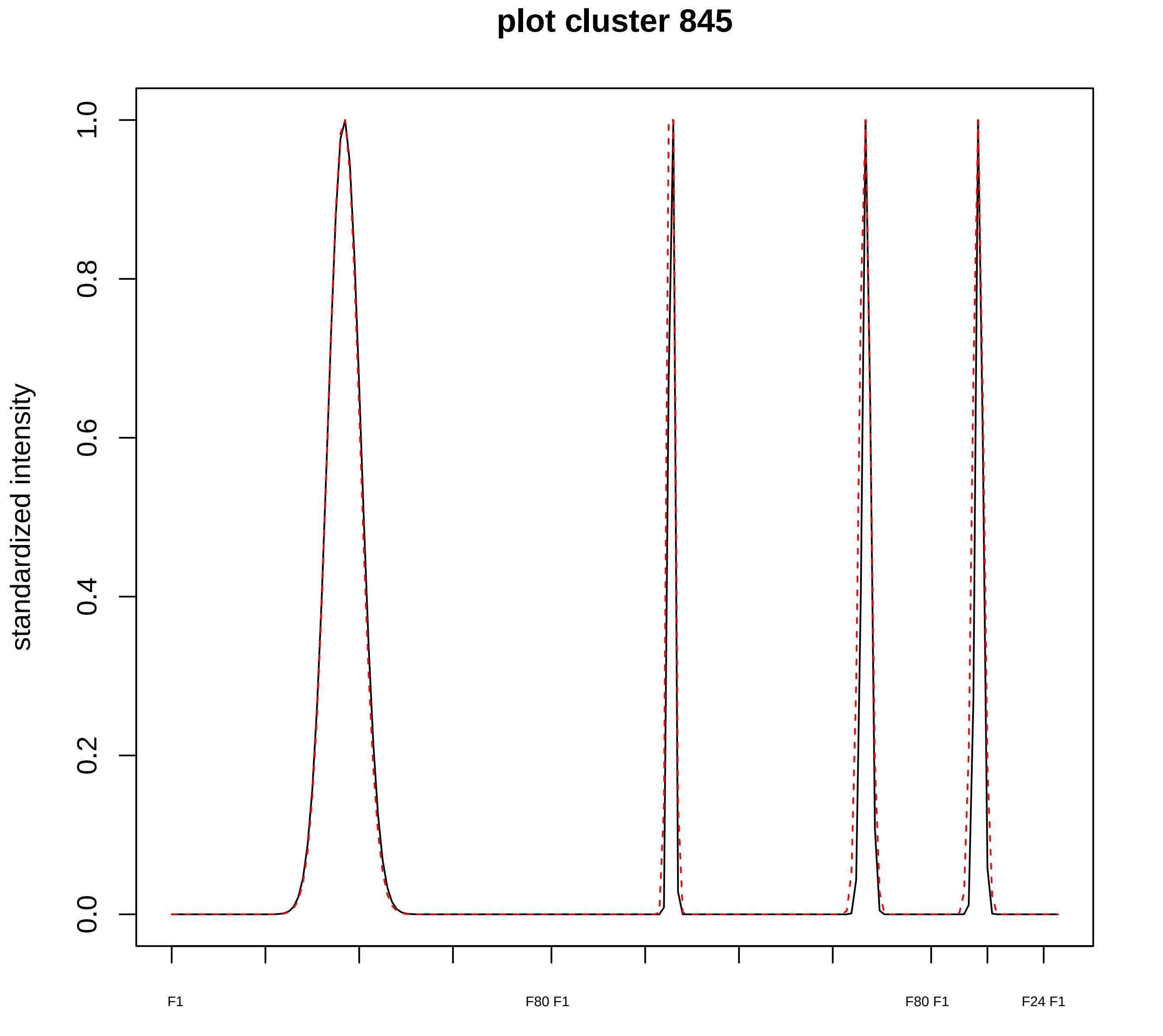

plot cluster 845
standardized intensity
0.0
0.2
0.4
0.6
0.8
1.0
F1
F80 F1
F80 F1
F24 F1

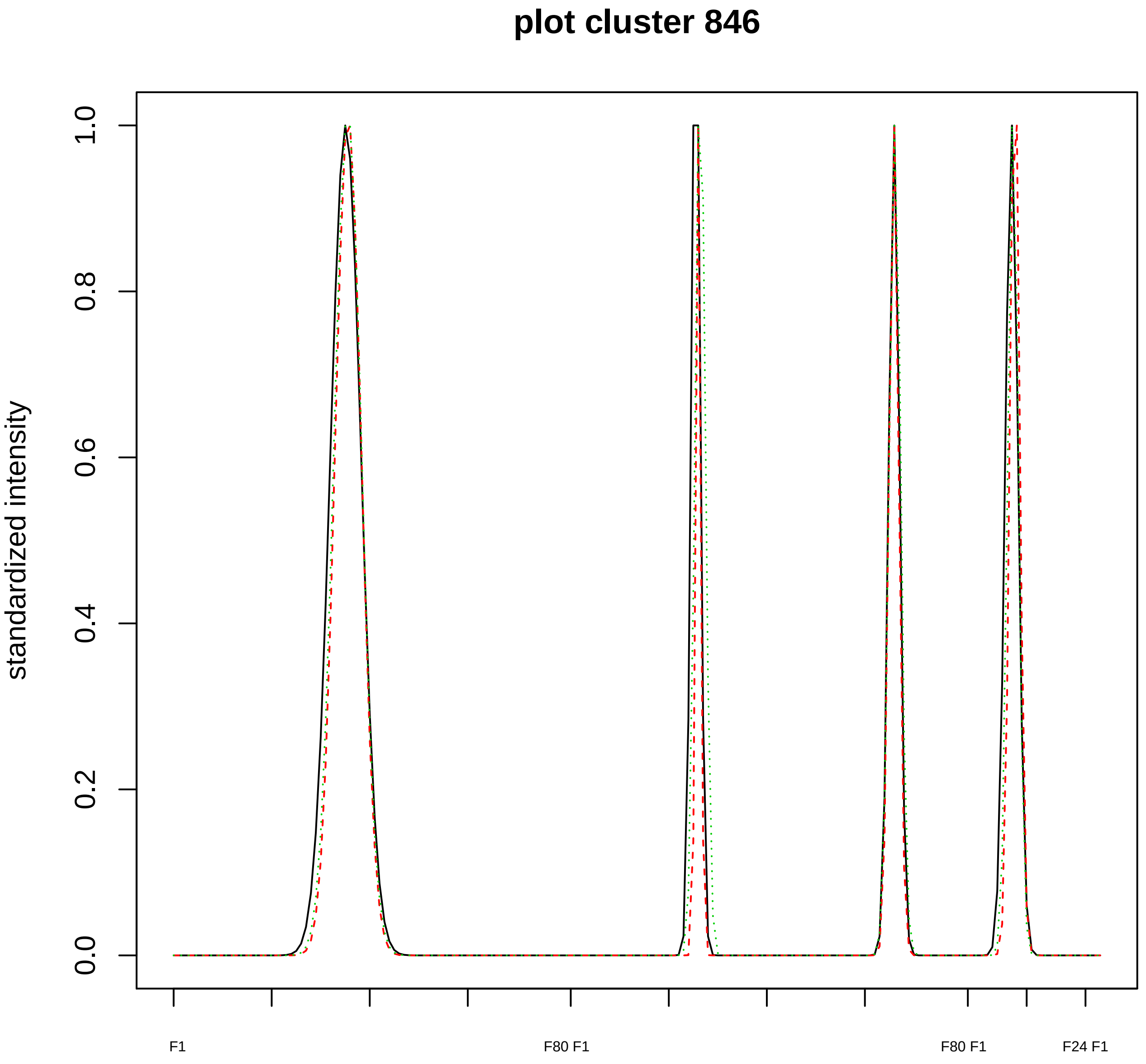
plot cluster 846
standardized intensity
1.0
0.8
0.6
0.4
0.2
0.0
F1
F80 F1
F80 F1
F24 F1

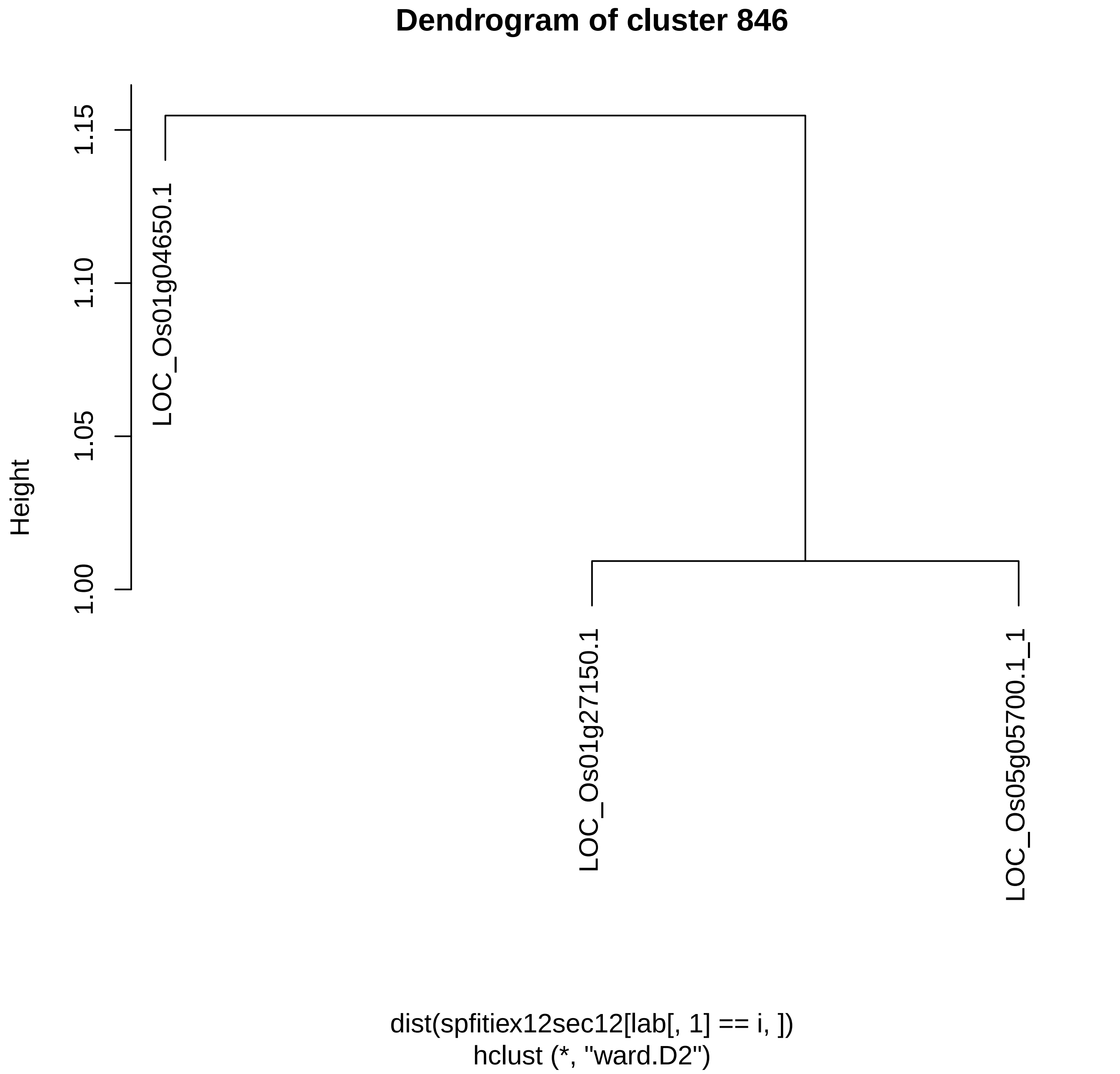
Dendrogram of cluster 846
Height
1.00
1.05
1.10
1.15
LOC_Os01g04650.1
LOC_Os01g27150.1
LOC_Os05g05700.1_1
dist(spfitiex12sec12[lab[, 1] == i, ])
hclust (*, "ward.D2")

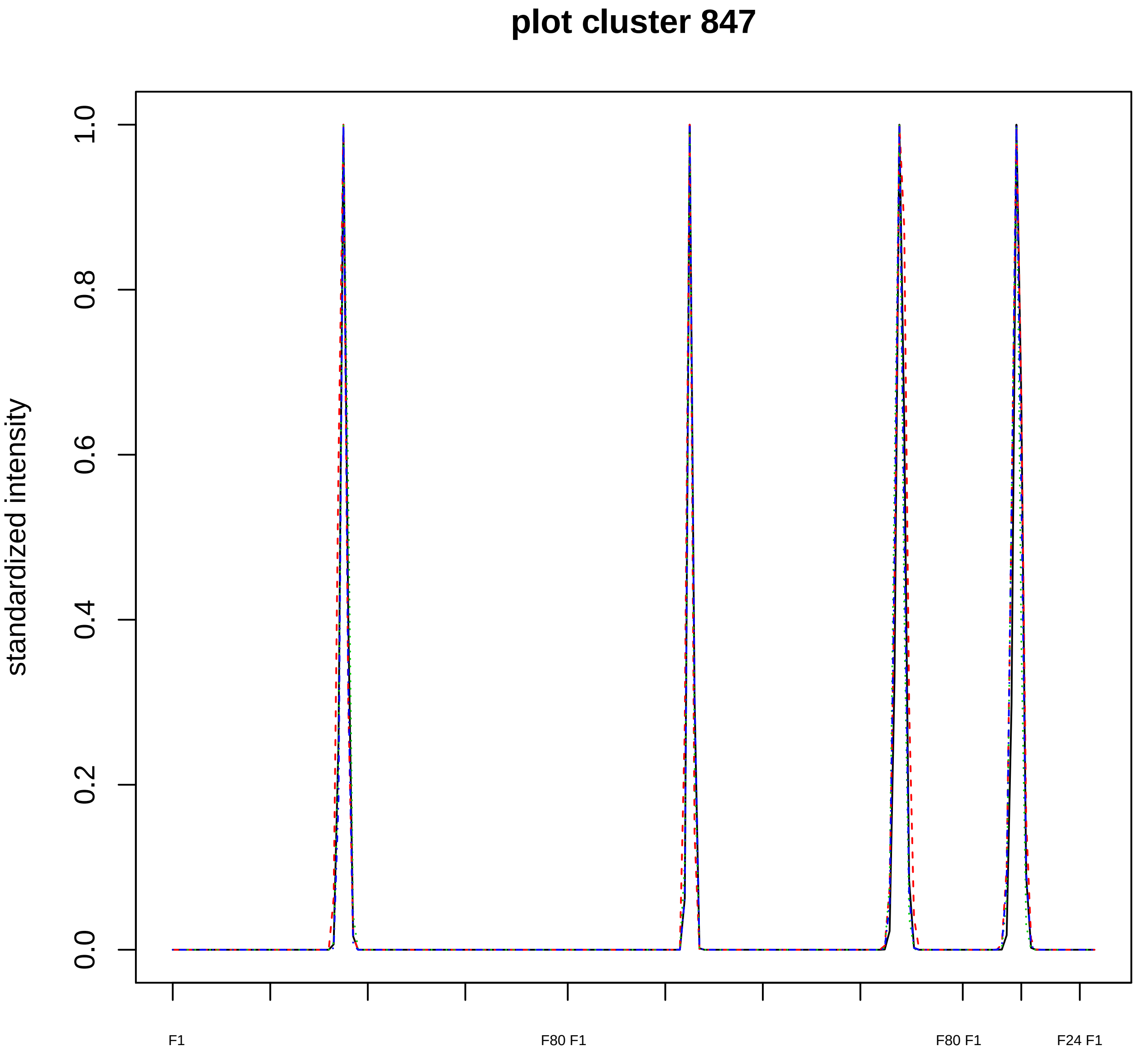

plot cluster 847
standardized intensity
1.0
0.8
0.6
0.4
0.2
0.0
F1
F80 F1
F80 F1
F24 F1

#### Dendrogram of cluster 847

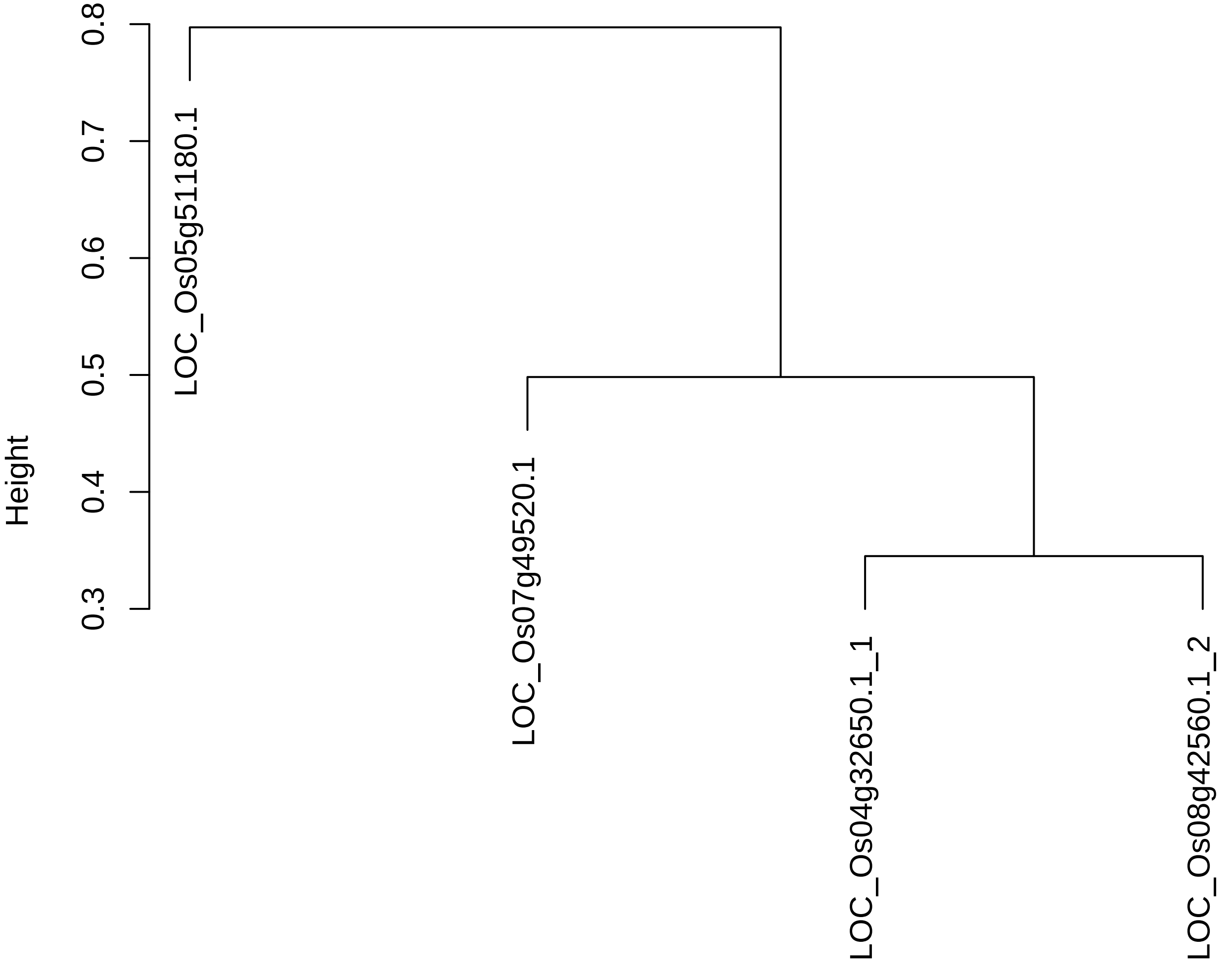


dist(spfitiex12sec12[lab[, 1] == i, ])
hclust (*, "ward.D2")

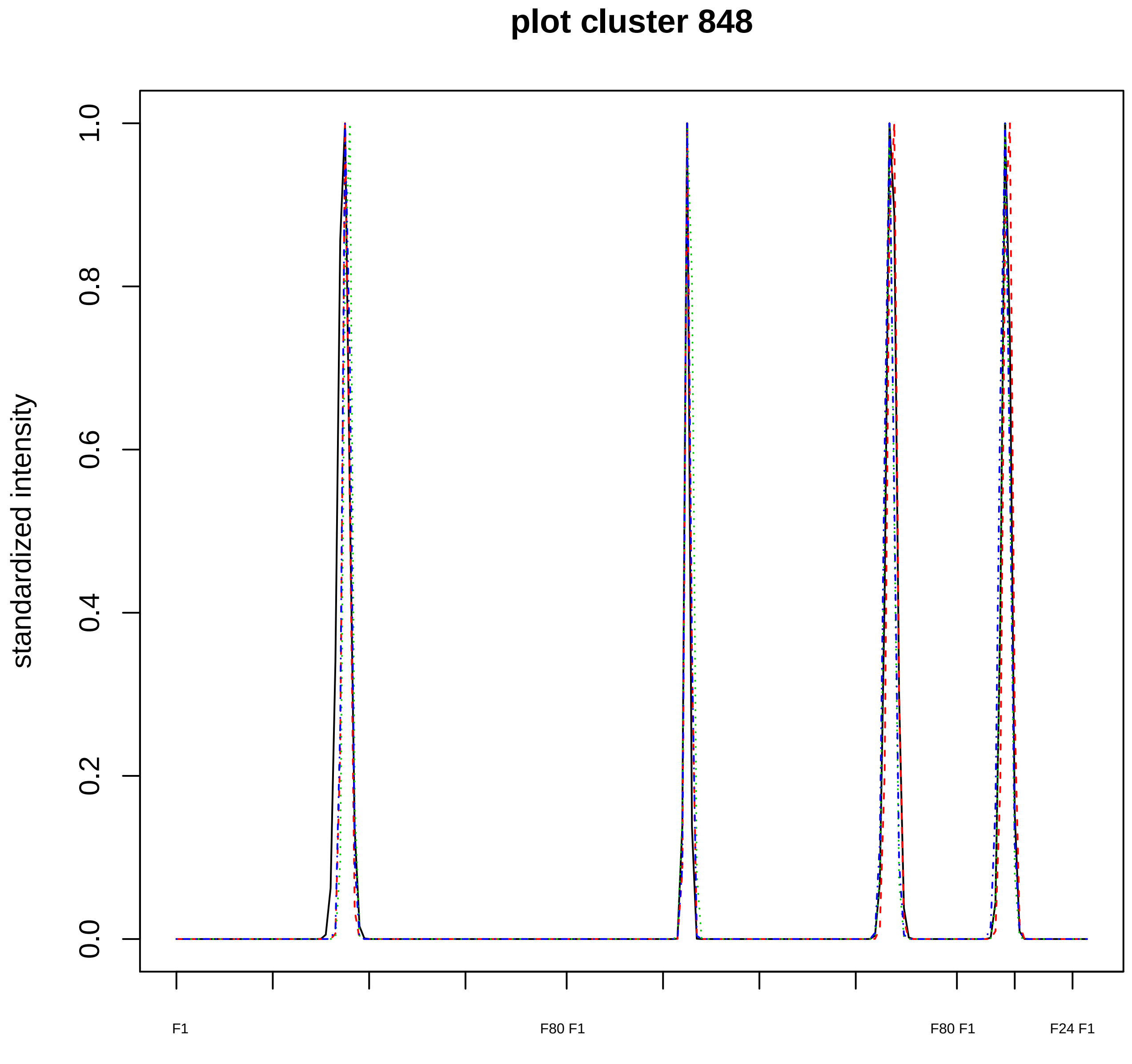

plot cluster 848
standardized intensity
0.0
0.2
0.4
0.6
0.8
1.0
F1
F80 F1
F80 F1
F24 F1

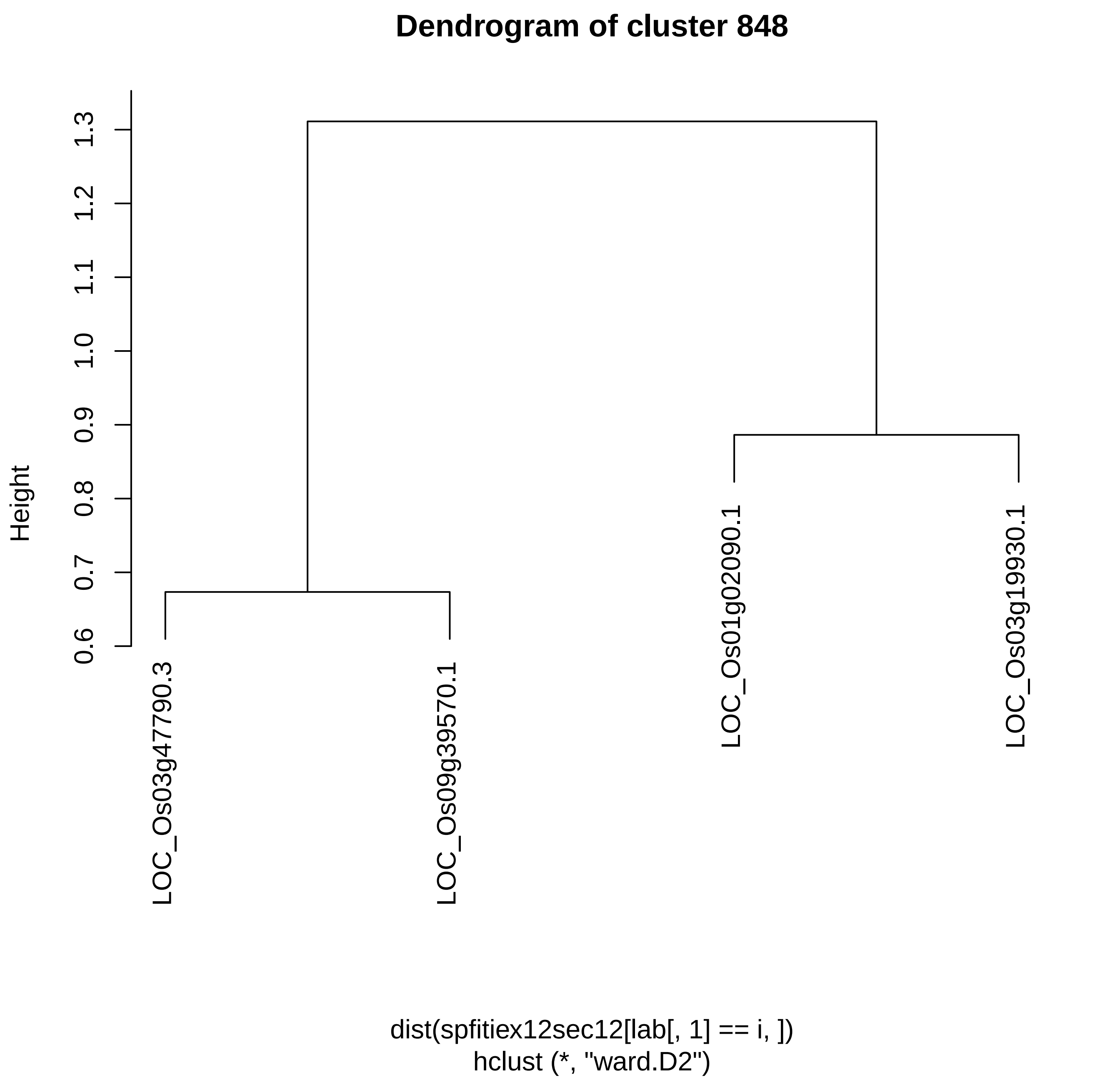

Dendrogram of cluster 848
Height
0.6
0.7
0.8
0.9
1.0
1.1
1.2
1.3
LOC_Os03g47790.3
LOC_Os09g39570.1
LOC_Os01g02090.1
LOC_Os03g19930.1
dist(spfitiex12sec12[lab[, 1] == i, ])
hclust (*, "ward.D2")

### plot cluster 849

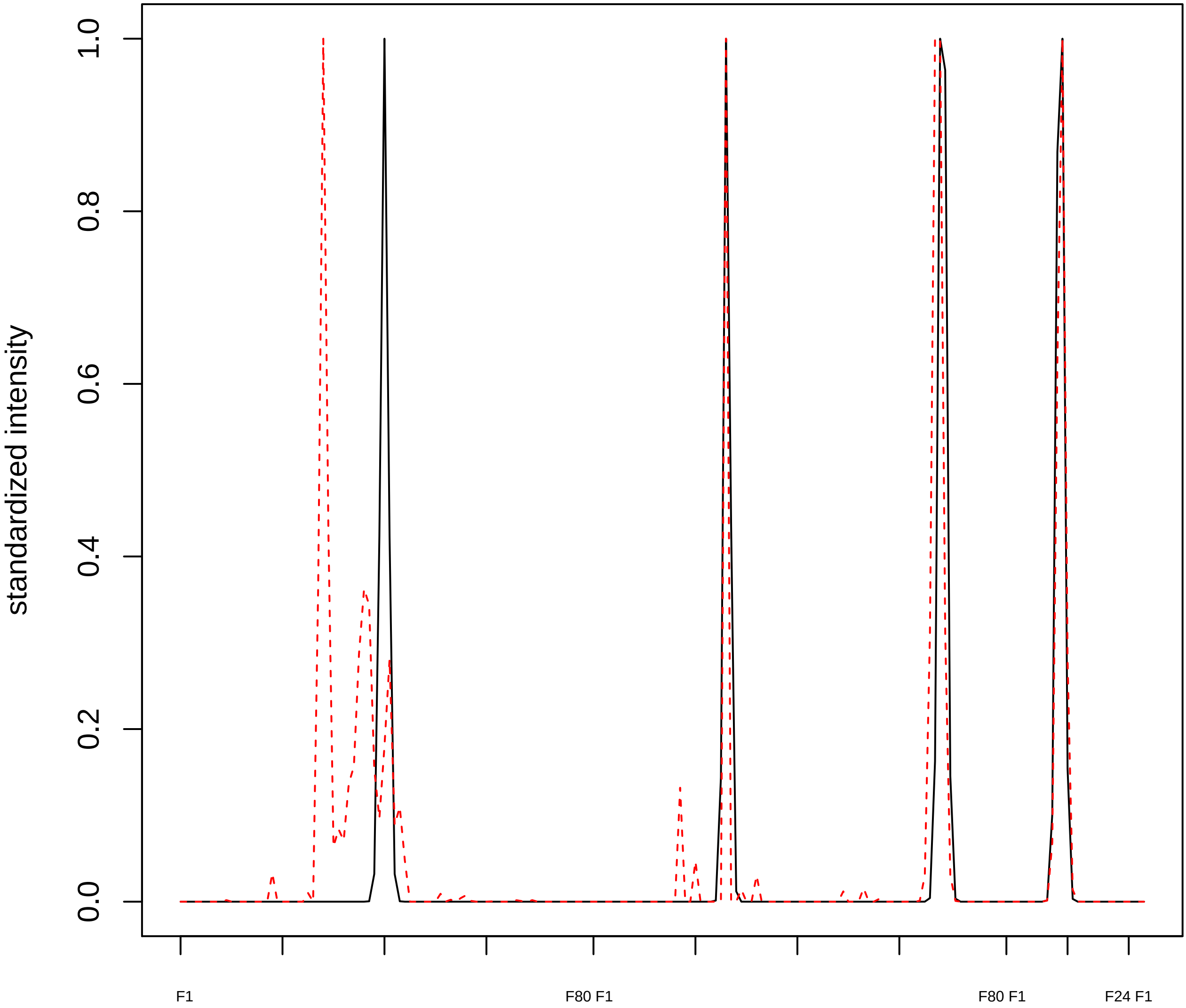

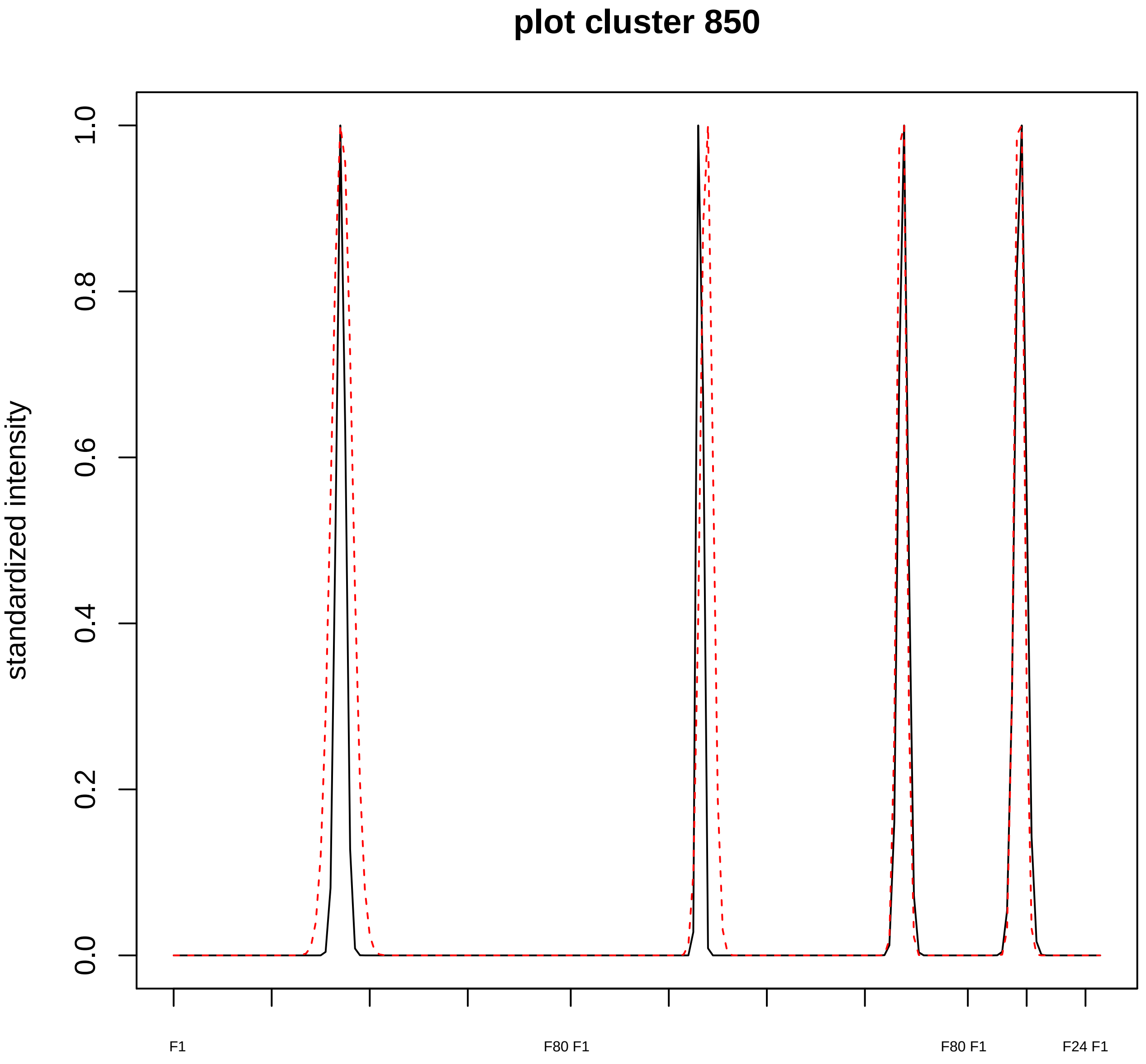

plot cluster 850
standardized intensity
0.0
0.2
0.4
0.6
0.8
1.0
F1
F80 F1
F80 F1
F24 F1

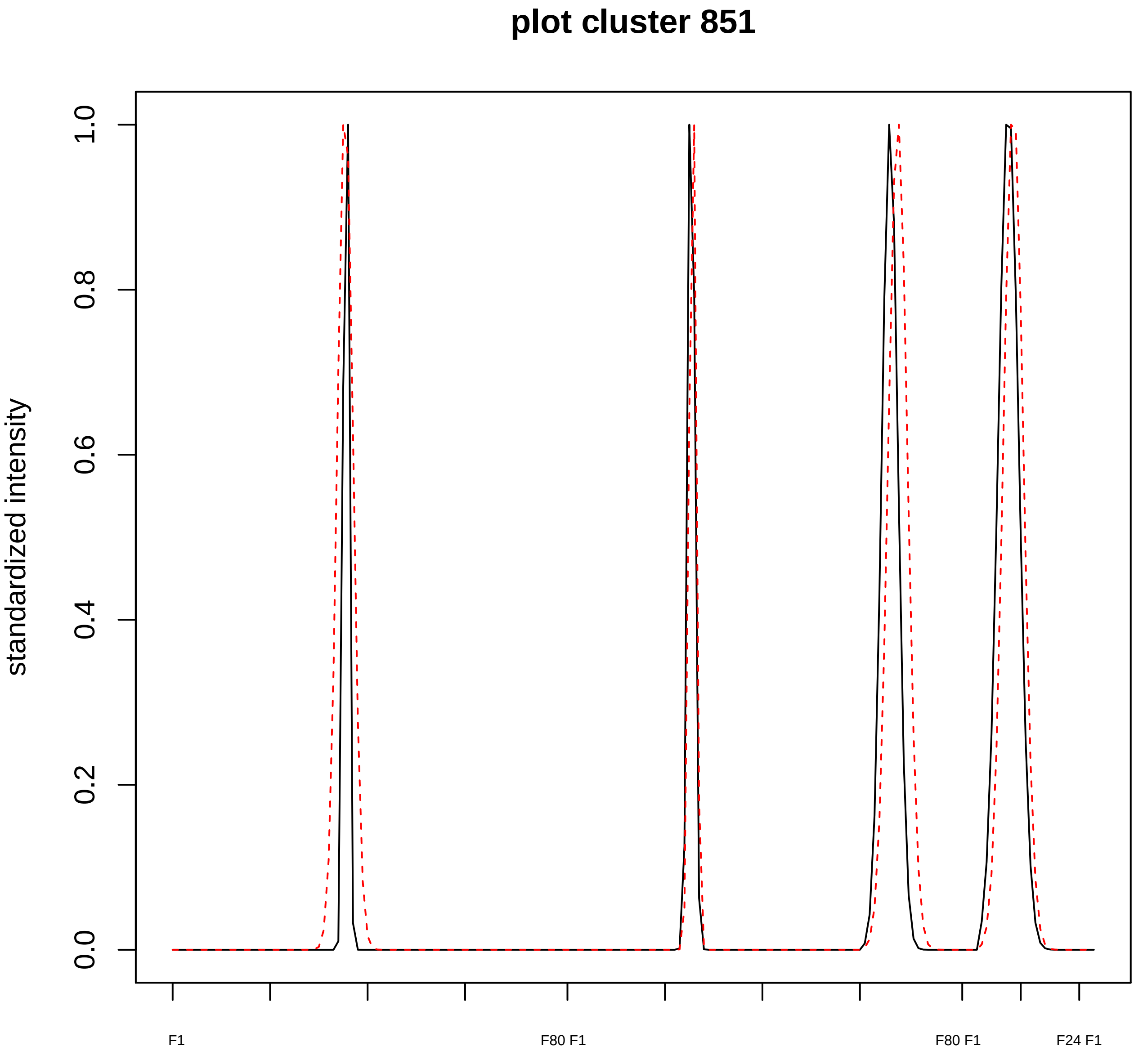

plot cluster 851
standardized intensity
0.0
0.2
0.4
0.6
0.8
1.0
F1
F80 F1
F80 F1
F24 F1

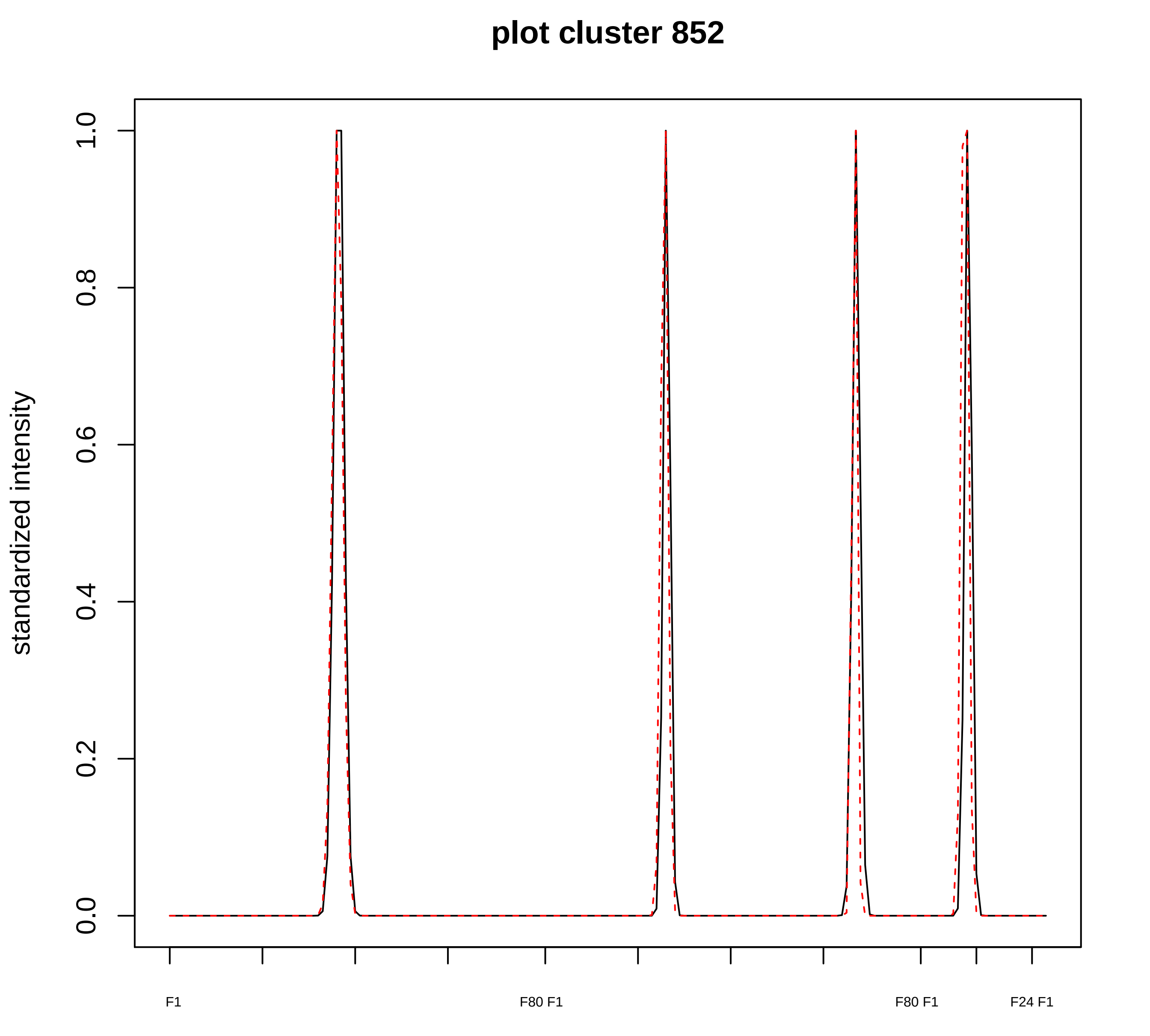
plot cluster 852
standardized intensity
0.0
0.2
0.4
0.6
0.8
1.0
F1
F80 F1
F80 F1
F24 F1

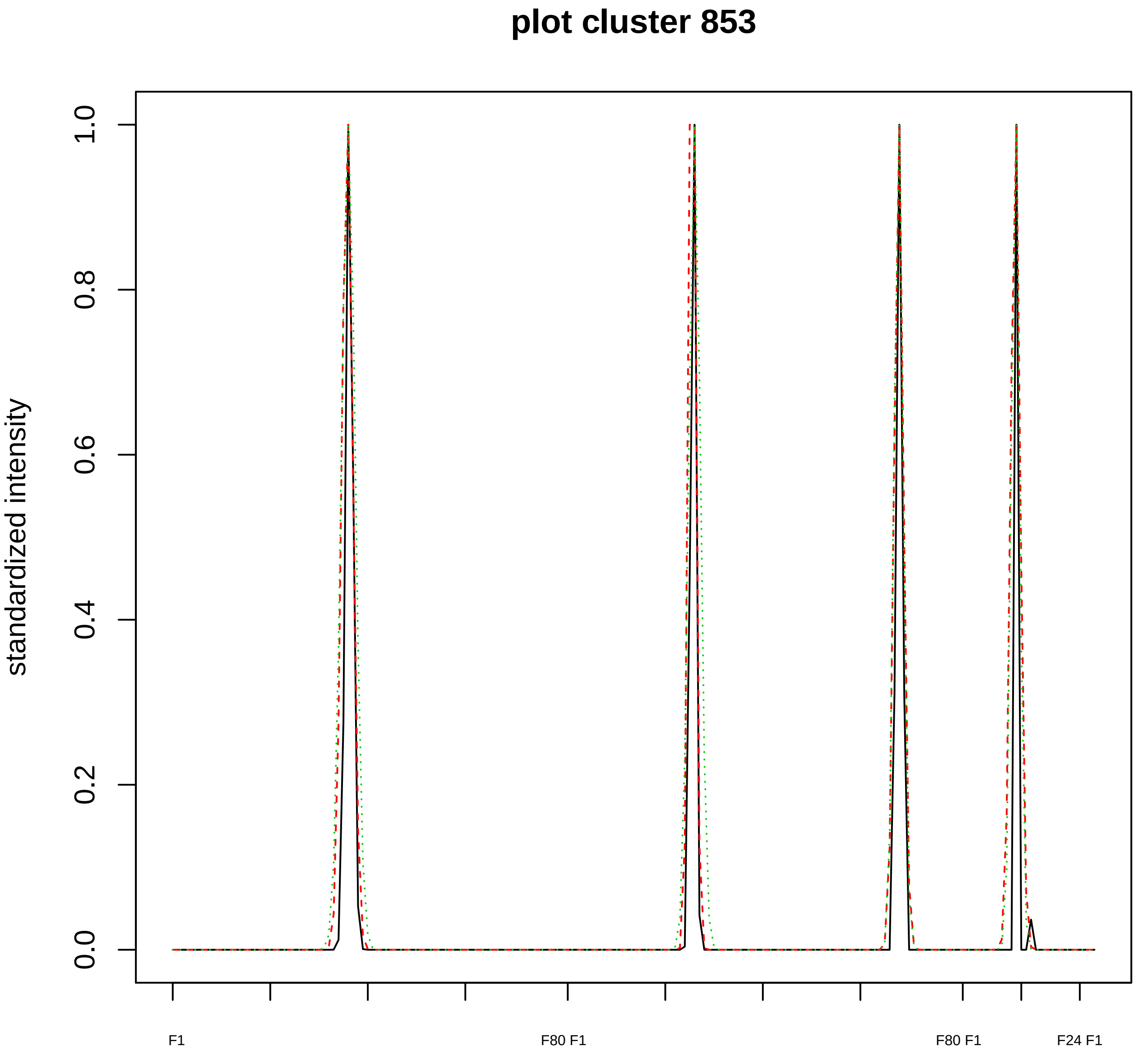

plot cluster 853
standardized intensity
0.0
0.2
0.4
0.6
0.8
1.0
F1
F80 F1
F80 F1
F24 F1

#### Dendrogram of cluster 853

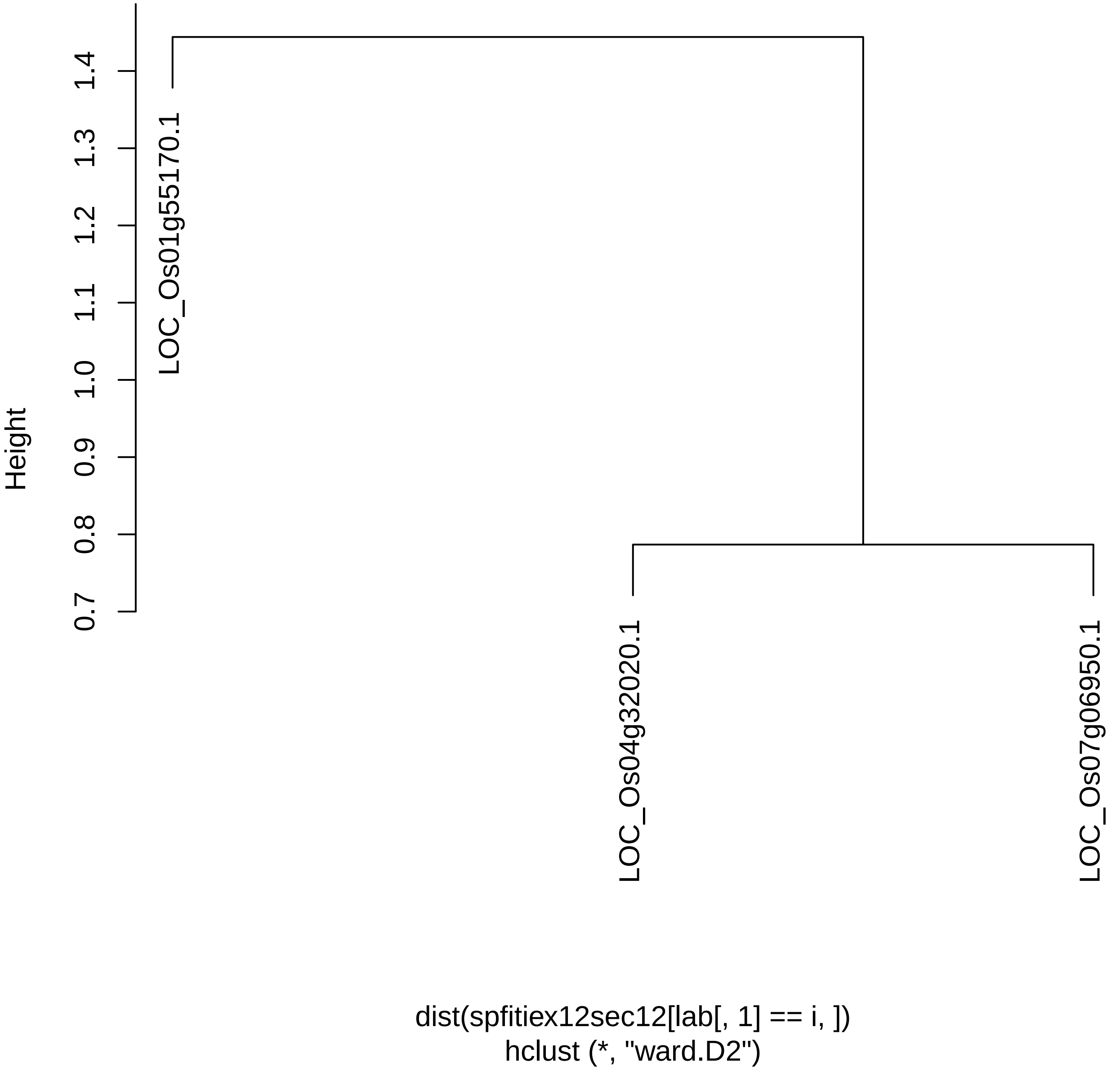

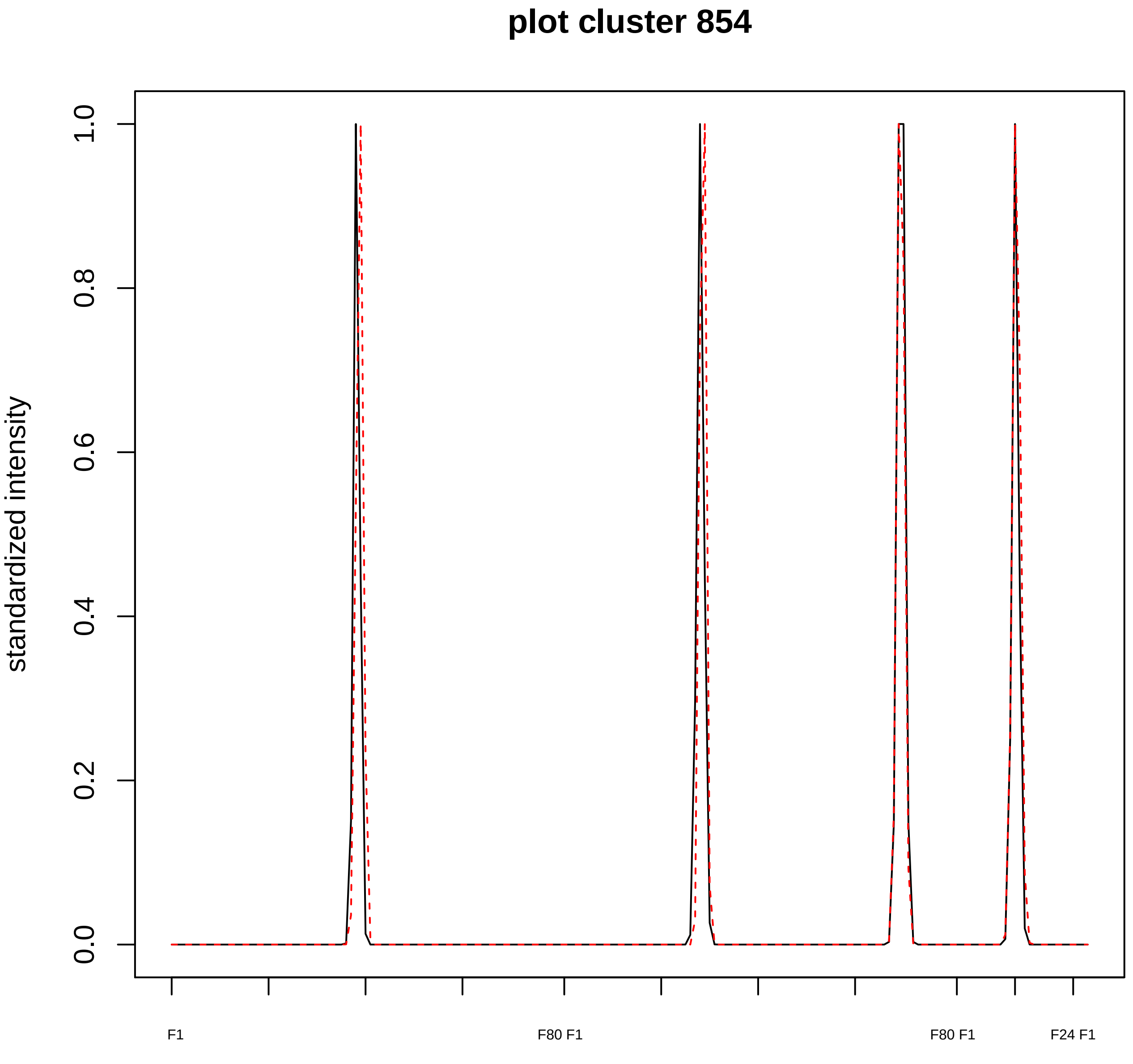
plot cluster 854
standardized intensity
1.0
0.8
0.6
0.4
0.2
0.0
F1
F80 F1
F80 F1
F24 F1

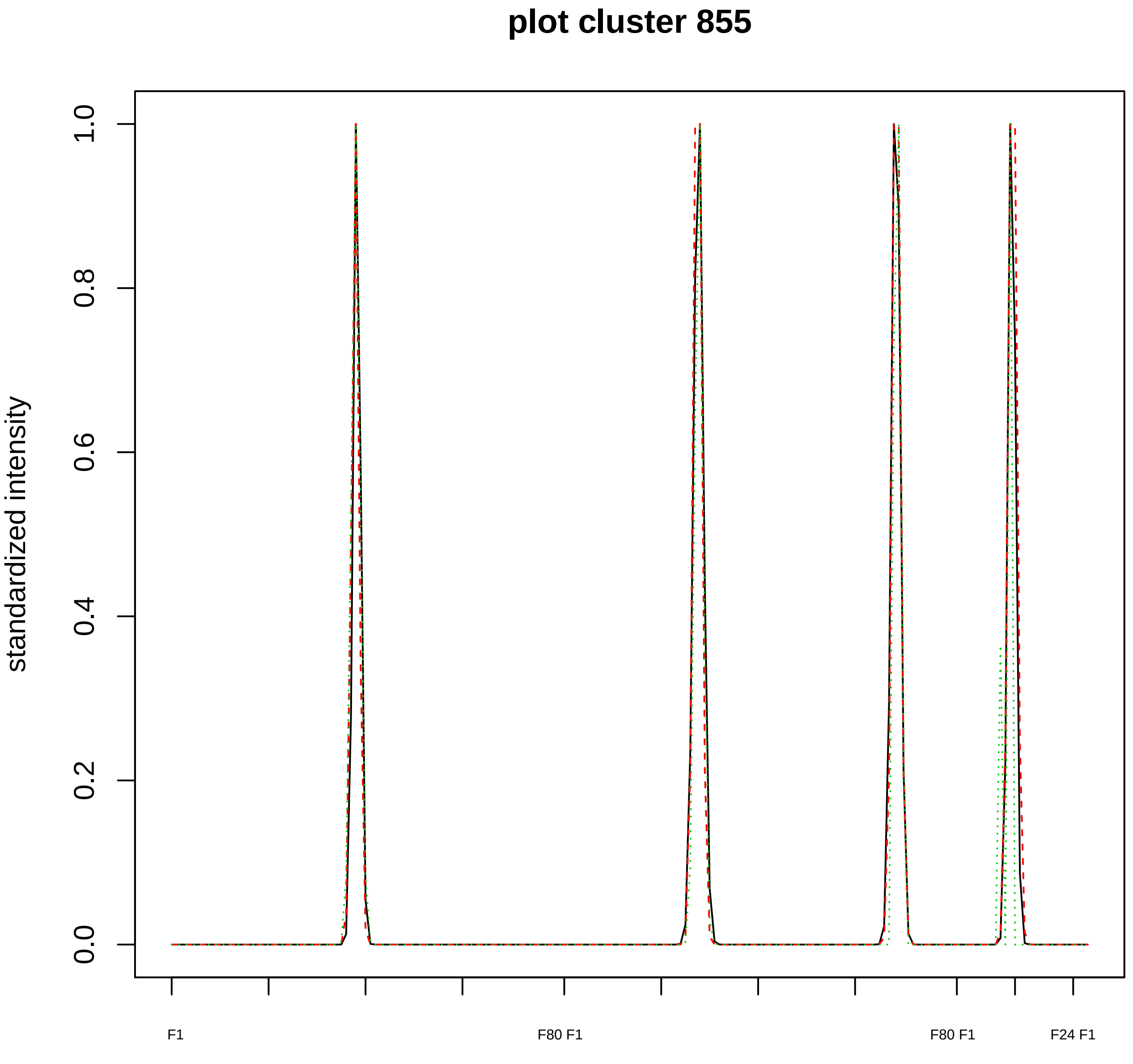
plot cluster 855
standardized intensity
0.0
0.2
0.4
0.6
0.8
1.0
F1
F80 F1
F80 F1
F24 F1

#### Dendrogram of cluster 855

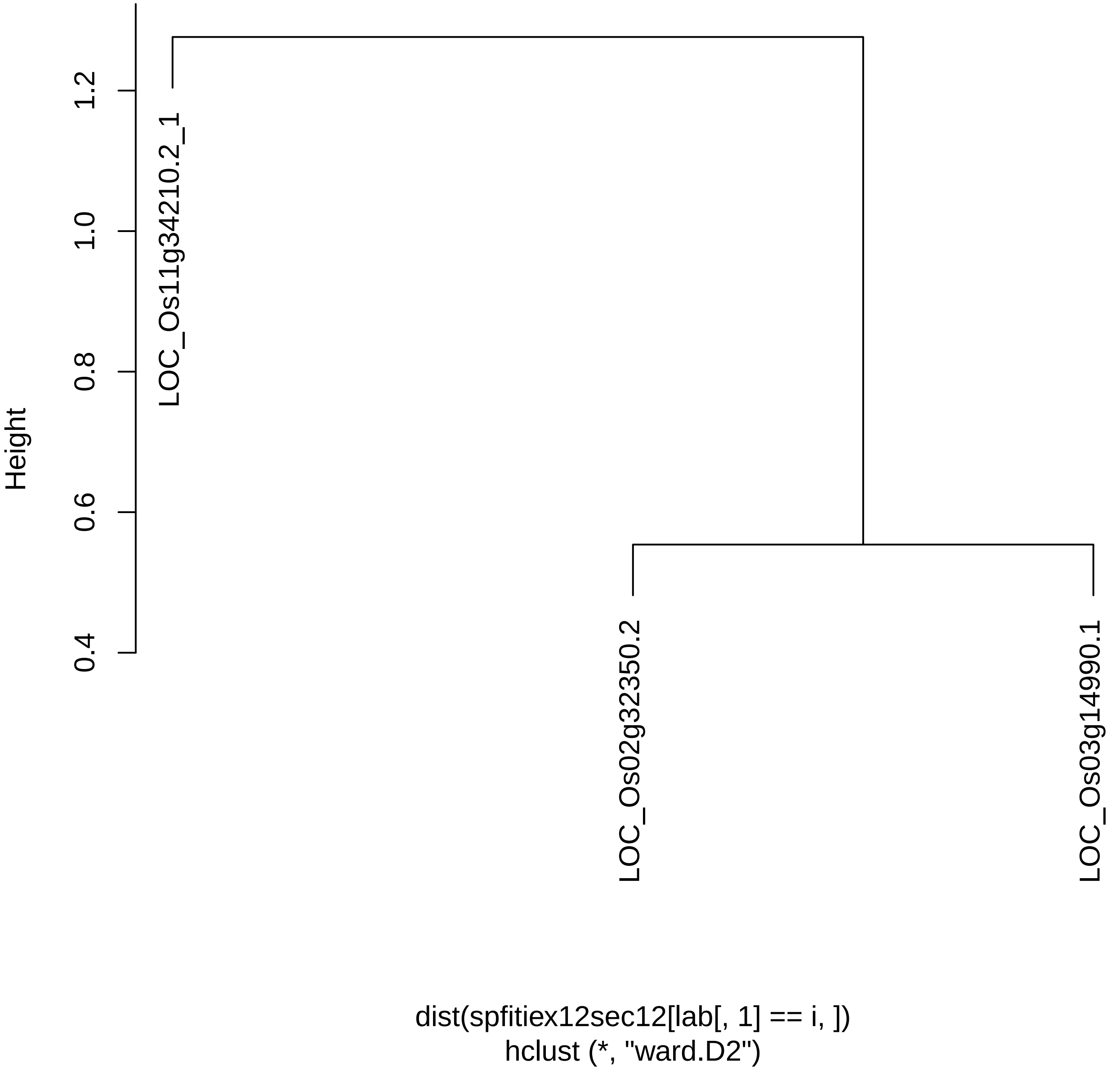

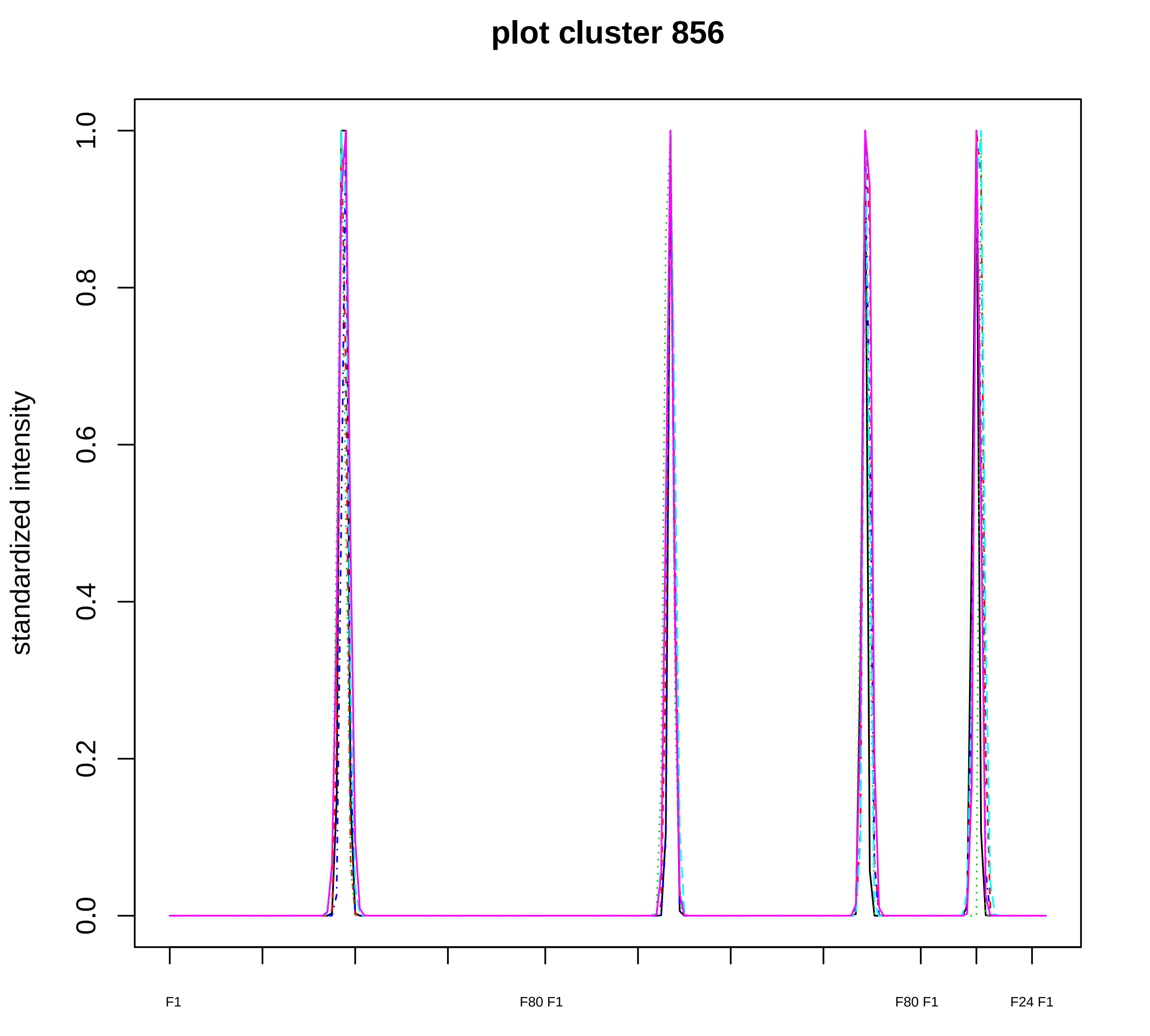
plot cluster 856
standardized intensity
0.0
0.2
0.4
0.6
0.8
1.0
F1
F80 F1
F80 F1
F24 F1

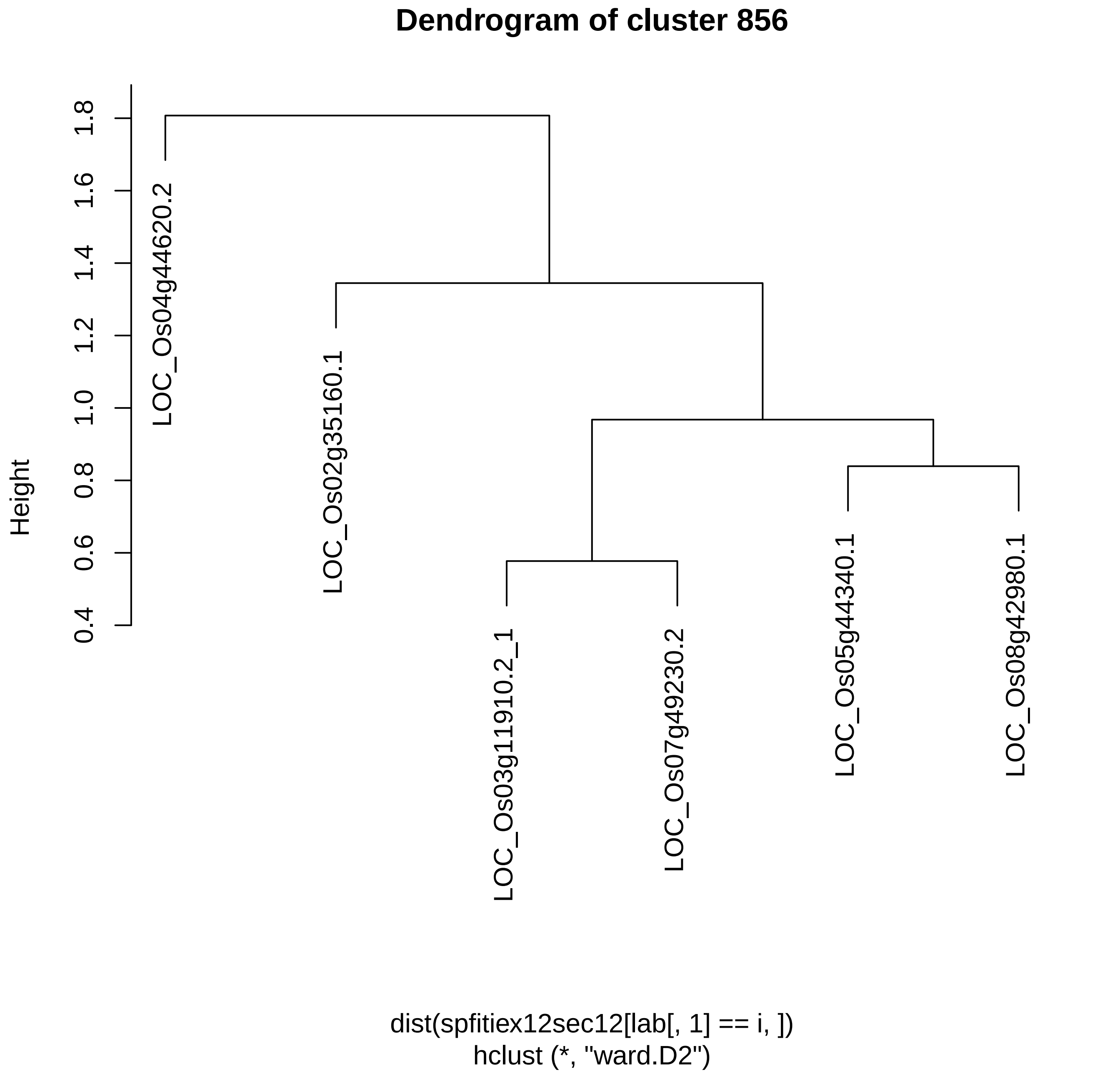

Dendrogram of cluster 856
Height
1.8
1.6
1.4
1.2
1.0
0.8
0.6
0.4
LOC_Os04g44620.2
LOC_Os02g35160.1
LOC_Os03g11910.2_1
LOC_Os07g49230.2
LOC_Os05g44340.1
LOC_Os08g42980.1
dist(spfitiex12sec12[lab[, 1] == i, ])
hclust (*, "ward.D2")

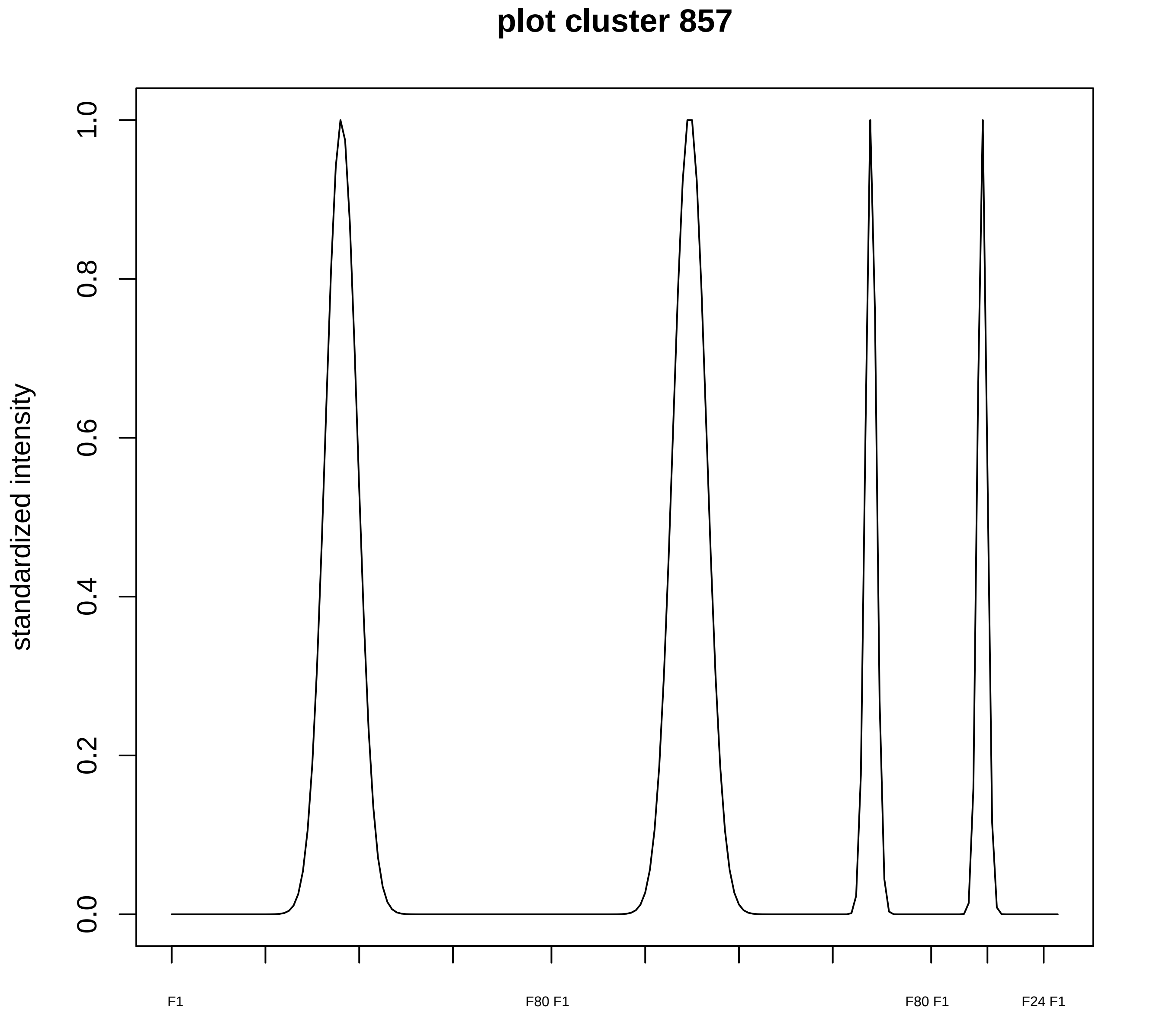
plot cluster 857
standardized intensity
0.0
0.2
0.4
0.6
0.8
1.0
F1
F80 F1
F80 F1
F24 F1

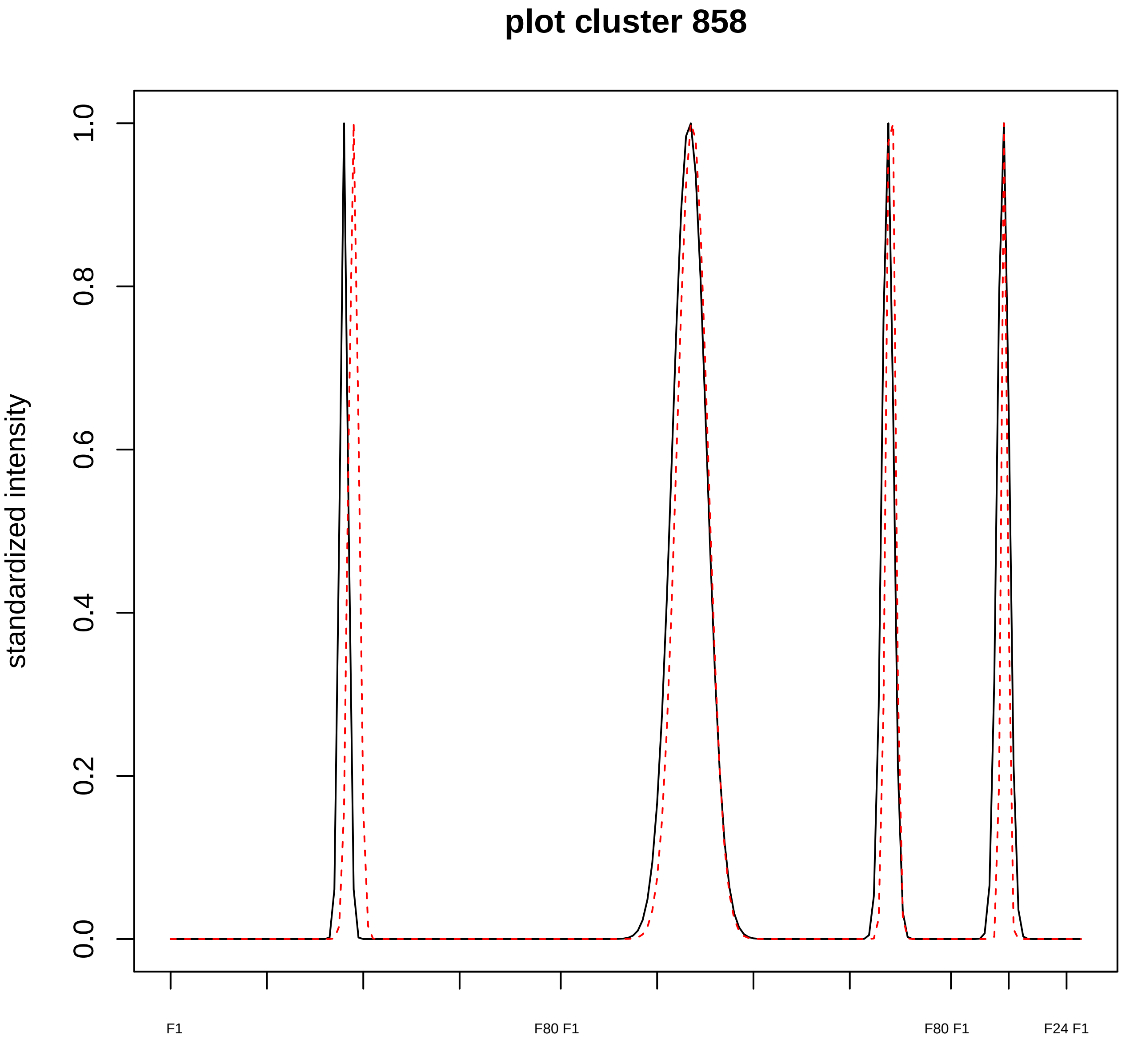

plot cluster 858
standardized intensity
0.0
0.2
0.4
0.6
0.8
1.0
F1
F80 F1
F80 F1
F24 F1

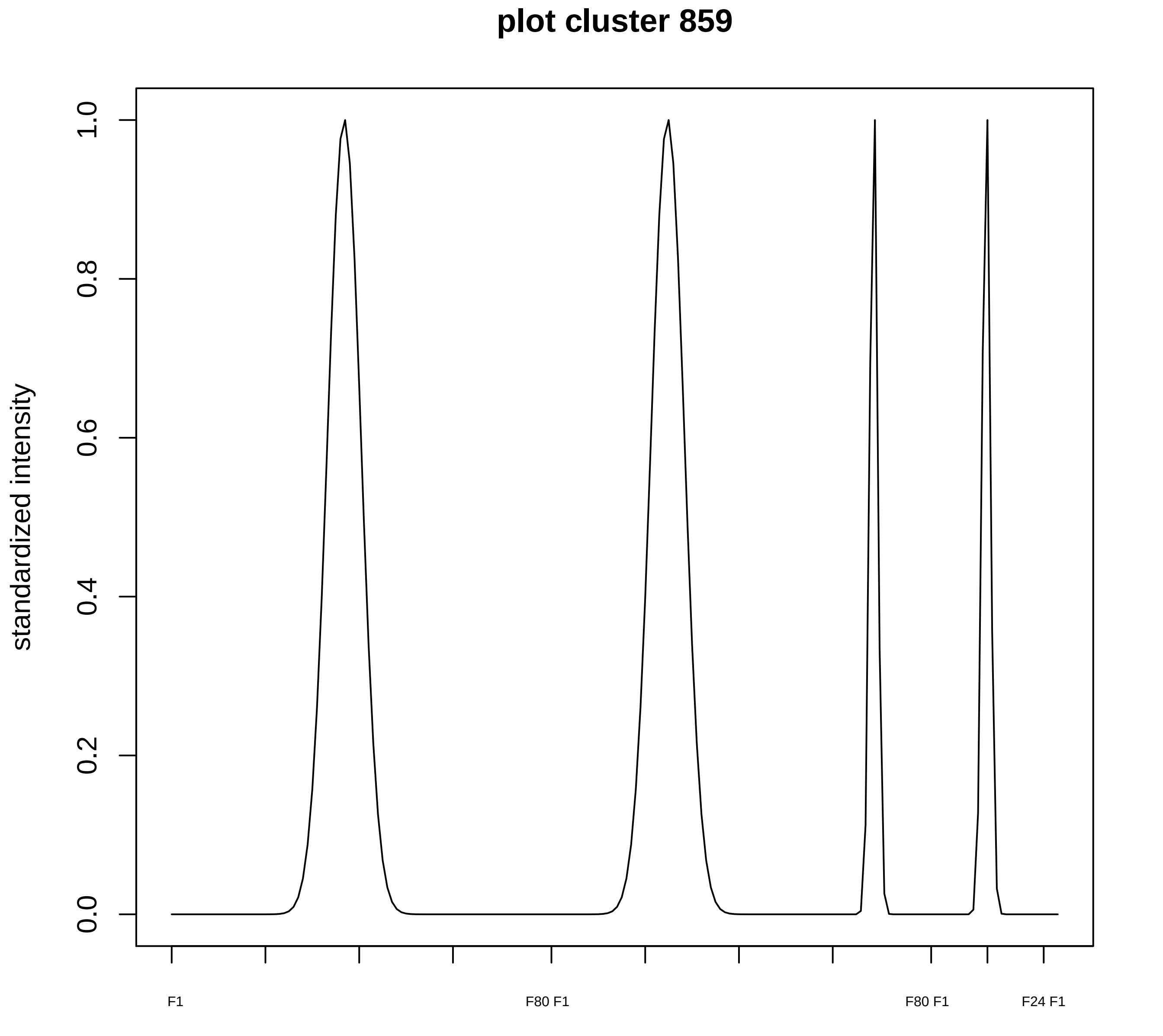

plot cluster 859
standardized intensity
0.0
0.2
0.4
0.6
0.8
1.0
F1
F80 F1
F80 F1
F24 F1

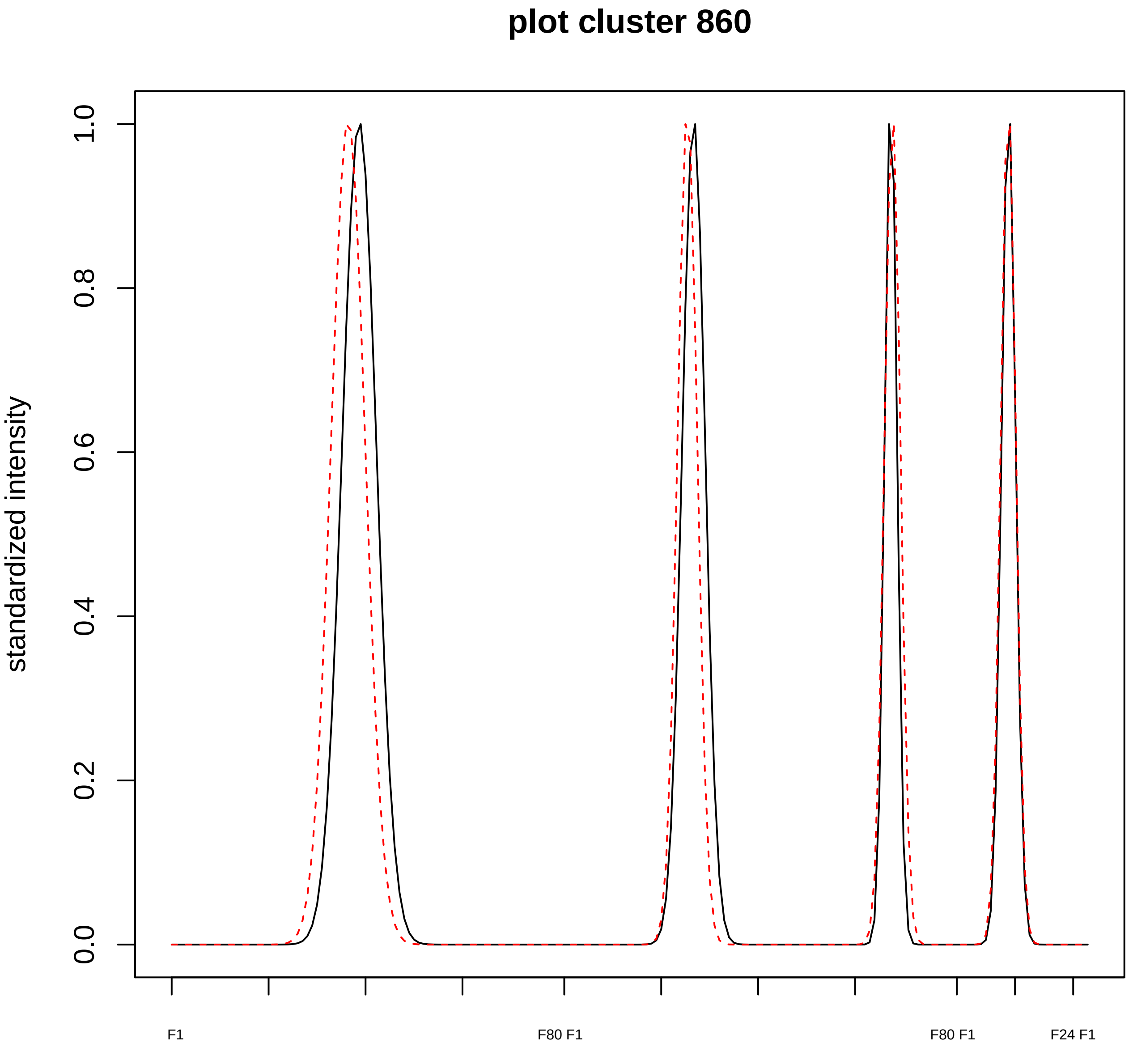
plot cluster 860
standardized intensity
0.0
0.2
0.4
0.6
0.8
1.0
F1
F80 F1
F80 F1
F24 F1

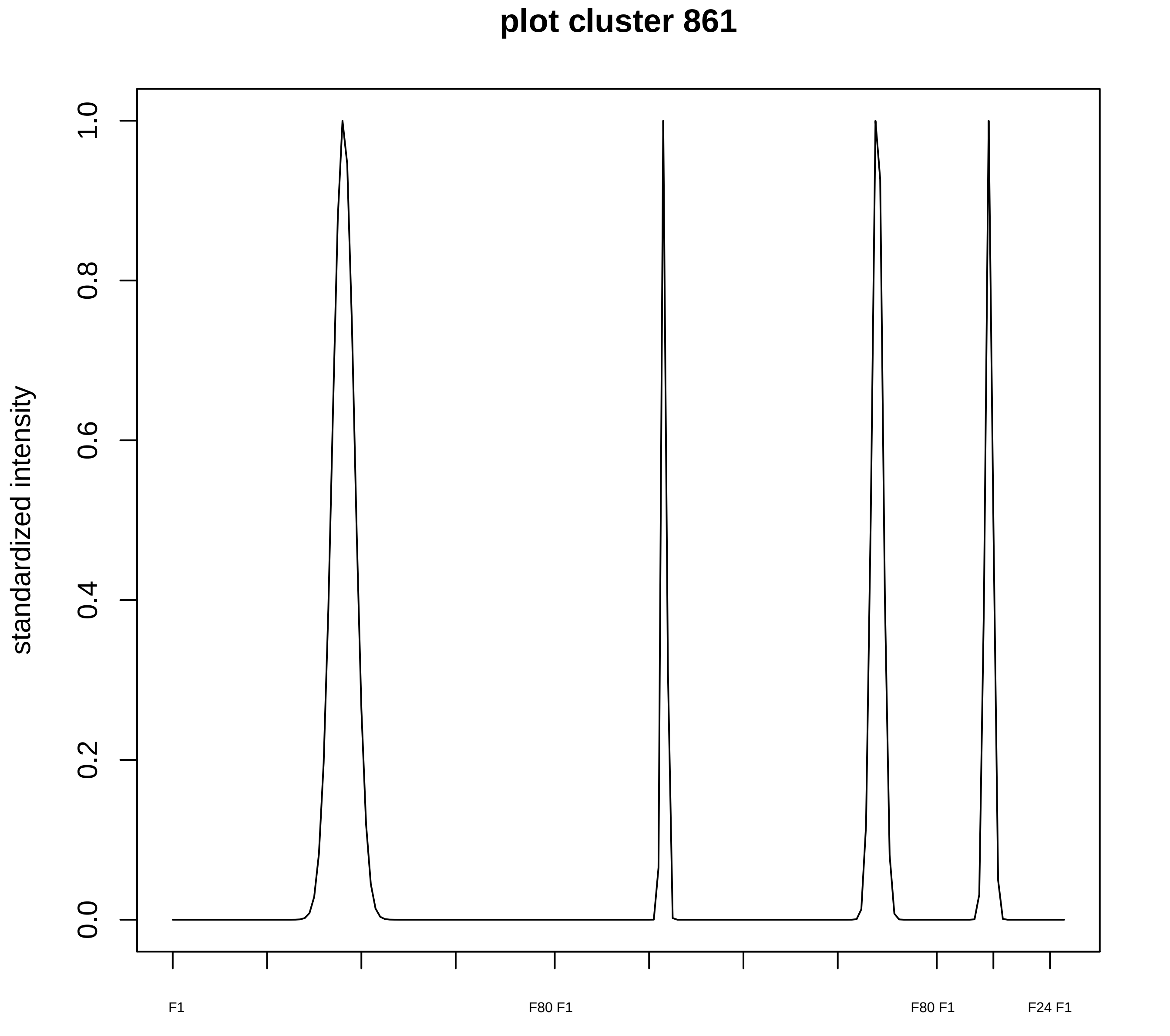
plot cluster 861
standardized intensity
0.0
0.2
0.4
0.6
0.8
1.0
F1
F80 F1
F80 F1
F24 F1

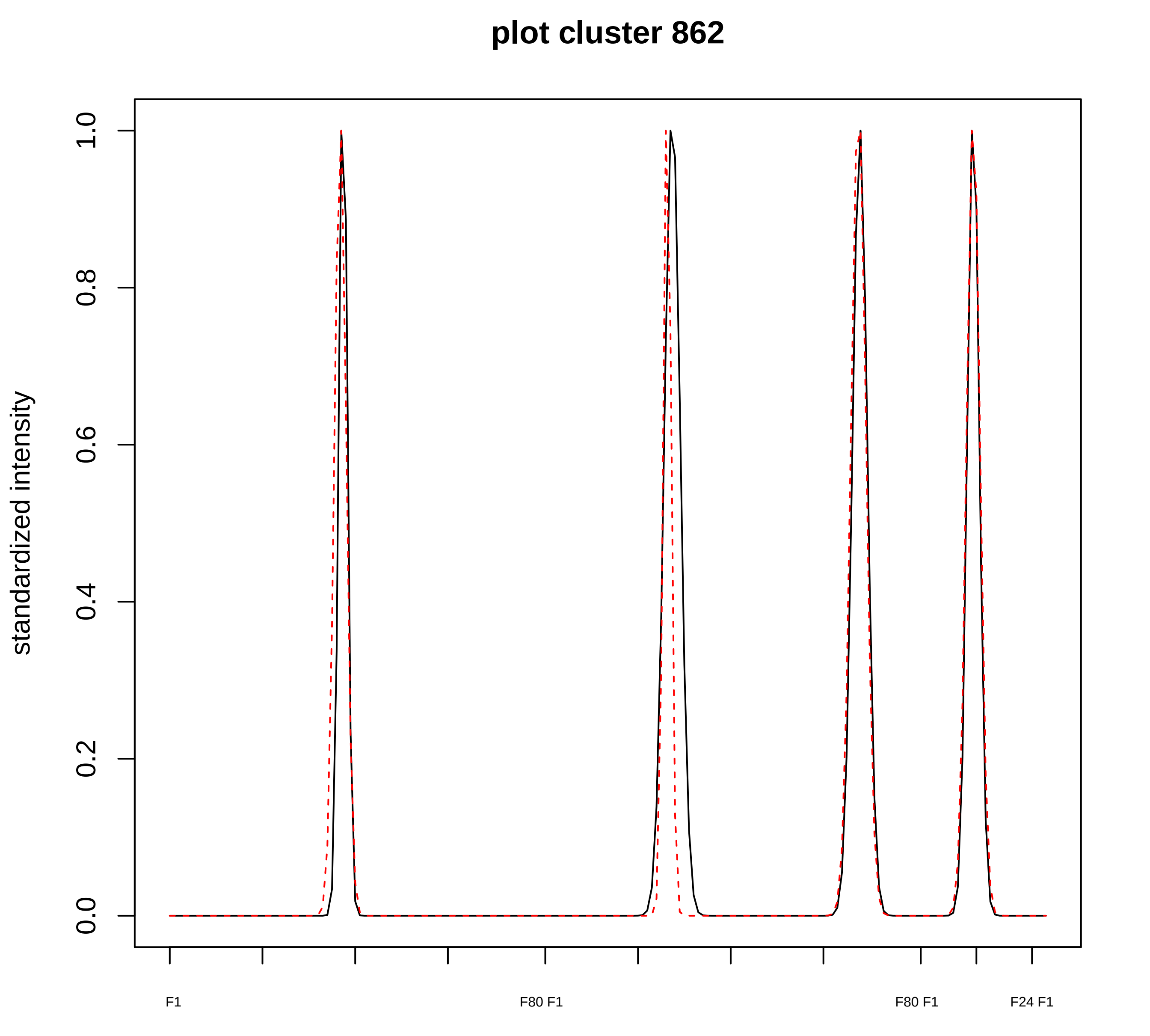
plot cluster 862
standardized intensity
0.0
0.2
0.4
0.6
0.8
1.0
F1
F80 F1
F80 F1
F24 F1

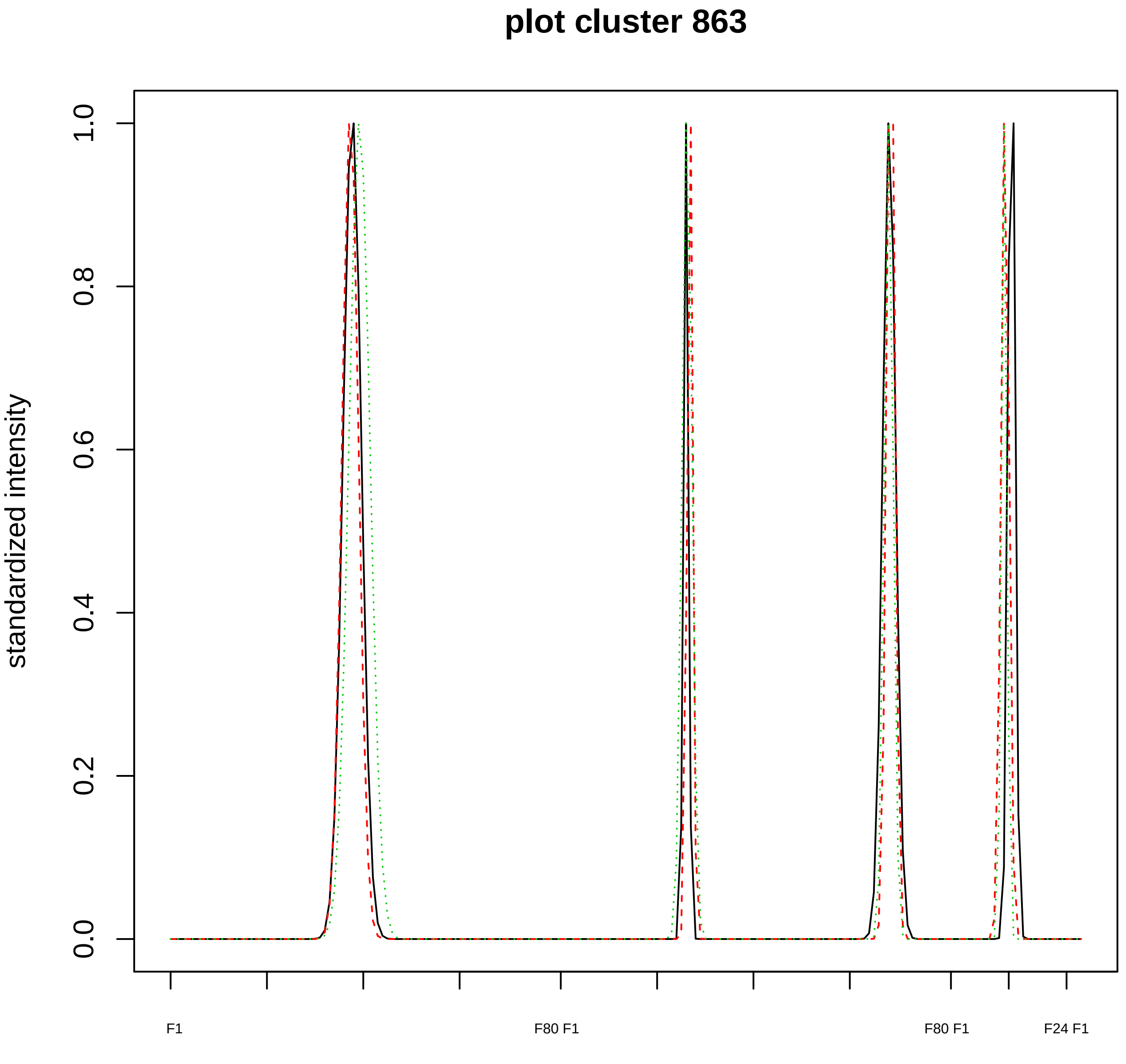

plot cluster 863
standardized intensity
0.0
0.2
0.4
0.6
0.8
1.0
F1
F80 F1
F80 F1
F24 F1

### Dendrogram of cluster 863

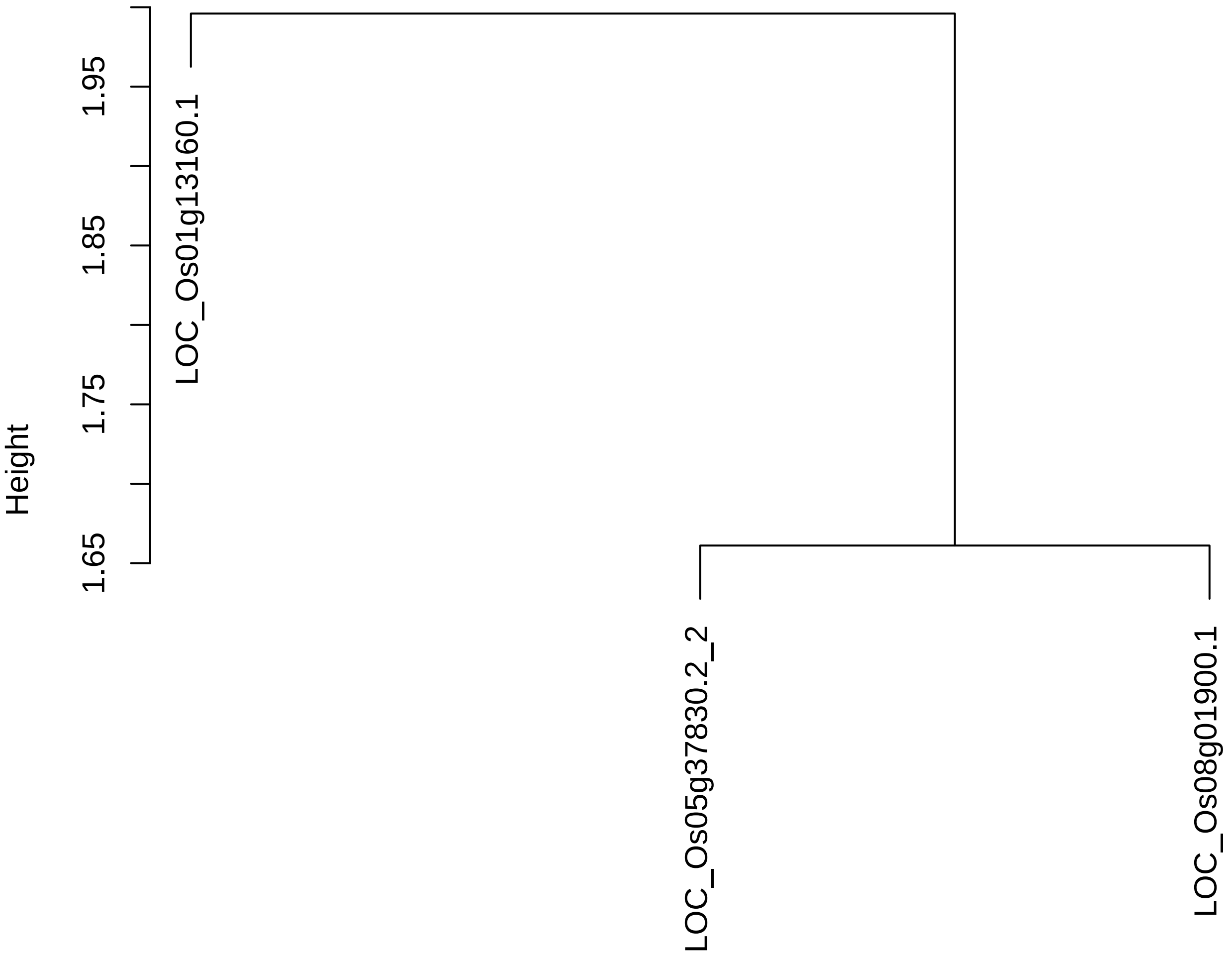


dist(spfitiex12sec12[lab[, 1] == i, ])
hclust (*, "ward.D2")

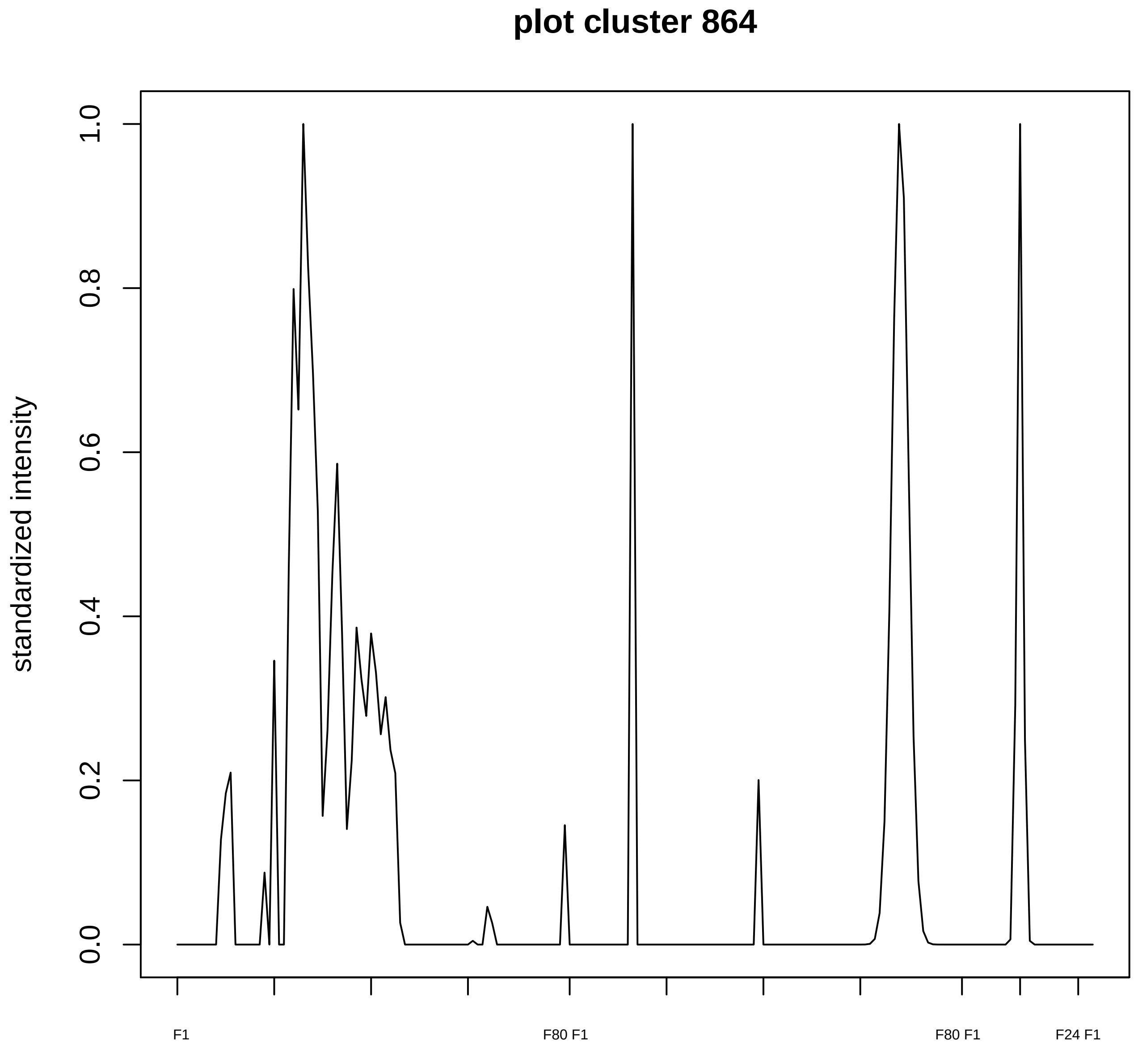
plot cluster 864
standardized intensity
0.0
0.2
0.4
0.6
0.8
1.0
F1
F80 F1
F80 F1
F24 F1

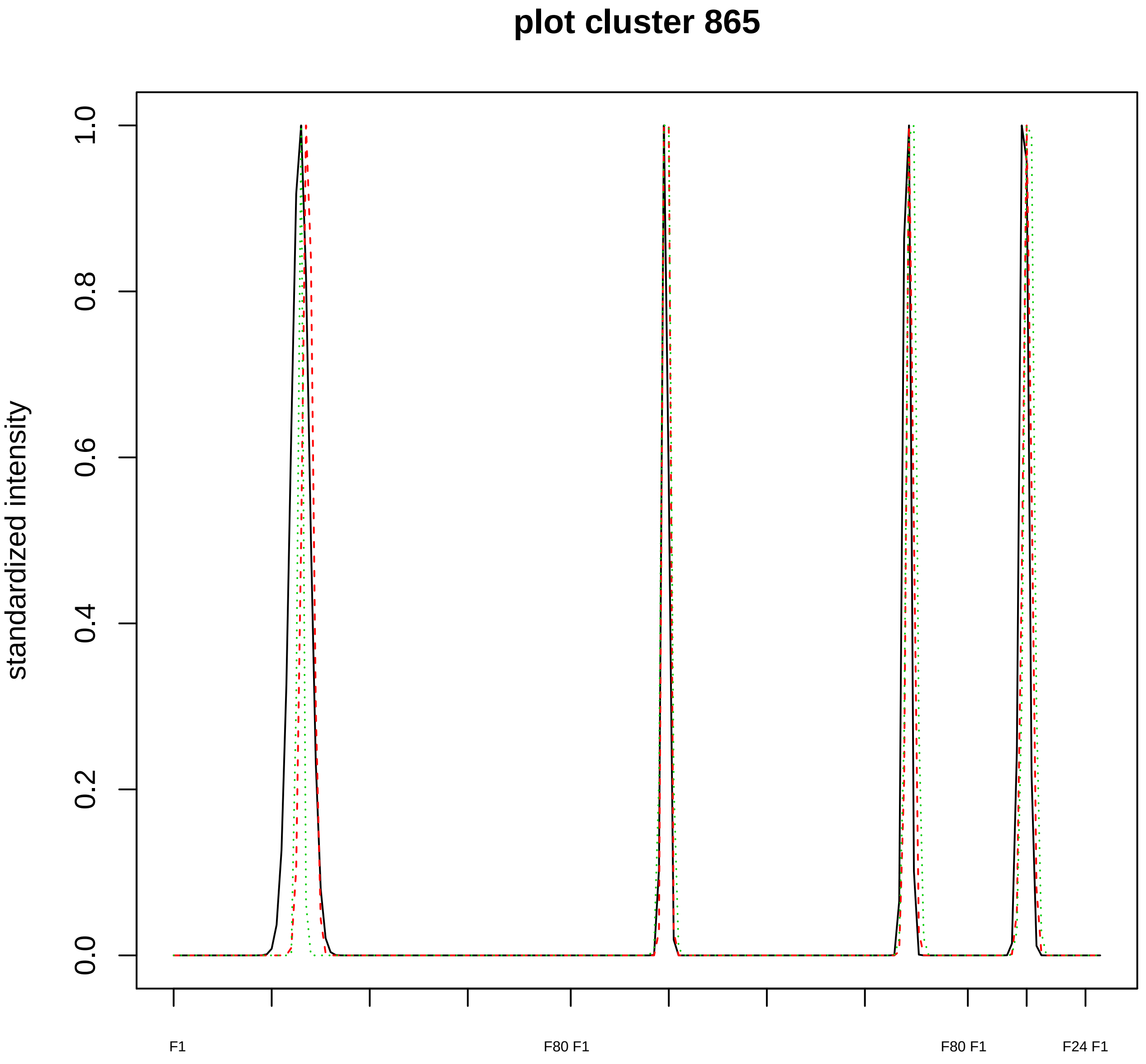

plot cluster 865
standardized intensity
0.0
0.2
0.4
0.6
0.8
1.0
F1
F80 F1
F80 F1
F24 F1

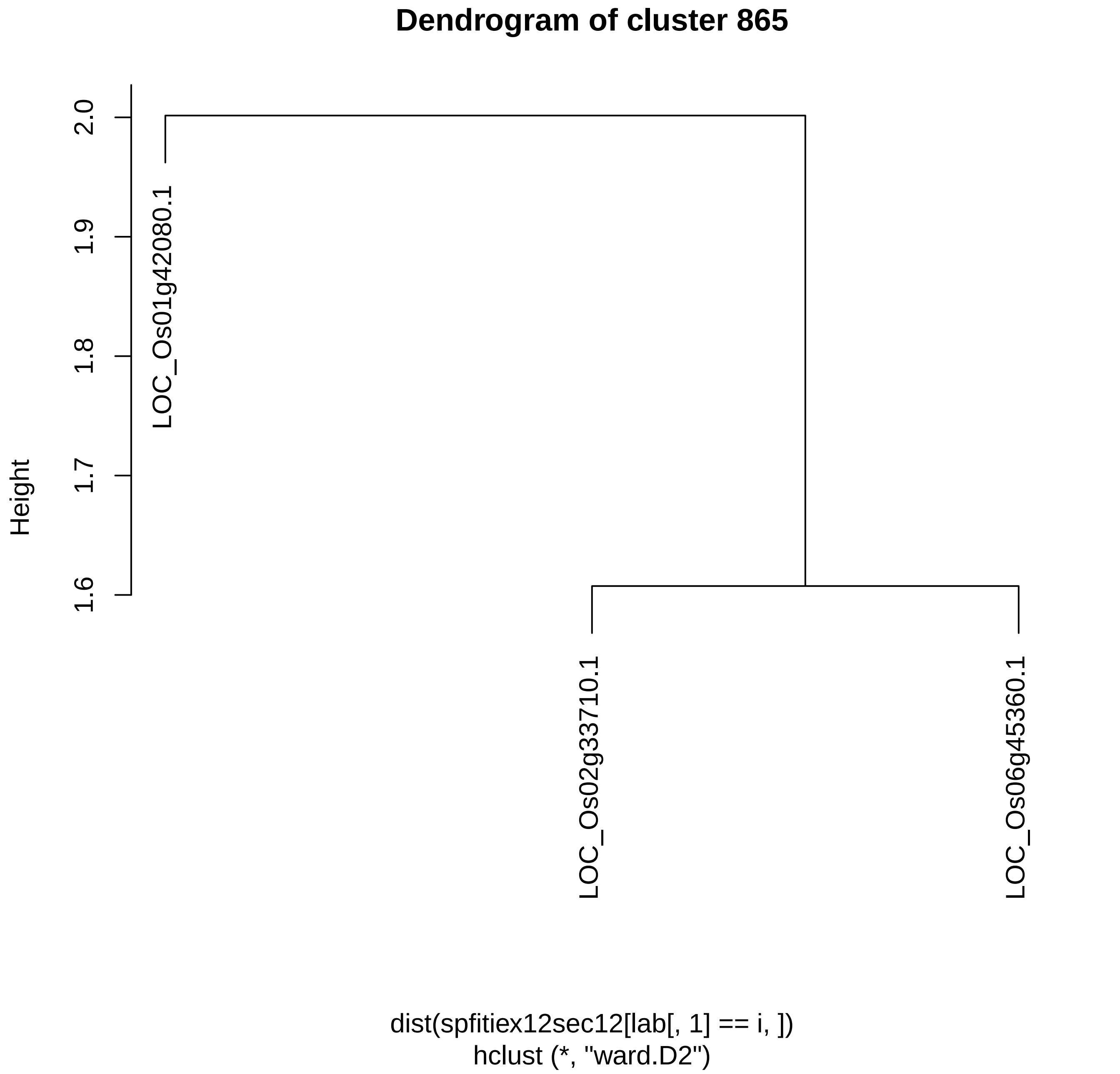
Dendrogram of cluster 865
Height
2.0
1.9
1.8
1.7
1.6
LOC_Os01g42080.1
LOC_Os02g33710.1
LOC_Os06g45360.1
dist(spfitiex12sec12[lab[, 1] == i, ])
hclust (*, "ward.D2")

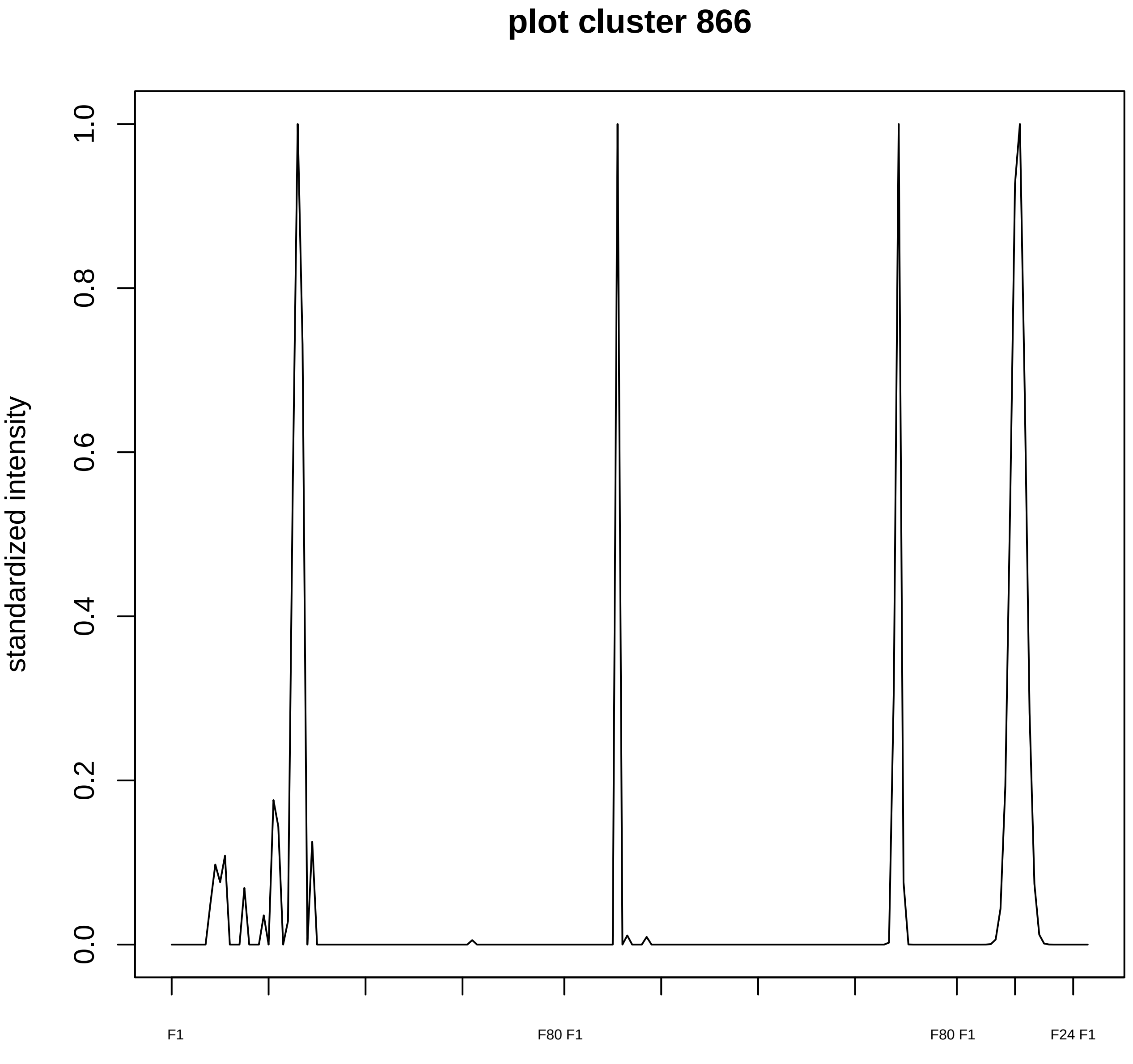

plot cluster 866
standardized intensity
0.0
0.2
0.4
0.6
0.8
1.0
F1
F80 F1
F80 F1
F24 F1

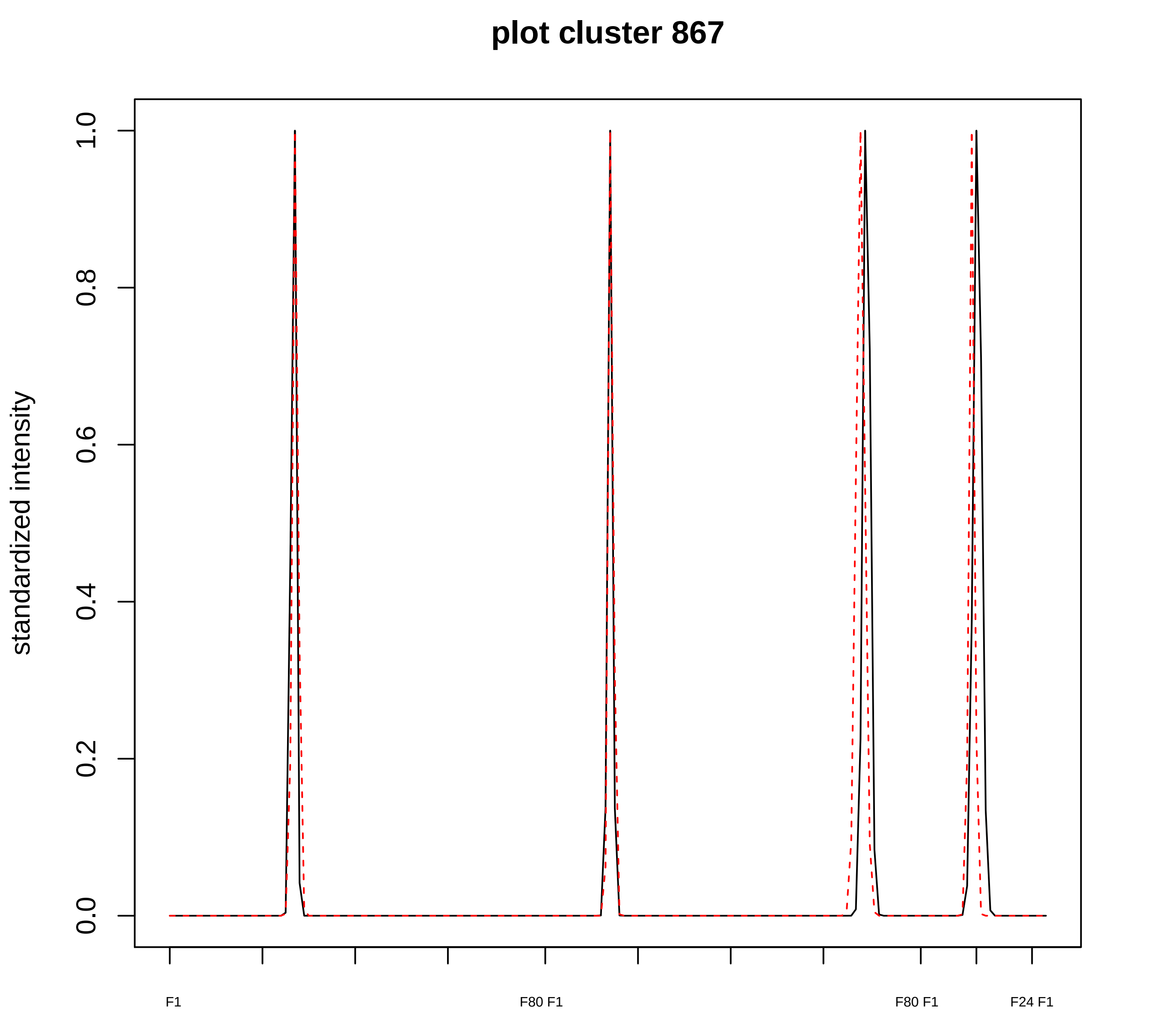

plot cluster 867
standardized intensity
0.0
0.2
0.4
0.6
0.8
1.0
F1
F80 F1
F80 F1
F24 F1

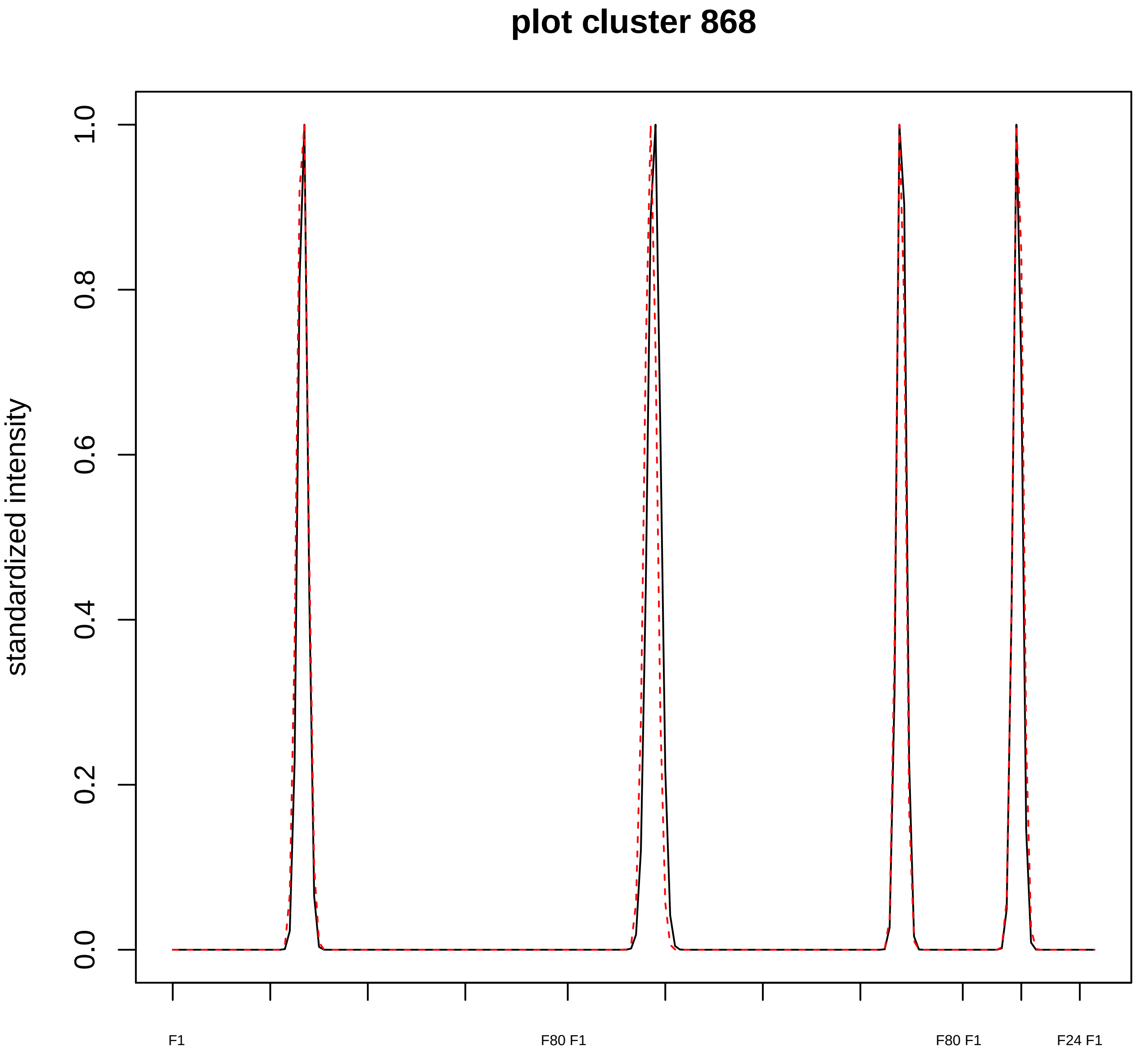

plot cluster 868
standardized intensity
0.0
0.2
0.4
0.6
0.8
1.0
F1
F80 F1
F80 F1
F24 F1

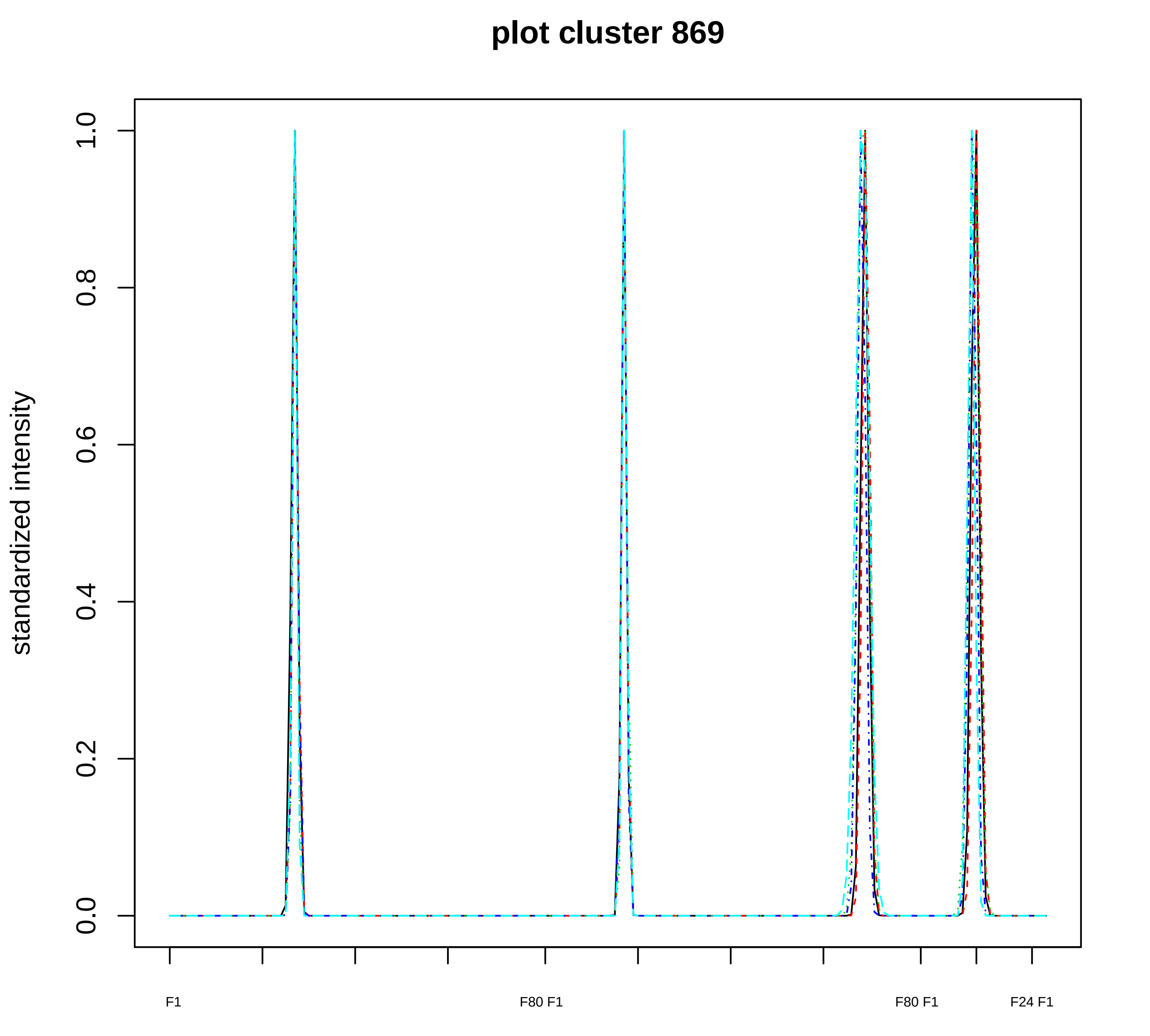

plot cluster 869
standardized intensity
0.0
0.2
0.4
0.6
0.8
1.0
F1
F80 F1
F80 F1
F24 F1

#### Dendrogram of cluster 869

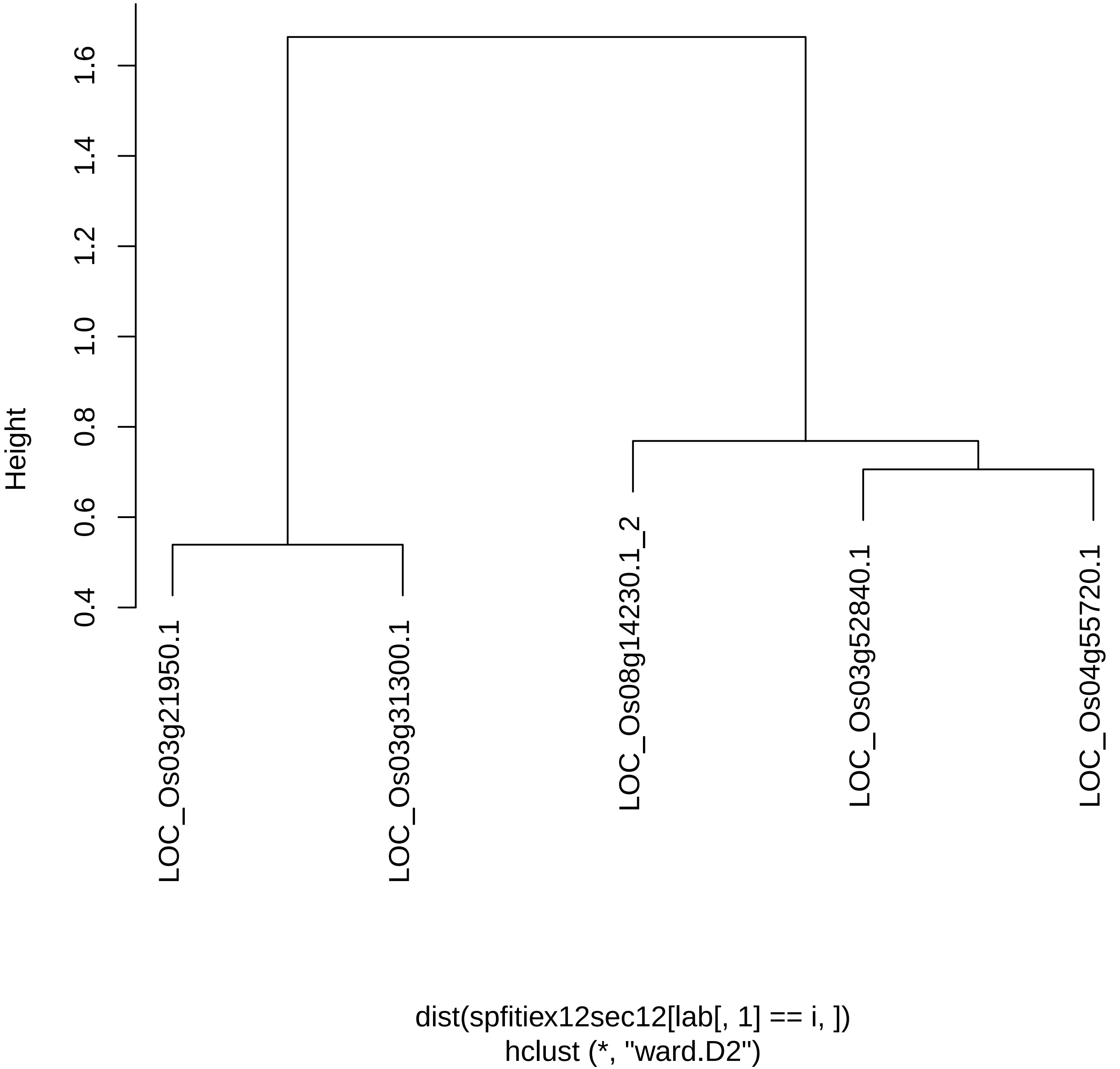

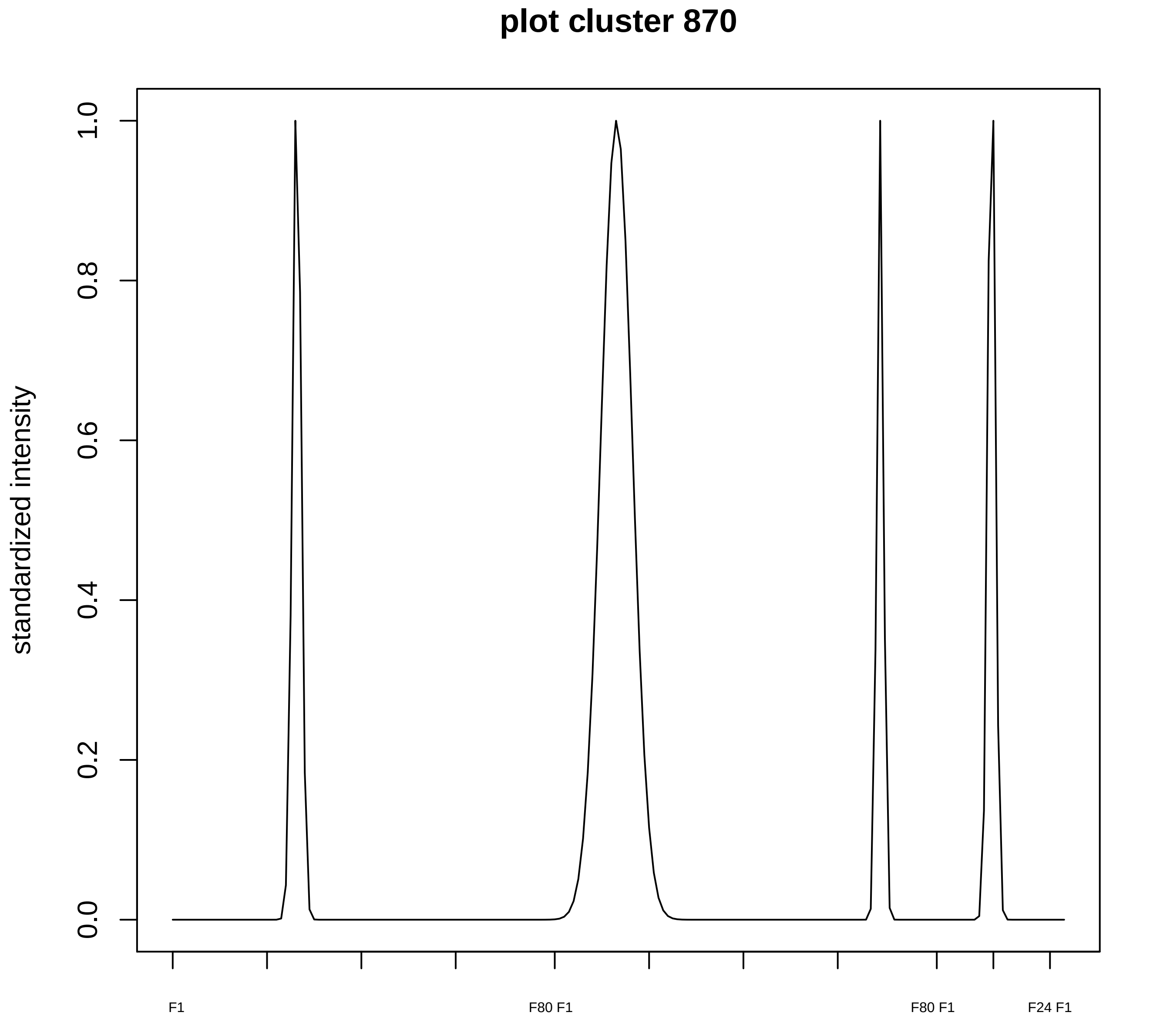
plot cluster 870
standardized intensity
0.0
0.2
0.4
0.6
0.8
1.0
F1
F80 F1
F80 F1
F24 F1

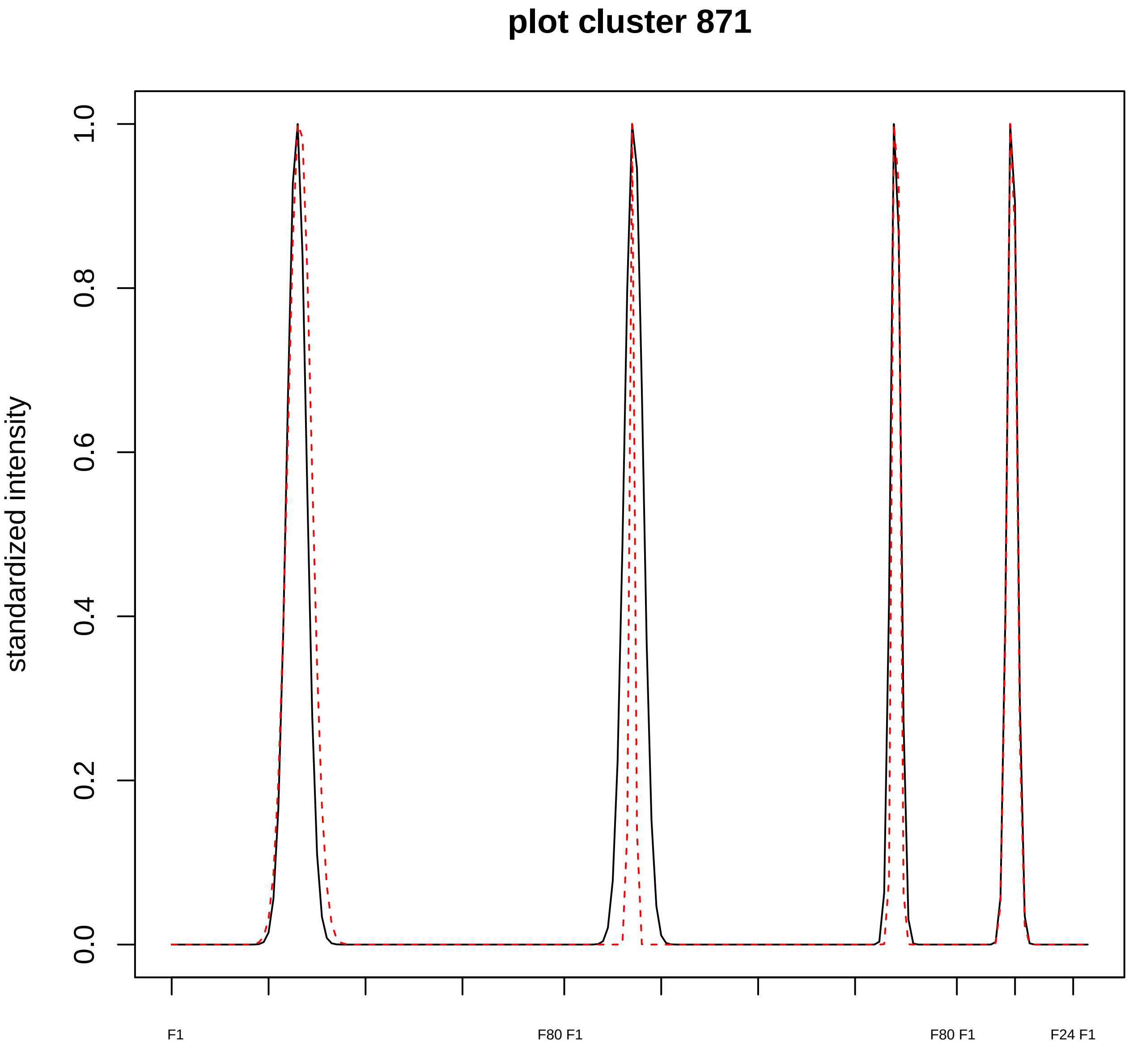

plot cluster 871
standardized intensity
0.0
0.2
0.4
0.6
0.8
1.0
F1
F80 F1
F80 F1
F24 F1

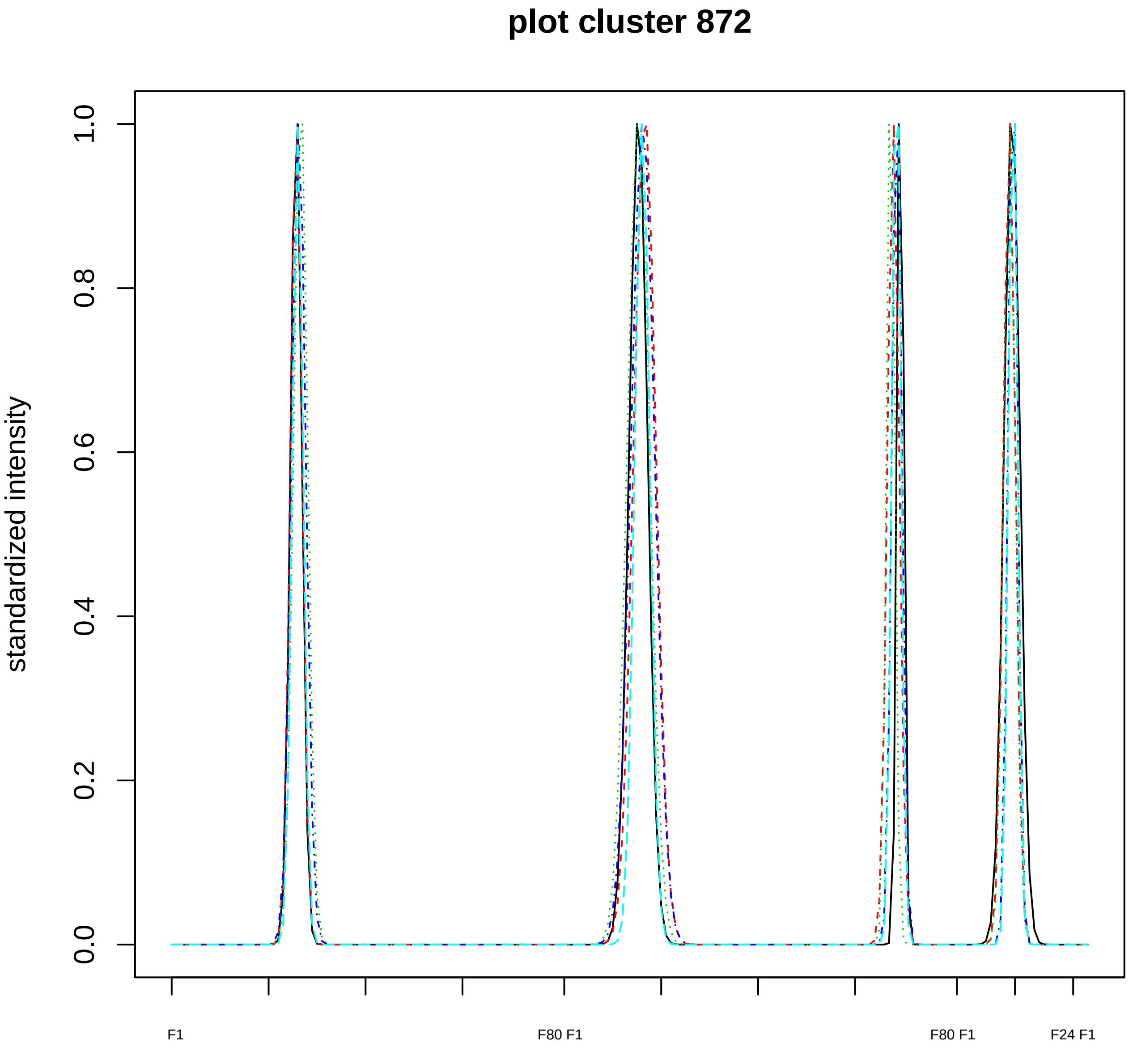
plot cluster 872
standardized intensity
0.0
0.2
0.4
0.6
0.8
1.0
F1
F80 F1
F80 F1
F24 F1

### Dendrogram of cluster 872

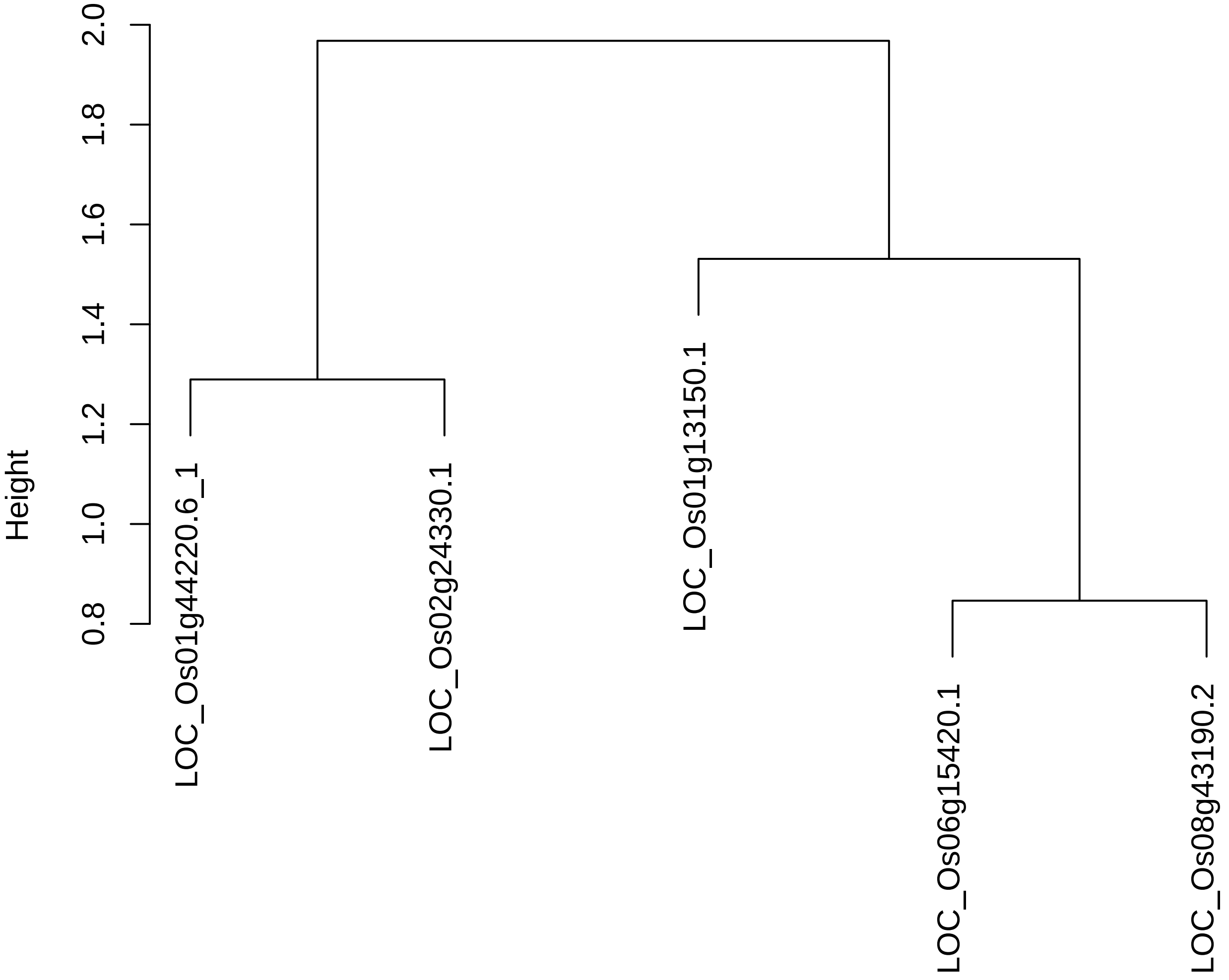


dist(spfitiex12sec12[lab[, 1] == i, ])
hclust (*, "ward.D2")

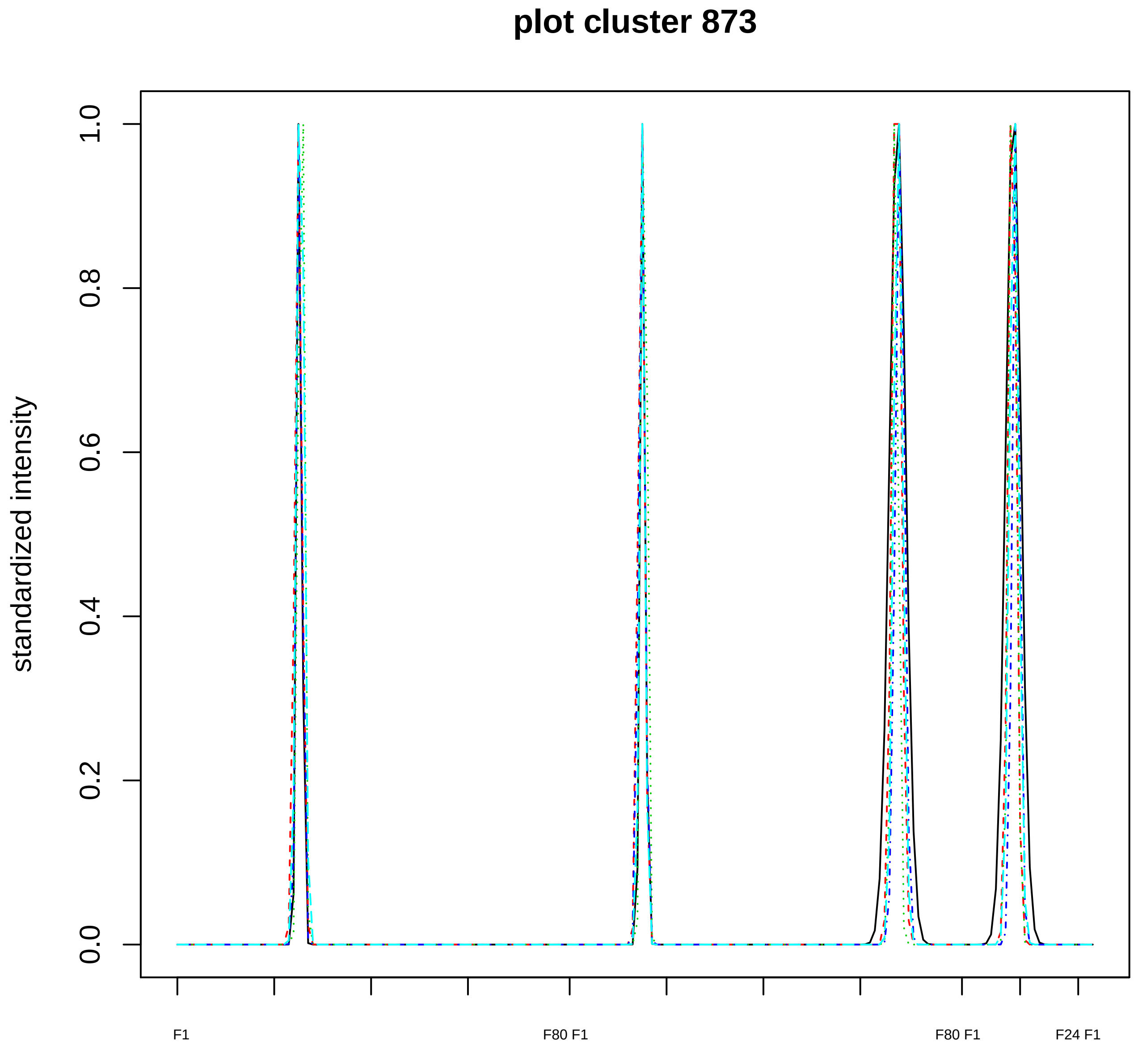

plot cluster 873
standardized intensity
1.0
0.8
0.6
0.4
0.2
0.0
F1
F80 F1
F80 F1
F24 F1

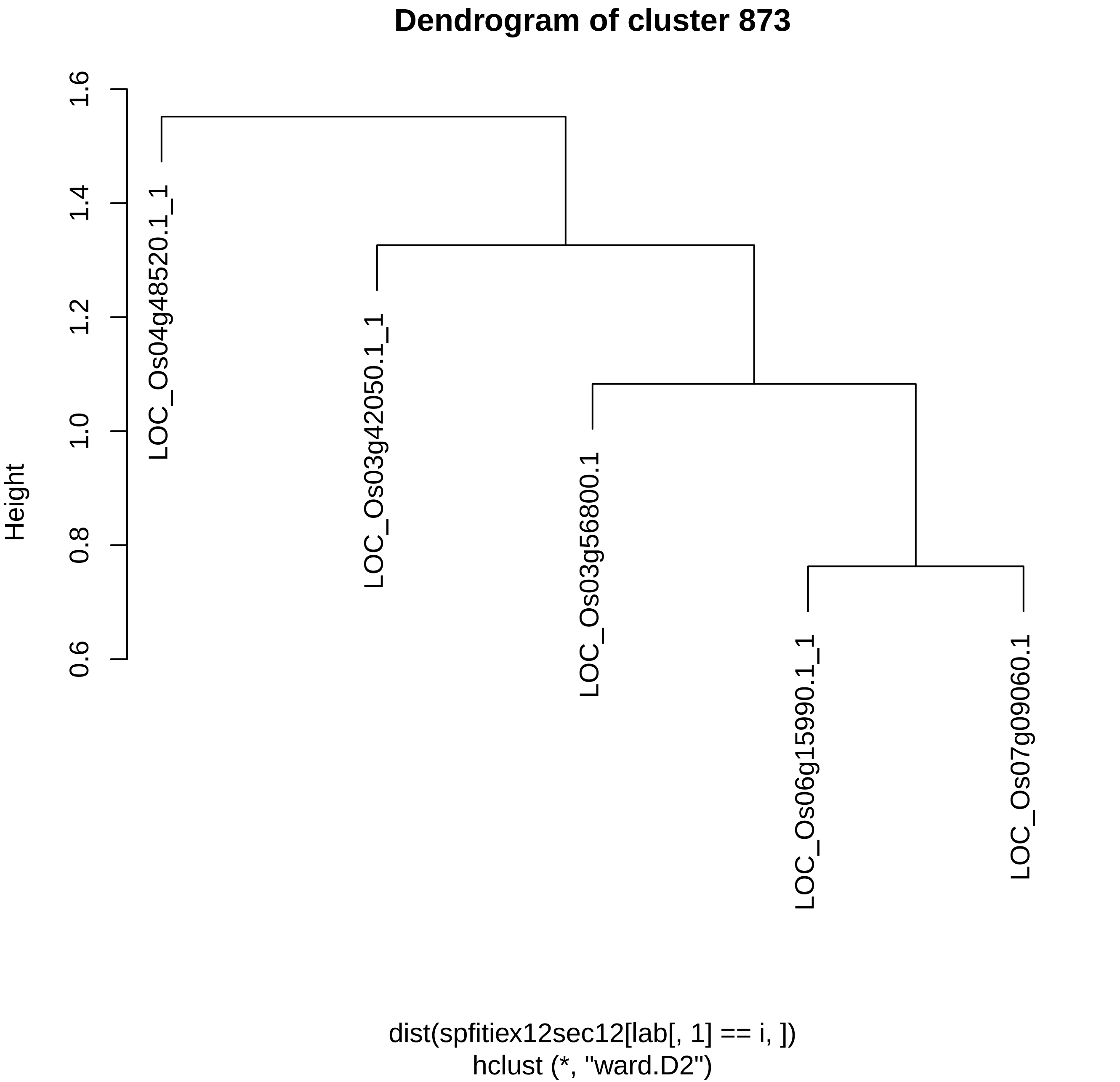

Dendrogram of cluster 873
Height
1.6
1.4
1.2
1.0
0.8
0.6
LOC_Os04g48520.1_1
LOC_Os03g42050.1_1
LOC_Os03g56800.1
LOC_Os06g15990.1_1
LOC_Os07g09060.1
dist(spfitiex12sec12[lab[, 1] == i, ])
hclust (*, "ward.D2")

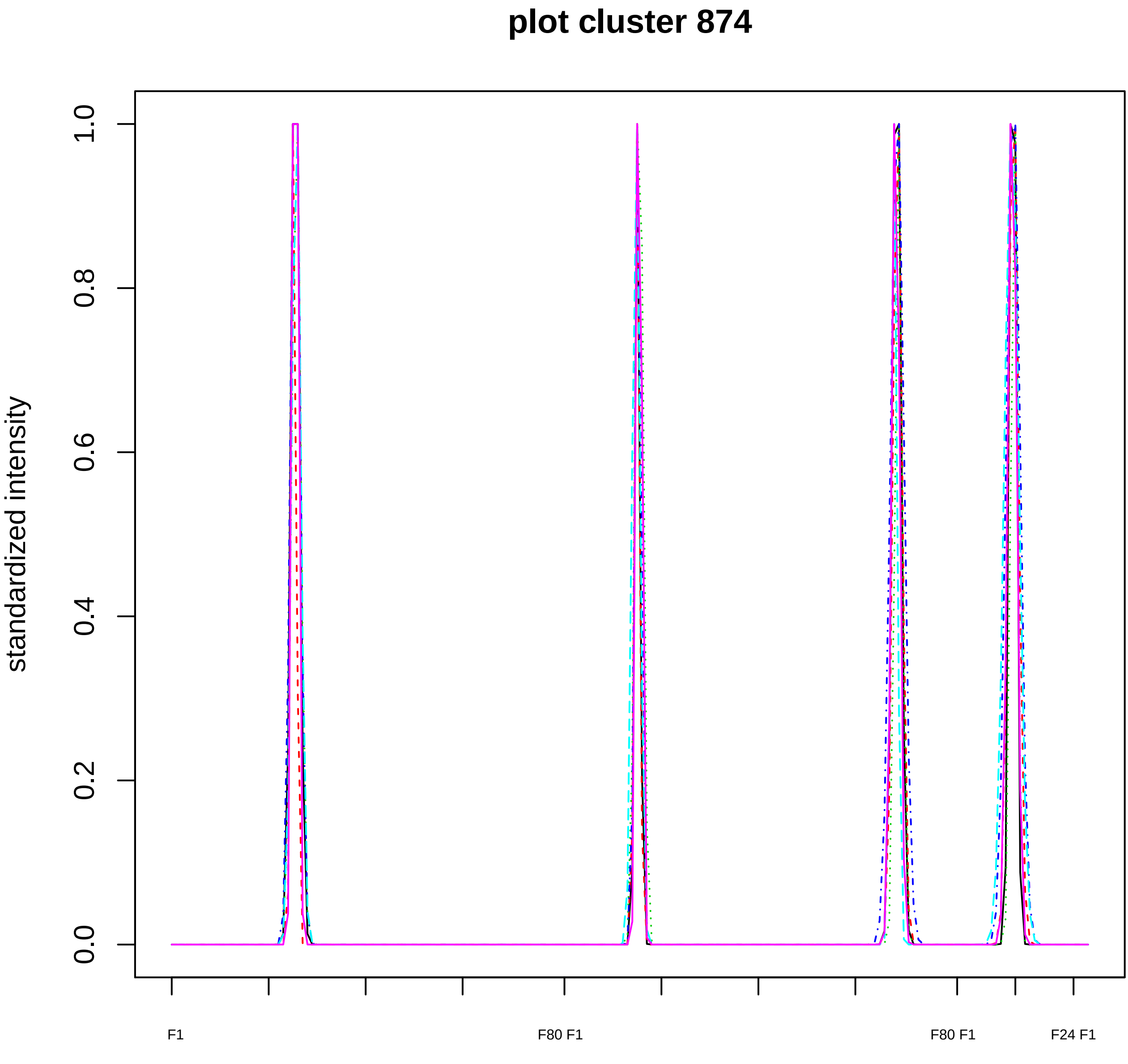

plot cluster 874
standardized intensity
0.0
0.2
0.4
0.6
0.8
1.0
F1
F80 F1
F80 F1
F24 F1

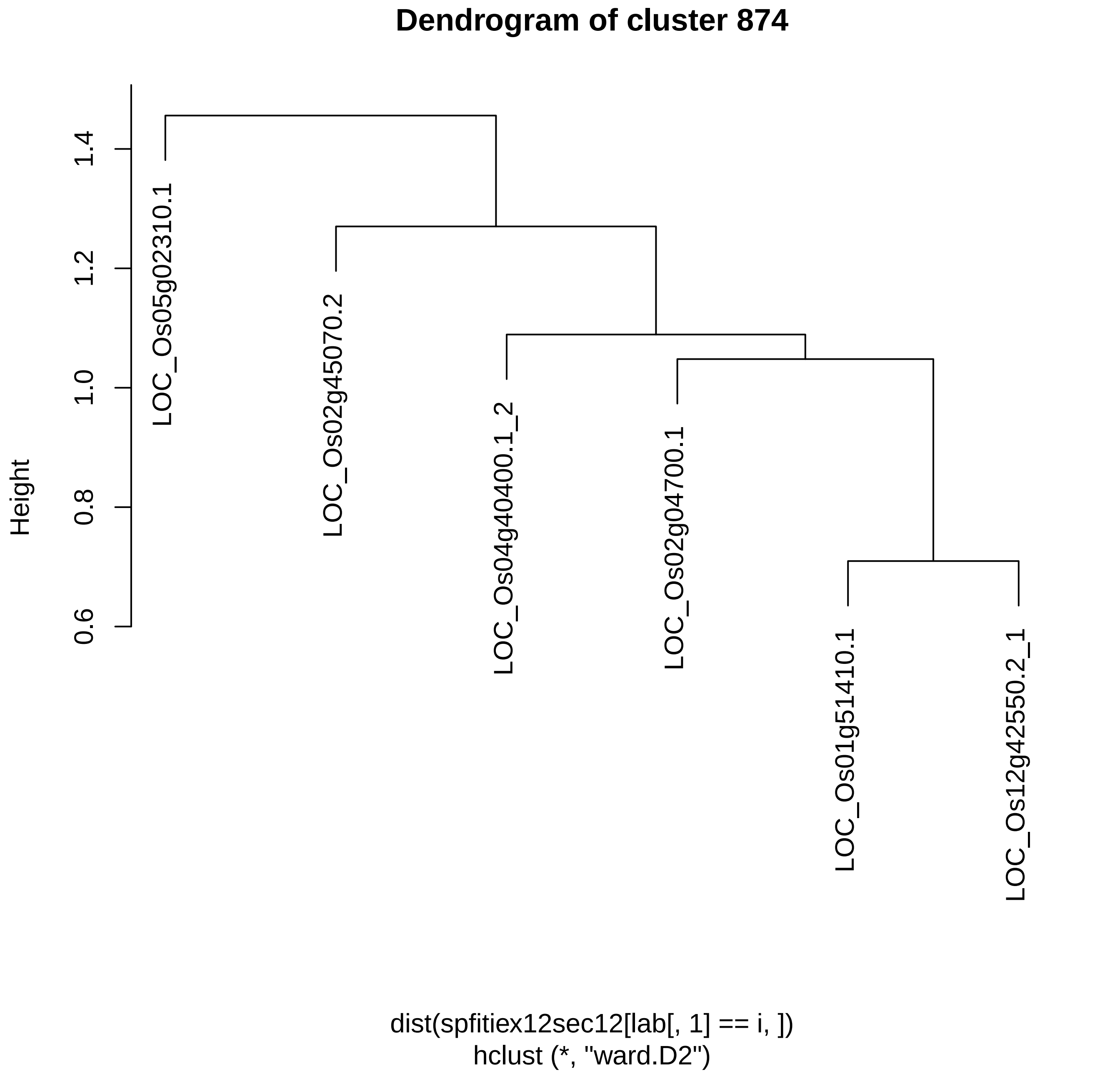

Dendrogram of cluster 874
Height
0.6
0.8
1.0
1.2
1.4
LOC_Os05g02310.1
LOC_Os02g45070.2
LOC_Os04g40400.1_2
LOC_Os02g04700.1
LOC_Os01g51410.1
LOC_Os12g42550.2_1
dist(spfitiex12sec12[lab[, 1] == i, ])
hclust (*, "ward.D2")

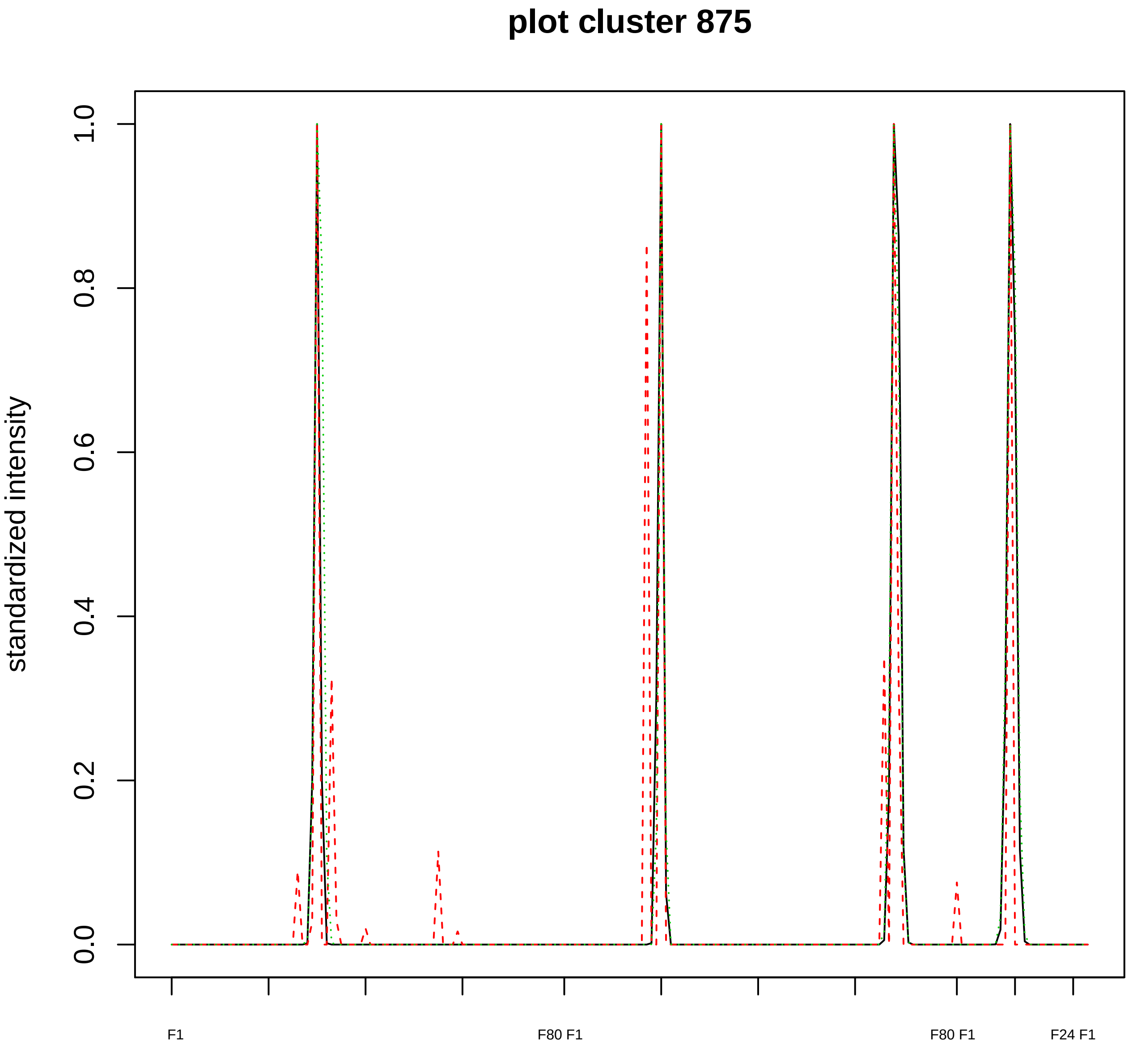

plot cluster 875
standardized intensity
0.0
0.2
0.4
0.6
0.8
1.0
F1
F80 F1
F80 F1
F24 F1

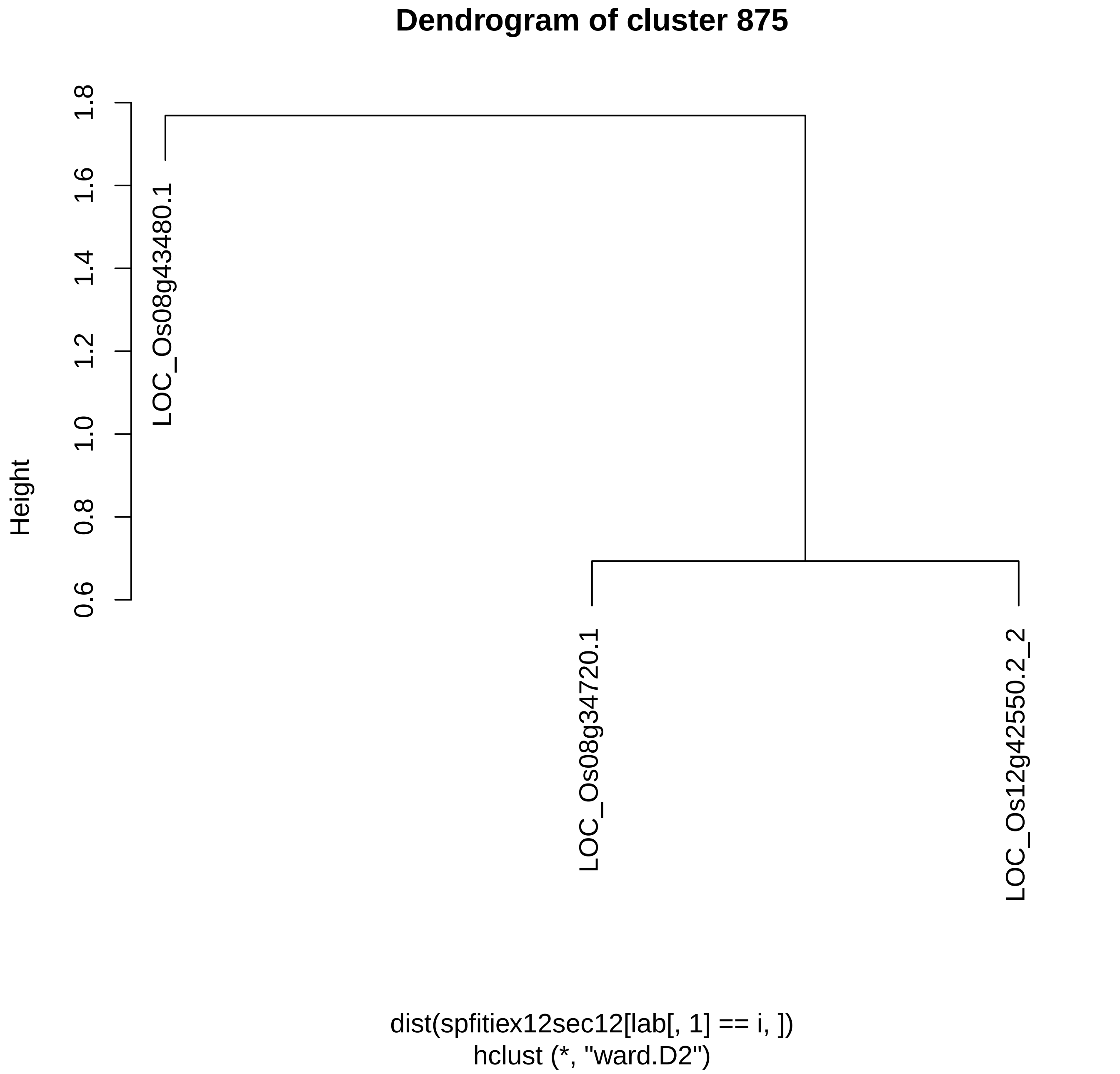

Dendrogram of cluster 875
Height
0.6
0.8
1.0
1.2
1.4
1.6
1.8
LOC_Os08g43480.1
LOC_Os08g34720.1
LOC_Os12g42550.2_2
dist(spfitiex12sec12[lab[, 1] == i, ])
hclust (*, "ward.D2")

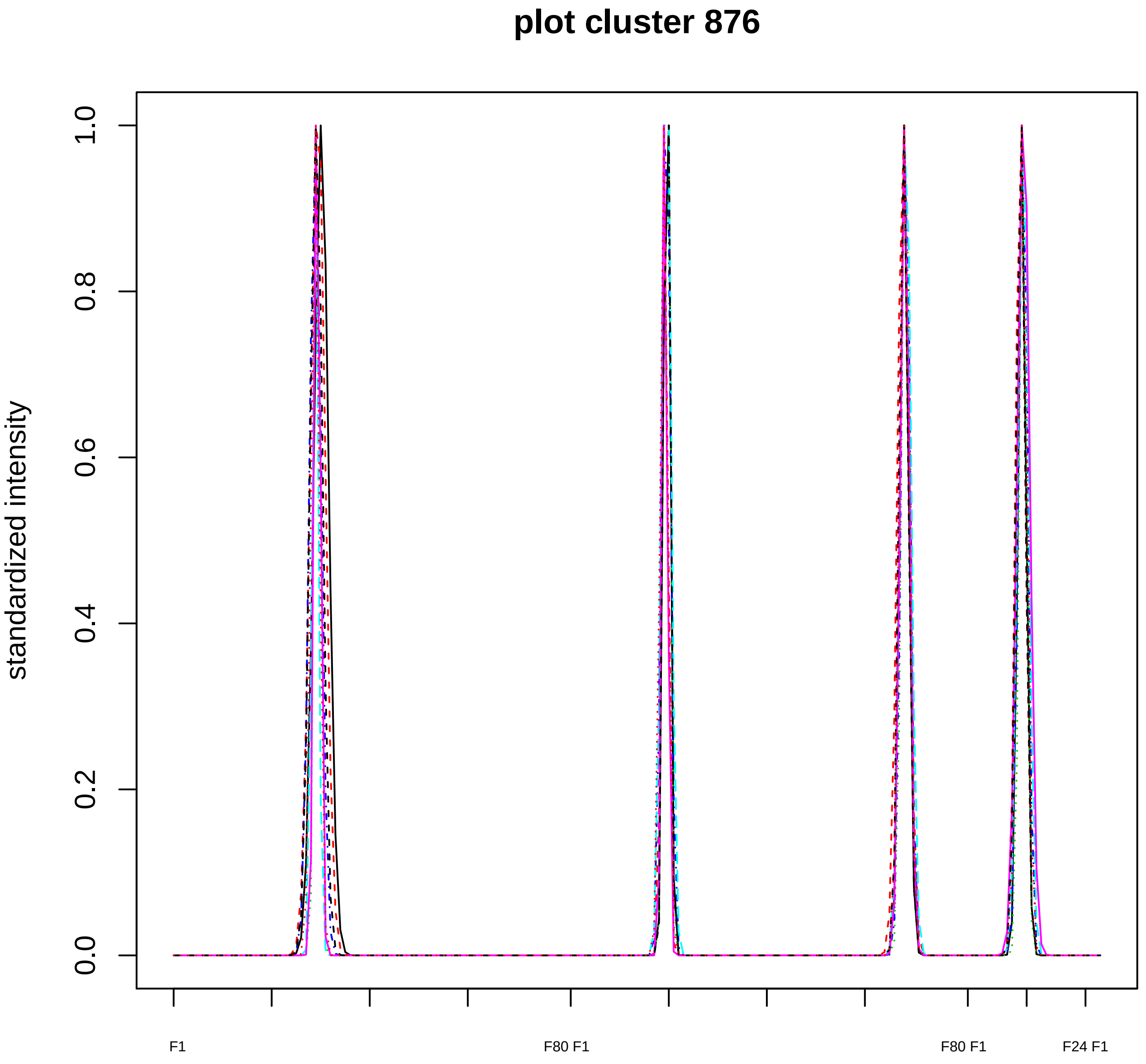
plot cluster 876
standardized intensity
0.0
0.2
0.4
0.6
0.8
1.0
F1
F80 F1
F80 F1
F24 F1

**Dendrogram of cluster 876**

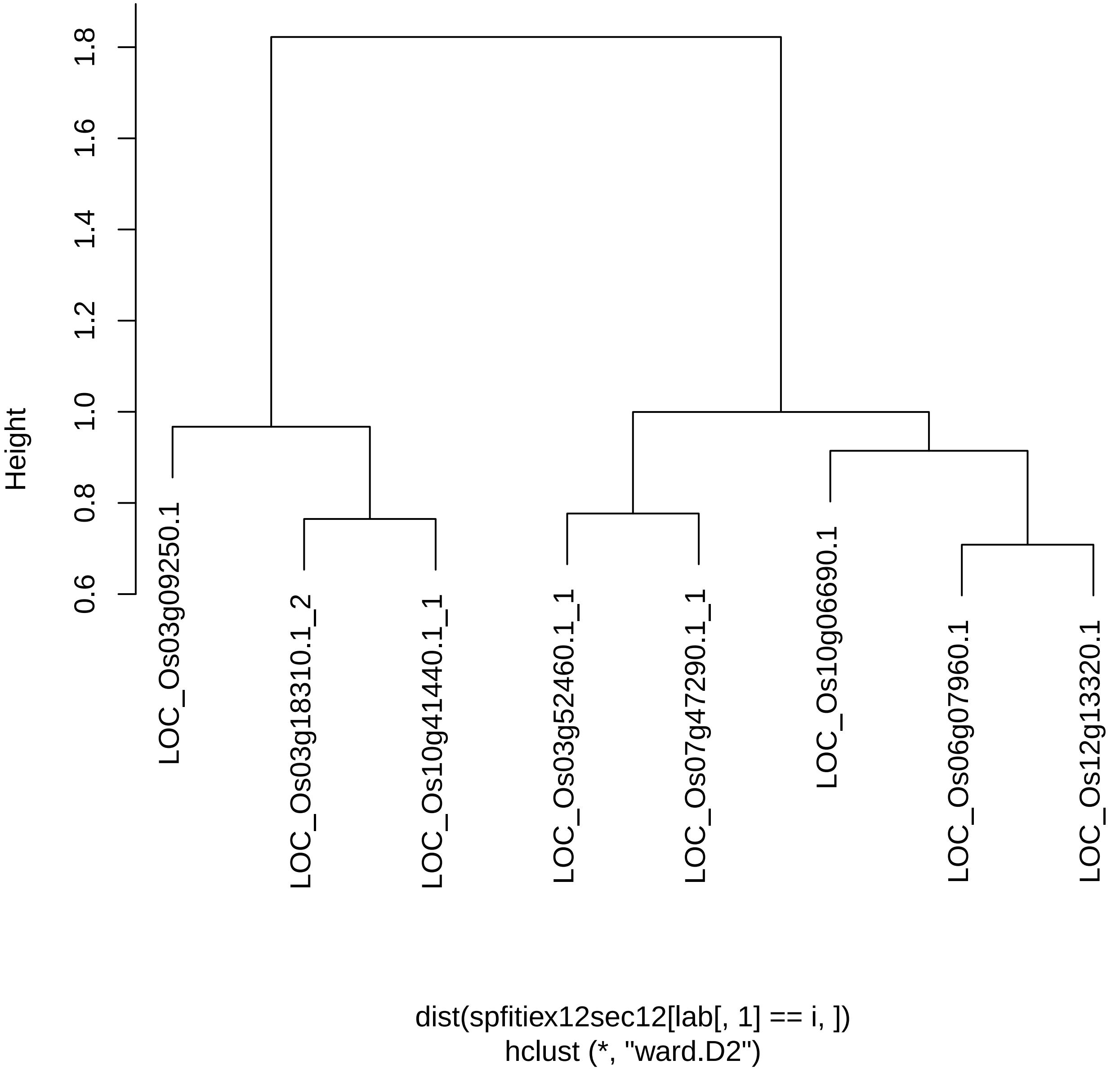

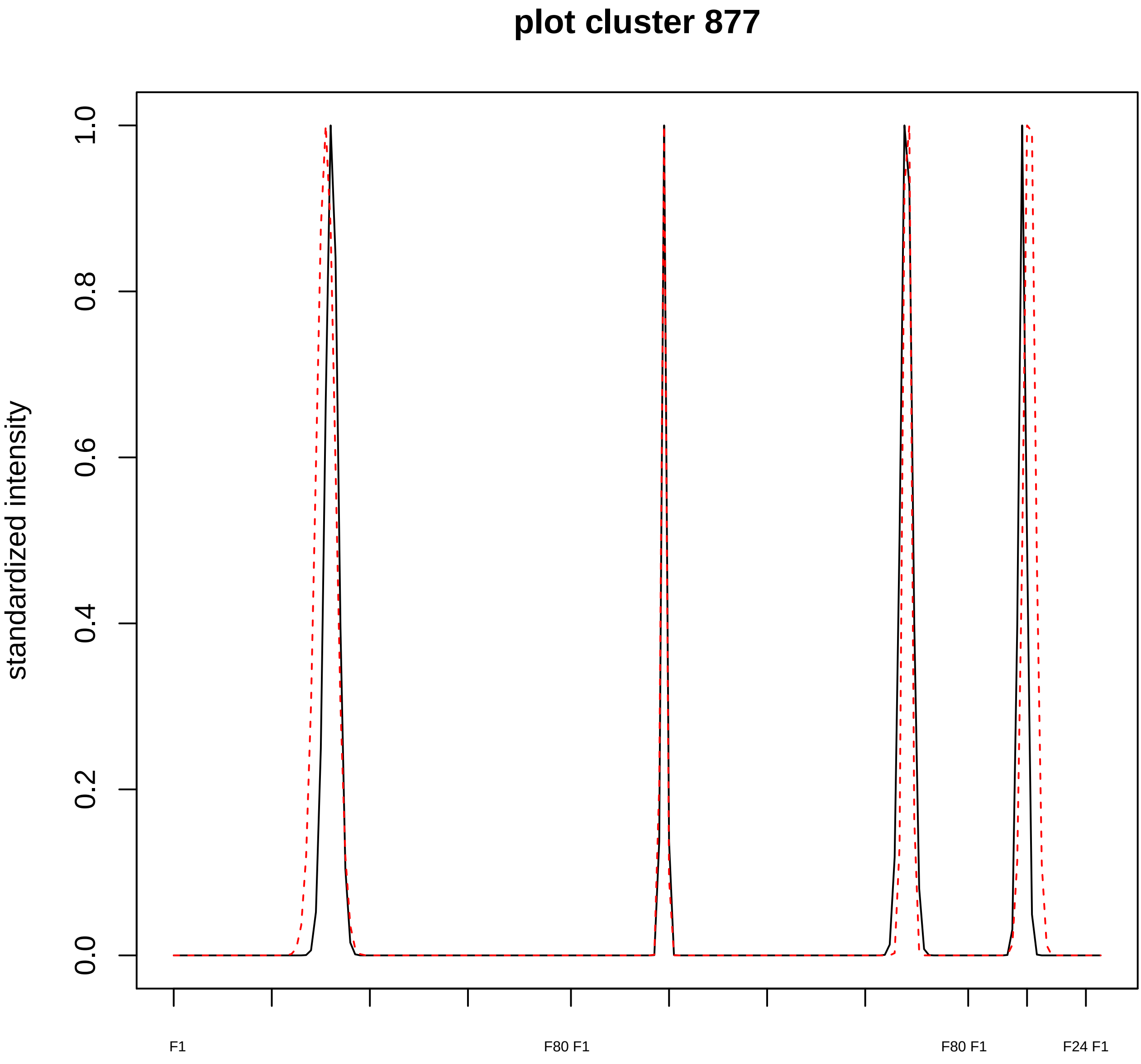

plot cluster 877
standardized intensity
0.0
0.2
0.4
0.6
0.8
1.0
F1
F80 F1
F80 F1
F24 F1

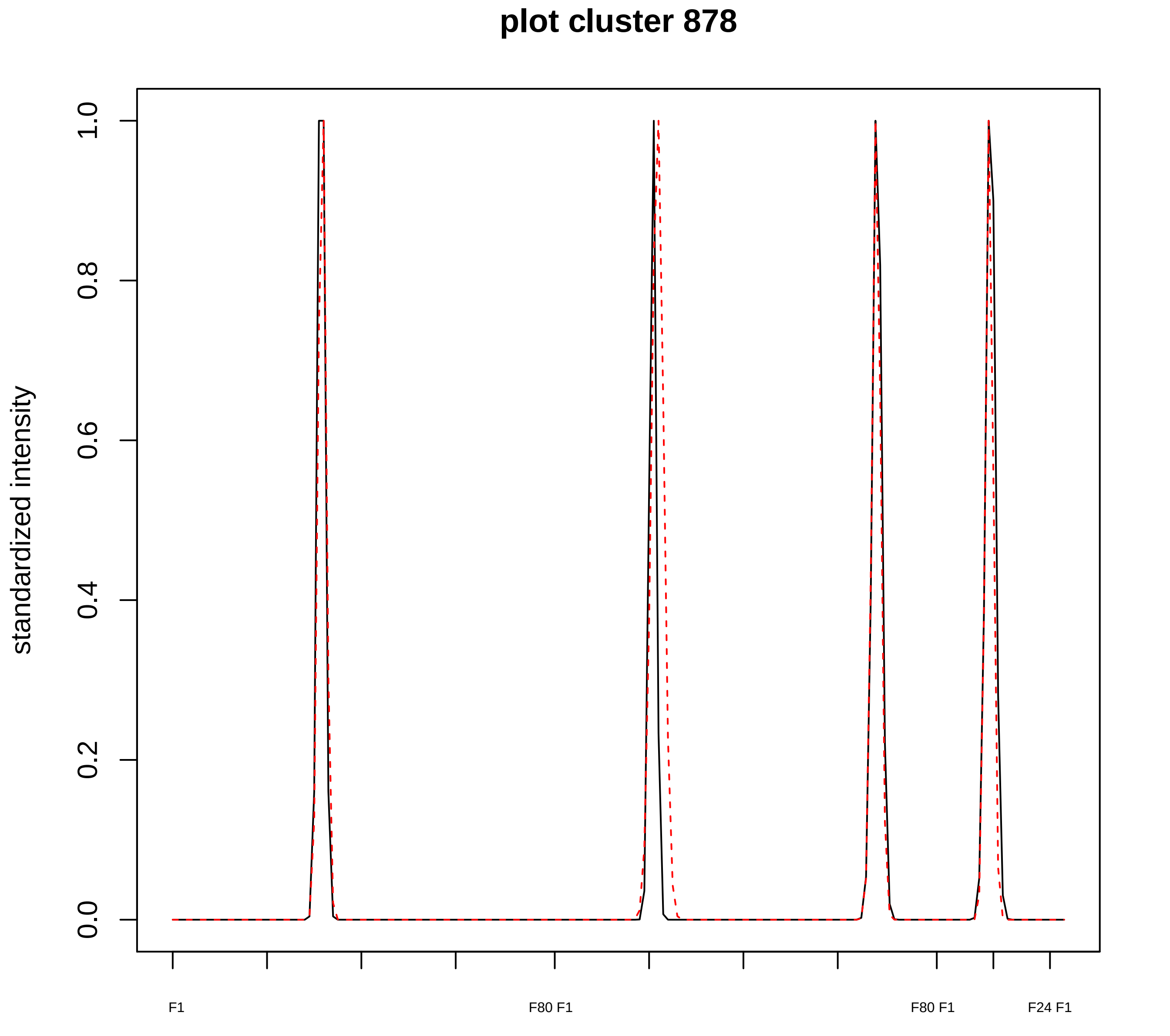

plot cluster 878
standardized intensity
0.0
0.2
0.4
0.6
0.8
1.0
F1
F80 F1
F80 F1
F24 F1

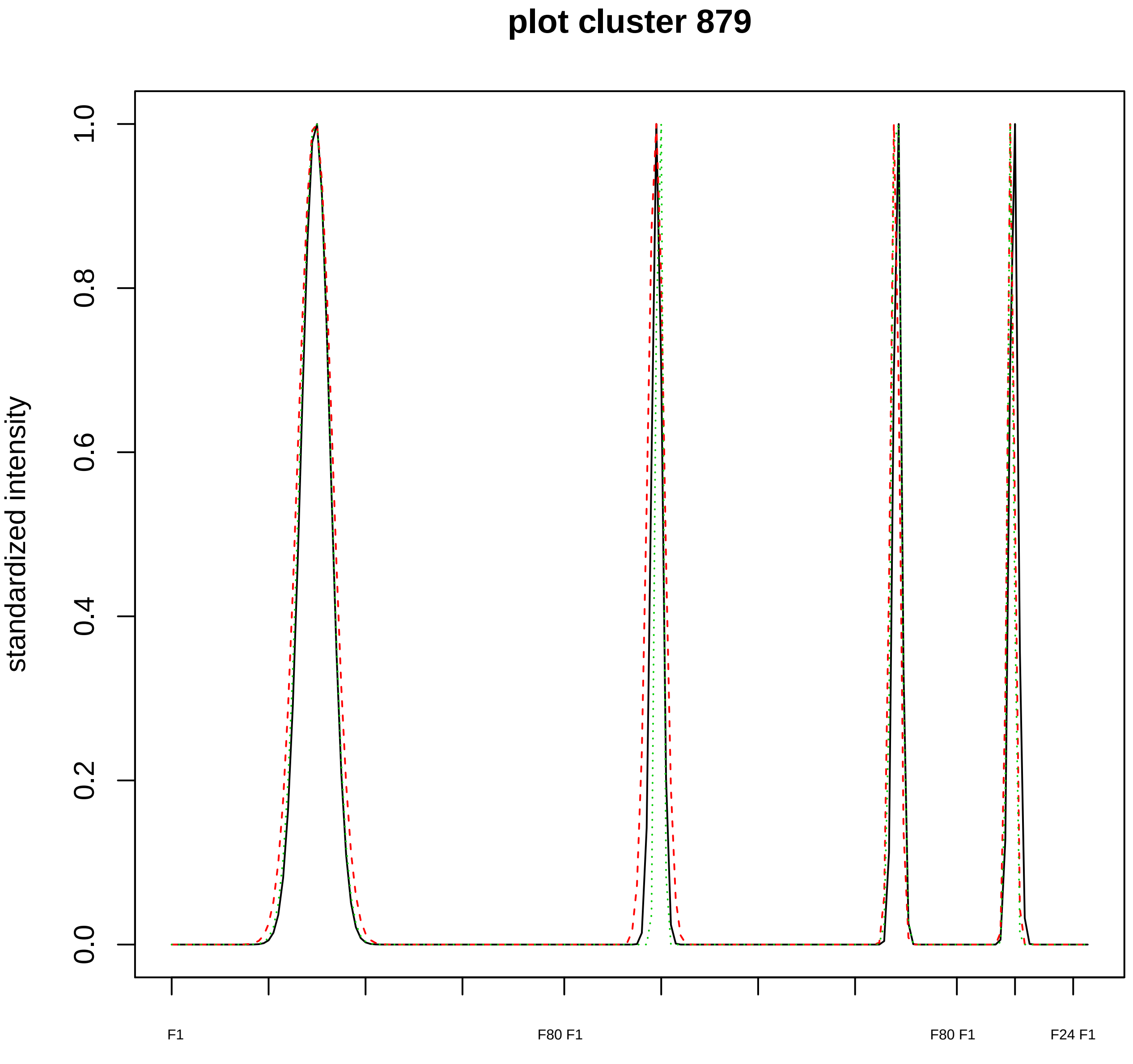

plot cluster 879
standardized intensity
0.0
0.2
0.4
0.6
0.8
1.0
F1
F80 F1
F80 F1
F24 F1

#### Dendrogram of cluster 879

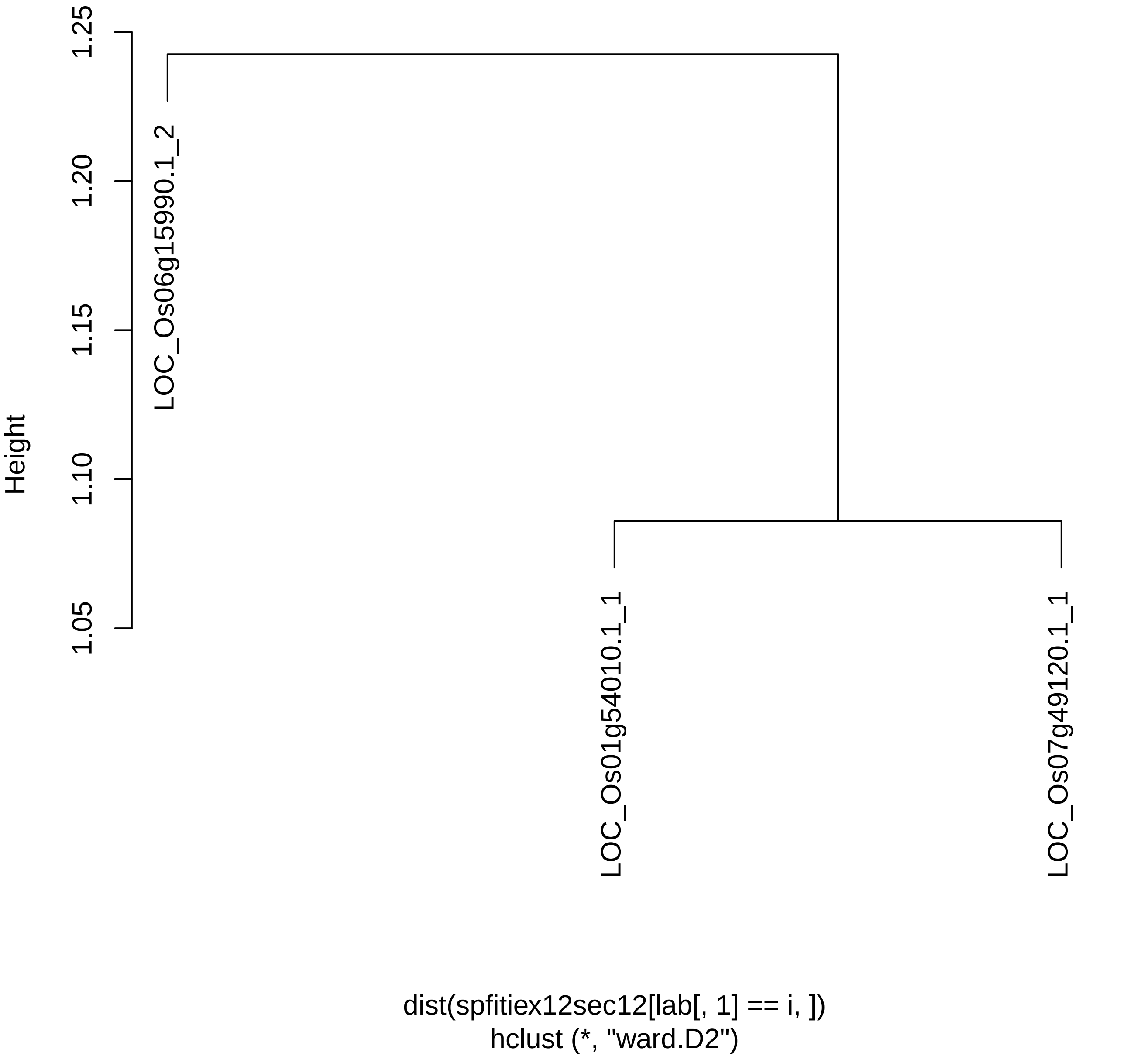

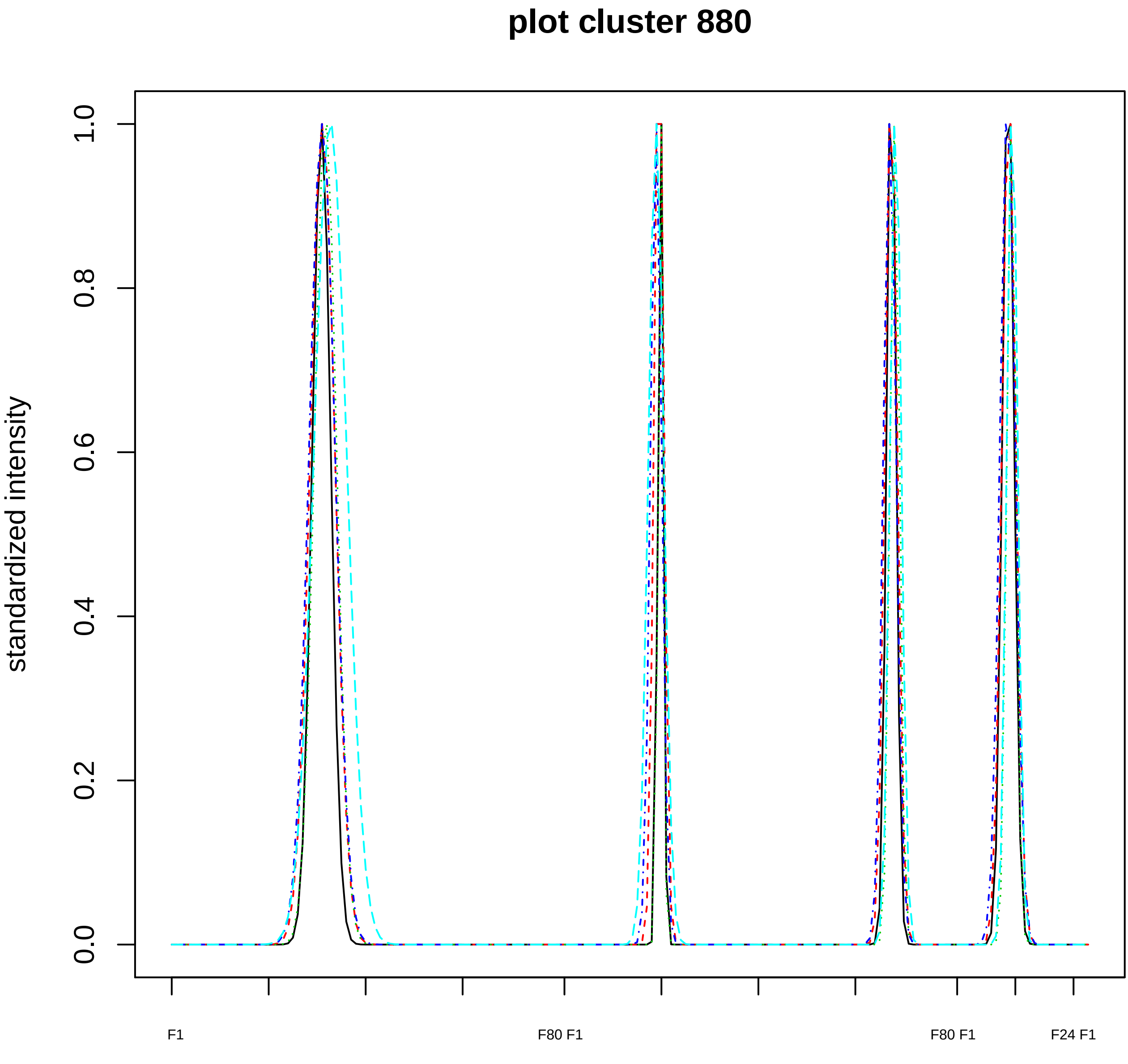
plot cluster 880
standardized intensity
0.0
0.2
0.4
0.6
0.8
1.0
F1
F80 F1
F80 F1
F24 F1

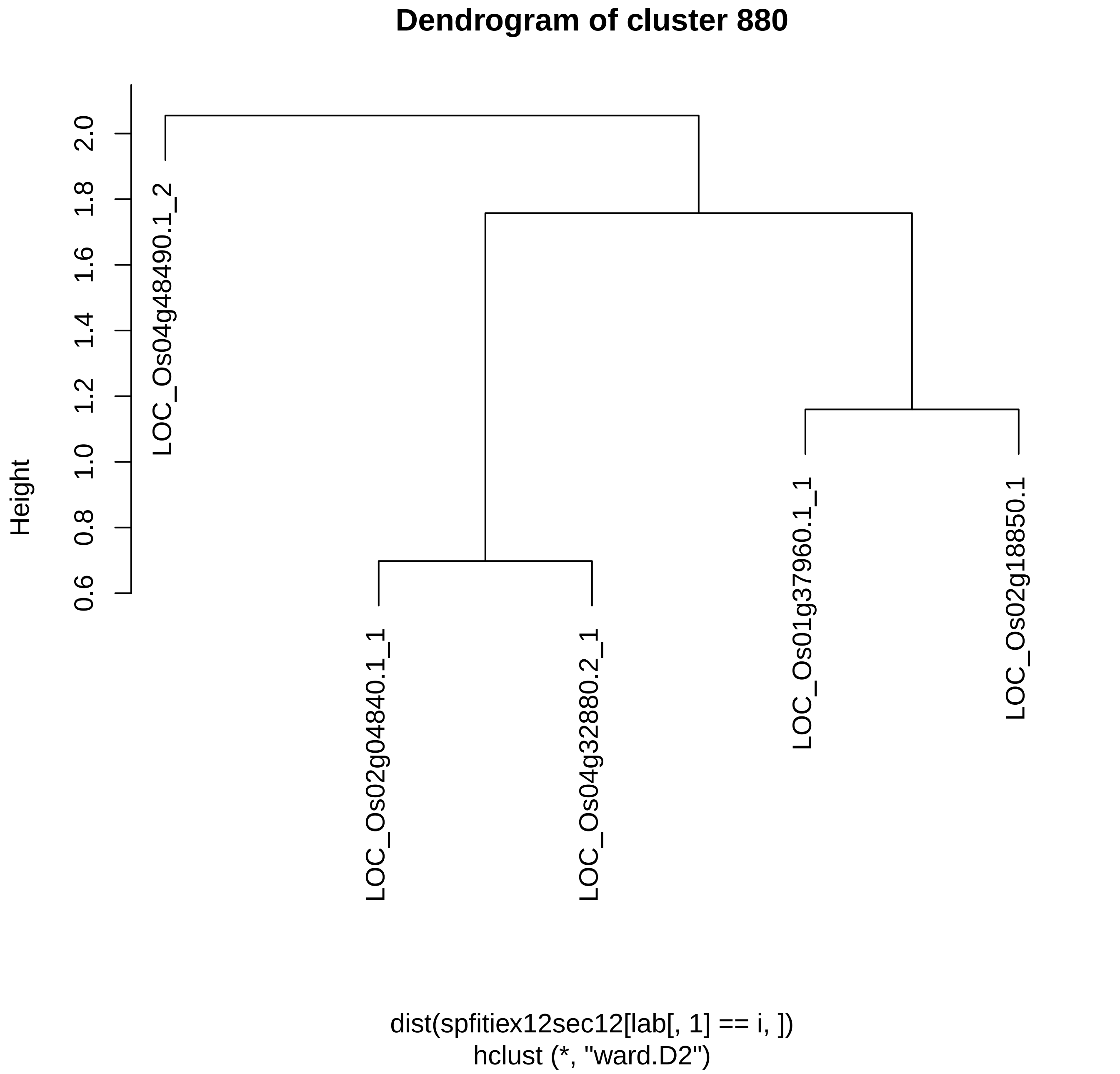
Dendrogram of cluster 880
Height
0.6
0.8
1.0
1.2
1.4
1.6
1.8
2.0
LOC_Os04g48490.1_2
LOC_Os02g04840.1_1
LOC_Os04g32880.2_1
LOC_Os01g37960.1_1
LOC_Os02g18850.1
dist(spfitiex12sec12[lab[, 1] == i, ])
hclust (*, "ward.D2")

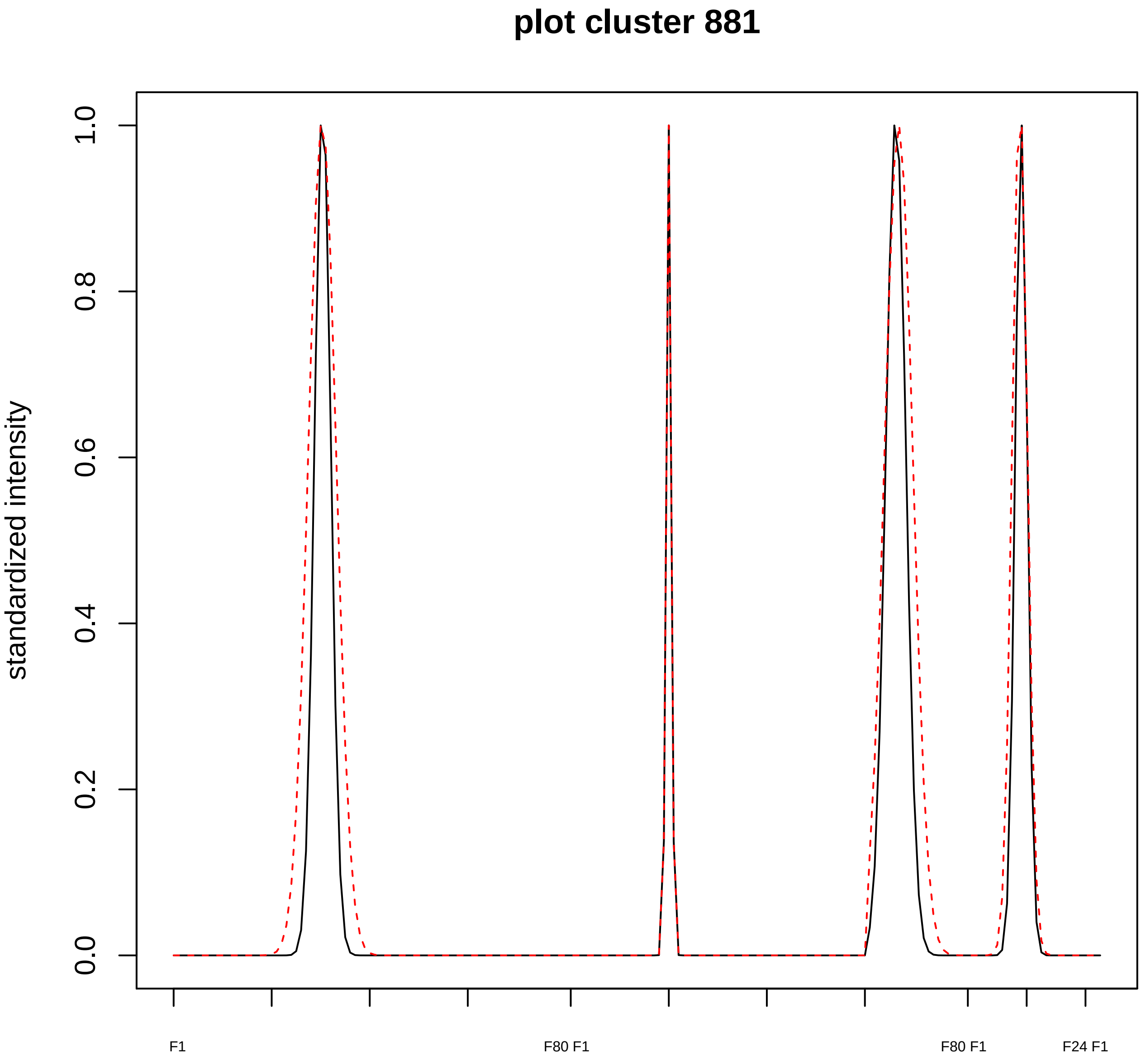
plot cluster 881
standardized intensity
0.0
0.2
0.4
0.6
0.8
1.0
F1
F80 F1
F80 F1
F24 F1

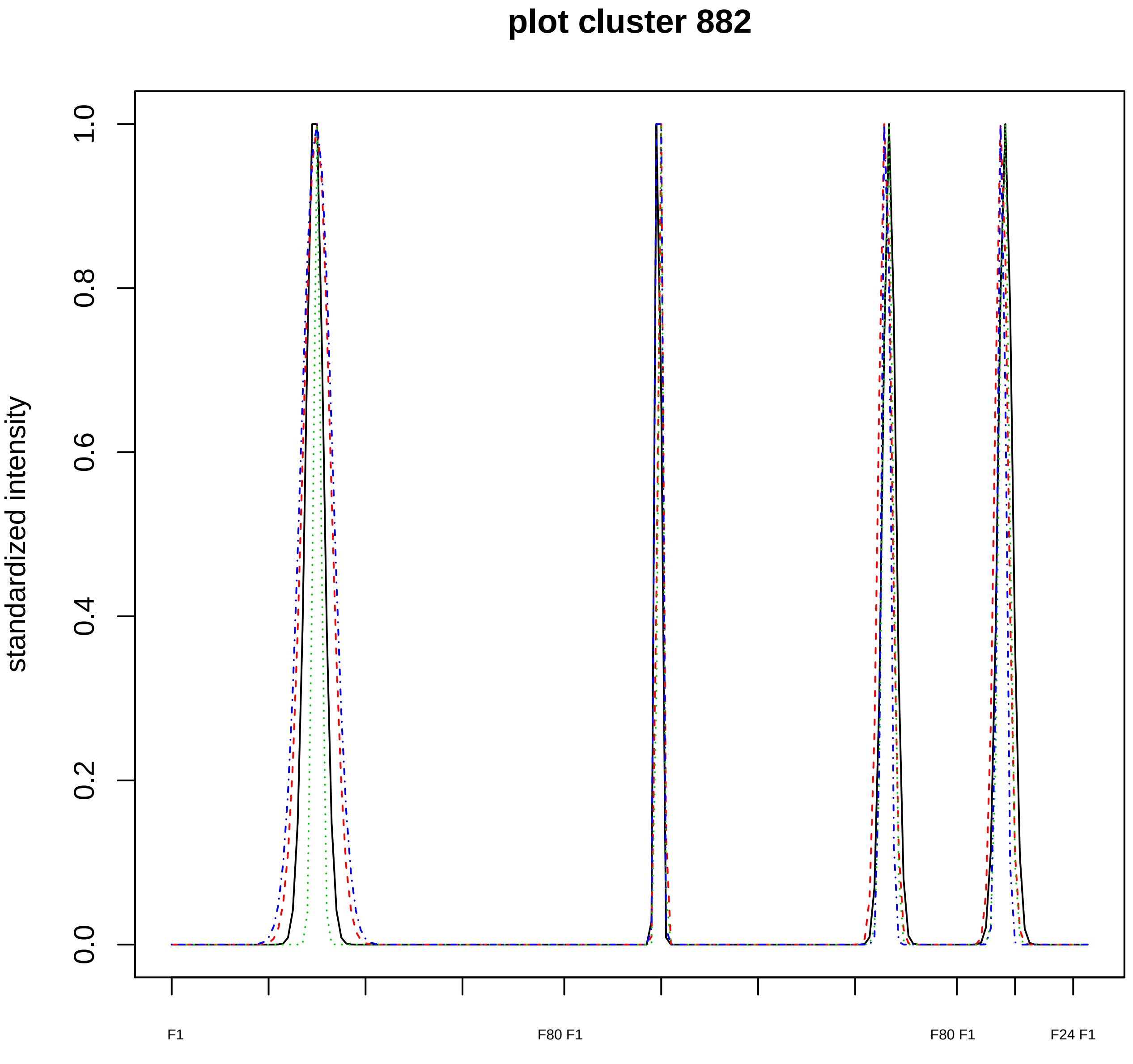

plot cluster 882
standardized intensity
0.0
0.2
0.4
0.6
0.8
1.0
F1
F80 F1
F80 F1
F24 F1

### Dendrogram of cluster 882

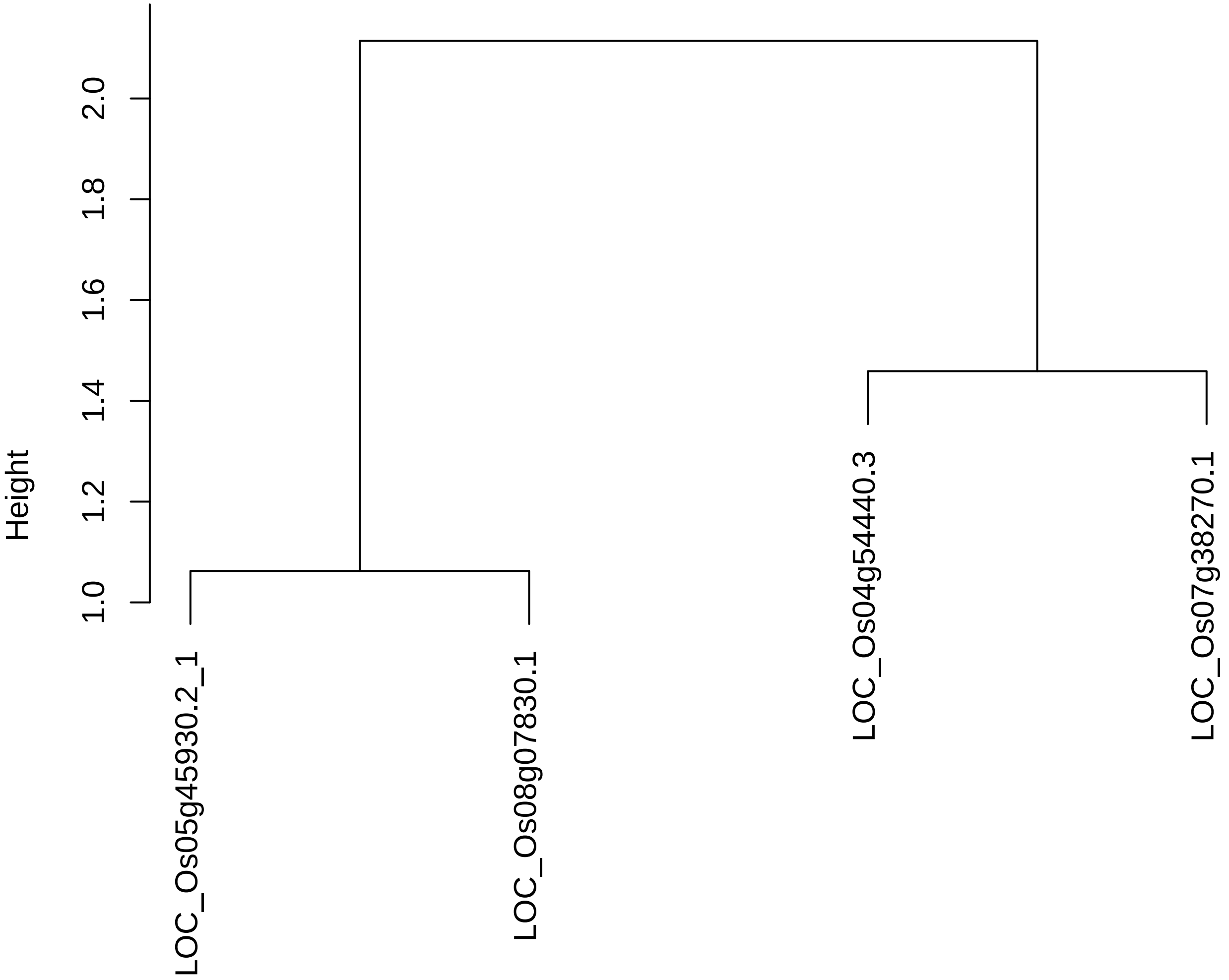


dist(spfitiex12sec12[lab[, 1] == i, ])
hclust (*, "ward.D2")

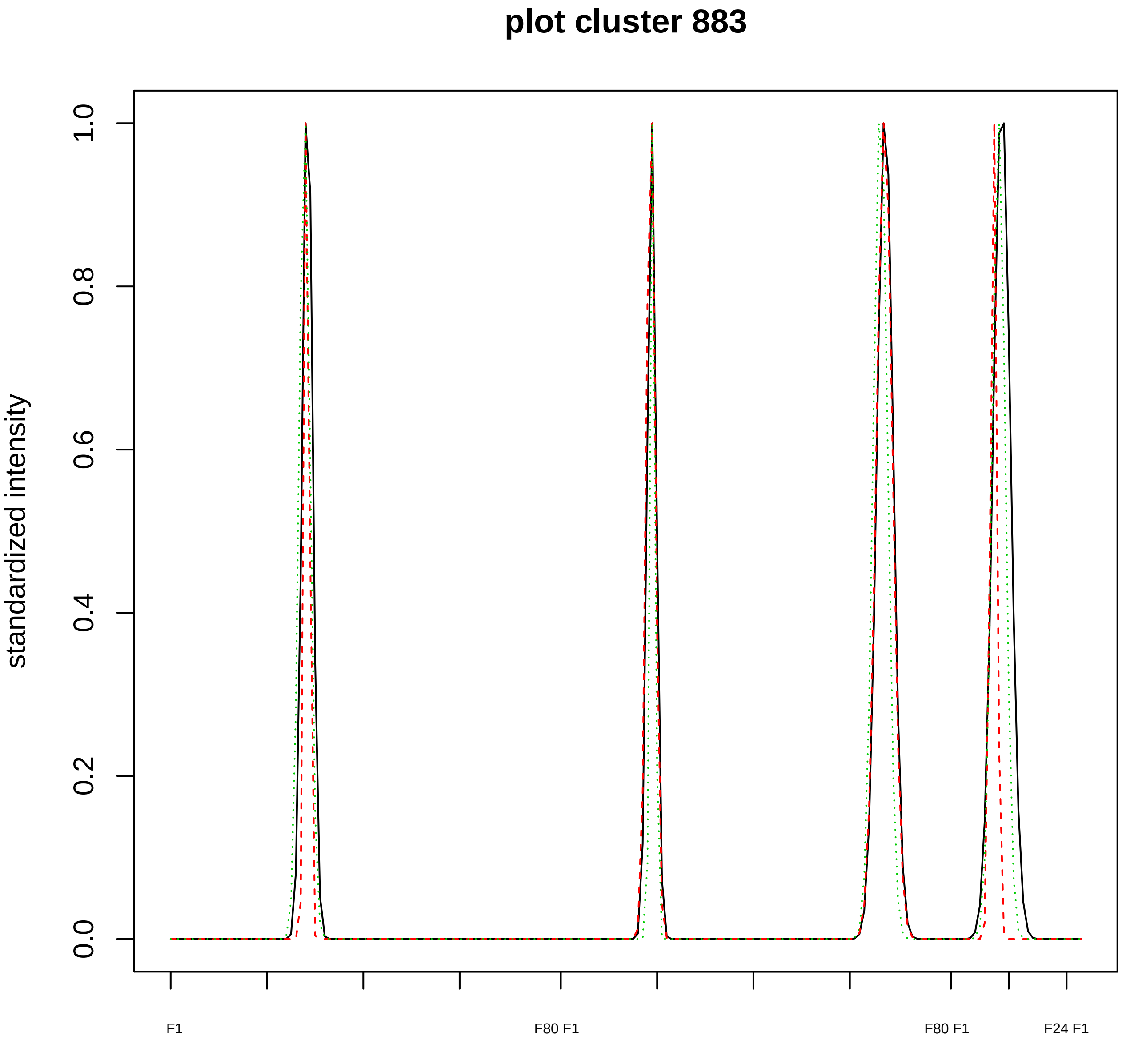

plot cluster 883
standardized intensity
0.0
0.2
0.4
0.6
0.8
1.0
F1
F80 F1
F80 F1
F24 F1

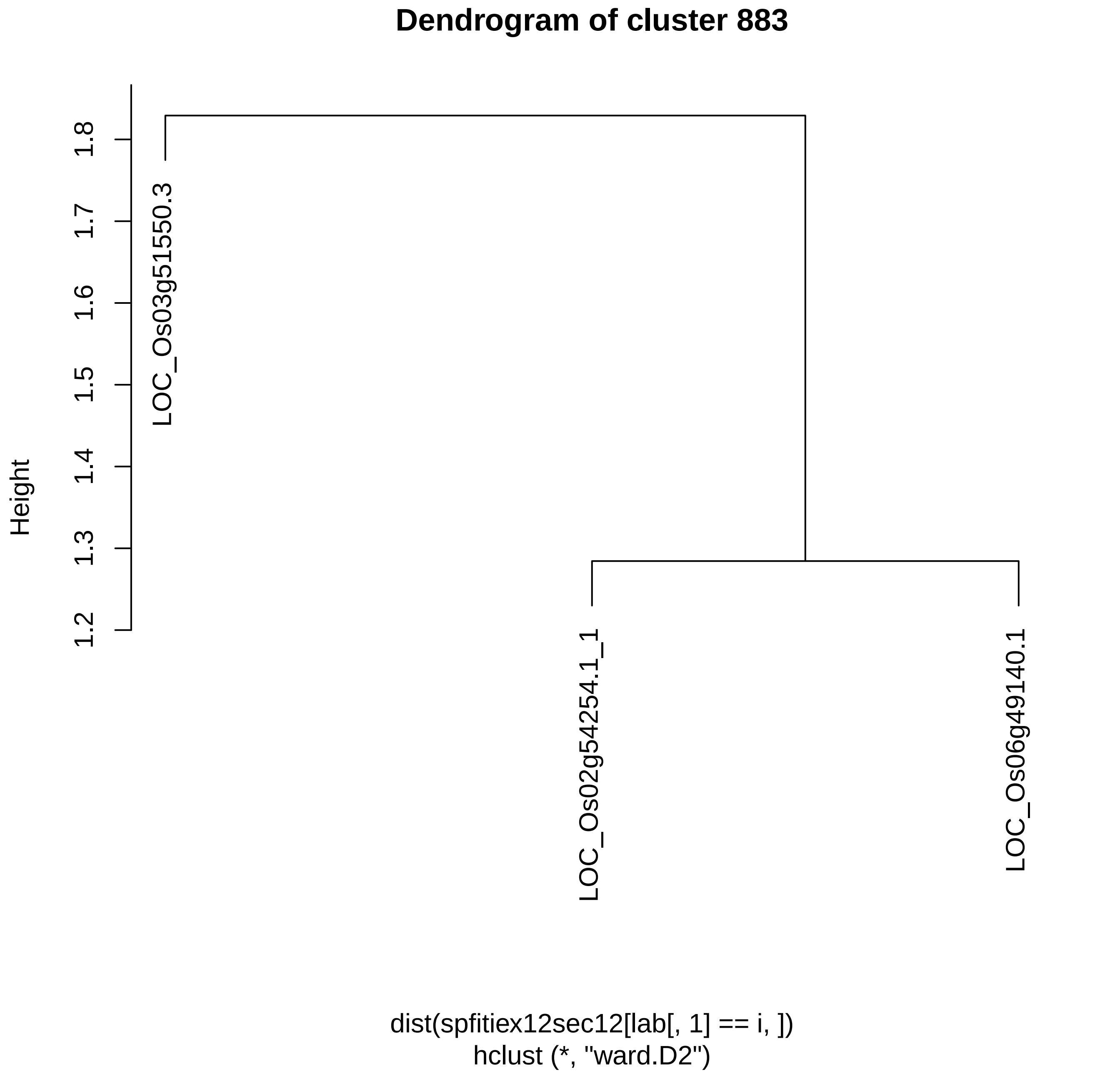
Dendrogram of cluster 883
Height
1.2
1.3
1.4
1.5
1.6
1.7
1.8
LOC_Os03g51550.3
LOC_Os02g54254.1_1
LOC_Os06g49140.1
dist(spfitiex12sec12[lab[, 1] == i, ])
hclust (*, "ward.D2")

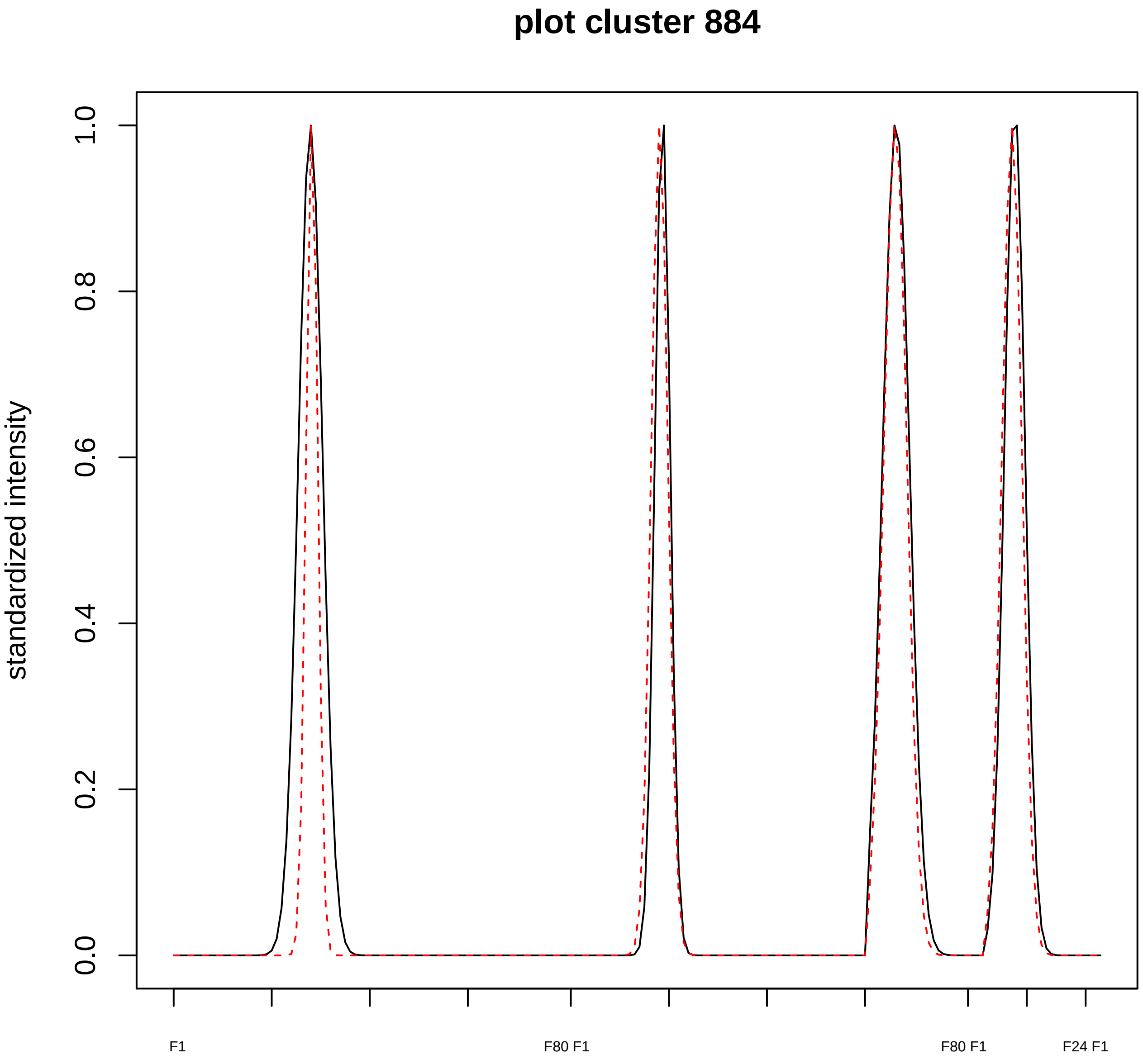
plot cluster 884
standardized intensity
1.0
0.8
0.6
0.4
0.2
0.0
F1
F80 F1
F80 F1
F24 F1

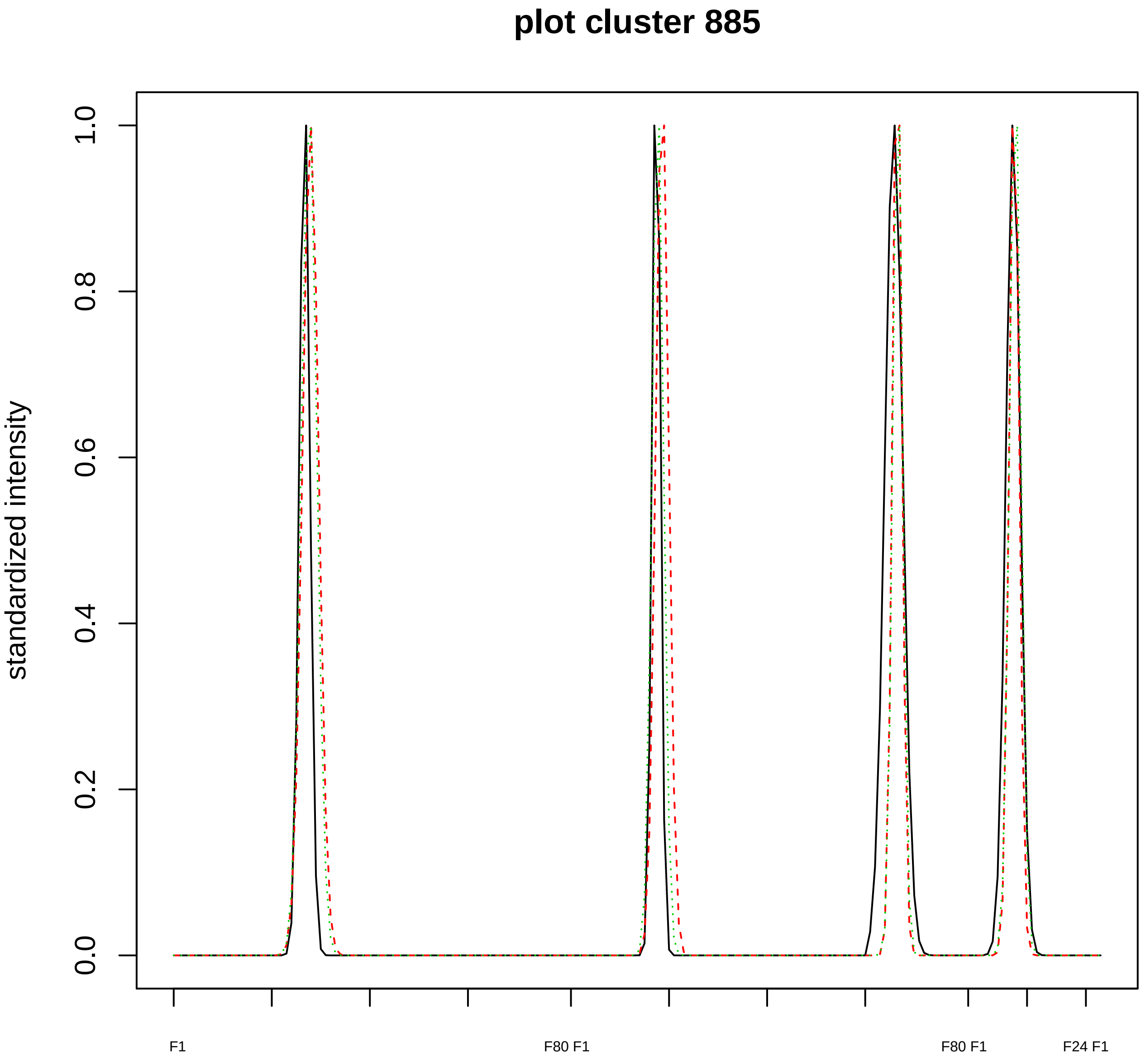

plot cluster 885
standardized intensity
0.0
0.2
0.4
0.6
0.8
1.0
F1
F80 F1
F80 F1
F24 F1

#### Dendrogram of cluster 885

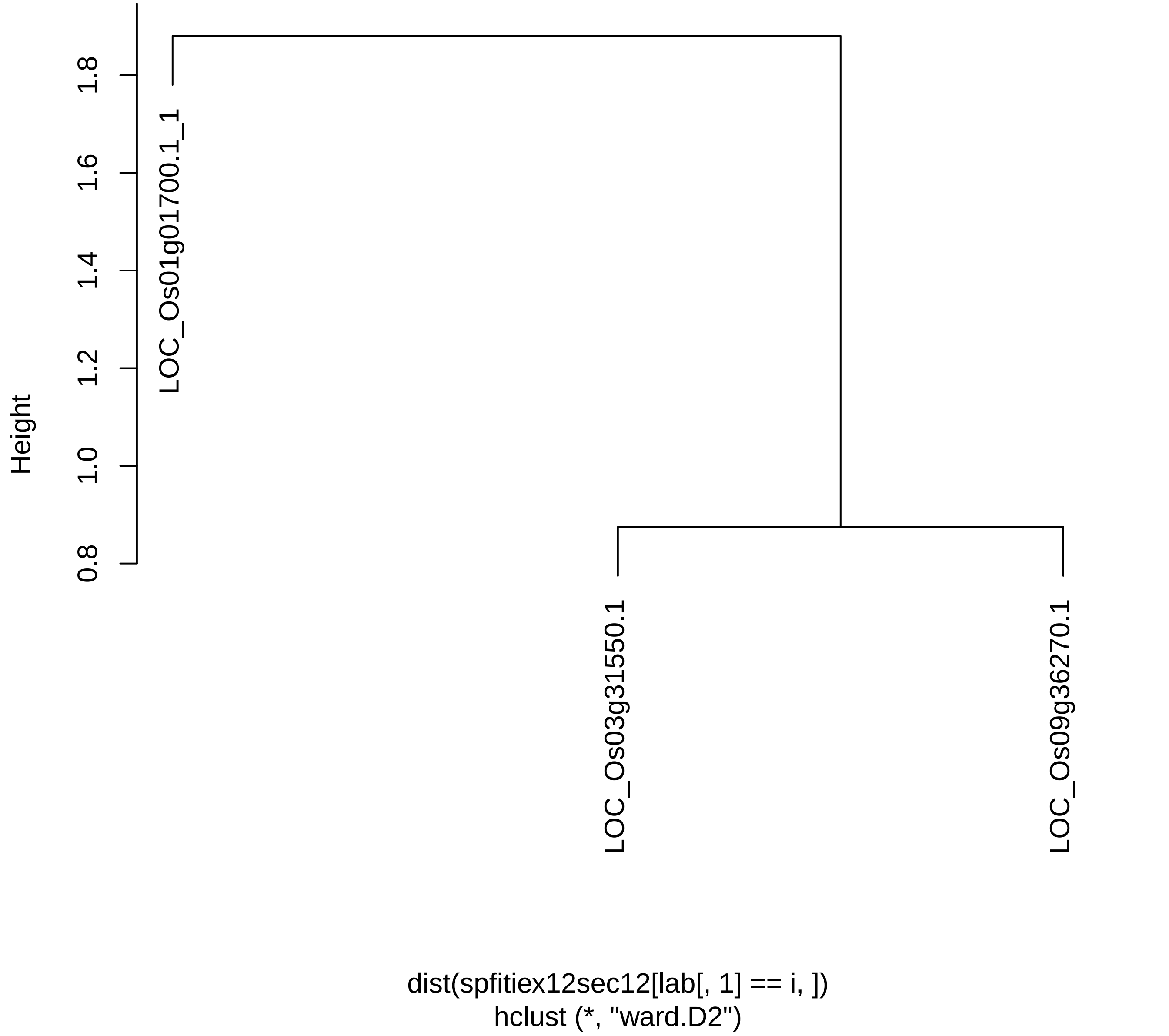

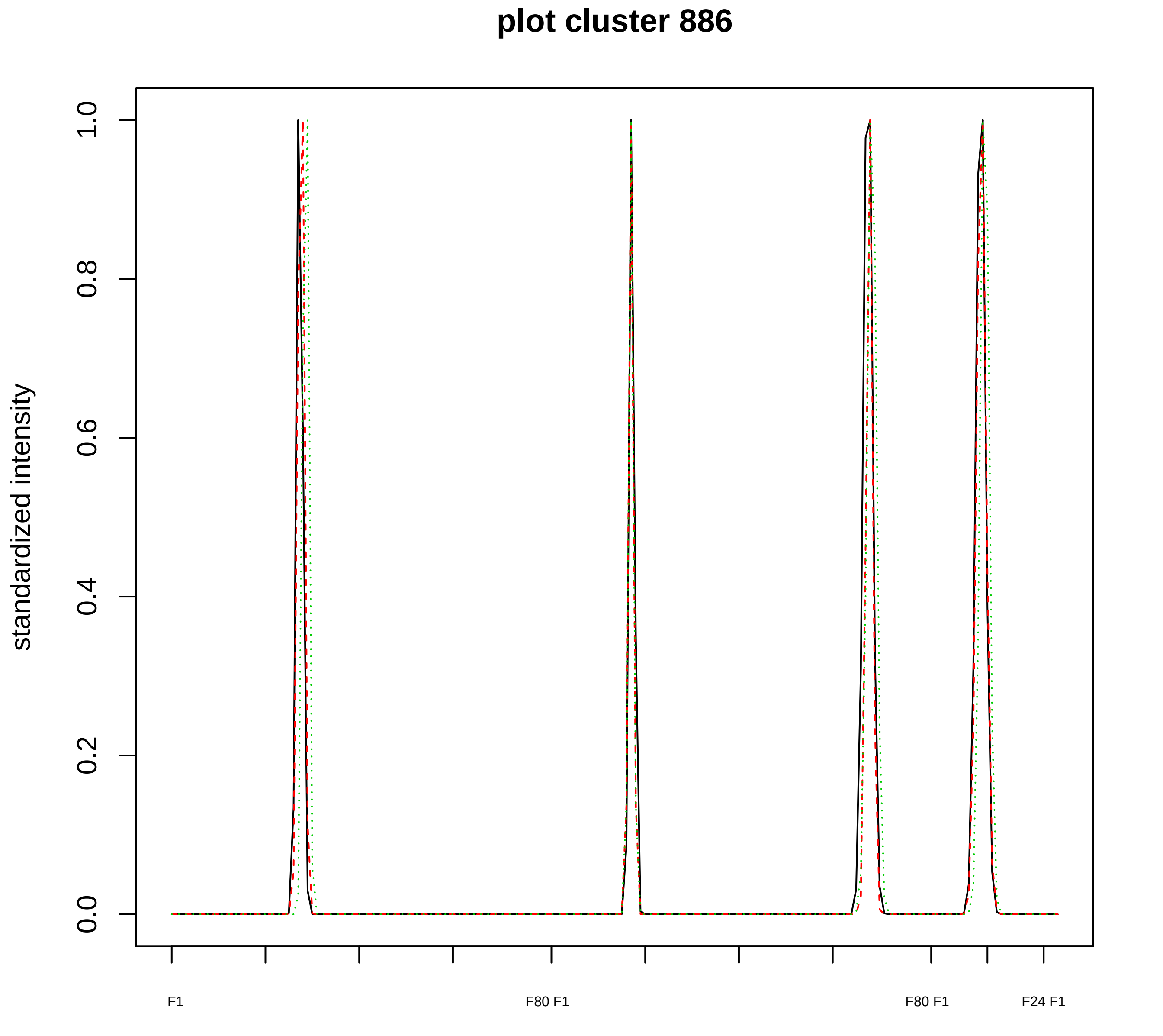
plot cluster 886
standardized intensity
0.0
0.2
0.4
0.6
0.8
1.0
F1
F80 F1
F80 F1
F24 F1

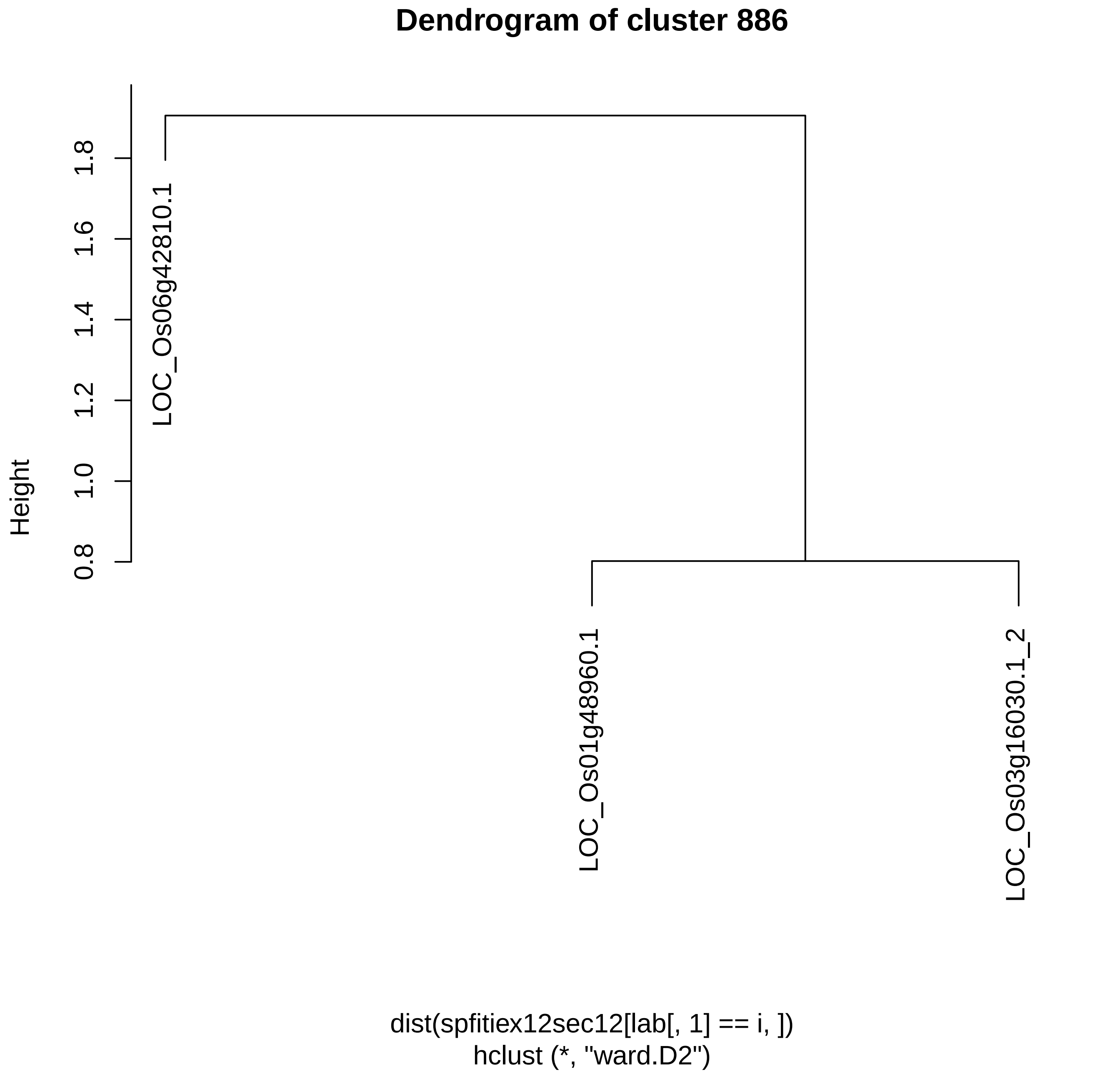

Dendrogram of cluster 886
Height
0.8
1.0
1.2
1.4
1.6
1.8
LOC_Os06g42810.1
LOC_Os01g48960.1
LOC_Os03g16030.1_2
dist(spfitiex12sec12[lab[, 1] == i, ])
hclust (*, "ward.D2")

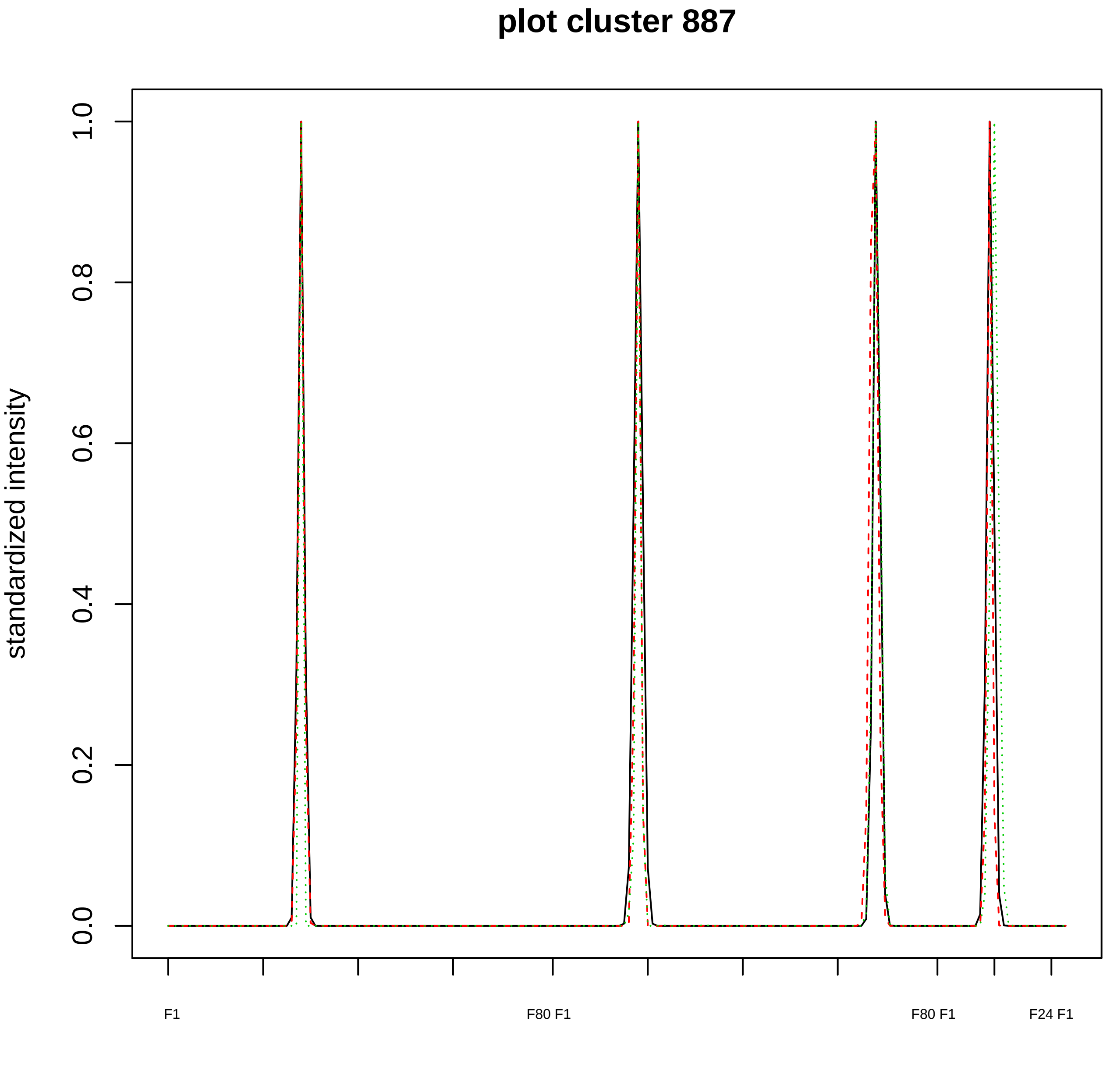

plot cluster 887
standardized intensity
0.0
0.2
0.4
0.6
0.8
1.0
F1
F80 F1
F80 F1
F24 F1

### Dendrogram of cluster 887

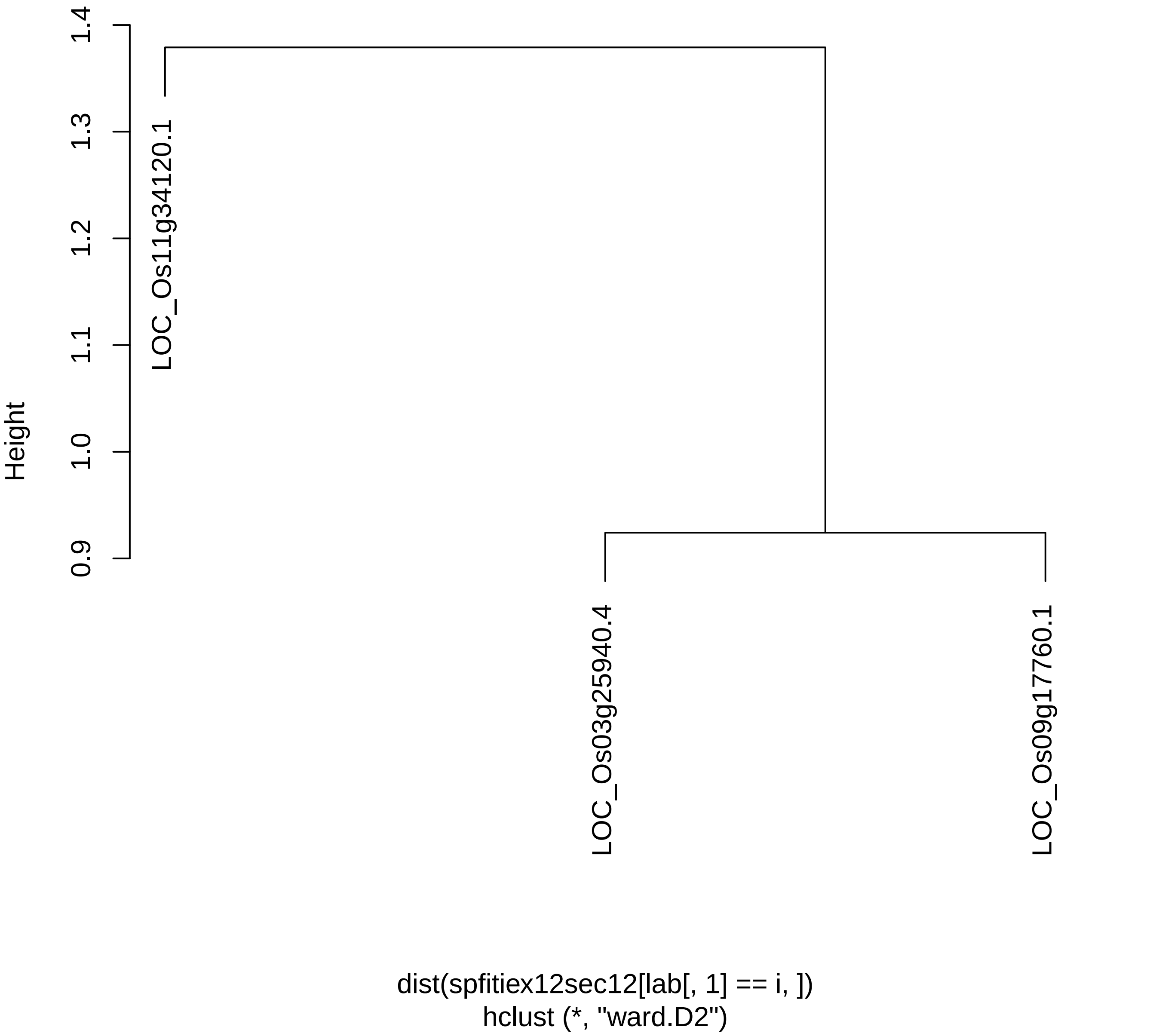

### plot cluster 888

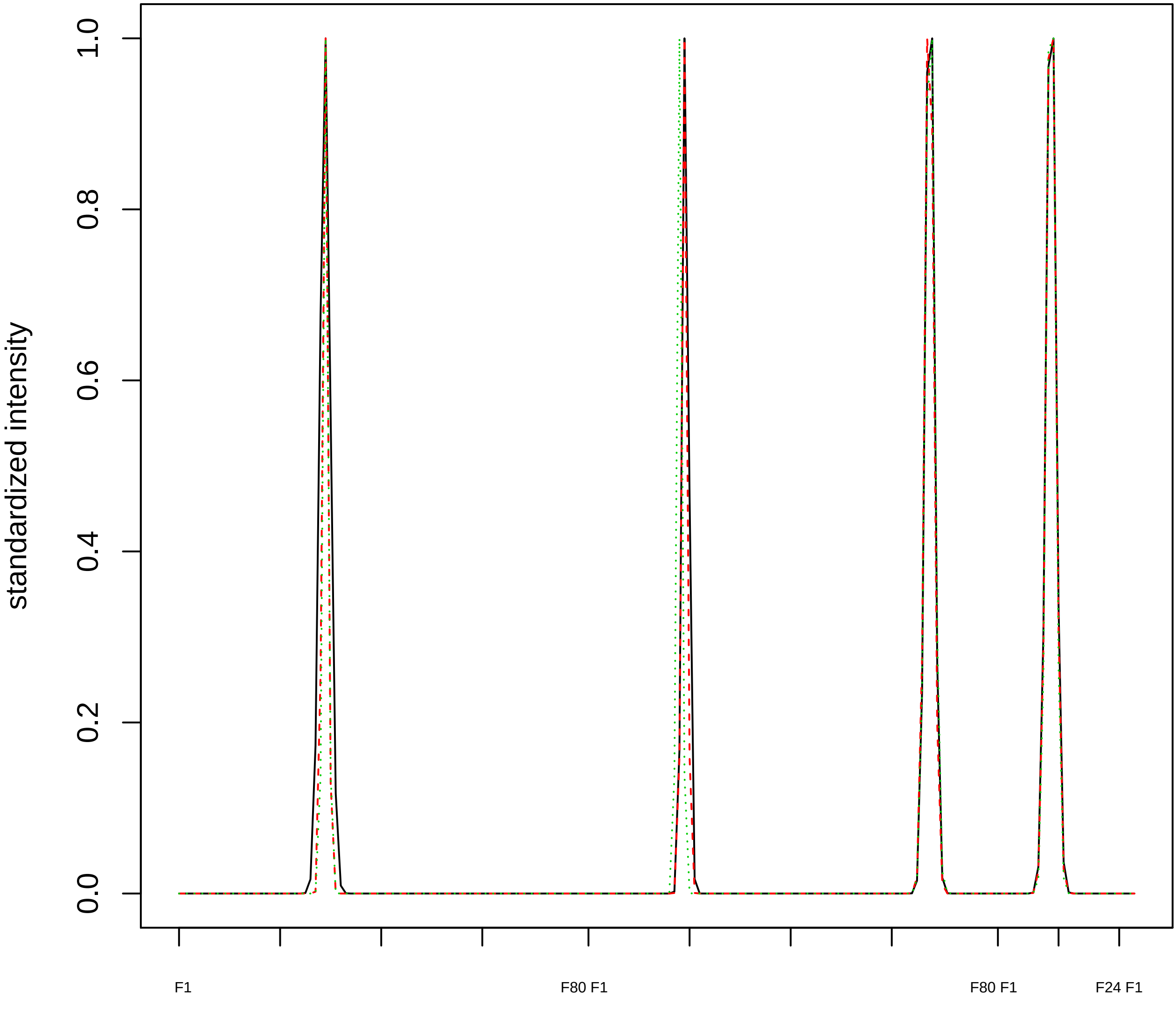

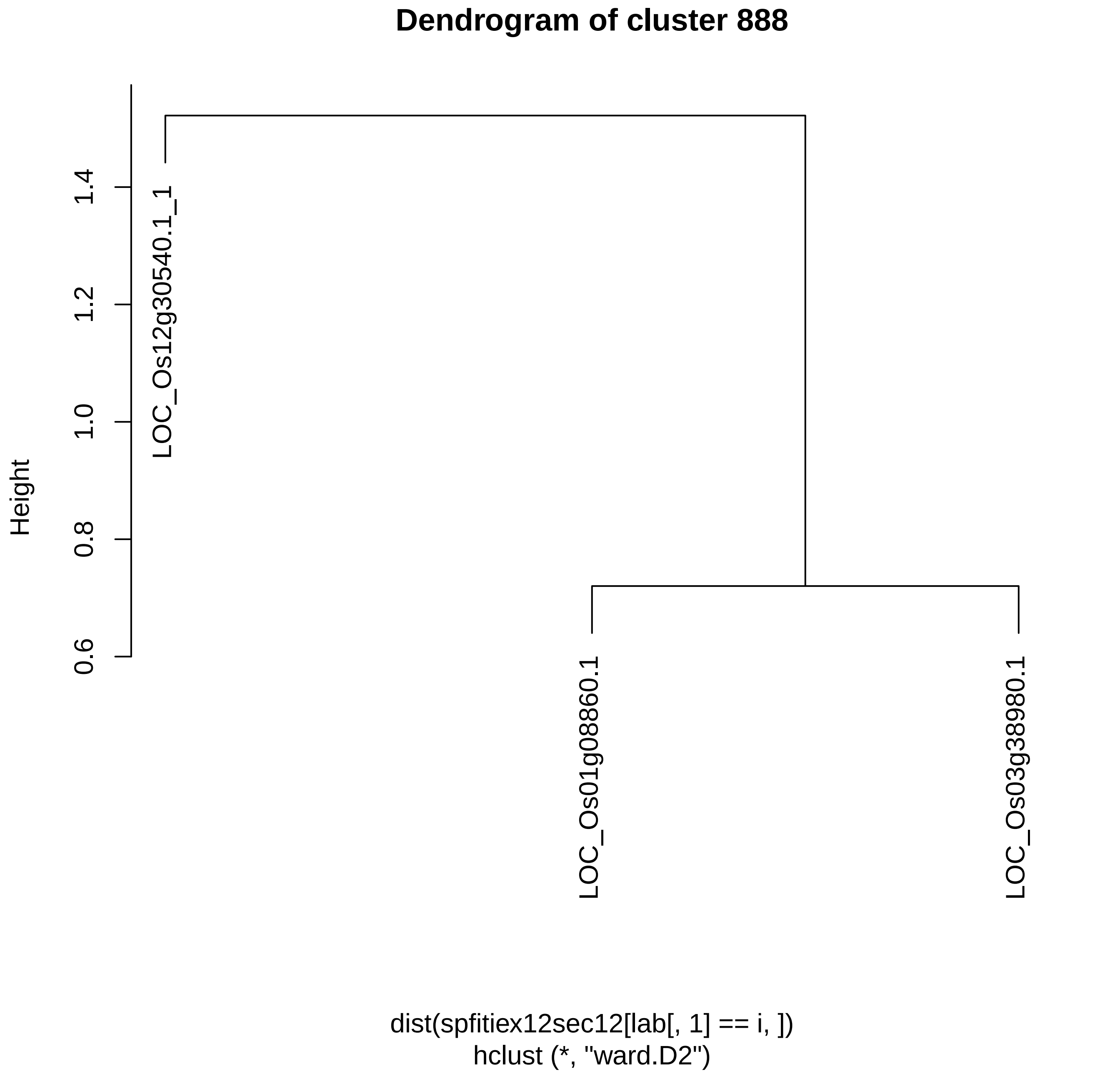
Dendrogram of cluster 888
Height
0.6
0.8
1.0
1.2
1.4
LOC_Os12g30540.1_1
LOC_Os01g08860.1
LOC_Os03g38980.1
dist(spfitiex12sec12[lab[, 1] == i, ])
hclust (*, "ward.D2")

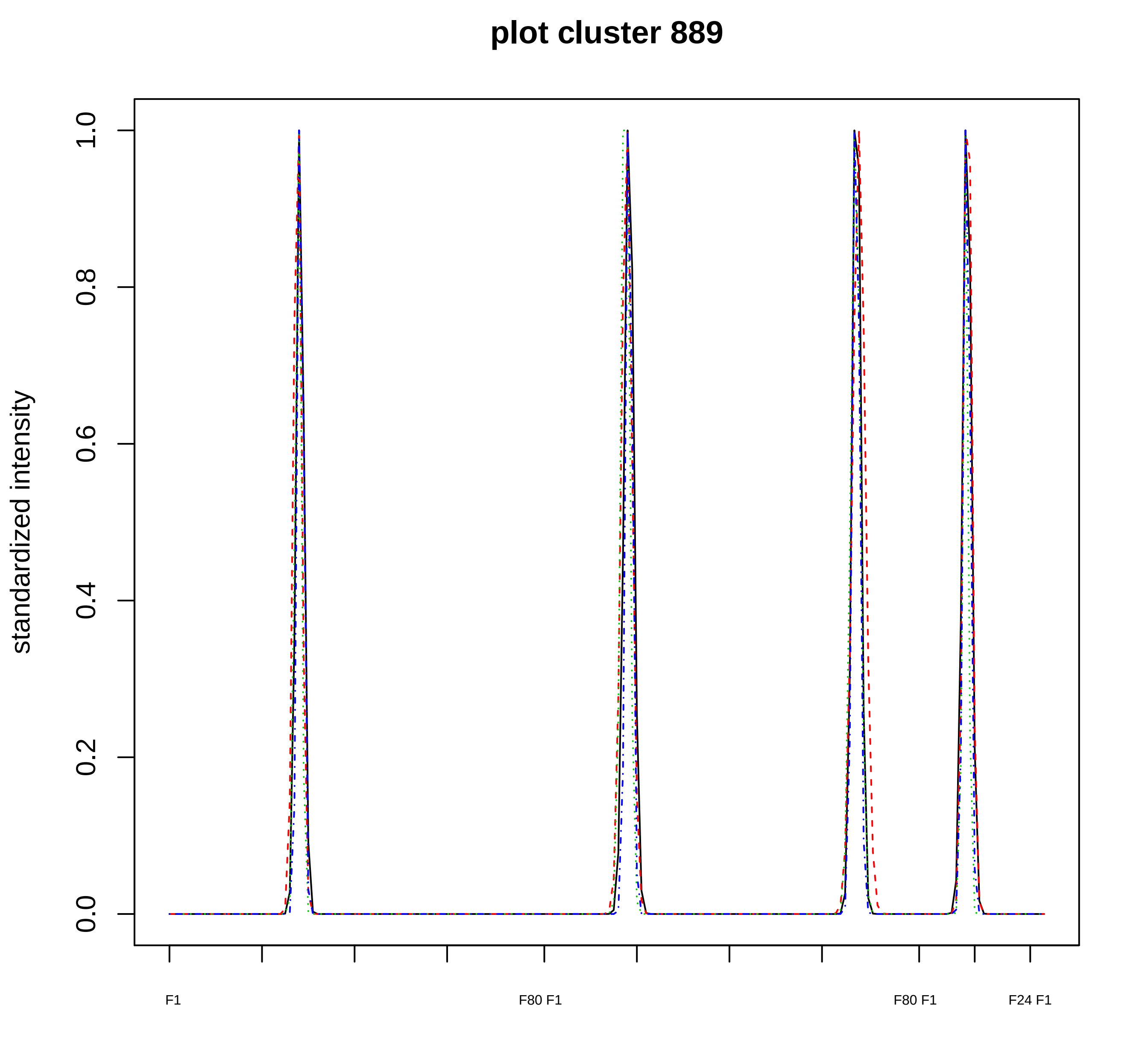

plot cluster 889
standardized intensity
0.0
0.2
0.4
0.6
0.8
1.0
F1
F80 F1
F80 F1
F24 F1

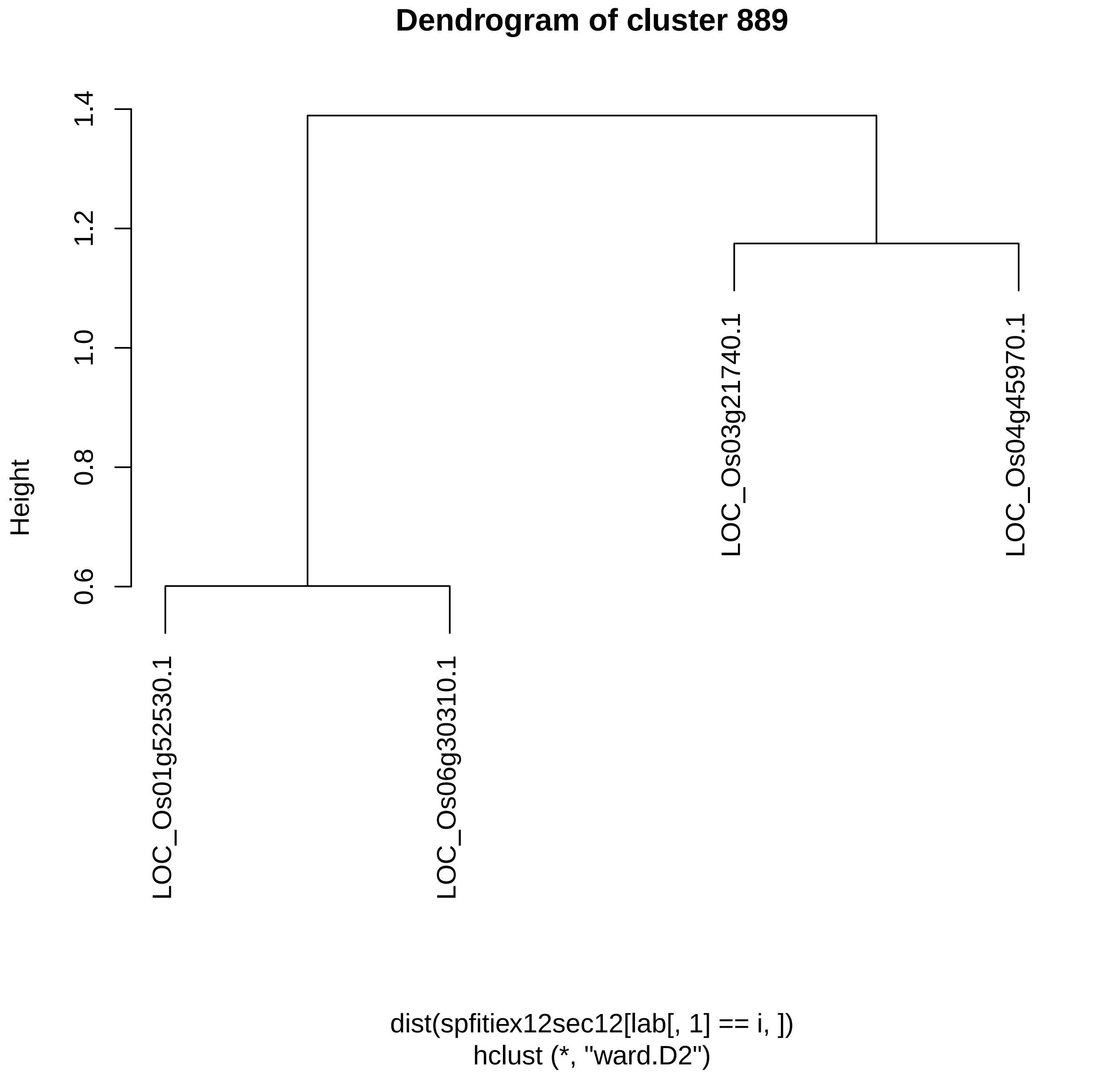
Dendrogram of cluster 889
Height
1.4
1.2
1.0
0.8
0.6
LOC_Os01g52530.1
LOC_Os06g30310.1
LOC_Os03g21740.1
LOC_Os04g45970.1
dist(spfitiex12sec12[lab[, 1] == i, ])
hclust (*, "ward.D2")

### plot cluster 890

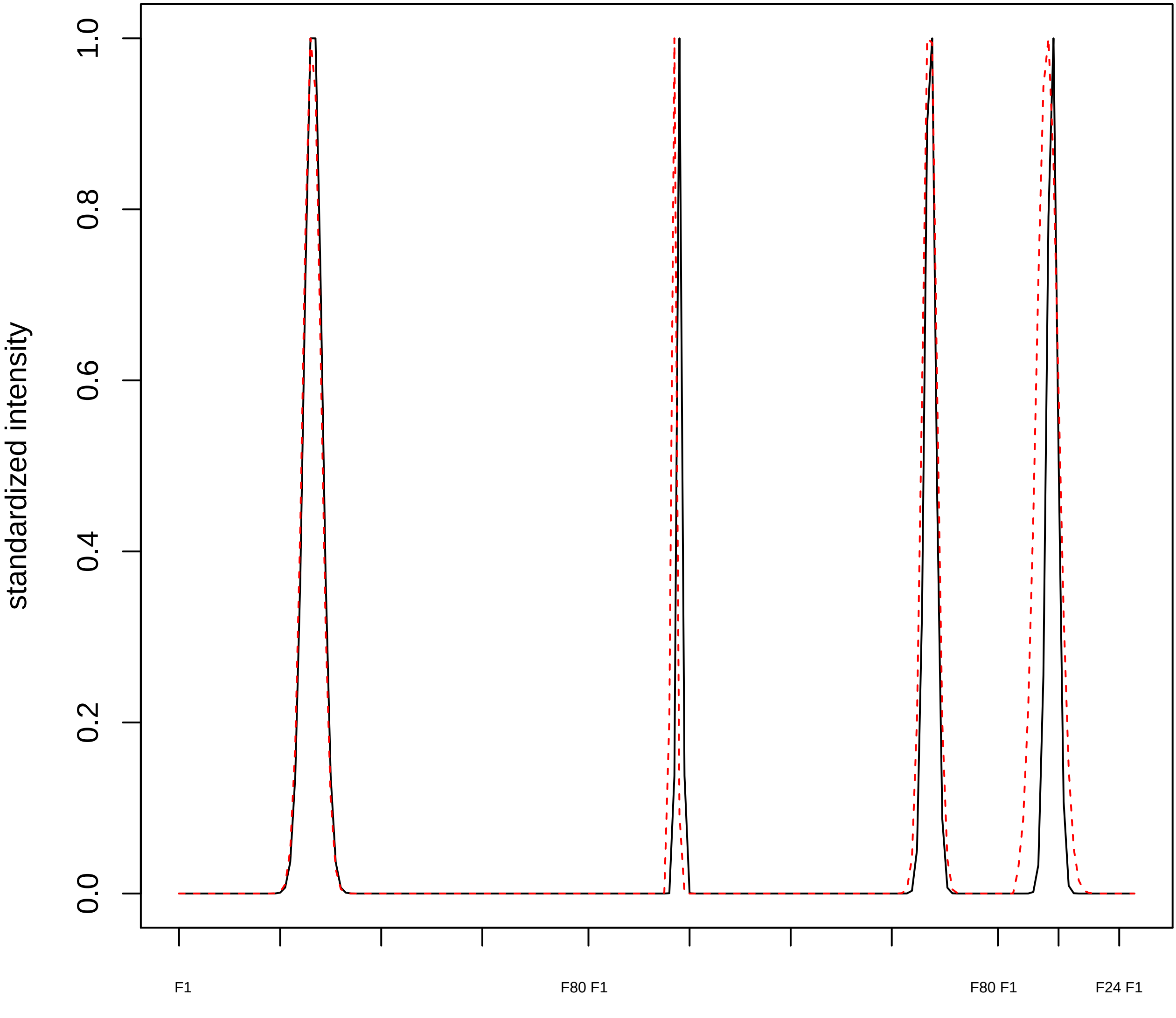

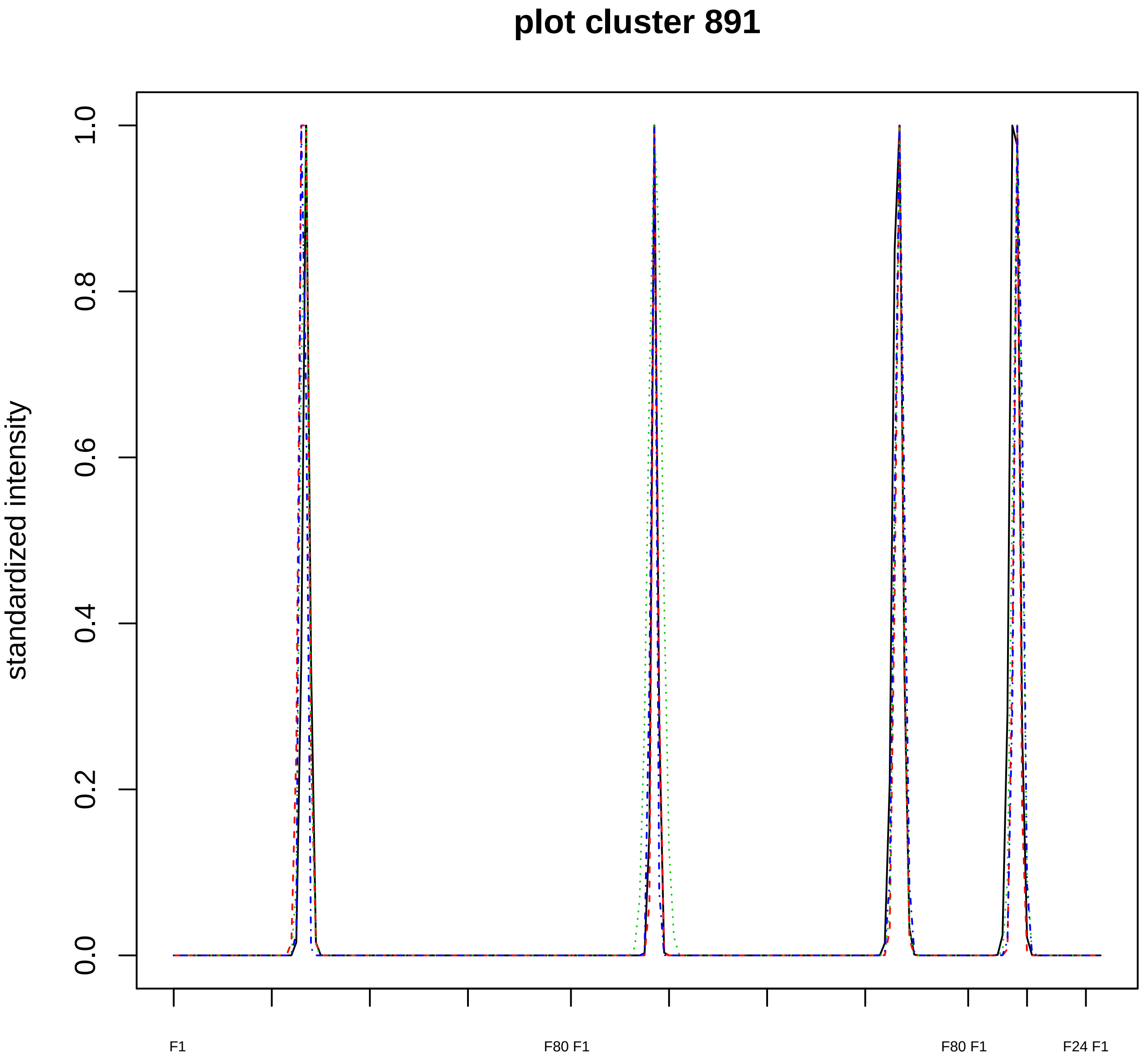
plot cluster 891
standardized intensity
0.0
0.2
0.4
0.6
0.8
1.0
F1
F80 F1
F80 F1
F24 F1

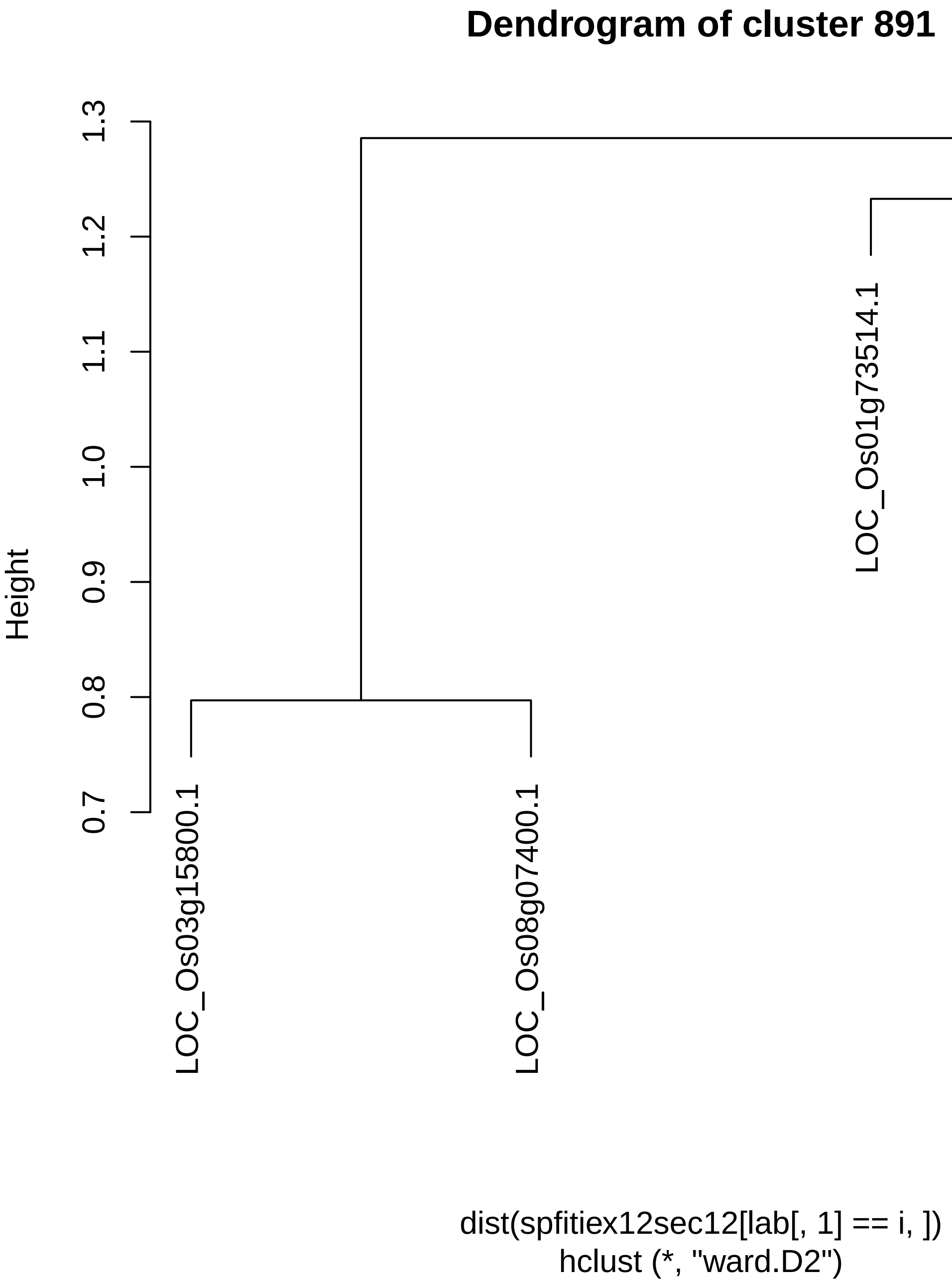
Dendrogram of cluster 891
Height
0.7
0.8
0.9
1.0
1.1
1.2
1.3
LOC_Os03g15800.1
LOC_Os08g07400.1
LOC_Os01g73514.1
LOC_Os07g23720.1
dist(spfitiex12sec12[lab[, 1] == i, ])
hclust (*, "ward.D2")

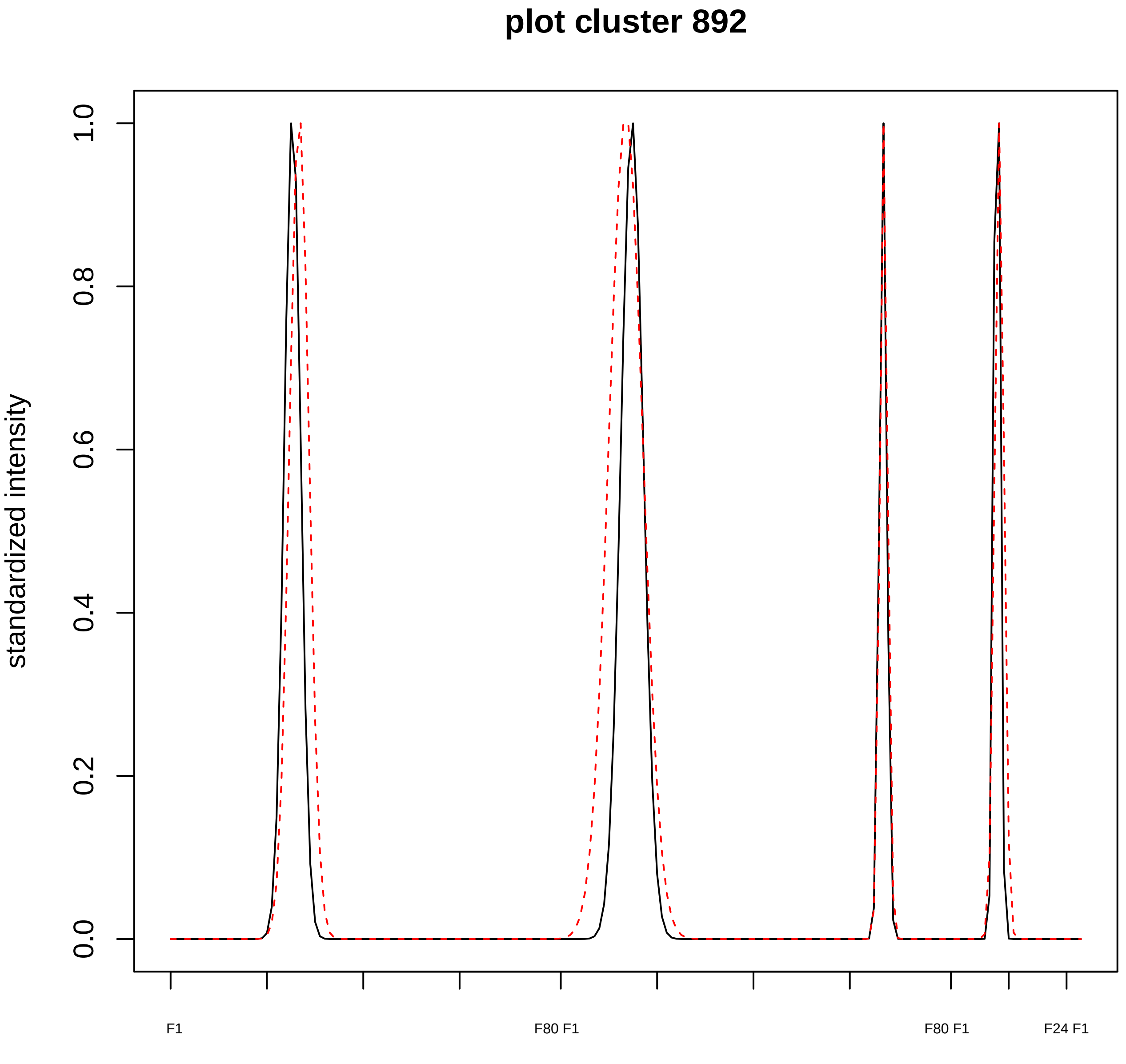

plot cluster 892
standardized intensity
0.0
0.2
0.4
0.6
0.8
1.0
F1
F80 F1
F80 F1
F24 F1

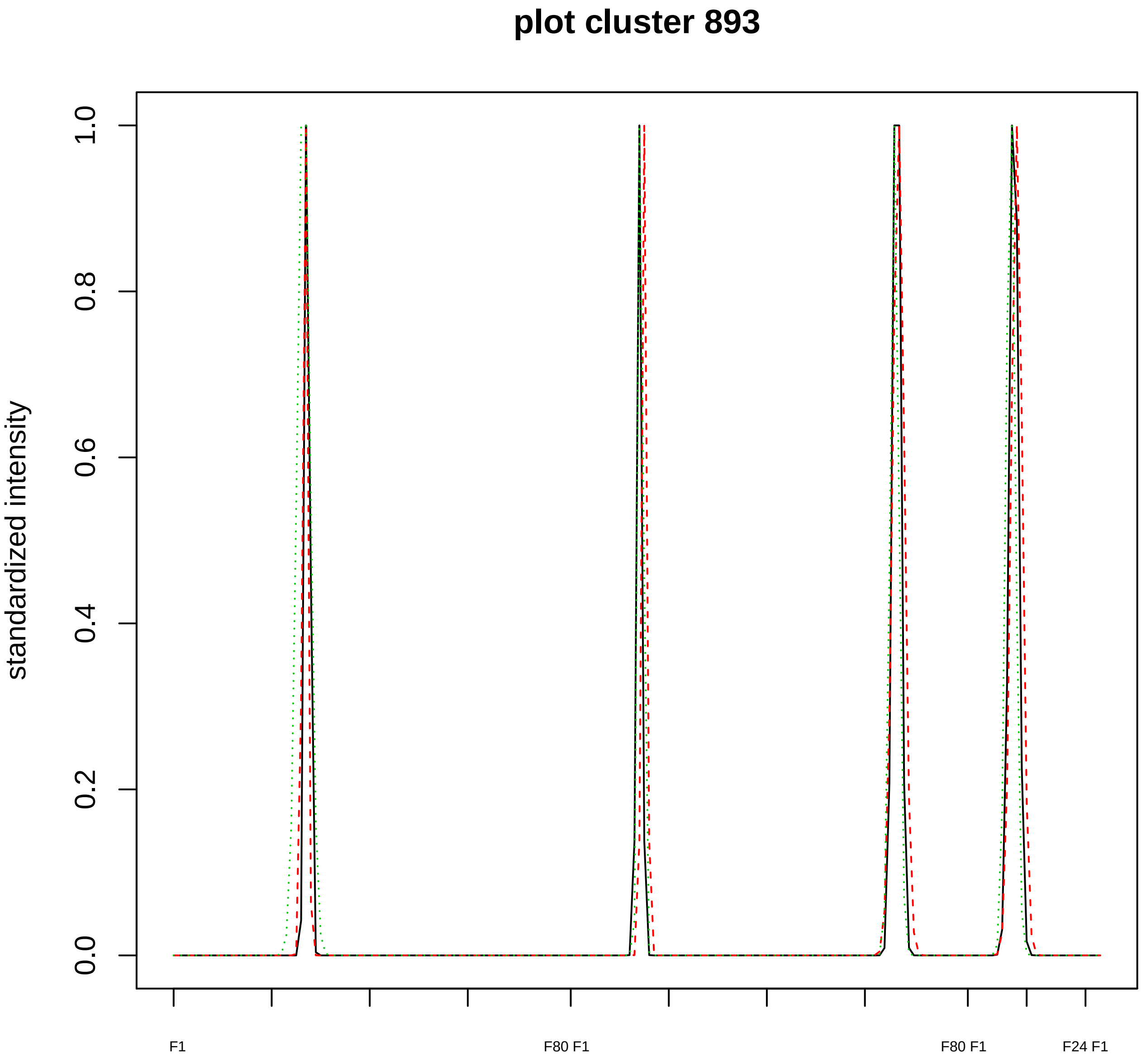
plot cluster 893
standardized intensity
0.0
0.2
0.4
0.6
0.8
1.0
F1
F80 F1
F80 F1
F24 F1

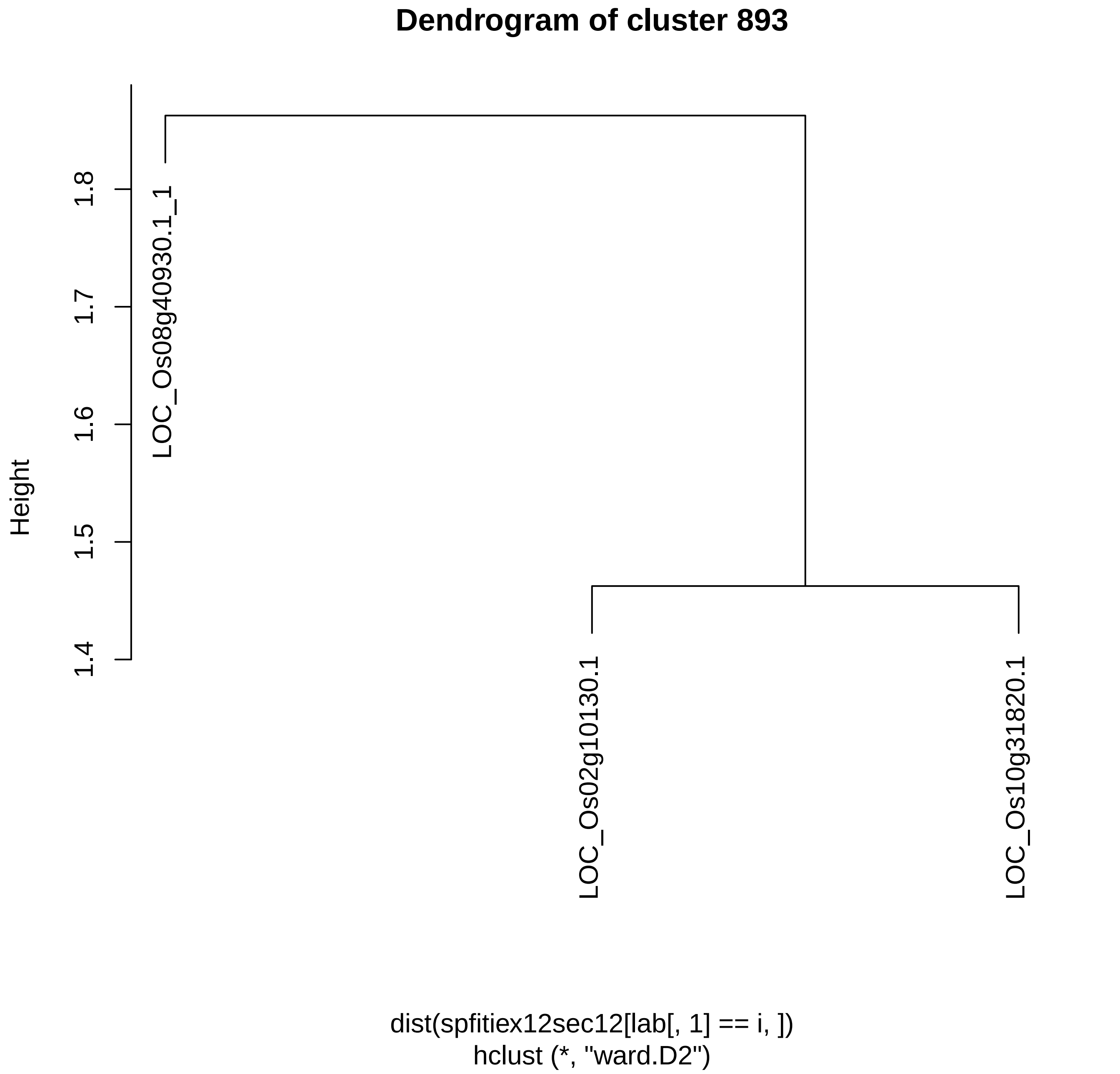

Dendrogram of cluster 893
Height
1.4
1.5
1.6
1.7
1.8
LOC_Os08g40930.1_1
LOC_Os02g10130.1
LOC_Os10g31820.1
dist(spfitiex12sec12[lab[, 1] == i, ])
hclust (*, "ward.D2")

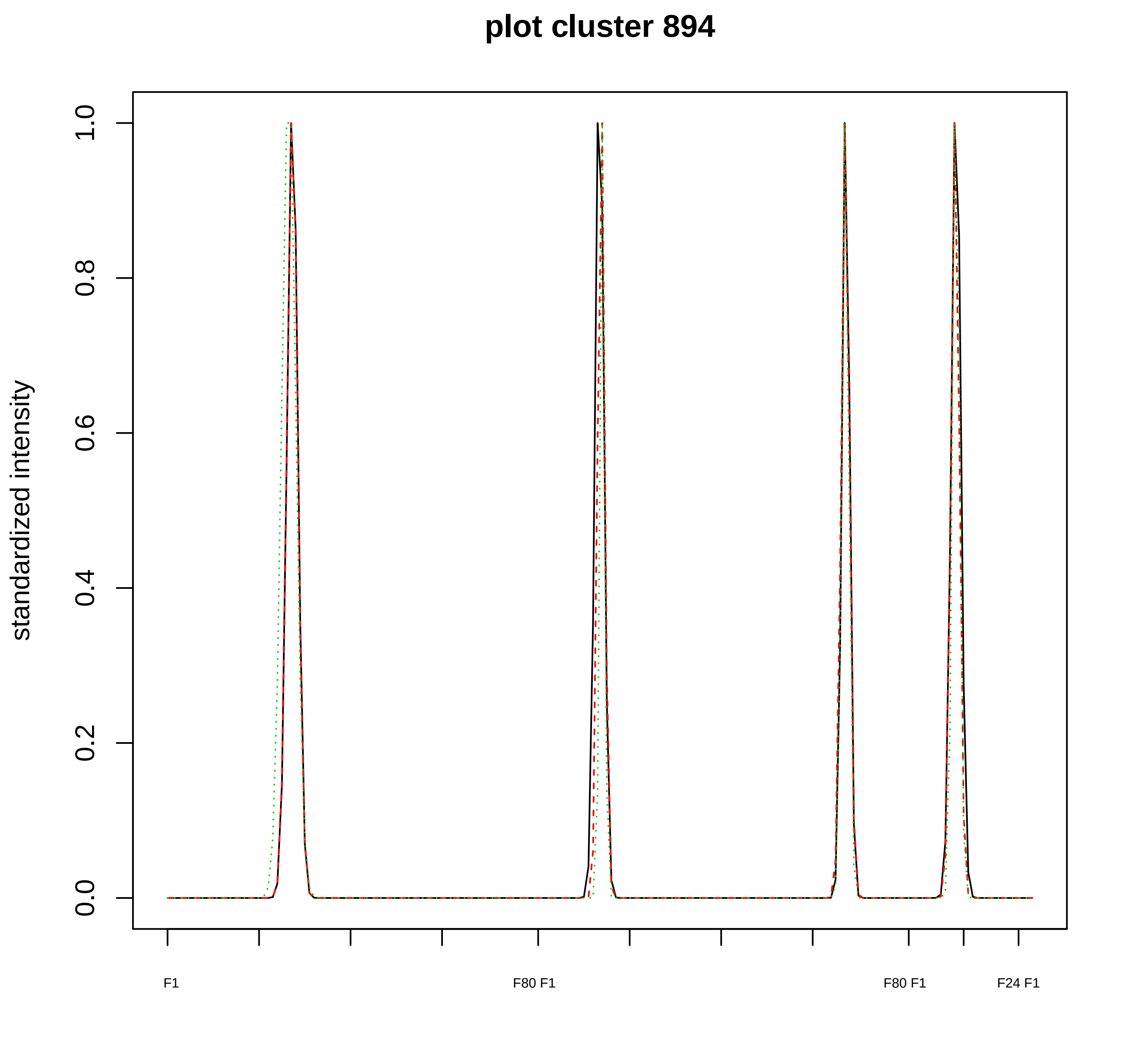

plot cluster 894
standardized intensity
0.0
0.2
0.4
0.6
0.8
1.0
F1
F80 F1
F80 F1
F24 F1

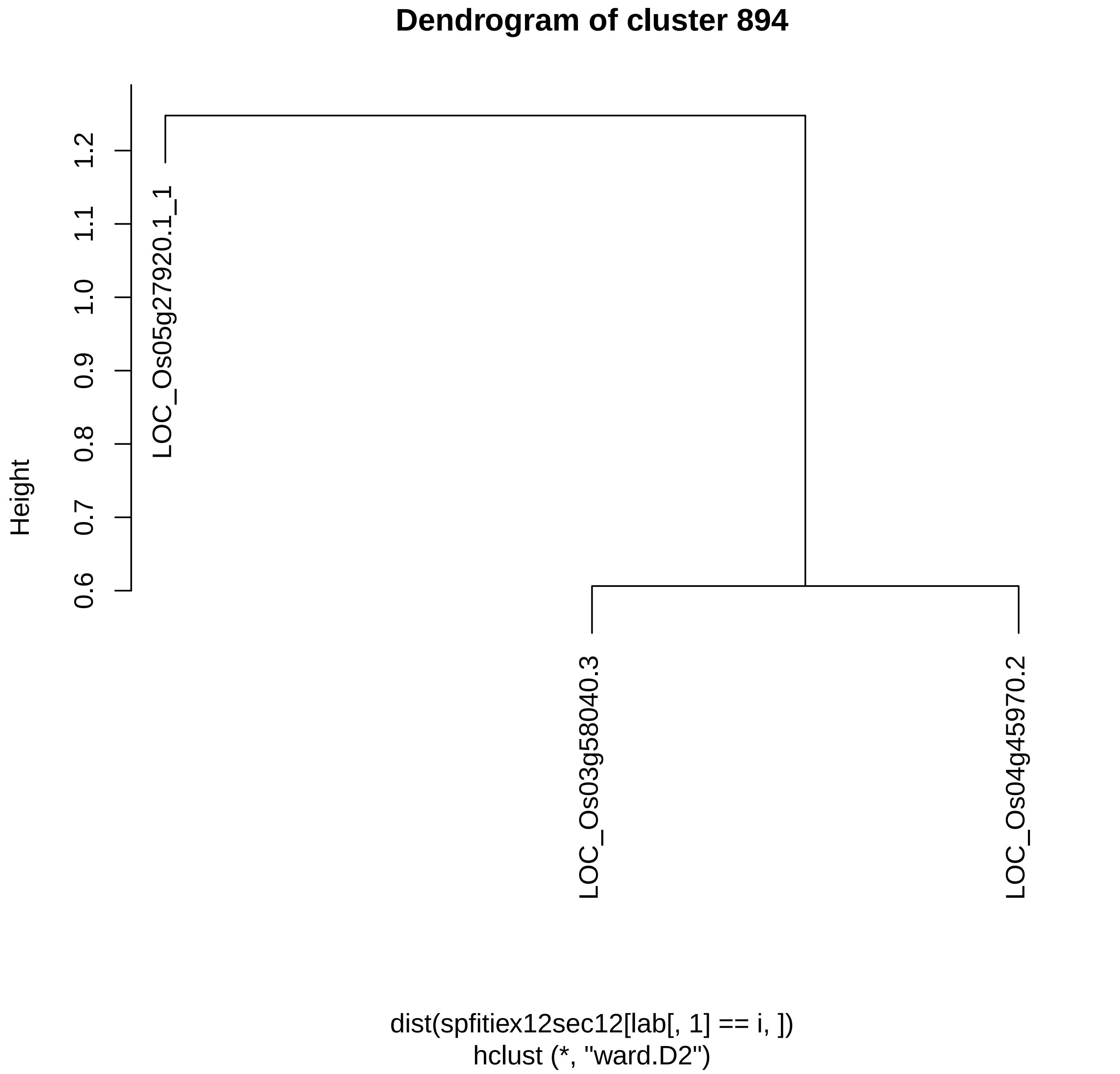
Dendrogram of cluster 894
Height
0.6
0.7
0.8
0.9
1.0
1.1
1.2
LOC_Os05g27920.1_1
LOC_Os03g58040.3
LOC_Os04g45970.2
dist(spfitiex12sec12[lab[, 1] == i, ])
hclust (*, "ward.D2")

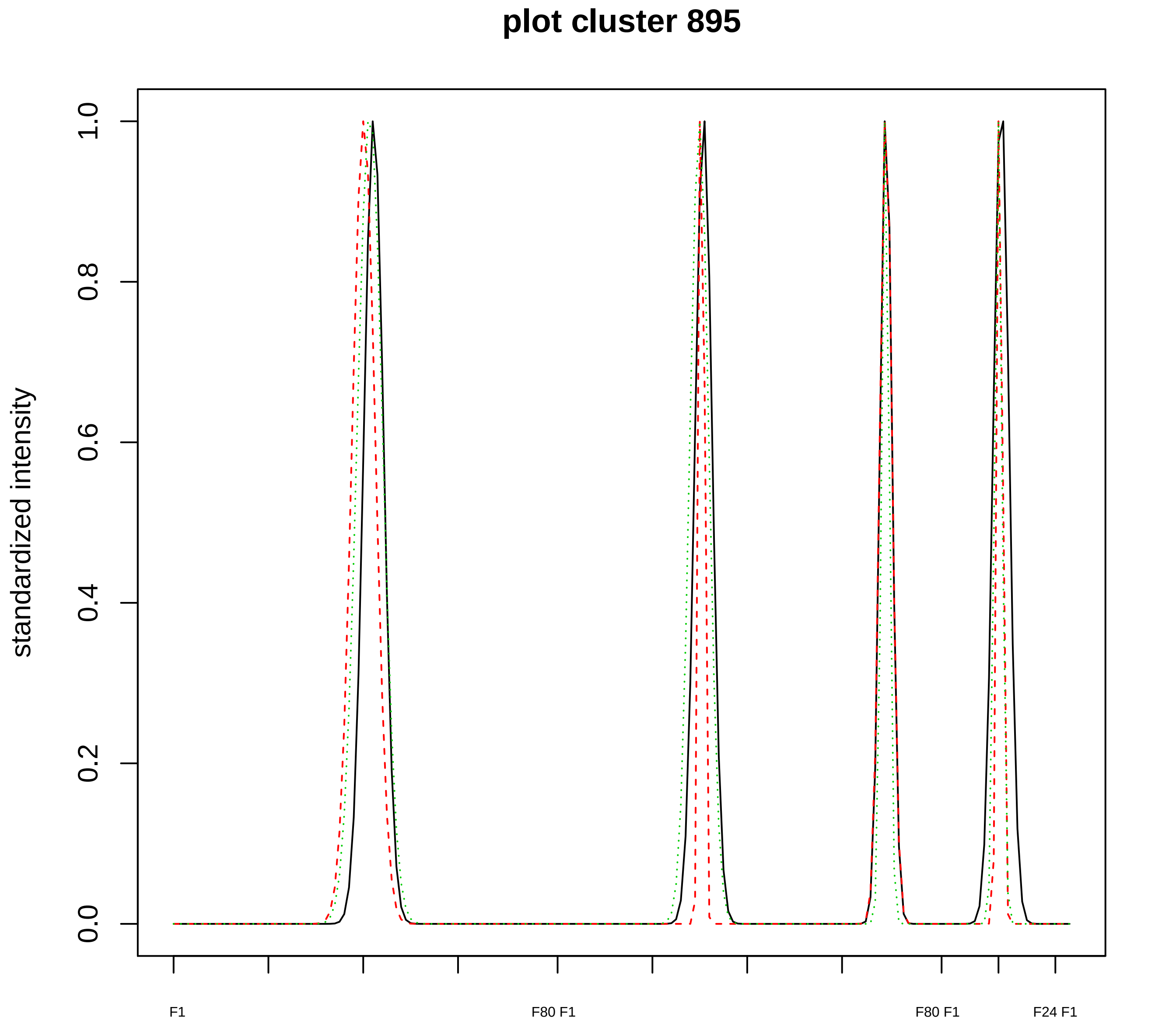

plot cluster 895
standardized intensity
0.0
0.2
0.4
0.6
0.8
1.0
F1
F80 F1
F80 F1
F24 F1

#### Dendrogram of cluster 895

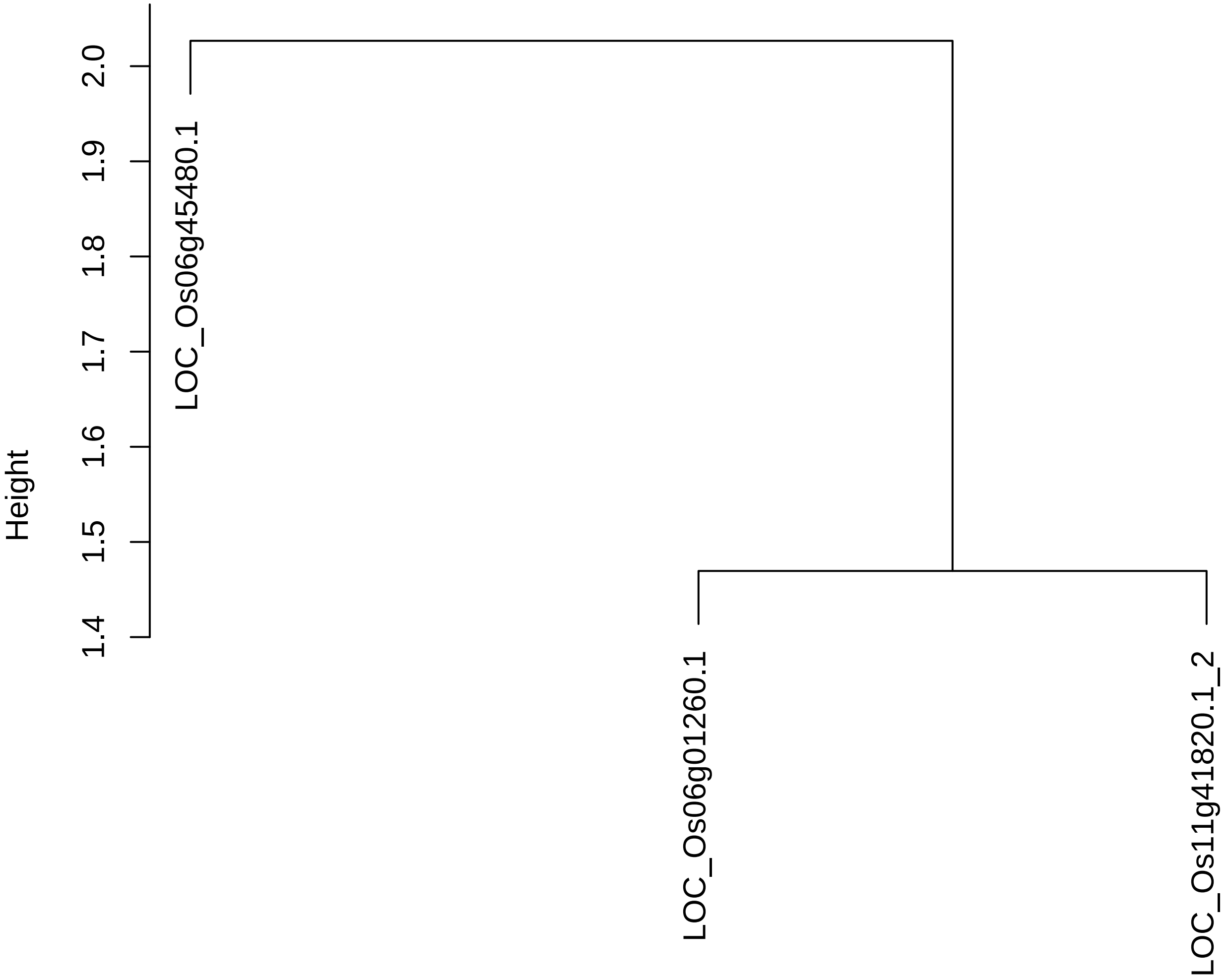


dist(spfitiex12sec12[lab[, 1] == i, ])
hclust (*, "ward.D2")

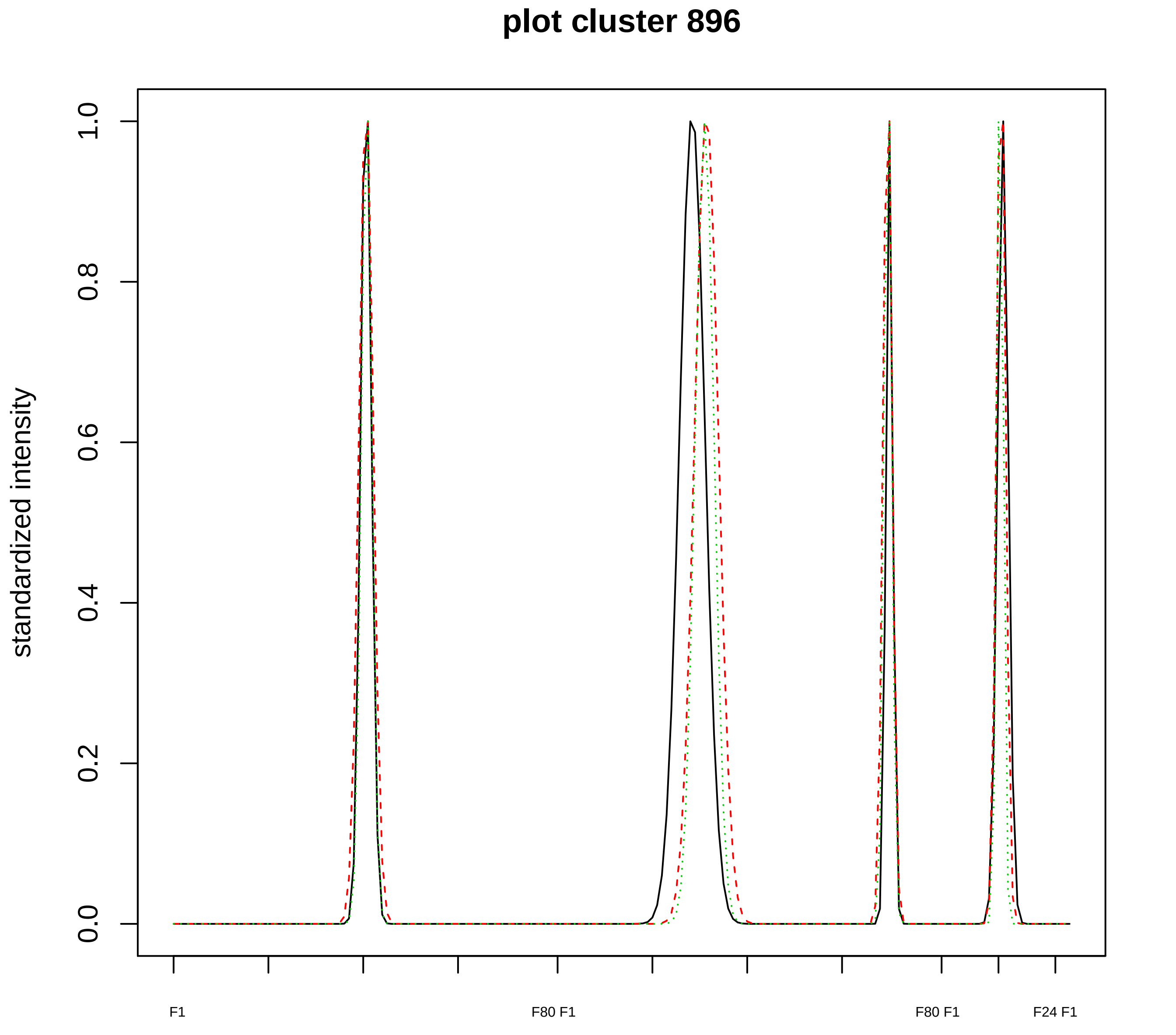
plot cluster 896
standardized intensity
0.0
0.2
0.4
0.6
0.8
1.0
F1
F80 F1
F80 F1
F24 F1

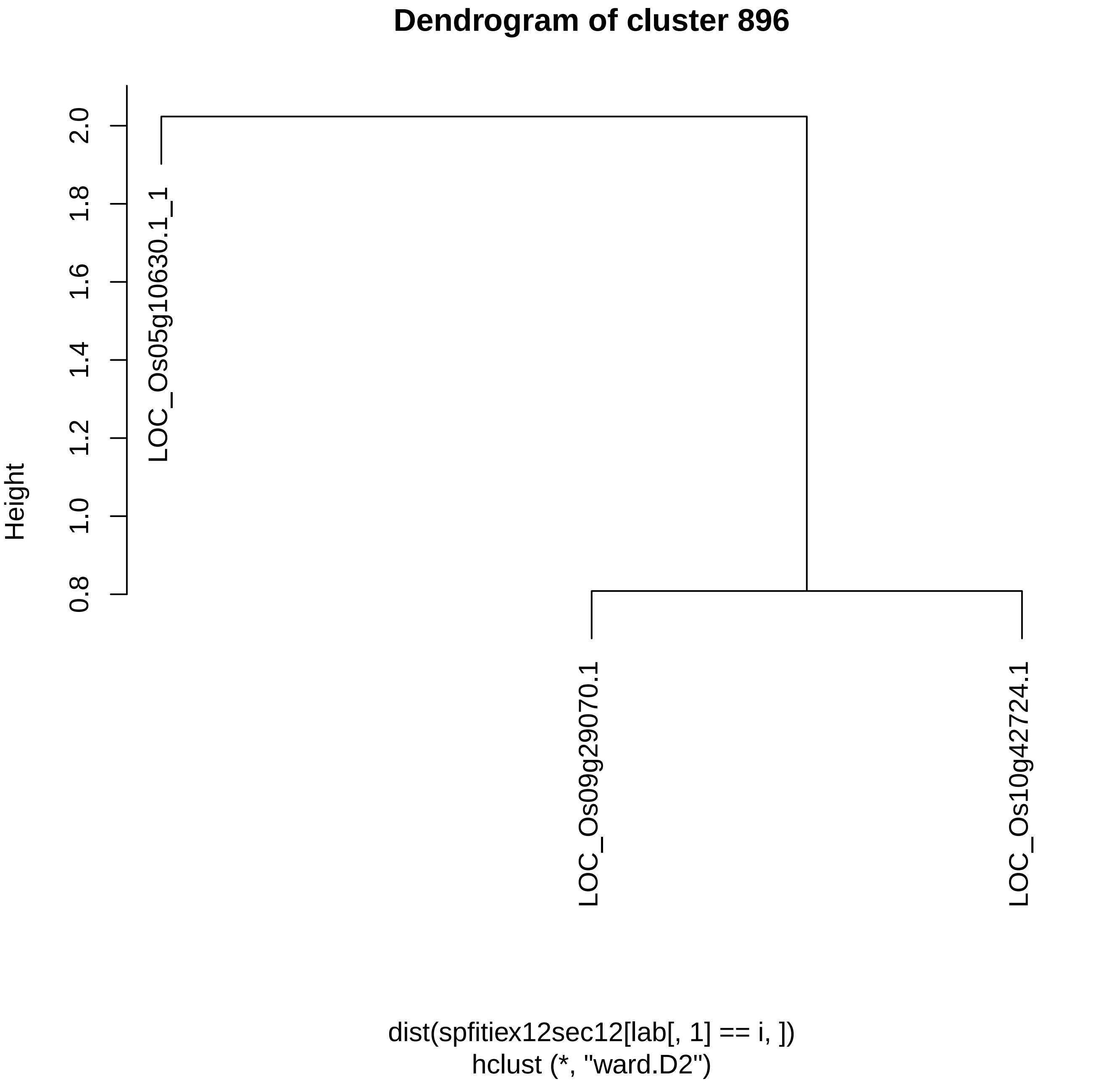
Dendrogram of cluster 896
Height
0.8
1.0
1.2
1.4
1.6
1.8
2.0
LOC_Os05g10630.1_1
LOC_Os09g29070.1
LOC_Os10g42724.1
dist(spfitiex12sec12[lab[, 1] == i, ])
hclust (*, "ward.D2")

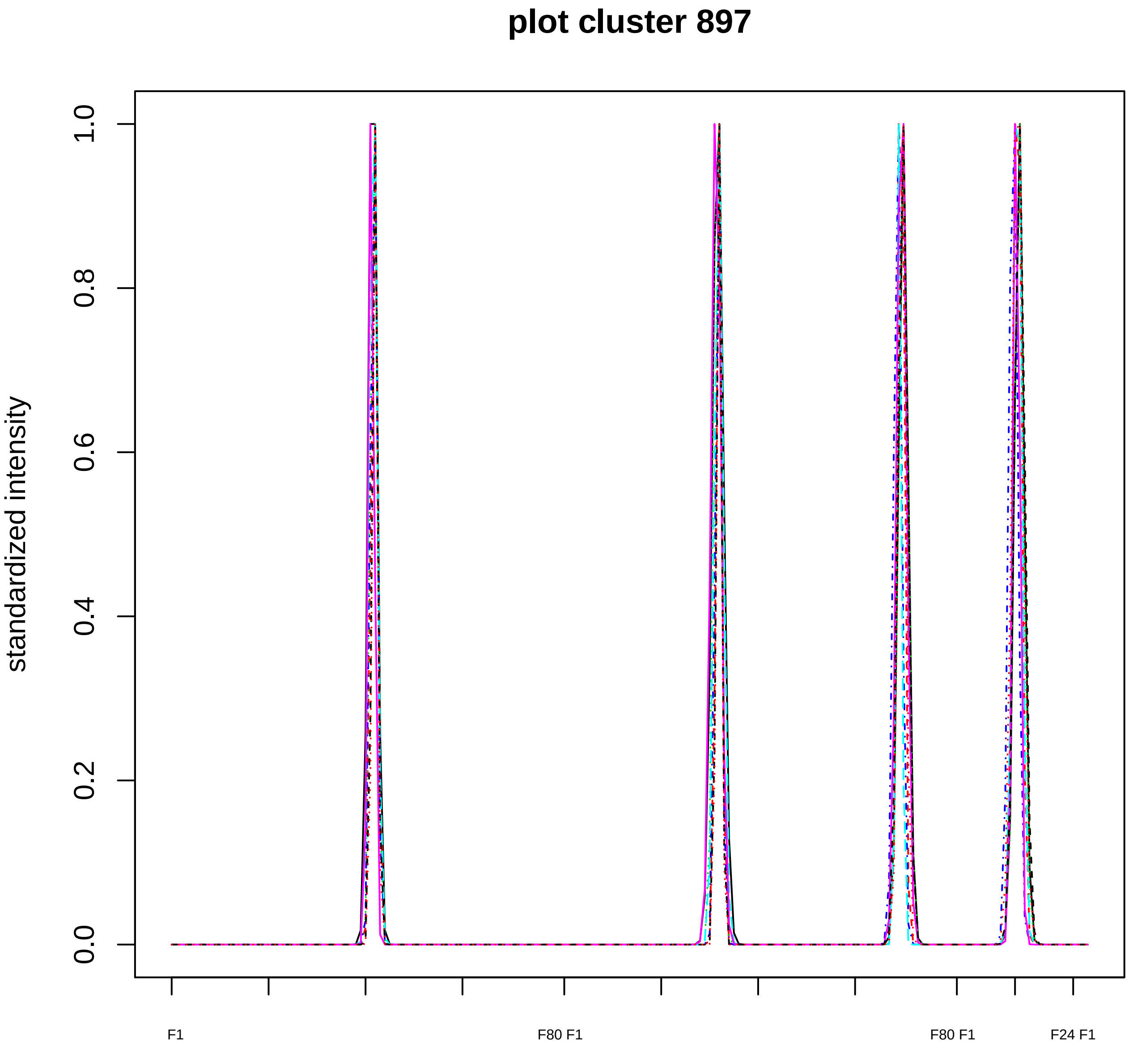
plot cluster 897
standardized intensity
0.0
0.2
0.4
0.6
0.8
1.0
F1
F80 F1
F80 F1
F24 F1

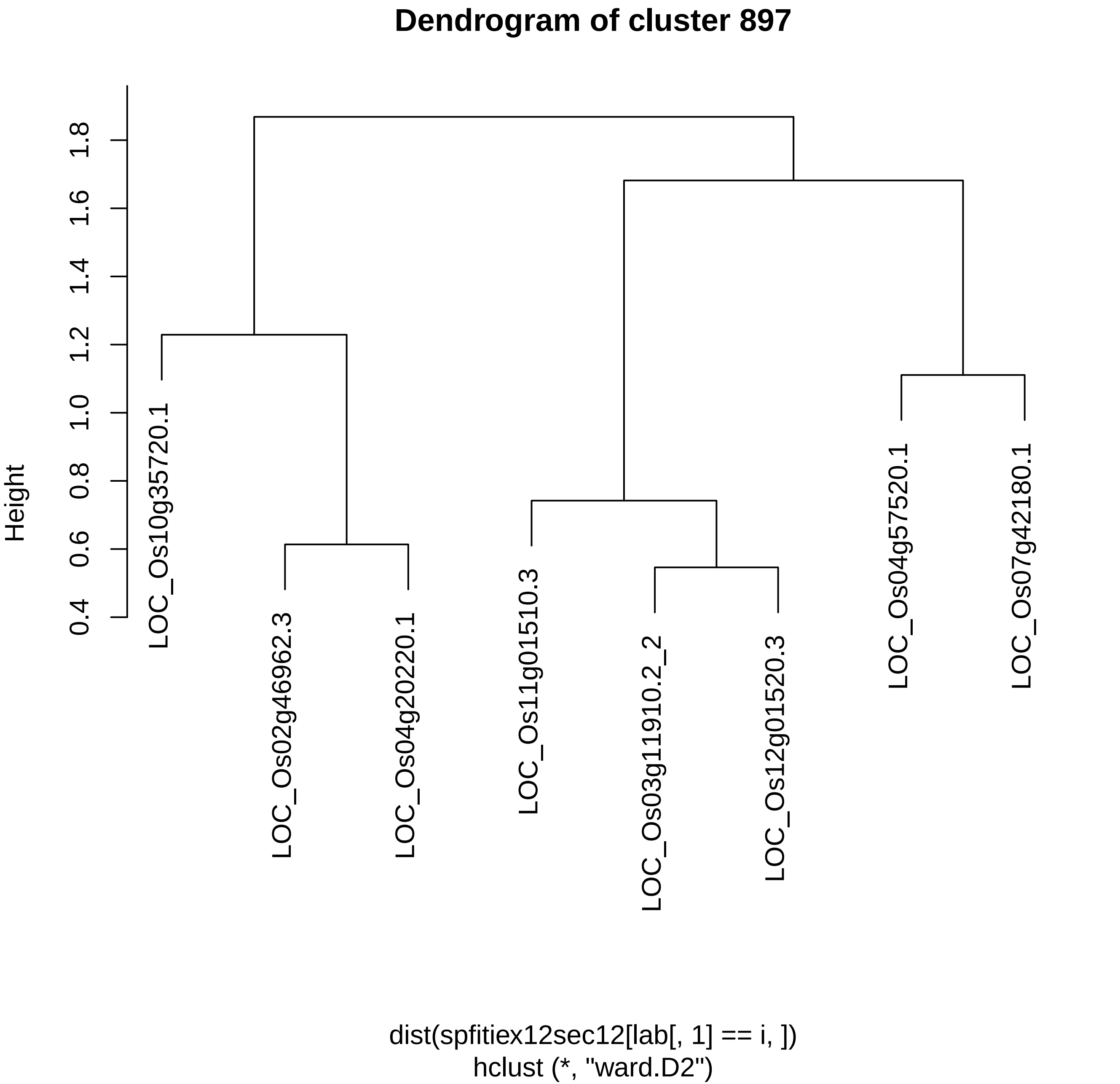

Dendrogram of cluster 897
Height
0.4
0.6
0.8
1.0
1.2
1.4
1.6
1.8
LOC_Os10g35720.1
LOC_Os02g46962.3
LOC_Os04g20220.1
LOC_Os11g01510.3
LOC_Os03g11910.2_2
LOC_Os12g01520.3
LOC_Os04g57520.1
LOC_Os07g42180.1
dist(spfitiex12sec12[lab[, 1] == i, ])
hclust (*, "ward.D2")

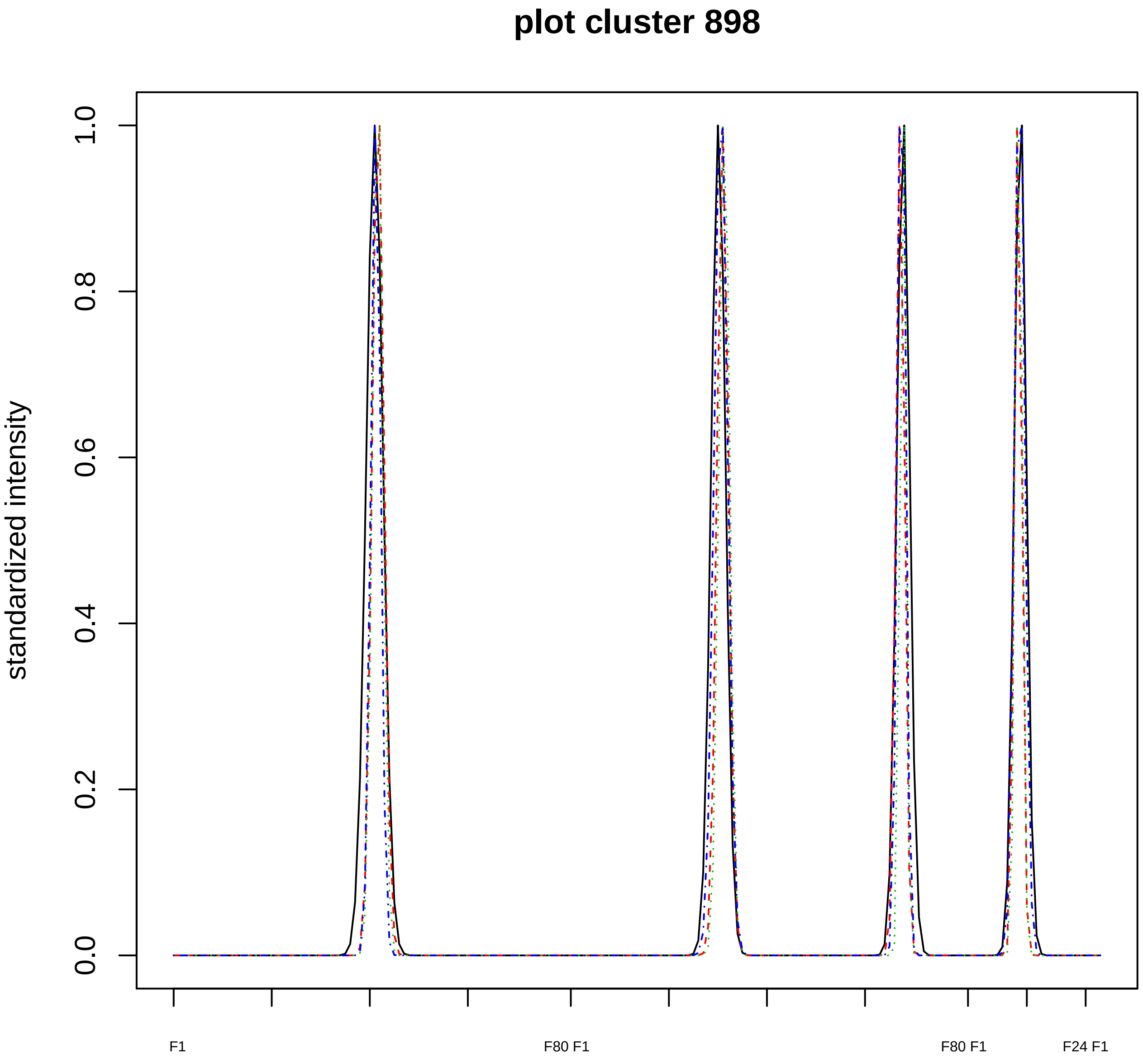
plot cluster 898
standardized intensity
0.0
0.2
0.4
0.6
0.8
1.0
F1
F80 F1
F80 F1
F24 F1

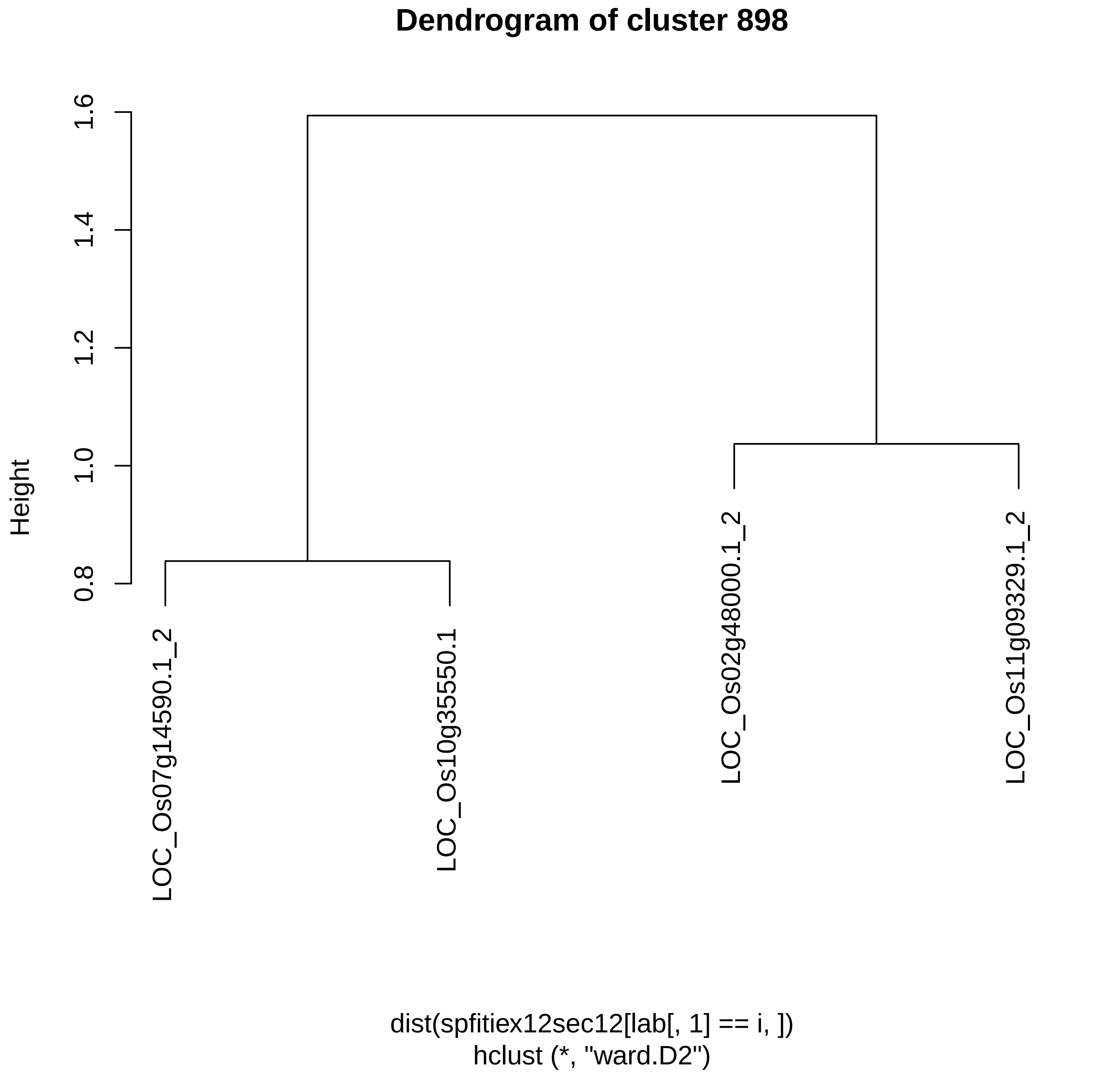
Dendrogram of cluster 898
Height
1.6
1.4
1.2
1.0
0.8
LOC_Os07g14590.1_2
LOC_Os10g35550.1
LOC_Os02g48000.1_2
LOC_Os11g09329.1_2
dist(spfitiex12sec12[lab[, 1] == i, ])
hclust (*, "ward.D2")

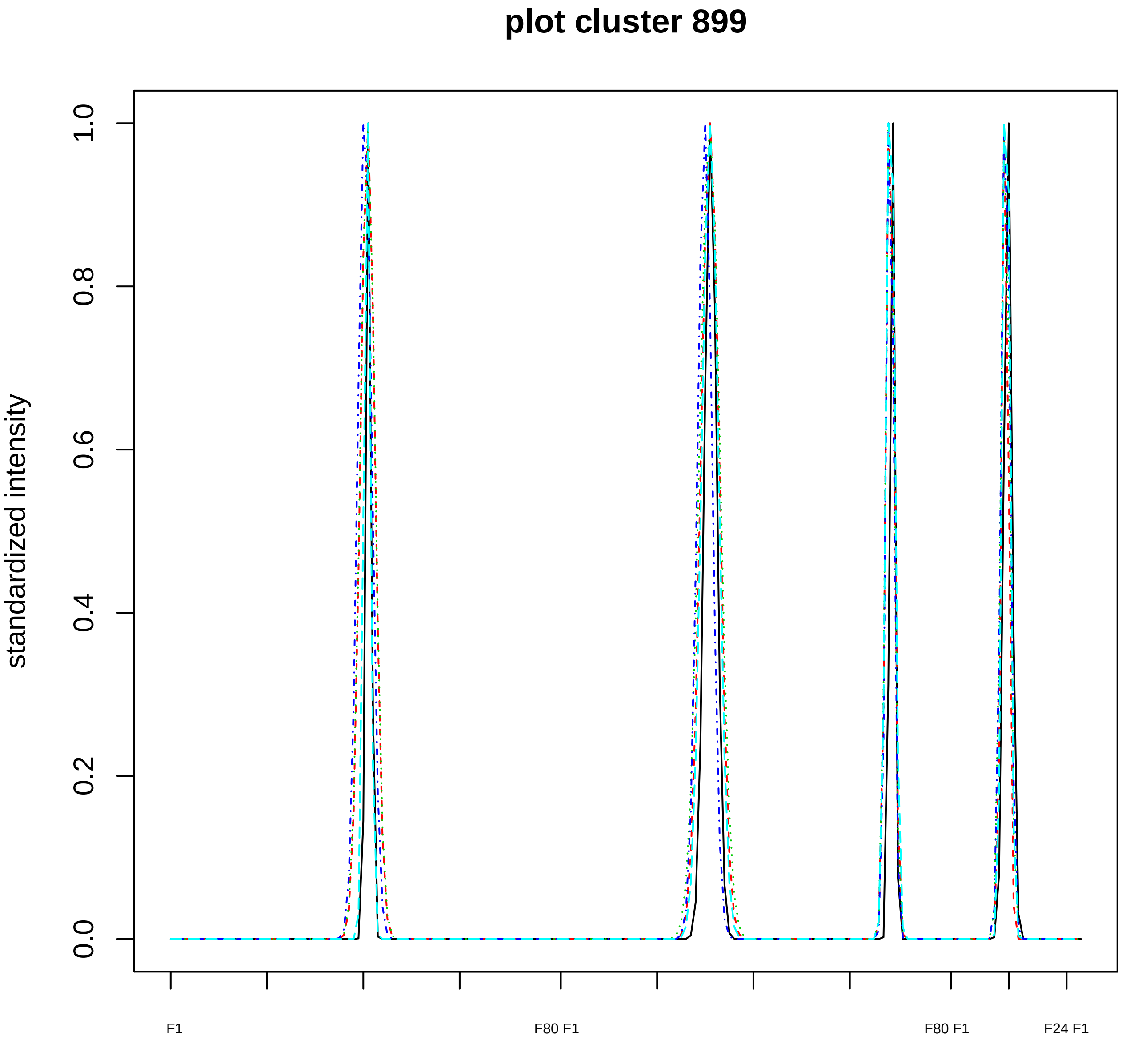

plot cluster 899
standardized intensity
0.0
0.2
0.4
0.6
0.8
1.0
F1
F80 F1
F80 F1
F24 F1

### Dendrogram of cluster 899

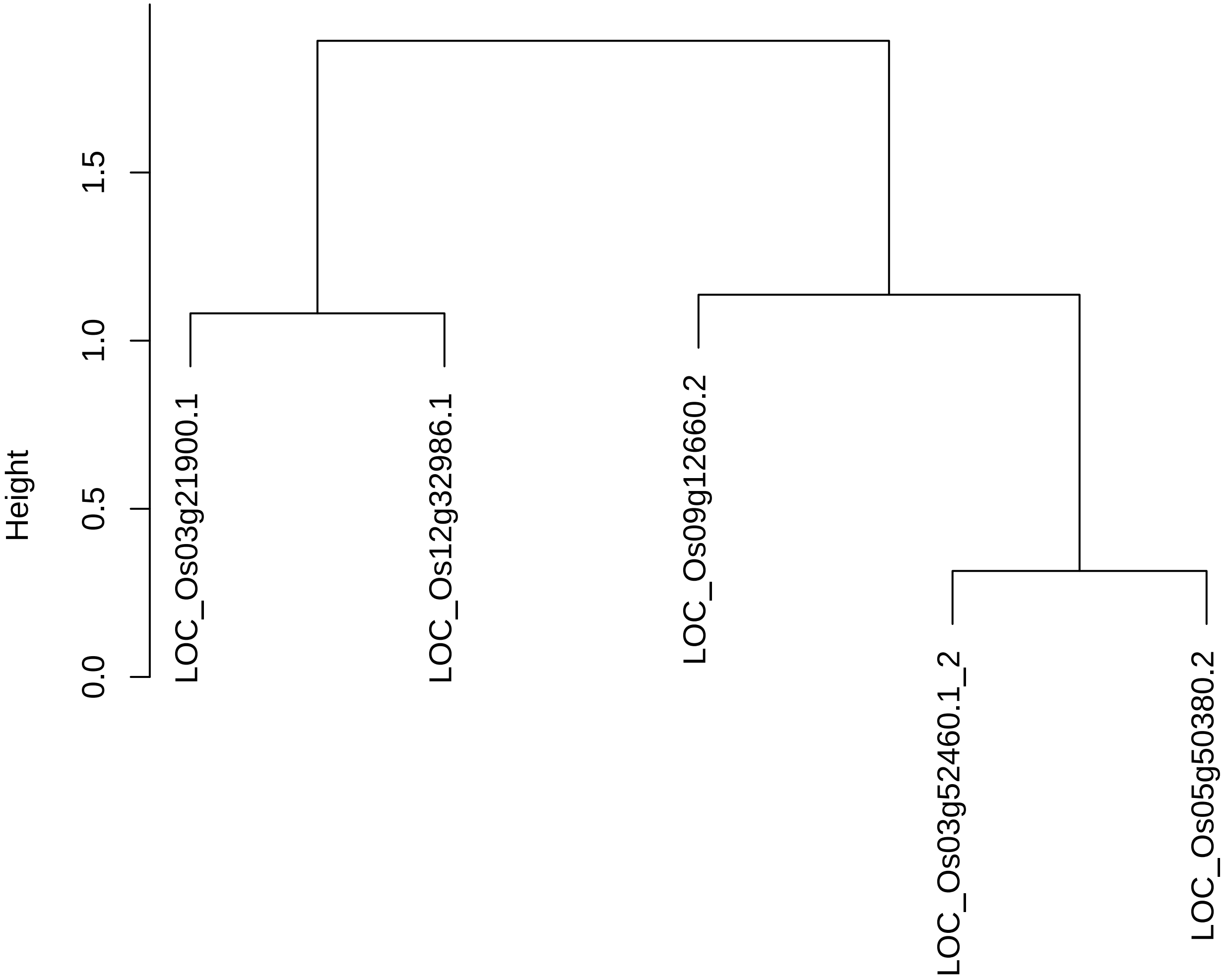


dist(spfitiex12sec12[lab[, 1] == i, ])
hclust (*, "ward.D2")

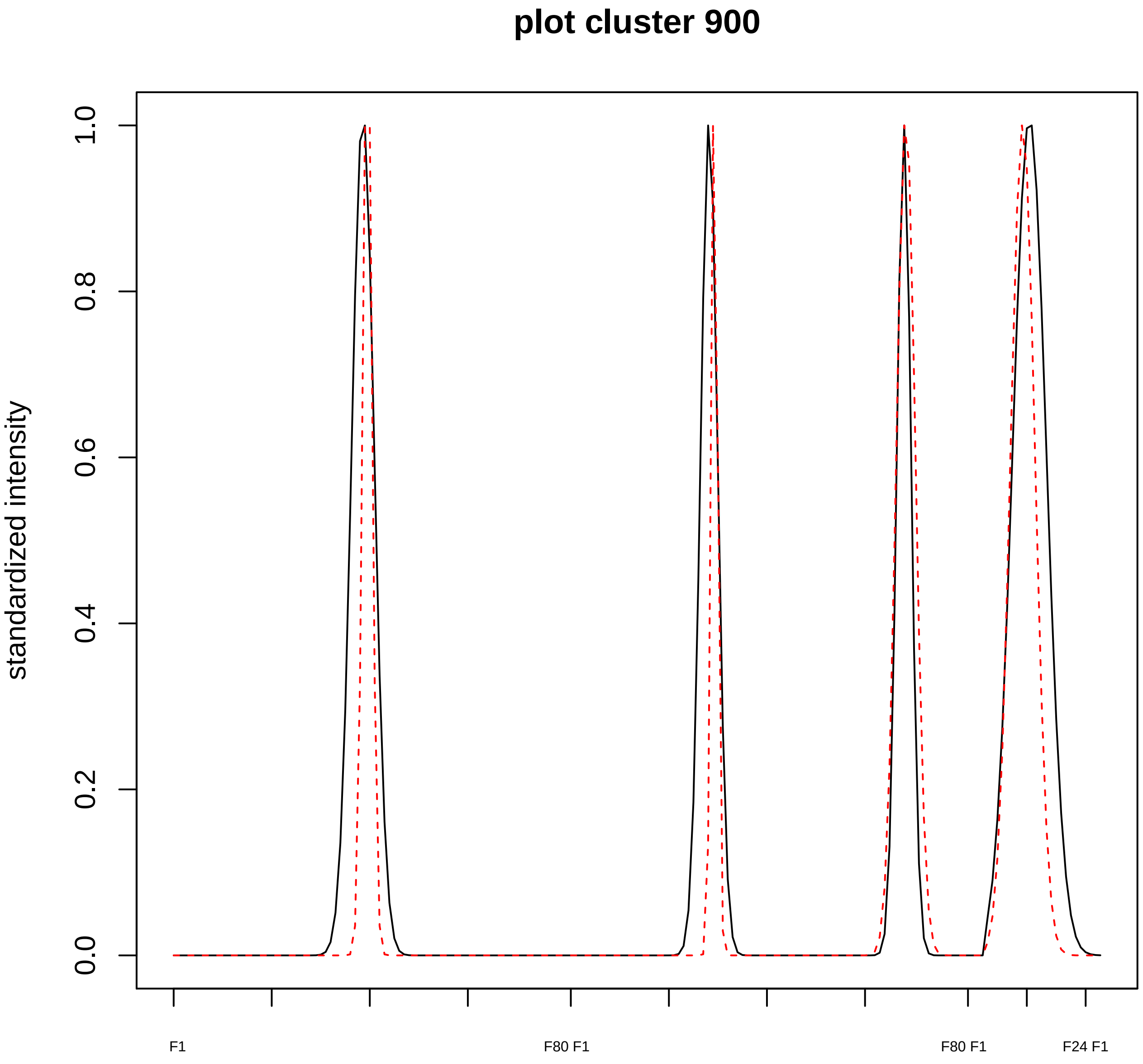
plot cluster 900
standardized intensity
0.0
0.2
0.4
0.6
0.8
1.0
F1
F80 F1
F80 F1
F24 F1

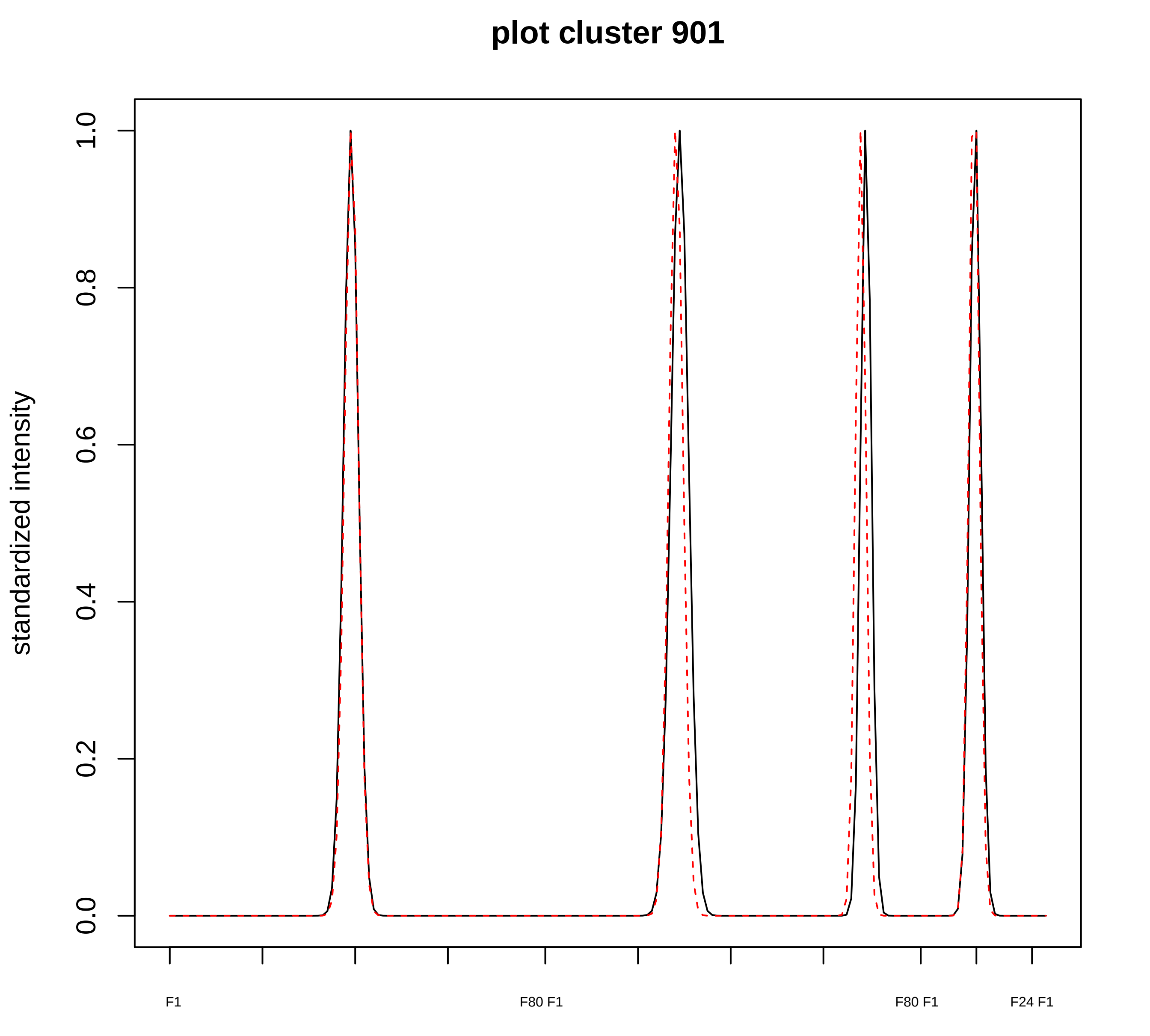
plot cluster 901
standardized intensity
0.0
0.2
0.4
0.6
0.8
1.0
F1
F80 F1
F80 F1
F24 F1

### plot cluster 902

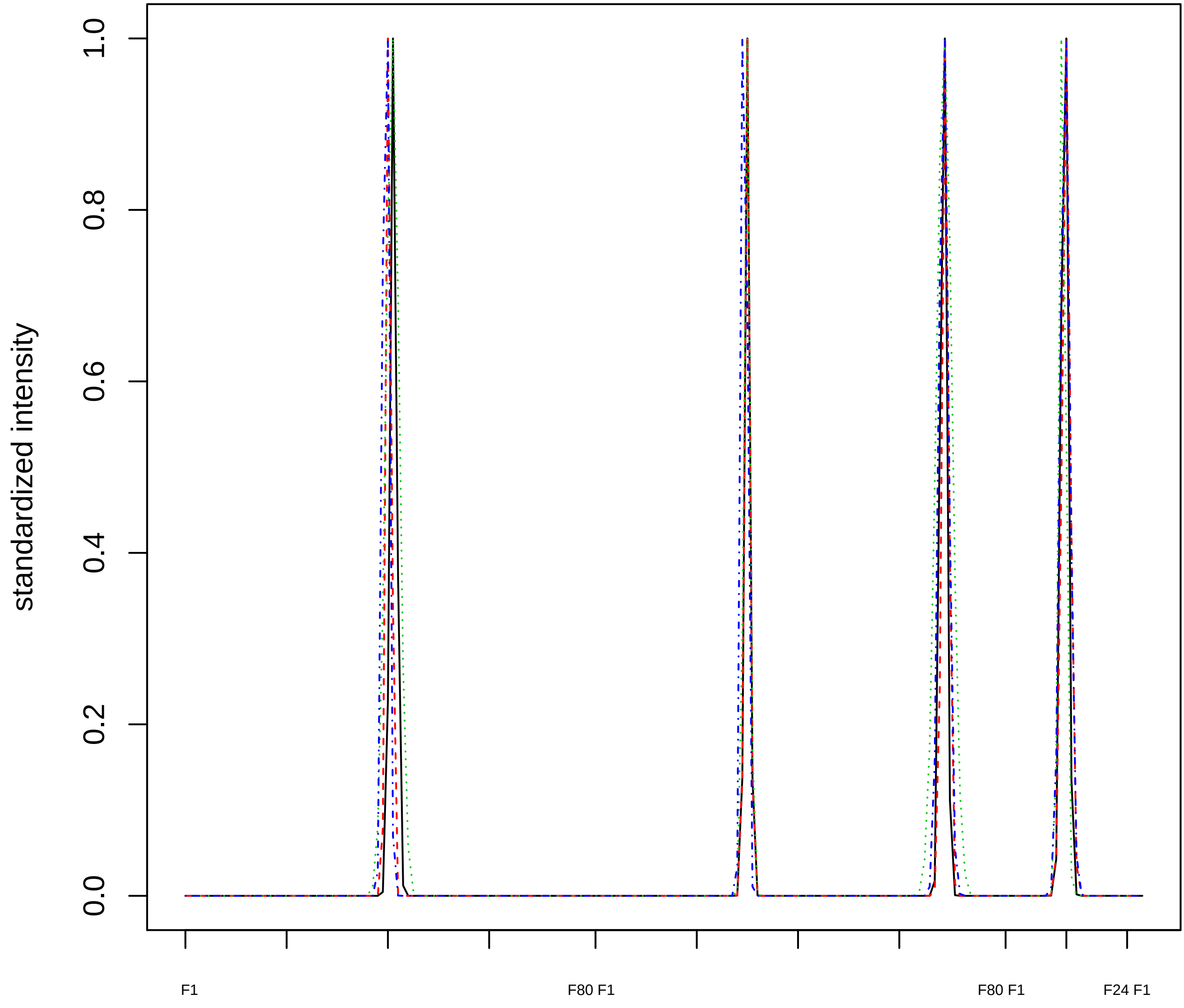

### Dendrogram of cluster 902

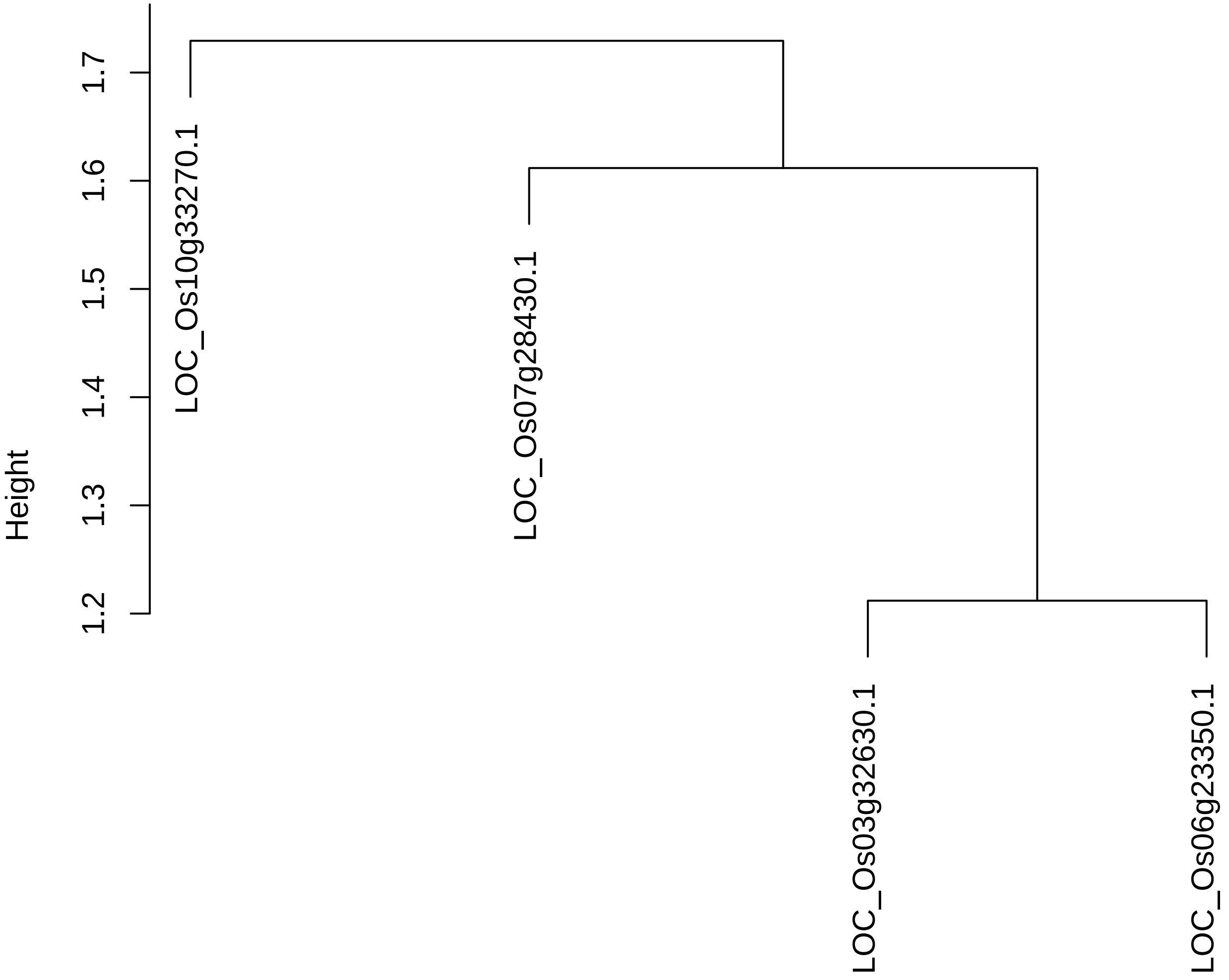


dist(spfitiex12sec12[lab[, 1] == i, ])
hclust (*, "ward.D2")

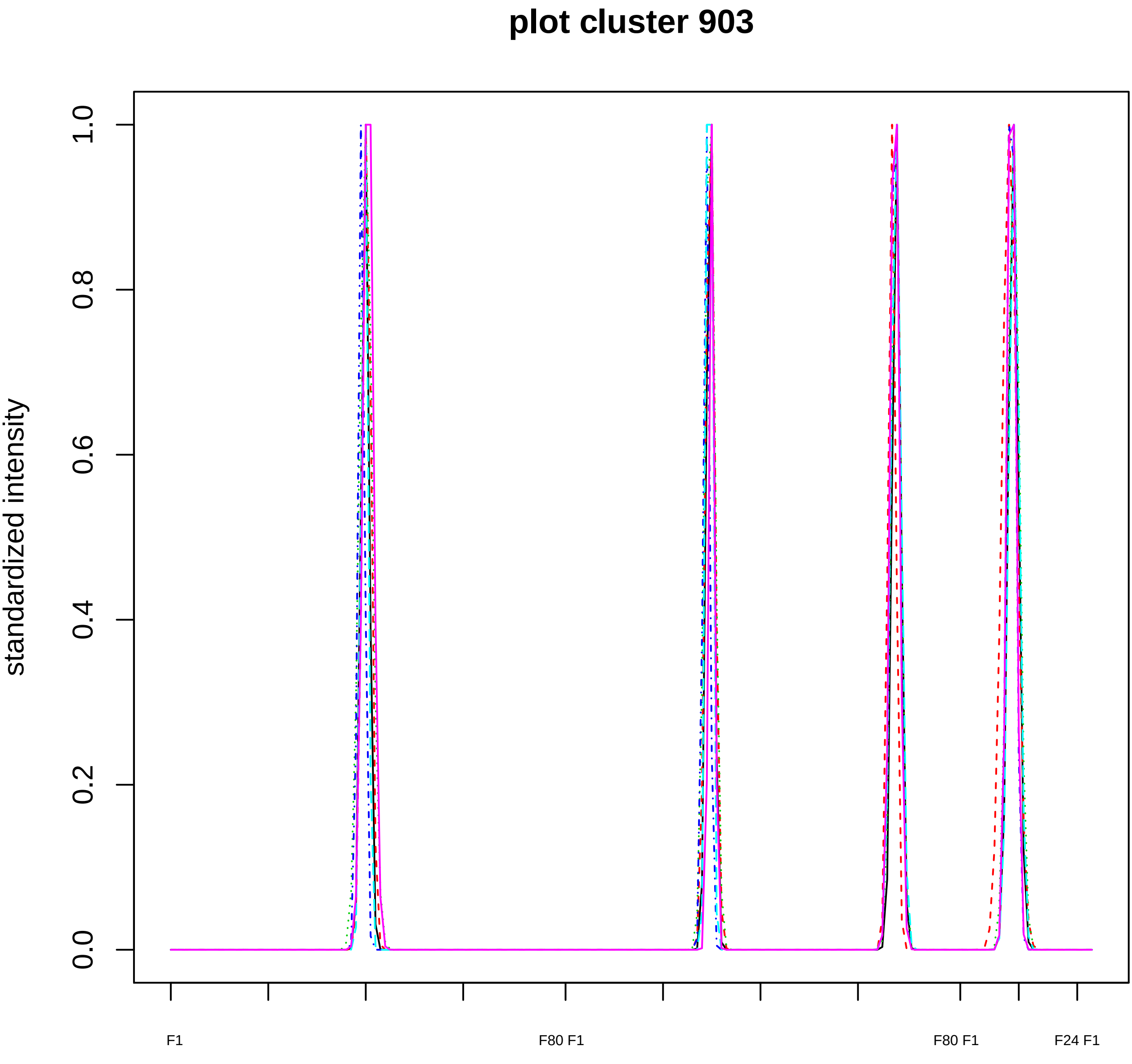

plot cluster 903
standardized intensity
1.0
0.8
0.6
0.4
0.2
0.0
F1
F80 F1
F80 F1
F24 F1

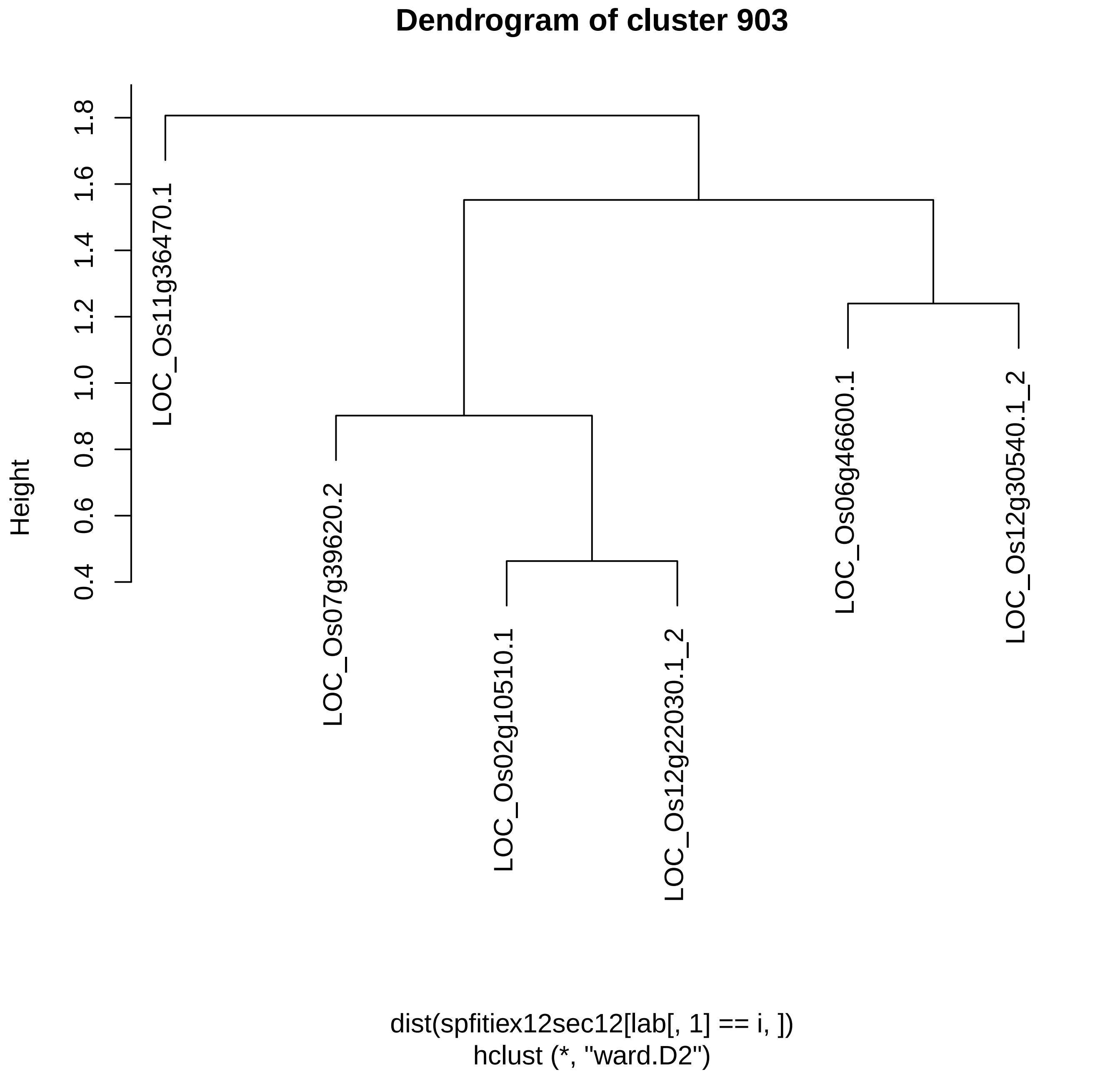
Dendrogram of cluster 903
Height
0.4
0.6
0.8
1.0
1.2
1.4
1.6
1.8
LOC_Os11g36470.1
LOC_Os07g39620.2
LOC_Os02g10510.1
LOC_Os12g22030.1_2
LOC_Os06g46600.1
LOC_Os12g30540.1_2
dist(spfitiex12sec12[lab[, 1] == i, ])
hclust (*, "ward.D2")

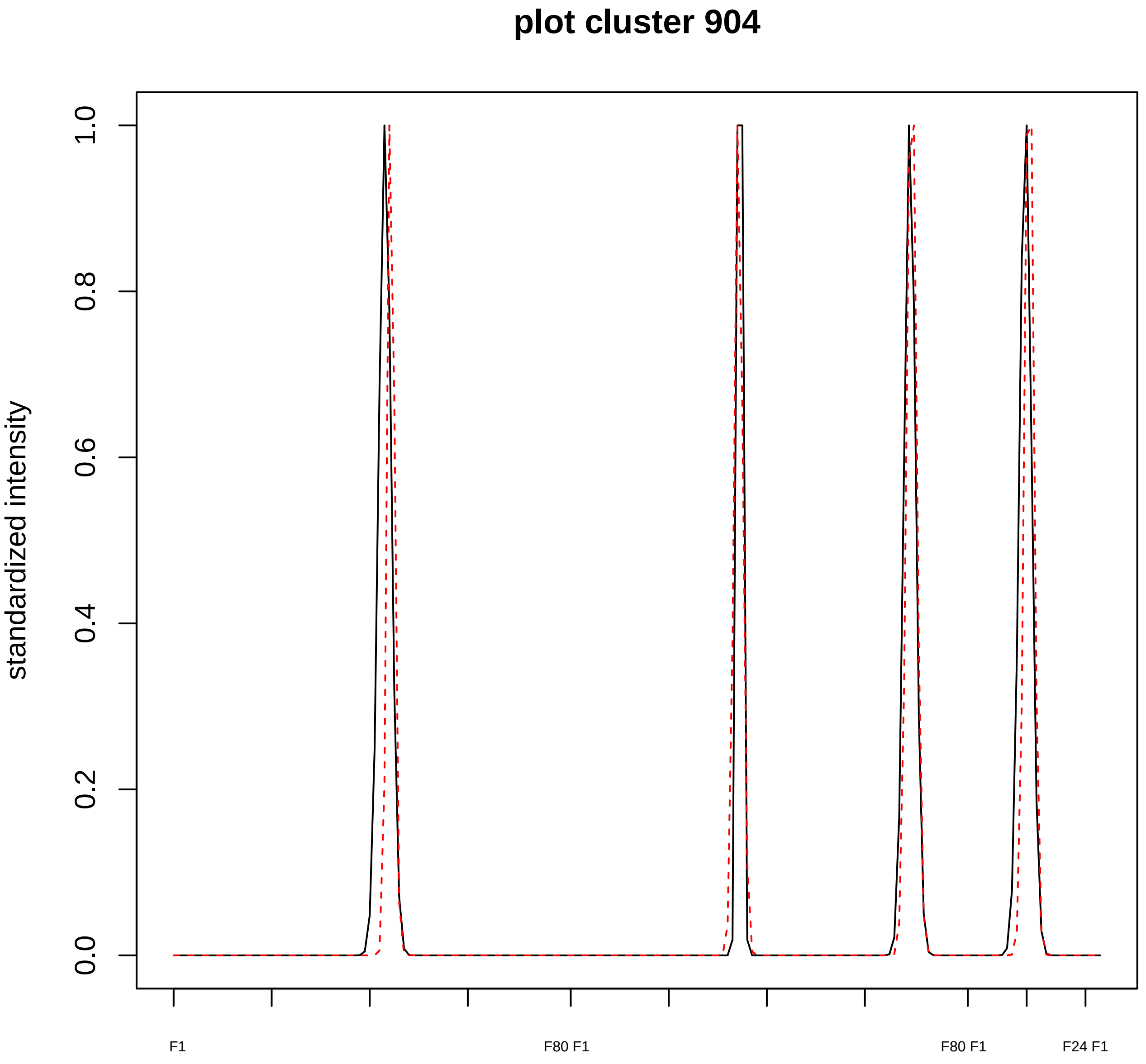

plot cluster 904
standardized intensity
0.0
0.2
0.4
0.6
0.8
1.0
F1
F80 F1
F80 F1
F24 F1

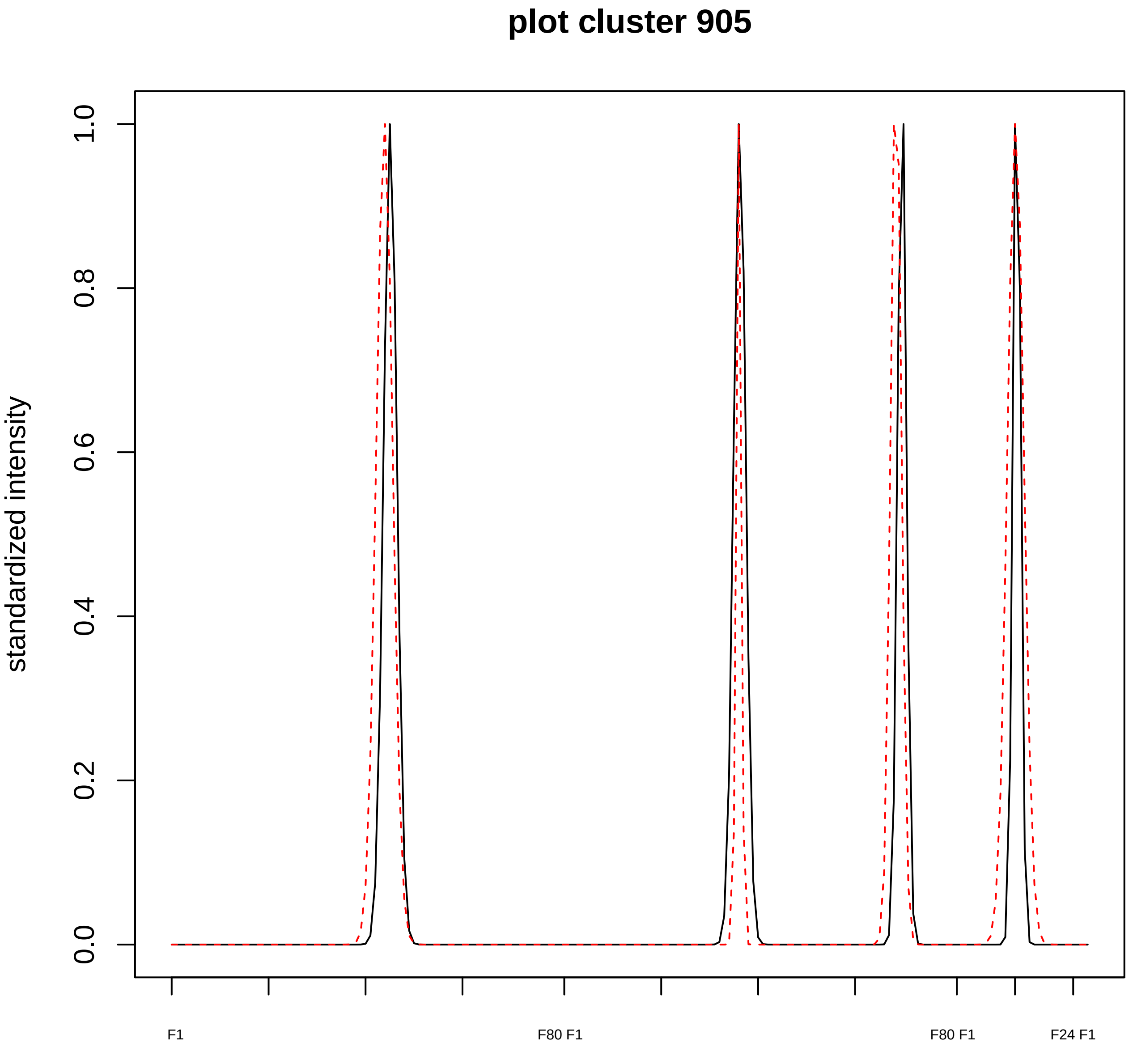

plot cluster 905
standardized intensity
0.0
0.2
0.4
0.6
0.8
1.0
F1
F80 F1
F80 F1
F24 F1

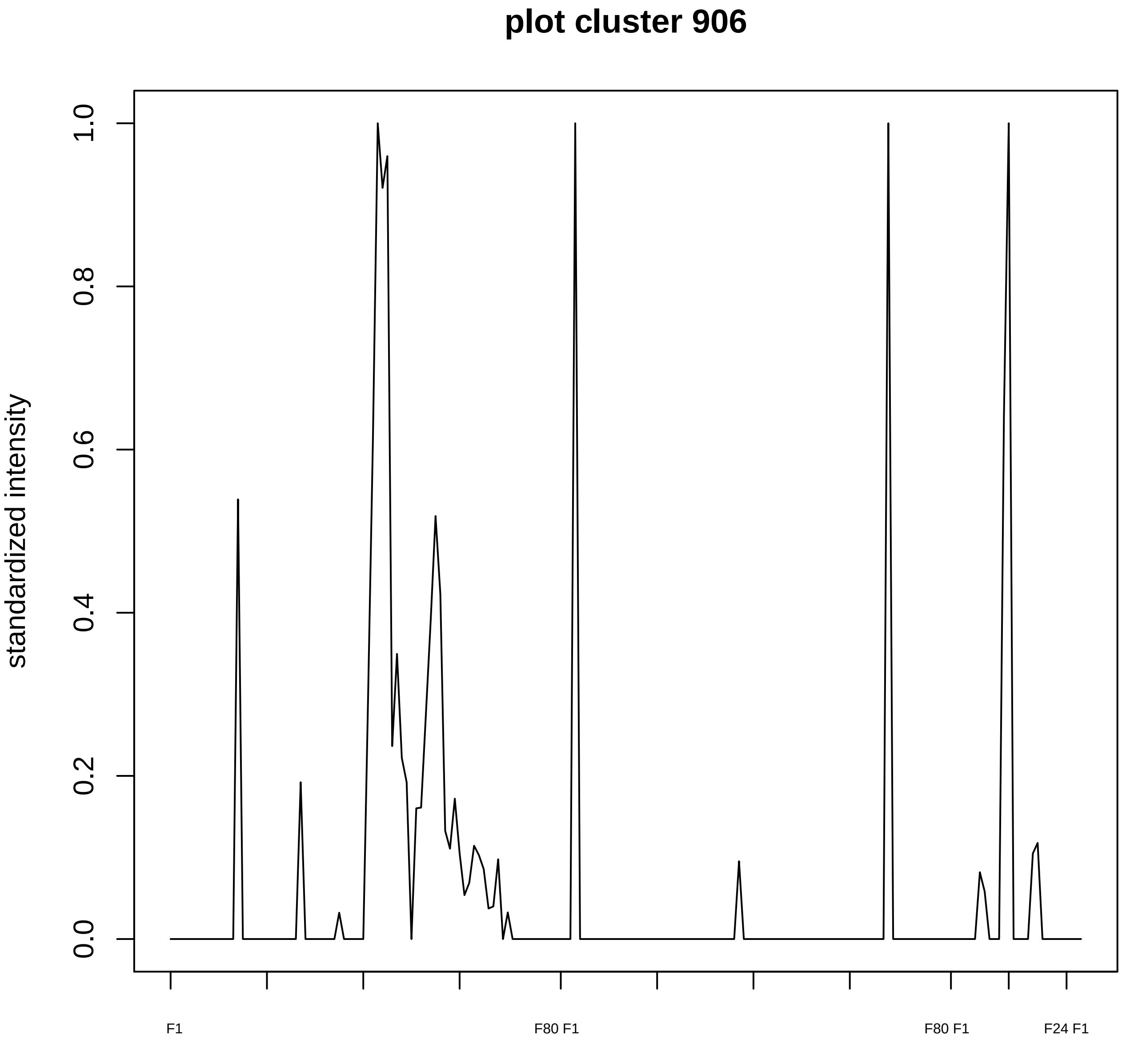

plot cluster 906
standardized intensity
0.0
0.2
0.4
0.6
0.8
1.0
F1
F80 F1
F80 F1
F24 F1

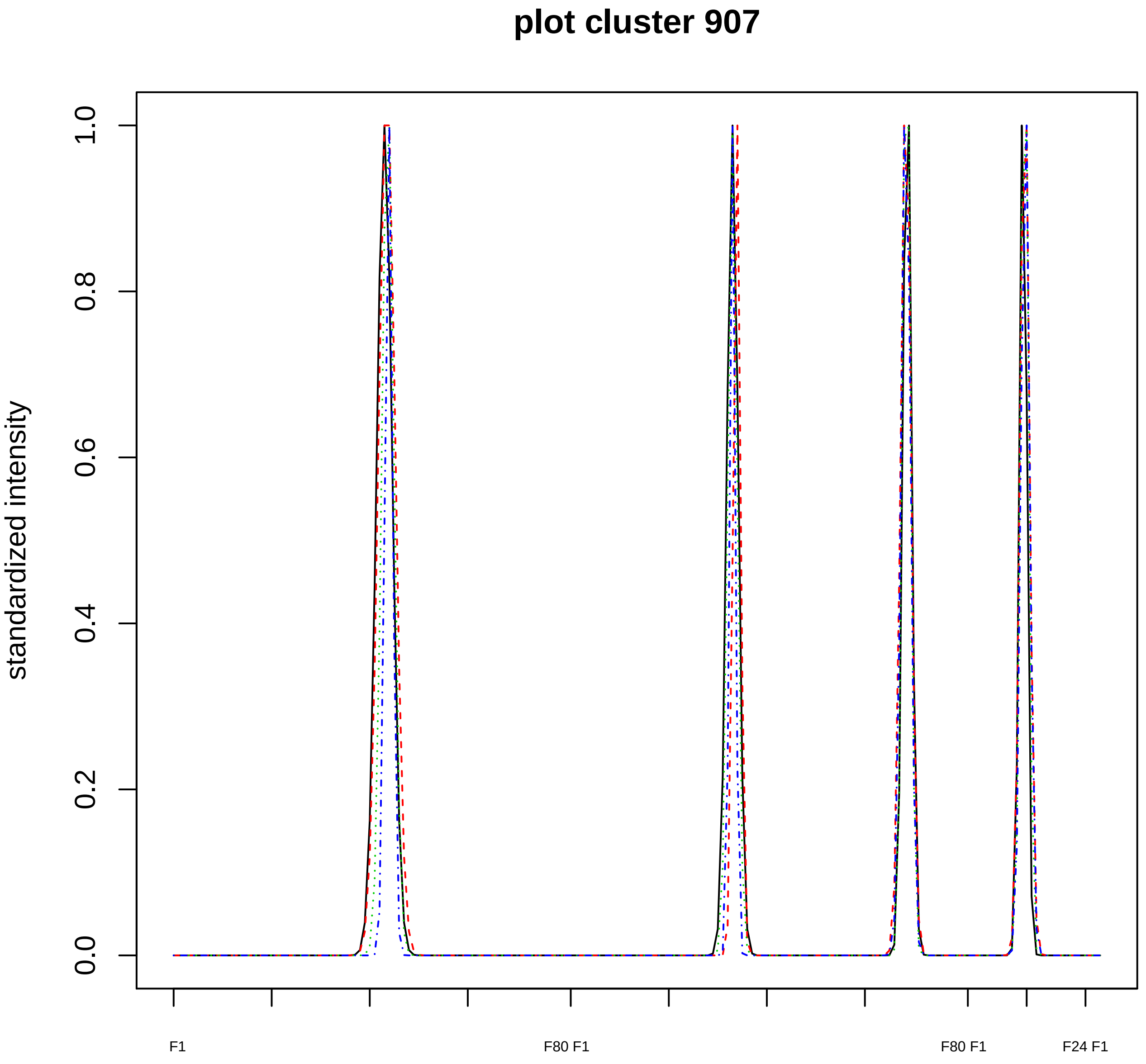

plot cluster 907
standardized intensity
0.0
0.2
0.4
0.6
0.8
1.0
F1
F80 F1
F80 F1
F24 F1

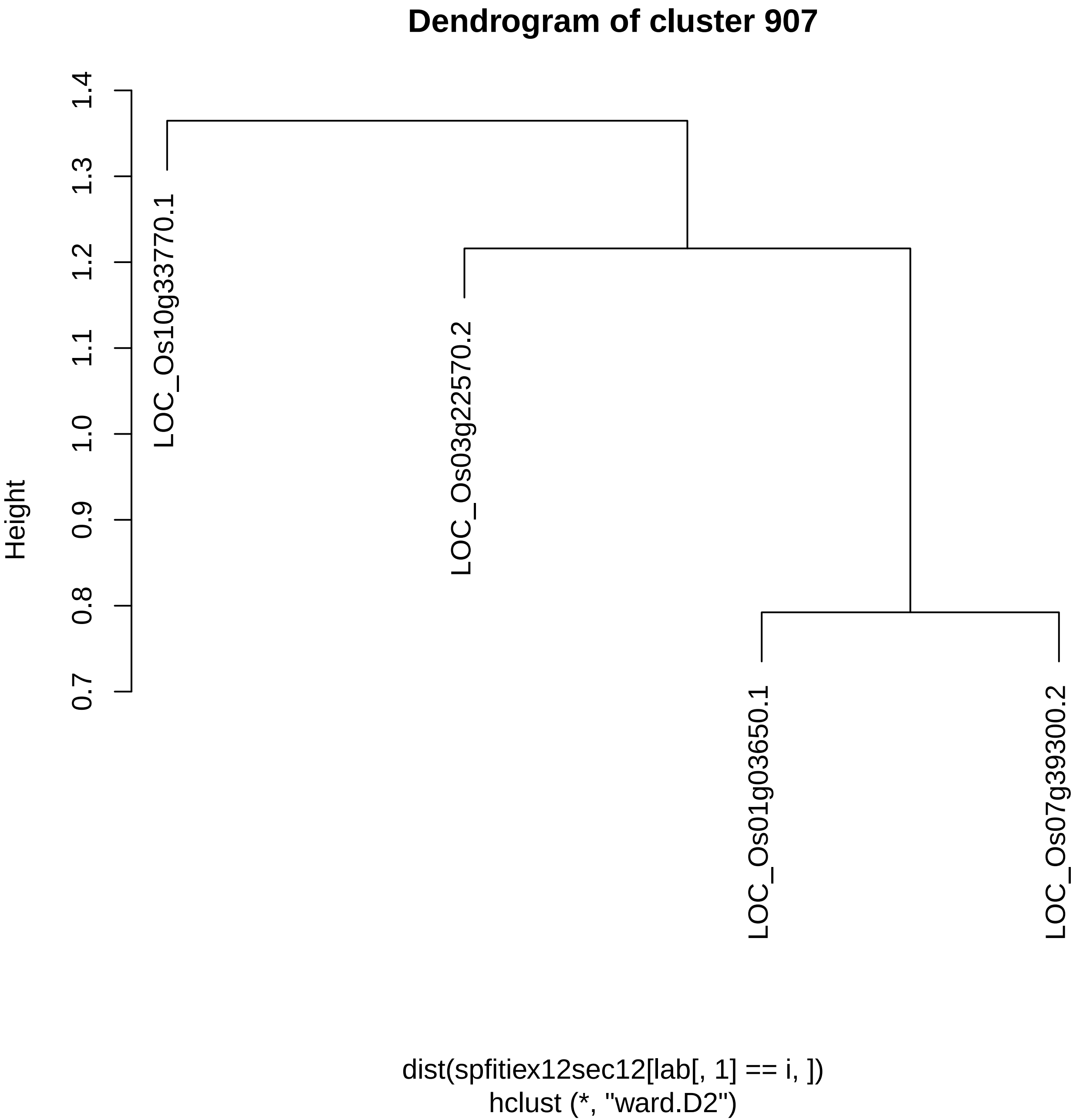

Dendrogram of cluster 907
Height
1.4
1.3
1.2
1.1
1.0
0.9
0.8
0.7
LOC_Os10g33770.1
LOC_Os03g22570.2
LOC_Os01g03650.1
LOC_Os07g39300.2
dist(spfitiex12sec12[lab[, 1] == i, ])
hclust (*, "ward.D2")

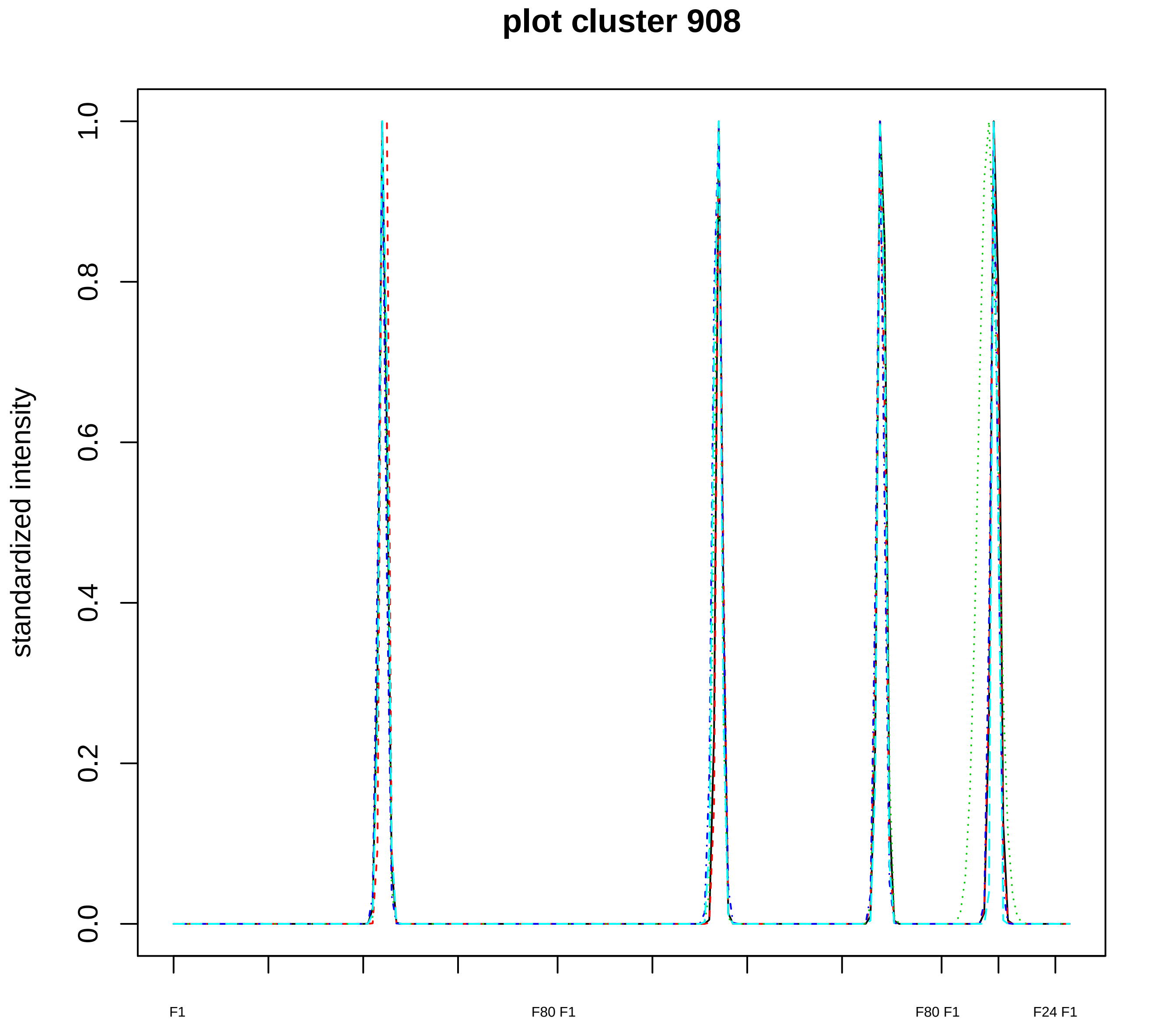

plot cluster 908
standardized intensity
1.0
0.8
0.6
0.4
0.2
0.0
F1
F80 F1
F80 F1
F24 F1

### Dendrogram of cluster 908

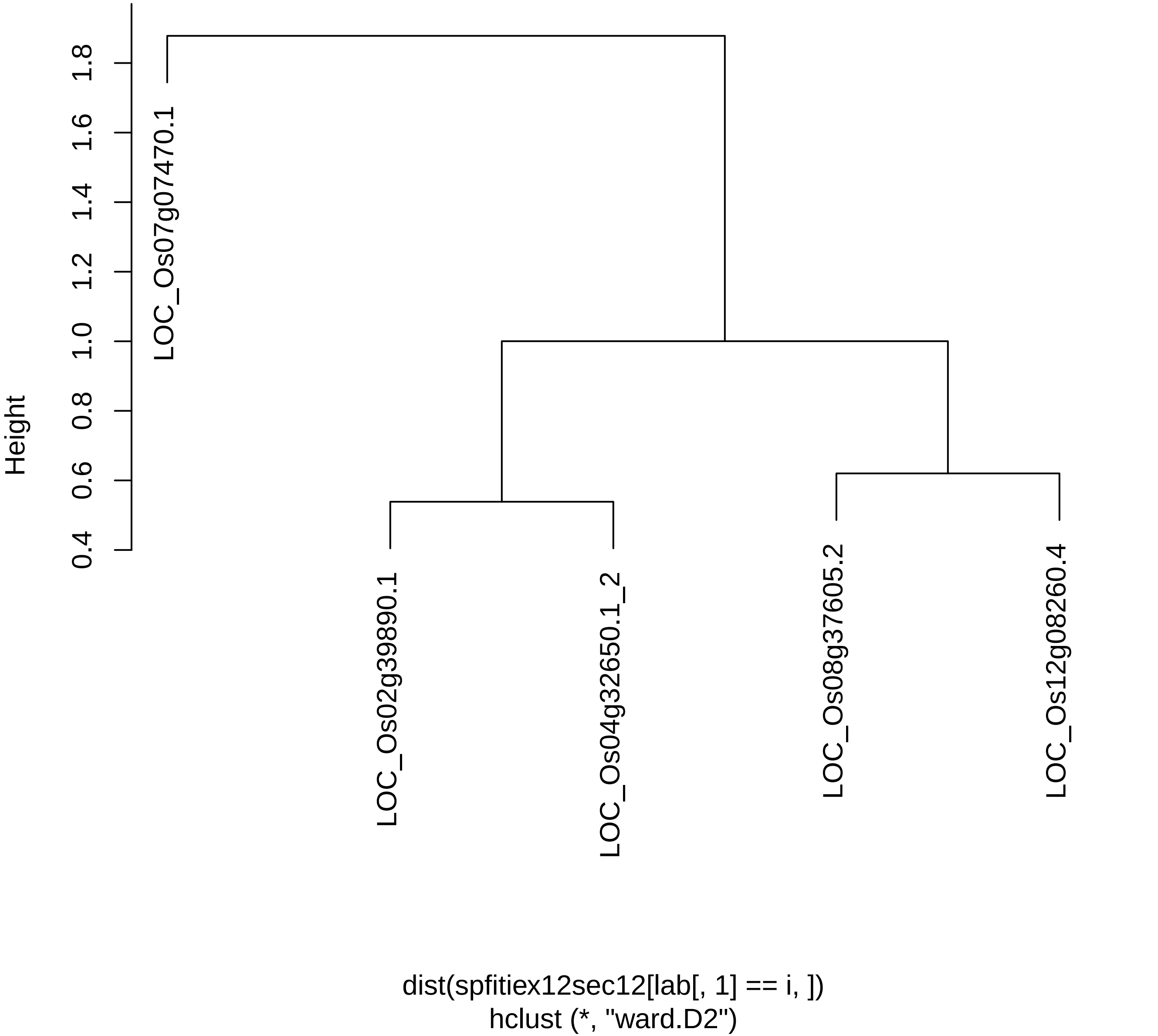

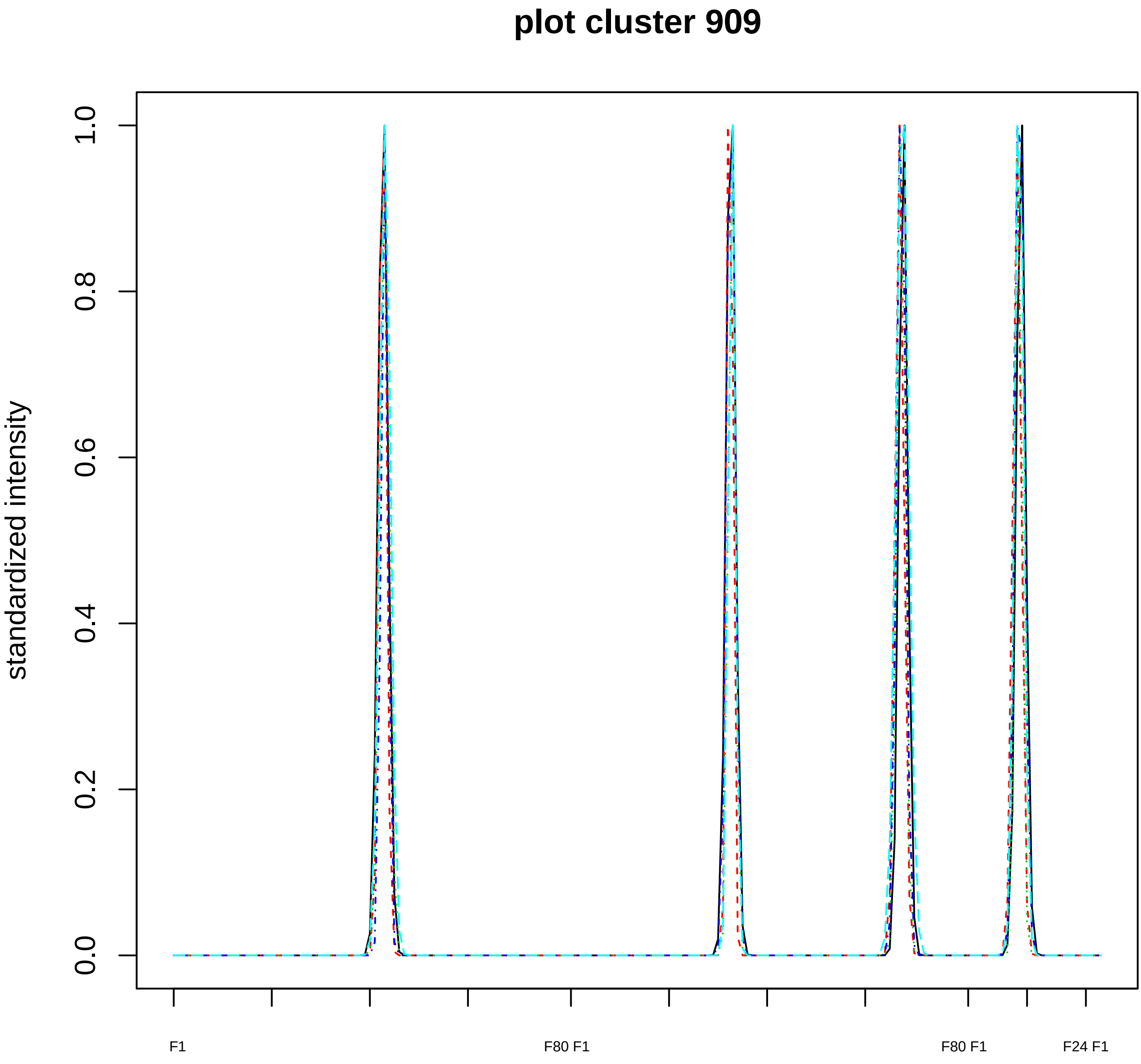

plot cluster 909
standardized intensity
0.0
0.2
0.4
0.6
0.8
1.0
F1
F80 F1
F80 F1
F24 F1

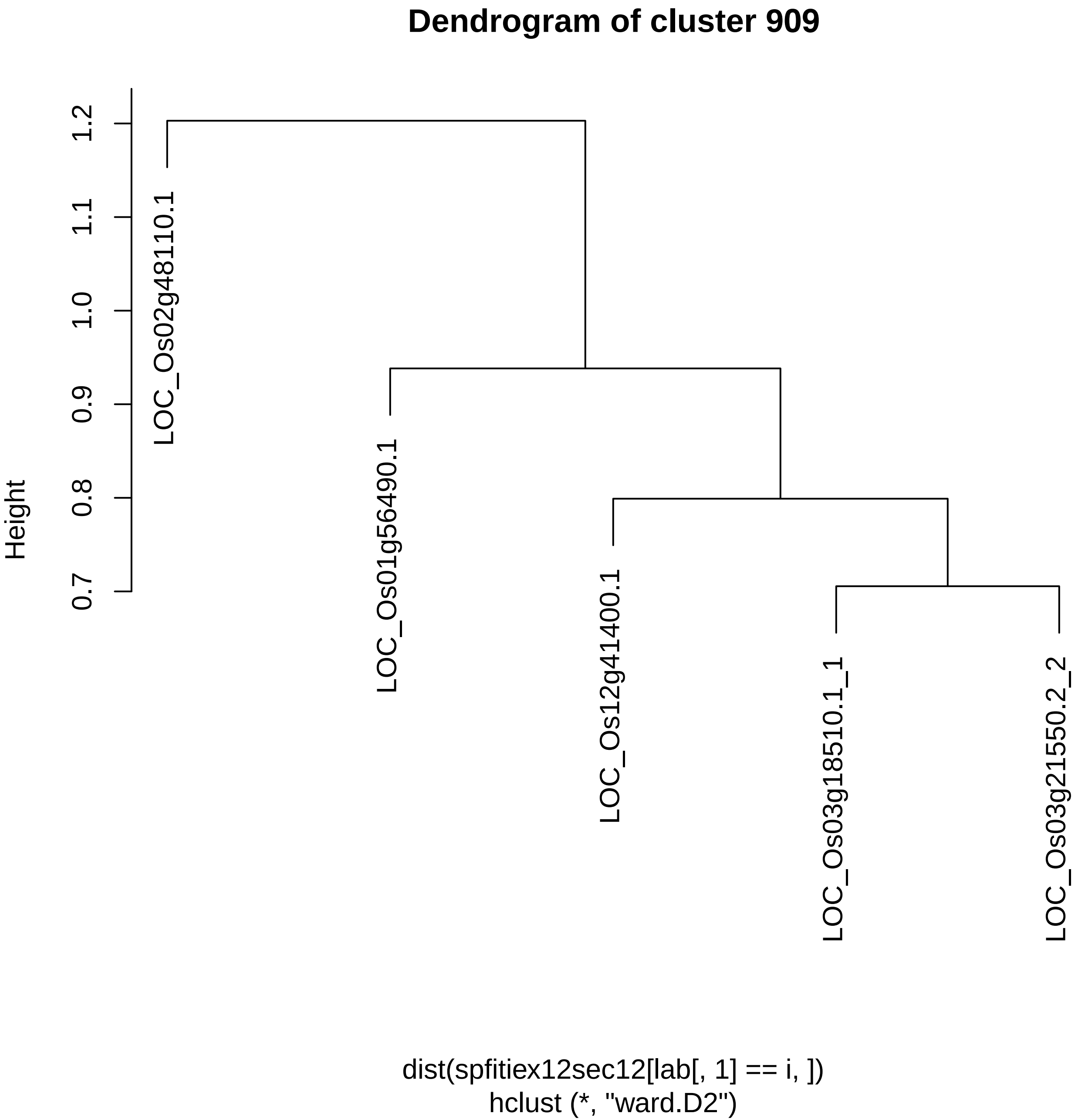

Dendrogram of cluster 909
Height
0.7
0.8
0.9
1.0
1.1
1.2
LOC_Os02g48110.1
LOC_Os01g56490.1
LOC_Os12g41400.1
LOC_Os03g18510.1_1
LOC_Os03g21550.2_2
dist(spfitiex12sec12[lab[, 1] == i, ])
hclust (*, "ward.D2")

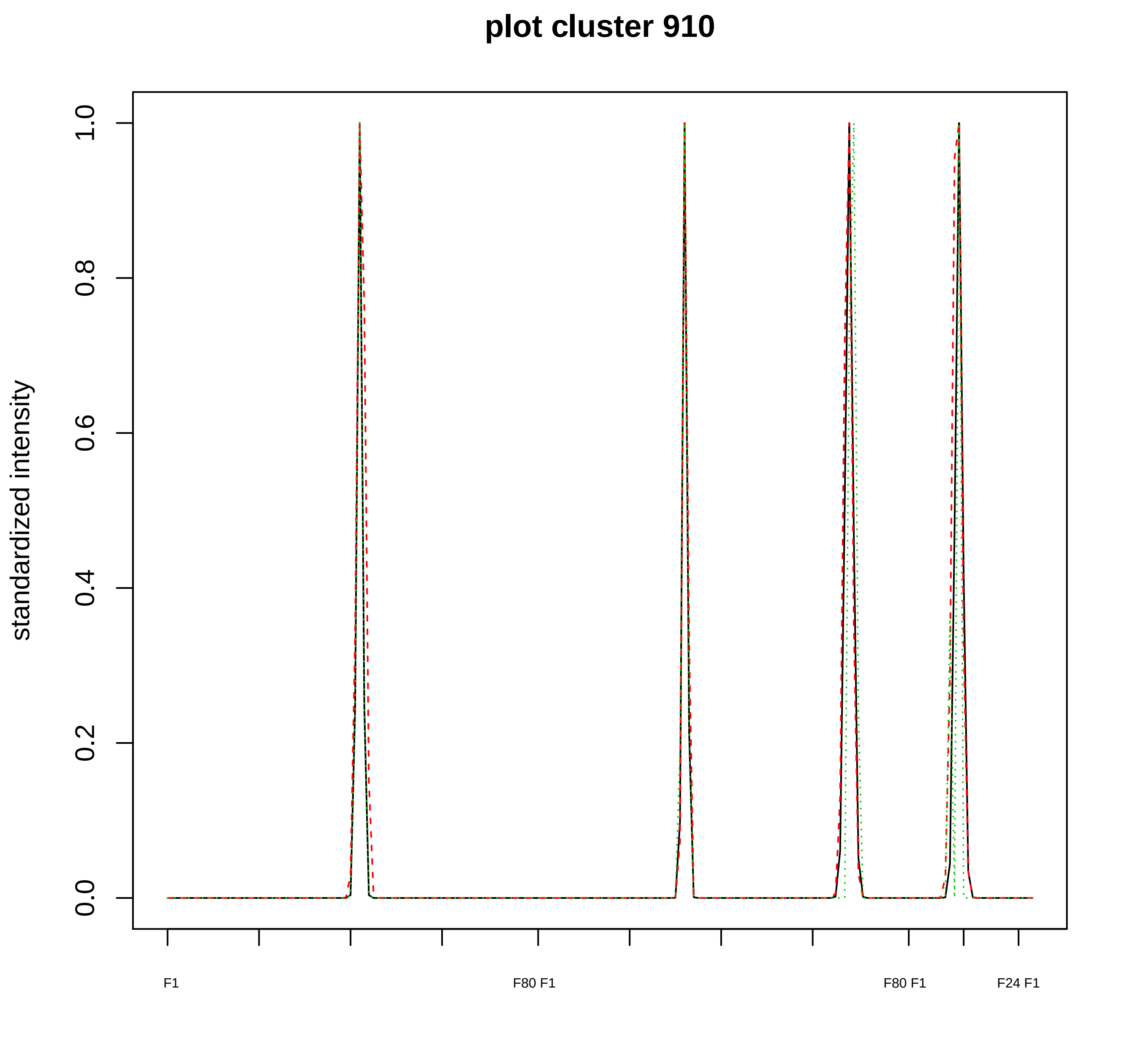

plot cluster 910
standardized intensity
0.0
0.2
0.4
0.6
0.8
1.0
F1
F80 F1
F80 F1
F24 F1

### Dendrogram of cluster 910

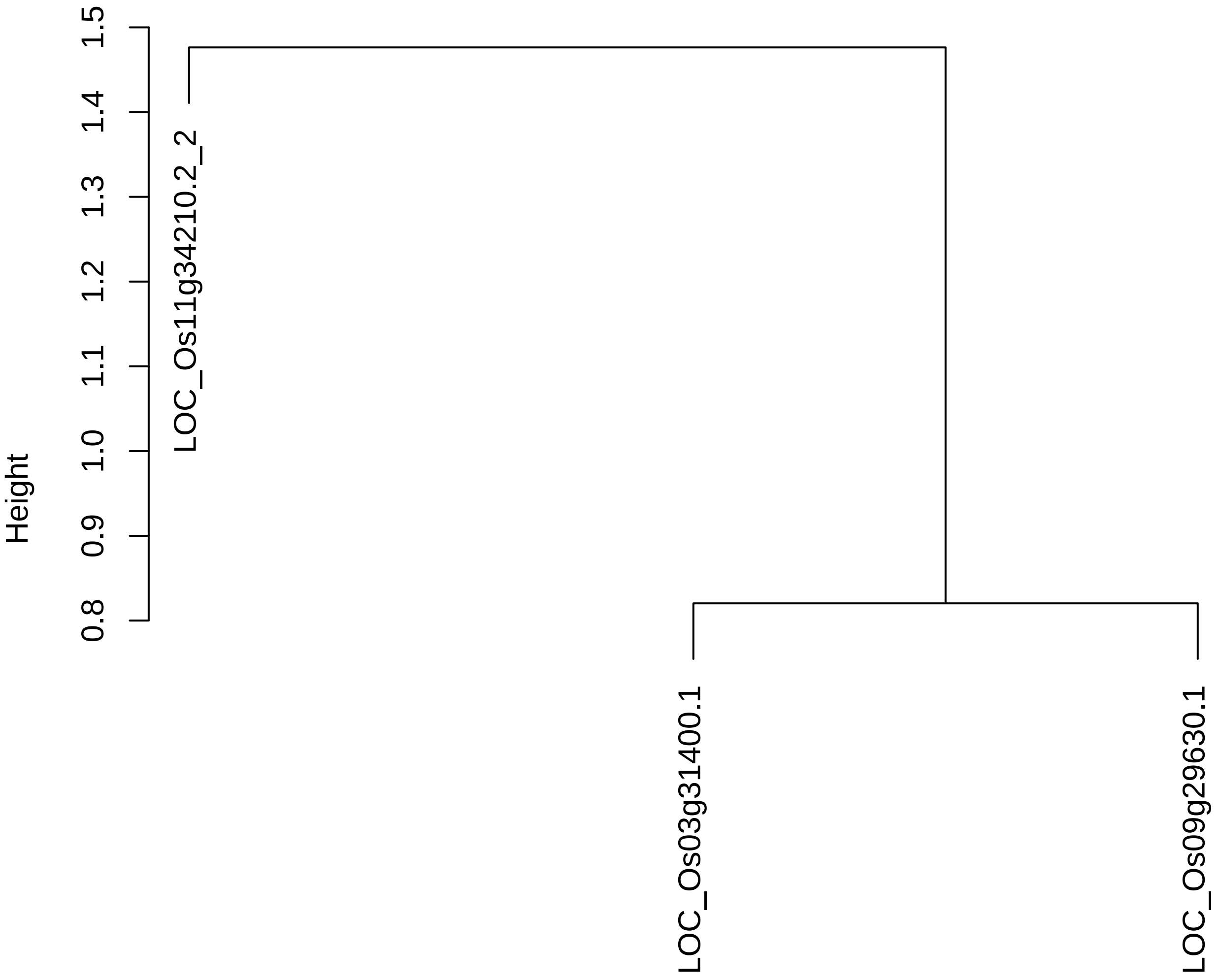


dist(spfitiex12sec12[lab[, 1] == i, ])
hclust (*, "ward.D2")

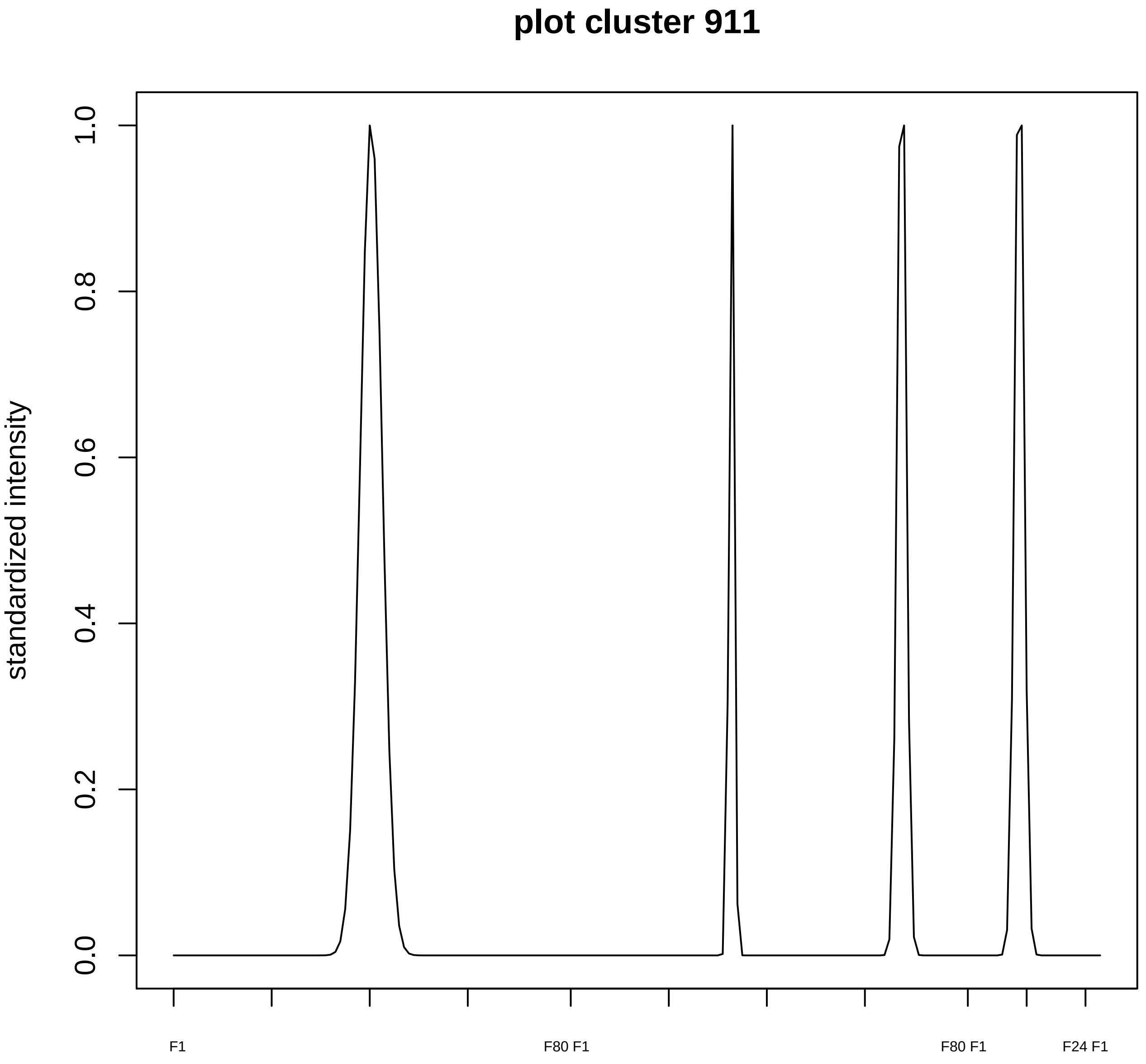

plot cluster 911
standardized intensity
0.0
0.2
0.4
0.6
0.8
1.0
F1
F80 F1
F80 F1
F24 F1

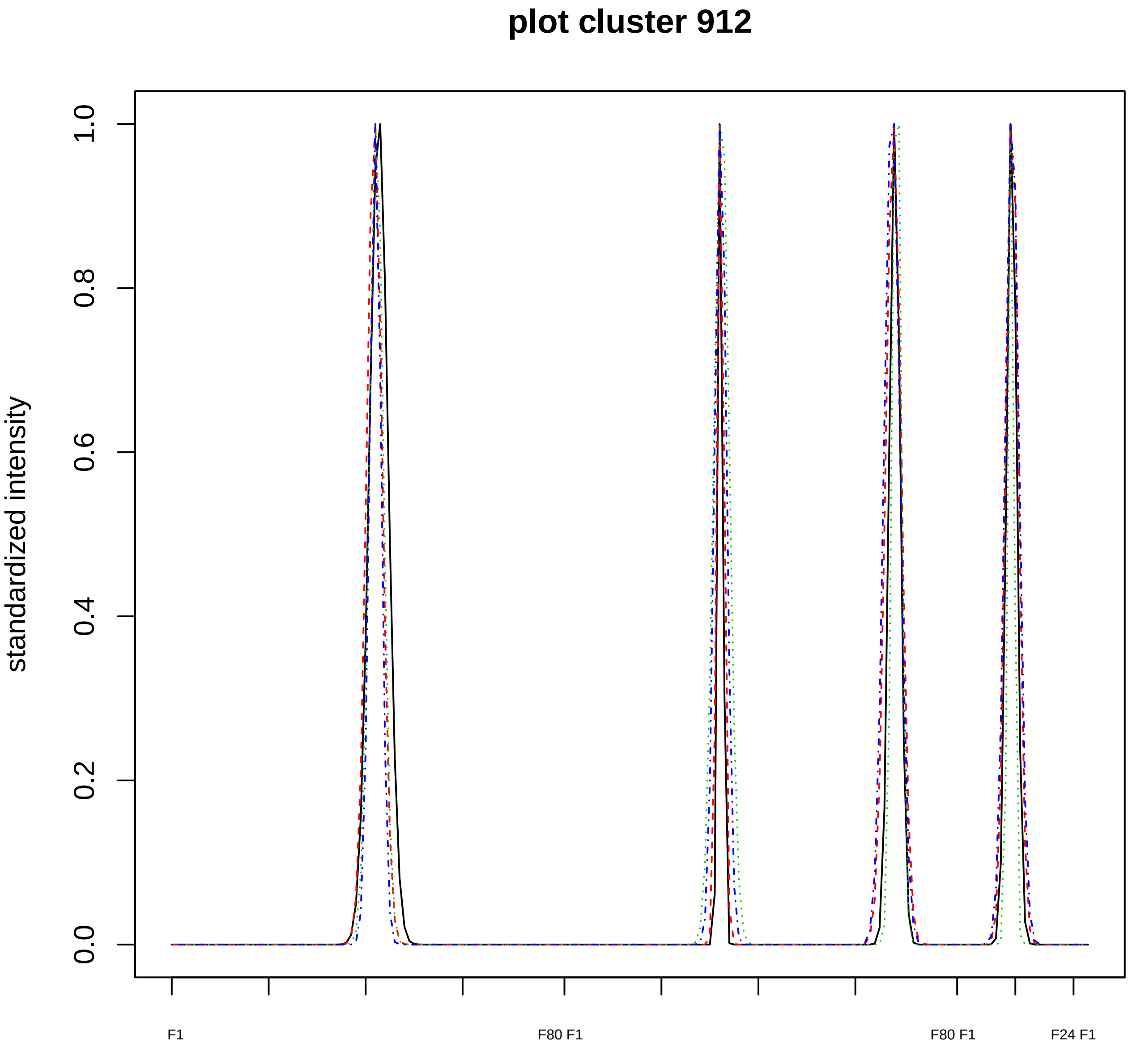
plot cluster 912
standardized intensity
0.0
0.2
0.4
0.6
0.8
1.0
F1
F80 F1
F80 F1
F24 F1

### Dendrogram of cluster 912

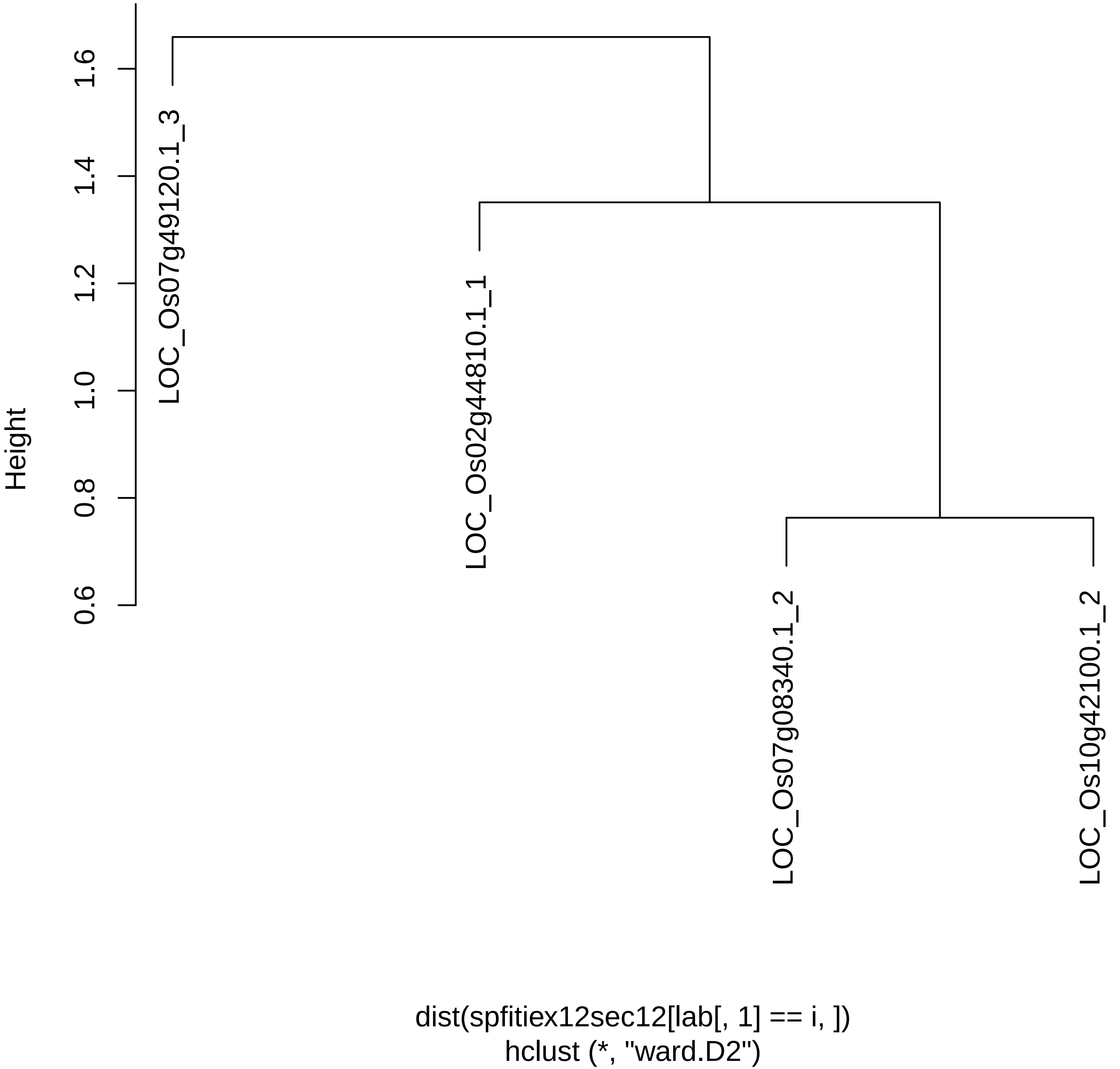

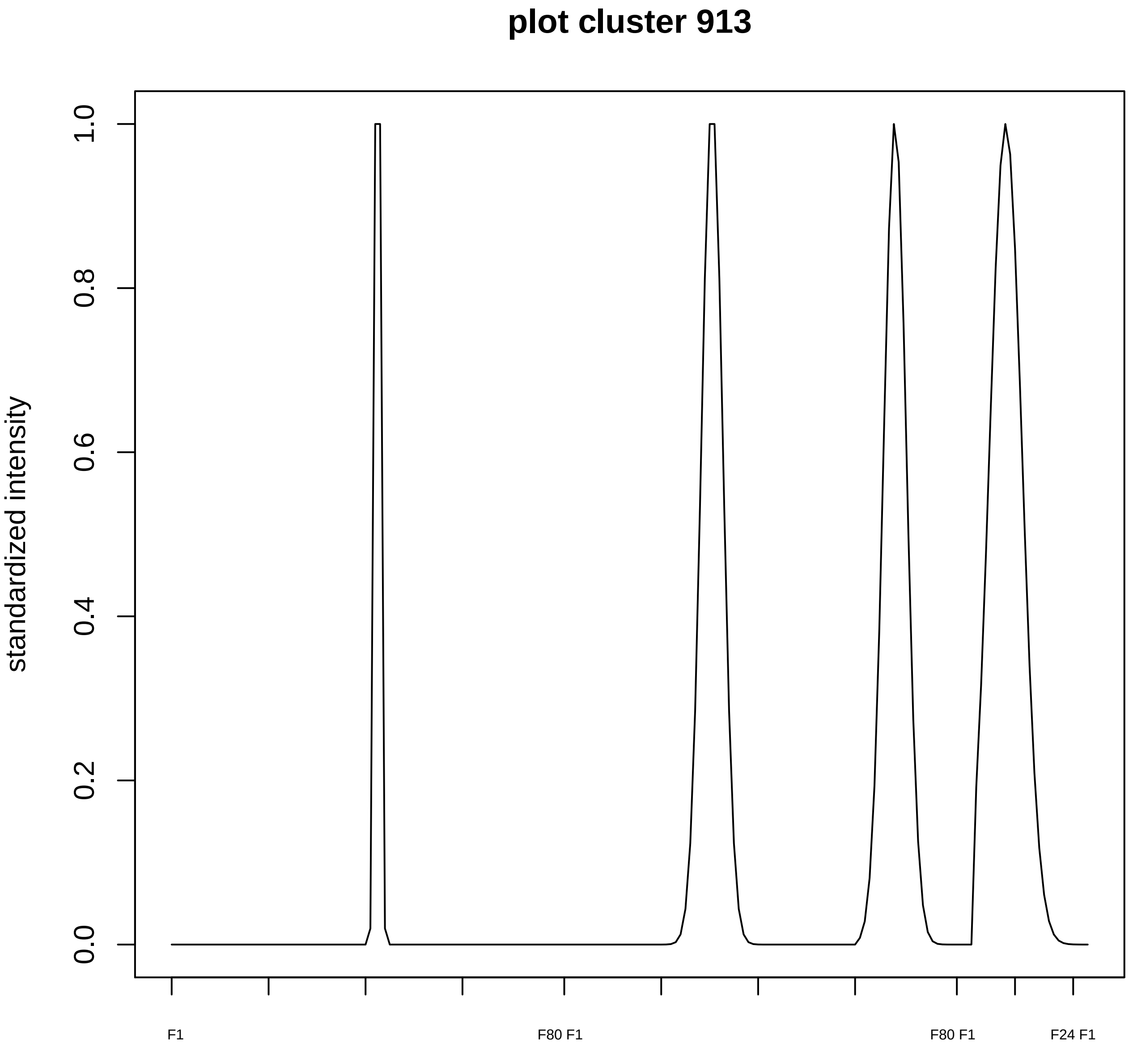

plot cluster 913
standardized intensity
0.0
0.2
0.4
0.6
0.8
1.0
F1
F80 F1
F80 F1
F24 F1

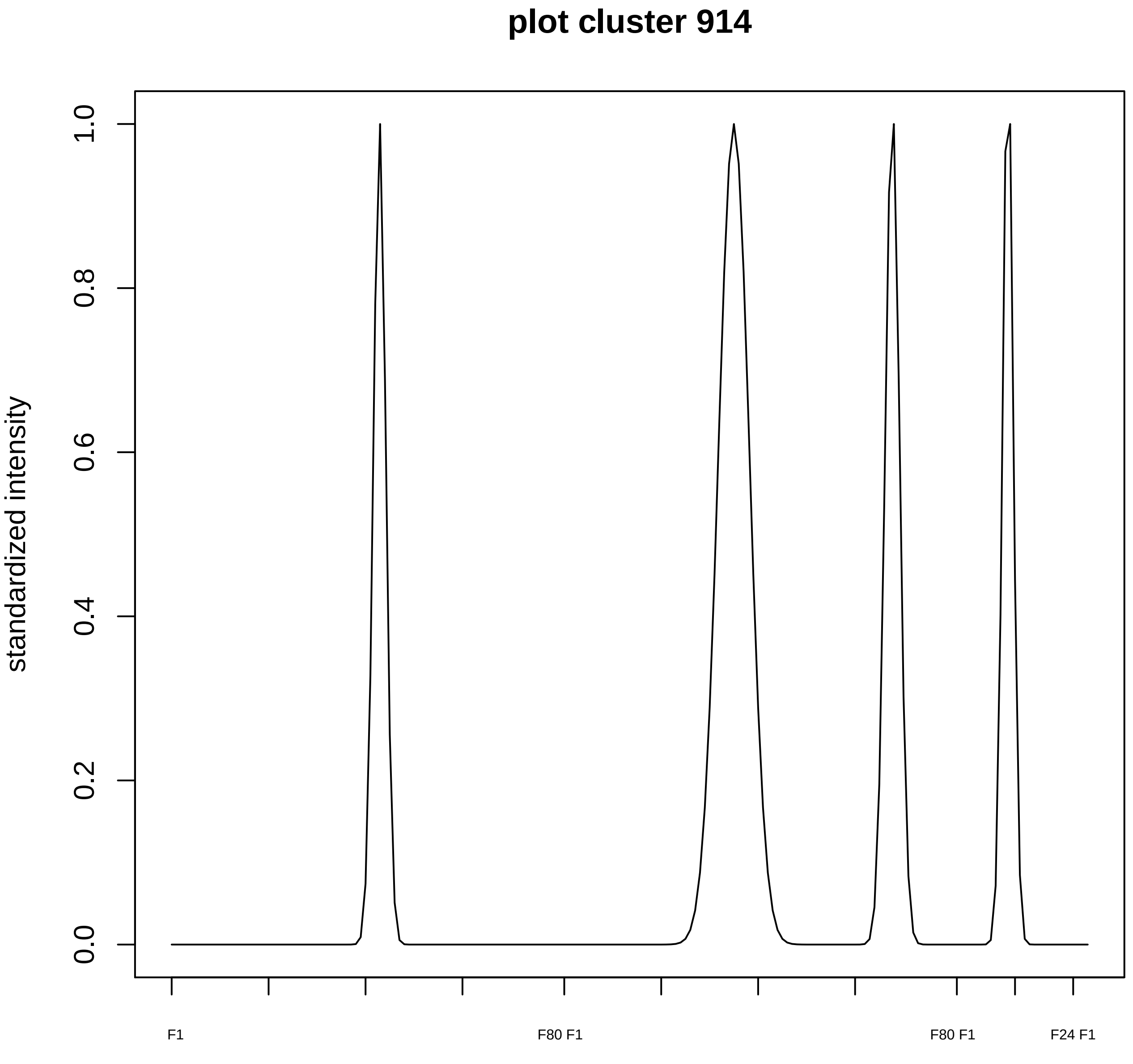
plot cluster 914
standardized intensity
0.0
0.2
0.4
0.6
0.8
1.0
F1
F80 F1
F80 F1
F24 F1

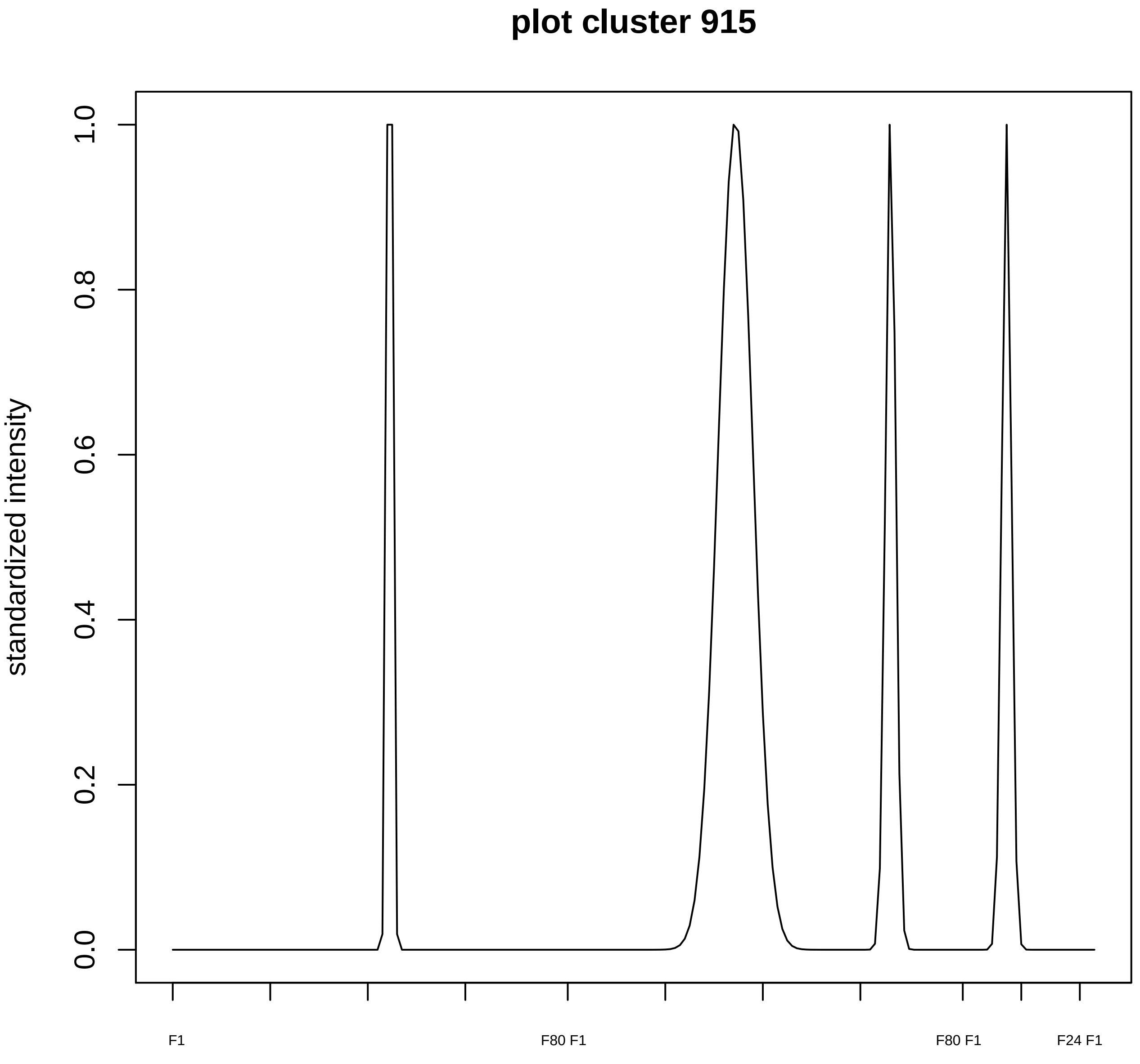

plot cluster 915
standardized intensity
1.0
0.8
0.6
0.4
0.2
0.0
F1
F80 F1
F80 F1
F24 F1

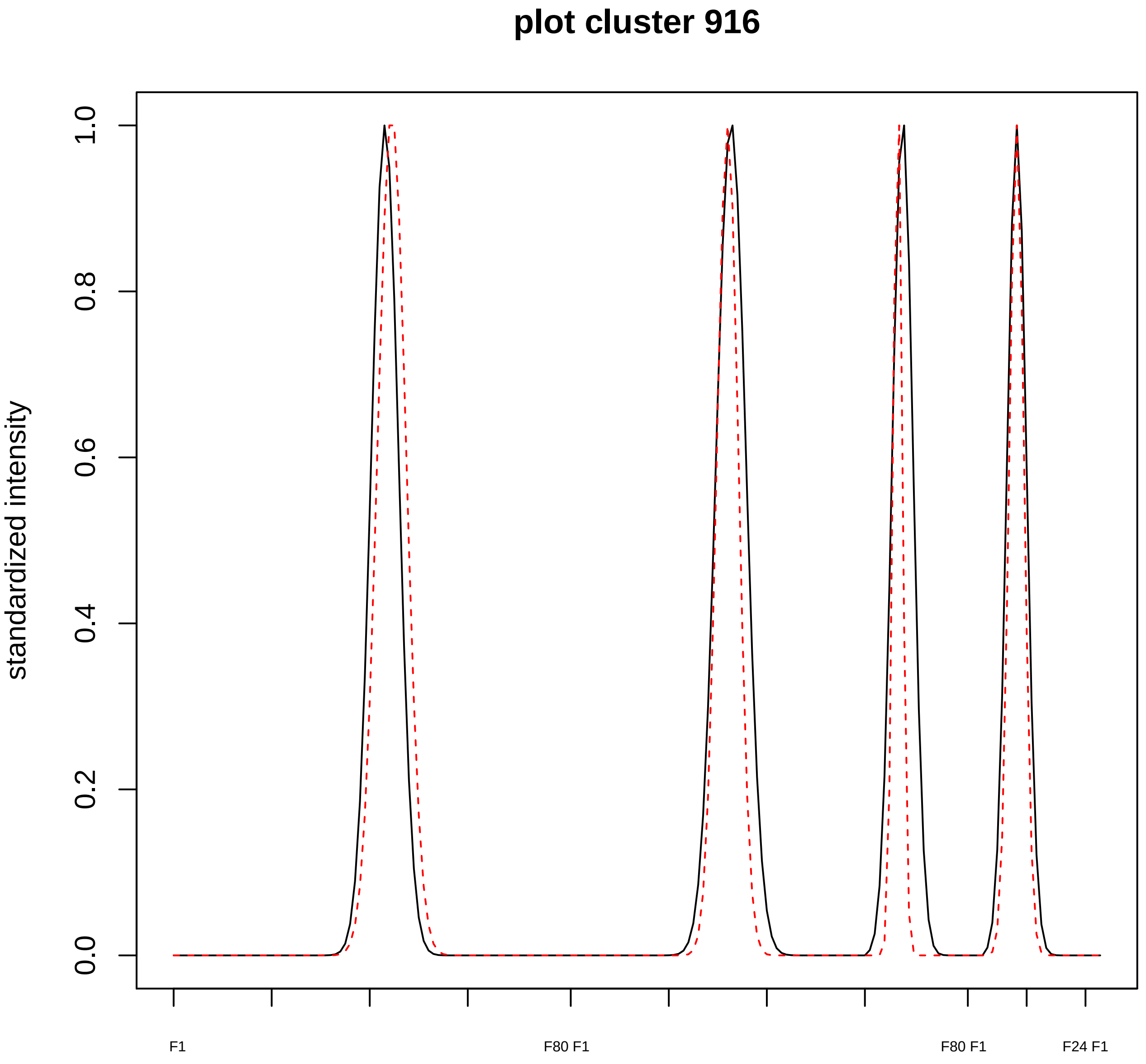

plot cluster 916
standardized intensity
0.0
0.2
0.4
0.6
0.8
1.0
F1
F80 F1
F80 F1
F24 F1

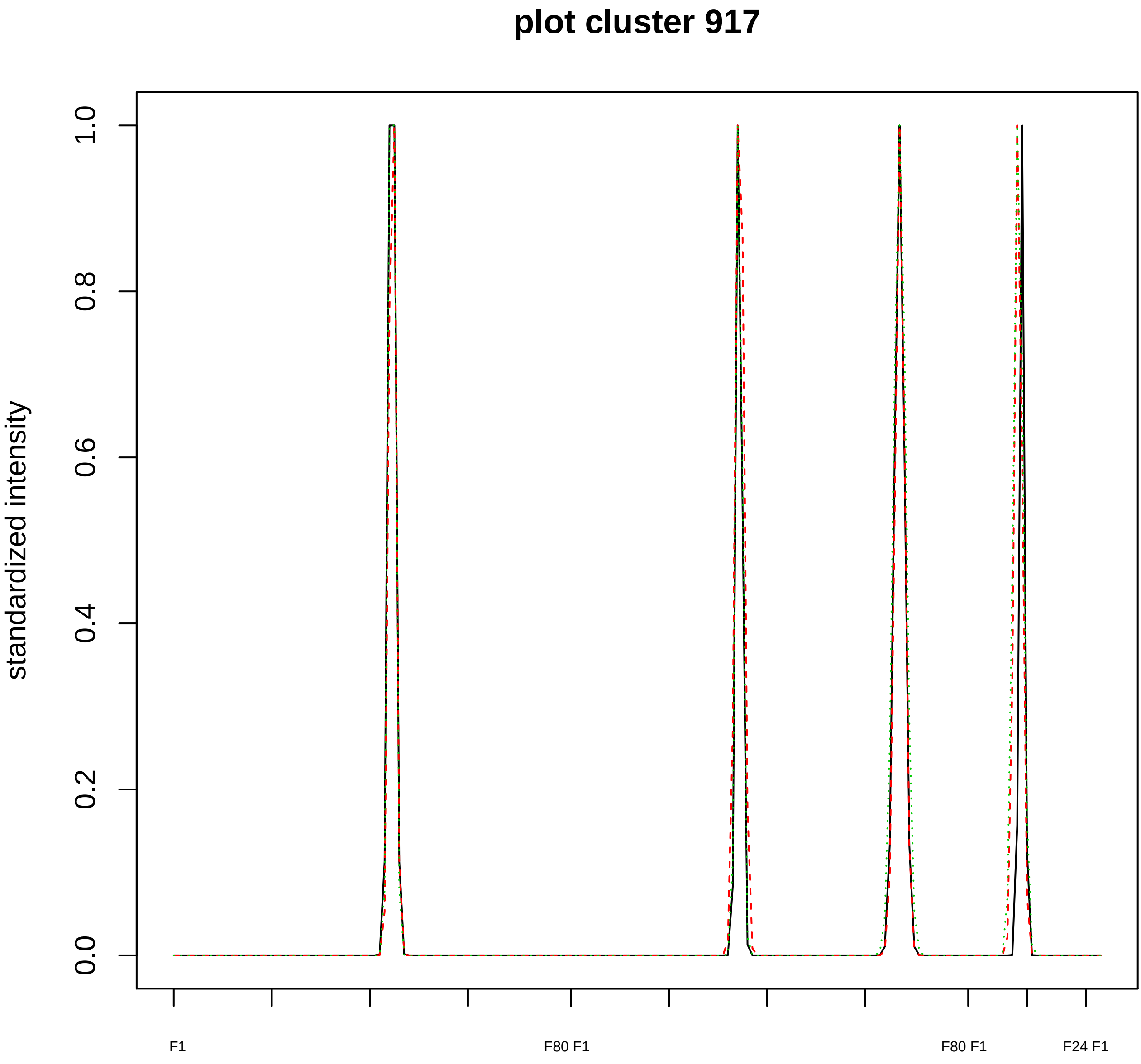

plot cluster 917
standardized intensity
0.0
0.2
0.4
0.6
0.8
1.0
F1
F80 F1
F80 F1
F24 F1

### Dendrogram of cluster 917

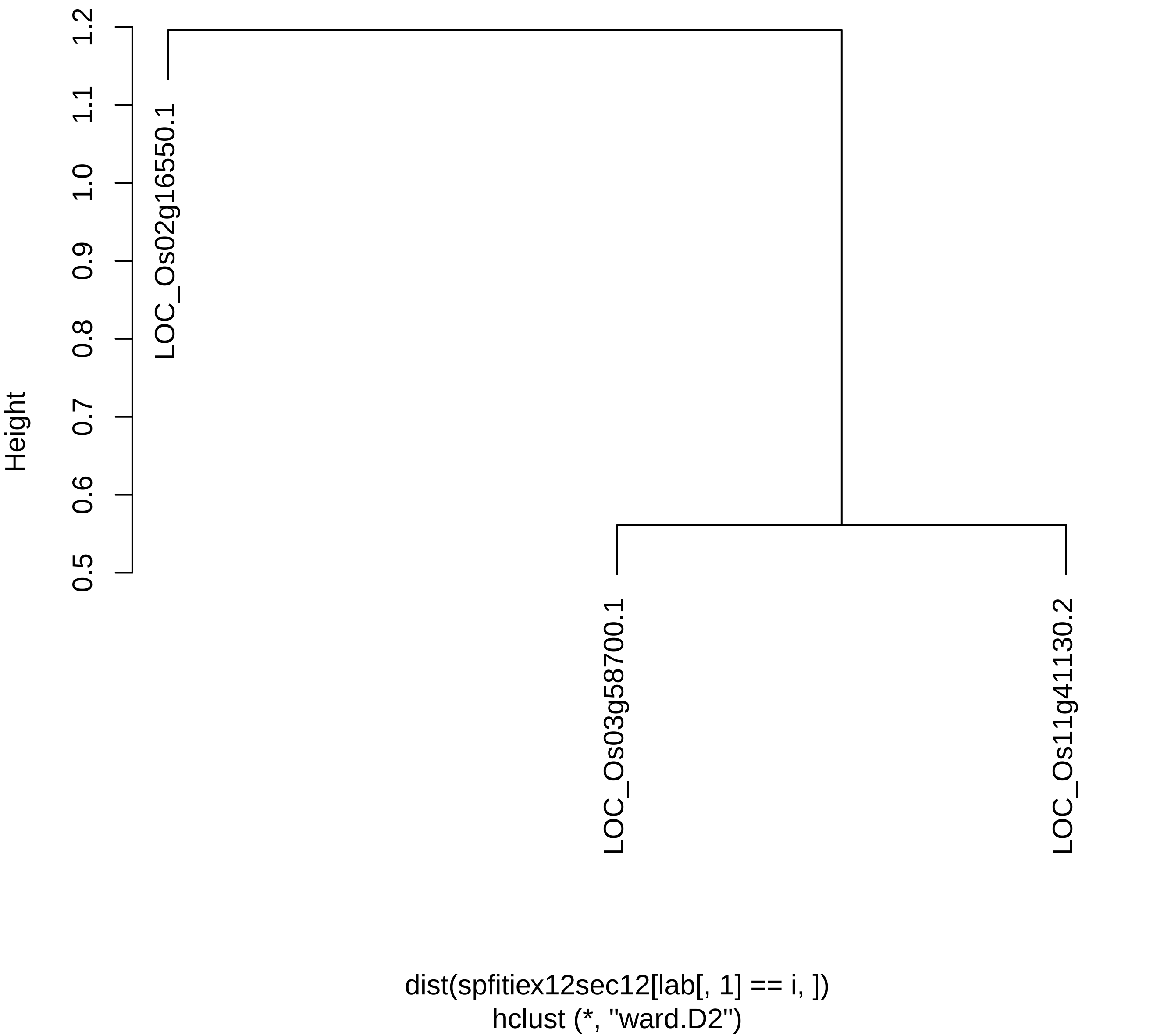

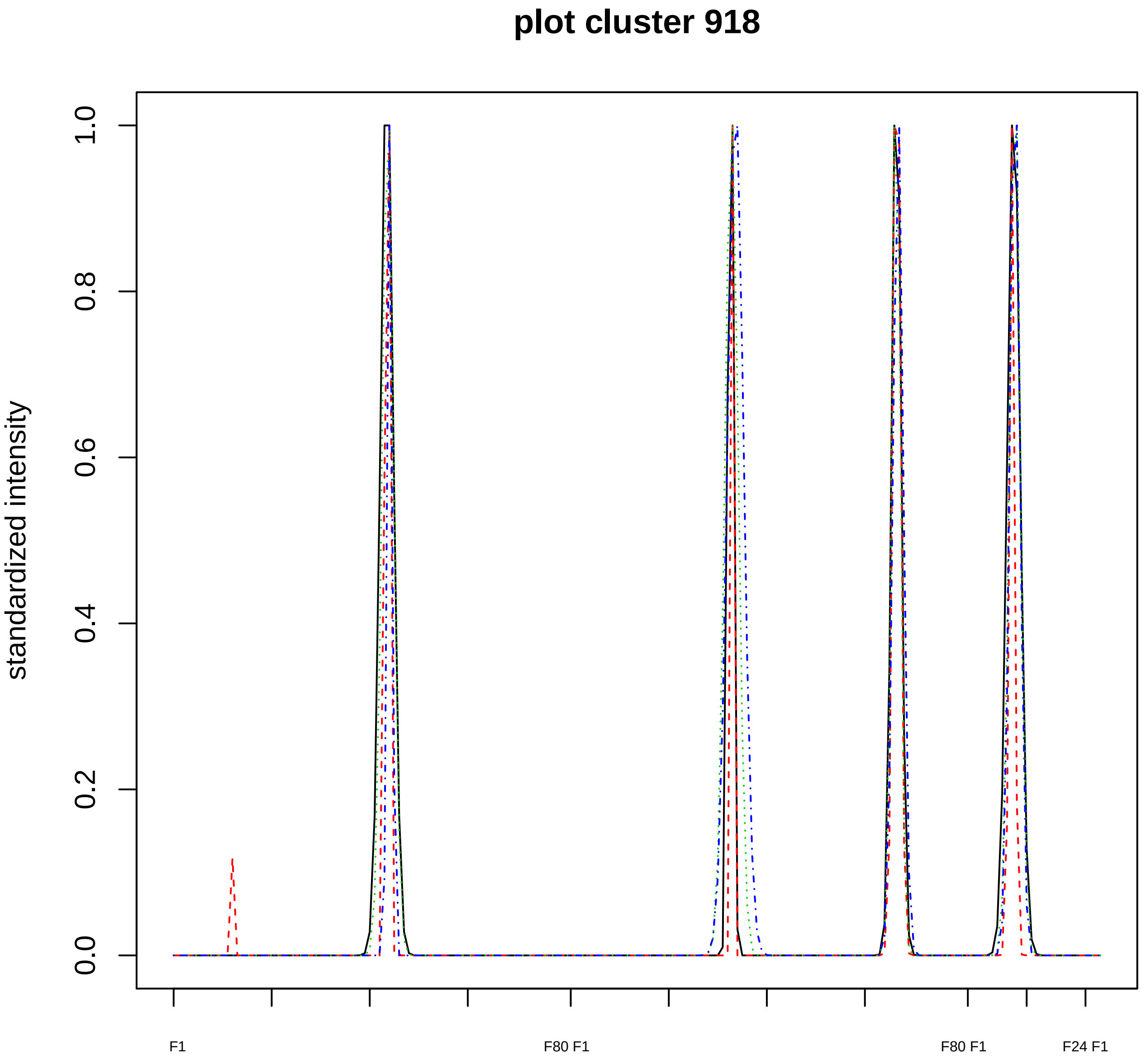

plot cluster 918
standardized intensity
0.0
0.2
0.4
0.6
0.8
1.0
F1
F80 F1
F80 F1
F24 F1

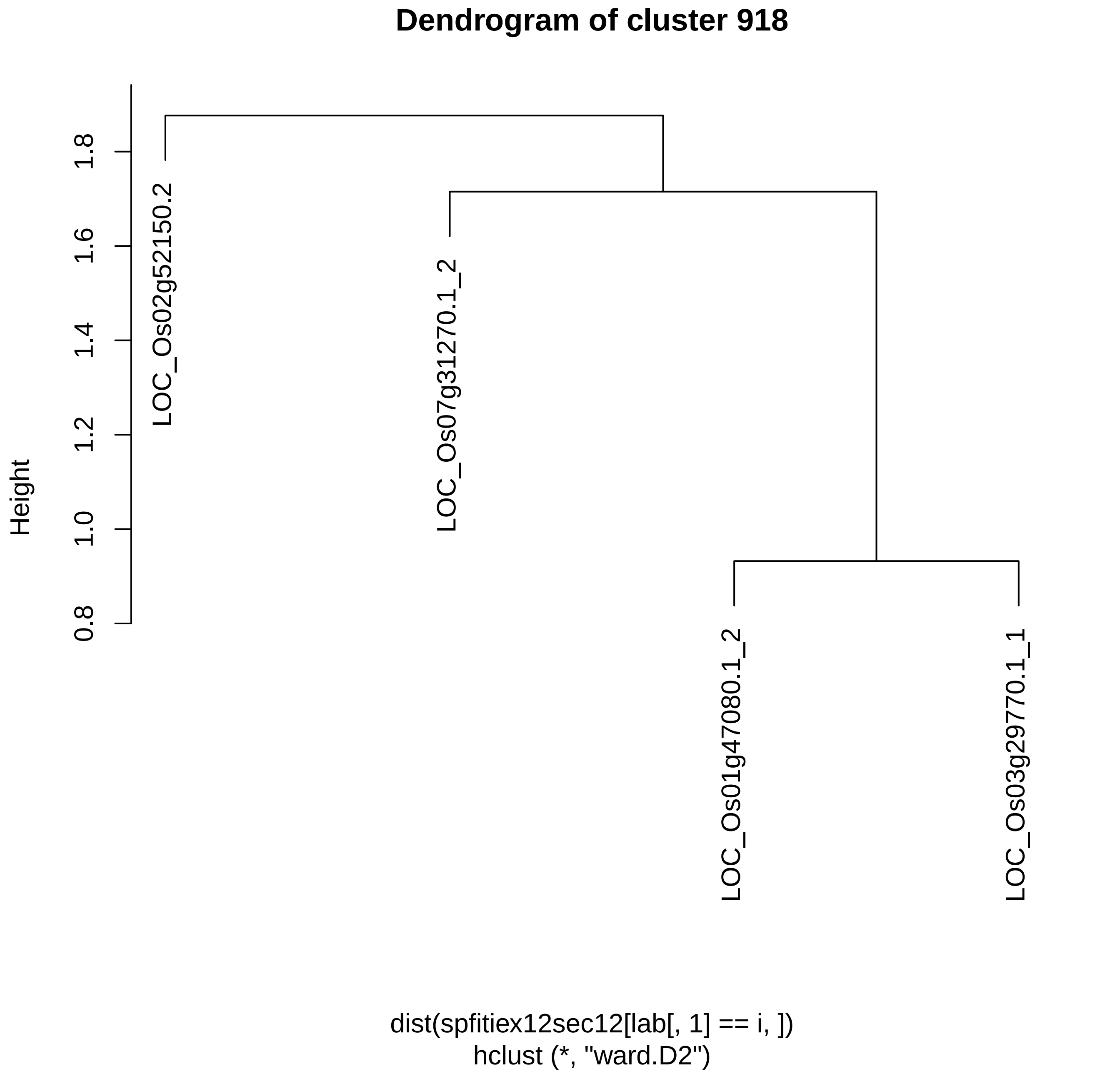
Dendrogram of cluster 918
Height
0.8
1.0
1.2
1.4
1.6
1.8
LOC_Os02g52150.2
LOC_Os07g31270.1_2
LOC_Os01g47080.1_2
LOC_Os03g29770.1_1
dist(spfitiex12sec12[lab[, 1] == i, ])
hclust (*, "ward.D2")

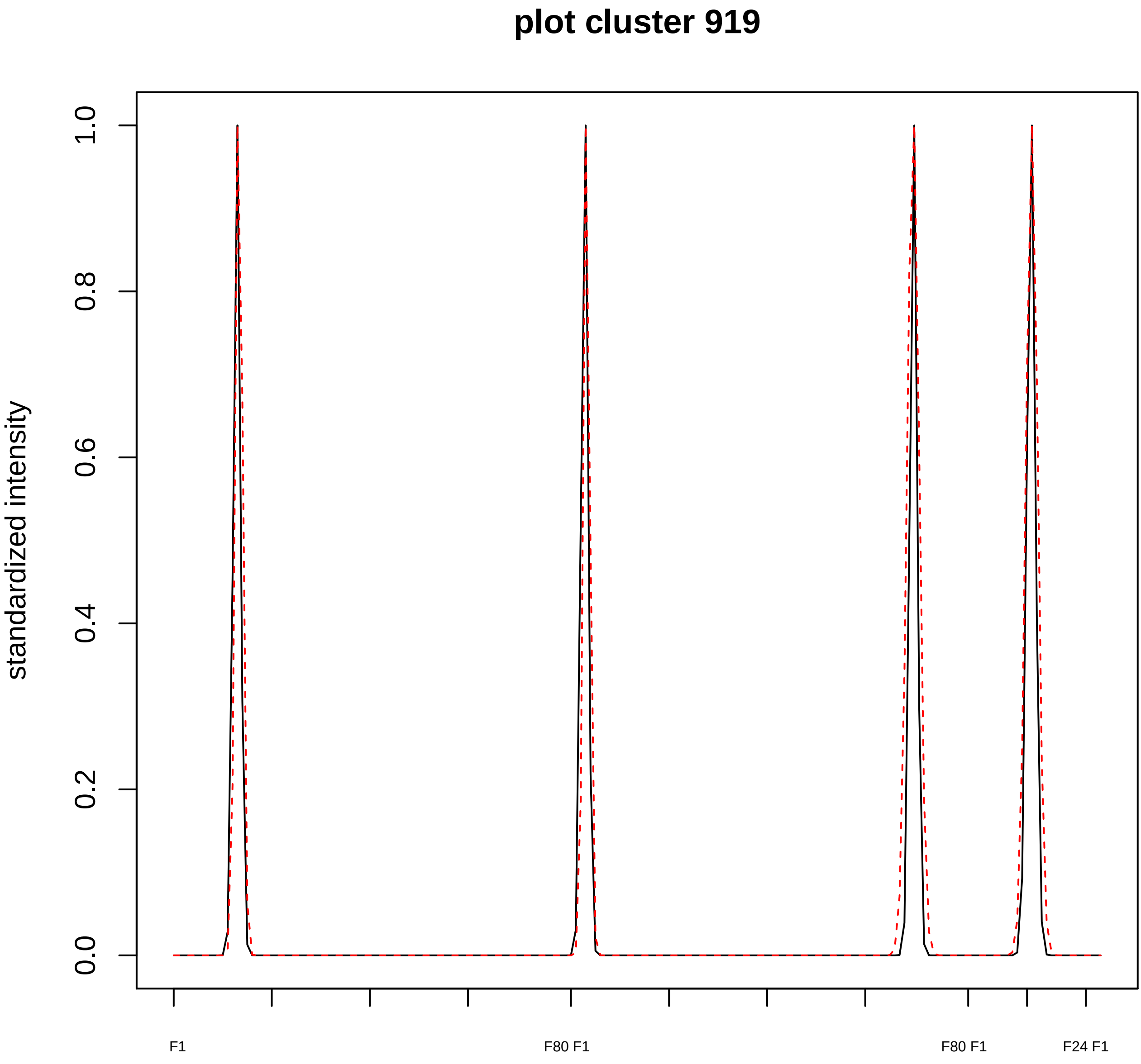

plot cluster 919
standardized intensity
1.0
0.8
0.6
0.4
0.2
0.0
F1
F80 F1
F80 F1
F24 F1

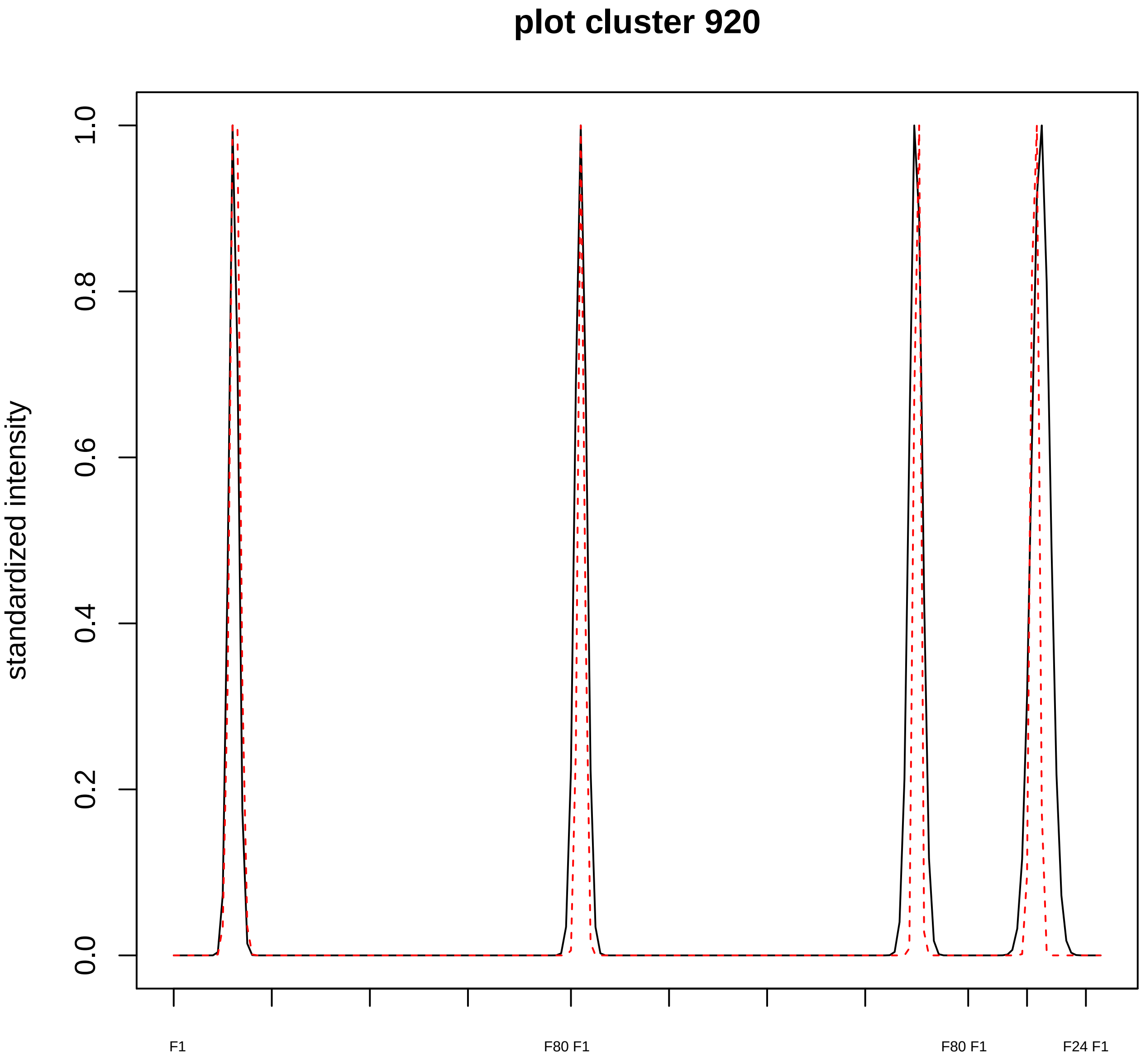

plot cluster 920
standardized intensity
0.0
0.2
0.4
0.6
0.8
1.0
F1
F80 F1
F80 F1
F24 F1

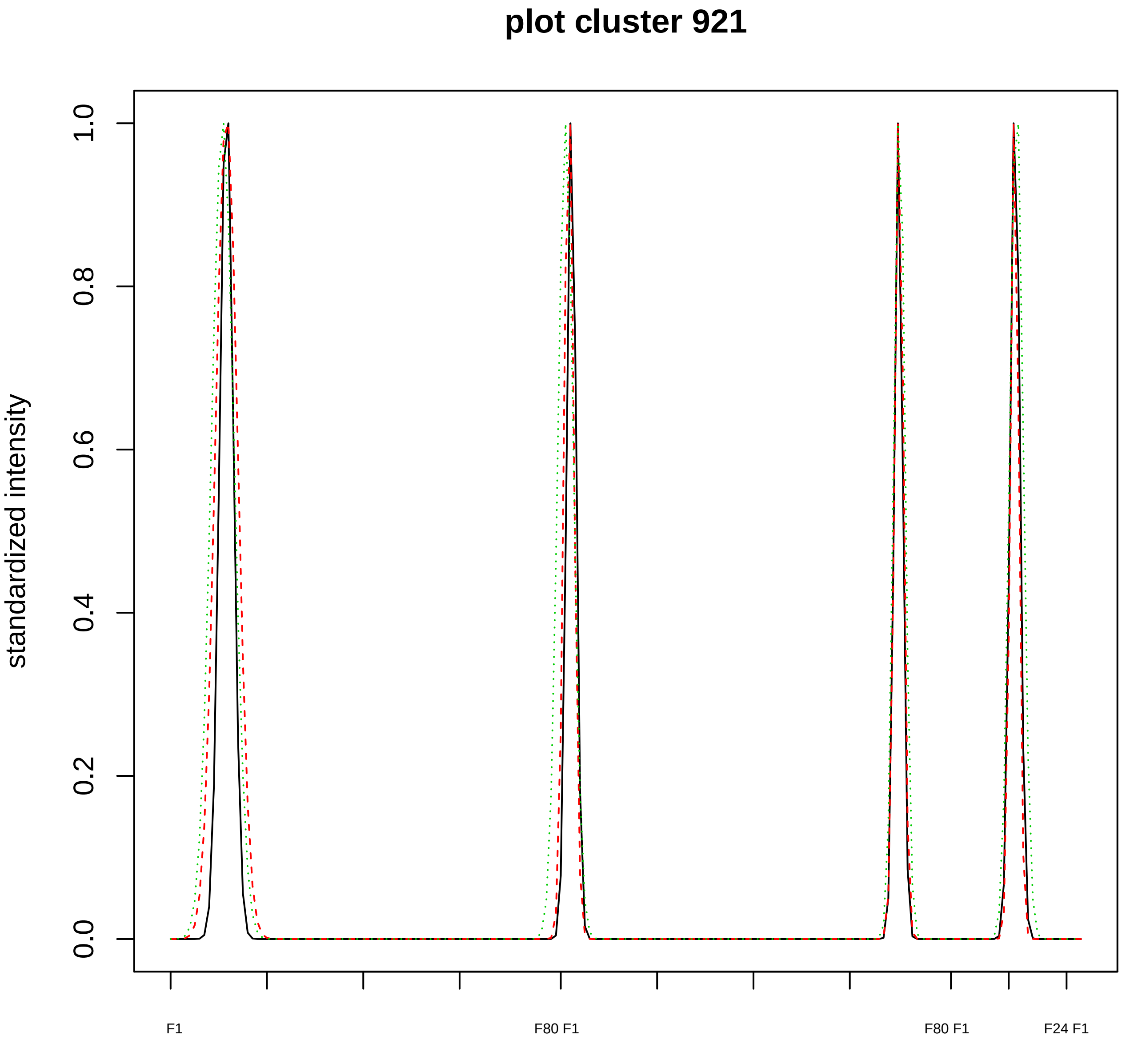
plot cluster 921
standardized intensity
0.0
0.2
0.4
0.6
0.8
1.0
F1
F80 F1
F80 F1
F24 F1

### Dendrogram of cluster 921

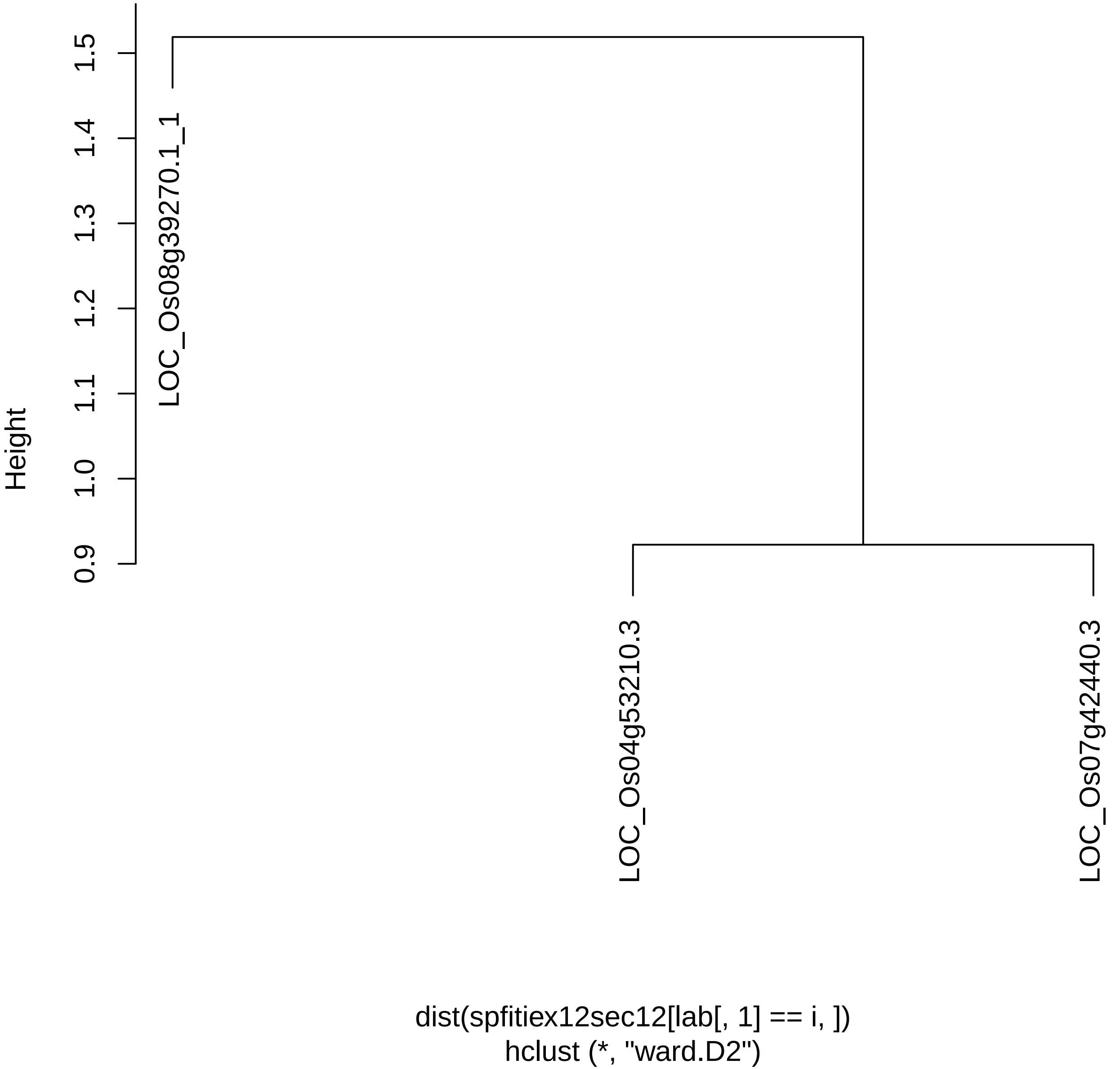

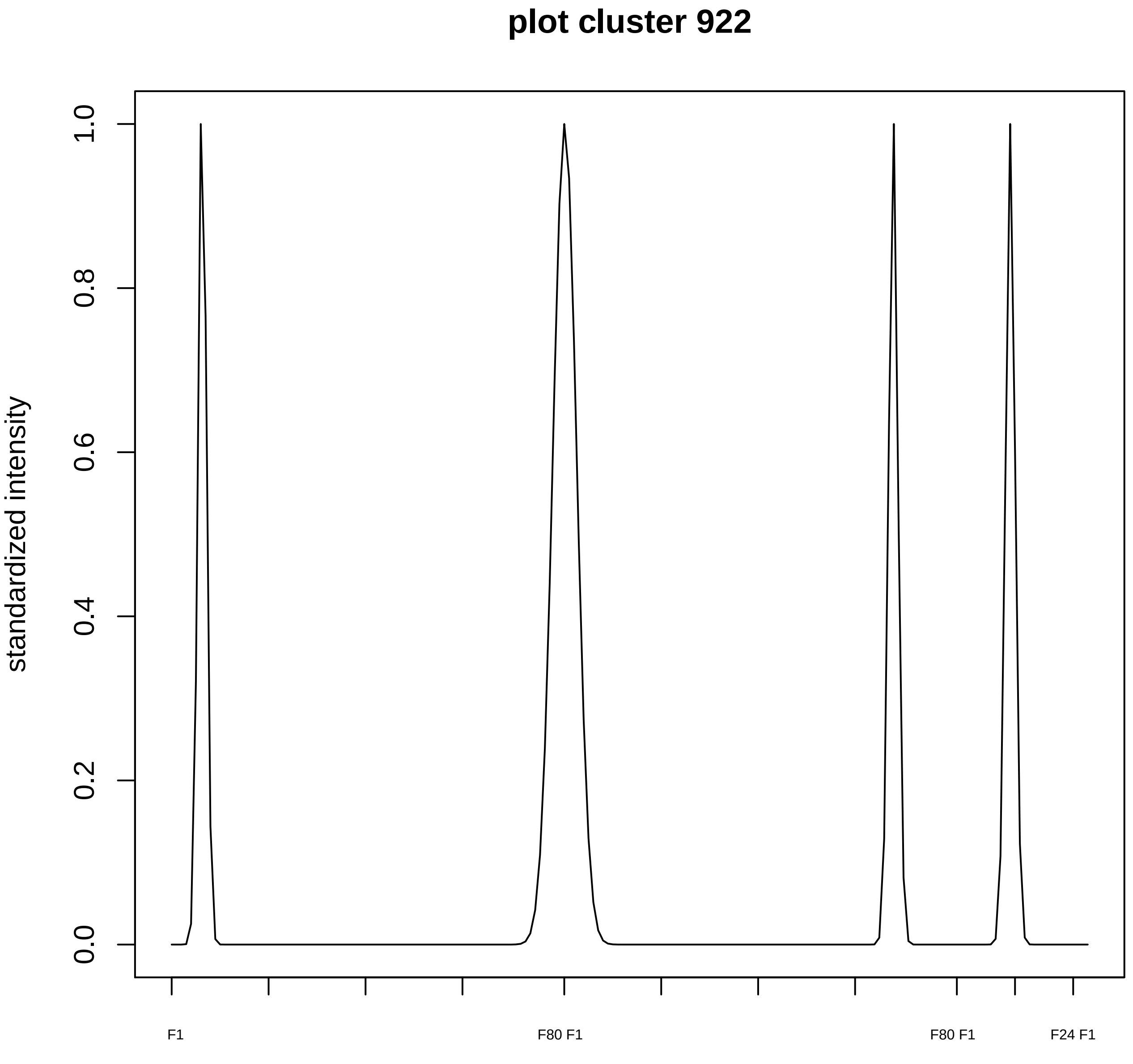
plot cluster 922
standardized intensity
0.0
0.2
0.4
0.6
0.8
1.0
F1
F80 F1
F80 F1
F24 F1

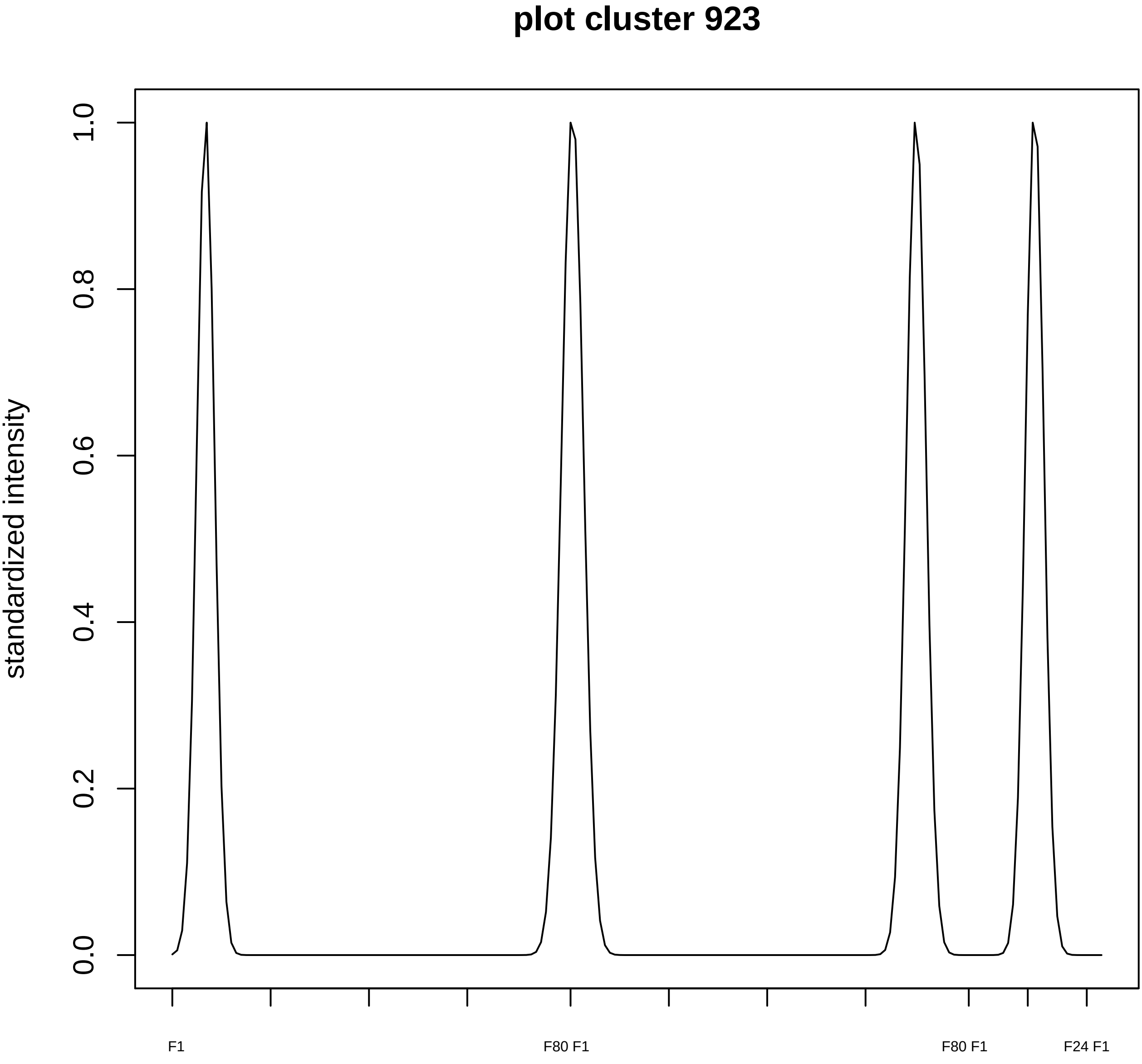
plot cluster 923
standardized intensity
1.0
0.8
0.6
0.4
0.2
0.0
F1
F80 F1
F80 F1
F24 F1

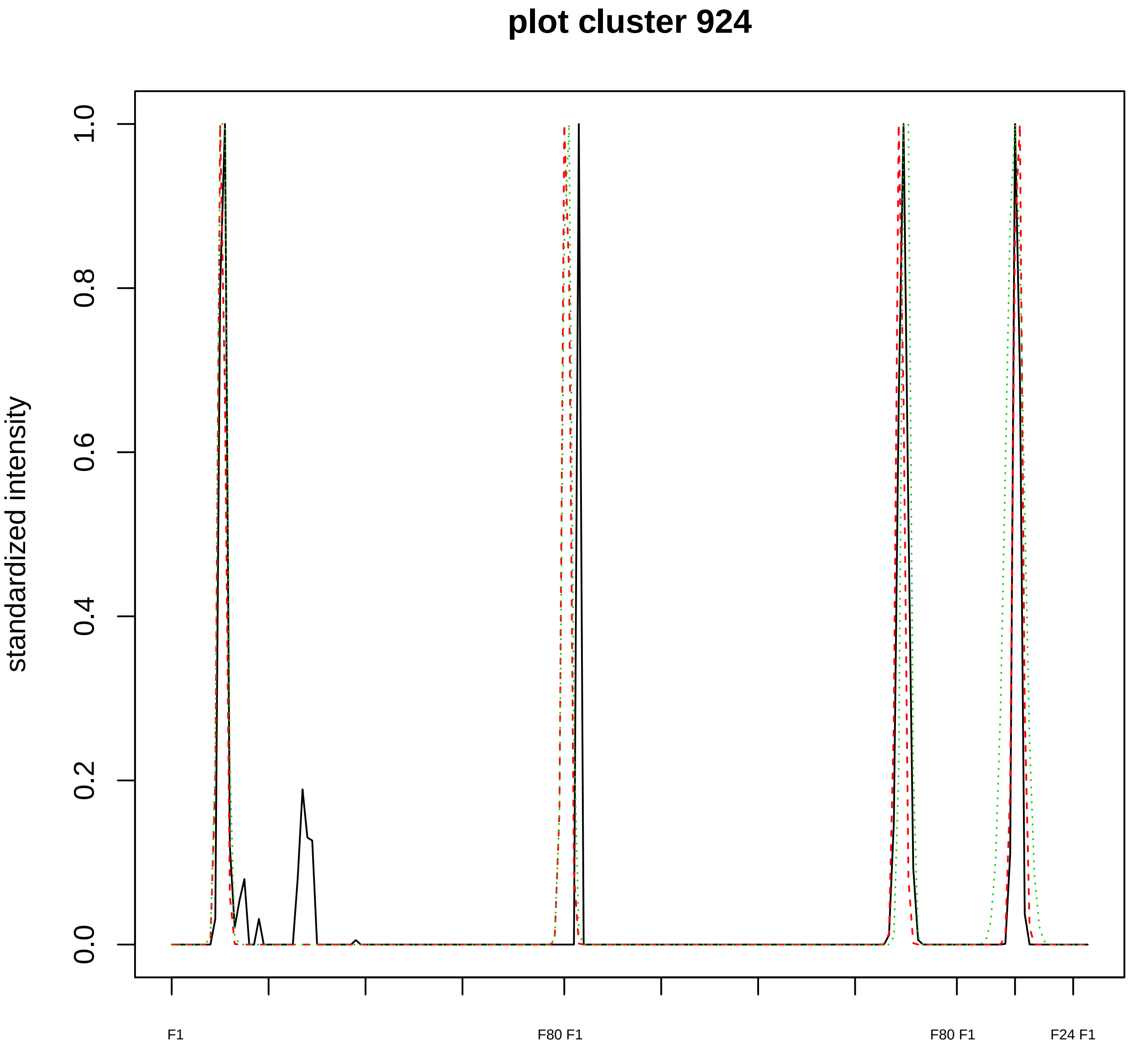

plot cluster 924
standardized intensity
0.0
0.2
0.4
0.6
0.8
1.0
F1
F80 F1
F80 F1
F24 F1

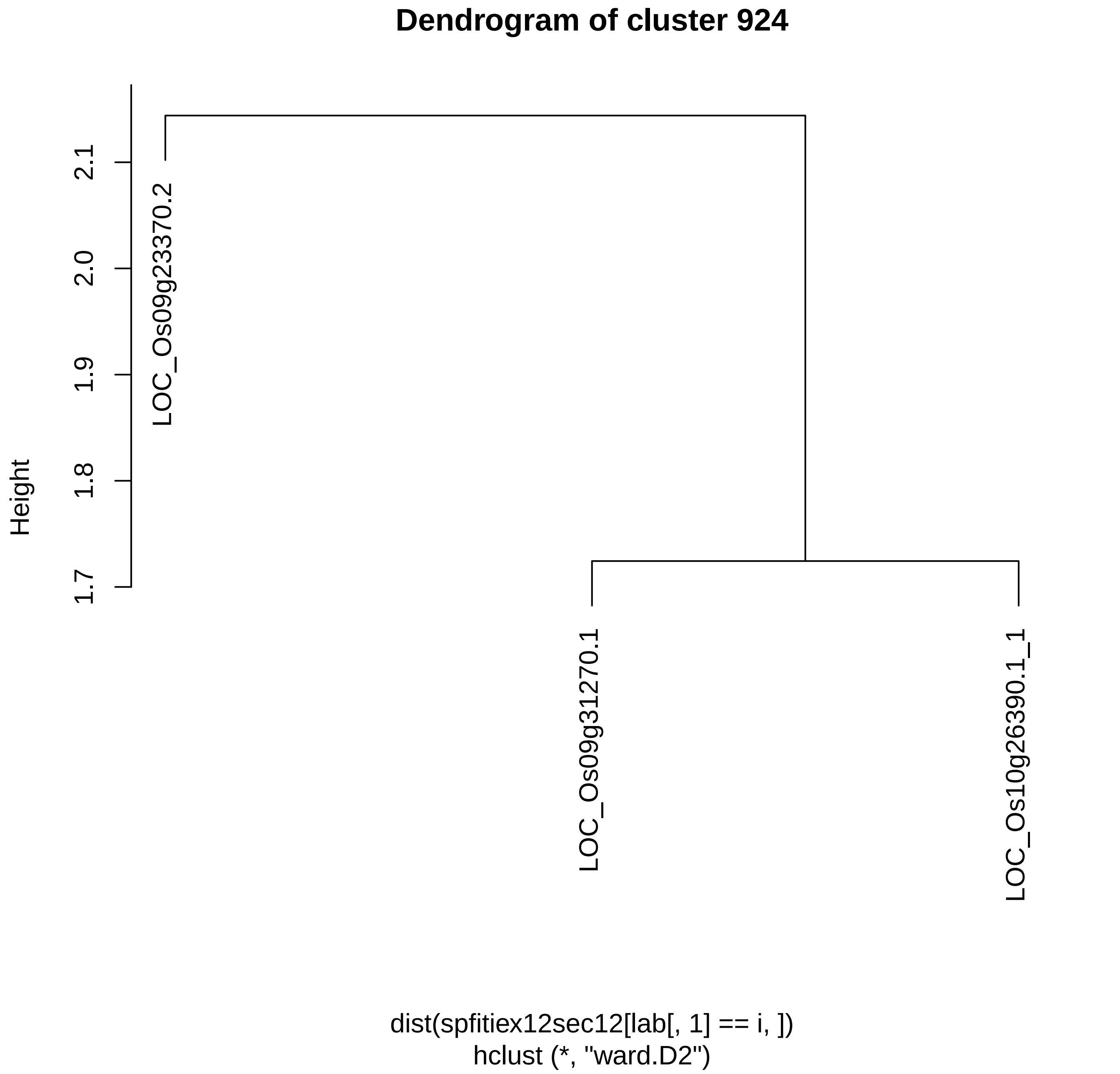
Dendrogram of cluster 924
Height
1.7
1.8
1.9
2.0
2.1
LOC_Os09g23370.2
LOC_Os09g31270.1
LOC_Os10g26390.1_1
dist(spfitiex12sec12[lab[, 1] == i, ])
hclust (*, "ward.D2")

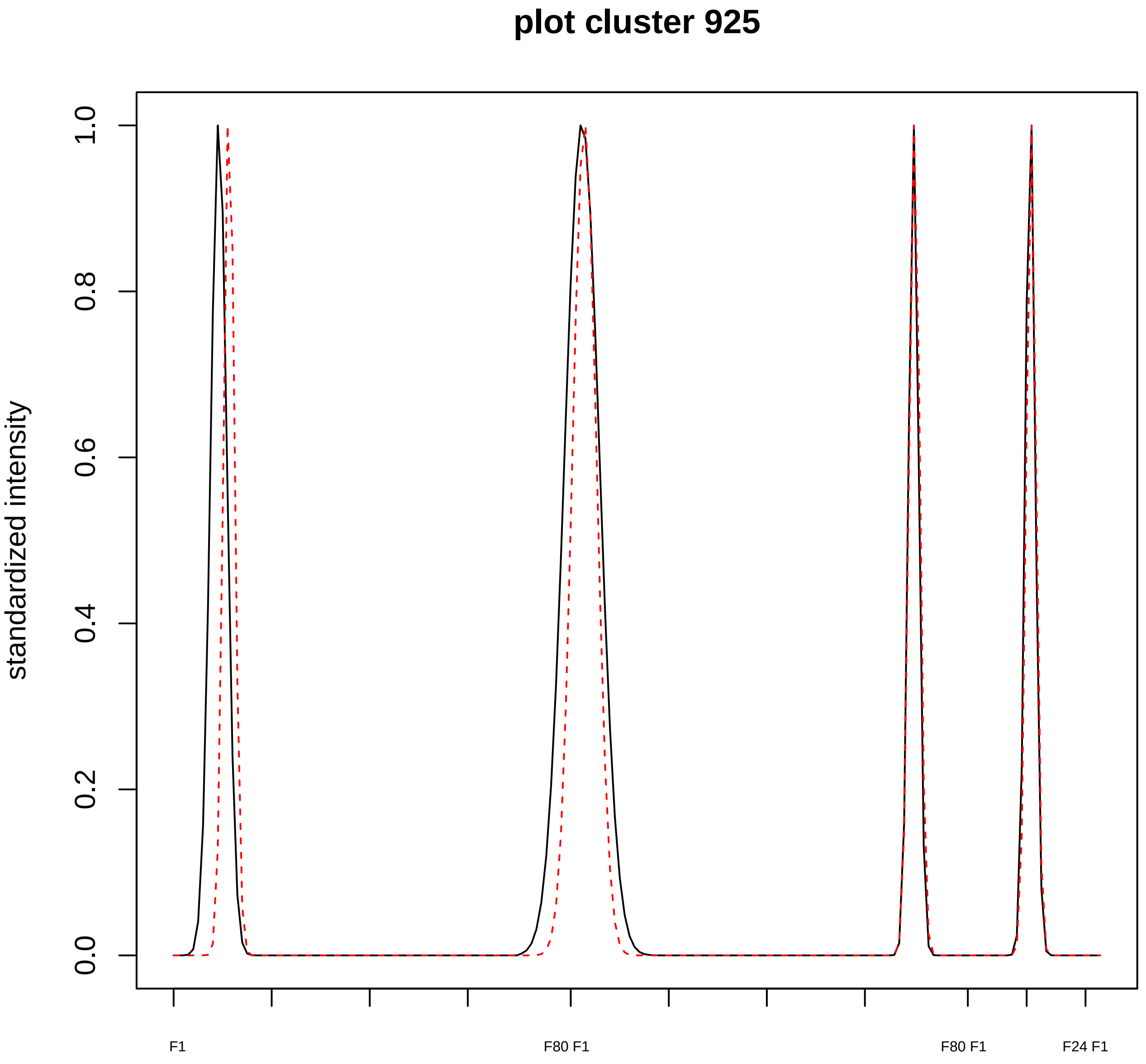
plot cluster 925
standardized intensity
0.0
0.2
0.4
0.6
0.8
1.0
F1
F80 F1
F80 F1
F24 F1

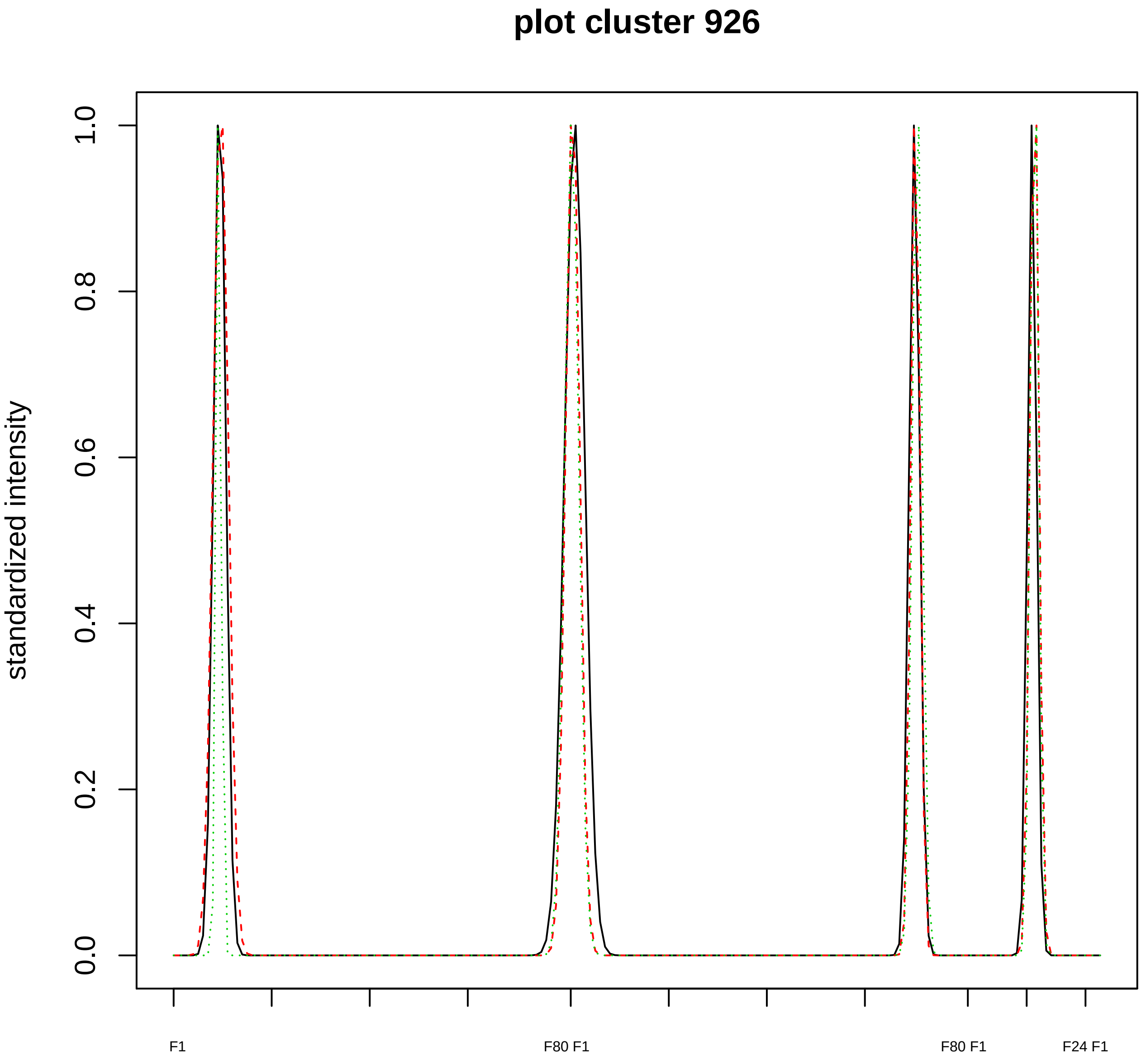
plot cluster 926
standardized intensity
1.0
0.8
0.6
0.4
0.2
0.0
F1
F80 F1
F80 F1
F24 F1

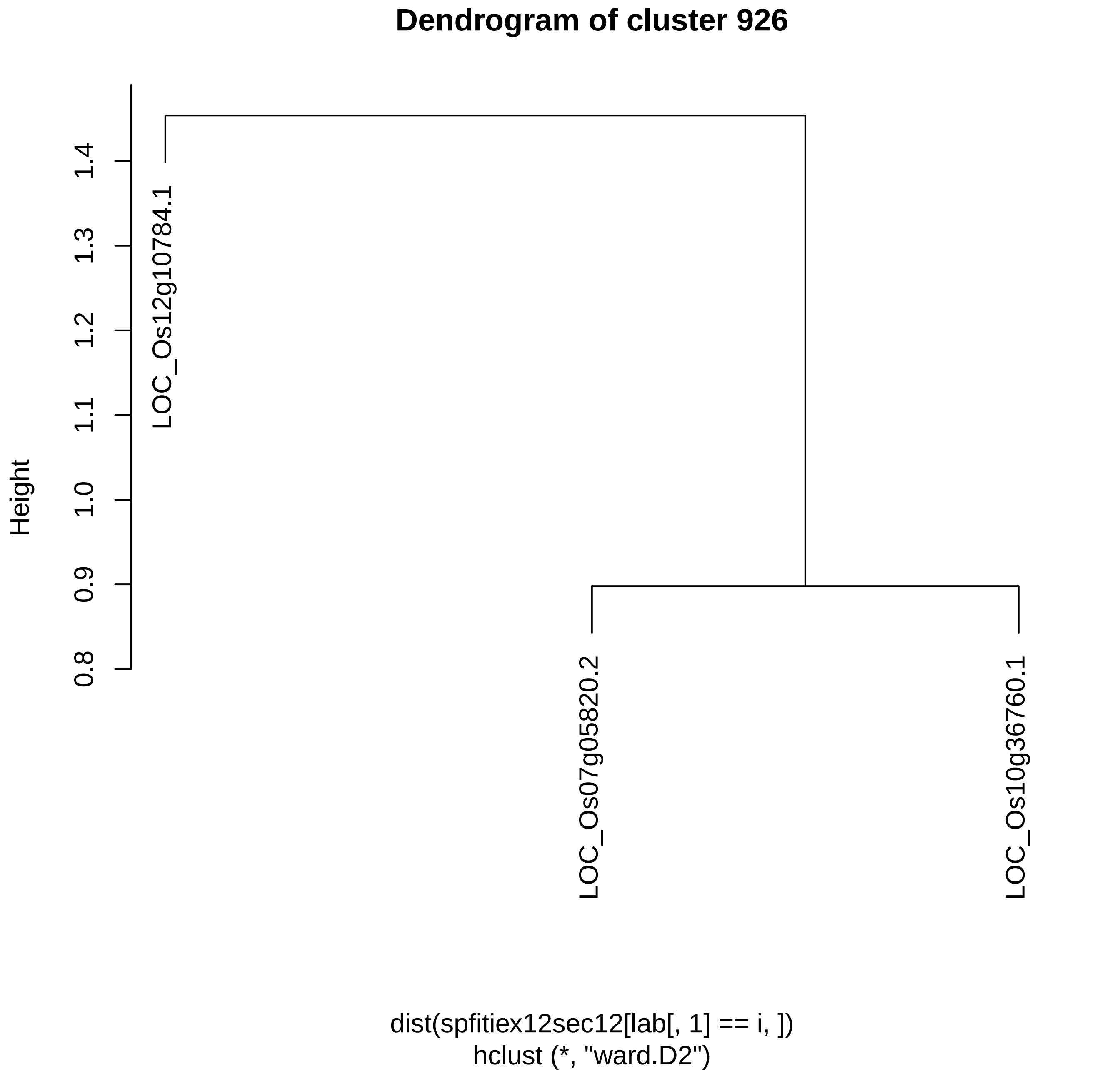

Dendrogram of cluster 926
Height
1.4
1.3
1.2
1.1
1.0
0.9
0.8
LOC_Os12g10784.1
LOC_Os07g05820.2
LOC_Os10g36760.1
dist(spfitiex12sec12[lab[, 1] == i, ])
hclust (*, "ward.D2")

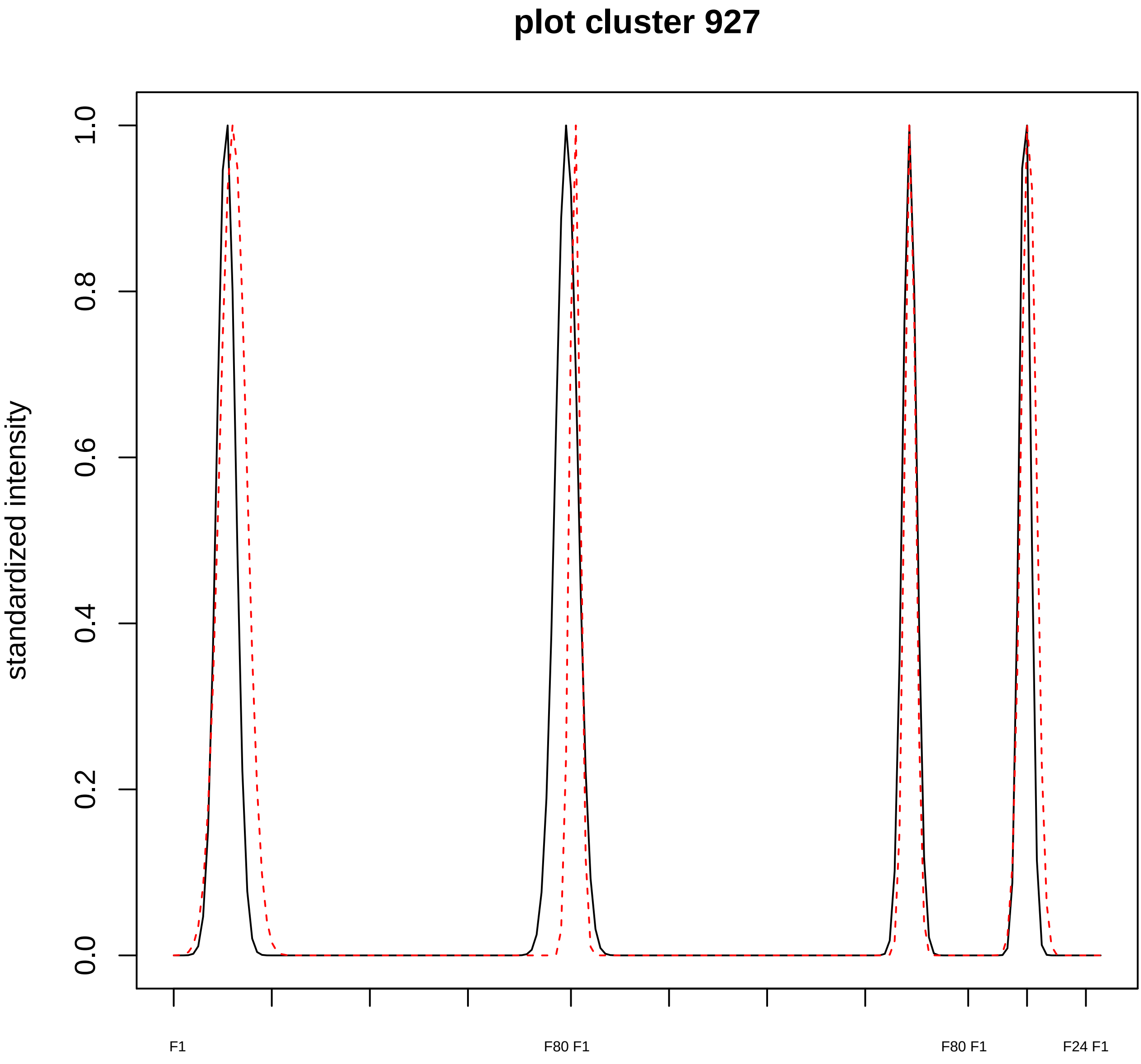

plot cluster 927
standardized intensity
0.0
0.2
0.4
0.6
0.8
1.0
F1
F80 F1
F80 F1
F24 F1

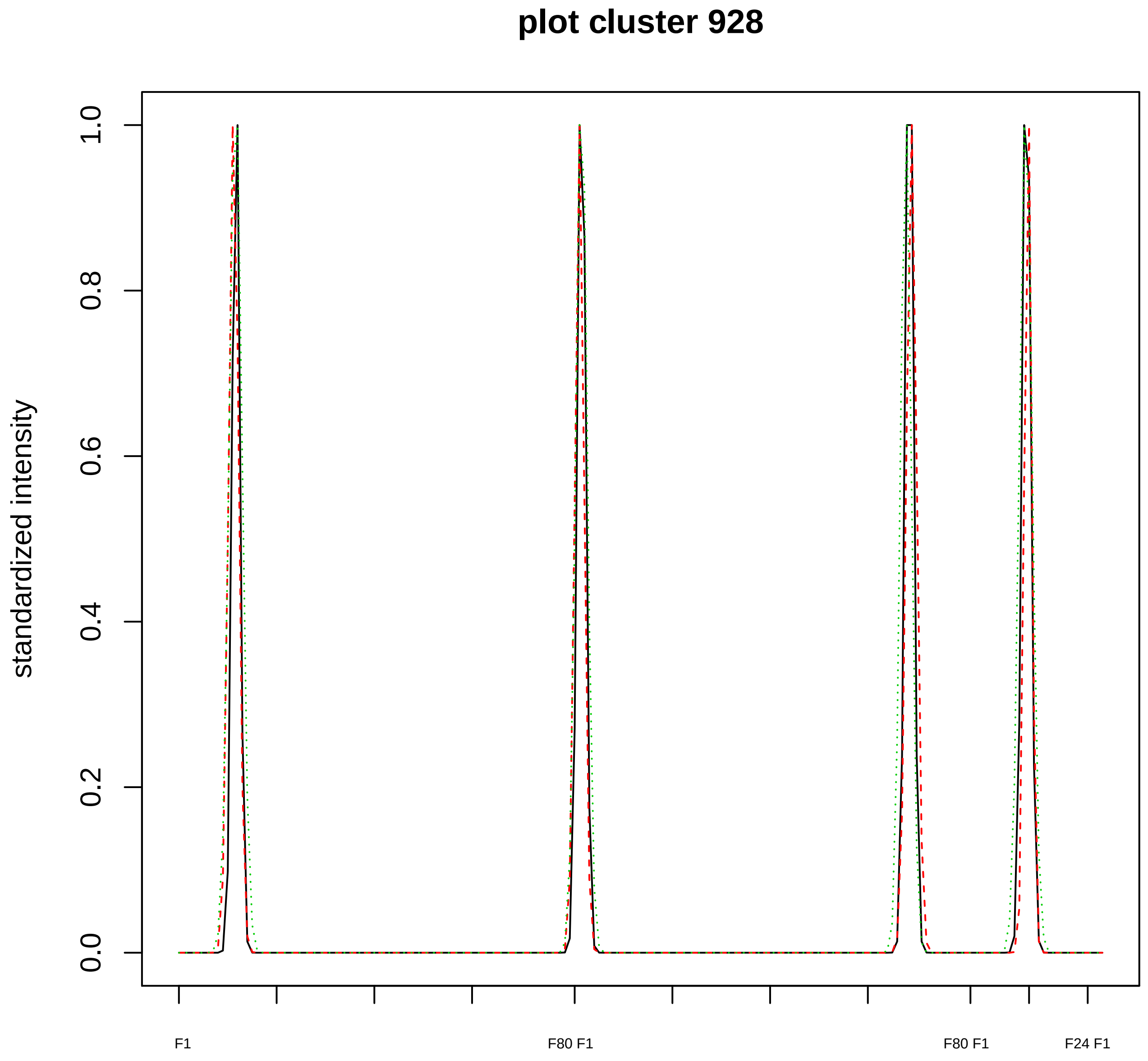
plot cluster 928
standardized intensity
0.0
0.2
0.4
0.6
0.8
1.0
F1
F80 F1
F80 F1
F24 F1

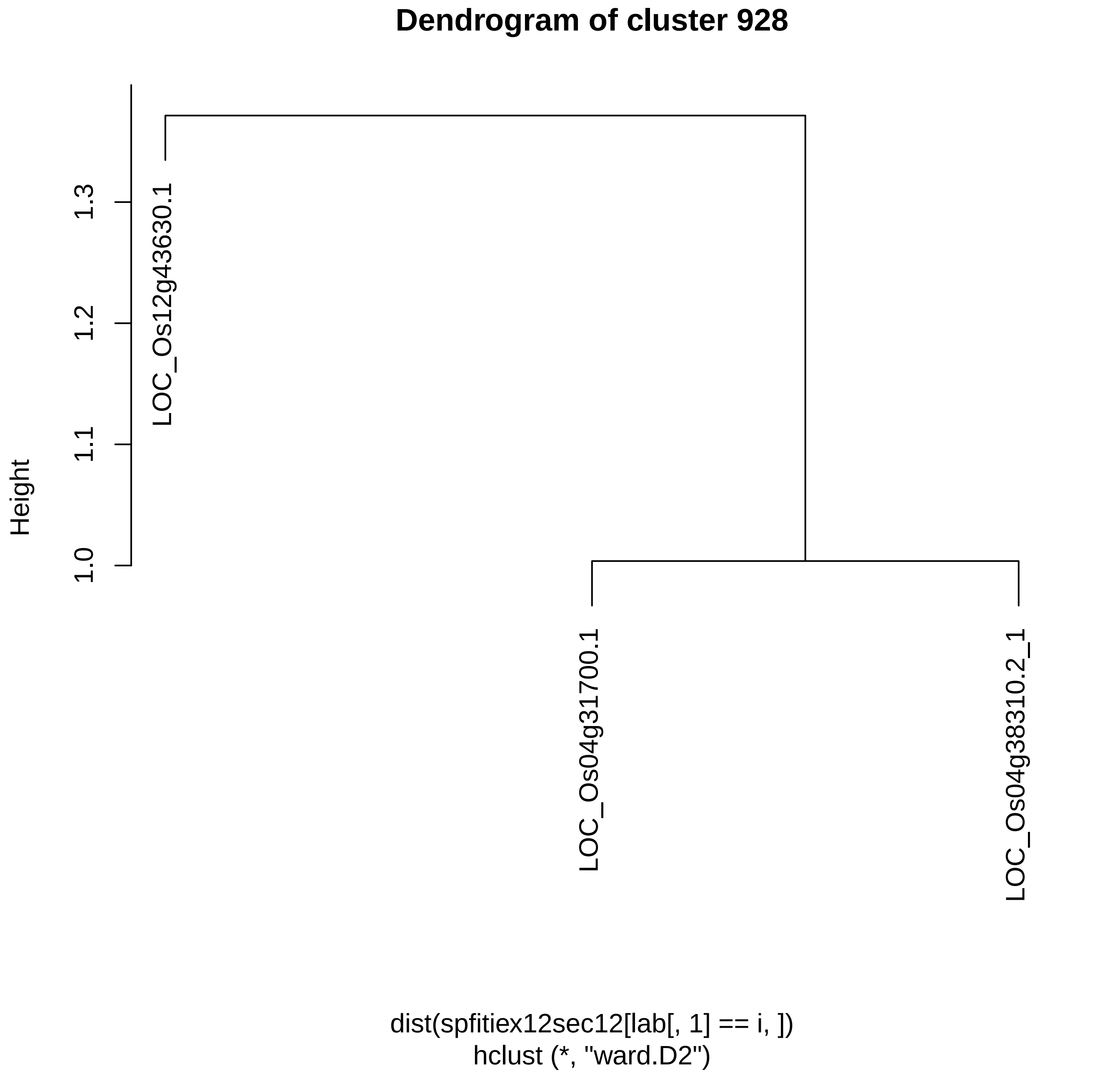
Dendrogram of cluster 928
Height
1.0
1.1
1.2
1.3
LOC_Os12g43630.1
LOC_Os04g31700.1
LOC_Os04g38310.2_1
dist(spfitiex12sec12[lab[, 1] == i, ])
hclust (*, "ward.D2")

#### plot cluster 929

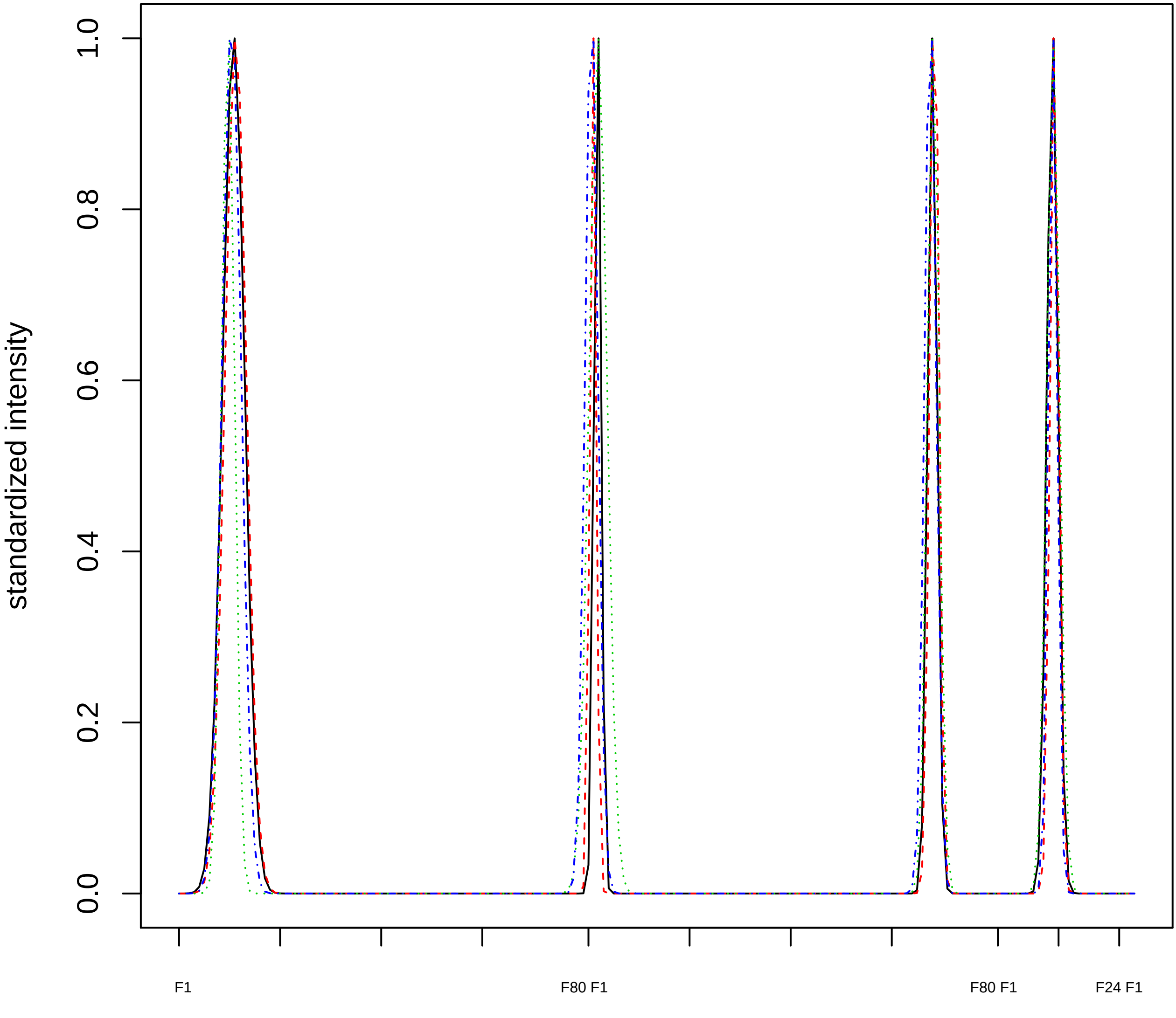

#### Dendrogram of cluster 929

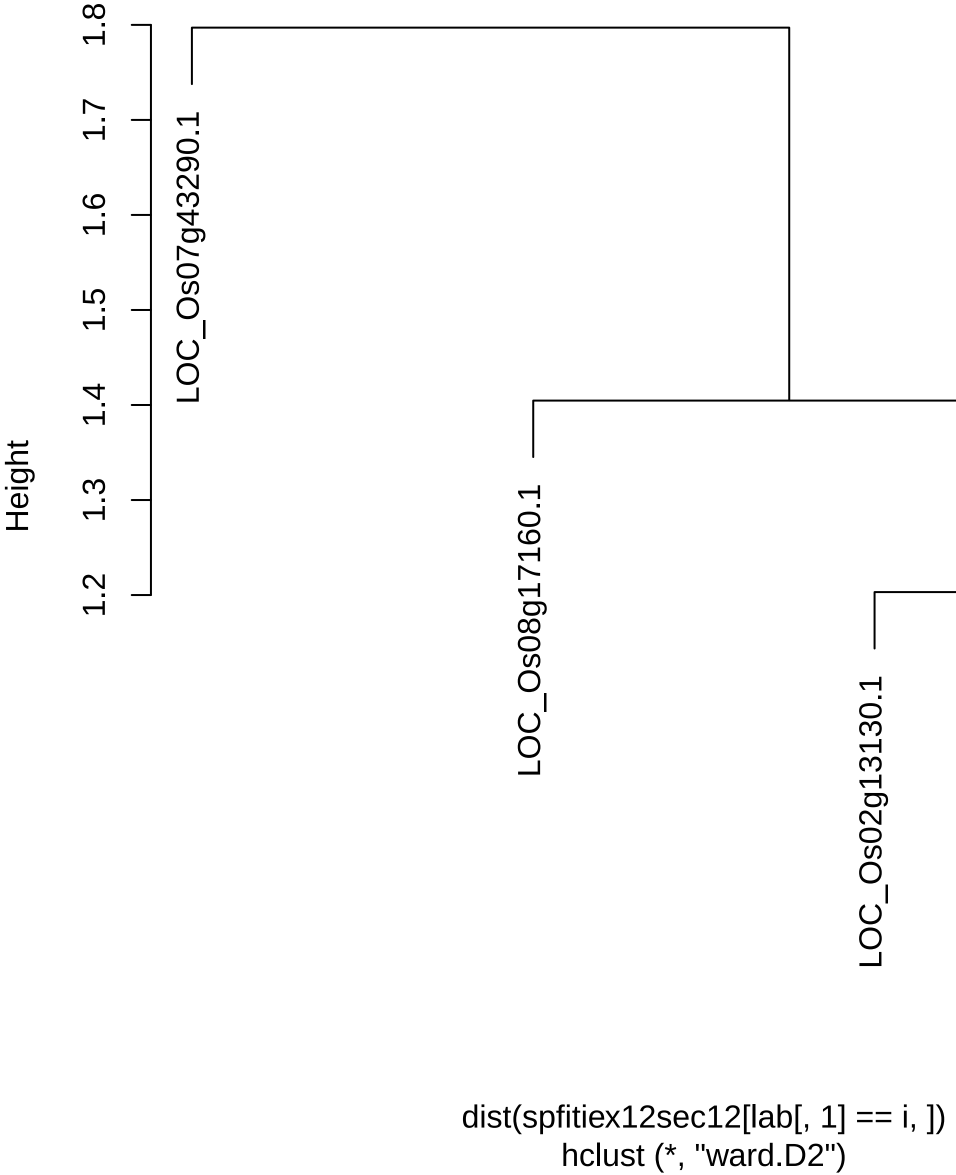

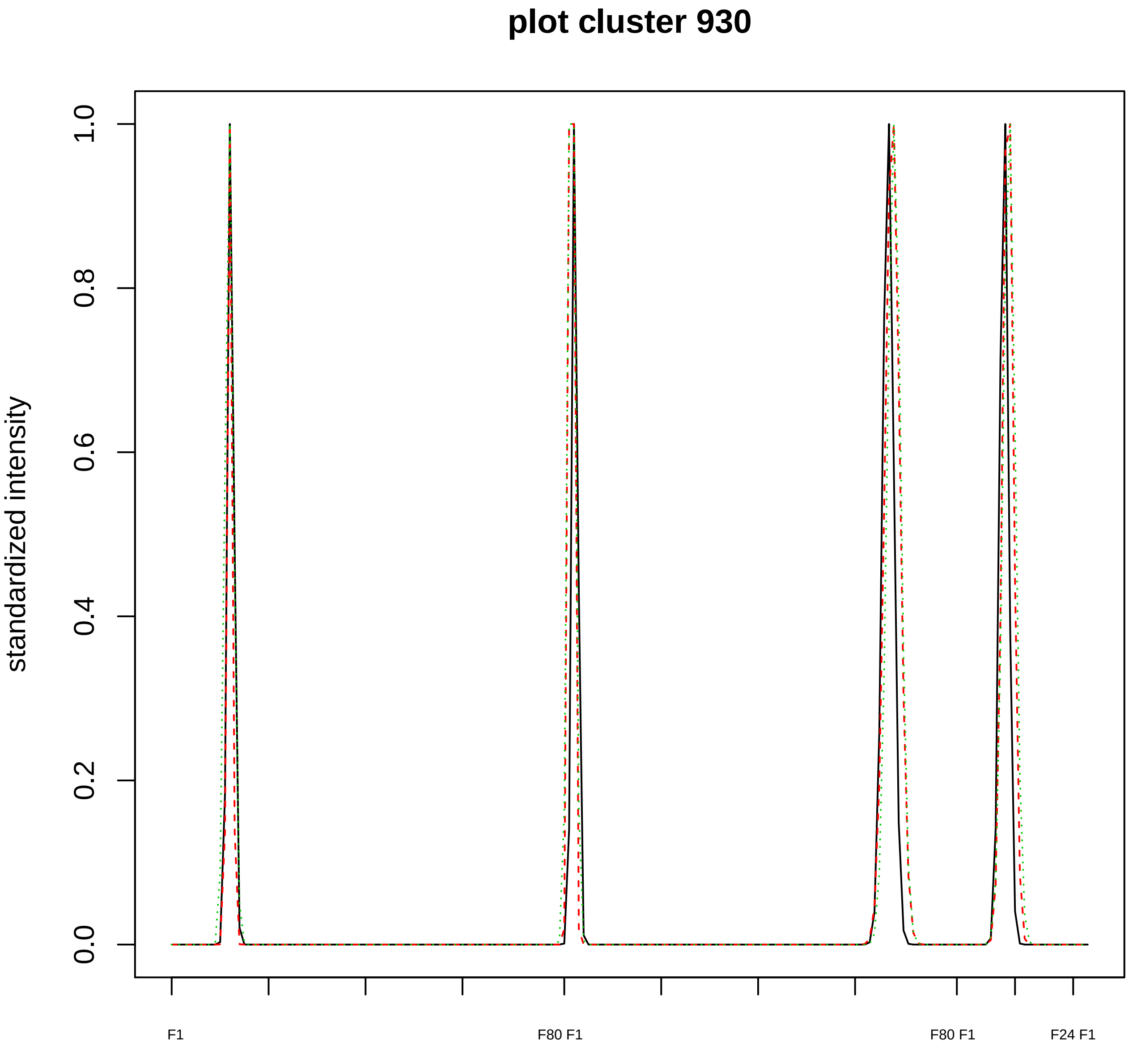

plot cluster 930
standardized intensity
0.0
0.2
0.4
0.6
0.8
1.0
F1
F80 F1
F80 F1
F24 F1

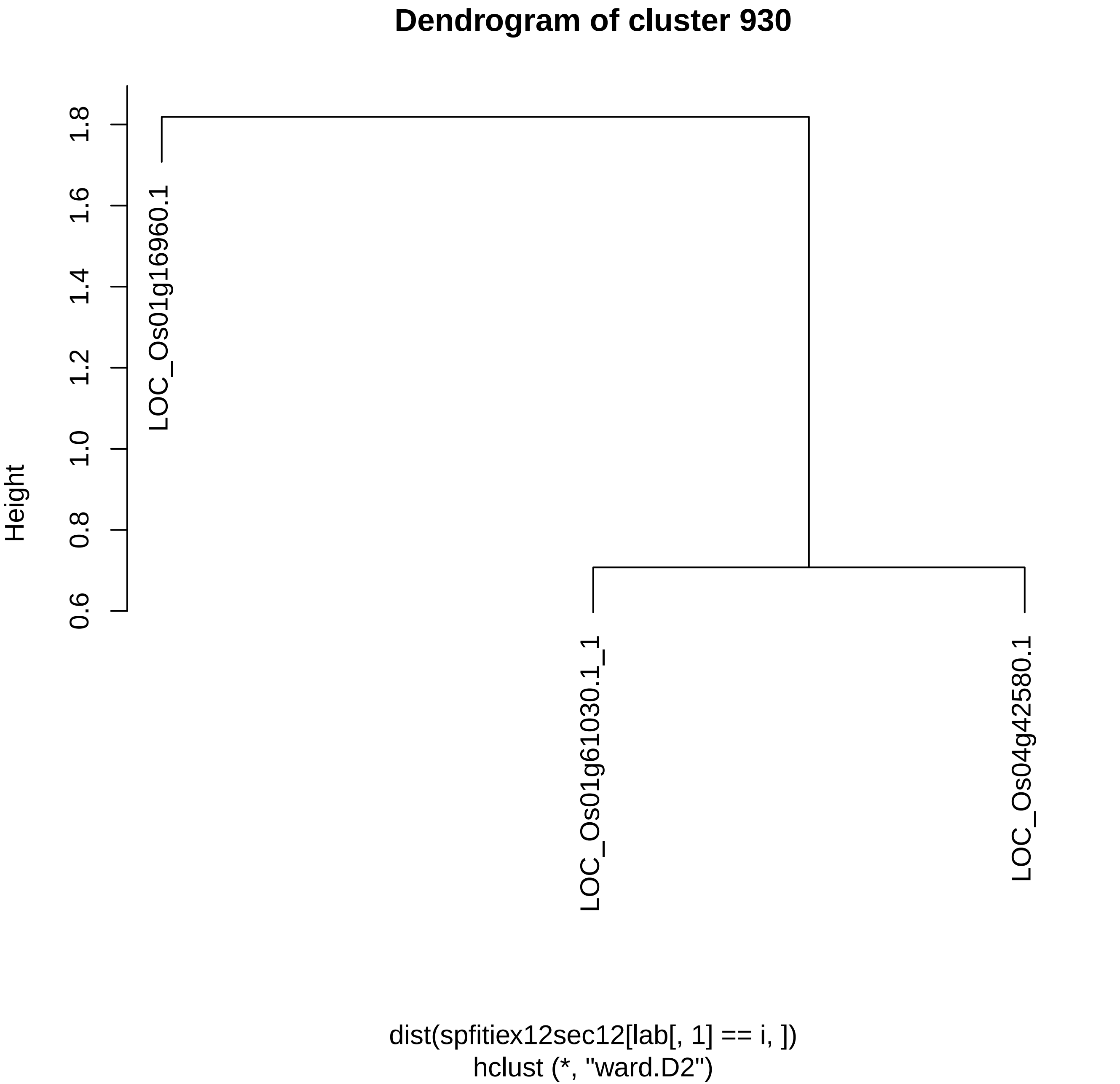
Dendrogram of cluster 930
Height
0.6
0.8
1.0
1.2
1.4
1.6
1.8
LOC_Os01g16960.1
LOC_Os01g61030.1_1
LOC_Os04g42580.1
dist(spfitiex12sec12[lab[, 1] == i, ])
hclust (*, "ward.D2")

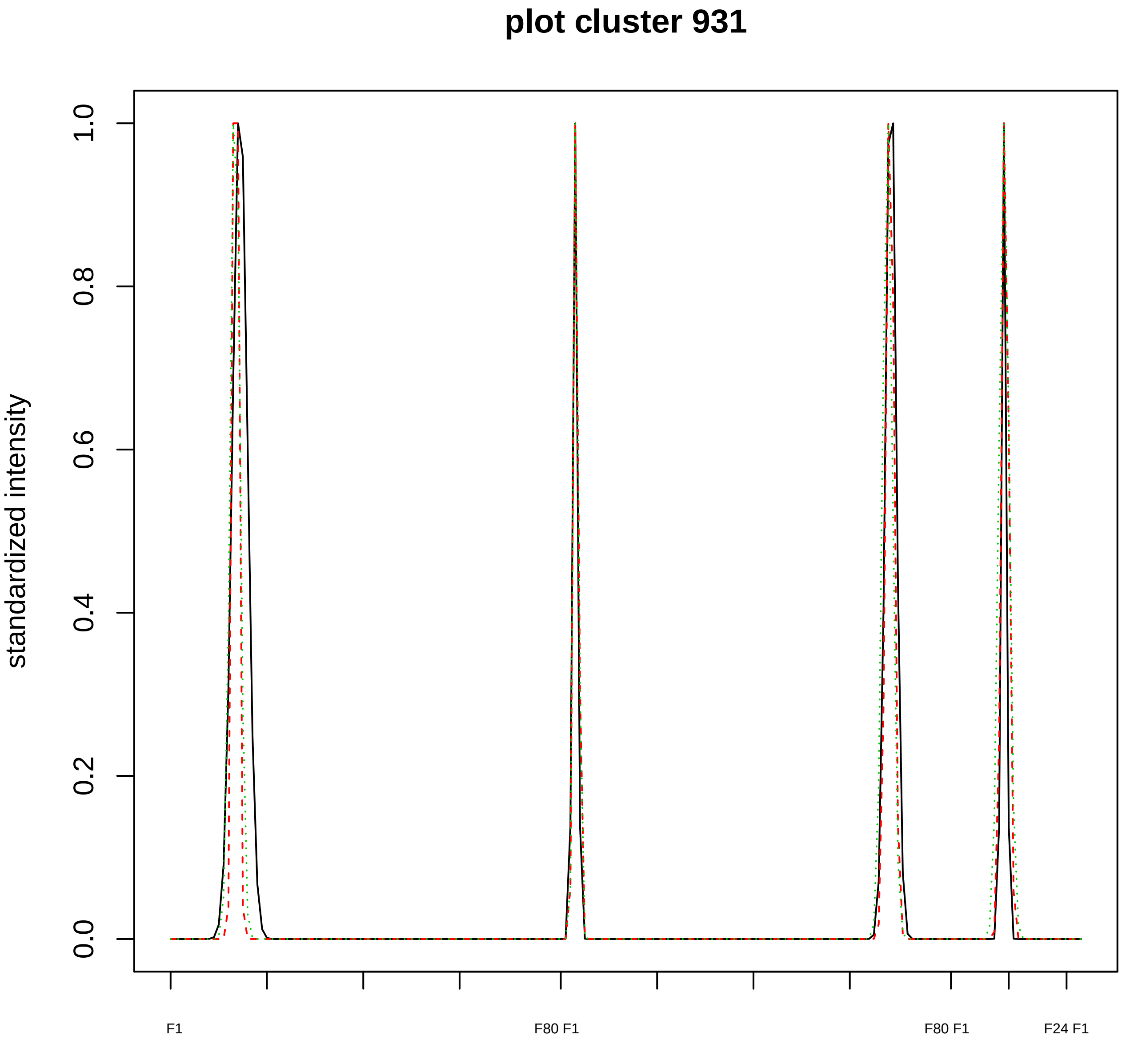

plot cluster 931
standardized intensity
0.0
0.2
0.4
0.6
0.8
1.0
F1
F80 F1
F80 F1
F24 F1

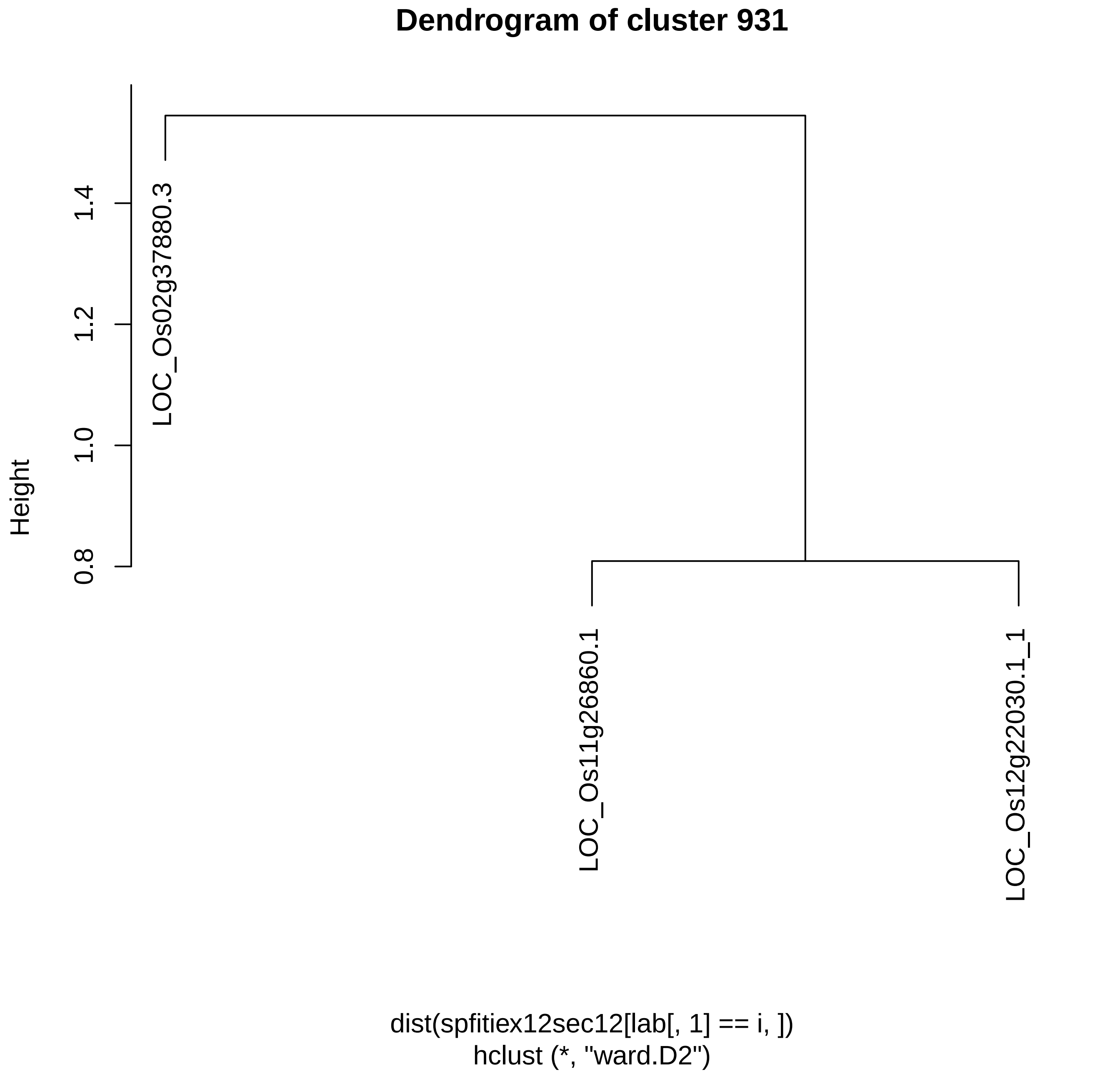

Dendrogram of cluster 931
Height
1.4
1.2
1.0
0.8
LOC_Os02g37880.3
LOC_Os11g26860.1
LOC_Os12g22030.1_1
dist(spfitiex12sec12[lab[, 1] == i, ])
hclust (*, "ward.D2")

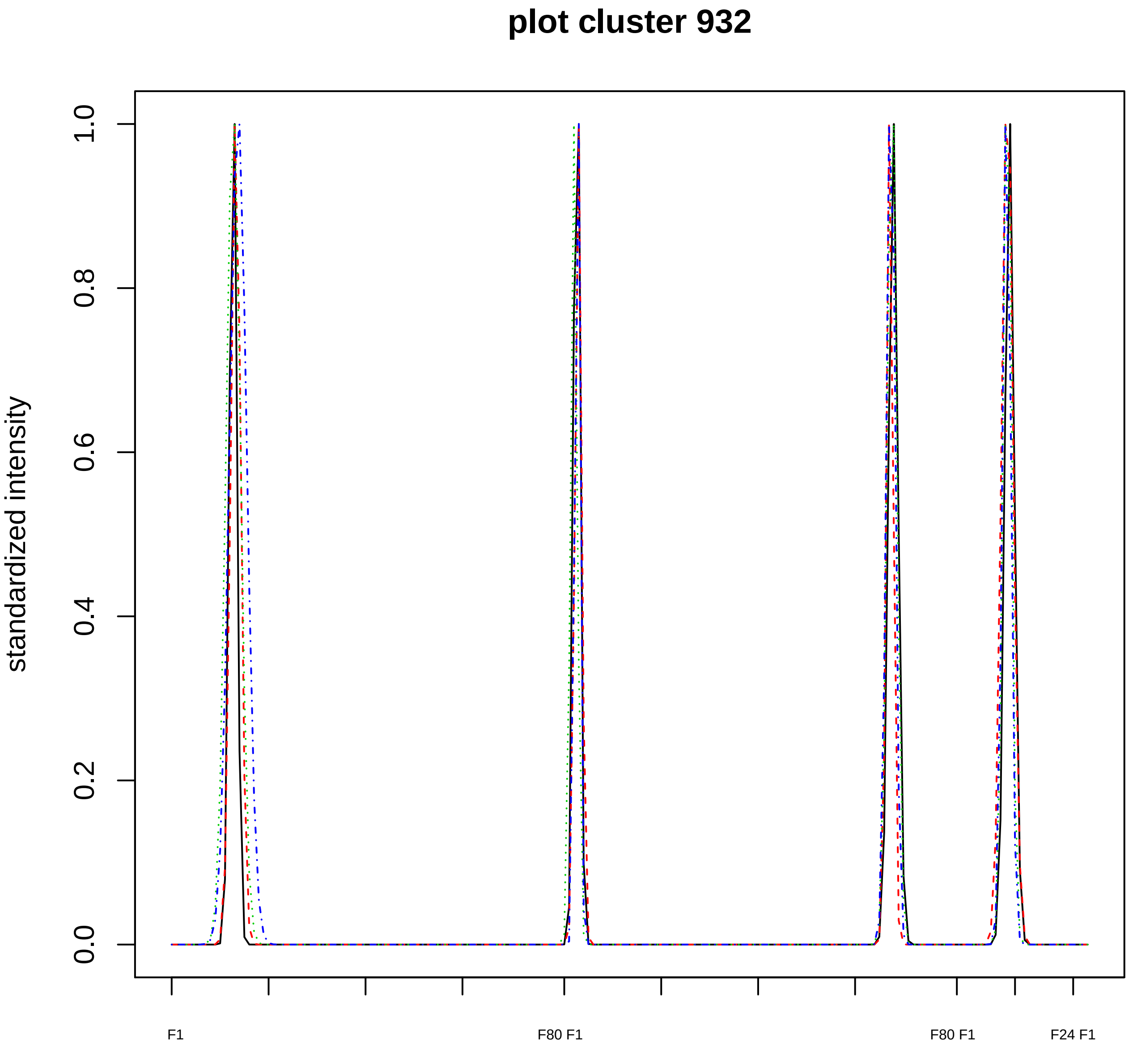
plot cluster 932
standardized intensity
0.0
0.2
0.4
0.6
0.8
1.0
F1
F80 F1
F80 F1
F24 F1

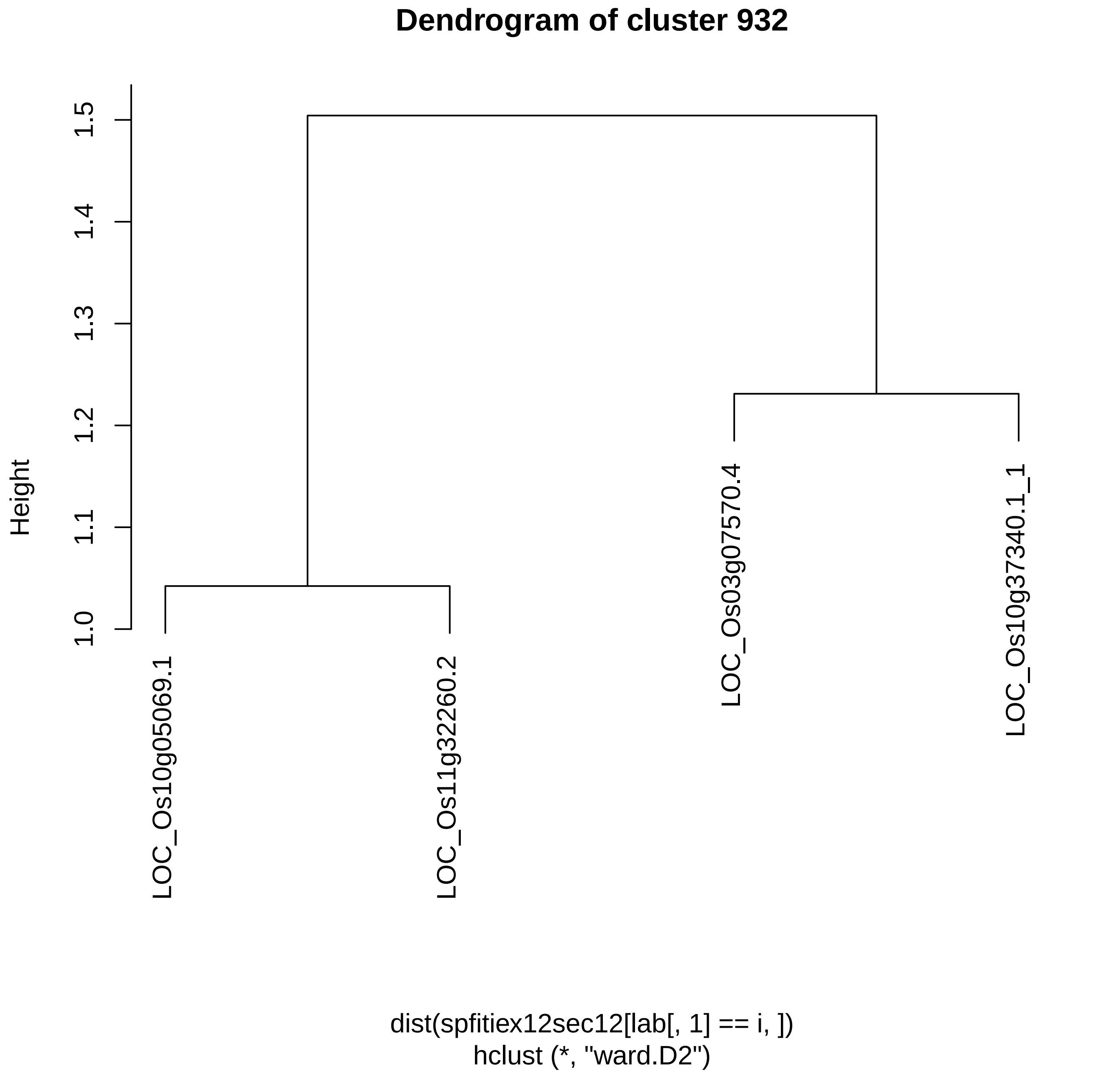

Dendrogram of cluster 932
Height
1.0
1.1
1.2
1.3
1.4
1.5
LOC_Os10g05069.1
LOC_Os11g32260.2
LOC_Os03g07570.4
LOC_Os10g37340.1_1
dist(spfitiex12sec12[lab[, 1] == i, ])
hclust (*, "ward.D2")

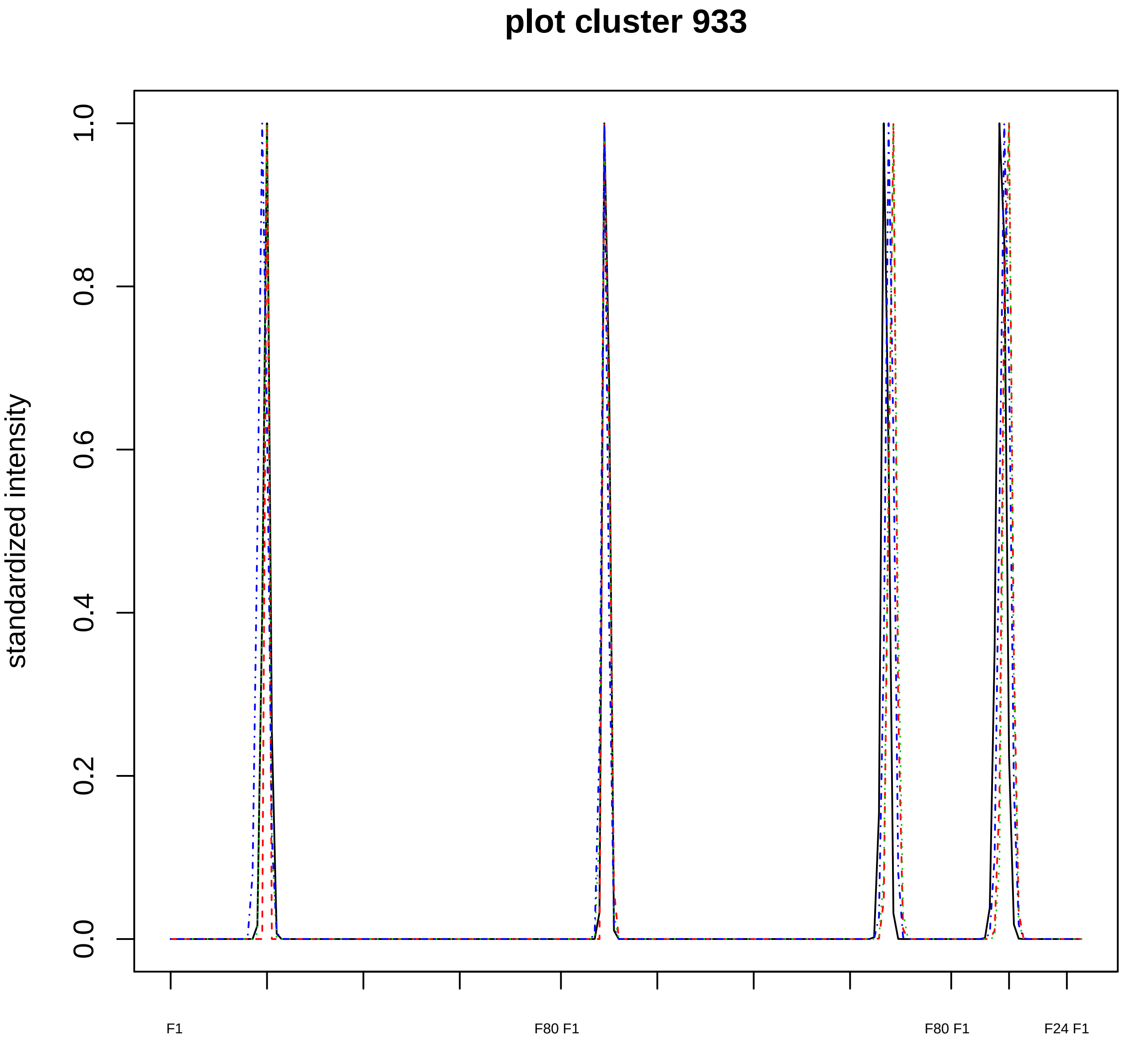
plot cluster 933
standardized intensity
0.0
0.2
0.4
0.6
0.8
1.0
F1
F80 F1
F80 F1
F24 F1

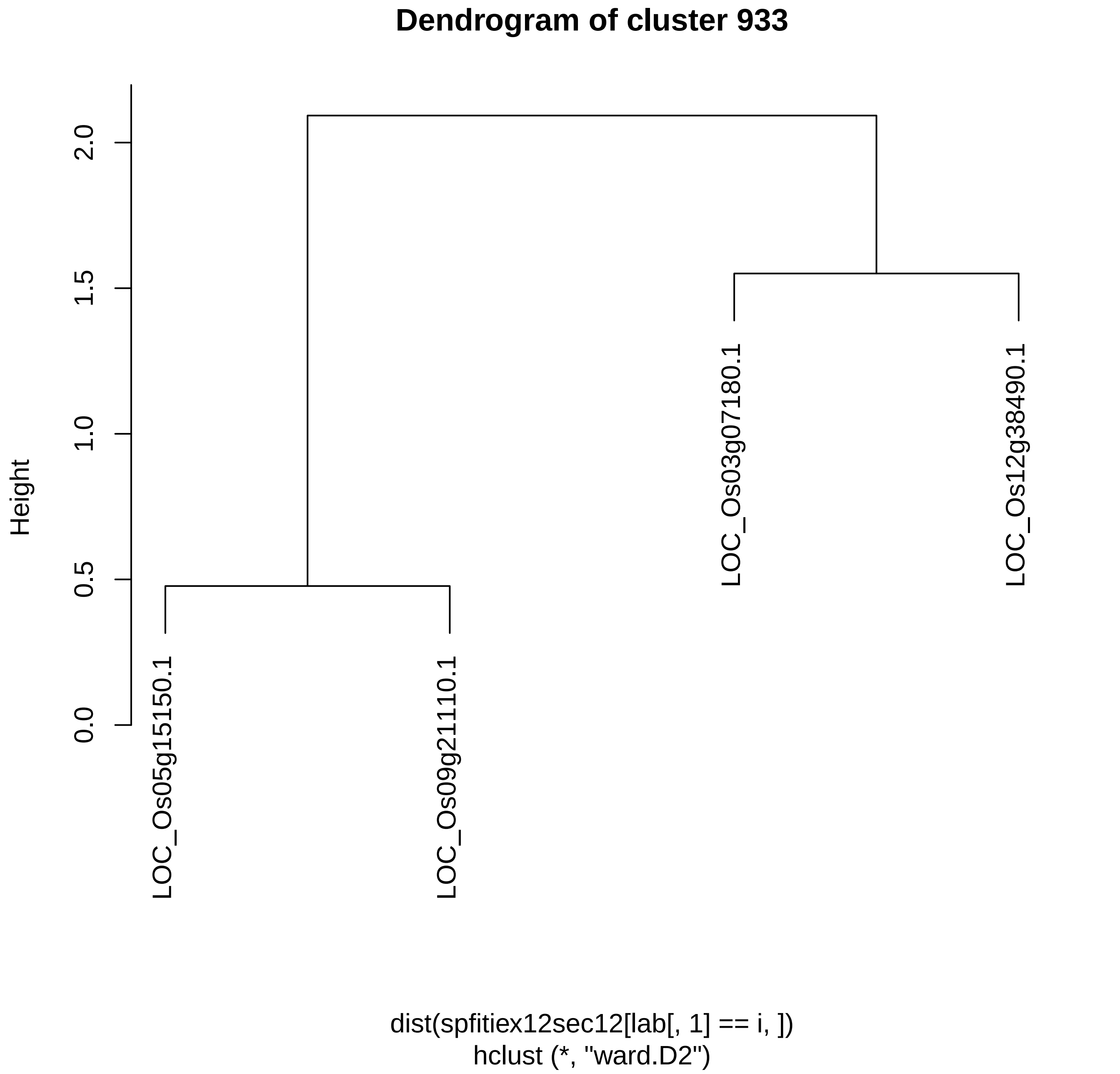
Dendrogram of cluster 933
Height
0.0
0.5
1.0
1.5
2.0
LOC_Os05g15150.1
LOC_Os09g21110.1
LOC_Os03g07180.1
LOC_Os12g38490.1
dist(spfitiex12sec12[lab[, 1] == i, ])
hclust (*, "ward.D2")

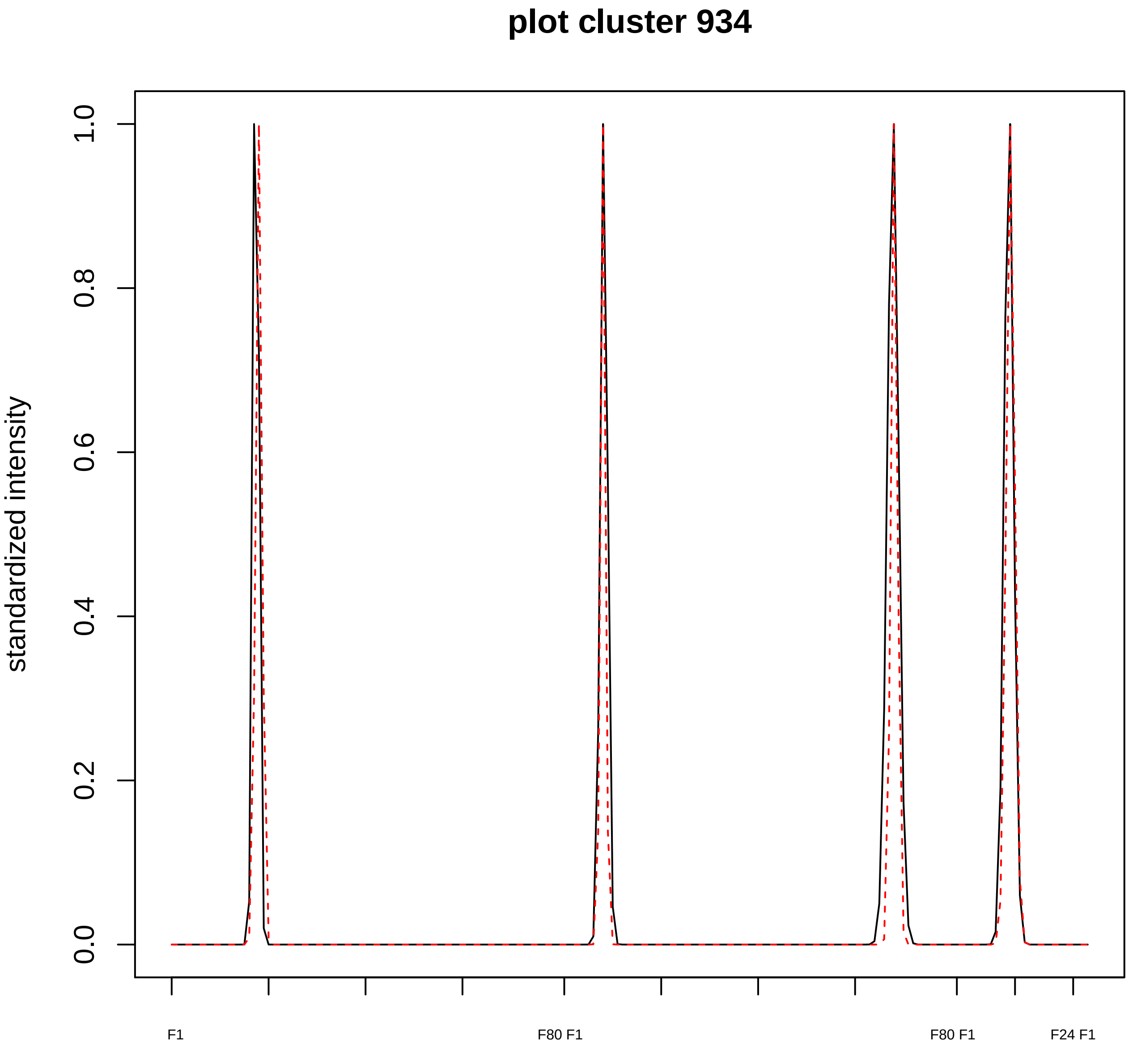

plot cluster 934
standardized intensity
0.0
0.2
0.4
0.6
0.8
1.0
F1
F80 F1
F80 F1
F24 F1

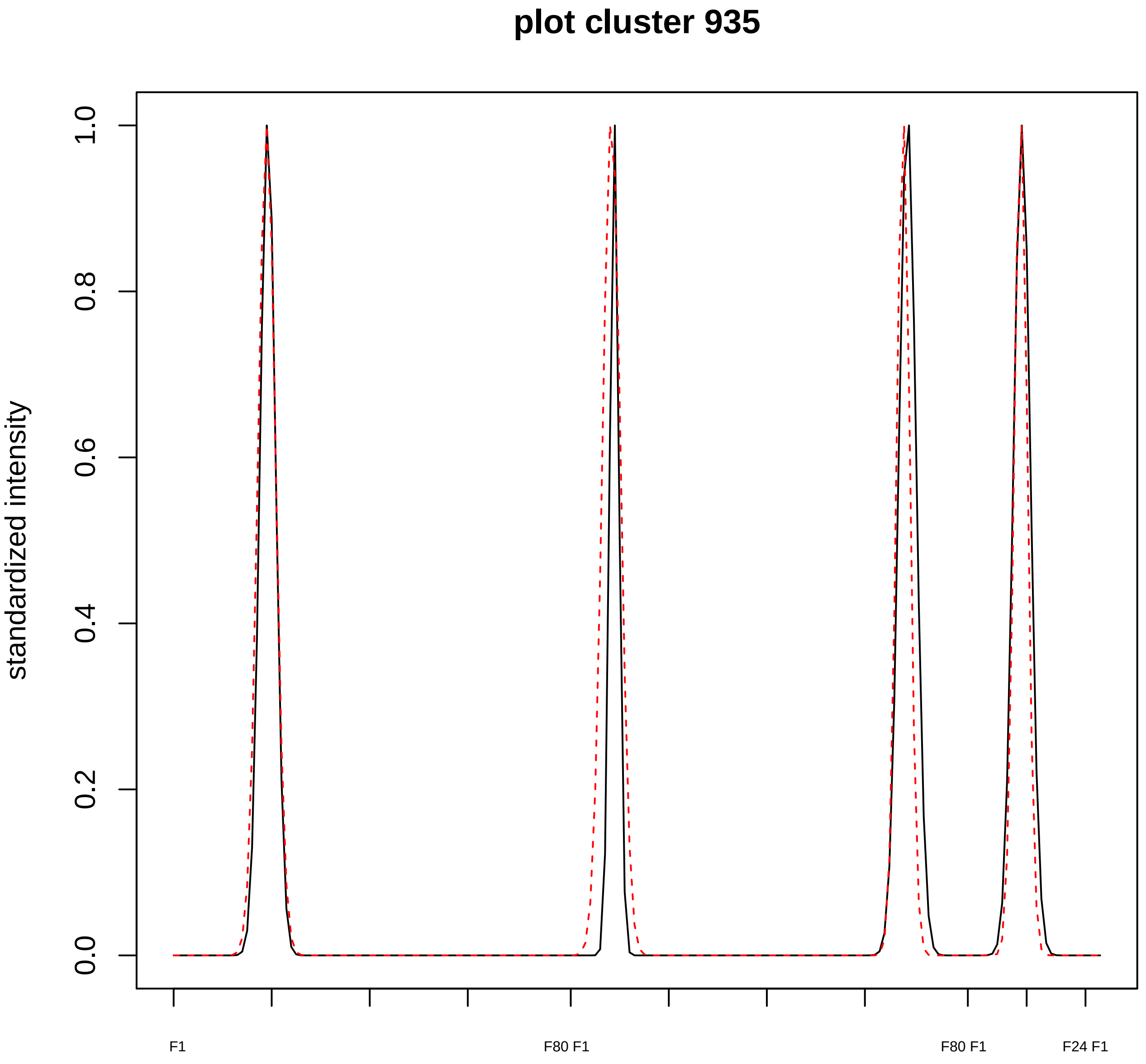

plot cluster 935
standardized intensity
0.0
0.2
0.4
0.6
0.8
1.0
F1
F80 F1
F80 F1
F24 F1

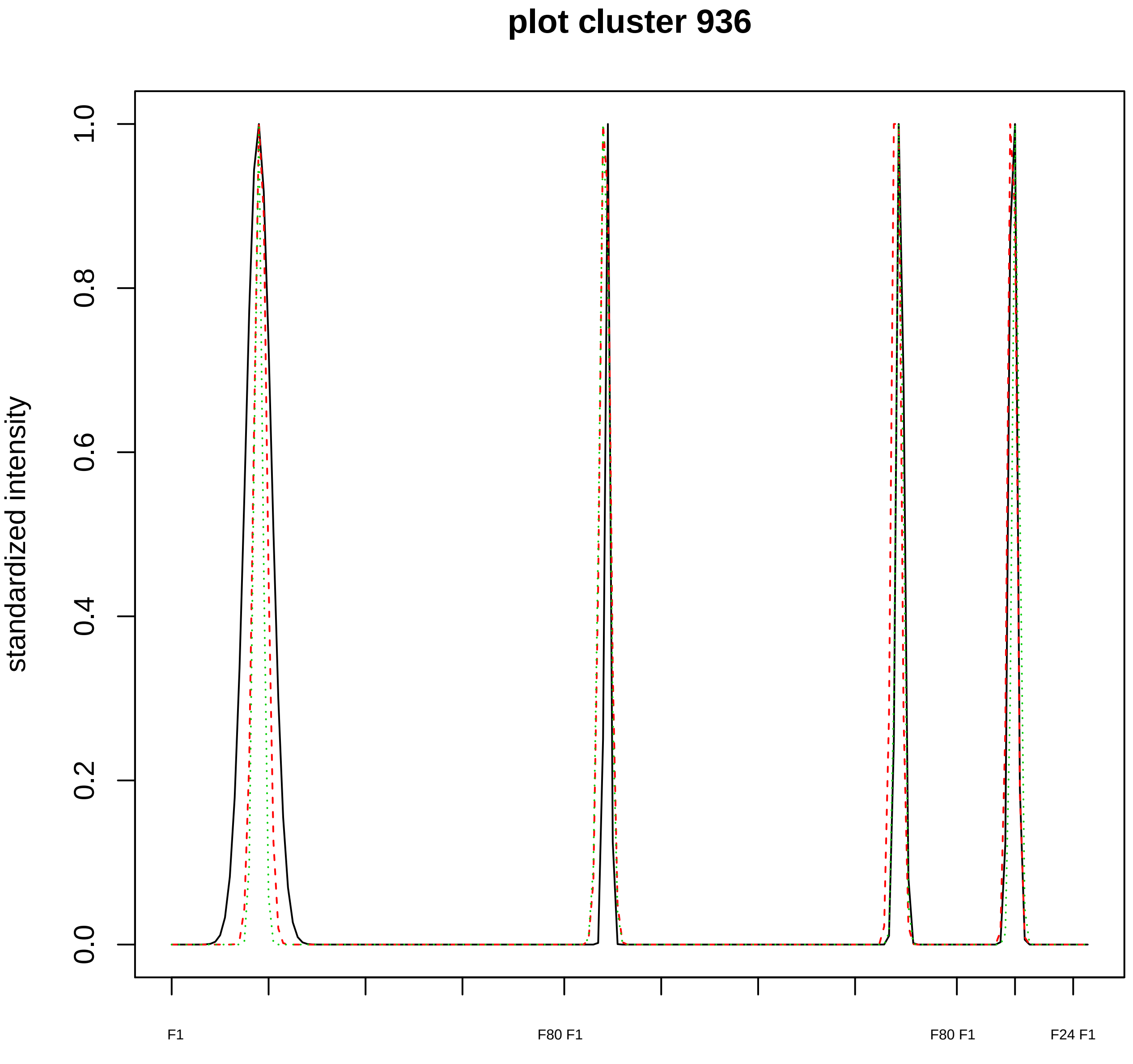

plot cluster 936
standardized intensity
0.0
0.2
0.4
0.6
0.8
1.0
F1
F80 F1
F80 F1
F24 F1

### Dendrogram of cluster 936

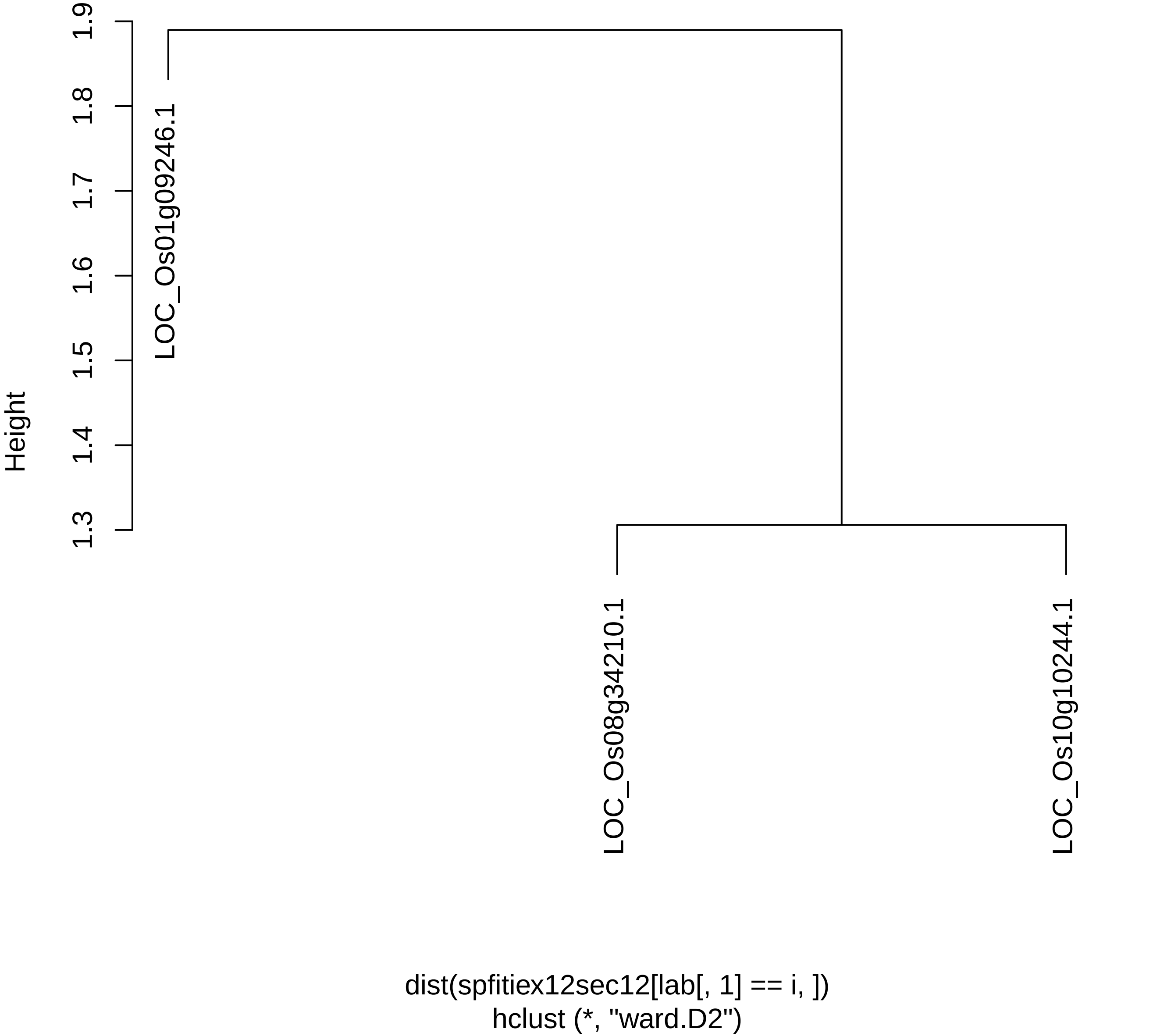

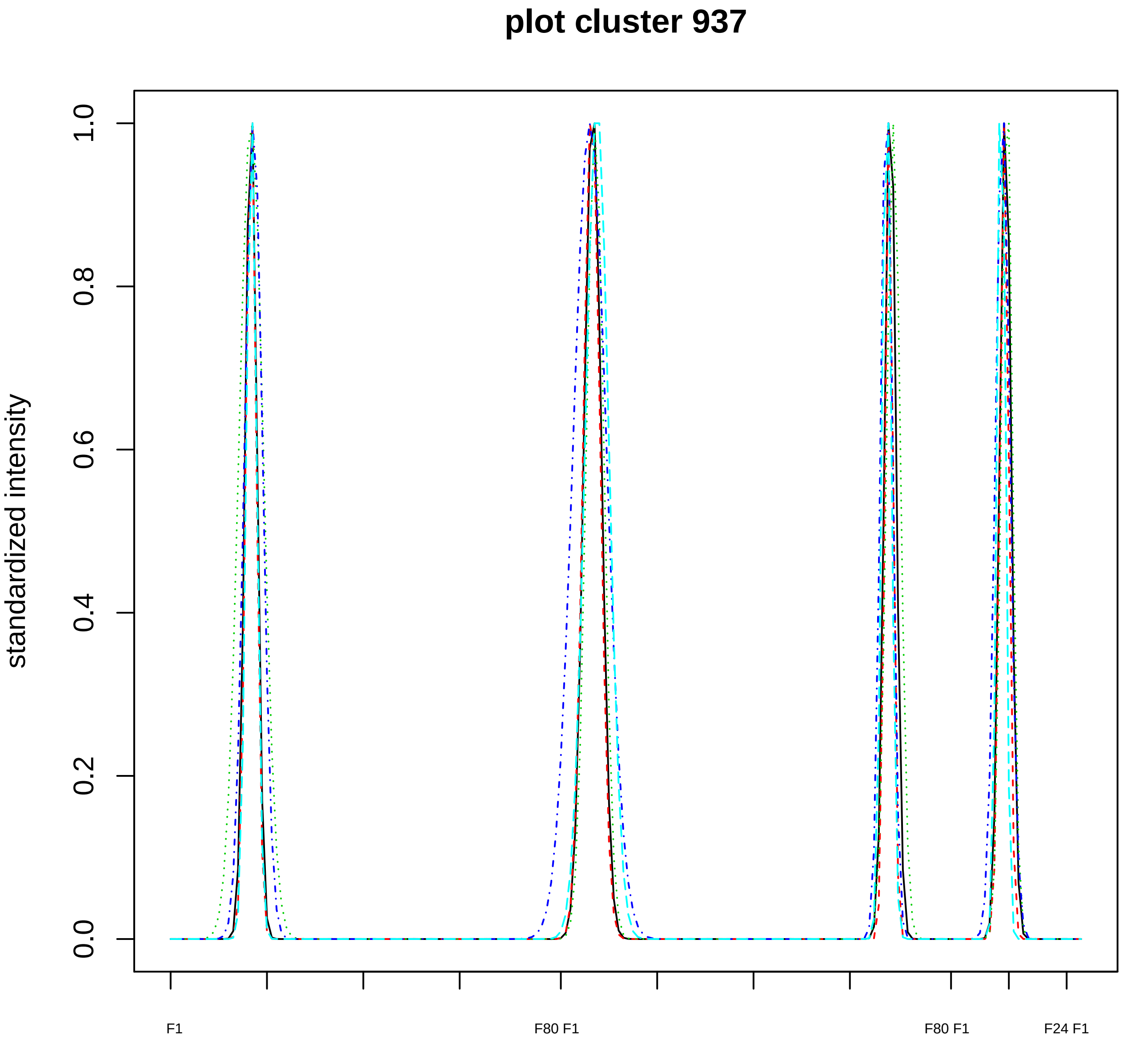
plot cluster 937
standardized intensity
0.0
0.2
0.4
0.6
0.8
1.0
F1
F80 F1
F80 F1
F24 F1

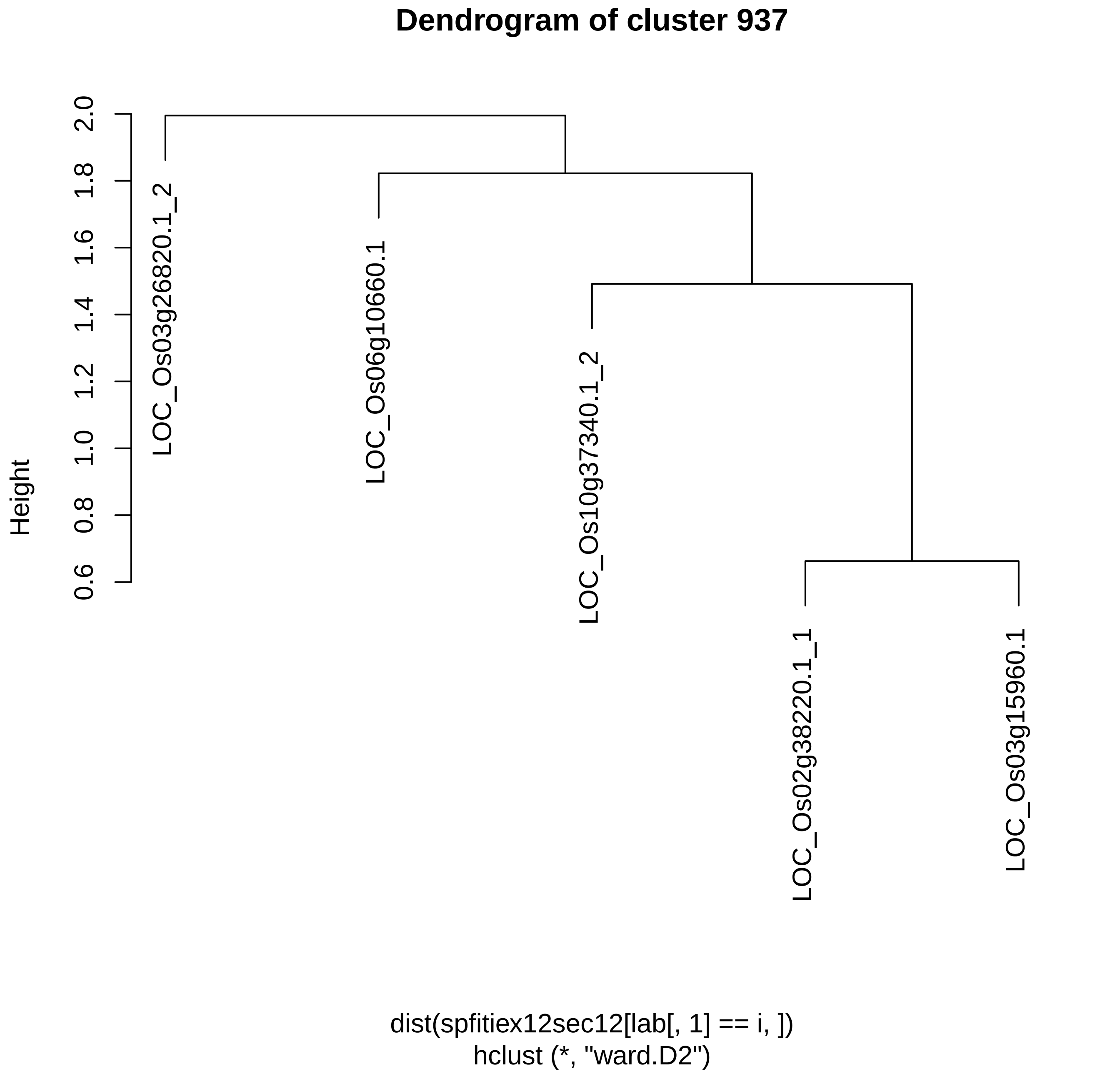
Dendrogram of cluster 937
Height
2.0
1.8
1.6
1.4
1.2
1.0
0.8
0.6
LOC_Os03g26820.1_2
LOC_Os06g10660.1
LOC_Os10g37340.1_2
LOC_Os02g38220.1_1
LOC_Os03g15960.1
dist(spfitiex12sec12[lab[, 1] == i, ])
hclust (*, "ward.D2")

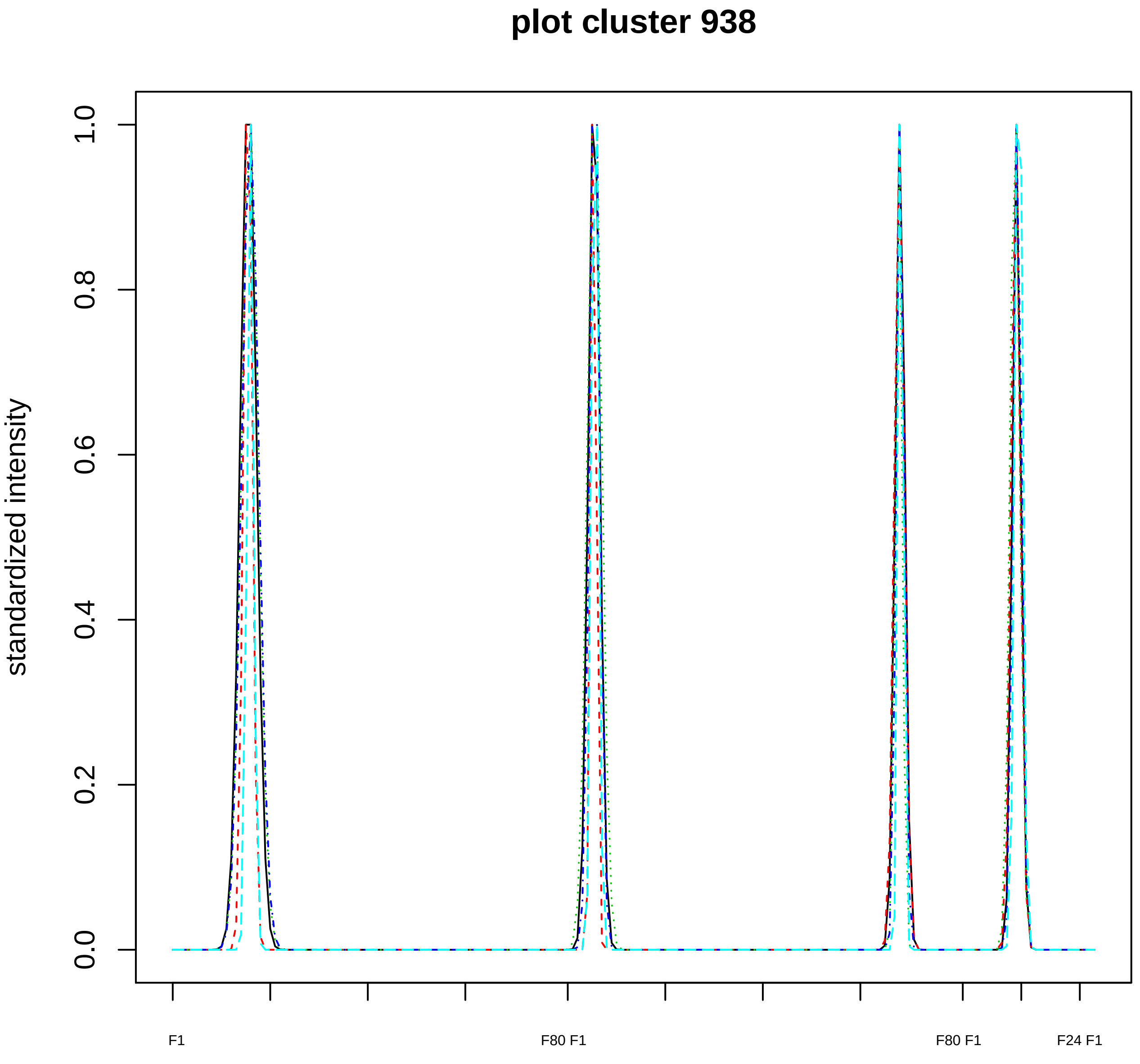
plot cluster 938
standardized intensity
0.0
0.2
0.4
0.6
0.8
1.0
F1
F80 F1
F80 F1
F24 F1

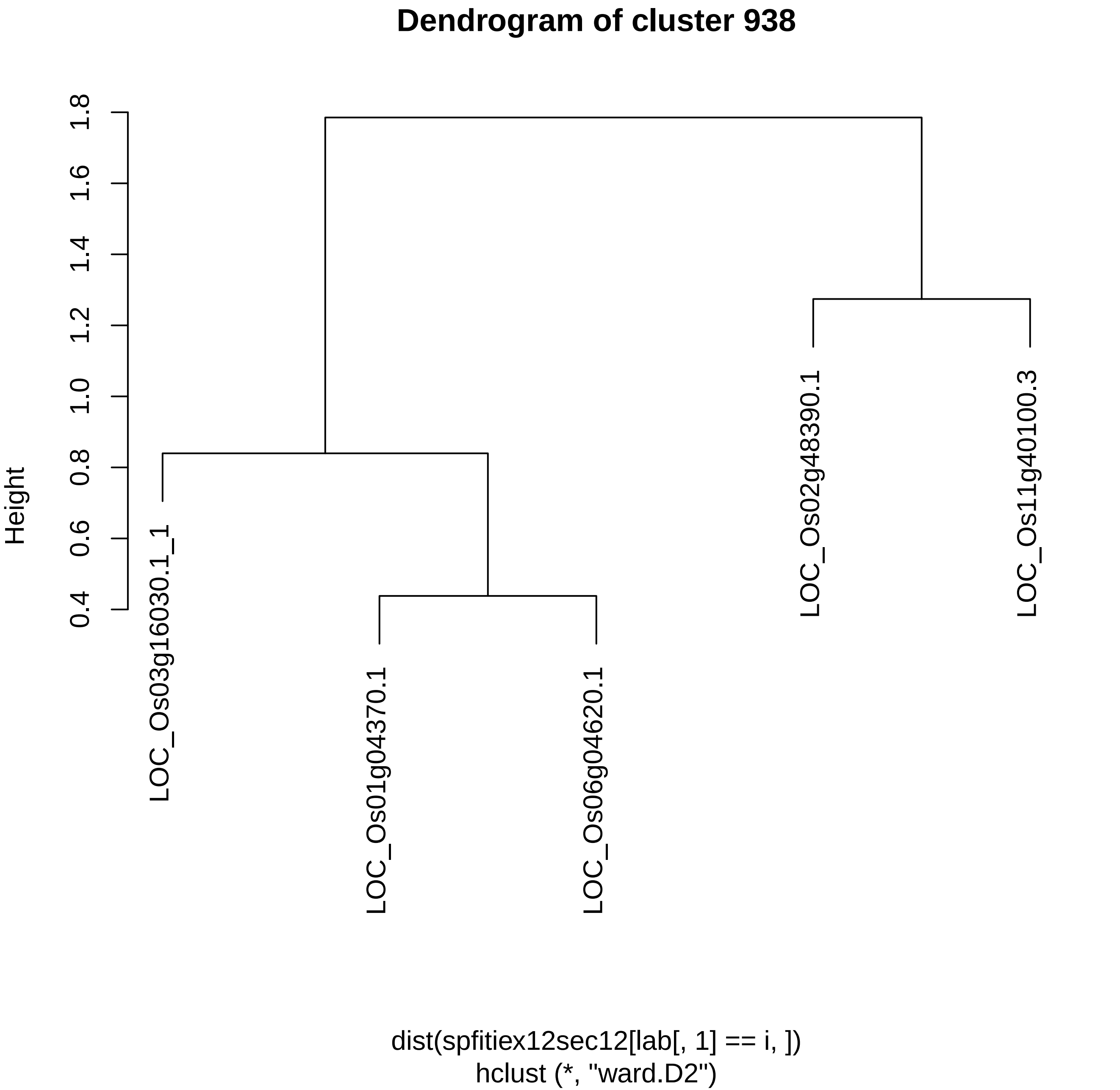

Dendrogram of cluster 938
Height
0.4
0.6
0.8
1.0
1.2
1.4
1.6
1.8
LOC_Os03g16030.1_1
LOC_Os01g04370.1
LOC_Os06g04620.1
LOC_Os02g48390.1
LOC_Os11g40100.3
dist(spfitiex12sec12[lab[, 1] == i, ])
hclust (*, "ward.D2")

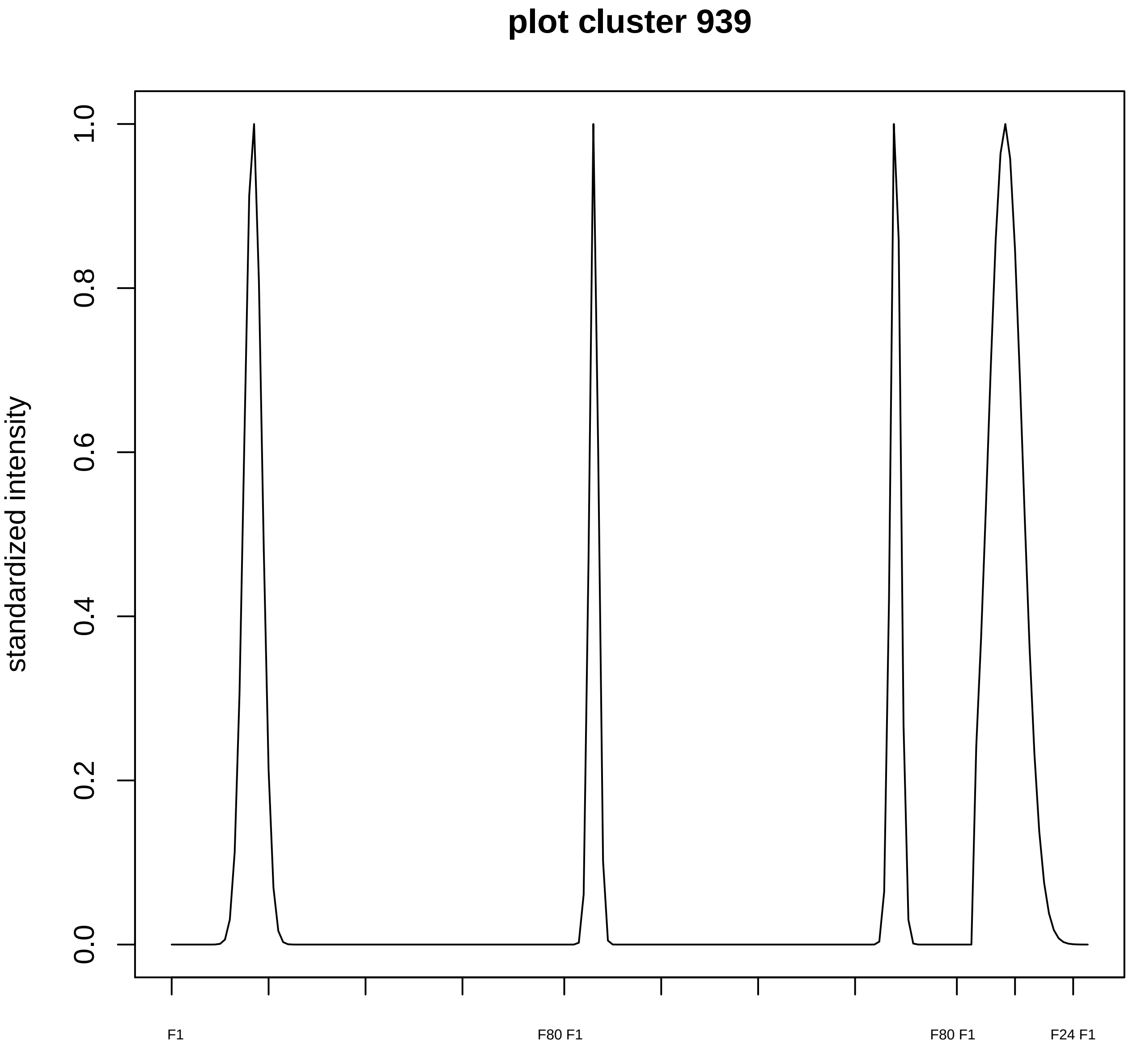
plot cluster 939
standardized intensity
0.0
0.2
0.4
0.6
0.8
1.0
F1
F80 F1
F80 F1
F24 F1

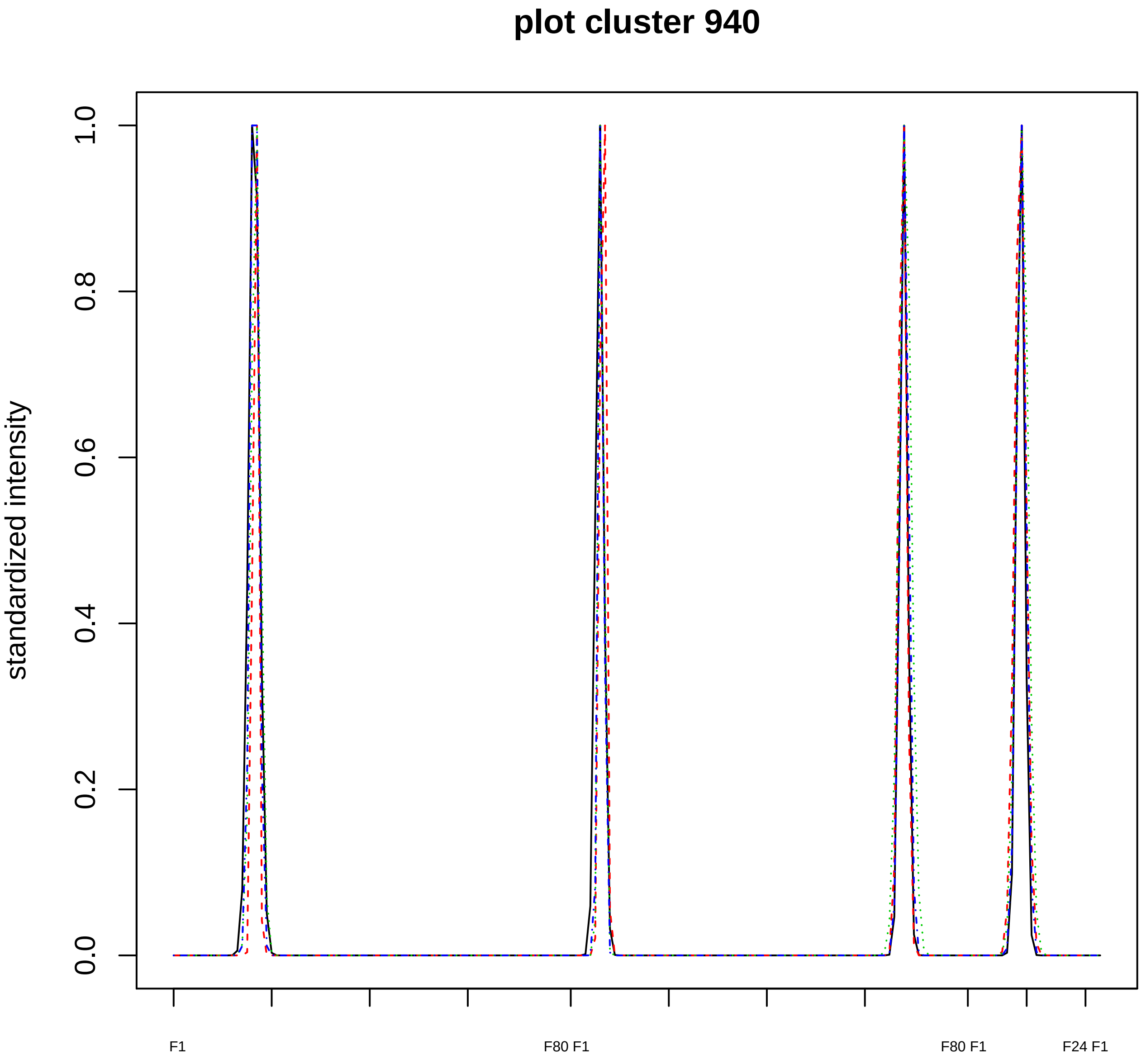
plot cluster 940
standardized intensity
0.0
0.2
0.4
0.6
0.8
1.0
F1
F80 F1
F80 F1
F24 F1

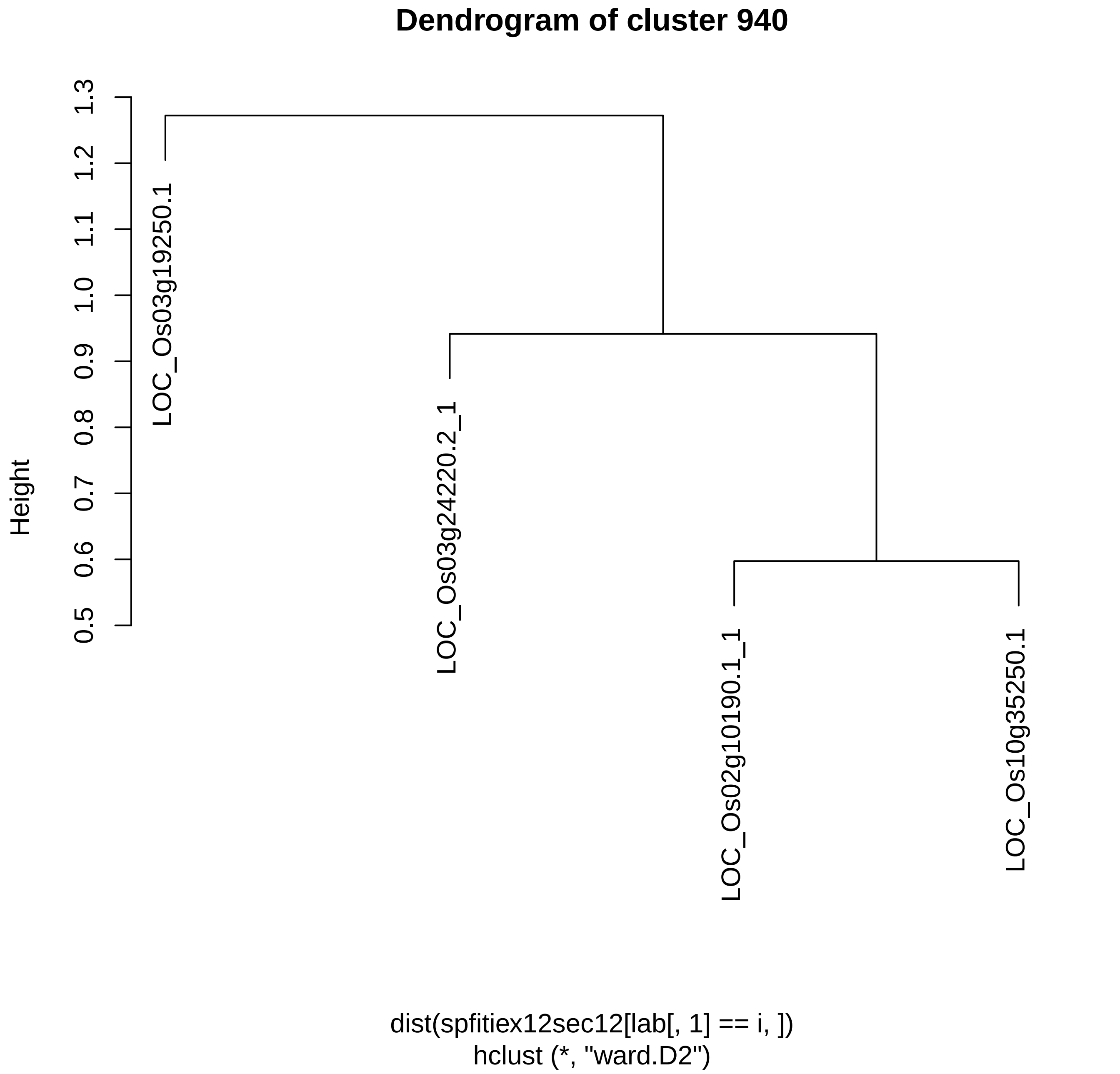
Dendrogram of cluster 940
Height
0.5
0.6
0.7
0.8
0.9
1.0
1.1
1.2
1.3
LOC_Os03g19250.1
LOC_Os03g24220.2_1
LOC_Os02g10190.1_1
LOC_Os10g35250.1
dist(spfitiex12sec12[lab[, 1] == i, ])
hclust (*, "ward.D2")

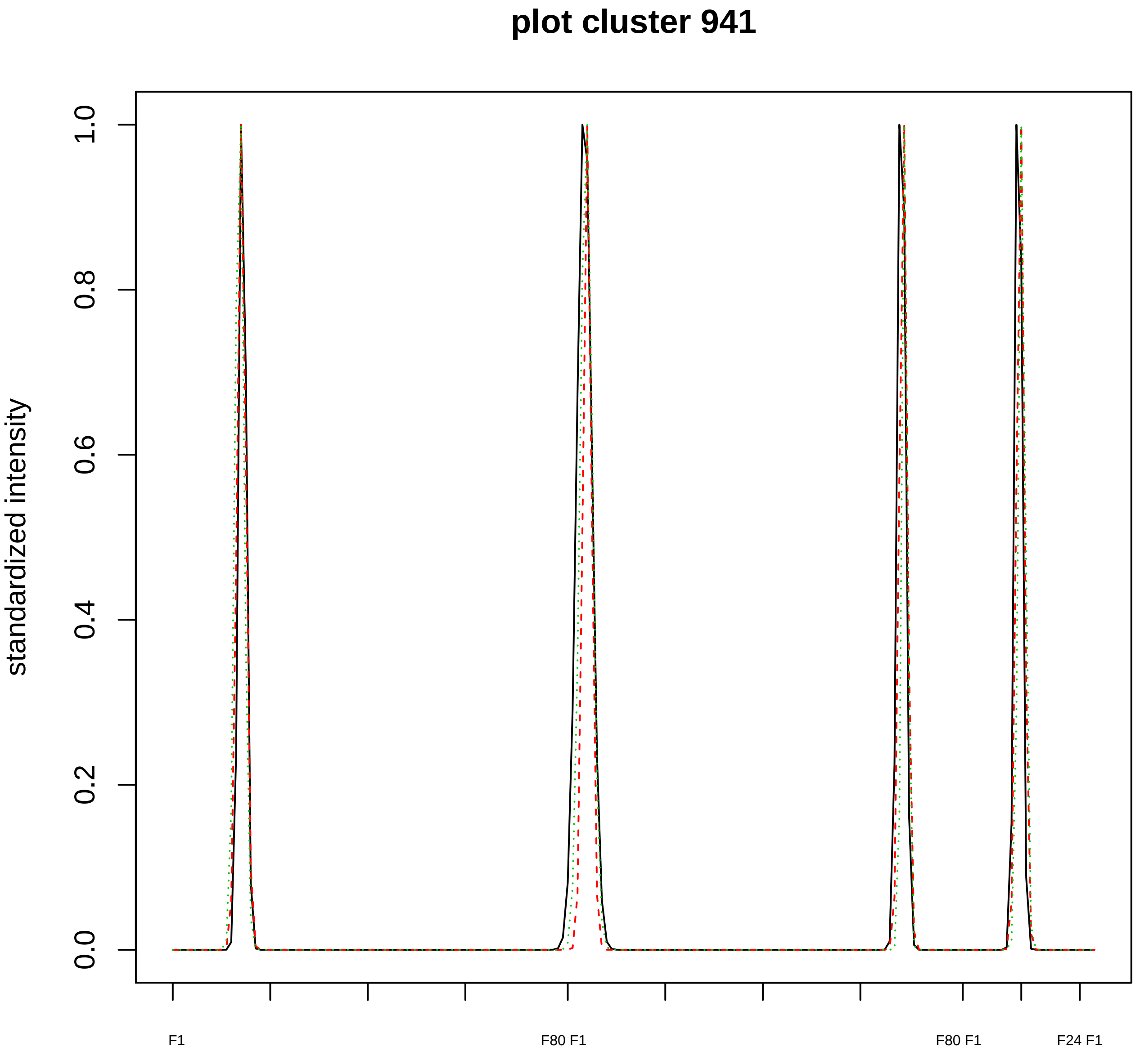
plot cluster 941
standardized intensity
0.0
0.2
0.4
0.6
0.8
1.0
F1
F80 F1
F80 F1
F24 F1

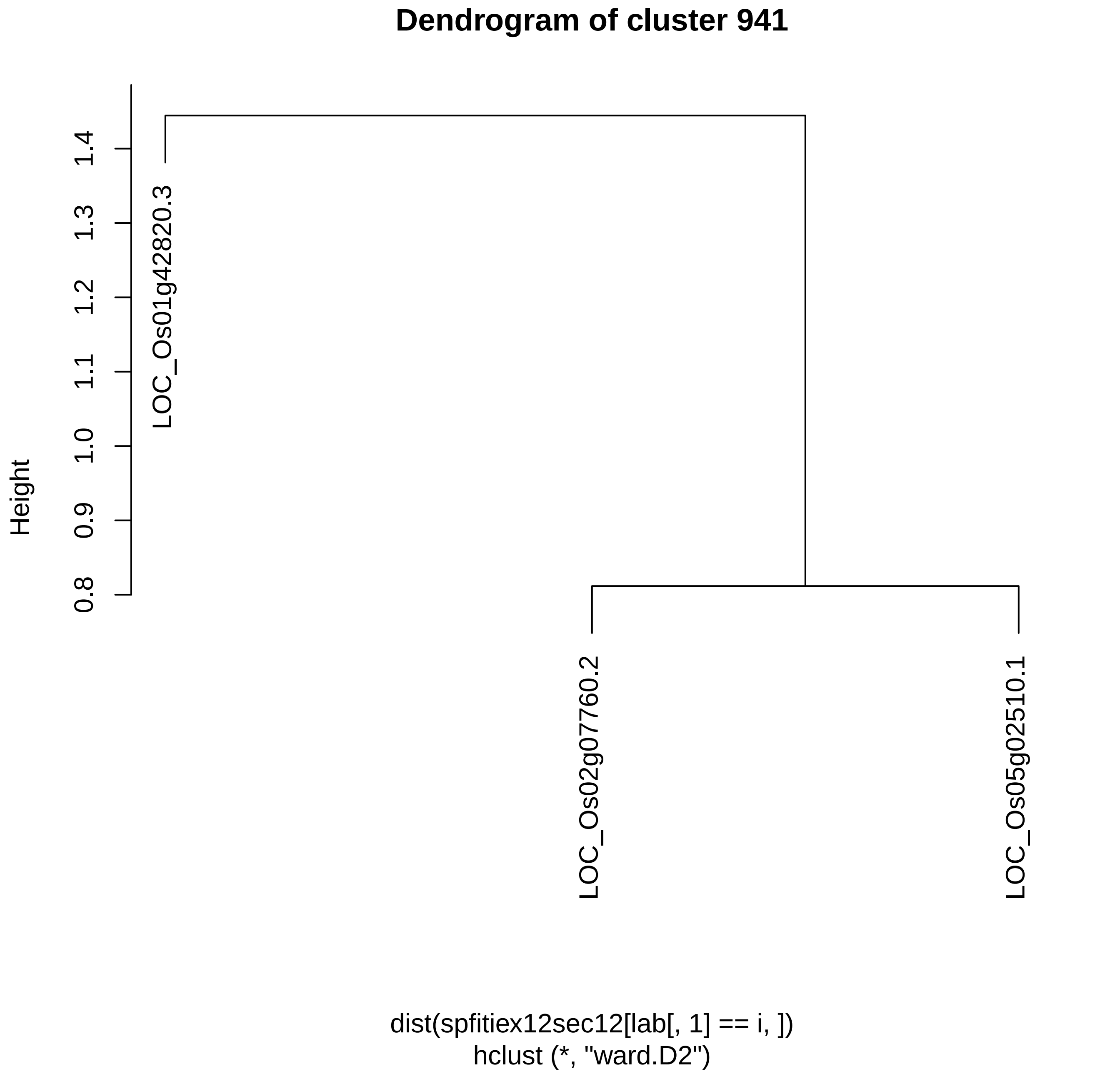
Dendrogram of cluster 941
Height
0.8
0.9
1.0
1.1
1.2
1.3
1.4
LOC_Os01g42820.3
LOC_Os02g07760.2
LOC_Os05g02510.1
dist(spfitiex12sec12[lab[, 1] == i, ])
hclust (*, "ward.D2")

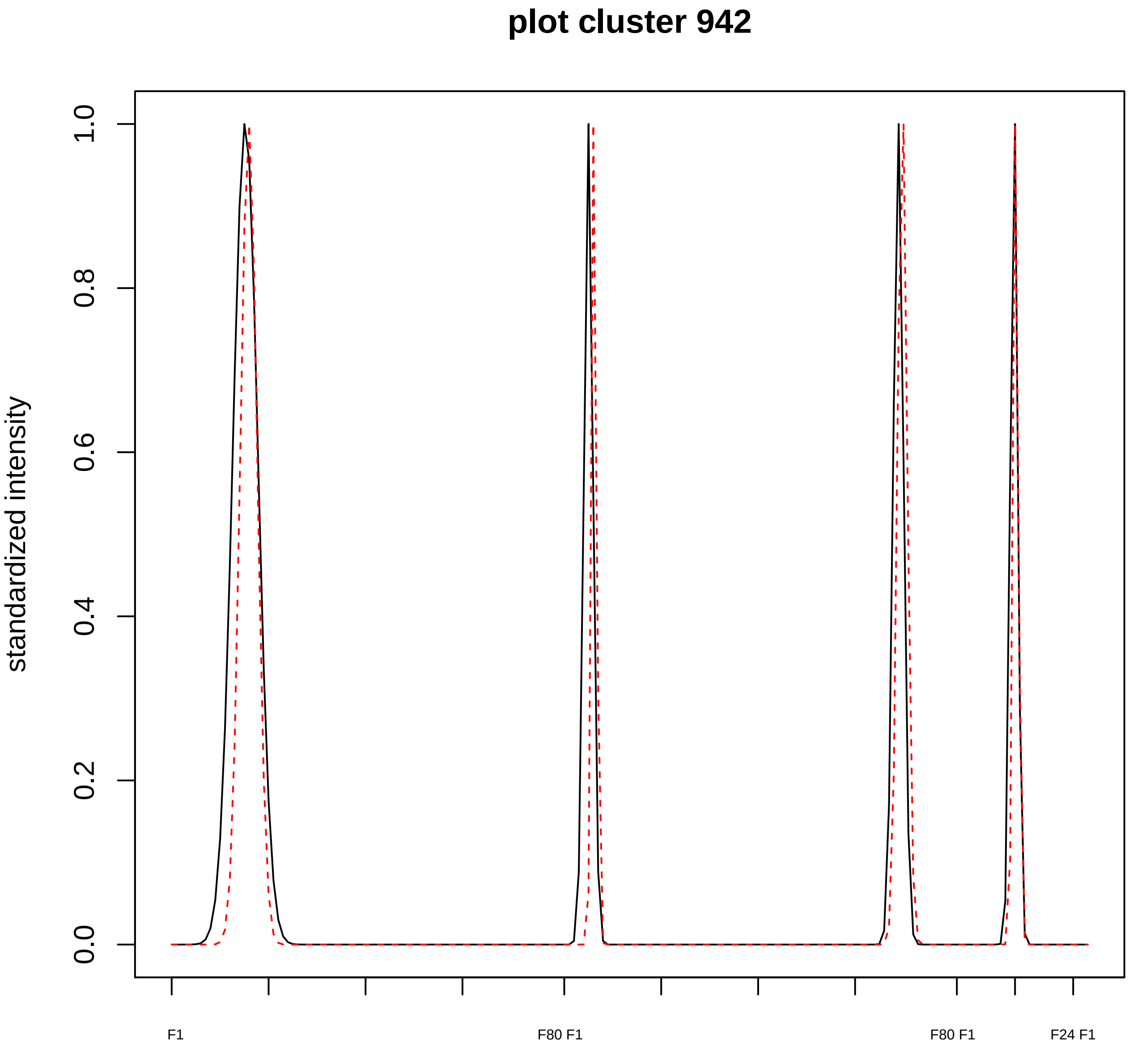

plot cluster 942
standardized intensity
0.0
0.2
0.4
0.6
0.8
1.0
F1
F80 F1
F80 F1
F24 F1

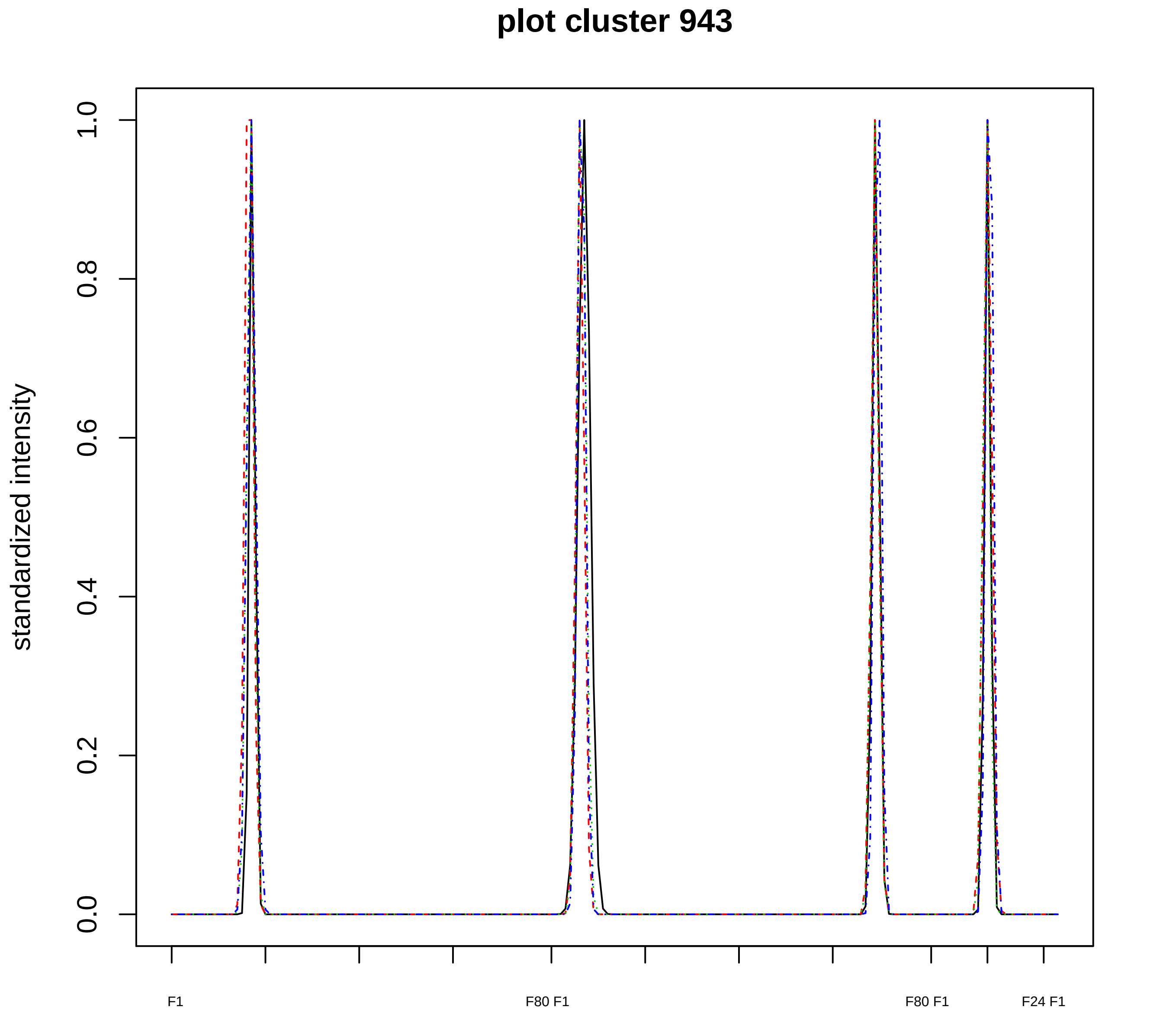
plot cluster 943
standardized intensity
0.0
0.2
0.4
0.6
0.8
1.0
F1
F80 F1
F80 F1
F24 F1

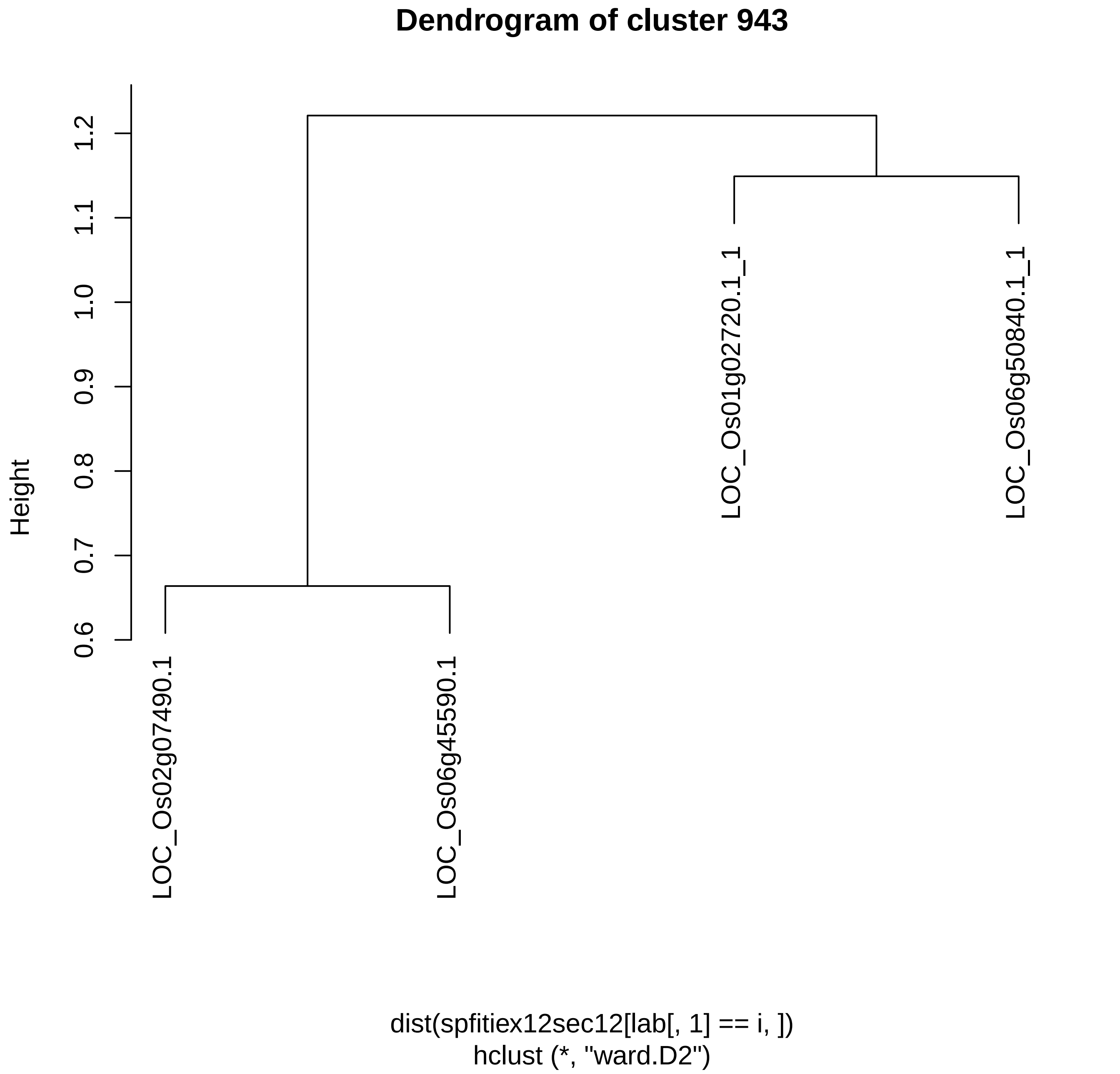
Dendrogram of cluster 943
Height
0.6
0.7
0.8
0.9
1.0
1.1
1.2
LOC_Os02g07490.1
LOC_Os06g45590.1
LOC_Os01g02720.1_1
LOC_Os06g50840.1_1
dist(spfitiex12sec12[lab[, 1] == i, ])
hclust (*, "ward.D2")

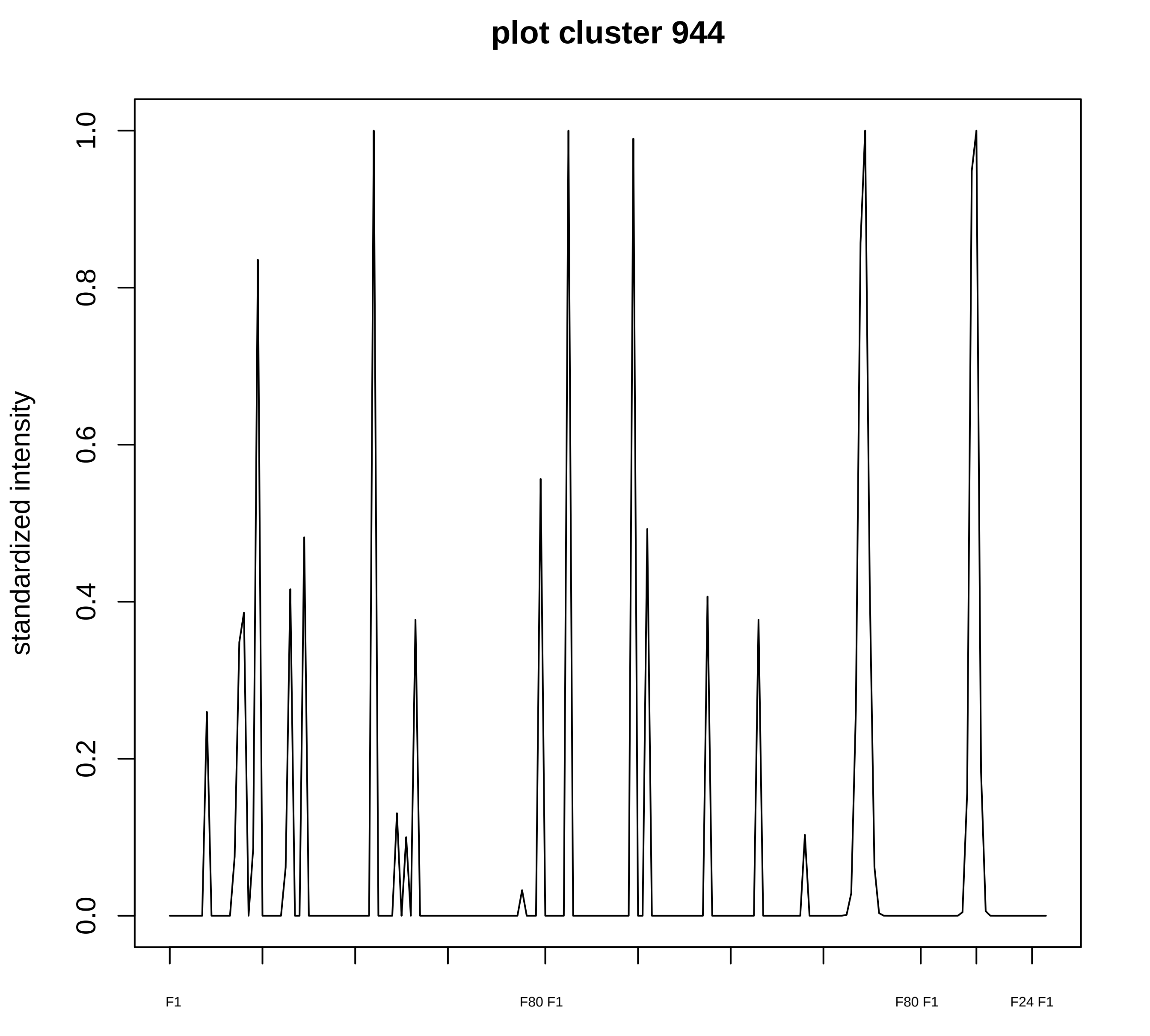

plot cluster 944
standardized intensity
0.0
0.2
0.4
0.6
0.8
1.0
F1
F80 F1
F80 F1
F24 F1

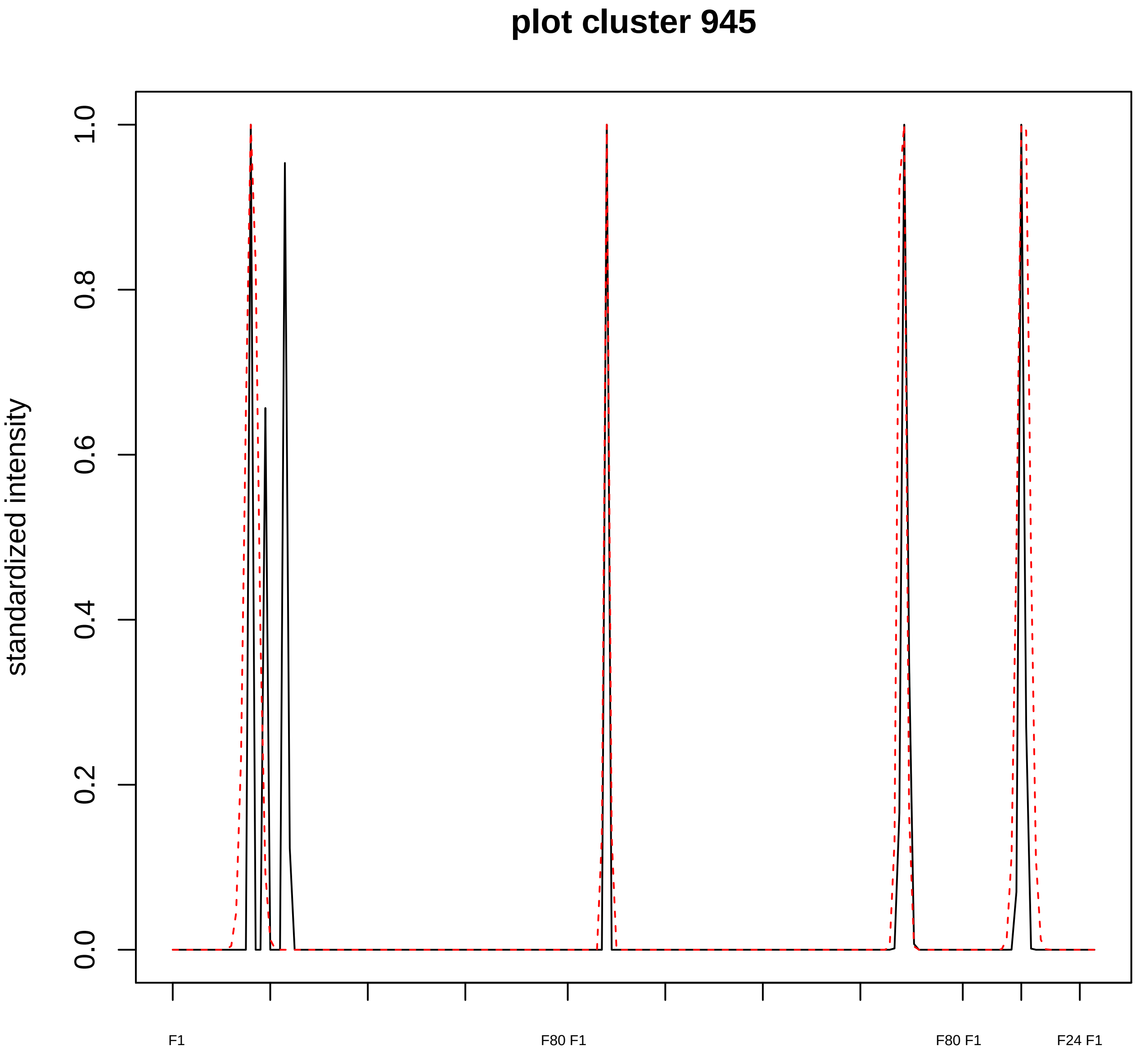
plot cluster 945
standardized intensity
0.0
0.2
0.4
0.6
0.8
1.0
F1
F80 F1
F80 F1
F24 F1

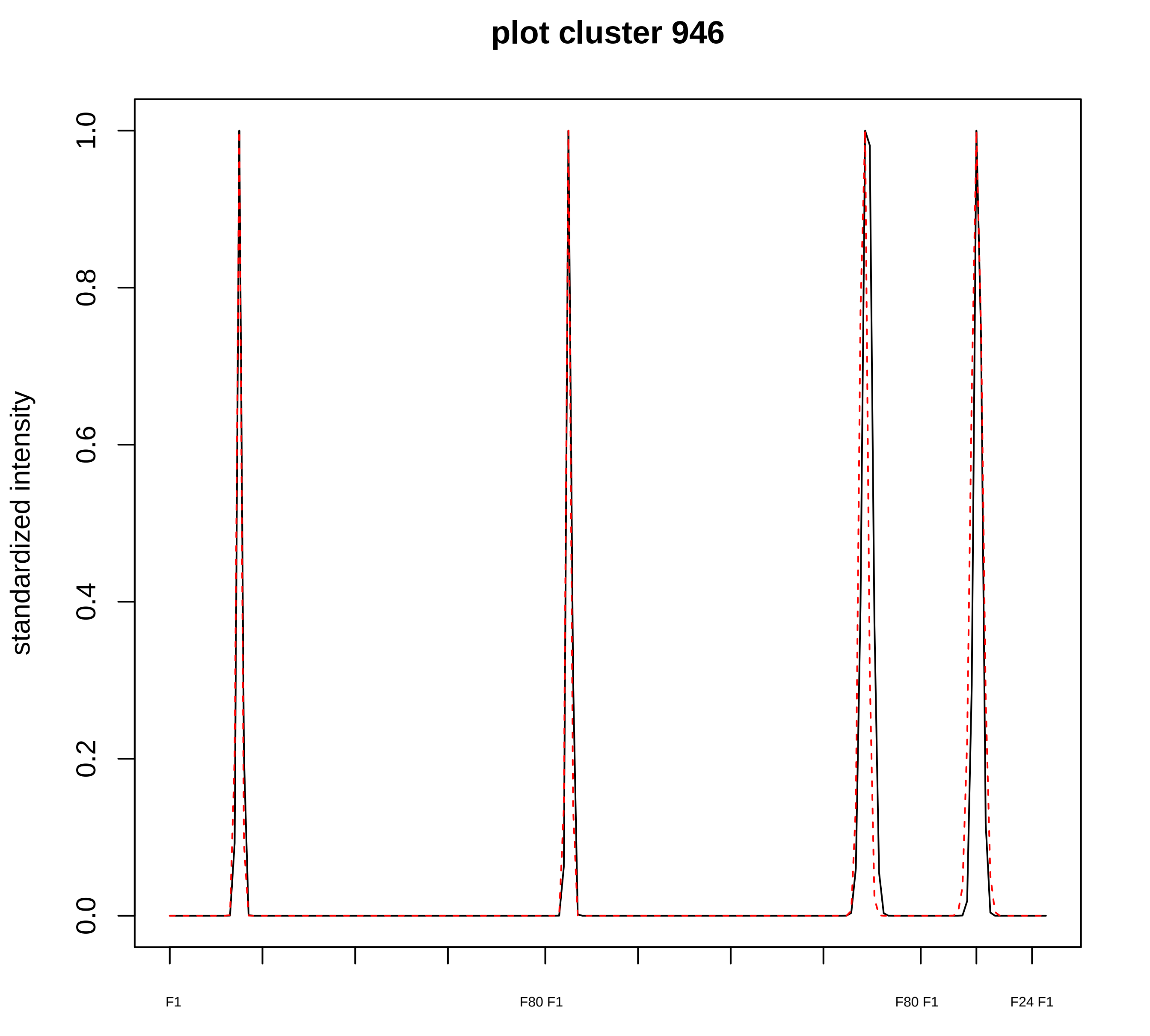

plot cluster 946
standardized intensity
0.0
0.2
0.4
0.6
0.8
1.0
F1
F80 F1
F80 F1
F24 F1

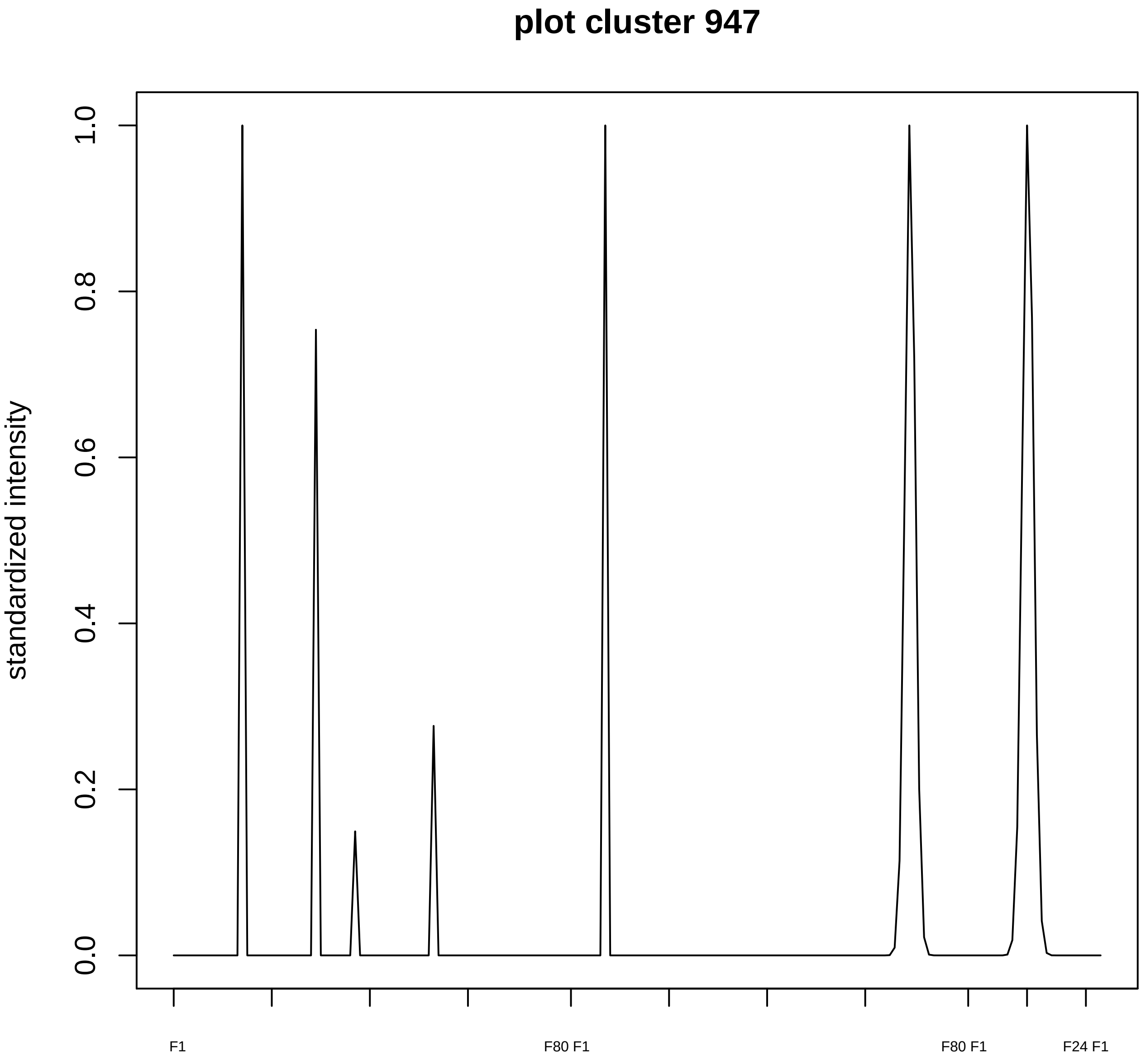

plot cluster 947
standardized intensity
0.0
0.2
0.4
0.6
0.8
1.0
F1
F80 F1
F80 F1
F24 F1

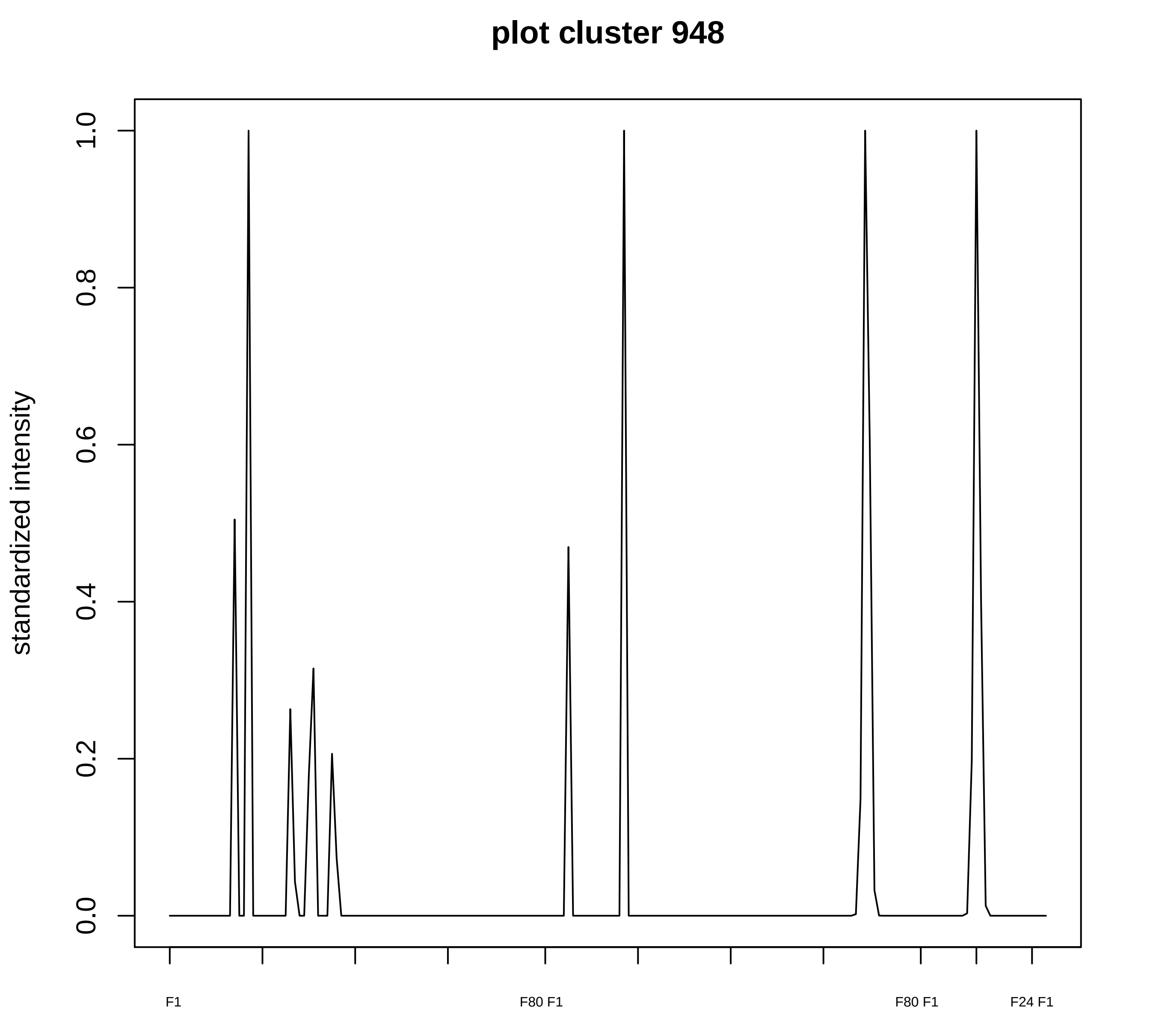
plot cluster 948
standardized intensity
0.0
0.2
0.4
0.6
0.8
1.0
F1
F80 F1
F80 F1
F24 F1

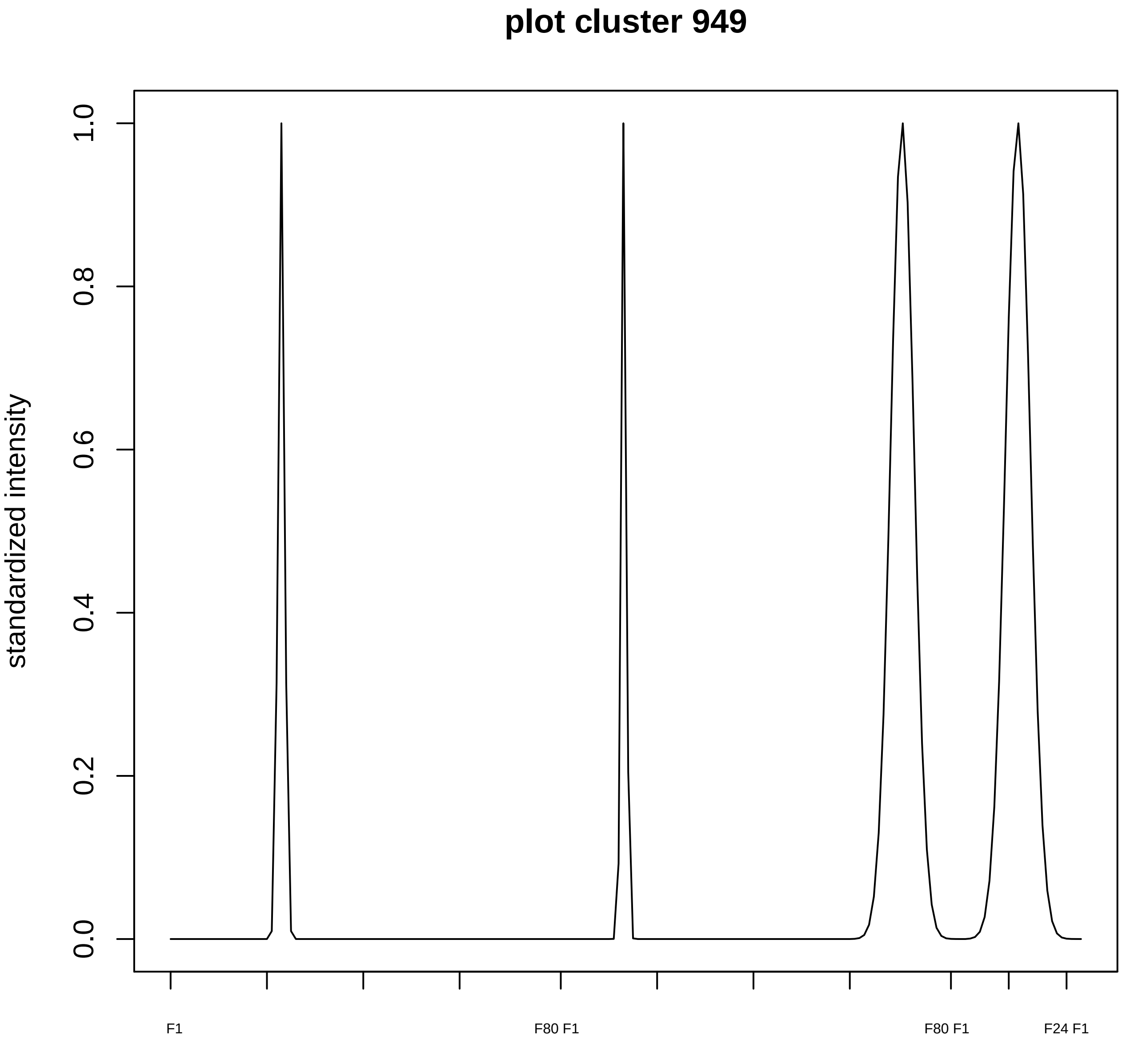

plot cluster 949
standardized intensity
0.0
0.2
0.4
0.6
0.8
1.0
F1
F80 F1
F80 F1
F24 F1

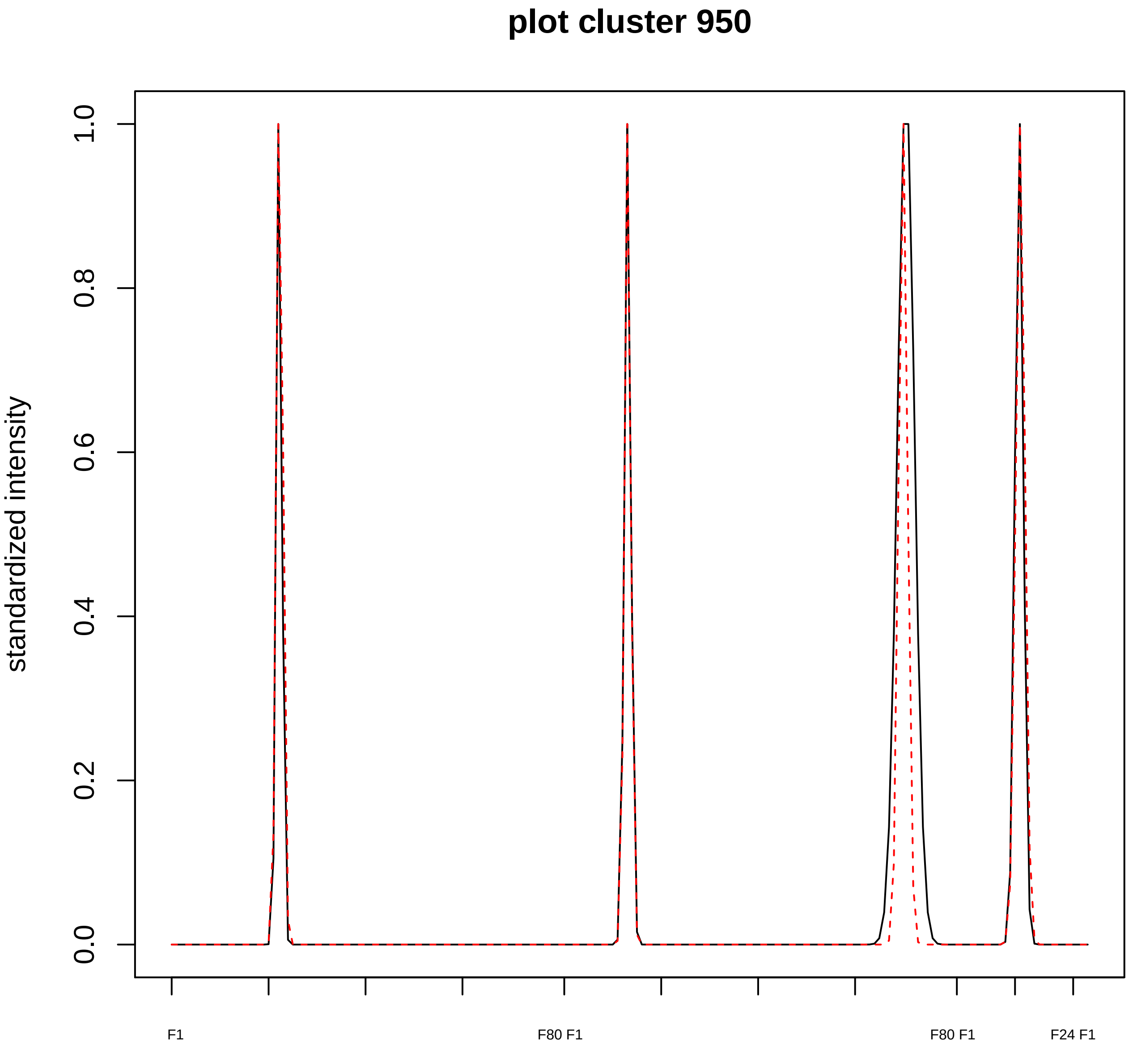

plot cluster 950
standardized intensity
0.0
0.2
0.4
0.6
0.8
1.0
F1
F80 F1
F80 F1
F24 F1

### plot cluster 951

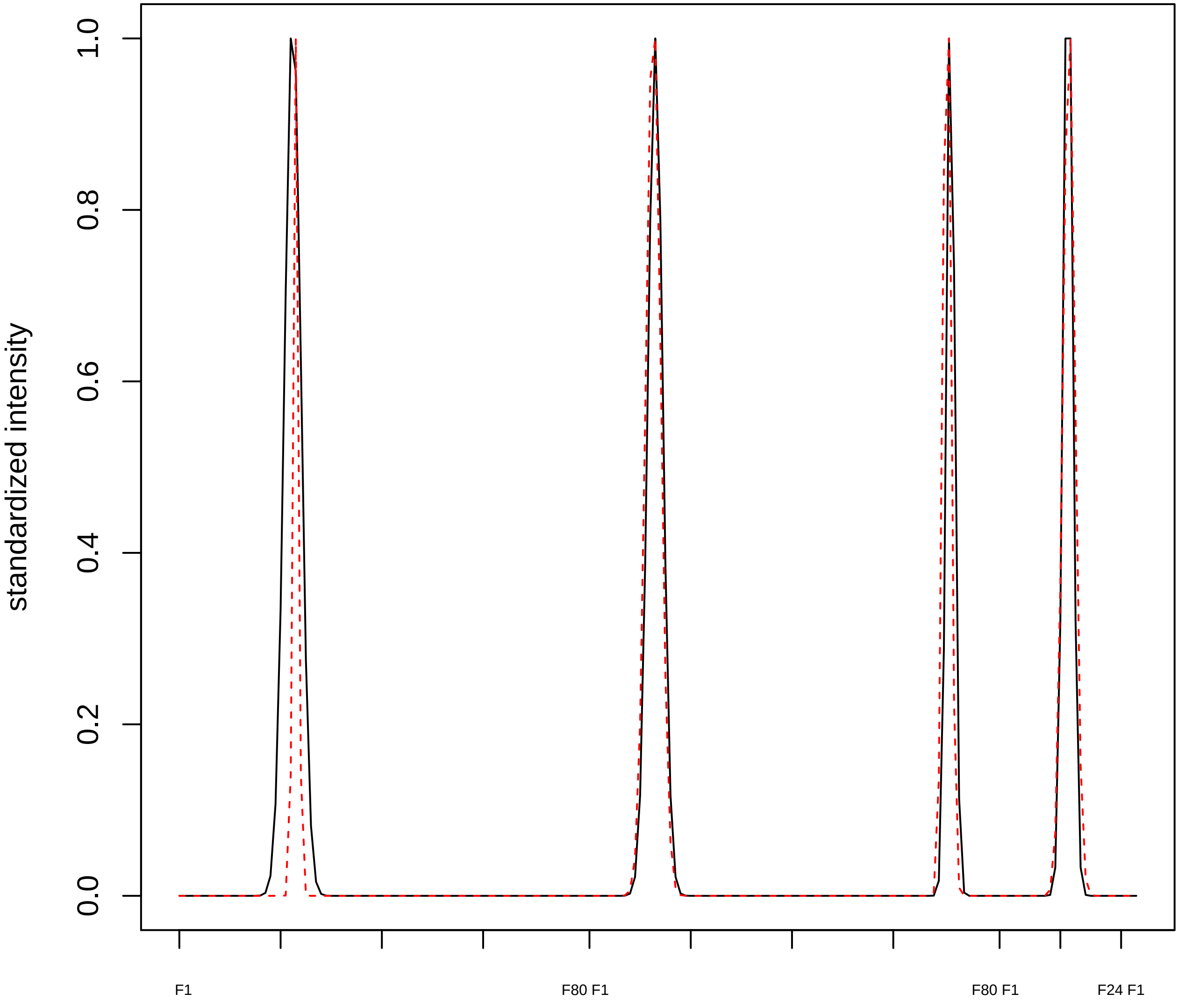

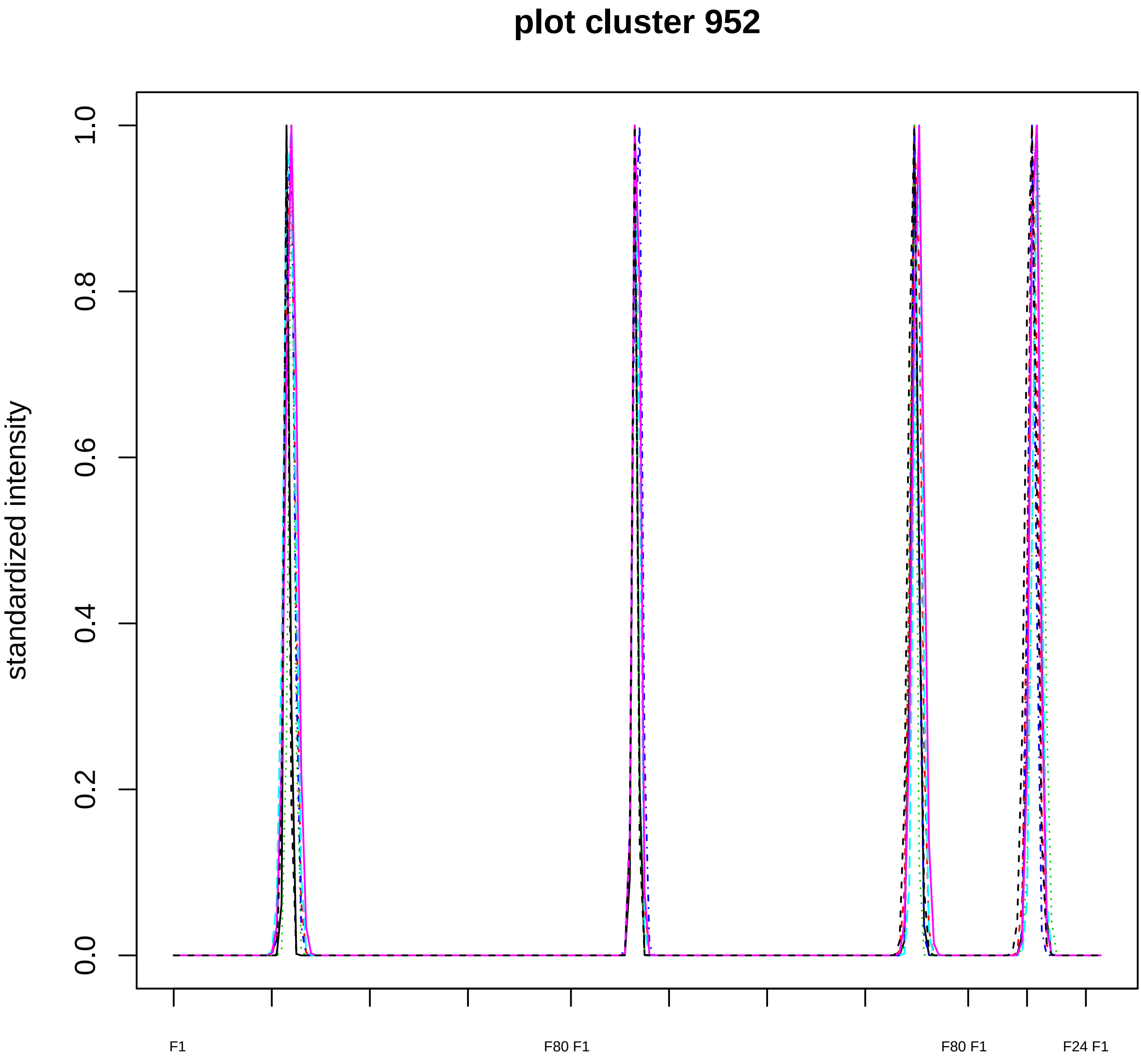

plot cluster 952
standardized intensity
0.0
0.2
0.4
0.6
0.8
1.0
F1
F80 F1
F80 F1
F24 F1

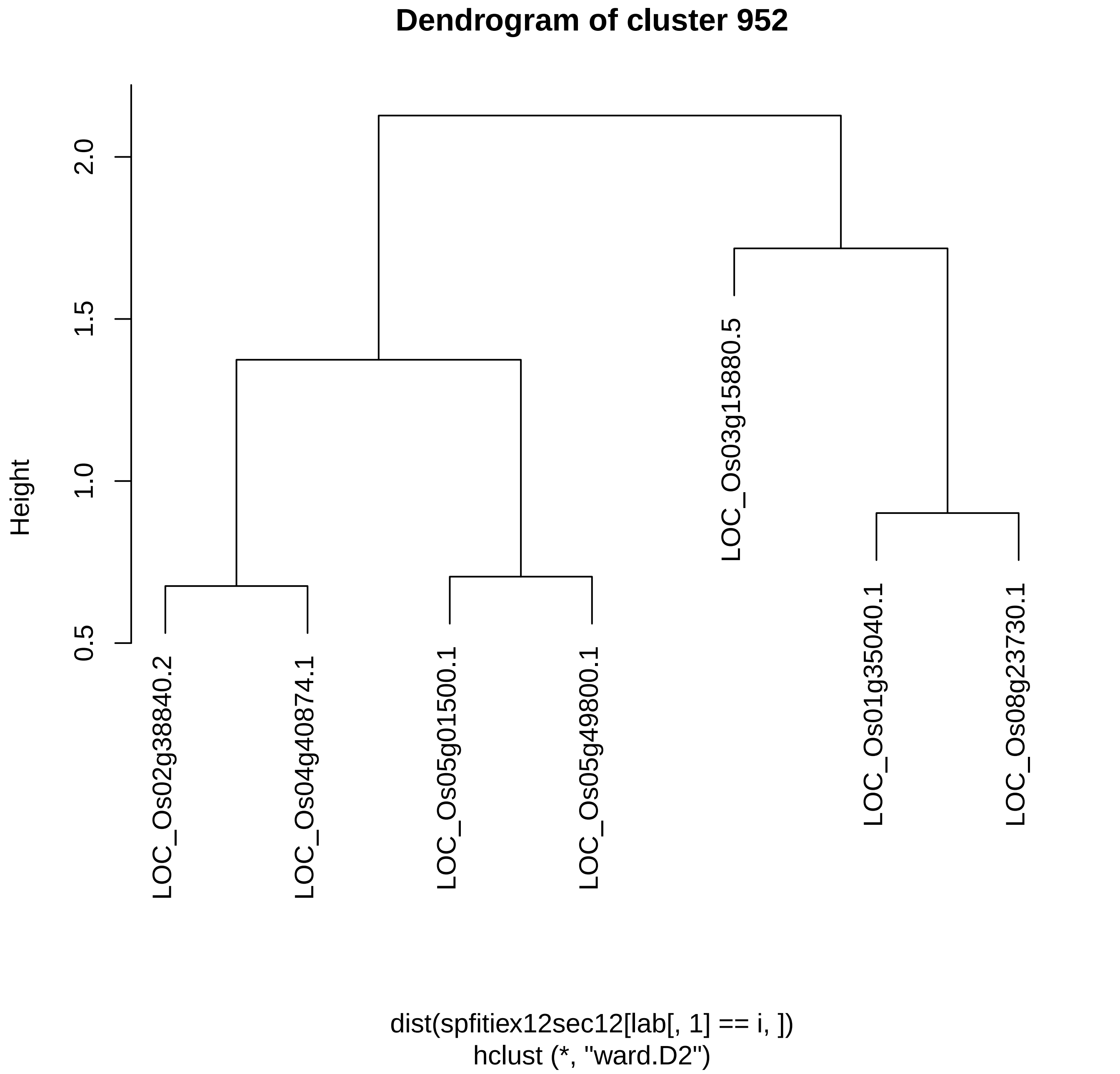
Dendrogram of cluster 952
Height
2.0
1.5
1.0
0.5
LOC_Os02g38840.2
LOC_Os04g40874.1
LOC_Os05g01500.1
LOC_Os05g49800.1
LOC_Os03g15880.5
LOC_Os01g35040.1
LOC_Os08g23730.1
dist(spfitiex12sec12[lab[, 1] == i, ])
hclust (*, "ward.D2")

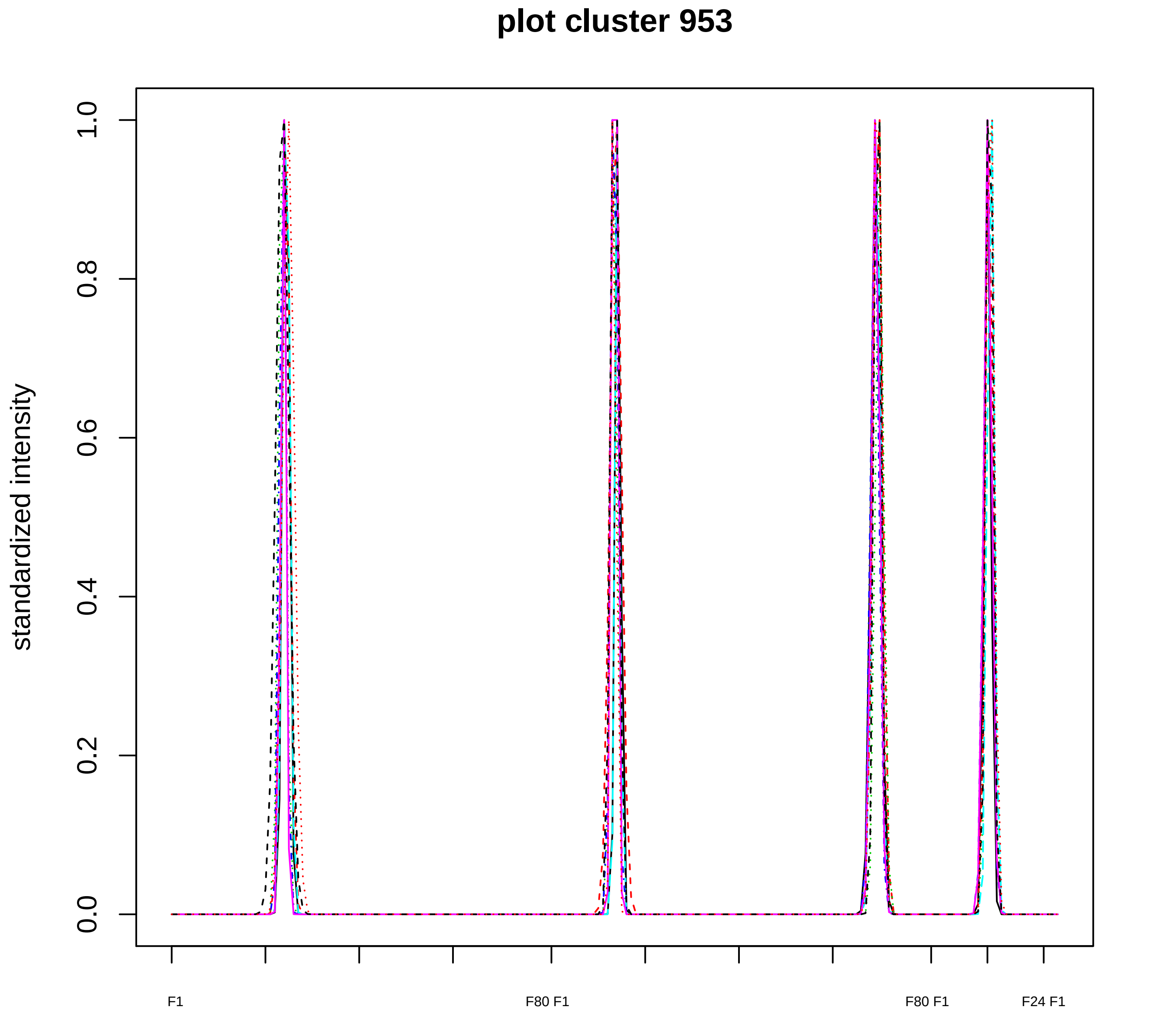
plot cluster 953
standardized intensity
0.0
0.2
0.4
0.6
0.8
1.0
F1
F80 F1
F80 F1
F24 F1

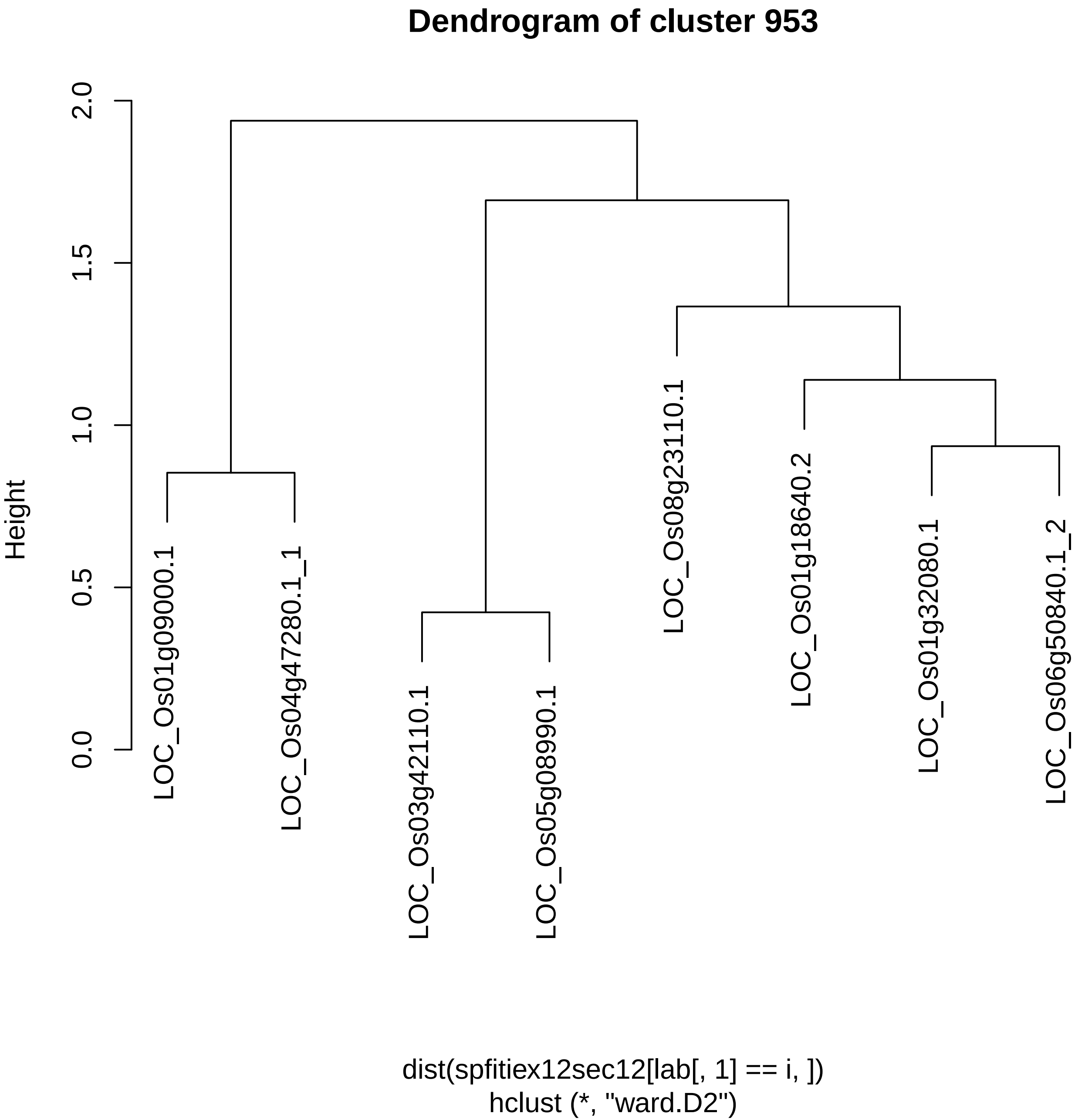

Dendrogram of cluster 953
Height
2.0
1.5
1.0
0.5
0.0
LOC_Os01g09000.1
LOC_Os04g47280.1_1
LOC_Os03g42110.1
LOC_Os05g08990.1
LOC_Os08g23110.1
LOC_Os01g18640.2
LOC_Os01g32080.1
LOC_Os06g50840.1_2
dist(spfitiex12sec12[lab[, 1] == i, ])
hclust (*, "ward.D2")

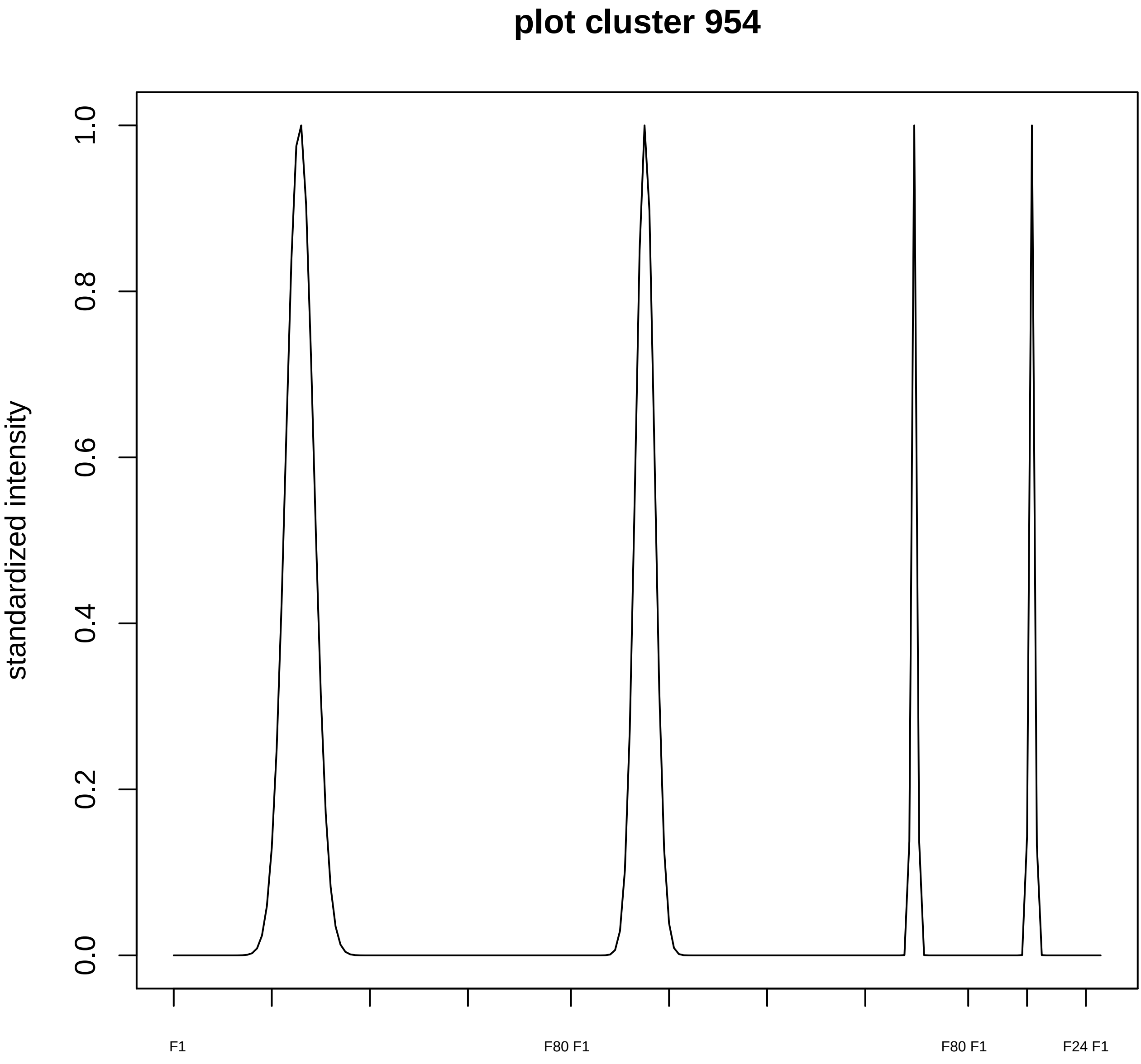

plot cluster 954
standardized intensity
0.0
0.2
0.4
0.6
0.8
1.0
F1
F80 F1
F80 F1
F24 F1

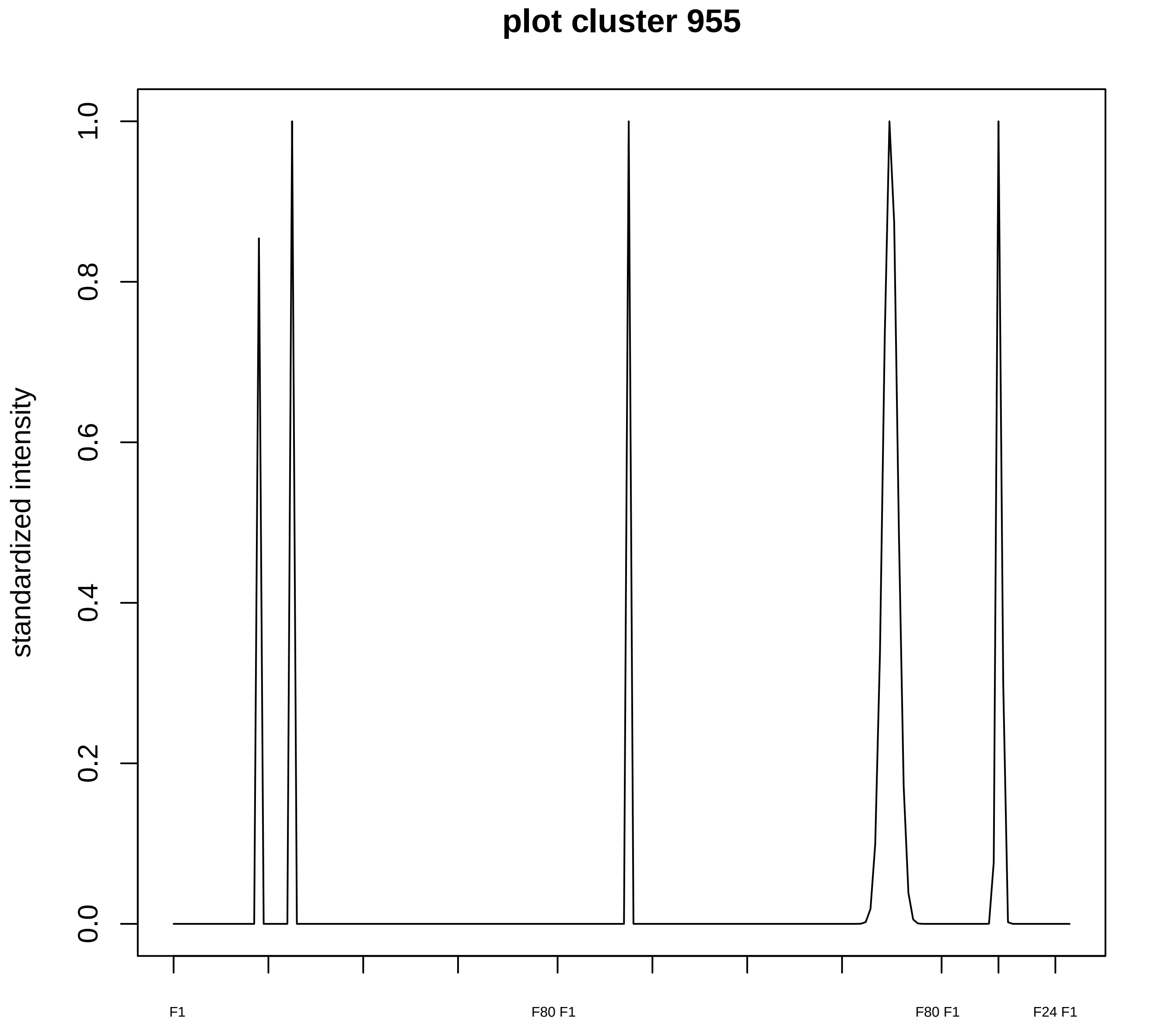

plot cluster 955
standardized intensity
0.0
0.2
0.4
0.6
0.8
1.0
F1
F80 F1
F80 F1
F24 F1

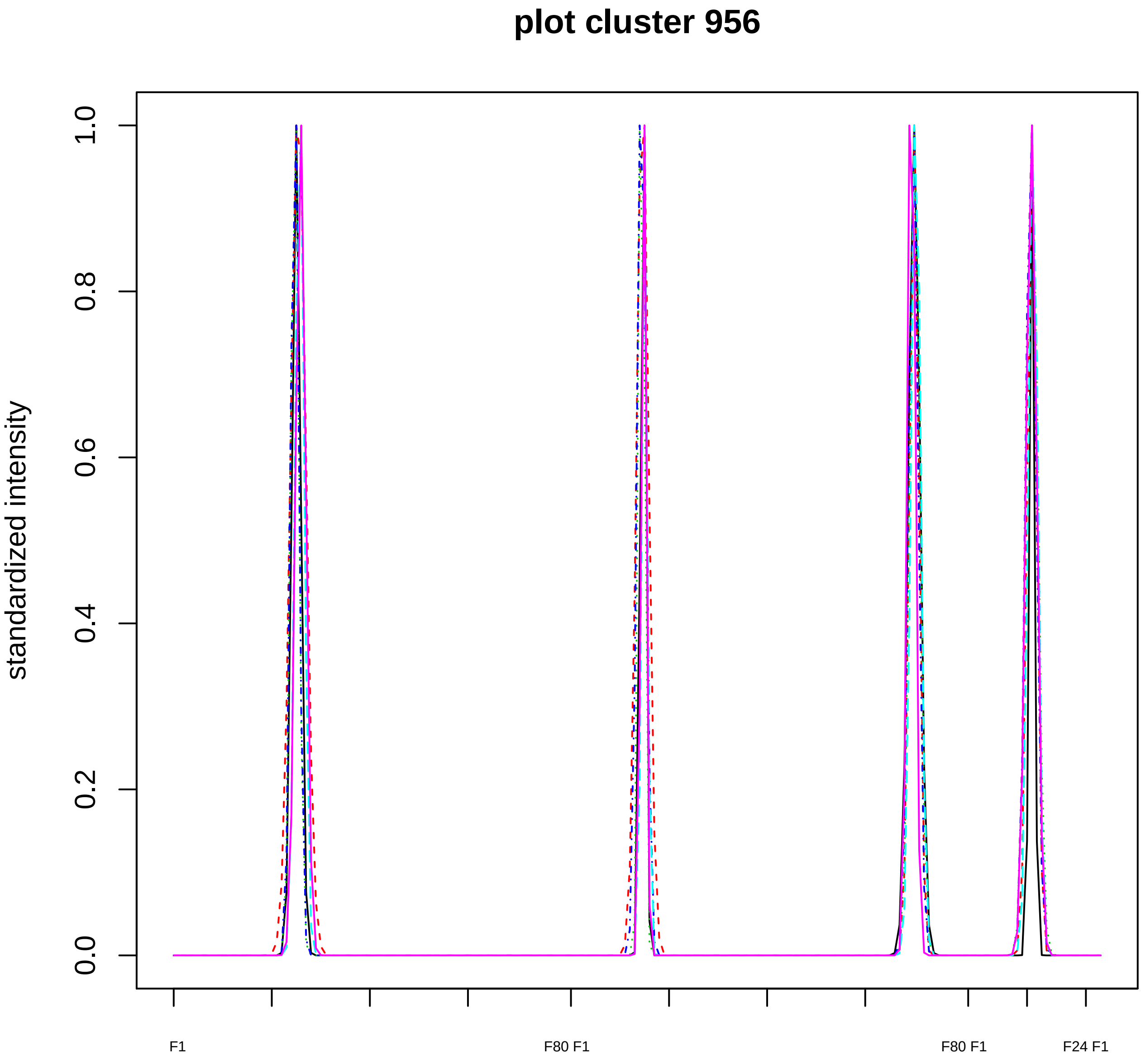

plot cluster 956
standardized intensity
0.0
0.2
0.4
0.6
0.8
1.0
F1
F80 F1
F80 F1
F24 F1

#### Dendrogram of cluster 956

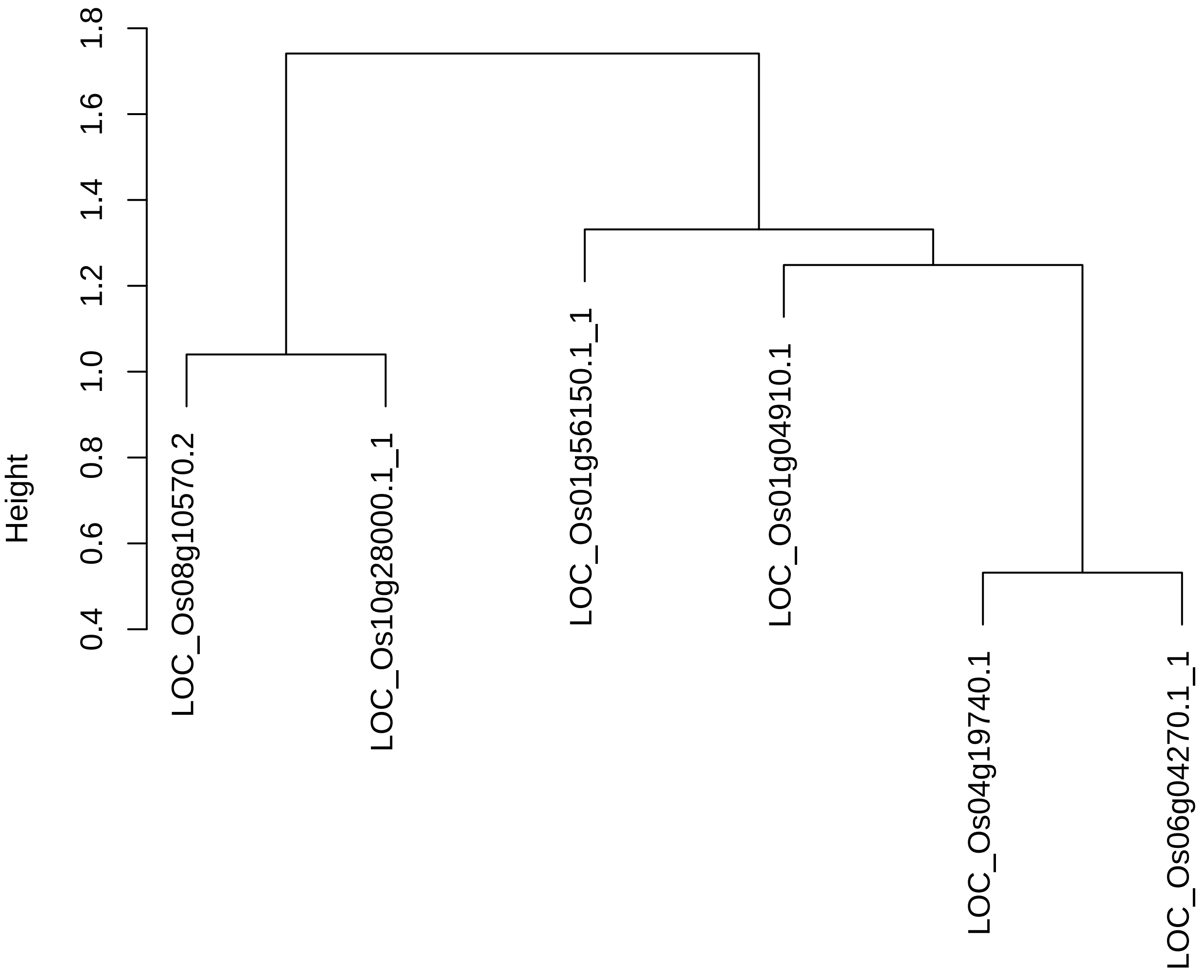


dist(spfitiex12sec12[lab[, 1] == i, ])
hclust (*, "ward.D2")

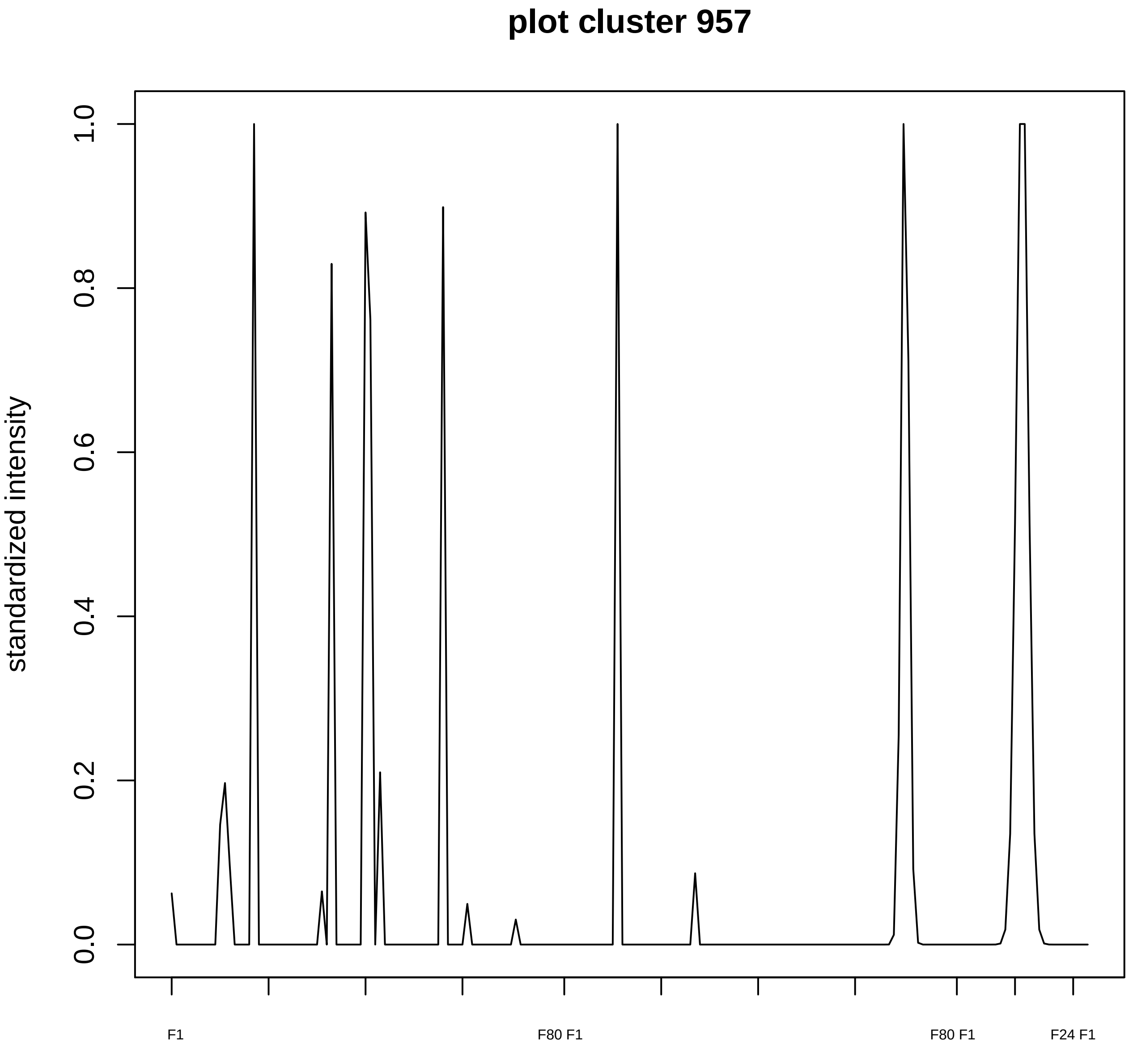

plot cluster 957
standardized intensity
0.0
0.2
0.4
0.6
0.8
1.0
F1
F80 F1
F80 F1
F24 F1

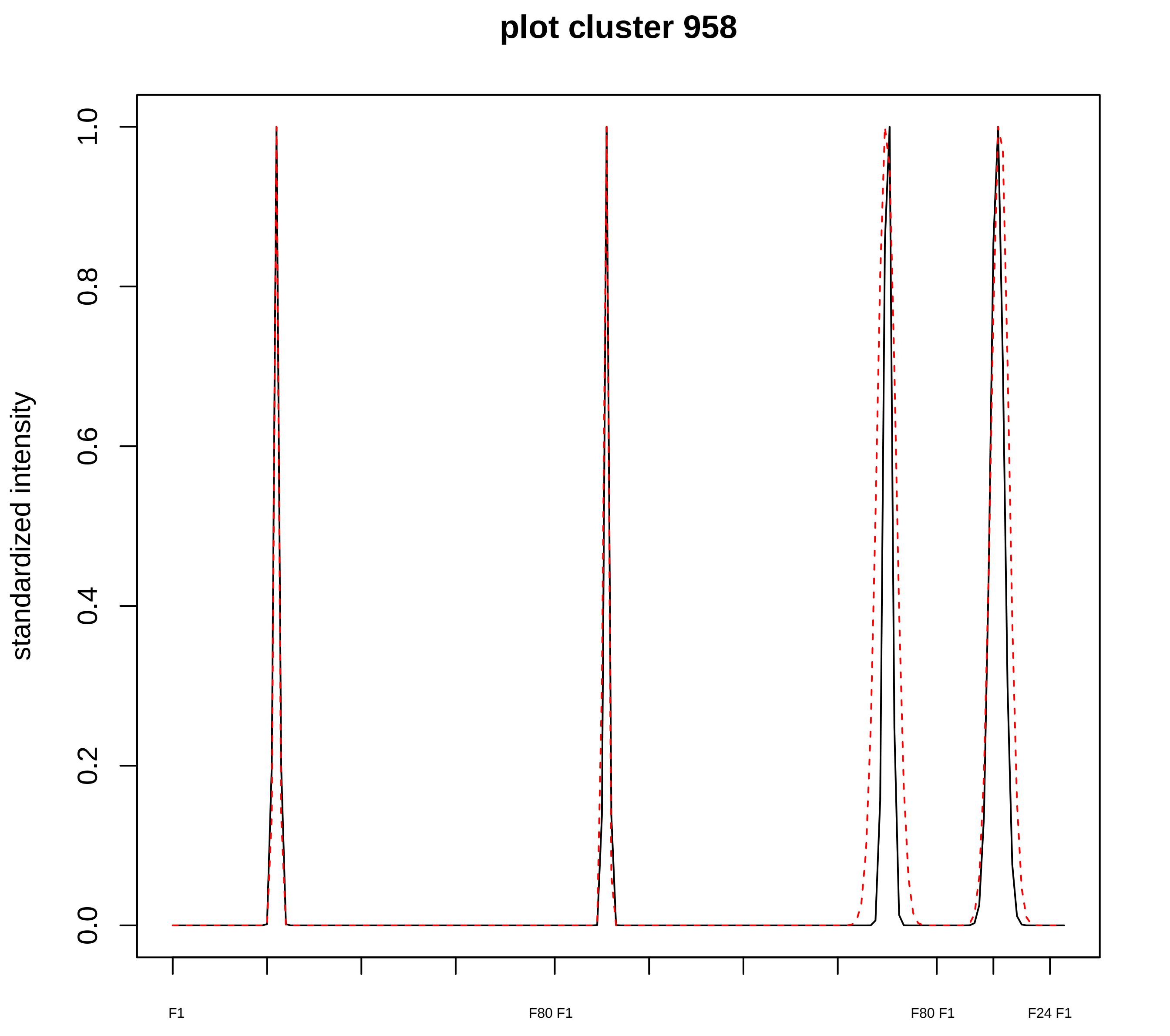
plot cluster 958
standardized intensity
0.0
0.2
0.4
0.6
0.8
1.0
F1
F80 F1
F80 F1
F24 F1

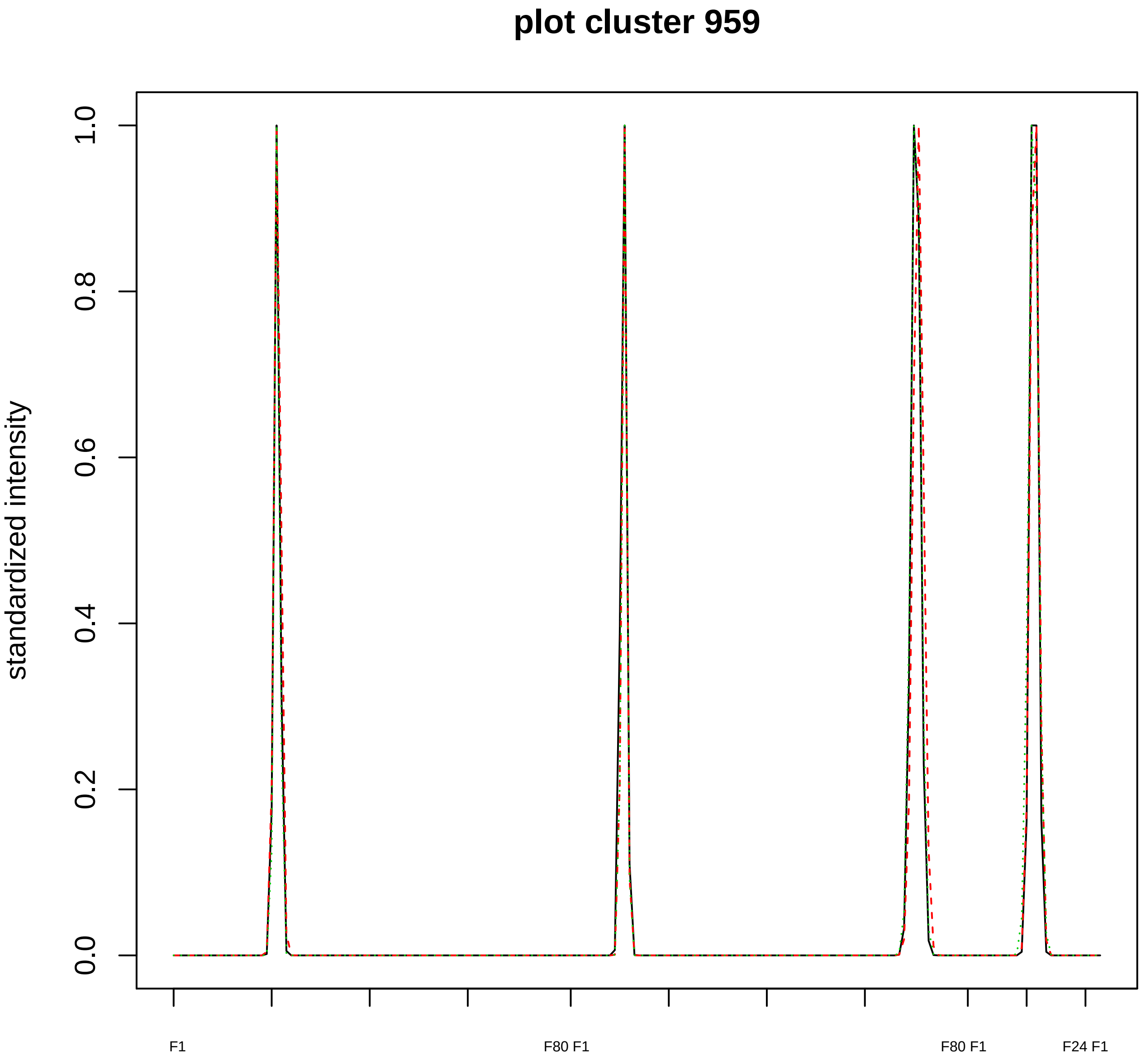

plot cluster 959
standardized intensity
0.0
0.2
0.4
0.6
0.8
1.0
F1
F80 F1
F80 F1
F24 F1

### Dendrogram of cluster 959

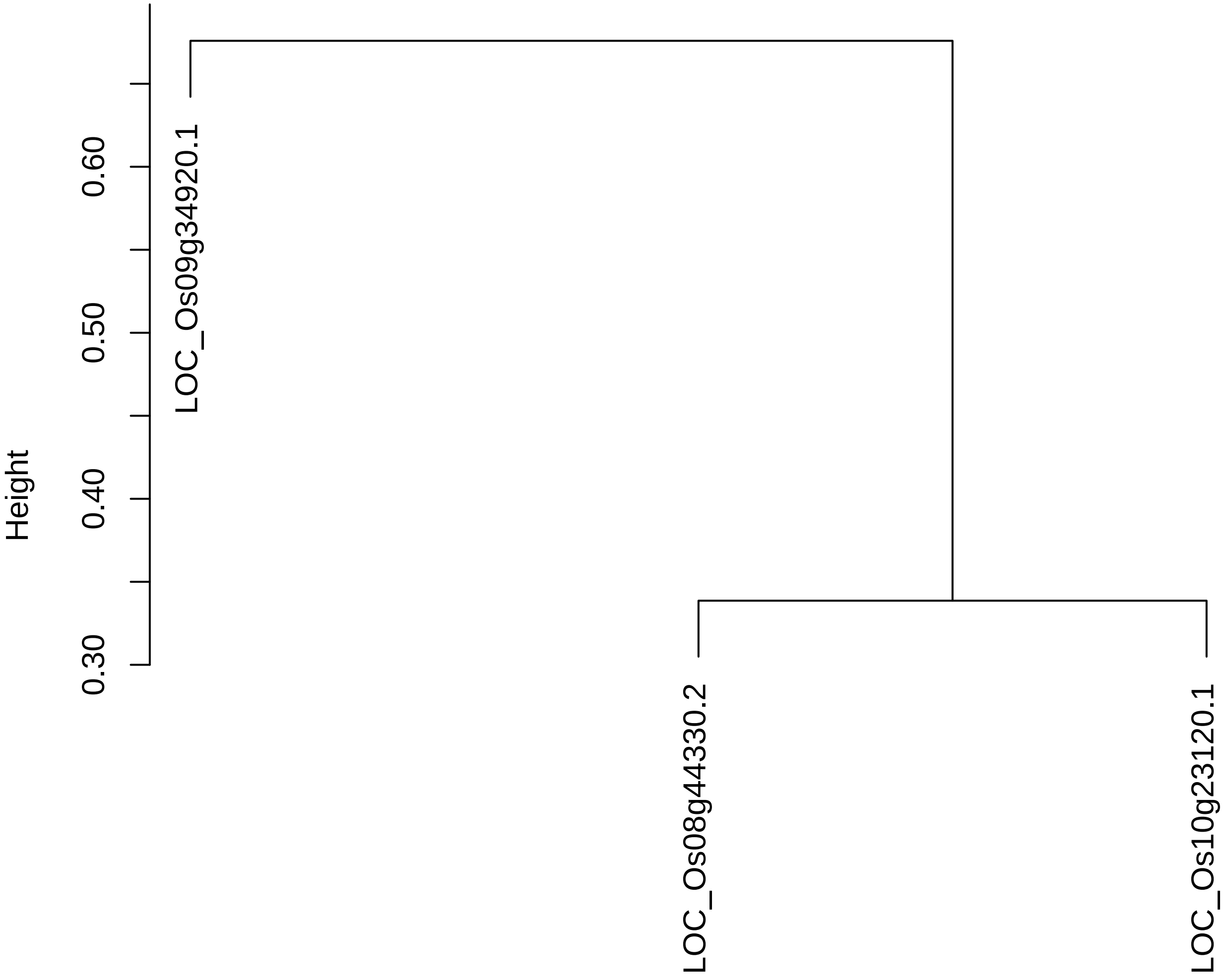


dist(spfitiex12sec12[lab[, 1] == i, ])
hclust (*, "ward.D2")

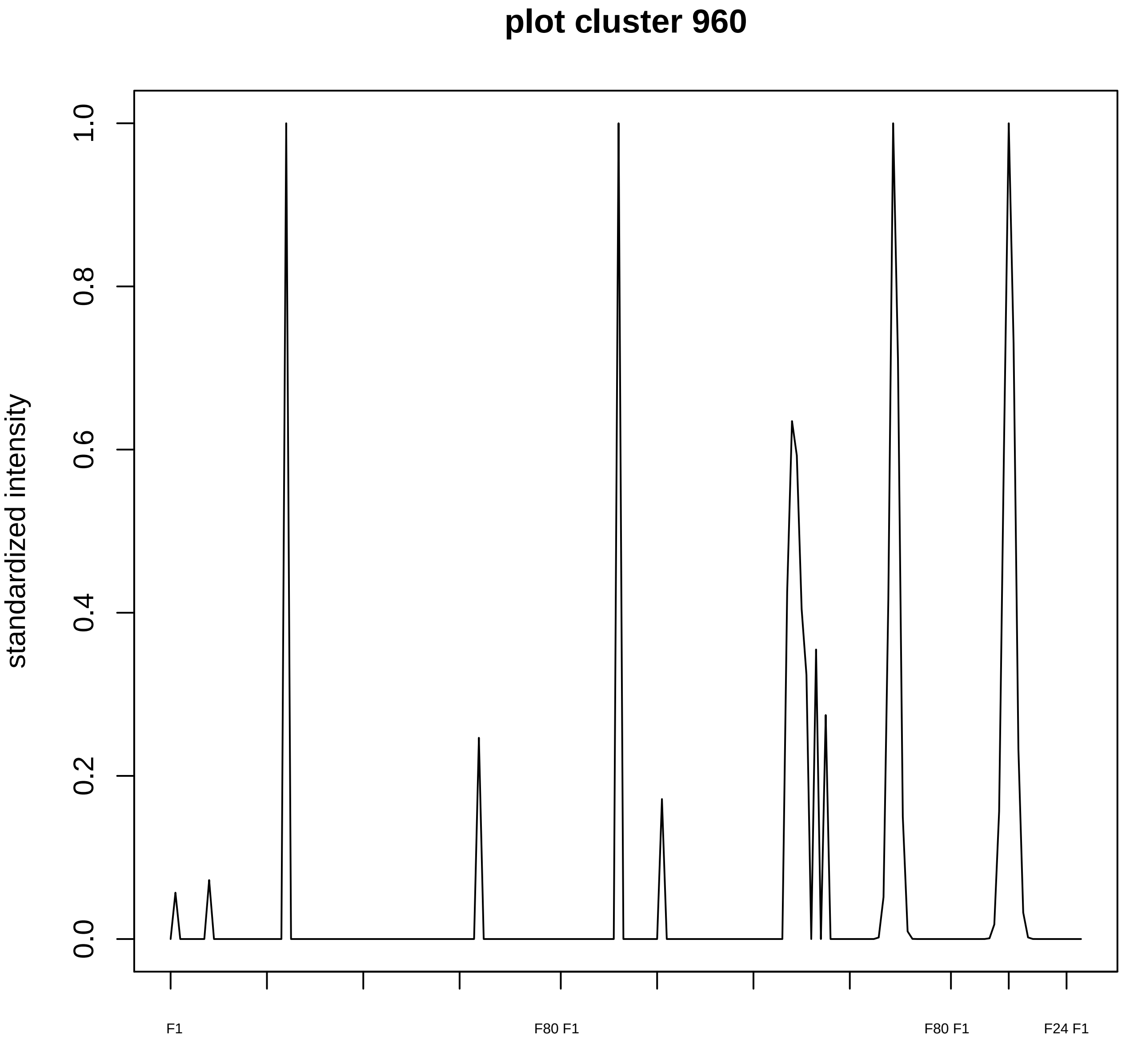

plot cluster 960
standardized intensity
0.0
0.2
0.4
0.6
0.8
1.0
F1
F80 F1
F80 F1
F24 F1

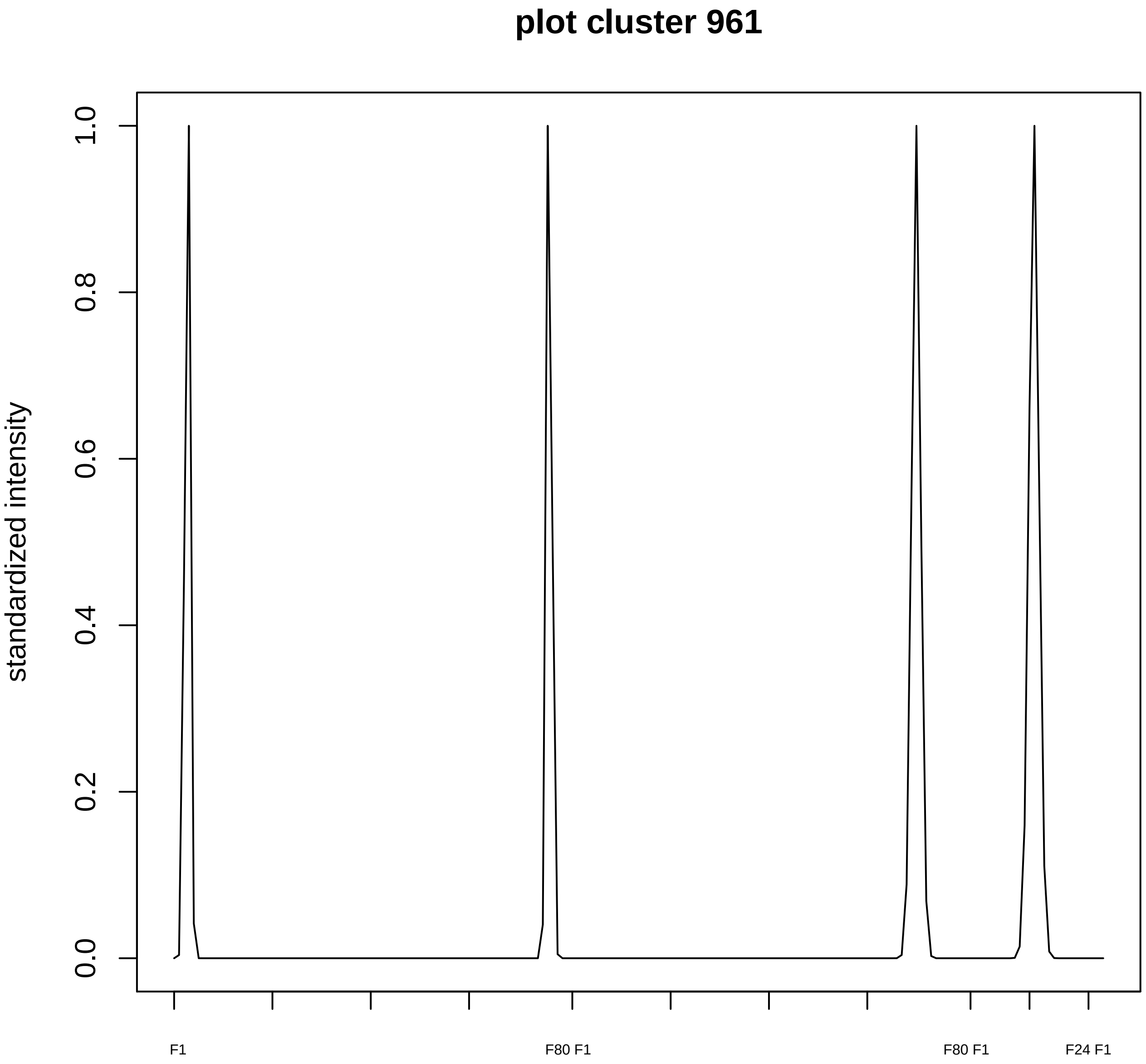
plot cluster 961
standardized intensity
0.0
0.2
0.4
0.6
0.8
1.0
F1
F80 F1
F80 F1
F24 F1

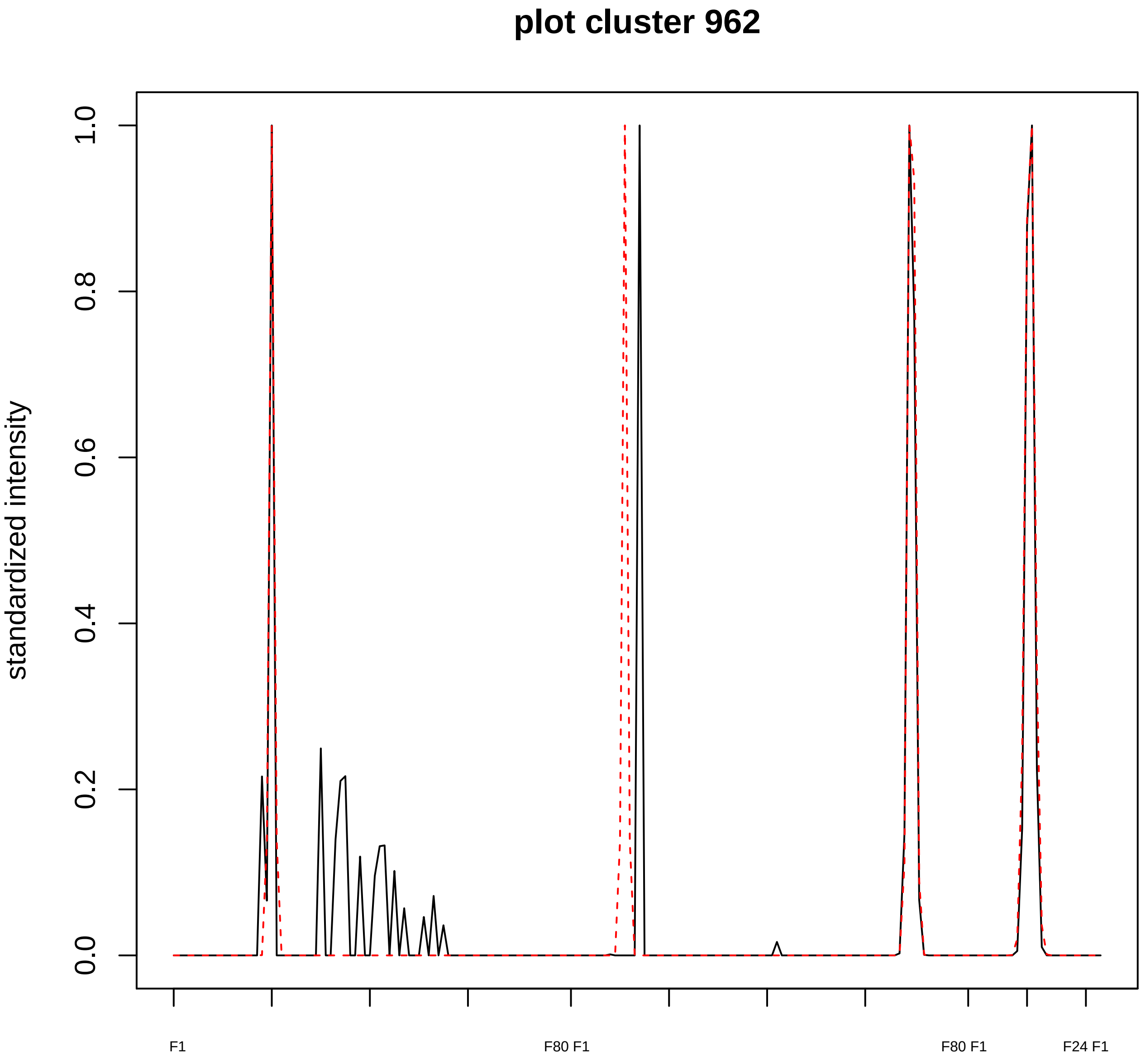

plot cluster 962
standardized intensity
0.0
0.2
0.4
0.6
0.8
1.0
F1
F80 F1
F80 F1
F24 F1

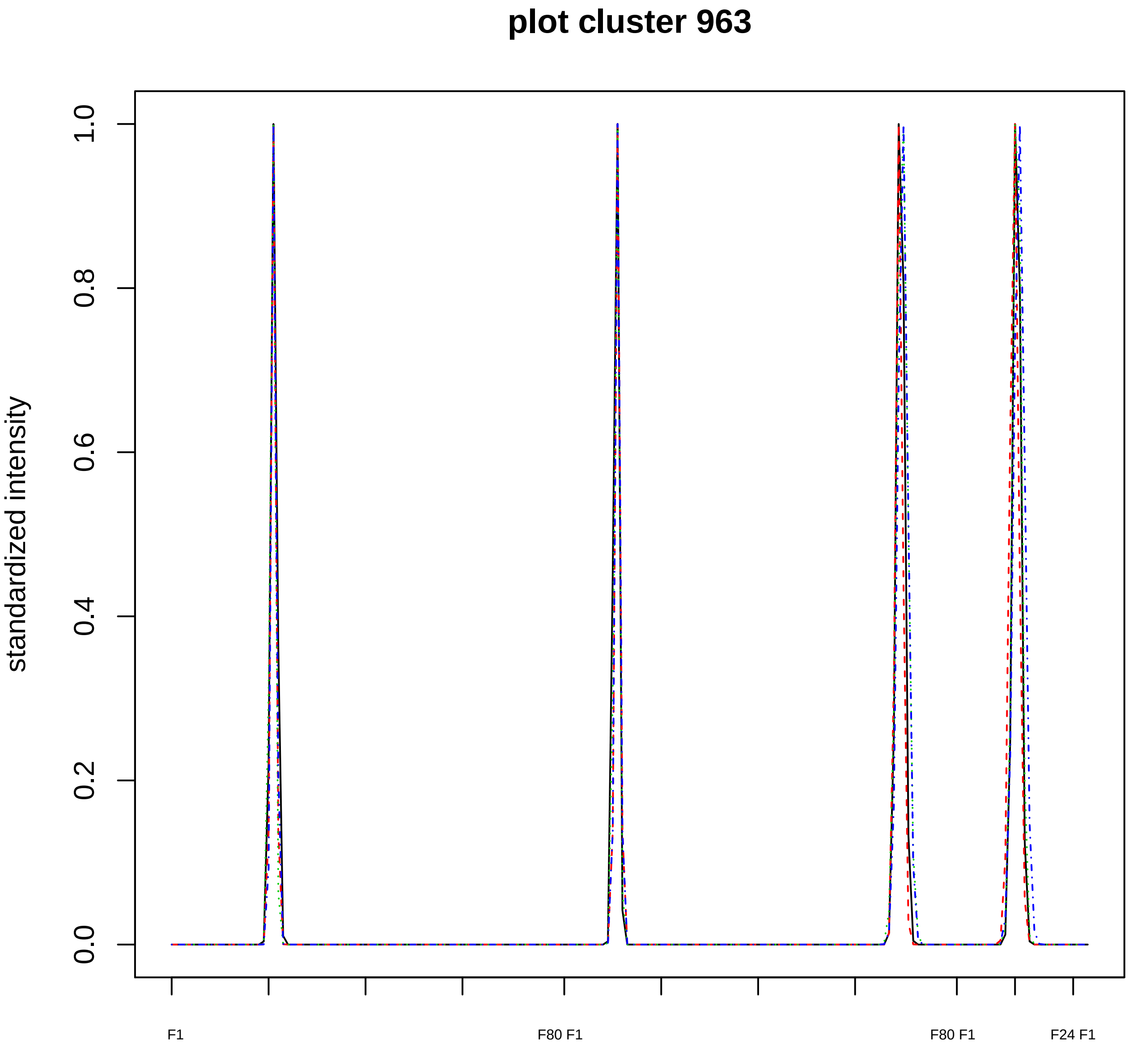

plot cluster 963
standardized intensity
0.0
0.2
0.4
0.6
0.8
1.0
F1
F80 F1
F80 F1
F24 F1

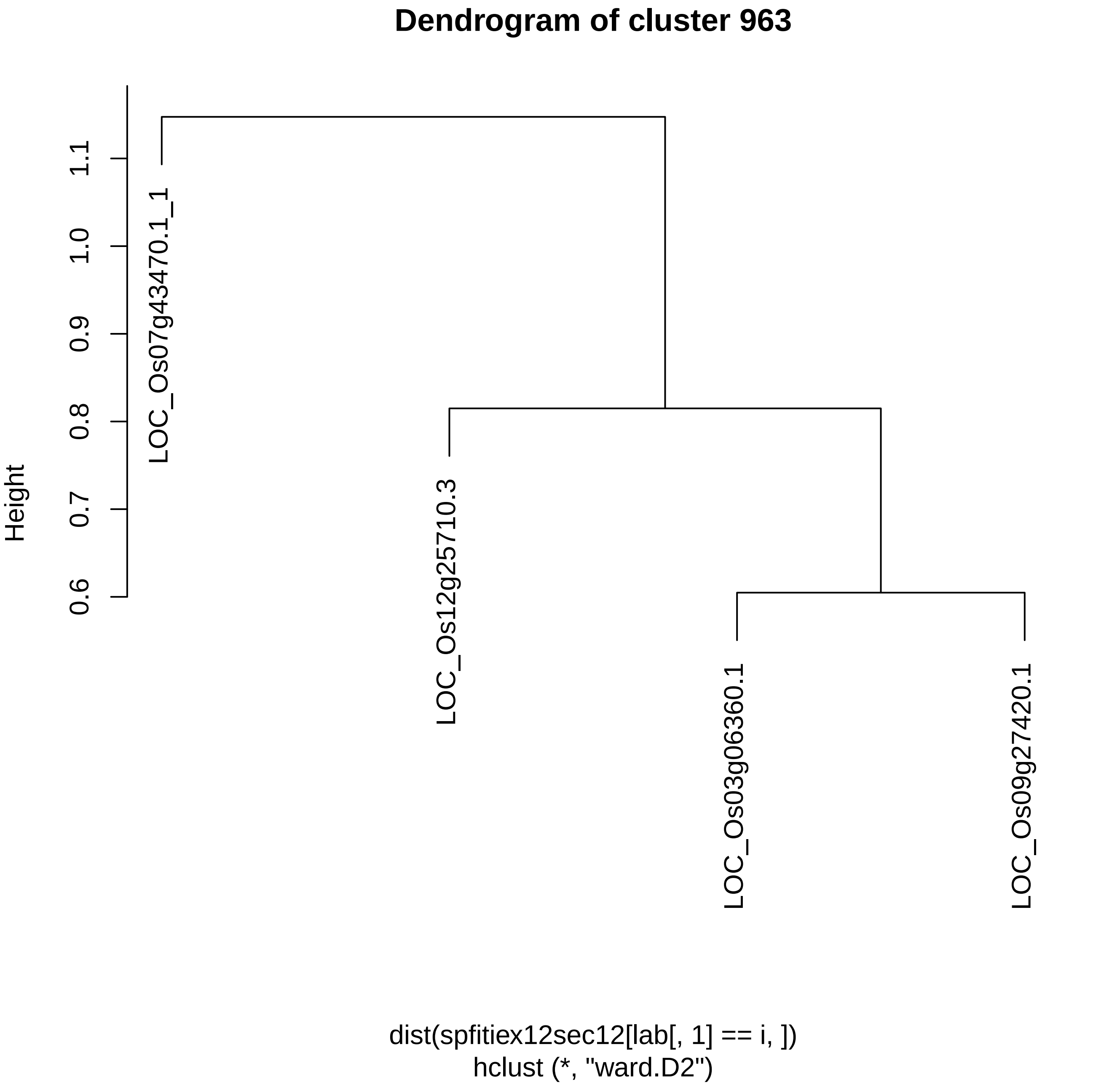

Dendrogram of cluster 963
Height
1.1
1.0
0.9
0.8
0.7
0.6
LOC_Os07g43470.1_1
LOC_Os12g25710.3
LOC_Os03g06360.1
LOC_Os09g27420.1
dist(spfitiex12sec12[lab[, 1] == i, ])
hclust (*, "ward.D2")

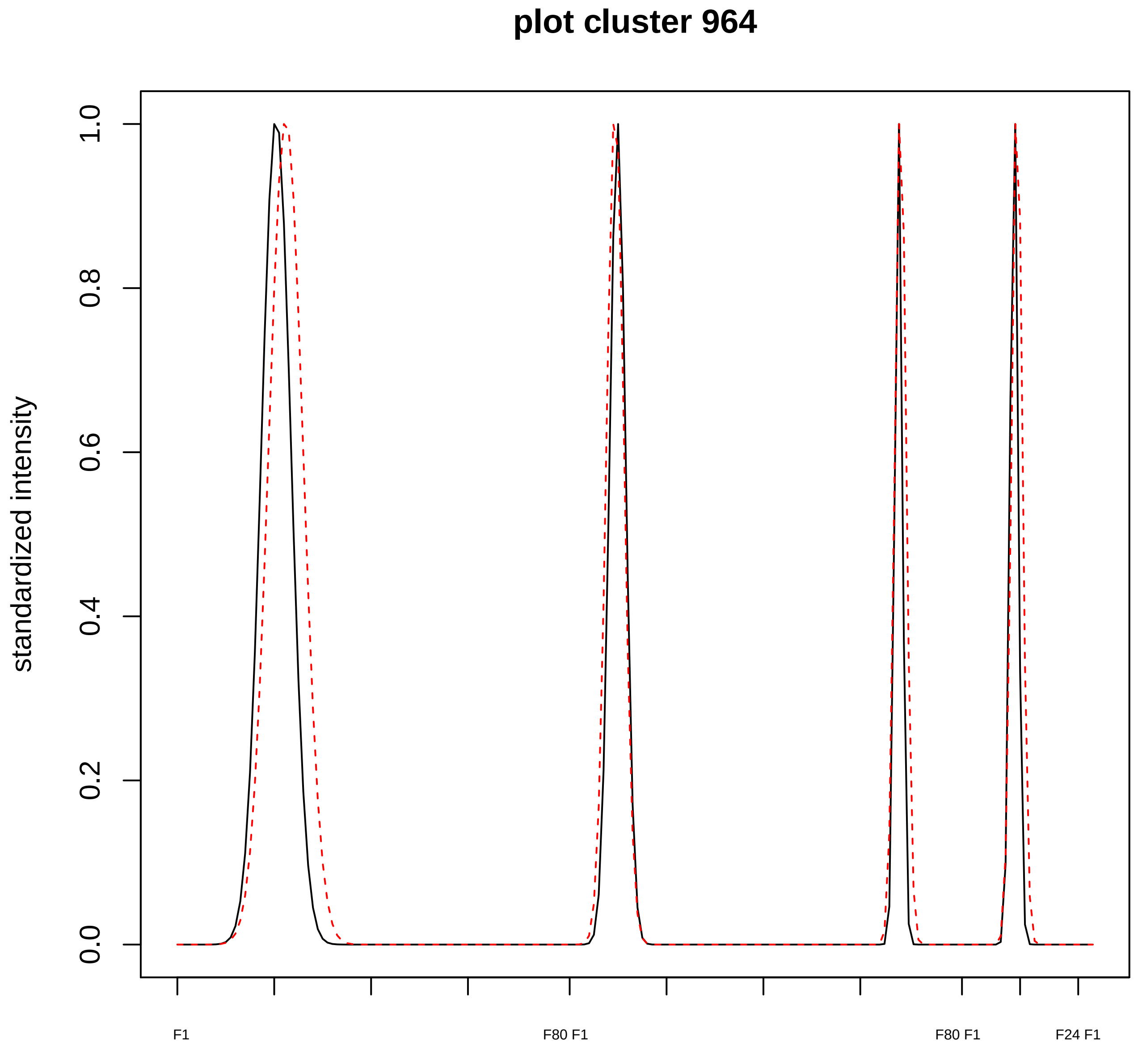

plot cluster 964
standardized intensity
0.0
0.2
0.4
0.6
0.8
1.0
F1
F80 F1
F80 F1
F24 F1

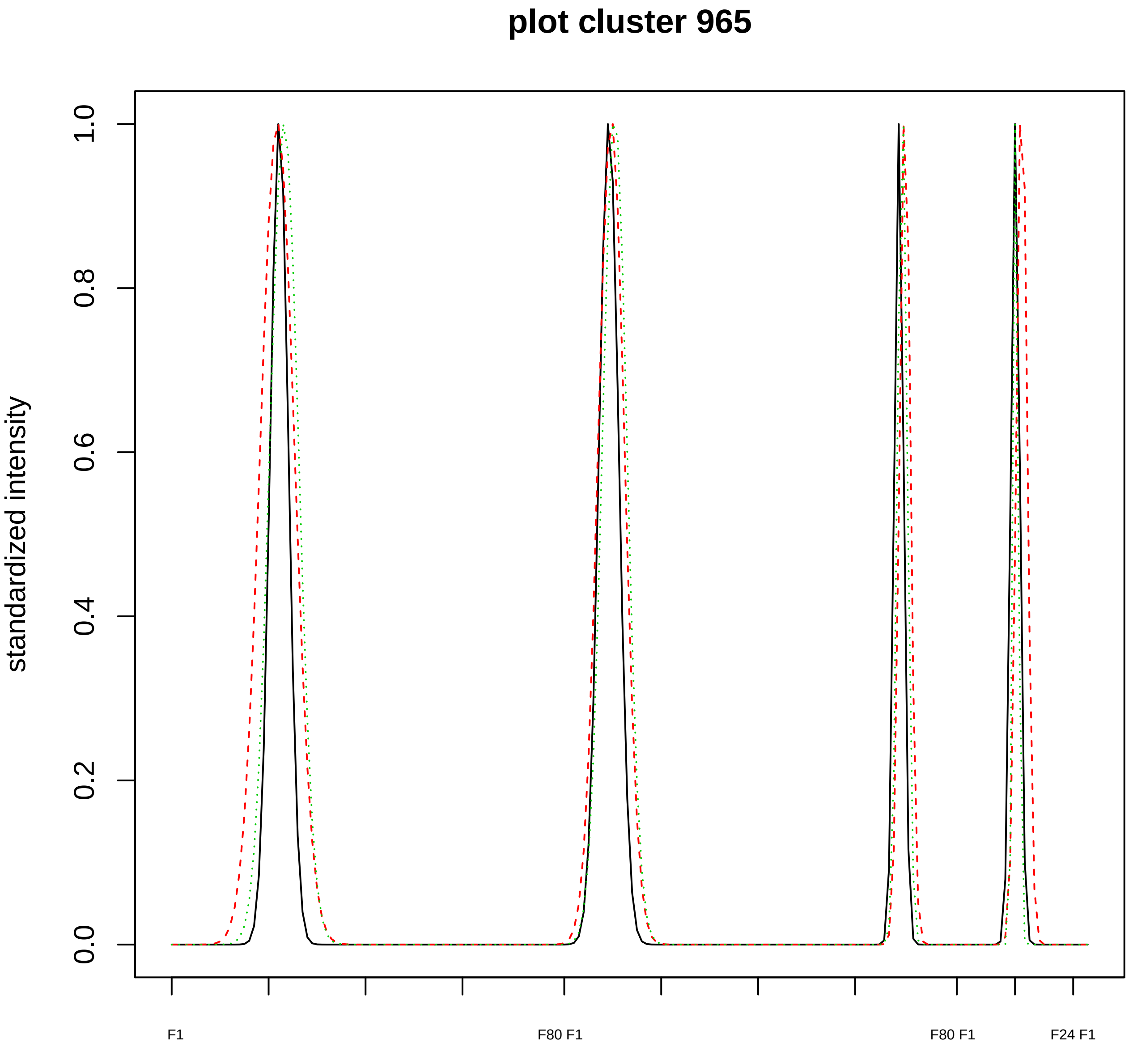

plot cluster 965
standardized intensity
0.0
0.2
0.4
0.6
0.8
1.0
F1
F80 F1
F80 F1
F24 F1

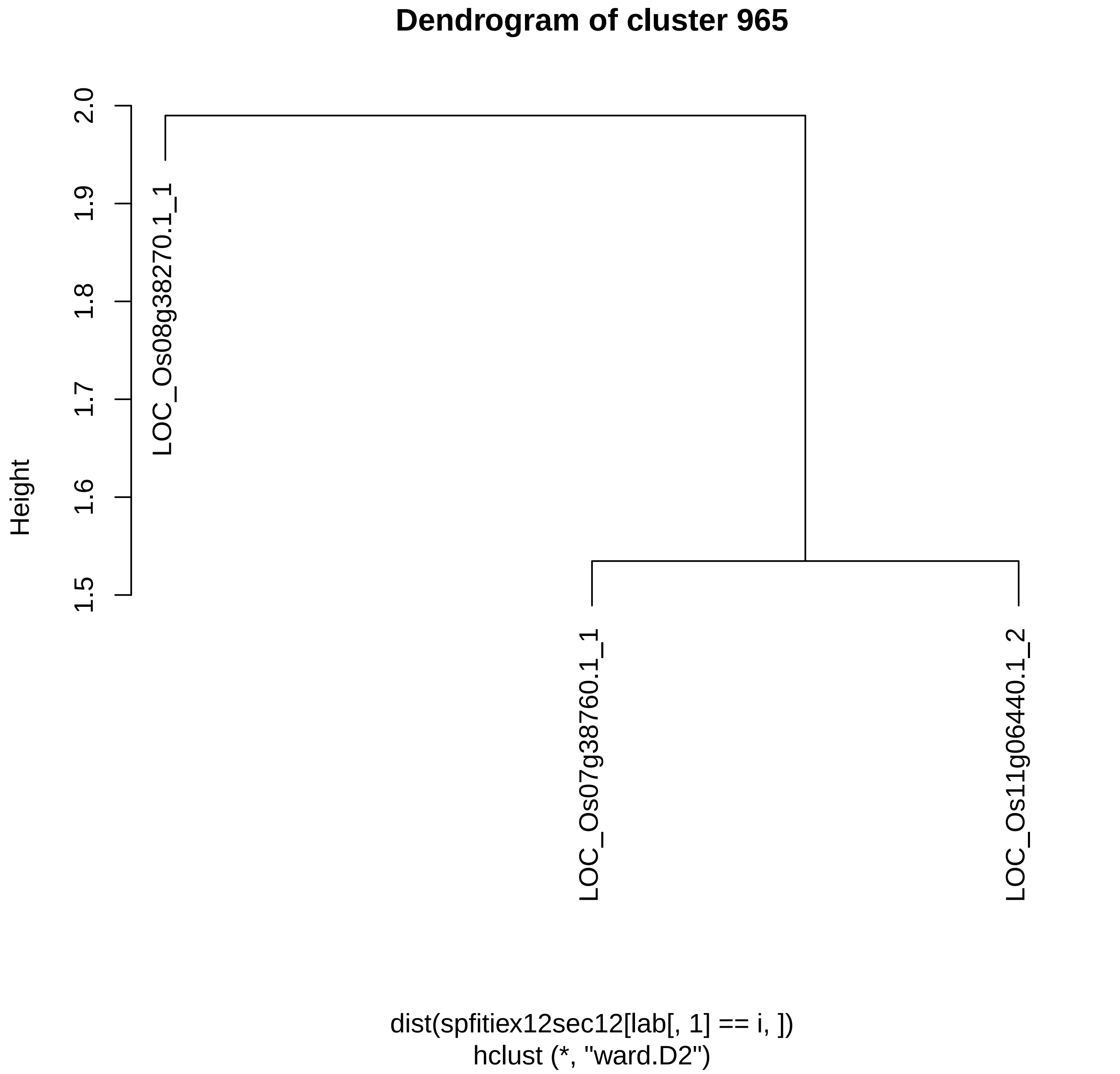
Dendrogram of cluster 965
Height
2.0
1.9
1.8
1.7
1.6
1.5
LOC_Os08g38270.1_1
LOC_Os07g38760.1_1
LOC_Os11g06440.1_2
dist(spfitiex12sec12[lab[, 1] == i, ])
hclust (*, "ward.D2")

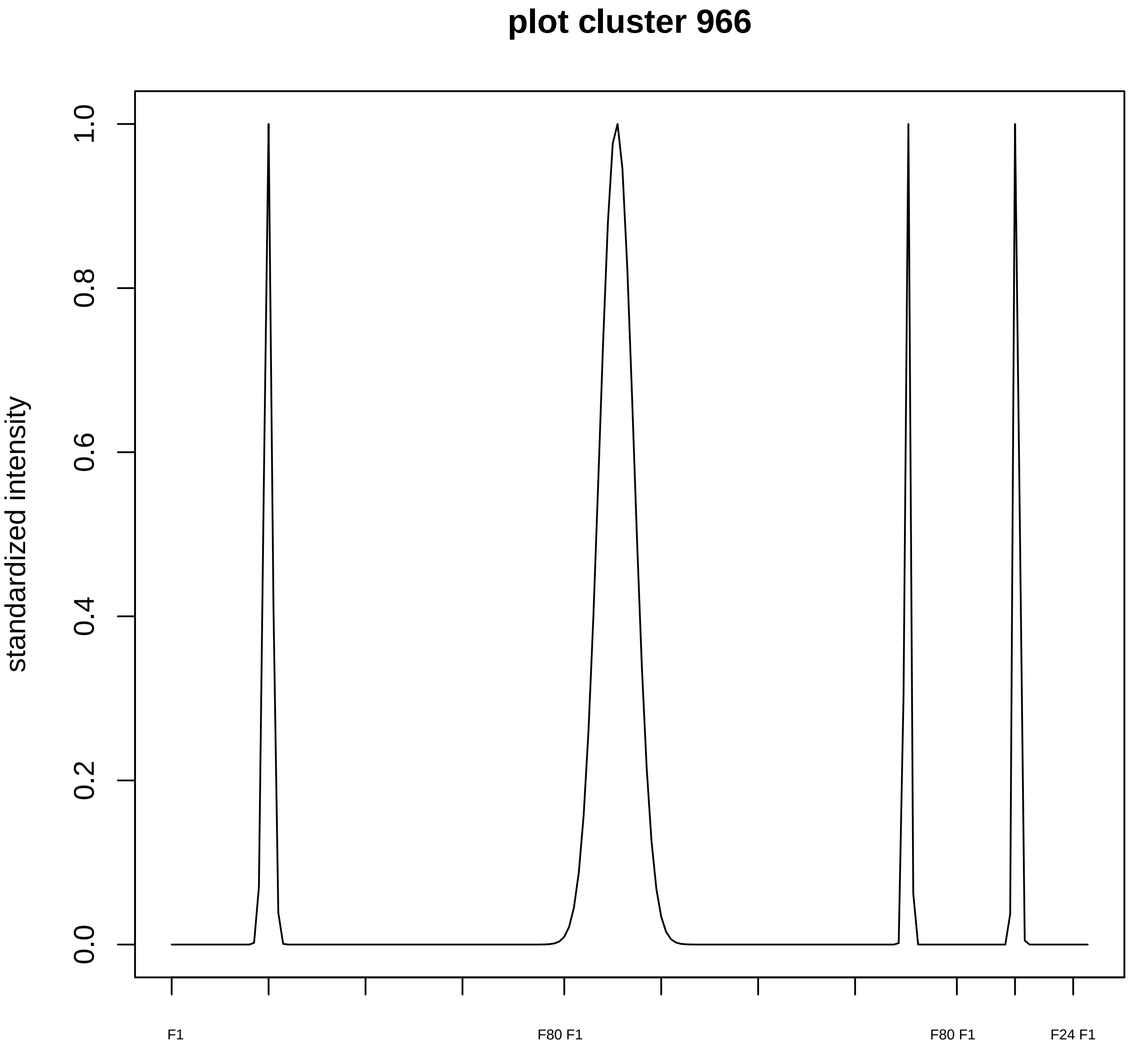
plot cluster 966
standardized intensity
0.0
0.2
0.4
0.6
0.8
1.0
F1
F80 F1
F80 F1
F24 F1

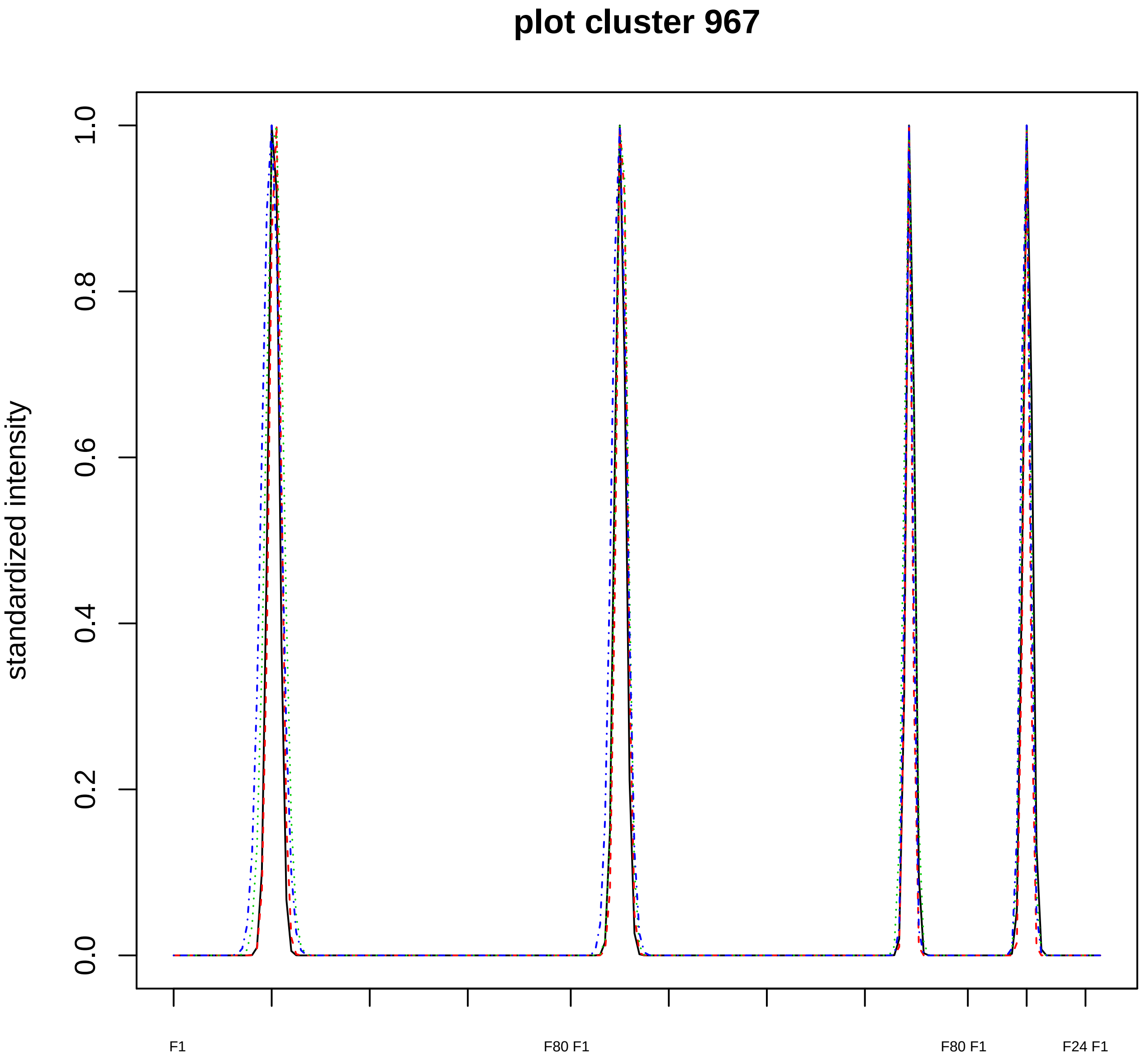
plot cluster 967
standardized intensity
0.0
0.2
0.4
0.6
0.8
1.0
F1
F80 F1
F80 F1
F24 F1

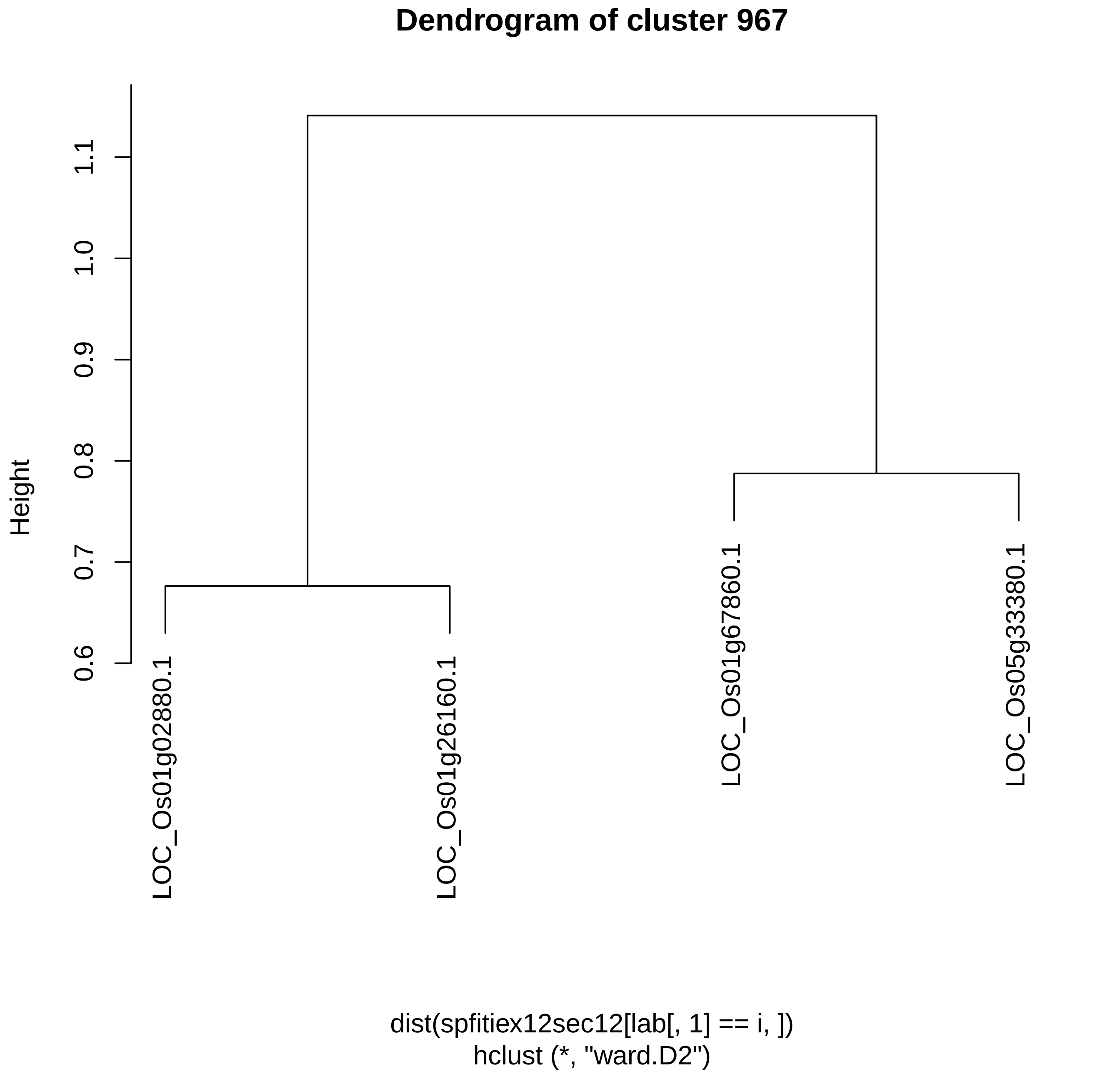

Dendrogram of cluster 967
Height
0.6
0.7
0.8
0.9
1.0
1.1
LOC_Os01g02880.1
LOC_Os01g26160.1
LOC_Os01g67860.1
LOC_Os05g33380.1
dist(spfitiex12sec12[lab[, 1] == i, ])
hclust (*, "ward.D2")

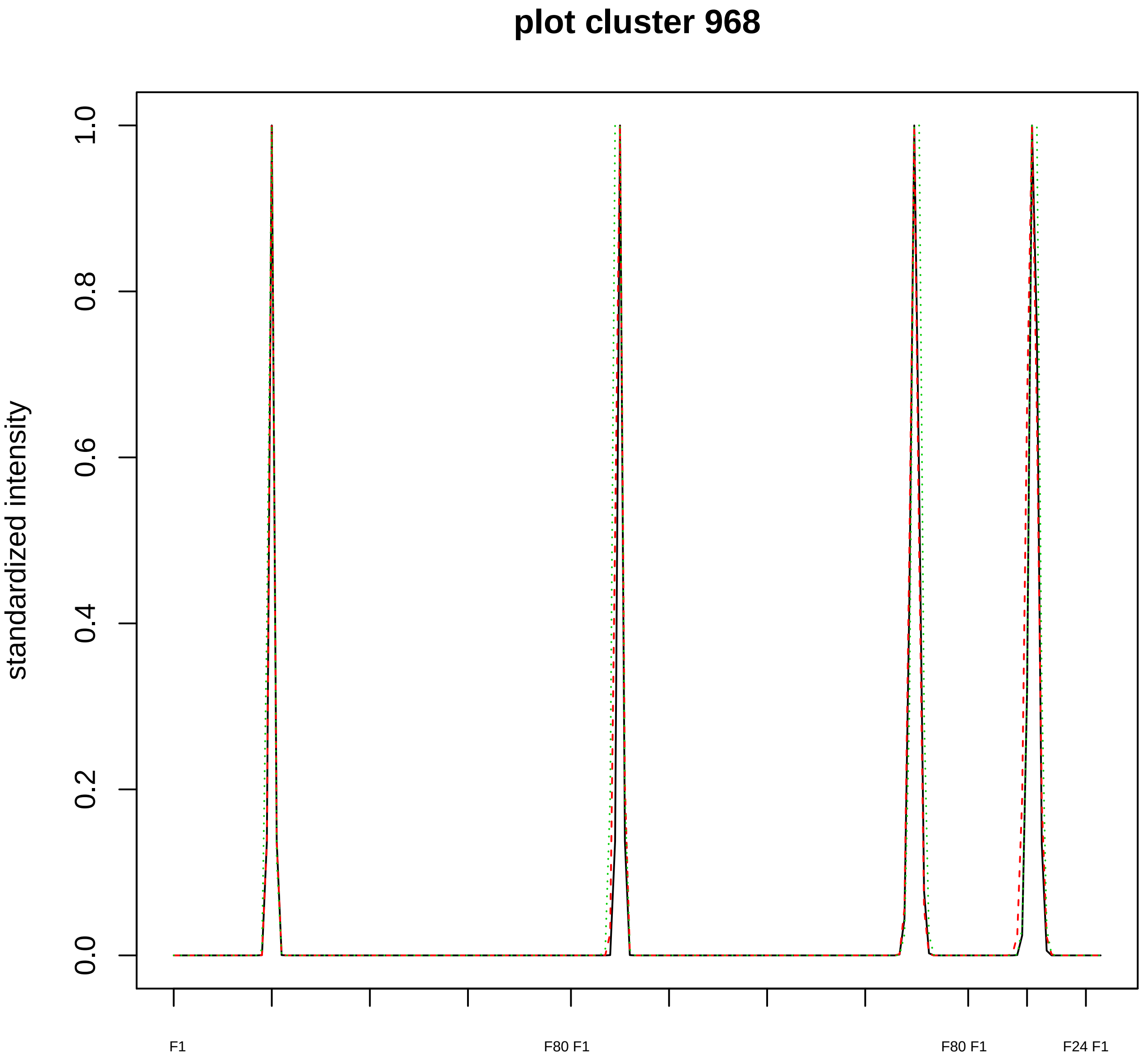

plot cluster 968
standardized intensity
0.0
0.2
0.4
0.6
0.8
1.0
F1
F80 F1
F80 F1
F24 F1

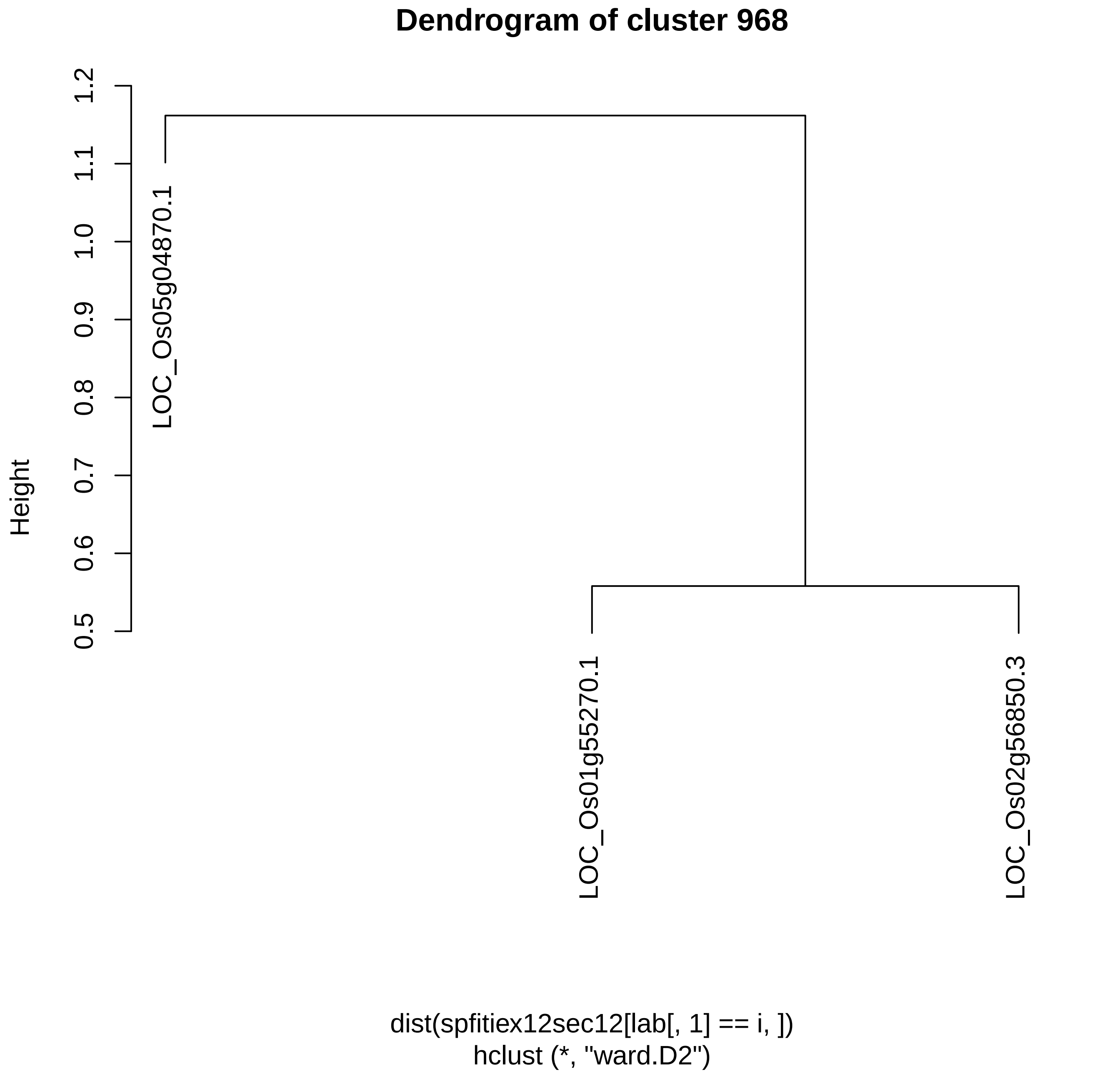

Dendrogram of cluster 968
Height
1.2
1.1
1.0
0.9
0.8
0.7
0.6
0.5
LOC_Os05g04870.1
LOC_Os01g55270.1
LOC_Os02g56850.3
dist(spfitiex12sec12[lab[, 1] == i, ])
hclust (*, "ward.D2")

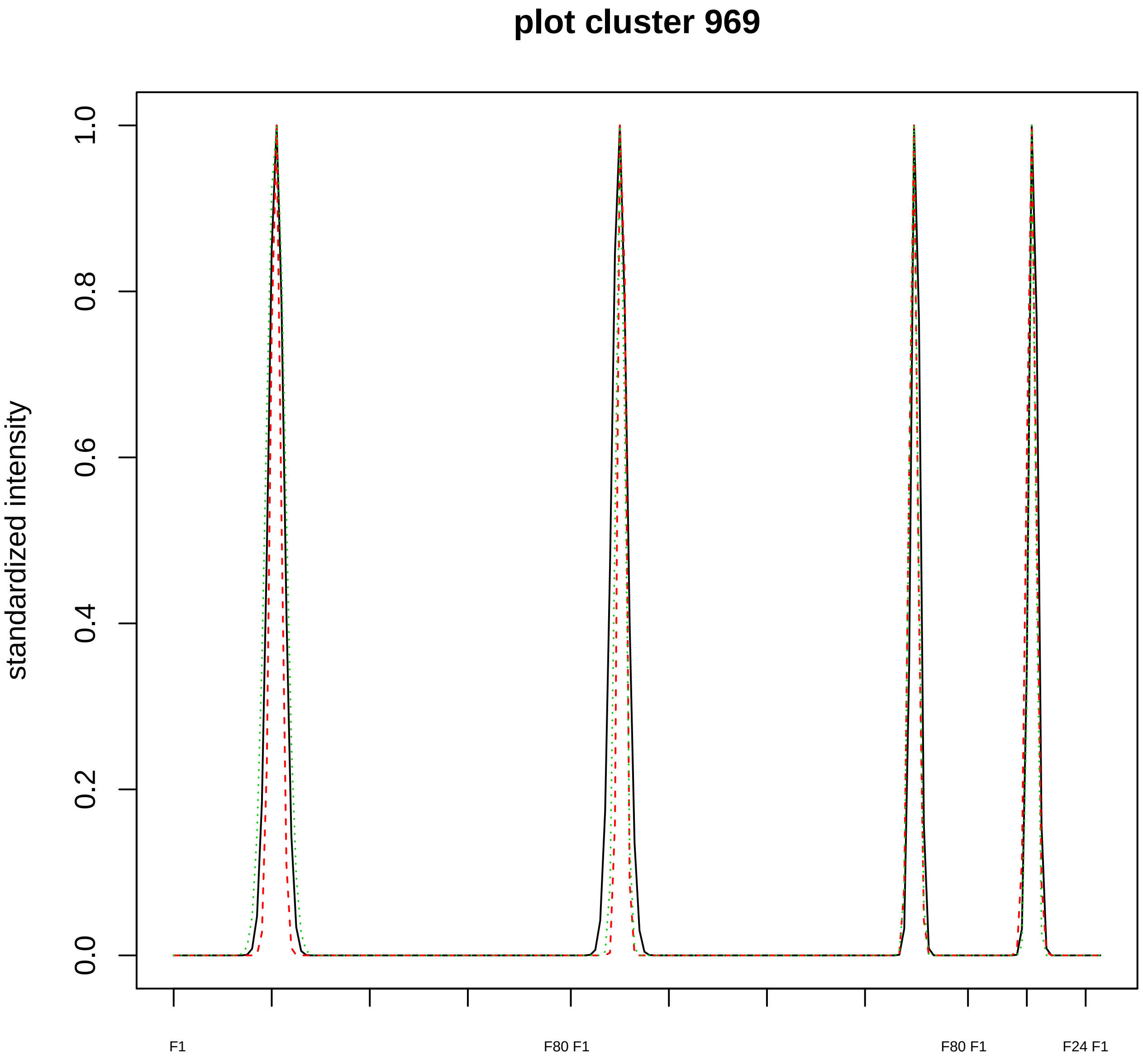

plot cluster 969
standardized intensity
0.0
0.2
0.4
0.6
0.8
1.0
F1
F80 F1
F80 F1
F24 F1

#### Dendrogram of cluster 969

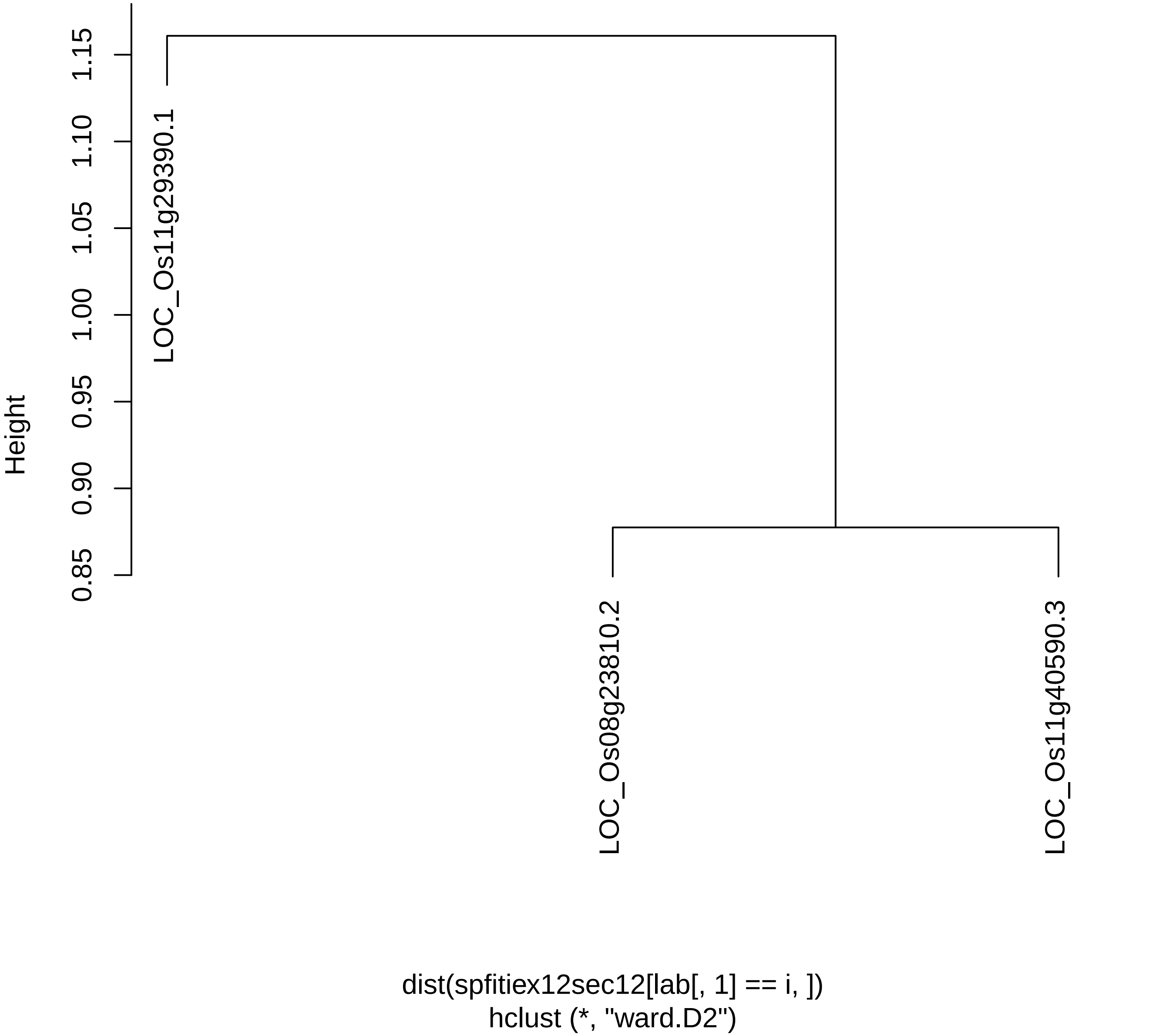

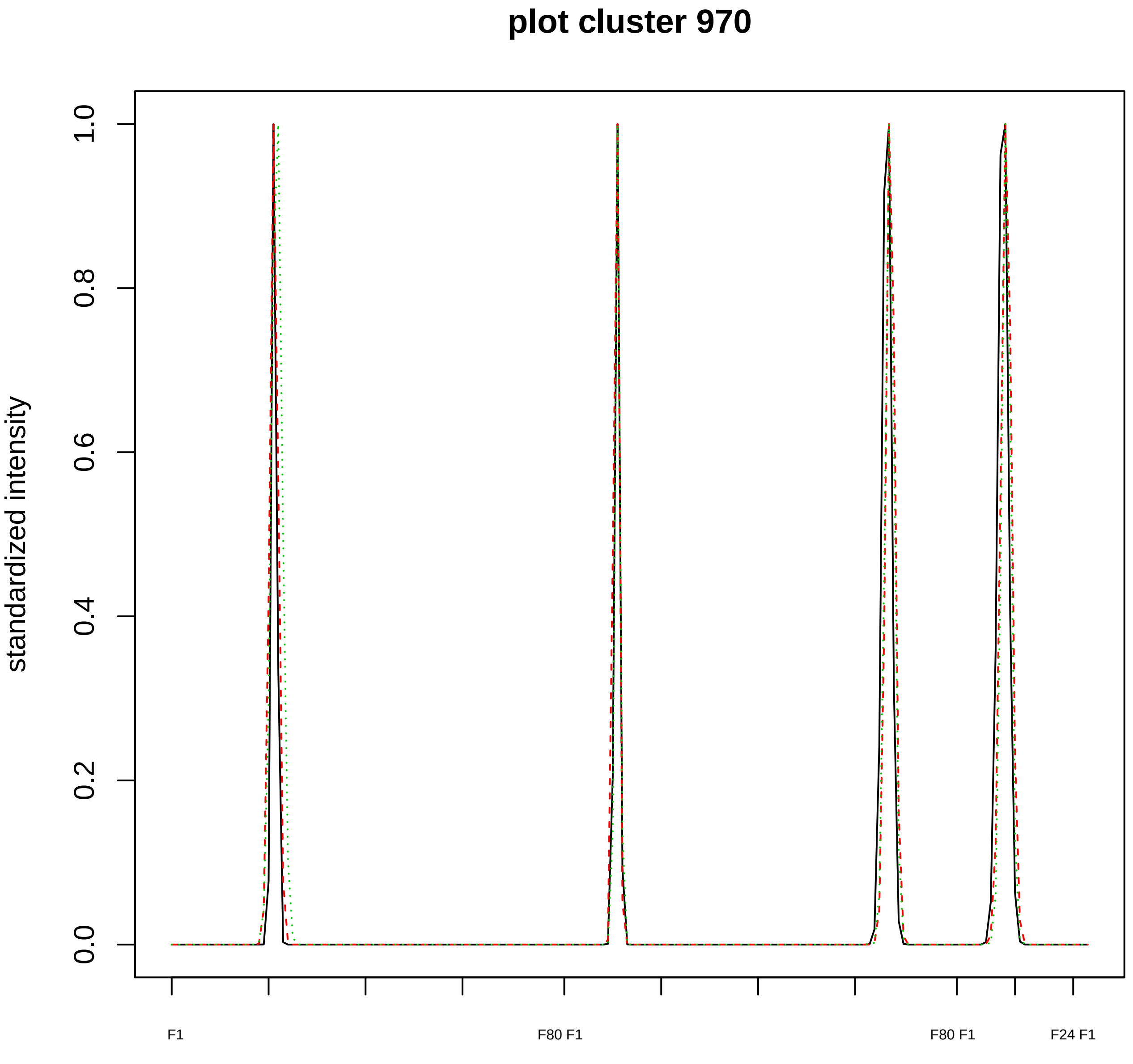
plot cluster 970
standardized intensity
0.0
0.2
0.4
0.6
0.8
1.0
F1
F80 F1
F80 F1
F24 F1

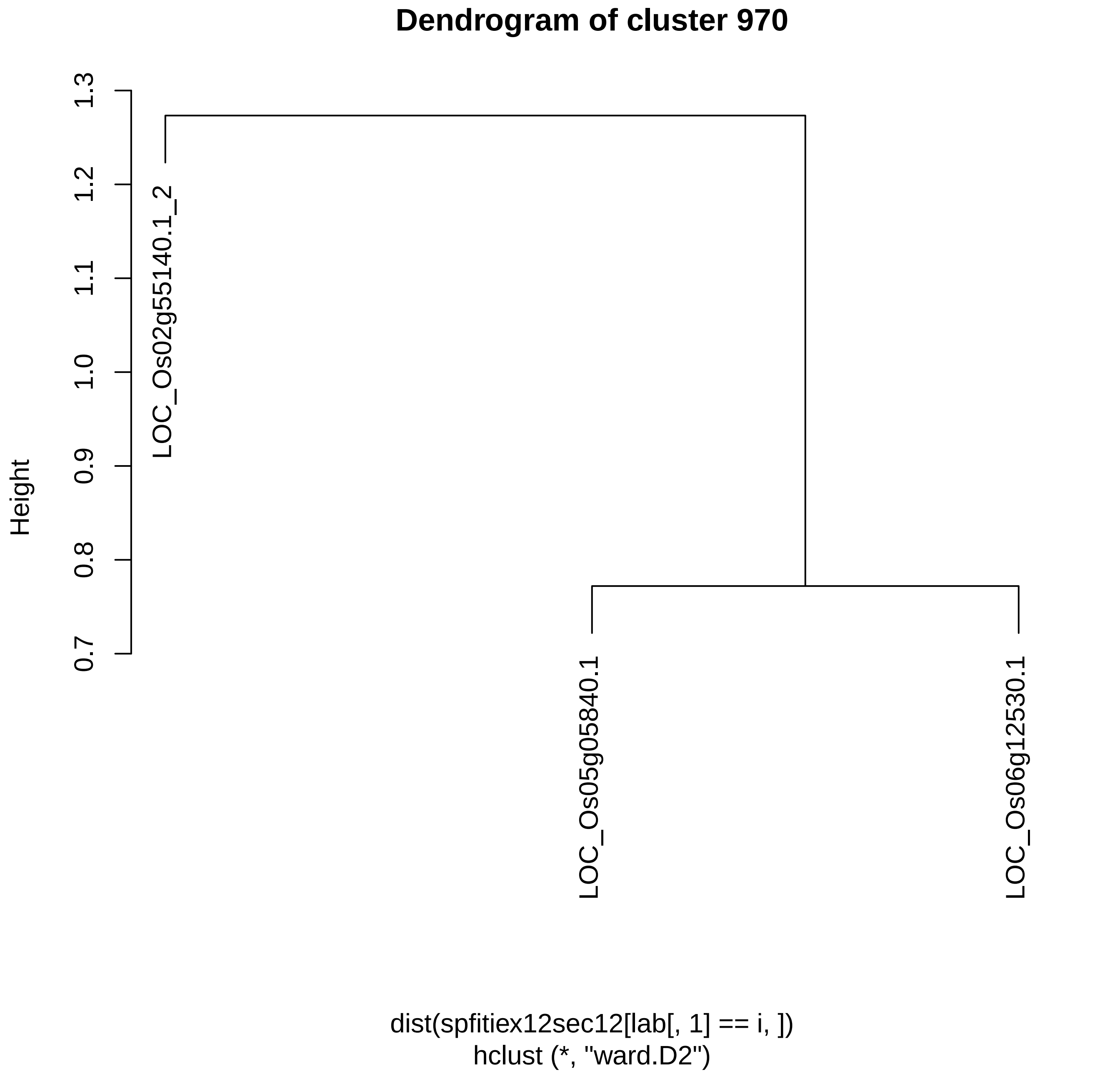

Dendrogram of cluster 970
Height
1.3
1.2
1.1
1.0
0.9
0.8
0.7
LOC_Os02g55140.1_2
LOC_Os05g05840.1
LOC_Os06g12530.1
dist(spfitiex12sec12[lab[, 1] == i, ])
hclust (*, "ward.D2")

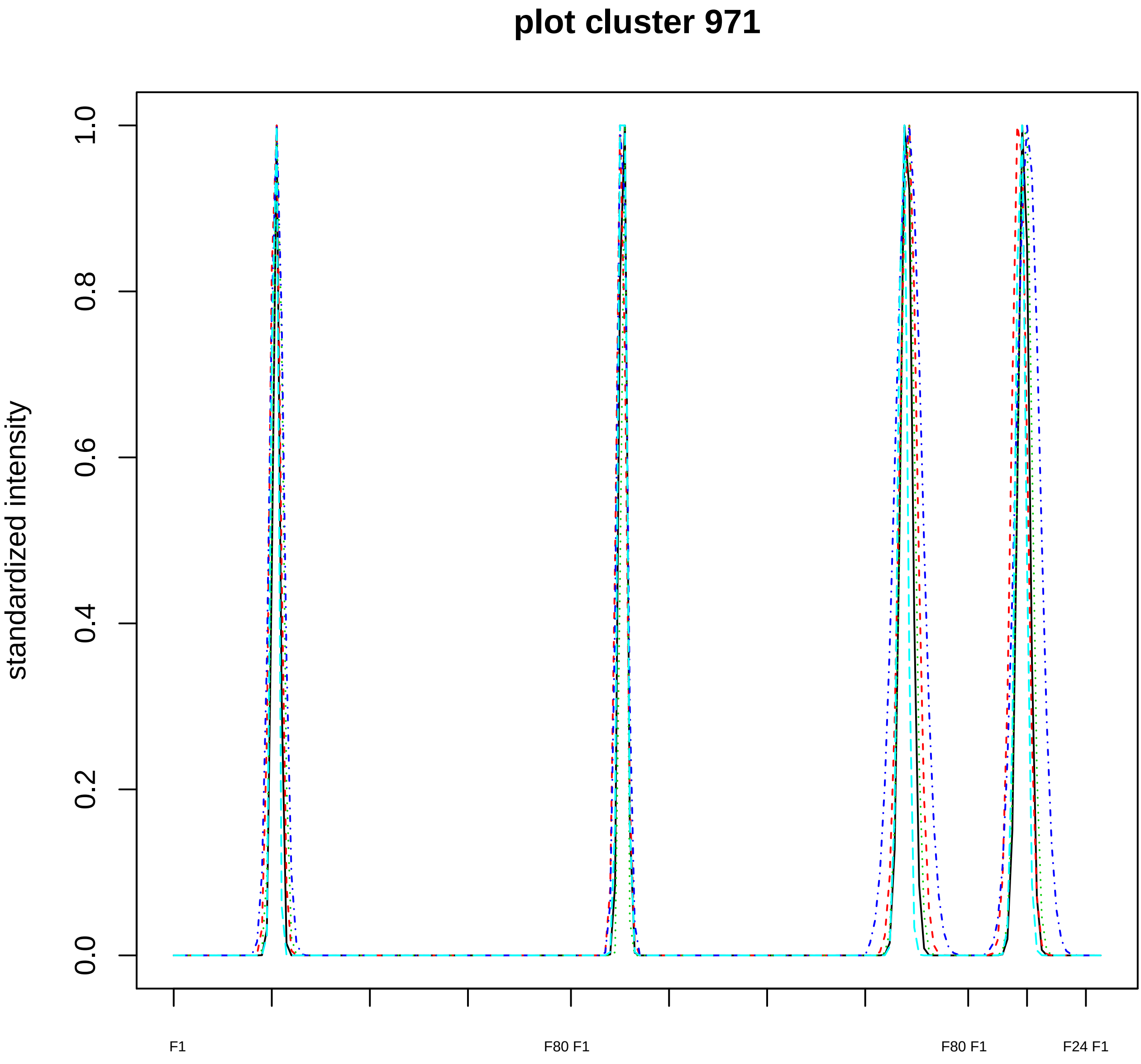

plot cluster 971
standardized intensity
0.0
0.2
0.4
0.6
0.8
1.0
F1
F80 F1
F80 F1
F24 F1

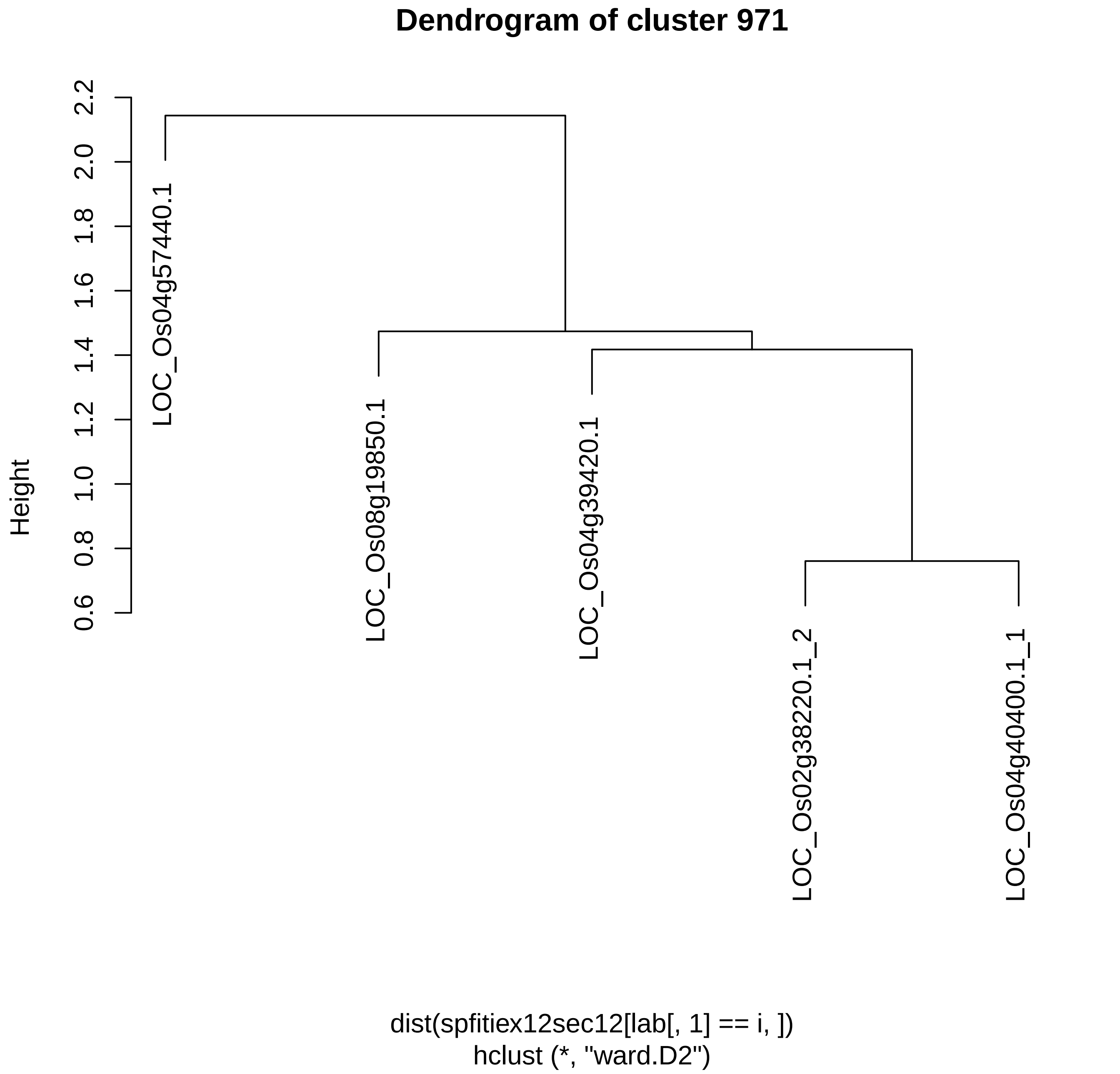
Dendrogram of cluster 971
Height
0.6
0.8
1.0
1.2
1.4
1.6
1.8
2.0
2.2
LOC_Os04g57440.1
LOC_Os08g19850.1
LOC_Os04g39420.1
LOC_Os02g38220.1_2
LOC_Os04g40400.1_1
dist(spfitiex12sec12[lab[, 1] == i, ])
hclust (*, "ward.D2")

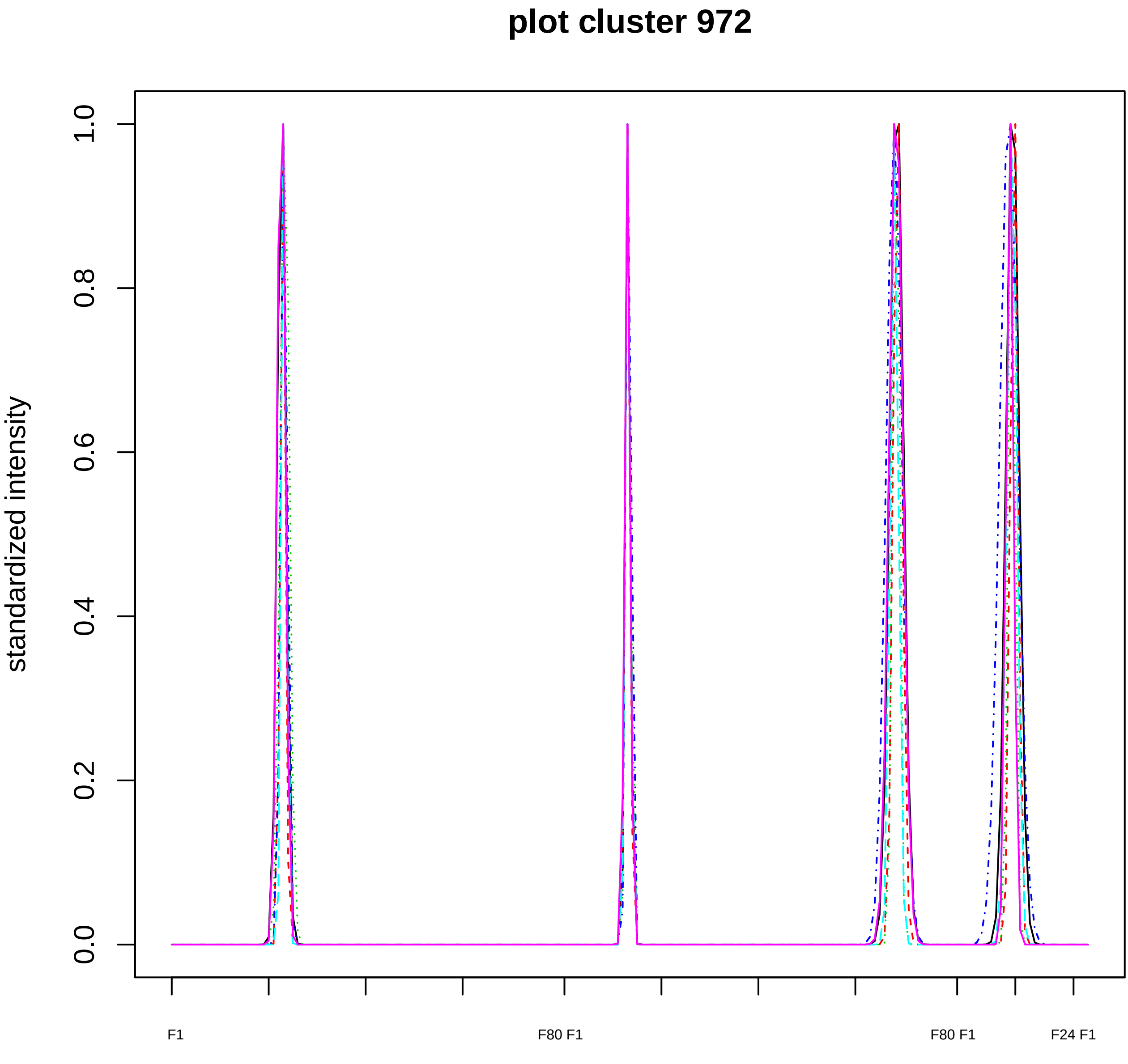
plot cluster 972
standardized intensity
0.0
0.2
0.4
0.6
0.8
1.0
F1
F80 F1
F80 F1
F24 F1

### Dendrogram of cluster 972

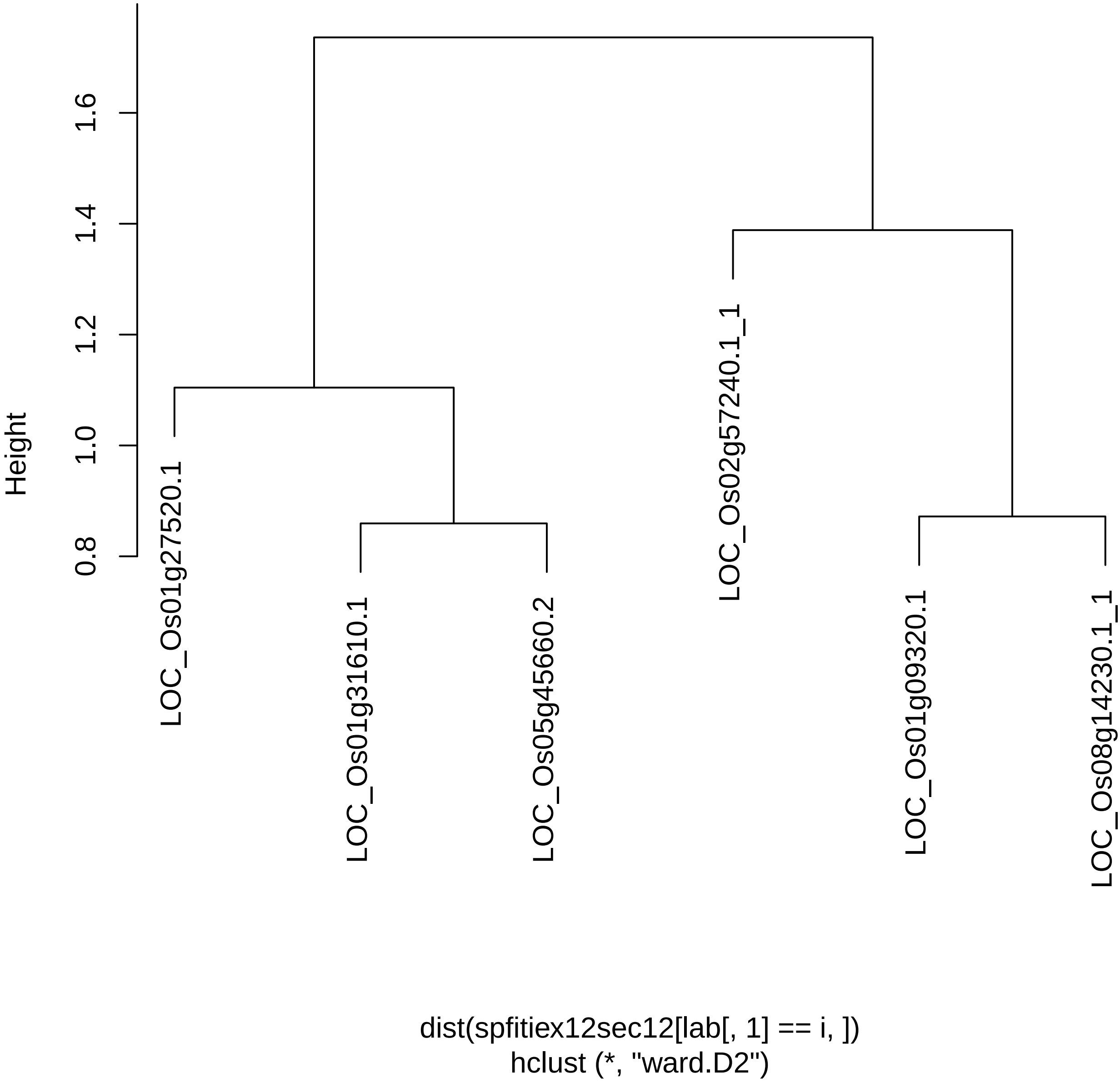

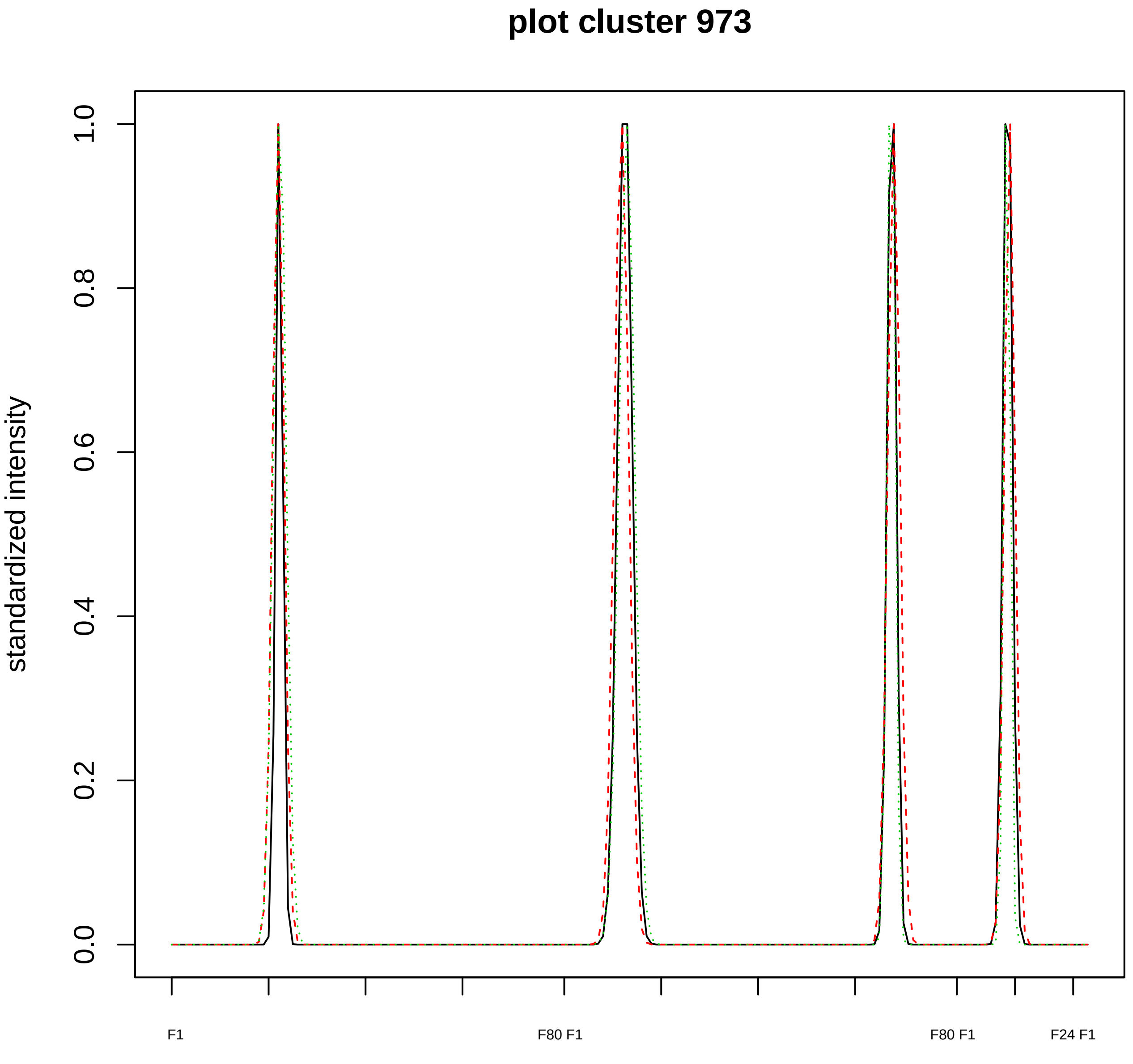
plot cluster 973
standardized intensity
0.0
0.2
0.4
0.6
0.8
1.0
F1
F80 F1
F80 F1
F24 F1

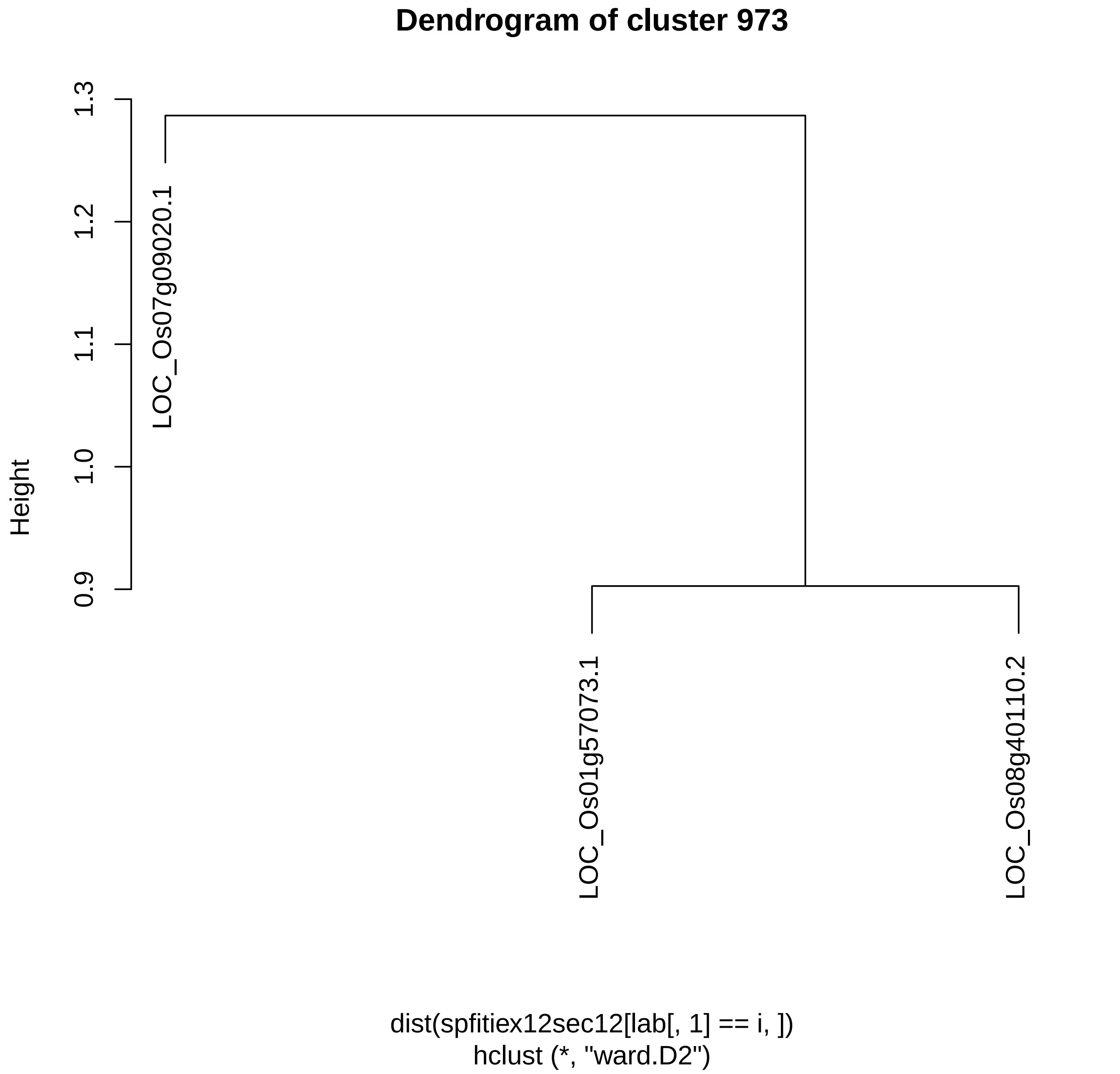

Dendrogram of cluster 973
Height
1.3
1.2
1.1
1.0
0.9
LOC_Os07g09020.1
LOC_Os01g57073.1
LOC_Os08g40110.2
dist(spfitiex12sec12[lab[, 1] == i, ])
hclust (*, "ward.D2")

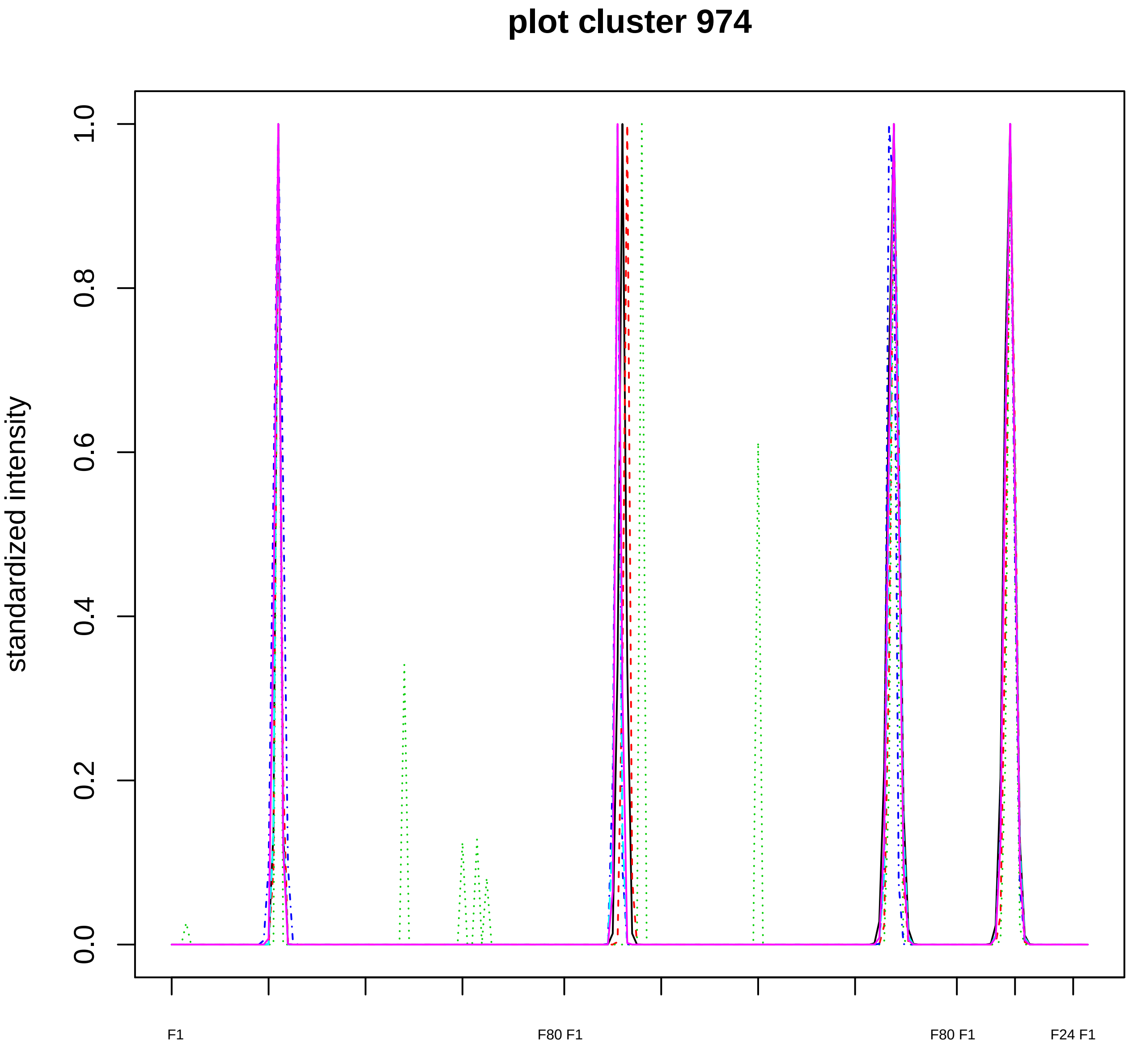

plot cluster 974
standardized intensity
0.0
0.2
0.4
0.6
0.8
1.0
F1
F80 F1
F80 F1
F24 F1

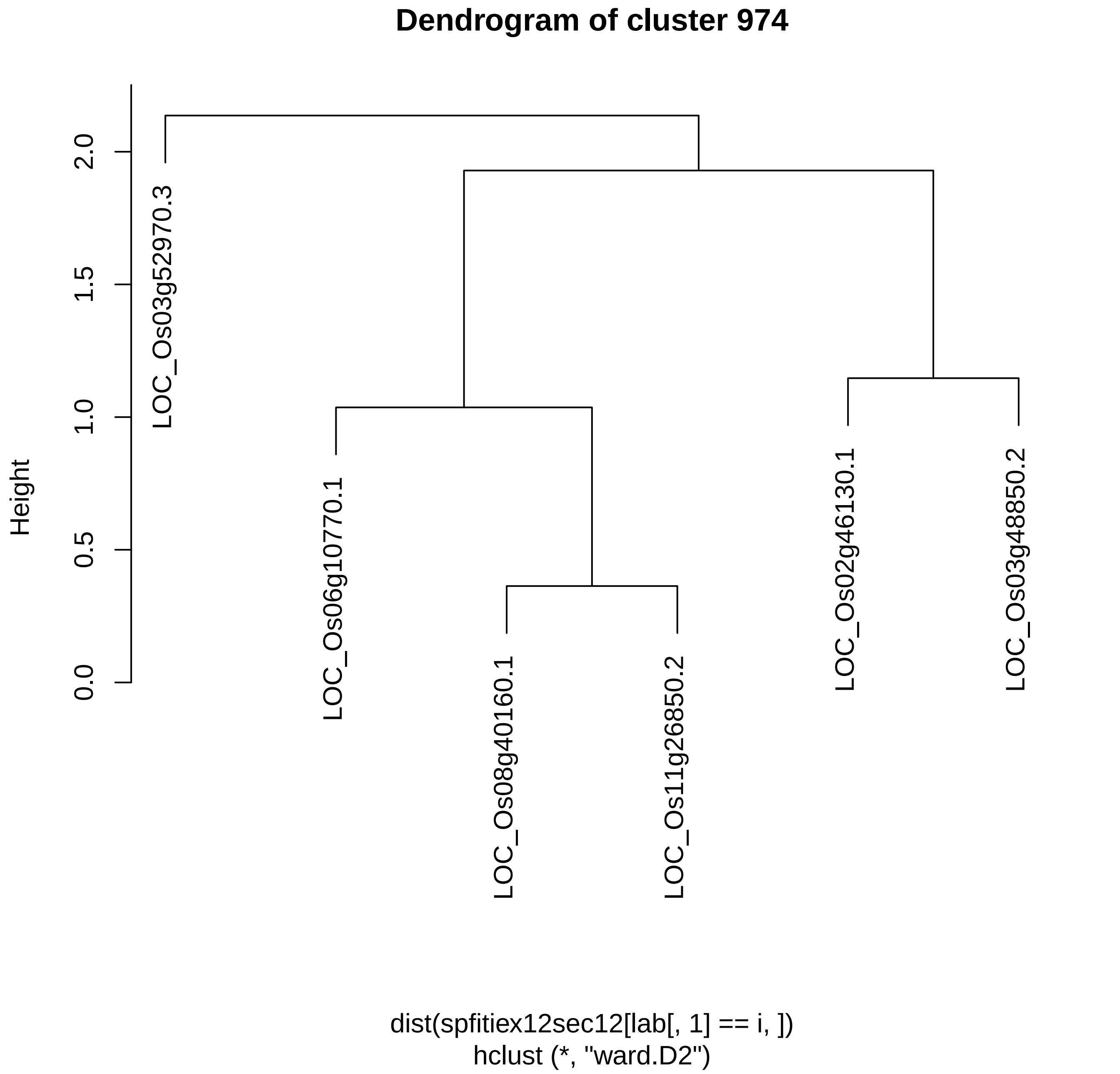
Dendrogram of cluster 974
Height
0.0
0.5
1.0
1.5
2.0
LOC_Os03g52970.3
LOC_Os06g10770.1
LOC_Os08g40160.1
LOC_Os11g26850.2
LOC_Os02g46130.1
LOC_Os03g48850.2
dist(spfitiex12sec12[lab[, 1] == i, ])
hclust (*, "ward.D2")

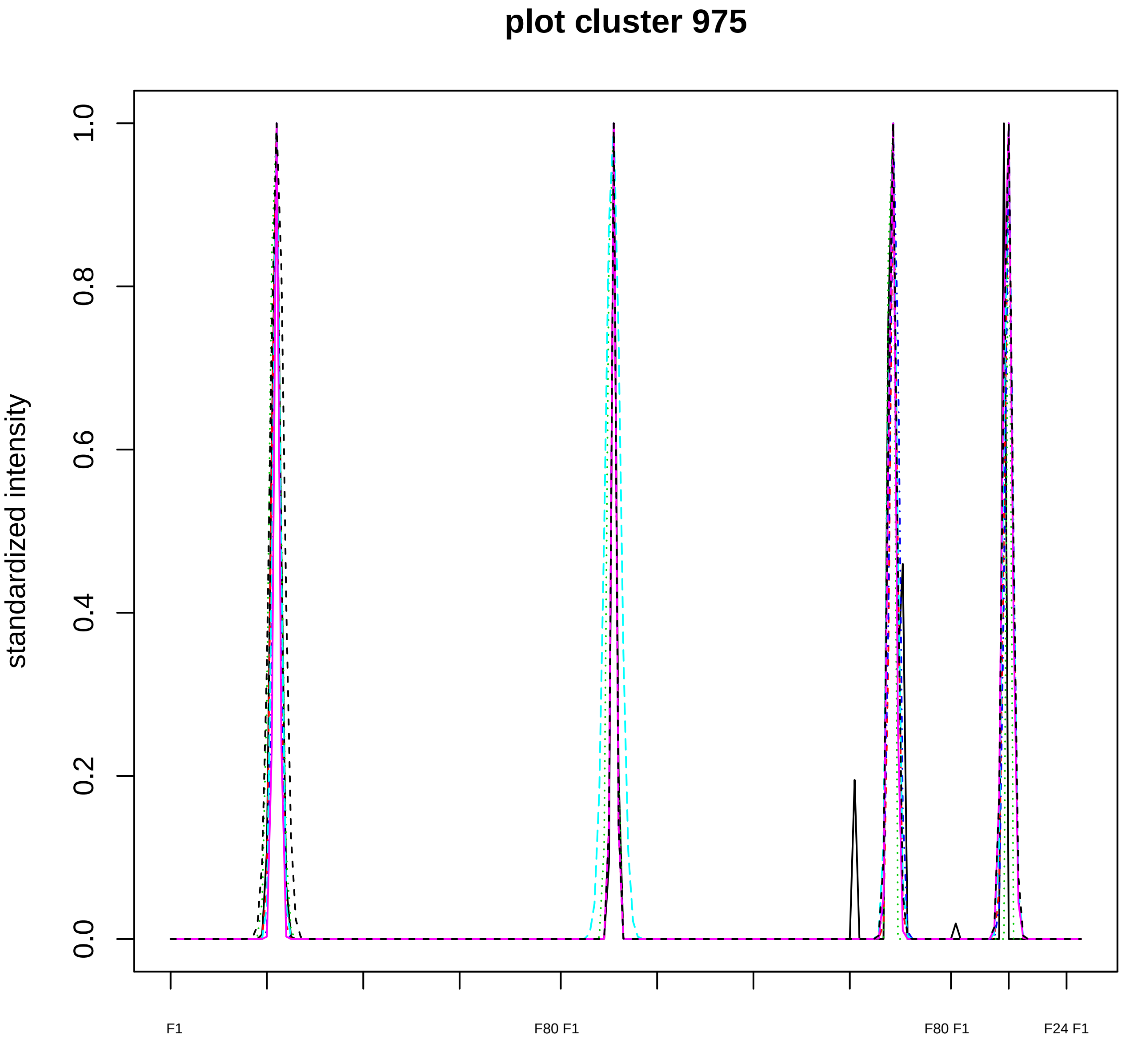

plot cluster 975
standardized intensity
0.0
0.2
0.4
0.6
0.8
1.0
F1
F80 F1
F80 F1
F24 F1

#### Dendrogram of cluster 975

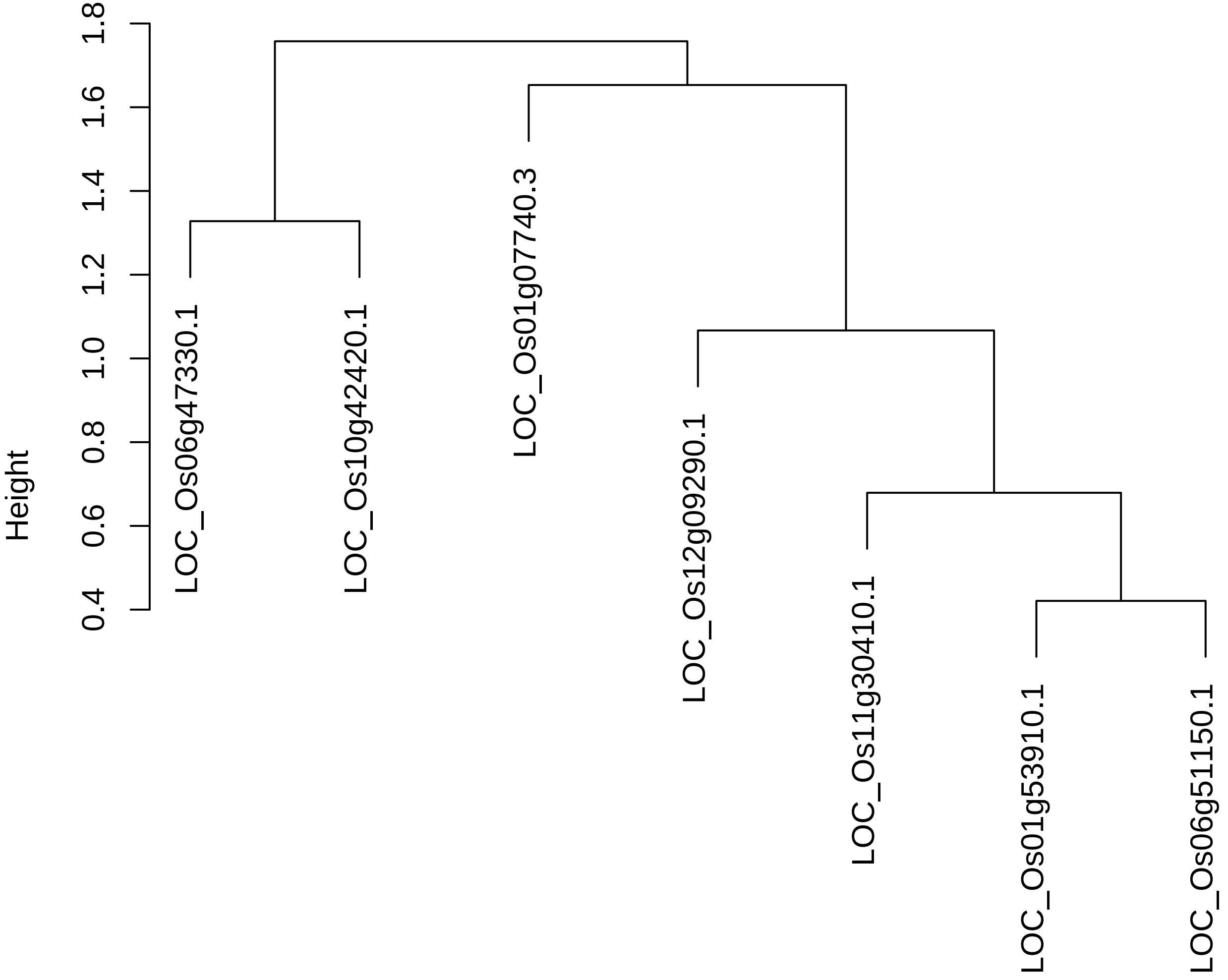


dist(spfitiex12sec12[lab[, 1] == i, ])
hclust (*, "ward.D2")

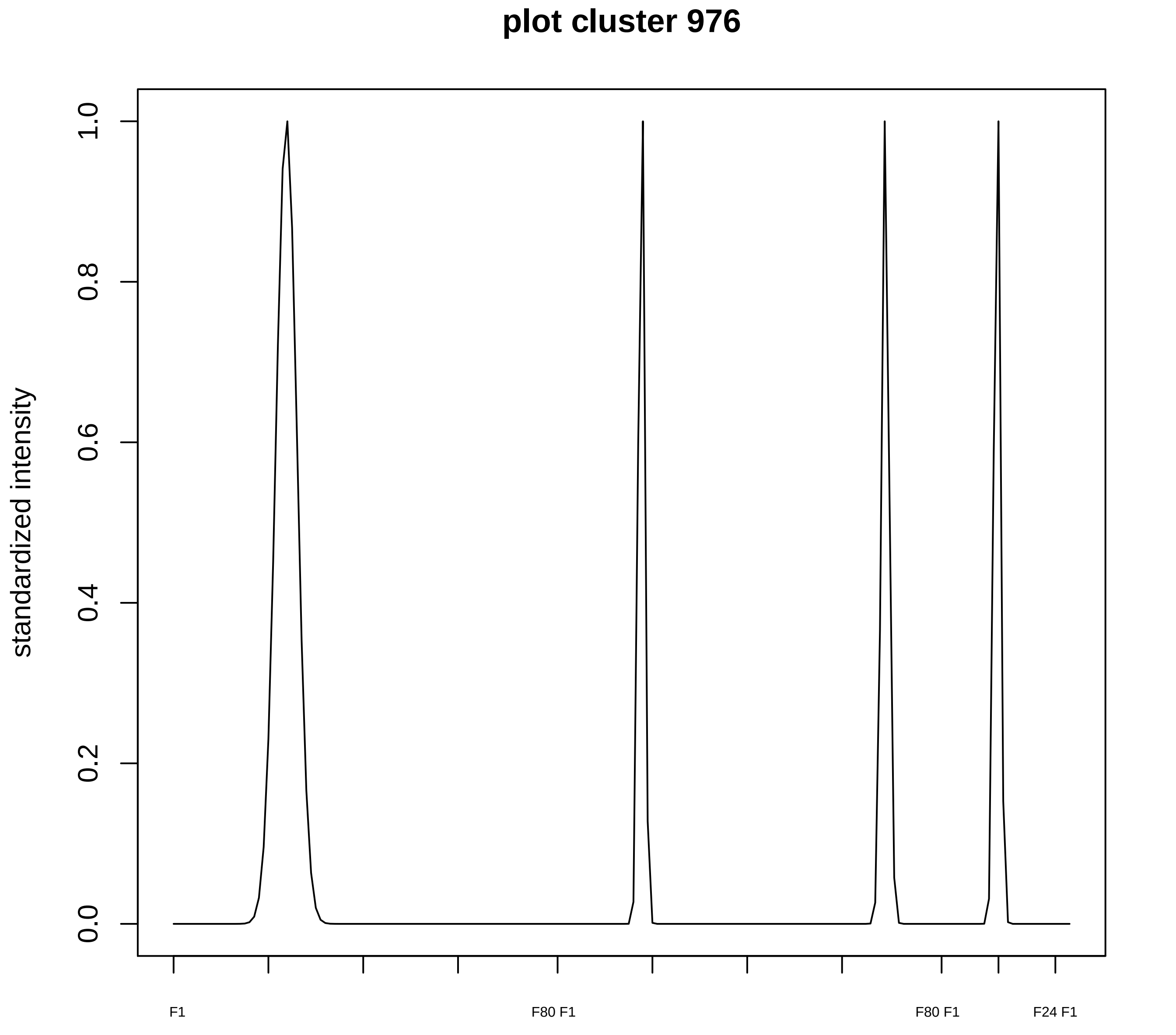

plot cluster 976
standardized intensity
0.0
0.2
0.4
0.6
0.8
1.0
F1
F80 F1
F80 F1
F24 F1

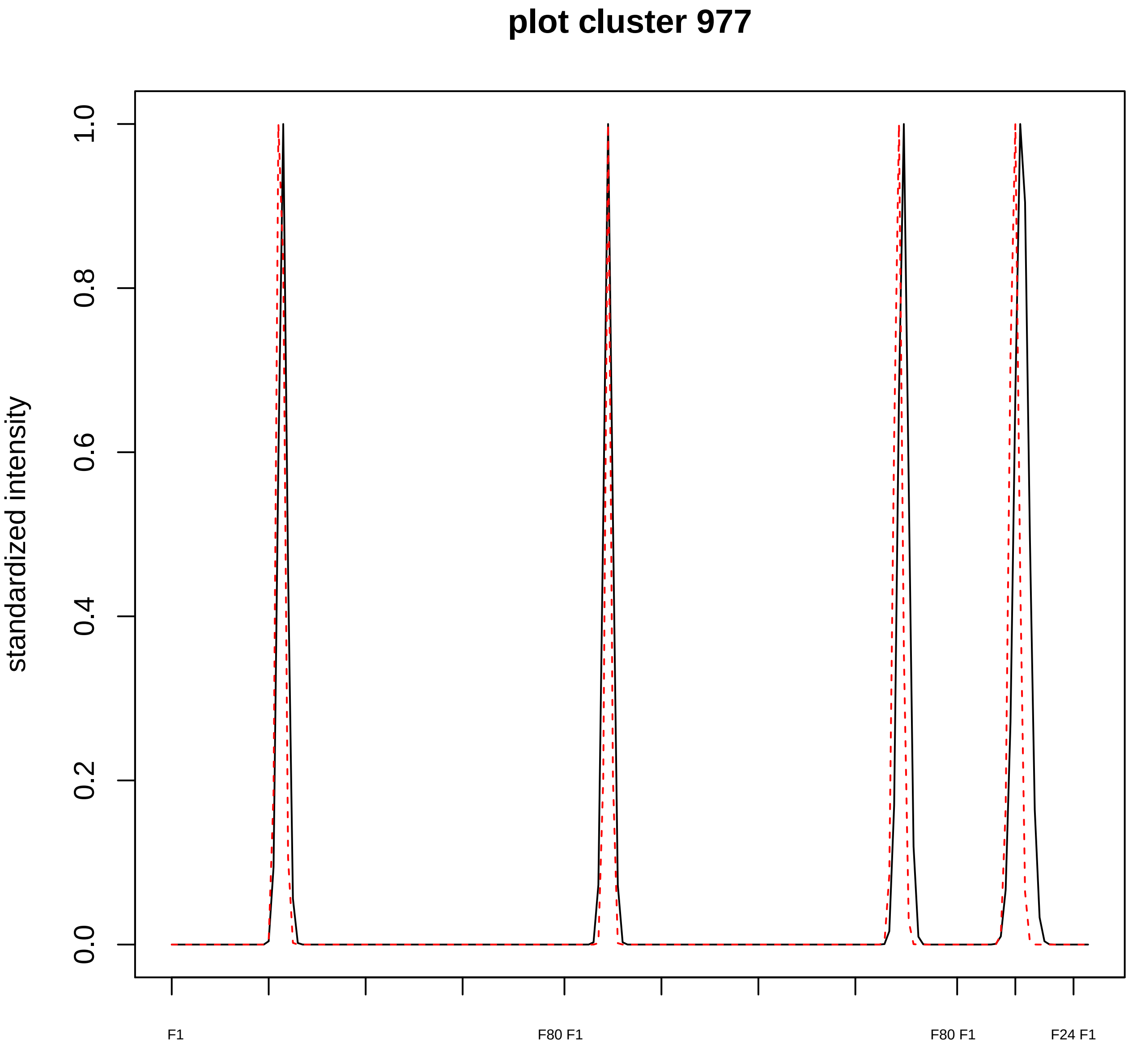

plot cluster 977
standardized intensity
0.0
0.2
0.4
0.6
0.8
1.0
F1
F80 F1
F80 F1
F24 F1

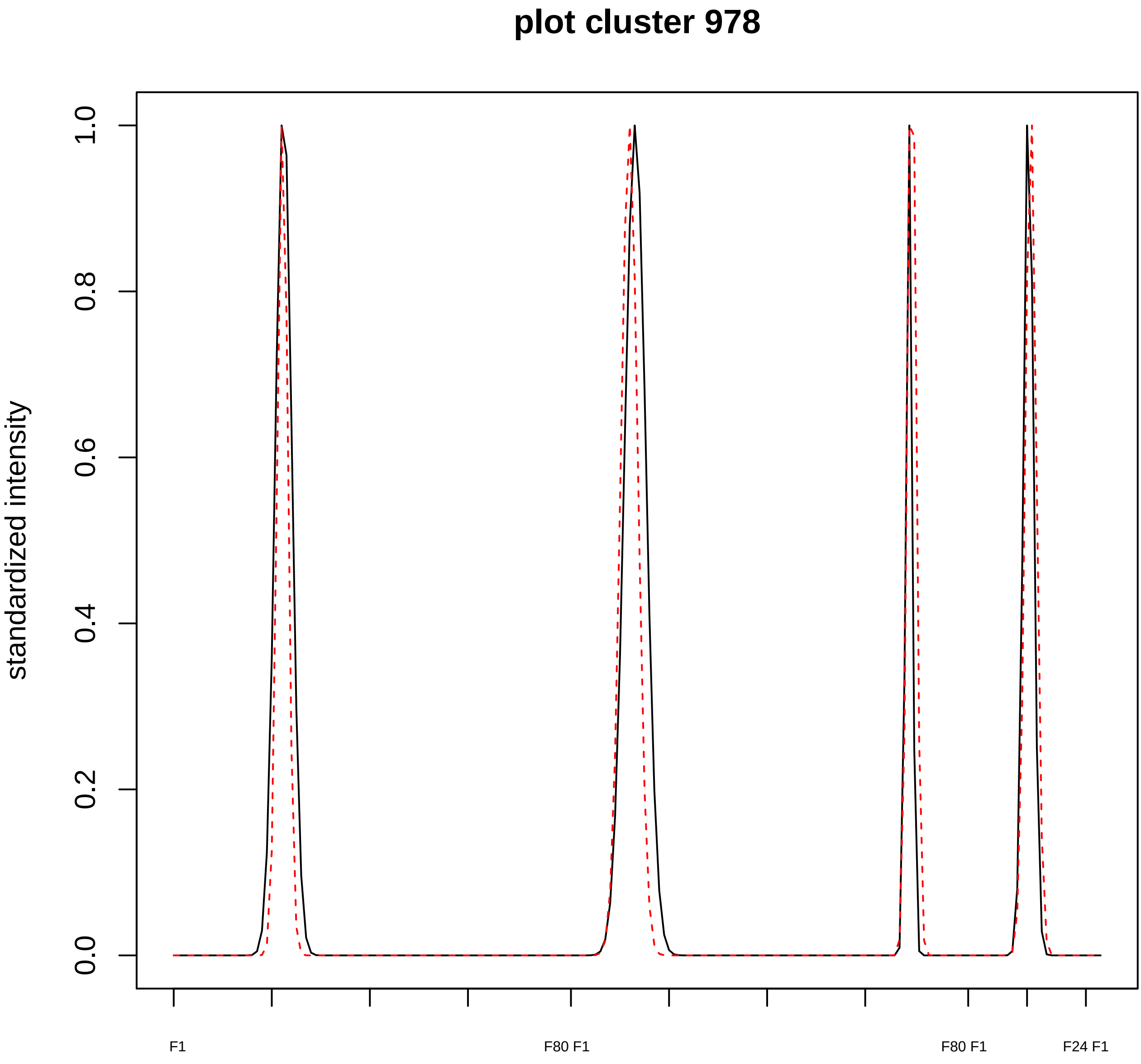
plot cluster 978
standardized intensity
0.0
0.2
0.4
0.6
0.8
1.0
F1
F80 F1
F80 F1
F24 F1

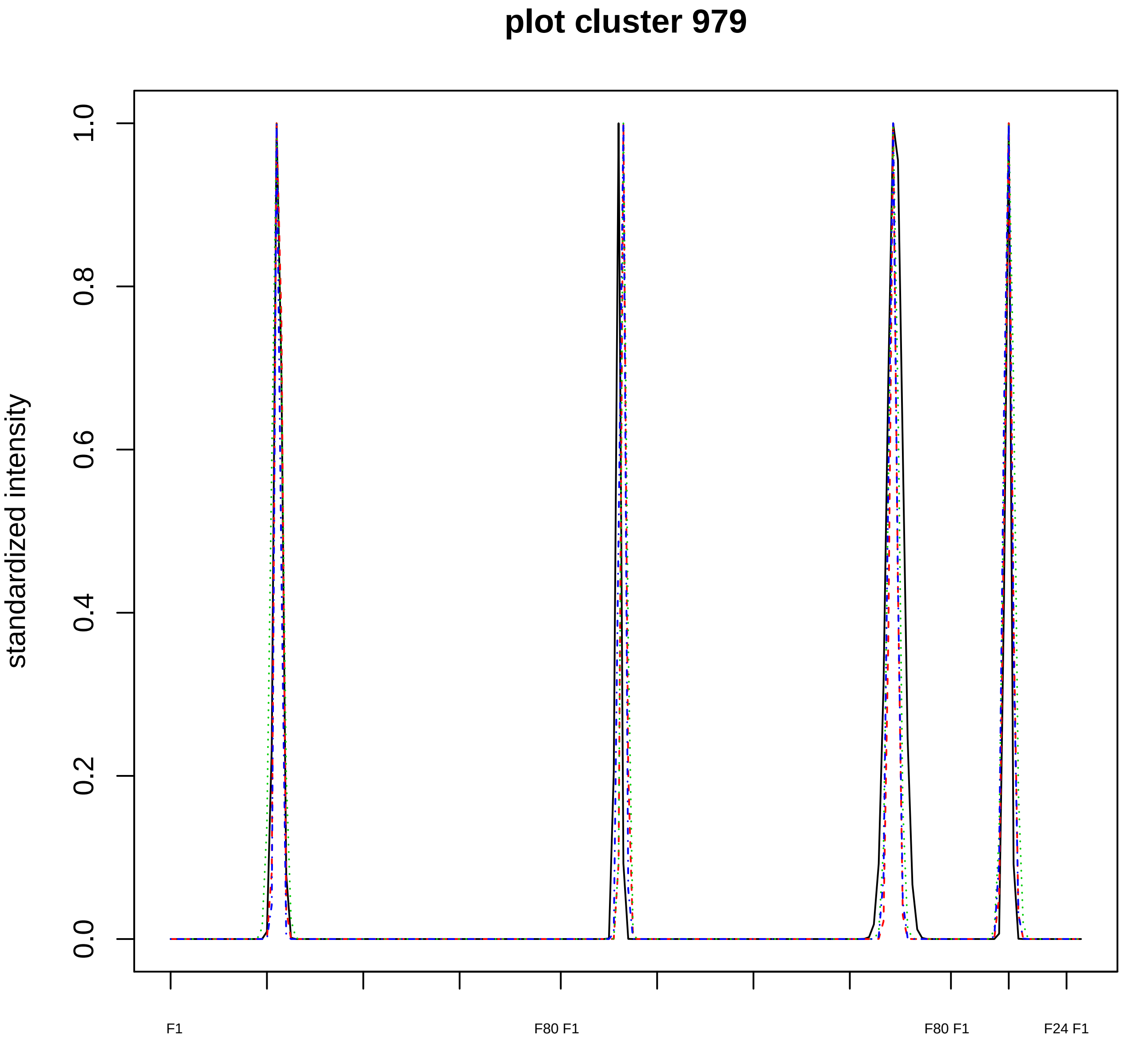
plot cluster 979
standardized intensity
0.0
0.2
0.4
0.6
0.8
1.0
F1
F80 F1
F80 F1
F24 F1

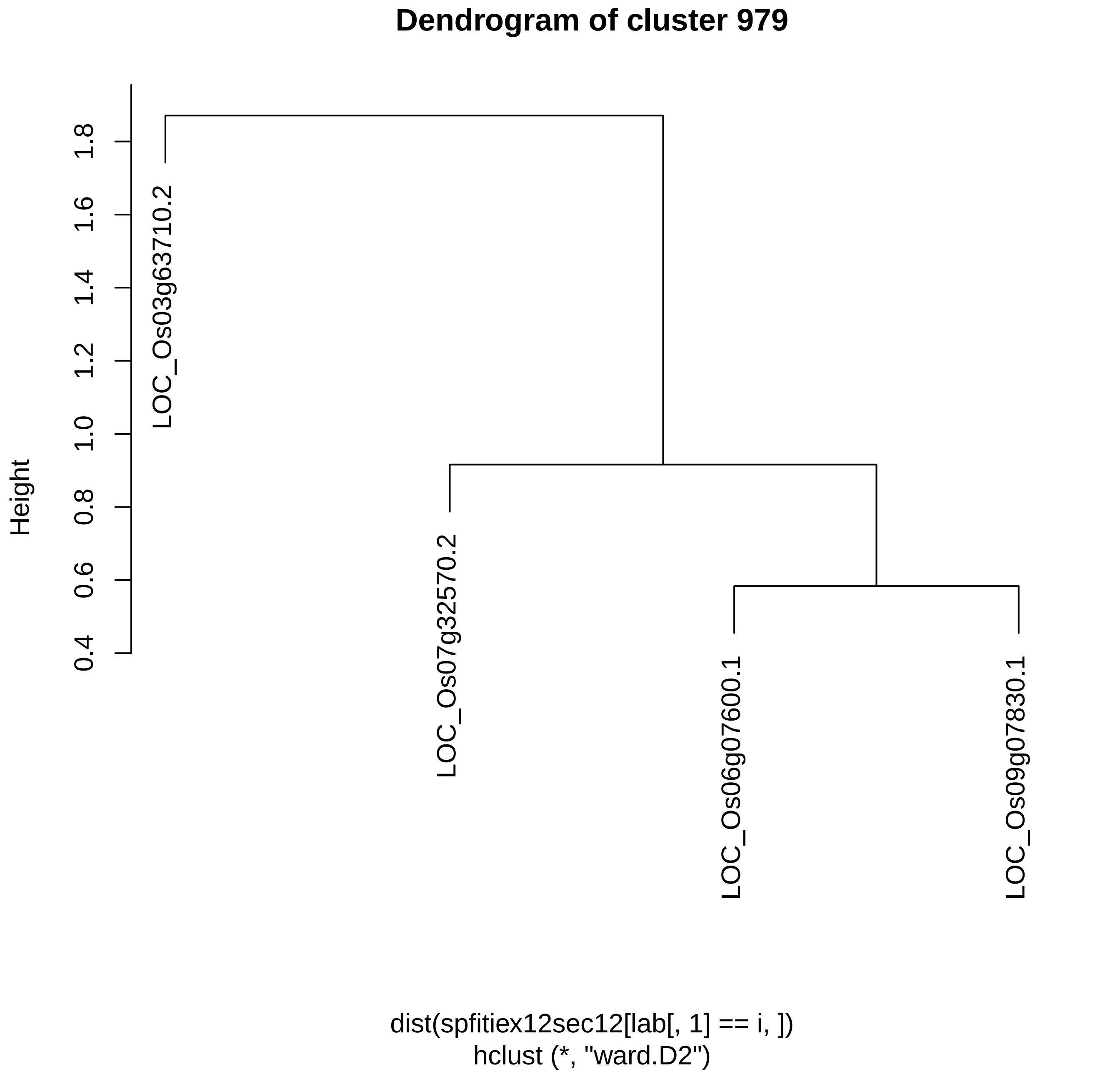
Dendrogram of cluster 979
Height
0.4
0.6
0.8
1.0
1.2
1.4
1.6
1.8
LOC_Os03g63710.2
LOC_Os07g32570.2
LOC_Os06g07600.1
LOC_Os09g07830.1
dist(spfitiex12sec12[lab[, 1] == i, ])
hclust (*, "ward.D2")

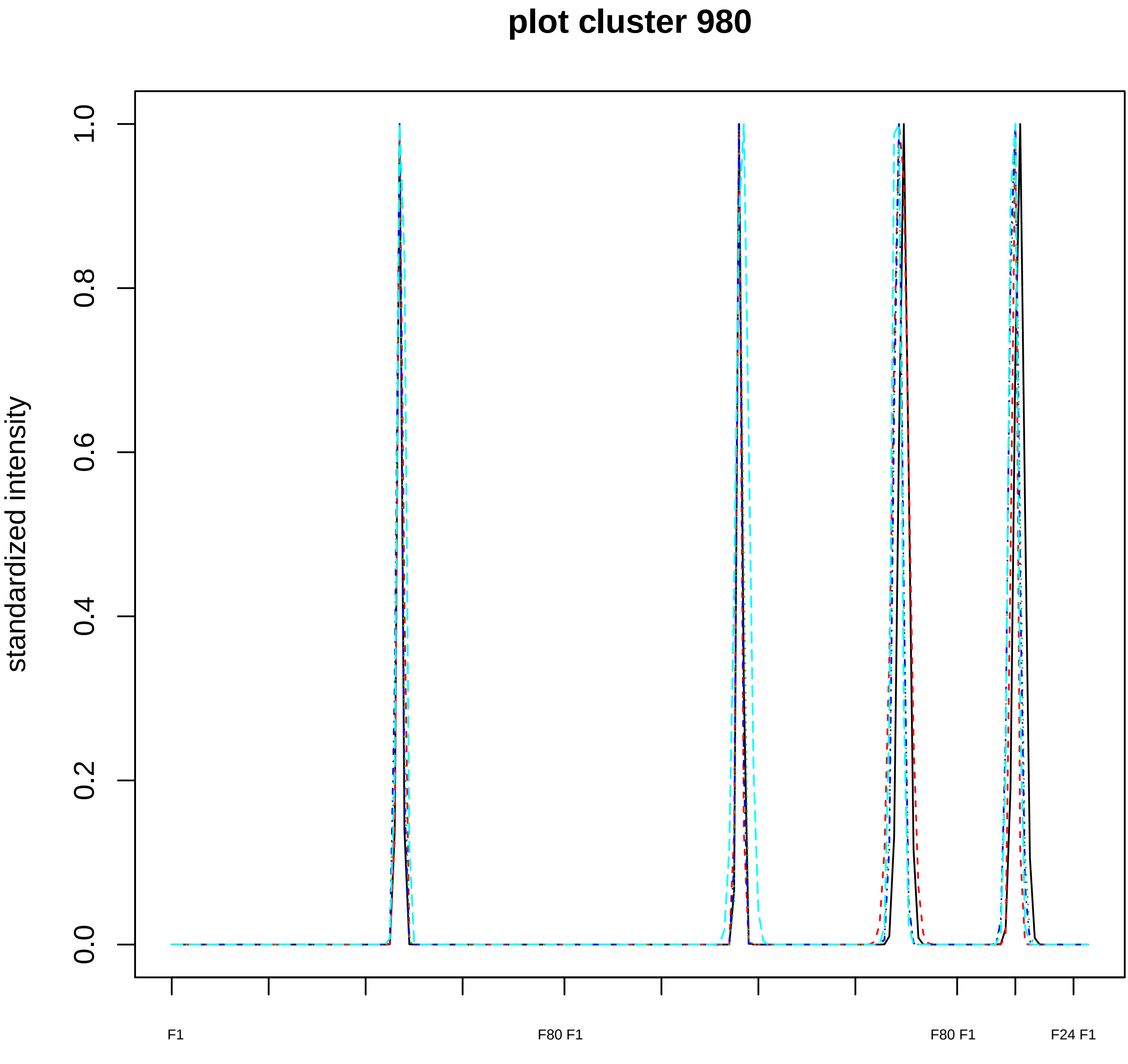
plot cluster 980
standardized intensity
0.0
0.2
0.4
0.6
0.8
1.0
F1
F80 F1
F80 F1
F24 F1

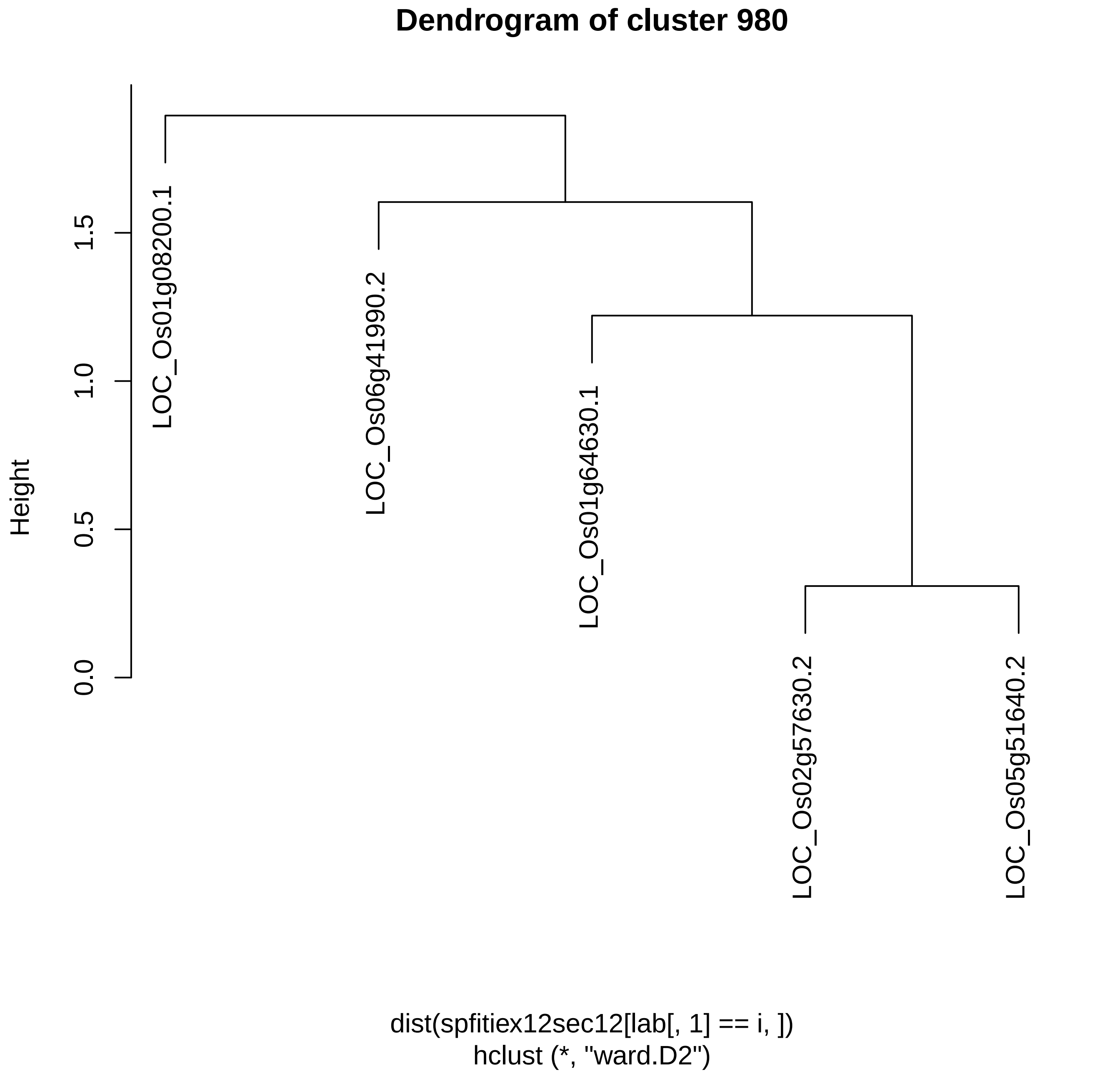

Dendrogram of cluster 980
Height
0.0
0.5
1.0
1.5
LOC_Os01g08200.1
LOC_Os06g41990.2
LOC_Os01g64630.1
LOC_Os02g57630.2
LOC_Os05g51640.2
dist(spfitiex12sec12[lab[, 1] == i, ])
hclust (*, "ward.D2")

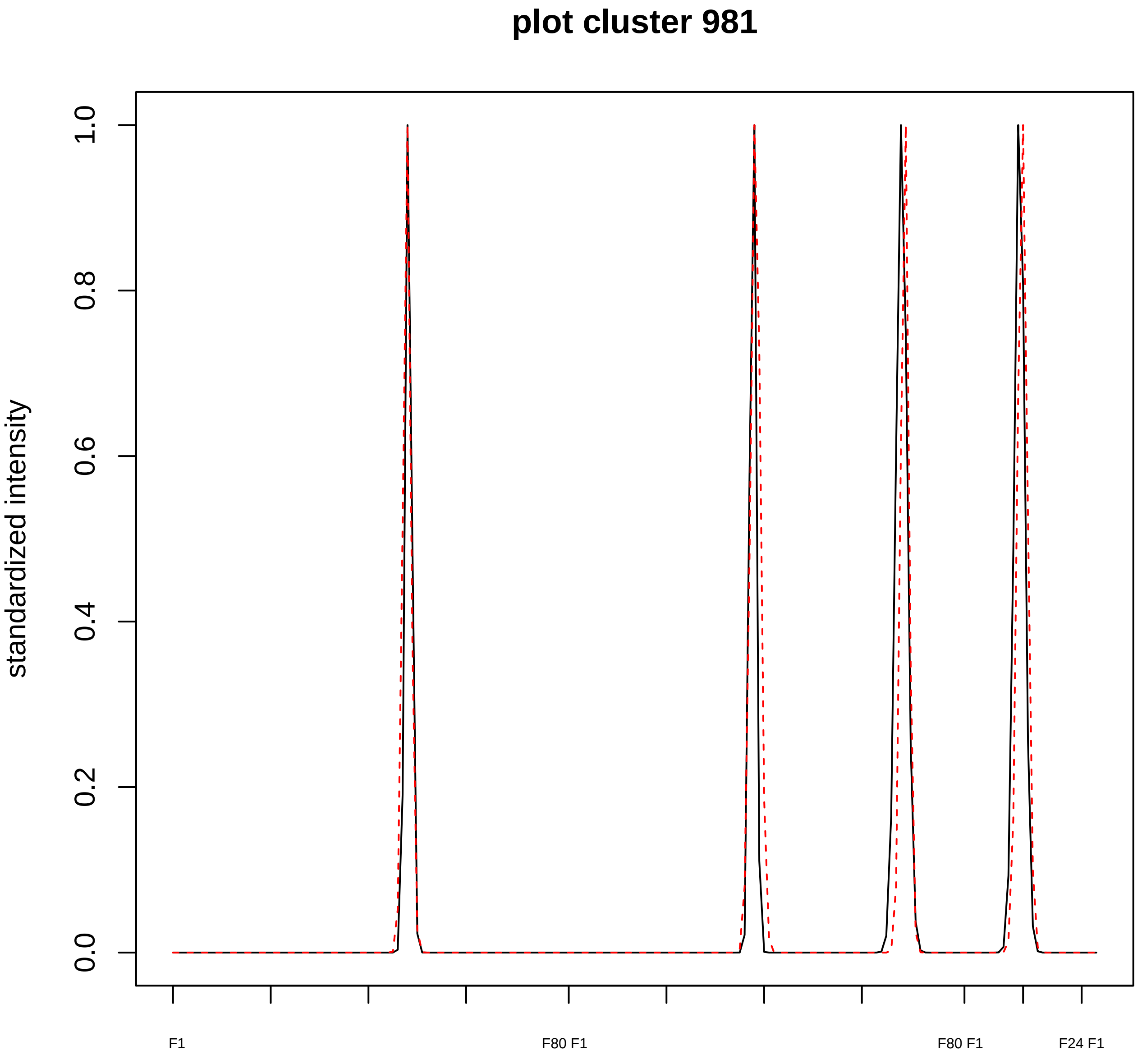

plot cluster 981
standardized intensity
0.0
0.2
0.4
0.6
0.8
1.0
F1
F80 F1
F80 F1
F24 F1

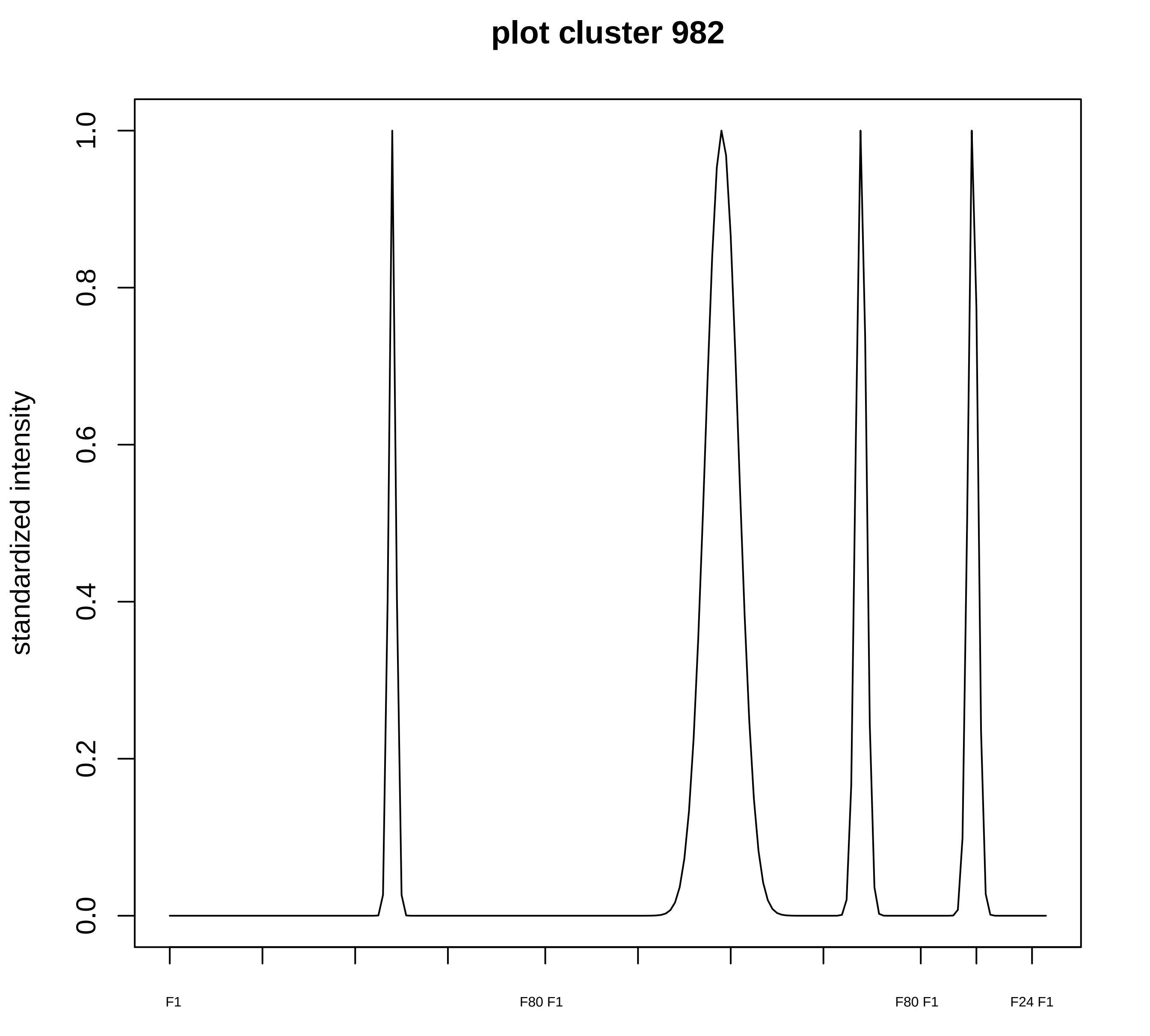
plot cluster 982
standardized intensity
0.0
0.2
0.4
0.6
0.8
1.0
F1
F80 F1
F80 F1
F24 F1

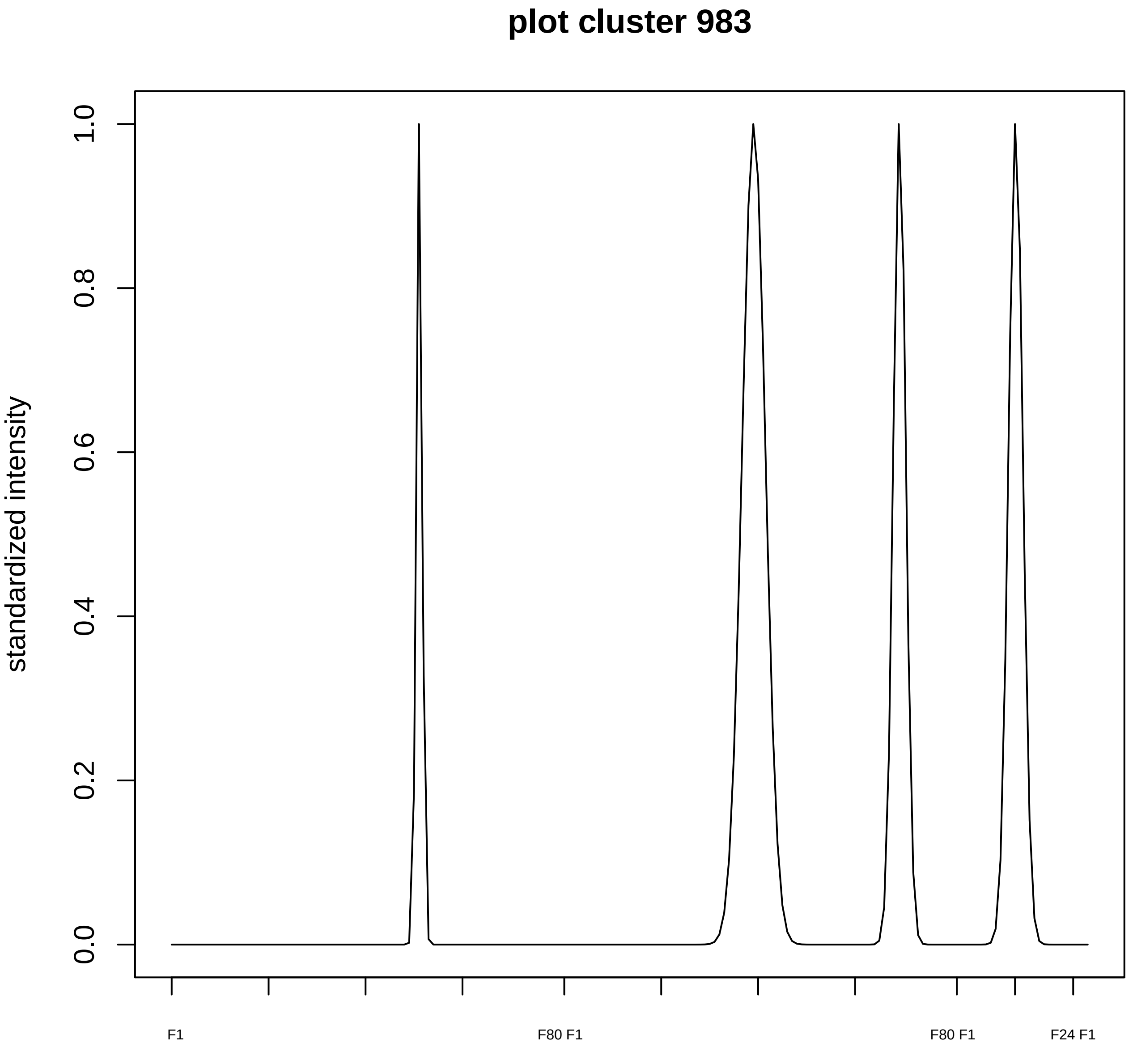
plot cluster 983
standardized intensity
1.0
0.8
0.6
0.4
0.2
0.0
F1
F80 F1
F80 F1
F24 F1

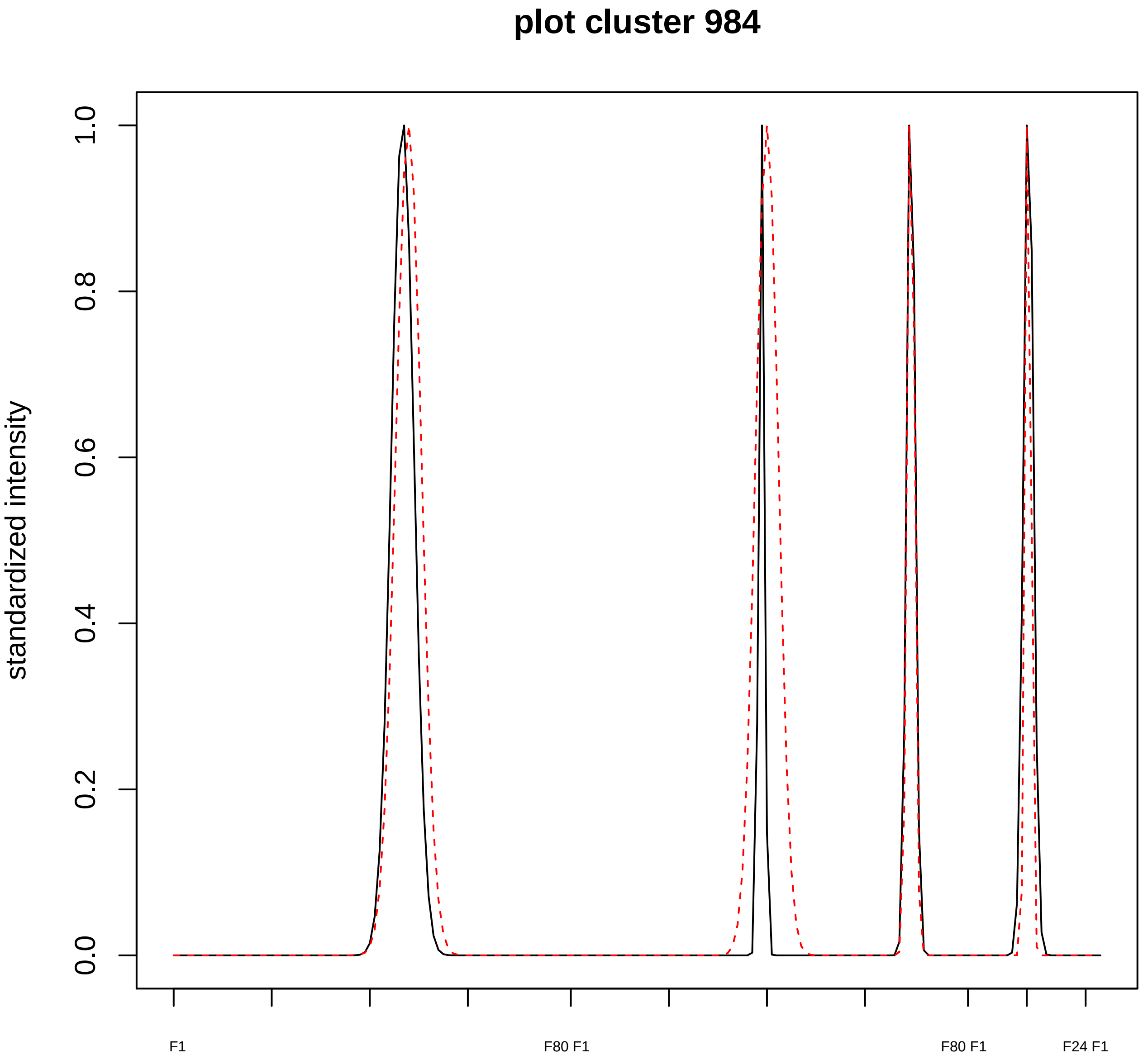

plot cluster 984
standardized intensity
0.0
0.2
0.4
0.6
0.8
1.0
F1
F80 F1
F80 F1
F24 F1

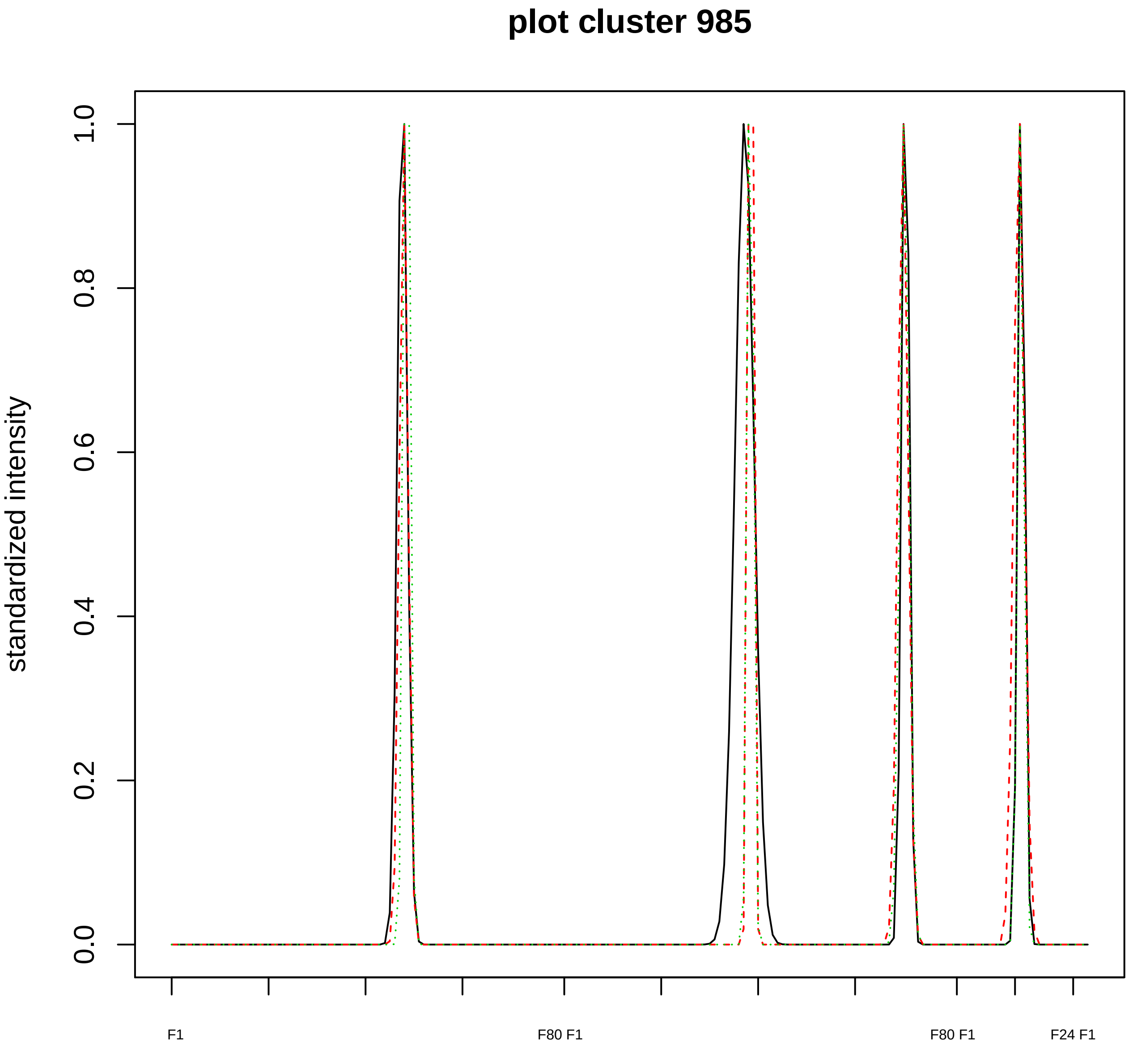

plot cluster 985
standardized intensity
0.0
0.2
0.4
0.6
0.8
1.0
F1
F80 F1
F80 F1
F24 F1

### Dendrogram of cluster 985

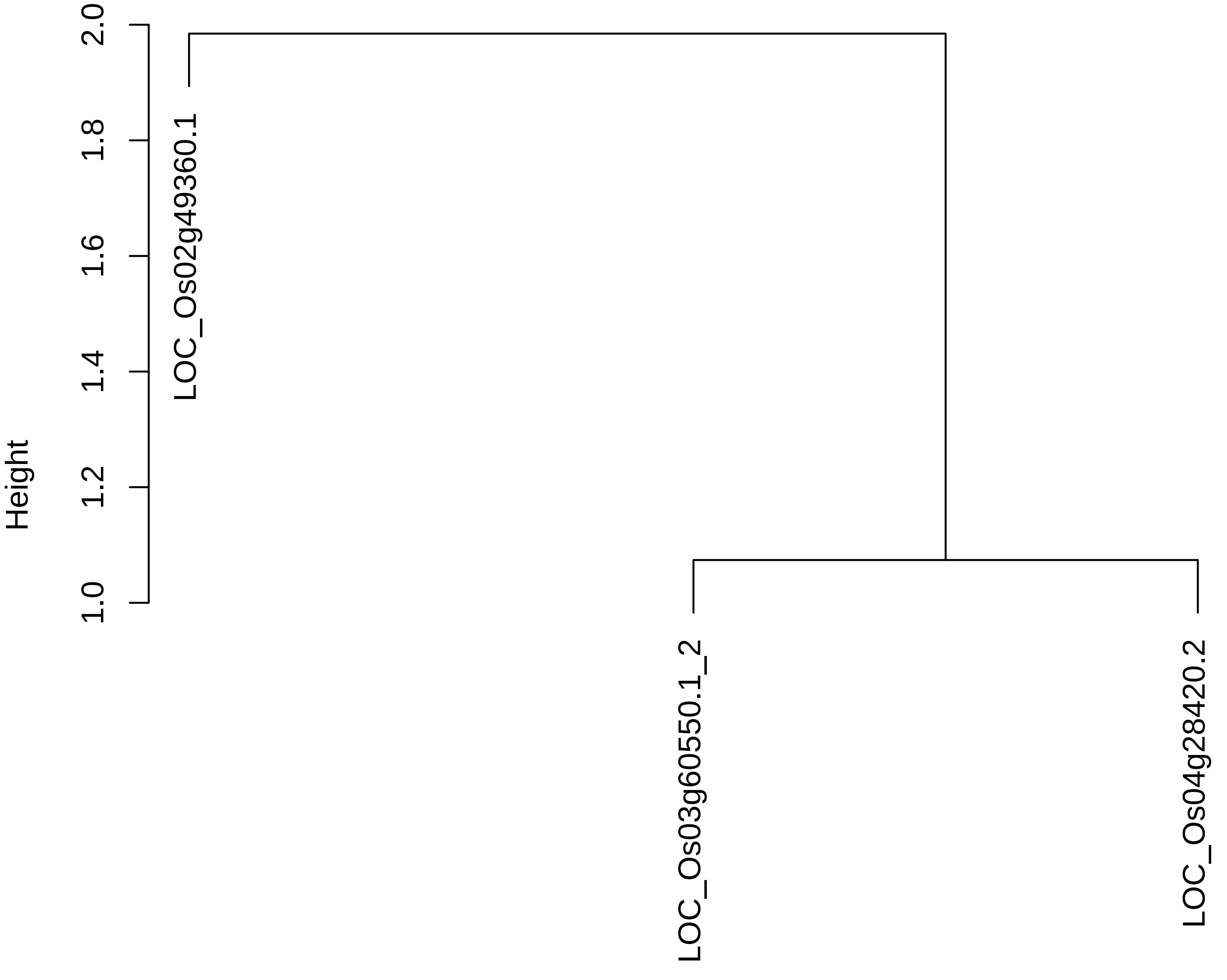


dist(spfitiex12sec12[lab[, 1] == i, ])
hclust (*, "ward.D2")

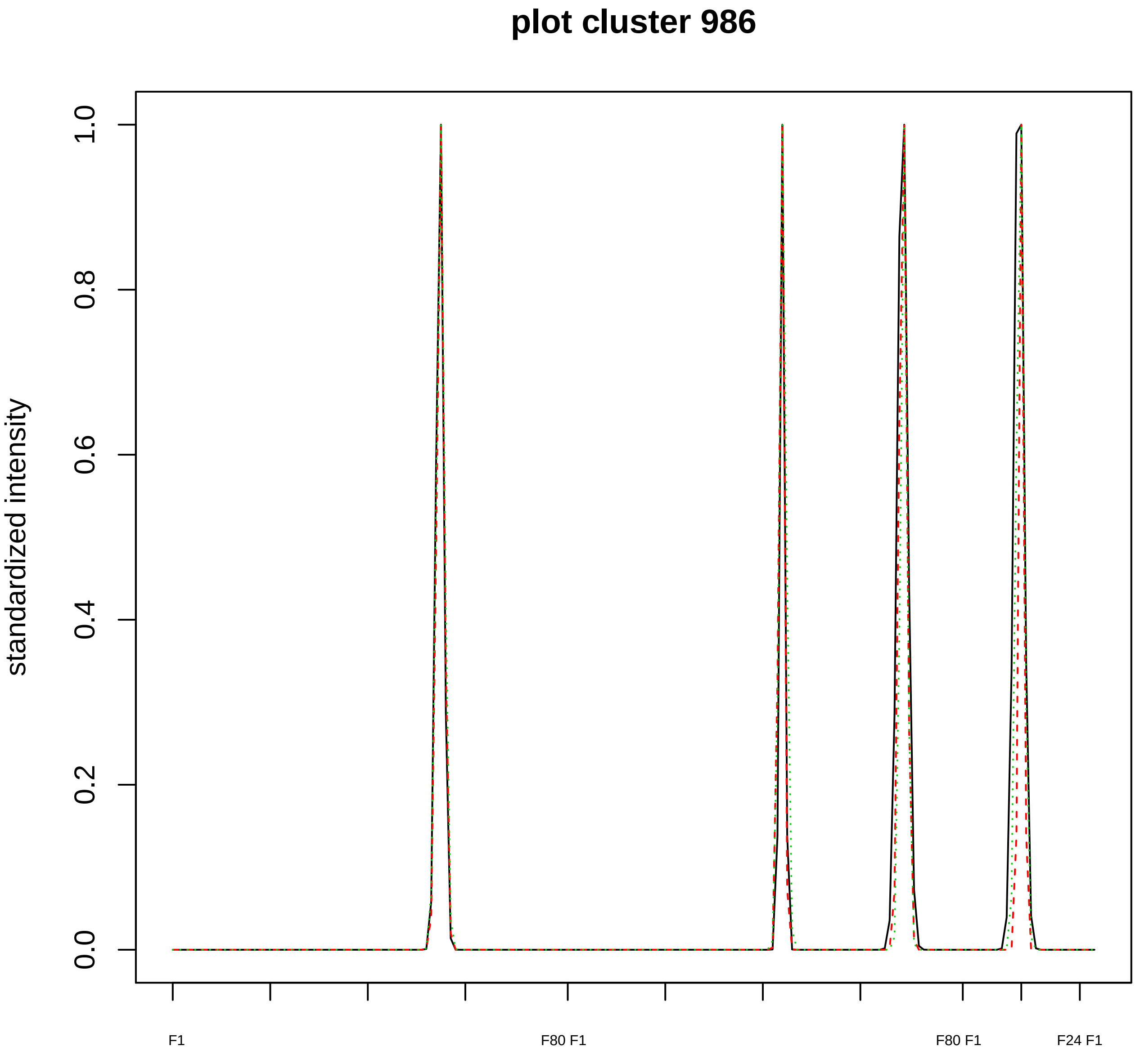

plot cluster 986
standardized intensity
0.0
0.2
0.4
0.6
0.8
1.0
F1
F80 F1
F80 F1
F24 F1

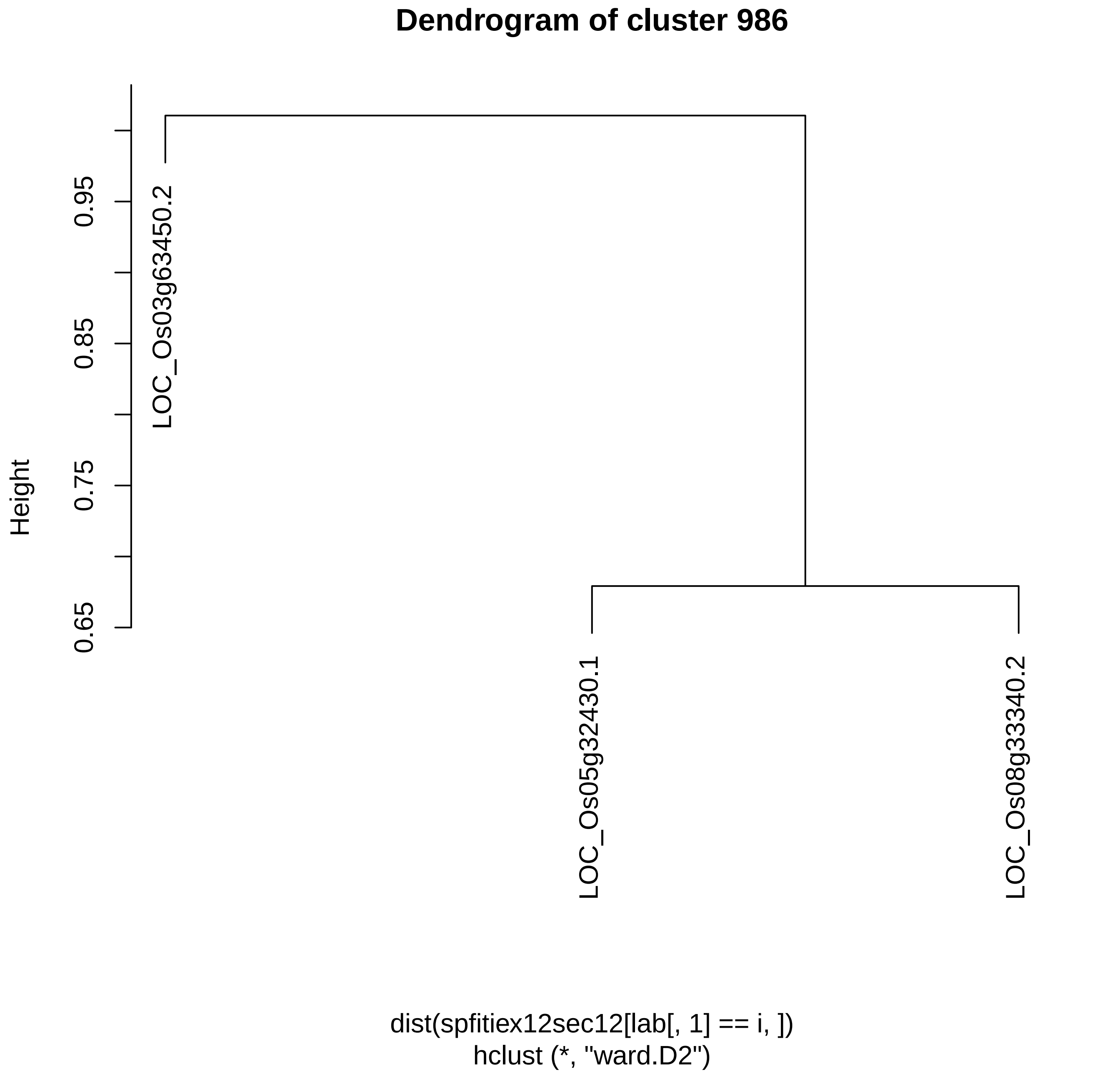

Dendrogram of cluster 986
Height
0.65
0.75
0.85
0.95
LOC_Os03g63450.2
LOC_Os05g32430.1
LOC_Os08g33340.2
dist(spfitiex12sec12[lab[, 1] == i, ])
hclust (*, "ward.D2")

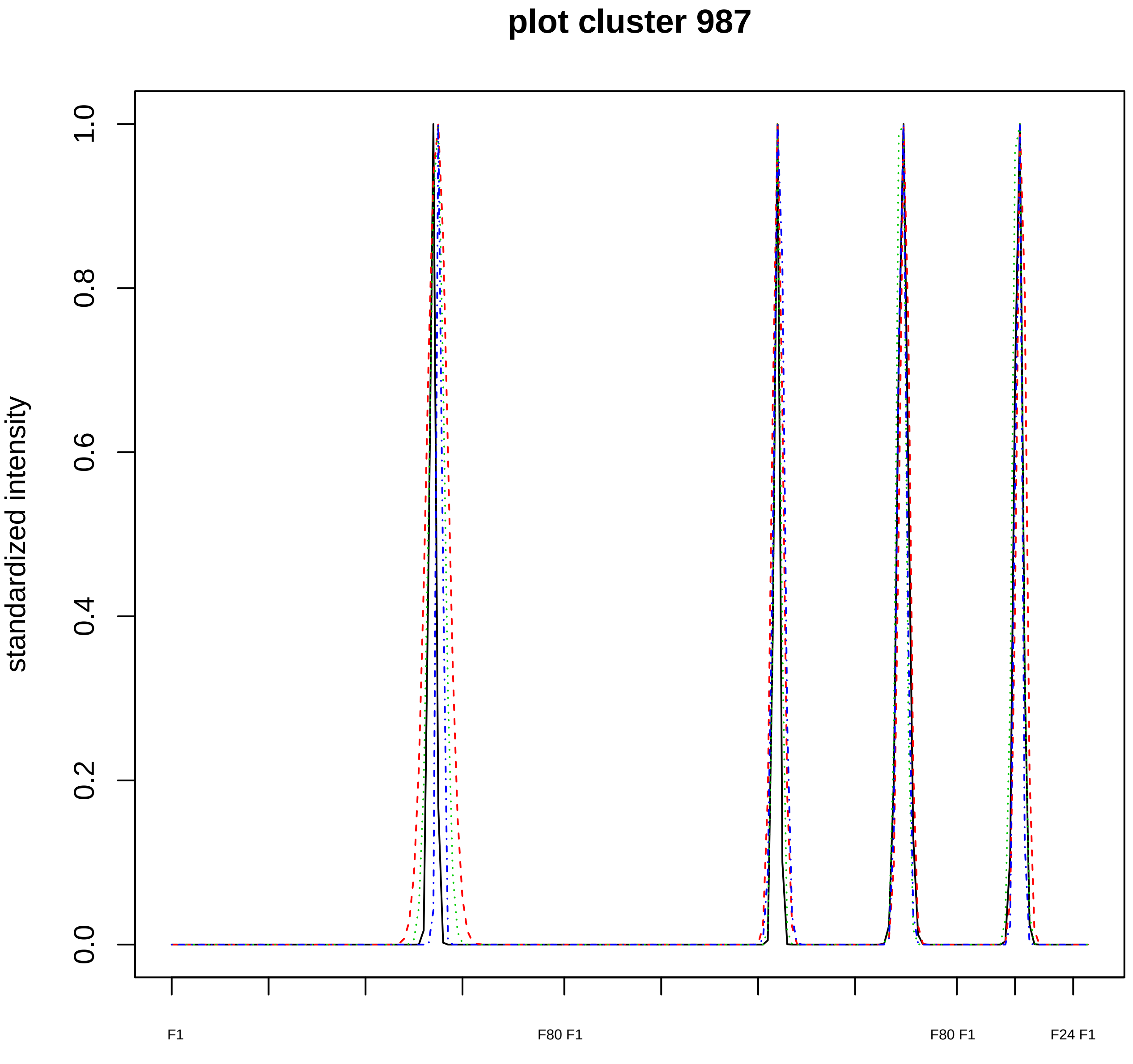

plot cluster 987
standardized intensity
0.0
0.2
0.4
0.6
0.8
1.0
F1
F80 F1
F80 F1
F24 F1

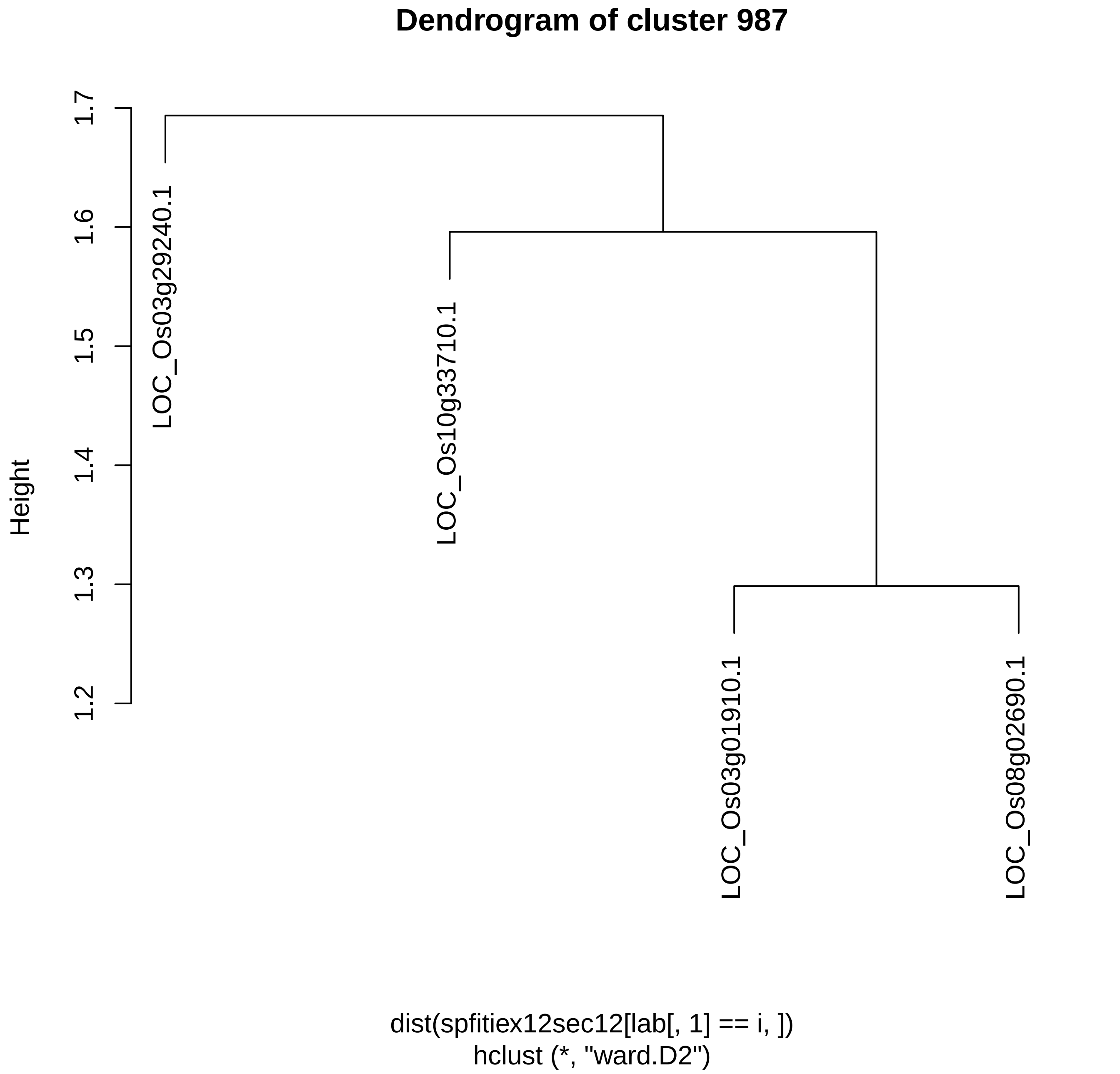
Dendrogram of cluster 987
Height
1.2
1.3
1.4
1.5
1.6
1.7
LOC_Os03g29240.1
LOC_Os10g33710.1
LOC_Os03g01910.1
LOC_Os08g02690.1
dist(spfitiex12sec12[lab[, 1] == i, ])
hclust (*, "ward.D2")

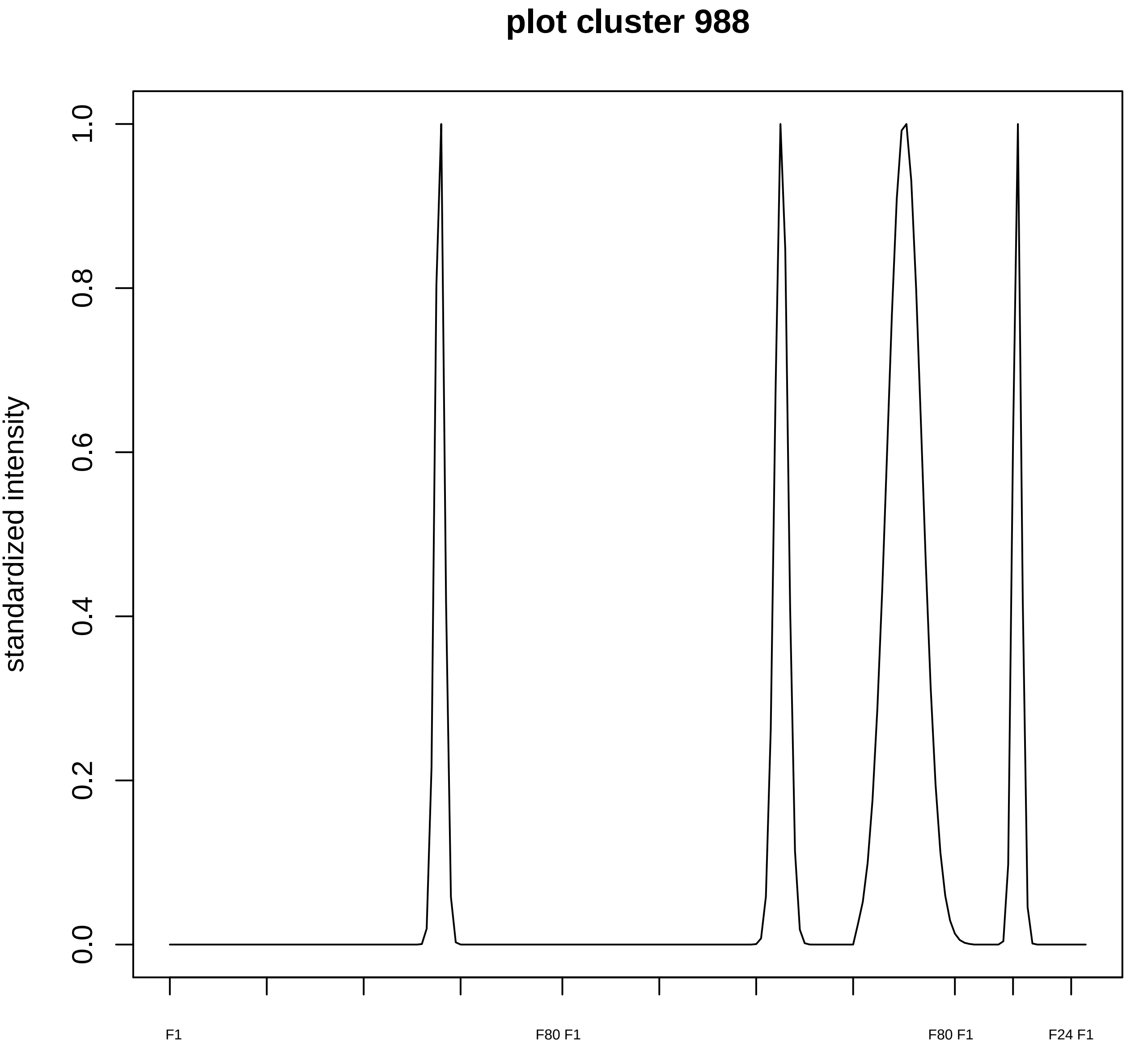
plot cluster 988
standardized intensity
0.0
0.2
0.4
0.6
0.8
1.0
F1
F80 F1
F80 F1
F24 F1

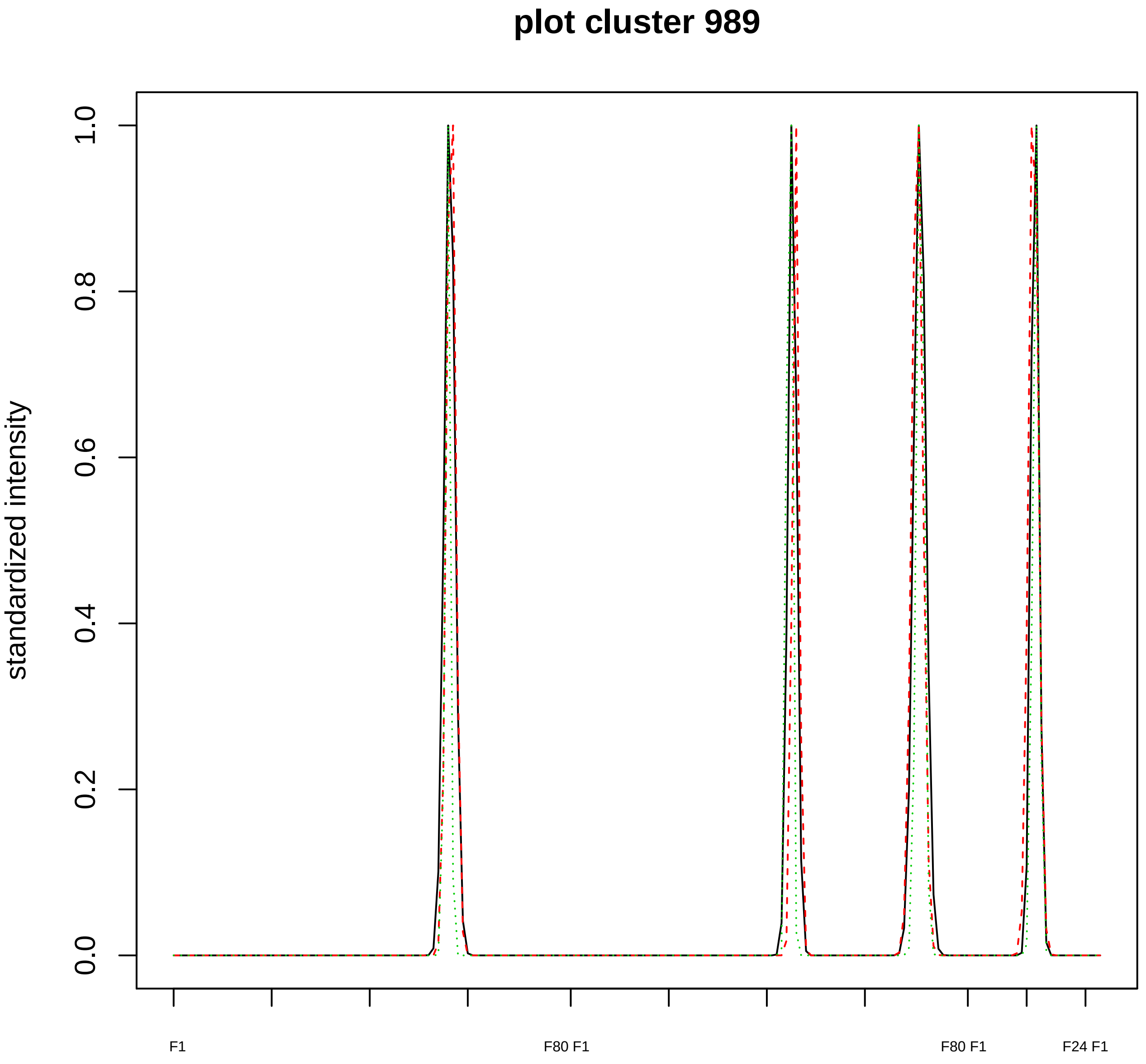
plot cluster 989
standardized intensity
0.0
0.2
0.4
0.6
0.8
1.0
F1
F80 F1
F80 F1
F24 F1

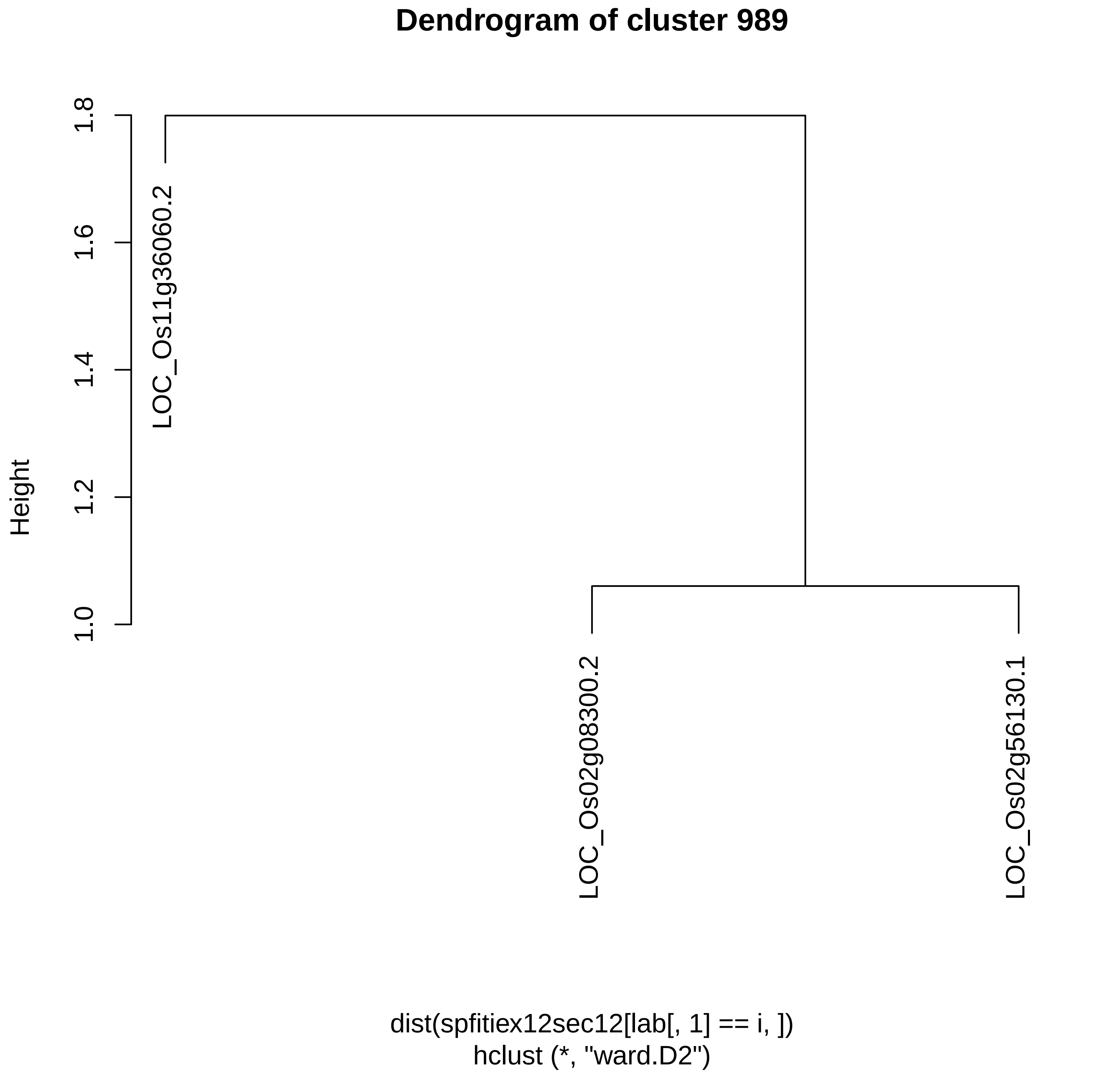
Dendrogram of cluster 989
Height
1.0
1.2
1.4
1.6
1.8
LOC_Os11g36060.2
LOC_Os02g08300.2
LOC_Os02g56130.1
dist(spfitiex12sec12[lab[, 1] == i, ])
hclust (*, "ward.D2")

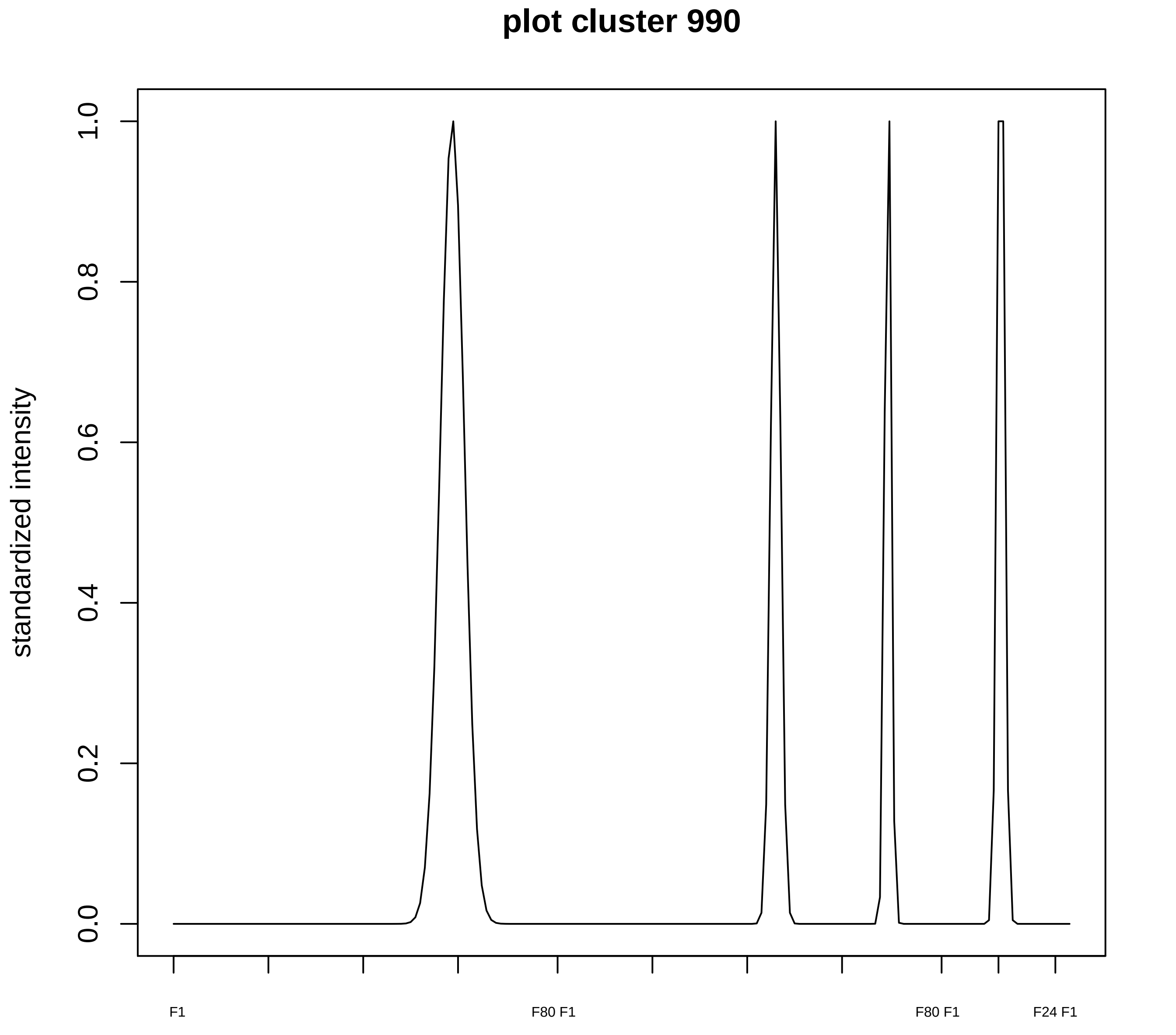

plot cluster 990
standardized intensity
0.0
0.2
0.4
0.6
0.8
1.0
F1
F80 F1
F80 F1
F24 F1

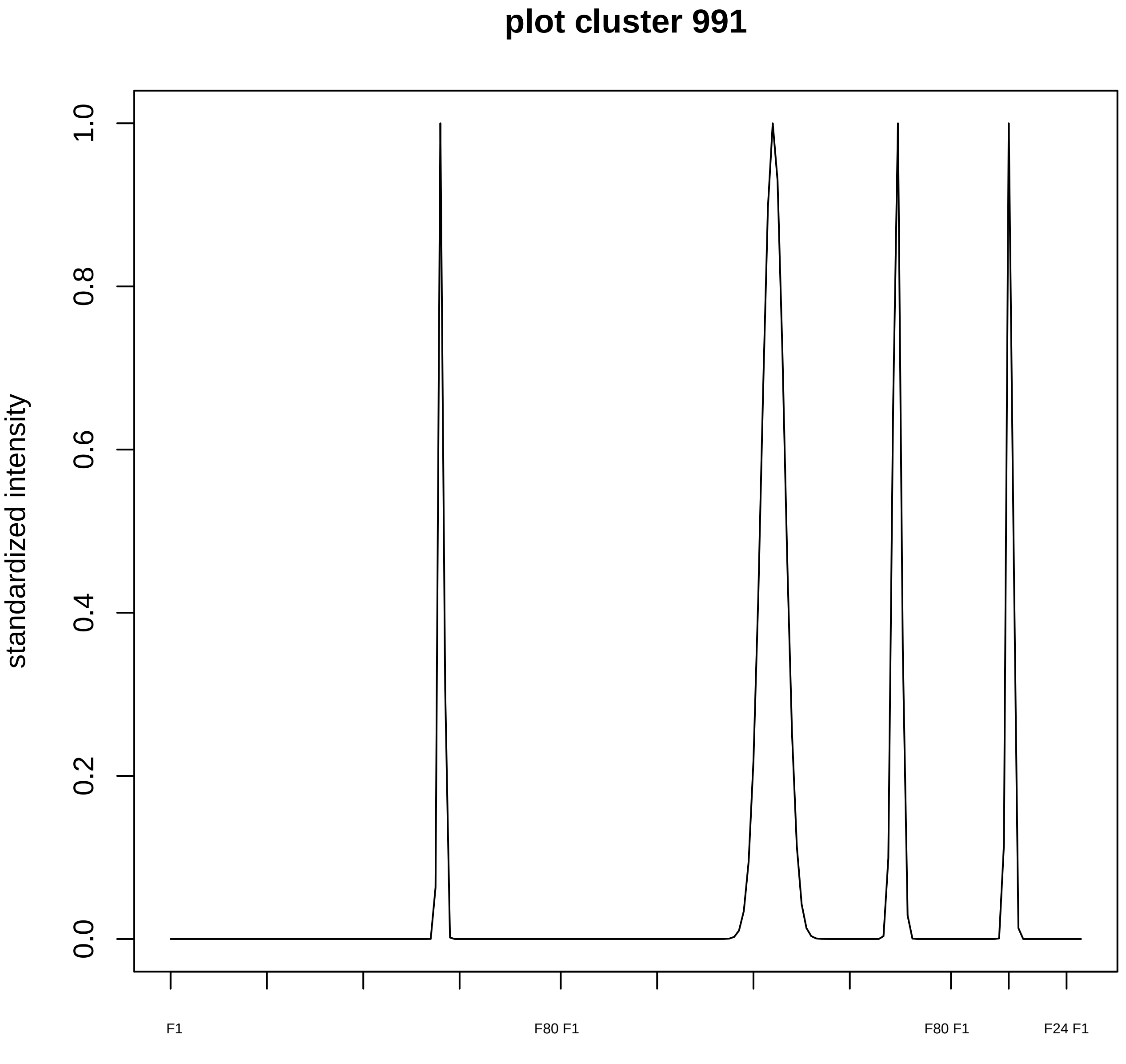

plot cluster 991
standardized intensity
0.0
0.2
0.4
0.6
0.8
1.0
F1
F80 F1
F80 F1
F24 F1

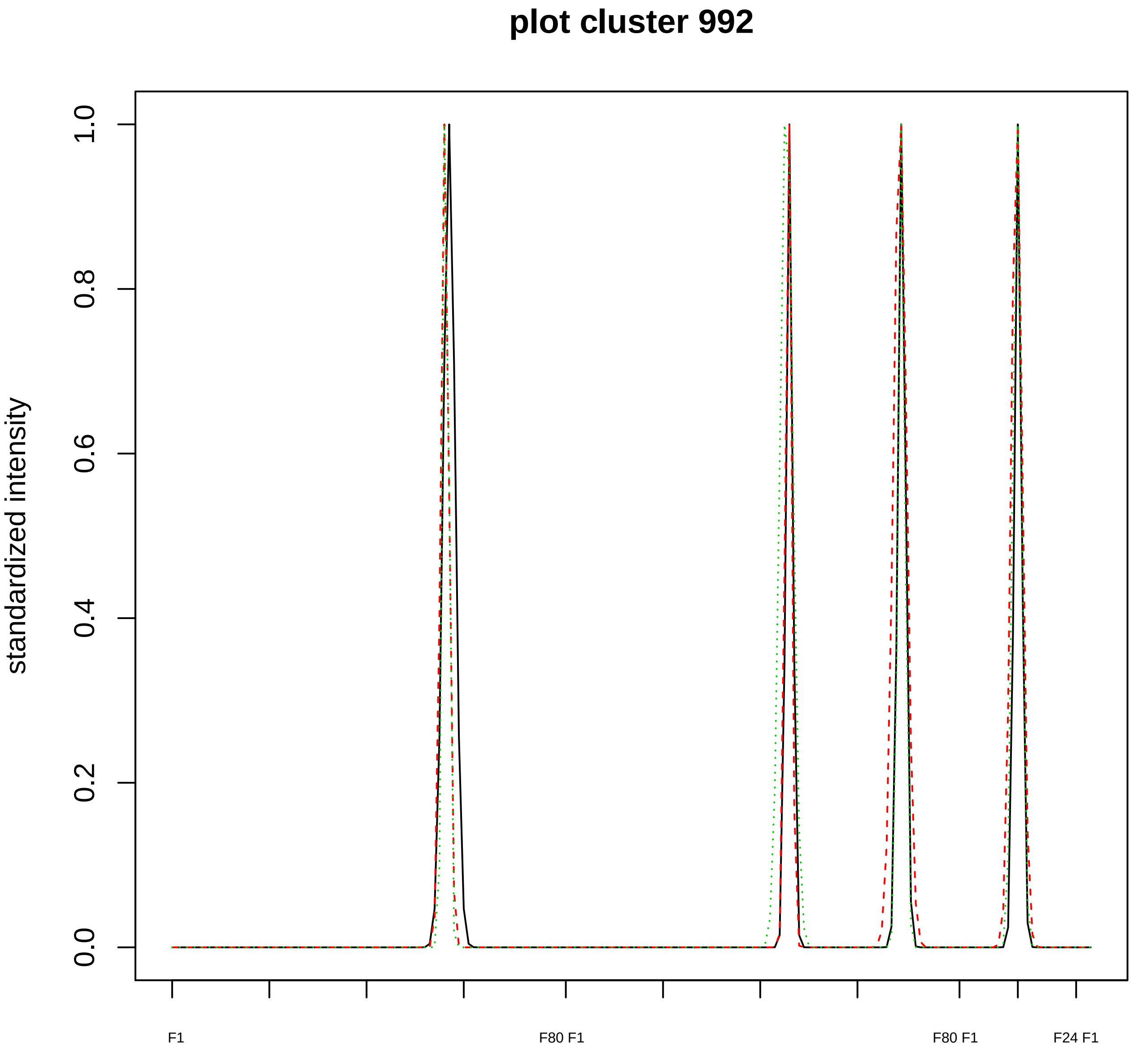
plot cluster 992
standardized intensity
0.0
0.2
0.4
0.6
0.8
1.0
F1
F80 F1
F80 F1
F24 F1

### Dendrogram of cluster 992

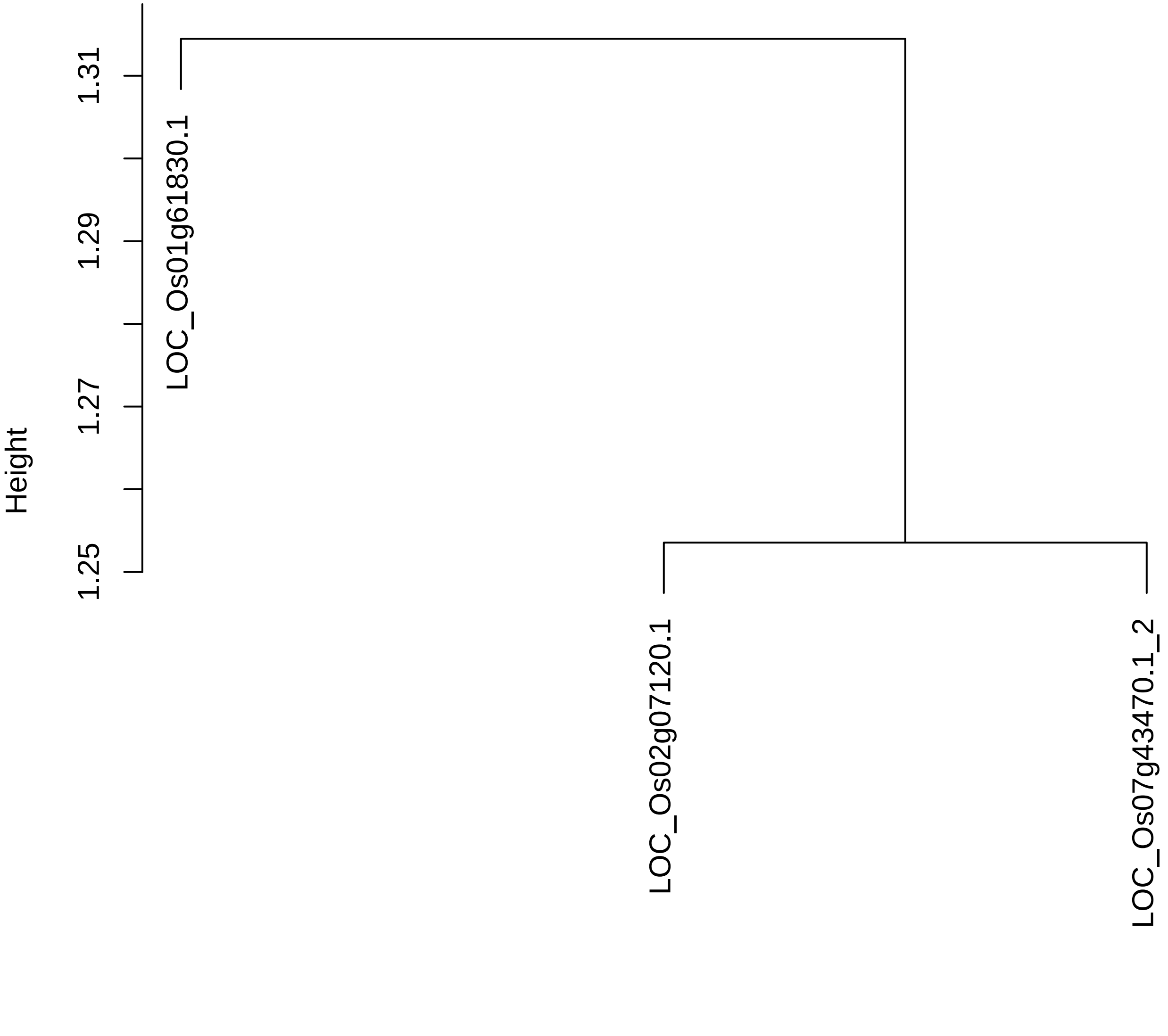


dist(spfitiex12sec12[lab[, 1] == i, ])
hclust (*, "ward.D2")

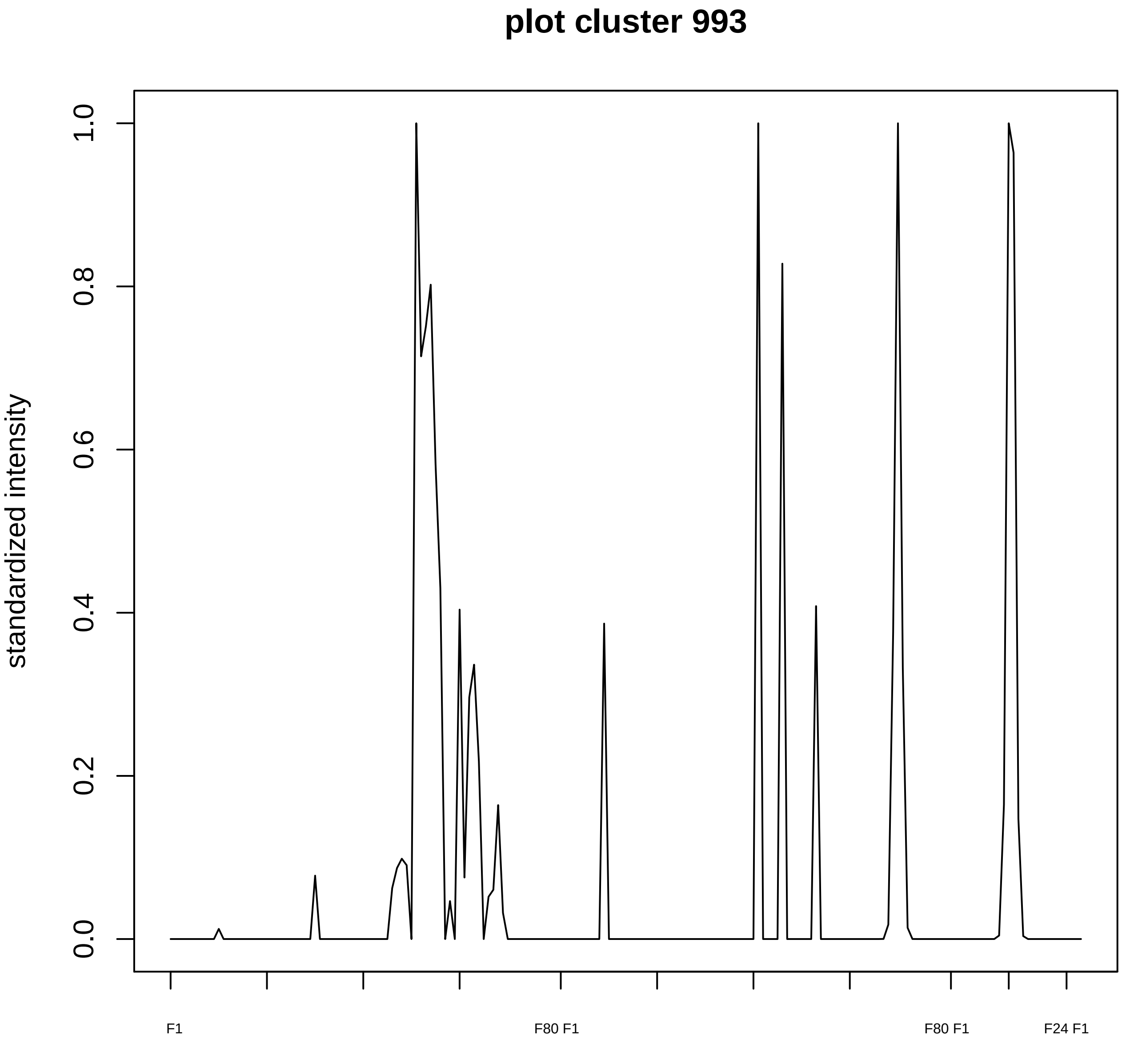
plot cluster 993
standardized intensity
0.0
0.2
0.4
0.6
0.8
1.0
F1
F80 F1
F80 F1
F24 F1

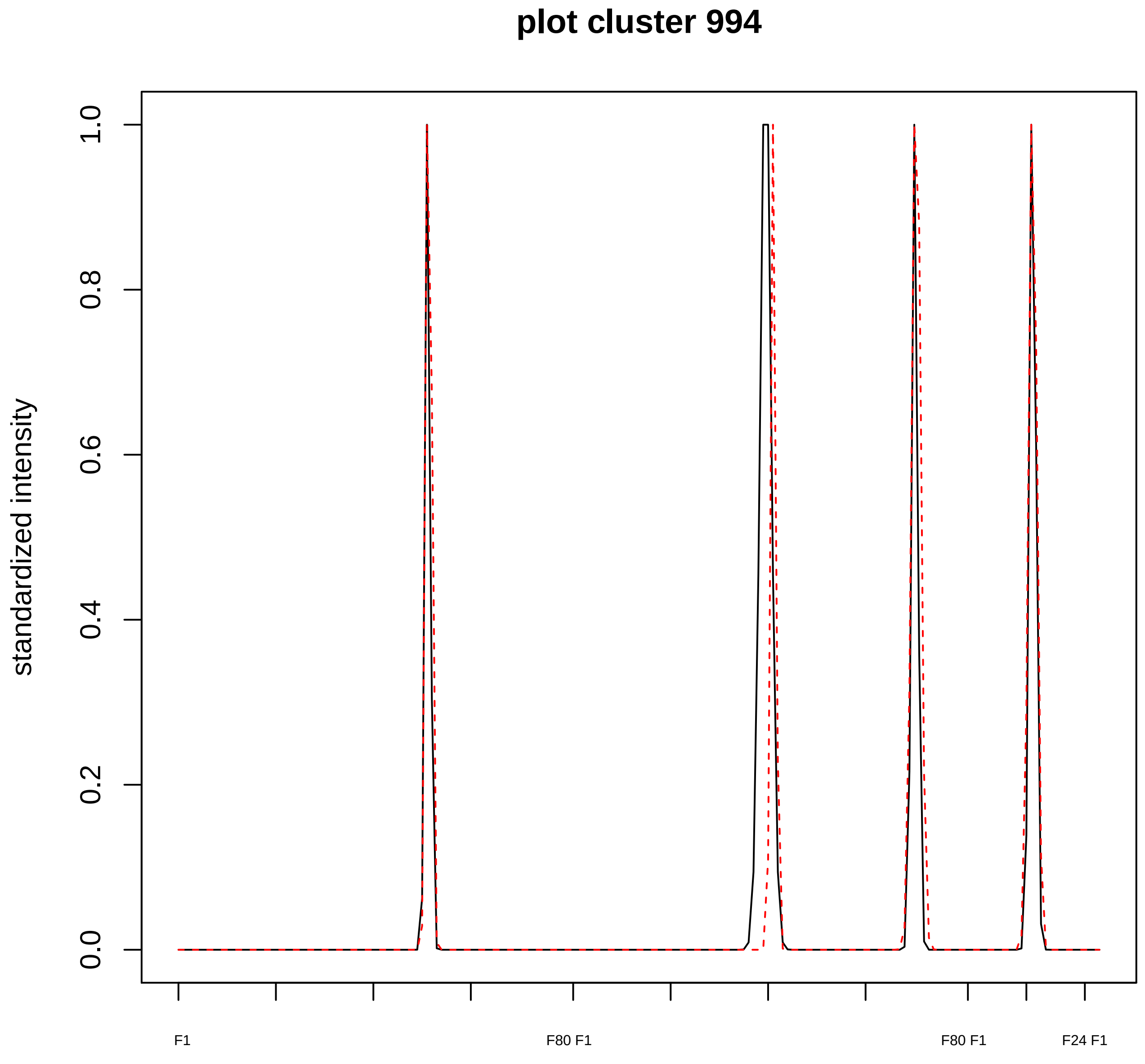

plot cluster 994
standardized intensity
0.0
0.2
0.4
0.6
0.8
1.0
F1
F80 F1
F80 F1
F24 F1

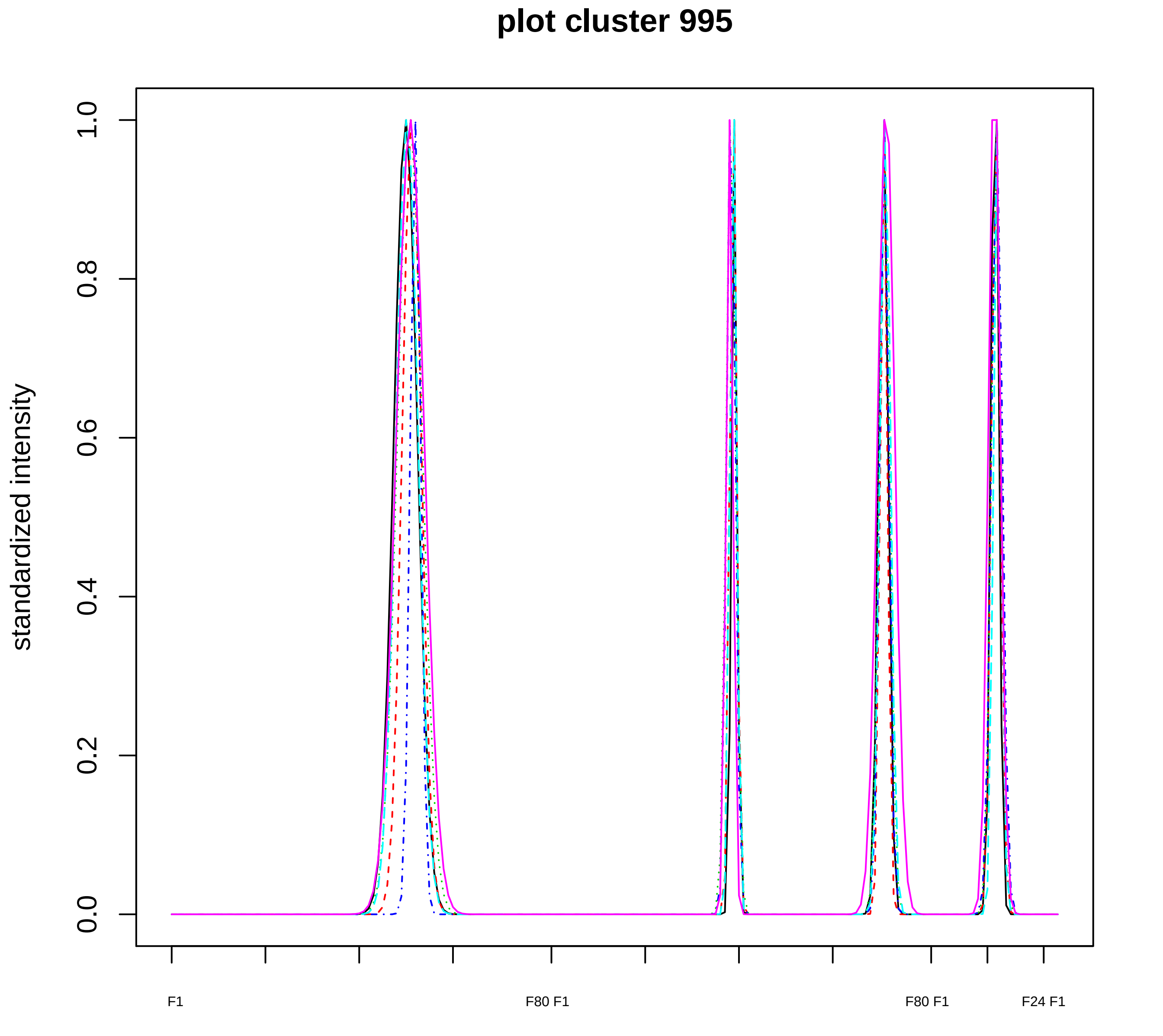

plot cluster 995
standardized intensity
1.0
0.8
0.6
0.4
0.2
0.0
F1
F80 F1
F80 F1
F24 F1

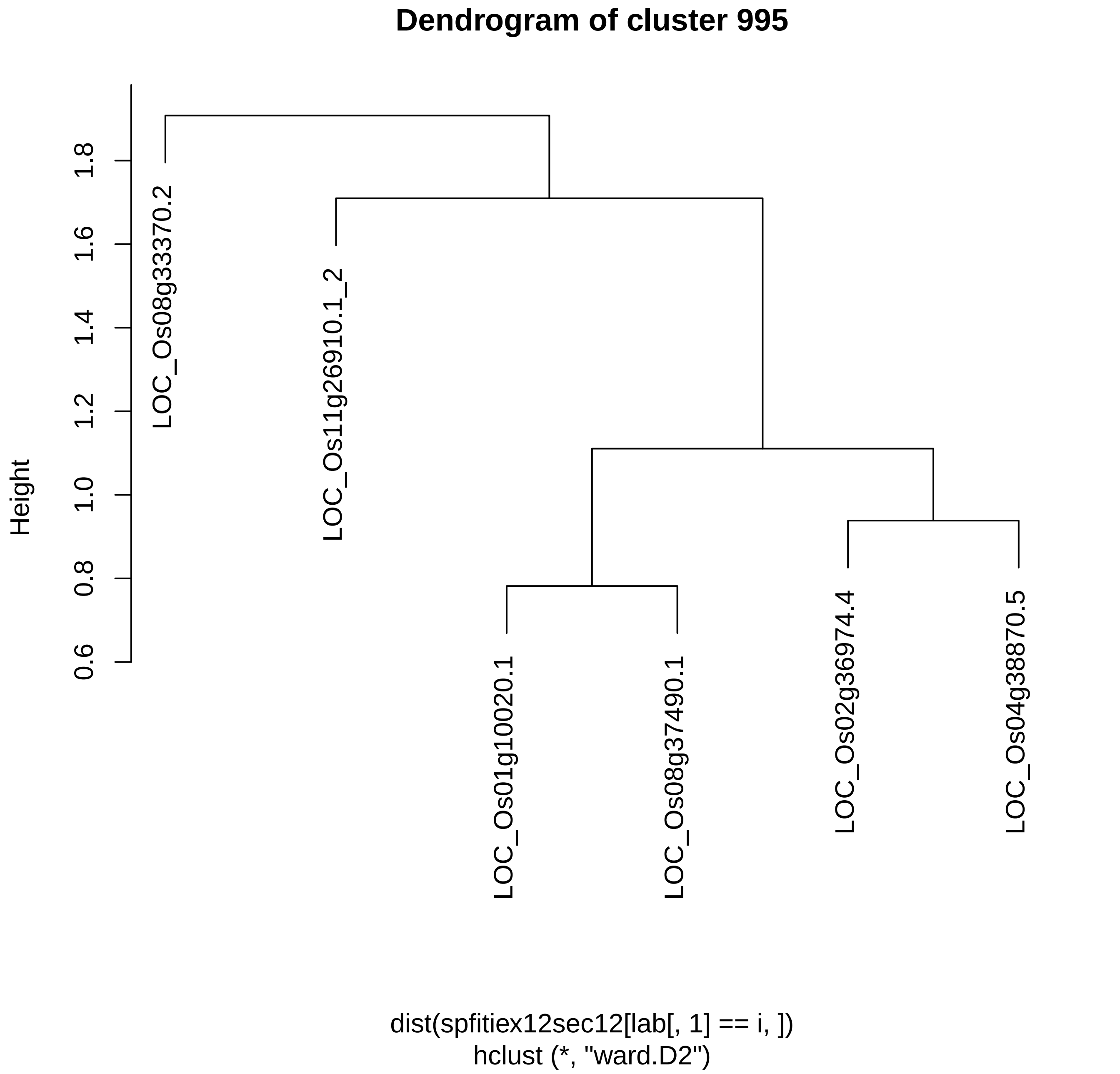

Dendrogram of cluster 995
Height
0.6
0.8
1.0
1.2
1.4
1.6
1.8
LOC_Os08g33370.2
LOC_Os11g26910.1_2
LOC_Os01g10020.1
LOC_Os08g37490.1
LOC_Os02g36974.4
LOC_Os04g38870.5
dist(spfitiex12sec12[lab[, 1] == i, ])
hclust (*, "ward.D2")

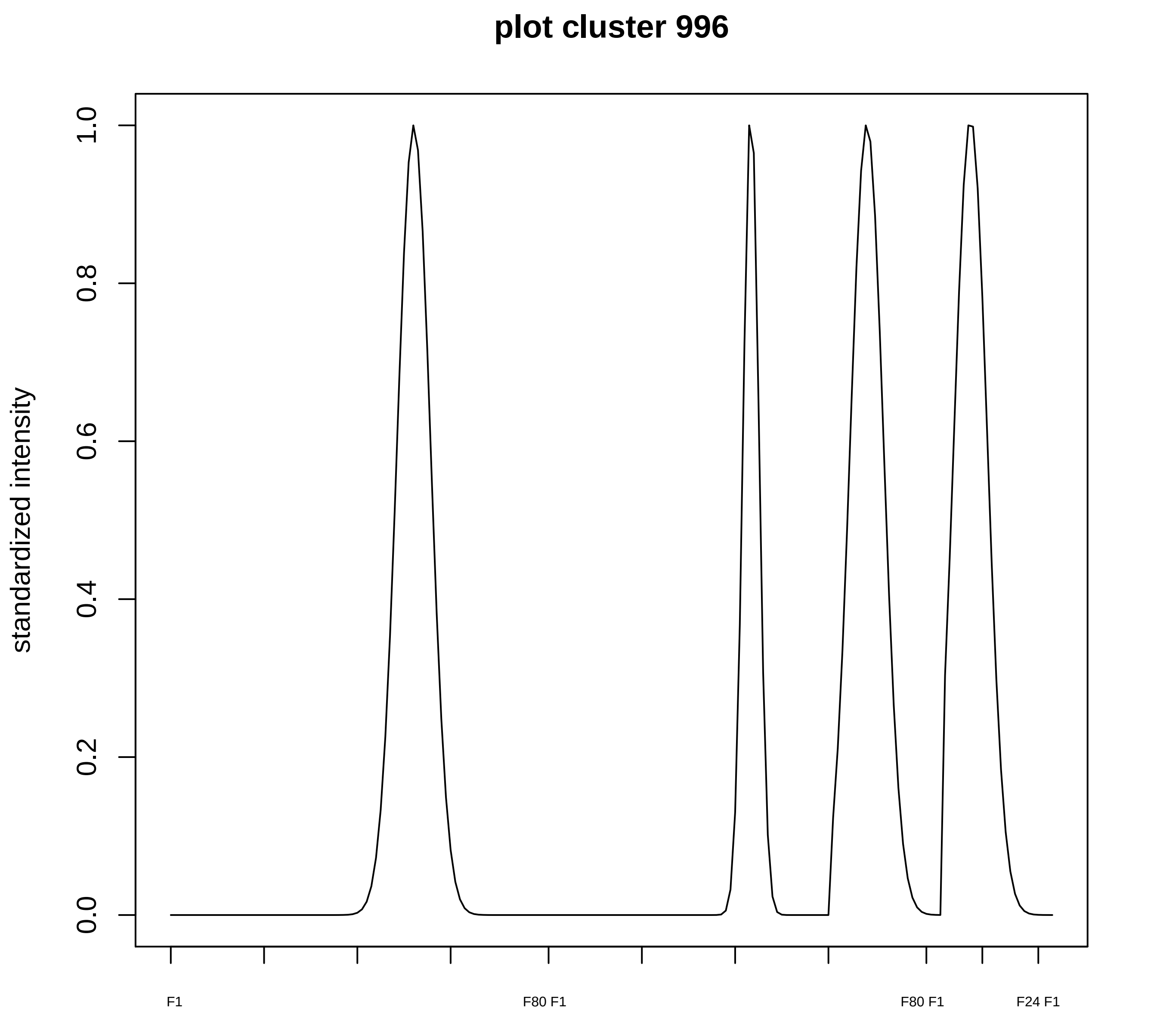

plot cluster 996
standardized intensity
0.0
0.2
0.4
0.6
0.8
1.0
F1
F80 F1
F80 F1
F24 F1

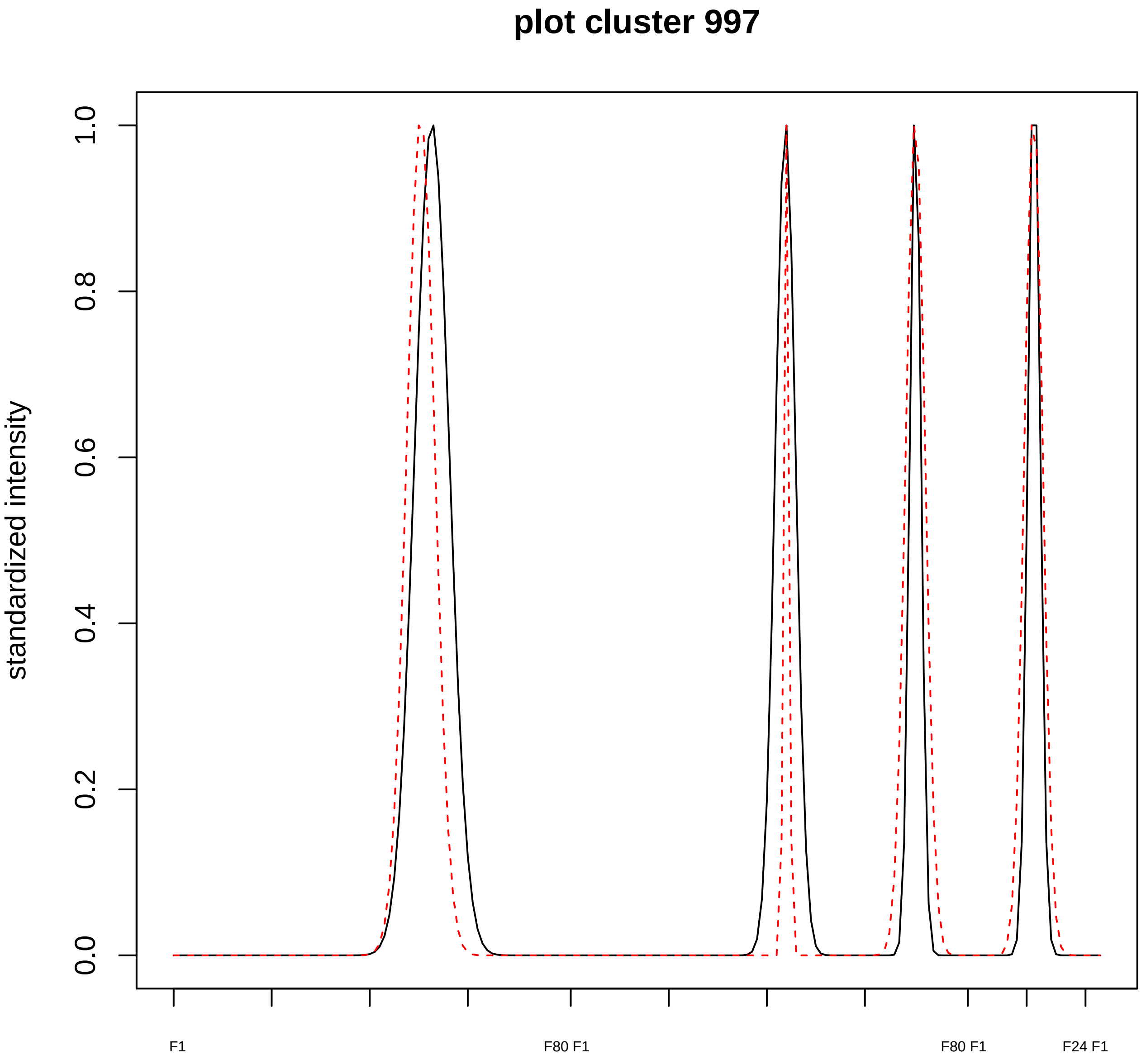

plot cluster 997
standardized intensity
1.0
0.8
0.6
0.4
0.2
0.0
F1
F80 F1
F80 F1
F24 F1

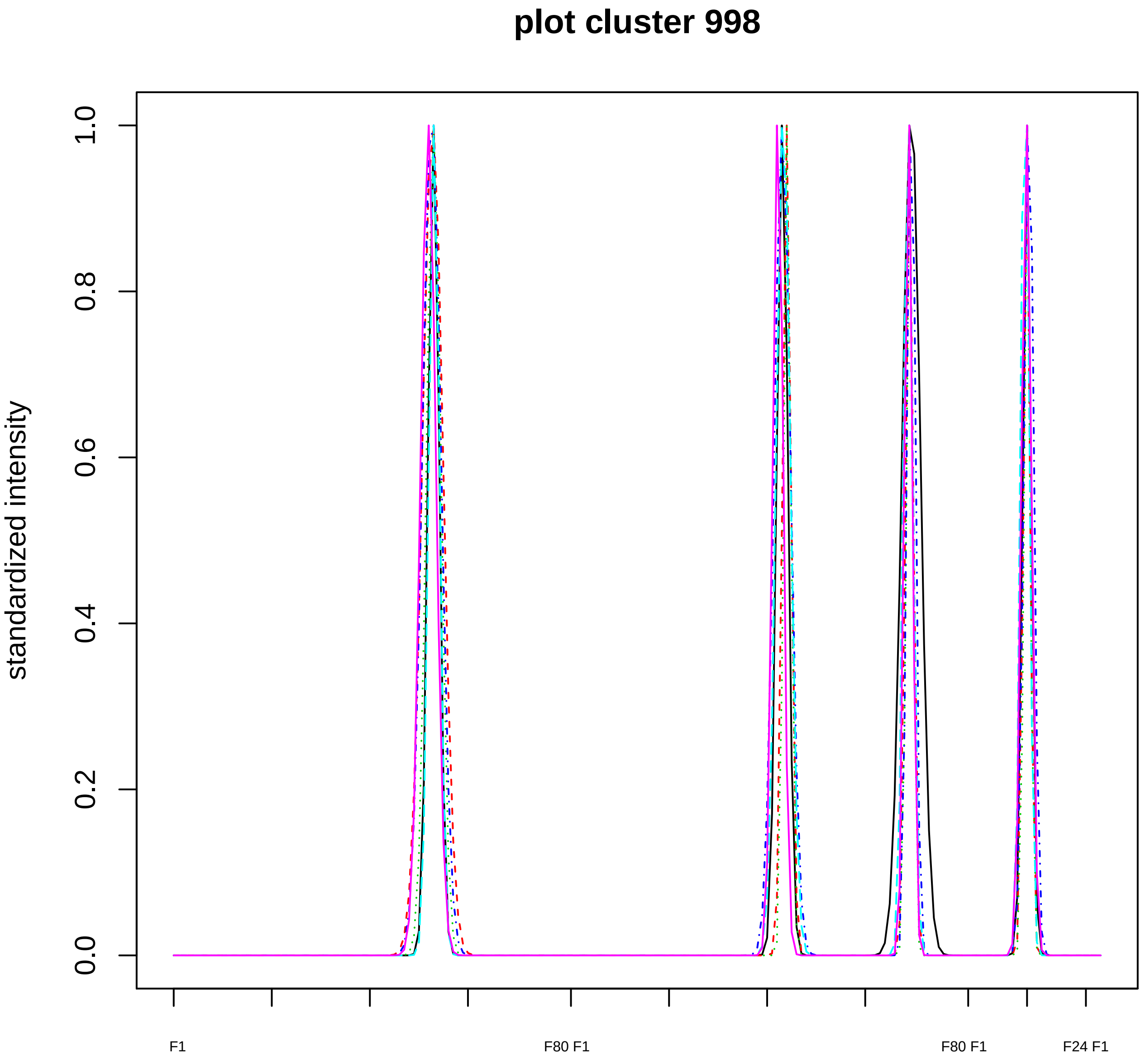

plot cluster 998
standardized intensity
0.0
0.2
0.4
0.6
0.8
1.0
F1
F80 F1
F80 F1
F24 F1

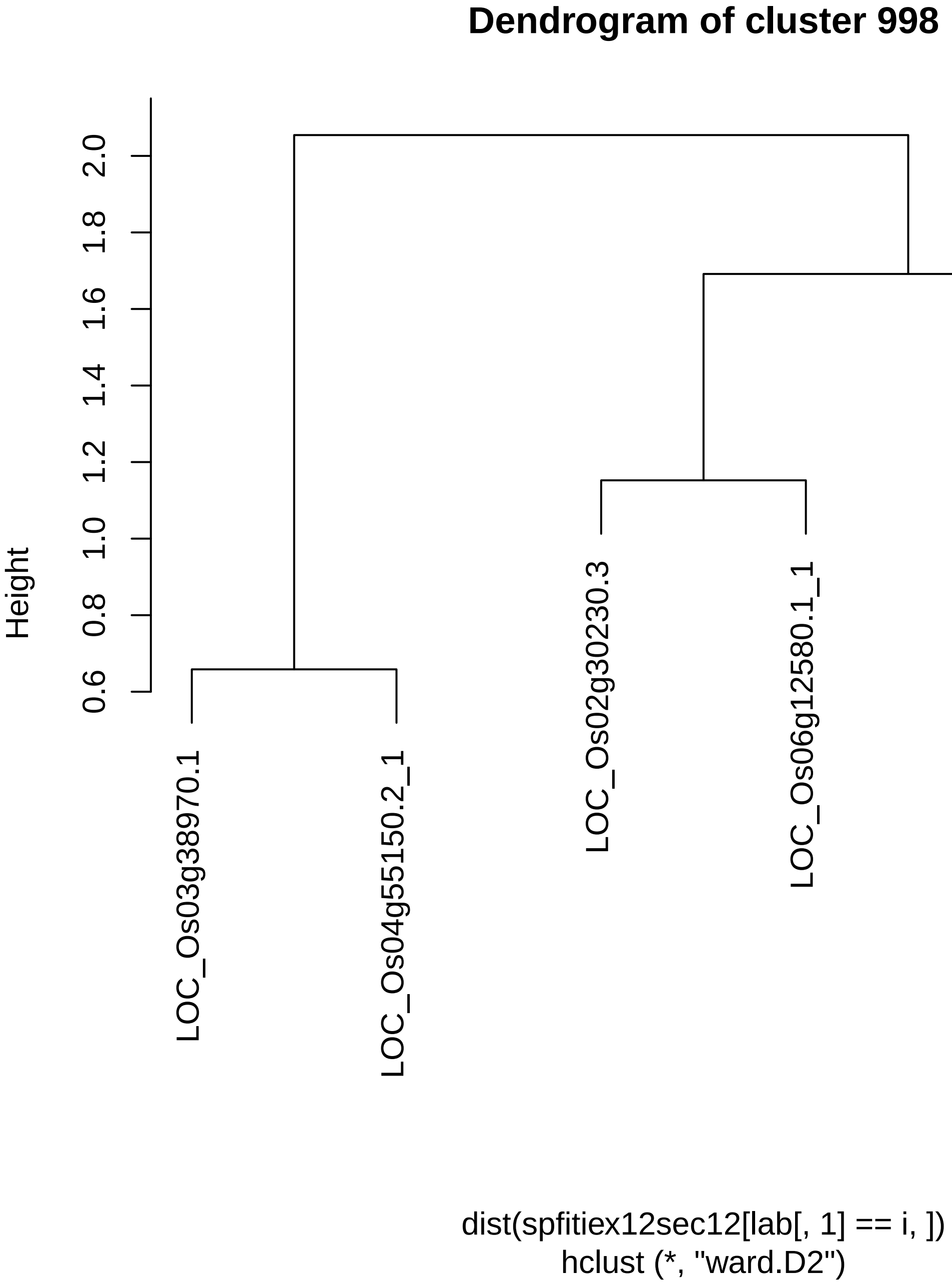
Dendrogram of cluster 998
Height
0.6
0.8
1.0
1.2
1.4
1.6
1.8
2.0
LOC_Os03g38970.1
LOC_Os04g55150.2_1
LOC_Os02g30230.3
LOC_Os06g12580.1_1
LOC_Os05g50970.1_2
LOC_Os07g39780.2_2
dist(spfitiex12sec12[lab[, 1] == i, ])
hclust (*, "ward.D2")

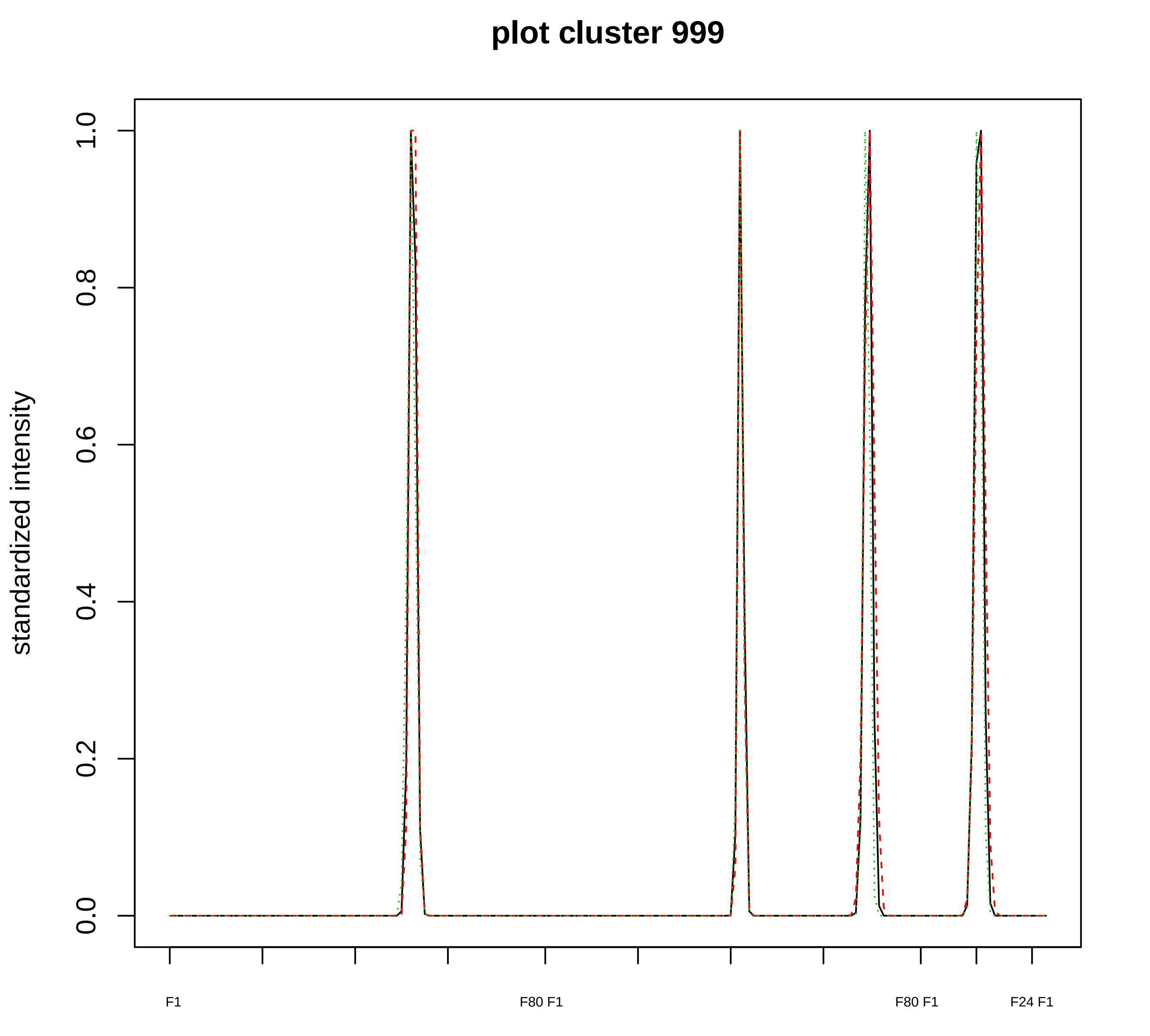

plot cluster 999
standardized intensity
0.0
0.2
0.4
0.6
0.8
1.0
F1
F80 F1
F80 F1
F24 F1

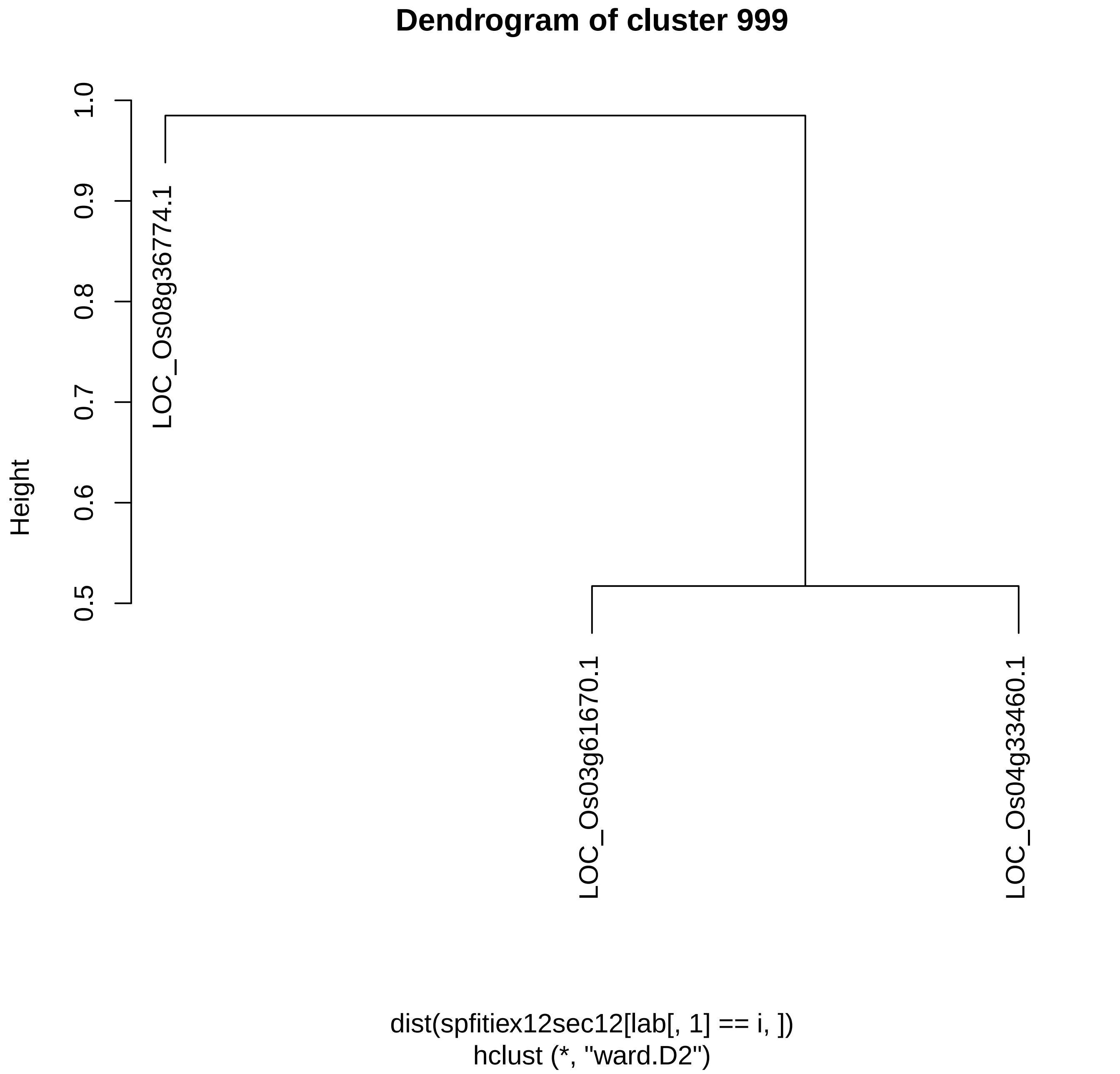
Dendrogram of cluster 999
Height
1.0
0.9
0.8
0.7
0.6
0.5
LOC_Os08g36774.1
LOC_Os03g61670.1
LOC_Os04g33460.1
dist(spfitiex12sec12[lab[, 1] == i, ])
hclust (*, "ward.D2")

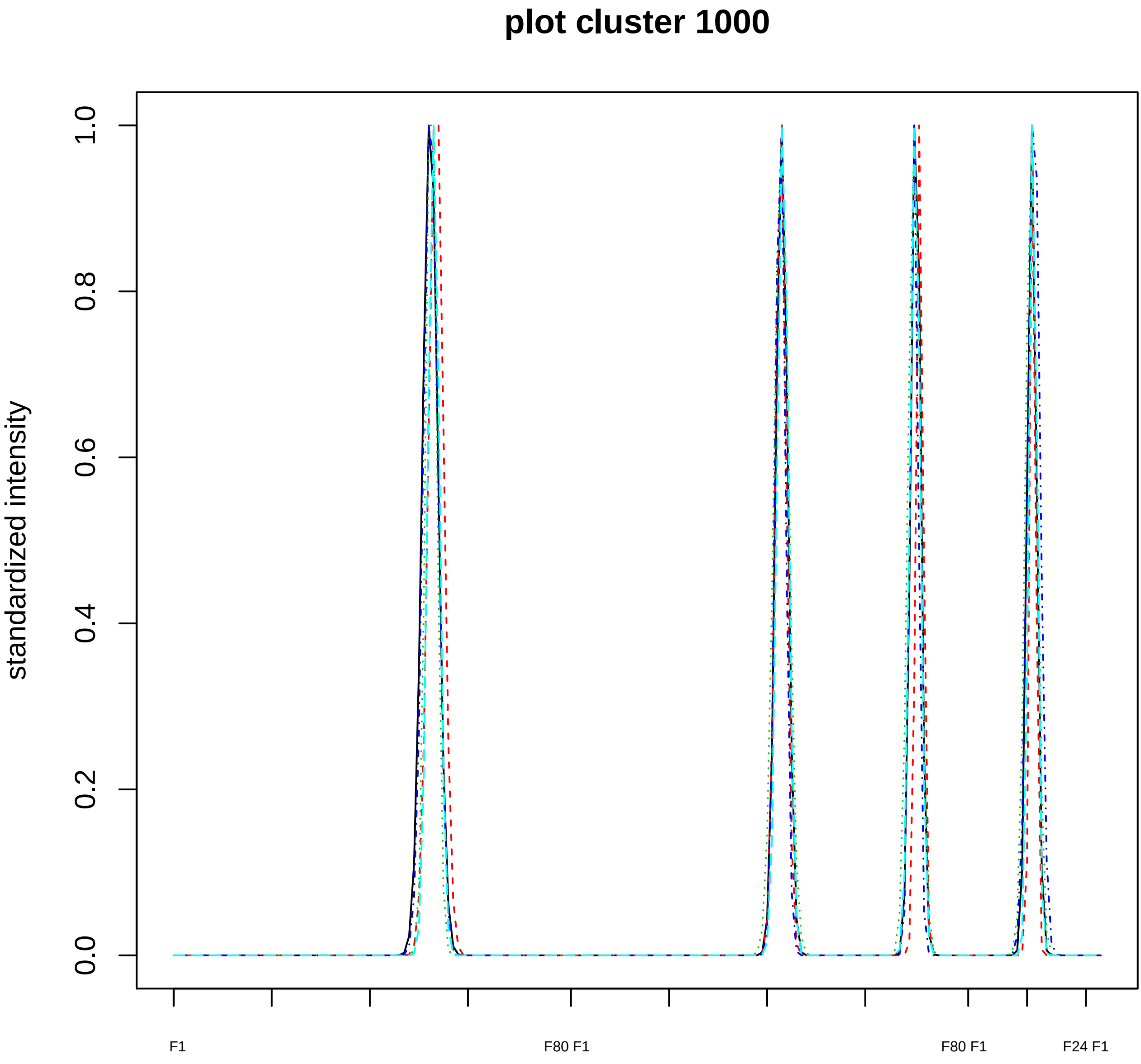
plot cluster 1000
standardized intensity
1.0
0.8
0.6
0.4
0.2
0.0
F1
F80 F1
F80 F1
F24 F1

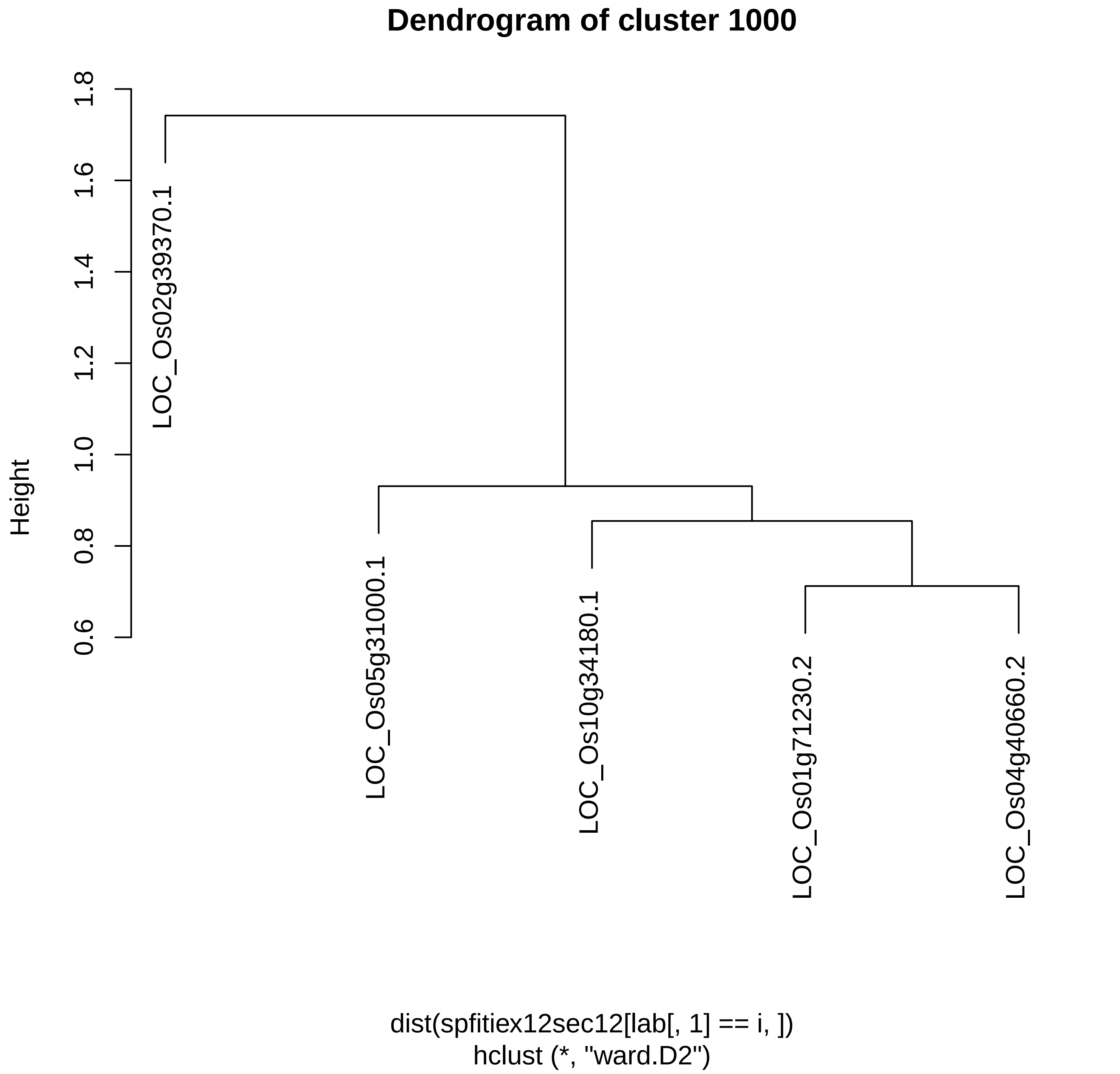

Dendrogram of cluster 1000
Height
1.8
1.6
1.4
1.2
1.0
0.8
0.6
LOC_Os02g39370.1
LOC_Os05g31000.1
LOC_Os10g34180.1
LOC_Os01g71230.2
LOC_Os04g40660.2
dist(spfitiex12sec12[lab[, 1] == i, ])
hclust (*, "ward.D2")
