## Supplemental Data Set 2 for "A Co-Fractionation Mass Spectrometry-based Prediction of Protein Complex Assemblies in the Developing Rice Aleurone-subaleurone"

### Supplemental Data Set 2. Supplemental texts for discussions on the predicted novel and known protein complexes

#### Supplemental text 1. Discussion on two known protein complexes: CCT and 19S proteasome complexes. This text supports Figure 3, D–E.

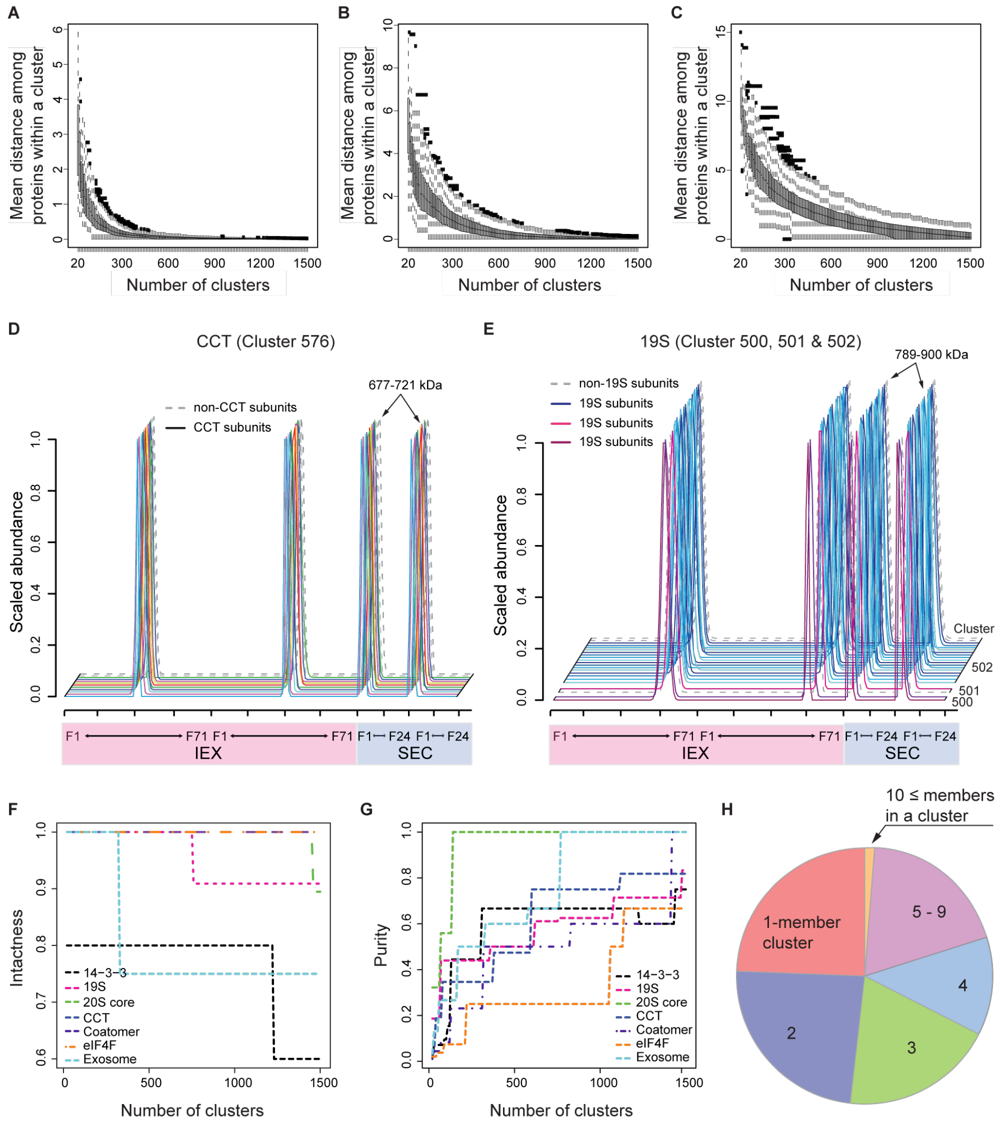


**Figure 3. The strong resolving power of the clustering analysis evaluated by the intrinsic and extrinsic tests.**
**A to C,** Intrinsic tests evaluated the resolving power for independent and concatenated protein profile datasets. Box-and-whisker plots visualize the distributions of average distances of elution profiles within the clusters in the SEC only (A), IEX only (B), and combined SEC and IEX (C) datasets as a function of ascending cluster number. **D and E,** The concatenated profiles of known protein complexes are visualized over the two IEX (pink) and two SEC (light blue) separations. Solid lines indicate elution profiles of CCT (D) and 19S proteasome complex (E) subunits. Dashed lines represent elution

profiles of proteins that are not subunits of the known complexes. Numbers shown above the SEC peaks show $M_{app}$ values of subunits in a given complex. Numbers on the right-handed side of profiles are cluster numbers, when multiple clusters are plotted. **F and G,** Extrinsic tests evaluated the resolving power for the independent and combined protein profile datasets as a function of ascending cluster number. (F) The intactness of known complexes was measured. (G) The purities of the known complexes were evaluated. Individual intactness and purity plots are provided in Supplemental Figure S3. **H,** The distribution of cluster size in the resulting clustering. The pie chart shows the distribution of predicted protein complexes by number of members in each cluster.

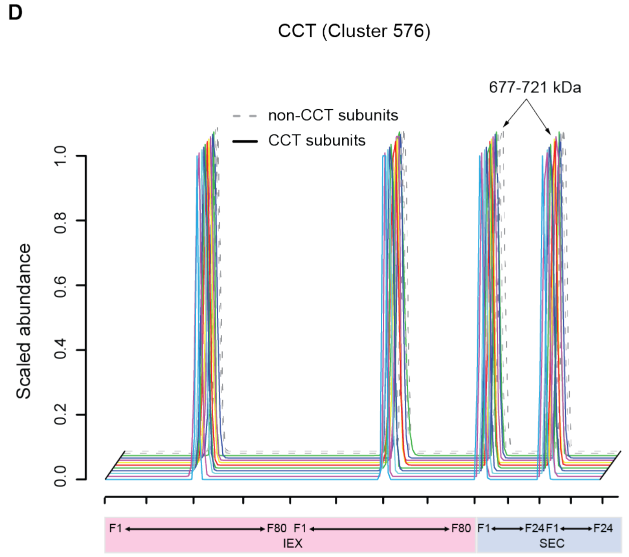


**Figure 3D. Elution profiles of CCT subunits.** The concatenated profiles of known protein complexes are visualized over the two IEX (pink) and two SEC (light blue) separations. Solid lines indicate elution profiles of CCT subunits. Dashed lines represent elution profiles of proteins that are not subunits of the known complexes. Numbers shown above the SEC peaks show $M_{app}$ values of subunits in a given complex. Numbers on the right-handed side of profiles are cluster numbers, when multiple clusters are plotted.

The chaperonin containing TCP1 folding complex (CCT) is a eukaryotic cytosolic chaperonin complex (Yébenes et al., 2011), which consists of two rings, each comprised of eight distinct subunits (Liou and Willison, 1997; Horwich et al., 2007). 12 known CCT subunits were assigned to the cluster 576 with $M_{app}$ of 677-721 kDa corresponding to the fully assembled complex (Figure 3D). The two additional members were present in this cluster. A vacuolar protein sorting-associated protein 52 (VPS52; LOC_Os03g30460.1) that had the most distinct elution profiles compared to the other cluster members. The VPS52 forms a tetrameric Golgi-associated retrograde protein (GARP) complex (Conibear et al., 2002). The other three subunits of the GARP complex, VPS51, VPS53, and VPS54, were found in the neighboring cluster 575, suggesting that VPS52 is a false positive in cluster 576. The other unexpected member of this cluster was a prefoldin subunit (LOC_Os12g17310.1_2).

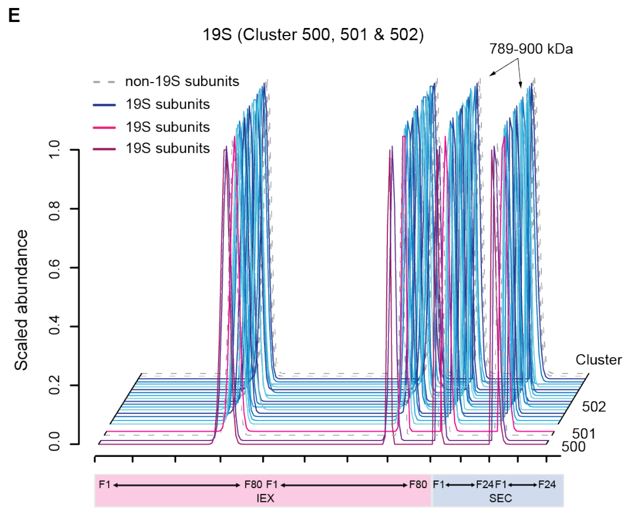


**Figure 3E. Elution profiles of 19S proteasome complex subunits.** The concatenated profiles of known protein complexes are visualized over the two IEX (pink) and two SEC (light blue) separations. Solid lines indicate elution profiles of 19S proteasome complex subunits. Dashed lines represent elution profiles of proteins that are not subunits of the known complexes. Numbers shown above the SEC peaks show $M_{app}$ values of subunits in a given complex. Numbers on the right-handed side of profiles are cluster numbers, when multiple clusters are plotted.

Besides, the 19S regulator, known as PA700, assembles 26S proteasome with the 20S proteolytic core. Seventeen subunits of the 19S form the 19S complex, functioning in regulation and localization of 26S proteasome complex (Ferrell et al., 2000). Six of the 17 subunits consist of the 19S regulator base, while 11 out of the 14 members associates into the 19S regulator lid. The seventeen different 19S subunits were also clustered into the cluster 502 (14 19S subunits / 16 cluster members) with $M_{app}$ of 789-900 kDa and neighboring clusters 500 (2 / 2) and 501 (1 / 2) (Figure 3E). Other subunits in the cluster 500 and 501 also had the similar $M_{app}$ and IEX fractions to those in the cluster 502, but a slight shift in the IEX fractions assigned these three subunits into the adjacent clusters. Nearby clusters for finding putative interactors in this clustering were searched (McBride et al., 2019).

**Supplemental text 2. Discussion on predicted homomers: One known homomer and three novel homomers are discussed. This text supports Figure 4, D, E, and G.**

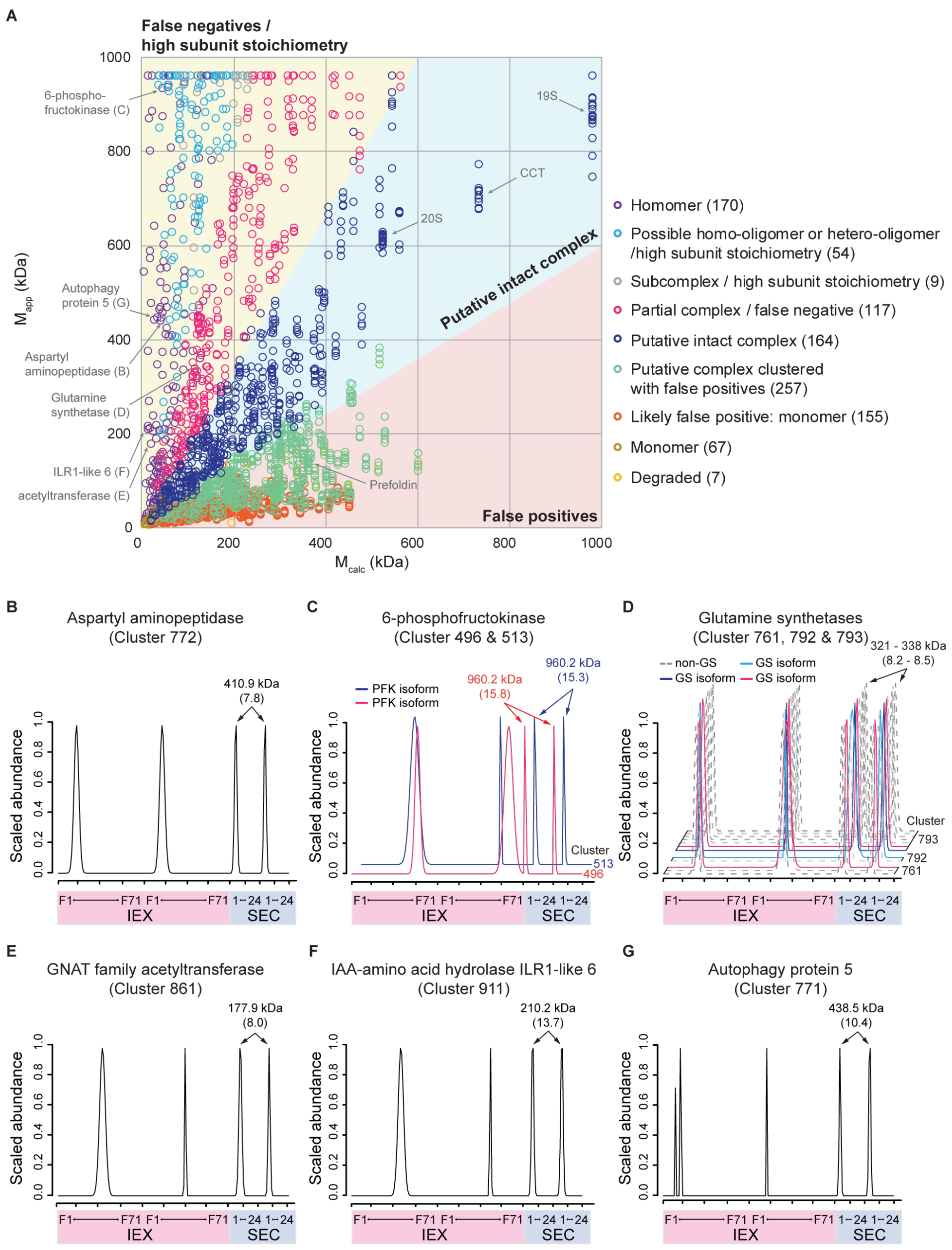


**Figure 4. The systematic classification of the resulting clustering analysis assessed the stoichiometry of subunits in the predicted protein complexes.** **A,** The clustering analysis was evaluated by comparing $M_{calc}$ and $M_{app}$. The most reliable predictions are present in the light blue area where there are less than 2-fold size differences between $M_{calc}$ and $M_{app}$. Another reliable predictions are on the light yellow area where $M_{app}$ is 2-fold greater than $M_{calc}$ including “false negatives” and “proteins with high subunit stoichiometry”. Proteins present in “putative complexes with false positives” in the clustering were saved from the false positives in the light pink

region where $M_{calc}$ is 2-fold greater than $M_{app}$. Number in parenthesis next to each category indicates number of clusters that belong to the category. **B to G,** Many of self-interacting homomers were assigned into single entry clusters and classified as homomers. B to D, The profiles of known homomers: Aspartyl aminopeptidase (B), 6-phosphofructokinase isoforms (C), and glutamine synthetase isoforms (D). Dashed lines represent elution profiles of proteins that are not subunits of the known complexes. E to G, The profiles of novel homomers: GNAT family acetyltransferase (E), IAA-amino acid hydrolase ILR1-like 6 (F), and ATG5 (G). Profiles of homomers are visualized over the combined two IEX (pink) and two SEC (light blue) separations. Numbers shown above the SEC peaks show $M_{app}$ values of homomers. Numbers in parenthesis indicate $R_{app}$ values of homomers. Numbers on the right-handed side of profiles are cluster numbers, when multiple clusters are plotted.

Lots of known homomers were identified from the single-entry clusters (Figure 4B to 4D and Supplemental Table 3A). The stoichiometries in the biological assembly of aspartyl aminopeptidase (LOC_Os12g13390.1) as homooctameric, 6-phosphofructokinases (LOC_Os05g44922.1 and LOC_Os06g05860.1) as homohexadecameric, and 3-dehydroquinate synthase (LOC_Os03g07420.1) as homododecameric, were annotated as homomers in our systematic classification result.

D

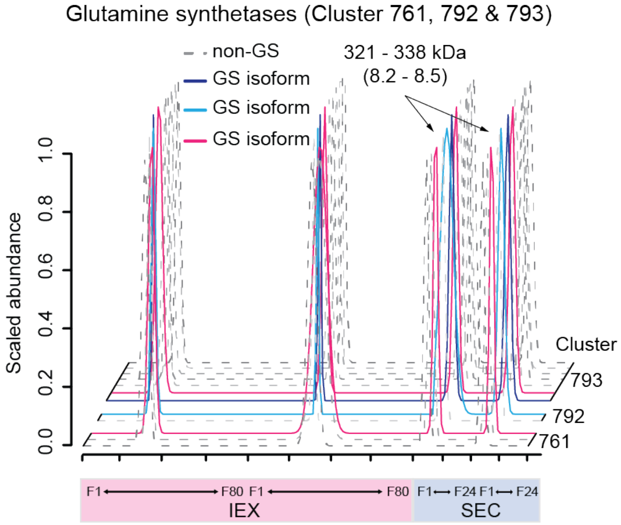


**Figure 4D. The profiles of known homomers - Glutamine synthetase isoforms.** Profiles of homomers are visualized over the combined two IEX (pink) and two SEC (light blue) separations. Numbers shown above the SEC peaks show $M_{\mathrm{app}}$ values of homomers. Numbers in parenthesis indicate $R_{\mathrm{app}}$ values of homomers. Numbers on the right-handed side of profiles are cluster numbers, when multiple clusters are plotted.

Glutamine synthetase (GS) is a critical enzyme that initiates ammonia fixation. Varied numbers (6 to 12) of monomers self-interact to form functional GS homomers (Almassy et al., 1986). There were three GS isoforms (LOC_Os02g50240.1, LOC_Os03g12290.1, and LOC_Os03g50490.1) identified in the clustering (Figure 4D and Supplemental Table S3A). LOC_Os03g12290.1 and LOC_Os03g50490.1 were assigned into the “putative intact complex” cluster 793 with several kinases, while LOC_Os02g50240.1 was clustered with a kinase into the “partial complex/false negative” cluster 792. Also, the second IEX peak of LOC_Os03g50490.1 was assigned with other two proteins into the “partial complex/false negative” cluster 761. Due to the high similarity of elution profiles, the clustering placed GS isoforms into the “partial complex/false negative” clusters which are neighboring.

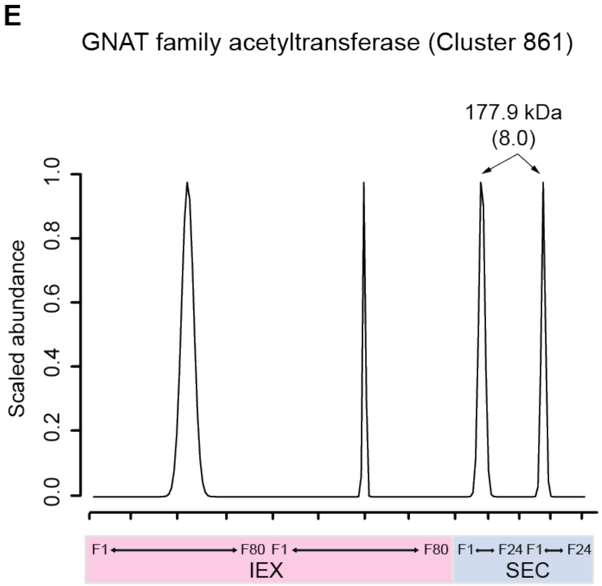


**Figure 4E. The profiles of novel homomers - GNAT family acetyltransferase.** Profiles of homomers are visualized over the combined two IEX (pink) and two SEC (light blue) separations. Numbers shown above the SEC peaks show $M_{app}$ values of homomers. Numbers in parenthesis indicate $R_{app}$ values of homomers.

An acetyltransferase (LOC_Os04g54330.3) was classified as a novel homomer with $R_{app}$ of 8.0 (Figure 4E). This enzyme belongs to a large superfamily of Gcn5-related N-acetyltransferases (GNAT) that transfer an acetyl group from acyl-CoAs. The dimer formation is common and is required for the active site formation for acetylation (Vetting et al., 2005). A tetrameric histone acetyltransferase complex was observed in *Saccharomyces cerevisiae* (Angus-Hill et al., 1999). Thus, the acetyltransferase showed an unexpected oligomeric state that differs from known homomeric states.

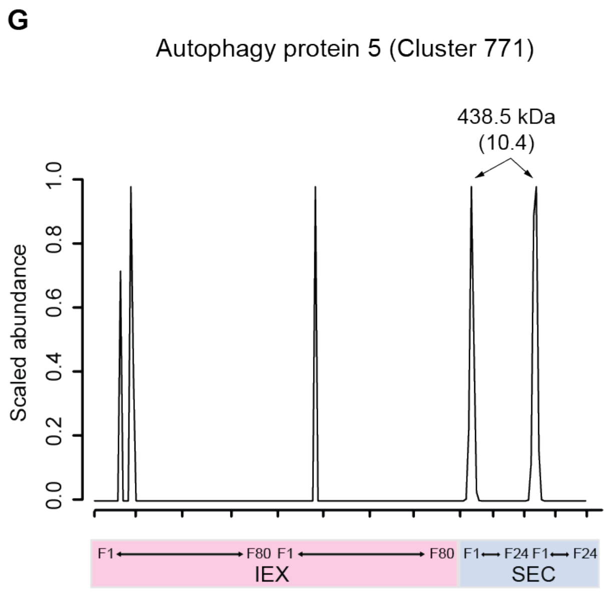


**Figure 4G. The profiles of novel homomers - ATG5.** Profiles of homomers are visualized over the combined two IEX (pink) and two SEC (light blue) separations. Numbers shown above the SEC peaks show $M_{app}$ values of homomers. Numbers in parenthesis indicate $R_{app}$ values of homomers.

Furthermore, autophagy protein (ATG) 5 (LOC_Os02g02570.1) was also classified as a homomer with $R_{app}$ of 10.4 in the systematic classification. Autophagy is a cell recycle process in which unnecessary or dysfunctional organelles are turned-over. This pathway is essential for maintaining cellular material homeostasis. ATGs were identified from yeast autophagy-defective mutants (Tsukada and Ohsumi, 1993), and 18 ATGs in yeast and 30 homologous ATGs in plants were characterized to be in the central autophagy machinery (Yoshimoto, 2012). Conjugation of human ATG5 and ATG12 associates with antiviral immune responses, degenerative neuro disease (Jounai et al., 2007), and developmental control (Kim et al., 2016). In plants, ATG5 has been shown to function in the control of plant lipid metabolism and catabolism via alleviating stress on ER where β-oxidation and synthesis of long chain fatty acids occur (Havé et al., 2019). Arabidopsis mutant *atg5* showed early browning and higher total protein concentrations at about 15 DAF, while earlier accumulation and low contents of 12S globulin (Di Berardino et al., 2018). Thus, these unexpected homomers could be essential for seed development.

**Supplemental text 3. Discussion on predicted heteromers: Two known and two novel heteromers are discussed. This text supports Figure 5, A–D.**

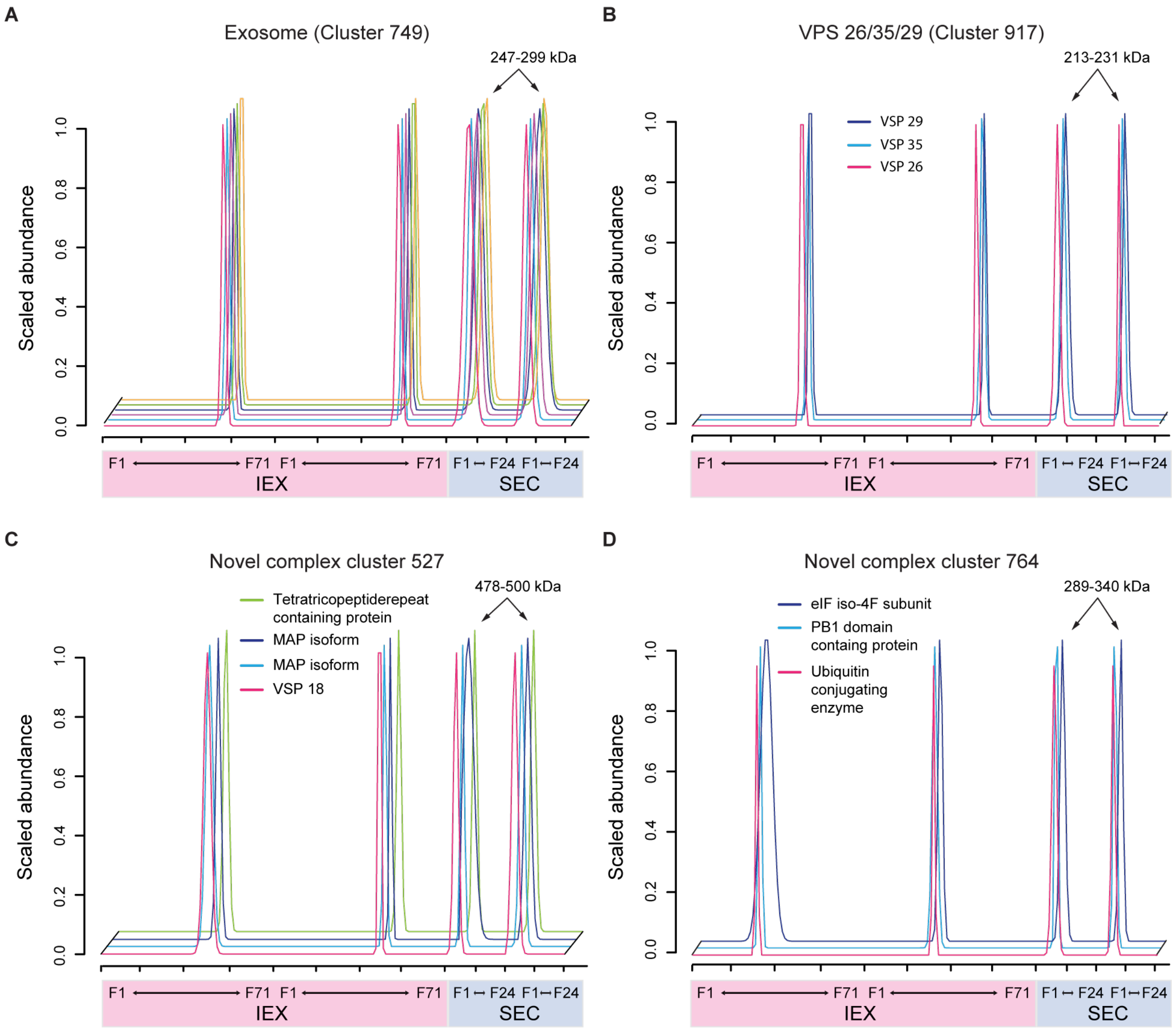


**Figure 5. The elution profiling-based clustering result placed known or novel protein complex subunits into the “putative intact complex” clusters.**
The profiles of known and novel protein complexes are overlapped each other across two IEX (pink) and two SEC (light blue) separations. The clusters were classified as “putative intact complexes”. **A and B,** Known protein complexes. Elution profiles of five exosome subunits (A) and a heterotrimeric VPS complex (B) are visualized. **C and D,** Novel protein complexes. Elution profiles of a MAP containing heterotetrameric complex (C) and a heterotrimeric novel complex (D). Numbers shown above the SEC peaks show $M_{app}$ values of subunits in a given complex.

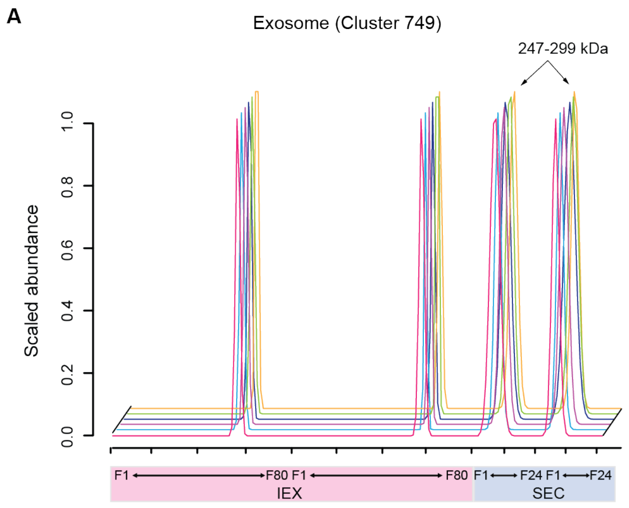


**Figure 5A. Known protein complex - Elution profiles of five exosome subunits.** The profiles of known protein complexes are overlapped each other across two IEX (pink) and two SEC (light blue) separations. The cluster was classified as “putative intact complex”. Numbers shown above the SEC peaks show $M_{app}$ values of subunits in a given complex.

Exosome complex is a conserved known protein complex that degrades various types of RNA molecules in eukaryote. The ring-shape core of exosome consists of six different proteins that possess RNase activity and three additional proteins form the cap structure (van Hoof and Parker, 1999). Six out of the nine exosome complex subunits coeluted with distinct elution patterns and were clustered into the cluster 749 (Figure 5A). $M_{calc}$ of this cluster was about 173 kDa and present within 40 % of their $M_{app}$, classifying the subunits into the “putative intact complex” (Supplemental Table S3A).

B

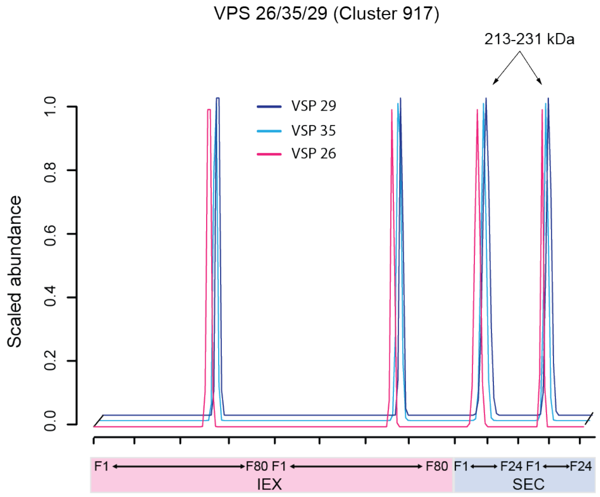


**Figure 5B. Known protein complex - Elution profiles of a heterotrimeric VPS complex.** The profiles of known protein complexes are overlapped each other across two IEX (pink) and two SEC (light blue) separations. The cluster was classified as "putative intact complex". Numbers shown above the SEC peaks show $M_{app}$ values of subunits in a given complex.

VPS35, VPS29, and VPS26 form a trimeric complex, a cargo recognition core complex that assembles into retromer complex with membrane-associated sorting nexin dimer subcomplex. The formation of retromer is required for retrograde transport from endosome to Golgi apparatus (Seaman et al., 1998). In the clustering analysis, the VPS 26/35/29 trimeric complex was assigned into cluster 917 with $M_{app}$ of 213-231 kDa (Figure 5B). $M_{calc}$ of the complex was present within 40 % of $M_{app}$ of the VPS cluster (Supplemental Table S3A). Thus the heteromeric complex was classified into the "putative intact complex" category

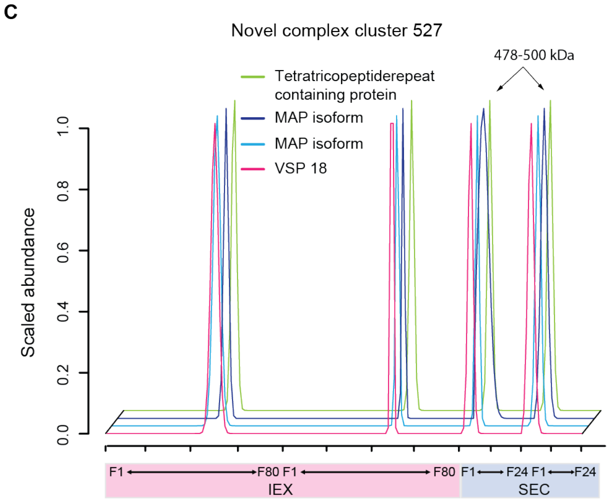


**Figure 5C. Novel protein complex - Elution profiles of a MAP containing heterotetrameric complex.** The profiles of novel protein complexes are overlapped each other across two IEX (pink) and two SEC (light blue) separations. The cluster was classified as “putative intact complex”. Numbers shown above the SEC peaks show $M_{\mathrm{app}}$ values of subunits in a given complex.

In cluster 527, VPS18 (LOC_Os08g08060.1), two isoforms of microtubule associated protein (MAP) (LOC_Os02g48830.1 and LOC_Os06g20370.1), and tetratricopeptide repeat (TPR)-containing protein (LOC_Os02g48620.2) were clustered with $M_{\mathrm{app}}$ of 478-500 kDa and $M_{\mathrm{calc}}$ of 397 kDa, classifying as a putative intact complex (Figure 5C). The VPS18 is a subunit of the shared core with VPS16, VPS11, and VPS33 in the class C core vacuole/endosome tethering complex (Peplowska et al., 2007) and/or homotypic fusion and protein sorting complex (Wurmser et al., 2000). Early endosome fusion and late endosome-lysosome fusion are regulated by the tethering complexes, respectively. MAPs belong to a family of plant microtubule-bundling proteins that control microtubule dynamics (Smertenko et al., 2004). Dimer formation is required to function in microtubule-bundling. The TPR-containing protein is an Arabidopsis homolog of *FRIENDLY*, a member of the CLUSTERED MITOCHONDRIA superfamily, that regulates mitochondrial fission and fusion on actin because mitochondrial clusters in a *friendly* mutant did not show tethering to microtubules (El Zawily et al., 2014). Damaged mitochondria could be purified via the formation of autophagosome in fibroblast cells (Twig et al., 2008). Autophagosomes are further delivered to lysosomes for hydrolysis (Shen and Mizushima, 2014). Presumably this novel complex might have a role in mediating autophagosome and endosome/lysosome fusion.

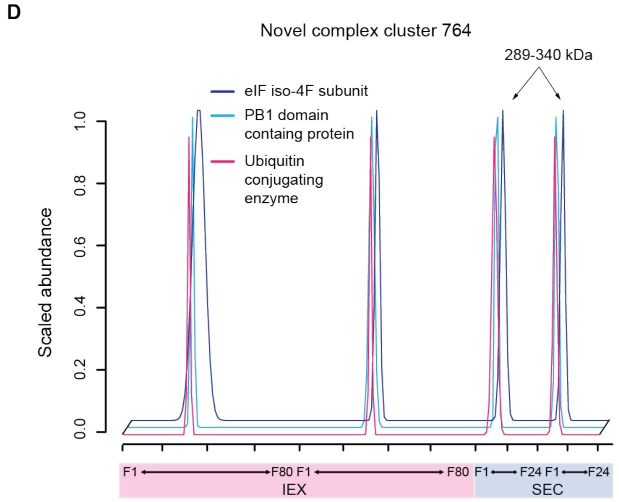


**Figure 5D. Novel protein complex - Elution profiles of a heterotrimeric VPS complex.** The profiles of novel protein complexes are overlapped each other across two IEX (pink) and two SEC (light blue) separations. The cluster was classified as "putative intact complex". Numbers shown above the SEC peaks show $M_{app}$ values of subunits in a given complex.

Another example of a putative intact complex predicted in the clustering is a novel complex cluster 764 with $M_{app}$ of 289-340 kDa that is similar to the cluster $M_{calc}$ of 256 kDa (Figure 5D). Eukaryotic initiation factor iso-4F subunit (eIFiso4G) (LOC_Os02g39840.1), PB1 domain containing protein (LOC_Os02g56480.1), and ubiquitin conjugating enzyme E2 34 (UBC 34) (LOC_Os01g03520.1_1) were assigned in cluster 764. The eIFiso4G is an isozyme of eIF4G that interacts with eIF4E to form an eIF4F cap-binding complex. The Arabidopsis double mutants showed abnormal phenotype, reduced growth, and decreased in weight and root growth in responses to dehydration and salinity, respectively (Lellis et al., 2010). In addition to the formation of the cap-binding complex, eIFiso4G can interact with Pokeweed antiviral protein to selectively depurinate uncapped viral RNA in plants (Wang and Hudak, 2006; Domashevskiy et al., 2017). UBCs transfer ubiquitin to E3 ligases that ubiquitinate substrate. The *OsUBC34* showed relatively higher expression levels during seed development than other developmental stages in other tissues. Expression of *OsUBC34* in 7-day-old seedlings was induced by ABA at 3-hours after treatment, while the expression level was continuous reduced after IAA, BA, and GA treatments (E et al., 2015). PB1 domain functions in high-order protein assembly (Ito et al., 2001; Noda et al., 2003). In plant domain III/IV on auxin response proteins Aux/IAAs were renamed to the PB1 domain (Mutte and Weijers, 2020). This domain is involved in auxin signaling (Hagen and Guilfoyle, 2002; Guilfoyle and Hagen, 2012) and selective autophagy (Svenning et al., 2011; Zientara-Rytter and Sirko, 2014). This novel protein complex might be involved in a mechanism of translational regulation via autophagy as a response to hormone signaling in the developing rice aleurone-subaleurone cells.

**Supplemental text 4. Discussion on predicted RPB-associated novel heteromeric complexes: Three novel RBP complexes are discussed. This text supports Figure 6, A, B, D and E.**

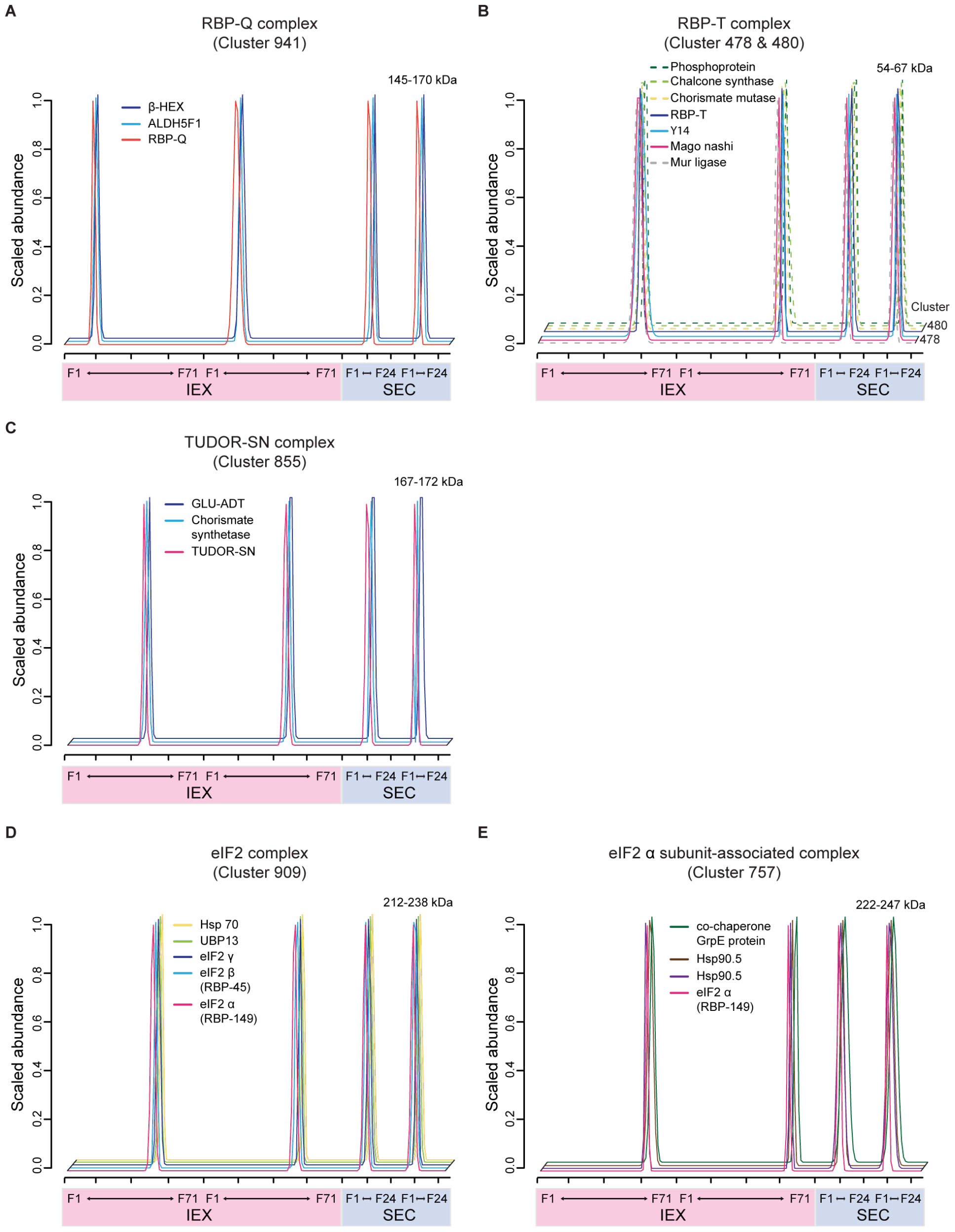


**Figure 6. Association of RBPs into novel protein complexes.**
The elution profiles of subunits in RBP-associated novel protein complexes: RBP-Q associated putative complex (A), RBP-T associated putative EJC (C), TUDOR-SN associated putative complex (C), eIF2 complex (D), and eIF2$\alpha$ associated complex (E). Dashed lines represent elution profiles of proteins that are not subunits of the known complexes. Numbers shown above the SEC peaks show $M_{app}$ values of subunits in a given complex. Numbers on the right-handed side of profiles are cluster numbers, when multiple clusters are plotted.

A

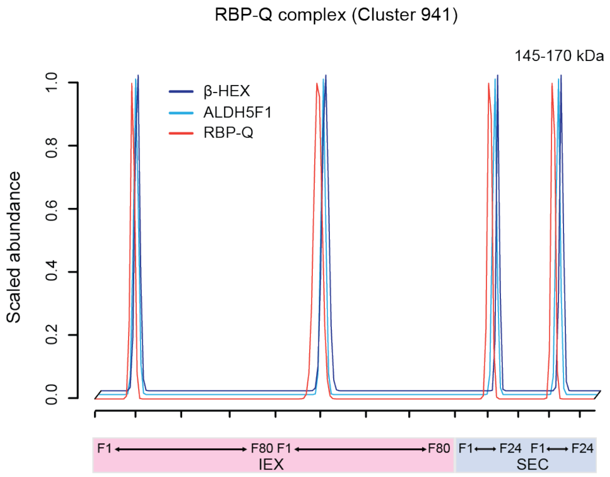


**Figure 6A. The elution profiles of subunits in RBP-Q associated putative complex.** Dashed lines represent elution profiles of proteins that are not subunits of the known complexes. Numbers shown above the SEC peaks show $M_{app}$ values of subunits in a given complex.

RBP-Q that recognizes the prolamine zipcode sequence involves in the transportation of prolamine mRNA to the PB-ER with the RBP A-J-K and I-J-K complexes. The RBP-Q assembles a third multiprotein complex with unknown other proteins (Yang et al., 2014). In our complex prediction, the RBP-Q (LOC_Os01g42820.3) was placed as a “putative intact complex” with two other proteins into cluster 941 (Figure 6A). RBP-Q is not detected by an immunoblotting method because it is blocked by its potential interactors in the third multiprotein complex when it consists of the prolamine mRNP complex in the nucleus. While its antibody detected the RBP-Q in the cytoplasm, the oligomerization state of the RBP-Q was not discovered (Yang et al., 2014). The $M_{app}$ and $R_{app}$ of RBP-Q were ~170 kDa and 3.8, respectively. The $M_{calc}$ of the cluster was 157 kDa similar to the $M_{app\text{-}avg}$ for the cluster. Our data suggest the presence of the RBP-Q associated multiprotein complex. The two subunits of this novel protein complex were *beta*-hexosaminidase (LOC_Os05g02510.1) and aldehyde dehydrogenase (LOC_Os02g07760.2). *Beta*-hexosaminidase hydrolyzes non-reducing N-acetyl-D-hexosamine terminal residues, and aldehyde dehydrogenase performs the oxidation of aldehydes. Considering the enzymatic functions of its binding partners, the RBP-Q associated novel protein complex might have a moonlighting function in regulation of enzymatic activity rather than RNA translocation.

B

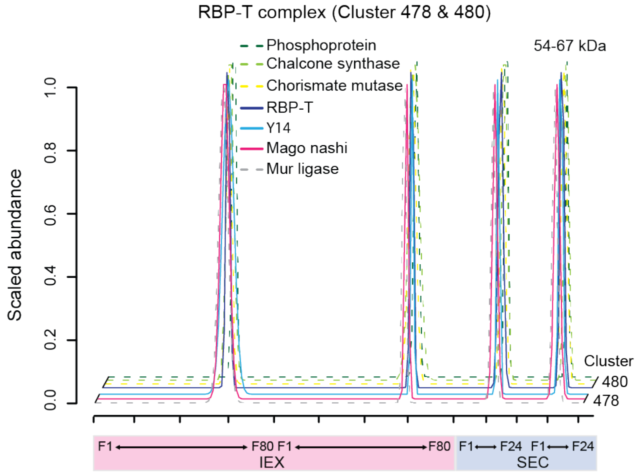


**Figure 6B. The elution profiles of subunits in RBP-T associated putative EJC.** Numbers shown above the SEC peaks show $M_{app}$ values of subunits in a given complex. Numbers on the right-handed side of profiles are cluster numbers, when multiple clusters are plotted.

Exon-junction complexes (EJC) are involved in mRNA decay, splicing, mRNA export, mRNA localization during expression of eukaryotic genes (Hir et al., 2001; Kim et al., 2001; Le Hir et al., 2001). The core EJC components are MAGO·Y14 heterodimer. Several peripheral proteins transiently associate into the EJC core during EJC assembly or subsequent mRNA metabolism (Tange et al., 2004). Most recently, highly conserved heterodimerization between paralogous MAGO-Y14 pairs was observed in rice (Gong and He, 2014). MAGO (LOC_Os12g18880.1) and Y14 (LOC_Os05g04850.2) were present in the cluster 478, and RBP-T (LOC_Os01g36920.2) was placed into the cluster 480 (Figure 6B). The RBP-T was annotated as a DEAD-box ATP-dependent RNA helicase, an ortholog of human DEAD-box helicase UAP56. The UAP56 participates in pre-mRNA splicing and was recently clarified to recruit REF/Aly to mRNAs for mRNA export (Gatfield et al., 2001; Luo et al., 2001). REF/Aly was not detected in our experiments. eIF4A can be recruited into the core EJC heterodimer for mRNA localization (Chan et al., 2004), and was present as a subunit of EJC heterotetramer in plants (McWhite et al., 2020). Two eIF4A paralogous proteins were identified as monomers and placed into the cluster 352 far from this EJC cluster. The $M_{app}$ and $R_{app}$ of RBP-T were ~65 kDa and 1.7, respectively. The $M_{calc}$ of the cluster was 74 kDa similar to the $M_{app\text{-}avg}$ of 61 kDa for the cluster. Presumably, the new heterotrimeric EJC (RBP-T·MAGO·Y14) might have a unique function in mRNA metabolism, rather mRNA splicing, exporting, and translocating.

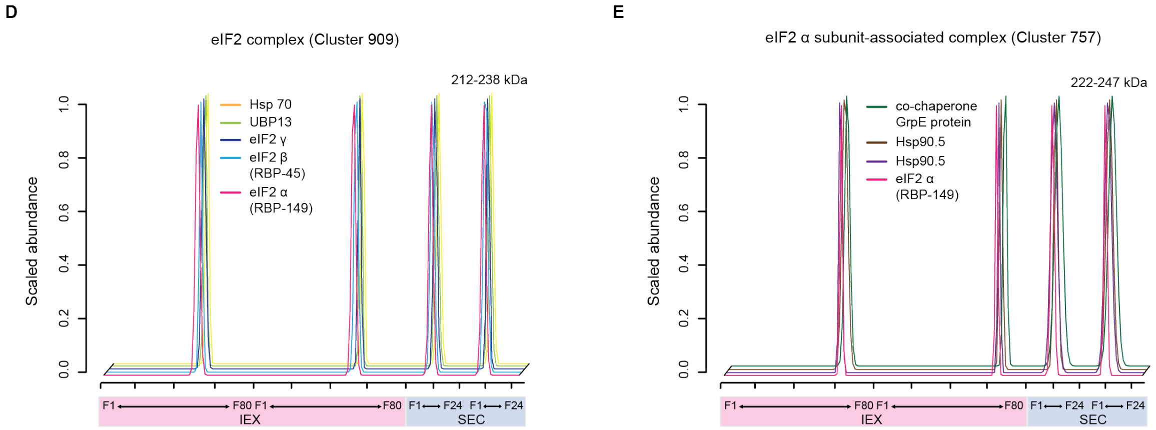


**Figure 6D and 6E. The elution profiles of subunits in eIF2 complex (D), and putative eIF2 α associated complex (E).** Dashed lines represent elution profiles of proteins that are not subunits of the known complexes. Numbers shown above the SEC peaks show $M_{\text{app}}$ values of subunits in a given complex.

We also detected the known translation initiation factor eIF2 complex where eIF2α (RBP-149), eIF2β (RBP-45) and eIF2γ are associated together at cluster 909 (Figure 6D). The heterotrimeric eIF2 binds to methionyl transfer RNA (Met-tRNAi$^{\text{Met}}$) and GTP to form the ternary complex. This ternary complex of eIF2·Met-tRNAi$^{\text{Met}}$·GTP serves to provide Met-tRNAi$^{\text{Met}}$ and GTP to the ribosome (Karen and Bailey-Serres, 2015). Interestingly, eIF2α (RBP-149) and eIF2β had two resolved peaks. Two heat shock proteins (LOC_Os08g38086.3 and LOC_Os09g29840.1) and co-chaperone GrpE protein (LOC_Os08g25090.2) were placed with the second peak of eIF2α (RBP-149) into the cluster 757 (Figure 6E). Recycling eIF2 ternary complex is required for initiating translation. In the animal system, eIF2B recycles eIF2·GDP to eIF2·GTP, but the presence of this recycle system is still questionable in plants (Karen and Bailey-Serres, 2015). Presumably, the interaction of eIF2α with the chaperone and co-chaperone proteins might be a step in eIF2 recycling. Protein complexes that function as translation machineries were annotated in Supplemental table S3A according to translational processes defined by Karen and Bailey-Serres (2015).
